# Cytokine priming of naïve CD8^+^ T lymphocytes modulates chromatin accessibility that partially overlaps with changes induced by antigen simulation

**DOI:** 10.1101/2020.08.11.246553

**Authors:** Akouavi Julite Quenum, Maryse Cloutier, Madanraj Appiya Santharam, Marian Mayhue, Sheela Ramanathan, Subburaj Ilangumaran

## Abstract

**Background:** Naïve CD8^+^ T lymphocytes undergo antigen non-specific proliferation following exposure to certain synergistic combination of inflammatory (IL-6, IL-21) and homeostatic (IL-7, IL-15) cytokines. Such cytokine-stimulated naïve CD8^+^ T cells display increased T cell antigen receptor (TCR) sensitivity, allowing them to respond to limiting concentrations of cognate antigenic peptides and altered peptide ligands of lower affinity towards the TCR. The purpose of this study is to gain insight into the molecular mechanisms of such ‘cytokine priming’.

**Methods:** Naïve CD8^+^ T lymphocytes expressing the PMEL-1 transgenic TCR were stimulated with IL-15 and IL-21, and chromatin accessibility was assessed using the assay for transposase-accessible chromatin (ATAC) sequencing. Cells stimulated by the cognate antigenic peptide mgp100_25-33_ were used as controls.

**Results:** Compared to naïve cells, cytokine-primed cells showed 212 opening and 484 closing peaks, whereas antigen-stimulated cells showed 12087 opening and 6982 closing peaks. However, a significant fraction of the opening (33%) and closing (63%) peaks of cytokine-primed cells overlapped with those of the antigenic stimulated cells. Chromatin accessibility peaks modulated in cytokine-primed cells were strongly represented in gene ontology pathways for T cell signaling, activation, regulation and effector functions. Many of the transcription factor binding motifs located close to the opening and closing peaks of cytokine-primed cells also occurred in antigen-stimulated cells.

**Conclusions:** Our data suggest that by modulating the gene expression programs involved in TCR signaling, cytokine priming induces a poised state that lowers the TCR signaling threshold in naïve CD8^+^ T cells and increases their antigen responsiveness.

## Background

CD8^+^ T lymphocytes confer immune protection against viral and bacterial pathogens, as well as prevent tumor development (1, 2). To carry out these functions, naïve CD8^+^ T cells must receive two essential signals delivered via the T cell antigen receptor (**TCR**) (signal 1) and the costimulatory receptors such as CD28 (signal 2) (3). Non-antigen (Ag) specific innate immune responses often precede Ag-specific adaptive immune responses mediated by T cells (4). Activation of innate immune cells via the pattern recognition receptors generates inflammatory cytokines, chemokines and other effector molecules (5-7). In addition to recruiting T cells and upregulating costimulatory ligands on antigen presenting cells (APC), the inflammatory mediators of the innate immune response may directly contribute to T cell activation and boost their effector functions. Molecular events that accompany the transition from innate to adaptive immune response continue to be an important area of investigation due to their relevance to vaccinology and autoimmunity (8-10).

Inflammatory cytokines such as type-I interferons (IFN-I), interleukin-12 (IL-12), IL-27 and IL-21 have been shown to provide a ‘third signal’ to activated CD8^+^ T cells, stimulating more efficient clonal expansion and effector functions (11-17). IFNγ produced by natural killer cells is also implicated in efficient expansion of antigen-stimulated CD8^+^ T cells, and autocrine IFNγ synergize with IFN-I to induce strong effector functions in Ag-stimulated CD8^+^ T cells (18-20). Distinct from the third signal, we and others have shown that inflammatory cytokines such as IL-6 and IL-21, produced during the innate immune response, can synergize with IL-7 or IL-15 that regulate T cell homeostasis to stimulate Ag non-specific proliferation of naïve CD8^+^ T cells independently of TCR and costimulatory signaling (21-23). IL-7 and IL-15 are also induced by innate immune stimulation (24, 25). Importantly, naïve CD8^+^ T cells pre-stimulated with IL-21 and IL-7 or IL-15 display increased sensitivity to Ag, proliferate strongly and exhibit potent Ag-specific cytolytic activity upon subsequent Ag stimulation (22). Moreover, cytokine-primed CD8^+^ T cells gain sensitivity toward altered peptide ligands of lower affinity than the cognate peptide (26). Indirect evidences suggest that such inflammatory cytokine driven increase in Ag responsiveness of CD8^+^ T cells also occurs *in vivo* (27, 28). Therefore, the ability of the inflammatory cytokines to boost Ag-specific responses of naïve CD8^+^ T cells, referred here as ‘cytokine priming’, can play a direct role in shaping the adaptive immune response mediated by CD8^+^ T cells, which may have implications for immunity, autoimmunity and antitumor immunity (29).

Molecular mechanisms underlying the heightened antigen responsiveness of cytokine-primed CD8^+^ T cells remains unclear. We have shown that cytokine priming is accompanied by profound changes in the expression of several cell surface molecules including many involved in TCR and costimulatory signaling such as CD5, CD8, CD44, CD132, CD134 (OX40) and GITR (TNFRSF18) (22, 30). However, all these changes were also induced by homeostatic cytokines alone even in the absence of synergistic stimulation by inflammatory cytokines. Besides, cytokine induced augmentation of TCR signaling can occur even without costimulation (22, 30). We have shown that inflammatory cytokines enhanced the activation of STAT5 induced by homeostatic cytokines and its DNA-binding activity (22). Moreover, virus-induced inflammation was shown to increase activation of proximal TCR signaling molecules ZAP70 and PLCγ, and ERK following TCR stimulation (28).

The above studies suggest that cytokine priming by inflammatory and homeostatic cytokines elicit a ‘poised state’ in naïve CD8^+^ T cells that allows them to respond robustly upon subsequent encounter with antigen. To gain molecular insight into this poised state, we compared the chromatin accessibility of naïve, cytokine-primed and Ag-stimulated CD8^+^ T cells.

## Methods

### Mice, peptides and cytokines

Pmel-1 TCR transgenic mice (31) were purchased from the Jackson Laboratory (Bar Harbor, ME, USA) and were used with the approval of the Université de Sherbrooke Ethics Committee for Animal Care and Use in accordance with guidelines established by the Canadian Council on Animal Care. The Pmel-1 melanoma antigen-derived peptide mgp100_25-33_ (EGSRNQDWL) (32) was custom synthesized by GenScript (Scotch Plains, NJ, USA) to more than 80% purity. Recombinant human IL-15 and mouse IL-21 were from R&D Systems (Minneapolis, MN).

### Cell purification and stimulation

Naïve CD8^+^ T cells were purified from the lymph node cells of Pmel-1 mice by negative selection using Invitrogen Magnisort CD8 Naïve T cell Enrichment kit (ThermoFisher, #8804-6825-74) following the manufacturer’s instructions. Cells were stimulated with IL-15 and IL-21 (both at 10 ng/ml) for 48 h or with the cognate antigenic peptide mgp100_25-33_ (1 μg/ml) for 36 h in the presence of irradiated splenocytes from C57BL/6 mice as APC. CD8^+^ T cells from Ag-stimulated cultures were purified again by negative selection.

### ATAC sequencing

The assay for transposase-accessible chromatin (ATAC) sequencing was performed following the methods described by Buenrostro et al., (33, 34). Unstimulated (naïve, N), cytokine stimulated (cytokine-primed, CytP) and Ag-stimulated (AgS) cells were washed in cold PBS, and 50,000 cells were suspended in lysis buffer (10 mM Tris-Cl, pH 7.4, 10 mM NaCl, 3 mM MgCl_2_ and 0.1% IGEPAL CA-630) at 4°C for 5 min. The nuclei were sedimented by centrifugation at 500g for 10 min, and DNA library was prepared using the Nextera DNA Library Prep kit (Illumina). Briefly, nuclei were resuspended in transposition reaction mix and incubated at 37°C for 30 min. DNA fragments were purified using Qiagen Mini-Elute kit (Qiagen Cat# 28004). The DNA fragments were PCR amplified using custom Nextera primers containing different barcodes for naïve, CytP and AgS cells. The amplified fragment libraries were sequenced at the Université Laval (Quebec, QC) sequencing facility and analyzed at the Bioinformatics service platform of the Université de Sherbrooke.

Quality assessment of the amplified TN5 transposition fragment libraries showed an enrichment of single nucleosomes (200 bp peak) in naïve and CytP cells, whereas the AgS cells showed an enrichment of nucleosome dimers (400 bp peak) (33) (Supplementary Fig. S1). Even though the latter may arise from annealing of the PCR products due to shortage of primers at the later PCR cycles that can be resolved by an additional round of PCR (https://dnatech.genomecenter.ucdavis.edu/faqs/), it contained libraries harboring sites for CCCTC-binding factor (CTCF), a highly conserved transcriptional regulator throughout the genome (35) again and sites encompassing transcriptional start sites (TSS) (33). Accordingly, fragment length distribution analysis showed enrichment of ATACseq reads around TSS spanning approximately 200 bp covering a mono-nucleosomal peak in all three samples (Fig. 1A).

**Figure 1.**
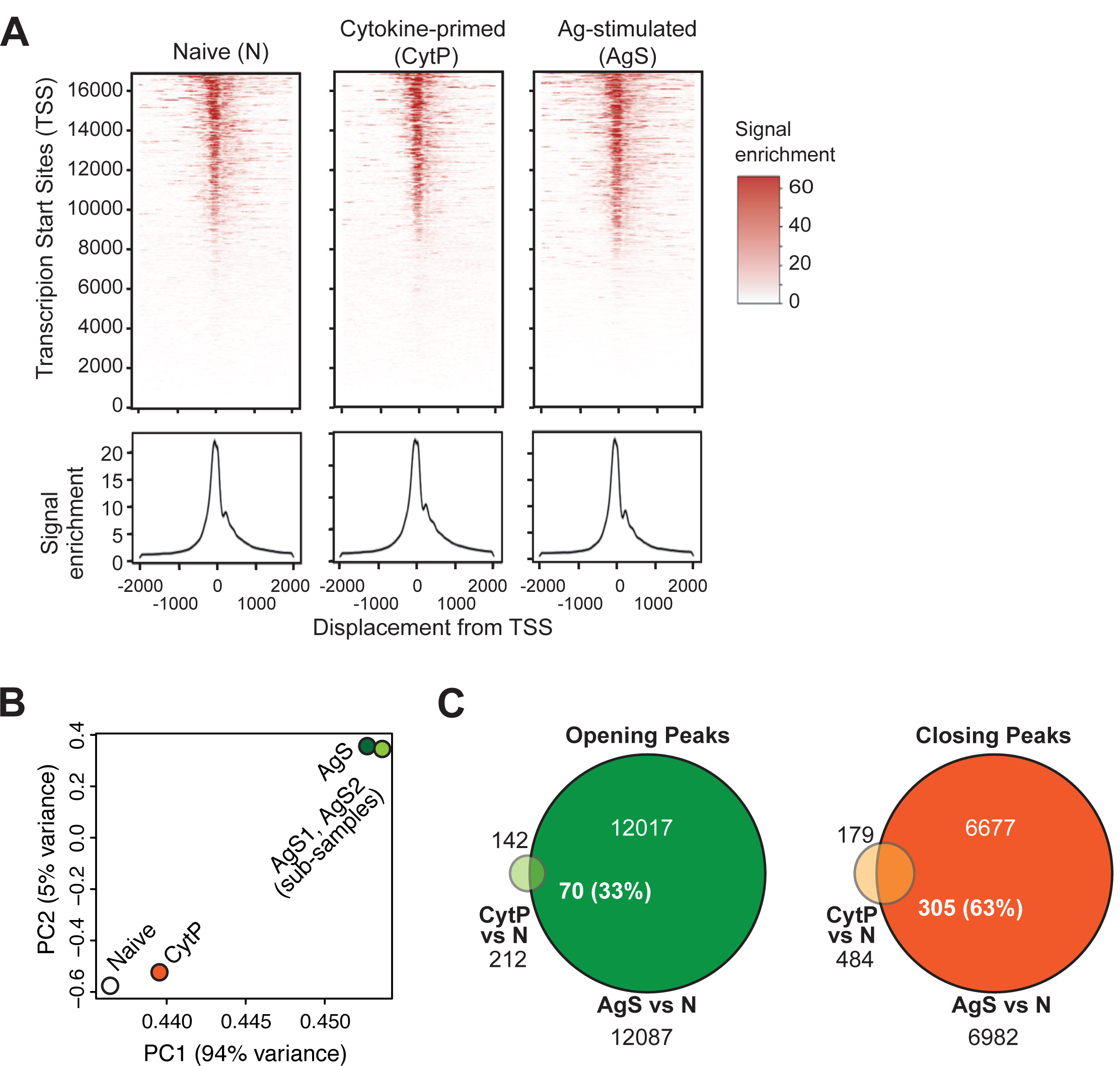
Cytokine-priming of CD8^+^ T cells modulates only limited number of ATACseq peaks compared to Ag-primed cells that show substantial overlap with the latter. (A) Fragment length distribution analysis of naïve (N), cytokine-primed (CytS) and Ag-stimulated (Ags) Pmel1 TCR transgenic CD8^+^ T cells. (B) Principal component analysis of the ATACseq reads of N, CytP and AgS cells. AgS1 and AgS2 represent random subgrouping of reads from AgS cells to ensure the random distribution of opening and closing ATACseq peaks (also see Supplementary Fig. S2B). (C) Overlap between the opening and closing peaks of CytP and AgS cells compared to naïve cells.

### ATACseq data analysis

The raw ATACseq data was processed using Trimmomatic (36) before analysis using the ENCODE ATAC-seq pipeline for non-replicated data to obtain signal and peak files. Whereas naïve and CytP cells generated 34M and 30M paired-end (PE) reads, AgS cells generated 134M reads. From these data, 50945, 53297 and 135836 peaks were identified in N, CytP and AgS cells respectively. A peak atlas was generated using Bedops (37) to concatenate peak files and iteratively merge >75% overlapping peaks. The coverage of the peaks was determined using the coverage tool in BEDTools (38).

To identify opening and closing peaks in stimulated cells compared to naïve cells, the DESeq2 package, developed to allow quantitative analysis based on strength rather than differential expression alone (39), was used. To avoid high fold change (FC) caused by low count, the regularized log transformation (rlog) was used to define meaningful change in chromatin accessibility. The distribution of rlogFC values for comparison between CytP versus N and AgS versus N cells are shown in Supplementary Fig. S2A. Because the number of peaks detected in AgS cells were more than double compared to those of naïve or CytP cells, they were randomly assigned to two subgroups (AgS1, AgS2) of 85640 and 86748 peaks using seqtk (https://github.com/lh3/seqtk) in order to determine if there was any skewing of opening and closing peaks. These two subgroups behaved similarly to AgS cells in the principal component analysis (Fig. 1B) and showed a similar enrichment of ATACseq reads around TSS and a similar pattern of opening and closing peaks (Supplementary Fig. S2B, S2C). Hence, all peaks in AgS cells were used for subsequent analysis. The rlogFC threshold was set to log2(1.5) as log2(2) was too stringent to identify opening and closing peaks in CytP and AgS cells. The peaks were subsequently analyzed using HOMER (40) to identify genes that are nearest to the peaks and the transcription factor binding motifs. Gene positions obtained from the Mouse Genome informatics database (http://www.informatics.jax.org/) was used to interrogate the UCSC Mouse Genome browser mm10 assembly (http://ucscbrowser.genap.ca) to visualize and capture snapshots of genome accessibility.

The STRING database (41) was used to study the interaction network analysis of proteins coded by genes in the opening and closing peaks and to identify gene ontology (GO) groups related to T cell activation, differentiation, effector functions and regulation. Only medium and high confidence interactions with a score above 0.40, supported by experiments, curated databases, protein homology, text mining and co-expression studies were considered for data interpretation.

## Results

### ATACseq peaks modulated by cytokine priming show substantial overlap with those altered by Ag stimulation

We have previously shown that the inflammatory cytokines IL-21 and IL-6, which stimulate STAT3, potentiate IL-7-induced STAT5 activation upon synergistic stimulation (22). Other groups have shown that IL-2-induced STAT5 activation stabilizes TCR signaling to promote differentiation of effector CD8^+^ T cells (42, 43). Similar role for STAT5 in promoting T helper cell differentiation via modulation of chromatin accessibility has also been documented (44-46). These reports raised the possibility that cytokine priming may induce changes in chromatin accessibility that would facilitate and strengthen TCR signaling, and thereby increase Ag sensitivity of CytP naïve CD8^+^ T cells. To test this hypothesis, we evaluated genome-wide transcription factor (TF) occupancy in PMEL1 TCR transgenic CD8^+^ T cells that were stimulated with IL-15 and IL-21, or with the cognate antigenic peptide using the ATACseq. Principal component (PC) analysis of ATACseq reads revealed that CytP cells were more closely related to naïve T cells than AgS cells (Fig. 1B). AgS cells showed 12087 opening and 6982 closing peaks with a rlog fold change value of 1.5 compared to naïve cells, whereas the CytP cells showed only 212 opening peaks and 484 closing peaks (Fig. 1C). Nevertheless, 70 of the opening peaks (33%) and 305 of the closing peaks (63%) in CytP cells were represented in AgS cells (Fig. 1C), suggesting that cytokine priming alters the accessibility of a subset of genes that are modulated by TCR signaling.

### Comparison of ATACseq peaks modulated by cytokine priming and Ag stimulation

The opening and closing chromatin accessibility peaks found in CytP and AgS cells were annotated using HOMER to associate the peaks with nearby genes. The open peaks found in CytP cells, in AgS cells and in both cells are listed in Supplementary Tables S1A to S1C, and the closing peaks in these three groups of cells are given in Supplementary Tables S2A to S2C. Protein encoded by genes nearest to the opening and closing peaks of CytP cells were analyzed using the STRING database to study their functional enrichment based on biological processes. This analysis identified several GO groups related to T cell activation, differentiation and functions that are modulated in CytP cells (Supplementary Table S3, Table 1A). Similar observations were made on opening and closing peaks found in both CytP cells and in AgS cells (Supplementary Table S4, Table 1B). Protein network analysis of all genes near the opening or closing peaks in CytP cells revealed a complex network (Supplementary Fig. S3). Therefore, we restricted this analysis to genes within the GO groups related to T lymphocyte activation to understand how cytokine priming might influence TCR signaling (Fig. 2A). When analyzed within the opening and closing peaks of both CytP and AgS cells, these GO enrichment groups showed significantly low false discovery rate (FDR; Fig. 2B). Among the genes that were modulated in both CytP and AgS cells, the opening peaks were found adjacent to *Rftn1* (Raftlin 1), *Dpp4* (Dipeptidylpeptidase 4; CD26), *Tnfsf8* (CD153), *Pkrcq* (PKCq) and *Zfp609*, whereas closing peaks were found near several other genes (*Cd28, Itk, Cblb, Bcl10, CD47, Tigit, Prkca* and *Dusp10*) (Fig. 2A). Functions of these genes related to T lymphocyte activation, regulation and functions are described in Table 2. Cytokine priming also modulated chromatin accessibility near a few other genes that fall within the GO terms of T lymphocyte functions but were not significantly affected in AgS cells (*Btla, Gadd45b, Cd300a, Jun, Xbp, Tnfsf11, Cd101*) (Fig. 2A and Table 2).

**Table 1.** GO analysis of genes related to T lymphocyte activation and functions that show changes in chromosome accessibility peaks.

| **A) in cytokine-primed cells** |  |  |  |  |
| --- | --- | --- | --- | --- |
| #term ID | term description | observed gene count | background gene count | FDR |
| GO:0002376 | immune system process | 63 | 1703 | 2.19E-07 |
| GO:1902105 | regulation of leukocyte differentiation | 22 | 284 | 7.52E-07 |
| GO:0006955 | immune response | 35 | 914 | 0.00015 |
| GO:0045582 | positive regulation of T cell differentiation | 9 | 90 | 0.00085 |
| GO:0046634 | regulation of alpha-beta T cell activation | 9 | 98 | 0.0014 |
| GO:0030098 | lymphocyte differentiation | 14 | 243 | 0.0016 |
| GO:0045619 | regulation of lymphocyte differentiation | 11 | 171 | 0.0033 |
| GO:0030217 | T cell differentiation | 10 | 154 | 0.0051 |
| GO:2000516 | positive regulation of CD4-positive, alpha-beta T cell activation | 5 | 34 | 0.0059 |
| GO:0046649 | lymphocyte activation | 16 | 378 | 0.009 |
| GO:0046632 | alpha-beta T cell differentiation | 6 | 60 | 0.0091 |
| GO:0042110 | T cell activation | 12 | 244 | 0.0118 |
| GO:0050670 | regulation of lymphocyte proliferation | 11 | 215 | 0.014 |
| GO:0070229 | negative regulation of lymphocyte apoptotic process | 4 | 39 | 0.0468 |
| **B) in both cytokine-primed and Ag-stimulated cells** |  |  |  |  |
| GO:0051251 | positive regulation of lymphocyte activation | 18 | 253 | 2.70E-05 |
| GO:0050870 | positive regulation of T cell activation | 16 | 180 | 1.00E-05 |
| GO:0051249 | regulation of lymphocyte activation | 23 | 396 | 2.13E-05 |
| GO:0050863 | regulation of T cell activation | 20 | 287 | 1.05E-05 |
| GO:0045621 | positive regulation of lymphocyte differentiation | 10 | 103 | 0.00045 |
| GO:0050852 | T cell receptor signaling pathway | 6 | 56 | 0.0069 |
| GO:0046635 | positive regulation of alpha-beta T cell activation | 6 | 64 | 0.0118 |
| GO:0050856 | regulation of T cell receptor signaling pathway | 5 | 39 | 0.0093 |
| GO:0042129 | regulation of T cell proliferation | 9 | 162 | 0.0207 |

FDR, false discovery rate.

**Table 2.**
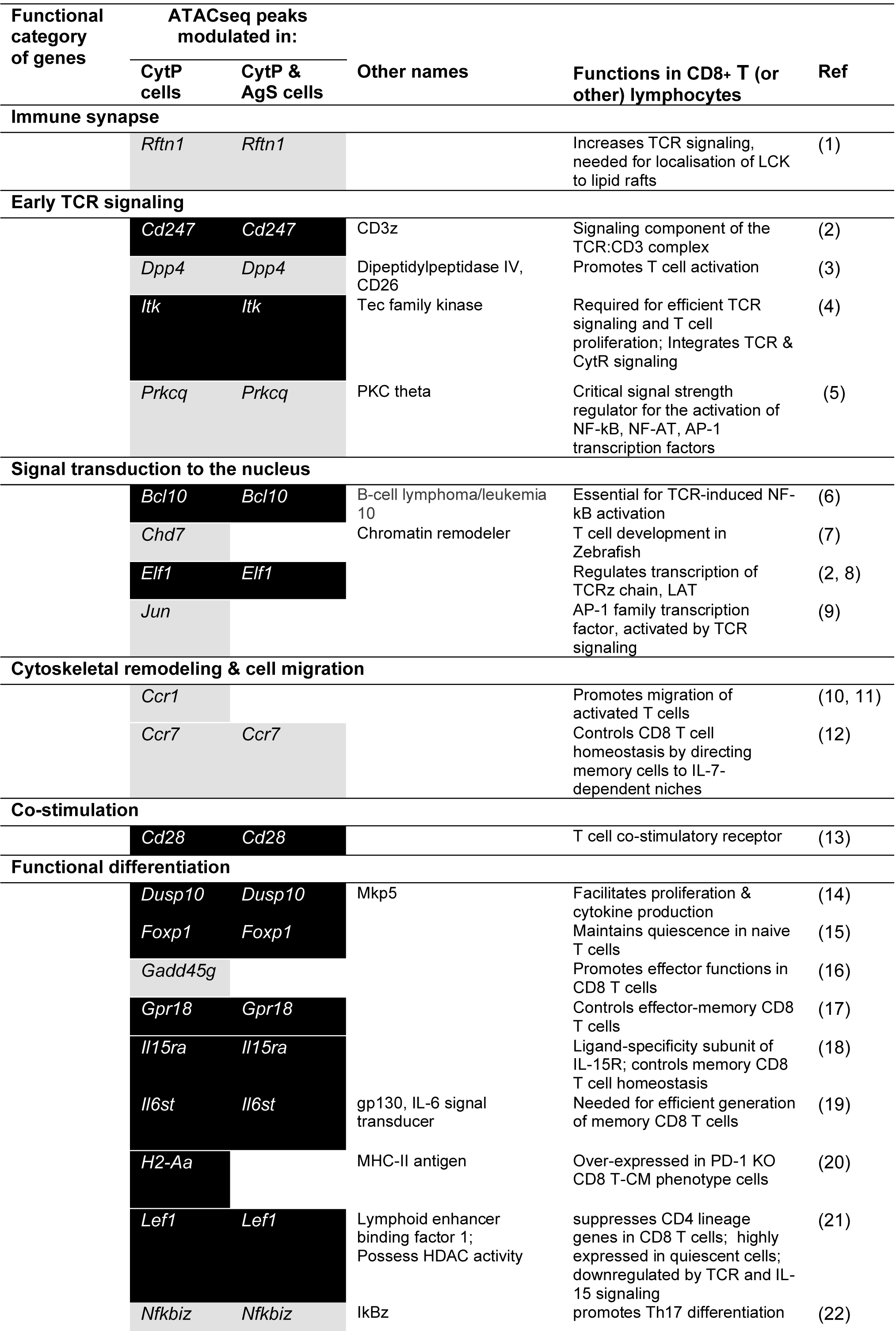
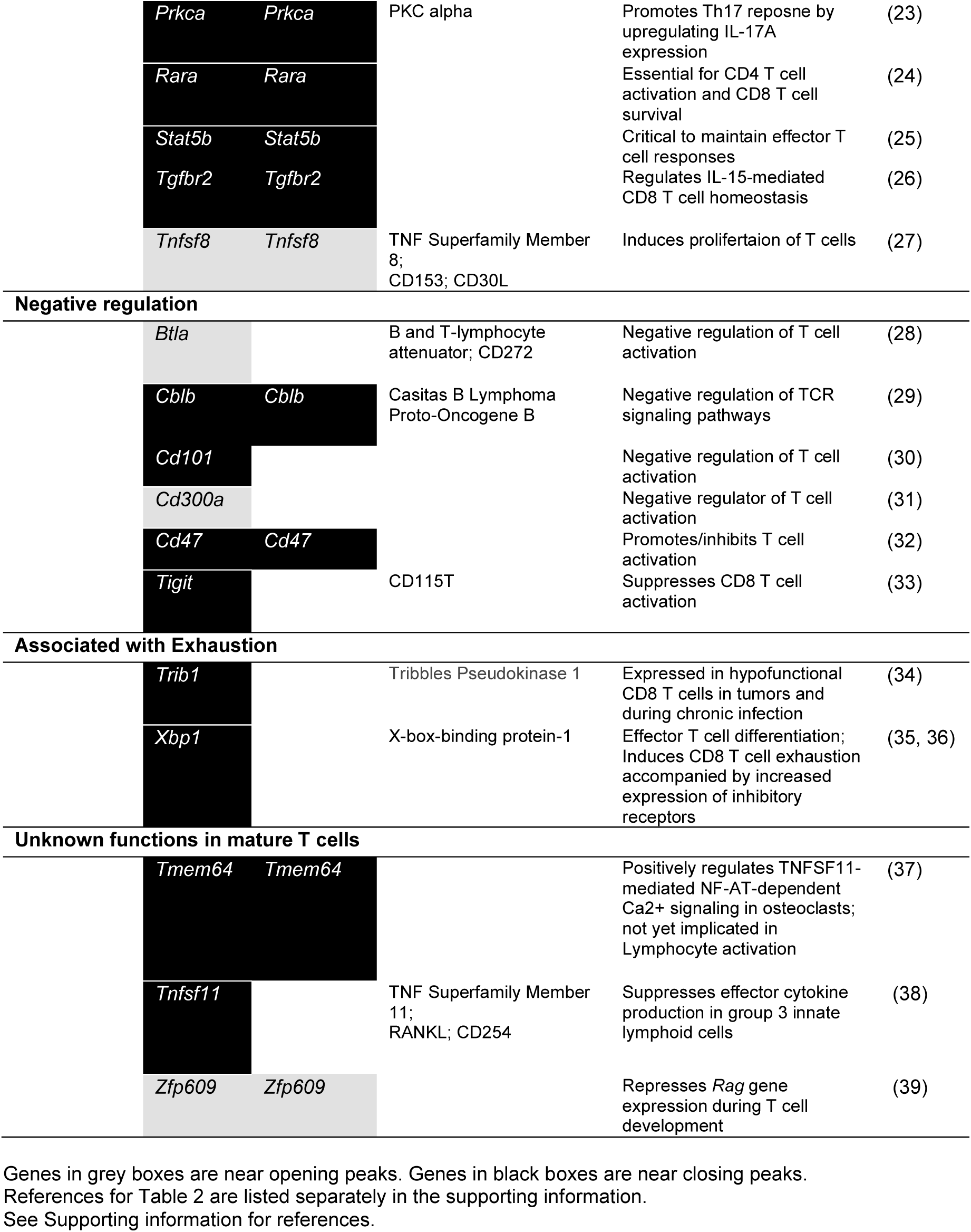
Genes related to T lymphocyte activation, regulation and functions that are adjacent to the opening and closing ATACseq peaks in CytP and in both CytP and AgS CD8_+_ T cells.

**Figure 2.**
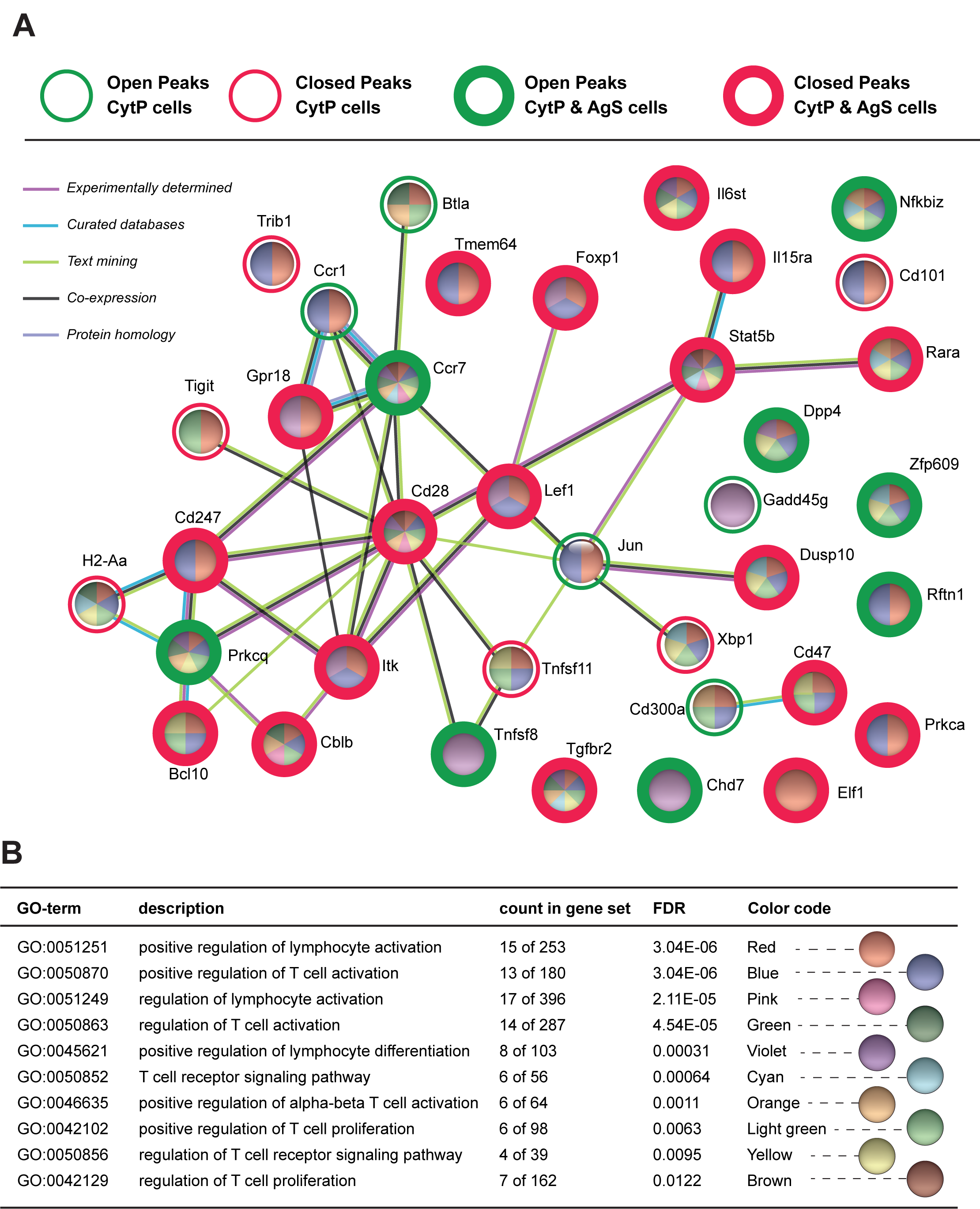
Chromatin accessibility of genes implicated in T cell activation, effector functions and regulation in cytokine-primed and Ag-stimulated cells. (A) Protein network analysis of genes within the GO groups related to T lymphocyte activation near the opening or closing peaks in CytP and AgS cells using the STRING database. Open and closed peaks are indicated by green and red colored circles, respectively. Peaks modulated in CytP cells alone are indicated by thin circles, and those modulated in both CytP and AgS cells by thick circles. The pie diagrams for individual genes are again color-coded based on their inclusion within the various gene ontology (GO) groups listed in (B). Only the GO groups that show significantly low FDR (false discovery rate) are shown.

### Chromatin accessibility in Ag-stimulated cells

To understand how cytokine priming modulates chromatin accessibility, we first analyzed the ATACseq peaks near the genes that are known to be modulated by Ag stimulation in CD8^+^ T cells. As expected, strong modulation of chromatin accessibility was observed near several genes such as *Il2, Ifng* and *Il2ra* (Fig. 3A-C), as indicated by the opening peaks that correspond to the binding sites of several TF activated by TCR signaling including NFATc, IRF4, Jun, Nur77 and RUNX (Fig. 3D). Genes coding for the checkpoint inhibitors PD1 (*Pdcd1*) and LAG3, which are strongly induced following TCR stimulation, also showed marked changes in chromatic accessibility with opening peaks for NFATc, NF-κB, IRF4, STAT5 and RUNX (Fig. 3E-F). Notably, almost none of these sites were accessible in CytP cells, indicating that cytokine priming does not induce many of the genes that promote effector cell differentiation. AgS cells also showed several other opening peaks (*Cd3e*), closing peaks (*Il7r, P2rx7*) or both (*Ctla4*), that were not affected in CytP cells (Supplementary Fig. S4).

**Figure 3.**
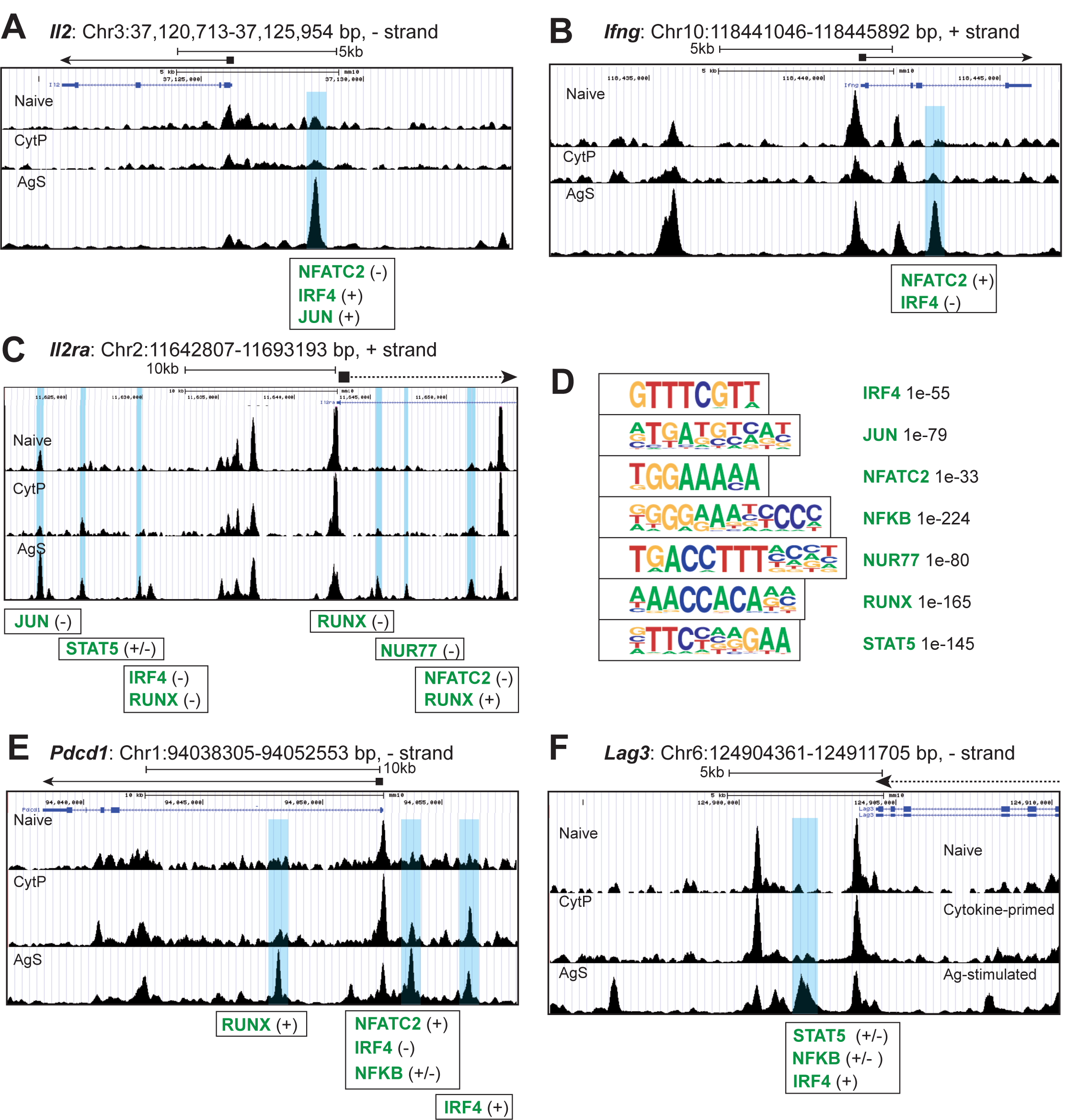
Chromatin accessibility in Ag-stimulated cells. Genome browser snapshots of chromatin accessibility signal at (A) *Il2*, (B) *Ifng* and (C) *Il2ra*, (D) *Pdcd1* and (E) *Lag3* genes. The chromosomal locations of the genes, accessibility peaks opening only in AgS cells (shaded blue) and the corresponding transcription factor bindings sites are indicated. Position of genes are indicated by solid lines for full genes and dotted lines for partially covered genes within the genome area shown. The peak heights (FC signal) are adjusted to the same value for all three tracks before taking the snapshots. (F) The transcription factors binding motifs significantly enriched in chromosome accessibility peaks opening in AgS cells.

### Chromatin accessibility in cytokine-primed cells

Next we compared the chromatin accessibility sites in CytP cells near genes implicated in T lymphocyte activation and functions, many of which are also modulated in a similar fashion in AgS cells (Table 2). These sites included many closing peaks and a fewer opening peaks. Analysis of chromatin accessibility near these genes revealed similar changes of comparable magnitude in the opening peaks near *Rftn1* (immune synapse component), *Dpp4* (modulator of TCR signaling), *Ccr7* (cytoskeletal remodeling and cell migration), *Tnfsf8* (T cell proliferation) and *Zfp609* (repressor of Rag genes) genes (Table 2, Fig. 4A). Whereas all the above opening peaks harbored the STAT5b binding motif, some of these opening peaks (Eg., *Dpp4, Tnfsf8*) also contained motifs for other TF such as NFAT, RUNX and IRF4 that are also found near the opening peaks of AgS cells. Strikingly, a majority of these genes also showed additional opening peaks only in AgS cells that contained motifs for various TF (Fig. 4A), suggesting that these genes are likely expressed in Ag-stimulated cells, whereas they are poised for expression in CytP cells.

**Figure 4.**
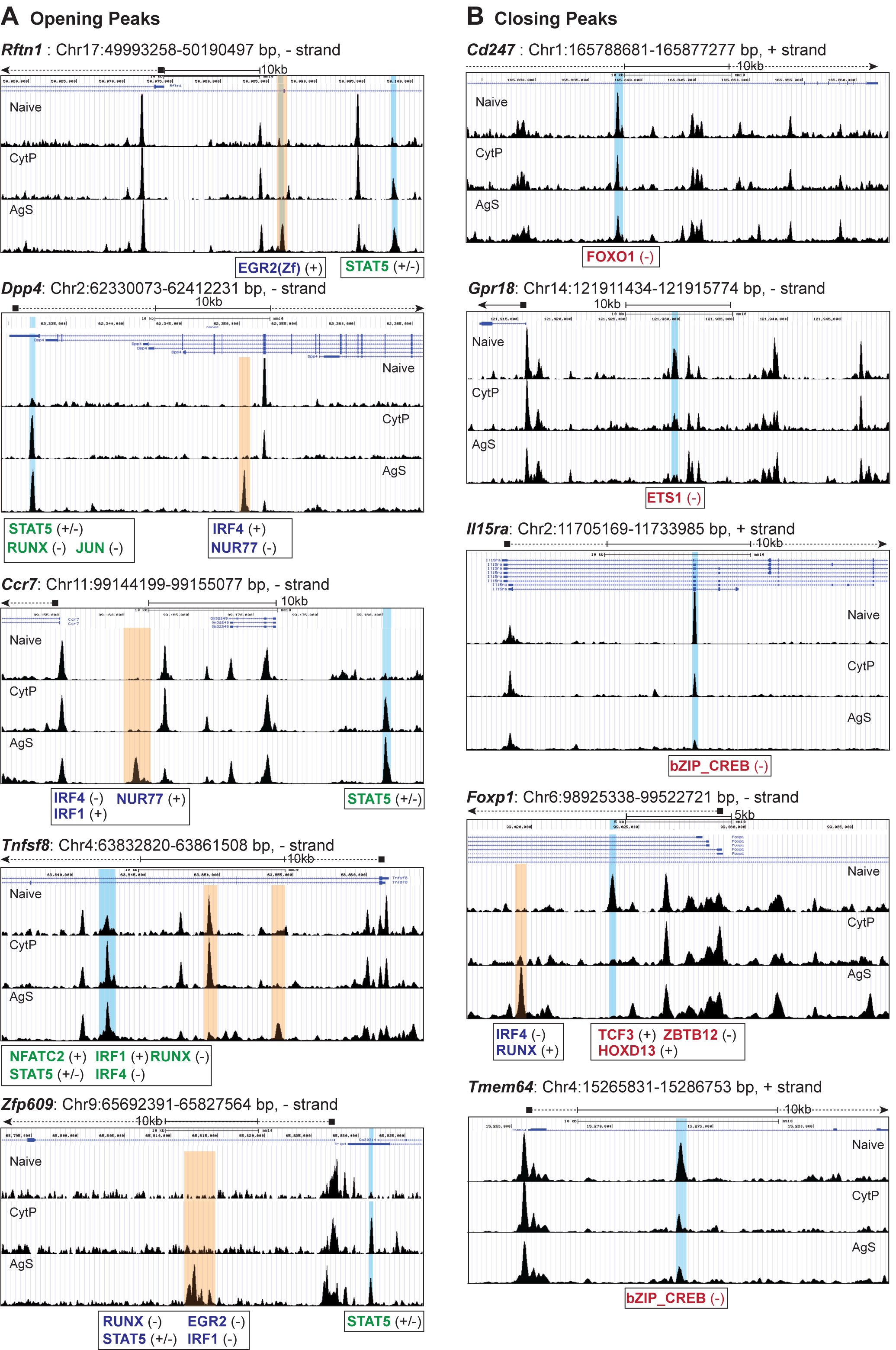
Chromatin accessibility in cytokine-primed cells. Genome browser snapshots of chromatin accessibility signal found in the opening (A) and closing (B) peaks of CytP cells that also occurred in AgS cells. Peaks modulated in both CytP and Ags cells are shaded blue, whereas those modulated only in AgS cells are shaded yellow. Color schemes for the transcription Factor binding motifs: green, in the opening peaks of CytP and AgS cells; red, in the closing peaks of CytP and AgS cells; blue, in the opening peaks of only AgS cells.

Analysis of the peaks closing in both CytP and AgS cells (Table 2, Fig. 4B) revealed similar changes near *Cd247* (CD3ζ, the signal transducing chain of the TCR), *Gpr18, Il15ra* (controls memory CD8 T cell generation and homeostasis), *Foxp1* (maintains quiescence in naïve T cells), *Tmem64* (unknown function in T cells; possibly involved in calcium signaling) genes. These peaks contained motifs for various TF such as FOXO1, ETS1, bZIP-CREB, TCF3 and HOXD13. Notably, the binding motifs for bZIP-CREB and ETS1 are found in several other closing peaks of both CytP and in AgS cells (Fig. S5A) or only in the latter (Fig. S5B). However, unlike for opening peaks, additional changes in chromatin accessibility near these closing peaks were uncommon.

### Transcription factor binding motifs enriched in CytP and AgS cells

Consistent with stimulation by IL-15 and IL-21, CytP cells showed an enrichment of binding sites for the STAT proteins STAT5, STAT1, STAT3 at the opening peaks (Fig. 5A; Supplementary Table S5A), although STAT4 and STAT6 binding was also observed. On the other hand, AgS cells predominantly showed motifs for the beta leucine zipper (bZIP) containing TF FRA1 (FOSL1), FOSL2, ATF3, BATF and AP1 (Fig. 5B; Supplementary table S5B) as previously reported (47). Several other TF motifs such as the ones for Tbet and Eomes, which are known to be activated by Ag, are also significantly enriched in AgS cells (Supplementary table S5B). The closing peaks of CytP and AgS cells remarkably differed in their accessibility to TF binding motifs, with the activating TF (ATF) family members ATF1, ATF7 and ATF2 dominating in CytP cells in contrast to the ETS (E26 transformation-specific or E-twenty-six) family members FLI1 and ETS1dominating in AgS cells (Fig. 5A-B; Supplementary tables S6A, S6B). However, analysis of the chromatin accessibility peaks present in both CytP and AgS cells showed a predominance of STAT and RUNX binding motifs in the opening peaks, almost in the same order as observed in CytP cells, and ATF family members in the closing peaks that occurred predominantly in CytP cells (Fig. 5C; Supplementary tables S5C, S6C). These findings indicate that chromatin accessibility to TF is markedly different in CytP and AgS cells while exhibiting a considerable degree of overlap.

**Figure 5.**
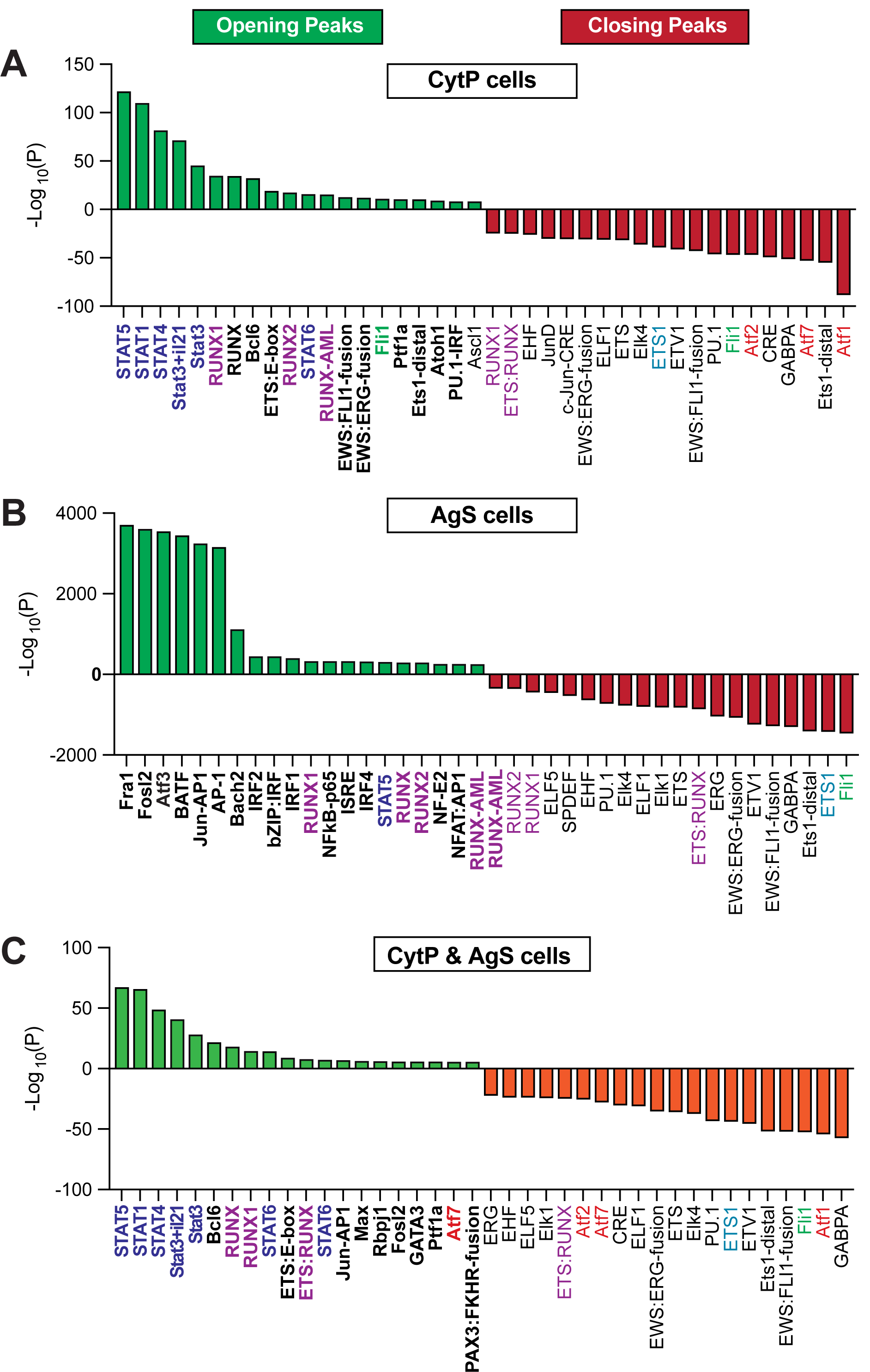
Enrichment of transcription factor binding motifs in cytokine-primed and Ag-stimulated cells. Top twenty transcription factor binding motifs found near the opening and closing peaks of CytP (A) and AgS (B) cells. Motifs found near the opening and closing peaks of both CytP and AgS cells are shown in (C).

### Chromatin accessibility near genes coding for proteins implicated in cytokine-priming, differentiation of AgS CD8^+^ T cells or differentially modulated by TCR affinity

Next we examined chromatin accessibility near genes coding for proteins that are mechanistically linked to the increased Ag sensitivity of CytP cells, conferring migratory potential to memory CD8^**+**^ T cells or modulated differentially depending on the TCR affinity towards the antigenic peptide. We have previously shown that CD5, a negative regulator of TCR signaling is downmodulated in CytP cells(30, 48). Analysis of chromatin accessibility at the *Cd5* locus revealed a closing peak upstream of the *Cd5* gene that is predicted to harbor motifs for ETS1 (FDR=1e-32) and FosL1 (FDR=1e-20) in CytP cells that was not observed in AgS cells (Fig. 6A). On the other hand, the locus containing *Gcnt1*, which is induced by IL-15 in memory CD8 T cells and codes for the enzyme that regulates 2-O-glycosylation of cell surface receptors to confer migratory potential (49), harbored an opening peak with binding motifs for NFATc, RUNX, IRF4 and STAT5 in AgS cells but not in CytP cells (Fig. 6B). Finally, IRF4, a key TF implicated in modulating the expression of many genes in AgS cells in a quantitative manner in proportion to the strength of TCR stimulation (50), revealed an opening peak harboring NFATc site in AgS cells but not in CytP cells, whereas a closing peak with TCF3 binding motif occurred in both AgS and CytP cells (Fig. 6C).

**Figure 6.**
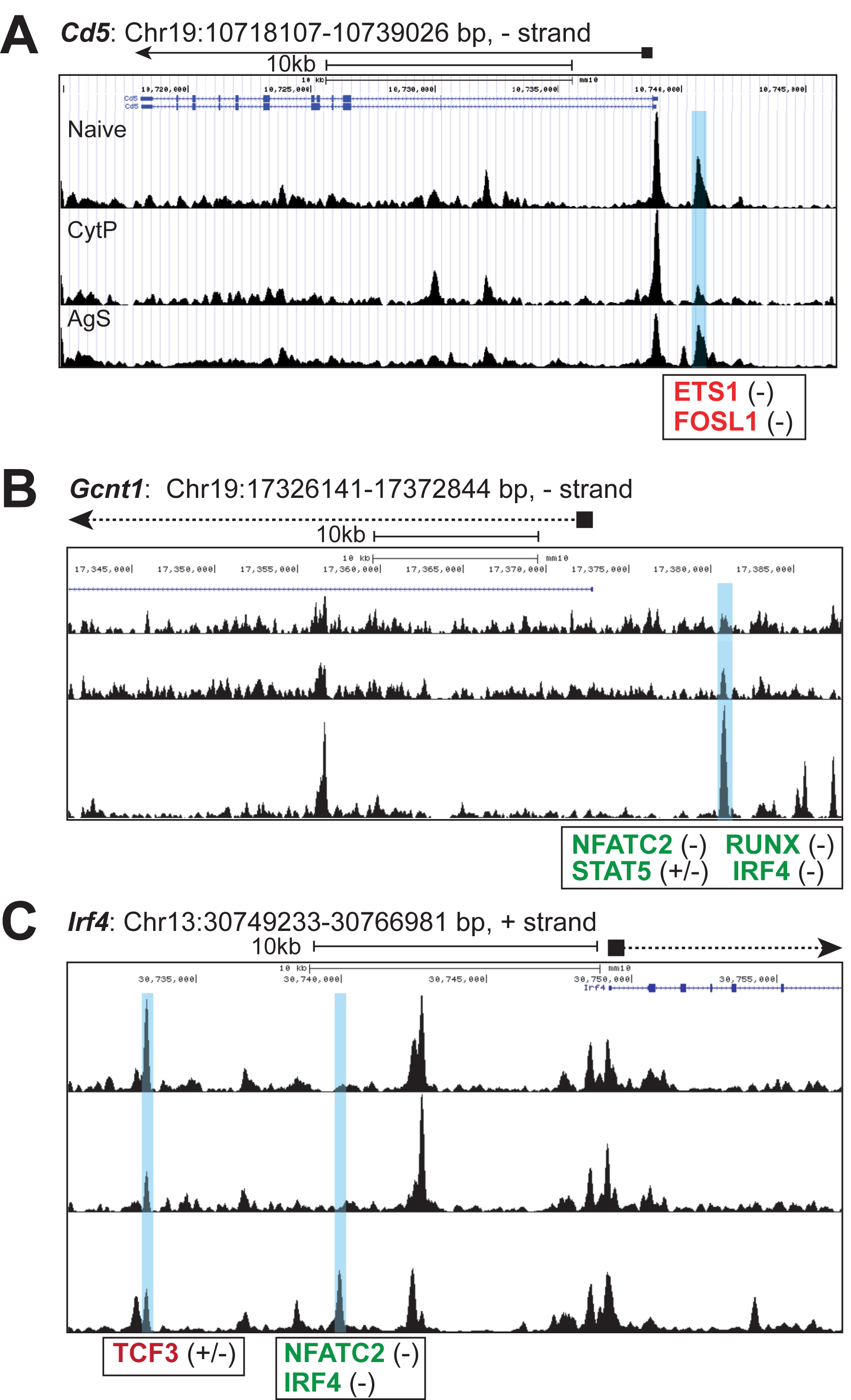
Chromatin accessibility of genes coding for proteins known to be modulated by cytokines in CD8^+^ T cells. Genome browser snapshots of chromatin accessibility signals near genes coding for proteins modulated by cytokine priming (A, *Cd5*), Ag stimulation that promotes migration potential of memory CD8^+^ T cells (B, *Gcnt1*), and the strength of pMHC-TCR interaction (C, *Irf4*) in CytP and AgS cells.

## Discussion

Modulation of CD8^+^ T cell responses by inflammatory cytokines produced by innate immune cells continues to be an area of intense scrutiny (11, 29, 51-53). At least three types of inflammatory cytokine-driven augmentation of CD8^+^ T cell responses are well recognized. First, the third signal cytokines IFN-I and IL-12, which promote efficient expansion and full activation of AgS CD8^+^ T cells, mediate these functions by sustaining the transcriptional program initiated by the TCR and costimulatory receptors via modulating histone acetylation at genetic loci that regulate cell survival, proliferation and effector functions (54). The second cytokine-mediated modulation of CD8^+^ T cell functions is the IL-15-dependent increase in Ag sensitivity of memory CD8^+^ T cells and their ability to migrate through tissues (28, 49, 55).

Whereas the latter function has been attributed to the increased expression of GCNT1, which promotes 2-O-glycosylation of cell surface molecules that interact with P- and E-selectins, the mechanistic basis for the IL-15-mediated increase in Ag sensitivity in memory cells remains to be elucidated. Third, the increase in Ag sensitivity of naïve CD8^+^ T cells mediated by inflammatory cytokines in synergy with homeostatic cytokines, which primes naïve T cells to respond to weak TCR agonists (22, 30, 56). In this study, we show that cytokine priming modulates chromatin accessibility at several key genetic loci, which are also associated with Ag stimulation, conferring a poised state for subsequent Ag encounter.

Indirect evidences suggest an important role for inflammatory cytokines in modulating the Ag responsiveness of naïve CD8^+^ T cells. One of them is the heterogeneity in the magnitude of OT-I TCR transgenic CD8^+^ T cell response towards the cognate peptide. Following Ag stimulation *in vivo*, naïve OT-I cells vary in the expression of effector molecules, the propensity to differentiate into various effector subsets and the ability to kill of virus-infected cells (57-59). As these studies used distinct viruses (influenza, vesicular stomatitis virus, murine cytomegalovirus) that expressed cognate OVA Ag, the heterogeneity of their responses could be explained at least partly by differential exposure to virus-induced cytokines and their impact on the strength of initial TCR activation. Such heterogeneity also occurs in the differentiation of CD4^+^ T cells with single TCR specificity *in vivo*, and were attributed to TCR signal strength as well as to environmental cues, particularly the cytokine milieu induced by adjuvants, although the effect of the latter can be nullified by quantitatively stronger TCR stimulation (60-62). Supporting this argument, CD8^+^ T cells in human peripheral blood that are specific to cytomegalovirus, Epstein-Barr virus and influenza virus antigenic epitopes were shown to vary in the expression of CTL effector molecules, cytokine and chemokines (63). CD8^+^ T cells specific to the viral epitopes in this study are unlikely to be monoclonal in origin and thus the TCR clonality could partially account for the variability. However, the cytokine context of their initial stimulation could also be a factor, which could account for the distinct response patterns observed against the Ag epitopes of the three different viruses. Therefore, there is a clear need to understand the molecular underpinnings that determine the role played by cytokines of the innate immune response in modulating the final outcome of T cell activation (60, 64).

The boosting of Ag-induced expansion and effector CD8^+^ T cell differentiation by the third signal cytokines IFN-I and IL-12, and the IL-15-mediated increase in the functional avidity of memory CD8^+^ T cells were demonstrated using cytokine or cytokine receptor knockout mice (14, 28, 49, 65). On the other hand, evidence for cytokine-induced increase in the functional avidity of the TCR in naïve CD8^+^ T cells mainly came from *in vitro* studies following stimulation with inflammatory cytokines in conjunction with IL-7 or IL-15 (22, 26). Obtaining genetic evidence for cytokine priming of naïve CD8^+^ T cells was complicated by the redundancy of cytokine combinations that could cause this effect. Either IL-6 or IL-21 (and possibly other inflammatory cytokines) along with IL-7 or IL-15 could achieve the cytokine priming effect of boosting TCR functional avidity *in vitro* (22, 26). Such *in vitro* cytokine-primed autoreactive CD8^+^ T cells, stimulated with weak agonists of the TCR, were able to cause disease in a mouse model of autoimmune diabetes (26). Moreover, PMEL-1 melanocyte Ag-specific TCR transgenic mice showed evidence of activation *in vivo* in the absence of SOCS1, the negative feedback regulator of IL-15 and IL-7 signaling, causing melanocyte destruction and autoreactive skin lesions (27). These findings lend support to the notion that cytokines play a key role in virus-induced and lymphopenia-associated triggering of autoreactive CD8^+^ T cells (66-68).

Cytokine priming may have role in activating CD8^+^ T cells bearing TCR with low affinity toward pathogen-derived Ag and these responses are known to be important in pathogen elimination (69-72). In fact, comparison of dodecamer and tetramer binding suggests that T cells expressing low-affinity TCRs are more abundant than those expressing high affinity TCR (73). While the latter are retained in the secondary lymphoid organs through stable interactions with DC, the former are released into circulation as the concentrations of the pMHC complexes decrease over time (74). Even though T cells bearing low-affinity TCR express effector molecules such as granzyme B and perforin and contribute to containing the infection, they fail to expand and persist in contrast to the high affinity clones during acute infections (70). However, during chronic infections, low affinity clones have been shown to predominate at later time points (75), and cytokine priming may have a role in these persistent immune responses.

Anti-tumor CD8^+^ T cells bearing low affinity TCR towards tumor Ag and neo-Ag may also benefit from cytokine priming. Due to central and peripheral tolerance mechanisms, anti-tumor CD8^+^ T cells necessarily bear low affinity TCR towards tumor Ag (76). Low affinity TCR clones are capable of containing a modest tumor burden (77, 78) and increasing the TCR affinity by genetic manipulation results in off-target effects and autoimmunity (76). We have shown that cytokine priming enables tumor Ag-specific CD8^+^ T cells to recognize and respond to an endogenously produced and presented tumor Ag peptide (79). Hence, cytokine priming could be used to expand antitumor CD8^+^ T cells bearing low affinity TCRs to circumvent the off-target toxicity associated with TCR engineering (80).

An essential step towards understanding and exploiting cytokine priming in protective immune responses against infections or cancers is to elucidate its mechanistic underpinnings. Even though our present study used one cytokine priming condition of IL-15 and IL-21 at a single time point on one TCR Tg CD8^+^ T cell model, our findings give important insights into how CD8+ T cell priming by inflammatory and homeostatic cytokines could impact on TCR signaling. Even though the transcriptional program activated by Ag as well as the TF involved have been well studied, the application of chromatin accessibility assays are beginning to broaden our understanding of these changes at the genome level (47, 81, 82). Consistent with these reports, the binding motifs for the bZIP containing TF were predominant in AgS cells.

Notably, the binding motifs for two bZIP factors Jun-AP1 and Fosl2 were also enriched in CytP cells. Many bZIP factors co-operate with IRF4, which is induced by TCR stimulation and plays a fundamental role in CD8 T cell differentiation in proportion to the TCR affinity and signal strength (50, 83). The IRF4 binding motif figured among the top TF motifs near the opening peaks of AgS cells and was found near key genes known to be induced by TCR stimulation (Fig. 3, 4). IL-15 was recently shown to enhance IRF4 expression in TCR-stimulated CD8 T cells (84). We did not find significant enrichment of IRF4 binding motif in CytP cells (Supplementary Table S5A). However, the *Irf4* locus of CytP cells contained one significant change among the many induced by Ag stimulation (Fig. 6), namely the loss of the binding peak for the transcriptional repressor TCF3 (E47A) (85). This suggests a poised state of the *Irf4* locus that could increase its induction in CytP cells following TCR stimulation, contributing to their increased Ag responsiveness.

Among the TF binding motifs shared by CytP cells with AgS cells, STAT and RUNX occurred predominantly near the common opening peaks (Fig. 5C). The enrichment of the binding motifs for STAT5, STAT3 and STAT1, which are activated by the IL-2 family cytokines (86), in AgS cells could be explained by autocrine IL-2 signaling that occurs within 36 hours of Ag stimulation. How the binding motifs for STAT4 and STAT6, which are activated during CD8^+^ T cell differentiation process induced by Ag stimulation (85), are enriched in CytP cells remains to be investigated. The Runt domain containing TF (RUNX) RUNX1, RUNX2 and RUNX3, which function as both transcriptional activators and repressors, are implicated in CD8^+^ T cell development and CTL differentiation (87-90). The bZIP domain containing ATF, which include several members including BATF, also function as both transcription activators and repressors and are implicated in the differentiation of activated CD8^+^ T cells (91, 92). Whereas RUNX binding motifs are shared between CytP and AgS cells near the opening peaks, the shared ATF motifs are found near the closing peaks (Fig. 5).

## Conclusions

In summary, even though the changes in chromatin accessibility induced by cytokine priming in naïve CD8^+^ T cells are >10-fold fewer than those induced by Ag stimulation, 30-50% of the changes induced in CytP cells also occur in AgS cells. Many of these changes happen near genes implicated in CD8^+^ T cell activation and differentiation and involve similar TF binding motifs modulated by Ag stimulation. Gene expression studies, especially RNAseq, would be needed to determine the proportion of chromatin accessibility changes in CytP cells that actually result in gene expression. Nevertheless, our findings strongly support our contention that inflammatory cytokines induced during innate immune responses have the potential to augment the CD8^+^ T cell responses that can be exploited for improving vaccine strategies and antitumor immunity.

## List of Abbreviations

Ag: antigen
AgS: antigen-stimulated
ATACseq: assay for transposase-accessible chromatin sequencing
bZIP: beta leucine zipper
CytP: cytokine-primed
FDR: false discovery rate
GO: gene ontology
TF: transcription factor
TSS: transcriptional start sites.

## Declarations

### Ethics approval and consent to participate

All experiments on mice were carried with the approval of the Université de Sherbrooke Ethics Committee for Animal Care and Use in accordance with guidelines established by the Canadian Council on Animal Care (Protocol # 212-17, 2017-2039).

### Consent for publication

Not applicable.

### Availability of data and materials

All data generated or analysed during this study are included in the supplementary information files.

## Authors’ contributions

SI and SR conceived the idea, analyzed data and wrote the manuscript; AJQ, MC, MAS and MM carried out the experiments and participated in data analysis, discussion and critical reading of the manuscript. All authors read and approved the final manuscript.

## Acknowledgements

We thank Dr. Frédérick Grenier and Mr. Jean-François Lucier (Bioinformatics Service platform, Université de Sherbrooke) for analyzing the ATACseq data. MAS received the ‘Abdenour Nabid, MD’ MSc scholarship from the Faculty of Medicine and Health Sciences, Université de Sherbrooke.

## Supporting information: References for Table 2

Cytokine priming of naïve CD8^+^ T lymphocytes modulates chromatin accessibility that partially overlaps with changes induced by antigen simulation MA Santharam et al.,

---

**Supplementary Fig. S1.**
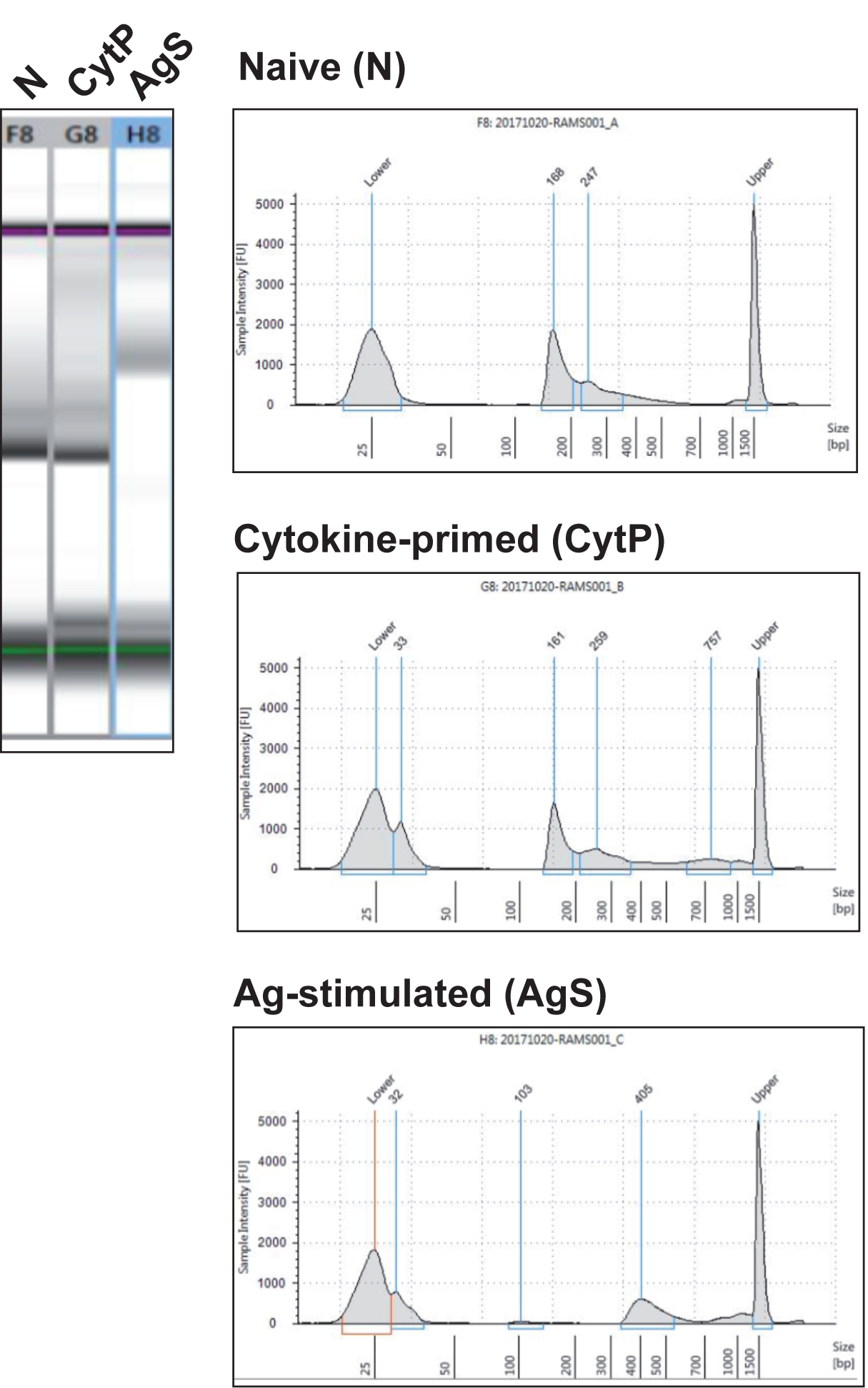
Bioanalyzer quality assessment of ATACseq libraries of naive, cytokine-primed and Ag-stimulated Pmel1 TCR transgenic CD8+ T ymphocytes..

**Supplementary Fig. S2.**
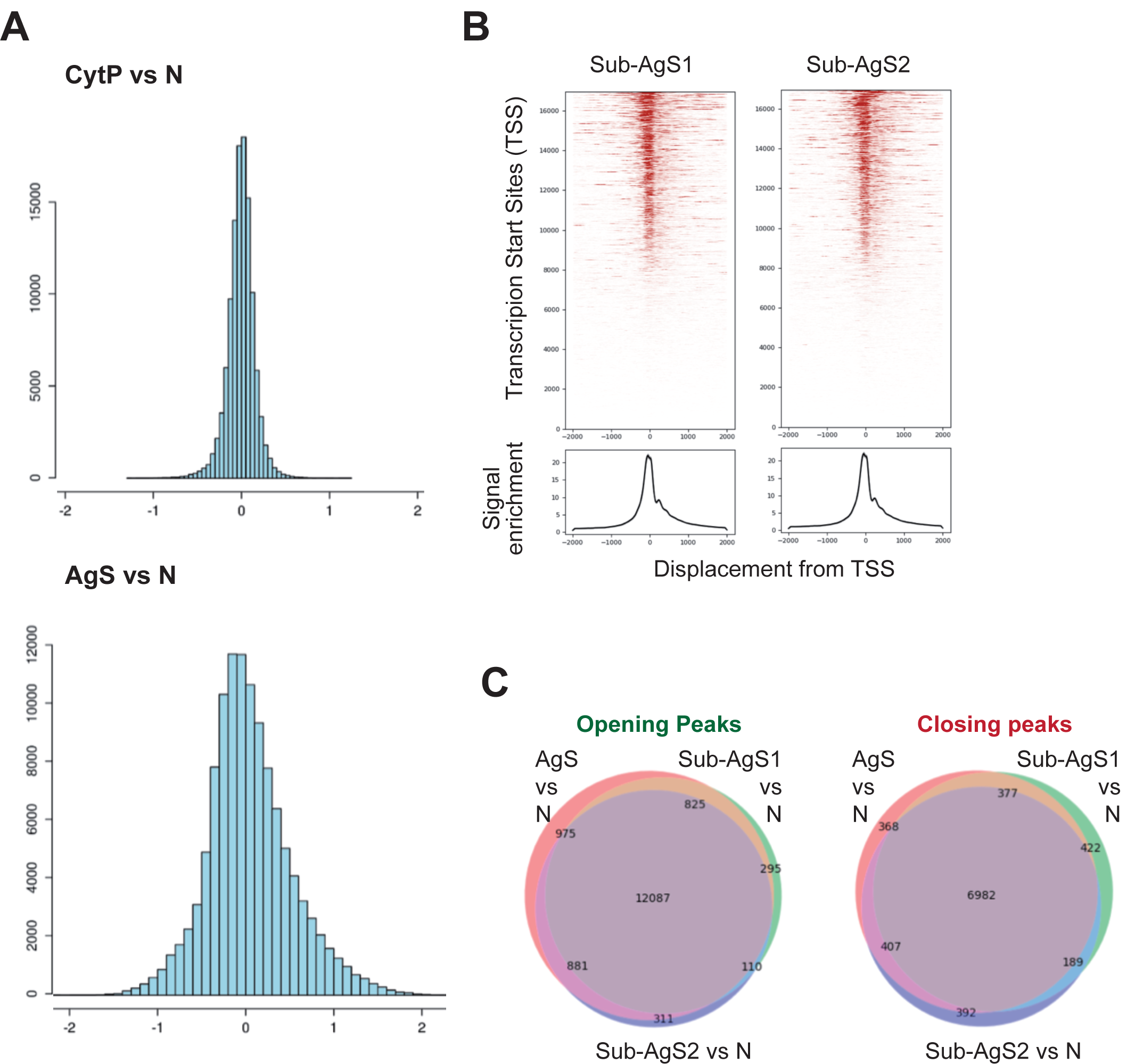
(A) Distribution of rlogFC values for comparison between CytP versus N and AgS) versus N cells. (B) Fragment length distribution analysis of ATACseq reads from AgS cells randomly subgrouped to sub-AgS1 and sub-AgS2 reads. (C) Comparison of the opening and closing peaks in sub-AgS1 and sub-AgS2 ATACseq reads compared to naive cells.

**Supplementary Fig. S3.**
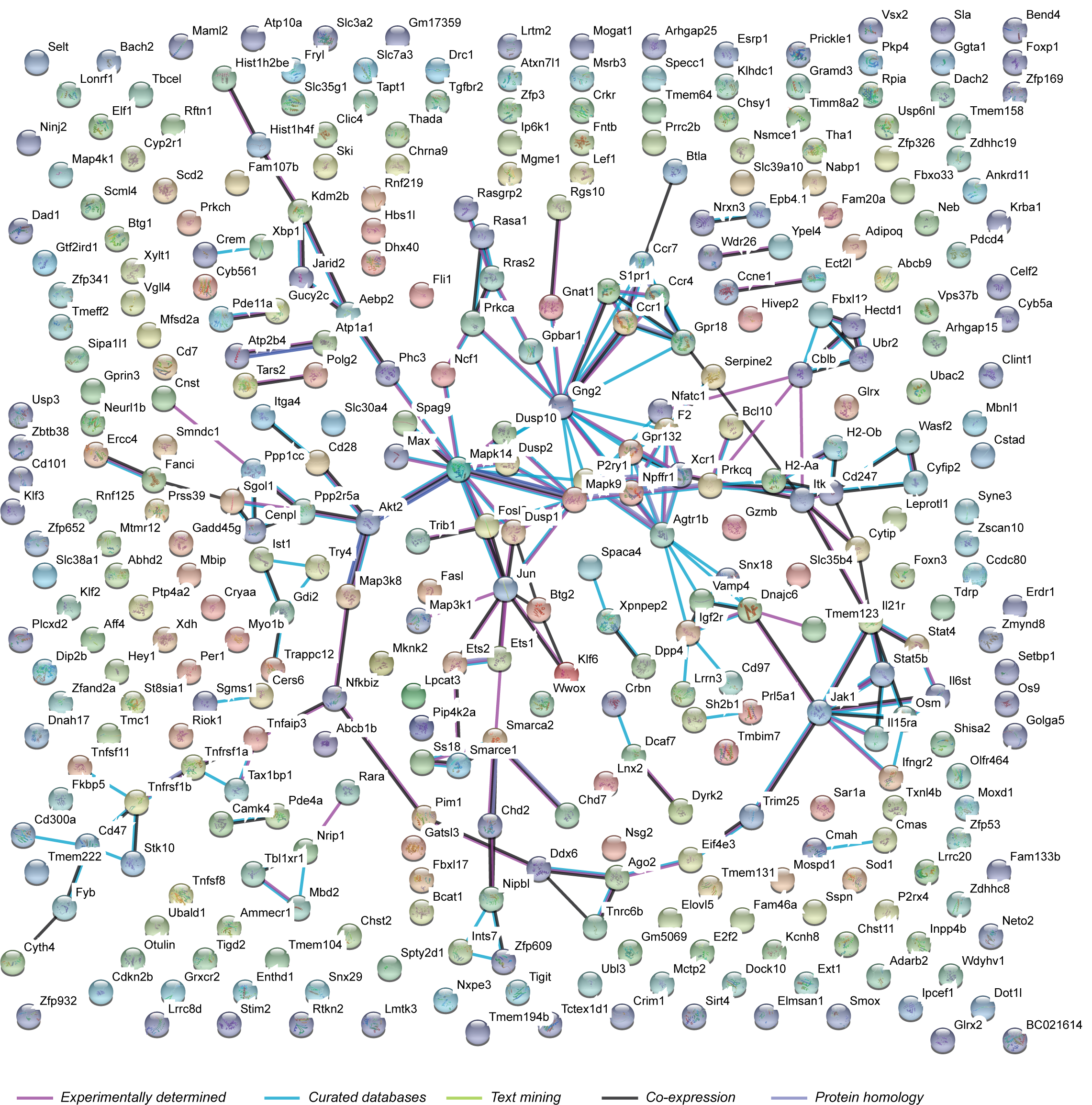
Protein interaction netwrok analsysis of genes in the vicinity of the ATACseq peaks modulated in cytokine-primed PMEL-1 cells compared to naive cells.

**Supplementary Fig. S4.**
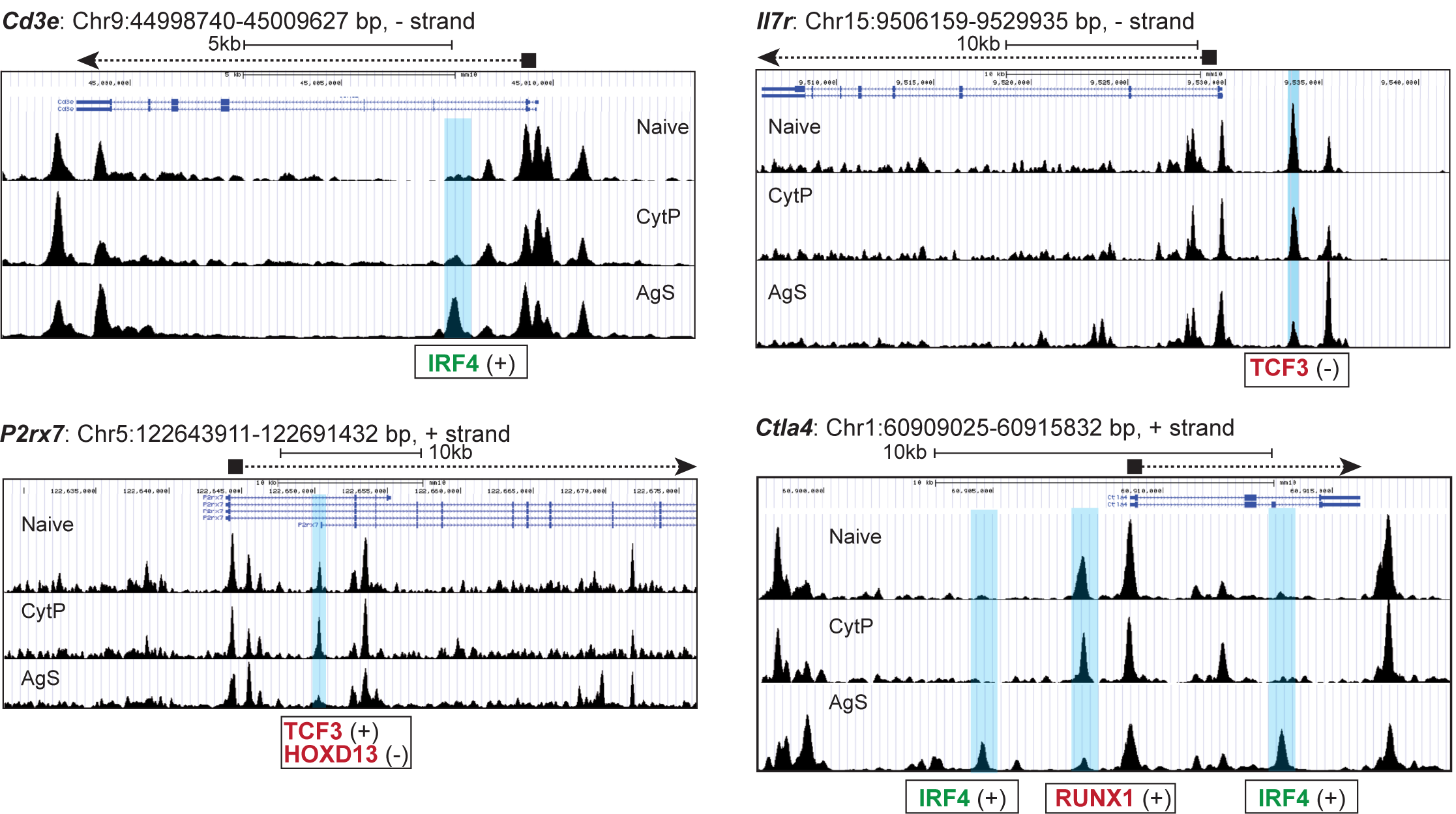
Other examples of chromosome accessibility peaks opening (*Cd3e*), closing (*Il7ra, P3rx7*) or both opening and closing (*Ctla4*) in Ag-stimulated but not in cytokine-primed cells.

**Supplementary Fig. S5.**
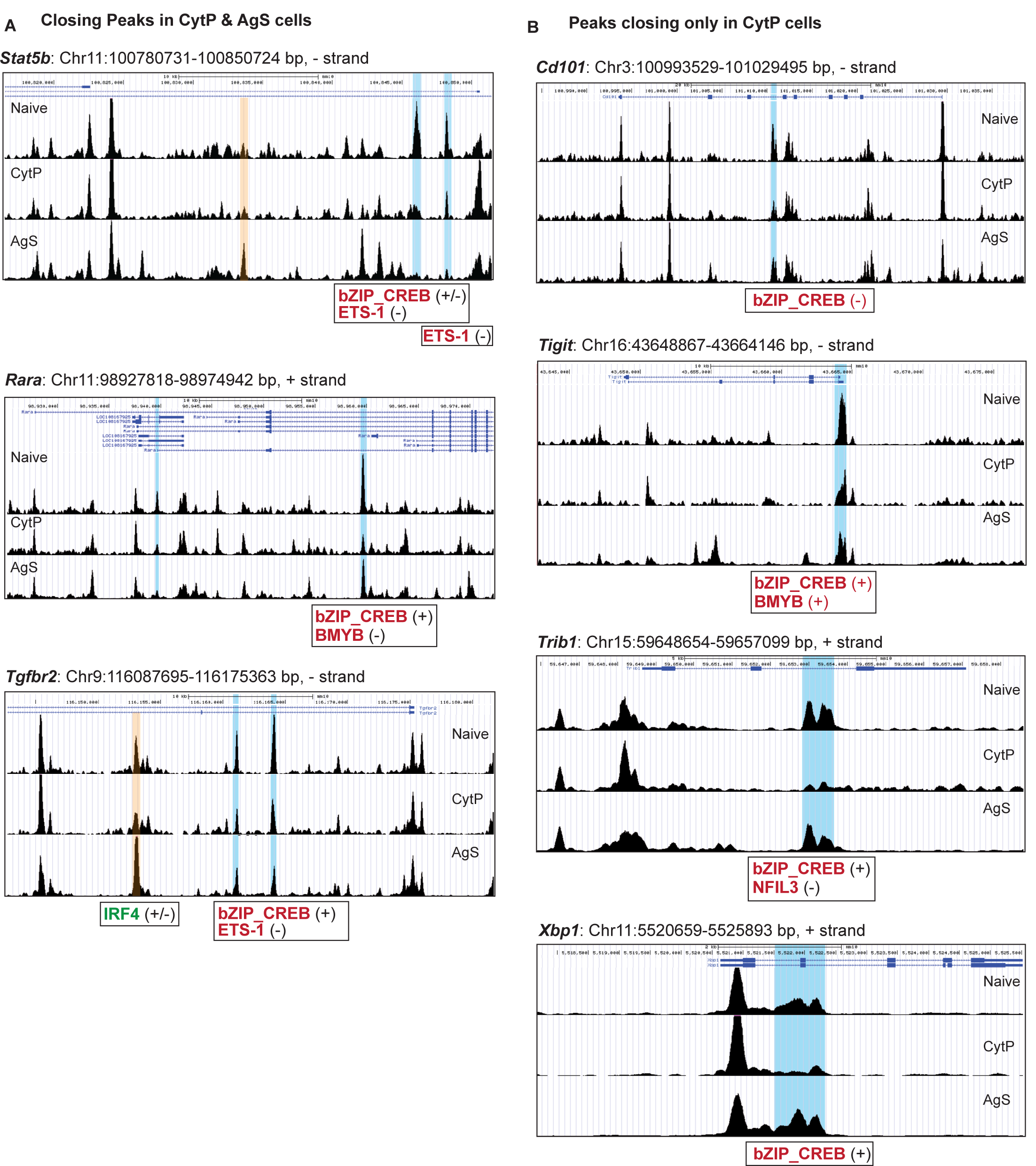
Chromosome accessibility peaks that are closing in both CytP and in AgS cells (A) or only in AgS cells (B).

## References

1. Harty JT, Tvinnereim AR, White DW. CD8+ T cell effector mechanisms in resistance to infection. Annu Rev Immunol. 2000;18:275–308.

2. Rosenberg SA, Restifo NP, Yang JC, Morgan RA, Dudley ME. Adoptive cell transfer: a clinical path to effective cancer immunotherapy. Nature reviews. 2008;8(4):299–308.

3. Sharpe AH. Mechanisms of costimulation. Immunol Rev. 2009;229(1):5–11.

4. Iwasaki A, Medzhitov R. Regulation of adaptive immunity by the innate immune system. Science. 2010;327(5963):291–5.

5. Luster AD. The role of chemokines in linking innate and adaptive immunity. Curr Opin Immunol. 2002;14(1):129–35.

6. Hoebe K, Janssen E, Beutler B. The interface between innate and adaptive immunity. Nature immunology. 2004;5(10):971–4.

7. Moretta A, Marcenaro E, Parolini S, Ferlazzo G, Moretta L. NK cells at the interface between innate and adaptive immunity. Cell death and differentiation. 2008;15(2):226–33.

8. Jain A, Pasare C. Innate Control of Adaptive Immunity: Beyond the Three-Signal Paradigm. J Immunol. 2017;198(10):3791–800.

9. Pulendran B, Ahmed R. Translating innate immunity into immunological memory: implications for vaccine development. Cell. 2006;124(4):849–63.

10. von Herrath MG, Fujinami RS, Whitton JL. Microorganisms and autoimmunity: making the barren field fertile? Nat Rev Microbiol. 2003;1(2):151–7.

11. Haring JS, Badovinac VP, Harty JT. Inflaming the CD8(+) T cell response. Immunity. 2006;25(1):19–29.

12. Curtsinger JM, Schmidt CS, Mondino A, Lins DC, Kedl RM, Jenkins MK, et al. Inflammatory cytokines provide a third signal for activation of naive CD4+ and CD8+ T cells. J Immunol. 1999;162(6):3256–62.

13. Curtsinger JM, Johnson CM, Mescher MF. CD8 T cell clonal expansion and development of effector function require prolonged exposure to antigen, costimulation, and signal 3 cytokine. J Immunol. 2003;171(10):5165–71.

14. Kolumam GA, Thomas S, Thompson LJ, Sprent J, Murali-Krishna K. Type I interferons act directly on CD8 T cells to allow clonal expansion and memory formation in response to viral infection. The Journal of experimental medicine. 2005;202(5):637–50.

15. Curtsinger JM, Valenzuela JO, Agarwal P, Lins D, Mescher MF. Type I IFNs provide a third signal to CD8 T cells to stimulate clonal expansion and differentiation. J Immunol. 2005;174(8):4465–9.

16. Morishima N, Owaki T, Asakawa M, Kamiya S, Mizuguchi J, Yoshimoto T. Augmentation of effector CD8+ T cell generation with enhanced granzyme B expression by IL-27. J Immunol. 2005;175(3):1686–93.

17. Le Bon A, Durand V, Kamphuis E, Thompson C, Bulfone-Paus S, Rossmann C, et al. Direct stimulation of T cells by type I IFN enhances the CD8+ T cell response during cross-priming. J Immunol. 2006;176(8):4682–9.

18. Whitmire JK, Tan JT, Whitton JL. Interferon-gamma acts directly on CD8+ T cells to increase their abundance during virus infection. The Journal of experimental medicine. 2005;201(7):1053–9.

19. Sercan O, Hammerling GJ, Arnold B, Schuler T. Innate immune cells contribute to the IFN-gamma-dependent regulation of antigen-specific CD8+ T cell homeostasis. J Immunol. 2006;176(2):735–9.

20. Curtsinger JM, Agarwal P, Lins DC, Mescher MF. Autocrine IFN-gamma promotes naive CD8 T cell differentiation and synergizes with IFN-alpha to stimulate strong function. J Immunol. 2012;189(2):659–68.

21. Gagnon J, Ramanathan S, Leblanc C, Ilangumaran S. Regulation of IL-21 signaling by suppressor of cytokine signaling-1 (SOCS1) in CD8(+) T lymphocytes. Cell Signal. 2007;19(4):806–16.

22. Gagnon J, Ramanathan S, Leblanc C, Cloutier A, McDonald PP, Ilangumaran S. IL-6, in Synergy with IL-7 or IL-15, Stimulates TCR-Independent Proliferation and Functional Differentiation of CD8+ T Lymphocytes. J Immunol. 2008;180(12):7958–68.

23. Zeng R, Spolski R, Finkelstein SE, Oh S, Kovanen PE, Hinrichs CS, et al. Synergy of IL-21 and IL-15 in regulating CD8+ T cell expansion and function. The Journal of experimental medicine. 2005;201(1):139–48.

24. Sawa Y, Arima Y, Ogura H, Kitabayashi C, Jiang JJ, Fukushima T, et al. Hepatic interleukin-7 expression regulates T cell responses. Immunity. 2009;30(3):447–57.

25. Mattei F, Schiavoni G, Belardelli F, Tough DF. IL-15 is expressed by dendritic cells in response to type I IFN, double-stranded RNA, or lipopolysaccharide and promotes dendritic cell activation. J Immunol. 2001;167(3):1179–87.

26. Ramanathan S, Dubois S, Chen XL, Leblanc C, Ohashi PS, Ilangumaran S. Exposure to IL-15 and IL-21 enables autoreactive CD8 T cells to respond to weak antigens and cause disease in a mouse model of autoimmune diabetes. J Immunol. 2011;186(9):5131–41.

27. Rodriguez GM, D’Urbano D, Bobbala D, Chen XL, Yeganeh M, Ramanathan S, et al. SOCS1 Prevents Potentially Skin-Reactive Cytotoxic T Lymphocytes from Gaining the Ability to Cause Inflammatory Lesions. The Journal of investigative dermatology. 2013;133(8):2013–22.

28. Richer MJ, Nolz JC, Harty JT. Pathogen-specific inflammatory milieux tune the antigen sensitivity of CD8(+) T cells by enhancing T cell receptor signaling. Immunity. 2013;38(1):140–52.

29. Ramanathan S, Gagnon J, Dubois S, Forand-Boulerice M, Richter MV, Ilangumaran S. Cytokine Synergy in Antigen-Independent Activation and Priming of Naive CD8+ T Lymphocytes. Critical reviews in immunology. 2009;29(3):219–39.

30. Gagnon J, Chen XL, Forand-Boulerice M, Leblanc C, Raman C, Ramanathan S, et al. Increased antigen responsiveness of naive CD8 T cells exposed to IL-7 and IL-21 is associated with decreased CD5 expression. Immunol Cell Biol. 2010;88(4):451–60.

31. Overwijk WW, Restifo NP. B16 as a mouse model for human melanoma. Current protocols in immunology / edited by John E Coligan [et al]. 2001;Chapter 20:Unit 201.

32. Overwijk WW, Tsung A, Irvine KR, Parkhurst MR, Goletz TJ, Tsung K, et al. gp100/pmel 17 is a murine tumor rejection antigen: induction of “self”-reactive, tumoricidal T cells using high-affinity, altered peptide ligand. The Journal of experimental medicine. 1998;188(2):277–86.

33. Buenrostro JD, Giresi PG, Zaba LC, Chang HY, Greenleaf WJ. Transposition of native chromatin for fast and sensitive epigenomic profiling of open chromatin, DNA-binding proteins and nucleosome position. Nat Methods. 2013;10(12):1213–8.

34. Buenrostro JD, Wu B, Chang HY, Greenleaf WJ. ATAC-seq: A Method for Assaying Chromatin Accessibility Genome-Wide. Curr Protoc Mol Biol. 2015;109:21 9 1–9.

35. Kim S, Yu NK, Kaang BK. CTCF as a multifunctional protein in genome regulation and gene expression. Exp Mol Med. 2015;47:e166.

36. Bolger AM, Lohse M, Usadel B. Trimmomatic: a flexible trimmer for Illumina sequence data. Bioinformatics. 2014;30(15):2114–20.

37. Neph S, Kuehn MS, Reynolds AP, Haugen E, Thurman RE, Johnson AK, et al. BEDOPS: high-performance genomic feature operations. Bioinformatics. 2012;28(14):1919–20.

38. Quinlan AR. BEDTools: The Swiss-Army Tool for Genome Feature Analysis. Curr Protoc Bioinformatics. 2014;47:11 2 1–34.

39. Love MI, Huber W, Anders S. Moderated estimation of fold change and dispersion for RNA-seq data with DESeq2. Genome Biol. 2014;15(12):550.

40. Heinz S, Benner C, Spann N, Bertolino E, Lin YC, Laslo P, et al. Simple combinations of lineage-determining transcription factors prime cis-regulatory elements required for macrophage and B cell identities. Mol Cell. 2010;38(4):576–89.

41. Szklarczyk D, Gable AL, Lyon D, Junge A, Wyder S, Huerta-Cepas J, et al. STRING v11: protein-protein association networks with increased coverage, supporting functional discovery in genome-wide experimental datasets. Nucleic Acids Res. 2019;47(D1):D607–D13.

42. Verdeil G, Chaix J, Schmitt-Verhulst AM, Auphan-Anezin N. Temporal cross-talk between TCR and STAT signals for CD8 T cell effector differentiation. European journal of immunology. 2006;36(12):3090–100.

43. Verdeil G, Puthier D, Nguyen C, Schmitt-Verhulst AM, Auphan-Anezin N. STAT5-mediated signals sustain a TCR-initiated gene expression program toward differentiation of CD8 T cell effectors. J Immunol. 2006;176(8):4834–42.

44. Bezbradica JS, Medzhitov R. Integration of cytokine and heterologous receptor signaling pathways. Nature immunology. 2009;10(4):333–9.

45. Shi M, Lin TH, Appell KC, Berg LJ. Janus-kinase-3-dependent signals induce chromatin remodeling at the Ifng locus during T helper 1 cell differentiation. Immunity. 2008;28(6):763–73.

46. Zhu J, Cote-Sierra J, Guo L, Paul WE. Stat5 activation plays a critical role in Th2 differentiation. Immunity. 2003;19(5):739–48.

47. Scott-Browne JP, Lopez-Moyado IF, Trifari S, Wong V, Chavez L, Rao A, et al. Dynamic Changes in Chromatin Accessibility Occur in CD8(+) T Cells Responding to Viral Infection. Immunity. 2016;45(6):1327–40.

48. Voisinne G, Gonzalez de Peredo A, Roncagalli R. CD5, an Undercover Regulator of TCR Signaling. Front Immunol. 2018;9:2900.

49. Nolz JC, Harty JT. IL-15 regulates memory CD8+ T cell O-glycan synthesis and affects trafficking. J Clin Invest. 2014;124(3):1013–26.

50. Conley JM, Gallagher MP, Berg LJ. T Cells and Gene Regulation: The Switching On and Turning Up of Genes after T Cell Receptor Stimulation in CD8 T Cells. Front Immunol. 2016;7:76.

51. Mescher MF, Curtsinger JM, Agarwal P, Casey KA, Gerner M, Hammerbeck CD, et al. Signals required for programming effector and memory development by CD8+ T cells. Immunol Rev. 2006;211:81–92.

52. Cox MA, Harrington LE, Zajac AJ. Cytokines and the inception of CD8 T cell responses. Trends Immunol. 2011;32(4):180–6.

53. Valbon SF, Condotta SA, Richer MJ. Regulation of effector and memory CD8(+) T cell function by inflammatory cytokines. Cytokine. 2016;82:16–23.

54. Agarwal P, Raghavan A, Nandiwada SL, Curtsinger JM, Bohjanen PR, Mueller DL, et al. Gene regulation and chromatin remodeling by IL-12 and type I IFN in programming for CD8 T cell effector function and memory. J Immunol. 2009;183(3):1695–704.

55. Raue HP, Beadling C, Haun J, Slifka MK. Cytokine-mediated programmed proliferation of virus-specific CD8(+) memory T cells. Immunity. 2013;38(1):131–9.

56. Ramanathan S, Dubois S, Gagnon J, Leblanc C, Mariathasan S, Ferbeyre G, et al. Regulation of cytokine-driven functional differentiation of CD8 T cells by suppressor of cytokine signaling 1 controls autoimmunity and preserves their proliferative capacity toward foreign antigens. J Immunol. 2010;185(1):357–66.

57. Jenkins MR, Mintern J, La Gruta NL, Kedzierska K, Doherty PC, Turner SJ. Cell cycle-related acquisition of cytotoxic mediators defines the progressive differentiation to effector status for virus-specific CD8+ T cells. J Immunol. 2008;181(6):3818–22.

58. Plumlee CR, Sheridan BS, Cicek BB, Lefrancois L. Environmental cues dictate the fate of individual CD8+ T cells responding to infection. Immunity. 2013;39(2):347–56.

59. Halle S, Keyser KA, Stahl FR, Busche A, Marquardt A, Zheng X, et al. In Vivo Killing Capacity of Cytotoxic T Cells Is Limited and Involves Dynamic Interactions and T Cell Cooperativity. Immunity. 2016;44(2):233–45.

60. Tubo NJ, Jenkins MK. TCR signal quantity and quality in CD4(+) T cell differentiation. Trends Immunol. 2014;35(12):591–6.

61. Tubo NJ, Pagan AJ, Taylor JJ, Nelson RW, Linehan JL, Ertelt JM, et al. Single naive CD4+ T cells from a diverse repertoire produce different effector cell types during infection. Cell. 2013;153(4):785–96.

62. van Panhuys N, Klauschen F, Germain RN. T-cell-receptor-dependent signal intensity dominantly controls CD4(+) T cell polarization In Vivo. Immunity. 2014;41(1):63–74.

63. Newell EW, Sigal N, Bendall SC, Nolan GP, Davis MM. Cytometry by time-of-flight shows combinatorial cytokine expression and virus-specific cell niches within a continuum of CD8+ T cell phenotypes. Immunity. 2012;36(1):142–52.

64. Wilson EB, Brooks DG. Inflammation makes T cells sensitive. Immunity. 2013;38(1):5–7.

65. Keppler SJ, Theil K, Vucikuja S, Aichele P. Effector T-cell differentiation during viral and bacterial infections: Role of direct IL-12 signals for cell fate decision of CD8(+) T cells. European journal of immunology. 2009;39(7):1774–83.

66. Krupica T, Jr., Fry TJ, Mackall CL. Autoimmunity during lymphopenia: a two-hit model. Clin Immunol. 2006;120(2):121–8.

67. Fujinami RS, von Herrath MG, Christen U, Whitton JL. Molecular mimicry, bystander activation, or viral persistence: infections and autoimmune disease. Clin Microbiol Rev. 2006;19(1):80–94.

68. Ellestad KK, Anderson CC. Two Strikes and You’re Out? The Pathogenic Interplay of Coinhibitor Deficiency and Lymphopenia-Induced Proliferation. J Immunol. 2017;198(7):2534–41.

69. Horwitz MS, Yanagi Y, Oldstone MB. T-cell receptors from virus-specific cytotoxic T lymphocytes recognizing a single immunodominant nine-amino-acid viral epitope show marked diversity. J Virol. 1994;68(1):352–7.

70. Zehn D, Lee SY, Bevan MJ. Complete but curtailed T-cell response to very low-affinity antigen. Nature. 2009;458(7235):211–4.

71. van Gisbergen KP, Klarenbeek PL, Kragten NA, Unger PP, Nieuwenhuis MB, Wensveen FM, et al. The costimulatory molecule CD27 maintains clonally diverse CD8(+) T cell responses of low antigen affinity to protect against viral variants. Immunity. 2011;35(1):97–108.

72. Martinez RJ, Evavold BD. Lower Affinity T Cells are Critical Components and Active Participants of the Immune Response. Front Immunol. 2015;6:468.

73. Huang J, Zeng X, Sigal N, Lund PJ, Su LF, Huang H, et al. Detection, phenotyping, and quantification of antigen-specific T cells using a peptide-MHC dodecamer. Proc Natl Acad Sci U S A. 2016;113(13):E1890–7.

74. Ozga AJ, Moalli F, Abe J, Swoger J, Sharpe J, Zehn D, et al. pMHC affinity controls duration of CD8+ T cell-DC interactions and imprints timing of effector differentiation versus expansion. The Journal of experimental medicine. 2016;213(12):2811–29.

75. Schober K, Voit F, Grassmann S, Muller TR, Eggert J, Jarosch S, et al. Reverse TCR repertoire evolution toward dominant low-affinity clones during chronic CMV infection. Nature immunology. 2020;21(4):434–41.

76. Hebeisen M, Allard M, Gannon PO, Schmidt J, Speiser DE, Rufer N. Identifying Individual T Cell Receptors of Optimal Avidity for Tumor Antigens. Front Immunol. 2015;6:582.

77. Miller AM, Bahmanof M, Zehn D, Cohen EEW, Schoenberger SP. Leveraging TCR Affinity in Adoptive Immunotherapy against Shared Tumor/Self-Antigens. Cancer Immunol Res. 2019;7(1):40–9.

78. Segal G, Prato S, Zehn D, Mintern JD, Villadangos JA. Target Density, Not Affinity or Avidity of Antigen Recognition, Determines Adoptive T Cell Therapy Outcomes in a Mouse Lymphoma Model. J Immunol. 2016;196(9):3935–42.

79. Rodriguez GM, Bobbala D, Serrano D, Mayhue M, Champagne A, Saucier C, et al. NLRC5 elicits antitumor immunity by enhancing processing and presentation of tumor antigens to CD8(+) T lymphocytes. Oncoimmunology. 2016;5(6):e1151593.

80. He Q, Jiang X, Zhou X, Weng J. Targeting cancers through TCR-peptide/MHC interactions. J Hematol Oncol. 2019;12(1):139.

81. Scharer CD, Bally AP, Gandham B, Boss JM. Cutting Edge: Chromatin Accessibility Programs CD8 T Cell Memory. J Immunol. 2017;198(6):2238–43.

82. Mognol GP, Spreafico R, Wong V, Scott-Browne JP, Togher S, Hoffmann A, et al. Exhaustion-associated regulatory regions in CD8(+) tumor-infiltrating T cells. Proc Natl Acad Sci U S A. 2017;114(13):E2776–E85.

83. Huber M, Lohoff M. IRF4 at the crossroads of effector T-cell fate decision. European journal of immunology. 2014;44(7):1886–95.

84. Huang S, Shen Y, Pham D, Jiang L, Wang Z, Kaplan MH, et al. IRF4 Modulates CD8(+) T Cell Sensitivity to IL-2 Family Cytokines. Immunohorizons. 2017;1(6):92–100.

85. Best JA, Blair DA, Knell J, Yang E, Mayya V, Doedens A, et al. Transcriptional insights into the CD8(+) T cell response to infection and memory T cell formation. Nature immunology. 2013;14(4):404–12.

86. Spolski R, Gromer D, Leonard WJ. The gamma c family of cytokines: fine-tuning signals from IL-2 and IL-21 in the regulation of the immune response. F1000Res. 2017;6:1872.

87. Shan Q, Zeng Z, Xing S, Li F, Hartwig SM, Gullicksrud JA, et al. The transcription factor Runx3 guards cytotoxic CD8(+) effector T cells against deviation towards follicular helper T cell lineage. Nature immunology. 2017;18(8):931–9.

88. Woolf E, Xiao C, Fainaru O, Lotem J, Rosen D, Negreanu V, et al. Runx3 and Runx1 are required for CD8 T cell development during thymopoiesis. Proc Natl Acad Sci U S A. 2003;100(13):7731–6.

89. Cruz-Guilloty F, Pipkin ME, Djuretic IM, Levanon D, Lotem J, Lichtenheld MG, et al. Runx3 and T-box proteins cooperate to establish the transcriptional program of effector CTLs. The Journal of experimental medicine. 2009;206(1):51–9.

90. Wang D, Diao H, Getzler AJ, Rogal W, Frederick MA, Milner J, et al. The Transcription Factor Runx3 Establishes Chromatin Accessibility of cis-Regulatory Landscapes that Drive Memory Cytotoxic T Lymphocyte Formation. Immunity. 2018;48(4):659–74 e6.

91. Hai T, Hartman MG. The molecular biology and nomenclature of the activating transcription factor/cAMP responsive element binding family of transcription factors: activating transcription factor proteins and homeostasis. Gene. 2001;273(1):1–11.

92. Kurachi M, Barnitz RA, Yosef N, Odorizzi PM, DiIorio MA, Lemieux ME, et al. The transcription factor BATF operates as an essential differentiation checkpoint in early effector CD8+ T cells. Nature immunology. 2014;15(4):373–83.

## References

1. Saeki K, Fukuyama S, Ayada T, Nakaya M, Aki D, Takaesu G, et al. A major lipid raft protein raftlin modulates T cell receptor signaling and enhances th17-mediated autoimmune responses. J Immunol. 2009;182(10):5929–37.

2. Rellahan BL, Jensen JP, Howcroft TK, Singer DS, Bonvini E, Weissman AM. Elf-1 regulates basal expression from the T cell antigen receptor zeta-chain gene promoter. J Immunol. 1998;160(6):2794–801.

3. Ohnuma K, Uchiyama M, Yamochi T, Nishibashi K, Hosono O, Takahashi N, et al. Caveolin-1 triggers T-cell activation via CD26 in association with CARMA1. J Biol Chem. 2007;282(13):10117–31.

4. Nayar R, Enos M, Prince A, Shin H, Hemmers S, Jiang JK, et al. TCR signaling via Tec kinase ITK and interferon regulatory factor 4 (IRF4) regulates CD8+ T-cell differentiation. Proc Natl Acad Sci U S A. 2012;109(41):E2794–802.

5. Pfeifhofer C, Kofler K, Gruber T, Tabrizi NG, Lutz C, Maly K, et al. Protein kinase C theta affects Ca2+ mobilization and NFAT cell activation in primary mouse T cells. The Journal of experimental medicine. 2003;197(11):1525–35.

6. Ruland J, Duncan GS, Elia A, del Barco Barrantes I, Nguyen L, Plyte S, et al. Bcl10 is a positive regulator of antigen receptor-induced activation of NF-kappaB and neural tube closure. Cell. 2001;104(1):33–42.

7. Liu ZZ, Wang ZL, Choi TI, Huang WT, Wang HT, Han YY, et al. Chd7 Is Critical for Early T-Cell Development and Thymus Organogenesis in Zebrafish. Am J Pathol. 2018;188(4):1043–58.

8. Finco TS, Justice-Healy GE, Patel SJ, Hamilton VE. Regulation of the human LAT gene by the Elf-1 transcription factor. BMC Mol Biol. 2006;7:4.

9. Riera-Sans L, Behrens A. Regulation of alphabeta/gammadelta T cell development by the activator protein 1 transcription factor c-Jun. J Immunol. 2007;178(9):5690–700.

10. Sallusto F, Kremmer E, Palermo B, Hoy A, Ponath P, Qin S, et al. Switch in chemokine receptor expression upon TCR stimulation reveals novel homing potential for recently activated T cells. European journal of immunology. 1999;29(6):2037–45.

11. Schaller MA, Kallal LE, Lukacs NW. A key role for CC chemokine receptor 1 in T-cell-mediated respiratory inflammation. Am J Pathol. 2008;172(2):386–94.

12. Jung YW, Kim HG, Perry CJ, Kaech SM. CCR7 expression alters memory CD8 T-cell homeostasis by regulating occupancy in IL-7- and IL-15-dependent niches. Proc Natl Acad Sci U S A. 2016;113(29):8278–83.

13. Acuto O, Michel F. CD28-mediated co-stimulation: a quantitative support for TCR signalling. Nat Rev Immunol. 2003;3(12):939–51.

14. Zhang Y, Blattman JN, Kennedy NJ, Duong J, Nguyen T, Wang Y, et al. Regulation of innate and adaptive immune responses by MAP kinase phosphatase 5. Nature. 2004;430(7001):793–7.

15. Wei H, Geng J, Shi B, Liu Z, Wang YH, Stevens AC, et al. Cutting Edge: Foxp1 Controls Naive CD8+ T Cell Quiescence by Simultaneously Repressing Key Pathways in Cellular Metabolism and Cell Cycle Progression. J Immunol. 2016;196(9):3537–41.

16. Ju S, Zhu Y, Liu L, Dai S, Li C, Chen E, et al. Gadd45b and Gadd45g are important for anti-tumor immune responses. European journal of immunology. 2009;39(11):3010–8.

17. Sumida H, Cyster JG. G-Protein Coupled Receptor 18 Contributes to Establishment of the CD8 Effector T Cell Compartment. Front Immunol. 2018;9:660.

18. Schluns KS, Williams K, Ma A, Zheng XX, Lefrancois L. Cutting edge: requirement for IL-15 in the generation of primary and memory antigen-specific CD8 T cells. J Immunol. 2002;168(10):4827–31.

19. Harker JA, Wong KA, Dolgoter A, Zuniga EI. Cell-Intrinsic gp130 Signaling on CD4+ T Cells Shapes Long-Lasting Antiviral Immunity. J Immunol. 2015;195(3):1071–81.

20. Charlton JJ, Chatzidakis I, Tsoukatou D, Boumpas DT, Garinis GA, Mamalaki C. Programmed death-1 shapes memory phenotype CD8 T cell subsets in a cell-intrinsic manner. J Immunol. 2013;190(12):6104–14.

21. Willinger T, Freeman T, Herbert M, Hasegawa H, McMichael AJ, Callan MF. Human naive CD8 T cells down-regulate expression of the WNT pathway transcription factors lymphoid enhancer binding factor 1 and transcription factor 7 (T cell factor-1) following antigen encounter in vitro and in vivo. J Immunol. 2006;176(3):1439–46.

22. MaruYama T. TGF-beta-induced IkappaB-zeta controls Foxp3 gene expression. Biochem Biophys Res Commun. 2015;464(2):586–9.

23. Meisel M, Hermann-Kleiter N, Hinterleitner R, Gruber T, Wachowicz K, Pfeifhofer-Obermair C, et al. The kinase PKCalpha selectively upregulates interleukin-17A during Th17 cell immune responses. Immunity. 2013;38(1):41–52.

24. Guo Y, Lee YC, Brown C, Zhang W, Usherwood E, Noelle RJ. Dissecting the role of retinoic acid receptor isoforms in the CD8 response to infection. J Immunol. 2014;192(7):3336–44.

25. Tripathi P, Kurtulus S, Wojciechowski S, Sholl A, Hoebe K, Morris SC, et al. STAT5 is critical to maintain effector CD8+ T cell responses. J Immunol. 2010;185(4):2116–24.

26. Ma C, Zhang N. Transforming growth factor-beta signaling is constantly shaping memory T-cell population. Proc Natl Acad Sci U S A. 2015;112(35):11013–7.

27. Nishimura H, Yajima T, Muta H, Podack ER, Tani K, Yoshikai Y. A novel role of CD30/CD30 ligand signaling in the generation of long-lived memory CD8+ T cells. J Immunol. 2005;175(7):4627–34.

28. Ritthipichai K, Haymaker CL, Martinez M, Aschenbrenner A, Yi X, Zhang M, et al. Multifaceted Role of BTLA in the Control of CD8(+) T-cell Fate after Antigen Encounter. Clin Cancer Res. 2017;23(20):6151–64.

29. Shamim M, Nanjappa SG, Singh A, Plisch EH, LeBlanc SE, Walent J, et al. Cbl-b regulates antigen-induced TCR down-regulation and IFN-gamma production by effector CD8 T cells without affecting functional avidity. J Immunol. 2007;179(11):7233–43.

30. Soares LR, Tsavaler L, Rivas A, Engleman EG. V7 (CD101) ligation inhibits TCR/CD3-induced IL-2 production by blocking Ca2+ flux and nuclear factor of activated T cell nuclear translocation. J Immunol. 1998;161(1):209–17.

31. DeBell KE, Simhadri VR, Mariano JL, Borrego F. Functional requirements for inhibitory signal transmission by the immunomodulatory receptor CD300a. BMC Immunol. 2012;13:23.

32. Bouguermouh S, Van VQ, Martel J, Gautier P, Rubio M, Sarfati M. CD47 expression on T cell is a self-control negative regulator of type 1 immune response. J Immunol. 2008;180(12):8073–82.

33. Chauvin JM, Pagliano O, Fourcade J, Sun Z, Wang H, Sander C, et al. TIGIT and PD-1 impair tumor antigen-specific CD8(+) T cells in melanoma patients. J Clin Invest. 2015;125(5):2046–58.

34. Waugh KA, Leach SM, Moore BL, Bruno TC, Buhrman JD, Slansky JE. Molecular Profile of Tumor-Specific CD8+ T Cell Hypofunction in a Transplantable Murine Cancer Model. J Immunol. 2016;197(4):1477–88.

35. Kamimura D, Bevan MJ. Endoplasmic reticulum stress regulator XBP-1 contributes to effector CD8+ T cell differentiation during acute infection. J Immunol. 2008;181(8):5433–41.

36. Ma X, Bi E, Lu Y, Su P, Huang C, Liu L, et al. Cholesterol Induces CD8(+) T Cell Exhaustion in the Tumor Microenvironment. Cell Metab. 2019;30(1):143–56 e5.

37. Kim H, Kim T, Jeong BC, Cho IT, Han D, Takegahara N, et al. Tmem64 modulates calcium signaling during RANKL-mediated osteoclast differentiation. Cell Metab. 2013;17(2):249–60.

38. Bando JK, Gilfillan S, Song C, McDonald KG, Huang SC, Newberry RD, et al. The Tumor Necrosis Factor Superfamily Member RANKL Suppresses Effector Cytokine Production in Group 3 Innate Lymphoid Cells. Immunity. 2018;48(6):1208–19 e4.

39. Reed NP, Henderson MA, Oltz EM, Aune TM. Reciprocal regulation of Rag expression in thymocytes by the zinc-finger proteins, Zfp608 and Zfp609. Genes Immun. 2013;14(1):7–12.

