## Supplementary Tables for "Cytokine priming of naïve CD8^+^ T lymphocytes modulates chromatin accessibility that partially overlaps with changes induced by antigen simulation"

**Table S1A. HOMER Opening peaks in Cytokine-primed cells**

Data is sorted alphabetically by the gene name; Only peaks above the peak score value of 0.5 are shown.

| PeakID (cmd=annotatePeaks.pl ../opening_peaks_BA.bed mm10 -go go_opening_peaks_BA) | Chr | Start | End | Strand | Peak Score | Annotation | Detailed Annotation | Distance to TSS | Nearest PromoterID | Entrez ID | Nearest Unigene | Nearest Refseq | Nearest Ensembl | Gene Name | Gene Alias | Gene Description | Gene Type |
|---|---|---|---|---|---|---|---|---|---|---|---|---|---|---|---|---|---|
| chr9_94562991_94563677 | chr9 | 94562992 | 94563677 | + | 0.630204 | Intergenic | Intergenic | -25253 | NM_001033145 | 68861 | Mm.258746 | NM_001033145 | ENSMUSG00000045414 | **1190002N15Rik** | GoPro49\|HASF | RIKEN cDNA 1190002N15 gene | protein-coding |
| chr13_5666242_5666569 | chr13 | 5666243 | 5666569 | + | 0.721066 | Intergenic | Intergenic | 191686 | NR_045891 | 71867 | Mm.179181 | NR_045891 | | **1700016G22Rik** | - | RIKEN cDNA 1700016G22 gene | ncRNA |
| chr2_46371045_46371404 | chr2 | 46371046 | 46371404 | + | 0.734234 | Intergenic | Intergenic | 674620 | NR_040497 | 69405 | Mm.396184 | NR_040497 | ENSMUSG00000086202 | **1700019E08Rik** | - | RIKEN cDNA 1700019E08 gene | ncRNA |
| chr2_46370821_46371458 | chr2 | 46370822 | 46371458 | + | 0.725918 | Intergenic | Intergenic | 674535 | NR_040497 | 69405 | Mm.396184 | NR_040497 | ENSMUSG00000086202 | **1700019E08Rik** | - | RIKEN cDNA 1700019E08 gene | ncRNA |
| chr1_183962301_183962948 | chr1 | 183962302 | 183962948 | + | 0.620688 | Intergenic | Intergenic | 71373 | NM_028516 | 73363 | Mm.50115 | NM_028516 | ENSMUSG00000044854 | **1700056E22Rik** | - | RIKEN cDNA 1700056E22 gene | protein-coding |
| chr14_54675212_54675674 | chr14 | 54675213 | 54675674 | + | 0.811176 | intron (NM_001085473, intron 4 of 18) | B1_Mur3\|SINE\|Alu | -10728 | NM_021437 | 58248 | Mm.197509 | NM_021437 | ENSMUSG00000040822 | **1700123O20Rik** | - | RIKEN cDNA 1700123O20 gene | protein-coding |
| chr12_12547387_12548252 | chr12 | 12547388 | 12548252 | + | 0.801045 | Intergenic | Intergenic | -155345 | NR_045829 | 70858 | Mm.410838 | NR_045829 | | **4921511I17Rik** | - | RIKEN cDNA 4921511I17 gene | ncRNA |
| chr16_64766690_64767290 | chr16 | 64766691 | 64767290 | + | 0.630606 | exon (NM_026273, exon 3 of 3) | exon (NM_026273, exon 3 of 3) | 3949 | NM_026273 | 67609 | Mm.212157 | NM_026273 | ENSMUSG00000059920 | **4930453N24Rik** | AI649104 | RIKEN cDNA 4930453N24 gene | protein-coding |
| chr10_96879557_96880067 | chr10 | 96879558 | 96880067 | + | 0.638227 | Intergenic | Intergenic | -155956 | NR_045358 | 75301 | Mm.108650 | NR_045358 | | **4930556N09Rik** | - | RIKEN cDNA 4930556N09 gene | ncRNA |
| chr2_52310704_52311206 | chr2 | 52310705 | 52311206 | + | 0.748639 | intron (NM_010889, intron 12 of 156) | intron (NM_010889, intron 12 of 156) | 19953 | NR_040620 | 78049 | Mm.243883 | NR_040620 | | **4930573O16Rik** | - | RIKEN cDNA 4930573O16 gene | ncRNA |
| chr9_118360838_118361565 | chr9 | 118360839 | 118361565 | + | 0.649248 | Intergenic | Intergenic | -68637 | NR_040716 | 71196 | Mm.73218 | NR_040716 | | **4933432G23Rik** | - | RIKEN cDNA 4933432G23 gene | ncRNA |
| chr9_118400636_118401020 | chr9 | 118400637 | 118401020 | + | 0.646094 | Intergenic | Intergenic | -29011 | NR_040716 | 71196 | Mm.73218 | NR_040716 | | **4933432G23Rik** | - | RIKEN cDNA 4933432G23 gene | ncRNA |
| chr2_167804760_167805647 | chr2 | 167804761 | 167805647 | + | 0.739567 | Intergenic | Intergenic | 114027 | NR_015500 | 329562 | Mm.123101 | NM_177851 | ENSMUSG00000006462 | **A530013C23Rik** | - | RIKEN cDNA A530013C23 gene | ncRNA |
| chr3_115857750_115858282 | chr3 | 115857751 | 115858282 | + | 0.695971 | Intergenic | Intergenic | 30043 | NR_027894 | 68161 | Mm.392043 | NR_015487 | | **A930005H10Rik** | - | RIKEN cDNA A930005H10 gene | ncRNA |
| chr3_115857603_115858461 | chr3 | 115857604 | 115858461 | + | 0.648701 | Intergenic | Intergenic | 30027 | NR_027894 | 68161 | Mm.392043 | NR_015487 | | **A930005H10Rik** | - | RIKEN cDNA A930005H10 gene | ncRNA |
| chr13_8457865_8458308 | chr13 | 8457866 | 8458308 | + | 0.720708 | intron (NM_001289530, intron 1 of 9) | intron (NM_001289530, intron 1 of 9) | 255221 | NM_052977 | 94191 | Mm.425794 | NM_052977 | ENSMUSG00000052551 | **Adarb2** | Adar3\|RED2 | adenosine deaminase, RNA-specific, B2 | protein-coding |
| chr3_20340469_20340968 | chr3 | 20340470 | 20340968 | + | 0.599635 | intron (NM_175086, intron 1 of 1) | L1MEc\|LINE\|L1 | 26458 | NM_175086 | 11608 | Mm.134863 | NM_175086 | ENSMUSG00000054988 | **Agtr1b** | AT1B\|AT2R1B\|Agtr-1b\|Angtr-1b | angiotensin II receptor, type 1b | protein-coding |
| chr3_107047867_107048534 | chr3 | 107047868 | 107048534 | + | 0.607763 | intron (NR_033498, intron 1 of 1) | intron (NR_033498, intron 1 of 1) | 8697 | NR_033498 | 229694 | Mm.347584 | NM_173787 | ENSMUSG00000056145 | **AI504432** | A230102G19 | expressed sequence AI504432 | ncRNA |
| chr7_27609565_27610156 | chr7 | 27609566 | 27610156 | + | 0.622262 | intron (NM_001110208, intron 2 of 13) | intron (NM_001110208, intron 2 of 13) | 4001 | NM_007434 | 11652 | Mm.177194 | NM_007434 | ENSMUSG00000004056 | **Akt2** | 2410016A19Rik\|AW554154\|PKB\|PKBbeta | thymoma viral proto-oncogene 2 | protein-coding |
| chr7_27609737_27610012 | chr7 | 27609738 | 27610012 | + | 0.618092 | intron (NM_001110208, intron 2 of 13) | intron (NM_001110208, intron 2 of 13) | 4015 | NM_007434 | 11652 | Mm.177194 | NM_007434 | ENSMUSG00000004056 | **Akt2** | 2410016A19Rik\|AW554154\|PKB\|PKBbeta | thymoma viral proto-oncogene 2 | protein-coding |
| chr7_27609367_27610166 | chr7 | 27609368 | 27610166 | + | 0.589094 | intron (NM_001110208, intron 2 of 13) | L2b\|LINE\|L2 | 3907 | NM_007434 | 11652 | Mm.177194 | NM_007434 | ENSMUSG00000004056 | **Akt2** | 2410016A19Rik\|AW554154\|PKB\|PKBbeta | thymoma viral proto-oncogene 2 | protein-coding |
| chrX_142907468_142908077 | chrX | 142907469 | 142908077 | + | 0.897741 | intron (NM_019496, intron 2 of 5) | intron (NM_019496, intron 2 of 5) | 58955 | NM_019496 | 56068 | Mm.143724 | NM_019496 | | **Ammecr1** | 6230420G18Rik | Alport syndrome, mental retardation, midface hypoplasia and elliptocytosis chromosomal region gene 1 | protein-coding |
| chr7_58475100_58475677 | chr7 | 58475101 | 58475677 | + | 0.856517 | Intergenic | Intergenic | -182813 | NM_009728 | 11982 | Mm.135129 | NM_009728 | ENSMUSG00000025324 | **Atp10a** | Atp10c\|pfatp | ATPase, class V, type 10A | protein-coding |
| chr3_101570263_101571144 | chr3 | 101570264 | 101571144 | + | 0.741426 | Intergenic | RLTR9A\|LTR\|ERVK | 34003 | NM_144900 | 11928 | Mm.280103 | NM_144900 | ENSMUSG00000033161 | **Atp1a1** | Atpa-1\|BC010319 | ATPase, Na+/K+ transporting, alpha 1 polypeptide | protein-coding |
| chr3_101570731_101571134 | chr3 | 101570732 | 101571134 | + | 0.604042 | Intergenic | ORR1B2\|LTR\|ERVL-MaLR | 33774 | NM_144900 | 11928 | Mm.280103 | NM_144900 | ENSMUSG00000033161 | **Atp1a1** | Atpa-1\|BC010319 | ATPase, Na+/K+ transporting, alpha 1 polypeptide | protein-coding |
| chr8_57592722_57592917 | chr8 | 57592723 | 57592917 | + | 0.590167 | intron (NM_001167981, intron 1 of 11) | B3A\|SINE\|B2 | -58933 | NR_040698 | 100502619 | Mm.393851 | NR_040698 | | **AW046200** | - | expressed sequence AW046200 | ncRNA |
| chr17_48318811_48319083 | chr17 | 48318812 | 48319083 | + | 0.771546 | intron (NM_177083, intron 2 of 5) | intron (NM_177083, intron 2 of 5) | 2785 | NM_177083 | 320148 | Mm.123455 | NM_177083 | ENSMUSG00000043740 | **B430306N03Rik** | Treml6 | RIKEN cDNA B430306N03 gene | protein-coding |
| chr17_48318698_48319311 | chr17 | 48318699 | 48319311 | + | 0.710582 | intron (NM_177083, intron 2 of 5) | intron (NM_177083, intron 2 of 5) | 2843 | NM_177083 | 320148 | Mm.123455 | NM_177083 | ENSMUSG00000043740 | **B430306N03Rik** | Treml6 | RIKEN cDNA B430306N03 gene | protein-coding |
| chr6_144922800_144923124 | chr6 | 144922801 | 144923124 | + | 0.595069 | Intergenic | Intergenic | 125850 | NM_001024468 | 12035 | Mm.4606 | NM_007532 | ENSMUSG00000030268 | **Bcat1** | BCATc\|Eca39 | branched chain aminotransferase 1, cytosolic | protein-coding |
| chr16_45261281_45261674 | chr16 | 45261282 | 45261674 | + | 0.686686 | Intergenic | ORR1B1\|LTR\|ERVL-MaLR | 37141 | NM_177584 | 208154 | Mm.38199 | NM_177584 | ENSMUSG00000052013 | **Btla** | A630002H24 | B and T lymphocyte associated | protein-coding |
| chr16_45261225_45261974 | chr16 | 45261226 | 45261974 | + | 0.625697 | Intergenic | Intergenic | 37263 | NM_177584 | 208154 | Mm.38199 | NM_177584 | ENSMUSG00000052013 | **Btla** | A630002H24 | B and T lymphocyte associated | protein-coding |
| chr1_151105181_151105611 | chr1 | 151105182 | 151105611 | + | 0.633366 | Intergenic | Intergenic | -32638 | NR_038011 | 402734 | Mm.337666 | NR_038011 | | **C730036E19Rik** | - | RIKEN cDNA C730036E19 gene | ncRNA |
| chr16_45038590_45039050 | chr16 | 45038591 | 45039050 | + | 0.701492 | Intergenic | Intergenic | -55233 | NM_026439 | 67896 | Mm.181074 | NM_026439 | ENSMUSG00000022665 | **Ccdc80** | 2610001E17Rik\|DRO1\|Ssg1\|Urb | coiled-coil domain containing 80 | protein-coding |
| chr16_45038390_45039051 | chr16 | 45038391 | 45039051 | + | 0.659909 | Intergenic | MLT2B2\|LTR\|ERVL | -55332 | NM_026439 | 67896 | Mm.181074 | NM_026439 | ENSMUSG00000022665 | **Ccdc80** | 2610001E17Rik\|DRO1\|Ssg1\|Urb | coiled-coil domain containing 80 | protein-coding |
| chr7_38090339_38090645 | chr7 | 38090340 | 38090645 | + | 0.716626 | Intergenic | Intergenic | 16998 | NM_007633 | 12447 | Mm.16110 | NM_007633 | ENSMUSG00000002068 | **Ccne1** | AW538188\|CycE1 | cyclin E1 | protein-coding |
| chr7_38090108_38090705 | chr7 | 38090109 | 38090705 | + | 0.642869 | Intergenic | Intergenic | 17083 | NM_007633 | 12447 | Mm.16110 | NM_007633 | ENSMUSG00000002068 | **Ccne1** | AW538188\|CycE1 | cyclin E1 | protein-coding |
| chr9_123942035_123942587 | chr9 | 123942036 | 123942587 | + | 0.805568 | Intergenic | Intergenic | 26381 | NM_009912 | 12768 | Mm.274927 | NM_009912 | ENSMUSG00000025804 | **Ccr1** | Cmkbr1\|Mip-1a-R | chemokine (C-C motif) receptor 1 | protein-coding |
| chr9_114488332_114488757 | chr9 | 114488333 | 114488757 | + | 0.878455 | Intergenic | Intergenic | 7999 | NM_009916 | 12773 | Mm.1337 | NM_009916 | ENSMUSG00000047898 | **Ccr4** | C-C CKR-4\|CHEMR1\|Cmkbr4\|LESTR\|Sdf1r | chemokine (C-C motif) receptor 4 | protein-coding |
| chr9_114488171_114488772 | chr9 | 114488172 | 114488772 | + | 0.854584 | Intergenic | Intergenic | 8072 | NM_009916 | 12773 | Mm.1337 | NM_009916 | ENSMUSG00000047898 | **Ccr4** | C-C CKR-4\|CHEMR1\|Cmkbr4\|LESTR\|Sdf1r | chemokine (C-C motif) receptor 4 | protein-coding |
| chr9_114487351_114488780 | chr9 | 114487352 | 114488780 | + | 0.718364 | Intergenic | LTR16A\|LTR\|ERVL | 8478 | NM_009916 | 12773 | Mm.1337 | NM_009916 | ENSMUSG00000047898 | **Ccr4** | C-C CKR-4\|CHEMR1\|Cmkbr4\|LESTR\|Sdf1r | chemokine (C-C motif) receptor 4 | protein-coding |
| chr11_99180054_99180723 | chr11 | 99180055 | 99180723 | + | 0.73048 | Intergenic | Intergenic | -25312 | NM_007719 | 12775 | Mm.2932 | NM_007719 | ENSMUSG00000037944 | **Ccr7** | CD197\|Cdw197\|Cmkbr7\|EBI1\|Ebi1h | chemokine (C-C motif) receptor 7 | protein-coding |
| chr11_114885070_114885538 | chr11 | 114885071 | 114885538 | + | 0.69734 | Intergenic | Intergenic | -4737 | NM_170758 | 217303 | Mm.294439 | NM_170758 | ENSMUSG00000034652 | **Cd300a** | B230315M08Rik\|Clm8\|LMIR1\|MAIR-I\|MAIR-Ia\|MMAC8\|Pigr4\|mcpir1 | CD300A antigen | protein-coding |
| chr11_121019446_121019875 | chr11 | 121019447 | 121019875 | + | 0.730161 | Intergenic | Intergenic | 19817 | NM_009854 | 12516 | Mm.4100 | NM_009854 | ENSMUSG00000025163 | **Cd7** | - | CD7 antigen | protein-coding |
| chr4_89388116_89388489 | chr4 | 89388117 | 89388489 | + | 0.595401 | Intergenic | Intergenic | -77271 | NM_007670 | 12579 | Mm.423094 | NM_007670 | ENSMUSG00000073802 | **Cdkn2b** | AV083695\|INK4b\|MTS2\|p15\|p15(INK4b)\|p15INK4b | cyclin-dependent kinase inhibitor 2B (p15, inhibits CDK4) | protein-coding |
| chr4_8733956_8734372 | chr4 | 8733957 | 8734372 | + | 0.679613 | intron (NM_001277149, intron 1 of 37) | intron (NM_001277149, intron 1 of 37) | 43758 | NM_001277149 | 320790 | Mm.138792 | NM_001033395 | ENSMUSG00000041235 | **Chd7** | A730019I05Rik\|Cycn\|Cyn\|Dz\|Edy\|Flo\|GENA 47\|GENA 60\|Gena 52\|Lda\|Mt\|Obt\|Todo\|WBE1\|Whi\|metis | chromodomain helicase DNA binding protein 7 | protein-coding |
| chr10_82969531_82969979 | chr10 | 82969532 | 82969979 | + | 0.591308 | Intergenic | RMER17D2\|LTR\|ERVK | -15742 | NM_021439 | 58250 | Mm.360747 | NM_021439 | ENSMUSG00000034612 | **Chst11** | 1110020P09Rik\|C4ST\|C4ST-1\|C4ST1\|C4s | carbohydrate sulfotransferase 11 | protein-coding |
| chr7_66183442_66183831 | chr7 | 66183443 | 66183831 | + | 0.674296 | Intergenic | Intergenic | 74122 | NM_001081163 | 269941 | Mm.333916 | NM_001081163 | ENSMUSG00000032640 | **Chsy1** | mKIAA0990 | chondroitin sulfate synthase 1 | protein-coding |
| chr11_45874235_45874640 | chr11 | 45874236 | 45874640 | + | 0.625044 | intron (NM_001045520, intron 1 of 11) | intron (NM_001045520, intron 1 of 11) | 22474 | NM_001045520 | 216705 | Mm.169673 | NM_001045520 | ENSMUSG00000006169 | **Clint1** | AI642036\|AW537616\|C530049I24Rik\|Enth\|Epn4\|F630206G17Rik\|mKIAA0171 | clathrin interactor 1 | protein-coding |
| chr17_78271345_78271924 | chr17 | 78271346 | 78271924 | + | 0.819302 | intron (NM_015800, intron 2 of 16) | intron (NM_015800, intron 2 of 16) | 71387 | NM_015800 | 50766 | Mm.311912 | NM_015800 | ENSMUSG00000024074 | **Crim1** | AU015004 | cysteine rich transmembrane BMP regulator 1 (chordin like) | protein-coding |
| chr18_84872385_84872663 | chr18 | 84872386 | 84872663 | + | 0.901509 | intron (NM_025797, intron 2 of 4) | intron (NM_025797, intron 2 of 4) | 21110 | NM_025797 | 109672 | Mm.31018 | NM_025797 | | **Cyb5a** | 0610009N12Rik\|Cyb5 | cytochrome b5 type A (microsomal) | protein-coding |
| chrX_113826837_113827153 | chrX | 113826838 | 113827153 | + | 0.679146 | intron (NM_001289734, intron 15 of 15) | B3A\|SINE\|B2 | 528739 | NM_001142570 | 93837 | Mm.79760 | NM_033605 | ENSMUSG00000025592 | **Dach2** | 9430028N04Rik | dachshund 2 (Drosophila) | protein-coding |
| chr2_62331849_62332356 | chr2 | 62331850 | 62332356 | + | 0.940502 | 3' UTR (NM_010074, exon 26 of 26) | 3' UTR (NM_010074, exon 26 of 26) | 80128 | NM_010074 | 13482 | Mm.1151 | NM_010074 | ENSMUSG00000035000 | **Dpp4** | Cd26\|Dpp-4\|THAM | dipeptidylpeptidase 4 | protein-coding |
| chr5_30353639_30353971 | chr5 | 30353640 | 30353971 | + | 0.926949 | intron (NM_001033460, intron 6 of 15) | intron (NM_001033460, intron 6 of 15) | 12142 | NM_001033460 | 381738 | Mm.389604 | NM_001033460 | ENSMUSG00000073102 | **Drc1** | Ccdc164\|Gm1060\|b2b1654Clo | dynein regulatory complex subunit 1 | protein-coding |
| chr7_90067893_90068425 | chr7 | 90067894 | 90068425 | + | 0.748751 | Intergenic | Intergenic | 39000 | NR_110364 | 319711 | Mm.31753 | NR_015614 | | **E230029C05Rik** | AI452102\|B230337J15 | RIKEN cDNA E230029C05 gene | ncRNA |
| chr7_90067998_90068378 | chr7 | 90067999 | 90068378 | + | 0.741466 | Intergenic | Intergenic | 39029 | NR_110364 | 319711 | Mm.31753 | NR_015614 | | **E230029C05Rik** | AI452102\|B230337J15 | RIKEN cDNA E230029C05 gene | ncRNA |

| | | | | | | | | | | | | | | | | | |
|---|---|---|---|---|---|---|---|---|---|---|---|---|---|---|---|---|---|
| chr7_90067888_90068928 | chr7 | 90067889 | 90068928 | + | 0.699109 | Intergenic | Intergenic | 39249 | NR_110364 | 319711 | Mm.31753 | NR_015614 | | **E230029C05Rik** | AI452102\|B230337J15 | RIKEN cDNA E230029C05 gene | ncRNA |
| chr4_136177541_136178310 | chr4 | 136177542 | 136178310 | + | 0.722056 | intron (NM_177733, intron 1 of 6) | intron (NM_177733, intron 1 of 6) | 5532 | NM_177733 | 242705 | Mm.307932 | NM_177733 | ENSMUSG00000018983 | **E2f2** | 9230110J10\|E130207A07 | E2F transcription factor 2 | protein-coding |
| chr4_136177250_136178359 | chr4 | 136177251 | 136178359 | + | 0.689281 | intron (NM_177733, intron 1 of 6) | intron (NM_177733, intron 1 of 6) | 5411 | NM_177733 | 242705 | Mm.307932 | NM_177733 | ENSMUSG00000018983 | **E2f2** | 9230110J10\|E130207A07 | E2F transcription factor 2 | protein-coding |
| chr9_77955595_77955870 | chr9 | 77955596 | 77955870 | + | 0.651188 | intron (NM_134255, intron 1 of 7) | intron (NM_134255, intron 1 of 7) | 38368 | NM_134255 | 68801 | Mm.430736 | NM_134255 | ENSMUSG00000032349 | **Elovl5** | 1110059L23Rik\|AI747313\|AU043003\|HELO1 | ELOVL family member 5, elongation of long chain fatty acids (yeast) | protein-coding |
| chr4_11419530_11420224 | chr4 | 11419531 | 11420224 | + | 0.664304 | Intergenic | Intergenic | -33094 | NM_194055 | 207920 | Mm.387719 | NM_194055 | ENSMUSG00000040728 | **Esrp1** | 2210008M09Rik\|A630065D16\|BC031468\|Rbm35a | epithelial splicing regulatory protein 1 | protein-coding |
| chr5_3460776_3461381 | chr5 | 3460777 | 3461381 | + | 0.929646 | intron (NM_009873, intron 3 of 6) | intron (NM_009873, intron 3 of 6) | -82754 | NM_001042501 | 68152 | Mm.240265 | NM_026583 | | **Fam133b** | 2900022K02Rik\|5830415L20Rik | family with sequence similarity 133, member B | protein-coding |
| chr5_3460214_3461370 | chr5 | 3460215 | 3461370 | + | 0.85972 | intron (NM_009873, intron 3 of 6) | intron (NM_009873, intron 3 of 6) | -83041 | NM_001042501 | 68152 | Mm.240265 | NM_026583 | | **Fam133b** | 2900022K02Rik\|5830415L20Rik | family with sequence similarity 133, member B | protein-coding |
| chr5_3477687_3478305 | chr5 | 3477688 | 3478305 | + | 0.629167 | intron (NM_009873, intron 4 of 6) | intron (NM_009873, intron 4 of 6) | -65837 | NM_001042501 | 68152 | Mm.240265 | NM_026583 | | **Fam133b** | 2900022K02Rik\|5830415L20Rik | family with sequence similarity 133, member B | protein-coding |
| chr5_3477606_3478593 | chr5 | 3477607 | 3478593 | + | 0.596764 | intron (NM_009873, intron 4 of 6) | intron (NM_009873, intron 4 of 6) | -65733 | NM_001042501 | 68152 | Mm.240265 | NM_026583 | | **Fam133b** | 2900022K02Rik\|5830415L20Rik | family with sequence similarity 133, member B | protein-coding |
| chr11_109758486_109759304 | chr11 | 109758487 | 109759304 | + | 0.763172 | Intergenic | Lx9\|LINE\|L1 | -36639 | NM_153782 | 208659 | Mm.208662 | NM_153782 | ENSMUSG00000020614 | **Fam20a** | AI606893 | family with sequence similarity 20, member A | protein-coding |
| chr9_85107251_85107927 | chr9 | 85107252 | 85107927 | + | 0.66836 | Intergenic | Intergenic | 219535 | NM_001160378 | 212943 | Mm.74610 | NM_001160378 | ENSMUSG00000032265 | **Fam46a** | D930050G01Rik | family with sequence similarity 46, member A | protein-coding |
| chr9_85107295_85107691 | chr9 | 85107296 | 85107691 | + | 0.632925 | Intergenic | Intergenic | 219631 | NM_001160378 | 212943 | Mm.74610 | NM_001160378 | ENSMUSG00000032265 | **Fam46a** | D930050G01Rik | family with sequence similarity 46, member A | protein-coding |
| chr17_63288405_63288776 | chr17 | 63288406 | 63288776 | + | 0.647626 | intron (NM_015794, intron 6 of 8) | intron (NM_015794, intron 6 of 8) | 211989 | NM_015794 | 50758 | Mm.136366 | NM_015794 | | **Fbxl17** | 6330576B01Rik\|AI452053\|BB073797\|C130023C01Rik\|Fbx13\|Fbxo13 | F-box and leucine-rich repeat protein 17 | protein-coding |
| chr17_63362109_63362576 | chr17 | 63362110 | 63362576 | + | 0.591702 | intron (NM_015794, intron 5 of 8) | RMER19A\|LTR\|ERVK | 138237 | NM_015794 | 50758 | Mm.136366 | NM_015794 | | **Fbxl17** | 6330576B01Rik\|AI452053\|BB073797\|C130023C01Rik\|Fbx13\|Fbxo13 | F-box and leucine-rich repeat protein 17 | protein-coding |
| chr12_59250509_59250901 | chr12 | 59250510 | 59250901 | + | 0.771916 | Intergenic | B3A\|SINE\|B2 | -31222 | NM_001033156 | 70611 | Mm.311026 | NM_001033156 | ENSMUSG00000035329 | **Fbxo33** | 5730501N20Rik\|AI642135 | F-box protein 33 | protein-coding |
| chr17_28488204_28488676 | chr17 | 28488205 | 28488676 | + | 0.634981 | Intergenic | Intergenic | -2291 | NM_010220 | 14229 | Mm.276405 | NM_010220 | ENSMUSG00000024222 | **Fkbp5** | D17Ertd592e\|Dit1\|FKBP-5\|FKBP51 | FK506 binding protein 5 | protein-coding |
| chr12_76848092_76848903 | chr12 | 76848093 | 76848903 | + | 0.586956 | intron (NM_145927, intron 1 of 11) | B4\|SINE\|B4 | 11031 | NM_145927 | 110606 | Mm.151174 | NM_145927 | ENSMUSG00000033373 | **Fntb** | 2010013E13Rik\|AA409500 | farnesyltransferase, CAAX box, beta | protein-coding |
| chr3_21965590_21966020 | chr3 | 21965591 | 21966020 | + | 0.873417 | Intergenic | Intergenic | -110847 | NM_030732 | 81004 | Mm.202966 | NM_030732 | ENSMUSG00000027630 | **Foxn3** | 8030499H02Rik\|A630076E03Rik\|AW539987\|C21\|C230089I12Rik\|DC42\|Ira1\|TBLR1 | transducin (beta)-like 1X-linked receptor 1 | protein-coding |
| chr12_99510119_99510476 | chr12 | 99510120 | 99510476 | + | 0.639924 | Intergenic | B3\|SINE\|B2 | -60224 | NM_183186 | 71375 | Mm.341972 | NM_183186 | ENSMUSG00000033713 | **Foxn3** | 5430426H20Rik\|AA589593\|AW556347\|Ches1\|Ches1l\|HTLFL1 | forkhead box N3 | protein-coding |
| chr12_99509971_99510622 | chr12 | 99509972 | 99510622 | + | 0.63291 | Intergenic | B3\|SINE\|B2 | -60223 | NM_183186 | 71375 | Mm.341972 | NM_183186 | ENSMUSG00000033713 | **Foxn3** | 5430426H20Rik\|AA589593\|AW556347\|Ches1\|Ches1l\|HTLFL1 | forkhead box N3 | protein-coding |
| chr13_51849256_51849628 | chr13 | 51849257 | 51849628 | + | 0.688273 | TTS (NM_011817) | TTS (NM_011817) | 2767 | NM_011817 | 23882 | Mm.281298 | NM_011817 | ENSMUSG00000021453 | **Gadd45g** | AI327420\|C86281\|CR6\|DDIT2\|OIG37 | growth arrest and DNA-damage-inducible 45 gamma | protein-coding |
| chr11_4223407_4223809 | chr11 | 4223408 | 4223809 | + | 0.886307 | Intergenic | Intergenic | 5357 | NM_028022 | 71962 | Mm.7852 | NM_028022 | ENSMUSG00000020424 | **Gatsl3** | 2410008K03Rik | GATS protein-like 3 | protein-coding |
| chr13_75843984_75844328 | chr13 | 75843985 | 75844328 | + | 0.633316 | intron (NM_053108, intron 1 of 2) | intron (NM_053108, intron 1 of 2) | 4270 | NM_053108 | 93692 | Mm.25844 | NM_053108 | ENSMUSG00000021591 | **Glrx** | C86710\|D13Wsu156e\|Glrx1\|Grx1\|TTase | glutaredoxin | protein-coding |
| chr13_75843969_75844577 | chr13 | 75843970 | 75844577 | + | 0.585179 | intron (NM_053108, intron 1 of 2) | intron (NM_053108, intron 1 of 2) | 4387 | NM_053108 | 93692 | Mm.25844 | NM_053108 | ENSMUSG00000021591 | **Glrx** | C86710\|D13Wsu156e\|Glrx1\|Grx1\|TTase | glutaredoxin | protein-coding |
| chr1_143749632_143750217 | chr1 | 143749633 | 143750217 | + | 0.685218 | TTS (NM_013835) | TTS (NM_013835) | 9478 | NM_001038594 | 69367 | Mm.272727 | NM_023505 | ENSMUSG00000018196 | **Glrx2** | 1700010P22Rik\|AI645710\|Grx2 | glutaredoxin 2 (thioltransferase) | protein-coding |
| chr3_79335990_79336510 | chr3 | 79335991 | 79336510 | + | 0.769416 | Intergenic | Intergenic | -9126 | NM_001142953 | 100233207 | Mm.373801 | NM_001142953 | ENSMUSG00000091685 | **Gm17359** | - | predicted gene, 17359 | protein-coding |
| chr1_88584472_88584923 | chr1 | 88584473 | 88584923 | + | 0.616397 | Intergenic | Intergenic | 33181 | NR_037971 | 100503197 | Mm.99352 | NR_037971 | | **Gm19589** | - | predicted gene, 19589 | ncRNA |
| chr7_106556503_106556813 | chr7 | 106556504 | 106556813 | + | 0.780708 | Intergenic | RLTR13D4\|LTR\|ERVK | 87987 | NM_001277179 | 434223 | Mm.276443 | NM_001277179 | | **Gm1966** | - | predicted gene 1966 | protein-coding |
| chr7_106556404_106556905 | chr7 | 106556405 | 106556905 | + | 0.779681 | Intergenic | RLTR13D4\|LTR\|ERVK | 87990 | NM_001277179 | 434223 | Mm.276443 | NM_001277179 | | **Gm1966** | - | predicted gene 1966 | protein-coding |
| chr3_143581740_143582323 | chr3 | 143581741 | 143582323 | + | 0.68765 | Intergenic | MER45B\|DNA\|hAT-Tip100 | 130111 | NR_040405 | 621809 | Mm.360134 | NR_040405 | | **Gm6260** | EG621809 | predicted gene 6260 | ncRNA |
| chr3_143581787_143582518 | chr3 | 143581788 | 143582518 | + | 0.671478 | Intergenic | MER45B\|DNA\|hAT-Tip100 | 129990 | NR_040405 | 621809 | Mm.360134 | NR_040405 | | **Gm6260** | EG621809 | predicted gene 6260 | ncRNA |
| chr3_143582028_143582518 | chr3 | 143582029 | 143582518 | + | 0.59783 | Intergenic | Intergenic | 129870 | NR_040405 | 621809 | Mm.360134 | NR_040405 | | **Gm6260** | EG621809 | predicted gene 6260 | ncRNA |
| chr14_19994101_19994474 | chr14 | 19994102 | 19994474 | + | 0.628 | Intergenic | ORR1D1\|LTR\|ERVL-MaLR | -17039 | NM_001285909 | 14702 | Mm.41737 | NM_010315 | ENSMUSG00000043004 | **Gng2** | 82 | guanine nucleotide binding protein (G protein), gamma 2 | protein-coding |
| chr14_19994176_19994423 | chr14 | 19994177 | 19994423 | + | 0.601404 | Intergenic | ORR1D1\|LTR\|ERVL-MaLR | -17051 | NM_001285909 | 14702 | Mm.41737 | NM_010315 | ENSMUSG00000043004 | **Gng2** | 82 | guanine nucleotide binding protein (G protein), gamma 2 | protein-coding |
| chr12_102504633_102505304 | chr12 | 102504634 | 102505304 | + | 0.62236 | Intergenic | Intergenic | 35059 | NM_013747 | 27277 | Mm.273370 | NM_013747 | ENSMUSG00000021192 | **Golga5** | Ret-II | golgi autoantigen, golgin subfamily a, 5 | protein-coding |
| chr12_102504839_102505137 | chr12 | 102504840 | 102505137 | + | 0.597133 | Intergenic | Intergenic | 35078 | NM_013747 | 27277 | Mm.273370 | NM_013747 | ENSMUSG00000021192 | **Golga5** | Ret-II | golgi autoantigen, golgin subfamily a, 5 | protein-coding |
| chr6_59324861_59325380 | chr6 | 59324862 | 59325380 | + | 0.633796 | Intergenic | Intergenic | 101169 | NM_183183 | 243385 | Mm.138080 | NM_183183 | ENSMUSG00000045441 | **Gprin3** | C030038J10Rik\|C730021L23 | GPRIN family member 3 | protein-coding |
| chr18_41982655_41983098 | chr18 | 41982656 | 41983098 | + | 1.13814 | Intergenic | Intergenic | 16172 | NM_001033426 | 332309 | Mm.240520 | NM_001033426 | ENSMUSG00000073574 | **Grxcr2** | Gm851 | glutaredoxin, cysteine rich 2 | protein-coding |
| chr18_41982765_41983002 | chr18 | 41982766 | 41983002 | + | 1.12434 | Intergenic | Intergenic | 16165 | NM_001033426 | 332309 | Mm.240520 | NM_001033426 | ENSMUSG00000073574 | **Grxcr2** | Gm851 | glutaredoxin, cysteine rich 2 | protein-coding |
| chr18_41979475_41980038 | chr18 | 41979476 | 41980038 | + | 0.964365 | Intergenic | Intergenic | 19292 | NM_001033426 | 332309 | Mm.240520 | NM_001033426 | ENSMUSG00000073574 | **Grxcr2** | Gm851 | glutaredoxin, cysteine rich 2 | protein-coding |
| chr18_41979205_41980041 | chr18 | 41979206 | 41980041 | + | 0.954428 | Intergenic | Intergenic | 19426 | NM_001033426 | 332309 | Mm.240520 | NM_001033426 | ENSMUSG00000073574 | **Grxcr2** | Gm851 | glutaredoxin, cysteine rich 2 | protein-coding |
| chr18_41978908_41980088 | chr18 | 41978909 | 41980088 | + | 0.936775 | Intergenic | Intergenic | 19551 | NM_001033426 | 332309 | Mm.240520 | NM_001033426 | ENSMUSG00000073574 | **Grxcr2** | Gm851 | glutaredoxin, cysteine rich 2 | protein-coding |
| chr18_41985622_41985997 | chr18 | 41985623 | 41985997 | + | 0.78381 | TTS (NM_001033426) | TTS (NM_001033426) | 13239 | NM_001033426 | 332309 | Mm.240520 | NM_001033426 | ENSMUSG00000073574 | **Grxcr2** | Gm851 | glutaredoxin, cysteine rich 2 | protein-coding |
| chr5_30353527_30354176 | chr5 | 30353528 | 30354176 | + | 0.928267 | intron (NM_001033460, intron 6 of 15) | intron (NM_001033460, intron 6 of 15) | 12189 | NM_001033460 | 381738 | Mm.389604 | NM_001033460 | ENSMUSG00000073102 | **Gzmb** | Ccdc164\|Gm1060\|b2b1654Clo | dynein regulatory complex subunit 1 | protein-coding |
| chr14_56266097_56266407 | chr14 | 56266098 | 56266407 | + | 0.706226 | Intergenic | Intergenic | -3992 | NM_013542 | 14939 | Mm.14874 | NM_013542 | ENSMUSG00000015437 | **Gzmb** | AI553453\|CCP-1/C11\|CCP1\|Ctla-1\|Ctla1\|GZB | granzyme B | protein-coding |
| chr14_56275673_56276521 | chr14 | 56275674 | 56276521 | + | 0.595582 | Intergenic | Intergenic | -13837 | NM_013542 | 14939 | Mm.14874 | NM_013542 | ENSMUSG00000015437 | **Gzmb** | AI553453\|CCP-1/C11\|CCP1\|Ctla-1\|Ctla1\|GZB | granzyme B | protein-coding |
| chr13_23593064_23593674 | chr13 | 23593065 | 23593674 | + | 0.607475 | intron (NM_001177653, intron 1 of 1) | intron (NM_001177653, intron 1 of 1) | -7399 | NM_001290530 | 319179 | Mm.440391 | NM_178194 | ENSMUSG00000047246 | **Hist1h2be** | - | histone cluster 1, H2be | protein-coding |
| chr10_14189156_14189577 | chr10 | 14189157 | 14189577 | + | 0.599711 | Intergenic | Intergenic | 222988 | NM_010437 | 15273 | Mm.42157 | NM_010437 | ENSMUSG00000015501 | **Hivep2** | MIBP1\|Schnurri-2\|Shn-2 | human immunodeficiency virus type I enhancer binding protein 2 | protein-coding |
| chr17_12759132_12759907 | chr17 | 12759133 | 12759907 | + | 0.61974 | intron (NM_010515, intron 1 of 47) | intron (NM_010515, intron 1 of 47) | 10186 | NM_010515 | 16004 | Mm.26553 | NM_010515 | ENSMUSG00000023830 | **Igf2r** | AI661837\|CD222\|CI-MPR\|M6P/IGF2R\|Mpr300 | insulin-like growth factor 2 receptor | protein-coding |
| chr10_6927887_6928280 | chr10 | 6927888 | 6928280 | + | 0.760251 | intron (NM_001033391, intron 6 of 7) | RSINE1\|SINE\|B4 | 52381 | NM_001170800 | 320495 | Mm.331674 | NM_001033391 | ENSMUSG00000064065 | **Ipcef1** | A130090K04Rik | interaction protein for cytohesin exchange factors 1 | protein-coding |
| chr2_79276211_79276736 | chr2 | 79276212 | 79276736 | + | 0.694665 | intron (NM_010576, intron 4 of 27) | intron (NM_010576, intron 4 of 27) | 21048 | NM_010576 | 16401 | Mm.31903 | NM_010576 | ENSMUSG00000027009 | **Itga4** | CD49D\|Itga4B | integrin alpha 4 | protein-coding |
| chr2_79269978_79270560 | chr2 | 79269979 | 79270560 | + | 0.600306 | intron (NM_010576, intron 2 of 27) | intron (NM_010576, intron 2 of 27) | 14843 | NM_010576 | 16401 | Mm.31903 | NM_010576 | ENSMUSG00000027009 | **Itga4** | CD49D\|Itga4B | integrin alpha 4 | protein-coding |
| chr13_44763075_44763492 | chr13 | 44763076 | 44763492 | + | 0.647714 | intron (NM_001205043, intron 1 of 17) | B3\|SINE\|B2 | 32013 | NM_001205043 | 16468 | Mm.25059 | NM_021878 | ENSMUSG00000038518 | **Jarid2** | Jmj\|jumonji | jumonji, AT rich interactive domain 2 | protein-coding |
| chr4_95199626_95200451 | chr4 | 95199627 | 95200451 | + | 0.717159 | Intergenic | L2\|LINE\|L2 | -147817 | NM_010591 | 16476 | Mm.275071 | NM_010591 | ENSMUSG00000052684 | **Jun** | AP-1\|Junc\|c-jun | jun proto-oncogene | protein-coding |
| chr4_95302749_95303854 | chr4 | 95302750 | 95303854 | + | 0.669742 | Intergenic | Intergenic | -251080 | NM_010591 | 16476 | Mm.275071 | NM_010591 | ENSMUSG00000052684 | **Jun** | AP-1\|Junc\|c-jun | jun proto-oncogene | protein-coding |
| chr6_48398457_48398789 | chr6 | 48398458 | 48398789 | + | 0.604403 | intron (NM_133922, intron 1 of 17) | ORR1E\|LTR\|ERVL-MaLR | 3037 | NM_133922 | 77827 | Mm.107646 | NM_133922 | ENSMUSG00000042810 | **Krba1** | A930040G15Rik\|AI448780 | KRAB-A domain containing 1 | protein-coding |
| chr8_34139899_34140213 | chr8 | 34139900 | 34140213 | + | 0.691345 | intron (NM_026609, intron 2 of 3) | intron (NM_026609, intron 2 of 3) | 6683 | NM_026609 | 68192 | Mm.34212 | NM_026609 | ENSMUSG00000031513 | **Leprotl1** | 1110067H13Rik\|1520402O14Rik\|AI854296 | leptin receptor overlapping transcript-like 1 | protein-coding |
| chr6_124683595_124684249 | chr6 | 124683596 | 124684249 | + | 0.720894 | intron (NM_145130, intron 1 of 12) | intron (NM_145130, intron 1 of 12) | 20818 | NM_145130 | 14792 | Mm.273915 | NM_145130 | ENSMUSG00000004270 | **Lpcat3** | C3f\|Grcc3f\|Lpcat\|Lpeat\|Lplat5\|Lpsat\|Mboat5\|Moact5\|Oact5\|PTG | lysophosphatidylcholine acyltransferase 3 | protein-coding |
| chr10_61534499_61535134 | chr10 | 61534500 | 61535134 | + | 0.725086 | intron (NM_153542, intron 3 of 4) | intron (NM_153542, intron 3 of 4) | 58984 | NM_153542 | 216011 | Mm.136116 | NM_153542 | ENSMUSG00000037151 | **Lrrc20** | BC036304 | leucine rich repeat containing 20 | protein-coding |

| | | | | | | | | | | | | | | | | | |
|---|---|---|---|---|---|---|---|---|---|---|---|---|---|---|---|---|---|
| chr12_41347710_41348344 | chr12 | 41347711 | 41348344 | + | 0.682573 | intron (NM_053122, intron 4 of 6) | RMER1B\|Other\|Other | 138030 | NM_001271708 | 16981 | Mm.404786 | NM_010733 | ENSMUSG00000036295 | **Lrrn3** | NLRR-3 | leucine rich repeat protein 3, neuronal | protein-coding |
| chr6_119343402_119344060 | chr6 | 119343403 | 119344060 | + | 0.625599 | intron (NM_001033382, intron 30 of 37) | intron (NM_001033382, intron 30 of 37) | -12965 | NM_172492 | 211187 | Mm.121498 | NM_172492 | ENSMUSG00000055003 | **Lrtm2** | A230084J22\|AI841794 | leucine-rich repeats and transmembrane domains 2 | protein-coding |
| chr6_119343614_119343911 | chr6 | 119343615 | 119343911 | + | 0.606952 | intron (NM_001033382, intron 30 of 37) | intron (NM_001033382, intron 30 of 37) | -12997 | NM_172492 | 211187 | Mm.121498 | NM_172492 | ENSMUSG00000055003 | **Lrtm2** | A230084J22\|AI841794 | leucine-rich repeats and transmembrane domains 2 | protein-coding |
| chr18_4278298_4278812 | chr18 | 4278299 | 4278812 | + | 0.850708 | Intergenic | Intergenic | 74398 | NM_007746 | 26410 | Mm.3275 | NM_007746 | ENSMUSG00000024235 | **Map3k8** | Cot\|Cot/Tpl2\|Est\|Estf\|Tpl-2\|Tpl2\|c-COT | mitogen-activated protein kinase kinase kinase 8 | protein-coding |
| chr7_29023407_29024014 | chr7 | 29023408 | 29024014 | + | 0.658333 | intron (NM_009109, intron 90 of 105) | intron (NM_009109, intron 90 of 105) | 40857 | NM_008279 | 26411 | Mm.148278 | NM_008279 | ENSMUSG00000037337 | **Map4k1** | Hpk1\|mHPK1 | mitogen-activated protein kinase kinase kinase kinase 1 | protein-coding |
| chr12_77056754_77057422 | chr12 | 77056755 | 77057422 | + | 0.682196 | Intergenic | Intergenic | -94840 | NM_001146176 | 17187 | Mm.268548 | NM_008558 | ENSMUSG00000059436 | **Max** | AA960152\|AI875693\|bHLHd4\|bHLHd5\|bHLHd6\|bHLHd7\|bHLHd8 | Max protein | protein-coding |
| chr12_56327445_56327955 | chr12 | 56327446 | 56327955 | + | 0.606611 | TTS (NM_145442) | TTS (NM_145442) | 18194 | NM_145442 | 217588 | Mm.25656 | NM_145442 | ENSMUSG00000021028 | **Mbip** | 4933408E06Rik | MAP3K12 binding inhibitory protein 1 | protein-coding |
| chr7_72283696_72284147 | chr7 | 72283697 | 72284147 | + | 0.619588 | intron (NM_001024703, intron 1 of 22) | intron (NM_001024703, intron 1 of 22) | 22673 | NM_001024703 | 244049 | Mm.217412 | NM_001024703 | ENSMUSG00000032776 | **Mctp2** | Gm489 | multiple C2 domains, transmembrane 2 | protein-coding |
| chr4_122930960_122931199 | chr4 | 122930961 | 122931199 | + | 0.631292 | Intergenic | B2_Mm2\|SINE\|B2 | 30108 | NM_029662 | 76574 | Mm.331842 | NM_029662 | ENSMUSG00000028655 | **Mfsd2a** | 1700018O18Rik\|Mfsd2\|NLS1 | major facilitator superfamily domain containing 2A | protein-coding |
| chr4_122930932_122931255 | chr4 | 122930933 | 122931255 | + | 0.611076 | Intergenic | B2_Mm2\|SINE\|B2 | 30094 | NM_029662 | 76574 | Mm.331842 | NM_029662 | ENSMUSG00000028655 | **Mfsd2a** | 1700018O18Rik\|Mfsd2\|NLS1 | major facilitator superfamily domain containing 2A | protein-coding |
| chr12_105022703_105023184 | chr12 | 105022704 | 105023184 | + | 0.60682 | Intergenic | Intergenic | 8197 | NR_037229 | 100526471 | | NR_037229 | | **Mir3069** | mmu-mir-3069 | microRNA 3069 | ncRNA |
| chr8_108890850_108891372 | chr8 | 108890851 | 108891372 | + | 0.626954 | intron (NM_007496, intron 4 of 9) | (CAGA)n\|Simple_repeat\|Simple_repeat | -45749 | NR_037295 | 100526511 | | NR_037295 | | **Mir3108** | mmu-mir-3108 | microRNA 3108 | ncRNA |
| chr14_102828474_102828924 | chr14 | 102828475 | 102828924 | + | 0.588744 | Intergenic | Intergenic | 153933 | NR_039592 | 100628599 | | NR_039592 | miRBase:MI0018042 | **Mir5130** | mmu-mir-5130\|mu-mir-5130 | microRNA 5130 | ncRNA |
| chr6_83392196_83392971 | chr6 | 83392197 | 83392971 | + | 0.766826 | intron (NM_183138, intron 3 of 10) | intron (NM_183138, intron 3 of 10) | 8518 | NR_105793 | 102465194 | | NR_105793 | | **Mir6374** | mmu-mir-6374 | microRNA mir-6374 | ncRNA |
| chrX_53379431_53379829 | chrX | 53379432 | 53379829 | + | 0.662563 | Intergenic | Intergenic | -9071 | NR_110964 | 70380 | Mm.491083 | NM_027409 | ENSMUSG00000023074 | **Mospd1** | 1810018L05Rik | motile sperm domain containing 1 | protein-coding |
| chr10_24291512_24292084 | chr10 | 24291513 | 24292084 | + | 0.588507 | intron (NM_021509, intron 10 of 11) | intron (NM_021509, intron 10 of 11) | 68281 | NM_021509 | 59012 | Mm.285934 | NM_021509 | ENSMUSG00000020000 | **Moxd1** | 3230402N08Rik\|MNCb-5203 | monooxygenase, DBH-like 1 | protein-coding |
| chr1_51514289_51515019 | chr1 | 51514290 | 51515019 | + | 0.79447 | Intergenic | Intergenic | -36255 | NM_028696 | 109019 | Mm.196290 | NM_028696 | ENSMUSG00000026107 | **Nabp1** | 4930434H03Rik\|4930442A21Rik\|4930488J04Rik\|4933440J18Rik\|5830411E10Rik\|AI852561\|Nbp1\|Obfc2a\|Ssb2 | nucleic acid binding protein 1 | protein-coding |
| chr2_52319180_52319426 | chr2 | 52319181 | 52319426 | + | 0.681932 | intron (NM_010889, intron 10 of 156) | intron (NM_010889, intron 10 of 156) | 19495 | NM_010889 | 17996 | Mm.389247 | NM_010889 | ENSMUSG00000026950 | **Neb** | AI595938 | nebulin | protein-coding |
| chr2_52319162_52319699 | chr2 | 52319163 | 52319699 | + | 0.659646 | intron (NM_010889, intron 10 of 156) | intron (NM_010889, intron 10 of 156) | 19367 | NM_010889 | 17996 | Mm.389247 | NM_010889 | ENSMUSG00000026950 | **Neb** | AI595938 | nebulin | protein-coding |
| chr2_52315754_52316218 | chr2 | 52315755 | 52316218 | + | 0.658419 | TTS (NR_040620) | TTS (NR_040620) | 22812 | NM_010889 | 17996 | Mm.389247 | NM_010889 | ENSMUSG00000026950 | **Neb** | AI595938 | nebulin | protein-coding |
| chr2_52318959_52319728 | chr2 | 52318960 | 52319728 | + | 0.633382 | intron (NM_010889, intron 10 of 156) | intron (NM_010889, intron 10 of 156) | 19454 | NM_010889 | 17996 | Mm.389247 | NM_010889 | ENSMUSG00000026950 | **Neb** | AI595938 | nebulin | protein-coding |
| chr16_55582865_55583434 | chr16 | 55582866 | 55583434 | + | 0.768457 | Intergenic | Intergenic | 238988 | NM_001159394 | 80859 | Mm.247272 | NM_030612 | ENSMUSG00000035356 | **Nfkbiz** | AA408868\|INAP\|Mail | nuclear factor of kappa light polypeptide gene enhancer in B cells inhibitor, zeta | protein-coding |
| chr16_55582470_55583434 | chr16 | 55582471 | 55583434 | + | 0.711834 | Intergenic | Intergenic | 239186 | NM_001159394 | 80859 | Mm.247272 | NM_030612 | ENSMUSG00000035356 | **Nfkbiz** | AA408868\|INAP\|Mail | nuclear factor of kappa light polypeptide gene enhancer in B cells inhibitor, zeta | protein-coding |
| chr6_120122826_120123273 | chr6 | 120122827 | 120123273 | + | 0.725501 | intron (NM_016718, intron 1 of 3) | L1M3\|LINE\|L1 | 29670 | NM_016718 | 29862 | Mm.103668 | NM_016718 | ENSMUSG00000041377 | **Ninj2** | - | ninjurin 2 | protein-coding |
| chr10_61538040_61538439 | chr10 | 61538041 | 61538439 | + | 0.682067 | intron (NM_153542, intron 3 of 4) | MTEb\|LTR\|ERVL-MaLR | -57244 | NM_001177511 | 237362 | Mm.131297 | NM_001177511 | ENSMUSG00000020090 | **Npffr1** | Gm236\|Gpr147\|NPFF1\|OT7T022 | neuropeptide FF receptor 1 | protein-coding |
| chr12_89950044_89950535 | chr12 | 89950045 | 89950535 | + | 0.682798 | intron (NM_001252074, intron 1 of 6) | intron (NM_001252074, intron 1 of 6) | 137807 | NM_001252074 | 18191 | Mm.425766 | NM_172544 | ENSMUSG00000066392 | **Nrxn3** | - | neurexin III | protein-coding |
| chr12_89954522_89954841 | chr12 | 89954523 | 89954841 | + | 0.60885 | intron (NM_001252074, intron 1 of 6) | intron (NM_001252074, intron 1 of 6) | 142199 | NM_001252074 | 18191 | Mm.425766 | NM_172544 | ENSMUSG00000066392 | **Nrxn3** | - | neurexin III | protein-coding |
| chr11_32013428_32014054 | chr11 | 32013429 | 32014054 | + | 0.78205 | intron (NM_001290681, intron 2 of 4) | intron (NM_001290681, intron 2 of 4) | 12464 | NM_001290681 | 18197 | Mm.3304 | NM_008741 | ENSMUSG00000020297 | **Nsg2** | - | neuron specific gene family member 2 | protein-coding |
| chr11_32013024_32013861 | chr11 | 32013025 | 32013861 | + | 0.660811 | intron (NM_001290681, intron 2 of 4) | intron (NM_001290681, intron 2 of 4) | 12166 | NM_001290681 | 18197 | Mm.3304 | NM_008741 | ENSMUSG00000020297 | **Nsg2** | - | neuron specific gene family member 2 | protein-coding |
| chr16_55891377_55891726 | chr16 | 55891378 | 55891726 | + | 0.613062 | intron (NM_001134457, intron 1 of 5) | intron (NM_001134457, intron 1 of 5) | 3727 | NM_001134457 | 385658 | Mm.310635 | NM_001134457 | ENSMUSG00000075033 | **Nxpe3** | Fam55c\|Gm1752\|Gm611 | neurexophilin and PC-esterase domain family, member 3 | protein-coding |
| chr11_87936381_87936730 | chr11 | 87936382 | 87936730 | + | 0.782289 | Intergenic | B1_Mur4\|SINE\|Alu | -21553 | NM_146412 | 258407 | Mm.271337 | NM_146412 | ENSMUSG00000060787 | **Olfr464** | MOR240-2 | olfactory receptor 464 | protein-coding |
| chr11_4235728_4237061 | chr11 | 4235729 | 4237061 | + | 0.671039 | promoter-TSS (NM_001013365) | promoter-TSS (NM_001013365) | -390 | NM_001013365 | 18413 | Mm.131422 | NM_001013365 | ENSMUSG00000058755 | **Osm** | OncoM | oncostatin M | protein-coding |
| chr11_4236070_4237055 | chr11 | 4236071 | 4237055 | + | 0.627282 | promoter-TSS (NM_001013365) | promoter-TSS (NM_001013365) | -222 | NM_001013365 | 18413 | Mm.131422 | NM_001013365 | ENSMUSG00000058755 | **Osm** | OncoM | oncostatin M | protein-coding |
| chr3_60821483_60821894 | chr3 | 60821484 | 60821894 | + | 0.624055 | Intergenic | MTD\|LTR\|ERVL-MaLR | -181096 | NM_008772 | 18441 | Mm.281452 | NM_008772 | ENSMUSG00000027765 | **P2ry1** | P2Y1 | purinergic receptor P2Y, G-protein coupled 1 | protein-coding |
| chr19_53791724_53792843 | chr19 | 53791725 | 53792843 | + | 0.605398 | intron (NM_001170847, intron 1 of 13) | intron (NM_001170847, intron 1 of 13) | -99947 | NM_011050 | 18569 | Mm.1605 | NM_011050 | ENSMUSG00000024975 | **Pdcd4** | D19Ucla1\|Ma3\|Tis | programmed cell death 4 | protein-coding |
| chr2_76219613_76219907 | chr2 | 76219614 | 76219907 | + | 0.684142 | intron (NM_001081033, intron 2 of 19) | intron (NM_001081033, intron 2 of 19) | 118893 | NM_001081033 | 241489 | Mm.246613 | NM_001081033 | | **Pde11a** | 6330414F14Rik\|A630086N24Rik\|Gm350\|PDE11A1 | phosphodiesterase 11A | protein-coding |
| chr2_76219418_76219942 | chr2 | 76219419 | 76219942 | + | 0.626267 | intron (NM_001081033, intron 2 of 19) | intron (NM_001081033, intron 2 of 19) | 118973 | NM_001081033 | 241489 | Mm.246613 | NM_001081033 | | **Pde11a** | 6330414F14Rik\|A630086N24Rik\|Gm350\|PDE11A1 | phosphodiesterase 11A | protein-coding |
| chr3_30965598_30966342 | chr3 | 30965599 | 30966342 | + | 0.608164 | intron (NM_001165954, intron 1 of 14) | intron (NM_001165954, intron 1 of 14) | 3445 | NM_001165954 | 241915 | Mm.233173 | NM_153421 | ENSMUSG00000037652 | **Phc3** | E030046K01Rik\|Edr3\|Hph3 | polyhomeotic-like 3 (Drosophila) | protein-coding |
| chr17_29500634_29501279 | chr17 | 29500635 | 29501279 | + | 0.700984 | Intergenic | Intergenic | 9912 | NM_008842 | 18712 | Mm.405293 | NM_008842 | | **Pim1** | Pim-1 | proviral integration site 1 | protein-coding |
| chr2_59171560_59172058 | chr2 | 59171561 | 59172058 | + | 0.664179 | intron (NM_026361, intron 1 of 21) | intron (NM_026361, intron 1 of 21) | 10959 | NM_175464 | 227937 | Mm.260938 | NM_026361 | ENSMUSG00000026991 | **Pkp4** | 5031422I09Rik\|9430019K17Rik\|Armrp\|p0071 | plakophilin 4 | protein-coding |
| chr2_59171568_59171921 | chr2 | 59171569 | 59171921 | + | 0.656482 | intron (NM_026361, intron 1 of 21) | intron (NM_026361, intron 1 of 21) | 10895 | NM_175464 | 227937 | Mm.260938 | NM_026361 | ENSMUSG00000026991 | **Pkp4** | 5031422I09Rik\|9430019K17Rik\|Armrp\|p0071 | plakophilin 4 | protein-coding |
| chr2_59171246_59172068 | chr2 | 59171247 | 59172068 | + | 0.634375 | intron (NM_026361, intron 1 of 21) | intron (NM_026361, intron 1 of 21) | 10807 | NM_175464 | 227937 | Mm.260938 | NM_026361 | ENSMUSG00000026991 | **Pkp4** | 5031422I09Rik\|9430019K17Rik\|Armrp\|p0071 | plakophilin 4 | protein-coding |
| chr2_59171380_59171950 | chr2 | 59171381 | 59171950 | + | 0.625394 | intron (NM_026361, intron 1 of 21) | intron (NM_026361, intron 1 of 21) | 10815 | NM_175464 | 227937 | Mm.260938 | NM_026361 | ENSMUSG00000026991 | **Pkp4** | 5031422I09Rik\|9430019K17Rik\|Armrp\|p0071 | plakophilin 4 | protein-coding |
| chr2_11173085_11173457 | chr2 | 11173086 | 11173457 | + | 0.659105 | intron (NM_008859, intron 1 of 17) | intron (NM_008859, intron 1 of 17) | 889 | NM_008859 | 18761 | Mm.329993 | NM_008859 | ENSMUSG00000026778 | **Prkcq** | A130035A12Rik\|AW494342\|PKC-0\|PKC-theta\|PKCtheta\|Pkcq | protein kinase C, theta | protein-coding |
| chr13_85498608_85499003 | chr13 | 85498609 | 85499003 | + | 0.590259 | Intergenic | Intergenic | -209320 | NM_145452 | 218397 | Mm.259653 | NM_145452 | ENSMUSG00000021549 | **Rasa1** | Gap\|RasGAP\|Rasa | RAS p21 protein activator 1 | protein-coding |
| chr17_50097790_50098343 | chr17 | 50097791 | 50098343 | + | 0.587408 | intron (NM_181397, intron 3 of 9) | intron (NM_181397, intron 3 of 9) | 92430 | NM_181397 | 76438 | Mm.41854 | NM_181397 | | **Rftn1** | 2310015N21Rik\|AI853526 | raftlin lipid raft linker 1 | protein-coding |
| chr14_104519057_104519487 | chr14 | 104519058 | 104519487 | + | 0.594683 | intron (NM_026047, intron 1 of 5) | ORR1D2-int\|LTR\|ERVL-MaLR | 3394 | NM_026047 | 72486 | Mm.291811 | NM_026047 | ENSMUSG00000022120 | **Rnf219** | 2610206B13Rik\|2810449K13Rik\|AI451544 | ring finger protein 219 | protein-coding |
| chr7_114171284_114172183 | chr7 | 114171285 | 114172183 | + | 0.639695 | Intergenic | Intergenic | -53953 | NM_025846 | 66922 | Mm.276572 | NM_025846 | ENSMUSG00000055723 | **Rras2** | 2610016H24Rik\|C86394\|TC21 | related RAS viral (r-ras) oncogene homolog 2 | protein-coding |
| chr10_61675995_61676335 | chr10 | 61675996 | 61676335 | + | 0.619957 | Intergenic | RLTR30E_MM\|LTR\|ERV1 | -4156 | NM_009120 | 20224 | Mm.391794 | NM_009120 | ENSMUSG00000020088 | **Sar1a** | 1600019H17Rik\|Sara\|Sara1 | SAR1 gene homolog A (S. cerevisiae) | protein-coding |

| | | | | | | | | | | | | | | | | | |
|---|---|---|---|---|---|---|---|---|---|---|---|---|---|---|---|---|---|
| chr1_80015311_80015841 | chr1 | 80015312 | 80015841 | + | 0.597272 | Intergenic | L2\|LINE\|L2 | -156911 | NM_009255 | 20720 | Mm.3093 | NM_009255 | ENSMUSG00000026249 | **Serpine2** | B230326M24Rik\|PAI-1\|PI-7\|PI7\|PN-1\|Spi4 | serine (or cysteine) peptidase inhibitor, clade E, member 2 | protein-coding |
| chr18_79459488_79460007 | chr18 | 79459489 | 79460007 | + | 0.955102 | Intergenic | Intergenic | -350357 | NM_053099 | 240427 | Mm.312871 | NM_053099 | ENSMUSG00000024548 | **Setbp1** | C130092E12\|Seb\|mKIAA0437 | SET binding protein 1 | protein-coding |
| chr18_79434525_79435241 | chr18 | 79434526 | 79435241 | + | 0.698228 | Intergenic | Intergenic | -325492 | NM_053099 | 240427 | Mm.312871 | NM_053099 | ENSMUSG00000024548 | **Setbp1** | C130092E12\|Seb\|mKIAA0437 | SET binding protein 1 | protein-coding |
| chr17_53650120_53650607 | chr17 | 53650121 | 53650607 | + | 0.673712 | intron (NM_020005, intron 9 of 17) | intron (NM_020005, intron 9 of 17) | 38951 | NM_028232 | 72415 | Mm.153202 | NM_028232 | ENSMUSG00000023940 | **Sgol1** | 3300001M08Rik\|C81037\|Sgo1 | shugoshin-like 1 (S. pombe) | protein-coding |
| chr14_59694817_59695881 | chr14 | 59694818 | 59695881 | + | 0.746441 | intron (NM_015803, intron 32 of 35) | intron (NM_015803, intron 32 of 35) | 70068 | NM_145463 | 219134 | Mm.275409 | NM_145463 | ENSMUSG00000044461 | **Shisa2** | 9430059P22Rik\|MAd2\|Tmem46\|mShisa\|shisa | shisa homolog 2 (Xenopus laevis) | protein-coding |
| chr14_59695192_59695831 | chr14 | 59695193 | 59695831 | + | 0.680527 | intron (NM_015803, intron 32 of 35) | PB1D7\|SINE\|Alu | 70231 | NM_145463 | 219134 | Mm.275409 | NM_145463 | ENSMUSG00000044461 | **Shisa2** | 9430059P22Rik\|MAd2\|Tmem46\|mShisa\|shisa | shisa homolog 2 (Xenopus laevis) | protein-coding |
| chr12_82294937_82295820 | chr12 | 82294938 | 82295820 | + | 0.750129 | intron (NM_001167983, intron 3 of 23) | intron (NM_001167983, intron 3 of 23) | -15970 | NM_172579 | 217692 | Mm.261333 | NM_172579 | ENSMUSG00000042700 | **Sipa1l1** | 4931426N11Rik\|AW213287\|Spar\|mKIAA0440 | signal-induced proliferation-associated 1 like 1 | protein-coding |
| chr12_82295326_82295841 | chr12 | 82295327 | 82295841 | + | 0.690842 | intron (NM_001167983, intron 3 of 23) | intron (NM_001167983, intron 3 of 23) | -15765 | NM_172579 | 217692 | Mm.261333 | NM_172579 | ENSMUSG00000042700 | **Sipa1l1** | 4931426N11Rik\|AW213287\|Spar\|mKIAA0440 | signal-induced proliferation-associated 1 like 1 | protein-coding |
| chr12_82322755_82323106 | chr12 | 82322756 | 82323106 | + | 0.613151 | intron (NM_172579, intron 1 of 20) | intron (NM_172579, intron 1 of 20) | 11582 | NM_172579 | 217692 | Mm.261333 | NM_172579 | ENSMUSG00000042700 | **Sipa1l1** | 4931426N11Rik\|AW213287\|Spar\|mKIAA0440 | signal-induced proliferation-associated 1 like 1 | protein-coding |
| chr12_82322824_82323437 | chr12 | 82322825 | 82323437 | + | 0.597632 | intron (NM_172579, intron 1 of 20) | intron (NM_172579, intron 1 of 20) | 11782 | NM_172579 | 217692 | Mm.261333 | NM_172579 | ENSMUSG00000042700 | **Sipa1l1** | 4931426N11Rik\|AW213287\|Spar\|mKIAA0440 | signal-induced proliferation-associated 1 like 1 | protein-coding |
| chr2_122679704_122680505 | chr2 | 122679705 | 122680505 | + | 1.01091 | Intergenic | Intergenic | 22558 | NM_011774 | 22785 | Mm.27801 | NM_011774 | ENSMUSG00000005802 | **Slc30a4** | Znt4\|lm\|znT-4 | solute carrier family 30 (zinc transporter), member 4 | protein-coding |
| chr2_122672669_122673588 | chr2 | 122672670 | 122673588 | + | 0.680528 | Intergenic | ORR1D2\|LTR\|ERVL-MaLR | 29534 | NM_011774 | 22785 | Mm.27801 | NM_011774 | ENSMUSG00000005802 | **Slc30a4** | Znt4\|lm\|znT-4 | solute carrier family 30 (zinc transporter), member 4 | protein-coding |
| chr19_38395118_38395504 | chr19 | 38395119 | 38395504 | + | 0.910601 | promoter-TSS (NM_175507) | promoter-TSS (NM_175507) | -669 | NM_175507 | 240660 | Mm.262085 | NM_175507 | ENSMUSG00000044026 | **Slc35g1** | AA590464\|D330039I19Rik\|Tmem20 | solute carrier family 35, member G1 | protein-coding |
| chr1_46864072_46864769 | chr1 | 46864073 | 46864769 | + | 0.740001 | Intergenic | Intergenic | -10912 | NM_172653 | 227059 | Mm.233889 | NM_172653 | ENSMUSG00000025986 | **Slc39a10** | 2900042E17Rik\|5430433I10\|mKIAA1265 | solute carrier family 39 (zinc transporter), member 10 | protein-coding |
| chr19_26641369_26641678 | chr19 | 26641370 | 26641678 | + | 0.853144 | intron (NM_011416, intron 5 of 32) | intron (NM_011416, intron 5 of 32) | 36364 | NM_011416 | 67155 | Mm.491146 | NM_011416 | ENSMUSG00000024921 | **Smarca2** | 2610209L14Rik\|SNF2alpha\|Snf2l2\|brahma\|brm | SWI/SNF related, matrix associated, actin dependent regulator of chromatin, subfamily a, member 2 | protein-coding |
| chr16_11577248_11578354 | chr16 | 11577249 | 11578354 | + | 0.586139 | intron (NM_001290148, intron 8 of 13) | intron (NM_001290148, intron 8 of 13) | 172153 | NM_001290148 | 74478 | Mm.325831 | NM_028964 | ENSMUSG00000071669 | **Snx29** | 4933437K13Rik\|ENSMUSG00000079738\|Gm11170\|Gm1737\|Gm930\|Rundc2a | sorting nexin 29 | protein-coding |
| chr11_62148181_62149063 | chr11 | 62148182 | 62149063 | + | 1.15396 | intron (NM_001281818, intron 9 of 14) | intron (NM_001281818, intron 9 of 14) | 71598 | NM_001029936 | 432572 | Mm.458025 | NM_001029936 | ENSMUSG00000042331 | **Specc1** | 2810012G08Rik\|B230396K10Rik\|Cytsb | sperm antigen with calponin homology and coiled-coil domains 1 | protein-coding |
| chr5_54070594_54070918 | chr5 | 54070595 | 54070918 | + | 0.716338 | intron (NM_001081103, intron 2 of 11) | intron (NM_001081103, intron 2 of 11) | 72233 | NM_001081103 | 116873 | Mm.265985 | NM_054083 | ENSMUSG00000039156 | **Stim2** | - | stromal interaction molecule 2 | protein-coding |
| chr5_54067631_54067938 | chr5 | 54067632 | 54067938 | + | 0.587571 | intron (NM_001081103, intron 2 of 11) | L1MEc\|LINE\|L1 | 69262 | NM_001081103 | 116873 | Mm.265985 | NM_054083 | ENSMUSG00000039156 | **Stim2** | - | stromal interaction molecule 2 | protein-coding |
| chr12_105019262_105019794 | chr12 | 105019263 | 105019794 | + | 0.655259 | Intergenic | Intergenic | -9719 | NM_172500 | 212073 | Mm.212688 | NM_172500 | ENSMUSG00000054150 | **Syne3** | nesprin-3 | spectrin repeat containing, nuclear envelope family member 3 | protein-coding |
| chr6_52801790_52802789 | chr6 | 52801791 | 52802789 | + | 0.617957 | intron (NM_173406, intron 3 of 4) | intron (NM_173406, intron 3 of 4) | 88561 | NM_025816 | 52440 | Mm.389757 | NM_025816 | ENSMUSG00000004535 | **Tax1bp1** | 1200003J11Rik\|1700069J21Rik\|AA930106\|D6Ertd404e\|D6Ertd772e\|T6bp\|TXBP151 | Tax1 (human T cell leukemia virus type I) binding protein 1 | protein-coding |
| chr3_21965407_21965959 | chr3 | 21965408 | 21965959 | + | 0.922694 | Intergenic | Intergenic | -110969 | NM_030732 | 81004 | Mm.202966 | NM_030732 | ENSMUSG00000027630 | **Tbl1xr1** | 8030499H02Rik\|A630076E03Rik\|AW539987\|C21\|C230089I12Rik\|DC42\|Ira1\|TBLR1 | transducin (beta)-like 1X-linked receptor 1 | protein-coding |
| chr11_117881853_117882231 | chr11 | 117881854 | 117882231 | + | 0.685261 | Intergenic | Intergenic | -8516 | NM_027919 | 71776 | Mm.3200 | NM_027919 | ENSMUSG00000017713 | **Tha1** | 1300017K07Rik\|GLY1 | threonine aldolase 1 | protein-coding |
| chr11_117879072_117879313 | chr11 | 117879073 | 117879313 | + | 0.659174 | Intergenic | Intergenic | -5667 | NM_027919 | 71776 | Mm.3200 | NM_027919 | ENSMUSG00000017713 | **Tha1** | 1300017K07Rik\|GLY1 | threonine aldolase 1 | protein-coding |
| chr6_59312928_59313891 | chr6 | 59312929 | 59313891 | + | 0.593323 | Intergenic | RLTR14_RN\|LTR\|ERV1 | 104540 | NM_001081145 | 68140 | Mm.25843 | NM_001081145 | ENSMUSG00000049232 | **Tigd2** | 3632410O17Rik | tigger transposable element derived 2 | protein-coding |
| chr5_3669510_3670721 | chr5 | 3669511 | 3670721 | + | 0.62201 | exon (NM_029154, exon 8 of 11) | exon (NM_029154, exon 8 of 11) | 13112 | NM_029154 | 75010 | Mm.85064 | NM_029141 | ENSMUSG00000014529 | **Tmbim7** | 4930403J02Rik\|4930500J03Rik\|4930511M11Rik\|Lgf5\|Tmbim1b | transmembrane BAX inhibitor motif containing 7 | protein-coding |
| chr19_21051101_21051368 | chr19 | 21051102 | 21051368 | + | 0.691023 | Intergenic | Intergenic | -97033 | NM_028953 | 13409 | Mm.219585 | NM_028953 | ENSMUSG00000024749 | **Tmc1** | 4933416G09Rik\|Beethoven\|Bth\|CWEA1\|dn | transmembrane channel-like gene family 1 | protein-coding |
| chr19_21051066_21051687 | chr19 | 21051067 | 21051687 | + | 0.658762 | Intergenic | Intergenic | -97175 | NM_028953 | 13409 | Mm.219585 | NM_028953 | ENSMUSG00000024749 | **Tmc1** | 4933416G09Rik\|Beethoven\|Bth\|CWEA1\|dn | transmembrane channel-like gene family 1 | protein-coding |
| chr1_51092371_51092706 | chr1 | 51092372 | 51092706 | + | 0.773696 | intron (NM_019790, intron 5 of 9) | intron (NM_019790, intron 5 of 9) | 165016 | NM_019790 | 56363 | Mm.245154 | NM_019790 | ENSMUSG00000026109 | **Tmeff2** | 4832418D20Rik\|7630402F16Rik\|TR-2 | transmembrane protein with EGF-like and two follistatin-like domains 2 | protein-coding |
| chr1_36892379_36892960 | chr1 | 36892380 | 36892960 | + | 0.640591 | intron (NM_018872, intron 1 of 40) | MTEa\|LTR\|ERVL-MaLR | 46857 | NM_018872 | 56030 | Mm.186943 | NM_018872 | ENSMUSG00000026116 | **Tmem131** | 2610524E03Rik\|CC28\|D1Bwg0491e\|Neg\|RW1\|YR-23\|mKIAA0257 | transmembrane protein 131 | protein-coding |
| chr9_123271591_123272147 | chr9 | 123271592 | 123272147 | + | 0.660123 | Intergenic | Lx2\|LINE\|L1 | -11080 | NM_001002267 | 72309 | Mm.8569 | NM_001002267 | ENSMUSG00000054871 | **Tmem158** | 2310037P21Rik\|Ris1 | transmembrane protein 158 | protein-coding |
| chr4_63841786_63842938 | chr4 | 63841787 | 63842938 | + | 0.598879 | intron (NM_009403, intron 2 of 3) | intron (NM_009403, intron 2 of 3) | 18922 | NM_009403 | 21949 | Mm.4664 | NM_009403 | | **Tnfsf8** | CD153\|CD30LG\|Cd30l | tumor necrosis factor (ligand) superfamily, member 8 | protein-coding |
| chr12_28735238_28736135 | chr12 | 28735239 | 28736135 | + | 0.58676 | intron (NM_178811, intron 3 of 11) | intron (NM_178811, intron 3 of 11) | 14707 | NM_001161410 | 217449 | Mm.132006 | NM_178811 | ENSMUSG00000020628 | **Trappc12** | CGI-87\|D930014A20Rik\|Ttc15 | trafficking protein particle complex 12 | protein-coding |
| chr6_41202893_41203503 | chr6 | 41202894 | 41203503 | + | 0.683062 | Intergenic | Intergenic | -99074 | NM_011646 | 22074 | Mm.383263 | NM_011646 | ENSMUSG00000054106 | **Try4** | 0910001B19Rik\|Td | trypsin 4 | protein-coding |
| chr9_66616813_66617168 | chr9 | 66616814 | 66617168 | + | 0.724345 | Intergenic | Intergenic | -24011 | NM_144937 | 235441 | Mm.38976 | NM_144937 | ENSMUSG00000032376 | **Usp3** | AA409661\|BC017156 | ubiquitin specific peptidase 3 | protein-coding |
| chr9_66616835_66617502 | chr9 | 66616836 | 66617502 | + | 0.720467 | Intergenic | Intergenic | -24189 | NM_144937 | 235441 | Mm.38976 | NM_144937 | ENSMUSG00000032376 | **Usp3** | AA409661\|BC017156 | ubiquitin specific peptidase 3 | protein-coding |
| chr9_66616872_66617303 | chr9 | 66616873 | 66617303 | + | 0.716338 | Intergenic | Intergenic | -24108 | NM_144937 | 235441 | Mm.38976 | NM_144937 | ENSMUSG00000032376 | **Usp3** | AA409661\|BC017156 | ubiquitin specific peptidase 3 | protein-coding |
| chr2_6338083_6338603 | chr2 | 6338084 | 6338603 | + | 0.669914 | intron (NM_001080548, intron 2 of 14) | intron (NM_001080548, intron 2 of 14) | -14393 | NM_181399 | 98910 | Mm.40673 | NM_181399 | ENSMUSG00000039046 | **Usp6nl** | AI316785\|RNTRE\|TRE2NL\|mKIAA0019 | USP6 N-terminal like | protein-coding |
| chr2_6337866_6338618 | chr2 | 6337867 | 6338618 | + | 0.662231 | intron (NM_001080548, intron 2 of 14) | intron (NM_001080548, intron 2 of 14) | -14494 | NM_181399 | 98910 | Mm.40673 | NM_181399 | ENSMUSG00000039046 | **Usp6nl** | AI316785\|RNTRE\|TRE2NL\|mKIAA0019 | USP6 N-terminal like | protein-coding |
| chr12_84525017_84525837 | chr12 | 84525018 | 84525837 | + | 0.743931 | intron (NM_173756, intron 5 of 5) | B4\|SINE\|B4 | -44425 | NM_007701 | 12677 | Mm.4405 | NM_007701 | ENSMUSG00000021239 | **Vsx2** | Chx10\|Hox-10\|Hox10\|or | visual system homeobox 2 | protein-coding |
| chr16_32520432_32520900 | chr16 | 32520433 | 32520900 | + | 0.713872 | Intergenic | Intergenic | 24385 | NM_199309 | 245308 | Mm.76859 | NM_199309 | ENSMUSG00000052363 | **Zdhhc19** | Gm1744\|Gm616 | zinc finger, DHHC domain containing 19 | protein-coding |
| chr16_32520117_32521039 | chr16 | 32520118 | 32521039 | + | 0.710355 | Intergenic | Intergenic | 24297 | NM_199309 | 245308 | Mm.76859 | NM_199309 | ENSMUSG00000052363 | **Zdhhc19** | Gm1744\|Gm616 | zinc finger, DHHC domain containing 19 | protein-coding |
| chr16_18233481_18233802 | chr16 | 18233482 | 18233802 | + | 0.676592 | intron (NM_172151, intron 1 of 10) | intron (NM_172151, intron 1 of 10) | 1494 | NM_172151 | 27801 | Mm.326263 | NM_172151 | ENSMUSG00000060166 | **Zdhhc8** | D16H22S1738E\|DHHC-8\|E330009O14Rik\|Op53c05\|ZDHHCL1 | zinc finger, DHHC domain containing 8 | protein-coding |
| chr5_105859149_105859701 | chr5 | 105859150 | 105859701 | + | 0.631 | Intergenic | Intergenic | -17143 | NR_030777 | 54367 | Mm.248876 | NM_018759 | ENSMUSG00000029290 | **Zfp326** | 5730470H14Rik\|ZAN75\|Znf326 | zinc finger protein 326 | protein-coding |
| chr9_65831225_65831684 | chr9 | 65831226 | 65831684 | + | 0.739415 | 3' UTR (NM_001170907, exon 12 of 12) | 3' UTR (NM_001170907, exon 12 of 12) | -3891 | NM_172536 | 214812 | Mm.139231 | NM_172536 | ENSMUSG00000040524 | **Zfp609** | 9830165N24\|BC016271\|Znf609\|mKIAA0295 | zinc finger protein 609 | protein-coding |
| chr11_95748121_95748792 | chr11 | 95748122 | 95748792 | + | 0.668623 | promoter-TSS (NM_201609) | promoter-TSS (NM_201609) | -610 | NM_201609 | 268469 | Mm.389038 | NM_201609 | ENSMUSG00000075595 | **Zfp652** | 9530033F24Rik\|ZNF652 | zinc finger protein 652 | protein-coding |
| chr5_109996131_109996814 | chr5 | 109996132 | 109996814 | + | 1.23067 | promoter-TSS (NM_145563) | promoter-TSS (NM_145563) | -54 | NM_145563 | 69504 | Mm.441438 | NM_145563 | ENSMUSG00000066613 | **Zfp932** | 2310001H12Rik\|Znf431 | zinc finger protein 932 | protein-coding |
| chr2_165877959_165878431 | chr2 | 165877960 | 165878431 | + | 0.632331 | 5' UTR (NM_001291158, exon 1 of 24) | 5' UTR (NM_001291158, exon 1 of 24) | 242 | NM_001291158 | 228880 | Mm.227598 | NM_027230 | ENSMUSG00000039671 | **Zmynd8** | 1110013E22Rik\|2010005I16Rik\|3632413B07Rik\|AI316811\|AL024039\|Prkcbp1\|RACK7 | zinc finger, MYND-type containing 8 | protein-coding |
| chr2_165877689_165878431 | chr2 | 165877690 | 165878431 | + | 0.599773 | 5' UTR (NM_001291158, exon 1 of 24) | 5' UTR (NM_001291158, exon 1 of 24) | 377 | NM_001291158 | 228880 | Mm.227598 | NM_027230 | ENSMUSG00000039671 | **Zmynd8** | 1110013E22Rik\|2010005I16Rik\|3632413B07Rik\|AI316811\|AL024039\|Prkcbp1\|RACK7 | zinc finger, MYND-type containing 8 | protein-coding |

**Table S1B. HOMER Opening peaks in Ag-stimulated cells**
Data is sorted alphabetically by the gene name; Only peaks above the peak score value of 0.5 are shown.

| PeakID (cmd=annotatePeaks.pl ../averagescore_opening_peaks_allCA.bed mm10 -go go_opening_peaks_allCA) | Chr | Start | End | Strand | Peak Score | Annotation | Detailed Annotation | Distance to TSS | Nearest PromoterID | Entrez ID | Nearest Unigene | Nearest Refseq | Nearest Ensembl | Gene Name | Gene Alias | Gene Description | Gene Type |
|---|---|---|---|---|---|---|---|---|---|---|---|---|---|---|---|---|---|
| chr12_85823551_85823931 | chr12 | 85823552 | 85823931 | + | 1.3124 | intron (NM_021446, intron 1 of 4) | intron (NM_021446, intron 1 of 4) | 804 | NM_021446 | 58520 | Mm.143795 | NM_021446 | ENSMUSG00000021252 | **0610007P14Rik** | 1190004E09Rik\|AU019315\|C77855\|ORF11 | RIKEN cDNA 0610007P14 gene | protein-coding |
| chr8_77435979_77436662 | chr8 | 77435980 | 77436662 | + | 1.17372 | intron (NM_030113, intron 3 of 22) | intron (NM_030113, intron 3 of 22) | -80735 | NR_028125 | 70345 | Mm.411813 | NR_028125 | ENSMUSG00000097882 | **0610038B21Rik** | - | RIKEN cDNA 0610038B21 gene | ncRNA |
| chr8_77210190_77210775 | chr8 | 77210191 | 77210775 | + | 1.09747 | intron (NM_001083906, intron 6 of 8) | intron (NM_001083906, intron 6 of 8) | -306573 | NR_028125 | 70345 | Mm.411813 | NR_028125 | ENSMUSG00000097882 | **0610038B21Rik** | - | RIKEN cDNA 0610038B21 gene | ncRNA |
| chr8_77307237_77307578 | chr8 | 77307238 | 77307578 | + | 0.904463 | intron (NM_030113, intron 19 of 22) | L1VL4\|LINE\|L1 | -209648 | NR_028125 | 70345 | Mm.411813 | NR_028125 | ENSMUSG00000097882 | **0610038B21Rik** | - | RIKEN cDNA 0610038B21 gene | ncRNA |
| chr8_77508935_77509462 | chr8 | 77508936 | 77509462 | + | 0.783472 | intron (NM_030113, intron 1 of 22) | intron (NM_030113, intron 1 of 22) | -7857 | NR_028125 | 70345 | Mm.411813 | NR_028125 | ENSMUSG00000097882 | **0610038B21Rik** | - | RIKEN cDNA 0610038B21 gene | ncRNA |
| chr8_77309029_77309646 | chr8 | 77309030 | 77309646 | + | 0.763863 | intron (NM_030113, intron 19 of 22) | intron (NM_030113, intron 19 of 22) | -207718 | NR_028125 | 70345 | Mm.411813 | NR_028125 | ENSMUSG00000097882 | **0610038B21Rik** | - | RIKEN cDNA 0610038B21 gene | ncRNA |
| chr8_77262364_77262819 | chr8 | 77262365 | 77262819 | + | 0.751566 | intron (NM_030113, intron 20 of 22) | ORR1D1\|LTR\|ERVL-MaLR | -254464 | NR_028125 | 70345 | Mm.411813 | NR_028125 | ENSMUSG00000097882 | **0610038B21Rik** | - | RIKEN cDNA 0610038B21 gene | ncRNA |
| chr8_77498195_77498682 | chr8 | 77498196 | 77498682 | + | 0.655832 | intron (NM_030113, intron 1 of 22) | intron (NM_030113, intron 1 of 22) | -18617 | NR_028125 | 70345 | Mm.411813 | NR_028125 | ENSMUSG00000097882 | **0610038B21Rik** | - | RIKEN cDNA 0610038B21 gene | ncRNA |
| chr8_77433259_77433686 | chr8 | 77433260 | 77433686 | + | 0.646763 | intron (NM_030113, intron 3 of 22) | intron (NM_030113, intron 3 of 22) | -83583 | NR_028125 | 70345 | Mm.411813 | NR_028125 | ENSMUSG00000097882 | **0610038B21Rik** | - | RIKEN cDNA 0610038B21 gene | ncRNA |
| chr8_77401201_77401495 | chr8 | 77401202 | 77401495 | + | 0.624122 | intron (NM_030113, intron 10 of 22) | intron (NM_030113, intron 10 of 22) | -115708 | NR_028125 | 70345 | Mm.411813 | NR_028125 | ENSMUSG00000097882 | **0610038B21Rik** | - | RIKEN cDNA 0610038B21 gene | ncRNA |
| chr2_163639791_163640028 | chr2 | 163639792 | 163640028 | + | 0.773356 | intron (NM_012032, intron 1 of 9) | B1_Mur1\|SINE\|Alu | -4940 | NR_028113 | 68386 | Mm.410649 | NR_028113 | ENSMUSG00000058812 | **0610039K10Rik** | - | RIKEN cDNA 0610039K10 gene | ncRNA |
| chr6_108665465_108666001 | chr6 | 108665466 | 108666001 | + | 1.67297 | 3' UTR (NM_011498, exon 5 of 5) | 3' UTR (NM_011498, exon 5 of 5) | -4799 | NR_104577 | 75394 | Mm.395061 | NR_040757 | ENSMUSG00000087341 | **0610040F04Rik** | - | RIKEN cDNA 0610040F04 gene | ncRNA |
| chr6_108672126_108672489 | chr6 | 108672127 | 108672489 | + | 1.29769 | Intergenic | L1MB5\|LINE\|L1 | -11374 | NR_104577 | 75394 | Mm.395061 | NR_040757 | ENSMUSG00000087341 | **0610040F04Rik** | - | RIKEN cDNA 0610040F04 gene | ncRNA |
| chr6_108688167_108688975 | chr6 | 108688168 | 108688975 | + | 1.27154 | Intergenic | Intergenic | -27637 | NR_104577 | 75394 | Mm.395061 | NR_040757 | ENSMUSG00000087341 | **0610040F04Rik** | - | RIKEN cDNA 0610040F04 gene | ncRNA |
| chr6_108569376_108569996 | chr6 | 108569377 | 108569996 | + | 1.09305 | Intergenic | Intergenic | 53481 | NR_040757 | 75394 | Mm.395061 | NR_040757 | ENSMUSG00000087341 | **0610040F04Rik** | - | RIKEN cDNA 0610040F04 gene | ncRNA |
| chr6_108706073_108706389 | chr6 | 108706074 | 108706389 | + | 0.965052 | Intergenic | Intergenic | -45297 | NR_104577 | 75394 | Mm.395061 | NR_040757 | ENSMUSG00000087341 | **0610040F04Rik** | - | RIKEN cDNA 0610040F04 gene | ncRNA |
| chr6_108710765_108710966 | chr6 | 108710766 | 108710966 | + | 0.76599 | Intergenic | B3A\|SINE\|B2 | -49932 | NR_104577 | 75394 | Mm.395061 | NR_040757 | ENSMUSG00000087341 | **0610040F04Rik** | - | RIKEN cDNA 0610040F04 gene | ncRNA |
| chr6_108558064_108558269 | chr6 | 108558065 | 108558269 | + | 0.746463 | Intergenic | ORR1C2\|LTR\|ERVL-MaLR | 65000 | NR_040757 | 75394 | Mm.395061 | NR_040757 | ENSMUSG00000087341 | **0610040F04Rik** | - | RIKEN cDNA 0610040F04 gene | ncRNA |
| chr6_108709415_108710337 | chr6 | 108709416 | 108710337 | + | 0.62445 | Intergenic | Intergenic | -48942 | NR_104577 | 75394 | Mm.395061 | NR_040757 | ENSMUSG00000087341 | **0610040F04Rik** | - | RIKEN cDNA 0610040F04 gene | ncRNA |
| chr5_115629713_115630043 | chr5 | 115629714 | 115630043 | + | 1.61598 | Intergenic | Intergenic | 1938 | NR_027810 | 66123 | Mm.133063 | NM_021417 | | **1110006O24Rik** | - | RIKEN cDNA 1110006O24 gene | ncRNA |
| chr5_115629107_115630043 | chr5 | 115629108 | 115630043 | + | 1.52384 | Intergenic | Intergenic | 2241 | NR_027810 | 66123 | Mm.133063 | NM_021417 | | **1110006O24Rik** | - | RIKEN cDNA 1110006O24 gene | ncRNA |
| chr3_68828604_68828964 | chr3 | 68828605 | 68828964 | + | 1.07632 | Intergenic | Intergenic | -40802 | NM_001167996 | 68725 | Mm.45741 | NM_001167996 | ENSMUSG00000046999 | **1110032F04Rik** | AI115547 | RIKEN cDNA 1110032F04 gene | protein-coding |
| chr3_68927654_68928126 | chr3 | 68927655 | 68928126 | + | 0.987194 | intron (NM_026641, intron 12 of 19) | intron (NM_026641, intron 12 of 19) | 58304 | NM_001167996 | 68725 | Mm.45741 | NM_001167996 | ENSMUSG00000046999 | **1110032F04Rik** | AI115547 | RIKEN cDNA 1110032F04 gene | protein-coding |
| chr3_68829794_68830510 | chr3 | 68829795 | 68830510 | + | 0.739682 | Intergenic | Intergenic | -39434 | NM_001167996 | 68725 | Mm.45741 | NM_001167996 | ENSMUSG00000046999 | **1110032F04Rik** | AI115547 | RIKEN cDNA 1110032F04 gene | protein-coding |
| chr3_68786280_68786582 | chr3 | 68786281 | 68786582 | + | 0.686288 | Intergenic | Intergenic | -83155 | NM_001167996 | 68725 | Mm.45741 | NM_001167996 | ENSMUSG00000046999 | **1110032F04Rik** | AI115547 | RIKEN cDNA 1110032F04 gene | protein-coding |
| chr2_132703647_132704434 | chr2 | 132703648 | 132704434 | + | 0.84092 | intron (NM_028637, intron 2 of 2) | intron (NM_028637, intron 2 of 2) | 13758 | NM_028637 | 73747 | Mm.284114 | NM_028637 | ENSMUSG00000044991 | **1110034G24Rik** | - | RIKEN cDNA 1110034G24 gene | protein-coding |
| chr2_132688638_132688911 | chr2 | 132688639 | 132688911 | + | 0.827393 | Intergenic | L1M4\|LINE\|L1 | -1508 | NM_028637 | 73747 | Mm.284114 | NM_028637 | ENSMUSG00000044991 | **1110034G24Rik** | - | RIKEN cDNA 1110034G24 gene | protein-coding |
| chr2_132645397_132645654 | chr2 | 132645398 | 132645654 | + | 0.688431 | intron (NR_033468, intron 4 of 5) | ORR1B1\|LTR\|ERVL-MaLR | -44757 | NM_028637 | 73747 | Mm.284114 | NM_028637 | ENSMUSG00000044991 | **1110034G24Rik** | - | RIKEN cDNA 1110034G24 gene | protein-coding |
| chr2_132688409_132689108 | chr2 | 132688410 | 132689108 | + | 0.676368 | Intergenic | L1M4\|LINE\|L1 | -1524 | NM_028637 | 73747 | Mm.284114 | NM_028637 | ENSMUSG00000044991 | **1110034G24Rik** | - | RIKEN cDNA 1110034G24 gene | protein-coding |
| chr2_132702198_132702523 | chr2 | 132702199 | 132702523 | + | 0.65461 | intron (NM_028637, intron 2 of 2) | B1F\|SINE\|Alu | 12078 | NM_028637 | 73747 | Mm.284114 | NM_028637 | ENSMUSG00000044991 | **1110034G24Rik** | - | RIKEN cDNA 1110034G24 gene | protein-coding |
| chr1_24045593_24046020 | chr1 | 24045594 | 24046020 | + | 0.86093 | intron (NM_026604, intron 10 of 20) | intron (NM_026604, intron 10 of 20) | -40167 | NM_026503 | 68002 | Mm.250424 | NM_026503 | ENSMUSG00000026154 | **1110058L19Rik** | 1700001E18Rik\|C6orf57 | RIKEN cDNA 1110058L19 gene | protein-coding |
| chr1_24045354_24046042 | chr1 | 24045355 | 24046042 | + | 0.755787 | intron (NM_026604, intron 10 of 20) | intron (NM_026604, intron 10 of 20) | -40058 | NM_026503 | 68002 | Mm.250424 | NM_026503 | ENSMUSG00000026154 | **1110058L19Rik** | 1700001E18Rik\|C6orf57 | RIKEN cDNA 1110058L19 gene | protein-coding |
| chr19_21643752_21644304 | chr19 | 21643753 | 21644304 | + | 0.857118 | intron (NM_025423, intron 1 of 3) | intron (NM_025423, intron 1 of 3) | 8763 | NM_025423 | 66206 | Mm.272430 | NM_025423 | ENSMUSG00000035171 | **1110059E24Rik** | - | RIKEN cDNA 1110059E24 gene | protein-coding |
| chr19_21643979_21644298 | chr19 | 21643980 | 21644298 | + | 0.774982 | intron (NM_025423, intron 1 of 3) | intron (NM_025423, intron 1 of 3) | 8652 | NM_025423 | 66206 | Mm.272430 | NM_025423 | ENSMUSG00000035171 | **1110059E24Rik** | - | RIKEN cDNA 1110059E24 gene | protein-coding |
| chr9_94411137_94412005 | chr9 | 94411138 | 94412005 | + | 0.961184 | Intergenic | Intergenic | 126510 | NM_001033145 | 68861 | Mm.258746 | NM_001033145 | ENSMUSG00000045414 | **1190002N15Rik** | GoPro49\|HASF | RIKEN cDNA 1190002N15 gene | protein-coding |
| chr11_73036935_73037293 | chr11 | 73036936 | 73037293 | + | 0.65271 | intron (NM_018883, intron 11 of 15) | intron (NM_018883, intron 11 of 15) | -10753 | NM_025818 | 66874 | Mm.41903 | NM_025818 | | **1200014J11Rik** | C130061O14Rik\|C78393 | RIKEN cDNA 1200014J11 gene | protein-coding |
| chr16_21793382_21793758 | chr16 | 21793383 | 21793758 | + | 0.697129 | promoter-TSS (NR_037958) | promoter-TSS (NR_037958) | -777 | NR_037958 | 100043489 | Mm.326535 | NR_037957 | ENSMUSG00000097290 | **1300002E11Rik** | 1500026P21Rik | RIKEN cDNA 1300002E11 gene | ncRNA |
| chr19_20424138_20424510 | chr19 | 20424139 | 20424510 | + | 0.78697 | Intergenic | Intergenic | 19036 | NR_045817 | 68994 | Mm.261028 | NM_183277 | ENSMUSG00000094732 | **1500015L24Rik** | - | RIKEN cDNA 1500015L24 gene | ncRNA |
| chr1_43766292_43766749 | chr1 | 43766293 | 43766749 | + | 1.80481 | intron (NM_026430, intron 9 of 14) | intron (NM_026430, intron 9 of 14) | 35919 | NM_024283 | 78896 | Mm.50109 | NM_024283 | ENSMUSG00000026051 | **1500015O10Rik** | Ecrg4 | RIKEN cDNA 1500015O10 gene | protein-coding |
| chr1_43675389_43676038 | chr1 | 43675390 | 43676038 | + | 0.866079 | Intergenic | Intergenic | -54888 | NM_024283 | 78896 | Mm.50109 | NM_024283 | ENSMUSG00000026051 | **1500015O10Rik** | Ecrg4 | RIKEN cDNA 1500015O10 gene | protein-coding |
| chr1_43767873_43768267 | chr1 | 43767874 | 43768267 | + | 0.775162 | intron (NM_026430, intron 9 of 14) | intron (NM_026430, intron 9 of 14) | 37468 | NM_024283 | 78896 | Mm.50109 | NM_024283 | ENSMUSG00000026051 | **1500015O10Rik** | Ecrg4 | RIKEN cDNA 1500015O10 gene | protein-coding |
| chr16_95897855_95898471 | chr16 | 95897856 | 95898471 | + | 1.42016 | intron (NR_040485, intron 1 of 3) | Tigger7\|DNA\|TcMar-Tigger | 550 | NR_040485 | 69776 | Mm.120470 | NR_040484 | ENSMUSG00000086899 | **1600002D24Rik** | - | RIKEN cDNA 1600002D24 gene | ncRNA |
| chr6_86547238_86547675 | chr6 | 86547239 | 86547675 | + | 1.16262 | intron (NR_037961, intron 3 of 4) | intron (NR_037961, intron 3 of 4) | 20056 | NR_037961 | 72012 | Mm.349174 | NR_037960 | ENSMUSG00000097048 | **1600020E01Rik** | - | RIKEN cDNA 1600020E01 gene | ncRNA |
| chr6_86547829_86548415 | chr6 | 86547830 | 86548415 | + | 0.76419 | intron (NR_037961, intron 3 of 4) | intron (NR_037961, intron 3 of 4) | 20721 | NR_037961 | 72012 | Mm.349174 | NR_037960 | ENSMUSG00000097048 | **1600020E01Rik** | - | RIKEN cDNA 1600020E01 gene | ncRNA |
| chr17_47462798_47463261 | chr17 | 47462799 | 47463261 | + | 0.781931 | intron (NM_145489, intron 2 of 4) | RSINE1\|SINE\|B4 | -25655 | NM_029296 | 75462 | Mm.46129 | NM_029296 | ENSMUSG00000047150 | **1700001C19Rik** | - | RIKEN cDNA 1700001C19 gene | protein-coding |
| chr8_61316378_61316744 | chr8 | 61316379 | 61316744 | + | 0.705055 | intron (NM_021506, intron 2 of 11) | intron (NM_021506, intron 2 of 11) | -27689 | NR_045475 | 69279 | Mm.417857 | NR_045475 | | **1700001D01Rik** | - | RIKEN cDNA 1700001D01 gene | ncRNA |
| chr19_53199336_53199947 | chr19 | 53199337 | 53199947 | + | 1.079 | intron (NM_013758, intron 1 of 15) | intron (NM_013758, intron 1 of 15) | 55563 | NR_036590 | 69319 | Mm.57452 | NR_036590 | | **1700001K23Rik** | - | RIKEN cDNA 1700001K23 gene | ncRNA |
| chr19_53219113_53219690 | chr19 | 53219114 | 53219690 | + | 1.07182 | intron (NM_001164100, intron 2 of 13) | MLT1F\|LTR\|ERVL-MaLR | 35803 | NR_036590 | 69319 | Mm.57452 | NR_036590 | | **1700001K23Rik** | - | RIKEN cDNA 1700001K23 gene | ncRNA |
| chr19_53217390_53217826 | chr19 | 53217391 | 53217826 | + | 0.771085 | intron (NM_001164100, intron 2 of 13) | URR1B\|DNA\|hAT-Charlie | 37597 | NR_036590 | 69319 | Mm.57452 | NR_036590 | | **1700001K23Rik** | - | RIKEN cDNA 1700001K23 gene | ncRNA |
| chrX_95778203_95778507 | chrX | 95778204 | 95778507 | + | 0.719085 | intron (NM_001034907, intron 2 of 8) | intron (NM_001034907, intron 2 of 8) | -45094 | NM_029590 | 76386 | Mm.443249 | NM_029590 | ENSMUSG00000071721 | **1700010D01Rik** | - | RIKEN cDNA 1700010D01 gene | protein-coding |
| chr17_9002395_9002803 | chr17 | 9002396 | 9002803 | + | 0.674459 | intron (NM_025851, intron 7 of 9) | intron (NM_025851, intron 7 of 9) | 14266 | NM_025851 | 66931 | Mm.67591 | NM_025851 | ENSMUSG00000023873 | **1700010I14Rik** | - | RIKEN cDNA 1700010I14 gene | protein-coding |
| chr8_79150540_79151126 | chr8 | 79150541 | 79151126 | + | 1.79638 | intron (NM_001294279, intron 8 of 14) | intron (NM_001294279, intron 8 of 14) | 97750 | NM_026315 | 67687 | Mm.178707 | NM_026315 | ENSMUSG00000031682 | **1700011L22Rik** | - | RIKEN cDNA 1700011L22 gene | protein-coding |
| chr8_79161212_79162135 | chr8 | 79161213 | 79162135 | + | 1.13587 | intron (NM_001294279, intron 8 of 14) | ORR1D2\|LTR\|ERVL-MaLR | 86909 | NM_026315 | 67687 | Mm.178707 | NM_026315 | ENSMUSG00000031682 | **1700011L22Rik** | - | RIKEN cDNA 1700011L22 gene | protein-coding |

| | | | | | | | | | | | | | | | | | |
|---|---|---|---|---|---|---|---|---|---|---|---|---|---|---|---|---|---|
| chr8_79162824_79163272 | chr8 | 79162825 | 79163272 | + | 1.07509 | intron (NM_001294279, intron 8 of 14) | intron (NM_001294279, intron 8 of 14) | 85535 | NM_026315 | 67687 | Mm.178707 | NM_026315 | ENSMUSG00000031682 | **1700011L22Rik** | - | RIKEN cDNA 1700011L22 gene | protein-coding |
| chr8_79243492_79243867 | chr8 | 79243493 | 79243867 | + | 0.8431 | intron (NM_026315, intron 1 of 5) | L1MC3\|LINE\|L1 | 4903 | NM_026315 | 67687 | Mm.178707 | NM_026315 | ENSMUSG00000031682 | **1700011L22Rik** | - | RIKEN cDNA 1700011L22 gene | protein-coding |
| chr8_79155510_79156022 | chr8 | 79155511 | 79156022 | + | 0.595707 | intron (NM_001294279, intron 8 of 14) | intron (NM_001294279, intron 8 of 14) | 92817 | NM_026315 | 67687 | Mm.178707 | NM_026315 | ENSMUSG00000031682 | **1700011L22Rik** | - | RIKEN cDNA 1700011L22 gene | protein-coding |
| chr9_14772106_14772444 | chr9 | 14772107 | 14772444 | + | 1.02346 | Intergenic | ORR1A4\|LTR\|ERVL-MaLR | -1245 | NM_029306 | 69325 | Mm.440453 | NM_029306 | ENSMUSG00000031927 | **1700012B09Rik** | 1700007P19Rik | RIKEN cDNA 1700012B09 gene | protein-coding |
| chr9_14775583_14775851 | chr9 | 14775584 | 14775851 | + | 0.620504 | Intergenic | L1MA10\|LINE\|L1 | -4687 | NM_029306 | 69325 | Mm.440453 | NM_029306 | ENSMUSG00000031927 | **1700012B09Rik** | 1700007P19Rik | RIKEN cDNA 1700012B09 gene | protein-coding |
| chr10_127670290_127670508 | chr10 | 127670291 | 127670508 | + | 0.782918 | Intergenic | B3\|SINE\|B2 | 3276 | NR_045171 | 72243 | Mm.482902 | NR_045171 | ENSMUSG00000040195 | **1700012D01Rik** | - | RIKEN cDNA 1700012D01 gene | ncRNA |
| chr15_67413567_67413981 | chr15 | 67413568 | 67413981 | + | 0.866147 | Intergenic | Intergenic | 187005 | NR_045140 | 69334 | Mm.63770 | NR_045140 | | **1700012I11Rik** | - | RIKEN cDNA 1700012I11 gene | ncRNA |
| chr15_67290896_67291239 | chr15 | 67290897 | 67291239 | + | 0.837983 | intron (NR_045140, intron 5 of 8) | intron (NR_045140, intron 5 of 8) | 64299 | NR_045140 | 69334 | Mm.63770 | NR_045140 | | **1700012I11Rik** | - | RIKEN cDNA 1700012I11 gene | ncRNA |
| chr15_67290717_67291241 | chr15 | 67290718 | 67291241 | + | 0.787448 | intron (NR_045140, intron 5 of 8) | intron (NR_045140, intron 5 of 8) | 64210 | NR_045140 | 69334 | Mm.63770 | NR_045140 | | **1700012I11Rik** | - | RIKEN cDNA 1700012I11 gene | ncRNA |
| chr15_67294761_67295090 | chr15 | 67294762 | 67295090 | + | 0.647409 | intron (NR_045140, intron 5 of 8) | intron (NR_045140, intron 5 of 8) | 68157 | NR_045140 | 69334 | Mm.63770 | NR_045140 | | **1700012I11Rik** | - | RIKEN cDNA 1700012I11 gene | ncRNA |
| chr13_5769585_5770105 | chr13 | 5769586 | 5770105 | + | 1.79815 | Intergenic | Intergenic | 88247 | NR_045891 | 71867 | Mm.179181 | NR_045891 | | **1700016G22Rik** | - | RIKEN cDNA 1700016G22 gene | ncRNA |
| chr13_5724326_5724725 | chr13 | 5724327 | 5724725 | + | 1.55443 | Intergenic | Intergenic | 133566 | NR_045891 | 71867 | Mm.179181 | NR_045891 | | **1700016G22Rik** | - | RIKEN cDNA 1700016G22 gene | ncRNA |
| chr13_5767841_5768304 | chr13 | 5767842 | 5768304 | + | 1.46619 | Intergenic | Intergenic | 90019 | NR_045891 | 71867 | Mm.179181 | NR_045891 | | **1700016G22Rik** | - | RIKEN cDNA 1700016G22 gene | ncRNA |
| chr13_5710986_5711373 | chr13 | 5710987 | 5711373 | + | 1.30101 | Intergenic | Intergenic | 146912 | NR_045891 | 71867 | Mm.179181 | NR_045891 | | **1700016G22Rik** | - | RIKEN cDNA 1700016G22 gene | ncRNA |
| chr13_5405923_5406463 | chr13 | 5405924 | 5406463 | + | 1.2952 | Intergenic | Intergenic | 451899 | NR_045891 | 71867 | Mm.179181 | NR_045891 | | **1700016G22Rik** | - | RIKEN cDNA 1700016G22 gene | ncRNA |
| chr13_5674447_5675070 | chr13 | 5674448 | 5675070 | + | 1.18734 | Intergenic | Intergenic | 183333 | NR_045891 | 71867 | Mm.179181 | NR_045891 | | **1700016G22Rik** | - | RIKEN cDNA 1700016G22 gene | ncRNA |
| chr13_5763555_5763910 | chr13 | 5763556 | 5763910 | + | 1.0524 | Intergenic | MLT1A\|LTR\|ERVL-MaLR | 94359 | NR_045891 | 71867 | Mm.179181 | NR_045891 | | **1700016G22Rik** | - | RIKEN cDNA 1700016G22 gene | ncRNA |
| chr13_5787228_5787449 | chr13 | 5787229 | 5787449 | + | 1.03313 | Intergenic | Intergenic | 70753 | NR_045891 | 71867 | Mm.179181 | NR_045891 | | **1700016G22Rik** | - | RIKEN cDNA 1700016G22 gene | ncRNA |
| chr13_5764464_5764788 | chr13 | 5764465 | 5764788 | + | 1.02471 | Intergenic | RMER6A\|LTR\|ERVK | 93466 | NR_045891 | 71867 | Mm.179181 | NR_045891 | | **1700016G22Rik** | - | RIKEN cDNA 1700016G22 gene | ncRNA |
| chr13_5785538_5786114 | chr13 | 5785539 | 5786114 | + | 1.01186 | Intergenic | Intergenic | 72266 | NR_045891 | 71867 | Mm.179181 | NR_045891 | | **1700016G22Rik** | - | RIKEN cDNA 1700016G22 gene | ncRNA |
| chr13_5717997_5718226 | chr13 | 5717998 | 5718226 | + | 0.963891 | Intergenic | Intergenic | 139980 | NR_045891 | 71867 | Mm.179181 | NR_045891 | | **1700016G22Rik** | - | RIKEN cDNA 1700016G22 gene | ncRNA |
| chr13_5837281_5837778 | chr13 | 5837282 | 5837778 | + | 0.927228 | Intergenic | MLT1F1\|LTR\|ERVL-MaLR | 20562 | NR_045891 | 71867 | Mm.179181 | NR_045891 | | **1700016G22Rik** | - | RIKEN cDNA 1700016G22 gene | ncRNA |
| chr13_5511393_5511834 | chr13 | 5511394 | 5511834 | + | 0.910733 | Intergenic | Intergenic | 346478 | NR_045891 | 71867 | Mm.179181 | NR_045891 | | **1700016G22Rik** | - | RIKEN cDNA 1700016G22 gene | ncRNA |
| chr13_5601633_5602020 | chr13 | 5601634 | 5602020 | + | 0.890798 | Intergenic | Intergenic | 256265 | NR_045891 | 71867 | Mm.179181 | NR_045891 | | **1700016G22Rik** | - | RIKEN cDNA 1700016G22 gene | ncRNA |
| chr13_5791861_5792398 | chr13 | 5791862 | 5792398 | + | 0.801024 | Intergenic | Intergenic | 65962 | NR_045891 | 71867 | Mm.179181 | NR_045891 | | **1700016G22Rik** | - | RIKEN cDNA 1700016G22 gene | ncRNA |
| chr13_5785561_5785768 | chr13 | 5785562 | 5785768 | + | 0.785438 | Intergenic | LTR40a\|LTR\|ERVL | 72427 | NR_045891 | 71867 | Mm.179181 | NR_045891 | | **1700016G22Rik** | - | RIKEN cDNA 1700016G22 gene | ncRNA |
| chr13_5773888_5774139 | chr13 | 5773889 | 5774139 | + | 0.727732 | Intergenic | Intergenic | 84078 | NR_045891 | 71867 | Mm.179181 | NR_045891 | | **1700016G22Rik** | - | RIKEN cDNA 1700016G22 gene | ncRNA |
| chr13_5310096_5310414 | chr13 | 5310097 | 5310414 | + | 0.700799 | Intergenic | Intergenic | 547837 | NR_045891 | 71867 | Mm.179181 | NR_045891 | | **1700016G22Rik** | - | RIKEN cDNA 1700016G22 gene | ncRNA |
| chr13_5782610_5782859 | chr13 | 5782611 | 5782859 | + | 0.697882 | Intergenic | Intergenic | 75357 | NR_045891 | 71867 | Mm.179181 | NR_045891 | | **1700016G22Rik** | - | RIKEN cDNA 1700016G22 gene | ncRNA |
| chr13_5511314_5512118 | chr13 | 5511315 | 5512118 | + | 0.684784 | Intergenic | Intergenic | 346376 | NR_045891 | 71867 | Mm.179181 | NR_045891 | | **1700016G22Rik** | - | RIKEN cDNA 1700016G22 gene | ncRNA |
| chr13_5789316_5789708 | chr13 | 5789317 | 5789708 | + | 0.670403 | Intergenic | Intergenic | 68580 | NR_045891 | 71867 | Mm.179181 | NR_045891 | | **1700016G22Rik** | - | RIKEN cDNA 1700016G22 gene | ncRNA |
| chr13_5846826_5847250 | chr13 | 5846827 | 5847250 | + | 0.64397 | Intergenic | Intergenic | 11054 | NR_045891 | 71867 | Mm.179181 | NR_045891 | | **1700016G22Rik** | - | RIKEN cDNA 1700016G22 gene | ncRNA |
| chr1_80404164_80404722 | chr1 | 80404165 | 80404722 | + | 1.08749 | Intergenic | Lx8\|LINE\|L1 | -41489 | NR_040460 | 72208 | Mm.396181 | NR_040460 | | **1700016L21Rik** | Jr3 | RIKEN cDNA 1700016L21 gene | ncRNA |
| chr1_80447599_80447902 | chr1 | 80447600 | 80447902 | + | 0.956167 | intron (NR_040460, intron 1 of 2) | intron (NR_040460, intron 1 of 2) | 1819 | NR_040460 | 72208 | Mm.396181 | NR_040460 | | **1700016L21Rik** | Jr3 | RIKEN cDNA 1700016L21 gene | ncRNA |
| chr1_80484492_80484812 | chr1 | 80484493 | 80484812 | + | 0.8556 | Intergenic | Intergenic | 38720 | NR_040460 | 72208 | Mm.396181 | NR_040460 | | **1700016L21Rik** | Jr3 | RIKEN cDNA 1700016L21 gene | ncRNA |
| chr1_80496374_80496830 | chr1 | 80496375 | 80496830 | + | 0.782955 | Intergenic | Intergenic | 50670 | NR_040460 | 72208 | Mm.396181 | NR_040460 | | **1700016L21Rik** | Jr3 | RIKEN cDNA 1700016L21 gene | ncRNA |
| chr1_80453924_80454356 | chr1 | 80453925 | 80454356 | + | 0.725771 | intron (NR_040460, intron 2 of 2) | ORR1E\|LTR\|ERVL-MaLR | 8208 | NR_040460 | 72208 | Mm.396181 | NR_040460 | | **1700016L21Rik** | Jr3 | RIKEN cDNA 1700016L21 gene | ncRNA |
| chr1_80418500_80419131 | chr1 | 80418501 | 80419131 | + | 0.681913 | Intergenic | Intergenic | -27116 | NR_040460 | 72208 | Mm.396181 | NR_040460 | | **1700016L21Rik** | Jr3 | RIKEN cDNA 1700016L21 gene | ncRNA |
| chr1_80418705_80419092 | chr1 | 80418706 | 80419092 | + | 0.680546 | Intergenic | MLT2B1\|LTR\|ERVL | -27033 | NR_040460 | 72208 | Mm.396181 | NR_040460 | | **1700016L21Rik** | Jr3 | RIKEN cDNA 1700016L21 gene | ncRNA |
| chr1_80447151_80447424 | chr1 | 80447152 | 80447424 | + | 0.666716 | intron (NR_040460, intron 1 of 2) | intron (NR_040460, intron 1 of 2) | 1356 | NR_040460 | 72208 | Mm.396181 | NR_040460 | | **1700016L21Rik** | Jr3 | RIKEN cDNA 1700016L21 gene | ncRNA |
| chr1_80405285_80405583 | chr1 | 80405286 | 80405583 | + | 0.657278 | Intergenic | RMER6C\|LTR\|ERVK | -40498 | NR_040460 | 72208 | Mm.396181 | NR_040460 | | **1700016L21Rik** | Jr3 | RIKEN cDNA 1700016L21 gene | ncRNA |
| chr19_11129736_11130378 | chr19 | 11129737 | 11130378 | + | 1.14127 | intron (NM_027058, intron 1 of 6) | intron (NM_027058, intron 1 of 6) | 821 | NM_027058 | 69369 | Mm.391200 | NM_027058 | ENSMUSG00000024729 | **1700017D01Rik** | - | RIKEN cDNA 1700017D01 gene | protein-coding |
| chr4_63643948_63644344 | chr4 | 63643949 | 63644344 | + | 1.59601 | Intergenic | Intergenic | -21717 | NM_029324 | 75524 | Mm.38543 | NM_029324 | ENSMUSG00000058935 | **1700018C11Rik** | bM230C2.3 | RIKEN cDNA 1700018C11 gene | protein-coding |
| chr4_63636116_63636637 | chr4 | 63636117 | 63636637 | + | 1.38872 | Intergenic | Intergenic | -13948 | NM_029324 | 75524 | Mm.38543 | NM_029324 | ENSMUSG00000058935 | **1700018C11Rik** | bM230C2.3 | RIKEN cDNA 1700018C11 gene | protein-coding |
| chr4_63639838_63640222 | chr4 | 63639839 | 63640222 | + | 1.30805 | Intergenic | Intergenic | -17601 | NM_029324 | 75524 | Mm.38543 | NM_029324 | ENSMUSG00000058935 | **1700018C11Rik** | bM230C2.3 | RIKEN cDNA 1700018C11 gene | protein-coding |
| chr4_63668274_63668670 | chr4 | 63668275 | 63668670 | + | 1.13505 | Intergenic | Intergenic | -46043 | NM_029324 | 75524 | Mm.38543 | NM_029324 | ENSMUSG00000058935 | **1700018C11Rik** | bM230C2.3 | RIKEN cDNA 1700018C11 gene | protein-coding |
| chr4_63668122_63668772 | chr4 | 63668123 | 63668772 | + | 1.0688 | Intergenic | Intergenic | -46018 | NM_029324 | 75524 | Mm.38543 | NM_029324 | ENSMUSG00000058935 | **1700018C11Rik** | bM230C2.3 | RIKEN cDNA 1700018C11 gene | protein-coding |
| chr4_63644418_63644809 | chr4 | 63644419 | 63644809 | + | 0.87485 | Intergenic | Intergenic | -22185 | NM_029324 | 75524 | Mm.38543 | NM_029324 | ENSMUSG00000058935 | **1700018C11Rik** | bM230C2.3 | RIKEN cDNA 1700018C11 gene | protein-coding |
| chr4_63627083_63627494 | chr4 | 63627084 | 63627494 | + | 0.694463 | Intergenic | L1MD2\|LINE\|L1 | -4860 | NM_029324 | 75524 | Mm.38543 | NM_029324 | ENSMUSG00000058935 | **1700018C11Rik** | bM230C2.3 | RIKEN cDNA 1700018C11 gene | protein-coding |
| chr2_46371045_46371404 | chr2 | 46371046 | 46371404 | + | 1.31434 | Intergenic | Intergenic | 674620 | NR_040497 | 69405 | Mm.396184 | NR_040497 | ENSMUSG00000086202 | **1700019E08Rik** | - | RIKEN cDNA 1700019E08 gene | ncRNA |
| chr2_46370821_46371458 | chr2 | 46370822 | 46371458 | + | 1.29154 | Intergenic | Intergenic | 674535 | NR_040497 | 69405 | Mm.396184 | NR_040497 | ENSMUSG00000086202 | **1700019E08Rik** | - | RIKEN cDNA 1700019E08 gene | ncRNA |
| chr2_46372845_46373148 | chr2 | 46372846 | 46373148 | + | 0.955095 | Intergenic | Lx9\|LINE\|L1 | 676392 | NR_040497 | 69405 | Mm.396184 | NR_040497 | ENSMUSG00000086202 | **1700019E08Rik** | - | RIKEN cDNA 1700019E08 gene | ncRNA |
| chr1_86439939_86440213 | chr1 | 86439940 | 86440213 | + | 0.839988 | Intergenic | Intergenic | 13747 | NM_027966 | 71863 | Mm.179627 | NM_027966 | ENSMUSG00000036574 | **1700019O17Rik** | - | RIKEN cDNA 1700019O17 gene | protein-coding |
| chr2_119596987_119597280 | chr2 | 119596988 | 119597280 | + | 1.361 | intron (NR_015473, intron 2 of 2) | ORR1B2\|LTR\|ERVL-MaLR | 2838 | NR_015473 | 66602 | Mm.379181 | NR_015473 | ENSMUSG00000085438 | **1700020I14Rik** | 1190006L01Rik\|4933437I03Rik\|AI451541\|AI585852\|AW538376\|AW558897 | RIKEN cDNA 1700020I14 gene | ncRNA |
| chr10_21535264_21536058 | chr10 | 21535265 | 21536058 | + | 0.956567 | Intergenic | Intergenic | -57484 | NR_027968 | 67692 | Mm.54306 | NM_183266 | | **1700020N01Rik** | - | RIKEN cDNA 1700020N01 gene | ncRNA |
| chr10_21504468_21504836 | chr10 | 21504469 | 21504836 | + | 0.76286 | Intergenic | Intergenic | -88493 | NR_027968 | 67692 | Mm.54306 | NM_183266 | | **1700020N01Rik** | - | RIKEN cDNA 1700020N01 gene | ncRNA |
| chr10_21504150_21504836 | chr10 | 21504151 | 21504836 | + | 0.757441 | Intergenic | Intergenic | -88652 | NR_027968 | 67692 | Mm.54306 | NM_183266 | | **1700020N01Rik** | - | RIKEN cDNA 1700020N01 gene | ncRNA |
| chr10_21611886_21612322 | chr10 | 21611887 | 21612322 | + | 0.740463 | intron (NR_027968, intron 2 of 2) | intron (NR_027968, intron 2 of 2) | 18959 | NR_027968 | 67692 | Mm.54306 | NM_183266 | | **1700020N01Rik** | - | RIKEN cDNA 1700020N01 gene | ncRNA |
| chr10_21504093_21505490 | chr10 | 21504094 | 21505490 | + | 0.728501 | Intergenic | Intergenic | -88353 | NR_027968 | 67692 | Mm.54306 | NM_183266 | | **1700020N01Rik** | - | RIKEN cDNA 1700020N01 gene | ncRNA |
| chr10_43519878_43520378 | chr10 | 43519879 | 43520378 | + | 0.780635 | Intergenic | Intergenic | 20866 | NM_026411 | 67851 | Mm.209953 | NM_026411 | ENSMUSG00000019797 | **1700021F05Rik** | - | RIKEN cDNA 1700021F05 gene | protein-coding |
| chr2_173448497_173449243 | chr2 | 173448498 | 173449243 | + | 0.630869 | Intergenic | ORR1D1-int\|LTR\|ERVL-MaLR | -73722 | NM_028158 | 72221 | Mm.46132 | NM_028158 | ENSMUSG00000027518 | **1700021F07Rik** | - | RIKEN cDNA 1700021F07 gene | protein-coding |
| chr16_32870985_32871213 | chr16 | 32870986 | 32871213 | + | 0.648062 | Intergenic | B3A\|SINE\|B2 | -2733 | NM_172615 | 100502698 | Mm.327319 | NM_172615 | ENSMUSG00000035629 | **1700021K19Rik** | 5330403K09\|Kiaa0226\|Rubicon\|mKIAA0226 | RIKEN cDNA 1700021K19 gene | protein-coding |
| chr5_24629574_24630212 | chr5 | 24629575 | 24630212 | + | 0.708712 | Intergenic | Intergenic | -15560 | NR_003953 | 72252 | Mm.87446 | NR_003953 | ENSMUSG00000038165 | **1700022A21Rik** | - | glycerol-3-phosphate dehydrogenase 1-like pseudogene | pseudo |
| chr16_59453977_59454308 | chr16 | 59453978 | 59454308 | + | 0.809207 | intron (NM_001081190, intron 8 of 8) | ORR1B1\|LTR\|ERVL-MaLR | 15095 | NR_040568 | 75636 | Mm.490522 | NR_040568 | | **1700022E09Rik** | - | RIKEN cDNA 1700022E09 gene | ncRNA |
| chr1_152015612_152015978 | chr1 | 152015613 | 152015978 | + | 0.655281 | intron (NM_197990, intron 1 of 5) | intron (NM_197990, intron 1 of 5) | 74525 | NM_197990 | 69399 | Mm.278031 | NM_197990 | ENSMUSG00000032666 | **1700025G04Rik** | 2610510E02Rik\|2900042O04Rik\|ENSMUSG00000073540 | RIKEN cDNA 1700025G04 gene | protein-coding |
| chr5_73285638_73285931 | chr5 | 73285639 | 73285931 | + | 0.992951 | Intergenic | RSINE1\|SINE\|B4 | -1601 | NR_040688 | 72276 | Mm.271215 | NR_040687 | | **1700025M24Rik** | - | RIKEN cDNA 1700025M24 gene | ncRNA |
| chr5_73285534_73286017 | chr5 | 73285535 | 73286017 | + | 0.909317 | Intergenic | RSINE1\|SINE\|B4 | -1592 | NR_040688 | 72276 | Mm.271215 | NR_040687 | | **1700025M24Rik** | - | RIKEN cDNA 1700025M24 gene | ncRNA |
| chr5_73271223_73271624 | chr5 | 73271224 | 73271624 | + | 0.870756 | intron (NR_040688, intron 3 of 3) | intron (NR_040688, intron 3 of 3) | 12760 | NR_040687 | 72276 | Mm.271215 | NR_040687 | | **1700025M24Rik** | - | RIKEN cDNA 1700025M24 gene | ncRNA |
| chr5_73286525_73287013 | chr5 | 73286526 | 73287013 | + | 0.811846 | Intergenic | Intergenic | -2585 | NR_040688 | 72276 | Mm.271215 | NR_040687 | | **1700025M24Rik** | - | RIKEN cDNA 1700025M24 gene | ncRNA |
| chr2_132524539_132525041 | chr2 | 132524540 | 132525041 | + | 0.95097 | TTS (NR_028286) | TTS (NR_028286) | 34062 | NR_028286 | 70371 | Mm.394667 | NR_028286 | ENSMUSG00000086379 | **1700026D11Rik** | - | RIKEN cDNA 1700026D11 gene | ncRNA |
| chr10_43746613_43746968 | chr10 | 43746614 | 43746968 | + | 0.770826 | intron (NR_040581, intron 1 of 3) | MTEa\|LTR\|ERVL-MaLR | 633 | NR_040581 | 69443 | Mm.484042 | NR_040581 | | **1700027J07Rik** | - | RIKEN cDNA 1700027J07 gene | ncRNA |
| chr7_28479993_28480565 | chr7 | 28479994 | 28480565 | + | 0.735058 | intron (NR_030567, intron 4 of 4) | intron (NR_030567, intron 4 of 4) | -16662 | NR_033605 | 70001 | Mm.417851 | NR_033605 | | **1700028B04Rik** | - | NADH dehydrogenase (ubiquinone) 1 beta subcomplex 8 pseudogene | pseudo |
| chr5_151374890_151375159 | chr5 | 151374891 | 151375159 | + | 1.52111 | intron (NR_045700, intron 1 of 3) | intron (NR_045700, intron 1 of 3) | 6350 | NR_045699 | 625175 | Mm.443286 | NR_045699 | ENSMUSG00000097321 | **1700028E10Rik** | A630054D14\|Gm6561 | RIKEN cDNA 1700028E10 gene | ncRNA |

| | | | | | | | | | | | | | | | | | |
|---|---|---|---|---|---|---|---|---|---|---|---|---|---|---|---|---|---|
| chr5_151376409_151376688 | chr5 | 151376410 | 151376688 | + | 1.24098 | intron (NR_045700, intron 1 of 3) | intron (NR_045700, intron 1 of 3) | 7874 | NR_045699 | 625175 | Mm.443286 | NR_045699 | ENSMUSG00000097321 | **1700028E10Rik** | A630054D14\|Gm6561 | RIKEN cDNA 1700028E10 gene | ncRNA |
| chr5_151412484_151413498 | chr5 | 151412485 | 151413498 | + | 1.01184 | intron (NR_045700, intron 3 of 3) | intron (NR_045700, intron 3 of 3) | 44316 | NR_045699 | 625175 | Mm.443286 | NR_045699 | ENSMUSG00000097321 | **1700028E10Rik** | A630054D14\|Gm6561 | RIKEN cDNA 1700028E10 gene | ncRNA |
| chr5_151376946_151377746 | chr5 | 151376947 | 151377746 | + | 0.952994 | intron (NR_045700, intron 1 of 3) | L2c\|LINE\|L2 | 8671 | NR_045699 | 625175 | Mm.443286 | NR_045699 | ENSMUSG00000097321 | **1700028E10Rik** | A630054D14\|Gm6561 | RIKEN cDNA 1700028E10 gene | ncRNA |
| chr10_82794991_82795694 | chr10 | 82794992 | 82795694 | + | 1.02531 | Intergenic | Intergenic | -16780 | NR_038042 | 70003 | Mm.48789 | NR_038042 | ENSMUSG00000056821 | **1700028I16Rik** | - | RIKEN cDNA 1700028I16 gene | ncRNA |
| chr10_82798556_82798912 | chr10 | 82798557 | 82798912 | + | 1.01396 | Intergenic | Intergenic | -13389 | NR_038042 | 70003 | Mm.48789 | NR_038042 | ENSMUSG00000056821 | **1700028I16Rik** | - | RIKEN cDNA 1700028I16 gene | ncRNA |
| chr10_82796632_82797487 | chr10 | 82796633 | 82797487 | + | 1.00858 | Intergenic | Intergenic | -15063 | NR_038042 | 70003 | Mm.48789 | NR_038042 | ENSMUSG00000056821 | **1700028I16Rik** | - | RIKEN cDNA 1700028I16 gene | ncRNA |
| chr10_82807262_82807545 | chr10 | 82807263 | 82807545 | + | 0.612813 | Intergenic | L2\|LINE\|L2 | -4719 | NR_038042 | 70003 | Mm.48789 | NR_038042 | ENSMUSG00000056821 | **1700028I16Rik** | - | RIKEN cDNA 1700028I16 gene | ncRNA |
| chr5_107528520_107529192 | chr5 | 107528521 | 107529192 | + | 0.994173 | Intergenic | MLTR18_MM\|LTR\|ERV K | -5855 | NM_182745 | 76421 | Mm.67037 | NM_175241 | ENSMUSG00000089798 | **1700028K03Rik** | - | RIKEN cDNA 1700028K03 gene | protein-coding |
| chr8_13602094_13602657 | chr8 | 13602095 | 13602657 | + | 0.741965 | intron (NM_009025, intron 4 of 23) | intron (NM_009025, intron 4 of 23) | -39915 | NM_001080781 | 66501 | Mm.46136 | NM_025601 | ENSMUSG00000031452 | **1700029H14Rik** | - | RIKEN cDNA 1700029H14 gene | protein-coding |
| chr2_92379842_92380206 | chr2 | 92379843 | 92380206 | + | 0.67891 | intron (NR_028114, intron 9 of 10) | intron (NR_028114, intron 9 of 10) | -2894 | NM_183112 | 75641 | Mm.330409 | NM_183112 | ENSMUSG00000044916 | **1700029I15Rik** | 1700021K07Rik | RIKEN cDNA 1700029I15 gene | protein-coding |
| chr13_44469614_44470201 | chr13 | 44469615 | 44470201 | + | 1.03336 | Intergenic | Intergenic | 30181 | NR_045489 | 69491 | Mm.392410 | NR_045489 | ENSMUSG00000098144 | **1700029N11Rik** | - | RIKEN cDNA 1700029N11 gene | ncRNA |
| chr3_136431305_136431836 | chr3 | 136431306 | 136431836 | + | 0.982427 | Intergenic | Intergenic | 17778 | NR_040592 | 70055 | Mm.261063 | NR_040592 | | **1700030L20Rik** | - | RIKEN cDNA 1700030L20 gene | ncRNA |
| chr3_40903031_40903805 | chr3 | 40903032 | 40903805 | + | 1.06129 | promoter-TSS (NR_045380) | promoter-TSS (NR_045380) | -875 | NR_045380 | 73297 | Mm.422673 | NM_028494 | ENSMUSG00000074619 | **1700034I23Rik** | - | FUN14 domain containing 2 pseudogene | pseudo |
| chr6_146959874_146960544 | chr6 | 146959875 | 146960544 | + | 0.798166 | intron (NM_001170433, intron 2 of 27) | intron (NM_001170433, intron 2 of 27) | -5788 | NM_028509 | 73344 | Mm.391543 | NM_028509 | ENSMUSG00000040163 | **1700034J05Rik** | - | RIKEN cDNA 1700034J05 gene | protein-coding |
| chr6_146963793_146964085 | chr6 | 146963794 | 146964085 | + | 0.704383 | intron (NM_001170433, intron 2 of 27) | intron (NM_001170433, intron 2 of 27) | -9518 | NM_028509 | 73344 | Mm.391543 | NM_028509 | ENSMUSG00000040163 | **1700034J05Rik** | - | RIKEN cDNA 1700034J05 gene | protein-coding |
| chr10_68558250_68558742 | chr10 | 68558251 | 68558742 | + | 1.07531 | Intergenic | Intergenic | -16621 | NM_028491 | 73287 | Mm.141830 | NM_028491 | ENSMUSG00000019945 | **1700040L02Rik** | - | RIKEN cDNA 1700040L02 gene | protein-coding |
| chr10_68512393_68512859 | chr10 | 68512394 | 68512859 | + | 0.629234 | intron (NM_028491, intron 4 of 7) | RMER15\|LTR\|ERVL | 29249 | NM_028491 | 73287 | Mm.141830 | NM_028491 | ENSMUSG00000019945 | **1700040L02Rik** | - | RIKEN cDNA 1700040L02 gene | protein-coding |
| chr16_76762761_76763284 | chr16 | 76762762 | 76763284 | + | 1.45361 | Intergenic | Intergenic | 247858 | NR_040573 | 76601 | Mm.396217 | NR_040573 | | **1700041M19Rik** | - | RIKEN cDNA 1700041M19 gene | ncRNA |
| chr16_76900948_76901483 | chr16 | 76900949 | 76901483 | + | 1.00762 | Intergenic | Intergenic | 109665 | NR_040573 | 76601 | Mm.396217 | NR_040573 | | **1700041M19Rik** | - | RIKEN cDNA 1700041M19 gene | ncRNA |
| chr16_76900841_76901688 | chr16 | 76900842 | 76901688 | + | 0.880205 | Intergenic | Intergenic | 109616 | NR_040573 | 76601 | Mm.396217 | NR_040573 | | **1700041M19Rik** | - | RIKEN cDNA 1700041M19 gene | ncRNA |
| chr16_76763777_76764057 | chr16 | 76763778 | 76764057 | + | 0.704915 | Intergenic | Intergenic | 246964 | NR_040573 | 76601 | Mm.396217 | NR_040573 | | **1700041M19Rik** | - | RIKEN cDNA 1700041M19 gene | ncRNA |
| chr4_150799503_150799792 | chr4 | 150799504 | 150799792 | + | 0.773772 | Intergenic | Intergenic | 55301 | NR_040649 | 73348 | Mm.396218 | NR_040649 | ENSMUSG00000078492 | **1700045H11Rik** | - | RIKEN cDNA 1700045H11 gene | ncRNA |
| chr4_150807996_150809672 | chr4 | 150807997 | 150809672 | + | 0.695487 | Intergenic | Intergenic | 46115 | NR_040649 | 73348 | Mm.396218 | NR_040649 | ENSMUSG00000078492 | **1700045H11Rik** | - | RIKEN cDNA 1700045H11 gene | ncRNA |
| chr4_150808376_150808897 | chr4 | 150808377 | 150808897 | + | 0.660288 | Intergenic | Intergenic | 46312 | NR_040649 | 73348 | Mm.396218 | NR_040649 | ENSMUSG00000078492 | **1700045H11Rik** | - | RIKEN cDNA 1700045H11 gene | ncRNA |
| chr6_100479356_100480438 | chr6 | 100479357 | 100480438 | + | 0.999459 | Intergenic | Intergenic | -47503 | NR_040525 | 73395 | Mm.69413 | NR_040525 | | **1700049E22Rik** | - | RIKEN cDNA 1700049E22 gene | ncRNA |
| chr19_53060175_53060600 | chr19 | 53060176 | 53060600 | + | 1.02575 | Intergenic | MTB_Mm\|LTR\|ERVL-MaLR | -15864 | NR_045320 | 78457 | Mm.76932 | NR_045320 | | **1700054A03Rik** | - | RIKEN cDNA 1700054A03 gene | ncRNA |
| chr1_183933879_183934278 | chr1 | 183933880 | 183934278 | + | 0.926651 | Intergenic | Intergenic | 99919 | NM_028516 | 73363 | Mm.50115 | NM_028516 | ENSMUSG00000044854 | **1700056E22Rik** | - | RIKEN cDNA 1700056E22 gene | protein-coding |
| chr1_183933805_183934381 | chr1 | 183933806 | 183934381 | + | 0.867421 | Intergenic | Intergenic | 99905 | NM_028516 | 73363 | Mm.50115 | NM_028516 | ENSMUSG00000044854 | **1700056E22Rik** | - | RIKEN cDNA 1700056E22 gene | protein-coding |
| chr1_183964171_183964526 | chr1 | 183964172 | 183964526 | + | 0.866707 | Intergenic | Intergenic | 69649 | NM_028516 | 73363 | Mm.50115 | NM_028516 | ENSMUSG00000044854 | **1700056E22Rik** | - | RIKEN cDNA 1700056E22 gene | protein-coding |
| chr1_183964010_183964536 | chr1 | 183964011 | 183964536 | + | 0.788451 | Intergenic | Intergenic | 69725 | NM_028516 | 73363 | Mm.50115 | NM_028516 | ENSMUSG00000044854 | **1700056E22Rik** | - | RIKEN cDNA 1700056E22 gene | protein-coding |
| chr1_183788771_183789085 | chr1 | 183788772 | 183789085 | + | 0.636868 | Intergenic | MER63A\|DNA\|hAT-Blackjack | 245070 | NM_028516 | 73363 | Mm.50115 | NM_028516 | ENSMUSG00000044854 | **1700056E22Rik** | - | RIKEN cDNA 1700056E22 gene | protein-coding |
| chr6_143871107_143871543 | chr6 | 143871108 | 143871543 | + | 0.912489 | intron (NM_011444, intron 10 of 14) | intron (NM_011444, intron 10 of 14) | -220037 | NR_045732 | 73425 | Mm.417844 | NR_045732 | ENSMUSG00000086887 | **1700060C16Rik** | - | RIKEN cDNA 1700060C16 gene | ncRNA |
| chr6_143816340_143816955 | chr6 | 143816341 | 143816955 | + | 0.902353 | Intergenic | Intergenic | -165360 | NR_045732 | 73425 | Mm.417844 | NR_045732 | ENSMUSG00000086887 | **1700060C16Rik** | - | RIKEN cDNA 1700060C16 gene | ncRNA |
| chr2_8865356_8865878 | chr2 | 8865357 | 8865878 | + | 1.68405 | Intergenic | Intergenic | -323898 | NR_038180 | 73438 | Mm.23558 | NR_038180 | ENSMUSG00000085580 | **1700061F12Rik** | - | RIKEN cDNA 1700061F12 gene | ncRNA |
| chr2_8870099_8870496 | chr2 | 8870100 | 8870496 | + | 1.46718 | Intergenic | L2\|LINE\|L2 | -319217 | NR_038180 | 73438 | Mm.23558 | NR_038180 | ENSMUSG00000085580 | **1700061F12Rik** | - | RIKEN cDNA 1700061F12 gene | ncRNA |
| chr2_9359928_9360472 | chr2 | 9359929 | 9360472 | + | 1.27185 | Intergenic | Intergenic | 170685 | NR_038180 | 73438 | Mm.23558 | NR_038180 | ENSMUSG00000085580 | **1700061F12Rik** | - | RIKEN cDNA 1700061F12 gene | ncRNA |
| chr2_8942399_8942963 | chr2 | 8942400 | 8942963 | + | 1.18578 | Intergenic | Intergenic | -246834 | NR_038180 | 73438 | Mm.23558 | NR_038180 | ENSMUSG00000085580 | **1700061F12Rik** | - | RIKEN cDNA 1700061F12 gene | ncRNA |
| chr2_9193058_9193598 | chr2 | 9193059 | 9193598 | + | 1.13196 | intron (NR_038180, intron 2 of 3) | intron (NR_038180, intron 2 of 3) | 3813 | NR_038180 | 73438 | Mm.23558 | NR_038180 | ENSMUSG00000085580 | **1700061F12Rik** | - | RIKEN cDNA 1700061F12 gene | ncRNA |
| chr2_9294434_9294937 | chr2 | 9294435 | 9294937 | + | 0.922482 | Intergenic | Intergenic | 105171 | NR_038180 | 73438 | Mm.23558 | NR_038180 | ENSMUSG00000085580 | **1700061F12Rik** | - | RIKEN cDNA 1700061F12 gene | ncRNA |
| chr2_8964908_8965610 | chr2 | 8964909 | 8965610 | + | 0.905597 | Intergenic | Intergenic | -224256 | NR_038180 | 73438 | Mm.23558 | NR_038180 | ENSMUSG00000085580 | **1700061F12Rik** | - | RIKEN cDNA 1700061F12 gene | ncRNA |
| chr2_9343171_9343630 | chr2 | 9343172 | 9343630 | + | 0.662692 | Intergenic | Intergenic | 153886 | NR_038180 | 73438 | Mm.23558 | NR_038180 | ENSMUSG00000085580 | **1700061F12Rik** | - | RIKEN cDNA 1700061F12 gene | ncRNA |
| chr9_41193529_41194195 | chr9 | 41193530 | 41194195 | + | 0.653343 | Intergenic | MTC\|LTR\|ERVL-MaLR | -12293 | NR_040392 | 78500 | Mm.275715 | NR_040392 | | **1700063D05Rik** | - | RIKEN cDNA 1700063D05 gene | ncRNA |
| chr10_119070582_119071354 | chr10 | 119070583 | 119071354 | + | 0.895986 | Intergenic | URR1A\|DNA\|hAT-Charlie | -21937 | NR_045348 | 73400 | Mm.416864 | NR_045348 | | **1700064J06Rik** | - | RIKEN cDNA 1700064J06 gene | ncRNA |
| chr10_119028514_119029061 | chr10 | 119028515 | 119029061 | + | 0.870103 | Intergenic | Intergenic | -64117 | NR_045348 | 73400 | Mm.416864 | NR_045348 | | **1700064J06Rik** | - | RIKEN cDNA 1700064J06 gene | ncRNA |
| chr12_99626146_99626522 | chr12 | 99626147 | 99626522 | + | 1.6143 | intron (NR_045288, intron 1 of 1) | intron (NR_045288, intron 1 of 1) | 1640 | NR_045288 | 73457 | Mm.383439 | NR_045288 | | **1700064M15Rik** | - | RIKEN cDNA 1700064M15 gene | ncRNA |
| chr12_99639153_99639507 | chr12 | 99639154 | 99639507 | + | 1.34069 | Intergenic | URR1B\|DNA\|hAT-Charlie | -11356 | NR_045288 | 73457 | Mm.383439 | NR_045288 | | **1700064M15Rik** | - | RIKEN cDNA 1700064M15 gene | ncRNA |
| chr12_99681500_99682042 | chr12 | 99681501 | 99682042 | + | 1.28003 | Intergenic | RLTR16B_MM\|LTR\|ER VK | -53797 | NR_045288 | 73457 | Mm.383439 | NR_045288 | | **1700064M15Rik** | - | RIKEN cDNA 1700064M15 gene | ncRNA |
| chr12_99686699_99687068 | chr12 | 99686700 | 99687068 | + | 1.17521 | Intergenic | Intergenic | -58910 | NR_045288 | 73457 | Mm.383439 | NR_045288 | | **1700064M15Rik** | - | RIKEN cDNA 1700064M15 gene | ncRNA |
| chr12_99610246_99611353 | chr12 | 99610247 | 99611353 | + | 0.909886 | Intergenic | Intergenic | 17174 | NR_045288 | 73457 | Mm.383439 | NR_045288 | | **1700064M15Rik** | - | RIKEN cDNA 1700064M15 gene | ncRNA |
| chr12_99686510_99687147 | chr12 | 99686511 | 99687147 | + | 0.902017 | Intergenic | Intergenic | -58855 | NR_045288 | 73457 | Mm.383439 | NR_045288 | | **1700064M15Rik** | - | RIKEN cDNA 1700064M15 gene | ncRNA |
| chr12_99587300_99587762 | chr12 | 99587301 | 99587762 | + | 0.862547 | Intergenic | Intergenic | 40443 | NR_045288 | 73457 | Mm.383439 | NR_045288 | | **1700064M15Rik** | - | RIKEN cDNA 1700064M15 gene | ncRNA |
| chr12_99622248_99622633 | chr12 | 99622249 | 99622633 | + | 0.727544 | Intergenic | URR1A\|DNA\|hAT-Charlie | 5533 | NR_045288 | 73457 | Mm.383439 | NR_045288 | | **1700064M15Rik** | - | RIKEN cDNA 1700064M15 gene | ncRNA |
| chr12_99633561_99634068 | chr12 | 99633562 | 99634068 | + | 0.621436 | Intergenic | Intergenic | -5841 | NR_045288 | 73457 | Mm.383439 | NR_045288 | | **1700064M15Rik** | - | RIKEN cDNA 1700064M15 gene | ncRNA |
| chr9_95869916_95870187 | chr9 | 95869917 | 95870187 | + | 0.87854 | exon (NM_019864, exon 7 of 47) | exon (NM_019864, exon 7 of 47) | -12170 | NM_001271569 | 73410 | Mm.443314 | NM_028533 | | **1700065D16Rik** | - | RIKEN cDNA 1700065D16 gene | protein-coding |
| chr1_89015396_89016318 | chr1 | 89015397 | 89016318 | + | 0.657942 | promoter-TSS (NR_040471) | promoter-TSS (NR_040471) | -256 | NR_040471 | 73475 | Mm.491040 | NR_040471 | | **1700067G17Rik** | - | RIKEN cDNA 1700067G17 gene | ncRNA |
| chr6_145371666_145371921 | chr6 | 145371667 | 145371921 | + | 0.779881 | Intergenic | RSINE1\|SINE\|B4 | -15831 | NR_003625 | 381827 | Mm.179075 | NM_001001987 | | **1700073E17Rik** | - | ribosomal protein L7 pseudogene | pseudo |
| chr13_119654783_119655130 | chr13 | 119654784 | 119655130 | + | 0.887248 | TTS (NM_020279) | TTS (NM_020279) | -25085 | NR_045296 | 73480 | Mm.71100 | NR_045296 | | **1700074H08Rik** | CH25-211A23.1 | RIKEN cDNA 1700074H08 gene | ncRNA |
| chr13_119681042_119681471 | chr13 | 119681043 | 119681471 | + | 0.662595 | intron (NR_045296, intron 2 of 2) | intron (NR_045296, intron 2 of 2) | 1215 | NR_045296 | 73480 | Mm.71100 | NR_045296 | | **1700074H08Rik** | CH25-211A23.1 | RIKEN cDNA 1700074H08 gene | ncRNA |
| chr1_169917848_169918460 | chr1 | 169917849 | 169918460 | + | 0.723833 | Intergenic | Intergenic | -10785 | NM_001033185 | 78465 | Mm.291078 | NM_001033185 | ENSMUSG00000070532 | **1700084C01Rik** | - | RIKEN cDNA 1700084C01 gene | protein-coding |
| chr1_169918033_169918431 | chr1 | 169918034 | 169918431 | + | 0.712455 | Intergenic | Intergenic | -10707 | NM_001033185 | 78465 | Mm.291078 | NM_001033185 | ENSMUSG00000070532 | **1700084C01Rik** | - | RIKEN cDNA 1700084C01 gene | protein-coding |
| chr11_23510110_23510784 | chr11 | 23510111 | 23510784 | + | 0.929012 | Intergenic | MTC-int\|LTR\|ERVL-MaLR | 8748 | NM_026105 | 67358 | Mm.87328 | NM_026105 | ENSMUSG00000020286 | **1700093K21Rik** | - | RIKEN cDNA 1700093K21 gene | protein-coding |
| chr17_30622908_30623591 | chr17 | 30622909 | 30623591 | + | 0.744648 | intron (NR_045287, intron 3 of 3) | intron (NR_045287, intron 3 of 3) | 3655 | NR_045287 | 67522 | Mm.254797 | NR_045287 | | **1700097N02Rik** | - | RIKEN cDNA 1700097N02 gene | ncRNA |
| chr6_129545087_129545594 | chr6 | 129545088 | 129545594 | + | 1.07061 | Intergenic | Intergenic | -12057 | NR_045270 | 100504641 | Mm.398286 | NR_045270 | ENSMUSG00000097675 | **1700101I11Rik** | - | RIKEN cDNA 1700101I11 gene | ncRNA |
| chr12_54299530_54299948 | chr12 | 54299531 | 54299948 | + | 1.22553 | Intergenic | Intergenic | 66124 | NR_108033 | 73592 | Mm.416874 | NR_108033 | | **1700104L18Rik** | - | RIKEN cDNA 1700104L18 gene | ncRNA |
| chr12_54299155_54299956 | chr12 | 54299156 | 54299956 | + | 1.11286 | Intergenic | Intergenic | 65941 | NR_108033 | 73592 | Mm.416874 | NR_108033 | | **1700104L18Rik** | - | RIKEN cDNA 1700104L18 gene | ncRNA |
| chr14_76760462_76760860 | chr14 | 76760463 | 76760860 | + | 0.891838 | Intergenic | Intergenic | 83093 | NR_015485 | 73272 | Mm.116804 | NR_015485 | | **1700108F19Rik** | 1700020I02Rik\|BB065030 | RIKEN cDNA 1700108F19 gene | ncRNA |
| chr14_76786987_76788135 | chr14 | 76786988 | 76788135 | + | 0.871827 | Intergenic | Intergenic | 109993 | NR_015485 | 73272 | Mm.116804 | NR_015485 | | **1700108F19Rik** | 1700020I02Rik\|BB065030 | RIKEN cDNA 1700108F19 gene | ncRNA |
| chr14_76787495_76788061 | chr14 | 76787496 | 76788061 | + | 0.815995 | Intergenic | Intergenic | 110210 | NR_015485 | 73272 | Mm.116804 | NR_015485 | | **1700108F19Rik** | 1700020I02Rik\|BB065030 | RIKEN cDNA 1700108F19 gene | ncRNA |
| chr14_76698857_76699366 | chr14 | 76698858 | 76699366 | + | 0.726942 | Intergenic | Intergenic | 21544 | NR_015485 | 73272 | Mm.116804 | NR_015485 | | **1700108F19Rik** | 1700020I02Rik\|BB065030 | RIKEN cDNA 1700108F19 gene | ncRNA |
| chr14_61290805_61291316 | chr14 | 61290806 | 61291316 | + | 1.07233 | Intergenic | Intergenic | 14273 | NR_033788 | 67355 | Mm.193728 | NR_033788 | | **1700109G14Rik** | 1700126G13Rik | RIKEN cDNA 1700109G14 gene | ncRNA |
| chr9_40348760_40349202 | chr9 | 40348761 | 40349202 | + | 1.72234 | intron (NM_172768, intron 2 of 19) | intron (NM_172768, intron 2 of 19) | 15509 | NR_040728 | 73558 | Mm.398307 | NR_040728 | | **1700110K17Rik** | - | RIKEN cDNA 1700110K17 gene | ncRNA |
| chr9_40342026_40342554 | chr9 | 40342027 | 40342554 | + | 0.938638 | non-coding (NR_040728, exon 2 of 3) | non-coding (NR_040728, exon 2 of 3) | 8818 | NR_040728 | 73558 | Mm.398307 | NR_040728 | | **1700110K17Rik** | - | RIKEN cDNA 1700110K17 gene | ncRNA |

| | | | | | | | | | | | | | | | | | |
|---|---|---|---|---|---|---|---|---|---|---|---|---|---|---|---|---|---|
| chr9_40349548_40349835 | chr9 | 40349549 | 40349835 | + | 0.819474 | intron (NM_172768, intron 2 of 19) | intron (NM_172768, intron 2 of 19) | 16220 | NR_040728 | 73558 | Mm.398307 | NR_040728 | | **1700110K17Rik** | - | RIKEN cDNA 1700110K17 gene | ncRNA |
| chr9_40320957_40321953 | chr9 | 40320958 | 40321953 | + | 0.667781 | intron (NM_172768, intron 4 of 19) | intron (NM_172768, intron 4 of 19) | -12017 | NR_040728 | 73558 | Mm.398307 | NR_040728 | | **1700110K17Rik** | - | RIKEN cDNA 1700110K17 gene | ncRNA |
| chr1_184534203_184535232 | chr1 | 184534204 | 184535232 | + | 0.896342 | intron (NR_040472, intron 2 of 5) | intron (NR_040472, intron 2 of 5) | 22973 | NR_040472 | 73590 | Mm.82792 | NR_040472 | | **1700112H15Rik** | - | RIKEN cDNA 1700112H15 gene | ncRNA |
| chr1_184526341_184526624 | chr1 | 184526342 | 184526624 | + | 0.791419 | Intergenic | L1MA5\|LINE\|L1 | 31208 | NR_040472 | 73590 | Mm.82792 | NR_040472 | | **1700112H15Rik** | - | RIKEN cDNA 1700112H15 gene | ncRNA |
| chr7_127607011_127607452 | chr7 | 127607012 | 127607452 | + | 0.920722 | 3' UTR (NM_177226, exon 3 of 3) | 3' UTR (NM_177226, exon 3 of 3) | -2787 | NR_027915 | 68232 | Mm.347960 | NM_026628 | | **1700120K04Rik** | - | RIKEN cDNA 1700120K04 gene | ncRNA |
| chr7_127607019_127607803 | chr7 | 127607020 | 127607803 | + | 0.784307 | 3' UTR (NM_177226, exon 3 of 3) | 3' UTR (NM_177226, exon 3 of 3) | -2966 | NR_027915 | 68232 | Mm.347960 | NM_026628 | | **1700120K04Rik** | - | RIKEN cDNA 1700120K04 gene | ncRNA |
| chrX_104760835_104761387 | chrX | 104760836 | 104761387 | + | 0.707417 | Intergenic | Intergenic | -55665 | NR_045453 | 76643 | Mm.439149 | NR_045453 | | **1700121L16Rik** | - | RIKEN cDNA 1700121L16 gene | ncRNA |
| chr17_48034352_48035109 | chr17 | 48034353 | 48035109 | + | 0.970466 | Intergenic | Intergenic | 3562 | NM_029689 | 76651 | Mm.76918 | NM_029689 | ENSMUSG00000094928 | **1700122O11Rik** | - | RIKEN cDNA 1700122O11 gene | protein-coding |
| chr4_11989920_11990629 | chr4 | 11989921 | 11990629 | + | 1.46423 | intron (NR_040577, intron 4 of 4) | intron (NR_040577, intron 4 of 4) | 23701 | NR_040577 | 100503636 | Mm.209078 | NR_040577 | | **1700123M08Rik** | 1600025K15Rik | RIKEN cDNA 1700123M08 gene | ncRNA |
| chr4_10357540_10357933 | chr4 | 10357541 | 10357933 | + | 0.631254 | Intergenic | L2c\|LINE\|L2 | 440065 | NR_045185 | 73624 | Mm.67392 | NR_045185 | ENSMUSG00000086080 | **1700123O12Rik** | - | RIKEN cDNA 1700123O12 gene | ncRNA |
| chr19_58629210_58629833 | chr19 | 58629211 | 58629833 | + | 0.896424 | non-coding (NR_038136, exon 3 of 3) | non-coding (NR_038136, exon 3 of 3) | 23193 | NR_038136 | 69055 | Mm.396318 | NR_038136 | | **1810007D17Rik** | - | RIKEN cDNA 1810007D17 gene | ncRNA |
| chr17_28252473_28252753 | chr17 | 28252474 | 28252753 | + | 0.705803 | intron (NM_011145, intron 2 of 7) | intron (NM_011145, intron 2 of 7) | -19095 | NR_045427 | 69050 | Mm.486938 | NR_045427 | ENSMUSG00000002250 | **1810013A23Rik** | - | RIKEN cDNA 1810013A23 gene | ncRNA |
| chr7_73576363_73576694 | chr7 | 73576364 | 73576694 | + | 1.1318 | Intergenic | RMER17C-int\|LTR\|ERVK | -18134 | NR_037569 | 69170 | Mm.34539 | NR_037569 | | **1810026B05Rik** | - | RIKEN cDNA 1810026B05 gene | ncRNA |
| chr7_73576067_73576722 | chr7 | 73576068 | 73576722 | + | 1.13153 | Intergenic | RMER17C-int\|LTR\|ERVK | -18000 | NR_037569 | 69170 | Mm.34539 | NR_037569 | | **1810026B05Rik** | - | RIKEN cDNA 1810026B05 gene | ncRNA |
| chr7_73608926_73609251 | chr7 | 73608927 | 73609251 | + | 0.71184 | Intergenic | Intergenic | -50694 | NR_037569 | 69170 | Mm.34539 | NR_037569 | | **1810026B05Rik** | - | RIKEN cDNA 1810026B05 gene | ncRNA |
| chr9_51398875_51399235 | chr9 | 51398876 | 51399235 | + | 0.820936 | Intergenic | Intergenic | -70138 | NM_027217 | 69809 | Mm.75315 | NM_027217 | ENSMUSG00000036027 | **1810046K07Rik** | - | RIKEN cDNA 1810046K07 gene | protein-coding |
| chr16_93190890_93191718 | chr16 | 93190891 | 93191718 | + | 0.859888 | Intergenic | Intergenic | 161885 | NR_040486 | 69857 | Mm.81192 | NR_040486 | | **1810053B23Rik** | 1810008L18Rik\|1810056K14Rik | RIKEN cDNA 1810053B23 gene | ncRNA |
| chr16_93192588_93192994 | chr16 | 93192589 | 93192994 | + | 0.688907 | Intergenic | Intergenic | 160398 | NR_040486 | 69857 | Mm.81192 | NR_040486 | | **1810053B23Rik** | 1810008L18Rik\|1810056K14Rik | RIKEN cDNA 1810053B23 gene | ncRNA |
| chr3_36489226_36489873 | chr3 | 36489227 | 36489873 | + | 0.721254 | Intergenic | MT2A\|LTR\|ERVL | 13613 | NM_028183 | 72282 | Mm.55317 | NM_028183 | ENSMUSG00000027713 | **1810062G17Rik** | - | RIKEN cDNA 1810062G17 gene | protein-coding |
| chr14_20543124_20543334 | chr14 | 20543125 | 20543334 | + | 1.03336 | intron (NM_008914, intron 1 of 13) | B3A\|SINE\|B2 | -3064 | NR_033571 | 75602 | Mm.437061 | NR_033571 | ENSMUSG00000084925 | **1810062O18Rik** | - | RIKEN cDNA 1810062O18 gene | ncRNA |
| chr9_22214585_22214984 | chr9 | 22214586 | 22214984 | + | 0.687121 | promoter-TSS (NR_027981) | promoter-TSS (NR_027981) | -925 | NR_027981 | 69862 | Mm.488068 | NM_001033151 | | **1810064F22Rik** | - | RIKEN cDNA 1810064F22 gene | ncRNA |
| chr9_22214564_22215142 | chr9 | 22214565 | 22215142 | + | 0.680761 | promoter-TSS (NR_027981) | promoter-TSS (NR_027981) | -993 | NR_027981 | 69862 | Mm.488068 | NM_001033151 | | **1810064F22Rik** | - | RIKEN cDNA 1810064F22 gene | ncRNA |
| chr18_75279124_75279568 | chr18 | 75279125 | 75279568 | + | 0.844334 | intron (NM_027727, intron 16 of 16) | intron (NM_027727, intron 16 of 16) | 17971 | NR_045394 | 100503117 | Mm.250517 | NR_045393 | | **2010010A06Rik** | 5033420E22Rik | RIKEN cDNA 2010010A06 gene | ncRNA |
| chr18_75263476_75263787 | chr18 | 75263477 | 75263787 | + | 0.730151 | intron (NM_027727, intron 16 of 16) | intron (NM_027727, intron 16 of 16) | 33685 | NR_045394 | 100503117 | Mm.250517 | NR_045393 | | **2010010A06Rik** | 5033420E22Rik | RIKEN cDNA 2010010A06 gene | ncRNA |
| chr18_75250695_75251011 | chr18 | 75250696 | 75251011 | + | 0.678087 | intron (NM_027727, intron 16 of 16) | intron (NM_027727, intron 16 of 16) | 46464 | NR_045394 | 100503117 | Mm.250517 | NR_045393 | | **2010010A06Rik** | 5033420E22Rik | RIKEN cDNA 2010010A06 gene | ncRNA |
| chr6_34971371_34971995 | chr6 | 34971372 | 34971995 | + | 0.994509 | intron (NM_001025573, intron 2 of 7) | MTB\|LTR\|ERVL-MaLR | 6316 | NM_001025573 | 243753 | Mm.440221 | NM_001025573 | ENSMUSG00000029847 | **2010107G12Rik** | Gm468 | RIKEN cDNA 2010107G12 gene | protein-coding |
| chr5_93207700_93208023 | chr5 | 93207701 | 93208023 | + | 0.744639 | intron (NM_029363, intron 1 of 2) | intron (NM_029363, intron 1 of 2) | 1344 | NM_029363 | 75610 | Mm.175502 | NM_029363 | ENSMUSG00000029384 | **2010109A12Rik** | - | RIKEN cDNA 2010109A12 gene | protein-coding |
| chr15_74906090_74906947 | chr15 | 74906091 | 74906947 | + | 0.76341 | Intergenic | MLT2B1\|LTR\|ERVL | -24815 | NM_025929 | 67038 | Mm.154566 | NM_025929 | ENSMUSG00000063522 | **2010109I03Rik** | - | RIKEN cDNA 2010109I03 gene | protein-coding |
| chr15_74897383_74898148 | chr15 | 74897384 | 74898148 | + | 0.640229 | Intergenic | Intergenic | -16062 | NM_025929 | 67038 | Mm.154566 | NM_025929 | ENSMUSG00000063522 | **2010109I03Rik** | - | RIKEN cDNA 2010109I03 gene | protein-coding |
| chr13_63180517_63181209 | chr13 | 63180518 | 63181209 | + | 1.49984 | intron (NM_001289926, intron 7 of 15) | intron (NM_001289926, intron 7 of 15) | -59166 | NR_110520 | 72061 | Mm.188105 | NM_028079 | ENSMUSG00000021458 | **2010111I01Rik** | 2300006M17Rik\|AP-O\|Aopep | RIKEN cDNA 2010111I01 gene | protein-coding |
| chr13_63187089_63188012 | chr13 | 63187090 | 63188012 | + | 1.22511 | intron (NM_001289926, intron 7 of 15) | intron (NM_001289926, intron 7 of 15) | -52478 | NR_110520 | 72061 | Mm.188105 | NM_028079 | ENSMUSG00000021458 | **2010111I01Rik** | 2300006M17Rik\|AP-O\|Aopep | RIKEN cDNA 2010111I01 gene | protein-coding |
| chr13_63243646_63243857 | chr13 | 63243647 | 63243857 | + | 0.937046 | intron (NR_110520, intron 2 of 5) | intron (NR_110520, intron 2 of 5) | 3723 | NR_110521 | 72061 | Mm.188105 | NM_028079 | ENSMUSG00000021458 | **2010111I01Rik** | 2300006M17Rik\|AP-O\|Aopep | RIKEN cDNA 2010111I01 gene | protein-coding |
| chr13_63253050_63253565 | chr13 | 63253051 | 63253565 | + | 0.87013 | intron (NR_110520, intron 2 of 5) | intron (NR_110520, intron 2 of 5) | 13279 | NR_110521 | 72061 | Mm.188105 | NM_028079 | ENSMUSG00000021458 | **2010111I01Rik** | 2300006M17Rik\|AP-O\|Aopep | RIKEN cDNA 2010111I01 gene | protein-coding |
| chr13_63245892_63246195 | chr13 | 63245893 | 63246195 | + | 0.865817 | intron (NR_110520, intron 2 of 5) | intron (NR_110520, intron 2 of 5) | 6015 | NR_110521 | 72061 | Mm.188105 | NM_028079 | ENSMUSG00000021458 | **2010111I01Rik** | 2300006M17Rik\|AP-O\|Aopep | RIKEN cDNA 2010111I01 gene | protein-coding |
| chr13_63243597_63243929 | chr13 | 63243598 | 63243929 | + | 0.850202 | intron (NR_110520, intron 2 of 5) | intron (NR_110520, intron 2 of 5) | 3734 | NR_110521 | 72061 | Mm.188105 | NM_028079 | ENSMUSG00000021458 | **2010111I01Rik** | 2300006M17Rik\|AP-O\|Aopep | RIKEN cDNA 2010111I01 gene | protein-coding |
| chr13_63253913_63254201 | chr13 | 63253914 | 63254201 | + | 0.817861 | intron (NR_110520, intron 2 of 5) | intron (NR_110520, intron 2 of 5) | 14028 | NR_110521 | 72061 | Mm.188105 | NM_028079 | ENSMUSG00000021458 | **2010111I01Rik** | 2300006M17Rik\|AP-O\|Aopep | RIKEN cDNA 2010111I01 gene | protein-coding |
| chr13_62949903_62950298 | chr13 | 62949904 | 62950298 | + | 0.804267 | Intergenic | Intergenic | -14792 | NM_001289926 | 72061 | Mm.188105 | NM_028079 | ENSMUSG00000021458 | **2010111I01Rik** | 2300006M17Rik\|AP-O\|Aopep | RIKEN cDNA 2010111I01 gene | protein-coding |
| chr13_63140851_63141911 | chr13 | 63140852 | 63141911 | + | 0.770481 | intron (NM_001289926, intron 5 of 15) | intron (NM_001289926, intron 5 of 15) | -98648 | NR_110520 | 72061 | Mm.188105 | NM_028079 | ENSMUSG00000021458 | **2010111I01Rik** | 2300006M17Rik\|AP-O\|Aopep | RIKEN cDNA 2010111I01 gene | protein-coding |
| chr13_63140870_63141294 | chr13 | 63140871 | 63141294 | + | 0.742363 | intron (NM_001289926, intron 5 of 15) | intron (NM_001289926, intron 5 of 15) | -98947 | NR_110520 | 72061 | Mm.188105 | NM_028079 | ENSMUSG00000021458 | **2010111I01Rik** | 2300006M17Rik\|AP-O\|Aopep | RIKEN cDNA 2010111I01 gene | protein-coding |
| chr13_63216520_63217399 | chr13 | 63216521 | 63217399 | + | 0.734494 | intron (NM_001289926, intron 11 of 15) | intron (NM_001289926, intron 11 of 15) | -23069 | NR_110520 | 72061 | Mm.188105 | NM_028079 | ENSMUSG00000021458 | **2010111I01Rik** | 2300006M17Rik\|AP-O\|Aopep | RIKEN cDNA 2010111I01 gene | protein-coding |
| chr13_63189729_63189995 | chr13 | 63189730 | 63189995 | + | 0.725094 | intron (NM_001289926, intron 7 of 15) | intron (NM_001289926, intron 7 of 15) | -50167 | NR_110520 | 72061 | Mm.188105 | NM_028079 | ENSMUSG00000021458 | **2010111I01Rik** | 2300006M17Rik\|AP-O\|Aopep | RIKEN cDNA 2010111I01 gene | protein-coding |
| chr13_63141315_63141911 | chr13 | 63141316 | 63141911 | + | 0.700439 | intron (NM_001289926, intron 5 of 15) | intron (NM_001289926, intron 5 of 15) | -98416 | NR_110520 | 72061 | Mm.188105 | NM_028079 | ENSMUSG00000021458 | **2010111I01Rik** | 2300006M17Rik\|AP-O\|Aopep | RIKEN cDNA 2010111I01 gene | protein-coding |
| chrX_7443915_7444383 | chrX | 7443916 | 7444383 | + | 0.748533 | Intergenic | Intergenic | -21161 | NR_027924 | 68355 | Mm.45111 | NR_027924 | | **2010204K13Rik** | 0610010M13Rik\|Cip7 | RIKEN cDNA 2010204K13 gene | ncRNA |
| chr9_124262832_124264137 | chr9 | 124262833 | 124264137 | + | 1.02003 | Intergenic | Intergenic | 49209 | NM_001243118 | 630836 | Mm.458553 | NM_001243117 | | **2010315B03Rik** | BC029716 | RIKEN cDNA 2010315B03 gene | protein-coding |
| chr9_124263324_124264121 | chr9 | 124263325 | 124264121 | + | 0.823158 | Intergenic | Intergenic | 48971 | NM_001243118 | 630836 | Mm.458553 | NM_001243117 | | **2010315B03Rik** | BC029716 | RIKEN cDNA 2010315B03 gene | protein-coding |
| chr9_124262837_124263245 | chr9 | 124262838 | 124263245 | + | 0.720761 | Intergenic | Intergenic | 49653 | NM_001243118 | 630836 | Mm.458553 | NM_001243117 | | **2010315B03Rik** | BC029716 | RIKEN cDNA 2010315B03 gene | protein-coding |
| chr7_30698743_30699053 | chr7 | 30698744 | 30699053 | + | 0.696303 | promoter-TSS (NM_026961) | promoter-TSS (NM_026961) | -28 | NM_026961 | 69147 | Mm.307035 | NM_026961 | | **2200002J24Rik** | - | RIKEN cDNA 2200002J24 gene | protein-coding |
| chr6_41115138_41115552 | chr6 | 41115139 | 41115552 | + | 1.51023 | Intergenic | Intergenic | -79836 | NM_023333 | 67373 | Mm.153729 | NM_023333 | ENSMUSG00000029882 | **2210010C04Rik** | AV072249 | RIKEN cDNA 2210010C04 gene | protein-coding |
| chr6_41078634_41079255 | chr6 | 41078635 | 41079255 | + | 0.98579 | Intergenic | Intergenic | -43436 | NM_023333 | 67373 | Mm.153729 | NM_023333 | ENSMUSG00000029882 | **2210010C04Rik** | AV072249 | RIKEN cDNA 2210010C04 gene | protein-coding |
| chr5_114943516_114943929 | chr5 | 114943517 | 114943929 | + | 0.66169 | intron (NM_028211, intron 1 of 5) | intron (NM_028211, intron 1 of 5) | 1521 | NM_028211 | 72357 | Mm.292672 | NM_028211 | | **2210016L21Rik** | - | RIKEN cDNA 2210016L21 gene | protein-coding |
| chr12_73566436_73566911 | chr12 | 73566437 | 73566911 | + | 1.35911 | Intergenic | Intergenic | 18191 | NR_044985 | 73644 | Mm.369885 | NR_044985 | | **2210039B01Rik** | - | RIKEN cDNA 2210039B01 gene | ncRNA |
| chr5_148789187_148789612 | chr5 | 148789188 | 148789612 | + | 0.791915 | Intergenic | Intergenic | 47560 | NR_028285 | 70138 | Mm.417834 | NR_028285 | ENSMUSG00000086494 | **2210417A02Rik** | AV081004 | RIKEN cDNA 2210417A02 gene | ncRNA |
| chr6_129224947_129225617 | chr6 | 129224948 | 129225617 | + | 1.05103 | Intergenic | B3A\|SINE\|B2 | 12763 | NR_040265 | 76432 | Mm.261738 | NR_040265 | ENSMUSG00000097354 | **2310001H17Rik** | AI646108 | RIKEN cDNA 2310001H17 gene | ncRNA |
| chr6_129230212_129230691 | chr6 | 129230213 | 129230691 | + | 0.912834 | Intergenic | Intergenic | 7593 | NR_040265 | 76432 | Mm.261738 | NR_040265 | ENSMUSG00000097354 | **2310001H17Rik** | AI646108 | RIKEN cDNA 2310001H17 gene | ncRNA |
| chr6_129236038_129236638 | chr6 | 129236039 | 129236638 | + | 0.903658 | intron (NR_040268, intron 2 of 3) | intron (NR_040268, intron 2 of 3) | 1707 | NR_040265 | 76432 | Mm.261738 | NR_040265 | ENSMUSG00000097354 | **2310001H17Rik** | AI646108 | RIKEN cDNA 2310001H17 gene | ncRNA |
| chr6_129230359_129230626 | chr6 | 129230360 | 129230626 | + | 0.820122 | Intergenic | Intergenic | 7552 | NR_040265 | 76432 | Mm.261738 | NR_040265 | ENSMUSG00000097354 | **2310001H17Rik** | AI646108 | RIKEN cDNA 2310001H17 gene | ncRNA |
| chr6_129211352_129211589 | chr6 | 129211353 | 129211589 | + | 0.64718 | Intergenic | Intergenic | 26574 | NR_040265 | 76432 | Mm.261738 | NR_040265 | ENSMUSG00000097354 | **2310001H17Rik** | AI646108 | RIKEN cDNA 2310001H17 gene | ncRNA |
| chr8_26813132_26813564 | chr8 | 26813133 | 26813564 | + | 0.658729 | Intergenic | Intergenic | 11075 | NR_045904 | 72316 | Mm.490534 | NR_045904 | | **2310008N11Rik** | - | RIKEN cDNA 2310008N11 gene | ncRNA |
| chr7_27574101_27574535 | chr7 | 27574102 | 27574535 | + | 1.36366 | intron (NM_175107, intron 7 of 9) | intron (NM_175107, intron 7 of 9) | 13842 | NM_001122767 | 66367 | Mm.29831 | NM_175107 | ENSMUSG00000049643 | **2310022A10Rik** | - | RIKEN cDNA 2310022A10 gene | protein-coding |

| | | | | | | | | | | | | | | | | | |
|---|---|---|---|---|---|---|---|---|---|---|---|---|---|---|---|---|---|
| chr7_27573378_27573652 | chr7 | 27573379 | 27573652 | + | 0.773699 | intron (NM_175107, intron 7 of 9) | intron (NM_175107, intron 7 of 9) | 13039 | NM_001122767 | 66367 | Mm.29831 | NM_175107 | ENSMUSG00000049643 | **2310022A10Rik** | - | RIKEN cDNA 2310022A10 gene | protein-coding |
| chr1_105700895_105701334 | chr1 | 105700896 | 105701334 | + | 0.7498 | intron (NM_173187, intron 10 of 28) | intron (NM_173187, intron 10 of 28) | 37254 | NM_173187 | 227446 | Mm.337339 | NM_029349 | ENSMUSG00000026319 | **2310035C23Rik** | 6430401N10\|Kiaa1468\|mKIAA1468 | RIKEN cDNA 2310035C23 gene | protein-coding |
| chr14_70818913_70819490 | chr14 | 70818914 | 70819490 | + | 0.893699 | Intergenic | Intergenic | -33731 | NR_045939 | 73648 | Mm.410851 | NR_045939 | | **2410012E07Rik** | - | RIKEN cDNA 2410012E07 gene | ncRNA |
| chr15_8234083_8234638 | chr15 | 8234084 | 8234638 | + | 0.920945 | intron (NM_001162906, intron 38 of 52) | LTRIS2\|LTR\|ERV1 | 65255 | NM_001162906 | 73692 | Mm.164436 | NM_001162906 | ENSMUSG00000039801 | **2410089E03Rik** | 4732468D17Rik\|b2b012Clo | RIKEN cDNA 2410089E03 gene | protein-coding |
| chr4_35006200_35006983 | chr4 | 35006201 | 35006983 | + | 1.15898 | intron (NM_178061, intron 2 of 3) | intron (NM_178061, intron 2 of 3) | -95111 | NR_040652 | 73696 | Mm.1361 | NR_040652 | ENSMUSG00000085824 | **2410114N07Rik** | - | RIKEN cDNA 2410114N07 gene | ncRNA |
| chr4_34966687_34966985 | chr4 | 34966688 | 34966985 | + | 0.78026 | intron (NM_178061, intron 3 of 3) | intron (NM_178061, intron 3 of 3) | -55355 | NR_040652 | 73696 | Mm.1361 | NR_040652 | ENSMUSG00000085824 | **2410114N07Rik** | - | RIKEN cDNA 2410114N07 gene | ncRNA |
| chr4_35009190_35009485 | chr4 | 35009191 | 35009485 | + | 0.779952 | intron (NM_178061, intron 2 of 3) | MTD\|LTR\|ERVL-MaLR | -97857 | NR_040652 | 73696 | Mm.1361 | NR_040652 | ENSMUSG00000085824 | **2410114N07Rik** | - | RIKEN cDNA 2410114N07 gene | ncRNA |
| chr4_35009562_35009779 | chr4 | 35009563 | 35009779 | + | 0.75348 | intron (NM_178061, intron 2 of 3) | intron (NM_178061, intron 2 of 3) | -98190 | NR_040652 | 73696 | Mm.1361 | NR_040652 | ENSMUSG00000085824 | **2410114N07Rik** | - | RIKEN cDNA 2410114N07 gene | ncRNA |
| chr4_34966646_34967070 | chr4 | 34966647 | 34967070 | + | 0.683023 | intron (NM_178061, intron 3 of 3) | intron (NM_178061, intron 3 of 3) | -55377 | NR_040652 | 73696 | Mm.1361 | NR_040652 | ENSMUSG00000085824 | **2410114N07Rik** | - | RIKEN cDNA 2410114N07 gene | ncRNA |
| chr5_118361762_118362476 | chr5 | 118361763 | 118362476 | + | 1.59026 | Intergenic | Intergenic | 116892 | NM_001081236 | 76792 | Mm.103428 | NM_001081236 | | **2410131K14Rik** | - | RIKEN cDNA 2410131K14 gene | protein-coding |
| chr5_118361446_118362504 | chr5 | 118361447 | 118362504 | + | 1.43575 | Intergenic | Intergenic | 116748 | NM_001081236 | 76792 | Mm.103428 | NM_001081236 | | **2410131K14Rik** | - | RIKEN cDNA 2410131K14 gene | protein-coding |
| chr5_118377442_118377951 | chr5 | 118377443 | 118377951 | + | 0.664851 | Intergenic | Intergenic | 132470 | NM_001081236 | 76792 | Mm.103428 | NM_001081236 | | **2410131K14Rik** | - | RIKEN cDNA 2410131K14 gene | protein-coding |
| chr5_118375649_118376449 | chr5 | 118375650 | 118376449 | + | 0.654919 | Intergenic | Intergenic | 130822 | NM_001081236 | 76792 | Mm.103428 | NM_001081236 | | **2410131K14Rik** | - | RIKEN cDNA 2410131K14 gene | protein-coding |
| chr5_118360730_118361033 | chr5 | 118360731 | 118361033 | + | 0.649137 | Intergenic | Intergenic | 115655 | NM_001081236 | 76792 | Mm.103428 | NM_001081236 | | **2410131K14Rik** | - | RIKEN cDNA 2410131K14 gene | protein-coding |
| chr7_119801017_119801473 | chr7 | 119801018 | 119801473 | + | 1.1075 | exon (NM_001004187, exon 3 of 13) | exon (NM_001004187, exon 3 of 13) | 7058 | NM_001004187 | 434234 | Mm.38237 | NM_028129 | ENSMUSG00000030924 | **2610020H08Rik** | - | RIKEN cDNA 2610020H08 gene | protein-coding |
| chr7_119800536_119801512 | chr7 | 119800537 | 119801512 | + | 0.723942 | intron (NM_001004187, intron 2 of 12) | intron (NM_001004187, intron 2 of 12) | 6837 | NM_001004187 | 434234 | Mm.38237 | NM_028129 | ENSMUSG00000030924 | **2610020H08Rik** | - | RIKEN cDNA 2610020H08 gene | protein-coding |
| chr11_113060829_113061480 | chr11 | 113060830 | 113061480 | + | 1.06389 | intron (NR_015556, intron 2 of 2) | intron (NR_015556, intron 2 of 2) | 140683 | NR_015556 | 72386 | Mm.266961 | NR_015556 | ENSMUSG00000087259 | **2610035D17Rik** | - | RIKEN cDNA 2610035D17 gene | ncRNA |
| chr7_84744462_84744986 | chr7 | 84744463 | 84744986 | + | 1.28912 | intron (NR_038175, intron 2 of 4) | intron (NR_038175, intron 2 of 4) | 55084 | NR_038175 | 72495 | Mm.382914 | NR_038175 | | **2610206C17Rik** | - | RIKEN cDNA 2610206C17 gene | ncRNA |
| chr7_84741750_84742069 | chr7 | 84741751 | 84742069 | + | 0.718626 | intron (NR_038175, intron 2 of 4) | L1MC1\|LINE\|L1 | 52270 | NR_038175 | 72495 | Mm.382914 | NR_038175 | | **2610206C17Rik** | - | RIKEN cDNA 2610206C17 gene | ncRNA |
| chr4_10915333_10915595 | chr4 | 10915334 | 10915595 | + | 0.791298 | Intergenic | Intergenic | 40966 | NM_026005 | 67157 | Mm.440552 | NM_026005 | ENSMUSG00000059482 | **2610301B20Rik** | AI428449 | RIKEN cDNA 2610301B20 gene | protein-coding |
| chr13_28855247_28855869 | chr13 | 28855248 | 28855869 | + | 0.780218 | intron (NR_045053, intron 2 of 8) | MER2\|DNA\|TcMar-Tigger | 29864 | NR_045054 | 72518 | Mm.37454 | NR_045053 | | **2610307P16Rik** | AI666457 | RIKEN cDNA 2610307P16 gene | ncRNA |
| chr3_45377660_45378480 | chr3 | 45377661 | 45378480 | + | 0.79853 | promoter-TSS (NM_001098172) | promoter-TSS (NM_001098172) | 190 | NR_045172 | 72511 | Mm.140700 | NR_045172 | ENSMUSG00000097040 | **2610316D01Rik** | - | RIKEN cDNA 2610316D01 gene | ncRNA |
| chr8_60915051_60915392 | chr8 | 60915052 | 60915392 | + | 1.20902 | intron (NM_173874, intron 12 of 13) | intron (NM_173874, intron 12 of 13) | 24771 | NM_028299 | 72612 | Mm.25263 | NM_028299 | ENSMUSG00000038005 | **2700029M09Rik** | C230006B22Rik | RIKEN cDNA 2700029M09 gene | protein-coding |
| chr8_60902122_60902658 | chr8 | 60902123 | 60902658 | + | 1.05712 | intron (NM_028299, intron 5 of 7) | intron (NM_028299, intron 5 of 7) | 11939 | NM_028299 | 72612 | Mm.25263 | NM_028299 | ENSMUSG00000038005 | **2700029M09Rik** | C230006B22Rik | RIKEN cDNA 2700029M09 gene | protein-coding |
| chr8_60889054_60889364 | chr8 | 60889055 | 60889364 | + | 0.924315 | Intergenic | Intergenic | -1242 | NM_028299 | 72612 | Mm.25263 | NM_028299 | ENSMUSG00000038005 | **2700029M09Rik** | C230006B22Rik | RIKEN cDNA 2700029M09 gene | protein-coding |
| chr8_60893802_60894071 | chr8 | 60893803 | 60894071 | + | 0.723457 | intron (NM_028299, intron 2 of 7) | intron (NM_028299, intron 2 of 7) | 3486 | NM_028299 | 72612 | Mm.25263 | NM_028299 | ENSMUSG00000038005 | **2700029M09Rik** | C230006B22Rik | RIKEN cDNA 2700029M09 gene | protein-coding |
| chr8_60859622_60860049 | chr8 | 60859623 | 60860049 | + | 0.671965 | Intergenic | Intergenic | -30615 | NM_028299 | 72612 | Mm.25263 | NM_028299 | ENSMUSG00000038005 | **2700029M09Rik** | C230006B22Rik | RIKEN cDNA 2700029M09 gene | protein-coding |
| chr8_60859619_60860212 | chr8 | 60859620 | 60860212 | + | 0.647742 | Intergenic | Intergenic | -30535 | NM_028299 | 72612 | Mm.25263 | NM_028299 | ENSMUSG00000038005 | **2700029M09Rik** | C230006B22Rik | RIKEN cDNA 2700029M09 gene | protein-coding |
| chr12_71144212_71145006 | chr12 | 71144213 | 71145006 | + | 1.03365 | intron (NM_029818, intron 4 of 27) | Lx9\|LINE\|L1 | 7761 | NM_001163378 | 76967 | Mm.87290 | NM_029818 | ENSMUSG00000034601 | **2700049A03Rik** | Talpid3\|mKIAA0586 | RIKEN cDNA 2700049A03 gene | protein-coding |
| chr12_71144221_71144769 | chr12 | 71144222 | 71144769 | + | 1.02712 | intron (NM_029818, intron 4 of 27) | Lx9\|LINE\|L1 | 7647 | NM_001163378 | 76967 | Mm.87290 | NM_029818 | ENSMUSG00000034601 | **2700049A03Rik** | Talpid3\|mKIAA0586 | RIKEN cDNA 2700049A03 gene | protein-coding |
| chr1_64656435_64656747 | chr1 | 64656436 | 64656747 | + | 0.734224 | Intergenic | ORR1A2\|LTR\|ERVL-MaLR | 34068 | NR_038009 | 69941 | Mm.69044 | NR_038009 | ENSMUSG00000087213 | **2810408I11Rik** | - | RIKEN cDNA 2810408I11 gene | ncRNA |
| chr1_64656409_64656927 | chr1 | 64656410 | 64656927 | + | 0.664694 | Intergenic | Intergenic | 33991 | NR_038009 | 69941 | Mm.69044 | NR_038009 | ENSMUSG00000087213 | **2810408I11Rik** | - | RIKEN cDNA 2810408I11 gene | ncRNA |
| chr6_134903430_134904449 | chr6 | 134903431 | 134904449 | + | 0.647218 | Intergenic | Intergenic | 5979 | NR_029441 | 72813 | Mm.409034 | NR_029441 | | **2810454H06Rik** | - | RIKEN cDNA 2810454H06 gene | ncRNA |
| chr6_149355008_149355698 | chr6 | 149355009 | 149355698 | + | 0.825837 | Intergenic | Intergenic | 45939 | NM_001289662 | 67246 | Mm.333515 | NM_026054 | ENSMUSG00000032712 | **2810474O19Rik** | 6720435I21Rik\|AA536804\|AU020969\|GET\|Kiaa1551 | RIKEN cDNA 2810474O19 gene | protein-coding |
| chr6_149355153_149355558 | chr6 | 149355154 | 149355558 | + | 0.793464 | Intergenic | Intergenic | 45942 | NM_001289662 | 67246 | Mm.333515 | NM_026054 | ENSMUSG00000032712 | **2810474O19Rik** | 6720435I21Rik\|AA536804\|AU020969\|GET\|Kiaa1551 | RIKEN cDNA 2810474O19 gene | protein-coding |
| chr6_149355811_149356217 | chr6 | 149355812 | 149356217 | + | 0.737153 | Intergenic | RLTR1F_Mm\|LTR\|ERVK | 46600 | NM_001289662 | 67246 | Mm.333515 | NM_026054 | ENSMUSG00000032712 | **2810474O19Rik** | 6720435I21Rik\|AA536804\|AU020969\|GET\|Kiaa1551 | RIKEN cDNA 2810474O19 gene | protein-coding |
| chr6_149330115_149330775 | chr6 | 149330116 | 149330775 | + | 0.685063 | intron (NM_026054, intron 3 of 4) | intron (NM_026054, intron 3 of 4) | 21031 | NM_001289662 | 67246 | Mm.333515 | NM_026054 | ENSMUSG00000032712 | **2810474O19Rik** | 6720435I21Rik\|AA536804\|AU020969\|GET\|Kiaa1551 | RIKEN cDNA 2810474O19 gene | protein-coding |
| chr6_149260655_149261274 | chr6 | 149260656 | 149261274 | + | 0.664654 | Intergenic | Intergenic | -48449 | NM_001289661 | 67246 | Mm.333515 | NM_026054 | ENSMUSG00000032712 | **2810474O19Rik** | 6720435I21Rik\|AA536804\|AU020969\|GET\|Kiaa1551 | RIKEN cDNA 2810474O19 gene | protein-coding |
| chrX_12124419_12125063 | chrX | 12124420 | 12125063 | + | 0.78107 | intron (NM_001168321, intron 1 of 13) | intron (NM_001168321, intron 1 of 13) | 35605 | NR_045434 | 72937 | Mm.440386 | NR_045434 | ENSMUSG00000087271 | **2900008C10Rik** | - | RIKEN cDNA 2900008C10 gene | ncRNA |
| chrX_12129845_12130383 | chrX | 12129846 | 12130383 | + | 0.627256 | intron (NM_001168321, intron 1 of 13) | intron (NM_001168321, intron 1 of 13) | 30232 | NR_045434 | 72937 | Mm.440386 | NR_045434 | ENSMUSG00000087271 | **2900008C10Rik** | - | RIKEN cDNA 2900008C10 gene | ncRNA |
| chr18_76034580_76034976 | chr18 | 76034581 | 76034976 | + | 0.66528 | intron (NM_145356, intron 3 of 4) | MIR\|SINE\|MIR | 63693 | NR_045365 | 73009 | Mm.443541 | NR_045365 | | **2900057B20Rik** | - | RIKEN cDNA 2900057B20 gene | ncRNA |
| chr1_118458314_118458560 | chr1 | 118458315 | 118458560 | + | 0.69046 | TTS (NR_027901) | TTS (NR_027901) | 828 | NR_027901 | 68204 | Mm.443536 | NR_027901 | | **2900060B14Rik** | - | RIKEN cDNA 2900060B14 gene | ncRNA |
| chr7_81516169_81517039 | chr7 | 81516170 | 81517039 | + | 0.883234 | Intergenic | Intergenic | -6946 | NR_045299 | 100504421 | Mm.217071 | NR_045299 | ENSMUSG00000097277 | **2900076A07Rik** | WI1-1616M8.2 | RIKEN cDNA 2900076A07 gene | ncRNA |
| chr9_112546208_112546637 | chr9 | 112546209 | 112546637 | + | 1.35716 | Intergenic | L1MA6\|LINE\|L1 | 289849 | NR_033433 | 620760 | Mm.259863 | NR_015468 | ENSMUSG00000087038 | **2900079G21Rik** | C130083B14\|Gm6179 | RIKEN cDNA 2900079G21 gene | ncRNA |
| chr1_120136024_120136819 | chr1 | 120136025 | 120136819 | + | 1.39841 | intron (NM_001172074, intron 1 of 4) | intron (NM_001172074, intron 1 of 4) | 15235 | NM_001172074 | 73103 | Mm.271602 | NM_028439 | ENSMUSG00000026388 | **3110009E18Rik** | - | RIKEN cDNA 3110009E18 gene | protein-coding |
| chr13_111186854_111187320 | chr13 | 111186855 | 111187320 | + | 1.18921 | Intergenic | Intergenic | 42344 | NR_045908 | 73129 | Mm.417826 | NR_045908 | | **3110015C05Rik** | - | RIKEN cDNA 3110015C05 gene | ncRNA |
| chr13_111186747_111187394 | chr13 | 111186748 | 111187394 | + | 1.00998 | Intergenic | Intergenic | 42360 | NR_045908 | 73129 | Mm.417826 | NR_045908 | | **3110015C05Rik** | - | RIKEN cDNA 3110015C05 gene | ncRNA |
| chr13_110960007_110960615 | chr13 | 110960008 | 110960615 | + | 0.818807 | Intergenic | Intergenic | 269120 | NR_045908 | 73129 | Mm.417826 | NR_045908 | | **3110015C05Rik** | - | RIKEN cDNA 3110015C05 gene | ncRNA |
| chr13_110960170_110960613 | chr13 | 110960171 | 110960613 | + | 0.804101 | Intergenic | Intergenic | 269039 | NR_045908 | 73129 | Mm.417826 | NR_045908 | | **3110015C05Rik** | - | RIKEN cDNA 3110015C05 gene | ncRNA |
| chr6_119855507_119855904 | chr6 | 119855508 | 119855904 | + | 0.863728 | Intergenic | Intergenic | 7513 | NR_030776 | 67289 | Mm.418299 | NR_030776 | ENSMUSG00000050604 | **3110021A11Rik** | - | RIKEN cDNA 3110021A11 gene | ncRNA |
| chr9_41439321_41439942 | chr9 | 41439322 | 41439942 | + | 0.959602 | Intergenic | Intergenic | 63071 | NR_040725 | 73144 | Mm.482850 | NR_040725 | | **3110039I08Rik** | - | RIKEN cDNA 3110039I08 gene | ncRNA |
| chr9_41426321_41427180 | chr9 | 41426322 | 41427180 | + | 0.881519 | intron (NR_040725, intron 4 of 4) | intron (NR_040725, intron 4 of 4) | 50190 | NR_040725 | 73144 | Mm.482850 | NR_040725 | | **3110039I08Rik** | - | RIKEN cDNA 3110039I08 gene | ncRNA |
| chr4_35224029_35224511 | chr4 | 35224030 | 35224511 | + | 1.36076 | intron (NM_001081343, intron 1 of 10) | MER44B\|DNA\|TcMar-Tigger | 1610 | NM_001081343 | 73205 | Mm.331544 | NM_001081343 | ENSMUSG00000028300 | **3110043O21Rik** | AI840585 | RIKEN cDNA 3110043O21 gene | protein-coding |
| chr17_21652525_21652973 | chr17 | 21652526 | 21652973 | + | 1.52332 | intron (NM_001166497, intron 2 of 5) | RLTR48B\|LTR\|ERV1 | 2138 | NM_001166497 | 73229 | Mm.296733 | NM_001166497 | | **3110052M02Rik** | mszf57\|mszf71 | RIKEN cDNA 3110052M02 gene | protein-coding |
| chr17_21652010_21652375 | chr17 | 21652011 | 21652375 | + | 0.883275 | intron (NM_001166497, intron 1 of 5) | intron (NM_001166497, intron 1 of 5) | 1582 | NM_001166497 | 73229 | Mm.296733 | NM_001166497 | | **3110052M02Rik** | mszf57\|mszf71 | RIKEN cDNA 3110052M02 gene | protein-coding |
| chr12_70995115_70995696 | chr12 | 70995116 | 70995696 | + | 1.01801 | TTS (NM_011184) | TTS (NM_011184) | 20417 | NR_045056 | 73204 | Mm.363770 | NR_045055 | ENSMUSG00000085622 | **3110056K07Rik** | 4921536K06Rik | RIKEN cDNA 3110056K07 gene | ncRNA |
| chr5_30126693_30127202 | chr5 | 30126694 | 30127202 | + | 1.15584 | intron (NM_178878, intron 12 of 19) | intron (NM_178878, intron 12 of 19) | -20862 | NM_001256263 | 433868 | Mm.450696 | NM_001256263 | | **3110082J24Rik** | ENSMUSG00000073106 | RIKEN cDNA 3110082J24 gene | protein-coding |
| chr17_5755775_5756511 | chr17 | 5755776 | 5756511 | + | 1.89487 | Intergenic | Intergenic | -42514 | NR_045081 | 78512 | Mm.23696 | NR_045079 | ENSMUSG00000096965 | **3300005D01Rik** | - | RIKEN cDNA 3300005D01 gene | ncRNA |

| | | | | | | | | | | | | | | | | | |
|---|---|---|---|---|---|---|---|---|---|---|---|---|---|---|---|---|---|
| chr17_5795378_5795763 | chr17 | 5795379 | 5795763 | + | 1.13708 | Intergenic | Intergenic | -3086 | NR_045081 | 78512 | Mm.23696 | NR_045079 | ENSMUSG00000096965 | **3300005D01Rik** | - | RIKEN cDNA 3300005D01 gene | ncRNA |
| chr17_5668244_5668927 | chr17 | 5668245 | 5668927 | + | 0.993679 | intron (NM_146073, intron 2 of 8) | intron (NM_146073, intron 2 of 8) | -130071 | NR_045081 | 78512 | Mm.23696 | NR_045079 | ENSMUSG00000096965 | **3300005D01Rik** | - | RIKEN cDNA 3300005D01 gene | ncRNA |
| chr17_5747655_5748474 | chr17 | 5747656 | 5748474 | + | 0.970643 | intron (NM_146073, intron 8 of 8) | (TG)n\|Simple_repeat\|Simple_repeat | -50592 | NR_045081 | 78512 | Mm.23696 | NR_045079 | ENSMUSG00000096965 | **3300005D01Rik** | - | RIKEN cDNA 3300005D01 gene | ncRNA |
| chr17_5729070_5729449 | chr17 | 5729071 | 5729449 | + | 0.636461 | intron (NM_146073, intron 7 of 8) | ORR1B1\|LTR\|ERVL-MaLR | -69397 | NR_045081 | 78512 | Mm.23696 | NR_045079 | ENSMUSG00000096965 | **3300005D01Rik** | - | RIKEN cDNA 3300005D01 gene | ncRNA |
| chr14_32696322_32697073 | chr14 | 32696323 | 32697073 | + | 1.18898 | Intergenic | Intergenic | -11426 | NM_001195097 | 100504518 | Mm.318319 | NM_001195097 | ENSMUSG00000071540 | **3425401B19Rik** | - | RIKEN cDNA 3425401B19 gene | protein-coding |
| chr14_12243398_12244301 | chr14 | 12243399 | 12244301 | + | 0.842678 | Intergenic | Intergenic | -40353 | NM_001284384 | 218734 | Mm.44068 | NM_146051 | ENSMUSG00000033111 | **3830406C13Rik** | - | RIKEN cDNA 3830406C13 gene | protein-coding |
| chr13_107133917_107134607 | chr13 | 107133918 | 107134607 | + | 1.63837 | Intergenic | Intergenic | 111692 | NR_015471 | 100040322 | Mm.133867 | NR_015471 | | **3830408C21Rik** | - | RIKEN cDNA 3830408C21 gene | ncRNA |
| chr13_107241685_107242223 | chr13 | 107241686 | 107242223 | + | 1.56063 | Intergenic | Intergenic | 219384 | NR_015471 | 100040322 | Mm.133867 | NR_015471 | | **3830408C21Rik** | - | RIKEN cDNA 3830408C21 gene | ncRNA |
| chr13_107096933_107097234 | chr13 | 107096934 | 107097234 | + | 1.11034 | Intergenic | Intergenic | 74514 | NR_015471 | 100040322 | Mm.133867 | NR_015471 | | **3830408C21Rik** | - | RIKEN cDNA 3830408C21 gene | ncRNA |
| chr13_107095191_107095997 | chr13 | 107095192 | 107095997 | + | 0.636972 | Intergenic | Intergenic | 73024 | NR_015471 | 100040322 | Mm.133867 | NR_015471 | | **3830408C21Rik** | - | RIKEN cDNA 3830408C21 gene | ncRNA |
| chr2_92994377_92995147 | chr2 | 92994378 | 92995147 | + | 1.0444 | intron (NM_001177536, intron 4 of 6) | intron (NM_001177536, intron 4 of 6) | -34586 | NR_110333 | 433456 | Mm.235231 | NM_001002794 | | **4631405J19Rik** | - | RIKEN cDNA 4631405J19 gene | ncRNA |
| chr7_67854164_67854998 | chr7 | 67854165 | 67854998 | + | 0.885899 | Intergenic | Intergenic | -51085 | NR_045954 | 73904 | Mm.70992 | NR_045954 | ENSMUSG00000097697 | **4833412C05Rik** | - | RIKEN cDNA 4833412C05 gene | ncRNA |
| chr7_67802642_67804039 | chr7 | 67802643 | 67804039 | + | 0.808068 | intron (NR_045954, intron 1 of 2) | intron (NR_045954, intron 1 of 2) | 155 | NR_045954 | 73904 | Mm.70992 | NR_045954 | ENSMUSG00000097697 | **4833412C05Rik** | - | RIKEN cDNA 4833412C05 gene | ncRNA |
| chr18_4354236_4354846 | chr18 | 4354237 | 4354846 | + | 0.654814 | intron (NR_040328, intron 2 of 3) | intron (NR_040328, intron 2 of 3) | 994 | NR_040328 | 73915 | Mm.482777 | NR_040328 | | **4833419F23Rik** | - | RIKEN cDNA 4833419F23 gene | ncRNA |
| chr13_91768658_91769607 | chr13 | 91768659 | 91769607 | + | 1.00945 | intron (NM_028790, intron 5 of 14) | intron (NM_028790, intron 5 of 14) | -27261 | NR_028313 | 373852 | Mm.466817 | NR_015501 | ENSMUSG00000074782 | **4833422C13Rik** | - | RIKEN cDNA 4833422C13 gene | ncRNA |
| chr13_91750156_91750487 | chr13 | 91750157 | 91750487 | + | 0.979131 | intron (NM_028790, intron 2 of 14) | intron (NM_028790, intron 2 of 14) | -8450 | NR_028313 | 373852 | Mm.466817 | NR_015501 | ENSMUSG00000074782 | **4833422C13Rik** | - | RIKEN cDNA 4833422C13 gene | ncRNA |
| chr13_91750078_91750612 | chr13 | 91750079 | 91750612 | + | 0.936562 | intron (NM_028790, intron 2 of 14) | intron (NM_028790, intron 2 of 14) | -8473 | NR_028313 | 373852 | Mm.466817 | NR_015501 | ENSMUSG00000074782 | **4833422C13Rik** | - | RIKEN cDNA 4833422C13 gene | ncRNA |
| chr2_19551922_19552437 | chr2 | 19551923 | 19552437 | + | 1.00511 | intron (NM_027600, intron 1 of 16) | intron (NM_027600, intron 1 of 16) | 1730 | NM_027600 | 70909 | Mm.158488 | NM_027600 | ENSMUSG00000026734 | **4921504E06Rik** | 4921522E24Rik | RIKEN cDNA 4921504E06 gene | protein-coding |
| chr2_19579551_19579884 | chr2 | 19579552 | 19579884 | + | 0.823749 | Intergenic | Intergenic | -25808 | NM_027600 | 70909 | Mm.158488 | NM_027600 | ENSMUSG00000026734 | **4921504E06Rik** | 4921522E24Rik | RIKEN cDNA 4921504E06 gene | protein-coding |
| chr2_19579447_19580140 | chr2 | 19579448 | 19580140 | + | 0.729073 | Intergenic | Intergenic | -25884 | NM_027600 | 70909 | Mm.158488 | NM_027600 | ENSMUSG00000026734 | **4921504E06Rik** | 4921522E24Rik | RIKEN cDNA 4921504E06 gene | protein-coding |
| chr2_19540758_19541308 | chr2 | 19540759 | 19541308 | + | 0.70024 | intron (NM_027600, intron 2 of 16) | intron (NM_027600, intron 2 of 16) | 12877 | NM_027600 | 70909 | Mm.158488 | NM_027600 | ENSMUSG00000026734 | **4921504E06Rik** | 4921522E24Rik | RIKEN cDNA 4921504E06 gene | protein-coding |
| chr2_93979743_93980137 | chr2 | 93979744 | 93980137 | + | 1.64723 | promoter-TSS (NR_045827) | promoter-TSS (NR_045827) | -94 | NR_045827 | 70852 | Mm.437014 | NR_045827 | ENSMUSG00000027196 | **4921507L20Rik** | - | RIKEN cDNA 4921507L20 gene | ncRNA |
| chr2_93978805_93979343 | chr2 | 93978806 | 93979343 | + | 1.04592 | promoter-TSS (NR_045827) | promoter-TSS (NR_045827) | -960 | NR_045827 | 70852 | Mm.437014 | NR_045827 | ENSMUSG00000027196 | **4921507L20Rik** | - | RIKEN cDNA 4921507L20 gene | ncRNA |
| chr13_113466520_113467068 | chr13 | 113466521 | 113467068 | + | 0.853155 | intron (NR_045501, intron 3 of 6) | intron (NR_045501, intron 3 of 6) | 36050 | NR_045501 | 70917 | Mm.491036 | NR_045501 | | **4921509O07Rik** | - | RIKEN cDNA 4921509O07 gene | ncRNA |
| chr3_79233693_79234034 | chr3 | 79233694 | 79234034 | + | 1.06091 | intron (NR_045502, intron 1 of 2) | intron (NR_045502, intron 1 of 2) | 20945 | NR_045502 | 70924 | Mm.444622 | NR_045502 | | **4921511C10Rik** | - | RIKEN cDNA 4921511C10 gene | ncRNA |
| chr3_79269860_79270257 | chr3 | 79269861 | 79270257 | + | 0.855733 | Intergenic | Intergenic | 57140 | NR_045502 | 70924 | Mm.444622 | NR_045502 | | **4921511C10Rik** | - | RIKEN cDNA 4921511C10 gene | ncRNA |
| chr3_79244445_79244676 | chr3 | 79244446 | 79244676 | + | 0.622214 | intron (NR_045502, intron 1 of 2) | Lx8\|LINE\|L1 | 31642 | NR_045502 | 70924 | Mm.444622 | NR_045502 | | **4921511C10Rik** | - | RIKEN cDNA 4921511C10 gene | ncRNA |
| chr12_12546340_12546715 | chr12 | 12546341 | 12546715 | + | 1.10071 | Intergenic | Intergenic | -154053 | NR_045829 | 70858 | Mm.410838 | NR_045829 | | **4921511I17Rik** | - | RIKEN cDNA 4921511I17 gene | ncRNA |
| chr12_12437432_12437849 | chr12 | 12437433 | 12437849 | + | 0.895773 | Intergenic | Intergenic | -45166 | NR_045829 | 70858 | Mm.410838 | NR_045829 | | **4921511I17Rik** | - | RIKEN cDNA 4921511I17 gene | ncRNA |
| chr12_12547387_12548252 | chr12 | 12547388 | 12548252 | + | 0.843281 | Intergenic | Intergenic | -155345 | NR_045829 | 70858 | Mm.410838 | NR_045829 | | **4921511I17Rik** | - | RIKEN cDNA 4921511I17 gene | ncRNA |
| chr12_12590812_12591295 | chr12 | 12590813 | 12591295 | + | 0.814113 | Intergenic | Intergenic | -198579 | NR_045829 | 70858 | Mm.410838 | NR_045829 | | **4921511I17Rik** | - | RIKEN cDNA 4921511I17 gene | ncRNA |
| chr12_12437169_12437881 | chr12 | 12437170 | 12437881 | + | 0.755503 | Intergenic | Intergenic | -45050 | NR_045829 | 70858 | Mm.410838 | NR_045829 | | **4921511I17Rik** | - | RIKEN cDNA 4921511I17 gene | ncRNA |
| chr12_12617631_12618095 | chr12 | 12617632 | 12618095 | + | 0.70952 | Intergenic | MTD\|LTR\|ERVL-MaLR | -225388 | NR_045829 | 70858 | Mm.410838 | NR_045829 | | **4921511I17Rik** | - | RIKEN cDNA 4921511I17 gene | ncRNA |
| chr8_85408127_85408653 | chr8 | 85408128 | 85408653 | + | 0.853056 | TTS (NM_025722) | TTS (NM_025722) | 24451 | NR_033144 | 66714 | Mm.272748 | NM_025722 | ENSMUSG00000036934 | **4921524J17Rik** | - | RIKEN cDNA 4921524J17 gene | protein-coding |
| chr13_52167876_52168404 | chr13 | 52167877 | 52168404 | + | 0.970005 | Intergenic | Intergenic | 71399 | NR_045061 | 74050 | Mm.84860 | NR_045061 | | **4921525O09Rik** | - | RIKEN cDNA 4921525O09 gene | ncRNA |
| chr13_52188499_52188901 | chr13 | 52188500 | 52188901 | + | 0.813808 | Intergenic | MLT1A1\|LTR\|ERVL-MaLR | 91959 | NR_045061 | 74050 | Mm.84860 | NR_045061 | | **4921525O09Rik** | - | RIKEN cDNA 4921525O09 gene | ncRNA |
| chr13_52188305_52188901 | chr13 | 52188306 | 52188901 | + | 0.686131 | Intergenic | Intergenic | 91862 | NR_045061 | 74050 | Mm.84860 | NR_045061 | | **4921525O09Rik** | - | RIKEN cDNA 4921525O09 gene | ncRNA |
| chr18_83465021_83465575 | chr18 | 83465022 | 83465575 | + | 0.667645 | Intergenic | MT2A\|LTR\|ERVL | -139414 | NR_045361 | 70960 | Mm.410834 | NR_045361 | | **4921531P14Rik** | 1700108N05Rik | RIKEN cDNA 4921531P14 gene | ncRNA |
| chr6_129312493_129313048 | chr6 | 129312494 | 129313048 | + | 1.71293 | Intergenic | L1M4\|LINE\|L1 | 19010 | NM_199034 | 381816 | Mm.46114 | NM_199034 | ENSMUSG00000047720 | **4922502D21Rik** | Clec2m\|Gm1074 | RIKEN cDNA 4922502D21 gene | protein-coding |
| chr6_129313974_129314351 | chr6 | 129313975 | 129314351 | + | 1.19526 | Intergenic | Intergenic | 17618 | NM_199034 | 381816 | Mm.46114 | NM_199034 | ENSMUSG00000047720 | **4922502D21Rik** | Clec2m\|Gm1074 | RIKEN cDNA 4922502D21 gene | protein-coding |
| chr6_129329323_129329733 | chr6 | 129329324 | 129329733 | + | 0.820796 | intron (NM_199034, intron 2 of 4) | Lx8\|LINE\|L1 | 2253 | NM_199034 | 381816 | Mm.46114 | NM_199034 | ENSMUSG00000047720 | **4922502D21Rik** | Clec2m\|Gm1074 | RIKEN cDNA 4922502D21 gene | protein-coding |
| chr6_129314768_129314971 | chr6 | 129314769 | 129314971 | + | 0.793324 | Intergenic | Intergenic | 16911 | NM_199034 | 381816 | Mm.46114 | NM_199034 | ENSMUSG00000047720 | **4922502D21Rik** | Clec2m\|Gm1074 | RIKEN cDNA 4922502D21 gene | protein-coding |
| chr6_129316401_129316801 | chr6 | 129316402 | 129316801 | + | 0.644251 | Intergenic | B1F\|SINE\|Alu | 15180 | NM_199034 | 381816 | Mm.46114 | NM_199034 | ENSMUSG00000047720 | **4922502D21Rik** | Clec2m\|Gm1074 | RIKEN cDNA 4922502D21 gene | protein-coding |
| chr13_101465422_101465875 | chr13 | 101465423 | 101465875 | + | 0.835527 | Intergenic | Intergenic | -68069 | NR_046187 | 78136 | Mm.399065 | NR_046187 | | **4922502H24Rik** | - | RIKEN cDNA 4922502H24 gene | ncRNA |
| chr13_101465466_101465781 | chr13 | 101465467 | 101465781 | + | 0.800765 | Intergenic | Intergenic | -68044 | NR_046187 | 78136 | Mm.399065 | NR_046187 | | **4922502H24Rik** | - | RIKEN cDNA 4922502H24 gene | ncRNA |
| chr2_35416029_35417044 | chr2 | 35416030 | 35417044 | + | 0.97979 | intron (NM_010283, intron 4 of 8) | intron (NM_010283, intron 4 of 8) | -19363 | NM_001272040 | 74854 | Mm.159338 | NM_001080709 | ENSMUSG00000079421 | **4930402F06Rik** | - | RIKEN cDNA 4930402F06 gene | protein-coding |
| chr2_35411535_35412832 | chr2 | 35411536 | 35412832 | + | 0.878251 | intron (NM_010283, intron 7 of 8) | intron (NM_010283, intron 7 of 8) | -15010 | NM_001272040 | 74854 | Mm.159338 | NM_001080709 | ENSMUSG00000079421 | **4930402F06Rik** | - | RIKEN cDNA 4930402F06 gene | protein-coding |
| chr2_35412258_35412805 | chr2 | 35412259 | 35412805 | + | 0.869266 | intron (NM_010283, intron 7 of 8) | B4\|SINE\|B4 | -15358 | NM_001272040 | 74854 | Mm.159338 | NM_001080709 | ENSMUSG00000079421 | **4930402F06Rik** | - | RIKEN cDNA 4930402F06 gene | protein-coding |
| chr2_35411581_35412202 | chr2 | 35411582 | 35412202 | + | 0.817069 | intron (NM_010283, intron 7 of 8) | intron (NM_010283, intron 7 of 8) | -14718 | NM_001272040 | 74854 | Mm.159338 | NM_001080709 | ENSMUSG00000079421 | **4930402F06Rik** | - | RIKEN cDNA 4930402F06 gene | protein-coding |
| chr2_130835559_130836008 | chr2 | 130835560 | 130836008 | + | 0.911434 | intron (NM_029432, intron 1 of 36) | PB1D11\|SINE\|Alu | 4326 | NM_029432 | 228602 | Mm.271990 | NM_029432 | ENSMUSG00000027309 | **4930402H24Rik** | 2810487F15Rik\|AI413763\|C78926 | RIKEN cDNA 4930402H24 gene | protein-coding |
| chrX_9139580_9140026 | chrX | 9139581 | 9140026 | + | 1.00335 | Intergenic | Intergenic | 35241 | NM_001270700 | 75775 | Mm.50374 | NM_001270700 | ENSMUSG00000045797 | **4930402K13Rik** | - | RIKEN cDNA 4930402K13 gene | protein-coding |
| chrX_9139401_9139959 | chrX | 9139402 | 9139959 | + | 0.806289 | Intergenic | Intergenic | 35118 | NM_001270700 | 75775 | Mm.50374 | NM_001270700 | ENSMUSG00000045797 | **4930402K13Rik** | - | RIKEN cDNA 4930402K13 gene | protein-coding |
| chrX_9139025_9140019 | chrX | 9139026 | 9140019 | + | 0.685852 | Intergenic | Lx4A\|LINE\|L1 | 34960 | NM_001270700 | 75775 | Mm.50374 | NM_001270700 | ENSMUSG00000045797 | **4930402K13Rik** | - | RIKEN cDNA 4930402K13 gene | protein-coding |
| chr16_52879494_52880004 | chr16 | 52879495 | 52880004 | + | 0.671887 | Intergenic | Intergenic | -64005 | NR_040495 | 73820 | Mm.60493 | NR_040495 | | **4930404A05Rik** | - | RIKEN cDNA 4930404A05 gene | ncRNA |
| chr2_156682876_156683391 | chr2 | 156682877 | 156683391 | + | 0.832401 | intron (NM_146128, intron 2 of 12) | intron (NM_146128, intron 2 of 12) | 37775 | NR_040505 | 74846 | Mm.443946 | NR_040505 | ENSMUSG00000086638 | **4930405A21Rik** | - | RIKEN cDNA 4930405A21 gene | ncRNA |
| chr3_100443615_100444482 | chr3 | 100443616 | 100444482 | + | 1.7725 | Intergenic | Intergenic | 17644 | NR_040543 | 73806 | Mm.396549 | NR_040543 | ENSMUSG00000086923 | **4930406D18Rik** | - | RIKEN cDNA 4930406D18 gene | ncRNA |
| chr3_100442517_100443268 | chr3 | 100442518 | 100443268 | + | 0.926103 | Intergenic | Intergenic | 16488 | NR_040543 | 73806 | Mm.396549 | NR_040543 | ENSMUSG00000086923 | **4930406D18Rik** | - | RIKEN cDNA 4930406D18 gene | ncRNA |
| chr3_100436596_100437293 | chr3 | 100436597 | 100437293 | + | 0.895305 | intron (NR_040543, intron 2 of 2) | intron (NR_040543, intron 2 of 2) | 10540 | NR_040543 | 73806 | Mm.396549 | NR_040543 | ENSMUSG00000086923 | **4930406D18Rik** | - | RIKEN cDNA 4930406D18 gene | ncRNA |
| chr3_100438476_100438733 | chr3 | 100438477 | 100438733 | + | 0.650189 | TTS (NR_040543) | TTS (NR_040543) | 12200 | NR_040543 | 73806 | Mm.396549 | NR_040543 | ENSMUSG00000086923 | **4930406D18Rik** | - | RIKEN cDNA 4930406D18 gene | ncRNA |
| chr7_122844858_122845219 | chr7 | 122844859 | 122845219 | + | 0.80852 | Intergenic | Intergenic | 125420 | NR_045614 | 73951 | Mm.389936 | NR_045614 | | **4930413G21Rik** | - | RIKEN cDNA 4930413G21 gene | ncRNA |
| chr7_122924158_122924608 | chr7 | 122924159 | 122924608 | + | 0.704041 | Intergenic | MLT1B\|LTR\|ERVL-MaLR | 46076 | NR_045614 | 73951 | Mm.389936 | NR_045614 | | **4930413G21Rik** | - | RIKEN cDNA 4930413G21 gene | ncRNA |
| chr7_122924003_122924630 | chr7 | 122924004 | 122924630 | + | 0.663207 | Intergenic | MLT1B\|LTR\|ERVL-MaLR | 46142 | NR_045614 | 73951 | Mm.389936 | NR_045614 | | **4930413G21Rik** | - | RIKEN cDNA 4930413G21 gene | ncRNA |
| chr11_11545898_11546240 | chr11 | 11545899 | 11546240 | + | 1.04369 | Intergenic | Intergenic | 56803 | NM_028669 | 73862 | Mm.159077 | NM_028669 | ENSMUSG00000020191 | **4930415F15Rik** | - | RIKEN cDNA 4930415F15 gene | protein-coding |
| chr11_11545765_11546248 | chr11 | 11545766 | 11546248 | + | 1.03306 | Intergenic | Intergenic | 56741 | NM_028669 | 73862 | Mm.159077 | NM_028669 | ENSMUSG00000020191 | **4930415F15Rik** | - | RIKEN cDNA 4930415F15 gene | protein-coding |
| chr11_11566416_11566769 | chr11 | 11566417 | 11566769 | + | 0.916604 | Intergenic | MTC\|LTR\|ERVL-MaLR | 77327 | NM_028669 | 73862 | Mm.159077 | NM_028669 | ENSMUSG00000020191 | **4930415F15Rik** | - | RIKEN cDNA 4930415F15 gene | protein-coding |
| chr11_11519342_11519713 | chr11 | 11519343 | 11519713 | + | 0.828127 | Intergenic | Intergenic | 30262 | NM_028669 | 73862 | Mm.159077 | NM_028669 | ENSMUSG00000020191 | **4930415F15Rik** | - | RIKEN cDNA 4930415F15 gene | protein-coding |
| chr11_11519303_11519923 | chr11 | 11519304 | 11519923 | + | 0.777479 | Intergenic | Intergenic | 30347 | NM_028669 | 73862 | Mm.159077 | NM_028669 | ENSMUSG00000020191 | **4930415F15Rik** | - | RIKEN cDNA 4930415F15 gene | protein-coding |
| chr11_11587311_11587819 | chr11 | 11587312 | 11587819 | + | 0.710859 | Intergenic | Intergenic | 98299 | NM_028669 | 73862 | Mm.159077 | NM_028669 | ENSMUSG00000020191 | **4930415F15Rik** | - | RIKEN cDNA 4930415F15 gene | protein-coding |
| chr11_11546861_11547262 | chr11 | 11546862 | 11547262 | + | 0.707521 | Intergenic | Intergenic | 57796 | NM_028669 | 73862 | Mm.159077 | NM_028669 | ENSMUSG00000020191 | **4930415F15Rik** | - | RIKEN cDNA 4930415F15 gene | protein-coding |
| chr11_11517850_11518080 | chr11 | 11517851 | 11518080 | + | 0.6527 | Intergenic | Lx9\|LINE\|L1 | 28699 | NM_028669 | 73862 | Mm.159077 | NM_028669 | ENSMUSG00000020191 | **4930415F15Rik** | - | RIKEN cDNA 4930415F15 gene | protein-coding |
| chr6_125256281_125256563 | chr6 | 125256282 | 125256563 | + | 0.772195 | Intergenic | Intergenic | -9103 | NR_015527 | 73870 | Mm.440681 | NR_015527 | | **4930417O13Rik** | - | RIKEN cDNA 4930417O13 gene | ncRNA |
| chr5_107677201_107677839 | chr5 | 107677202 | 107677839 | + | 1.48705 | Intergenic | Intergenic | 25960 | NR_045871 | 654809 | Mm.399159 | NR_045871 | | **4930428O21Rik** | - | RIKEN cDNA 4930428O21 gene | ncRNA |
| chr5_107678399_107678831 | chr5 | 107678400 | 107678831 | + | 1.40201 | Intergenic | Intergenic | 24865 | NR_045871 | 654809 | Mm.399159 | NR_045871 | | **4930428O21Rik** | - | RIKEN cDNA 4930428O21 gene | ncRNA |
| chr5_107677191_107678236 | chr5 | 107677192 | 107678236 | + | 1.40025 | Intergenic | B1F\|SINE\|Alu | 25766 | NR_045871 | 654809 | Mm.399159 | NR_045871 | | **4930428O21Rik** | - | RIKEN cDNA 4930428O21 gene | ncRNA |

| | | | | | | | | | | | | | | | | |
|---|---|---|---|---|---|---|---|---|---|---|---|---|---|---|---|---|
| chr5_107694272_107694702 | chr5 | 107694273 | 107694702 | + | 0.794611 | Intergenic | Intergenic | 8993 | NR_045871 | 654809 | Mm.399159 | NR_045871 | | **4930428O21Rik** | - | RIKEN cDNA 4930428O21 gene | ncRNA |
| chr17_79455182_79455641 | chr17 | 79455183 | 79455641 | + | 0.646396 | Intergenic | Intergenic | 22473 | NR_033463 | 74622 | Mm.311921 | NR_033463 | | **4930429F11Rik** | - | RIKEN cDNA 4930429F11 gene | ncRNA |
| chr9_79802230_79802675 | chr9 | 79802231 | 79802675 | + | 1.06993 | non-coding (NR_040734, exon 3 of 3) | non-coding (NR_040734, exon 3 of 3) | 8812 | NR_040734 | 74633 | Mm.262175 | NR_040734 | ENSMUSG00000086918 | **4930429F24Rik** | - | RIKEN cDNA 4930429F24 gene | ncRNA |
| chr9_79806370_79806701 | chr9 | 79806371 | 79806701 | + | 0.663449 | Intergenic | Intergenic | 12895 | NR_040734 | 74633 | Mm.262175 | NR_040734 | ENSMUSG00000086918 | **4930429F24Rik** | - | RIKEN cDNA 4930429F24 gene | ncRNA |
| chr10_100542755_100543387 | chr10 | 100542756 | 100543387 | + | 0.642384 | intron (NM_146009, intron 35 of 52) | intron (NM_146009, intron 35 of 52) | 46188 | NM_175128 | 68281 | Mm.268474 | NM_175128 | ENSMUSG00000046567 | **4930430F08Rik** | 4930571E09Rik\|A730088G13Rik\|AW551239\|C430008C19Rik | RIKEN cDNA 4930430F08 gene | protein-coding |
| chr15_31125849_31126373 | chr15 | 31125850 | 31126373 | + | 0.864524 | Intergenic | Intergenic | 86985 | NR_045186 | 73784 | Mm.66947 | NR_045186 | | **4930430F21Rik** | - | RIKEN cDNA 4930430F21 gene | ncRNA |
| chr3_121686605_121687167 | chr3 | 121686606 | 121687167 | + | 0.930466 | Intergenic | Intergenic | 16123 | NM_001033814 | 619318 | Mm.371258 | NM_001033814 | | **4930432M17Rik** | - | RIKEN cDNA 4930432M17 gene | protein-coding |
| chr1_94827581_94828109 | chr1 | 94827582 | 94828109 | + | 1.20351 | Intergenic | RLTR14\|LTR\|ERV1 | 30403 | NR_040473 | 74689 | Mm.399187 | NR_040473 | | **4930440C22Rik** | - | RIKEN cDNA 4930440C22 gene | ncRNA |
| chr1_94456502_94456838 | chr1 | 94456503 | 94456838 | + | 1.14666 | Intergenic | ORR1B1\|LTR\|ERVL-MaLR | -340772 | NR_040473 | 74689 | Mm.399187 | NR_040473 | | **4930440C22Rik** | - | RIKEN cDNA 4930440C22 gene | ncRNA |
| chr1_94464736_94465238 | chr1 | 94464737 | 94465238 | + | 0.794923 | Intergenic | Intergenic | -332455 | NR_040473 | 74689 | Mm.399187 | NR_040473 | | **4930440C22Rik** | - | RIKEN cDNA 4930440C22 gene | ncRNA |
| chr2_78049218_78049762 | chr2 | 78049219 | 78049762 | + | 0.829419 | Intergenic | Intergenic | -1687 | NR_108098 | 78158 | Mm.78473 | NR_108098 | ENSMUSG00000085431 | **4930440I19Rik** | - | RIKEN cDNA 4930440I19 gene | ncRNA |
| chr2_78049145_78049882 | chr2 | 78049146 | 78049882 | + | 0.704864 | Intergenic | Intergenic | -1663 | NR_108098 | 78158 | Mm.78473 | NR_108098 | ENSMUSG00000085431 | **4930440I19Rik** | - | RIKEN cDNA 4930440I19 gene | ncRNA |
| chr16_64739451_64739872 | chr16 | 64739452 | 64739872 | + | 0.717963 | Intergenic | Intergenic | 31277 | NM_026273 | 67609 | Mm.212157 | NM_026273 | ENSMUSG00000059920 | **4930453N24Rik** | AI649104 | RIKEN cDNA 4930453N24 gene | protein-coding |
| chr16_64739480_64739763 | chr16 | 64739481 | 64739763 | + | 0.677158 | Intergenic | MER5A\|DNA\|hAT-Charlie | 31317 | NM_026273 | 67609 | Mm.212157 | NM_026273 | ENSMUSG00000059920 | **4930453N24Rik** | AI649104 | RIKEN cDNA 4930453N24 gene | protein-coding |
| chr10_21319446_21319960 | chr10 | 21319447 | 21319960 | + | 1.16301 | intron (NR_045352, intron 3 of 3) | B1F2\|SINE\|Alu | 22352 | NR_045352 | 73990 | Mm.399240 | NR_045352 | | **4930455C13Rik** | - | RIKEN cDNA 4930455C13 gene | ncRNA |
| chr10_21358028_21358538 | chr10 | 21358029 | 21358538 | + | 1.07204 | intron (NM_019702, intron 13 of 17) | intron (NM_019702, intron 13 of 17) | -16228 | NR_045352 | 73990 | Mm.399240 | NR_045352 | | **4930455C13Rik** | - | RIKEN cDNA 4930455C13 gene | ncRNA |
| chr10_21336667_21336936 | chr10 | 21336668 | 21336936 | + | 0.849082 | intron (NR_045352, intron 1 of 3) | intron (NR_045352, intron 1 of 3) | 5253 | NR_045352 | 73990 | Mm.399240 | NR_045352 | | **4930455C13Rik** | - | RIKEN cDNA 4930455C13 gene | ncRNA |
| chr10_96237649_96238074 | chr10 | 96237650 | 96238074 | + | 1.60975 | intron (NR_110448, intron 1 of 2) | intron (NR_110448, intron 1 of 2) | 11655 | NR_110448 | 74881 | Mm.410819 | NR_110448 | | **4930459C07Rik** | - | RIKEN cDNA 4930459C07 gene | ncRNA |
| chr10_96231132_96231791 | chr10 | 96231133 | 96231791 | + | 1.40758 | intron (NR_110448, intron 1 of 2) | intron (NR_110448, intron 1 of 2) | 5255 | NR_110448 | 74881 | Mm.410819 | NR_110448 | | **4930459C07Rik** | - | RIKEN cDNA 4930459C07 gene | ncRNA |
| chr10_96337804_96338983 | chr10 | 96337805 | 96338983 | + | 1.28477 | Intergenic | Intergenic | 112187 | NR_110448 | 74881 | Mm.410819 | NR_110448 | | **4930459C07Rik** | - | RIKEN cDNA 4930459C07 gene | ncRNA |
| chr10_96342388_96342838 | chr10 | 96342389 | 96342838 | + | 1.23008 | Intergenic | Intergenic | 116406 | NR_110448 | 74881 | Mm.410819 | NR_110448 | | **4930459C07Rik** | - | RIKEN cDNA 4930459C07 gene | ncRNA |
| chr10_96238822_96239280 | chr10 | 96238823 | 96239280 | + | 1.04471 | intron (NR_110448, intron 1 of 2) | intron (NR_110448, intron 1 of 2) | 12844 | NR_110448 | 74881 | Mm.410819 | NR_110448 | | **4930459C07Rik** | - | RIKEN cDNA 4930459C07 gene | ncRNA |
| chr10_96104240_96105465 | chr10 | 96104241 | 96105465 | + | 0.903455 | Intergenic | Intergenic | -121354 | NR_110448 | 74881 | Mm.410819 | NR_110448 | | **4930459C07Rik** | - | RIKEN cDNA 4930459C07 gene | ncRNA |
| chr10_96104242_96105122 | chr10 | 96104243 | 96105122 | + | 0.878612 | Intergenic | MTD\|LTR\|ERVL-MaLR | -121525 | NR_110448 | 74881 | Mm.410819 | NR_110448 | | **4930459C07Rik** | - | RIKEN cDNA 4930459C07 gene | ncRNA |
| chr10_96262923_96263249 | chr10 | 96262924 | 96263249 | + | 0.800013 | Intergenic | Intergenic | 36879 | NR_110448 | 74881 | Mm.410819 | NR_110448 | | **4930459C07Rik** | - | RIKEN cDNA 4930459C07 gene | ncRNA |
| chr10_96237095_96237341 | chr10 | 96237096 | 96237341 | + | 0.720536 | intron (NR_110448, intron 1 of 2) | intron (NR_110448, intron 1 of 2) | 11011 | NR_110448 | 74881 | Mm.410819 | NR_110448 | | **4930459C07Rik** | - | RIKEN cDNA 4930459C07 gene | ncRNA |
| chr10_96236125_96236326 | chr10 | 96236126 | 96236326 | + | 0.706981 | intron (NR_110448, intron 1 of 2) | RMER17C\|LTR\|ERVK | 10019 | NR_110448 | 74881 | Mm.410819 | NR_110448 | | **4930459C07Rik** | - | RIKEN cDNA 4930459C07 gene | ncRNA |
| chr10_96263659_96264628 | chr10 | 96263660 | 96264628 | + | 0.701169 | Intergenic | (TCTCTG)n\|Simple_repeat\|Simple_repeat | 37937 | NR_110448 | 74881 | Mm.410819 | NR_110448 | | **4930459C07Rik** | - | RIKEN cDNA 4930459C07 gene | ncRNA |
| chr10_96263782_96264320 | chr10 | 96263783 | 96264320 | + | 0.628285 | Intergenic | Intergenic | 37844 | NR_110448 | 74881 | Mm.410819 | NR_110448 | | **4930459C07Rik** | - | RIKEN cDNA 4930459C07 gene | ncRNA |
| chr5_58599876_58600249 | chr5 | 58599877 | 58600249 | + | 0.904419 | Intergenic | Lx7\|LINE\|L1 | -147065 | NR_046190 | 78163 | Mm.243828 | NR_046190 | | **4930459L07Rik** | - | RIKEN cDNA 4930459L07 gene | ncRNA |
| chr12_107641573_107642083 | chr12 | 107641574 | 107642083 | + | 1.27065 | Intergenic | Intergenic | 93660 | NR_045973 | 74968 | Mm.359821 | NR_045973 | ENSMUSG00000085641 | **4930465M20Rik** | 4930431H15Rik | RIKEN cDNA 4930465M20 gene | ncRNA |
| chr12_107540564_107541025 | chr12 | 107540565 | 107541025 | + | 0.808503 | Intergenic | ORR1F\|LTR\|ERVL-MaLR | 194693 | NR_045973 | 74968 | Mm.359821 | NR_045973 | ENSMUSG00000085641 | **4930465M20Rik** | 4930431H15Rik | RIKEN cDNA 4930465M20 gene | ncRNA |
| chr12_107640471_107640988 | chr12 | 107640472 | 107640988 | + | 0.757122 | Intergenic | MTD\|LTR\|ERVL-MaLR | 94758 | NR_045973 | 74968 | Mm.359821 | NR_045973 | ENSMUSG00000085641 | **4930465M20Rik** | 4930431H15Rik | RIKEN cDNA 4930465M20 gene | ncRNA |
| chr14_36317270_36317643 | chr14 | 36317271 | 36317643 | + | 0.941938 | Intergenic | Intergenic | 222492 | NM_175008 | 218921 | Mm.212464 | NM_175008 | ENSMUSG00000096405 | **4930474N05Rik** | - | RIKEN cDNA 4930474N05 gene | protein-coding |
| chr12_99183309_99183799 | chr12 | 99183310 | 99183799 | + | 0.671244 | Intergenic | Intergenic | -20746 | NR_038130 | 67639 | Mm.87374 | NR_038130 | | **4930474N09Rik** | - | RIKEN cDNA 4930474N09 gene | ncRNA |
| chr16_78502479_78503434 | chr16 | 78502480 | 78503434 | + | 1.00101 | Intergenic | Intergenic | -57359 | NR_040569 | 75818 | Mm.410817 | NR_040569 | | **4930478L05Rik** | - | RIKEN cDNA 4930478L05 gene | ncRNA |
| chr16_78518658_78519125 | chr16 | 78518659 | 78519125 | + | 0.820772 | Intergenic | Intergenic | -41424 | NR_040569 | 75818 | Mm.410817 | NR_040569 | | **4930478L05Rik** | - | RIKEN cDNA 4930478L05 gene | ncRNA |
| chr16_78547416_78547649 | chr16 | 78547417 | 78547649 | + | 0.72158 | intron (NM_001252438, intron 3 of 4) | intron (NM_001252438, intron 3 of 4) | -12783 | NR_040569 | 75818 | Mm.410817 | NR_040569 | | **4930478L05Rik** | - | RIKEN cDNA 4930478L05 gene | ncRNA |
| chr16_78547433_78547737 | chr16 | 78547434 | 78547737 | + | 0.700009 | exon (NM_025967, exon 3 of 5) | exon (NM_025967, exon 3 of 5) | -12731 | NR_040569 | 75818 | Mm.410817 | NR_040569 | | **4930478L05Rik** | - | RIKEN cDNA 4930478L05 gene | ncRNA |
| chr16_78547374_78547817 | chr16 | 78547375 | 78547817 | + | 0.692171 | exon (NM_025967, exon 3 of 5) | exon (NM_025967, exon 3 of 5) | -12720 | NR_040569 | 75818 | Mm.410817 | NR_040569 | | **4930478L05Rik** | - | RIKEN cDNA 4930478L05 gene | ncRNA |
| chr6_146674399_146675614 | chr6 | 146674400 | 146675614 | + | 1.3587 | Intergenic | Intergenic | 17789 | NR_046277 | 74925 | Mm.386304 | NR_046277 | ENSMUSG00000085238 | **4930479D17Rik** | - | RIKEN cDNA 4930479D17 gene | ncRNA |
| chr17_90797588_90798132 | chr17 | 90797589 | 90798132 | + | 0.84165 | intron (NM_020252, intron 4 of 22) | intron (NM_020252, intron 4 of 22) | -63309 | NR_045463 | 78800 | Mm.410772 | NR_045463 | ENSMUSG00000090160 | **4930480K15Rik** | - | RIKEN cDNA 4930480K15 gene | ncRNA |
| chr12_26059070_26059533 | chr12 | 26059071 | 26059533 | + | 1.69501 | Intergenic | Intergenic | -151965 | NR_046278 | 74930 | Mm.398932 | NR_046278 | | **4930480M12Rik** | - | RIKEN cDNA 4930480M12 gene | ncRNA |
| chr12_25989474_25990105 | chr12 | 25989475 | 25990105 | + | 1.58158 | Intergenic | Intergenic | -221477 | NR_046278 | 74930 | Mm.398932 | NR_046278 | | **4930480M12Rik** | - | RIKEN cDNA 4930480M12 gene | ncRNA |
| chr12_26062369_26062834 | chr12 | 26062370 | 26062834 | + | 1.07448 | Intergenic | Intergenic | -148665 | NR_046278 | 74930 | Mm.398932 | NR_046278 | | **4930480M12Rik** | - | RIKEN cDNA 4930480M12 gene | ncRNA |
| chr12_26087678_26088109 | chr12 | 26087679 | 26088109 | + | 0.953484 | Intergenic | Intergenic | -123373 | NR_046278 | 74930 | Mm.398932 | NR_046278 | | **4930480M12Rik** | - | RIKEN cDNA 4930480M12 gene | ncRNA |
| chr12_26206748_26207185 | chr12 | 26206749 | 26207185 | + | 0.773701 | Intergenic | L2\|LINE\|L2 | -4300 | NR_046278 | 74930 | Mm.398932 | NR_046278 | | **4930480M12Rik** | - | RIKEN cDNA 4930480M12 gene | ncRNA |
| chr12_26087497_26088081 | chr12 | 26087498 | 26088081 | + | 0.660504 | Intergenic | Intergenic | -123478 | NR_046278 | 74930 | Mm.398932 | NR_046278 | | **4930480M12Rik** | - | RIKEN cDNA 4930480M12 gene | ncRNA |
| chr3_153189995_153190592 | chr3 | 153189996 | 153190592 | + | 0.775072 | Intergenic | ORR1D2\|LTR\|ERVL-MaLR | -38186 | NR_108044 | 75001 | | NR_108044 | | **4930482G09Rik** | - | RIKEN cDNA 4930482G09 gene | ncRNA |
| chr3_153190048_153190420 | chr3 | 153190049 | 153190420 | + | 0.747909 | Intergenic | ORR1D2\|LTR\|ERVL-MaLR | -38126 | NR_108044 | 75001 | | NR_108044 | | **4930482G09Rik** | - | RIKEN cDNA 4930482G09 gene | ncRNA |
| chr2_104561386_104561875 | chr2 | 104561387 | 104561875 | + | 1.67769 | intron (NR_040322, intron 2 of 3) | MTEa\|LTR\|ERVL-MaLR | 2172 | NR_040323 | 74959 | Mm.417819 | NR_040322 | ENSMUSG00000086454 | **4930500J02Rik** | - | RIKEN cDNA 4930500J02 gene | ncRNA |
| chr2_104562112_104562471 | chr2 | 104562113 | 104562471 | + | 1.34358 | intron (NR_040322, intron 2 of 3) | intron (NR_040322, intron 2 of 3) | 2833 | NR_040323 | 74959 | Mm.417819 | NR_040322 | ENSMUSG00000086454 | **4930500J02Rik** | - | RIKEN cDNA 4930500J02 gene | ncRNA |
| chr2_104560164_104560558 | chr2 | 104560165 | 104560558 | + | 1.0676 | intron (NR_040322, intron 2 of 3) | intron (NR_040322, intron 2 of 3) | 902 | NR_040323 | 74959 | Mm.417819 | NR_040322 | ENSMUSG00000086454 | **4930500J02Rik** | - | RIKEN cDNA 4930500J02 gene | ncRNA |
| chr5_139506065_139506588 | chr5 | 139506066 | 139506588 | + | 0.980047 | Intergenic | ZP3AR\|Satellite\|Satellite | -17424 | NR_040701 | 74973 | Mm.141282 | NR_040701 | | **4930500L23Rik** | - | RIKEN cDNA 4930500L23 gene | ncRNA |
| chr3_146705133_146705830 | chr3 | 146705134 | 146705830 | + | 0.930382 | Intergenic | MER67B\|LTR\|ERV1 | -54165 | NM_029144 | 75015 | Mm.250432 | NM_029144 | | **4930503B20Rik** | 1700013A07Rik\|1700095G21Rik\|4930483N21Rik | RIKEN cDNA 4930503B20 gene | protein-coding |
| chr10_124138446_124138861 | chr10 | 124138447 | 124138861 | + | 1.16167 | Intergenic | Intergenic | -330420 | NR_028310 | 75030 | Mm.159403 | NR_028310 | | **4930503E24Rik** | - | eukaryotic translation initiation factor 3, subunit I pseudogene | pseudo |
| chr9_72520415_72521227 | chr9 | 72520416 | 72521227 | + | 1.36528 | intron (NR_045735, intron 2 of 2) | intron (NR_045735, intron 2 of 2) | 11037 | NR_045735 | 75115 | Mm.280755 | NR_045735 | ENSMUSG00000096986 | **4930509E16Rik** | - | RIKEN cDNA 4930509E16 gene | ncRNA |
| chr9_72520698_72521211 | chr9 | 72520699 | 72521211 | + | 1.32976 | intron (NR_045735, intron 2 of 2) | intron (NR_045735, intron 2 of 2) | 10903 | NR_045735 | 75115 | Mm.280755 | NR_045735 | ENSMUSG00000096986 | **4930509E16Rik** | - | RIKEN cDNA 4930509E16 gene | ncRNA |
| chr6_94929009_94929409 | chr6 | 94929010 | 94929409 | + | 1.03195 | Intergenic | Intergenic | -14625 | NR_040761 | 78773 | Mm.67065 | NR_040761 | | **4930511E03Rik** | - | RIKEN cDNA 4930511E03 gene | ncRNA |
| chr6_94915851_94916997 | chr6 | 94915852 | 94916997 | + | 0.950633 | Intergenic | Intergenic | -27410 | NR_040761 | 78773 | Mm.67065 | NR_040761 | | **4930511E03Rik** | - | RIKEN cDNA 4930511E03 gene | ncRNA |
| chr6_94935128_94935459 | chr6 | 94935129 | 94935459 | + | 0.743842 | Intergenic | Intergenic | -8540 | NR_040761 | 78773 | Mm.67065 | NR_040761 | | **4930511E03Rik** | - | RIKEN cDNA 4930511E03 gene | ncRNA |
| chr6_94935127_94936012 | chr6 | 94935128 | 94936012 | + | 0.630224 | Intergenic | MER113\|DNA\|hAT-Charlie | -8264 | NR_040761 | 78773 | Mm.67065 | NR_040761 | | **4930511E03Rik** | - | RIKEN cDNA 4930511E03 gene | ncRNA |
| chr5_39338794_39339117 | chr5 | 39338795 | 39339117 | + | 1.26493 | Intergenic | Intergenic | 264618 | NR_045416 | 75095 | Mm.332354 | NR_045416 | | **4930513D17Rik** | - | RIKEN cDNA 4930513D17 gene | ncRNA |
| chr5_39337971_39338430 | chr5 | 39337972 | 39338430 | + | 1.16543 | Intergenic | Intergenic | 265373 | NR_045416 | 75095 | Mm.332354 | NR_045416 | | **4930513D17Rik** | - | RIKEN cDNA 4930513D17 gene | ncRNA |
| chr5_39335410_39335845 | chr5 | 39335411 | 39335845 | + | 0.902963 | Intergenic | Lx\|LINE\|L1 | 267946 | NR_045416 | 75095 | Mm.332354 | NR_045416 | | **4930513D17Rik** | - | RIKEN cDNA 4930513D17 gene | ncRNA |
| chr5_39395208_39395842 | chr5 | 39395209 | 39395842 | + | 0.883307 | Intergenic | Intergenic | 208049 | NR_045416 | 75095 | Mm.332354 | NR_045416 | | **4930513D17Rik** | - | RIKEN cDNA 4930513D17 gene | ncRNA |
| chr13_30619725_30620074 | chr13 | 30619726 | 30620074 | + | 1.56757 | intron (NR_045975, intron 2 of 3) | intron (NR_045975, intron 2 of 3) | 4857 | NR_045975 | 75090 | Mm.292232 | NR_045975 | ENSMUSG00000086263 | **4930519D14Rik** | - | RIKEN cDNA 4930519D14 gene | ncRNA |
| chr13_30613968_30614525 | chr13 | 30613969 | 30614525 | + | 1.34917 | promoter-TSS (NR_045975) | promoter-TSS (NR_045975) | -796 | NR_045975 | 75090 | Mm.292232 | NR_045975 | ENSMUSG00000086263 | **4930519D14Rik** | - | RIKEN cDNA 4930519D14 gene | ncRNA |
| chr13_30585792_30586448 | chr13 | 30585793 | 30586448 | + | 1.25006 | Intergenic | Intergenic | -28923 | NR_045975 | 75090 | Mm.292232 | NR_045975 | ENSMUSG00000086263 | **4930519D14Rik** | - | RIKEN cDNA 4930519D14 gene | ncRNA |

| | | | | | | | | | | | | | | | | | |
|---|---|---|---|---|---|---|---|---|---|---|---|---|---|---|---|---|---|
| chr13_30628317_30628871 | chr13 | 30628318 | 30628871 | + | 1.16638 | Intergenic | Intergenic | 13551 | NR_045975 | 75090 | Mm.292232 | NR_045975 | ENSMUSG00000086263 | **4930519D14Rik** | - | RIKEN cDNA 4930519D14 gene | ncRNA |
| chr13_30609784_30610144 | chr13 | 30609785 | 30610144 | + | 1.05559 | Intergenic | Intergenic | -5079 | NR_045975 | 75090 | Mm.292232 | NR_045975 | ENSMUSG00000086263 | **4930519D14Rik** | - | RIKEN cDNA 4930519D14 gene | ncRNA |
| chr13_30603868_30604314 | chr13 | 30603869 | 30604314 | + | 1.0258 | Intergenic | Intergenic | -10952 | NR_045975 | 75090 | Mm.292232 | NR_045975 | ENSMUSG00000086263 | **4930519D14Rik** | - | RIKEN cDNA 4930519D14 gene | ncRNA |
| chr13_30607269_30607738 | chr13 | 30607270 | 30607738 | + | 1.02549 | Intergenic | Intergenic | -7539 | NR_045975 | 75090 | Mm.292232 | NR_045975 | ENSMUSG00000086263 | **4930519D14Rik** | - | RIKEN cDNA 4930519D14 gene | ncRNA |
| chr13_30627008_30627391 | chr13 | 30627009 | 30627391 | + | 0.860617 | Intergenic | Intergenic | 12157 | NR_045975 | 75090 | Mm.292232 | NR_045975 | ENSMUSG00000086263 | **4930519D14Rik** | - | RIKEN cDNA 4930519D14 gene | ncRNA |
| chr13_30600723_30601359 | chr13 | 30600724 | 30601359 | + | 0.857139 | Intergenic | Intergenic | -14002 | NR_045975 | 75090 | Mm.292232 | NR_045975 | ENSMUSG00000086263 | **4930519D14Rik** | - | RIKEN cDNA 4930519D14 gene | ncRNA |
| chr13_30634025_30634269 | chr13 | 30634026 | 30634269 | + | 0.845284 | Intergenic | B1F\|SINE\|Alu | 19104 | NR_045975 | 75090 | Mm.292232 | NR_045975 | ENSMUSG00000086263 | **4930519D14Rik** | - | RIKEN cDNA 4930519D14 gene | ncRNA |
| chr13_30619316_30619654 | chr13 | 30619317 | 30619654 | + | 0.819294 | intron (NR_045975, intron 2 of 3) | intron (NR_045975, intron 2 of 3) | 4442 | NR_045975 | 75090 | Mm.292232 | NR_045975 | ENSMUSG00000086263 | **4930519D14Rik** | - | RIKEN cDNA 4930519D14 gene | ncRNA |
| chr13_30614704_30614992 | chr13 | 30614705 | 30614992 | + | 0.699774 | promoter-TSS (NR_045975) | promoter-TSS (NR_045975) | -195 | NR_045975 | 75090 | Mm.292232 | NR_045975 | ENSMUSG00000086263 | **4930519D14Rik** | - | RIKEN cDNA 4930519D14 gene | ncRNA |
| chr13_30601579_30602162 | chr13 | 30601580 | 30602162 | + | 0.689881 | Intergenic | MTD\|LTR\|ERVL-MaLR | -13172 | NR_045975 | 75090 | Mm.292232 | NR_045975 | ENSMUSG00000086263 | **4930519D14Rik** | - | RIKEN cDNA 4930519D14 gene | ncRNA |
| chr13_30601081_30601363 | chr13 | 30601082 | 30601363 | + | 0.673377 | Intergenic | Intergenic | -13821 | NR_045975 | 75090 | Mm.292232 | NR_045975 | ENSMUSG00000086263 | **4930519D14Rik** | - | RIKEN cDNA 4930519D14 gene | ncRNA |
| chr10_28807310_28808025 | chr10 | 28807311 | 28808025 | + | 1.1643 | intron (NM_178666, intron 4 of 5) | intron (NM_178666, intron 4 of 5) | 115839 | NR_033601 | 114676 | Mm.390292 | NR_033601 | | **4930519F09Rik** | - | pyruvate dehydrogenase E1 alpha 1 pseudogene | pseudo |
| chr10_28860157_28860508 | chr10 | 28860158 | 28860508 | + | 1.00807 | intron (NM_178666, intron 4 of 5) | intron (NM_178666, intron 4 of 5) | 63174 | NR_033601 | 114676 | Mm.390292 | NR_033601 | | **4930519F09Rik** | - | pyruvate dehydrogenase E1 alpha 1 pseudogene | pseudo |
| chr10_28816095_28816584 | chr10 | 28816096 | 28816584 | + | 0.853203 | intron (NM_178666, intron 4 of 5) | ORR1B1\|LTR\|ERVL-MaLR | 107167 | NR_033601 | 114676 | Mm.390292 | NR_033601 | | **4930519F09Rik** | - | pyruvate dehydrogenase E1 alpha 1 pseudogene | pseudo |
| chr10_28816253_28816521 | chr10 | 28816254 | 28816521 | + | 0.833455 | intron (NM_178666, intron 4 of 5) | ORR1B1\|LTR\|ERVL-MaLR | 107120 | NR_033601 | 114676 | Mm.390292 | NR_033601 | | **4930519F09Rik** | - | pyruvate dehydrogenase E1 alpha 1 pseudogene | pseudo |
| chr10_28815859_28816593 | chr10 | 28815860 | 28816593 | + | 0.724092 | intron (NM_178666, intron 4 of 5) | intron (NM_178666, intron 4 of 5) | 107281 | NR_033601 | 114676 | Mm.390292 | NR_033601 | | **4930519F09Rik** | - | pyruvate dehydrogenase E1 alpha 1 pseudogene | pseudo |
| chr1_159999171_159999722 | chr1 | 159999172 | 159999722 | + | 0.987532 | Intergenic | Intergenic | -44933 | NR_028111 | 67647 | Mm.171518 | NM_001024470 | ENSMUSG00000090394 | **4930523C07Rik** | - | RIKEN cDNA 4930523C07 gene | protein-coding |
| chr1_160060573_160061023 | chr1 | 160060574 | 160061023 | + | 0.796233 | intron (NR_028111, intron 1 of 2) | intron (NR_028111, intron 1 of 2) | 16418 | NM_001162896 | 67647 | Mm.171518 | NM_001024470 | ENSMUSG00000090394 | **4930523C07Rik** | - | RIKEN cDNA 4930523C07 gene | protein-coding |
| chr1_160052607_160052869 | chr1 | 160052608 | 160052869 | + | 0.792064 | intron (NR_028111, intron 1 of 2) | intron (NR_028111, intron 1 of 2) | 8358 | NM_001162896 | 67647 | Mm.171518 | NM_001024470 | ENSMUSG00000090394 | **4930523C07Rik** | - | RIKEN cDNA 4930523C07 gene | protein-coding |
| chr1_160006002_160006511 | chr1 | 160006003 | 160006511 | + | 0.695146 | Intergenic | Intergenic | -38123 | NR_028111 | 67647 | Mm.171518 | NM_001024470 | ENSMUSG00000090394 | **4930523C07Rik** | - | RIKEN cDNA 4930523C07 gene | protein-coding |
| chr16_84682007_84682533 | chr16 | 84682008 | 84682533 | + | 2.07245 | intron (NR_040537, intron 2 of 2) | intron (NR_040537, intron 2 of 2) | 2481 | NR_040537 | 78023 | Mm.243853 | NR_040537 | | **4930529L06Rik** | - | RIKEN cDNA 4930529L06 gene | ncRNA |
| chr16_84681996_84682707 | chr16 | 84681997 | 84682707 | + | 2.06971 | intron (NR_040537, intron 2 of 2) | intron (NR_040537, intron 2 of 2) | 2563 | NR_040537 | 78023 | Mm.243853 | NR_040537 | | **4930529L06Rik** | - | RIKEN cDNA 4930529L06 gene | ncRNA |
| chr16_84627050_84627548 | chr16 | 84627051 | 84627548 | + | 1.84787 | Intergenic | Intergenic | -52490 | NR_040537 | 78023 | Mm.243853 | NR_040537 | | **4930529L06Rik** | - | RIKEN cDNA 4930529L06 gene | ncRNA |
| chr16_84646062_84646724 | chr16 | 84646063 | 84646724 | + | 1.7017 | Intergenic | Intergenic | -33396 | NR_040537 | 78023 | Mm.243853 | NR_040537 | | **4930529L06Rik** | - | RIKEN cDNA 4930529L06 gene | ncRNA |
| chr16_84610371_84611140 | chr16 | 84610372 | 84611140 | + | 1.40802 | Intergenic | Intergenic | -69033 | NR_040537 | 78023 | Mm.243853 | NR_040537 | | **4930529L06Rik** | - | RIKEN cDNA 4930529L06 gene | ncRNA |
| chr16_84680364_84681149 | chr16 | 84680365 | 84681149 | + | 1.33407 | intron (NR_040537, intron 1 of 2) | intron (NR_040537, intron 1 of 2) | 968 | NR_040537 | 78023 | Mm.243853 | NR_040537 | | **4930529L06Rik** | - | RIKEN cDNA 4930529L06 gene | ncRNA |
| chr16_84619956_84620204 | chr16 | 84619957 | 84620204 | + | 1.09377 | Intergenic | 7SK\|RNA\|RNA | -59709 | NR_040537 | 78023 | Mm.243853 | NR_040537 | | **4930529L06Rik** | - | RIKEN cDNA 4930529L06 gene | ncRNA |
| chr16_84603570_84604379 | chr16 | 84603571 | 84604379 | + | 0.833522 | Intergenic | Intergenic | -75814 | NR_040537 | 78023 | Mm.243853 | NR_040537 | | **4930529L06Rik** | - | RIKEN cDNA 4930529L06 gene | ncRNA |
| chr16_84616214_84616464 | chr16 | 84616215 | 84616464 | + | 0.806902 | Intergenic | LTR89\|LTR\|ERVL? | -63450 | NR_040537 | 78023 | Mm.243853 | NR_040537 | | **4930529L06Rik** | - | RIKEN cDNA 4930529L06 gene | ncRNA |
| chr16_84696147_84696474 | chr16 | 84696148 | 84696474 | + | 0.788322 | Intergenic | RMER1B\|Other\|Other | 16522 | NR_040537 | 78023 | Mm.243853 | NR_040537 | | **4930529L06Rik** | - | RIKEN cDNA 4930529L06 gene | ncRNA |
| chr16_84622463_84622687 | chr16 | 84622464 | 84622687 | + | 0.72477 | Intergenic | Intergenic | -57214 | NR_040537 | 78023 | Mm.243853 | NR_040537 | | **4930529L06Rik** | - | RIKEN cDNA 4930529L06 gene | ncRNA |
| chr16_84696044_84696596 | chr16 | 84696045 | 84696596 | + | 0.662888 | Intergenic | RMER1B\|Other\|Other | 16531 | NR_040537 | 78023 | Mm.243853 | NR_040537 | | **4930529L06Rik** | - | RIKEN cDNA 4930529L06 gene | ncRNA |
| chr4_119223700_119224522 | chr4 | 119223701 | 119224522 | + | 1.08616 | Intergenic | B1_Mus1\|SINE\|Alu | -5894 | NM_029198 | 75180 | Mm.179335 | NM_029198 | ENSMUSG00000028642 | **4930538K18Rik** | - | RIKEN cDNA 4930538K18 gene | protein-coding |
| chr4_119223744_119223954 | chr4 | 119223745 | 119223954 | + | 0.749314 | Intergenic | Intergenic | -5632 | NM_029198 | 75180 | Mm.179335 | NM_029198 | ENSMUSG00000028642 | **4930538K18Rik** | - | RIKEN cDNA 4930538K18 gene | protein-coding |
| chr3_9151601_9151948 | chr3 | 9151602 | 9151948 | + | 0.939786 | Intergenic | Intergenic | 89811 | NR_040597 | 75149 | Mm.399324 | NR_040597 | | **4930539M17Rik** | - | RIKEN cDNA 4930539M17 gene | ncRNA |
| chr16_38016889_38017475 | chr16 | 38016890 | 38017475 | + | 0.69049 | promoter-TSS (NR_040565) | promoter-TSS (NR_040565) | -141 | NR_040565 | 75175 | Mm.399335 | NR_040565 | | **4930542C21Rik** | - | RIKEN cDNA 4930542C21 gene | ncRNA |
| chr7_113139892_113140709 | chr7 | 113139893 | 113140709 | + | 0.77368 | intron (NR_045978, intron 2 of 5) | intron (NR_045978, intron 2 of 5) | 1714 | NR_045978 | 75239 | Mm.381688 | NR_045978 | | **4930543E12Rik** | 4930545H14Rik | RIKEN cDNA 4930543E12 gene | ncRNA |
| chr10_68356136_68356476 | chr10 | 68356137 | 68356476 | + | 0.894513 | Intergenic | Intergenic | -35164 | NR_045357 | 75148 | Mm.23505 | NR_045357 | | **4930545H06Rik** | - | RIKEN cDNA 4930545H06 gene | ncRNA |
| chr9_42209575_42210210 | chr9 | 42209576 | 42210210 | + | 1.53876 | promoter-TSS (NR_040754) | promoter-TSS (NR_040754) | -662 | NR_040754 | 78194 | Mm.251457 | NR_040754 | | **4930546K05Rik** | - | RIKEN cDNA 4930546K05 gene | ncRNA |
| chr9_42214157_42214505 | chr9 | 42214158 | 42214505 | + | 1.49879 | Intergenic | MLT1A\|LTR\|ERVL-MaLR | -5100 | NR_040754 | 78194 | Mm.251457 | NR_040754 | | **4930546K05Rik** | - | RIKEN cDNA 4930546K05 gene | ncRNA |
| chr9_42208515_42209474 | chr9 | 42208516 | 42209474 | + | 0.854279 | intron (NR_040754, intron 1 of 6) | MIRb\|SINE\|MIR | 236 | NR_040754 | 78194 | Mm.251457 | NR_040754 | | **4930546K05Rik** | - | RIKEN cDNA 4930546K05 gene | ncRNA |
| chr16_59605449_59605785 | chr16 | 59605450 | 59605785 | + | 1.1652 | Intergenic | Intergenic | -31328 | NR_040564 | 75145 | Mm.399348 | NR_040564 | | **4930547E14Rik** | - | RIKEN cDNA 4930547E14 gene | ncRNA |
| chr16_59624364_59624848 | chr16 | 59624365 | 59624848 | + | 1.00982 | intron (NM_019665, intron 2 of 7) | intron (NM_019665, intron 2 of 7) | -12339 | NR_040564 | 75145 | Mm.399348 | NR_040564 | | **4930547E14Rik** | - | RIKEN cDNA 4930547E14 gene | ncRNA |
| chr9_53452125_53452778 | chr9 | 53452126 | 53452778 | + | 1.93864 | intron (NM_007499, intron 55 of 63) | intron (NM_007499, intron 55 of 63) | 47166 | NM_029247 | 75311 | Mm.26078 | NM_029247 | ENSMUSG00000005131 | **4930550C14Rik** | AU017961 | RIKEN cDNA 4930550C14 gene | protein-coding |
| chr9_53435909_53436559 | chr9 | 53435910 | 53436559 | + | 0.830792 | TTS (NM_007499) | TTS (NM_007499) | 30948 | NM_029247 | 75311 | Mm.26078 | NM_029247 | ENSMUSG00000005131 | **4930550C14Rik** | AU017961 | RIKEN cDNA 4930550C14 gene | protein-coding |
| chr9_53435482_53436497 | chr9 | 53435483 | 53436497 | + | 0.830531 | Intergenic | URR1B\|DNA\|hAT-Charlie | 30704 | NM_029247 | 75311 | Mm.26078 | NM_029247 | ENSMUSG00000005131 | **4930550C14Rik** | AU017961 | RIKEN cDNA 4930550C14 gene | protein-coding |
| chr9_53405843_53406132 | chr9 | 53405844 | 53406132 | + | 0.616255 | intron (NM_029247, intron 1 of 8) | intron (NM_029247, intron 1 of 8) | 702 | NM_029247 | 75311 | Mm.26078 | NM_029247 | ENSMUSG00000005131 | **4930550C14Rik** | AU017961 | RIKEN cDNA 4930550C14 gene | protein-coding |
| chr19_55712750_55713081 | chr19 | 55712751 | 55713081 | + | 0.877712 | Intergenic | Intergenic | -10712 | NR_045318 | 77594 | Mm.329924 | NR_045318 | ENSMUSG00000097632 | **4930552P12Rik** | - | RIKEN cDNA 4930552P12 gene | ncRNA |
| chr16_84477433_84478350 | chr16 | 84477434 | 84478350 | + | 0.920034 | Intergenic | Intergenic | 106749 | NR_040567 | 75247 | Mm.417813 | NR_040567 | | **4930553E22Rik** | - | RIKEN cDNA 4930553E22 gene | ncRNA |
| chr16_82789214_82789876 | chr16 | 82789215 | 82789876 | + | 0.907114 | Intergenic | MTD\|LTR\|ERVL-MaLR | -1581598 | NR_040567 | 75247 | Mm.417813 | NR_040567 | | **4930553E22Rik** | - | RIKEN cDNA 4930553E22 gene | ncRNA |
| chr2_57184733_57185357 | chr2 | 57184734 | 57185357 | + | 1.13461 | intron (NR_040633, intron 1 of 3) | intron (NR_040633, intron 1 of 3) | 3214 | NR_040633 | 75279 | Mm.401987 | NR_040633 | ENSMUSG00000086396 | **4930555B11Rik** | - | RIKEN cDNA 4930555B11 gene | ncRNA |
| chr3_95405623_95405987 | chr3 | 95405624 | 95405987 | + | 0.653263 | Intergenic | ORR1C2\|LTR\|ERVL-MaLR | -2964 | NR_015490 | 67654 | Mm.75066 | NR_015490 | | **4930558C23Rik** | - | RIKEN cDNA 4930558C23 gene | ncRNA |
| chr8_116799471_116800021 | chr8 | 116799472 | 116800021 | + | 1.09152 | intron (NR_046193, intron 2 of 3) | MTD\|LTR\|ERVL-MaLR | 1666 | NR_046193 | 78213 | Mm.402039 | NR_046193 | | **4930563M20Rik** | - | RIKEN cDNA 4930563M20 gene | ncRNA |
| chr3_105059880_105060376 | chr3 | 105059881 | 105060376 | + | 0.833312 | Intergenic | Intergenic | -6716 | NM_029228 | 75269 | Mm.307994 | NM_029228 | ENSMUSG00000051788 | **4930564D02Rik** | - | RIKEN cDNA 4930564D02 gene | protein-coding |
| chr3_105090995_105091524 | chr3 | 105090996 | 105091524 | + | 0.791165 | Intergenic | Intergenic | 24416 | NM_029228 | 75269 | Mm.307994 | NM_029228 | ENSMUSG00000051788 | **4930564D02Rik** | - | RIKEN cDNA 4930564D02 gene | protein-coding |
| chr3_105090997_105091276 | chr3 | 105090998 | 105091276 | + | 0.781812 | Intergenic | Intergenic | 24293 | NM_029228 | 75269 | Mm.307994 | NM_029228 | ENSMUSG00000051788 | **4930564D02Rik** | - | RIKEN cDNA 4930564D02 gene | protein-coding |
| chr3_84388258_84389288 | chr3 | 84388259 | 84389288 | + | 1.70185 | Intergenic | L1ME3\|LINE\|L1 | -8852 | NR_040752 | 75309 | Mm.159497 | NR_040752 | | **4930565D16Rik** | - | RIKEN cDNA 4930565D16 gene | ncRNA |
| chr3_84355851_84356275 | chr3 | 84355852 | 84356275 | + | 1.09913 | intron (NR_040752, intron 3 of 3) | MER20\|DNA\|hAT-Charlie | 23858 | NR_040752 | 75309 | Mm.159497 | NR_040752 | | **4930565D16Rik** | - | RIKEN cDNA 4930565D16 gene | ncRNA |
| chr3_84427760_84428472 | chr3 | 84427761 | 84428472 | + | 0.971076 | Intergenic | Intergenic | -48195 | NR_040752 | 75309 | Mm.159497 | NR_040752 | | **4930565D16Rik** | - | RIKEN cDNA 4930565D16 gene | ncRNA |
| chr2_35381727_35382472 | chr2 | 35381728 | 35382472 | + | 1.47745 | intron (NM_001080709, intron 4 of 5) | L1VL2\|LINE\|L1 | -14371 | NM_029463 | 75859 | Mm.159604 | NM_029463 | ENSMUSG00000026882 | **4930568D16Rik** | Ggta1l1 | RIKEN cDNA 4930568D16 gene | protein-coding |
| chr14_25189071_25189866 | chr14 | 25189072 | 25189866 | + | 1.69401 | Intergenic | MIRb\|SINE\|MIR | -46228 | NR_045718 | 75862 | Mm.109191 | NR_045718 | | **4930572O13Rik** | - | RIKEN cDNA 4930572O13 gene | ncRNA |
| chr2_52189005_52189435 | chr2 | 52189006 | 52189435 | + | 0.837971 | intron (NM_010889, intron 120 of 156) | intron (NM_010889, intron 120 of 156) | -101782 | NR_040620 | 78049 | Mm.243883 | NR_040620 | | **4930573O16Rik** | - | RIKEN cDNA 4930573O16 gene | ncRNA |
| chr3_51254817_51255262 | chr3 | 51254818 | 51255262 | + | 0.693973 | 3' UTR (NM_001291059, exon 10 of 10) | 3' UTR (NM_001291059, exon 10 of 10) | -21721 | NR_073429 | 67746 | Mm.441177 | NR_073429 | | **4930577N17Rik** | - | RIKEN cDNA 4930577N17 gene | ncRNA |
| chr15_98986296_98986591 | chr15 | 98986297 | 98986591 | + | 0.953251 | intron (NR_045991, intron 1 of 3) | intron (NR_045991, intron 1 of 3) | 479 | NR_045991 | 75951 | Mm.441657 | NR_045991 | ENSMUSG00000097587 | **4930578M01Rik** | 3010001K12Rik | RIKEN cDNA 4930578M01 gene | ncRNA |
| chr16_76183573_76183920 | chr16 | 76183574 | 76183920 | + | 1.22712 | Intergenic | Intergenic | 61134 | NR_040575 | 78215 | Mm.243885 | NR_040575 | | **4930578N18Rik** | - | RIKEN cDNA 4930578N18 gene | ncRNA |
| chr16_76118670_76119161 | chr16 | 76118671 | 76119161 | + | 0.815904 | Intergenic | MTD\|LTR\|ERVL-MaLR | -3697 | NR_040575 | 78215 | Mm.243885 | NR_040575 | | **4930578N18Rik** | - | RIKEN cDNA 4930578N18 gene | ncRNA |
| chr3_58696883_58697197 | chr3 | 58696884 | 58697197 | + | 0.877159 | intron (NR_045167, intron 2 of 7) | intron (NR_045167, intron 2 of 7) | 4451 | NR_045167 | 71301 | Mm.254475 | NR_045167 | ENSMUSG00000097725 | **4930593A02Rik** | 4933440I01Rik | RIKEN cDNA 4930593A02 gene | ncRNA |
| chr1_139009556_139010162 | chr1 | 139009557 | 139010162 | + | 0.716412 | intron (NM_181347, intron 1 of 20) | intron (NM_181347, intron 1 of 20) | 32202 | NR_108103 | 78229 | Mm.437838 | NR_108103 | | **4930596I21Rik** | - | RIKEN cDNA 4930596I21 gene | ncRNA |

| | | | | | | | | | | | | | | | | | |
|---|---|---|---|---|---|---|---|---|---|---|---|---|---|---|---|---|---|
| chr1_138994932_138995436 | chr1 | 138994933 | 138995436 | + | 0.67654 | intron (NM_181347, intron 1 of 20) | ORR1B1\|LTR\|ERVL-MaLR | 17527 | NR_108103 | 78229 | Mm.437838 | NR_108103 | | **4930596I21Rik** | - | RIKEN cDNA 4930596I21 gene | ncRNA |
| chr7_34321733_34322453 | chr7 | 34321734 | 34322453 | + | 1.8219 | Intergenic | Intergenic | -36484 | NM_172741 | 233103 | Mm.491097 | NM_172741 | ENSMUSG00000066571 | **4931406P16Rik** | AA553327\|mKIAA0355 | RIKEN cDNA 4931406P16 gene | protein-coding |
| chr6_73368054_73368687 | chr6 | 73368055 | 73368687 | + | 0.851069 | Intergenic | RSINE1\|SINE\|B4 | 101296 | NM_025737 | 66740 | Mm.440760 | NM_025737 | ENSMUSG00000056197 | **4931417E11Rik** | - | RIKEN cDNA 4931417E11 gene | protein-coding |
| chr9_40920047_40920567 | chr9 | 40920048 | 40920567 | + | 1.62196 | intron (NM_001081121, intron 4 of 8) | intron (NM_001081121, intron 4 of 8) | 43805 | NM_001081121 | 70989 | Mm.335087 | NM_001081121 | ENSMUSG00000032023 | **4931429I11Rik** | BB239335\|jhy | RIKEN cDNA 4931429I11 gene | protein-coding |
| chr9_40933370_40933695 | chr9 | 40933371 | 40933695 | + | 0.891174 | intron (NM_001081121, intron 3 of 8) | intron (NM_001081121, intron 3 of 8) | 30579 | NM_001081121 | 70989 | Mm.335087 | NM_001081121 | ENSMUSG00000032023 | **4931429I11Rik** | BB239335\|jhy | RIKEN cDNA 4931429I11 gene | protein-coding |
| chr5_35609432_35610237 | chr5 | 35609433 | 35610237 | + | 0.748405 | intron (NM_030721, intron 18 of 18) | intron (NM_030721, intron 18 of 18) | -21072 | NR_045807 | 74364 | Mm.473123 | NM_175208 | ENSMUSG00000055061 | **4931431C16Rik** | - | RIKEN cDNA 4931431C16 gene | ncRNA |
| chr11_29810378_29810722 | chr11 | 29810379 | 29810722 | + | 1.24794 | intron (NM_146016, intron 16 of 41) | intron (NM_146016, intron 16 of 41) | 15118 | NM_176829 | 216622 | Mm.443260 | NM_176829 | ENSMUSG00000078157 | **4931440F15Rik** | Fem1ab | RIKEN cDNA 4931440F15 gene | protein-coding |
| chr11_29799061_29800330 | chr11 | 29799062 | 29800330 | + | 1.1061 | intron (NM_146016, intron 21 of 41) | MuLV-int\|LTR\|ERV1 | 25972 | NM_176829 | 216622 | Mm.443260 | NM_176829 | ENSMUSG00000078157 | **4931440F15Rik** | Fem1ab | RIKEN cDNA 4931440F15 gene | protein-coding |
| chr11_29807379_29807702 | chr11 | 29807380 | 29807702 | + | 0.74236 | intron (NM_146016, intron 17 of 41) | intron (NM_146016, intron 17 of 41) | 18127 | NM_176829 | 216622 | Mm.443260 | NM_176829 | ENSMUSG00000078157 | **4931440F15Rik** | Fem1ab | RIKEN cDNA 4931440F15 gene | protein-coding |
| chr3_65534487_65534968 | chr3 | 65534488 | 65534968 | + | 0.660434 | intron (NM_178892, intron 2 of 5) | intron (NM_178892, intron 2 of 5) | -5314 | NR_027955 | 71004 | Mm.475465 | NM_001033527 | ENSMUSG00000074580 | **4931440P22Rik** | - | RIKEN cDNA 4931440P22 gene | ncRNA |
| chr3_36956589_36956820 | chr3 | 36956590 | 36956820 | + | 0.668363 | exon (NM_172679, exon 34 of 86) | exon (NM_172679, exon 34 of 86) | 93599 | NM_172679 | 229227 | Mm.207907 | NM_172679 | ENSMUSG00000037270 | **4932438A13Rik** | 4732443H21\|B830039D19Rik\|D630029K19Rik\|FSA\|Kiaa1109\|Tweek | RIKEN cDNA 4932438A13 gene | protein-coding |
| chr13_23298512_23299622 | chr13 | 23298513 | 23299622 | + | 1.28903 | Intergenic | tRNA-Ala-GCY\|tRNA\|tRNA | 20717 | NR_046038 | 74061 | Mm.399501 | NR_046038 | ENSMUSG00000085122 | **4933404K08Rik** | - | RIKEN cDNA 4933404K08 gene | ncRNA |
| chr13_23307060_23307900 | chr13 | 23307061 | 23307900 | + | 0.956356 | Intergenic | Intergenic | 29130 | NR_046038 | 74061 | Mm.399501 | NR_046038 | ENSMUSG00000085122 | **4933404K08Rik** | - | RIKEN cDNA 4933404K08 gene | ncRNA |
| chr13_23270315_23270832 | chr13 | 23270316 | 23270832 | + | 0.866727 | Intergenic | Intergenic | -7776 | NR_046038 | 74061 | Mm.399501 | NR_046038 | ENSMUSG00000085122 | **4933404K08Rik** | - | RIKEN cDNA 4933404K08 gene | ncRNA |
| chr13_23285122_23286373 | chr13 | 23285123 | 23286373 | + | 0.796948 | Intergenic | Intergenic | 7398 | NR_046038 | 74061 | Mm.399501 | NR_046038 | ENSMUSG00000085122 | **4933404K08Rik** | - | RIKEN cDNA 4933404K08 gene | ncRNA |
| chr11_54187827_54188316 | chr11 | 54187828 | 54188316 | + | 1.0139 | intron (NR_045506, intron 2 of 2) | L1MC2\|LINE\|L1 | 25216 | NR_045506 | 71056 | Mm.56332 | NR_045506 | ENSMUSG00000086209 | **4933405E24Rik** | - | RIKEN cDNA 4933405E24 gene | ncRNA |
| chr11_54217399_54217867 | chr11 | 54217400 | 54217867 | + | 0.760104 | Intergenic | Intergenic | -4345 | NR_045506 | 71056 | Mm.56332 | NR_045506 | ENSMUSG00000086209 | **4933405E24Rik** | - | RIKEN cDNA 4933405E24 gene | ncRNA |
| chr2_146628815_146629300 | chr2 | 146628816 | 146629300 | + | 0.725016 | Intergenic | Intergenic | 86127 | NR_040502 | 74065 | Mm.399508 | NR_040502 | ENSMUSG00000085796 | **4933406D12Rik** | - | RIKEN cDNA 4933406D12 gene | ncRNA |
| chr12_107127546_107128135 | chr12 | 107127547 | 107128135 | + | 1.37899 | Intergenic | Intergenic | -411517 | NR_015512 | 71033 | Mm.159404 | NR_015512 | ENSMUSG00000085820 | **4933406K04Rik** | 4930426M03Rik\|4930485I22Rik\|4933427G14Rik | RIKEN cDNA 4933406K04 gene | ncRNA |
| chr12_107180499_107180847 | chr12 | 107180500 | 107180847 | + | 1.29379 | Intergenic | Intergenic | -464349 | NR_015512 | 71033 | Mm.159404 | NR_015512 | ENSMUSG00000085820 | **4933406K04Rik** | 4930426M03Rik\|4930485I22Rik\|4933427G14Rik | RIKEN cDNA 4933406K04 gene | ncRNA |
| chr12_106649424_106649831 | chr12 | 106649425 | 106649831 | + | 1.10636 | Intergenic | B3\|SINE\|B2 | 66696 | NR_015512 | 71033 | Mm.159404 | NR_015512 | ENSMUSG00000085820 | **4933406K04Rik** | 4930426M03Rik\|4930485I22Rik\|4933427G14Rik | RIKEN cDNA 4933406K04 gene | ncRNA |
| chr12_106649433_106650016 | chr12 | 106649434 | 106650016 | + | 1.04867 | Intergenic | L1ME3E\|LINE\|L1 | 66599 | NR_015512 | 71033 | Mm.159404 | NR_015512 | ENSMUSG00000085820 | **4933406K04Rik** | 4930426M03Rik\|4930485I22Rik\|4933427G14Rik | RIKEN cDNA 4933406K04 gene | ncRNA |
| chr12_107134992_107135337 | chr12 | 107134993 | 107135337 | + | 0.900873 | Intergenic | Intergenic | -418841 | NR_015512 | 71033 | Mm.159404 | NR_015512 | ENSMUSG00000085820 | **4933406K04Rik** | 4930426M03Rik\|4930485I22Rik\|4933427G14Rik | RIKEN cDNA 4933406K04 gene | ncRNA |
| chr12_107132534_107132848 | chr12 | 107132535 | 107132848 | + | 0.78373 | Intergenic | MIRc\|SINE\|MIR | -416367 | NR_015512 | 71033 | Mm.159404 | NR_015512 | ENSMUSG00000085820 | **4933406K04Rik** | 4930426M03Rik\|4930485I22Rik\|4933427G14Rik | RIKEN cDNA 4933406K04 gene | ncRNA |
| chr12_107145410_107145835 | chr12 | 107145411 | 107145835 | + | 0.745551 | Intergenic | Intergenic | -429299 | NR_015512 | 71033 | Mm.159404 | NR_015512 | ENSMUSG00000085820 | **4933406K04Rik** | 4930426M03Rik\|4930485I22Rik\|4933427G14Rik | RIKEN cDNA 4933406K04 gene | ncRNA |
| chr12_106649423_106650235 | chr12 | 106649424 | 106650235 | + | 0.730232 | Intergenic | Intergenic | 66495 | NR_015512 | 71033 | Mm.159404 | NR_015512 | ENSMUSG00000085820 | **4933406K04Rik** | 4930426M03Rik\|4930485I22Rik\|4933427G14Rik | RIKEN cDNA 4933406K04 gene | ncRNA |
| chr12_106769006_106770027 | chr12 | 106769007 | 106770027 | + | 0.681817 | Intergenic | Intergenic | -53193 | NR_015512 | 71033 | Mm.159404 | NR_015512 | ENSMUSG00000085820 | **4933406K04Rik** | 4930426M03Rik\|4930485I22Rik\|4933427G14Rik | RIKEN cDNA 4933406K04 gene | ncRNA |
| chr12_107146578_107147718 | chr12 | 107146579 | 107147718 | + | 0.629317 | Intergenic | Intergenic | -430824 | NR_015512 | 71033 | Mm.159404 | NR_015512 | ENSMUSG00000085820 | **4933406K04Rik** | 4930426M03Rik\|4930485I22Rik\|4933427G14Rik | RIKEN cDNA 4933406K04 gene | ncRNA |
| chr10_75433289_75433725 | chr10 | 75433290 | 75433725 | + | 1.06785 | intron (NM_133995, intron 6 of 9) | intron (NM_133995, intron 6 of 9) | -16003 | NR_045841 | 71125 | Mm.399516 | NR_045841 | | **4933407G14Rik** | - | RIKEN cDNA 4933407G14 gene | ncRNA |
| chr10_75434404_75434931 | chr10 | 75434405 | 75434931 | + | 0.806154 | intron (NM_133995, intron 6 of 9) | intron (NM_133995, intron 6 of 9) | -14842 | NR_045841 | 71125 | Mm.399516 | NR_045841 | | **4933407G14Rik** | - | RIKEN cDNA 4933407G14 gene | ncRNA |
| chrX_75711870_75712386 | chrX | 75711871 | 75712386 | + | 0.886286 | Intergenic | Intergenic | 52571 | NR_029443 | 74396 | Mm.162493 | NR_029443 | ENSMUSG00000087396 | **4933407K13Rik** | A130080L05Rik | RIKEN cDNA 4933407K13 gene | ncRNA |
| chr1_85936801_85937980 | chr1 | 85936802 | 85937980 | + | 1.29497 | Intergenic | Intergenic | 8908 | NR_037692 | 71141 | Mm.314965 | NR_037692 | ENSMUSG00000026224 | **4933407L21Rik** | - | RIKEN cDNA 4933407L21 gene | ncRNA |
| chr1_85936727_85937242 | chr1 | 85936728 | 85937242 | + | 0.997327 | Intergenic | Intergenic | 8502 | NR_037692 | 71141 | Mm.314965 | NR_037692 | ENSMUSG00000026224 | **4933407L21Rik** | - | RIKEN cDNA 4933407L21 gene | ncRNA |
| chr1_85933249_85933534 | chr1 | 85933250 | 85933534 | + | 0.943348 | Intergenic | LTR48B\|LTR\|ERV1 | 4909 | NR_037692 | 71141 | Mm.314965 | NR_037692 | ENSMUSG00000026224 | **4933407L21Rik** | - | RIKEN cDNA 4933407L21 gene | ncRNA |
| chr8_114351728_114352446 | chr8 | 114351729 | 114352446 | + | 1.02818 | promoter-TSS (NR_045831) | promoter-TSS (NR_045831) | -609 | NR_045831 | 71122 | Mm.437539 | NR_045831 | | **4933408N05Rik** | - | RIKEN cDNA 4933408N05 gene | ncRNA |
| chr2_68530241_68530589 | chr2 | 68530242 | 68530589 | + | 0.81479 | Intergenic | MTEb\|LTR\|ERVL-MaLR | -51998 | NM_177651 | 227998 | Mm.67562 | NM_177651 | ENSMUSG00000053896 | **4933409G03Rik** | - | RIKEN cDNA 4933409G03 gene | protein-coding |
| chr10_117910381_117910932 | chr10 | 117910382 | 117910932 | + | 1.51815 | Intergenic | MER44B\|DNA\|TcMar-Tigger | -14802 | NR_045342 | 71067 | Mm.410793 | NR_045342 | | **4933411E08Rik** | - | RIKEN cDNA 4933411E08 gene | ncRNA |
| chr10_117987579_117988516 | chr10 | 117987580 | 117988516 | + | 1.37378 | Intergenic | Intergenic | 62589 | NR_045342 | 71067 | Mm.410793 | NR_045342 | | **4933411E08Rik** | - | RIKEN cDNA 4933411E08 gene | ncRNA |
| chr10_117938486_117939285 | chr10 | 117938487 | 117939285 | + | 1.13244 | intron (NR_045342, intron 3 of 3) | intron (NR_045342, intron 3 of 3) | 13427 | NR_045342 | 71067 | Mm.410793 | NR_045342 | | **4933411E08Rik** | - | RIKEN cDNA 4933411E08 gene | ncRNA |
| chr10_117985026_117985597 | chr10 | 117985027 | 117985597 | + | 1.08018 | Intergenic | Intergenic | 59853 | NR_045342 | 71067 | Mm.410793 | NR_045342 | | **4933411E08Rik** | - | RIKEN cDNA 4933411E08 gene | ncRNA |
| chr10_117984770_117985632 | chr10 | 117984771 | 117985632 | + | 1.03003 | Intergenic | Intergenic | 59742 | NR_045342 | 71067 | Mm.410793 | NR_045342 | | **4933411E08Rik** | - | RIKEN cDNA 4933411E08 gene | ncRNA |
| chr10_117909220_117909527 | chr10 | 117909221 | 117909527 | + | 1.01038 | Intergenic | Intergenic | -16085 | NR_045342 | 71067 | Mm.410793 | NR_045342 | | **4933411E08Rik** | - | RIKEN cDNA 4933411E08 gene | ncRNA |
| chr10_117916341_117916599 | chr10 | 117916342 | 117916599 | + | 0.824441 | Intergenic | Intergenic | -8989 | NR_045342 | 71067 | Mm.410793 | NR_045342 | | **4933411E08Rik** | - | RIKEN cDNA 4933411E08 gene | ncRNA |
| chr10_117909690_117910333 | chr10 | 117909691 | 117910333 | + | 0.805399 | Intergenic | Intergenic | -15447 | NR_045342 | 71067 | Mm.410793 | NR_045342 | | **4933411E08Rik** | - | RIKEN cDNA 4933411E08 gene | ncRNA |
| chr10_117940523_117941303 | chr10 | 117940524 | 117941303 | + | 0.801107 | intron (NR_045342, intron 3 of 3) | intron (NR_045342, intron 3 of 3) | 15454 | NR_045342 | 71067 | Mm.410793 | NR_045342 | | **4933411E08Rik** | - | RIKEN cDNA 4933411E08 gene | ncRNA |
| chr10_117976178_117976664 | chr10 | 117976179 | 117976664 | + | 0.791561 | Intergenic | MER4B-int\|LTR\|ERV1 | 50962 | NR_045342 | 71067 | Mm.410793 | NR_045342 | | **4933411E08Rik** | - | RIKEN cDNA 4933411E08 gene | ncRNA |
| chr10_117976345_117976664 | chr10 | 117976346 | 117976664 | + | 0.70182 | Intergenic | MER4B-int\|LTR\|ERV1 | 51046 | NR_045342 | 71067 | Mm.410793 | NR_045342 | | **4933411E08Rik** | - | RIKEN cDNA 4933411E08 gene | ncRNA |
| chr15_60110984_60111580 | chr15 | 60110985 | 60111580 | + | 0.948075 | Intergenic | Intergenic | -94664 | NM_027668 | 71088 | Mm.425950 | NM_027668 | ENSMUSG00000071749 | **4933412E24Rik** | - | RIKEN cDNA 4933412E24 gene | protein-coding |
| chr15_60101294_60101887 | chr15 | 60101295 | 60101887 | + | 0.655315 | Intergenic | B4\|SINE\|B4 | -84973 | NM_027668 | 71088 | Mm.425950 | NM_027668 | ENSMUSG00000071749 | **4933412E24Rik** | - | RIKEN cDNA 4933412E24 gene | protein-coding |
| chr15_60101571_60101824 | chr15 | 60101572 | 60101824 | + | 0.641124 | Intergenic | B4\|SINE\|B4 | -85080 | NM_027668 | 71088 | Mm.425950 | NM_027668 | ENSMUSG00000071749 | **4933412E24Rik** | - | RIKEN cDNA 4933412E24 gene | protein-coding |
| chr13_15777605_15778218 | chr13 | 15777606 | 15778218 | + | 1.16392 | Intergenic | Intergenic | 24718 | NR_045507 | 71071 | Mm.380845 | NR_045507 | | **4933412O06Rik** | - | RIKEN cDNA 4933412O06 gene | ncRNA |
| chr9_35323516_35324291 | chr9 | 35323517 | 35324291 | + | 1.11061 | Intergenic | ORR1E\|LTR\|ERVL-MaLR | 42799 | NR_040715 | 71133 | Mm.158577 | NR_040715 | | **4933422A05Rik** | - | RIKEN cDNA 4933422A05 gene | ncRNA |
| chr1_36722477_36723100 | chr1 | 36722478 | 36723100 | + | 1.07843 | intron (NR_040290, intron 2 of 8) | intron (NR_040290, intron 2 of 8) | 8251 | NR_040290 | 71166 | Mm.157049 | NR_040290 | | **4933424G06Rik** | 1700104I15Rik | RIKEN cDNA 4933424G06 gene | ncRNA |
| chr14_46671056_46671466 | chr14 | 46671057 | 46671466 | + | 0.76328 | Intergenic | Intergenic | -33838 | NR_046460 | 74458 | Mm.390708 | NR_046460 | | **4933425B07Rik** | - | RIKEN cDNA 4933425B07 gene | ncRNA |
| chr6_89068269_89068683 | chr6 | 89068270 | 89068683 | + | 0.772374 | Intergenic | Intergenic | -27639 | NM_175017 | 232217 | Mm.281744 | NM_175017 | ENSMUSG00000055403 | **4933427D06Rik** | - | RIKEN cDNA 4933427D06 gene | protein-coding |
| chr14_48804112_48804415 | chr14 | 48804113 | 48804415 | + | 0.716602 | Intergenic | Intergenic | -74609 | NR_033783 | 66770 | Mm.136954 | NR_033783 | | **4933429O19Rik** | 1700031B11Rik | RIKEN cDNA 4933429O19 gene | ncRNA |
| chr7_82976612_82977050 | chr7 | 82976613 | 82977050 | + | 0.794402 | Intergenic | URR1A\|DNA\|hAT-Charlie | -5440 | NR_045855 | 71285 | Mm.76818 | NR_045855 | | **4933430H16Rik** | - | RIKEN cDNA 4933430H16 gene | ncRNA |
| chr7_82976603_82977299 | chr7 | 82976604 | 82977299 | + | 0.78548 | Intergenic | L1M5\|LINE\|L1 | -5560 | NR_045855 | 71285 | Mm.76818 | NR_045855 | | **4933430H16Rik** | - | RIKEN cDNA 4933430H16 gene | ncRNA |
| chr7_83261539_83261860 | chr7 | 83261540 | 83261860 | + | 0.695638 | Intergenic | MLT1H\|LTR\|ERVL-MaLR | -290309 | NR_045855 | 71285 | Mm.76818 | NR_045855 | | **4933430H16Rik** | - | RIKEN cDNA 4933430H16 gene | ncRNA |
| chr6_72140235_72140844 | chr6 | 72140236 | 72140844 | + | 1.67334 | intron (NM_001035228, intron 3 of 6) | intron (NM_001035228, intron 3 of 6) | -17343 | NR_045165 | 71265 | Mm.442035 | NR_045163 | | **4933431G14Rik** | 4930565C19Rik | RIKEN cDNA 4933431G14 gene | ncRNA |
| chr6_72140227_72141195 | chr6 | 72140228 | 72141195 | + | 1.59169 | intron (NM_001035228, intron 3 of 6) | intron (NM_001035228, intron 3 of 6) | -17514 | NR_045165 | 71265 | Mm.442035 | NR_045163 | | **4933431G14Rik** | 4930565C19Rik | RIKEN cDNA 4933431G14 gene | ncRNA |
| chr6_72113293_72113663 | chr6 | 72113294 | 72113663 | + | 1.16627 | intron (NM_011375, intron 1 of 6) | MTC\|LTR\|ERVL-MaLR | 9462 | NR_045163 | 71265 | Mm.442035 | NR_045163 | | **4933431G14Rik** | 4930565C19Rik | RIKEN cDNA 4933431G14 gene | ncRNA |

| | | | | | | | | | | | | | | | | | |
|---|---|---|---|---|---|---|---|---|---|---|---|---|---|---|---|---|---|
| chr9_118381929_118382403 | chr9 | 118381930 | 118382403 | + | 1.00649 | Intergenic | L1MB5\|LINE\|L1 | -47673 | NR_040716 | 71196 | Mm.73218 | NR_040716 | | **4933432G23Rik** | - | RIKEN cDNA 4933432G23 gene | ncRNA |
| chr9_118424144_118424803 | chr9 | 118424145 | 118424803 | + | 0.952359 | Intergenic | Intergenic | -5365 | NR_040716 | 71196 | Mm.73218 | NR_040716 | | **4933432G23Rik** | - | RIKEN cDNA 4933432G23 gene | ncRNA |
| chr9_118420907_118421436 | chr9 | 118420908 | 118421436 | + | 0.771789 | Intergenic | Intergenic | -8667 | NR_040716 | 71196 | Mm.73218 | NR_040716 | | **4933432G23Rik** | - | RIKEN cDNA 4933432G23 gene | ncRNA |
| chr9_118360838_118361565 | chr9 | 118360839 | 118361565 | + | 0.715456 | Intergenic | Intergenic | -68637 | NR_040716 | 71196 | Mm.73218 | NR_040716 | | **4933432G23Rik** | - | RIKEN cDNA 4933432G23 gene | ncRNA |
| chr9_118374509_118375238 | chr9 | 118374510 | 118375238 | + | 0.681529 | Intergenic | Intergenic | -54965 | NR_040716 | 71196 | Mm.73218 | NR_040716 | | **4933432G23Rik** | - | RIKEN cDNA 4933432G23 gene | ncRNA |
| chr14_103033285_103033934 | chr14 | 103033286 | 103033934 | + | 0.973981 | intron (NR_045657, intron 1 of 2) | intron (NR_045657, intron 1 of 2) | 193 | NR_045657 | 71264 | Mm.399572 | NR_045657 | | **4933432I03Rik** | - | RIKEN cDNA 4933432I03 gene | ncRNA |
| chr14_103037028_103037398 | chr14 | 103037029 | 103037398 | + | 0.971001 | Intergenic | Intergenic | -3410 | NR_045657 | 71264 | Mm.399572 | NR_045657 | | **4933432I03Rik** | - | RIKEN cDNA 4933432I03 gene | ncRNA |
| chr16_18194974_18195482 | chr16 | 18194975 | 18195482 | + | 0.65676 | Intergenic | Intergenic | 18305 | NR_027843 | 66779 | Mm.387525 | NR_015575 | | **4933432I09Rik** | - | RIKEN cDNA 4933432I09 gene | ncRNA |
| chr13_67028319_67028646 | chr13 | 67028320 | 67028646 | + | 0.94648 | intron (NR_045851, intron 1 of 8) | L1_Rod\|LINE\|L1 | 3922 | NR_045851 | 71226 | Mm.399574 | NR_045851 | | **4933433G19Rik** | - | RIKEN cDNA 4933433G19 gene | ncRNA |
| chr17_84010571_84010831 | chr17 | 84010572 | 84010831 | + | 0.750174 | Intergenic | B3\|SINE\|B2 | -67959 | NR_045458 | 74473 | Mm.399575 | NR_045458 | | **4933433H22Rik** | - | RIKEN cDNA 4933433H22 gene | ncRNA |
| chr17_84010576_84010958 | chr17 | 84010577 | 84010958 | + | 0.726288 | Intergenic | B3\|SINE\|B2 | -67893 | NR_045458 | 74473 | Mm.399575 | NR_045458 | | **4933433H22Rik** | - | RIKEN cDNA 4933433H22 gene | ncRNA |
| chr7_68478678_68479380 | chr7 | 68478679 | 68479380 | + | 0.871057 | Intergenic | Intergenic | 61605 | NR_046208 | 71244 | Mm.410787 | NR_046208 | | **4933436H12Rik** | - | RIKEN cDNA 4933436H12 gene | ncRNA |
| chr10_111619482_111620483 | chr10 | 111619483 | 111620483 | + | 1.04168 | Intergenic | L2\|LINE\|L2 | -25710 | NR_045344 | 71321 | Mm.410785 | NR_045344 | ENSMUSG00000097082 | **4933440J02Rik** | - | RIKEN cDNA 4933440J02 gene | ncRNA |
| chr10_111571091_111571417 | chr10 | 111571092 | 111571417 | + | 0.786745 | Intergenic | ORR1F\|LTR\|ERVL-MaLR | 23019 | NR_045344 | 71321 | Mm.410785 | NR_045344 | ENSMUSG00000097082 | **4933440J02Rik** | - | RIKEN cDNA 4933440J02 gene | ncRNA |
| chr10_111575978_111576303 | chr10 | 111575979 | 111576303 | + | 0.68673 | Intergenic | Intergenic | 18132 | NR_045344 | 71321 | Mm.410785 | NR_045344 | ENSMUSG00000097082 | **4933440J02Rik** | - | RIKEN cDNA 4933440J02 gene | ncRNA |
| chr7_125189767_125190537 | chr7 | 125189768 | 125190537 | + | 0.988414 | Intergenic | MLTR18B_MM\|LTR\|ERVK | 159634 | NR_045804 | 71208 | Mm.159465 | NM_172408 | | **4933440M02Rik** | - | RIKEN cDNA 4933440M02 gene | ncRNA |
| chr7_125191823_125192509 | chr7 | 125191824 | 125192509 | + | 0.966061 | Intergenic | L1_Rod\|LINE\|L1 | 157620 | NR_045804 | 71208 | Mm.159465 | NM_172408 | | **4933440M02Rik** | - | RIKEN cDNA 4933440M02 gene | ncRNA |
| chr7_125190005_125190452 | chr7 | 125190006 | 125190452 | + | 0.940209 | Intergenic | MLTR18B_MM\|LTR\|ERVK | 159557 | NR_045804 | 71208 | Mm.159465 | NM_172408 | | **4933440M02Rik** | - | RIKEN cDNA 4933440M02 gene | ncRNA |
| chr7_125191354_125191670 | chr7 | 125191355 | 125191670 | + | 0.874879 | Intergenic | L1_Rod\|LINE\|L1 | 158274 | NR_045804 | 71208 | Mm.159465 | NM_172408 | | **4933440M02Rik** | - | RIKEN cDNA 4933440M02 gene | ncRNA |
| chr14_74996294_74997075 | chr14 | 74996295 | 74997075 | + | 0.955744 | Intergenic | Intergenic | -19342 | NM_198642 | 271221 | Mm.320280 | NM_198642 | ENSMUSG00000034959 | **5031414D18Rik** | Gm744\|Gm911\|Kiaa0226l | RIKEN cDNA 5031414D18 gene | protein-coding |
| chr14_75045001_75045793 | chr14 | 75045002 | 75045793 | + | 0.947841 | intron (NM_198642, intron 9 of 13) | intron (NM_198642, intron 9 of 13) | 29370 | NM_198642 | 271221 | Mm.320280 | NM_198642 | ENSMUSG00000034959 | **5031414D18Rik** | Gm744\|Gm911\|Kiaa0226l | RIKEN cDNA 5031414D18 gene | protein-coding |
| chr14_74996438_74997451 | chr14 | 74996439 | 74997451 | + | 0.721324 | Intergenic | Intergenic | -19082 | NM_198642 | 271221 | Mm.320280 | NM_198642 | ENSMUSG00000034959 | **5031414D18Rik** | Gm744\|Gm911\|Kiaa0226l | RIKEN cDNA 5031414D18 gene | protein-coding |
| chr14_75005743_75006167 | chr14 | 75005744 | 75006167 | + | 0.71206 | intron (NR_030568, intron 3 of 3) | intron (NR_030568, intron 3 of 3) | -10072 | NM_198642 | 271221 | Mm.320280 | NM_198642 | ENSMUSG00000034959 | **5031414D18Rik** | Gm744\|Gm911\|Kiaa0226l | RIKEN cDNA 5031414D18 gene | protein-coding |
| chr5_23401977_23402383 | chr5 | 23401978 | 23402383 | + | 0.782378 | Intergenic | Intergenic | 32173 | NR_040469 | 269630 | Mm.425231 | NR_040469 | ENSMUSG00000073147 | **5031425E22Rik** | - | RIKEN cDNA 5031425E22 gene | ncRNA |
| chr5_23413746_23413972 | chr5 | 23413747 | 23413972 | + | 0.778054 | Intergenic | Intergenic | 20494 | NR_040469 | 269630 | Mm.425231 | NR_040469 | ENSMUSG00000073147 | **5031425E22Rik** | - | RIKEN cDNA 5031425E22 gene | ncRNA |
| chr5_23416145_23416559 | chr5 | 23416146 | 23416559 | + | 0.687881 | Intergenic | Intergenic | 18001 | NR_040469 | 269630 | Mm.425231 | NR_040469 | ENSMUSG00000073147 | **5031425E22Rik** | - | RIKEN cDNA 5031425E22 gene | ncRNA |
| chr2_166398969_166400061 | chr2 | 166398970 | 166400061 | + | 1.14733 | Intergenic | Intergenic | -47936 | NR_015558 | 319684 | Mm.399746 | NR_015558 | | **5031425F14Rik** | - | RIKEN cDNA 5031425F14 gene | ncRNA |
| chr2_131840494_131841676 | chr2 | 131840495 | 131841676 | + | 1.07241 | intron (NR_029381, intron 1 of 2) | intron (NR_029381, intron 1 of 2) | 6571 | NR_029381 | 414081 | Mm.246905 | NR_029381 | ENSMUSG00000086822 | **5330413P13Rik** | - | RIKEN cDNA 5330413P13 gene | ncRNA |
| chr2_131840947_131841650 | chr2 | 131840948 | 131841650 | + | 1.00926 | intron (NR_029381, intron 1 of 2) | MER21C\|LTR\|ERVL | 6357 | NR_029381 | 414081 | Mm.246905 | NR_029381 | ENSMUSG00000086822 | **5330413P13Rik** | - | RIKEN cDNA 5330413P13 gene | ncRNA |
| chr2_131841136_131841658 | chr2 | 131841137 | 131841658 | + | 0.913871 | intron (NR_029381, intron 1 of 2) | MER21C\|LTR\|ERVL | 6259 | NR_029381 | 414081 | Mm.246905 | NR_029381 | ENSMUSG00000086822 | **5330413P13Rik** | - | RIKEN cDNA 5330413P13 gene | ncRNA |
| chr16_51299903_51300311 | chr16 | 51299904 | 51300311 | + | 1.37416 | Intergenic | Intergenic | -567334 | NR_028300 | 68190 | Mm.139089 | NR_028300 | | **5330426P16Rik** | - | RIKEN cDNA 5330426P16 gene | ncRNA |
| chr16_51312892_51313337 | chr16 | 51312893 | 51313337 | + | 1.17477 | Intergenic | Intergenic | -580342 | NR_028300 | 68190 | Mm.139089 | NR_028300 | | **5330426P16Rik** | - | RIKEN cDNA 5330426P16 gene | ncRNA |
| chr16_51299042_51300311 | chr16 | 51299043 | 51300311 | + | 0.826161 | Intergenic | MLT1E\|LTR\|ERVL-MaLR | -566904 | NR_028300 | 68190 | Mm.139089 | NR_028300 | | **5330426P16Rik** | - | RIKEN cDNA 5330426P16 gene | ncRNA |
| chr16_50839572_50840077 | chr16 | 50839573 | 50840077 | + | 0.813839 | Intergenic | RLTR17\|LTR\|ERVK | -107052 | NR_028300 | 68190 | Mm.139089 | NR_028300 | | **5330426P16Rik** | - | RIKEN cDNA 5330426P16 gene | ncRNA |
| chr16_51316298_51316596 | chr16 | 51316299 | 51316596 | + | 0.729556 | Intergenic | Intergenic | -583674 | NR_028300 | 68190 | Mm.139089 | NR_028300 | | **5330426P16Rik** | - | RIKEN cDNA 5330426P16 gene | ncRNA |
| chr8_35881641_35882143 | chr8 | 35881642 | 35882143 | + | 0.692709 | intron (NR_046181, intron 1 of 2) | intron (NR_046181, intron 1 of 2) | 1065 | NR_046181 | 330737 | Mm.381743 | NR_046181 | | **5430403N17Rik** | 5430403N17 | RIKEN cDNA 5430403N17 gene | ncRNA |
| chr2_156857462_156857726 | chr2 | 156857463 | 156857726 | + | 0.619854 | intron (NR_015591, intron 1 of 2) | ID_B1\|SINE\|B4 | 5351 | NR_015591 | 74487 | Mm.490465 | NR_015591 | ENSMUSG00000085741 | **5430405H02Rik** | 2610307N17Rik | RIKEN cDNA 5430405H02 gene | ncRNA |
| chr8_25270533_25270822 | chr8 | 25270534 | 25270822 | + | 0.680566 | Intergenic | Intergenic | 35592 | NR_040352 | 74489 | Mm.256468 | NR_040352 | | **5430421F17Rik** | - | RIKEN cDNA 5430421F17 gene | ncRNA |
| chrX_85873823_85874472 | chrX | 85873824 | 85874472 | + | 1.24416 | intron (NM_001163539, intron 1 of 1) | L1M2\|LINE\|L1 | 3875 | NM_001163539 | 71398 | Mm.185296 | NM_001163539 | ENSMUSG00000025058 | **5430427O19Rik** | - | RIKEN cDNA 5430427O19 gene | protein-coding |
| chr1_131703486_131703841 | chr1 | 131703487 | 131703841 | + | 1.00485 | intron (NM_145509, intron 4 of 5) | intron (NM_145509, intron 4 of 5) | 14969 | NM_145509 | 226421 | Mm.44508 | NM_145509 | ENSMUSG00000052688 | **5430435G22Rik** | Rab7b | RIKEN cDNA 5430435G22 gene | protein-coding |
| chr1_131703366_131703938 | chr1 | 131703367 | 131703938 | + | 0.83694 | intron (NM_145509, intron 4 of 5) | intron (NM_145509, intron 4 of 5) | 14957 | NM_145509 | 226421 | Mm.44508 | NM_145509 | ENSMUSG00000052688 | **5430435G22Rik** | Rab7b | RIKEN cDNA 5430435G22 gene | protein-coding |
| chr1_131698835_131699223 | chr1 | 131698836 | 131699223 | + | 0.786248 | intron (NM_145509, intron 4 of 5) | intron (NM_145509, intron 4 of 5) | 10334 | NM_145509 | 226421 | Mm.44508 | NM_145509 | ENSMUSG00000052688 | **5430435G22Rik** | Rab7b | RIKEN cDNA 5430435G22 gene | protein-coding |
| chr15_5758254_5759335 | chr15 | 5758255 | 5759335 | + | 1.75121 | Intergenic | Intergenic | 262477 | NR_045274 | 71432 | Mm.291087 | NR_045274 | | **5430437J10Rik** | - | RIKEN cDNA 5430437J10 gene | ncRNA |
| chr15_5601929_5602730 | chr15 | 5601930 | 5602730 | + | 1.39967 | Intergenic | Intergenic | 106012 | NR_045274 | 71432 | Mm.291087 | NR_045274 | | **5430437J10Rik** | - | RIKEN cDNA 5430437J10 gene | ncRNA |
| chr15_5432067_5432670 | chr15 | 5432068 | 5432670 | + | 1.2919 | Intergenic | Intergenic | -63949 | NR_045274 | 71432 | Mm.291087 | NR_045274 | | **5430437J10Rik** | - | RIKEN cDNA 5430437J10 gene | ncRNA |
| chr15_5446178_5447267 | chr15 | 5446179 | 5447267 | + | 1.16725 | Intergenic | Intergenic | -49595 | NR_045274 | 71432 | Mm.291087 | NR_045274 | | **5430437J10Rik** | - | RIKEN cDNA 5430437J10 gene | ncRNA |
| chr15_5431237_5431618 | chr15 | 5431238 | 5431618 | + | 0.858835 | Intergenic | Intergenic | -64890 | NR_045274 | 71432 | Mm.291087 | NR_045274 | | **5430437J10Rik** | - | RIKEN cDNA 5430437J10 gene | ncRNA |
| chr15_5513227_5513570 | chr15 | 5513228 | 5513570 | + | 0.785194 | intron (NR_045274, intron 1 of 4) | intron (NR_045274, intron 1 of 4) | 17081 | NR_045274 | 71432 | Mm.291087 | NR_045274 | | **5430437J10Rik** | - | RIKEN cDNA 5430437J10 gene | ncRNA |
| chr15_5512579_5512832 | chr15 | 5512580 | 5512832 | + | 0.752761 | intron (NR_045274, intron 1 of 4) | intron (NR_045274, intron 1 of 4) | 16388 | NR_045274 | 71432 | Mm.291087 | NR_045274 | | **5430437J10Rik** | - | RIKEN cDNA 5430437J10 gene | ncRNA |
| chr15_5507661_5507961 | chr15 | 5507662 | 5507961 | + | 0.669641 | intron (NR_045274, intron 1 of 4) | intron (NR_045274, intron 1 of 4) | 11493 | NR_045274 | 71432 | Mm.291087 | NR_045274 | | **5430437J10Rik** | - | RIKEN cDNA 5430437J10 gene | ncRNA |
| chr15_5464682_5465066 | chr15 | 5464683 | 5465066 | + | 0.611514 | Intergenic | Intergenic | -31444 | NR_045274 | 71432 | Mm.291087 | NR_045274 | | **5430437J10Rik** | - | RIKEN cDNA 5430437J10 gene | ncRNA |
| chr4_44624276_44625355 | chr4 | 44624277 | 44625355 | + | 1.0187 | intron (NM_008782, intron 6 of 9) | intron (NM_008782, intron 6 of 9) | -77984 | NR_073462 | 70602 | Mm.491142 | NR_073462 | | **5730488B01Rik** | - | RIKEN cDNA 5730488B01 gene | ncRNA |
| chr3_127887514_127888272 | chr3 | 127887515 | 127888272 | + | 0.979799 | intron (NM_027482, intron 1 of 1) | intron (NM_027482, intron 1 of 1) | 8430 | NM_027482 | 70617 | Mm.317049 | NM_027482 | ENSMUSG00000050549 | **5730508B09Rik** | 2700063A17Rik\|C80506 | RIKEN cDNA 5730508B09 gene | protein-coding |
| chr3_127887565_127887963 | chr3 | 127887566 | 127887963 | + | 0.94824 | intron (NM_027482, intron 1 of 1) | RSINE1\|SINE\|B4 | 8559 | NM_027482 | 70617 | Mm.317049 | NM_027482 | ENSMUSG00000050549 | **5730508B09Rik** | 2700063A17Rik\|C80506 | RIKEN cDNA 5730508B09 gene | protein-coding |
| chr3_127888013_127888259 | chr3 | 127888014 | 127888259 | + | 0.834928 | intron (NM_027482, intron 1 of 1) | intron (NM_027482, intron 1 of 1) | 8187 | NM_027482 | 70617 | Mm.317049 | NM_027482 | ENSMUSG00000050549 | **5730508B09Rik** | 2700063A17Rik\|C80506 | RIKEN cDNA 5730508B09 gene | protein-coding |
| chr3_127890986_127891316 | chr3 | 127890987 | 127891316 | + | 0.704088 | intron (NM_027482, intron 1 of 1) | intron (NM_027482, intron 1 of 1) | 5172 | NM_027482 | 70617 | Mm.317049 | NM_027482 | ENSMUSG00000050549 | **5730508B09Rik** | 2700063A17Rik\|C80506 | RIKEN cDNA 5730508B09 gene | protein-coding |
| chr3_127891979_127892255 | chr3 | 127891980 | 127892255 | + | 0.645722 | intron (NM_027482, intron 1 of 1) | intron (NM_027482, intron 1 of 1) | 4206 | NM_027482 | 70617 | Mm.317049 | NM_027482 | ENSMUSG00000050549 | **5730508B09Rik** | 2700063A17Rik\|C80506 | RIKEN cDNA 5730508B09 gene | protein-coding |
| chr19_53490940_53492434 | chr19 | 53490941 | 53492434 | + | 1.09087 | Intergenic | Intergenic | -26891 | NR_028427 | 381232 | Mm.248624 | NR_028427 | ENSMUSG00000097636 | **5830416P10Rik** | 4833407H14Rik\|5430437C04Rik\|ENSMUSG00000074788\|Gm10758 | RIKEN cDNA 5830416P10 gene | ncRNA |
| chr19_53433888_53434603 | chr19 | 53433889 | 53434603 | + | 0.786833 | Intergenic | Intergenic | 30550 | NR_028427 | 381232 | Mm.248624 | NR_028427 | ENSMUSG00000097636 | **5830416P10Rik** | 4833407H14Rik\|5430437C04Rik\|ENSMUSG00000074788\|Gm10758 | RIKEN cDNA 5830416P10 gene | ncRNA |
| chr19_53454436_53454986 | chr19 | 53454437 | 53454986 | + | 0.694005 | intron (NR_028427, intron 2 of 9) | intron (NR_028427, intron 2 of 9) | 10085 | NR_028427 | 381232 | Mm.248624 | NR_028427 | ENSMUSG00000097636 | **5830416P10Rik** | 4833407H14Rik\|5430437C04Rik\|ENSMUSG00000074788\|Gm10758 | RIKEN cDNA 5830416P10 gene | ncRNA |
| chr12_69510718_69511429 | chr12 | 69510719 | 69511429 | + | 1.17836 | Intergenic | Intergenic | -35524 | NR_038061 | 76062 | Mm.380755 | NR_038060 | | **5830428M24Rik** | 100040862\|Gm3007 | RIKEN cDNA 5830428M24 gene | ncRNA |
| chr12_69510718_69511246 | chr12 | 69510719 | 69511246 | + | 1.17813 | Intergenic | ORR1B1\|LTR\|ERVL-MaLR | -35432 | NR_038061 | 76062 | Mm.380755 | NR_038060 | | **5830428M24Rik** | 100040862\|Gm3007 | RIKEN cDNA 5830428M24 gene | ncRNA |
| chr12_69456646_69456928 | chr12 | 69456647 | 69456928 | + | 0.967589 | Intergenic | Intergenic | 18763 | NR_038060 | 76062 | Mm.380755 | NR_038060 | | **5830428M24Rik** | 100040862\|Gm3007 | RIKEN cDNA 5830428M24 gene | ncRNA |
| chr12_69456596_69457010 | chr12 | 69456597 | 69457010 | + | 0.940661 | Intergenic | Intergenic | 18747 | NR_038060 | 76062 | Mm.380755 | NR_038060 | | **5830428M24Rik** | 100040862\|Gm3007 | RIKEN cDNA 5830428M24 gene | ncRNA |
| chr12_69437146_69437421 | chr12 | 69437147 | 69437421 | + | 0.828771 | Intergenic | Intergenic | 38266 | NR_038060 | 76062 | Mm.380755 | NR_038060 | | **5830428M24Rik** | 100040862\|Gm3007 | RIKEN cDNA 5830428M24 gene | ncRNA |
| chr12_69436725_69437434 | chr12 | 69436726 | 69437434 | + | 0.679864 | Intergenic | Intergenic | 38470 | NR_038060 | 76062 | Mm.380755 | NR_038060 | | **5830428M24Rik** | 100040862\|Gm3007 | RIKEN cDNA 5830428M24 gene | ncRNA |
| chr7_135644600_135644905 | chr7 | 135644601 | 135644905 | + | 0.708378 | intron (NM_011212, intron 3 of 20) | RMER15\|LTR\|ERVL | 7561 | NR_015548 | 67765 | Mm.445272 | NR_015548 | ENSMUSG00000097471 | **5830432E09Rik** | - | RIKEN cDNA 5830432E09 gene | ncRNA |

| | | | | | | | | | | | | | | | | | |
|---|---|---|---|---|---|---|---|---|---|---|---|---|---|---|---|---|---|
| chr13_101229836_101230305 | chr13 | 101229837 | 101230305 | + | 1.00566 | Intergenic | MLT1J1\|LTR\|ERVL-MaLR | -2056 | NR_046158 | 328324 | Mm.402274 | NR_046158 | | **5930438M14Rik** | 5930438M14 | RIKEN cDNA 5930438M14 gene | ncRNA |
| chr13_101182413_101182821 | chr13 | 101182414 | 101182821 | + | 0.894788 | Intergenic | Intergenic | -49510 | NR_046158 | 328324 | Mm.402274 | NR_046158 | | **5930438M14Rik** | 5930438M14 | RIKEN cDNA 5930438M14 gene | ncRNA |
| chr1_73936671_73937008 | chr1 | 73936672 | 73937008 | + | 1.03162 | intron (NM_027884, intron 22 of 34) | intron (NM_027884, intron 22 of 34) | -72374 | NR_045311 | 109198 | Mm.359721 | NR_045311 | | **6030407O03Rik** | - | RIKEN cDNA 6030407O03 gene | ncRNA |
| chr1_73947285_73948343 | chr1 | 73947286 | 73948343 | + | 0.980797 | intron (NM_001289895, intron 19 of 32) | L2\|LINE\|L2 | -83348 | NR_045311 | 109198 | Mm.359721 | NR_045311 | | **6030407O03Rik** | - | RIKEN cDNA 6030407O03 gene | ncRNA |
| chr1_73940474_73941226 | chr1 | 73940475 | 73941226 | + | 0.916086 | intron (NM_001289895, intron 21 of 32) | intron (NM_001289895, intron 21 of 32) | -76384 | NR_045311 | 109198 | Mm.359721 | NR_045311 | | **6030407O03Rik** | - | RIKEN cDNA 6030407O03 gene | ncRNA |
| chr1_73940890_73941158 | chr1 | 73940891 | 73941158 | + | 0.911618 | exon (NM_027884, exon 21 of 35) | exon (NM_027884, exon 21 of 35) | -76558 | NR_045311 | 109198 | Mm.359721 | NR_045311 | | **6030407O03Rik** | - | RIKEN cDNA 6030407O03 gene | ncRNA |
| chr1_73936663_73937131 | chr1 | 73936664 | 73937131 | + | 0.885957 | intron (NM_027884, intron 22 of 34) | intron (NM_027884, intron 22 of 34) | -72431 | NR_045311 | 109198 | Mm.359721 | NR_045311 | | **6030407O03Rik** | - | RIKEN cDNA 6030407O03 gene | ncRNA |
| chr1_73936644_73937437 | chr1 | 73936645 | 73937437 | + | 0.618848 | intron (NM_027884, intron 22 of 34) | intron (NM_027884, intron 22 of 34) | -72575 | NR_045311 | 109198 | Mm.359721 | NR_045311 | | **6030407O03Rik** | - | RIKEN cDNA 6030407O03 gene | ncRNA |
| chr3_129656600_129657588 | chr3 | 129656601 | 129657588 | + | 1.20252 | Intergenic | RMER1B\|Other\|Other | -96664 | NR_040589 | 319224 | Mm.477035 | NR_040589 | | **6330410L21Rik** | - | RIKEN cDNA 6330410L21 gene | ncRNA |
| chr7_139971467_139971867 | chr7 | 139971468 | 139971867 | + | 1.03726 | TTS (NM_001033465) | TTS (NM_001033465) | 7088 | NM_001033465 | 381933 | Mm.389985 | NM_001033465 | ENSMUSG00000073795 | **6430531B16Rik** | Gm1093 | RIKEN cDNA 6430531B16 gene | protein-coding |
| chr13_99410407_99411467 | chr13 | 99410408 | 99411467 | + | 0.610967 | non-coding (NR_015515, exon 2 of 2) | non-coding (NR_015515, exon 2 of 2) | 1889 | NR_015515 | 320893 | Mm.470842 | NR_015515 | | **6430562O15Rik** | - | RIKEN cDNA 6430562O15 gene | ncRNA |
| chr8_36452742_36453412 | chr8 | 36452743 | 36453412 | + | 1.01304 | Intergenic | Intergenic | -36114 | NM_176952 | 319582 | Mm.180924 | NM_176952 | ENSMUSG00000039620 | **6430573F11Rik** | Kiaa1456 | RIKEN cDNA 6430573F11 gene | protein-coding |
| chr2_72868177_72868731 | chr2 | 72868178 | 72868731 | + | 1.61397 | Intergenic | LTR48\|LTR\|ERV1 | -54737 | NR_102349 | 76218 | Mm.75260 | NR_102348 | ENSMUSG00000085427 | **6430710C18Rik** | 5033406J01Rik | RIKEN cDNA 6430710C18 gene | ncRNA |
| chr2_72867796_72868736 | chr2 | 72867797 | 72868736 | + | 1.36213 | Intergenic | Intergenic | -54549 | NR_102349 | 76218 | Mm.75260 | NR_102348 | ENSMUSG00000085427 | **6430710C18Rik** | 5033406J01Rik | RIKEN cDNA 6430710C18 gene | ncRNA |
| chr2_72841255_72841843 | chr2 | 72841256 | 72841843 | + | 1.31879 | Intergenic | Intergenic | -27832 | NR_102349 | 76218 | Mm.75260 | NR_102348 | ENSMUSG00000085427 | **6430710C18Rik** | 5033406J01Rik | RIKEN cDNA 6430710C18 gene | ncRNA |
| chr2_72865436_72865852 | chr2 | 72865437 | 72865852 | + | 0.787949 | Intergenic | Intergenic | -51927 | NR_102349 | 76218 | Mm.75260 | NR_102348 | ENSMUSG00000085427 | **6430710C18Rik** | 5033406J01Rik | RIKEN cDNA 6430710C18 gene | ncRNA |
| chr2_11429569_11430049 | chr2 | 11429570 | 11430049 | + | 1.34063 | Intergenic | Intergenic | -31309 | NR_040615 | 77543 | Mm.490670 | NR_040615 | | **8030442B05Rik** | - | RIKEN cDNA 8030442B05 gene | ncRNA |
| chr18_77652513_77652904 | chr18 | 77652514 | 77652904 | + | 1.06734 | intron (NM_178670, intron 3 of 4) | intron (NM_178670, intron 3 of 4) | 61301 | NM_178670 | 212163 | Mm.23293 | NM_178670 | ENSMUSG00000047466 | **8030462N17Rik** | Akd2\|Ark2N | RIKEN cDNA 8030462N17 gene | protein-coding |
| chr18_77681714_77682004 | chr18 | 77681715 | 77682004 | + | 0.718025 | intron (NM_178670, intron 1 of 4) | intron (NM_178670, intron 1 of 4) | 32151 | NM_178670 | 212163 | Mm.23293 | NM_178670 | ENSMUSG00000047466 | **8030462N17Rik** | Akd2\|Ark2N | RIKEN cDNA 8030462N17 gene | protein-coding |
| chr11_3442587_3443651 | chr11 | 3442588 | 3443651 | + | 0.811313 | intron (NM_145355, intron 1 of 7) | MIRb\|SINE\|MIR | -9318 | NR_028317 | 71523 | Mm.72795 | NM_027820 | ENSMUSG00000034587 | **8430429K09Rik** | - | RIKEN cDNA 8430429K09 gene | ncRNA |
| chr11_3442645_3443049 | chr11 | 3442646 | 3443049 | + | 0.734149 | intron (NM_145355, intron 1 of 7) | intron (NM_145355, intron 1 of 7) | -9590 | NR_028317 | 71523 | Mm.72795 | NM_027820 | ENSMUSG00000034587 | **8430429K09Rik** | - | RIKEN cDNA 8430429K09 gene | ncRNA |
| chr19_31321787_31322198 | chr19 | 31321788 | 31322198 | + | 0.837129 | intron (NM_001013833, intron 2 of 17) | intron (NM_001013833, intron 2 of 17) | -105597 | NR_002849 | 78103 | Mm.445198 | NR_002849 | | **8430431K14Rik** | - | RIKEN cDNA 8430431K14 gene | ncRNA |
| chr4_7689479_7689918 | chr4 | 7689480 | 7689918 | + | 1.2939 | Intergenic | Intergenic | 129011 | NR_040645 | 71510 | Mm.410775 | NR_040645 | ENSMUSG00000086070 | **8430436N08Rik** | - | RIKEN cDNA 8430436N08 gene | ncRNA |
| chr4_7490890_7491176 | chr4 | 7490891 | 7491176 | + | 0.69393 | Intergenic | Intergenic | -69655 | NR_040645 | 71510 | Mm.410775 | NR_040645 | ENSMUSG00000086070 | **8430436N08Rik** | - | RIKEN cDNA 8430436N08 gene | ncRNA |
| chr2_72729184_72729703 | chr2 | 72729185 | 72729703 | + | 0.899817 | TTS (NR_040503) | TTS (NR_040503) | 25934 | NR_040503 | 74541 | Mm.410774 | NR_040503 | ENSMUSG00000085252 | **8430437L04Rik** | - | RIKEN cDNA 8430437L04 gene | ncRNA |
| chr2_35881907_35882277 | chr2 | 35881908 | 35882277 | + | 0.69119 | intron (NM_029774, intron 6 of 9) | intron (NM_029774, intron 6 of 9) | 62902 | NR_040618 | 77665 | Mm.400558 | NR_040618 | ENSMUSG00000086449 | **9030204H09Rik** | - | RIKEN cDNA 9030204H09 gene | ncRNA |
| chr16_30024500_30025297 | chr16 | 30024501 | 30025297 | + | 0.768737 | intron (NR_045878, intron 3 of 3) | intron (NR_045878, intron 3 of 3) | 14580 | NR_045878 | 71528 | Mm.297640 | NR_045878 | | **9030404E10Rik** | - | RIKEN cDNA 9030404E10 gene | ncRNA |
| chr16_30024651_30024985 | chr16 | 30024652 | 30024985 | + | 0.753935 | intron (NR_045878, intron 3 of 3) | intron (NR_045878, intron 3 of 3) | 14661 | NR_045878 | 71528 | Mm.297640 | NR_045878 | | **9030404E10Rik** | - | RIKEN cDNA 9030404E10 gene | ncRNA |
| chr16_30024647_30025139 | chr16 | 30024648 | 30025139 | + | 0.733595 | intron (NR_045878, intron 3 of 3) | intron (NR_045878, intron 3 of 3) | 14586 | NR_045878 | 71528 | Mm.297640 | NR_045878 | | **9030404E10Rik** | - | RIKEN cDNA 9030404E10 gene | ncRNA |
| chr12_35640856_35641416 | chr12 | 35640857 | 35641416 | + | 1.20116 | Intergenic | MIR3\|SINE\|MIR | 45168 | NR_045051 | 71581 | Mm.100110 | NR_045050 | | **9130015A21Rik** | - | RIKEN cDNA 9130015A21 gene | ncRNA |
| chr12_35712607_35713115 | chr12 | 35712608 | 35713115 | + | 0.862251 | Intergenic | Intergenic | -17743 | NR_045050 | 71581 | Mm.100110 | NR_045050 | | **9130015A21Rik** | - | RIKEN cDNA 9130015A21 gene | ncRNA |
| chr12_35687584_35688062 | chr12 | 35687585 | 35688062 | + | 0.786786 | intron (NR_045050, intron 3 of 4) | intron (NR_045050, intron 3 of 4) | -1519 | NR_045051 | 71581 | Mm.100110 | NR_045050 | | **9130015A21Rik** | - | RIKEN cDNA 9130015A21 gene | ncRNA |
| chr6_72056505_72057091 | chr6 | 72056506 | 72057091 | + | 1.04372 | Intergenic | Intergenic | -14270 | NR_046180 | 330350 | Mm.246787 | NR_046180 | | **9130221F21Rik** | 9130221F21 | RIKEN cDNA 9130221F21 gene | ncRNA |
| chr5_65991155_65991628 | chr5 | 65991156 | 65991628 | + | 0.826697 | intron (NR_027961, intron 4 of 6) | ORR1B1\|LTR\|ERVL-MaLR | 12893 | NR_027961 | 231253 | Mm.436542 | NM_177679 | ENSMUSG00000054598 | **9130230L23Rik** | - | RIKEN cDNA 9130230L23 gene | ncRNA |
| chr5_65986457_65986874 | chr5 | 65986458 | 65986874 | + | 0.808242 | Intergenic | Lx8\|LINE\|L1 | 17619 | NR_027961 | 231253 | Mm.436542 | NM_177679 | ENSMUSG00000054598 | **9130230L23Rik** | - | RIKEN cDNA 9130230L23 gene | ncRNA |
| chr13_81169982_81170668 | chr13 | 81169983 | 81170668 | + | 1.29831 | intron (NM_054053, intron 86 of 89) | intron (NM_054053, intron 86 of 89) | 206251 | NR_015587 | 319983 | Mm.343027 | NR_015587 | ENSMUSG00000097098 | **9330111N05Rik** | - | RIKEN cDNA 9330111N05 gene | ncRNA |
| chr13_81175439_81175781 | chr13 | 81175440 | 81175781 | + | 0.80832 | intron (NM_054053, intron 86 of 89) | intron (NM_054053, intron 86 of 89) | 211536 | NR_015587 | 319983 | Mm.343027 | NR_015587 | ENSMUSG00000097098 | **9330111N05Rik** | - | RIKEN cDNA 9330111N05 gene | ncRNA |
| chr10_29253828_29254188 | chr10 | 29253829 | 29254188 | + | 0.657105 | Intergenic | ORR1D1\|LTR\|ERVL-MaLR | 42365 | NM_001162537 | 212448 | Mm.132789 | NM_001162537 | ENSMUSG00000004360 | **9330159F19Rik** | - | RIKEN cDNA 9330159F19 gene | protein-coding |
| chr8_94447136_94448081 | chr8 | 94447137 | 94448081 | + | 1.14655 | Intergenic | Intergenic | -12506 | NR_015514 | 320377 | Mm.486022 | NR_015514 | ENSMUSG00000097194 | **9330175E14Rik** | A630085K21 | RIKEN cDNA 9330175E14 gene | ncRNA |
| chr8_94437001_94437282 | chr8 | 94437002 | 94437282 | + | 1.1032 | Intergenic | Intergenic | -2039 | NR_015514 | 320377 | Mm.486022 | NR_015514 | ENSMUSG00000097194 | **9330175E14Rik** | A630085K21 | RIKEN cDNA 9330175E14 gene | ncRNA |
| chr8_94441616_94442044 | chr8 | 94441617 | 94442044 | + | 0.865194 | Intergenic | MER113\|DNA\|hAT-Charlie | -6727 | NR_015514 | 320377 | Mm.486022 | NR_015514 | ENSMUSG00000097194 | **9330175E14Rik** | A630085K21 | RIKEN cDNA 9330175E14 gene | ncRNA |
| chr6_127143104_127143767 | chr6 | 127143105 | 127143767 | + | 0.864525 | intron (NM_009829, intron 3 of 4) | intron (NM_009829, intron 3 of 4) | -5953 | NR_040274 | 77558 | Mm.358918 | NR_040273 | ENSMUSG00000097166 | **9330179D12Rik** | - | RIKEN cDNA 9330179D12 gene | ncRNA |
| chr6_127149559_127150063 | chr6 | 127149560 | 127150063 | + | 0.611568 | non-coding (NR_040275, exon 1 of 2) | non-coding (NR_040275, exon 1 of 2) | 185 | NR_040273 | 77558 | Mm.358918 | NR_040273 | ENSMUSG00000097166 | **9330179D12Rik** | - | RIKEN cDNA 9330179D12 gene | ncRNA |
| chr2_158354397_158354661 | chr2 | 158354398 | 158354661 | + | 0.754451 | non-coding (NR_015463, exon 6 of 6) | non-coding (NR_015463, exon 6 of 6) | 6980 | NR_015463 | 68108 | Mm.119198 | NR_015463 | | **9430008C03Rik** | - | RIKEN cDNA 9430008C03 gene | ncRNA |
| chr15_93948969_93949262 | chr15 | 93948970 | 93949262 | + | 0.653816 | Intergenic | Intergenic | -22005 | NR_045737 | 77248 | Mm.410768 | NR_045737 | | **9430014N10Rik** | - | RIKEN cDNA 9430014N10 gene | ncRNA |
| chr6_43382827_43383328 | chr6 | 43382828 | 43383328 | + | 1.36714 | intron (NM_013861, intron 8 of 8) | intron (NM_013861, intron 8 of 8) | -59351 | NR_045988 | 319812 | Mm.446393 | NR_045988 | | **9430018G01Rik** | - | RIKEN cDNA 9430018G01 gene | ncRNA |
| chr6_43420166_43420889 | chr6 | 43420167 | 43420889 | + | 1.1553 | intron (NM_013861, intron 8 of 8) | intron (NM_013861, intron 8 of 8) | -21901 | NR_045988 | 319812 | Mm.446393 | NR_045988 | | **9430018G01Rik** | - | RIKEN cDNA 9430018G01 gene | ncRNA |
| chr6_43382294_43382656 | chr6 | 43382295 | 43382656 | + | 0.953025 | intron (NM_013861, intron 8 of 8) | ORR1D1\|LTR\|ERVL-MaLR | -59954 | NR_045988 | 319812 | Mm.446393 | NR_045988 | | **9430018G01Rik** | - | RIKEN cDNA 9430018G01 gene | ncRNA |
| chr2_75286348_75286730 | chr2 | 75286349 | 75286730 | + | 1.07087 | Intergenic | Intergenic | 157640 | NR_040635 | 77269 | Mm.389278 | NR_040635 | | **9430019J16Rik** | - | RIKEN cDNA 9430019J16 gene | ncRNA |
| chr2_75285000_75285552 | chr2 | 75285001 | 75285552 | + | 0.817378 | Intergenic | Intergenic | 158903 | NR_040635 | 77269 | Mm.389278 | NR_040635 | | **9430019J16Rik** | - | RIKEN cDNA 9430019J16 gene | ncRNA |
| chr2_75547925_75548231 | chr2 | 75547926 | 75548231 | + | 0.626467 | Intergenic | Intergenic | -103899 | NR_040635 | 77269 | Mm.389278 | NR_040635 | | **9430019J16Rik** | - | RIKEN cDNA 9430019J16 gene | ncRNA |
| chr15_34336208_34336742 | chr15 | 34336209 | 34336742 | + | 0.791489 | intron (NM_016762, intron 2 of 18) | intron (NM_016762, intron 2 of 18) | 21397 | NM_001256161 | 77358 | Mm.402759 | NM_001256161 | | **9430069I07Rik** | - | RIKEN cDNA 9430069I07 gene | protein-coding |
| chr13_50958063_50958745 | chr13 | 50958064 | 50958745 | + | 0.843102 | Intergenic | Intergenic | 142323 | NR_029463 | 77428 | Mm.405630 | NR_029463 | | **9430083A17Rik** | - | RIKEN cDNA 9430083A17 gene | ncRNA |
| chr1_61423253_61423697 | chr1 | 61423254 | 61423697 | + | 0.74355 | Intergenic | Intergenic | 45043 | NR_040483 | 77341 | Mm.402964 | NR_040483 | ENSMUSG00000085447 | **9530026F06Rik** | - | RIKEN cDNA 9530026F06 gene | ncRNA |
| chr1_61423307_61423620 | chr1 | 61423308 | 61423620 | + | 0.711123 | Intergenic | Intergenic | 45032 | NR_040483 | 77341 | Mm.402964 | NR_040483 | ENSMUSG00000085447 | **9530026F06Rik** | - | RIKEN cDNA 9530026F06 gene | ncRNA |
| chr5_28397404_28397680 | chr5 | 28397405 | 28397680 | + | 0.608943 | intron (NM_028234, intron 15 of 18) | intron (NM_028234, intron 15 of 18) | -69442 | NR_015562 | 654796 | Mm.57202 | NR_015562 | ENSMUSG00000097726 | **9530036O11Rik** | - | RIKEN cDNA 9530036O11Rik | ncRNA |
| chr8_10993545_10994256 | chr8 | 10993546 | 10994256 | + | 1.44615 | intron (NM_001081212, intron 1 of 1) | intron (NM_001081212, intron 1 of 1) | -13949 | NR_046017 | 619321 | Mm.403007 | NR_046017 | ENSMUSG00000096938 | **9530052E02Rik** | - | RIKEN cDNA 9530052E02 gene | ncRNA |
| chr8_10995800_10996123 | chr8 | 10995801 | 10996123 | + | 1.00311 | intron (NM_001081212, intron 1 of 1) | intron (NM_001081212, intron 1 of 1) | -11888 | NR_046017 | 619321 | Mm.403007 | NR_046017 | ENSMUSG00000096938 | **9530052E02Rik** | - | RIKEN cDNA 9530052E02 gene | ncRNA |
| chr9_122525870_122526092 | chr9 | 122525871 | 122526092 | + | 0.686191 | Intergenic | MER121\|DNA\|TcMar? | -46520 | NR_015610 | 319626 | Mm.186748 | NR_015610 | ENSMUSG00000097736 | **9530059O14Rik** | - | RIKEN cDNA 9530059O14 gene | ncRNA |
| chr9_22429033_22429324 | chr9 | 22429034 | 22429324 | + | 0.901619 | intron (NM_026739, intron 2 of 6) | B3\|SINE\|B2 | 17602 | NM_026739 | 68283 | Mm.280669 | NM_026739 | ENSMUSG00000036411 | **9530077C05Rik** | 1110003N12Rik\|Kiaa0895\|mKIAA0895 | RIKEN cDNA 9530077C05 gene | protein-coding |

| | | | | | | | | | | | | | | | | | |
|---|---|---|---|---|---|---|---|---|---|---|---|---|---|---|---|---|---|
| chr9_22428900_22429316 | chr9 | 22428901 | 22429316 | + | 0.898723 | intron (NM_026739, intron 2 of 6) | B3\|SINE\|B2 | 17531 | NM_026739 | 68283 | Mm.280669 | NM_026739 | ENSMUSG00000036411 | **9530077C05Rik** | 1110003N12Rik\|Kiaa0895\|mKIAA0895 | RIKEN cDNA 9530077C05 gene | protein-coding |
| chr9_69025905_69026571 | chr9 | 69025906 | 69026571 | + | 1.54551 | intron (NM_013646, intron 1 of 10) | intron (NM_013646, intron 1 of 10) | 260892 | NR_033299 | 320440 | Mm.483129 | NM_177159 | | **9530091C08Rik** | - | RIKEN cDNA 9530091C08 gene | ncRNA |
| chr9_68970150_68970513 | chr9 | 68970151 | 68970513 | + | 0.683325 | intron (NM_013646, intron 1 of 10) | MER119\|DNA\|hAT-Charlie | 204986 | NR_033299 | 320440 | Mm.483129 | NM_177159 | | **9530091C08Rik** | - | RIKEN cDNA 9530091C08 gene | ncRNA |
| chr3_127914017_127914621 | chr3 | 127914018 | 127914621 | + | 0.8796 | Intergenic | Intergenic | -1852 | NR_040552 | 329753 | Mm.119720 | NR_040552 | | **9830132P13Rik** | 9830132P13 | RIKEN cDNA 9830132P13 gene | ncRNA |
| chr3_127932079_127932888 | chr3 | 127932080 | 127932888 | + | 0.803251 | intron (NR_040552, intron 3 of 4) | intron (NR_040552, intron 3 of 4) | 16313 | NR_040552 | 329753 | Mm.119720 | NR_040552 | | **9830132P13Rik** | 9830132P13 | RIKEN cDNA 9830132P13 gene | ncRNA |
| chr15_60739387_60740188 | chr15 | 60739388 | 60740188 | + | 0.93122 | Intergenic | Intergenic | -83177 | NR_030696 | 320469 | Mm.230853 | NM_177166 | ENSMUSG00000097493 | **9930014A18Rik** | - | RIKEN cDNA 9930014A18 gene | ncRNA |
| chr2_155038331_155038724 | chr2 | 155038332 | 155038724 | + | 0.912697 | intron (NM_015770, intron 1 of 3) | intron (NM_015770, intron 1 of 3) | 24958 | NM_015770 | 50518 | Mm.315593 | NM_015770 | ENSMUSG00000027596 | **a** | A<y>\|ASIP\|ASP\|As | nonagouti | protein-coding |
| chr1_193178701_193179230 | chr1 | 193178702 | 193179230 | + | 1.00491 | intron (NM_153137, intron 14 of 15) | intron (NM_153137, intron 14 of 15) | 5389 | NM_181048 | 319266 | Mm.482460 | NM_181048 | ENSMUSG00000079144 | **A130010J15Rik** | - | RIKEN cDNA A130010J15 gene | protein-coding |
| chr10_122603947_122604319 | chr10 | 122603948 | 122604319 | + | 0.666265 | Intergenic | Intergenic | -34242 | NR_040616 | 319272 | Mm.411763 | NR_040616 | | **A130077B15Rik** | - | RIKEN cDNA A130077B15 gene | ncRNA |
| chr16_24952851_24953349 | chr16 | 24952852 | 24953349 | + | 1.10835 | intron (NM_001145952, intron 9 of 11) | intron (NM_001145952, intron 9 of 11) | -106539 | NR_040374 | 319487 | Mm.137341 | NR_040374 | ENSMUSG00000096960 | **A230028O05Rik** | - | RIKEN cDNA A230028O05 gene | ncRNA |
| chr16_24952689_24953363 | chr16 | 24952690 | 24953363 | + | 1.02132 | intron (NM_001145952, intron 9 of 11) | intron (NM_001145952, intron 9 of 11) | -106613 | NR_040374 | 319487 | Mm.137341 | NR_040374 | ENSMUSG00000096960 | **A230028O05Rik** | - | RIKEN cDNA A230028O05 gene | ncRNA |
| chr9_20869311_20869731 | chr9 | 20869312 | 20869731 | + | 0.662514 | intron (NM_175687, intron 2 of 7) | intron (NM_175687, intron 2 of 7) | 879 | NM_175687 | 319278 | Mm.249932 | NM_175687 | ENSMUSG00000038884 | **A230050P20Rik** | - | RIKEN cDNA A230050P20 gene | protein-coding |
| chr14_122074278_122074627 | chr14 | 122074279 | 122074627 | + | 0.816035 | Intergenic | Intergenic | 32521 | NR_015586 | 319727 | Mm.215552 | NR_015586 | | **A330035P11Rik** | - | RIKEN cDNA A330035P11 gene | ncRNA |
| chr2_91242487_91243251 | chr2 | 91242488 | 91243251 | + | 1.20037 | Intergenic | Intergenic | 5723 | NR_015464 | 606735 | Mm.246808 | NR_015464 | ENSMUSG00000097868 | **A330069E16Rik** | - | RIKEN cDNA A330069E16 gene | ncRNA |
| chr2_91242423_91242991 | chr2 | 91242424 | 91242991 | + | 1.19682 | Intergenic | Intergenic | 5561 | NR_015464 | 606735 | Mm.246808 | NR_015464 | ENSMUSG00000097868 | **A330069E16Rik** | - | RIKEN cDNA A330069E16 gene | ncRNA |
| chr2_91242338_91243729 | chr2 | 91242339 | 91243729 | + | 1.05144 | Intergenic | (GA)n\|Simple_repeat\|Simple_repeat | 5888 | NR_015464 | 606735 | Mm.246808 | NR_015464 | ENSMUSG00000097868 | **A330069E16Rik** | - | RIKEN cDNA A330069E16 gene | ncRNA |
| chr8_120332435_120333187 | chr8 | 120332436 | 120333187 | + | 1.1811 | Intergenic | Intergenic | -104581 | NR_110496 | 434353 | Mm.411793 | NR_110496 | | **A330074K22Rik** | - | RIKEN cDNA A330074K22 gene | ncRNA |
| chr8_120225681_120226134 | chr8 | 120225682 | 120226134 | + | 0.864473 | intron (NR_110496, intron 1 of 2) | intron (NR_110496, intron 1 of 2) | 2322 | NR_110496 | 434353 | Mm.411793 | NR_110496 | | **A330074K22Rik** | - | RIKEN cDNA A330074K22 gene | ncRNA |
| chr8_120228100_120228542 | chr8 | 120228101 | 120228542 | + | 0.6768 | promoter-TSS (NR_110496) | promoter-TSS (NR_110496) | -91 | NR_110496 | 434353 | Mm.411793 | NR_110496 | | **A330074K22Rik** | - | RIKEN cDNA A330074K22 gene | ncRNA |
| chr13_44320849_44321676 | chr13 | 44320850 | 44321676 | + | 1.43678 | Intergenic | Intergenic | -104776 | NR_045088 | 100503592 | Mm.184925 | NR_045088 | | **A330076C08Rik** | - | RIKEN cDNA A330076C08 gene | ncRNA |
| chr13_113976469_113976823 | chr13 | 113976470 | 113976823 | + | 0.712554 | intron (NM_172595, intron 4 of 4) | intron (NM_172595, intron 4 of 4) | 174946 | NR_045836 | 319803 | Mm.386896 | NM_177004 | | **A430090L17Rik** | - | RIKEN cDNA A430090L17 gene | ncRNA |
| chr19_10781547_10782042 | chr19 | 10781548 | 10782042 | + | 1.17163 | non-coding (NR_027805, exon 2 of 4) | non-coding (NR_027805, exon 2 of 4) | 40848 | NR_027805 | 403202 | Mm.34482 | NM_207529 | ENSMUSG00000067577 | **A430093F15Rik** | Gm674 | RIKEN cDNA A430093F15 gene | ncRNA |
| chr19_10757076_10757610 | chr19 | 10757077 | 10757610 | + | 0.875884 | intron (NR_027805, intron 1 of 3) | intron (NR_027805, intron 1 of 3) | 16396 | NR_027805 | 403202 | Mm.34482 | NM_207529 | ENSMUSG00000067577 | **A430093F15Rik** | Gm674 | RIKEN cDNA A430093F15 gene | ncRNA |
| chr19_10757066_10757998 | chr19 | 10757067 | 10757998 | + | 0.834278 | intron (NR_027805, intron 1 of 3) | intron (NR_027805, intron 1 of 3) | 16585 | NR_027805 | 403202 | Mm.34482 | NM_207529 | ENSMUSG00000067577 | **A430093F15Rik** | Gm674 | RIKEN cDNA A430093F15 gene | ncRNA |
| chr19_10763325_10764423 | chr19 | 10763326 | 10764423 | + | 0.778823 | intron (NR_027805, intron 1 of 3) | intron (NR_027805, intron 1 of 3) | 22927 | NR_027805 | 403202 | Mm.34482 | NM_207529 | ENSMUSG00000067577 | **A430093F15Rik** | Gm674 | RIKEN cDNA A430093F15 gene | ncRNA |
| chr19_10739979_10740219 | chr19 | 10739980 | 10740219 | + | 0.755821 | promoter-TSS (NR_027805) | promoter-TSS (NR_027805) | -848 | NR_027805 | 403202 | Mm.34482 | NM_207529 | ENSMUSG00000067577 | **A430093F15Rik** | Gm674 | RIKEN cDNA A430093F15 gene | ncRNA |
| chr19_10763340_10764127 | chr19 | 10763341 | 10764127 | + | 0.738396 | intron (NR_027805, intron 1 of 3) | intron (NR_027805, intron 1 of 3) | 22787 | NR_027805 | 403202 | Mm.34482 | NM_207529 | ENSMUSG00000067577 | **A430093F15Rik** | Gm674 | RIKEN cDNA A430093F15 gene | ncRNA |
| chr19_10783951_10784605 | chr19 | 10783952 | 10784605 | + | 0.720323 | intron (NR_027805, intron 2 of 3) | intron (NR_027805, intron 2 of 3) | 43331 | NR_027805 | 403202 | Mm.34482 | NM_207529 | ENSMUSG00000067577 | **A430093F15Rik** | Gm674 | RIKEN cDNA A430093F15 gene | ncRNA |
| chr19_10763588_10764097 | chr19 | 10763589 | 10764097 | + | 0.670509 | intron (NR_027805, intron 1 of 3) | intron (NR_027805, intron 1 of 3) | 22896 | NR_027805 | 403202 | Mm.34482 | NM_207529 | ENSMUSG00000067577 | **A430093F15Rik** | Gm674 | RIKEN cDNA A430093F15 gene | ncRNA |
| chr14_53943341_53943740 | chr14 | 53943342 | 53943740 | + | 1.35249 | Intergenic | Intergenic | 277601 | NM_001205242 | 100568452 | Mm.213248 | NM_001205242 | | **A430107P09Rik** | Va10 | RIKEN cDNA A430107P09 gene | protein-coding |
| chr14_53736269_53736722 | chr14 | 53736270 | 53736722 | + | 1.03253 | Intergenic | Intergenic | 70556 | NM_001205242 | 100568452 | Mm.213248 | NM_001205242 | | **A430107P09Rik** | Va10 | RIKEN cDNA A430107P09 gene | protein-coding |
| chr14_53643640_53644414 | chr14 | 53643641 | 53644414 | + | 0.953182 | Intergenic | LTRIS2\|LTR\|ERV1 | -21913 | NM_001205242 | 100568452 | Mm.213248 | NM_001205242 | | **A430107P09Rik** | Va10 | RIKEN cDNA A430107P09 gene | protein-coding |
| chr14_53832040_53832501 | chr14 | 53832041 | 53832501 | + | 0.905968 | Intergenic | Intergenic | 166331 | NM_001205242 | 100568452 | Mm.213248 | NM_001205242 | | **A430107P09Rik** | Va10 | RIKEN cDNA A430107P09 gene | protein-coding |
| chr14_53559955_53560589 | chr14 | 53559956 | 53560589 | + | 0.713986 | Intergenic | Intergenic | -105668 | NM_001205242 | 100568452 | Mm.213248 | NM_001205242 | | **A430107P09Rik** | Va10 | RIKEN cDNA A430107P09 gene | protein-coding |
| chr13_15848032_15848464 | chr13 | 15848033 | 15848464 | + | 0.997741 | Intergenic | Intergenic | 41005 | NR_046131 | 328190 | Mm.446957 | NR_046131 | | **A530046M15Rik** | A530046M15 | RIKEN cDNA A530046M15 gene | ncRNA |
| chr18_61481577_61482048 | chr18 | 61481578 | 61482048 | + | 1.07359 | intron (NR_045419, intron 1 of 3) | MLT1K\|LTR\|ERVL-MaLR | 2794 | NR_045419 | 442846 | Mm.417648 | NR_045419 | | **A530050N04Rik** | - | RIKEN cDNA A530050N04 gene | ncRNA |
| chr4_16211422_16211941 | chr4 | 16211423 | 16211941 | + | 1.30545 | intron (NR_045765, intron 3 of 6) | Lx7\|LINE\|L1 | 47572 | NR_045765 | 100415915 | Mm.387180 | NR_045765 | ENSMUSG00000085112 | **A530072M11Rik** | - | RIKEN cDNA gene A530072M11 | ncRNA |
| chr1_85706082_85706446 | chr1 | 85706083 | 85706446 | + | 0.851656 | intron (NM_013673, intron 17 of 19) | MTD\|LTR\|ERVL-MaLR | 30342 | NM_177055 | 319997 | Mm.116691 | NM_177055 | ENSMUSG00000052760 | **A630001G21Rik** | - | RIKEN cDNA A630001G21 gene | protein-coding |
| chr13_93092942_93093254 | chr13 | 93092943 | 93093254 | + | 0.879985 | exon (NM_023821, exon 2 of 13) | exon (NM_023821, exon 2 of 13) | -1735 | NR_046182 | 408254 | Mm.474857 | NR_046182 | | **A630019I02Rik** | - | RIKEN cDNA A630019I02 gene | ncRNA |
| chr5_20929880_20930384 | chr5 | 20929881 | 20930384 | + | 0.78552 | intron (NM_001080977, intron 1 of 7) | B1_Mur2\|SINE\|Alu | -20857 | NR_030699 | 320770 | Mm.491217 | NR_030699 | | **A630072M18Rik** | - | RIKEN cDNA A630072M18 gene | ncRNA |
| chr6_61225361_61226006 | chr6 | 61225362 | 61226006 | + | 1.71113 | intron (NM_183310, intron 1 of 10) | intron (NM_183310, intron 1 of 10) | -44874 | NR_040287 | 100504262 | Mm.438066 | NR_040287 | | **A730020E08Rik** | - | RIKEN cDNA A730020E08 gene | ncRNA |
| chr6_61266504_61266875 | chr6 | 61266505 | 61266875 | + | 1.06326 | intron (NM_183310, intron 1 of 10) | intron (NM_183310, intron 1 of 10) | -85880 | NR_040287 | 100504262 | Mm.438066 | NR_040287 | | **A730020E08Rik** | - | RIKEN cDNA A730020E08 gene | ncRNA |
| chr6_61266326_61266888 | chr6 | 61266327 | 61266888 | + | 0.993253 | intron (NM_183310, intron 1 of 10) | intron (NM_183310, intron 1 of 10) | -85797 | NR_040287 | 100504262 | Mm.438066 | NR_040287 | | **A730020E08Rik** | - | RIKEN cDNA A730020E08 gene | ncRNA |
| chr6_62014364_62014963 | chr6 | 62014365 | 62014963 | + | 0.824473 | intron (NM_183310, intron 10 of 10) | intron (NM_183310, intron 10 of 10) | -833854 | NR_040287 | 100504262 | Mm.438066 | NR_040287 | | **A730020E08Rik** | - | RIKEN cDNA A730020E08 gene | ncRNA |
| chr6_61263257_61263874 | chr6 | 61263258 | 61263874 | + | 0.757874 | intron (NM_183310, intron 1 of 10) | intron (NM_183310, intron 1 of 10) | -82756 | NR_040287 | 100504262 | Mm.438066 | NR_040287 | | **A730020E08Rik** | - | RIKEN cDNA A730020E08 gene | ncRNA |
| chr6_61430715_61430967 | chr6 | 61430716 | 61430967 | + | 0.718443 | intron (NM_183310, intron 5 of 10) | intron (NM_183310, intron 5 of 10) | -250031 | NR_040287 | 100504262 | Mm.438066 | NR_040287 | | **A730020E08Rik** | - | RIKEN cDNA A730020E08 gene | ncRNA |
| chr6_61430689_61431088 | chr6 | 61430690 | 61431088 | + | 0.708091 | intron (NM_183310, intron 5 of 10) | intron (NM_183310, intron 5 of 10) | -250079 | NR_040287 | 100504262 | Mm.438066 | NR_040287 | | **A730020E08Rik** | - | RIKEN cDNA A730020E08 gene | ncRNA |
| chr6_61300631_61301375 | chr6 | 61300632 | 61301375 | + | 0.603712 | intron (NM_183310, intron 1 of 10) | intron (NM_183310, intron 1 of 10) | -120193 | NR_040287 | 100504262 | Mm.438066 | NR_040287 | | **A730020E08Rik** | - | RIKEN cDNA A730020E08 gene | ncRNA |
| chr3_121592911_121594042 | chr3 | 121592912 | 121594042 | + | 0.999761 | Intergenic | Intergenic | 50008 | NR_036456 | 100503044 | Mm.65320 | NM_176992 | ENSMUSG00000044522 | **A730020M07Rik** | - | RIKEN cDNA A730020M07 gene | ncRNA |
| chr9_62222115_62222901 | chr9 | 62222116 | 62222901 | + | 0.800918 | Intergenic | Intergenic | 21592 | NR_040769 | 330958 | Mm.411888 | NR_040769 | | **A730043L09Rik** | A730043L09 | RIKEN cDNA A730043L09 gene | ncRNA |
| chr9_62222449_62222888 | chr9 | 62222450 | 62222888 | + | 0.79057 | Intergenic | Intergenic | 21431 | NR_040769 | 330958 | Mm.411888 | NR_040769 | | **A730043L09Rik** | A730043L09 | RIKEN cDNA A730043L09 gene | ncRNA |
| chr3_65319321_65320108 | chr3 | 65319322 | 65320108 | + | 0.766451 | intron (NM_001289450, intron 9 of 14) | CT-rich\|Low_complexity\|Low_complexity | 29714 | NR_040390 | 319798 | Mm.445698 | NR_040390 | | **A730090N16Rik** | - | RIKEN cDNA A730090N16 gene | ncRNA |
| chr16_24506624_24507275 | chr16 | 24506625 | 24507275 | + | 1.51152 | intron (NM_001145952, intron 2 of 11) | intron (NM_001145952, intron 2 of 11) | -22364 | NR_038091 | 624582 | Mm.439775 | NR_038091 | | **A730098P11Rik** | - | mortality factor 4 like 1 pseudogene | pseudo |
| chr16_24506314_24507276 | chr16 | 24506315 | 24507276 | + | 1.4407 | intron (NM_001145952, intron 2 of 11) | intron (NM_001145952, intron 2 of 11) | -22519 | NR_038091 | 624582 | Mm.439775 | NR_038091 | | **A730098P11Rik** | - | mortality factor 4 like 1 pseudogene | pseudo |

| | | | | | | | | | | | | | | | | | |
|---|---|---|---|---|---|---|---|---|---|---|---|---|---|---|---|---|---|
| chr16_24507905_24508233 | chr16 | 24507906 | 24508233 | + | 1.10727 | intron (NM_001145952, intron 2 of 11) | intron (NM_001145952, intron 2 of 11) | -21245 | NR_038091 | 624582 | Mm.439775 | NR_038091 | | **A730098P11Rik** | - | mortality factor 4 like 1 pseudogene | pseudo |
| chr16_24526753_24528066 | chr16 | 24526754 | 24528066 | + | 0.992391 | intron (NM_001145952, intron 2 of 11) | intron (NM_001145952, intron 2 of 11) | -1904 | NR_038091 | 624582 | Mm.439775 | NR_038091 | | **A730098P11Rik** | - | mortality factor 4 like 1 pseudogene | pseudo |
| chr16_24594499_24594960 | chr16 | 24594500 | 24594960 | + | 0.842692 | intron (NM_001145952, intron 3 of 11) | intron (NM_001145952, intron 3 of 11) | 65416 | NR_038091 | 624582 | Mm.439775 | NR_038091 | | **A730098P11Rik** | - | mortality factor 4 like 1 pseudogene | pseudo |
| chr16_24594246_24594960 | chr16 | 24594247 | 24594960 | + | 0.834634 | intron (NM_001145952, intron 3 of 11) | intron (NM_001145952, intron 3 of 11) | 65289 | NR_038091 | 624582 | Mm.439775 | NR_038091 | | **A730098P11Rik** | - | mortality factor 4 like 1 pseudogene | pseudo |
| chr16_24603176_24603751 | chr16 | 24603177 | 24603751 | + | 0.685744 | intron (NM_001145952, intron 3 of 11) | intron (NM_001145952, intron 3 of 11) | 74150 | NR_038091 | 624582 | Mm.439775 | NR_038091 | | **A730098P11Rik** | - | mortality factor 4 like 1 pseudogene | pseudo |
| chr16_24608506_24609027 | chr16 | 24608507 | 24609027 | + | 0.677592 | intron (NM_001145952, intron 4 of 11) | intron (NM_001145952, intron 4 of 11) | 79453 | NR_038091 | 624582 | Mm.439775 | NR_038091 | | **A730098P11Rik** | - | mortality factor 4 like 1 pseudogene | pseudo |
| chr16_24608501_24608766 | chr16 | 24608502 | 24608766 | + | 0.621162 | intron (NM_001145952, intron 4 of 11) | intron (NM_001145952, intron 4 of 11) | 79320 | NR_038091 | 624582 | Mm.439775 | NR_038091 | | **A730098P11Rik** | - | mortality factor 4 like 1 pseudogene | pseudo |
| chr13_78161914_78162500 | chr13 | 78161915 | 78162500 | + | 0.858185 | intron (NR_028109, intron 10 of 10) | intron (NR_028109, intron 10 of 10) | -35810 | NR_045195 | 320174 | Mm.393122 | NR_045195 | | **A830082K12Rik** | - | RIKEN cDNA A830082K12 gene | ncRNA |
| chr3_15139528_15139929 | chr3 | 15139529 | 15139929 | + | 1.42955 | Intergenic | Intergenic | 168528 | NR_040550 | 77127 | Mm.195792 | NR_040549 | ENSMUSG00000098008 | **A930001A20Rik** | - | RIKEN cDNA A930001A20 gene | ncRNA |
| chr3_115858534_115859253 | chr3 | 115858535 | 115859253 | + | 1.03767 | Intergenic | RLTR30B_MM\|LTR\|ERV1 | 29165 | NR_027894 | 68161 | Mm.392043 | NR_015487 | | **A930005H10Rik** | - | RIKEN cDNA A930005H10 gene | ncRNA |
| chr3_115857750_115858282 | chr3 | 115857751 | 115858282 | + | 1.02382 | Intergenic | Intergenic | 30043 | NR_027894 | 68161 | Mm.392043 | NR_015487 | | **A930005H10Rik** | - | RIKEN cDNA A930005H10 gene | ncRNA |
| chr3_115857603_115858461 | chr3 | 115857604 | 115858461 | + | 0.97766 | Intergenic | Intergenic | 30027 | NR_027894 | 68161 | Mm.392043 | NR_015487 | | **A930005H10Rik** | - | RIKEN cDNA A930005H10 gene | ncRNA |
| chr3_115861077_115861320 | chr3 | 115861078 | 115861320 | + | 0.756187 | Intergenic | RMER6C\|LTR\|ERVK | 26860 | NR_027894 | 68161 | Mm.392043 | NR_015487 | | **A930005H10Rik** | - | RIKEN cDNA A930005H10 gene | ncRNA |
| chr16_91459786_91460215 | chr16 | 91459787 | 91460215 | + | 0.908414 | Intergenic | Intergenic | -5103 | NR_077219 | 100503120 | Mm.387130 | NR_077219 | ENSMUSG00000084866 | **A930006K02Rik** | - | RIKEN cDNA A930006K02 gene | ncRNA |
| chr16_91444917_91445118 | chr16 | 91444918 | 91445118 | + | 0.807579 | Intergenic | Intergenic | -20086 | NR_077219 | 100503120 | Mm.387130 | NR_077219 | ENSMUSG00000084866 | **A930006K02Rik** | - | RIKEN cDNA A930006K02 gene | ncRNA |
| chr16_91459472_91459739 | chr16 | 91459473 | 91459739 | + | 0.634228 | Intergenic | B3\|SINE\|B2 | -5498 | NR_077219 | 100503120 | Mm.387130 | NR_077219 | ENSMUSG00000084866 | **A930006K02Rik** | - | RIKEN cDNA A930006K02 gene | ncRNA |
| chr19_29477017_29477548 | chr19 | 29477018 | 29477548 | + | 1.73939 | Intergenic | Intergenic | 44704 | NR_015567 | 77779 | Mm.458738 | NR_015567 | ENSMUSG00000097855 | **A930007I19Rik** | - | RIKEN cDNA A930007I19 gene | ncRNA |
| chr19_29518965_29519303 | chr19 | 29518966 | 29519303 | + | 1.15461 | intron (NR_015567, intron 2 of 3) | intron (NR_015567, intron 2 of 3) | 2853 | NR_015567 | 77779 | Mm.458738 | NR_015567 | ENSMUSG00000097855 | **A930007I19Rik** | - | RIKEN cDNA A930007I19 gene | ncRNA |
| chr17_35998637_35998860 | chr17 | 35998638 | 35998860 | + | 0.979624 | intron (NR_015618, intron 1 of 1) | intron (NR_015618, intron 1 of 1) | 4245 | NR_015618 | 77810 | Mm.483143 | NR_015618 | | **A930015D03Rik** | - | RIKEN cDNA A930015D03 gene | ncRNA |
| chr17_36001156_36001493 | chr17 | 36001157 | 36001493 | + | 0.698329 | intron (NR_015618, intron 1 of 1) | intron (NR_015618, intron 1 of 1) | 6821 | NR_015618 | 77810 | Mm.483143 | NR_015618 | | **A930015D03Rik** | - | RIKEN cDNA A930015D03 gene | ncRNA |
| chr2_104176669_104177154 | chr2 | 104176670 | 104177154 | + | 1.01899 | intron (NM_001033347, intron 16 of 20) | intron (NM_001033347, intron 16 of 20) | 54143 | NM_026634 | 68243 | Mm.158191 | NM_026634 | ENSMUSG00000032671 | **A930018P22Rik** | - | RIKEN cDNA A930018P22 gene | protein-coding |
| chr2_104207924_104208367 | chr2 | 104207925 | 104208367 | + | 0.802936 | exon (NM_001033347, exon 13 of 21) | exon (NM_001033347, exon 13 of 21) | 85377 | NM_026634 | 68243 | Mm.158191 | NM_026634 | ENSMUSG00000032671 | **A930018P22Rik** | - | RIKEN cDNA A930018P22 gene | protein-coding |
| chr2_104207928_104208714 | chr2 | 104207929 | 104208714 | + | 0.747764 | intron (NM_001033347, intron 12 of 20) | intron (NM_001033347, intron 12 of 20) | 85552 | NM_026634 | 68243 | Mm.158191 | NM_026634 | ENSMUSG00000032671 | **A930018P22Rik** | - | RIKEN cDNA A930018P22 gene | protein-coding |
| chr2_146254181_146254886 | chr2 | 146254182 | 146254886 | + | 1.58322 | intron (NM_001033348, intron 39 of 40) | intron (NM_001033348, intron 39 of 40) | -4752 | NR_040619 | 77806 | Mm.392819 | NR_040619 | ENSMUSG00000085958 | **A930019D19Rik** | - | RIKEN cDNA A930019D19 gene | ncRNA |
| chr5_107648731_107649103 | chr5 | 107648732 | 107649103 | + | 0.673183 | intron (NM_144911, intron 11 of 12) | intron (NM_144911, intron 11 of 12) | -15067 | NR_046195 | 78443 | Mm.490703 | NR_046195 | ENSMUSG00000085900 | **A930041C12Rik** | - | RIKEN cDNA A930041C12 gene | ncRNA |
| chr5_107647644_107648013 | chr5 | 107647645 | 107648013 | + | 0.662832 | intron (NM_144911, intron 11 of 12) | intron (NM_144911, intron 11 of 12) | -13979 | NR_046195 | 78443 | Mm.490703 | NR_046195 | ENSMUSG00000085900 | **A930041C12Rik** | - | RIKEN cDNA A930041C12 gene | ncRNA |
| chr5_107645989_107646351 | chr5 | 107645990 | 107646351 | + | 0.611165 | intron (NM_144911, intron 11 of 12) | B1_Mur1\|SINE\|Alu | -12320 | NR_046195 | 78443 | Mm.490703 | NR_046195 | ENSMUSG00000085900 | **A930041C12Rik** | - | RIKEN cDNA A930041C12 gene | ncRNA |
| chrX_12959948_12960250 | chrX | 12959949 | 12960250 | + | 0.805968 | Intergenic | Intergenic | 23226 | NM_001272033 | 245350 | Mm.302695 | NM_001272033 | | **AA414768** | OTTMUSG00000017000 | expressed sequence AA414768 | protein-coding |
| chr1_130761079_130761682 | chr1 | 130761080 | 130761682 | + | 0.686046 | Intergenic | Intergenic | 29405 | NM_177604 | 212439 | Mm.268547 | NM_177604 | ENSMUSG00000042510 | **AA986860** | E030025M07\|Sarg | expressed sequence AA986860 | protein-coding |
| chr9_63678089_63678560 | chr9 | 63678090 | 63678560 | + | 0.695159 | intron (NM_016769, intron 1 of 8) | intron (NM_016769, intron 1 of 8) | 75670 | NM_025857 | 66939 | Mm.34338 | NM_025857 | ENSMUSG00000037257 | **Aagab** | 2310007F21Rik\|p34 | alpha- and gamma-adaptin binding protein | protein-coding |
| chr9_63674585_63674985 | chr9 | 63674586 | 63674985 | + | 0.686674 | intron (NM_016769, intron 1 of 8) | intron (NM_016769, intron 1 of 8) | 72130 | NM_025857 | 66939 | Mm.34338 | NM_025857 | ENSMUSG00000037257 | **Aagab** | 2310007F21Rik\|p34 | alpha- and gamma-adaptin binding protein | protein-coding |
| chr2_156539130_156539639 | chr2 | 156539131 | 156539639 | + | 0.791214 | intron (NM_001291122, intron 20 of 20) | intron (NM_001291122, intron 20 of 20) | -8191 | NM_026661 | 68295 | Mm.253648 | NM_026661 | ENSMUSG00000027628 | **Aar2** | 0610011L14Rik | AAR2 splicing factor homolog (S. cerevisiae) | protein-coding |
| chr15_52039530_52040636 | chr15 | 52039531 | 52040636 | + | 0.925402 | promoter-TSS (NM_175503) | promoter-TSS (NM_175503) | -24 | NM_175503 | 239435 | Mm.24204 | NM_175503 | ENSMUSG00000068522 | **Aard** | A5D3\|AV328152 | alanine and arginine rich domain containing protein | protein-coding |
| chr4_53118517_53119095 | chr4 | 53118518 | 53119095 | + | 0.681545 | intron (NM_013454, intron 5 of 49) | MIR\|SINE\|MIR | 41089 | NM_013454 | 11303 | Mm.277376 | NM_013454 | ENSMUSG00000015243 | **Abca1** | ABC-1\|Abc1 | ATP-binding cassette, sub-family A (ABC1), member 1 | protein-coding |
| chr5_8722250_8723196 | chr5 | 8722251 | 8723196 | + | 0.776475 | intron (NM_011076, intron 19 of 27) | intron (NM_011076, intron 19 of 27) | 62631 | NM_011076 | 18671 | Mm.207354 | NM_011076 | ENSMUSG00000040584 | **Abcb1a** | Abcb4\|Evi32\|Mdr1a\|Mdr3\|P-gp\|Pgp\|Pgy-3\|Pgy3\|mdr-3 | ATP-binding cassette, sub-family B (MDR/TAP), member 1A | protein-coding |
| chr5_8668346_8668928 | chr5 | 8668347 | 8668928 | + | 0.766801 | intron (NM_011076, intron 3 of 27) | intron (NM_011076, intron 3 of 27) | 8545 | NM_011076 | 18671 | Mm.207354 | NM_011076 | ENSMUSG00000040584 | **Abcb1a** | Abcb4\|Evi32\|Mdr1a\|Mdr3\|P-gp\|Pgp\|Pgy-3\|Pgy3\|mdr-3 | ATP-binding cassette, sub-family B (MDR/TAP), member 1A | protein-coding |
| chr5_8761651_8762211 | chr5 | 8761652 | 8762211 | + | 1.25077 | Intergenic | L1VL2\|LINE\|L1 | -36216 | NM_011075 | 18669 | Mm.146649 | NM_011075 | ENSMUSG00000028970 | **Abcb1b** | Abcb1\|Mdr1\|Mdr1b\|Pgy-1\|Pgy1\|mdr | ATP-binding cassette, sub-family B (MDR/TAP), member 1B | protein-coding |
| chr5_8826886_8827355 | chr5 | 8826887 | 8827355 | + | 0.699054 | intron (NM_011075, intron 15 of 27) | intron (NM_011075, intron 15 of 27) | 28974 | NM_011075 | 18669 | Mm.146649 | NM_011075 | ENSMUSG00000028970 | **Abcb1b** | Abcb1\|Mdr1\|Mdr1b\|Pgy-1\|Pgy1\|mdr | ATP-binding cassette, sub-family B (MDR/TAP), member 1B | protein-coding |
| chr5_8805766_8806115 | chr5 | 8805767 | 8806115 | + | 0.638817 | intron (NM_011075, intron 3 of 27) | intron (NM_011075, intron 3 of 27) | 7794 | NM_011075 | 18669 | Mm.146649 | NM_011075 | ENSMUSG00000028970 | **Abcb1b** | Abcb1\|Mdr1\|Mdr1b\|Pgy-1\|Pgy1\|mdr | ATP-binding cassette, sub-family B (MDR/TAP), member 1B | protein-coding |
| chrX_104375547_104376199 | chrX | 104375548 | 104376199 | + | 0.800652 | intron (NM_009592, intron 1 of 15) | ORR1E\|LTR\|ERVL-MaLR | 37973 | NM_009592 | 11306 | Mm.426128 | NM_009592 | ENSMUSG00000031333 | **Abcb7** | AA517758\|AU019072\|Abc7 | ATP-binding cassette, sub-family B (MDR/TAP), member 7 | protein-coding |
| chrX_104375783_104376174 | chrX | 104375784 | 104376174 | + | 0.768097 | intron (NM_009592, intron 1 of 15) | Tigger7\|DNA\|TcMar-Tigger | 37867 | NM_009592 | 11306 | Mm.426128 | NM_009592 | ENSMUSG00000031333 | **Abcb7** | AA517758\|AU019072\|Abc7 | ATP-binding cassette, sub-family B (MDR/TAP), member 7 | protein-coding |
| chr16_14350312_14351232 | chr16 | 14350313 | 14351232 | + | 0.881153 | Intergenic | L1MB3\|LINE\|L1 | -10786 | NM_008576 | 17250 | Mm.196634 | NM_008576 | ENSMUSG00000023088 | **Abcc1** | Abcc1a\|Abcc1b\|MRP\|Mdrap\|Mrp1 | ATP-binding cassette, sub-family C (CFTR/MRP), member 1 | protein-coding |
| chr16_14350629_14351212 | chr16 | 14350630 | 14351212 | + | 0.867891 | Intergenic | L1MB3\|LINE\|L1 | -10637 | NM_008576 | 17250 | Mm.196634 | NM_008576 | ENSMUSG00000023088 | **Abcc1** | Abcc1a\|Abcc1b\|MRP\|Mdrap\|Mrp1 | ATP-binding cassette, sub-family C (CFTR/MRP), member 1 | protein-coding |
| chr19_43771490_43771926 | chr19 | 43771491 | 43771926 | + | 1.04225 | Intergenic | L1M4\|LINE\|L1 | -10600 | NM_013806 | 12780 | Mm.39054 | NM_013806 | ENSMUSG00000025194 | **Abcc2** | AI173996\|Abc30\|Cmoat\|Mrp2\|cMRP | ATP-binding cassette, sub-family C (CFTR/MRP), member 2 | protein-coding |
| chr14_118657960_118658775 | chr14 | 118657961 | 118658775 | + | 0.943807 | intron (NM_001163675, intron 3 of 30) | intron (NM_001163675, intron 3 of 30) | 18563 | NM_001163675 | 239273 | Mm.40537 | NM_001033336 | ENSMUSG00000032849 | **Abcc4** | D630049P08Rik\|MOATB\|MRP4 | ATP-binding cassette, sub-family C (CFTR/MRP), member 4 | protein-coding |
| chr3_121796871_121797687 | chr3 | 121796872 | 121797687 | + | 1.17286 | intron (NM_008991, intron 1 of 22) | intron (NM_008991, intron 1 of 22) | 17936 | NM_008991 | 19299 | Mm.399042 | NM_008991 | ENSMUSG00000028127 | **Abcd3** | AI313901\|AU018866\|AW146054\|PMP68\|PMP70\|Pxmp1 | ATP-binding cassette, sub-family D (ALD), member 3 | protein-coding |
| chr3_121872690_121873174 | chr3 | 121872691 | 121873174 | + | 0.989593 | Intergenic | Intergenic | -57717 | NM_008991 | 19299 | Mm.399042 | NM_008991 | ENSMUSG00000028127 | **Abcd3** | AI313901\|AU018866\|AW146054\|PMP68\|PMP70\|Pxmp1 | ATP-binding cassette, sub-family D (ALD), member 3 | protein-coding |
| chr3_121874415_121874936 | chr3 | 121874416 | 121874936 | + | 0.984107 | Intergenic | Intergenic | -59461 | NM_008991 | 19299 | Mm.399042 | NM_008991 | ENSMUSG00000028127 | **Abcd3** | AI313901\|AU018866\|AW146054\|PMP68\|PMP70\|Pxmp1 | ATP-binding cassette, sub-family D (ALD), member 3 | protein-coding |
| chr3_121796866_121798265 | chr3 | 121796867 | 121798265 | + | 0.830606 | intron (NM_008991, intron 1 of 22) | RLTR11A\|LTR\|ERVK | 17649 | NM_008991 | 19299 | Mm.399042 | NM_008991 | ENSMUSG00000028127 | **Abcd3** | AI313901\|AU018866\|AW146054\|PMP68\|PMP70\|Pxmp1 | ATP-binding cassette, sub-family D (ALD), member 3 | protein-coding |

| | | | | | | | | | | | | | | | | | |
|---|---|---|---|---|---|---|---|---|---|---|---|---|---|---|---|---|---|
| chr3_121844012_121844605 | chr3 | 121844013 | 121844605 | + | 0.727783 | Intergenic | MIRb\|SINE\|MIR | -29094 | NM_008991 | 19299 | Mm.399042 | NM_008991 | ENSMUSG00000028127 | **Abcd3** | AI313901\|AU018866\|AW146054\|PMP68\|PMP70\|Pxmp1 | ATP-binding cassette, sub-family D (ALD), member 3 | protein-coding |
| chr8_79678921_79679368 | chr8 | 79678922 | 79679368 | + | 1.72763 | Intergenic | Intergenic | 32595 | NM_015751 | 24015 | Mm.5831 | NM_015751 | ENSMUSG00000058355 | **Abce1** | C79080\|Oabp\|RLI\|RNS41\|Rnaseli | ATP-binding cassette, sub-family E (OABP), member 1 | protein-coding |
| chr6_58624176_58625054 | chr6 | 58624177 | 58625054 | + | 0.679221 | intron (NM_011920, intron 1 of 15) | ORR1E\|LTR\|ERVL-MaLR | 27943 | NM_011920 | 26357 | Mm.333096 | NM_011920 | ENSMUSG00000029802 | **Abcg2** | ABC15\|ABCP\|AI428558\|BCRP\|Bcrp1\|MXR\|MXR1 | ATP-binding cassette, sub-family G (WHITE), member 2 | protein-coding |
| chr5_105004383_105004786 | chr5 | 105004384 | 105004786 | + | 1.28647 | Intergenic | Intergenic | -21868 | NM_030239 | 27405 | Mm.389663 | NM_030239 | ENSMUSG00000029299 | **Abcg3** | Abcp2\|Mxr2 | ATP-binding cassette, sub-family G (WHITE), member 3 | protein-coding |
| chr5_104946589_104947086 | chr5 | 104946590 | 104947086 | + | 1.00727 | intron (NM_030239, intron 13 of 15) | Lx8b\|LINE\|L1 | 35879 | NM_030239 | 27405 | Mm.389663 | NM_030239 | ENSMUSG00000029299 | **Abcg3** | Abcp2\|Mxr2 | ATP-binding cassette, sub-family G (WHITE), member 3 | protein-coding |
| chr5_105010390_105010815 | chr5 | 105010391 | 105010815 | + | 0.934303 | Intergenic | Intergenic | -27886 | NM_030239 | 27405 | Mm.389663 | NM_030239 | ENSMUSG00000029299 | **Abcg3** | Abcp2\|Mxr2 | ATP-binding cassette, sub-family G (WHITE), member 3 | protein-coding |
| chr5_105010154_105010913 | chr5 | 105010155 | 105010913 | + | 0.926529 | Intergenic | Intergenic | -27817 | NM_030239 | 27405 | Mm.389663 | NM_030239 | ENSMUSG00000029299 | **Abcg3** | Abcp2\|Mxr2 | ATP-binding cassette, sub-family G (WHITE), member 3 | protein-coding |
| chr5_105005088_105005495 | chr5 | 105005089 | 105005495 | + | 0.710655 | Intergenic | Intergenic | -22575 | NM_030239 | 27405 | Mm.389663 | NM_030239 | ENSMUSG00000029299 | **Abcg3** | Abcp2\|Mxr2 | ATP-binding cassette, sub-family G (WHITE), member 3 | protein-coding |
| chr5_104938639_104939044 | chr5 | 104938640 | 104939044 | + | 0.702663 | intron (NM_030239, intron 14 of 15) | intron (NM_030239, intron 14 of 15) | 43875 | NM_030239 | 27405 | Mm.389663 | NM_030239 | ENSMUSG00000029299 | **Abcg3** | Abcp2\|Mxr2 | ATP-binding cassette, sub-family G (WHITE), member 3 | protein-coding |
| chr5_104960312_104960797 | chr5 | 104960313 | 104960797 | + | 0.675605 | intron (NM_030239, intron 10 of 15) | L1MB3\|LINE\|L1 | 22162 | NM_030239 | 27405 | Mm.389663 | NM_030239 | ENSMUSG00000029299 | **Abcg3** | Abcp2\|Mxr2 | ATP-binding cassette, sub-family G (WHITE), member 3 | protein-coding |
| chr16_45758441_45759189 | chr16 | 45758442 | 45759189 | + | 0.753372 | intron (NM_153412, intron 13 of 17) | intron (NM_153412, intron 13 of 17) | -15860 | NM_172511 | 213012 | Mm.247453 | NM_172511 | ENSMUSG00000033157 | **Abhd10** | - | abhydrolase domain containing 10 | protein-coding |
| chr11_77531980_77532754 | chr11 | 77531981 | 77532754 | + | 0.701507 | 3' UTR (NM_144825, exon 20 of 20) | 3' UTR (NM_144825, exon 20 of 20) | 17250 | NM_026185 | 67477 | Mm.432526 | NM_026185 | ENSMUSG00000000686 | **Abhd15** | 1300007F04Rik | abhydrolase domain containing 15 | protein-coding |
| chr7_84148879_84149456 | chr7 | 84148880 | 84149456 | + | 1.11961 | intron (NM_133722, intron 1 of 2) | intron (NM_133722, intron 1 of 2) | 2725 | NM_133722 | 70178 | Mm.23914 | NM_133722 | ENSMUSG00000038459 | **Abhd17c** | 2210412D01Rik\|AL023007\|Fam108c\|Fam108c1 | abhydrolase domain containing 17C | protein-coding |
| chr7_84158760_84159118 | chr7 | 84158761 | 84159118 | + | 0.754213 | Intergenic | Intergenic | -7046 | NM_133722 | 70178 | Mm.23914 | NM_133722 | ENSMUSG00000038459 | **Abhd17c** | 2210412D01Rik\|AL023007\|Fam108c\|Fam108c1 | abhydrolase domain containing 17C | protein-coding |
| chr2_23048002_23048534 | chr2 | 23048003 | 23048534 | + | 1.25742 | Intergenic | ORR1B1\|LTR\|ERVL-MaLR | -8027 | NM_145994 | 11308 | Mm.205647 | NM_007380 | ENSMUSG00000058835 | **Abi1** | E3B1\|NAP1\|Ssh3bp1 | abl-interactor 1 | protein-coding |
| chr2_23047993_23048859 | chr2 | 23047994 | 23048859 | + | 1.00969 | Intergenic | (TC)n\|Simple_repeat\|Simple_repeat | -8185 | NM_145994 | 11308 | Mm.205647 | NM_007380 | ENSMUSG00000058835 | **Abi1** | E3B1\|NAP1\|Ssh3bp1 | abl-interactor 1 | protein-coding |
| chr2_23005336_23005655 | chr2 | 23005337 | 23005655 | + | 0.865465 | intron (NM_145994, intron 1 of 9) | MLTR25A\|LTR\|ERVK | 34745 | NM_001077192 | 11308 | Mm.205647 | NM_007380 | ENSMUSG00000058835 | **Abi1** | E3B1\|NAP1\|Ssh3bp1 | abl-interactor 1 | protein-coding |
| chr2_23005436_23005655 | chr2 | 23005437 | 23005655 | + | 0.736719 | intron (NM_145994, intron 1 of 9) | MLTR25A\|LTR\|ERVK | 34695 | NM_001077192 | 11308 | Mm.205647 | NM_007380 | ENSMUSG00000058835 | **Abi1** | E3B1\|NAP1\|Ssh3bp1 | abl-interactor 1 | protein-coding |
| chr2_23006128_23006817 | chr2 | 23006129 | 23006817 | + | 0.649551 | intron (NM_145994, intron 1 of 9) | Lx8\|LINE\|L1 | 33768 | NM_001077192 | 11308 | Mm.205647 | NM_007380 | ENSMUSG00000058835 | **Abi1** | E3B1\|NAP1\|Ssh3bp1 | abl-interactor 1 | protein-coding |
| chr16_56556139_56556528 | chr16 | 56556140 | 56556528 | + | 0.617384 | intron (NM_001014424, intron 2 of 36) | intron (NM_001014424, intron 2 of 36) | 78488 | NM_001014424 | 320712 | Mm.209236 | NM_178790 | ENSMUSG00000035258 | **Abi3bp** | 5033411B22Rik\|AI506287\|BG172926\|D930038M13Rik\|TARSH\|eratin | ABI gene family, member 3 (NESH) binding protein | protein-coding |
| chr19_57167996_57168571 | chr19 | 57167997 | 57168571 | + | 0.948892 | intron (NM_001290813, intron 2 of 20) | intron (NM_001290813, intron 2 of 20) | 29347 | NM_001103177 | 226251 | Mm.217161 | NM_178688 | ENSMUSG00000025085 | **Ablim1** | 2210411C18Rik\|2610209L21Rik\|4833406P10Rik\|9330196J19Rik\|AV079770\|AW060987\|AW215784\|Limab1\|mKIAA0059 | actin-binding LIM protein 1 | protein-coding |
| chr19_57167610_57168575 | chr19 | 57167611 | 57168575 | + | 0.808724 | intron (NM_001290813, intron 2 of 20) | intron (NM_001290813, intron 2 of 20) | 29538 | NM_001103177 | 226251 | Mm.217161 | NM_178688 | ENSMUSG00000025085 | **Ablim1** | 2210411C18Rik\|2610209L21Rik\|4833406P10Rik\|9330196J19Rik\|AV079770\|AW060987\|AW215784\|Limab1\|mKIAA0059 | actin-binding LIM protein 1 | protein-coding |
| chr19_57082309_57082785 | chr19 | 57082310 | 57082785 | + | 0.724404 | intron (NM_001290813, intron 7 of 20) | intron (NM_001290813, intron 7 of 20) | 36477 | NM_001103178 | 226251 | Mm.217161 | NM_178688 | ENSMUSG00000025085 | **Ablim1** | 2210411C18Rik\|2610209L21Rik\|4833406P10Rik\|9330196J19Rik\|AV079770\|AW060987\|AW215784\|Limab1\|mKIAA0059 | actin-binding LIM protein 1 | protein-coding |
| chr5_35817283_35817678 | chr5 | 35817284 | 35817678 | + | 1.05339 | intron (NM_001177700, intron 6 of 16) | intron (NM_001177700, intron 6 of 16) | 59601 | NM_177678 | 231148 | Mm.254446 | NM_177678 | ENSMUSG00000029095 | **Ablim2** | AI606905\|C230091L11 | actin-binding LIM protein 2 | protein-coding |
| chr5_35817896_35818308 | chr5 | 35817897 | 35818308 | + | 0.775459 | intron (NM_001177700, intron 6 of 16) | intron (NM_001177700, intron 6 of 16) | 60222 | NM_177678 | 231148 | Mm.254446 | NM_177678 | ENSMUSG00000029095 | **Ablim2** | AI606905\|C230091L11 | actin-binding LIM protein 2 | protein-coding |
| chr5_35817882_35818434 | chr5 | 35817883 | 35818434 | + | 0.758953 | intron (NM_001177700, intron 6 of 16) | intron (NM_001177700, intron 6 of 16) | 60278 | NM_177678 | 231148 | Mm.254446 | NM_177678 | ENSMUSG00000029095 | **Ablim2** | AI606905\|C230091L11 | actin-binding LIM protein 2 | protein-coding |
| chr11_76572810_76573289 | chr11 | 76572811 | 76573289 | + | 0.797275 | intron (NM_198894, intron 1 of 22) | intron (NM_198894, intron 1 of 22) | -1666 | NM_198895 | 109934 | Mm.258939 | NM_198018 | ENSMUSG00000017631 | **Abr** | - | active BCR-related gene | protein-coding |
| chr11_76483967_76484483 | chr11 | 76483968 | 76484483 | + | 0.643925 | intron (NM_198894, intron 3 of 22) | MIR\|SINE\|MIR | -15760 | NM_001291186 | 109934 | Mm.258939 | NM_198018 | ENSMUSG00000017631 | **Abr** | - | active BCR-related gene | protein-coding |
| chr15_41914471_41915122 | chr15 | 41914472 | 41915122 | + | 1.20885 | Intergenic | RLTR19\|LTR\|ERVK | -45077 | NM_175456 | 223513 | Mm.148913 | NM_175456 | ENSMUSG00000042895 | **Abra** | C130068O12Rik\|STARS | actin-binding Rho activating protein | protein-coding |
| chr15_42167429_42167849 | chr15 | 42167430 | 42167849 | + | 1.04366 | Intergenic | Intergenic | -297919 | NM_175456 | 223513 | Mm.148913 | NM_175456 | ENSMUSG00000042895 | **Abra** | C130068O12Rik\|STARS | actin-binding Rho activating protein | protein-coding |
| chr15_41905030_41905403 | chr15 | 41905031 | 41905403 | + | 0.954614 | Intergenic | Intergenic | -35497 | NM_175456 | 223513 | Mm.148913 | NM_175456 | ENSMUSG00000042895 | **Abra** | C130068O12Rik\|STARS | actin-binding Rho activating protein | protein-coding |
| chr13_23426902_23427219 | chr13 | 23426903 | 23427219 | + | 1.08561 | TTS (NR_015492) | TTS (NR_015492) | -3195 | NM_013924 | 30946 | Mm.292094 | NM_013924 | | **Abt1** | - | activator of basal transcription 1 | protein-coding |
| chr13_23412353_23413162 | chr13 | 23412354 | 23413162 | + | 1.03556 | Intergenic | Intergenic | 11108 | NM_013924 | 30946 | Mm.292094 | NM_013924 | | **Abt1** | - | activator of basal transcription 1 | protein-coding |
| chr13_23396992_23397269 | chr13 | 23396993 | 23397269 | + | 0.859247 | Intergenic | tRNA-Ile-ATT\|tRNA\|tRNA | 26735 | NM_013924 | 30946 | Mm.292094 | NM_013924 | | **Abt1** | - | activator of basal transcription 1 | protein-coding |
| chr13_23425505_23425792 | chr13 | 23425506 | 23425792 | + | 0.810445 | Intergenic | tRNA-Tyr-TAC\|tRNA\|tRNA | -1783 | NM_013924 | 30946 | Mm.292094 | NM_013924 | | **Abt1** | - | activator of basal transcription 1 | protein-coding |
| chr13_23426463_23426698 | chr13 | 23426464 | 23426698 | + | 0.806457 | Intergenic | tRNA-Tyr-TAC\|tRNA\|tRNA | -2715 | NM_013924 | 30946 | Mm.292094 | NM_013924 | | **Abt1** | - | activator of basal transcription 1 | protein-coding |
| chr2_103657592_103658154 | chr2 | 103657593 | 103658154 | + | 1.52674 | intron (NM_178890, intron 1 of 16) | intron (NM_178890, intron 1 of 16) | 91563 | NM_178890 | 99382 | Mm.35850 | NM_178890 | ENSMUSG00000032724 | **Abtb2** | AW539457\|BPOZ-2 | ankyrin repeat and BTB (POZ) domain containing 2 | protein-coding |
| chr2_103568774_103569127 | chr2 | 103568775 | 103569127 | + | 1.45879 | intron (NM_178890, intron 1 of 16) | MER21B\|LTR\|ERVL | 2641 | NM_178890 | 99382 | Mm.35850 | NM_178890 | ENSMUSG00000032724 | **Abtb2** | AW539457\|BPOZ-2 | ankyrin repeat and BTB (POZ) domain containing 2 | protein-coding |
| chr2_103655531_103656357 | chr2 | 103655532 | 103656357 | + | 0.997735 | intron (NM_178890, intron 1 of 16) | intron (NM_178890, intron 1 of 16) | 89634 | NM_178890 | 99382 | Mm.35850 | NM_178890 | ENSMUSG00000032724 | **Abtb2** | AW539457\|BPOZ-2 | ankyrin repeat and BTB (POZ) domain containing 2 | protein-coding |
| chr2_103642428_103642932 | chr2 | 103642429 | 103642932 | + | 0.912381 | intron (NM_178890, intron 1 of 16) | Tigger7\|DNA\|TcMar-Tigger | 76370 | NM_178890 | 99382 | Mm.35850 | NM_178890 | ENSMUSG00000032724 | **Abtb2** | AW539457\|BPOZ-2 | ankyrin repeat and BTB (POZ) domain containing 2 | protein-coding |
| chr2_103569725_103570135 | chr2 | 103569726 | 103570135 | + | 0.901172 | intron (NM_178890, intron 1 of 16) | intron (NM_178890, intron 1 of 16) | 3620 | NM_178890 | 99382 | Mm.35850 | NM_178890 | ENSMUSG00000032724 | **Abtb2** | AW539457\|BPOZ-2 | ankyrin repeat and BTB (POZ) domain containing 2 | protein-coding |
| chr2_103561014_103561305 | chr2 | 103561015 | 103561305 | + | 0.884779 | Intergenic | RLTR53_Mm\|LTR\|ERVK | -5150 | NM_178890 | 99382 | Mm.35850 | NM_178890 | ENSMUSG00000032724 | **Abtb2** | AW539457\|BPOZ-2 | ankyrin repeat and BTB (POZ) domain containing 2 | protein-coding |
| chr2_103595008_103595323 | chr2 | 103595009 | 103595323 | + | 0.754242 | intron (NM_178890, intron 1 of 16) | intron (NM_178890, intron 1 of 16) | 28856 | NM_178890 | 99382 | Mm.35850 | NM_178890 | ENSMUSG00000032724 | **Abtb2** | AW539457\|BPOZ-2 | ankyrin repeat and BTB (POZ) domain containing 2 | protein-coding |
| chr2_103640910_103641132 | chr2 | 103640911 | 103641132 | + | 0.652806 | intron (NM_178890, intron 1 of 16) | intron (NM_178890, intron 1 of 16) | 74711 | NM_178890 | 99382 | Mm.35850 | NM_178890 | ENSMUSG00000032724 | **Abtb2** | AW539457\|BPOZ-2 | ankyrin repeat and BTB (POZ) domain containing 2 | protein-coding |
| chr2_103607094_103607378 | chr2 | 103607095 | 103607378 | + | 0.624717 | intron (NM_178890, intron 1 of 16) | intron (NM_178890, intron 1 of 16) | 40926 | NM_178890 | 99382 | Mm.35850 | NM_178890 | ENSMUSG00000032724 | **Abtb2** | AW539457\|BPOZ-2 | ankyrin repeat and BTB (POZ) domain containing 2 | protein-coding |
| chr18_74786239_74786704 | chr18 | 74786240 | 74786704 | + | 1.20222 | intron (NM_177470, intron 1 of 9) | intron (NM_177470, intron 1 of 9) | 7260 | NM_177470 | 52538 | Mm.245724 | NM_177470 | ENSMUSG00000036880 | **Acaa2** | 0610011L04Rik\|AI255831\|AI265397\|D18Ertd240e | acetyl-Coenzyme A acyltransferase 2 (mitochondrial 3-oxoacyl-Coenzyme A thiolase) | protein-coding |
| chr18_74787138_74787424 | chr18 | 74787139 | 74787424 | + | 0.827372 | exon (NM_177470, exon 2 of 10) | exon (NM_177470, exon 2 of 10) | 8069 | NM_177470 | 52538 | Mm.245724 | NM_177470 | ENSMUSG00000036880 | **Acaa2** | 0610011L04Rik\|AI255831\|AI265397\|D18Ertd240e | acetyl-Coenzyme A acyltransferase 2 (mitochondrial 3-oxoacyl-Coenzyme A thiolase) | protein-coding |
| chr5_114175514_114176245 | chr5 | 114175515 | 114176245 | + | 0.766199 | intron (NM_133904, intron 1 of 51) | intron (NM_133904, intron 1 of 51) | 10362 | NM_133904 | 100705 | Mm.81793 | NM_133904 | ENSMUSG00000042010 | **Acacb** | AI597064\|AW743042\|Acc2\|Accb | acetyl-Coenzyme A carboxylase beta | protein-coding |

| | | | | | | | | | | | | | | | | | |
|---|---|---|---|---|---|---|---|---|---|---|---|---|---|---|---|---|---|
| chr5_121623248_121623627 | chr5 | 121623249 | 121623627 | + | 0.72914 | intron (NM_028037, intron 18 of 20) | PB1D9\|SINE\|Alu | -4500 | NM_178799 | 338350 | Mm.482918 | NM_178799 | ENSMUSG00000042647 | **Acad12** | 9330129D05Rik | acyl-Coenzyme A dehydrogenase family, member 12 | protein-coding |
| chr1_66878402_66878755 | chr1 | 66878403 | 66878755 | + | 0.61856 | Intergenic | ORR1E\|LTR\|ERVL-MaLR | -15270 | NM_007381 | 11363 | Mm.2445 | NM_007381 | ENSMUSG00000026003 | **Acadl** | AA960361\|AU018452\|C79855\|LCAD | acyl-Coenzyme A dehydrogenase, long-chain | protein-coding |
| chr3_153957669_153958396 | chr3 | 153957670 | 153958396 | + | 1.22213 | Intergenic | RLTR48A\|LTR\|ERV1 | -13390 | NM_007382 | 11364 | Mm.10530 | NM_007382 | ENSMUSG00000062908 | **Acadm** | AU018656\|MCAD | acyl-Coenzyme A dehydrogenase, medium chain | protein-coding |
| chr16_31233572_31234467 | chr16 | 31233573 | 31234467 | + | 0.938693 | intron (NR_035427, intron 1 of 2) | intron (NR_035427, intron 1 of 2) | -32782 | NM_030138 | 78618 | Mm.274646 | NM_030138 | ENSMUSG00000049076 | **Acap2** | 4832442G16\|9530039J15Rik\|CNT-B2\|Centb2\|mKIAA0041 | ArfGAP with coiled-coil, ankyrin repeat and PH domains 2 | protein-coding |
| chr16_31235422_31235855 | chr16 | 31235423 | 31235855 | + | 0.83729 | intron (NR_035427, intron 1 of 2) | intron (NR_035427, intron 1 of 2) | -34401 | NM_030138 | 78618 | Mm.274646 | NM_030138 | ENSMUSG00000049076 | **Acap2** | 4832442G16\|9530039J15Rik\|CNT-B2\|Centb2\|mKIAA0041 | ArfGAP with coiled-coil, ankyrin repeat and PH domains 2 | protein-coding |
| chr16_31235488_31235766 | chr16 | 31235489 | 31235766 | + | 0.836983 | intron (NR_035427, intron 1 of 2) | intron (NR_035427, intron 1 of 2) | -34389 | NM_030138 | 78618 | Mm.274646 | NM_030138 | ENSMUSG00000049076 | **Acap2** | 4832442G16\|9530039J15Rik\|CNT-B2\|Centb2\|mKIAA0041 | ArfGAP with coiled-coil, ankyrin repeat and PH domains 2 | protein-coding |
| chr1_180765443_180765799 | chr1 | 180765444 | 180765799 | + | 0.937782 | Intergenic | Intergenic | 39578 | NM_133225 | 170760 | Mm.272981 | NM_133225 | ENSMUSG00000026499 | **Acbd3** | 60kDa\|8430407O11Rik\|D1Ertd10e\|GCP60\|GOLPH1\|Gocap1\|Pap7 | acyl-Coenzyme A binding domain containing 3 | protein-coding |
| chr2_23088744_23089229 | chr2 | 23088745 | 23089229 | + | 1.31136 | intron (NM_028793, intron 6 of 12) | intron (NM_028793, intron 6 of 12) | 19915 | NM_001102438 | 74159 | Mm.181973 | NM_028793 | ENSMUSG00000026781 | **Acbd5** | 1300014E15Rik | acyl-Coenzyme A binding domain containing 5 | protein-coding |
| chr1_155510215_155510739 | chr1 | 155510216 | 155510739 | + | 0.996413 | Intergenic | Intergenic | -47643 | NM_001145781 | 72482 | Mm.288249 | NM_026683 | ENSMUSG00000033701 | **Acbd6** | 0610010G04Rik\|2610100E10Rik | acyl-Coenzyme A binding domain containing 6 | protein-coding |
| chr1_155647605_155647882 | chr1 | 155647606 | 155647882 | + | 0.895602 | intron (NM_028250, intron 6 of 7) | intron (NM_028250, intron 6 of 7) | 89624 | NM_026683 | 72482 | Mm.288249 | NM_026683 | ENSMUSG00000033701 | **Acbd6** | 0610010G04Rik\|2610100E10Rik | acyl-Coenzyme A binding domain containing 6 | protein-coding |
| chr2_3339021_3339559 | chr2 | 3339022 | 3339559 | + | 1.07223 | intron (NM_030063, intron 1 of 3) | intron (NM_030063, intron 1 of 3) | 3122 | NM_030063 | 78245 | Mm.99733 | NM_030063 | ENSMUSG00000026644 | **Acbd7** | 9230116B18Rik | acyl-Coenzyme A binding domain containing 7 | protein-coding |
| chr11_100524356_100524681 | chr11 | 100524357 | 100524681 | + | 0.955775 | intron (NM_134037, intron 1 of 27) | intron (NM_134037, intron 1 of 27) | 3481 | NM_134037 | 104112 | Mm.282039 | NM_134037 | ENSMUSG00000020917 | **Acly** | A730098H14Rik\|AW538652 | ATP citrate lyase | protein-coding |
| chr4_40171244_40171794 | chr4 | 40171245 | 40171794 | + | 0.865132 | intron (NM_007386, intron 4 of 20) | URR1A\|DNA\|hAT-Charlie | 28254 | NM_007386 | 11428 | Mm.331547 | NM_007386 | ENSMUSG00000028405 | **Aco1** | Aco-1\|Irebp\|Irp1 | aconitase 1 | protein-coding |
| chr15_81875625_81876013 | chr15 | 81875626 | 81876013 | + | 0.858926 | intron (NM_080633, intron 1 of 17) | intron (NM_080633, intron 1 of 17) | 3356 | NM_080633 | 11429 | Mm.154581 | NM_080633 | ENSMUSG00000022477 | **Aco2** | Aco-2\|Aco3\|D10Wsu183e | aconitase 2, mitochondrial | protein-coding |
| chr15_19858681_19859021 | chr15 | 19858682 | 19859021 | + | 0.873031 | Intergenic | Intergenic | 807899 | NM_022816 | 64833 | Mm.352763 | NM_022816 | ENSMUSG00000047565 | **Acot10** | Acate3\|MT-ACT48\|p48 | acyl-CoA thioesterase 10 | protein-coding |
| chr13_91663343_91663887 | chr13 | 91663344 | 91663887 | + | 1.42706 | intron (NM_024272, intron 5 of 16) | intron (NM_024272, intron 5 of 16) | -77906 | NM_028790 | 74156 | Mm.275963 | NM_028790 | ENSMUSG00000021620 | **Acot12** | 1300004O04Rik\|4930449F15Rik\|AV027244\|Cach\|mCACH-1 | acyl-CoA thioesterase 12 | protein-coding |
| chr13_91681108_91681416 | chr13 | 91681109 | 91681416 | + | 0.767227 | intron (NM_024272, intron 10 of 16) | intron (NM_024272, intron 10 of 16) | -60259 | NM_028790 | 74156 | Mm.275963 | NM_028790 | ENSMUSG00000021620 | **Acot12** | 1300004O04Rik\|4930449F15Rik\|AV027244\|Cach\|mCACH-1 | acyl-CoA thioesterase 12 | protein-coding |
| chr13_91680919_91681423 | chr13 | 91680920 | 91681423 | + | 0.643844 | intron (NM_024272, intron 10 of 16) | intron (NM_024272, intron 10 of 16) | -60350 | NM_028790 | 74156 | Mm.275963 | NM_028790 | ENSMUSG00000021620 | **Acot12** | 1300004O04Rik\|4930449F15Rik\|AV027244\|Cach\|mCACH-1 | acyl-CoA thioesterase 12 | protein-coding |
| chr13_24829015_24829284 | chr13 | 24829016 | 24829284 | + | 1.22283 | intron (NM_025790, intron 1 of 2) | intron (NM_025790, intron 1 of 2) | 2339 | NM_025790 | 66834 | Mm.2125 | NM_025790 | ENSMUSG00000006717 | **Acot13** | 0610006O17Rik\|Them2 | acyl-CoA thioesterase 13 | protein-coding |
| chr4_152185123_152186185 | chr4 | 152185124 | 152186185 | + | 0.822974 | promoter-TSS (NM_001146057) | promoter-TSS (NM_001146057) | -480 | NM_001146057 | 70025 | Mm.296191 | NM_133348 | ENSMUSG00000028937 | **Acot7** | 2410041A17Rik\|Ach1\|Act\|Bach\|CTE-IIa\|Cte-II\|Lach1 | acyl-CoA thioesterase 7 | protein-coding |
| chrX_155278442_155278852 | chrX | 155278443 | 155278852 | + | 1.07259 | intron (NM_019736, intron 5 of 14) | intron (NM_019736, intron 5 of 14) | 16147 | NM_019736 | 56360 | Mm.268710 | NM_019736 | ENSMUSG00000025287 | **Acot9** | 0610041P13Rik\|Acate2\|C76421\|MT-ACT48\|U8\|p48 | acyl-CoA thioesterase 9 | protein-coding |
| chr2_127929859_127930232 | chr2 | 127929860 | 127930232 | + | 0.743633 | intron (NM_028765, intron 9 of 17) | intron (NM_028765, intron 9 of 17) | 75418 | NM_028765 | 74121 | Mm.244835 | NM_028765 | ENSMUSG00000027380 | **Acoxl** | 1200014P05Rik\|AV025673 | acyl-Coenzyme A oxidase-like | protein-coding |
| chr8_46469245_46469445 | chr8 | 46469246 | 46469445 | + | 0.784036 | Intergenic | Intergenic | -1697 | NM_007981 | 14081 | Mm.210323 | NM_007981 | ENSMUSG00000018796 | **Acsl1** | Acas\|Acas1\|Acs\|FACS\|Facl2 | acyl-CoA synthetase long-chain family member 1 | protein-coding |
| chr8_46511101_46511710 | chr8 | 46511102 | 46511710 | + | 0.769154 | exon (NM_007981, exon 5 of 21) | exon (NM_007981, exon 5 of 21) | 40364 | NM_007981 | 14081 | Mm.210323 | NM_007981 | ENSMUSG00000018796 | **Acsl1** | Acas\|Acas1\|Acs\|FACS\|Facl2 | acyl-CoA synthetase long-chain family member 1 | protein-coding |
| chr8_46511313_46511668 | chr8 | 46511314 | 46511668 | + | 0.709688 | intron (NM_007981, intron 5 of 20) | intron (NM_007981, intron 5 of 20) | 40449 | NM_007981 | 14081 | Mm.210323 | NM_007981 | ENSMUSG00000018796 | **Acsl1** | Acas\|Acas1\|Acs\|FACS\|Facl2 | acyl-CoA synthetase long-chain family member 1 | protein-coding |
| chrX_142489667_142490738 | chrX | 142489668 | 142490738 | + | 1.3564 | Intergenic | Intergenic | -99668 | NM_019477 | 50790 | Mm.391337 | NM_019477 | ENSMUSG00000031278 | **Acsl4** | 9430020A05Rik\|ACS4\|AU018108\|Facl4\|Lacs4 | acyl-CoA synthetase long-chain family member 4 | protein-coding |
| chrX_142472954_142473276 | chrX | 142472955 | 142473276 | + | 0.930736 | Intergenic | Intergenic | -82580 | NM_019477 | 50790 | Mm.391337 | NM_019477 | ENSMUSG00000031278 | **Acsl4** | 9430020A05Rik\|ACS4\|AU018108\|Facl4\|Lacs4 | acyl-CoA synthetase long-chain family member 4 | protein-coding |
| chrX_142488906_142489231 | chrX | 142488907 | 142489231 | + | 0.775998 | Intergenic | Intergenic | -98534 | NM_019477 | 50790 | Mm.391337 | NM_019477 | ENSMUSG00000031278 | **Acsl4** | 9430020A05Rik\|ACS4\|AU018108\|Facl4\|Lacs4 | acyl-CoA synthetase long-chain family member 4 | protein-coding |
| chr11_54293993_54294964 | chr11 | 54293994 | 54294964 | + | 1.29572 | Intergenic | Intergenic | -9345 | NM_001033598 | 216739 | Mm.267478 | NM_144823 | ENSMUSG00000020333 | **Acsl6** | A330035H04Rik\|AW050338\|Facl6\|LACS\|Lacsl\|mKIAA0837 | acyl-CoA synthetase long-chain family member 6 | protein-coding |
| chr11_54287771_54288723 | chr11 | 54287772 | 54288723 | + | 0.82927 | Intergenic | Intergenic | -15577 | NM_001033598 | 216739 | Mm.267478 | NM_144823 | ENSMUSG00000020333 | **Acsl6** | A330035H04Rik\|AW050338\|Facl6\|LACS\|Lacsl\|mKIAA0837 | acyl-CoA synthetase long-chain family member 6 | protein-coding |
| chr11_54287828_54288253 | chr11 | 54287829 | 54288253 | + | 0.666788 | Intergenic | Intergenic | -15783 | NM_001033598 | 216739 | Mm.267478 | NM_144823 | ENSMUSG00000020333 | **Acsl6** | A330035H04Rik\|AW050338\|Facl6\|LACS\|Lacsl\|mKIAA0837 | acyl-CoA synthetase long-chain family member 6 | protein-coding |
| chr2_155562834_155563347 | chr2 | 155562835 | 155563347 | + | 0.973643 | TTS (NM_019811) | TTS (NM_019811) | -6161 | NR_040613 | 74945 | Mm.401956 | NR_040613 | ENSMUSG00000086529 | **Acss2os** | 4930471I20Rik | acyl-CoA synthetase short-chain family member 2, opposite strand | ncRNA |
| chr2_155570697_155571097 | chr2 | 155570698 | 155571097 | + | 0.864958 | intron (NM_008180, intron 9 of 12) | intron (NM_008180, intron 9 of 12) | -13967 | NR_040613 | 74945 | Mm.401956 | NR_040613 | ENSMUSG00000086529 | **Acss2os** | 4930471I20Rik | acyl-CoA synthetase short-chain family member 2, opposite strand | ncRNA |
| chr2_155562824_155563727 | chr2 | 155562825 | 155563727 | + | 0.863848 | TTS (NM_019811) | TTS (NM_019811) | -6346 | NR_040613 | 74945 | Mm.401956 | NR_040613 | ENSMUSG00000086529 | **Acss2os** | 4930471I20Rik | acyl-CoA synthetase short-chain family member 2, opposite strand | ncRNA |
| chr5_142930928_142931231 | chr5 | 142930929 | 142931231 | + | 1.22666 | Intergenic | Intergenic | -24356 | NM_007393 | 11461 | Mm.328431 | NM_007393 | ENSMUSG00000029580 | **Actb** | Actx\|E430023M04Rik\|beta-actin | actin, beta | protein-coding |
| chr5_142932075_142932637 | chr5 | 142932076 | 142932637 | + | 0.902946 | Intergenic | Intergenic | -25632 | NM_007393 | 11461 | Mm.328431 | NM_007393 | ENSMUSG00000029580 | **Actb** | Actx\|E430023M04Rik\|beta-actin | actin, beta | protein-coding |
| chr5_142931790_142932652 | chr5 | 142931791 | 142932652 | + | 0.890933 | Intergenic | B3\|SINE\|B2 | -25497 | NM_007393 | 11461 | Mm.328431 | NM_007393 | ENSMUSG00000029580 | **Actb** | Actx\|E430023M04Rik\|beta-actin | actin, beta | protein-coding |
| chr13_111248973_111249667 | chr13 | 111248974 | 111249667 | + | 1.97568 | Intergenic | Intergenic | -5693 | NM_175497 | 238880 | Mm.107293 | NM_175497 | ENSMUSG00000055194 | **Actbl2** | 4732495G21Rik | actin, beta-like 2 | protein-coding |
| chr13_111265620_111266048 | chr13 | 111265621 | 111266048 | + | 1.64477 | Intergenic | Intergenic | 10821 | NM_175497 | 238880 | Mm.107293 | NM_175497 | ENSMUSG00000055194 | **Actbl2** | 4732495G21Rik | actin, beta-like 2 | protein-coding |
| chr13_111265283_111266085 | chr13 | 111265284 | 111266085 | + | 1.61607 | Intergenic | Intergenic | 10671 | NM_175497 | 238880 | Mm.107293 | NM_175497 | ENSMUSG00000055194 | **Actbl2** | 4732495G21Rik | actin, beta-like 2 | protein-coding |
| chr13_111247535_111247831 | chr13 | 111247536 | 111247831 | + | 0.966244 | Intergenic | Intergenic | -7330 | NM_175497 | 238880 | Mm.107293 | NM_175497 | ENSMUSG00000055194 | **Actbl2** | 4732495G21Rik | actin, beta-like 2 | protein-coding |
| chr13_111258046_111259188 | chr13 | 111258047 | 111259188 | + | 0.900289 | TTS (NM_175497) | TTS (NM_175497) | 3604 | NM_175497 | 238880 | Mm.107293 | NM_175497 | ENSMUSG00000055194 | **Actbl2** | 4732495G21Rik | actin, beta-like 2 | protein-coding |
| chr13_111269656_111269969 | chr13 | 111269657 | 111269969 | + | 0.772844 | Intergenic | Intergenic | 14800 | NM_175497 | 238880 | Mm.107293 | NM_175497 | ENSMUSG00000055194 | **Actbl2** | 4732495G21Rik | actin, beta-like 2 | protein-coding |
| chr4_56673892_56674222 | chr4 | 56673893 | 56674222 | + | 0.705624 | Intergenic | Intergenic | 67368 | NM_025271 | 11471 | Mm.251434 | NM_025271 | ENSMUSG00000070980 | **Actl7b** | Tact1 | actin-like 7b | protein-coding |
| chr12_80269251_80269656 | chr12 | 80269252 | 80269656 | + | 1.1385 | Intergenic | ID_B1\|SINE\|B4 | -9083 | NM_134156 | 109711 | Mm.253564 | NM_134156 | ENSMUSG00000015143 | **Actn1** | 3110023F10Rik\|Actn1a | actinin, alpha 1 | protein-coding |
| chr12_80239951_80240315 | chr12 | 80239952 | 80240315 | + | 0.912545 | intron (NM_134156, intron 1 of 20) | intron (NM_134156, intron 1 of 20) | 20238 | NM_134156 | 109711 | Mm.253564 | NM_134156 | ENSMUSG00000015143 | **Actn1** | 3110023F10Rik\|Actn1a | actinin, alpha 1 | protein-coding |
| chr12_80269743_80270037 | chr12 | 80269744 | 80270037 | + | 0.852227 | Intergenic | MTD\|LTR\|ERVL-MaLR | -9519 | NM_134156 | 109711 | Mm.253564 | NM_134156 | ENSMUSG00000015143 | **Actn1** | 3110023F10Rik\|Actn1a | actinin, alpha 1 | protein-coding |
| chr7_28930660_28931623 | chr7 | 28930661 | 28931623 | + | 1.43394 | intron (NM_021895, intron 1 of 20) | intron (NM_021895, intron 1 of 20) | 31138 | NM_021895 | 60595 | Mm.81144 | NM_021895 | ENSMUSG00000054808 | **Actn4** | C77391 | actinin alpha 4 | protein-coding |
| chr7_28949112_28949770 | chr7 | 28949113 | 28949770 | + | 1.36356 | intron (NM_021895, intron 1 of 20) | intron (NM_021895, intron 1 of 20) | 12839 | NM_021895 | 60595 | Mm.81144 | NM_021895 | ENSMUSG00000054808 | **Actn4** | C77391 | actinin alpha 4 | protein-coding |
| chr7_28930611_28931996 | chr7 | 28930612 | 28931996 | + | 1.30361 | intron (NM_021895, intron 1 of 20) | intron (NM_021895, intron 1 of 20) | 30976 | NM_021895 | 60595 | Mm.81144 | NM_021895 | ENSMUSG00000054808 | **Actn4** | C77391 | actinin alpha 4 | protein-coding |
| chr7_28945415_28945756 | chr7 | 28945416 | 28945756 | + | 0.724715 | intron (NM_021895, intron 1 of 20) | intron (NM_021895, intron 1 of 20) | 16694 | NM_021895 | 60595 | Mm.81144 | NM_021895 | ENSMUSG00000054808 | **Actn4** | C77391 | actinin alpha 4 | protein-coding |
| chr1_125389625_125390412 | chr1 | 125389626 | 125390412 | + | 0.789266 | Intergenic | Intergenic | 45708 | NM_023735 | 74117 | Mm.183102 | NM_023735 | ENSMUSG00000026341 | **Actr3** | 1200003A09Rik\|Arp3 | ARP3 actin-related protein 3 | protein-coding |
| chr1_125389625_125390713 | chr1 | 125389626 | 125390713 | + | 0.757472 | Intergenic | Intergenic | 45558 | NM_023735 | 74117 | Mm.183102 | NM_023735 | ENSMUSG00000026341 | **Actr3** | 1200003A09Rik\|Arp3 | ARP3 actin-related protein 3 | protein-coding |
| chr1_125450273_125450799 | chr1 | 125450274 | 125450799 | + | 0.64386 | Intergenic | Intergenic | -14809 | NM_001205386 | 74117 | Mm.183102 | NM_023735 | ENSMUSG00000026341 | **Actr3** | 1200003A09Rik\|Arp3 | ARP3 actin-related protein 3 | protein-coding |
| chr2_48327386_48327975 | chr2 | 48327387 | 48327975 | + | 0.947252 | Intergenic | Intergenic | -486428 | NM_007396 | 11480 | Mm.314338 | NM_007396 | ENSMUSG00000052155 | **Acvr2a** | ActrIIa\|Acvr2\|TactrII | activin receptor IIA | protein-coding |
| chr11_30676114_30676543 | chr11 | 30676115 | 30676543 | + | 0.787511 | Intergenic | MamRep1527\|LTR\|LTR | -26933 | NM_029344 | 75572 | Mm.390319 | NM_029344 | | **Acyp2** | 2310004B09Rik | acylphosphatase 2, muscle type | protein-coding |
| chr9_70688278_70688605 | chr9 | 70688279 | 70688605 | + | 0.939351 | intron (NM_007399, intron 1 of 15) | B1_Mur4\|SINE\|Alu | 9441 | NM_007399 | 11487 | Mm.3037 | NM_007399 | ENSMUSG00000054693 | **Adam10** | 1700031C13Rik\|MADM\|kuz\|kuzbanian | a disintegrin and metallopeptidase domain 10 | protein-coding |
| chr12_21370353_21370625 | chr12 | 21370354 | 21370625 | + | 1.01181 | intron (NR_120376, intron 1 of 18) | intron (NR_120376, intron 1 of 18) | 3143 | NR_102380 | 11491 | Mm.27681 | NM_009615 | ENSMUSG00000052593 | **Adam17** | CD156b\|Tace | a disintegrin and metallopeptidase domain 17 | protein-coding |
| chr11_46073492_46074133 | chr11 | 46073493 | 46074133 | + | 0.828233 | intron (NM_009616, intron 3 of 22) | MLT1A0\|LTR\|ERVL-MaLR | 17829 | NM_001291891 | 11492 | Mm.89940 | NM_009616 | ENSMUSG00000011256 | **Adam19** | AL024287\|Mltnb | a disintegrin and metallopeptidase domain 19 (meltrin beta) | protein-coding |
| chr5_8374245_8374802 | chr5 | 8374246 | 8374802 | + | 0.894187 | Intergenic | L1MC\|LINE\|L1 | -6443 | NM_001098225 | 11496 | Mm.275895 | NM_001007220 | ENSMUSG00000040537 | **Adam22** | 2900022I03Rik\|AI854032\|MDC2 | a disintegrin and metallopeptidase domain 22 | protein-coding |
| chr3_98187331_98188218 | chr3 | 98187332 | 98188218 | + | 1.24598 | Intergenic | Lx8\|LINE\|L1 | 26963 | NM_027665 | 71078 | Mm.50711 | NM_027665 | ENSMUSG00000043468 | **Adam30** | 4933424D07Rik\|svph4 | a disintegrin and metallopeptidase domain 30 | protein-coding |

| | | | | | | | | | | | | | | | | | |
|---|---|---|---|---|---|---|---|---|---|---|---|---|---|---|---|---|---|
| chr3_98187314_98188537 | chr3 | 98187315 | 98188537 | + | 1.15457 | Intergenic | MER21B\|LTR\|ERVL | 27114 | NM_027665 | 71078 | Mm.50711 | NM_027665 | ENSMUSG00000043468 | **Adam30** | 4933424D07Rik\|svph4 | a disintegrin and metallopeptidase domain 30 | protein-coding |
| chr12_113362283_113363079 | chr12 | 113362284 | 113363079 | + | 1.06124 | Intergenic | Intergenic | -126884 | NM_001009545 | 238405 | Mm.442216 | NM_001009545 | ENSMUSG00000051804 | **Adam6b** | 4930523C11Rik | a disintegrin and metallopeptidase domain 6B | protein-coding |
| chr12_113462956_113463277 | chr12 | 113462957 | 113463277 | + | 0.747041 | Intergenic | Intergenic | -26448 | NM_001009545 | 238405 | Mm.442216 | NM_001009545 | ENSMUSG00000051804 | **Adam6b** | 4930523C11Rik | a disintegrin and metallopeptidase domain 6B | protein-coding |
| chr17_33537326_33538049 | chr17 | 33537327 | 33538049 | + | 0.746808 | intron (NR_037707, intron 7 of 24) | intron (NR_037707, intron 7 of 24) | 13492 | NR_037708 | 224697 | Mm.29304 | NM_172619 | ENSMUSG00000024299 | **Adamts10** | 9430006O18\|AW045948\|Adam-ts10\|Adamts-10\|Znmp | a disintegrin-like and metallopeptidase (reprolysin type) with thrombospondin type 1 motif, 10 | protein-coding |
| chr10_61281106_61281593 | chr10 | 61281107 | 61281593 | + | 0.649544 | Intergenic | Intergenic | -7912 | NM_001081127 | 237360 | Mm.333252 | NM_001081127 | ENSMUSG00000059901 | **Adamts14** | Adamts-14\|TS14 | a disintegrin-like and metallopeptidase (reprolysin type) with thrombospondin type 1 motif, 14 | protein-coding |
| chr5_89812497_89813094 | chr5 | 89812498 | 89813094 | + | 0.820409 | intron (NM_177872, intron 3 of 21) | Lx7\|LINE\|L1 | 70538 | NM_177872 | 330119 | Mm.133062 | NM_177872 | ENSMUSG00000043635 | **Adamts3** | 1100001H14Rik\|6330442E02Rik | a disintegrin-like and metallopeptidase (reprolysin type) with thrombospondin type 1 motif, 3 | protein-coding |
| chr13_104259001_104259310 | chr13 | 104259002 | 104259310 | + | 0.741582 | Intergenic | MTE-int\|LTR\|ERVL-MaLR | -28717 | NM_001081020 | 108154 | Mm.208125 | NM_175496 | ENSMUSG00000046169 | **Adamts6** | 5031426K13\|A930019D11Rik\|ADAM-TS6\|b2b1879.1Clo\|b2b2029Clo\|b2b2182Clo\|b2b2187.1Clo\|b2b2228Clo | a disintegrin-like and metallopeptidase (reprolysin type) with thrombospondin type 1 motif, 6 | protein-coding |
| chr9_90165118_90165609 | chr9 | 90165119 | 90165609 | + | 0.799972 | intron (NM_001003911, intron 1 of 23) | intron (NM_001003911, intron 1 of 23) | 2386 | NM_001003911 | 108153 | Mm.102141 | NM_001003911 | ENSMUSG00000032363 | **Adamts7** | ADAM-TS7\|ADAMTS7B | a disintegrin-like and metallopeptidase (reprolysin type) with thrombospondin type 1 motif, 7 | protein-coding |
| chr5_139300379_139300897 | chr5 | 139300380 | 139300897 | + | 1.00651 | intron (NM_172723, intron 3 of 10) | intron (NM_172723, intron 3 of 10) | 24826 | NM_172723 | 231821 | Mm.297819 | NM_172723 | ENSMUSG00000056413 | **Adap1** | 4930431P11Rik\|Centa1\|GC1L\|p42IP4 | ArfGAP with dual PH domains 1 | protein-coding |
| chr3_89686009_89686504 | chr3 | 89686010 | 89686504 | + | 1.07894 | Intergenic | Intergenic | -28765 | NM_001038587 | 56417 | Mm.316628 | NM_019655 | ENSMUSG00000027951 | **Adar** | AV242451\|Adar1\|Adar1p110\|Adar1p150\|mZaADAR | adenosine deaminase, RNA-specific | protein-coding |
| chr10_77387273_77387979 | chr10 | 77387274 | 77387979 | + | 1.03928 | intron (NR_004429, intron 1 of 10) | intron (NR_004429, intron 1 of 10) | 30647 | NM_130895 | 110532 | Mm.276815 | NM_130895 | ENSMUSG00000020262 | **Adarb1** | 1700057H01Rik\|AW124433\|AW558573\|Adar2\|BB220382\|D10Bwg0447e\|Red1 | adenosine deaminase, RNA-specific, B1 | protein-coding |
| chr10_77366842_77367640 | chr10 | 77366843 | 77367640 | + | 0.743264 | intron (NR_004429, intron 1 of 10) | intron (NR_004429, intron 1 of 10) | 51032 | NM_130895 | 110532 | Mm.276815 | NM_130895 | ENSMUSG00000020262 | **Adarb1** | 1700057H01Rik\|AW124433\|AW558573\|Adar2\|BB220382\|D10Bwg0447e\|Red1 | adenosine deaminase, RNA-specific, B1 | protein-coding |
| chr10_77366540_77366777 | chr10 | 77366541 | 77366777 | + | 0.738887 | intron (NR_004429, intron 1 of 10) | (TGG)n\|Simple_repeat\|Simple_repeat | 51614 | NM_130895 | 110532 | Mm.276815 | NM_130895 | ENSMUSG00000020262 | **Adarb1** | 1700057H01Rik\|AW124433\|AW558573\|Adar2\|BB220382\|D10Bwg0447e\|Red1 | adenosine deaminase, RNA-specific, B1 | protein-coding |
| chr10_77415890_77416103 | chr10 | 77415891 | 77416103 | + | 0.734715 | intron (NR_004429, intron 1 of 10) | intron (NR_004429, intron 1 of 10) | 2276 | NM_130895 | 110532 | Mm.276815 | NM_130895 | ENSMUSG00000020262 | **Adarb1** | 1700057H01Rik\|AW124433\|AW558573\|Adar2\|BB220382\|D10Bwg0447e\|Red1 | adenosine deaminase, RNA-specific, B1 | protein-coding |
| chr13_8479024_8479611 | chr13 | 8479025 | 8479611 | + | 1.04308 | intron (NM_001289530, intron 1 of 9) | intron (NM_001289530, intron 1 of 9) | 276452 | NM_052977 | 94191 | Mm.425794 | NM_052977 | ENSMUSG00000052551 | **Adarb2** | Adar3\|RED2 | adenosine deaminase, RNA-specific, B2 | protein-coding |
| chr13_8480294_8480761 | chr13 | 8480295 | 8480761 | + | 0.998946 | intron (NM_001289530, intron 1 of 9) | intron (NM_001289530, intron 1 of 9) | 277662 | NM_052977 | 94191 | Mm.425794 | NM_052977 | ENSMUSG00000052551 | **Adarb2** | Adar3\|RED2 | adenosine deaminase, RNA-specific, B2 | protein-coding |
| chr13_8479877_8480767 | chr13 | 8479878 | 8480767 | + | 0.974543 | intron (NM_001289530, intron 1 of 9) | intron (NM_001289530, intron 1 of 9) | 277456 | NM_052977 | 94191 | Mm.425794 | NM_052977 | ENSMUSG00000052551 | **Adarb2** | Adar3\|RED2 | adenosine deaminase, RNA-specific, B2 | protein-coding |
| chr1_180199153_180199902 | chr1 | 180199154 | 180199902 | + | 0.917231 | Intergenic | L2a\|LINE\|L2 | -3508 | NM_023341 | 67426 | Mm.38330 | NM_023341 | ENSMUSG00000026489 | **Adck3** | 4632432J16Rik\|AI462003\|Cabc1\|mKIAA0451 | aarF domain containing kinase 3 | protein-coding |
| chr15_98617183_98617937 | chr15 | 98617184 | 98617937 | + | 0.841162 | Intergenic | Intergenic | -9927 | NM_007405 | 11512 | Mm.157091 | NM_007405 | ENSMUSG00000022994 | **Adcy6** | AC6\|mKIAA0422 | adenylate cyclase 6 | protein-coding |
| chr8_88297912_88298417 | chr8 | 88297913 | 88298417 | + | 0.885008 | intron (NM_001037723, intron 3 of 27) | intron (NM_001037723, intron 3 of 27) | 3961 | NM_001037724 | 11513 | Mm.288206 | NM_007406 | ENSMUSG00000031659 | **Adcy7** | AA407758 | adenylate cyclase 7 | protein-coding |
| chr8_88297898_88298592 | chr8 | 88297899 | 88298592 | + | 0.855069 | intron (NM_001037723, intron 3 of 27) | intron (NM_001037723, intron 3 of 27) | 4041 | NM_001037724 | 11513 | Mm.288206 | NM_007406 | ENSMUSG00000031659 | **Adcy7** | AA407758 | adenylate cyclase 7 | protein-coding |
| chr8_88268396_88268714 | chr8 | 88268397 | 88268714 | + | 0.74691 | Intergenic | MTEa\|LTR\|ERVL-MaLR | -3848 | NM_007406 | 11513 | Mm.288206 | NM_007406 | ENSMUSG00000031659 | **Adcy7** | AA407758 | adenylate cyclase 7 | protein-coding |
| chr16_4469625_4470300 | chr16 | 4469626 | 4470300 | + | 0.631395 | Intergenic | Intergenic | -49465 | NM_009624 | 11515 | Mm.439750 | NM_009624 | ENSMUSG00000005580 | **Adcy9** | ACtp10\|AW125421\|D16Wsu65e\|mKIAA0520 | adenylate cyclase 9 | protein-coding |
| chr19_53199336_53199756 | chr19 | 53199337 | 53199756 | + | 1.06548 | intron (NM_013758, intron 1 of 15) | intron (NM_013758, intron 1 of 15) | 55549 | NM_013758 | 27360 | Mm.426080 | NM_013758 | ENSMUSG00000025026 | **Add3** | AI463285\|R75380 | adducin 3 (gamma) | protein-coding |
| chr19_53192232_53192416 | chr19 | 53192233 | 53192416 | + | 0.889327 | intron (NM_013758, intron 1 of 15) | intron (NM_013758, intron 1 of 15) | 48327 | NM_013758 | 27360 | Mm.426080 | NM_013758 | ENSMUSG00000025026 | **Add3** | AI463285\|R75380 | adducin 3 (gamma) | protein-coding |
| chr10_10224674_10224888 | chr10 | 10224675 | 10224888 | + | 0.937673 | Intergenic | RMER3D-int\|LTR\|ERVK | 247533 | NM_001127353 | 215772 | Mm.246573 | NM_172547 | ENSMUSG00000050994 | **Adgb** | 9130014G24Rik | androglobin | protein-coding |
| chr10_10224617_10225144 | chr10 | 10224618 | 10225144 | + | 0.849463 | Intergenic | RMER3D-int\|LTR\|ERVK | 247433 | NM_001127353 | 215772 | Mm.246573 | NM_172547 | ENSMUSG00000050994 | **Adgb** | 9130014G24Rik | androglobin | protein-coding |
| chr3_138420457_138420973 | chr3 | 138420458 | 138420973 | + | 2.09405 | intron (NM_011996, intron 4 of 8) | intron (NM_011996, intron 4 of 8) | 5218 | NM_011996 | 26876 | Mm.158750 | NM_011996 | ENSMUSG00000037797 | **Adh4** | Adh2 | alcohol dehydrogenase 4 (class II), pi polypeptide | protein-coding |
| chr3_138423633_138423923 | chr3 | 138423634 | 138423923 | + | 1.67859 | intron (NM_011996, intron 5 of 8) | intron (NM_011996, intron 5 of 8) | 8281 | NM_011996 | 26876 | Mm.158750 | NM_011996 | ENSMUSG00000037797 | **Adh4** | Adh2 | alcohol dehydrogenase 4 (class II), pi polypeptide | protein-coding |
| chr3_138423606_138424048 | chr3 | 138423607 | 138424048 | + | 1.43865 | intron (NM_011996, intron 5 of 8) | intron (NM_011996, intron 5 of 8) | 8330 | NM_011996 | 26876 | Mm.158750 | NM_011996 | ENSMUSG00000037797 | **Adh4** | Adh2 | alcohol dehydrogenase 4 (class II), pi polypeptide | protein-coding |
| chr3_138420127_138420389 | chr3 | 138420128 | 138420389 | + | 1.36205 | intron (NM_011996, intron 3 of 8) | intron (NM_011996, intron 3 of 8) | 4761 | NM_011996 | 26876 | Mm.158750 | NM_011996 | ENSMUSG00000037797 | **Adh4** | Adh2 | alcohol dehydrogenase 4 (class II), pi polypeptide | protein-coding |
| chr3_138419506_138419894 | chr3 | 138419507 | 138419894 | + | 1.03332 | intron (NM_011996, intron 3 of 8) | intron (NM_011996, intron 3 of 8) | 4203 | NM_011996 | 26876 | Mm.158750 | NM_011996 | ENSMUSG00000037797 | **Adh4** | Adh2 | alcohol dehydrogenase 4 (class II), pi polypeptide | protein-coding |
| chr1_9548718_9548970 | chr1 | 9548719 | 9548970 | + | 1.07305 | intron (NR_027664, intron 1 of 12) | intron (NR_027664, intron 1 of 12) | 798 | NR_027664 | 76187 | Mm.28514 | NM_175236 | ENSMUSG00000025911 | **Adhfe1** | 6330565B14Rik\|AI043035\|Adh8 | alcohol dehydrogenase, iron containing, 1 | protein-coding |
| chr6_119390692_119391011 | chr6 | 119390693 | 119391011 | + | 0.963076 | intron (NM_197985, intron 1 of 7) | intron (NM_197985, intron 1 of 7) | 26631 | NM_197985 | 68465 | Mm.291826 | NM_197985 | ENSMUSG00000030168 | **Adipor2** | 1110001I14Rik\|ADCR2\|AI115388\|AW554121\|D6Ucla1e\|Paqr2 | adiponectin receptor 2 | protein-coding |
| chr6_119384809_119385108 | chr6 | 119384810 | 119385108 | + | 0.737811 | intron (NM_197985, intron 1 of 7) | B4\|SINE\|B4 | 32524 | NM_197985 | 68465 | Mm.291826 | NM_197985 | ENSMUSG00000030168 | **Adipor2** | 1110001I14Rik\|ADCR2\|AI115388\|AW554121\|D6Ucla1e\|Paqr2 | adiponectin receptor 2 | protein-coding |
| chr2_168197501_168197958 | chr2 | 168197502 | 168197958 | + | 1.22734 | intron (NM_009628, intron 1 of 3) | intron (NM_009628, intron 1 of 3) | 9332 | NM_009628 | 11538 | Mm.201322 | NM_009628 | ENSMUSG00000051149 | **Adnp** | AA589558\|mKIAA0784 | activity-dependent neuroprotective protein | protein-coding |
| chr2_168191858_168192183 | chr2 | 168191859 | 168192183 | + | 0.860031 | intron (NM_009628, intron 1 of 3) | B1_Mur1\|SINE\|Alu | 15041 | NM_009628 | 11538 | Mm.201322 | NM_009628 | ENSMUSG00000051149 | **Adnp** | AA589558\|mKIAA0784 | activity-dependent neuroprotective protein | protein-coding |
| chr10_67655539_67656143 | chr10 | 67655540 | 67656143 | + | 1.32273 | Intergenic | Intergenic | -106886 | NM_001005419 | 211488 | Mm.297250 | NM_001005419 | ENSMUSG00000057134 | **Ado** | Gm237 | 2-aminoethanethiol (cysteamine) dioxygenase | protein-coding |
| chr10_67717411_67718224 | chr10 | 67717412 | 67718224 | + | 1.14773 | Intergenic | Intergenic | -168863 | NM_001005419 | 211488 | Mm.297250 | NM_001005419 | ENSMUSG00000057134 | **Ado** | Gm237 | 2-aminoethanethiol (cysteamine) dioxygenase | protein-coding |
| chr10_67646635_67647026 | chr10 | 67646636 | 67647026 | + | 1.11469 | Intergenic | Intergenic | -97876 | NM_001005419 | 211488 | Mm.297250 | NM_001005419 | ENSMUSG00000057134 | **Ado** | Gm237 | 2-aminoethanethiol (cysteamine) dioxygenase | protein-coding |
| chr10_67651431_67651949 | chr10 | 67651432 | 67651949 | + | 1.10325 | Intergenic | Intergenic | -102735 | NM_001005419 | 211488 | Mm.297250 | NM_001005419 | ENSMUSG00000057134 | **Ado** | Gm237 | 2-aminoethanethiol (cysteamine) dioxygenase | protein-coding |
| chr10_67657734_67658508 | chr10 | 67657735 | 67658508 | + | 1.05672 | Intergenic | Intergenic | -109166 | NM_001005419 | 211488 | Mm.297250 | NM_001005419 | ENSMUSG00000057134 | **Ado** | Gm237 | 2-aminoethanethiol (cysteamine) dioxygenase | protein-coding |
| chr10_67598487_67599251 | chr10 | 67598488 | 67599251 | + | 1.04669 | Intergenic | Intergenic | -49914 | NM_001005419 | 211488 | Mm.297250 | NM_001005419 | ENSMUSG00000057134 | **Ado** | Gm237 | 2-aminoethanethiol (cysteamine) dioxygenase | protein-coding |
| chr10_67696351_67696818 | chr10 | 67696352 | 67696818 | + | 1.02746 | Intergenic | Intergenic | -147630 | NM_001005419 | 211488 | Mm.297250 | NM_001005419 | ENSMUSG00000057134 | **Ado** | Gm237 | 2-aminoethanethiol (cysteamine) dioxygenase | protein-coding |
| chr10_67651646_67651942 | chr10 | 67651647 | 67651942 | + | 1.02308 | Intergenic | Intergenic | -102839 | NM_001005419 | 211488 | Mm.297250 | NM_001005419 | ENSMUSG00000057134 | **Ado** | Gm237 | 2-aminoethanethiol (cysteamine) dioxygenase | protein-coding |
| chr10_67696028_67696847 | chr10 | 67696029 | 67696847 | + | 1.0007 | Intergenic | Intergenic | -147483 | NM_001005419 | 211488 | Mm.297250 | NM_001005419 | ENSMUSG00000057134 | **Ado** | Gm237 | 2-aminoethanethiol (cysteamine) dioxygenase | protein-coding |
| chr10_67606852_67607064 | chr10 | 67606853 | 67607064 | + | 0.643012 | Intergenic | Intergenic | -58003 | NM_001005419 | 211488 | Mm.297250 | NM_001005419 | ENSMUSG00000057134 | **Ado** | Gm237 | 2-aminoethanethiol (cysteamine) dioxygenase | protein-coding |
| chr10_75289778_75290222 | chr10 | 75289779 | 75290222 | + | 0.857796 | intron (NM_153406, intron 13 of 15) | intron (NM_153406, intron 13 of 15) | -26877 | NM_009630 | 11540 | Mm.333734 | NM_009630 | ENSMUSG00000020178 | **Adora2a** | A2AAR\|A2aR\|AA2AR | adenosine A2a receptor | protein-coding |
| chr11_62184925_62185486 | chr11 | 62184926 | 62185486 | + | 1.66923 | intron (NM_001281818, intron 12 of 14) | intron (NM_001281818, intron 12 of 14) | -63778 | NM_007413 | 11541 | Mm.40740 | NM_007413 | ENSMUSG00000018500 | **Adora2b** | A2BAR\|A2BR\|A2b\|AA2BR\|AI480866\|AI605384 | adenosine A2b receptor | protein-coding |

| | | | | | | | | | | | | | | | | | |
|---|---|---|---|---|---|---|---|---|---|---|---|---|---|---|---|---|---|
| chr11_62223798_62224479 | chr11 | 62223799 | 62224479 | + | 1.01939 | Intergenic | MLT1A0\|LTR\|ERVL-MaLR | -24845 | NM_007413 | 11541 | Mm.40740 | NM_007413 | ENSMUSG00000018500 | **Adora2b** | A2BAR\|A2BR\|A2b\|AA2BR\|AI480866\|AI605384 | adenosine A2b receptor | protein-coding |
| chr11_62186633_62187574 | chr11 | 62186634 | 62187574 | + | 0.942622 | intron (NM_001281818, intron 12 of 14) | intron (NM_001281818, intron 12 of 14) | -61880 | NM_007413 | 11541 | Mm.40740 | NM_007413 | ENSMUSG00000018500 | **Adora2b** | A2BAR\|A2BR\|A2b\|AA2BR\|AI480866\|AI605384 | adenosine A2b receptor | protein-coding |
| chr11_62217585_62218160 | chr11 | 62217586 | 62218160 | + | 0.824 | intron (NM_001281818, intron 14 of 14) | intron (NM_001281818, intron 14 of 14) | -31111 | NM_007413 | 11541 | Mm.40740 | NM_007413 | ENSMUSG00000018500 | **Adora2b** | A2BAR\|A2BR\|A2b\|AA2BR\|AI480866\|AI605384 | adenosine A2b receptor | protein-coding |
| chr9_59308276_59309044 | chr9 | 59308277 | 59309044 | + | 0.703216 | intron (NM_028121, intron 3 of 6) | intron (NM_028121, intron 3 of 6) | 17088 | NM_028121 | 72141 | Mm.22456 | NM_028121 | ENSMUSG00000025236 | **Adpgk** | 2610017G09Rik\|Adp-gk\|C77577 | ADP-dependent glucokinase | protein-coding |
| chr16_38449788_38450143 | chr16 | 38449789 | 38450143 | + | 1.24962 | intron (NM_007414, intron 2 of 3) | PB1D9\|SINE\|Alu | 2723 | NM_007414 | 11544 | Mm.20047 | NM_007414 | ENSMUSG00000002844 | **Adprh** | Arh1 | ADP-ribosylarginine hydrolase | protein-coding |
| chr8_13263231_13263775 | chr8 | 13263232 | 13263775 | + | 1.20481 | intron (NM_001024504, intron 4 of 6) | intron (NM_001024504, intron 4 of 6) | -9341 | NM_172750 | 234072 | Mm.132785 | NM_172750 | ENSMUSG00000031448 | **Adprhl1** | Arh2\|D330008N11Rik | ADP-ribosylhydrolase like 1 | protein-coding |
| chr11_67048522_67049098 | chr11 | 67048523 | 67049098 | + | 1.50573 | intron (NM_025510, intron 2 of 4) | intron (NM_025510, intron 2 of 4) | 3808 | NM_025510 | 66358 | Mm.22511 | NM_025510 | ENSMUSG00000020910 | **Adprm** | 2310004I24Rik\|MDS006 | ADP-ribose/CDP-alcohol diphosphatase, manganese dependent | protein-coding |
| chr11_67049277_67049574 | chr11 | 67049278 | 67049574 | + | 0.856823 | intron (NM_025510, intron 2 of 4) | B3A\|SINE\|B2 | 3192 | NM_025510 | 66358 | Mm.22511 | NM_025510 | ENSMUSG00000020910 | **Adprm** | 2310004I24Rik\|MDS006 | ADP-ribose/CDP-alcohol diphosphatase, manganese dependent | protein-coding |
| chr18_62162972_62163435 | chr18 | 62162973 | 62163435 | + | 1.15998 | Intergenic | Intergenic | 16777 | NM_007420 | 11555 | Mm.5598 | NM_007420 | ENSMUSG00000045730 | **Adrb2** | Adrb-2\|Badm\|Gpcr7 | adrenergic receptor, beta 2 | protein-coding |
| chr18_62169160_62169562 | chr18 | 62169161 | 62169562 | + | 1.08526 | Intergenic | Intergenic | 10620 | NM_007420 | 11555 | Mm.5598 | NM_007420 | ENSMUSG00000045730 | **Adrb2** | Adrb-2\|Badm\|Gpcr7 | adrenergic receptor, beta 2 | protein-coding |
| chr18_62169022_62169737 | chr18 | 62169023 | 62169737 | + | 1.02332 | Intergenic | Intergenic | 10601 | NM_007420 | 11555 | Mm.5598 | NM_007420 | ENSMUSG00000045730 | **Adrb2** | Adrb-2\|Badm\|Gpcr7 | adrenergic receptor, beta 2 | protein-coding |
| chr18_62122275_62122752 | chr18 | 62122276 | 62122752 | + | 0.646261 | Intergenic | L1ME3A\|LINE\|L1 | 57467 | NM_007420 | 11555 | Mm.5598 | NM_007420 | ENSMUSG00000045730 | **Adrb2** | Adrb-2\|Badm\|Gpcr7 | adrenergic receptor, beta 2 | protein-coding |
| chr19_4300884_4301058 | chr19 | 4300885 | 4301058 | + | 0.902639 | intron (NM_001290818, intron 1 of 20) | intron (NM_001290818, intron 1 of 20) | 5251 | NM_130863 | 110355 | Mm.254144 | NM_130863 | ENSMUSG00000024858 | **Adrbk1** | Adrbk-1\|Bark-1\|GRK2\|beta ARK\|betaARK1 | adrenergic receptor kinase, beta 1 | protein-coding |
| chr1_177792915_177793531 | chr1 | 177792916 | 177793531 | + | 1.05755 | intron (NM_007422, intron 1 of 12) | intron (NM_007422, intron 1 of 12) | 3286 | NM_007422 | 11566 | Mm.338021 | NM_007422 | ENSMUSG00000015961 | **Adss** | AI314886\|AS\|Adss2 | adenylosuccinate synthetase, non muscle | protein-coding |
| chr12_112617190_112617598 | chr12 | 112617191 | 112617598 | + | 0.760164 | Intergenic | Intergenic | -2653 | NM_007421 | 11565 | Mm.3440 | NM_007421 | ENSMUSG00000011148 | **Adssl1** | AI528595\|Adss\|Adss1 | adenylosuccinate synthetase like 1 | protein-coding |
| chr13_41761911_41762355 | chr13 | 41761912 | 41762355 | + | 0.894739 | Intergenic | Intergenic | 85355 | NM_175417 | 109254 | Mm.25195 | NM_175417 | ENSMUSG00000058022 | **Adtrp** | 9530008L14Rik | androgen dependent TFPI regulating protein | protein-coding |
| chr13_41824101_41824766 | chr13 | 41824102 | 41824766 | + | 0.814854 | intron (NM_175417, intron 2 of 6) | intron (NM_175417, intron 2 of 6) | 23054 | NM_175417 | 109254 | Mm.25195 | NM_175417 | ENSMUSG00000058022 | **Adtrp** | 9530008L14Rik | androgen dependent TFPI regulating protein | protein-coding |
| chr13_41803913_41804271 | chr13 | 41803914 | 41804271 | + | 0.687544 | intron (NM_175417, intron 4 of 6) | intron (NM_175417, intron 4 of 6) | 43396 | NM_175417 | 109254 | Mm.25195 | NM_175417 | ENSMUSG00000058022 | **Adtrp** | 9530008L14Rik | androgen dependent TFPI regulating protein | protein-coding |
| chr6_140620575_140621259 | chr6 | 140620576 | 140621259 | + | 1.4027 | Intergenic | LTR88b\|LTR\|Gypsy? | -1746 | NM_178803 | 11569 | Mm.86453 | NM_009637 | ENSMUSG00000030232 | **Aebp2** | AU023766\|B230313N05Rik | AE binding protein 2 | protein-coding |
| chr6_140613614_140614177 | chr6 | 140613615 | 140614177 | + | 1.30885 | Intergenic | RMER15\|LTR\|ERVL | -8767 | NM_178803 | 11569 | Mm.86453 | NM_009637 | ENSMUSG00000030232 | **Aebp2** | AU023766\|B230313N05Rik | AE binding protein 2 | protein-coding |
| chr6_140722553_140723300 | chr6 | 140722554 | 140723300 | + | 0.888875 | Intergenic | Intergenic | 98629 | NM_009637 | 11569 | Mm.86453 | NM_009637 | ENSMUSG00000030232 | **Aebp2** | AU023766\|B230313N05Rik | AE binding protein 2 | protein-coding |
| chr6_140801073_140801486 | chr6 | 140801074 | 140801486 | + | 0.825666 | Intergenic | Intergenic | 176982 | NM_009637 | 11569 | Mm.86453 | NM_009637 | ENSMUSG00000030232 | **Aebp2** | AU023766\|B230313N05Rik | AE binding protein 2 | protein-coding |
| chr6_140750307_140750722 | chr6 | 140750308 | 140750722 | + | 0.811289 | Intergenic | Intergenic | 126217 | NM_009637 | 11569 | Mm.86453 | NM_009637 | ENSMUSG00000030232 | **Aebp2** | AU023766\|B230313N05Rik | AE binding protein 2 | protein-coding |
| chr6_140661920_140662318 | chr6 | 140661921 | 140662318 | + | 0.705202 | intron (NM_009637, intron 8 of 8) | MTEa\|LTR\|ERVL-MaLR | 37821 | NM_009637 | 11569 | Mm.86453 | NM_009637 | ENSMUSG00000030232 | **Aebp2** | AU023766\|B230313N05Rik | AE binding protein 2 | protein-coding |
| chr6_140598517_140599330 | chr6 | 140598518 | 140599330 | + | 0.663854 | Intergenic | B3A\|SINE\|B2 | -23739 | NM_178803 | 11569 | Mm.86453 | NM_009637 | ENSMUSG00000030232 | **Aebp2** | AU023766\|B230313N05Rik | AE binding protein 2 | protein-coding |
| chr6_140598523_140598960 | chr6 | 140598524 | 140598960 | + | 0.641514 | Intergenic | Intergenic | -23921 | NM_178803 | 11569 | Mm.86453 | NM_009637 | ENSMUSG00000030232 | **Aebp2** | AU023766\|B230313N05Rik | AE binding protein 2 | protein-coding |
| chr18_61834690_61835343 | chr18 | 61834691 | 61835343 | + | 0.914505 | intron (NM_001164491, intron 8 of 23) | intron (NM_001164491, intron 8 of 23) | -48355 | NM_178928 | 106877 | Mm.26632 | NM_178928 | ENSMUSG00000033032 | **Afap1l1** | AI173486 | actin filament associated protein 1-like 1 | protein-coding |
| chr19_57014828_57015150 | chr19 | 57014829 | 57015150 | + | 1.09335 | Intergenic | Intergenic | -6414 | NM_001177797 | 226250 | Mm.226284 | NM_146102 | ENSMUSG00000025083 | **Afap1l2** | AU041783\|C86904\|mKIAA1914 | actin filament associated protein 1-like 2 | protein-coding |
| chr19_57055556_57056168 | chr19 | 57055557 | 57056168 | + | 0.847805 | intron (NM_001290813, intron 12 of 20) | URR1B\|DNA\|hAT-Charlie | -47287 | NM_001177797 | 226250 | Mm.226284 | NM_146102 | ENSMUSG00000025083 | **Afap1l2** | AU041783\|C86904\|mKIAA1914 | actin filament associated protein 1-like 2 | protein-coding |
| chr19_57013196_57013818 | chr19 | 57013197 | 57013818 | + | 0.82227 | Intergenic | Intergenic | -4932 | NM_001177797 | 226250 | Mm.226284 | NM_146102 | ENSMUSG00000025083 | **Afap1l2** | AU041783\|C86904\|mKIAA1914 | actin filament associated protein 1-like 2 | protein-coding |
| chr5_103806114_103806688 | chr5 | 103806115 | 103806688 | + | 0.753475 | intron (NM_133919, intron 3 of 19) | intron (NM_133919, intron 3 of 19) | 51828 | NM_001080798 | 17355 | Mm.6949 | NM_133919 | ENSMUSG00000029313 | **Aff1** | 9630032B01Rik\|AW319193\|Af4\|Mllt2h\|Rob | AF4/FMR2 family, member 1 | protein-coding |
| chr1_38647267_38647879 | chr1 | 38647268 | 38647879 | + | 1.43828 | Intergenic | Intergenic | -20329 | NM_001290814 | 16764 | Mm.336679 | NM_010678 | ENSMUSG00000037138 | **Aff3** | 3222402O04Rik\|A730046J16\|LAF-4\|Laf4 | AF4/FMR2 family, member 3 | protein-coding |
| chr1_38723306_38724277 | chr1 | 38723307 | 38724277 | + | 1.39468 | Intergenic | MER74A\|LTR\|ERVL | -96548 | NM_001290814 | 16764 | Mm.336679 | NM_010678 | ENSMUSG00000037138 | **Aff3** | 3222402O04Rik\|A730046J16\|LAF-4\|Laf4 | AF4/FMR2 family, member 3 | protein-coding |
| chr1_38658395_38658731 | chr1 | 38658396 | 38658731 | + | 1.25292 | Intergenic | Intergenic | -31319 | NM_001290814 | 16764 | Mm.336679 | NM_010678 | ENSMUSG00000037138 | **Aff3** | 3222402O04Rik\|A730046J16\|LAF-4\|Laf4 | AF4/FMR2 family, member 3 | protein-coding |
| chr1_38437729_38438153 | chr1 | 38437730 | 38438153 | + | 1.17532 | intron (NM_001290814, intron 4 of 21) | intron (NM_001290814, intron 4 of 21) | 189303 | NM_001290814 | 16764 | Mm.336679 | NM_010678 | ENSMUSG00000037138 | **Aff3** | 3222402O04Rik\|A730046J16\|LAF-4\|Laf4 | AF4/FMR2 family, member 3 | protein-coding |
| chr1_38597507_38598140 | chr1 | 38597508 | 38598140 | + | 0.776099 | intron (NM_001290814, intron 1 of 21) | intron (NM_001290814, intron 1 of 21) | 29420 | NM_001290814 | 16764 | Mm.336679 | NM_010678 | ENSMUSG00000037138 | **Aff3** | 3222402O04Rik\|A730046J16\|LAF-4\|Laf4 | AF4/FMR2 family, member 3 | protein-coding |
| chr8_123478626_123479279 | chr8 | 123478627 | 123479279 | + | 0.856835 | intron (NM_054070, intron 1 of 16) | intron (NM_054070, intron 1 of 16) | 1091 | NR_028260 | 114896 | Mm.287475 | NM_054070 | ENSMUSG00000031967 | **Afg3l1** | 1700047G05Rik\|3110061K15Rik | AFG3-like AAA ATPase 1 | protein-coding |
| chr8_123478875_123479273 | chr8 | 123478876 | 123479273 | + | 0.829504 | intron (NM_054070, intron 1 of 16) | intron (NM_054070, intron 1 of 16) | 1212 | NR_028260 | 114896 | Mm.287475 | NM_054070 | ENSMUSG00000031967 | **Afg3l1** | 1700047G05Rik\|3110061K15Rik | AFG3-like AAA ATPase 1 | protein-coding |
| chr5_90504687_90505226 | chr5 | 90504688 | 90505226 | + | 0.822644 | exon (NM_007423, exon 11 of 15) | exon (NM_007423, exon 11 of 15) | -13992 | NM_145146 | 280662 | Mm.348786 | NM_145146 | ENSMUSG00000029369 | **Afm** | Alf\|alpha-Alb | afamin | protein-coding |
| chr5_90512332_90512596 | chr5 | 90512333 | 90512596 | + | 0.760095 | Intergenic | Intergenic | -6485 | NM_145146 | 280662 | Mm.348786 | NM_145146 | ENSMUSG00000029369 | **Afm** | Alf\|alpha-Alb | afamin | protein-coding |
| chr5_90512318_90512793 | chr5 | 90512319 | 90512793 | + | 0.733975 | Intergenic | Intergenic | -6393 | NM_145146 | 280662 | Mm.348786 | NM_145146 | ENSMUSG00000029369 | **Afm** | Alf\|alpha-Alb | afamin | protein-coding |
| chr11_117825360_117826013 | chr11 | 117825361 | 117826013 | + | 0.684907 | promoter-TSS (NM_027827) | promoter-TSS (NM_027827) | -232 | NM_027827 | 71562 | Mm.273958 | NM_027827 | ENSMUSG00000017718 | **Afmid** | 9030621K19Rik\|Ammd\|FKF\|KFA\|Kf\|formylase | arylformamidase | protein-coding |
| chr11_20743400_20743672 | chr11 | 20743401 | 20743672 | + | 0.658906 | Intergenic | Intergenic | -1980 | NM_181411 | 216549 | Mm.356580 | NM_181411 | ENSMUSG00000049659 | **Aftph** | - | aftiphilin | protein-coding |
| chr7_76407884_76408603 | chr7 | 76407885 | 76408603 | + | 1.35595 | intron (NM_001199224, intron 6 of 22) | RMER15\|LTR\|ERVL | 178357 | NM_001199224 | 244071 | Mm.322829 | NM_172902 | ENSMUSG00000025754 | **Agbl1** | Ccp4\|D430020F16\|EG244071\|Nna1-l1 | ATP/GTP binding protein-like 1 | protein-coding |
| chr7_76146277_76146771 | chr7 | 76146278 | 76146771 | + | 1.19941 | Intergenic | Intergenic | -83363 | NM_001199224 | 244071 | Mm.322829 | NM_172902 | ENSMUSG00000025754 | **Agbl1** | Ccp4\|D430020F16\|EG244071\|Nna1-l1 | ATP/GTP binding protein-like 1 | protein-coding |
| chr7_76142346_76142858 | chr7 | 76142347 | 76142858 | + | 0.910025 | Intergenic | Intergenic | -87285 | NM_001199224 | 244071 | Mm.322829 | NM_172902 | ENSMUSG00000025754 | **Agbl1** | Ccp4\|D430020F16\|EG244071\|Nna1-l1 | ATP/GTP binding protein-like 1 | protein-coding |
| chr7_76047694_76048286 | chr7 | 76047695 | 76048286 | + | 0.861773 | Intergenic | Intergenic | -181897 | NM_001199224 | 244071 | Mm.322829 | NM_172902 | ENSMUSG00000025754 | **Agbl1** | Ccp4\|D430020F16\|EG244071\|Nna1-l1 | ATP/GTP binding protein-like 1 | protein-coding |
| chr1_82879187_82879600 | chr1 | 82879188 | 82879600 | + | 1.452 | intron (NM_010472, intron 5 of 12) | B4A\|SINE\|B4 | 39911 | NM_010472 | 15463 | Mm.392569 | NM_010472 | ENSMUSG00000026159 | **Agfg1** | AU045498\|C130049H11Rik\|C85612\|D730048C23Rik\|Hrb\|RAB\|Rip | ArfGAP with FG repeats 1 | protein-coding |
| chr1_82869235_82870086 | chr1 | 82869236 | 82870086 | + | 1.36006 | intron (NM_010472, intron 2 of 12) | L1M5\|LINE\|L1 | 30178 | NM_010472 | 15463 | Mm.392569 | NM_010472 | ENSMUSG00000026159 | **Agfg1** | AU045498\|C130049H11Rik\|C85612\|D730048C23Rik\|Hrb\|RAB\|Rip | ArfGAP with FG repeats 1 | protein-coding |
| chr1_82831527_82832251 | chr1 | 82831528 | 82832251 | + | 1.01169 | Intergenic | Intergenic | -7594 | NM_010472 | 15463 | Mm.392569 | NM_010472 | ENSMUSG00000026159 | **Agfg1** | AU045498\|C130049H11Rik\|C85612\|D730048C23Rik\|Hrb\|RAB\|Rip | ArfGAP with FG repeats 1 | protein-coding |
| chr1_82834525_82834910 | chr1 | 82834526 | 82834910 | + | 0.856004 | Intergenic | Intergenic | -4765 | NM_010472 | 15463 | Mm.392569 | NM_010472 | ENSMUSG00000026159 | **Agfg1** | AU045498\|C130049H11Rik\|C85612\|D730048C23Rik\|Hrb\|RAB\|Rip | ArfGAP with FG repeats 1 | protein-coding |
| chr1_82849799_82850092 | chr1 | 82849800 | 82850092 | + | 0.833629 | intron (NM_010472, intron 1 of 12) | B2_Mm2\|SINE\|B2 | 10463 | NM_010472 | 15463 | Mm.392569 | NM_010472 | ENSMUSG00000026159 | **Agfg1** | AU045498\|C130049H11Rik\|C85612\|D730048C23Rik\|Hrb\|RAB\|Rip | ArfGAP with FG repeats 1 | protein-coding |
| chr1_82820174_82820473 | chr1 | 82820175 | 82820473 | + | 0.645604 | Intergenic | Intergenic | -19159 | NM_010472 | 15463 | Mm.392569 | NM_010472 | ENSMUSG00000026159 | **Agfg1** | AU045498\|C130049H11Rik\|C85612\|D730048C23Rik\|Hrb\|RAB\|Rip | ArfGAP with FG repeats 1 | protein-coding |
| chr5_137671687_137672340 | chr5 | 137671688 | 137672340 | + | 0.940997 | intron (NM_145566, intron 1 of 4) | intron (NM_145566, intron 1 of 4) | 12603 | NM_178162 | 231801 | Mm.260869 | NM_145566 | ENSMUSG00000029722 | **Agfg2** | A630095P14Rik\|Hrbl\|RABR | ArfGAP with FG repeats 2 | protein-coding |
| chr4_141766075_141766751 | chr4 | 141766076 | 141766751 | + | 0.74366 | intron (NM_172338, intron 13 of 14) | intron (NM_172338, intron 13 of 14) | 19738 | NM_001081408 | 75986 | Mm.30032 | NM_001081408 | ENSMUSG00000040706 | **Agmat** | 5033405N08Rik | agmatine ureohydrolase (agmatinase) | protein-coding |

| | | | | | | | | | | | | | | | | | |
|---|---|---|---|---|---|---|---|---|---|---|---|---|---|---|---|---|---|
| chr15_73155365_73155701 | chr15 | 73155366 | 73155701 | + | 1.1142 | intron (NM_153178, intron 1 of 18) | intron (NM_153178, intron 1 of 18) | 29414 | NM_153178 | 239528 | Mm.23095 | NM_153178 | | **Ago2** | 1110029L17Rik\|2310051F07Rik\|AI225898\|AL022874\|AW546247\|ENSMUSG00000072493\|Eif2c2\|Gerp95\|Gm10365\|mKIAA4215 | argonaute RISC catalytic subunit 2 | protein-coding |
| chr15_73169766_73170340 | chr15 | 73169767 | 73170340 | + | 1.00957 | intron (NM_153178, intron 1 of 18) | ID\|SINE\|ID | 14894 | NM_153178 | 239528 | Mm.23095 | NM_153178 | | **Ago2** | 1110029L17Rik\|2310051F07Rik\|AI225898\|AL022874\|AW546247\|ENSMUSG00000072493\|Eif2c2\|Gerp95\|Gm10365\|mKIAA4215 | argonaute RISC catalytic subunit 2 | protein-coding |
| chr15_73265342_73265657 | chr15 | 73265343 | 73265657 | + | 0.957772 | intron (NM_007982, intron 18 of 31) | intron (NM_007982, intron 18 of 31) | -80553 | NM_153178 | 239528 | Mm.23095 | NM_153178 | | **Ago2** | 1110029L17Rik\|2310051F07Rik\|AI225898\|AL022874\|AW546247\|ENSMUSG00000072493\|Eif2c2\|Gerp95\|Gm10365\|mKIAA4215 | argonaute RISC catalytic subunit 2 | protein-coding |
| chr15_73169756_73170917 | chr15 | 73169757 | 73170917 | + | 0.871338 | intron (NM_153178, intron 1 of 18) | B1_Mm\|SINE\|Alu | 14610 | NM_153178 | 239528 | Mm.23095 | NM_153178 | | **Ago2** | 1110029L17Rik\|2310051F07Rik\|AI225898\|AL022874\|AW546247\|ENSMUSG00000072493\|Eif2c2\|Gerp95\|Gm10365\|mKIAA4215 | argonaute RISC catalytic subunit 2 | protein-coding |
| chr15_73170536_73170816 | chr15 | 73170537 | 73170816 | + | 0.731265 | intron (NM_153178, intron 1 of 18) | intron (NM_153178, intron 1 of 18) | 14271 | NM_153178 | 239528 | Mm.23095 | NM_153178 | | **Ago2** | 1110029L17Rik\|2310051F07Rik\|AI225898\|AL022874\|AW546247\|ENSMUSG00000072493\|Eif2c2\|Gerp95\|Gm10365\|mKIAA4215 | argonaute RISC catalytic subunit 2 | protein-coding |
| chr15_73170393_73170917 | chr15 | 73170394 | 73170917 | + | 0.682286 | intron (NM_153178, intron 1 of 18) | intron (NM_153178, intron 1 of 18) | 14292 | NM_153178 | 239528 | Mm.23095 | NM_153178 | | **Ago2** | 1110029L17Rik\|2310051F07Rik\|AI225898\|AL022874\|AW546247\|ENSMUSG00000072493\|Eif2c2\|Gerp95\|Gm10365\|mKIAA4215 | argonaute RISC catalytic subunit 2 | protein-coding |
| chr15_73148680_73149051 | chr15 | 73148681 | 73149051 | + | 0.601533 | intron (NM_153178, intron 1 of 18) | intron (NM_153178, intron 1 of 18) | 36081 | NM_153178 | 239528 | Mm.23095 | NM_153178 | | **Ago2** | 1110029L17Rik\|2310051F07Rik\|AI225898\|AL022874\|AW546247\|ENSMUSG00000072493\|Eif2c2\|Gerp95\|Gm10365\|mKIAA4215 | argonaute RISC catalytic subunit 2 | protein-coding |
| chr4_126415740_126416166 | chr4 | 126415741 | 126416166 | + | 1.12648 | intron (NM_153402, intron 2 of 18) | intron (NM_153402, intron 2 of 18) | 13589 | NM_153402 | 214150 | Mm.440915 | NM_153402 | | **Ago3** | AW048688\|C130014L07Rik\|Eif2c3 | argonaute RISC catalytic subunit 3 | protein-coding |
| chr10_78300139_78300488 | chr10 | 78300140 | 78300488 | + | 1.25916 | intron (NM_053014, intron 2 of 9) | intron (NM_053014, intron 2 of 9) | 51386 | NM_053014 | 28169 | Mm.141230 | NM_053014 | ENSMUSG00000001211 | **Agpat3** | AW061257\|AW493985\|D10Jhu12e\|Ipaat3 | 1-acylglycerol-3-phosphate O-acyltransferase 3 | protein-coding |
| chr10_78360141_78360814 | chr10 | 78360142 | 78360814 | + | 0.693594 | Intergenic | Intergenic | -8778 | NM_053014 | 28169 | Mm.141230 | NM_053014 | ENSMUSG00000001211 | **Agpat3** | AW061257\|AW493985\|D10Jhu12e\|Ipaat3 | 1-acylglycerol-3-phosphate O-acyltransferase 3 | protein-coding |
| chr17_12165090_12165931 | chr17 | 12165091 | 12165931 | + | 1.4145 | intron (NM_026644, intron 2 of 8) | intron (NM_026644, intron 2 of 8) | 46227 | NM_026644 | 68262 | Mm.258300 | NM_026644 | ENSMUSG00000023827 | **Agpat4** | 1500003P24Rik | 1-acylglycerol-3-phosphate O-acyltransferase 4 (lysophosphatidic acid acyltransferase, delta) | protein-coding |
| chr17_12165418_12165708 | chr17 | 12165419 | 12165708 | + | 1.23881 | intron (NM_026644, intron 2 of 8) | intron (NM_026644, intron 2 of 8) | 46279 | NM_026644 | 68262 | Mm.258300 | NM_026644 | ENSMUSG00000023827 | **Agpat4** | 1500003P24Rik | 1-acylglycerol-3-phosphate O-acyltransferase 4 (lysophosphatidic acid acyltransferase, delta) | protein-coding |
| chr17_12121434_12121788 | chr17 | 12121435 | 12121788 | + | 1.22214 | intron (NM_026644, intron 1 of 8) | intron (NM_026644, intron 1 of 8) | 2327 | NM_026644 | 68262 | Mm.258300 | NM_026644 | ENSMUSG00000023827 | **Agpat4** | 1500003P24Rik | 1-acylglycerol-3-phosphate O-acyltransferase 4 (lysophosphatidic acid acyltransferase, delta) | protein-coding |
| chr17_12164681_12164950 | chr17 | 12164682 | 12164950 | + | 0.843593 | intron (NM_026644, intron 2 of 8) | intron (NM_026644, intron 2 of 8) | 45532 | NM_026644 | 68262 | Mm.258300 | NM_026644 | ENSMUSG00000023827 | **Agpat4** | 1500003P24Rik | 1-acylglycerol-3-phosphate O-acyltransferase 4 (lysophosphatidic acid acyltransferase, delta) | protein-coding |
| chr17_12205419_12205960 | chr17 | 12205420 | 12205960 | + | 0.805603 | intron (NM_026644, intron 3 of 8) | L2a\|LINE\|L2 | 86406 | NM_026644 | 68262 | Mm.258300 | NM_026644 | ENSMUSG00000023827 | **Agpat4** | 1500003P24Rik | 1-acylglycerol-3-phosphate O-acyltransferase 4 (lysophosphatidic acid acyltransferase, delta) | protein-coding |
| chr17_12205128_12206449 | chr17 | 12205129 | 12206449 | + | 0.777427 | intron (NM_026644, intron 3 of 8) | intron (NM_026644, intron 3 of 8) | 86505 | NM_026644 | 68262 | Mm.258300 | NM_026644 | ENSMUSG00000023827 | **Agpat4** | 1500003P24Rik | 1-acylglycerol-3-phosphate O-acyltransferase 4 (lysophosphatidic acid acyltransferase, delta) | protein-coding |
| chr17_12171194_12171582 | chr17 | 12171195 | 12171582 | + | 0.722789 | intron (NM_026644, intron 2 of 8) | intron (NM_026644, intron 2 of 8) | 52104 | NM_026644 | 68262 | Mm.258300 | NM_026644 | ENSMUSG00000023827 | **Agpat4** | 1500003P24Rik | 1-acylglycerol-3-phosphate O-acyltransferase 4 (lysophosphatidic acid acyltransferase, delta) | protein-coding |
| chr17_12195146_12195547 | chr17 | 12195147 | 12195547 | + | 0.658271 | intron (NM_026644, intron 2 of 8) | intron (NM_026644, intron 2 of 8) | 76063 | NM_026644 | 68262 | Mm.258300 | NM_026644 | ENSMUSG00000023827 | **Agpat4** | 1500003P24Rik | 1-acylglycerol-3-phosphate O-acyltransferase 4 (lysophosphatidic acid acyltransferase, delta) | protein-coding |
| chr17_12162880_12163187 | chr17 | 12162881 | 12163187 | + | 0.645373 | intron (NM_026644, intron 2 of 8) | intron (NM_026644, intron 2 of 8) | 43750 | NM_026644 | 68262 | Mm.258300 | NM_026644 | ENSMUSG00000023827 | **Agpat4** | 1500003P24Rik | 1-acylglycerol-3-phosphate O-acyltransferase 4 (lysophosphatidic acid acyltransferase, delta) | protein-coding |
| chr5_101240925_101241548 | chr5 | 101240926 | 101241548 | + | 1.39521 | Intergenic | Intergenic | 395008 | NM_172715 | 231510 | Mm.271911 | NM_172715 | ENSMUSG00000029314 | **Agpat9** | 1-AGPAT\|1-AGPAT 9\|4933407I02Rik\|4933408F15\|9\|A230097K15Rik\|GPAT-3\|mGPAT3 | 1-acylglycerol-3-phosphate O-acyltransferase 9 | protein-coding |
| chr5_100861806_100862234 | chr5 | 100861807 | 100862234 | + | 0.816283 | intron (NM_172715, intron 3 of 12) | intron (NM_172715, intron 3 of 12) | 15791 | NM_172715 | 231510 | Mm.271911 | NM_172715 | ENSMUSG00000029314 | **Agpat9** | 1-AGPAT\|1-AGPAT 9\|4933407I02Rik\|4933408F15\|9\|A230097K15Rik\|GPAT-3\|mGPAT3 | 1-acylglycerol-3-phosphate O-acyltransferase 9 | protein-coding |
| chr5_101233327_101233853 | chr5 | 101233328 | 101233853 | + | 0.750959 | Intergenic | Intergenic | 387361 | NM_172715 | 231510 | Mm.271911 | NM_172715 | ENSMUSG00000029314 | **Agpat9** | 1-AGPAT\|1-AGPAT 9\|4933407I02Rik\|4933408F15\|9\|A230097K15Rik\|GPAT-3\|mGPAT3 | 1-acylglycerol-3-phosphate O-acyltransferase 9 | protein-coding |
| chr2_75880391_75880905 | chr2 | 75880392 | 75880905 | + | 0.907733 | intron (NM_172666, intron 11 of 19) | B2_Mm2\|SINE\|B2 | 48471 | NM_172666 | 228061 | Mm.31227 | NM_172666 | ENSMUSG00000042410 | **Agps** | 5832437L22\|9930035G10Rik\|AW123847\|Adaps\|Adas\|Adhaps\|Adps\|Aldhpsy\|bs2 | alkylglycerone phosphate synthase | protein-coding |
| chr12_35896695_35897697 | chr12 | 35896696 | 35897697 | + | 1.1238 | Intergenic | Intergenic | -28425 | NM_207531 | 403205 | Mm.190508 | NM_207531 | ENSMUSG00000036231 | **Agr3** | BCMP11\|E030025L21Rik\|Gm888 | anterior gradient 3 | protein-coding |
| chr12_35934924_35935222 | chr12 | 35934925 | 35935222 | + | 0.869262 | intron (NM_207531, intron 3 of 7) | intron (NM_207531, intron 3 of 7) | 9452 | NM_207531 | 403205 | Mm.190508 | NM_207531 | ENSMUSG00000036231 | **Agr3** | BCMP11\|E030025L21Rik\|Gm888 | anterior gradient 3 | protein-coding |
| chr1_179828780_179829424 | chr1 | 179828781 | 179829424 | + | 0.758986 | Intergenic | Intergenic | -25087 | NM_026375 | 226747 | Mm.128165 | NM_026375 | ENSMUSG00000026491 | **Ahctf1** | 6230412P20Rik\|AV011447\|Elys | AT hook containing transcription factor 1 | protein-coding |
| chr2_155046873_155047161 | chr2 | 155046874 | 155047161 | + | 1.24911 | intron (NM_015770, intron 2 of 3) | B3\|SINE\|B2 | 27480 | NM_016661 | 269378 | Mm.330692 | NM_016661 | ENSMUSG00000027597 | **Ahcy** | AA987153\|AL024110\|CuBP\|SAHH | S-adenosylhomocysteine hydrolase | protein-coding |
| chr3_107683680_107684440 | chr3 | 107683681 | 107684440 | + | 1.05857 | intron (NM_145542, intron 1 of 16) | intron (NM_145542, intron 1 of 16) | 12488 | NM_145542 | 229709 | Mm.220328 | NM_145542 | ENSMUSG00000027893 | **Ahcyl1** | 1110034F20Rik\|AA409031\|AA414901\|Ahcy-rs3\|DCAL\|Irbit | S-adenosylhomocysteine hydrolase-like 1 | protein-coding |
| chr6_29843075_29843420 | chr6 | 29843076 | 29843420 | + | 1.41741 | intron (NM_001171000, intron 1 of 16) | ORR1D1\|LTR\|ERVL-MaLR | -16348 | NM_001171001 | 74340 | Mm.210899 | NM_021414 | ENSMUSG00000029772 | **Ahcyl2** | 4631427C17Rik\|AI227036\|Adohcyase3\|mKIAA0828 | S-adenosylhomocysteine hydrolase-like 2 | protein-coding |
| chr6_29826884_29827828 | chr6 | 29826885 | 29827828 | + | 1.01484 | intron (NM_001171000, intron 1 of 16) | intron (NM_001171000, intron 1 of 16) | -32240 | NM_001171001 | 74340 | Mm.210899 | NM_021414 | ENSMUSG00000029772 | **Ahcyl2** | 4631427C17Rik\|AI227036\|Adohcyase3\|mKIAA0828 | S-adenosylhomocysteine hydrolase-like 2 | protein-coding |
| chr6_29837620_29837923 | chr6 | 29837621 | 29837923 | + | 0.89347 | intron (NM_001171000, intron 1 of 16) | intron (NM_001171000, intron 1 of 16) | -21824 | NM_001171001 | 74340 | Mm.210899 | NM_021414 | ENSMUSG00000029772 | **Ahcyl2** | 4631427C17Rik\|AI227036\|Adohcyase3\|mKIAA0828 | S-adenosylhomocysteine hydrolase-like 2 | protein-coding |
| chr6_29824705_29824961 | chr6 | 29824706 | 29824961 | + | 0.718078 | intron (NM_001171000, intron 1 of 16) | intron (NM_001171000, intron 1 of 16) | -34763 | NM_001171001 | 74340 | Mm.210899 | NM_021414 | ENSMUSG00000029772 | **Ahcyl2** | 4631427C17Rik\|AI227036\|Adohcyase3\|mKIAA0828 | S-adenosylhomocysteine hydrolase-like 2 | protein-coding |
| chr10_21028871_21029494 | chr10 | 21028872 | 21029494 | + | 1.24482 | intron (NM_026203, intron 18 of 23) | intron (NM_026203, intron 18 of 23) | -11637 | NM_001177776 | 52906 | Mm.253280 | NM_026203 | ENSMUSG00000019986 | **Ahi1** | 1700015F03Rik\|Ahi-1\|D10Bwg0629e | Abelson helper integration site 1 | protein-coding |
| chr10_21099171_21100029 | chr10 | 21099172 | 21100029 | + | 1.2169 | Intergenic | Intergenic | 58780 | NM_001177776 | 52906 | Mm.253280 | NM_026203 | ENSMUSG00000019986 | **Ahi1** | 1700015F03Rik\|Ahi-1\|D10Bwg0629e | Abelson helper integration site 1 | protein-coding |
| chr10_21029179_21029487 | chr10 | 21029180 | 21029487 | + | 1.14481 | intron (NM_026203, intron 18 of 23) | intron (NM_026203, intron 18 of 23) | -11487 | NM_001177776 | 52906 | Mm.253280 | NM_026203 | ENSMUSG00000019986 | **Ahi1** | 1700015F03Rik\|Ahi-1\|D10Bwg0629e | Abelson helper integration site 1 | protein-coding |

| | | | | | | | | | | | | | | | | | |
|---|---|---|---|---|---|---|---|---|---|---|---|---|---|---|---|---|---|
| chr10_20991780_20992209 | chr10 | 20991781 | 20992209 | + | 1.03142 | intron (NM_026203, intron 15 of 23) | intron (NM_026203, intron 15 of 23) | 39448 | NM_026203 | 52906 | Mm.253280 | NM_026203 | ENSMUSG00000019986 | **Ahi1** | 1700015F03Rik\|Ahi-1\|D10Bwg0629e | Abelson helper integration site 1 | protein-coding |
| chr10_20984429_20984863 | chr10 | 20984430 | 20984863 | + | 0.773222 | intron (NM_026203, intron 13 of 23) | MER20\|DNA\|hAT-Charlie | 32099 | NM_026203 | 52906 | Mm.253280 | NM_026203 | ENSMUSG00000019986 | **Ahi1** | 1700015F03Rik\|Ahi-1\|D10Bwg0629e | Abelson helper integration site 1 | protein-coding |
| chr10_20864943_20865525 | chr10 | 20864944 | 20865525 | + | 0.613205 | Intergenic | MT2A\|LTR\|ERVL | -87313 | NM_026203 | 52906 | Mm.253280 | NM_026203 | ENSMUSG00000019986 | **Ahi1** | 1700015F03Rik\|Ahi-1\|D10Bwg0629e | Abelson helper integration site 1 | protein-coding |
| chr19_8982732_8983042 | chr19 | 8982733 | 8983042 | + | 0.864205 | Intergenic | Intergenic | -6397 | NR_104460 | 66395 | Mm.203866 | NM_009643 | ENSMUSG00000069833 | **Ahnak** | DY6 | AHNAK nucleoprotein (desmoyokin) | protein-coding |
| chr19_8983303_8983601 | chr19 | 8983304 | 8983601 | + | 0.7272 | Intergenic | Intergenic | -5832 | NR_104460 | 66395 | Mm.203866 | NM_009643 | ENSMUSG00000069833 | **Ahnak** | DY6 | AHNAK nucleoprotein (desmoyokin) | protein-coding |
| chr12_35562798_35563364 | chr12 | 35562799 | 35563364 | + | 1.51205 | Intergenic | Intergenic | -28092 | NM_013464 | 11622 | Mm.341377 | NM_013464 | ENSMUSG00000019256 | **Ahr** | Ah\|Ahh\|Ahre\|In\|bHLHe76 | aryl-hydrocarbon receptor | protein-coding |
| chr12_35553102_35553582 | chr12 | 35553103 | 35553582 | + | 0.769829 | Intergenic | Intergenic | -18353 | NM_013464 | 11622 | Mm.341377 | NM_013464 | ENSMUSG00000019256 | **Ahr** | Ah\|Ahh\|Ahre\|In\|bHLHe76 | aryl-hydrocarbon receptor | protein-coding |
| chr9_55378924_55379242 | chr9 | 55378925 | 55379242 | + | 1.1455 | intron (NM_172923, intron 2 of 10) | intron (NM_172923, intron 2 of 10) | 52134 | NM_172923 | 244886 | Mm.24361 | NM_172923 | ENSMUSG00000032313 | **AI118078** | 9630029F15 | expressed sequence AI118078 | protein-coding |
| chr9_55378560_55379628 | chr9 | 55378561 | 55379628 | + | 1.01449 | intron (NM_172923, intron 2 of 10) | intron (NM_172923, intron 2 of 10) | 52145 | NM_172923 | 244886 | Mm.24361 | NM_172923 | ENSMUSG00000032313 | **AI118078** | 9630029F15 | expressed sequence AI118078 | protein-coding |
| chr9_55378586_55378877 | chr9 | 55378587 | 55378877 | + | 0.673901 | intron (NM_172923, intron 2 of 10) | intron (NM_172923, intron 2 of 10) | 51783 | NM_172923 | 244886 | Mm.24361 | NM_172923 | ENSMUSG00000032313 | **AI118078** | 9630029F15 | expressed sequence AI118078 | protein-coding |
| chr13_107485499_107486156 | chr13 | 107485500 | 107486156 | + | 1.17249 | Intergenic | Intergenic | 16006 | NR_045084 | 105120 | Mm.417700 | NR_045083 | | **AI197445** | - | expressed sequence AI197445 | ncRNA |
| chr13_107574774_107575401 | chr13 | 107574775 | 107575401 | + | 1.08681 | Intergenic | Intergenic | 105266 | NR_045084 | 105120 | Mm.417700 | NR_045083 | | **AI197445** | - | expressed sequence AI197445 | ncRNA |
| chr13_107573535_107574017 | chr13 | 107573536 | 107574017 | + | 1.03183 | Intergenic | Intergenic | 103954 | NR_045084 | 105120 | Mm.417700 | NR_045083 | | **AI197445** | - | expressed sequence AI197445 | ncRNA |
| chr13_107574482_107575392 | chr13 | 107574483 | 107575392 | + | 0.934796 | Intergenic | Intergenic | 105115 | NR_045084 | 105120 | Mm.417700 | NR_045083 | | **AI197445** | - | expressed sequence AI197445 | ncRNA |
| chr13_107400276_107400773 | chr13 | 107400277 | 107400773 | + | 0.848457 | Intergenic | MusHAL1\|LINE\|L1 | -69297 | NR_045083 | 105120 | Mm.417700 | NR_045083 | | **AI197445** | - | expressed sequence AI197445 | ncRNA |
| chr4_58901940_58902311 | chr4 | 58901941 | 58902311 | + | 0.766665 | intron (NM_172381, intron 1 of 48) | intron (NM_172381, intron 1 of 48) | 10599 | NM_172381 | 230249 | Mm.234801 | NM_172381 | ENSMUSG00000050812 | **AI314180** | AW558785\|BB181316\|mKIAA0368 | expressed sequence AI314180 | protein-coding |
| chr4_53322820_53323650 | chr4 | 53322821 | 53323650 | + | 0.637496 | Intergenic | Intergenic | -53003 | NR_033139 | 381524 | Mm.23351 | NM_001033454 | | **AI427809** | Gm1659 | expressed sequence AI427809 | ncRNA |
| chr4_41502993_41503430 | chr4 | 41502994 | 41503430 | + | 0.656653 | promoter-TSS (NM_001085515) | promoter-TSS (NM_001085515) | -137 | NM_001085515 | 329828 | Mm.27054 | NM_001085515 | ENSMUSG00000046312 | **AI464131** | AA408153\|AI604836\|Gm762\|Kiaa1161\|NET37 | expressed sequence AI464131 | protein-coding |
| chr4_41497491_41498171 | chr4 | 41497492 | 41498171 | + | 0.619714 | exon (NM_001085515, exon 2 of 2) | exon (NM_001085515, exon 2 of 2) | 5244 | NM_001085515 | 329828 | Mm.27054 | NM_001085515 | ENSMUSG00000046312 | **AI464131** | AA408153\|AI604836\|Gm762\|Kiaa1161\|NET37 | expressed sequence AI464131 | protein-coding |
| chr3_107054884_107055267 | chr3 | 107054885 | 107055267 | + | 1.0138 | TTS (NR_033498) | TTS (NR_033498) | 15572 | NR_033498 | 229694 | Mm.347584 | NM_173787 | ENSMUSG00000056145 | **AI504432** | A230102G19 | expressed sequence AI504432 | ncRNA |
| chr5_23713573_23713980 | chr5 | 23713574 | 23713980 | + | 0.930927 | Intergenic | Intergenic | -1110 | NR_015554 | 433855 | Mm.205125 | NR_015554 | | **AI506816** | - | expressed sequence AI506816 | ncRNA |
| chr11_83239596_83240138 | chr11 | 83239597 | 83240138 | + | 1.22391 | Intergenic | RMER17C-int\|LTR\|ERVK | 16291 | NR_015519 | 100043636 | Mm.295569 | NR_015519 | ENSMUSG00000087107 | **AI662270** | AW496474 | expressed sequence AI662270 | ncRNA |
| chr19_5431787_5432413 | chr19 | 5431788 | 5432413 | + | 0.789696 | Intergenic | Intergenic | 6956 | NM_001256515 | 107242 | Mm.329658 | NM_134149 | ENSMUSG00000047423 | **AI837181** | Bles03\|N28173 | expressed sequence AI837181 | protein-coding |
| chr19_5432128_5432415 | chr19 | 5432129 | 5432415 | + | 0.785226 | Intergenic | Intergenic | 7128 | NM_001256515 | 107242 | Mm.329658 | NM_134149 | ENSMUSG00000047423 | **AI837181** | Bles03\|N28173 | expressed sequence AI837181 | protein-coding |
| chr5_31576451_31577133 | chr5 | 31576452 | 31577133 | + | 0.632687 | Intergenic | Intergenic | -5395 | NR_102275 | 100740 | Mm.23370 | NR_102275 | | **AI839979** | - | expressed sequence AI839979 | ncRNA |
| chr7_79563657_79564192 | chr7 | 79563658 | 79564192 | + | 0.728729 | Intergenic | Intergenic | 48490 | NR_040312 | 101694 | Mm.41661 | NR_040311 | | **AI854517** | - | expressed sequence AI854517 | ncRNA |
| chr6_122575718_122576547 | chr6 | 122575719 | 122576547 | + | 1.07108 | Intergenic | Intergenic | 22324 | NM_009645 | 11628 | Mm.391503 | NM_009645 | ENSMUSG00000040627 | **Aicda** | Aid\|Arp2 | activation-induced cytidine deaminase | protein-coding |
| chr10_13832480_13833435 | chr10 | 13832481 | 13833435 | + | 1.71308 | intron (NM_025446, intron 1 of 5) | intron (NM_025446, intron 1 of 5) | 35872 | NM_025446 | 66253 | Mm.45755 | NM_025446 | ENSMUSG00000019806 | **Aig1** | 1500031O19Rik\|AV064870\|AW413422\|CGI-103 | androgen-induced 1 | protein-coding |
| chr10_13775863_13776619 | chr10 | 13775864 | 13776619 | + | 1.49487 | intron (NM_025446, intron 3 of 5) | intron (NM_025446, intron 3 of 5) | 92589 | NM_025446 | 66253 | Mm.45755 | NM_025446 | ENSMUSG00000019806 | **Aig1** | 1500031O19Rik\|AV064870\|AW413422\|CGI-103 | androgen-induced 1 | protein-coding |
| chr10_13730837_13731485 | chr10 | 13730838 | 13731485 | + | 1.43118 | intron (NM_025446, intron 3 of 5) | intron (NM_025446, intron 3 of 5) | 137669 | NM_025446 | 66253 | Mm.45755 | NM_025446 | ENSMUSG00000019806 | **Aig1** | 1500031O19Rik\|AV064870\|AW413422\|CGI-103 | androgen-induced 1 | protein-coding |
| chr10_13875062_13875360 | chr10 | 13875063 | 13875360 | + | 0.948443 | Intergenic | Intergenic | -6381 | NM_025446 | 66253 | Mm.45755 | NM_025446 | ENSMUSG00000019806 | **Aig1** | 1500031O19Rik\|AV064870\|AW413422\|CGI-103 | androgen-induced 1 | protein-coding |
| chr10_13874398_13875495 | chr10 | 13874399 | 13875495 | + | 0.807438 | Intergenic | Intergenic | -6117 | NM_025446 | 66253 | Mm.45755 | NM_025446 | ENSMUSG00000019806 | **Aig1** | 1500031O19Rik\|AV064870\|AW413422\|CGI-103 | androgen-induced 1 | protein-coding |
| chr10_13874725_13875475 | chr10 | 13874726 | 13875475 | + | 0.806355 | Intergenic | Intergenic | -6270 | NM_025446 | 66253 | Mm.45755 | NM_025446 | ENSMUSG00000019806 | **Aig1** | 1500031O19Rik\|AV064870\|AW413422\|CGI-103 | androgen-induced 1 | protein-coding |
| chr10_13869688_13870158 | chr10 | 13869689 | 13870158 | + | 0.736041 | Intergenic | Intergenic | -1093 | NM_025446 | 66253 | Mm.45755 | NM_025446 | ENSMUSG00000019806 | **Aig1** | 1500031O19Rik\|AV064870\|AW413422\|CGI-103 | androgen-induced 1 | protein-coding |
| chr10_43998496_43998895 | chr10 | 43998497 | 43998895 | + | 1.11078 | exon (NM_172393, exon 2 of 20) | exon (NM_172393, exon 2 of 20) | 6150 | NM_172393 | 11630 | Mm.292082 | NM_172393 | ENSMUSG00000019866 | **Aim1** | AI462491\|AI463325 | absent in melanoma 1 | protein-coding |
| chr4_134059830_134060194 | chr4 | 134059831 | 134060194 | + | 1.00895 | Intergenic | Intergenic | -8440 | NM_001162970 | 230806 | Mm.240080 | NM_001162970 | ENSMUSG00000012123 | **Aim1l** | - | absent in melanoma 1-like | protein-coding |
| chr17_12750616_12751481 | chr17 | 12750617 | 12751481 | + | 1.08648 | intron (NM_010515, intron 1 of 47) | intron (NM_010515, intron 1 of 47) | 9738 | NR_027784 | 104103 | Mm.26553 | NR_002853 | ENSMUSG00000078247 | **Airn** | 2810051F02Rik\|2810434M15Rik\|AI256653\|AI597500\|AW049873\|Air\|B930018I07Rik\|D17Ertd663e\|IGF2RAS | antisense Igf2r RNA | ncRNA |
| chr17_12750854_12751488 | chr17 | 12750855 | 12751488 | + | 1.08361 | intron (NM_010515, intron 1 of 47) | intron (NM_010515, intron 1 of 47) | 9860 | NR_027784 | 104103 | Mm.26553 | NR_002853 | ENSMUSG00000078247 | **Airn** | 2810051F02Rik\|2810434M15Rik\|AI256653\|AI597500\|AW049873\|Air\|B930018I07Rik\|D17Ertd663e\|IGF2RAS | antisense Igf2r RNA | ncRNA |
| chr17_12754280_12754859 | chr17 | 12754281 | 12754859 | + | 0.847271 | intron (NM_010515, intron 1 of 47) | intron (NM_010515, intron 1 of 47) | 13259 | NR_027784 | 104103 | Mm.26553 | NR_002853 | ENSMUSG00000078247 | **Airn** | 2810051F02Rik\|2810434M15Rik\|AI256653\|AI597500\|AW049873\|Air\|B930018I07Rik\|D17Ertd663e\|IGF2RAS | antisense Igf2r RNA | ncRNA |
| chr17_12714995_12715478 | chr17 | 12714996 | 12715478 | + | 0.709213 | intron (NM_010515, intron 17 of 47) | intron (NM_010515, intron 17 of 47) | -26074 | NR_002853 | 104103 | Mm.26553 | NR_002853 | ENSMUSG00000078247 | **Airn** | 2810051F02Rik\|2810434M15Rik\|AI256653\|AI597500\|AW049873\|Air\|B930018I07Rik\|D17Ertd663e\|IGF2RAS | antisense Igf2r RNA | ncRNA |
| chr4_129017081_129017720 | chr4 | 129017082 | 129017720 | + | 0.803991 | Intergenic | Intergenic | 24177 | NM_001033966 | 11637 | Mm.29460 | NM_016895 | ENSMUSG00000028792 | **Ak2** | Ak-2\|D4Ertd220e | adenylate kinase 2 | protein-coding |
| chr14_78473655_78474594 | chr14 | 78473656 | 78474594 | + | 1.39647 | Intergenic | L1MA6\|LINE\|L1 | 62735 | NM_001164503 | 219181 | Mm.89413 | NM_001164503 | ENSMUSG00000022016 | **Akap11** | 6330501D17Rik\|AKAP220\|Gm80\|PRKA11\|mKIAA0629 | A kinase (PRKA) anchor protein 11 | protein-coding |
| chr14_78458138_78458477 | chr14 | 78458139 | 78458477 | + | 0.77176 | Intergenic | Intergenic | 78552 | NM_001164503 | 219181 | Mm.89413 | NM_001164503 | ENSMUSG00000022016 | **Akap11** | 6330501D17Rik\|AKAP220\|Gm80\|PRKA11\|mKIAA0629 | A kinase (PRKA) anchor protein 11 | protein-coding |
| chr14_78454003_78454459 | chr14 | 78454004 | 78454459 | + | 0.691414 | Intergenic | Charlie1a\|DNA\|hAT-Charlie | 82629 | NM_001164503 | 219181 | Mm.89413 | NM_001164503 | ENSMUSG00000022016 | **Akap11** | 6330501D17Rik\|AKAP220\|Gm80\|PRKA11\|mKIAA0629 | A kinase (PRKA) anchor protein 11 | protein-coding |
| chr7_75397513_75398063 | chr7 | 75397514 | 75398063 | + | 1.22765 | Intergenic | MER5A1\|DNA\|hAT-Charlie | -57746 | NM_029332 | 75547 | Mm.216107 | NM_029332 | ENSMUSG00000066406 | **Akap13** | 1700026G02Rik\|5730522G15Rik\|5830460E08Rik\|AKAP-Lbc\|BRX\|Ht31\|LBC\|PROTO-LB\|PROTO-LBC | A kinase (PRKA) anchor protein 13 | protein-coding |
| chr7_75387788_75388490 | chr7 | 75387789 | 75388490 | + | 1.21117 | Intergenic | Intergenic | -67395 | NM_029332 | 75547 | Mm.216107 | NM_029332 | ENSMUSG00000066406 | **Akap13** | 1700026G02Rik\|5730522G15Rik\|5830460E08Rik\|AKAP-Lbc\|BRX\|Ht31\|LBC\|PROTO-LB\|PROTO-LBC | A kinase (PRKA) anchor protein 13 | protein-coding |
| chr7_75633863_75634457 | chr7 | 75633864 | 75634457 | + | 0.864272 | intron (NM_029332, intron 8 of 36) | intron (NM_029332, intron 8 of 36) | 178626 | NM_029332 | 75547 | Mm.216107 | NM_029332 | ENSMUSG00000066406 | **Akap13** | 1700026G02Rik\|5730522G15Rik\|5830460E08Rik\|AKAP-Lbc\|BRX\|Ht31\|LBC\|PROTO-LB\|PROTO-LBC | A kinase (PRKA) anchor protein 13 | protein-coding |
| chr7_75642253_75642740 | chr7 | 75642254 | 75642740 | + | 0.776023 | intron (NM_029332, intron 8 of 36) | intron (NM_029332, intron 8 of 36) | 186963 | NM_029332 | 75547 | Mm.216107 | NM_029332 | ENSMUSG00000066406 | **Akap13** | 1700026G02Rik\|5730522G15Rik\|5830460E08Rik\|AKAP-Lbc\|BRX\|Ht31\|LBC\|PROTO-LB\|PROTO-LBC | A kinase (PRKA) anchor protein 13 | protein-coding |
| chr7_75598563_75599009 | chr7 | 75598564 | 75599009 | + | 0.768203 | intron (NM_029332, intron 5 of 36) | Lx2\|LINE\|L1 | 143252 | NM_029332 | 75547 | Mm.216107 | NM_029332 | ENSMUSG00000066406 | **Akap13** | 1700026G02Rik\|5730522G15Rik\|5830460E08Rik\|AKAP-Lbc\|BRX\|Ht31\|LBC\|PROTO-LB\|PROTO-LBC | A kinase (PRKA) anchor protein 13 | protein-coding |
| chr7_75649226_75649657 | chr7 | 75649227 | 75649657 | + | 0.672159 | intron (NM_029332, intron 8 of 36) | intron (NM_029332, intron 8 of 36) | 193908 | NM_029332 | 75547 | Mm.216107 | NM_029332 | ENSMUSG00000066406 | **Akap13** | 1700026G02Rik\|5730522G15Rik\|5830460E08Rik\|AKAP-Lbc\|BRX\|Ht31\|LBC\|PROTO-LB\|PROTO-LBC | A kinase (PRKA) anchor protein 13 | protein-coding |

| | | | | | | | | | | | | | | | | | |
|---|---|---|---|---|---|---|---|---|---|---|---|---|---|---|---|---|---|
| chr4_57725442_57726333 | chr4 | 57725443 | 57726333 | + | 1.06226 | Intergenic | Intergenic | -119360 | NM_009649 | 11641 | Mm.467752 | NM_009649 | ENSMUSG00000038729 | **Akap2** | AA959716\|AI649048\|Akap-2\|Akap-kl\|B230340M18Rik\|Prka2 | A kinase (PRKA) anchor protein 2 | protein-coding |
| chr4_57800403_57800709 | chr4 | 57800404 | 57800709 | + | 0.745221 | Intergenic | Intergenic | -44692 | NM_009649 | 11641 | Mm.467752 | NM_009649 | ENSMUSG00000038729 | **Akap2** | AA959716\|AI649048\|Akap-2\|Akap-kl\|B230340M18Rik\|Prka2 | A kinase (PRKA) anchor protein 2 | protein-coding |
| chr4_57786272_57786484 | chr4 | 57786273 | 57786484 | + | 0.62357 | Intergenic | RMER1B\|Other\|Other | -58870 | NM_009649 | 11641 | Mm.467752 | NM_009649 | ENSMUSG00000038729 | **Akap2** | AA959716\|AI649048\|Akap-2\|Akap-kl\|B230340M18Rik\|Prka2 | A kinase (PRKA) anchor protein 2 | protein-coding |
| chr12_52853540_52853885 | chr12 | 52853541 | 52853885 | + | 0.671869 | intron (NM_198111, intron 2 of 13) | intron (NM_198111, intron 2 of 13) | 154330 | NM_198111 | 238161 | Mm.310822 | NM_198111 | ENSMUSG00000061603 | **Akap6** | AI482140\|Akapalpha\|Akapbeta\|PRKA | A kinase (PRKA) anchor protein 6 | protein-coding |
| chr10_25200676_25201121 | chr10 | 25200677 | 25201121 | + | 1.16002 | intron (NM_018747, intron 7 of 7) | intron (NM_018747, intron 7 of 7) | 98264 | NM_018747 | 432442 | Mm.485937 | NM_018747 | ENSMUSG00000039166 | **Akap7** | 6430401D08\|AI662165\|AKAP15\|Akap18\|BB170514 | A kinase (PRKA) anchor protein 7 | protein-coding |
| chr10_25220629_25221438 | chr10 | 25220630 | 25221438 | + | 0.776644 | intron (NM_018747, intron 6 of 7) | intron (NM_018747, intron 6 of 7) | 78129 | NM_018747 | 432442 | Mm.485937 | NM_018747 | ENSMUSG00000039166 | **Akap7** | 6430401D08\|AI662165\|AKAP15\|Akap18\|BB170514 | A kinase (PRKA) anchor protein 7 | protein-coding |
| chr10_25128396_25128932 | chr10 | 25128397 | 25128932 | + | 0.616783 | Intergenic | RMER15\|LTR\|ERVL | 170499 | NM_018747 | 432442 | Mm.485937 | NM_018747 | ENSMUSG00000039166 | **Akap7** | 6430401D08\|AI662165\|AKAP15\|Akap18\|BB170514 | A kinase (PRKA) anchor protein 7 | protein-coding |
| chr5_3952445_3953254 | chr5 | 3952446 | 3953254 | + | 1.4669 | intron (NM_194462, intron 3 of 47) | L1M2\|LINE\|L1 | 24664 | NM_194462 | 100986 | Mm.46044 | NM_194462 | ENSMUSG00000040407 | **Akap9** | 5730481H23Rik\|AKAP-9\|AKAP450\|AW545847\|C79026\|G1-448-15\|PRKA9\|mKIAA0803\|mei2-5\|repro12 | A kinase (PRKA) anchor protein (yotiao) 9 | protein-coding |
| chr5_3992905_3993630 | chr5 | 3992906 | 3993630 | + | 1.34398 | intron (NM_194462, intron 15 of 47) | MIRb\|SINE\|MIR | 65082 | NM_194462 | 100986 | Mm.46044 | NM_194462 | ENSMUSG00000040407 | **Akap9** | 5730481H23Rik\|AKAP-9\|AKAP450\|AW545847\|C79026\|G1-448-15\|PRKA9\|mKIAA0803\|mei2-5\|repro12 | A kinase (PRKA) anchor protein (yotiao) 9 | protein-coding |
| chr5_3968357_3968773 | chr5 | 3968358 | 3968773 | + | 0.973145 | intron (NM_194462, intron 7 of 47) | intron (NM_194462, intron 7 of 47) | 40379 | NM_194462 | 100986 | Mm.46044 | NM_194462 | ENSMUSG00000040407 | **Akap9** | 5730481H23Rik\|AKAP-9\|AKAP450\|AW545847\|C79026\|G1-448-15\|PRKA9\|mKIAA0803\|mei2-5\|repro12 | A kinase (PRKA) anchor protein (yotiao) 9 | protein-coding |
| chr5_3992554_3992846 | chr5 | 3992555 | 3992846 | + | 0.723641 | intron (NM_194462, intron 15 of 47) | intron (NM_194462, intron 15 of 47) | 64514 | NM_194462 | 100986 | Mm.46044 | NM_194462 | ENSMUSG00000040407 | **Akap9** | 5730481H23Rik\|AKAP-9\|AKAP450\|AW545847\|C79026\|G1-448-15\|PRKA9\|mKIAA0803\|mei2-5\|repro12 | A kinase (PRKA) anchor protein (yotiao) 9 | protein-coding |
| chr4_123752402_123752627 | chr4 | 123752403 | 123752627 | + | 0.713132 | Intergenic | Intergenic | -2216 | NM_023423 | 68050 | Mm.27903 | NM_023423 | | **Akirin1** | 6330407G11Rik | akirin 1 | protein-coding |
| chr4_34581301_34581958 | chr4 | 34581302 | 34581958 | + | 0.857013 | intron (NM_015824, intron 13 of 19) | ORR1D2\|LTR\|ERVL-MaLR | 31015 | NM_001007589 | 433693 | Mm.389532 | NM_026019 | ENSMUSG00000028291 | **Akirin2** | 2700059D21Rik\|AA114675\|AA522011\|AU019887\|Akirin-2 | akirin 2 | protein-coding |
| chr4_34580930_34581953 | chr4 | 34580931 | 34581953 | + | 0.84742 | intron (NM_015824, intron 13 of 19) | PB1D7\|SINE\|Alu | 30827 | NM_001007589 | 433693 | Mm.389532 | NM_026019 | ENSMUSG00000028291 | **Akirin2** | 2700059D21Rik\|AA114675\|AA522011\|AU019887\|Akirin-2 | akirin 2 | protein-coding |
| chr4_63385284_63385632 | chr4 | 63385285 | 63385632 | + | 1.13459 | intron (NM_001045514, intron 6 of 21) | intron (NM_001045514, intron 6 of 21) | 17987 | NM_001045514 | 100182 | Mm.387725 | NM_001045514 | ENSMUSG00000039158 | **Akna** | AI597013\|mKIAA1968 | AT-hook transcription factor | protein-coding |
| chr4_63383448_63383775 | chr4 | 63383449 | 63383775 | + | 0.818194 | intron (NM_001045514, intron 7 of 21) | MIRb\|SINE\|MIR | 19833 | NM_001045514 | 100182 | Mm.387725 | NM_001045514 | ENSMUSG00000039158 | **Akna** | AI597013\|mKIAA1968 | AT-hook transcription factor | protein-coding |
| chr4_63409243_63409759 | chr4 | 63409244 | 63409759 | + | 0.61374 | Intergenic | Intergenic | -6056 | NM_001045514 | 100182 | Mm.387725 | NM_001045514 | ENSMUSG00000039158 | **Akna** | AI597013\|mKIAA1968 | AT-hook transcription factor | protein-coding |
| chr3_108781448_108781880 | chr3 | 108781449 | 108781880 | + | 1.62488 | intron (NM_177859, intron 11 of 11) | intron (NM_177859, intron 11 of 11) | 42006 | NM_177859 | 329738 | Mm.72322 | NM_177859 | ENSMUSG00000049565 | **Aknad1** | 4921525H12Rik | AKNA domain containing 1 | protein-coding |
| chr3_108781393_108782007 | chr3 | 108781394 | 108782007 | + | 1.41882 | intron (NM_177859, intron 11 of 11) | intron (NM_177859, intron 11 of 11) | 42042 | NM_177859 | 329738 | Mm.72322 | NM_177859 | ENSMUSG00000049565 | **Aknad1** | 4921525H12Rik | AKNA domain containing 1 | protein-coding |
| chr3_108767319_108767787 | chr3 | 108767320 | 108767787 | + | 1.30582 | intron (NM_177859, intron 6 of 11) | intron (NM_177859, intron 6 of 11) | 27895 | NM_177859 | 329738 | Mm.72322 | NM_177859 | ENSMUSG00000049565 | **Aknad1** | 4921525H12Rik | AKNA domain containing 1 | protein-coding |
| chr3_108767113_108767793 | chr3 | 108767114 | 108767793 | + | 1.27464 | intron (NM_177859, intron 6 of 11) | intron (NM_177859, intron 6 of 11) | 27795 | NM_177859 | 329738 | Mm.72322 | NM_177859 | ENSMUSG00000049565 | **Aknad1** | 4921525H12Rik | AKNA domain containing 1 | protein-coding |
| chr3_108737349_108738090 | chr3 | 108737350 | 108738090 | + | 1.00169 | Intergenic | ID_B1\|SINE\|B4 | -1938 | NM_177859 | 329738 | Mm.72322 | NM_177859 | ENSMUSG00000049565 | **Aknad1** | 4921525H12Rik | AKNA domain containing 1 | protein-coding |
| chr4_116668021_116668609 | chr4 | 116668022 | 116668609 | + | 0.947336 | Intergenic | Intergenic | -16641 | NM_021473 | 58810 | Mm.30085 | NM_021473 | ENSMUSG00000028692 | **Akr1a1** | 2610201A18Rik\|Akr1a4 | aldo-keto reductase family 1, member A1 (aldehyde reductase) | protein-coding |
| chr6_34395930_34396553 | chr6 | 34395931 | 34396553 | + | 1.57267 | intron (NM_172398, intron 9 of 9) | MIRb\|SINE\|MIR | 11995 | NM_172398 | 67861 | Mm.70371 | NM_172398 | ENSMUSG00000061758 | **Akr1b10** | 2310005E10Rik\|Akr1b16 | aldo-keto reductase family 1, member B10 (aldose reductase) | protein-coding |
| chr6_37563884_37564450 | chr6 | 37563885 | 37564450 | + | 1.96654 | intron (NM_145364, intron 7 of 8) | intron (NM_145364, intron 7 of 8) | 33994 | NM_145364 | 208665 | Mm.262635 | NM_145364 | ENSMUSG00000038641 | **Akr1d1** | - | aldo-keto reductase family 1, member D1 | protein-coding |
| chr6_37523260_37523607 | chr6 | 37523261 | 37523607 | + | 1.24471 | Intergenic | RLTR25B\|LTR\|ERVK | -6739 | NM_145364 | 208665 | Mm.262635 | NM_145364 | ENSMUSG00000038641 | **Akr1d1** | - | aldo-keto reductase family 1, member D1 | protein-coding |
| chr13_4712699_4713150 | chr13 | 4712700 | 4713150 | + | 0.807488 | Intergenic | Intergenic | -103761 | NM_018859 | 56043 | Mm.251908 | NM_018859 | ENSMUSG00000045410 | **Akr1e1** | 1810061I10Rik\|Akr1e2 | aldo-keto reductase family 1, member E1 | protein-coding |
| chr13_4707357_4707709 | chr13 | 4707358 | 4707709 | + | 0.729832 | Intergenic | Intergenic | -98369 | NM_018859 | 56043 | Mm.251908 | NM_018859 | ENSMUSG00000045410 | **Akr1e1** | 1810061I10Rik\|Akr1e2 | aldo-keto reductase family 1, member E1 | protein-coding |
| chr1_177260398_177260802 | chr1 | 177260399 | 177260802 | + | 1.13261 | Intergenic | Intergenic | -11833 | NM_011785 | 23797 | Mm.235194 | NM_011785 | ENSMUSG00000019699 | **Akt3** | AI851531\|D930002M15Rik\|Nmf350 | thymoma viral proto-oncogene 3 | protein-coding |
| chr5_90433540_90434229 | chr5 | 90433541 | 90434229 | + | 0.940414 | Intergenic | MIR\|SINE\|MIR | -27004 | NM_009654 | 11657 | Mm.16773 | NM_009654 | ENSMUSG00000029368 | **Alb** | Alb-1\|Alb1 | albumin | protein-coding |
| chr16_52292337_52293116 | chr16 | 52292338 | 52293116 | + | 2.07366 | intron (NM_009655, intron 6 of 15) | intron (NM_009655, intron 6 of 15) | 160270 | NM_009655 | 11658 | Mm.288282 | NM_009655 | ENSMUSG00000022636 | **Alcam** | AI853494\|BEN\|CD166\|DM-GRASP\|MuSC\|SC1 | activated leukocyte cell adhesion molecule | protein-coding |
| chr16_52442808_52443475 | chr16 | 52442809 | 52443475 | + | 1.09861 | intron (NM_009655, intron 1 of 15) | (TCTCTG)n\|Simple_repeat\|Simple_repeat | 9855 | NM_009655 | 11658 | Mm.288282 | NM_009655 | ENSMUSG00000022636 | **Alcam** | AI853494\|BEN\|CD166\|DM-GRASP\|MuSC\|SC1 | activated leukocyte cell adhesion molecule | protein-coding |
| chr16_52332034_52332526 | chr16 | 52332035 | 52332526 | + | 0.953223 | intron (NM_009655, intron 1 of 15) | Lx7\|LINE\|L1 | 120717 | NM_009655 | 11658 | Mm.288282 | NM_009655 | ENSMUSG00000022636 | **Alcam** | AI853494\|BEN\|CD166\|DM-GRASP\|MuSC\|SC1 | activated leukocyte cell adhesion molecule | protein-coding |
| chr16_52470092_52470833 | chr16 | 52470093 | 52470833 | + | 0.943341 | Intergenic | Intergenic | -17466 | NM_009655 | 11658 | Mm.288282 | NM_009655 | ENSMUSG00000022636 | **Alcam** | AI853494\|BEN\|CD166\|DM-GRASP\|MuSC\|SC1 | activated leukocyte cell adhesion molecule | protein-coding |
| chr16_52443654_52444390 | chr16 | 52443655 | 52444390 | + | 0.940917 | intron (NM_009655, intron 1 of 15) | intron (NM_009655, intron 1 of 15) | 8975 | NM_009655 | 11658 | Mm.288282 | NM_009655 | ENSMUSG00000022636 | **Alcam** | AI853494\|BEN\|CD166\|DM-GRASP\|MuSC\|SC1 | activated leukocyte cell adhesion molecule | protein-coding |
| chr16_52472815_52473180 | chr16 | 52472816 | 52473180 | + | 0.846639 | Intergenic | L1M5\|LINE\|L1 | -20001 | NM_009655 | 11658 | Mm.288282 | NM_009655 | ENSMUSG00000022636 | **Alcam** | AI853494\|BEN\|CD166\|DM-GRASP\|MuSC\|SC1 | activated leukocyte cell adhesion molecule | protein-coding |
| chr16_52470268_52470562 | chr16 | 52470269 | 52470562 | + | 0.829014 | Intergenic | Intergenic | -17418 | NM_009655 | 11658 | Mm.288282 | NM_009655 | ENSMUSG00000022636 | **Alcam** | AI853494\|BEN\|CD166\|DM-GRASP\|MuSC\|SC1 | activated leukocyte cell adhesion molecule | protein-coding |
| chr4_45804634_45805021 | chr4 | 45804635 | 45805021 | + | 0.735511 | TTS (NM_028270) | TTS (NM_028270) | 5806 | NM_028270 | 72535 | Mm.331583 | NM_028270 | ENSMUSG00000035561 | **Aldh1b1** | 2700007F14Rik | aldehyde dehydrogenase 1 family, member B1 | protein-coding |
| chr19_3975832_3976247 | chr19 | 3975833 | 3976247 | + | 0.824763 | intron (NM_001177438, intron 1 of 9) | RMER17C2\|LTR\|ERVK | 3712 | NM_001177438 | 621603 | Mm.34137 | NM_001177438 | ENSMUSG00000075296 | **Aldh3b2** | AI848594\|C130048D07Rik | aldehyde dehydrogenase 3 family, member B2 | protein-coding |
| chr19_3975547_3976267 | chr19 | 3975548 | 3976267 | + | 0.76811 | intron (NM_001177438, intron 1 of 9) | RMER17C2\|LTR\|ERVK | 3579 | NM_001177438 | 621603 | Mm.34137 | NM_001177438 | ENSMUSG00000075296 | **Aldh3b2** | AI848594\|C130048D07Rik | aldehyde dehydrogenase 3 family, member B2 | protein-coding |
| chr10_21365613_21366312 | chr10 | 21365614 | 21366312 | + | 0.729202 | intron (NM_019702, intron 16 of 17) | intron (NM_019702, intron 16 of 17) | -11337 | NM_178713 | 237320 | Mm.90181 | NM_178713 | ENSMUSG00000037542 | **Aldh8a1** | Raldh4 | aldehyde dehydrogenase 8 family, member A1 | protein-coding |
| chr3_121327383_121327715 | chr3 | 121327384 | 121327715 | + | 1.16733 | intron (NM_024178, intron 3 of 3) | intron (NM_024178, intron 3 of 3) | 35732 | NM_024178 | 66789 | Mm.269881 | NM_024178 | ENSMUSG00000039887 | **Alg14** | 5430428G01Rik\|AI854024 | asparagine-linked glycosylation 14 | protein-coding |
| chr9_50774006_50774488 | chr9 | 50774007 | 50774488 | + | 0.820493 | promoter-TSS (NM_133981) | promoter-TSS (NM_133981) | -978 | NM_133981 | 102580 | Mm.160035 | NM_133981 | ENSMUSG00000032059 | **Alg9** | 8230402H15Rik\|AI747665\|B430313H07Rik\|Dibd1 | asparagine-linked glycosylation 9 (alpha 1,2 mannosyltransferase) | protein-coding |
| chr17_72604518_72604945 | chr17 | 72604519 | 72604945 | + | 0.651825 | promoter-TSS (NM_007439) | promoter-TSS (NM_007439) | -425 | NM_007439 | 11682 | Mm.311854 | NM_007439 | ENSMUSG00000055471 | **Alk** | CD246\|Tcrz | anaplastic lymphoma kinase | protein-coding |
| chr2_94051922_94052493 | chr2 | 94051923 | 94052493 | + | 1.97266 | intron (NM_019657, intron 7 of 10) | intron (NM_019657, intron 7 of 10) | -41478 | NM_026944 | 69113 | Mm.272498 | NM_026944 | ENSMUSG00000040174 | **Alkbh3** | 1700108H04Rik\|1810020C19Rik\|Abh3\|mABH3 | alkB, alkylation repair homolog 3 (E. coli) | protein-coding |
| chr2_94051892_94052742 | chr2 | 94051893 | 94052742 | + | 1.79383 | intron (NM_019657, intron 6 of 10) | intron (NM_019657, intron 6 of 10) | -41587 | NM_026944 | 69113 | Mm.272498 | NM_026944 | ENSMUSG00000040174 | **Alkbh3** | 1700108H04Rik\|1810020C19Rik\|Abh3\|mABH3 | alkB, alkylation repair homolog 3 (E. coli) | protein-coding |
| chr5_136143310_136143820 | chr5 | 136143311 | 136143820 | + | 0.687219 | Intergenic | Intergenic | 4784 | NM_028070 | 72041 | Mm.486605 | NM_028070 | ENSMUSG00000039754 | **Alkbh4** | 2010004B12Rik\|Abj4 | alkB, alkylation repair homolog 4 (E. coli) | protein-coding |

| | | | | | | | | | | | | | | | | | |
|---|---|---|---|---|---|---|---|---|---|---|---|---|---|---|---|---|---|
| chr9_3344317_3344698 | chr9 | 3344318 | 3344698 | + | 0.790294 | intron (NM_026303, intron 3 of 11) | intron (NM_026303, intron 3 of 11) | 9277 | NM_026303 | 67667 | Mm.116968 | NM_026303 | ENSMUSG00000025899 | **Alkbh8** | 4930562C03Rik\|8030431D03Rik\|9430088N01Rik\|Abh8 | alkB, alkylation repair homolog 8 (E. coli) | protein-coding |
| chr11_69172240_69173167 | chr11 | 69172241 | 69173167 | + | 1.03502 | Intergenic | Intergenic | 15632 | NM_009659 | 11686 | Mm.340329 | NM_009659 | ENSMUSG00000032807 | **Alox12b** | Aloxe2\|e-LOX2 | arachidonate 12-lipoxygenase, 12R type | protein-coding |
| chr18_65388916_65389517 | chr18 | 65388917 | 65389517 | + | 1.36752 | intron (NM_001037294, intron 1 of 12) | intron (NM_001037294, intron 1 of 12) | 4671 | NM_001037294 | 225638 | Mm.296528 | NM_001037294 | ENSMUSG00000032845 | **Alpk2** | Gm549\|Hak | alpha-kinase 2 | protein-coding |
| chr18_65321573_65322739 | chr18 | 65321574 | 65322739 | + | 0.908713 | intron (NM_001037294, intron 4 of 12) | intron (NM_001037294, intron 4 of 12) | 71732 | NM_001037294 | 225638 | Mm.296528 | NM_001037294 | ENSMUSG00000032845 | **Alpk2** | Gm549\|Hak | alpha-kinase 2 | protein-coding |
| chr18_65321967_65322739 | chr18 | 65321968 | 65322739 | + | 0.859034 | intron (NM_001037294, intron 4 of 12) | intron (NM_001037294, intron 4 of 12) | 71535 | NM_001037294 | 225638 | Mm.296528 | NM_001037294 | ENSMUSG00000032845 | **Alpk2** | Gm549\|Hak | alpha-kinase 2 | protein-coding |
| chr18_65321573_65321804 | chr18 | 65321574 | 65321804 | + | 0.844602 | intron (NM_001037294, intron 4 of 12) | MLT1C\|LTR\|ERVL-MaLR | 72199 | NM_001037294 | 225638 | Mm.296528 | NM_001037294 | ENSMUSG00000032845 | **Alpk2** | Gm549\|Hak | alpha-kinase 2 | protein-coding |
| chr18_65325870_65326179 | chr18 | 65325871 | 65326179 | + | 0.794612 | intron (NM_001037294, intron 4 of 12) | intron (NM_001037294, intron 4 of 12) | 67863 | NM_001037294 | 225638 | Mm.296528 | NM_001037294 | ENSMUSG00000032845 | **Alpk2** | Gm549\|Hak | alpha-kinase 2 | protein-coding |
| chr18_65328639_65329043 | chr18 | 65328640 | 65329043 | + | 0.718378 | intron (NM_001037294, intron 4 of 12) | intron (NM_001037294, intron 4 of 12) | 65047 | NM_001037294 | 225638 | Mm.296528 | NM_001037294 | ENSMUSG00000032845 | **Alpk2** | Gm549\|Hak | alpha-kinase 2 | protein-coding |
| chr18_65327086_65327502 | chr18 | 65327087 | 65327502 | + | 0.66377 | intron (NM_001037294, intron 4 of 12) | intron (NM_001037294, intron 4 of 12) | 66594 | NM_001037294 | 225638 | Mm.296528 | NM_001037294 | ENSMUSG00000032845 | **Alpk2** | Gm549\|Hak | alpha-kinase 2 | protein-coding |
| chr18_65395311_65395985 | chr18 | 65395312 | 65395985 | + | 0.66186 | intron (NR_105742, intron 3 of 6) | intron (NR_105742, intron 3 of 6) | -1760 | NM_001037294 | 225638 | Mm.296528 | NM_001037294 | ENSMUSG00000032845 | **Alpk2** | Gm549\|Hak | alpha-kinase 2 | protein-coding |
| chr9_110880979_110881252 | chr9 | 110880980 | 110881252 | + | 0.951809 | intron (NM_001146060, intron 1 of 26) | intron (NM_001146060, intron 1 of 26) | 794 | NM_001146060 | 235633 | Mm.86338 | NM_146228 | ENSMUSG00000044037 | **Als2cl** | 79H19D\|79H19E\|BC022645\|D930044G19Rik\|mRn.49018 | ALS2 C-terminal like | protein-coding |
| chr9_110880980_110881350 | chr9 | 110880981 | 110881350 | + | 0.922266 | intron (NM_001146060, intron 1 of 26) | intron (NM_001146060, intron 1 of 26) | 843 | NM_001146060 | 235633 | Mm.86338 | NM_146228 | ENSMUSG00000044037 | **Als2cl** | 79H19D\|79H19E\|BC022645\|D930044G19Rik\|mRn.49018 | ALS2 C-terminal like | protein-coding |
| chr1_171504729_171504966 | chr1 | 171504730 | 171504966 | + | 0.95157 | TTS (NM_019484) | TTS (NM_019484) | 1370 | NM_019484 | 56009 | Mm.389208 | NM_019484 | ENSMUSG00000060244 | **Alyref2** | C130042O11Rik\|REF2\|Refbp2 | Aly/REF export factor 2 | protein-coding |
| chr10_40309815_40310843 | chr10 | 40309816 | 40310843 | + | 0.811002 | Intergenic | ORR1B1\|LTR\|ERVL-MaLR | -8141 | NM_009665 | 11702 | Mm.253533 | NM_009665 | ENSMUSG00000075232 | **Amd1** | 1\|AdoMetDC\|Amd-1\|SAMDC\|SAMDC 1\|adoMetDC1 | S-adenosylmethionine decarboxylase 1 | protein-coding |
| chr8_93981532_93982264 | chr8 | 93981533 | 93982264 | + | 1.48882 | intron (NM_011787, intron 9 of 13) | RLTR15\|LTR\|ERVK | 30742 | NM_011787 | 23802 | Mm.34641 | NM_011787 | ENSMUSG00000031751 | **Amfr** | gp78 | autocrine motility factor receptor | protein-coding |
| chr8_94010067_94010499 | chr8 | 94010068 | 94010499 | + | 1.14215 | intron (NM_011787, intron 1 of 13) | intron (NM_011787, intron 1 of 13) | 2357 | NM_011787 | 23802 | Mm.34641 | NM_011787 | ENSMUSG00000031751 | **Amfr** | gp78 | autocrine motility factor receptor | protein-coding |
| chr8_94010050_94010726 | chr8 | 94010051 | 94010726 | + | 1.09621 | intron (NM_011787, intron 1 of 13) | intron (NM_011787, intron 1 of 13) | 2252 | NM_011787 | 23802 | Mm.34641 | NM_011787 | ENSMUSG00000031751 | **Amfr** | gp78 | autocrine motility factor receptor | protein-coding |
| chr9_45082828_45083556 | chr9 | 45082829 | 45083556 | + | 1.02738 | intron (NM_001005421, intron 2 of 10) | intron (NM_001005421, intron 2 of 10) | 4009 | NM_001005421 | 270152 | Mm.190461 | NM_001005421 | ENSMUSG00000048534 | **Amica1** | AMICA\|Crea7\|Gm638 | adhesion molecule, interacts with CXADR antigen 1 | protein-coding |
| chr15_97258817_97260233 | chr15 | 97258818 | 97260233 | + | 0.64272 | intron (NM_172293, intron 1 of 5) | intron (NM_172293, intron 1 of 5) | -12238 | NM_001164602 | 105827 | Mm.24005 | NM_178114 | ENSMUSG00000048218 | **Amigo2** | AI415330\|AMIGO-2\|AW208913\|Ali1 | adhesion molecule with Ig like domain 2 | protein-coding |
| chr15_97259286_97260002 | chr15 | 97259287 | 97260002 | + | 0.641692 | intron (NM_172293, intron 1 of 5) | intron (NM_172293, intron 1 of 5) | -12357 | NM_001164602 | 105827 | Mm.24005 | NM_178114 | ENSMUSG00000048218 | **Amigo2** | AI415330\|AMIGO-2\|AW208913\|Ali1 | adhesion molecule with Ig like domain 2 | protein-coding |
| chrX_142883371_142884211 | chrX | 142883372 | 142884211 | + | 1.6121 | intron (NM_019496, intron 2 of 5) | intron (NM_019496, intron 2 of 5) | 82937 | NM_019496 | 56068 | Mm.143724 | NM_019496 | | **Ammecr1** | 6230420G18Rik | Alport syndrome, mental retardation, midface hypoplasia and elliptocytosis chromosomal region gene 1 | protein-coding |
| chrX_142898184_142898907 | chrX | 142898185 | 142898907 | + | 1.14589 | intron (NM_019496, intron 2 of 5) | intron (NM_019496, intron 2 of 5) | 68182 | NM_019496 | 56068 | Mm.143724 | NM_019496 | | **Ammecr1** | 6230420G18Rik | Alport syndrome, mental retardation, midface hypoplasia and elliptocytosis chromosomal region gene 1 | protein-coding |
| chrX_142907468_142908077 | chrX | 142907469 | 142908077 | + | 1.10156 | intron (NM_019496, intron 2 of 5) | intron (NM_019496, intron 2 of 5) | 58955 | NM_019496 | 56068 | Mm.143724 | NM_019496 | | **Ammecr1** | 6230420G18Rik | Alport syndrome, mental retardation, midface hypoplasia and elliptocytosis chromosomal region gene 1 | protein-coding |
| chrX_142991780_142992099 | chrX | 142991781 | 142992099 | + | 1.08482 | Intergenic | Intergenic | -25212 | NM_019496 | 56068 | Mm.143724 | NM_019496 | | **Ammecr1** | 6230420G18Rik | Alport syndrome, mental retardation, midface hypoplasia and elliptocytosis chromosomal region gene 1 | protein-coding |
| chrX_142914170_142914547 | chrX | 142914171 | 142914547 | + | 1.01073 | intron (NM_019496, intron 1 of 5) | RMER1B\|Other\|Other | 52369 | NM_019496 | 56068 | Mm.143724 | NM_019496 | | **Ammecr1** | 6230420G18Rik | Alport syndrome, mental retardation, midface hypoplasia and elliptocytosis chromosomal region gene 1 | protein-coding |
| chrX_142914162_142914935 | chrX | 142914163 | 142914935 | + | 0.999536 | intron (NM_019496, intron 1 of 5) | RMER1B\|Other\|Other | 52179 | NM_019496 | 56068 | Mm.143724 | NM_019496 | | **Ammecr1** | 6230420G18Rik | Alport syndrome, mental retardation, midface hypoplasia and elliptocytosis chromosomal region gene 1 | protein-coding |
| chrX_142991778_142992245 | chrX | 142991779 | 142992245 | + | 0.951687 | Intergenic | Intergenic | -25284 | NM_019496 | 56068 | Mm.143724 | NM_019496 | | **Ammecr1** | 6230420G18Rik | Alport syndrome, mental retardation, midface hypoplasia and elliptocytosis chromosomal region gene 1 | protein-coding |
| chr18_31706023_31706547 | chr18 | 31706024 | 31706547 | + | 1.17061 | intron (NM_172965, intron 16 of 20) | RLTR42-int\|LTR\|ERVK | -53539 | NM_153515 | 225339 | Mm.290804 | NM_153515 | ENSMUSG00000041915 | **Ammecr1l** | 5430429D03Rik\|AU022236\|AU040755\|AW111353\|E230022H04Rik | AMME chromosomal region gene 1-like | protein-coding |
| chr18_31706246_31706547 | chr18 | 31706247 | 31706547 | + | 1.11298 | intron (NM_172965, intron 16 of 20) | intron (NM_172965, intron 16 of 20) | -53427 | NM_153515 | 225339 | Mm.290804 | NM_153515 | ENSMUSG00000041915 | **Ammecr1l** | 5430429D03Rik\|AU022236\|AU040755\|AW111353\|E230022H04Rik | AMME chromosomal region gene 1-like | protein-coding |
| chr6_149189263_149189490 | chr6 | 149189264 | 149189490 | + | 0.961068 | promoter-TSS (NM_001113424) | promoter-TSS (NM_001113424) | -665 | NM_001113424 | 232566 | Mm.209147 | NM_001113424 | ENSMUSG00000068250 | **Amn1** | 5830467E07Rik\|A230083D09\|C730024G19Rik | antagonist of mitotic exit network 1 | protein-coding |
| chr6_149193437_149194006 | chr6 | 149193438 | 149194006 | + | 0.710969 | Intergenic | RLTR30B_MM\|LTR\|ERV1 | -5010 | NM_001113424 | 232566 | Mm.209147 | NM_001113424 | ENSMUSG00000068250 | **Amn1** | 5830467E07Rik\|A230083D09\|C730024G19Rik | antagonist of mitotic exit network 1 | protein-coding |
| chr7_110797505_110798026 | chr7 | 110797506 | 110798026 | + | 1.20533 | intron (NM_001276301, intron 6 of 14) | intron (NM_001276301, intron 6 of 14) | 25162 | NM_009667 | 11717 | Mm.3238 | NM_009667 | ENSMUSG00000005686 | **Ampd3** | - | adenosine monophosphate deaminase 3 | protein-coding |
| chr11_109413126_109413519 | chr11 | 109413127 | 109413519 | + | 1.10073 | Intergenic | MTE-int\|LTR\|ERVL-MaLR | -12623 | NM_025275 | 13929 | Mm.211850 | NM_025275 | ENSMUSG00000020610 | **Amz2** | AA408420\|ESTM12\|X83328 | archaelysin family metallopeptidase 2 | protein-coding |
| chr11_109417433_109417907 | chr11 | 109417434 | 109417907 | + | 0.951699 | Intergenic | Intergenic | -8276 | NM_025275 | 13929 | Mm.211850 | NM_025275 | ENSMUSG00000020610 | **Amz2** | AA408420\|ESTM12\|X83328 | archaelysin family metallopeptidase 2 | protein-coding |
| chr11_109417415_109418134 | chr11 | 109417416 | 109418134 | + | 0.84544 | Intergenic | Intergenic | -8171 | NM_025275 | 13929 | Mm.211850 | NM_025275 | ENSMUSG00000020610 | **Amz2** | AA408420\|ESTM12\|X83328 | archaelysin family metallopeptidase 2 | protein-coding |
| chr11_109413659_109414026 | chr11 | 109413660 | 109414026 | + | 0.638735 | Intergenic | Intergenic | -12103 | NM_025275 | 13929 | Mm.211850 | NM_025275 | ENSMUSG00000020610 | **Amz2** | AA408420\|ESTM12\|X83328 | archaelysin family metallopeptidase 2 | protein-coding |
| chr5_52855713_52856092 | chr5 | 52855714 | 52856092 | + | 1.35209 | intron (NM_024213, intron 17 of 27) | intron (NM_024213, intron 17 of 27) | 21768 | NM_024213 | 52206 | Mm.272568 | NM_024213 | | **Anapc4** | 2610306D21Rik\|APC4\|D5Ertd249e | anaphase promoting complex subunit 4 | protein-coding |
| chr5_52908206_52908804 | chr5 | 52908207 | 52908804 | + | 1.25607 | Intergenic | Intergenic | 74370 | NM_024213 | 52206 | Mm.272568 | NM_024213 | | **Anapc4** | 2610306D21Rik\|APC4\|D5Ertd249e | anaphase promoting complex subunit 4 | protein-coding |
| chr5_52847377_52847860 | chr5 | 52847378 | 52847860 | + | 0.686472 | intron (NM_024213, intron 10 of 27) | ORR1F\|LTR\|ERVL-MaLR | 13484 | NM_024213 | 52206 | Mm.272568 | NM_024213 | | **Anapc4** | 2610306D21Rik\|APC4\|D5Ertd249e | anaphase promoting complex subunit 4 | protein-coding |
| chr2_33176484_33176891 | chr2 | 33176485 | 33176891 | + | 0.729962 | intron (NM_001290570, intron 7 of 16) | MIR3\|SINE\|MIR | -39273 | NM_011923 | 26360 | Mm.208919 | NM_011923 | ENSMUSG00000004105 | **Angptl2** | AI593246\|AW260363\|Arp2 | angiopoietin-like 2 | protein-coding |
| chr8_23038376_23038753 | chr8 | 23038377 | 23038753 | + | 1.25713 | intron (NM_001277284, intron 1 of 42) | intron (NM_001277284, intron 1 of 42) | 3466 | NM_001277284 | 11733 | Mm.334444 | NM_031158 | ENSMUSG00000031543 | **Ank1** | Ank-1\|nb\|pale | ankyrin 1, erythroid | protein-coding |
| chr8_23038003_23038816 | chr8 | 23038004 | 23038816 | + | 0.795849 | intron (NM_001277284, intron 1 of 42) | intron (NM_001277284, intron 1 of 42) | 3311 | NM_001277284 | 11733 | Mm.334444 | NM_031158 | ENSMUSG00000031543 | **Ank1** | Ank-1\|nb\|pale | ankyrin 1, erythroid | protein-coding |
| chr3_126920280_126920914 | chr3 | 126920281 | 126920914 | + | 1.73745 | Intergenic | Intergenic | 22789 | NM_001034168 | 109676 | Mm.220242 | NM_178655 | ENSMUSG00000032826 | **Ank2** | AI835472\|AW491075\|Ank-2 | ankyrin 2, brain | protein-coding |

| | | | | | | | | | | | | | | | | | |
|---|---|---|---|---|---|---|---|---|---|---|---|---|---|---|---|---|---|
| chr3_126831366_126831901 | chr3 | 126831367 | 126831901 | + | 1.65848 | intron (NM_023813, intron 14 of 17) | intron (NM_023813, intron 14 of 17) | 111752 | NM_001034168 | 109676 | Mm.220242 | NM_178655 | ENSMUSG00000032826 | **Ank2** | AI835472\|AW491075\|Ank-2 | ankyrin 2, brain | protein-coding |
| chr3_126827899_126828258 | chr3 | 126827900 | 126828258 | + | 1.07248 | intron (NM_023813, intron 14 of 17) | intron (NM_023813, intron 14 of 17) | 115307 | NM_001034168 | 109676 | Mm.220242 | NM_178655 | ENSMUSG00000032826 | **Ank2** | AI835472\|AW491075\|Ank-2 | ankyrin 2, brain | protein-coding |
| chr3_126827904_126828393 | chr3 | 126827905 | 126828393 | + | 1.02821 | intron (NM_023813, intron 14 of 17) | intron (NM_023813, intron 14 of 17) | 115237 | NM_001034168 | 109676 | Mm.220242 | NM_178655 | ENSMUSG00000032826 | **Ank2** | AI835472\|AW491075\|Ank-2 | ankyrin 2, brain | protein-coding |
| chr3_126829857_126830141 | chr3 | 126829858 | 126830141 | + | 0.887972 | intron (NM_023813, intron 14 of 17) | intron (NM_023813, intron 14 of 17) | 113387 | NM_001034168 | 109676 | Mm.220242 | NM_178655 | ENSMUSG00000032826 | **Ank2** | AI835472\|AW491075\|Ank-2 | ankyrin 2, brain | protein-coding |
| chr13_96487175_96487845 | chr13 | 96487176 | 96487845 | + | 1.565 | intron (NM_012048, intron 11 of 14) | intron (NM_012048, intron 11 of 14) | -16350 | NM_001042714 | 271144 | Mm.470057 | NM_001042714 | ENSMUSG00000047117 | **Ankdd1b** | 9330128J19Rik | ankyrin repeat and death domain containing 1B | protein-coding |
| chr18_36548397_36548844 | chr18 | 36548398 | 36548844 | + | 1.00604 | intron (NM_001271547, intron 19 of 19) | MTD\|LTR\|ERVL-MaLR | -11982 | NM_175375 | 108857 | Mm.482287 | NM_175375 | ENSMUSG00000024483 | **Ankhd1** | 1110004O12Rik\|4933432B13Rik\|9130019P20Rik\|A530027J04Rik\|A630021B20Rik\|AA571404\|Mask\|mFLJ00246\|mKIAA1085 | ankyrin repeat and KH domain containing 1 | protein-coding |
| chr5_3747605_3748137 | chr5 | 3747606 | 3748137 | + | 1.31141 | intron (NM_001289527, intron 4 of 19) | intron (NM_001289527, intron 4 of 19) | 24916 | NM_001289529 | 70797 | Mm.130148 | NM_001003909 | ENSMUSG00000040351 | **Ankib1** | 2310061P20Rik\|4631416I11Rik\|AW494740\|C80642\|mKIAA1386 | ankyrin repeat and IBR domain containing 1 | protein-coding |
| chr5_3799947_3800215 | chr5 | 3799948 | 3800215 | + | 0.898691 | intron (NM_001289527, intron 1 of 19) | intron (NM_001289527, intron 1 of 19) | 3043 | NM_001289528 | 70797 | Mm.130148 | NM_001003909 | ENSMUSG00000040351 | **Ankib1** | 2310061P20Rik\|4631416I11Rik\|AW494740\|C80642\|mKIAA1386 | ankyrin repeat and IBR domain containing 1 | protein-coding |
| chr19_36184343_36184740 | chr19 | 36184344 | 36184740 | + | 0.720852 | Intergenic | Intergenic | -64698 | NM_013468 | 107765 | Mm.10279 | NM_013468 | ENSMUSG00000024803 | **Ankrd1** | Alrp\|CARP\|Crap\|MARP1 | ankyrin repeat domain 1 (cardiac muscle) | protein-coding |
| chr8_122962081_122962338 | chr8 | 122962082 | 122962338 | + | 0.743049 | intron (NR_037865, intron 2 of 13) | intron (NR_037865, intron 2 of 13) | 80074 | NR_037865 | 77087 | Mm.31567 | NM_001081379 | ENSMUSG00000035569 | **Ankrd11** | 2410104C19Rik\|3010027A04Rik\|6330578C09Rik\|9530048I21Rik\|AA930108\|Gm176\|Yod | ankyrin repeat domain 11 | protein-coding |
| chr8_122962005_122962478 | chr8 | 122962006 | 122962478 | + | 0.739793 | intron (NR_037865, intron 2 of 13) | intron (NR_037865, intron 2 of 13) | 80042 | NR_037865 | 77087 | Mm.31567 | NM_001081379 | ENSMUSG00000035569 | **Ankrd11** | 2410104C19Rik\|3010027A04Rik\|6330578C09Rik\|9530048I21Rik\|AA930108\|Gm176\|Yod | ankyrin repeat domain 11 | protein-coding |
| chr17_66065109_66065488 | chr17 | 66065110 | 66065488 | + | 1.04618 | intron (NM_001025572, intron 1 of 12) | intron (NM_001025572, intron 1 of 12) | 11747 | NM_001025572 | 106585 | Mm.34706 | NM_001025572 | ENSMUSG00000034647 | **Ankrd12** | 2900001A12Rik\|AI447928\|ANCO-2\|AV347965\|GAC-1\|mKIAA0874 | ankyrin repeat domain 12 | protein-coding |
| chr17_66048685_66049332 | chr17 | 66048686 | 66049332 | + | 0.884507 | intron (NM_001025572, intron 3 of 12) | ORR1C1\|LTR\|ERVL-MaLR | 28037 | NM_001025572 | 106585 | Mm.34706 | NM_001025572 | ENSMUSG00000034647 | **Ankrd12** | 2900001A12Rik\|AI447928\|ANCO-2\|AV347965\|GAC-1\|mKIAA0874 | ankyrin repeat domain 12 | protein-coding |
| chr17_66065008_66065910 | chr17 | 66065009 | 66065910 | + | 0.810551 | intron (NM_001025572, intron 1 of 12) | intron (NM_001025572, intron 1 of 12) | 11587 | NM_001025572 | 106585 | Mm.34706 | NM_001025572 | ENSMUSG00000034647 | **Ankrd12** | 2900001A12Rik\|AI447928\|ANCO-2\|AV347965\|GAC-1\|mKIAA0874 | ankyrin repeat domain 12 | protein-coding |
| chr3_157971225_157971545 | chr3 | 157971226 | 157971545 | + | 0.797112 | intron (NM_001013806, intron 3 of 12) | B3A\|SINE\|B2 | 23919 | NM_001013806 | 433667 | Mm.265806 | NM_001013806 | ENSMUSG00000039988 | **Ankrd13c** | AI505652\|AU022220 | ankyrin repeat domain 13c | protein-coding |
| chr5_90344178_90344557 | chr5 | 90344179 | 90344557 | + | 0.976451 | intron (NM_198010, intron 1 of 32) | intron (NM_198010, intron 1 of 32) | 21817 | NM_030886 | 81702 | Mm.245522 | NM_030886 | ENSMUSG00000055204 | **Ankrd17** | 4933425K22Rik\|A130069E23Rik\|A930008M01\|AA407558\|AA516750\|AU040470\|Foe\|Gtar\|Mask\|mKIAA0697 | ankyrin repeat domain 17 | protein-coding |
| chr5_90363973_90364210 | chr5 | 90363974 | 90364210 | + | 0.738288 | intron (NM_198010, intron 1 of 32) | intron (NM_198010, intron 1 of 32) | 2093 | NM_030886 | 81702 | Mm.245522 | NM_030886 | ENSMUSG00000055204 | **Ankrd17** | 4933425K22Rik\|A130069E23Rik\|A930008M01\|AA407558\|AA516750\|AU040470\|Foe\|Gtar\|Mask\|mKIAA0697 | ankyrin repeat domain 17 | protein-coding |
| chr5_90364032_90364198 | chr5 | 90364033 | 90364198 | + | 0.700039 | intron (NM_198010, intron 1 of 32) | intron (NM_198010, intron 1 of 32) | 2070 | NM_030886 | 81702 | Mm.245522 | NM_030886 | ENSMUSG00000055204 | **Ankrd17** | 4933425K22Rik\|A130069E23Rik\|A930008M01\|AA407558\|AA516750\|AU040470\|Foe\|Gtar\|Mask\|mKIAA0697 | ankyrin repeat domain 17 | protein-coding |
| chr19_42017891_42018209 | chr19 | 42017892 | 42018209 | + | 0.686389 | intron (NM_145500, intron 1 of 2) | intron (NM_145500, intron 1 of 2) | -17988 | NM_020033 | 56642 | Mm.143737 | NM_020033 | ENSMUSG00000025172 | **Ankrd2** | Arpp\|mArpp | ankyrin repeat domain 2 (stretch responsive muscle) | protein-coding |
| chr19_34171264_34171707 | chr19 | 34171265 | 34171707 | + | 0.847715 | Intergenic | Intergenic | -5445 | NM_024204 | 52024 | Mm.183030 | NM_024204 | ENSMUSG00000024774 | **Ankrd22** | 5430429D21Rik\|D19Ertd675e | ankyrin repeat domain 22 | protein-coding |
| chr19_34171248_34171864 | chr19 | 34171249 | 34171864 | + | 0.847473 | Intergenic | Intergenic | -5515 | NM_024204 | 52024 | Mm.183030 | NM_024204 | ENSMUSG00000024774 | **Ankrd22** | 5430429D21Rik\|D19Ertd675e | ankyrin repeat domain 22 | protein-coding |
| chr19_34166470_34166757 | chr19 | 34166471 | 34166757 | + | 0.644538 | promoter-TSS (NM_024204) | promoter-TSS (NM_024204) | -573 | NM_024204 | 52024 | Mm.183030 | NM_024204 | ENSMUSG00000024774 | **Ankrd22** | 5430429D21Rik\|D19Ertd675e | ankyrin repeat domain 22 | protein-coding |
| chr14_31768602_31768877 | chr14 | 31768603 | 31768877 | + | 1.0305 | intron (NM_001024604, intron 2 of 27) | intron (NM_001024604, intron 2 of 27) | 61675 | NM_001024604 | 105522 | Mm.37660 | NM_001024604 | ENSMUSG00000014496 | **Ankrd28** | AI465466\|AI604979\|E430019N21Rik\|mKIAA0379 | ankyrin repeat domain 28 | protein-coding |
| chr14_31785654_31786056 | chr14 | 31785655 | 31786056 | + | 0.654637 | intron (NM_001024604, intron 1 of 27) | MLT1L\|LTR\|ERVL-MaLR | 44560 | NM_001024604 | 105522 | Mm.37660 | NM_001024604 | ENSMUSG00000014496 | **Ankrd28** | AI465466\|AI604979\|E430019N21Rik\|mKIAA0379 | ankyrin repeat domain 28 | protein-coding |
| chr14_31779957_31780322 | chr14 | 31779958 | 31780322 | + | 0.599105 | intron (NM_001024604, intron 1 of 27) | intron (NM_001024604, intron 1 of 27) | 50275 | NM_001024604 | 105522 | Mm.37660 | NM_001024604 | ENSMUSG00000014496 | **Ankrd28** | AI465466\|AI604979\|E430019N21Rik\|mKIAA0379 | ankyrin repeat domain 28 | protein-coding |
| chr15_31365677_31366162 | chr15 | 31365678 | 31366162 | + | 0.932526 | intron (NM_027496, intron 1 of 3) | intron (NM_027496, intron 1 of 3) | 1839 | NM_001164441 | 67434 | Mm.102470 | NM_026153 | ENSMUSG00000022237 | **Ankrd33b** | 0610012A05Rik\|3021401C12Rik\|5730557B15Rik | ankyrin repeat domain 33B | protein-coding |
| chr15_31365483_31366164 | chr15 | 31365484 | 31366164 | + | 0.798699 | intron (NM_027496, intron 1 of 3) | intron (NM_027496, intron 1 of 3) | 1935 | NM_001164441 | 67434 | Mm.102470 | NM_026153 | ENSMUSG00000022237 | **Ankrd33b** | 0610012A05Rik\|3021401C12Rik\|5730557B15Rik | ankyrin repeat domain 33B | protein-coding |
| chr15_31356687_31357339 | chr15 | 31356688 | 31357339 | + | 0.72915 | intron (NM_027496, intron 1 of 3) | intron (NM_027496, intron 1 of 3) | 10746 | NM_001164441 | 67434 | Mm.102470 | NM_026153 | ENSMUSG00000022237 | **Ankrd33b** | 0610012A05Rik\|3021401C12Rik\|5730557B15Rik | ankyrin repeat domain 33B | protein-coding |
| chr15_31356903_31357312 | chr15 | 31356904 | 31357312 | + | 0.682155 | intron (NM_027496, intron 1 of 3) | intron (NM_027496, intron 1 of 3) | 10651 | NM_001164441 | 67434 | Mm.102470 | NM_026153 | ENSMUSG00000022237 | **Ankrd33b** | 0610012A05Rik\|3021401C12Rik\|5730557B15Rik | ankyrin repeat domain 33B | protein-coding |
| chr13_92416696_92417315 | chr13 | 92416697 | 92417315 | + | 0.777775 | Intergenic | Intergenic | -8963 | NM_175455 | 218440 | Mm.213623 | NM_175455 | ENSMUSG00000045034 | **Ankrd34b** | 6430502M16Rik\|DP58 | ankyrin repeat domain 34B | protein-coding |
| chr1_54935462_54935991 | chr1 | 54935463 | 54935991 | + | 1.31138 | Intergenic | URR1B\|DNA\|hAT-Charlie | -9340 | NM_001081433 | 329154 | Mm.397910 | NM_001081433 | ENSMUSG00000052331 | **Ankrd44** | 4930444A19Rik\|A130096K20\|E130014H08Rik\|PP6-ARS-B | ankyrin repeat domain 44 | protein-coding |
| chr1_54934897_54936002 | chr1 | 54934898 | 54936002 | + | 1.06065 | Intergenic | Lx10\|LINE\|L1 | -9063 | NM_001081433 | 329154 | Mm.397910 | NM_001081433 | ENSMUSG00000052331 | **Ankrd44** | 4930444A19Rik\|A130096K20\|E130014H08Rik\|PP6-ARS-B | ankyrin repeat domain 44 | protein-coding |
| chr1_54870876_54871355 | chr1 | 54870877 | 54871355 | + | 1.00146 | intron (NM_001081433, intron 1 of 27) | intron (NM_001081433, intron 1 of 27) | 55271 | NM_001081433 | 329154 | Mm.397910 | NM_001081433 | ENSMUSG00000052331 | **Ankrd44** | 4930444A19Rik\|A130096K20\|E130014H08Rik\|PP6-ARS-B | ankyrin repeat domain 44 | protein-coding |
| chr1_54821871_54822471 | chr1 | 54821872 | 54822471 | + | 0.916779 | intron (NM_001081433, intron 2 of 27) | intron (NM_001081433, intron 2 of 27) | 104216 | NM_001081433 | 329154 | Mm.397910 | NM_001081433 | ENSMUSG00000052331 | **Ankrd44** | 4930444A19Rik\|A130096K20\|E130014H08Rik\|PP6-ARS-B | ankyrin repeat domain 44 | protein-coding |
| chr1_54839757_54840015 | chr1 | 54839758 | 54840015 | + | 0.82838 | intron (NM_001081433, intron 1 of 27) | intron (NM_001081433, intron 1 of 27) | 86501 | NM_001081433 | 329154 | Mm.397910 | NM_001081433 | ENSMUSG00000052331 | **Ankrd44** | 4930444A19Rik\|A130096K20\|E130014H08Rik\|PP6-ARS-B | ankyrin repeat domain 44 | protein-coding |
| chr1_54934907_54935358 | chr1 | 54934908 | 54935358 | + | 0.751954 | Intergenic | ERVL-E-int\|LTR\|ERVL | -8746 | NM_001081433 | 329154 | Mm.397910 | NM_001081433 | ENSMUSG00000052331 | **Ankrd44** | 4930444A19Rik\|A130096K20\|E130014H08Rik\|PP6-ARS-B | ankyrin repeat domain 44 | protein-coding |
| chr1_54904146_54904506 | chr1 | 54904147 | 54904506 | + | 0.67887 | intron (NM_001081433, intron 1 of 27) | Lx5c\|LINE\|L1 | 22061 | NM_001081433 | 329154 | Mm.397910 | NM_001081433 | ENSMUSG00000052331 | **Ankrd44** | 4930444A19Rik\|A130096K20\|E130014H08Rik\|PP6-ARS-B | ankyrin repeat domain 44 | protein-coding |
| chr1_54823524_54823824 | chr1 | 54823525 | 54823824 | + | 0.67685 | intron (NM_001081433, intron 2 of 27) | intron (NM_001081433, intron 2 of 27) | 102713 | NM_001081433 | 329154 | Mm.397910 | NM_001081433 | ENSMUSG00000052331 | **Ankrd44** | 4930444A19Rik\|A130096K20\|E130014H08Rik\|PP6-ARS-B | ankyrin repeat domain 44 | protein-coding |
| chr1_54922296_54922660 | chr1 | 54922297 | 54922660 | + | 0.654299 | intron (NM_001081433, intron 1 of 27) | MIR\|SINE\|MIR | 3909 | NM_001081433 | 329154 | Mm.397910 | NM_001081433 | ENSMUSG00000052331 | **Ankrd44** | 4930444A19Rik\|A130096K20\|E130014H08Rik\|PP6-ARS-B | ankyrin repeat domain 44 | protein-coding |
| chr1_161140091_161140686 | chr1 | 161140092 | 161140686 | + | 1.01201 | Intergenic | B1F\|SINE\|Alu | -2323 | NM_028664 | 73844 | Mm.38830 | NM_028664 | | **Ankrd45** | 4933409K03Rik | ankyrin repeat domain 45 | protein-coding |
| chr15_36491334_36492419 | chr15 | 36491335 | 36492419 | + | 1.74935 | intron (NM_175134, intron 2 of 4) | intron (NM_175134, intron 2 of 4) | 4914 | NM_175134 | 68839 | Mm.396537 | NM_175134 | ENSMUSG00000048307 | **Ankrd46** | 1110054N06Rik\|AI314978\|AI987733 | ankyrin repeat domain 46 | protein-coding |

| | | | | | | | | | | | | | | | | | |
|---|---|---|---|---|---|---|---|---|---|---|---|---|---|---|---|---|---|
| chr15_36397341_36397572 | chr15 | 36397342 | 36397572 | + | 0.879848 | Intergenic | MLT1J2\|LTR\|ERVL-MaLR | 99334 | NM_175134 | 68839 | Mm.396537 | NM_175134 | ENSMUSG00000048307 | **Ankrd46** | 1110054N06Rik\|AI314978\|AI987733 | ankyrin repeat domain 46 | protein-coding |
| chr15_36423890_36424328 | chr15 | 36423891 | 36424328 | + | 0.840485 | Intergenic | Intergenic | 72682 | NM_175134 | 68839 | Mm.396537 | NM_175134 | ENSMUSG00000048307 | **Ankrd46** | 1110054N06Rik\|AI314978\|AI987733 | ankyrin repeat domain 46 | protein-coding |
| chr15_36423921_36424471 | chr15 | 36423922 | 36424471 | + | 0.832665 | Intergenic | Intergenic | 72595 | NM_175134 | 68839 | Mm.396537 | NM_175134 | ENSMUSG00000048307 | **Ankrd46** | 1110054N06Rik\|AI314978\|AI987733 | ankyrin repeat domain 46 | protein-coding |
| chr3_38491511_38492148 | chr3 | 38491512 | 38492148 | + | 0.941932 | Intergenic | MLT1E2\|LTR\|ERVL-MaLR | -7014 | NM_001167883 | 99696 | Mm.290937 | NM_001033198 | ENSMUSG00000044864 | **Ankrd50** | AI662170\|E430012K20Rik\|mKIAA1223\|mKIAA3019 | ankyrin repeat domain 50 | protein-coding |
| chr3_38491311_38492188 | chr3 | 38491312 | 38492188 | + | 0.899092 | Intergenic | MLT1E2\|LTR\|ERVL-MaLR | -6934 | NM_001167883 | 99696 | Mm.290937 | NM_001033198 | ENSMUSG00000044864 | **Ankrd50** | AI662170\|E430012K20Rik\|mKIAA1223\|mKIAA3019 | ankyrin repeat domain 50 | protein-coding |
| chr13_112318160_112318492 | chr13 | 112318161 | 112318492 | + | 1.41746 | intron (NM_001168403, intron 2 of 11) | intron (NM_001168403, intron 2 of 11) | 29808 | NM_001168404 | 77318 | Mm.159895 | NM_029898 | ENSMUSG00000049985 | **Ankrd55** | C030011J08Rik | ankyrin repeat domain 55 | protein-coding |
| chr13_112317953_112318601 | chr13 | 112317954 | 112318601 | + | 1.22464 | intron (NM_001168403, intron 2 of 11) | intron (NM_001168403, intron 2 of 11) | 29759 | NM_001168404 | 77318 | Mm.159895 | NM_029898 | ENSMUSG00000049985 | **Ankrd55** | C030011J08Rik | ankyrin repeat domain 55 | protein-coding |
| chr13_112258446_112258953 | chr13 | 112258447 | 112258953 | + | 1.11468 | Intergenic | ORR1F\|LTR\|ERVL-MaLR | -29751 | NM_001168403 | 77318 | Mm.159895 | NM_029898 | ENSMUSG00000049985 | **Ankrd55** | C030011J08Rik | ankyrin repeat domain 55 | protein-coding |
| chr13_112271944_112272576 | chr13 | 112271945 | 112272576 | + | 0.754714 | Intergenic | Intergenic | -16191 | NM_001168403 | 77318 | Mm.159895 | NM_029898 | ENSMUSG00000049985 | **Ankrd55** | C030011J08Rik | ankyrin repeat domain 55 | protein-coding |
| chr4_32905396_32905767 | chr4 | 32905397 | 32905767 | + | 0.891365 | intron (NM_001012450, intron 1 of 16) | intron (NM_001012450, intron 1 of 16) | 17923 | NM_080471 | 140577 | Mm.214766 | NM_080471 | ENSMUSG00000040183 | **Ankrd6** | - | ankyrin repeat domain 6 | protein-coding |
| chr12_110939395_110939979 | chr12 | 110939396 | 110939979 | + | 0.920207 | intron (NM_001081057, intron 12 of 20) | intron (NM_001081057, intron 12 of 20) | 39334 | NM_175207 | 74251 | Mm.250989 | NM_175207 | ENSMUSG00000037904 | **Ankrd9** | 2500003O20Rik | ankyrin repeat domain 9 | protein-coding |
| chr15_95802621_95803463 | chr15 | 95802622 | 95803463 | + | 0.916145 | intron (NM_175344, intron 1 of 19) | intron (NM_175344, intron 1 of 19) | 12199 | NM_001253813 | 105722 | Mm.38087 | NM_175344 | ENSMUSG00000064210 | **Ano6** | 2900059G15Rik\|AA407480\|AW554778\|F730003B03Rik\|Tmem16f | anoctamin 6 | protein-coding |
| chr15_95805112_95805493 | chr15 | 95805113 | 95805493 | + | 0.808074 | intron (NM_175344, intron 1 of 19) | intron (NM_175344, intron 1 of 19) | 14460 | NM_001253813 | 105722 | Mm.38087 | NM_175344 | ENSMUSG00000064210 | **Ano6** | 2900059G15Rik\|AA407480\|AW554778\|F730003B03Rik\|Tmem16f | anoctamin 6 | protein-coding |
| chr1_93391433_93392020 | chr1 | 93391434 | 93392020 | + | 1.01829 | intron (NM_207031, intron 10 of 23) | intron (NM_207031, intron 10 of 23) | 17831 | NM_001271884 | 404545 | Mm.297411 | NM_207031 | ENSMUSG00000034107 | **Ano7** | Ngep\|Ngep-L\|Pcanap5\|Tmem16g | anoctamin 7 | protein-coding |
| chr1_93388981_93389322 | chr1 | 93388982 | 93389322 | + | 0.829208 | intron (NM_207031, intron 8 of 23) | intron (NM_207031, intron 8 of 23) | 15256 | NM_001271884 | 404545 | Mm.297411 | NM_207031 | ENSMUSG00000034107 | **Ano7** | Ngep\|Ngep-L\|Pcanap5\|Tmem16g | anoctamin 7 | protein-coding |
| chr9_62372337_62372825 | chr9 | 62372338 | 62372825 | + | 1.36307 | intron (NM_009672, intron 3 of 6) | intron (NM_009672, intron 3 of 6) | 31238 | NM_009672 | 11737 | Mm.269088 | NM_009672 | ENSMUSG00000032249 | **Anp32a** | Anp32\|I1PP2A\|LANP\|PHAP1\|W91701\|pp32 | acidic (leucine-rich) nuclear phosphoprotein 32 family, member A | protein-coding |
| chr4_46458068_46458750 | chr4 | 46458069 | 46458750 | + | 0.722752 | intron (NM_130889, intron 1 of 6) | PB1D9\|SINE\|Alu | 7292 | NM_130889 | 67628 | Mm.263913 | NM_130889 | ENSMUSG00000028333 | **Anp32b** | 2410015B15Rik\|PAL31\|PHAPI2a\|Ssp29 | acidic (leucine-rich) nuclear phosphoprotein 32 family, member B | protein-coding |
| chr5_98034982_98035270 | chr5 | 98034983 | 98035270 | + | 1.50437 | Intergenic | Intergenic | -4164 | NM_133738 | 71914 | Mm.24842 | NM_133738 | | **Antxr2** | 2310046B19Rik\|AW561899\|CMG-2\|CMG2\|cl-35 | anthrax toxin receptor 2 | protein-coding |
| chr5_98081223_98081632 | chr5 | 98081224 | 98081632 | + | 1.14325 | Intergenic | Intergenic | -50466 | NM_133738 | 71914 | Mm.24842 | NM_133738 | | **Antxr2** | 2310046B19Rik\|AW561899\|CMG-2\|CMG2\|cl-35 | anthrax toxin receptor 2 | protein-coding |
| chr5_97999307_98000097 | chr5 | 97999308 | 98000097 | + | 1.115 | intron (NM_133738, intron 7 of 16) | intron (NM_133738, intron 7 of 16) | 31260 | NM_133738 | 71914 | Mm.24842 | NM_133738 | | **Antxr2** | 2310046B19Rik\|AW561899\|CMG-2\|CMG2\|cl-35 | anthrax toxin receptor 2 | protein-coding |
| chr5_97849120_97849800 | chr5 | 97849121 | 97849800 | + | 0.841712 | Intergenic | Intergenic | 181502 | NM_133738 | 71914 | Mm.24842 | NM_133738 | | **Antxr2** | 2310046B19Rik\|AW561899\|CMG-2\|CMG2\|cl-35 | anthrax toxin receptor 2 | protein-coding |
| chr5_97846516_97846856 | chr5 | 97846517 | 97846856 | + | 0.767994 | Intergenic | MTE2b\|LTR\|ERVL-MaLR | 184276 | NM_133738 | 71914 | Mm.24842 | NM_133738 | | **Antxr2** | 2310046B19Rik\|AW561899\|CMG-2\|CMG2\|cl-35 | anthrax toxin receptor 2 | protein-coding |
| chr5_97890668_97891024 | chr5 | 97890669 | 97891024 | + | 0.662594 | intron (NM_133738, intron 16 of 16) | intron (NM_133738, intron 16 of 16) | 140116 | NM_133738 | 71914 | Mm.24842 | NM_133738 | | **Antxr2** | 2310046B19Rik\|AW561899\|CMG-2\|CMG2\|cl-35 | anthrax toxin receptor 2 | protein-coding |
| chr19_20393598_20394110 | chr19 | 20393599 | 20394110 | + | 0.91249 | Intergenic | Intergenic | -3183 | NM_010730 | 16952 | Mm.248360 | NM_010730 | ENSMUSG00000024659 | **Anxa1** | Anx-1\|Anx-A1\|C430014K04Rik\|Lpc-1\|Lpc1 | annexin A1 | protein-coding |
| chr14_25839638_25840136 | chr14 | 25839639 | 25840136 | + | 1.1213 | Intergenic | URR1A\|DNA\|hAT-Charlie | -2268 | NM_013469 | 11744 | Mm.294083 | NM_013469 | ENSMUSG00000021866 | **Anxa11** | A830099O17Rik\|Anx11 | annexin A11 | protein-coding |
| chr14_25861771_25862138 | chr14 | 25861772 | 25862138 | + | 0.955298 | intron (NM_013469, intron 1 of 14) | intron (NM_013469, intron 1 of 14) | 19800 | NM_013469 | 11744 | Mm.294083 | NM_013469 | ENSMUSG00000021866 | **Anxa11** | A830099O17Rik\|Anx11 | annexin A11 | protein-coding |
| chr14_25857399_25857730 | chr14 | 25857400 | 25857730 | + | 0.769054 | intron (NM_013469, intron 1 of 14) | intron (NM_013469, intron 1 of 14) | 15410 | NM_013469 | 11744 | Mm.294083 | NM_013469 | ENSMUSG00000021866 | **Anxa11** | A830099O17Rik\|Anx11 | annexin A11 | protein-coding |
| chr5_96857018_96857679 | chr5 | 96857019 | 96857679 | + | 1.03017 | Intergenic | Intergenic | 63964 | NM_013470 | 11745 | Mm.7214 | NM_013470 | ENSMUSG00000029484 | **Anxa3** | Anx3 | annexin A3 | protein-coding |
| chr5_96824665_96825303 | chr5 | 96824666 | 96825303 | + | 0.860725 | intron (NM_013470, intron 7 of 12) | intron (NM_013470, intron 7 of 12) | 31599 | NM_013470 | 11745 | Mm.7214 | NM_013470 | ENSMUSG00000029484 | **Anxa3** | Anx3 | annexin A3 | protein-coding |
| chr5_96856132_96856933 | chr5 | 96856133 | 96856933 | + | 0.82006 | Intergenic | RMER1B\|Other\|Other | 63148 | NM_013470 | 11745 | Mm.7214 | NM_013470 | ENSMUSG00000029484 | **Anxa3** | Anx3 | annexin A3 | protein-coding |
| chr6_86784698_86785275 | chr6 | 86784699 | 86785275 | + | 0.792205 | intron (NM_013471, intron 1 of 12) | B4A\|SINE\|B4 | 8597 | NM_013471 | 11746 | Mm.259702 | NM_013471 | ENSMUSG00000029994 | **Anxa4** | AI265406\|AIV\|AW106930\|Anx4\|Xanx-4 | annexin A4 | protein-coding |
| chr6_86777279_86778244 | chr6 | 86777280 | 86778244 | + | 0.788134 | intron (NM_013471, intron 1 of 12) | intron (NM_013471, intron 1 of 12) | 15822 | NM_013471 | 11746 | Mm.259702 | NM_013471 | ENSMUSG00000029994 | **Anxa4** | AI265406\|AIV\|AW106930\|Anx4\|Xanx-4 | annexin A4 | protein-coding |
| chr1_58169827_58170507 | chr1 | 58169828 | 58170507 | + | 0.719052 | intron (NM_023617, intron 23 of 34) | intron (NM_023617, intron 23 of 34) | -40230 | NM_023631 | 71872 | Mm.244525 | NM_023631 | ENSMUSG00000038242 | **Aox4** | 2310003G12Rik\|Aoh2 | aldehyde oxidase 4 | protein-coding |
| chr3_127860619_127861275 | chr3 | 127860620 | 127861275 | + | 1.20287 | Intergenic | Intergenic | -23455 | NM_145964 | 211556 | Mm.403486 | NM_145964 | ENSMUSG00000074238 | **Ap1ar** | AA407621\|C78909\|Gadkin | adaptor-related protein complex 1 associated regulatory protein | protein-coding |
| chr11_5000135_5000582 | chr11 | 5000136 | 5000582 | + | 1.07443 | intron (NM_007454, intron 1 of 20) | intron (NM_007454, intron 1 of 20) | 13535 | NM_007454 | 11764 | Mm.274816 | NM_007454 | ENSMUSG00000009090 | **Ap1b1** | Adtb1\|b2b1660Clo | adaptor protein complex AP-1, beta 1 subunit | protein-coding |
| chr11_4999899_5000603 | chr11 | 4999900 | 5000603 | + | 0.973791 | intron (NM_007454, intron 1 of 20) | intron (NM_007454, intron 1 of 20) | 13427 | NM_007454 | 11764 | Mm.274816 | NM_007454 | ENSMUSG00000009090 | **Ap1b1** | Adtb1\|b2b1660Clo | adaptor protein complex AP-1, beta 1 subunit | protein-coding |
| chr8_109773164_109773634 | chr8 | 109773165 | 109773634 | + | 0.894529 | Intergenic | Intergenic | -5184 | NM_009677 | 11765 | Mm.37210 | NM_009677 | | **Ap1g1** | AA409002\|AU041323\|AW551707\|Adtg\|D8Ertd374e | adaptor protein complex AP-1, gamma 1 subunit | protein-coding |
| chr1_79701550_79702118 | chr1 | 79701551 | 79702118 | + | 1.54738 | TTS (NM_001111279) | TTS (NM_001111279) | -29862 | NM_183027 | 252903 | Mm.32409 | NM_183027 | ENSMUSG00000054702 | **Ap1s3** | 1190009B22\|Jr2\|[s]3A | adaptor-related protein complex AP-1, sigma 3 | protein-coding |
| chr1_79559220_79559939 | chr1 | 79559221 | 79559939 | + | 1.15958 | Intergenic | Intergenic | 112392 | NM_183027 | 252903 | Mm.32409 | NM_183027 | ENSMUSG00000054702 | **Ap1s3** | 1190009B22\|Jr2\|[s]3A | adaptor-related protein complex AP-1, sigma 3 | protein-coding |
| chr1_79559233_79559731 | chr1 | 79559234 | 79559731 | + | 1.08228 | Intergenic | Intergenic | 112490 | NM_183027 | 252903 | Mm.32409 | NM_183027 | ENSMUSG00000054702 | **Ap1s3** | 1190009B22\|Jr2\|[s]3A | adaptor-related protein complex AP-1, sigma 3 | protein-coding |
| chr1_79581422_79581745 | chr1 | 79581423 | 79581745 | + | 0.921403 | Intergenic | Intergenic | 90388 | NM_183027 | 252903 | Mm.32409 | NM_183027 | ENSMUSG00000054702 | **Ap1s3** | 1190009B22\|Jr2\|[s]3A | adaptor-related protein complex AP-1, sigma 3 | protein-coding |
| chr1_79581147_79582027 | chr1 | 79581148 | 79582027 | + | 0.869849 | Intergenic | Intergenic | 90385 | NM_183027 | 252903 | Mm.32409 | NM_183027 | ENSMUSG00000054702 | **Ap1s3** | 1190009B22\|Jr2\|[s]3A | adaptor-related protein complex AP-1, sigma 3 | protein-coding |
| chr7_141580088_141580511 | chr7 | 141580089 | 141580511 | + | 1.10636 | intron (NM_007459, intron 1 of 21) | intron (NM_007459, intron 1 of 21) | 18120 | NM_007459 | 11772 | Mm.253090 | NM_007459 | ENSMUSG00000002957 | **Ap2a2** | 2410074K14Rik\|AF006990\|AW146353\|Adtab\|C78001\|L25 | adaptor-related protein complex 2, alpha 2 subunit | protein-coding |
| chr7_141564563_141564969 | chr7 | 141564564 | 141564969 | + | 1.01586 | intron (NM_007459, intron 1 of 21) | intron (NM_007459, intron 1 of 21) | 2586 | NM_007459 | 11772 | Mm.253090 | NM_007459 | ENSMUSG00000002957 | **Ap2a2** | 2410074K14Rik\|AF006990\|AW146353\|Adtab\|C78001\|L25 | adaptor-related protein complex 2, alpha 2 subunit | protein-coding |
| chr10_91028859_91029488 | chr10 | 91028860 | 91029488 | + | 1.60269 | intron (NM_001282947, intron 17 of 26) | L1MB4\|LINE\|L1 | 53569 | NM_001042558 | 11783 | Mm.220289 | NM_009684 | ENSMUSG00000019979 | **Apaf1** | 6230400I06Rik\|Apaf-1\|Apaf1l\|fog\|mKIAA0413 | apoptotic peptidase activating factor 1 | protein-coding |
| chr10_91076636_91077088 | chr10 | 91076637 | 91077088 | + | 0.894919 | intron (NM_001282947, intron 4 of 26) | ID4\|SINE\|ID | 5881 | NM_001042558 | 11783 | Mm.220289 | NM_009684 | ENSMUSG00000019979 | **Apaf1** | 6230400I06Rik\|Apaf-1\|Apaf1l\|fog\|mKIAA0413 | apoptotic peptidase activating factor 1 | protein-coding |
| chr10_91076680_91077315 | chr10 | 91076681 | 91077315 | + | 0.7689 | intron (NM_001282947, intron 4 of 26) | MLT1D\|LTR\|ERVL-MaLR | 5745 | NM_001042558 | 11783 | Mm.220289 | NM_009684 | ENSMUSG00000019979 | **Apaf1** | 6230400I06Rik\|Apaf-1\|Apaf1l\|fog\|mKIAA0413 | apoptotic peptidase activating factor 1 | protein-coding |
| chr2_22723990_22724655 | chr2 | 22723991 | 22724655 | + | 1.31404 | Intergenic | Lx8\|LINE\|L1 | -50004 | NM_019456 | 54519 | Mm.14255 | NM_019456 | ENSMUSG00000026786 | **Apbb1ip** | 9930118P07Rik\|Prp48 | amyloid beta (A4) precursor protein-binding, family B, member 1 interacting protein | protein-coding |
| chr2_22724233_22724637 | chr2 | 22724234 | 22724637 | + | 1.21653 | Intergenic | ORR1B2\|LTR\|ERVL-MaLR | -49892 | NM_019456 | 54519 | Mm.14255 | NM_019456 | ENSMUSG00000026786 | **Apbb1ip** | 9930118P07Rik\|Prp48 | amyloid beta (A4) precursor protein-binding, family B, member 1 interacting protein | protein-coding |
| chr2_22761216_22761670 | chr2 | 22761217 | 22761670 | + | 1.18771 | Intergenic | Intergenic | -12884 | NM_019456 | 54519 | Mm.14255 | NM_019456 | ENSMUSG00000026786 | **Apbb1ip** | 9930118P07Rik\|Prp48 | amyloid beta (A4) precursor protein-binding, family B, member 1 interacting protein | protein-coding |
| chr2_22761067_22761687 | chr2 | 22761068 | 22761687 | + | 1.16292 | Intergenic | Intergenic | -12950 | NM_019456 | 54519 | Mm.14255 | NM_019456 | ENSMUSG00000026786 | **Apbb1ip** | 9930118P07Rik\|Prp48 | amyloid beta (A4) precursor protein-binding, family B, member 1 interacting protein | protein-coding |

| | | | | | | | | | | | | | | | | | |
|---|---|---|---|---|---|---|---|---|---|---|---|---|---|---|---|---|---|
| chr2_22799333_22799742 | chr2 | 22799334 | 22799742 | + | 1.08792 | intron (NM_019456, intron 1 of 13) | MTD\|LTR\|ERVL-MaLR | 25211 | NM_019456 | 54519 | Mm.14255 | NM_019456 | ENSMUSG00000026786 | **Apbb1ip** | 9930118P07Rik\|Prp48 | amyloid beta (A4) precursor protein-binding, family B, member 1 interacting protein | protein-coding |
| chr2_22793684_22794175 | chr2 | 22793685 | 22794175 | + | 0.936202 | intron (NM_019456, intron 1 of 13) | intron (NM_019456, intron 1 of 13) | 19603 | NM_019456 | 54519 | Mm.14255 | NM_019456 | ENSMUSG00000026786 | **Apbb1ip** | 9930118P07Rik\|Prp48 | amyloid beta (A4) precursor protein-binding, family B, member 1 interacting protein | protein-coding |
| chr2_22783404_22783661 | chr2 | 22783405 | 22783661 | + | 0.926516 | intron (NM_019456, intron 1 of 13) | intron (NM_019456, intron 1 of 13) | 9206 | NM_019456 | 54519 | Mm.14255 | NM_019456 | ENSMUSG00000026786 | **Apbb1ip** | 9930118P07Rik\|Prp48 | amyloid beta (A4) precursor protein-binding, family B, member 1 interacting protein | protein-coding |
| chr2_22769267_22769552 | chr2 | 22769268 | 22769552 | + | 0.720035 | Intergenic | ORR1F\|LTR\|ERVL-MaLR | -4917 | NM_019456 | 54519 | Mm.14255 | NM_019456 | ENSMUSG00000026786 | **Apbb1ip** | 9930118P07Rik\|Prp48 | amyloid beta (A4) precursor protein-binding, family B, member 1 interacting protein | protein-coding |
| chr2_22781408_22781677 | chr2 | 22781409 | 22781677 | + | 0.686999 | intron (NM_019456, intron 1 of 13) | L1_Mus4\|LINE\|L1 | 7216 | NM_019456 | 54519 | Mm.14255 | NM_019456 | ENSMUSG00000026786 | **Apbb1ip** | 9930118P07Rik\|Prp48 | amyloid beta (A4) precursor protein-binding, family B, member 1 interacting protein | protein-coding |
| chr2_22792292_22792787 | chr2 | 22792293 | 22792787 | + | 0.645273 | intron (NM_019456, intron 1 of 13) | intron (NM_019456, intron 1 of 13) | 18213 | NM_019456 | 54519 | Mm.14255 | NM_019456 | ENSMUSG00000026786 | **Apbb1ip** | 9930118P07Rik\|Prp48 | amyloid beta (A4) precursor protein-binding, family B, member 1 interacting protein | protein-coding |
| chr9_108120651_108121100 | chr9 | 108120652 | 108121100 | + | 1.17942 | intron (NM_007567, intron 3 of 11) | intron (NM_007567, intron 3 of 11) | -26396 | NM_146226 | 235606 | Mm.219639 | NM_146226 | ENSMUSG00000032590 | **Apeh** | - | acylpeptide hydrolase | protein-coding |
| chr9_108120198_108121100 | chr9 | 108120199 | 108121100 | + | 1.04609 | intron (NM_007567, intron 3 of 11) | intron (NM_007567, intron 3 of 11) | -26169 | NM_146226 | 235606 | Mm.219639 | NM_146226 | ENSMUSG00000032590 | **Apeh** | - | acylpeptide hydrolase | protein-coding |
| chr9_66802447_66803371 | chr9 | 66802448 | 66803371 | + | 0.621183 | Intergenic | Intergenic | -7486 | NM_177583 | 208117 | Mm.41291 | NM_177583 | ENSMUSG00000032375 | **Aph1b** | 2310057K14Rik\|Aph1c | anterior pharynx defective 1b homolog (C. elegans) | protein-coding |
| chrX_48046057_48046680 | chrX | 48046058 | 48046680 | + | 1.62207 | Intergenic | ORR1E\|LTR\|ERVL-MaLR | -11517 | NM_013912 | 30878 | Mm.29262 | NM_013912 | ENSMUSG00000037010 | **Apln** | 6030430G11Rik\|Apel | apelin | protein-coding |
| chr9_46168511_46169247 | chr9 | 46168512 | 46169247 | + | 1.53488 | intron (NM_027498, intron 4 of 24) | intron (NM_027498, intron 4 of 24) | -59751 | NM_009692 | 11806 | Mm.26743 | NM_009692 | ENSMUSG00000032083 | **Apoa1** | Alp-1\|Apoa-1\|Brp-14\|Ltw-1\|Lvtw-1\|Sep-1\|Sep-2\|Sep2\|apo-AI\|apoA-I | apolipoprotein A-I | protein-coding |
| chr9_46172284_46172589 | chr9 | 46172285 | 46172589 | + | 1.16821 | intron (NM_027498, intron 4 of 24) | intron (NM_027498, intron 4 of 24) | -56193 | NM_009692 | 11806 | Mm.26743 | NM_009692 | ENSMUSG00000032083 | **Apoa1** | Alp-1\|Apoa-1\|Brp-14\|Ltw-1\|Lvtw-1\|Sep-1\|Sep-2\|Sep2\|apo-AI\|apoA-I | apolipoprotein A-I | protein-coding |
| chr9_46163494_46163966 | chr9 | 46163495 | 46163966 | + | 0.921869 | intron (NM_027498, intron 4 of 24) | intron (NM_027498, intron 4 of 24) | -64900 | NM_009692 | 11806 | Mm.26743 | NM_009692 | ENSMUSG00000032083 | **Apoa1** | Alp-1\|Apoa-1\|Brp-14\|Ltw-1\|Lvtw-1\|Sep-1\|Sep-2\|Sep2\|apo-AI\|apoA-I | apolipoprotein A-I | protein-coding |
| chr9_46120942_46121185 | chr9 | 46120943 | 46121185 | + | 0.715716 | intron (NM_027498, intron 1 of 24) | MER44C\|DNA\|TcMar-Tigger | -107566 | NM_009692 | 11806 | Mm.26743 | NM_009692 | ENSMUSG00000032083 | **Apoa1** | Alp-1\|Apoa-1\|Brp-14\|Ltw-1\|Lvtw-1\|Sep-1\|Sep-2\|Sep2\|apo-AI\|apoA-I | apolipoprotein A-I | protein-coding |
| chr9_46173001_46173328 | chr9 | 46173002 | 46173328 | + | 0.667214 | intron (NM_027498, intron 4 of 24) | intron (NM_027498, intron 4 of 24) | -55465 | NM_009692 | 11806 | Mm.26743 | NM_009692 | ENSMUSG00000032083 | **Apoa1** | Alp-1\|Apoa-1\|Brp-14\|Ltw-1\|Lvtw-1\|Sep-1\|Sep-2\|Sep2\|apo-AI\|apoA-I | apolipoprotein A-I | protein-coding |
| chr16_31352816_31353355 | chr16 | 31352817 | 31353355 | + | 0.680695 | Intergenic | Intergenic | -38278 | NM_001301354 | 11815 | Mm.2082 | NM_007470 | ENSMUSG00000022548 | **Apod** | - | apolipoprotein D | protein-coding |
| chr15_77785745_77786219 | chr15 | 77785746 | 77786219 | + | 0.879343 | intron (NM_022410, intron 13 of 40) | intron (NM_022410, intron 13 of 40) | -30753 | NM_001081970 | 239552 | Mm.125650 | NM_001081970 | ENSMUSG00000056656 | **Apol8** | 9830006J20Rik\|APOL-II\|Apol2 | apolipoprotein L 8 | protein-coding |
| chr15_77785523_77786207 | chr15 | 77785524 | 77786207 | + | 0.819368 | intron (NM_022410, intron 13 of 40) | intron (NM_022410, intron 13 of 40) | -30636 | NM_001081970 | 239552 | Mm.125650 | NM_001081970 | ENSMUSG00000056656 | **Apol8** | 9830006J20Rik\|APOL-II\|Apol2 | apolipoprotein L 8 | protein-coding |
| chr10_128256332_128256729 | chr10 | 128256333 | 128256729 | + | 1.23935 | TTS (NM_133996) | TTS (NM_133996) | 2400 | NM_133996 | 28194 | Mm.439846 | NM_133996 | ENSMUSG00000051716 | **Apon** | AI195275\|D10Ucla2 | apolipoprotein N | protein-coding |
| chr10_128258953_128259318 | chr10 | 128258954 | 128259318 | + | 0.829362 | Intergenic | Lx8\|LINE\|L1 | 5005 | NM_133996 | 28194 | Mm.439846 | NM_133996 | ENSMUSG00000051716 | **Apon** | AI195275\|D10Ucla2 | apolipoprotein N | protein-coding |
| chrX_112347985_112348379 | chrX | 112347986 | 112348379 | + | 0.913247 | intron (NM_026565, intron 3 of 8) | intron (NM_026565, intron 3 of 8) | 36774 | NM_026565 | 68117 | Mm.161014 | NM_026565 | ENSMUSG00000025525 | **Apool** | 6720473G16Rik\|9430083G14Rik\|E130106L15Rik | apolipoprotein O-like | protein-coding |
| chr4_40704477_40704724 | chr4 | 40704478 | 40704724 | + | 0.706936 | Intergenic | L1MB1\|LINE\|L1 | -1407 | NM_001025445 | 66408 | Mm.430710 | NM_025545 | ENSMUSG00000028411 | **Aptx** | 2410016G21Rik\|AA388047\|FHA-HIT | aprataxin | protein-coding |
| chrX_20816970_20817876 | chrX | 20816971 | 20817876 | + | 1.22244 | Intergenic | Intergenic | -31120 | NM_009703 | 11836 | Mm.220946 | NM_009703 | ENSMUSG00000001127 | **Araf** | 1200013E08Rik\|A-Raf\|AW495444\|Araf1 | v-raf murine sarcoma 3611 viral oncogene homolog | protein-coding |
| chrX_20816991_20817556 | chrX | 20816992 | 20817556 | + | 1.18493 | Intergenic | Intergenic | -31269 | NM_009703 | 11836 | Mm.220946 | NM_009703 | ENSMUSG00000001127 | **Araf** | 1200013E08Rik\|A-Raf\|AW495444\|Araf1 | v-raf murine sarcoma 3611 viral oncogene homolog | protein-coding |
| chrX_20816993_20817269 | chrX | 20816994 | 20817269 | + | 1.06633 | Intergenic | Intergenic | -31412 | NM_009703 | 11836 | Mm.220946 | NM_009703 | ENSMUSG00000001127 | **Araf** | 1200013E08Rik\|A-Raf\|AW495444\|Araf1 | v-raf murine sarcoma 3611 viral oncogene homolog | protein-coding |
| chr7_101361676_101362108 | chr7 | 101361677 | 101362108 | + | 0.828572 | intron (NM_001040111, intron 1 of 33) | intron (NM_001040111, intron 1 of 33) | 13823 | NM_001040111 | 69710 | Mm.277687 | NM_027180 | ENSMUSG00000032812 | **Arap1** | 2410002L19Rik\|Centd2\|mKIAA0782 | ArfGAP with RhoGAP domain, ankyrin repeat and PH domain 1 | protein-coding |
| chr7_101361794_101362108 | chr7 | 101361795 | 101362108 | + | 0.823153 | intron (NM_001040111, intron 1 of 33) | intron (NM_001040111, intron 1 of 33) | 13882 | NM_001040111 | 69710 | Mm.277687 | NM_027180 | ENSMUSG00000032812 | **Arap1** | 2410002L19Rik\|Centd2\|mKIAA0782 | ArfGAP with RhoGAP domain, ankyrin repeat and PH domain 1 | protein-coding |
| chr5_62780550_62780872 | chr5 | 62780551 | 62780872 | + | 1.27912 | Intergenic | Intergenic | -14534 | NM_178407 | 212285 | Mm.244403 | NM_178407 | ENSMUSG00000037999 | **Arap2** | Centd1\|Gm148\|mKIAA0580 | ArfGAP with RhoGAP domain, ankyrin repeat and PH domain 2 | protein-coding |
| chr5_62786806_62787108 | chr5 | 62786807 | 62787108 | + | 0.890162 | Intergenic | Lx8\|LINE\|L1 | -20780 | NM_178407 | 212285 | Mm.244403 | NM_178407 | ENSMUSG00000037999 | **Arap2** | Centd1\|Gm148\|mKIAA0580 | ArfGAP with RhoGAP domain, ankyrin repeat and PH domain 2 | protein-coding |
| chr5_62788477_62788701 | chr5 | 62788478 | 62788701 | + | 0.8746 | Intergenic | Intergenic | -22412 | NM_178407 | 212285 | Mm.244403 | NM_178407 | ENSMUSG00000037999 | **Arap2** | Centd1\|Gm148\|mKIAA0580 | ArfGAP with RhoGAP domain, ankyrin repeat and PH domain 2 | protein-coding |
| chr15_74687314_74688053 | chr15 | 74687315 | 74688053 | + | 0.872847 | Intergenic | Intergenic | -15114 | NM_018790 | 11838 | Mm.491310 | NM_018790 | ENSMUSG00000022602 | **Arc** | Arc3.1\|C86064\|arg3.1\|mArc | activity regulated cytoskeletal-associated protein | protein-coding |
| chr12_84958219_84958712 | chr12 | 84958220 | 84958712 | + | 1.15024 | intron (NM_178065, intron 1 of 19) | intron (NM_178065, intron 1 of 19) | 12420 | NM_178065 | 68497 | Mm.24446 | NM_178065 | ENSMUSG00000042350 | **Arel1** | 1110018G07Rik\|AI649076\|AW610792\|Kiaa0317\|mKIAA0317 | apoptosis resistant E3 ubiquitin protein ligase 1 | protein-coding |
| chr12_84968897_84969752 | chr12 | 84968898 | 84969752 | + | 0.911991 | intron (NM_178065, intron 1 of 19) | intron (NM_178065, intron 1 of 19) | 1561 | NM_178065 | 68497 | Mm.24446 | NM_178065 | ENSMUSG00000042350 | **Arel1** | 1110018G07Rik\|AI649076\|AW610792\|Kiaa0317\|mKIAA0317 | apoptosis resistant E3 ubiquitin protein ligase 1 | protein-coding |
| chr12_84956324_84956719 | chr12 | 84956325 | 84956719 | + | 0.85647 | intron (NM_178065, intron 1 of 19) | L3b\|LINE\|CR1 | 14364 | NM_178065 | 68497 | Mm.24446 | NM_178065 | ENSMUSG00000042350 | **Arel1** | 1110018G07Rik\|AI649076\|AW610792\|Kiaa0317\|mKIAA0317 | apoptosis resistant E3 ubiquitin protein ligase 1 | protein-coding |
| chr11_59226285_59226999 | chr11 | 59226286 | 59226999 | + | 0.909041 | intron (NM_007476, intron 1 of 4) | intron (NM_007476, intron 1 of 4) | 1520 | NM_001130408 | 11840 | Mm.371546 | NM_007476 | ENSMUSG00000048076 | **Arf1** | - | ADP-ribosylation factor 1 | protein-coding |
| chr11_59219680_59220004 | chr11 | 59219681 | 59220004 | + | 0.682986 | intron (NM_007476, intron 1 of 4) | intron (NM_007476, intron 1 of 4) | 8320 | NM_001130408 | 11840 | Mm.371546 | NM_007476 | ENSMUSG00000048076 | **Arf1** | - | ADP-ribosylation factor 1 | protein-coding |
| chr11_103964002_103964406 | chr11 | 103964003 | 103964406 | + | 1.31088 | Intergenic | MTD\|LTR\|ERVL-MaLR | -2667 | NM_007477 | 11841 | Mm.5061 | NM_007477 | ENSMUSG00000062421 | **Arf2** | - | ADP-ribosylation factor 2 | protein-coding |
| chr12_69416804_69417128 | chr12 | 69416805 | 69417128 | + | 1.53177 | Intergenic | Intergenic | 44816 | NM_007481 | 11845 | Mm.27308 | NM_007481 | ENSMUSG00000044147 | **Arf6** | AI788669\|AW496366 | ADP-ribosylation factor 6 | protein-coding |
| chr12_69416784_69417315 | chr12 | 69416785 | 69417315 | + | 1.39885 | Intergenic | Intergenic | 44900 | NM_007481 | 11845 | Mm.27308 | NM_007481 | ENSMUSG00000044147 | **Arf6** | AI788669\|AW496366 | ADP-ribosylation factor 6 | protein-coding |
| chr1_10219913_10220551 | chr1 | 10219914 | 10220551 | + | 0.732337 | intron (NM_001102430, intron 1 of 37) | Lx9\|LINE\|L1 | 12438 | NM_001102430 | 211673 | Mm.229141 | NM_001102430 | ENSMUSG00000067851 | **Arfgef1** | ARFGEP1\|BIG1\|D130059B05Rik\|D730028O18Rik\|P200 | ADP-ribosylation factor guanine nucleotide-exchange factor 1(brefeldin A-inhibited) | protein-coding |
| chr3_84579959_84580247 | chr3 | 84579960 | 84580247 | + | 1.00012 | intron (NM_001293801, intron 1 of 7) | URR1B\|DNA\|hAT-Charlie | 2522 | NM_001081093 | 99889 | Mm.316536 | NM_001081093 | ENSMUSG00000074513 | **Arfip1** | 2310030I10Rik\|Arfaptin\|D3Ertd598e | ADP-ribosylation factor interacting protein 1 | protein-coding |
| chr3_84570515_84571209 | chr3 | 84570516 | 84571209 | + | 0.683494 | intron (NM_001293801, intron 1 of 7) | intron (NM_001293801, intron 1 of 7) | 11763 | NM_001081093 | 99889 | Mm.316536 | NM_001081093 | ENSMUSG00000074513 | **Arfip1** | 2310030I10Rik\|Arfaptin\|D3Ertd598e | ADP-ribosylation factor interacting protein 1 | protein-coding |
| chr2_113886077_113886451 | chr2 | 113886078 | 113886451 | + | 0.694736 | Intergenic | Intergenic | -37603 | NM_181416 | 228482 | Mm.45753 | NM_181416 | ENSMUSG00000041219 | **Arhgap11a** | 6530401L14Rik\|mKIAA0013 | Rho GTPase activating protein 11A | protein-coding |
| chr2_43856747_43857196 | chr2 | 43856748 | 43857196 | + | 1.07754 | intron (NM_153820, intron 5 of 13) | intron (NM_153820, intron 5 of 13) | 108145 | NM_001025377 | 76117 | Mm.217350 | NM_153820 | ENSMUSG00000049744 | **Arhgap15** | 5830480G12Rik | Rho GTPase activating protein 15 | protein-coding |
| chr2_43753312_43753553 | chr2 | 43753313 | 43753553 | + | 0.866415 | intron (NM_153820, intron 1 of 13) | intron (NM_153820, intron 1 of 13) | 4606 | NM_001025377 | 76117 | Mm.217350 | NM_153820 | ENSMUSG00000049744 | **Arhgap15** | 5830480G12Rik | Rho GTPase activating protein 15 | protein-coding |
| chr2_43720286_43720791 | chr2 | 43720287 | 43720791 | + | 0.759923 | Intergenic | ID4_\|SINE\|ID | -28285 | NM_153820 | 76117 | Mm.217350 | NM_153820 | ENSMUSG00000049744 | **Arhgap15** | 5830480G12Rik | Rho GTPase activating protein 15 | protein-coding |
| chr2_43752985_43753622 | chr2 | 43752986 | 43753622 | + | 0.642116 | intron (NM_153820, intron 1 of 13) | intron (NM_153820, intron 1 of 13) | 4477 | NM_001025377 | 76117 | Mm.217350 | NM_153820 | ENSMUSG00000049744 | **Arhgap15** | 5830480G12Rik | Rho GTPase activating protein 15 | protein-coding |
| chr2_44345308_44345736 | chr2 | 44345309 | 44345736 | + | 1.32552 | intron (NM_153820, intron 13 of 13) | intron (NM_153820, intron 13 of 13) | -280198 | NR_040622 | 320872 | Mm.445023 | NR_040622 | ENSMUSG00000085089 | **Arhgap15os** | A430092G05Rik | Rho GTPase activating protein 15, opposite strand | ncRNA |
| chr2_44166723_44167088 | chr2 | 44166724 | 44167088 | + | 1.23255 | intron (NM_153820, intron 10 of 13) | intron (NM_153820, intron 10 of 13) | -101582 | NR_040622 | 320872 | Mm.445023 | NR_040622 | ENSMUSG00000085089 | **Arhgap15os** | A430092G05Rik | Rho GTPase activating protein 15, opposite strand | ncRNA |
| chr2_44344867_44345242 | chr2 | 44344868 | 44345242 | + | 1.11013 | intron (NM_153820, intron 13 of 13) | intron (NM_153820, intron 13 of 13) | -279731 | NR_040622 | 320872 | Mm.445023 | NR_040622 | ENSMUSG00000085089 | **Arhgap15os** | A430092G05Rik | Rho GTPase activating protein 15, opposite strand | ncRNA |

| | | | | | | | | | | | | | | | | | |
|---|---|---|---|---|---|---|---|---|---|---|---|---|---|---|---|---|---|
| chr2_44165759_44166235 | chr2 | 44165760 | 44166235 | + | 0.90882 | intron (NM_153820, intron 10 of 13) | intron (NM_153820, intron 10 of 13) | -100673 | NR_040622 | 320872 | Mm.445023 | NR_040622 | ENSMUSG00000085089 | **Arhgap15os** | A430092G05Rik | Rho GTPase activating protein 15, opposite strand | ncRNA |
| chr2_44165696_44166459 | chr2 | 44165697 | 44166459 | + | 0.837368 | intron (NM_153820, intron 10 of 13) | intron (NM_153820, intron 10 of 13) | -100754 | NR_040622 | 320872 | Mm.445023 | NR_040622 | ENSMUSG00000085089 | **Arhgap15os** | A430092G05Rik | Rho GTPase activating protein 15, opposite strand | ncRNA |
| chr7_123361707_123362007 | chr7 | 123361708 | 123362007 | + | 1.35118 | intron (NM_001122643, intron 1 of 19) | intron (NM_001122643, intron 1 of 19) | 8058 | NM_001122643 | 70497 | Mm.403318 | NM_144529 | ENSMUSG00000030766 | **Arhgap17** | 5730403H17Rik\|Nadrin\|Nadrin2\|Rich1\|Wbp15 | Rho GTPase activating protein 17 | protein-coding |
| chr7_123322130_123323065 | chr7 | 123322131 | 123323065 | + | 1.2769 | intron (NM_001122643, intron 4 of 19) | intron (NM_001122643, intron 4 of 19) | 47317 | NM_001122643 | 70497 | Mm.403318 | NM_144529 | ENSMUSG00000030766 | **Arhgap17** | 5730403H17Rik\|Nadrin\|Nadrin2\|Rich1\|Wbp15 | Rho GTPase activating protein 17 | protein-coding |
| chr7_123346612_123347270 | chr7 | 123346613 | 123347270 | + | 1.2638 | intron (NM_001122643, intron 1 of 19) | intron (NM_001122643, intron 1 of 19) | 22974 | NM_001122643 | 70497 | Mm.403318 | NM_144529 | ENSMUSG00000030766 | **Arhgap17** | 5730403H17Rik\|Nadrin\|Nadrin2\|Rich1\|Wbp15 | Rho GTPase activating protein 17 | protein-coding |
| chr7_123361223_123362017 | chr7 | 123361224 | 123362017 | + | 1.25629 | intron (NM_001122643, intron 1 of 19) | intron (NM_001122643, intron 1 of 19) | 8295 | NM_001122643 | 70497 | Mm.403318 | NM_144529 | ENSMUSG00000030766 | **Arhgap17** | 5730403H17Rik\|Nadrin\|Nadrin2\|Rich1\|Wbp15 | Rho GTPase activating protein 17 | protein-coding |
| chr7_123361223_123361662 | chr7 | 123361224 | 123361662 | + | 0.76697 | intron (NM_001122643, intron 1 of 19) | intron (NM_001122643, intron 1 of 19) | 8472 | NM_001122643 | 70497 | Mm.403318 | NM_144529 | ENSMUSG00000030766 | **Arhgap17** | 5730403H17Rik\|Nadrin\|Nadrin2\|Rich1\|Wbp15 | Rho GTPase activating protein 17 | protein-coding |
| chr10_26878931_26879581 | chr10 | 26878932 | 26879581 | + | 1.14467 | intron (NM_176837, intron 9 of 14) | intron (NM_176837, intron 9 of 14) | 106744 | NM_176837 | 73910 | Mm.356496 | NM_176837 | ENSMUSG00000039031 | **Arhgap18** | 4833419J07Rik\|AI314906\|AI843501 | Rho GTPase activating protein 18 | protein-coding |
| chr10_26996008_26996467 | chr10 | 26996009 | 26996467 | + | 1.07109 | intron (NM_008481, intron 57 of 63) | intron (NM_008481, intron 57 of 63) | 223726 | NM_176837 | 73910 | Mm.356496 | NM_176837 | ENSMUSG00000039031 | **Arhgap18** | 4833419J07Rik\|AI314906\|AI843501 | Rho GTPase activating protein 18 | protein-coding |
| chr10_26995632_26996545 | chr10 | 26995633 | 26996545 | + | 1.006 | intron (NM_008481, intron 57 of 63) | intron (NM_008481, intron 57 of 63) | 223577 | NM_176837 | 73910 | Mm.356496 | NM_176837 | ENSMUSG00000039031 | **Arhgap18** | 4833419J07Rik\|AI314906\|AI843501 | Rho GTPase activating protein 18 | protein-coding |
| chr10_26995632_26995944 | chr10 | 26995633 | 26995944 | + | 0.630954 | intron (NM_008481, intron 57 of 63) | intron (NM_008481, intron 57 of 63) | 223276 | NM_176837 | 73910 | Mm.356496 | NM_176837 | ENSMUSG00000039031 | **Arhgap18** | 4833419J07Rik\|AI314906\|AI843501 | Rho GTPase activating protein 18 | protein-coding |
| chr5_102481099_102482343 | chr5 | 102481100 | 102482343 | + | 1.30872 | 5' UTR (NM_029270, exon 1 of 10) | 5' UTR (NM_029270, exon 1 of 10) | 330 | NM_029270 | 231532 | Mm.440191 | NM_029270 | ENSMUSG00000057315 | **Arhgap24** | 0610025G21Rik | Rho GTPase activating protein 24 | protein-coding |
| chr5_102451687_102452245 | chr5 | 102451688 | 102452245 | + | 0.701163 | Intergenic | Intergenic | -29425 | NM_146161 | 231532 | Mm.440191 | NM_029270 | ENSMUSG00000057315 | **Arhgap24** | 0610025G21Rik | Rho GTPase activating protein 24 | protein-coding |
| chr6_87536168_87536469 | chr6 | 87536169 | 87536469 | + | 0.896702 | Intergenic | Intergenic | -3060 | NM_001286610 | 232201 | Mm.119564 | NM_175476 | | **Arhgap25** | A130039I20Rik | Rho GTPase activating protein 25 | protein-coding |
| chr18_39019851_39020247 | chr18 | 39019852 | 39020247 | + | 0.82231 | intron (NM_175164, intron 1 of 22) | intron (NM_175164, intron 1 of 22) | 26904 | NM_175164 | 71302 | Mm.329396 | NM_175164 | ENSMUSG00000036452 | **Arhgap26** | 1810044B20Rik\|2610010G17Rik\|4933432P15Rik\|AI853435\|mKIAA0621 | Rho GTPase activating protein 26 | protein-coding |
| chr18_39137440_39137805 | chr18 | 39137441 | 39137805 | + | 0.675196 | intron (NM_175164, intron 10 of 22) | intron (NM_175164, intron 10 of 22) | 144478 | NM_175164 | 71302 | Mm.329396 | NM_175164 | ENSMUSG00000036452 | **Arhgap26** | 1810044B20Rik\|2610010G17Rik\|4933432P15Rik\|AI853435\|mKIAA0621 | Rho GTPase activating protein 26 | protein-coding |
| chr18_39010301_39011080 | chr18 | 39010302 | 39011080 | + | 0.611057 | intron (NM_175164, intron 1 of 22) | intron (NM_175164, intron 1 of 22) | 17546 | NM_175164 | 71302 | Mm.329396 | NM_175164 | ENSMUSG00000036452 | **Arhgap26** | 1810044B20Rik\|2610010G17Rik\|4933432P15Rik\|AI853435\|mKIAA0621 | Rho GTPase activating protein 26 | protein-coding |
| chr16_38678767_38679223 | chr16 | 38678768 | 38679223 | + | 1.12768 | intron (NM_020260, intron 1 of 11) | RCHARR1\|DNA\|hAT-Charlie | 34040 | NM_020260 | 12549 | Mm.370848 | NM_020260 | ENSMUSG00000022799 | **Arhgap31** | 5830477L08Rik\|AU041750\|Cdgap\|D330026I07Rik | Rho GTPase activating protein 31 | protein-coding |
| chr16_38679317_38680051 | chr16 | 38679318 | 38680051 | + | 0.654484 | intron (NM_020260, intron 1 of 11) | intron (NM_020260, intron 1 of 11) | 33351 | NM_020260 | 12549 | Mm.370848 | NM_020260 | ENSMUSG00000022799 | **Arhgap31** | 5830477L08Rik\|AU041750\|Cdgap\|D330026I07Rik | Rho GTPase activating protein 31 | protein-coding |
| chr16_38679311_38680480 | chr16 | 38679312 | 38680480 | + | 0.618462 | intron (NM_020260, intron 1 of 11) | intron (NM_020260, intron 1 of 11) | 33139 | NM_020260 | 12549 | Mm.370848 | NM_020260 | ENSMUSG00000022799 | **Arhgap31** | 5830477L08Rik\|AU041750\|Cdgap\|D330026I07Rik | Rho GTPase activating protein 31 | protein-coding |
| chr9_32089876_32090288 | chr9 | 32089877 | 32090288 | + | 1.14725 | Intergenic | Intergenic | -26054 | NM_001195632 | 330914 | Mm.46683 | NM_177379 | ENSMUSG00000041444 | **Arhgap32** | 3426406O18Rik\|Gc-gap\|Grit\|Px-rics\|Rics\|mKIAA0712\|p200Rhogap\|p250Gap | Rho GTPase activating protein 32 | protein-coding |
| chrX_49407709_49408029 | chrX | 49407710 | 49408029 | + | 0.725983 | Intergenic | B3A\|SINE\|B2 | -62581 | NM_001081123 | 75404 | Mm.44305 | NM_001081123 | ENSMUSG00000036198 | **Arhgap36** | 1100001E04Rik\|rho-gap | Rho GTPase activating protein 36 | protein-coding |
| chr12_52382864_52383480 | chr12 | 52382865 | 52383480 | + | 0.668193 | Intergenic | Intergenic | -132905 | NM_009706 | 11855 | Mm.35059 | NM_009706 | ENSMUSG00000035133 | **Arhgap5** | p190-B\|p190B | Rho GTPase activating protein 5 | protein-coding |
| chrX_169054441_169054851 | chrX | 169054442 | 169054851 | + | 0.777629 | intron (NR_109847, intron 1 of 14) | intron (NR_109847, intron 1 of 14) | 18035 | NR_109847 | 11856 | Mm.441810 | NM_009707 | ENSMUSG00000031355 | **Arhgap6** | AI504284\|RhoGAPX-1 | Rho GTPase activating protein 6 | protein-coding |
| chr8_3408998_3409450 | chr8 | 3408999 | 3409450 | + | 0.702332 | intron (NM_133962, intron 1 of 19) | RSINE1\|SINE\|B4 | 16216 | NM_133962 | 102098 | Mm.170461 | NM_133962 | ENSMUSG00000004568 | **Arhgef18** | AI467246\|D030053O22Rik | rho/rac guanine nucleotide exchange factor (GEF) 18 | protein-coding |
| chr3_88625051_88625386 | chr3 | 88625052 | 88625386 | + | 0.965667 | intron (NM_001198913, intron 1 of 21) | intron (NM_001198913, intron 1 of 21) | 3825 | NM_001198912 | 16800 | Mm.239329 | NM_008487 | ENSMUSG00000028059 | **Arhgef2** | AA408978\|GEF\|GEF-H1\|GEFH1\|LFP40\|Lbcl1\|Lfc\|P40\|mKIAA0651 | rho/rac guanine nucleotide exchange factor (GEF) 2 | protein-coding |
| chr3_88619744_88620891 | chr3 | 88619745 | 88620891 | + | 0.757571 | promoter-TSS (NM_008487) | promoter-TSS (NM_008487) | -788 | NM_008487 | 16800 | Mm.239329 | NM_008487 | ENSMUSG00000028059 | **Arhgef2** | AA408978\|GEF\|GEF-H1\|GEFH1\|LFP40\|Lbcl1\|Lfc\|P40\|mKIAA0651 | rho/rac guanine nucleotide exchange factor (GEF) 2 | protein-coding |
| chr13_97903196_97903985 | chr13 | 97903197 | 97903985 | + | 0.711903 | intron (NM_012026, intron 35 of 35) | intron (NM_012026, intron 35 of 35) | 302574 | NM_012026 | 110596 | Mm.252718 | NM_012026 | ENSMUSG00000021662 | **Arhgef28** | 9230110L08Rik\|AI323540\|D13Bwg1089e\|RIP2\|Rgnef\|RhoGEF\|Rhoip2\|p190RhoGEF | Rho guanine nucleotide exchange factor (GEF) 28 | protein-coding |
| chr14_27376063_27376726 | chr14 | 27376064 | 27376726 | + | 1.44089 | intron (NM_027871, intron 3 of 10) | intron (NM_027871, intron 3 of 10) | 15006 | NM_001289688 | 71704 | Mm.248606 | NM_027871 | ENSMUSG00000021895 | **Arhgef3** | 1200004I24Rik\|9830169H03Rik\|C76747\|Xpln | Rho guanine nucleotide exchange factor (GEF) 3 | protein-coding |
| chr14_27303776_27304094 | chr14 | 27303777 | 27304094 | + | 1.2152 | intron (NM_027871, intron 2 of 10) | PB1D10\|SINE\|Alu | -32102 | NM_001289687 | 71704 | Mm.248606 | NM_027871 | ENSMUSG00000021895 | **Arhgef3** | 1200004I24Rik\|9830169H03Rik\|C76747\|Xpln | Rho guanine nucleotide exchange factor (GEF) 3 | protein-coding |
| chr14_27308483_27309155 | chr14 | 27308484 | 27309155 | + | 1.0789 | intron (NM_027871, intron 2 of 10) | intron (NM_027871, intron 2 of 10) | -27218 | NM_001289687 | 71704 | Mm.248606 | NM_027871 | ENSMUSG00000021895 | **Arhgef3** | 1200004I24Rik\|9830169H03Rik\|C76747\|Xpln | Rho guanine nucleotide exchange factor (GEF) 3 | protein-coding |
| chr14_27234223_27234644 | chr14 | 27234224 | 27234644 | + | 0.943131 | intron (NM_001289686, intron 3 of 12) | intron (NM_001289686, intron 3 of 12) | -3576 | NM_027871 | 71704 | Mm.248606 | NM_027871 | ENSMUSG00000021895 | **Arhgef3** | 1200004I24Rik\|9830169H03Rik\|C76747\|Xpln | Rho guanine nucleotide exchange factor (GEF) 3 | protein-coding |
| chr14_27198443_27199168 | chr14 | 27198444 | 27199168 | + | 0.938251 | intron (NM_001289686, intron 2 of 12) | B1_Mur4\|SINE\|Alu | -39204 | NM_027871 | 71704 | Mm.248606 | NM_027871 | ENSMUSG00000021895 | **Arhgef3** | 1200004I24Rik\|9830169H03Rik\|C76747\|Xpln | Rho guanine nucleotide exchange factor (GEF) 3 | protein-coding |
| chr14_27198447_27199846 | chr14 | 27198448 | 27199846 | + | 0.886026 | intron (NM_001289686, intron 2 of 12) | intron (NM_001289686, intron 2 of 12) | -38863 | NM_027871 | 71704 | Mm.248606 | NM_027871 | ENSMUSG00000021895 | **Arhgef3** | 1200004I24Rik\|9830169H03Rik\|C76747\|Xpln | Rho guanine nucleotide exchange factor (GEF) 3 | protein-coding |
| chr14_27123464_27123984 | chr14 | 27123465 | 27123984 | + | 0.869866 | Intergenic | Intergenic | -20269 | NM_001289686 | 71704 | Mm.248606 | NM_027871 | ENSMUSG00000021895 | **Arhgef3** | 1200004I24Rik\|9830169H03Rik\|C76747\|Xpln | Rho guanine nucleotide exchange factor (GEF) 3 | protein-coding |
| chr14_27303329_27303696 | chr14 | 27303330 | 27303696 | + | 0.868175 | intron (NM_027871, intron 2 of 10) | intron (NM_027871, intron 2 of 10) | -32524 | NM_001289687 | 71704 | Mm.248606 | NM_027871 | ENSMUSG00000021895 | **Arhgef3** | 1200004I24Rik\|9830169H03Rik\|C76747\|Xpln | Rho guanine nucleotide exchange factor (GEF) 3 | protein-coding |
| chr14_27304499_27304746 | chr14 | 27304500 | 27304746 | + | 0.781592 | intron (NM_027871, intron 2 of 10) | intron (NM_027871, intron 2 of 10) | -31414 | NM_001289687 | 71704 | Mm.248606 | NM_027871 | ENSMUSG00000021895 | **Arhgef3** | 1200004I24Rik\|9830169H03Rik\|C76747\|Xpln | Rho guanine nucleotide exchange factor (GEF) 3 | protein-coding |
| chr14_27304396_27304760 | chr14 | 27304397 | 27304760 | + | 0.743032 | intron (NM_027871, intron 2 of 10) | intron (NM_027871, intron 2 of 10) | -31459 | NM_001289687 | 71704 | Mm.248606 | NM_027871 | ENSMUSG00000021895 | **Arhgef3** | 1200004I24Rik\|9830169H03Rik\|C76747\|Xpln | Rho guanine nucleotide exchange factor (GEF) 3 | protein-coding |
| chr14_27344948_27345560 | chr14 | 27344949 | 27345560 | + | 0.688268 | intron (NM_001289687, intron 1 of 9) | intron (NM_001289687, intron 1 of 9) | 9217 | NM_001289687 | 71704 | Mm.248606 | NM_027871 | ENSMUSG00000021895 | **Arhgef3** | 1200004I24Rik\|9830169H03Rik\|C76747\|Xpln | Rho guanine nucleotide exchange factor (GEF) 3 | protein-coding |
| chr14_27236531_27236801 | chr14 | 27236532 | 27236801 | + | 0.687183 | intron (NM_001289686, intron 3 of 12) | intron (NM_001289686, intron 3 of 12) | -1344 | NM_027871 | 71704 | Mm.248606 | NM_027871 | ENSMUSG00000021895 | **Arhgef3** | 1200004I24Rik\|9830169H03Rik\|C76747\|Xpln | Rho guanine nucleotide exchange factor (GEF) 3 | protein-coding |
| chr14_27268105_27268729 | chr14 | 27268106 | 27268729 | + | 0.640237 | intron (NM_027871, intron 2 of 10) | intron (NM_027871, intron 2 of 10) | 30407 | NM_027871 | 71704 | Mm.248606 | NM_027871 | ENSMUSG00000021895 | **Arhgef3** | 1200004I24Rik\|9830169H03Rik\|C76747\|Xpln | Rho guanine nucleotide exchange factor (GEF) 3 | protein-coding |
| chrX_57282216_57282632 | chrX | 57282217 | 57282632 | + | 0.699884 | intron (NM_152801, intron 6 of 21) | intron (NM_152801, intron 6 of 21) | 56305 | NM_152801 | 73341 | Mm.261443 | NM_152801 | ENSMUSG00000031133 | **Arhgef6** | 1600028C08Rik\|1700038J06Rik\|4930592P22Rik\|alpha-PIX | Rac/Cdc42 guanine nucleotide exchange factor (GEF) 6 | protein-coding |
| chrX_95161004_95161287 | chrX | 95161005 | 95161287 | + | 1.14909 | intron (NM_001290384, intron 1 of 9) | intron (NM_001290384, intron 1 of 9) | 5393 | NM_001290385 | 236915 | Mm.44841 | NM_001033329 | ENSMUSG00000025656 | **Arhgef9** | 9630036L12Rik\|A230067K14\|mKIAA0424 | CDC42 guanine nucleotide exchange factor (GEF) 9 | protein-coding |

| | | | | | | | | | | | | | | | | | |
|---|---|---|---|---|---|---|---|---|---|---|---|---|---|---|---|---|---|
| chrX_95164181_95164470 | chrX | 95164182 | 95164470 | + | 0.778179 | intron (NM_001290384, intron 1 of 9) | intron (NM_001290384, intron 1 of 9) | 2213 | NM_001290385 | 236915 | Mm.44841 | NM_001033329 | ENSMUSG00000025656 | **Arhgef9** | 9630036L12Rik\|A230067K14\|mKIAA0424 | CDC42 guanine nucleotide exchange factor (GEF) 9 | protein-coding |
| chr17_4991705_4992271 | chr17 | 4991706 | 4992271 | + | 1.31536 | Intergenic | Intergenic | -3086 | NM_001085355 | 239985 | Mm.133401 | NM_001085355 | ENSMUSG00000069729 | **Arid1b** | 8030481M12\|9330189K18Rik\|AI836955\|Ardi1b\|B230217J03Rik\|mKIAA1235 | AT rich interactive domain 1B (SWI-like) | protein-coding |
| chr17_5094651_5095085 | chr17 | 5094652 | 5095085 | + | 0.960362 | intron (NM_001085355, intron 2 of 19) | intron (NM_001085355, intron 2 of 19) | 99794 | NM_001085355 | 239985 | Mm.133401 | NM_001085355 | ENSMUSG00000069729 | **Arid1b** | 8030481M12\|9330189K18Rik\|AI836955\|Ardi1b\|B230217J03Rik\|mKIAA1235 | AT rich interactive domain 1B (SWI-like) | protein-coding |
| chr17_4767854_4768348 | chr17 | 4767855 | 4768348 | + | 0.775277 | Intergenic | MTC\|LTR\|ERVL-MaLR | -226973 | NM_001085355 | 239985 | Mm.133401 | NM_001085355 | ENSMUSG00000069729 | **Arid1b** | 8030481M12\|9330189K18Rik\|AI836955\|Ardi1b\|B230217J03Rik\|mKIAA1235 | AT rich interactive domain 1B (SWI-like) | protein-coding |
| chr17_5030874_5031221 | chr17 | 5030875 | 5031221 | + | 0.686312 | intron (NM_001085355, intron 1 of 19) | intron (NM_001085355, intron 1 of 19) | 35974 | NM_001085355 | 239985 | Mm.133401 | NM_001085355 | ENSMUSG00000069729 | **Arid1b** | 8030481M12\|9330189K18Rik\|AI836955\|Ardi1b\|B230217J03Rik\|mKIAA1235 | AT rich interactive domain 1B (SWI-like) | protein-coding |
| chr15_96331534_96331833 | chr15 | 96331535 | 96331833 | + | 0.739769 | intron (NM_175251, intron 3 of 20) | intron (NM_175251, intron 3 of 20) | 44162 | NM_175251 | 77044 | Mm.17166 | NM_175251 | ENSMUSG00000033237 | **Arid2** | 1700124K17Rik\|4432409D24Rik\|zipzap/p200 | AT rich interactive domain 2 (ARID, RFX-like) | protein-coding |
| chr9_57813262_57813626 | chr9 | 57813263 | 57813626 | + | 0.632459 | intron (NM_019689, intron 1 of 7) | intron (NM_019689, intron 1 of 7) | 20790 | NM_019689 | 56380 | Mm.425588 | NM_019689 | ENSMUSG00000004661 | **Arid3b** | Bdp\|Dri2 | AT rich interactive domain 3B (BRIGHT-like) | protein-coding |
| chr12_71045024_71045286 | chr12 | 71045025 | 71045286 | + | 0.980686 | intron (NM_001081195, intron 9 of 23) | intron (NM_001081195, intron 9 of 23) | 29188 | NM_001081195 | 238247 | Mm.241601 | NM_001081195 | ENSMUSG00000048118 | **Arid4a** | A630009N03\|A630067N03Rik\|MmRBBP1\|Rbbp1 | AT rich interactive domain 4A (RBP1-like) | protein-coding |
| chr12_71049603_71049919 | chr12 | 71049604 | 71049919 | + | 0.801671 | intron (NM_001081195, intron 11 of 23) | intron (NM_001081195, intron 11 of 23) | 33794 | NM_001081195 | 238247 | Mm.241601 | NM_001081195 | ENSMUSG00000048118 | **Arid4a** | A630009N03\|A630067N03Rik\|MmRBBP1\|Rbbp1 | AT rich interactive domain 4A (RBP1-like) | protein-coding |
| chr12_71020797_71021155 | chr12 | 71020798 | 71021155 | + | 0.780474 | intron (NM_001081195, intron 3 of 23) | intron (NM_001081195, intron 3 of 23) | 5009 | NM_001081195 | 238247 | Mm.241601 | NM_001081195 | ENSMUSG00000048118 | **Arid4a** | A630009N03\|A630067N03Rik\|MmRBBP1\|Rbbp1 | AT rich interactive domain 4A (RBP1-like) | protein-coding |
| chr1_36325717_36326458 | chr1 | 36325718 | 36326458 | + | 1.37263 | Intergenic | Intergenic | 18355 | NM_001172205 | 214855 | Mm.34316 | NM_145996 | ENSMUSG00000037447 | **Arid5a** | D430024K22Rik\|Mrf1 | AT rich interactive domain 5A (MRF1-like) | protein-coding |
| chr1_36325351_36326491 | chr1 | 36325352 | 36326491 | + | 1.22292 | Intergenic | Intergenic | 18188 | NM_001172205 | 214855 | Mm.34316 | NM_145996 | ENSMUSG00000037447 | **Arid5a** | D430024K22Rik\|Mrf1 | AT rich interactive domain 5A (MRF1-like) | protein-coding |
| chr1_36293923_36294167 | chr1 | 36293924 | 36294167 | + | 0.887218 | Intergenic | L1MEd\|LINE\|L1 | -13688 | NM_001290727 | 214855 | Mm.34316 | NM_145996 | ENSMUSG00000037447 | **Arid5a** | D430024K22Rik\|Mrf1 | AT rich interactive domain 5A (MRF1-like) | protein-coding |
| chr10_68136510_68137913 | chr10 | 68136511 | 68137913 | + | 0.921369 | intron (NM_023598, intron 4 of 9) | intron (NM_023598, intron 4 of 9) | 141514 | NM_023598 | 71371 | Mm.440357 | NM_023598 | ENSMUSG00000019947 | **Arid5b** | 4930580B11\|5430435G07Rik\|AI467247\|Desrt\|Mrf2\|Mrf2alpha\|Mrf2beta | AT rich interactive domain 5B (MRF1-like) | protein-coding |
| chr9_108643308_108643742 | chr9 | 108643309 | 108643742 | + | 0.860611 | intron (NM_011790, intron 3 of 15) | intron (NM_011790, intron 3 of 15) | 5855 | NM_011790 | 23807 | Mm.290447 | NM_011790 | ENSMUSG00000064145 | **Arih2** | AI843547\|ARI2\|TRIAD1\|UIP48 | ariadne homolog 2 (Drosophila) | protein-coding |
| chr13_54570190_54570449 | chr13 | 54570191 | 54570449 | + | 0.918542 | Intergenic | Intergenic | -4693 | NM_019968 | 56795 | Mm.89484 | NM_019968 | ENSMUSG00000025870 | **Arl10** | Arl10a\|Arm1 | ADP-ribosylation factor-like 10 | protein-coding |
| chr3_69182338_69182751 | chr3 | 69182339 | 69182751 | + | 1.45695 | Intergenic | Intergenic | -39874 | NM_027843 | 71619 | Mm.98122 | NM_027843 | ENSMUSG00000098207 | **Arl14** | 9130014L17Rik\|Arf7 | ADP-ribosylation factor-like 14 | protein-coding |
| chr2_107005602_107006246 | chr2 | 107005603 | 107006246 | + | 0.682508 | Intergenic | ERVB4_3-LTR_MM\|LTR\|ERVK | -31527 | NM_001025102 | 212772 | Mm.381231 | NM_173750 | ENSMUSG00000027122 | **Arl14ep** | 2700007P21Rik\|4930448O08Rik\|ARF7EP | ADP-ribosylation factor-like 14 effector protein | protein-coding |
| chr13_113829088_113829417 | chr13 | 113829089 | 113829417 | + | 0.766295 | intron (NM_172595, intron 1 of 4) | LTR16B1\|LTR\|ERVL | 34745 | NM_172595 | 218639 | Mm.79837 | NM_172595 | ENSMUSG00000042348 | **Arl15** | A430036I03\|Arfrp2\|C230032K13Rik | ADP-ribosylation factor-like 15 | protein-coding |
| chr13_113828663_113829539 | chr13 | 113828664 | 113829539 | + | 0.727642 | intron (NM_172595, intron 1 of 4) | intron (NM_172595, intron 1 of 4) | 34593 | NM_172595 | 218639 | Mm.79837 | NM_172595 | ENSMUSG00000042348 | **Arl15** | A430036I03\|Arfrp2\|C230032K13Rik | ADP-ribosylation factor-like 15 | protein-coding |
| chr12_39648944_39649351 | chr12 | 39648945 | 39649351 | + | 1.24214 | Intergenic | Intergenic | 388228 | NM_007487 | 11861 | Mm.12723 | NM_007487 | ENSMUSG00000047446 | **Arl4a** | AI467555\|Arl4 | ADP-ribosylation factor-like 4A | protein-coding |
| chr12_40009868_40010305 | chr12 | 40009869 | 40010305 | + | 1.13031 | Intergenic | Intergenic | 27289 | NM_007487 | 11861 | Mm.12723 | NM_007487 | ENSMUSG00000047446 | **Arl4a** | AI467555\|Arl4 | ADP-ribosylation factor-like 4A | protein-coding |
| chr12_40008253_40008826 | chr12 | 40008254 | 40008826 | + | 1.08173 | Intergenic | RCHARR1\|DNA\|hAT-Charlie | 28836 | NM_007487 | 11861 | Mm.12723 | NM_007487 | ENSMUSG00000047446 | **Arl4a** | AI467555\|Arl4 | ADP-ribosylation factor-like 4A | protein-coding |
| chr1_88639761_88640206 | chr1 | 88639762 | 88640206 | + | 0.848523 | Intergenic | Intergenic | 62207 | NM_177305 | 320982 | Mm.27968 | NM_177305 | ENSMUSG00000049866 | **Arl4c** | A630084M22Rik\|Arl7\|LAK | ADP-ribosylation factor-like 4C | protein-coding |
| chr2_52420032_52420305 | chr2 | 52420033 | 52420305 | + | 0.764902 | intron (NM_182994, intron 1 of 5) | intron (NM_182994, intron 1 of 5) | 4705 | NM_182994 | 75423 | Mm.314378 | NM_182994 | ENSMUSG00000036093 | **Arl5a** | 2410015N24Rik\|2810410P22Rik\|AA408731\|AW610751\|Arl5\|T25534 | ADP-ribosylation factor-like 5A | protein-coding |
| chr7_118135319_118136026 | chr7 | 118135320 | 118136026 | + | 1.85726 | intron (NM_001031814, intron 62 of 62) | intron (NM_001031814, intron 62 of 62) | -6048 | NM_019419 | 54208 | Mm.29924 | NM_019419 | ENSMUSG00000030654 | **Arl6ip1** | AIP-6\|AL022945\|ARMER\|AU042858\|Aip-1\|Arl6ip\|C85138\|mKIAA0069 | ADP-ribosylation factor-like 6 interacting protein 1 | protein-coding |
| chr7_118136090_118136582 | chr7 | 118136091 | 118136582 | + | 1.35054 | intron (NM_001031814, intron 62 of 62) | intron (NM_001031814, intron 62 of 62) | -6711 | NM_019419 | 54208 | Mm.29924 | NM_019419 | ENSMUSG00000030654 | **Arl6ip1** | AIP-6\|AL022945\|ARMER\|AU042858\|Aip-1\|Arl6ip\|C85138\|mKIAA0069 | ADP-ribosylation factor-like 6 interacting protein 1 | protein-coding |
| chr7_118141217_118141922 | chr7 | 118141218 | 118141922 | + | 0.980539 | intron (NM_001031814, intron 56 of 62) | intron (NM_001031814, intron 56 of 62) | -11945 | NM_019419 | 54208 | Mm.29924 | NM_019419 | ENSMUSG00000030654 | **Arl6ip1** | AIP-6\|AL022945\|ARMER\|AU042858\|Aip-1\|Arl6ip\|C85138\|mKIAA0069 | ADP-ribosylation factor-like 6 interacting protein 1 | protein-coding |
| chr7_118185954_118186400 | chr7 | 118185955 | 118186400 | + | 0.815585 | intron (NM_001031814, intron 18 of 62) | intron (NM_001031814, intron 18 of 62) | -56552 | NM_019419 | 54208 | Mm.29924 | NM_019419 | ENSMUSG00000030654 | **Arl6ip1** | AIP-6\|AL022945\|ARMER\|AU042858\|Aip-1\|Arl6ip\|C85138\|mKIAA0069 | ADP-ribosylation factor-like 6 interacting protein 1 | protein-coding |
| chr7_118185950_118186217 | chr7 | 118185951 | 118186217 | + | 0.796634 | exon (NM_001031814, exon 19 of 63) | exon (NM_001031814, exon 19 of 63) | -56459 | NM_019419 | 54208 | Mm.29924 | NM_019419 | ENSMUSG00000030654 | **Arl6ip1** | AIP-6\|AL022945\|ARMER\|AU042858\|Aip-1\|Arl6ip\|C85138\|mKIAA0069 | ADP-ribosylation factor-like 6 interacting protein 1 | protein-coding |
| chr6_97209325_97209535 | chr6 | 97209326 | 97209535 | + | 0.754272 | Intergenic | B3\|SINE\|B2 | -1362 | NM_022992 | 65106 | Mm.291014 | NM_022992 | ENSMUSG00000035199 | **Arl6ip5** | 5930404D22Rik\|AV001879\|Aip-5\|Gtrap3-18\|addiscin | ADP-ribosylation factor-like 6 interacting protein 5 | protein-coding |
| chr10_41978160_41979234 | chr10 | 41978161 | 41979234 | + | 1.36607 | intron (NM_001034858, intron 6 of 17) | intron (NM_001034858, intron 6 of 17) | 39685 | NM_001034858 | 213402 | Mm.211320 | NM_001034858 | ENSMUSG00000071324 | **Armc2** | 2610018I05Rik | armadillo repeat containing 2 | protein-coding |
| chr10_42033604_42034139 | chr10 | 42033605 | 42034139 | + | 0.973107 | Intergenic | MLT1L\|LTR\|ERVL-MaLR | -15490 | NM_001034858 | 213402 | Mm.211320 | NM_001034858 | ENSMUSG00000071324 | **Armc2** | 2610018I05Rik | armadillo repeat containing 2 | protein-coding |
| chr10_41984057_41984697 | chr10 | 41984058 | 41984697 | + | 0.968212 | intron (NM_001034858, intron 5 of 17) | L3\|LINE\|CR1 | 34005 | NM_001034858 | 213402 | Mm.211320 | NM_001034858 | ENSMUSG00000071324 | **Armc2** | 2610018I05Rik | armadillo repeat containing 2 | protein-coding |
| chr10_42130489_42130833 | chr10 | 42130490 | 42130833 | + | 0.71444 | Intergenic | Intergenic | -112279 | NM_001034858 | 213402 | Mm.211320 | NM_001034858 | ENSMUSG00000071324 | **Armc2** | 2610018I05Rik | armadillo repeat containing 2 | protein-coding |
| chr7_84420669_84420943 | chr7 | 84420670 | 84420943 | + | 0.633861 | Intergenic | Intergenic | -10768 | NM_007488 | 11864 | Mm.38594 | NM_007488 | ENSMUSG00000015709 | **Arnt2** | Hif-2b\|bHLHe1\|mKIAA0307 | aryl hydrocarbon receptor nuclear translocator 2 | protein-coding |
| chr7_113183775_113184338 | chr7 | 113183776 | 113184338 | + | 1.05076 | Intergenic | B3A\|SINE\|B2 | -23408 | NM_001243048 | 11865 | Mm.440371 | NM_007489 | ENSMUSG00000055116 | **Arntl** | Arnt3\|BMAL1b\|Bmal1\|MOP3\|bHLHe5\|bmal1b' | aryl hydrocarbon receptor nuclear translocator-like | protein-coding |
| chr7_113183775_113184565 | chr7 | 113183776 | 113184565 | + | 0.884382 | Intergenic | B3A\|SINE\|B2 | -23295 | NM_001243048 | 11865 | Mm.440371 | NM_007489 | ENSMUSG00000055116 | **Arntl** | Arnt3\|BMAL1b\|Bmal1\|MOP3\|bHLHe5\|bmal1b' | aryl hydrocarbon receptor nuclear translocator-like | protein-coding |
| chr6_146813007_146813689 | chr6 | 146813008 | 146813689 | + | 1.5724 | intron (NM_001289681, intron 5 of 14) | intron (NM_001289681, intron 5 of 14) | 7818 | NM_001289681 | 272322 | Mm.333500 | NM_172309 | ENSMUSG00000040187 | **Arntl2** | 4632430A05Rik\|BMAL2\|CLIF\|MOP9\|bHLHe6 | aryl hydrocarbon receptor nuclear translocator-like 2 | protein-coding |
| chr6_146776691_146776926 | chr6 | 146776692 | 146776926 | + | 0.99405 | 3' UTR (NM_172734, exon 14 of 14) | 3' UTR (NM_172734, exon 14 of 14) | -28721 | NM_172309 | 272322 | Mm.333500 | NM_172309 | ENSMUSG00000040187 | **Arntl2** | 4632430A05Rik\|BMAL2\|CLIF\|MOP9\|bHLHe6 | aryl hydrocarbon receptor nuclear translocator-like 2 | protein-coding |
| chr6_146776546_146777276 | chr6 | 146776547 | 146777276 | + | 0.946841 | 3' UTR (NM_172734, exon 14 of 14) | 3' UTR (NM_172734, exon 14 of 14) | -28619 | NM_172309 | 272322 | Mm.333500 | NM_172309 | ENSMUSG00000040187 | **Arntl2** | 4632430A05Rik\|BMAL2\|CLIF\|MOP9\|bHLHe6 | aryl hydrocarbon receptor nuclear translocator-like 2 | protein-coding |
| chr6_146793587_146793908 | chr6 | 146793588 | 146793908 | + | 0.782121 | Intergenic | Intergenic | -11782 | NM_172309 | 272322 | Mm.333500 | NM_172309 | ENSMUSG00000040187 | **Arntl2** | 4632430A05Rik\|BMAL2\|CLIF\|MOP9\|bHLHe6 | aryl hydrocarbon receptor nuclear translocator-like 2 | protein-coding |
| chr13_80864230_80864545 | chr13 | 80864231 | 80864545 | + | 0.749252 | Intergenic | Intergenic | -19034 | NM_001042591 | 105171 | Mm.423137 | NM_001042591 | ENSMUSG00000074794 | **Arrdc3** | AI450344\|mKIAA1376 | arrestin domain containing 3 | protein-coding |
| chr5_92336830_92337400 | chr5 | 92336831 | 92337400 | + | 0.926625 | intron (NM_181728, intron 2 of 9) | L1M4c\|LINE\|L1 | 5274 | NM_181728 | 109979 | Mm.263514 | NM_181728 | ENSMUSG00000034842 | **Art3** | 4930569O04Rik\|ARTC3 | ADP-ribosyltransferase 3 | protein-coding |
| chr6_136852132_136852567 | chr6 | 136852133 | 136852567 | + | 0.930568 | intron (NM_026639, intron 2 of 2) | L2a\|LINE\|L2 | 5250 | NM_026639 | 109978 | Mm.31840 | NM_026639 | ENSMUSG00000030217 | **Art4** | 4432404K01Rik\|ARTC4\|DO\|DOK1 | ADP-ribosyltransferase 4 | protein-coding |
| chr4_117981148_117981894 | chr4 | 117981149 | 117981894 | + | 0.780477 | intron (NM_001285521, intron 4 of 10) | intron (NM_001285521, intron 4 of 10) | -51758 | NM_001284192 | 11876 | Mm.56897 | NM_009711 | ENSMUSG00000028539 | **Artn** | neublastin | artemin | protein-coding |

| | | | | | | | | | | | | | | | | | |
|---|---|---|---|---|---|---|---|---|---|---|---|---|---|---|---|---|---|
| chr4_117926459_117927375 | chr4 | 117926460 | 117927375 | + | 0.610914 | exon (NM_001284193, exon 5 of 5) | exon (NM_001284193, exon 5 of 5) | 2846 | NM_001284191 | 11876 | Mm.56897 | NM_009711 | ENSMUSG00000028539 | **Artn** | neublastin | artemin | protein-coding |
| chr19_32120532_32121195 | chr19 | 32120533 | 32121195 | + | 1.04507 | Intergenic | Intergenic | -17724 | NM_018830 | 54447 | Mm.491229 | NM_018830 | ENSMUSG00000024887 | **Asah2** | AI585898 | N-acylsphingosine amidohydrolase 2 | protein-coding |
| chr19_32113001_32113402 | chr19 | 32113002 | 32113402 | + | 0.729525 | Intergenic | Intergenic | -10062 | NM_018830 | 54447 | Mm.491229 | NM_018830 | ENSMUSG00000024887 | **Asah2** | AI585898 | N-acylsphingosine amidohydrolase 2 | protein-coding |
| chr19_32147522_32148033 | chr19 | 32147523 | 32148033 | + | 0.722623 | intron (NM_001168526, intron 2 of 5) | MTC\|LTR\|ERVL-MaLR | -44638 | NM_018830 | 54447 | Mm.491229 | NM_018830 | ENSMUSG00000024887 | **Asah2** | AI585898 | N-acylsphingosine amidohydrolase 2 | protein-coding |
| chr15_64235588_64236153 | chr15 | 64235589 | 64236153 | + | 1.34086 | intron (NM_001276462, intron 1 of 27) | intron (NM_001276462, intron 1 of 27) | 76802 | NM_001276462 | 13196 | Mm.277236 | NM_010026 | ENSMUSG00000022377 | **Asap1** | AV239055\|DEF-1\|Ddef1\|PAP\|mKIAA1249\|s19 | ArfGAP with SH3 domain, ankyrin repeat and PH domain1 | protein-coding |
| chr15_64211433_64211849 | chr15 | 64211434 | 64211849 | + | 1.10293 | intron (NM_001276462, intron 1 of 27) | intron (NM_001276462, intron 1 of 27) | 101032 | NM_001276462 | 13196 | Mm.277236 | NM_010026 | ENSMUSG00000022377 | **Asap1** | AV239055\|DEF-1\|Ddef1\|PAP\|mKIAA1249\|s19 | ArfGAP with SH3 domain, ankyrin repeat and PH domain1 | protein-coding |
| chr15_64477093_64477774 | chr15 | 64477094 | 64477774 | + | 0.897425 | Intergenic | L1MB8\|LINE\|L1 | -94515 | NM_010026 | 13196 | Mm.277236 | NM_010026 | ENSMUSG00000022377 | **Asap1** | AV239055\|DEF-1\|Ddef1\|PAP\|mKIAA1249\|s19 | ArfGAP with SH3 domain, ankyrin repeat and PH domain1 | protein-coding |
| chr15_64250153_64250603 | chr15 | 64250154 | 64250603 | + | 0.820982 | intron (NM_001276462, intron 1 of 27) | PB1D10\|SINE\|Alu | 62295 | NM_001276462 | 13196 | Mm.277236 | NM_010026 | ENSMUSG00000022377 | **Asap1** | AV239055\|DEF-1\|Ddef1\|PAP\|mKIAA1249\|s19 | ArfGAP with SH3 domain, ankyrin repeat and PH domain1 | protein-coding |
| chr15_64192641_64193223 | chr15 | 64192642 | 64193223 | + | 0.789017 | intron (NM_001276462, intron 2 of 27) | intron (NM_001276462, intron 2 of 27) | 119741 | NM_001276462 | 13196 | Mm.277236 | NM_010026 | ENSMUSG00000022377 | **Asap1** | AV239055\|DEF-1\|Ddef1\|PAP\|mKIAA1249\|s19 | ArfGAP with SH3 domain, ankyrin repeat and PH domain1 | protein-coding |
| chr15_64197056_64197374 | chr15 | 64197057 | 64197374 | + | 0.776868 | intron (NM_001276462, intron 2 of 27) | intron (NM_001276462, intron 2 of 27) | 115458 | NM_001276462 | 13196 | Mm.277236 | NM_010026 | ENSMUSG00000022377 | **Asap1** | AV239055\|DEF-1\|Ddef1\|PAP\|mKIAA1249\|s19 | ArfGAP with SH3 domain, ankyrin repeat and PH domain1 | protein-coding |
| chr15_64284502_64284914 | chr15 | 64284503 | 64284914 | + | 0.764941 | intron (NM_001276462, intron 1 of 27) | intron (NM_001276462, intron 1 of 27) | 27965 | NM_001276462 | 13196 | Mm.277236 | NM_010026 | ENSMUSG00000022377 | **Asap1** | AV239055\|DEF-1\|Ddef1\|PAP\|mKIAA1249\|s19 | ArfGAP with SH3 domain, ankyrin repeat and PH domain1 | protein-coding |
| chr15_64194071_64194348 | chr15 | 64194072 | 64194348 | + | 0.727075 | intron (NM_001276462, intron 2 of 27) | intron (NM_001276462, intron 2 of 27) | 118463 | NM_001276462 | 13196 | Mm.277236 | NM_010026 | ENSMUSG00000022377 | **Asap1** | AV239055\|DEF-1\|Ddef1\|PAP\|mKIAA1249\|s19 | ArfGAP with SH3 domain, ankyrin repeat and PH domain1 | protein-coding |
| chr15_64284312_64284924 | chr15 | 64284313 | 64284924 | + | 0.698487 | intron (NM_001276462, intron 1 of 27) | intron (NM_001276462, intron 1 of 27) | 28055 | NM_001276462 | 13196 | Mm.277236 | NM_010026 | ENSMUSG00000022377 | **Asap1** | AV239055\|DEF-1\|Ddef1\|PAP\|mKIAA1249\|s19 | ArfGAP with SH3 domain, ankyrin repeat and PH domain1 | protein-coding |
| chr14_26919260_26920006 | chr14 | 26919261 | 26920006 | + | 1.36349 | 3' UTR (NM_145221, exon 22 of 22) | 3' UTR (NM_145221, exon 22 of 22) | 25029 | NM_080856 | 142687 | Mm.138259 | NM_080856 | ENSMUSG00000021898 | **Asb14** | Asb-14 | ankyrin repeat and SOCS box-containing 14 | protein-coding |
| chr14_26904406_26904771 | chr14 | 26904407 | 26904771 | + | 1.00837 | intron (NM_080856, intron 6 of 8) | L1Md_F2\|LINE\|L1 | 9985 | NM_080856 | 142687 | Mm.138259 | NM_080856 | ENSMUSG00000021898 | **Asb14** | Asb-14 | ankyrin repeat and SOCS box-containing 14 | protein-coding |
| chr12_103346957_103347334 | chr12 | 103346958 | 103347334 | + | 0.941427 | intron (NM_023049, intron 1 of 9) | intron (NM_023049, intron 1 of 9) | 8855 | NM_023049 | 65256 | Mm.27159 | NM_023049 | ENSMUSG00000021200 | **Asb2** | 1110008E15Rik | ankyrin repeat and SOCS box-containing 2 | protein-coding |
| chr10_60015752_60016577 | chr10 | 60015753 | 60016577 | + | 1.14108 | intron (NM_001199187, intron 5 of 10) | B1_Mur4\|SINE\|Alu | 12838 | NM_001199187 | 69090 | Mm.155839 | NM_026937 | ENSMUSG00000044475 | **Ascc1** | 1810015P09Rik\|AI550520\|ASC1p50\|CGI-18 | activating signal cointegrator 1 complex subunit 1 | protein-coding |
| chr10_50586586_50586900 | chr10 | 50586587 | 50586900 | + | 0.975928 | Intergenic | Intergenic | -5926 | NM_198007 | 77987 | Mm.222497 | NM_198007 | ENSMUSG00000038774 | **Ascc3** | ASC1p200\|B630009I04Rik\|BC023451\|D430001L07Rik\|D630041L21\|Helic1\|RNAH | activating signal cointegrator 1 complex subunit 3 | protein-coding |
| chr10_53598761_53599175 | chr10 | 53598762 | 53599175 | + | 0.704078 | intron (NM_025541, intron 1 of 3) | intron (NM_025541, intron 1 of 3) | 2007 | NM_025541 | 66403 | Mm.272989 | NM_025541 | ENSMUSG00000019857 | **Asf1a** | 2310079C17Rik\|C79052 | anti-silencing function 1A histone chaperone | protein-coding |
| chr5_24404740_24405407 | chr5 | 24404741 | 24405407 | + | 0.998769 | intron (NM_029020, intron 9 of 15) | intron (NM_029020, intron 9 of 15) | -8377 | NM_183000 | 171209 | Mm.299636 | NM_183000 | ENSMUSG00000038276 | **Asic3** | AW742291\|Accn3\|DRASIC\|SLNAC1\|TNAC1 | acid-sensing (proton-gated) ion channel 3 | protein-coding |
| chr3_81992508_81992983 | chr3 | 81992509 | 81992983 | + | 0.695898 | Intergenic | X6B_LINE\|LINE\|CR1 | -4176 | NM_021370 | 58170 | Mm.88839 | NM_021370 | ENSMUSG00000028008 | **Asic5** | Accn5\|Blinac\|Inac | acid-sensing (proton-gated) ion channel family member 5 | protein-coding |
| chr6_7692796_7693519 | chr6 | 7692797 | 7693519 | + | 1.3604 | promoter-TSS (NM_012055) | promoter-TSS (NM_012055) | 24 | NM_012055 | 27053 | Mm.2942 | NM_012055 | ENSMUSG00000029752 | **Asns** | - | asparagine synthetase | protein-coding |
| chr6_7692545_7693548 | chr6 | 7692546 | 7693548 | + | 1.34716 | intron (NM_012055, intron 1 of 11) | CpG | 135 | NM_012055 | 27053 | Mm.2942 | NM_012055 | ENSMUSG00000029752 | **Asns** | - | asparagine synthetase | protein-coding |
| chr11_73323998_73324404 | chr11 | 73323999 | 73324404 | + | 0.689688 | intron (NM_023113, intron 1 of 5) | intron (NM_023113, intron 1 of 5) | 436 | NM_023113 | 11484 | Mm.293574 | NM_023113 | ENSMUSG00000020774 | **Aspa** | Acy-2\|Acy2\|nur7 | aspartoacylase | protein-coding |
| chr7_44462736_44463033 | chr7 | 44462737 | 44463033 | + | 0.823799 | 3' UTR (NM_198250, exon 3 of 3) | 3' UTR (NM_198250, exon 3 of 3) | -2550 | NM_026690 | 68352 | Mm.88478 | NM_026690 | ENSMUSG00000038704 | **Aspdh** | 0610012D14Rik | aspartate dehydrogenase domain containing | protein-coding |
| chr4_9605318_9605878 | chr4 | 9605319 | 9605878 | + | 1.81246 | intron (NM_001177854, intron 3 of 11) | Tigger7\|DNA\|TcMar-Tigger | 38119 | NM_001177853 | 65973 | Mm.222206 | NM_023066 | ENSMUSG00000028207 | **Asph** | 2310005F16Rik\|3110001L23Rik\|AI848629\|AW261690\|AW561901\|BAH\|C79816\|cI-37 | aspartate-beta-hydroxylase | protein-coding |
| chr4_9640682_9641398 | chr4 | 9640683 | 9641398 | + | 1.68915 | intron (NM_001290367, intron 1 of 22) | intron (NM_001290367, intron 1 of 22) | 2677 | NM_001177853 | 65973 | Mm.222206 | NM_023066 | ENSMUSG00000028207 | **Asph** | 2310005F16Rik\|3110001L23Rik\|AI848629\|AW261690\|AW561901\|BAH\|C79816\|cI-37 | aspartate-beta-hydroxylase | protein-coding |
| chr4_9674312_9675068 | chr4 | 9674313 | 9675068 | + | 0.775268 | Intergenic | Intergenic | -5346 | NM_001290367 | 65973 | Mm.222206 | NM_023066 | ENSMUSG00000028207 | **Asph** | 2310005F16Rik\|3110001L23Rik\|AI848629\|AW261690\|AW561901\|BAH\|C79816\|cI-37 | aspartate-beta-hydroxylase | protein-coding |
| chr4_9593099_9593609 | chr4 | 9593100 | 9593609 | + | 0.745956 | intron (NM_001290367, intron 11 of 22) | intron (NM_001290367, intron 11 of 22) | 50363 | NM_001177853 | 65973 | Mm.222206 | NM_023066 | ENSMUSG00000028207 | **Asph** | 2310005F16Rik\|3110001L23Rik\|AI848629\|AW261690\|AW561901\|BAH\|C79816\|cI-37 | aspartate-beta-hydroxylase | protein-coding |
| chr4_9642154_9642415 | chr4 | 9642155 | 9642415 | + | 0.699622 | intron (NM_001290367, intron 1 of 22) | intron (NM_001290367, intron 1 of 22) | 1432 | NM_001177853 | 65973 | Mm.222206 | NM_023066 | ENSMUSG00000028207 | **Asph** | 2310005F16Rik\|3110001L23Rik\|AI848629\|AW261690\|AW561901\|BAH\|C79816\|cI-37 | aspartate-beta-hydroxylase | protein-coding |
| chr4_9592902_9593879 | chr4 | 9592903 | 9593879 | + | 0.677183 | intron (NM_001290367, intron 11 of 22) | intron (NM_001290367, intron 11 of 22) | 50326 | NM_001177853 | 65973 | Mm.222206 | NM_023066 | ENSMUSG00000028207 | **Asph** | 2310005F16Rik\|3110001L23Rik\|AI848629\|AW261690\|AW561901\|BAH\|C79816\|cI-37 | aspartate-beta-hydroxylase | protein-coding |
| chr13_49556921_49557341 | chr13 | 49556922 | 49557341 | + | 1.71391 | intron (NM_001172481, intron 3 of 7) | URR1B\|DNA\|hAT-Charlie | 12688 | NM_025711 | 66695 | Mm.383216 | NM_025711 | ENSMUSG00000021388 | **Aspn** | 4631401G09Rik\|AA986886\|Plap1\|Slrr1c | asporin | protein-coding |
| chr13_49556157_49556633 | chr13 | 49556158 | 49556633 | + | 0.834257 | intron (NM_001172481, intron 3 of 7) | intron (NM_001172481, intron 3 of 7) | 11952 | NM_025711 | 66695 | Mm.383216 | NM_025711 | ENSMUSG00000021388 | **Aspn** | 4631401G09Rik\|AA986886\|Plap1\|Slrr1c | asporin | protein-coding |
| chr13_49559245_49559533 | chr13 | 49559246 | 49559533 | + | 0.746612 | intron (NM_001172481, intron 5 of 7) | intron (NM_001172481, intron 5 of 7) | 14946 | NM_025711 | 66695 | Mm.383216 | NM_025711 | ENSMUSG00000021388 | **Aspn** | 4631401G09Rik\|AA986886\|Plap1\|Slrr1c | asporin | protein-coding |
| chr13_49547384_49548014 | chr13 | 49547385 | 49548014 | + | 0.737308 | intron (NM_001172481, intron 1 of 7) | intron (NM_001172481, intron 1 of 7) | 3256 | NM_025711 | 66695 | Mm.383216 | NM_025711 | ENSMUSG00000021388 | **Aspn** | 4631401G09Rik\|AA986886\|Plap1\|Slrr1c | asporin | protein-coding |
| chr6_86605245_86605768 | chr6 | 86605246 | 86605768 | + | 0.619468 | Intergenic | Intergenic | -22667 | NM_026414 | 67855 | Mm.183043 | NM_026414 | ENSMUSG00000033508 | **Asprv1** | 2300003P22Rik\|AA986851\|SASP\|SASPase\|Taps | aspartic peptidase, retroviral-like 1 | protein-coding |
| chr19_9124083_9124398 | chr19 | 9124084 | 9124398 | + | 0.816388 | intron (NM_025610, intron 2 of 6) | L1VL2\|LINE\|L1 | 11325 | NM_025610 | 66514 | Mm.272847 | NM_025610 | | **Asrgl1** | 2410004D18Rik\|ALP\|ALP1\|AU040643\|AW060375 | asparaginase like 1 | protein-coding |
| chr2_31447531_31447866 | chr2 | 31447532 | 31447866 | + | 0.619465 | Intergenic | Intergenic | -22571 | NM_007494 | 11898 | Mm.3217 | NM_007494 | ENSMUSG00000076441 | **Ass1** | AA408052\|ASS\|Ass-1\|fold | argininosuccinate synthetase 1 | protein-coding |
| chr9_105401798_105402153 | chr9 | 105401799 | 105402153 | + | 0.805081 | intron (NM_025651, intron 3 of 4) | intron (NM_025651, intron 3 of 4) | 1188 | NM_001164828 | 66595 | Mm.84941 | NM_025651 | ENSMUSG00000032567 | **Aste1** | 1100001A21Rik\|AV006403 | asteroid homolog 1 (Drosophila) | protein-coding |
| chr2_153350496_153351273 | chr2 | 153350497 | 153351273 | + | 0.617961 | intron (NM_001039939, intron 1 of 11) | intron (NM_001039939, intron 1 of 11) | 4746 | NM_001039939 | 228790 | Mm.330677 | NM_001039939 | ENSMUSG00000042548 | **Asxl1** | mKIAA0978 | additional sex combs like 1 | protein-coding |

| | | | | | | | | | | | | | | | | | |
|---|---|---|---|---|---|---|---|---|---|---|---|---|---|---|---|---|---|
| chr12_4929624_4930107 | chr12 | 4929625 | 4930107 | + | 1.07984 | intron (NM_001099628, intron 1 of 30) | intron (NM_001099628, intron 1 of 30) | 12513 | NM_001099628 | 320817 | Mm.210966 | NM_001099628 | | **Atad2b** | 1110014E10Rik\|BC032887\|C79189\|D530031C13Rik | ATPase family, AAA domain containing 2B | protein-coding |
| chr15_100236920_100237615 | chr15 | 100236921 | 100237615 | + | 1.08495 | intron (NM_007497, intron 2 of 6) | intron (NM_007497, intron 2 of 6) | 9409 | NM_007497 | 11908 | Mm.676 | NM_007497 | ENSMUSG00000023027 | **Atf1** | - | activating transcription factor 1 | protein-coding |
| chr15_100242602_100243080 | chr15 | 100242603 | 100243080 | + | 0.697402 | intron (NM_007497, intron 2 of 6) | intron (NM_007497, intron 2 of 6) | 14982 | NM_007497 | 11908 | Mm.676 | NM_007497 | ENSMUSG00000023027 | **Atf1** | - | activating transcription factor 1 | protein-coding |
| chr1_191216096_191216523 | chr1 | 191216097 | 191216523 | + | 1.35765 | Intergenic | Intergenic | -32977 | NM_007498 | 11910 | Mm.2706 | NM_007498 | ENSMUSG00000026628 | **Atf3** | LRG-21 | activating transcription factor 3 | protein-coding |
| chr1_191215917_191216813 | chr1 | 191215918 | 191216813 | + | 1.1621 | Intergenic | Intergenic | -33032 | NM_007498 | 11910 | Mm.2706 | NM_007498 | ENSMUSG00000026628 | **Atf3** | LRG-21 | activating transcription factor 3 | protein-coding |
| chr1_191189947_191190207 | chr1 | 191189948 | 191190207 | + | 0.95832 | Intergenic | Intergenic | -6744 | NM_007498 | 11910 | Mm.2706 | NM_007498 | ENSMUSG00000026628 | **Atf3** | LRG-21 | activating transcription factor 3 | protein-coding |
| chr1_191182375_191183777 | chr1 | 191182376 | 191183777 | + | 0.890475 | intron (NM_007498, intron 1 of 3) | CpG | 257 | NM_007498 | 11910 | Mm.2706 | NM_007498 | ENSMUSG00000026628 | **Atf3** | LRG-21 | activating transcription factor 3 | protein-coding |
| chr1_191183004_191183707 | chr1 | 191183005 | 191183707 | + | 0.862324 | promoter-TSS (NM_007498) | promoter-TSS (NM_007498) | -23 | NM_007498 | 11910 | Mm.2706 | NM_007498 | ENSMUSG00000026628 | **Atf3** | LRG-21 | activating transcription factor 3 | protein-coding |
| chr1_191197996_191198197 | chr1 | 191197997 | 191198197 | + | 0.732554 | Intergenic | Intergenic | -14764 | NM_007498 | 11910 | Mm.2706 | NM_007498 | ENSMUSG00000026628 | **Atf3** | LRG-21 | activating transcription factor 3 | protein-coding |
| chr1_170853523_170854125 | chr1 | 170853524 | 170854125 | + | 1.60358 | intron (NM_001081304, intron 2 of 15) | intron (NM_001081304, intron 2 of 15) | 13947 | NM_001081304 | 226641 | Mm.377046 | NM_001081304 | ENSMUSG00000026663 | **Atf6** | 9130025P16Rik\|9630036G24\|AA789574\|Atf6alpha\|ESTM49 | activating transcription factor 6 | protein-coding |
| chr1_170853343_170854176 | chr1 | 170853344 | 170854176 | + | 1.56357 | intron (NM_001081304, intron 2 of 15) | intron (NM_001081304, intron 2 of 15) | 14011 | NM_001081304 | 226641 | Mm.377046 | NM_001081304 | ENSMUSG00000026663 | **Atf6** | 9130025P16Rik\|9630036G24\|AA789574\|Atf6alpha\|ESTM49 | activating transcription factor 6 | protein-coding |
| chr1_170845702_170846055 | chr1 | 170845703 | 170846055 | + | 0.740439 | intron (NM_001081304, intron 4 of 15) | RMER4B\|LTR\|ERVK | 21892 | NM_001081304 | 226641 | Mm.377046 | NM_001081304 | ENSMUSG00000026663 | **Atf6** | 9130025P16Rik\|9630036G24\|AA789574\|Atf6alpha\|ESTM49 | activating transcription factor 6 | protein-coding |
| chr1_170845691_170846267 | chr1 | 170845692 | 170846267 | + | 0.696935 | intron (NM_001081304, intron 4 of 15) | RMER4B\|LTR\|ERVK | 21792 | NM_001081304 | 226641 | Mm.377046 | NM_001081304 | ENSMUSG00000026663 | **Atf6** | 9130025P16Rik\|9630036G24\|AA789574\|Atf6alpha\|ESTM49 | activating transcription factor 6 | protein-coding |
| chr6_136499282_136499581 | chr6 | 136499283 | 136499581 | + | 0.937167 | Intergenic | Intergenic | -19419 | NM_019426 | 54343 | Mm.173271 | NM_019426 | ENSMUSG00000030213 | **Atf7ip** | 2610204M12Rik\|AM\|Mcaf1 | activating transcription factor 7 interacting protein | protein-coding |
| chr6_136499192_136499729 | chr6 | 136499193 | 136499729 | + | 0.88879 | Intergenic | Intergenic | -19390 | NM_019426 | 54343 | Mm.173271 | NM_019426 | ENSMUSG00000030213 | **Atf7ip** | 2610204M12Rik\|AM\|Mcaf1 | activating transcription factor 7 interacting protein | protein-coding |
| chr6_136492208_136492679 | chr6 | 136492209 | 136492679 | + | 0.671259 | Intergenic | Intergenic | -26407 | NM_019426 | 54343 | Mm.173271 | NM_019426 | ENSMUSG00000030213 | **Atf7ip** | 2610204M12Rik\|AM\|Mcaf1 | activating transcription factor 7 interacting protein | protein-coding |
| chr13_91206200_91206643 | chr13 | 91206201 | 91206643 | + | 1.7581 | intron (NM_025770, intron 2 of 7) | intron (NM_025770, intron 2 of 7) | 17565 | NM_025770 | 66795 | Mm.235385 | NM_025770 | ENSMUSG00000021619 | **Atg10** | 5330424L23Rik\|5430428K15Rik\|AI852123\|APG10\|Agp10\|Apg10l\|Apg10p\|Atg10l | autophagy related 10 | protein-coding |
| chr13_91134229_91134839 | chr13 | 91134230 | 91134839 | + | 0.893526 | intron (NM_025770, intron 3 of 7) | intron (NM_025770, intron 3 of 7) | 89453 | NM_025770 | 66795 | Mm.235385 | NM_025770 | ENSMUSG00000021619 | **Atg10** | 5330424L23Rik\|5430428K15Rik\|AI852123\|APG10\|Agp10\|Apg10l\|Apg10p\|Atg10l | autophagy related 10 | protein-coding |
| chr13_91137764_91138098 | chr13 | 91137765 | 91138098 | + | 0.705722 | intron (NM_025770, intron 3 of 7) | intron (NM_025770, intron 3 of 7) | 86056 | NM_025770 | 66795 | Mm.235385 | NM_025770 | ENSMUSG00000021619 | **Atg10** | 5330424L23Rik\|5430428K15Rik\|AI852123\|APG10\|Agp10\|Apg10l\|Apg10p\|Atg10l | autophagy related 10 | protein-coding |
| chr13_91147502_91147973 | chr13 | 91147503 | 91147973 | + | 0.652978 | intron (NM_025770, intron 3 of 7) | L1VL2\|LINE\|L1 | 76249 | NM_025770 | 66795 | Mm.235385 | NM_025770 | ENSMUSG00000021619 | **Atg10** | 5330424L23Rik\|5430428K15Rik\|AI852123\|APG10\|Agp10\|Apg10l\|Apg10p\|Atg10l | autophagy related 10 | protein-coding |
| chr15_101282423_101282823 | chr15 | 101282424 | 101282823 | + | 1.03967 | Intergenic | Intergenic | -1678 | NM_026566 | 68118 | Mm.482360 | NM_026566 | ENSMUSG00000037204 | **Atg101** | 9430023L20Rik\|AA407441\|AU022967 | autophagy related 101 | protein-coding |
| chr2_91695747_91696711 | chr2 | 91695748 | 91696711 | + | 1.08677 | intron (NM_145528, intron 1 of 15) | intron (NM_145528, intron 1 of 15) | 14363 | NM_145528 | 51897 | Mm.28492 | NM_145528 | ENSMUSG00000027244 | **Atg13** | 1110053A20Rik\|D2Ertd391e\|Harbi1 | autophagy related 13 | protein-coding |
| chr7_101287631_101288756 | chr7 | 101287632 | 101288756 | + | 1.34482 | Intergenic | B3\|SINE\|B2 | 13894 | NM_001111111 | 73683 | Mm.275744 | NM_001111111 | ENSMUSG00000047767 | **Atg16l2** | 2410118P20Rik | autophagy related 16-like 2 (S. cerevisiae) | protein-coding |
| chr7_101275397_101275747 | chr7 | 101275398 | 101275747 | + | 0.872767 | intron (NM_001146010, intron 14 of 19) | intron (NM_001146010, intron 14 of 19) | 26516 | NM_001111111 | 73683 | Mm.275744 | NM_001111111 | ENSMUSG00000047767 | **Atg16l2** | 2410118P20Rik | autophagy related 16-like 2 (S. cerevisiae) | protein-coding |
| chr7_101275399_101275973 | chr7 | 101275400 | 101275973 | + | 0.790076 | intron (NM_001146010, intron 14 of 19) | ID_B1\|SINE\|B4 | 26402 | NM_001111111 | 73683 | Mm.275744 | NM_001111111 | ENSMUSG00000047767 | **Atg16l2** | 2410118P20Rik | autophagy related 16-like 2 (S. cerevisiae) | protein-coding |
| chr7_101275168_101276027 | chr7 | 101275169 | 101276027 | + | 0.637074 | intron (NM_001146010, intron 14 of 19) | ID_B1\|SINE\|B4 | 26490 | NM_001111111 | 73683 | Mm.275744 | NM_001111111 | ENSMUSG00000047767 | **Atg16l2** | 2410118P20Rik | autophagy related 16-like 2 (S. cerevisiae) | protein-coding |
| chr3_103639321_103639905 | chr3 | 103639322 | 103639905 | + | 1.82387 | intron (NM_001276680, intron 4 of 4) | intron (NM_001276680, intron 4 of 4) | 6455 | NM_174875 | 666468 | Mm.102230 | NM_174875 | ENSMUSG00000079418 | **Atg4a** | AI627006\|AV169859\|Apg4a\|Atg4al\|Autl2 | autophagy related 4A, cysteine peptidase | protein-coding |
| chr3_103640077_103640633 | chr3 | 103640078 | 103640633 | + | 1.37209 | intron (NM_001276680, intron 4 of 4) | intron (NM_001276680, intron 4 of 4) | 5713 | NM_174875 | 666468 | Mm.102230 | NM_174875 | ENSMUSG00000079418 | **Atg4a** | AI627006\|AV169859\|Apg4a\|Atg4al\|Autl2 | autophagy related 4A, cysteine peptidase | protein-coding |
| chr3_103682419_103683032 | chr3 | 103682420 | 103683032 | + | 1.28031 | intron (NR_106164, intron 2 of 4) | intron (NR_106164, intron 2 of 4) | -36658 | NM_174875 | 666468 | Mm.102230 | NM_174875 | ENSMUSG00000079418 | **Atg4a** | AI627006\|AV169859\|Apg4a\|Atg4al\|Autl2 | autophagy related 4A, cysteine peptidase | protein-coding |
| chr3_103683732_103684047 | chr3 | 103683733 | 103684047 | + | 0.761112 | intron (NR_106164, intron 2 of 4) | intron (NR_106164, intron 2 of 4) | -37822 | NM_174875 | 666468 | Mm.102230 | NM_174875 | ENSMUSG00000079418 | **Atg4a** | AI627006\|AV169859\|Apg4a\|Atg4al\|Autl2 | autophagy related 4A, cysteine peptidase | protein-coding |
| chr4_99336072_99336580 | chr4 | 99336073 | 99336580 | + | 0.794866 | Intergenic | RCHARR1\|DNA\|hAT-Charlie | 142179 | NM_001145967 | 242557 | Mm.241663 | NM_175029 | ENSMUSG00000028550 | **Atg4c** | Apg4-C\|Apg4c\|Atg4cl\|Autl1 | autophagy related 4C, cysteine peptidase | protein-coding |
| chr6_114680569_114681507 | chr6 | 114680570 | 114681507 | + | 1.21408 | intron (NM_001253717, intron 6 of 19) | intron (NM_001253717, intron 6 of 19) | 37941 | NM_001253717 | 74244 | Mm.275332 | NM_028835 | ENSMUSG00000030314 | **Atg7** | 1810013K23Rik\|Agp7\|Apg7l\|Atg7l\|Gm21553 | autophagy related 7 | protein-coding |
| chr1_71594236_71594580 | chr1 | 71594237 | 71594580 | + | 0.685175 | intron (NM_001276408, intron 40 of 44) | intron (NM_001276408, intron 40 of 44) | 37252 | NM_026195 | 108147 | Mm.38010 | NM_026195 | ENSMUSG00000026192 | **Atic** | 2610509C24Rik\|AA536954\|AW212393 | 5-aminoimidazole-4-carboxamide ribonucleotide formyltransferase/IMP cyclohydrolase | protein-coding |
| chr12_69884878_69885792 | chr12 | 69884879 | 69885792 | + | 1.39923 | intron (NM_201519, intron 1 of 30) | intron (NM_201519, intron 1 of 30) | -7770 | NM_178628 | 73991 | Mm.474462 | NM_178628 | ENSMUSG00000021066 | **Atl1** | 4930435M24Rik\|Adfsp\|Fsp1\|Spg3\|Spg3a | atlastin GTPase 1 | protein-coding |
| chr12_69879144_69879481 | chr12 | 69879145 | 69879481 | + | 0.73435 | intron (NM_201519, intron 1 of 30) | intron (NM_201519, intron 1 of 30) | -13792 | NM_178628 | 73991 | Mm.474462 | NM_178628 | ENSMUSG00000021066 | **Atl1** | 4930435M24Rik\|Adfsp\|Fsp1\|Spg3\|Spg3a | atlastin GTPase 1 | protein-coding |
| chr17_79939051_79939588 | chr17 | 79939052 | 79939588 | + | 1.06789 | Intergenic | L2c\|LINE\|L2 | -43197 | NM_001286647 | 56298 | Mm.175403 | NM_019717 | ENSMUSG00000059811 | **Atl2** | 2010110I21Rik\|AA407293\|AV334690\|Aip-2\|Arl6ip2 | atlastin GTPase 2 | protein-coding |
| chr19_7498945_7499243 | chr19 | 7498946 | 7499243 | + | 1.18585 | intron (NM_146091, intron 1 of 12) | intron (NM_146091, intron 1 of 12) | 4656 | NM_146091 | 109168 | Mm.264016 | NM_146091 | ENSMUSG00000024759 | **Atl3** | 4633402C03Rik\|5730596K20Rik\|AI465397\|AW228836 | atlastin GTPase 3 | protein-coding |
| chr7_58722278_58722860 | chr7 | 58722279 | 58722860 | + | 1.34216 | intron (NM_009728, intron 1 of 20) | intron (NM_009728, intron 1 of 20) | 64367 | NM_009728 | 11982 | Mm.135129 | NM_009728 | ENSMUSG00000025324 | **Atp10a** | Atp10c\|pfatp | ATPase, class V, type 10A | protein-coding |
| chr7_58216619_58217257 | chr7 | 58216620 | 58217257 | + | 1.33586 | Intergenic | RMER17B2\|LTR\|ERVK | -441264 | NM_009728 | 11982 | Mm.135129 | NM_009728 | ENSMUSG00000025324 | **Atp10a** | Atp10c\|pfatp | ATPase, class V, type 10A | protein-coding |
| chr7_58822343_58823153 | chr7 | 58822344 | 58823153 | + | 1.32528 | intron (NM_009728, intron 16 of 20) | intron (NM_009728, intron 16 of 20) | 164546 | NM_009728 | 11982 | Mm.135129 | NM_009728 | ENSMUSG00000025324 | **Atp10a** | Atp10c\|pfatp | ATPase, class V, type 10A | protein-coding |
| chr7_58388398_58388827 | chr7 | 58388399 | 58388827 | + | 1.29662 | Intergenic | Intergenic | -269589 | NM_009728 | 11982 | Mm.135129 | NM_009728 | ENSMUSG00000025324 | **Atp10a** | Atp10c\|pfatp | ATPase, class V, type 10A | protein-coding |
| chr7_58671946_58672328 | chr7 | 58671947 | 58672328 | + | 1.25324 | intron (NM_009728, intron 1 of 20) | Lx8b\|LINE\|L1 | 13935 | NM_009728 | 11982 | Mm.135129 | NM_009728 | ENSMUSG00000025324 | **Atp10a** | Atp10c\|pfatp | ATPase, class V, type 10A | protein-coding |
| chr7_58824668_58825023 | chr7 | 58824669 | 58825023 | + | 1.16202 | intron (NM_009728, intron 17 of 20) | intron (NM_009728, intron 17 of 20) | 166644 | NM_009728 | 11982 | Mm.135129 | NM_009728 | ENSMUSG00000025324 | **Atp10a** | Atp10c\|pfatp | ATPase, class V, type 10A | protein-coding |
| chr7_58216614_58217626 | chr7 | 58216615 | 58217626 | + | 1.15872 | Intergenic | RLTR41\|LTR\|ERV1 | -441082 | NM_009728 | 11982 | Mm.135129 | NM_009728 | ENSMUSG00000025324 | **Atp10a** | Atp10c\|pfatp | ATPase, class V, type 10A | protein-coding |
| chr7_58475100_58475677 | chr7 | 58475101 | 58475677 | + | 0.970329 | Intergenic | Intergenic | -182813 | NM_009728 | 11982 | Mm.135129 | NM_009728 | ENSMUSG00000025324 | **Atp10a** | Atp10c\|pfatp | ATPase, class V, type 10A | protein-coding |
| chr7_58474482_58474899 | chr7 | 58474483 | 58474899 | + | 0.867266 | Intergenic | Intergenic | -183511 | NM_009728 | 11982 | Mm.135129 | NM_009728 | ENSMUSG00000025324 | **Atp10a** | Atp10c\|pfatp | ATPase, class V, type 10A | protein-coding |
| chr7_58709401_58709708 | chr7 | 58709402 | 58709708 | + | 0.803705 | intron (NM_009728, intron 1 of 20) | intron (NM_009728, intron 1 of 20) | 51353 | NM_009728 | 11982 | Mm.135129 | NM_009728 | ENSMUSG00000025324 | **Atp10a** | Atp10c\|pfatp | ATPase, class V, type 10A | protein-coding |
| chr7_58483515_58483861 | chr7 | 58483516 | 58483861 | + | 0.731662 | Intergenic | Intergenic | -174514 | NM_009728 | 11982 | Mm.135129 | NM_009728 | ENSMUSG00000025324 | **Atp10a** | Atp10c\|pfatp | ATPase, class V, type 10A | protein-coding |

| | | | | | | | | | | | | | | | | | |
|---|---|---|---|---|---|---|---|---|---|---|---|---|---|---|---|---|---|
| chr7_58481388_58481808 | chr7 | 58481389 | 58481808 | + | 0.701651 | Intergenic | Lx\|LINE\|L1 | -176604 | NM_009728 | 11982 | Mm.135129 | NM_009728 | ENSMUSG00000025324 | **Atp10a** | Atp10c\|pfatp | ATPase, class V, type 10A | protein-coding |
| chr7_58829100_58829557 | chr7 | 58829101 | 58829557 | + | 0.674414 | TTS (NM_009728) | TTS (NM_009728) | 171127 | NM_009728 | 11982 | Mm.135129 | NM_009728 | ENSMUSG00000025324 | **Atp10a** | Atp10c\|pfatp | ATPase, class V, type 10A | protein-coding |
| chr5_72250025_72250688 | chr5 | 72250026 | 72250688 | + | 1.83798 | intron (NR_003966, intron 9 of 22) | intron (NR_003966, intron 9 of 22) | 47014 | NM_153389 | 231287 | Mm.32775 | NM_153389 | ENSMUSG00000046808 | **Atp10d** | 9830145H18Rik\|ATPVD\|D5Buc24e | ATPase, class V, type 10D | protein-coding |
| chr5_72212444_72213128 | chr5 | 72212445 | 72213128 | + | 0.695199 | intron (NR_003966, intron 1 of 22) | L2\|LINE\|L2 | 9443 | NM_153389 | 231287 | Mm.32775 | NM_153389 | ENSMUSG00000046808 | **Atp10d** | 9830145H18Rik\|ATPVD\|D5Buc24e | ATPase, class V, type 10D | protein-coding |
| chr3_35749529_35749907 | chr3 | 35749530 | 35749907 | + | 1.64872 | Intergenic | Intergenic | -4420 | NM_029570 | 76295 | Mm.239416 | NM_029570 | ENSMUSG00000037400 | **Atp11b** | 1110019I14Rik\|ATPIF\|ATPIR\|mKIAA0956 | ATPase, class VI, type 11B | protein-coding |
| chr3_35763160_35763550 | chr3 | 35763161 | 35763550 | + | 1.36576 | intron (NM_029570, intron 1 of 29) | L1ME3G\|LINE\|L1 | 9217 | NM_029570 | 76295 | Mm.239416 | NM_029570 | ENSMUSG00000037400 | **Atp11b** | 1110019I14Rik\|ATPIF\|ATPIR\|mKIAA0956 | ATPase, class VI, type 11B | protein-coding |
| chr3_35764645_35765483 | chr3 | 35764646 | 35765483 | + | 0.747217 | intron (NM_029570, intron 1 of 29) | RMER17C2\|LTR\|ERVK | 10926 | NM_029570 | 76295 | Mm.239416 | NM_029570 | ENSMUSG00000037400 | **Atp11b** | 1110019I14Rik\|ATPIF\|ATPIR\|mKIAA0956 | ATPase, class VI, type 11B | protein-coding |
| chr14_56325521_56325862 | chr14 | 56325522 | 56325862 | + | 0.801407 | Intergenic | Intergenic | -39376 | NM_138652 | 192113 | Mm.273271 | NM_138652 | ENSMUSG00000022229 | **Atp12a** | Atp1al1\|HKalpha2\|cHKA | ATPase, H+/K+ transporting, nongastric, alpha polypeptide | protein-coding |
| chr8_69792758_69793096 | chr8 | 69792759 | 69793096 | + | 0.697332 | intron (NM_133224, intron 1 of 25) | intron (NM_133224, intron 1 of 25) | 1764 | NM_133224 | 170759 | Mm.186066 | NM_133224 | ENSMUSG00000031862 | **Atp13a1** | Atp13a\|Cgi152\|catp | ATPase type 13A1 | protein-coding |
| chr3_101606848_101607286 | chr3 | 101606849 | 101607286 | + | 1.41983 | Intergenic | Intergenic | -2360 | NM_144900 | 11928 | Mm.280103 | NM_144900 | ENSMUSG00000033161 | **Atp1a1** | Atpa-1\|BC010319 | ATPase, Na+/K+ transporting, alpha 1 polypeptide | protein-coding |
| chr3_101650348_101650988 | chr3 | 101650349 | 101650988 | + | 1.00429 | Intergenic | Intergenic | -45961 | NM_144900 | 11928 | Mm.280103 | NM_144900 | ENSMUSG00000033161 | **Atp1a1** | Atpa-1\|BC010319 | ATPase, Na+/K+ transporting, alpha 1 polypeptide | protein-coding |
| chr3_101650348_101650771 | chr3 | 101650349 | 101650771 | + | 0.9871 | Intergenic | Intergenic | -45853 | NM_144900 | 11928 | Mm.280103 | NM_144900 | ENSMUSG00000033161 | **Atp1a1** | Atpa-1\|BC010319 | ATPase, Na+/K+ transporting, alpha 1 polypeptide | protein-coding |
| chr3_101600307_101600627 | chr3 | 101600308 | 101600627 | + | 0.973238 | intron (NM_144900, intron 1 of 22) | intron (NM_144900, intron 1 of 22) | 4240 | NM_144900 | 11928 | Mm.280103 | NM_144900 | ENSMUSG00000033161 | **Atp1a1** | Atpa-1\|BC010319 | ATPase, Na+/K+ transporting, alpha 1 polypeptide | protein-coding |
| chr3_101651087_101651809 | chr3 | 101651088 | 101651809 | + | 0.696636 | Intergenic | Intergenic | -46741 | NM_144900 | 11928 | Mm.280103 | NM_144900 | ENSMUSG00000033161 | **Atp1a1** | Atpa-1\|BC010319 | ATPase, Na+/K+ transporting, alpha 1 polypeptide | protein-coding |
| chr9_96393201_96393462 | chr9 | 96393202 | 96393462 | + | 0.636977 | Intergenic | Lx8\|LINE\|L1 | -29033 | NM_007502 | 11933 | Mm.424 | NM_007502 | ENSMUSG00000032412 | **Atp1b3** | AA409958\|AI664000\|AW212096 | ATPase, Na+/K+ transporting, beta 3 polypeptide | protein-coding |
| chr11_72948311_72948624 | chr11 | 72948312 | 72948624 | + | 1.28017 | Intergenic | Intergenic | -12701 | NM_016745 | 53313 | Mm.6306 | NM_016745 | ENSMUSG00000020788 | **Atp2a3** | SERCA3b\|Serca3 | ATPase, Ca++ transporting, ubiquitous | protein-coding |
| chr11_72953230_72954456 | chr11 | 72953231 | 72954456 | + | 0.967683 | Intergenic | Intergenic | -7326 | NM_016745 | 53313 | Mm.6306 | NM_016745 | ENSMUSG00000020788 | **Atp2a3** | SERCA3b\|Serca3 | ATPase, Ca++ transporting, ubiquitous | protein-coding |
| chr11_72946131_72946907 | chr11 | 72946132 | 72946907 | + | 0.657476 | Intergenic | Intergenic | -14650 | NM_016745 | 53313 | Mm.6306 | NM_016745 | ENSMUSG00000020788 | **Atp2a3** | SERCA3b\|Serca3 | ATPase, Ca++ transporting, ubiquitous | protein-coding |
| chr10_98880100_98880484 | chr10 | 98880101 | 98880484 | + | 1.28527 | Intergenic | Intergenic | -34860 | NM_026482 | 67972 | Mm.166944 | NM_026482 | ENSMUSG00000019943 | **Atp2b1** | 2810442I22Rik\|E130111D10Rik\|Pmca1 | ATPase, Ca++ transporting, plasma membrane 1 | protein-coding |
| chr10_98879896_98880488 | chr10 | 98879897 | 98880488 | + | 1.24188 | Intergenic | Intergenic | -34960 | NM_026482 | 67972 | Mm.166944 | NM_026482 | ENSMUSG00000019943 | **Atp2b1** | 2810442I22Rik\|E130111D10Rik\|Pmca1 | ATPase, Ca++ transporting, plasma membrane 1 | protein-coding |
| chr10_98708575_98709033 | chr10 | 98708576 | 98709033 | + | 1.02282 | Intergenic | Intergenic | -206348 | NM_026482 | 67972 | Mm.166944 | NM_026482 | ENSMUSG00000019943 | **Atp2b1** | 2810442I22Rik\|E130111D10Rik\|Pmca1 | ATPase, Ca++ transporting, plasma membrane 1 | protein-coding |
| chr10_98861396_98861922 | chr10 | 98861397 | 98861922 | + | 0.871488 | Intergenic | Intergenic | -53493 | NM_026482 | 67972 | Mm.166944 | NM_026482 | ENSMUSG00000019943 | **Atp2b1** | 2810442I22Rik\|E130111D10Rik\|Pmca1 | ATPase, Ca++ transporting, plasma membrane 1 | protein-coding |
| chr10_98798994_98799433 | chr10 | 98798995 | 98799433 | + | 0.759434 | Intergenic | Intergenic | -115938 | NM_026482 | 67972 | Mm.166944 | NM_026482 | ENSMUSG00000019943 | **Atp2b1** | 2810442I22Rik\|E130111D10Rik\|Pmca1 | ATPase, Ca++ transporting, plasma membrane 1 | protein-coding |
| chr10_98962100_98962572 | chr10 | 98962101 | 98962572 | + | 0.738942 | intron (NM_026482, intron 1 of 20) | intron (NM_026482, intron 1 of 20) | 47184 | NM_026482 | 67972 | Mm.166944 | NM_026482 | ENSMUSG00000019943 | **Atp2b1** | 2810442I22Rik\|E130111D10Rik\|Pmca1 | ATPase, Ca++ transporting, plasma membrane 1 | protein-coding |
| chr10_98786080_98786605 | chr10 | 98786081 | 98786605 | + | 0.709726 | Intergenic | Intergenic | -128809 | NM_026482 | 67972 | Mm.166944 | NM_026482 | ENSMUSG00000019943 | **Atp2b1** | 2810442I22Rik\|E130111D10Rik\|Pmca1 | ATPase, Ca++ transporting, plasma membrane 1 | protein-coding |
| chr10_98976902_98977374 | chr10 | 98976903 | 98977374 | + | 0.644731 | intron (NM_026482, intron 2 of 20) | intron (NM_026482, intron 2 of 20) | 61986 | NM_026482 | 67972 | Mm.166944 | NM_026482 | ENSMUSG00000019943 | **Atp2b1** | 2810442I22Rik\|E130111D10Rik\|Pmca1 | ATPase, Ca++ transporting, plasma membrane 1 | protein-coding |
| chrX_73567280_73567784 | chrX | 73567281 | 73567784 | + | 0.998741 | intron (NM_177236, intron 20 of 20) | intron (NM_177236, intron 20 of 20) | 64446 | NM_177236 | 320707 | Mm.210095 | NM_177236 | ENSMUSG00000031376 | **Atp2b3** | 6430519O13Rik\|Pmca3 | ATPase, Ca++ transporting, plasma membrane 3 | protein-coding |
| chrX_73567065_73567753 | chrX | 73567066 | 73567753 | + | 0.804664 | intron (NM_177236, intron 20 of 20) | intron (NM_177236, intron 20 of 20) | 64323 | NM_177236 | 320707 | Mm.210095 | NM_177236 | ENSMUSG00000031376 | **Atp2b3** | 6430519O13Rik\|Pmca3 | ATPase, Ca++ transporting, plasma membrane 3 | protein-coding |
| chr1_133797663_133798409 | chr1 | 133797664 | 133798409 | + | 1.34354 | Intergenic | Intergenic | -44289 | NM_213616 | 381290 | Mm.188617 | NM_213616 | ENSMUSG00000026463 | **Atp2b4** | Pmca4 | ATPase, Ca++ transporting, plasma membrane 4 | protein-coding |
| chr9_105483847_105484234 | chr9 | 105483848 | 105484234 | + | 0.839015 | intron (NM_001253831, intron 1 of 26) | L1MB7\|LINE\|L1 | 11092 | NM_175025 | 235574 | Mm.326247 | NM_175025 | ENSMUSG00000032570 | **Atp2c1** | 1700121J11Rik\|ATP2C1A\|AW061228\|BCPM\|D930003G21Rik\|HHD\|SPCA\|pmr1 | ATPase, Ca++-sequestering | protein-coding |
| chr8_13382608_13383343 | chr8 | 13382609 | 13383343 | + | 0.73373 | Intergenic | L1MB3\|LINE\|L1 | 13802 | NM_009724 | 11945 | Mm.154039 | NM_009724 | ENSMUSG00000031449 | **Atp4b** | AV080843 | ATPase, H+/K+ exchanging, beta polypeptide | protein-coding |
| chr2_74044424_74044833 | chr2 | 74044425 | 74044833 | + | 0.695244 | Intergenic | Intergenic | -133335 | NM_175015 | 228033 | Mm.2966 | NM_175015 | ENSMUSG00000018770 | **Atp5g3** | 6030447M23 | ATP synthase, H+ transporting, mitochondrial F0 complex, subunit C3 (subunit 9) | protein-coding |
| chr16_91962269_91962836 | chr16 | 91962270 | 91962836 | + | 1.50466 | Intergenic | (T)n\|Simple_repeat\|Simple_repeat | -30923 | NM_138597 | 28080 | Mm.41 | NM_138597 | ENSMUSG00000022956 | **Atp5o** | ATPO\|D12Wsu28e\|OSCP | ATP synthase, H+ transporting, mitochondrial F1 complex, O subunit | protein-coding |
| chr16_91870175_91870556 | chr16 | 91870176 | 91870556 | + | 1.29724 | 3' UTR (NM_001110275, exon 29 of 29) | 3' UTR (NM_001110275, exon 29 of 29) | 61264 | NM_138597 | 28080 | Mm.41 | NM_138597 | ENSMUSG00000022956 | **Atp5o** | ATPO\|D12Wsu28e\|OSCP | ATP synthase, H+ transporting, mitochondrial F1 complex, O subunit | protein-coding |
| chr11_101002962_101003355 | chr11 | 101002963 | 101003355 | + | 0.736842 | Intergenic | Intergenic | -6293 | NM_016920 | 11975 | Mm.340818 | NM_016920 | ENSMUSG00000019302 | **Atp6v0a1** | AA959968\|ATP6a1\|Atp6n1\|Atp6n1a\|Atpv0a1\|Vpp-1\|Vpp1 | ATPase, H+ transporting, lysosomal V0 subunit A1 | protein-coding |
| chr5_124626166_124626391 | chr5 | 124626167 | 124626391 | + | 0.719322 | intron (NM_026486, intron 17 of 17) | MLTR18D_MM\|LTR\|ERVK | -2773 | NM_011596 | 21871 | Mm.1158 | NM_011596 | | **Atp6v0a2** | 8430408C20Rik\|AI385560\|ATP6a2\|AW489264\|Atp6n1d\|Atp6n2\|C76904\|ISF\|J6B7\|SHIF\|Stv1\|TJ6M\|TJ6s\|Tj6\|V-ATPase 116 kDa\|V-ATPase a2 | ATPase, H+ transporting, lysosomal V0 subunit A2 | protein-coding |
| chr4_19922335_19922801 | chr4 | 19922336 | 19922801 | + | 1.20667 | promoter-TSS (NM_175406) | promoter-TSS (NM_175406) | -2 | NM_175406 | 242341 | Mm.19298 | NM_175406 | ENSMUSG00000028238 | **Atp6v0d2** | 1620401A02Rik\|AI324824\|V-ATPase | ATPase, H+ transporting, lysosomal V0 subunit D2 | protein-coding |
| chr15_38709411_38710594 | chr15 | 38709412 | 38710594 | + | 1.12172 | Intergenic | Intergenic | 48099 | NM_025494 | 66335 | Mm.276618 | NM_025494 | ENSMUSG00000022295 | **Atp6v1c1** | 1700025B18Rik\|U13839 | ATPase, H+ transporting, lysosomal V1 subunit C1 | protein-coding |
| chr6_120798082_120798452 | chr6 | 120798083 | 120798452 | + | 1.02678 | intron (NM_007510, intron 7 of 8) | intron (NM_007510, intron 7 of 8) | 24418 | NM_007510 | 11973 | Mm.29045 | NM_007510 | ENSMUSG00000019210 | **Atp6v1e1** | 2410029D23Rik\|Atp6e\|Atp6e2\|Atp6v1e\|D6Ertd385e\|E2\|P31\|Vma4 | ATPase, H+ transporting, lysosomal V1 subunit E1 | protein-coding |
| chr4_63530936_63531828 | chr4 | 63530937 | 63531828 | + | 0.907874 | Intergenic | Intergenic | -13383 | NM_024173 | 66290 | Mm.371615 | NM_024173 | ENSMUSG00000039105 | **Atp6v1g1** | 1810024D14Rik\|AA960677\|ATP6J\|Atp6g1\|VAG1\|Vma10 | ATPase, H+ transporting, lysosomal V1 subunit G1 | protein-coding |
| chr4_63530934_63531495 | chr4 | 63530935 | 63531495 | + | 0.897848 | Intergenic | Intergenic | -13550 | NM_024173 | 66290 | Mm.371615 | NM_024173 | ENSMUSG00000039105 | **Atp6v1g1** | 1810024D14Rik\|AA960677\|ATP6J\|Atp6g1\|VAG1\|Vma10 | ATPase, H+ transporting, lysosomal V1 subunit G1 | protein-coding |
| chr1_138418631_138419167 | chr1 | 138418632 | 138419167 | + | 1.61331 | Intergenic | B4A\|SINE\|B4 | 145161 | NM_177397 | 338375 | Mm.114424 | NM_177397 | ENSMUSG00000026394 | **Atp6v1g3** | - | ATPase, H+ transporting, lysosomal V1 subunit G3 | protein-coding |
| chr1_138322694_138323521 | chr1 | 138322695 | 138323521 | + | 0.970876 | Intergenic | Intergenic | 49370 | NM_177397 | 338375 | Mm.114424 | NM_177397 | ENSMUSG00000026394 | **Atp6v1g3** | - | ATPase, H+ transporting, lysosomal V1 subunit G3 | protein-coding |
| chr1_138248729_138249150 | chr1 | 138248730 | 138249150 | + | 0.964386 | Intergenic | RMER15\|LTR\|ERVL | -24798 | NM_177397 | 338375 | Mm.114424 | NM_177397 | ENSMUSG00000026394 | **Atp6v1g3** | - | ATPase, H+ transporting, lysosomal V1 subunit G3 | protein-coding |
| chr1_138323087_138323501 | chr1 | 138323088 | 138323501 | + | 0.950184 | Intergenic | Intergenic | 49556 | NM_177397 | 338375 | Mm.114424 | NM_177397 | ENSMUSG00000026394 | **Atp6v1g3** | - | ATPase, H+ transporting, lysosomal V1 subunit G3 | protein-coding |
| chr1_138293338_138293676 | chr1 | 138293339 | 138293676 | + | 0.817763 | Intergenic | Intergenic | 19769 | NM_177397 | 338375 | Mm.114424 | NM_177397 | ENSMUSG00000026394 | **Atp6v1g3** | - | ATPase, H+ transporting, lysosomal V1 subunit G3 | protein-coding |
| chr1_138289837_138290463 | chr1 | 138289838 | 138290463 | + | 0.812856 | TTS (NM_177397) | TTS (NM_177397) | 16412 | NM_177397 | 338375 | Mm.114424 | NM_177397 | ENSMUSG00000026394 | **Atp6v1g3** | - | ATPase, H+ transporting, lysosomal V1 subunit G3 | protein-coding |
| chr1_138361438_138361761 | chr1 | 138361439 | 138361761 | + | 0.800431 | Intergenic | L1VL2\|LINE\|L1 | 87862 | NM_177397 | 338375 | Mm.114424 | NM_177397 | ENSMUSG00000026394 | **Atp6v1g3** | - | ATPase, H+ transporting, lysosomal V1 subunit G3 | protein-coding |
| chr1_138322696_138323038 | chr1 | 138322697 | 138323038 | + | 0.791392 | Intergenic | Intergenic | 49129 | NM_177397 | 338375 | Mm.114424 | NM_177397 | ENSMUSG00000026394 | **Atp6v1g3** | - | ATPase, H+ transporting, lysosomal V1 subunit G3 | protein-coding |
| chr1_138298761_138299192 | chr1 | 138298762 | 138299192 | + | 0.705285 | Intergenic | Intergenic | 25239 | NM_177397 | 338375 | Mm.114424 | NM_177397 | ENSMUSG00000026394 | **Atp6v1g3** | - | ATPase, H+ transporting, lysosomal V1 subunit G3 | protein-coding |
| chr1_138343394_138343797 | chr1 | 138343395 | 138343797 | + | 0.663406 | Intergenic | MTE2b\|LTR\|ERVL-MaLR | 69858 | NM_177397 | 338375 | Mm.114424 | NM_177397 | ENSMUSG00000026394 | **Atp6v1g3** | - | ATPase, H+ transporting, lysosomal V1 subunit G3 | protein-coding |

| | | | | | | | | | | | | | | | | | |
|---|---|---|---|---|---|---|---|---|---|---|---|---|---|---|---|---|---|
| chr8_22055559_22055818 | chr8 | 22055560 | 22055818 | + | 1.05849 | intron (NM_007511, intron 1 of 20) | intron (NM_007511, intron 1 of 20) | 4385 | NM_007511 | 11979 | Mm.87854 | NM_007511 | ENSMUSG00000006567 | **Atp7b** | Atp7a\|WND\|tx | ATPase, Cu++ transporting, beta polypeptide | protein-coding |
| chr18_64749462_64749789 | chr18 | 64749463 | 64749789 | + | 0.700419 | Intergenic | ORR1F\|LTR\|ERVL-MaLR | -88626 | NM_001001488 | 54670 | Mm.270043 | NM_001001488 | ENSMUSG00000039529 | **Atp8b1** | AI451886\|FIC1\|Ic | ATPase, class I, type 8B, member 1 | protein-coding |
| chr3_89957802_89958149 | chr3 | 89957803 | 89958149 | + | 0.990122 | intron (NM_001081182, intron 6 of 27) | intron (NM_001081182, intron 6 of 27) | 5356 | NM_001081182 | 54667 | Mm.30455 | NM_001081182 | ENSMUSG00000060671 | **Atp8b2** | Id | ATPase, class I, type 8B, member 2 | protein-coding |
| chr3_89959564_89959790 | chr3 | 89959565 | 89959790 | + | 0.626317 | intron (NM_001081182, intron 2 of 27) | intron (NM_001081182, intron 2 of 27) | 3655 | NM_001081182 | 54667 | Mm.30455 | NM_001081182 | ENSMUSG00000060671 | **Atp8b2** | Id | ATPase, class I, type 8B, member 2 | protein-coding |
| chr2_126515101_126515951 | chr2 | 126515102 | 126515951 | + | 1.46018 | Intergenic | Lx6\|LINE\|L1 | -23973 | NM_001080944 | 241633 | Mm.222235 | NM_001080944 | ENSMUSG00000060131 | **Atp8b4** | A530043E15\|Im | ATPase, class I, type 8B, member 4 | protein-coding |
| chr2_126331749_126332211 | chr2 | 126331750 | 126332211 | + | 1.22158 | intron (NM_001080944, intron 25 of 27) | intron (NM_001080944, intron 25 of 27) | 159573 | NM_001080944 | 241633 | Mm.222235 | NM_001080944 | ENSMUSG00000060131 | **Atp8b4** | A530043E15\|Im | ATPase, class I, type 8B, member 4 | protein-coding |
| chr2_126464390_126464811 | chr2 | 126464391 | 126464811 | + | 0.96169 | intron (NM_001080944, intron 3 of 27) | MIR\|SINE\|MIR | 26952 | NM_001080944 | 241633 | Mm.222235 | NM_001080944 | ENSMUSG00000060131 | **Atp8b4** | A530043E15\|Im | ATPase, class I, type 8B, member 4 | protein-coding |
| chr2_126481598_126482055 | chr2 | 126481599 | 126482055 | + | 0.654619 | intron (NM_001080944, intron 2 of 27) | intron (NM_001080944, intron 2 of 27) | 9726 | NM_001080944 | 241633 | Mm.222235 | NM_001080944 | ENSMUSG00000060131 | **Atp8b4** | A530043E15\|Im | ATPase, class I, type 8B, member 4 | protein-coding |
| chr19_57825236_57825745 | chr19 | 57825237 | 57825745 | + | 0.837937 | intron (NM_181415, intron 26 of 28) | intron (NM_181415, intron 26 of 28) | 214457 | NM_181415 | 226255 | Mm.245340 | NM_181415 | ENSMUSG00000054843 | **Atrnl1** | AI504415\|AW555641\|Alp\|Atrnl\|Atrnlp | attractin like 1 | protein-coding |
| chr13_45837486_45838223 | chr13 | 45837487 | 45838223 | + | 1.53942 | intron (NM_001199305, intron 3 of 7) | intron (NM_001199305, intron 3 of 7) | 127136 | NM_001199304 | 20238 | Mm.342683 | NM_009124 | ENSMUSG00000046876 | **Atxn1** | 2900016G23Rik\|Atx1\|C85907\|ENSMUSG00000074917\|Gm10786\|Sca1 | ataxin 1 | protein-coding |
| chr13_45872956_45873493 | chr13 | 45872957 | 45873493 | + | 1.43949 | intron (NM_001199305, intron 2 of 7) | intron (NM_001199305, intron 2 of 7) | 91766 | NM_001199304 | 20238 | Mm.342683 | NM_009124 | ENSMUSG00000046876 | **Atxn1** | 2900016G23Rik\|Atx1\|C85907\|ENSMUSG00000074917\|Gm10786\|Sca1 | ataxin 1 | protein-coding |
| chr13_45911905_45912367 | chr13 | 45911906 | 45912367 | + | 1.28142 | intron (NM_001199305, intron 2 of 7) | intron (NM_001199305, intron 2 of 7) | 52855 | NM_001199304 | 20238 | Mm.342683 | NM_009124 | ENSMUSG00000046876 | **Atxn1** | 2900016G23Rik\|Atx1\|C85907\|ENSMUSG00000074917\|Gm10786\|Sca1 | ataxin 1 | protein-coding |
| chr13_45839837_45840292 | chr13 | 45839838 | 45840292 | + | 1.25826 | intron (NM_001199305, intron 3 of 7) | intron (NM_001199305, intron 3 of 7) | 124926 | NM_001199304 | 20238 | Mm.342683 | NM_009124 | ENSMUSG00000046876 | **Atxn1** | 2900016G23Rik\|Atx1\|C85907\|ENSMUSG00000074917\|Gm10786\|Sca1 | ataxin 1 | protein-coding |
| chr13_45856124_45856509 | chr13 | 45856125 | 45856509 | + | 1.0875 | intron (NM_001199305, intron 2 of 7) | intron (NM_001199305, intron 2 of 7) | 108674 | NM_001199304 | 20238 | Mm.342683 | NM_009124 | ENSMUSG00000046876 | **Atxn1** | 2900016G23Rik\|Atx1\|C85907\|ENSMUSG00000074917\|Gm10786\|Sca1 | ataxin 1 | protein-coding |
| chr13_45932349_45932770 | chr13 | 45932350 | 45932770 | + | 1.06836 | intron (NM_001199305, intron 2 of 7) | intron (NM_001199305, intron 2 of 7) | 32431 | NM_001199304 | 20238 | Mm.342683 | NM_009124 | ENSMUSG00000046876 | **Atxn1** | 2900016G23Rik\|Atx1\|C85907\|ENSMUSG00000074917\|Gm10786\|Sca1 | ataxin 1 | protein-coding |
| chr13_45866631_45866995 | chr13 | 45866632 | 45866995 | + | 0.857069 | intron (NM_001199305, intron 2 of 7) | Tigger7\|DNA\|TcMar-Tigger | 98178 | NM_001199304 | 20238 | Mm.342683 | NM_009124 | ENSMUSG00000046876 | **Atxn1** | 2900016G23Rik\|Atx1\|C85907\|ENSMUSG00000074917\|Gm10786\|Sca1 | ataxin 1 | protein-coding |
| chr13_45827305_45827862 | chr13 | 45827306 | 45827862 | + | 0.856193 | intron (NM_001199305, intron 3 of 7) | intron (NM_001199305, intron 3 of 7) | 137407 | NM_001199304 | 20238 | Mm.342683 | NM_009124 | ENSMUSG00000046876 | **Atxn1** | 2900016G23Rik\|Atx1\|C85907\|ENSMUSG00000074917\|Gm10786\|Sca1 | ataxin 1 | protein-coding |
| chr13_45861870_45862237 | chr13 | 45861871 | 45862237 | + | 0.828927 | intron (NM_001199305, intron 2 of 7) | MER2\|DNA\|TcMar-Tigger | 102937 | NM_001199304 | 20238 | Mm.342683 | NM_009124 | ENSMUSG00000046876 | **Atxn1** | 2900016G23Rik\|Atx1\|C85907\|ENSMUSG00000074917\|Gm10786\|Sca1 | ataxin 1 | protein-coding |
| chr13_45884845_45885376 | chr13 | 45884846 | 45885376 | + | 0.719526 | intron (NM_001199305, intron 2 of 7) | intron (NM_001199305, intron 2 of 7) | 79880 | NM_001199304 | 20238 | Mm.342683 | NM_009124 | ENSMUSG00000046876 | **Atxn1** | 2900016G23Rik\|Atx1\|C85907\|ENSMUSG00000074917\|Gm10786\|Sca1 | ataxin 1 | protein-coding |
| chr15_85319914_85320231 | chr15 | 85319915 | 85320231 | + | 0.685742 | Intergenic | Intergenic | -16308 | NM_016843 | 54138 | Mm.248906 | NM_016843 | ENSMUSG00000016541 | **Atxn10** | AI325283\|C77170\|E46\|Sca10\|TEG-169\|Tex169 | ataxin 10 | protein-coding |
| chr15_85346988_85347518 | chr15 | 85346989 | 85347518 | + | 0.624879 | intron (NM_016843, intron 1 of 11) | intron (NM_016843, intron 1 of 11) | 10872 | NM_016843 | 54138 | Mm.248906 | NM_016843 | ENSMUSG00000016541 | **Atxn10** | AI325283\|C77170\|E46\|Sca10\|TEG-169\|Tex169 | ataxin 10 | protein-coding |
| chr14_14024532_14025299 | chr14 | 14024533 | 14025299 | + | 1.09944 | intron (NM_139227, intron 3 of 11) | intron (NM_139227, intron 3 of 11) | 12425 | NM_139227 | 246103 | Mm.133625 | NM_139227 | ENSMUSG00000021738 | **Atxn7** | A430107N12Rik\|AI627028\|Sca7\|ataxin-7 | ataxin 7 | protein-coding |
| chr14_13993820_13994515 | chr14 | 13993821 | 13994515 | + | 1.07137 | Intergenic | B1_Mur2\|SINE\|Alu | -18323 | NM_139227 | 246103 | Mm.133625 | NM_139227 | ENSMUSG00000021738 | **Atxn7** | A430107N12Rik\|AI627028\|Sca7\|ataxin-7 | ataxin 7 | protein-coding |
| chr14_14059855_14060292 | chr14 | 14059856 | 14060292 | + | 0.836809 | intron (NM_139227, intron 4 of 11) | intron (NM_139227, intron 4 of 11) | 47583 | NM_139227 | 246103 | Mm.133625 | NM_139227 | ENSMUSG00000021738 | **Atxn7** | A430107N12Rik\|AI627028\|Sca7\|ataxin-7 | ataxin 7 | protein-coding |
| chr14_14025893_14026340 | chr14 | 14025894 | 14026340 | + | 0.791236 | intron (NM_139227, intron 3 of 11) | Lx9\|LINE\|L1 | 13626 | NM_139227 | 246103 | Mm.133625 | NM_139227 | ENSMUSG00000021738 | **Atxn7** | A430107N12Rik\|AI627028\|Sca7\|ataxin-7 | ataxin 7 | protein-coding |
| chr14_14059640_14060317 | chr14 | 14059641 | 14060317 | + | 0.751979 | intron (NM_139227, intron 4 of 11) | intron (NM_139227, intron 4 of 11) | 47488 | NM_139227 | 246103 | Mm.133625 | NM_139227 | ENSMUSG00000021738 | **Atxn7** | A430107N12Rik\|AI627028\|Sca7\|ataxin-7 | ataxin 7 | protein-coding |
| chr12_33170836_33171329 | chr12 | 33170837 | 33171329 | + | 1.85963 | intron (NM_028139, intron 2 of 3) | MLT1G3\|LTR\|ERVL-MaLR | 23397 | NM_028139 | 380753 | Mm.41540 | NM_028139 | ENSMUSG00000020564 | **Atxn7l1** | 2810423G08Rik\|Atxn7l4 | ataxin 7-like 1 | protein-coding |
| chr12_33170040_33170518 | chr12 | 33170041 | 33170518 | + | 0.855952 | intron (NM_028139, intron 2 of 3) | L1MB7\|LINE\|L1 | 22593 | NM_028139 | 380753 | Mm.41540 | NM_028139 | ENSMUSG00000020564 | **Atxn7l1** | 2810423G08Rik\|Atxn7l4 | ataxin 7-like 1 | protein-coding |
| chr12_33235701_33236066 | chr12 | 33235702 | 33236066 | + | 0.750826 | intron (NM_028139, intron 3 of 3) | intron (NM_028139, intron 3 of 3) | -66631 | NM_001033436 | 380753 | Mm.41540 | NM_028139 | ENSMUSG00000020564 | **Atxn7l1** | 2810423G08Rik\|Atxn7l4 | ataxin 7-like 1 | protein-coding |
| chr12_33175949_33176396 | chr12 | 33175950 | 33176396 | + | 0.671707 | intron (NM_028139, intron 2 of 3) | intron (NM_028139, intron 2 of 3) | 28487 | NM_028139 | 380753 | Mm.41540 | NM_028139 | ENSMUSG00000020564 | **Atxn7l1** | 2810423G08Rik\|Atxn7l4 | ataxin 7-like 1 | protein-coding |
| chr12_33284369_33284928 | chr12 | 33284370 | 33284928 | + | 0.606133 | Intergenic | RLTR53_Mm\|LTR\|ERVK | -17866 | NM_001033436 | 380753 | Mm.41540 | NM_028139 | ENSMUSG00000020564 | **Atxn7l1** | 2810423G08Rik\|Atxn7l4 | ataxin 7-like 1 | protein-coding |
| chr2_132613566_132614081 | chr2 | 132613567 | 132614081 | + | 0.827352 | intron (NR_033469, intron 2 of 3) | intron (NR_033469, intron 2 of 3) | 15628 | NR_033469 | 767814 | Mm.371752 | NR_033468 | ENSMUSG00000074783 | **AU019990** | - | expressed sequence AU019990 | ncRNA |
| chr2_132593742_132594172 | chr2 | 132593743 | 132594172 | + | 0.816595 | Intergenic | Intergenic | -4239 | NR_033468 | 767814 | Mm.371752 | NR_033468 | ENSMUSG00000074783 | **AU019990** | - | expressed sequence AU019990 | ncRNA |
| chr4_119226825_119227145 | chr4 | 119226826 | 119227145 | + | 0.711038 | intron (NM_001012400, intron 3 of 4) | intron (NM_001012400, intron 3 of 4) | 5739 | NM_001012400 | 230696 | Mm.473722 | NM_001012400 | | **AU022252** | AI450326 | expressed sequence AU022252 | protein-coding |
| chr11_79413181_79413822 | chr11 | 79413182 | 79413822 | + | 1.01879 | intron (NM_010897, intron 11 of 57) | intron (NM_010897, intron 11 of 57) | 70529 | NR_045305 | 104522 | Mm.436987 | NR_045305 | ENSMUSG00000091523 | **AU040972** | AU040954 | expressed sequence AU040972 | ncRNA |
| chr11_79413231_79413585 | chr11 | 79413232 | 79413585 | + | 0.967525 | intron (NM_010897, intron 11 of 57) | intron (NM_010897, intron 11 of 57) | 70623 | NR_045305 | 104522 | Mm.436987 | NR_045305 | ENSMUSG00000091523 | **AU040972** | AU040954 | expressed sequence AU040972 | ncRNA |
| chr13_52914474_52915101 | chr13 | 52914475 | 52915101 | + | 1.48783 | intron (NM_016709, intron 3 of 9) | intron (NM_016709, intron 3 of 9) | 14889 | NM_016709 | 11992 | Mm.252034 | NM_016709 | ENSMUSG00000021460 | **Auh** | C77140\|W91705 | AU RNA binding protein/enoyl-coenzyme A hydratase | protein-coding |
| chr13_52915546_52915819 | chr13 | 52915547 | 52915819 | + | 1.10134 | intron (NM_016709, intron 3 of 9) | intron (NM_016709, intron 3 of 9) | 13994 | NM_016709 | 11992 | Mm.252034 | NM_016709 | ENSMUSG00000021460 | **Auh** | C77140\|W91705 | AU RNA binding protein/enoyl-coenzyme A hydratase | protein-coding |
| chr13_52903544_52904172 | chr13 | 52903545 | 52904172 | + | 0.782393 | intron (NM_016709, intron 3 of 9) | intron (NM_016709, intron 3 of 9) | 25819 | NM_016709 | 11992 | Mm.252034 | NM_016709 | ENSMUSG00000021460 | **Auh** | C77140\|W91705 | AU RNA binding protein/enoyl-coenzyme A hydratase | protein-coding |
| chr13_52913771_52914282 | chr13 | 52913772 | 52914282 | + | 0.764546 | intron (NM_016709, intron 3 of 9) | ORR1F\|LTR\|ERVL-MaLR | 15650 | NM_016709 | 11992 | Mm.252034 | NM_016709 | ENSMUSG00000021460 | **Auh** | C77140\|W91705 | AU RNA binding protein/enoyl-coenzyme A hydratase | protein-coding |
| chr13_52860039_52860743 | chr13 | 52860040 | 52860743 | + | 0.730665 | intron (NM_016709, intron 6 of 9) | L1MC\|LINE\|L1 | 69286 | NM_016709 | 11992 | Mm.252034 | NM_016709 | ENSMUSG00000021460 | **Auh** | C77140\|W91705 | AU RNA binding protein/enoyl-coenzyme A hydratase | protein-coding |
| chr13_52883147_52883496 | chr13 | 52883148 | 52883496 | + | 0.62041 | intron (NM_016709, intron 6 of 9) | intron (NM_016709, intron 6 of 9) | 46355 | NM_016709 | 11992 | Mm.252034 | NM_016709 | ENSMUSG00000021460 | **Auh** | C77140\|W91705 | AU RNA binding protein/enoyl-coenzyme A hydratase | protein-coding |
| chr2_112596484_112597374 | chr2 | 112596485 | 112597374 | + | 0.768324 | intron (NM_001165935, intron 1 of 4) | Lx9\|LINE\|L1 | 37247 | NM_001165935 | 74268 | Mm.315125 | NM_028844 | | **Aven** | 1700013A01Rik\|1700056A21Rik\|mAven-L\|mAven-S | apoptosis, caspase activation inhibitor | protein-coding |

| | | | | | | | | | | | | | | | | | |
|---|---|---|---|---|---|---|---|---|---|---|---|---|---|---|---|---|---|
| chr19_42131113_42131404 | chr19 | 42131114 | 42131404 | + | 0.991795 | Intergenic | Intergenic | -2266 | NM_027106 | 69534 | Mm.30060 | NM_027106 | ENSMUSG00000018821 | **Avpi1** | 2310008N12Rik\|AA410041\|VIT32\|esu\|mVIT32 | arginine vasopressin-induced 1 | protein-coding |
| chr8_57589037_57589802 | chr8 | 57589038 | 57589802 | + | 1.62854 | intron (NM_001167981, intron 1 of 11) | intron (NM_001167981, intron 1 of 11) | -62333 | NR_040698 | 100502619 | Mm.393851 | NR_040698 | | **AW046200** | - | expressed sequence AW046200 | ncRNA |
| chr8_57591420_57591637 | chr8 | 57591421 | 57591637 | + | 1.04775 | intron (NM_001167981, intron 1 of 11) | intron (NM_001167981, intron 1 of 11) | -60224 | NR_040698 | 100502619 | Mm.393851 | NR_040698 | | **AW046200** | - | expressed sequence AW046200 | ncRNA |
| chr8_57624006_57624535 | chr8 | 57624007 | 57624535 | + | 0.978862 | intron (NM_001167981, intron 1 of 11) | intron (NM_001167981, intron 1 of 11) | -27482 | NR_040698 | 100502619 | Mm.393851 | NR_040698 | | **AW046200** | - | expressed sequence AW046200 | ncRNA |
| chr8_57627938_57628755 | chr8 | 57627939 | 57628755 | + | 0.865254 | intron (NM_001167981, intron 1 of 11) | B1_Mur4\|SINE\|Alu | -23406 | NR_040698 | 100502619 | Mm.393851 | NR_040698 | | **AW046200** | - | expressed sequence AW046200 | ncRNA |
| chr8_57627956_57629115 | chr8 | 57627957 | 57629115 | + | 0.700389 | intron (NM_001167981, intron 1 of 11) | intron (NM_001167981, intron 1 of 11) | -23217 | NR_040698 | 100502619 | Mm.393851 | NR_040698 | | **AW046200** | - | expressed sequence AW046200 | ncRNA |
| chr7_25781348_25782004 | chr7 | 25781349 | 25782004 | + | 0.983319 | intron (NM_009465, intron 4 of 19) | intron (NM_009465, intron 4 of 19) | 7057 | NM_001190974 | 26362 | Mm.4128 | NM_009465 | ENSMUSG00000002602 | **Axl** | AI323647\|Ark\|Tyro7\|Ufo | AXL receptor tyrosine kinase | protein-coding |
| chr7_25780028_25780323 | chr7 | 25780029 | 25780323 | + | 0.89506 | intron (NM_009465, intron 4 of 19) | intron (NM_009465, intron 4 of 19) | 8557 | NM_001190974 | 26362 | Mm.4128 | NM_009465 | ENSMUSG00000002602 | **Axl** | AI323647\|Ark\|Tyro7\|Ufo | AXL receptor tyrosine kinase | protein-coding |
| chr7_25781271_25782526 | chr7 | 25781272 | 25782526 | + | 0.809005 | intron (NM_009465, intron 4 of 19) | intron (NM_009465, intron 4 of 19) | 6834 | NM_001190974 | 26362 | Mm.4128 | NM_009465 | ENSMUSG00000002602 | **Axl** | AI323647\|Ark\|Tyro7\|Ufo | AXL receptor tyrosine kinase | protein-coding |
| chr6_95519359_95519911 | chr6 | 95519360 | 95519911 | + | 1.03558 | intron (NM_011507, intron 9 of 10) | intron (NM_011507, intron 9 of 10) | -185642 | NR_033559 | 414067 | Mm.355990 | NR_033559 | ENSMUSG00000068180 | **AY512915** | - | cDNA sequence AY512915 | ncRNA |
| chr9_117910322_117910689 | chr9 | 117910323 | 117910689 | + | 0.847416 | Intergenic | Lx8b\|LINE\|L1 | -130016 | NM_001286508 | 27215 | Mm.92705 | NM_013727 | ENSMUSG00000039285 | **Azi2** | AA410145\|AZ2 | 5-azacytidine induced gene 2 | protein-coding |
| chr9_117910176_117910707 | chr9 | 117910177 | 117910707 | + | 0.800784 | Intergenic | Lx8b\|LINE\|L1 | -130080 | NM_001286508 | 27215 | Mm.92705 | NM_013727 | ENSMUSG00000039285 | **Azi2** | AA410145\|AZ2 | 5-azacytidine induced gene 2 | protein-coding |
| chr11_95734357_95735096 | chr11 | 95734358 | 95735096 | + | 0.783114 | Intergenic | Intergenic | 11141 | NR_028263 | 320616 | Mm.428140 | NM_177209 | ENSMUSG00000075596 | **B130006D01Rik** | - | RIKEN cDNA B130006D01 gene | ncRNA |
| chr16_87524334_87524586 | chr16 | 87524335 | 87524586 | + | 0.612879 | Intergenic | B4A\|SINE\|B4 | 28387 | NR_040375 | 414095 | Mm.166361 | NR_040375 | ENSMUSG00000093606 | **B130034C11Rik** | - | RIKEN cDNA B130034C11 gene | ncRNA |
| chr10_12893430_12893834 | chr10 | 12893431 | 12893834 | + | 0.65127 | Intergenic | (TG)n\|Simple_repeat\|Simple_repeat | 29495 | NR_038027 | 320273 | Mm.417706 | NR_038027 | ENSMUSG00000097378 | **B230208H11Rik** | - | RIKEN cDNA B230208H11 gene | ncRNA |
| chr6_142412883_142413414 | chr6 | 142412884 | 142413414 | + | 1.19751 | promoter-TSS (NM_001242345) | promoter-TSS (NM_001242345) | -282 | NM_001242345 | 319552 | Mm.39649 | NM_001242345 | ENSMUSG00000071112 | **B230216G23Rik** | Npq\|Spx | RIKEN cDNA B230216G23 gene | protein-coding |
| chr6_142412216_142412745 | chr6 | 142412217 | 142412745 | + | 1.08809 | promoter-TSS (NM_001242345) | promoter-TSS (NM_001242345) | -950 | NM_001242345 | 319552 | Mm.39649 | NM_001242345 | ENSMUSG00000071112 | **B230216G23Rik** | Npq\|Spx | RIKEN cDNA B230216G23 gene | protein-coding |
| chr6_83442570_83442977 | chr6 | 83442571 | 83442977 | + | 1.45621 | intron (NR_028382, intron 2 of 2) | MIR3\|SINE\|MIR | 1019 | NR_028382 | 320775 | Mm.434334 | NR_028382 | ENSMUSG00000051844 | **B230319C09Rik** | - | RIKEN cDNA B230319C09 gene | ncRNA |
| chr9_69870302_69870825 | chr9 | 69870303 | 69870825 | + | 0.8203 | Intergenic | Intergenic | 109418 | NR_040765 | 319970 | Mm.186434 | NR_040765 | ENSMUSG00000097623 | **B230323A14Rik** | - | RIKEN cDNA B230323A14 gene | ncRNA |
| chr4_141547448_141547708 | chr4 | 141547449 | 141547708 | + | 0.998198 | non-coding (NR_030695, exon 1 of 1) | non-coding (NR_030695, exon 1 of 1) | 1416 | NR_030695 | 320456 | Mm.32859 | NM_177163 | | **B330016D10Rik** | - | RIKEN cDNA B330016D10 gene | ncRNA |
| chr3_69524788_69525437 | chr3 | 69524789 | 69525437 | + | 0.662061 | intron (NM_178726, intron 2 of 3) | intron (NM_178726, intron 2 of 3) | 73847 | NM_020026 | 26879 | Mm.153710 | NM_020026 | ENSMUSG00000043300 | **B3galnt1** | B3galt3\|b3GT3 | UDP-GalNAc:betaGlcNAc beta 1,3-galactosaminyltransferase, polypeptide 1 | protein-coding |
| chr2_68219856_68220194 | chr2 | 68219857 | 68220194 | + | 0.854206 | intron (NM_016866, intron 17 of 17) | intron (NM_016866, intron 17 of 17) | 115353 | NM_020283 | 26877 | Mm.226435 | NM_020283 | ENSMUSG00000034780 | **B3galt1** | beta3Gal-T1 | UDP-Gal:betaGlcNAc beta 1,3-galactosyltransferase, polypeptide 1 | protein-coding |
| chr2_68219132_68220227 | chr2 | 68219133 | 68220227 | + | 0.74984 | intron (NM_016866, intron 17 of 17) | intron (NM_016866, intron 17 of 17) | 115008 | NM_020283 | 26877 | Mm.226435 | NM_020283 | ENSMUSG00000034780 | **B3galt1** | beta3Gal-T1 | UDP-Gal:betaGlcNAc beta 1,3-galactosyltransferase, polypeptide 1 | protein-coding |
| chr2_68162093_68162349 | chr2 | 68162094 | 68162349 | + | 0.624064 | Intergenic | Intergenic | 57549 | NM_020283 | 26877 | Mm.226435 | NM_020283 | ENSMUSG00000034780 | **B3galt1** | beta3Gal-T1 | UDP-Gal:betaGlcNAc beta 1,3-galactosyltransferase, polypeptide 1 | protein-coding |
| chr1_143100454_143100801 | chr1 | 143100455 | 143100801 | + | 0.709342 | Intergenic | Intergenic | -540069 | NM_020025 | 26878 | Mm.285580 | NM_020025 | ENSMUSG00000033849 | **B3galt2** | - | UDP-Gal:betaGlcNAc beta 1,3-galactosyltransferase, polypeptide 2 | protein-coding |
| chr16_96301525_96301930 | chr16 | 96301526 | 96301930 | + | 0.680686 | intron (NM_033149, intron 1 of 2) | ORR1A1\|LTR\|ERVL-MaLR | 20696 | NM_033149 | 93961 | Mm.154783 | NM_033149 | ENSMUSG00000074892 | **B3galt5** | 1190002B21Rik\|AU045265\|b3Galt-V | UDP-Gal:betaGlcNAc beta 1,3-galactosyltransferase, polypeptide 5 | protein-coding |
| chr16_96298141_96298518 | chr16 | 96298142 | 96298518 | + | 0.667076 | intron (NM_033149, intron 1 of 2) | intron (NM_033149, intron 1 of 2) | 17298 | NM_033149 | 93961 | Mm.154783 | NM_033149 | ENSMUSG00000074892 | **B3galt5** | 1190002B21Rik\|AU045265\|b3Galt-V | UDP-Gal:betaGlcNAc beta 1,3-galactosyltransferase, polypeptide 5 | protein-coding |
| chr5_149677288_149677510 | chr5 | 149677289 | 149677510 | + | 0.648111 | promoter-TSS (NM_001081204) | promoter-TSS (NM_001081204) | -858 | NM_001081204 | 381694 | Mm.45060 | NM_001081204 | ENSMUSG00000051950 | **B3glct** | B3galtl\|Gm1057 | beta-3-glucosyltransferase | protein-coding |
| chr11_22845259_22845793 | chr11 | 22845260 | 22845793 | + | 1.00059 | intron (NM_001169114, intron 1 of 1) | intron (NM_001169114, intron 1 of 1) | 14209 | NM_016888 | 53625 | Mm.395495 | NM_016888 | ENSMUSG00000051650 | **B3gnt2** | AA408337\|AA409404\|AW260308\|B3Galt6\|B3gnt1\|BGnT-1\|BGnT-2\|BGnT1\|BGnT2\|Beta3gnt | UDP-GlcNAc:betaGal beta-1,3-N-acetylglucosaminyltransferase 2 | protein-coding |
| chr11_22811513_22811725 | chr11 | 22811514 | 22811725 | + | 0.929156 | Intergenic | Intergenic | 48116 | NM_016888 | 53625 | Mm.395495 | NM_016888 | ENSMUSG00000051650 | **B3gnt2** | AA408337\|AA409404\|AW260308\|B3Galt6\|B3gnt1\|BGnT-1\|BGnT-2\|BGnT1\|BGnT2\|Beta3gnt | UDP-GlcNAc:betaGal beta-1,3-N-acetylglucosaminyltransferase 2 | protein-coding |
| chr11_22815904_22816223 | chr11 | 22815905 | 22816223 | + | 0.865789 | Intergenic | Intergenic | 43671 | NM_016888 | 53625 | Mm.395495 | NM_016888 | ENSMUSG00000051650 | **B3gnt2** | AA408337\|AA409404\|AW260308\|B3Galt6\|B3gnt1\|BGnT-1\|BGnT-2\|BGnT1\|BGnT2\|Beta3gnt | UDP-GlcNAc:betaGal beta-1,3-N-acetylglucosaminyltransferase 2 | protein-coding |
| chr11_22813483_22814787 | chr11 | 22813484 | 22814787 | + | 0.746123 | Intergenic | Intergenic | 45600 | NM_016888 | 53625 | Mm.395495 | NM_016888 | ENSMUSG00000051650 | **B3gnt2** | AA408337\|AA409404\|AW260308\|B3Galt6\|B3gnt1\|BGnT-1\|BGnT-2\|BGnT1\|BGnT2\|Beta3gnt | UDP-GlcNAc:betaGal beta-1,3-N-acetylglucosaminyltransferase 2 | protein-coding |
| chr11_22865066_22865452 | chr11 | 22865067 | 22865452 | + | 0.691122 | Intergenic | Intergenic | -4923 | NM_001169114 | 53625 | Mm.395495 | NM_016888 | ENSMUSG00000051650 | **B3gnt2** | AA408337\|AA409404\|AW260308\|B3Galt6\|B3gnt1\|BGnT-1\|BGnT-2\|BGnT1\|BGnT2\|Beta3gnt | UDP-GlcNAc:betaGal beta-1,3-N-acetylglucosaminyltransferase 2 | protein-coding |
| chr11_22839410_22839722 | chr11 | 22839411 | 22839722 | + | 0.645173 | intron (NM_001169114, intron 1 of 1) | B3\|SINE\|B2 | 20169 | NM_016888 | 53625 | Mm.395495 | NM_016888 | ENSMUSG00000051650 | **B3gnt2** | AA408337\|AA409404\|AW260308\|B3Galt6\|B3gnt1\|BGnT-1\|BGnT-2\|BGnT1\|BGnT2\|Beta3gnt | UDP-GlcNAc:betaGal beta-1,3-N-acetylglucosaminyltransferase 2 | protein-coding |
| chr10_127144081_127144290 | chr10 | 127144082 | 127144290 | + | 0.84179 | Intergenic | Intergenic | -20970 | NR_045166 | 14421 | Mm.386762 | NM_008080 | ENSMUSG00000006731 | **B4galnt1** | 4933429D13Rik\|Gal-NAc-T\|GalNAc-T\|GalNAcT\|Galgt1\|Ggm-2\|Ggm2 | beta-1,4-N-acetyl-galactosaminyl transferase 1 | protein-coding |
| chr6_120279631_120280722 | chr6 | 120279632 | 120280722 | + | 0.919646 | intron (NM_198884, intron 1 of 19) | intron (NM_198884, intron 1 of 19) | 14382 | NM_198884 | 330406 | Mm.321840 | NM_198884 | ENSMUSG00000041372 | **B4galnt3** | AB114826\|B4GalNAcT3\|C330047A21 | beta-1,4-N-acetyl-galactosaminyl transferase 3 | protein-coding |
| chr6_120197554_120197850 | chr6 | 120197555 | 120197850 | + | 0.721093 | intron (NM_016718, intron 1 of 3) | intron (NM_016718, intron 1 of 3) | 96857 | NM_198884 | 330406 | Mm.321840 | NM_198884 | ENSMUSG00000041372 | **B4galnt3** | AB114826\|B4GalNAcT3\|C330047A21 | beta-1,4-N-acetyl-galactosaminyl transferase 3 | protein-coding |
| chr2_167362967_167363926 | chr2 | 167362968 | 167363926 | + | 1.43137 | Intergenic | MLT-int\|LTR\|ERVL-MaLR | -14269 | NM_019835 | 56336 | Mm.200886 | NM_019835 | ENSMUSG00000017929 | **B4galt5** | 9430078I07Rik\|AW049941\|AW539721 | UDP-Gal:betaGlcNAc beta 1,4-galactosyltransferase, polypeptide 5 | protein-coding |
| chr6_13664926_13665273 | chr6 | 13664927 | 13665273 | + | 0.63371 | intron (NM_175312, intron 1 of 4) | intron (NM_175312, intron 1 of 4) | 12866 | NM_175312 | 101148 | Mm.354863 | NM_175312 | ENSMUSG00000042742 | **B630005N14Rik** | AI666701 | RIKEN cDNA B630005N14 gene | protein-coding |
| chr8_34523719_34524370 | chr8 | 34523720 | 34524370 | + | 1.67501 | Intergenic | Intergenic | 116271 | NR_040706 | 330734 | Mm.149998 | NR_040706 | | **B930018H19Rik** | B930018H19 | RIKEN cDNA B930018H19 gene | ncRNA |
| chr8_34672381_34673025 | chr8 | 34672382 | 34673025 | + | 1.53033 | Intergenic | Intergenic | -32387 | NR_040706 | 330734 | Mm.149998 | NR_040706 | | **B930018H19Rik** | B930018H19 | RIKEN cDNA B930018H19 gene | ncRNA |
| chr8_34672759_34672989 | chr8 | 34672760 | 34672989 | + | 1.38121 | Intergenic | Lx9\|LINE\|L1 | -32558 | NR_040706 | 330734 | Mm.149998 | NR_040706 | | **B930018H19Rik** | B930018H19 | RIKEN cDNA B930018H19 gene | ncRNA |
| chr8_34590914_34591814 | chr8 | 34590915 | 34591814 | + | 1.33599 | intron (NR_040706, intron 4 of 4) | intron (NR_040706, intron 4 of 4) | 48952 | NR_040706 | 330734 | Mm.149998 | NR_040706 | | **B930018H19Rik** | B930018H19 | RIKEN cDNA B930018H19 gene | ncRNA |
| chr8_34671431_34672116 | chr8 | 34671432 | 34672116 | + | 1.27535 | Intergenic | Intergenic | -31458 | NR_040706 | 330734 | Mm.149998 | NR_040706 | | **B930018H19Rik** | B930018H19 | RIKEN cDNA B930018H19 gene | ncRNA |
| chr8_34671001_34672119 | chr8 | 34671002 | 34672119 | + | 1.23664 | Intergenic | Intergenic | -31244 | NR_040706 | 330734 | Mm.149998 | NR_040706 | | **B930018H19Rik** | B930018H19 | RIKEN cDNA B930018H19 gene | ncRNA |
| chr8_34672389_34672690 | chr8 | 34672390 | 34672690 | + | 1.09506 | Intergenic | Intergenic | -32224 | NR_040706 | 330734 | Mm.149998 | NR_040706 | | **B930018H19Rik** | B930018H19 | RIKEN cDNA B930018H19 gene | ncRNA |
| chr8_34530015_34530496 | chr8 | 34530016 | 34530496 | + | 1.04201 | Intergenic | Intergenic | 110060 | NR_040706 | 330734 | Mm.149998 | NR_040706 | | **B930018H19Rik** | B930018H19 | RIKEN cDNA B930018H19 gene | ncRNA |
| chr8_34622401_34623089 | chr8 | 34622402 | 34623089 | + | 0.989407 | intron (NR_040706, intron 1 of 4) | intron (NR_040706, intron 1 of 4) | 17571 | NR_040706 | 330734 | Mm.149998 | NR_040706 | | **B930018H19Rik** | B930018H19 | RIKEN cDNA B930018H19 gene | ncRNA |
| chr8_34683234_34683534 | chr8 | 34683235 | 34683534 | + | 0.907043 | Intergenic | Intergenic | -43068 | NR_040706 | 330734 | Mm.149998 | NR_040706 | | **B930018H19Rik** | B930018H19 | RIKEN cDNA B930018H19 gene | ncRNA |
| chr8_34682278_34682655 | chr8 | 34682279 | 34682655 | + | 0.893058 | Intergenic | MLT1E2\|LTR\|ERVL-MaLR | -42151 | NR_040706 | 330734 | Mm.149998 | NR_040706 | | **B930018H19Rik** | B930018H19 | RIKEN cDNA B930018H19 gene | ncRNA |
| chr8_34613020_34613605 | chr8 | 34613021 | 34613605 | + | 0.834663 | intron (NR_040706, intron 2 of 4) | intron (NR_040706, intron 2 of 4) | 27003 | NR_040706 | 330734 | Mm.149998 | NR_040706 | | **B930018H19Rik** | B930018H19 | RIKEN cDNA B930018H19 gene | ncRNA |

| | | | | | | | | | | | | | | | | | |
|---|---|---|---|---|---|---|---|---|---|---|---|---|---|---|---|---|---|
| chr8_34607439_34608294 | chr8 | 34607440 | 34608294 | + | 0.812384 | intron (NR_040706, intron 2 of 4) | intron (NR_040706, intron 2 of 4) | 32449 | NR_040706 | 330734 | Mm.149998 | NR_040706 | | **B930018H19Rik** | B930018H19 | RIKEN cDNA B930018H19 gene | ncRNA |
| chr8_34522876_34523227 | chr8 | 34522877 | 34523227 | + | 0.810127 | Intergenic | Intergenic | 117264 | NR_040706 | 330734 | Mm.149998 | NR_040706 | | **B930018H19Rik** | B930018H19 | RIKEN cDNA B930018H19 gene | ncRNA |
| chr8_34529527_34530608 | chr8 | 34529528 | 34530608 | + | 0.763468 | Intergenic | CYRA11_Mm\|Unknown\|Y-chromosome | 110248 | NR_040706 | 330734 | Mm.149998 | NR_040706 | | **B930018H19Rik** | B930018H19 | RIKEN cDNA B930018H19 gene | ncRNA |
| chr8_10534718_10535356 | chr8 | 10534719 | 10535356 | + | 1.23535 | intron (NM_001081397, intron 28 of 35) | intron (NM_001081397, intron 28 of 35) | 347417 | NR_040705 | 320014 | Mm.403203 | NR_040705 | | **B930025P03Rik** | - | RIKEN cDNA B930025P03 gene | ncRNA |
| chr12_110558173_110558541 | chr12 | 110558174 | 110558541 | + | 0.867015 | intron (NM_001135001, intron 10 of 13) | intron (NM_001135001, intron 10 of 13) | -33017 | NR_033340 | 319786 | Mm.129691 | NR_033340 | | **B930059L03Rik** | - | RIKEN cDNA B930059L03 gene | ncRNA |
| chr9_61243388_61244076 | chr9 | 61243389 | 61244076 | + | 0.617887 | Intergenic | MLT1K\|LTR\|ERVL-MaLR | 50077 | NR_045334 | 622037 | Mm.126243 | NR_045334 | | **B930092H01Rik** | - | RIKEN cDNA B930092H01 gene | ncRNA |
| chr9_45847077_45847511 | chr9 | 45847078 | 45847511 | + | 0.640299 | intron (NM_001145947, intron 1 of 7) | intron (NM_001145947, intron 1 of 7) | 8765 | NM_011792 | 23821 | Mm.24044 | NM_011792 | ENSMUSG00000032086 | **Bace1** | C76936 | beta-site APP cleaving enzyme 1 | protein-coding |
| chr16_97401364_97402151 | chr16 | 97401365 | 97402151 | + | 0.942391 | intron (NM_019517, intron 2 of 8) | intron (NM_019517, intron 2 of 8) | 45030 | NM_019517 | 56175 | Mm.97885 | NM_019517 | ENSMUSG00000040605 | **Bace2** | 1110059C24Rik\|AEPLC\|AI850424\|ALP56\|ARP1\|ASP21\|BAE2\|CDA13\|CEAP1\|DRAP | beta-site APP-cleaving enzyme 2 | protein-coding |
| chr16_97402388_97402710 | chr16 | 97402389 | 97402710 | + | 0.63961 | intron (NM_019517, intron 2 of 8) | intron (NM_019517, intron 2 of 8) | 45821 | NM_019517 | 56175 | Mm.97885 | NM_019517 | ENSMUSG00000040605 | **Bace2** | 1110059C24Rik\|AEPLC\|AI850424\|ALP56\|ARP1\|ASP21\|BAE2\|CDA13\|CEAP1\|DRAP | beta-site APP-cleaving enzyme 2 | protein-coding |
| chr16_87642107_87642624 | chr16 | 87642108 | 87642624 | + | 1.68189 | Intergenic | L1VL2\|LINE\|L1 | -56588 | NM_007520 | 12013 | Mm.26147 | NM_007520 | ENSMUSG00000025612 | **Bach1** | 6230421P05Rik\|AI323795 | BTB and CNC homology 1 | protein-coding |
| chr16_87724804_87725148 | chr16 | 87724805 | 87725148 | + | 1.36226 | intron (NM_007520, intron 4 of 4) | intron (NM_007520, intron 4 of 4) | 26022 | NM_007520 | 12013 | Mm.26147 | NM_007520 | ENSMUSG00000025612 | **Bach1** | 6230421P05Rik\|AI323795 | BTB and CNC homology 1 | protein-coding |
| chr16_87724818_87725317 | chr16 | 87724819 | 87725317 | + | 1.34544 | intron (NM_007520, intron 4 of 4) | intron (NM_007520, intron 4 of 4) | 26114 | NM_007520 | 12013 | Mm.26147 | NM_007520 | ENSMUSG00000025612 | **Bach1** | 6230421P05Rik\|AI323795 | BTB and CNC homology 1 | protein-coding |
| chr16_87724804_87725553 | chr16 | 87724805 | 87725553 | + | 1.26343 | intron (NM_007520, intron 4 of 4) | RSINE1\|SINE\|B4 | 26225 | NM_007520 | 12013 | Mm.26147 | NM_007520 | ENSMUSG00000025612 | **Bach1** | 6230421P05Rik\|AI323795 | BTB and CNC homology 1 | protein-coding |
| chr16_87665271_87665600 | chr16 | 87665272 | 87665600 | + | 1.19054 | Intergenic | RLTR30D_MM\|LTR\|ERV1 | -33518 | NM_007520 | 12013 | Mm.26147 | NM_007520 | ENSMUSG00000025612 | **Bach1** | 6230421P05Rik\|AI323795 | BTB and CNC homology 1 | protein-coding |
| chr16_87658422_87658808 | chr16 | 87658423 | 87658808 | + | 1.06753 | Intergenic | Intergenic | -40339 | NM_007520 | 12013 | Mm.26147 | NM_007520 | ENSMUSG00000025612 | **Bach1** | 6230421P05Rik\|AI323795 | BTB and CNC homology 1 | protein-coding |
| chr16_87652865_87653610 | chr16 | 87652866 | 87653610 | + | 0.778317 | Intergenic | Intergenic | -45716 | NM_007520 | 12013 | Mm.26147 | NM_007520 | ENSMUSG00000025612 | **Bach1** | 6230421P05Rik\|AI323795 | BTB and CNC homology 1 | protein-coding |
| chr16_87652563_87653641 | chr16 | 87652564 | 87653641 | + | 0.657858 | Intergenic | Intergenic | -45852 | NM_007520 | 12013 | Mm.26147 | NM_007520 | ENSMUSG00000025612 | **Bach1** | 6230421P05Rik\|AI323795 | BTB and CNC homology 1 | protein-coding |
| chr4_32405492_32405931 | chr4 | 32405493 | 32405931 | + | 1.46221 | Intergenic | Intergenic | -11723 | NM_001109661 | 12014 | Mm.491223 | NM_007521 | ENSMUSG00000040270 | **Bach2** | E030004N02Rik | BTB and CNC homology 2 | protein-coding |
| chr4_32391213_32391672 | chr4 | 32391214 | 32391672 | + | 1.42317 | Intergenic | Intergenic | -25992 | NM_001109661 | 12014 | Mm.491223 | NM_007521 | ENSMUSG00000040270 | **Bach2** | E030004N02Rik | BTB and CNC homology 2 | protein-coding |
| chr4_32424234_32424977 | chr4 | 32424235 | 32424977 | + | 1.41414 | intron (NM_001109661, intron 1 of 4) | intron (NM_001109661, intron 1 of 4) | 7171 | NM_001109661 | 12014 | Mm.491223 | NM_007521 | ENSMUSG00000040270 | **Bach2** | E030004N02Rik | BTB and CNC homology 2 | protein-coding |
| chr4_32405491_32406353 | chr4 | 32405492 | 32406353 | + | 1.38822 | Intergenic | Intergenic | -11513 | NM_001109661 | 12014 | Mm.491223 | NM_007521 | ENSMUSG00000040270 | **Bach2** | E030004N02Rik | BTB and CNC homology 2 | protein-coding |
| chr4_32292273_32292670 | chr4 | 32292274 | 32292670 | + | 1.31923 | Intergenic | Intergenic | -124963 | NM_001109661 | 12014 | Mm.491223 | NM_007521 | ENSMUSG00000040270 | **Bach2** | E030004N02Rik | BTB and CNC homology 2 | protein-coding |
| chr4_32405997_32406352 | chr4 | 32405998 | 32406352 | + | 1.1691 | Intergenic | Intergenic | -11260 | NM_001109661 | 12014 | Mm.491223 | NM_007521 | ENSMUSG00000040270 | **Bach2** | E030004N02Rik | BTB and CNC homology 2 | protein-coding |
| chr4_32425569_32426017 | chr4 | 32425570 | 32426017 | + | 1.12202 | intron (NM_001109661, intron 1 of 4) | intron (NM_001109661, intron 1 of 4) | 8358 | NM_001109661 | 12014 | Mm.491223 | NM_007521 | ENSMUSG00000040270 | **Bach2** | E030004N02Rik | BTB and CNC homology 2 | protein-coding |
| chr4_32319365_32320409 | chr4 | 32319366 | 32320409 | + | 1.0402 | Intergenic | Intergenic | -97548 | NM_001109661 | 12014 | Mm.491223 | NM_007521 | ENSMUSG00000040270 | **Bach2** | E030004N02Rik | BTB and CNC homology 2 | protein-coding |
| chr4_32366303_32366530 | chr4 | 32366304 | 32366530 | + | 1.00382 | Intergenic | ORR1F\|LTR\|ERVL-MaLR | -51018 | NM_001109661 | 12014 | Mm.491223 | NM_007521 | ENSMUSG00000040270 | **Bach2** | E030004N02Rik | BTB and CNC homology 2 | protein-coding |
| chr4_32401463_32401812 | chr4 | 32401464 | 32401812 | + | 0.930627 | Intergenic | Intergenic | -15797 | NM_001109661 | 12014 | Mm.491223 | NM_007521 | ENSMUSG00000040270 | **Bach2** | E030004N02Rik | BTB and CNC homology 2 | protein-coding |
| chr4_32217557_32217902 | chr4 | 32217558 | 32217902 | + | 0.890145 | Intergenic | Intergenic | -199705 | NM_001109661 | 12014 | Mm.491223 | NM_007521 | ENSMUSG00000040270 | **Bach2** | E030004N02Rik | BTB and CNC homology 2 | protein-coding |
| chr4_32330614_32331851 | chr4 | 32330615 | 32331851 | + | 0.863431 | Intergenic | Intergenic | -86202 | NM_001109661 | 12014 | Mm.491223 | NM_007521 | ENSMUSG00000040270 | **Bach2** | E030004N02Rik | BTB and CNC homology 2 | protein-coding |
| chr4_32366242_32366555 | chr4 | 32366243 | 32366555 | + | 0.854832 | Intergenic | ORR1F\|LTR\|ERVL-MaLR | -51036 | NM_001109661 | 12014 | Mm.491223 | NM_007521 | ENSMUSG00000040270 | **Bach2** | E030004N02Rik | BTB and CNC homology 2 | protein-coding |
| chr4_32376086_32376341 | chr4 | 32376087 | 32376341 | + | 0.844954 | Intergenic | Intergenic | -41221 | NM_001109661 | 12014 | Mm.491223 | NM_007521 | ENSMUSG00000040270 | **Bach2** | E030004N02Rik | BTB and CNC homology 2 | protein-coding |
| chr4_32375266_32375539 | chr4 | 32375267 | 32375539 | + | 0.811691 | Intergenic | Intergenic | -42032 | NM_001109661 | 12014 | Mm.491223 | NM_007521 | ENSMUSG00000040270 | **Bach2** | E030004N02Rik | BTB and CNC homology 2 | protein-coding |
| chr4_32283884_32284219 | chr4 | 32283885 | 32284219 | + | 0.810204 | Intergenic | Intergenic | -133383 | NM_001109661 | 12014 | Mm.491223 | NM_007521 | ENSMUSG00000040270 | **Bach2** | E030004N02Rik | BTB and CNC homology 2 | protein-coding |
| chr4_32268034_32268426 | chr4 | 32268035 | 32268426 | + | 0.808504 | Intergenic | Intergenic | -149205 | NM_001109661 | 12014 | Mm.491223 | NM_007521 | ENSMUSG00000040270 | **Bach2** | E030004N02Rik | BTB and CNC homology 2 | protein-coding |
| chr4_32319426_32320069 | chr4 | 32319427 | 32320069 | + | 0.803113 | Intergenic | Intergenic | -97687 | NM_001109661 | 12014 | Mm.491223 | NM_007521 | ENSMUSG00000040270 | **Bach2** | E030004N02Rik | BTB and CNC homology 2 | protein-coding |
| chr4_32374806_32376629 | chr4 | 32374807 | 32376629 | + | 0.786833 | Intergenic | Intergenic | -41717 | NM_001109661 | 12014 | Mm.491223 | NM_007521 | ENSMUSG00000040270 | **Bach2** | E030004N02Rik | BTB and CNC homology 2 | protein-coding |
| chr4_32330990_32331231 | chr4 | 32330991 | 32331231 | + | 0.764795 | Intergenic | Intergenic | -86324 | NM_001109661 | 12014 | Mm.491223 | NM_007521 | ENSMUSG00000040270 | **Bach2** | E030004N02Rik | BTB and CNC homology 2 | protein-coding |
| chr4_32426493_32426781 | chr4 | 32426494 | 32426781 | + | 0.747854 | intron (NM_001109661, intron 1 of 4) | intron (NM_001109661, intron 1 of 4) | 9202 | NM_001109661 | 12014 | Mm.491223 | NM_007521 | ENSMUSG00000040270 | **Bach2** | E030004N02Rik | BTB and CNC homology 2 | protein-coding |
| chr4_32330636_32330846 | chr4 | 32330637 | 32330846 | + | 0.722789 | Intergenic | Intergenic | -86694 | NM_001109661 | 12014 | Mm.491223 | NM_007521 | ENSMUSG00000040270 | **Bach2** | E030004N02Rik | BTB and CNC homology 2 | protein-coding |
| chr4_32256131_32256310 | chr4 | 32256132 | 32256310 | + | 0.717503 | Intergenic | Lx\|LINE\|L1 | -161214 | NM_001109661 | 12014 | Mm.491223 | NM_007521 | ENSMUSG00000040270 | **Bach2** | E030004N02Rik | BTB and CNC homology 2 | protein-coding |
| chr4_32315258_32315490 | chr4 | 32315259 | 32315490 | + | 0.672003 | Intergenic | Intergenic | -102061 | NM_001109661 | 12014 | Mm.491223 | NM_007521 | ENSMUSG00000040270 | **Bach2** | E030004N02Rik | BTB and CNC homology 2 | protein-coding |
| chr4_32405202_32405395 | chr4 | 32405203 | 32405395 | + | 0.670048 | Intergenic | Intergenic | -12136 | NM_001109661 | 12014 | Mm.491223 | NM_007521 | ENSMUSG00000040270 | **Bach2** | E030004N02Rik | BTB and CNC homology 2 | protein-coding |
| chr4_32544546_32545744 | chr4 | 32544547 | 32545744 | + | 0.997305 | intron (NM_001109661, intron 2 of 4) | intron (NM_001109661, intron 2 of 4) | 26517 | NR_026843 | 414068 | Mm.389550 | NR_026843 | ENSMUSG00000086150 | **Bach2os** | - | BTB and CNC homology 2, opposite strand | ncRNA |
| chr4_32540974_32541293 | chr4 | 32540975 | 32541293 | + | 0.775612 | intron (NM_001109661, intron 2 of 4) | intron (NM_001109661, intron 2 of 4) | 30528 | NR_026843 | 414068 | Mm.389550 | NR_026843 | ENSMUSG00000086150 | **Bach2os** | - | BTB and CNC homology 2, opposite strand | ncRNA |
| chr7_128554390_128555009 | chr7 | 128554391 | 128555009 | + | 1.63983 | Intergenic | Intergenic | 31117 | NM_013863 | 29810 | Mm.84073 | NM_013863 | ENSMUSG00000030847 | **Bag3** | AA407278\|Bis\|mg638 | BCL2-associated athanogene 3 | protein-coding |
| chr11_120256900_120257296 | chr11 | 120256901 | 120257296 | + | 0.868127 | intron (NM_198423, intron 2 of 27) | intron (NM_198423, intron 2 of 27) | 24151 | NM_198423 | 268515 | Mm.309498 | NM_198423 | ENSMUSG00000039741 | **Bahcc1** | B930044J06\|BC060615 | BAH domain and coiled-coil containing 1 | protein-coding |
| chr11_120256643_120257414 | chr11 | 120256644 | 120257414 | + | 0.674512 | intron (NM_198423, intron 2 of 27) | intron (NM_198423, intron 2 of 27) | 24082 | NM_198423 | 268515 | Mm.309498 | NM_198423 | ENSMUSG00000039741 | **Bahcc1** | B930044J06\|BC060615 | BAH domain and coiled-coil containing 1 | protein-coding |
| chr2_118905815_118906353 | chr2 | 118905816 | 118906353 | + | 0.956228 | intron (NM_001045523, intron 1 of 6) | intron (NM_001045523, intron 1 of 6) | 4469 | NM_001045523 | 228536 | Mm.330487 | NM_001045523 | ENSMUSG00000040007 | **Bahd1** | AL022997\|AW541238\|Gm117\|mKIAA0945 | bromo adjacent homology domain containing 1 | protein-coding |
| chr11_119975773_119976565 | chr11 | 119975774 | 119976565 | + | 1.38816 | intron (NM_001037754, intron 3 of 13) | intron (NM_001037754, intron 3 of 13) | 33077 | NM_130862 | 108100 | Mm.197534 | NM_130862 | ENSMUSG00000025372 | **Baiap2** | IRSp53\|R75030 | brain-specific angiogenesis inhibitor 1-associated protein 2 | protein-coding |
| chr3_136101999_136102279 | chr3 | 136102000 | 136102279 | + | 1.00578 | intron (NM_001033350, intron 8 of 16) | intron (NM_001033350, intron 8 of 16) | 223907 | NM_001033350 | 242248 | Mm.30832 | NM_001033350 | ENSMUSG00000037922 | **Bank1** | A530094C12Rik\|AI451642\|AVIEF\|BANK | B cell scaffold protein with ankyrin repeats 1 | protein-coding |
| chr3_136086896_136087238 | chr3 | 136086897 | 136087238 | + | 0.847175 | intron (NM_001033350, intron 10 of 16) | intron (NM_001033350, intron 10 of 16) | 238979 | NM_001033350 | 242248 | Mm.30832 | NM_001033350 | ENSMUSG00000037922 | **Bank1** | A530094C12Rik\|AI451642\|AVIEF\|BANK | B cell scaffold protein with ankyrin repeats 1 | protein-coding |
| chr8_121955703_121956235 | chr8 | 121955704 | 121956235 | + | 0.706858 | intron (NM_001285981, intron 1 of 12) | intron (NM_001285981, intron 1 of 12) | 5477 | NM_001285983 | 53325 | Mm.287163 | NM_016812 | ENSMUSG00000025316 | **Banp** | AA408158\|SMAR1 | BTG3 associated nuclear protein | protein-coding |
| chr15_25401945_25402400 | chr15 | 25401946 | 25402400 | + | 0.837105 | intron (NM_027395, intron 1 of 1) | intron (NM_027395, intron 1 of 1) | 11591 | NM_027395 | 70350 | Mm.29586 | NM_027395 | ENSMUSG00000045763 | **Basp1** | 2610024P12Rik\|CAP-23\|CAP23\|Ckap3\|NAP-22\|NAP22 | brain abundant, membrane attached signal protein 1 | protein-coding |
| chr15_25271875_25272301 | chr15 | 25271876 | 25272301 | + | 0.714634 | Intergenic | Intergenic | 141676 | NM_027395 | 70350 | Mm.29586 | NM_027395 | ENSMUSG00000045763 | **Basp1** | 2610024P12Rik\|CAP-23\|CAP23\|Ckap3\|NAP-22\|NAP22 | brain abundant, membrane attached signal protein 1 | protein-coding |
| chr12_85664265_85665070 | chr12 | 85664266 | 85665070 | + | 0.719978 | Intergenic | Intergenic | -22052 | NM_016767 | 53314 | Mm.6672 | NM_016767 | ENSMUSG00000034266 | **Batf** | B-ATF\|SFA-2 | basic leucine zipper transcription factor, ATF-like | protein-coding |
| chr1_191123778_191124717 | chr1 | 191123779 | 191124717 | + | 2.01233 | Intergenic | Intergenic | 25834 | NM_030060 | 381319 | Mm.6922 | NM_030060 | ENSMUSG00000026630 | **Batf3** | 9130211I03Rik\|Snft | basic leucine zipper transcription factor, ATF-like 3 | protein-coding |

| | | | | | | | | | | | | | | | | | |
|---|---|---|---|---|---|---|---|---|---|---|---|---|---|---|---|---|---|
| chr1_191123036_191123539 | chr1 | 191123037 | 191123539 | + | 1.57454 | Intergenic | B1_Mur1\|SINE\|Alu | 24874 | NM_030060 | 381319 | Mm.6922 | NM_030060 | ENSMUSG00000026630 | **Batf3** | 9130211I03Rik\|Snft | basic leucine zipper transcription factor, ATF-like 3 | protein-coding |
| chr1_191095086_191095750 | chr1 | 191095087 | 191095750 | + | 1.31862 | Intergenic | Intergenic | -2996 | NM_030060 | 381319 | Mm.6922 | NM_030060 | ENSMUSG00000026630 | **Batf3** | 9130211I03Rik\|Snft | basic leucine zipper transcription factor, ATF-like 3 | protein-coding |
| chr1_191099993_191100389 | chr1 | 191099994 | 191100389 | + | 1.24383 | intron (NM_030060, intron 1 of 2) | intron (NM_030060, intron 1 of 2) | 1777 | NM_030060 | 381319 | Mm.6922 | NM_030060 | ENSMUSG00000026630 | **Batf3** | 9130211I03Rik\|Snft | basic leucine zipper transcription factor, ATF-like 3 | protein-coding |
| chr1_191114852_191115510 | chr1 | 191114853 | 191115510 | + | 1.0573 | Intergenic | Intergenic | 16767 | NM_030060 | 381319 | Mm.6922 | NM_030060 | ENSMUSG00000026630 | **Batf3** | 9130211I03Rik\|Snft | basic leucine zipper transcription factor, ATF-like 3 | protein-coding |
| chr1_191114902_191115260 | chr1 | 191114903 | 191115260 | + | 1.05049 | Intergenic | Intergenic | 16667 | NM_030060 | 381319 | Mm.6922 | NM_030060 | ENSMUSG00000026630 | **Batf3** | 9130211I03Rik\|Snft | basic leucine zipper transcription factor, ATF-like 3 | protein-coding |
| chr1_191110996_191111841 | chr1 | 191110997 | 191111841 | + | 0.96855 | Intergenic | Intergenic | 13005 | NM_030060 | 381319 | Mm.6922 | NM_030060 | ENSMUSG00000026630 | **Batf3** | 9130211I03Rik\|Snft | basic leucine zipper transcription factor, ATF-like 3 | protein-coding |
| chr1_191097241_191098023 | chr1 | 191097242 | 191098023 | + | 0.961193 | promoter-TSS (NM_030060) | promoter-TSS (NM_030060) | -782 | NM_030060 | 381319 | Mm.6922 | NM_030060 | ENSMUSG00000026630 | **Batf3** | 9130211I03Rik\|Snft | basic leucine zipper transcription factor, ATF-like 3 | protein-coding |
| chr1_191100539_191100949 | chr1 | 191100540 | 191100949 | + | 0.867052 | exon (NM_030060, exon 2 of 3) | exon (NM_030060, exon 2 of 3) | 2330 | NM_030060 | 381319 | Mm.6922 | NM_030060 | ENSMUSG00000026630 | **Batf3** | 9130211I03Rik\|Snft | basic leucine zipper transcription factor, ATF-like 3 | protein-coding |
| chr1_191118780_191119263 | chr1 | 191118781 | 191119263 | + | 0.682055 | Intergenic | Intergenic | 20608 | NM_030060 | 381319 | Mm.6922 | NM_030060 | ENSMUSG00000026630 | **Batf3** | 9130211I03Rik\|Snft | basic leucine zipper transcription factor, ATF-like 3 | protein-coding |
| chr1_191098241_191098703 | chr1 | 191098242 | 191098703 | + | 0.623042 | promoter-TSS (NM_030060) | promoter-TSS (NM_030060) | 58 | NM_030060 | 381319 | Mm.6922 | NM_030060 | ENSMUSG00000026630 | **Batf3** | 9130211I03Rik\|Snft | basic leucine zipper transcription factor, ATF-like 3 | protein-coding |
| chr1_191118977_191119263 | chr1 | 191118978 | 191119263 | + | 0.613769 | Intergenic | Intergenic | 20706 | NM_030060 | 381319 | Mm.6922 | NM_030060 | ENSMUSG00000026630 | **Batf3** | 9130211I03Rik\|Snft | basic leucine zipper transcription factor, ATF-like 3 | protein-coding |
| chr12_55016304_55016939 | chr12 | 55016305 | 55016939 | + | 1.09343 | Intergenic | Intergenic | -30286 | NM_013815 | 217578 | Mm.479583 | NM_013815 | ENSMUSG00000035021 | **Baz1a** | Acf1\|B930060C03\|BC065123\|Gtl5\|Wcrf180\|cbp146 | bromodomain adjacent to zinc finger domain 1A | protein-coding |
| chr12_55016304_55017215 | chr12 | 55016305 | 55017215 | + | 0.896139 | Intergenic | B3\|SINE\|B2 | -30424 | NM_013815 | 217578 | Mm.479583 | NM_013815 | ENSMUSG00000035021 | **Baz1a** | Acf1\|B930060C03\|BC065123\|Gtl5\|Wcrf180\|cbp146 | bromodomain adjacent to zinc finger domain 1A | protein-coding |
| chr2_60080009_60080395 | chr2 | 60080010 | 60080395 | + | 1.2775 | intron (NM_001001182, intron 1 of 35) | intron (NM_001001182, intron 1 of 35) | 45538 | NM_001001182 | 407823 | Mm.228071 | NM_001001182 | ENSMUSG00000026987 | **Baz2b** | 5830435C13Rik\|BC053917\|D2Ertd794e | bromodomain adjacent to zinc finger domain, 2B | protein-coding |
| chr10_58713001_58713538 | chr10 | 58713002 | 58713538 | + | 0.645238 | Intergenic | Intergenic | 16525 | NR_033565 | 103505 | Mm.50261 | NR_033565 | | **BB019430** | - | expressed sequence BB019430 | ncRNA |
| chr13_52828920_52829602 | chr13 | 52828921 | 52829602 | + | 0.68316 | Intergenic | G-rich\|Low_complexity\|Low_complexity | -72056 | NR_027893 | 105404 | Mm.184173 | NR_027893 | | **BB123696** | - | expressed sequence BB123696 | ncRNA |
| chr13_52821827_52822522 | chr13 | 52821828 | 52822522 | + | 0.676588 | Intergenic | Intergenic | -64970 | NR_027893 | 105404 | Mm.184173 | NR_027893 | | **BB123696** | - | expressed sequence BB123696 | ncRNA |
| chr13_52793387_52794027 | chr13 | 52793388 | 52794027 | + | 0.659916 | Intergenic | Intergenic | -36502 | NR_027893 | 105404 | Mm.184173 | NR_027893 | | **BB123696** | - | expressed sequence BB123696 | ncRNA |
| chr7_16343250_16344249 | chr7 | 16343251 | 16344249 | + | 0.864224 | intron (NM_019748, intron 6 of 8) | intron (NM_019748, intron 6 of 8) | 34167 | NM_133234 | 170770 | Mm.7660 | NM_133234 | ENSMUSG00000002083 | **Bbc3** | PUMA\|PUMA/JFY1 | BCL2 binding component 3 | protein-coding |
| chr10_111234331_111235011 | chr10 | 111234332 | 111235011 | + | 1.59502 | intron (NM_175489, intron 4 of 23) | Charlie1a\|DNA\|hAT-Charlie | -64008 | NM_027914 | 71769 | Mm.45256 | NM_027914 | ENSMUSG00000035759 | **Bbs10** | 1300007O09Rik\|AI452285 | Bardet-Biedl syndrome 10 (human) | protein-coding |
| chr10_111234353_111234848 | chr10 | 111234354 | 111234848 | + | 1.56518 | intron (NM_175489, intron 4 of 23) | Charlie1a\|DNA\|hAT-Charlie | -64078 | NM_027914 | 71769 | Mm.45256 | NM_027914 | ENSMUSG00000035759 | **Bbs10** | 1300007O09Rik\|AI452285 | Bardet-Biedl syndrome 10 (human) | protein-coding |
| chr10_111233504_111234171 | chr10 | 111233505 | 111234171 | + | 1.27849 | intron (NM_175489, intron 4 of 23) | intron (NM_175489, intron 4 of 23) | -64841 | NM_027914 | 71769 | Mm.45256 | NM_027914 | ENSMUSG00000035759 | **Bbs10** | 1300007O09Rik\|AI452285 | Bardet-Biedl syndrome 10 (human) | protein-coding |
| chr10_111368759_111369677 | chr10 | 111368760 | 111369677 | + | 1.27333 | Intergenic | LTRIS2\|LTR\|ERV1 | 70539 | NM_027914 | 71769 | Mm.45256 | NM_027914 | ENSMUSG00000035759 | **Bbs10** | 1300007O09Rik\|AI452285 | Bardet-Biedl syndrome 10 (human) | protein-coding |
| chr10_111233837_111234168 | chr10 | 111233838 | 111234168 | + | 1.2052 | intron (NM_175489, intron 4 of 23) | intron (NM_175489, intron 4 of 23) | -64676 | NM_027914 | 71769 | Mm.45256 | NM_027914 | ENSMUSG00000035759 | **Bbs10** | 1300007O09Rik\|AI452285 | Bardet-Biedl syndrome 10 (human) | protein-coding |
| chr10_111233502_111233767 | chr10 | 111233503 | 111233767 | + | 0.989483 | exon (NM_001003717, exon 3 of 23) | exon (NM_001003717, exon 3 of 23) | -65044 | NM_027914 | 71769 | Mm.45256 | NM_027914 | ENSMUSG00000035759 | **Bbs10** | 1300007O09Rik\|AI452285 | Bardet-Biedl syndrome 10 (human) | protein-coding |
| chr10_111240389_111240730 | chr10 | 111240390 | 111240730 | + | 0.728703 | intron (NM_175489, intron 4 of 23) | intron (NM_175489, intron 4 of 23) | -58119 | NM_027914 | 71769 | Mm.45256 | NM_027914 | ENSMUSG00000035759 | **Bbs10** | 1300007O09Rik\|AI452285 | Bardet-Biedl syndrome 10 (human) | protein-coding |
| chr10_111235067_111235406 | chr10 | 111235068 | 111235406 | + | 0.698615 | intron (NM_175489, intron 4 of 23) | intron (NM_175489, intron 4 of 23) | -63442 | NM_027914 | 71769 | Mm.45256 | NM_027914 | ENSMUSG00000035759 | **Bbs10** | 1300007O09Rik\|AI452285 | Bardet-Biedl syndrome 10 (human) | protein-coding |
| chr3_37313391_37314017 | chr3 | 37313392 | 37314017 | + | 0.76516 | intron (NM_001255992, intron 1 of 1) | intron (NM_001255992, intron 1 of 1) | 1150 | NM_001255992 | 241950 | Mm.32171 | NM_001008502 | ENSMUSG00000051444 | **Bbs12** | Gm1805\|Gm407\|Gm721 | Bardet-Biedl syndrome 12 (human) | protein-coding |
| chr9_59387533_59388141 | chr9 | 59387534 | 59388141 | + | 1.03172 | TTS (NM_019927) | TTS (NM_019927) | -34329 | NM_175325 | 102774 | Mm.200714 | NM_175325 | ENSMUSG00000025235 | **Bbs4** | AW537059\|AW742241\|D9Ertd464e | Bardet-Biedl syndrome 4 (human) | protein-coding |
| chr3_36646906_36647231 | chr3 | 36646907 | 36647231 | + | 0.820818 | intron (NM_019510, intron 6 of 11) | L1VL2\|LINE\|L1 | -33680 | NM_027810 | 71492 | Mm.286187 | NM_027810 | ENSMUSG00000037325 | **Bbs7** | 8430406N16Rik | Bardet-Biedl syndrome 7 (human) | protein-coding |
| chr9_22483845_22484894 | chr9 | 22483846 | 22484894 | + | 1.06538 | intron (NM_178415, intron 2 of 22) | intron (NM_178415, intron 2 of 22) | 8655 | NM_178415 | 319845 | Mm.176725 | NM_178415 | ENSMUSG00000035919 | **Bbs9** | E130103I17Rik | Bardet-Biedl syndrome 9 (human) | protein-coding |
| chr16_50302553_50302824 | chr16 | 50302554 | 50302824 | + | 0.869969 | intron (NM_027444, intron 3 of 16) | intron (NM_027444, intron 3 of 16) | 129700 | NM_027444 | 70508 | Mm.28940 | NM_027444 | ENSMUSG00000022641 | **Bbx** | 5530401J07Rik\|5730403O13Rik | bobby sox homolog (Drosophila) | protein-coding |
| chr16_45647033_45647721 | chr16 | 45647034 | 45647721 | + | 0.989515 | intron (NM_145389, intron 2 of 5) | intron (NM_145389, intron 2 of 5) | 6741 | NM_145389 | 212998 | Mm.97511 | NM_145389 | ENSMUSG00000033187 | **BC016579** | - | cDNA sequence, BC016579 | protein-coding |
| chr16_45648718_45649130 | chr16 | 45648719 | 45649130 | + | 0.825898 | exon (NM_145389, exon 2 of 6) | exon (NM_145389, exon 2 of 6) | 5194 | NM_145389 | 212998 | Mm.97511 | NM_145389 | ENSMUSG00000033187 | **BC016579** | - | cDNA sequence, BC016579 | protein-coding |
| chr16_45640849_45641288 | chr16 | 45640850 | 45641288 | + | 0.797406 | intron (NM_145389, intron 2 of 5) | intron (NM_145389, intron 2 of 5) | 13049 | NM_145389 | 212998 | Mm.97511 | NM_145389 | ENSMUSG00000033187 | **BC016579** | - | cDNA sequence, BC016579 | protein-coding |
| chr19_4052060_4053183 | chr19 | 4052061 | 4053183 | + | 0.710717 | Intergenic | B1_Mur1\|SINE\|Alu | 6672 | NM_144869 | 225884 | Mm.440201 | NM_144869 | ENSMUSG00000058216 | **BC021614** | - | cDNA sequence BC021614 | protein-coding |
| chr12_112797683_112798464 | chr12 | 112797684 | 112798464 | + | 0.720471 | Intergenic | Intergenic | -10901 | NM_145450 | 217887 | Mm.186678 | NM_145450 | ENSMUSG00000037594 | **BC022687** | AW050009\|C130001I08Rik\|Flj20080 | cDNA sequence BC022687 | protein-coding |
| chr16_44734107_44734424 | chr16 | 44734108 | 44734424 | + | 0.822388 | intron (NM_145972, intron 7 of 8) | intron (NM_145972, intron 7 of 8) | 9965 | NM_145972 | 212547 | Mm.310573 | NM_145972 | ENSMUSG00000036208 | **BC027231** | Nepro | cDNA sequence BC027231 | protein-coding |
| chr16_38181004_38181579 | chr16 | 38181005 | 38181579 | + | 1.1609 | intron (NM_019827, intron 3 of 10) | intron (NM_019827, intron 3 of 10) | -92032 | NR_033221 | 414072 | Mm.483602 | NR_033221 | | **BC031361** | - | cDNA sequence BC031361 | ncRNA |
| chr6_128901952_128902584 | chr6 | 128901953 | 128902584 | + | 1.22865 | Intergenic | BGLII_Mus\|LTR\|ERVK | -11144 | NM_001254947 | 232406 | Mm.30745 | NM_001254946 | ENSMUSG00000090164 | **BC035044** | - | cDNA sequence BC035044 | protein-coding |
| chr10_121694369_121695017 | chr10 | 121694370 | 121695017 | + | 0.761034 | TTS (NM_177335) | TTS (NM_177335) | -45244 | NM_173022 | 270802 | Mm.209172 | NM_173022 | ENSMUSG00000053684 | **BC048403** | E030027L10 | cDNA sequence BC048403 | protein-coding |
| chr17_34106809_34107562 | chr17 | 34106810 | 34107562 | + | 0.914658 | Intergenic | Intergenic | -11877 | NR_046183 | 414076 | Mm.473092 | NR_046183 | ENSMUSG00000098197 | **BC051537** | - | cDNA sequence BC051537 | ncRNA |
| chr2_18802365_18802607 | chr2 | 18802366 | 18802607 | + | 0.747829 | Intergenic | Intergenic | 103464 | NM_001001334 | 381350 | Mm.330024 | NM_001001334 | | **BC061194** | - | cDNA sequence BC061194 | protein-coding |
| chr1_173520740_173521366 | chr1 | 173520741 | 173521366 | + | 1.24651 | TTS (NM_001024721) | TTS (NM_001024721) | 14904 | NM_001024721 | 545384 | Mm.371698 | NM_001024721 | | **BC094916** | Ifi214\|p214 | cDNA sequence BC094916 | protein-coding |
| chr1_173521492_173521813 | chr1 | 173521493 | 173521813 | + | 0.822691 | exon (NM_001024721, exon 5 of 5) | exon (NM_001024721, exon 5 of 5) | 14304 | NM_001024721 | 545384 | Mm.371698 | NM_001024721 | | **BC094916** | Ifi214\|p214 | cDNA sequence BC094916 | protein-coding |
| chr1_173510580_173511172 | chr1 | 173510581 | 173511172 | + | 0.657886 | Intergenic | Lx2\|LINE\|L1 | 25081 | NM_001024721 | 545384 | Mm.371698 | NM_001024721 | | **BC094916** | Ifi214\|p214 | cDNA sequence BC094916 | protein-coding |
| chr1_173489376_173489864 | chr1 | 173489377 | 173489864 | + | 0.610938 | Intergenic | Intergenic | 46337 | NM_001024721 | 545384 | Mm.371698 | NM_001024721 | | **BC094916** | Ifi214\|p214 | cDNA sequence BC094916 | protein-coding |
| chr11_118332778_118333154 | chr11 | 118332779 | 118333154 | + | 1.31857 | exon (NM_021440, exon 3 of 3) | exon (NM_021440, exon 3 of 3) | 9568 | NM_021440 | 58251 | Mm.439815 | NM_021440 | ENSMUSG00000076433 | **BC100451** | Ddc8\|FS39 | cDNA sequence BC100451 | protein-coding |
| chr3_96486041_96486388 | chr3 | 96486042 | 96486388 | + | 0.893714 | Intergenic | Intergenic | -33909 | NM_001256180 | 329716 | Mm.275223 | NM_183223 | ENSMUSG00000046317 | **BC107364** | D730018G16 | cDNA sequence BC107364 | protein-coding |
| chr3_96479342_96479918 | chr3 | 96479343 | 96479918 | + | 0.807803 | Intergenic | Intergenic | -27324 | NM_001256180 | 329716 | Mm.275223 | NM_183223 | ENSMUSG00000046317 | **BC107364** | D730018G16 | cDNA sequence BC107364 | protein-coding |
| chr8_111747930_111748241 | chr8 | 111747931 | 111748241 | + | 0.733859 | Intergenic | Intergenic | -4237 | NM_009954 | 12927 | Mm.3758 | NM_009954 | ENSMUSG00000031955 | **Bcar1** | AI385681\|Cas\|Crkas | breast cancer anti-estrogen resistance 1 | protein-coding |
| chr2_170321626_170322549 | chr2 | 170321627 | 170322549 | + | 1.72502 | Intergenic | Intergenic | -34273 | NR_040610 | 73303 | Mm.443297 | NR_040610 | ENSMUSG00000086999 | **Bcas1os2** | 1700040N02Rik | breast carcinoma amplified sequence 1, opposite strand 2 | ncRNA |
| chr2_170310717_170311069 | chr2 | 170310718 | 170311069 | + | 1.07854 | Intergenic | Intergenic | -45468 | NR_040610 | 73303 | Mm.443297 | NR_040610 | ENSMUSG00000086999 | **Bcas1os2** | 1700040N02Rik | breast carcinoma amplified sequence 1, opposite strand 2 | ncRNA |
| chr2_170311723_170312140 | chr2 | 170311724 | 170312140 | + | 1.06385 | Intergenic | Intergenic | -44429 | NR_040610 | 73303 | Mm.443297 | NR_040610 | ENSMUSG00000086999 | **Bcas1os2** | 1700040N02Rik | breast carcinoma amplified sequence 1, opposite strand 2 | ncRNA |
| chr2_170316448_170316714 | chr2 | 170316449 | 170316714 | + | 0.981813 | Intergenic | Intergenic | -39780 | NR_040610 | 73303 | Mm.443297 | NR_040610 | ENSMUSG00000086999 | **Bcas1os2** | 1700040N02Rik | breast carcinoma amplified sequence 1, opposite strand 2 | ncRNA |
| chr2_170307343_170307907 | chr2 | 170307344 | 170307907 | + | 0.788648 | Intergenic | B4A\|SINE\|B4 | -48736 | NR_040610 | 73303 | Mm.443297 | NR_040610 | ENSMUSG00000086999 | **Bcas1os2** | 1700040N02Rik | breast carcinoma amplified sequence 1, opposite strand 2 | ncRNA |

| | | | | | | | | | | | | | | | | | |
|---|---|---|---|---|---|---|---|---|---|---|---|---|---|---|---|---|---|
| chr2_170306654_170307953 | chr2 | 170306655 | 170307953 | + | 0.613666 | Intergenic | Intergenic | -49057 | NR_040610 | 73303 | Mm.443297 | NR_040610 | ENSMUSG00000086999 | **Bcas1os2** | 1700040N02Rik | breast carcinoma amplified sequence 1, opposite strand 2 | ncRNA |
| chr11_85407624_85408353 | chr11 | 85407625 | 85408353 | + | 1.1197 | intron (NM_001166643, intron 6 of 23) | intron (NM_001166643, intron 6 of 23) | 54825 | NM_001166643 | 192197 | Mm.287663 | NM_138681 | ENSMUSG00000059439 | **Bcas3** | 1500019F07Rik\|2610028P08Rik\|AU021018\|BC028339\|K20D4\|rudhira | breast carcinoma amplified sequence 3 | protein-coding |
| chr6_145001495_145002631 | chr6 | 145001496 | 145002631 | + | 1.57539 | intron (NM_007532, intron 10 of 10) | intron (NM_007532, intron 10 of 10) | 46749 | NM_001024468 | 12035 | Mm.4606 | NM_007532 | ENSMUSG00000030268 | **Bcat1** | BCATc\|Eca39 | branched chain aminotransferase 1, cytosolic | protein-coding |
| chr6_145001434_145003012 | chr6 | 145001435 | 145003012 | + | 1.49675 | intron (NM_007532, intron 10 of 10) | intron (NM_007532, intron 10 of 10) | 46589 | NM_001024468 | 12035 | Mm.4606 | NM_007532 | ENSMUSG00000030268 | **Bcat1** | BCATc\|Eca39 | branched chain aminotransferase 1, cytosolic | protein-coding |
| chr6_144942550_144943455 | chr6 | 144942551 | 144943455 | + | 1.15958 | Intergenic | Intergenic | 105809 | NM_001024468 | 12035 | Mm.4606 | NM_007532 | ENSMUSG00000030268 | **Bcat1** | BCATc\|Eca39 | branched chain aminotransferase 1, cytosolic | protein-coding |
| chr6_145015015_145015449 | chr6 | 145015016 | 145015449 | + | 0.682444 | intron (NM_007532, intron 7 of 10) | B3A\|SINE\|B2 | 33580 | NM_001024468 | 12035 | Mm.4606 | NM_007532 | ENSMUSG00000030268 | **Bcat1** | BCATc\|Eca39 | branched chain aminotransferase 1, cytosolic | protein-coding |
| chr7_25647741_25648198 | chr7 | 25647742 | 25648198 | + | 0.712822 | intron (NM_007533, intron 1 of 8) | intron (NM_007533, intron 1 of 8) | 10791 | NM_007533 | 12039 | Mm.25848 | NM_007533 | ENSMUSG00000060376 | **Bckdha** | - | branched chain ketoacid dehydrogenase E1, alpha polypeptide | protein-coding |
| chr7_25647570_25648308 | chr7 | 25647571 | 25648308 | + | 0.623974 | intron (NM_007533, intron 1 of 8) | intron (NM_007533, intron 1 of 8) | 10822 | NM_007533 | 12039 | Mm.25848 | NM_007533 | ENSMUSG00000060376 | **Bckdha** | - | branched chain ketoacid dehydrogenase E1, alpha polypeptide | protein-coding |
| chr9_83967390_83968209 | chr9 | 83967391 | 83968209 | + | 0.765199 | intron (NM_199195, intron 3 of 10) | intron (NM_199195, intron 3 of 10) | 19019 | NM_199195 | 12040 | Mm.12819 | NM_199195 | ENSMUSG00000032263 | **Bckdhb** | - | branched chain ketoacid dehydrogenase E1, beta polypeptide | protein-coding |
| chr12_107969875_107970308 | chr12 | 107969876 | 107970308 | + | 1.0917 | intron (NM_001079883, intron 2 of 3) | intron (NM_001079883, intron 2 of 3) | 33322 | NM_001079883 | 58208 | Mm.392694 | NM_021399 | ENSMUSG00000048251 | **Bcl11b** | 9130430L19Rik\|AI604821\|B630002E05Rik\|BCL-11B\|Ctip2\|Rit1 | B cell leukemia/lymphoma 11B | protein-coding |
| chr12_108010790_108011013 | chr12 | 108010791 | 108011013 | + | 0.764431 | Intergenic | Intergenic | -7488 | NM_021399 | 58208 | Mm.392694 | NM_021399 | ENSMUSG00000048251 | **Bcl11b** | 9130430L19Rik\|AI604821\|B630002E05Rik\|BCL-11B\|Ctip2\|Rit1 | B cell leukemia/lymphoma 11B | protein-coding |
| chr12_107972978_107973245 | chr12 | 107972979 | 107973245 | + | 0.745606 | intron (NM_001079883, intron 2 of 3) | intron (NM_001079883, intron 2 of 3) | 30302 | NM_001079883 | 58208 | Mm.392694 | NM_021399 | ENSMUSG00000048251 | **Bcl11b** | 9130430L19Rik\|AI604821\|B630002E05Rik\|BCL-11B\|Ctip2\|Rit1 | B cell leukemia/lymphoma 11B | protein-coding |
| chr12_108009240_108009497 | chr12 | 108009241 | 108009497 | + | 0.69192 | Intergenic | Intergenic | -5955 | NM_021399 | 58208 | Mm.392694 | NM_021399 | ENSMUSG00000048251 | **Bcl11b** | 9130430L19Rik\|AI604821\|B630002E05Rik\|BCL-11B\|Ctip2\|Rit1 | B cell leukemia/lymphoma 11B | protein-coding |
| chr12_107993621_107993855 | chr12 | 107993622 | 107993855 | + | 0.659037 | intron (NM_001079883, intron 1 of 3) | intron (NM_001079883, intron 1 of 3) | 9676 | NM_001079883 | 58208 | Mm.392694 | NM_021399 | ENSMUSG00000048251 | **Bcl11b** | 9130430L19Rik\|AI604821\|B630002E05Rik\|BCL-11B\|Ctip2\|Rit1 | B cell leukemia/lymphoma 11B | protein-coding |
| chr12_108005099_108005743 | chr12 | 108005100 | 108005743 | + | 0.652638 | Intergenic | Intergenic | -2007 | NM_021399 | 58208 | Mm.392694 | NM_021399 | ENSMUSG00000048251 | **Bcl11b** | 9130430L19Rik\|AI604821\|B630002E05Rik\|BCL-11B\|Ctip2\|Rit1 | B cell leukemia/lymphoma 11B | protein-coding |
| chr12_107961913_107962276 | chr12 | 107961914 | 107962276 | + | 0.620791 | intron (NM_001286343, intron 1 of 1) | intron (NM_001286343, intron 1 of 1) | 41319 | NM_001079883 | 58208 | Mm.392694 | NM_021399 | ENSMUSG00000048251 | **Bcl11b** | 9130430L19Rik\|AI604821\|B630002E05Rik\|BCL-11B\|Ctip2\|Rit1 | B cell leukemia/lymphoma 11B | protein-coding |
| chr1_106570259_106570729 | chr1 | 106570260 | 106570729 | + | 1.67179 | intron (NM_009741, intron 1 of 1) | intron (NM_009741, intron 1 of 1) | 143796 | NM_009741 | 12043 | Mm.257460 | NM_009741 | ENSMUSG00000057329 | **Bcl2** | AW986256\|Bcl-2\|C430015F12Rik\|D630044D05Rik\|D830018M01Rik | B cell leukemia/lymphoma 2 | protein-coding |
| chr1_106567493_106567852 | chr1 | 106567494 | 106567852 | + | 1.62141 | intron (NM_009741, intron 1 of 1) | intron (NM_009741, intron 1 of 1) | 146617 | NM_009741 | 12043 | Mm.257460 | NM_009741 | ENSMUSG00000057329 | **Bcl2** | AW986256\|Bcl-2\|C430015F12Rik\|D630044D05Rik\|D830018M01Rik | B cell leukemia/lymphoma 2 | protein-coding |
| chr1_106569362_106570737 | chr1 | 106569363 | 106570737 | + | 1.50675 | intron (NM_009741, intron 1 of 1) | PB1D10\|SINE\|Alu | 144240 | NM_009741 | 12043 | Mm.257460 | NM_009741 | ENSMUSG00000057329 | **Bcl2** | AW986256\|Bcl-2\|C430015F12Rik\|D630044D05Rik\|D830018M01Rik | B cell leukemia/lymphoma 2 | protein-coding |
| chr1_106616907_106617401 | chr1 | 106616908 | 106617401 | + | 1.49093 | intron (NM_009741, intron 1 of 1) | intron (NM_009741, intron 1 of 1) | 97136 | NM_009741 | 12043 | Mm.257460 | NM_009741 | ENSMUSG00000057329 | **Bcl2** | AW986256\|Bcl-2\|C430015F12Rik\|D630044D05Rik\|D830018M01Rik | B cell leukemia/lymphoma 2 | protein-coding |
| chr1_106569429_106569946 | chr1 | 106569430 | 106569946 | + | 1.30633 | intron (NM_009741, intron 1 of 1) | intron (NM_009741, intron 1 of 1) | 144602 | NM_009741 | 12043 | Mm.257460 | NM_009741 | ENSMUSG00000057329 | **Bcl2** | AW986256\|Bcl-2\|C430015F12Rik\|D630044D05Rik\|D830018M01Rik | B cell leukemia/lymphoma 2 | protein-coding |
| chr1_106599408_106599767 | chr1 | 106599409 | 106599767 | + | 1.28296 | intron (NM_009741, intron 1 of 1) | intron (NM_009741, intron 1 of 1) | 114702 | NM_009741 | 12043 | Mm.257460 | NM_009741 | ENSMUSG00000057329 | **Bcl2** | AW986256\|Bcl-2\|C430015F12Rik\|D630044D05Rik\|D830018M01Rik | B cell leukemia/lymphoma 2 | protein-coding |
| chr1_106561040_106561715 | chr1 | 106561041 | 106561715 | + | 1.25215 | intron (NM_009741, intron 1 of 1) | intron (NM_009741, intron 1 of 1) | 152912 | NM_009741 | 12043 | Mm.257460 | NM_009741 | ENSMUSG00000057329 | **Bcl2** | AW986256\|Bcl-2\|C430015F12Rik\|D630044D05Rik\|D830018M01Rik | B cell leukemia/lymphoma 2 | protein-coding |
| chr1_106585266_106585895 | chr1 | 106585267 | 106585895 | + | 1.20276 | intron (NM_009741, intron 1 of 1) | intron (NM_009741, intron 1 of 1) | 128709 | NM_009741 | 12043 | Mm.257460 | NM_009741 | ENSMUSG00000057329 | **Bcl2** | AW986256\|Bcl-2\|C430015F12Rik\|D630044D05Rik\|D830018M01Rik | B cell leukemia/lymphoma 2 | protein-coding |
| chr1_106618951_106619352 | chr1 | 106618952 | 106619352 | + | 1.09809 | intron (NM_009741, intron 1 of 1) | intron (NM_009741, intron 1 of 1) | 95138 | NM_009741 | 12043 | Mm.257460 | NM_009741 | ENSMUSG00000057329 | **Bcl2** | AW986256\|Bcl-2\|C430015F12Rik\|D630044D05Rik\|D830018M01Rik | B cell leukemia/lymphoma 2 | protein-coding |
| chr1_106640926_106641309 | chr1 | 106640927 | 106641309 | + | 0.937308 | intron (NM_009741, intron 1 of 1) | intron (NM_009741, intron 1 of 1) | 73172 | NM_009741 | 12043 | Mm.257460 | NM_009741 | ENSMUSG00000057329 | **Bcl2** | AW986256\|Bcl-2\|C430015F12Rik\|D630044D05Rik\|D830018M01Rik | B cell leukemia/lymphoma 2 | protein-coding |
| chr1_106620471_106620908 | chr1 | 106620472 | 106620908 | + | 0.862188 | intron (NM_009741, intron 1 of 1) | B4A\|SINE\|B4 | 93600 | NM_009741 | 12043 | Mm.257460 | NM_009741 | ENSMUSG00000057329 | **Bcl2** | AW986256\|Bcl-2\|C430015F12Rik\|D630044D05Rik\|D830018M01Rik | B cell leukemia/lymphoma 2 | protein-coding |
| chr1_106586372_106586629 | chr1 | 106586373 | 106586629 | + | 0.751453 | intron (NM_009741, intron 1 of 1) | intron (NM_009741, intron 1 of 1) | 127789 | NM_009741 | 12043 | Mm.257460 | NM_009741 | ENSMUSG00000057329 | **Bcl2** | AW986256\|Bcl-2\|C430015F12Rik\|D630044D05Rik\|D830018M01Rik | B cell leukemia/lymphoma 2 | protein-coding |
| chr1_106588463_106588862 | chr1 | 106588464 | 106588862 | + | 0.74003 | intron (NM_009741, intron 1 of 1) | MTEa\|LTR\|ERVL-MaLR | 125627 | NM_009741 | 12043 | Mm.257460 | NM_009741 | ENSMUSG00000057329 | **Bcl2** | AW986256\|Bcl-2\|C430015F12Rik\|D630044D05Rik\|D830018M01Rik | B cell leukemia/lymphoma 2 | protein-coding |
| chr1_106627078_106627580 | chr1 | 106627079 | 106627580 | + | 0.696962 | intron (NM_009741, intron 1 of 1) | intron (NM_009741, intron 1 of 1) | 86961 | NM_009741 | 12043 | Mm.257460 | NM_009741 | ENSMUSG00000057329 | **Bcl2** | AW986256\|Bcl-2\|C430015F12Rik\|D630044D05Rik\|D830018M01Rik | B cell leukemia/lymphoma 2 | protein-coding |
| chr1_106707127_106707600 | chr1 | 106707128 | 106707600 | + | 0.687559 | intron (NM_009741, intron 1 of 1) | intron (NM_009741, intron 1 of 1) | 6926 | NM_009741 | 12043 | Mm.257460 | NM_009741 | ENSMUSG00000057329 | **Bcl2** | AW986256\|Bcl-2\|C430015F12Rik\|D630044D05Rik\|D830018M01Rik | B cell leukemia/lymphoma 2 | protein-coding |
| chr1_106588455_106589108 | chr1 | 106588456 | 106589108 | + | 0.652853 | intron (NM_009741, intron 1 of 1) | intron (NM_009741, intron 1 of 1) | 125508 | NM_009741 | 12043 | Mm.257460 | NM_009741 | ENSMUSG00000057329 | **Bcl2** | AW986256\|Bcl-2\|C430015F12Rik\|D630044D05Rik\|D830018M01Rik | B cell leukemia/lymphoma 2 | protein-coding |
| chr1_106560209_106560487 | chr1 | 106560210 | 106560487 | + | 0.611255 | intron (NM_009741, intron 1 of 1) | intron (NM_009741, intron 1 of 1) | 153942 | NM_009741 | 12043 | Mm.257460 | NM_009741 | ENSMUSG00000057329 | **Bcl2** | AW986256\|Bcl-2\|C430015F12Rik\|D630044D05Rik\|D830018M01Rik | B cell leukemia/lymphoma 2 | protein-coding |
| chr2_152825398_152826239 | chr2 | 152825399 | 152826239 | + | 0.828316 | intron (NM_001289717, intron 2 of 2) | intron (NM_001289717, intron 2 of 2) | 4898 | NM_001289716 | 12048 | Mm.238213 | NM_009743 | ENSMUSG00000007659 | **Bcl2l1** | Bcl(X)L\|Bcl-XL\|Bcl2l\|BclX\|bcl-x\|bcl2-L-1 | BCL2-like 1 | protein-coding |
| chr2_152825753_152826105 | chr2 | 152825754 | 152826105 | + | 0.737966 | intron (NM_001289717, intron 2 of 2) | intron (NM_001289717, intron 2 of 2) | 4788 | NM_001289716 | 12048 | Mm.238213 | NM_009743 | ENSMUSG00000007659 | **Bcl2l1** | Bcl(X)L\|Bcl-XL\|Bcl2l\|BclX\|bcl-x\|bcl2-L-1 | BCL2-like 1 | protein-coding |
| chr2_128073585_128073980 | chr2 | 128073586 | 128073980 | + | 1.33744 | intron (NM_028765, intron 14 of 17) | Lx8\|LINE\|L1 | -52255 | NM_207681 | 12125 | Mm.141083 | NM_009754 | ENSMUSG00000027381 | **Bcl2l11** | 1500006F24Rik\|Bim\|Bod\|bcl2-L-11 | BCL2-like 11 (apoptosis facilitator) | protein-coding |
| chr2_128072768_128073523 | chr2 | 128072769 | 128073523 | + | 1.15491 | intron (NM_028765, intron 14 of 17) | intron (NM_028765, intron 14 of 17) | -52892 | NM_207681 | 12125 | Mm.141083 | NM_009754 | ENSMUSG00000027381 | **Bcl2l11** | 1500006F24Rik\|Bim\|Bod\|bcl2-L-11 | BCL2-like 11 (apoptosis facilitator) | protein-coding |
| chr2_128087742_128088110 | chr2 | 128087743 | 128088110 | + | 1.12965 | intron (NM_028765, intron 16 of 17) | intron (NM_028765, intron 16 of 17) | -38112 | NM_207681 | 12125 | Mm.141083 | NM_009754 | ENSMUSG00000027381 | **Bcl2l11** | 1500006F24Rik\|Bim\|Bod\|bcl2-L-11 | BCL2-like 11 (apoptosis facilitator) | protein-coding |
| chr2_128072857_128073185 | chr2 | 128072858 | 128073185 | + | 1.09766 | intron (NM_028765, intron 14 of 17) | intron (NM_028765, intron 14 of 17) | -53017 | NM_207681 | 12125 | Mm.141083 | NM_009754 | ENSMUSG00000027381 | **Bcl2l11** | 1500006F24Rik\|Bim\|Bod\|bcl2-L-11 | BCL2-like 11 (apoptosis facilitator) | protein-coding |

| | | | | | | | | | | | | | | | | | |
|---|---|---|---|---|---|---|---|---|---|---|---|---|---|---|---|---|---|
| chr2_128117397_128118057 | chr2 | 128117398 | 128118057 | + | 1.0847 | intron (NM_028765, intron 17 of 17) | intron (NM_028765, intron 17 of 17) | -8311 | NM_207681 | 12125 | Mm.141083 | NM_009754 | ENSMUSG00000027381 | **Bcl2l11** | 1500006F24Rik\|Bim\|Bod\|bcl2-L-11 | BCL2-like 11 (apoptosis facilitator) | protein-coding |
| chr2_128092018_128092534 | chr2 | 128092019 | 128092534 | + | 0.904619 | intron (NM_028765, intron 16 of 17) | intron (NM_028765, intron 16 of 17) | -33762 | NM_207681 | 12125 | Mm.141083 | NM_009754 | ENSMUSG00000027381 | **Bcl2l11** | 1500006F24Rik\|Bim\|Bod\|bcl2-L-11 | BCL2-like 11 (apoptosis facilitator) | protein-coding |
| chr2_128094539_128094925 | chr2 | 128094540 | 128094925 | + | 0.861995 | intron (NM_028765, intron 16 of 17) | intron (NM_028765, intron 16 of 17) | -31306 | NM_207681 | 12125 | Mm.141083 | NM_009754 | ENSMUSG00000027381 | **Bcl2l11** | 1500006F24Rik\|Bim\|Bod\|bcl2-L-11 | BCL2-like 11 (apoptosis facilitator) | protein-coding |
| chr2_128229777_128230493 | chr2 | 128229778 | 128230493 | + | 0.810745 | Intergenic | Intergenic | 102560 | NM_001284410 | 12125 | Mm.141083 | NM_009754 | ENSMUSG00000027381 | **Bcl2l11** | 1500006F24Rik\|Bim\|Bod\|bcl2-L-11 | BCL2-like 11 (apoptosis facilitator) | protein-coding |
| chr2_128230950_128231530 | chr2 | 128230951 | 128231530 | + | 0.746048 | Intergenic | L1M4\|LINE\|L1 | 103665 | NM_001284410 | 12125 | Mm.141083 | NM_009754 | ENSMUSG00000027381 | **Bcl2l11** | 1500006F24Rik\|Bim\|Bod\|bcl2-L-11 | BCL2-like 11 (apoptosis facilitator) | protein-coding |
| chr2_128230955_128231298 | chr2 | 128230956 | 128231298 | + | 0.735587 | Intergenic | L1M4\|LINE\|L1 | 103552 | NM_001284410 | 12125 | Mm.141083 | NM_009754 | ENSMUSG00000027381 | **Bcl2l11** | 1500006F24Rik\|Bim\|Bod\|bcl2-L-11 | BCL2-like 11 (apoptosis facilitator) | protein-coding |
| chr2_128229785_128230233 | chr2 | 128229786 | 128230233 | + | 0.692406 | Intergenic | L1M4\|LINE\|L1 | 102434 | NM_001284410 | 12125 | Mm.141083 | NM_009754 | ENSMUSG00000027381 | **Bcl2l11** | 1500006F24Rik\|Bim\|Bod\|bcl2-L-11 | BCL2-like 11 (apoptosis facilitator) | protein-coding |
| chr2_128221325_128222083 | chr2 | 128221326 | 128222083 | + | 0.659208 | Intergenic | Intergenic | 94129 | NM_001284410 | 12125 | Mm.141083 | NM_009754 | ENSMUSG00000027381 | **Bcl2l11** | 1500006F24Rik\|Bim\|Bod\|bcl2-L-11 | BCL2-like 11 (apoptosis facilitator) | protein-coding |
| chr2_128060814_128061862 | chr2 | 128060815 | 128061862 | + | 0.622697 | intron (NM_028765, intron 14 of 17) | (GGGAGA)n\|Simple_repeat\|Simple_repeat | -64700 | NM_207681 | 12125 | Mm.141083 | NM_009754 | ENSMUSG00000027381 | **Bcl2l11** | 1500006F24Rik\|Bim\|Bod\|bcl2-L-11 | BCL2-like 11 (apoptosis facilitator) | protein-coding |
| chr6_134281930_134282658 | chr6 | 134281931 | 134282658 | + | 2.09947 | Intergenic | Intergenic | -114035 | NM_025778 | 66813 | Mm.474472 | NM_025778 | ENSMUSG00000030200 | **Bcl2l14** | 4930452K23Rik\|4933405K19Rik\|9030625M01Rik\|AI429211\|Bclg | BCL2-like 14 (apoptosis facilitator) | protein-coding |
| chr6_134251855_134252651 | chr6 | 134251856 | 134252651 | + | 1.84 | intron (NM_007961, intron 5 of 7) | intron (NM_007961, intron 5 of 7) | -144076 | NM_025778 | 66813 | Mm.474472 | NM_025778 | ENSMUSG00000030200 | **Bcl2l14** | 4930452K23Rik\|4933405K19Rik\|9030625M01Rik\|AI429211\|Bclg | BCL2-like 14 (apoptosis facilitator) | protein-coding |
| chr6_134225053_134225981 | chr6 | 134225054 | 134225981 | + | 1.77398 | intron (NM_007961, intron 3 of 7) | Lx5\|LINE\|L1 | -170812 | NM_025778 | 66813 | Mm.474472 | NM_025778 | ENSMUSG00000030200 | **Bcl2l14** | 4930452K23Rik\|4933405K19Rik\|9030625M01Rik\|AI429211\|Bclg | BCL2-like 14 (apoptosis facilitator) | protein-coding |
| chr6_134220067_134220862 | chr6 | 134220068 | 134220862 | + | 1.51927 | intron (NM_007961, intron 2 of 7) | intron (NM_007961, intron 2 of 7) | -175864 | NM_025778 | 66813 | Mm.474472 | NM_025778 | ENSMUSG00000030200 | **Bcl2l14** | 4930452K23Rik\|4933405K19Rik\|9030625M01Rik\|AI429211\|Bclg | BCL2-like 14 (apoptosis facilitator) | protein-coding |
| chr6_134372670_134373406 | chr6 | 134372671 | 134373406 | + | 1.34623 | Intergenic | Intergenic | -23291 | NM_025778 | 66813 | Mm.474472 | NM_025778 | ENSMUSG00000030200 | **Bcl2l14** | 4930452K23Rik\|4933405K19Rik\|9030625M01Rik\|AI429211\|Bclg | BCL2-like 14 (apoptosis facilitator) | protein-coding |
| chr6_134261875_134262855 | chr6 | 134261876 | 134262855 | + | 1.23811 | intron (NM_007961, intron 6 of 7) | intron (NM_007961, intron 6 of 7) | -133964 | NM_025778 | 66813 | Mm.474472 | NM_025778 | ENSMUSG00000030200 | **Bcl2l14** | 4930452K23Rik\|4933405K19Rik\|9030625M01Rik\|AI429211\|Bclg | BCL2-like 14 (apoptosis facilitator) | protein-coding |
| chr6_134450669_134451173 | chr6 | 134450670 | 134451173 | + | 0.995044 | exon (NM_008514, exon 23 of 23) | exon (NM_008514, exon 23 of 23) | 54592 | NM_025778 | 66813 | Mm.474472 | NM_025778 | ENSMUSG00000030200 | **Bcl2l14** | 4930452K23Rik\|4933405K19Rik\|9030625M01Rik\|AI429211\|Bclg | BCL2-like 14 (apoptosis facilitator) | protein-coding |
| chr6_134366321_134366683 | chr6 | 134366322 | 134366683 | + | 0.821225 | Intergenic | Intergenic | -29827 | NM_025778 | 66813 | Mm.474472 | NM_025778 | ENSMUSG00000030200 | **Bcl2l14** | 4930452K23Rik\|4933405K19Rik\|9030625M01Rik\|AI429211\|Bclg | BCL2-like 14 (apoptosis facilitator) | protein-coding |
| chr6_134291784_134292266 | chr6 | 134291785 | 134292266 | + | 0.798848 | Intergenic | Intergenic | -104304 | NM_025778 | 66813 | Mm.474472 | NM_025778 | ENSMUSG00000030200 | **Bcl2l14** | 4930452K23Rik\|4933405K19Rik\|9030625M01Rik\|AI429211\|Bclg | BCL2-like 14 (apoptosis facilitator) | protein-coding |
| chr6_134302000_134302419 | chr6 | 134302001 | 134302419 | + | 0.784218 | Intergenic | ORR1A2\|LTR\|ERVL-MaLR | -94119 | NM_025778 | 66813 | Mm.474472 | NM_025778 | ENSMUSG00000030200 | **Bcl2l14** | 4930452K23Rik\|4933405K19Rik\|9030625M01Rik\|AI429211\|Bclg | BCL2-like 14 (apoptosis facilitator) | protein-coding |
| chr6_134342865_134343581 | chr6 | 134342866 | 134343581 | + | 0.755992 | Intergenic | Intergenic | -53106 | NM_025778 | 66813 | Mm.474472 | NM_025778 | ENSMUSG00000030200 | **Bcl2l14** | 4930452K23Rik\|4933405K19Rik\|9030625M01Rik\|AI429211\|Bclg | BCL2-like 14 (apoptosis facilitator) | protein-coding |
| chr6_134301972_134302619 | chr6 | 134301973 | 134302619 | + | 0.735685 | Intergenic | ORR1A2\|LTR\|ERVL-MaLR | -94033 | NM_025778 | 66813 | Mm.474472 | NM_025778 | ENSMUSG00000030200 | **Bcl2l14** | 4930452K23Rik\|4933405K19Rik\|9030625M01Rik\|AI429211\|Bclg | BCL2-like 14 (apoptosis facilitator) | protein-coding |
| chr6_134342993_134343354 | chr6 | 134342994 | 134343354 | + | 0.733061 | Intergenic | Intergenic | -53155 | NM_025778 | 66813 | Mm.474472 | NM_025778 | ENSMUSG00000030200 | **Bcl2l14** | 4930452K23Rik\|4933405K19Rik\|9030625M01Rik\|AI429211\|Bclg | BCL2-like 14 (apoptosis facilitator) | protein-coding |
| chr6_134365953_134368036 | chr6 | 134365954 | 134368036 | + | 0.614468 | Intergenic | Intergenic | -29334 | NM_025778 | 66813 | Mm.474472 | NM_025778 | ENSMUSG00000030200 | **Bcl2l14** | 4930452K23Rik\|4933405K19Rik\|9030625M01Rik\|AI429211\|Bclg | BCL2-like 14 (apoptosis facilitator) | protein-coding |
| chr3_103828876_103829552 | chr3 | 103828877 | 103829552 | + | 1.44751 | intron (NR_106164, intron 4 of 4) | intron (NR_106164, intron 4 of 4) | -3482 | NM_001142959 | 229672 | Mm.297245 | NM_001142959 | ENSMUSG00000044165 | **Bcl2l15** | Bfk\|Gm566 | BCLl2-like 15 | protein-coding |
| chr3_103830689_103831055 | chr3 | 103830690 | 103831055 | + | 0.658255 | intron (NR_106164, intron 4 of 4) | RLTR13D2\|LTR\|ERVK | -1824 | NM_001142959 | 229672 | Mm.297245 | NM_001142959 | ENSMUSG00000044165 | **Bcl2l15** | Bfk\|Gm566 | BCLl2-like 15 | protein-coding |
| chr16_24095629_24095984 | chr16 | 24095630 | 24095984 | + | 1.18514 | Intergenic | Intergenic | -107195 | NM_009744 | 12053 | Mm.347398 | NM_009744 | ENSMUSG00000022508 | **Bcl6** | Bcl5 | B cell leukemia/lymphoma 6 | protein-coding |
| chr16_24182415_24182787 | chr16 | 24182416 | 24182787 | + | 0.925434 | Intergenic | URR1A\|DNA\|hAT-Charlie | -193989 | NM_009744 | 12053 | Mm.347398 | NM_009744 | ENSMUSG00000022508 | **Bcl6** | Bcl5 | B cell leukemia/lymphoma 6 | protein-coding |
| chr16_24037537_24037982 | chr16 | 24037538 | 24037982 | + | 0.745246 | Intergenic | Intergenic | -49148 | NM_009744 | 12053 | Mm.347398 | NM_009744 | ENSMUSG00000022508 | **Bcl6** | Bcl5 | B cell leukemia/lymphoma 6 | protein-coding |
| chr16_23997083_23997573 | chr16 | 23997084 | 23997573 | + | 0.660598 | Intergenic | Intergenic | -8716 | NM_009744 | 12053 | Mm.347398 | NM_009744 | ENSMUSG00000022508 | **Bcl6** | Bcl5 | B cell leukemia/lymphoma 6 | protein-coding |
| chr5_123346393_123346728 | chr5 | 123346394 | 123346728 | + | 0.63965 | intron (NM_029850, intron 1 of 5) | intron (NM_029850, intron 1 of 5) | 2113 | NM_029850 | 77045 | Mm.151960 | NM_029850 | ENSMUSG00000029438 | **Bcl7a** | 4432415N06Rik\|AI448316 | B cell CLL/lymphoma 7A | protein-coding |
| chr8_117125988_117126629 | chr8 | 117125989 | 117126629 | + | 0.878916 | intron (NM_021486, intron 7 of 10) | MER44B\|DNA\|TcMar-Tigger | 30444 | NM_001163028 | 63857 | Mm.174133 | NM_021486 | ENSMUSG00000031845 | **Bco1** | Bcdo\|Bcdo1\|Bcmo1\|CMO1\|Cmoi\|beta-CD\|betaCMOOX | beta-carotene oxygenase 1 | protein-coding |
| chr9_50547739_50548469 | chr9 | 50547740 | 50548469 | + | 1.01451 | intron (NM_133217, intron 2 of 11) | MER34\|LTR\|ERV1 | 7034 | NM_133217 | 170752 | Mm.390149 | NM_133217 | ENSMUSG00000032066 | **Bco2** | B-diox-II\|Bcdo2\|Bcmo2\|CMO2 | beta-carotene oxygenase 2 | protein-coding |
| chrX_11839583_11840275 | chrX | 11839584 | 11840275 | + | 0.9797 | Intergenic | Intergenic | 240624 | NM_175045 | 71458 | Mm.196328 | NM_029510 | ENSMUSG00000040363 | **Bcor** | 5830466J11Rik\|8430401K06Rik\|BcorR\|D930024N20Rik\|mKIAA1575 | BCL6 interacting corepressor | protein-coding |
| chrX_11845459_11846235 | chrX | 11845460 | 11846235 | + | 0.96584 | Intergenic | Intergenic | 234706 | NM_175045 | 71458 | Mm.196328 | NM_029510 | ENSMUSG00000040363 | **Bcor** | 5830466J11Rik\|8430401K06Rik\|BcorR\|D930024N20Rik\|mKIAA1575 | BCL6 interacting corepressor | protein-coding |
| chrX_12060032_12060571 | chrX | 12060033 | 12060571 | + | 0.718218 | intron (NM_175046, intron 2 of 13) | intron (NM_175046, intron 2 of 13) | 20251 | NM_175045 | 71458 | Mm.196328 | NM_029510 | ENSMUSG00000040363 | **Bcor** | 5830466J11Rik\|8430401K06Rik\|BcorR\|D930024N20Rik\|mKIAA1575 | BCL6 interacting corepressor | protein-coding |
| chrX_12060003_12060775 | chrX | 12060004 | 12060775 | + | 0.637364 | intron (NM_175046, intron 2 of 13) | intron (NM_175046, intron 2 of 13) | 20164 | NM_175045 | 71458 | Mm.196328 | NM_029510 | ENSMUSG00000040363 | **Bcor** | 5830466J11Rik\|8430401K06Rik\|BcorR\|D930024N20Rik\|mKIAA1575 | BCL6 interacting corepressor | protein-coding |
| chrX_12007147_12007629 | chrX | 12007148 | 12007629 | + | 0.610246 | Intergenic | CpG | 73165 | NM_175045 | 71458 | Mm.196328 | NM_029510 | ENSMUSG00000040363 | **Bcor** | 5830466J11Rik\|8430401K06Rik\|BcorR\|D930024N20Rik\|mKIAA1575 | BCL6 interacting corepressor | protein-coding |
| chrX_48398392_48399037 | chrX | 48398393 | 48399037 | + | 1.04958 | intron (NM_178782, intron 12 of 14) | intron (NM_178782, intron 12 of 14) | 57357 | NM_178782 | 320376 | Mm.210146 | NM_178782 | ENSMUSG00000036959 | **Bcorl1** | 6720425J07Rik\|B930011H20Rik | BCL6 co-repressor-like 1 | protein-coding |
| chr10_75062608_75062970 | chr10 | 75062609 | 75062970 | + | 0.852758 | intron (NM_001081412, intron 1 of 22) | intron (NM_001081412, intron 1 of 22) | 1893 | NM_001081412 | 110279 | Mm.333722 | NM_001081412 | ENSMUSG00000009681 | **Bcr** | 5133400C09Rik\|AI561783\|AI853148\|mKIAA3017 | breakpoint cluster region | protein-coding |
| chr10_75091465_75091858 | chr10 | 75091466 | 75091858 | + | 0.745459 | intron (NM_001081412, intron 1 of 22) | intron (NM_001081412, intron 1 of 22) | 30766 | NM_001081412 | 110279 | Mm.333722 | NM_001081412 | ENSMUSG00000009681 | **Bcr** | 5133400C09Rik\|AI561783\|AI853148\|mKIAA3017 | breakpoint cluster region | protein-coding |
| chr16_31424325_31424843 | chr16 | 31424326 | 31424843 | + | 1.07421 | intron (NM_175177, intron 1 of 6) | intron (NM_175177, intron 1 of 6) | 2287 | NM_175177 | 71911 | Mm.293470 | NM_175177 | ENSMUSG00000046598 | **Bdh1** | 2310032J20Rik\|AI327223\|Bdh | 3-hydroxybutyrate dehydrogenase, type 1 | protein-coding |
| chr12_109055912_109056390 | chr12 | 109055913 | 109056390 | + | 0.753213 | intron (NM_001163175, intron 1 of 6) | intron (NM_001163175, intron 1 of 6) | 12066 | NM_001163175 | 380785 | Mm.342085 | NM_001163175 | ENSMUSG00000040867 | **Begain** | BM948371\|Gm897 | brain-enriched guanylate kinase-associated | protein-coding |
| chr16_32181857_32182185 | chr16 | 32181858 | 32182185 | + | 0.75315 | intron (NM_001033539, intron 1 of 2) | L1MB8\|LINE\|L1 | 2221 | NM_001033539 | 328660 | Mm.441006 | NM_001033539 | | **Bex6** | B020003O03Rik | brain expressed gene 6 | protein-coding |
| chr9_103503647_103504340 | chr9 | 103503648 | 103504340 | + | 1.01376 | intron (NM_178638, intron 3 of 5) | RLTR23\|LTR\|ERV1 | -23666 | NM_001002896 | 107993 | Mm.335403 | NM_001002896 | ENSMUSG00000032556 | **Bfsp2** | AI448593\|CP49 | beaded filament structural protein 2, phakinin | protein-coding |
| chr9_103494377_103494675 | chr9 | 103494378 | 103494675 | + | 0.84582 | intron (NM_178638, intron 4 of 5) | intron (NM_178638, intron 4 of 5) | -14198 | NM_001002896 | 107993 | Mm.335403 | NM_001002896 | ENSMUSG00000032556 | **Bfsp2** | AI448593\|CP49 | beaded filament structural protein 2, phakinin | protein-coding |
| chr9_103501827_103502443 | chr9 | 103501828 | 103502443 | + | 0.758245 | intron (NM_178638, intron 3 of 5) | intron (NM_178638, intron 3 of 5) | -21807 | NM_001002896 | 107993 | Mm.335403 | NM_001002896 | ENSMUSG00000032556 | **Bfsp2** | AI448593\|CP49 | beaded filament structural protein 2, phakinin | protein-coding |
| chr6_108658109_108659026 | chr6 | 108658110 | 108659026 | + | 1.13335 | intron (NR_104577, intron 1 of 6) | intron (NR_104577, intron 1 of 6) | -2061 | NM_011498 | 20893 | Mm.2436 | NM_011498 | ENSMUSG00000030103 | **Bhlhe40** | Bhlhb2\|C130042M06Rik\|CR8\|Clast5\|Dec1\|Sharp2\|Stra13\|Stra14 | basic helix-loop-helix family, member e40 | protein-coding |
| chr6_108659336_108659888 | chr6 | 108659337 | 108659888 | + | 0.603259 | intron (NR_104577, intron 1 of 6) | CpG | -1017 | NM_011498 | 20893 | Mm.2436 | NM_011498 | ENSMUSG00000030103 | **Bhlhe40** | Bhlhb2\|C130042M06Rik\|CR8\|Clast5\|Dec1\|Sharp2\|Stra13\|Stra14 | basic helix-loop-helix family, member e40 | protein-coding |

| | | | | | | | | | | | | | | | | | |
|---|---|---|---|---|---|---|---|---|---|---|---|---|---|---|---|---|---|
| chr6_145882976_145883387 | chr6 | 145882977 | 145883387 | + | 1.30855 | Intergenic | L1MB7\|LINE\|L1 | -17762 | NM_024469 | 79362 | Mm.154529 | NM_024469 | ENSMUSG00000030256 | **Bhlhe41** | 6430520M22Rik\|Bhlhb2l\|Bhlhb3\|DEC2\|Sharp1 | basic helix-loop-helix family, member e41 | protein-coding |
| chr6_145883678_145884038 | chr6 | 145883679 | 145884038 | + | 0.854888 | Intergenic | L1MB7\|LINE\|L1 | -18438 | NM_024469 | 79362 | Mm.154529 | NM_024469 | ENSMUSG00000030256 | **Bhlhe41** | 6430520M22Rik\|Bhlhb2l\|Bhlhb3\|DEC2\|Sharp1 | basic helix-loop-helix family, member e41 | protein-coding |
| chr6_120926763_120927260 | chr6 | 120926764 | 120927260 | + | 1.25746 | Intergenic | Intergenic | -10192 | NM_007544 | 12122 | Mm.235081 | NM_007544 | ENSMUSG00000004446 | **Bid** | 2700049M22Rik\|AI875481\|AU022477 | BH3 interacting domain death agonist | protein-coding |
| chr6_120994283_120994913 | chr6 | 120994284 | 120994913 | + | 1.08167 | intron (NM_001270475, intron 19 of 31) | intron (NM_001270475, intron 19 of 31) | -77778 | NM_007544 | 12122 | Mm.235081 | NM_007544 | ENSMUSG00000004446 | **Bid** | 2700049M22Rik\|AI875481\|AU022477 | BH3 interacting domain death agonist | protein-coding |
| chr18_32335504_32336003 | chr18 | 32335505 | 32336003 | + | 1.75273 | Intergenic | L1M4a2\|LINE\|L1 | -41463 | NM_001083334 | 30948 | Mm.4383 | NM_009668 | ENSMUSG00000024381 | **Bin1** | ALP-1\|Amphl\|BRAMP-2\|SH3P9 | bridging integrator 1 | protein-coding |
| chr18_32358916_32359481 | chr18 | 32358917 | 32359481 | + | 0.976332 | non-coding (NR_045403, exon 3 of 3) | non-coding (NR_045403, exon 3 of 3) | -18018 | NM_001083334 | 30948 | Mm.4383 | NM_009668 | ENSMUSG00000024381 | **Bin1** | ALP-1\|Amphl\|BRAMP-2\|SH3P9 | bridging integrator 1 | protein-coding |
| chr15_100669945_100670337 | chr15 | 100669946 | 100670337 | + | 0.64597 | promoter-TSS (NM_001270537) | promoter-TSS (NM_001270537) | -641 | NM_001270537 | 668218 | Mm.451225 | NM_001270537 | ENSMUSG00000098112 | **Bin2** | - | bridging integrator 2 | protein-coding |
| chr14_70113986_70114305 | chr14 | 70113987 | 70114305 | + | 0.656862 | intron (NM_021328, intron 1 of 8) | L1MA4\|LINE\|L1 | 14001 | NM_021328 | 57784 | Mm.24186 | NM_021328 | ENSMUSG00000022089 | **Bin3** | 1700015F07Rik | bridging integrator 3 | protein-coding |
| chr9_7862605_7863512 | chr9 | 7862606 | 7863512 | + | 0.870885 | intron (NM_007464, intron 1 of 8) | RLTR30B_MM\|LTR\|ERV1 | 10111 | NM_007464 | 11796 | Mm.2026 | NM_007464 | ENSMUSG00000032000 | **Birc3** | AW107670\|Api1\|Api2\|Birc2\|HIAP2\|IAP1\|IAP2\|MIAP1\|MIAP2\|MIHB\|MIHC\|RNF49\|cIAP-1\|cIAP-2\|cIAP1\|cIAP2 | baculoviral IAP repeat-containing 3 | protein-coding |
| chr5_36724278_36724663 | chr5 | 36724279 | 36724663 | + | 1.15292 | Intergenic | Intergenic | 24208 | NM_133724 | 117197 | Mm.46744 | NM_133724 | ENSMUSG00000060708 | **Bloc1s4** | 2610101N07Rik\|Blos4\|Cno | biogenesis of lysosomal organelles complex-1, subunit 4, cappuccino | protein-coding |
| chr5_36722385_36723009 | chr5 | 36722386 | 36723009 | + | 0.768597 | Intergenic | Intergenic | 25982 | NM_133724 | 117197 | Mm.46744 | NM_133724 | ENSMUSG00000060708 | **Bloc1s4** | 2610101N07Rik\|Blos4\|Cno | biogenesis of lysosomal organelles complex-1, subunit 4, cappuccino | protein-coding |
| chr5_36726441_36726816 | chr5 | 36726442 | 36726816 | + | 0.666533 | Intergenic | Intergenic | 22050 | NM_133724 | 117197 | Mm.46744 | NM_133724 | ENSMUSG00000060708 | **Bloc1s4** | 2610101N07Rik\|Blos4\|Cno | biogenesis of lysosomal organelles complex-1, subunit 4, cappuccino | protein-coding |
| chr5_36722447_36722675 | chr5 | 36722448 | 36722675 | + | 0.603319 | Intergenic | Intergenic | 26118 | NM_133724 | 117197 | Mm.46744 | NM_133724 | ENSMUSG00000060708 | **Bloc1s4** | 2610101N07Rik\|Blos4\|Cno | biogenesis of lysosomal organelles complex-1, subunit 4, cappuccino | protein-coding |
| chr2_122721041_122721394 | chr2 | 122721042 | 122721394 | + | 0.690949 | Intergenic | ORR1E\|LTR\|ERVL-MaLR | -17287 | NM_019788 | 18457 | Mm.268921 | NM_019788 | ENSMUSG00000005804 | **Bloc1s6** | BLOC-1\|Pldn\|Stx13bp1\|pa | biogenesis of lysosomal organelles complex-1, subunit 6, pallidin | protein-coding |
| chr2_122731249_122731905 | chr2 | 122731250 | 122731905 | + | 0.644709 | Intergenic | Intergenic | -6928 | NM_019788 | 18457 | Mm.268921 | NM_019788 | ENSMUSG00000005804 | **Bloc1s6** | BLOC-1\|Pldn\|Stx13bp1\|pa | biogenesis of lysosomal organelles complex-1, subunit 6, pallidin | protein-coding |
| chr2_118540887_118541158 | chr2 | 118540888 | 118541158 | + | 1.13847 | intron (NM_138313, intron 4 of 4) | intron (NM_138313, intron 4 of 4) | 8655 | NM_138313 | 171543 | Mm.210125 | NM_138313 | ENSMUSG00000040093 | **Bmf** | AW260063\|AW411872 | BCL2 modifying factor | protein-coding |
| chr2_118540681_118541314 | chr2 | 118540682 | 118541314 | + | 1.11831 | intron (NM_138313, intron 4 of 4) | intron (NM_138313, intron 4 of 4) | 8680 | NM_138313 | 171543 | Mm.210125 | NM_138313 | ENSMUSG00000040093 | **Bmf** | AW260063\|AW411872 | BCL2 modifying factor | protein-coding |
| chr6_87447135_87448087 | chr6 | 87447136 | 87448087 | + | 1.30049 | Intergenic | Intergenic | 18609 | NM_009756 | 12154 | Mm.483321 | NM_009756 | ENSMUSG00000030046 | **Bmp10** | - | bone morphogenetic protein 10 | protein-coding |
| chr6_87436584_87437236 | chr6 | 87436585 | 87437236 | + | 0.889397 | Intergenic | Intergenic | 7908 | NM_009756 | 12154 | Mm.483321 | NM_009756 | ENSMUSG00000030046 | **Bmp10** | - | bone morphogenetic protein 10 | protein-coding |
| chr6_87435817_87436107 | chr6 | 87435818 | 87436107 | + | 0.701532 | Intergenic | Intergenic | 6960 | NM_009756 | 12154 | Mm.483321 | NM_009756 | ENSMUSG00000030046 | **Bmp10** | - | bone morphogenetic protein 10 | protein-coding |
| chr5_97001241_97001620 | chr5 | 97001242 | 97001620 | + | 0.945276 | intron (NM_080708, intron 1 of 15) | intron (NM_080708, intron 1 of 15) | 3742 | NM_080708 | 140780 | Mm.281490 | NM_080708 | ENSMUSG00000034663 | **Bmp2k** | 4933417M22Rik\|AA673486\|AV128808\|BIKE | BMP2 inducible kinase | protein-coding |
| chr5_96962224_96962630 | chr5 | 96962225 | 96962630 | + | 0.763121 | Intergenic | MER58A\|DNA\|hAT-Charlie | -35262 | NM_080708 | 140780 | Mm.281490 | NM_080708 | ENSMUSG00000034663 | **Bmp2k** | 4933417M22Rik\|AA673486\|AV128808\|BIKE | BMP2 inducible kinase | protein-coding |
| chr5_96921066_96921368 | chr5 | 96921067 | 96921368 | + | 0.706243 | Intergenic | Intergenic | -76472 | NM_080708 | 140780 | Mm.281490 | NM_080708 | ENSMUSG00000034663 | **Bmp2k** | 4933417M22Rik\|AA673486\|AV128808\|BIKE | BMP2 inducible kinase | protein-coding |
| chr5_96948669_96949674 | chr5 | 96948670 | 96949674 | + | 0.70334 | Intergenic | Intergenic | -48517 | NM_080708 | 140780 | Mm.281490 | NM_080708 | ENSMUSG00000034663 | **Bmp2k** | 4933417M22Rik\|AA673486\|AV128808\|BIKE | BMP2 inducible kinase | protein-coding |
| chr5_97000578_97000833 | chr5 | 97000579 | 97000833 | + | 0.695281 | intron (NM_080708, intron 1 of 15) | Tigger17a\|DNA\|TcMar-Tigger | 3017 | NM_080708 | 140780 | Mm.281490 | NM_080708 | ENSMUSG00000034663 | **Bmp2k** | 4933417M22Rik\|AA673486\|AV128808\|BIKE | BMP2 inducible kinase | protein-coding |
| chr5_96961962_96963276 | chr5 | 96961963 | 96963276 | + | 0.649765 | Intergenic | Intergenic | -35070 | NM_080708 | 140780 | Mm.281490 | NM_080708 | ENSMUSG00000034663 | **Bmp2k** | 4933417M22Rik\|AA673486\|AV128808\|BIKE | BMP2 inducible kinase | protein-coding |
| chr5_96901271_96901750 | chr5 | 96901272 | 96901750 | + | 0.612095 | Intergenic | L1MA6\|LINE\|L1 | -96178 | NM_080708 | 140780 | Mm.281490 | NM_080708 | ENSMUSG00000034663 | **Bmp2k** | 4933417M22Rik\|AA673486\|AV128808\|BIKE | BMP2 inducible kinase | protein-coding |
| chr9_23123586_23124295 | chr9 | 23123587 | 23124295 | + | 0.715359 | Intergenic | Intergenic | -99135 | NM_028472 | 73230 | Mm.335020 | NM_028472 | ENSMUSG00000031963 | **Bmper** | 3110056H04Rik\|CV-2\|Crim3\|Cv2 | BMP-binding endothelial regulator | protein-coding |
| chr3_142222479_142222931 | chr3 | 142222480 | 142222931 | + | 1.19375 | Intergenic | Intergenic | -53477 | NM_001277216 | 12167 | Mm.39089 | NM_007560 | ENSMUSG00000052430 | **Bmpr1b** | AI385617\|ALK-6\|AV355320\|Acvrlk6\|Alk6\|BMPR-1B\|BMPR-IB\|CFK-43a\|SKR6 | bone morphogenetic protein receptor, type 1B | protein-coding |
| chr3_142249349_142249835 | chr3 | 142249350 | 142249835 | + | 0.605788 | intron (NM_019808, intron 9 of 12) | intron (NM_019808, intron 9 of 12) | -80364 | NM_001277216 | 12167 | Mm.39089 | NM_007560 | ENSMUSG00000052430 | **Bmpr1b** | AI385617\|ALK-6\|AV355320\|Acvrlk6\|Alk6\|BMPR-1B\|BMPR-IB\|CFK-43a\|SKR6 | bone morphogenetic protein receptor, type 1B | protein-coding |
| chr17_26773169_26773643 | chr17 | 26773170 | 26773643 | + | 0.845717 | 3' UTR (NM_029870, exon 9 of 9) | 3' UTR (NM_029870, exon 9 of 9) | -7673 | NM_172149 | 224630 | Mm.21795 | NM_172149 | ENSMUSG00000024191 | **Bnip1** | 2010005M06Rik\|5930429G21Rik\|Sec20 | BCL2/adenovirus E1B interacting protein 1 | protein-coding |
| chr14_67019774_67020355 | chr14 | 67019775 | 67020355 | + | 0.993209 | intron (NM_028032, intron 8 of 9) | ORR1C1\|LTR\|ERVL-MaLR | -11188 | NM_009761 | 12177 | Mm.29820 | NM_009761 | ENSMUSG00000022051 | **Bnip3l** | C86132\|D14Ertd719e\|Nip3L\|Nix | BCL2/adenovirus E1B interacting protein 3-like | protein-coding |
| chr11_31737873_31738422 | chr11 | 31737874 | 31738422 | + | 1.05132 | Intergenic | Intergenic | -66286 | NM_001024919 | 69556 | Mm.34183 | NM_001024919 | ENSMUSG00000044502 | **Bod1** | 2310022M17Rik\|BOD2.1\|Fam44b | biorientation of chromosomes in cell division 1 | protein-coding |
| chr7_126697479_126697971 | chr7 | 126697480 | 126697971 | + | 0.98527 | Intergenic | Intergenic | 1725 | NM_175103 | 66162 | Mm.358692 | NM_175103 | | **Bola2** | 1110025L05Rik\|BolA-2 | bolA-like 2 (E. coli) | protein-coding |
| chr13_34050471_34050880 | chr13 | 34050472 | 34050880 | + | 0.79299 | intron (NM_026512, intron 4 of 6) | intron (NM_026512, intron 4 of 6) | 13035 | NM_026512 | 68021 | Mm.140243 | NM_026512 | ENSMUSG00000038286 | **Bphl** | 2010012D11Rik\|5730533B08Rik\|AI115341 | biphenyl hydrolase-like (serine hydrolase, breast epithelial mucin-associated antigen) | protein-coding |
| chr11_107119826_107120292 | chr11 | 107119827 | 107120292 | + | 0.789754 | intron (NM_176850, intron 1 of 27) | intron (NM_176850, intron 1 of 27) | 11863 | NM_176850 | 207165 | Mm.343986 | NM_176850 | ENSMUSG00000040481 | **Bptf** | 9430093H17Rik\|Falz | bromodomain PHD finger transcription factor | protein-coding |
| chr6_39714280_39714887 | chr6 | 39714281 | 39714887 | + | 1.0755 | intron (NM_139294, intron 1 of 21) | intron (NM_139294, intron 1 of 21) | 10879 | NM_139294 | 109880 | Mm.245513 | NM_139294 | ENSMUSG00000002413 | **Braf** | 9930012E13Rik\|AA120551\|AA387315\|AA473386\|B-raf\|Braf-2\|Braf2\|C230098H17\|C87398\|D6Ertd631e | Braf transforming gene | protein-coding |
| chr6_39722752_39722984 | chr6 | 39722753 | 39722984 | + | 0.796508 | intron (NM_139294, intron 1 of 21) | intron (NM_139294, intron 1 of 21) | 2595 | NM_139294 | 109880 | Mm.245513 | NM_139294 | ENSMUSG00000002413 | **Braf** | 9930012E13Rik\|AA120551\|AA387315\|AA473386\|B-raf\|Braf-2\|Braf2\|C230098H17\|C87398\|D6Ertd631e | Braf transforming gene | protein-coding |
| chr6_39681872_39682155 | chr6 | 39681873 | 39682155 | + | 0.77547 | intron (NM_139294, intron 2 of 21) | intron (NM_139294, intron 2 of 21) | 43449 | NM_139294 | 109880 | Mm.245513 | NM_139294 | ENSMUSG00000002413 | **Braf** | 9930012E13Rik\|AA120551\|AA387315\|AA473386\|B-raf\|Braf-2\|Braf2\|C230098H17\|C87398\|D6Ertd631e | Braf transforming gene | protein-coding |
| chr11_101544045_101544455 | chr11 | 101544046 | 101544455 | + | 0.668882 | intron (NM_009764, intron 2 of 22) | intron (NM_009764, intron 2 of 22) | 7705 | NM_009764 | 12189 | Mm.244975 | NM_009764 | ENSMUSG00000017146 | **Brca1** | - | breast cancer 1 | protein-coding |
| chr5_150538746_150539116 | chr5 | 150538747 | 150539116 | + | 0.736471 | exon (NM_009765, exon 11 of 27) | exon (NM_009765, exon 11 of 27) | 16310 | NM_001081001 | 12190 | Mm.236256 | NM_009765 | ENSMUSG00000041147 | **Brca2** | Fancd1\|RAB163 | breast cancer 2 | protein-coding |
| chr5_107355821_107356454 | chr5 | 107355822 | 107356454 | + | 0.993803 | intron (NM_054054, intron 7 of 18) | intron (NM_054054, intron 7 of 18) | 24944 | NM_054054 | 114642 | Mm.182836 | NM_054054 | ENSMUSG00000029279 | **Brdt** | 7420412D09Rik\|Brd6\|Fsrg3 | bromodomain, testis-specific | protein-coding |
| chr6_113612215_113613104 | chr6 | 113612216 | 113613104 | + | 1.0623 | intron (NM_133937, intron 1 of 2) | intron (NM_133937, intron 1 of 2) | 7888 | NM_133937 | 101314 | Mm.485793 | NM_133937 | ENSMUSG00000033940 | **Brk1** | 6720456B07Rik\|AW011779 | BRICK1, SCAR/WAVE actin-nucleating complex subunit | protein-coding |
| chr6_113612632_113613006 | chr6 | 113612633 | 113613006 | + | 0.985174 | intron (NM_133937, intron 1 of 2) | intron (NM_133937, intron 1 of 2) | 8047 | NM_133937 | 101314 | Mm.485793 | NM_133937 | ENSMUSG00000033940 | **Brk1** | 6720456B07Rik\|AW011779 | BRICK1, SCAR/WAVE actin-nucleating complex subunit | protein-coding |
| chr7_4694559_4695571 | chr7 | 4694560 | 4695571 | + | 1.09986 | intron (NM_001003920, intron 4 of 18) | (TG)n\|Simple_repeat\|Simple_repeat | 4137 | NM_001168572 | 381979 | Mm.297064 | NM_001003920 | ENSMUSG00000035390 | **Brsk1** | Gm1100\|SAD-B\|SADB | BR serine/threonine kinase 1 | protein-coding |
| chr9_108154685_108155255 | chr9 | 108154686 | 108155255 | + | 1.63785 | intron (NM_007567, intron 1 of 11) | L1MB4\|LINE\|L1 | 35413 | NM_007567 | 12217 | Mm.20425 | NM_007567 | ENSMUSG00000032589 | **Bsn** | - | bassoon | protein-coding |
| chr7_130800703_130801504 | chr7 | 130800704 | 130801504 | + | 1.06689 | intron (NM_001081038, intron 9 of 15) | intron (NM_001081038, intron 9 of 15) | 27035 | NM_001081038 | 330660 | Mm.271550 | NM_001081038 | ENSMUSG00000040298 | **Btbd16** | E330040A16Rik | BTB (POZ) domain containing 16 | protein-coding |

| | | | | | | | | | | | | | | | | | |
|---|---|---|---|---|---|---|---|---|---|---|---|---|---|---|---|---|---|
| chr7_130800428_130801464 | chr7 | 130800429 | 130801464 | + | 0.962089 | intron (NM_001081038, intron 9 of 15) | intron (NM_001081038, intron 9 of 15) | 26877 | NM_001081038 | 330660 | Mm.271550 | NM_001081038 | ENSMUSG00000040298 | **Btbd16** | E330040A16Rik | BTB (POZ) domain containing 16 | protein-coding |
| chr12_112975600_112975907 | chr12 | 112975601 | 112975907 | + | 0.758098 | promoter-TSS (NM_001145900) | promoter-TSS (NM_001145900) | -728 | NM_001145900 | 399566 | Mm.440140 | NM_201646 | ENSMUSG00000002803 | **Btbd6** | - | BTB (POZ) domain containing 6 | protein-coding |
| chr5_107449313_107449958 | chr5 | 107449314 | 107449958 | + | 1.46373 | intron (NM_001255991, intron 2 of 8) | intron (NM_001255991, intron 2 of 8) | 11639 | NM_001255991 | 100503185 | Mm.366650 | NM_001255991 | ENSMUSG00000070632 | **Btbd8** | EG627196\|Gm16115\|OTTMUSG00000030610 | BTB (POZ) domain containing 8 | protein-coding |
| chr5_107448754_107449238 | chr5 | 107448755 | 107449238 | + | 1.24371 | intron (NM_001255991, intron 2 of 8) | intron (NM_001255991, intron 2 of 8) | 10999 | NM_001255991 | 100503185 | Mm.366650 | NM_001255991 | ENSMUSG00000070632 | **Btbd8** | EG627196\|Gm16115\|OTTMUSG00000030610 | BTB (POZ) domain containing 8 | protein-coding |
| chr5_107450287_107450704 | chr5 | 107450288 | 107450704 | + | 0.92675 | intron (NM_001255991, intron 2 of 8) | MER74A\|LTR\|ERVL | 12499 | NM_001255991 | 100503185 | Mm.366650 | NM_001255991 | ENSMUSG00000070632 | **Btbd8** | EG627196\|Gm16115\|OTTMUSG00000030610 | BTB (POZ) domain containing 8 | protein-coding |
| chr5_107466990_107467458 | chr5 | 107466991 | 107467458 | + | 0.67225 | intron (NM_001255991, intron 4 of 8) | intron (NM_001255991, intron 4 of 8) | 29227 | NM_001255991 | 100503185 | Mm.366650 | NM_001255991 | ENSMUSG00000070632 | **Btbd8** | EG627196\|Gm16115\|OTTMUSG00000030610 | BTB (POZ) domain containing 8 | protein-coding |
| chr17_30530148_30530578 | chr17 | 30530149 | 30530578 | + | 1.28558 | 5' UTR (NM_172618, exon 2 of 12) | 5' UTR (NM_172618, exon 2 of 12) | 45924 | NM_172618 | 224671 | Mm.41220 | NM_027060 | ENSMUSG00000062202 | **Btbd9** | 1700023F20Rik\|4930402L05\|AI464421 | BTB (POZ) domain containing 9 | protein-coding |
| chr17_30530154_30530736 | chr17 | 30530155 | 30530736 | + | 1.23065 | intron (NM_027060, intron 1 of 10) | intron (NM_027060, intron 1 of 10) | 45842 | NM_172618 | 224671 | Mm.41220 | NM_027060 | ENSMUSG00000062202 | **Btbd9** | 1700023F20Rik\|4930402L05\|AI464421 | BTB (POZ) domain containing 9 | protein-coding |
| chr14_31680636_31681170 | chr14 | 31680637 | 31681170 | + | 0.63759 | Intergenic | Intergenic | 39846 | NM_025295 | 26363 | Mm.282679 | NM_025295 | ENSMUSG00000021900 | **Btd** | - | biotinidase | protein-coding |
| chr13_98307871_98308380 | chr13 | 98307872 | 98308380 | + | 0.641711 | Intergenic | Intergenic | 8841 | NM_145455 | 218490 | Mm.371688 | NM_145455 | ENSMUSG00000021660 | **Btf3** | 1700054E11Rik | basic transcription factor 3 | protein-coding |
| chr13_98307704_98308774 | chr13 | 98307705 | 98308774 | + | 0.636837 | Intergenic | Intergenic | 8728 | NM_145455 | 218490 | Mm.371688 | NM_145455 | ENSMUSG00000021660 | **Btf3** | 1700054E11Rik | basic transcription factor 3 | protein-coding |
| chr4_108782357_108783178 | chr4 | 108782358 | 108783178 | + | 0.932973 | TTS (NM_001254730) | TTS (NM_001254730) | 50816 | NM_027453 | 70533 | Mm.379178 | NM_027453 | ENSMUSG00000028568 | **Btf3l4** | 4632412E09Rik\|5730434I03Rik\|AI316863\|AU022207\|AW552446 | basic transcription factor 3-like 4 | protein-coding |
| chr10_96688735_96688962 | chr10 | 96688736 | 96688962 | + | 1.23258 | Intergenic | Intergenic | 71848 | NM_007569 | 12226 | Mm.23811 | NM_007569 | ENSMUSG00000036478 | **Btg1** | AI426953\|AW546738 | B cell translocation gene 1, anti-proliferative | protein-coding |
| chr10_96421492_96422257 | chr10 | 96421493 | 96422257 | + | 1.02279 | Intergenic | L1VL2\|LINE\|L1 | -195126 | NM_007569 | 12226 | Mm.23811 | NM_007569 | ENSMUSG00000036478 | **Btg1** | AI426953\|AW546738 | B cell translocation gene 1, anti-proliferative | protein-coding |
| chr10_96459074_96459568 | chr10 | 96459075 | 96459568 | + | 0.990397 | Intergenic | Intergenic | -157680 | NM_007569 | 12226 | Mm.23811 | NM_007569 | ENSMUSG00000036478 | **Btg1** | AI426953\|AW546738 | B cell translocation gene 1, anti-proliferative | protein-coding |
| chr10_96463849_96464091 | chr10 | 96463850 | 96464091 | + | 0.780551 | Intergenic | Intergenic | -153031 | NM_007569 | 12226 | Mm.23811 | NM_007569 | ENSMUSG00000036478 | **Btg1** | AI426953\|AW546738 | B cell translocation gene 1, anti-proliferative | protein-coding |
| chr10_96727542_96727785 | chr10 | 96727543 | 96727785 | + | 0.634066 | Intergenic | Intergenic | 110663 | NM_007569 | 12226 | Mm.23811 | NM_007569 | ENSMUSG00000036478 | **Btg1** | AI426953\|AW546738 | B cell translocation gene 1, anti-proliferative | protein-coding |
| chr17_50817441_50817825 | chr17 | 50817442 | 50817825 | + | 0.763699 | intron (NM_001285993, intron 15 of 21) | intron (NM_001285993, intron 15 of 21) | 118952 | NM_001297747 | 12228 | Mm.392910 | NM_009770 | ENSMUSG00000022863 | **Btg3** | ANA\|tob5 | B cell translocation gene 3 | protein-coding |
| chr16_45310420_45310891 | chr16 | 45310421 | 45310891 | + | 1.41132 | Intergenic | Lx7\|LINE\|L1 | 86319 | NM_177584 | 208154 | Mm.38199 | NM_177584 | ENSMUSG00000052013 | **Btla** | A630002H24 | B and T lymphocyte associated | protein-coding |
| chr16_45310398_45311080 | chr16 | 45310399 | 45311080 | + | 1.31707 | Intergenic | Lx7\|LINE\|L1 | 86402 | NM_177584 | 208154 | Mm.38199 | NM_177584 | ENSMUSG00000052013 | **Btla** | A630002H24 | B and T lymphocyte associated | protein-coding |
| chr16_45289498_45290516 | chr16 | 45289499 | 45290516 | + | 0.969739 | Intergenic | Intergenic | 65670 | NM_177584 | 208154 | Mm.38199 | NM_177584 | ENSMUSG00000052013 | **Btla** | A630002H24 | B and T lymphocyte associated | protein-coding |
| chr16_45294213_45294656 | chr16 | 45294214 | 45294656 | + | 0.821917 | Intergenic | Intergenic | 70098 | NM_177584 | 208154 | Mm.38199 | NM_177584 | ENSMUSG00000052013 | **Btla** | A630002H24 | B and T lymphocyte associated | protein-coding |
| chr16_45305512_45306083 | chr16 | 45305513 | 45306083 | + | 0.710613 | Intergenic | Intergenic | 81461 | NM_177584 | 208154 | Mm.38199 | NM_177584 | ENSMUSG00000052013 | **Btla** | A630002H24 | B and T lymphocyte associated | protein-coding |
| chr16_45286339_45286699 | chr16 | 45286340 | 45286699 | + | 0.668422 | Intergenic | Intergenic | 62182 | NM_177584 | 208154 | Mm.38199 | NM_177584 | ENSMUSG00000052013 | **Btla** | A630002H24 | B and T lymphocyte associated | protein-coding |
| chr13_23463426_23463774 | chr13 | 23463427 | 23463774 | + | 0.676644 | intron (NM_013483, intron 3 of 7) | tRNA-Glu-GAG_\|tRNA\|tRNA | 2301 | NM_013483 | 12231 | Mm.1420 | NM_013483 | ENSMUSG00000000706 | **Btn1a1** | Btn | butyrophilin, subfamily 1, member A1 | protein-coding |
| chr13_23449178_23449500 | chr13 | 23449179 | 23449500 | + | 0.627994 | Intergenic | B3A\|SINE\|B2 | 16562 | NM_013483 | 12231 | Mm.1420 | NM_013483 | ENSMUSG00000000706 | **Btn1a1** | Btn | butyrophilin, subfamily 1, member A1 | protein-coding |
| chr13_23500461_23500775 | chr13 | 23500462 | 23500775 | + | 0.684399 | Intergenic | Intergenic | -11761 | NM_001289614 | 238555 | Mm.129668 | NM_175938 | | **Btn2a2** | Btn2\|D030063K05 | butyrophilin, subfamily 2, member A2 | protein-coding |
| chr13_23500452_23501177 | chr13 | 23500453 | 23501177 | + | 0.654974 | Intergenic | Intergenic | -11958 | NM_001289614 | 238555 | Mm.129668 | NM_175938 | | **Btn2a2** | Btn2\|D030063K05 | butyrophilin, subfamily 2, member A2 | protein-coding |
| chr11_49177526_49178141 | chr11 | 49177527 | 49178141 | + | 1.10778 | intron (NM_172793, intron 4 of 9) | Lx5c\|LINE\|L1 | 9255 | NM_172793 | 237754 | Mm.130787 | NM_172793 | | **Btnl9** | B430208I01\|Btn3\|D330012D11Rik | butyrophilin-like 9 | protein-coding |
| chr2_118620662_118620927 | chr2 | 118620663 | 118620927 | + | 0.854178 | intron (NM_009773, intron 9 of 22) | intron (NM_009773, intron 9 of 22) | 22584 | NM_009773 | 12236 | Mm.29133 | NM_009773 | ENSMUSG00000040084 | **Bub1b** | AU045529\|BUBR1 | budding uninhibited by benzimidazoles 1 homolog, beta (S. cerevisiae) | protein-coding |
| chr2_118620452_118621114 | chr2 | 118620453 | 118621114 | + | 0.833624 | intron (NM_009773, intron 9 of 22) | intron (NM_009773, intron 9 of 22) | 22572 | NM_009773 | 12236 | Mm.29133 | NM_009773 | ENSMUSG00000040084 | **Bub1b** | AU045529\|BUBR1 | budding uninhibited by benzimidazoles 1 homolog, beta (S. cerevisiae) | protein-coding |
| chr10_45376845_45377346 | chr10 | 45376846 | 45377346 | + | 0.895646 | 3' UTR (NM_001031772, exon 5 of 5) | 3' UTR (NM_001031772, exon 5 of 5) | 41334 | NM_024285 | 23828 | Mm.332793 | NM_024285 | ENSMUSG00000071317 | **Bves** | Pop1\|Popdc1\|mBVES | blood vessel epicardial substance | protein-coding |
| chr1_58359577_58359871 | chr1 | 58359578 | 58359871 | + | 0.648965 | intron (NM_001008419, intron 34 of 35) | intron (NM_001008419, intron 34 of 35) | -33412 | NM_025824 | 66882 | Mm.21848 | NM_025824 | ENSMUSG00000051223 | **Bzw1** | 1200015E15Rik\|mKIAA0005 | basic leucine zipper and W2 domains 1 | protein-coding |
| chr12_36134142_36134493 | chr12 | 36134143 | 36134493 | + | 0.981185 | intron (NM_025840, intron 2 of 11) | intron (NM_025840, intron 2 of 11) | 22507 | NM_025840 | 66912 | Mm.288669 | NM_025840 | ENSMUSG00000020547 | **Bzw2** | 1110001I24Rik\|Bdm2\|HSPC028 | basic leucine zipper and W2 domains 2 | protein-coding |
| chr7_136269496_136270834 | chr7 | 136269497 | 136270834 | + | 0.719416 | intron (NR_102277, intron 1 of 3) | intron (NR_102277, intron 1 of 3) | 1841 | NR_102277 | 77383 | Mm.227745 | NR_102277 | | **C030029H02Rik** | - | RIKEN cDNA C030029H02 gene | ncRNA |
| chr7_136164411_136164836 | chr7 | 136164412 | 136164836 | + | 0.711859 | Intergenic | Intergenic | -103700 | NR_102277 | 77383 | Mm.227745 | NR_102277 | | **C030029H02Rik** | - | RIKEN cDNA C030029H02 gene | ncRNA |
| chr3_9345846_9346317 | chr3 | 9345847 | 9346317 | + | 0.878308 | Intergenic | Intergenic | -56996 | NR_015499 | 320088 | Mm.212908 | NR_015499 | ENSMUSG00000097365 | **C030034L19Rik** | A130044O20Rik | RIKEN cDNA C030034L19 gene | ncRNA |
| chr6_65472402_65473028 | chr6 | 65472403 | 65473028 | + | 0.790405 | Intergenic | Intergenic | 91421 | NM_175524 | 243407 | Mm.389758 | NM_175524 | ENSMUSG00000029917 | **C130060K24Rik** | - | RIKEN cDNA C130060K24 gene | protein-coding |
| chr2_114059732_114059999 | chr2 | 114059733 | 114059999 | + | 0.824378 | intron (NR_028422, intron 2 of 2) | intron (NR_028422, intron 2 of 2) | 5470 | NR_028422 | 100303644 | Mm.330469 | NR_028422 | ENSMUSG00000092397 | **C130080G10Rik** | - | RIKEN cDNA C130080G10 gene | ncRNA |
| chr6_7853508_7853972 | chr6 | 7853509 | 7853972 | + | 0.694168 | intron (NM_052993, intron 1 of 3) | L1M4c\|LINE\|L1 | 8516 | NM_052993 | 94192 | Mm.102752 | NM_052993 | ENSMUSG00000042460 | **C1galt1** | 2210410E06Rik\|AV284120\|T-synthase | core 1 synthase, glycoprotein-N-acetylgalactosamine 3-beta-galactosyltransferase, 1 | protein-coding |
| chr6_7848269_7848615 | chr6 | 7848270 | 7848615 | + | 0.632695 | intron (NM_052993, intron 1 of 3) | MER20\|DNA\|hAT-Charlie | 3218 | NM_052993 | 94192 | Mm.102752 | NM_052993 | ENSMUSG00000042460 | **C1galt1** | 2210410E06Rik\|AV284120\|T-synthase | core 1 synthase, glycoprotein-N-acetylgalactosamine 3-beta-galactosyltransferase, 1 | protein-coding |
| chrX_38622695_38623245 | chrX | 38622696 | 38623245 | + | 0.770776 | Intergenic | Intergenic | 12173 | NM_021550 | 59048 | Mm.441410 | NM_021550 | ENSMUSG00000048970 | **C1galt1c1** | 1500002I11Rik\|AB030184\|C1GalT2\|C81205\|Cosmc | C1GALT1-specific chaperone 1 | protein-coding |
| chr2_12977012_12977766 | chr2 | 12977013 | 12977766 | + | 1.58945 | intron (NM_008961, intron 1 of 4) | MTE2a\|LTR\|ERVL-MaLR | 33475 | NM_153155 | 227580 | Mm.229322 | NM_153155 | | **C1ql3** | 1110065A22Rik\|AI661623\|C1ql\|C1qtnf13\|CTRP13\|K100 | C1q-like 3 | protein-coding |
| chr2_12977899_12978175 | chr2 | 12977900 | 12978175 | + | 1.32729 | intron (NM_008961, intron 1 of 4) | intron (NM_008961, intron 1 of 4) | 32827 | NM_153155 | 227580 | Mm.229322 | NM_153155 | | **C1ql3** | 1110065A22Rik\|AI661623\|C1ql\|C1qtnf13\|CTRP13\|K100 | C1q-like 3 | protein-coding |
| chr2_12977897_12978610 | chr2 | 12977898 | 12978610 | + | 1.24517 | exon (NM_008961, exon 2 of 5) | exon (NM_008961, exon 2 of 5) | 32610 | NM_153155 | 227580 | Mm.229322 | NM_153155 | | **C1ql3** | 1110065A22Rik\|AI661623\|C1ql\|C1qtnf13\|CTRP13\|K100 | C1q-like 3 | protein-coding |
| chr2_12975826_12976014 | chr2 | 12975827 | 12976014 | + | 0.712104 | intron (NM_008961, intron 1 of 4) | intron (NM_008961, intron 1 of 4) | 34944 | NM_153155 | 227580 | Mm.229322 | NM_153155 | | **C1ql3** | 1110065A22Rik\|AI661623\|C1ql\|C1qtnf13\|CTRP13\|K100 | C1q-like 3 | protein-coding |
| chr11_118427294_118427717 | chr11 | 118427295 | 118427717 | + | 0.720655 | promoter-TSS (NM_001204129) | promoter-TSS (NM_001204129) | -951 | NM_001204129 | 56745 | Mm.23845 | NM_019959 | | **C1qtnf1** | 1600017K21Rik\|CTRP1\|Zsig37 | C1q and tumor necrosis factor related protein 1 | protein-coding |
| chr2_90894329_90894606 | chr2 | 90894330 | 90894606 | + | 0.679527 | TTS (NM_026688) | TTS (NM_026688) | 8682 | NM_026161 | 67445 | Mm.258993 | NM_026161 | ENSMUSG00000040794 | **C1qtnf4** | 0710001E10Rik\|9430004J15Rik\|CTRP4 | C1q and tumor necrosis factor related protein 4 | protein-coding |
| chr9_44105422_44106411 | chr9 | 44105423 | 44106411 | + | 1.37398 | intron (NM_145613, intron 11 of 13) | intron (NM_145613, intron 11 of 13) | -1328 | NM_001040631 | 235312 | Mm.227110 | NM_145613 | ENSMUSG00000079592 | **C1qtnf5** | Ctrp5 | C1q and tumor necrosis factor related protein 5 | protein-coding |
| chr14_60726084_60726461 | chr14 | 60726085 | 60726461 | + | 0.885788 | intron (NM_001033272, intron 4 of 12) | intron (NM_001033272, intron 4 of 12) | -41861 | NM_183175 | 239126 | Mm.319636 | NM_183175 | ENSMUSG00000071347 | **C1qtnf9** | 9130217G22Rik\|CTRP9\|Ciqtnf9 | C1q and tumor necrosis factor related protein 9 | protein-coding |
| chr6_124631194_124631854 | chr6 | 124631195 | 124631854 | + | 1.16369 | intron (NM_173864, intron 4 of 10) | intron (NM_173864, intron 4 of 10) | 4561 | NM_173864 | 317677 | Mm.483270 | NM_173864 | ENSMUSG00000079343 | **C1s2** | C1sb\|EG317677\|Gm5077 | complement component 1, s subcomponent 2 | protein-coding |
| chr6_124637100_124637510 | chr6 | 124637101 | 124637510 | + | 0.752781 | Intergenic | Intergenic | -1220 | NM_173864 | 317677 | Mm.483270 | NM_173864 | ENSMUSG00000079343 | **C1s2** | C1sb\|EG317677\|Gm5077 | complement component 1, s subcomponent 2 | protein-coding |
| chr7_100390880_100391307 | chr7 | 100390881 | 100391307 | + | 0.92091 | exon (NM_001017985, exon 5 of 33) | exon (NM_001017985, exon 5 of 33) | 18861 | NM_001017985 | 277939 | Mm.276374 | NM_001017985 | ENSMUSG00000047248 | **C2cd3** | AU020772 | C2 calcium-dependent domain containing 3 | protein-coding |
| chr9_67813606_67814410 | chr9 | 67813607 | 67814410 | + | 1.48272 | Intergenic | Intergenic | 18322 | NM_001163143 | 244911 | Mm.325849 | NM_001163143 | ENSMUSG00000047990 | **C2cd4a** | EG244911\|Fam148a | C2 calcium-dependent domain containing 4A | protein-coding |
| chrX_36867716_36867979 | chrX | 36867717 | 36867979 | + | 0.782224 | Intergenic | Intergenic | -3602 | NM_029951 | 77644 | Mm.290704 | NM_029951 | ENSMUSG00000006423 | **C330007P06Rik** | - | RIKEN cDNA C330007P06 gene | protein-coding |
| chr7_16254076_16254675 | chr7 | 16254077 | 16254675 | + | 0.87262 | intron (NM_001173550, intron 1 of 1) | intron (NM_001173550, intron 1 of 1) | 4962 | NM_001173550 | 12273 | Mm.247623 | NM_007577 | ENSMUSG00000049130 | **C5ar1** | C5aR\|C5r1\|Cd88\|D7Msu1 | complement component 5a receptor 1 | protein-coding |

| | | | | | | | | | | | | | | | | | |
|---|---|---|---|---|---|---|---|---|---|---|---|---|---|---|---|---|---|
| chr15_5082986_5083398 | chr15 | 5082987 | 5083398 | + | 0.819008 | Intergenic | RSINE1\|SINE\|B4 | -19419 | NM_001243837 | 109828 | Mm.208034 | NM_001243837 | ENSMUSG00000079105 | **C7** | A530026G17\|Gm1238 | complement component 7 | protein-coding |
| chr1_151236439_151237047 | chr1 | 151236440 | 151237047 | + | 0.903131 | Intergenic | Intergenic | 98709 | NR_038011 | 402734 | Mm.337666 | NR_038011 | | **C730036E19Rik** | - | RIKEN cDNA C730036E19 gene | ncRNA |
| chr15_6492319_6492843 | chr15 | 6492320 | 6492843 | + | 1.67357 | intron (NM_013485, intron 9 of 10) | intron (NM_013485, intron 9 of 10) | 47248 | NM_013485 | 12279 | Mm.29095 | NM_013485 | ENSMUSG00000022149 | **C9** | - | complement component 9 | protein-coding |
| chr9_78149606_78150148 | chr9 | 78149607 | 78150148 | + | 0.685615 | intron (NM_019987, intron 5 of 13) | intron (NM_019987, intron 5 of 13) | -26037 | NR_040401 | 320295 | Mm.131285 | NR_040401 | ENSMUSG00000097574 | **C920006O11Rik** | - | RIKEN cDNA C920006O11 gene | ncRNA |
| chr10_22298108_22298490 | chr10 | 22298109 | 22298490 | + | 0.642813 | intron (NM_009018, intron 5 of 5) | intron (NM_009018, intron 5 of 5) | -8421 | NR_015465 | 606736 | Mm.387042 | NR_015465 | ENSMUSG00000097352 | **C920009B18Rik** | - | RIKEN cDNA C920009B18 gene | ncRNA |
| chr4_94542865_94543357 | chr4 | 94542866 | 94543357 | + | 1.05004 | intron (NM_026368, intron 4 of 5) | intron (NM_026368, intron 4 of 5) | 13685 | NM_026368 | 67770 | Mm.35170 | NM_026368 | ENSMUSG00000028578 | **Caap1** | 5830433M19Rik\|AI536248\|CAAP | caspase activity and apoptosis inhibitor 1 | protein-coding |
| chr4_94466690_94467003 | chr4 | 94466691 | 94467003 | + | 0.79607 | Intergenic | Intergenic | 89949 | NM_026368 | 67770 | Mm.35170 | NM_026368 | ENSMUSG00000028578 | **Caap1** | 5830433M19Rik\|AI536248\|CAAP | caspase activity and apoptosis inhibitor 1 | protein-coding |
| chr4_94522591_94523210 | chr4 | 94522592 | 94523210 | + | 0.686559 | intron (NM_026368, intron 4 of 5) | L1MC4\|LINE\|L1 | 33895 | NM_026368 | 67770 | Mm.35170 | NM_026368 | ENSMUSG00000028578 | **Caap1** | 5830433M19Rik\|AI536248\|CAAP | caspase activity and apoptosis inhibitor 1 | protein-coding |
| chr1_85828925_85829378 | chr1 | 85828926 | 85829378 | + | 0.81379 | intron (NM_133781, intron 2 of 8) | intron (NM_133781, intron 2 of 8) | 35705 | NM_133781 | 12283 | Mm.26135 | NM_133781 | ENSMUSG00000036707 | **Cab39** | AA408805\|AA960512\|C78372\|MO25\|MO25alpha | calcium binding protein 39 | protein-coding |
| chr18_11791773_11792136 | chr18 | 11791774 | 11792136 | + | 0.735203 | Intergenic | RLTR23\|LTR\|ERV1 | -47319 | NM_022021 | 63955 | Mm.40717 | NM_022021 | ENSMUSG00000040957 | **Cables1** | Cables\|ik3-1 | CDK5 and Abl enzyme substrate 1 | protein-coding |
| chr5_115186667_115187048 | chr5 | 115186668 | 115187048 | + | 0.672785 | promoter-TSS (NM_013879) | promoter-TSS (NM_013879) | -737 | NM_013879 | 29867 | Mm.38551 | NM_013879 | ENSMUSG00000029544 | **Cabp1** | caldendrin | calcium binding protein 1 | protein-coding |
| chr2_24776206_24776718 | chr2 | 24776207 | 24776718 | + | 1.13695 | Intergenic | Intergenic | -13310 | NM_007579 | 12287 | Mm.4424 | NM_007579 | ENSMUSG00000004113 | **Cacna1b** | AW050276\|AW060892\|AW822256\|Cav2.2\|Cchn1a\|alpha(1B) | calcium channel, voltage-dependent, N type, alpha 1B subunit | protein-coding |
| chr2_24642489_24643290 | chr2 | 24642490 | 24643290 | + | 0.979534 | intron (NM_007579, intron 31 of 45) | intron (NM_007579, intron 31 of 45) | 120262 | NM_001042528 | 12287 | Mm.4424 | NM_007579 | ENSMUSG00000004113 | **Cacna1b** | AW050276\|AW060892\|AW822256\|Cav2.2\|Cchn1a\|alpha(1B) | calcium channel, voltage-dependent, N type, alpha 1B subunit | protein-coding |
| chr6_119091127_119091750 | chr6 | 119091128 | 119091750 | + | 1.44156 | intron (NM_001255999, intron 1 of 46) | B4A\|SINE\|B4 | 17056 | NM_001159534 | 12288 | Mm.41628 | NM_009781 | ENSMUSG00000051331 | **Cacna1c** | Cav1.2\|Cchl1a1\|D930026N18Rik\|MBC\|MELC-CC | calcium channel, voltage-dependent, L type, alpha 1C subunit | protein-coding |
| chr6_119090280_119090639 | chr6 | 119090281 | 119090639 | + | 0.87618 | intron (NM_001255999, intron 1 of 46) | L1MEd\|LINE\|L1 | 18035 | NM_001159534 | 12288 | Mm.41628 | NM_009781 | ENSMUSG00000051331 | **Cacna1c** | Cav1.2\|Cchl1a1\|D930026N18Rik\|MBC\|MELC-CC | calcium channel, voltage-dependent, L type, alpha 1C subunit | protein-coding |
| chr6_119084608_119085200 | chr6 | 119084609 | 119085200 | + | 0.834353 | intron (NM_001255999, intron 1 of 46) | intron (NM_001255999, intron 1 of 46) | 23591 | NM_001159534 | 12288 | Mm.41628 | NM_009781 | ENSMUSG00000051331 | **Cacna1c** | Cav1.2\|Cchl1a1\|D930026N18Rik\|MBC\|MELC-CC | calcium channel, voltage-dependent, L type, alpha 1C subunit | protein-coding |
| chr15_78119524_78120231 | chr15 | 78119525 | 78120231 | + | 0.952025 | promoter-TSS (NM_007583) | promoter-TSS (NM_007583) | -598 | NM_007583 | 12300 | Mm.277338 | NM_007583 | | **Cacng2** | AW060990\|B230105C07Rik\|B930041E13Rik\|stargazer\|stargazin\|stg\|wag\|waggler | calcium channel, voltage-dependent, gamma subunit 2 | protein-coding |
| chr7_3451091_3451548 | chr7 | 3451092 | 3451548 | + | 1.17863 | Intergenic | Intergenic | 26658 | NM_133183 | 54378 | Mm.424047 | NM_133183 | | **Cacng6** | 2310033H20Rik\|AW050150 | calcium channel, voltage-dependent, gamma subunit 6 | protein-coding |
| chr7_3450578_3451553 | chr7 | 3450579 | 3451553 | + | 1.02312 | Intergenic | Intergenic | 26404 | NM_133183 | 54378 | Mm.424047 | NM_133183 | | **Cacng6** | 2310033H20Rik\|AW050150 | calcium channel, voltage-dependent, gamma subunit 6 | protein-coding |
| chr1_160251408_160251925 | chr1 | 160251409 | 160251925 | + | 0.644135 | intron (NM_001038621, intron 3 of 8) | intron (NM_001038621, intron 3 of 8) | -38775 | NM_009786 | 12301 | Mm.10702 | NM_009786 | ENSMUSG00000014226 | **Cacybp** | SIP | calcyclin binding protein | protein-coding |
| chr13_38027926_38028344 | chr13 | 38027927 | 38028344 | + | 0.798616 | exon (NM_027724, exon 3 of 13) | exon (NM_027724, exon 3 of 13) | 8802 | NM_027724 | 71213 | Mm.314688 | NM_027724 | ENSMUSG00000044566 | **Cage1** | 4933427I01Rik\|Ctag3 | cancer antigen 1 | protein-coding |
| chr13_38027921_38028748 | chr13 | 38027922 | 38028748 | + | 0.649996 | intron (NM_027724, intron 2 of 12) | intron (NM_027724, intron 2 of 12) | 8602 | NM_027724 | 71213 | Mm.314688 | NM_027724 | ENSMUSG00000044566 | **Cage1** | 4933427I01Rik\|Ctag3 | cancer antigen 1 | protein-coding |
| chr7_114671408_114671805 | chr7 | 114671409 | 114671805 | + | 1.10313 | Intergenic | Intergenic | -35250 | NM_001033954 | 12310 | Mm.4361 | NM_007587 | ENSMUSG00000030669 | **Calca** | CA\|CGRP-1\|CGRP1\|Calc\|Calc1\|Cgrp\|Ct\|Ctn | calcitonin/calcitonin-related polypeptide, alpha | protein-coding |
| chr7_114670435_114670788 | chr7 | 114670436 | 114670788 | + | 0.854042 | Intergenic | Intergenic | -34255 | NM_001033954 | 12310 | Mm.4361 | NM_007587 | ENSMUSG00000030669 | **Calca** | CA\|CGRP-1\|CGRP1\|Calc\|Calc1\|Cgrp\|Ct\|Ctn | calcitonin/calcitonin-related polypeptide, alpha | protein-coding |
| chr7_114717016_114717393 | chr7 | 114717017 | 114717393 | + | 0.998176 | Intergenic | Intergenic | -1438 | NM_054084 | 116903 | Mm.273838 | NM_054084 | ENSMUSG00000030666 | **Calcb** | CGRP2\|Calc2 | calcitonin-related polypeptide, beta | protein-coding |
| chr7_114728778_114729250 | chr7 | 114728779 | 114729250 | + | 0.748354 | Intergenic | MTD\|LTR\|ERVL-MaLR | 10371 | NM_054084 | 116903 | Mm.273838 | NM_054084 | ENSMUSG00000030666 | **Calcb** | CGRP2\|Calc2 | calcitonin-related polypeptide, beta | protein-coding |
| chr7_114719800_114720047 | chr7 | 114719801 | 114720047 | + | 0.714471 | intron (NM_054084, intron 2 of 4) | intron (NM_054084, intron 2 of 4) | 1281 | NM_054084 | 116903 | Mm.273838 | NM_054084 | ENSMUSG00000030666 | **Calcb** | CGRP2\|Calc2 | calcitonin-related polypeptide, beta | protein-coding |
| chr7_114719379_114720060 | chr7 | 114719380 | 114720060 | + | 0.66367 | intron (NM_054084, intron 1 of 4) | intron (NM_054084, intron 1 of 4) | 1077 | NM_054084 | 116903 | Mm.273838 | NM_054084 | ENSMUSG00000030666 | **Calcb** | CGRP2\|Calc2 | calcitonin-related polypeptide, beta | protein-coding |
| chr2_84221992_84222400 | chr2 | 84221993 | 84222400 | + | 1.10696 | Intergenic | Intergenic | 203070 | NM_018782 | 54598 | Mm.75467 | NM_018782 | ENSMUSG00000059588 | **Calcrl** | AV071593\|CRLR | calcitonin receptor-like | protein-coding |
| chr2_84221838_84222684 | chr2 | 84221839 | 84222684 | + | 1.00739 | Intergenic | Intergenic | 203005 | NM_018782 | 54598 | Mm.75467 | NM_018782 | ENSMUSG00000059588 | **Calcrl** | AV071593\|CRLR | calcitonin receptor-like | protein-coding |
| chr2_84212007_84212284 | chr2 | 84212008 | 84212284 | + | 0.751403 | Intergenic | L1_Mus3\|LINE\|L1 | 213120 | NM_018782 | 54598 | Mm.75467 | NM_018782 | ENSMUSG00000059588 | **Calcrl** | AV071593\|CRLR | calcitonin receptor-like | protein-coding |
| chr19_47142518_47142802 | chr19 | 47142519 | 47142802 | + | 0.799048 | intron (NM_001081271, intron 1 of 1) | MER102b\|DNA\|hAT-Charlie | 1514 | NM_001081271 | 546729 | Mm.439624 | NM_001081271 | ENSMUSG00000079258 | **Calhm1** | EG546729 | calcium homeostasis modulator 1 | protein-coding |
| chr19_47142506_47143045 | chr19 | 47142507 | 47143045 | + | 0.745825 | intron (NM_001081271, intron 1 of 1) | MER102b\|DNA\|hAT-Charlie | 1398 | NM_001081271 | 546729 | Mm.439624 | NM_001081271 | ENSMUSG00000079258 | **Calhm1** | EG546729 | calcium homeostasis modulator 1 | protein-coding |
| chr19_47148223_47148977 | chr19 | 47148224 | 47148977 | + | 0.744784 | Intergenic | Intergenic | -4426 | NM_001081271 | 546729 | Mm.439624 | NM_001081271 | ENSMUSG00000079258 | **Calhm1** | EG546729 | calcium homeostasis modulator 1 | protein-coding |
| chr17_87463006_87463519 | chr17 | 87463007 | 87463519 | + | 1.12993 | Intergenic | RLTR24\|LTR\|ERV1 | -16328 | NM_007589 | 12314 | Mm.329243 | NM_007589 | ENSMUSG00000036438 | **Calm2** | 1500001E21Rik\|AL024017 | calmodulin 2 | protein-coding |
| chr7_16924933_16925199 | chr7 | 16924934 | 16925199 | + | 1.06211 | Intergenic | Intergenic | -1034 | NM_007590 | 12315 | Mm.288630 | NM_007590 | ENSMUSG00000019370 | **Calm3** | CaMA\|R75142 | calmodulin 3 | protein-coding |
| chr13_3865602_3866099 | chr13 | 3865603 | 3866099 | + | 0.632488 | Intergenic | Intergenic | 11678 | NM_001008706 | 494124 | Mm.483992 | NM_001008706 | | **Calm5** | AY314008\|Scarf2 | calmodulin 5 | protein-coding |
| chr9_62848823_62849241 | chr9 | 62848824 | 62849241 | + | 0.634602 | intron (NM_001033175, intron 5 of 6) | intron (NM_001033175, intron 5 of 6) | -9072 | NM_001102468 | 75600 | Mm.440576 | NM_138304 | ENSMUSG00000032246 | **Calml4** | 2010002G05Rik | calmodulin-like 4 | protein-coding |
| chr7_140080806_140081128 | chr7 | 140080807 | 140081128 | + | 0.698359 | intron (NM_026769, intron 1 of 5) | intron (NM_026769, intron 1 of 5) | 1334 | NM_001190386 | 68566 | Mm.44241 | NM_026769 | ENSMUSG00000025468 | **Caly** | 0710001P07Rik\|1110004A22Rik\|Calcyon\|Drd1ip | calcyon neuron-specific vesicular protein | protein-coding |
| chr2_5650940_5651853 | chr2 | 5650941 | 5651853 | + | 0.750605 | intron (NM_177343, intron 1 of 10) | intron (NM_177343, intron 1 of 10) | 24649 | NM_001290376 | 227541 | Mm.191949 | NM_177343 | ENSMUSG00000039145 | **Camk1d** | A630059D12Rik\|CKLiK\|CaMKIdelta\|E030025C11Rik\|mCKLiK | calcium/calmodulin-dependent protein kinase ID | protein-coding |
| chr2_5651412_5651822 | chr2 | 5651413 | 5651822 | + | 0.720029 | intron (NM_177343, intron 1 of 10) | intron (NM_177343, intron 1 of 10) | 24429 | NM_001290376 | 227541 | Mm.191949 | NM_177343 | ENSMUSG00000039145 | **Camk1d** | A630059D12Rik\|CKLiK\|CaMKIdelta\|E030025C11Rik\|mCKLiK | calcium/calmodulin-dependent protein kinase ID | protein-coding |
| chr1_193365736_193366297 | chr1 | 193365737 | 193366297 | + | 0.881852 | intron (NM_144817, intron 1 of 12) | intron (NM_144817, intron 1 of 12) | 4265 | NM_144817 | 215303 | Mm.40329 | NM_144817 | ENSMUSG00000016179 | **Camk1g** | CLICK-III\|CaMKIgamma | calcium/calmodulin-dependent protein kinase I gamma | protein-coding |
| chr18_60925393_60926355 | chr18 | 60925394 | 60926355 | + | 1.11614 | intron (NM_177407, intron 1 of 17) | intron (NM_177407, intron 1 of 17) | 242 | NM_177407 | 12322 | Mm.131530 | NM_009792 | ENSMUSG00000024617 | **Camk2a** | CaMKII\|R74975\|mKIAA0968 | calcium/calmodulin-dependent protein kinase II alpha | protein-coding |
| chr3_126652408_126652724 | chr3 | 126652409 | 126652724 | + | 1.07013 | intron (NM_023813, intron 2 of 17) | intron (NM_023813, intron 2 of 17) | 55615 | NM_023813 | 108058 | Mm.255822 | NM_023813 | ENSMUSG00000053819 | **Camk2d** | 2810011D23Rik\|8030469K03Rik\|CaMK II\|[d]-CaMKII | calcium/calmodulin-dependent protein kinase II, delta | protein-coding |
| chr3_126661140_126661342 | chr3 | 126661141 | 126661342 | + | 0.866321 | intron (NM_023813, intron 2 of 17) | intron (NM_023813, intron 2 of 17) | 64290 | NM_023813 | 108058 | Mm.255822 | NM_023813 | ENSMUSG00000053819 | **Camk2d** | 2810011D23Rik\|8030469K03Rik\|CaMK II\|[d]-CaMKII | calcium/calmodulin-dependent protein kinase II, delta | protein-coding |
| chr3_126659864_126661387 | chr3 | 126659865 | 126661387 | + | 0.674062 | intron (NM_023813, intron 2 of 17) | intron (NM_023813, intron 2 of 17) | 63675 | NM_023813 | 108058 | Mm.255822 | NM_023813 | ENSMUSG00000053819 | **Camk2d** | 2810011D23Rik\|8030469K03Rik\|CaMK II\|[d]-CaMKII | calcium/calmodulin-dependent protein kinase II, delta | protein-coding |
| chr3_126570664_126571178 | chr3 | 126570665 | 126571178 | + | 0.647934 | Intergenic | Intergenic | -26030 | NM_001293663 | 108058 | Mm.255822 | NM_023813 | ENSMUSG00000053819 | **Camk2d** | 2810011D23Rik\|8030469K03Rik\|CaMK II\|[d]-CaMKII | calcium/calmodulin-dependent protein kinase II, delta | protein-coding |
| chr14_20790930_20791275 | chr14 | 20790931 | 20791275 | + | 0.829311 | intron (NM_178597, intron 2 of 20) | intron (NM_178597, intron 2 of 20) | 2985 | NM_001039138 | 12325 | Mm.235182 | NM_178597 | ENSMUSG00000021820 | **Camk2g** | Camkg | calcium/calmodulin-dependent protein kinase II gamma | protein-coding |
| chr18_32992520_32993308 | chr18 | 32992521 | 32993308 | + | 1.09566 | intron (NM_009793, intron 1 of 10) | intron (NM_009793, intron 1 of 10) | 53873 | NM_009793 | 12326 | Mm.222329 | NM_009793 | ENSMUSG00000038128 | **Camk4** | A430110E23Rik\|AI666733\|CaMKIV\|CaMKIV/Gr\|D18Bwg0362e | calcium/calmodulin-dependent protein kinase IV | protein-coding |
| chr18_32992539_32993014 | chr18 | 32992540 | 32993014 | + | 1.03798 | intron (NM_009793, intron 1 of 10) | intron (NM_009793, intron 1 of 10) | 53736 | NM_009793 | 12326 | Mm.222329 | NM_009793 | ENSMUSG00000038128 | **Camk4** | A430110E23Rik\|AI666733\|CaMKIV\|CaMKIV/Gr\|D18Bwg0362e | calcium/calmodulin-dependent protein kinase IV | protein-coding |
| chr4_151827139_151827910 | chr4 | 151827140 | 151827910 | + | 0.776359 | intron (NM_001195565, intron 3 of 3) | intron (NM_001195565, intron 3 of 3) | 34243 | NM_001081557 | 100072 | Mm.318846 | NM_001081557 | ENSMUSG00000014592 | **Camta1** | 1810059M14Rik\|2310058O09Rik\|AI316882\|AW987475\|mKIAA0833 | calmodulin binding transcription activator 1 | protein-coding |
| chr10_119199695_119200137 | chr10 | 119199696 | 119200137 | + | 0.659997 | 3' UTR (NM_027994, exon 15 of 15) | 3' UTR (NM_027994, exon 15 of 15) | 40139 | NM_027994 | 71902 | Mm.203965 | NM_027994 | ENSMUSG00000020114 | **Cand1** | 2310038O07Rik\|6330512O03Rik\|AI195005\|AI846556\|D10Ertd516e\|Tp120a\|mKIAA0829 | cullin associated and neddylation disassociated 1 | protein-coding |

| | | | | | | | | | | | | | | | | | |
|---|---|---|---|---|---|---|---|---|---|---|---|---|---|---|---|---|---|
| chr4_122868079_122868446 | chr4 | 122868080 | 122868446 | + | 1.56941 | intron (NM_007598, intron 4 of 12) | intron (NM_007598, intron 4 of 12) | 17794 | NM_007598 | 12331 | Mm.8687 | NM_007598 | ENSMUSG00000028656 | **Cap1** | - | CAP, adenylate cyclase-associated protein 1 (yeast) | protein-coding |
| chr4_122868073_122868636 | chr4 | 122868074 | 122868636 | + | 1.51479 | intron (NM_007598, intron 4 of 12) | intron (NM_007598, intron 4 of 12) | 17702 | NM_007598 | 12331 | Mm.8687 | NM_007598 | ENSMUSG00000028656 | **Cap1** | - | CAP, adenylate cyclase-associated protein 1 (yeast) | protein-coding |
| chr4_122915841_122916267 | chr4 | 122915842 | 122916267 | + | 0.659926 | Intergenic | B3\|SINE\|B2 | -29997 | NM_001301067 | 12331 | Mm.8687 | NM_007598 | ENSMUSG00000028656 | **Cap1** | - | CAP, adenylate cyclase-associated protein 1 (yeast) | protein-coding |
| chr13_46469565_46469887 | chr13 | 46469566 | 46469887 | + | 0.852672 | Intergenic | Intergenic | -32177 | NM_026056 | 67252 | Mm.44529 | NM_026056 | ENSMUSG00000021373 | **Cap2** | 2810452G09Rik\|AV045235 | CAP, adenylate cyclase-associated protein, 2 (yeast) | protein-coding |
| chr6_72511097_72511359 | chr6 | 72511098 | 72511359 | + | 0.755072 | Intergenic | ID_B1\|SINE\|B4 | -33163 | NM_007599 | 12332 | Mm.18626 | NM_007599 | ENSMUSG00000056737 | **Capg** | gCap39\|mbh1 | capping protein (actin filament), gelsolin-like | protein-coding |
| chr17_45637895_45638345 | chr17 | 45637896 | 45638345 | + | 0.800952 | intron (NM_001013767, intron 11 of 21) | intron (NM_001013767, intron 11 of 21) | 21189 | NM_001013767 | 268958 | Mm.426026 | NM_001013767 | ENSMUSG00000058626 | **Capn11** | - | calpain 11 | protein-coding |
| chr17_73386543_73387001 | chr17 | 73386544 | 73387001 | + | 0.998404 | intron (NM_001033444, intron 1 of 22) | intron (NM_001033444, intron 1 of 22) | 12524 | NM_001033444 | 381122 | Mm.81496 | NM_001033444 | ENSMUSG00000043705 | **Capn13** | CANP 13\|Gm943 | calpain 13 | protein-coding |
| chr17_73386358_73386985 | chr17 | 73386359 | 73386985 | + | 0.981966 | intron (NM_001033444, intron 1 of 22) | intron (NM_001033444, intron 1 of 22) | 12624 | NM_001033444 | 381122 | Mm.81496 | NM_001033444 | ENSMUSG00000043705 | **Capn13** | CANP 13\|Gm943 | calpain 13 | protein-coding |
| chr17_73271507_73272284 | chr17 | 73271508 | 73272284 | + | 0.731944 | Intergenic | Intergenic | 127400 | NM_001033444 | 381122 | Mm.81496 | NM_001033444 | ENSMUSG00000043705 | **Capn13** | CANP 13\|Gm943 | calpain 13 | protein-coding |
| chr17_73271741_73272274 | chr17 | 73271742 | 73272274 | + | 0.728548 | Intergenic | Intergenic | 127288 | NM_001033444 | 381122 | Mm.81496 | NM_001033444 | ENSMUSG00000043705 | **Capn13** | CANP 13\|Gm943 | calpain 13 | protein-coding |
| chr1_182527923_182528364 | chr1 | 182527924 | 182528364 | + | 0.761907 | Intergenic | Intergenic | -10661 | NM_009794 | 12334 | Mm.19306 | NM_009794 | ENSMUSG00000026509 | **Capn2** | AI326419\|Capa-2\|Capa2\|m-calpain\|m-calpin | calpain 2 | protein-coding |
| chr1_182527923_182528252 | chr1 | 182527924 | 182528252 | + | 0.75768 | Intergenic | Intergenic | -10605 | NM_009794 | 12334 | Mm.19306 | NM_009794 | ENSMUSG00000026509 | **Capn2** | AI326419\|Capa-2\|Capa2\|m-calpain\|m-calpin | calpain 2 | protein-coding |
| chr1_182527885_182528601 | chr1 | 182527886 | 182528601 | + | 0.73953 | Intergenic | Intergenic | -10760 | NM_009794 | 12334 | Mm.19306 | NM_009794 | ENSMUSG00000026509 | **Capn2** | AI326419\|Capa-2\|Capa2\|m-calpain\|m-calpin | calpain 2 | protein-coding |
| chr14_31367598_31368388 | chr14 | 31367599 | 31368388 | + | 1.76057 | intron (NM_009796, intron 16 of 20) | intron (NM_009796, intron 16 of 20) | 31269 | NM_009796 | 12339 | Mm.201535 | NM_009796 | ENSMUSG00000021893 | **Capn7** | AU022319\|PalBH | calpain 7 | protein-coding |
| chr1_182543121_182543641 | chr1 | 182543122 | 182543641 | + | 1.08841 | Intergenic | Intergenic | -21626 | NM_001145806 | 170725 | Mm.189201 | NM_130890 | ENSMUSG00000038599 | **Capn8** | nCL-2\|nCL-2' | calpain 8 | protein-coding |
| chr2_103801074_103801425 | chr2 | 103801075 | 103801425 | + | 1.35767 | Intergenic | Intergenic | -3610 | NM_016739 | 53872 | Mm.427589 | NM_016739 | ENSMUSG00000027184 | **Caprin1** | AL022980\|Caprin-1\|Gpiap\|Gpiap1\|Mmgpip137\|P137\|rng105 | cell cycle associated protein 1 | protein-coding |
| chr15_9462325_9462578 | chr15 | 9462326 | 9462578 | + | 1.00232 | intron (NM_029341, intron 3 of 4) | intron (NM_029341, intron 3 of 4) | 26424 | NM_029341 | 75568 | Mm.82143 | NM_029341 | ENSMUSG00000039676 | **Capsl** | 1700028N11Rik\|AW125205 | calcyphosine-like | protein-coding |
| chr15_9443608_9444153 | chr15 | 9443609 | 9444153 | + | 0.990752 | intron (NM_029341, intron 1 of 4) | intron (NM_029341, intron 1 of 4) | 7853 | NM_029341 | 75568 | Mm.82143 | NM_029341 | ENSMUSG00000039676 | **Capsl** | 1700028N11Rik\|AW125205 | calcyphosine-like | protein-coding |
| chr15_9462873_9464052 | chr15 | 9462874 | 9464052 | + | 0.892144 | intron (NM_029341, intron 4 of 4) | intron (NM_029341, intron 4 of 4) | 27435 | NM_029341 | 75568 | Mm.82143 | NM_029341 | ENSMUSG00000039676 | **Capsl** | 1700028N11Rik\|AW125205 | calcyphosine-like | protein-coding |
| chr15_9463448_9464041 | chr15 | 9463449 | 9464041 | + | 0.760453 | intron (NM_029341, intron 4 of 4) | intron (NM_029341, intron 4 of 4) | 27717 | NM_029341 | 75568 | Mm.82143 | NM_029341 | ENSMUSG00000039676 | **Capsl** | 1700028N11Rik\|AW125205 | calcyphosine-like | protein-coding |
| chr15_9469739_9470045 | chr15 | 9469740 | 9470045 | + | 0.73155 | Intergenic | Intergenic | 33864 | NM_029341 | 75568 | Mm.82143 | NM_029341 | ENSMUSG00000039676 | **Capsl** | 1700028N11Rik\|AW125205 | calcyphosine-like | protein-coding |
| chr6_17652108_17652535 | chr6 | 17652109 | 17652535 | + | 1.58384 | intron (NM_007604, intron 3 of 9) | ORR1E\|LTR\|ERVL-MaLR | 15224 | NM_007604 | 12343 | Mm.392504 | NM_007604 | | **Capza2** | 1110053K06Rik\|AW208808\|Cappa2 | capping protein (actin filament) muscle Z-line, alpha 2 | protein-coding |
| chr6_17634963_17635366 | chr6 | 17634964 | 17635366 | + | 1.33999 | Intergenic | Intergenic | -1933 | NM_007604 | 12343 | Mm.392504 | NM_007604 | | **Capza2** | 1110053K06Rik\|AW208808\|Cappa2 | capping protein (actin filament) muscle Z-line, alpha 2 | protein-coding |
| chr6_17594017_17594369 | chr6 | 17594018 | 17594369 | + | 1.02705 | Intergenic | RMER6C\|LTR\|ERVK | -42905 | NM_007604 | 12343 | Mm.392504 | NM_007604 | | **Capza2** | 1110053K06Rik\|AW208808\|Cappa2 | capping protein (actin filament) muscle Z-line, alpha 2 | protein-coding |
| chr6_17652697_17653165 | chr6 | 17652698 | 17653165 | + | 1.01718 | intron (NM_007604, intron 3 of 9) | intron (NM_007604, intron 3 of 9) | 15833 | NM_007604 | 12343 | Mm.392504 | NM_007604 | | **Capza2** | 1110053K06Rik\|AW208808\|Cappa2 | capping protein (actin filament) muscle Z-line, alpha 2 | protein-coding |
| chr6_17633813_17634202 | chr6 | 17633814 | 17634202 | + | 1.00616 | Intergenic | RMER15\|LTR\|ERVL | -3090 | NM_007604 | 12343 | Mm.392504 | NM_007604 | | **Capza2** | 1110053K06Rik\|AW208808\|Cappa2 | capping protein (actin filament) muscle Z-line, alpha 2 | protein-coding |
| chr6_17639502_17639771 | chr6 | 17639503 | 17639771 | + | 0.672774 | intron (NM_007604, intron 1 of 9) | PB1D10\|SINE\|Alu | 2539 | NM_007604 | 12343 | Mm.392504 | NM_007604 | | **Capza2** | 1110053K06Rik\|AW208808\|Cappa2 | capping protein (actin filament) muscle Z-line, alpha 2 | protein-coding |
| chr6_17645316_17645649 | chr6 | 17645317 | 17645649 | + | 0.644287 | intron (NM_007604, intron 1 of 9) | ORR1D2\|LTR\|ERVL-MaLR | 8385 | NM_007604 | 12343 | Mm.392504 | NM_007604 | | **Capza2** | 1110053K06Rik\|AW208808\|Cappa2 | capping protein (actin filament) muscle Z-line, alpha 2 | protein-coding |
| chr9_66721970_66722751 | chr9 | 66721971 | 66722751 | + | 1.15913 | intron (NM_178396, intron 2 of 10) | B4\|SINE\|B4 | 8675 | NM_178396 | 76459 | Mm.277921 | NM_178396 | ENSMUSG00000032373 | **Car12** | 2310047E01Rik\|AI314958\|Ca12 | carbonic anyhydrase 12 | protein-coding |
| chr3_95909258_95909830 | chr3 | 95909259 | 95909830 | + | 0.863994 | Intergenic | Intergenic | -4905 | NM_011797 | 23831 | Mm.489647 | NM_011797 | ENSMUSG00000038526 | **Car14** | AW536446\|Ca14 | carbonic anhydrase 14 | protein-coding |
| chr11_85003496_85003926 | chr11 | 85003497 | 85003926 | + | 1.03313 | intron (NM_001029934, intron 29 of 33) | intron (NM_001029934, intron 29 of 33) | 45957 | NM_007607 | 12351 | Mm.1641 | NM_007607 | ENSMUSG00000000805 | **Car4** | AW456718\|Ca4 | carbonic anhydrase 4 | protein-coding |
| chrX_164013124_164013950 | chrX | 164013125 | 164013950 | + | 1.33987 | intron (NM_181315, intron 2 of 7) | intron (NM_181315, intron 2 of 7) | 14473 | NM_181315 | 56078 | Mm.180953 | NM_181315 | ENSMUSG00000031373 | **Car5b** | 7330410H16Rik\|CAVB\|Ca5b\|CarVb\|D730005F19Rik | carbonic anhydrase 5b, mitochondrial | protein-coding |
| chr4_8033132_8033808 | chr4 | 8033133 | 8033808 | + | 1.10747 | Intergenic | MERV1_LTR\|LTR\|ERV1 | 205571 | NM_007592 | 12319 | Mm.119320 | NM_007592 | | **Car8** | AW546993\|Ca8\|Cals\|Cals1\|Carp\|wdl | carbonic anhydrase 8 | protein-coding |
| chr4_7912647_7913011 | chr4 | 7912648 | 7913011 | + | 0.910234 | Intergenic | Intergenic | 326212 | NM_007592 | 12319 | Mm.119320 | NM_007592 | | **Car8** | AW546993\|Ca8\|Cals\|Cals1\|Carp\|wdl | carbonic anhydrase 8 | protein-coding |
| chr4_7982759_7983051 | chr4 | 7982760 | 7983051 | + | 0.747358 | Intergenic | Intergenic | 256136 | NM_007592 | 12319 | Mm.119320 | NM_007592 | | **Car8** | AW546993\|Ca8\|Cals\|Cals1\|Carp\|wdl | carbonic anhydrase 8 | protein-coding |
| chr5_140994444_140994695 | chr5 | 140994445 | 140994695 | + | 0.919102 | intron (NM_175362, intron 1 of 24) | intron (NM_175362, intron 1 of 24) | 6026 | NM_175362 | 108723 | Mm.46187 | NM_175362 | ENSMUSG00000036526 | **Card11** | 0610008L17Rik\|2410011D02Rik\|BIMP3\|CARMA1 | caspase recruitment domain family, member 11 | protein-coding |
| chr5_140994837_140995148 | chr5 | 140994838 | 140995148 | + | 0.653773 | intron (NM_175362, intron 1 of 24) | B1F\|SINE\|Alu | 5603 | NM_175362 | 108723 | Mm.46187 | NM_175362 | ENSMUSG00000036526 | **Card11** | 0610008L17Rik\|2410011D02Rik\|BIMP3\|CARMA1 | caspase recruitment domain family, member 11 | protein-coding |
| chr16_8666524_8666894 | chr16 | 8666525 | 8666894 | + | 0.978114 | intron (NM_025821, intron 1 of 3) | intron (NM_025821, intron 1 of 3) | 5444 | NM_025821 | 52502 | Mm.142095 | NM_025821 | ENSMUSG00000008393 | **Carhsp1** | 1200011K09Rik\|Crhsp-24\|D16Ertd465e | calcium regulated heat stable protein 1 | protein-coding |
| chr16_8666193_8666908 | chr16 | 8666194 | 8666908 | + | 0.74261 | intron (NM_025821, intron 1 of 3) | intron (NM_025821, intron 1 of 3) | 5602 | NM_025821 | 52502 | Mm.142095 | NM_025821 | ENSMUSG00000008393 | **Carhsp1** | 1200011K09Rik\|Crhsp-24\|D16Ertd465e | calcium regulated heat stable protein 1 | protein-coding |
| chr8_46655677_46656451 | chr8 | 46655678 | 46656451 | + | 1.5077 | Intergenic | Intergenic | 38773 | NM_009810 | 12367 | Mm.34405 | NM_009810 | | **Casp3** | A830040C14Rik\|AC-3\|Apopain\|CASP-3\|CC3\|CPP-32\|CPP32\|Caspase-3\|Lice\|SCA-1\|Yama\|mldy | caspase 3 | protein-coding |
| chr8_46647900_46648506 | chr8 | 46647901 | 46648506 | + | 1.49175 | Intergenic | Intergenic | 30912 | NM_009810 | 12367 | Mm.34405 | NM_009810 | | **Casp3** | A830040C14Rik\|AC-3\|Apopain\|CASP-3\|CC3\|CPP-32\|CPP32\|Caspase-3\|Lice\|SCA-1\|Yama\|mldy | caspase 3 | protein-coding |
| chr8_46644998_46645340 | chr8 | 46644999 | 46645340 | + | 1.2751 | Intergenic | Intergenic | 27878 | NM_009810 | 12367 | Mm.34405 | NM_009810 | | **Casp3** | A830040C14Rik\|AC-3\|Apopain\|CASP-3\|CC3\|CPP-32\|CPP32\|Caspase-3\|Lice\|SCA-1\|Yama\|mldy | caspase 3 | protein-coding |
| chr8_46639921_46640146 | chr8 | 46639922 | 46640146 | + | 1.00143 | TTS (NM_001284409) | TTS (NM_001284409) | 22743 | NM_009810 | 12367 | Mm.34405 | NM_009810 | | **Casp3** | A830040C14Rik\|AC-3\|Apopain\|CASP-3\|CC3\|CPP-32\|CPP32\|Caspase-3\|Lice\|SCA-1\|Yama\|mldy | caspase 3 | protein-coding |
| chr8_46641216_46642005 | chr8 | 46641217 | 46642005 | + | 0.885833 | Intergenic | Intergenic | 24320 | NM_009810 | 12367 | Mm.34405 | NM_009810 | | **Casp3** | A830040C14Rik\|AC-3\|Apopain\|CASP-3\|CC3\|CPP-32\|CPP32\|Caspase-3\|Lice\|SCA-1\|Yama\|mldy | caspase 3 | protein-coding |
| chr8_46651905_46652158 | chr8 | 46651906 | 46652158 | + | 0.687714 | Intergenic | ORR1E\|LTR\|ERVL-MaLR | 34741 | NM_009810 | 12367 | Mm.34405 | NM_009810 | | **Casp3** | A830040C14Rik\|AC-3\|Apopain\|CASP-3\|CC3\|CPP-32\|CPP32\|Caspase-3\|Lice\|SCA-1\|Yama\|mldy | caspase 3 | protein-coding |

| | | | | | | | | | | | | | | | | | |
|---|---|---|---|---|---|---|---|---|---|---|---|---|---|---|---|---|---|
| chr8_46663208_46663652 | chr8 | 46663209 | 46663652 | + | 0.65477 | Intergenic | MTB\|LTR\|ERVL-MaLR | 46139 | NM_009810 | 12367 | Mm.34405 | NM_009810 | | **Casp3** | A830040C14Rik\|AC-3\|Apopain\|CASP-3\|CC3\|CPP-32\|CPP32\|Caspase-3\|Lice\|SCA-1\|Yama\|mldy | caspase 3 | protein-coding |
| chr3_129892840_129893464 | chr3 | 129892841 | 129893464 | + | 1.69676 | intron (NM_183423, intron 3 of 3) | intron (NM_183423, intron 3 of 3) | -8273 | NM_009811 | 12368 | Mm.281379 | NM_009811 | ENSMUSG00000027997 | **Casp6** | Mch2\|mCASP-6 | caspase 6 | protein-coding |
| chr3_129904500_129905109 | chr3 | 129904501 | 129905109 | + | 0.99771 | intron (NM_009811, intron 1 of 6) | intron (NM_009811, intron 1 of 6) | 3380 | NM_009811 | 12368 | Mm.281379 | NM_009811 | ENSMUSG00000027997 | **Casp6** | Mch2\|mCASP-6 | caspase 6 | protein-coding |
| chr1_58856823_58857172 | chr1 | 58856824 | 58857172 | + | 0.752596 | Intergenic | Intergenic | 54496 | NM_001080126 | 12370 | Mm.336851 | NM_009812 | ENSMUSG00000026029 | **Casp8** | CASP-8\|FLICE\|MACH\|Mch5 | caspase 8 | protein-coding |
| chr16_36602455_36603074 | chr16 | 36602456 | 36603074 | + | 1.61899 | Intergenic | Intergenic | -40631 | NM_013803 | 12374 | Mm.103619 | NM_013803 | ENSMUSG00000051980 | **Casr** | CaR\|Gprc2a | calcium-sensing receptor | protein-coding |
| chr16_36598139_36598755 | chr16 | 36598140 | 36598755 | + | 0.677097 | Intergenic | Intergenic | -36313 | NM_013803 | 12374 | Mm.103619 | NM_013803 | ENSMUSG00000051980 | **Casr** | CaR\|Gprc2a | calcium-sensing receptor | protein-coding |
| chr16_36540728_36541276 | chr16 | 36540729 | 36541276 | + | 0.658022 | intron (NM_013803, intron 1 of 6) | intron (NM_013803, intron 1 of 6) | 21132 | NM_013803 | 12374 | Mm.103619 | NM_013803 | ENSMUSG00000051980 | **Casr** | CaR\|Gprc2a | calcium-sensing receptor | protein-coding |
| chr2_172408965_172409299 | chr2 | 172408966 | 172409299 | + | 1.32766 | intron (NM_001276422, intron 1 of 5) | intron (NM_001276422, intron 1 of 5) | 15338 | NM_001276423 | 320664 | Mm.210195 | NM_001033538 | ENSMUSG00000074570 | **Cass4** | F730031O20Rik | Cas scaffolding protein family member 4 | protein-coding |
| chr2_172405021_172405372 | chr2 | 172405022 | 172405372 | + | 1.27105 | intron (NM_001276422, intron 1 of 5) | intron (NM_001276422, intron 1 of 5) | 11403 | NM_001276423 | 320664 | Mm.210195 | NM_001033538 | ENSMUSG00000074570 | **Cass4** | F730031O20Rik | Cas scaffolding protein family member 4 | protein-coding |
| chr2_172411204_172411491 | chr2 | 172411205 | 172411491 | + | 0.833578 | intron (NM_001276422, intron 1 of 5) | intron (NM_001276422, intron 1 of 5) | 17554 | NM_001276423 | 320664 | Mm.210195 | NM_001033538 | ENSMUSG00000074570 | **Cass4** | F730031O20Rik | Cas scaffolding protein family member 4 | protein-coding |
| chr2_172401045_172401819 | chr2 | 172401046 | 172401819 | + | 0.763964 | intron (NM_001276422, intron 1 of 5) | intron (NM_001276422, intron 1 of 5) | 7638 | NM_001276423 | 320664 | Mm.210195 | NM_001033538 | ENSMUSG00000074570 | **Cass4** | F730031O20Rik | Cas scaffolding protein family member 4 | protein-coding |
| chr2_172401103_172401560 | chr2 | 172401104 | 172401560 | + | 0.700116 | intron (NM_001276422, intron 1 of 5) | intron (NM_001276422, intron 1 of 5) | 7538 | NM_001276423 | 320664 | Mm.210195 | NM_001033538 | ENSMUSG00000074570 | **Cass4** | F730031O20Rik | Cas scaffolding protein family member 4 | protein-coding |
| chr2_172401121_172401363 | chr2 | 172401122 | 172401363 | + | 0.678089 | intron (NM_001276422, intron 1 of 5) | intron (NM_001276422, intron 1 of 5) | 7448 | NM_001276423 | 320664 | Mm.210195 | NM_001033538 | ENSMUSG00000074570 | **Cass4** | F730031O20Rik | Cas scaffolding protein family member 4 | protein-coding |
| chr2_172392012_172392595 | chr2 | 172392013 | 172392595 | + | 0.674737 | Intergenic | Intergenic | -1490 | NM_001080820 | 320664 | Mm.210195 | NM_001033538 | ENSMUSG00000074570 | **Cass4** | F730031O20Rik | Cas scaffolding protein family member 4 | protein-coding |
| chr13_74771322_74771658 | chr13 | 74771323 | 74771658 | + | 1.17154 | promoter-TSS (NM_001301155) | promoter-TSS (NM_001301155) | -754 | NM_001301155 | 12380 | Mm.29163 | NM_009817 | ENSMUSG00000021585 | **Cast** | - | calpastatin | protein-coding |
| chr13_74772245_74772479 | chr13 | 74772246 | 74772479 | + | 1.13974 | intron (NM_001301153, intron 3 of 31) | intron (NM_001301153, intron 3 of 31) | -1626 | NM_001301155 | 12380 | Mm.29163 | NM_009817 | ENSMUSG00000021585 | **Cast** | - | calpastatin | protein-coding |
| chr13_74822086_74822444 | chr13 | 74822087 | 74822444 | + | 1.03892 | Intergenic | Intergenic | -13505 | NM_001301153 | 12380 | Mm.29163 | NM_009817 | ENSMUSG00000021585 | **Cast** | - | calpastatin | protein-coding |
| chr13_74903717_74904028 | chr13 | 74903718 | 74904028 | + | 0.90278 | Intergenic | Intergenic | -95113 | NM_001301153 | 12380 | Mm.29163 | NM_009817 | ENSMUSG00000021585 | **Cast** | - | calpastatin | protein-coding |
| chr4_148865208_148865519 | chr4 | 148865209 | 148865519 | + | 0.685907 | intron (NM_027195, intron 2 of 15) | Charlie1\|DNA\|hAT-Charlie | 60972 | NM_027195 | 69743 | Mm.233879 | NM_027195 | ENSMUSG00000028977 | **Casz1** | 2410019P08Rik\|AV096627\|BC040081\|Cst\|D4Ertd432e | castor zinc finger 1 | protein-coding |
| chr4_148901688_148902077 | chr4 | 148901689 | 148902077 | + | 0.672963 | intron (NM_027195, intron 4 of 15) | intron (NM_027195, intron 4 of 15) | 97491 | NM_027195 | 69743 | Mm.233879 | NM_027195 | ENSMUSG00000028977 | **Casz1** | 2410019P08Rik\|AV096627\|BC040081\|Cst\|D4Ertd432e | castor zinc finger 1 | protein-coding |
| chr1_74351155_74351948 | chr1 | 74351156 | 74351948 | + | 0.778099 | exon (NM_019999, exon 8 of 9) | exon (NM_019999, exon 8 of 9) | -10556 | NM_001033345 | 241112 | Mm.310460 | NM_001033345 | ENSMUSG00000073650 | **Catip** | Gm216 | ciliogenesis associated TTC17 interacting protein | protein-coding |
| chr6_112465012_112465859 | chr6 | 112465013 | 112465859 | + | 1.05543 | intron (NM_007617, intron 1 of 1) | intron (NM_007617, intron 1 of 1) | 5931 | NM_007617 | 12391 | Mm.3924 | NM_007617 | ENSMUSG00000062694 | **Cav3** | AI385751\|Cav-3\|M-cav | caveolin 3 | protein-coding |
| chr9_44197961_44198314 | chr9 | 44197962 | 44198314 | + | 0.732796 | intron (NM_007619, intron 2 of 15) | B3\|SINE\|B2 | 35908 | NM_007619 | 12402 | Mm.266871 | NM_007619 | ENSMUSG00000034342 | **Cbl** | 4732447J05Rik\|Cbl-2\|c-Cbl\|cbl | Casitas B-lineage lymphoma | protein-coding |
| chr9_44208967_44209548 | chr9 | 44208968 | 44209548 | + | 0.642113 | intron (NM_007619, intron 1 of 15) | intron (NM_007619, intron 1 of 15) | 24788 | NM_007619 | 12402 | Mm.266871 | NM_007619 | ENSMUSG00000034342 | **Cbl** | 4732447J05Rik\|Cbl-2\|c-Cbl\|cbl | Casitas B-lineage lymphoma | protein-coding |
| chr9_44192835_44193175 | chr9 | 44192836 | 44193175 | + | 0.63247 | intron (NM_007619, intron 2 of 15) | L1MB4\|LINE\|L1 | 41041 | NM_007619 | 12402 | Mm.266871 | NM_007619 | ENSMUSG00000034342 | **Cbl** | 4732447J05Rik\|Cbl-2\|c-Cbl\|cbl | Casitas B-lineage lymphoma | protein-coding |
| chr16_51747037_51747565 | chr16 | 51747038 | 51747565 | + | 2.03557 | Intergenic | MER44B\|DNA\|TcMar-Tigger | -284248 | NM_001033238 | 208650 | Mm.328206 | NM_001033238 | ENSMUSG00000022637 | **Cblb** | AI429560\|AI851073\|Cbl-b | Casitas B-lineage lymphoma b | protein-coding |
| chr16_51878080_51878745 | chr16 | 51878081 | 51878745 | + | 1.96072 | Intergenic | Intergenic | -153136 | NM_001033238 | 208650 | Mm.328206 | NM_001033238 | ENSMUSG00000022637 | **Cblb** | AI429560\|AI851073\|Cbl-b | Casitas B-lineage lymphoma b | protein-coding |
| chr16_51767570_51768170 | chr16 | 51767571 | 51768170 | + | 1.45316 | Intergenic | Intergenic | -263679 | NM_001033238 | 208650 | Mm.328206 | NM_001033238 | ENSMUSG00000022637 | **Cblb** | AI429560\|AI851073\|Cbl-b | Casitas B-lineage lymphoma b | protein-coding |
| chr16_51813736_51814403 | chr16 | 51813737 | 51814403 | + | 1.43719 | Intergenic | RMER15\|LTR\|ERVL | -217479 | NM_001033238 | 208650 | Mm.328206 | NM_001033238 | ENSMUSG00000022637 | **Cblb** | AI429560\|AI851073\|Cbl-b | Casitas B-lineage lymphoma b | protein-coding |
| chr16_51701684_51702150 | chr16 | 51701685 | 51702150 | + | 1.4031 | Intergenic | MTE2a\|LTR\|ERVL-MaLR | -329632 | NM_001033238 | 208650 | Mm.328206 | NM_001033238 | ENSMUSG00000022637 | **Cblb** | AI429560\|AI851073\|Cbl-b | Casitas B-lineage lymphoma b | protein-coding |
| chr16_52229723_52230384 | chr16 | 52229724 | 52230384 | + | 1.38184 | Intergenic | Intergenic | 198505 | NM_001033238 | 208650 | Mm.328206 | NM_001033238 | ENSMUSG00000022637 | **Cblb** | AI429560\|AI851073\|Cbl-b | Casitas B-lineage lymphoma b | protein-coding |
| chr16_52184974_52185541 | chr16 | 52184975 | 52185541 | + | 1.31202 | intron (NM_001033238, intron 14 of 17) | intron (NM_001033238, intron 14 of 17) | 153709 | NM_001033238 | 208650 | Mm.328206 | NM_001033238 | ENSMUSG00000022637 | **Cblb** | AI429560\|AI851073\|Cbl-b | Casitas B-lineage lymphoma b | protein-coding |
| chr16_52107734_52108262 | chr16 | 52107735 | 52108262 | + | 1.09984 | intron (NM_001033238, intron 3 of 17) | intron (NM_001033238, intron 3 of 17) | 76449 | NM_001033238 | 208650 | Mm.328206 | NM_001033238 | ENSMUSG00000022637 | **Cblb** | AI429560\|AI851073\|Cbl-b | Casitas B-lineage lymphoma b | protein-coding |
| chr16_51766551_51767207 | chr16 | 51766552 | 51767207 | + | 1.07387 | Intergenic | MTD\|LTR\|ERVL-MaLR | -264670 | NM_001033238 | 208650 | Mm.328206 | NM_001033238 | ENSMUSG00000022637 | **Cblb** | AI429560\|AI851073\|Cbl-b | Casitas B-lineage lymphoma b | protein-coding |
| chr16_52067642_52068549 | chr16 | 52067643 | 52068549 | + | 1.04781 | intron (NM_001033238, intron 3 of 17) | intron (NM_001033238, intron 3 of 17) | 36547 | NM_001033238 | 208650 | Mm.328206 | NM_001033238 | ENSMUSG00000022637 | **Cblb** | AI429560\|AI851073\|Cbl-b | Casitas B-lineage lymphoma b | protein-coding |
| chr16_51683243_51683803 | chr16 | 51683244 | 51683803 | + | 1.03478 | Intergenic | Intergenic | -348026 | NM_001033238 | 208650 | Mm.328206 | NM_001033238 | ENSMUSG00000022637 | **Cblb** | AI429560\|AI851073\|Cbl-b | Casitas B-lineage lymphoma b | protein-coding |
| chr16_51909990_51910452 | chr16 | 51909991 | 51910452 | + | 1.00732 | Intergenic | Intergenic | -121328 | NM_001033238 | 208650 | Mm.328206 | NM_001033238 | ENSMUSG00000022637 | **Cblb** | AI429560\|AI851073\|Cbl-b | Casitas B-lineage lymphoma b | protein-coding |
| chr16_51650124_51650631 | chr16 | 51650125 | 51650631 | + | 0.970448 | Intergenic | Intergenic | -381171 | NM_001033238 | 208650 | Mm.328206 | NM_001033238 | ENSMUSG00000022637 | **Cblb** | AI429560\|AI851073\|Cbl-b | Casitas B-lineage lymphoma b | protein-coding |
| chr16_51766560_51766819 | chr16 | 51766561 | 51766819 | + | 0.861636 | Intergenic | Intergenic | -264859 | NM_001033238 | 208650 | Mm.328206 | NM_001033238 | ENSMUSG00000022637 | **Cblb** | AI429560\|AI851073\|Cbl-b | Casitas B-lineage lymphoma b | protein-coding |
| chr16_51766877_51767199 | chr16 | 51766878 | 51767199 | + | 0.858573 | Intergenic | MTD\|LTR\|ERVL-MaLR | -264511 | NM_001033238 | 208650 | Mm.328206 | NM_001033238 | ENSMUSG00000022637 | **Cblb** | AI429560\|AI851073\|Cbl-b | Casitas B-lineage lymphoma b | protein-coding |
| chr16_51750430_51750685 | chr16 | 51750431 | 51750685 | + | 0.852253 | Intergenic | MTE2b\|LTR\|ERVL-MaLR | -280991 | NM_001033238 | 208650 | Mm.328206 | NM_001033238 | ENSMUSG00000022637 | **Cblb** | AI429560\|AI851073\|Cbl-b | Casitas B-lineage lymphoma b | protein-coding |
| chr16_51496002_51496421 | chr16 | 51496003 | 51496421 | + | 0.822751 | Intergenic | ORR1E\|LTR\|ERVL-MaLR | -535337 | NM_001033238 | 208650 | Mm.328206 | NM_001033238 | ENSMUSG00000022637 | **Cblb** | AI429560\|AI851073\|Cbl-b | Casitas B-lineage lymphoma b | protein-coding |
| chr16_51594090_51594556 | chr16 | 51594091 | 51594556 | + | 0.800792 | Intergenic | Lx8b\|LINE\|L1 | -437226 | NM_001033238 | 208650 | Mm.328206 | NM_001033238 | ENSMUSG00000022637 | **Cblb** | AI429560\|AI851073\|Cbl-b | Casitas B-lineage lymphoma b | protein-coding |
| chr16_52179737_52180384 | chr16 | 52179738 | 52180384 | + | 0.752763 | intron (NM_001033238, intron 13 of 17) | intron (NM_001033238, intron 13 of 17) | 148512 | NM_001033238 | 208650 | Mm.328206 | NM_001033238 | ENSMUSG00000022637 | **Cblb** | AI429560\|AI851073\|Cbl-b | Casitas B-lineage lymphoma b | protein-coding |
| chr16_52179737_52180148 | chr16 | 52179738 | 52180148 | + | 0.729485 | intron (NM_001033238, intron 13 of 17) | intron (NM_001033238, intron 13 of 17) | 148394 | NM_001033238 | 208650 | Mm.328206 | NM_001033238 | ENSMUSG00000022637 | **Cblb** | AI429560\|AI851073\|Cbl-b | Casitas B-lineage lymphoma b | protein-coding |
| chr16_51918986_51919449 | chr16 | 51918987 | 51919449 | + | 0.728586 | Intergenic | Intergenic | -112331 | NM_001033238 | 208650 | Mm.328206 | NM_001033238 | ENSMUSG00000022637 | **Cblb** | AI429560\|AI851073\|Cbl-b | Casitas B-lineage lymphoma b | protein-coding |
| chr16_51686748_51686984 | chr16 | 51686749 | 51686984 | + | 0.706438 | Intergenic | Intergenic | -344683 | NM_001033238 | 208650 | Mm.328206 | NM_001033238 | ENSMUSG00000022637 | **Cblb** | AI429560\|AI851073\|Cbl-b | Casitas B-lineage lymphoma b | protein-coding |
| chr16_52066025_52066334 | chr16 | 52066026 | 52066334 | + | 0.630719 | intron (NM_001033238, intron 3 of 17) | intron (NM_001033238, intron 3 of 17) | 34631 | NM_001033238 | 208650 | Mm.328206 | NM_001033238 | ENSMUSG00000022637 | **Cblb** | AI429560\|AI851073\|Cbl-b | Casitas B-lineage lymphoma b | protein-coding |
| chr8_61475374_61475695 | chr8 | 61475375 | 61475695 | + | 1.31874 | Intergenic | RMER15\|LTR\|ERVL | -12199 | NM_145595 | 234309 | Mm.29059 | NM_145595 | ENSMUSG00000031641 | **Cbr4** | A730083J17Rik\|BC009118 | carbonyl reductase 4 | protein-coding |
| chr8_61475317_61475904 | chr8 | 61475318 | 61475904 | + | 1.30533 | Intergenic | Intergenic | -12123 | NM_145595 | 234309 | Mm.29059 | NM_145595 | ENSMUSG00000031641 | **Cbr4** | A730083J17Rik\|BC009118 | carbonyl reductase 4 | protein-coding |
| chr8_61440373_61440984 | chr8 | 61440374 | 61440984 | + | 1.07036 | Intergenic | Intergenic | -47055 | NM_145595 | 234309 | Mm.29059 | NM_145595 | ENSMUSG00000031641 | **Cbr4** | A730083J17Rik\|BC009118 | carbonyl reductase 4 | protein-coding |
| chr8_61461952_61462225 | chr8 | 61461953 | 61462225 | + | 0.859453 | Intergenic | RMER1A\|Other\|Other | -25645 | NM_145595 | 234309 | Mm.29059 | NM_145595 | ENSMUSG00000031641 | **Cbr4** | A730083J17Rik\|BC009118 | carbonyl reductase 4 | protein-coding |
| chr8_61517490_61518187 | chr8 | 61517491 | 61518187 | + | 0.747098 | intron (NM_001293772, intron 18 of 21) | intron (NM_001293772, intron 18 of 21) | 30105 | NM_145595 | 234309 | Mm.29059 | NM_145595 | ENSMUSG00000031641 | **Cbr4** | A730083J17Rik\|BC009118 | carbonyl reductase 4 | protein-coding |
| chr8_61440011_61440286 | chr8 | 61440012 | 61440286 | + | 0.673158 | Intergenic | Intergenic | -47585 | NM_145595 | 234309 | Mm.29059 | NM_145595 | ENSMUSG00000031641 | **Cbr4** | A730083J17Rik\|BC009118 | carbonyl reductase 4 | protein-coding |
| chr8_61425207_61427034 | chr8 | 61425208 | 61427034 | + | 0.659508 | Intergenic | Intergenic | -61613 | NM_145595 | 234309 | Mm.29059 | NM_145595 | ENSMUSG00000031641 | **Cbr4** | A730083J17Rik\|BC009118 | carbonyl reductase 4 | protein-coding |
| chr8_61476248_61476847 | chr8 | 61476249 | 61476847 | + | 0.630996 | Intergenic | Intergenic | -11186 | NM_145595 | 234309 | Mm.29059 | NM_145595 | ENSMUSG00000031641 | **Cbr4** | A730083J17Rik\|BC009118 | carbonyl reductase 4 | protein-coding |

| | | | | | | | | | | | | | | | | | |
|---|---|---|---|---|---|---|---|---|---|---|---|---|---|---|---|---|---|
| chr11_119109377_119109906 | chr11 | 119109378 | 119109906 | + | 1.06584 | Intergenic | Intergenic | -23405 | NM_007625 | 12418 | Mm.268070 | NM_007625 | ENSMUSG00000039989 | **Cbx4** | MPc2\|PC2 | chromobox 4 | protein-coding |
| chr15_79933731_79934419 | chr15 | 79933732 | 79934419 | + | 0.774575 | Intergenic | Intergenic | -1429 | NM_144811 | 52609 | Mm.323442 | NM_144811 | ENSMUSG00000053411 | **Cbx7** | 1600014J01Rik\|AI851678\|D15Ertd417e | chromobox 7 | protein-coding |
| chr15_79933736_79934126 | chr15 | 79933737 | 79934126 | + | 0.657676 | Intergenic | Intergenic | -1285 | NM_144811 | 52609 | Mm.323442 | NM_144811 | ENSMUSG00000053411 | **Cbx7** | 1600014J01Rik\|AI851678\|D15Ertd417e | chromobox 7 | protein-coding |
| chr15_79933441_79934421 | chr15 | 79933442 | 79934421 | + | 0.617548 | Intergenic | Intergenic | -1285 | NM_144811 | 52609 | Mm.323442 | NM_144811 | ENSMUSG00000053411 | **Cbx7** | 1600014J01Rik\|AI851678\|D15Ertd417e | chromobox 7 | protein-coding |
| chr3_142706632_142707185 | chr3 | 142706633 | 142707185 | + | 0.692652 | intron (NM_173763, intron 2 of 13) | intron (NM_173763, intron 2 of 13) | 5861 | NM_001293560 | 229905 | Mm.289643 | NM_173763 | ENSMUSG00000040213 | **Ccbl2** | KATIII\|Kat3 | cysteine conjugate-beta lyase 2 | protein-coding |
| chr3_129940795_129941147 | chr3 | 129940796 | 129941147 | + | 1.24441 | intron (NM_001293644, intron 1 of 6) | intron (NM_001293644, intron 1 of 6) | 29235 | NM_025779 | 66815 | Mm.31056 | NM_025779 | ENSMUSG00000027994 | **Ccdc109b** | 9030408N13Rik\|MCUb | coiled-coil domain containing 109B | protein-coding |
| chr3_129940804_129941322 | chr3 | 129940805 | 129941322 | + | 1.0125 | intron (NM_001293644, intron 1 of 6) | intron (NM_001293644, intron 1 of 6) | 29143 | NM_025779 | 66815 | Mm.31056 | NM_025779 | ENSMUSG00000027994 | **Ccdc109b** | 9030408N13Rik\|MCUb | coiled-coil domain containing 109B | protein-coding |
| chr18_74287714_74288683 | chr18 | 74287715 | 74288683 | + | 0.813356 | intron (NR_029424, intron 2 of 8) | intron (NR_029424, intron 2 of 8) | 5099 | NM_028948 | 74453 | Mm.182293 | NM_028948 | ENSMUSG00000035394 | **Ccdc11** | 4933415I03Rik | coiled-coil domain containing 11 | protein-coding |
| chr18_74285965_74286409 | chr18 | 74285966 | 74286409 | + | 0.690829 | intron (NR_029424, intron 2 of 8) | intron (NR_029424, intron 2 of 8) | 3087 | NM_028948 | 74453 | Mm.182293 | NM_028948 | ENSMUSG00000035394 | **Ccdc11** | 4933415I03Rik | coiled-coil domain containing 11 | protein-coding |
| chr18_46380756_46381516 | chr18 | 46380757 | 46381516 | + | 0.603984 | Intergenic | Intergenic | -69208 | NM_001160399 | 240261 | Mm.329416 | NM_001160399 | ENSMUSG00000071855 | **Ccdc112** | 8430438M01Rik\|AW108467 | coiled-coil domain containing 112 | protein-coding |
| chr1_181446739_181447846 | chr1 | 181446740 | 181447846 | + | 1.34149 | intron (NM_028408, intron 3 of 5) | intron (NM_028408, intron 3 of 5) | 64158 | NM_207280 | 403180 | Mm.220754 | NM_207280 | ENSMUSG00000050625 | **Ccdc121** | 6530421E24Rik\|Gm755 | coiled-coil domain containing 121 | protein-coding |
| chr1_181615976_181616496 | chr1 | 181615977 | 181616496 | + | 0.639412 | Intergenic | Intergenic | -104785 | NM_207280 | 403180 | Mm.220754 | NM_207280 | ENSMUSG00000050625 | **Ccdc121** | 6530421E24Rik\|Gm755 | coiled-coil domain containing 121 | protein-coding |
| chr2_77206922_77207371 | chr2 | 77206923 | 77207371 | + | 1.3564 | intron (NM_175465, intron 9 of 17) | intron (NM_175465, intron 9 of 17) | -36512 | NM_001025576 | 545428 | Mm.26579 | NM_001025576 | ENSMUSG00000044033 | **Ccdc141** | 2610301F02Rik\|CAMDI\|ENSMUSG00000075261 | coiled-coil domain containing 141 | protein-coding |
| chr2_77166309_77166717 | chr2 | 77166310 | 77166717 | + | 1.30196 | intron (NM_001025576, intron 2 of 23) | intron (NM_001025576, intron 2 of 23) | 4122 | NM_001025576 | 545428 | Mm.26579 | NM_001025576 | ENSMUSG00000044033 | **Ccdc141** | 2610301F02Rik\|CAMDI\|ENSMUSG00000075261 | coiled-coil domain containing 141 | protein-coding |
| chr2_77180652_77181078 | chr2 | 77180653 | 77181078 | + | 0.9093 | 3' UTR (NM_175465, exon 18 of 18) | 3' UTR (NM_175465, exon 18 of 18) | -10230 | NM_001025576 | 545428 | Mm.26579 | NM_001025576 | ENSMUSG00000044033 | **Ccdc141** | 2610301F02Rik\|CAMDI\|ENSMUSG00000075261 | coiled-coil domain containing 141 | protein-coding |
| chr2_77102344_77103040 | chr2 | 77102345 | 77103040 | + | 0.885378 | intron (NM_001025576, intron 6 of 23) | intron (NM_001025576, intron 6 of 23) | 67943 | NM_001025576 | 545428 | Mm.26579 | NM_001025576 | ENSMUSG00000044033 | **Ccdc141** | 2610301F02Rik\|CAMDI\|ENSMUSG00000075261 | coiled-coil domain containing 141 | protein-coding |
| chr2_77187314_77188268 | chr2 | 77187315 | 77188268 | + | 0.832622 | intron (NM_175465, intron 17 of 17) | intron (NM_175465, intron 17 of 17) | -17156 | NM_001025576 | 545428 | Mm.26579 | NM_001025576 | ENSMUSG00000044033 | **Ccdc141** | 2610301F02Rik\|CAMDI\|ENSMUSG00000075261 | coiled-coil domain containing 141 | protein-coding |
| chr2_59042363_59043138 | chr2 | 59042364 | 59043138 | + | 1.6215 | Intergenic | Intergenic | -24145 | NM_001001178 | 227933 | Mm.330234 | NM_177650 | ENSMUSG00000036641 | **Ccdc148** | 5830402J09 | coiled-coil domain containing 148 | protein-coding |
| chr2_59055677_59056340 | chr2 | 59055678 | 59056340 | + | 1.15492 | Intergenic | L1MA9\|LINE\|L1 | -37403 | NM_001001178 | 227933 | Mm.330234 | NM_177650 | ENSMUSG00000036641 | **Ccdc148** | 5830402J09 | coiled-coil domain containing 148 | protein-coding |
| chr2_59053446_59054000 | chr2 | 59053447 | 59054000 | + | 0.739254 | Intergenic | Intergenic | -35117 | NM_001001178 | 227933 | Mm.330234 | NM_177650 | ENSMUSG00000036641 | **Ccdc148** | 5830402J09 | coiled-coil domain containing 148 | protein-coding |
| chrX_52819373_52819772 | chrX | 52819374 | 52819772 | + | 0.697355 | Intergenic | Tigger5\|DNA\|TcMar-Tigger | 28373 | NM_001034059 | 434778 | Mm.23411 | NM_001034059 | ENSMUSG00000073207 | **Ccdc160** | 6330534C20Rik | coiled-coil domain containing 160 | protein-coding |
| chr4_83514540_83515032 | chr4 | 83514541 | 83515032 | + | 0.843989 | Intergenic | Intergenic | -10759 | NM_001081012 | 320226 | Mm.180032 | NM_029117 | ENSMUSG00000052407 | **Ccdc171** | 4930418J05Rik\|4930473A06Rik\|A330015D16Rik | coiled-coil domain containing 171 | protein-coding |
| chr4_83944483_83945120 | chr4 | 83944484 | 83945120 | + | 0.692146 | Intergenic | MIR3\|SINE\|MIR | 419257 | NM_001081012 | 320226 | Mm.180032 | NM_029117 | ENSMUSG00000052407 | **Ccdc171** | 4930418J05Rik\|4930473A06Rik\|A330015D16Rik | coiled-coil domain containing 171 | protein-coding |
| chr2_69775432_69776035 | chr2 | 69775433 | 69776035 | + | 1.02436 | intron (NM_001077684, intron 6 of 8) | intron (NM_001077684, intron 6 of 8) | 13752 | NM_001077684 | 75051 | Mm.195726 | NM_001077684 | ENSMUSG00000070883 | **Ccdc173** | 4930525K21Rik\|4930578N16Rik | coiled-coil domain containing 173 | protein-coding |
| chr5_108160483_108161288 | chr5 | 108160484 | 108161288 | + | 1.63471 | intron (NM_028481, intron 8 of 28) | intron (NM_028481, intron 8 of 28) | 27972 | NM_028481 | 73254 | Mm.441231 | NM_028481 | ENSMUSG00000056531 | **Ccdc18** | 1700021E15Rik\|4932411G06Rik\|NY-SAR-41 | coiled-coil domain containing 18 | protein-coding |
| chr5_108189600_108190046 | chr5 | 108189601 | 108190046 | + | 0.811365 | intron (NM_028481, intron 17 of 28) | ORR1D1\|LTR\|ERVL-MaLR | 56909 | NM_028481 | 73254 | Mm.441231 | NM_028481 | ENSMUSG00000056531 | **Ccdc18** | 1700021E15Rik\|4932411G06Rik\|NY-SAR-41 | coiled-coil domain containing 18 | protein-coding |
| chr15_98170343_98170766 | chr15 | 98170344 | 98170766 | + | 1.03908 | TTS (NM_177716) | TTS (NM_177716) | 2749 | NM_177716 | 239650 | Mm.54140 | NM_177716 | ENSMUSG00000029875 | **Ccdc184** | 5630400A08\|AI836003 | coiled-coil domain containing 184 | protein-coding |
| chr2_118999250_118999954 | chr2 | 118999251 | 118999954 | + | 1.50099 | Intergenic | Intergenic | 29791 | NM_199310 | 269336 | Mm.292465 | NM_199310 | ENSMUSG00000039983 | **Ccdc32** | Gm631 | coiled-coil domain containing 32 | protein-coding |
| chr2_118983799_118984281 | chr2 | 118983800 | 118984281 | + | 0.901248 | Intergenic | ORR1F\|LTR\|ERVL-MaLR | 45353 | NM_199310 | 269336 | Mm.292465 | NM_199310 | ENSMUSG00000039983 | **Ccdc32** | Gm631 | coiled-coil domain containing 32 | protein-coding |
| chr2_118995535_118995827 | chr2 | 118995536 | 118995827 | + | 0.728451 | Intergenic | Intergenic | 33712 | NM_199310 | 269336 | Mm.292465 | NM_199310 | ENSMUSG00000039983 | **Ccdc32** | Gm631 | coiled-coil domain containing 32 | protein-coding |
| chr2_118997968_118998461 | chr2 | 118997969 | 118998461 | + | 0.670752 | Intergenic | Intergenic | 31178 | NM_199310 | 269336 | Mm.292465 | NM_199310 | ENSMUSG00000039983 | **Ccdc32** | Gm631 | coiled-coil domain containing 32 | protein-coding |
| chr6_90428308_90428695 | chr6 | 90428309 | 90428695 | + | 0.619615 | promoter-TSS (NM_173775) | promoter-TSS (NM_173775) | -22 | NM_173775 | 243538 | Mm.386876 | NM_173775 | ENSMUSG00000048794 | **Ccdc37** | AW048498\|C030041G11Rik\|C230069K22Rik | coiled-coil domain containing 37 | protein-coding |
| chr16_27401573_27401968 | chr16 | 27401574 | 27401968 | + | 1.27874 | intron (NM_026202, intron 1 of 10) | intron (NM_026202, intron 1 of 10) | 12815 | NM_001289436 | 67501 | Mm.258985 | NM_026202 | ENSMUSG00000038127 | **Ccdc50** | C3orf6 | coiled-coil domain containing 50 | protein-coding |
| chr16_27399851_27400254 | chr16 | 27399852 | 27400254 | + | 1.24717 | intron (NM_026202, intron 1 of 10) | intron (NM_026202, intron 1 of 10) | 11097 | NM_001289436 | 67501 | Mm.258985 | NM_026202 | ENSMUSG00000038127 | **Ccdc50** | C3orf6 | coiled-coil domain containing 50 | protein-coding |
| chr16_27401071_27401968 | chr16 | 27401072 | 27401968 | + | 1.21873 | intron (NM_026202, intron 1 of 10) | intron (NM_026202, intron 1 of 10) | 12564 | NM_001289436 | 67501 | Mm.258985 | NM_026202 | ENSMUSG00000038127 | **Ccdc50** | C3orf6 | coiled-coil domain containing 50 | protein-coding |
| chr16_27407327_27407658 | chr16 | 27407328 | 27407658 | + | 0.835128 | intron (NM_001289436, intron 4 of 11) | intron (NM_001289436, intron 4 of 11) | 18537 | NM_001289436 | 67501 | Mm.258985 | NM_026202 | ENSMUSG00000038127 | **Ccdc50** | C3orf6 | coiled-coil domain containing 50 | protein-coding |
| chr16_27387722_27388568 | chr16 | 27387723 | 27388568 | + | 0.771549 | promoter-TSS (NM_001025615) | promoter-TSS (NM_001025615) | -811 | NM_001025615 | 67501 | Mm.258985 | NM_026202 | ENSMUSG00000038127 | **Ccdc50** | C3orf6 | coiled-coil domain containing 50 | protein-coding |
| chr10_88205991_88206236 | chr10 | 88205992 | 88206236 | + | 0.620899 | intron (NM_027487, intron 2 of 5) | intron (NM_027487, intron 2 of 5) | 5017 | NM_027487 | 67282 | Mm.208465 | NM_026070 | ENSMUSG00000020056 | **Ccdc53** | 2900091E11Rik\|5730495F03Rik | coiled-coil domain containing 53 | protein-coding |
| chr16_50596504_50597169 | chr16 | 50596505 | 50597169 | + | 0.804499 | intron (NR_040566, intron 3 of 4) | intron (NR_040566, intron 3 of 4) | -5683 | NM_027046 | 69339 | Mm.422869 | NM_027046 | ENSMUSG00000050685 | **Ccdc54** | 1700007N18Rik\|AI644412\|BB013989\|NYD-SP17 | coiled-coil domain containing 54 | protein-coding |
| chr11_120888555_120889341 | chr11 | 120888556 | 120889341 | + | 0.650062 | intron (NM_027745, intron 10 of 18) | intron (NM_027745, intron 10 of 18) | 43924 | NM_027745 | 71276 | Mm.275751 | NM_027745 | ENSMUSG00000048445 | **Ccdc57** | 4933434G05Rik | coiled-coil domain containing 57 | protein-coding |
| chr16_36097687_36098569 | chr16 | 36097688 | 36098569 | + | 1.56303 | Intergenic | RLTR1B\|LTR\|ERV1 | 16853 | NM_001159422 | 381045 | Mm.310434 | NM_198645 | ENSMUSG00000075229 | **Ccdc58** | A930007B11Rik\|AI413631 | coiled-coil domain containing 58 | protein-coding |
| chr10_70116319_70116821 | chr10 | 70116320 | 70116821 | + | 0.86447 | intron (NM_001111121, intron 1 of 8) | Lx7\|LINE\|L1 | 19449 | NM_001111121 | 76551 | Mm.281741 | NM_001111121 | ENSMUSG00000048701 | **Ccdc6** | 2810012H18Rik\|AA536681\|AA960498\|AW061011 | coiled-coil domain containing 6 | protein-coding |
| chr10_70112431_70112714 | chr10 | 70112432 | 70112714 | + | 0.61493 | intron (NM_001111121, intron 1 of 8) | intron (NM_001111121, intron 1 of 8) | 15452 | NM_001111121 | 76551 | Mm.281741 | NM_001111121 | ENSMUSG00000048701 | **Ccdc6** | 2810012H18Rik\|AA536681\|AA960498\|AW061011 | coiled-coil domain containing 6 | protein-coding |
| chr5_115702000_115702409 | chr5 | 115702001 | 115702409 | + | 1.13923 | intron (NM_001080808, intron 2 of 8) | intron (NM_001080808, intron 2 of 8) | 29354 | NM_001080808 | 75665 | Mm.233914 | NM_001080808 | ENSMUSG00000041609 | **Ccdc64** | 2210403N09Rik\|BICDR-1 | coiled-coil domain containing 64 | protein-coding |
| chr5_115714696_115714938 | chr5 | 115714697 | 115714938 | + | 0.9589 | intron (NM_001080808, intron 2 of 8) | intron (NM_001080808, intron 2 of 8) | 16742 | NM_001080808 | 75665 | Mm.233914 | NM_001080808 | ENSMUSG00000041609 | **Ccdc64** | 2210403N09Rik\|BICDR-1 | coiled-coil domain containing 64 | protein-coding |
| chr5_115710803_115711056 | chr5 | 115710804 | 115711056 | + | 0.829409 | intron (NM_001080808, intron 2 of 8) | intron (NM_001080808, intron 2 of 8) | 20629 | NM_001080808 | 75665 | Mm.233914 | NM_001080808 | ENSMUSG00000041609 | **Ccdc64** | 2210403N09Rik\|BICDR-1 | coiled-coil domain containing 64 | protein-coding |
| chr17_23661569_23661871 | chr17 | 23661570 | 23661871 | + | 0.715076 | intron (NM_153784, intron 1 of 9) | intron (NM_153784, intron 1 of 9) | 1197 | NM_153784 | 212733 | Mm.59997 | NM_153784 | ENSMUSG00000043782 | **Ccdc64b** | BICDR-2 | coiled-coil domain containing 64B | protein-coding |
| chr14_27521158_27521508 | chr14 | 27521159 | 27521508 | + | 0.855653 | Intergenic | RLTR19A\|LTR\|ERVK | -12873 | NM_177111 | 320234 | Mm.247767 | NM_177111 | ENSMUSG00000046753 | **Ccdc66** | E230015L20Rik | coiled-coil domain containing 66 | protein-coding |
| chr14_27521097_27521596 | chr14 | 27521098 | 27521596 | + | 0.843405 | Intergenic | RLTR19A\|LTR\|ERVK | -12887 | NM_177111 | 320234 | Mm.247767 | NM_177111 | ENSMUSG00000046753 | **Ccdc66** | E230015L20Rik | coiled-coil domain containing 66 | protein-coding |
| chr18_69644441_69645204 | chr18 | 69644442 | 69645204 | + | 0.837648 | intron (NM_001083967, intron 11 of 19) | intron (NM_001083967, intron 11 of 19) | -280736 | NM_201362 | 381175 | Mm.266831 | NM_201362 | ENSMUSG00000038903 | **Ccdc68** | BC046343 | coiled-coil domain containing 68 | protein-coding |
| chr18_69642948_69643600 | chr18 | 69642949 | 69643600 | + | 0.762609 | intron (NM_001083967, intron 11 of 19) | intron (NM_001083967, intron 11 of 19) | -282285 | NM_201362 | 381175 | Mm.266831 | NM_201362 | ENSMUSG00000038903 | **Ccdc68** | BC046343 | coiled-coil domain containing 68 | protein-coding |

| | | | | | | | | | | | | | | | | | |
|---|---|---|---|---|---|---|---|---|---|---|---|---|---|---|---|---|---|
| chr12_32343499_32343881 | chr12 | 32343500 | 32343881 | + | 1.84589 | Intergenic | MERVL_2A-int\|LTR\|ERVL | -35099 | NM_001162903 | 72123 | Mm.341352 | NM_028111 | ENSMUSG00000090946 | **Ccdc71l** | 2010109K11Rik\|AV375874 | coiled-coil domain containing 71 like | protein-coding |
| chr12_32336777_32337211 | chr12 | 32336778 | 32337211 | + | 1.18582 | Intergenic | Intergenic | -41795 | NM_001162903 | 72123 | Mm.341352 | NM_028111 | ENSMUSG00000090946 | **Ccdc71l** | 2010109K11Rik\|AV375874 | coiled-coil domain containing 71 like | protein-coding |
| chr12_32325273_32325885 | chr12 | 32325274 | 32325885 | + | 1.12458 | Intergenic | L1MC1\|LINE\|L1 | -53210 | NM_001162903 | 72123 | Mm.341352 | NM_028111 | ENSMUSG00000090946 | **Ccdc71l** | 2010109K11Rik\|AV375874 | coiled-coil domain containing 71 like | protein-coding |
| chr16_45124249_45125423 | chr16 | 45124250 | 45125423 | + | 1.94267 | intron (NM_026439, intron 6 of 7) | intron (NM_026439, intron 6 of 7) | 30783 | NM_026439 | 67896 | Mm.181074 | NM_026439 | ENSMUSG00000022665 | **Ccdc80** | 2610001E17Rik\|DRO1\|Ssg1\|Urb | coiled-coil domain containing 80 | protein-coding |
| chr16_45038590_45039050 | chr16 | 45038591 | 45039050 | + | 1.51786 | Intergenic | Intergenic | -55233 | NM_026439 | 67896 | Mm.181074 | NM_026439 | ENSMUSG00000022665 | **Ccdc80** | 2610001E17Rik\|DRO1\|Ssg1\|Urb | coiled-coil domain containing 80 | protein-coding |
| chr16_45038390_45039051 | chr16 | 45038391 | 45039051 | + | 1.46744 | Intergenic | MLT2B2\|LTR\|ERVL | -55332 | NM_026439 | 67896 | Mm.181074 | NM_026439 | ENSMUSG00000022665 | **Ccdc80** | 2610001E17Rik\|DRO1\|Ssg1\|Urb | coiled-coil domain containing 80 | protein-coding |
| chr16_45093159_45093730 | chr16 | 45093160 | 45093730 | + | 0.751703 | promoter-TSS (NM_026439) | promoter-TSS (NM_026439) | -608 | NM_026439 | 67896 | Mm.181074 | NM_026439 | ENSMUSG00000022665 | **Ccdc80** | 2610001E17Rik\|DRO1\|Ssg1\|Urb | coiled-coil domain containing 80 | protein-coding |
| chr9_13287434_13287917 | chr9 | 13287435 | 13287917 | + | 1.45 | intron (NM_025534, intron 8 of 8) | intron (NM_025534, intron 8 of 8) | 40697 | NM_025534 | 66396 | Mm.290620 | NM_025534 | | **Ccdc82** | 2310043N13Rik | coiled-coil domain containing 82 | protein-coding |
| chr9_13304528_13304860 | chr9 | 13304529 | 13304860 | + | 1.13432 | Intergenic | Intergenic | 57715 | NM_025534 | 66396 | Mm.290620 | NM_025534 | | **Ccdc82** | 2310043N13Rik | coiled-coil domain containing 82 | protein-coding |
| chr9_13320805_13321312 | chr9 | 13320806 | 13321312 | + | 0.82707 | Intergenic | Intergenic | 74080 | NM_025534 | 66396 | Mm.290620 | NM_025534 | | **Ccdc82** | 2310043N13Rik | coiled-coil domain containing 82 | protein-coding |
| chr9_13320791_13321514 | chr9 | 13320792 | 13321514 | + | 0.719858 | Intergenic | Intergenic | 74174 | NM_025534 | 66396 | Mm.290620 | NM_025534 | | **Ccdc82** | 2310043N13Rik | coiled-coil domain containing 82 | protein-coding |
| chr7_90218164_90218453 | chr7 | 90218165 | 90218453 | + | 0.662558 | Intergenic | Intergenic | 47123 | NM_029256 | 75338 | Mm.487222 | NM_029256 | ENSMUSG00000030617 | **Ccdc83** | 4930549K11Rik\|4930554C01Rik\|4932423M01Rik | coiled-coil domain containing 83 | protein-coding |
| chr7_90214657_90214948 | chr7 | 90214658 | 90214948 | + | 0.647401 | Intergenic | Intergenic | 50629 | NM_029256 | 75338 | Mm.487222 | NM_029256 | ENSMUSG00000030617 | **Ccdc83** | 4930549K11Rik\|4930554C01Rik\|4932423M01Rik | coiled-coil domain containing 83 | protein-coding |
| chr7_25707532_25708233 | chr7 | 25707533 | 25708233 | + | 0.762665 | Intergenic | Intergenic | 11170 | NM_028771 | 52132 | Mm.278798 | NM_028771 | ENSMUSG00000002608 | **Ccdc97** | 1200014H14Rik\|2810446P04Rik\|AA545218\|D7Ertd462e | coiled-coil domain containing 97 | protein-coding |
| chr7_25707501_25708532 | chr7 | 25707502 | 25708532 | + | 0.703818 | Intergenic | Intergenic | 11036 | NM_028771 | 52132 | Mm.278798 | NM_028771 | ENSMUSG00000002608 | **Ccdc97** | 1200014H14Rik\|2810446P04Rik\|AA545218\|D7Ertd462e | coiled-coil domain containing 97 | protein-coding |
| chr11_82194936_82195453 | chr11 | 82194937 | 82195453 | + | 1.07954 | Intergenic | Intergenic | -15383 | NM_011329 | 20290 | Mm.1283 | NM_011329 | ENSMUSG00000020702 | **Ccl1** | BF534335\|I-309\|Scya1\|Tca-3 | chemokine (C-C motif) ligand 1 | protein-coding |
| chr8_94802200_94802732 | chr8 | 94802201 | 94802732 | + | 0.847106 | Intergenic | ORR1E\|LTR\|ERVL-MaLR | -7987 | NM_011332 | 20295 | Mm.41988 | NM_011332 | ENSMUSG00000031780 | **Ccl17** | Abcd-2\|Scya17\|Scya17l\|Tarc | chemokine (C-C motif) ligand 17 | protein-coding |
| chr11_83646544_83646883 | chr11 | 83646545 | 83646883 | + | 0.862389 | Intergenic | Intergenic | 2664 | NM_011337 | 20302 | Mm.1282 | NM_011337 | ENSMUSG00000000982 | **Ccl3** | AI323804\|G0S19-1\|LD78alpha\|MIP-1alpha\|MIP1-(a)\|MIP1-alpha\|Mip1a\|Scya3 | chemokine (C-C motif) ligand 3 | protein-coding |
| chr11_83648850_83649926 | chr11 | 83648851 | 83649926 | + | 0.828252 | promoter-TSS (NM_011337) | promoter-TSS (NM_011337) | -10 | NM_011337 | 20302 | Mm.1282 | NM_011337 | ENSMUSG00000000982 | **Ccl3** | AI323804\|G0S19-1\|LD78alpha\|MIP-1alpha\|MIP1-(a)\|MIP1-alpha\|Mip1a\|Scya3 | chemokine (C-C motif) ligand 3 | protein-coding |
| chr11_83528010_83528328 | chr11 | 83528011 | 83528328 | + | 0.922351 | intron (NM_013653, intron 2 of 2) | intron (NM_013653, intron 2 of 2) | 2349 | NM_013653 | 20304 | Mm.284248 | NM_013653 | ENSMUSG00000035042 | **Ccl5** | MuRantes\|RANTES\|SISd\|Scya5\|TCP228 | chemokine (C-C motif) ligand 5 | protein-coding |
| chr11_83527128_83527455 | chr11 | 83527129 | 83527455 | + | 0.915877 | intron (NM_013653, intron 2 of 2) | intron (NM_013653, intron 2 of 2) | 3226 | NM_013653 | 20304 | Mm.284248 | NM_013653 | ENSMUSG00000035042 | **Ccl5** | MuRantes\|RANTES\|SISd\|Scya5\|TCP228 | chemokine (C-C motif) ligand 5 | protein-coding |
| chr11_83535062_83535422 | chr11 | 83535063 | 83535422 | + | 0.687776 | Intergenic | Intergenic | -4724 | NM_013653 | 20304 | Mm.284248 | NM_013653 | ENSMUSG00000035042 | **Ccl5** | MuRantes\|RANTES\|SISd\|Scya5\|TCP228 | chemokine (C-C motif) ligand 5 | protein-coding |
| chr13_100790093_100790662 | chr13 | 100790094 | 100790662 | + | 1.01446 | Intergenic | Intergenic | -3892 | NM_172301 | 268697 | Mm.260114 | NM_172301 | ENSMUSG00000041431 | **Ccnb1** | Ccnb1-rs1\|Ccnb1-rs13\|CycB1\|Cycb-4\|Cycb-5\|Cycb1-rs1 | cyclin B1 | protein-coding |
| chr6_127207323_127207832 | chr6 | 127207324 | 127207832 | + | 0.939943 | intron (NR_040273, intron 2 of 2) | intron (NR_040273, intron 2 of 2) | -56530 | NM_009829 | 12444 | Mm.333406 | NM_009829 | ENSMUSG00000000184 | **Ccnd2** | 2600016F06Rik\|AI256817\|BF642806\|C86853\|Vin-1\|Vin1\|cD2 | cyclin D2 | protein-coding |
| chr6_127219181_127219768 | chr6 | 127219182 | 127219768 | + | 0.915979 | Intergenic | Intergenic | -68427 | NM_009829 | 12444 | Mm.333406 | NM_009829 | ENSMUSG00000000184 | **Ccnd2** | 2600016F06Rik\|AI256817\|BF642806\|C86853\|Vin-1\|Vin1\|cD2 | cyclin D2 | protein-coding |
| chr6_127157296_127157868 | chr6 | 127157297 | 127157868 | + | 0.885412 | intron (NR_040275, intron 1 of 1) | intron (NR_040275, intron 1 of 1) | -6534 | NM_009829 | 12444 | Mm.333406 | NM_009829 | ENSMUSG00000000184 | **Ccnd2** | 2600016F06Rik\|AI256817\|BF642806\|C86853\|Vin-1\|Vin1\|cD2 | cyclin D2 | protein-coding |
| chr17_47515237_47516117 | chr17 | 47515238 | 47516117 | + | 1.25805 | intron (NM_001081636, intron 1 of 6) | intron (NM_001081636, intron 1 of 6) | 10626 | NM_001081635 | 12445 | Mm.27291 | NM_007632 | ENSMUSG00000034165 | **Ccnd3** | 9230106B05Rik\|AA682053\|AL024085\|AW146355\|C78795 | cyclin D3 | protein-coding |
| chr17_47554514_47554999 | chr17 | 47554515 | 47554999 | + | 1.21534 | intron (NM_001081636, intron 1 of 6) | intron (NM_001081636, intron 1 of 6) | -38710 | NM_007632 | 12445 | Mm.27291 | NM_007632 | ENSMUSG00000034165 | **Ccnd3** | 9230106B05Rik\|AA682053\|AL024085\|AW146355\|C78795 | cyclin D3 | protein-coding |
| chr17_47554309_47554997 | chr17 | 47554310 | 47554997 | + | 1.17202 | intron (NM_001081636, intron 1 of 6) | intron (NM_001081636, intron 1 of 6) | -38814 | NM_007632 | 12445 | Mm.27291 | NM_007632 | ENSMUSG00000034165 | **Ccnd3** | 9230106B05Rik\|AA682053\|AL024085\|AW146355\|C78795 | cyclin D3 | protein-coding |
| chr17_47564529_47565073 | chr17 | 47564530 | 47565073 | + | 0.760927 | intron (NM_001081636, intron 1 of 6) | Arthur1\|DNA\|hAT-Tip100 | -28666 | NM_007632 | 12445 | Mm.27291 | NM_007632 | ENSMUSG00000034165 | **Ccnd3** | 9230106B05Rik\|AA682053\|AL024085\|AW146355\|C78795 | cyclin D3 | protein-coding |
| chr7_38077119_38077777 | chr7 | 38077120 | 38077777 | + | 1.87807 | Intergenic | Intergenic | 30042 | NM_007633 | 12447 | Mm.16110 | NM_007633 | ENSMUSG00000002068 | **Ccne1** | AW538188\|CycE1 | cyclin E1 | protein-coding |
| chr7_38071009_38071696 | chr7 | 38071010 | 38071696 | + | 1.20746 | Intergenic | Intergenic | 36137 | NM_007633 | 12447 | Mm.16110 | NM_007633 | ENSMUSG00000002068 | **Ccne1** | AW538188\|CycE1 | cyclin E1 | protein-coding |
| chr7_38070666_38071728 | chr7 | 38070667 | 38071728 | + | 1.14398 | Intergenic | Intergenic | 36293 | NM_007633 | 12447 | Mm.16110 | NM_007633 | ENSMUSG00000002068 | **Ccne1** | AW538188\|CycE1 | cyclin E1 | protein-coding |
| chr7_38088492_38088816 | chr7 | 38088493 | 38088816 | + | 0.791645 | Intergenic | PB1D10\|SINE\|Alu | 18836 | NM_007633 | 12447 | Mm.16110 | NM_007633 | ENSMUSG00000002068 | **Ccne1** | AW538188\|CycE1 | cyclin E1 | protein-coding |
| chr18_9393754_9394613 | chr18 | 9393755 | 9394613 | + | 1.48466 | intron (NM_026484, intron 1 of 9) | intron (NM_026484, intron 1 of 9) | 55966 | NM_026484 | 67974 | Mm.86523 | NM_026484 | ENSMUSG00000024286 | **Ccny** | 1700025H17Rik\|3110050L10Rik\|4631402G10Rik\|5730405I09Rik | cyclin Y | protein-coding |
| chr18_9408923_9409756 | chr18 | 9408924 | 9409756 | + | 1.47542 | intron (NM_026484, intron 1 of 9) | PB1D7\|SINE\|Alu | 40810 | NM_026484 | 67974 | Mm.86523 | NM_026484 | ENSMUSG00000024286 | **Ccny** | 1700025H17Rik\|3110050L10Rik\|4631402G10Rik\|5730405I09Rik | cyclin Y | protein-coding |
| chr9_123936527_123937015 | chr9 | 123936528 | 123937015 | + | 1.34878 | Intergenic | Intergenic | 31921 | NM_009912 | 12768 | Mm.274927 | NM_009912 | ENSMUSG00000025804 | **Ccr1** | Cmkbr1\|Mip-1a-R | chemokine (C-C motif) receptor 1 | protein-coding |
| chr9_124099082_124099579 | chr9 | 124099083 | 124099579 | + | 0.811965 | Intergenic | Intergenic | -2852 | NM_009915 | 12772 | Mm.6272 | NM_009915 | ENSMUSG00000049103 | **Ccr2** | Cc-ckr-2\|Ccr2a\|Ccr2b\|Ckr2\|Ckr2a\|Ckr2b\|Cmkbr2\|mJe-r | chemokine (C-C motif) receptor 2 | protein-coding |
| chr9_124028309_124028994 | chr9 | 124028310 | 124028994 | + | 0.811006 | exon (NM_009914, exon 2 of 2) | exon (NM_009914, exon 2 of 2) | 6681 | NM_009914 | 12771 | Mm.57050 | NM_009914 | ENSMUSG00000035448 | **Ccr3** | CC-CKR3\|CKR3\|Cmkbr1l2\|Cmkbr3 | chemokine (C-C motif) receptor 3 | protein-coding |
| chr9_124050965_124051411 | chr9 | 124050966 | 124051411 | + | 0.784624 | Intergenic | Intergenic | 29217 | NM_009914 | 12771 | Mm.57050 | NM_009914 | ENSMUSG00000035448 | **Ccr3** | CC-CKR3\|CKR3\|Cmkbr1l2\|Cmkbr3 | chemokine (C-C motif) receptor 3 | protein-coding |
| chr9_124028568_124028928 | chr9 | 124028569 | 124028928 | + | 0.680691 | exon (NM_009914, exon 2 of 2) | exon (NM_009914, exon 2 of 2) | 6777 | NM_009914 | 12771 | Mm.57050 | NM_009914 | ENSMUSG00000035448 | **Ccr3** | CC-CKR3\|CKR3\|Cmkbr1l2\|Cmkbr3 | chemokine (C-C motif) receptor 3 | protein-coding |
| chr9_114487383_114488109 | chr9 | 114487384 | 114488109 | + | 1.57755 | Intergenic | Intergenic | 8798 | NM_009916 | 12773 | Mm.1337 | NM_009916 | ENSMUSG00000047898 | **Ccr4** | C-C CKR-4\|CHEMR1\|Cmkbr4\|LESTR\|Sdf1r | chemokine (C-C motif) receptor 4 | protein-coding |
| chr9_114487351_114488780 | chr9 | 114487352 | 114488780 | + | 1.54732 | Intergenic | LTR16A\|LTR\|ERVL | 8478 | NM_009916 | 12773 | Mm.1337 | NM_009916 | ENSMUSG00000047898 | **Ccr4** | C-C CKR-4\|CHEMR1\|Cmkbr4\|LESTR\|Sdf1r | chemokine (C-C motif) receptor 4 | protein-coding |
| chr9_114488332_114488757 | chr9 | 114488333 | 114488757 | + | 1.23265 | Intergenic | Intergenic | 7999 | NM_009916 | 12773 | Mm.1337 | NM_009916 | ENSMUSG00000047898 | **Ccr4** | C-C CKR-4\|CHEMR1\|Cmkbr4\|LESTR\|Sdf1r | chemokine (C-C motif) receptor 4 | protein-coding |
| chr9_114488171_114488772 | chr9 | 114488172 | 114488772 | + | 1.227 | Intergenic | Intergenic | 8072 | NM_009916 | 12773 | Mm.1337 | NM_009916 | ENSMUSG00000047898 | **Ccr4** | C-C CKR-4\|CHEMR1\|Cmkbr4\|LESTR\|Sdf1r | chemokine (C-C motif) receptor 4 | protein-coding |
| chr9_114468227_114469012 | chr9 | 114468228 | 114469012 | + | 0.802543 | intron (NR_108101, intron 14 of 14) | intron (NR_108101, intron 14 of 14) | 27924 | NM_009916 | 12773 | Mm.1337 | NM_009916 | ENSMUSG00000047898 | **Ccr4** | C-C CKR-4\|CHEMR1\|Cmkbr4\|LESTR\|Sdf1r | chemokine (C-C motif) receptor 4 | protein-coding |
| chr9_114480271_114480583 | chr9 | 114480272 | 114480583 | + | 0.739048 | Intergenic | Intergenic | 16117 | NM_009916 | 12773 | Mm.1337 | NM_009916 | ENSMUSG00000047898 | **Ccr4** | C-C CKR-4\|CHEMR1\|Cmkbr4\|LESTR\|Sdf1r | chemokine (C-C motif) receptor 4 | protein-coding |
| chr9_114471833_114472429 | chr9 | 114471834 | 114472429 | + | 0.726348 | intron (NR_108101, intron 14 of 14) | intron (NR_108101, intron 14 of 14) | 24413 | NM_009916 | 12773 | Mm.1337 | NM_009916 | ENSMUSG00000047898 | **Ccr4** | C-C CKR-4\|CHEMR1\|Cmkbr4\|LESTR\|Sdf1r | chemokine (C-C motif) receptor 4 | protein-coding |
| chr9_114476358_114476998 | chr9 | 114476359 | 114476998 | + | 0.709307 | Intergenic | Intergenic | 19866 | NM_009916 | 12773 | Mm.1337 | NM_009916 | ENSMUSG00000047898 | **Ccr4** | C-C CKR-4\|CHEMR1\|Cmkbr4\|LESTR\|Sdf1r | chemokine (C-C motif) receptor 4 | protein-coding |
| chr9_124145519_124146961 | chr9 | 124145520 | 124146961 | + | 1.29315 | Intergenic | Intergenic | 24697 | NM_009917 | 12774 | Mm.14302 | NM_009917 | ENSMUSG00000079227 | **Ccr5** | AM4-7\|CD195\|Cmkbr5 | chemokine (C-C motif) receptor 5 | protein-coding |
| chr9_124146321_124146693 | chr9 | 124146322 | 124146693 | + | 1.26801 | Intergenic | RMER15\|LTR\|ERVL | 24964 | NM_009917 | 12774 | Mm.14302 | NM_009917 | ENSMUSG00000079227 | **Ccr5** | AM4-7\|CD195\|Cmkbr5 | chemokine (C-C motif) receptor 5 | protein-coding |
| chr9_124145564_124146276 | chr9 | 124145565 | 124146276 | + | 1.00055 | Intergenic | Intergenic | 24377 | NM_009917 | 12774 | Mm.14302 | NM_009917 | ENSMUSG00000079227 | **Ccr5** | AM4-7\|CD195\|Cmkbr5 | chemokine (C-C motif) receptor 5 | protein-coding |
| chr17_8251703_8252120 | chr17 | 8251704 | 8252120 | + | 1.01297 | intron (NM_001190336, intron 2 of 2) | intron (NM_001190336, intron 2 of 2) | 6854 | NM_001190338 | 12458 | Mm.8007 | NM_009835 | ENSMUSG00000040899 | **Ccr6** | CC-CKR-6\|CCR-6\|Cmkbr6\|KY411 | chemokine (C-C motif) receptor 6 | protein-coding |
| chr17_8234506_8234964 | chr17 | 8234507 | 8234964 | + | 0.843494 | Intergenic | Intergenic | -1308 | NM_001190333 | 12458 | Mm.8007 | NM_009835 | ENSMUSG00000040899 | **Ccr6** | CC-CKR-6\|CCR-6\|Cmkbr6\|KY411 | chemokine (C-C motif) receptor 6 | protein-coding |
| chr17_8234420_8235137 | chr17 | 8234421 | 8235137 | + | 0.773468 | Intergenic | Intergenic | -1264 | NM_001190333 | 12458 | Mm.8007 | NM_009835 | ENSMUSG00000040899 | **Ccr6** | CC-CKR-6\|CCR-6\|Cmkbr6\|KY411 | chemokine (C-C motif) receptor 6 | protein-coding |
| chr11_99160661_99161246 | chr11 | 99160662 | 99161246 | + | 1.68298 | Intergenic | Intergenic | -5877 | NM_007719 | 12775 | Mm.2932 | NM_007719 | ENSMUSG00000037944 | **Ccr7** | CD197\|Cdw197\|Cmkbr7\|EBI1\|Ebi1h | chemokine (C-C motif) receptor 7 | protein-coding |

| | | | | | | | | | | | | | | | | | |
|---|---|---|---|---|---|---|---|---|---|---|---|---|---|---|---|---|---|
| chr11_99180054_99180723 | chr11 | 99180055 | 99180723 | + | 1.28808 | Intergenic | Intergenic | -25312 | NM_007719 | 12775 | Mm.2932 | NM_007719 | ENSMUSG00000037944 | **Ccr7** | CD197\|Cdw197\|Cmkbr7\|EBI1\|Ebi1h | chemokine (C-C motif) receptor 7 | protein-coding |
| chr11_99129367_99129660 | chr11 | 99129368 | 99129660 | + | 0.732506 | Intergenic | Intergenic | 25563 | NM_007719 | 12775 | Mm.2932 | NM_007719 | ENSMUSG00000037944 | **Ccr7** | CD197\|Cdw197\|Cmkbr7\|EBI1\|Ebi1h | chemokine (C-C motif) receptor 7 | protein-coding |
| chr9_123765954_123766316 | chr9 | 123765955 | 123766316 | + | 0.960555 | Intergenic | Intergenic | -1076 | NM_009913 | 12769 | Mm.440604 | NM_009913 | | **Ccr9** | A130091K22Rik\|Cmkbr10\|GPR-9-6 | chemokine (C-C motif) receptor 9 | protein-coding |
| chr9_123768440_123768924 | chr9 | 123768441 | 123768924 | + | 0.78369 | intron (NM_009913, intron 1 of 2) | intron (NM_009913, intron 1 of 2) | 1471 | NM_001166625 | 12769 | Mm.440604 | NM_009913 | | **Ccr9** | A130091K22Rik\|Cmkbr10\|GPR-9-6 | chemokine (C-C motif) receptor 9 | protein-coding |
| chr9_123762901_123763275 | chr9 | 123762902 | 123763275 | + | 0.68646 | Intergenic | Intergenic | -4123 | NM_009913 | 12769 | Mm.440604 | NM_009913 | | **Ccr9** | A130091K22Rik\|Cmkbr10\|GPR-9-6 | chemokine (C-C motif) receptor 9 | protein-coding |
| chr9_111086876_111087490 | chr9 | 111086877 | 111087490 | + | 0.695904 | Intergenic | Intergenic | -29913 | NM_017466 | 54199 | Mm.7336 | NM_017466 | ENSMUSG00000043953 | **Ccrl2** | 1810047I05Rik\|Ackr5\|CCR11\|Cmkbr1l2\|E01\|L-CCR | chemokine (C-C motif) receptor-like 2 | protein-coding |
| chr9_111086166_111086681 | chr9 | 111086167 | 111086681 | + | 0.621897 | Intergenic | Intergenic | -29154 | NM_017466 | 54199 | Mm.7336 | NM_017466 | ENSMUSG00000043953 | **Ccrl2** | 1810047I05Rik\|Ackr5\|CCR11\|Cmkbr1l2\|E01\|L-CCR | chemokine (C-C motif) receptor-like 2 | protein-coding |
| chr3_51195174_51195754 | chr3 | 51195175 | 51195754 | + | 1.60401 | Intergenic | Intergenic | -28983 | NM_009834 | 12457 | Mm.86541 | NM_009834 | ENSMUSG00000023087 | **Ccrn4l** | AU043840\|Ccr4\|nocturnin | CCR4 carbon catabolite repression 4-like (S. cerevisiae) | protein-coding |
| chr6_61065753_61066327 | chr6 | 61065754 | 61066327 | + | 1.57811 | Intergenic | Intergenic | -114285 | NM_001164316 | 232035 | Mm.150324 | NM_183310 | ENSMUSG00000039578 | **Ccser1** | 6230405M12Rik\|C130092O11Rik\|Fam190a | coiled-coil serine rich 1 | protein-coding |
| chr6_61065753_61066560 | chr6 | 61065754 | 61066560 | + | 1.53866 | Intergenic | Intergenic | -114168 | NM_001164316 | 232035 | Mm.150324 | NM_183310 | ENSMUSG00000039578 | **Ccser1** | 6230405M12Rik\|C130092O11Rik\|Fam190a | coiled-coil serine rich 1 | protein-coding |
| chr6_61065441_61065707 | chr6 | 61065442 | 61065707 | + | 0.971439 | Intergenic | Intergenic | -114751 | NM_001164316 | 232035 | Mm.150324 | NM_183310 | ENSMUSG00000039578 | **Ccser1** | 6230405M12Rik\|C130092O11Rik\|Fam190a | coiled-coil serine rich 1 | protein-coding |
| chr6_61068150_61068404 | chr6 | 61068151 | 61068404 | + | 0.636224 | Intergenic | RLTR14\|LTR\|ERV1 | -112048 | NM_001164316 | 232035 | Mm.150324 | NM_183310 | ENSMUSG00000039578 | **Ccser1** | 6230405M12Rik\|C130092O11Rik\|Fam190a | coiled-coil serine rich 1 | protein-coding |
| chr14_36956754_36956991 | chr14 | 36956755 | 36956991 | + | 0.684765 | intron (NM_027045, intron 1 of 10) | intron (NM_027045, intron 1 of 10) | 11891 | NM_027045 | 72972 | Mm.27621 | NM_027045 | ENSMUSG00000058690 | **Ccser2** | 1700012P13Rik\|2900054P12Rik\|AI035535\|AW108503\|Fam190b\|Gcap14 | coiled-coil serine rich 2 | protein-coding |
| chr10_117103911_117104544 | chr10 | 117103912 | 117104544 | + | 0.777371 | intron (NM_177798, intron 1 of 9) | RMER15\|LTR\|ERVL | -40414 | NM_007636 | 12461 | Mm.247788 | NM_007636 | ENSMUSG00000034024 | **Cct2** | Cctb | chaperonin containing Tcp1, subunit 2 (beta) | protein-coding |
| chr10_117094960_117095713 | chr10 | 117094961 | 117095713 | + | 0.73614 | intron (NM_177798, intron 4 of 9) | intron (NM_177798, intron 4 of 9) | -31523 | NM_007636 | 12461 | Mm.247788 | NM_007636 | ENSMUSG00000034024 | **Cct2** | Cctb | chaperonin containing Tcp1, subunit 2 (beta) | protein-coding |
| chr10_117105053_117105640 | chr10 | 117105054 | 117105640 | + | 0.66768 | intron (NM_177798, intron 1 of 9) | intron (NM_177798, intron 1 of 9) | -41533 | NM_007636 | 12461 | Mm.247788 | NM_007636 | ENSMUSG00000034024 | **Cct2** | Cctb | chaperonin containing Tcp1, subunit 2 (beta) | protein-coding |
| chr10_117105033_117105951 | chr10 | 117105034 | 117105951 | + | 0.664032 | intron (NM_177798, intron 1 of 9) | L2a\|LINE\|L2 | -41678 | NM_007636 | 12461 | Mm.247788 | NM_007636 | ENSMUSG00000034024 | **Cct2** | Cctb | chaperonin containing Tcp1, subunit 2 (beta) | protein-coding |
| chr3_101047433_101048219 | chr3 | 101047434 | 101048219 | + | 0.745695 | intron (NM_011197, intron 7 of 8) | intron (NM_011197, intron 7 of 8) | -18331 | NM_001167906 | 630146 | Mm.190915 | NM_001099332 | ENSMUSG00000086564 | **Cd101** | Gm1016\|Gm734\|Igsf2 | CD101 antigen | protein-coding |
| chr3_101047608_101047929 | chr3 | 101047609 | 101047929 | + | 0.70867 | intron (NM_011197, intron 7 of 8) | intron (NM_011197, intron 7 of 8) | -18274 | NM_001167906 | 630146 | Mm.190915 | NM_001099332 | ENSMUSG00000086564 | **Cd101** | Gm1016\|Gm734\|Igsf2 | CD101 antigen | protein-coding |
| chr3_101051313_101051651 | chr3 | 101051314 | 101051651 | + | 0.634458 | intron (NM_011197, intron 6 of 8) | intron (NM_011197, intron 6 of 8) | -21987 | NM_001167906 | 630146 | Mm.190915 | NM_001099332 | ENSMUSG00000086564 | **Cd101** | Gm1016\|Gm734\|Igsf2 | CD101 antigen | protein-coding |
| chr3_101021483_101021954 | chr3 | 101021484 | 101021954 | + | 0.624403 | intron (NM_001167906, intron 1 of 8) | intron (NM_001167906, intron 1 of 8) | 457 | NM_001099332 | 630146 | Mm.190915 | NM_001099332 | ENSMUSG00000086564 | **Cd101** | Gm1016\|Gm734\|Igsf2 | CD101 antigen | protein-coding |
| chr9_78623254_78623833 | chr9 | 78623255 | 78623833 | + | 1.31536 | intron (NM_153098, intron 2 of 32) | intron (NM_153098, intron 2 of 32) | 7998 | NM_153098 | 235505 | Mm.32955 | NM_153098 | ENSMUSG00000046186 | **Cd109** | 9930012E15Rik\|AI480638\|GARP | CD109 antigen | protein-coding |
| chr3_96778296_96778600 | chr3 | 96778297 | 96778600 | + | 0.835818 | intron (NM_026406, intron 4 of 8) | MTA_Mm\|LTR\|ERVL-MaLR | 36073 | NM_018767 | 54215 | Mm.34693 | NM_018767 | ENSMUSG00000038304 | **Cd160** | AU045688\|By55 | CD160 antigen | protein-coding |
| chr3_96774355_96774840 | chr3 | 96774356 | 96774840 | + | 0.747723 | intron (NM_026406, intron 3 of 8) | MTEa\|LTR\|ERVL-MaLR | 39923 | NM_018767 | 54215 | Mm.34693 | NM_018767 | ENSMUSG00000038304 | **Cd160** | AU045688\|By55 | CD160 antigen | protein-coding |
| chr13_102705909_102706263 | chr13 | 102705910 | 102706263 | + | 1.10944 | exon (NM_008533, exon 3 of 3) | exon (NM_008533, exon 3 of 3) | 12528 | NM_008533 | 17079 | Mm.373974 | NM_008533 | ENSMUSG00000021624 | **Cd180** | F630107B15\|Ly78\|RP105 | CD180 antigen | protein-coding |
| chr13_102679757_102680112 | chr13 | 102679758 | 102680112 | + | 0.773791 | Intergenic | Lx\|LINE\|L1 | -13623 | NM_008533 | 17079 | Mm.373974 | NM_008533 | ENSMUSG00000021624 | **Cd180** | F630107B15\|Ly78\|RP105 | CD180 antigen | protein-coding |
| chr3_101235328_101235692 | chr3 | 101235329 | 101235692 | + | 1.78743 | Intergenic | Intergenic | 52429 | NM_013486 | 12481 | Mm.22842 | NM_013486 | ENSMUSG00000027863 | **Cd2** | LFA-2\|Ly-37\|Ly37 | CD2 antigen | protein-coding |
| chr3_101272508_101273381 | chr3 | 101272509 | 101273381 | + | 1.31983 | Intergenic | Intergenic | 14994 | NM_013486 | 12481 | Mm.22842 | NM_013486 | ENSMUSG00000027863 | **Cd2** | LFA-2\|Ly-37\|Ly37 | CD2 antigen | protein-coding |
| chr3_101228855_101229241 | chr3 | 101228856 | 101229241 | + | 1.19508 | Intergenic | Intergenic | 58891 | NM_013486 | 12481 | Mm.22842 | NM_013486 | ENSMUSG00000027863 | **Cd2** | LFA-2\|Ly-37\|Ly37 | CD2 antigen | protein-coding |
| chr3_101228675_101229246 | chr3 | 101228676 | 101229246 | + | 1.14101 | Intergenic | Intergenic | 58978 | NM_013486 | 12481 | Mm.22842 | NM_013486 | ENSMUSG00000027863 | **Cd2** | LFA-2\|Ly-37\|Ly37 | CD2 antigen | protein-coding |
| chr3_101293499_101294045 | chr3 | 101293500 | 101294045 | + | 1.09709 | Intergenic | Intergenic | -5833 | NM_013486 | 12481 | Mm.22842 | NM_013486 | ENSMUSG00000027863 | **Cd2** | LFA-2\|Ly-37\|Ly37 | CD2 antigen | protein-coding |
| chr3_101237285_101237600 | chr3 | 101237286 | 101237600 | + | 1.04162 | Intergenic | Intergenic | 50496 | NM_013486 | 12481 | Mm.22842 | NM_013486 | ENSMUSG00000027863 | **Cd2** | LFA-2\|Ly-37\|Ly37 | CD2 antigen | protein-coding |
| chr3_101293509_101293782 | chr3 | 101293510 | 101293782 | + | 1.02981 | Intergenic | Intergenic | -5707 | NM_013486 | 12481 | Mm.22842 | NM_013486 | ENSMUSG00000027863 | **Cd2** | LFA-2\|Ly-37\|Ly37 | CD2 antigen | protein-coding |
| chr3_101290083_101290339 | chr3 | 101290084 | 101290339 | + | 0.942484 | Intergenic | Intergenic | -2272 | NM_013486 | 12481 | Mm.22842 | NM_013486 | ENSMUSG00000027863 | **Cd2** | LFA-2\|Ly-37\|Ly37 | CD2 antigen | protein-coding |
| chr3_101279564_101279874 | chr3 | 101279565 | 101279874 | + | 0.641937 | intron (NM_013486, intron 4 of 4) | intron (NM_013486, intron 4 of 4) | 8220 | NM_013486 | 12481 | Mm.22842 | NM_013486 | ENSMUSG00000027863 | **Cd2** | LFA-2\|Ly-37\|Ly37 | CD2 antigen | protein-coding |
| chr16_45332278_45332930 | chr16 | 45332279 | 45332930 | + | 1.1026 | Intergenic | Intergenic | 76449 | NM_010818 | 17470 | Mm.245851 | NM_010818 | ENSMUSG00000022661 | **Cd200** | Mox2\|OX2 | CD200 antigen | protein-coding |
| chr16_45332630_45332925 | chr16 | 45332631 | 45332925 | + | 1.10038 | Intergenic | Intergenic | 76275 | NM_010818 | 17470 | Mm.245851 | NM_010818 | ENSMUSG00000022661 | **Cd200** | Mox2\|OX2 | CD200 antigen | protein-coding |
| chr16_45379594_45380108 | chr16 | 45379595 | 45380108 | + | 0.749184 | Intergenic | Intergenic | 29202 | NM_010818 | 17470 | Mm.245851 | NM_010818 | ENSMUSG00000022661 | **Cd200** | Mox2\|OX2 | CD200 antigen | protein-coding |
| chr16_45327086_45327582 | chr16 | 45327087 | 45327582 | + | 0.656838 | Intergenic | Intergenic | 81719 | NM_010818 | 17470 | Mm.245851 | NM_010818 | ENSMUSG00000022661 | **Cd200** | Mox2\|OX2 | CD200 antigen | protein-coding |
| chr16_45007513_45008145 | chr16 | 45007514 | 45008145 | + | 1.04367 | Intergenic | MLT1C\|LTR\|ERVL-MaLR | 64151 | NM_001128132 | 74603 | Mm.297337 | NM_027578 | ENSMUSG00000036172 | **Cd200r3** | 4733401I18Rik\|4833409J19Rik\|AI505817\|BB106877\|mCD200RLb | CD200 receptor 3 | protein-coding |
| chr16_45013439_45014239 | chr16 | 45013440 | 45014239 | + | 0.89336 | Intergenic | Lx8\|LINE\|L1 | 70161 | NM_001128132 | 74603 | Mm.297337 | NM_027578 | ENSMUSG00000036172 | **Cd200r3** | 4733401I18Rik\|4833409J19Rik\|AI505817\|BB106877\|mCD200RLb | CD200 receptor 3 | protein-coding |
| chr16_44976105_44976505 | chr16 | 44976106 | 44976505 | + | 0.86583 | intron (NM_029018, intron 5 of 5) | intron (NM_029018, intron 5 of 5) | 32627 | NM_001128132 | 74603 | Mm.297337 | NM_027578 | ENSMUSG00000036172 | **Cd200r3** | 4733401I18Rik\|4833409J19Rik\|AI505817\|BB106877\|mCD200RLb | CD200 receptor 3 | protein-coding |
| chr6_83689366_83689795 | chr6 | 83689367 | 83689795 | + | 0.641314 | Intergenic | Intergenic | -11724 | NM_144943 | 246278 | Mm.136079 | NM_144943 | ENSMUSG00000034783 | **Cd207** | - | CD207 antigen | protein-coding |
| chr18_89433068_89433628 | chr18 | 89433069 | 89433628 | + | 1.3597 | intron (NM_001039173, intron 5 of 7) | RMER15\|LTR\|ERVL | 235921 | NM_001039149 | 225825 | Mm.139293 | NM_178687 | ENSMUSG00000034028 | **Cd226** | BC051526\|DNAM-1\|DNAM1\|Pta1\|TLiSA1 | CD226 antigen | protein-coding |
| chr18_89332543_89332905 | chr18 | 89332544 | 89332905 | + | 1.25459 | intron (NM_001039173, intron 7 of 7) | intron (NM_001039173, intron 7 of 7) | 135297 | NM_001039149 | 225825 | Mm.139293 | NM_178687 | ENSMUSG00000034028 | **Cd226** | BC051526\|DNAM-1\|DNAM1\|Pta1\|TLiSA1 | CD226 antigen | protein-coding |
| chr18_89253929_89254391 | chr18 | 89253930 | 89254391 | + | 1.13452 | intron (NM_178687, intron 4 of 6) | intron (NM_178687, intron 4 of 6) | 56733 | NM_001039149 | 225825 | Mm.139293 | NM_178687 | ENSMUSG00000034028 | **Cd226** | BC051526\|DNAM-1\|DNAM1\|Pta1\|TLiSA1 | CD226 antigen | protein-coding |
| chr18_89435304_89435809 | chr18 | 89435305 | 89435809 | + | 1.01509 | intron (NM_001039173, intron 5 of 7) | RMER15\|LTR\|ERVL | 238130 | NM_001039149 | 225825 | Mm.139293 | NM_178687 | ENSMUSG00000034028 | **Cd226** | BC051526\|DNAM-1\|DNAM1\|Pta1\|TLiSA1 | CD226 antigen | protein-coding |
| chr18_89222594_89223308 | chr18 | 89222595 | 89223308 | + | 0.931697 | intron (NM_178687, intron 3 of 6) | MTE-int\|LTR\|ERVL-MaLR | 25524 | NM_001039149 | 225825 | Mm.139293 | NM_178687 | ENSMUSG00000034028 | **Cd226** | BC051526\|DNAM-1\|DNAM1\|Pta1\|TLiSA1 | CD226 antigen | protein-coding |
| chr18_89272326_89272856 | chr18 | 89272327 | 89272856 | + | 0.923105 | Intergenic | MTC\|LTR\|ERVL-MaLR | 75164 | NM_001039149 | 225825 | Mm.139293 | NM_178687 | ENSMUSG00000034028 | **Cd226** | BC051526\|DNAM-1\|DNAM1\|Pta1\|TLiSA1 | CD226 antigen | protein-coding |
| chr18_89222781_89223247 | chr18 | 89222782 | 89223247 | + | 0.916654 | intron (NM_178687, intron 3 of 6) | MTE-int\|LTR\|ERVL-MaLR | 25587 | NM_001039149 | 225825 | Mm.139293 | NM_178687 | ENSMUSG00000034028 | **Cd226** | BC051526\|DNAM-1\|DNAM1\|Pta1\|TLiSA1 | CD226 antigen | protein-coding |
| chr18_89459093_89459780 | chr18 | 89459094 | 89459780 | + | 0.868182 | intron (NM_001039173, intron 5 of 7) | intron (NM_001039173, intron 5 of 7) | 262010 | NM_001039149 | 225825 | Mm.139293 | NM_178687 | ENSMUSG00000034028 | **Cd226** | BC051526\|DNAM-1\|DNAM1\|Pta1\|TLiSA1 | CD226 antigen | protein-coding |
| chr18_89256740_89257635 | chr18 | 89256741 | 89257635 | + | 0.837908 | intron (NM_178687, intron 4 of 6) | MIR\|SINE\|MIR | 59761 | NM_001039149 | 225825 | Mm.139293 | NM_178687 | ENSMUSG00000034028 | **Cd226** | BC051526\|DNAM-1\|DNAM1\|Pta1\|TLiSA1 | CD226 antigen | protein-coding |
| chr18_89198538_89198847 | chr18 | 89198539 | 89198847 | + | 0.830422 | intron (NM_178687, intron 1 of 6) | intron (NM_178687, intron 1 of 6) | 1266 | NM_001039149 | 225825 | Mm.139293 | NM_178687 | ENSMUSG00000034028 | **Cd226** | BC051526\|DNAM-1\|DNAM1\|Pta1\|TLiSA1 | CD226 antigen | protein-coding |
| chr18_89219574_89220114 | chr18 | 89219575 | 89220114 | + | 0.788753 | intron (NM_178687, intron 3 of 6) | intron (NM_178687, intron 3 of 6) | 22417 | NM_001039149 | 225825 | Mm.139293 | NM_178687 | ENSMUSG00000034028 | **Cd226** | BC051526\|DNAM-1\|DNAM1\|Pta1\|TLiSA1 | CD226 antigen | protein-coding |
| chr18_89159017_89159657 | chr18 | 89159018 | 89159657 | + | 0.786869 | Intergenic | Intergenic | -38090 | NM_178687 | 225825 | Mm.139293 | NM_178687 | ENSMUSG00000034028 | **Cd226** | BC051526\|DNAM-1\|DNAM1\|Pta1\|TLiSA1 | CD226 antigen | protein-coding |

| | | | | | | | | | | | | | | | | | |
|---|---|---|---|---|---|---|---|---|---|---|---|---|---|---|---|---|---|
| chr18_89159293_89159632 | chr18 | 89159294 | 89159632 | + | 0.738304 | Intergenic | Intergenic | -37964 | NM_178687 | 225825 | Mm.139293 | NM_178687 | ENSMUSG00000034028 | **Cd226** | BC051526\|DNAM-1\|DNAM1\|Pta1\|TLiSA1 | CD226 antigen | protein-coding |
| chr18_89218989_89219461 | chr18 | 89218990 | 89219461 | + | 0.728989 | intron (NM_178687, intron 3 of 6) | intron (NM_178687, intron 3 of 6) | 21798 | NM_001039149 | 225825 | Mm.139293 | NM_178687 | ENSMUSG00000034028 | **Cd226** | BC051526\|DNAM-1\|DNAM1\|Pta1\|TLiSA1 | CD226 antigen | protein-coding |
| chr18_89272324_89273438 | chr18 | 89272325 | 89273438 | + | 0.726623 | Intergenic | Lx5\|LINE\|L1 | 75454 | NM_001039149 | 225825 | Mm.139293 | NM_178687 | ENSMUSG00000034028 | **Cd226** | BC051526\|DNAM-1\|DNAM1\|Pta1\|TLiSA1 | CD226 antigen | protein-coding |
| chr18_89331483_89331972 | chr18 | 89331484 | 89331972 | + | 0.714898 | intron (NM_001039173, intron 7 of 7) | intron (NM_001039173, intron 7 of 7) | 134301 | NM_001039149 | 225825 | Mm.139293 | NM_178687 | ENSMUSG00000034028 | **Cd226** | BC051526\|DNAM-1\|DNAM1\|Pta1\|TLiSA1 | CD226 antigen | protein-coding |
| chr18_89325329_89325592 | chr18 | 89325330 | 89325592 | + | 0.697912 | intron (NM_001039173, intron 7 of 7) | intron (NM_001039173, intron 7 of 7) | 128034 | NM_001039149 | 225825 | Mm.139293 | NM_178687 | ENSMUSG00000034028 | **Cd226** | BC051526\|DNAM-1\|DNAM1\|Pta1\|TLiSA1 | CD226 antigen | protein-coding |
| chr18_89219567_89219859 | chr18 | 89219568 | 89219859 | + | 0.666849 | intron (NM_178687, intron 3 of 6) | intron (NM_178687, intron 3 of 6) | 22286 | NM_001039149 | 225825 | Mm.139293 | NM_178687 | ENSMUSG00000034028 | **Cd226** | BC051526\|DNAM-1\|DNAM1\|Pta1\|TLiSA1 | CD226 antigen | protein-coding |
| chr1_165801543_165802785 | chr1 | 165801544 | 165802785 | + | 1.2553 | intron (NM_031162, intron 1 of 7) | intron (NM_031162, intron 1 of 7) | 13483 | NM_001113392 | 12503 | Mm.217308 | NM_031162 | ENSMUSG00000005763 | **Cd247** | 4930549J05Rik\|A430104F18Rik\|AW552088\|Cd3\|Cd3-eta\|Cd3-zeta\|Cd3h\|Cd3z\|Cd3zeta\|T3z\|Tcrk\|Tcrz | CD247 antigen | protein-coding |
| chr1_165801546_165802208 | chr1 | 165801547 | 165802208 | + | 1.24727 | intron (NM_031162, intron 1 of 7) | intron (NM_031162, intron 1 of 7) | 13196 | NM_001113392 | 12503 | Mm.217308 | NM_031162 | ENSMUSG00000005763 | **Cd247** | 4930549J05Rik\|A430104F18Rik\|AW552088\|Cd3\|Cd3-eta\|Cd3-zeta\|Cd3h\|Cd3z\|Cd3zeta\|T3z\|Tcrk\|Tcrz | CD247 antigen | protein-coding |
| chr1_165802249_165802785 | chr1 | 165802250 | 165802785 | + | 1.11056 | intron (NM_031162, intron 1 of 7) | intron (NM_031162, intron 1 of 7) | 13836 | NM_001113392 | 12503 | Mm.217308 | NM_031162 | ENSMUSG00000005763 | **Cd247** | 4930549J05Rik\|A430104F18Rik\|AW552088\|Cd3\|Cd3-eta\|Cd3-zeta\|Cd3h\|Cd3z\|Cd3zeta\|T3z\|Tcrk\|Tcrz | CD247 antigen | protein-coding |
| chr1_165843592_165843993 | chr1 | 165843593 | 165843993 | + | 0.70174 | intron (NM_031162, intron 1 of 7) | intron (NM_031162, intron 1 of 7) | 55112 | NM_001113392 | 12503 | Mm.217308 | NM_031162 | ENSMUSG00000005763 | **Cd247** | 4930549J05Rik\|A430104F18Rik\|AW552088\|Cd3\|Cd3-eta\|Cd3-zeta\|Cd3h\|Cd3z\|Cd3zeta\|T3z\|Tcrk\|Tcrz | CD247 antigen | protein-coding |
| chr1_165850785_165851123 | chr1 | 165850786 | 165851123 | + | 0.644559 | intron (NM_031162, intron 1 of 7) | intron (NM_031162, intron 1 of 7) | 62273 | NM_001113392 | 12503 | Mm.217308 | NM_031162 | ENSMUSG00000005763 | **Cd247** | 4930549J05Rik\|A430104F18Rik\|AW552088\|Cd3\|Cd3-eta\|Cd3-zeta\|Cd3h\|Cd3z\|Cd3zeta\|T3z\|Tcrk\|Tcrz | CD247 antigen | protein-coding |
| chr10_43577709_43578054 | chr10 | 43577710 | 43578054 | + | 0.892239 | Intergenic | Intergenic | -1287 | NM_009846 | 12484 | Mm.29742 | NM_009846 | ENSMUSG00000047139 | **Cd24a** | Cd24\|HSA\|Ly-52\|nectadrin | CD24a antigen | protein-coding |
| chr6_125240627_125240988 | chr6 | 125240628 | 125240988 | + | 0.910649 | TTS (NR_015604) | TTS (NR_015604) | -3781 | NM_001286753 | 21940 | Mm.367714 | NM_001033126 | ENSMUSG00000030336 | **Cd27** | S152\|Tnfrsf7\|Tp55 | CD27 antigen | protein-coding |
| chr6_125238391_125238926 | chr6 | 125238392 | 125238926 | + | 0.798379 | intron (NR_015604, intron 1 of 1) | intron (NR_015604, intron 1 of 1) | -1632 | NM_001286753 | 21940 | Mm.367714 | NM_001033126 | ENSMUSG00000030336 | **Cd27** | S152\|Tnfrsf7\|Tp55 | CD27 antigen | protein-coding |
| chr19_29374911_29375377 | chr19 | 29374912 | 29375377 | + | 1.1649 | intron (NM_021893, intron 3 of 6) | intron (NM_021893, intron 3 of 6) | 7706 | NM_021893 | 60533 | Mm.245363 | NM_021893 | ENSMUSG00000016496 | **Cd274** | A530045L16Rik\|B7h1\|Pdcd1l1\|Pdcd1lg1\|Pdl1 | CD274 antigen | protein-coding |
| chr19_29365241_29365453 | chr19 | 29365242 | 29365453 | + | 0.934715 | Intergenic | Intergenic | -2091 | NM_021893 | 60533 | Mm.245363 | NM_021893 | ENSMUSG00000016496 | **Cd274** | A530045L16Rik\|B7h1\|Pdcd1l1\|Pdcd1lg1\|Pdl1 | CD274 antigen | protein-coding |
| chr1_60677300_60677796 | chr1 | 60677301 | 60677796 | + | 1.60937 | Intergenic | Lx5\|LINE\|L1 | -68840 | NM_007642 | 12487 | Mm.255003 | NM_007642 | ENSMUSG00000026012 | **Cd28** | - | CD28 antigen | protein-coding |
| chr1_60771799_60772416 | chr1 | 60771800 | 60772416 | + | 1.02953 | 3' UTR (NM_007642, exon 4 of 4) | 3' UTR (NM_007642, exon 4 of 4) | 25720 | NM_007642 | 12487 | Mm.255003 | NM_007642 | ENSMUSG00000026012 | **Cd28** | - | CD28 antigen | protein-coding |
| chr7_127176947_127177315 | chr7 | 127176948 | 127177315 | + | 0.758968 | Intergenic | Intergenic | 18946 | NM_001285907 | 70233 | Mm.28050 | NM_027353 | ENSMUSG00000042502 | **Cd2bp2** | 1500011B02Rik\|2410024K20Rik\|AW048183 | CD2 antigen (cytoplasmic tail) binding protein 2 | protein-coding |
| chr7_127176916_127177497 | chr7 | 127176917 | 127177497 | + | 0.67341 | Intergenic | ORR1F\|LTR\|ERVL-MaLR | 18870 | NM_001285907 | 70233 | Mm.28050 | NM_027353 | ENSMUSG00000042502 | **Cd2bp2** | 1500011B02Rik\|2410024K20Rik\|AW048183 | CD2 antigen (cytoplasmic tail) binding protein 2 | protein-coding |
| chr11_115131762_115132192 | chr11 | 115131763 | 115132192 | + | 1.13478 | intron (NM_001169153, intron 1 of 6) | L2\|LINE\|L2 | 2015 | NM_001169153 | 246746 | Mm.277387 | NM_145634 | ENSMUSG00000047798 | **Cd300lf** | CLIM1\|CLM-1\|CLM1\|Digr2\|F730004D16Rik\|IREM1\|IgSF13\|LMIR3\|Pigr3 | CD300 antigen like family member F | protein-coding |
| chr5_17854227_17855007 | chr5 | 17854228 | 17855007 | + | 0.701467 | intron (NM_001159558, intron 1 of 15) | intron (NM_001159558, intron 1 of 15) | -4825 | NM_001159557 | 12491 | Mm.18628 | NM_007643 | ENSMUSG00000002944 | **Cd36** | FAT\|GPIV\|Scarb3 | CD36 antigen | protein-coding |
| chr5_43915201_43915723 | chr5 | 43915202 | 43915723 | + | 0.659926 | Intergenic | URR1A\|DNA\|hAT-Charlie | 46635 | NM_007646 | 12494 | Mm.249873 | NM_007646 | ENSMUSG00000029084 | **Cd38** | ADPRC 1\|Cd38-rs1 | CD38 antigen | protein-coding |
| chr9_45007463_45007871 | chr9 | 45007464 | 45007871 | + | 1.02519 | intron (NM_007648, intron 2 of 7) | intron (NM_007648, intron 2 of 7) | 1923 | NM_007648 | 12501 | Mm.210361 | NM_007648 | ENSMUSG00000032093 | **Cd3e** | AI504783\|CD3\|CD3epsilon\|T3e | CD3 antigen, epsilon polypeptide | protein-coding |
| chr9_45018965_45019255 | chr9 | 45018966 | 45019255 | + | 0.818937 | Intergenic | L1M4\|LINE\|L1 | -9520 | NM_007648 | 12501 | Mm.210361 | NM_007648 | ENSMUSG00000032093 | **Cd3e** | AI504783\|CD3\|CD3epsilon\|T3e | CD3 antigen, epsilon polypeptide | protein-coding |
| chr9_45018853_45019264 | chr9 | 45018854 | 45019264 | + | 0.652055 | Intergenic | L1M4\|LINE\|L1 | -9469 | NM_007648 | 12501 | Mm.210361 | NM_007648 | ENSMUSG00000032093 | **Cd3e** | AI504783\|CD3\|CD3epsilon\|T3e | CD3 antigen, epsilon polypeptide | protein-coding |
| chr2_102956876_102957475 | chr2 | 102956877 | 102957475 | + | 1.91228 | Intergenic | MTEb\|LTR\|ERVL-MaLR | -55511 | NM_001039150 | 12505 | Mm.423621 | NM_009851 | ENSMUSG00000005087 | **Cd44** | AU023126\|AW121933\|AW146109\|HERMES\|Ly-24\|Pgp-1 | CD44 antigen | protein-coding |
| chr2_102955129_102955557 | chr2 | 102955130 | 102955557 | + | 1.4215 | Intergenic | Intergenic | -53678 | NM_001039150 | 12505 | Mm.423621 | NM_009851 | ENSMUSG00000005087 | **Cd44** | AU023126\|AW121933\|AW146109\|HERMES\|Ly-24\|Pgp-1 | CD44 antigen | protein-coding |
| chr2_102980899_102982212 | chr2 | 102980900 | 102982212 | + | 1.3182 | Intergenic | Intergenic | -79891 | NM_001039150 | 12505 | Mm.423621 | NM_009851 | ENSMUSG00000005087 | **Cd44** | AU023126\|AW121933\|AW146109\|HERMES\|Ly-24\|Pgp-1 | CD44 antigen | protein-coding |
| chr2_102912466_102912820 | chr2 | 102912467 | 102912820 | + | 1.00774 | Intergenic | Intergenic | -10978 | NM_001039150 | 12505 | Mm.423621 | NM_009851 | ENSMUSG00000005087 | **Cd44** | AU023126\|AW121933\|AW146109\|HERMES\|Ly-24\|Pgp-1 | CD44 antigen | protein-coding |
| chr2_102812667_102813383 | chr2 | 102812668 | 102813383 | + | 1.0057 | 3' UTR (NM_001039150, exon 14 of 14) | 3' UTR (NM_001039150, exon 14 of 14) | 88640 | NM_001177785 | 12505 | Mm.423621 | NM_009851 | ENSMUSG00000005087 | **Cd44** | AU023126\|AW121933\|AW146109\|HERMES\|Ly-24\|Pgp-1 | CD44 antigen | protein-coding |
| chr2_102853052_102853682 | chr2 | 102853053 | 102853682 | + | 0.891306 | intron (NM_001177787, intron 4 of 10) | intron (NM_001177787, intron 4 of 10) | 48298 | NM_001177785 | 12505 | Mm.423621 | NM_009851 | ENSMUSG00000005087 | **Cd44** | AU023126\|AW121933\|AW146109\|HERMES\|Ly-24\|Pgp-1 | CD44 antigen | protein-coding |
| chr2_102948506_102948819 | chr2 | 102948507 | 102948819 | + | 0.878531 | Intergenic | Intergenic | -46998 | NM_001039150 | 12505 | Mm.423621 | NM_009851 | ENSMUSG00000005087 | **Cd44** | AU023126\|AW121933\|AW146109\|HERMES\|Ly-24\|Pgp-1 | CD44 antigen | protein-coding |
| chr2_102986536_102986835 | chr2 | 102986537 | 102986835 | + | 0.75258 | Intergenic | Intergenic | -85021 | NM_001039150 | 12505 | Mm.423621 | NM_009851 | ENSMUSG00000005087 | **Cd44** | AU023126\|AW121933\|AW146109\|HERMES\|Ly-24\|Pgp-1 | CD44 antigen | protein-coding |
| chr2_102947975_102948906 | chr2 | 102947976 | 102948906 | + | 0.693823 | Intergenic | Intergenic | -46776 | NM_001039150 | 12505 | Mm.423621 | NM_009851 | ENSMUSG00000005087 | **Cd44** | AU023126\|AW121933\|AW146109\|HERMES\|Ly-24\|Pgp-1 | CD44 antigen | protein-coding |
| chr16_49847370_49847945 | chr16 | 49847371 | 49847945 | + | 0.954367 | Intergenic | MER5A\|DNA\|hAT-Charlie | -7996 | NM_010581 | 16423 | Mm.31752 | NM_010581 | ENSMUSG00000055447 | **Cd47** | 9130415E20Rik\|AA407862\|AI848868\|AW108519\|B430305P08Rik\|IAP\|Itgp | CD47 antigen (Rh-related antigen, integrin-associated signal transducer) | protein-coding |
| chr16_49892045_49892446 | chr16 | 49892046 | 49892446 | + | 0.839038 | intron (NM_010581, intron 4 of 9) | intron (NM_010581, intron 4 of 9) | 36592 | NM_010581 | 16423 | Mm.31752 | NM_010581 | ENSMUSG00000055447 | **Cd47** | 9130415E20Rik\|AA407862\|AI848868\|AW108519\|B430305P08Rik\|IAP\|Itgp | CD47 antigen (Rh-related antigen, integrin-associated signal transducer) | protein-coding |
| chr16_49857547_49857904 | chr16 | 49857548 | 49857904 | + | 0.766758 | intron (NM_010581, intron 1 of 9) | intron (NM_010581, intron 1 of 9) | 2072 | NM_010581 | 16423 | Mm.31752 | NM_010581 | ENSMUSG00000055447 | **Cd47** | 9130415E20Rik\|AA407862\|AI848868\|AW108519\|B430305P08Rik\|IAP\|Itgp | CD47 antigen (Rh-related antigen, integrin-associated signal transducer) | protein-coding |
| chr16_49796970_49797468 | chr16 | 49796971 | 49797468 | + | 0.691792 | Intergenic | ERV3-16A3_LTR\|LTR\|ERVL | -58435 | NM_010581 | 16423 | Mm.31752 | NM_010581 | ENSMUSG00000055447 | **Cd47** | 9130415E20Rik\|AA407862\|AI848868\|AW108519\|B430305P08Rik\|IAP\|Itgp | CD47 antigen (Rh-related antigen, integrin-associated signal transducer) | protein-coding |
| chr16_49799719_49800597 | chr16 | 49799720 | 49800597 | + | 0.66056 | Intergenic | Intergenic | -55496 | NM_010581 | 16423 | Mm.31752 | NM_010581 | ENSMUSG00000055447 | **Cd47** | 9130415E20Rik\|AA407862\|AI848868\|AW108519\|B430305P08Rik\|IAP\|Itgp | CD47 antigen (Rh-related antigen, integrin-associated signal transducer) | protein-coding |
| chr16_49796992_49797242 | chr16 | 49796993 | 49797242 | + | 0.648454 | Intergenic | ERV3-16A3_LTR\|LTR\|ERVL | -58537 | NM_010581 | 16423 | Mm.31752 | NM_010581 | ENSMUSG00000055447 | **Cd47** | 9130415E20Rik\|AA407862\|AI848868\|AW108519\|B430305P08Rik\|IAP\|Itgp | CD47 antigen (Rh-related antigen, integrin-associated signal transducer) | protein-coding |
| chr1_171684625_171685474 | chr1 | 171684626 | 171685474 | + | 1.14721 | intron (NM_007649, intron 1 of 3) | intron (NM_007649, intron 1 of 3) | 2995 | NM_007649 | 12506 | Mm.1738 | NM_007649 | ENSMUSG00000015355 | **Cd48** | AI449234\|AW610730\|BCM1\|BLAST\|BLAST-1\|BLAST1\|Bcm-1\|MEM-102\|SLAMF2\|Sgp-60 | CD48 antigen | protein-coding |

| | | | | | | | | | | | | | | | | | |
|---|---|---|---|---|---|---|---|---|---|---|---|---|---|---|---|---|---|
| chr3_106768508_106768921 | chr3 | 106768509 | 106768921 | + | 0.935132 | intron (NM_007651, intron 3 of 7) | intron (NM_007651, intron 3 of 7) | 21434 | NM_007651 | 12508 | Mm.316861 | NM_007651 | ENSMUSG00000040747 | **Cd53** | AI323659\|Ox-44\|Tspan25 | CD53 antigen | protein-coding |
| chr19_10795191_10795489 | chr19 | 10795192 | 10795489 | + | 0.862919 | intron (NM_009852, intron 6 of 12) | intron (NM_009852, intron 6 of 12) | 34718 | NM_001037801 | 12511 | Mm.290897 | NM_009852 | ENSMUSG00000024670 | **Cd6** | - | CD6 antigen | protein-coding |
| chr19_10795160_10795937 | chr19 | 10795161 | 10795937 | + | 0.848655 | intron (NM_009852, intron 6 of 12) | intron (NM_009852, intron 6 of 12) | 34509 | NM_001037801 | 12511 | Mm.290897 | NM_009852 | ENSMUSG00000024670 | **Cd6** | - | CD6 antigen | protein-coding |
| chr6_129265930_129267357 | chr6 | 129265931 | 129267357 | + | 1.04746 | TTS (NM_001033122) | TTS (NM_001033122) | 8725 | NM_001033122 | 12515 | Mm.74745 | NM_001033122 | ENSMUSG00000030156 | **Cd69** | 5830438K24Rik\|AI452015\|AIM\|VEA | CD69 antigen | protein-coding |
| chr6_129267416_129267704 | chr6 | 129267417 | 129267704 | + | 0.723535 | 3' UTR (NM_001033122, exon 5 of 5) | 3' UTR (NM_001033122, exon 5 of 5) | 7809 | NM_001033122 | 12515 | Mm.74745 | NM_001033122 | ENSMUSG00000030156 | **Cd69** | 5830438K24Rik\|AI452015\|AIM\|VEA | CD69 antigen | protein-coding |
| chr11_121042589_121042872 | chr11 | 121042590 | 121042872 | + | 0.670901 | Intergenic | Intergenic | -3253 | NM_009854 | 12516 | Mm.4100 | NM_009854 | ENSMUSG00000025163 | **Cd7** | - | CD7 antigen | protein-coding |
| chr18_60804258_60804494 | chr18 | 60804259 | 60804494 | + | 0.942824 | intron (NM_001042605, intron 1 of 8) | intron (NM_001042605, intron 1 of 8) | 527 | NM_010545 | 16149 | Mm.439737 | NM_010545 | ENSMUSG00000024610 | **Cd74** | CLIP\|DHLAG\|HLADG\|Ia-GAMMA\|Ii | CD74 antigen (invariant polypeptide of major histocompatibility complex, class II antigen-associated) | protein-coding |
| chr16_38458627_38459184 | chr16 | 38458628 | 38459184 | + | 1.31501 | promoter-TSS (NM_009855) | promoter-TSS (NM_009855) | -27 | NM_009855 | 12519 | Mm.89474 | NM_009855 | ENSMUSG00000075122 | **Cd80** | B71\|Cd28l\|Ly-53\|Ly53\|MIC17\|TSA1 | CD80 antigen | protein-coding |
| chr2_93460872_93461532 | chr2 | 93460873 | 93461532 | + | 0.685367 | intron (NR_040532, intron 2 of 4) | intron (NR_040532, intron 2 of 4) | 649 | NM_001271461 | 12521 | Mm.4261 | NM_007656 | ENSMUSG00000027215 | **Cd82** | AA682076\|AL023070\|C33\|IA4\|Kai1\|Tspan27 | CD82 antigen | protein-coding |
| chr13_43778010_43778646 | chr13 | 43778011 | 43778646 | + | 1.27436 | Intergenic | Intergenic | -6784 | NM_009856 | 12522 | Mm.57175 | NM_009856 | ENSMUSG00000015396 | **Cd83** | - | CD83 antigen | protein-coding |
| chr13_43866445_43866799 | chr13 | 43866446 | 43866799 | + | 1.1957 | Intergenic | MLT1A0\|LTR\|ERVL-MaLR | 81510 | NM_001289915 | 12522 | Mm.57175 | NM_009856 | ENSMUSG00000015396 | **Cd83** | - | CD83 antigen | protein-coding |
| chr13_43777414_43777899 | chr13 | 43777415 | 43777899 | + | 1.05066 | Intergenic | Intergenic | -7455 | NM_009856 | 12522 | Mm.57175 | NM_009856 | ENSMUSG00000015396 | **Cd83** | - | CD83 antigen | protein-coding |
| chr13_43774060_43774577 | chr13 | 43774061 | 43774577 | + | 0.919928 | Intergenic | Intergenic | -10793 | NM_009856 | 12522 | Mm.57175 | NM_009856 | ENSMUSG00000015396 | **Cd83** | - | CD83 antigen | protein-coding |
| chr13_43774779_43775643 | chr13 | 43774780 | 43775643 | + | 0.894262 | Intergenic | Charlie8\|DNA\|hAT-Charlie | -9901 | NM_009856 | 12522 | Mm.57175 | NM_009856 | ENSMUSG00000015396 | **Cd83** | - | CD83 antigen | protein-coding |
| chr6_71387069_71387914 | chr6 | 71387070 | 71387914 | + | 0.816137 | Intergenic | MLTR73\|LTR\|ERV1 | 14065 | NM_001081110 | 12525 | Mm.1858 | NM_009857 | ENSMUSG00000053977 | **Cd8a** | BB154331\|Ly-2\|Ly-35\|Ly-B\|Lyt-2 | CD8 antigen, alpha chain | protein-coding |
| chr6_71358752_71359118 | chr6 | 71358753 | 71359118 | + | 0.731021 | Intergenic | Intergenic | -14492 | NM_009857 | 12525 | Mm.1858 | NM_009857 | ENSMUSG00000053977 | **Cd8a** | BB154331\|Ly-2\|Ly-35\|Ly-B\|Lyt-2 | CD8 antigen, alpha chain | protein-coding |
| chr6_71361264_71361715 | chr6 | 71361265 | 71361715 | + | 0.716684 | Intergenic | Intergenic | -11937 | NM_009857 | 12525 | Mm.1858 | NM_009857 | ENSMUSG00000053977 | **Cd8a** | BB154331\|Ly-2\|Ly-35\|Ly-B\|Lyt-2 | CD8 antigen, alpha chain | protein-coding |
| chr6_71324008_71324232 | chr6 | 71324009 | 71324232 | + | 1.07012 | intron (NM_009858, intron 1 of 5) | intron (NM_009858, intron 1 of 5) | 1308 | NM_009858 | 12526 | Mm.333148 | NM_009858 | ENSMUSG00000053044 | **Cd8b1** | Cd8b\|Ly-3\|Ly-C\|Lyt-3 | CD8 antigen, beta chain 1 | protein-coding |
| chr6_71323947_71324260 | chr6 | 71323948 | 71324260 | + | 0.95004 | intron (NM_009858, intron 1 of 5) | intron (NM_009858, intron 1 of 5) | 1292 | NM_009858 | 12526 | Mm.333148 | NM_009858 | ENSMUSG00000053044 | **Cd8b1** | Cd8b\|Ly-3\|Ly-C\|Lyt-3 | CD8 antigen, beta chain 1 | protein-coding |
| chr6_71308263_71308655 | chr6 | 71308264 | 71308655 | + | 0.895309 | Intergenic | B3\|SINE\|B2 | -14353 | NM_009858 | 12526 | Mm.333148 | NM_009858 | ENSMUSG00000053044 | **Cd8b1** | Cd8b\|Ly-3\|Ly-C\|Lyt-3 | CD8 antigen, beta chain 1 | protein-coding |
| chr6_71313517_71313908 | chr6 | 71313518 | 71313908 | + | 0.860635 | Intergenic | Intergenic | -9099 | NM_009858 | 12526 | Mm.333148 | NM_009858 | ENSMUSG00000053044 | **Cd8b1** | Cd8b\|Ly-3\|Ly-C\|Lyt-3 | CD8 antigen, beta chain 1 | protein-coding |
| chr6_125483473_125483941 | chr6 | 125483474 | 125483941 | + | 1.08604 | intron (NM_007657, intron 1 of 7) | intron (NM_007657, intron 1 of 7) | 11048 | NM_007657 | 12527 | Mm.210676 | NM_007657 | ENSMUSG00000030342 | **Cd9** | Tspan29 | CD9 antigen | protein-coding |
| chr6_125483469_125484327 | chr6 | 125483470 | 125484327 | + | 0.929361 | intron (NM_007657, intron 1 of 7) | URR1A\|DNA\|hAT-Charlie | 10857 | NM_007657 | 12527 | Mm.210676 | NM_007657 | ENSMUSG00000030342 | **Cd9** | Tspan29 | CD9 antigen | protein-coding |
| chr16_46117530_46118157 | chr16 | 46117531 | 46118157 | + | 1.56821 | exon (NM_032465, exon 2 of 15) | exon (NM_032465, exon 2 of 15) | 2404 | NM_032465 | 84544 | Mm.29204 | NM_032465 | ENSMUSG00000022657 | **Cd96** | 1700109I12Rik\|Tactile | CD96 antigen | protein-coding |
| chr8_83756957_83757307 | chr8 | 83756958 | 83757307 | + | 0.715723 | Intergenic | Intergenic | -15821 | NM_001163031 | 26364 | Mm.334648 | NM_011925 | ENSMUSG00000002885 | **Cd97** | AA409984\|TM7LN1 | CD97 antigen | protein-coding |
| chrX_71479307_71479739 | chrX | 71479308 | 71479739 | + | 1.17291 | intron (NM_138309, intron 1 of 8) | intron (NM_138309, intron 1 of 8) | 13326 | NM_138309 | 171486 | Mm.283879 | NM_138309 | ENSMUSG00000035776 | **Cd99l2** | AW548191\|Mic2l1\|Xap89 | CD99 antigen-like 2 | protein-coding |
| chrX_71475155_71476313 | chrX | 71475156 | 71476313 | + | 0.977526 | intron (NM_138309, intron 1 of 8) | intron (NM_138309, intron 1 of 8) | 17115 | NM_138309 | 171486 | Mm.283879 | NM_138309 | ENSMUSG00000035776 | **Cd99l2** | AW548191\|Mic2l1\|Xap89 | CD99 antigen-like 2 | protein-coding |
| chrX_71475529_71476287 | chrX | 71475530 | 71476287 | + | 0.96562 | intron (NM_138309, intron 1 of 8) | intron (NM_138309, intron 1 of 8) | 16941 | NM_138309 | 171486 | Mm.283879 | NM_138309 | ENSMUSG00000035776 | **Cd99l2** | AW548191\|Mic2l1\|Xap89 | CD99 antigen-like 2 | protein-coding |
| chrX_71464168_71464758 | chrX | 71464169 | 71464758 | + | 0.660902 | intron (NM_138309, intron 1 of 8) | intron (NM_138309, intron 1 of 8) | 28386 | NM_138309 | 171486 | Mm.283879 | NM_138309 | ENSMUSG00000035776 | **Cd99l2** | AW548191\|Mic2l1\|Xap89 | CD99 antigen-like 2 | protein-coding |
| chr3_116380277_116381210 | chr3 | 116380278 | 116381210 | + | 1.05368 | intron (NM_001080818, intron 4 of 15) | intron (NM_001080818, intron 4 of 15) | 43288 | NM_001080818 | 229776 | Mm.17647 | NM_001080818 | ENSMUSG00000033502 | **Cdc14a** | A830059A17Rik\|CDC14A2\|CDC14a1\|Cdc14 | CDC14 cell division cycle 14A | protein-coding |
| chr3_116380313_116380787 | chr3 | 116380314 | 116380787 | + | 1.02617 | intron (NM_001080818, intron 4 of 15) | intron (NM_001080818, intron 4 of 15) | 43482 | NM_001080818 | 229776 | Mm.17647 | NM_001080818 | ENSMUSG00000033502 | **Cdc14a** | A830059A17Rik\|CDC14A2\|CDC14a1\|Cdc14 | CDC14 cell division cycle 14A | protein-coding |
| chr3_116368976_116369558 | chr3 | 116368977 | 116369558 | + | 0.96929 | intron (NM_001080818, intron 4 of 15) | intron (NM_001080818, intron 4 of 15) | 54765 | NM_001080818 | 229776 | Mm.17647 | NM_001080818 | ENSMUSG00000033502 | **Cdc14a** | A830059A17Rik\|CDC14A2\|CDC14a1\|Cdc14 | CDC14 cell division cycle 14A | protein-coding |
| chr3_116465658_116466121 | chr3 | 116465659 | 116466121 | + | 0.703444 | Intergenic | L1MEc\|LINE\|L1 | -41858 | NM_001173553 | 229776 | Mm.17647 | NM_001080818 | ENSMUSG00000033502 | **Cdc14a** | A830059A17Rik\|CDC14A2\|CDC14a1\|Cdc14 | CDC14 cell division cycle 14A | protein-coding |
| chr13_113031323_113031659 | chr13 | 113031324 | 113031659 | + | 0.731412 | Intergenic | B4A\|SINE\|B4 | -3888 | NM_001281487 | 238896 | | NM_001281487 | ENSMUSG00000078926 | **Cdc20b** | EG238896\|EG622422 | cell division cycle 20B | protein-coding |
| chr13_113033925_113034137 | chr13 | 113033926 | 113034137 | + | 0.691864 | Intergenic | Intergenic | -1348 | NM_001281487 | 238896 | | NM_001281487 | ENSMUSG00000078926 | **Cdc20b** | EG238896\|EG622422 | cell division cycle 20B | protein-coding |
| chr9_109876863_109877281 | chr9 | 109876864 | 109877281 | + | 0.945223 | intron (NM_007658, intron 1 of 14) | intron (NM_007658, intron 1 of 14) | 1493 | NM_007658 | 12530 | Mm.307103 | NM_007658 | ENSMUSG00000032477 | **Cdc25a** | D9Ertd393e | cell division cycle 25A | protein-coding |
| chr4_137371680_137371955 | chr4 | 137371681 | 137371955 | + | 0.920727 | Intergenic | MER20\|DNA\|hAT-Charlie | -14059 | NM_001243769 | 12540 | Mm.1022 | NM_009861 | ENSMUSG00000006699 | **Cdc42** | AI747189\|AU018915 | cell division cycle 42 | protein-coding |
| chr19_6297466_6298043 | chr19 | 6297467 | 6298043 | + | 1.2435 | intron (NM_010119, intron 4 of 4) | intron (NM_010119, intron 4 of 4) | -8702 | NM_001033342 | 240505 | Mm.329622 | NM_001033342 | ENSMUSG00000024769 | **Cdc42bpg** | BC046418\|MRCKgamma | CDC42 binding protein kinase gamma (DMPK-like) | protein-coding |
| chr19_6296589_6296876 | chr19 | 6296590 | 6296876 | + | 1.00311 | intron (NM_010119, intron 3 of 4) | intron (NM_010119, intron 3 of 4) | -9724 | NM_001033342 | 240505 | Mm.329622 | NM_001033342 | ENSMUSG00000024769 | **Cdc42bpg** | BC046418\|MRCKgamma | CDC42 binding protein kinase gamma (DMPK-like) | protein-coding |
| chr19_6296391_6296879 | chr19 | 6296392 | 6296879 | + | 0.940197 | intron (NM_010119, intron 3 of 4) | intron (NM_010119, intron 3 of 4) | -9822 | NM_001033342 | 240505 | Mm.329622 | NM_001033342 | ENSMUSG00000024769 | **Cdc42bpg** | BC046418\|MRCKgamma | CDC42 binding protein kinase gamma (DMPK-like) | protein-coding |
| chr17_79298630_79299373 | chr17 | 79298631 | 79299373 | + | 1.1846 | Intergenic | Intergenic | 56089 | NM_026514 | 260409 | Mm.140601 | NM_026514 | ENSMUSG00000036533 | **Cdc42ep3** | 3200001F04Rik\|AA986861\|Borg2\|Cep3\|UB1 | CDC42 effector protein (Rho GTPase binding) 3 | protein-coding |
| chr17_79282758_79283273 | chr17 | 79282759 | 79283273 | + | 1.04737 | Intergenic | Intergenic | 72075 | NM_026514 | 260409 | Mm.140601 | NM_026514 | ENSMUSG00000036533 | **Cdc42ep3** | 3200001F04Rik\|AA986861\|Borg2\|Cep3\|UB1 | CDC42 effector protein (Rho GTPase binding) 3 | protein-coding |
| chr17_79300027_79300301 | chr17 | 79300028 | 79300301 | + | 0.980801 | Intergenic | MTEb\|LTR\|ERVL-MaLR | 54927 | NM_026514 | 260409 | Mm.140601 | NM_026514 | ENSMUSG00000036533 | **Cdc42ep3** | 3200001F04Rik\|AA986861\|Borg2\|Cep3\|UB1 | CDC42 effector protein (Rho GTPase binding) 3 | protein-coding |
| chr11_54742850_54743374 | chr11 | 54742851 | 54743374 | + | 1.2971 | intron (NM_178626, intron 2 of 5) | intron (NM_178626, intron 2 of 5) | 44591 | NM_178626 | 72729 | Mm.29476 | NM_178626 | ENSMUSG00000052298 | **Cdc42se2** | 2810404F18Rik\|AA408783\|AA536669\|SPEC2 | CDC42 small effector 2 | protein-coding |
| chr11_54748860_54749315 | chr11 | 54748861 | 54749315 | + | 1.05285 | intron (NM_178626, intron 2 of 5) | B3A\|SINE\|B2 | 38615 | NM_178626 | 72729 | Mm.29476 | NM_178626 | ENSMUSG00000052298 | **Cdc42se2** | 2810404F18Rik\|AA408783\|AA536669\|SPEC2 | CDC42 small effector 2 | protein-coding |
| chr5_107013361_107013872 | chr5 | 107013362 | 107013872 | + | 1.19417 | Intergenic | B4A\|SINE\|B4 | 49056 | NM_001271566 | 12545 | Mm.20842 | NM_009863 | ENSMUSG00000029283 | **Cdc7** | AI597260\|Cdc7l1\|muCdc7 | cell division cycle 7 (S. cerevisiae) | protein-coding |
| chr5_106959277_106959771 | chr5 | 106959278 | 106959771 | + | 1.01433 | Intergenic | Intergenic | -4798 | NM_009863 | 12545 | Mm.20842 | NM_009863 | ENSMUSG00000029283 | **Cdc7** | AI597260\|Cdc7l1\|muCdc7 | cell division cycle 7 (S. cerevisiae) | protein-coding |
| chr19_6085471_6085973 | chr19 | 6085472 | 6085973 | + | 0.651342 | promoter-TSS (NM_024231) | promoter-TSS (NM_024231) | 625 | NM_026410 | 67849 | Mm.23526 | NM_026410 | ENSMUSG00000024791 | **Cdca5** | 2610036L13Rik\|AL024086\|AW536684\|C85404 | cell division cycle associated 5 | protein-coding |
| chr19_6085578_6085789 | chr19 | 6085579 | 6085789 | + | 0.602439 | promoter-TSS (NM_024231) | promoter-TSS (NM_024231) | 587 | NM_026410 | 67849 | Mm.23526 | NM_026410 | ENSMUSG00000024791 | **Cdca5** | 2610036L13Rik\|AL024086\|AW536684\|C85404 | cell division cycle associated 5 | protein-coding |
| chr2_72535836_72536275 | chr2 | 72535837 | 72536275 | + | 1.30661 | Intergenic | Intergenic | 59837 | NM_025866 | 66953 | Mm.270676 | NM_025866 | ENSMUSG00000055612 | **Cdca7** | 2310021G01Rik\|AU044109\|JPO1 | cell division cycle associated 7 | protein-coding |
| chr2_72535836_72536438 | chr2 | 72535837 | 72536438 | + | 1.24912 | Intergenic | Intergenic | 59918 | NM_025866 | 66953 | Mm.270676 | NM_025866 | ENSMUSG00000055612 | **Cdca7** | 2310021G01Rik\|AU044109\|JPO1 | cell division cycle associated 7 | protein-coding |
| chr2_72500344_72500854 | chr2 | 72500345 | 72500854 | + | 0.771568 | Intergenic | Intergenic | 24380 | NM_025866 | 66953 | Mm.270676 | NM_025866 | ENSMUSG00000055612 | **Cdca7** | 2310021G01Rik\|AU044109\|JPO1 | cell division cycle associated 7 | protein-coding |
| chr2_72500107_72500880 | chr2 | 72500108 | 72500880 | + | 0.727247 | Intergenic | Intergenic | 24275 | NM_025866 | 66953 | Mm.270676 | NM_025866 | ENSMUSG00000055612 | **Cdca7** | 2310021G01Rik\|AU044109\|JPO1 | cell division cycle associated 7 | protein-coding |
| chr12_117810053_117810861 | chr12 | 117810054 | 117810861 | + | 0.800653 | Intergenic | Intergenic | -33404 | NM_146040 | 217946 | Mm.281149 | NM_146040 | ENSMUSG00000021175 | **Cdca7l** | BC006933\|JPO2 | cell division cycle associated 7 like | protein-coding |
| chr12_117810058_117810556 | chr12 | 117810059 | 117810556 | + | 0.798612 | Intergenic | Intergenic | -33554 | NM_146040 | 217946 | Mm.281149 | NM_146040 | ENSMUSG00000021175 | **Cdca7l** | BC006933\|JPO2 | cell division cycle associated 7 like | protein-coding |
| chr4_124927898_124928613 | chr4 | 124927899 | 124928613 | + | 1.24632 | intron (NM_026560, intron 3 of 9) | Charlie19a\|DNA\|hAT-Charlie | 8661 | NM_026560 | 52276 | Mm.28038 | NM_026560 | ENSMUSG00000028873 | **Cdca8** | 4831429J16Rik\|AU044747\|BOR\|Borealin\|D4Ertd421e\|DasraB\|MESRGP | cell division cycle associated 8 | protein-coding |
| chr4_11734214_11734641 | chr4 | 11734215 | 11734641 | + | 0.767953 | Intergenic | Intergenic | -23729 | NM_019753 | 12557 | Mm.33402 | NM_019753 | ENSMUSG00000028217 | **Cdh17** | HPT-1\|HPT-1/LI | cadherin 17 | protein-coding |
| chr12_33040747_33041115 | chr12 | 33040748 | 33041115 | + | 0.717958 | intron (NM_001024478, intron 15 of 19) | intron (NM_001024478, intron 15 of 19) | 51944 | NM_001024478 | 68764 | Mm.60083 | NM_001024478 | ENSMUSG00000035860 | **Cdhr3** | 1110049B09Rik\|BB172694\|Cdh28 | cadherin-related family member 3 | protein-coding |

| | | | | | | | | | | | | | | | | |
|---|---|---|---|---|---|---|---|---|---|---|---|---|---|---|---|---|
| chr13_17783720_17784715 | chr13 | 17783721 | 17784715 | + | 1.08127 | intron (NM_001081058, intron 1 of 13) | MLTR11B\|LTR\|ERVK | 20879 | NM_001081058 | 69562 | Mm.193924 | NM_027118 | ENSMUSG00000041297 | **Cdk13** | 2310015O17Rik\|Cdc2l5 | cyclin-dependent kinase 13 | protein-coding |
| chr10_93049377_93049991 | chr10 | 93049378 | 93049991 | + | 1.31752 | Intergenic | Intergenic | -111192 | NM_146239 | 237459 | Mm.217404 | NM_146239 | ENSMUSG00000020015 | **Cdk17** | 6430598J10Rik\|Pctk2 | cyclin-dependent kinase 17 | protein-coding |
| chr10_93083240_93083794 | chr10 | 93083241 | 93083794 | + | 1.1494 | Intergenic | Intergenic | -77359 | NM_146239 | 237459 | Mm.217404 | NM_146239 | ENSMUSG00000020015 | **Cdk17** | 6430598J10Rik\|Pctk2 | cyclin-dependent kinase 17 | protein-coding |
| chr10_93029528_93030103 | chr10 | 93029529 | 93030103 | + | 1.08107 | Intergenic | Intergenic | -131060 | NM_146239 | 237459 | Mm.217404 | NM_146239 | ENSMUSG00000020015 | **Cdk17** | 6430598J10Rik\|Pctk2 | cyclin-dependent kinase 17 | protein-coding |
| chr10_93138294_93138701 | chr10 | 93138295 | 93138701 | + | 0.967396 | Intergenic | L1MB3\|LINE\|L1 | -22378 | NM_146239 | 237459 | Mm.217404 | NM_146239 | ENSMUSG00000020015 | **Cdk17** | 6430598J10Rik\|Pctk2 | cyclin-dependent kinase 17 | protein-coding |
| chr10_93029525_93030417 | chr10 | 93029526 | 93030417 | + | 0.903936 | Intergenic | Intergenic | -130905 | NM_146239 | 237459 | Mm.217404 | NM_146239 | ENSMUSG00000020015 | **Cdk17** | 6430598J10Rik\|Pctk2 | cyclin-dependent kinase 17 | protein-coding |
| chr10_93084363_93084808 | chr10 | 93084364 | 93084808 | + | 0.672965 | Intergenic | Tigger5\|DNA\|TcMar-Tigger | -76290 | NM_146239 | 237459 | Mm.217404 | NM_146239 | ENSMUSG00000020015 | **Cdk17** | 6430598J10Rik\|Pctk2 | cyclin-dependent kinase 17 | protein-coding |
| chr1_132145137_132145485 | chr1 | 132145138 | 132145485 | + | 0.75779 | Intergenic | Intergenic | -5626 | NM_008795 | 18557 | Mm.28130 | NM_008795 | ENSMUSG00000026437 | **Cdk18** | AA682070\|Ctk18\|Pctk3 | cyclin-dependent kinase 18 | protein-coding |
| chr10_40372263_40372620 | chr10 | 40372264 | 40372620 | + | 0.694387 | intron (NM_001291816, intron 1 of 11) | intron (NM_001291816, intron 1 of 11) | 23134 | NM_001291816 | 78334 | Mm.200924 | NM_198164 | ENSMUSG00000038481 | **Cdk19** | 2700084L06Rik\|AI845279\|AW228747\|CDK11\|Cdc2l6\|mKIAA1028 | cyclin-dependent kinase 19 | protein-coding |
| chr10_40372208_40372720 | chr10 | 40372209 | 40372720 | + | 0.625272 | intron (NM_001291816, intron 1 of 11) | intron (NM_001291816, intron 1 of 11) | 23156 | NM_001291816 | 78334 | Mm.200924 | NM_198164 | ENSMUSG00000038481 | **Cdk19** | 2700084L06Rik\|AI845279\|AW228747\|CDK11\|Cdc2l6\|mKIAA1028 | cyclin-dependent kinase 19 | protein-coding |
| chr11_80496631_80497104 | chr11 | 80496632 | 80497104 | + | 1.05354 | intron (NM_177390, intron 21 of 21) | intron (NM_177390, intron 21 of 21) | 19822 | NM_009871 | 12569 | Mm.142275 | NM_009871 | ENSMUSG00000048895 | **Cdk5r1** | Cdk5r\|D11Bwg0379e\|p25\|p35 | cyclin-dependent kinase 5, regulatory subunit 1 (p35) | protein-coding |
| chr11_80504889_80505388 | chr11 | 80504890 | 80505388 | + | 0.778357 | intron (NM_177390, intron 21 of 21) | intron (NM_177390, intron 21 of 21) | 28093 | NM_009871 | 12569 | Mm.142275 | NM_009871 | ENSMUSG00000048895 | **Cdk5r1** | Cdk5r\|D11Bwg0379e\|p25\|p35 | cyclin-dependent kinase 5, regulatory subunit 1 (p35) | protein-coding |
| chr5_3151921_3152797 | chr5 | 3151922 | 3152797 | + | 1.39015 | Intergenic | Intergenic | -191953 | NM_009873 | 12571 | Mm.31672 | NM_009873 | ENSMUSG00000040274 | **Cdk6** | 5830411I20\|AI504062\|Crk2 | cyclin-dependent kinase 6 | protein-coding |
| chr5_3355908_3356200 | chr5 | 3355909 | 3356200 | + | 1.32815 | intron (NM_009873, intron 1 of 6) | intron (NM_009873, intron 1 of 6) | 11742 | NM_009873 | 12571 | Mm.31672 | NM_009873 | ENSMUSG00000040274 | **Cdk6** | 5830411I20\|AI504062\|Crk2 | cyclin-dependent kinase 6 | protein-coding |
| chr5_3300762_3301529 | chr5 | 3300763 | 3301529 | + | 1.14809 | Intergenic | Intergenic | -43166 | NM_009873 | 12571 | Mm.31672 | NM_009873 | ENSMUSG00000040274 | **Cdk6** | 5830411I20\|AI504062\|Crk2 | cyclin-dependent kinase 6 | protein-coding |
| chr5_3355502_3356573 | chr5 | 3355503 | 3356573 | + | 1.10734 | intron (NM_009873, intron 1 of 6) | intron (NM_009873, intron 1 of 6) | 11726 | NM_009873 | 12571 | Mm.31672 | NM_009873 | ENSMUSG00000040274 | **Cdk6** | 5830411I20\|AI504062\|Crk2 | cyclin-dependent kinase 6 | protein-coding |
| chr5_3301185_3301490 | chr5 | 3301186 | 3301490 | + | 1.04067 | Intergenic | Intergenic | -42974 | NM_009873 | 12571 | Mm.31672 | NM_009873 | ENSMUSG00000040274 | **Cdk6** | 5830411I20\|AI504062\|Crk2 | cyclin-dependent kinase 6 | protein-coding |
| chr5_3355960_3356508 | chr5 | 3355961 | 3356508 | + | 1.02569 | intron (NM_009873, intron 1 of 6) | intron (NM_009873, intron 1 of 6) | 11922 | NM_009873 | 12571 | Mm.31672 | NM_009873 | ENSMUSG00000040274 | **Cdk6** | 5830411I20\|AI504062\|Crk2 | cyclin-dependent kinase 6 | protein-coding |
| chr5_3219766_3220151 | chr5 | 3219767 | 3220151 | + | 0.798585 | Intergenic | Intergenic | -124353 | NM_009873 | 12571 | Mm.31672 | NM_009873 | ENSMUSG00000040274 | **Cdk6** | 5830411I20\|AI504062\|Crk2 | cyclin-dependent kinase 6 | protein-coding |
| chr5_3351771_3352097 | chr5 | 3351772 | 3352097 | + | 0.758837 | intron (NM_009873, intron 1 of 6) | intron (NM_009873, intron 1 of 6) | 7622 | NM_009873 | 12571 | Mm.31672 | NM_009873 | ENSMUSG00000040274 | **Cdk6** | 5830411I20\|AI504062\|Crk2 | cyclin-dependent kinase 6 | protein-coding |
| chr5_146238076_146238630 | chr5 | 146238077 | 146238630 | + | 0.894277 | intron (NM_153599, intron 1 of 12) | intron (NM_153599, intron 1 of 12) | 6678 | NM_153599 | 264064 | Mm.260576 | NM_153599 | ENSMUSG00000029635 | **Cdk8** | - | cyclin-dependent kinase 8 | protein-coding |
| chr5_146286974_146287459 | chr5 | 146286975 | 146287459 | + | 0.818584 | intron (NM_153599, intron 6 of 12) | intron (NM_153599, intron 6 of 12) | 55542 | NM_153599 | 264064 | Mm.260576 | NM_153599 | ENSMUSG00000029635 | **Cdk8** | - | cyclin-dependent kinase 8 | protein-coding |
| chr5_146282216_146282526 | chr5 | 146282217 | 146282526 | + | 0.706554 | intron (NM_153599, intron 4 of 12) | intron (NM_153599, intron 4 of 12) | 50696 | NM_153599 | 264064 | Mm.260576 | NM_153599 | ENSMUSG00000029635 | **Cdk8** | - | cyclin-dependent kinase 8 | protein-coding |
| chr5_146281964_146282552 | chr5 | 146281965 | 146282552 | + | 0.647773 | intron (NM_153599, intron 4 of 12) | intron (NM_153599, intron 4 of 12) | 50583 | NM_153599 | 264064 | Mm.260576 | NM_153599 | ENSMUSG00000029635 | **Cdk8** | - | cyclin-dependent kinase 8 | protein-coding |
| chr13_29640244_29640633 | chr13 | 29640245 | 29640633 | + | 0.762061 | intron (NM_144536, intron 8 of 15) | intron (NM_144536, intron 8 of 15) | 215149 | NM_144536 | 68916 | Mm.212227 | NM_144536 | ENSMUSG00000006191 | **Cdkal1** | 1190005B03Rik\|6620401C13Rik | CDK5 regulatory subunit associated protein 1-like 1 | protein-coding |
| chr11_52035125_52035687 | chr11 | 52035126 | 52035687 | + | 0.989973 | intron (NM_001166654, intron 12 of 13) | RSINE1\|SINE\|B4 | 31185 | NM_001166653 | 213084 | Mm.280557 | NM_153785 | ENSMUSG00000020389 | **Cdkl3** | AI852479\|B230379H01Rik | cyclin-dependent kinase-like 3 | protein-coding |
| chr17_80632394_80632783 | chr17 | 80632395 | 80632783 | + | 0.640214 | intron (NM_001290345, intron 13 of 33) | intron (NM_001290345, intron 13 of 33) | -68755 | NM_001033443 | 381113 | Mm.329216 | NM_001033443 | ENSMUSG00000033966 | **Cdkl4** | AU067824\|Gm942 | cyclin-dependent kinase-like 4 | protein-coding |
| chr4_89296531_89296842 | chr4 | 89296532 | 89296842 | + | 1.18452 | Intergenic | Intergenic | -2068 | NM_009877 | 12578 | Mm.4733 | NM_009877 | ENSMUSG00000044303 | **Cdkn2a** | ARF-INK4a\|Arf\|INK4a-ARF\|Ink4a/Arf\|MTS1\|Pctr1\|p16\|p16(INK4a)\|p16INK4a\|p19<ARF>\|p19ARF | cyclin-dependent kinase inhibitor 2A | protein-coding |
| chr4_89257775_89258155 | chr4 | 89257776 | 89258155 | + | 0.881108 | Intergenic | RMER15\|LTR\|ERVL | 24227 | NM_001040654 | 12578 | Mm.4733 | NM_009877 | ENSMUSG00000044303 | **Cdkn2a** | ARF-INK4a\|Arf\|INK4a-ARF\|Ink4a/Arf\|MTS1\|Pctr1\|p16\|p16(INK4a)\|p16INK4a\|p19<ARF>\|p19ARF | cyclin-dependent kinase inhibitor 2A | protein-coding |
| chr4_89302100_89302497 | chr4 | 89302101 | 89302497 | + | 0.836609 | Intergenic | Lx7\|LINE\|L1 | -7680 | NM_009877 | 12578 | Mm.4733 | NM_009877 | ENSMUSG00000044303 | **Cdkn2a** | ARF-INK4a\|Arf\|INK4a-ARF\|Ink4a/Arf\|MTS1\|Pctr1\|p16\|p16(INK4a)\|p16INK4a\|p19<ARF>\|p19ARF | cyclin-dependent kinase inhibitor 2A | protein-coding |
| chr11_51915288_51915989 | chr11 | 51915289 | 51915989 | + | 1.11964 | Intergenic | Intergenic | -51992 | NM_029976 | 52626 | Mm.289456 | NM_029976 | ENSMUSG00000020392 | **Cdkn2aipnl** | A430101B06Rik\|AA409260\|D11Ertd497e\|MNCb-1520 | CDKN2A interacting protein N-terminal like | protein-coding |
| chr4_89395359_89395757 | chr4 | 89395360 | 89395757 | + | 1.60484 | Intergenic | Intergenic | -84526 | NM_007670 | 12579 | Mm.423094 | NM_007670 | ENSMUSG00000073802 | **Cdkn2b** | AV083695\|INK4b\|MTS2\|p15\|p15(INK4b)\|p15INK4b | cyclin-dependent kinase inhibitor 2B (p15, inhibits CDK4) | protein-coding |
| chr4_89393162_89393844 | chr4 | 89393163 | 89393844 | + | 1.53376 | Intergenic | Intergenic | -82471 | NM_007670 | 12579 | Mm.423094 | NM_007670 | ENSMUSG00000073802 | **Cdkn2b** | AV083695\|INK4b\|MTS2\|p15\|p15(INK4b)\|p15INK4b | cyclin-dependent kinase inhibitor 2B (p15, inhibits CDK4) | protein-coding |
| chr4_89388116_89388489 | chr4 | 89388117 | 89388489 | + | 1.51541 | Intergenic | Intergenic | -77271 | NM_007670 | 12579 | Mm.423094 | NM_007670 | ENSMUSG00000073802 | **Cdkn2b** | AV083695\|INK4b\|MTS2\|p15\|p15(INK4b)\|p15INK4b | cyclin-dependent kinase inhibitor 2B (p15, inhibits CDK4) | protein-coding |
| chr4_89420838_89421287 | chr4 | 89420839 | 89421287 | + | 1.09406 | Intergenic | Intergenic | -110031 | NM_007670 | 12579 | Mm.423094 | NM_007670 | ENSMUSG00000073802 | **Cdkn2b** | AV083695\|INK4b\|MTS2\|p15\|p15(INK4b)\|p15INK4b | cyclin-dependent kinase inhibitor 2B (p15, inhibits CDK4) | protein-coding |
| chr4_89359476_89359998 | chr4 | 89359477 | 89359998 | + | 1.01234 | Intergenic | Intergenic | -48705 | NM_007670 | 12579 | Mm.423094 | NM_007670 | ENSMUSG00000073802 | **Cdkn2b** | AV083695\|INK4b\|MTS2\|p15\|p15(INK4b)\|p15INK4b | cyclin-dependent kinase inhibitor 2B (p15, inhibits CDK4) | protein-coding |
| chr4_89351395_89351870 | chr4 | 89351396 | 89351870 | + | 0.81616 | Intergenic | Intergenic | -40601 | NM_007670 | 12579 | Mm.423094 | NM_007670 | ENSMUSG00000073802 | **Cdkn2b** | AV083695\|INK4b\|MTS2\|p15\|p15(INK4b)\|p15INK4b | cyclin-dependent kinase inhibitor 2B (p15, inhibits CDK4) | protein-coding |
| chr4_89387181_89387586 | chr4 | 89387182 | 89387586 | + | 0.777342 | Intergenic | Intergenic | -76352 | NM_007670 | 12579 | Mm.423094 | NM_007670 | ENSMUSG00000073802 | **Cdkn2b** | AV083695\|INK4b\|MTS2\|p15\|p15(INK4b)\|p15INK4b | cyclin-dependent kinase inhibitor 2B (p15, inhibits CDK4) | protein-coding |
| chr4_89350874_89351870 | chr4 | 89350875 | 89351870 | + | 0.638943 | Intergenic | Intergenic | -40340 | NM_007670 | 12579 | Mm.423094 | NM_007670 | ENSMUSG00000073802 | **Cdkn2b** | AV083695\|INK4b\|MTS2\|p15\|p15(INK4b)\|p15INK4b | cyclin-dependent kinase inhibitor 2B (p15, inhibits CDK4) | protein-coding |
| chr9_35483537_35484465 | chr9 | 35483538 | 35484465 | + | 1.2342 | intron (NM_021339, intron 14 of 18) | intron (NM_021339, intron 14 of 18) | 31925 | NM_021339 | 57810 | Mm.80509 | NM_021339 | ENSMUSG00000038119 | **Cdon** | CDO\|Cd0\|Orcam | cell adhesion molecule-related/down-regulated by oncogenes | protein-coding |
| chr9_35432908_35433309 | chr9 | 35432909 | 35433309 | + | 0.790007 | Intergenic | Lx\|LINE\|L1 | -18967 | NM_021339 | 57810 | Mm.80509 | NM_021339 | ENSMUSG00000038119 | **Cdon** | CDO\|Cd0\|Orcam | cell adhesion molecule-related/down-regulated by oncogenes | protein-coding |
| chr11_62957760_62958138 | chr11 | 62957761 | 62958138 | + | 1.53046 | intron (NM_025496, intron 1 of 1) | L1M5\|LINE\|L1 | 6756 | NM_025496 | 66338 | Mm.46166 | NM_025496 | ENSMUSG00000042200 | **Cdrt4** | 1700019I23Rik | CMT1A duplicated region transcript 4 | protein-coding |
| chr11_62963281_62963777 | chr11 | 62963282 | 62963777 | + | 1.04591 | intron (NR_105772, intron 5 of 6) | intron (NR_105772, intron 5 of 6) | 12336 | NM_025496 | 66338 | Mm.46166 | NM_025496 | ENSMUSG00000042200 | **Cdrt4** | 1700019I23Rik | CMT1A duplicated region transcript 4 | protein-coding |
| chr2_132286486_132286703 | chr2 | 132286487 | 132286703 | + | 0.629207 | intron (NM_001291039, intron 1 of 12) | RMER3D-int\|LTR\|ERVK | 656 | NM_001291039 | 110911 | Mm.284503 | NM_138651 | ENSMUSG00000058793 | **Cds2** | 5730450N06Rik\|5730460C18Rik\|AI854580\|D2Wsu127e | CDP-diacylglycerol synthase (phosphatidate cytidylyltransferase) 2 | protein-coding |
| chr2_132292052_132292652 | chr2 | 132292053 | 132292652 | + | 0.6092 | intron (NM_001291039, intron 1 of 12) | intron (NM_001291039, intron 1 of 12) | 6413 | NM_001291039 | 110911 | Mm.284503 | NM_138651 | ENSMUSG00000058793 | **Cds2** | 5730450N06Rik\|5730460C18Rik\|AI854580\|D2Wsu127e | CDP-diacylglycerol synthase (phosphatidate cytidylyltransferase) 2 | protein-coding |
| chr13_35742870_35743288 | chr13 | 35742871 | 35743288 | + | 0.907439 | intron (NM_001123386, intron 1 of 6) | intron (NM_001123386, intron 1 of 6) | 1677 | NM_001123386 | 12593 | Mm.29002 | NM_009881 | ENSMUSG00000059288 | **Cdyl** | AI325931 | chromodomain protein, Y chromosome-like | protein-coding |
| chr8_116735346_116736008 | chr8 | 116735347 | 116736008 | + | 1.2279 | Intergenic | Intergenic | -2686 | NM_029441 | 75796 | Mm.159575 | NM_029441 | ENSMUSG00000031758 | **Cdyl2** | 1700029M19Rik\|4930453I21Rik\|Gm20194 | chromodomain protein, Y chromosome-like 2 | protein-coding |
| chr8_116676381_116676710 | chr8 | 116676382 | 116676710 | + | 1.04999 | intron (NM_029441, intron 1 of 6) | intron (NM_029441, intron 1 of 6) | 56445 | NM_029441 | 75796 | Mm.159575 | NM_029441 | ENSMUSG00000031758 | **Cdyl2** | 1700029M19Rik\|4930453I21Rik\|Gm20194 | chromodomain protein, Y chromosome-like 2 | protein-coding |
| chr8_116673975_116674678 | chr8 | 116673976 | 116674678 | + | 0.853042 | intron (NM_029441, intron 1 of 6) | (CAGGG)n\|Simple_repeat\|Simple_repeat | 58664 | NM_029441 | 75796 | Mm.159575 | NM_029441 | ENSMUSG00000031758 | **Cdyl2** | 1700029M19Rik\|4930453I21Rik\|Gm20194 | chromodomain protein, Y chromosome-like 2 | protein-coding |

| | | | | | | | | | | | | | | | | | |
|---|---|---|---|---|---|---|---|---|---|---|---|---|---|---|---|---|---|
| chr8_116673975_116674418 | chr8 | 116673976 | 116674418 | + | 0.842494 | intron (NM_029441, intron 1 of 6) | intron (NM_029441, intron 1 of 6) | 58794 | NM_029441 | 75796 | Mm.159575 | NM_029441 | ENSMUSG00000031758 | **Cdyl2** | 1700029M19Rik\|4930453I21Rik\|Gm20194 | chromodomain protein, Y chromosome-like 2 | protein-coding |
| chr8_116673811_116674398 | chr8 | 116673812 | 116674398 | + | 0.81838 | intron (NM_029441, intron 1 of 6) | intron (NM_029441, intron 1 of 6) | 58886 | NM_029441 | 75796 | Mm.159575 | NM_029441 | ENSMUSG00000031758 | **Cdyl2** | 1700029M19Rik\|4930453I21Rik\|Gm20194 | chromodomain protein, Y chromosome-like 2 | protein-coding |
| chr8_116633857_116634181 | chr8 | 116633858 | 116634181 | + | 0.761152 | intron (NM_029441, intron 1 of 6) | MTD\|LTR\|ERVL-MaLR | 98972 | NM_029441 | 75796 | Mm.159575 | NM_029441 | ENSMUSG00000031758 | **Cdyl2** | 1700029M19Rik\|4930453I21Rik\|Gm20194 | chromodomain protein, Y chromosome-like 2 | protein-coding |
| chr8_116621702_116622104 | chr8 | 116621703 | 116622104 | + | 0.734751 | intron (NM_029441, intron 2 of 6) | intron (NM_029441, intron 2 of 6) | 111088 | NM_029441 | 75796 | Mm.159575 | NM_029441 | ENSMUSG00000031758 | **Cdyl2** | 1700029M19Rik\|4930453I21Rik\|Gm20194 | chromodomain protein, Y chromosome-like 2 | protein-coding |
| chr8_116716858_116717674 | chr8 | 116716859 | 116717674 | + | 0.656347 | intron (NM_029441, intron 1 of 6) | MTC-int\|LTR\|ERVL-MaLR | 15725 | NM_029441 | 75796 | Mm.159575 | NM_029441 | ENSMUSG00000031758 | **Cdyl2** | 1700029M19Rik\|4930453I21Rik\|Gm20194 | chromodomain protein, Y chromosome-like 2 | protein-coding |
| chr7_16681538_16682098 | chr7 | 16681539 | 16682098 | + | 0.791627 | Intergenic | RLTR48A\|LTR\|ERV1 | -6113 | NM_175315 | 101434 | Mm.102767 | NM_175315 | ENSMUSG00000078795 | **Ceacam15** | AA407838\|C430002N04Rik | carcinoembryonic antigen-related cell adhesion molecule 15 | protein-coding |
| chr7_16680986_16681960 | chr7 | 16680987 | 16681960 | + | 0.727886 | Intergenic | RLTR48A\|LTR\|ERV1 | -5768 | NM_175315 | 101434 | Mm.102767 | NM_175315 | ENSMUSG00000078795 | **Ceacam15** | AA407838\|C430002N04Rik | carcinoembryonic antigen-related cell adhesion molecule 15 | protein-coding |
| chr7_17135325_17135676 | chr7 | 17135326 | 17135676 | + | 0.652342 | Intergenic | B3\|SINE\|B2 | -14899 | NM_054059 | 384557 | Mm.486145 | NM_054059 | ENSMUSG00000053228 | **Ceacam3** | EG384557\|Psg24\|cea12 | carcinoembryonic antigen-related cell adhesion molecule 3 | protein-coding |
| chr16_15973784_15974149 | chr16 | 15973785 | 15974149 | + | 1.01739 | intron (NM_146068, intron 8 of 19) | Lx2A1\|LINE\|L1 | 86681 | NM_007679 | 12609 | Mm.347407 | NM_007679 | | **Cebpd** | - | CCAAT/enhancer binding protein (C/EBP), delta | protein-coding |
| chr16_15972616_15972986 | chr16 | 15972617 | 15972986 | + | 0.678189 | intron (NM_146068, intron 8 of 19) | Lx2A1\|LINE\|L1 | 85515 | NM_007679 | 12609 | Mm.347407 | NM_007679 | | **Cebpd** | - | CCAAT/enhancer binding protein (C/EBP), delta | protein-coding |
| chr14_54727542_54727940 | chr14 | 54727543 | 54727940 | + | 0.876506 | intron (NM_016972, intron 8 of 10) | intron (NM_016972, intron 8 of 10) | -15567 | NM_207131 | 110794 | Mm.236223 | NM_207131 | ENSMUSG00000052435 | **Cebpe** | C/EBPe\|CRP1\|Gm294 | CCAAT/enhancer binding protein (C/EBP), epsilon | protein-coding |
| chr15_100687010_100687933 | chr15 | 100687011 | 100687933 | + | 0.702421 | intron (NM_033612, intron 1 of 7) | intron (NM_033612, intron 1 of 7) | 448 | NM_033612 | 109901 | Mm.2131 | NM_033612 | ENSMUSG00000023031 | **Cela1** | 1810009A17Rik\|1810062B19Rik\|Ela-1\|Ela1\|PC-TsF | chymotrypsin-like elastase family, member 1 | protein-coding |
| chr2_90975064_90975804 | chr2 | 90975065 | 90975804 | + | 1.60408 | intron (NM_001244891, intron 2 of 14) | intron (NM_001244891, intron 2 of 14) | 35037 | NM_017368 | 13046 | Mm.29495 | NM_017368 | ENSMUSG00000005506 | **Celf1** | 1600010O03Rik\|AA407467\|Brunol2\|CUG-BP\|CUG-BP1\|CUGBP\|Cugbp1\|D2Wsu101e\|HNAB50\|NAB50 | CUGBP, Elav-like family member 1 | protein-coding |
| chr2_6646455_6647337 | chr2 | 6646456 | 6647337 | + | 1.20338 | intron (NM_001110232, intron 5 of 15) | intron (NM_001110232, intron 5 of 15) | 238100 | NM_010160 | 14007 | Mm.398543 | NM_010160 | ENSMUSG00000002107 | **Celf2** | B230218O03\|B230345P09Rik\|C88023\|CELF-2\|CUG-BP2\|Cugbp2\|D230046B21Rik\|Etr-3\|Napor\|Napor-2\|mETR-3 | CUGBP, Elav-like family member 2 | protein-coding |
| chr2_6704660_6705340 | chr2 | 6704661 | 6705340 | + | 1.04557 | intron (NM_001110232, intron 4 of 15) | intron (NM_001110232, intron 4 of 15) | 179996 | NM_010160 | 14007 | Mm.398543 | NM_010160 | ENSMUSG00000002107 | **Celf2** | B230218O03\|B230345P09Rik\|C88023\|CELF-2\|CUG-BP2\|Cugbp2\|D230046B21Rik\|Etr-3\|Napor\|Napor-2\|mETR-3 | CUGBP, Elav-like family member 2 | protein-coding |
| chr2_6695421_6696208 | chr2 | 6695422 | 6696208 | + | 0.901773 | intron (NM_001110232, intron 4 of 15) | intron (NM_001110232, intron 4 of 15) | 189181 | NM_010160 | 14007 | Mm.398543 | NM_010160 | ENSMUSG00000002107 | **Celf2** | B230218O03\|B230345P09Rik\|C88023\|CELF-2\|CUG-BP2\|Cugbp2\|D230046B21Rik\|Etr-3\|Napor\|Napor-2\|mETR-3 | CUGBP, Elav-like family member 2 | protein-coding |
| chr2_6733243_6733950 | chr2 | 6733244 | 6733950 | + | 0.629478 | intron (NM_001110232, intron 3 of 15) | intron (NM_001110232, intron 3 of 15) | 151399 | NM_010160 | 14007 | Mm.398543 | NM_010160 | ENSMUSG00000002107 | **Celf2** | B230218O03\|B230345P09Rik\|C88023\|CELF-2\|CUG-BP2\|Cugbp2\|D230046B21Rik\|Etr-3\|Napor\|Napor-2\|mETR-3 | CUGBP, Elav-like family member 2 | protein-coding |
| chr2_6695406_6696634 | chr2 | 6695407 | 6696634 | + | 0.61729 | intron (NM_001110232, intron 4 of 15) | intron (NM_001110232, intron 4 of 15) | 188976 | NM_010160 | 14007 | Mm.398543 | NM_010160 | ENSMUSG00000002107 | **Celf2** | B230218O03\|B230345P09Rik\|C88023\|CELF-2\|CUG-BP2\|Cugbp2\|D230046B21Rik\|Etr-3\|Napor\|Napor-2\|mETR-3 | CUGBP, Elav-like family member 2 | protein-coding |
| chr3_94482852_94483137 | chr3 | 94482853 | 94483137 | + | 0.733427 | intron (NR_110353, intron 1 of 11) | intron (NR_110353, intron 1 of 11) | 273 | NR_110353 | 78784 | Mm.44292 | NM_172434 | ENSMUSG00000028137 | **Celf3** | 4930415M08Rik\|BRUNOL1\|CAGH4\|ERDA4\|Tnrc4 | CUGBP, Elav-like family member 3 | protein-coding |
| chr15_86020022_86020402 | chr15 | 86020023 | 86020402 | + | 0.931828 | intron (NM_009886, intron 1 of 34) | intron (NM_009886, intron 1 of 34) | 13565 | NM_009886 | 12614 | Mm.22680 | NM_009886 | ENSMUSG00000016028 | **Celsr1** | Crsh\|Scy\|crash | cadherin, EGF LAG seven-pass G-type receptor 1 (flamingo homolog, Drosophila) | protein-coding |
| chr15_86034800_86035108 | chr15 | 86034801 | 86035108 | + | 0.621563 | Intergenic | Intergenic | -1177 | NM_009886 | 12614 | Mm.22680 | NM_009886 | ENSMUSG00000016028 | **Celsr1** | Crsh\|Scy\|crash | cadherin, EGF LAG seven-pass G-type receptor 1 (flamingo homolog, Drosophila) | protein-coding |
| chr5_30679272_30680006 | chr5 | 30679273 | 30680006 | + | 1.19145 | Intergenic | RLTR19\|LTR\|ERVK | 12737 | NM_007681 | 12615 | Mm.290563 | NM_007681 | ENSMUSG00000029177 | **Cenpa** | Cenp-A | centromere protein A | protein-coding |
| chr5_30678990_30680033 | chr5 | 30678991 | 30680033 | + | 1.04915 | Intergenic | PB1D10\|SINE\|Alu | 12610 | NM_007681 | 12615 | Mm.290563 | NM_007681 | ENSMUSG00000029177 | **Cenpa** | Cenp-A | centromere protein A | protein-coding |
| chr5_86068504_86068712 | chr5 | 86068505 | 86068712 | + | 1.1943 | Intergenic | Intergenic | -3025 | NM_007683 | 12617 | Mm.4649 | NM_007683 | ENSMUSG00000029253 | **Cenpc1** | CENP-C\|Cenpc | centromere protein C1 | protein-coding |
| chr1_189615268_189615729 | chr1 | 189615269 | 189615729 | + | 1.35896 | Intergenic | Intergenic | 72587 | NM_001081363 | 108000 | Mm.129746 | NM_001081363 | ENSMUSG00000026605 | **Cenpf** | 6530404A22Rik\|AI325968\|CENF\|Lek1\|PRO1779\|mitosin | centromere protein F | protein-coding |
| chr1_189703169_189704043 | chr1 | 189703170 | 189704043 | + | 0.67665 | Intergenic | Intergenic | -15520 | NM_001081363 | 108000 | Mm.129746 | NM_001081363 | ENSMUSG00000026605 | **Cenpf** | 6530404A22Rik\|AI325968\|CENF\|Lek1\|PRO1779\|mitosin | centromere protein F | protein-coding |
| chr12_4293835_4294392 | chr12 | 4293836 | 4294392 | + | 0.886312 | intron (NM_010881, intron 12 of 22) | intron (NM_010881, intron 12 of 22) | -59820 | NM_134046 | 52504 | Mm.70546 | NM_134046 | ENSMUSG00000020652 | **Cenpo** | 2810429O05Rik\|8430427C03Rik\|D12Ertd482e | centromere protein O | protein-coding |
| chr12_4279270_4280124 | chr12 | 4279271 | 4280124 | + | 0.625358 | intron (NM_010881, intron 13 of 22) | intron (NM_010881, intron 13 of 22) | -45403 | NM_134046 | 52504 | Mm.70546 | NM_134046 | ENSMUSG00000020652 | **Cenpo** | 2810429O05Rik\|8430427C03Rik\|D12Ertd482e | centromere protein O | protein-coding |
| chr13_49649933_49650125 | chr13 | 49649934 | 49650125 | + | 0.855331 | intron (NM_025495, intron 2 of 7) | intron (NM_025495, intron 2 of 7) | 2702 | NM_025495 | 66336 | Mm.440323 | NM_025495 | ENSMUSG00000021391 | **Cenpp** | 1700022C02Rik\|4921518G09Rik | centromere protein P | protein-coding |
| chr13_49644580_49645193 | chr13 | 49644581 | 49645193 | + | 0.834951 | intron (NM_025495, intron 3 of 7) | MTC\|LTR\|ERVL-MaLR | 7844 | NM_025495 | 66336 | Mm.440323 | NM_025495 | ENSMUSG00000021391 | **Cenpp** | 1700022C02Rik\|4921518G09Rik | centromere protein P | protein-coding |
| chr18_53801386_53801808 | chr18 | 53801387 | 53801808 | + | 0.927104 | Intergenic | MTE2a\|LTR\|ERVL-MaLR | -57050 | NM_178686 | 225523 | Mm.193678 | NM_178686 | ENSMUSG00000048799 | **Cep120** | A230075C01\|AU016693\|Ccdc100 | centrosomal protein 120 | protein-coding |
| chr12_91268721_91269524 | chr12 | 91268722 | 91269524 | + | 1.09513 | intron (NM_181815, intron 14 of 25) | intron (NM_181815, intron 14 of 25) | 115286 | NM_181815 | 75216 | Mm.312204 | NM_181815 | ENSMUSG00000061533 | **Cep128** | 4930534B04Rik\|5430424K18Rik\|AI449708\|AI480494\|E230024F20 | centrosomal protein 128 | protein-coding |
| chr12_91298536_91299304 | chr12 | 91298537 | 91299304 | + | 0.868383 | intron (NM_181815, intron 11 of 25) | intron (NM_181815, intron 11 of 25) | 85489 | NM_181815 | 75216 | Mm.312204 | NM_181815 | ENSMUSG00000061533 | **Cep128** | 4930534B04Rik\|5430424K18Rik\|AI449708\|AI480494\|E230024F20 | centrosomal protein 128 | protein-coding |
| chr12_91298749_91299290 | chr12 | 91298750 | 91299290 | + | 0.863857 | exon (NM_181815, exon 11 of 26) | exon (NM_181815, exon 11 of 26) | 85389 | NM_181815 | 75216 | Mm.312204 | NM_181815 | ENSMUSG00000061533 | **Cep128** | 4930534B04Rik\|5430424K18Rik\|AI449708\|AI480494\|E230024F20 | centrosomal protein 128 | protein-coding |
| chr12_91284264_91284638 | chr12 | 91284265 | 91284638 | + | 0.670392 | intron (NM_181815, intron 14 of 25) | intron (NM_181815, intron 14 of 25) | 99958 | NM_181815 | 75216 | Mm.312204 | NM_181815 | ENSMUSG00000061533 | **Cep128** | 4930534B04Rik\|5430424K18Rik\|AI449708\|AI480494\|E230024F20 | centrosomal protein 128 | protein-coding |
| chr12_112702417_112702721 | chr12 | 112702418 | 112702721 | + | 0.861107 | Intergenic | MTD\|LTR\|ERVL-MaLR | -19605 | NM_001024602 | 217882 | Mm.23689 | NM_001024602 | ENSMUSG00000072825 | **Cep170b** | AI466840\|AW555464\|Kiaa0284\|mKIAA0284 | centrosomal protein 170B | protein-coding |
| chr16_32124272_32125069 | chr16 | 32124273 | 32125069 | + | 1.21647 | Intergenic | Intergenic | 24869 | NM_025892 | 66994 | Mm.22990 | NM_025892 | ENSMUSG00000035790 | **Cep19** | 1500031L02Rik\|AI428934\|AL022620 | centrosomal protein 19 | protein-coding |
| chr18_67808874_67809481 | chr18 | 67808875 | 67809481 | + | 1.15816 | intron (NM_027556, intron 5 of 45) | B3\|SINE\|B2 | 9071 | NM_027556 | 70799 | Mm.30857 | NM_027556 | ENSMUSG00000024542 | **Cep192** | 4631422C13Rik\|D430014P18Rik | centrosomal protein 192 | protein-coding |
| chr1_155961061_155961437 | chr1 | 155961062 | 155961437 | + | 0.740586 | intron (NM_001039184, intron 2 of 37) | intron (NM_001039184, intron 2 of 37) | 12006 | NM_001039184 | 74081 | Mm.260247 | NM_001039184 | ENSMUSG00000033671 | **Cep350** | 4933409L06Rik\|6430546F08Rik | centrosomal protein 350 | protein-coding |
| chr1_155960802_155961459 | chr1 | 155960803 | 155961459 | + | 0.652959 | intron (NM_001039184, intron 3 of 37) | intron (NM_001039184, intron 3 of 37) | 12124 | NM_001039184 | 74081 | Mm.260247 | NM_001039184 | ENSMUSG00000033671 | **Cep350** | 4933409L06Rik\|6430546F08Rik | centrosomal protein 350 | protein-coding |
| chr1_156002054_156003010 | chr1 | 156002055 | 156003010 | + | 0.641322 | Intergenic | Intergenic | -29277 | NM_001039184 | 74081 | Mm.260247 | NM_001039184 | ENSMUSG00000033671 | **Cep350** | 4933409L06Rik\|6430546F08Rik | centrosomal protein 350 | protein-coding |
| chr6_30690041_30690457 | chr6 | 30690042 | 30690457 | + | 1.04354 | intron (NM_031998, intron 1 of 10) | L1MD1\|LINE\|L1 | 3433 | NM_031998 | 83922 | Mm.65389 | NM_031998 | ENSMUSG00000029790 | **Cep41** | 1700017E11Rik\|2810431D15Rik\|AI449046\|AI503610\|Tsga14 | centrosomal protein 41 | protein-coding |
| chr6_30687757_30688186 | chr6 | 30687758 | 30688186 | + | 0.822732 | intron (NM_031998, intron 1 of 10) | intron (NM_031998, intron 1 of 10) | 5710 | NM_031998 | 83922 | Mm.65389 | NM_031998 | ENSMUSG00000029790 | **Cep41** | 1700017E11Rik\|2810431D15Rik\|AI449046\|AI503610\|Tsga14 | centrosomal protein 41 | protein-coding |
| chr8_56443525_56444116 | chr8 | 56443526 | 56444116 | + | 0.733939 | Intergenic | Intergenic | 106745 | NM_001009951 | 382010 | Mm.425534 | NM_001009951 | ENSMUSG00000038215 | **Cep44** | 4933440G23Rik\|mKIAA1712 | centrosomal protein 44 | protein-coding |
| chr10_53395314_53395976 | chr10 | 53395315 | 53395976 | + | 1.01247 | Intergenic | Intergenic | -15794 | NM_001204983 | 100038725 | Mm.441403 | NM_001204983 | ENSMUSG00000038594 | **Cep85l** | ENSMUSG00000038594\|Gm9766 | centrosomal protein 85-like | protein-coding |
| chr10_53419112_53419759 | chr10 | 53419113 | 53419759 | + | 0.73668 | Intergenic | B1F\|SINE\|Alu | -39585 | NM_001204983 | 100038725 | Mm.441403 | NM_001204983 | ENSMUSG00000038594 | **Cep85l** | ENSMUSG00000038594\|Gm9766 | centrosomal protein 85-like | protein-coding |
| chr10_53394459_53395170 | chr10 | 53394460 | 53395170 | + | 0.696772 | Intergenic | MIRc\|SINE\|MIR | -14964 | NM_001204983 | 100038725 | Mm.441403 | NM_001204983 | ENSMUSG00000038594 | **Cep85l** | ENSMUSG00000038594\|Gm9766 | centrosomal protein 85-like | protein-coding |
| chr10_53412864_53413349 | chr10 | 53412865 | 53413349 | + | 0.672562 | Intergenic | ORR1B1\|LTR\|ERVL-MaLR | -33256 | NM_001204983 | 100038725 | Mm.441403 | NM_001204983 | ENSMUSG00000038594 | **Cep85l** | ENSMUSG00000038594\|Gm9766 | centrosomal protein 85-like | protein-coding |

| | | | | | | | | | | | | | | | | | |
|---|---|---|---|---|---|---|---|---|---|---|---|---|---|---|---|---|---|
| chr16_55946375_55946678 | chr16 | 55946376 | 55946678 | + | 0.915354 | Intergenic | MTC\|LTR\|ERVL-MaLR | -11679 | NM_001159365 | 74201 | Mm.178427 | NM_028815 | ENSMUSG00000022604 | **Cep97** | 2810403B08Rik\|4932439K18Rik\|AA437768\|E130116N02Rik\|Lrriq2 | centrosomal protein 97 | protein-coding |
| chr15_99754192_99755135 | chr15 | 99754193 | 99755135 | + | 0.904824 | intron (NM_028015, intron 1 of 9) | B1_Mur1\|SINE\|Alu | 17851 | NM_028015 | 71949 | Mm.390821 | NM_028015 | | **Cers5** | 2310081H14Rik\|AU045339\|AW544927\|Lass5\|Trh4 | ceramide synthase 5 | protein-coding |
| chr2_68884560_68885034 | chr2 | 68884561 | 68885034 | + | 1.27273 | intron (NM_172856, intron 1 of 9) | intron (NM_172856, intron 1 of 9) | 23240 | NM_172856 | 241447 | Mm.222222 | NM_172856 | ENSMUSG00000027035 | **Cers6** | 4732462C07Rik\|AW544719\|Lass6\|T1L | ceramide synthase 6 | protein-coding |
| chr2_68915661_68916083 | chr2 | 68915662 | 68916083 | + | 1.13977 | intron (NM_172856, intron 1 of 9) | RLTR42\|LTR\|ERVK | 54315 | NM_172856 | 241447 | Mm.222222 | NM_172856 | ENSMUSG00000027035 | **Cers6** | 4732462C07Rik\|AW544719\|Lass6\|T1L | ceramide synthase 6 | protein-coding |
| chr2_68915822_68916050 | chr2 | 68915823 | 68916050 | + | 1.13757 | intron (NM_172856, intron 1 of 9) | RLTR42\|LTR\|ERVK | 54379 | NM_172856 | 241447 | Mm.222222 | NM_172856 | ENSMUSG00000027035 | **Cers6** | 4732462C07Rik\|AW544719\|Lass6\|T1L | ceramide synthase 6 | protein-coding |
| chr2_68989101_68989760 | chr2 | 68989102 | 68989760 | + | 0.883725 | intron (NM_172856, intron 3 of 9) | intron (NM_172856, intron 3 of 9) | 127874 | NM_172856 | 241447 | Mm.222222 | NM_172856 | ENSMUSG00000027035 | **Cers6** | 4732462C07Rik\|AW544719\|Lass6\|T1L | ceramide synthase 6 | protein-coding |
| chr3_37292677_37293077 | chr3 | 37292678 | 37293077 | + | 2.08484 | Intergenic | Intergenic | 19569 | NM_145825 | 207175 | Mm.344708 | NM_145825 | ENSMUSG00000045031 | **Cetn4** | - | centrin 4 | protein-coding |
| chr3_37299219_37300149 | chr3 | 37299220 | 37300149 | + | 1.71273 | Intergenic | Intergenic | 12762 | NM_145825 | 207175 | Mm.344708 | NM_145825 | ENSMUSG00000045031 | **Cetn4** | - | centrin 4 | protein-coding |
| chr3_37272375_37272940 | chr3 | 37272376 | 37272940 | + | 1.60078 | Intergenic | Intergenic | 39788 | NM_145825 | 207175 | Mm.344708 | NM_145825 | ENSMUSG00000045031 | **Cetn4** | - | centrin 4 | protein-coding |
| chr3_37280871_37281576 | chr3 | 37280872 | 37281576 | + | 1.59786 | Intergenic | Intergenic | 31222 | NM_145825 | 207175 | Mm.344708 | NM_145825 | ENSMUSG00000045031 | **Cetn4** | - | centrin 4 | protein-coding |
| chr3_37280279_37281576 | chr3 | 37280280 | 37281576 | + | 1.57142 | Intergenic | Intergenic | 31518 | NM_145825 | 207175 | Mm.344708 | NM_145825 | ENSMUSG00000045031 | **Cetn4** | - | centrin 4 | protein-coding |
| chr3_37272196_37272975 | chr3 | 37272197 | 37272975 | + | 1.56359 | Intergenic | Intergenic | 39860 | NM_145825 | 207175 | Mm.344708 | NM_145825 | ENSMUSG00000045031 | **Cetn4** | - | centrin 4 | protein-coding |
| chr3_37280289_37280530 | chr3 | 37280290 | 37280530 | + | 1.0847 | Intergenic | Intergenic | 32036 | NM_145825 | 207175 | Mm.344708 | NM_145825 | ENSMUSG00000045031 | **Cetn4** | - | centrin 4 | protein-coding |
| chr3_37280273_37280795 | chr3 | 37280274 | 37280795 | + | 1.02096 | Intergenic | Intergenic | 31912 | NM_145825 | 207175 | Mm.344708 | NM_145825 | ENSMUSG00000045031 | **Cetn4** | - | centrin 4 | protein-coding |
| chr3_37291867_37292286 | chr3 | 37291868 | 37292286 | + | 0.84917 | Intergenic | MER70A\|LTR\|ERVL | 20369 | NM_145825 | 207175 | Mm.344708 | NM_145825 | ENSMUSG00000045031 | **Cetn4** | - | centrin 4 | protein-coding |
| chr3_37283350_37283606 | chr3 | 37283351 | 37283606 | + | 0.821292 | Intergenic | Intergenic | 28968 | NM_145825 | 207175 | Mm.344708 | NM_145825 | ENSMUSG00000045031 | **Cetn4** | - | centrin 4 | protein-coding |
| chr8_95403727_95404201 | chr8 | 95403728 | 95404201 | + | 1.00416 | Intergenic | Intergenic | 30905 | NM_008187 | 14894 | Mm.2080 | NM_008187 | ENSMUSG00000031796 | **Cfap20** | 2600014O15Rik\|AL024127\|Gtl3\|T10-2A2 | cilia and flagella associated protein 20 | protein-coding |
| chr16_44454341_44455029 | chr16 | 44454342 | 44455029 | + | 0.691033 | intron (NM_001033247, intron 26 of 34) | ORR1B1\|LTR\|ERVL-MaLR | 59886 | NM_001033247 | 212517 | Mm.327888 | NM_001033247 | ENSMUSG00000071550 | **Cfap44** | 6330444M21Rik\|D16Ertd642e\|Gm1750\|Wdr52 | cilia and flagella associated protein 44 | protein-coding |
| chr8_46172507_46173335 | chr8 | 46172508 | 46173335 | + | 0.728217 | intron (NM_025747, intron 1 of 3) | intron (NM_025747, intron 1 of 3) | 3365 | NM_025747 | 66756 | Mm.133623 | NM_025747 | ENSMUSG00000031631 | **Cfap97** | 1110068E21Rik\|4933411K20Rik\|AW049273\|Kiaa1430\|mKIAA1430 | cilia and flagella associated protein 97 | protein-coding |
| chr8_46172507_46172958 | chr8 | 46172508 | 46172958 | + | 0.727927 | intron (NM_025747, intron 1 of 3) | intron (NM_025747, intron 1 of 3) | 3177 | NM_025747 | 66756 | Mm.133623 | NM_025747 | ENSMUSG00000031631 | **Cfap97** | 1110068E21Rik\|4933411K20Rik\|AW049273\|Kiaa1430\|mKIAA1430 | cilia and flagella associated protein 97 | protein-coding |
| chr12_54834895_54835538 | chr12 | 54834896 | 54835538 | + | 0.624494 | Intergenic | Intergenic | 27660 | NM_007688 | 12632 | Mm.276826 | NM_007688 | ENSMUSG00000062929 | **Cfl2** | - | cofilin 2, muscle | protein-coding |
| chr1_58722089_58722982 | chr1 | 58722090 | 58722982 | + | 1.26194 | intron (NM_207653, intron 1 of 9) | intron (NM_207653, intron 1 of 9) | -8016 | NM_001293805 | 12633 | Mm.336848 | NM_009805 | ENSMUSG00000026031 | **Cflar** | 2310024N18Rik\|A430105C05Rik\|CLARP\|Cash\|Casper\|ENSMUSG00000072980\|FLAME\|FLAME-1\|Flip\|Gm9845\|I-FLICE\|MRIT\|c-Flip | CASP8 and FADD-like apoptosis regulator | protein-coding |
| chr1_58722032_58722659 | chr1 | 58722033 | 58722659 | + | 1.17464 | intron (NM_207653, intron 1 of 9) | intron (NM_207653, intron 1 of 9) | -8206 | NM_001293805 | 12633 | Mm.336848 | NM_009805 | ENSMUSG00000026031 | **Cflar** | 2310024N18Rik\|A430105C05Rik\|CLARP\|Cash\|Casper\|ENSMUSG00000072980\|FLAME\|FLAME-1\|Flip\|Gm9845\|I-FLICE\|MRIT\|c-Flip | CASP8 and FADD-like apoptosis regulator | protein-coding |
| chr1_58742532_58742892 | chr1 | 58742533 | 58742892 | + | 0.952017 | intron (NM_207653, intron 8 of 9) | intron (NM_207653, intron 8 of 9) | 12160 | NM_001293805 | 12633 | Mm.336848 | NM_009805 | ENSMUSG00000026031 | **Cflar** | 2310024N18Rik\|A430105C05Rik\|CLARP\|Cash\|Casper\|ENSMUSG00000072980\|FLAME\|FLAME-1\|Flip\|Gm9845\|I-FLICE\|MRIT\|c-Flip | CASP8 and FADD-like apoptosis regulator | protein-coding |
| chr1_58742444_58743070 | chr1 | 58742445 | 58743070 | + | 0.913469 | intron (NM_207653, intron 8 of 9) | intron (NM_207653, intron 8 of 9) | 12205 | NM_001293805 | 12633 | Mm.336848 | NM_009805 | ENSMUSG00000026031 | **Cflar** | 2310024N18Rik\|A430105C05Rik\|CLARP\|Cash\|Casper\|ENSMUSG00000072980\|FLAME\|FLAME-1\|Flip\|Gm9845\|I-FLICE\|MRIT\|c-Flip | CASP8 and FADD-like apoptosis regulator | protein-coding |
| chr1_58725785_58726119 | chr1 | 58725786 | 58726119 | + | 0.891807 | intron (NM_207653, intron 1 of 9) | intron (NM_207653, intron 1 of 9) | -4600 | NM_001293805 | 12633 | Mm.336848 | NM_009805 | ENSMUSG00000026031 | **Cflar** | 2310024N18Rik\|A430105C05Rik\|CLARP\|Cash\|Casper\|ENSMUSG00000072980\|FLAME\|FLAME-1\|Flip\|Gm9845\|I-FLICE\|MRIT\|c-Flip | CASP8 and FADD-like apoptosis regulator | protein-coding |
| chr1_58742213_58743072 | chr1 | 58742214 | 58743072 | + | 0.798045 | intron (NM_207653, intron 8 of 9) | intron (NM_207653, intron 8 of 9) | 12091 | NM_001293805 | 12633 | Mm.336848 | NM_009805 | ENSMUSG00000026031 | **Cflar** | 2310024N18Rik\|A430105C05Rik\|CLARP\|Cash\|Casper\|ENSMUSG00000072980\|FLAME\|FLAME-1\|Flip\|Gm9845\|I-FLICE\|MRIT\|c-Flip | CASP8 and FADD-like apoptosis regulator | protein-coding |
| chr1_58728222_58728783 | chr1 | 58728223 | 58728783 | + | 0.726955 | intron (NM_001293804, intron 3 of 9) | intron (NM_001293804, intron 3 of 9) | -2049 | NM_001293805 | 12633 | Mm.336848 | NM_009805 | ENSMUSG00000026031 | **Cflar** | 2310024N18Rik\|A430105C05Rik\|CLARP\|Cash\|Casper\|ENSMUSG00000072980\|FLAME\|FLAME-1\|Flip\|Gm9845\|I-FLICE\|MRIT\|c-Flip | CASP8 and FADD-like apoptosis regulator | protein-coding |
| chr1_58739545_58739837 | chr1 | 58739546 | 58739837 | + | 0.700816 | intron (NM_207653, intron 6 of 9) | intron (NM_207653, intron 6 of 9) | 9139 | NM_001293805 | 12633 | Mm.336848 | NM_009805 | ENSMUSG00000026031 | **Cflar** | 2310024N18Rik\|A430105C05Rik\|CLARP\|Cash\|Casper\|ENSMUSG00000072980\|FLAME\|FLAME-1\|Flip\|Gm9845\|I-FLICE\|MRIT\|c-Flip | CASP8 and FADD-like apoptosis regulator | protein-coding |
| chr1_58727026_58727374 | chr1 | 58727027 | 58727374 | + | 0.690885 | intron (NM_001293804, intron 3 of 9) | intron (NM_001293804, intron 3 of 9) | -3352 | NM_001293805 | 12633 | Mm.336848 | NM_009805 | ENSMUSG00000026031 | **Cflar** | 2310024N18Rik\|A430105C05Rik\|CLARP\|Cash\|Casper\|ENSMUSG00000072980\|FLAME\|FLAME-1\|Flip\|Gm9845\|I-FLICE\|MRIT\|c-Flip | CASP8 and FADD-like apoptosis regulator | protein-coding |
| chr16_64825653_64826028 | chr16 | 64825654 | 64826028 | + | 0.910175 | Intergenic | ORR1A4\|LTR\|ERVL-MaLR | -26244 | NM_178647 | 106143 | Mm.10160 | NM_178647 | ENSMUSG00000054604 | **Cggbp1** | AA960172\|AL023996\|AW045841 | CGG triplet repeat binding protein 1 | protein-coding |
| chr9_71696714_71697147 | chr9 | 71696715 | 71697147 | + | 0.673363 | intron (NM_026599, intron 8 of 18) | intron (NM_026599, intron 8 of 18) | 74671 | NM_026599 | 68178 | Mm.99961 | NM_026599 | ENSMUSG00000032232 | **Cgnl1** | 4933421H10Rik\|9930020M10Rik\|AI503810\|Jacop | cingulin-like 1 | protein-coding |
| chr5_30945175_30945896 | chr5 | 30945176 | 30945896 | + | 0.684375 | promoter-TSS (NM_001160149) | promoter-TSS (NM_001160149) | -56 | NM_026770 | 68567 | Mm.45127 | NM_026770 | ENSMUSG00000029161 | **Cgref1** | 1110004G24Rik\|AW045201\|Cgr11 | cell growth regulator with EF hand domain 1 | protein-coding |
| chr19_34450881_34451368 | chr19 | 34450882 | 34451368 | + | 1.14248 | Intergenic | Intergenic | 24010 | NM_009890 | 12642 | Mm.30824 | NM_009890 | ENSMUSG00000050370 | **Ch25h** | AI462618\|m25OH | cholesterol 25-hydroxylase | protein-coding |
| chr19_34410845_34411426 | chr19 | 34410846 | 34411426 | + | 0.803457 | Intergenic | Intergenic | 63999 | NM_009890 | 12642 | Mm.30824 | NM_009890 | ENSMUSG00000050370 | **Ch25h** | AI462618\|m25OH | cholesterol 25-hydroxylase | protein-coding |
| chr2_119350907_119351499 | chr2 | 119350908 | 119351499 | + | 1.44507 | promoter-TSS (NM_026929) | promoter-TSS (NM_026929) | -39 | NM_026929 | 69065 | Mm.35083 | NM_026929 | ENSMUSG00000027313 | **Chac1** | 1810008K03Rik | ChaC, cation transport regulator 1 | protein-coding |
| chr2_119350409_119351527 | chr2 | 119350410 | 119351527 | + | 1.40964 | promoter-TSS (NM_026929) | promoter-TSS (NM_026929) | -274 | NM_026929 | 69065 | Mm.35083 | NM_026929 | ENSMUSG00000027313 | **Chac1** | 1810008K03Rik | ChaC, cation transport regulator 1 | protein-coding |
| chr2_119355726_119355956 | chr2 | 119355727 | 119355956 | + | 0.724187 | Intergenic | Intergenic | 4599 | NM_026929 | 69065 | Mm.35083 | NM_026929 | ENSMUSG00000027313 | **Chac1** | 1810008K03Rik | ChaC, cation transport regulator 1 | protein-coding |
| chr10_75937250_75937654 | chr10 | 75937251 | 75937654 | + | 0.700856 | TTS (NM_001277132) | TTS (NM_001277132) | 1879 | NM_175329 | 103172 | Mm.333849 | NM_175329 | ENSMUSG00000049422 | **Chchd10** | 1620401E04Rik\|AI267078\|Ndg2 | coiled-coil-helix-coiled-coil-helix domain containing 10 | protein-coding |
| chr4_3952332_3953108 | chr4 | 3952333 | 3953108 | + | 0.92813 | Intergenic | Intergenic | 13296 | NR_104356 | 66433 | Mm.27406 | NM_181391 | ENSMUSG00000042198 | **Chchd7** | 1810049H20Rik | coiled-coil-helix-coiled-coil-helix domain containing 7 | protein-coding |
| chr7_73503202_73503785 | chr7 | 73503203 | 73503785 | + | 1.56657 | exon (NM_001081345, exon 8 of 39) | exon (NM_001081345, exon 8 of 39) | 38252 | NM_001081345 | 244059 | Mm.34539 | NM_001081345 | ENSMUSG00000078671 | **Chd2** | 2810013C04Rik\|2810040A01Rik\|5630401D06Rik\|AI851092\|BC029703\|CHD-2 | chromodomain helicase DNA binding protein 2 | protein-coding |
| chr4_152360869_152361156 | chr4 | 152360870 | 152361156 | + | 0.657938 | exon (NM_029216, exon 9 of 41) | exon (NM_029216, exon 9 of 41) | 22362 | NM_001081376 | 269610 | Mm.40192 | NM_029216 | ENSMUSG00000005045 | **Chd5** | 4930532L22Rik\|AW060752\|B230399N07Rik\|CHD-5 | chromodomain helicase DNA binding protein 5 | protein-coding |
| chr4_8935964_8936338 | chr4 | 8935965 | 8936338 | + | 1.38558 | Intergenic | Intergenic | 245745 | NM_001277149 | 320790 | Mm.138792 | NM_001033395 | ENSMUSG00000041235 | **Chd7** | A730019I05Rik\|Cycn\|Cyn\|Dz\|Edy\|Flo\|GENA 47\|GENA 60\|Gena 52\|Lda\|Mt\|Obt\|Todo\|WBE1\|Whi\|metis | chromodomain helicase DNA binding protein 7 | protein-coding |
| chr4_8906630_8907381 | chr4 | 8906631 | 8907381 | + | 1.22555 | Intergenic | Intergenic | 216600 | NM_001277149 | 320790 | Mm.138792 | NM_001033395 | ENSMUSG00000041235 | **Chd7** | A730019I05Rik\|Cycn\|Cyn\|Dz\|Edy\|Flo\|GENA 47\|GENA 60\|Gena 52\|Lda\|Mt\|Obt\|Todo\|WBE1\|Whi\|metis | chromodomain helicase DNA binding protein 7 | protein-coding |
| chr4_8755338_8755783 | chr4 | 8755339 | 8755783 | + | 1.18783 | intron (NM_001277149, intron 2 of 37) | intron (NM_001277149, intron 2 of 37) | 65155 | NM_001277149 | 320790 | Mm.138792 | NM_001033395 | ENSMUSG00000041235 | **Chd7** | A730019I05Rik\|Cycn\|Cyn\|Dz\|Edy\|Flo\|GENA 47\|GENA 60\|Gena 52\|Lda\|Mt\|Obt\|Todo\|WBE1\|Whi\|metis | chromodomain helicase DNA binding protein 7 | protein-coding |

| | | | | | | | | | | | | | | | | | |
|---|---|---|---|---|---|---|---|---|---|---|---|---|---|---|---|---|---|
| chr4_8928414_8929131 | chr4 | 8928415 | 8929131 | + | 0.777279 | Intergenic | Intergenic | 238367 | NM_001277149 | 320790 | Mm.138792 | NM_001033395 | ENSMUSG00000041235 | **Chd7** | A730019I05Rik\|Cycn\|Cyn\|Dz\|Edy\|Flo\|GENA 47\|GENA 60\|Gena 52\|Lda\|Mt\|Obt\|Todo\|WBE1\|Whi\|metis | chromodomain helicase DNA binding protein 7 | protein-coding |
| chr4_8857548_8857933 | chr4 | 8857549 | 8857933 | + | 0.627339 | intron (NM_001277149, intron 32 of 37) | intron (NM_001277149, intron 32 of 37) | 167335 | NM_001277149 | 320790 | Mm.138792 | NM_001033395 | ENSMUSG00000041235 | **Chd7** | A730019I05Rik\|Cycn\|Cyn\|Dz\|Edy\|Flo\|GENA 47\|GENA 60\|Gena 52\|Lda\|Mt\|Obt\|Todo\|WBE1\|Whi\|metis | chromodomain helicase DNA binding protein 7 | protein-coding |
| chr14_52246210_52246478 | chr14 | 52246211 | 52246478 | + | 0.92824 | Intergenic | B2_Mm2\|SINE\|B2 | -8772 | NM_201637 | 67772 | Mm.289934 | NM_201637 | ENSMUSG00000053754 | **Chd8** | 5830451P18Rik\|AU015341\|Duplin\|HELSNF1\|mKIAA1564 | chromodomain helicase DNA binding protein 8 | protein-coding |
| chr8_90871341_90872017 | chr8 | 90871342 | 90872017 | + | 1.14779 | intron (NM_177224, intron 2 of 39) | RSINE1\|SINE\|B4 | 42844 | NM_177224 | 109151 | Mm.100615 | NM_177224 | ENSMUSG00000056608 | **Chd9** | 1810014J18Rik\|9030205D12Rik\|A330063D19Rik\|AD013\|PRIC320\|mKIAA0308 | chromodomain helicase DNA binding protein 9 | protein-coding |
| chr8_90870410_90870844 | chr8 | 90870411 | 90870844 | + | 1.0707 | intron (NM_177224, intron 2 of 39) | intron (NM_177224, intron 2 of 39) | 41792 | NM_177224 | 109151 | Mm.100615 | NM_177224 | ENSMUSG00000056608 | **Chd9** | 1810014J18Rik\|9030205D12Rik\|A330063D19Rik\|AD013\|PRIC320\|mKIAA0308 | chromodomain helicase DNA binding protein 9 | protein-coding |
| chr8_90797714_90798099 | chr8 | 90797715 | 90798099 | + | 0.756164 | Intergenic | ORR1F\|LTR\|ERVL-MaLR | -30928 | NM_177224 | 109151 | Mm.100615 | NM_177224 | ENSMUSG00000056608 | **Chd9** | 1810014J18Rik\|9030205D12Rik\|A330063D19Rik\|AD013\|PRIC320\|mKIAA0308 | chromodomain helicase DNA binding protein 9 | protein-coding |
| chr8_90825390_90825763 | chr8 | 90825391 | 90825763 | + | 0.719568 | Intergenic | Intergenic | -3258 | NM_177224 | 109151 | Mm.100615 | NM_177224 | ENSMUSG00000056608 | **Chd9** | 1810014J18Rik\|9030205D12Rik\|A330063D19Rik\|AD013\|PRIC320\|mKIAA0308 | chromodomain helicase DNA binding protein 9 | protein-coding |
| chr8_90825390_90825894 | chr8 | 90825391 | 90825894 | + | 0.674907 | Intergenic | Intergenic | -3193 | NM_177224 | 109151 | Mm.100615 | NM_177224 | ENSMUSG00000056608 | **Chd9** | 1810014J18Rik\|9030205D12Rik\|A330063D19Rik\|AD013\|PRIC320\|mKIAA0308 | chromodomain helicase DNA binding protein 9 | protein-coding |
| chr9_36727384_36727693 | chr9 | 36727385 | 36727693 | + | 0.715129 | promoter-TSS (NM_007691) | promoter-TSS (NM_007691) | -881 | NM_007691 | 12649 | Mm.16753 | NM_007691 | ENSMUSG00000032113 | **Chek1** | C85740\|Chk1\|rad27 | checkpoint kinase 1 | protein-coding |
| chr9_36727235_36727697 | chr9 | 36727236 | 36727697 | + | 0.670964 | promoter-TSS (NM_007691) | promoter-TSS (NM_007691) | -808 | NM_007691 | 12649 | Mm.16753 | NM_007691 | ENSMUSG00000032113 | **Chek1** | C85740\|Chk1\|rad27 | checkpoint kinase 1 | protein-coding |
| chr2_132781522_132781883 | chr2 | 132781523 | 132781883 | + | 0.92536 | 5' UTR (NM_007694, exon 1 of 5) | 5' UTR (NM_007694, exon 1 of 5) | 425 | NM_007694 | 12653 | Mm.255241 | NM_007694 | ENSMUSG00000027350 | **Chgb** | Scg-1 | chromogranin B | protein-coding |
| chr2_132781352_132782109 | chr2 | 132781353 | 132782109 | + | 0.869461 | 5' UTR (NM_007694, exon 1 of 5) | 5' UTR (NM_007694, exon 1 of 5) | 453 | NM_007694 | 12653 | Mm.255241 | NM_007694 | ENSMUSG00000027350 | **Chgb** | Scg-1 | chromogranin B | protein-coding |
| chr1_134110959_134111384 | chr1 | 134110960 | 134111384 | + | 0.869684 | promoter-TSS (NM_001284525) | promoter-TSS (NM_001284525) | -70 | NM_001284524 | 71884 | Mm.328268 | NM_027979 | ENSMUSG00000026450 | **Chit1** | 2300002L19Rik | chitinase 1 (chitotriosidase) | protein-coding |
| chr19_3810528_3811186 | chr19 | 3810529 | 3811186 | + | 0.725315 | intron (NM_144871, intron 11 of 11) | intron (NM_144871, intron 11 of 11) | -40916 | NM_001271496 | 12660 | Mm.225505 | NM_013490 | ENSMUSG00000024843 | **Chka** | CK\|CK/EK-alpha\|Chetk-alpha\|Chk\|ChoK\|EK\|EtnK-alpha | choline kinase alpha | protein-coding |
| chr6_103345035_103345540 | chr6 | 103345036 | 103345540 | + | 1.17687 | Intergenic | Intergenic | -165588 | NM_007697 | 12661 | Mm.251288 | NM_007697 | ENSMUSG00000030077 | **Chl1** | A530023M13Rik\|AI465420\|CALL\|LICAM2 | cell adhesion molecule with homology to L1CAM | protein-coding |
| chr6_103379892_103380378 | chr6 | 103379893 | 103380378 | + | 1.06021 | Intergenic | Intergenic | -130741 | NM_007697 | 12661 | Mm.251288 | NM_007697 | ENSMUSG00000030077 | **Chl1** | A530023M13Rik\|AI465420\|CALL\|LICAM2 | cell adhesion molecule with homology to L1CAM | protein-coding |
| chr6_103339614_103339867 | chr6 | 103339615 | 103339867 | + | 0.920675 | Intergenic | B1F\|SINE\|Alu | -171135 | NM_007697 | 12661 | Mm.251288 | NM_007697 | ENSMUSG00000030077 | **Chl1** | A530023M13Rik\|AI465420\|CALL\|LICAM2 | cell adhesion molecule with homology to L1CAM | protein-coding |
| chr6_103308632_103309283 | chr6 | 103308633 | 103309283 | + | 0.734048 | Intergenic | CT-rich\|Low_complexity\|Low_complexity | -201918 | NM_007697 | 12661 | Mm.251288 | NM_007697 | ENSMUSG00000030077 | **Chl1** | A530023M13Rik\|AI465420\|CALL\|LICAM2 | cell adhesion molecule with homology to L1CAM | protein-coding |
| chr6_103336675_103336939 | chr6 | 103336676 | 103336939 | + | 0.701916 | Intergenic | Intergenic | -174069 | NM_007697 | 12661 | Mm.251288 | NM_007697 | ENSMUSG00000030077 | **Chl1** | A530023M13Rik\|AI465420\|CALL\|LICAM2 | cell adhesion molecule with homology to L1CAM | protein-coding |
| chrX_113074703_113075218 | chrX | 113074704 | 113075218 | + | 1.56958 | intron (NM_018818, intron 9 of 14) | intron (NM_018818, intron 9 of 14) | 110554 | NM_018818 | 12662 | Mm.257316 | NM_018818 | ENSMUSG00000025531 | **Chm** | Rep-1 | choroidermia | protein-coding |
| chrX_113081357_113082001 | chrX | 113081358 | 113082001 | + | 1.44224 | intron (NM_018818, intron 8 of 14) | intron (NM_018818, intron 8 of 14) | 103836 | NM_018818 | 12662 | Mm.257316 | NM_018818 | ENSMUSG00000025531 | **Chm** | Rep-1 | choroidermia | protein-coding |
| chrX_113002085_113002743 | chrX | 113002086 | 113002743 | + | 1.38075 | Intergenic | ORR1C2\|LTR\|ERVL-MaLR | 183101 | NM_018818 | 12662 | Mm.257316 | NM_018818 | ENSMUSG00000025531 | **Chm** | Rep-1 | choroidermia | protein-coding |
| chrX_113178332_113179074 | chrX | 113178333 | 113179074 | + | 1.18692 | intron (NM_018818, intron 1 of 14) | intron (NM_018818, intron 1 of 14) | 6812 | NM_018818 | 12662 | Mm.257316 | NM_018818 | ENSMUSG00000025531 | **Chm** | Rep-1 | choroidermia | protein-coding |
| chrX_113056855_113057288 | chrX | 113056856 | 113057288 | + | 0.790794 | intron (NM_018818, intron 12 of 14) | intron (NM_018818, intron 12 of 14) | 128443 | NM_018818 | 12662 | Mm.257316 | NM_018818 | ENSMUSG00000025531 | **Chm** | Rep-1 | choroidermia | protein-coding |
| chrX_113177593_113178009 | chrX | 113177594 | 113178009 | + | 0.693104 | intron (NM_018818, intron 1 of 14) | L1MA6\|LINE\|L1 | 7714 | NM_018818 | 12662 | Mm.257316 | NM_018818 | ENSMUSG00000025531 | **Chm** | Rep-1 | choroidermia | protein-coding |
| chr6_54092873_54093272 | chr6 | 54092874 | 54093272 | + | 0.934098 | intron (NM_001163640, intron 1 of 12) | intron (NM_001163640, intron 1 of 12) | 53141 | NM_001163640 | 69993 | Mm.321147 | NM_023543 | ENSMUSG00000004633 | **Chn2** | 1700026N20Rik\|4930557O16Rik\|ARHGAP3\|Bch | chimerin 2 | protein-coding |
| chr13_10210931_10211454 | chr13 | 10210932 | 10211454 | + | 0.760194 | intron (NM_033269, intron 2 of 4) | B4A\|SINE\|B4 | 149610 | NM_033269 | 12671 | Mm.342315 | NM_033269 | | **Chrm3** | Chrm-3\|M3\|M3R | cholinergic receptor, muscarinic 3, cardiac | protein-coding |
| chr2_91880324_91880633 | chr2 | 91880325 | 91880633 | + | 0.995676 | intron (NM_172669, intron 12 of 17) | intron (NM_172669, intron 12 of 17) | -41710 | NM_007699 | 12672 | Mm.330405 | NM_007699 | ENSMUSG00000040495 | **Chrm4** | Chrm-4\|M4 | cholinergic receptor, muscarinic 4 | protein-coding |
| chr2_91878263_91878505 | chr2 | 91878264 | 91878505 | + | 0.654371 | intron (NM_172669, intron 12 of 17) | L1ME4b\|LINE\|L1 | -43805 | NM_007699 | 12672 | Mm.330405 | NM_007699 | ENSMUSG00000040495 | **Chrm4** | Chrm-4\|M4 | cholinergic receptor, muscarinic 4 | protein-coding |
| chr9_54985739_54986304 | chr9 | 54985740 | 54986304 | + | 0.69773 | intron (NM_176844, intron 1 of 5) | Lx7\|LINE\|L1 | 5142 | NM_176844 | 110835 | Mm.103778 | NM_176844 | ENSMUSG00000035594 | **Chrna5** | Acra-5\|Acra5 | cholinergic receptor, nicotinic, alpha polypeptide 5 | protein-coding |
| chr9_54990407_54990792 | chr9 | 54990408 | 54990792 | + | 0.640637 | intron (NM_176844, intron 1 of 5) | intron (NM_176844, intron 1 of 5) | 9720 | NM_176844 | 110835 | Mm.103778 | NM_176844 | ENSMUSG00000035594 | **Chrna5** | Acra-5\|Acra5 | cholinergic receptor, nicotinic, alpha polypeptide 5 | protein-coding |
| chr9_54990412_54990668 | chr9 | 54990413 | 54990668 | + | 0.634788 | intron (NM_176844, intron 1 of 5) | intron (NM_176844, intron 1 of 5) | 9660 | NM_176844 | 110835 | Mm.103778 | NM_176844 | ENSMUSG00000035594 | **Chrna5** | Acra-5\|Acra5 | cholinergic receptor, nicotinic, alpha polypeptide 5 | protein-coding |
| chr5_65936779_65937249 | chr5 | 65936780 | 65937249 | + | 0.757445 | Intergenic | Intergenic | -30110 | NM_001081104 | 231252 | Mm.486671 | NM_001081104 | ENSMUSG00000029205 | **Chrna9** | 2410015I05Rik\|Acra9\|EG666827\|Gm8311 | cholinergic receptor, nicotinic, alpha polypeptide 9 | protein-coding |
| chr2_92627426_92628108 | chr2 | 92627427 | 92628108 | + | 1.03649 | Intergenic | Intergenic | 28060 | NM_023850 | 76969 | Mm.38021 | NM_023850 | ENSMUSG00000027221 | **Chst1** | 2610008E20Rik\|AW125896\|C6ST\|GST-1\|Gst1\|KSGAL6ST | carbohydrate (keratan sulfate Gal-6) sulfotransferase 1 | protein-coding |
| chr10_83093685_83094633 | chr10 | 83093686 | 83094633 | + | 0.964004 | intron (NM_021439, intron 2 of 2) | intron (NM_021439, intron 2 of 2) | 108662 | NM_021439 | 58250 | Mm.360747 | NM_021439 | ENSMUSG00000034612 | **Chst11** | 1110020P09Rik\|C4ST\|C4ST-1\|C4ST1\|C4s | carbohydrate sulfotransferase 11 | protein-coding |
| chr10_82991935_82992627 | chr10 | 82991936 | 82992627 | + | 0.621488 | intron (NM_021439, intron 1 of 2) | intron (NM_021439, intron 1 of 2) | 6784 | NM_021439 | 58250 | Mm.360747 | NM_021439 | ENSMUSG00000034612 | **Chst11** | 1110020P09Rik\|C4ST\|C4ST-1\|C4ST1\|C4s | carbohydrate sulfotransferase 11 | protein-coding |
| chr5_140507838_140508160 | chr5 | 140507839 | 140508160 | + | 0.90361 | intron (NM_021528, intron 1 of 1) | intron (NM_021528, intron 1 of 1) | 2390 | NM_021528 | 59031 | Mm.28934 | NM_021528 | | **Chst12** | AI595374\|C4S-2\|C4ST2 | carbohydrate sulfotransferase 12 | protein-coding |
| chr5_140517698_140518175 | chr5 | 140517699 | 140518175 | + | 0.780324 | intron (NM_021528, intron 1 of 1) | intron (NM_021528, intron 1 of 1) | 12328 | NM_021528 | 59031 | Mm.28934 | NM_021528 | | **Chst12** | AI595374\|C4S-2\|C4ST2 | carbohydrate sulfotransferase 12 | protein-coding |
| chr5_140517512_140518220 | chr5 | 140517513 | 140518220 | + | 0.625321 | intron (NM_021528, intron 1 of 1) | intron (NM_021528, intron 1 of 1) | 12257 | NM_021528 | 59031 | Mm.28934 | NM_021528 | | **Chst12** | AI595374\|C4S-2\|C4ST2 | carbohydrate sulfotransferase 12 | protein-coding |
| chr2_118935820_118936258 | chr2 | 118935821 | 118936258 | + | 1.57153 | Intergenic | Intergenic | 9542 | NM_028117 | 72136 | Mm.486238 | NM_028117 | ENSMUSG00000074916 | **Chst14** | 2600016L03Rik\|D4ST-1\|D4st1 | carbohydrate (N-acetylgalactosamine 4-0) sulfotransferase 14 | protein-coding |
| chr2_118961470_118961936 | chr2 | 118961471 | 118961936 | + | 1.50329 | Intergenic | Intergenic | 35206 | NM_028117 | 72136 | Mm.486238 | NM_028117 | ENSMUSG00000074916 | **Chst14** | 2600016L03Rik\|D4ST-1\|D4st1 | carbohydrate (N-acetylgalactosamine 4-0) sulfotransferase 14 | protein-coding |
| chr7_132336649_132337381 | chr7 | 132336650 | 132337381 | + | 1.13599 | Intergenic | Intergenic | -19860 | NM_029935 | 77590 | Mm.213582 | NM_029935 | ENSMUSG00000030930 | **Chst15** | 4631426J05Rik\|BRAG\|GalNAcS-6ST\|MAd5\|mKIAA0598 | carbohydrate (N-acetylgalactosamine 4-sulfate 6-O) sulfotransferase 15 | protein-coding |
| chr7_132336101_132336555 | chr7 | 132336102 | 132336555 | + | 1.05841 | Intergenic | Intergenic | -19173 | NM_029935 | 77590 | Mm.213582 | NM_029935 | ENSMUSG00000030930 | **Chst15** | 4631426J05Rik\|BRAG\|GalNAcS-6ST\|MAd5\|mKIAA0598 | carbohydrate (N-acetylgalactosamine 4-sulfate 6-O) sulfotransferase 15 | protein-coding |
| chr7_132428283_132428831 | chr7 | 132428284 | 132428831 | + | 0.879656 | Intergenic | MLT1J2\|LTR\|ERVL-MaLR | -111402 | NM_029935 | 77590 | Mm.213582 | NM_029935 | ENSMUSG00000030930 | **Chst15** | 4631426J05Rik\|BRAG\|GalNAcS-6ST\|MAd5\|mKIAA0598 | carbohydrate (N-acetylgalactosamine 4-sulfate 6-O) sulfotransferase 15 | protein-coding |
| chr9_95278259_95278923 | chr9 | 95278260 | 95278923 | + | 1.85318 | Intergenic | Intergenic | 128679 | NM_018763 | 54371 | Mm.212446 | NM_018763 | ENSMUSG00000033350 | **Chst2** | AI428561\|AW121776\|C130041E03Rik\|Chts2\|GST-2\|GlcNAc6ST\|Gn6st | carbohydrate sulfotransferase 2 | protein-coding |

| | | | | | | | | | | | | | | | | | |
|---|---|---|---|---|---|---|---|---|---|---|---|---|---|---|---|---|---|
| chr9_95312716_95313131 | chr9 | 95312717 | 95313131 | + | 1.45933 | Intergenic | Intergenic | 94346 | NM_018763 | 54371 | Mm.212446 | NM_018763 | ENSMUSG00000033350 | **Chst2** | AI428561\|AW121776\|C130041E03Rik\|Chts2\|GST-2\|GlcNAc6ST\|Gn6st | carbohydrate sulfotransferase 2 | protein-coding |
| chr9_95151574_95152252 | chr9 | 95151575 | 95152252 | + | 1.04279 | intron (NM_177909, intron 14 of 15) | intron (NM_177909, intron 14 of 15) | 255357 | NM_018763 | 54371 | Mm.212446 | NM_018763 | ENSMUSG00000033350 | **Chst2** | AI428561\|AW121776\|C130041E03Rik\|Chts2\|GST-2\|GlcNAc6ST\|Gn6st | carbohydrate sulfotransferase 2 | protein-coding |
| chr9_95151849_95152220 | chr9 | 95151850 | 95152220 | + | 1.03565 | intron (NM_177909, intron 14 of 15) | MIRb\|SINE\|MIR | 255235 | NM_018763 | 54371 | Mm.212446 | NM_018763 | ENSMUSG00000033350 | **Chst2** | AI428561\|AW121776\|C130041E03Rik\|Chts2\|GST-2\|GlcNAc6ST\|Gn6st | carbohydrate sulfotransferase 2 | protein-coding |
| chr9_95311598_95312094 | chr9 | 95311599 | 95312094 | + | 1.0128 | Intergenic | Intergenic | 95424 | NM_018763 | 54371 | Mm.212446 | NM_018763 | ENSMUSG00000033350 | **Chst2** | AI428561\|AW121776\|C130041E03Rik\|Chts2\|GST-2\|GlcNAc6ST\|Gn6st | carbohydrate sulfotransferase 2 | protein-coding |
| chr9_95311749_95312004 | chr9 | 95311750 | 95312004 | + | 0.972827 | Intergenic | Intergenic | 95393 | NM_018763 | 54371 | Mm.212446 | NM_018763 | ENSMUSG00000033350 | **Chst2** | AI428561\|AW121776\|C130041E03Rik\|Chts2\|GST-2\|GlcNAc6ST\|Gn6st | carbohydrate sulfotransferase 2 | protein-coding |
| chr9_95265876_95266509 | chr9 | 95265877 | 95266509 | + | 0.94344 | Intergenic | Intergenic | 141077 | NM_018763 | 54371 | Mm.212446 | NM_018763 | ENSMUSG00000033350 | **Chst2** | AI428561\|AW121776\|C130041E03Rik\|Chts2\|GST-2\|GlcNAc6ST\|Gn6st | carbohydrate sulfotransferase 2 | protein-coding |
| chr9_95408176_95408409 | chr9 | 95408177 | 95408409 | + | 0.887036 | Intergenic | Intergenic | -1023 | NM_018763 | 54371 | Mm.212446 | NM_018763 | ENSMUSG00000033350 | **Chst2** | AI428561\|AW121776\|C130041E03Rik\|Chts2\|GST-2\|GlcNAc6ST\|Gn6st | carbohydrate sulfotransferase 2 | protein-coding |
| chr9_95407645_95408112 | chr9 | 95407646 | 95408112 | + | 0.856865 | promoter-TSS (NM_018763) | promoter-TSS (NM_018763) | -609 | NM_018763 | 54371 | Mm.212446 | NM_018763 | ENSMUSG00000033350 | **Chst2** | AI428561\|AW121776\|C130041E03Rik\|Chts2\|GST-2\|GlcNAc6ST\|Gn6st | carbohydrate sulfotransferase 2 | protein-coding |
| chr9_95391999_95392472 | chr9 | 95392000 | 95392472 | + | 0.73973 | Intergenic | Intergenic | 15034 | NM_018763 | 54371 | Mm.212446 | NM_018763 | ENSMUSG00000033350 | **Chst2** | AI428561\|AW121776\|C130041E03Rik\|Chts2\|GST-2\|GlcNAc6ST\|Gn6st | carbohydrate sulfotransferase 2 | protein-coding |
| chr10_60225181_60225775 | chr10 | 60225182 | 60225775 | + | 0.876481 | Intergenic | Intergenic | -6218 | NM_016803 | 53374 | Mm.12866 | NM_016803 | ENSMUSG00000057337 | **Chst3** | C6ST\|C6ST-1\|GST-0 | carbohydrate (chondroitin 6/keratan) sulfotransferase 3 | protein-coding |
| chr10_60193549_60194162 | chr10 | 60193550 | 60194162 | + | 0.75913 | intron (NM_016803, intron 2 of 3) | intron (NM_016803, intron 2 of 3) | 25404 | NM_016803 | 53374 | Mm.12866 | NM_016803 | ENSMUSG00000057337 | **Chst3** | C6ST\|C6ST-1\|GST-0 | carbohydrate (chondroitin 6/keratan) sulfotransferase 3 | protein-coding |
| chr7_66118098_66118586 | chr7 | 66118099 | 66118586 | + | 0.997793 | intron (NM_001081163, intron 1 of 2) | intron (NM_001081163, intron 1 of 2) | 8827 | NM_001081163 | 269941 | Mm.333916 | NM_001081163 | ENSMUSG00000032640 | **Chsy1** | mKIAA0990 | chondroitin sulfate synthase 1 | protein-coding |
| chr7_66219166_66219734 | chr7 | 66219167 | 66219734 | + | 0.960241 | Intergenic | Intergenic | 109935 | NM_001081163 | 269941 | Mm.333916 | NM_001081163 | ENSMUSG00000032640 | **Chsy1** | mKIAA0990 | chondroitin sulfate synthase 1 | protein-coding |
| chr7_66118034_66118735 | chr7 | 66118035 | 66118735 | + | 0.924927 | intron (NM_001081163, intron 1 of 2) | intron (NM_001081163, intron 1 of 2) | 8870 | NM_001081163 | 269941 | Mm.333916 | NM_001081163 | ENSMUSG00000032640 | **Chsy1** | mKIAA0990 | chondroitin sulfate synthase 1 | protein-coding |
| chr7_66180828_66181405 | chr7 | 66180829 | 66181405 | + | 0.729227 | Intergenic | MIRb\|SINE\|MIR | 71602 | NM_001081163 | 269941 | Mm.333916 | NM_001081163 | ENSMUSG00000032640 | **Chsy1** | mKIAA0990 | chondroitin sulfate synthase 1 | protein-coding |
| chr7_66147038_66147559 | chr7 | 66147039 | 66147559 | + | 0.689001 | intron (NM_001081163, intron 2 of 2) | intron (NM_001081163, intron 2 of 2) | 37784 | NM_001081163 | 269941 | Mm.333916 | NM_001081163 | ENSMUSG00000032640 | **Chsy1** | mKIAA0990 | chondroitin sulfate synthase 1 | protein-coding |
| chr7_66147248_66147514 | chr7 | 66147249 | 66147514 | + | 0.657068 | intron (NM_001081163, intron 2 of 2) | intron (NM_001081163, intron 2 of 2) | 37866 | NM_001081163 | 269941 | Mm.333916 | NM_001081163 | ENSMUSG00000032640 | **Chsy1** | mKIAA0990 | chondroitin sulfate synthase 1 | protein-coding |
| chr8_106888855_106889164 | chr8 | 106888856 | 106889164 | + | 1.52876 | intron (NM_145412, intron 1 of 3) | intron (NM_145412, intron 1 of 3) | 4583 | NM_145412 | 214987 | Mm.490308 | NM_145412 | ENSMUSG00000046691 | **Chtf8** | 5830457O10Rik\|Ctf8\|Derpc | CTF8, chromosome transmission fidelity factor 8 | protein-coding |
| chr8_106888633_106889165 | chr8 | 106888634 | 106889165 | + | 1.47964 | intron (NM_145412, intron 1 of 3) | intron (NM_145412, intron 1 of 3) | 4694 | NM_145412 | 214987 | Mm.490308 | NM_145412 | ENSMUSG00000046691 | **Chtf8** | 5830457O10Rik\|Ctf8\|Derpc | CTF8, chromosome transmission fidelity factor 8 | protein-coding |
| chr19_44111711_44112170 | chr19 | 44111712 | 44112170 | + | 0.754797 | intron (NM_001081077, intron 13 of 13) | intron (NM_001081077, intron 13 of 13) | -4464 | NM_001162410 | 12675 | Mm.3996 | NM_007700 | ENSMUSG00000025199 | **Chuk** | AI256658\|Chuk1\|Fbx24\|Fbxo24\|IKBKA\|IKK1\|Ikka\|NFKBIKA | conserved helix-loop-helix ubiquitous kinase | protein-coding |
| chr8_94827909_94828552 | chr8 | 94827910 | 94828552 | + | 1.22151 | intron (NM_134141, intron 5 of 8) | intron (NM_134141, intron 5 of 8) | 10109 | NM_134141 | 109006 | Mm.27582 | NM_134141 | ENSMUSG00000031781 | **Ciapin1** | 2810413N20Rik\|AA617265\|AU021794\|anamorsin | cytokine induced apoptosis inhibitor 1 | protein-coding |
| chr8_72208168_72209104 | chr8 | 72208169 | 72209104 | + | 1.41014 | intron (NM_001080812, intron 2 of 4) | ORR1C1\|LTR\|ERVL-MaLR | 4201 | NM_001080812 | 234421 | Mm.211247 | NM_001080812 | ENSMUSG00000074240 | **Cib3** | C730014M21Rik\|Gm1107\|KIP3 | calcium and integrin binding family member 3 | protein-coding |
| chr7_25279672_25280159 | chr7 | 25279673 | 25280159 | + | 0.718032 | Intergenic | Intergenic | -2263 | NM_001110132 | 71722 | Mm.28833 | NM_027882 | ENSMUSG00000005442 | **Cic** | 1200010B10Rik\|mKIAA0306 | capicua homolog (Drosophila) | protein-coding |
| chr7_25279854_25280130 | chr7 | 25279855 | 25280130 | + | 0.660245 | Intergenic | Intergenic | -2187 | NM_001110132 | 71722 | Mm.28833 | NM_027882 | ENSMUSG00000005442 | **Cic** | 1200010B10Rik\|mKIAA0306 | capicua homolog (Drosophila) | protein-coding |
| chr9_107303021_107303737 | chr9 | 107303022 | 107303737 | + | 1.79586 | Intergenic | Intergenic | 6690 | NM_009895 | 12700 | Mm.4592 | NM_009895 | | **Cish** | AI385595\|CIS1\|Cis\|F17\|F23 | cytokine inducible SH2-containing protein | protein-coding |
| chr9_107303021_107303546 | chr9 | 107303022 | 107303546 | + | 1.78086 | Intergenic | Intergenic | 6595 | NM_009895 | 12700 | Mm.4592 | NM_009895 | | **Cish** | AI385595\|CIS1\|Cis\|F17\|F23 | cytokine inducible SH2-containing protein | protein-coding |
| chr9_107307516_107308448 | chr9 | 107307517 | 107308448 | + | 1.72035 | Intergenic | Intergenic | 11293 | NM_009895 | 12700 | Mm.4592 | NM_009895 | | **Cish** | AI385595\|CIS1\|Cis\|F17\|F23 | cytokine inducible SH2-containing protein | protein-coding |
| chr9_107309172_107309538 | chr9 | 107309173 | 107309538 | + | 1.21885 | Intergenic | Intergenic | 12666 | NM_009895 | 12700 | Mm.4592 | NM_009895 | | **Cish** | AI385595\|CIS1\|Cis\|F17\|F23 | cytokine inducible SH2-containing protein | protein-coding |
| chr9_107306025_107306522 | chr9 | 107306026 | 107306522 | + | 0.838893 | Intergenic | Intergenic | 9585 | NM_009895 | 12700 | Mm.4592 | NM_009895 | | **Cish** | AI385595\|CIS1\|Cis\|F17\|F23 | cytokine inducible SH2-containing protein | protein-coding |
| chr9_107295377_107297017 | chr9 | 107295378 | 107297017 | + | 0.683956 | promoter-TSS (NM_009895) | promoter-TSS (NM_009895) | -492 | NM_009895 | 12700 | Mm.4592 | NM_009895 | | **Cish** | AI385595\|CIS1\|Cis\|F17\|F23 | cytokine inducible SH2-containing protein | protein-coding |
| chr9_107298175_107298375 | chr9 | 107298176 | 107298375 | + | 0.672444 | intron (NM_009895, intron 1 of 2) | intron (NM_009895, intron 1 of 2) | 1586 | NM_009895 | 12700 | Mm.4592 | NM_009895 | | **Cish** | AI385595\|CIS1\|Cis\|F17\|F23 | cytokine inducible SH2-containing protein | protein-coding |
| chr9_107295644_107296031 | chr9 | 107295645 | 107296031 | + | 0.621888 | promoter-TSS (NM_009895) | promoter-TSS (NM_009895) | -851 | NM_009895 | 12700 | Mm.4592 | NM_009895 | | **Cish** | AI385595\|CIS1\|Cis\|F17\|F23 | cytokine inducible SH2-containing protein | protein-coding |
| chr15_102743156_102743640 | chr15 | 102743157 | 102743640 | + | 1.55523 | Intergenic | Intergenic | 3511 | NR_104334 | 102216269 | | NR_104334 | | **Cistr-act** | Cistr | chondrogenic regulator lncRNA | ncRNA |
| chr15_102746130_102746435 | chr15 | 102746131 | 102746435 | + | 0.744473 | TTS (NR_104334) | TTS (NR_104334) | 626 | NR_104334 | 102216269 | | NR_104334 | | **Cistr-act** | Cistr | chondrogenic regulator lncRNA | ncRNA |
| chr10_17753566_17754196 | chr10 | 17753567 | 17754196 | + | 1.2402 | Intergenic | Intergenic | 30653 | NM_010828 | 17684 | Mm.272321 | NM_010828 | ENSMUSG00000039910 | **Cited2** | AI835299\|ER154-like\|Mrg1\|Msg2\|p35srj | Cbp/p300-interacting transactivator, with Glu/Asp-rich carboxy-terminal domain, 2 | protein-coding |
| chr10_17747964_17748363 | chr10 | 17747965 | 17748363 | + | 1.22671 | Intergenic | Intergenic | 24936 | NM_010828 | 17684 | Mm.272321 | NM_010828 | ENSMUSG00000039910 | **Cited2** | AI835299\|ER154-like\|Mrg1\|Msg2\|p35srj | Cbp/p300-interacting transactivator, with Glu/Asp-rich carboxy-terminal domain, 2 | protein-coding |
| chr10_17753119_17754196 | chr10 | 17753120 | 17754196 | + | 1.13243 | Intergenic | Intergenic | 30430 | NM_010828 | 17684 | Mm.272321 | NM_010828 | ENSMUSG00000039910 | **Cited2** | AI835299\|ER154-like\|Mrg1\|Msg2\|p35srj | Cbp/p300-interacting transactivator, with Glu/Asp-rich carboxy-terminal domain, 2 | protein-coding |
| chr10_17538805_17539821 | chr10 | 17538806 | 17539821 | + | 1.09262 | Intergenic | Intergenic | -183915 | NM_010828 | 17684 | Mm.272321 | NM_010828 | ENSMUSG00000039910 | **Cited2** | AI835299\|ER154-like\|Mrg1\|Msg2\|p35srj | Cbp/p300-interacting transactivator, with Glu/Asp-rich carboxy-terminal domain, 2 | protein-coding |
| chr10_17753106_17753455 | chr10 | 17753107 | 17753455 | + | 0.834673 | Intergenic | Intergenic | 30053 | NM_010828 | 17684 | Mm.272321 | NM_010828 | ENSMUSG00000039910 | **Cited2** | AI835299\|ER154-like\|Mrg1\|Msg2\|p35srj | Cbp/p300-interacting transactivator, with Glu/Asp-rich carboxy-terminal domain, 2 | protein-coding |
| chr10_17548803_17549262 | chr10 | 17548804 | 17549262 | + | 0.780011 | Intergenic | Intergenic | -174195 | NM_010828 | 17684 | Mm.272321 | NM_010828 | ENSMUSG00000039910 | **Cited2** | AI835299\|ER154-like\|Mrg1\|Msg2\|p35srj | Cbp/p300-interacting transactivator, with Glu/Asp-rich carboxy-terminal domain, 2 | protein-coding |
| chr10_17584909_17585191 | chr10 | 17584910 | 17585191 | + | 0.738415 | Intergenic | Tigger7\|DNA\|TcMar-Tigger | -138178 | NM_010828 | 17684 | Mm.272321 | NM_010828 | ENSMUSG00000039910 | **Cited2** | AI835299\|ER154-like\|Mrg1\|Msg2\|p35srj | Cbp/p300-interacting transactivator, with Glu/Asp-rich carboxy-terminal domain, 2 | protein-coding |
| chr10_17521910_17522365 | chr10 | 17521911 | 17522365 | + | 0.635124 | Intergenic | Intergenic | -201090 | NM_010828 | 17684 | Mm.272321 | NM_010828 | ENSMUSG00000039910 | **Cited2** | AI835299\|ER154-like\|Mrg1\|Msg2\|p35srj | Cbp/p300-interacting transactivator, with Glu/Asp-rich carboxy-terminal domain, 2 | protein-coding |
| chr4_120678172_120678665 | chr4 | 120678173 | 120678665 | + | 0.741339 | Intergenic | Intergenic | 11856 | NM_019563 | 56222 | Mm.272646 | NM_019563 | ENSMUSG00000070803 | **Cited4** | MRG-2\|Mrg2 | Cbp/p300-interacting transactivator, with Glu/Asp-rich carboxy-terminal domain, 4 | protein-coding |
| chr8_104255036_104255367 | chr8 | 104255037 | 104255367 | + | 0.68548 | intron (NM_029295, intron 1 of 3) | L2b\|LINE\|L2 | 4341 | NM_001286383 | 75458 | Mm.269219 | NM_029295 | ENSMUSG00000054400 | **Cklf** | 1700001C14Rik\|1810018M11Rik\|C32\|CKLF1\|CKLF3\|CKLF4\|CKLF5\|Cklf2\|Cklf6\|HSPC224\|UCK-1 | chemokine-like factor | protein-coding |
| chr1_118399557_118400247 | chr1 | 118399558 | 118400247 | + | 1.23804 | intron (NM_001293301, intron 1 of 35) | intron (NM_001293301, intron 1 of 35) | 10844 | NM_001293301 | 76707 | Mm.138740 | NM_029709 | ENSMUSG00000064302 | **Clasp1** | 1700030C23Rik\|5730583A19Rik\|B130045P17Rik\|CLASP1alpha | CLIP associating protein 1 | protein-coding |
| chr1_118398681_118399342 | chr1 | 118398682 | 118399342 | + | 0.818364 | intron (NM_001293301, intron 1 of 35) | intron (NM_001293301, intron 1 of 35) | 9954 | NM_001293301 | 76707 | Mm.138740 | NM_029709 | ENSMUSG00000064302 | **Clasp1** | 1700030C23Rik\|5730583A19Rik\|B130045P17Rik\|CLASP1alpha | CLIP associating protein 1 | protein-coding |
| chr1_118403459_118404152 | chr1 | 118403460 | 118404152 | + | 0.775786 | intron (NM_001293301, intron 1 of 35) | intron (NM_001293301, intron 1 of 35) | 14748 | NM_001293301 | 76707 | Mm.138740 | NM_029709 | ENSMUSG00000064302 | **Clasp1** | 1700030C23Rik\|5730583A19Rik\|B130045P17Rik\|CLASP1alpha | CLIP associating protein 1 | protein-coding |

<table>
<tr><td>chr8_60954637_60954992</td><td>chr8</td><td>60954638</td><td>60954992</td><td>+</td><td>0.838388</td><td>promoter-TSS (NM_173876)</td><td>promoter-TSS (NM_173876)</td><td>-16</td><td>NM_173876</td><td>12725</td><td>Mm.25263</td><td>NM_007711</td><td>ENSMUSG00000004319</td><td>**Clcn3**</td><td>Clc3</td><td>chloride channel 3</td><td>protein-coding</td></tr>
<tr><td>chr8_60960241_60960646</td><td>chr8</td><td>60960242</td><td>60960646</td><td>+</td><td>0.710983</td><td>intron (NM_007711, intron 1 of 11)</td><td>intron (NM_007711, intron 1 of 11)</td><td>-5645</td><td>NM_173876</td><td>12725</td><td>Mm.25263</td><td>NM_007711</td><td>ENSMUSG00000004319</td><td>**Clcn3**</td><td>Clc3</td><td>chloride channel 3</td><td>protein-coding</td></tr>
<tr><td>chrX_7308173_7308695</td><td>chrX</td><td>7308174</td><td>7308695</td><td>+</td><td>0.707795</td><td>intron (NM_001243762, intron 2 of 13)</td><td>intron (NM_001243762, intron 2 of 13)</td><td>10924</td><td>NM_001243762</td><td>12728</td><td>Mm.486257</td><td>NM_016691</td><td>ENSMUSG00000004317</td><td>**Clcn5**</td><td>5430408K11Rik|ClC-5|Clc5|D930009B12Rik|DXImx42e|Sfc13|T25545</td><td>chloride channel 5</td><td>protein-coding</td></tr>
<tr><td>chr4_148033866_148034218</td><td>chr4</td><td>148033867</td><td>148034218</td><td>+</td><td>0.796613</td><td>intron (NM_011929, intron 2 of 22)</td><td>intron (NM_011929, intron 2 of 22)</td><td>4725</td><td>NM_011929</td><td>26372</td><td>Mm.89987</td><td>NM_011929</td><td>ENSMUSG00000029016</td><td>**Clcn6**</td><td>AI850629</td><td>chloride channel 6</td><td>protein-coding</td></tr>
<tr><td>chr14_118836870_118837138</td><td>chr14</td><td>118836871</td><td>118837138</td><td>+</td><td>0.809061</td><td>intron (NM_001160097, intron 1 of 3)</td><td>intron (NM_001160097, intron 1 of 3)</td><td>-17946</td><td>NM_021386</td><td>58187</td><td>Mm.390755</td><td>NM_021386</td><td>ENSMUSG00000022132</td><td>**Cldn10**</td><td>6720456I16Rik|Cldn10a|Cldn10b|D14Ertd728e</td><td>claudin 10</td><td>protein-coding</td></tr>
<tr><td>chr8_35716498_35717112</td><td>chr8</td><td>35716499</td><td>35717112</td><td>+</td><td>1.16835</td><td>Intergenic</td><td>Intergenic</td><td>109754</td><td>NM_027998</td><td>71908</td><td>Mm.37817</td><td>NM_027998</td><td>ENSMUSG00000055976</td><td>**Cldn23**</td><td>2310014B08Rik</td><td>claudin 23</td><td>protein-coding</td></tr>
<tr><td>chr8_35728112_35728807</td><td>chr8</td><td>35728113</td><td>35728807</td><td>+</td><td>0.715487</td><td>Intergenic</td><td>MTD|LTR|ERVL-MaLR</td><td>98099</td><td>NM_027998</td><td>71908</td><td>Mm.37817</td><td>NM_027998</td><td>ENSMUSG00000055976</td><td>**Cldn23**</td><td>2310014B08Rik</td><td>claudin 23</td><td>protein-coding</td></tr>
<tr><td>chr16_18791165_18792089</td><td>chr16</td><td>18791166</td><td>18792089</td><td>+</td><td>0.83937</td><td>intron (NM_009862, intron 14 of 19)</td><td>L1MB8|LINE|L1</td><td>14780</td><td>NM_013805</td><td>12741</td><td>Mm.22768</td><td>NM_013805</td><td>ENSMUSG00000041378</td><td>**Cldn5**</td><td>AI854493|MBEC1|Tmvcf</td><td>claudin 5</td><td>protein-coding</td></tr>
<tr><td>chr16_18733039_18733969</td><td>chr16</td><td>18733040</td><td>18733969</td><td>+</td><td>0.715328</td><td>Intergenic</td><td>Intergenic</td><td>-43343</td><td>NM_013805</td><td>12741</td><td>Mm.22768</td><td>NM_013805</td><td>ENSMUSG00000041378</td><td>**Cldn5**</td><td>AI854493|MBEC1|Tmvcf</td><td>claudin 5</td><td>protein-coding</td></tr>
<tr><td>chr11_70151140_70151446</td><td>chr11</td><td>70151141</td><td>70151446</td><td>+</td><td>0.79913</td><td>Intergenic</td><td>B3|SINE|B2</td><td>-15330</td><td>NM_001204252</td><td>17312</td><td>Mm.252405</td><td>NM_010796</td><td>ENSMUSG00000000318</td><td>**Clec10a**</td><td>CD301a|M-ASGP-BP-1|Mgl|Mgl1</td><td>C-type lectin domain family 10, member A</td><td>protein-coding</td></tr>
<tr><td>chr16_10566517_10567086</td><td>chr16</td><td>10566518</td><td>10567086</td><td>+</td><td>1.11923</td><td>intron (NM_177562, intron 3 of 22)</td><td>intron (NM_177562, intron 3 of 22)</td><td>21463</td><td>NM_001204229</td><td>74374</td><td>Mm.159176</td><td>NM_177562</td><td>ENSMUSG00000068663</td><td>**Clec16a**</td><td>4932416N17Rik|curt|mKIAA0350</td><td>C-type lectin domain family 16, member A</td><td>protein-coding</td></tr>
<tr><td>chr16_10660605_10660998</td><td>chr16</td><td>10660606</td><td>10660998</td><td>+</td><td>1.09634</td><td>intron (NM_177562, intron 18 of 22)</td><td>intron (NM_177562, intron 18 of 22)</td><td>115463</td><td>NM_001204229</td><td>74374</td><td>Mm.159176</td><td>NM_177562</td><td>ENSMUSG00000068663</td><td>**Clec16a**</td><td>4932416N17Rik|curt|mKIAA0350</td><td>C-type lectin domain family 16, member A</td><td>protein-coding</td></tr>
<tr><td>chr16_10653970_10654443</td><td>chr16</td><td>10653971</td><td>10654443</td><td>+</td><td>0.787535</td><td>intron (NM_177562, intron 18 of 22)</td><td>intron (NM_177562, intron 18 of 22)</td><td>108868</td><td>NM_001204229</td><td>74374</td><td>Mm.159176</td><td>NM_177562</td><td>ENSMUSG00000068663</td><td>**Clec16a**</td><td>4932416N17Rik|curt|mKIAA0350</td><td>C-type lectin domain family 16, member A</td><td>protein-coding</td></tr>
<tr><td>chr16_10638981_10639475</td><td>chr16</td><td>10638982</td><td>10639475</td><td>+</td><td>0.725804</td><td>intron (NM_177562, intron 17 of 22)</td><td>intron (NM_177562, intron 17 of 22)</td><td>93889</td><td>NM_001204229</td><td>74374</td><td>Mm.159176</td><td>NM_177562</td><td>ENSMUSG00000068663</td><td>**Clec16a**</td><td>4932416N17Rik|curt|mKIAA0350</td><td>C-type lectin domain family 16, member A</td><td>protein-coding</td></tr>
<tr><td>chr16_10640674_10640875</td><td>chr16</td><td>10640675</td><td>10640875</td><td>+</td><td>0.700065</td><td>intron (NM_177562, intron 17 of 22)</td><td>intron (NM_177562, intron 17 of 22)</td><td>95436</td><td>NM_001204229</td><td>74374</td><td>Mm.159176</td><td>NM_177562</td><td>ENSMUSG00000068663</td><td>**Clec16a**</td><td>4932416N17Rik|curt|mKIAA0350</td><td>C-type lectin domain family 16, member A</td><td>protein-coding</td></tr>
<tr><td>chr16_10641229_10641506</td><td>chr16</td><td>10641230</td><td>10641506</td><td>+</td><td>0.697409</td><td>intron (NM_177562, intron 17 of 22)</td><td>intron (NM_177562, intron 17 of 22)</td><td>96029</td><td>NM_001204229</td><td>74374</td><td>Mm.159176</td><td>NM_177562</td><td>ENSMUSG00000068663</td><td>**Clec16a**</td><td>4932416N17Rik|curt|mKIAA0350</td><td>C-type lectin domain family 16, member A</td><td>protein-coding</td></tr>
<tr><td>chr16_10638897_10639852</td><td>chr16</td><td>10638898</td><td>10639852</td><td>+</td><td>0.64277</td><td>intron (NM_177562, intron 17 of 22)</td><td>intron (NM_177562, intron 17 of 22)</td><td>94036</td><td>NM_001204229</td><td>74374</td><td>Mm.159176</td><td>NM_177562</td><td>ENSMUSG00000068663</td><td>**Clec16a**</td><td>4932416N17Rik|curt|mKIAA0350</td><td>C-type lectin domain family 16, member A</td><td>protein-coding</td></tr>
<tr><td>chr6_129434220_129434674</td><td>chr6</td><td>129434221</td><td>129434674</td><td>+</td><td>0.746916</td><td>intron (NM_175526, intron 3 of 5)</td><td>intron (NM_175526, intron 3 of 5)</td><td>17553</td><td>NM_175526</td><td>243653</td><td>Mm.131289</td><td>NM_175526</td><td>ENSMUSG00000033082</td><td>**Clec1a**</td><td>5930406N14Rik</td><td>C-type lectin domain family 1, member a</td><td>protein-coding</td></tr>
<tr><td>chr6_129208835_129209536</td><td>chr6</td><td>129208836</td><td>129209536</td><td>+</td><td>1.40798</td><td>Intergenic</td><td>Intergenic</td><td>28571</td><td>NM_053109</td><td>93694</td><td>Mm.197536</td><td>NM_053109</td><td>ENSMUSG00000030157</td><td>**Clec2d**</td><td>Clr-b|Clrb|Ocil</td><td>C-type lectin domain family 2, member d</td><td>protein-coding</td></tr>
<tr><td>chr6_38669617_38670165</td><td>chr6</td><td>38669618</td><td>38670165</td><td>+</td><td>0.886018</td><td>intron (NM_001101507, intron 1 of 4)</td><td>HAL1|LINE|L1</td><td>6822</td><td>NM_001101507</td><td>665180</td><td>Mm.349066</td><td>NM_001101507</td><td>ENSMUSG00000079598</td><td>**Clec2l**</td><td>AI414054|AW122724|Gm1064</td><td>C-type lectin domain family 2, member L</td><td>protein-coding</td></tr>
<tr><td>chr6_129427269_129427694</td><td>chr6</td><td>129427270</td><td>129427694</td><td>+</td><td>1.1353</td><td>3' UTR (NM_175526, exon 6 of 6)</td><td>3' UTR (NM_175526, exon 6 of 6)</td><td>18620</td><td>NM_172732</td><td>232414</td><td>Mm.391518</td><td>NM_172732</td><td>ENSMUSG00000046080</td><td>**Clec9a**</td><td>9830005G06Rik|DNGR-1</td><td>C-type lectin domain family 9, member a</td><td>protein-coding</td></tr>
<tr><td>chr8_83389582_83390349</td><td>chr8</td><td>83389583</td><td>83390349</td><td>+</td><td>0.716932</td><td>promoter-TSS (NM_009904)</td><td>promoter-TSS (NM_009904)</td><td>75</td><td>NM_009904</td><td>12745</td><td>Mm.358581</td><td>NM_009904</td><td>ENSMUSG00000002190</td><td>**Clgn**</td><td>4930459O04Rik|AI528775|Cln</td><td>calmegin</td><td>protein-coding</td></tr>
<tr><td>chr4_135242382_135242849</td><td>chr4</td><td>135242383</td><td>135242849</td><td>+</td><td>0.671731</td><td>intron (NM_013885, intron 1 of 5)</td><td>intron (NM_013885, intron 1 of 5)</td><td>30144</td><td>NM_013885</td><td>29876</td><td>Mm.257765</td><td>NM_013885</td><td>ENSMUSG00000037242</td><td>**Clic4**</td><td>D0Jmb3|TU-74|mc3s5|mtCLIC</td><td>chloride intracellular channel 4 (mitochondrial)</td><td>protein-coding</td></tr>
<tr><td>chr4_135242372_135243109</td><td>chr4</td><td>135242373</td><td>135243109</td><td>+</td><td>0.64317</td><td>intron (NM_013885, intron 1 of 5)</td><td>B1_Mur2|SINE|Alu</td><td>30019</td><td>NM_013885</td><td>29876</td><td>Mm.257765</td><td>NM_013885</td><td>ENSMUSG00000037242</td><td>**Clic4**</td><td>D0Jmb3|TU-74|mc3s5|mtCLIC</td><td>chloride intracellular channel 4 (mitochondrial)</td><td>protein-coding</td></tr>
<tr><td>chr17_44458981_44459401</td><td>chr17</td><td>44458982</td><td>44459401</td><td>+</td><td>1.19457</td><td>Intergenic</td><td>Intergenic</td><td>270619</td><td>NM_172621</td><td>224796</td><td>Mm.37666</td><td>NM_172621</td><td>ENSMUSG00000023959</td><td>**Clic5**</td><td>5730531E12Rik|B330005L24|Gm322|jbg</td><td>chloride intracellular channel 5</td><td>protein-coding</td></tr>
<tr><td>chr11_45855856_45856426</td><td>chr11</td><td>45855857</td><td>45856426</td><td>+</td><td>1.02107</td><td>intron (NM_001045520, intron 1 of 11)</td><td>intron (NM_001045520, intron 1 of 11)</td><td>4177</td><td>NM_001045520</td><td>216705</td><td>Mm.169673</td><td>NM_001045520</td><td>ENSMUSG00000006169</td><td>**Clint1**</td><td>AI642036|AW537616|C530049I24Rik|Enth|Epn4|F630206G17Rik|mKIAA0171</td><td>clathrin interactor 1</td><td>protein-coding</td></tr>
<tr><td>chr11_45850376_45850589</td><td>chr11</td><td>45850377</td><td>45850589</td><td>+</td><td>0.742385</td><td>Intergenic</td><td>Intergenic</td><td>-1481</td><td>NM_001045520</td><td>216705</td><td>Mm.169673</td><td>NM_001045520</td><td>ENSMUSG00000006169</td><td>**Clint1**</td><td>AI642036|AW537616|C530049I24Rik|Enth|Epn4|F630206G17Rik|mKIAA0171</td><td>clathrin interactor 1</td><td>protein-coding</td></tr>
<tr><td>chr5_123661623_123662095</td><td>chr5</td><td>123661624</td><td>123662095</td><td>+</td><td>0.855943</td><td>intron (NM_001291229, intron 1 of 23)</td><td>PB1D7|SINE|Alu</td><td>22502</td><td>NM_001291229</td><td>56430</td><td>Mm.241109</td><td>NM_019765</td><td>ENSMUSG00000049550</td><td>**Clip1**</td><td>1110007I12Rik|4631429H07Rik|AV017631|C81039|CLIP-170|CLIP170|CYLN1|Clip50|Rsn|mKIAA4046|restin</td><td>CAP-GLY domain containing linker protein 1</td><td>protein-coding</td></tr>
<tr><td>chr9_40698736_40699654</td><td>chr9</td><td>40698737</td><td>40699654</td><td>+</td><td>0.958042</td><td>intron (NM_133733, intron 1 of 6)</td><td>intron (NM_133733, intron 1 of 6)</td><td>13231</td><td>NM_133733</td><td>71566</td><td>Mm.280560</td><td>NM_133733</td><td>ENSMUSG00000032024</td><td>**Clmp**</td><td>9030425E11Rik|ACAM|ASP5|AW557819</td><td>CXADR-like membrane protein</td><td>protein-coding</td></tr>
<tr><td>chr9_40612265_40612770</td><td>chr9</td><td>40612266</td><td>40612770</td><td>+</td><td>0.858935</td><td>Intergenic</td><td>ORR1E|LTR|ERVL-MaLR</td><td>-73446</td><td>NM_133733</td><td>71566</td><td>Mm.280560</td><td>NM_133733</td><td>ENSMUSG00000032024</td><td>**Clmp**</td><td>9030425E11Rik|ACAM|ASP5|AW557819</td><td>CXADR-like membrane protein</td><td>protein-coding</td></tr>
<tr><td>chr8_14890307_14890640</td><td>chr8</td><td>14890308</td><td>14890640</td><td>+</td><td>0.648953</td><td>intron (NM_012000, intron 1 of 2)</td><td>intron (NM_012000, intron 1 of 2)</td><td>1938</td><td>NM_012000</td><td>26889</td><td>Mm.254027</td><td>NM_012000</td><td>ENSMUSG00000026317</td><td>**Cln8**</td><td>mnd</td><td>ceroid-lipofuscinosis, neuronal 8</td><td>protein-coding</td></tr>
<tr><td>chr5_39155749_39156206</td><td>chr5</td><td>39155750</td><td>39156206</td><td>+</td><td>1.09872</td><td>Intergenic</td><td>Intergenic</td><td>-279285</td><td>NM_013748</td><td>27278</td><td>Mm.440845</td><td>NM_013748</td><td>ENSMUSG00000039315</td><td>**Clnk**</td><td>MIST</td><td>cytokine-dependent hematopoietic cell linker</td><td>protein-coding</td></tr>
<tr><td>chr7_101726790_101727355</td><td>chr7</td><td>101726791</td><td>101727355</td><td>+</td><td>0.762921</td><td>intron (NM_009191, intron 5 of 15)</td><td>intron (NM_009191, intron 5 of 15)</td><td>63305</td><td>NM_009191</td><td>20480</td><td>Mm.3990</td><td>NM_009191</td><td>ENSMUSG00000001829</td><td>**Clpb**</td><td>AL118244|Skd3</td><td>ClpB caseinolytic peptidase B</td><td>protein-coding</td></tr>
<tr><td>chr7_101726770_101727592</td><td>chr7</td><td>101726771</td><td>101727592</td><td>+</td><td>0.752161</td><td>intron (NM_009191, intron 5 of 15)</td><td>intron (NM_009191, intron 5 of 15)</td><td>63413</td><td>NM_009191</td><td>20480</td><td>Mm.3990</td><td>NM_009191</td><td>ENSMUSG00000001829</td><td>**Clpb**</td><td>AL118244|Skd3</td><td>ClpB caseinolytic peptidase B</td><td>protein-coding</td></tr>
<tr><td>chr9_65288103_65288574</td><td>chr9</td><td>65288104</td><td>65288574</td><td>+</td><td>0.76081</td><td>Intergenic</td><td>RLTR45|LTR|ERVK</td><td>-5921</td><td>NM_001044389</td><td>270166</td><td>Mm.30088</td><td>NM_011802</td><td>ENSMUSG00000015357</td><td>**Clpx**</td><td>AU014732|E330029I21</td><td>caseinolytic mitochondrial matrix peptidase chaperone subunit</td><td>protein-coding</td></tr>
<tr><td>chr7_135489642_135490804</td><td>chr7</td><td>135489643</td><td>135490804</td><td>+</td><td>0.763323</td><td>Intergenic</td><td>Intergenic</td><td>38431</td><td>NM_178669</td><td>212070</td><td>Mm.39721</td><td>NM_178669</td><td>ENSMUSG00000050866</td><td>**Clrn3**</td><td>AI649392|Tmem12</td><td>clarin 3</td><td>protein-coding</td></tr>
<tr><td>chr7_135489641_135490426</td><td>chr7</td><td>135489642</td><td>135490426</td><td>+</td><td>0.734944</td><td>Intergenic</td><td>Intergenic</td><td>38620</td><td>NM_178669</td><td>212070</td><td>Mm.39721</td><td>NM_178669</td><td>ENSMUSG00000050866</td><td>**Clrn3**</td><td>AI649392|Tmem12</td><td>clarin 3</td><td>protein-coding</td></tr>
<tr><td>chr11_86752236_86752776</td><td>chr11</td><td>86752237</td><td>86752776</td><td>+</td><td>0.838406</td><td>intron (NM_001003908, intron 1 of 31)</td><td>intron (NM_001003908, intron 1 of 31)</td><td>4986</td><td>NM_001003908</td><td>67300</td><td>Mm.254588</td><td>NM_001003908</td><td>ENSMUSG00000047126</td><td>**Cltc**</td><td>3110065L21Rik|CHC|R74732</td><td>clathrin, heavy polypeptide (Hc)</td><td>protein-coding</td></tr>
<tr><td>chr11_86760479_86760728</td><td>chr11</td><td>86760480</td><td>86760728</td><td>+</td><td>0.791655</td><td>intron (NR_106195, intron 2 of 8)</td><td>PB1D7|SINE|Alu</td><td>-3112</td><td>NM_001003908</td><td>67300</td><td>Mm.254588</td><td>NM_001003908</td><td>ENSMUSG00000047126</td><td>**Cltc**</td><td>3110065L21Rik|CHC|R74732</td><td>clathrin, heavy polypeptide (Hc)</td><td>protein-coding</td></tr>
<tr><td>chr14_65968299_65968882</td><td>chr14</td><td>65968300</td><td>65968882</td><td>+</td><td>0.987157</td><td>5' UTR (NM_013492, exon 1 of 9)</td><td>5' UTR (NM_013492, exon 1 of 9)</td><td>108</td><td>NM_013492</td><td>12759</td><td>Mm.200608</td><td>NM_013492</td><td>ENSMUSG00000022037</td><td>**Clu**</td><td>AI893575|ApoJ|Cli|D14Ucla3|SP-40|Sgp-2|Sgp2|Sugp-2</td><td>clusterin</td><td>protein-coding</td></tr>
<tr><td>chr16_3934693_3935429</td><td>chr16</td><td>3934694</td><td>3935429</td><td>+</td><td>0.861803</td><td>intron (NM_029738, intron 10 of 11)</td><td>RMER1A|Other|Other</td><td>26052</td><td>NM_029738</td><td>76779</td><td>Mm.458921</td><td>NM_029738</td><td>ENSMUSG00000014232</td><td>**Cluap1**</td><td>2310030D15Rik|2610111M03Rik</td><td>clusterin associated protein 1</td><td>protein-coding</td></tr>
<tr><td>chr16_3933585_3934006</td><td>chr16</td><td>3933586</td><td>3934006</td><td>+</td><td>0.781068</td><td>exon (NM_029738, exon 10 of 12)</td><td>exon (NM_029738, exon 10 of 12)</td><td>24787</td><td>NM_029738</td><td>76779</td><td>Mm.458921</td><td>NM_029738</td><td>ENSMUSG00000014232</td><td>**Cluap1**</td><td>2310030D15Rik|2610111M03Rik</td><td>clusterin associated protein 1</td><td>protein-coding</td></tr>
<tr><td>chr11_74655596_74656478</td><td>chr11</td><td>74655597</td><td>74656478</td><td>+</td><td>1.4455</td><td>intron (NM_001081158, intron 1 of 25)</td><td>intron (NM_001081158, intron 1 of 25)</td><td>6542</td><td>NM_001081158</td><td>74148</td><td>Mm.214574</td><td>NM_001081158</td><td>ENSMUSG00000020741</td><td>**Cluh**</td><td>1300001I01Rik|Kiaa0664|mKIAA0664</td><td>clustered mitochondria (cluA/CLU1) homolog</td><td>protein-coding</td></tr>
<tr><td>chr4_9133323_9133627</td><td>chr4</td><td>9133324</td><td>9133627</td><td>+</td><td>0.851538</td><td>Intergenic</td><td>Intergenic</td><td>-135842</td><td>NM_028940</td><td>74438</td><td>Mm.473368</td><td>NM_028940</td><td>ENSMUSG00000041216</td><td>**Clvs1**</td><td>4933402J24Rik|Clvl1|Rlbp1l1</td><td>clavesin 1</td><td>protein-coding</td></tr>
<tr><td>chr4_9133306_9134015</td><td>chr4</td><td>9133307</td><td>9134015</td><td>+</td><td>0.665692</td><td>Intergenic</td><td>Intergenic</td><td>-135656</td><td>NM_028940</td><td>74438</td><td>Mm.473368</td><td>NM_028940</td><td>ENSMUSG00000041216</td><td>**Clvs1**</td><td>4933402J24Rik|Clvl1|Rlbp1l1</td><td>clavesin 1</td><td>protein-coding</td></tr>
<tr><td>chr14_122193730_122194287</td><td>chr14</td><td>122193731</td><td>122194287</td><td>+</td><td>0.629205</td><td>intron (NM_029556, intron 1 of 7)</td><td>intron (NM_029556, intron 1 of 7)</td><td>12315</td><td>NM_029556</td><td>69634</td><td>Mm.34608</td><td>NM_029556</td><td>ENSMUSG00000025545</td><td>**Clybl**</td><td>0610033J05Rik|2310014M14Rik|AI256068|Clb</td><td>citrate lyase beta like</td><td>protein-coding</td></tr>
<tr><td>chr13_24467669_24468349</td><td>chr13</td><td>24467670</td><td>24468349</td><td>+</td><td>1.50273</td><td>intron (NM_001284520, intron 13 of 14)</td><td>intron (NM_001284520, intron 13 of 14)</td><td>52887</td><td>NM_001284520</td><td>12763</td><td>Mm.8396</td><td>NM_007717</td><td>ENSMUSG00000016756</td><td>**Cmah**</td><td>-</td><td>cytidine monophospho-N-acetylneuraminic acid hydroxylase</td><td>protein-coding</td></tr>
<tr><td>chr6_142747001_142747569</td><td>chr6</td><td>142747002</td><td>142747569</td><td>+</td><td>1.03997</td><td>Intergenic</td><td>(CA)n|Simple_repeat|Simple_repeat</td><td>-9401</td><td>NM_009908</td><td>12764</td><td>Mm.3820</td><td>NM_009908</td><td>ENSMUSG00000030282</td><td>**Cmas**</td><td>AW208911|CMPNeu5Ac|D6Bwg0250e</td><td>cytidine monophospho-N-acetylneuraminic acid synthetase</td><td>protein-coding</td></tr>
<tr><td>chr6_142746794_142747579</td><td>chr6</td><td>142746795</td><td>142747579</td><td>+</td><td>1.02127</td><td>Intergenic</td><td>Lx8b|LINE|L1</td><td>-9499</td><td>NM_009908</td><td>12764</td><td>Mm.3820</td><td>NM_009908</td><td>ENSMUSG00000030282</td><td>**Cmas**</td><td>AW208911|CMPNeu5Ac|D6Bwg0250e</td><td>cytidine monophospho-N-acetylneuraminic acid synthetase</td><td>protein-coding</td></tr>
<tr><td>chr9_118271483_118272166</td><td>chr9</td><td>118271484</td><td>118272166</td><td>+</td><td>1.43714</td><td>Intergenic</td><td>Intergenic</td><td>-121629</td><td>NM_026442</td><td>67899</td><td>Mm.9953</td><td>NM_026442</td><td>ENSMUSG00000039163</td><td>**Cmc1**</td><td>2010110K16Rik|2010312P05Rik|AI596001</td><td>COX assembly mitochondrial protein 1</td><td>protein-coding</td></tr>
<tr><td>chr9_118282576_118283070</td><td>chr9</td><td>118282577</td><td>118283070</td><td>+</td><td>1.2774</td><td>Intergenic</td><td>Intergenic</td><td>-132627</td><td>NM_026442</td><td>67899</td><td>Mm.9953</td><td>NM_026442</td><td>ENSMUSG00000039163</td><td>**Cmc1**</td><td>2010110K16Rik|2010312P05Rik|AI596001</td><td>COX assembly mitochondrial protein 1</td><td>protein-coding</td></tr>
</table>

| | | | | | | | | | | | | | | | | | |
|---|---|---|---|---|---|---|---|---|---|---|---|---|---|---|---|---|---|
| chr9_118109612_118110668 | chr9 | 118109613 | 118110668 | + | 0.64343 | intron (NM_026442, intron 2 of 3) | intron (NM_026442, intron 2 of 3) | 40056 | NM_026442 | 67899 | Mm.9953 | NM_026442 | ENSMUSG00000039163 | **Cmc1** | 2010110K16Rik\|2010312P05Rik\|AI596001 | COX assembly mitochondrial protein 1 | protein-coding |
| chr8_117284924_117285581 | chr8 | 117284925 | 117285581 | + | 0.802225 | intron (NM_001163262, intron 1 of 20) | intron (NM_001163262, intron 1 of 20) | 28234 | NM_001163262 | 74440 | Mm.233181 | NM_028941 | ENSMUSG00000034390 | **Cmip** | 4933407C03Rik\|5830471E12Rik\|AI314065\|AI464126 | c-Maf inducing protein | protein-coding |
| chr8_117285114_117285493 | chr8 | 117285115 | 117285493 | + | 0.788402 | intron (NM_001163262, intron 1 of 20) | intron (NM_001163262, intron 1 of 20) | 28285 | NM_001163262 | 74440 | Mm.233181 | NM_028941 | ENSMUSG00000034390 | **Cmip** | 4933407C03Rik\|5830471E12Rik\|AI314065\|AI464126 | c-Maf inducing protein | protein-coding |
| chr8_117394974_117395654 | chr8 | 117394975 | 117395654 | + | 0.622061 | intron (NM_001163262, intron 3 of 20) | intron (NM_001163262, intron 3 of 20) | 46144 | NM_028941 | 74440 | Mm.233181 | NM_028941 | ENSMUSG00000034390 | **Cmip** | 4933407C03Rik\|5830471E12Rik\|AI314065\|AI464126 | c-Maf inducing protein | protein-coding |
| chr5_113671606_113671916 | chr5 | 113671607 | 113671916 | + | 0.672349 | Intergenic | Intergenic | -21371 | NM_008153 | 14747 | Mm.5196 | NM_008153 | ENSMUSG00000042190 | **Cmklr1** | ChemR23\|DEZ\|Gpcr27\|mcmklr1 | chemokine-like receptor 1 | protein-coding |
| chr16_57585011_57586097 | chr16 | 57585012 | 57586097 | + | 1.44491 | intron (NM_025599, intron 1 of 9) | intron (NM_025599, intron 1 of 9) | 21313 | NM_025599 | 66497 | Mm.425976 | NM_025599 | | **Cmss1** | 1110001A06Rik\|2610528E23Rik\|4930572F24Rik\|AV099812\|AV122629 | cms small ribosomal subunit 1 | protein-coding |
| chr16_57608147_57608404 | chr16 | 57608148 | 57608404 | + | 0.820072 | Intergenic | RLTR48A\|LTR\|ERV1 | -1409 | NM_025599 | 66497 | Mm.425976 | NM_025599 | | **Cmss1** | 1110001A06Rik\|2610528E23Rik\|4930572F24Rik\|AV099812\|AV122629 | cms small ribosomal subunit 1 | protein-coding |
| chr8_104373661_104374272 | chr8 | 104373662 | 104374272 | + | 0.679468 | intron (NM_153582, intron 1 of 3) | intron (NM_153582, intron 1 of 3) | 21840 | NM_153582 | 97487 | Mm.29658 | NM_153582 | ENSMUSG00000096188 | **Cmtm4** | Cklfsf4\|D19397\|ENSMUSG00000051554\|Gm9853 | CKLF-like MARVEL transmembrane domain containing 4 | protein-coding |
| chr9_114760224_114760804 | chr9 | 114760225 | 114760804 | + | 0.991053 | intron (NM_001252479, intron 2 of 3) | intron (NM_001252479, intron 2 of 3) | 21479 | NM_133978 | 102545 | Mm.35600 | NM_133978 | ENSMUSG00000032436 | **Cmtm7** | AI481279\|Cklfsf7\|LNV | CKLF-like MARVEL transmembrane domain containing 7 | protein-coding |
| chr9_114760227_114761071 | chr9 | 114760228 | 114761071 | + | 0.914378 | intron (NM_001252479, intron 2 of 3) | intron (NM_001252479, intron 2 of 3) | 21344 | NM_133978 | 102545 | Mm.35600 | NM_133978 | ENSMUSG00000032436 | **Cmtm7** | AI481279\|Cklfsf7\|LNV | CKLF-like MARVEL transmembrane domain containing 7 | protein-coding |
| chr13_93133525_93133883 | chr13 | 93133526 | 93133883 | + | 0.730795 | intron (NM_023821, intron 1 of 12) | intron (NM_023821, intron 1 of 12) | 11020 | NM_023821 | 76469 | Mm.33857 | NM_023821 | ENSMUSG00000047419 | **Cmya5** | 2310076E16Rik\|2310076E21Rik\|AA420382\|AI504003\|Srfsd\|Tims\|sr553 | cardiomyopathy associated 5 | protein-coding |
| chr18_84683996_84684515 | chr18 | 84683997 | 84684515 | + | 1.02733 | intron (NM_023149, intron 1 of 11) | intron (NM_023149, intron 1 of 11) | 1446 | NM_023149 | 66054 | Mm.29646 | NM_023149 | ENSMUSG00000024644 | **Cndp2** | 0610010E05Rik\|C76600\|Cn2\|Dip-2\|Pep-1\|Pep1 | CNDP dipeptidase 2 (metallopeptidase M20 family) | protein-coding |
| chr8_88098870_88099259 | chr8 | 88098871 | 88099259 | + | 0.621802 | Intergenic | Intergenic | -19694 | NM_029074 | 382030 | Mm.216574 | NM_029074 | ENSMUSG00000036810 | **Cnep1r1** | 5033428A16Rik\|NEP1-R1\|Tmem188 | CTD nuclear envelope phosphatase 1 regulatory subunit 1 | protein-coding |
| chr1_181157065_181157520 | chr1 | 181157066 | 181157520 | + | 0.812506 | intron (NM_030131, intron 3 of 4) | intron (NM_030131, intron 3 of 4) | 6362 | NM_030131 | 98417 | Mm.38229 | NM_030131 | ENSMUSG00000062169 | **Cnih4** | AI647760\|AW050376\|CNIH-4\|D530030D03Rik\|E430023H19Rik\|HSPC163 | cornichon homolog 4 (Drosophila) | protein-coding |
| chr10_7117867_7118975 | chr10 | 7117868 | 7118975 | + | 1.45013 | TTS (NM_172546) | TTS (NM_172546) | 93816 | NM_172546 | 215748 | Mm.37984 | NM_172546 | ENSMUSG00000015202 | **Cnksr3** | 6820402C05\|BC024086\|Magi1 | Cnksr family member 3 | protein-coding |
| chr10_7145639_7146174 | chr10 | 7145640 | 7146174 | + | 0.912911 | intron (NM_172546, intron 4 of 12) | intron (NM_172546, intron 4 of 12) | 66330 | NM_172546 | 215748 | Mm.37984 | NM_172546 | ENSMUSG00000015202 | **Cnksr3** | 6820402C05\|BC024086\|Magi1 | Cnksr family member 3 | protein-coding |
| chr10_7212541_7213014 | chr10 | 7212542 | 7213014 | + | 0.707462 | promoter-TSS (NM_172546) | promoter-TSS (NM_172546) | -541 | NM_172546 | 215748 | Mm.37984 | NM_172546 | ENSMUSG00000015202 | **Cnksr3** | 6820402C05\|BC024086\|Magi1 | Cnksr family member 3 | protein-coding |
| chr3_121371917_121372202 | chr3 | 121371918 | 121372202 | + | 0.68152 | Intergenic | MLT1G\|LTR\|ERVL-MaLR | -54481 | NM_028044 | 71994 | Mm.275555 | NM_028044 | ENSMUSG00000053931 | **Cnn3** | 1600014M03Rik\|C85854\|Calpo3 | calponin 3, acidic | protein-coding |
| chr19_46804760_46805144 | chr19 | 46804761 | 46805144 | + | 0.650696 | intron (NM_033569, intron 1 of 7) | B3\|SINE\|B2 | 43343 | NM_001102471 | 94219 | Mm.306903 | NM_033569 | ENSMUSG00000064105 | **Cnnm2** | AU015877\|AW048635\|Acdp2\|Clp2 | cyclin M2 | protein-coding |
| chr19_46819172_46819655 | chr19 | 46819173 | 46819655 | + | 0.626044 | intron (NM_033569, intron 1 of 7) | ORR1B2\|LTR\|ERVL-MaLR | 57805 | NM_001102471 | 94219 | Mm.306903 | NM_033569 | ENSMUSG00000064105 | **Cnnm2** | AU015877\|AW048635\|Acdp2\|Clp2 | cyclin M2 | protein-coding |
| chr10_116548005_116548343 | chr10 | 116548006 | 116548343 | + | 0.797463 | intron (NM_001037847, intron 2 of 15) | intron (NM_001037847, intron 2 of 15) | 33337 | NM_001037848 | 72068 | Mm.351553 | NM_028082 | ENSMUSG00000020166 | **Cnot2** | 2600016M12Rik\|2810470K03Rik\|AA537049\|AA959607\|AW557563\|C79650 | CCR4-NOT transcription complex, subunit 2 | protein-coding |
| chr10_116547786_116548338 | chr10 | 116547787 | 116548338 | + | 0.794773 | intron (NM_001037847, intron 2 of 15) | intron (NM_001037847, intron 2 of 15) | 33449 | NM_001037848 | 72068 | Mm.351553 | NM_028082 | ENSMUSG00000020166 | **Cnot2** | 2600016M12Rik\|2810470K03Rik\|AA537049\|AA959607\|AW557563\|C79650 | CCR4-NOT transcription complex, subunit 2 | protein-coding |
| chr6_35100044_35100764 | chr6 | 35100045 | 35100764 | + | 0.638228 | intron (NM_001164411, intron 1 of 10) | Lx8\|LINE\|L1 | 33333 | NM_001164412 | 53621 | Mm.214525 | NM_016877 | ENSMUSG00000038784 | **Cnot4** | Not4\|Not4h\|Not4hp | CCR4-NOT transcription complex, subunit 4 | protein-coding |
| chr11_49694953_49695447 | chr11 | 49694954 | 49695447 | + | 0.986047 | intron (NM_001290741, intron 1 of 10) | intron (NM_001290741, intron 1 of 10) | 7454 | NM_001290741 | 104625 | Mm.247113 | NM_212484 | ENSMUSG00000020362 | **Cnot6** | CCR4 | CCR4-NOT transcription complex, subunit 6 | protein-coding |
| chr11_49746656_49747110 | chr11 | 49746657 | 49747110 | + | 0.715292 | Intergenic | Intergenic | -34161 | NM_212484 | 104625 | Mm.247113 | NM_212484 | ENSMUSG00000020362 | **Cnot6** | CCR4 | CCR4-NOT transcription complex, subunit 6 | protein-coding |
| chr5_96171584_96172303 | chr5 | 96171585 | 96172303 | + | 0.844185 | Intergenic | Intergenic | -9954 | NM_144910 | 231464 | Mm.28374 | NM_144910 | ENSMUSG00000034724 | **Cnot6l** | 4932442K20Rik | CCR4-NOT transcription complex, subunit 6-like | protein-coding |
| chr17_46749361_46749613 | chr17 | 46749362 | 46749613 | + | 0.94759 | intron (NM_028065, intron 1 of 5) | intron (NM_028065, intron 1 of 5) | 2725 | NM_028065 | 72029 | Mm.240325 | NM_028065 | ENSMUSG00000023973 | **Cnpy3** | 1600025D17Rik\|2410050O22Rik\|AI413153\|CAG4A\|ERDA5\|PRAT4A\|Tnrc5 | canopy 3 homolog (zebrafish) | protein-coding |
| chr1_179570531_179570947 | chr1 | 179570532 | 179570947 | + | 1.14194 | intron (NM_146105, intron 1 of 10) | Charlie5\|DNA\|hAT-Charlie | 24210 | NM_146105 | 226744 | Mm.87662 | NM_146105 | ENSMUSG00000038949 | **Cnst** | 9630058J23Rik | consortin, connexin sorting protein | protein-coding |
| chr1_179575634_179576127 | chr1 | 179575635 | 179576127 | + | 0.886599 | intron (NM_146105, intron 1 of 10) | B4\|SINE\|B4 | 29352 | NM_146105 | 226744 | Mm.87662 | NM_146105 | ENSMUSG00000038949 | **Cnst** | 9630058J23Rik | consortin, connexin sorting protein | protein-coding |
| chr9_11328809_11329378 | chr9 | 11328810 | 11329378 | + | 1.35328 | Intergenic | MMVL30-int\|LTR\|ERV1 | -424319 | NM_001170787 | 244682 | Mm.187021 | NM_001033359 | ENSMUSG00000039488 | **Cntn5** | 6720426O10Rik\|A830025P08Rik\|Gm507\|NB-2 | contactin 5 | protein-coding |
| chr9_11380419_11380859 | chr9 | 11380420 | 11380859 | + | 0.903787 | Intergenic | RLTR1F_Mm\|LTR\|ERVK | -475864 | NM_001170787 | 244682 | Mm.187021 | NM_001033359 | ENSMUSG00000039488 | **Cntn5** | 6720426O10Rik\|A830025P08Rik\|Gm507\|NB-2 | contactin 5 | protein-coding |
| chr9_11671191_11671595 | chr9 | 11671192 | 11671595 | + | 0.772773 | Intergenic | Intergenic | -766618 | NM_001170787 | 244682 | Mm.187021 | NM_001033359 | ENSMUSG00000039488 | **Cntn5** | 6720426O10Rik\|A830025P08Rik\|Gm507\|NB-2 | contactin 5 | protein-coding |
| chr9_11366512_11366864 | chr9 | 11366513 | 11366864 | + | 0.745306 | Intergenic | Intergenic | -461913 | NM_001170787 | 244682 | Mm.187021 | NM_001033359 | ENSMUSG00000039488 | **Cntn5** | 6720426O10Rik\|A830025P08Rik\|Gm507\|NB-2 | contactin 5 | protein-coding |
| chr11_101194573_101195215 | chr11 | 101194574 | 101195215 | + | 1.26713 | exon (NM_007970, exon 17 of 21) | exon (NM_007970, exon 17 of 21) | 18777 | NM_016782 | 53321 | Mm.474527 | NM_016782 | ENSMUSG00000017167 | **Cntnap1** | AI841080\|Caspr\|NCP1\|Nrxn4\|p190\|shm | contactin associated protein-like 1 | protein-coding |
| chr11_101194373_101195400 | chr11 | 101194374 | 101195400 | + | 1.17436 | exon (NM_007970, exon 17 of 21) | exon (NM_007970, exon 17 of 21) | 18770 | NM_016782 | 53321 | Mm.474527 | NM_016782 | ENSMUSG00000017167 | **Cntnap1** | AI841080\|Caspr\|NCP1\|Nrxn4\|p190\|shm | contactin associated protein-like 1 | protein-coding |
| chr6_47344538_47345641 | chr6 | 47344539 | 47345641 | + | 0.945636 | Intergenic | Intergenic | 100694 | NM_025771 | 66797 | Mm.440084 | NM_025771 | ENSMUSG00000039419 | **Cntnap2** | 5430425M22Rik\|Caspr2\|mKIAA0868 | contactin associated protein-like 2 | protein-coding |
| chr2_35143548_35143853 | chr2 | 35143549 | 35143853 | + | 0.729582 | exon (NM_001290635, exon 6 of 9) | exon (NM_001290635, exon 6 of 9) | 11467 | NM_001290635 | 26920 | Mm.231332 | NM_012018 | ENSMUSG00000057110 | **Cntrl** | 6720467O09Rik\|AA968343\|Cep1\|Cep110\|IB3/5\|Ma2a8\|b2b1468Clo | centriolin | protein-coding |
| chr2_35153869_35155276 | chr2 | 35153870 | 35155276 | + | 0.673253 | intron (NM_030000, intron 14 of 33) | intron (NM_030000, intron 14 of 33) | 22339 | NM_001290635 | 26920 | Mm.231332 | NM_012018 | ENSMUSG00000057110 | **Cntrl** | 6720467O09Rik\|AA968343\|Cep1\|Cep110\|IB3/5\|Ma2a8\|b2b1468Clo | centriolin | protein-coding |
| chr2_35153898_35154510 | chr2 | 35153899 | 35154510 | + | 0.629985 | intron (NM_030000, intron 14 of 33) | RMER15\|LTR\|ERVL | 21970 | NM_001290635 | 26920 | Mm.231332 | NM_012018 | ENSMUSG00000057110 | **Cntrl** | 6720467O09Rik\|AA968343\|Cep1\|Cep110\|IB3/5\|Ma2a8\|b2b1468Clo | centriolin | protein-coding |
| chr2_65135558_65135976 | chr2 | 65135559 | 65135976 | + | 1.2162 | intron (NM_177025, intron 3 of 13) | intron (NM_177025, intron 3 of 13) | 102859 | NM_177025 | 319876 | Mm.34159 | NM_027225 | ENSMUSG00000034903 | **Cobll1** | 1810047P18Rik\|Coblr1\|D430044D16Rik | Cobl-like 1 | protein-coding |
| chr2_65240450_65240748 | chr2 | 65240451 | 65240748 | + | 0.996531 | Intergenic | Intergenic | -1973 | NM_027225 | 319876 | Mm.34159 | NM_027225 | ENSMUSG00000034903 | **Cobll1** | 1810047P18Rik\|Coblr1\|D430044D16Rik | Cobl-like 1 | protein-coding |
| chr2_65161572_65162143 | chr2 | 65161573 | 65162143 | + | 0.93316 | intron (NM_177025, intron 1 of 13) | intron (NM_177025, intron 1 of 13) | 76768 | NM_177025 | 319876 | Mm.34159 | NM_027225 | ENSMUSG00000034903 | **Cobll1** | 1810047P18Rik\|Coblr1\|D430044D16Rik | Cobl-like 1 | protein-coding |
| chr2_65195215_65196423 | chr2 | 65195216 | 65196423 | + | 0.733933 | intron (NM_177025, intron 1 of 13) | intron (NM_177025, intron 1 of 13) | 42807 | NM_177025 | 319876 | Mm.34159 | NM_027225 | ENSMUSG00000034903 | **Cobll1** | 1810047P18Rik\|Coblr1\|D430044D16Rik | Cobl-like 1 | protein-coding |
| chr12_51629805_51630105 | chr12 | 51629806 | 51630105 | + | 0.639854 | intron (NM_001172098, intron 7 of 15) | intron (NM_001172098, intron 7 of 15) | 36614 | NM_001198835 | 12810 | Mm.21325 | NM_007728 | ENSMUSG00000020953 | **Coch** | AW122937\|Coch-5B2\|D12H14S564E | coagulation factor C homolog (Limulus polyphemus) | protein-coding |

| | | | | | | | | | | | | | | | | | |
|---|---|---|---|---|---|---|---|---|---|---|---|---|---|---|---|---|---|
| chr14_75747848_75748636 | chr14 | 75747849 | 75748636 | + | 0.77882 | intron (NM_177381, intron 1 of 22) | intron (NM_177381, intron 1 of 22) | 6251 | NM_177381 | 338337 | Mm.86507 | NM_177381 | ENSMUSG00000034893 | **Cog3** | E430004N23 | component of oligomeric golgi complex 3 | protein-coding |
| chr14_75747570_75748649 | chr14 | 75747571 | 75748649 | + | 0.70502 | intron (NM_177381, intron 1 of 22) | intron (NM_177381, intron 1 of 22) | 6383 | NM_177381 | 338337 | Mm.86507 | NM_177381 | ENSMUSG00000034893 | **Cog3** | E430004N23 | component of oligomeric golgi complex 3 | protein-coding |
| chr8_110864081_110864457 | chr8 | 110864082 | 110864457 | + | 0.713274 | intron (NM_133973, intron 9 of 18) | B1_Mur1\|SINE\|Alu | 17245 | NM_133973 | 102339 | Mm.272930 | NM_133973 | ENSMUSG00000031753 | **Cog4** | AW554810\|D8Ertd515e | component of oligomeric golgi complex 4 | protein-coding |
| chr4_47273657_47274812 | chr4 | 47273658 | 47274812 | + | 1.15032 | intron (NM_009928, intron 14 of 39) | intron (NM_009928, intron 14 of 39) | 66223 | NM_009928 | 12819 | Mm.233547 | NM_009928 | ENSMUSG00000028339 | **Col15a1** | - | collagen, type XV, alpha 1 | protein-coding |
| chr4_47274425_47274802 | chr4 | 47274426 | 47274802 | + | 1.11377 | intron (NM_009928, intron 14 of 39) | MTEb\|LTR\|ERVL-MaLR | 66602 | NM_009928 | 12819 | Mm.233547 | NM_009928 | ENSMUSG00000028339 | **Col15a1** | - | collagen, type XV, alpha 1 | protein-coding |
| chr4_47273826_47274271 | chr4 | 47273827 | 47274271 | + | 0.967416 | intron (NM_009928, intron 14 of 39) | intron (NM_009928, intron 14 of 39) | 66037 | NM_009928 | 12819 | Mm.233547 | NM_009928 | ENSMUSG00000028339 | **Col15a1** | - | collagen, type XV, alpha 1 | protein-coding |
| chr4_47279951_47280621 | chr4 | 47279952 | 47280621 | + | 0.61405 | intron (NM_009928, intron 17 of 39) | intron (NM_009928, intron 17 of 39) | 72274 | NM_009928 | 12819 | Mm.233547 | NM_009928 | ENSMUSG00000028339 | **Col15a1** | - | collagen, type XV, alpha 1 | protein-coding |
| chr10_77104299_77104906 | chr10 | 77104300 | 77104906 | + | 0.74074 | intron (NM_009929, intron 2 of 41) | MLTR14\|LTR\|ERV1 | 9102 | NM_001109991 | 12822 | Mm.4352 | NM_009929 | ENSMUSG00000001435 | **Col18a1** | - | collagen, type XVIII, alpha 1 | protein-coding |
| chr8_11203236_11203773 | chr8 | 11203237 | 11203773 | + | 0.858908 | intron (NM_009931, intron 49 of 51) | intron (NM_009931, intron 49 of 51) | 109321 | NM_009931 | 12826 | Mm.738 | NM_009931 | ENSMUSG00000031502 | **Col4a1** | Bru\|Col4a-1\|Del(8)44H\|Raw\|Svc | collagen, type IV, alpha 1 | protein-coding |
| chr9_20812167_20813299 | chr9 | 20812168 | 20813299 | + | 1.29956 | intron (NM_016919, intron 1 of 66) | intron (NM_016919, intron 1 of 66) | 2301 | NM_016919 | 53867 | Mm.334994 | NM_016919 | ENSMUSG00000004098 | **Col5a3** | - | collagen, type V, alpha 3 | protein-coding |
| chr10_76736847_76737172 | chr10 | 76736848 | 76737172 | + | 0.787047 | Intergenic | LTR16A1\|LTR\|ERVL | -10966 | NM_009933 | 12833 | Mm.2509 | NM_009933 | ENSMUSG00000001119 | **Col6a1** | AI747156\|Col6a-1 | collagen, type VI, alpha 1 | protein-coding |
| chr1_152422198_152422670 | chr1 | 152422199 | 152422670 | + | 0.803968 | intron (NM_177756, intron 1 of 11) | intron (NM_177756, intron 1 of 11) | 22567 | NM_177756 | 269132 | Mm.23782 | NM_177756 | ENSMUSG00000032649 | **Colgalt2** | AI427652\|D730024P12\|Glt25d2 | collagen beta(1-O)galactosyltransferase 2 | protein-coding |
| chr1_152422226_152422567 | chr1 | 152422227 | 152422567 | + | 0.801974 | intron (NM_177756, intron 1 of 11) | intron (NM_177756, intron 1 of 11) | 22530 | NM_177756 | 269132 | Mm.23782 | NM_177756 | ENSMUSG00000032649 | **Colgalt2** | AI427652\|D730024P12\|Glt25d2 | collagen beta(1-O)galactosyltransferase 2 | protein-coding |
| chr14_31597398_31597947 | chr14 | 31597399 | 31597947 | + | 1.18206 | Intergenic | Intergenic | -20290 | NM_009937 | 382864 | Mm.239821 | NM_009937 | ENSMUSG00000057606 | **Colq** | A130034K24Rik | collagen-like tail subunit (single strand of homotrimer) of asymmetric acetylcholinesterase | protein-coding |
| chr18_47040607_47041049 | chr18 | 47040608 | 47041049 | + | 1.8269 | intron (NM_178377, intron 5 of 6) | intron (NM_178377, intron 5 of 6) | 81952 | NM_178377 | 69456 | Mm.210734 | NM_178377 | ENSMUSG00000042705 | **Commd10** | 2310003A05Rik\|AU019438\|DRWMS2 | COMM domain containing 10 | protein-coding |
| chr18_47035925_47036429 | chr18 | 47035926 | 47036429 | + | 1.16841 | intron (NM_178377, intron 5 of 6) | intron (NM_178377, intron 5 of 6) | 77301 | NM_178377 | 69456 | Mm.210734 | NM_178377 | ENSMUSG00000042705 | **Commd10** | 2310003A05Rik\|AU019438\|DRWMS2 | COMM domain containing 10 | protein-coding |
| chr18_47044670_47045145 | chr18 | 47044671 | 47045145 | + | 0.856542 | intron (NM_178377, intron 5 of 6) | intron (NM_178377, intron 5 of 6) | 86032 | NM_178377 | 69456 | Mm.210734 | NM_178377 | ENSMUSG00000042705 | **Commd10** | 2310003A05Rik\|AU019438\|DRWMS2 | COMM domain containing 10 | protein-coding |
| chr18_47033600_47034071 | chr18 | 47033601 | 47034071 | + | 0.828609 | intron (NM_178377, intron 5 of 6) | intron (NM_178377, intron 5 of 6) | 74960 | NM_178377 | 69456 | Mm.210734 | NM_178377 | ENSMUSG00000042705 | **Commd10** | 2310003A05Rik\|AU019438\|DRWMS2 | COMM domain containing 10 | protein-coding |
| chr2_18664003_18664358 | chr2 | 18664004 | 18664358 | + | 0.896747 | Intergenic | Intergenic | -8281 | NM_147778 | 12238 | Mm.249586 | NM_147778 | ENSMUSG00000051154 | **Commd3** | AW550818\|Bup\|D2Ertd542e | COMM domain containing 3 | protein-coding |
| chr2_18604374_18604644 | chr2 | 18604375 | 18604644 | + | 0.819464 | Intergenic | Intergenic | -67953 | NM_147778 | 12238 | Mm.249586 | NM_147778 | ENSMUSG00000051154 | **Commd3** | AW550818\|Bup\|D2Ertd542e | COMM domain containing 3 | protein-coding |
| chr2_102034382_102034844 | chr2 | 102034383 | 102034844 | + | 1.55132 | intron (NM_178886, intron 4 of 5) | MIRc\|SINE\|MIR | 148351 | NM_029635 | 76501 | Mm.330424 | NM_029635 | ENSMUSG00000027163 | **Commd9** | 1810029F08Rik\|AA409293 | COMM domain containing 9 | protein-coding |
| chr2_102031860_102032090 | chr2 | 102031861 | 102032090 | + | 0.661503 | intron (NM_178886, intron 4 of 5) | intron (NM_178886, intron 4 of 5) | 145713 | NM_029635 | 76501 | Mm.330424 | NM_029635 | ENSMUSG00000027163 | **Commd9** | 1810029F08Rik\|AA409293 | COMM domain containing 9 | protein-coding |
| chr16_18452238_18452754 | chr16 | 18452239 | 18452754 | + | 0.894905 | intron (NM_013711, intron 8 of 17) | intron (NM_013711, intron 8 of 17) | -25780 | NM_001111062 | 12846 | Mm.100940 | NM_007744 | ENSMUSG00000000326 | **Comt** | Comt1\|D16Wsu103e\|D330014B15Rik | catechol-O-methyltransferase | protein-coding |
| chr7_114231781_114232274 | chr7 | 114231782 | 114232274 | + | 0.907681 | intron (NM_033370, intron 14 of 21) | intron (NM_033370, intron 14 of 21) | 22652 | NM_033370 | 70349 | Mm.277024 | NM_033370 | ENSMUSG00000030754 | **Copb1** | 2610019B04Rik | coatomer protein complex, subunit beta 1 | protein-coding |
| chr5_100535528_100536322 | chr5 | 100535529 | 100536322 | + | 1.82513 | intron (NM_012001, intron 6 of 9) | MTD\|LTR\|ERVL-MaLR | 17616 | NM_012001 | 26891 | Mm.957 | NM_012001 | ENSMUSG00000035297 | **Cops4** | AW208976\|D5Ertd774e | COP9 (constitutive photomorphogenic) homolog, subunit 4 (Arabidopsis thaliana) | protein-coding |
| chr5_100533878_100534113 | chr5 | 100533879 | 100534113 | + | 1.09361 | intron (NM_012001, intron 6 of 9) | RSINE1\|SINE\|B4 | 15687 | NM_012001 | 26891 | Mm.957 | NM_012001 | ENSMUSG00000035297 | **Cops4** | AW208976\|D5Ertd774e | COP9 (constitutive photomorphogenic) homolog, subunit 4 (Arabidopsis thaliana) | protein-coding |
| chr6_124950819_124951202 | chr6 | 124950820 | 124951202 | + | 0.672597 | Intergenic | LTRIS2\|LTR\|ERV1 | 14386 | NM_012003 | 26894 | Mm.244536 | NM_012003 | ENSMUSG00000030127 | **Cops7a** | D6Ertd35e | COP9 (constitutive photomorphogenic) homolog, subunit 7a (Arabidopsis thaliana) | protein-coding |
| chr1_90677596_90678247 | chr1 | 90677597 | 90678247 | + | 1.77601 | Intergenic | ORR1F\|LTR\|ERVL-MaLR | 74497 | NM_133805 | 108679 | Mm.119177 | NM_133805 | ENSMUSG00000034432 | **Cops8** | 9430009J09Rik\|AA408242\|Csn8\|Sgn8 | COP9 (constitutive photomorphogenic) homolog, subunit 8 (Arabidopsis thaliana) | protein-coding |
| chr1_90676617_90677008 | chr1 | 90676618 | 90677008 | + | 0.824802 | Intergenic | Intergenic | 73388 | NM_133805 | 108679 | Mm.119177 | NM_133805 | ENSMUSG00000034432 | **Cops8** | 9430009J09Rik\|AA408242\|Csn8\|Sgn8 | COP9 (constitutive photomorphogenic) homolog, subunit 8 (Arabidopsis thaliana) | protein-coding |
| chr1_55049171_55049511 | chr1 | 55049172 | 55049511 | + | 1.35371 | Intergenic | Intergenic | -3429 | NM_026424 | 67876 | Mm.281019 | NM_026424 | ENSMUSG00000025981 | **Coq10b** | 1500041J02Rik\|1700030I21Rik\|9530077A17Rik\|AV002237\|AW742641 | coenzyme Q10 homolog B (S. cerevisiae) | protein-coding |
| chr1_55051810_55051979 | chr1 | 55051811 | 55051979 | + | 0.66739 | promoter-TSS (NM_001039710) | promoter-TSS (NM_001039710) | -875 | NM_026424 | 67876 | Mm.281019 | NM_026424 | ENSMUSG00000025981 | **Coq10b** | 1500041J02Rik\|1700030I21Rik\|9530077A17Rik\|AV002237\|AW742641 | coenzyme Q10 homolog B (S. cerevisiae) | protein-coding |
| chr5_100635221_100635786 | chr5 | 100635222 | 100635786 | + | 0.861735 | Intergenic | Intergenic | 38752 | NM_027978 | 71883 | Mm.260661 | NM_027978 | ENSMUSG00000029319 | **Coq2** | 2310002F18Rik | coenzyme Q2 homolog, prenyltransferase (yeast) | protein-coding |
| chr5_100627950_100628499 | chr5 | 100627951 | 100628499 | + | 0.761322 | Intergenic | Intergenic | 46031 | NM_027978 | 71883 | Mm.260661 | NM_027978 | ENSMUSG00000029319 | **Coq2** | 2310002F18Rik | coenzyme Q2 homolog, prenyltransferase (yeast) | protein-coding |
| chr4_21877808_21878151 | chr4 | 21877809 | 21878151 | + | 1.1194 | Intergenic | B3A\|SINE\|B2 | -1695 | NM_172687 | 230027 | Mm.5662 | NM_172687 | ENSMUSG00000028247 | **Coq3** | 4732433J24\|C77934 | coenzyme Q3 homolog, methyltransferase (yeast) | protein-coding |
| chr7_126708237_126708514 | chr7 | 126708238 | 126708514 | + | 0.930639 | Intergenic | MLT1K\|LTR\|ERVL-MaLR | -3560 | NM_009898 | 12721 | Mm.290482 | NM_009898 | ENSMUSG00000030707 | **Coro1a** | Clabp\|Lmb3\|TACO\|p57 | coronin, actin binding protein 1A | protein-coding |
| chr9_62509176_62509620 | chr9 | 62509177 | 62509620 | + | 0.936186 | intron (NM_175484, intron 1 of 11) | intron (NM_175484, intron 1 of 11) | 27646 | NM_175484 | 235431 | Mm.335229 | NM_175484 | ENSMUSG00000041729 | **Coro2b** | CLIPINC\|E130012P22Rik | coronin, actin binding protein, 2B | protein-coding |
| chr8_119817038_119817820 | chr8 | 119817039 | 119817820 | + | 0.600877 | intron (NM_028071, intron 3 of 3) | intron (NM_028071, intron 3 of 3) | 23150 | NM_028071 | 72042 | Mm.393405 | NM_028071 | ENSMUSG00000031827 | **Cotl1** | 1810074P22Rik\|2010004C08Rik\|Clp | coactosin-like 1 (Dictyostelium) | protein-coding |
| chr5_90271271_90271725 | chr5 | 90271272 | 90271725 | + | 1.11038 | intron (NM_198010, intron 15 of 32) | intron (NM_198010, intron 15 of 32) | -47502 | NM_001163456 | 231430 | Mm.319697 | NM_001033310 | ENSMUSG00000035505 | **Cox18** | BC038311 | cytochrome c oxidase assembly protein 18 | protein-coding |
| chr8_120699616_120699872 | chr8 | 120699617 | 120699872 | + | 0.667521 | Intergenic | Intergenic | 31424 | NM_001293559 | 12857 | Mm.386758 | NM_009941 | ENSMUSG00000031818 | **Cox4i1** | AL024441\|COX\|COX IV-1\|COXIV\|Cox4\|Cox4a\|IV-1 | cytochrome c oxidase subunit IV isoform 1 | protein-coding |
| chr2_152787591_152787839 | chr2 | 152787592 | 152787839 | + | 0.698513 | intron (NM_001289717, intron 2 of 2) | intron (NM_001289717, intron 2 of 2) | 33542 | NM_053091 | 84682 | Mm.196668 | NM_053091 | ENSMUSG00000009876 | **Cox4i2** | Cox4b\|CoxIV-2 | cytochrome c oxidase subunit IV isoform 2 | protein-coding |
| chr2_152787581_152788019 | chr2 | 152787582 | 152788019 | + | 0.695299 | intron (NM_001289717, intron 2 of 2) | intron (NM_001289717, intron 2 of 2) | 33627 | NM_053091 | 84682 | Mm.196668 | NM_053091 | ENSMUSG00000009876 | **Cox4i2** | Cox4b\|CoxIV-2 | cytochrome c oxidase subunit IV isoform 2 | protein-coding |
| chr9_57525632_57525877 | chr9 | 57525633 | 57525877 | + | 0.871913 | intron (NM_007747, intron 1 of 4) | ID_B1\|SINE\|B4 | 4523 | NM_007747 | 12858 | Mm.273403 | NM_007747 | ENSMUSG00000000088 | **Cox5a** | AA959768\|CcOX | cytochrome c oxidase subunit Va | protein-coding |
| chr9_79772086_79772564 | chr9 | 79772087 | 79772564 | + | 1.51855 | intron (NM_133718, intron 6 of 6) | intron (NM_133718, intron 6 of 6) | -12472 | NM_009945 | 12866 | Mm.152627 | NM_009945 | ENSMUSG00000032330 | **Cox7a2** | COX7AL\|Cox7a3\|CoxVIIa-L | cytochrome c oxidase subunit VIIa 2 | protein-coding |
| chr17_83483689_83484505 | chr17 | 83483690 | 83484505 | + | 1.53969 | Intergenic | Intergenic | 30236 | NM_009187 | 20463 | Mm.30072 | NM_009187 | ENSMUSG00000024248 | **Cox7a2l** | COX7AR\|COX7RP\|EB1\|SIG-81\|SIG81\|Silg81 | cytochrome c oxidase subunit VIIa polypeptide 2-like | protein-coding |
| chr17_83456286_83456848 | chr17 | 83456287 | 83456848 | + | 0.752523 | intron (NM_001286567, intron 17 of 23) | intron (NM_001286567, intron 17 of 23) | 57766 | NM_009187 | 20463 | Mm.30072 | NM_009187 | ENSMUSG00000024248 | **Cox7a2l** | COX7AR\|COX7RP\|EB1\|SIG-81\|SIG81\|Silg81 | cytochrome c oxidase subunit VIIa polypeptide 2-like | protein-coding |
| chr13_85724686_85724996 | chr13 | 85724687 | 85724996 | + | 1.02255 | Intergenic | MLTR31FA_MM\|LTR\|ERVK | 321954 | NM_007749 | 12867 | Mm.378898 | NM_007749 | | **Cox7c** | COXVIIc\|Cox7c1 | cytochrome c oxidase subunit VIIc | protein-coding |
| chr13_85724651_85725114 | chr13 | 85724652 | 85725114 | + | 1.00695 | Intergenic | MusHAL1\|LINE\|L1 | 321912 | NM_007749 | 12867 | Mm.378898 | NM_007749 | | **Cox7c** | COXVIIc\|Cox7c1 | cytochrome c oxidase subunit VIIc | protein-coding |
| chr13_85724120_85725120 | chr13 | 85724121 | 85725120 | + | 0.94924 | Intergenic | MLTR31FA_MM\|LTR\|ERVK | 322175 | NM_007749 | 12867 | Mm.378898 | NM_007749 | | **Cox7c** | COXVIIc\|Cox7c1 | cytochrome c oxidase subunit VIIc | protein-coding |
| chr6_30657133_30657771 | chr6 | 30657134 | 30657771 | + | 1.09309 | intron (NM_031998, intron 8 of 10) | intron (NM_031998, intron 8 of 10) | 18231 | NM_025350 | 109697 | Mm.25377 | NM_025350 | ENSMUSG00000054446 | **Cpa1** | 0910001L12Rik\|Cpa | carboxypeptidase A1, pancreatic | protein-coding |

| | | | | | | | | | | | | | | | | | |
|---|---|---|---|---|---|---|---|---|---|---|---|---|---|---|---|---|---|
| chr6_30550993_30551507 | chr6 | 30550994 | 30551507 | + | 0.885891 | intron (NM_001024698, intron 7 of 10) | intron (NM_001024698, intron 7 of 10) | 9608 | NM_001024698 | 232680 | Mm.268855 | NM_001024698 | ENSMUSG00000071553 | **Cpa2** | - | carboxypeptidase A2, pancreatic | protein-coding |
| chr14_75190677_75191138 | chr14 | 75190678 | 75191138 | + | 1.23978 | intron (NM_008879, intron 2 of 16) | intron (NM_008879, intron 2 of 16) | -51379 | NM_019775 | 56373 | Mm.24242 | NM_019775 | ENSMUSG00000021999 | **Cpb2** | 1110032P04Rik\|4930405E17Rik\|AI255929\|CPR\|Cpu\|TAFI | carboxypeptidase B2 (plasma) | protein-coding |
| chr11_76823626_76824074 | chr11 | 76823627 | 76824074 | + | 0.989222 | intron (NM_007754, intron 2 of 20) | MER44B\|DNA\|TcMar-Tigger | 23158 | NM_007754 | 12874 | Mm.276736 | NM_007754 | ENSMUSG00000020841 | **Cpd** | AA960140\|AW322530\|D830034L15Rik | carboxypeptidase D | protein-coding |
| chr11_76815465_76816015 | chr11 | 76815466 | 76816015 | + | 0.794926 | intron (NM_007754, intron 2 of 20) | MTEb\|LTR\|ERVL-MaLR | 31268 | NM_007754 | 12874 | Mm.276736 | NM_007754 | ENSMUSG00000020841 | **Cpd** | AA960140\|AW322530\|D830034L15Rik | carboxypeptidase D | protein-coding |
| chr10_117647700_117648070 | chr10 | 117647701 | 117648070 | + | 0.935159 | intron (NM_027468, intron 2 of 8) | intron (NM_027468, intron 2 of 8) | 18385 | NM_027468 | 70574 | Mm.339332 | NM_027468 | ENSMUSG00000020183 | **Cpm** | 1110060I01Rik\|5730456K23Rik\|AA589379\|E030045M14Rik | carboxypeptidase M | protein-coding |
| chr10_117624166_117624466 | chr10 | 117624167 | 117624466 | + | 0.736413 | Intergenic | L2\|LINE\|L2 | -5184 | NM_027468 | 70574 | Mm.339332 | NM_027468 | ENSMUSG00000020183 | **Cpm** | 1110060I01Rik\|5730456K23Rik\|AA589379\|E030045M14Rik | carboxypeptidase M | protein-coding |
| chr16_30211930_30212484 | chr16 | 30211931 | 30212484 | + | 0.937194 | Intergenic | RLTR24\|LTR\|ERV1 | 55325 | NM_027904 | 71756 | Mm.290614 | NM_027904 | ENSMUSG00000023176 | **Cpn2** | 1300018K11Rik | carboxypeptidase N, polypeptide 2 | protein-coding |
| chr16_30211708_30212493 | chr16 | 30211709 | 30212493 | + | 0.730305 | Intergenic | RLTR24\|LTR\|ERV1 | 55431 | NM_027904 | 71756 | Mm.290614 | NM_027904 | ENSMUSG00000023176 | **Cpn2** | 1300018K11Rik | carboxypeptidase N, polypeptide 2 | protein-coding |
| chr16_58661601_58662295 | chr16 | 58661602 | 58662295 | + | 1.2479 | Intergenic | ORR1E\|LTR\|ERVL-MaLR | -8260 | NM_007757 | 12892 | Mm.291519 | NM_007757 | ENSMUSG00000022742 | **Cpox** | CPX\|Cpo\|HCP\|cac\|nct | coproporphyrinogen oxidase | protein-coding |
| chr15_33126898_33127253 | chr15 | 33126899 | 33127253 | + | 0.620942 | intron (NM_176073, intron 1 of 7) | intron (NM_176073, intron 1 of 7) | 43947 | NM_176073 | 54381 | Mm.158251 | NM_018755 | ENSMUSG00000039007 | **Cpq** | 1190003P12Rik\|2610034C17Rik\|His2\|Lal-1\|Pgcp | carboxypeptidase Q | protein-coding |
| chr4_155867128_155867672 | chr4 | 155867129 | 155867672 | + | 0.703594 | intron (NM_024472, intron 1 of 2) | intron (NM_024472, intron 1 of 2) | 2040 | NM_024472 | 79554 | Mm.25556 | NM_024472 | ENSMUSG00000029073 | **Cptp** | Gltpd1 | ceramide-1-phosphate transfer protein | protein-coding |
| chr3_87959338_87959686 | chr3 | 87959339 | 87959686 | + | 0.769521 | Intergenic | Intergenic | 10819 | NM_007759 | 12904 | Mm.4757 | NM_007759 | ENSMUSG00000004885 | **Crabp2** | AI893628\|Crabp-2\|CrabpII | cellular retinoic acid binding protein II | protein-coding |
| chr10_95142261_95143236 | chr10 | 95142262 | 95143236 | + | 1.16732 | Intergenic | Intergenic | 181348 | NM_009950 | 12905 | Mm.473739 | NM_009950 | ENSMUSG00000045867 | **Cradd** | RAIDD | CASP2 and RIPK1 domain containing adaptor with death domain | protein-coding |
| chr10_95146576_95146984 | chr10 | 95146577 | 95146984 | + | 0.997596 | Intergenic | Intergenic | 177317 | NM_009950 | 12905 | Mm.473739 | NM_009950 | ENSMUSG00000045867 | **Cradd** | RAIDD | CASP2 and RIPK1 domain containing adaptor with death domain | protein-coding |
| chr10_95299434_95300191 | chr10 | 95299435 | 95300191 | + | 0.966811 | intron (NM_009950, intron 2 of 2) | intron (NM_009950, intron 2 of 2) | 24284 | NM_009950 | 12905 | Mm.473739 | NM_009950 | ENSMUSG00000045867 | **Cradd** | RAIDD | CASP2 and RIPK1 domain containing adaptor with death domain | protein-coding |
| chr1_139179474_139180292 | chr1 | 139179475 | 139180292 | + | 1.35655 | Intergenic | Lx7\|LINE\|L1 | 197193 | NM_133239 | 170788 | Mm.95700 | NM_133239 | ENSMUSG00000063681 | **Crb1** | 7530426H14Rik\|A930008G09Rik | crumbs homolog 1 (Drosophila) | protein-coding |
| chr1_139340560_139340812 | chr1 | 139340561 | 139340812 | + | 0.71453 | intron (NM_133239, intron 1 of 11) | intron (NM_133239, intron 1 of 11) | 36390 | NM_133239 | 170788 | Mm.95700 | NM_133239 | ENSMUSG00000063681 | **Crb1** | 7530426H14Rik\|A930008G09Rik | crumbs homolog 1 (Drosophila) | protein-coding |
| chr2_37716298_37716721 | chr2 | 37716299 | 37716721 | + | 0.659481 | Intergenic | Intergenic | -59739 | NM_001163566 | 241324 | Mm.392369 | NM_001163566 | ENSMUSG00000035403 | **Crb2** | 5930402A21\|BC043114 | crumbs homolog 2 (Drosophila) | protein-coding |
| chr6_106985195_106985588 | chr6 | 106985196 | 106985588 | + | 0.730681 | Intergenic | Intergenic | -185305 | NM_175357 | 58799 | Mm.290085 | NM_021449 | | **Crbn** | 2610203G15Rik\|2900045O07Rik\|AF229032\|AW108261\|piL | cereblon | protein-coding |
| chr6_106984032_106984632 | chr6 | 106984033 | 106984632 | + | 0.67865 | Intergenic | MTD\|LTR\|ERVL-MaLR | -184245 | NM_175357 | 58799 | Mm.290085 | NM_021449 | | **Crbn** | 2610203G15Rik\|2900045O07Rik\|AF229032\|AW108261\|piL | cereblon | protein-coding |
| chr1_64549461_64549750 | chr1 | 64549462 | 64549750 | + | 1.19929 | intron (NM_001037726, intron 1 of 6) | intron (NM_001037726, intron 1 of 6) | 16802 | NM_001037726 | 12912 | Mm.422634 | NM_009952 | ENSMUSG00000025958 | **Creb1** | 2310001E10Rik\|3526402H21Rik\|AV083133\|Creb\|Creb-1 | cAMP responsive element binding protein 1 | protein-coding |
| chr1_64507579_64507955 | chr1 | 64507580 | 64507955 | + | 1.06892 | Intergenic | MTD\|LTR\|ERVL-MaLR | -25037 | NM_133828 | 12912 | Mm.422634 | NM_009952 | ENSMUSG00000025958 | **Creb1** | 2310001E10Rik\|3526402H21Rik\|AV083133\|Creb\|Creb-1 | cAMP responsive element binding protein 1 | protein-coding |
| chr1_64546653_64547223 | chr1 | 64546654 | 64547223 | + | 0.738518 | intron (NM_001037726, intron 1 of 6) | intron (NM_001037726, intron 1 of 6) | 14134 | NM_001037726 | 12912 | Mm.422634 | NM_009952 | ENSMUSG00000025958 | **Creb1** | 2310001E10Rik\|3526402H21Rik\|AV083133\|Creb\|Creb-1 | cAMP responsive element binding protein 1 | protein-coding |
| chr1_64531137_64531312 | chr1 | 64531138 | 64531312 | + | 0.627236 | Intergenic | Intergenic | -1579 | NM_133828 | 12912 | Mm.422634 | NM_009952 | ENSMUSG00000025958 | **Creb1** | 2310001E10Rik\|3526402H21Rik\|AV083133\|Creb\|Creb-1 | cAMP responsive element binding protein 1 | protein-coding |
| chr6_37455535_37456264 | chr6 | 37455536 | 37456264 | + | 0.822284 | Intergenic | Intergenic | -13752 | NM_178661 | 208647 | Mm.169929 | NM_178661 | | **Creb3l2** | BBF2H7\|C530025K05Rik\|SCIRR69 | cAMP responsive element binding protein 3-like 2 | protein-coding |
| chr6_37388584_37389157 | chr6 | 37388585 | 37389157 | + | 0.603217 | intron (NM_178661, intron 1 of 11) | intron (NM_178661, intron 1 of 11) | 53277 | NM_178661 | 208647 | Mm.169929 | NM_178661 | | **Creb3l2** | BBF2H7\|C530025K05Rik\|SCIRR69 | cAMP responsive element binding protein 3-like 2 | protein-coding |
| chr6_53497815_53498404 | chr6 | 53497816 | 53498404 | + | 1.39484 | Intergenic | Intergenic | -75264 | NM_172728 | 231991 | Mm.321138 | NM_172728 | ENSMUSG00000053007 | **Creb5** | Crebpa\|D430026C09Rik | cAMP responsive element binding protein 5 | protein-coding |
| chr6_53455409_53455844 | chr6 | 53455410 | 53455844 | + | 0.695451 | Intergenic | Intergenic | -117747 | NM_172728 | 231991 | Mm.321138 | NM_172728 | ENSMUSG00000053007 | **Creb5** | Crebpa\|D430026C09Rik | cAMP responsive element binding protein 5 | protein-coding |
| chr16_4151164_4151692 | chr16 | 4151165 | 4151692 | + | 0.729141 | intron (NM_001025432, intron 3 of 30) | intron (NM_001025432, intron 3 of 30) | 61976 | NM_001025432 | 12914 | Mm.132238 | NM_001025432 | ENSMUSG00000022521 | **Crebbp** | AW558298\|CBP\|CBP/p300\|KAT3A\|p300/CBP | CREB binding protein | protein-coding |
| chr6_134843459_134843774 | chr6 | 134843460 | 134843774 | + | 0.671787 | intron (NM_177687, intron 1 of 4) | intron (NM_177687, intron 1 of 4) | 13418 | NM_177687 | 232430 | Mm.267010 | NM_177687 | | **Crebl2** | AI046348\|B230205M03 | cAMP responsive element binding protein-like 2 | protein-coding |
| chr1_39680574_39680903 | chr1 | 39680575 | 39680903 | + | 0.608272 | intron (NM_001145660, intron 11 of 14) | intron (NM_001145660, intron 11 of 14) | -29557 | NM_170597 | 263764 | Mm.122091 | NM_170597 | ENSMUSG00000050967 | **Creg2** | A830098L22Rik | cellular repressor of E1A-stimulated genes 2 | protein-coding |
| chr13_95427492_95427945 | chr13 | 95427493 | 95427945 | + | 1.24046 | Intergenic | ORR1E\|LTR\|ERVL-MaLR | 17112 | NM_198408 | 12919 | Mm.316614 | NM_198408 | ENSMUSG00000021680 | **Crhbp** | CRH-BP | corticotropin releasing hormone binding protein | protein-coding |
| chr13_95427308_95427962 | chr13 | 95427309 | 95427962 | + | 1.19524 | Intergenic | ORR1E\|LTR\|ERVL-MaLR | 17196 | NM_198408 | 12919 | Mm.316614 | NM_198408 | ENSMUSG00000021680 | **Crhbp** | CRH-BP | corticotropin releasing hormone binding protein | protein-coding |
| chr11_104083679_104084158 | chr11 | 104083680 | 104084158 | + | 1.30876 | Intergenic | Intergenic | -48963 | NM_007762 | 12921 | Mm.1892 | NM_007762 | ENSMUSG00000018634 | **Crhr1** | CRFR1\|Crhr | corticotropin releasing hormone receptor 1 | protein-coding |
| chr17_78271345_78271924 | chr17 | 78271346 | 78271924 | + | 1.80871 | intron (NM_015800, intron 2 of 16) | intron (NM_015800, intron 2 of 16) | 71387 | NM_015800 | 50766 | Mm.311912 | NM_015800 | ENSMUSG00000024074 | **Crim1** | AU015004 | cysteine rich transmembrane BMP regulator 1 (chordin like) | protein-coding |
| chr17_78268731_78269112 | chr17 | 78268732 | 78269112 | + | 1.50668 | intron (NM_015800, intron 2 of 16) | intron (NM_015800, intron 2 of 16) | 68674 | NM_015800 | 50766 | Mm.311912 | NM_015800 | ENSMUSG00000024074 | **Crim1** | AU015004 | cysteine rich transmembrane BMP regulator 1 (chordin like) | protein-coding |
| chr17_78271002_78271286 | chr17 | 78271003 | 78271286 | + | 0.924781 | intron (NM_015800, intron 2 of 16) | intron (NM_015800, intron 2 of 16) | 70896 | NM_015800 | 50766 | Mm.311912 | NM_015800 | ENSMUSG00000024074 | **Crim1** | AU015004 | cysteine rich transmembrane BMP regulator 1 (chordin like) | protein-coding |
| chr17_78165327_78165569 | chr17 | 78165328 | 78165569 | + | 0.880426 | Intergenic | Intergenic | -34800 | NM_015800 | 50766 | Mm.311912 | NM_015800 | ENSMUSG00000024074 | **Crim1** | AU015004 | cysteine rich transmembrane BMP regulator 1 (chordin like) | protein-coding |
| chr17_78165653_78165902 | chr17 | 78165654 | 78165902 | + | 0.7143 | Intergenic | Intergenic | -34470 | NM_015800 | 50766 | Mm.311912 | NM_015800 | ENSMUSG00000024074 | **Crim1** | AU015004 | cysteine rich transmembrane BMP regulator 1 (chordin like) | protein-coding |
| chr17_87052365_87052890 | chr17 | 87052366 | 87052890 | + | 1.73192 | Intergenic | ORR1C2\|LTR\|ERVL-MaLR | 27067 | NM_019936 | 56724 | Mm.109329 | NM_019936 | ENSMUSG00000024146 | **Cript** | 1200020A08Rik\|AA589469 | cysteine-rich PDZ-binding protein | protein-coding |
| chr17_40763725_40763955 | chr17 | 40763726 | 40763955 | + | 0.690251 | TTS (NM_009420) | TTS (NM_009420) | 30306 | NM_001204071 | 22024 | Mm.1296 | NM_009420 | ENSMUSG00000023930 | **Crisp2** | CRISP-2\|GAPDL5\|Tpx-1\|Tpx1 | cysteine-rich secretory protein 2 | protein-coding |
| chr8_119992208_119992766 | chr8 | 119992209 | 119992766 | + | 0.753947 | promoter-TSS (NM_030209) | promoter-TSS (NM_030209) | 17 | NM_030209 | 78892 | Mm.264680 | NM_030209 | ENSMUSG00000031825 | **Crispld2** | 1810049K24Rik\|Lcrisp2\|Lgl1\|coffeecrisp | cysteine-rich secretory protein LCCL domain containing 2 | protein-coding |
| chr5_109544576_109544838 | chr5 | 109544577 | 109544838 | + | 0.795294 | Intergenic | RMER1C\|Other\|Other | 14286 | NM_016715 | 57914 | Mm.35771 | NM_016715 | ENSMUSG00000033467 | **Crlf2** | CRLM2\|Ly114\|Tpte2\|Tslpr | cytokine receptor-like factor 2 | protein-coding |
| chr5_37172299_37172886 | chr5 | 37172300 | 37172886 | + | 1.0224 | Intergenic | Intergenic | -69465 | NM_001136058 | 12933 | Mm.290995 | NM_007765 | ENSMUSG00000029121 | **Crmp1** | CRMP-1\|DRP-1\|Dpysl1\|ULIP-3\|Ulip3 | collapsin response mediator protein 1 | protein-coding |
| chr5_9043614_9044458 | chr5 | 9043615 | 9044458 | + | 1.25411 | Intergenic | Intergenic | -46890 | NM_023733 | 74114 | Mm.28197 | NM_023733 | ENSMUSG00000003623 | **Crot** | 1200003H03Rik | carnitine O-octanoyltransferase | protein-coding |
| chr5_8989076_8989849 | chr5 | 8989077 | 8989849 | + | 1.20033 | intron (NM_023733, intron 4 of 17) | intron (NM_023733, intron 4 of 17) | 7683 | NM_023733 | 74114 | Mm.28197 | NM_023733 | ENSMUSG00000003623 | **Crot** | 1200003H03Rik | carnitine O-octanoyltransferase | protein-coding |
| chr5_8995526_8995781 | chr5 | 8995527 | 8995781 | + | 1.04532 | intron (NM_023733, intron 2 of 17) | intron (NM_023733, intron 2 of 17) | 1492 | NM_023733 | 74114 | Mm.28197 | NM_023733 | ENSMUSG00000003623 | **Crot** | 1200003H03Rik | carnitine O-octanoyltransferase | protein-coding |
| chr5_8946175_8946469 | chr5 | 8946176 | 8946469 | + | 0.718569 | intron (NM_008830, intron 22 of 27) | intron (NM_008830, intron 22 of 27) | 50824 | NM_023733 | 74114 | Mm.28197 | NM_023733 | ENSMUSG00000003623 | **Crot** | 1200003H03Rik | carnitine O-octanoyltransferase | protein-coding |
| chr5_8946214_8946652 | chr5 | 8946215 | 8946652 | + | 0.691582 | intron (NM_008830, intron 22 of 27) | intron (NM_008830, intron 22 of 27) | 50713 | NM_023733 | 74114 | Mm.28197 | NM_023733 | ENSMUSG00000003623 | **Crot** | 1200003H03Rik | carnitine O-octanoyltransferase | protein-coding |
| chr9_40995108_40995437 | chr9 | 40995109 | 40995437 | + | 0.974535 | intron (NM_019465, intron 1 of 9) | intron (NM_019465, intron 1 of 9) | 9355 | NM_001281954 | 54698 | Mm.42791 | NM_019465 | ENSMUSG00000032021 | **Crtam** | - | cytotoxic and regulatory T cell molecule | protein-coding |
| chr9_41011731_41012316 | chr9 | 41011732 | 41012316 | + | 0.79745 | Intergenic | Intergenic | -7396 | NM_019465 | 54698 | Mm.42791 | NM_019465 | ENSMUSG00000032021 | **Crtam** | - | cytotoxic and regulatory T cell molecule | protein-coding |
| chr9_40994078_40994484 | chr9 | 40994079 | 40994484 | + | 0.726064 | intron (NM_019465, intron 2 of 9) | intron (NM_019465, intron 2 of 9) | 10347 | NM_001281954 | 54698 | Mm.42791 | NM_019465 | ENSMUSG00000032021 | **Crtam** | - | cytotoxic and regulatory T cell molecule | protein-coding |
| chr9_40993949_40994927 | chr9 | 40993950 | 40994927 | + | 0.70324 | intron (NM_019465, intron 2 of 9) | intron (NM_019465, intron 2 of 9) | 10190 | NM_001281954 | 54698 | Mm.42791 | NM_019465 | ENSMUSG00000032021 | **Crtam** | - | cytotoxic and regulatory T cell molecule | protein-coding |
| chr7_80715116_80715834 | chr7 | 80715117 | 80715834 | + | 0.942214 | intron (NM_016721, intron 37 of 37) | intron (NM_016721, intron 37 of 37) | -26598 | NM_173863 | 70461 | Mm.392175 | NM_173863 | ENSMUSG00000030527 | **Crtc3** | 2610312F20Rik\|6332415K15Rik\|AI429792\|BC011210\|TORC3 | CREB regulated transcription coactivator 3 | protein-coding |

| | | | | | | | | | | | | | | | | |
|---|---|---|---|---|---|---|---|---|---|---|---|---|---|---|---|---|
| chr7_80742662_80743030 | chr7 | 80742663 | 80743030 | + | 0.791989 | intron (NM_016721, intron 18 of 37) | intron (NM_016721, intron 18 of 37) | -53969 | NM_173863 | 70461 | Mm.392175 | NM_173863 | ENSMUSG00000030527 | **Crtc3** | 2610312F20Rik\|6332415K15Rik\|AI429792\|BC011210\|TORC3 | CREB regulated transcription coactivator 3 | protein-coding |
| chr2_92445905_92446232 | chr2 | 92445906 | 92446232 | + | 1.00446 | Intergenic | Intergenic | -12000 | NM_009963 | 12953 | Mm.254181 | NM_009963 | ENSMUSG00000068742 | **Cry2** | AV006279\|D130054K12Rik | cryptochrome 2 (photolyase-like) | protein-coding |
| chr2_92445905_92446398 | chr2 | 92445906 | 92446398 | + | 0.933538 | Intergenic | Intergenic | -12083 | NM_009963 | 12953 | Mm.254181 | NM_009963 | ENSMUSG00000068742 | **Cry2** | AV006279\|D130054K12Rik | cryptochrome 2 (photolyase-like) | protein-coding |
| chr17_31753271_31754145 | chr17 | 31753272 | 31754145 | + | 1.09897 | Intergenic | Intergenic | 75777 | NM_001278570 | 12954 | Mm.1228 | NM_013501 | ENSMUSG00000024041 | **Cryaa** | Acry-1\|Crya-1\|Crya1\|DAcry-1\|lop18 | crystallin, alpha A | protein-coding |
| chr5_112264082_112264501 | chr5 | 112264083 | 112264501 | + | 0.888025 | intron (NM_023695, intron 4 of 5) | intron (NM_023695, intron 4 of 5) | 8471 | NM_023695 | 12960 | Mm.29488 | NM_023695 | ENSMUSG00000029343 | **Crybb1** | 3110006K12Rik\|BB1CRY | crystallin, beta B1 | protein-coding |
| chr5_113108343_113108868 | chr5 | 113108344 | 113108868 | + | 0.958005 | intron (NM_172884, intron 1 of 7) | intron (NM_172884, intron 1 of 7) | -27022 | NM_021352 | 12962 | Mm.378899 | NM_021352 | ENSMUSG00000029352 | **Crybb3** | AI852419 | crystallin, beta B3 | protein-coding |
| chr16_59521955_59522480 | chr16 | 59521956 | 59522480 | + | 1.59315 | intron (NM_174848, intron 13 of 18) | intron (NM_174848, intron 13 of 18) | 33534 | NM_174848 | 224273 | Mm.246424 | NM_174848 | ENSMUSG00000022723 | **Crybg3** | 5031404N19\|AU022790 | beta-gamma crystallin domain containing 3 | protein-coding |
| chr16_59558904_59559306 | chr16 | 59558905 | 59559306 | + | 0.964865 | Intergenic | Intergenic | -3353 | NM_174848 | 224273 | Mm.246424 | NM_174848 | ENSMUSG00000022723 | **Crybg3** | 5031404N19\|AU022790 | beta-gamma crystallin domain containing 3 | protein-coding |
| chr16_59558816_59559512 | chr16 | 59558817 | 59559512 | + | 0.92137 | Intergenic | Intergenic | -3412 | NM_174848 | 224273 | Mm.246424 | NM_174848 | ENSMUSG00000022723 | **Crybg3** | 5031404N19\|AU022790 | beta-gamma crystallin domain containing 3 | protein-coding |
| chr5_24759243_24760228 | chr5 | 24759244 | 24760228 | + | 0.836474 | Intergenic | RLTR19\|LTR\|ERVK | -1893 | NM_153076 | 214301 | Mm.63484 | NM_153076 | | **Crygn** | - | crystallin, gamma N | protein-coding |
| chr16_22750796_22751208 | chr16 | 22750797 | 22751208 | + | 0.725892 | Intergenic | MER31-int\|LTR\|ERV1 | 60408 | NM_009967 | 12970 | Mm.6253 | NM_009967 | ENSMUSG00000033501 | **Crygs** | AI327013\|Opj\|rncat | crystallin, gamma S | protein-coding |
| chr10_128349008_128349248 | chr10 | 128349009 | 128349248 | + | 0.760406 | intron (NM_026444, intron 1 of 10) | B3\|SINE\|B2 | 11296 | NM_026444 | 12974 | Mm.58836 | NM_026444 | ENSMUSG00000005683 | **Cs** | 2610511A05Rik\|9030605P22Rik\|BB234005\|Cis\|ahl4 | citrate synthase | protein-coding |
| chr15_102172527_102173023 | chr15 | 102172528 | 102173023 | + | 0.997005 | Intergenic | ID_B1\|SINE\|B4 | 16268 | NM_144942 | 246277 | Mm.296382 | NM_144942 | ENSMUSG00000023044 | **Csad** | Csd | cysteine sulfinic acid decarboxylase | protein-coding |
| chr15_81943324_81943751 | chr15 | 81943325 | 81943751 | + | 0.642498 | intron (NM_145473, intron 1 of 3) | intron (NM_145473, intron 1 of 3) | 6779 | NM_145473 | 105859 | Mm.22524 | NM_145473 | ENSMUSG00000042109 | **Csdc2** | AI415250\|AI481750\|Pippin | cold shock domain containing C2, RNA binding | protein-coding |
| chr2_166910602_166910882 | chr2 | 166910603 | 166910882 | + | 1.20704 | intron (NM_023565, intron 1 of 24) | intron (NM_023565, intron 1 of 24) | 4646 | NM_023565 | 110750 | Mm.22417 | NM_023565 | ENSMUSG00000002718 | **Cse1l** | 2610100P18Rik\|AA407533\|Capts\|Cas\|Xpo2 | chromosome segregation 1-like (S. cerevisiae) | protein-coding |
| chr2_166909780_166909960 | chr2 | 166909781 | 166909960 | + | 0.698923 | intron (NM_023565, intron 1 of 24) | intron (NM_023565, intron 1 of 24) | 3774 | NM_023565 | 110750 | Mm.22417 | NM_023565 | ENSMUSG00000002718 | **Cse1l** | 2610100P18Rik\|AA407533\|Capts\|Cas\|Xpo2 | chromosome segregation 1-like (S. cerevisiae) | protein-coding |
| chr3_107815121_107815413 | chr3 | 107815122 | 107815413 | + | 0.803667 | Intergenic | Intergenic | -54798 | NM_001113529 | 12977 | Mm.795 | NM_007778 | ENSMUSG00000014599 | **Csf1** | C87615\|Csfm\|MCSF\|op | colony stimulating factor 1 (macrophage) | protein-coding |
| chr3_107761500_107761769 | chr3 | 107761501 | 107761769 | + | 0.648854 | Intergenic | URR1A\|DNA\|hAT-Charlie | -1166 | NM_001113529 | 12977 | Mm.795 | NM_007778 | ENSMUSG00000014599 | **Csf1** | C87615\|Csfm\|MCSF\|op | colony stimulating factor 1 (macrophage) | protein-coding |
| chr9_57637337_57637578 | chr9 | 57637338 | 57637578 | + | 0.693343 | intron (NM_007783, intron 1 of 12) | intron (NM_007783, intron 1 of 12) | 7722 | NM_007783 | 12988 | Mm.21974 | NM_007783 | ENSMUSG00000032312 | **Csk** | AW212630 | c-src tyrosine kinase | protein-coding |
| chr9_65929496_65930244 | chr9 | 65929497 | 65930244 | + | 1.29239 | intron (NM_173185, intron 1 of 13) | ORR1A4\|LTR\|ERVL-MaLR | 20860 | NM_173185 | 214897 | Mm.325816 | NM_173185 | ENSMUSG00000032384 | **Csnk1g1** | 9130020E21Rik | casein kinase 1, gamma 1 | protein-coding |
| chr18_53926701_53927273 | chr18 | 53926702 | 53927273 | + | 1.01649 | intron (NM_152809, intron 6 of 12) | RMER15\|LTR\|ERVL | 64874 | NM_152809 | 70425 | Mm.368668 | NM_152809 | ENSMUSG00000073563 | **Csnk1g3** | 3300002K07Rik\|C330049O21Rik | casein kinase 1, gamma 3 | protein-coding |
| chr1_10077791_10078150 | chr1 | 10077792 | 10078150 | + | 0.807624 | intron (NM_026493, intron 9 of 29) | ORR1C2\|LTR\|ERVL-MaLR | 39753 | NM_026493 | 211660 | Mm.244046 | NM_026493 | ENSMUSG00000056763 | **Cspp1** | 2310020J12Rik\|4930413O22Rik\|AA387550\|BC055915\|Cspp2\|Cspp3 | centrosome and spindle pole associated protein 1 | protein-coding |
| chr15_100508781_100509153 | chr15 | 100508782 | 100509153 | + | 0.959813 | intron (NM_033476, intron 14 of 15) | intron (NM_033476, intron 14 of 15) | -13728 | NM_153407 | 207785 | Mm.294969 | NM_153407 | ENSMUSG00000044636 | **Csrnp2** | CSRNP-2\|Csnrp2\|taip-12 | cysteine-serine-rich nuclear protein 2 | protein-coding |
| chr15_100489063_100489547 | chr15 | 100489064 | 100489547 | + | 0.789614 | intron (NM_153407, intron 2 of 4) | intron (NM_153407, intron 2 of 4) | 5934 | NM_153407 | 207785 | Mm.294969 | NM_153407 | ENSMUSG00000044636 | **Csrnp2** | CSRNP-2\|Csnrp2\|taip-12 | cysteine-serine-rich nuclear protein 2 | protein-coding |
| chr2_65826745_65827247 | chr2 | 65826746 | 65827247 | + | 0.707683 | Intergenic | UCON28a\|Unknown\|Unknown | -18771 | NM_001290666 | 77771 | Mm.102025 | NM_153409 | ENSMUSG00000044647 | **Csrnp3** | A330102K23Rik\|CSRNP-3\|Csnrp3\|Mbu1\|taip-2 | cysteine-serine-rich nuclear protein 3 | protein-coding |
| chr2_65818402_65818884 | chr2 | 65818403 | 65818884 | + | 0.664132 | Intergenic | Intergenic | -27124 | NM_001290666 | 77771 | Mm.102025 | NM_153409 | ENSMUSG00000044647 | **Csrnp3** | A330102K23Rik\|CSRNP-3\|Csnrp3\|Mbu1\|taip-2 | cysteine-serine-rich nuclear protein 3 | protein-coding |
| chr2_65818185_65818910 | chr2 | 65818186 | 65818910 | + | 0.607992 | Intergenic | Intergenic | -27219 | NM_001290666 | 77771 | Mm.102025 | NM_153409 | ENSMUSG00000044647 | **Csrnp3** | A330102K23Rik\|CSRNP-3\|Csnrp3\|Mbu1\|taip-2 | cysteine-serine-rich nuclear protein 3 | protein-coding |
| chr2_150567469_150567826 | chr2 | 150567470 | 150567826 | + | 0.658779 | Intergenic | Intergenic | -2767 | NM_009977 | 13011 | Mm.12965 | NM_009977 | ENSMUSG00000068129 | **Cst7** | Cmap | cystatin F (leukocystatin) | protein-coding |
| chr19_31062361_31062716 | chr19 | 31062362 | 31062716 | + | 0.849216 | intron (NM_001013833, intron 3 of 17) | intron (NM_001013833, intron 3 of 17) | -20302 | NM_031249 | 83410 | Mm.22031 | NM_031249 | ENSMUSG00000053536 | **Cstf2t** | 64kDa\|C77975\|tCstF-64\|tauCstF-64 | cleavage stimulation factor, 3' pre-RNA subunit 2, tau | protein-coding |
| chr2_104607457_104607977 | chr2 | 104607458 | 104607977 | + | 0.944511 | intron (NM_177253, intron 1 of 2) | B3\|SINE\|B2 | 17233 | NM_001037326 | 228410 | Mm.259876 | NM_145529 | ENSMUSG00000027176 | **Cstf3** | 4732468G05Rik\|C79532\|CstF-77 | cleavage stimulation factor, 3' pre-RNA, subunit 3 | protein-coding |
| chr2_104607067_104607977 | chr2 | 104607068 | 104607977 | + | 0.694396 | intron (NM_177253, intron 1 of 2) | intron (NM_177253, intron 1 of 2) | 17038 | NM_001037326 | 228410 | Mm.259876 | NM_145529 | ENSMUSG00000027176 | **Cstf3** | 4732468G05Rik\|C79532\|CstF-77 | cleavage stimulation factor, 3' pre-RNA, subunit 3 | protein-coding |
| chr12_59162828_59163245 | chr12 | 59162829 | 59163245 | + | 0.858879 | intron (NM_001165254, intron 12 of 22) | intron (NM_001165254, intron 12 of 22) | 31584 | NM_001165253 | 217615 | Mm.244118 | NM_146034 | ENSMUSG00000021000 | **Ctage5** | D12Bwg0579e\|Mea6\|Mgea\|Mgea6 | CTAGE family, member 5 | protein-coding |
| chr5_33265588_33266148 | chr5 | 33265589 | 33266148 | + | 1.16112 | intron (NM_013502, intron 2 of 8) | intron (NM_013502, intron 2 of 8) | 9136 | NM_001198861 | 13016 | Mm.7286 | NM_013502 | ENSMUSG00000037373 | **Ctbp1** | BARS\|CtBP1-L\|CtBP1-S\|CtBP3/BARS\|D4S115h\|D5H4S115\|D5H4S115E | C-terminal binding protein 1 | protein-coding |
| chr5_33266394_33266897 | chr5 | 33266395 | 33266897 | + | 0.886537 | intron (NM_013502, intron 2 of 8) | intron (NM_013502, intron 2 of 8) | 8358 | NM_001198861 | 13016 | Mm.7286 | NM_013502 | ENSMUSG00000037373 | **Ctbp1** | BARS\|CtBP1-L\|CtBP1-S\|CtBP3/BARS\|D4S115h\|D5H4S115\|D5H4S115E | C-terminal binding protein 1 | protein-coding |
| chr8_105629203_105629774 | chr8 | 105629204 | 105629774 | + | 1.15957 | Intergenic | Intergenic | -7049 | NM_181322 | 13018 | Mm.269474 | NM_181322 | ENSMUSG00000005698 | **Ctcf** | AW108038 | CCCTC-binding factor | protein-coding |
| chr2_173138527_173138982 | chr2 | 173138528 | 173138982 | + | 0.744988 | Intergenic | Intergenic | 14006 | NR_040321 | 74161 | Mm.208098 | NR_040321 | ENSMUSG00000087382 | **Ctcflos** | 1300015D01Rik\|A530094K13 | CCCTC-binding factor (zinc finger protein)-like, opposite strand | ncRNA |
| chr9_118843518_118844462 | chr9 | 118843519 | 118844462 | + | 0.751224 | intron (NM_133721, intron 22 of 27) | intron (NM_133721, intron 22 of 27) | -82546 | NM_133710 | 69274 | Mm.439928 | NM_133710 | ENSMUSG00000047409 | **Ctdspl** | 2810418J22Rik\|AI426263\|HYA22\|SCP3 | CTD (carboxy-terminal domain, RNA polymerase II, polypeptide A) small phosphatase-like | protein-coding |
| chr9_118843520_118844033 | chr9 | 118843521 | 118844033 | + | 0.681227 | intron (NM_133721, intron 22 of 27) | intron (NM_133721, intron 22 of 27) | -82759 | NM_133710 | 69274 | Mm.439928 | NM_133710 | ENSMUSG00000047409 | **Ctdspl** | 2810418J22Rik\|AI426263\|HYA22\|SCP3 | CTD (carboxy-terminal domain, RNA polymerase II, polypeptide A) small phosphatase-like | protein-coding |
| chr3_157883874_157884484 | chr3 | 157883875 | 157884484 | + | 1.83813 | Intergenic | Intergenic | 40884 | NM_145953 | 107869 | Mm.28301 | NM_145953 | ENSMUSG00000028179 | **Cth** | 0610010I13Rik\|AI314617\|BC019483\|CSE\|Cys3 | cystathionase (cystathionine gamma-lyase) | protein-coding |
| chr3_157883284_157883777 | chr3 | 157883285 | 157883777 | + | 0.922244 | Intergenic | L1_Mus3\|LINE\|L1 | 41532 | NM_145953 | 107869 | Mm.28301 | NM_145953 | ENSMUSG00000028179 | **Cth** | 0610010I13Rik\|AI314617\|BC019483\|CSE\|Cys3 | cystathionase (cystathionine gamma-lyase) | protein-coding |
| chr3_157924668_157925474 | chr3 | 157924669 | 157925474 | + | 0.853237 | promoter-TSS (NM_145953) | promoter-TSS (NM_145953) | -8 | NM_145953 | 107869 | Mm.28301 | NM_145953 | ENSMUSG00000028179 | **Cth** | 0610010I13Rik\|AI314617\|BC019483\|CSE\|Cys3 | cystathionase (cystathionine gamma-lyase) | protein-coding |
| chr18_75694798_75695172 | chr18 | 75694799 | 75695172 | + | 0.956537 | intron (NM_201354, intron 1 of 11) | intron (NM_201354, intron 1 of 11) | 2711 | NM_201354 | 269037 | Mm.36745 | NM_201354 | ENSMUSG00000052928 | **Ctif** | Gm672 | CBP80/20-dependent translation initiation factor | protein-coding |
| chr1_60880055_60880664 | chr1 | 60880056 | 60880664 | + | 1.30978 | Intergenic | Intergenic | -28665 | NM_009843 | 12477 | Mm.390 | NM_009843 | ENSMUSG00000026011 | **Ctla4** | Cd152\|Ctla-4\|Ly-56 | cytotoxic T-lymphocyte-associated protein 4 | protein-coding |
| chr1_60875886_60876180 | chr1 | 60875887 | 60876180 | + | 1.30622 | Intergenic | Intergenic | -32992 | NM_009843 | 12477 | Mm.390 | NM_009843 | ENSMUSG00000026011 | **Ctla4** | Cd152\|Ctla-4\|Ly-56 | cytotoxic T-lymphocyte-associated protein 4 | protein-coding |
| chr1_60913307_60913709 | chr1 | 60913308 | 60913709 | + | 1.06833 | intron (NM_009843, intron 3 of 3) | intron (NM_009843, intron 3 of 3) | 4483 | NM_001281976 | 12477 | Mm.390 | NM_009843 | ENSMUSG00000026011 | **Ctla4** | Cd152\|Ctla-4\|Ly-56 | cytotoxic T-lymphocyte-associated protein 4 | protein-coding |
| chr1_60937030_60937343 | chr1 | 60937031 | 60937343 | + | 1.03617 | Intergenic | MamGypLTR1d\|LTR\|Gypsy | 28162 | NM_001281976 | 12477 | Mm.390 | NM_009843 | ENSMUSG00000026011 | **Ctla4** | Cd152\|Ctla-4\|Ly-56 | cytotoxic T-lymphocyte-associated protein 4 | protein-coding |
| chr1_60833533_60834052 | chr1 | 60833534 | 60834052 | + | 0.99864 | Intergenic | Intergenic | -75232 | NM_009843 | 12477 | Mm.390 | NM_009843 | ENSMUSG00000026011 | **Ctla4** | Cd152\|Ctla-4\|Ly-56 | cytotoxic T-lymphocyte-associated protein 4 | protein-coding |
| chr1_60845692_60846101 | chr1 | 60845693 | 60846101 | + | 0.942301 | Intergenic | Intergenic | -63128 | NM_009843 | 12477 | Mm.390 | NM_009843 | ENSMUSG00000026011 | **Ctla4** | Cd152\|Ctla-4\|Ly-56 | cytotoxic T-lymphocyte-associated protein 4 | protein-coding |
| chr1_60854079_60854392 | chr1 | 60854080 | 60854392 | + | 0.93959 | Intergenic | Intergenic | -54789 | NM_009843 | 12477 | Mm.390 | NM_009843 | ENSMUSG00000026011 | **Ctla4** | Cd152\|Ctla-4\|Ly-56 | cytotoxic T-lymphocyte-associated protein 4 | protein-coding |
| chr1_60879230_60879499 | chr1 | 60879231 | 60879499 | + | 0.936057 | Intergenic | Intergenic | -29660 | NM_009843 | 12477 | Mm.390 | NM_009843 | ENSMUSG00000026011 | **Ctla4** | Cd152\|Ctla-4\|Ly-56 | cytotoxic T-lymphocyte-associated protein 4 | protein-coding |
| chr1_60854100_60854623 | chr1 | 60854101 | 60854623 | + | 0.930943 | Intergenic | Intergenic | -54663 | NM_009843 | 12477 | Mm.390 | NM_009843 | ENSMUSG00000026011 | **Ctla4** | Cd152\|Ctla-4\|Ly-56 | cytotoxic T-lymphocyte-associated protein 4 | protein-coding |
| chr1_60875203_60875679 | chr1 | 60875204 | 60875679 | + | 0.916766 | Intergenic | Intergenic | -33584 | NM_009843 | 12477 | Mm.390 | NM_009843 | ENSMUSG00000026011 | **Ctla4** | Cd152\|Ctla-4\|Ly-56 | cytotoxic T-lymphocyte-associated protein 4 | protein-coding |
| chr1_60859883_60860213 | chr1 | 60859884 | 60860213 | + | 0.904371 | Intergenic | Intergenic | -48977 | NM_009843 | 12477 | Mm.390 | NM_009843 | ENSMUSG00000026011 | **Ctla4** | Cd152\|Ctla-4\|Ly-56 | cytotoxic T-lymphocyte-associated protein 4 | protein-coding |
| chr1_60874889_60875679 | chr1 | 60874890 | 60875679 | + | 0.894556 | Intergenic | Intergenic | -33741 | NM_009843 | 12477 | Mm.390 | NM_009843 | ENSMUSG00000026011 | **Ctla4** | Cd152\|Ctla-4\|Ly-56 | cytotoxic T-lymphocyte-associated protein 4 | protein-coding |
| chr1_60903139_60903491 | chr1 | 60903140 | 60903491 | + | 0.872691 | Intergenic | Intergenic | -5710 | NM_009843 | 12477 | Mm.390 | NM_009843 | ENSMUSG00000026011 | **Ctla4** | Cd152\|Ctla-4\|Ly-56 | cytotoxic T-lymphocyte-associated protein 4 | protein-coding |
| chr1_60846902_60847340 | chr1 | 60846903 | 60847340 | + | 0.854409 | Intergenic | Intergenic | -61904 | NM_009843 | 12477 | Mm.390 | NM_009843 | ENSMUSG00000026011 | **Ctla4** | Cd152\|Ctla-4\|Ly-56 | cytotoxic T-lymphocyte-associated protein 4 | protein-coding |
| chr1_60904518_60904834 | chr1 | 60904519 | 60904834 | + | 0.830087 | Intergenic | Intergenic | -4349 | NM_009843 | 12477 | Mm.390 | NM_009843 | ENSMUSG00000026011 | **Ctla4** | Cd152\|Ctla-4\|Ly-56 | cytotoxic T-lymphocyte-associated protein 4 | protein-coding |
| chr1_60899117_60899751 | chr1 | 60899118 | 60899751 | + | 0.816668 | Intergenic | Intergenic | -9591 | NM_009843 | 12477 | Mm.390 | NM_009843 | ENSMUSG00000026011 | **Ctla4** | Cd152\|Ctla-4\|Ly-56 | cytotoxic T-lymphocyte-associated protein 4 | protein-coding |
| chr1_60941575_60942101 | chr1 | 60941576 | 60942101 | + | 0.81013 | Intergenic | Intergenic | 32813 | NM_001281976 | 12477 | Mm.390 | NM_009843 | ENSMUSG00000026011 | **Ctla4** | Cd152\|Ctla-4\|Ly-56 | cytotoxic T-lymphocyte-associated protein 4 | protein-coding |
| chr1_60888889_60889213 | chr1 | 60888890 | 60889213 | + | 0.80771 | Intergenic | B1_Mur1\|SINE\|Alu | -19974 | NM_009843 | 12477 | Mm.390 | NM_009843 | ENSMUSG00000026011 | **Ctla4** | Cd152\|Ctla-4\|Ly-56 | cytotoxic T-lymphocyte-associated protein 4 | protein-coding |
| chr1_60828196_60828802 | chr1 | 60828197 | 60828802 | + | 0.65387 | Intergenic | MTD\|LTR\|ERVL-MaLR | -80526 | NM_009843 | 12477 | Mm.390 | NM_009843 | ENSMUSG00000026011 | **Ctla4** | Cd152\|Ctla-4\|Ly-56 | cytotoxic T-lymphocyte-associated protein 4 | protein-coding |

| | | | | | | | | | | | | | | | | | |
|---|---|---|---|---|---|---|---|---|---|---|---|---|---|---|---|---|---|
| chr2_157807848_157809169 | chr2 | 157807849 | 157809169 | + | 0.723644 | intron (NM_025680, intron 6 of 15) | intron (NM_025680, intron 6 of 15) | 71108 | NM_025680 | 66642 | Mm.45193 | NM_025680 | ENSMUSG00000027649 | **Ctnnbl1** | 5730471K09Rik\|NAP\|NYD-SP19\|P14L | catenin, beta like 1 | protein-coding |
| chrX_162921438_162922548 | chrX | 162921439 | 162922548 | + | 0.631665 | intron (NR_033143, intron 6 of 18) | intron (NR_033143, intron 6 of 18) | 19822 | NR_033143 | 55936 | Mm.2065 | NM_018737 | ENSMUSG00000031360 | **Ctps2** | A830031M15Rik\|AI326475\|Ctpsh | cytidine 5'-triphosphate synthase 2 | protein-coding |
| chr14_63132100_63132794 | chr14 | 63132101 | 63132794 | + | 1.1046 | intron (NM_007798, intron 1 of 9) | intron (NM_007798, intron 1 of 9) | 9985 | NM_007798 | 13030 | Mm.236553 | NM_007798 | ENSMUSG00000021939 | **Ctsb** | CB | cathepsin B | protein-coding |
| chr7_88259203_88259583 | chr7 | 88259204 | 88259583 | + | 1.43623 | Intergenic | Intergenic | -18700 | NM_009982 | 13032 | Mm.322945 | NM_009982 | ENSMUSG00000030560 | **Ctsc** | AI047818\|DPP1\|DPPI | cathepsin C | protein-coding |
| chr7_88259206_88259915 | chr7 | 88259207 | 88259915 | + | 1.2243 | Intergenic | Intergenic | -18532 | NM_009982 | 13032 | Mm.322945 | NM_009982 | ENSMUSG00000030560 | **Ctsc** | AI047818\|DPP1\|DPPI | cathepsin C | protein-coding |
| chr7_88258712_88259927 | chr7 | 88258713 | 88259927 | + | 1.17329 | Intergenic | Intergenic | -18773 | NM_009982 | 13032 | Mm.322945 | NM_009982 | ENSMUSG00000030560 | **Ctsc** | AI047818\|DPP1\|DPPI | cathepsin C | protein-coding |
| chr7_88200214_88200794 | chr7 | 88200215 | 88200794 | + | 0.984005 | Intergenic | Intergenic | -77589 | NM_009982 | 13032 | Mm.322945 | NM_009982 | ENSMUSG00000030560 | **Ctsc** | AI047818\|DPP1\|DPPI | cathepsin C | protein-coding |
| chr7_88200219_88200638 | chr7 | 88200220 | 88200638 | + | 0.94799 | Intergenic | Intergenic | -77664 | NM_009982 | 13032 | Mm.322945 | NM_009982 | ENSMUSG00000030560 | **Ctsc** | AI047818\|DPP1\|DPPI | cathepsin C | protein-coding |
| chr7_88258756_88259141 | chr7 | 88258757 | 88259141 | + | 0.756263 | Intergenic | ORR1A2\|LTR\|ERVL-MaLR | -19144 | NM_009982 | 13032 | Mm.322945 | NM_009982 | ENSMUSG00000030560 | **Ctsc** | AI047818\|DPP1\|DPPI | cathepsin C | protein-coding |
| chr7_88164836_88165230 | chr7 | 88164837 | 88165230 | + | 0.686071 | Intergenic | B3\|SINE\|B2 | -113060 | NM_009982 | 13032 | Mm.322945 | NM_009982 | ENSMUSG00000030560 | **Ctsc** | AI047818\|DPP1\|DPPI | cathepsin C | protein-coding |
| chr3_81918233_81918697 | chr3 | 81918234 | 81918697 | + | 1.48719 | Intergenic | Intergenic | -14151 | NM_177662 | 229445 | Mm.254642 | NM_177662 | ENSMUSG00000028015 | **Ctso** | A330105D01Rik\|A430056C07\|AI118514 | cathepsin O | protein-coding |
| chr3_81873878_81874240 | chr3 | 81873879 | 81874240 | + | 1.05092 | Intergenic | Lx2\|LINE\|L1 | -58557 | NM_177662 | 229445 | Mm.254642 | NM_177662 | ENSMUSG00000028015 | **Ctso** | A330105D01Rik\|A430056C07\|AI118514 | cathepsin O | protein-coding |
| chr3_81873870_81874380 | chr3 | 81873871 | 81874380 | + | 0.999736 | Intergenic | Lx2\|LINE\|L1 | -58491 | NM_177662 | 229445 | Mm.254642 | NM_177662 | ENSMUSG00000028015 | **Ctso** | A330105D01Rik\|A430056C07\|AI118514 | cathepsin O | protein-coding |
| chr3_81917594_81918082 | chr3 | 81917595 | 81918082 | + | 0.952177 | Intergenic | Intergenic | -14778 | NM_177662 | 229445 | Mm.254642 | NM_177662 | ENSMUSG00000028015 | **Ctso** | A330105D01Rik\|A430056C07\|AI118514 | cathepsin O | protein-coding |
| chr2_174442430_174442850 | chr2 | 174442431 | 174442850 | + | 0.818214 | Intergenic | Intergenic | -3648 | NM_022325 | 64138 | Mm.156919 | NM_022325 | ENSMUSG00000016256 | **Ctsz** | AI787083\|AU019819\|CTSX\|D2Wsu143e | cathepsin Z | protein-coding |
| chr2_174437356_174438274 | chr2 | 174437357 | 174438274 | + | 0.758478 | intron (NM_022325, intron 2 of 5) | intron (NM_022325, intron 2 of 5) | 1177 | NM_022325 | 64138 | Mm.156919 | NM_022325 | ENSMUSG00000016256 | **Ctsz** | AI787083\|AU019819\|CTSX\|D2Wsu143e | cathepsin Z | protein-coding |
| chr2_13416797_13417108 | chr2 | 13416798 | 13417108 | + | 1.2174 | intron (NM_001081084, intron 27 of 66) | intron (NM_001081084, intron 27 of 66) | 74923 | NM_001081084 | 65969 | Mm.313915 | NM_001081084 | ENSMUSG00000026726 | **Cubn** | AA408369\|AL022750\|D2Wsu88e | cubilin (intrinsic factor-cobalamin receptor) | protein-coding |
| chr2_13410667_13411057 | chr2 | 13410668 | 13411057 | + | 1.03244 | intron (NM_001081084, intron 27 of 66) | intron (NM_001081084, intron 27 of 66) | 81014 | NM_001081084 | 65969 | Mm.313915 | NM_001081084 | ENSMUSG00000026726 | **Cubn** | AA408369\|AL022750\|D2Wsu88e | cubilin (intrinsic factor-cobalamin receptor) | protein-coding |
| chr2_13409668_13410507 | chr2 | 13409669 | 13410507 | + | 1.00477 | intron (NM_001081084, intron 27 of 66) | intron (NM_001081084, intron 27 of 66) | 81788 | NM_001081084 | 65969 | Mm.313915 | NM_001081084 | ENSMUSG00000026726 | **Cubn** | AA408369\|AL022750\|D2Wsu88e | cubilin (intrinsic factor-cobalamin receptor) | protein-coding |
| chr2_13492795_13493612 | chr2 | 13492796 | 13493612 | + | 0.856501 | Intergenic | Intergenic | -1328 | NM_001081084 | 65969 | Mm.313915 | NM_001081084 | ENSMUSG00000026726 | **Cubn** | AA408369\|AL022750\|D2Wsu88e | cubilin (intrinsic factor-cobalamin receptor) | protein-coding |
| chr2_13429825_13430420 | chr2 | 13429826 | 13430420 | + | 0.800575 | intron (NM_001081084, intron 23 of 66) | MTE2b\|LTR\|ERVL-MaLR | 61753 | NM_001081084 | 65969 | Mm.313915 | NM_001081084 | ENSMUSG00000026726 | **Cubn** | AA408369\|AL022750\|D2Wsu88e | cubilin (intrinsic factor-cobalamin receptor) | protein-coding |
| chr11_88103237_88103670 | chr11 | 88103238 | 88103670 | + | 1.03743 | intron (NM_198013, intron 1 of 10) | intron (NM_198013, intron 1 of 10) | 4308 | NM_198013 | 103841 | Mm.374846 | NM_198013 | ENSMUSG00000018378 | **Cuedc1** | AI841487\|C330016O16Rik | CUE domain containing 1 | protein-coding |
| chr11_88105776_88106195 | chr11 | 88105777 | 88106195 | + | 1.01485 | intron (NM_198013, intron 1 of 10) | intron (NM_198013, intron 1 of 10) | 6840 | NM_198013 | 103841 | Mm.374846 | NM_198013 | ENSMUSG00000018378 | **Cuedc1** | AI841487\|C330016O16Rik | CUE domain containing 1 | protein-coding |
| chr11_88105875_88106183 | chr11 | 88105876 | 88106183 | + | 0.977424 | intron (NM_198013, intron 1 of 10) | intron (NM_198013, intron 1 of 10) | 6883 | NM_198013 | 103841 | Mm.374846 | NM_198013 | ENSMUSG00000018378 | **Cuedc1** | AI841487\|C330016O16Rik | CUE domain containing 1 | protein-coding |
| chr11_88103335_88103634 | chr11 | 88103336 | 88103634 | + | 0.975872 | intron (NM_198013, intron 1 of 10) | intron (NM_198013, intron 1 of 10) | 4339 | NM_198013 | 103841 | Mm.374846 | NM_198013 | ENSMUSG00000018378 | **Cuedc1** | AI841487\|C330016O16Rik | CUE domain containing 1 | protein-coding |
| chr11_88110257_88110514 | chr11 | 88110258 | 88110514 | + | 0.628722 | intron (NM_198013, intron 1 of 10) | intron (NM_198013, intron 1 of 10) | 11240 | NM_198013 | 103841 | Mm.374846 | NM_198013 | ENSMUSG00000018378 | **Cuedc1** | AI841487\|C330016O16Rik | CUE domain containing 1 | protein-coding |
| chr6_47480762_47481572 | chr6 | 47480763 | 47481572 | + | 0.690624 | intron (NM_012042, intron 1 of 21) | intron (NM_012042, intron 1 of 21) | 26843 | NM_012042 | 26965 | Mm.87611 | NM_012042 | ENSMUSG00000029686 | **Cul1** | - | cullin 1 | protein-coding |
| chr6_47524119_47524581 | chr6 | 47524120 | 47524581 | + | 0.653709 | intron (NM_012042, intron 20 of 21) | intron (NM_012042, intron 20 of 21) | 70026 | NM_012042 | 26965 | Mm.87611 | NM_012042 | ENSMUSG00000029686 | **Cul1** | - | cullin 1 | protein-coding |
| chr1_80321186_80321997 | chr1 | 80321187 | 80321997 | + | 0.788781 | intron (NM_016716, intron 2 of 15) | intron (NM_016716, intron 2 of 15) | 18838 | NM_016716 | 26554 | Mm.12665 | NM_016716 | ENSMUSG00000004364 | **Cul3** | AI467304\|AW146203\|mKIAA0617 | cullin 3 | protein-coding |
| chr1_80362384_80362897 | chr1 | 80362385 | 80362897 | + | 0.719043 | Intergenic | ORR1F\|LTR\|ERVL-MaLR | -22211 | NM_016716 | 26554 | Mm.12665 | NM_016716 | ENSMUSG00000004364 | **Cul3** | AI467304\|AW146203\|mKIAA0617 | cullin 3 | protein-coding |
| chrX_38553260_38553682 | chrX | 38553261 | 38553682 | + | 0.662595 | intron (NM_001110142, intron 5 of 22) | intron (NM_001110142, intron 5 of 22) | 22725 | NM_028288 | 72584 | Mm.327675 | NM_028288 | | **Cul4b** | 2700050M05Rik\|AA409770\|CUL-4B\|mKIAA0695 | cullin 4B | protein-coding |
| chr5_136543012_136543547 | chr5 | 136543013 | 136543547 | + | 0.828988 | intron (NM_001291234, intron 1 of 22) | L1M4\|LINE\|L1 | 22701 | NM_001291239 | 13047 | Mm.320317 | NM_009986 | ENSMUSG00000029705 | **Cux1** | CDP\|Cutl1\|Cux\|Cux-1 | cut-like homeobox 1 | protein-coding |
| chr13_104762911_104763815 | chr13 | 104762912 | 104763815 | + | 1.96854 | intron (NM_026072, intron 10 of 13) | intron (NM_026072, intron 10 of 13) | 53590 | NM_026072 | 67285 | Mm.235398 | NM_026072 | ENSMUSG00000021715 | **Cwc27** | 3110009E13Rik\|NY-CO-10\|Sdccag10 | CWC27 spliceosome-associated protein homolog (S. cerevisiae) | protein-coding |
| chr13_104619809_104620275 | chr13 | 104619810 | 104620275 | + | 1.81863 | Intergenic | Intergenic | 196911 | NM_026072 | 67285 | Mm.235398 | NM_026072 | ENSMUSG00000021715 | **Cwc27** | 3110009E13Rik\|NY-CO-10\|Sdccag10 | CWC27 spliceosome-associated protein homolog (S. cerevisiae) | protein-coding |
| chr13_104625039_104625345 | chr13 | 104625040 | 104625345 | + | 1.19212 | Intergenic | B3\|SINE\|B2 | 191761 | NM_026072 | 67285 | Mm.235398 | NM_026072 | ENSMUSG00000021715 | **Cwc27** | 3110009E13Rik\|NY-CO-10\|Sdccag10 | CWC27 spliceosome-associated protein homolog (S. cerevisiae) | protein-coding |
| chr13_104553290_104553797 | chr13 | 104553291 | 104553797 | + | 1.12707 | Intergenic | Intergenic | 263409 | NM_026072 | 67285 | Mm.235398 | NM_026072 | ENSMUSG00000021715 | **Cwc27** | 3110009E13Rik\|NY-CO-10\|Sdccag10 | CWC27 spliceosome-associated protein homolog (S. cerevisiae) | protein-coding |
| chr13_104776784_104777277 | chr13 | 104776785 | 104777277 | + | 0.993024 | intron (NM_026072, intron 10 of 13) | RMER6D\|LTR\|ERVK | 39922 | NM_026072 | 67285 | Mm.235398 | NM_026072 | ENSMUSG00000021715 | **Cwc27** | 3110009E13Rik\|NY-CO-10\|Sdccag10 | CWC27 spliceosome-associated protein homolog (S. cerevisiae) | protein-coding |
| chr13_104610536_104611290 | chr13 | 104610537 | 104611290 | + | 0.960805 | Intergenic | Intergenic | 206040 | NM_026072 | 67285 | Mm.235398 | NM_026072 | ENSMUSG00000021715 | **Cwc27** | 3110009E13Rik\|NY-CO-10\|Sdccag10 | CWC27 spliceosome-associated protein homolog (S. cerevisiae) | protein-coding |
| chr13_104622430_104622800 | chr13 | 104622431 | 104622800 | + | 0.828138 | Intergenic | ORR1F\|LTR\|ERVL-MaLR | 194338 | NM_026072 | 67285 | Mm.235398 | NM_026072 | ENSMUSG00000021715 | **Cwc27** | 3110009E13Rik\|NY-CO-10\|Sdccag10 | CWC27 spliceosome-associated protein homolog (S. cerevisiae) | protein-coding |
| chr13_104617259_104617558 | chr13 | 104617260 | 104617558 | + | 0.819814 | Intergenic | Intergenic | 199544 | NM_026072 | 67285 | Mm.235398 | NM_026072 | ENSMUSG00000021715 | **Cwc27** | 3110009E13Rik\|NY-CO-10\|Sdccag10 | CWC27 spliceosome-associated protein homolog (S. cerevisiae) | protein-coding |
| chr13_104758751_104758947 | chr13 | 104758752 | 104758947 | + | 0.787565 | intron (NM_026072, intron 10 of 13) | RLTR12BD_Mm\|LTR\|ERVK | 58104 | NM_026072 | 67285 | Mm.235398 | NM_026072 | ENSMUSG00000021715 | **Cwc27** | 3110009E13Rik\|NY-CO-10\|Sdccag10 | CWC27 spliceosome-associated protein homolog (S. cerevisiae) | protein-coding |
| chr13_104625421_104625684 | chr13 | 104625422 | 104625684 | + | 0.696923 | Intergenic | L2a\|LINE\|L2 | 191400 | NM_026072 | 67285 | Mm.235398 | NM_026072 | ENSMUSG00000021715 | **Cwc27** | 3110009E13Rik\|NY-CO-10\|Sdccag10 | CWC27 spliceosome-associated protein homolog (S. cerevisiae) | protein-coding |
| chr13_104610528_104610786 | chr13 | 104610529 | 104610786 | + | 0.646349 | Intergenic | Intergenic | 206296 | NM_026072 | 67285 | Mm.235398 | NM_026072 | ENSMUSG00000021715 | **Cwc27** | 3110009E13Rik\|NY-CO-10\|Sdccag10 | CWC27 spliceosome-associated protein homolog (S. cerevisiae) | protein-coding |
| chr9_3405629_3405875 | chr9 | 3405630 | 3405875 | + | 0.701977 | intron (NM_027545, intron 1 of 17) | Lx2B\|LINE\|L1 | 1667 | NM_027545 | 244672 | Mm.245482 | NM_027545 | ENSMUSG00000025898 | **Cwf19l2** | 3230401L03Rik\|9030209E10 | CWF19-like 2, cell cycle control (S. pombe) | protein-coding |
| chr5_73406883_73407299 | chr5 | 73406884 | 73407299 | + | 1.17465 | intron (NM_181323, intron 1 of 15) | intron (NM_181323, intron 1 of 15) | 1013 | NM_181323 | 231293 | Mm.280211 | NM_181323 | ENSMUSG00000029154 | **Cwh43** | C130090K23Rik\|imds-60 | cell wall biogenesis 43 C-terminal homolog (S. cerevisiae) | protein-coding |
| chr5_73381982_73382569 | chr5 | 73381983 | 73382569 | + | 0.854365 | Intergenic | Intergenic | -23802 | NM_181323 | 231293 | Mm.280211 | NM_181323 | ENSMUSG00000029154 | **Cwh43** | C130090K23Rik\|imds-60 | cell wall biogenesis 43 C-terminal homolog (S. cerevisiae) | protein-coding |
| chr8_94779261_94779704 | chr8 | 94779262 | 94779704 | + | 0.787686 | intron (NM_009142, intron 2 of 2) | intron (NM_009142, intron 2 of 2) | 7303 | NM_009142 | 20312 | Mm.103711 | NM_009142 | ENSMUSG00000031778 | **Cx3cl1** | AB030188\|ABCD-3\|AI848747\|CX3C\|Cxc3\|D8Bwg0439e\|Scyd1 | chemokine (C-X3-C motif) ligand 1 | protein-coding |
| chr9_120049381_120049765 | chr9 | 120049382 | 120049765 | + | 1.12865 | 3' UTR (NM_009987, exon 2 of 2) | 3' UTR (NM_009987, exon 2 of 2) | 18723 | NM_009987 | 13051 | Mm.44065 | NM_009987 | ENSMUSG00000052336 | **Cx3cr1** | - | chemokine (C-X3-C motif) receptor 1 | protein-coding |
| chr5_92373130_92373536 | chr5 | 92373131 | 92373536 | + | 0.84347 | intron (NM_181728, intron 2 of 9) | intron (NM_181728, intron 2 of 9) | -10037 | NR_038116 | 56066 | Mm.131723 | NM_019494 | ENSMUSG00000060183 | **Cxcl11** | Cxc11\|H174\|I-tac\|Ip9\|Itac\|Scyb11\|Scyb9b\|b-R1\|betaR1 | chemokine (C-X-C motif) ligand 11 | protein-coding |
| chr5_92363105_92363597 | chr5 | 92363106 | 92363597 | + | 0.775689 | promoter-TSS (NM_019494) | promoter-TSS (NM_019494) | -55 | NR_038116 | 56066 | Mm.131723 | NM_019494 | ENSMUSG00000060183 | **Cxcl11** | Cxc11\|H174\|I-tac\|Ip9\|Itac\|Scyb11\|Scyb9b\|b-R1\|betaR1 | chemokine (C-X-C motif) ligand 11 | protein-coding |

| | | | | | | | | | | | | | | | | | |
|---|---|---|---|---|---|---|---|---|---|---|---|---|---|---|---|---|---|
| chr5_95965002_95965311 | chr5 | 95965003 | 95965311 | + | 0.816279 | Intergenic | Intergenic | 8218 | NM_018866 | 55985 | Mm.10116 | NM_018866 | ENSMUSG00000023078 | **Cxcl13** | 4631412M08Rik\|ANGIE2\|Angie\|BCA-1\|BLC\|BLR1L\|Scyb13 | chemokine (C-X-C motif) ligand 13 | protein-coding |
| chr7_25429774_25430220 | chr7 | 25429775 | 25430220 | + | 1.37316 | intron (NR_015569, intron 4 of 6) | intron (NR_015569, intron 4 of 6) | -17111 | NM_153576 | 232983 | Mm.10545 | NM_153576 | ENSMUSG00000060188 | **Cxcl17** | VCC-1\|Vcc1 | chemokine (C-X-C motif) ligand 17 | protein-coding |
| chr7_25411090_25411735 | chr7 | 25411091 | 25411735 | + | 0.741555 | intron (NM_153576, intron 1 of 3) | intron (NM_153576, intron 1 of 3) | 1473 | NM_153576 | 232983 | Mm.10545 | NM_153576 | ENSMUSG00000060188 | **Cxcl17** | VCC-1\|Vcc1 | chemokine (C-X-C motif) ligand 17 | protein-coding |
| chr7_25411092_25412085 | chr7 | 25411093 | 25412085 | + | 0.633413 | intron (NM_153576, intron 1 of 3) | intron (NM_153576, intron 1 of 3) | 1297 | NM_153576 | 232983 | Mm.10545 | NM_153576 | ENSMUSG00000060188 | **Cxcl17** | VCC-1\|Vcc1 | chemokine (C-X-C motif) ligand 17 | protein-coding |
| chr1_74201060_74201796 | chr1 | 74201061 | 74201796 | + | 0.723663 | Intergenic | (TTTTG)n\|Simple_repeat\|Simple_repeat | -6797 | NM_178241 | 227288 | Mm.337035 | NM_178241 | ENSMUSG00000048480 | **Cxcr1** | Il8ra | chemokine (C-X-C motif) receptor 1 | protein-coding |
| chr1_74200793_74201798 | chr1 | 74200794 | 74201798 | + | 0.691091 | Intergenic | L1MA9\|LINE\|L1 | -6665 | NM_178241 | 227288 | Mm.337035 | NM_178241 | ENSMUSG00000048480 | **Cxcr1** | Il8ra | chemokine (C-X-C motif) receptor 1 | protein-coding |
| chrX_101730068_101730527 | chrX | 101730069 | 101730527 | + | 1.33174 | Intergenic | Intergenic | 3849 | NM_009910 | 12766 | Mm.12876 | NM_009910 | ENSMUSG00000050232 | **Cxcr3** | Cd183\|Cmkar3 | chemokine (C-X-C motif) receptor 3 | protein-coding |
| chrX_101765051_101765611 | chrX | 101765052 | 101765611 | + | 1.21184 | Intergenic | Intergenic | -31184 | NM_009910 | 12766 | Mm.12876 | NM_009910 | ENSMUSG00000050232 | **Cxcr3** | Cd183\|Cmkar3 | chemokine (C-X-C motif) receptor 3 | protein-coding |
| chrX_101764616_101764989 | chrX | 101764617 | 101764989 | + | 0.756152 | Intergenic | Intergenic | -30656 | NM_009910 | 12766 | Mm.12876 | NM_009910 | ENSMUSG00000050232 | **Cxcr3** | Cd183\|Cmkar3 | chemokine (C-X-C motif) receptor 3 | protein-coding |
| chrX_101737017_101737775 | chrX | 101737018 | 101737775 | + | 0.659899 | Intergenic | Intergenic | -3249 | NM_009910 | 12766 | Mm.12876 | NM_009910 | ENSMUSG00000050232 | **Cxcr3** | Cd183\|Cmkar3 | chemokine (C-X-C motif) receptor 3 | protein-coding |
| chrX_101737196_101737578 | chrX | 101737197 | 101737578 | + | 0.653095 | Intergenic | Intergenic | -3240 | NM_009910 | 12766 | Mm.12876 | NM_009910 | ENSMUSG00000050232 | **Cxcr3** | Cd183\|Cmkar3 | chemokine (C-X-C motif) receptor 3 | protein-coding |
| chr1_128785176_128785422 | chr1 | 128785177 | 128785422 | + | 1.02731 | Intergenic | Intergenic | -193000 | NM_009911 | 12767 | Mm.1401 | NM_009911 | ENSMUSG00000045382 | **Cxcr4** | CD184\|Cmkar4\|LESTR\|PB-CKR\|PBSF/SDF-1\|Sdf1r\|b2b220Clo | chemokine (C-X-C motif) receptor 4 | protein-coding |
| chr1_128672316_128672723 | chr1 | 128672317 | 128672723 | + | 0.951306 | Intergenic | Intergenic | -80221 | NM_009911 | 12767 | Mm.1401 | NM_009911 | ENSMUSG00000045382 | **Cxcr4** | CD184\|Cmkar4\|LESTR\|PB-CKR\|PBSF/SDF-1\|Sdf1r\|b2b220Clo | chemokine (C-X-C motif) receptor 4 | protein-coding |
| chr1_128770871_128771178 | chr1 | 128770872 | 128771178 | + | 0.928736 | Intergenic | Intergenic | -178726 | NM_009911 | 12767 | Mm.1401 | NM_009911 | ENSMUSG00000045382 | **Cxcr4** | CD184\|Cmkar4\|LESTR\|PB-CKR\|PBSF/SDF-1\|Sdf1r\|b2b220Clo | chemokine (C-X-C motif) receptor 4 | protein-coding |
| chr1_128765637_128766001 | chr1 | 128765638 | 128766001 | + | 0.87515 | Intergenic | Intergenic | -173520 | NM_009911 | 12767 | Mm.1401 | NM_009911 | ENSMUSG00000045382 | **Cxcr4** | CD184\|Cmkar4\|LESTR\|PB-CKR\|PBSF/SDF-1\|Sdf1r\|b2b220Clo | chemokine (C-X-C motif) receptor 4 | protein-coding |
| chr1_128519265_128519572 | chr1 | 128519266 | 128519572 | + | 0.737341 | Intergenic | Intergenic | 72880 | NM_009911 | 12767 | Mm.1401 | NM_009911 | ENSMUSG00000045382 | **Cxcr4** | CD184\|Cmkar4\|LESTR\|PB-CKR\|PBSF/SDF-1\|Sdf1r\|b2b220Clo | chemokine (C-X-C motif) receptor 4 | protein-coding |
| chr1_128767955_128768212 | chr1 | 128767956 | 128768212 | + | 0.703844 | Intergenic | Intergenic | -175785 | NM_009911 | 12767 | Mm.1401 | NM_009911 | ENSMUSG00000045382 | **Cxcr4** | CD184\|Cmkar4\|LESTR\|PB-CKR\|PBSF/SDF-1\|Sdf1r\|b2b220Clo | chemokine (C-X-C motif) receptor 4 | protein-coding |
| chr9_44540009_44540396 | chr9 | 44540010 | 44540396 | + | 1.19192 | Intergenic | Intergenic | -13782 | NM_007551 | 12145 | Mm.6246 | NM_007551 | ENSMUSG00000047880 | **Cxcr5** | Blr1\|CXC-R5\|CXCR-5\|Gpcr6\|MDR15 | chemokine (C-X-C motif) receptor 5 | protein-coding |
| chr9_44537656_44538314 | chr9 | 44537657 | 44538314 | + | 0.600616 | Intergenic | Intergenic | -11564 | NM_007551 | 12145 | Mm.6246 | NM_007551 | ENSMUSG00000047880 | **Cxcr5** | Blr1\|CXC-R5\|CXCR-5\|Gpcr6\|MDR15 | chemokine (C-X-C motif) receptor 5 | protein-coding |
| chr18_35847903_35848788 | chr18 | 35847904 | 35848788 | + | 0.762779 | intron (NM_133687, intron 1 of 2) | intron (NM_133687, intron 1 of 2) | 18528 | NM_133687 | 67393 | Mm.94560 | NM_133687 | ENSMUSG00000046668 | **Cxxc5** | 4930415K17Rik | CXXC finger 5 | protein-coding |
| chr18_35847899_35849185 | chr18 | 35847900 | 35849185 | + | 0.745721 | intron (NM_133687, intron 1 of 2) | intron (NM_133687, intron 1 of 2) | 18724 | NM_133687 | 67393 | Mm.94560 | NM_133687 | ENSMUSG00000046668 | **Cxxc5** | 4930415K17Rik | CXXC finger 5 | protein-coding |
| chr18_35813539_35813959 | chr18 | 35813540 | 35813959 | + | 0.660774 | Intergenic | Intergenic | -16069 | NM_133687 | 67393 | Mm.94560 | NM_133687 | ENSMUSG00000046668 | **Cxxc5** | 4930415K17Rik | CXXC finger 5 | protein-coding |
| chr18_84867142_84867648 | chr18 | 84867143 | 84867648 | + | 1.01337 | intron (NM_025797, intron 1 of 4) | intron (NM_025797, intron 1 of 4) | 15981 | NM_025797 | 109672 | Mm.31018 | NM_025797 | | **Cyb5a** | 0610009N12Rik\|Cyb5 | cytochrome b5 type A (microsomal) | protein-coding |
| chr18_84872385_84872663 | chr18 | 84872386 | 84872663 | + | 0.737658 | intron (NM_025797, intron 2 of 4) | intron (NM_025797, intron 2 of 4) | 21110 | NM_025797 | 109672 | Mm.31018 | NM_025797 | | **Cyb5a** | 0610009N12Rik\|Cyb5 | cytochrome b5 type A (microsomal) | protein-coding |
| chr11_72782144_72782493 | chr11 | 72782145 | 72782493 | + | 1.07918 | intron (NM_001024926, intron 3 of 3) | ORR1E\|LTR\|ERVL-MaLR | 13520 | NM_001024926 | 192986 | Mm.194755 | NM_001024926 | ENSMUSG00000057778 | **Cyb5d2** | 9330151E16Rik\|Gm2 | cytochrome b5 domain containing 2 | protein-coding |
| chr7_107720622_107721574 | chr7 | 107720623 | 107721574 | + | 1.01654 | intron (NM_008905, intron 11 of 24) | intron (NM_008905, intron 11 of 24) | 36934 | NM_001205227 | 320635 | Mm.132828 | NM_177216 | ENSMUSG00000048065 | **Cyb5r2** | D630003K02Rik\|b5R.2 | cytochrome b5 reductase 2 | protein-coding |
| chr8_122436155_122436428 | chr8 | 122436156 | 122436428 | + | 0.676198 | intron (NM_138656, intron 8 of 9) | intron (NM_138656, intron 8 of 9) | -3352 | NM_007806 | 13057 | Mm.271671 | NM_007806 | ENSMUSG00000006519 | **Cyba** | b558\|nmf333\|p22-phox\|p22phox | cytochrome b-245, alpha polypeptide | protein-coding |
| chrX_9447268_9447770 | chrX | 9447269 | 9447770 | + | 0.680406 | intron (NM_007807, intron 8 of 12) | intron (NM_007807, intron 8 of 12) | 21805 | NM_007807 | 13058 | Mm.200362 | NM_007807 | ENSMUSG00000015340 | **Cybb** | C88302\|CGD91-phox\|Cgd\|Cyd\|Nox2\|gp91-1\|gp91phox | cytochrome b-245, beta polypeptide | protein-coding |
| chrX_9447296_9447643 | chrX | 9447297 | 9447643 | + | 0.66973 | intron (NM_007807, intron 8 of 12) | intron (NM_007807, intron 8 of 12) | 21854 | NM_007807 | 13058 | Mm.200362 | NM_007807 | ENSMUSG00000015340 | **Cybb** | C88302\|CGD91-phox\|Cgd\|Cyd\|Nox2\|gp91-1\|gp91phox | cytochrome b-245, beta polypeptide | protein-coding |
| chr11_46322427_46322825 | chr11 | 46322428 | 46322825 | + | 0.980489 | intron (NR_105955, intron 2 of 4) | intron (NR_105955, intron 2 of 4) | -9767 | NM_001252459 | 76884 | Mm.154358 | NM_133769 | ENSMUSG00000020340 | **Cyfip2** | 1500004I01Rik\|6430511D02Rik\|AA930218\|AU022376\|Pir121\|mKIAA1168 | cytoplasmic FMR1 interacting protein 2 | protein-coding |
| chr11_46349078_46349927 | chr11 | 46349079 | 46349927 | + | 0.863538 | intron (NM_001281966, intron 7 of 16) | MER44B\|DNA\|TcMar-Tigger | -36644 | NM_001252459 | 76884 | Mm.154358 | NM_133769 | ENSMUSG00000020340 | **Cyfip2** | 1500004I01Rik\|6430511D02Rik\|AA930218\|AU022376\|Pir121\|mKIAA1168 | cytoplasmic FMR1 interacting protein 2 | protein-coding |
| chr11_46304059_46304287 | chr11 | 46304060 | 46304287 | + | 0.609178 | intron (NM_001252459, intron 1 of 30) | intron (NM_001252459, intron 1 of 30) | 8185 | NM_001252460 | 76884 | Mm.154358 | NM_133769 | ENSMUSG00000020340 | **Cyfip2** | 1500004I01Rik\|6430511D02Rik\|AA930218\|AU022376\|Pir121\|mKIAA1168 | cytoplasmic FMR1 interacting protein 2 | protein-coding |
| chr9_57989201_57989711 | chr9 | 57989202 | 57989711 | + | 0.661496 | Intergenic | Intergenic | -25561 | NM_019779 | 13070 | Mm.302865 | NM_019779 | ENSMUSG00000032323 | **Cyp11a1** | Cyp11a\|Cypxia1\|D9Ertd411e\|P450scc\|Scc\|cscc | cytochrome P450, family 11, subfamily a, polypeptide 1 | protein-coding |
| chr6_84798697_84799123 | chr6 | 84798698 | 84799123 | + | 0.786834 | intron (NM_177077, intron 18 of 21) | Lx4A\|LINE\|L1 | -205002 | NM_001177713 | 232174 | Mm.255246 | NM_175475 | ENSMUSG00000063415 | **Cyp26b1** | CP26\|P450RAI-2 | cytochrome P450, family 26, subfamily b, polypeptide 1 | protein-coding |
| chr19_37673409_37673905 | chr19 | 37673410 | 37673905 | + | 0.677265 | intron (NM_175353, intron 20 of 21) | intron (NM_175353, intron 20 of 21) | -12023 | NM_001105201 | 546726 | Mm.457750 | NM_001105201 | ENSMUSG00000062432 | **Cyp26c1** | EG546726 | cytochrome P450, family 26, subfamily c, polypeptide 1 | protein-coding |
| chr7_114520142_114520678 | chr7 | 114520143 | 114520678 | + | 1.0612 | intron (NM_011055, intron 9 of 15) | RMER6C\|LTR\|ERVK | 42562 | NR_110407 | 244209 | Mm.108037 | NM_177382 | ENSMUSG00000030670 | **Cyp2r1** | - | cytochrome P450, family 2, subfamily r, polypeptide 1 | protein-coding |
| chr17_32470778_32471320 | chr17 | 32470779 | 32471320 | + | 0.980205 | intron (NM_177307, intron 4 of 13) | intron (NM_177307, intron 4 of 13) | 18326 | NM_177307 | 320997 | Mm.130900 | NM_177307 | | **Cyp4f39** | 4732474A20Rik | cytochrome P450, family 4, subfamily f, polypeptide 39 | protein-coding |
| chr17_32470415_32471339 | chr17 | 32470416 | 32471339 | + | 0.877411 | exon (NM_177307, exon 4 of 14) | exon (NM_177307, exon 4 of 14) | 18154 | NM_177307 | 320997 | Mm.130900 | NM_177307 | | **Cyp4f39** | 4732474A20Rik | cytochrome P450, family 4, subfamily f, polypeptide 39 | protein-coding |
| chr9_121924932_121925484 | chr9 | 121924933 | 121925484 | + | 0.99255 | Intergenic | Lx8\|LINE\|L1 | -8903 | NM_010012 | 13124 | Mm.20889 | NM_010012 | ENSMUSG00000050445 | **Cyp8b1** | - | cytochrome P450, family 8, subfamily b, polypeptide 1 | protein-coding |
| chr3_145617979_145618948 | chr3 | 145617980 | 145618948 | + | 1.47593 | Intergenic | Intergenic | 31521 | NM_010516 | 16007 | Mm.1231 | NM_010516 | ENSMUSG00000028195 | **Cyr61** | AI325051\|CCN1\|Igfbp10 | cysteine rich protein 61 | protein-coding |
| chr3_145613051_145613552 | chr3 | 145613052 | 145613552 | + | 0.99832 | Intergenic | Intergenic | 36683 | NM_010516 | 16007 | Mm.1231 | NM_010516 | ENSMUSG00000028195 | **Cyr61** | AI325051\|CCN1\|Igfbp10 | cysteine rich protein 61 | protein-coding |
| chr3_145627884_145628170 | chr3 | 145627885 | 145628170 | + | 0.907737 | Intergenic | Intergenic | 21958 | NM_010516 | 16007 | Mm.1231 | NM_010516 | ENSMUSG00000028195 | **Cyr61** | AI325051\|CCN1\|Igfbp10 | cysteine rich protein 61 | protein-coding |
| chr3_145612766_145613557 | chr3 | 145612767 | 145613557 | + | 0.855918 | Intergenic | Intergenic | 36823 | NM_010516 | 16007 | Mm.1231 | NM_010516 | ENSMUSG00000028195 | **Cyr61** | AI325051\|CCN1\|Igfbp10 | cysteine rich protein 61 | protein-coding |
| chr18_36371366_36371760 | chr18 | 36371367 | 36371760 | + | 0.829863 | intron (NM_001081365, intron 2 of 2) | PB1D10\|SINE\|Alu | 22939 | NM_001081365 | 66060 | Mm.272527 | NM_001081365 | | **Cystm1** | 0610010O12Rik | cysteine-rich transmembrane module containing 1 | protein-coding |
| chr18_36399687_36400140 | chr18 | 36399688 | 36400140 | + | 0.73564 | Intergenic | Intergenic | 51290 | NM_001081365 | 66060 | Mm.272527 | NM_001081365 | | **Cystm1** | 0610010O12Rik | cysteine-rich transmembrane module containing 1 | protein-coding |
| chr11_118215896_118216515 | chr11 | 118215897 | 118216515 | + | 0.824737 | intron (NM_001112699, intron 1 of 12) | MTD\|LTR\|ERVL-MaLR | -3464 | NM_001112700 | 19157 | Mm.86413 | NM_011180 | ENSMUSG00000017132 | **Cyth1** | CLM1\|CTH-1\|CYTIP\|Pscd1\|mKIAA4240 | cytohesin 1 | protein-coding |
| chr5_143683566_143684600 | chr5 | 143683567 | 143684600 | + | 1.18262 | intron (NM_001163548, intron 1 of 12) | intron (NM_001163548, intron 1 of 12) | 32842 | NM_001163548 | 19159 | Mm.281003 | NM_011182 | ENSMUSG00000018001 | **Cyth3** | AI648983\|ARNO3\|CLM3\|Grp1\|Pscd3 | cytohesin 3 | protein-coding |
| chr15_78610803_78611052 | chr15 | 78610804 | 78611052 | + | 0.661911 | intron (NM_028195, intron 7 of 12) | MIR\|SINE\|MIR | 13881 | NM_028195 | 72318 | Mm.32911 | NM_028195 | ENSMUSG00000018008 | **Cyth4** | 2510004M07Rik\|5830469K17Rik\|AI467541\|Pscd4\|mFLJ00017 | cytohesin 4 | protein-coding |
| chr15_78607363_78607857 | chr15 | 78607364 | 78607857 | + | 0.653933 | intron (NM_028195, intron 5 of 12) | intron (NM_028195, intron 5 of 12) | 10563 | NM_028195 | 72318 | Mm.32911 | NM_028195 | ENSMUSG00000018008 | **Cyth4** | 2510004M07Rik\|5830469K17Rik\|AI467541\|Pscd4\|mFLJ00017 | cytohesin 4 | protein-coding |

| | | | | | | | | | | | | | | | | | |
|---|---|---|---|---|---|---|---|---|---|---|---|---|---|---|---|---|---|
| chr15_78591748_78591988 | chr15 | 78591749 | 78591988 | + | 0.623133 | Intergenic | Intergenic | -5179 | NM_028195 | 72318 | Mm.32911 | NM_028195 | ENSMUSG00000018008 | **Cyth4** | 2510004M07Rik\|5830469K17Rik\|AI467541\|Pscd4\|mFLJ00017 | cytohesin 4 | protein-coding |
| chr15_78591689_78592672 | chr15 | 78591690 | 78592672 | + | 0.603265 | Intergenic | Intergenic | -4866 | NM_028195 | 72318 | Mm.32911 | NM_028195 | ENSMUSG00000018008 | **Cyth4** | 2510004M07Rik\|5830469K17Rik\|AI467541\|Pscd4\|mFLJ00017 | cytohesin 4 | protein-coding |
| chr2_58124741_58125131 | chr2 | 58124742 | 58125131 | + | 1.00458 | Intergenic | RMER15\|LTR\|ERVL | 35186 | NM_139200 | 227929 | Mm.273905 | NM_139200 | ENSMUSG00000026832 | **Cytip** | A130053M09Rik\|AI462064\|C80816\|Cbp\|Cybr\|Pscdbp | cytohesin 1 interacting protein | protein-coding |
| chr2_58161643_58161934 | chr2 | 58161644 | 58161934 | + | 0.813792 | Intergenic | Intergenic | -1667 | NM_139200 | 227929 | Mm.273905 | NM_139200 | ENSMUSG00000026832 | **Cytip** | A130053M09Rik\|AI462064\|C80816\|Cbp\|Cybr\|Pscdbp | cytohesin 1 interacting protein | protein-coding |
| chr2_58129489_58130401 | chr2 | 58129490 | 58130401 | + | 0.631325 | 3' UTR (NM_139200, exon 8 of 8) | 3' UTR (NM_139200, exon 8 of 8) | 30177 | NM_139200 | 227929 | Mm.273905 | NM_139200 | ENSMUSG00000026832 | **Cytip** | A130053M09Rik\|AI462064\|C80816\|Cbp\|Cybr\|Pscdbp | cytohesin 1 interacting protein | protein-coding |
| chr15_61454871_61455422 | chr15 | 61454872 | 61455422 | + | 1.60628 | Intergenic | Intergenic | 319304 | NR_040350 | 432956 | Mm.404918 | NR_040350 | | **D030024E09Rik** | - | RIKEN cDNA D030024E09 gene | ncRNA |
| chr15_61396991_61397419 | chr15 | 61396992 | 61397419 | + | 1.19903 | Intergenic | Intergenic | 377246 | NR_040350 | 432956 | Mm.404918 | NR_040350 | | **D030024E09Rik** | - | RIKEN cDNA D030024E09 gene | ncRNA |
| chr15_61396727_61397437 | chr15 | 61396728 | 61397437 | + | 1.16019 | Intergenic | Intergenic | 377369 | NR_040350 | 432956 | Mm.404918 | NR_040350 | | **D030024E09Rik** | - | RIKEN cDNA D030024E09 gene | ncRNA |
| chr15_61548882_61549134 | chr15 | 61548883 | 61549134 | + | 1.12584 | Intergenic | Intergenic | 225443 | NR_040350 | 432956 | Mm.404918 | NR_040350 | | **D030024E09Rik** | - | RIKEN cDNA D030024E09 gene | ncRNA |
| chr15_61448402_61449059 | chr15 | 61448403 | 61449059 | + | 1.03181 | Intergenic | MER20\|DNA\|hAT-Charlie | 325720 | NR_040350 | 432956 | Mm.404918 | NR_040350 | | **D030024E09Rik** | - | RIKEN cDNA D030024E09 gene | ncRNA |
| chr15_61407903_61408627 | chr15 | 61407904 | 61408627 | + | 1.02634 | Intergenic | L1MCa\|LINE\|L1 | 366186 | NR_040350 | 432956 | Mm.404918 | NR_040350 | | **D030024E09Rik** | - | RIKEN cDNA D030024E09 gene | ncRNA |
| chr15_61482643_61483465 | chr15 | 61482644 | 61483465 | + | 0.983003 | Intergenic | Intergenic | 291397 | NR_040350 | 432956 | Mm.404918 | NR_040350 | | **D030024E09Rik** | - | RIKEN cDNA D030024E09 gene | ncRNA |
| chr15_61545334_61545568 | chr15 | 61545335 | 61545568 | + | 0.948109 | Intergenic | B2_Mm2\|SINE\|B2 | 229000 | NR_040350 | 432956 | Mm.404918 | NR_040350 | | **D030024E09Rik** | - | RIKEN cDNA D030024E09 gene | ncRNA |
| chr15_61548388_61548757 | chr15 | 61548389 | 61548757 | + | 0.945262 | Intergenic | Intergenic | 225878 | NR_040350 | 432956 | Mm.404918 | NR_040350 | | **D030024E09Rik** | - | RIKEN cDNA D030024E09 gene | ncRNA |
| chr15_61441480_61442122 | chr15 | 61441481 | 61442122 | + | 0.944081 | Intergenic | Intergenic | 332650 | NR_040350 | 432956 | Mm.404918 | NR_040350 | | **D030024E09Rik** | - | RIKEN cDNA D030024E09 gene | ncRNA |
| chr15_61441263_61442122 | chr15 | 61441264 | 61442122 | + | 0.91826 | Intergenic | Intergenic | 332758 | NR_040350 | 432956 | Mm.404918 | NR_040350 | | **D030024E09Rik** | - | RIKEN cDNA D030024E09 gene | ncRNA |
| chr15_61682569_61683356 | chr15 | 61682570 | 61683356 | + | 0.848451 | Intergenic | Intergenic | 91488 | NR_040350 | 432956 | Mm.404918 | NR_040350 | | **D030024E09Rik** | - | RIKEN cDNA D030024E09 gene | ncRNA |
| chr15_61454133_61454823 | chr15 | 61454134 | 61454823 | + | 0.782411 | Intergenic | Intergenic | 319973 | NR_040350 | 432956 | Mm.404918 | NR_040350 | | **D030024E09Rik** | - | RIKEN cDNA D030024E09 gene | ncRNA |
| chr15_61430304_61430780 | chr15 | 61430305 | 61430780 | + | 0.750282 | Intergenic | Intergenic | 343909 | NR_040350 | 432956 | Mm.404918 | NR_040350 | | **D030024E09Rik** | - | RIKEN cDNA D030024E09 gene | ncRNA |
| chr15_61413193_61413373 | chr15 | 61413194 | 61413373 | + | 0.677733 | Intergenic | Intergenic | 361168 | NR_040350 | 432956 | Mm.404918 | NR_040350 | | **D030024E09Rik** | - | RIKEN cDNA D030024E09 gene | ncRNA |
| chr15_61622369_61623089 | chr15 | 61622370 | 61623089 | + | 0.668734 | Intergenic | ORR1E\|LTR\|ERVL-MaLR | 151722 | NR_040350 | 432956 | Mm.404918 | NR_040350 | | **D030024E09Rik** | - | RIKEN cDNA D030024E09 gene | ncRNA |
| chr3_128080885_128081691 | chr3 | 128080886 | 128081691 | + | 1.42465 | Intergenic | Intergenic | 150317 | NR_045704 | 402774 | Mm.209170 | NR_045704 | | **D030025E07Rik** | - | RIKEN cDNA D030025E07 gene | ncRNA |
| chr3_128106449_128106856 | chr3 | 128106450 | 128106856 | + | 0.657733 | Intergenic | Intergenic | 124952 | NR_045704 | 402774 | Mm.209170 | NR_045704 | | **D030025E07Rik** | - | RIKEN cDNA D030025E07 gene | ncRNA |
| chr10_45710692_45711356 | chr10 | 45710693 | 45711356 | + | 1.03734 | intron (NM_172473, intron 23 of 23) | intron (NM_172473, intron 23 of 23) | -96680 | NR_040624 | 320363 | Mm.123343 | NR_040624 | | **D030045P18Rik** | - | RIKEN cDNA D030045P18 gene | ncRNA |
| chr10_45710334_45711360 | chr10 | 45710335 | 45711360 | + | 0.970817 | intron (NM_172473, intron 23 of 23) | intron (NM_172473, intron 23 of 23) | -96857 | NR_040624 | 320363 | Mm.123343 | NR_040624 | | **D030045P18Rik** | - | RIKEN cDNA D030045P18 gene | ncRNA |
| chr10_45778231_45778726 | chr10 | 45778232 | 45778726 | + | 0.68107 | Intergenic | Intergenic | -29225 | NR_040624 | 320363 | Mm.123343 | NR_040624 | | **D030045P18Rik** | - | RIKEN cDNA D030045P18 gene | ncRNA |
| chr13_24866227_24867380 | chr13 | 24866228 | 24867380 | + | 0.713968 | intron (NM_001081051, intron 7 of 20) | intron (NM_001081051, intron 7 of 20) | 21673 | NM_001081051 | 210108 | Mm.261514 | NM_001081051 | ENSMUSG00000006711 | **D130043K22Rik** | 4930451E12Rik\|Kiaa0319 | RIKEN cDNA D130043K22 gene | protein-coding |
| chr6_127243097_127243402 | chr6 | 127243098 | 127243402 | + | 1.34313 | Intergenic | Intergenic | 64717 | NR_073373 | 232386 | Mm.257367 | NR_073373 | | **D130058E03** | - | uncharacterized D130058E03 | ncRNA |
| chr6_127293790_127294220 | chr6 | 127293791 | 127294220 | + | 1.09516 | Intergenic | Intergenic | 13962 | NR_073373 | 232386 | Mm.257367 | NR_073373 | | **D130058E03** | - | uncharacterized D130058E03 | ncRNA |
| chr6_127239656_127241137 | chr6 | 127239657 | 127241137 | + | 1.03815 | Intergenic | Intergenic | 67570 | NR_073373 | 232386 | Mm.257367 | NR_073373 | | **D130058E03** | - | uncharacterized D130058E03 | ncRNA |
| chr6_127371639_127372214 | chr6 | 127371640 | 127372214 | + | 1.00633 | Intergenic | Intergenic | -63960 | NR_073373 | 232386 | Mm.257367 | NR_073373 | | **D130058E03** | - | uncharacterized D130058E03 | ncRNA |
| chr6_127293389_127294220 | chr6 | 127293390 | 127294220 | + | 0.939046 | Intergenic | Intergenic | 14162 | NR_073373 | 232386 | Mm.257367 | NR_073373 | | **D130058E03** | - | uncharacterized D130058E03 | ncRNA |
| chr6_127239695_127240076 | chr6 | 127239696 | 127240076 | + | 0.911463 | Intergenic | Intergenic | 68081 | NR_073373 | 232386 | Mm.257367 | NR_073373 | | **D130058E03** | - | uncharacterized D130058E03 | ncRNA |
| chr6_127263865_127264185 | chr6 | 127263866 | 127264185 | + | 0.866967 | Intergenic | Intergenic | 43942 | NR_073373 | 232386 | Mm.257367 | NR_073373 | | **D130058E03** | - | uncharacterized D130058E03 | ncRNA |
| chr6_127236508_127236919 | chr6 | 127236509 | 127236919 | + | 0.754986 | Intergenic | Intergenic | 71253 | NR_073373 | 232386 | Mm.257367 | NR_073373 | | **D130058E03** | - | uncharacterized D130058E03 | ncRNA |
| chr6_127264964_127265700 | chr6 | 127264965 | 127265700 | + | 0.696011 | Intergenic | (TC)n\|Simple_repeat\|Simple_repeat | 42635 | NR_073373 | 232386 | Mm.257367 | NR_073373 | | **D130058E03** | - | uncharacterized D130058E03 | ncRNA |
| chr6_127351505_127352039 | chr6 | 127351506 | 127352039 | + | 0.661889 | Intergenic | MLTR11B\|LTR\|ERVK | -43805 | NR_073373 | 232386 | Mm.257367 | NR_073373 | | **D130058E03** | - | uncharacterized D130058E03 | ncRNA |
| chr15_58585320_58586411 | chr15 | 58585321 | 58586411 | + | 1.52585 | Intergenic | Intergenic | 170398 | NM_145959 | 210998 | Mm.3224 | NM_145959 | ENSMUSG00000037119 | **D15Ertd621e** | AV220772\|BC033609\|Fam91a1\|SKIN\|mKIAA0493 | DNA segment, Chr 15, ERATO Doi 621, expressed | protein-coding |
| chr15_58417079_58417403 | chr15 | 58417080 | 58417403 | + | 0.777645 | intron (NM_145959, intron 1 of 23) | intron (NM_145959, intron 1 of 23) | 1773 | NM_145959 | 210998 | Mm.3224 | NM_145959 | ENSMUSG00000037119 | **D15Ertd621e** | AV220772\|BC033609\|Fam91a1\|SKIN\|mKIAA0493 | DNA segment, Chr 15, ERATO Doi 621, expressed | protein-coding |
| chr16_78619472_78619706 | chr16 | 78619473 | 78619706 | + | 1.1646 | Intergenic | RMER15\|LTR\|ERVL | -42901 | NM_025967 | 67102 | Mm.37332 | NM_025967 | | **D16Ertd472e** | 1700010I10Rik\|2310009O17Rik\|E330003K22Rik\|Eurl | DNA segment, Chr 16, ERATO Doi 472, expressed | protein-coding |
| chr16_78619315_78619761 | chr16 | 78619316 | 78619761 | + | 1.13226 | Intergenic | RMER15\|LTR\|ERVL | -42850 | NM_025967 | 67102 | Mm.37332 | NM_025967 | | **D16Ertd472e** | 1700010I10Rik\|2310009O17Rik\|E330003K22Rik\|Eurl | DNA segment, Chr 16, ERATO Doi 472, expressed | protein-coding |
| chr16_78632631_78632960 | chr16 | 78632632 | 78632960 | + | 0.991809 | Intergenic | Intergenic | -56108 | NM_025967 | 67102 | Mm.37332 | NM_025967 | | **D16Ertd472e** | 1700010I10Rik\|2310009O17Rik\|E330003K22Rik\|Eurl | DNA segment, Chr 16, ERATO Doi 472, expressed | protein-coding |
| chr16_78575260_78575438 | chr16 | 78575261 | 78575438 | + | 0.68552 | intron (NM_001252438, intron 2 of 4) | intron (NM_001252438, intron 2 of 4) | 1339 | NM_001252438 | 67102 | Mm.37332 | NM_025967 | | **D16Ertd472e** | 1700010I10Rik\|2310009O17Rik\|E330003K22Rik\|Eurl | DNA segment, Chr 16, ERATO Doi 472, expressed | protein-coding |
| chr16_78688781_78689876 | chr16 | 78688782 | 78689876 | + | 0.653802 | Intergenic | MTE2b\|LTR\|ERVL-MaLR | -112641 | NM_025967 | 67102 | Mm.37332 | NM_025967 | | **D16Ertd472e** | 1700010I10Rik\|2310009O17Rik\|E330003K22Rik\|Eurl | DNA segment, Chr 16, ERATO Doi 472, expressed | protein-coding |
| chr17_27795249_27795567 | chr17 | 27795250 | 27795567 | + | 0.825327 | intron (NM_001033279, intron 1 of 4) | intron (NM_001033279, intron 1 of 4) | 25134 | NM_001033279 | 224647 | Mm.196577 | NM_001033279 | ENSMUSG00000056692 | **D17Wsu92e** | AU020189\|C030010A15\|C80239 | DNA segment, Chr 17, Wayne State University 92, expressed | protein-coding |
| chr17_27789930_27790526 | chr17 | 27789931 | 27790526 | + | 0.666424 | intron (NM_001033279, intron 2 of 4) | intron (NM_001033279, intron 2 of 4) | 30314 | NM_001033279 | 224647 | Mm.196577 | NM_001033279 | ENSMUSG00000056692 | **D17Wsu92e** | AU020189\|C030010A15\|C80239 | DNA segment, Chr 17, Wayne State University 92, expressed | protein-coding |
| chr1_97577166_97577412 | chr1 | 97577167 | 97577412 | + | 0.856119 | Intergenic | Intergenic | 84729 | NM_133825 | 52392 | Mm.12309 | NM_133825 | ENSMUSG00000044768 | **D1Ertd622e** | AI987691\|AW047450 | DNA segment, Chr 1, ERATO Doi 622, expressed | protein-coding |
| chr2_116972049_116972539 | chr2 | 116972050 | 116972539 | + | 0.979557 | Intergenic | Intergenic | 72142 | NR_040335 | 320975 | Mm.132509 | NR_040335 | ENSMUSG00000085316 | **D330050G23Rik** | - | RIKEN cDNA D330050G23 gene | ncRNA |
| chr2_116972201_116972541 | chr2 | 116972202 | 116972541 | + | 0.904664 | Intergenic | Intergenic | 72219 | NR_040335 | 320975 | Mm.132509 | NR_040335 | ENSMUSG00000085316 | **D330050G23Rik** | - | RIKEN cDNA D330050G23 gene | ncRNA |
| chr9_81624931_81626564 | chr9 | 81624932 | 81626564 | + | 1.3512 | Intergenic | Intergenic | -6182 | NR_040394 | 319405 | Mm.246494 | NR_040393 | ENSMUSG00000097466 | **D430036J16Rik** | - | RIKEN cDNA D430036J16 gene | ncRNA |
| chr9_81593840_81594144 | chr9 | 81593841 | 81594144 | + | 0.744677 | Intergenic | Intergenic | -37938 | NR_040394 | 319405 | Mm.246494 | NR_040393 | ENSMUSG00000097466 | **D430036J16Rik** | - | RIKEN cDNA D430036J16 gene | ncRNA |
| chr7_125832752_125833956 | chr7 | 125832753 | 125833956 | + | 1.2494 | intron (NM_001081022, intron 14 of 27) | intron (NM_001081022, intron 14 of 27) | 125478 | NM_001081022 | 233865 | Mm.274389 | NM_001081022 | ENSMUSG00000032743 | **D430042O09Rik** | Kiaa0556 | RIKEN cDNA D430042O09 gene | protein-coding |
| chr5_36720844_36721281 | chr5 | 36720845 | 36721281 | + | 1.51658 | Intergenic | Intergenic | -25042 | NM_001081232 | 320661 | Mm.26480 | NM_177221 | ENSMUSG00000029190 | **D5Ertd579e** | 9030221A05Rik\|A930018H20Rik\|Kiaa0232\|mKIAA0232 | DNA segment, Chr 5, ERATO Doi 579, expressed | protein-coding |
| chr5_36679867_36680783 | chr5 | 36679868 | 36680783 | + | 0.701396 | intron (NM_001081232, intron 1 of 8) | intron (NM_001081232, intron 1 of 8) | 15696 | NM_001081232 | 320661 | Mm.26480 | NM_177221 | ENSMUSG00000029190 | **D5Ertd579e** | 9030221A05Rik\|A930018H20Rik\|Kiaa0232\|mKIAA0232 | DNA segment, Chr 5, ERATO Doi 579, expressed | protein-coding |
| chr2_158233045_158233560 | chr2 | 158233046 | 158233560 | + | 0.67164 | Intergenic | Intergenic | -4079 | NM_177657 | 228846 | Mm.187574 | NM_177657 | ENSMUSG00000037813 | **D630003M21Rik** | - | RIKEN cDNA D630003M21 gene | protein-coding |
| chr15_94090321_94091230 | chr15 | 94090322 | 94091230 | + | 1.37539 | Intergenic | Intergenic | -156426 | NR_045629 | 328594 | Mm.135523 | NR_045629 | | **D630010B17Rik** | Gm811 | RIKEN cDNA D630010B17 gene | ncRNA |
| chr15_94089551_94091234 | chr15 | 94089552 | 94091234 | + | 1.34842 | Intergenic | Intergenic | -156809 | NR_045629 | 328594 | Mm.135523 | NR_045629 | | **D630010B17Rik** | Gm811 | RIKEN cDNA D630010B17 gene | ncRNA |
| chr15_94089585_94090273 | chr15 | 94089586 | 94090273 | + | 1.21704 | Intergenic | RLTR17\|LTR\|ERVK | -157273 | NR_045629 | 328594 | Mm.135523 | NR_045629 | | **D630010B17Rik** | Gm811 | RIKEN cDNA D630010B17 gene | ncRNA |
| chr15_94227242_94227743 | chr15 | 94227243 | 94227743 | + | 0.785865 | Intergenic | RMER15\|LTR\|ERVL | -19709 | NR_045629 | 328594 | Mm.135523 | NR_045629 | | **D630010B17Rik** | Gm811 | RIKEN cDNA D630010B17 gene | ncRNA |
| chr15_94206420_94206796 | chr15 | 94206421 | 94206796 | + | 0.684705 | Intergenic | Intergenic | -40594 | NR_045629 | 328594 | Mm.135523 | NR_045629 | | **D630010B17Rik** | Gm811 | RIKEN cDNA D630010B17 gene | ncRNA |
| chr11_31828733_31829039 | chr11 | 31828734 | 31829039 | + | 0.901528 | Intergenic | Intergenic | -4339 | NR_102312 | 414116 | Mm.150222 | NR_102310 | ENSMUSG00000085772 | **D630024D03Rik** | - | RIKEN cDNA D630024D03 gene | ncRNA |
| chr11_31833875_31834584 | chr11 | 31833876 | 31834584 | + | 0.810247 | Intergenic | Intergenic | -9683 | NR_102312 | 414116 | Mm.150222 | NR_102310 | ENSMUSG00000085772 | **D630024D03Rik** | - | RIKEN cDNA D630024D03 gene | ncRNA |
| chr11_31828664_31829108 | chr11 | 31828665 | 31829108 | + | 0.771405 | Intergenic | Intergenic | -4339 | NR_102312 | 414116 | Mm.150222 | NR_102310 | ENSMUSG00000085772 | **D630024D03Rik** | - | RIKEN cDNA D630024D03 gene | ncRNA |
| chr11_31807860_31808202 | chr11 | 31807861 | 31808202 | + | 0.714018 | intron (NR_102312, intron 1 of 4) | intron (NR_102312, intron 1 of 4) | 16516 | NR_102310 | 414116 | Mm.150222 | NR_102310 | ENSMUSG00000085772 | **D630024D03Rik** | - | RIKEN cDNA D630024D03 gene | ncRNA |
| chr9_43261743_43262378 | chr9 | 43261744 | 43262378 | + | 1.10522 | intron (NM_001243261, intron 1 of 2) | intron (NM_001243261, intron 1 of 2) | 2182 | NM_001243261 | 235302 | Mm.135745 | NM_001243261 | ENSMUSG00000095385 | **D630033O11Rik** | Gm588 | RIKEN cDNA D630033O11 gene | protein-coding |
| chr6_38255270_38255545 | chr6 | 38255271 | 38255545 | + | 0.91852 | Intergenic | Intergenic | -1399 | NM_194061 | 330286 | Mm.277163 | NM_194061 | ENSMUSG00000063455 | **D630045J12Rik** | Kiaa1549\|mKIAA1549 | RIKEN cDNA D630045J12 gene | protein-coding |

| | | | | | | | | | | | | | | | | | |
|---|---|---|---|---|---|---|---|---|---|---|---|---|---|---|---|---|---|
| chr6_38255239_38255851 | chr6 | 38255240 | 38255851 | + | 0.869783 | Intergenic | Intergenic | -1536 | NM_194061 | 330286 | Mm.277163 | NM_194061 | ENSMUSG00000063455 | **D630045J12Rik** | Kiaa1549\|mKIAA1549 | RIKEN cDNA D630045J12 gene | protein-coding |
| chr6_87145932_87146400 | chr6 | 87145933 | 87146400 | + | 1.50284 | intron (NM_054041, intron 17 of 17) | intron (NM_054041, intron 17 of 17) | 41420 | NM_001167937 | 52372 | Mm.282851 | NM_001167937 | | **D6Ertd527e** | AU043841 | DNA segment, Chr 6, ERATO Doi 527, expressed | protein-coding |
| chr15_79894713_79895340 | chr15 | 79894714 | 79895340 | + | 0.721097 | intron (NM_030255, intron 1 of 7) | B4A\|SINE\|B4 | -1889 | NR_030675 | 109361 | Mm.394656 | NR_030675 | | **D730005E14Rik** | - | RIKEN cDNA D730005E14 gene | ncRNA |
| chr8_36164528_36165610 | chr8 | 36164529 | 36165610 | + | 1.8611 | Intergenic | Intergenic | 70241 | NM_172911 | 244418 | Mm.244971 | NM_172911 | ENSMUSG00000050271 | **D8Ertd82e** | 9830148H23\|Sgk223\|mFLJ00269 | DNA segment, Chr 8, ERATO Doi 82, expressed | protein-coding |
| chr8_36059657_36060441 | chr8 | 36059658 | 36060441 | + | 1.50663 | Intergenic | MTA_Mm\|LTR\|ERVL-MaLR | -34779 | NM_172911 | 244418 | Mm.244971 | NM_172911 | ENSMUSG00000050271 | **D8Ertd82e** | 9830148H23\|Sgk223\|mFLJ00269 | DNA segment, Chr 8, ERATO Doi 82, expressed | protein-coding |
| chr15_103152776_103153307 | chr15 | 103152777 | 103153307 | + | 1.03811 | TTS (NM_027885) | TTS (NM_027885) | -6214 | NR_045743 | 432989 | Mm.383979 | NR_045743 | ENSMUSG00000097003 | **D930007P13Rik** | - | Riken cDNA D930007P13 gene | ncRNA |
| chr15_103152973_103153260 | chr15 | 103152974 | 103153260 | + | 1.03461 | TTS (NM_027885) | TTS (NM_027885) | -6289 | NR_045743 | 432989 | Mm.383979 | NR_045743 | ENSMUSG00000097003 | **D930007P13Rik** | - | Riken cDNA D930007P13 gene | ncRNA |
| chr3_83987764_83988073 | chr3 | 83987765 | 83988073 | + | 1.13276 | intron (NM_172681, intron 3 of 34) | intron (NM_172681, intron 3 of 34) | 52242 | NM_172681 | 229473 | Mm.28838 | NM_172681 | ENSMUSG00000033767 | **D930015E06Rik** | Kiaa0922\|mKIAA0922 | RIKEN cDNA D930015E06 gene | protein-coding |
| chr3_83987572_83988746 | chr3 | 83987573 | 83988746 | + | 1.11695 | intron (NM_172681, intron 3 of 34) | intron (NM_172681, intron 3 of 34) | 52002 | NM_172681 | 229473 | Mm.28838 | NM_172681 | ENSMUSG00000033767 | **D930015E06Rik** | Kiaa0922\|mKIAA0922 | RIKEN cDNA D930015E06 gene | protein-coding |
| chr3_84061896_84062517 | chr3 | 84061897 | 84062517 | + | 1.10807 | Intergenic | L2a\|LINE\|L2 | -22046 | NM_172681 | 229473 | Mm.28838 | NM_172681 | ENSMUSG00000033767 | **D930015E06Rik** | Kiaa0922\|mKIAA0922 | RIKEN cDNA D930015E06 gene | protein-coding |
| chr3_83987610_83988188 | chr3 | 83987611 | 83988188 | + | 1.0497 | intron (NM_172681, intron 3 of 34) | intron (NM_172681, intron 3 of 34) | 52262 | NM_172681 | 229473 | Mm.28838 | NM_172681 | ENSMUSG00000033767 | **D930015E06Rik** | Kiaa0922\|mKIAA0922 | RIKEN cDNA D930015E06 gene | protein-coding |
| chr3_84004403_84004967 | chr3 | 84004404 | 84004967 | + | 0.983103 | intron (NM_172681, intron 3 of 34) | intron (NM_172681, intron 3 of 34) | 35476 | NM_172681 | 229473 | Mm.28838 | NM_172681 | ENSMUSG00000033767 | **D930015E06Rik** | Kiaa0922\|mKIAA0922 | RIKEN cDNA D930015E06 gene | protein-coding |
| chr3_84022632_84023341 | chr3 | 84022633 | 84023341 | + | 0.92521 | intron (NM_172681, intron 3 of 34) | intron (NM_172681, intron 3 of 34) | 17174 | NM_172681 | 229473 | Mm.28838 | NM_172681 | ENSMUSG00000033767 | **D930015E06Rik** | Kiaa0922\|mKIAA0922 | RIKEN cDNA D930015E06 gene | protein-coding |
| chr3_84068953_84069781 | chr3 | 84068954 | 84069781 | + | 0.791931 | Intergenic | Intergenic | -29206 | NM_172681 | 229473 | Mm.28838 | NM_172681 | ENSMUSG00000033767 | **D930015E06Rik** | Kiaa0922\|mKIAA0922 | RIKEN cDNA D930015E06 gene | protein-coding |
| chr3_84068909_84070097 | chr3 | 84068910 | 84070097 | + | 0.784005 | Intergenic | Intergenic | -29342 | NM_172681 | 229473 | Mm.28838 | NM_172681 | ENSMUSG00000033767 | **D930015E06Rik** | Kiaa0922\|mKIAA0922 | RIKEN cDNA D930015E06 gene | protein-coding |
| chr3_84008668_84008918 | chr3 | 84008669 | 84008918 | + | 0.761545 | intron (NM_172681, intron 3 of 34) | intron (NM_172681, intron 3 of 34) | 31368 | NM_172681 | 229473 | Mm.28838 | NM_172681 | ENSMUSG00000033767 | **D930015E06Rik** | Kiaa0922\|mKIAA0922 | RIKEN cDNA D930015E06 gene | protein-coding |
| chr10_121640000_121640785 | chr10 | 121640001 | 121640785 | + | 0.91308 | Intergenic | Intergenic | -1308 | NM_177335 | 216393 | Mm.119759 | NM_177335 | | **D930020B18Rik** | 1500017I17Rik | RIKEN cDNA D930020B18 gene | protein-coding |
| chr15_5929093_5929896 | chr15 | 5929094 | 5929896 | + | 1.00306 | Intergenic | Intergenic | -370294 | NM_001008702 | 13132 | Mm.240830 | NM_023118 | ENSMUSG00000022150 | **Dab2** | 5730435J12Rik\|AA960054\|AI957090\|D15Wsu122e\|D630005B22Rik\|Doc-2\|Doc2\|p96 | disabled 2, mitogen-responsive phosphoprotein | protein-coding |
| chr15_5927822_5928317 | chr15 | 5927823 | 5928317 | + | 0.708433 | Intergenic | Intergenic | -371719 | NM_001008702 | 13132 | Mm.240830 | NM_023118 | ENSMUSG00000022150 | **Dab2** | 5730435J12Rik\|AA960054\|AI957090\|D15Wsu122e\|D630005B22Rik\|Doc-2\|Doc2\|p96 | disabled 2, mitogen-responsive phosphoprotein | protein-coding |
| chr14_97356968_97357362 | chr14 | 97356969 | 97357362 | + | 0.836935 | Intergenic | RMER6C\|LTR\|ERVK | 812600 | NM_001038610 | 13134 | Mm.320593 | NM_007826 | ENSMUSG00000055639 | **Dach1** | Dac\|Dach | dachshund 1 (Drosophila) | protein-coding |
| chrX_113732589_113733014 | chrX | 113732590 | 113733014 | + | 0.881099 | intron (NM_001289734, intron 8 of 15) | RMER15-int\|LTR\|ERVL | 434546 | NM_001142570 | 93837 | Mm.79760 | NM_033605 | ENSMUSG00000025592 | **Dach2** | 9430028N04Rik | dachshund 2 (Drosophila) | protein-coding |
| chrX_113682353_113682761 | chrX | 113682354 | 113682761 | + | 0.706008 | intron (NM_001289734, intron 6 of 15) | ORR1A1\|LTR\|ERVL-MaLR | 384301 | NM_001142570 | 93837 | Mm.79760 | NM_033605 | ENSMUSG00000025592 | **Dach2** | 9430028N04Rik | dachshund 2 (Drosophila) | protein-coding |
| chr14_54179337_54179609 | chr14 | 54179338 | 54179609 | + | 1.08341 | Intergenic | Intergenic | 74456 | NM_001113358 | 13135 | Mm.319038 | NM_010015 | ENSMUSG00000022174 | **Dad1** | AI323713 | defender against cell death 1 | protein-coding |
| chr14_54043848_54044213 | chr14 | 54043849 | 54044213 | + | 0.795402 | Intergenic | Intergenic | 209898 | NM_001113358 | 13135 | Mm.319038 | NM_010015 | ENSMUSG00000022174 | **Dad1** | AI323713 | defender against cell death 1 | protein-coding |
| chr14_53973429_53973964 | chr14 | 53973430 | 53973964 | + | 0.719117 | Intergenic | Intergenic | 280232 | NM_001113358 | 13135 | Mm.319038 | NM_010015 | ENSMUSG00000022174 | **Dad1** | AI323713 | defender against cell death 1 | protein-coding |
| chr14_54178946_54179609 | chr14 | 54178947 | 54179609 | + | 0.64013 | Intergenic | Intergenic | 74651 | NM_001113358 | 13135 | Mm.319038 | NM_010015 | ENSMUSG00000022174 | **Dad1** | AI323713 | defender against cell death 1 | protein-coding |
| chr15_31239719_31240462 | chr15 | 31239720 | 31240462 | + | 0.828415 | intron (NM_146057, intron 2 of 3) | intron (NM_146057, intron 2 of 3) | 15706 | NM_146057 | 223453 | Mm.222867 | NM_146057 | ENSMUSG00000039168 | **Dap** | 4921531N22Rik\|AI789753\|AU040360 | death-associated protein | protein-coding |
| chr15_31187393_31187865 | chr15 | 31187394 | 31187865 | + | 0.810361 | Intergenic | Intergenic | -36756 | NM_146057 | 223453 | Mm.222867 | NM_146057 | ENSMUSG00000039168 | **Dap** | 4921531N22Rik\|AI789753\|AU040360 | death-associated protein | protein-coding |
| chr15_31226601_31226782 | chr15 | 31226602 | 31226782 | + | 0.703415 | intron (NM_146057, intron 1 of 3) | intron (NM_146057, intron 1 of 3) | 2307 | NM_146057 | 223453 | Mm.222867 | NM_146057 | ENSMUSG00000039168 | **Dap** | 4921531N22Rik\|AI789753\|AU040360 | death-associated protein | protein-coding |
| chr9_66142159_66142839 | chr9 | 66142160 | 66142839 | + | 1.14052 | Intergenic | MTC\|LTR\|ERVL-MaLR | -15727 | NM_010019 | 13143 | Mm.304472 | NM_010019 | ENSMUSG00000032380 | **Dapk2** | - | death-associated protein kinase 2 | protein-coding |
| chr2_59489818_59490351 | chr2 | 59489819 | 59490351 | + | 1.27124 | intron (NM_029723, intron 1 of 3) | intron (NM_029723, intron 1 of 3) | 5432 | NM_029723 | 76747 | Mm.275769 | NM_029723 | ENSMUSG00000026989 | **Dapl1** | 2310032F03Rik\|EEDA | death associated protein-like 1 | protein-coding |
| chr2_59473721_59474306 | chr2 | 59473722 | 59474306 | + | 0.919884 | Intergenic | Intergenic | -10639 | NM_029723 | 76747 | Mm.275769 | NM_029723 | ENSMUSG00000026989 | **Dapl1** | 2310032F03Rik\|EEDA | death associated protein-like 1 | protein-coding |
| chr2_59489806_59490668 | chr2 | 59489807 | 59490668 | + | 0.909326 | intron (NM_029723, intron 1 of 3) | intron (NM_029723, intron 1 of 3) | 5584 | NM_029723 | 76747 | Mm.275769 | NM_029723 | ENSMUSG00000026989 | **Dapl1** | 2310032F03Rik\|EEDA | death associated protein-like 1 | protein-coding |
| chr2_59513579_59513914 | chr2 | 59513580 | 59513914 | + | 0.796412 | Intergenic | Intergenic | 29094 | NM_029723 | 76747 | Mm.275769 | NM_029723 | ENSMUSG00000026989 | **Dapl1** | 2310032F03Rik\|EEDA | death associated protein-like 1 | protein-coding |
| chr2_59513541_59514098 | chr2 | 59513542 | 59514098 | + | 0.76279 | Intergenic | Intergenic | 29167 | NM_029723 | 76747 | Mm.275769 | NM_029723 | ENSMUSG00000026989 | **Dapl1** | 2310032F03Rik\|EEDA | death associated protein-like 1 | protein-coding |
| chr2_59534195_59534839 | chr2 | 59534196 | 59534839 | + | 0.734062 | Intergenic | Intergenic | 49864 | NM_029723 | 76747 | Mm.275769 | NM_029723 | ENSMUSG00000026989 | **Dapl1** | 2310032F03Rik\|EEDA | death associated protein-like 1 | protein-coding |
| chr3_137959536_137959831 | chr3 | 137959537 | 137959831 | + | 1.48751 | intron (NM_011932, intron 3 of 8) | intron (NM_011932, intron 3 of 8) | 21865 | NM_011932 | 26377 | Mm.254835 | NM_011932 | ENSMUSG00000028159 | **Dapp1** | Bam32 | dual adaptor for phosphotyrosine and 3-phosphoinositides 1 | protein-coding |
| chr3_137954229_137954778 | chr3 | 137954230 | 137954778 | + | 1.05556 | intron (NM_011932, intron 3 of 8) | MIR\|SINE\|MIR | 27045 | NM_011932 | 26377 | Mm.254835 | NM_011932 | ENSMUSG00000028159 | **Dapp1** | Bam32 | dual adaptor for phosphotyrosine and 3-phosphoinositides 1 | protein-coding |
| chr3_137952446_137952722 | chr3 | 137952447 | 137952722 | + | 0.997709 | intron (NM_011932, intron 3 of 8) | B3A\|SINE\|B2 | 28965 | NM_011932 | 26377 | Mm.254835 | NM_011932 | ENSMUSG00000028159 | **Dapp1** | Bam32 | dual adaptor for phosphotyrosine and 3-phosphoinositides 1 | protein-coding |
| chr3_137973407_137973648 | chr3 | 137973408 | 137973648 | + | 0.945506 | intron (NM_011932, intron 1 of 8) | intron (NM_011932, intron 1 of 8) | 8021 | NM_011932 | 26377 | Mm.254835 | NM_011932 | ENSMUSG00000028159 | **Dapp1** | Bam32 | dual adaptor for phosphotyrosine and 3-phosphoinositides 1 | protein-coding |
| chr3_138024100_138024561 | chr3 | 138024101 | 138024561 | + | 0.702909 | Intergenic | MTC\|LTR\|ERVL-MaLR | -42782 | NM_011932 | 26377 | Mm.254835 | NM_011932 | ENSMUSG00000028159 | **Dapp1** | Bam32 | dual adaptor for phosphotyrosine and 3-phosphoinositides 1 | protein-coding |
| chr3_138001640_138002201 | chr3 | 138001641 | 138002201 | + | 0.685249 | Intergenic | Intergenic | -20372 | NM_011932 | 26377 | Mm.254835 | NM_011932 | ENSMUSG00000028159 | **Dapp1** | Bam32 | dual adaptor for phosphotyrosine and 3-phosphoinositides 1 | protein-coding |
| chr12_74333058_74333348 | chr12 | 74333059 | 74333348 | + | 0.691559 | Intergenic | Intergenic | 35729 | NM_198866 | 386753 | Mm.341686 | NM_198866 | ENSMUSG00000029878 | **Dbpht2** | Spot2 | DNA binding protein with his-thr domain | protein-coding |
| chr3_116512091_116512435 | chr3 | 116512092 | 116512435 | + | 1.33233 | promoter-TSS (NM_010022) | promoter-TSS (NM_010022) | -816 | NM_010022 | 13171 | Mm.3636 | NM_010022 | ENSMUSG00000000340 | **Dbt** | D3Wsu60e | dihydrolipoamide branched chain transacylase E2 | protein-coding |
| chr7_49674961_49675569 | chr7 | 49674962 | 49675569 | + | 1.12831 | Intergenic | Intergenic | -38430 | NM_001005232 | 13172 | Mm.307176 | NM_001005232 | ENSMUSG00000030507 | **Dbx1** | AI426026\|Dbx | developing brain homeobox 1 | protein-coding |
| chr7_49472965_49473357 | chr7 | 49472966 | 49473357 | + | 0.693805 | intron (NM_001111016, intron 11 of 38) | intron (NM_001111016, intron 11 of 38) | 163674 | NM_001005232 | 13172 | Mm.307176 | NM_001005232 | ENSMUSG00000030507 | **Dbx1** | AI426026\|Dbx | developing brain homeobox 1 | protein-coding |
| chr4_41307435_41307828 | chr4 | 41307436 | 41307828 | + | 0.952202 | intron (NM_026893, intron 2 of 8) | B1_Mm\|SINE\|Alu | 7269 | NM_026893 | 68970 | Mm.28891 | NM_026893 | ENSMUSG00000028436 | **Dcaf12** | 1500001L20Rik\|5830424K06Rik\|AA420338\|AI851081\|Wdr40a | DDB1 and CUL4 associated factor 12 | protein-coding |
| chr8_84106686_84106951 | chr8 | 84106687 | 84106951 | + | 0.645138 | Intergenic | Intergenic | -2057 | NM_172502 | 212123 | Mm.260557 | NM_172502 | ENSMUSG00000037103 | **Dcaf15** | 6720484B16 | DDB1 and CUL4 associated factor 15 | protein-coding |
| chr1_165432945_165433228 | chr1 | 165432946 | 165433228 | + | 1.13163 | intron (NM_028759, intron 1 of 18) | RMER3D-int\|LTR\|ERVK | 27376 | NM_028759 | 74106 | Mm.227605 | NM_028759 | ENSMUSG00000026571 | **Dcaf6** | 1200006M05Rik\|Iqwd1\|PC326 | DDB1 and CUL4 associated factor 6 | protein-coding |
| chr1_172158876_172159298 | chr1 | 172158877 | 172159298 | + | 1.06851 | intron (NM_153555, intron 2 of 13) | L1ME1\|LINE\|L1 | 11072 | NM_153555 | 98193 | Mm.159915 | NM_153555 | ENSMUSG00000026554 | **Dcaf8** | AA408877\|AA433120\|AA792214\|AU018973\|C76055\|D1Dau35e\|D1Ucla4\|H326\|Wdr42a | DDB1 and CUL4 associated factor 8 | protein-coding |
| chr3_55461545_55462224 | chr3 | 55461546 | 55462224 | + | 1.44903 | 5' UTR (NM_001195540, exon 1 of 3) | 5' UTR (NM_001195540, exon 1 of 3) | 127 | NM_001195540 | 13175 | Mm.393242 | NM_019978 | ENSMUSG00000027797 | **Dclk1** | 1700113D08Rik\|2810480F11Rik\|AI836758\|Click-I\|Cpg16\|Dcamkl1\|Dcl\|Dclk\|mKIAA0369 | doublecortin-like kinase 1 | protein-coding |
| chr3_55438299_55439489 | chr3 | 55438300 | 55439489 | + | 1.25571 | intron (NM_019978, intron 4 of 16) | intron (NM_019978, intron 4 of 16) | -22864 | NM_001195539 | 13175 | Mm.393242 | NM_019978 | ENSMUSG00000027797 | **Dclk1** | 1700113D08Rik\|2810480F11Rik\|AI836758\|Click-I\|Cpg16\|Dcamkl1\|Dcl\|Dclk\|mKIAA0369 | doublecortin-like kinase 1 | protein-coding |
| chr3_55419248_55419845 | chr3 | 55419249 | 55419845 | + | 0.90967 | intron (NM_019978, intron 4 of 16) | intron (NM_019978, intron 4 of 16) | -42211 | NM_001195539 | 13175 | Mm.393242 | NM_019978 | ENSMUSG00000027797 | **Dclk1** | 1700113D08Rik\|2810480F11Rik\|AI836758\|Click-I\|Cpg16\|Dcamkl1\|Dcl\|Dclk\|mKIAA0369 | doublecortin-like kinase 1 | protein-coding |
| chr3_86756377_86757027 | chr3 | 86756378 | 86757027 | + | 1.1572 | intron (NM_030695, intron 52 of 56) | intron (NM_030695, intron 52 of 56) | 164182 | NM_001195496 | 70762 | Mm.44490 | NM_027539 | ENSMUSG00000028078 | **Dclk2** | 6330415M09Rik\|AU044875\|CL2\|CLICK2\|Click-II\|Dcamkl2 | doublecortin-like kinase 2 | protein-coding |

| | | | | | | | | | | | | | | | | | |
|---|---|---|---|---|---|---|---|---|---|---|---|---|---|---|---|---|---|
| chr3_86837089_86837851 | chr3 | 86837090 | 86837851 | + | 1.11779 | intron (NM_001195497, intron 2 of 15) | intron (NM_001195497, intron 2 of 15) | 83414 | NM_001195496 | 70762 | Mm.44490 | NM_027539 | ENSMUSG00000028078 | **Dclk2** | 6330415M09Rik\|AU044875\|CL2\|CLICK2\|Click-II\|Dcamkl2 | doublecortin-like kinase 2 | protein-coding |
| chr3_86839602_86839985 | chr3 | 86839603 | 86839985 | + | 0.960435 | intron (NM_001195497, intron 2 of 15) | intron (NM_001195497, intron 2 of 15) | 81090 | NM_001195496 | 70762 | Mm.44490 | NM_027539 | ENSMUSG00000028078 | **Dclk2** | 6330415M09Rik\|AU044875\|CL2\|CLICK2\|Click-II\|Dcamkl2 | doublecortin-like kinase 2 | protein-coding |
| chr3_86868606_86869143 | chr3 | 86868607 | 86869143 | + | 0.778059 | intron (NM_001195497, intron 2 of 15) | intron (NM_001195497, intron 2 of 15) | 52009 | NM_001195496 | 70762 | Mm.44490 | NM_027539 | ENSMUSG00000028078 | **Dclk2** | 6330415M09Rik\|AU044875\|CL2\|CLICK2\|Click-II\|Dcamkl2 | doublecortin-like kinase 2 | protein-coding |
| chr9_111383546_111384363 | chr9 | 111383547 | 111384363 | + | 1.49827 | intron (NM_001164659, intron 16 of 21) | intron (NM_001164659, intron 16 of 21) | -55126 | NM_172928 | 245038 | Mm.26361 | NM_172928 | ENSMUSG00000032500 | **Dclk3** | BC056929\|C730036H08\|Dcamkl3 | doublecortin-like kinase 3 | protein-coding |
| chr14_30467157_30468008 | chr14 | 30467158 | 30468008 | + | 1.44682 | intron (NM_028981, intron 1 of 46) | intron (NM_028981, intron 1 of 46) | -11982 | NM_133761 | 75901 | Mm.28733 | NM_133761 | ENSMUSG00000021962 | **Dcp1a** | 1110066A22Rik\|4930568L04Rik\|AU019772\|D14Ertd817e\|Mitc1\|SMIF | DCP1 decapping enzyme homolog A (S. cerevisiae) | protein-coding |
| chr9_35173154_35173802 | chr9 | 35173155 | 35173802 | + | 1.11255 | intron (NM_027030, intron 1 of 5) | B4A\|SINE\|B4 | 2509 | NM_027030 | 69305 | Mm.229110 | NM_027030 | ENSMUSG00000032040 | **Dcps** | 1700001E16Rik\|AA408441 | decapping enzyme, scavenger | protein-coding |
| chr9_35153002_35153390 | chr9 | 35153003 | 35153390 | + | 0.636661 | intron (NM_027030, intron 2 of 5) | Lx5c\|LINE\|L1 | 22791 | NM_027030 | 69305 | Mm.229110 | NM_027030 | ENSMUSG00000032040 | **Dcps** | 1700001E16Rik\|AA408441 | decapping enzyme, scavenger | protein-coding |
| chr3_35898705_35899303 | chr3 | 35898706 | 35899303 | + | 0.720092 | intron (NM_033623, intron 4 of 6) | intron (NM_033623, intron 4 of 6) | 31070 | NM_033623 | 114893 | Mm.374810 | NM_033623 | ENSMUSG00000027708 | **Dcun1d1** | Rp42\|SCCRO\|Tes3\|pTes3 | DCN1, defective in cullin neddylation 1, domain containing 1 (S. cerevisiae) | protein-coding |
| chr8_13293555_13294041 | chr8 | 13293556 | 13294041 | + | 0.902032 | intron (NM_172282, intron 3 of 12) | intron (NM_172282, intron 3 of 12) | -5672 | NM_001042650 | 102323 | Mm.368890 | NM_001024504 | ENSMUSG00000038506 | **Dcun1d2** | AW495713\|mFLJ10704 | DCN1, defective in cullin neddylation 1, domain containing 2 (S. cerevisiae) | protein-coding |
| chr7_119871636_119872582 | chr7 | 119871637 | 119872582 | + | 1.37099 | intron (NM_001163703, intron 1 of 2) | intron (NM_001163703, intron 1 of 2) | 23349 | NM_173408 | 233805 | Mm.31539 | NM_173408 | ENSMUSG00000048787 | **Dcun1d3** | 1700020A13Rik\|BC030335 | DCN1, defective in cullin neddylation 1, domain containing 3 (S. cerevisiae) | protein-coding |
| chr7_119854805_119855364 | chr7 | 119854806 | 119855364 | + | 1.21638 | 3' UTR (NM_173408, exon 3 of 3) | 3' UTR (NM_173408, exon 3 of 3) | 40373 | NM_173408 | 233805 | Mm.31539 | NM_173408 | ENSMUSG00000048787 | **Dcun1d3** | 1700020A13Rik\|BC030335 | DCN1, defective in cullin neddylation 1, domain containing 3 (S. cerevisiae) | protein-coding |
| chr7_119873912_119874219 | chr7 | 119873913 | 119874219 | + | 1.20208 | intron (NM_001163703, intron 1 of 2) | ORR1D1\|LTR\|ERVL-MaLR | 21392 | NM_173408 | 233805 | Mm.31539 | NM_173408 | ENSMUSG00000048787 | **Dcun1d3** | 1700020A13Rik\|BC030335 | DCN1, defective in cullin neddylation 1, domain containing 3 (S. cerevisiae) | protein-coding |
| chr9_7220928_7222096 | chr9 | 7220929 | 7222096 | + | 0.667999 | Intergenic | Intergenic | 36946 | NM_029775 | 76863 | Mm.27293 | NM_029775 | ENSMUSG00000032002 | **Dcun1d5** | 3110001A18Rik\|4833420K19Rik\|AW060460\|D430047L21Rik | DCN1, defective in cullin neddylation 1, domain containing 5 (S. cerevisiae) | protein-coding |
| chr3_145742747_145743430 | chr3 | 145742748 | 145743430 | + | 0.89108 | Intergenic | Intergenic | -15603 | NM_026993 | 69219 | Mm.234247 | NM_026993 | ENSMUSG00000028194 | **Ddah1** | 2410006N07Rik\|2510015N06Rik\|AI987801\|AW050362 | dimethylarginine dimethylaminohydrolase 1 | protein-coding |
| chr11_11960841_11961307 | chr11 | 11960842 | 11961307 | + | 0.910952 | intron (NM_010345, intron 4 of 16) | intron (NM_010345, intron 4 of 16) | -62930 | NM_001190448 | 13195 | Mm.12906 | NM_016672 | ENSMUSG00000020182 | **Ddc** | Aadc | dopa decarboxylase | protein-coding |
| chr14_45764335_45764655 | chr14 | 45764336 | 45764655 | + | 0.890964 | Intergenic | RLTR13D6\|LTR\|ERVK | -106352 | NM_001042719 | 114874 | Mm.121918 | NM_176845 | ENSMUSG00000037697 | **Ddhd1** | PA-PLA1 | DDHD domain containing 1 | protein-coding |
| chr7_92837253_92837746 | chr7 | 92837254 | 92837746 | + | 0.64557 | TTS (NM_029494) | TTS (NM_029494) | 36732 | NM_001080995 | 74041 | Mm.26468 | NM_028729 | ENSMUSG00000030641 | **Ddias** | 4632434I11Rik\|noxin | DNA damage-induced apoptosis suppressor | protein-coding |
| chr10_127294245_127294463 | chr10 | 127294246 | 127294463 | + | 0.894718 | intron (NM_007837, intron 2 of 3) | intron (NM_007837, intron 2 of 3) | 3561 | NM_001290183 | 13198 | Mm.110220 | NM_007837 | ENSMUSG00000025408 | **Ddit3** | CHOP-10\|CHOP10\|chop\|gadd153 | DNA-damage inducible transcript 3 | protein-coding |
| chr17_35707840_35708312 | chr17 | 35707841 | 35708312 | + | 1.06301 | Intergenic | Charlie29a\|DNA\|hAT-Charlie | -3937 | NM_172962 | 12305 | Mm.5021 | NM_007584 | ENSMUSG00000003534 | **Ddr1** | 6030432F18\|AI323681\|CD167a\|Cak\|Nep\|PTK3A | discoidin domain receptor family, member 1 | protein-coding |
| chr17_35705930_35706376 | chr17 | 35705931 | 35706376 | + | 0.713417 | Intergenic | Intergenic | -2014 | NM_172962 | 12305 | Mm.5021 | NM_007584 | ENSMUSG00000003534 | **Ddr1** | 6030432F18\|AI323681\|CD167a\|Cak\|Nep\|PTK3A | discoidin domain receptor family, member 1 | protein-coding |
| chr17_35697980_35698342 | chr17 | 35697981 | 35698342 | + | 0.687947 | intron (NM_007584, intron 1 of 17) | intron (NM_007584, intron 1 of 17) | 2429 | NM_001198833 | 12305 | Mm.5021 | NM_007584 | ENSMUSG00000003534 | **Ddr1** | 6030432F18\|AI323681\|CD167a\|Cak\|Nep\|PTK3A | discoidin domain receptor family, member 1 | protein-coding |
| chr9_53154428_53154881 | chr9 | 53154429 | 53154881 | + | 1.02003 | intron (NM_029936, intron 15 of 17) | intron (NM_029936, intron 15 of 17) | 93457 | NM_029936 | 77591 | Mm.133560 | NM_029936 | ENSMUSG00000053289 | **Ddx10** | 4632415A01Rik\|AI646054 | DEAD (Asp-Glu-Ala-Asp) box polypeptide 10 | protein-coding |
| chr9_53240560_53241180 | chr9 | 53240561 | 53241180 | + | 1.00572 | intron (NM_029936, intron 1 of 17) | RMER17D2\|LTR\|ERVK | 7242 | NM_029936 | 77591 | Mm.133560 | NM_029936 | ENSMUSG00000053289 | **Ddx10** | 4632415A01Rik\|AI646054 | DEAD (Asp-Glu-Ala-Asp) box polypeptide 10 | protein-coding |
| chr9_53132871_53133295 | chr9 | 53132872 | 53133295 | + | 0.661253 | intron (NM_029936, intron 16 of 17) | intron (NM_029936, intron 16 of 17) | 115029 | NM_029936 | 77591 | Mm.133560 | NM_029936 | ENSMUSG00000053289 | **Ddx10** | 4632415A01Rik\|AI646054 | DEAD (Asp-Glu-Ala-Asp) box polypeptide 10 | protein-coding |
| chr17_66136714_66137428 | chr17 | 66136715 | 66137428 | + | 1.31212 | intron (NM_001003919, intron 9 of 26) | intron (NM_001003919, intron 9 of 26) | 13551 | NM_001003919 | 320209 | Mm.259605 | NM_001003919 | ENSMUSG00000035842 | **Ddx11** | 4732462I11Rik\|CHL1\|CHLR1\|KRG2\|essa15a | DEAD/H (Asp-Glu-Ala-Asp/His) box helicase 11 | protein-coding |
| chr3_105627188_105628091 | chr3 | 105627189 | 105628091 | + | 1.25371 | intron (NM_001039347, intron 2 of 6) | ORR1E\|LTR\|ERVL-MaLR | 59931 | NM_017397 | 53975 | Mm.272826 | NM_017397 | ENSMUSG00000027905 | **Ddx20** | GEMIN3\|dp103 | DEAD (Asp-Glu-Ala-Asp) box polypeptide 20 | protein-coding |
| chr3_105672077_105672694 | chr3 | 105672078 | 105672694 | + | 0.866809 | 3' UTR (NM_001039347, exon 7 of 7) | 3' UTR (NM_001039347, exon 7 of 7) | 15185 | NM_017397 | 53975 | Mm.272826 | NM_017397 | ENSMUSG00000027905 | **Ddx20** | GEMIN3\|dp103 | DEAD (Asp-Glu-Ala-Asp) box polypeptide 20 | protein-coding |
| chr3_105671692_105672696 | chr3 | 105671693 | 105672696 | + | 0.840188 | 3' UTR (NM_001039347, exon 7 of 7) | 3' UTR (NM_001039347, exon 7 of 7) | 15377 | NM_017397 | 53975 | Mm.272826 | NM_017397 | ENSMUSG00000027905 | **Ddx20** | GEMIN3\|dp103 | DEAD (Asp-Glu-Ala-Asp) box polypeptide 20 | protein-coding |
| chr3_85022766_85023342 | chr3 | 85022767 | 85023342 | + | 0.910659 | Intergenic | Intergenic | -57474 | NM_001040461 | 654362 | Mm.466598 | NM_001040461 | | **Dear1** | Dear\|Despr | dual endothelin 1/angiotensin II receptor 1 | protein-coding |
| chr17_28192573_28193210 | chr17 | 28192574 | 28193210 | + | 1.00389 | intron (NM_172617, intron 3 of 14) | intron (NM_172617, intron 3 of 14) | -14886 | NM_027185 | 23853 | Mm.204731 | NM_027185 | ENSMUSG00000002257 | **Def6** | 2410003F05Rik\|6430538D02Rik\|AV094905\|Ibp\|Slat\|Slat2\|Slat6 | differentially expressed in FDCP 6 | protein-coding |
| chr17_28193529_28194033 | chr17 | 28193530 | 28194033 | + | 0.801012 | intron (NM_172617, intron 3 of 14) | B4\|SINE\|B4 | -13997 | NM_027185 | 23853 | Mm.204731 | NM_027185 | ENSMUSG00000002257 | **Def6** | 2410003F05Rik\|6430538D02Rik\|AV094905\|Ibp\|Slat\|Slat2\|Slat6 | differentially expressed in FDCP 6 | protein-coding |
| chr17_28193611_28193877 | chr17 | 28193612 | 28193877 | + | 0.752956 | intron (NM_172617, intron 3 of 14) | intron (NM_172617, intron 3 of 14) | -14034 | NM_027185 | 23853 | Mm.204731 | NM_027185 | ENSMUSG00000002257 | **Def6** | 2410003F05Rik\|6430538D02Rik\|AV094905\|Ibp\|Slat\|Slat2\|Slat6 | differentially expressed in FDCP 6 | protein-coding |
| chr2_152480645_152481264 | chr2 | 152480646 | 152481264 | + | 0.822309 | Intergenic | Intergenic | 3892 | NM_176950 | 319579 | Mm.190454 | NM_176950 | ENSMUSG00000049560 | **Defb20** | 9230107O10Rik\|BD-20\|mBD-20 | defensin beta 20 | protein-coding |
| chr2_152480837_152481237 | chr2 | 152480838 | 152481237 | + | 0.815726 | Intergenic | Intergenic | 3974 | NM_176950 | 319579 | Mm.190454 | NM_176950 | ENSMUSG00000049560 | **Defb20** | 9230107O10Rik\|BD-20\|mBD-20 | defensin beta 20 | protein-coding |
| chr2_152598626_152598959 | chr2 | 152598627 | 152598959 | + | 0.832078 | Intergenic | Intergenic | -2308 | NM_001037752 | 433490 | Mm.23779 | NM_001037752 | ENSMUSG00000062124 | **Defb45** | Defb27\|OTTMUSG00000015859 | defensin beta 45 | protein-coding |
| chr13_47104825_47105303 | chr13 | 47104826 | 47105303 | + | 0.976413 | intron (NM_025900, intron 2 of 10) | intron (NM_025900, intron 2 of 10) | 1157 | NM_025900 | 110052 | Mm.131150 | NM_025900 | ENSMUSG00000021377 | **Dek** | 1810019E15Rik\|D13H6S231E | DEK oncogene (DNA binding) | protein-coding |
| chr2_38282143_38282494 | chr2 | 38282144 | 38282494 | + | 0.995829 | intron (NM_146122, intron 1 of 21) | L2a\|LINE\|L2 | 5065 | NM_146122 | 227801 | Mm.314245 | NM_146122 | ENSMUSG00000035392 | **Dennd1a** | 6030446I19Rik\|connecdenn | DENN/MADD domain containing 1A | protein-coding |
| chr2_38154496_38154848 | chr2 | 38154497 | 38154848 | + | 0.883401 | intron (NM_146122, intron 3 of 21) | intron (NM_146122, intron 3 of 21) | 132712 | NM_146122 | 227801 | Mm.314245 | NM_146122 | ENSMUSG00000035392 | **Dennd1a** | 6030446I19Rik\|connecdenn | DENN/MADD domain containing 1A | protein-coding |
| chr2_38293361_38294015 | chr2 | 38293362 | 38294015 | + | 0.723924 | Intergenic | Intergenic | -6304 | NM_146122 | 227801 | Mm.314245 | NM_146122 | ENSMUSG00000035392 | **Dennd1a** | 6030446I19Rik\|connecdenn | DENN/MADD domain containing 1A | protein-coding |
| chr2_38191265_38191686 | chr2 | 38191266 | 38191686 | + | 0.675193 | intron (NM_146122, intron 2 of 21) | intron (NM_146122, intron 2 of 21) | 95908 | NM_146122 | 227801 | Mm.314245 | NM_146122 | ENSMUSG00000035392 | **Dennd1a** | 6030446I19Rik\|connecdenn | DENN/MADD domain containing 1A | protein-coding |
| chr2_38090812_38091234 | chr2 | 38090813 | 38091234 | + | 0.63401 | intron (NM_146122, intron 5 of 21) | intron (NM_146122, intron 5 of 21) | 196361 | NM_146122 | 227801 | Mm.314245 | NM_146122 | ENSMUSG00000035392 | **Dennd1a** | 6030446I19Rik\|connecdenn | DENN/MADD domain containing 1A | protein-coding |
| chr1_138939683_138940164 | chr1 | 138939684 | 138940164 | + | 1.51361 | Intergenic | MIRb\|SINE\|MIR | -23785 | NM_001166501 | 329260 | Mm.337604 | NM_181347 | ENSMUSG00000056268 | **Dennd1b** | 4632404N19Rik\|4930467M19Rik\|6820401H01Rik\|F730008N07Rik | DENN/MADD domain containing 1B | protein-coding |
| chr1_138933361_138933693 | chr1 | 138933362 | 138933693 | + | 1.40641 | Intergenic | Intergenic | -30182 | NM_001166501 | 329260 | Mm.337604 | NM_181347 | ENSMUSG00000056268 | **Dennd1b** | 4632404N19Rik\|4930467M19Rik\|6820401H01Rik\|F730008N07Rik | DENN/MADD domain containing 1B | protein-coding |
| chr1_138930786_138931244 | chr1 | 138930787 | 138931244 | + | 0.942639 | Intergenic | ORR1F\|LTR\|ERVL-MaLR | -32694 | NM_001166501 | 329260 | Mm.337604 | NM_181347 | ENSMUSG00000056268 | **Dennd1b** | 4632404N19Rik\|4930467M19Rik\|6820401H01Rik\|F730008N07Rik | DENN/MADD domain containing 1B | protein-coding |
| chr17_57079848_57080368 | chr17 | 57079849 | 57080368 | + | 0.87351 | TTS (NM_009451) | TTS (NM_009451) | -1598 | NM_153551 | 70785 | Mm.284447 | NM_153551 | ENSMUSG00000002668 | **Dennd1c** | 4432409M07Rik | DENN/MADD domain containing 1C | protein-coding |
| chr3_103146995_103147390 | chr3 | 103146996 | 103147390 | + | 0.787557 | intron (NM_177857, intron 7 of 18) | intron (NM_177857, intron 7 of 18) | 19637 | NM_177857 | 329727 | Mm.293641 | NM_177857 | ENSMUSG00000007379 | **Dennd2c** | A930010I20Rik | DENN/MADD domain containing 2C | protein-coding |
| chr15_73514055_73514304 | chr15 | 73514056 | 73514304 | + | 0.897764 | intron (NM_001081066, intron 1 of 22) | intron (NM_001081066, intron 1 of 22) | 1620 | NM_001081066 | 105841 | Mm.322557 | NM_001081066 | ENSMUSG00000036661 | **Dennd3** | AI447457\|E030003N15Rik | DENN/MADD domain containing 3 | protein-coding |
| chr9_64805850_64806157 | chr9 | 64805851 | 64806157 | + | 0.769944 | Intergenic | Intergenic | -5007 | NM_001162917 | 102442 | Mm.222473 | NM_176935 | ENSMUSG00000053641 | **Dennd4a** | AI115600\|AI851943\|F730015K02Rik | DENN/MADD domain containing 4A | protein-coding |

| | | | | | | | | | | | | | | | | | |
|---|---|---|---|---|---|---|---|---|---|---|---|---|---|---|---|---|---|
| chr7_109936668_109937292 | chr7 | 109936669 | 109937292 | + | 0.77932 | intron (NM_021494, intron 1 of 22) | intron (NM_021494, intron 1 of 22) | 23442 | NM_021494 | 19347 | Mm.21904 | NM_021494 | ENSMUSG00000035901 | **Dennd5a** | 1500012B19Rik\|AJ245569\|AW496491\|ORF37\|Rab6ip1\|mKIAA1091 | DENN/MADD domain containing 5A | protein-coding |
| chr7_109936145_109936599 | chr7 | 109936146 | 109936599 | + | 0.746531 | intron (NM_021494, intron 1 of 22) | B3\|SINE\|B2 | 24050 | NM_021494 | 19347 | Mm.21904 | NM_021494 | ENSMUSG00000035901 | **Dennd5a** | 1500012B19Rik\|AJ245569\|AW496491\|ORF37\|Rab6ip1\|mKIAA1091 | DENN/MADD domain containing 5A | protein-coding |
| chr5_32930778_32931260 | chr5 | 32930779 | 32931260 | + | 1.14789 | intron (NM_001025426, intron 25 of 42) | intron (NM_001025426, intron 25 of 42) | 67298 | NM_001025426 | 277854 | Mm.101524 | NM_177786 | ENSMUSG00000037426 | **Depdc5** | AV016528 | DEP domain containing 5 | protein-coding |
| chr2_104763007_104763703 | chr2 | 104763008 | 104763703 | + | 1.26248 | intron (NM_001123327, intron 9 of 11) | intron (NM_001123327, intron 9 of 11) | -20554 | NM_144804 | 211896 | Mm.288805 | NM_144804 | ENSMUSG00000027173 | **Depdc7** | AV216087 | DEP domain containing 7 | protein-coding |
| chr2_104740284_104740818 | chr2 | 104740285 | 104740818 | + | 0.787517 | intron (NM_144804, intron 1 of 8) | B3\|SINE\|B2 | 2250 | NM_144804 | 211896 | Mm.288805 | NM_144804 | ENSMUSG00000027173 | **Depdc7** | AV216087 | DEP domain containing 7 | protein-coding |
| chr15_57886586_57887099 | chr15 | 57886587 | 57887099 | + | 0.961991 | intron (NM_024207, intron 1 of 7) | intron (NM_024207, intron 1 of 7) | 5575 | NM_024207 | 67819 | Mm.289387 | NM_024207 | ENSMUSG00000022365 | **Derl1** | 1110021N07Rik\|AI195141\|AW551338\|Derlin-1 | Der1-like domain family, member 1 | protein-coding |
| chr15_57853460_57853710 | chr15 | 57853461 | 57853710 | + | 0.690138 | Intergenic | Intergenic | 38833 | NM_024207 | 67819 | Mm.289387 | NM_024207 | ENSMUSG00000022365 | **Derl1** | 1110021N07Rik\|AI195141\|AW551338\|Derlin-1 | Der1-like domain family, member 1 | protein-coding |
| chr15_57842202_57842429 | chr15 | 57842203 | 57842429 | + | 0.656564 | Intergenic | Intergenic | 50102 | NM_024207 | 67819 | Mm.289387 | NM_024207 | ENSMUSG00000022365 | **Derl1** | 1110021N07Rik\|AI195141\|AW551338\|Derlin-1 | Der1-like domain family, member 1 | protein-coding |
| chr10_75895641_75896579 | chr10 | 75895642 | 75896579 | + | 1.36119 | TTS (NM_024440) | TTS (NM_024440) | 2712 | NM_024440 | 70377 | Mm.275878 | NM_024440 | ENSMUSG00000009092 | **Derl3** | 1810006I20Rik\|1810063P04Rik\|IZP6\|derlin-3 | Der1-like domain family, member 3 | protein-coding |
| chr10_75895646_75896310 | chr10 | 75895647 | 75896310 | + | 1.35413 | TTS (NM_024440) | TTS (NM_024440) | 2580 | NM_024440 | 70377 | Mm.275878 | NM_024440 | ENSMUSG00000009092 | **Derl3** | 1810006I20Rik\|1810063P04Rik\|IZP6\|derlin-3 | Der1-like domain family, member 3 | protein-coding |
| chr1_178181999_178182717 | chr1 | 178182000 | 178182717 | + | 1.56037 | Intergenic | ORR1E\|LTR\|ERVL-MaLR | -5059 | NM_024282 | 78825 | Mm.440132 | NM_024282 | ENSMUSG00000026502 | **Desi2** | 5830417C01Rik\|DeSI-2\|Fam152a\|Pppde1 | desumoylating isopeptidase 2 | protein-coding |
| chr1_178181976_178182975 | chr1 | 178181977 | 178182975 | + | 1.51352 | Intergenic | ORR1E\|LTR\|ERVL-MaLR | -4941 | NM_024282 | 78825 | Mm.440132 | NM_024282 | ENSMUSG00000026502 | **Desi2** | 5830417C01Rik\|DeSI-2\|Fam152a\|Pppde1 | desumoylating isopeptidase 2 | protein-coding |
| chr1_178218287_178218590 | chr1 | 178218288 | 178218590 | + | 0.881254 | intron (NM_024282, intron 1 of 4) | intron (NM_024282, intron 1 of 4) | 31022 | NM_024282 | 78825 | Mm.440132 | NM_024282 | ENSMUSG00000026502 | **Desi2** | 5830417C01Rik\|DeSI-2\|Fam152a\|Pppde1 | desumoylating isopeptidase 2 | protein-coding |
| chr1_178218289_178218812 | chr1 | 178218290 | 178218812 | + | 0.834869 | intron (NM_024282, intron 1 of 4) | intron (NM_024282, intron 1 of 4) | 31134 | NM_024282 | 78825 | Mm.440132 | NM_024282 | ENSMUSG00000026502 | **Desi2** | 5830417C01Rik\|DeSI-2\|Fam152a\|Pppde1 | desumoylating isopeptidase 2 | protein-coding |
| chr6_50254956_50255336 | chr6 | 50254957 | 50255336 | + | 0.710917 | intron (NM_018769, intron 1 of 9) | intron (NM_018769, intron 1 of 9) | 6623 | NM_018769 | 54722 | Mm.248361 | NM_018769 | ENSMUSG00000029821 | **Dfna5** | 2310037D07Rik\|4932441K13Rik\|Dfna5h\|EG14210\|Fin15 | deafness, autosomal dominant 5 (human) | protein-coding |
| chr6_50192874_50193220 | chr6 | 50192875 | 50193220 | + | 0.665595 | intron (NM_019939, intron 10 of 12) | intron (NM_019939, intron 10 of 12) | 68722 | NM_018769 | 54722 | Mm.248361 | NM_018769 | ENSMUSG00000029821 | **Dfna5** | 2310037D07Rik\|4932441K13Rik\|Dfna5h\|EG14210\|Fin15 | deafness, autosomal dominant 5 (human) | protein-coding |
| chr10_128731839_128732384 | chr10 | 128731840 | 128732384 | + | 0.92018 | intron (NM_016811, intron 9 of 23) | intron (NM_016811, intron 9 of 23) | 11944 | NM_016811 | 13139 | Mm.291235 | NM_016811 | ENSMUSG00000025357 | **Dgka** | 80kDa\|AW146112\|Dagk1 | diacylglycerol kinase, alpha | protein-coding |
| chr10_128731627_128732386 | chr10 | 128731628 | 128732386 | + | 0.821527 | intron (NM_016811, intron 9 of 23) | B1F2\|SINE\|Alu | 12049 | NM_016811 | 13139 | Mm.291235 | NM_016811 | ENSMUSG00000025357 | **Dgka** | 80kDa\|AW146112\|Dagk1 | diacylglycerol kinase, alpha | protein-coding |
| chr1_87885388_87885923 | chr1 | 87885389 | 87885923 | + | 1.41442 | intron (NM_177646, intron 3 of 29) | ORR1C2\|LTR\|ERVL-MaLR | 32369 | NM_177646 | 227333 | Mm.277217 | NM_177646 | ENSMUSG00000070738 | **Dgkd** | AI841987\|D330025K09\|DGKdelta\|dgkd-2 | diacylglycerol kinase, delta | protein-coding |
| chr1_87861932_87862611 | chr1 | 87861933 | 87862611 | + | 0.776674 | intron (NM_177646, intron 1 of 29) | intron (NM_177646, intron 1 of 29) | 8985 | NM_177646 | 227333 | Mm.277217 | NM_177646 | ENSMUSG00000070738 | **Dgkd** | AI841987\|D330025K09\|DGKdelta\|dgkd-2 | diacylglycerol kinase, delta | protein-coding |
| chr11_89031141_89031538 | chr11 | 89031142 | 89031538 | + | 0.698799 | Intergenic | Intergenic | 29408 | NM_019505 | 56077 | Mm.153695 | NM_019505 | ENSMUSG00000000276 | **Dgke** | C87606\|DAGK6\|DGK | diacylglycerol kinase, epsilon | protein-coding |
| chr16_22632387_22633453 | chr16 | 22632388 | 22633453 | + | 1.33367 | intron (NM_138650, intron 1 of 24) | MLT1G\|LTR\|ERVL-MaLR | 24311 | NM_138650 | 110197 | Mm.194986 | NM_138650 | ENSMUSG00000022861 | **Dgkg** | 2900055E17Rik\|90kDa\|AI854428\|Dagk3\|E430001K23Rik\|mKIAA4131 | diacylglycerol kinase, gamma | protein-coding |
| chr16_22660980_22661481 | chr16 | 22660981 | 22661481 | + | 1.22687 | Intergenic | Intergenic | -4000 | NM_138650 | 110197 | Mm.194986 | NM_138650 | ENSMUSG00000022861 | **Dgkg** | 2900055E17Rik\|90kDa\|AI854428\|Dagk3\|E430001K23Rik\|mKIAA4131 | diacylglycerol kinase, gamma | protein-coding |
| chr14_78761206_78761513 | chr14 | 78761207 | 78761513 | + | 1.27065 | Intergenic | ID_B1\|SINE\|B4 | -36271 | NM_001081336 | 380921 | Mm.89591 | NM_001081336 | ENSMUSG00000034731 | **Dgkh** | 5930402B05Rik\|D130015C16 | diacylglycerol kinase, eta | protein-coding |
| chr14_78710682_78711134 | chr14 | 78710683 | 78711134 | + | 0.947475 | intron (NM_001081336, intron 1 of 28) | intron (NM_001081336, intron 1 of 28) | 14181 | NM_001081336 | 380921 | Mm.89591 | NM_001081336 | ENSMUSG00000034731 | **Dgkh** | 5930402B05Rik\|D130015C16 | diacylglycerol kinase, eta | protein-coding |
| chr14_78612205_78612763 | chr14 | 78612206 | 78612763 | + | 0.788515 | intron (NR_104044, intron 10 of 27) | intron (NR_104044, intron 10 of 27) | 16806 | NM_001253766 | 380921 | Mm.89591 | NM_001081336 | ENSMUSG00000034731 | **Dgkh** | 5930402B05Rik\|D130015C16 | diacylglycerol kinase, eta | protein-coding |
| chr14_78611960_78612756 | chr14 | 78611961 | 78612756 | + | 0.767476 | intron (NR_104044, intron 10 of 27) | intron (NR_104044, intron 10 of 27) | 16932 | NM_001253766 | 380921 | Mm.89591 | NM_001081336 | ENSMUSG00000034731 | **Dgkh** | 5930402B05Rik\|D130015C16 | diacylglycerol kinase, eta | protein-coding |
| chr14_78610892_78611186 | chr14 | 78610893 | 78611186 | + | 0.624715 | intron (NR_104044, intron 10 of 27) | URR1A\|DNA\|hAT-Charlie | 18251 | NM_001253766 | 380921 | Mm.89591 | NM_001081336 | ENSMUSG00000034731 | **Dgkh** | 5930402B05Rik\|D130015C16 | diacylglycerol kinase, eta | protein-coding |
| chr6_37370481_37371145 | chr6 | 37370482 | 37371145 | + | 1.08042 | intron (NM_178661, intron 2 of 11) | intron (NM_178661, intron 2 of 11) | -70837 | NM_001081206 | 320127 | Mm.119469 | NM_001081206 | ENSMUSG00000038665 | **Dgki** | C130010K08Rik | diacylglycerol kinase, iota | protein-coding |
| chr2_91947715_91948094 | chr2 | 91947716 | 91948094 | + | 1.09965 | intron (NM_001166597, intron 1 of 31) | intron (NM_001166597, intron 1 of 31) | 2443 | NM_001166597 | 104418 | Mm.314923 | NM_138306 | ENSMUSG00000040479 | **Dgkz** | E130307B02Rik\|ENSMUSG00000075035\|F730209L11Rik\|mDGK[z] | diacylglycerol kinase zeta | protein-coding |
| chr2_91947534_91948158 | chr2 | 91947535 | 91948158 | + | 1.02509 | intron (NM_001166597, intron 1 of 31) | intron (NM_001166597, intron 1 of 31) | 2502 | NM_001166597 | 104418 | Mm.314923 | NM_138306 | ENSMUSG00000040479 | **Dgkz** | E130307B02Rik\|ENSMUSG00000075035\|F730209L11Rik\|mDGK[z] | diacylglycerol kinase zeta | protein-coding |
| chr11_78021605_78022060 | chr11 | 78021606 | 78022060 | + | 0.696387 | intron (NM_174852, intron 7 of 14) | intron (NM_174852, intron 7 of 14) | -10480 | NM_183286 | 70451 | Mm.390342 | NM_183286 | ENSMUSG00000020834 | **Dhrs13** | 2610209N15Rik | dehydrogenase/reductase (SDR family) member 13 | protein-coding |
| chr14_55207963_55208882 | chr14 | 55207964 | 55208882 | + | 0.698524 | Intergenic | Intergenic | -13584 | NM_027790 | 71412 | Mm.85484 | NM_027790 | ENSMUSG00000022209 | **Dhrs2** | 5430405K24Rik\|Hep27 | dehydrogenase/reductase member 2 | protein-coding |
| chr4_144897986_144898540 | chr4 | 144897987 | 144898540 | + | 1.04188 | intron (NM_011303, intron 1 of 5) | intron (NM_011303, intron 1 of 5) | 5126 | NM_011303 | 20148 | Mm.14063 | NM_011303 | ENSMUSG00000066026 | **Dhrs3** | Rsdr1\|retSDR1 | dehydrogenase/reductase (SDR family) member 3 | protein-coding |
| chr4_144979933_144980473 | chr4 | 144979934 | 144980473 | + | 1.0206 | intron (NM_001276465, intron 68 of 69) | intron (NM_001276465, intron 68 of 69) | 87066 | NM_011303 | 20148 | Mm.14063 | NM_011303 | ENSMUSG00000066026 | **Dhrs3** | Rsdr1\|retSDR1 | dehydrogenase/reductase (SDR family) member 3 | protein-coding |
| chr4_144981102_144981497 | chr4 | 144981103 | 144981497 | + | 0.663419 | intron (NM_001276465, intron 68 of 69) | intron (NM_001276465, intron 68 of 69) | 88163 | NM_011303 | 20148 | Mm.14063 | NM_011303 | ENSMUSG00000066026 | **Dhrs3** | Rsdr1\|retSDR1 | dehydrogenase/reductase (SDR family) member 3 | protein-coding |
| chr4_144978836_144980478 | chr4 | 144978837 | 144980478 | + | 0.65984 | intron (NM_001276465, intron 68 of 69) | intron (NM_001276465, intron 68 of 69) | 86520 | NM_011303 | 20148 | Mm.14063 | NM_011303 | ENSMUSG00000066026 | **Dhrs3** | Rsdr1\|retSDR1 | dehydrogenase/reductase (SDR family) member 3 | protein-coding |
| chr4_144901507_144901942 | chr4 | 144901508 | 144901942 | + | 0.631701 | intron (NM_011303, intron 1 of 5) | intron (NM_011303, intron 1 of 5) | 8588 | NM_011303 | 20148 | Mm.14063 | NM_011303 | ENSMUSG00000066026 | **Dhrs3** | Rsdr1\|retSDR1 | dehydrogenase/reductase (SDR family) member 3 | protein-coding |
| chr2_69380259_69381051 | chr2 | 69380260 | 69381051 | + | 0.955514 | intron (NM_175512, intron 1 of 4) | intron (NM_175512, intron 1 of 4) | 193 | NM_175512 | 241452 | Mm.211655 | NM_175512 | | **Dhrs9** | C730025I08Rik\|Rdh15 | dehydrogenase/reductase (SDR family) member 9 | protein-coding |
| chr2_69401075_69401709 | chr2 | 69401076 | 69401709 | + | 0.804196 | exon (NM_175512, exon 5 of 5) | exon (NM_175512, exon 5 of 5) | 20930 | NM_175512 | 241452 | Mm.211655 | NM_175512 | | **Dhrs9** | C730025I08Rik\|Rdh15 | dehydrogenase/reductase (SDR family) member 9 | protein-coding |
| chr9_110108057_110108430 | chr9 | 110108058 | 110108430 | + | 0.872423 | intron (NM_001252682, intron 5 of 22) | intron (NM_001252682, intron 5 of 22) | -7157 | NM_001252683 | 72831 | Mm.276305 | NM_133347 | ENSMUSG00000032480 | **Dhx30** | 2810477H02Rik\|C130058C04Rik\|Ddx30\|HELG\|Ret-CoR | DEAH (Asp-Glu-Ala-His) box polypeptide 30 | protein-coding |
| chr7_133793443_133793850 | chr7 | 133793444 | 133793850 | + | 0.893903 | intron (NM_025850, intron 1 of 10) | intron (NM_025850, intron 1 of 10) | -10862 | NM_001286031 | 101437 | Mm.199223 | NM_133941 | ENSMUSG00000030986 | **Dhx32** | 3110079L04Rik\|4732469F02Rik\|AA408140\|Ddx32 | DEAH (Asp-Glu-Ala-His) box polypeptide 32 | protein-coding |
| chr7_16198512_16198962 | chr7 | 16198513 | 16198962 | + | 0.972313 | intron (NM_001285931, intron 15 of 16) | intron (NM_001285931, intron 15 of 16) | 23256 | NM_001285931 | 71723 | Mm.75259 | NM_027883 | ENSMUSG00000006019 | **Dhx34** | 1200013B07Rik\|1810012L18Rik\|Ddx34\|mKIAA0134 | DEAH (Asp-Glu-Ala-His) box polypeptide 34 | protein-coding |
| chr7_16207057_16207926 | chr7 | 16207058 | 16207926 | + | 0.679023 | intron (NM_001285931, intron 7 of 16) | intron (NM_001285931, intron 7 of 16) | 14501 | NM_001285931 | 71723 | Mm.75259 | NM_027883 | ENSMUSG00000006019 | **Dhx34** | 1200013B07Rik\|1810012L18Rik\|Ddx34\|mKIAA0134 | DEAH (Asp-Glu-Ala-His) box polypeptide 34 | protein-coding |
| chr7_16198487_16199258 | chr7 | 16198488 | 16199258 | + | 0.675063 | exon (NM_027883, exon 15 of 17) | exon (NM_027883, exon 15 of 17) | 23120 | NM_001285931 | 71723 | Mm.75259 | NM_027883 | ENSMUSG00000006019 | **Dhx34** | 1200013B07Rik\|1810012L18Rik\|Ddx34\|mKIAA0134 | DEAH (Asp-Glu-Ala-His) box polypeptide 34 | protein-coding |

| | | | | | | | | | | | | | | | | | |
|---|---|---|---|---|---|---|---|---|---|---|---|---|---|---|---|---|---|
| chr2_159133876_159134560 | chr2 | 159133877 | 159134560 | + | 0.699986 | Intergenic | Intergenic | 339411 | NM_001291144 | 71715 | Mm.315652 | NM_145742 | ENSMUSG00000027655 | **Dhx35** | 1200009D07Rik\|Ddx35 | DEAH (Asp-Glu-Ala-His) box polypeptide 35 | protein-coding |
| chr11_86844312_86844999 | chr11 | 86844313 | 86844999 | + | 0.776468 | intron (NR_106195, intron 4 of 8) | URR1B\|DNA\|hAT-Charlie | -36996 | NM_026191 | 67487 | Mm.260627 | NM_026191 | ENSMUSG00000018425 | **Dhx40** | 2410016C14Rik\|ARG147\|DDX40\|PAD | DEAH (Asp-Glu-Ala-His) box polypeptide 40 | protein-coding |
| chr11_86851529_86852076 | chr11 | 86851530 | 86852076 | + | 0.698394 | intron (NR_106195, intron 4 of 8) | intron (NR_106195, intron 4 of 8) | -44143 | NM_026191 | 67487 | Mm.260627 | NM_026191 | ENSMUSG00000018425 | **Dhx40** | 2410016C14Rik\|ARG147\|DDX40\|PAD | DEAH (Asp-Glu-Ala-His) box polypeptide 40 | protein-coding |
| chr18_37910206_37910718 | chr18 | 37910207 | 37910718 | + | 0.860248 | intron (NM_007858, intron 1 of 26) | intron (NM_007858, intron 1 of 26) | 24949 | NM_007858 | 13367 | Mm.195916 | NM_007858 | ENSMUSG00000024456 | **Diap1** | D18Wsu154e\|Dia1\|Diaph1\|Drf1\|mKIAA4062\|p140mDia | diaphanous homolog 1 (Drosophila) | protein-coding |
| chrX_129480499_129480920 | chrX | 129480500 | 129480920 | + | 1.33686 | Intergenic | B3A\|SINE\|B2 | -269032 | NM_172493 | 54004 | Mm.10763 | NM_017398 | ENSMUSG00000034480 | **Diap2** | Dia3\|Diaph2\|Drf2\|E430022I22Rik | diaphanous homolog 2 (Drosophila) | protein-coding |
| chrX_130768516_130769297 | chrX | 130768517 | 130769297 | + | 1.18025 | Intergenic | RMER15\|LTR\|ERVL | 1019165 | NM_172493 | 54004 | Mm.10763 | NM_017398 | ENSMUSG00000034480 | **Diap2** | Dia3\|Diaph2\|Drf2\|E430022I22Rik | diaphanous homolog 2 (Drosophila) | protein-coding |
| chrX_130744242_130744543 | chrX | 130744243 | 130744543 | + | 0.846251 | Intergenic | Intergenic | 994651 | NM_172493 | 54004 | Mm.10763 | NM_017398 | ENSMUSG00000034480 | **Diap2** | Dia3\|Diaph2\|Drf2\|E430022I22Rik | diaphanous homolog 2 (Drosophila) | protein-coding |
| chrX_130743984_130744770 | chrX | 130743985 | 130744770 | + | 0.785549 | Intergenic | Intergenic | 994635 | NM_172493 | 54004 | Mm.10763 | NM_017398 | ENSMUSG00000034480 | **Diap2** | Dia3\|Diaph2\|Drf2\|E430022I22Rik | diaphanous homolog 2 (Drosophila) | protein-coding |
| chrX_130473342_130473844 | chrX | 130473343 | 130473844 | + | 0.707791 | Intergenic | ORR1D2\|LTR\|ERVL-MaLR | 723851 | NM_172493 | 54004 | Mm.10763 | NM_017398 | ENSMUSG00000034480 | **Diap2** | Dia3\|Diaph2\|Drf2\|E430022I22Rik | diaphanous homolog 2 (Drosophila) | protein-coding |
| chrX_130533508_130533744 | chrX | 130533509 | 130533744 | + | 0.673528 | Intergenic | Intergenic | 783884 | NM_172493 | 54004 | Mm.10763 | NM_017398 | ENSMUSG00000034480 | **Diap2** | Dia3\|Diaph2\|Drf2\|E430022I22Rik | diaphanous homolog 2 (Drosophila) | protein-coding |
| chrX_130480160_130480908 | chrX | 130480161 | 130480908 | + | 0.672519 | Intergenic | Intergenic | 730792 | NM_172493 | 54004 | Mm.10763 | NM_017398 | ENSMUSG00000034480 | **Diap2** | Dia3\|Diaph2\|Drf2\|E430022I22Rik | diaphanous homolog 2 (Drosophila) | protein-coding |
| chrX_131140600_131140935 | chrX | 131140601 | 131140935 | + | 0.658935 | Intergenic | Intergenic | 1391026 | NM_172493 | 54004 | Mm.10763 | NM_017398 | ENSMUSG00000034480 | **Diap2** | Dia3\|Diaph2\|Drf2\|E430022I22Rik | diaphanous homolog 2 (Drosophila) | protein-coding |
| chrX_130533355_130533852 | chrX | 130533356 | 130533852 | + | 0.649549 | Intergenic | Intergenic | 783862 | NM_172493 | 54004 | Mm.10763 | NM_017398 | ENSMUSG00000034480 | **Diap2** | Dia3\|Diaph2\|Drf2\|E430022I22Rik | diaphanous homolog 2 (Drosophila) | protein-coding |
| chr14_87051146_87051707 | chr14 | 87051147 | 87051707 | + | 1.63723 | intron (NM_019670, intron 4 of 27) | intron (NM_019670, intron 4 of 27) | 89687 | NM_019670 | 56419 | Mm.440585 | NM_019670 | ENSMUSG00000022021 | **Diap3** | 4930417P13Rik\|Dia2\|Diaph3\|Drf3\|mDia2\|mKIAA4117\|p134MDia2 | diaphanous homolog 3 (Drosophila) | protein-coding |
| chr14_86861520_86861935 | chr14 | 86861521 | 86861935 | + | 1.06927 | intron (NM_019670, intron 21 of 27) | intron (NM_019670, intron 21 of 27) | 279386 | NM_019670 | 56419 | Mm.440585 | NM_019670 | ENSMUSG00000022021 | **Diap3** | 4930417P13Rik\|Dia2\|Diaph3\|Drf3\|mDia2\|mKIAA4117\|p134MDia2 | diaphanous homolog 3 (Drosophila) | protein-coding |
| chr14_86861380_86862184 | chr14 | 86861381 | 86862184 | + | 0.981051 | intron (NM_019670, intron 21 of 27) | intron (NM_019670, intron 21 of 27) | 279332 | NM_019670 | 56419 | Mm.440585 | NM_019670 | ENSMUSG00000022021 | **Diap3** | 4930417P13Rik\|Dia2\|Diaph3\|Drf3\|mDia2\|mKIAA4117\|p134MDia2 | diaphanous homolog 3 (Drosophila) | protein-coding |
| chr2_180695248_180696030 | chr2 | 180695249 | 180696030 | + | 1.22935 | intron (NM_001291432, intron 1 of 5) | Charlie1\|DNA\|hAT-Charlie | -3591 | NR_111965 | 23856 | Mm.253836 | NM_011805 | ENSMUSG00000038914 | **Dido1** | 6720461J16Rik\|D130048F08Rik\|DATF-1\|DIO-1\|Datf1 | death inducer-obliterator 1 | protein-coding |
| chr2_180681703_180682196 | chr2 | 180681704 | 180682196 | + | 0.928654 | 3' UTR (NM_011805, exon 6 of 6) | 3' UTR (NM_011805, exon 6 of 6) | 10098 | NR_111965 | 23856 | Mm.253836 | NM_011805 | ENSMUSG00000038914 | **Dido1** | 6720461J16Rik\|D130048F08Rik\|DATF-1\|DIO-1\|Datf1 | death inducer-obliterator 1 | protein-coding |
| chr1_193130546_193131183 | chr1 | 193130547 | 193131183 | + | 0.77304 | promoter-TSS (NM_145415) | promoter-TSS (NM_145415) | -614 | NM_145415 | 215193 | Mm.173758 | NM_145415 | ENSMUSG00000016181 | **Diexf** | AA408296\|AW045375\|Def | digestive organ expansion factor homolog (zebrafish) | protein-coding |
| chr4_107285539_107285847 | chr4 | 107285540 | 107285847 | + | 0.616255 | Intergenic | Intergenic | 21450 | NM_007860 | 13370 | Mm.148342 | NM_007860 | ENSMUSG00000034785 | **Dio1** | 5DI\|D1\|ITDI1\|TXDI1 | deiodinase, iodothyronine, type I | protein-coding |
| chr12_90575965_90576522 | chr12 | 90575966 | 90576522 | + | 0.979986 | Intergenic | Intergenic | 162194 | NM_010050 | 13371 | Mm.21389 | NM_010050 | ENSMUSG00000007682 | **Dio2** | 5DII\|AI324267\|DIOII | deiodinase, iodothyronine, type II | protein-coding |
| chr12_110332524_110333266 | chr12 | 110332525 | 110333266 | + | 1.04892 | Intergenic | L1MC\|LINE\|L1 | 53665 | NM_172119 | 107585 | Mm.154427 | NM_172119 | ENSMUSG00000075707 | **Dio3** | - | deiodinase, iodothyronine type III | protein-coding |
| chr15_100018274_100019047 | chr15 | 100018275 | 100019047 | + | 0.870866 | Intergenic | RLTR18B\|LTR\|ERVK | -20003 | NM_001159361 | 239667 | Mm.243658 | NM_172819 | ENSMUSG00000023026 | **Dip2b** | 4932422C22\|AI317237\|AI854602\|mKIAA1463 | DIP2 disco-interacting protein 2 homolog B (Drosophila) | protein-coding |
| chr13_9353797_9354306 | chr13 | 9353798 | 9354306 | + | 1.55782 | intron (NM_001081426, intron 1 of 36) | intron (NM_001081426, intron 1 of 36) | 77527 | NM_001081426 | 208440 | Mm.217326 | NM_172459 | ENSMUSG00000048264 | **Dip2c** | 2900024P20Rik\|9630044M06\|mKIAA0934 | DIP2 disco-interacting protein 2 homolog C (Drosophila) | protein-coding |
| chr13_9304734_9305296 | chr13 | 9304735 | 9305296 | + | 1.10555 | intron (NM_001081426, intron 1 of 36) | intron (NM_001081426, intron 1 of 36) | 28490 | NM_001081426 | 208440 | Mm.217326 | NM_172459 | ENSMUSG00000048264 | **Dip2c** | 2900024P20Rik\|9630044M06\|mKIAA0934 | DIP2 disco-interacting protein 2 homolog C (Drosophila) | protein-coding |
| chr13_9380697_9381311 | chr13 | 9380698 | 9381311 | + | 1.05312 | intron (NM_001081426, intron 1 of 36) | intron (NM_001081426, intron 1 of 36) | 104479 | NM_001081426 | 208440 | Mm.217326 | NM_172459 | ENSMUSG00000048264 | **Dip2c** | 2900024P20Rik\|9630044M06\|mKIAA0934 | DIP2 disco-interacting protein 2 homolog C (Drosophila) | protein-coding |
| chr13_9304523_9305271 | chr13 | 9304524 | 9305271 | + | 1.00408 | intron (NM_001081426, intron 1 of 36) | intron (NM_001081426, intron 1 of 36) | 28372 | NM_001081426 | 208440 | Mm.217326 | NM_172459 | ENSMUSG00000048264 | **Dip2c** | 2900024P20Rik\|9630044M06\|mKIAA0934 | DIP2 disco-interacting protein 2 homolog C (Drosophila) | protein-coding |
| chr13_9427808_9428321 | chr13 | 9427809 | 9428321 | + | 0.98762 | intron (NM_001081426, intron 1 of 36) | intron (NM_001081426, intron 1 of 36) | 151540 | NM_001081426 | 208440 | Mm.217326 | NM_172459 | ENSMUSG00000048264 | **Dip2c** | 2900024P20Rik\|9630044M06\|mKIAA0934 | DIP2 disco-interacting protein 2 homolog C (Drosophila) | protein-coding |
| chr13_9380526_9381365 | chr13 | 9380527 | 9381365 | + | 0.864204 | intron (NM_001081426, intron 1 of 36) | intron (NM_001081426, intron 1 of 36) | 104421 | NM_001081426 | 208440 | Mm.217326 | NM_172459 | ENSMUSG00000048264 | **Dip2c** | 2900024P20Rik\|9630044M06\|mKIAA0934 | DIP2 disco-interacting protein 2 homolog C (Drosophila) | protein-coding |
| chr13_9191147_9191601 | chr13 | 9191148 | 9191601 | + | 0.70422 | Intergenic | Intergenic | -85151 | NM_001081426 | 208440 | Mm.217326 | NM_172459 | ENSMUSG00000048264 | **Dip2c** | 2900024P20Rik\|9630044M06\|mKIAA0934 | DIP2 disco-interacting protein 2 homolog C (Drosophila) | protein-coding |
| chr13_9190947_9191601 | chr13 | 9190948 | 9191601 | + | 0.660002 | Intergenic | Intergenic | -85251 | NM_001081426 | 208440 | Mm.217326 | NM_172459 | ENSMUSG00000048264 | **Dip2c** | 2900024P20Rik\|9630044M06\|mKIAA0934 | DIP2 disco-interacting protein 2 homolog C (Drosophila) | protein-coding |
| chr13_9431830_9432389 | chr13 | 9431831 | 9432389 | + | 0.639602 | intron (NM_001081426, intron 1 of 36) | intron (NM_001081426, intron 1 of 36) | 155585 | NM_001081426 | 208440 | Mm.217326 | NM_172459 | ENSMUSG00000048264 | **Dip2c** | 2900024P20Rik\|9630044M06\|mKIAA0934 | DIP2 disco-interacting protein 2 homolog C (Drosophila) | protein-coding |
| chr16_35751869_35752231 | chr16 | 35751870 | 35752231 | + | 0.730429 | intron (NM_153550, intron 1 of 8) | RLTR13D\|LTR\|ERVK | 17306 | NM_153550 | 224132 | Mm.24094 | NM_153550 | ENSMUSG00000022848 | **Dirc2** | RCC4 | disrupted in renal carcinoma 2 (human) | protein-coding |
| chr16_35768348_35768721 | chr16 | 35768349 | 35768721 | + | 0.633218 | intron (NM_153550, intron 1 of 8) | intron (NM_153550, intron 1 of 8) | 821 | NM_153550 | 224132 | Mm.24094 | NM_153550 | ENSMUSG00000022848 | **Dirc2** | RCC4 | disrupted in renal carcinoma 2 (human) | protein-coding |
| chr1_86886858_86887609 | chr1 | 86886859 | 86887609 | + | 1.08575 | intron (NM_153530, intron 9 of 20) | RLTR14\|LTR\|ERV1 | 183430 | NM_001172157 | 208718 | Mm.389152 | NM_153530 | ENSMUSG00000053333 | **Dis3l2** | 4930429A22Rik\|8030493P09Rik | DIS3 mitotic control homolog (S. cerevisiae)-like 2 | protein-coding |
| chr1_86884421_86885010 | chr1 | 86884422 | 86885010 | + | 1.07106 | intron (NM_153530, intron 9 of 20) | intron (NM_153530, intron 9 of 20) | 180912 | NM_001172157 | 208718 | Mm.389152 | NM_153530 | ENSMUSG00000053333 | **Dis3l2** | 4930429A22Rik\|8030493P09Rik | DIS3 mitotic control homolog (S. cerevisiae)-like 2 | protein-coding |
| chr1_183154442_183155504 | chr1 | 183154443 | 183155504 | + | 0.969688 | intron (NM_001278219, intron 2 of 8) | intron (NM_001278219, intron 2 of 8) | -7416 | NM_001278220 | 68897 | Mm.327216 | NM_026866 | ENSMUSG00000030768 | **Disp1** | 1190008H24Rik\|DispA | dispatched homolog 1 (Drosophila) | protein-coding |
| chr1_183138596_183138897 | chr1 | 183138597 | 183138897 | + | 0.6129 | intron (NM_001278219, intron 2 of 8) | intron (NM_001278219, intron 2 of 8) | 8810 | NM_001278220 | 68897 | Mm.327216 | NM_026866 | ENSMUSG00000030768 | **Disp1** | 1190008H24Rik\|DispA | dispatched homolog 1 (Drosophila) | protein-coding |
| chr9_50732774_50733509 | chr9 | 50732775 | 50733509 | + | 1.40957 | Intergenic | L2a\|LINE\|L2 | -5158 | NM_178118 | 330938 | Mm.82598 | NM_178118 | ENSMUSG00000032064 | **Dixdc1** | 4930563F16Rik\|BC048182\|Ccd1\|Ccd1Aalpha1L\|Ccd1Abeta1L\|Ccd1BalphaL\|Ccd1BbetaL\|Ccd1CL\|mKIAA1735 | DIX domain containing 1 | protein-coding |
| chr9_50686479_50687240 | chr9 | 50686480 | 50687240 | + | 1.02553 | intron (NM_178118, intron 10 of 19) | intron (NM_178118, intron 10 of 19) | -27080 | NM_145614 | 235339 | Mm.285076 | NM_145614 | ENSMUSG00000000168 | **Dlat** | 6332404G05Rik\|DLTA\|PDC-E2 | dihydrolipoamide S-acetyltransferase (E2 component of pyruvate dehydrogenase complex) | protein-coding |
| chr14_61819228_61819577 | chr14 | 61819229 | 61819577 | + | 1.50579 | Intergenic | Intergenic | -137030 | NR_028264 | 668253 | Mm.32886 | NR_028264 | ENSMUSG00000097589 | **Dleu2** | 1810047A16Rik\|4932411A06Rik\|7730401J12Rik\|AI197277\|AI788824\|Alt1\|Gm9069\|Leu2 | deleted in lymphocytic leukemia, 2 | ncRNA |
| chr14_61819213_61819846 | chr14 | 61819214 | 61819846 | + | 1.40664 | Intergenic | Intergenic | -137157 | NR_028264 | 668253 | Mm.32886 | NR_028264 | ENSMUSG00000097589 | **Dleu2** | 1810047A16Rik\|4932411A06Rik\|7730401J12Rik\|AI197277\|AI788824\|Alt1\|Gm9069\|Leu2 | deleted in lymphocytic leukemia, 2 | ncRNA |
| chr14_61661574_61661995 | chr14 | 61661575 | 61661995 | + | 1.3231 | intron (NR_028264, intron 1 of 6) | intron (NR_028264, intron 1 of 6) | 20588 | NR_028264 | 668253 | Mm.32886 | NR_028264 | ENSMUSG00000097589 | **Dleu2** | 1810047A16Rik\|4932411A06Rik\|7730401J12Rik\|AI197277\|AI788824\|Alt1\|Gm9069\|Leu2 | deleted in lymphocytic leukemia, 2 | ncRNA |
| chr14_61661574_61662425 | chr14 | 61661575 | 61662425 | + | 1.22148 | intron (NR_028264, intron 1 of 6) | PB1D10\|SINE\|Alu | 20373 | NR_028264 | 668253 | Mm.32886 | NR_028264 | ENSMUSG00000097589 | **Dleu2** | 1810047A16Rik\|4932411A06Rik\|7730401J12Rik\|AI197277\|AI788824\|Alt1\|Gm9069\|Leu2 | deleted in lymphocytic leukemia, 2 | ncRNA |
| chr14_61769757_61770279 | chr14 | 61769758 | 61770279 | + | 1.14693 | Intergenic | Intergenic | -87645 | NR_028264 | 668253 | Mm.32886 | NR_028264 | ENSMUSG00000097589 | **Dleu2** | 1810047A16Rik\|4932411A06Rik\|7730401J12Rik\|AI197277\|AI788824\|Alt1\|Gm9069\|Leu2 | deleted in lymphocytic leukemia, 2 | ncRNA |
| chr14_61818388_61818800 | chr14 | 61818389 | 61818800 | + | 0.838378 | Intergenic | Intergenic | -136221 | NR_028264 | 668253 | Mm.32886 | NR_028264 | ENSMUSG00000097589 | **Dleu2** | 1810047A16Rik\|4932411A06Rik\|7730401J12Rik\|AI197277\|AI788824\|Alt1\|Gm9069\|Leu2 | deleted in lymphocytic leukemia, 2 | ncRNA |
| chr14_61953043_61953495 | chr14 | 61953044 | 61953495 | + | 0.827046 | Intergenic | Intergenic | -270896 | NR_028264 | 668253 | Mm.32886 | NR_028264 | ENSMUSG00000097589 | **Dleu2** | 1810047A16Rik\|4932411A06Rik\|7730401J12Rik\|AI197277\|AI788824\|Alt1\|Gm9069\|Leu2 | deleted in lymphocytic leukemia, 2 | ncRNA |

| | | | | | | | | | | | | | | | | | |
|---|---|---|---|---|---|---|---|---|---|---|---|---|---|---|---|---|---|
| chr14_61815758_61816236 | chr14 | 61815759 | 61816236 | + | 0.808683 | Intergenic | MTEa\|LTR\|ERVL-MaLR | -133624 | NR_028264 | 668253 | Mm.32886 | NR_028264 | ENSMUSG00000097589 | **Dleu2** | 1810047A16Rik\|4932411A06Rik\|7730401J12Rik\|AI197277\|AI788824\|Alt1\|Gm9069\|Leu2 | deleted in lymphocytic leukemia, 2 | ncRNA |
| chr14_61739535_61739860 | chr14 | 61739536 | 61739860 | + | 0.7217 | Intergenic | RLTR40\|LTR\|ERVK | -57325 | NR_028264 | 668253 | Mm.32886 | NR_028264 | ENSMUSG00000097589 | **Dleu2** | 1810047A16Rik\|4932411A06Rik\|7730401J12Rik\|AI197277\|AI788824\|Alt1\|Gm9069\|Leu2 | deleted in lymphocytic leukemia, 2 | ncRNA |
| chr14_62022202_62022555 | chr14 | 62022203 | 62022555 | + | 1.15129 | Intergenic | L1MB3\|LINE\|L1 | 270600 | NM_173419 | 239133 | Mm.127330 | NM_173419 | ENSMUSG00000048281 | **Dleu7** | BC038059 | deleted in lymphocytic leukemia, 7 | protein-coding |
| chr14_62022802_62023465 | chr14 | 62022803 | 62023465 | + | 0.764206 | Intergenic | RMER15\|LTR\|ERVL | 269845 | NM_173419 | 239133 | Mm.127330 | NM_173419 | ENSMUSG00000048281 | **Dleu7** | BC038059 | deleted in lymphocytic leukemia, 7 | protein-coding |
| chr14_62023086_62023465 | chr14 | 62023087 | 62023465 | + | 0.753774 | Intergenic | RMER15\|LTR\|ERVL | 269703 | NM_173419 | 239133 | Mm.127330 | NM_173419 | ENSMUSG00000048281 | **Dleu7** | BC038059 | deleted in lymphocytic leukemia, 7 | protein-coding |
| chr16_31570639_31571239 | chr16 | 31570640 | 31571239 | + | 1.0596 | Intergenic | Intergenic | -92504 | NM_001252435 | 13383 | Mm.382 | NM_007862 | ENSMUSG00000022770 | **Dlg1** | B130052P05Rik\|Dlgh1\|E-dlg/SAP97\|SAP-97\|SAP97\|mKIAA4187 | discs, large homolog 1 (Drosophila) | protein-coding |
| chr16_31788876_31789185 | chr16 | 31788877 | 31789185 | + | 0.933951 | intron (NM_001252434, intron 7 of 24) | intron (NM_001252434, intron 7 of 24) | 34681 | NM_001252436 | 13383 | Mm.382 | NM_007862 | ENSMUSG00000022770 | **Dlg1** | B130052P05Rik\|Dlgh1\|E-dlg/SAP97\|SAP-97\|SAP97\|mKIAA4187 | discs, large homolog 1 (Drosophila) | protein-coding |
| chr7_92134148_92134796 | chr7 | 92134149 | 92134796 | + | 1.22656 | intron (NM_011807, intron 13 of 22) | intron (NM_011807, intron 13 of 22) | 72078 | NM_001243046 | 23859 | Mm.147213 | NM_011807 | | **Dlg2** | A330103J02Rik\|B230218P12Rik\|B330007M19Rik\|Dlgh2\|Gm1197\|PSD93 | discs, large homolog 2 (Drosophila) | protein-coding |
| chr7_92165003_92165508 | chr7 | 92165004 | 92165508 | + | 0.792258 | intron (NM_011807, intron 13 of 22) | intron (NM_011807, intron 13 of 22) | -69714 | NM_001243047 | 23859 | Mm.147213 | NM_011807 | | **Dlg2** | A330103J02Rik\|B230218P12Rik\|B330007M19Rik\|Dlgh2\|Gm1197\|PSD93 | discs, large homolog 2 (Drosophila) | protein-coding |
| chr7_92087497_92088091 | chr7 | 92087498 | 92088091 | + | 0.717603 | intron (NM_001243046, intron 2 of 14) | intron (NM_001243046, intron 2 of 14) | 25400 | NM_001243046 | 23859 | Mm.147213 | NM_011807 | | **Dlg2** | A330103J02Rik\|B230218P12Rik\|B330007M19Rik\|Dlgh2\|Gm1197\|PSD93 | discs, large homolog 2 (Drosophila) | protein-coding |
| chrX_100787786_100788029 | chrX | 100787787 | 100788029 | + | 0.742156 | intron (NM_001177780, intron 1 of 11) | intron (NM_001177780, intron 1 of 11) | 10637 | NM_001177780 | 53310 | Mm.4615 | NM_016747 | ENSMUSG00000000881 | **Dlg3** | Dlgh3\|SAP102\|mKIAA1232 | discs, large homolog 3 (Drosophila) | protein-coding |
| chr17_46302804_46303588 | chr17 | 46302805 | 46303588 | + | 0.650427 | TTS (NM_001286028) | TTS (NM_001286028) | 5268 | NM_001286013 | 106565 | Mm.23633 | NM_134120 | ENSMUSG00000047428 | **Dlk2** | AI413481\|DLK-2\|Egfl9 | delta-like 2 homolog (Drosophila) | protein-coding |
| chr2_71543033_71543322 | chr2 | 71543034 | 71543322 | + | 0.632002 | TTS (NM_010054) | TTS (NM_010054) | 3576 | NM_010054 | 13392 | Mm.3896 | NM_010054 | ENSMUSG00000023391 | **Dlx2** | AW121999\|Dlx-2\|Tes-1 | distal-less homeobox 2 | protein-coding |
| chr15_79600871_79601380 | chr15 | 79600872 | 79601380 | + | 0.865996 | intron (NM_001278226, intron 3 of 11) | intron (NM_001278226, intron 3 of 11) | 3983 | NR_103477 | 13404 | Mm.2524 | NM_010059 | ENSMUSG00000022429 | **Dmc1** | Dmc1h\|Lim15\|Mei11\|sgdp | DMC1 dosage suppressor of mck1 homolog, meiosis-specific homologous recombination | protein-coding |
| chr4_89530007_89530572 | chr4 | 89530008 | 89530572 | + | 1.42806 | Intergenic | Intergenic | -157908 | NM_175647 | 242523 | Mm.130167 | NM_175647 | ENSMUSG00000043753 | **Dmrta1** | Dmrt4 | doublesex and mab-3 related transcription factor like family A1 | protein-coding |
| chr4_89505214_89505864 | chr4 | 89505215 | 89505864 | + | 0.834675 | Intergenic | Intergenic | -182659 | NM_175647 | 242523 | Mm.130167 | NM_175647 | ENSMUSG00000043753 | **Dmrta1** | Dmrt4 | doublesex and mab-3 related transcription factor like family A1 | protein-coding |
| chr4_89523412_89523886 | chr4 | 89523413 | 89523886 | + | 0.680215 | Intergenic | Intergenic | -164549 | NM_175647 | 242523 | Mm.130167 | NM_175647 | ENSMUSG00000043753 | **Dmrta1** | Dmrt4 | doublesex and mab-3 related transcription factor like family A1 | protein-coding |
| chr4_89523524_89523876 | chr4 | 89523525 | 89523876 | + | 0.65358 | Intergenic | Intergenic | -164498 | NM_175647 | 242523 | Mm.130167 | NM_175647 | ENSMUSG00000043753 | **Dmrta1** | Dmrt4 | doublesex and mab-3 related transcription factor like family A1 | protein-coding |
| chr4_109829798_109830850 | chr4 | 109829799 | 109830850 | + | 0.994176 | intron (NM_007983, intron 9 of 18) | intron (NM_007983, intron 9 of 18) | -147701 | NM_172296 | 242620 | Mm.32825 | NM_172296 | ENSMUSG00000047143 | **Dmrta2** | Dmrt5 | doublesex and mab-3 related transcription factor like family A2 | protein-coding |
| chr4_109829803_109830149 | chr4 | 109829804 | 109830149 | + | 0.824681 | intron (NM_007983, intron 9 of 18) | intron (NM_007983, intron 9 of 18) | -148049 | NM_172296 | 242620 | Mm.32825 | NM_172296 | ENSMUSG00000047143 | **Dmrta2** | Dmrt5 | doublesex and mab-3 related transcription factor like family A2 | protein-coding |
| chr4_107605954_107606521 | chr4 | 107605955 | 107606521 | + | 0.745437 | intron (NM_147221, intron 4 of 10) | intron (NM_147221, intron 4 of 10) | 77924 | NM_019872 | 56296 | Mm.141324 | NM_019872 | ENSMUSG00000028610 | **Dmrtb1** | Dmrt6\|Prp13 | DMRT-like family B with proline-rich C-terminal, 1 | protein-coding |
| chr5_9141738_9142783 | chr5 | 9141739 | 9142783 | + | 1.04657 | intron (NM_011806, intron 4 of 17) | L2\|LINE\|L2 | 19515 | NM_001110327 | 23857 | Mm.22480 | NM_011806 | ENSMUSG00000042508 | **Dmtf1** | Dimp\|Dmp1 | cyclin D binding myb-like transcription factor 1 | protein-coding |
| chr7_19075879_19076375 | chr7 | 19075880 | 19076375 | + | 0.821879 | promoter-TSS (NM_010058) | promoter-TSS (NM_010058) | -73 | NM_010058 | 13401 | Mm.2852 | NM_010058 | ENSMUSG00000030410 | **Dmwd** | DMR-N9\|Dm9 | dystrophia myotonica-containing WD repeat motif | protein-coding |
| chr18_49850105_49850791 | chr18 | 49850106 | 49850791 | + | 1.30186 | intron (NM_001081371, intron 5 of 42) | intron (NM_001081371, intron 5 of 42) | 17451 | NM_001081371 | 240283 | Mm.257790 | NM_001081371 | ENSMUSG00000037416 | **Dmxl1** | C630007L23Rik\|D030030C22 | Dmx-like 1 | protein-coding |
| chr12_118071520_118072021 | chr12 | 118071521 | 118072021 | + | 0.970867 | intron (NM_010060, intron 30 of 81) | intron (NM_010060, intron 30 of 81) | 127272 | NM_010060 | 13411 | Mm.390540 | NM_010060 | ENSMUSG00000018581 | **Dnah11** | Dnahc11\|b2b1203Clo\|b2b1279Clo\|b2b1289Clo\|b2b1727Clo\|b2b598Clo\|iv\|lrd | dynein, axonemal, heavy chain 11 | protein-coding |
| chr12_118071064_118072035 | chr12 | 118071065 | 118072035 | + | 0.946372 | intron (NM_010060, intron 30 of 81) | L1M3\|LINE\|L1 | 127493 | NM_010060 | 13411 | Mm.390540 | NM_010060 | ENSMUSG00000018581 | **Dnah11** | Dnahc11\|b2b1203Clo\|b2b1279Clo\|b2b1289Clo\|b2b1727Clo\|b2b598Clo\|iv\|lrd | dynein, axonemal, heavy chain 11 | protein-coding |
| chr6_73139340_73139817 | chr6 | 73139341 | 73139817 | + | 0.958017 | intron (NM_001164669, intron 30 of 76) | ORR1E\|LTR\|ERVL-MaLR | 82052 | NM_001164669 | 330355 | Mm.333166 | NM_001164669 | ENSMUSG00000052861 | **Dnah6** | 9830168K20\|A730004I20Rik\|Dnahc6\|mKIAA1697\|mdhc6 | dynein, axonemal, heavy chain 6 | protein-coding |
| chr8_85532854_85533300 | chr8 | 85532855 | 85533300 | + | 1.09992 | Intergenic | Intergenic | 22194 | NM_019794 | 56445 | Mm.475573 | NM_019794 | ENSMUSG00000031701 | **Dnaja2** | 1500017M13Rik\|2010206B19Rik\|DNAJ\|DNJ3\|Dnaj3\|HIRIP4\|PRO3015\|mDj3 | DnaJ (Hsp40) homolog, subfamily A, member 2 | protein-coding |
| chr16_22860759_22860994 | chr16 | 22860760 | 22860994 | + | 0.863835 | intron (NM_001190804, intron 1 of 9) | intron (NM_001190804, intron 1 of 9) | 3032 | NM_001190804 | 67838 | Mm.37516 | NM_026400 | ENSMUSG00000004460 | **Dnajb11** | 1810031F23Rik\|ABBP-2\|AL024055\|Dj9\|ERdj3\|ERj3p | DnaJ (Hsp40) homolog, subfamily B, member 11 | protein-coding |
| chr10_59895947_59896264 | chr10 | 59895948 | 59896264 | + | 1.13874 | intron (NM_019965, intron 7 of 8) | intron (NM_019965, intron 7 of 8) | 16515 | NM_019965 | 56709 | Mm.103610 | NM_019965 | ENSMUSG00000020109 | **Dnajb12** | Dj10\|mDj10 | DnaJ (Hsp40) homolog, subfamily B, member 12 | protein-coding |
| chr10_59895944_59896482 | chr10 | 59895945 | 59896482 | + | 1.02562 | intron (NM_019965, intron 7 of 8) | intron (NM_019965, intron 7 of 8) | 16622 | NM_019965 | 56709 | Mm.103610 | NM_019965 | ENSMUSG00000020109 | **Dnajb12** | Dj10\|mDj10 | DnaJ (Hsp40) homolog, subfamily B, member 12 | protein-coding |
| chr3_152188752_152189109 | chr3 | 152188753 | 152189109 | + | 0.92172 | intron (NM_025926, intron 1 of 2) | intron (NM_025926, intron 1 of 2) | 4914 | NM_025926 | 67035 | Mm.46746 | NM_025926 | ENSMUSG00000028035 | **Dnajb4** | 1700029A20Rik\|2010306G19Rik\|5730460G06Rik | DnaJ (Hsp40) homolog, subfamily B, member 4 | protein-coding |
| chr4_42952334_42952665 | chr4 | 42952335 | 42952665 | + | 1.58867 | promoter-TSS (NM_019874) | promoter-TSS (NM_019874) | -594 | NM_019874 | 56323 | Mm.20437 | NM_019874 | ENSMUSG00000036052 | **Dnajb5** | 1110058L06Rik\|AI462558\|Hsc40\|Hsp40-3 | DnaJ (Hsp40) homolog, subfamily B, member 5 | protein-coding |
| chr4_42951645_42952100 | chr4 | 42951646 | 42952100 | + | 1.47982 | Intergenic | Intergenic | -1221 | NM_019874 | 56323 | Mm.20437 | NM_019874 | ENSMUSG00000036052 | **Dnajb5** | 1110058L06Rik\|AI462558\|Hsc40\|Hsp40-3 | DnaJ (Hsp40) homolog, subfamily B, member 5 | protein-coding |
| chr4_42941271_42941946 | chr4 | 42941272 | 42941946 | + | 1.07306 | TTS (NM_172690) | TTS (NM_172690) | -11485 | NM_019874 | 56323 | Mm.20437 | NM_019874 | ENSMUSG00000036052 | **Dnajb5** | 1110058L06Rik\|AI462558\|Hsc40\|Hsp40-3 | DnaJ (Hsp40) homolog, subfamily B, member 5 | protein-coding |
| chr2_18415263_18415603 | chr2 | 18415264 | 18415603 | + | 1.16083 | Intergenic | RSINE1\|SINE\|B4 | -22603 | NM_007869 | 13418 | Mm.246674 | NM_007869 | ENSMUSG00000026740 | **Dnajc1** | 4733401K02Rik\|AA960110\|D230036H06Rik\|Dnajl1\|ERdj1\|ERj1p\|MTJ1 | DnaJ (Hsp40) homolog, subfamily C, member 1 | protein-coding |
| chr2_18415283_18415724 | chr2 | 18415284 | 18415724 | + | 1.1301 | Intergenic | RSINE1\|SINE\|B4 | -22674 | NM_007869 | 13418 | Mm.246674 | NM_007869 | ENSMUSG00000026740 | **Dnajc1** | 4733401K02Rik\|AA960110\|D230036H06Rik\|Dnajl1\|ERdj1\|ERj1p\|MTJ1 | DnaJ (Hsp40) homolog, subfamily C, member 1 | protein-coding |
| chr2_18415256_18415936 | chr2 | 18415257 | 18415936 | + | 1.03861 | Intergenic | (CA)n\|Simple_repeat\|Simple_repeat | -22766 | NM_007869 | 13418 | Mm.246674 | NM_007869 | ENSMUSG00000026740 | **Dnajc1** | 4733401K02Rik\|AA960110\|D230036H06Rik\|Dnajl1\|ERdj1\|ERj1p\|MTJ1 | DnaJ (Hsp40) homolog, subfamily C, member 1 | protein-coding |
| chr2_18406841_18407153 | chr2 | 18406842 | 18407153 | + | 0.808204 | Intergenic | L1VL4\|LINE\|L1 | -14167 | NM_007869 | 13418 | Mm.246674 | NM_007869 | ENSMUSG00000026740 | **Dnajc1** | 4733401K02Rik\|AA960110\|D230036H06Rik\|Dnajl1\|ERdj1\|ERj1p\|MTJ1 | DnaJ (Hsp40) homolog, subfamily C, member 1 | protein-coding |
| chr2_18436173_18436618 | chr2 | 18436174 | 18436618 | + | 0.6989 | Intergenic | Intergenic | -43566 | NM_007869 | 13418 | Mm.246674 | NM_007869 | ENSMUSG00000026740 | **Dnajc1** | 4733401K02Rik\|AA960110\|D230036H06Rik\|Dnajl1\|ERdj1\|ERj1p\|MTJ1 | DnaJ (Hsp40) homolog, subfamily C, member 1 | protein-coding |
| chr2_80326153_80326565 | chr2 | 80326154 | 80326565 | + | 1.08521 | intron (NM_024181, intron 7 of 22) | intron (NM_024181, intron 7 of 22) | 10893 | NM_024181 | 66861 | Mm.21762 | NM_024181 | ENSMUSG00000027006 | **Dnajc10** | 1200006L06Rik\|D2Ertd706e\|ERdj5\|JPDI | DnaJ (Hsp40) homolog, subfamily C, member 10 | protein-coding |
| chr2_80304390_80305182 | chr2 | 80304391 | 80305182 | + | 1.06303 | Intergenic | MERV1_LTR\|LTR\|ERV1 | -10680 | NM_024181 | 66861 | Mm.21762 | NM_024181 | ENSMUSG00000027006 | **Dnajc10** | 1200006L06Rik\|D2Ertd706e\|ERdj5\|JPDI | DnaJ (Hsp40) homolog, subfamily C, member 10 | protein-coding |
| chr2_80321858_80322153 | chr2 | 80321859 | 80322153 | + | 0.946811 | intron (NM_024181, intron 4 of 22) | intron (NM_024181, intron 4 of 22) | 6540 | NM_024181 | 66861 | Mm.21762 | NM_024181 | ENSMUSG00000027006 | **Dnajc10** | 1200006L06Rik\|D2Ertd706e\|ERdj5\|JPDI | DnaJ (Hsp40) homolog, subfamily C, member 10 | protein-coding |

| | | | | | | | | | | | | | | | | | |
|---|---|---|---|---|---|---|---|---|---|---|---|---|---|---|---|---|---|
| chr14_118959696_118960249 | chr14 | 118959697 | 118960249 | + | 1.30479 | intron (NM_008929, intron 3 of 11) | intron (NM_008929, intron 3 of 11) | 22041 | NM_008929 | 100037258 | Mm.12616 | NM_008929 | ENSMUSG00000022136 | **Dnajc3** | AA408985\|AU067833\|Dnajc3a\|Dnajc3b\|Prkri\|mp58\|p58\|p58IPK | DnaJ (Hsp40) homolog, subfamily C, member 3 | protein-coding |
| chr14_118948948_118949481 | chr14 | 118948949 | 118949481 | + | 1.19741 | intron (NM_008929, intron 1 of 11) | intron (NM_008929, intron 1 of 11) | 11283 | NM_008929 | 100037258 | Mm.12616 | NM_008929 | ENSMUSG00000022136 | **Dnajc3** | AA408985\|AU067833\|Dnajc3a\|Dnajc3b\|Prkri\|mp58\|p58\|p58IPK | DnaJ (Hsp40) homolog, subfamily C, member 3 | protein-coding |
| chr14_118965577_118965910 | chr14 | 118965578 | 118965910 | + | 1.19667 | intron (NM_008929, intron 4 of 11) | L1M2\|LINE\|L1 | 27812 | NM_008929 | 100037258 | Mm.12616 | NM_008929 | ENSMUSG00000022136 | **Dnajc3** | AA408985\|AU067833\|Dnajc3a\|Dnajc3b\|Prkri\|mp58\|p58\|p58IPK | DnaJ (Hsp40) homolog, subfamily C, member 3 | protein-coding |
| chr14_118965261_118966242 | chr14 | 118965262 | 118966242 | + | 1.07403 | intron (NM_008929, intron 4 of 11) | L1M2\|LINE\|L1 | 27820 | NM_008929 | 100037258 | Mm.12616 | NM_008929 | ENSMUSG00000022136 | **Dnajc3** | AA408985\|AU067833\|Dnajc3a\|Dnajc3b\|Prkri\|mp58\|p58\|p58IPK | DnaJ (Hsp40) homolog, subfamily C, member 3 | protein-coding |
| chr14_118979762_118980119 | chr14 | 118979763 | 118980119 | + | 0.751434 | 3' UTR (NM_008929, exon 12 of 12) | 3' UTR (NM_008929, exon 12 of 12) | 42009 | NM_008929 | 100037258 | Mm.12616 | NM_008929 | ENSMUSG00000022136 | **Dnajc3** | AA408985\|AU067833\|Dnajc3a\|Dnajc3b\|Prkri\|mp58\|p58\|p58IPK | DnaJ (Hsp40) homolog, subfamily C, member 3 | protein-coding |
| chr3_19458474_19459395 | chr3 | 19458475 | 19459395 | + | 1.52245 | Intergenic | Intergenic | -49660 | NM_001163536 | 66326 | Mm.45615 | NM_025489 | ENSMUSG00000027606 | **Dnajc5b** | 1700008A05Rik | DnaJ (Hsp40) homolog, subfamily C, member 5 beta | protein-coding |
| chr3_19419523_19420521 | chr3 | 19419524 | 19420521 | + | 1.17424 | Intergenic | Intergenic | -88573 | NM_001163536 | 66326 | Mm.45615 | NM_025489 | ENSMUSG00000027606 | **Dnajc5b** | 1700008A05Rik | DnaJ (Hsp40) homolog, subfamily C, member 5 beta | protein-coding |
| chr3_19421288_19421798 | chr3 | 19421289 | 19421798 | + | 1.01314 | Intergenic | Intergenic | -87052 | NM_001163536 | 66326 | Mm.45615 | NM_025489 | ENSMUSG00000027606 | **Dnajc5b** | 1700008A05Rik | DnaJ (Hsp40) homolog, subfamily C, member 5 beta | protein-coding |
| chr3_19480505_19481009 | chr3 | 19480506 | 19481009 | + | 0.969824 | Intergenic | RMER17C-int\|LTR\|ERVK | -27838 | NM_001163536 | 66326 | Mm.45615 | NM_025489 | ENSMUSG00000027606 | **Dnajc5b** | 1700008A05Rik | DnaJ (Hsp40) homolog, subfamily C, member 5 beta | protein-coding |
| chr3_19421421_19421783 | chr3 | 19421422 | 19421783 | + | 0.966231 | Intergenic | Intergenic | -86993 | NM_001163536 | 66326 | Mm.45615 | NM_025489 | ENSMUSG00000027606 | **Dnajc5b** | 1700008A05Rik | DnaJ (Hsp40) homolog, subfamily C, member 5 beta | protein-coding |
| chr3_19456259_19456646 | chr3 | 19456260 | 19456646 | + | 0.873484 | Intergenic | Intergenic | -52142 | NM_001163536 | 66326 | Mm.45615 | NM_025489 | ENSMUSG00000027606 | **Dnajc5b** | 1700008A05Rik | DnaJ (Hsp40) homolog, subfamily C, member 5 beta | protein-coding |
| chr3_19426059_19426359 | chr3 | 19426060 | 19426359 | + | 0.702757 | Intergenic | Intergenic | -82386 | NM_001163536 | 66326 | Mm.45615 | NM_025489 | ENSMUSG00000027606 | **Dnajc5b** | 1700008A05Rik | DnaJ (Hsp40) homolog, subfamily C, member 5 beta | protein-coding |
| chr4_101568429_101568990 | chr4 | 101568430 | 101568990 | + | 0.752775 | intron (NM_001164583, intron 1 of 18) | intron (NM_001164583, intron 1 of 18) | 18116 | NM_001164583 | 72685 | Mm.76494 | NM_198412 | ENSMUSG00000028528 | **Dnajc6** | 2810027M23Rik\|mKIAA0473 | DnaJ (Hsp40) homolog, subfamily C, member 6 | protein-coding |
| chr4_101585743_101586358 | chr4 | 101585744 | 101586358 | + | 0.636961 | intron (NM_001164583, intron 1 of 18) | intron (NM_001164583, intron 1 of 18) | 35457 | NM_001164583 | 72685 | Mm.76494 | NM_198412 | ENSMUSG00000028528 | **Dnajc6** | 2810027M23Rik\|mKIAA0473 | DnaJ (Hsp40) homolog, subfamily C, member 6 | protein-coding |
| chr14_7989326_7989869 | chr14 | 7989327 | 7989869 | + | 1.8875 | intron (NM_007870, intron 1 of 7) | B1_Mur1\|SINE\|Alu | 4584 | NM_007870 | 13421 | Mm.272258 | NM_007870 | | **Dnase1l3** | DNasegamma\|Dhp2\|Lsd | deoxyribonuclease 1-like 3 | protein-coding |
| chr14_7928782_7929815 | chr14 | 7928783 | 7929815 | + | 1.15769 | intron (NM_001081427, intron 35 of 45) | intron (NM_001081427, intron 35 of 45) | 64883 | NM_007870 | 13421 | Mm.272258 | NM_007870 | | **Dnase1l3** | DNasegamma\|Dhp2\|Lsd | deoxyribonuclease 1-like 3 | protein-coding |
| chr14_7986868_7987332 | chr14 | 7986869 | 7987332 | + | 0.741239 | intron (NM_007870, intron 2 of 7) | intron (NM_007870, intron 2 of 7) | 7082 | NM_007870 | 13421 | Mm.272258 | NM_007870 | | **Dnase1l3** | DNasegamma\|Dhp2\|Lsd | deoxyribonuclease 1-like 3 | protein-coding |
| chr9_21439265_21439788 | chr9 | 21439266 | 21439788 | + | 0.888245 | intron (NM_007871, intron 1 of 19) | intron (NM_007871, intron 1 of 19) | 14619 | NM_001039520 | 13430 | Mm.433257 | NM_007871 | ENSMUSG00000033335 | **Dnm2** | Dyn2\|Udnm\|b2b2159Clo | dynamin 2 | protein-coding |
| chr1_162509323_162509970 | chr1 | 162509324 | 162509970 | + | 1.54237 | Intergenic | Intergenic | -31613 | NM_172646 | 103967 | Mm.441620 | NM_172646 | ENSMUSG00000040265 | **Dnm3** | 9630020E24Rik\|AW061159\|B230343F03Rik\|C530045C17\|mKIAA0820 | dynamin 3 | protein-coding |
| chr1_162509325_162510189 | chr1 | 162509326 | 162510189 | + | 1.44333 | Intergenic | Intergenic | -31723 | NM_172646 | 103967 | Mm.441620 | NM_172646 | ENSMUSG00000040265 | **Dnm3** | 9630020E24Rik\|AW061159\|B230343F03Rik\|C530045C17\|mKIAA0820 | dynamin 3 | protein-coding |
| chr12_3729590_3730046 | chr12 | 3729591 | 3730046 | + | 1.62292 | intron (NM_001162465, intron 10 of 17) | intron (NM_001162465, intron 10 of 17) | -77162 | NM_007872 | 13435 | Mm.5001 | NM_007872 | ENSMUSG00000020661 | **Dnmt3a** | MmuIIIA | DNA methyltransferase 3A | protein-coding |
| chr12_3886231_3886691 | chr12 | 3886232 | 3886691 | + | 0.652958 | intron (NM_007872, intron 6 of 22) | intron (NM_007872, intron 6 of 22) | -5283 | NM_153743 | 13435 | Mm.5001 | NM_007872 | ENSMUSG00000020661 | **Dnmt3a** | MmuIIIA | DNA methyltransferase 3A | protein-coding |
| chr12_3886265_3886567 | chr12 | 3886266 | 3886567 | + | 0.644765 | intron (NM_007872, intron 6 of 22) | intron (NM_007872, intron 6 of 22) | -5328 | NM_153743 | 13435 | Mm.5001 | NM_007872 | ENSMUSG00000020661 | **Dnmt3a** | MmuIIIA | DNA methyltransferase 3A | protein-coding |
| chr12_3845769_3846683 | chr12 | 3845770 | 3846683 | + | 1.45194 | intron (NM_007872, intron 2 of 22) | intron (NM_007872, intron 2 of 22) | 16018 | NR_045884 | 100038522 | Mm.417270 | NR_045884 | ENSMUSG00000073242 | **Dnmt3aos** | ENSMUSG00000073242\|Gm10485 | DNA methyltransferase 3A, opposite strand | ncRNA |
| chr12_3835566_3835976 | chr12 | 3835567 | 3835976 | + | 1.02822 | intron (NM_007872, intron 2 of 22) | intron (NM_007872, intron 2 of 22) | 26473 | NR_045884 | 100038522 | Mm.417270 | NR_045884 | ENSMUSG00000073242 | **Dnmt3aos** | ENSMUSG00000073242\|Gm10485 | DNA methyltransferase 3A, opposite strand | ncRNA |
| chr12_3861360_3862739 | chr12 | 3861361 | 3862739 | + | 0.699248 | non-coding (NR_045884, exon 1 of 4) | non-coding (NR_045884, exon 1 of 4) | 194 | NR_045884 | 100038522 | Mm.417270 | NR_045884 | ENSMUSG00000073242 | **Dnmt3aos** | ENSMUSG00000073242\|Gm10485 | DNA methyltransferase 3A, opposite strand | ncRNA |
| chr12_3834130_3835421 | chr12 | 3834131 | 3835421 | + | 0.645642 | intron (NM_001271753, intron 1 of 22) | intron (NM_001271753, intron 1 of 22) | 27468 | NR_045884 | 100038522 | Mm.417270 | NR_045884 | ENSMUSG00000073242 | **Dnmt3aos** | ENSMUSG00000073242\|Gm10485 | DNA methyltransferase 3A, opposite strand | ncRNA |
| chr17_46496548_46497267 | chr17 | 46496549 | 46497267 | + | 0.650518 | exon (NM_207161, exon 1 of 4) | exon (NM_207161, exon 1 of 4) | 119 | NM_207161 | 381101 | Mm.35801 | NM_207161 | ENSMUSG00000040658 | **Dnph1** | C76683\|Rcl | 2'-deoxynucleoside 5'-phosphate N-hydrolase 1 | protein-coding |
| chr1_80855070_80855704 | chr1 | 80855071 | 80855704 | + | 2.04325 | Intergenic | Intergenic | -96834 | NM_175291 | 210293 | Mm.133473 | NM_175291 | ENSMUSG00000038608 | **Dock10** | 9330153B10Rik\|A630054M16Rik\|Jr4\|Jr5\|R75174\|ZIZ3\|Zizimin3\|mKIAA0694 | dedicator of cytokinesis 10 | protein-coding |
| chr1_80603050_80603563 | chr1 | 80603051 | 80603563 | + | 1.5544 | intron (NM_001285927, intron 7 of 55) | intron (NM_001285927, intron 7 of 55) | 62045 | NM_001285927 | 210293 | Mm.133473 | NM_175291 | ENSMUSG00000038608 | **Dock10** | 9330153B10Rik\|A630054M16Rik\|Jr4\|Jr5\|R75174\|ZIZ3\|Zizimin3\|mKIAA0694 | dedicator of cytokinesis 10 | protein-coding |
| chr1_80655218_80655719 | chr1 | 80655219 | 80655719 | + | 1.43967 | intron (NM_175291, intron 1 of 55) | intron (NM_175291, intron 1 of 55) | 9883 | NM_001285927 | 210293 | Mm.133473 | NM_175291 | ENSMUSG00000038608 | **Dock10** | 9330153B10Rik\|A630054M16Rik\|Jr4\|Jr5\|R75174\|ZIZ3\|Zizimin3\|mKIAA0694 | dedicator of cytokinesis 10 | protein-coding |
| chr1_80749308_80749649 | chr1 | 80749309 | 80749649 | + | 1.37677 | intron (NM_175291, intron 1 of 55) | intron (NM_175291, intron 1 of 55) | 9074 | NM_175291 | 210293 | Mm.133473 | NM_175291 | ENSMUSG00000038608 | **Dock10** | 9330153B10Rik\|A630054M16Rik\|Jr4\|Jr5\|R75174\|ZIZ3\|Zizimin3\|mKIAA0694 | dedicator of cytokinesis 10 | protein-coding |
| chr1_80749084_80749657 | chr1 | 80749085 | 80749657 | + | 1.3562 | intron (NM_175291, intron 1 of 55) | MTA_Mm\|LTR\|ERVL-MaLR | 9182 | NM_175291 | 210293 | Mm.133473 | NM_175291 | ENSMUSG00000038608 | **Dock10** | 9330153B10Rik\|A630054M16Rik\|Jr4\|Jr5\|R75174\|ZIZ3\|Zizimin3\|mKIAA0694 | dedicator of cytokinesis 10 | protein-coding |
| chr1_80619085_80619426 | chr1 | 80619086 | 80619426 | + | 1.2296 | intron (NM_001285927, intron 3 of 55) | B3\|SINE\|B2 | 46096 | NM_001285927 | 210293 | Mm.133473 | NM_175291 | ENSMUSG00000038608 | **Dock10** | 9330153B10Rik\|A630054M16Rik\|Jr4\|Jr5\|R75174\|ZIZ3\|Zizimin3\|mKIAA0694 | dedicator of cytokinesis 10 | protein-coding |
| chr1_80737711_80738418 | chr1 | 80737712 | 80738418 | + | 1.21865 | intron (NM_175291, intron 1 of 55) | intron (NM_175291, intron 1 of 55) | 20488 | NM_175291 | 210293 | Mm.133473 | NM_175291 | ENSMUSG00000038608 | **Dock10** | 9330153B10Rik\|A630054M16Rik\|Jr4\|Jr5\|R75174\|ZIZ3\|Zizimin3\|mKIAA0694 | dedicator of cytokinesis 10 | protein-coding |
| chr1_80639648_80640044 | chr1 | 80639649 | 80640044 | + | 1.0971 | intron (NM_001285927, intron 2 of 55) | L1MB7\|LINE\|L1 | 25506 | NM_001285927 | 210293 | Mm.133473 | NM_175291 | ENSMUSG00000038608 | **Dock10** | 9330153B10Rik\|A630054M16Rik\|Jr4\|Jr5\|R75174\|ZIZ3\|Zizimin3\|mKIAA0694 | dedicator of cytokinesis 10 | protein-coding |
| chr1_80641842_80642166 | chr1 | 80641843 | 80642166 | + | 0.896533 | intron (NM_001285927, intron 2 of 55) | intron (NM_001285927, intron 2 of 55) | 23348 | NM_001285927 | 210293 | Mm.133473 | NM_175291 | ENSMUSG00000038608 | **Dock10** | 9330153B10Rik\|A630054M16Rik\|Jr4\|Jr5\|R75174\|ZIZ3\|Zizimin3\|mKIAA0694 | dedicator of cytokinesis 10 | protein-coding |
| chr1_80613083_80613955 | chr1 | 80613084 | 80613955 | + | 0.861776 | intron (NM_001285927, intron 4 of 55) | intron (NM_001285927, intron 4 of 55) | 51833 | NM_001285927 | 210293 | Mm.133473 | NM_175291 | ENSMUSG00000038608 | **Dock10** | 9330153B10Rik\|A630054M16Rik\|Jr4\|Jr5\|R75174\|ZIZ3\|Zizimin3\|mKIAA0694 | dedicator of cytokinesis 10 | protein-coding |
| chr1_80746764_80747049 | chr1 | 80746765 | 80747049 | + | 0.825761 | intron (NM_175291, intron 1 of 55) | intron (NM_175291, intron 1 of 55) | 11646 | NM_175291 | 210293 | Mm.133473 | NM_175291 | ENSMUSG00000038608 | **Dock10** | 9330153B10Rik\|A630054M16Rik\|Jr4\|Jr5\|R75174\|ZIZ3\|Zizimin3\|mKIAA0694 | dedicator of cytokinesis 10 | protein-coding |
| chr1_80630192_80631035 | chr1 | 80630193 | 80631035 | + | 0.760339 | intron (NM_001285927, intron 3 of 55) | intron (NM_001285927, intron 3 of 55) | 34738 | NM_001285927 | 210293 | Mm.133473 | NM_175291 | ENSMUSG00000038608 | **Dock10** | 9330153B10Rik\|A630054M16Rik\|Jr4\|Jr5\|R75174\|ZIZ3\|Zizimin3\|mKIAA0694 | dedicator of cytokinesis 10 | protein-coding |

| | | | | | | | | | | | | | | | | | |
|---|---|---|---|---|---|---|---|---|---|---|---|---|---|---|---|---|---|
| chr1_80614730_80615119 | chr1 | 80614731 | 80615119 | + | 0.676916 | intron (NM_001285927, intron 4 of 55) | intron (NM_001285927, intron 4 of 55) | 50427 | NM_001285927 | 210293 | Mm.133473 | NM_175291 | ENSMUSG00000038608 | **Dock10** | 9330153B10Rik\|A630054M16Rik\|Jr4\|Jr5\|R75174\|ZIZ3\|Zizimin3\|mKIAA0694 | dedicator of cytokinesis 10 | protein-coding |
| chrX_35930191_35931358 | chrX | 35930192 | 35931358 | + | 1.06521 | intron (NM_001009947, intron 1 of 52) | intron (NM_001009947, intron 1 of 52) | 41943 | NM_001009947 | 75974 | Mm.32873 | NM_177745 | ENSMUSG00000031093 | **Dock11** | 5033414A21Rik\|8030476J24\|Zizimin2\|sph238 | dedicator of cytokinesis 11 | protein-coding |
| chr11_34763754_34764131 | chr11 | 34763755 | 34764131 | + | 1.18897 | intron (NM_033374, intron 1 of 51) | ORR1F\|LTR\|ERVL-MaLR | 19962 | NM_033374 | 94176 | Mm.380679 | NM_033374 | ENSMUSG00000020143 | **Dock2** | AI662014\|AW122239\|CED-5\|Hch\|MBC | dedicator of cyto-kinesis 2 | protein-coding |
| chr11_34762143_34762496 | chr11 | 34762144 | 34762496 | + | 0.834564 | intron (NM_033374, intron 1 of 51) | L1MB5\|LINE\|L1 | 21585 | NM_033374 | 94176 | Mm.380679 | NM_033374 | ENSMUSG00000020143 | **Dock2** | AI662014\|AW122239\|CED-5\|Hch\|MBC | dedicator of cyto-kinesis 2 | protein-coding |
| chr11_34737147_34737620 | chr11 | 34737148 | 34737620 | + | 0.819956 | intron (NM_033374, intron 2 of 51) | L1M3\|LINE\|L1 | 46521 | NM_033374 | 94176 | Mm.380679 | NM_033374 | ENSMUSG00000020143 | **Dock2** | AI662014\|AW122239\|CED-5\|Hch\|MBC | dedicator of cyto-kinesis 2 | protein-coding |
| chr11_34776716_34777239 | chr11 | 34776717 | 34777239 | + | 0.644165 | intron (NM_033374, intron 1 of 51) | intron (NM_033374, intron 1 of 51) | 6927 | NM_033374 | 94176 | Mm.380679 | NM_033374 | ENSMUSG00000020143 | **Dock2** | AI662014\|AW122239\|CED-5\|Hch\|MBC | dedicator of cyto-kinesis 2 | protein-coding |
| chr11_34782280_34782830 | chr11 | 34782281 | 34782830 | + | 0.629168 | intron (NM_033374, intron 1 of 51) | intron (NM_033374, intron 1 of 51) | 1350 | NM_033374 | 94176 | Mm.380679 | NM_033374 | ENSMUSG00000020143 | **Dock2** | AI662014\|AW122239\|CED-5\|Hch\|MBC | dedicator of cyto-kinesis 2 | protein-coding |
| chr9_107238727_107239278 | chr9 | 107238728 | 107239278 | + | 0.643467 | Intergenic | Intergenic | -7094 | NM_153413 | 208869 | Mm.150259 | NM_153413 | ENSMUSG00000039716 | **Dock3** | MOCA\|PBP\|mKIAA0299 | dedicator of cyto-kinesis 3 | protein-coding |
| chr9_21854040_21854321 | chr9 | 21854041 | 21854321 | + | 0.641173 | Intergenic | Intergenic | -1546 | NM_177030 | 319899 | Mm.128153 | NM_177030 | ENSMUSG00000032198 | **Dock6** | 2410095B20Rik\|4931431C02Rik\|C330023D02Rik\|mKIAA1395 | dedicator of cytokinesis 6 | protein-coding |
| chr19_25078103_25078700 | chr19 | 25078104 | 25078700 | + | 1.42012 | intron (NM_028785, intron 5 of 47) | intron (NM_028785, intron 5 of 47) | 78873 | NM_028785 | 76088 | Mm.102485 | NM_028785 | ENSMUSG00000052085 | **Dock8** | 1200017A24Rik\|5830472H07Rik\|A130095G14Rik\|AI461977 | dedicator of cytokinesis 8 | protein-coding |
| chr19_24984993_24985394 | chr19 | 24984994 | 24985394 | + | 0.965769 | Intergenic | Lx8\|LINE\|L1 | -14335 | NM_028785 | 76088 | Mm.102485 | NM_028785 | ENSMUSG00000052085 | **Dock8** | 1200017A24Rik\|5830472H07Rik\|A130095G14Rik\|AI461977 | dedicator of cytokinesis 8 | protein-coding |
| chr19_24984793_24985410 | chr19 | 24984794 | 24985410 | + | 0.945544 | Intergenic | Lx8\|LINE\|L1 | -14427 | NM_028785 | 76088 | Mm.102485 | NM_028785 | ENSMUSG00000052085 | **Dock8** | 1200017A24Rik\|5830472H07Rik\|A130095G14Rik\|AI461977 | dedicator of cytokinesis 8 | protein-coding |
| chr19_25074839_25075188 | chr19 | 25074840 | 25075188 | + | 0.877505 | intron (NM_028785, intron 4 of 47) | intron (NM_028785, intron 4 of 47) | 75485 | NM_028785 | 76088 | Mm.102485 | NM_028785 | ENSMUSG00000052085 | **Dock8** | 1200017A24Rik\|5830472H07Rik\|A130095G14Rik\|AI461977 | dedicator of cytokinesis 8 | protein-coding |
| chr19_25062203_25063163 | chr19 | 25062204 | 25063163 | + | 0.857242 | intron (NM_028785, intron 3 of 47) | intron (NM_028785, intron 3 of 47) | 63154 | NM_028785 | 76088 | Mm.102485 | NM_028785 | ENSMUSG00000052085 | **Dock8** | 1200017A24Rik\|5830472H07Rik\|A130095G14Rik\|AI461977 | dedicator of cytokinesis 8 | protein-coding |
| chr19_24983855_24984392 | chr19 | 24983856 | 24984392 | + | 0.729329 | Intergenic | Intergenic | -15405 | NM_028785 | 76088 | Mm.102485 | NM_028785 | ENSMUSG00000052085 | **Dock8** | 1200017A24Rik\|5830472H07Rik\|A130095G14Rik\|AI461977 | dedicator of cytokinesis 8 | protein-coding |
| chr14_121823068_121823368 | chr14 | 121823069 | 121823368 | + | 1.40922 | Intergenic | Intergenic | -25484 | NM_134074 | 105445 | Mm.441054 | NM_134074 | ENSMUSG00000025558 | **Dock9** | AA959601\|AW538057\|B230309H04Rik\|D14Wsu89e\|Zizimin1\|mKIAA1058 | dedicator of cytokinesis 9 | protein-coding |
| chr14_121710065_121710585 | chr14 | 121710066 | 121710585 | + | 1.38216 | intron (NM_134074, intron 1 of 52) | intron (NM_134074, intron 1 of 52) | -11908 | NM_001081039 | 105445 | Mm.441054 | NM_134074 | ENSMUSG00000025558 | **Dock9** | AA959601\|AW538057\|B230309H04Rik\|D14Wsu89e\|Zizimin1\|mKIAA1058 | dedicator of cytokinesis 9 | protein-coding |
| chr14_121822958_121823525 | chr14 | 121822959 | 121823525 | + | 1.37797 | Intergenic | Intergenic | -25508 | NM_134074 | 105445 | Mm.441054 | NM_134074 | ENSMUSG00000025558 | **Dock9** | AA959601\|AW538057\|B230309H04Rik\|D14Wsu89e\|Zizimin1\|mKIAA1058 | dedicator of cytokinesis 9 | protein-coding |
| chr14_121826161_121826429 | chr14 | 121826162 | 121826429 | + | 1.14211 | Intergenic | MIRb\|SINE\|MIR | -28561 | NM_134074 | 105445 | Mm.441054 | NM_134074 | ENSMUSG00000025558 | **Dock9** | AA959601\|AW538057\|B230309H04Rik\|D14Wsu89e\|Zizimin1\|mKIAA1058 | dedicator of cytokinesis 9 | protein-coding |
| chr14_121658184_121659352 | chr14 | 121658185 | 121659352 | + | 1.08511 | exon (NM_001128307, exon 6 of 56) | exon (NM_001128307, exon 6 of 56) | 39649 | NM_001081039 | 105445 | Mm.441054 | NM_134074 | ENSMUSG00000025558 | **Dock9** | AA959601\|AW538057\|B230309H04Rik\|D14Wsu89e\|Zizimin1\|mKIAA1058 | dedicator of cytokinesis 9 | protein-coding |
| chr14_121824481_121825131 | chr14 | 121824482 | 121825131 | + | 0.95562 | Intergenic | Intergenic | -27072 | NM_134074 | 105445 | Mm.441054 | NM_134074 | ENSMUSG00000025558 | **Dock9** | AA959601\|AW538057\|B230309H04Rik\|D14Wsu89e\|Zizimin1\|mKIAA1058 | dedicator of cytokinesis 9 | protein-coding |
| chr14_121666651_121666949 | chr14 | 121666652 | 121666949 | + | 0.680916 | intron (NM_001081039, intron 3 of 53) | intron (NM_001081039, intron 3 of 53) | 31617 | NM_001081039 | 105445 | Mm.441054 | NM_134074 | ENSMUSG00000025558 | **Dock9** | AA959601\|AW538057\|B230309H04Rik\|D14Wsu89e\|Zizimin1\|mKIAA1058 | dedicator of cytokinesis 9 | protein-coding |
| chr6_83027906_83028877 | chr6 | 83027907 | 83028877 | + | 0.942114 | exon (NM_033079, exon 9 of 10) | exon (NM_033079, exon 9 of 10) | 5079 | NM_010070 | 13448 | Mm.156 | NM_010070 | ENSMUSG00000068335 | **Dok1** | AW557123\|p62DOK | docking protein 1 | protein-coding |
| chr6_83017149_83017737 | chr6 | 83017150 | 83017737 | + | 0.871132 | intron (NM_033079, intron 7 of 9) | intron (NM_033079, intron 7 of 9) | 16028 | NM_010070 | 13448 | Mm.156 | NM_010070 | ENSMUSG00000068335 | **Dok1** | AW557123\|p62DOK | docking protein 1 | protein-coding |
| chr6_83017154_83017396 | chr6 | 83017155 | 83017396 | + | 0.861891 | intron (NM_033079, intron 7 of 9) | L3\|LINE\|CR1 | 16196 | NM_010070 | 13448 | Mm.156 | NM_010070 | ENSMUSG00000068335 | **Dok1** | AW557123\|p62DOK | docking protein 1 | protein-coding |
| chr14_70778060_70778382 | chr14 | 70778061 | 70778382 | + | 1.23482 | 3' UTR (NM_010071, exon 5 of 5) | 3' UTR (NM_010071, exon 5 of 5) | 3840 | NM_010071 | 13449 | Mm.243323 | NM_010071 | ENSMUSG00000022102 | **Dok2** | DokR\|Frip | docking protein 2 | protein-coding |
| chr14_70798327_70798978 | chr14 | 70798328 | 70798978 | + | 0.758625 | Intergenic | L2\|LINE\|L2 | 24272 | NM_010071 | 13449 | Mm.243323 | NM_010071 | ENSMUSG00000022102 | **Dok2** | DokR\|Frip | docking protein 2 | protein-coding |
| chr14_70798625_70798933 | chr14 | 70798626 | 70798933 | + | 0.725508 | Intergenic | L2\|LINE\|L2 | 24398 | NM_010071 | 13449 | Mm.243323 | NM_010071 | ENSMUSG00000022102 | **Dok2** | DokR\|Frip | docking protein 2 | protein-coding |
| chr18_89502325_89502872 | chr18 | 89502326 | 89502872 | + | 0.800727 | intron (NM_001039173, intron 3 of 7) | MTE2b\|LTR\|ERVL-MaLR | 266537 | NM_001039173 | 623279 | Mm.447164 | NM_001039173 | | **Dok6** | Dok-6 | docking protein 6 | protein-coding |
| chr18_89606147_89606504 | chr18 | 89606148 | 89606504 | + | 0.764162 | intron (NM_001039173, intron 1 of 7) | B3A\|SINE\|B2 | 162810 | NM_001039173 | 623279 | Mm.447164 | NM_001039173 | | **Dok6** | Dok-6 | docking protein 6 | protein-coding |
| chr16_93746381_93746789 | chr16 | 93746382 | 93746789 | + | 1.18326 | intron (NM_027293, intron 4 of 36) | intron (NM_027293, intron 4 of 36) | 34678 | NM_027293 | 70028 | Mm.23230 | NM_026700 | ENSMUSG00000022946 | **Dopey2** | 0610038M01Rik\|2610510B01Rik\|AI503246\|C21orf5\|mKIAA0933 | dopey family member 2 | protein-coding |
| chr16_93746241_93746807 | chr16 | 93746242 | 93746807 | + | 1.02768 | intron (NM_027293, intron 4 of 36) | intron (NM_027293, intron 4 of 36) | 34617 | NM_027293 | 70028 | Mm.23230 | NM_026700 | ENSMUSG00000022946 | **Dopey2** | 0610038M01Rik\|2610510B01Rik\|AI503246\|C21orf5\|mKIAA0933 | dopey family member 2 | protein-coding |
| chr10_80758695_80759079 | chr10 | 80758696 | 80759079 | + | 1.15376 | intron (NM_199322, intron 1 of 27) | intron (NM_199322, intron 1 of 27) | 3681 | NM_199322 | 208266 | Mm.152371 | NM_199322 | ENSMUSG00000061589 | **Dot1l** | A630076O07\|AW907654\|Dot1\|KMT4\|mDot1 | DOT1-like, histone H3 methyltransferase (S. cerevisiae) | protein-coding |
| chr10_80752060_80752623 | chr10 | 80752061 | 80752623 | + | 0.813741 | Intergenic | Intergenic | -2864 | NM_199322 | 208266 | Mm.152371 | NM_199322 | ENSMUSG00000061589 | **Dot1l** | A630076O07\|AW907654\|Dot1\|KMT4\|mDot1 | DOT1-like, histone H3 methyltransferase (S. cerevisiae) | protein-coding |
| chr10_80751198_80751933 | chr10 | 80751199 | 80751933 | + | 0.69669 | Intergenic | Intergenic | -3640 | NM_199322 | 208266 | Mm.152371 | NM_199322 | ENSMUSG00000061589 | **Dot1l** | A630076O07\|AW907654\|Dot1\|KMT4\|mDot1 | DOT1-like, histone H3 methyltransferase (S. cerevisiae) | protein-coding |
| chr8_123197069_123197416 | chr8 | 123197070 | 123197416 | + | 0.7901 | intron (NM_007876, intron 2 of 10) | ORR1A2\|LTR\|ERVL-MaLR | 11008 | NM_007876 | 13479 | Mm.20388 | NM_007876 | ENSMUSG00000019278 | **Dpep1** | AI327012\|MBD | dipeptidase 1 (renal) | protein-coding |
| chr11_75186159_75186470 | chr11 | 75186160 | 75186470 | + | 1.02316 | intron (NM_144491, intron 1 of 12) | intron (NM_144491, intron 1 of 12) | 4168 | NM_144491 | 116905 | Mm.41496 | NM_144491 | ENSMUSG00000078789 | **Dph1** | 2310011M22Rik\|4930488F09Rik\|AW551873\|Dph2l1\|Ovca1 | DPH1 homolog (S. cerevisiae) | protein-coding |
| chr2_114658762_114659337 | chr2 | 114658763 | 114659337 | + | 1.56186 | Intergenic | Intergenic | -4122 | NM_025675 | 66632 | Mm.117809 | NM_025675 | ENSMUSG00000057147 | **Dph6** | 5730421E18Rik\|Atpbd4 | diphthamine biosynthesis 6 | protein-coding |
| chr2_62350188_62350831 | chr2 | 62350189 | 62350831 | + | 1.40129 | intron (NM_001159543, intron 20 of 24) | intron (NM_001159543, intron 20 of 24) | 61721 | NM_010074 | 13482 | Mm.1151 | NM_010074 | ENSMUSG00000035000 | **Dpp4** | Cd26\|Dpp-4\|THAM | dipeptidylpeptidase 4 | protein-coding |
| chr2_62331849_62332356 | chr2 | 62331850 | 62332356 | + | 1.16439 | 3' UTR (NM_010074, exon 26 of 26) | 3' UTR (NM_010074, exon 26 of 26) | 80128 | NM_010074 | 13482 | Mm.1151 | NM_010074 | ENSMUSG00000035000 | **Dpp4** | Cd26\|Dpp-4\|THAM | dipeptidylpeptidase 4 | protein-coding |
| chr2_62398538_62398811 | chr2 | 62398539 | 62398811 | + | 1.14646 | intron (NM_001159543, intron 2 of 24) | intron (NM_001159543, intron 2 of 24) | 13556 | NM_010074 | 13482 | Mm.1151 | NM_010074 | ENSMUSG00000035000 | **Dpp4** | Cd26\|Dpp-4\|THAM | dipeptidylpeptidase 4 | protein-coding |
| chr2_62399456_62399718 | chr2 | 62399457 | 62399718 | + | 0.819947 | intron (NM_001159543, intron 2 of 24) | intron (NM_001159543, intron 2 of 24) | 12644 | NM_010074 | 13482 | Mm.1151 | NM_010074 | ENSMUSG00000035000 | **Dpp4** | Cd26\|Dpp-4\|THAM | dipeptidylpeptidase 4 | protein-coding |
| chr2_62410314_62410501 | chr2 | 62410315 | 62410501 | + | 0.782009 | intron (NM_001159543, intron 2 of 24) | intron (NM_001159543, intron 2 of 24) | 1823 | NM_010074 | 13482 | Mm.1151 | NM_010074 | ENSMUSG00000035000 | **Dpp4** | Cd26\|Dpp-4\|THAM | dipeptidylpeptidase 4 | protein-coding |
| chr7_35745106_35745714 | chr7 | 35745107 | 35745714 | + | 1.35678 | intron (NM_178704, intron 3 of 18) | intron (NM_178704, intron 3 of 18) | 9044 | NM_178704 | 233115 | Mm.257808 | NM_178704 | ENSMUSG00000043671 | **Dpy19l3** | 6030410G08\|9330164H19Rik | dpy-19-like 3 (C. elegans) | protein-coding |
| chr7_35742298_35742709 | chr7 | 35742299 | 35742709 | + | 1.28315 | intron (NM_178704, intron 3 of 18) | intron (NM_178704, intron 3 of 18) | 11950 | NM_178704 | 233115 | Mm.257808 | NM_178704 | ENSMUSG00000043671 | **Dpy19l3** | 6030410G08\|9330164H19Rik | dpy-19-like 3 (C. elegans) | protein-coding |

| | | | | | | | | | | | | | | | | | |
|---|---|---|---|---|---|---|---|---|---|---|---|---|---|---|---|---|---|
| chr7_35715858_35716442 | chr7 | 35715859 | 35716442 | + | 1.10215 | intron (NM_178704, intron 9 of 18) | intron (NM_178704, intron 9 of 18) | 38304 | NM_178704 | 233115 | Mm.257808 | NM_178704 | ENSMUSG00000043671 | **Dpy19l3** | 6030410G08\|9330164H19Rik | dpy-19-like 3 (C. elegans) | protein-coding |
| chr7_35728422_35728740 | chr7 | 35728423 | 35728740 | + | 0.782111 | intron (NM_178704, intron 4 of 18) | intron (NM_178704, intron 4 of 18) | 25873 | NM_178704 | 233115 | Mm.257808 | NM_178704 | ENSMUSG00000043671 | **Dpy19l3** | 6030410G08\|9330164H19Rik | dpy-19-like 3 (C. elegans) | protein-coding |
| chr7_35718449_35718876 | chr7 | 35718450 | 35718876 | + | 0.670308 | intron (NM_178704, intron 8 of 18) | Lx8\|LINE\|L1 | 35791 | NM_178704 | 233115 | Mm.257808 | NM_178704 | ENSMUSG00000043671 | **Dpy19l3** | 6030410G08\|9330164H19Rik | dpy-19-like 3 (C. elegans) | protein-coding |
| chr5_108245986_108246918 | chr5 | 108245987 | 108246918 | + | 0.764651 | Intergenic | Intergenic | -22445 | NM_026106 | 13486 | Mm.303534 | NM_026106 | ENSMUSG00000029265 | **Dr1** | 1700121L09Rik\|Dr1l\|NC2\|NC2beta | down-regulator of transcription 1 | protein-coding |
| chr16_43805406_43806076 | chr16 | 43805407 | 43806076 | + | 1.24363 | intron (NM_007877, intron 2 of 6) | intron (NM_007877, intron 2 of 6) | 43499 | NM_007877 | 13490 | Mm.439735 | NM_007877 | ENSMUSG00000022705 | **Drd3** | D3R | dopamine receptor D3 | protein-coding |
| chr17_64740066_64740584 | chr17 | 64740067 | 64740584 | + | 1.13935 | intron (NM_008549, intron 19 of 21) | intron (NM_008549, intron 19 of 21) | -26786 | NR_105051 | 101202683 | Mm.393309 | NR_105051 | | **Dreh** | - | down-regulated in hepatocellular carcinoma | ncRNA |
| chr17_64740066_64740818 | chr17 | 64740067 | 64740818 | + | 0.969896 | intron (NM_008549, intron 19 of 21) | intron (NM_008549, intron 19 of 21) | -26669 | NR_105051 | 101202683 | Mm.393309 | NR_105051 | | **Dreh** | - | down-regulated in hepatocellular carcinoma | ncRNA |
| chr17_64833685_64834114 | chr17 | 64833686 | 64834114 | + | 0.95111 | Intergenic | Intergenic | 66789 | NR_105051 | 101202683 | Mm.393309 | NR_105051 | | **Dreh** | - | down-regulated in hepatocellular carcinoma | ncRNA |
| chr17_64826632_64827883 | chr17 | 64826633 | 64827883 | + | 0.692433 | Intergenic | Intergenic | 60147 | NR_105051 | 101202683 | Mm.393309 | NR_105051 | | **Dreh** | - | down-regulated in hepatocellular carcinoma | ncRNA |
| chr17_64826814_64827267 | chr17 | 64826815 | 64827267 | + | 0.665697 | Intergenic | Intergenic | 59930 | NR_105051 | 101202683 | Mm.393309 | NR_105051 | | **Dreh** | - | down-regulated in hepatocellular carcinoma | ncRNA |
| chr17_64826627_64827392 | chr17 | 64826628 | 64827392 | + | 0.641733 | Intergenic | Intergenic | 59899 | NR_105051 | 101202683 | Mm.393309 | NR_105051 | | **Dreh** | - | down-regulated in hepatocellular carcinoma | ncRNA |
| chr11_3246901_3247729 | chr11 | 3246902 | 3247729 | + | 1.66033 | intron (NR_037301, intron 1 of 1) | intron (NR_037301, intron 1 of 1) | 19071 | NM_007879 | 13494 | Mm.3250 | NM_007879 | ENSMUSG00000020457 | **Drg1** | AA408859\|AI132520\|Nedd3 | developmentally regulated GTP binding protein 1 | protein-coding |
| chrX_134374095_134374571 | chrX | 134374096 | 134374571 | + | 1.01021 | Intergenic | RLTR28\|LTR\|ERVL | -30447 | NM_010078 | 13497 | Mm.121662 | NM_010078 | ENSMUSG00000000223 | **Drp2** | AW495265\|DRP-2 | dystrophin related protein 2 | protein-coding |
| chr16_97204052_97204633 | chr16 | 97204053 | 97204633 | + | 0.784906 | Intergenic | Intergenic | -33608 | NM_031174 | 13508 | Mm.328737 | NM_031174 | ENSMUSG00000050272 | **Dscam** | 4932410A21Rik | Down syndrome cell adhesion molecule | protein-coding |
| chr16_96819655_96820075 | chr16 | 96819656 | 96820075 | + | 0.675941 | exon (NM_031174, exon 8 of 33) | exon (NM_031174, exon 8 of 33) | 350870 | NM_031174 | 13508 | Mm.328737 | NM_031174 | ENSMUSG00000050272 | **Dscam** | 4932410A21Rik | Down syndrome cell adhesion molecule | protein-coding |
| chr16_94524771_94525026 | chr16 | 94524772 | 94525026 | + | 0.826088 | intron (NM_007834, intron 1 of 7) | intron (NM_007834, intron 1 of 7) | 1730 | NM_007834 | 13185 | Mm.7472 | NM_007834 | ENSMUSG00000022898 | **Dscr3** | AW538125\|Dcra | Down syndrome critical region gene 3 | protein-coding |
| chr12_52002830_52003146 | chr12 | 52002831 | 52003146 | + | 0.80836 | intron (NM_029545, intron 2 of 2) | intron (NM_029545, intron 2 of 2) | 3513 | NM_029545 | 328092 | Mm.2821 | NM_029545 | ENSMUSG00000020956 | **Dtd2** | 4930578F06Rik\|6530401N04Rik\|B830049N13Rik | D-tyrosyl-tRNA deacylase 2 | protein-coding |
| chr5_62892841_62893663 | chr5 | 62892842 | 62893663 | + | 1.28418 | Intergenic | Intergenic | 79429 | NM_001170705 | 100322896 | Mm.479821 | NM_001170705 | ENSMUSG00000090326 | **Dthd1** | Gm17384 | death domain containing 1 | protein-coding |
| chr5_62854130_62854872 | chr5 | 62854131 | 62854872 | + | 0.801271 | intron (NM_001170705, intron 7 of 8) | intron (NM_001170705, intron 7 of 8) | 40678 | NM_001170705 | 100322896 | Mm.479821 | NM_001170705 | ENSMUSG00000090326 | **Dthd1** | Gm17384 | death domain containing 1 | protein-coding |
| chr5_62814155_62814472 | chr5 | 62814156 | 62814472 | + | 0.657981 | exon (NM_001170705, exon 1 of 9) | exon (NM_001170705, exon 1 of 9) | 491 | NM_001170705 | 100322896 | Mm.479821 | NM_001170705 | ENSMUSG00000090326 | **Dthd1** | Gm17384 | death domain containing 1 | protein-coding |
| chr5_62820755_62821233 | chr5 | 62820756 | 62821233 | + | 0.636519 | intron (NM_001170705, intron 2 of 8) | intron (NM_001170705, intron 2 of 8) | 7171 | NM_001170705 | 100322896 | Mm.479821 | NM_001170705 | ENSMUSG00000090326 | **Dthd1** | Gm17384 | death domain containing 1 | protein-coding |
| chr1_191569682_191570155 | chr1 | 191569683 | 191570155 | + | 0.790271 | intron (NM_029766, intron 2 of 14) | B1_Mm\|SINE\|Alu | 5615 | NM_029766 | 76843 | Mm.189102 | NM_029766 | ENSMUSG00000037474 | **Dtl** | 2810047L02Rik\|5730564G15Rik\|Ramp | denticleless homolog (Drosophila) | protein-coding |
| chr12_3610821_3611228 | chr12 | 3610822 | 3611228 | + | 1.06197 | intron (NM_001162465, intron 4 of 17) | MER21C\|LTR\|ERVL | 37984 | NM_007886 | 13528 | Mm.286202 | NM_007886 | ENSMUSG00000071454 | **Dtnb** | - | dystrobrevin, beta | protein-coding |
| chr12_3610682_3611630 | chr12 | 3610683 | 3611630 | + | 1.04111 | intron (NM_001162465, intron 4 of 17) | intron (NM_001162465, intron 4 of 17) | 38115 | NM_007886 | 13528 | Mm.286202 | NM_007886 | ENSMUSG00000071454 | **Dtnb** | - | dystrobrevin, beta | protein-coding |
| chr12_3549278_3549871 | chr12 | 3549279 | 3549871 | + | 0.994302 | Intergenic | Intergenic | -22816 | NM_001162465 | 13528 | Mm.286202 | NM_007886 | ENSMUSG00000071454 | **Dtnb** | - | dystrobrevin, beta | protein-coding |
| chr12_3555978_3556370 | chr12 | 3555979 | 3556370 | + | 0.78776 | Intergenic | B4A\|SINE\|B4 | -16217 | NM_001162465 | 13528 | Mm.286202 | NM_007886 | ENSMUSG00000071454 | **Dtnb** | - | dystrobrevin, beta | protein-coding |
| chr2_126310027_126310688 | chr2 | 126310028 | 126310688 | + | 0.756828 | Intergenic | Intergenic | 158217 | NM_026981 | 69185 | Mm.29019 | NM_026981 | ENSMUSG00000023330 | **Dtwd1** | 1810033A06Rik | DTW domain containing 1 | protein-coding |
| chr18_49784112_49784685 | chr18 | 49784113 | 49784685 | + | 1.40863 | Intergenic | Intergenic | -28798 | NM_001170960 | 68857 | Mm.96010 | NM_026854 | ENSMUSG00000024505 | **Dtwd2** | 1190002H09Rik\|8030470C17Rik\|AI414673\|BB115449 | DTW domain containing 2 | protein-coding |
| chr18_49721673_49722176 | chr18 | 49721674 | 49722176 | + | 0.652362 | intron (NM_001170960, intron 2 of 3) | intron (NM_001170960, intron 2 of 3) | 33676 | NM_026854 | 68857 | Mm.96010 | NM_026854 | ENSMUSG00000024505 | **Dtwd2** | 1190002H09Rik\|8030470C17Rik\|AI414673\|BB115449 | DTW domain containing 2 | protein-coding |
| chr18_49721456_49722219 | chr18 | 49721457 | 49722219 | + | 0.63983 | intron (NM_001170960, intron 2 of 3) | intron (NM_001170960, intron 2 of 3) | 33763 | NM_026854 | 68857 | Mm.96010 | NM_026854 | ENSMUSG00000024505 | **Dtwd2** | 1190002H09Rik\|8030470C17Rik\|AI414673\|BB115449 | DTW domain containing 2 | protein-coding |
| chr10_127199701_127200071 | chr10 | 127199702 | 127200071 | + | 0.835622 | intron (NM_054097, intron 7 of 9) | intron (NM_054097, intron 7 of 9) | -4177 | NM_030714 | 80904 | Mm.271724 | NM_030714 | ENSMUSG00000040415 | **Dtx3** | Deltex3\|mg310 | deltex 3 homolog (Drosophila) | protein-coding |
| chr8_106007004_106007762 | chr8 | 106007005 | 106007762 | + | 1.46476 | Intergenic | RLTR19\|LTR\|ERVK | -4095 | NM_025518 | 66369 | Mm.287500 | NM_025518 | ENSMUSG00000031901 | **Dus2** | 2310016K04Rik\|Dus2l | dihydrouridine synthase 2 | protein-coding |
| chr17_26518331_26518696 | chr17 | 26518332 | 26518696 | + | 1.06499 | Intergenic | Intergenic | -10042 | NM_013642 | 19252 | Mm.239041 | NM_013642 | ENSMUSG00000024190 | **Dusp1** | 3CH134\|MKP1\|Ptpn16\|U19515\|erp\|mkp-1 | dual specificity phosphatase 1 | protein-coding |
| chr1_184196988_184197801 | chr1 | 184196989 | 184197801 | + | 0.908762 | Intergenic | MER21B\|LTR\|ERVL | 162934 | NM_022019 | 63953 | Mm.404024 | NM_022019 | ENSMUSG00000039384 | **Dusp10** | 2610306G15Rik\|AI158871\|MKP-5\|MKP5 | dual specificity phosphatase 10 | protein-coding |
| chr1_184084836_184085324 | chr1 | 184084837 | 184085324 | + | 0.756237 | Intergenic | Intergenic | 50619 | NM_022019 | 63953 | Mm.404024 | NM_022019 | ENSMUSG00000039384 | **Dusp10** | 2610306G15Rik\|AI158871\|MKP-5\|MKP5 | dual specificity phosphatase 10 | protein-coding |
| chr1_184182154_184182524 | chr1 | 184182155 | 184182524 | + | 0.669602 | Intergenic | RSINE1\|SINE\|B4 | 147878 | NM_022019 | 63953 | Mm.404024 | NM_022019 | ENSMUSG00000039384 | **Dusp10** | 2610306G15Rik\|AI158871\|MKP-5\|MKP5 | dual specificity phosphatase 10 | protein-coding |
| chr6_85969943_85970450 | chr6 | 85969944 | 85970450 | + | 0.958118 | Intergenic | Intergenic | -8530 | NM_028099 | 72102 | Mm.24007 | NM_028099 | ENSMUSG00000030002 | **Dusp11** | 2010300F21Rik\|AI481497 | dual specificity phosphatase 11 (RNA/RNP complex 1-interacting) | protein-coding |
| chr11_84066947_84067294 | chr11 | 84066948 | 84067294 | + | 0.817165 | intron (NM_019819, intron 1 of 2) | intron (NM_019819, intron 1 of 2) | 1236 | NM_019819 | 56405 | Mm.240885 | NM_019819 | ENSMUSG00000018648 | **Dusp14** | 1110014C10Rik\|2310042C07Rik\|D11Ertd395e\|MKP-L\|Mkp6 | dual specificity phosphatase 14 | protein-coding |
| chr11_84069067_84069387 | chr11 | 84069068 | 84069387 | + | 0.65149 | promoter-TSS (NM_019819) | promoter-TSS (NM_019819) | -870 | NM_019819 | 56405 | Mm.240885 | NM_019819 | ENSMUSG00000018648 | **Dusp14** | 1110014C10Rik\|2310042C07Rik\|D11Ertd395e\|MKP-L\|Mkp6 | dual specificity phosphatase 14 | protein-coding |
| chr6_134807074_134807446 | chr6 | 134807075 | 134807446 | + | 0.857724 | Intergenic | Intergenic | -14632 | NM_001048054 | 70686 | Mm.491182 | NM_130447 | ENSMUSG00000030203 | **Dusp16** | 3830417M17Rik\|AW558566\|D6Ertd213e\|MKP-7\|MKP7\|Mkpm | dual specificity phosphatase 16 | protein-coding |
| chr2_127332753_127332942 | chr2 | 127332754 | 127332942 | + | 1.29775 | Intergenic | L2b\|LINE\|L2 | -3311 | NM_010090 | 13537 | Mm.4729 | NM_010090 | ENSMUSG00000027368 | **Dusp2** | PAC1 | dual specificity phosphatase 2 | protein-coding |
| chr2_127323622_127324088 | chr2 | 127323623 | 127324088 | + | 0.670438 | Intergenic | Intergenic | -12304 | NM_010090 | 13537 | Mm.4729 | NM_010090 | ENSMUSG00000027368 | **Dusp2** | PAC1 | dual specificity phosphatase 2 | protein-coding |
| chr2_127333251_127333578 | chr2 | 127333252 | 127333578 | + | 0.641677 | Intergenic | Intergenic | -2744 | NM_010090 | 13537 | Mm.4729 | NM_010090 | ENSMUSG00000027368 | **Dusp2** | PAC1 | dual specificity phosphatase 2 | protein-coding |
| chr13_30657746_30658148 | chr13 | 30657747 | 30658148 | + | 1.21709 | Intergenic | Intergenic | -2112 | NM_001037955 | 105352 | Mm.289646 | NM_134068 | ENSMUSG00000069255 | **Dusp22** | JKAP\|JSP1\|LMW-DSP2\|MKPX | dual specificity phosphatase 22 | protein-coding |
| chr13_30638375_30639370 | chr13 | 30638376 | 30639370 | + | 1.11555 | Intergenic | L1MB4\|LINE\|L1 | -21186 | NM_001037955 | 105352 | Mm.289646 | NM_134068 | ENSMUSG00000069255 | **Dusp22** | JKAP\|JSP1\|LMW-DSP2\|MKPX | dual specificity phosphatase 22 | protein-coding |
| chr13_30702379_30702948 | chr13 | 30702380 | 30702948 | + | 0.829587 | intron (NM_001037955, intron 4 of 6) | intron (NM_001037955, intron 4 of 6) | 42605 | NM_134068 | 105352 | Mm.289646 | NM_134068 | ENSMUSG00000069255 | **Dusp22** | JKAP\|JSP1\|LMW-DSP2\|MKPX | dual specificity phosphatase 22 | protein-coding |
| chr13_30692578_30693446 | chr13 | 30692579 | 30693446 | + | 0.77556 | intron (NM_001037955, intron 3 of 6) | intron (NM_001037955, intron 3 of 6) | 32953 | NM_134068 | 105352 | Mm.289646 | NM_134068 | ENSMUSG00000069255 | **Dusp22** | JKAP\|JSP1\|LMW-DSP2\|MKPX | dual specificity phosphatase 22 | protein-coding |
| chr13_30642041_30642415 | chr13 | 30642042 | 30642415 | + | 0.751985 | Intergenic | L1MC3\|LINE\|L1 | -17831 | NM_001037955 | 105352 | Mm.289646 | NM_134068 | ENSMUSG00000069255 | **Dusp22** | JKAP\|JSP1\|LMW-DSP2\|MKPX | dual specificity phosphatase 22 | protein-coding |
| chr13_30650571_30651126 | chr13 | 30650572 | 30651126 | + | 0.751629 | Intergenic | Intergenic | -9210 | NM_001037955 | 105352 | Mm.289646 | NM_134068 | ENSMUSG00000069255 | **Dusp22** | JKAP\|JSP1\|LMW-DSP2\|MKPX | dual specificity phosphatase 22 | protein-coding |
| chr13_30702137_30702997 | chr13 | 30702138 | 30702997 | + | 0.747946 | intron (NM_001037955, intron 4 of 6) | intron (NM_001037955, intron 4 of 6) | 42508 | NM_134068 | 105352 | Mm.289646 | NM_134068 | ENSMUSG00000069255 | **Dusp22** | JKAP\|JSP1\|LMW-DSP2\|MKPX | dual specificity phosphatase 22 | protein-coding |
| chr1_92905864_92906149 | chr1 | 92905865 | 92906149 | + | 0.65767 | promoter-TSS (NM_175118) | promoter-TSS (NM_175118) | -982 | NM_175118 | 67446 | Mm.46179 | NM_175118 | ENSMUSG00000047067 | **Dusp28** | 0710001B24Rik\|AV005521 | dual specificity phosphatase 28 | protein-coding |
| chr8_34755390_34756109 | chr8 | 34755391 | 34756109 | + | 1.63701 | Intergenic | Intergenic | -51860 | NM_176933 | 319520 | Mm.170276 | NM_176933 | ENSMUSG00000031530 | **Dusp4** | 2700078F24Rik\|AI844617\|BB104621\|E130306H24Rik\|MKP2 | dual specificity phosphatase 4 | protein-coding |
| chr8_34759056_34759374 | chr8 | 34759057 | 34759374 | + | 1.12328 | Intergenic | Intergenic | -48395 | NM_176933 | 319520 | Mm.170276 | NM_176933 | ENSMUSG00000031530 | **Dusp4** | 2700078F24Rik\|AI844617\|BB104621\|E130306H24Rik\|MKP2 | dual specificity phosphatase 4 | protein-coding |
| chr8_34756248_34756561 | chr8 | 34756249 | 34756561 | + | 1.07098 | Intergenic | Intergenic | -51205 | NM_176933 | 319520 | Mm.170276 | NM_176933 | ENSMUSG00000031530 | **Dusp4** | 2700078F24Rik\|AI844617\|BB104621\|E130306H24Rik\|MKP2 | dual specificity phosphatase 4 | protein-coding |
| chr8_34756228_34756782 | chr8 | 34756229 | 34756782 | + | 0.978746 | Intergenic | MIR\|SINE\|MIR | -51105 | NM_176933 | 319520 | Mm.170276 | NM_176933 | ENSMUSG00000031530 | **Dusp4** | 2700078F24Rik\|AI844617\|BB104621\|E130306H24Rik\|MKP2 | dual specificity phosphatase 4 | protein-coding |

| | | | | | | | | | | | | | | | | | |
|---|---|---|---|---|---|---|---|---|---|---|---|---|---|---|---|---|---|
| chr8_34756978_34757291 | chr8 | 34756979 | 34757291 | + | 0.941364 | Intergenic | Intergenic | -50475 | NM_176933 | 319520 | Mm.170276 | NM_176933 | ENSMUSG00000031530 | **Dusp4** | 2700078F24Rik\|AI844617\|BB104621\|E130306H24Rik\|MKP2 | dual specificity phosphatase 4 | protein-coding |
| chr8_34781721_34782423 | chr8 | 34781722 | 34782423 | + | 0.916418 | Intergenic | Intergenic | -25538 | NM_176933 | 319520 | Mm.170276 | NM_176933 | ENSMUSG00000031530 | **Dusp4** | 2700078F24Rik\|AI844617\|BB104621\|E130306H24Rik\|MKP2 | dual specificity phosphatase 4 | protein-coding |
| chr8_34743196_34743758 | chr8 | 34743197 | 34743758 | + | 0.713396 | Intergenic | Intergenic | -64133 | NM_176933 | 319520 | Mm.170276 | NM_176933 | ENSMUSG00000031530 | **Dusp4** | 2700078F24Rik\|AI844617\|BB104621\|E130306H24Rik\|MKP2 | dual specificity phosphatase 4 | protein-coding |
| chr19_53544230_53545128 | chr19 | 53544231 | 53545128 | + | 0.840215 | Intergenic | Intergenic | 15361 | NM_001085390 | 240672 | Mm.482424 | NM_001085390 | | **Dusp5** | Gm337 | dual specificity phosphatase 5 | protein-coding |
| chr19_53536116_53537433 | chr19 | 53536117 | 53537433 | + | 0.714817 | intron (NM_001085390, intron 2 of 3) | intron (NM_001085390, intron 2 of 3) | 7457 | NM_001085390 | 240672 | Mm.482424 | NM_001085390 | | **Dusp5** | Gm337 | dual specificity phosphatase 5 | protein-coding |
| chr19_53536730_53537230 | chr19 | 53536731 | 53537230 | + | 0.646156 | intron (NM_001085390, intron 2 of 3) | intron (NM_001085390, intron 2 of 3) | 7662 | NM_001085390 | 240672 | Mm.482424 | NM_001085390 | | **Dusp5** | Gm337 | dual specificity phosphatase 5 | protein-coding |
| chr10_99379210_99379748 | chr10 | 99379211 | 99379748 | + | 1.26328 | Intergenic | Intergenic | 116248 | NM_026268 | 67603 | Mm.1791 | NM_026268 | ENSMUSG00000019960 | **Dusp6** | 1300019I03Rik\|MKP-3\|MKP3\|PYST1 | dual specificity phosphatase 6 | protein-coding |
| chr10_99379233_99379940 | chr10 | 99379234 | 99379940 | + | 1.14678 | Intergenic | Intergenic | 116356 | NM_026268 | 67603 | Mm.1791 | NM_026268 | ENSMUSG00000019960 | **Dusp6** | 1300019I03Rik\|MKP-3\|MKP3\|PYST1 | dual specificity phosphatase 6 | protein-coding |
| chr10_99396583_99396981 | chr10 | 99396584 | 99396981 | + | 0.809958 | Intergenic | Intergenic | 133551 | NM_026268 | 67603 | Mm.1791 | NM_026268 | ENSMUSG00000019960 | **Dusp6** | 1300019I03Rik\|MKP-3\|MKP3\|PYST1 | dual specificity phosphatase 6 | protein-coding |
| chr10_99350547_99350880 | chr10 | 99350548 | 99350880 | + | 0.772497 | Intergenic | Intergenic | 87483 | NM_026268 | 67603 | Mm.1791 | NM_026268 | ENSMUSG00000019960 | **Dusp6** | 1300019I03Rik\|MKP-3\|MKP3\|PYST1 | dual specificity phosphatase 6 | protein-coding |
| chr10_99376209_99376691 | chr10 | 99376210 | 99376691 | + | 0.768157 | Intergenic | Intergenic | 113219 | NM_026268 | 67603 | Mm.1791 | NM_026268 | ENSMUSG00000019960 | **Dusp6** | 1300019I03Rik\|MKP-3\|MKP3\|PYST1 | dual specificity phosphatase 6 | protein-coding |
| chr10_99210879_99211838 | chr10 | 99210880 | 99211838 | + | 0.697052 | Intergenic | L1MC4\|LINE\|L1 | -51872 | NM_026268 | 67603 | Mm.1791 | NM_026268 | ENSMUSG00000019960 | **Dusp6** | 1300019I03Rik\|MKP-3\|MKP3\|PYST1 | dual specificity phosphatase 6 | protein-coding |
| chr7_142115131_142115501 | chr7 | 142115132 | 142115501 | + | 1.03534 | Intergenic | Intergenic | -20032 | NM_008748 | 18218 | Mm.39725 | NM_008748 | ENSMUSG00000037887 | **Dusp8** | 5530400B01Rik\|AI593498\|Nttp1 | dual specificity phosphatase 8 | protein-coding |
| chr7_142113784_142114232 | chr7 | 142113785 | 142114232 | + | 0.740138 | Intergenic | PB1D10\|SINE\|Alu | -18724 | NM_008748 | 18218 | Mm.39725 | NM_008748 | ENSMUSG00000037887 | **Dusp8** | 5530400B01Rik\|AI593498\|Nttp1 | dual specificity phosphatase 8 | protein-coding |
| chr7_142113567_142114405 | chr7 | 142113568 | 142114405 | + | 0.656724 | Intergenic | PB1D10\|SINE\|Alu | -18702 | NM_008748 | 18218 | Mm.39725 | NM_008748 | ENSMUSG00000037887 | **Dusp8** | 5530400B01Rik\|AI593498\|Nttp1 | dual specificity phosphatase 8 | protein-coding |
| chr2_125245995_125246219 | chr2 | 125245996 | 125246219 | + | 1.1373 | Intergenic | B3A\|SINE\|B2 | -1141 | NM_001159646 | 110074 | Mm.282499 | NM_023595 | ENSMUSG00000027203 | **Dut** | 5031412I06Rik\|5133400F09Rik\|D2Bwg0749e\|Dutp\|dUTPase | deoxyuridine triphosphatase | protein-coding |
| chr2_125217215_125217650 | chr2 | 125217216 | 125217650 | + | 0.826261 | intron (NM_183354, intron 21 of 26) | ORR1E\|LTR\|ERVL-MaLR | -29815 | NM_001159646 | 110074 | Mm.282499 | NM_023595 | ENSMUSG00000027203 | **Dut** | 5031412I06Rik\|5133400F09Rik\|D2Bwg0749e\|Dutp\|dUTPase | deoxyuridine triphosphatase | protein-coding |
| chr2_71274400_71274985 | chr2 | 71274401 | 71274985 | + | 1.46559 | 3' UTR (NM_172436, exon 18 of 18) | 3' UTR (NM_172436, exon 18 of 18) | 62747 | NM_010064 | 13427 | Mm.249479 | NM_010064 | ENSMUSG00000027012 | **Dync1i2** | 3110079H08Rik\|AW554389\|Dncic2 | dynein cytoplasmic 1 intermediate chain 2 | protein-coding |
| chr2_71274359_71275252 | chr2 | 71274360 | 71275252 | + | 1.25472 | 3' UTR (NM_172436, exon 18 of 18) | 3' UTR (NM_172436, exon 18 of 18) | 62860 | NM_010064 | 13427 | Mm.249479 | NM_010064 | ENSMUSG00000027012 | **Dync1i2** | 3110079H08Rik\|AW554389\|Dncic2 | dynein cytoplasmic 1 intermediate chain 2 | protein-coding |
| chr9_7158665_7158971 | chr9 | 7158666 | 7158971 | + | 0.66653 | intron (NM_029851, intron 15 of 88) | intron (NM_029851, intron 15 of 88) | 18228 | NM_029851 | 110350 | Mm.237596 | NM_029851 | ENSMUSG00000047193 | **Dync2h1** | 4432416O06Rik\|AI448217\|D030010H02Rik\|D330044F14Rik\|DHC1b\|DHC2\|Dnchc2\|b2b414Clo\|mKIAA1997 | dynein cytoplasmic 2 heavy chain 1 | protein-coding |
| chr17_84616488_84617218 | chr17 | 84616489 | 84617218 | + | 1.20902 | intron (NM_177606, intron 28 of 29) | intron (NM_177606, intron 28 of 29) | -9646 | NM_172256 | 213575 | Mm.246119 | NM_172256 | ENSMUSG00000024253 | **Dync2li1** | 4933404O11Rik\|CGI-60\|D2lic\|LIC3\|mD2LIC | dynein cytoplasmic 2 light intermediate chain 1 | protein-coding |
| chr2_155242303_155242625 | chr2 | 155242304 | 155242625 | + | 0.660598 | intron (NR_111890, intron 1 of 3) | intron (NR_111890, intron 1 of 3) | 5856 | NM_001291108 | 67068 | Mm.23693 | NM_025947 | ENSMUSG00000047459 | **Dynlrb1** | 2010012N15Rik\|2010320M17Rik\|9430076K19Rik\|AV124457\|DNLC2A\|Dncl2a\|km23-1 | dynein light chain roadblock-type 1 | protein-coding |
| chr8_116618656_116619268 | chr8 | 116618657 | 116619268 | + | 1.41249 | intron (NM_029441, intron 2 of 6) | intron (NM_029441, intron 2 of 6) | 113947 | NM_029297 | 75465 | Mm.23114 | NM_029297 | ENSMUSG00000034467 | **Dynlrb2** | 1700009A04Rik\|DNLC2B\|Dncl2b | dynein light chain roadblock-type 2 | protein-coding |
| chr8_116618452_116619308 | chr8 | 116618453 | 116619308 | + | 1.1967 | intron (NM_029441, intron 2 of 6) | intron (NM_029441, intron 2 of 6) | 113865 | NM_029297 | 75465 | Mm.23114 | NM_029297 | ENSMUSG00000034467 | **Dynlrb2** | 1700009A04Rik\|DNLC2B\|Dncl2b | dynein light chain roadblock-type 2 | protein-coding |
| chr8_116552117_116552607 | chr8 | 116552118 | 116552607 | + | 1.14998 | Intergenic | Intergenic | 47347 | NM_029297 | 75465 | Mm.23114 | NM_029297 | ENSMUSG00000034467 | **Dynlrb2** | 1700009A04Rik\|DNLC2B\|Dncl2b | dynein light chain roadblock-type 2 | protein-coding |
| chr8_116552825_116553271 | chr8 | 116552826 | 116553271 | + | 0.732 | Intergenic | Intergenic | 48033 | NM_029297 | 75465 | Mm.23114 | NM_029297 | ENSMUSG00000034467 | **Dynlrb2** | 1700009A04Rik\|DNLC2B\|Dncl2b | dynein light chain roadblock-type 2 | protein-coding |
| chr16_94611370_94611764 | chr16 | 94611371 | 94611764 | + | 1.52461 | intron (NM_007890, intron 2 of 11) | intron (NM_007890, intron 2 of 11) | 40660 | NM_001113389 | 13548 | Mm.310973 | NM_007890 | ENSMUSG00000022897 | **Dyrk1a** | 2310043O08Rik\|D16Ertd272e\|D16Ertd493e\|Dyrk\|ENSMUSG00000074897\|Gm10783\|Mnbh\|Mp86\|mmb | dual-specificity tyrosine-(Y)-phosphorylation regulated kinase 1a | protein-coding |
| chr16_94647788_94648304 | chr16 | 94647789 | 94648304 | + | 1.32989 | intron (NM_007890, intron 2 of 11) | intron (NM_007890, intron 2 of 11) | 77139 | NM_001113389 | 13548 | Mm.310973 | NM_007890 | ENSMUSG00000022897 | **Dyrk1a** | 2310043O08Rik\|D16Ertd272e\|D16Ertd493e\|Dyrk\|ENSMUSG00000074897\|Gm10783\|Mnbh\|Mp86\|mmb | dual-specificity tyrosine-(Y)-phosphorylation regulated kinase 1a | protein-coding |
| chr16_94586396_94586692 | chr16 | 94586397 | 94586692 | + | 0.889563 | intron (NM_001113389, intron 1 of 11) | MLT1A1\|LTR\|ERVL-MaLR | 15637 | NM_001113389 | 13548 | Mm.310973 | NM_007890 | ENSMUSG00000022897 | **Dyrk1a** | 2310043O08Rik\|D16Ertd272e\|D16Ertd493e\|Dyrk\|ENSMUSG00000074897\|Gm10783\|Mnbh\|Mp86\|mmb | dual-specificity tyrosine-(Y)-phosphorylation regulated kinase 1a | protein-coding |
| chr10_118872443_118872746 | chr10 | 118872444 | 118872746 | + | 0.948813 | Intergenic | Tigger5\|DNA\|TcMar-Tigger | -3692 | NM_001014390 | 69181 | Mm.45565 | NM_001014390 | ENSMUSG00000028630 | **Dyrk2** | 1810038L18Rik | dual-specificity tyrosine-(Y)-phosphorylation regulated kinase 2 | protein-coding |
| chr10_118870314_118870647 | chr10 | 118870315 | 118870647 | + | 0.649293 | Intergenic | Intergenic | -1578 | NM_001014390 | 69181 | Mm.45565 | NM_001014390 | ENSMUSG00000028630 | **Dyrk2** | 1810038L18Rik | dual-specificity tyrosine-(Y)-phosphorylation regulated kinase 2 | protein-coding |
| chr1_131142522_131143043 | chr1 | 131142523 | 131143043 | + | 1.63726 | Intergenic | ORR1B2\|LTR\|ERVL-MaLR | -4549 | NM_145508 | 226419 | Mm.39299 | NM_145508 | ENSMUSG00000016526 | **Dyrk3** | BC006704 | dual-specificity tyrosine-(Y)-phosphorylation regulated kinase 3 | protein-coding |
| chr1_131135183_131135480 | chr1 | 131135184 | 131135480 | + | 0.676779 | intron (NM_145508, intron 2 of 2) | intron (NM_145508, intron 2 of 2) | 2902 | NM_145508 | 226419 | Mm.39299 | NM_145508 | ENSMUSG00000016526 | **Dyrk3** | BC006704 | dual-specificity tyrosine-(Y)-phosphorylation regulated kinase 3 | protein-coding |
| chr6_83993838_83994169 | chr6 | 83993839 | 83994169 | + | 0.94373 | Intergenic | B3\|SINE\|B2 | -14594 | NM_001077694 | 26903 | Mm.220982 | NM_021469 | ENSMUSG00000033788 | **Dysf** | 2310004N10Rik\|AI604795\|D6Pas3 | dysferlin | protein-coding |
| chr14_118912628_118913684 | chr14 | 118912629 | 118913684 | + | 1.85364 | intron (NM_025943, intron 3 of 19) | intron (NM_025943, intron 3 of 19) | 10011 | NM_025943 | 66573 | Mm.87456 | NM_025943 | ENSMUSG00000042156 | **Dzip1** | 2510025K24Rik\|2810422M04Rik\|mKIAA0996 | DAZ interacting protein 1 | protein-coding |
| chr9_96756784_96757116 | chr9 | 96756785 | 96757116 | + | 0.639507 | Intergenic | L1MB8\|LINE\|L1 | -4119 | NR_015511 | 319859 | Mm.259909 | NM_177020 | | **E030011O05Rik** | - | RIKEN cDNA E030011O05 gene | ncRNA |
| chr19_16204783_16205568 | chr19 | 16204784 | 16205568 | + | 1.35596 | intron (NM_008139, intron 1 of 6) | intron (NM_008139, intron 1 of 6) | 41078 | NR_033228 | 595139 | Mm.405662 | NR_033228 | | **E030024N20Rik** | 9030417F11Rik\|Ppia-ps8 | peptidylprolyl isomerase A pseudogene 8 | pseudo |
| chr11_109149369_109149936 | chr11 | 109149370 | 109149936 | + | 1.12893 | Intergenic | Intergenic | -5284 | NR_037978 | 268498 | Mm.227970 | NM_001085512 | ENSMUSG00000078605 | **E030025P04Rik** | Gm664 | RIKEN cDNA E030025P04 gene | ncRNA |
| chr2_168150095_168150533 | chr2 | 168150096 | 168150533 | + | 0.799948 | Intergenic | Intergenic | 4199 | NR_040327 | 77017 | Mm.407822 | NR_040327 | ENSMUSG00000087648 | **E130018N17Rik** | - | RIKEN cDNA E130018N17 gene | ncRNA |
| chr2_150645204_150645635 | chr2 | 150645205 | 150645635 | + | 0.927008 | intron (NM_080575, intron 2 of 13) | intron (NM_080575, intron 2 of 13) | -22074 | NR_040331 | 109344 | Mm.440012 | NR_040331 | ENSMUSG00000087406 | **E130215H24Rik** | - | RIKEN cDNA E130215H24 gene | ncRNA |
| chr4_59682469_59683189 | chr4 | 59682470 | 59683189 | + | 1.89084 | intron (NM_153158, intron 1 of 2) | intron (NM_153158, intron 1 of 2) | 56618 | NM_153158 | 230259 | Mm.45688 | NM_153158 | ENSMUSG00000045071 | **E130308A19Rik** | - | RIKEN cDNA E130308A19 gene | protein-coding |
| chr3_63941196_63942083 | chr3 | 63941197 | 63942083 | + | 0.772639 | TTS (NM_015728) | TTS (NM_015728) | -12255 | NM_177856 | 329659 | Mm.39342 | NM_177856 | ENSMUSG00000048581 | **E130311K13Rik** | - | RIKEN cDNA E130311K13 gene | protein-coding |
| chr3_63941208_63941870 | chr3 | 63941209 | 63941870 | + | 0.766097 | TTS (NM_015728) | TTS (NM_015728) | -12154 | NM_177856 | 329659 | Mm.39342 | NM_177856 | ENSMUSG00000048581 | **E130311K13Rik** | - | RIKEN cDNA E130311K13 gene | protein-coding |
| chr3_63905126_63905715 | chr3 | 63905127 | 63905715 | + | 0.721786 | Intergenic | CpG | 23964 | NM_177856 | 329659 | Mm.39342 | NM_177856 | ENSMUSG00000048581 | **E130311K13Rik** | - | RIKEN cDNA E130311K13 gene | protein-coding |
| chr11_83580839_83581232 | chr11 | 83580840 | 83581232 | + | 0.81897 | Intergenic | Intergenic | -1025 | NR_036453 | 100504464 | Mm.341706 | NR_036452 | ENSMUSG00000051832 | **E230016K23Rik** | - | RIKEN cDNA E230016K23 gene | ncRNA |
| chr6_67014238_67015053 | chr6 | 67014239 | 67015053 | + | 0.81395 | intron (NM_001177560, intron 3 of 4) | intron (NM_001177560, intron 3 of 4) | -21953 | NR_040278 | 320172 | Mm.449039 | NR_040278 | ENSMUSG00000087231 | **E230016M11Rik** | - | RIKEN cDNA E230016M11 gene | ncRNA |
| chr18_36708614_36709312 | chr18 | 36708615 | 36709312 | + | 1.53056 | Intergenic | MER4B-int\|LTR\|ERV1 | -13038 | NM_172831 | 240216 | Mm.252323 | NM_172831 | ENSMUSG00000044719 | **E230025N22Rik** | EG240216 | Riken cDNA E230025N22 gene | protein-coding |
| chr18_36708325_36709317 | chr18 | 36708326 | 36709317 | + | 1.50198 | Intergenic | MER4B-int\|LTR\|ERV1 | -12896 | NM_172831 | 240216 | Mm.252323 | NM_172831 | ENSMUSG00000044719 | **E230025N22Rik** | EG240216 | Riken cDNA E230025N22 gene | protein-coding |
| chr7_90007068_90007818 | chr7 | 90007069 | 90007818 | + | 1.12525 | Intergenic | Intergenic | -21716 | NR_110365 | 319711 | Mm.31753 | NR_015614 | | **E230029C05Rik** | AI452102\|B230337J15 | RIKEN cDNA E230029C05 gene | ncRNA |
| chr7_90067888_90068928 | chr7 | 90067889 | 90068928 | + | 0.784514 | Intergenic | Intergenic | 39249 | NR_110364 | 319711 | Mm.31753 | NR_015614 | | **E230029C05Rik** | AI452102\|B230337J15 | RIKEN cDNA E230029C05 gene | ncRNA |
| chr7_90067893_90068425 | chr7 | 90067894 | 90068425 | + | 0.698103 | Intergenic | Intergenic | 39000 | NR_110364 | 319711 | Mm.31753 | NR_015614 | | **E230029C05Rik** | AI452102\|B230337J15 | RIKEN cDNA E230029C05 gene | ncRNA |
| chr7_90067998_90068378 | chr7 | 90067999 | 90068378 | + | 0.672306 | Intergenic | Intergenic | 39029 | NR_110364 | 319711 | Mm.31753 | NR_015614 | | **E230029C05Rik** | AI452102\|B230337J15 | RIKEN cDNA E230029C05 gene | ncRNA |
| chr13_30022506_30022827 | chr13 | 30022507 | 30022827 | + | 0.770412 | Intergenic | Intergenic | -36604 | NM_010093 | 13557 | Mm.268356 | NM_010093 | ENSMUSG00000016477 | **E2f3** | E2F3b\|E2f3a | E2F transcription factor 3 | protein-coding |
| chr13_30022369_30023024 | chr13 | 30022370 | 30023024 | + | 0.761609 | Intergenic | Intergenic | -36634 | NM_010093 | 13557 | Mm.268356 | NM_010093 | ENSMUSG00000016477 | **E2f3** | E2F3b\|E2f3a | E2F transcription factor 3 | protein-coding |
| chr8_105298904_105299148 | chr8 | 105298905 | 105299148 | + | 0.861366 | exon (NM_148952, exon 4 of 10) | exon (NM_148952, exon 4 of 10) | 1363 | NM_148952 | 104394 | Mm.34554 | NM_148952 | ENSMUSG00000014859 | **E2f4** | 2010111M04Rik\|AI427446 | E2F transcription factor 4 | protein-coding |

| | | | | | | | | | | | | | | | | | |
|---|---|---|---|---|---|---|---|---|---|---|---|---|---|---|---|---|---|
| chr7_48919100_48919858 | chr7 | 48919101 | 48919858 | + | 0.89637 | Intergenic | Intergenic | -38438 | NM_001013368 | 108961 | Mm.240566 | NM_001013368 | ENSMUSG00000046179 | **E2f8** | 4432406C08Rik\|AA410048 | E2F transcription factor 8 | protein-coding |
| chr16_78200397_78200709 | chr16 | 78200398 | 78200709 | + | 0.617854 | Intergenic | Intergenic | 54901 | NR_045698 | 100503619 | Mm.387531 | NR_045698 | | **E330011O21Rik** | - | RIKEN cDNA E330011O21 gene | ncRNA |
| chr19_60182083_60182621 | chr19 | 60182084 | 60182621 | + | 1.01065 | Intergenic | ORR1B1\|LTR\|ERVL-MaLR | 37677 | NR_026942 | 107376 | Mm.245813 | NR_026942 | ENSMUSG00000053117 | **E330013P04Rik** | 9930018I18Rik\|AW492152 | RIKEN cDNA E330013P04 gene | ncRNA |
| chr4_129894892_129895183 | chr4 | 129894893 | 129895183 | + | 0.652549 | Intergenic | B3A\|SINE\|B2 | -11188 | NR_045190 | 319894 | Mm.59196 | NR_045190 | ENSMUSG00000086369 | **E330017L17Rik** | - | RIKEN cDNA E330017L17 gene | ncRNA |
| chr4_129894853_129895604 | chr4 | 129894854 | 129895604 | + | 0.642873 | Intergenic | Intergenic | -10997 | NR_045190 | 319894 | Mm.59196 | NR_045190 | ENSMUSG00000086369 | **E330017L17Rik** | - | RIKEN cDNA E330017L17 gene | ncRNA |
| chr15_96265941_96266380 | chr15 | 96265942 | 96266380 | + | 0.964604 | Intergenic | L4_A_Mam\|LINE\|RTE-X | 9114 | NR_030690 | 319722 | Mm.103823 | NR_030690 | | **E330033B04Rik** | - | RIKEN cDNA E330033B04 gene | ncRNA |
| chr15_96276806_96277565 | chr15 | 96276807 | 96277565 | + | 0.688148 | Intergenic | Intergenic | -1911 | NR_030690 | 319722 | Mm.103823 | NR_030690 | | **E330033B04Rik** | - | RIKEN cDNA E330033B04 gene | ncRNA |
| chr7_78723452_78723902 | chr7 | 78723453 | 78723902 | + | 1.34831 | Intergenic | Intergenic | -5457 | NR_015542 | 414121 | Mm.482537 | NR_015542 | | **E430016F16Rik** | - | RIKEN cDNA E430016F16 gene | ncRNA |
| chr7_78725879_78726197 | chr7 | 78725880 | 78726197 | + | 1.13921 | Intergenic | L1MEf\|LINE\|L1 | -7818 | NR_015542 | 414121 | Mm.482537 | NR_015542 | | **E430016F16Rik** | - | RIKEN cDNA E430016F16 gene | ncRNA |
| chr7_78690795_78691376 | chr7 | 78690796 | 78691376 | + | 1.10319 | intron (NR_015542, intron 5 of 10) | intron (NR_015542, intron 5 of 10) | 27134 | NR_015542 | 414121 | Mm.482537 | NR_015542 | | **E430016F16Rik** | - | RIKEN cDNA E430016F16 gene | ncRNA |
| chr17_24453390_24453758 | chr17 | 24453391 | 24453758 | + | 0.702693 | intron (NM_007893, intron 1 of 13) | intron (NM_007893, intron 1 of 13) | 1737 | NM_007893 | 13560 | Mm.163132 | NM_007893 | ENSMUSG00000024137 | **E4f1** | - | E4F transcription factor 1 | protein-coding |
| chr9_121741108_121741526 | chr9 | 121741109 | 121741526 | + | 0.774459 | intron (NM_010918, intron 7 of 16) | intron (NM_010918, intron 7 of 16) | 18626 | NR_033503 | 320301 | Mm.373883 | NR_033503 | | **E530011L22Rik** | - | RIKEN cDNA E530011L22 gene | ncRNA |
| chr16_36821507_36822231 | chr16 | 36821508 | 36822231 | + | 1.31667 | intron (NM_001113401, intron 2 of 5) | intron (NM_001113401, intron 2 of 5) | 6390 | NM_001113401 | 106389 | Mm.432078 | NM_134111 | ENSMUSG00000022838 | **Eaf2** | AW048865\|FESTA-L\|FESTA-S\|Festa\|Traits\|U19 | ELL associated factor 2 | protein-coding |
| chr16_36821507_36821938 | chr16 | 36821508 | 36821938 | + | 1.28128 | intron (NM_001113401, intron 2 of 5) | intron (NM_001113401, intron 2 of 5) | 6536 | NM_001113401 | 106389 | Mm.432078 | NM_134111 | ENSMUSG00000022838 | **Eaf2** | AW048865\|FESTA-L\|FESTA-S\|Festa\|Traits\|U19 | ELL associated factor 2 | protein-coding |
| chr15_44597534_44598517 | chr15 | 44597535 | 44598517 | + | 0.892513 | TTS (NM_138674) | TTS (NM_138674) | -21615 | NM_019480 | 55960 | Mm.287896 | NM_019480 | ENSMUSG00000022339 | **Ebag9** | AI835379\|Rcas1 | estrogen receptor-binding fragment-associated gene 9 | protein-coding |
| chr15_44597895_44598510 | chr15 | 44597896 | 44598510 | + | 0.870618 | Intergenic | Intergenic | -21438 | NM_019480 | 55960 | Mm.287896 | NM_019480 | ENSMUSG00000022339 | **Ebag9** | AI835379\|Rcas1 | estrogen receptor-binding fragment-associated gene 9 | protein-coding |
| chr11_44636145_44636639 | chr11 | 44636146 | 44636639 | + | 1.15122 | intron (NM_001290709, intron 5 of 15) | MIRb\|SINE\|MIR | 18292 | NM_001290710 | 13591 | Mm.439662 | NM_007897 | | **Ebf1** | Ebf\|O/E-1\|OE-1\|Olf-1\|Olf1 | early B cell factor 1 | protein-coding |
| chr11_44636912_44637438 | chr11 | 44636913 | 44637438 | + | 0.885912 | intron (NM_001290709, intron 5 of 15) | intron (NM_001290709, intron 5 of 15) | 19075 | NM_001290710 | 13591 | Mm.439662 | NM_007897 | | **Ebf1** | Ebf\|O/E-1\|OE-1\|Olf-1\|Olf1 | early B cell factor 1 | protein-coding |
| chr11_44606913_44607234 | chr11 | 44606914 | 44607234 | + | 0.721792 | Intergenic | Intergenic | -11026 | NM_001290711 | 13591 | Mm.439662 | NM_007897 | | **Ebf1** | Ebf\|O/E-1\|OE-1\|Olf-1\|Olf1 | early B cell factor 1 | protein-coding |
| chr14_61376104_61376758 | chr14 | 61376105 | 61376758 | + | 0.726588 | intron (NM_008466, intron 10 of 16) | ORR1C1\|LTR\|ERVL-MaLR | -15986 | NM_026598 | 68177 | Mm.22758 | NM_026598 | ENSMUSG00000021928 | **Ebpl** | 5730442K12Rik\|EBRP | emopamil binding protein-like | protein-coding |
| chr4_137860004_137860370 | chr4 | 137860005 | 137860370 | + | 1.31348 | Intergenic | Intergenic | -2050 | NM_199307 | 230857 | Mm.401062 | NM_199307 | ENSMUSG00000057530 | **Ece1** | AW322500\|ECE-1\|ECE-1a\|ECE-1b\|ECE-1d | endothelin converting enzyme 1 | protein-coding |
| chr4_137875317_137875806 | chr4 | 137875318 | 137875806 | + | 1.28314 | intron (NM_199307, intron 1 of 19) | intron (NM_199307, intron 1 of 19) | 13325 | NM_199307 | 230857 | Mm.401062 | NM_199307 | ENSMUSG00000057530 | **Ece1** | AW322500\|ECE-1\|ECE-1a\|ECE-1b\|ECE-1d | endothelin converting enzyme 1 | protein-coding |
| chr4_137874684_137876011 | chr4 | 137874685 | 137876011 | + | 1.06923 | intron (NM_199307, intron 1 of 19) | intron (NM_199307, intron 1 of 19) | 13111 | NM_199307 | 230857 | Mm.401062 | NM_199307 | ENSMUSG00000057530 | **Ece1** | AW322500\|ECE-1\|ECE-1a\|ECE-1b\|ECE-1d | endothelin converting enzyme 1 | protein-coding |
| chr4_137879456_137880115 | chr4 | 137879457 | 137880115 | + | 0.85876 | intron (NM_199307, intron 1 of 19) | intron (NM_199307, intron 1 of 19) | 17549 | NM_199307 | 230857 | Mm.401062 | NM_199307 | ENSMUSG00000057530 | **Ece1** | AW322500\|ECE-1\|ECE-1a\|ECE-1b\|ECE-1d | endothelin converting enzyme 1 | protein-coding |
| chr4_137853645_137854227 | chr4 | 137853646 | 137854227 | + | 0.633848 | Intergenic | Intergenic | -8301 | NM_199307 | 230857 | Mm.401062 | NM_199307 | ENSMUSG00000057530 | **Ece1** | AW322500\|ECE-1\|ECE-1a\|ECE-1b\|ECE-1d | endothelin converting enzyme 1 | protein-coding |
| chr7_28821552_28821837 | chr7 | 28821553 | 28821837 | + | 0.844302 | intron (NM_177301, intron 12 of 12) | intron (NM_177301, intron 12 of 12) | -3643 | NM_016772 | 51798 | Mm.291776 | NM_016772 | ENSMUSG00000053898 | **Ech1** | AA617331 | enoyl coenzyme A hydratase 1, peroxisomal | protein-coding |
| chr7_28820581_28821487 | chr7 | 28820582 | 28821487 | + | 0.609391 | intron (NM_177301, intron 10 of 12) | intron (NM_177301, intron 10 of 12) | -4304 | NM_016772 | 51798 | Mm.291776 | NM_016772 | ENSMUSG00000053898 | **Ech1** | AA617331 | enoyl coenzyme A hydratase 1, peroxisomal | protein-coding |
| chr13_49513328_49514000 | chr13 | 49513329 | 49514000 | + | 1.03014 | intron (NM_001012324, intron 1 of 9) | intron (NM_001012324, intron 1 of 9) | 8854 | NM_001012324 | 407800 | Mm.386931 | NM_001012324 | ENSMUSG00000043631 | **Ecm2** | 9030618O22Rik\|BC065151\|tenonectin | extracellular matrix protein 2, female organ and adipocyte specific | protein-coding |
| chr13_49513338_49514389 | chr13 | 49513339 | 49514389 | + | 0.882785 | intron (NM_001012324, intron 1 of 9) | B4\|SINE\|B4 | 9054 | NM_001012324 | 407800 | Mm.386931 | NM_001012324 | ENSMUSG00000043631 | **Ecm2** | 9030618O22Rik\|BC065151\|tenonectin | extracellular matrix protein 2, female organ and adipocyte specific | protein-coding |
| chr6_108928927_108929362 | chr6 | 108928928 | 108929362 | + | 0.996365 | Intergenic | Intergenic | 100504 | NM_138677 | 192193 | Mm.21596 | NM_138677 | ENSMUSG00000030104 | **Edem1** | A130059K23Rik\|EDEM\|mKIAA0212 | ER degradation enhancer, mannosidase alpha-like 1 | protein-coding |
| chr6_108894996_108895859 | chr6 | 108894997 | 108895859 | + | 0.812902 | Intergenic | Intergenic | 66787 | NM_138677 | 192193 | Mm.21596 | NM_138677 | ENSMUSG00000030104 | **Edem1** | A130059K23Rik\|EDEM\|mKIAA0212 | ER degradation enhancer, mannosidase alpha-like 1 | protein-coding |
| chr2_155722886_155723553 | chr2 | 155722887 | 155723553 | + | 1.33511 | intron (NM_145537, intron 3 of 10) | B3\|SINE\|B2 | 6255 | NM_145537 | 108687 | Mm.273706 | NM_145537 | ENSMUSG00000038312 | **Edem2** | 9530090G24Rik\|AI327354 | ER degradation enhancer, mannosidase alpha-like 2 | protein-coding |
| chr1_151697464_151698585 | chr1 | 151697465 | 151698585 | + | 1.52741 | intron (NM_022018, intron 8 of 13) | intron (NM_022018, intron 8 of 13) | -57349 | NM_001039644 | 66967 | Mm.337691 | NM_001039644 | ENSMUSG00000043019 | **Edem3** | 2310050N11Rik | ER degradation enhancer, mannosidase alpha-like 3 | protein-coding |
| chr1_151766247_151766489 | chr1 | 151766248 | 151766489 | + | 0.695544 | intron (NM_001039644, intron 2 of 19) | intron (NM_001039644, intron 2 of 19) | 10994 | NM_001039644 | 66967 | Mm.337691 | NM_001039644 | ENSMUSG00000043019 | **Edem3** | 2310050N11Rik | ER degradation enhancer, mannosidase alpha-like 3 | protein-coding |
| chr1_151766106_151766506 | chr1 | 151766107 | 151766506 | + | 0.632564 | intron (NM_001039644, intron 2 of 19) | intron (NM_001039644, intron 2 of 19) | 10932 | NM_001039644 | 66967 | Mm.337691 | NM_001039644 | ENSMUSG00000043019 | **Edem3** | 2310050N11Rik | ER degradation enhancer, mannosidase alpha-like 3 | protein-coding |
| chr4_120027687_120028010 | chr4 | 120027688 | 120028010 | + | 1.36767 | intron (NM_010657, intron 2 of 8) | intron (NM_010657, intron 2 of 8) | -133575 | NM_007902 | 13615 | Mm.284855 | NM_007902 | ENSMUSG00000028635 | **Edn2** | VIC | endothelin 2 | protein-coding |
| chr4_120062210_120062717 | chr4 | 120062211 | 120062717 | + | 1.22038 | intron (NM_010657, intron 2 of 8) | intron (NM_010657, intron 2 of 8) | -98960 | NM_007902 | 13615 | Mm.284855 | NM_007902 | ENSMUSG00000028635 | **Edn2** | VIC | endothelin 2 | protein-coding |
| chr4_120059281_120060048 | chr4 | 120059282 | 120060048 | + | 1.21898 | intron (NM_010657, intron 2 of 8) | URR1B\|DNA\|hAT-Charlie | -101759 | NM_007902 | 13615 | Mm.284855 | NM_007902 | ENSMUSG00000028635 | **Edn2** | VIC | endothelin 2 | protein-coding |
| chr4_120058634_120059196 | chr4 | 120058635 | 120059196 | + | 0.866983 | intron (NM_010657, intron 2 of 8) | intron (NM_010657, intron 2 of 8) | -102509 | NM_007902 | 13615 | Mm.284855 | NM_007902 | ENSMUSG00000028635 | **Edn2** | VIC | endothelin 2 | protein-coding |
| chr4_120148419_120149510 | chr4 | 120148420 | 120149510 | + | 0.851165 | Intergenic | CpG | -12459 | NM_007902 | 13615 | Mm.284855 | NM_007902 | ENSMUSG00000028635 | **Edn2** | VIC | endothelin 2 | protein-coding |
| chr4_120058913_120059186 | chr4 | 120058914 | 120059186 | + | 0.817981 | intron (NM_010657, intron 2 of 8) | intron (NM_010657, intron 2 of 8) | -102374 | NM_007902 | 13615 | Mm.284855 | NM_007902 | ENSMUSG00000028635 | **Edn2** | VIC | endothelin 2 | protein-coding |
| chr10_95914452_95915377 | chr10 | 95914453 | 95915377 | + | 1.59092 | Intergenic | Intergenic | -25748 | NM_001001932 | 216238 | Mm.210035 | NM_001001932 | ENSMUSG00000036499 | **Eea1** | A430109M19Rik\|B230358H09Rik\|ZFYVE2 | early endosome antigen 1 | protein-coding |
| chr10_95914129_95915427 | chr10 | 95914130 | 95915427 | + | 1.4989 | Intergenic | MLT2B2\|LTR\|ERVL | -25885 | NM_001001932 | 216238 | Mm.210035 | NM_001001932 | ENSMUSG00000036499 | **Eea1** | A430109M19Rik\|B230358H09Rik\|ZFYVE2 | early endosome antigen 1 | protein-coding |
| chr10_95964336_95964707 | chr10 | 95964337 | 95964707 | + | 0.955765 | intron (NM_001001932, intron 1 of 28) | B3\|SINE\|B2 | 23859 | NM_001001932 | 216238 | Mm.210035 | NM_001001932 | ENSMUSG00000036499 | **Eea1** | A430109M19Rik\|B230358H09Rik\|ZFYVE2 | early endosome antigen 1 | protein-coding |
| chr10_96080516_96081582 | chr10 | 96080517 | 96081582 | + | 0.919618 | Intergenic | Intergenic | 140386 | NM_001001932 | 216238 | Mm.210035 | NM_001001932 | ENSMUSG00000036499 | **Eea1** | A430109M19Rik\|B230358H09Rik\|ZFYVE2 | early endosome antigen 1 | protein-coding |
| chr10_95964326_95965144 | chr10 | 95964327 | 95965144 | + | 0.793575 | intron (NM_001001932, intron 1 of 28) | intron (NM_001001932, intron 1 of 28) | 24072 | NM_001001932 | 216238 | Mm.210035 | NM_001001932 | ENSMUSG00000036499 | **Eea1** | A430109M19Rik\|B230358H09Rik\|ZFYVE2 | early endosome antigen 1 | protein-coding |
| chr9_78474292_78474678 | chr9 | 78474293 | 78474678 | + | 1.1574 | TTS (NM_026658) | TTS (NM_026658) | 7239 | NM_010106 | 13627 | Mm.138471 | NM_010106 | ENSMUSG00000037742 | **Eef1a1** | - | eukaryotic translation elongation factor 1 alpha 1 | protein-coding |
| chr9_78474264_78474954 | chr9 | 78474265 | 78474954 | + | 1.09805 | TTS (NM_026658) | TTS (NM_026658) | 7115 | NM_010106 | 13627 | Mm.138471 | NM_010106 | ENSMUSG00000037742 | **Eef1a1** | - | eukaryotic translation elongation factor 1 alpha 1 | protein-coding |
| chr9_78475985_78477003 | chr9 | 78475986 | 78477003 | + | 1.02405 | Intergenic | Intergenic | 5230 | NM_010106 | 13627 | Mm.138471 | NM_010106 | ENSMUSG00000037742 | **Eef1a1** | - | eukaryotic translation elongation factor 1 alpha 1 | protein-coding |
| chr9_78476568_78477167 | chr9 | 78476569 | 78477167 | + | 1.02189 | Intergenic | Intergenic | 4856 | NM_010106 | 13627 | Mm.138471 | NM_010106 | ENSMUSG00000037742 | **Eef1a1** | - | eukaryotic translation elongation factor 1 alpha 1 | protein-coding |

| | | | | | | | | | | | | | | | | | |
|---|---|---|---|---|---|---|---|---|---|---|---|---|---|---|---|---|---|
| chr9_78513938_78514438 | chr9 | 78513939 | 78514438 | + | 0.721027 | Intergenic | Intergenic | -32464 | NM_010106 | 13627 | Mm.138471 | NM_010106 | ENSMUSG00000037742 | **Eef1a1** | - | eukaryotic translation elongation factor 1 alpha 1 | protein-coding |
| chr9_78475987_78477471 | chr9 | 78475988 | 78477471 | + | 0.661984 | Intergenic | Intergenic | 4995 | NM_010106 | 13627 | Mm.138471 | NM_010106 | ENSMUSG00000037742 | **Eef1a1** | - | eukaryotic translation elongation factor 1 alpha 1 | protein-coding |
| chr7_120848795_120849081 | chr7 | 120848796 | 120849081 | + | 0.749559 | intron (NM_001267710, intron 1 of 17) | intron (NM_001267710, intron 1 of 17) | -2251 | NM_001267711 | 13631 | Mm.25997 | NM_007908 | ENSMUSG00000035064 | **Eef2k** | C86191\|eEF-2K | eukaryotic elongation factor-2 kinase | protein-coding |
| chr9_25539155_25539821 | chr9 | 25539156 | 25539821 | + | 1.34589 | intron (NM_026189, intron 2 of 7) | intron (NM_026189, intron 2 of 7) | 57891 | NM_026189 | 67484 | Mm.112977 | NM_026189 | ENSMUSG00000036611 | **Eepd1** | 2310005P05Rik\|AI481005 | endonuclease/exonuclease/phosphatase family domain containing 1 | protein-coding |
| chr16_14885011_14885876 | chr16 | 14885012 | 14885876 | + | 1.23803 | Intergenic | Intergenic | -21202 | NM_025769 | 66793 | Mm.100270 | NM_025769 | | **Efcab1** | 4930449A16Rik\|5430404L10Rik | EF hand calcium binding domain 1 | protein-coding |
| chr12_99874201_99875251 | chr12 | 99874202 | 99875251 | + | 1.3539 | intron (NM_030172, intron 3 of 5) | intron (NM_030172, intron 3 of 5) | 8716 | NM_030172 | 78767 | Mm.235920 | NM_030172 | ENSMUSG00000021176 | **Efcab11** | 2610021K21Rik\|2610110A13Rik | EF-hand calcium binding domain 11 | protein-coding |
| chr6_87741959_87742427 | chr6 | 87741960 | 87742427 | + | 1.41877 | intron (NM_001159697, intron 2 of 7) | MER44B\|DNA\|TcMar-Tigger | 11324 | NM_001159697 | 58229 | Mm.333229 | NM_021418 | ENSMUSG00000068263 | **Efcc1** | Ccdc48 | EF hand and coiled-coil domain containing 1 | protein-coding |
| chr1_87255136_87255644 | chr1 | 87255137 | 87255644 | + | 1.72362 | Intergenic | ORR1F\|LTR\|ERVL-MaLR | -8974 | NM_028889 | 98363 | Mm.247951 | NM_028889 | ENSMUSG00000026255 | **Efhd1** | 4931430I01Rik\|AI452351\|PP3051\|mitocalcin | EF hand domain containing 1 | protein-coding |
| chr1_87259147_87259691 | chr1 | 87259148 | 87259691 | + | 0.797506 | Intergenic | Intergenic | -4945 | NM_028889 | 98363 | Mm.247951 | NM_028889 | ENSMUSG00000026255 | **Efhd1** | 4931430I01Rik\|AI452351\|PP3051\|mitocalcin | EF hand domain containing 1 | protein-coding |
| chr3_89297166_89297427 | chr3 | 89297167 | 89297427 | + | 0.717042 | Intergenic | HAL1\|LINE\|L1 | -16346 | NM_001162425 | 13636 | Mm.15675 | NM_010107 | ENSMUSG00000027954 | **Efna1** | AI325262\|B61\|Efl1\|Epl1\|Eplg1\|Lerk1 | ephrin A1 | protein-coding |
| chr3_89299470_89300008 | chr3 | 89299471 | 89300008 | + | 0.677757 | Intergenic | Intergenic | -18788 | NM_001162425 | 13636 | Mm.15675 | NM_010107 | ENSMUSG00000027954 | **Efna1** | AI325262\|B61\|Efl1\|Epl1\|Eplg1\|Lerk1 | ephrin A1 | protein-coding |
| chr7_82649085_82649410 | chr7 | 82649086 | 82649410 | + | 0.803651 | promoter-TSS (NM_177894) | promoter-TSS (NM_177894) | 634 | NM_175317 | 101592 | Mm.238020 | NM_175317 | ENSMUSG00000038563 | **Eftud1** | 4932434J20Rik\|6030468D11Rik\|AI451340\|AU019507\|AU022896\|D7Ertd791e | elongation factor Tu GTP binding domain containing 1 | protein-coding |
| chr8_124962007_124962391 | chr8 | 124962008 | 124962391 | + | 1.16043 | Intergenic | PB1D10\|SINE\|Alu | -12945 | NM_053207 | 112405 | Mm.140619 | NM_053207 | ENSMUSG00000031987 | **Egln1** | AI503754\|C1orf12\|HIF-PH2\|HPH-2\|Hif-p4h-2\|ORF13\|Phd2\|SM-20 | egl-9 family hypoxia-inducible factor 1 | protein-coding |
| chr7_27161852_27162065 | chr7 | 27161853 | 27162065 | + | 0.91944 | intron (NM_053208, intron 2 of 5) | intron (NM_053208, intron 2 of 5) | 4843 | NM_053208 | 112406 | Mm.29978 | NM_053208 | ENSMUSG00000058709 | **Egln2** | 0610011A13Rik\|C85656\|Hif-p4h-1\|Ier4\|Phd1\|SM-20 | egl-9 family hypoxia-inducible factor 2 | protein-coding |
| chr12_54210633_54211183 | chr12 | 54210634 | 54211183 | + | 1.76648 | Intergenic | MLT1L\|LTR\|ERVL-MaLR | -7034 | NM_028133 | 112407 | Mm.133037 | NM_028133 | ENSMUSG00000035105 | **Egln3** | 2610021G09Rik\|AI505553\|AI648162\|Hif-p4h-3\|Phd3\|SM-20 | egl-9 family hypoxia-inducible factor 3 | protein-coding |
| chr12_54198353_54198737 | chr12 | 54198354 | 54198737 | + | 0.63519 | intron (NM_028133, intron 1 of 4) | intron (NM_028133, intron 1 of 4) | 5329 | NM_028133 | 112407 | Mm.133037 | NM_028133 | ENSMUSG00000035105 | **Egln3** | 2610021G09Rik\|AI505553\|AI648162\|Hif-p4h-3\|Phd3\|SM-20 | egl-9 family hypoxia-inducible factor 3 | protein-coding |
| chr10_67493534_67493940 | chr10 | 67493535 | 67493940 | + | 1.423 | Intergenic | Intergenic | -44132 | NM_010118 | 13654 | Mm.290421 | NM_010118 | ENSMUSG00000037868 | **Egr2** | Egr-2\|Krox-20\|Krox20\|NGF1-B\|Zfp-25\|Zfp-6 | early growth response 2 | protein-coding |
| chr10_67484083_67484397 | chr10 | 67484084 | 67484397 | + | 1.09997 | Intergenic | IAPLTR2a\|LTR\|ERVK | -53629 | NM_010118 | 13654 | Mm.290421 | NM_010118 | ENSMUSG00000037868 | **Egr2** | Egr-2\|Krox-20\|Krox20\|NGF1-B\|Zfp-25\|Zfp-6 | early growth response 2 | protein-coding |
| chr10_67542257_67542670 | chr10 | 67542258 | 67542670 | + | 0.977593 | TTS (NM_010118) | TTS (NM_010118) | 4595 | NM_010118 | 13654 | Mm.290421 | NM_010118 | ENSMUSG00000037868 | **Egr2** | Egr-2\|Krox-20\|Krox20\|NGF1-B\|Zfp-25\|Zfp-6 | early growth response 2 | protein-coding |
| chr10_67488533_67488763 | chr10 | 67488534 | 67488763 | + | 0.887993 | Intergenic | Intergenic | -49221 | NM_010118 | 13654 | Mm.290421 | NM_010118 | ENSMUSG00000037868 | **Egr2** | Egr-2\|Krox-20\|Krox20\|NGF1-B\|Zfp-25\|Zfp-6 | early growth response 2 | protein-coding |
| chr10_67480022_67480301 | chr10 | 67480023 | 67480301 | + | 0.870471 | Intergenic | L1MB4\|LINE\|L1 | -57707 | NM_010118 | 13654 | Mm.290421 | NM_010118 | ENSMUSG00000037868 | **Egr2** | Egr-2\|Krox-20\|Krox20\|NGF1-B\|Zfp-25\|Zfp-6 | early growth response 2 | protein-coding |
| chr10_67497132_67498084 | chr10 | 67497133 | 67498084 | + | 0.786069 | Intergenic | Intergenic | -40261 | NM_010118 | 13654 | Mm.290421 | NM_010118 | ENSMUSG00000037868 | **Egr2** | Egr-2\|Krox-20\|Krox20\|NGF1-B\|Zfp-25\|Zfp-6 | early growth response 2 | protein-coding |
| chr10_67497098_67497827 | chr10 | 67497099 | 67497827 | + | 0.779099 | Intergenic | Intergenic | -40406 | NM_010118 | 13654 | Mm.290421 | NM_010118 | ENSMUSG00000037868 | **Egr2** | Egr-2\|Krox-20\|Krox20\|NGF1-B\|Zfp-25\|Zfp-6 | early growth response 2 | protein-coding |
| chr10_67487499_67487841 | chr10 | 67487500 | 67487841 | + | 0.74333 | Intergenic | MIR\|SINE\|MIR | -50199 | NM_010118 | 13654 | Mm.290421 | NM_010118 | ENSMUSG00000037868 | **Egr2** | Egr-2\|Krox-20\|Krox20\|NGF1-B\|Zfp-25\|Zfp-6 | early growth response 2 | protein-coding |
| chr10_67529527_67530174 | chr10 | 67529528 | 67530174 | + | 0.636759 | Intergenic | Intergenic | -8018 | NM_010118 | 13654 | Mm.290421 | NM_010118 | ENSMUSG00000037868 | **Egr2** | Egr-2\|Krox-20\|Krox20\|NGF1-B\|Zfp-25\|Zfp-6 | early growth response 2 | protein-coding |
| chr14_70061686_70061968 | chr14 | 70061687 | 70061968 | + | 0.897214 | Intergenic | Intergenic | -15618 | NM_018781 | 13655 | Mm.103737 | NM_018781 | ENSMUSG00000033730 | **Egr3** | Pilot | early growth response 3 | protein-coding |
| chr11_22163691_22164160 | chr11 | 22163692 | 22164160 | + | 1.26933 | intron (NM_153078, intron 7 of 24) | Lx8\|LINE\|L1 | 122869 | NM_153078 | 216565 | Mm.491265 | NM_153078 | ENSMUSG00000042302 | **Ehbp1** | AF424697\|Flj21950\|NACSIN | EH domain binding protein 1 | protein-coding |
| chr11_22255286_22255896 | chr11 | 22255287 | 22255896 | + | 0.856475 | intron (NM_153078, intron 2 of 24) | intron (NM_153078, intron 2 of 24) | 31204 | NM_153078 | 216565 | Mm.491265 | NM_153078 | ENSMUSG00000042302 | **Ehbp1** | AF424697\|Flj21950\|NACSIN | EH domain binding protein 1 | protein-coding |
| chr17_73853389_73854207 | chr17 | 73853390 | 73854207 | + | 1.39058 | Intergenic | Intergenic | 48957 | NM_020578 | 57440 | Mm.18526 | NM_020578 | ENSMUSG00000024065 | **Ehd3** | Ehd2 | EH-domain containing 3 | protein-coding |
| chr17_73806284_73806683 | chr17 | 73806285 | 73806683 | + | 0.884626 | intron (NM_020578, intron 1 of 5) | intron (NM_020578, intron 1 of 5) | 1643 | NM_020578 | 57440 | Mm.18526 | NM_020578 | ENSMUSG00000024065 | **Ehd3** | Ehd2 | EH-domain containing 3 | protein-coding |
| chr17_73818064_73818331 | chr17 | 73818065 | 73818331 | + | 0.75611 | intron (NM_020578, intron 2 of 5) | intron (NM_020578, intron 2 of 5) | 13357 | NM_020578 | 57440 | Mm.18526 | NM_020578 | ENSMUSG00000024065 | **Ehd3** | Ehd2 | EH-domain containing 3 | protein-coding |
| chr9_36800690_36801327 | chr9 | 36800691 | 36801327 | + | 1.3549 | Intergenic | Intergenic | -3616 | NM_001199494 | 13663 | Mm.4337 | NM_007915 | ENSMUSG00000062762 | **Ei24** | AA536736\|AI115355\|PIG8 | etoposide induced 2.4 mRNA | protein-coding |
| chr9_36804067_36804523 | chr9 | 36804068 | 36804523 | + | 0.675024 | Intergenic | Intergenic | -6902 | NM_001199494 | 13663 | Mm.4337 | NM_007915 | ENSMUSG00000062762 | **Ei24** | AA536736\|AI115355\|PIG8 | etoposide induced 2.4 mRNA | protein-coding |
| chr18_46598465_46599402 | chr18 | 46598466 | 46599402 | + | 0.831633 | intron (NM_010120, intron 2 of 2) | intron (NM_010120, intron 2 of 2) | 1230 | NM_010120 | 13664 | Mm.262037 | NM_010120 | ENSMUSG00000057561 | **Eif1a** | C76390\|Ef1a\|Eftu\|Eif4c\|eIF-1A\|eIF-4C | eukaryotic translation initiation factor 1A | protein-coding |
| chr6_70872192_70872493 | chr6 | 70872193 | 70872493 | + | 0.946531 | intron (NM_010121, intron 2 of 16) | intron (NM_010121, intron 2 of 16) | 27816 | NM_010121 | 13666 | Mm.247167 | NM_010121 | ENSMUSG00000031668 | **Eif2ak3** | AI427929\|Pek\|Perk | eukaryotic translation initiation factor 2 alpha kinase 3 | protein-coding |
| chr6_70872110_70872631 | chr6 | 70872111 | 70872631 | + | 0.8985 | intron (NM_010121, intron 2 of 16) | intron (NM_010121, intron 2 of 16) | 27844 | NM_010121 | 13666 | Mm.247167 | NM_010121 | ENSMUSG00000031668 | **Eif2ak3** | AI427929\|Pek\|Perk | eukaryotic translation initiation factor 2 alpha kinase 3 | protein-coding |
| chr2_118390086_118390352 | chr2 | 118390087 | 118390352 | + | 1.04487 | intron (NM_001177806, intron 1 of 37) | intron (NM_001177806, intron 1 of 37) | 1107 | NM_001177806 | 27103 | Mm.217616 | NM_013719 | ENSMUSG00000005102 | **Eif2ak4** | 2610011M03\|GCN2\|MGCN2 | eukaryotic translation initiation factor 2 alpha kinase 4 | protein-coding |
| chr16_20464722_20465591 | chr16 | 20464723 | 20465591 | + | 1.6072 | Intergenic | MTB\|LTR\|ERVL-MaLR | -33660 | NM_172265 | 224045 | Mm.233855 | NM_172265 | ENSMUSG00000003235 | **Eif2b5** | C81315 | eukaryotic translation initiation factor 2B, subunit 5 epsilon | protein-coding |
| chr1_131184631_131184961 | chr1 | 131184632 | 131184961 | + | 1.22068 | intron (NM_018750, intron 2 of 5) | intron (NM_018750, intron 2 of 5) | 31589 | NM_010709 | 16865 | Mm.276294 | NM_010709 | ENSMUSG00000026427 | **Eif2d** | D1Ertd5e\|Lgtn | eukaryotic translation initiation factor 2D | protein-coding |
| chr1_131186184_131186464 | chr1 | 131186185 | 131186464 | + | 0.925669 | intron (NM_018750, intron 2 of 5) | intron (NM_018750, intron 2 of 5) | 33117 | NM_010709 | 16865 | Mm.276294 | NM_010709 | ENSMUSG00000026427 | **Eif2d** | D1Ertd5e\|Lgtn | eukaryotic translation initiation factor 2D | protein-coding |
| chr1_131184371_131184982 | chr1 | 131184372 | 131184982 | + | 0.876859 | intron (NM_018750, intron 2 of 5) | intron (NM_018750, intron 2 of 5) | 31470 | NM_010709 | 16865 | Mm.276294 | NM_010709 | ENSMUSG00000026427 | **Eif2d** | D1Ertd5e\|Lgtn | eukaryotic translation initiation factor 2D | protein-coding |
| chr2_154901690_154902454 | chr2 | 154901691 | 154902454 | + | 1.08288 | Intergenic | MTC\|LTR\|ERVL-MaLR | -9166 | NM_026030 | 67204 | Mm.377134 | NM_026030 | ENSMUSG00000074656 | **Eif2s2** | 2810026E11Rik\|38kDa\|AA408636\|AA571381\|AA986487\|AW822225\|D2Ertd303e\|EIF2\|EIF2B | eukaryotic translation initiation factor 2, subunit 2 (beta) | protein-coding |
| chr7_126545583_126545967 | chr7 | 126545584 | 126545967 | + | 0.733585 | Intergenic | RLTR11A2\|LTR\|ERVK | 20591 | NM_146200 | 56347 | Mm.22776 | NM_146200 | ENSMUSG00000030738 | **Eif3c** | 110kDa\|3230401O13Rik\|Eif3s8\|NIPIL(A3)\|NipilA3\|Xs\|Xsl | eukaryotic translation initiation factor 3, subunit C | protein-coding |
| chr7_108954200_108954675 | chr7 | 108954201 | 108954675 | + | 0.796329 | Intergenic | L2a\|LINE\|L2 | 20023 | NM_025344 | 66085 | Mm.182962 | NM_025344 | | **Eif3f** | 0610037M02Rik\|AA409853\|Eif3s5 | eukaryotic translation initiation factor 3, subunit F | protein-coding |
| chr15_51845089_51845439 | chr15 | 51845090 | 51845439 | + | 1.33515 | intron (NM_080635, intron 1 of 7) | intron (NM_080635, intron 1 of 7) | 20197 | NM_080635 | 68135 | Mm.289800 | NM_080635 | ENSMUSG00000022312 | **Eif3h** | 1110008A16Rik\|40kD\|9430017H16Rik\|EIF3-P40\|EIF3-gamma\|Eif3s3 | eukaryotic translation initiation factor 3, subunit H | protein-coding |
| chr15_51730705_51731323 | chr15 | 51730706 | 51731323 | + | 1.02843 | Intergenic | Intergenic | 134447 | NM_080635 | 68135 | Mm.289800 | NM_080635 | ENSMUSG00000022312 | **Eif3h** | 1110008A16Rik\|40kD\|9430017H16Rik\|EIF3-P40\|EIF3-gamma\|Eif3s3 | eukaryotic translation initiation factor 3, subunit H | protein-coding |
| chr15_51838392_51838823 | chr15 | 51838393 | 51838823 | + | 0.871328 | intron (NM_080635, intron 2 of 7) | intron (NM_080635, intron 2 of 7) | 26853 | NM_080635 | 68135 | Mm.289800 | NM_080635 | ENSMUSG00000022312 | **Eif3h** | 1110008A16Rik\|40kD\|9430017H16Rik\|EIF3-P40\|EIF3-gamma\|Eif3s3 | eukaryotic translation initiation factor 3, subunit H | protein-coding |
| chr15_51730143_51730499 | chr15 | 51730144 | 51730499 | + | 0.727467 | Intergenic | Intergenic | 135140 | NM_080635 | 68135 | Mm.289800 | NM_080635 | ENSMUSG00000022312 | **Eif3h** | 1110008A16Rik\|40kD\|9430017H16Rik\|EIF3-P40\|EIF3-gamma\|Eif3s3 | eukaryotic translation initiation factor 3, subunit H | protein-coding |

| | | | | | | | | | | | | | | | | | |
|---|---|---|---|---|---|---|---|---|---|---|---|---|---|---|---|---|---|
| chr15_51726864_51727326 | chr15 | 51726865 | 51727326 | + | 0.713769 | Intergenic | Intergenic | 138366 | NM_080635 | 68135 | Mm.289800 | NM_080635 | ENSMUSG00000022312 | **Eif3h** | 1110008A16Rik\|40kD\|9430017H16Rik\|EIF3-P40\|EIF3-gamma\|Eif3s3 | eukaryotic translation initiation factor 3, subunit H | protein-coding |
| chr15_51726955_51727298 | chr15 | 51726956 | 51727298 | + | 0.686031 | Intergenic | Intergenic | 138334 | NM_080635 | 68135 | Mm.289800 | NM_080635 | ENSMUSG00000022312 | **Eif3h** | 1110008A16Rik\|40kD\|9430017H16Rik\|EIF3-P40\|EIF3-gamma\|Eif3s3 | eukaryotic translation initiation factor 3, subunit H | protein-coding |
| chr2_122020476_122021064 | chr2 | 122020477 | 122021064 | + | 1.67186 | Intergenic | URR1A\|DNA\|hAT-Charlie | -7810 | NM_144545 | 78655 | Mm.458184 | NM_144545 | | **Eif3j1** | 1810016I04Rik\|2700079K05Rik\|AA409117\|AA409446\|C78575\|Eif3j\|Eif3j-1\|Eif3j-2\|Eif3j2\|Eif3s1\|eIF-3-alpha-A\|eIF-3-alpha-B\|eIF3-alpha\|eIF3-p35\|eIF3j-A\|eIF3j-B | eukaryotic translation initiation factor 3, subunit J1 | protein-coding |
| chr2_122069393_122069898 | chr2 | 122069394 | 122069898 | + | 0.775211 | intron (NM_145531, intron 27 of 39) | intron (NM_145531, intron 27 of 39) | 41063 | NM_001256055 | 100042807 | Mm.329403 | NM_001256055 | ENSMUSG00000043424 | **Eif3j2** | ENSMUSG00000043424\|Eif3j-2\|Gm9781\|eIF3 p35\|eIF3j-B | eukaryotic translation initiation factor 3, subunit J2 | protein-coding |
| chr2_104991877_104992225 | chr2 | 104991878 | 104992225 | + | 0.640321 | exon (NM_177600, exon 16 of 18) | exon (NM_177600, exon 16 of 18) | 24976 | NM_145380 | 98221 | Mm.379278 | NM_145380 | ENSMUSG00000027170 | **Eif3m** | Ga17\|Pcid1\|Tango7 | eukaryotic translation initiation factor 3, subunit M | protein-coding |
| chr3_138514831_138515101 | chr3 | 138514832 | 138515101 | + | 0.678471 | Intergenic | Intergenic | -11225 | NM_007917 | 13684 | Mm.3941 | NM_007917 | ENSMUSG00000028156 | **Eif4e** | EG668879\|Eif4e-ps\|If4e\|eIF-4E | eukaryotic translation initiation factor 4E | protein-coding |
| chr6_99663259_99663568 | chr6 | 99663260 | 99663568 | + | 1.02235 | intron (NM_025829, intron 1 of 6) | intron (NM_025829, intron 1 of 6) | 3357 | NM_025829 | 66892 | Mm.32889 | NM_025829 | ENSMUSG00000093661 | **Eif4e3** | 1300018P11Rik\|AI451927\|eIF4E-3 | eukaryotic translation initiation factor 4E member 3 | protein-coding |
| chr6_99631466_99631953 | chr6 | 99631467 | 99631953 | + | 0.837049 | intron (NM_025829, intron 6 of 6) | intron (NM_025829, intron 6 of 6) | 35061 | NM_025829 | 66892 | Mm.32889 | NM_025829 | ENSMUSG00000093661 | **Eif4e3** | 1300018P11Rik\|AI451927\|eIF4E-3 | eukaryotic translation initiation factor 4E member 3 | protein-coding |
| chr8_27260982_27261671 | chr8 | 27260983 | 27261671 | + | 0.75041 | intron (NM_007918, intron 1 of 2) | intron (NM_007918, intron 1 of 2) | 1000 | NM_007918 | 13685 | Mm.6700 | NM_007918 | ENSMUSG00000031490 | **Eif4ebp1** | 4e-bp1\|AA959816\|PHAS-I | eukaryotic translation initiation factor 4E binding protein 1 | protein-coding |
| chr11_3203650_3204052 | chr11 | 3203651 | 3204052 | + | 1.3633 | intron (NM_001166547, intron 2 of 18) | intron (NM_001166547, intron 2 of 18) | 832 | NM_001166549 | 74203 | Mm.255649 | NM_023743 | ENSMUSG00000020454 | **Eif4enif1** | 2610509L04Rik\|A930019J01Rik\|AA410001\|AU021239\|Clast4\|D11Ertd166e | eukaryotic translation initiation factor 4E nuclear import factor 1 | protein-coding |
| chr11_3206487_3207286 | chr11 | 3206488 | 3207286 | + | 1.36169 | intron (NM_001166547, intron 2 of 18) | intron (NM_001166547, intron 2 of 18) | 3868 | NM_001166549 | 74203 | Mm.255649 | NM_023743 | ENSMUSG00000020454 | **Eif4enif1** | 2610509L04Rik\|A930019J01Rik\|AA410001\|AU021239\|Clast4\|D11Ertd166e | eukaryotic translation initiation factor 4E nuclear import factor 1 | protein-coding |
| chr11_3206890_3207258 | chr11 | 3206891 | 3207258 | + | 1.29979 | intron (NM_001166547, intron 2 of 18) | intron (NM_001166547, intron 2 of 18) | 4055 | NM_001166549 | 74203 | Mm.255649 | NM_023743 | ENSMUSG00000020454 | **Eif4enif1** | 2610509L04Rik\|A930019J01Rik\|AA410001\|AU021239\|Clast4\|D11Ertd166e | eukaryotic translation initiation factor 4E nuclear import factor 1 | protein-coding |
| chr11_3208833_3209040 | chr11 | 3208834 | 3209040 | + | 0.811358 | intron (NM_001166547, intron 2 of 18) | intron (NM_001166547, intron 2 of 18) | 5918 | NM_001166549 | 74203 | Mm.255649 | NM_023743 | ENSMUSG00000020454 | **Eif4enif1** | 2610509L04Rik\|A930019J01Rik\|AA410001\|AU021239\|Clast4\|D11Ertd166e | eukaryotic translation initiation factor 4E nuclear import factor 1 | protein-coding |
| chr7_111095234_111095613 | chr7 | 111095235 | 111095613 | + | 0.843484 | Intergenic | URR1B\|DNA\|hAT-Charlie | -12394 | NM_013507 | 13690 | Mm.185453 | NM_013507 | ENSMUSG00000005610 | **Eif4g2** | AA589388\|DAP-5\|E130105L11Rik\|Nat1\|Natm1\|p97 | eukaryotic translation initiation factor 4, gamma 2 | protein-coding |
| chr1_38038970_38039272 | chr1 | 38038971 | 38039272 | + | 0.969576 | intron (NM_198303, intron 12 of 23) | B3\|SINE\|B2 | 41111 | NM_198303 | 226982 | Mm.260943 | NM_198303 | ENSMUSG00000026083 | **Eif5b** | A030003E17Rik\|BC018347\|IF2 | eukaryotic translation initiation factor 5B | protein-coding |
| chr1_38038862_38039329 | chr1 | 38038863 | 38039329 | + | 0.962435 | intron (NM_198303, intron 12 of 23) | B3\|SINE\|B2 | 41086 | NM_198303 | 226982 | Mm.260943 | NM_198303 | ENSMUSG00000026083 | **Eif5b** | A030003E17Rik\|BC018347\|IF2 | eukaryotic translation initiation factor 5B | protein-coding |
| chr18_73748909_73749174 | chr18 | 73748910 | 73749174 | + | 0.870826 | intron (NM_053255, intron 1 of 3) | intron (NM_053255, intron 1 of 3) | 5437 | NM_053255 | 114615 | Mm.50642 | NM_053255 | ENSMUSG00000036941 | **Elac1** | 2610018O07Rik\|8430417G19Rik | elaC homolog 1 (E. coli) | protein-coding |
| chr11_16950136_16950376 | chr11 | 16950137 | 16950376 | + | 0.769369 | intron (NR_110421, intron 2 of 5) | intron (NR_110421, intron 2 of 5) | 1026 | NR_110421 | 72735 | Mm.268598 | NR_015469 | | **Eldr** | 2810442I21Rik\|4930500E12Rik\|Fabl | Egfr long non-coding downstream RNA | ncRNA |
| chr14_79530813_79531021 | chr14 | 79530814 | 79531021 | + | 0.720009 | intron (NM_001286411, intron 1 of 8) | intron (NM_001286411, intron 1 of 8) | 15243 | NM_001286411 | 13709 | Mm.20938 | NM_007920 | ENSMUSG00000036461 | **Elf1** | Elf-1\|Sts1\|mElf-1\|p70 | E74-like factor 1 | protein-coding |
| chr14_79519701_79520002 | chr14 | 79519702 | 79520002 | + | 0.685524 | intron (NM_001286411, intron 1 of 8) | ID4_\|SINE\|ID | 4178 | NM_001286411 | 13709 | Mm.20938 | NM_007920 | ENSMUSG00000036461 | **Elf1** | Elf-1\|Sts1\|mElf-1\|p70 | E74-like factor 1 | protein-coding |
| chr14_79519441_79520028 | chr14 | 79519442 | 79520028 | + | 0.68122 | intron (NM_001286411, intron 1 of 8) | intron (NM_001286411, intron 1 of 8) | 4061 | NM_001286411 | 13709 | Mm.20938 | NM_007920 | ENSMUSG00000036461 | **Elf1** | Elf-1\|Sts1\|mElf-1\|p70 | E74-like factor 1 | protein-coding |
| chr14_79548377_79548613 | chr14 | 79548378 | 79548613 | + | 0.677614 | intron (NM_007920, intron 2 of 8) | Lx5\|LINE\|L1 | 32821 | NM_001286411 | 13709 | Mm.20938 | NM_007920 | ENSMUSG00000036461 | **Elf1** | Elf-1\|Sts1\|mElf-1\|p70 | E74-like factor 1 | protein-coding |
| chr14_79547513_79547828 | chr14 | 79547514 | 79547828 | + | 0.674924 | intron (NM_007920, intron 2 of 8) | Lx5\|LINE\|L1 | 31997 | NM_001286411 | 13709 | Mm.20938 | NM_007920 | ENSMUSG00000036461 | **Elf1** | Elf-1\|Sts1\|mElf-1\|p70 | E74-like factor 1 | protein-coding |
| chr14_79533379_79533724 | chr14 | 79533380 | 79533724 | + | 0.659672 | intron (NM_001286411, intron 1 of 8) | intron (NM_001286411, intron 1 of 8) | 17878 | NM_001286411 | 13709 | Mm.20938 | NM_007920 | ENSMUSG00000036461 | **Elf1** | Elf-1\|Sts1\|mElf-1\|p70 | E74-like factor 1 | protein-coding |
| chr3_51327385_51327746 | chr3 | 51327386 | 51327746 | + | 0.672506 | intron (NM_023502, intron 1 of 9) | B3\|SINE\|B2 | 13078 | NM_001291059 | 69257 | Mm.131038 | NM_023502 | ENSMUSG00000037174 | **Elf2** | 2610036A20Rik\|A230104O07Rik\|AW111824\|BB183398\|EU32\|NERF\|NERF-1A\|NERF-1B\|NERF-2 | E74-like factor 2 | protein-coding |
| chr10_93272251_93272594 | chr10 | 93272252 | 93272594 | + | 1.10096 | intron (NM_013508, intron 2 of 4) | intron (NM_013508, intron 2 of 4) | 38736 | NM_013508 | 13713 | Mm.4454 | NM_013508 | ENSMUSG00000008398 | **Elk3** | D430049E23Rik\|Erp\|Etrp\|Net\|Sap-2 | ELK3, member of ETS oncogene family | protein-coding |
| chr10_93271921_93272646 | chr10 | 93271922 | 93272646 | + | 1.08706 | intron (NM_013508, intron 2 of 4) | intron (NM_013508, intron 2 of 4) | 38875 | NM_013508 | 13713 | Mm.4454 | NM_013508 | ENSMUSG00000008398 | **Elk3** | D430049E23Rik\|Erp\|Etrp\|Net\|Sap-2 | ELK3, member of ETS oncogene family | protein-coding |
| chr10_93264129_93264649 | chr10 | 93264130 | 93264649 | + | 0.740055 | intron (NM_001282967, intron 2 of 3) | intron (NM_001282967, intron 2 of 3) | 46770 | NM_013508 | 13713 | Mm.4454 | NM_013508 | ENSMUSG00000008398 | **Elk3** | D430049E23Rik\|Erp\|Etrp\|Net\|Sap-2 | ELK3, member of ETS oncogene family | protein-coding |
| chr10_93275218_93275421 | chr10 | 93275219 | 93275421 | + | 0.611501 | intron (NM_013508, intron 2 of 4) | intron (NM_013508, intron 2 of 4) | 35839 | NM_013508 | 13713 | Mm.4454 | NM_013508 | ENSMUSG00000008398 | **Elk3** | D430049E23Rik\|Erp\|Etrp\|Net\|Sap-2 | ELK3, member of ETS oncogene family | protein-coding |
| chr1_131993159_131993514 | chr1 | 131993160 | 131993514 | + | 0.826326 | Intergenic | MER44D\|DNA\|TcMar-Tigger | -14268 | NM_007923 | 13714 | Mm.254233 | NM_007923 | ENSMUSG00000026436 | **Elk4** | 2310011G17Rik\|A130026I01Rik\|BB162516\|SAP-1\|Sap1 | ELK4, member of ETS oncogene family | protein-coding |
| chr13_75691646_75692021 | chr13 | 75691647 | 75692021 | + | 0.76812 | Intergenic | Intergenic | -15650 | NM_138953 | 192657 | Mm.21288 | NM_138953 | ENSMUSG00000001542 | **Ell2** | - | elongation factor RNA polymerase II 2 | protein-coding |
| chr13_20111298_20111999 | chr13 | 20111299 | 20111999 | + | 1.56726 | intron (NM_080288, intron 1 of 21) | ORR1F\|LTR\|ERVL-MaLR | 21142 | NM_080288 | 140580 | Mm.342392 | NM_080288 | ENSMUSG00000041112 | **Elmo1** | 6330578D22Rik\|C230095H21Rik\|CED-12 | engulfment and cell motility 1 | protein-coding |
| chr13_20163456_20164152 | chr13 | 20163457 | 20164152 | + | 1.45741 | intron (NM_080288, intron 1 of 21) | intron (NM_080288, intron 1 of 21) | 73297 | NM_080288 | 140580 | Mm.342392 | NM_080288 | ENSMUSG00000041112 | **Elmo1** | 6330578D22Rik\|C230095H21Rik\|CED-12 | engulfment and cell motility 1 | protein-coding |
| chr13_20136690_20137165 | chr13 | 20136691 | 20137165 | + | 1.44591 | intron (NM_080288, intron 1 of 21) | intron (NM_080288, intron 1 of 21) | 46421 | NM_080288 | 140580 | Mm.342392 | NM_080288 | ENSMUSG00000041112 | **Elmo1** | 6330578D22Rik\|C230095H21Rik\|CED-12 | engulfment and cell motility 1 | protein-coding |
| chr13_20048075_20048773 | chr13 | 20048076 | 20048773 | + | 1.36316 | Intergenic | Intergenic | -42083 | NM_080288 | 140580 | Mm.342392 | NM_080288 | ENSMUSG00000041112 | **Elmo1** | 6330578D22Rik\|C230095H21Rik\|CED-12 | engulfment and cell motility 1 | protein-coding |
| chr13_20161985_20162353 | chr13 | 20161986 | 20162353 | + | 1.16679 | intron (NM_080288, intron 1 of 21) | intron (NM_080288, intron 1 of 21) | 71662 | NM_080288 | 140580 | Mm.342392 | NM_080288 | ENSMUSG00000041112 | **Elmo1** | 6330578D22Rik\|C230095H21Rik\|CED-12 | engulfment and cell motility 1 | protein-coding |
| chr13_20162556_20162988 | chr13 | 20162557 | 20162988 | + | 1.15042 | intron (NM_080288, intron 1 of 21) | intron (NM_080288, intron 1 of 21) | 72265 | NM_080288 | 140580 | Mm.342392 | NM_080288 | ENSMUSG00000041112 | **Elmo1** | 6330578D22Rik\|C230095H21Rik\|CED-12 | engulfment and cell motility 1 | protein-coding |
| chr13_20162693_20162974 | chr13 | 20162694 | 20162974 | + | 1.09072 | intron (NM_080288, intron 1 of 21) | intron (NM_080288, intron 1 of 21) | 72327 | NM_080288 | 140580 | Mm.342392 | NM_080288 | ENSMUSG00000041112 | **Elmo1** | 6330578D22Rik\|C230095H21Rik\|CED-12 | engulfment and cell motility 1 | protein-coding |
| chr13_20306174_20306812 | chr13 | 20306175 | 20306812 | + | 1.08192 | intron (NM_080288, intron 13 of 21) | intron (NM_080288, intron 13 of 21) | -166497 | NM_198093 | 140580 | Mm.342392 | NM_080288 | ENSMUSG00000041112 | **Elmo1** | 6330578D22Rik\|C230095H21Rik\|CED-12 | engulfment and cell motility 1 | protein-coding |
| chr13_20305502_20306019 | chr13 | 20305503 | 20306019 | + | 1.06583 | intron (NM_080288, intron 13 of 21) | intron (NM_080288, intron 13 of 21) | -167229 | NM_198093 | 140580 | Mm.342392 | NM_080288 | ENSMUSG00000041112 | **Elmo1** | 6330578D22Rik\|C230095H21Rik\|CED-12 | engulfment and cell motility 1 | protein-coding |
| chr13_20306180_20307150 | chr13 | 20306181 | 20307150 | + | 0.974762 | intron (NM_080288, intron 13 of 21) | intron (NM_080288, intron 13 of 21) | -166325 | NM_198093 | 140580 | Mm.342392 | NM_080288 | ENSMUSG00000041112 | **Elmo1** | 6330578D22Rik\|C230095H21Rik\|CED-12 | engulfment and cell motility 1 | protein-coding |
| chr13_20256605_20257004 | chr13 | 20256606 | 20257004 | + | 0.863051 | intron (NM_080288, intron 6 of 21) | intron (NM_080288, intron 6 of 21) | 166298 | NM_080288 | 140580 | Mm.342392 | NM_080288 | ENSMUSG00000041112 | **Elmo1** | 6330578D22Rik\|C230095H21Rik\|CED-12 | engulfment and cell motility 1 | protein-coding |
| chr13_20133669_20134226 | chr13 | 20133670 | 20134226 | + | 0.839456 | intron (NM_080288, intron 1 of 21) | intron (NM_080288, intron 1 of 21) | 43441 | NM_080288 | 140580 | Mm.342392 | NM_080288 | ENSMUSG00000041112 | **Elmo1** | 6330578D22Rik\|C230095H21Rik\|CED-12 | engulfment and cell motility 1 | protein-coding |
| chr13_20256615_20257253 | chr13 | 20256616 | 20257253 | + | 0.782572 | intron (NM_080288, intron 6 of 21) | (TG)n\|Simple_repeat\|Simple_repeat | 166427 | NM_080288 | 140580 | Mm.342392 | NM_080288 | ENSMUSG00000041112 | **Elmo1** | 6330578D22Rik\|C230095H21Rik\|CED-12 | engulfment and cell motility 1 | protein-coding |

| | | | | | | | | | | | | | | | | | |
|---|---|---|---|---|---|---|---|---|---|---|---|---|---|---|---|---|---|
| chr6_72596840_72597448 | chr6 | 72596841 | 72597448 | + | 0.653404 | intron (NM_144917, intron 1 of 12) | intron (NM_144917, intron 1 of 12) | 1269 | NM_144917 | 232089 | Mm.167232 | NM_144917 | ENSMUSG00000056698 | **Elmod3** | AI844780\|C330008I15Rik\|RBM29\|Rbed1 | ELMO/CED-12 domain containing 3 | protein-coding |
| chr9_77955595_77955870 | chr9 | 77955596 | 77955870 | + | 0.717671 | intron (NM_134255, intron 1 of 7) | intron (NM_134255, intron 1 of 7) | 38368 | NM_134255 | 68801 | Mm.430736 | NM_134255 | ENSMUSG00000032349 | **Elovl5** | 1110059L23Rik\|AI747313\|AU043003\|HELO1 | ELOVL family member 5, elongation of long chain fatty acids (yeast) | protein-coding |
| chr9_77954440_77954844 | chr9 | 77954441 | 77954844 | + | 0.635135 | intron (NM_134255, intron 1 of 7) | intron (NM_134255, intron 1 of 7) | 37277 | NM_134255 | 68801 | Mm.430736 | NM_134255 | ENSMUSG00000032349 | **Elovl5** | 1110059L23Rik\|AI747313\|AU043003\|HELO1 | ELOVL family member 5, elongation of long chain fatty acids (yeast) | protein-coding |
| chr9_77954440_77955020 | chr9 | 77954441 | 77955020 | + | 0.624142 | intron (NM_134255, intron 1 of 7) | intron (NM_134255, intron 1 of 7) | 37365 | NM_134255 | 68801 | Mm.430736 | NM_134255 | ENSMUSG00000032349 | **Elovl5** | 1110059L23Rik\|AI747313\|AU043003\|HELO1 | ELOVL family member 5, elongation of long chain fatty acids (yeast) | protein-coding |
| chr3_129588610_129589207 | chr3 | 129588611 | 129589207 | + | 1.65956 | intron (NM_130450, intron 2 of 4) | intron (NM_130450, intron 2 of 4) | 56523 | NM_130450 | 170439 | Mm.314113 | NM_130450 | ENSMUSG00000041220 | **Elovl6** | C77826\|FAE\|LCE | ELOVL family member 6, elongation of long chain fatty acids (yeast) | protein-coding |
| chr3_129565936_129566360 | chr3 | 129565937 | 129566360 | + | 1.13977 | intron (NM_130450, intron 2 of 4) | intron (NM_130450, intron 2 of 4) | 33762 | NM_130450 | 170439 | Mm.314113 | NM_130450 | ENSMUSG00000041220 | **Elovl6** | C77826\|FAE\|LCE | ELOVL family member 6, elongation of long chain fatty acids (yeast) | protein-coding |
| chr3_129565601_129566365 | chr3 | 129565602 | 129566365 | + | 1.09446 | intron (NM_130450, intron 2 of 4) | intron (NM_130450, intron 2 of 4) | 33597 | NM_130450 | 170439 | Mm.314113 | NM_130450 | ENSMUSG00000041220 | **Elovl6** | C77826\|FAE\|LCE | ELOVL family member 6, elongation of long chain fatty acids (yeast) | protein-coding |
| chr13_108228642_108229693 | chr13 | 108228643 | 108229693 | + | 0.629575 | intron (NM_029001, intron 1 of 7) | intron (NM_029001, intron 1 of 7) | 14764 | NM_029001 | 74559 | Mm.286127 | NM_029001 | ENSMUSG00000021696 | **Elovl7** | 9130013K24Rik\|AI840082 | ELOVL family member 7, elongation of long chain fatty acids (yeast) | protein-coding |
| chr18_24619355_24619826 | chr18 | 24619356 | 24619826 | + | 0.646194 | intron (NM_021448, intron 11 of 21) | intron (NM_021448, intron 11 of 21) | 15630 | NM_021448 | 58523 | Mm.25298 | NM_021448 | ENSMUSG00000024271 | **Elp2** | AU023723\|Epl2\|StIP1\|Statip1 | elongator acetyltransferase complex subunit 2 | protein-coding |
| chr12_108395993_108396468 | chr12 | 108395994 | 108396468 | + | 1.26991 | intron (NM_001286346, intron 1 of 21) | intron (NM_001286346, intron 1 of 21) | -14402 | NM_001043336 | 68519 | Mm.236645 | NM_001043335 | ENSMUSG00000058070 | **Eml1** | 1110008N23Rik\|A930030P13Rik\|AA171013\|AI847476\|AI853955\|ELP79\|EMAP\|EMAPL | echinoderm microtubule associated protein like 1 | protein-coding |
| chr12_108453422_108454203 | chr12 | 108453423 | 108454203 | + | 0.962026 | intron (NM_001043335, intron 1 of 21) | intron (NM_001043335, intron 1 of 21) | 30997 | NM_001043335 | 68519 | Mm.236645 | NM_001043335 | ENSMUSG00000058070 | **Eml1** | 1110008N23Rik\|A930030P13Rik\|AA171013\|AI847476\|AI853955\|ELP79\|EMAP\|EMAPL | echinoderm microtubule associated protein like 1 | protein-coding |
| chr17_83379313_83380162 | chr17 | 83379314 | 83380162 | + | 1.24965 | intron (NM_001114361, intron 1 of 23) | intron (NM_001114361, intron 1 of 23) | 28807 | NM_001114362 | 78798 | Mm.295565 | NM_199466 | ENSMUSG00000032624 | **Eml4** | 4930443C24Rik\|AI644019 | echinoderm microtubule associated protein like 4 | protein-coding |
| chr17_83379799_83380161 | chr17 | 83379800 | 83380161 | + | 1.16811 | intron (NM_001114361, intron 1 of 23) | L1MA8\|LINE\|L1 | 29049 | NM_001114362 | 78798 | Mm.295565 | NM_199466 | ENSMUSG00000032624 | **Eml4** | 4930443C24Rik\|AI644019 | echinoderm microtubule associated protein like 4 | protein-coding |
| chr17_83367995_83368514 | chr17 | 83367996 | 83368514 | + | 1.10957 | intron (NM_001114361, intron 1 of 23) | intron (NM_001114361, intron 1 of 23) | 17324 | NM_001114362 | 78798 | Mm.295565 | NM_199466 | ENSMUSG00000032624 | **Eml4** | 4930443C24Rik\|AI644019 | echinoderm microtubule associated protein like 4 | protein-coding |
| chr17_83379313_83379726 | chr17 | 83379314 | 83379726 | + | 0.857635 | intron (NM_001114361, intron 1 of 23) | RSINE1\|SINE\|B4 | 28589 | NM_001114362 | 78798 | Mm.295565 | NM_199466 | ENSMUSG00000032624 | **Eml4** | 4930443C24Rik\|AI644019 | echinoderm microtubule associated protein like 4 | protein-coding |
| chr6_135362615_135363050 | chr6 | 135362616 | 135363050 | + | 1.1938 | promoter-TSS (NM_010128) | promoter-TSS (NM_010128) | -98 | NM_010128 | 13730 | Mm.182785 | NM_010128 | ENSMUSG00000030208 | **Emp1** | I-8-09\|TMP | epithelial membrane protein 1 | protein-coding |
| chr6_135362403_135363209 | chr6 | 135362404 | 135363209 | + | 1.172 | promoter-TSS (NM_010128) | promoter-TSS (NM_010128) | -125 | NM_010128 | 13730 | Mm.182785 | NM_010128 | ENSMUSG00000030208 | **Emp1** | I-8-09\|TMP | epithelial membrane protein 1 | protein-coding |
| chr6_135362610_135362911 | chr6 | 135362611 | 135362911 | + | 1.1547 | promoter-TSS (NM_010128) | promoter-TSS (NM_010128) | -170 | NM_010128 | 13730 | Mm.182785 | NM_010128 | ENSMUSG00000030208 | **Emp1** | I-8-09\|TMP | epithelial membrane protein 1 | protein-coding |
| chr19_59402152_59402871 | chr19 | 59402153 | 59402871 | + | 0.644079 | Intergenic | Intergenic | 56123 | NR_002863 | 329078 | Mm.80382 | NR_002863 | ENSMUSG00000087095 | **Emx2os** | - | Emx2 opposite strand/antisense transcript (non-protein coding) | ncRNA |
| chr5_88499750_88500201 | chr5 | 88499751 | 88500201 | + | 0.679411 | intron (NM_017468, intron 9 of 9) | intron (NM_017468, intron 9 of 9) | 12001 | NM_017468 | 13801 | Mm.8014 | NM_017468 | ENSMUSG00000029286 | **Enam** | abte | enamelin | protein-coding |
| chr13_97295850_97296182 | chr13 | 97295851 | 97296182 | + | 0.634566 | Intergenic | Intergenic | 54911 | NM_007930 | 13803 | Mm.241073 | NM_007930 | ENSMUSG00000041773 | **Enc1** | Nrpb\|PIG10 | ectodermal-neural cortex 1 | protein-coding |
| chr9_14385653_14386028 | chr9 | 14385654 | 14386028 | + | 1.26348 | Intergenic | Intergenic | -4599 | NM_028013 | 71946 | Mm.41423 | NM_028013 | ENSMUSG00000037419 | **Endod1** | 2210414F18Rik\|2310067E08Rik\|C85344 | endonuclease domain containing 1 | protein-coding |
| chr9_14385640_14386156 | chr9 | 14385641 | 14386156 | + | 1.18643 | Intergenic | Intergenic | -4656 | NM_028013 | 71946 | Mm.41423 | NM_028013 | ENSMUSG00000037419 | **Endod1** | 2210414F18Rik\|2310067E08Rik\|C85344 | endonuclease domain containing 1 | protein-coding |
| chr9_14376170_14376540 | chr9 | 14376171 | 14376540 | + | 0.984999 | intron (NM_028013, intron 1 of 1) | intron (NM_028013, intron 1 of 1) | 4887 | NM_028013 | 71946 | Mm.41423 | NM_028013 | ENSMUSG00000037419 | **Endod1** | 2210414F18Rik\|2310067E08Rik\|C85344 | endonuclease domain containing 1 | protein-coding |
| chr9_14375749_14376056 | chr9 | 14375750 | 14376056 | + | 0.784825 | intron (NM_028013, intron 1 of 1) | intron (NM_028013, intron 1 of 1) | 5339 | NM_028013 | 71946 | Mm.41423 | NM_028013 | ENSMUSG00000037419 | **Endod1** | 2210414F18Rik\|2310067E08Rik\|C85344 | endonuclease domain containing 1 | protein-coding |
| chr9_14375623_14376070 | chr9 | 14375624 | 14376070 | + | 0.756059 | intron (NM_028013, intron 1 of 1) | intron (NM_028013, intron 1 of 1) | 5395 | NM_028013 | 71946 | Mm.41423 | NM_028013 | ENSMUSG00000037419 | **Endod1** | 2210414F18Rik\|2310067E08Rik\|C85344 | endonuclease domain containing 1 | protein-coding |
| chr9_14352365_14352779 | chr9 | 14352366 | 14352779 | + | 0.745943 | Intergenic | Intergenic | 28670 | NM_028013 | 71946 | Mm.41423 | NM_028013 | ENSMUSG00000037419 | **Endod1** | 2210414F18Rik\|2310067E08Rik\|C85344 | endonuclease domain containing 1 | protein-coding |
| chr9_14352365_14353005 | chr9 | 14352366 | 14353005 | + | 0.654631 | Intergenic | Intergenic | 28557 | NM_028013 | 71946 | Mm.41423 | NM_028013 | ENSMUSG00000037419 | **Endod1** | 2210414F18Rik\|2310067E08Rik\|C85344 | endonuclease domain containing 1 | protein-coding |
| chr11_118483269_118483688 | chr11 | 118483270 | 118483688 | + | 0.734418 | intron (NM_172573, intron 7 of 13) | intron (NM_172573, intron 7 of 13) | 6519 | NM_172573 | 217364 | Mm.473963 | NM_172573 | ENSMUSG00000033857 | **Engase** | C130099A03\|D230014K01Rik | endo-beta-N-acetylglucosaminidase | protein-coding |
| chr3_129313665_129314345 | chr3 | 129313666 | 129314345 | + | 1.12543 | intron (NM_007934, intron 3 of 19) | intron (NM_007934, intron 3 of 19) | 18744 | NM_007934 | 13809 | Mm.1193 | NM_007934 | ENSMUSG00000028024 | **Enpep** | 6030431M22Rik\|APA\|Bp-1/6C3\|Ly-51\|Ly51 | glutamyl aminopeptidase | protein-coding |
| chr3_129313674_129314056 | chr3 | 129313675 | 129314056 | + | 1.10443 | intron (NM_007934, intron 3 of 19) | intron (NM_007934, intron 3 of 19) | 18884 | NM_007934 | 13809 | Mm.1193 | NM_007934 | ENSMUSG00000028024 | **Enpep** | 6030431M22Rik\|APA\|Bp-1/6C3\|Ly-51\|Ly51 | glutamyl aminopeptidase | protein-coding |
| chr8_46943259_46943846 | chr8 | 46943260 | 46943846 | + | 0.995358 | Intergenic | Intergenic | -43372 | NM_177304 | 320981 | Mm.211429 | NM_177304 | ENSMUSG00000038173 | **Enpp6** | 4833421B01Rik\|B830047L21Rik\|D8Ertd514e\|E-NPP 6\|Npp6 | ectonucleotide pyrophosphatase/phosphodiesterase 6 | protein-coding |
| chr8_46914396_46914947 | chr8 | 46914397 | 46914947 | + | 0.840209 | Intergenic | L1MB5\|LINE\|L1 | -72253 | NM_177304 | 320981 | Mm.211429 | NM_177304 | ENSMUSG00000038173 | **Enpp6** | 4833421B01Rik\|B830047L21Rik\|D8Ertd514e\|E-NPP 6\|Npp6 | ectonucleotide pyrophosphatase/phosphodiesterase 6 | protein-coding |
| chr8_46921013_46921518 | chr8 | 46921014 | 46921518 | + | 0.730931 | Intergenic | Intergenic | -65659 | NM_177304 | 320981 | Mm.211429 | NM_177304 | ENSMUSG00000038173 | **Enpp6** | 4833421B01Rik\|B830047L21Rik\|D8Ertd514e\|E-NPP 6\|Npp6 | ectonucleotide pyrophosphatase/phosphodiesterase 6 | protein-coding |
| chr8_46990934_46991529 | chr8 | 46990935 | 46991529 | + | 0.634058 | intron (NM_177304, intron 1 of 7) | intron (NM_177304, intron 1 of 7) | 4307 | NM_177304 | 320981 | Mm.211429 | NM_177304 | ENSMUSG00000038173 | **Enpp6** | 4833421B01Rik\|B830047L21Rik\|D8Ertd514e\|E-NPP 6\|Npp6 | ectonucleotide pyrophosphatase/phosphodiesterase 6 | protein-coding |
| chr3_95637931_95638572 | chr3 | 95637932 | 95638572 | + | 1.2163 | Intergenic | Intergenic | 13272 | NM_001026212 | 56205 | Mm.241365 | NM_019561 | ENSMUSG00000038619 | **Ensa** | 1700020C18Rik\|2610007F17Rik\|AI451924 | endosulfine alpha | protein-coding |
| chr3_95637593_95638746 | chr3 | 95637594 | 95638746 | + | 0.961332 | Intergenic | Intergenic | 13190 | NM_001026212 | 56205 | Mm.241365 | NM_019561 | ENSMUSG00000038619 | **Ensa** | 1700020C18Rik\|2610007F17Rik\|AI451924 | endosulfine alpha | protein-coding |
| chr19_40639761_40640625 | chr19 | 40639762 | 40640625 | + | 0.703503 | Intergenic | Intergenic | -19601 | NM_009848 | 12495 | Mm.2824 | NM_009848 | ENSMUSG00000048120 | **Entpd1** | 2610206B08Rik\|AA408691\|Cd39\|NTPDase-1 | ectonucleoside triphosphate diphosphohydrolase 1 | protein-coding |
| chr19_40639789_40640305 | chr19 | 40639790 | 40640305 | + | 0.663573 | Intergenic | Intergenic | -19747 | NM_009848 | 12495 | Mm.2824 | NM_009848 | ENSMUSG00000048120 | **Entpd1** | 2610206B08Rik\|AA408691\|Cd39\|NTPDase-1 | ectonucleoside triphosphate diphosphohydrolase 1 | protein-coding |
| chr15_44390425_44390687 | chr15 | 44390426 | 44390687 | + | 0.725405 | intron (NM_001113554, intron 7 of 9) | L1_Mus2\|LINE\|L1 | -37555 | NM_175009 | 223527 | Mm.291828 | NM_175009 | ENSMUSG00000022338 | **Eny2** | 1810057B09Rik\|6720481I12\|DC6\|Ey2 | enhancer of yellow 2 homolog (Drosophila) | protein-coding |
| chr15_81534473_81534992 | chr15 | 81534474 | 81534992 | + | 1.64174 | Intergenic | RMER15\|LTR\|ERVL | -51481 | NM_177821 | 328572 | Mm.258397 | NM_177821 | ENSMUSG00000055024 | **Ep300** | A430090G16\|A730011L11\|KAT3B\|p300\|p300 HAT | E1A binding protein p300 | protein-coding |
| chr4_132076772_132077021 | chr4 | 132076773 | 132077021 | + | 0.652736 | Intergenic | Intergenic | -1584 | NM_001128607 | 269587 | Mm.30038 | NM_183428 | ENSMUSG00000028906 | **Epb4.1** | 4.1R\|AI415518\|D4Ertd442e\|Elp-1\|Elp1\|Epb41\|mKIAA4056 | erythrocyte protein band 4.1 | protein-coding |
| chr10_25432994_25433389 | chr10 | 25432995 | 25433389 | + | 1.07022 | intron (NM_013511, intron 1 of 19) | intron (NM_013511, intron 1 of 19) | 24824 | NM_013511 | 13822 | Mm.306026 | NM_013511 | ENSMUSG00000019978 | **Epb4.1l2** | 4.1G\|AW555191\|D10Ertd398e\|Epb41l2\|NBL2 | erythrocyte protein band 4.1-like 2 | protein-coding |
| chr10_25331633_25332104 | chr10 | 25331634 | 25332104 | + | 0.70044 | Intergenic | Intergenic | -27929 | NM_001199265 | 13822 | Mm.306026 | NM_013511 | ENSMUSG00000019978 | **Epb4.1l2** | 4.1G\|AW555191\|D10Ertd398e\|Epb41l2\|NBL2 | erythrocyte protein band 4.1-like 2 | protein-coding |
| chr18_34006869_34007591 | chr18 | 34006870 | 34007591 | + | 0.70624 | promoter-TSS (NM_013512) | promoter-TSS (NM_013512) | -24 | NM_013512 | 13824 | Mm.3465 | NM_013512 | ENSMUSG00000024376 | **Epb4.1l4a** | Epb4.1l4\|Epb41l4a\|NBL4 | erythrocyte protein band 4.1-like 4a | protein-coding |
| chr2_49483998_49484454 | chr2 | 49483999 | 49484454 | + | 0.979588 | intron (NM_172663, intron 1 of 13) | intron (NM_172663, intron 1 of 13) | 32740 | NM_172663 | 227867 | Mm.29167 | NM_172663 | ENSMUSG00000069495 | **Epc2** | 5830499L14\|D2Ertd694e | enhancer of polycomb homolog 2 (Drosophila) | protein-coding |
| chr17_87609729_87610171 | chr17 | 87609730 | 87610171 | + | 0.78064 | Intergenic | Intergenic | -26029 | NM_008532 | 17075 | Mm.4259 | NM_008532 | ENSMUSG00000045394 | **Epcam** | CD326\|EGP\|EGP-2\|Egp314\|Ep-CAM\|EpCAM1\|GA733-2\|Ly74\|TROP1\|Tacsd1\|Tacstd1\|gp40 | epithelial cell adhesion molecule | protein-coding |

| | | | | | | | | | | | | | | | | | |
|---|---|---|---|---|---|---|---|---|---|---|---|---|---|---|---|---|---|
| chr11_61597422_61598129 | chr11 | 61597423 | 61598129 | + | 1.18872 | Intergenic | Intergenic | -18089 | NM_001252189 | 13855 | Mm.139695 | NM_010148 | ENSMUSG00000001036 | **Epn2** | 9530051D10Rik\|AA536924\|Ibp2 | epsin 2 | protein-coding |
| chr11_61563668_61564394 | chr11 | 61563669 | 61564394 | + | 0.728532 | intron (NM_001252189, intron 1 of 8) | intron (NM_001252189, intron 1 of 8) | 15656 | NM_010148 | 13855 | Mm.139695 | NM_010148 | ENSMUSG00000001036 | **Epn2** | 9530051D10Rik\|AA536924\|Ibp2 | epsin 2 | protein-coding |
| chr4_109281985_109282450 | chr4 | 109281986 | 109282450 | + | 1.10946 | intron (NM_007943, intron 1 of 24) | intron (NM_007943, intron 1 of 24) | 1943 | NM_007943 | 13858 | Mm.318250 | NM_007943 | ENSMUSG00000028552 | **Eps15** | 2410112D09Rik | epidermal growth factor receptor pathway substrate 15 | protein-coding |
| chr4_109281160_109281513 | chr4 | 109281161 | 109281513 | + | 0.838953 | intron (NM_007943, intron 1 of 24) | intron (NM_007943, intron 1 of 24) | 1062 | NM_007943 | 13858 | Mm.318250 | NM_007943 | ENSMUSG00000028552 | **Eps15** | 2410112D09Rik | epidermal growth factor receptor pathway substrate 15 | protein-coding |
| chr6_137558674_137559267 | chr6 | 137558675 | 137559267 | + | 0.952502 | intron (NM_001271588, intron 1 of 20) | intron (NM_001271588, intron 1 of 20) | 12038 | NM_001271595 | 13860 | Mm.235346 | NM_007945 | ENSMUSG00000015766 | **Eps8** | AW261790 | epidermal growth factor receptor pathway substrate 8 | protein-coding |
| chr6_137558427_137559573 | chr6 | 137558428 | 137559573 | + | 0.805539 | intron (NM_001271588, intron 1 of 20) | intron (NM_001271588, intron 1 of 20) | 12009 | NM_001271595 | 13860 | Mm.235346 | NM_007945 | ENSMUSG00000015766 | **Eps8** | AW261790 | epidermal growth factor receptor pathway substrate 8 | protein-coding |
| chr6_137647239_137647872 | chr6 | 137647240 | 137647872 | + | 0.665419 | intron (NM_007945, intron 1 of 21) | (TC)n\|Simple_repeat\|Simple_repeat | 1729 | NM_001271588 | 13860 | Mm.235346 | NM_007945 | ENSMUSG00000015766 | **Eps8** | AW261790 | epidermal growth factor receptor pathway substrate 8 | protein-coding |
| chr3_107819182_107819601 | chr3 | 107819183 | 107819601 | + | 0.968813 | Intergenic | MER5C1\|DNA\|hAT-Charlie | -57838 | NM_133867 | 99662 | Mm.108491 | NM_133867 | ENSMUSG00000040600 | **Eps8l3** | AI504489\|Eps8r3 | EPS8-like 3 | protein-coding |
| chr14_77932166_77933019 | chr14 | 77932167 | 77933019 | + | 1.41925 | intron (NM_178825, intron 5 of 6) | intron (NM_178825, intron 5 of 6) | 28354 | NM_029495 | 108670 | Mm.68134 | NM_029495 | ENSMUSG00000022014 | **Epsti1** | 2310046K10Rik\|5033415K03Rik\|BRESI1 | epithelial stromal interaction 1 (breast) | protein-coding |
| chr5_30240585_30240988 | chr5 | 30240586 | 30240988 | + | 1.19273 | intron (NM_027652, intron 1 of 9) | intron (NM_027652, intron 1 of 9) | 8169 | NM_027652 | 28042 | Mm.168854 | NM_027652 | ENSMUSG00000075703 | **Ept1** | 4933402G07Rik\|AI448296\|AI452230\|C79563\|D5Wsu178e\|SELI\|mKIAA1724 | ethanolaminephosphotransferase 1 (CDP-ethanolamine-specific) | protein-coding |
| chr5_30234729_30234955 | chr5 | 30234730 | 30234955 | + | 0.826718 | intron (NM_027652, intron 1 of 9) | intron (NM_027652, intron 1 of 9) | 2224 | NM_027652 | 28042 | Mm.168854 | NM_027652 | ENSMUSG00000075703 | **Ept1** | 4933402G07Rik\|AI448296\|AI452230\|C79563\|D5Wsu178e\|SELI\|mKIAA1724 | ethanolaminephosphotransferase 1 (CDP-ethanolamine-specific) | protein-coding |
| chr13_74647830_74648379 | chr13 | 74647831 | 74648379 | + | 1.43392 | intron (NM_030711, intron 2 of 18) | intron (NM_030711, intron 2 of 18) | 8233 | NM_030711 | 80898 | Mm.83526 | NM_030711 | ENSMUSG00000021583 | **Erap1** | Arts1\|ERAAP\|PILSA\|PILSAP | endoplasmic reticulum aminopeptidase 1 | protein-coding |
| chr13_74647893_74648213 | chr13 | 74647894 | 74648213 | + | 1.41739 | intron (NM_030711, intron 2 of 18) | intron (NM_030711, intron 2 of 18) | 8181 | NM_030711 | 80898 | Mm.83526 | NM_030711 | ENSMUSG00000021583 | **Erap1** | Arts1\|ERAAP\|PILSA\|PILSAP | endoplasmic reticulum aminopeptidase 1 | protein-coding |
| chr13_103955426_103956066 | chr13 | 103955427 | 103956066 | + | 0.877772 | Intergenic | ORR1F\|LTR\|ERVL-MaLR | -35160 | NM_001005868 | 59079 | Mm.277354 | NM_021563 | ENSMUSG00000021709 | **Erbb2ip** | 1700028E05Rik\|Erbin\|mKIAA1225 | Erbb2 interacting protein | protein-coding |
| chr1_69286315_69286781 | chr1 | 69286316 | 69286781 | + | 1.56148 | Intergenic | Intergenic | -178489 | NM_010154 | 13869 | Mm.442420 | NM_010154 | ENSMUSG00000062209 | **Erbb4** | Her4\|c-erbB-4 | v-erb-a erythroblastic leukemia viral oncogene homolog 4 (avian) | protein-coding |
| chr1_69343567_69343963 | chr1 | 69343568 | 69343963 | + | 0.726183 | Intergenic | Intergenic | -235706 | NM_010154 | 13869 | Mm.442420 | NM_010154 | ENSMUSG00000062209 | **Erbb4** | Her4\|c-erbB-4 | v-erb-a erythroblastic leukemia viral oncogene homolog 4 (avian) | protein-coding |
| chr1_44177001_44177452 | chr1 | 44177002 | 44177452 | + | 0.687792 | intron (NM_011729, intron 12 of 14) | intron (NM_011729, intron 12 of 14) | 29483 | NM_011729 | 22592 | Mm.2213 | NM_011729 | ENSMUSG00000026048 | **Ercc5** | Xpg | excision repair cross-complementing rodent repair deficiency, complementation group 5 | protein-coding |
| chr14_32493380_32493732 | chr14 | 32493381 | 32493732 | + | 0.989583 | Intergenic | Intergenic | -19965 | NM_001081221 | 319955 | Mm.318310 | NM_177043 | ENSMUSG00000054051 | **Ercc6** | 4732403I04\|C130058G22Rik\|CSB | excision repair cross-complementing rodent repair deficiency, complementation group 6 | protein-coding |
| chr14_32516112_32516656 | chr14 | 32516113 | 32516656 | + | 0.824457 | intron (NM_001081221, intron 1 of 20) | intron (NM_001081221, intron 1 of 20) | 2863 | NM_001081221 | 319955 | Mm.318310 | NM_177043 | ENSMUSG00000054051 | **Ercc6** | 4732403I04\|C130058G22Rik\|CSB | excision repair cross-complementing rodent repair deficiency, complementation group 6 | protein-coding |
| chr14_32493199_32493877 | chr14 | 32493200 | 32493877 | + | 0.697914 | Intergenic | Intergenic | -19983 | NM_001081221 | 319955 | Mm.318310 | NM_177043 | ENSMUSG00000054051 | **Ercc6** | 4732403I04\|C130058G22Rik\|CSB | excision repair cross-complementing rodent repair deficiency, complementation group 6 | protein-coding |
| chr14_32496098_32496447 | chr14 | 32496099 | 32496447 | + | 0.619521 | Intergenic | Intergenic | -17248 | NM_001081221 | 319955 | Mm.318310 | NM_177043 | ENSMUSG00000054051 | **Ercc6** | 4732403I04\|C130058G22Rik\|CSB | excision repair cross-complementing rodent repair deficiency, complementation group 6 | protein-coding |
| chr13_63799763_63800538 | chr13 | 63799764 | 63800538 | + | 1.11629 | Intergenic | Intergenic | -15169 | NM_001013608 | 76251 | Mm.186610 | NM_023507 | ENSMUSG00000021470 | **Ercc6l2** | 0610007P08Rik\|1700019D06Rik\|9330134C04Rik\|Rad26l\|Sr278 | excision repair cross-complementing rodent repair deficiency, complementation group 6 like 2 | protein-coding |
| chr13_63793149_63793692 | chr13 | 63793150 | 63793692 | + | 0.912902 | Intergenic | Intergenic | -21899 | NM_001013608 | 76251 | Mm.186610 | NM_023507 | ENSMUSG00000021470 | **Ercc6l2** | 0610007P08Rik\|1700019D06Rik\|9330134C04Rik\|Rad26l\|Sr278 | excision repair cross-complementing rodent repair deficiency, complementation group 6 like 2 | protein-coding |
| chr13_63864858_63865725 | chr13 | 63864859 | 63865725 | + | 0.900638 | intron (NM_001013608, intron 12 of 18) | B4\|SINE\|B4 | 49958 | NM_023507 | 76251 | Mm.186610 | NM_023507 | ENSMUSG00000021470 | **Ercc6l2** | 0610007P08Rik\|1700019D06Rik\|9330134C04Rik\|Rad26l\|Sr278 | excision repair cross-complementing rodent repair deficiency, complementation group 6 like 2 | protein-coding |
| chr13_63793040_63794117 | chr13 | 63793041 | 63794117 | + | 0.839254 | Intergenic | Intergenic | -21741 | NM_001013608 | 76251 | Mm.186610 | NM_023507 | ENSMUSG00000021470 | **Ercc6l2** | 0610007P08Rik\|1700019D06Rik\|9330134C04Rik\|Rad26l\|Sr278 | excision repair cross-complementing rodent repair deficiency, complementation group 6 like 2 | protein-coding |
| chr13_63845776_63846176 | chr13 | 63845777 | 63846176 | + | 0.812937 | intron (NM_001013608, intron 7 of 18) | MLTR14\|LTR\|ERV1 | 30642 | NM_023507 | 76251 | Mm.186610 | NM_023507 | ENSMUSG00000021470 | **Ercc6l2** | 0610007P08Rik\|1700019D06Rik\|9330134C04Rik\|Rad26l\|Sr278 | excision repair cross-complementing rodent repair deficiency, complementation group 6 like 2 | protein-coding |
| chr16_95415943_95416611 | chr16 | 95415944 | 95416611 | + | 0.783701 | intron (NM_133659, intron 2 of 10) | intron (NM_133659, intron 2 of 10) | 114088 | NM_133659 | 13876 | Mm.164531 | NM_133659 | ENSMUSG00000040732 | **Erg** | D030036I24Rik | avian erythroblastosis virus E-26 (v-ets) oncogene related | protein-coding |
| chr17_26550941_26551723 | chr17 | 26550942 | 26551723 | + | 1.19907 | Intergenic | Intergenic | -10180 | NM_026170 | 67458 | Mm.30496 | NM_026170 | ENSMUSG00000001576 | **Ergic1** | 1200007D18Rik\|maa-136 | endoplasmic reticulum-golgi intermediate compartment (ERGIC) 1 | protein-coding |
| chr17_26588853_26589484 | chr17 | 26588854 | 26589484 | + | 1.17983 | intron (NM_026170, intron 1 of 9) | intron (NM_026170, intron 1 of 9) | 27657 | NM_026170 | 67458 | Mm.30496 | NM_026170 | ENSMUSG00000001576 | **Ergic1** | 1200007D18Rik\|maa-136 | endoplasmic reticulum-golgi intermediate compartment (ERGIC) 1 | protein-coding |
| chr17_26605564_26606013 | chr17 | 26605565 | 26606013 | + | 0.830876 | intron (NM_026170, intron 1 of 9) | intron (NM_026170, intron 1 of 9) | 44277 | NM_026170 | 67458 | Mm.30496 | NM_026170 | ENSMUSG00000001576 | **Ergic1** | 1200007D18Rik\|maa-136 | endoplasmic reticulum-golgi intermediate compartment (ERGIC) 1 | protein-coding |
| chr6_148238918_148239707 | chr6 | 148238919 | 148239707 | + | 0.991597 | intron (NM_198967, intron 19 of 19) | intron (NM_198967, intron 19 of 19) | -26939 | NM_001286560 | 67456 | Mm.59812 | NM_026168 | ENSMUSG00000030304 | **Ergic2** | 1200009B18Rik\|4930572C01Rik\|AA408438\|Ptx1 | ERGIC and golgi 2 | protein-coding |
| chr6_148279850_148280594 | chr6 | 148279851 | 148280594 | + | 0.914836 | intron (NM_198967, intron 11 of 19) | ORR1D1\|LTR\|ERVL-MaLR | -67848 | NM_001286560 | 67456 | Mm.59812 | NM_026168 | ENSMUSG00000030304 | **Ergic2** | 1200009B18Rik\|4930572C01Rik\|AA408438\|Ptx1 | ERGIC and golgi 2 | protein-coding |
| chr8_35481964_35482760 | chr8 | 35481965 | 35482760 | + | 1.07711 | intron (NM_026067, intron 3 of 6) | intron (NM_026067, intron 3 of 6) | 13171 | NM_026067 | 67276 | Mm.207534 | NM_026067 | ENSMUSG00000031527 | **Eri1** | 3'hexo\|3110010F15Rik\|Thex1\|eri-1 | exoribonuclease 1 | protein-coding |
| chr8_14051359_14052447 | chr8 | 14051360 | 14052447 | + | 1.63778 | intron (NM_001034862, intron 3 of 5) | intron (NM_001034862, intron 3 of 5) | 38424 | NM_001034862 | 234086 | Mm.268896 | NM_001034862 | ENSMUSG00000051978 | **Erich1** | - | glutamate rich 1 | protein-coding |
| chr8_14091651_14091953 | chr8 | 14091652 | 14091953 | + | 1.42744 | Intergenic | L2\|LINE\|L2 | -1475 | NM_001034862 | 234086 | Mm.268896 | NM_001034862 | ENSMUSG00000051978 | **Erich1** | - | glutamate rich 1 | protein-coding |
| chr8_14088689_14088973 | chr8 | 14088690 | 14088973 | + | 0.733777 | intron (NM_001034862, intron 1 of 5) | intron (NM_001034862, intron 1 of 5) | 1496 | NM_001034862 | 234086 | Mm.268896 | NM_001034862 | ENSMUSG00000051978 | **Erich1** | - | glutamate rich 1 | protein-coding |
| chr8_14088642_14089077 | chr8 | 14088643 | 14089077 | + | 0.714717 | intron (NM_001034862, intron 1 of 5) | intron (NM_001034862, intron 1 of 5) | 1467 | NM_001034862 | 234086 | Mm.268896 | NM_001034862 | ENSMUSG00000051978 | **Erich1** | - | glutamate rich 1 | protein-coding |
| chr2_58096818_58097430 | chr2 | 58096819 | 58097430 | + | 1.70739 | Intergenic | MT2A\|LTR\|ERVL | -44372 | NM_029972 | 77767 | Mm.91712 | NM_029972 | ENSMUSG00000026830 | **Ermn** | A330104H05Rik\|AI854460\|ermin\|mKIAA1189 | ermin, ERM-like protein | protein-coding |
| chr6_136927269_136927557 | chr6 | 136927270 | 136927557 | + | 0.809048 | intron (NM_001301309, intron 3 of 4) | intron (NM_001301309, intron 3 of 4) | -5233 | NM_026983 | 69187 | Mm.33692 | NM_026983 | ENSMUSG00000030219 | **Erp27** | 1810033M07Rik\|1810047B09Rik | endoplasmic reticulum protein 27 | protein-coding |
| chr5_121449209_121449626 | chr5 | 121449210 | 121449626 | + | 1.13064 | intron (NM_026129, intron 1 of 2) | intron (NM_026129, intron 1 of 2) | 3056 | NM_026129 | 67397 | Mm.154570 | NM_026129 | | **Erp29** | 1200015M03Rik\|2810446M09Rik\|AW209030\|Erp28\|Erp31\|PDI-Db | endoplasmic reticulum protein 29 | protein-coding |
| chr4_48267672_48268530 | chr4 | 48267673 | 48268530 | + | 0.952221 | intron (NM_029572, intron 1 of 11) | intron (NM_029572, intron 1 of 11) | 11488 | NM_029572 | 76299 | Mm.317701 | NM_029572 | ENSMUSG00000028343 | **Erp44** | 1110001E24Rik\|AI849526\|AL033348\|Txndc4 | endoplasmic reticulum protein 44 | protein-coding |
| chr4_48241931_48242378 | chr4 | 48241932 | 48242378 | + | 0.75533 | intron (NM_029572, intron 2 of 11) | Lx4A\|LINE\|L1 | 37434 | NM_029572 | 76299 | Mm.317701 | NM_029572 | ENSMUSG00000028343 | **Erp44** | 1110001E24Rik\|AI849526\|AL033348\|Txndc4 | endoplasmic reticulum protein 44 | protein-coding |
| chr14_74827574_74827934 | chr14 | 74827575 | 74827934 | + | 1.31905 | intron (NM_001252132, intron 3 of 18) | intron (NM_001252132, intron 3 of 18) | 95457 | NM_016903 | 13885 | Mm.38055 | NM_016903 | ENSMUSG00000021996 | **Esd** | Es-10\|Es10\|FGH\|sid478 | esterase D/formylglutathione hydrolase | protein-coding |

| | | | | | | | | | | | | | | | | | |
|---|---|---|---|---|---|---|---|---|---|---|---|---|---|---|---|---|---|
| chr14_74827381_74827985 | chr14 | 74827382 | 74827985 | + | 1.27821 | intron (NM_001252132, intron 3 of 18) | intron (NM_001252132, intron 3 of 18) | 95386 | NM_016903 | 13885 | Mm.38055 | NM_016903 | ENSMUSG00000021996 | **Esd** | Es-10\|Es10\|FGH\|sid478 | esterase D/formylglutathione hydrolase | protein-coding |
| chr14_74823371_74823754 | chr14 | 74823372 | 74823754 | + | 0.810865 | intron (NM_001252132, intron 4 of 18) | Lx8b\|LINE\|L1 | 91266 | NM_016903 | 13885 | Mm.38055 | NM_016903 | ENSMUSG00000021996 | **Esd** | Es-10\|Es10\|FGH\|sid478 | esterase D/formylglutathione hydrolase | protein-coding |
| chr14_74767850_74768318 | chr14 | 74767851 | 74768318 | + | 0.756933 | intron (NM_001252132, intron 18 of 18) | intron (NM_001252132, intron 18 of 18) | 35787 | NM_016903 | 13885 | Mm.38055 | NM_016903 | ENSMUSG00000021996 | **Esd** | Es-10\|Es10\|FGH\|sid478 | esterase D/formylglutathione hydrolase | protein-coding |
| chr10_4670970_4671810 | chr10 | 4670971 | 4671810 | + | 1.52551 | intron (NM_007956, intron 1 of 8) | intron (NM_007956, intron 1 of 8) | 59344 | NM_007956 | 13982 | Mm.9213 | NM_007956 | ENSMUSG00000019768 | **Esr1** | AA420328\|AU041214\|ER-alpha\|ER[a]\|ERa\|ERalpha\|ESR\|Estr\|Estra\|Nr3a1 | estrogen receptor 1 (alpha) | protein-coding |
| chr12_76042936_76043807 | chr12 | 76042937 | 76043807 | + | 1.24322 | intron (NM_001005510, intron 82 of 115) | intron (NM_001005510, intron 82 of 115) | 133887 | NM_010157 | 13983 | Mm.2561 | NM_010157 | ENSMUSG00000021055 | **Esr2** | ER[b]\|ERbeta\|Estrb | estrogen receptor 2 (beta) | protein-coding |
| chr6_143174249_143174564 | chr6 | 143174250 | 143174564 | + | 0.911713 | intron (NM_029250, intron 1 of 7) | B3\|SINE\|B2 | 7177 | NM_029250 | 75320 | Mm.272548 | NM_029250 | ENSMUSG00000030275 | **Etnk1** | 1110061E11Rik\|4930555L11Rik\|AI195356\|AI841245\|C80956\|D6Ertd3e\|EKI1 | ethanolamine kinase 1 | protein-coding |
| chr9_33000252_33000668 | chr9 | 33000253 | 33000668 | + | 1.42225 | Intergenic | Intergenic | 304418 | NM_001038642 | 23871 | Mm.292415 | NM_011808 | ENSMUSG00000032035 | **Ets1** | AI196000\|AI448617\|D230050P06\|Ets-1\|Tpl1 | E26 avian leukemia oncogene 1, 5' domain | protein-coding |
| chr9_32661677_32662138 | chr9 | 32661678 | 32662138 | + | 1.40504 | Intergenic | Intergenic | -34134 | NM_011808 | 23871 | Mm.292415 | NM_011808 | ENSMUSG00000032035 | **Ets1** | AI196000\|AI448617\|D230050P06\|Ets-1\|Tpl1 | E26 avian leukemia oncogene 1, 5' domain | protein-coding |
| chr9_32661392_32662185 | chr9 | 32661393 | 32662185 | + | 1.32364 | Intergenic | Intergenic | -34253 | NM_011808 | 23871 | Mm.292415 | NM_011808 | ENSMUSG00000032035 | **Ets1** | AI196000\|AI448617\|D230050P06\|Ets-1\|Tpl1 | E26 avian leukemia oncogene 1, 5' domain | protein-coding |
| chr9_32999889_33000676 | chr9 | 32999890 | 33000676 | + | 1.2582 | Intergenic | Intergenic | 304241 | NM_001038642 | 23871 | Mm.292415 | NM_011808 | ENSMUSG00000032035 | **Ets1** | AI196000\|AI448617\|D230050P06\|Ets-1\|Tpl1 | E26 avian leukemia oncogene 1, 5' domain | protein-coding |
| chr9_33010977_33011605 | chr9 | 33010978 | 33011605 | + | 1.23372 | Intergenic | Intergenic | 315249 | NM_001038642 | 23871 | Mm.292415 | NM_011808 | ENSMUSG00000032035 | **Ets1** | AI196000\|AI448617\|D230050P06\|Ets-1\|Tpl1 | E26 avian leukemia oncogene 1, 5' domain | protein-coding |
| chr9_33024672_33025227 | chr9 | 33024673 | 33025227 | + | 1.16372 | Intergenic | Intergenic | 328908 | NM_001038642 | 23871 | Mm.292415 | NM_011808 | ENSMUSG00000032035 | **Ets1** | AI196000\|AI448617\|D230050P06\|Ets-1\|Tpl1 | E26 avian leukemia oncogene 1, 5' domain | protein-coding |
| chr9_33011217_33011604 | chr9 | 33011218 | 33011604 | + | 1.15597 | Intergenic | Intergenic | 315369 | NM_001038642 | 23871 | Mm.292415 | NM_011808 | ENSMUSG00000032035 | **Ets1** | AI196000\|AI448617\|D230050P06\|Ets-1\|Tpl1 | E26 avian leukemia oncogene 1, 5' domain | protein-coding |
| chr9_33186265_33186803 | chr9 | 33186266 | 33186803 | + | 1.04414 | Intergenic | MTE2a-int\|LTR\|ERVL-MaLR | 490492 | NM_001038642 | 23871 | Mm.292415 | NM_011808 | ENSMUSG00000032035 | **Ets1** | AI196000\|AI448617\|D230050P06\|Ets-1\|Tpl1 | E26 avian leukemia oncogene 1, 5' domain | protein-coding |
| chr9_32744884_32745728 | chr9 | 32744885 | 32745728 | + | 0.981727 | intron (NM_011808, intron 6 of 7) | intron (NM_011808, intron 6 of 7) | 49264 | NM_001038642 | 23871 | Mm.292415 | NM_011808 | ENSMUSG00000032035 | **Ets1** | AI196000\|AI448617\|D230050P06\|Ets-1\|Tpl1 | E26 avian leukemia oncogene 1, 5' domain | protein-coding |
| chr9_32778913_32779275 | chr9 | 32778914 | 32779275 | + | 0.885036 | Intergenic | Intergenic | 83052 | NM_001038642 | 23871 | Mm.292415 | NM_011808 | ENSMUSG00000032035 | **Ets1** | AI196000\|AI448617\|D230050P06\|Ets-1\|Tpl1 | E26 avian leukemia oncogene 1, 5' domain | protein-coding |
| chr9_32875345_32876119 | chr9 | 32875346 | 32876119 | + | 0.867091 | Intergenic | MER44A\|DNA\|TcMar-Tigger | 179690 | NM_001038642 | 23871 | Mm.292415 | NM_011808 | ENSMUSG00000032035 | **Ets1** | AI196000\|AI448617\|D230050P06\|Ets-1\|Tpl1 | E26 avian leukemia oncogene 1, 5' domain | protein-coding |
| chr9_32669416_32669792 | chr9 | 32669417 | 32669792 | + | 0.865758 | Intergenic | Intergenic | -26438 | NM_011808 | 23871 | Mm.292415 | NM_011808 | ENSMUSG00000032035 | **Ets1** | AI196000\|AI448617\|D230050P06\|Ets-1\|Tpl1 | E26 avian leukemia oncogene 1, 5' domain | protein-coding |
| chr9_32738476_32738904 | chr9 | 32738477 | 32738904 | + | 0.854675 | intron (NM_011808, intron 6 of 7) | intron (NM_011808, intron 6 of 7) | 42648 | NM_001038642 | 23871 | Mm.292415 | NM_011808 | ENSMUSG00000032035 | **Ets1** | AI196000\|AI448617\|D230050P06\|Ets-1\|Tpl1 | E26 avian leukemia oncogene 1, 5' domain | protein-coding |
| chr9_33036180_33036632 | chr9 | 33036181 | 33036632 | + | 0.821454 | Intergenic | Intergenic | 340364 | NM_001038642 | 23871 | Mm.292415 | NM_011808 | ENSMUSG00000032035 | **Ets1** | AI196000\|AI448617\|D230050P06\|Ets-1\|Tpl1 | E26 avian leukemia oncogene 1, 5' domain | protein-coding |
| chr9_33187104_33187564 | chr9 | 33187105 | 33187564 | + | 0.797727 | Intergenic | MTE2a\|LTR\|ERVL-MaLR | 491292 | NM_001038642 | 23871 | Mm.292415 | NM_011808 | ENSMUSG00000032035 | **Ets1** | AI196000\|AI448617\|D230050P06\|Ets-1\|Tpl1 | E26 avian leukemia oncogene 1, 5' domain | protein-coding |
| chr9_32738117_32738904 | chr9 | 32738118 | 32738904 | + | 0.675295 | intron (NM_011808, intron 6 of 7) | intron (NM_011808, intron 6 of 7) | 42469 | NM_001038642 | 23871 | Mm.292415 | NM_011808 | ENSMUSG00000032035 | **Ets1** | AI196000\|AI448617\|D230050P06\|Ets-1\|Tpl1 | E26 avian leukemia oncogene 1, 5' domain | protein-coding |
| chr16_95665725_95666176 | chr16 | 95665726 | 95666176 | + | 0.813577 | Intergenic | Intergenic | -36456 | NM_011809 | 23872 | Mm.290207 | NM_011809 | ENSMUSG00000022895 | **Ets2** | AU022856\|Ets-2 | E26 avian leukemia oncogene 2, 3' domain | protein-coding |
| chr16_95682142_95683031 | chr16 | 95682143 | 95683031 | + | 0.801591 | Intergenic | Intergenic | -19820 | NM_011809 | 23872 | Mm.290207 | NM_011809 | ENSMUSG00000022895 | **Ets2** | AU022856\|Ets-2 | E26 avian leukemia oncogene 2, 3' domain | protein-coding |
| chr16_95682202_95682658 | chr16 | 95682203 | 95682658 | + | 0.745461 | Intergenic | Intergenic | -19977 | NM_011809 | 23872 | Mm.290207 | NM_011809 | ENSMUSG00000022895 | **Ets2** | AU022856\|Ets-2 | E26 avian leukemia oncogene 2, 3' domain | protein-coding |
| chr16_95665526_95666360 | chr16 | 95665527 | 95666360 | + | 0.6427 | Intergenic | Intergenic | -36464 | NM_011809 | 23872 | Mm.290207 | NM_011809 | ENSMUSG00000022895 | **Ets2** | AU022856\|Ets-2 | E26 avian leukemia oncogene 2, 3' domain | protein-coding |
| chr3_87487881_87488601 | chr3 | 87487882 | 87488601 | + | 1.15819 | intron (NM_001113238, intron 11 of 11) | intron (NM_001113238, intron 11 of 11) | -37337 | NM_001083318 | 27049 | Mm.219460 | NM_012051 | ENSMUSG00000003382 | **Etv3** | AI414410\|METS\|Pe1 | ets variant 3 | protein-coding |
| chr3_87487495_87488633 | chr3 | 87487496 | 87488633 | + | 1.1521 | intron (NM_001113238, intron 11 of 11) | intron (NM_001113238, intron 11 of 11) | -37514 | NM_001083318 | 27049 | Mm.219460 | NM_012051 | ENSMUSG00000003382 | **Etv3** | AI414410\|METS\|Pe1 | ets variant 3 | protein-coding |
| chr3_87521900_87522167 | chr3 | 87521901 | 87522167 | + | 0.934746 | Intergenic | Intergenic | -3544 | NM_001083318 | 27049 | Mm.219460 | NM_012051 | ENSMUSG00000003382 | **Etv3** | AI414410\|METS\|Pe1 | ets variant 3 | protein-coding |
| chr3_87512349_87512859 | chr3 | 87512350 | 87512859 | + | 0.731346 | Intergenic | Intergenic | -12974 | NM_001083318 | 27049 | Mm.219460 | NM_012051 | ENSMUSG00000003382 | **Etv3** | AI414410\|METS\|Pe1 | ets variant 3 | protein-coding |
| chr16_22460935_22461404 | chr16 | 22460936 | 22461404 | + | 1.32218 | Intergenic | Intergenic | -21600 | NM_023794 | 104156 | Mm.155708 | NM_023794 | ENSMUSG00000013089 | **Etv5** | 1110005E01Rik\|8430401F14Rik\|ERM | ets variant 5 | protein-coding |
| chr6_134156203_134156799 | chr6 | 134156204 | 134156799 | + | 1.09058 | intron (NM_007961, intron 2 of 7) | intron (NM_007961, intron 2 of 7) | 120794 | NM_007961 | 14011 | Mm.269995 | NM_007961 | ENSMUSG00000030199 | **Etv6** | AW123102\|AW557856\|Tel | ets variant 6 | protein-coding |
| chr6_134124365_134124792 | chr6 | 134124366 | 134124792 | + | 0.966055 | intron (NM_007961, intron 1 of 7) | RMER15\|LTR\|ERVL | 88872 | NM_007961 | 14011 | Mm.269995 | NM_007961 | ENSMUSG00000030199 | **Etv6** | AW123102\|AW557856\|Tel | ets variant 6 | protein-coding |
| chr6_133985995_133986711 | chr6 | 133985996 | 133986711 | + | 0.82989 | Intergenic | ORR1E\|LTR\|ERVL-MaLR | -49354 | NM_007961 | 14011 | Mm.269995 | NM_007961 | ENSMUSG00000030199 | **Etv6** | AW123102\|AW557856\|Tel | ets variant 6 | protein-coding |
| chr6_134066937_134067907 | chr6 | 134066938 | 134067907 | + | 0.819403 | intron (NM_007961, intron 1 of 7) | MTD\|LTR\|ERVL-MaLR | 31715 | NM_007961 | 14011 | Mm.269995 | NM_007961 | ENSMUSG00000030199 | **Etv6** | AW123102\|AW557856\|Tel | ets variant 6 | protein-coding |
| chr6_134066964_134067365 | chr6 | 134066965 | 134067365 | + | 0.784481 | intron (NM_007961, intron 1 of 7) | MTD\|LTR\|ERVL-MaLR | 31458 | NM_007961 | 14011 | Mm.269995 | NM_007961 | ENSMUSG00000030199 | **Etv6** | AW123102\|AW557856\|Tel | ets variant 6 | protein-coding |
| chr6_134122951_134123291 | chr6 | 134122952 | 134123291 | + | 0.748289 | intron (NM_007961, intron 1 of 7) | intron (NM_007961, intron 1 of 7) | 87414 | NM_007961 | 14011 | Mm.269995 | NM_007961 | ENSMUSG00000030199 | **Etv6** | AW123102\|AW557856\|Tel | ets variant 6 | protein-coding |
| chr6_134059537_134060005 | chr6 | 134059538 | 134060005 | + | 0.736351 | intron (NM_007961, intron 1 of 7) | intron (NM_007961, intron 1 of 7) | 24064 | NM_007961 | 14011 | Mm.269995 | NM_007961 | ENSMUSG00000030199 | **Etv6** | AW123102\|AW557856\|Tel | ets variant 6 | protein-coding |
| chr6_134069951_134070365 | chr6 | 134069952 | 134070365 | + | 0.707238 | intron (NM_007961, intron 1 of 7) | intron (NM_007961, intron 1 of 7) | 34451 | NM_007961 | 14011 | Mm.269995 | NM_007961 | ENSMUSG00000030199 | **Etv6** | AW123102\|AW557856\|Tel | ets variant 6 | protein-coding |
| chr6_134020556_134021501 | chr6 | 134020557 | 134021501 | + | 0.691455 | Intergenic | Intergenic | -14678 | NM_007961 | 14011 | Mm.269995 | NM_007961 | ENSMUSG00000030199 | **Etv6** | AW123102\|AW557856\|Tel | ets variant 6 | protein-coding |
| chr6_134067433_134067905 | chr6 | 134067434 | 134067905 | + | 0.680956 | intron (NM_007961, intron 1 of 7) | intron (NM_007961, intron 1 of 7) | 31962 | NM_007961 | 14011 | Mm.269995 | NM_007961 | ENSMUSG00000030199 | **Etv6** | AW123102\|AW557856\|Tel | ets variant 6 | protein-coding |
| chr6_134059292_134060022 | chr6 | 134059293 | 134060022 | + | 0.675159 | intron (NM_007961, intron 1 of 7) | intron (NM_007961, intron 1 of 7) | 23950 | NM_007961 | 14011 | Mm.269995 | NM_007961 | ENSMUSG00000030199 | **Etv6** | AW123102\|AW557856\|Tel | ets variant 6 | protein-coding |
| chr4_126144738_126145171 | chr4 | 126144739 | 126145171 | + | 1.13956 | Intergenic | Intergenic | -3048 | NM_172145 | 230752 | Mm.41569 | NM_172145 | ENSMUSG00000050212 | **Eva1b** | 2610027C15Rik\|Fam176b | eva-1 homolog B (C. elegans) | protein-coding |
| chr8_4165171_4165457 | chr8 | 4165172 | 4165457 | + | 1.28146 | Intergenic | Intergenic | -1253 | NM_001039578 | 213027 | Mm.437106 | NM_153536 | ENSMUSG00000011832 | **Evi5l** | 1700084G18Rik\|2310039H16Rik\|3110007G05Rik\|B130050I23Rik | ecotropic viral integration site 5 like | protein-coding |
| chr12_108646669_108647172 | chr12 | 108646670 | 108647172 | + | 1.42437 | intron (NM_001163396, intron 2 of 14) | intron (NM_001163396, intron 2 of 14) | 11168 | NM_001163396 | 14026 | Mm.238841 | NM_007965 | ENSMUSG00000021262 | **Evl** | AI528774 | Ena-vasodilator stimulated phosphoprotein | protein-coding |
| chr12_108641104_108641355 | chr12 | 108641105 | 108641355 | + | 1.07578 | intron (NM_001163396, intron 2 of 14) | intron (NM_001163396, intron 2 of 14) | 5477 | NM_001163396 | 14026 | Mm.238841 | NM_007965 | ENSMUSG00000021262 | **Evl** | AI528774 | Ena-vasodilator stimulated phosphoprotein | protein-coding |
| chr12_108642070_108642294 | chr12 | 108642071 | 108642294 | + | 0.915079 | intron (NM_001163396, intron 2 of 14) | intron (NM_001163396, intron 2 of 14) | 6429 | NM_001163396 | 14026 | Mm.238841 | NM_007965 | ENSMUSG00000021262 | **Evl** | AI528774 | Ena-vasodilator stimulated phosphoprotein | protein-coding |
| chr12_108646129_108647173 | chr12 | 108646130 | 108647173 | + | 0.908598 | intron (NM_001163396, intron 2 of 14) | intron (NM_001163396, intron 2 of 14) | 10898 | NM_001163396 | 14026 | Mm.238841 | NM_007965 | ENSMUSG00000021262 | **Evl** | AI528774 | Ena-vasodilator stimulated phosphoprotein | protein-coding |

| | | | | | | | | | | | | | | | | | |
|---|---|---|---|---|---|---|---|---|---|---|---|---|---|---|---|---|---|
| chr12_108645531_108646018 | chr12 | 108645532 | 108646018 | + | 0.765743 | intron (NM_001163396, intron 2 of 14) | intron (NM_001163396, intron 2 of 14) | 10022 | NM_001163396 | 14026 | Mm.238841 | NM_007965 | ENSMUSG00000021262 | **Evl** | AI528774 | Ena-vasodilator stimulated phosphoprotein | protein-coding |
| chr12_108639078_108639527 | chr12 | 108639079 | 108639527 | + | 0.729661 | intron (NM_001163396, intron 2 of 14) | intron (NM_001163396, intron 2 of 14) | 3550 | NM_001163396 | 14026 | Mm.238841 | NM_007965 | ENSMUSG00000021262 | **Evl** | AI528774 | Ena-vasodilator stimulated phosphoprotein | protein-coding |
| chr12_108617572_108618051 | chr12 | 108617573 | 108618051 | + | 0.679036 | intron (NM_001163395, intron 1 of 12) | intron (NM_001163395, intron 1 of 12) | 12046 | NM_001163395 | 14026 | Mm.238841 | NM_007965 | ENSMUSG00000021262 | **Evl** | AI528774 | Ena-vasodilator stimulated phosphoprotein | protein-coding |
| chr14_49089545_49090050 | chr14 | 49089546 | 49090050 | + | 1.11199 | Intergenic | Intergenic | -23131 | NM_207214 | 105504 | Mm.31607 | NM_207214 | ENSMUSG00000061244 | **Exoc5** | AI447711\|AI448003\|Gm76\|PRO1912\|SEC10\|Sec10l1 | exocyst complex component 5 | protein-coding |
| chr19_37526833_37527200 | chr19 | 37526834 | 37527200 | + | 0.73633 | Intergenic | Intergenic | -23401 | NM_175353 | 107371 | Mm.24865 | NM_175353 | ENSMUSG00000053799 | **Exoc6** | 4833405E05Rik\|AW413330\|C430002C19\|Sec15\|Sec15l1\|hbd\|msec15 | exocyst complex component 6 | protein-coding |
| chr19_37553829_37554217 | chr19 | 37553830 | 37554217 | + | 0.680419 | intron (NM_175353, intron 1 of 21) | intron (NM_175353, intron 1 of 21) | 3605 | NM_175353 | 107371 | Mm.24865 | NM_175353 | ENSMUSG00000053799 | **Exoc6** | 4833405E05Rik\|AW413330\|C430002C19\|Sec15\|Sec15l1\|hbd\|msec15 | exocyst complex component 6 | protein-coding |
| chr6_84962600_84963067 | chr6 | 84962601 | 84963067 | + | 1.80855 | intron (NM_177077, intron 6 of 21) | ORR1D1\|LTR\|ERVL-MaLR | 106679 | NM_177077 | 75914 | Mm.159621 | NM_177077 | ENSMUSG00000033769 | **Exoc6b** | 4930569O18Rik\|G430127E12Rik\|Sec15b\|Sec15l2\|mKIAA0919 | exocyst complex component 6B | protein-coding |
| chr6_84962585_84963249 | chr6 | 84962586 | 84963249 | + | 1.64956 | intron (NM_177077, intron 6 of 21) | ORR1D1\|LTR\|ERVL-MaLR | 106596 | NM_177077 | 75914 | Mm.159621 | NM_177077 | ENSMUSG00000033769 | **Exoc6b** | 4930569O18Rik\|G430127E12Rik\|Sec15b\|Sec15l2\|mKIAA0919 | exocyst complex component 6B | protein-coding |
| chr6_84961719_84962115 | chr6 | 84961720 | 84962115 | + | 1.00827 | intron (NM_177077, intron 6 of 21) | intron (NM_177077, intron 6 of 21) | 107596 | NM_177077 | 75914 | Mm.159621 | NM_177077 | ENSMUSG00000033769 | **Exoc6b** | 4930569O18Rik\|G430127E12Rik\|Sec15b\|Sec15l2\|mKIAA0919 | exocyst complex component 6B | protein-coding |
| chr6_84920908_84921494 | chr6 | 84920909 | 84921494 | + | 0.983465 | intron (NM_177077, intron 6 of 21) | intron (NM_177077, intron 6 of 21) | 148312 | NM_177077 | 75914 | Mm.159621 | NM_177077 | ENSMUSG00000033769 | **Exoc6b** | 4930569O18Rik\|G430127E12Rik\|Sec15b\|Sec15l2\|mKIAA0919 | exocyst complex component 6B | protein-coding |
| chr9_119455974_119456385 | chr9 | 119455975 | 119456385 | + | 1.14881 | intron (NM_172456, intron 5 of 5) | B1_Mur4\|SINE\|Alu | 11257 | NM_172456 | 208194 | Mm.101578 | NM_172456 | ENSMUSG00000042787 | **Exog** | AW557704\|ENGL-B\|ENGL-a\|Endogl1\|Endogl2\|Engl\|Engla\|Englb | endo/exonuclease (5'-3'), endonuclease G-like | protein-coding |
| chr9_53328763_53329697 | chr9 | 53328764 | 53329697 | + | 1.03832 | intron (NM_176846, intron 1 of 5) | MER2\|DNA\|TcMar-Tigger | 27560 | NM_176846 | 320051 | Mm.277540 | NM_176846 | | **Exph5** | AC079869.22gm5\|B130009M24Rik\|E030050P12\|Kiaa0624\|Slac2b\|slac2-b | exophilin 5 | protein-coding |
| chr9_53328785_53329187 | chr9 | 53328786 | 53329187 | + | 1.00339 | intron (NM_176846, intron 1 of 5) | intron (NM_176846, intron 1 of 5) | 27316 | NM_176846 | 320051 | Mm.277540 | NM_176846 | | **Exph5** | AC079869.22gm5\|B130009M24Rik\|E030050P12\|Kiaa0624\|Slac2b\|slac2-b | exophilin 5 | protein-coding |
| chr9_53329231_53329615 | chr9 | 53329232 | 53329615 | + | 0.785992 | intron (NM_176846, intron 1 of 5) | MER2\|DNA\|TcMar-Tigger | 27753 | NM_176846 | 320051 | Mm.277540 | NM_176846 | | **Exph5** | AC079869.22gm5\|B130009M24Rik\|E030050P12\|Kiaa0624\|Slac2b\|slac2-b | exophilin 5 | protein-coding |
| chr9_53277541_53278342 | chr9 | 53277542 | 53278342 | + | 0.655218 | Intergenic | Intergenic | -23728 | NM_176846 | 320051 | Mm.277540 | NM_176846 | | **Exph5** | AC079869.22gm5\|B130009M24Rik\|E030050P12\|Kiaa0624\|Slac2b\|slac2-b | exophilin 5 | protein-coding |
| chr15_53492065_53492563 | chr15 | 53492066 | 53492563 | + | 1.438 | intron (NM_177225, intron 4 of 4) | intron (NM_177225, intron 4 of 4) | -146131 | NM_010162 | 14042 | Mm.309395 | NM_010162 | | **Ext1** | AA409028 | exostoses (multiple) 1 | protein-coding |
| chr15_53222322_53222824 | chr15 | 53222323 | 53222824 | + | 1.20486 | intron (NM_010162, intron 1 of 10) | intron (NM_010162, intron 1 of 10) | 123610 | NM_010162 | 14042 | Mm.309395 | NM_010162 | | **Ext1** | AA409028 | exostoses (multiple) 1 | protein-coding |
| chr15_53142851_53143359 | chr15 | 53142852 | 53143359 | + | 1.09161 | intron (NM_010162, intron 1 of 10) | intron (NM_010162, intron 1 of 10) | 203078 | NM_010162 | 14042 | Mm.309395 | NM_010162 | | **Ext1** | AA409028 | exostoses (multiple) 1 | protein-coding |
| chr15_53258772_53259637 | chr15 | 53258773 | 53259637 | + | 0.90411 | intron (NM_010162, intron 1 of 10) | MLT1C\|LTR\|ERVL-MaLR | 86978 | NM_010162 | 14042 | Mm.309395 | NM_010162 | | **Ext1** | AA409028 | exostoses (multiple) 1 | protein-coding |
| chr15_53563166_53563640 | chr15 | 53563167 | 53563640 | + | 0.874951 | intron (NM_177225, intron 4 of 4) | intron (NM_177225, intron 4 of 4) | -217220 | NM_010162 | 14042 | Mm.309395 | NM_010162 | | **Ext1** | AA409028 | exostoses (multiple) 1 | protein-coding |
| chr15_53562817_53563640 | chr15 | 53562818 | 53563640 | + | 0.848708 | intron (NM_177225, intron 4 of 4) | intron (NM_177225, intron 4 of 4) | -217046 | NM_010162 | 14042 | Mm.309395 | NM_010162 | | **Ext1** | AA409028 | exostoses (multiple) 1 | protein-coding |
| chr15_53259168_53259524 | chr15 | 53259169 | 53259524 | + | 0.814812 | intron (NM_010162, intron 1 of 10) | intron (NM_010162, intron 1 of 10) | 86837 | NM_010162 | 14042 | Mm.309395 | NM_010162 | | **Ext1** | AA409028 | exostoses (multiple) 1 | protein-coding |
| chr14_65106605_65107041 | chr14 | 65106606 | 65107041 | + | 1.36245 | Intergenic | Lx7\|LINE\|L1 | -8717 | NM_018788 | 54616 | Mm.103748 | NM_018788 | ENSMUSG00000021978 | **Extl3** | 2900009G18Rik\|AU040105\|Ext1l\|mKIAA0519 | exostoses (multiple)-like 3 | protein-coding |
| chr14_65091031_65091590 | chr14 | 65091032 | 65091590 | + | 0.921112 | intron (NM_018788, intron 1 of 6) | intron (NM_018788, intron 1 of 6) | 6795 | NM_018788 | 54616 | Mm.103748 | NM_018788 | ENSMUSG00000021978 | **Extl3** | 2900009G18Rik\|AU040105\|Ext1l\|mKIAA0519 | exostoses (multiple)-like 3 | protein-coding |
| chr14_65113589_65114344 | chr14 | 65113590 | 65114344 | + | 0.634473 | Intergenic | GA-rich\|Low_complexity\|Low_complexity | -15861 | NM_018788 | 54616 | Mm.103748 | NM_018788 | ENSMUSG00000021978 | **Extl3** | 2900009G18Rik\|AU040105\|Ext1l\|mKIAA0519 | exostoses (multiple)-like 3 | protein-coding |
| chr2_165644116_165644854 | chr2 | 165644117 | 165644854 | + | 1.01921 | intron (NM_001271962, intron 1 of 15) | intron (NM_001271962, intron 1 of 15) | -10813 | NM_010165 | 14049 | Mm.282719 | NM_010165 | ENSMUSG00000017897 | **Eya2** | Eab1 | eyes absent 2 homolog (Drosophila) | protein-coding |
| chr6_47620983_47621612 | chr6 | 47620984 | 47621612 | + | 1.76832 | Intergenic | Intergenic | -26268 | NM_001146689 | 14056 | Mm.246688 | NM_007971 | ENSMUSG00000029687 | **Ezh2** | Enx-1\|Enx1h\|KMT6\|mKIAA4065 | enhancer of zeste homolog 2 (Drosophila) | protein-coding |
| chr6_47618905_47619145 | chr6 | 47618906 | 47619145 | + | 1.03188 | Intergenic | URR1A\|DNA\|hAT-Charlie | -23995 | NM_001146689 | 14056 | Mm.246688 | NM_007971 | ENSMUSG00000029687 | **Ezh2** | Enx-1\|Enx1h\|KMT6\|mKIAA4065 | enhancer of zeste homolog 2 (Drosophila) | protein-coding |
| chr17_6834961_6835977 | chr17 | 6834962 | 6835977 | + | 1.82372 | Intergenic | LTR40a\|LTR\|ERVL | -52689 | NM_009510 | 22350 | Mm.277812 | NM_009510 | ENSMUSG00000052397 | **Ezr** | AW146364\|R75297\|Vil2\|p81 | ezrin | protein-coding |
| chr17_6758757_6759281 | chr17 | 6758758 | 6759281 | + | 0.924566 | intron (NM_009510, intron 2 of 12) | intron (NM_009510, intron 2 of 12) | 23761 | NM_009510 | 22350 | Mm.277812 | NM_009510 | ENSMUSG00000052397 | **Ezr** | AW146364\|R75297\|Vil2\|p81 | ezrin | protein-coding |
| chr17_6803585_6804119 | chr17 | 6803586 | 6804119 | + | 0.914171 | Intergenic | Intergenic | -21072 | NM_009510 | 22350 | Mm.277812 | NM_009510 | ENSMUSG00000052397 | **Ezr** | AW146364\|R75297\|Vil2\|p81 | ezrin | protein-coding |
| chr17_6749046_6749576 | chr17 | 6749047 | 6749576 | + | 0.770017 | intron (NM_009510, intron 6 of 12) | intron (NM_009510, intron 6 of 12) | 33469 | NM_009510 | 22350 | Mm.277812 | NM_009510 | ENSMUSG00000052397 | **Ezr** | AW146364\|R75297\|Vil2\|p81 | ezrin | protein-coding |
| chr17_6793754_6794864 | chr17 | 6793755 | 6794864 | + | 0.755119 | Intergenic | Intergenic | -11529 | NM_009510 | 22350 | Mm.277812 | NM_009510 | ENSMUSG00000052397 | **Ezr** | AW146364\|R75297\|Vil2\|p81 | ezrin | protein-coding |
| chr17_6757221_6757623 | chr17 | 6757222 | 6757623 | + | 0.640556 | intron (NM_009510, intron 2 of 12) | intron (NM_009510, intron 2 of 12) | 25358 | NM_009510 | 22350 | Mm.277812 | NM_009510 | ENSMUSG00000052397 | **Ezr** | AW146364\|R75297\|Vil2\|p81 | ezrin | protein-coding |
| chr13_37008899_37009360 | chr13 | 37008900 | 37009360 | + | 1.73573 | intron (NM_028784, intron 3 of 14) | intron (NM_028784, intron 3 of 14) | 40083 | NM_028784 | 74145 | Mm.235105 | NM_028784 | ENSMUSG00000039109 | **F13a1** | 1200014I03Rik\|AI462306\|F13a | coagulation factor XIII, A1 subunit | protein-coding |
| chr13_37071300_37071983 | chr13 | 37071301 | 37071983 | + | 1.41062 | Intergenic | Intergenic | -21398 | NM_001166391 | 74145 | Mm.235105 | NM_028784 | ENSMUSG00000039109 | **F13a1** | 1200014I03Rik\|AI462306\|F13a | coagulation factor XIII, A1 subunit | protein-coding |
| chr13_36962208_36962718 | chr13 | 36962209 | 36962718 | + | 1.10082 | intron (NM_028784, intron 6 of 14) | (CACCAT)n\|Simple_repeat\|Simple_repeat | 86750 | NM_028784 | 74145 | Mm.235105 | NM_028784 | ENSMUSG00000039109 | **F13a1** | 1200014I03Rik\|AI462306\|F13a | coagulation factor XIII, A1 subunit | protein-coding |
| chr13_37010528_37011096 | chr13 | 37010529 | 37011096 | + | 1.09577 | intron (NM_028784, intron 3 of 14) | intron (NM_028784, intron 3 of 14) | 38401 | NM_028784 | 74145 | Mm.235105 | NM_028784 | ENSMUSG00000039109 | **F13a1** | 1200014I03Rik\|AI462306\|F13a | coagulation factor XIII, A1 subunit | protein-coding |
| chr13_37010822_37011081 | chr13 | 37010823 | 37011081 | + | 0.994151 | intron (NM_028784, intron 3 of 14) | intron (NM_028784, intron 3 of 14) | 38261 | NM_028784 | 74145 | Mm.235105 | NM_028784 | ENSMUSG00000039109 | **F13a1** | 1200014I03Rik\|AI462306\|F13a | coagulation factor XIII, A1 subunit | protein-coding |
| chr13_37007322_37007909 | chr13 | 37007323 | 37007909 | + | 0.911619 | intron (NM_028784, intron 3 of 14) | intron (NM_028784, intron 3 of 14) | 41597 | NM_028784 | 74145 | Mm.235105 | NM_028784 | ENSMUSG00000039109 | **F13a1** | 1200014I03Rik\|AI462306\|F13a | coagulation factor XIII, A1 subunit | protein-coding |
| chr13_37140596_37141477 | chr13 | 37140597 | 37141477 | + | 0.895866 | Intergenic | RMER15\|LTR\|ERVL | -90793 | NM_001166391 | 74145 | Mm.235105 | NM_028784 | ENSMUSG00000039109 | **F13a1** | 1200014I03Rik\|AI462306\|F13a | coagulation factor XIII, A1 subunit | protein-coding |
| chr13_36965647_36966164 | chr13 | 36965648 | 36966164 | + | 0.807098 | intron (NM_028784, intron 6 of 14) | intron (NM_028784, intron 6 of 14) | 83307 | NM_028784 | 74145 | Mm.235105 | NM_028784 | ENSMUSG00000039109 | **F13a1** | 1200014I03Rik\|AI462306\|F13a | coagulation factor XIII, A1 subunit | protein-coding |
| chr13_37010542_37010771 | chr13 | 37010543 | 37010771 | + | 0.781686 | intron (NM_028784, intron 3 of 14) | intron (NM_028784, intron 3 of 14) | 38556 | NM_028784 | 74145 | Mm.235105 | NM_028784 | ENSMUSG00000039109 | **F13a1** | 1200014I03Rik\|AI462306\|F13a | coagulation factor XIII, A1 subunit | protein-coding |
| chr13_95566907_95567559 | chr13 | 95566908 | 95567559 | + | 0.752895 | Intergenic | Intergenic | -41993 | NM_007974 | 14063 | Mm.1614 | NM_007974 | ENSMUSG00000021678 | **F2rl1** | Gpcr11\|PAR-2\|Par2 | coagulation factor II (thrombin) receptor-like 1 | protein-coding |
| chr13_95548297_95548863 | chr13 | 95548298 | 95548863 | + | 0.680285 | Intergenic | Intergenic | -23340 | NM_007974 | 14063 | Mm.1614 | NM_007974 | ENSMUSG00000021678 | **F2rl1** | Gpcr11\|PAR-2\|Par2 | coagulation factor II (thrombin) receptor-like 1 | protein-coding |
| chr1_164136653_164136919 | chr1 | 164136654 | 164136919 | + | 0.939205 | intron (NM_011347, intron 8 of 15) | intron (NM_011347, intron 8 of 15) | -15073 | NM_007976 | 14067 | Mm.12900 | NM_007976 | ENSMUSG00000026579 | **F5** | AI173222\|Cf-5\|Cf5 | coagulation factor V | protein-coding |
| chr1_164136616_164137076 | chr1 | 164136617 | 164137076 | + | 0.928499 | intron (NM_011347, intron 8 of 15) | intron (NM_011347, intron 8 of 15) | -15013 | NM_007976 | 14067 | Mm.12900 | NM_007976 | ENSMUSG00000026579 | **F5** | AI173222\|Cf-5\|Cf5 | coagulation factor V | protein-coding |
| chr4_116008474_116008964 | chr4 | 116008475 | 116008964 | + | 0.676095 | intron (NM_010173, intron 1 of 14) | intron (NM_010173, intron 1 of 14) | 9183 | NM_010173 | 14073 | Mm.256025 | NM_010173 | ENSMUSG00000034171 | **Faah** | AW412498 | fatty acid amide hydrolase | protein-coding |

| | | | | | | | | | | | | | | | | | |
|---|---|---|---|---|---|---|---|---|---|---|---|---|---|---|---|---|---|
| chr6_71224223_71224624 | chr6 | 71224224 | 71224624 | + | 0.762071 | intron (NM_001160127, intron 6 of 9) | intron (NM_001160127, intron 6 of 9) | 24536 | NM_017399 | 14080 | Mm.22126 | NM_017399 | ENSMUSG00000054422 | **Fabp1** | Fabpl\|L-FABP | fatty acid binding protein 1, liver | protein-coding |
| chr3_10259277_10259883 | chr3 | 10259278 | 10259883 | + | 1.01495 | intron (NM_029310, intron 2 of 5) | L1MD3\|LINE\|L1 | 41603 | NM_029310 | 75497 | Mm.47052 | NM_029310 | ENSMUSG00000027530 | **Fabp12** | 1700008G05Rik | fatty acid binding protein 12 | protein-coding |
| chr3_10243499_10244157 | chr3 | 10243500 | 10244157 | + | 1.3822 | TTS (NM_029310) | TTS (NM_029310) | -35252 | NM_024406 | 11770 | Mm.582 | NM_024406 | ENSMUSG00000062515 | **Fabp4** | 422/aP2\|ALBP/Ap2\|Ap2\|Lbpl | fatty acid binding protein 4, adipocyte | protein-coding |
| chr3_10204795_10206000 | chr3 | 10204796 | 10206000 | + | 0.833417 | intron (NM_024406, intron 2 of 3) | intron (NM_024406, intron 2 of 3) | 3178 | NM_024406 | 11770 | Mm.582 | NM_024406 | ENSMUSG00000062515 | **Fabp4** | 422/aP2\|ALBP/Ap2\|Ap2\|Lbpl | fatty acid binding protein 4, adipocyte | protein-coding |
| chr3_10204800_10205461 | chr3 | 10204801 | 10205461 | + | 0.741026 | intron (NM_024406, intron 3 of 3) | intron (NM_024406, intron 3 of 3) | 3445 | NM_024406 | 11770 | Mm.582 | NM_024406 | ENSMUSG00000062515 | **Fabp4** | 422/aP2\|ALBP/Ap2\|Ap2\|Lbpl | fatty acid binding protein 4, adipocyte | protein-coding |
| chr3_10211879_10212310 | chr3 | 10211880 | 10212310 | + | 0.735768 | Intergenic | Intergenic | -3519 | NM_024406 | 11770 | Mm.582 | NM_024406 | ENSMUSG00000062515 | **Fabp4** | 422/aP2\|ALBP/Ap2\|Ap2\|Lbpl | fatty acid binding protein 4, adipocyte | protein-coding |
| chr3_10212944_10213374 | chr3 | 10212945 | 10213374 | + | 0.689024 | Intergenic | Intergenic | -4583 | NM_024406 | 11770 | Mm.582 | NM_024406 | ENSMUSG00000062515 | **Fabp4** | 422/aP2\|ALBP/Ap2\|Ap2\|Lbpl | fatty acid binding protein 4, adipocyte | protein-coding |
| chr3_10212977_10213714 | chr3 | 10212978 | 10213714 | + | 0.659252 | Intergenic | Intergenic | -4770 | NM_024406 | 11770 | Mm.582 | NM_024406 | ENSMUSG00000062515 | **Fabp4** | 422/aP2\|ALBP/Ap2\|Ap2\|Lbpl | fatty acid binding protein 4, adipocyte | protein-coding |
| chr3_10043348_10044373 | chr3 | 10043349 | 10044373 | + | 0.777641 | Intergenic | RMER15\|LTR\|ERVL | 31276 | NM_001272098 | 16592 | Mm.741 | NM_010634 | ENSMUSG00000027533 | **Fabp5** | E-FABP\|Fabpe\|Klbp\|PA-FABP\|mal1 | fatty acid binding protein 5, epidermal | protein-coding |
| chr19_10100735_10102286 | chr19 | 10100736 | 10102286 | + | 0.63494 | promoter-TSS (NM_019699) | promoter-TSS (NM_019699) | -8 | NM_019699 | 56473 | Mm.38901 | NM_019699 | ENSMUSG00000024665 | **Fads2** | 2900042M13Rik\|Fadsd2 | fatty acid desaturase 2 | protein-coding |
| chr4_109726308_109726760 | chr4 | 109726309 | 109726760 | + | 1.82309 | intron (NM_007983, intron 3 of 18) | intron (NM_007983, intron 3 of 18) | 49907 | NM_007983 | 14084 | Mm.318259 | NM_007983 | ENSMUSG00000010517 | **Faf1** | AA408698\|Dffrx\|Fam | Fas-associated factor 1 | protein-coding |
| chr4_109726299_109727156 | chr4 | 109726300 | 109727156 | + | 1.80168 | intron (NM_007983, intron 3 of 18) | intron (NM_007983, intron 3 of 18) | 50101 | NM_007983 | 14084 | Mm.318259 | NM_007983 | ENSMUSG00000010517 | **Faf1** | AA408698\|Dffrx\|Fam | Fas-associated factor 1 | protein-coding |
| chr4_109705217_109705734 | chr4 | 109705218 | 109705734 | + | 1.45365 | intron (NM_007983, intron 1 of 18) | RMER15\|LTR\|ERVL | 28849 | NM_007983 | 14084 | Mm.318259 | NM_007983 | ENSMUSG00000010517 | **Faf1** | AA408698\|Dffrx\|Fam | Fas-associated factor 1 | protein-coding |
| chr4_109705006_109705770 | chr4 | 109705007 | 109705770 | + | 1.37602 | intron (NM_007983, intron 1 of 18) | RMER15\|LTR\|ERVL | 28761 | NM_007983 | 14084 | Mm.318259 | NM_007983 | ENSMUSG00000010517 | **Faf1** | AA408698\|Dffrx\|Fam | Fas-associated factor 1 | protein-coding |
| chr4_109705382_109705883 | chr4 | 109705383 | 109705883 | + | 1.369 | intron (NM_007983, intron 1 of 18) | RMER15\|LTR\|ERVL | 29006 | NM_007983 | 14084 | Mm.318259 | NM_007983 | ENSMUSG00000010517 | **Faf1** | AA408698\|Dffrx\|Fam | Fas-associated factor 1 | protein-coding |
| chr4_109726845_109727145 | chr4 | 109726846 | 109727145 | + | 1.10768 | intron (NM_007983, intron 3 of 18) | RSINE1\|SINE\|B4 | 50368 | NM_007983 | 14084 | Mm.318259 | NM_007983 | ENSMUSG00000010517 | **Faf1** | AA408698\|Dffrx\|Fam | Fas-associated factor 1 | protein-coding |
| chr7_84638610_84639339 | chr7 | 84638611 | 84639339 | + | 0.861498 | intron (NM_022985, intron 2 of 7) | intron (NM_022985, intron 2 of 7) | -33033 | NM_010176 | 14085 | Mm.3798 | NM_010176 | ENSMUSG00000030630 | **Fah** | - | fumarylacetoacetate hydrolase | protein-coding |
| chr7_84591406_84591675 | chr7 | 84591407 | 84591675 | + | 0.723529 | intron (NM_010176, intron 11 of 13) | L1ME3A\|LINE\|L1 | 14401 | NM_010176 | 14085 | Mm.3798 | NM_010176 | ENSMUSG00000030630 | **Fah** | - | fumarylacetoacetate hydrolase | protein-coding |
| chr7_84624503_84624812 | chr7 | 84624504 | 84624812 | + | 0.653264 | intron (NM_022985, intron 5 of 7) | intron (NM_022985, intron 5 of 7) | -18716 | NM_010176 | 14085 | Mm.3798 | NM_010176 | ENSMUSG00000030630 | **Fah** | - | fumarylacetoacetate hydrolase | protein-coding |
| chr5_124953003_124953419 | chr5 | 124953004 | 124953419 | + | 0.886283 | Intergenic | URR1B\|DNA\|hAT-Charlie | -50264 | NM_028443 | 73121 | Mm.296138 | NM_028443 | ENSMUSG00000037962 | **Fam101a** | 3110032G18Rik\|cfm\|cfm2 | family with sequence similarity 101, member A | protein-coding |
| chr3_109010884_109011199 | chr3 | 109010885 | 109011199 | + | 0.858301 | intron (NM_001163567, intron 1 of 10) | intron (NM_001163567, intron 1 of 10) | 16565 | NM_001163567 | 329739 | Mm.69013 | NM_001163567 | ENSMUSG00000040339 | **Fam102b** | 1600010D10Rik\|B430201A12Rik | family with sequence similarity 102, member B | protein-coding |
| chr3_109010174_109010522 | chr3 | 109010175 | 109010522 | + | 0.807388 | intron (NM_001163567, intron 1 of 10) | B1_Mur4\|SINE\|Alu | 17259 | NM_001163567 | 329739 | Mm.69013 | NM_001163567 | ENSMUSG00000040339 | **Fam102b** | 1600010D10Rik\|B430201A12Rik | family with sequence similarity 102, member B | protein-coding |
| chr15_27849694_27850496 | chr15 | 27849695 | 27850496 | + | 0.900541 | intron (NM_001081302, intron 15 of 56) | intron (NM_001081302, intron 15 of 56) | -168553 | NM_001242423 | 223433 | Mm.266485 | NM_198301 | ENSMUSG00000056069 | **Fam105a** | 9830126M18 | family with sequence similarity 105, member A | protein-coding |
| chr15_27703379_27703777 | chr15 | 27703380 | 27703777 | + | 0.801831 | Intergenic | Intergenic | -22036 | NM_001242423 | 223433 | Mm.266485 | NM_198301 | ENSMUSG00000056069 | **Fam105a** | 9830126M18 | family with sequence similarity 105, member A | protein-coding |
| chr2_3679802_3680488 | chr2 | 3679803 | 3680488 | + | 1.13042 | Intergenic | Intergenic | -33313 | NM_025626 | 66540 | Mm.277864 | NM_025626 | ENSMUSG00000026655 | **Fam107b** | 3110001A13Rik\|AA589595\|AA959793\|AI132312 | family with sequence similarity 107, member B | protein-coding |
| chr2_3659341_3659821 | chr2 | 3659342 | 3659821 | + | 1.01145 | Intergenic | Intergenic | -53877 | NM_025626 | 66540 | Mm.277864 | NM_025626 | ENSMUSG00000026655 | **Fam107b** | 3110001A13Rik\|AA589595\|AA959793\|AI132312 | family with sequence similarity 107, member B | protein-coding |
| chr2_3709394_3709623 | chr2 | 3709395 | 3709623 | + | 0.807879 | Intergenic | Intergenic | -3949 | NM_025626 | 66540 | Mm.277864 | NM_025626 | ENSMUSG00000026655 | **Fam107b** | 3110001A13Rik\|AA589595\|AA959793\|AI132312 | family with sequence similarity 107, member B | protein-coding |
| chr11_95339392_95339928 | chr11 | 95339393 | 95339928 | + | 0.749152 | intron (NM_172543, intron 1 of 7) | intron (NM_172543, intron 1 of 7) | 2642 | NM_172543 | 215512 | Mm.262113 | NM_172543 | ENSMUSG00000038893 | **Fam117a** | 5730593F17Rik\|AW548096 | family with sequence similarity 117, member A | protein-coding |
| chr11_95355105_95356036 | chr11 | 95355106 | 95356036 | + | 0.687019 | intron (NM_172543, intron 1 of 7) | intron (NM_172543, intron 1 of 7) | 18553 | NM_172543 | 215512 | Mm.262113 | NM_172543 | ENSMUSG00000038893 | **Fam117a** | 5730593F17Rik\|AW548096 | family with sequence similarity 117, member A | protein-coding |
| chr1_59886876_59887662 | chr1 | 59886877 | 59887662 | + | 1.75682 | Intergenic | Intergenic | -25737 | NM_001037725 | 72750 | Mm.258250 | NM_001037725 | ENSMUSG00000041040 | **Fam117b** | 2810425F24Rik\|6330416D14Rik\|Als2cr13 | family with sequence similarity 117, member B | protein-coding |
| chr13_48974709_48975068 | chr13 | 48974710 | 48975068 | + | 1.34607 | Intergenic | Intergenic | 6777 | NR_015601 | 68128 | Mm.426304 | NR_015601 | | **Fam120aos** | C030044B11Rik | family with sequence similarity 120A, opposite strand | ncRNA |
| chr13_48990202_48990771 | chr13 | 48990203 | 48990771 | + | 0.85183 | Intergenic | Intergenic | 22375 | NR_015601 | 68128 | Mm.426304 | NR_015601 | | **Fam120aos** | C030044B11Rik | family with sequence similarity 120A, opposite strand | ncRNA |
| chr17_15416438_15417502 | chr17 | 15416439 | 15417502 | + | 0.624678 | intron (NR_033586, intron 5 of 10) | MTEa\|LTR\|ERVL-MaLR | 20724 | NM_024203 | 67544 | Mm.35039 | NM_024203 | ENSMUSG00000014763 | **Fam120b** | 4932442K08Rik\|AW049283\|AW413338\|CCPG\|Pgcc1\|mKIAA1838 | family with sequence similarity 120, member B | protein-coding |
| chr1_80177540_80177975 | chr1 | 80177541 | 80177975 | + | 0.650609 | Intergenic | Intergenic | 36186 | NM_173425 | 241128 | Mm.145511 | NM_173425 | ENSMUSG00000043230 | **Fam124b** | A830043J08Rik | family with sequence similarity 124, member B | protein-coding |
| chr1_80177683_80177959 | chr1 | 80177684 | 80177959 | + | 0.636954 | Intergenic | Intergenic | 36123 | NM_173425 | 241128 | Mm.145511 | NM_173425 | ENSMUSG00000043230 | **Fam124b** | A830043J08Rik | family with sequence similarity 124, member B | protein-coding |
| chr1_151644035_151644657 | chr1 | 151644036 | 151644657 | + | 1.29366 | intron (NM_022018, intron 3 of 13) | G-rich\|Low_complexity\|Low_complexity | 72973 | NM_022018 | 63913 | Mm.157700 | NM_022018 | | **Fam129a** | AI256368\|AU019833\|Niban | family with sequence similarity 129, member A | protein-coding |
| chr1_151610554_151611037 | chr1 | 151610555 | 151611037 | + | 0.903616 | intron (NM_022018, intron 1 of 13) | MTC\|LTR\|ERVL-MaLR | 39423 | NM_022018 | 63913 | Mm.157700 | NM_022018 | | **Fam129a** | AI256368\|AU019833\|Niban | family with sequence similarity 129, member A | protein-coding |
| chr1_151572648_151572948 | chr1 | 151572649 | 151572948 | + | 0.903134 | intron (NM_022018, intron 1 of 13) | Lx4A\|LINE\|L1 | 1425 | NM_022018 | 63913 | Mm.157700 | NM_022018 | | **Fam129a** | AI256368\|AU019833\|Niban | family with sequence similarity 129, member A | protein-coding |
| chr1_151571111_151572495 | chr1 | 151571112 | 151572495 | + | 0.711585 | intron (NM_022018, intron 1 of 13) | intron (NM_022018, intron 1 of 13) | 430 | NM_022018 | 63913 | Mm.157700 | NM_022018 | | **Fam129a** | AI256368\|AU019833\|Niban | family with sequence similarity 129, member A | protein-coding |
| chr2_32867383_32867934 | chr2 | 32867384 | 32867934 | + | 0.609703 | Intergenic | Intergenic | -8475 | NM_146119 | 227737 | Mm.301646 | NM_146119 | ENSMUSG00000026796 | **Fam129b** | 9130404D14Rik | family with sequence similarity 129, member B | protein-coding |
| chr5_3477687_3478305 | chr5 | 3477688 | 3478305 | + | 1.74764 | intron (NM_009873, intron 4 of 6) | intron (NM_009873, intron 4 of 6) | -65837 | NM_001042501 | 68152 | Mm.240265 | NM_026583 | | **Fam133b** | 2900022K02Rik\|5830415L20Rik | family with sequence similarity 133, member B | protein-coding |
| chr5_3477606_3478593 | chr5 | 3477607 | 3478593 | + | 1.68231 | intron (NM_009873, intron 4 of 6) | intron (NM_009873, intron 4 of 6) | -65733 | NM_001042501 | 68152 | Mm.240265 | NM_026583 | | **Fam133b** | 2900022K02Rik\|5830415L20Rik | family with sequence similarity 133, member B | protein-coding |
| chr5_3494381_3495171 | chr5 | 3494382 | 3495171 | + | 1.58381 | intron (NM_009873, intron 4 of 6) | intron (NM_009873, intron 4 of 6) | -49057 | NM_001042501 | 68152 | Mm.240265 | NM_026583 | | **Fam133b** | 2900022K02Rik\|5830415L20Rik | family with sequence similarity 133, member B | protein-coding |
| chr5_3509472_3509872 | chr5 | 3509473 | 3509872 | + | 1.37419 | intron (NM_009873, intron 4 of 6) | MIRb\|SINE\|MIR | -34161 | NM_001042501 | 68152 | Mm.240265 | NM_026583 | | **Fam133b** | 2900022K02Rik\|5830415L20Rik | family with sequence similarity 133, member B | protein-coding |
| chr5_3463120_3463537 | chr5 | 3463121 | 3463537 | + | 1.22361 | intron (NM_009873, intron 3 of 6) | intron (NM_009873, intron 3 of 6) | -80504 | NM_001042501 | 68152 | Mm.240265 | NM_026583 | | **Fam133b** | 2900022K02Rik\|5830415L20Rik | family with sequence similarity 133, member B | protein-coding |
| chr5_3460214_3461370 | chr5 | 3460215 | 3461370 | + | 1.13489 | intron (NM_009873, intron 3 of 6) | intron (NM_009873, intron 3 of 6) | -83041 | NM_001042501 | 68152 | Mm.240265 | NM_026583 | | **Fam133b** | 2900022K02Rik\|5830415L20Rik | family with sequence similarity 133, member B | protein-coding |
| chr5_3501506_3502099 | chr5 | 3501507 | 3502099 | + | 1.12337 | intron (NM_009873, intron 4 of 6) | intron (NM_009873, intron 4 of 6) | -42030 | NM_001042501 | 68152 | Mm.240265 | NM_026583 | | **Fam133b** | 2900022K02Rik\|5830415L20Rik | family with sequence similarity 133, member B | protein-coding |
| chr5_3460776_3461381 | chr5 | 3460777 | 3461381 | + | 1.10391 | intron (NM_009873, intron 3 of 6) | intron (NM_009873, intron 3 of 6) | -82754 | NM_001042501 | 68152 | Mm.240265 | NM_026583 | | **Fam133b** | 2900022K02Rik\|5830415L20Rik | family with sequence similarity 133, member B | protein-coding |
| chr5_3501072_3502099 | chr5 | 3501073 | 3502099 | + | 1.0795 | intron (NM_009873, intron 4 of 6) | intron (NM_009873, intron 4 of 6) | -42247 | NM_001042501 | 68152 | Mm.240265 | NM_026583 | | **Fam133b** | 2900022K02Rik\|5830415L20Rik | family with sequence similarity 133, member B | protein-coding |
| chr5_3538855_3539125 | chr5 | 3538856 | 3539125 | + | 0.951251 | Intergenic | Intergenic | -4843 | NM_001042501 | 68152 | Mm.240265 | NM_026583 | | **Fam133b** | 2900022K02Rik\|5830415L20Rik | family with sequence similarity 133, member B | protein-coding |
| chr5_3502570_3503033 | chr5 | 3502571 | 3503033 | + | 0.939407 | intron (NM_009873, intron 4 of 6) | intron (NM_009873, intron 4 of 6) | -41031 | NM_001042501 | 68152 | Mm.240265 | NM_026583 | | **Fam133b** | 2900022K02Rik\|5830415L20Rik | family with sequence similarity 133, member B | protein-coding |
| chr5_3538494_3539131 | chr5 | 3538495 | 3539131 | + | 0.866757 | Intergenic | L1ME1\|LINE\|L1 | -5020 | NM_001042501 | 68152 | Mm.240265 | NM_026583 | | **Fam133b** | 2900022K02Rik\|5830415L20Rik | family with sequence similarity 133, member B | protein-coding |
| chr5_3466142_3466570 | chr5 | 3466143 | 3466570 | + | 0.807404 | intron (NM_009873, intron 3 of 6) | intron (NM_009873, intron 3 of 6) | -77477 | NM_001042501 | 68152 | Mm.240265 | NM_026583 | | **Fam133b** | 2900022K02Rik\|5830415L20Rik | family with sequence similarity 133, member B | protein-coding |

| | | | | | | | | | | | | | | | | | |
|---|---|---|---|---|---|---|---|---|---|---|---|---|---|---|---|---|---|
| chr5_3508464_3508779 | chr5 | 3508465 | 3508779 | + | 0.792143 | intron (NM_009873, intron 4 of 6) | intron (NM_009873, intron 4 of 6) | -35211 | NM_001042501 | 68152 | Mm.240265 | NM_026583 | | **Fam133b** | 2900022K02Rik\|5830415L20Rik | family with sequence similarity 133, member B | protein-coding |
| chr5_3497762_3498153 | chr5 | 3497763 | 3498153 | + | 0.766536 | intron (NM_009873, intron 4 of 6) | intron (NM_009873, intron 4 of 6) | -45875 | NM_001042501 | 68152 | Mm.240265 | NM_026583 | | **Fam133b** | 2900022K02Rik\|5830415L20Rik | family with sequence similarity 133, member B | protein-coding |
| chr5_3512590_3512945 | chr5 | 3512591 | 3512945 | + | 0.710764 | intron (NM_009873, intron 4 of 6) | intron (NM_009873, intron 4 of 6) | -31065 | NM_001042501 | 68152 | Mm.240265 | NM_026583 | | **Fam133b** | 2900022K02Rik\|5830415L20Rik | family with sequence similarity 133, member B | protein-coding |
| chr15_25884218_25884952 | chr15 | 25884219 | 25884952 | + | 1.44431 | intron (NM_001034851, intron 1 of 8) | intron (NM_001034851, intron 1 of 8) | 41287 | NM_001034851 | 66270 | Mm.25311 | NM_025459 | ENSMUSG00000022270 | **Fam134b** | 1810015C04Rik\|AU015349 | family with sequence similarity 134, member B | protein-coding |
| chr15_25884210_25885278 | chr15 | 25884211 | 25885278 | + | 1.2181 | intron (NM_001034851, intron 1 of 8) | RSINE1\|SINE\|B4 | 41446 | NM_001034851 | 66270 | Mm.25311 | NM_025459 | ENSMUSG00000022270 | **Fam134b** | 1810015C04Rik\|AU015349 | family with sequence similarity 134, member B | protein-coding |
| chr15_25916224_25916593 | chr15 | 25916225 | 25916593 | + | 0.955532 | intron (NM_001034851, intron 3 of 8) | intron (NM_001034851, intron 3 of 8) | -17143 | NM_001277318 | 66270 | Mm.25311 | NM_025459 | ENSMUSG00000022270 | **Fam134b** | 1810015C04Rik\|AU015349 | family with sequence similarity 134, member B | protein-coding |
| chr15_25915357_25916599 | chr15 | 25915358 | 25916599 | + | 0.884806 | intron (NM_001034851, intron 3 of 8) | intron (NM_001034851, intron 3 of 8) | -17574 | NM_001277318 | 66270 | Mm.25311 | NM_025459 | ENSMUSG00000022270 | **Fam134b** | 1810015C04Rik\|AU015349 | family with sequence similarity 134, member B | protein-coding |
| chr15_25879152_25879589 | chr15 | 25879153 | 25879589 | + | 0.71222 | intron (NM_001034851, intron 1 of 8) | intron (NM_001034851, intron 1 of 8) | 36073 | NM_001034851 | 66270 | Mm.25311 | NM_025459 | ENSMUSG00000022270 | **Fam134b** | 1810015C04Rik\|AU015349 | family with sequence similarity 134, member B | protein-coding |
| chr15_25915403_25915974 | chr15 | 25915404 | 25915974 | + | 0.695376 | intron (NM_001034851, intron 3 of 8) | intron (NM_001034851, intron 3 of 8) | -17863 | NM_001277318 | 66270 | Mm.25311 | NM_025459 | ENSMUSG00000022270 | **Fam134b** | 1810015C04Rik\|AU015349 | family with sequence similarity 134, member B | protein-coding |
| chr6_86318649_86319258 | chr6 | 86318650 | 86319258 | + | 0.941579 | Intergenic | Intergenic | -46729 | NM_025591 | 66488 | Mm.422972 | NM_025591 | ENSMUSG00000057497 | **Fam136a** | 2010309E21Rik\|AW124904\|Hyccin | family with sequence similarity 136, member A | protein-coding |
| chr6_86363896_86364154 | chr6 | 86363897 | 86364154 | + | 0.906096 | Intergenic | Intergenic | -1658 | NM_025591 | 66488 | Mm.422972 | NM_025591 | ENSMUSG00000057497 | **Fam136a** | 2010309E21Rik\|AW124904\|Hyccin | family with sequence similarity 136, member A | protein-coding |
| chr6_59020393_59020666 | chr6 | 59020394 | 59020666 | + | 0.841326 | intron (NM_153574, intron 1 of 17) | AmnSINE1\|SINE\|Deu | 3972 | NM_153574 | 58909 | Mm.28813 | NM_153574 | | **Fam13a** | AI462012\|D430015B01Rik\|FAM13A1\|Precm1 | family with sequence similarity 13, member A | protein-coding |
| chr4_108371319_108371791 | chr4 | 108371320 | 108371791 | + | 1.18013 | intron (NM_001099303, intron 2 of 2) | intron (NM_001099303, intron 2 of 2) | 12100 | NM_001099303 | 545667 | Mm.380025 | NM_001099303 | | **Fam159a** | OTTMUSG00000008243 | family with sequence similarity 159, member A | protein-coding |
| chr4_108370425_108370756 | chr4 | 108370426 | 108370756 | + | 0.897897 | intron (NM_001099303, intron 2 of 2) | intron (NM_001099303, intron 2 of 2) | 13064 | NM_001099303 | 545667 | Mm.380025 | NM_001099303 | | **Fam159a** | OTTMUSG00000008243 | family with sequence similarity 159, member A | protein-coding |
| chr13_104896247_104896627 | chr13 | 104896248 | 104896627 | + | 0.790795 | Intergenic | Lx9\|LINE\|L1 | -32544 | NM_029984 | 77803 | Mm.159995 | NM_029984 | ENSMUSG00000042655 | **Fam159b** | A930021C24Rik | family with sequence similarity 159, member B | protein-coding |
| chr19_57398671_57399292 | chr19 | 57398672 | 57399292 | + | 0.906608 | Intergenic | RLTR19\|LTR\|ERVK | 37973 | NM_145505 | 226252 | Mm.358870 | NM_145505 | ENSMUSG00000033478 | **Fam160b1** | AI450540\|mKIAA1600 | family with sequence similarity 160, member B1 | protein-coding |
| chr16_36057449_36057892 | chr16 | 36057450 | 36057892 | + | 1.1466 | intron (NM_027342, intron 1 of 4) | intron (NM_027342, intron 1 of 4) | 13844 | NM_027342 | 70186 | Mm.212991 | NM_027342 | ENSMUSG00000003955 | **Fam162a** | 2310056P07Rik\|HGTD-P | family with sequence similarity 162, member A | protein-coding |
| chr16_36058376_36058908 | chr16 | 36058377 | 36058908 | + | 0.902863 | intron (NM_027342, intron 1 of 4) | intron (NM_027342, intron 1 of 4) | 12873 | NM_027342 | 70186 | Mm.212991 | NM_027342 | ENSMUSG00000003955 | **Fam162a** | 2310056P07Rik\|HGTD-P | family with sequence similarity 162, member A | protein-coding |
| chr16_36058180_36058918 | chr16 | 36058181 | 36058918 | + | 0.66817 | intron (NM_027342, intron 1 of 4) | intron (NM_027342, intron 1 of 4) | 12966 | NM_027342 | 70186 | Mm.212991 | NM_027342 | ENSMUSG00000003955 | **Fam162a** | 2310056P07Rik\|HGTD-P | family with sequence similarity 162, member A | protein-coding |
| chr1_156206046_156206434 | chr1 | 156206047 | 156206434 | + | 0.896062 | Intergenic | Intergenic | -1214 | NM_177838 | 329274 | Mm.150857 | NM_177838 | ENSMUSG00000015484 | **Fam163a** | A230106N23Rik | family with sequence similarity 163, member A | protein-coding |
| chr1_156205907_156206656 | chr1 | 156205908 | 156206656 | + | 0.700513 | Intergenic | Intergenic | -1256 | NM_177838 | 329274 | Mm.150857 | NM_177838 | ENSMUSG00000015484 | **Fam163a** | A230106N23Rik | family with sequence similarity 163, member A | protein-coding |
| chr1_156205892_156206432 | chr1 | 156205893 | 156206432 | + | 0.690071 | Intergenic | Intergenic | -1136 | NM_177838 | 329274 | Mm.150857 | NM_177838 | ENSMUSG00000015484 | **Fam163a** | A230106N23Rik | family with sequence similarity 163, member A | protein-coding |
| chr4_129580304_129580832 | chr4 | 129580305 | 129580832 | + | 0.732294 | Intergenic | Intergenic | -1988 | NM_182783 | 230766 | Mm.20377 | NM_182783 | ENSMUSG00000050493 | **Fam167b** | SEC | family with sequence similarity 167, member B | protein-coding |
| chr7_100693033_100693673 | chr7 | 100693034 | 100693673 | + | 1.20692 | Intergenic | Intergenic | -13349 | NM_178764 | 319604 | Mm.260362 | NM_178764 | ENSMUSG00000029461 | **Fam168a** | 2610030B18Rik\|B930006L02Rik\|mKIAA0280 | family with sequence similarity 168, member A | protein-coding |
| chr2_83813589_83813885 | chr2 | 83813590 | 83813885 | + | 1.01993 | intron (NM_175514, intron 1 of 7) | intron (NM_175514, intron 1 of 7) | 1009 | NM_175514 | 241520 | Mm.330351 | NM_175514 | ENSMUSG00000048388 | **Fam171b** | D430039N05Rik | family with sequence similarity 171, member B | protein-coding |
| chr1_95267870_95268914 | chr1 | 95267871 | 95268914 | + | 0.805172 | Intergenic | Intergenic | -45236 | NM_026321 | 67698 | Mm.272705 | NM_026321 | ENSMUSG00000051185 | **Fam174a** | 2310044D20Rik\|Fam174\|Naa6\|Tmem157 | family with sequence similarity 174, member A | protein-coding |
| chr1_95367668_95368104 | chr1 | 95367669 | 95368104 | + | 0.7711 | Intergenic | Intergenic | 54258 | NM_026321 | 67698 | Mm.272705 | NM_026321 | ENSMUSG00000051185 | **Fam174a** | 2310044D20Rik\|Fam174\|Naa6\|Tmem157 | family with sequence similarity 174, member A | protein-coding |
| chr7_73771356_73771971 | chr7 | 73771357 | 73771971 | + | 0.914627 | intron (NM_001162532, intron 2 of 2) | intron (NM_001162532, intron 2 of 2) | 31357 | NM_001162532 | 100038347 | Mm.425464 | NM_001162532 | ENSMUSG00000078670 | **Fam174b** | - | family with sequence similarity 174, member B | protein-coding |
| chr1_36646165_36646916 | chr1 | 36646166 | 36646916 | + | 1.25502 | intron (NM_001126046, intron 3 of 14) | intron (NM_001126046, intron 3 of 14) | 36642 | NM_001126046 | 381337 | Mm.17413 | NM_027957 | ENSMUSG00000046337 | **Fam178b** | 1700024G10Rik\|Gm990 | family with sequence similarity 178, member B | protein-coding |
| chr12_64970632_64970901 | chr12 | 64970633 | 64970901 | + | 1.21378 | intron (NM_177805, intron 1 of 19) | B1_Mur2\|SINE\|Alu | 5025 | NM_177805 | 328108 | Mm.138465 | NM_177805 | ENSMUSG00000035614 | **Fam179b** | A430041B07Rik\|BC062107\|C130034K06\|mKIAA0423 | family with sequence similarity 179, member B | protein-coding |
| chr12_64976194_64976565 | chr12 | 64976195 | 64976565 | + | 0.953429 | intron (NM_177805, intron 1 of 19) | intron (NM_177805, intron 1 of 19) | 10638 | NM_177805 | 328108 | Mm.138465 | NM_177805 | ENSMUSG00000035614 | **Fam179b** | A430041B07Rik\|BC062107\|C130034K06\|mKIAA0423 | family with sequence similarity 179, member B | protein-coding |
| chr10_53848267_53849071 | chr10 | 53848268 | 53849071 | + | 1.31073 | Intergenic | L2b\|LINE\|L2 | -97760 | NM_001081428 | 75906 | Mm.436785 | NM_001081428 | ENSMUSG00000019856 | **Fam184a** | 3110012E06Rik\|4930438C08Rik\|4930589M24Rik | family with sequence similarity 184, member A | protein-coding |
| chr10_53812900_53813226 | chr10 | 53812901 | 53813226 | + | 0.743032 | Intergenic | Intergenic | -62154 | NM_001081428 | 75906 | Mm.436785 | NM_001081428 | ENSMUSG00000019856 | **Fam184a** | 3110012E06Rik\|4930438C08Rik\|4930589M24Rik | family with sequence similarity 184, member A | protein-coding |
| chr2_12424293_12424824 | chr2 | 12424294 | 12424824 | + | 2.12788 | Intergenic | Lx9\|LINE\|L1 | -5099 | NM_024185 | 66960 | Mm.34059 | NM_024185 | ENSMUSG00000026767 | **Fam188a** | 1810041E18Rik\|2310047O13Rik\|5830410F13Rik\|AI447827\|AW111958 | family with sequence similarity 188, member A | protein-coding |
| chr2_12406717_12407184 | chr2 | 12406718 | 12407184 | + | 0.999889 | intron (NM_024185, intron 2 of 14) | intron (NM_024185, intron 2 of 14) | 12509 | NM_024185 | 66960 | Mm.34059 | NM_024185 | ENSMUSG00000026767 | **Fam188a** | 1810041E18Rik\|2310047O13Rik\|5830410F13Rik\|AI447827\|AW111958 | family with sequence similarity 188, member A | protein-coding |
| chr5_34430831_34431097 | chr5 | 34430832 | 34431097 | + | 1.28335 | intron (NM_001243123, intron 5 of 20) | intron (NM_001243123, intron 5 of 20) | 61031 | NM_001243123 | 231128 | Mm.286885 | NM_001030306 | ENSMUSG00000037210 | **Fam193a** | - | family with sequence similarity 193, member A | protein-coding |
| chr5_34430334_34431166 | chr5 | 34430335 | 34431166 | + | 1.162 | intron (NM_001243123, intron 5 of 20) | RSINE1\|SINE\|B4 | 60817 | NM_001243123 | 231128 | Mm.286885 | NM_001030306 | ENSMUSG00000037210 | **Fam193a** | - | family with sequence similarity 193, member A | protein-coding |
| chr5_34407366_34407717 | chr5 | 34407367 | 34407717 | + | 0.970958 | intron (NM_001243123, intron 1 of 20) | intron (NM_001243123, intron 1 of 20) | 37609 | NM_001243123 | 231128 | Mm.286885 | NM_001030306 | ENSMUSG00000037210 | **Fam193a** | - | family with sequence similarity 193, member A | protein-coding |
| chr5_34407242_34407923 | chr5 | 34407243 | 34407923 | + | 0.910387 | intron (NM_001243123, intron 1 of 20) | intron (NM_001243123, intron 1 of 20) | 37650 | NM_001243123 | 231128 | Mm.286885 | NM_001030306 | ENSMUSG00000037210 | **Fam193a** | - | family with sequence similarity 193, member A | protein-coding |
| chr5_34407210_34408211 | chr5 | 34407211 | 34408211 | + | 0.854845 | intron (NM_001243123, intron 1 of 20) | PB1\|SINE\|Alu | 37778 | NM_001243123 | 231128 | Mm.286885 | NM_001030306 | ENSMUSG00000037210 | **Fam193a** | - | family with sequence similarity 193, member A | protein-coding |
| chr5_34378864_34379591 | chr5 | 34378865 | 34379591 | + | 0.730261 | intron (NM_001243123, intron 1 of 20) | RSINE1\|SINE\|B4 | 9295 | NM_001243123 | 231128 | Mm.286885 | NM_001030306 | ENSMUSG00000037210 | **Fam193a** | - | family with sequence similarity 193, member A | protein-coding |
| chr11_34429458_34430044 | chr11 | 34429459 | 34430044 | + | 1.13732 | intron (NM_033374, intron 27 of 51) | intron (NM_033374, intron 27 of 51) | 114929 | NM_001025382 | 574403 | Mm.444796 | NM_001025382 | ENSMUSG00000069911 | **Fam196b** | EG574403\|Gm6041 | family with sequence similarity 196, member B | protein-coding |
| chr11_34423057_34423646 | chr11 | 34423058 | 34423646 | + | 0.730912 | TTS (NM_001025382) | TTS (NM_001025382) | 108530 | NM_001025382 | 574403 | Mm.444796 | NM_001025382 | ENSMUSG00000069911 | **Fam196b** | EG574403\|Gm6041 | family with sequence similarity 196, member B | protein-coding |
| chr11_34423058_34423355 | chr11 | 34423059 | 34423355 | + | 0.699733 | TTS (NM_001025382) | TTS (NM_001025382) | 108385 | NM_001025382 | 574403 | Mm.444796 | NM_001025382 | ENSMUSG00000069911 | **Fam196b** | EG574403\|Gm6041 | family with sequence similarity 196, member B | protein-coding |
| chr11_34372204_34372679 | chr11 | 34372205 | 34372679 | + | 0.693067 | intron (NM_001025382, intron 1 of 3) | MIR\|SINE\|MIR | 57620 | NM_001025382 | 574403 | Mm.444796 | NM_001025382 | ENSMUSG00000069911 | **Fam196b** | EG574403\|Gm6041 | family with sequence similarity 196, member B | protein-coding |

| | | | | | | | | | | | | | | | | | |
|---|---|---|---|---|---|---|---|---|---|---|---|---|---|---|---|---|---|
| chr9_121961796_121962261 | chr9 | 121961797 | 121962261 | + | 0.919155 | intron (NM_001199927, intron 1 of 4) | intron (NM_001199927, intron 1 of 4) | 11009 | NM_001199927 | 245050 | Mm.71062 | NM_177743 | ENSMUSG00000038233 | **Fam198a** | C730027P07Rik | family with sequence similarity 198, member A | protein-coding |
| chr9_121961551_121962389 | chr9 | 121961552 | 121962389 | + | 0.768628 | intron (NM_001199927, intron 1 of 4) | intron (NM_001199927, intron 1 of 4) | 10950 | NM_001199927 | 245050 | Mm.71062 | NM_177743 | ENSMUSG00000038233 | **Fam198a** | C730027P07Rik | family with sequence similarity 198, member A | protein-coding |
| chr3_80083179_80083525 | chr3 | 80083180 | 80083525 | + | 0.854616 | Intergenic | Intergenic | 197422 | NM_133187 | 68659 | Mm.427274 | NM_133187 | ENSMUSG00000027955 | **Fam198b** | 1110032E23Rik\|2210419I08Rik\|AV011458\|Ened | family with sequence similarity 198, member B | protein-coding |
| chr6_97012432_97012968 | chr6 | 97012433 | 97012968 | + | 0.668346 | intron (NM_177233, intron 3 of 5) | intron (NM_177233, intron 3 of 5) | 47711 | NM_177233 | 320701 | Mm.131700 | NM_177233 | ENSMUSG00000046500 | **Fam19a4** | C130034I18Rik\|Fam19a4 Tafa-4\|Tafa-4 | family with sequence similarity 19, member A4 | protein-coding |
| chr6_97012634_97012929 | chr6 | 97012635 | 97012929 | + | 0.642637 | intron (NM_177233, intron 3 of 5) | intron (NM_177233, intron 3 of 5) | 47629 | NM_177233 | 320701 | Mm.131700 | NM_177233 | ENSMUSG00000046500 | **Fam19a4** | C130034I18Rik\|Fam19a4 Tafa-4\|Tafa-4 | family with sequence similarity 19, member A4 | protein-coding |
| chr11_109743897_109744341 | chr11 | 109743898 | 109744341 | + | 1.2413 | Intergenic | Intergenic | -21863 | NM_153782 | 208659 | Mm.208662 | NM_153782 | ENSMUSG00000020614 | **Fam20a** | AI606893 | family with sequence similarity 20, member A | protein-coding |
| chr11_109766752_109767540 | chr11 | 109766753 | 109767540 | + | 1.02432 | Intergenic | Intergenic | -44890 | NM_153782 | 208659 | Mm.208662 | NM_153782 | ENSMUSG00000020614 | **Fam20a** | AI606893 | family with sequence similarity 20, member A | protein-coding |
| chr11_109758486_109759304 | chr11 | 109758487 | 109759304 | + | 0.688818 | Intergenic | Lx9\|LINE\|L1 | -36639 | NM_153782 | 208659 | Mm.208662 | NM_153782 | ENSMUSG00000020614 | **Fam20a** | AI606893 | family with sequence similarity 20, member A | protein-coding |
| chr11_109757873_109758221 | chr11 | 109757874 | 109758221 | + | 0.64291 | Intergenic | Lx8\|LINE\|L1 | -35791 | NM_153782 | 208659 | Mm.208662 | NM_153782 | ENSMUSG00000020614 | **Fam20a** | AI606893 | family with sequence similarity 20, member A | protein-coding |
| chr3_105721844_105722294 | chr3 | 105721845 | 105722294 | + | 0.64522 | Intergenic | Intergenic | 16611 | NM_001163356 | 109050 | Mm.212591 | NM_175398 | ENSMUSG00000048458 | **Fam212b** | 6530418L21Rik | family with sequence similarity 212, member B | protein-coding |
| chr14_40999870_41000579 | chr14 | 40999871 | 41000579 | + | 1.43988 | intron (NM_027464, intron 3 of 5) | intron (NM_027464, intron 3 of 5) | 13550 | NM_027464 | 70564 | Mm.27227 | NM_027464 | ENSMUSG00000021792 | **Fam213a** | 5730469M10Rik\|AU040822\|Pamm | family with sequence similarity 213, member A | protein-coding |
| chr14_41005542_41005901 | chr14 | 41005543 | 41005901 | + | 0.757733 | intron (NM_027464, intron 1 of 5) | intron (NM_027464, intron 1 of 5) | 8053 | NM_027464 | 70564 | Mm.27227 | NM_027464 | ENSMUSG00000021792 | **Fam213a** | 5730469M10Rik\|AU040822\|Pamm | family with sequence similarity 213, member A | protein-coding |
| chr4_43040967_43041165 | chr4 | 43040968 | 43041165 | + | 0.841499 | promoter-TSS (NM_001253354) | promoter-TSS (NM_001253354) | -765 | NM_001253354 | 230088 | Mm.260162 | NM_172691 | ENSMUSG00000036002 | **Fam214b** | B230312A22Rik | family with sequence similarity 214, member B | protein-coding |
| chr14_78040678_78041120 | chr14 | 78040679 | 78041120 | + | 0.982866 | Intergenic | Intergenic | 48108 | NM_177629 | 219170 | Mm.320428 | NM_177629 | ENSMUSG00000045655 | **Fam216b** | 9330158N17\|AU021034 | family with sequence similarity 216, member B | protein-coding |
| chr5_143561531_143562009 | chr5 | 143561532 | 143562009 | + | 1.06523 | intron (NM_026050, intron 2 of 2) | intron (NM_026050, intron 2 of 2) | 13064 | NM_026050 | 67238 | Mm.383219 | NM_026050 | ENSMUSG00000048910 | **Fam220a** | 1500026F15Rik\|2810453I06Rik\|AW047776\|Acpin1\|Sipar | family with sequence similarity 220, member A | protein-coding |
| chr5_143579826_143580150 | chr5 | 143579827 | 143580150 | + | 0.683703 | Intergenic | Intergenic | 31282 | NM_026050 | 67238 | Mm.383219 | NM_026050 | ENSMUSG00000048910 | **Fam220a** | 1500026F15Rik\|2810453I06Rik\|AW047776\|Acpin1\|Sipar | family with sequence similarity 220, member A | protein-coding |
| chr14_34363444_34364087 | chr14 | 34363445 | 34364087 | + | 0.679705 | Intergenic | Intergenic | -8373 | NM_183278 | 69134 | Mm.27156 | NM_183278 | ENSMUSG00000043681 | **Fam25c** | 2200001I15Rik\|Fam25 | family with sequence similarity 25, member C | protein-coding |
| chr6_22378419_22379107 | chr6 | 22378420 | 22379107 | + | 2.02444 | Intergenic | Intergenic | -22682 | NM_138587 | 27999 | Mm.490307 | NM_138587 | ENSMUSG00000029672 | **Fam3c** | D6Wsu176e\|Ilei | family with sequence similarity 3, member C | protein-coding |
| chr6_22385004_22385815 | chr6 | 22385005 | 22385815 | + | 1.50317 | Intergenic | RLTR23\|LTR\|ERV1 | -29329 | NM_138587 | 27999 | Mm.490307 | NM_138587 | ENSMUSG00000029672 | **Fam3c** | D6Wsu176e\|Ilei | family with sequence similarity 3, member C | protein-coding |
| chr6_22373451_22373836 | chr6 | 22373452 | 22373836 | + | 1.34776 | Intergenic | MLT1A\|LTR\|ERVL-MaLR | -17563 | NM_138587 | 27999 | Mm.490307 | NM_138587 | ENSMUSG00000029672 | **Fam3c** | D6Wsu176e\|Ilei | family with sequence similarity 3, member C | protein-coding |
| chr6_22409993_22410272 | chr6 | 22409994 | 22410272 | + | 1.14636 | Intergenic | PB1D9\|SINE\|Alu | -54052 | NM_138587 | 27999 | Mm.490307 | NM_138587 | ENSMUSG00000029672 | **Fam3c** | D6Wsu176e\|Ilei | family with sequence similarity 3, member C | protein-coding |
| chr6_22339747_22340503 | chr6 | 22339748 | 22340503 | + | 1.04995 | intron (NM_138587, intron 2 of 9) | intron (NM_138587, intron 2 of 9) | 15956 | NM_138587 | 27999 | Mm.490307 | NM_138587 | ENSMUSG00000029672 | **Fam3c** | D6Wsu176e\|Ilei | family with sequence similarity 3, member C | protein-coding |
| chr6_22379327_22379629 | chr6 | 22379328 | 22379629 | + | 0.81936 | Intergenic | MTD\|LTR\|ERVL-MaLR | -23397 | NM_138587 | 27999 | Mm.490307 | NM_138587 | ENSMUSG00000029672 | **Fam3c** | D6Wsu176e\|Ilei | family with sequence similarity 3, member C | protein-coding |
| chr6_22331754_22332620 | chr6 | 22331755 | 22332620 | + | 0.730435 | intron (NM_138587, intron 3 of 9) | B3\|SINE\|B2 | 23894 | NM_138587 | 27999 | Mm.490307 | NM_138587 | ENSMUSG00000029672 | **Fam3c** | D6Wsu176e\|Ilei | family with sequence similarity 3, member C | protein-coding |
| chr9_85107251_85107927 | chr9 | 85107252 | 85107927 | + | 0.982071 | Intergenic | Intergenic | 219535 | NM_001160378 | 212943 | Mm.74610 | NM_001160378 | ENSMUSG00000032265 | **Fam46a** | D930050G01Rik | family with sequence similarity 46, member A | protein-coding |
| chr9_85107295_85107691 | chr9 | 85107296 | 85107691 | + | 0.904623 | Intergenic | Intergenic | 219631 | NM_001160378 | 212943 | Mm.74610 | NM_001160378 | ENSMUSG00000032265 | **Fam46a** | D930050G01Rik | family with sequence similarity 46, member A | protein-coding |
| chr12_12172066_12172853 | chr12 | 12172067 | 12172853 | + | 1.45518 | Intergenic | Intergenic | -89679 | NM_029758 | 76820 | Mm.440802 | NM_029758 | ENSMUSG00000020589 | **Fam49a** | 2410157M17Rik\|9630047E15\|D12Ertd553e | family with sequence similarity 49, member A | protein-coding |
| chr12_12264967_12265460 | chr12 | 12264968 | 12265460 | + | 1.21425 | intron (NM_001146119, intron 1 of 11) | intron (NM_001146119, intron 1 of 11) | 3075 | NM_001146119 | 76820 | Mm.440802 | NM_029758 | ENSMUSG00000020589 | **Fam49a** | 2410157M17Rik\|9630047E15\|D12Ertd553e | family with sequence similarity 49, member A | protein-coding |
| chr12_12172938_12173447 | chr12 | 12172939 | 12173447 | + | 1.0894 | Intergenic | Intergenic | -88946 | NM_029758 | 76820 | Mm.440802 | NM_029758 | ENSMUSG00000020589 | **Fam49a** | 2410157M17Rik\|9630047E15\|D12Ertd553e | family with sequence similarity 49, member A | protein-coding |
| chr12_12274304_12274662 | chr12 | 12274305 | 12274662 | + | 0.736773 | intron (NM_001146119, intron 1 of 11) | intron (NM_001146119, intron 1 of 11) | 12344 | NM_001146119 | 76820 | Mm.440802 | NM_029758 | ENSMUSG00000020589 | **Fam49a** | 2410157M17Rik\|9630047E15\|D12Ertd553e | family with sequence similarity 49, member A | protein-coding |
| chr12_12274416_12274682 | chr12 | 12274417 | 12274682 | + | 0.692692 | intron (NM_001146119, intron 1 of 11) | intron (NM_001146119, intron 1 of 11) | 12410 | NM_001146119 | 76820 | Mm.440802 | NM_029758 | ENSMUSG00000020589 | **Fam49a** | 2410157M17Rik\|9630047E15\|D12Ertd553e | family with sequence similarity 49, member A | protein-coding |
| chr15_64018443_64018841 | chr15 | 64018444 | 64018841 | + | 0.99814 | intron (NM_144846, intron 1 of 11) | intron (NM_144846, intron 1 of 11) | 41806 | NM_144846 | 223601 | Mm.246108 | NM_144846 | ENSMUSG00000022378 | **Fam49b** | 0910001A06Rik\|AW122079 | family with sequence similarity 49, member B | protein-coding |
| chr15_64184915_64185549 | chr15 | 64184916 | 64185549 | + | 0.963635 | intron (NM_001276462, intron 3 of 27) | Lx9\|LINE\|L1 | -124784 | NM_144846 | 223601 | Mm.246108 | NM_144846 | ENSMUSG00000022378 | **Fam49b** | 0910001A06Rik\|AW122079 | family with sequence similarity 49, member B | protein-coding |
| chr15_64018414_64018973 | chr15 | 64018415 | 64018973 | + | 0.929915 | intron (NM_144846, intron 1 of 11) | intron (NM_144846, intron 1 of 11) | 41754 | NM_144846 | 223601 | Mm.246108 | NM_144846 | ENSMUSG00000022378 | **Fam49b** | 0910001A06Rik\|AW122079 | family with sequence similarity 49, member B | protein-coding |
| chr15_64155726_64156202 | chr15 | 64155727 | 64156202 | + | 0.879767 | intron (NM_010026, intron 11 of 30) | intron (NM_010026, intron 11 of 30) | -95516 | NM_144846 | 223601 | Mm.246108 | NM_144846 | ENSMUSG00000022378 | **Fam49b** | 0910001A06Rik\|AW122079 | family with sequence similarity 49, member B | protein-coding |
| chr15_64173378_64173804 | chr15 | 64173379 | 64173804 | + | 0.72209 | intron (NM_001276462, intron 3 of 27) | intron (NM_001276462, intron 3 of 27) | -113143 | NM_144846 | 223601 | Mm.246108 | NM_144846 | ENSMUSG00000022378 | **Fam49b** | 0910001A06Rik\|AW122079 | family with sequence similarity 49, member B | protein-coding |
| chr15_64075082_64076169 | chr15 | 64075083 | 64076169 | + | 0.721288 | Intergenic | Intergenic | -15178 | NM_144846 | 223601 | Mm.246108 | NM_144846 | ENSMUSG00000022378 | **Fam49b** | 0910001A06Rik\|AW122079 | family with sequence similarity 49, member B | protein-coding |
| chr15_64176396_64176810 | chr15 | 64176397 | 64176810 | + | 0.70857 | intron (NM_001276462, intron 3 of 27) | intron (NM_001276462, intron 3 of 27) | -116155 | NM_144846 | 223601 | Mm.246108 | NM_144846 | ENSMUSG00000022378 | **Fam49b** | 0910001A06Rik\|AW122079 | family with sequence similarity 49, member B | protein-coding |
| chr7_132756825_132757412 | chr7 | 132756826 | 132757412 | + | 1.13208 | intron (NM_175268, intron 4 of 4) | B3\|SINE\|B2 | 29816 | NM_175268 | 77938 | Mm.225649 | NM_175268 | ENSMUSG00000030956 | **Fam53b** | A930008G19Rik\|mKIAA0140 | family with sequence similarity 53, member B | protein-coding |
| chr7_132806863_132807185 | chr7 | 132806864 | 132807185 | + | 1.10958 | intron (NM_212473, intron 1 of 4) | intron (NM_212473, intron 1 of 4) | 6172 | NM_212473 | 77938 | Mm.225649 | NM_175268 | ENSMUSG00000030956 | **Fam53b** | A930008G19Rik\|mKIAA0140 | family with sequence similarity 53, member B | protein-coding |
| chr7_132719239_132720054 | chr7 | 132719240 | 132720054 | + | 1.09136 | intron (NM_175268, intron 4 of 4) | intron (NM_175268, intron 4 of 4) | 67288 | NM_175268 | 77938 | Mm.225649 | NM_175268 | ENSMUSG00000030956 | **Fam53b** | A930008G19Rik\|mKIAA0140 | family with sequence similarity 53, member B | protein-coding |
| chr7_132764863_132765553 | chr7 | 132764864 | 132765553 | + | 0.994833 | intron (NM_175268, intron 3 of 4) | intron (NM_175268, intron 3 of 4) | 21727 | NM_175268 | 77938 | Mm.225649 | NM_175268 | ENSMUSG00000030956 | **Fam53b** | A930008G19Rik\|mKIAA0140 | family with sequence similarity 53, member B | protein-coding |
| chr7_132765126_132765522 | chr7 | 132765127 | 132765522 | + | 0.966485 | intron (NM_175268, intron 3 of 4) | intron (NM_175268, intron 3 of 4) | 21611 | NM_175268 | 77938 | Mm.225649 | NM_175268 | ENSMUSG00000030956 | **Fam53b** | A930008G19Rik\|mKIAA0140 | family with sequence similarity 53, member B | protein-coding |
| chr7_132770271_132770446 | chr7 | 132770272 | 132770446 | + | 0.938048 | intron (NM_175268, intron 3 of 4) | intron (NM_175268, intron 3 of 4) | 16576 | NM_175268 | 77938 | Mm.225649 | NM_175268 | ENSMUSG00000030956 | **Fam53b** | A930008G19Rik\|mKIAA0140 | family with sequence similarity 53, member B | protein-coding |
| chr7_132768043_132768380 | chr7 | 132768044 | 132768380 | + | 0.792385 | intron (NM_175268, intron 3 of 4) | intron (NM_175268, intron 3 of 4) | 18723 | NM_175268 | 77938 | Mm.225649 | NM_175268 | ENSMUSG00000030956 | **Fam53b** | A930008G19Rik\|mKIAA0140 | family with sequence similarity 53, member B | protein-coding |
| chr7_132756399_132757419 | chr7 | 132756400 | 132757419 | + | 0.778251 | intron (NM_175268, intron 4 of 4) | intron (NM_175268, intron 4 of 4) | 30026 | NM_175268 | 77938 | Mm.225649 | NM_175268 | ENSMUSG00000030956 | **Fam53b** | A930008G19Rik\|mKIAA0140 | family with sequence similarity 53, member B | protein-coding |
| chr7_132800657_132801273 | chr7 | 132800658 | 132801273 | + | 0.774359 | intron (NM_212473, intron 1 of 4) | intron (NM_212473, intron 1 of 4) | 12231 | NM_212473 | 77938 | Mm.225649 | NM_175268 | ENSMUSG00000030956 | **Fam53b** | A930008G19Rik\|mKIAA0140 | family with sequence similarity 53, member B | protein-coding |
| chr7_132762993_132763303 | chr7 | 132762994 | 132763303 | + | 0.763361 | intron (NM_175268, intron 3 of 4) | intron (NM_175268, intron 3 of 4) | 23787 | NM_175268 | 77938 | Mm.225649 | NM_175268 | ENSMUSG00000030956 | **Fam53b** | A930008G19Rik\|mKIAA0140 | family with sequence similarity 53, member B | protein-coding |
| chr6_148962545_148962853 | chr6 | 148962546 | 148962853 | + | 0.987368 | Intergenic | Intergenic | -16267 | NM_019643 | 56306 | Mm.18637 | NM_019643 | ENSMUSG00000039985 | **Fam60a** | Ppcs1\|Pptcs1\|Tera | family with sequence similarity 60, member A | protein-coding |
| chr9_70636789_70637084 | chr9 | 70636790 | 70637084 | + | 0.867056 | intron (NM_172772, intron 2 of 8) | URR1A\|DNA\|hAT-Charlie | 20237 | NM_172772 | 235461 | Mm.170855 | NM_172772 | ENSMUSG00000042444 | **Fam63b** | AI465301\|B230380D07Rik\|mKIAA1164 | family with sequence similarity 63, member B | protein-coding |
| chr13_24536641_24537145 | chr13 | 24536642 | 24537145 | + | 1.2418 | Intergenic | MTEa\|LTR\|ERVL-MaLR | -45296 | NM_001286100 | 193385 | Mm.217319 | NM_029679 | ENSMUSG00000036006 | **Fam65b** | - | family with sequence similarity 65, member B | protein-coding |
| chr13_24524542_24525223 | chr13 | 24524543 | 24525223 | + | 0.996882 | Intergenic | L1MB7\|LINE\|L1 | -57306 | NM_001286100 | 193385 | Mm.217319 | NM_029679 | ENSMUSG00000036006 | **Fam65b** | - | family with sequence similarity 65, member B | protein-coding |
| chr13_24524297_24525231 | chr13 | 24524298 | 24525231 | + | 0.823501 | Intergenic | L1MB7\|LINE\|L1 | -57425 | NM_001286100 | 193385 | Mm.217319 | NM_029679 | ENSMUSG00000036006 | **Fam65b** | - | family with sequence similarity 65, member B | protein-coding |
| chr13_24646335_24646777 | chr13 | 24646336 | 24646777 | + | 0.735486 | intron (NM_029679, intron 1 of 22) | intron (NM_029679, intron 1 of 22) | 7908 | NM_029679 | 193385 | Mm.217319 | NM_029679 | ENSMUSG00000036006 | **Fam65b** | - | family with sequence similarity 65, member B | protein-coding |
| chr13_24509011_24509750 | chr13 | 24509012 | 24509750 | + | 0.701238 | Intergenic | MTEb\|LTR\|ERVL-MaLR | -72808 | NM_001286100 | 193385 | Mm.217319 | NM_029679 | ENSMUSG00000036006 | **Fam65b** | - | family with sequence similarity 65, member B | protein-coding |
| chr13_24509964_24510388 | chr13 | 24509965 | 24510388 | + | 0.654328 | Intergenic | Intergenic | -72013 | NM_001286100 | 193385 | Mm.217319 | NM_029679 | ENSMUSG00000036006 | **Fam65b** | - | family with sequence similarity 65, member B | protein-coding |

| | | | | | | | | | | | | | | | | | |
|---|---|---|---|---|---|---|---|---|---|---|---|---|---|---|---|---|---|
| chr5_107952932_107953345 | chr5 | 107952933 | 107953345 | + | 1.14478 | intron (NM_026062, intron 1 of 4) | intron (NM_026062, intron 1 of 4) | 33938 | NM_026062 | 67266 | Mm.277897 | NM_026062 | ENSMUSG00000029270 | **Fam69a** | 2900024C23Rik\|AI315274 | family with sequence similarity 69, member A | protein-coding |
| chr5_107980817_107981123 | chr5 | 107980818 | 107981123 | + | 0.726469 | intron (NM_026062, intron 1 of 4) | intron (NM_026062, intron 1 of 4) | 6107 | NM_026062 | 67266 | Mm.277897 | NM_026062 | ENSMUSG00000029270 | **Fam69a** | 2900024C23Rik\|AI315274 | family with sequence similarity 69, member A | protein-coding |
| chr1_191172557_191172853 | chr1 | 191172558 | 191172853 | + | 0.649625 | intron (NM_007498, intron 2 of 3) | B3\|SINE\|B2 | -7888 | NM_001109759 | 619288 | Mm.61148 | NM_001109759 | ENSMUSG00000091017 | **Fam71a** | 4933417M04Rik\|GARI-L4 | family with sequence similarity 71, member A | protein-coding |
| chr11_46403585_46404127 | chr11 | 46403586 | 46404127 | + | 1.47403 | promoter-TSS (NM_001013783) | promoter-TSS (NM_001013783) | -872 | NM_001013783 | 432552 | Mm.443244 | NM_001013783 | ENSMUSG00000020401 | **Fam71b** | Gari-l3\|OTTMUSG00000005491 | family with sequence similarity 71, member B | protein-coding |
| chr6_29313721_29313990 | chr6 | 29313722 | 29313990 | + | 0.684843 | Intergenic | Intergenic | -5284 | NM_001289663 | 330277 | Mm.87624 | NM_207258 | ENSMUSG00000039742 | **Fam71f1** | GARI-L1\|Gm764\|NYD-SP18 | family with sequence similarity 71, member F1 | protein-coding |
| chr1_131533644_131534313 | chr1 | 131533645 | 131534313 | + | 0.951034 | intron (NM_175382, intron 3 of 3) | intron (NM_175382, intron 3 of 3) | 5990 | NM_175382 | 108900 | Mm.31278 | NM_175382 | ENSMUSG00000055184 | **Fam72a** | 2700049P18Rik\|P17 | family with sequence similarity 72, member A | protein-coding |
| chr3_152342954_152343170 | chr3 | 152342955 | 152343170 | + | 0.643415 | Intergenic | ORR1D2\|LTR\|ERVL-MaLR | -2655 | NM_174868 | 215708 | Mm.135018 | NM_174868 | ENSMUSG00000054942 | **Fam73a** | A930038D06\|AI852168\|C030011O14Rik | family with sequence similarity 73, member A | protein-coding |
| chr4_132940569_132940800 | chr4 | 132940570 | 132940800 | + | 0.608115 | Intergenic | Intergenic | -18134 | NM_001163792 | 230789 | Mm.11473 | NM_145553 | ENSMUSG00000028878 | **Fam76a** | - | family with sequence similarity 76, member A | protein-coding |
| chr9_13904048_13904732 | chr9 | 13904049 | 13904732 | + | 1.18082 | Intergenic | Intergenic | 76663 | NM_176836 | 72826 | Mm.246174 | NM_176836 | ENSMUSG00000037808 | **Fam76b** | 2810485I05Rik\|C78303 | family with sequence similarity 76, member B | protein-coding |
| chr9_14010221_14011017 | chr9 | 14010222 | 14011017 | + | 1.15171 | Intergenic | Intergenic | 182892 | NM_176836 | 72826 | Mm.246174 | NM_176836 | ENSMUSG00000037808 | **Fam76b** | 2810485I05Rik\|C78303 | family with sequence similarity 76, member B | protein-coding |
| chr9_13894291_13895275 | chr9 | 13894292 | 13895275 | + | 0.686788 | Intergenic | Intergenic | 67056 | NM_176836 | 72826 | Mm.246174 | NM_176836 | ENSMUSG00000037808 | **Fam76b** | 2810485I05Rik\|C78303 | family with sequence similarity 76, member B | protein-coding |
| chr2_158718071_158718279 | chr2 | 158718072 | 158718279 | + | 0.800387 | intron (NM_153089, intron 2 of 10) | intron (NM_153089, intron 2 of 10) | -49924 | NM_027975 | 71878 | Mm.85162 | NM_027975 | ENSMUSG00000027654 | **Fam83d** | 2310007D09Rik\|BB104611 | family with sequence similarity 83, member D | protein-coding |
| chr11_61698669_61699386 | chr11 | 61698670 | 61699386 | + | 1.20765 | intron (NM_178618, intron 2 of 4) | intron (NM_178618, intron 2 of 4) | 14618 | NM_178618 | 69640 | Mm.458559 | NM_178618 | ENSMUSG00000042377 | **Fam83g** | 2310040C09Rik\|AI506990\|wly | family with sequence similarity 83, member G | protein-coding |
| chr16_5520147_5520826 | chr16 | 5520148 | 5520826 | + | 1.18793 | Intergenic | Intergenic | -264531 | NM_027446 | 70511 | Mm.45780 | NM_027446 | ENSMUSG00000022544 | **Fam86** | 5730409G15Rik\|Fam86a | family with sequence similarity 86 | protein-coding |
| chr16_5520480_5520824 | chr16 | 5520481 | 5520824 | + | 0.959164 | Intergenic | Intergenic | -264696 | NM_027446 | 70511 | Mm.45780 | NM_027446 | ENSMUSG00000022544 | **Fam86** | 5730409G15Rik\|Fam86a | family with sequence similarity 86 | protein-coding |
| chr8_124756631_124757145 | chr8 | 124756632 | 124757145 | + | 1.33215 | Intergenic | Intergenic | -5079 | NM_001081120 | 69627 | Mm.271600 | NM_001081120 | ENSMUSG00000043068 | **Fam89a** | 2310031A18Rik | family with sequence similarity 89, member A | protein-coding |
| chr8_124756631_124757004 | chr8 | 124756632 | 124757004 | + | 1.32934 | Intergenic | Intergenic | -5009 | NM_001081120 | 69627 | Mm.271600 | NM_001081120 | ENSMUSG00000043068 | **Fam89a** | 2310031A18Rik | family with sequence similarity 89, member A | protein-coding |
| chr17_28311823_28312042 | chr17 | 28311824 | 28312042 | + | 0.874648 | Intergenic | Intergenic | -1597 | NR_028297 | 72775 | Mm.201859 | NM_028348 | | **Fance** | 2810451D06Rik\|AI415634\|AW209126 | Fanconi anemia, complementation group E | protein-coding |
| chr7_51865440_51865762 | chr7 | 51865441 | 51865762 | + | 0.755062 | Intergenic | Intergenic | -3334 | NM_001115087 | 100040608 | Mm.322643 | NM_001115087 | ENSMUSG00000092118 | **Fancf** | A730016A17 | Fanconi anemia, complementation group F | protein-coding |
| chr7_51865522_51865747 | chr7 | 51865523 | 51865747 | + | 0.676876 | Intergenic | Intergenic | -3368 | NM_001115087 | 100040608 | Mm.322643 | NM_001115087 | ENSMUSG00000092118 | **Fancf** | A730016A17 | Fanconi anemia, complementation group F | protein-coding |
| chr7_113610238_113610605 | chr7 | 113610239 | 113610605 | + | 0.7093 | Intergenic | Intergenic | 96588 | NM_027379 | 67420 | Mm.206919 | NM_026143 | ENSMUSG00000030759 | **Far1** | 2600011M19Rik\|2900034E22Rik\|3732409C05Rik\|AI850429\|Mlstd2 | fatty acyl CoA reductase 1 | protein-coding |
| chr14_121102550_121103145 | chr14 | 121102551 | 121103145 | + | 0.839586 | intron (NM_134082, intron 2 of 26) | intron (NM_134082, intron 2 of 26) | 67274 | NM_134082 | 223254 | Mm.223980 | NM_134082 | ENSMUSG00000025555 | **Farp1** | AW228844\|BC030329\|Cdep | FERM, RhoGEF (Arhgef) and pleckstrin domain protein 1 (chondrocyte-derived) | protein-coding |
| chr13_36405915_36406367 | chr13 | 36405916 | 36406367 | + | 1.35231 | intron (NM_001039189, intron 4 of 5) | B4A\|SINE\|B4 | 288498 | NM_001039189 | 69955 | Mm.70690 | NM_024274 | ENSMUSG00000021420 | **Fars2** | Fars1\|pheRS | phenylalanine-tRNA synthetase 2 (mitochondrial) | protein-coding |
| chr13_36287638_36288542 | chr13 | 36287639 | 36288542 | + | 1.27757 | intron (NM_001039189, intron 3 of 5) | intron (NM_001039189, intron 3 of 5) | 170447 | NM_001039189 | 69955 | Mm.70690 | NM_024274 | ENSMUSG00000021420 | **Fars2** | Fars1\|pheRS | phenylalanine-tRNA synthetase 2 (mitochondrial) | protein-coding |
| chr13_36214026_36214583 | chr13 | 36214027 | 36214583 | + | 1.01935 | intron (NM_024274, intron 2 of 6) | intron (NM_024274, intron 2 of 6) | 96662 | NM_001039189 | 69955 | Mm.70690 | NM_024274 | ENSMUSG00000021420 | **Fars2** | Fars1\|pheRS | phenylalanine-tRNA synthetase 2 (mitochondrial) | protein-coding |
| chr13_36369445_36369765 | chr13 | 36369446 | 36369765 | + | 0.970463 | intron (NM_001039189, intron 4 of 5) | L1M3\|LINE\|L1 | 251962 | NM_001039189 | 69955 | Mm.70690 | NM_024274 | ENSMUSG00000021420 | **Fars2** | Fars1\|pheRS | phenylalanine-tRNA synthetase 2 (mitochondrial) | protein-coding |
| chr13_36211031_36211352 | chr13 | 36211032 | 36211352 | + | 0.842778 | intron (NM_024274, intron 2 of 6) | intron (NM_024274, intron 2 of 6) | 93549 | NM_001039189 | 69955 | Mm.70690 | NM_024274 | ENSMUSG00000021420 | **Fars2** | Fars1\|pheRS | phenylalanine-tRNA synthetase 2 (mitochondrial) | protein-coding |
| chr13_36309017_36309427 | chr13 | 36309018 | 36309427 | + | 0.70506 | intron (NM_001039189, intron 3 of 5) | intron (NM_001039189, intron 3 of 5) | 191579 | NM_001039189 | 69955 | Mm.70690 | NM_024274 | ENSMUSG00000021420 | **Fars2** | Fars1\|pheRS | phenylalanine-tRNA synthetase 2 (mitochondrial) | protein-coding |
| chr19_34275117_34275478 | chr19 | 34275118 | 34275478 | + | 1.09194 | Intergenic | Intergenic | -15361 | NM_007987 | 14102 | Mm.1626 | NM_007987 | ENSMUSG00000024778 | **Fas** | AI196731\|APO1\|APT1\|CD95\|TNFR6\|Tnfrsf6\|lpr | Fas (TNF receptor superfamily member 6) | protein-coding |
| chr19_34284782_34285129 | chr19 | 34284783 | 34285129 | + | 1.01994 | Intergenic | Intergenic | -5703 | NM_007987 | 14102 | Mm.1626 | NM_007987 | ENSMUSG00000024778 | **Fas** | AI196731\|APO1\|APT1\|CD95\|TNFR6\|Tnfrsf6\|lpr | Fas (TNF receptor superfamily member 6) | protein-coding |
| chr19_34284508_34285131 | chr19 | 34284509 | 34285131 | + | 0.639777 | Intergenic | Intergenic | -5839 | NM_007987 | 14102 | Mm.1626 | NM_007987 | ENSMUSG00000024778 | **Fas** | AI196731\|APO1\|APT1\|CD95\|TNFR6\|Tnfrsf6\|lpr | Fas (TNF receptor superfamily member 6) | protein-coding |
| chr1_161693254_161694427 | chr1 | 161693255 | 161694427 | + | 1.0399 | Intergenic | Intergenic | 94654 | NM_001205243 | 14103 | Mm.3355 | NM_010177 | ENSMUSG00000000817 | **Fasl** | APT1LG1\|CD178\|CD95-L\|CD95L\|Fas-L\|Faslg\|Tnfsf6\|gld | Fas ligand (TNF superfamily, member 6) | protein-coding |
| chr1_161653556_161654187 | chr1 | 161653557 | 161654187 | + | 0.971812 | Intergenic | Intergenic | 134623 | NM_001205243 | 14103 | Mm.3355 | NM_010177 | ENSMUSG00000000817 | **Fasl** | APT1LG1\|CD178\|CD95-L\|CD95L\|Fas-L\|Faslg\|Tnfsf6\|gld | Fas ligand (TNF superfamily, member 6) | protein-coding |
| chr1_161653882_161654178 | chr1 | 161653883 | 161654178 | + | 0.909839 | Intergenic | Intergenic | 134465 | NM_001205243 | 14103 | Mm.3355 | NM_010177 | ENSMUSG00000000817 | **Fasl** | APT1LG1\|CD178\|CD95-L\|CD95L\|Fas-L\|Faslg\|Tnfsf6\|gld | Fas ligand (TNF superfamily, member 6) | protein-coding |
| chr1_161714413_161714735 | chr1 | 161714414 | 161714735 | + | 0.780876 | Intergenic | Intergenic | 73921 | NM_001205243 | 14103 | Mm.3355 | NM_010177 | ENSMUSG00000000817 | **Fasl** | APT1LG1\|CD178\|CD95-L\|CD95L\|Fas-L\|Faslg\|Tnfsf6\|gld | Fas ligand (TNF superfamily, member 6) | protein-coding |
| chr13_68586708_68587130 | chr13 | 68586709 | 68587130 | + | 0.845399 | intron (NM_027123, intron 3 of 6) | intron (NM_027123, intron 3 of 6) | 4671 | NM_027123 | 69577 | Mm.23670 | NM_027123 | ENSMUSG00000021532 | **Fastkd3** | 2310010B21Rik\|AW048587 | FAST kinase domains 3 | protein-coding |
| chr3_38887416_38888446 | chr3 | 38887417 | 38888446 | + | 0.678159 | exon (NM_183221, exon 1 of 17) | exon (NM_183221, exon 1 of 17) | 991 | NM_183221 | 329628 | Mm.316210 | NM_183221 | ENSMUSG00000046743 | **Fat4** | 6030410K14Rik\|9430004M15 | FAT tumor suppressor homolog 4 (Drosophila) | protein-coding |
| chr4_21957595_21957929 | chr4 | 21957596 | 21957929 | + | 0.776285 | intron (NM_175234, intron 3 of 5) | intron (NM_175234, intron 3 of 5) | 26436 | NM_175234 | 76132 | Mm.159689 | NM_175234 | | **Faxc** | 6230409E13Rik\|AW742931 | failed axon connections homolog (Drosophila) | protein-coding |
| chr2_125454596_125454993 | chr2 | 125454597 | 125454993 | + | 1.04526 | intron (NM_007993, intron 6 of 65) | intron (NM_007993, intron 6 of 65) | 51643 | NM_007993 | 14118 | Mm.271644 | NM_007993 | ENSMUSG00000027204 | **Fbn1** | AI536462\|B430209H23\|Fib-1\|Tsk | fibrillin 1 | protein-coding |
| chr17_25803181_25803505 | chr17 | 25803182 | 25803505 | + | 0.932157 | Intergenic | Intergenic | -5742 | NM_001164225 | 214931 | Mm.152289 | NM_001164225 | ENSMUSG00000025738 | **Fbxl16** | BC042620\|Scirr1 | F-box and leucine-rich repeat protein 16 | protein-coding |
| chr17_63288002_63288322 | chr17 | 63288003 | 63288322 | + | 0.851679 | intron (NM_015794, intron 6 of 8) | intron (NM_015794, intron 6 of 8) | 212418 | NM_015794 | 50758 | Mm.136366 | NM_015794 | | **Fbxl17** | 6330576B01Rik\|AI452053\|BB073797\|C130023C01Rik\|Fbx13\|Fbxo13 | F-box and leucine-rich repeat protein 17 | protein-coding |
| chr17_63282044_63282371 | chr17 | 63282045 | 63282371 | + | 0.663467 | intron (NM_015794, intron 6 of 8) | intron (NM_015794, intron 6 of 8) | 218372 | NM_015794 | 50758 | Mm.136366 | NM_015794 | | **Fbxl17** | 6330576B01Rik\|AI452053\|BB073797\|C130023C01Rik\|Fbx13\|Fbxo13 | F-box and leucine-rich repeat protein 17 | protein-coding |
| chr5_142858966_142859672 | chr5 | 142858967 | 142859672 | + | 0.77449 | Intergenic | Intergenic | 35919 | NM_001033312 | 231863 | Mm.89753 | NM_001033312 | ENSMUSG00000066640 | **Fbxl18** | AI427077\|B130019G13Rik\|C330021B20Rik | F-box and leucine-rich repeat protein 18 | protein-coding |
| chr9_66520289_66520667 | chr9 | 66520290 | 66520667 | + | 1.12669 | intron (NM_144937, intron 12 of 14) | intron (NM_144937, intron 12 of 14) | -5885 | NM_175206 | 74165 | Mm.297998 | NM_175206 | ENSMUSG00000050503 | **Fbxl22** | 1110004B15Rik | F-box and leucine-rich repeat protein 22 | protein-coding |
| chr9_66520072_66520753 | chr9 | 66520073 | 66520753 | + | 1.11315 | intron (NM_144937, intron 12 of 14) | intron (NM_144937, intron 12 of 14) | -5820 | NM_175206 | 74165 | Mm.297998 | NM_175206 | ENSMUSG00000050503 | **Fbxl22** | 1110004B15Rik | F-box and leucine-rich repeat protein 22 | protein-coding |
| chr5_43784099_43784328 | chr5 | 43784100 | 43784328 | + | 0.812369 | Intergenic | Intergenic | -2065 | NM_178729 | 242960 | Mm.25794 | NM_178729 | ENSMUSG00000039753 | **Fbxl5** | Fbl4\|Fir4 | F-box and leucine-rich repeat protein 5 | protein-coding |
| chr5_117921871_117922511 | chr5 | 117921872 | 117922511 | + | 1.21683 | intron (NM_008712, intron 16 of 27) | intron (NM_008712, intron 16 of 27) | -54579 | NM_145564 | 231670 | Mm.21912 | NM_145564 | ENSMUSG00000032898 | **Fbxo21** | 2810425J22Rik\|AU016673\|mKIAA0875 | F-box protein 21 | protein-coding |
| chr5_117922108_117922505 | chr5 | 117922109 | 117922505 | + | 1.17565 | intron (NM_008712, intron 16 of 27) | intron (NM_008712, intron 16 of 27) | -54463 | NM_145564 | 231670 | Mm.21912 | NM_145564 | ENSMUSG00000032898 | **Fbxo21** | 2810425J22Rik\|AU016673\|mKIAA0875 | F-box protein 21 | protein-coding |
| chr1_182361851_182362220 | chr1 | 182361852 | 182362220 | + | 1.11818 | Intergenic | Intergenic | -20430 | NM_175127 | 67948 | Mm.30737 | NM_175127 | ENSMUSG00000047539 | **Fbxo28** | 4833428J17Rik\|5730505P19Rik\|D1Ertd578e\|Fbx28\|mKIAA0483 | F-box protein 28 | protein-coding |
| chr15_58194427_58195375 | chr15 | 58194428 | 58195375 | + | 0.757058 | intron (NM_026346, intron 4 of 8) | intron (NM_026346, intron 4 of 8) | 19991 | NM_026346 | 67731 | Mm.25683 | NM_026346 | ENSMUSG00000022358 | **Fbxo32** | 4833442G10Rik\|AI430017\|ATROGIN1\|Gm20361\|MAFbx | F-box protein 32 | protein-coding |
| chr12_59212553_59212922 | chr12 | 59212554 | 59212922 | + | 0.91412 | intron (NM_001033156, intron 1 of 3) | L1VL4\|LINE\|L1 | 6745 | NM_001033156 | 70611 | Mm.311026 | NM_001033156 | ENSMUSG00000035329 | **Fbxo33** | 5730501N20Rik\|AI642135 | F-box protein 33 | protein-coding |
| chr14_47474634_47474925 | chr14 | 47474635 | 47474925 | + | 0.864195 | intron (NM_030236, intron 1 of 1) | MLT-int\|LTR\|ERVL-MaLR | 2219 | NM_001146085 | 78938 | Mm.11935 | NM_030236 | ENSMUSG00000037536 | **Fbxo34** | 2900057B08Rik\|5830426G16Rik | F-box protein 34 | protein-coding |
| chr1_84881548_84882054 | chr1 | 84881549 | 84882054 | + | 0.914066 | intron (NM_025386, intron 2 of 3) | intron (NM_025386, intron 2 of 3) | 41960 | NM_025386 | 66153 | Mm.381192 | NM_025386 | ENSMUSG00000073633 | **Fbxo36** | 0610008D19Rik\|1110020F21Rik\|2410002G19Rik\|D1Ertd757e | F-box protein 36 | protein-coding |
| chr1_84881357_84882079 | chr1 | 84881358 | 84882079 | + | 0.71384 | intron (NM_025386, intron 2 of 3) | RMER15\|LTR\|ERVL | 41877 | NM_025386 | 66153 | Mm.381192 | NM_025386 | ENSMUSG00000073633 | **Fbxo36** | 0610008D19Rik\|1110020F21Rik\|2410002G19Rik\|D1Ertd757e | F-box protein 36 | protein-coding |

| | | | | | | | | | | | | | | | | | |
|---|---|---|---|---|---|---|---|---|---|---|---|---|---|---|---|---|---|
| chr11_72311120_72311915 | chr11 | 72311121 | 72311915 | + | 0.848465 | intron (NM_001291153, intron 3 of 3) | intron (NM_001291153, intron 3 of 3) | -2926 | NM_001099688 | 628100 | Mm.390365 | NM_001099688 | ENSMUSG00000070388 | **Fbxo39** | 1700010H23Rik | F-box protein 39 | protein-coding |
| chr16_36983769_36984159 | chr16 | 36983770 | 36984159 | + | 1.23094 | Intergenic | Lx9\|LINE\|L1 | -4828 | NM_001037321 | 207215 | Mm.211899 | NM_001037321 | ENSMUSG00000047746 | **Fbxo40** | 9830003A13Rik\|mKIAA1195 | F-box protein 40 | protein-coding |
| chr16_36989362_36989933 | chr16 | 36989363 | 36989933 | + | 0.674183 | Intergenic | Intergenic | -10512 | NM_001037321 | 207215 | Mm.211899 | NM_001037321 | ENSMUSG00000047746 | **Fbxo40** | 9830003A13Rik\|mKIAA1195 | F-box protein 40 | protein-coding |
| chr16_32214943_32215659 | chr16 | 32214944 | 32215659 | + | 0.889497 | Intergenic | B4\|SINE\|B4 | 31724 | NM_173439 | 268882 | Mm.256137 | NM_173439 | ENSMUSG00000035764 | **Fbxo45** | 2610017J04Rik\|AI463119\|BC026799 | F-box protein 45 | protein-coding |
| chr11_97908439_97909038 | chr11 | 97908440 | 97909038 | + | 1.14716 | Intergenic | Intergenic | -24585 | NM_001081435 | 72973 | Mm.245939 | NM_001081435 | | **Fbxo47** | 2900052P03Rik\|Gm1562 | F-box protein 47 | protein-coding |
| chr11_97908425_97909224 | chr11 | 97908426 | 97909224 | + | 0.965904 | Intergenic | MIR\|SINE\|MIR | -24671 | NM_001081435 | 72973 | Mm.245939 | NM_001081435 | | **Fbxo47** | 2900052P03Rik\|Gm1562 | F-box protein 47 | protein-coding |
| chr11_16952087_16952467 | chr11 | 16952088 | 16952467 | + | 0.737381 | promoter-TSS (NR_110425) | promoter-TSS (NR_110425) | 867 | NM_176982 | 319701 | Mm.120672 | NM_176982 | ENSMUSG00000044966 | **Fbxo48** | A630050E13Rik | F-box protein 48 | protein-coding |
| chr10_86032214_86032536 | chr10 | 86032215 | 86032536 | + | 0.705258 | intron (NM_153195, intron 4 of 8) | intron (NM_153195, intron 4 of 8) | 10446 | NM_153195 | 69754 | Mm.44236 | NM_153195 | ENSMUSG00000001786 | **Fbxo7** | 2410015K21Rik\|A230052G17Rik\|AI117679\|AW260462\|C85118 | F-box protein 7 | protein-coding |
| chr11_32661695_32662046 | chr11 | 32661696 | 32662046 | + | 0.885516 | intron (NM_001271349, intron 1 of 11) | PB1D10\|SINE\|Alu | 19316 | NM_001271347 | 103583 | Mm.28017 | NM_134015 | ENSMUSG00000020271 | **Fbxw11** | 2310065A07Rik\|AA536858\|BTRC2\|BTRCP2\|Fbxw1b\|HOS | F-box and WD-40 domain protein 11 | protein-coding |
| chr11_32693513_32694127 | chr11 | 32693514 | 32694127 | + | 0.875173 | intron (NM_001271347, intron 2 of 12) | B1_Mus2\|SINE\|Alu | 51265 | NM_001271347 | 103583 | Mm.28017 | NM_134015 | ENSMUSG00000020271 | **Fbxw11** | 2310065A07Rik\|AA536858\|BTRC2\|BTRCP2\|Fbxw1b\|HOS | F-box and WD-40 domain protein 11 | protein-coding |
| chr11_32693571_32693970 | chr11 | 32693572 | 32693970 | + | 0.858206 | intron (NM_001271347, intron 2 of 12) | B1_Mus2\|SINE\|Alu | 51216 | NM_001271347 | 103583 | Mm.28017 | NM_134015 | ENSMUSG00000020271 | **Fbxw11** | 2310065A07Rik\|AA536858\|BTRC2\|BTRCP2\|Fbxw1b\|HOS | F-box and WD-40 domain protein 11 | protein-coding |
| chr11_32705123_32705424 | chr11 | 32705124 | 32705424 | + | 0.778185 | intron (NM_001271347, intron 2 of 12) | intron (NM_001271347, intron 2 of 12) | 62719 | NM_001271347 | 103583 | Mm.28017 | NM_134015 | ENSMUSG00000020271 | **Fbxw11** | 2310065A07Rik\|AA536858\|BTRC2\|BTRCP2\|Fbxw1b\|HOS | F-box and WD-40 domain protein 11 | protein-coding |
| chr11_32708872_32709357 | chr11 | 32708873 | 32709357 | + | 0.757295 | intron (NM_001271348, intron 2 of 12) | intron (NM_001271348, intron 2 of 12) | 66560 | NM_001271347 | 103583 | Mm.28017 | NM_134015 | ENSMUSG00000020271 | **Fbxw11** | 2310065A07Rik\|AA536858\|BTRC2\|BTRCP2\|Fbxw1b\|HOS | F-box and WD-40 domain protein 11 | protein-coding |
| chr11_32644060_32644361 | chr11 | 32644061 | 32644361 | + | 0.75209 | intron (NM_001271349, intron 1 of 11) | intron (NM_001271349, intron 1 of 11) | 1656 | NM_001271347 | 103583 | Mm.28017 | NM_134015 | ENSMUSG00000020271 | **Fbxw11** | 2310065A07Rik\|AA536858\|BTRC2\|BTRCP2\|Fbxw1b\|HOS | F-box and WD-40 domain protein 11 | protein-coding |
| chr11_32708521_32709394 | chr11 | 32708522 | 32709394 | + | 0.65226 | intron (NM_001271348, intron 2 of 12) | intron (NM_001271348, intron 2 of 12) | 66403 | NM_001271347 | 103583 | Mm.28017 | NM_134015 | ENSMUSG00000020271 | **Fbxw11** | 2310065A07Rik\|AA536858\|BTRC2\|BTRCP2\|Fbxw1b\|HOS | F-box and WD-40 domain protein 11 | protein-coding |
| chr3_84808901_84809401 | chr3 | 84808902 | 84809401 | + | 1.15612 | Intergenic | Intergenic | -6426 | NM_001177773 | 50754 | Mm.196475 | NM_080428 | ENSMUSG00000028086 | **Fbxw7** | 1110001A17Rik\|AGO\|Cdc4\|Fbw7\|Fbwd6\|Fbx30\|Fbxo30\|Fbxw6\|SEL-10 | F-box and WD-40 domain protein 7 | protein-coding |
| chr3_84835360_84835775 | chr3 | 84835361 | 84835775 | + | 0.975851 | intron (NM_001177773, intron 1 of 13) | intron (NM_001177773, intron 1 of 13) | 19991 | NM_001177773 | 50754 | Mm.196475 | NM_080428 | ENSMUSG00000028086 | **Fbxw7** | 1110001A17Rik\|AGO\|Cdc4\|Fbw7\|Fbwd6\|Fbx30\|Fbxo30\|Fbxw6\|SEL-10 | F-box and WD-40 domain protein 7 | protein-coding |
| chr3_84771883_84772400 | chr3 | 84771884 | 84772400 | + | 0.709854 | Intergenic | Intergenic | -43435 | NM_001177773 | 50754 | Mm.196475 | NM_080428 | ENSMUSG00000028086 | **Fbxw7** | 1110001A17Rik\|AGO\|Cdc4\|Fbw7\|Fbwd6\|Fbx30\|Fbxo30\|Fbxw6\|SEL-10 | F-box and WD-40 domain protein 7 | protein-coding |
| chr1_130789638_130790500 | chr1 | 130789639 | 130790500 | + | 1.34197 | Intergenic | RMER1A\|Other\|Other | -10833 | NM_001170632 | 64435 | Mm.31821 | NM_144960 | ENSMUSG00000026415 | **Fcamr** | - | Fc receptor, IgA, IgM, high affinity | protein-coding |
| chr1_130795271_130795910 | chr1 | 130795272 | 130795910 | + | 1.15601 | Intergenic | Lx8\|LINE\|L1 | -5311 | NM_001170632 | 64435 | Mm.31821 | NM_144960 | ENSMUSG00000026415 | **Fcamr** | - | Fc receptor, IgA, IgM, high affinity | protein-coding |
| chr3_96340866_96341291 | chr3 | 96340867 | 96341291 | + | 0.764174 | Intergenic | Intergenic | -47110 | NM_010186 | 14129 | Mm.150 | NM_010186 | ENSMUSG00000015947 | **Fcgr1** | AI323638\|AV092959\|CD64\|FcgammaRI\|IGGHAFC | Fc receptor, IgG, high affinity I | protein-coding |
| chr3_96328298_96329275 | chr3 | 96328299 | 96329275 | + | 0.733508 | Intergenic | B2_Mm2\|SINE\|B2 | -34818 | NM_010186 | 14129 | Mm.150 | NM_010186 | ENSMUSG00000015947 | **Fcgr1** | AI323638\|AV092959\|CD64\|FcgammaRI\|IGGHAFC | Fc receptor, IgG, high affinity I | protein-coding |
| chr3_96332616_96333441 | chr3 | 96332617 | 96333441 | + | 0.675323 | Intergenic | Intergenic | -39060 | NM_010186 | 14129 | Mm.150 | NM_010186 | ENSMUSG00000015947 | **Fcgr1** | AI323638\|AV092959\|CD64\|FcgammaRI\|IGGHAFC | Fc receptor, IgG, high affinity I | protein-coding |
| chr7_45098589_45099205 | chr7 | 45098590 | 45099205 | + | 0.694925 | intron (NM_010189, intron 4 of 6) | PB1D9\|SINE\|Alu | 4925 | NM_010189 | 14132 | Mm.3303 | NM_010189 | ENSMUSG00000003420 | **Fcgrt** | FcRn | Fc receptor, IgG, alpha chain transporter | protein-coding |
| chr3_87383268_87383697 | chr3 | 87383269 | 87383697 | + | 1.01375 | intron (NM_153090, intron 3 of 10) | intron (NM_153090, intron 3 of 10) | 7096 | NM_001136236 | 229499 | Mm.255203 | NM_153090 | ENSMUSG00000059994 | **Fcrl1** | A230020G22Rik\|Bxmas1\|Bxmh1\|Bxmh1b\|Fcrh1\|Fcrh1l\|Fcrh1s\|Ifgp1 | Fc receptor-like 1 | protein-coding |
| chr9_51961168_51961546 | chr9 | 51961169 | 51961546 | + | 1.01071 | intron (NM_007996, intron 1 of 3) | intron (NM_007996, intron 1 of 3) | 2176 | NM_007996 | 14148 | Mm.1061 | NM_007996 | ENSMUSG00000032051 | **Fdx1** | - | ferredoxin 1 | protein-coding |
| chr17_56265984_56266889 | chr17 | 56265985 | 56266889 | + | 1.36504 | Intergenic | Intergenic | 9644 | NM_010192 | 14154 | Mm.290813 | NM_010192 | ENSMUSG00000043683 | **Fem1a** | AW611390\|Fem1aa | feminization 1 homolog a (C. elegans) | protein-coding |
| chr18_46548918_46549765 | chr18 | 46548919 | 46549765 | + | 0.834669 | Intergenic | Intergenic | -23371 | NM_173423 | 240263 | Mm.38249 | NM_173423 | | **Fem1c** | 2610312A07Rik\|3632443A22Rik\|AI646567 | fem-1 homolog c (C.elegans) | protein-coding |
| chr8_120999877_121000177 | chr8 | 120999878 | 121000177 | + | 0.914804 | Intergenic | Lx8\|LINE\|L1 | 83005 | NR_045471 | 68790 | Mm.246916 | NR_045471 | ENSMUSG00000097336 | **Fendrr** | 1110050K14Rik\|linc-Foxf1a | Foxf1 adjacent non-coding developmental regulatory RNA | ncRNA |
| chr8_121003042_121003559 | chr8 | 121003043 | 121003559 | + | 0.772996 | Intergenic | Intergenic | 79731 | NR_045471 | 68790 | Mm.246916 | NR_045471 | ENSMUSG00000097336 | **Fendrr** | 1110050K14Rik\|linc-Foxf1a | Foxf1 adjacent non-coding developmental regulatory RNA | ncRNA |
| chr8_121001573_121001888 | chr8 | 121001574 | 121001888 | + | 0.713997 | Intergenic | Intergenic | 81301 | NR_045471 | 68790 | Mm.246916 | NR_045471 | ENSMUSG00000097336 | **Fendrr** | 1110050K14Rik\|linc-Foxf1a | Foxf1 adjacent non-coding developmental regulatory RNA | ncRNA |
| chr17_78363479_78364150 | chr17 | 78363480 | 78364150 | + | 1.06911 | intron (NM_015800, intron 13 of 16) | intron (NM_015800, intron 13 of 16) | 54337 | NM_001285949 | 225020 | Mm.236835 | NM_199448 | ENSMUSG00000056121 | **Fez2** | 9030616F16\|D17Ertd315e | fasciculation and elongation protein zeta 2 (zygin II) | protein-coding |
| chr17_78419197_78419492 | chr17 | 78419198 | 78419492 | + | 0.969592 | Intergenic | Intergenic | -1193 | NM_001285940 | 225020 | Mm.236835 | NM_199448 | ENSMUSG00000056121 | **Fez2** | 9030616F16\|D17Ertd315e | fasciculation and elongation protein zeta 2 (zygin II) | protein-coding |
| chr17_29395319_29395819 | chr17 | 29395320 | 29395819 | + | 1.07568 | Intergenic | Intergenic | 34655 | NM_001159538 | 26382 | Mm.279187 | NM_013710 | ENSMUSG00000024013 | **Fgd2** | Tcd-2\|Tcd2\|ZFYVE4\|tcs-2\|tcs2 | FYVE, RhoGEF and PH domain containing 2 | protein-coding |
| chr17_29397861_29398391 | chr17 | 29397862 | 29398391 | + | 0.658737 | Intergenic | Intergenic | 37212 | NM_001159538 | 26382 | Mm.279187 | NM_013710 | ENSMUSG00000024013 | **Fgd2** | Tcd-2\|Tcd2\|ZFYVE4\|tcs-2\|tcs2 | FYVE, RhoGEF and PH domain containing 2 | protein-coding |
| chr17_29393364_29394072 | chr17 | 29393365 | 29394072 | + | 0.629181 | Intergenic | Intergenic | 32804 | NM_001159538 | 26382 | Mm.279187 | NM_013710 | ENSMUSG00000024013 | **Fgd2** | Tcd-2\|Tcd2\|ZFYVE4\|tcs-2\|tcs2 | FYVE, RhoGEF and PH domain containing 2 | protein-coding |
| chrX_59251631_59251933 | chrX | 59251632 | 59251933 | + | 0.782987 | intron (NM_001290414, intron 2 of 6) | intron (NM_001290414, intron 2 of 6) | -85694 | NM_001290415 | 14168 | Mm.7995 | NM_010200 | ENSMUSG00000031137 | **Fgf13** | Fhf2 | fibroblast growth factor 13 | protein-coding |
| chrX_105715800_105716348 | chrX | 105715801 | 105716348 | + | 0.916029 | Intergenic | Intergenic | -48403 | NM_030614 | 80903 | Mm.154768 | NM_030614 | | **Fgf16** | - | fibroblast growth factor 16 | protein-coding |
| chr11_33149517_33149767 | chr11 | 33149518 | 33149767 | + | 0.924199 | Intergenic | ID_B1\|SINE\|B4 | -2242 | NR_102395 | 14172 | Mm.339812 | NM_008005 | | **Fgf18** | D130055P09Rik\|FGF-18 | fibroblast growth factor 18 | protein-coding |
| chr11_33151576_33151752 | chr11 | 33151577 | 33151752 | + | 0.712512 | TTS (NM_008722) | TTS (NM_008722) | -4264 | NR_102395 | 14172 | Mm.339812 | NM_008005 | | **Fgf18** | D130055P09Rik\|FGF-18 | fibroblast growth factor 18 | protein-coding |
| chr3_37370337_37370942 | chr3 | 37370338 | 37370942 | + | 1.50596 | intron (NM_008006, intron 1 of 2) | ORR1F\|LTR\|ERVL-MaLR | 21987 | NM_008006 | 14173 | Mm.473689 | NM_008006 | ENSMUSG00000037225 | **Fgf2** | Fgf-2\|Fgfb\|bFGF | fibroblast growth factor 2 | protein-coding |
| chr3_37365223_37365791 | chr3 | 37365224 | 37365791 | + | 0.969777 | intron (NM_008006, intron 1 of 2) | MTE-int\|LTR\|ERVL-MaLR | 16854 | NM_008006 | 14173 | Mm.473689 | NM_008006 | ENSMUSG00000037225 | **Fgf2** | Fgf-2\|Fgfb\|bFGF | fibroblast growth factor 2 | protein-coding |
| chr3_37365008_37365796 | chr3 | 37365009 | 37365796 | + | 0.692977 | intron (NM_008006, intron 1 of 2) | MTE-int\|LTR\|ERVL-MaLR | 16749 | NM_008006 | 14173 | Mm.473689 | NM_008006 | ENSMUSG00000037225 | **Fgf2** | Fgf-2\|Fgfb\|bFGF | fibroblast growth factor 2 | protein-coding |
| chr6_126995533_126997022 | chr6 | 126995534 | 126997022 | + | 0.868907 | Intergenic | Intergenic | -19264 | NM_010204 | 14177 | Mm.3403 | NM_010204 | ENSMUSG00000000183 | **Fgf6** | Fgf-6\|HSTF-2 | fibroblast growth factor 6 | protein-coding |
| chr6_126996093_126997014 | chr6 | 126996094 | 126997014 | + | 0.80401 | Intergenic | Intergenic | -18988 | NM_010204 | 14177 | Mm.3403 | NM_010204 | ENSMUSG00000000183 | **Fgf6** | Fgf-6\|HSTF-2 | fibroblast growth factor 6 | protein-coding |
| chr6_126995541_126995991 | chr6 | 126995542 | 126995991 | + | 0.689484 | Intergenic | Intergenic | -19776 | NM_010204 | 14177 | Mm.3403 | NM_010204 | ENSMUSG00000000183 | **Fgf6** | Fgf-6\|HSTF-2 | fibroblast growth factor 6 | protein-coding |
| chr2_125994264_125994791 | chr2 | 125994265 | 125994791 | + | 0.815759 | intron (NM_029455, intron 13 of 16) | intron (NM_029455, intron 13 of 16) | -40130 | NM_008008 | 14178 | Mm.330557 | NM_008008 | ENSMUSG00000027208 | **Fgf7** | Kgf | fibroblast growth factor 7 | protein-coding |
| chr5_43948800_43949358 | chr5 | 43948801 | 43949358 | + | 0.822955 | Intergenic | Intergenic | 32720 | NM_008009 | 14181 | Mm.46053 | NM_008009 | ENSMUSG00000048373 | **Fgfbp1** | FGF-BP\|FGF-BP1\|FGFBP-1 | fibroblast growth factor binding protein 1 | protein-coding |
| chr5_43947299_43947737 | chr5 | 43947300 | 43947737 | + | 0.746492 | Intergenic | Intergenic | 34281 | NM_008009 | 14181 | Mm.46053 | NM_008009 | ENSMUSG00000048373 | **Fgfbp1** | FGF-BP\|FGF-BP1\|FGFBP-1 | fibroblast growth factor binding protein 1 | protein-coding |
| chr8_25465471_25465953 | chr8 | 25465472 | 25465953 | + | 0.745003 | Intergenic | Intergenic | -53047 | NM_001079909 | 14182 | Mm.265716 | NM_010206 | ENSMUSG00000031565 | **Fgfr1** | AW208770\|Eask\|FLG\|Fgfr-1\|Flt-2\|Hspy\|MFR\|bFGF-R-1\|c-fgr | fibroblast growth factor receptor 1 | protein-coding |
| chr17_8190450_8191185 | chr17 | 8190451 | 8191185 | + | 0.682038 | intron (NM_201230, intron 9 of 11) | ID4\|SINE\|ID | 25300 | NM_201230 | 75296 | Mm.227250 | NM_201230 | ENSMUSG00000069135 | **Fgfr1op** | 4930553O10Rik\|FOP | Fgfr1 oncogene partner | protein-coding |
| chr17_8190716_8191197 | chr17 | 8190717 | 8191197 | + | 0.668614 | intron (NM_201230, intron 9 of 11) | intron (NM_201230, intron 9 of 11) | 25439 | NM_201230 | 75296 | Mm.227250 | NM_201230 | ENSMUSG00000069135 | **Fgfr1op** | 4930553O10Rik\|FOP | Fgfr1 oncogene partner | protein-coding |
| chr4_95707860_95708539 | chr4 | 95707861 | 95708539 | + | 1.94216 | intron (NM_029347, intron 6 of 10) | intron (NM_029347, intron 6 of 10) | 128779 | NM_029347 | 75578 | Mm.276098 | NM_029347 | ENSMUSG00000028573 | **Fggy** | 2310009E04Rik | FGGY carbohydrate kinase domain containing | protein-coding |
| chr4_95439792_95440233 | chr4 | 95439793 | 95440233 | + | 1.07686 | Intergenic | Intergenic | -117494 | NM_001113412 | 75578 | Mm.276098 | NM_029347 | ENSMUSG00000028573 | **Fggy** | 2310009E04Rik | FGGY carbohydrate kinase domain containing | protein-coding |

| | | | | | | | | | | | | | | | | | |
|---|---|---|---|---|---|---|---|---|---|---|---|---|---|---|---|---|---|
| chr4_95686647_95687151 | chr4 | 95686648 | 95687151 | + | 0.949966 | intron (NM_029347, intron 5 of 10) | L1M5\|LINE\|L1 | 107478 | NM_029347 | 75578 | Mm.276098 | NM_029347 | ENSMUSG00000028573 | **Fggy** | 2310009E04Rik | FGGY carbohydrate kinase domain containing | protein-coding |
| chr4_95448764_95449324 | chr4 | 95448765 | 95449324 | + | 0.889475 | Intergenic | RMER13B\|LTR\|ERVK | -108463 | NM_001113412 | 75578 | Mm.276098 | NM_029347 | ENSMUSG00000028573 | **Fggy** | 2310009E04Rik | FGGY carbohydrate kinase domain containing | protein-coding |
| chr4_95482236_95482660 | chr4 | 95482237 | 95482660 | + | 0.79159 | Intergenic | Intergenic | -75059 | NM_001113412 | 75578 | Mm.276098 | NM_029347 | ENSMUSG00000028573 | **Fggy** | 2310009E04Rik | FGGY carbohydrate kinase domain containing | protein-coding |
| chr4_95482210_95482873 | chr4 | 95482211 | 95482873 | + | 0.760624 | Intergenic | Intergenic | -74965 | NM_001113412 | 75578 | Mm.276098 | NM_029347 | ENSMUSG00000028573 | **Fggy** | 2310009E04Rik | FGGY carbohydrate kinase domain containing | protein-coding |
| chr4_95440438_95440718 | chr4 | 95440439 | 95440718 | + | 0.748129 | Intergenic | Lx9\|LINE\|L1 | -116929 | NM_001113412 | 75578 | Mm.276098 | NM_029347 | ENSMUSG00000028573 | **Fggy** | 2310009E04Rik | FGGY carbohydrate kinase domain containing | protein-coding |
| chr4_95481931_95482880 | chr4 | 95481932 | 95482880 | + | 0.719567 | Intergenic | Intergenic | -75101 | NM_001113412 | 75578 | Mm.276098 | NM_029347 | ENSMUSG00000028573 | **Fggy** | 2310009E04Rik | FGGY carbohydrate kinase domain containing | protein-coding |
| chr5_21342293_21343080 | chr5 | 21342294 | 21343080 | + | 1.20341 | intron (NM_029195, intron 2 of 18) | L1_Mus3\|LINE\|L1 | -29986 | NM_008013 | 14190 | Mm.292100 | NM_008013 | ENSMUSG00000039899 | **Fgl2** | AI385601\|musfiblp | fibrinogen-like protein 2 | protein-coding |
| chr5_21342304_21342697 | chr5 | 21342305 | 21342697 | + | 1.09426 | intron (NM_029195, intron 2 of 18) | L1_Mus3\|LINE\|L1 | -30172 | NM_008013 | 14190 | Mm.292100 | NM_008013 | ENSMUSG00000039899 | **Fgl2** | AI385601\|musfiblp | fibrinogen-like protein 2 | protein-coding |
| chr1_175586768_175587186 | chr1 | 175586769 | 175587186 | + | 1.33132 | Intergenic | Intergenic | 38658 | NM_010209 | 14194 | Mm.41502 | NM_010209 | | **Fh1** | Fh\|Fh-1 | fumarate hydratase 1 | protein-coding |
| chr4_142014714_142015270 | chr4 | 142014715 | 142015270 | + | 0.616277 | Intergenic | Intergenic | -3341 | NM_177868 | 329977 | Mm.252019 | NM_028429 | ENSMUSG00000051435 | **Fhad1** | 2900090M10Rik\|B230311F11 | forkhead-associated (FHA) phosphopeptide binding domain 1 | protein-coding |
| chr4_141951996_141953239 | chr4 | 141951997 | 141953239 | + | 1.35472 | intron (NM_177868, intron 12 of 31) | (TG)n\|Simple_repeat\|Simple_repeat | -30374 | NR_040672 | 545691 | Mm.332015 | NR_040672 | ENSMUSG00000085424 | **Fhad1os1** | Gm13037\|OTTMUSG00000010113 | forkhead-associated (FHA) phosphopeptide binding domain 1, opposite strand 1 | ncRNA |
| chr4_141928797_141929288 | chr4 | 141928798 | 141929288 | + | 1.33717 | intron (NM_177868, intron 17 of 31) | intron (NM_177868, intron 17 of 31) | -53949 | NR_040672 | 545691 | Mm.332015 | NR_040672 | ENSMUSG00000085424 | **Fhad1os1** | Gm13037\|OTTMUSG00000010113 | forkhead-associated (FHA) phosphopeptide binding domain 1, opposite strand 1 | ncRNA |
| chr3_84477427_84477785 | chr3 | 84477428 | 84477785 | + | 0.934331 | intron (NM_001033301, intron 1 of 11) | intron (NM_001033301, intron 1 of 11) | 1885 | NM_001205355 | 229474 | Mm.331107 | NM_001033301 | | **Fhdc1** | 6330505N24Rik\|Gm126 | FH2 domain containing 1 | protein-coding |
| chr1_43144321_43144944 | chr1 | 43144322 | 43144944 | + | 1.38895 | intron (NM_001289533, intron 2 of 5) | intron (NM_001289533, intron 2 of 5) | 19328 | NM_010212 | 14200 | Mm.6799 | NM_010212 | ENSMUSG00000008136 | **Fhl2** | FHL-2\|SLIM-3\|SLIM3 | four and a half LIM domains 2 | protein-coding |
| chr1_43191066_43191601 | chr1 | 43191067 | 43191601 | + | 1.05135 | intron (NM_001289533, intron 1 of 5) | intron (NM_001289533, intron 1 of 5) | 5427 | NM_001289533 | 14200 | Mm.6799 | NM_010212 | ENSMUSG00000008136 | **Fhl2** | FHL-2\|SLIM-3\|SLIM3 | four and a half LIM domains 2 | protein-coding |
| chr1_43168589_43169426 | chr1 | 43168590 | 43169426 | + | 1.0239 | intron (NM_001289533, intron 1 of 5) | intron (NM_001289533, intron 1 of 5) | -5047 | NM_010212 | 14200 | Mm.6799 | NM_010212 | ENSMUSG00000008136 | **Fhl2** | FHL-2\|SLIM-3\|SLIM3 | four and a half LIM domains 2 | protein-coding |
| chr1_43302639_43303173 | chr1 | 43302640 | 43303173 | + | 0.885928 | Intergenic | Intergenic | -106145 | NM_001289533 | 14200 | Mm.6799 | NM_010212 | ENSMUSG00000008136 | **Fhl2** | FHL-2\|SLIM-3\|SLIM3 | four and a half LIM domains 2 | protein-coding |
| chr1_43143339_43143800 | chr1 | 43143340 | 43143800 | + | 0.804084 | intron (NM_001289533, intron 2 of 5) | intron (NM_001289533, intron 2 of 5) | 20391 | NM_010212 | 14200 | Mm.6799 | NM_010212 | ENSMUSG00000008136 | **Fhl2** | FHL-2\|SLIM-3\|SLIM3 | four and a half LIM domains 2 | protein-coding |
| chr1_43225439_43226045 | chr1 | 43225440 | 43226045 | + | 0.651405 | Intergenic | L1MB3\|LINE\|L1 | -28981 | NM_001289533 | 14200 | Mm.6799 | NM_010212 | ENSMUSG00000008136 | **Fhl2** | FHL-2\|SLIM-3\|SLIM3 | four and a half LIM domains 2 | protein-coding |
| chr11_11825480_11826375 | chr11 | 11825481 | 11826375 | + | 1.42103 | intron (NM_001190448, intron 10 of 14) | intron (NM_001190448, intron 10 of 14) | -16966 | NM_001163360 | 60530 | Mm.236114 | NM_021891 | ENSMUSG00000035455 | **Fignl1** | - | fidgetin-like 1 | protein-coding |
| chr11_11820684_11821093 | chr11 | 11820685 | 11821093 | + | 1.32739 | intron (NM_001190448, intron 12 of 14) | intron (NM_001190448, intron 12 of 14) | -11927 | NM_001163360 | 60530 | Mm.236114 | NM_021891 | ENSMUSG00000035455 | **Fignl1** | - | fidgetin-like 1 | protein-coding |
| chr11_11820260_11821156 | chr11 | 11820261 | 11821156 | + | 0.96566 | intron (NM_001190448, intron 12 of 14) | intron (NM_001190448, intron 12 of 14) | -11746 | NM_001163360 | 60530 | Mm.236114 | NM_021891 | ENSMUSG00000035455 | **Fignl1** | - | fidgetin-like 1 | protein-coding |
| chr11_11837631_11838265 | chr11 | 11837632 | 11838265 | + | 0.833448 | intron (NM_001190448, intron 8 of 14) | intron (NM_001190448, intron 8 of 14) | -28986 | NM_001163360 | 60530 | Mm.236114 | NM_021891 | ENSMUSG00000035455 | **Fignl1** | - | fidgetin-like 1 | protein-coding |
| chr11_11837657_11837909 | chr11 | 11837658 | 11837909 | + | 0.746443 | intron (NM_001190448, intron 8 of 14) | intron (NM_001190448, intron 8 of 14) | -28821 | NM_001163360 | 60530 | Mm.236114 | NM_021891 | ENSMUSG00000035455 | **Fignl1** | - | fidgetin-like 1 | protein-coding |
| chr9_79989640_79990170 | chr9 | 79989641 | 79990170 | + | 1.05364 | Intergenic | Intergenic | -12023 | NM_001081243 | 70598 | Mm.23049 | NM_001081243 | ENSMUSG00000034898 | **Filip1** | 5730485H21Rik\|AI843021 | filamin A interacting protein 1 | protein-coding |
| chr9_79947181_79947876 | chr9 | 79947182 | 79947876 | + | 0.915689 | intron (NM_001081243, intron 1 of 5) | intron (NM_001081243, intron 1 of 5) | 30353 | NM_001081243 | 70598 | Mm.23049 | NM_001081243 | ENSMUSG00000034898 | **Filip1** | 5730485H21Rik\|AI843021 | filamin A interacting protein 1 | protein-coding |
| chr16_57571629_57572344 | chr16 | 57571630 | 57572344 | + | 1.69646 | exon (NM_001040397, exon 5 of 5) | exon (NM_001040397, exon 5 of 5) | 22745 | NM_001177871 | 78749 | Mm.328360 | NM_001040397 | ENSMUSG00000043336 | **Filip1l** | 4631422O05Rik\|Doc1 | filamin A interacting protein 1-like | protein-coding |
| chr16_57569013_57569400 | chr16 | 57569014 | 57569400 | + | 1.63371 | intron (NM_001177871, intron 1 of 1) | MIR\|SINE\|MIR | 19965 | NM_001177871 | 78749 | Mm.328360 | NM_001040397 | ENSMUSG00000043336 | **Filip1l** | 4631422O05Rik\|Doc1 | filamin A interacting protein 1-like | protein-coding |
| chr16_57570640_57571153 | chr16 | 57570641 | 57571153 | + | 1.39807 | exon (NM_001040397, exon 5 of 5) | exon (NM_001040397, exon 5 of 5) | 21655 | NM_001177871 | 78749 | Mm.328360 | NM_001040397 | ENSMUSG00000043336 | **Filip1l** | 4631422O05Rik\|Doc1 | filamin A interacting protein 1-like | protein-coding |
| chr16_57558159_57558578 | chr16 | 57558160 | 57558578 | + | 1.17599 | intron (NM_001177871, intron 1 of 1) | intron (NM_001177871, intron 1 of 1) | 9127 | NM_001177871 | 78749 | Mm.328360 | NM_001040397 | ENSMUSG00000043336 | **Filip1l** | 4631422O05Rik\|Doc1 | filamin A interacting protein 1-like | protein-coding |
| chr16_57574903_57575163 | chr16 | 57574904 | 57575163 | + | 1.09055 | intron (NM_025599, intron 1 of 9) | intron (NM_025599, intron 1 of 9) | 25791 | NM_001177871 | 78749 | Mm.328360 | NM_001040397 | ENSMUSG00000043336 | **Filip1l** | 4631422O05Rik\|Doc1 | filamin A interacting protein 1-like | protein-coding |
| chr16_57431324_57431726 | chr16 | 57431325 | 57431726 | + | 0.975474 | intron (NM_001040397, intron 1 of 4) | intron (NM_001040397, intron 1 of 4) | 78248 | NM_001040397 | 78749 | Mm.328360 | NM_001040397 | ENSMUSG00000043336 | **Filip1l** | 4631422O05Rik\|Doc1 | filamin A interacting protein 1-like | protein-coding |
| chr16_57431270_57431881 | chr16 | 57431271 | 57431881 | + | 0.952645 | intron (NM_001040397, intron 1 of 4) | intron (NM_001040397, intron 1 of 4) | 78299 | NM_001040397 | 78749 | Mm.328360 | NM_001040397 | ENSMUSG00000043336 | **Filip1l** | 4631422O05Rik\|Doc1 | filamin A interacting protein 1-like | protein-coding |
| chr16_57557010_57557374 | chr16 | 57557011 | 57557374 | + | 0.894881 | intron (NM_001177871, intron 1 of 1) | intron (NM_001177871, intron 1 of 1) | 7950 | NM_001177871 | 78749 | Mm.328360 | NM_001040397 | ENSMUSG00000043336 | **Filip1l** | 4631422O05Rik\|Doc1 | filamin A interacting protein 1-like | protein-coding |
| chr16_57486303_57486698 | chr16 | 57486304 | 57486698 | + | 0.844671 | intron (NM_001040397, intron 1 of 4) | intron (NM_001040397, intron 1 of 4) | -62741 | NM_001177871 | 78749 | Mm.328360 | NM_001040397 | ENSMUSG00000043336 | **Filip1l** | 4631422O05Rik\|Doc1 | filamin A interacting protein 1-like | protein-coding |
| chr16_57500813_57501230 | chr16 | 57500814 | 57501230 | + | 0.781782 | intron (NM_001040397, intron 1 of 4) | intron (NM_001040397, intron 1 of 4) | -48220 | NM_001177871 | 78749 | Mm.328360 | NM_001040397 | ENSMUSG00000043336 | **Filip1l** | 4631422O05Rik\|Doc1 | filamin A interacting protein 1-like | protein-coding |
| chr16_57556338_57556824 | chr16 | 57556339 | 57556824 | + | 0.717035 | intron (NM_001177871, intron 1 of 1) | intron (NM_001177871, intron 1 of 1) | 7339 | NM_001177871 | 78749 | Mm.328360 | NM_001040397 | ENSMUSG00000043336 | **Filip1l** | 4631422O05Rik\|Doc1 | filamin A interacting protein 1-like | protein-coding |
| chr5_74533991_74534188 | chr5 | 74533992 | 74534188 | + | 0.87126 | Intergenic | Intergenic | -1392 | NM_001159573 | 66899 | Mm.272468 | NM_024183 | ENSMUSG00000029227 | **Fip1l1** | 1300019H17Rik\|Rje | FIP1 like 1 (S. cerevisiae) | protein-coding |
| chrX_50632030_50632610 | chrX | 50632031 | 50632610 | + | 0.795583 | intron (NR_026976, intron 1 of 23) | intron (NR_026976, intron 1 of 23) | 2938 | NR_015505 | 103012 | Mm.482523 | NR_015505 | | **Firre** | 5830467J12Rik\|6720401G13Rik\|AW048145 | functional intergenic repeating RNA element | ncRNA |
| chr2_151540444_151540768 | chr2 | 151540445 | 151540768 | + | 1.319 | Intergenic | RLTR24\|LTR\|ERV1 | -1893 | NM_008019 | 14225 | Mm.278458 | NM_008019 | ENSMUSG00000032966 | **Fkbp1a** | 12kDa\|FKBP12\|FKBP12-T1\|FKBP12-T2\|Fkbp\|Fkbp1\|mFKBP1\|mFKBP12 | FK506 binding protein 1a | protein-coding |
| chr2_151533657_151534073 | chr2 | 151533658 | 151534073 | + | 1.23529 | Intergenic | Intergenic | -8634 | NM_008019 | 14225 | Mm.278458 | NM_008019 | ENSMUSG00000032966 | **Fkbp1a** | 12kDa\|FKBP12\|FKBP12-T1\|FKBP12-T2\|Fkbp\|Fkbp1\|mFKBP1\|mFKBP12 | FK506 binding protein 1a | protein-coding |
| chr2_151549719_151549982 | chr2 | 151549720 | 151549982 | + | 1.04474 | intron (NM_008019, intron 2 of 4) | intron (NM_008019, intron 2 of 4) | 7352 | NM_008019 | 14225 | Mm.278458 | NM_008019 | ENSMUSG00000032966 | **Fkbp1a** | 12kDa\|FKBP12\|FKBP12-T1\|FKBP12-T2\|Fkbp\|Fkbp1\|mFKBP1\|mFKBP12 | FK506 binding protein 1a | protein-coding |

| | | | | | | | | | | | | | | | | | |
|---|---|---|---|---|---|---|---|---|---|---|---|---|---|---|---|---|---|
| chr2_151549719_151550170 | chr2 | 151549720 | 151550170 | + | 0.996272 | intron (NM_008019, intron 2 of 4) | intron (NM_008019, intron 2 of 4) | 7446 | NM_008019 | 14225 | Mm.278458 | NM_008019 | ENSMUSG00000032966 | **Fkbp1a** | 12kDa\|FKBP12\|FKBP12-T1\|FKBP12-T2\|Fkbp\|Fkbp1\|mFKBP1\|mFKBP12 | FK506 binding protein 1a | protein-coding |
| chr2_151549515_151550174 | chr2 | 151549516 | 151550174 | + | 0.956663 | intron (NM_008019, intron 2 of 4) | intron (NM_008019, intron 2 of 4) | 7346 | NM_008019 | 14225 | Mm.278458 | NM_008019 | ENSMUSG00000032966 | **Fkbp1a** | 12kDa\|FKBP12\|FKBP12-T1\|FKBP12-T2\|Fkbp\|Fkbp1\|mFKBP1\|mFKBP12 | FK506 binding protein 1a | protein-coding |
| chr17_28500196_28500792 | chr17 | 28500197 | 28500792 | + | 1.01504 | Intergenic | RSINE1\|SINE\|B4 | -14345 | NM_010220 | 14229 | Mm.276405 | NM_010220 | ENSMUSG00000024222 | **Fkbp5** | D17Ertd592e\|Dit1\|FKBP-5\|FKBP51 | FK506 binding protein 5 | protein-coding |
| chr17_28488204_28488676 | chr17 | 28488205 | 28488676 | + | 0.812832 | Intergenic | Intergenic | -2291 | NM_010220 | 14229 | Mm.276405 | NM_010220 | ENSMUSG00000024222 | **Fkbp5** | D17Ertd592e\|Dit1\|FKBP-5\|FKBP51 | FK506 binding protein 5 | protein-coding |
| chr4_53718131_53718427 | chr4 | 53718132 | 53718427 | + | 1.35272 | intron (NM_139309, intron 1 of 9) | B3A\|SINE\|B2 | 4097 | NM_139309 | 246179 | Mm.247210 | NM_139309 | ENSMUSG00000028414 | **Fktn** | D830030O17Rik\|Fcmd | fukutin | protein-coding |
| chr9_32533853_32534416 | chr9 | 32533854 | 32534416 | + | 1.37666 | intron (NM_008026, intron 1 of 8) | intron (NM_008026, intron 1 of 8) | 7317 | NM_008026 | 14247 | Mm.258908 | NM_008026 | ENSMUSG00000016087 | **Fli1** | EWSR2\|Fli-1\|SIC-1\|Sic1 | Friend leukemia integration 1 | protein-coding |
| chr9_32510470_32510898 | chr9 | 32510471 | 32510898 | + | 0.917744 | intron (NM_008026, intron 1 of 8) | intron (NM_008026, intron 1 of 8) | 30768 | NM_008026 | 14247 | Mm.258908 | NM_008026 | ENSMUSG00000016087 | **Fli1** | EWSR2\|Fli-1\|SIC-1\|Sic1 | Friend leukemia integration 1 | protein-coding |
| chr9_32487835_32488221 | chr9 | 32487836 | 32488221 | + | 0.86122 | intron (NM_008026, intron 1 of 8) | B4A\|SINE\|B4 | 53424 | NM_008026 | 14247 | Mm.258908 | NM_008026 | ENSMUSG00000016087 | **Fli1** | EWSR2\|Fli-1\|SIC-1\|Sic1 | Friend leukemia integration 1 | protein-coding |
| chr9_32521901_32522358 | chr9 | 32521902 | 32522358 | + | 0.854782 | intron (NM_008026, intron 1 of 8) | intron (NM_008026, intron 1 of 8) | 19322 | NM_008026 | 14247 | Mm.258908 | NM_008026 | ENSMUSG00000016087 | **Fli1** | EWSR2\|Fli-1\|SIC-1\|Sic1 | Friend leukemia integration 1 | protein-coding |
| chr9_32486735_32487080 | chr9 | 32486736 | 32487080 | + | 0.798495 | intron (NM_008026, intron 1 of 8) | intron (NM_008026, intron 1 of 8) | 54544 | NM_008026 | 14247 | Mm.258908 | NM_008026 | ENSMUSG00000016087 | **Fli1** | EWSR2\|Fli-1\|SIC-1\|Sic1 | Friend leukemia integration 1 | protein-coding |
| chr9_32494692_32495136 | chr9 | 32494693 | 32495136 | + | 0.70569 | intron (NM_008026, intron 1 of 8) | intron (NM_008026, intron 1 of 8) | 46538 | NM_008026 | 14247 | Mm.258908 | NM_008026 | ENSMUSG00000016087 | **Fli1** | EWSR2\|Fli-1\|SIC-1\|Sic1 | Friend leukemia integration 1 | protein-coding |
| chr9_32494813_32495130 | chr9 | 32494814 | 32495130 | + | 0.697002 | intron (NM_008026, intron 1 of 8) | intron (NM_008026, intron 1 of 8) | 46480 | NM_008026 | 14247 | Mm.258908 | NM_008026 | ENSMUSG00000016087 | **Fli1** | EWSR2\|Fli-1\|SIC-1\|Sic1 | Friend leukemia integration 1 | protein-coding |
| chr14_7857814_7858212 | chr14 | 7857815 | 7858212 | + | 0.851172 | intron (NM_001081427, intron 1 of 45) | intron (NM_001081427, intron 1 of 45) | 40056 | NM_134080 | 286940 | Mm.489652 | NM_134080 | ENSMUSG00000025278 | **Flnb** | AL024016\|Fln-b | filamin, beta | protein-coding |
| chr6_29449923_29450248 | chr6 | 29449924 | 29450248 | + | 0.814342 | intron (NM_001081185, intron 24 of 47) | MLTR31C_MM\|LTR\|ERVK | 16933 | NM_001081185 | 68794 | Mm.39046 | NM_001081185 | ENSMUSG00000068699 | **Flnc** | 1110055E19Rik\|ABP-280\|ABPL\|Fln2 | filamin C, gamma | protein-coding |
| chr7_45134477_45135172 | chr7 | 45134478 | 45135172 | + | 0.590247 | intron (NR_102725, intron 4 of 7) | intron (NR_102725, intron 4 of 7) | 1607 | NM_013520 | 14256 | Mm.1658 | NM_013520 | ENSMUSG00000089989 | **Flt3l** | Flt3lg\|Ly72L | FMS-like tyrosine kinase 3 ligand | protein-coding |
| chr2_52938237_52938672 | chr2 | 52938238 | 52938672 | + | 1.14779 | intron (NM_172409, intron 1 of 26) | intron (NM_172409, intron 1 of 26) | 80587 | NM_172409 | 71409 | Mm.394427 | NM_172409 | ENSMUSG00000036053 | **Fmnl2** | 5430425K04Rik\|Man | formin-like 2 | protein-coding |
| chr2_52938071_52938684 | chr2 | 52938072 | 52938684 | + | 1.07514 | intron (NM_172409, intron 1 of 26) | intron (NM_172409, intron 1 of 26) | 80510 | NM_172409 | 71409 | Mm.394427 | NM_172409 | ENSMUSG00000036053 | **Fmnl2** | 5430425K04Rik\|Man | formin-like 2 | protein-coding |
| chr2_52861811_52862356 | chr2 | 52861812 | 52862356 | + | 0.907836 | intron (NM_172409, intron 1 of 26) | intron (NM_172409, intron 1 of 26) | 4216 | NM_172409 | 71409 | Mm.394427 | NM_172409 | ENSMUSG00000036053 | **Fmnl2** | 5430425K04Rik\|Man | formin-like 2 | protein-coding |
| chr2_52891832_52892141 | chr2 | 52891833 | 52892141 | + | 0.907076 | intron (NM_172409, intron 1 of 26) | intron (NM_172409, intron 1 of 26) | 34119 | NM_172409 | 71409 | Mm.394427 | NM_172409 | ENSMUSG00000036053 | **Fmnl2** | 5430425K04Rik\|Man | formin-like 2 | protein-coding |
| chr2_52885023_52885581 | chr2 | 52885024 | 52885581 | + | 0.786792 | intron (NM_172409, intron 1 of 26) | intron (NM_172409, intron 1 of 26) | 27434 | NM_172409 | 71409 | Mm.394427 | NM_172409 | ENSMUSG00000036053 | **Fmnl2** | 5430425K04Rik\|Man | formin-like 2 | protein-coding |
| chr2_31060748_31061034 | chr2 | 31060749 | 31061034 | + | 1.07235 | intron (NM_001038700, intron 6 of 14) | intron (NM_001038700, intron 6 of 14) | 35609 | NM_001177650 | 14269 | Mm.52297 | NM_019406 | ENSMUSG00000075415 | **Fnbp1** | 1110057E06Rik\|2210010H06Rik\|FBP1\|Fbp17 | formin binding protein 1 | protein-coding |
| chr2_31068815_31069302 | chr2 | 31068816 | 31069302 | + | 0.879657 | intron (NM_001038700, intron 6 of 14) | intron (NM_001038700, intron 6 of 14) | 27441 | NM_001177650 | 14269 | Mm.52297 | NM_019406 | ENSMUSG00000075415 | **Fnbp1** | 1110057E06Rik\|2210010H06Rik\|FBP1\|Fbp17 | formin binding protein 1 | protein-coding |
| chr2_31042637_31043405 | chr2 | 31042638 | 31043405 | + | 0.73998 | intron (NM_001038700, intron 9 of 14) | intron (NM_001038700, intron 9 of 14) | 53479 | NM_001177650 | 14269 | Mm.52297 | NM_019406 | ENSMUSG00000075415 | **Fnbp1** | 1110057E06Rik\|2210010H06Rik\|FBP1\|Fbp17 | formin binding protein 1 | protein-coding |
| chr14_72807162_72807692 | chr14 | 72807163 | 72807692 | + | 1.81535 | Intergenic | Intergenic | -97424 | NM_207636 | 319448 | Mm.205421 | NM_207636 | ENSMUSG00000033487 | **Fndc3a** | 1700094E19Rik\|D14Ertd453e\|F730017H24Rik\|Fndc3\|sys | fibronectin type III domain containing 3A | protein-coding |
| chr14_72752619_72753367 | chr14 | 72752620 | 72753367 | + | 1.49275 | Intergenic | Intergenic | -42990 | NM_207636 | 319448 | Mm.205421 | NM_207636 | ENSMUSG00000033487 | **Fndc3a** | 1700094E19Rik\|D14Ertd453e\|F730017H24Rik\|Fndc3\|sys | fibronectin type III domain containing 3A | protein-coding |
| chr14_72741696_72742346 | chr14 | 72741697 | 72742346 | + | 1.40296 | Intergenic | Intergenic | -32018 | NM_207636 | 319448 | Mm.205421 | NM_207636 | ENSMUSG00000033487 | **Fndc3a** | 1700094E19Rik\|D14Ertd453e\|F730017H24Rik\|Fndc3\|sys | fibronectin type III domain containing 3A | protein-coding |
| chr14_72777785_72778473 | chr14 | 72777786 | 72778473 | + | 1.11137 | Intergenic | Intergenic | -68126 | NM_207636 | 319448 | Mm.205421 | NM_207636 | ENSMUSG00000033487 | **Fndc3a** | 1700094E19Rik\|D14Ertd453e\|F730017H24Rik\|Fndc3\|sys | fibronectin type III domain containing 3A | protein-coding |
| chr14_72777940_72778300 | chr14 | 72777941 | 72778300 | + | 1.03378 | Intergenic | Intergenic | -68117 | NM_207636 | 319448 | Mm.205421 | NM_207636 | ENSMUSG00000033487 | **Fndc3a** | 1700094E19Rik\|D14Ertd453e\|F730017H24Rik\|Fndc3\|sys | fibronectin type III domain containing 3A | protein-coding |
| chr14_72740790_72741369 | chr14 | 72740791 | 72741369 | + | 0.889708 | Intergenic | Intergenic | -31077 | NM_207636 | 319448 | Mm.205421 | NM_207636 | ENSMUSG00000033487 | **Fndc3a** | 1700094E19Rik\|D14Ertd453e\|F730017H24Rik\|Fndc3\|sys | fibronectin type III domain containing 3A | protein-coding |
| chr14_72779717_72779982 | chr14 | 72779718 | 72779982 | + | 0.882399 | Intergenic | Intergenic | -69847 | NM_207636 | 319448 | Mm.205421 | NM_207636 | ENSMUSG00000033487 | **Fndc3a** | 1700094E19Rik\|D14Ertd453e\|F730017H24Rik\|Fndc3\|sys | fibronectin type III domain containing 3A | protein-coding |
| chr14_72809216_72809462 | chr14 | 72809217 | 72809462 | + | 0.873215 | Intergenic | Intergenic | -99336 | NM_207636 | 319448 | Mm.205421 | NM_207636 | ENSMUSG00000033487 | **Fndc3a** | 1700094E19Rik\|D14Ertd453e\|F730017H24Rik\|Fndc3\|sys | fibronectin type III domain containing 3A | protein-coding |
| chr14_72737668_72737972 | chr14 | 72737669 | 72737972 | + | 0.863793 | Intergenic | RMER4A\|LTR\|ERVK | -27817 | NM_207636 | 319448 | Mm.205421 | NM_207636 | ENSMUSG00000033487 | **Fndc3a** | 1700094E19Rik\|D14Ertd453e\|F730017H24Rik\|Fndc3\|sys | fibronectin type III domain containing 3A | protein-coding |
| chr14_72862957_72863186 | chr14 | 72862958 | 72863186 | + | 0.692131 | Intergenic | Intergenic | -153069 | NM_207636 | 319448 | Mm.205421 | NM_207636 | ENSMUSG00000033487 | **Fndc3a** | 1700094E19Rik\|D14Ertd453e\|F730017H24Rik\|Fndc3\|sys | fibronectin type III domain containing 3A | protein-coding |
| chr3_108886833_108887507 | chr3 | 108886834 | 108887507 | + | 1.74657 | intron (NM_177091, intron 2 of 12) | URR1B\|DNA\|hAT-Charlie | 2838 | NM_177091 | 320181 | Mm.244233 | NM_177091 | ENSMUSG00000045326 | **Fndc7** | E230011A21Rik | fibronectin type III domain containing 7 | protein-coding |
| chr3_108881958_108882224 | chr3 | 108881959 | 108882224 | + | 0.84254 | intron (NM_177091, intron 3 of 12) | intron (NM_177091, intron 3 of 12) | 7917 | NM_177091 | 320181 | Mm.244233 | NM_177091 | ENSMUSG00000045326 | **Fndc7** | E230011A21Rik | fibronectin type III domain containing 7 | protein-coding |
| chr11_54427813_54428675 | chr11 | 54427814 | 54428675 | + | 1.10856 | Intergenic | B4A\|SINE\|B4 | -9935 | NM_173753 | 216742 | Mm.34087 | NM_173753 | ENSMUSG00000035992 | **Fnip1** | A730024A03Rik\|AI838773\|AW557298 | folliculin interacting protein 1 | protein-coding |
| chr11_54464469_54465260 | chr11 | 54464470 | 54465260 | + | 0.637685 | intron (NM_173753, intron 1 of 17) | intron (NM_173753, intron 1 of 17) | 26686 | NM_173753 | 216742 | Mm.34087 | NM_173753 | ENSMUSG00000035992 | **Fnip1** | A730024A03Rik\|AI838773\|AW557298 | folliculin interacting protein 1 | protein-coding |
| chr3_79542602_79543022 | chr3 | 79542603 | 79543022 | + | 0.708424 | intron (NM_001162999, intron 1 of 16) | intron (NM_001162999, intron 1 of 16) | 24867 | NM_001162999 | 329679 | Mm.244005 | NM_001162999 | ENSMUSG00000061175 | **Fnip2** | D630023B12Rik\|mKIAA1450 | folliculin interacting protein 2 | protein-coding |
| chr4_88066572_88066943 | chr4 | 88066573 | 88066943 | + | 0.71827 | Intergenic | MTD\|LTR\|ERVL-MaLR | -27872 | NM_001081184 | 230393 | Mm.287210 | NM_001081184 | ENSMUSG00000038368 | **Focad** | Kiaa1797\|mKIAA1797 | focadhesin | protein-coding |
| chr9_14921206_14921713 | chr9 | 14921207 | 14921713 | + | 1.08132 | Intergenic | (ACTG)n\|Simple_repeat\|Simple_repeat | -17509 | NM_022888 | 64931 | Mm.86738 | NM_022888 | ENSMUSG00000031933 | **Folr4** | 0910001L11Rik\|C86255\|Folbp3\|Juno | folate receptor 4 (delta) | protein-coding |
| chr9_14965380_14965813 | chr9 | 14965381 | 14965813 | + | 0.998004 | Intergenic | L1ME3A\|LINE\|L1 | -61646 | NM_022888 | 64931 | Mm.86738 | NM_022888 | ENSMUSG00000031933 | **Folr4** | 0910001L11Rik\|C86255\|Folbp3\|Juno | folate receptor 4 (delta) | protein-coding |
| chr9_14965231_14965998 | chr9 | 14965232 | 14965998 | + | 0.67165 | Intergenic | L1ME3A\|LINE\|L1 | -61664 | NM_022888 | 64931 | Mm.86738 | NM_022888 | ENSMUSG00000031933 | **Folr4** | 0910001L11Rik\|C86255\|Folbp3\|Juno | folate receptor 4 (delta) | protein-coding |
| chr7_19325727_19326162 | chr7 | 19325728 | 19326162 | + | 1.0474 | Intergenic | Intergenic | -15900 | NM_008036 | 14282 | Mm.248335 | NM_008036 | ENSMUSG00000003545 | **Fosb** | - | FBJ osteosarcoma oncogene B | protein-coding |
| chr7_19321442_19321827 | chr7 | 19321443 | 19321827 | + | 0.732902 | Intergenic | Intergenic | -11590 | NM_008036 | 14282 | Mm.248335 | NM_008036 | ENSMUSG00000003545 | **Fosb** | - | FBJ osteosarcoma oncogene B | protein-coding |
| chr7_19321546_19321818 | chr7 | 19321547 | 19321818 | + | 0.712348 | Intergenic | Intergenic | -11637 | NM_008036 | 14282 | Mm.248335 | NM_008036 | ENSMUSG00000003545 | **Fosb** | - | FBJ osteosarcoma oncogene B | protein-coding |
| chr5_32055689_32056061 | chr5 | 32055690 | 32056061 | + | 1.77604 | intron (NM_144541, intron 10 of 11) | intron (NM_144541, intron 10 of 11) | -80597 | NM_008037 | 14284 | Mm.24684 | NM_008037 | ENSMUSG00000029135 | **Fosl2** | Fra-2 | fos-like antigen 2 | protein-coding |
| chr5_32055655_32056166 | chr5 | 32055656 | 32056166 | + | 1.68504 | intron (NM_144541, intron 10 of 11) | intron (NM_144541, intron 10 of 11) | -80561 | NM_008037 | 14284 | Mm.24684 | NM_008037 | ENSMUSG00000029135 | **Fosl2** | Fra-2 | fos-like antigen 2 | protein-coding |
| chr5_32013076_32014111 | chr5 | 32013077 | 32014111 | + | 1.23852 | intron (NM_144541, intron 10 of 11) | intron (NM_144541, intron 10 of 11) | -122878 | NM_008037 | 14284 | Mm.24684 | NM_008037 | ENSMUSG00000029135 | **Fosl2** | Fra-2 | fos-like antigen 2 | protein-coding |

| | | | | | | | | | | | | | | | | | |
|---|---|---|---|---|---|---|---|---|---|---|---|---|---|---|---|---|---|
| chr5_32035661_32036249 | chr5 | 32035662 | 32036249 | + | 0.984279 | intron (NM_144541, intron 10 of 11) | intron (NM_144541, intron 10 of 11) | -100517 | NM_008037 | 14284 | Mm.24684 | NM_008037 | ENSMUSG00000029135 | **Fosl2** | Fra-2 | fos-like antigen 2 | protein-coding |
| chr5_32052393_32052915 | chr5 | 32052394 | 32052915 | + | 0.820321 | intron (NM_144541, intron 10 of 11) | ID4_\|SINE\|ID | -83818 | NM_008037 | 14284 | Mm.24684 | NM_008037 | ENSMUSG00000029135 | **Fosl2** | Fra-2 | fos-like antigen 2 | protein-coding |
| chr12_57546393_57547201 | chr12 | 57546394 | 57547201 | + | 0.627729 | promoter-TSS (NM_008259) | promoter-TSS (NM_008259) | -676 | NM_008259 | 15375 | Mm.4578 | NM_008259 | ENSMUSG00000035451 | **Foxa1** | Hnf-3a\|Hnf3a\|Tcf-3a\|Tcf3a | forkhead box A1 | protein-coding |
| chr9_69666689_69667416 | chr9 | 69666690 | 69667416 | + | 1.69693 | Intergenic | Intergenic | 93887 | NM_022378 | 64290 | Mm.440890 | NM_022378 | ENSMUSG00000059246 | **Foxb1** | C43\|Fkh5\|Foxb1a\|Foxb1b\|Hfh-e5.1\|Mf3\|Twh | forkhead box B1 | protein-coding |
| chr9_69702059_69702835 | chr9 | 69702060 | 69702835 | + | 1.18333 | Intergenic | Intergenic | 58493 | NM_022378 | 64290 | Mm.440890 | NM_022378 | ENSMUSG00000059246 | **Foxb1** | C43\|Fkh5\|Foxb1a\|Foxb1b\|Hfh-e5.1\|Mf3\|Twh | forkhead box B1 | protein-coding |
| chr9_69651627_69652157 | chr9 | 69651628 | 69652157 | + | 0.932221 | Intergenic | Intergenic | 109048 | NM_022378 | 64290 | Mm.440890 | NM_022378 | ENSMUSG00000059246 | **Foxb1** | C43\|Fkh5\|Foxb1a\|Foxb1b\|Hfh-e5.1\|Mf3\|Twh | forkhead box B1 | protein-coding |
| chr9_69638528_69639222 | chr9 | 69638529 | 69639222 | + | 0.917785 | Intergenic | Intergenic | 122065 | NM_022378 | 64290 | Mm.440890 | NM_022378 | ENSMUSG00000059246 | **Foxb1** | C43\|Fkh5\|Foxb1a\|Foxb1b\|Hfh-e5.1\|Mf3\|Twh | forkhead box B1 | protein-coding |
| chr19_16898365_16899126 | chr19 | 16898366 | 16899126 | + | 1.27167 | Intergenic | Intergenic | -24916 | NM_008023 | 14240 | Mm.7971 | NM_008023 | ENSMUSG00000056829 | **Foxb2** | Fkh4 | forkhead box B2 | protein-coding |
| chr19_16908873_16909153 | chr19 | 16908874 | 16909153 | + | 0.707515 | Intergenic | MLT1-int\|LTR\|ERVL-MaLR | -35183 | NM_008023 | 14240 | Mm.7971 | NM_008023 | ENSMUSG00000056829 | **Foxb2** | Fkh4 | forkhead box B2 | protein-coding |
| chr13_32071561_32072149 | chr13 | 32071562 | 32072149 | + | 0.646637 | intron (NM_146041, intron 7 of 10) | intron (NM_146041, intron 7 of 10) | 265209 | NM_008592 | 17300 | Mm.12949 | NM_008592 | ENSMUSG00000050295 | **Foxc1** | FREAC3\|Fkh1\|Mf1\|Mf4\|ch\|fkh-1\|frkhda | forkhead box C1 | protein-coding |
| chr5_142407378_142407737 | chr5 | 142407379 | 142407737 | + | 0.893974 | intron (NM_199068, intron 1 of 8) | B1F1\|SINE\|Alu | 6061 | NM_199068 | 17425 | Mm.24214 | NM_010812 | ENSMUSG00000056493 | **Foxk1** | A630048H08Rik\|AI463295\|ENSMUSG00000075577\|Gm10868\|Mnf | forkhead box K1 | protein-coding |
| chr12_99325931_99326877 | chr12 | 99325932 | 99326877 | + | 1.78469 | intron (NM_183186, intron 3 of 5) | MER112\|DNA\|hAT-Charlie | 123670 | NM_183186 | 71375 | Mm.341972 | NM_183186 | ENSMUSG00000033713 | **Foxn3** | 5430426H20Rik\|AA589593\|AW556347\|Ches1\|Ches1l\|HTLFL1 | forkhead box N3 | protein-coding |
| chr12_99510119_99510476 | chr12 | 99510120 | 99510476 | + | 1.75708 | Intergenic | B3\|SINE\|B2 | -60224 | NM_183186 | 71375 | Mm.341972 | NM_183186 | ENSMUSG00000033713 | **Foxn3** | 5430426H20Rik\|AA589593\|AW556347\|Ches1\|Ches1l\|HTLFL1 | forkhead box N3 | protein-coding |
| chr12_99509971_99510622 | chr12 | 99509972 | 99510622 | + | 1.7381 | Intergenic | B3\|SINE\|B2 | -60223 | NM_183186 | 71375 | Mm.341972 | NM_183186 | ENSMUSG00000033713 | **Foxn3** | 5430426H20Rik\|AA589593\|AW556347\|Ches1\|Ches1l\|HTLFL1 | forkhead box N3 | protein-coding |
| chr12_99394174_99394616 | chr12 | 99394175 | 99394616 | + | 1.00829 | intron (NM_183186, intron 1 of 5) | intron (NM_183186, intron 1 of 5) | 55679 | NM_183186 | 71375 | Mm.341972 | NM_183186 | ENSMUSG00000033713 | **Foxn3** | 5430426H20Rik\|AA589593\|AW556347\|Ches1\|Ches1l\|HTLFL1 | forkhead box N3 | protein-coding |
| chr12_99340491_99341116 | chr12 | 99340492 | 99341116 | + | 0.678635 | intron (NM_183186, intron 3 of 5) | intron (NM_183186, intron 3 of 5) | 109270 | NM_183186 | 71375 | Mm.341972 | NM_183186 | ENSMUSG00000033713 | **Foxn3** | 5430426H20Rik\|AA589593\|AW556347\|Ches1\|Ches1l\|HTLFL1 | forkhead box N3 | protein-coding |
| chr12_99380784_99381640 | chr12 | 99380785 | 99381640 | + | 0.677892 | intron (NM_183186, intron 2 of 5) | Lx4A\|LINE\|L1 | 68862 | NM_183186 | 71375 | Mm.341972 | NM_183186 | ENSMUSG00000033713 | **Foxn3** | 5430426H20Rik\|AA589593\|AW556347\|Ches1\|Ches1l\|HTLFL1 | forkhead box N3 | protein-coding |
| chr12_99380780_99381349 | chr12 | 99380781 | 99381349 | + | 0.634295 | intron (NM_183186, intron 2 of 5) | Lx4A\|LINE\|L1 | 69009 | NM_183186 | 71375 | Mm.341972 | NM_183186 | ENSMUSG00000033713 | **Foxn3** | 5430426H20Rik\|AA589593\|AW556347\|Ches1\|Ches1l\|HTLFL1 | forkhead box N3 | protein-coding |
| chr12_99350293_99350614 | chr12 | 99350294 | 99350614 | + | 0.6266 | intron (NM_183186, intron 2 of 5) | intron (NM_183186, intron 2 of 5) | 99620 | NM_183186 | 71375 | Mm.341972 | NM_183186 | ENSMUSG00000033713 | **Foxn3** | 5430426H20Rik\|AA589593\|AW556347\|Ches1\|Ches1l\|HTLFL1 | forkhead box N3 | protein-coding |
| chr3_52369119_52369605 | chr3 | 52369120 | 52369605 | + | 1.27067 | Intergenic | Intergenic | 101025 | NM_019739 | 56458 | Mm.29891 | NM_019739 | ENSMUSG00000044167 | **Foxo1** | AI876417\|Afxh\|FKHR\|Fkhr1\|Foxo1a | forkhead box O1 | protein-coding |
| chr3_52271412_52271819 | chr3 | 52271413 | 52271819 | + | 1.23473 | intron (NM_019739, intron 1 of 2) | intron (NM_019739, intron 1 of 2) | 3279 | NM_019739 | 56458 | Mm.29891 | NM_019739 | ENSMUSG00000044167 | **Foxo1** | AI876417\|Afxh\|FKHR\|Fkhr1\|Foxo1a | forkhead box O1 | protein-coding |
| chr3_52270053_52270926 | chr3 | 52270054 | 52270926 | + | 0.76343 | intron (NM_019739, intron 1 of 2) | intron (NM_019739, intron 1 of 2) | 2153 | NM_019739 | 56458 | Mm.29891 | NM_019739 | ENSMUSG00000044167 | **Foxo1** | AI876417\|Afxh\|FKHR\|Fkhr1\|Foxo1a | forkhead box O1 | protein-coding |
| chr3_52443820_52444293 | chr3 | 52443821 | 52444293 | + | 0.640022 | Intergenic | Intergenic | 175720 | NM_019739 | 56458 | Mm.29891 | NM_019739 | ENSMUSG00000044167 | **Foxo1** | AI876417\|Afxh\|FKHR\|Fkhr1\|Foxo1a | forkhead box O1 | protein-coding |
| chr10_42216557_42216938 | chr10 | 42216558 | 42216938 | + | 0.840318 | intron (NM_019740, intron 2 of 3) | intron (NM_019740, intron 2 of 3) | 59994 | NM_019740 | 56484 | Mm.338613 | NM_019740 | ENSMUSG00000048756 | **Foxo3** | 1110048B16Rik\|2010203A17Rik\|C76856\|FKHRL1\|Fkhr2\|Foxo3a | forkhead box O3 | protein-coding |
| chr10_42261502_42262307 | chr10 | 42261503 | 42262307 | + | 0.736523 | intron (NM_019740, intron 2 of 3) | intron (NM_019740, intron 2 of 3) | 14837 | NM_019740 | 56484 | Mm.338613 | NM_019740 | ENSMUSG00000048756 | **Foxo3** | 1110048B16Rik\|2010203A17Rik\|C76856\|FKHRL1\|Fkhr2\|Foxo3a | forkhead box O3 | protein-coding |
| chr6_99405904_99406677 | chr6 | 99405905 | 99406677 | + | 1.7683 | intron (NM_053202, intron 1 of 18) | intron (NM_053202, intron 1 of 18) | 29054 | NM_053202 | 108655 | Mm.234965 | NM_053202 | ENSMUSG00000030067 | **Foxp1** | 3110052D19Rik\|4932443N09Rik\|AI461938\|AW494214 | forkhead box P1 | protein-coding |
| chr6_99045813_99046203 | chr6 | 99045814 | 99046203 | + | 1.7481 | intron (NM_053202, intron 5 of 18) | MIR3\|SINE\|MIR | -17710 | NM_001197322 | 108655 | Mm.234965 | NM_053202 | ENSMUSG00000030067 | **Foxp1** | 3110052D19Rik\|4932443N09Rik\|AI461938\|AW494214 | forkhead box P1 | protein-coding |
| chr6_99340975_99341315 | chr6 | 99340976 | 99341315 | + | 1.47722 | intron (NM_053202, intron 1 of 18) | intron (NM_053202, intron 1 of 18) | -74630 | NM_001197321 | 108655 | Mm.234965 | NM_053202 | ENSMUSG00000030067 | **Foxp1** | 3110052D19Rik\|4932443N09Rik\|AI461938\|AW494214 | forkhead box P1 | protein-coding |
| chr6_99274080_99275007 | chr6 | 99274081 | 99275007 | + | 1.38378 | intron (NM_053202, intron 2 of 18) | intron (NM_053202, intron 2 of 18) | -8029 | NM_001197321 | 108655 | Mm.234965 | NM_053202 | ENSMUSG00000030067 | **Foxp1** | 3110052D19Rik\|4932443N09Rik\|AI461938\|AW494214 | forkhead box P1 | protein-coding |
| chr6_99259740_99260101 | chr6 | 99259741 | 99260101 | + | 1.29612 | intron (NM_053202, intron 3 of 18) | intron (NM_053202, intron 3 of 18) | 6594 | NM_001197321 | 108655 | Mm.234965 | NM_053202 | ENSMUSG00000030067 | **Foxp1** | 3110052D19Rik\|4932443N09Rik\|AI461938\|AW494214 | forkhead box P1 | protein-coding |
| chr6_99344861_99345358 | chr6 | 99344862 | 99345358 | + | 1.27838 | intron (NM_053202, intron 1 of 18) | intron (NM_053202, intron 1 of 18) | -78595 | NM_001197321 | 108655 | Mm.234965 | NM_053202 | ENSMUSG00000030067 | **Foxp1** | 3110052D19Rik\|4932443N09Rik\|AI461938\|AW494214 | forkhead box P1 | protein-coding |
| chr6_99187976_99189109 | chr6 | 99187977 | 99189109 | + | 1.25391 | intron (NM_053202, intron 3 of 18) | intron (NM_053202, intron 3 of 18) | 77972 | NM_001197321 | 108655 | Mm.234965 | NM_053202 | ENSMUSG00000030067 | **Foxp1** | 3110052D19Rik\|4932443N09Rik\|AI461938\|AW494214 | forkhead box P1 | protein-coding |
| chr6_99274716_99274990 | chr6 | 99274717 | 99274990 | + | 1.13824 | intron (NM_053202, intron 2 of 18) | intron (NM_053202, intron 2 of 18) | -8338 | NM_001197321 | 108655 | Mm.234965 | NM_053202 | ENSMUSG00000030067 | **Foxp1** | 3110052D19Rik\|4932443N09Rik\|AI461938\|AW494214 | forkhead box P1 | protein-coding |
| chr6_99050233_99051136 | chr6 | 99050234 | 99051136 | + | 1.1148 | intron (NM_053202, intron 5 of 18) | intron (NM_053202, intron 5 of 18) | -22387 | NM_001197322 | 108655 | Mm.234965 | NM_053202 | ENSMUSG00000030067 | **Foxp1** | 3110052D19Rik\|4932443N09Rik\|AI461938\|AW494214 | forkhead box P1 | protein-coding |
| chr6_99056483_99056868 | chr6 | 99056484 | 99056868 | + | 1.08027 | intron (NM_053202, intron 5 of 18) | Lx5c\|LINE\|L1 | -28378 | NM_001197322 | 108655 | Mm.234965 | NM_053202 | ENSMUSG00000030067 | **Foxp1** | 3110052D19Rik\|4932443N09Rik\|AI461938\|AW494214 | forkhead box P1 | protein-coding |
| chr6_99050581_99051109 | chr6 | 99050582 | 99051109 | + | 1.07427 | intron (NM_053202, intron 5 of 18) | intron (NM_053202, intron 5 of 18) | -22547 | NM_001197322 | 108655 | Mm.234965 | NM_053202 | ENSMUSG00000030067 | **Foxp1** | 3110052D19Rik\|4932443N09Rik\|AI461938\|AW494214 | forkhead box P1 | protein-coding |
| chr6_99413631_99414205 | chr6 | 99413632 | 99414205 | + | 1.06921 | intron (NM_053202, intron 1 of 18) | ORR1D1\|LTR\|ERVL-MaLR | 21427 | NM_053202 | 108655 | Mm.234965 | NM_053202 | ENSMUSG00000030067 | **Foxp1** | 3110052D19Rik\|4932443N09Rik\|AI461938\|AW494214 | forkhead box P1 | protein-coding |
| chr6_99419651_99419904 | chr6 | 99419652 | 99419904 | + | 1.02182 | intron (NM_053202, intron 1 of 18) | intron (NM_053202, intron 1 of 18) | 15567 | NM_053202 | 108655 | Mm.234965 | NM_053202 | ENSMUSG00000030067 | **Foxp1** | 3110052D19Rik\|4932443N09Rik\|AI461938\|AW494214 | forkhead box P1 | protein-coding |
| chr6_99060973_99061250 | chr6 | 99060974 | 99061250 | + | 0.976674 | intron (NM_053202, intron 5 of 18) | intron (NM_053202, intron 5 of 18) | -32814 | NM_001197322 | 108655 | Mm.234965 | NM_053202 | ENSMUSG00000030067 | **Foxp1** | 3110052D19Rik\|4932443N09Rik\|AI461938\|AW494214 | forkhead box P1 | protein-coding |
| chr6_99415379_99415790 | chr6 | 99415380 | 99415790 | + | 0.955182 | intron (NM_053202, intron 1 of 18) | intron (NM_053202, intron 1 of 18) | 19760 | NM_053202 | 108655 | Mm.234965 | NM_053202 | ENSMUSG00000030067 | **Foxp1** | 3110052D19Rik\|4932443N09Rik\|AI461938\|AW494214 | forkhead box P1 | protein-coding |
| chr6_99035523_99035998 | chr6 | 99035524 | 99035998 | + | 0.858686 | intron (NM_053202, intron 5 of 18) | intron (NM_053202, intron 5 of 18) | -7463 | NM_001197322 | 108655 | Mm.234965 | NM_053202 | ENSMUSG00000030067 | **Foxp1** | 3110052D19Rik\|4932443N09Rik\|AI461938\|AW494214 | forkhead box P1 | protein-coding |
| chr6_99056482_99057713 | chr6 | 99056483 | 99057713 | + | 0.841319 | intron (NM_053202, intron 5 of 18) | intron (NM_053202, intron 5 of 18) | -28800 | NM_001197322 | 108655 | Mm.234965 | NM_053202 | ENSMUSG00000030067 | **Foxp1** | 3110052D19Rik\|4932443N09Rik\|AI461938\|AW494214 | forkhead box P1 | protein-coding |
| chr6_99035639_99035964 | chr6 | 99035640 | 99035964 | + | 0.836892 | intron (NM_053202, intron 5 of 18) | intron (NM_053202, intron 5 of 18) | -7504 | NM_001197322 | 108655 | Mm.234965 | NM_053202 | ENSMUSG00000030067 | **Foxp1** | 3110052D19Rik\|4932443N09Rik\|AI461938\|AW494214 | forkhead box P1 | protein-coding |
| chr6_98951564_98952130 | chr6 | 98951565 | 98952130 | + | 0.813018 | intron (NM_053202, intron 12 of 18) | intron (NM_053202, intron 12 of 18) | 76451 | NM_001197322 | 108655 | Mm.234965 | NM_053202 | ENSMUSG00000030067 | **Foxp1** | 3110052D19Rik\|4932443N09Rik\|AI461938\|AW494214 | forkhead box P1 | protein-coding |
| chr6_99019340_99019710 | chr6 | 99019341 | 99019710 | + | 0.810276 | intron (NM_053202, intron 5 of 18) | intron (NM_053202, intron 5 of 18) | 8773 | NM_001197322 | 108655 | Mm.234965 | NM_053202 | ENSMUSG00000030067 | **Foxp1** | 3110052D19Rik\|4932443N09Rik\|AI461938\|AW494214 | forkhead box P1 | protein-coding |
| chr6_99418554_99419236 | chr6 | 99418555 | 99419236 | + | 0.795396 | intron (NM_053202, intron 1 of 18) | intron (NM_053202, intron 1 of 18) | 16450 | NM_053202 | 108655 | Mm.234965 | NM_053202 | ENSMUSG00000030067 | **Foxp1** | 3110052D19Rik\|4932443N09Rik\|AI461938\|AW494214 | forkhead box P1 | protein-coding |
| chr6_98932953_98933507 | chr6 | 98932954 | 98933507 | + | 0.782732 | intron (NM_053202, intron 18 of 18) | intron (NM_053202, intron 18 of 18) | 95068 | NM_001197322 | 108655 | Mm.234965 | NM_053202 | ENSMUSG00000030067 | **Foxp1** | 3110052D19Rik\|4932443N09Rik\|AI461938\|AW494214 | forkhead box P1 | protein-coding |
| chr6_99056953_99057669 | chr6 | 99056954 | 99057669 | + | 0.776506 | intron (NM_053202, intron 5 of 18) | intron (NM_053202, intron 5 of 18) | -29013 | NM_001197322 | 108655 | Mm.234965 | NM_053202 | ENSMUSG00000030067 | **Foxp1** | 3110052D19Rik\|4932443N09Rik\|AI461938\|AW494214 | forkhead box P1 | protein-coding |
| chr6_99418554_99419009 | chr6 | 99418555 | 99419009 | + | 0.762603 | intron (NM_053202, intron 1 of 18) | intron (NM_053202, intron 1 of 18) | 16563 | NM_053202 | 108655 | Mm.234965 | NM_053202 | ENSMUSG00000030067 | **Foxp1** | 3110052D19Rik\|4932443N09Rik\|AI461938\|AW494214 | forkhead box P1 | protein-coding |
| chr6_99338601_99338966 | chr6 | 99338602 | 99338966 | + | 0.759052 | intron (NM_053202, intron 1 of 18) | intron (NM_053202, intron 1 of 18) | -72269 | NM_001197321 | 108655 | Mm.234965 | NM_053202 | ENSMUSG00000030067 | **Foxp1** | 3110052D19Rik\|4932443N09Rik\|AI461938\|AW494214 | forkhead box P1 | protein-coding |
| chr6_99519177_99519432 | chr6 | 99519178 | 99519432 | + | 0.736 | Intergenic | Intergenic | -83960 | NM_053202 | 108655 | Mm.234965 | NM_053202 | ENSMUSG00000030067 | **Foxp1** | 3110052D19Rik\|4932443N09Rik\|AI461938\|AW494214 | forkhead box P1 | protein-coding |

| | | | | | | | | | | | | | | | | | |
|---|---|---|---|---|---|---|---|---|---|---|---|---|---|---|---|---|---|
| chr6_98932939_98933730 | chr6 | 98932940 | 98933730 | + | 0.730226 | intron (NM_053202, intron 18 of 18) | intron (NM_053202, intron 18 of 18) | 94963 | NM_001197322 | 108655 | Mm.234965 | NM_053202 | ENSMUSG00000030067 | **Foxp1** | 3110052D19Rik\|4932443N09Rik\|AI461938\|AW494214 | forkhead box P1 | protein-coding |
| chr6_99336471_99336878 | chr6 | 99336472 | 99336878 | + | 0.723344 | intron (NM_053202, intron 1 of 18) | intron (NM_053202, intron 1 of 18) | -70160 | NM_001197321 | 108655 | Mm.234965 | NM_053202 | ENSMUSG00000030067 | **Foxp1** | 3110052D19Rik\|4932443N09Rik\|AI461938\|AW494214 | forkhead box P1 | protein-coding |
| chr6_99432614_99433326 | chr6 | 99432615 | 99433326 | + | 0.721507 | intron (NM_053202, intron 1 of 18) | intron (NM_053202, intron 1 of 18) | 2375 | NM_053202 | 108655 | Mm.234965 | NM_053202 | ENSMUSG00000030067 | **Foxp1** | 3110052D19Rik\|4932443N09Rik\|AI461938\|AW494214 | forkhead box P1 | protein-coding |
| chr6_99122078_99122479 | chr6 | 99122079 | 99122479 | + | 0.721138 | intron (NM_053202, intron 4 of 18) | MIRb\|SINE\|MIR | -93981 | NM_001197322 | 108655 | Mm.234965 | NM_053202 | ENSMUSG00000030067 | **Foxp1** | 3110052D19Rik\|4932443N09Rik\|AI461938\|AW494214 | forkhead box P1 | protein-coding |
| chr6_98992763_98993479 | chr6 | 98992764 | 98993479 | + | 0.7168 | intron (NM_053202, intron 9 of 18) | intron (NM_053202, intron 9 of 18) | 35177 | NM_001197322 | 108655 | Mm.234965 | NM_053202 | ENSMUSG00000030067 | **Foxp1** | 3110052D19Rik\|4932443N09Rik\|AI461938\|AW494214 | forkhead box P1 | protein-coding |
| chr6_99426850_99427432 | chr6 | 99426851 | 99427432 | + | 0.693929 | intron (NM_053202, intron 1 of 18) | intron (NM_053202, intron 1 of 18) | 8204 | NM_053202 | 108655 | Mm.234965 | NM_053202 | ENSMUSG00000030067 | **Foxp1** | 3110052D19Rik\|4932443N09Rik\|AI461938\|AW494214 | forkhead box P1 | protein-coding |
| chr6_99426801_99427831 | chr6 | 99426802 | 99427831 | + | 0.678451 | intron (NM_053202, intron 1 of 18) | intron (NM_053202, intron 1 of 18) | 8029 | NM_053202 | 108655 | Mm.234965 | NM_053202 | ENSMUSG00000030067 | **Foxp1** | 3110052D19Rik\|4932443N09Rik\|AI461938\|AW494214 | forkhead box P1 | protein-coding |
| chr6_99429547_99430045 | chr6 | 99429548 | 99430045 | + | 0.619854 | intron (NM_053202, intron 1 of 18) | intron (NM_053202, intron 1 of 18) | 5549 | NM_053202 | 108655 | Mm.234965 | NM_053202 | ENSMUSG00000030067 | **Foxp1** | 3110052D19Rik\|4932443N09Rik\|AI461938\|AW494214 | forkhead box P1 | protein-coding |
| chr17_47943000_47943688 | chr17 | 47943001 | 47943688 | + | 1.20634 | Intergenic | Intergenic | -18712 | NM_001110825 | 74123 | Mm.240062 | NM_028767 | ENSMUSG00000023991 | **Foxp4** | 1200010K03Rik\|2310007G05Rik\|mFKHLA | forkhead box P4 | protein-coding |
| chr13_31567698_31568019 | chr13 | 31567699 | 31568019 | + | 0.652184 | intron (NR_029439, intron 3 of 5) | intron (NR_029439, intron 3 of 5) | 9689 | NM_008239 | 15220 | Mm.44235 | NM_008239 | ENSMUSG00000038415 | **Foxq1** | HFH-1\|Hfh1\|Hfh1l\|sa | forkhead box Q1 | protein-coding |
| chr17_17904563_17905036 | chr17 | 17904564 | 17905036 | + | 1.16024 | Intergenic | Intergenic | 16976 | NM_008039 | 14289 | Mm.378918 | NM_008039 | ENSMUSG00000052270 | **Fpr2** | E330010I07Rik\|Fpr-rs2 | formyl peptide receptor 2 | protein-coding |
| chr17_17904361_17905051 | chr17 | 17904362 | 17905051 | + | 0.900975 | Intergenic | PB1D10\|SINE\|Alu | 16882 | NM_008039 | 14289 | Mm.378918 | NM_008039 | ENSMUSG00000052270 | **Fpr2** | E330010I07Rik\|Fpr-rs2 | formyl peptide receptor 2 | protein-coding |
| chr3_53580331_53581123 | chr3 | 53580332 | 53581123 | + | 0.69583 | intron (NM_172862, intron 4 of 23) | intron (NM_172862, intron 4 of 23) | 76628 | NM_172862 | 242022 | Mm.38378 | NM_172862 | ENSMUSG00000037016 | **Frem2** | 6030440P17Rik\|8430406N05Rik\|Gm409\|b2b1562Clo\|my\|ne\|nv1 | Fras1 related extracellular matrix protein 2 | protein-coding |
| chr2_4380394_4380765 | chr2 | 4380395 | 4380765 | + | 0.768777 | intron (NM_172475, intron 1 of 22) | intron (NM_172475, intron 1 of 22) | -20396 | NM_001177843 | 209630 | Mm.37932 | NM_172475 | ENSMUSG00000026657 | **Frmd4a** | 2700017I06Rik\|5330420J21\|AI594879\|C230040M21Rik\|Frmd4\|Gm13190\|OTTMUSG00000010806\|mKIAA1294 | FERM domain containing 4A | protein-coding |
| chr12_70698525_70699201 | chr12 | 70698526 | 70699201 | + | 1.42218 | Intergenic | Intergenic | -126651 | NM_028127 | 319710 | Mm.2962 | NM_028127 | ENSMUSG00000048285 | **Frmd6** | 2610019M19Rik\|4930488L10Rik\|AW212977 | FERM domain containing 6 | protein-coding |
| chr12_70698041_70698431 | chr12 | 70698042 | 70698431 | + | 1.29408 | Intergenic | Intergenic | -127278 | NM_028127 | 319710 | Mm.2962 | NM_028127 | ENSMUSG00000048285 | **Frmd6** | 2610019M19Rik\|4930488L10Rik\|AW212977 | FERM domain containing 6 | protein-coding |
| chr12_70850549_70851049 | chr12 | 70850550 | 70851049 | + | 1.03281 | intron (NM_028127, intron 1 of 13) | intron (NM_028127, intron 1 of 13) | 25285 | NM_028127 | 319710 | Mm.2962 | NM_028127 | ENSMUSG00000048285 | **Frmd6** | 2610019M19Rik\|4930488L10Rik\|AW212977 | FERM domain containing 6 | protein-coding |
| chr12_70826593_70826818 | chr12 | 70826594 | 70826818 | + | 0.705001 | intron (NM_028127, intron 1 of 13) | intron (NM_028127, intron 1 of 13) | 1192 | NM_028127 | 319710 | Mm.2962 | NM_028127 | ENSMUSG00000048285 | **Frmd6** | 2610019M19Rik\|4930488L10Rik\|AW212977 | FERM domain containing 6 | protein-coding |
| chr12_70720424_70720905 | chr12 | 70720425 | 70720905 | + | 0.653137 | Intergenic | Intergenic | -104849 | NM_028127 | 319710 | Mm.2962 | NM_028127 | ENSMUSG00000048285 | **Frmd6** | 2610019M19Rik\|4930488L10Rik\|AW212977 | FERM domain containing 6 | protein-coding |
| chr12_70707047_70707378 | chr12 | 70707048 | 70707378 | + | 0.647117 | Intergenic | Intergenic | -118301 | NM_028127 | 319710 | Mm.2962 | NM_028127 | ENSMUSG00000048285 | **Frmd6** | 2610019M19Rik\|4930488L10Rik\|AW212977 | FERM domain containing 6 | protein-coding |
| chr12_70681483_70682125 | chr12 | 70681484 | 70682125 | + | 0.646214 | Intergenic | ORR1G\|LTR\|ERVL-MaLR | -143710 | NM_028127 | 319710 | Mm.2962 | NM_028127 | ENSMUSG00000048285 | **Frmd6** | 2610019M19Rik\|4930488L10Rik\|AW212977 | FERM domain containing 6 | protein-coding |
| chr12_70849070_70849391 | chr12 | 70849071 | 70849391 | + | 0.640236 | intron (NM_028127, intron 1 of 13) | intron (NM_028127, intron 1 of 13) | 23717 | NM_028127 | 319710 | Mm.2962 | NM_028127 | ENSMUSG00000048285 | **Frmd6** | 2610019M19Rik\|4930488L10Rik\|AW212977 | FERM domain containing 6 | protein-coding |
| chrX_167650441_167650872 | chrX | 167650442 | 167650872 | + | 0.887839 | intron (NM_001033330, intron 5 of 19) | intron (NM_001033330, intron 5 of 19) | 204420 | NM_001290428 | 333605 | Mm.426168 | NM_001033330 | ENSMUSG00000049176 | **Frmpd4** | Gm196\|PKAP1\|Pdzd10\|Pdzk10\|Preso1\|preso | FERM and PDZ domain containing 4 | protein-coding |
| chr3_116865188_116865639 | chr3 | 116865189 | 116865639 | + | 0.687986 | intron (NM_009146, intron 1 of 16) | MTB_Mm\|LTR\|ERVL-MaLR | 3734 | NM_009146 | 20321 | Mm.66293 | NM_009146 | ENSMUSG00000033386 | **Frrs1** | AI131732\|Sdfr2 | ferric-chelate reductase 1 | protein-coding |
| chr4_56977238_56978356 | chr4 | 56977239 | 56978356 | + | 0.748451 | intron (NM_001142965, intron 1 of 4) | intron (NM_001142965, intron 1 of 4) | 12594 | NM_001142965 | 230235 | Mm.207654 | NM_001142965 | | **Frrs1l** | 6430704M03Rik | ferric-chelate reductase 1 like | protein-coding |
| chr10_117110933_117111524 | chr10 | 117110934 | 117111524 | + | 0.785947 | intron (NM_177798, intron 1 of 9) | intron (NM_177798, intron 1 of 9) | 37245 | NM_177798 | 327826 | Mm.135965 | NM_177798 | ENSMUSG00000020170 | **Frs2** | 4732458E18\|C330018A15Rik\|Frs2alpha\|SNT1 | fibroblast growth factor receptor substrate 2 | protein-coding |
| chr5_73210662_73211148 | chr5 | 73210663 | 73211148 | + | 0.86793 | intron (NM_028194, intron 2 of 62) | intron (NM_028194, intron 2 of 62) | 45713 | NM_028194 | 72313 | Mm.260594 | NM_028194 | ENSMUSG00000070733 | **Fryl** | 2010313D22Rik\|2310004H21Rik\|2510002A14Rik\|9030227G01Rik\|mKIAA0826 | furry homolog-like (Drosophila) | protein-coding |
| chr5_73223027_73223560 | chr5 | 73223028 | 73223560 | + | 0.776121 | intron (NM_028194, intron 1 of 62) | Lx8b\|LINE\|L1 | 33324 | NM_028194 | 72313 | Mm.260594 | NM_028194 | ENSMUSG00000070733 | **Fryl** | 2010313D22Rik\|2310004H21Rik\|2510002A14Rik\|9030227G01Rik\|mKIAA0826 | furry homolog-like (Drosophila) | protein-coding |
| chr5_73174274_73174741 | chr5 | 73174275 | 73174741 | + | 0.770144 | intron (NM_028194, intron 3 of 62) | intron (NM_028194, intron 3 of 62) | 82110 | NM_028194 | 72313 | Mm.260594 | NM_028194 | ENSMUSG00000070733 | **Fryl** | 2010313D22Rik\|2310004H21Rik\|2510002A14Rik\|9030227G01Rik\|mKIAA0826 | furry homolog-like (Drosophila) | protein-coding |
| chr2_80461863_80462436 | chr2 | 80461864 | 80462436 | + | 0.808975 | Intergenic | RMER5\|LTR\|ERV1 | -14754 | NM_011356 | 20378 | Mm.427436 | NM_011356 | ENSMUSG00000027004 | **Frzb** | Frp\|Sfrp3\|frezzled\|fritz\|frzb-1 | frizzled-related protein | protein-coding |
| chr5_142963957_142964553 | chr5 | 142963958 | 142964553 | + | 0.791113 | intron (NM_007984, intron 1 of 4) | intron (NM_007984, intron 1 of 4) | 3900 | NM_007984 | 14086 | Mm.289707 | NM_007984 | ENSMUSG00000029581 | **Fscn1** | AI663989\|Fan1\|fascin-1 | fascin homolog 1, actin bundling protein (Strongylocentrotus purpuratus) | protein-coding |
| chr11_52919673_52920227 | chr11 | 52919674 | 52920227 | + | 1.06962 | intron (NM_177059, intron 3 of 15) | intron (NM_177059, intron 3 of 15) | 155244 | NM_177059 | 320027 | Mm.44518 | NM_177059 | | **Fstl4** | B230374F23Rik\|SPIG1\|mKIAA1061 | follistatin-like 4 | protein-coding |
| chr11_52919145_52920224 | chr11 | 52919146 | 52920224 | + | 0.95017 | intron (NM_177059, intron 3 of 15) | intron (NM_177059, intron 3 of 15) | 154979 | NM_177059 | 320027 | Mm.44518 | NM_177059 | | **Fstl4** | B230374F23Rik\|SPIG1\|mKIAA1061 | follistatin-like 4 | protein-coding |
| chr8_91395465_91396280 | chr8 | 91395466 | 91396280 | + | 1.33358 | intron (NM_011936, intron 1 of 8) | intron (NM_011936, intron 1 of 8) | 82348 | NM_011936 | 26383 | Mm.4375 | NM_011936 | ENSMUSG00000055932 | **Fto** | AW743446\|mKIAA1752 | fat mass and obesity associated | protein-coding |
| chr8_91545566_91546015 | chr8 | 91545567 | 91546015 | + | 1.07324 | intron (NM_011936, intron 8 of 8) | intron (NM_011936, intron 8 of 8) | 232266 | NM_011936 | 26383 | Mm.4375 | NM_011936 | ENSMUSG00000055932 | **Fto** | AW743446\|mKIAA1752 | fat mass and obesity associated | protein-coding |
| chr8_91555315_91555967 | chr8 | 91555316 | 91555967 | + | 0.955741 | intron (NM_011936, intron 8 of 8) | intron (NM_011936, intron 8 of 8) | 242116 | NM_011936 | 26383 | Mm.4375 | NM_011936 | ENSMUSG00000055932 | **Fto** | AW743446\|mKIAA1752 | fat mass and obesity associated | protein-coding |
| chr8_91507410_91508145 | chr8 | 91507411 | 91508145 | + | 0.730077 | intron (NM_011936, intron 7 of 8) | intron (NM_011936, intron 7 of 8) | 194253 | NM_011936 | 26383 | Mm.4375 | NM_011936 | ENSMUSG00000055932 | **Fto** | AW743446\|mKIAA1752 | fat mass and obesity associated | protein-coding |
| chr8_91455636_91456051 | chr8 | 91455637 | 91456051 | + | 0.681978 | intron (NM_011936, intron 4 of 8) | intron (NM_011936, intron 4 of 8) | 142319 | NM_011936 | 26383 | Mm.4375 | NM_011936 | ENSMUSG00000055932 | **Fto** | AW743446\|mKIAA1752 | fat mass and obesity associated | protein-coding |
| chr8_91396475_91396648 | chr8 | 91396476 | 91396648 | + | 0.651769 | intron (NM_011936, intron 1 of 8) | B3\|SINE\|B2 | 83037 | NM_011936 | 26383 | Mm.4375 | NM_011936 | ENSMUSG00000055932 | **Fto** | AW743446\|mKIAA1752 | fat mass and obesity associated | protein-coding |
| chrX_17519989_17520469 | chrX | 17519990 | 17520469 | + | 0.952488 | Intergenic | RMER17A2\|LTR\|ERVK | 52068 | NM_028058 | 72018 | Mm.41558 | NM_028058 | ENSMUSG00000025040 | **Fundc1** | 1500005J14Rik\|1810033P05Rik | FUN14 domain containing 1 | protein-coding |
| chrX_17547896_17548361 | chrX | 17547897 | 17548361 | + | 0.940886 | Intergenic | RMER17C-int\|LTR\|ERVK | 24168 | NM_028058 | 72018 | Mm.41558 | NM_028058 | ENSMUSG00000025040 | **Fundc1** | 1500005J14Rik\|1810033P05Rik | FUN14 domain containing 1 | protein-coding |
| chr7_80416918_80417626 | chr7 | 80416919 | 80417626 | + | 0.902024 | Intergenic | Intergenic | -11841 | NM_001081454 | 18550 | Mm.5241 | NM_011046 | ENSMUSG00000030530 | **Furin** | 9130404I01Rik\|Fur\|PACE\|Pcsk3\|SPC1 | furin (paired basic amino acid cleaving enzyme) | protein-coding |
| chr7_80416918_80417942 | chr7 | 80416919 | 80417942 | + | 0.882316 | Intergenic | Intergenic | -11999 | NM_001081454 | 18550 | Mm.5241 | NM_011046 | ENSMUSG00000030530 | **Furin** | 9130404I01Rik\|Fur\|PACE\|Pcsk3\|SPC1 | furin (paired basic amino acid cleaving enzyme) | protein-coding |
| chr7_80405383_80405580 | chr7 | 80405384 | 80405580 | + | 0.847557 | promoter-TSS (NM_001081454) | promoter-TSS (NM_001081454) | -51 | NM_001081454 | 18550 | Mm.5241 | NM_011046 | ENSMUSG00000030530 | **Furin** | 9130404I01Rik\|Fur\|PACE\|Pcsk3\|SPC1 | furin (paired basic amino acid cleaving enzyme) | protein-coding |
| chr7_80410204_80410876 | chr7 | 80410205 | 80410876 | + | 0.759902 | Intergenic | Intergenic | -5109 | NM_001081454 | 18550 | Mm.5241 | NM_011046 | ENSMUSG00000030530 | **Furin** | 9130404I01Rik\|Fur\|PACE\|Pcsk3\|SPC1 | furin (paired basic amino acid cleaving enzyme) | protein-coding |
| chr7_80412309_80412612 | chr7 | 80412310 | 80412612 | + | 0.639642 | Intergenic | Charlie29a\|DNA\|hAT-Charlie | -7030 | NM_001081454 | 18550 | Mm.5241 | NM_011046 | ENSMUSG00000030530 | **Furin** | 9130404I01Rik\|Fur\|PACE\|Pcsk3\|SPC1 | furin (paired basic amino acid cleaving enzyme) | protein-coding |
| chr9_14756124_14756423 | chr9 | 14756125 | 14756423 | + | 0.847839 | Intergenic | Intergenic | -4152 | NM_010242 | 14345 | Mm.63450 | NM_010242 | ENSMUSG00000049307 | **Fut4** | AI451562\|CD15\|FAL\|FucT-IV\|LeX\|SSEA-1\|Ssea1 | fucosyltransferase 4 | protein-coding |
| chr12_77235856_77236394 | chr12 | 77235857 | 77236394 | + | 1.88196 | Intergenic | Intergenic | -1979 | NM_001252614 | 53618 | Mm.35628 | NM_016893 | ENSMUSG00000021065 | **Fut8** | - | fucosyltransferase 8 | protein-coding |
| chr12_77290934_77291456 | chr12 | 77290935 | 77291456 | + | 1.03535 | intron (NM_001252614, intron 3 of 12) | intron (NM_001252614, intron 3 of 12) | 52456 | NR_045554 | 53618 | Mm.35628 | NM_016893 | ENSMUSG00000021065 | **Fut8** | - | fucosyltransferase 8 | protein-coding |

| | | | | | | | | | | | | | | | | | |
|---|---|---|---|---|---|---|---|---|---|---|---|---|---|---|---|---|---|
| chr12_77302140_77302422 | chr12 | 77302141 | 77302422 | + | 0.706212 | intron (NM_001252614, intron 3 of 12) | PB1D11\|SINE\|Alu | 63542 | NR_045554 | 53618 | Mm.35628 | NM_016893 | ENSMUSG00000021065 | **Fut8** | - | fucosyltransferase 8 | protein-coding |
| chr7_44898364_44898698 | chr7 | 44898365 | 44898698 | + | 0.63547 | exon (NM_027376, exon 6 of 11) | exon (NM_027376, exon 6 of 11) | 2452 | NM_027376 | 70300 | Mm.276298 | NM_027376 | ENSMUSG00000011658 | **Fuz** | 2600013E07Rik\|b2b1273Clo | fuzzy homolog (Drosophila) | protein-coding |
| chr19_24280951_24281194 | chr19 | 24280952 | 24281194 | + | 0.806736 | promoter-TSS (NM_008044) | promoter-TSS (NM_008044) | -487 | NM_008044 | 14297 | Mm.7319 | NM_008044 | ENSMUSG00000059363 | **Fxn** | FA\|FARR\|Frda\|X25 | frataxin | protein-coding |
| chr19_24280881_24281262 | chr19 | 24280882 | 24281262 | + | 0.784162 | promoter-TSS (NM_008044) | promoter-TSS (NM_008044) | -486 | NM_008044 | 14297 | Mm.7319 | NM_008044 | ENSMUSG00000059363 | **Fxn** | FA\|FARR\|Frda\|X25 | frataxin | protein-coding |
| chr15_6555540_6556010 | chr15 | 6555541 | 6556010 | + | 1.06788 | Intergenic | Intergenic | -24072 | NM_001278269 | 23880 | Mm.170905 | NM_011815 | | **Fyb** | ADAP\|B630013F22Rik\|FYB-120/130\|Slap130\|p120/130\|p120/p130 | FYN binding protein | protein-coding |
| chr15_6603539_6604232 | chr15 | 6603540 | 6604232 | + | 0.99237 | intron (NM_011815, intron 1 of 17) | intron (NM_011815, intron 1 of 17) | 24039 | NM_011815 | 23880 | Mm.170905 | NM_011815 | | **Fyb** | ADAP\|B630013F22Rik\|FYB-120/130\|Slap130\|p120/130\|p120/p130 | FYN binding protein | protein-coding |
| chr15_6603458_6603996 | chr15 | 6603459 | 6603996 | + | 0.895276 | intron (NM_011815, intron 1 of 17) | intron (NM_011815, intron 1 of 17) | 23880 | NM_011815 | 23880 | Mm.170905 | NM_011815 | | **Fyb** | ADAP\|B630013F22Rik\|FYB-120/130\|Slap130\|p120/130\|p120/p130 | FYN binding protein | protein-coding |
| chr15_6523580_6524151 | chr15 | 6523581 | 6524151 | + | 0.817204 | Intergenic | Intergenic | -55981 | NM_001278269 | 23880 | Mm.170905 | NM_011815 | | **Fyb** | ADAP\|B630013F22Rik\|FYB-120/130\|Slap130\|p120/130\|p120/p130 | FYN binding protein | protein-coding |
| chr15_6539983_6540655 | chr15 | 6539984 | 6540655 | + | 0.801978 | Intergenic | Intergenic | -39528 | NM_001278269 | 23880 | Mm.170905 | NM_011815 | | **Fyb** | ADAP\|B630013F22Rik\|FYB-120/130\|Slap130\|p120/130\|p120/p130 | FYN binding protein | protein-coding |
| chr15_6584012_6584293 | chr15 | 6584013 | 6584293 | + | 0.650333 | intron (NM_011815, intron 1 of 17) | intron (NM_011815, intron 1 of 17) | 4306 | NM_011815 | 23880 | Mm.170905 | NM_011815 | | **Fyb** | ADAP\|B630013F22Rik\|FYB-120/130\|Slap130\|p120/130\|p120/p130 | FYN binding protein | protein-coding |
| chr15_6548745_6549422 | chr15 | 6548746 | 6549422 | + | 0.643048 | Intergenic | Intergenic | -30763 | NM_001278269 | 23880 | Mm.170905 | NM_011815 | | **Fyb** | ADAP\|B630013F22Rik\|FYB-120/130\|Slap130\|p120/130\|p120/p130 | FYN binding protein | protein-coding |
| chr9_123840217_123840550 | chr9 | 123840218 | 123840550 | + | 0.950139 | intron (NM_148925, intron 4 of 18) | intron (NM_148925, intron 4 of 18) | 11277 | NM_148925 | 17281 | Mm.252941 | NM_148925 | ENSMUSG00000025241 | **Fyco1** | 2810409M01Rik\|Mem2\|RUFY3\|ZFYVE7 | FYVE and coiled-coil domain containing 1 | protein-coding |
| chr10_39457393_39457942 | chr10 | 39457394 | 39457942 | + | 1.33051 | intron (NM_008054, intron 3 of 13) | intron (NM_008054, intron 3 of 13) | 87869 | NM_001122893 | 14360 | Mm.4848 | NM_008054 | ENSMUSG00000019843 | **Fyn** | AI448320\|AW552119 | Fyn proto-oncogene | protein-coding |
| chr10_39415586_39415951 | chr10 | 39415587 | 39415951 | + | 1.21513 | intron (NM_001122892, intron 2 of 12) | MIR3\|SINE\|MIR | 45970 | NM_001122893 | 14360 | Mm.4848 | NM_008054 | ENSMUSG00000019843 | **Fyn** | AI448320\|AW552119 | Fyn proto-oncogene | protein-coding |
| chr10_39430895_39431141 | chr10 | 39430896 | 39431141 | + | 1.09891 | intron (NM_001122892, intron 2 of 12) | intron (NM_001122892, intron 2 of 12) | 61219 | NM_001122893 | 14360 | Mm.4848 | NM_008054 | ENSMUSG00000019843 | **Fyn** | AI448320\|AW552119 | Fyn proto-oncogene | protein-coding |
| chr10_39456760_39458126 | chr10 | 39456761 | 39458126 | + | 1.08713 | intron (NM_008054, intron 3 of 13) | intron (NM_008054, intron 3 of 13) | 87644 | NM_001122893 | 14360 | Mm.4848 | NM_008054 | ENSMUSG00000019843 | **Fyn** | AI448320\|AW552119 | Fyn proto-oncogene | protein-coding |
| chr10_39434805_39435514 | chr10 | 39434806 | 39435514 | + | 1.0803 | intron (NM_001122892, intron 2 of 12) | intron (NM_001122892, intron 2 of 12) | 65361 | NM_001122893 | 14360 | Mm.4848 | NM_008054 | ENSMUSG00000019843 | **Fyn** | AI448320\|AW552119 | Fyn proto-oncogene | protein-coding |
| chr10_39377988_39378489 | chr10 | 39377989 | 39378489 | + | 1.05364 | intron (NM_008054, intron 1 of 13) | intron (NM_008054, intron 1 of 13) | 8440 | NM_001122893 | 14360 | Mm.4848 | NM_008054 | ENSMUSG00000019843 | **Fyn** | AI448320\|AW552119 | Fyn proto-oncogene | protein-coding |
| chr10_39479125_39479713 | chr10 | 39479126 | 39479713 | + | 0.900676 | intron (NM_008054, intron 3 of 13) | intron (NM_008054, intron 3 of 13) | 109620 | NM_001122893 | 14360 | Mm.4848 | NM_008054 | ENSMUSG00000019843 | **Fyn** | AI448320\|AW552119 | Fyn proto-oncogene | protein-coding |
| chr10_39478893_39479965 | chr10 | 39478894 | 39479965 | + | 0.893656 | intron (NM_008054, intron 3 of 13) | intron (NM_008054, intron 3 of 13) | 109630 | NM_001122893 | 14360 | Mm.4848 | NM_008054 | ENSMUSG00000019843 | **Fyn** | AI448320\|AW552119 | Fyn proto-oncogene | protein-coding |
| chr10_39430396_39430705 | chr10 | 39430397 | 39430705 | + | 0.821408 | intron (NM_001122892, intron 2 of 12) | intron (NM_001122892, intron 2 of 12) | 60752 | NM_001122893 | 14360 | Mm.4848 | NM_008054 | ENSMUSG00000019843 | **Fyn** | AI448320\|AW552119 | Fyn proto-oncogene | protein-coding |
| chr10_39456805_39457124 | chr10 | 39456806 | 39457124 | + | 0.771733 | intron (NM_008054, intron 3 of 13) | intron (NM_008054, intron 3 of 13) | 87166 | NM_001122893 | 14360 | Mm.4848 | NM_008054 | ENSMUSG00000019843 | **Fyn** | AI448320\|AW552119 | Fyn proto-oncogene | protein-coding |
| chr10_39314479_39314965 | chr10 | 39314480 | 39314965 | + | 0.735523 | Intergenic | URR1A\|DNA\|hAT-Charlie | -55077 | NM_001122892 | 14360 | Mm.4848 | NM_008054 | ENSMUSG00000019843 | **Fyn** | AI448320\|AW552119 | Fyn proto-oncogene | protein-coding |
| chr10_39433983_39434229 | chr10 | 39433984 | 39434229 | + | 0.721954 | intron (NM_001122892, intron 2 of 12) | MTC\|LTR\|ERVL-MaLR | 64307 | NM_001122893 | 14360 | Mm.4848 | NM_008054 | ENSMUSG00000019843 | **Fyn** | AI448320\|AW552119 | Fyn proto-oncogene | protein-coding |
| chr10_39435991_39436442 | chr10 | 39435992 | 39436442 | + | 0.699898 | intron (NM_001122892, intron 2 of 12) | intron (NM_001122892, intron 2 of 12) | 66418 | NM_001122893 | 14360 | Mm.4848 | NM_008054 | ENSMUSG00000019843 | **Fyn** | AI448320\|AW552119 | Fyn proto-oncogene | protein-coding |
| chr7_97041864_97042520 | chr7 | 97041865 | 97042520 | + | 0.867633 | intron (NM_153591, intron 11 of 13) | B1_Mur1\|SINE\|Alu | -39559 | NM_001162477 | 14389 | Mm.42033 | NM_010248 | ENSMUSG00000004508 | **Gab2** | AI463667\|D130058I17Rik\|p97 | growth factor receptor bound protein 2-associated protein 2 | protein-coding |
| chr8_111964318_111964839 | chr8 | 111964319 | 111964839 | + | 0.684431 | Intergenic | Intergenic | 23876 | NM_026693 | 93739 | Mm.371666 | NM_026693 | | **Gabarapl2** | 0610012F20Rik\|2900019O08Rik\|AI173605\|GATE-16\|Gef2 | gamma-aminobutyric acid (GABA) A receptor-associated protein-like 2 | protein-coding |
| chr7_57918208_57918650 | chr7 | 57918209 | 57918650 | + | 0.920972 | Intergenic | L1M2\|LINE\|L1 | 327256 | NM_008071 | 14402 | Mm.8004 | NM_008071 | ENSMUSG00000033676 | **Gabrb3** | A230092K12Rik\|AW049585\|Cp1\|Gabrb-3\|beta3 | gamma-aminobutyric acid (GABA) A receptor, subunit beta 3 | protein-coding |
| chr6_67104628_67104893 | chr6 | 67104629 | 67104893 | + | 1.00567 | Intergenic | Intergenic | -67354 | NM_007836 | 13197 | Mm.72235 | NM_007836 | ENSMUSG00000036390 | **Gadd45a** | AA545191\|Ddit1\|GADD45 | growth arrest and DNA-damage-inducible 45 alpha | protein-coding |
| chr6_67059934_67060325 | chr6 | 67059935 | 67060325 | + | 1.00466 | intron (NR_040278, intron 1 of 6) | intron (NR_040278, intron 1 of 6) | -22723 | NM_007836 | 13197 | Mm.72235 | NM_007836 | ENSMUSG00000036390 | **Gadd45a** | AA545191\|Ddit1\|GADD45 | growth arrest and DNA-damage-inducible 45 alpha | protein-coding |
| chr6_67105281_67106011 | chr6 | 67105282 | 67106011 | + | 0.974172 | Intergenic | MLT1C\|LTR\|ERVL-MaLR | -68239 | NM_007836 | 13197 | Mm.72235 | NM_007836 | ENSMUSG00000036390 | **Gadd45a** | AA545191\|Ddit1\|GADD45 | growth arrest and DNA-damage-inducible 45 alpha | protein-coding |
| chr6_67058326_67058656 | chr6 | 67058327 | 67058656 | + | 0.831236 | intron (NR_040278, intron 1 of 6) | intron (NR_040278, intron 1 of 6) | -21084 | NM_007836 | 13197 | Mm.72235 | NM_007836 | ENSMUSG00000036390 | **Gadd45a** | AA545191\|Ddit1\|GADD45 | growth arrest and DNA-damage-inducible 45 alpha | protein-coding |
| chr6_67059814_67060357 | chr6 | 67059815 | 67060357 | + | 0.830347 | intron (NR_040278, intron 1 of 6) | intron (NR_040278, intron 1 of 6) | -22679 | NM_007836 | 13197 | Mm.72235 | NM_007836 | ENSMUSG00000036390 | **Gadd45a** | AA545191\|Ddit1\|GADD45 | growth arrest and DNA-damage-inducible 45 alpha | protein-coding |
| chr10_80930229_80930936 | chr10 | 80930230 | 80930936 | + | 0.69018 | intron (NM_008655, intron 2 of 3) | CpG-1325 | 492 | NM_008655 | 17873 | Mm.1360 | NM_008655 | ENSMUSG00000015312 | **Gadd45b** | AI323528\|Myd118 | growth arrest and DNA-damage-inducible 45 beta | protein-coding |
| chr10_80939211_80939493 | chr10 | 80939212 | 80939493 | + | 0.659218 | Intergenic | Intergenic | 9261 | NM_008655 | 17873 | Mm.1360 | NM_008655 | ENSMUSG00000015312 | **Gadd45b** | AI323528\|Myd118 | growth arrest and DNA-damage-inducible 45 beta | protein-coding |
| chr13_51969960_51970457 | chr13 | 51969961 | 51970457 | + | 1.24307 | Intergenic | Intergenic | 123534 | NM_011817 | 23882 | Mm.281298 | NM_011817 | ENSMUSG00000021453 | **Gadd45g** | AI327420\|C86281\|CR6\|DDIT2\|OIG37 | growth arrest and DNA-damage-inducible 45 gamma | protein-coding |
| chr13_51969833_51970649 | chr13 | 51969834 | 51970649 | + | 1.21604 | Intergenic | Intergenic | 123566 | NM_011817 | 23882 | Mm.281298 | NM_011817 | ENSMUSG00000021453 | **Gadd45g** | AI327420\|C86281\|CR6\|DDIT2\|OIG37 | growth arrest and DNA-damage-inducible 45 gamma | protein-coding |
| chr13_51935789_51936976 | chr13 | 51935790 | 51936976 | + | 1.04661 | Intergenic | Intergenic | 89708 | NM_011817 | 23882 | Mm.281298 | NM_011817 | ENSMUSG00000021453 | **Gadd45g** | AI327420\|C86281\|CR6\|DDIT2\|OIG37 | growth arrest and DNA-damage-inducible 45 gamma | protein-coding |
| chr13_51933333_51933615 | chr13 | 51933334 | 51933615 | + | 0.915 | Intergenic | Intergenic | 86799 | NM_011817 | 23882 | Mm.281298 | NM_011817 | ENSMUSG00000021453 | **Gadd45g** | AI327420\|C86281\|CR6\|DDIT2\|OIG37 | growth arrest and DNA-damage-inducible 45 gamma | protein-coding |
| chr13_51848661_51849505 | chr13 | 51848662 | 51849505 | + | 0.762691 | TTS (NM_011817) | TTS (NM_011817) | 2408 | NM_011817 | 23882 | Mm.281298 | NM_011817 | ENSMUSG00000021453 | **Gadd45g** | AI327420\|C86281\|CR6\|DDIT2\|OIG37 | growth arrest and DNA-damage-inducible 45 gamma | protein-coding |
| chr13_51958433_51958751 | chr13 | 51958434 | 51958751 | + | 0.759532 | Intergenic | Intergenic | 111917 | NM_011817 | 23882 | Mm.281298 | NM_011817 | ENSMUSG00000021453 | **Gadd45g** | AI327420\|C86281\|CR6\|DDIT2\|OIG37 | growth arrest and DNA-damage-inducible 45 gamma | protein-coding |
| chr1_93859740_93860073 | chr1 | 93859741 | 93860073 | + | 0.644203 | Intergenic | ORR1D1\|LTR\|ERVL-MaLR | -1437 | NM_199366 | 381334 | Mm.482878 | NM_199366 | ENSMUSG00000094651 | **Gal3st2** | GP3ST\|Gm988 | galactose-3-O-sulfotransferase 2 | protein-coding |
| chr12_98244201_98245046 | chr12 | 98244202 | 98245046 | + | 1.62712 | intron (NM_008079, intron 6 of 16) | RMER5\|LTR\|ERV1 | 14713 | NM_008079 | 14420 | Mm.5120 | NM_008079 | ENSMUSG00000021003 | **Galc** | 2310068B06Rik\|A930008M05Rik\|AW212969\|AW413532\|Gacy\|twi\|twitcher | galactosylceramidase | protein-coding |

| | | | | | | | | | | | | | | | | | |
|---|---|---|---|---|---|---|---|---|---|---|---|---|---|---|---|---|---|
| chr12_98244201_98245390 | chr12 | 98244202 | 98245390 | + | 1.58187 | intron (NM_008079, intron 6 of 16) | RMER5\|LTR\|ERV1 | 14541 | NM_008079 | 14420 | Mm.5120 | NM_008079 | ENSMUSG00000021003 | **Galc** | 2310068B06Rik\|A930008M05Rik\|AW212969\|AW413532\|Gacy\|twi\|twitcher | galactosylceramidase | protein-coding |
| chr12_98249681_98250047 | chr12 | 98249682 | 98250047 | + | 1.19989 | intron (NM_008079, intron 5 of 16) | intron (NM_008079, intron 5 of 16) | 9473 | NM_008079 | 14420 | Mm.5120 | NM_008079 | ENSMUSG00000021003 | **Galc** | 2310068B06Rik\|A930008M05Rik\|AW212969\|AW413532\|Gacy\|twi\|twitcher | galactosylceramidase | protein-coding |
| chr12_98213412_98214082 | chr12 | 98213413 | 98214082 | + | 0.667315 | intron (NM_008079, intron 13 of 16) | intron (NM_008079, intron 13 of 16) | 45590 | NM_008079 | 14420 | Mm.5120 | NM_008079 | ENSMUSG00000021003 | **Galc** | 2310068B06Rik\|A930008M05Rik\|AW212969\|AW413532\|Gacy\|twi\|twitcher | galactosylceramidase | protein-coding |
| chr2_125896301_125897033 | chr2 | 125896302 | 125897033 | + | 0.977651 | exon (NM_001291002, exon 5 of 10) | exon (NM_001291002, exon 5 of 10) | 30444 | NM_001291002 | 69976 | Mm.20216 | NM_175154 | ENSMUSG00000027207 | **Galk2** | 2810017M24Rik\|AI327030\|Gk2 | galactokinase 2 | protein-coding |
| chr17_80165853_80166226 | chr17 | 80165854 | 80166226 | + | 0.9228 | intron (NM_176963, intron 4 of 6) | intron (NM_176963, intron 4 of 6) | 38569 | NM_176963 | 319625 | Mm.29098 | NM_176963 | ENSMUSG00000035473 | **Galm** | A530057M15Rik\|AU015645\|AU020959 | galactose mutarotase | protein-coding |
| chr14_31934814_31935268 | chr14 | 31934815 | 31935268 | + | 0.660859 | Intergenic | (TG)n\|Simple_repeat\|Simple_repeat | -94062 | NM_030166 | 78754 | Mm.40681 | NM_030166 | | **Galnt15** | 4631401E18Rik\|Galntl2\|mpp-GalNAc-T15 | UDP-N-acetyl-alpha-D-galactosamine:polypeptide N-acetylgalactosaminyltransferase 15 | protein-coding |
| chr8_124278213_124278916 | chr8 | 124278214 | 124278916 | + | 1.54893 | intron (NM_139272, intron 1 of 15) | intron (NM_139272, intron 1 of 15) | 47171 | NM_139272 | 108148 | Mm.33808 | NM_139272 | ENSMUSG00000089704 | **Galnt2** | AI480629 | UDP-N-acetyl-alpha-D-galactosamine:polypeptide N-acetylgalactosaminyltransferase 2 | protein-coding |
| chr8_124229897_124230190 | chr8 | 124229898 | 124230190 | + | 0.669396 | Intergenic | Intergenic | -1350 | NM_139272 | 108148 | Mm.33808 | NM_139272 | ENSMUSG00000089704 | **Galnt2** | AI480629 | UDP-N-acetyl-alpha-D-galactosamine:polypeptide N-acetylgalactosaminyltransferase 2 | protein-coding |
| chr8_124259172_124259575 | chr8 | 124259173 | 124259575 | + | 0.654514 | intron (NM_139272, intron 1 of 15) | intron (NM_139272, intron 1 of 15) | 27980 | NM_139272 | 108148 | Mm.33808 | NM_139272 | ENSMUSG00000089704 | **Galnt2** | AI480629 | UDP-N-acetyl-alpha-D-galactosamine:polypeptide N-acetylgalactosaminyltransferase 2 | protein-coding |
| chr2_66101755_66102408 | chr2 | 66101756 | 66102408 | + | 0.943498 | intron (NM_015736, intron 3 of 10) | MIR\|SINE\|MIR | 22711 | NM_015736 | 14425 | Mm.439760 | NM_015736 | ENSMUSG00000026994 | **Galnt3** | - | UDP-N-acetyl-alpha-D-galactosamine:polypeptide N-acetylgalactosaminyltransferase 3 | protein-coding |
| chr2_66184506_66185164 | chr2 | 66184507 | 66185164 | + | 0.874278 | exon (NM_001047604, exon 29 of 29) | exon (NM_001047604, exon 29 of 29) | -60042 | NM_015736 | 14425 | Mm.439760 | NM_015736 | ENSMUSG00000026994 | **Galnt3** | - | UDP-N-acetyl-alpha-D-galactosamine:polypeptide N-acetylgalactosaminyltransferase 3 | protein-coding |
| chr2_66101780_66102789 | chr2 | 66101781 | 66102789 | + | 0.80392 | intron (NM_015736, intron 3 of 10) | intron (NM_015736, intron 3 of 10) | 22508 | NM_015736 | 14425 | Mm.439760 | NM_015736 | ENSMUSG00000026994 | **Galnt3** | - | UDP-N-acetyl-alpha-D-galactosamine:polypeptide N-acetylgalactosaminyltransferase 3 | protein-coding |
| chr2_66120035_66120345 | chr2 | 66120036 | 66120345 | + | 0.746329 | intron (NM_015736, intron 1 of 10) | intron (NM_015736, intron 1 of 10) | 4603 | NM_015736 | 14425 | Mm.439760 | NM_015736 | ENSMUSG00000026994 | **Galnt3** | - | UDP-N-acetyl-alpha-D-galactosamine:polypeptide N-acetylgalactosaminyltransferase 3 | protein-coding |
| chr8_57741984_57742361 | chr8 | 57741985 | 57742361 | + | 1.35006 | Intergenic | Intergenic | -89118 | NM_144731 | 108150 | Mm.62886 | NM_144731 | ENSMUSG00000031608 | **Galnt7** | AI225872 | UDP-N-acetyl-alpha-D-galactosamine: polypeptide N-acetylgalactosaminyltransferase 7 | protein-coding |
| chr8_57791852_57792658 | chr8 | 57791853 | 57792658 | + | 1.25659 | intron (NM_175032, intron 11 of 11) | intron (NM_175032, intron 11 of 11) | -139200 | NM_144731 | 108150 | Mm.62886 | NM_144731 | ENSMUSG00000031608 | **Galnt7** | AI225872 | UDP-N-acetyl-alpha-D-galactosamine: polypeptide N-acetylgalactosaminyltransferase 7 | protein-coding |
| chr8_57809637_57810036 | chr8 | 57809638 | 57810036 | + | 0.696501 | intron (NM_175032, intron 9 of 11) | intron (NM_175032, intron 9 of 11) | -156782 | NM_144731 | 108150 | Mm.62886 | NM_144731 | ENSMUSG00000031608 | **Galnt7** | AI225872 | UDP-N-acetyl-alpha-D-galactosamine: polypeptide N-acetylgalactosaminyltransferase 7 | protein-coding |
| chr8_117127120_117127518 | chr8 | 117127121 | 117127518 | + | 0.6699 | intron (NM_021486, intron 7 of 10) | intron (NM_021486, intron 7 of 10) | -30816 | NM_001081151 | 209239 | Mm.132992 | NM_001081151 | ENSMUSG00000052557 | **Gan** | A330045G18\|gigaxonin | giant axonal neuropathy | protein-coding |
| chr2_34742439_34743357 | chr2 | 34742440 | 34743357 | + | 1.15322 | intron (NM_025709, intron 2 of 26) | RSINE1\|SINE\|B4 | 12334 | NM_025709 | 66691 | Mm.156452 | NM_025709 | ENSMUSG00000026867 | **Gapvd1** | 2010005B09Rik\|4432404J10Rik\|AW108497\|Gapex-5\|RAP6\|RME-6\|mKIAA1521 | GTPase activating protein and VPS9 domains 1 | protein-coding |
| chr2_34742468_34743009 | chr2 | 34742469 | 34743009 | + | 1.1469 | intron (NM_025709, intron 2 of 26) | intron (NM_025709, intron 2 of 26) | 12493 | NM_025709 | 66691 | Mm.156452 | NM_025709 | ENSMUSG00000026867 | **Gapvd1** | 2010005B09Rik\|4432404J10Rik\|AW108497\|Gapex-5\|RAP6\|RME-6\|mKIAA1521 | GTPase activating protein and VPS9 domains 1 | protein-coding |
| chr6_55015603_55016098 | chr6 | 55015604 | 55016098 | + | 0.946945 | Intergenic | MTC\|LTR\|ERVL-MaLR | -22150 | NM_180678 | 353172 | Mm.250004 | NM_180678 | ENSMUSG00000029777 | **Gars** | GENA202\|Gena201\|Nmf249\|Sgrp23 | glycyl-tRNA synthetase | protein-coding |
| chr7_51929211_51929594 | chr7 | 51929212 | 51929594 | + | 0.754428 | intron (NM_008087, intron 4 of 7) | intron (NM_008087, intron 4 of 7) | 50258 | NM_008087 | 14453 | Mm.207360 | NM_008087 | ENSMUSG00000030498 | **Gas2** | Gas-2 | growth arrest specific 2 | protein-coding |
| chr11_67563049_67563685 | chr11 | 67563050 | 67563685 | + | 0.738169 | intron (NM_008088, intron 1 of 13) | intron (NM_008088, intron 1 of 13) | -23133 | NM_001109657 | 14457 | Mm.40338 | NM_008088 | ENSMUSG00000033066 | **Gas7** | AW124766\|B230343A10Rik\|Gas7-cb | growth arrest specific 7 | protein-coding |
| chr2_9577810_9578449 | chr2 | 9577811 | 9578449 | + | 1.41708 | Intergenic | Intergenic | 300470 | NM_008091 | 14462 | Mm.313866 | NM_008091 | ENSMUSG00000015619 | **Gata3** | Gata-3\|jal | GATA binding protein 3 | protein-coding |
| chr2_9802862_9803424 | chr2 | 9802863 | 9803424 | + | 1.37694 | Intergenic | Intergenic | 75457 | NM_008091 | 14462 | Mm.313866 | NM_008091 | ENSMUSG00000015619 | **Gata3** | Gata-3\|jal | GATA binding protein 3 | protein-coding |
| chr2_9574151_9574551 | chr2 | 9574152 | 9574551 | + | 1.28094 | Intergenic | Intergenic | 304249 | NM_008091 | 14462 | Mm.313866 | NM_008091 | ENSMUSG00000015619 | **Gata3** | Gata-3\|jal | GATA binding protein 3 | protein-coding |
| chr2_9615976_9616615 | chr2 | 9615977 | 9616615 | + | 1.22709 | Intergenic | Intergenic | 262304 | NM_008091 | 14462 | Mm.313866 | NM_008091 | ENSMUSG00000015619 | **Gata3** | Gata-3\|jal | GATA binding protein 3 | protein-coding |
| chr2_9717880_9718415 | chr2 | 9717881 | 9718415 | + | 1.01179 | Intergenic | Intergenic | 160452 | NM_008091 | 14462 | Mm.313866 | NM_008091 | ENSMUSG00000015619 | **Gata3** | Gata-3\|jal | GATA binding protein 3 | protein-coding |
| chr2_9801326_9801945 | chr2 | 9801327 | 9801945 | + | 0.885368 | Intergenic | Intergenic | 76964 | NM_008091 | 14462 | Mm.313866 | NM_008091 | ENSMUSG00000015619 | **Gata3** | Gata-3\|jal | GATA binding protein 3 | protein-coding |
| chr2_9748191_9748866 | chr2 | 9748192 | 9748866 | + | 0.843036 | Intergenic | Intergenic | 130071 | NM_008091 | 14462 | Mm.313866 | NM_008091 | ENSMUSG00000015619 | **Gata3** | Gata-3\|jal | GATA binding protein 3 | protein-coding |
| chr2_9764411_9764707 | chr2 | 9764412 | 9764707 | + | 0.746224 | Intergenic | RMER15\|LTR\|ERVL | 114041 | NM_008091 | 14462 | Mm.313866 | NM_008091 | ENSMUSG00000015619 | **Gata3** | Gata-3\|jal | GATA binding protein 3 | protein-coding |
| chr2_9683742_9684062 | chr2 | 9683743 | 9684062 | + | 0.701187 | Intergenic | Intergenic | 194698 | NM_008091 | 14462 | Mm.313866 | NM_008091 | ENSMUSG00000015619 | **Gata3** | Gata-3\|jal | GATA binding protein 3 | protein-coding |
| chr2_9751063_9751523 | chr2 | 9751064 | 9751523 | + | 0.679707 | Intergenic | RSINE1\|SINE\|B4 | 127307 | NM_008091 | 14462 | Mm.313866 | NM_008091 | ENSMUSG00000015619 | **Gata3** | Gata-3\|jal | GATA binding protein 3 | protein-coding |
| chr2_9683733_9684440 | chr2 | 9683734 | 9684440 | + | 0.677294 | Intergenic | Intergenic | 194513 | NM_008091 | 14462 | Mm.313866 | NM_008091 | ENSMUSG00000015619 | **Gata3** | Gata-3\|jal | GATA binding protein 3 | protein-coding |
| chr5_134102718_134103201 | chr5 | 134102719 | 134103201 | + | 1.21837 | intron (NM_030719, intron 1 of 8) | intron (NM_030719, intron 1 of 8) | 3212 | NM_030719 | 80909 | Mm.45218 | NM_030719 | | **Gatsl2** | 7530428J21Rik\|AW742413\|Gats\|Gatsl1 | GATS protein-like 2 | protein-coding |
| chr5_134105748_134106221 | chr5 | 134105749 | 134106221 | + | 0.795702 | intron (NM_030719, intron 1 of 8) | ORR1E\|LTR\|ERVL-MaLR | 6237 | NM_030719 | 80909 | Mm.45218 | NM_030719 | | **Gatsl2** | 7530428J21Rik\|AW742413\|Gats\|Gatsl1 | GATS protein-like 2 | protein-coding |
| chr11_4219992_4220254 | chr11 | 4219993 | 4220254 | + | 0.704932 | exon (NM_028022, exon 3 of 9) | exon (NM_028022, exon 3 of 9) | 1872 | NM_028022 | 71962 | Mm.7852 | NM_028022 | ENSMUSG00000020424 | **Gatsl3** | 2410008K03Rik | GATS protein-like 3 | protein-coding |
| chr19_46198863_46199244 | chr19 | 46198864 | 46199244 | + | 0.889906 | intron (NM_178930, intron 3 of 39) | B4A\|SINE\|B4 | 46496 | NM_178930 | 107338 | Mm.271620 | NM_178930 | ENSMUSG00000025224 | **Gbf1** | 1700083E03Rik\|AI035702\|AW049900\|mKIAA0248 | golgi-specific brefeldin A-resistance factor 1 | protein-coding |
| chr5_105262720_105263846 | chr5 | 105262721 | 105263846 | + | 0.917815 | Intergenic | Intergenic | -23750 | NM_001039646 | 626578 | Mm.275893 | NM_001039646 | ENSMUSG00000054588 | **Gbp10** | - | guanylate-binding protein 10 | protein-coding |
| chr5_105353441_105353985 | chr5 | 105353442 | 105353985 | + | 0.846745 | Intergenic | Intergenic | -7241 | NM_001039647 | 634650 | Mm.458435 | NM_001039647 | ENSMUSG00000092021 | **Gbp11** | EG634650\|Gm7141 | guanylate binding protein 11 | protein-coding |
| chr5_105353418_105354417 | chr5 | 105353419 | 105354417 | + | 0.723092 | Intergenic | Intergenic | -7446 | NM_001039647 | 634650 | Mm.458435 | NM_001039647 | ENSMUSG00000092021 | **Gbp11** | EG634650\|Gm7141 | guanylate binding protein 11 | protein-coding |
| chr3_142618630_142618939 | chr3 | 142618631 | 142618939 | + | 0.7545 | 3' UTR (NM_010259, exon 11 of 11) | 3' UTR (NM_010259, exon 11 of 11) | -1878 | NM_010260 | 14469 | Mm.24038 | NM_010260 | ENSMUSG00000028270 | **Gbp2** | - | guanylate binding protein 2 | protein-coding |
| chr3_142578340_142578872 | chr3 | 142578341 | 142578872 | + | 0.864727 | Intergenic | Intergenic | -16241 | NM_010259 | 14468 | Mm.457978 | NM_010259 | ENSMUSG00000028269 | **Gbp2b** | Gbp-1\|Gbp1\|Mag-1\|Mpa-1\|Mpa1 | guanylate binding protein 2b | protein-coding |
| chr3_142577481_142578188 | chr3 | 142577482 | 142578188 | + | 0.649533 | Intergenic | IMPB_01\|Satellite\|Satellite | -17012 | NM_010259 | 14468 | Mm.457978 | NM_010259 | ENSMUSG00000028269 | **Gbp2b** | Gbp-1\|Gbp1\|Mag-1\|Mpa-1\|Mpa1 | guanylate binding protein 2b | protein-coding |
| chr3_142550117_142550799 | chr3 | 142550118 | 142550799 | + | 1.04533 | TTS (NM_001083312) | TTS (NM_001083312) | -9594 | NM_018734 | 55932 | Mm.439718 | NM_018734 | ENSMUSG00000028268 | **Gbp3** | AW228655\|Gbp4 | guanylate binding protein 3 | protein-coding |
| chr5_105141439_105141821 | chr5 | 105141440 | 105141821 | + | 0.735846 | Intergenic | Intergenic | -2044 | NM_001256005 | 17472 | Mm.440525 | NM_008620 | ENSMUSG00000079363 | **Gbp4** | AW228052\|Mag-2\|Mpa-2\|Mpa2\|mKIAA4245 | guanylate binding protein 4 | protein-coding |
| chr3_142495232_142495481 | chr3 | 142495233 | 142495481 | + | 0.98465 | Intergenic | Intergenic | -1577 | NM_153564 | 229898 | Mm.459245 | NM_153564 | ENSMUSG00000040264 | **Gbp5** | 5330409J06Rik\|Gbp5a | guanylate binding protein 5 | protein-coding |
| chr3_142499155_142499369 | chr3 | 142499156 | 142499369 | + | 0.834842 | intron (NM_153564, intron 1 of 10) | intron (NM_153564, intron 1 of 10) | 2328 | NM_153564 | 229898 | Mm.459245 | NM_153564 | ENSMUSG00000040264 | **Gbp5** | 5330409J06Rik\|Gbp5a | guanylate binding protein 5 | protein-coding |
| chr3_142498962_142499386 | chr3 | 142498963 | 142499386 | + | 0.746384 | intron (NM_153564, intron 1 of 10) | intron (NM_153564, intron 1 of 10) | 2240 | NM_153564 | 229898 | Mm.459245 | NM_153564 | ENSMUSG00000040264 | **Gbp5** | 5330409J06Rik\|Gbp5a | guanylate binding protein 5 | protein-coding |
| chr3_142496136_142497072 | chr3 | 142496137 | 142497072 | + | 0.681267 | promoter-TSS (NM_153564) | promoter-TSS (NM_153564) | -330 | NM_153564 | 229898 | Mm.459245 | NM_153564 | ENSMUSG00000040264 | **Gbp5** | 5330409J06Rik\|Gbp5a | guanylate binding protein 5 | protein-coding |
| chr3_142496487_142496987 | chr3 | 142496488 | 142496987 | + | 0.636253 | promoter-TSS (NM_153564) | promoter-TSS (NM_153564) | -197 | NM_153564 | 229898 | Mm.459245 | NM_153564 | ENSMUSG00000040264 | **Gbp5** | 5330409J06Rik\|Gbp5a | guanylate binding protein 5 | protein-coding |
| chr5_105305528_105306633 | chr5 | 105305529 | 105306633 | + | 1.27491 | Intergenic | Intergenic | -12382 | NM_194336 | 100702 | Mm.491210 | NM_194336 | ENSMUSG00000079362 | **Gbp6** | AI595338\|Mpa2l | guanylate binding protein 6 | protein-coding |
| chr5_105313501_105313815 | chr5 | 105313502 | 105313815 | + | 1.13202 | Intergenic | Intergenic | -19959 | NM_194336 | 100702 | Mm.491210 | NM_194336 | ENSMUSG00000079362 | **Gbp6** | AI595338\|Mpa2l | guanylate binding protein 6 | protein-coding |

| | | | | | | | | | | | | | | | | | |
|---|---|---|---|---|---|---|---|---|---|---|---|---|---|---|---|---|---|
| chr5_105313261_105314007 | chr5 | 105313262 | 105314007 | + | 1.01105 | Intergenic | Intergenic | -19935 | NM_194336 | 100702 | Mm.491210 | NM_194336 | ENSMUSG00000079362 | **Gbp6** | AI595338\|Mpa2l | guanylate binding protein 6 | protein-coding |
| chr5_105305517_105305984 | chr5 | 105305518 | 105305984 | + | 0.86608 | Intergenic | B4A\|SINE\|B4 | -12052 | NM_194336 | 100702 | Mm.491210 | NM_194336 | ENSMUSG00000079362 | **Gbp6** | AI595338\|Mpa2l | guanylate binding protein 6 | protein-coding |
| chr5_105307469_105307862 | chr5 | 105307470 | 105307862 | + | 0.83748 | Intergenic | Intergenic | -13967 | NM_194336 | 100702 | Mm.491210 | NM_194336 | ENSMUSG00000079362 | **Gbp6** | AI595338\|Mpa2l | guanylate binding protein 6 | protein-coding |
| chr3_142524764_142525144 | chr3 | 142524765 | 142525144 | + | 0.657043 | Intergenic | Intergenic | -5382 | NM_001083312 | 229900 | Mm.45740 | NM_145545 | ENSMUSG00000040253 | **Gbp7** | 9830147J24Rik | guanylate binding protein 7 | protein-coding |
| chr5_105056465_105056880 | chr5 | 105056466 | 105056880 | + | 1.15969 | Intergenic | Intergenic | -3112 | NM_029509 | 76074 | Mm.458519 | NM_029509 | ENSMUSG00000034438 | **Gbp8** | 5830443L24Rik\|mGBP8 | guanylate-binding protein 8 | protein-coding |
| chr6_81939535_81940198 | chr6 | 81939536 | 81940198 | + | 0.950624 | intron (NM_177884, intron 6 of 16) | intron (NM_177884, intron 6 of 16) | 16198 | NM_177884 | 330361 | Mm.474487 | NM_177884 | ENSMUSG00000035125 | **Gcfc2** | A130099G21\|AW146020\|GCF2\|Tcf9 | GC-rich sequence DNA binding factor 2 | protein-coding |
| chr2_62463345_62463701 | chr2 | 62463346 | 62463701 | + | 1.28543 | Intergenic | Intergenic | 20130 | NM_008100 | 14526 | Mm.45494 | NM_008100 | ENSMUSG00000000394 | **Gcg** | GLP-1\|Glu\|PPG | glucagon | protein-coding |
| chr2_62526305_62526797 | chr2 | 62526306 | 62526797 | + | 1.22471 | intron (NM_007986, intron 16 of 25) | intron (NM_007986, intron 16 of 25) | -42898 | NM_008100 | 14526 | Mm.45494 | NM_008100 | ENSMUSG00000000394 | **Gcg** | GLP-1\|Glu\|PPG | glucagon | protein-coding |
| chr2_62526875_62527385 | chr2 | 62526876 | 62527385 | + | 0.985325 | intron (NM_007986, intron 16 of 25) | intron (NM_007986, intron 16 of 25) | -43477 | NM_008100 | 14526 | Mm.45494 | NM_008100 | ENSMUSG00000000394 | **Gcg** | GLP-1\|Glu\|PPG | glucagon | protein-coding |
| chr2_62459819_62460672 | chr2 | 62459820 | 62460672 | + | 0.829442 | Intergenic | Intergenic | 23407 | NM_008100 | 14526 | Mm.45494 | NM_008100 | ENSMUSG00000000394 | **Gcg** | GLP-1\|Glu\|PPG | glucagon | protein-coding |
| chr14_47107428_47107899 | chr14 | 47107429 | 47107899 | + | 1.14228 | Intergenic | Intergenic | 81738 | NM_008102 | 14528 | Mm.10651 | NM_008102 | ENSMUSG00000037580 | **Gch1** | GTP-CH\|GTPCH\|Gch | GTP cyclohydrolase 1 | protein-coding |
| chr14_47165092_47165573 | chr14 | 47165093 | 47165573 | + | 0.678195 | intron (NM_008102, intron 3 of 5) | intron (NM_008102, intron 3 of 5) | 24069 | NM_008102 | 14528 | Mm.10651 | NM_008102 | ENSMUSG00000037580 | **Gch1** | GTP-CH\|GTPCH\|Gch | GTP cyclohydrolase 1 | protein-coding |
| chr3_122252944_122253620 | chr3 | 122252945 | 122253620 | + | 0.816629 | intron (NM_008129, intron 1 of 6) | intron (NM_008129, intron 1 of 6) | 7725 | NM_008129 | 14630 | Mm.292676 | NM_008129 | | **Gclm** | AI649393\|Gcmc\|Glclr | glutamate-cysteine ligase, modifier subunit | protein-coding |
| chr9_78022178_78022656 | chr9 | 78022179 | 78022656 | + | 1.31689 | Intergenic | Tigger7\|DNA\|TcMar-Tigger | -29541 | NM_008103 | 14531 | Mm.1400 | NM_008103 | ENSMUSG00000023333 | **Gcm1** | GCMa\|Gcm-rs2\|Gcm1-rs1\|glide | glial cells missing homolog 1 (Drosophila) | protein-coding |
| chr9_78022041_78022790 | chr9 | 78022042 | 78022790 | + | 1.20854 | Intergenic | Tigger7\|DNA\|TcMar-Tigger | -29542 | NM_008103 | 14531 | Mm.1400 | NM_008103 | ENSMUSG00000023333 | **Gcm1** | GCMa\|Gcm-rs2\|Gcm1-rs1\|glide | glial cells missing homolog 1 (Drosophila) | protein-coding |
| chr5_115563876_115564189 | chr5 | 115563877 | 115564189 | + | 0.694211 | TTS (NM_007475) | TTS (NM_007475) | -1230 | NM_172719 | 231659 | Mm.153014 | NM_172719 | ENSMUSG00000041638 | **Gcn1l1** | 4932409G22\|AL022764\|G431004K08Rik\|GCN1L\|mKIAA0219 | GCN1 general control of amino-acid synthesis 1-like 1 (yeast) | protein-coding |
| chr19_17264573_17265304 | chr19 | 17264574 | 17265304 | + | 1.57167 | Intergenic | Intergenic | 74566 | NM_001136484 | 14537 | Mm.244825 | NM_010265 | ENSMUSG00000038843 | **Gcnt1** | 5630400D21Rik\|B130048E03\|C2GlcNAcT\|C2GNT\|IGnT | glucosaminyl (N-acetyl) transferase 1, core 2 | protein-coding |
| chr19_17380490_17381043 | chr19 | 17380491 | 17381043 | + | 1.20344 | Intergenic | Intergenic | -7923 | NM_010265 | 14537 | Mm.244825 | NM_010265 | ENSMUSG00000038843 | **Gcnt1** | 5630400D21Rik\|B130048E03\|C2GlcNAcT\|C2GNT\|IGnT | glucosaminyl (N-acetyl) transferase 1, core 2 | protein-coding |
| chr19_17268722_17269032 | chr19 | 17268723 | 17269032 | + | 0.916384 | Intergenic | MLT1D\|LTR\|ERVL-MaLR | 70628 | NM_001136484 | 14537 | Mm.244825 | NM_010265 | ENSMUSG00000038843 | **Gcnt1** | 5630400D21Rik\|B130048E03\|C2GlcNAcT\|C2GNT\|IGnT | glucosaminyl (N-acetyl) transferase 1, core 2 | protein-coding |
| chr19_17263164_17263486 | chr19 | 17263165 | 17263486 | + | 0.909541 | Intergenic | Intergenic | 76180 | NM_001136484 | 14537 | Mm.244825 | NM_010265 | ENSMUSG00000038843 | **Gcnt1** | 5630400D21Rik\|B130048E03\|C2GlcNAcT\|C2GNT\|IGnT | glucosaminyl (N-acetyl) transferase 1, core 2 | protein-coding |
| chr13_96968642_96969014 | chr13 | 96968643 | 96969014 | + | 0.988035 | Intergenic | ORR1E-int\|LTR\|ERVL-MaLR | 44139 | NM_001166065 | 218476 | Mm.483490 | NM_001166065 | ENSMUSG00000091387 | **Gcnt4** | C2gnt3\|Gm279\|Gm73 | glucosaminyl (N-acetyl) transferase 4, core 2 (beta-1,6-N-acetylglucosaminyltransferase) | protein-coding |
| chr13_96869694_96870185 | chr13 | 96869695 | 96870185 | + | 0.922496 | Intergenic | RLTR17\|LTR\|ERVK | -54749 | NM_001166065 | 218476 | Mm.483490 | NM_001166065 | ENSMUSG00000091387 | **Gcnt4** | C2gnt3\|Gm279\|Gm73 | glucosaminyl (N-acetyl) transferase 4, core 2 (beta-1,6-N-acetylglucosaminyltransferase) | protein-coding |
| chr16_45582589_45582881 | chr16 | 45582590 | 45582881 | + | 0.705918 | intron (NM_198106, intron 18 of 27) | B3\|SINE\|B2 | -27705 | NM_008099 | 14525 | Mm.4799 | NM_008099 | ENSMUSG00000022659 | **Gcsam** | Gcet\|Gcet2\|M17\|M17-L | germinal center associated, signaling and motility | protein-coding |
| chr19_21507243_21508114 | chr19 | 21507244 | 21508114 | + | 0.711869 | Intergenic | Intergenic | -35018 | NM_010266 | 14544 | Mm.45054 | NM_010266 | ENSMUSG00000058624 | **Gda** | AU015411\|AW047581 | guanine deaminase | protein-coding |
| chr10_128870347_128870789 | chr10 | 128870348 | 128870789 | + | 0.948012 | intron (NM_025364, intron 9 of 10) | intron (NM_025364, intron 9 of 10) | 21150 | NM_010272 | 14561 | Mm.299218 | NM_010272 | | **Gdf11** | BMP-11\|Bmp11 | growth differentiation factor 11 | protein-coding |
| chr10_128871657_128872218 | chr10 | 128871658 | 128872218 | + | 0.926553 | intron (NM_025364, intron 9 of 10) | intron (NM_025364, intron 9 of 10) | 19780 | NM_010272 | 14561 | Mm.299218 | NM_010272 | | **Gdf11** | BMP-11\|Bmp11 | growth differentiation factor 11 | protein-coding |
| chr10_128870347_128871086 | chr10 | 128870348 | 128871086 | + | 0.891003 | intron (NM_025364, intron 9 of 10) | intron (NM_025364, intron 9 of 10) | 21001 | NM_010272 | 14561 | Mm.299218 | NM_010272 | | **Gdf11** | BMP-11\|Bmp11 | growth differentiation factor 11 | protein-coding |
| chr13_3541327_3541625 | chr13 | 3541328 | 3541625 | + | 0.843323 | intron (NM_008112, intron 1 of 10) | intron (NM_008112, intron 1 of 10) | 3401 | NM_008112 | 14569 | Mm.153226 | NM_008112 | ENSMUSG00000021218 | **Gdi2** | GDI-B\|GDIB\|Gdi3 | guanosine diphosphate (GDP) dissociation inhibitor 2 | protein-coding |
| chr7_99393752_99394363 | chr7 | 99393753 | 99394363 | + | 0.814985 | intron (NM_201352, intron 2 of 17) | intron (NM_201352, intron 2 of 17) | 12509 | NM_201352 | 233552 | Mm.286317 | NM_201352 | ENSMUSG00000035314 | **Gdpd5** | BC024955\|Gde2 | glycerophosphodiester phosphodiesterase domain containing 5 | protein-coding |
| chr17_80236725_80237341 | chr17 | 80236726 | 80237341 | + | 0.775813 | Intergenic | B3\|SINE\|B2 | 12544 | NM_026053 | 67242 | Mm.279985 | NM_026053 | ENSMUSG00000055760 | **Gemin6** | 2610019B15Rik\|2810470M17Rik | gem (nuclear organelle) associated protein 6 | protein-coding |
| chrX_166185082_166186132 | chrX | 166185083 | 166186132 | + | 1.01559 | intron (NM_146238, intron 3 of 3) | MTEa\|LTR\|ERVL-MaLR | 15127 | NM_146238 | 237221 | Mm.209490 | NM_146238 | ENSMUSG00000040621 | **Gemin8** | B130034M20Rik\|B930082C09Rik\|BC023488\|gemin-8 | gem (nuclear organelle) associated protein 8 | protein-coding |
| chr12_11241893_11242422 | chr12 | 11241894 | 11242422 | + | 1.5392 | exon (NM_177331, exon 14 of 14) | exon (NM_177331, exon 14 of 14) | 23629 | NM_177331 | 209334 | Mm.215295 | NM_177331 | ENSMUSG00000051235 | **Gen1** | 5830483C08Rik | Gen homolog 1, endonuclease (Drosophila) | protein-coding |
| chr5_107734996_107735268 | chr5 | 107734997 | 107735268 | + | 1.15394 | Intergenic | Intergenic | -9327 | NM_001267621 | 14581 | Mm.2078 | NM_010278 | ENSMUSG00000029275 | **Gfi1** | AW495828\|Gfi-1\|Pal-1\|Pal1 | growth factor independent 1 | protein-coding |
| chr5_107731357_107732104 | chr5 | 107731358 | 107732104 | + | 1.04451 | Intergenic | Intergenic | -5926 | NM_001267621 | 14581 | Mm.2078 | NM_010278 | ENSMUSG00000029275 | **Gfi1** | AW495828\|Gfi-1\|Pal-1\|Pal1 | growth factor independent 1 | protein-coding |
| chr5_107756041_107756631 | chr5 | 107756042 | 107756631 | + | 0.900612 | intron (NM_007964, intron 17 of 17) | intron (NM_007964, intron 17 of 17) | -30531 | NM_001267621 | 14581 | Mm.2078 | NM_010278 | ENSMUSG00000029275 | **Gfi1** | AW495828\|Gfi-1\|Pal-1\|Pal1 | growth factor independent 1 | protein-coding |
| chr5_107761713_107762195 | chr5 | 107761714 | 107762195 | + | 0.849604 | intron (NM_007964, intron 17 of 17) | L1_Mus3\|LINE\|L1 | -36149 | NM_001267621 | 14581 | Mm.2078 | NM_010278 | ENSMUSG00000029275 | **Gfi1** | AW495828\|Gfi-1\|Pal-1\|Pal1 | growth factor independent 1 | protein-coding |
| chr5_107777306_107777768 | chr5 | 107777307 | 107777768 | + | 0.843496 | intron (NM_007964, intron 16 of 17) | intron (NM_007964, intron 16 of 17) | -51732 | NM_001267621 | 14581 | Mm.2078 | NM_010278 | ENSMUSG00000029275 | **Gfi1** | AW495828\|Gfi-1\|Pal-1\|Pal1 | growth factor independent 1 | protein-coding |
| chr5_107756041_107756376 | chr5 | 107756042 | 107756376 | + | 0.808797 | intron (NM_007964, intron 17 of 17) | intron (NM_007964, intron 17 of 17) | -30404 | NM_001267621 | 14581 | Mm.2078 | NM_010278 | ENSMUSG00000029275 | **Gfi1** | AW495828\|Gfi-1\|Pal-1\|Pal1 | growth factor independent 1 | protein-coding |
| chr5_107773769_107774362 | chr5 | 107773770 | 107774362 | + | 0.684634 | intron (NM_007964, intron 16 of 17) | intron (NM_007964, intron 16 of 17) | -48261 | NM_001267621 | 14581 | Mm.2078 | NM_010278 | ENSMUSG00000029275 | **Gfi1** | AW495828\|Gfi-1\|Pal-1\|Pal1 | growth factor independent 1 | protein-coding |
| chr13_43261828_43262650 | chr13 | 43261829 | 43262650 | + | 1.54224 | intron (NM_001033399, intron 1 of 1) | Tigger7\|DNA\|TcMar-Tigger | 41933 | NM_001033399 | 328232 | Mm.441399 | NM_001033399 | ENSMUSG00000051335 | **Gfod1** | 9630032O13\|AI850995 | glucose-fructose oxidoreductase domain containing 1 | protein-coding |
| chr13_43261274_43262660 | chr13 | 43261275 | 43262660 | + | 1.08498 | intron (NM_001033399, intron 1 of 1) | (GGA)n\|Simple_repeat\|Simple_repeat | 42205 | NM_001033399 | 328232 | Mm.441399 | NM_001033399 | ENSMUSG00000051335 | **Gfod1** | 9630032O13\|AI850995 | glucose-fructose oxidoreductase domain containing 1 | protein-coding |
| chr13_43328981_43329360 | chr13 | 43328982 | 43329360 | + | 0.787311 | Intergenic | Intergenic | -24999 | NM_001033399 | 328232 | Mm.441399 | NM_001033399 | ENSMUSG00000051335 | **Gfod1** | 9630032O13\|AI850995 | glucose-fructose oxidoreductase domain containing 1 | protein-coding |
| chr13_43241847_43242073 | chr13 | 43241848 | 43242073 | + | 0.701698 | intron (NM_001033399, intron 1 of 1) | B1_Mur1\|SINE\|Alu | 62212 | NM_001033399 | 328232 | Mm.441399 | NM_001033399 | ENSMUSG00000051335 | **Gfod1** | 9630032O13\|AI850995 | glucose-fructose oxidoreductase domain containing 1 | protein-coding |
| chr13_43261276_43261748 | chr13 | 43261277 | 43261748 | + | 0.701329 | intron (NM_001033399, intron 1 of 1) | intron (NM_001033399, intron 1 of 1) | 42660 | NM_001033399 | 328232 | Mm.441399 | NM_001033399 | ENSMUSG00000051335 | **Gfod1** | 9630032O13\|AI850995 | glucose-fructose oxidoreductase domain containing 1 | protein-coding |
| chr13_43306597_43306886 | chr13 | 43306598 | 43306886 | + | 0.683454 | Intergenic | L2a\|LINE\|L2 | -2570 | NM_001033399 | 328232 | Mm.441399 | NM_001033399 | ENSMUSG00000051335 | **Gfod1** | 9630032O13\|AI850995 | glucose-fructose oxidoreductase domain containing 1 | protein-coding |
| chr13_43261272_43261582 | chr13 | 43261273 | 43261582 | + | 0.653734 | intron (NM_001033399, intron 1 of 1) | intron (NM_001033399, intron 1 of 1) | 42745 | NM_001033399 | 328232 | Mm.441399 | NM_001033399 | ENSMUSG00000051335 | **Gfod1** | 9630032O13\|AI850995 | glucose-fructose oxidoreductase domain containing 1 | protein-coding |
| chr11_49770698_49771593 | chr11 | 49770699 | 49771593 | + | 0.823913 | Intergenic | Intergenic | -23009 | NM_013529 | 14584 | Mm.24402 | NM_013529 | ENSMUSG00000020363 | **Gfpt2** | AI480523\|GFAT2 | glutamine fructose-6-phosphate transaminase 2 | protein-coding |
| chr11_49798395_49798736 | chr11 | 49798396 | 49798736 | + | 0.813093 | intron (NM_013529, intron 1 of 18) | MER63A\|DNA\|hAT-Blackjack | 4411 | NM_013529 | 14584 | Mm.24402 | NM_013529 | ENSMUSG00000020363 | **Gfpt2** | AI480523\|GFAT2 | glutamine fructose-6-phosphate transaminase 2 | protein-coding |
| chr11_115588864_115589399 | chr11 | 115588865 | 115589399 | + | 1.42844 | exon (NM_173048, exon 9 of 17) | exon (NM_173048, exon 9 of 17) | 14784 | NM_001252067 | 260302 | Mm.119479 | NM_173048 | ENSMUSG00000020740 | **Gga3** | C230037M19Rik\|mKIAA0154 | golgi associated, gamma adaptin ear containing, ARF binding protein 3 | protein-coding |
| chr14_122830219_122830866 | chr14 | 122830220 | 122830866 | + | 1.70535 | intron (NM_144844, intron 21 of 23) | intron (NM_144844, intron 21 of 23) | 82622 | NM_145466 | 223267 | Mm.379358 | NM_145466 | ENSMUSG00000041625 | **Ggact** | A2ld1 | gamma-glutamylamine cyclotransferase | protein-coding |
| chr14_122725870_122726198 | chr14 | 122725871 | 122726198 | + | 0.87402 | intron (NM_144844, intron 18 of 23) | intron (NM_144844, intron 18 of 23) | 187131 | NM_145466 | 223267 | Mm.379358 | NM_145466 | ENSMUSG00000041625 | **Ggact** | A2ld1 | gamma-glutamylamine cyclotransferase | protein-coding |
| chr14_122725873_122726386 | chr14 | 122725874 | 122726386 | + | 0.870396 | intron (NM_144844, intron 18 of 23) | L1MEf\|LINE\|L1 | 187035 | NM_145466 | 223267 | Mm.379358 | NM_145466 | ENSMUSG00000041625 | **Ggact** | A2ld1 | gamma-glutamylamine cyclotransferase | protein-coding |
| chr14_122747646_122748391 | chr14 | 122747647 | 122748391 | + | 0.750122 | intron (NM_144844, intron 19 of 23) | intron (NM_144844, intron 19 of 23) | 165146 | NM_145466 | 223267 | Mm.379358 | NM_145466 | ENSMUSG00000041625 | **Ggact** | A2ld1 | gamma-glutamylamine cyclotransferase | protein-coding |
| chr14_122725840_122726578 | chr14 | 122725841 | 122726578 | + | 0.68486 | intron (NM_144844, intron 18 of 23) | L1MEf\|LINE\|L1 | 186956 | NM_145466 | 223267 | Mm.379358 | NM_145466 | ENSMUSG00000041625 | **Ggact** | A2ld1 | gamma-glutamylamine cyclotransferase | protein-coding |

| | | | | | | | | | | | | | | | | | |
|---|---|---|---|---|---|---|---|---|---|---|---|---|---|---|---|---|---|
| chr14_122911989_122912253 | chr14 | 122911990 | 122912253 | + | 0.649784 | intron (NM_145466, intron 1 of 2) | intron (NM_145466, intron 1 of 2) | 1044 | NM_145466 | 223267 | Mm.379358 | NM_145466 | ENSMUSG00000041625 | **Ggact** | A2ld1 | gamma-glutamylamine cyclotransferase | protein-coding |
| chr4_20044463_20044876 | chr4 | 20044464 | 20044876 | + | 0.800839 | intron (NM_010281, intron 1 of 8) | intron (NM_010281, intron 1 of 8) | 2618 | NM_010281 | 14590 | Mm.20461 | NM_010281 | | **Ggh** | gamma-GH | gamma-glutamyl hydrolase | protein-coding |
| chr4_20044492_20044746 | chr4 | 20044493 | 20044746 | + | 0.702346 | intron (NM_010281, intron 1 of 8) | intron (NM_010281, intron 1 of 8) | 2567 | NM_010281 | 14590 | Mm.20461 | NM_010281 | | **Ggh** | gamma-GH | gamma-glutamyl hydrolase | protein-coding |
| chr11_84866339_84866528 | chr11 | 84866340 | 84866528 | + | 0.795197 | intron (NM_153144, intron 2 of 13) | intron (NM_153144, intron 2 of 13) | 4304 | NM_153144 | 217039 | Mm.356653 | NM_153144 | ENSMUSG00000020530 | **Ggnbp2** | AI451896\|C75939\|D330017P12Rik\|DIF-3\|Zfp403 | gametogenetin binding protein 2 | protein-coding |
| chr13_14062572_14062833 | chr13 | 14062573 | 14062833 | + | 0.609364 | intron (NM_010282, intron 1 of 3) | intron (NM_010282, intron 1 of 3) | 698 | NM_010282 | 14593 | Mm.148039 | NM_010282 | ENSMUSG00000021302 | **Ggps1** | 1810026C22Rik\|9530089B04Rik\|AI843169\|C79210 | geranylgeranyl diphosphate synthase 1 | protein-coding |
| chr2_35447459_35447913 | chr2 | 35447460 | 35447913 | + | 1.04312 | intron (NM_010283, intron 1 of 8) | intron (NM_010283, intron 1 of 8) | 13739 | NM_010283 | 14594 | Mm.281124 | NM_010283 | ENSMUSG00000035778 | **Ggta1** | AW108479\|GALT\|Gal\|Ggta\|Ggta-1\|alpha3GalT | glycoprotein galactosyltransferase alpha 1, 3 | protein-coding |
| chr2_35465757_35466011 | chr2 | 35465758 | 35466011 | + | 0.830937 | Intergenic | ORR1C2\|LTR\|ERVL-MaLR | -4459 | NM_001145821 | 14594 | Mm.281124 | NM_010283 | ENSMUSG00000035778 | **Ggta1** | AW108479\|GALT\|Gal\|Ggta\|Ggta-1\|alpha3GalT | glycoprotein galactosyltransferase alpha 1, 3 | protein-coding |
| chr1_87315873_87316198 | chr1 | 87315874 | 87316198 | + | 0.758894 | Intergenic | MLT1F\|LTR\|ERVL-MaLR | -10962 | NM_146112 | 227331 | Mm.23065 | NM_146112 | ENSMUSG00000048000 | **Gigyf2** | 2610016F01Rik\|A830080H02Rik\|AI852361\|AW259676\|BC006835\|Tnrc15\|mKIAA0642 | GRB10 interacting GYF protein 2 | protein-coding |
| chr1_97781574_97782219 | chr1 | 97781575 | 97782219 | + | 1.10732 | intron (NM_026250, intron 3 of 7) | intron (NM_026250, intron 3 of 7) | 11718 | NM_026250 | 252876 | Mm.86090 | NM_026250 | ENSMUSG00000026333 | **Gin1** | 4930429M06Rik\|8430411H10Rik\|Gin-1\|Zh2c2 | gypsy retrotransposon integrase 1 | protein-coding |
| chr2_150907170_150907344 | chr2 | 150907171 | 150907344 | + | 1.08713 | Intergenic | Intergenic | -2337 | NM_027014 | 69270 | Mm.271603 | NM_027014 | ENSMUSG00000027454 | **Gins1** | 2810418N01Rik\|Gins4\|PSF1\|mKIAA0186 | GINS complex subunit 1 (Psf1 homolog) | protein-coding |
| chr8_120563531_120564558 | chr8 | 120563532 | 120564558 | + | 0.638374 | intron (NM_001145897, intron 3 of 15) | intron (NM_001145897, intron 3 of 15) | 25030 | NM_178856 | 272551 | Mm.275910 | NM_178856 | | **Gins2** | 2210013I18Rik\|4833427B12Rik\|AI323585\|Pfs2 | GINS complex subunit 2 (Psf2 homolog) | protein-coding |
| chr8_120563766_120564097 | chr8 | 120563767 | 120564097 | + | 0.6273 | intron (NM_001145897, intron 3 of 15) | intron (NM_001145897, intron 3 of 15) | 25143 | NM_178856 | 272551 | Mm.275910 | NM_178856 | | **Gins2** | 2210013I18Rik\|4833427B12Rik\|AI323585\|Pfs2 | GINS complex subunit 2 (Psf2 homolog) | protein-coding |
| chr11_96017781_96018414 | chr11 | 96017782 | 96018414 | + | 0.808723 | Intergenic | Intergenic | -6447 | NM_008119 | 14607 | Mm.248452 | NM_008119 | ENSMUSG00000014351 | **Gip** | - | gastric inhibitory polypeptide | protein-coding |
| chr3_152127159_152127872 | chr3 | 152127160 | 152127872 | + | 1.23711 | intron (NM_016867, intron 3 of 5) | intron (NM_016867, intron 3 of 5) | 38384 | NM_016867 | 54120 | Mm.97 | NM_016867 | | **Gipc2** | 2200002N01Rik\|AU021850\|Semcap2 | GIPC PDZ domain containing family, member 2 | protein-coding |
| chr3_152151937_152152704 | chr3 | 152151938 | 152152704 | + | 1.21 | intron (NM_016867, intron 1 of 5) | B4\|SINE\|B4 | 13579 | NM_016867 | 54120 | Mm.97 | NM_016867 | | **Gipc2** | 2200002N01Rik\|AU021850\|Semcap2 | GIPC PDZ domain containing family, member 2 | protein-coding |
| chr3_152149290_152149901 | chr3 | 152149291 | 152149901 | + | 0.704364 | intron (NM_016867, intron 1 of 5) | intron (NM_016867, intron 1 of 5) | 16304 | NM_016867 | 54120 | Mm.97 | NM_016867 | | **Gipc2** | 2200002N01Rik\|AU021850\|Semcap2 | GIPC PDZ domain containing family, member 2 | protein-coding |
| chr3_152154181_152154392 | chr3 | 152154182 | 152154392 | + | 0.68295 | intron (NM_016867, intron 1 of 5) | ID_B1\|SINE\|B4 | 11613 | NM_016867 | 54120 | Mm.97 | NM_016867 | | **Gipc2** | 2200002N01Rik\|AU021850\|Semcap2 | GIPC PDZ domain containing family, member 2 | protein-coding |
| chr3_152149447_152149782 | chr3 | 152149448 | 152149782 | + | 0.680718 | intron (NM_016867, intron 1 of 5) | intron (NM_016867, intron 1 of 5) | 16285 | NM_016867 | 54120 | Mm.97 | NM_016867 | | **Gipc2** | 2200002N01Rik\|AU021850\|Semcap2 | GIPC PDZ domain containing family, member 2 | protein-coding |
| chr5_114746618_114747268 | chr5 | 114746619 | 114747268 | + | 0.738127 | intron (NM_001077360, intron 12 of 17) | Tigger17a\|DNA\|TcMar-Tigger | 26548 | NM_019834 | 26431 | Mm.195632 | NM_019834 | ENSMUSG00000041890 | **Git2** | 1500036H07Rik\|5830420E16Rik\|6430510B20Rik\|9630056M03Rik\|B230104M05Rik\|CAT2\|mKIAA0148 | G protein-coupled receptor kinase-interactor 2 | protein-coding |
| chr10_56486366_56486944 | chr10 | 56486367 | 56486944 | + | 1.32429 | Intergenic | MLT2F\|LTR\|ERVL | 109355 | NM_010288 | 14609 | Mm.378921 | NM_010288 | ENSMUSG00000050953 | **Gja1** | AU042049\|AW546267\|Cnx43\|Cx43\|Cx43alpha1\|Gja-1\|Npm1\|connexin43 | gap junction protein, alpha 1 | protein-coding |
| chr10_56484959_56485531 | chr10 | 56484960 | 56485531 | + | 1.02682 | Intergenic | Intergenic | 107945 | NM_010288 | 14609 | Mm.378921 | NM_010288 | ENSMUSG00000050953 | **Gja1** | AU042049\|AW546267\|Cnx43\|Cx43\|Cx43alpha1\|Gja-1\|Npm1\|connexin43 | gap junction protein, alpha 1 | protein-coding |
| chr10_56393325_56393752 | chr10 | 56393326 | 56393752 | + | 0.660079 | Intergenic | (TTTTG)n\|Simple_repeat\|Simple_repeat | 16239 | NM_010288 | 14609 | Mm.378921 | NM_010288 | ENSMUSG00000050953 | **Gja1** | AU042049\|AW546267\|Cnx43\|Cx43\|Cx43alpha1\|Gja-1\|Npm1\|connexin43 | gap junction protein, alpha 1 | protein-coding |
| chr3_97005840_97006431 | chr3 | 97005841 | 97006431 | + | 0.825735 | Intergenic | MLT1A1\|LTR\|ERVL-MaLR | -26266 | NM_001271628 | 14613 | Mm.281816 | NM_008121 | ENSMUSG00000057123 | **Gja5** | 5730555N10Rik\|Cnx40\|Cx40\|Gja-5 | gap junction protein, alpha 5 | protein-coding |
| chr3_97004400_97005320 | chr3 | 97004401 | 97005320 | + | 0.627689 | Intergenic | Intergenic | -27542 | NM_001271628 | 14613 | Mm.281816 | NM_008121 | ENSMUSG00000057123 | **Gja5** | 5730555N10Rik\|Cnx40\|Cx40\|Gja-5 | gap junction protein, alpha 5 | protein-coding |
| chrX_160941068_160941676 | chrX | 160941069 | 160941676 | + | 0.793939 | intron (NM_001024624, intron 1 of 18) | MER21B\|LTR\|ERVL | -34320 | NM_001001496 | 414089 | Mm.123113 | NM_001001496 | ENSMUSG00000055691 | **Gja6** | A530015O12Rik\|Cnx33\|Cx33\|Gja-6 | gap junction protein, alpha 6 | protein-coding |
| chrX_101368993_101369496 | chrX | 101368994 | 101369496 | + | 1.41842 | Intergenic | Intergenic | -8092 | NM_008124 | 14618 | Mm.21198 | NM_008124 | ENSMUSG00000047797 | **Gjb1** | AI118175\|Cnx32\|Cx32\|Gjb-1 | gap junction protein, beta 1 | protein-coding |
| chrX_101338728_101339504 | chrX | 101338729 | 101339504 | + | 0.68658 | Intergenic | Intergenic | -38221 | NM_008124 | 14618 | Mm.21198 | NM_008124 | ENSMUSG00000047797 | **Gjb1** | AI118175\|Cnx32\|Cx32\|Gjb-1 | gap junction protein, beta 1 | protein-coding |
| chr6_8638180_8639410 | chr6 | 8638181 | 8639410 | + | 1.23494 | intron (NM_001252266, intron 11 of 11) | B1F\|SINE\|Alu | 118737 | NM_001286729 | 170772 | Mm.210787 | NM_133236 | ENSMUSG00000029638 | **Glcci1** | 2310047L21Rik\|A130036A18Rik\|Fam117c\|Gig18\|Tssn1 | glucocorticoid induced transcript 1 | protein-coding |
| chr6_8638167_8639989 | chr6 | 8638168 | 8639989 | + | 1.07575 | intron (NM_001252266, intron 11 of 11) | intron (NM_001252266, intron 11 of 11) | 119020 | NM_001286729 | 170772 | Mm.210787 | NM_133236 | ENSMUSG00000029638 | **Glcci1** | 2310047L21Rik\|A130036A18Rik\|Fam117c\|Gig18\|Tssn1 | glucocorticoid induced transcript 1 | protein-coding |
| chr9_62054334_62054659 | chr9 | 62054335 | 62054659 | + | 0.716892 | Intergenic | Lx5b\|LINE\|L1 | 16109 | NM_033320 | 93683 | Mm.24411 | NM_033320 | ENSMUSG00000032252 | **Glce** | 1110017N23Rik\|AI747411\|C130034A12Rik\|Hsepi | glucuronyl C5-epimerase | protein-coding |
| chr19_30125196_30125654 | chr19 | 30125197 | 30125654 | + | 1.09127 | intron (NM_138595, intron 15 of 24) | intron (NM_138595, intron 15 of 24) | 50016 | NM_138595 | 104174 | Mm.274852 | NM_138595 | ENSMUSG00000024827 | **Gldc** | D030049L12Rik\|D19Wsu57e | glycine decarboxylase | protein-coding |
| chr19_30118679_30119208 | chr19 | 30118680 | 30119208 | + | 1.0711 | intron (NM_138595, intron 16 of 24) | intron (NM_138595, intron 16 of 24) | 56497 | NM_138595 | 104174 | Mm.274852 | NM_138595 | ENSMUSG00000024827 | **Gldc** | D030049L12Rik\|D19Wsu57e | glycine decarboxylase | protein-coding |
| chr19_30125171_30125957 | chr19 | 30125172 | 30125957 | + | 0.909533 | intron (NM_138595, intron 15 of 24) | intron (NM_138595, intron 15 of 24) | 49877 | NM_138595 | 104174 | Mm.274852 | NM_138595 | ENSMUSG00000024827 | **Gldc** | D030049L12Rik\|D19Wsu57e | glycine decarboxylase | protein-coding |
| chr9_54355632_54356165 | chr9 | 54355633 | 54356165 | + | 0.861503 | Intergenic | L1_Rod\|LINE\|L1 | -54329 | NR_045805 | 73567 | Mm.211290 | NM_001033169 | ENSMUSG00000074343 | **Gldnos** | 1700104A03Rik | gliomedin, opposite strand | ncRNA |
| chr8_111233781_111234315 | chr8 | 111233782 | 111234315 | + | 1.17966 | intron (NM_009149, intron 1 of 25) | intron (NM_009149, intron 1 of 25) | 25154 | NM_009149 | 20340 | Mm.276271 | NM_009149 | ENSMUSG00000003316 | **Glg1** | AI593353\|AW537898\|CFR\|CFR-1\|ESL-1\|MG-160\|MG160\|Selel | golgi apparatus protein 1 | protein-coding |
| chr4_43968697_43968988 | chr4 | 43968698 | 43968988 | + | 1.05539 | intron (NM_027450, intron 3 of 4) | intron (NM_027450, intron 3 of 4) | 11141 | NM_027450 | 384009 | Mm.22213 | NM_027450 | ENSMUSG00000028480 | **Glipr2** | 5730414A08Rik\|C77180\|GAPR-1 | GLI pathogenesis-related 2 | protein-coding |
| chr3_80867361_80867829 | chr3 | 80867362 | 80867829 | + | 0.800085 | intron (NM_001281969, intron 4 of 9) | intron (NM_001281969, intron 4 of 9) | 46019 | NM_001281969 | 14658 | Mm.275639 | NM_010298 | ENSMUSG00000028020 | **Glrb** | AI853901\|Glyrb\|spa\|spastic | glycine receptor, beta subunit | protein-coding |
| chr13_75832441_75833119 | chr13 | 75832442 | 75833119 | + | 1.40239 | Intergenic | Intergenic | -7106 | NM_053108 | 93692 | Mm.25844 | NM_053108 | ENSMUSG00000021591 | **Glrx** | C86710\|D13Wsu156e\|Glrx1\|Grx1\|TTase | glutaredoxin | protein-coding |
| chr13_75832725_75833113 | chr13 | 75832726 | 75833113 | + | 1.40087 | Intergenic | Intergenic | -6967 | NM_053108 | 93692 | Mm.25844 | NM_053108 | ENSMUSG00000021591 | **Glrx** | C86710\|D13Wsu156e\|Glrx1\|Grx1\|TTase | glutaredoxin | protein-coding |
| chr1_52366234_52366921 | chr1 | 52366235 | 52366921 | + | 1.19751 | Intergenic | Intergenic | -133346 | NM_001113383 | 14660 | Mm.398608 | NM_001081081 | ENSMUSG00000026103 | **Gls** | 6330442B14\|AI314027\|B230365M23Rik | glutaminase | protein-coding |
| chr1_52331482_52331771 | chr1 | 52331483 | 52331771 | + | 0.637126 | Intergenic | Intergenic | -98395 | NM_001113383 | 14660 | Mm.398608 | NM_001081081 | ENSMUSG00000026103 | **Gls** | 6330442B14\|AI314027\|B230365M23Rik | glutaminase | protein-coding |
| chr7_15947994_15949093 | chr7 | 15947995 | 15949093 | + | 0.66875 | TTS (NM_153068) | TTS (NM_153068) | -2436 | NM_133831 | 68077 | Mm.277634 | NM_133831 | ENSMUSG00000041560 | **Gltscr2** | 5330430H08Rik\|9430097C02Rik\|AU041936\|AW536441\|PICT-1\|R74911 | glioma tumor suppressor candidate region gene 2 | protein-coding |
| chr14_34317222_34317800 | chr14 | 34317223 | 34317800 | + | 0.654197 | intron (NM_008133, intron 1 of 12) | intron (NM_008133, intron 1 of 12) | 6784 | NM_008133 | 14661 | Mm.10600 | NM_008133 | ENSMUSG00000021794 | **Glud1** | AI118167\|Gdh-X\|Glud\|Gludl | glutamate dehydrogenase 1 | protein-coding |
| chr14_34314049_34314333 | chr14 | 34314050 | 34314333 | + | 0.627004 | intron (NM_008133, intron 1 of 12) | B3\|SINE\|B2 | 3464 | NM_008133 | 14661 | Mm.10600 | NM_008133 | ENSMUSG00000021794 | **Glud1** | AI118167\|Gdh-X\|Glud\|Gludl | glutamate dehydrogenase 1 | protein-coding |
| chr19_58357942_58359096 | chr19 | 58357943 | 58359096 | + | 0.830077 | intron (NM_001285457, intron 3 of 8) | intron (NM_001285457, intron 3 of 8) | 62315 | NR_040449 | 791347 | Mm.452208 | NR_040449 | | **Gm10007** | ENSMUSG00000056739 | predicted gene 10007 | ncRNA |

| | | | | | | | | | | | | | | | | | |
|---|---|---|---|---|---|---|---|---|---|---|---|---|---|---|---|---|---|
| chr19_58357964_58358503 | chr19 | 58357965 | 58358503 | + | 0.68002 | intron (NM_001285457, intron 3 of 8) | intron (NM_001285457, intron 3 of 8) | 62030 | NR_040449 | 791347 | Mm.452208 | NR_040449 | | **Gm10007** | ENSMUSG00000056739 | predicted gene 10007 | ncRNA |
| chr18_5740238_5740847 | chr18 | 5740239 | 5740847 | + | 1.04313 | intron (NM_011546, intron 2 of 7) | L1MD2\|LINE\|L1 | -148106 | NR_033552 | 791318 | Mm.84339 | NR_033552 | ENSMUSG00000063087 | **Gm10125** | ENSMUSG00000063087\|Zeb1os | predicted gene 10125 | ncRNA |
| chr18_5694222_5694830 | chr18 | 5694223 | 5694830 | + | 0.792683 | intron (NM_011546, intron 1 of 7) | intron (NM_011546, intron 1 of 7) | -102089 | NR_033552 | 791318 | Mm.84339 | NR_033552 | ENSMUSG00000063087 | **Gm10125** | ENSMUSG00000063087\|Zeb1os | predicted gene 10125 | ncRNA |
| chr17_80385324_80385694 | chr17 | 80385325 | 80385694 | + | 0.656594 | intron (NM_001145452, intron 16 of 17) | intron (NM_001145452, intron 16 of 17) | -11967 | NR_028385 | 791338 | Mm.204722 | NR_028385 | | **Gm10190** | ENSMUSG00000066938 | predicted gene 10190 | ncRNA |
| chr14_23458483_23458905 | chr14 | 23458484 | 23458905 | + | 0.836787 | intron (NM_001253369, intron 15 of 29) | intron (NM_001253369, intron 15 of 29) | -364123 | NR_033550 | 791307 | Mm.364384 | NR_033550 | ENSMUSG00000068680 | **Gm10248** | ENSMUSG00000068680 | predicted gene 10248 | ncRNA |
| chr5_120130158_120130488 | chr5 | 120130159 | 120130488 | + | 0.719826 | intron (NM_028762, intron 11 of 23) | ORR1D1\|LTR\|ERVL-MaLR | 8940 | NR_045793 | 100503427 | Mm.446834 | NR_045793 | ENSMUSG00000097368 | **Gm10390** | ENSMUSG00000072671 | predicted gene 10390 | ncRNA |
| chr5_120129953_120130665 | chr5 | 120129954 | 120130665 | + | 0.65433 | intron (NM_028762, intron 11 of 23) | ORR1D1\|LTR\|ERVL-MaLR | 8954 | NR_045793 | 100503427 | Mm.446834 | NR_045793 | ENSMUSG00000097368 | **Gm10390** | ENSMUSG00000072671 | predicted gene 10390 | ncRNA |
| chr5_109830232_109831142 | chr5 | 109830233 | 109831142 | + | 1.28866 | Intergenic | Intergenic | 3455 | NR_027964 | 667213 | Mm.454429 | NR_027964 | | **Gm10416** | EG667213\|ENSMUSG00000072761 | zinc finger protein pseudogene | pseudo |
| chr5_109778428_109778909 | chr5 | 109778429 | 109778909 | + | 0.868066 | Intergenic | Intergenic | 55473 | NR_027964 | 667213 | Mm.454429 | NR_027964 | | **Gm10416** | EG667213\|ENSMUSG00000072761 | zinc finger protein pseudogene | pseudo |
| chr18_75466807_75467242 | chr18 | 75466808 | 75467242 | + | 0.772702 | intron (NM_201354, intron 10 of 11) | L1MB1\|LINE\|L1 | -47620 | NR_045879 | 100038353 | Mm.440724 | NR_045879 | ENSMUSG00000097712 | **Gm10532** | ENSMUSG00000073534 | predicted gene 10532 | ncRNA |
| chr7_132312444_132312672 | chr7 | 132312445 | 132312672 | + | 1.06379 | intron (NM_029935, intron 1 of 7) | intron (NM_029935, intron 1 of 7) | -3106 | NR_028578 | 100043682 | Mm.482794 | NR_028578 | | **Gm10584** | ENSMUSG00000073814 | predicted gene 10584 | ncRNA |
| chr7_132283690_132284297 | chr7 | 132283691 | 132284297 | + | 0.683772 | intron (NM_029935, intron 1 of 7) | (GA)n\|Simple_repeat\|Simple_repeat | -31670 | NR_028578 | 100043682 | Mm.482794 | NR_028578 | | **Gm10584** | ENSMUSG00000073814 | predicted gene 10584 | ncRNA |
| chr3_146327267_146327499 | chr3 | 146327268 | 146327499 | + | 0.739022 | Intergenic | Intergenic | 51561 | NR_033542 | 100038607 | Mm.365888 | NR_033542 | | **Gm10636** | ENSMUSG00000074175 | predicted gene 10636 | ncRNA |
| chr8_86704521_86704761 | chr8 | 86704522 | 86704761 | + | 0.964537 | intron (NM_001168591, intron 5 of 8) | intron (NM_001168591, intron 5 of 8) | -41058 | NR_027829 | 666945 | Mm.391806 | NR_027829 | | **Gm10638** | ENSMUSG00000074178 | predicted gene 10638 | ncRNA |
| chr9_47733677_47734650 | chr9 | 47733678 | 47734650 | + | 0.700614 | intron (NM_001025600, intron 1 of 8) | intron (NM_001025600, intron 1 of 8) | 84044 | NR_046048 | 100038460 | Mm.447649 | NR_046048 | | **Gm10677** | ENSMUSG00000074363 | predicted gene 10677 | ncRNA |
| chr19_43880389_43880792 | chr19 | 43880390 | 43880792 | + | 0.741015 | intron (NM_028029, intron 4 of 15) | intron (NM_028029, intron 4 of 15) | 41788 | NR_033472 | 100038628 | Mm.212585 | NR_033472 | ENSMUSG00000074828 | **Gm10768** | ENSMUSG00000074828 | predicted gene 10768 | ncRNA |
| chr13_41900090_41900549 | chr13 | 41900091 | 41900549 | + | 0.922156 | intron (NR_033545, intron 3 of 6) | MTEa\|LTR\|ERVL-MaLR | 34405 | NR_033545 | 100038577 | Mm.387093 | NR_033545 | | **Gm10790** | ENSMUSG00000074927 | predicted gene 10790 | ncRNA |
| chr10_22391680_22392159 | chr10 | 22391681 | 22392159 | + | 0.915418 | Intergenic | Intergenic | -10893 | NR_028580 | 100038752 | Mm.392859 | NR_028580 | ENSMUSG00000096937 | **Gm10825** | ENSMUSG00000075295 | predicted gene 10825 | ncRNA |
| chr10_22391680_22392393 | chr10 | 22391681 | 22392393 | + | 0.876592 | Intergenic | Intergenic | -10776 | NR_028580 | 100038752 | Mm.392859 | NR_028580 | ENSMUSG00000096937 | **Gm10825** | ENSMUSG00000075295 | predicted gene 10825 | ncRNA |
| chr15_79176570_79177317 | chr15 | 79176571 | 79177317 | + | 1.15439 | intron (NR_029470, intron 2 of 4) | intron (NR_029470, intron 2 of 4) | 10878 | NR_029470 | 100041655 | Mm.483310 | NR_029470 | ENSMUSG00000075555 | **Gm10863** | ENSMUSG00000075555 | predicted gene 10863 | ncRNA |
| chr17_57108703_57109269 | chr17 | 57108704 | 57109269 | + | 1.85087 | Intergenic | Intergenic | -3044 | NR_033508 | 100169874 | Mm.76700 | NR_033508 | ENSMUSG00000079414 | **Gm11110** | ENSMUSG00000079414 | predicted gene 11110 | ncRNA |
| chr17_57125883_57126582 | chr17 | 57125884 | 57126582 | + | 1.03985 | Intergenic | MLTR14\|LTR\|ERV1 | -20291 | NR_033508 | 100169874 | Mm.76700 | NR_033508 | ENSMUSG00000079414 | **Gm11110** | ENSMUSG00000079414 | predicted gene 11110 | ncRNA |
| chr11_79059656_79060335 | chr11 | 79059657 | 79060335 | + | 1.48081 | intron (NM_013571, intron 2 of 17) | intron (NM_013571, intron 2 of 17) | 36399 | NR_045873 | 790910 | Mm.440410 | NR_045873 | | **Gm11201** | ENSMUSG00000072637\|OTTMUSG00000000169 | predicted gene 11201 | ncRNA |
| chr11_79056960_79057255 | chr11 | 79056961 | 79057255 | + | 0.664672 | intron (NM_013571, intron 2 of 17) | intron (NM_013571, intron 2 of 17) | 33511 | NR_045873 | 790910 | Mm.440410 | NR_045873 | | **Gm11201** | ENSMUSG00000072637\|OTTMUSG00000000169 | predicted gene 11201 | ncRNA |
| chr4_63590628_63591017 | chr4 | 63590629 | 63591017 | + | 0.759336 | Intergenic | Intergenic | 9941 | NR_028584 | 670833 | Mm.32213 | NR_028583 | ENSMUSG00000084782 | **Gm11213** | OTTMUSG00000000252 | predicted gene 11213 | ncRNA |
| chr13_24608919_24609180 | chr13 | 24608920 | 24609180 | + | 0.804121 | intron (NM_001286100, intron 1 of 22) | intron (NM_001286100, intron 1 of 22) | -4283 | NR_024599 | 76024 | Mm.440710 | NR_024599 | | **Gm11346** | 5830431A10Rik\|AI447617\|Gm11347\|OTTMUSG00000000601\|OTTMUSG00000000602 | X-linked lymphocyte-regulated 5 pseudogene | pseudo |
| chr4_83375613_83376115 | chr4 | 83375614 | 83376115 | + | 0.943905 | Intergenic | Intergenic | 14804 | NR_045450 | 790913 | Mm.406752 | NR_045450 | ENSMUSG00000086321 | **Gm11413** | OTTMUSG00000000829 | predicted gene 11413 | ncRNA |
| chr4_83380336_83380866 | chr4 | 83380337 | 83380866 | + | 0.710063 | intron (NR_045450, intron 2 of 2) | intron (NR_045450, intron 2 of 2) | 10067 | NR_045450 | 790913 | Mm.406752 | NR_045450 | ENSMUSG00000086321 | **Gm11413** | OTTMUSG00000000829 | predicted gene 11413 | ncRNA |
| chr3_36531933_36532387 | chr3 | 36531934 | 36532387 | + | 0.687251 | Intergenic | Intergenic | -10715 | NR_040411 | 100503068 | Mm.389423 | NR_040411 | | **Gm11549** | OTTMUSG00000002111 | predicted gene 11549 | ncRNA |
| chr11_98595936_98596353 | chr11 | 98595937 | 98596353 | + | 1.63041 | Intergenic | Intergenic | -2146 | NM_001195544 | 192852 | Mm.342821 | NM_001195544 | | **Gm12** | - | predicted gene 12 | protein-coding |
| chr11_98595491_98595850 | chr11 | 98595492 | 98595850 | + | 0.971265 | Intergenic | Intergenic | -2620 | NM_001195544 | 192852 | Mm.342821 | NM_001195544 | | **Gm12** | - | predicted gene 12 | protein-coding |
| chr11_23570000_23570574 | chr11 | 23570001 | 23570574 | + | 0.776751 | Intergenic | Intergenic | -11445 | NR_004857 | 100134990 | Mm.247328 | NR_004857 | ENSMUSG00000085415 | **Gm12060** | OTTMUSG00000005275 | selenoprotein K pseudogene | pseudo |
| chr11_46509786_46510485 | chr11 | 46509787 | 46510485 | + | 1.36931 | intron (NR_105955, intron 2 of 4) | intron (NR_105955, intron 2 of 4) | -14115 | NM_001163357 | 210535 | Mm.173153 | NM_001163357 | ENSMUSG00000078924 | **Gm12169** | OTTMUSG00000005497 | predicted gene 12169 | protein-coding |
| chr11_46506959_46507577 | chr11 | 46506960 | 46507577 | + | 1.31444 | intron (NR_105955, intron 2 of 4) | intron (NR_105955, intron 2 of 4) | -16983 | NM_001163357 | 210535 | Mm.173153 | NM_001163357 | ENSMUSG00000078924 | **Gm12169** | OTTMUSG00000005497 | predicted gene 12169 | protein-coding |
| chr11_53861001_53861313 | chr11 | 53861002 | 53861313 | + | 0.642615 | Intergenic | Intergenic | -1901 | NR_033332 | 622459 | Mm.479656 | NR_033332 | ENSMUSG00000081769 | **Gm12216** | 602-19-115\|OTTMUSG00000005650 | predicted gene 12216 | ncRNA |
| chr4_55443450_55444086 | chr4 | 55443451 | 55444086 | + | 1.21152 | Intergenic | Intergenic | -24926 | NR_040674 | 100415914 | Mm.261765 | NR_040674 | ENSMUSG00000087070 | **Gm12505** | OTTMUSG00000007226 | predicted gene 12505 | ncRNA |
| chr4_55451038_55451368 | chr4 | 55451039 | 55451368 | + | 0.727714 | Intergenic | B3\|SINE\|B2 | -32361 | NR_040674 | 100415914 | Mm.261765 | NR_040674 | ENSMUSG00000087070 | **Gm12505** | OTTMUSG00000007226 | predicted gene 12505 | ncRNA |
| chr4_55415473_55416101 | chr4 | 55415474 | 55416101 | + | 0.701226 | intron (NR_040674, intron 1 of 3) | intron (NR_040674, intron 1 of 3) | 3055 | NR_040674 | 100415914 | Mm.261765 | NR_040674 | ENSMUSG00000087070 | **Gm12505** | OTTMUSG00000007226 | predicted gene 12505 | ncRNA |
| chr4_55442135_55442664 | chr4 | 55442136 | 55442664 | + | 0.625144 | Intergenic | L2\|LINE\|L2 | -23558 | NR_040674 | 100415914 | Mm.261765 | NR_040674 | ENSMUSG00000087070 | **Gm12505** | OTTMUSG00000007226 | predicted gene 12505 | ncRNA |
| chr4_89002971_89003594 | chr4 | 89002972 | 89003594 | + | 1.86662 | Intergenic | L1MA4\|LINE\|L1 | -47024 | NR_033533 | 100040617 | Mm.171720 | NR_033533 | ENSMUSG00000085183 | **Gm12603** | OTTMUSG00000007693 | predicted gene 12603 | ncRNA |
| chr4_89008481_89009055 | chr4 | 89008482 | 89009055 | + | 1.27028 | Intergenic | Intergenic | -41539 | NR_033533 | 100040617 | Mm.171720 | NR_033533 | ENSMUSG00000085183 | **Gm12603** | OTTMUSG00000007693 | predicted gene 12603 | ncRNA |
| chr4_89008457_89009267 | chr4 | 89008458 | 89009267 | + | 1.22143 | Intergenic | Intergenic | -41445 | NR_033533 | 100040617 | Mm.171720 | NR_033533 | ENSMUSG00000085183 | **Gm12603** | OTTMUSG00000007693 | predicted gene 12603 | ncRNA |
| chr4_89006790_89007036 | chr4 | 89006791 | 89007036 | + | 1.21098 | Intergenic | RMER5\|LTR\|ERV1 | -43394 | NR_033533 | 100040617 | Mm.171720 | NR_033533 | ENSMUSG00000085183 | **Gm12603** | OTTMUSG00000007693 | predicted gene 12603 | ncRNA |
| chr4_89043937_89044444 | chr4 | 89043938 | 89044444 | + | 0.981131 | Intergenic | Intergenic | -6116 | NR_033533 | 100040617 | Mm.171720 | NR_033533 | ENSMUSG00000085183 | **Gm12603** | OTTMUSG00000007693 | predicted gene 12603 | ncRNA |
| chr4_88998354_88998623 | chr4 | 88998355 | 88998623 | + | 0.960301 | Intergenic | Intergenic | -51818 | NR_033533 | 100040617 | Mm.171720 | NR_033533 | ENSMUSG00000085183 | **Gm12603** | OTTMUSG00000007693 | predicted gene 12603 | ncRNA |
| chr4_89001441_89001877 | chr4 | 89001442 | 89001877 | + | 0.814182 | Intergenic | Intergenic | -48648 | NR_033533 | 100040617 | Mm.171720 | NR_033533 | ENSMUSG00000085183 | **Gm12603** | OTTMUSG00000007693 | predicted gene 12603 | ncRNA |
| chr4_89090861_89091293 | chr4 | 89090862 | 89091293 | + | 0.741439 | Intergenic | URR1B\|DNA\|hAT-Charlie | 40770 | NR_033533 | 100040617 | Mm.171720 | NR_033533 | ENSMUSG00000085183 | **Gm12603** | OTTMUSG00000007693 | predicted gene 12603 | ncRNA |
| chr4_89038785_89039241 | chr4 | 89038786 | 89039241 | + | 0.673808 | Intergenic | L1MB2\|LINE\|L1 | -11294 | NR_033533 | 100040617 | Mm.171720 | NR_033533 | ENSMUSG00000085183 | **Gm12603** | OTTMUSG00000007693 | predicted gene 12603 | ncRNA |
| chr4_102990103_102990401 | chr4 | 102990104 | 102990401 | + | 0.650789 | promoter-TSS (NR_040444) | promoter-TSS (NR_040444) | -497 | NR_040444 | 100504717 | Mm.394445 | NR_040444 | ENSMUSG00000085181 | **Gm12709** | OTTMUSG00000008056 | predicted gene 12709 | ncRNA |
| chr4_102104444_102105010 | chr4 | 102104445 | 102105010 | + | 0.990966 | Intergenic | Intergenic | 117887 | NM_001085520 | 381536 | Mm.297685 | NM_001085520 | | **Gm12789** | OTTMUSG00000008445 | predicted gene 12789 | protein-coding |
| chr4_127114573_127115745 | chr4 | 127114574 | 127115745 | + | 1.1531 | intron (NM_001285887, intron 12 of 13) | MER44D\|DNA\|TcMar-Tigger | -10884 | NM_001099319 | 100039968 | Mm.392655 | NM_001099319 | ENSMUSG00000070737 | **Gm12942** | OTTMUSG00000009332\|Zmym6nb | predicted gene 12942 | protein-coding |
| chr4_147808372_147808937 | chr4 | 147808373 | 147808937 | + | 1.36917 | intron (NM_001127189, intron 1 of 5) | LTRIS2\|LTR\|ERV1 | 1133 | NM_001127189 | 100041677 | Mm.327194 | NM_001127189 | ENSMUSG00000078495 | **Gm13157** | OTTMUSG00000010679 | predicted gene 13157 | protein-coding |
| chr4_145553618_145554027 | chr4 | 145553619 | 145554027 | + | 1.17223 | Intergenic | RMER15\|LTR\|ERVL | -63323 | NM_001205101 | 433801 | Mm.276881 | NM_001013808 | | **Gm13212** | OTTMUSG00000010965 | predicted gene 13212 | protein-coding |
| chr4_145585906_145586456 | chr4 | 145585907 | 145586456 | + | 0.943566 | Intergenic | LTRIS2\|LTR\|ERV1 | -30965 | NM_001205101 | 433801 | Mm.276881 | NM_001013808 | | **Gm13212** | OTTMUSG00000010965 | predicted gene 13212 | protein-coding |
| chr4_88747693_88748043 | chr4 | 88747694 | 88748043 | + | 0.652799 | Intergenic | RMER2\|LTR\|ERV1 | -7000 | NM_001085528 | 435791 | Mm.473673 | NM_001085528 | ENSMUSG00000094618 | **Gm13271** | Ifnz\|OTTMUSG00000011262 | predicted gene 13271 | protein-coding |
| chr2_11315726_11316106 | chr2 | 11315727 | 11316106 | + | 1.12497 | Intergenic | Intergenic | -23573 | NR_040369 | 100503147 | Mm.91961 | NR_040369 | | **Gm13293** | OTTMUSG00000011311 | predicted gene 13293 | ncRNA |
| chr2_11312237_11313570 | chr2 | 11312238 | 11313570 | + | 0.894141 | Intergenic | Intergenic | -26585 | NR_040369 | 100503147 | Mm.91961 | NR_040369 | | **Gm13293** | OTTMUSG00000011311 | predicted gene 13293 | ncRNA |
| chr2_11282891_11283747 | chr2 | 11282892 | 11283747 | + | 0.62265 | intron (NM_008859, intron 14 of 17) | intron (NM_008859, intron 14 of 17) | -56170 | NR_040369 | 100503147 | Mm.91961 | NR_040369 | | **Gm13293** | OTTMUSG00000011311 | predicted gene 13293 | ncRNA |
| chr2_14770480_14771475 | chr2 | 14770481 | 14771475 | + | 0.818718 | intron (NM_023116, intron 2 of 13) | intron (NM_023116, intron 2 of 13) | 50336 | NR_028497 | 625342 | Mm.484784 | NR_028497 | | **Gm13315** | OTTMUSG00000011384 | lactate dehydrogenase A pseudogene | pseudo |
| chr2_21054508_21054948 | chr2 | 21054509 | 21054948 | + | 0.685701 | Intergenic | Intergenic | 85854 | NR_033225 | 433408 | Mm.386941 | NR_033225 | | **Gm13375** | OTTMUSG00000011664 | predicted gene 13375 | ncRNA |
| chr2_50415874_50416694 | chr2 | 50415875 | 50416694 | + | 0.996484 | intron (NR_040362, intron 1 of 2) | intron (NR_040362, intron 1 of 2) | 119474 | NR_040362 | 100502624 | Mm.382825 | NR_040361 | | **Gm13483** | OTTMUSG00000012382 | predicted gene 13483 | ncRNA |

| | | | | | | | | | | | | | | | | | |
|---|---|---|---|---|---|---|---|---|---|---|---|---|---|---|---|---|---|
| chr2_50416091_50416683 | chr2 | 50416092 | 50416683 | + | 0.985383 | intron (NR_040362, intron 1 of 2) | intron (NR_040362, intron 1 of 2) | 119577 | NR_040362 | 100502624 | Mm.382825 | NR_040361 | | **Gm13483** | OTTMUSG00000012382 | predicted gene 13483 | ncRNA |
| chr2_58187660_58188168 | chr2 | 58187661 | 58188168 | + | 1.43869 | Intergenic | Intergenic | 23940 | NR_045895 | 100042926 | Mm.446515 | NR_045895 | ENSMUSG00000086862 | **Gm13546** | OTTMUSG00000012675 | predicted gene 13546 | ncRNA |
| chr2_58189546_58190009 | chr2 | 58189547 | 58190009 | + | 1.05169 | Intergenic | Intergenic | 25804 | NR_045895 | 100042926 | Mm.446515 | NR_045895 | ENSMUSG00000086862 | **Gm13546** | OTTMUSG00000012675 | predicted gene 13546 | ncRNA |
| chr2_58169116_58169605 | chr2 | 58169117 | 58169605 | + | 0.829374 | intron (NR_045895, intron 2 of 3) | intron (NR_045895, intron 2 of 3) | 5387 | NR_045895 | 100042926 | Mm.446515 | NR_045895 | ENSMUSG00000086862 | **Gm13546** | OTTMUSG00000012675 | predicted gene 13546 | ncRNA |
| chr2_58169121_58169867 | chr2 | 58169122 | 58169867 | + | 0.738402 | intron (NR_045895, intron 2 of 3) | intron (NR_045895, intron 2 of 3) | 5520 | NR_045895 | 100042926 | Mm.446515 | NR_045895 | ENSMUSG00000086862 | **Gm13546** | OTTMUSG00000012675 | predicted gene 13546 | ncRNA |
| chr2_58183739_58184038 | chr2 | 58183740 | 58184038 | + | 0.663855 | Intergenic | Intergenic | 19915 | NR_045895 | 100042926 | Mm.446515 | NR_045895 | ENSMUSG00000086862 | **Gm13546** | OTTMUSG00000012675 | predicted gene 13546 | ncRNA |
| chr2_29693912_29695207 | chr2 | 29693913 | 29695207 | + | 0.963144 | intron (NM_001039086, intron 4 of 25) | intron (NM_001039086, intron 4 of 25) | -66968 | NM_001177392 | 433416 | Mm.330112 | NM_001034887 | ENSMUSG00000075425 | **Gm13547** | OTTMUSG00000012686 | predicted gene 13547 | protein-coding |
| chr2_80374857_80375204 | chr2 | 80374858 | 80375204 | + | 1.23671 | Intergenic | Intergenic | -16333 | NR_040370 | 100503167 | Mm.265892 | NR_040370 | | **Gm13752** | OTTMUSG00000013925 | predicted gene 13752 | ncRNA |
| chr2_80374688_80375190 | chr2 | 80374689 | 80375190 | + | 1.13361 | Intergenic | Intergenic | -16425 | NR_040370 | 100503167 | Mm.265892 | NR_040370 | | **Gm13752** | OTTMUSG00000013925 | predicted gene 13752 | ncRNA |
| chr2_80374465_80375256 | chr2 | 80374466 | 80375256 | + | 1.05737 | Intergenic | Intergenic | -16503 | NR_040370 | 100503167 | Mm.265892 | NR_040370 | | **Gm13752** | OTTMUSG00000013925 | predicted gene 13752 | ncRNA |
| chr2_93411157_93411620 | chr2 | 93411158 | 93411620 | + | 0.765114 | intron (NR_040529, intron 3 of 5) | intron (NR_040529, intron 3 of 5) | 7842 | NR_040529 | 639658 | Mm.66880 | NR_040529 | ENSMUSG00000085852 | **Gm13807** | OTTMUSG00000014546 | predicted gene 13807 | ncRNA |
| chr5_115068691_115069331 | chr5 | 115068692 | 115069331 | + | 1.07068 | intron (NM_029012, intron 2 of 10) | intron (NM_029012, intron 2 of 10) | 38951 | NM_001271591 | 100502825 | Mm.361657 | NM_001271590 | | **Gm13826** | OTTMUSG00000014624 | predicted gene 13826 | protein-coding |
| chr2_128457467_128458367 | chr2 | 128457468 | 128458367 | + | 1.5065 | Intergenic | Intergenic | -28566 | NR_028590 | 100043424 | Mm.297222 | NR_028589 | | **Gm14005** | OTTMUSG00000015351 | predicted gene 14005 | ncRNA |
| chr2_128544339_128545199 | chr2 | 128544340 | 128545199 | + | 1.3667 | Intergenic | Intergenic | -115418 | NR_028590 | 100043424 | Mm.297222 | NR_028589 | | **Gm14005** | OTTMUSG00000015351 | predicted gene 14005 | ncRNA |
| chr2_128379630_128379911 | chr2 | 128379631 | 128379911 | + | 1.22877 | intron (NR_028591, intron 1 of 1) | intron (NR_028591, intron 1 of 1) | 49580 | NR_028591 | 100043424 | Mm.297222 | NR_028589 | | **Gm14005** | OTTMUSG00000015351 | predicted gene 14005 | ncRNA |
| chr2_128465291_128465995 | chr2 | 128465292 | 128465995 | + | 1.14532 | Intergenic | Intergenic | -36292 | NR_028590 | 100043424 | Mm.297222 | NR_028589 | | **Gm14005** | OTTMUSG00000015351 | predicted gene 14005 | ncRNA |
| chr2_128333411_128334019 | chr2 | 128333412 | 128334019 | + | 1.093 | intron (NR_028589, intron 1 of 2) | intron (NR_028589, intron 1 of 2) | 95636 | NR_028591 | 100043424 | Mm.297222 | NR_028589 | | **Gm14005** | OTTMUSG00000015351 | predicted gene 14005 | ncRNA |
| chr2_128377027_128378007 | chr2 | 128377028 | 128378007 | + | 0.989899 | intron (NR_028591, intron 1 of 1) | intron (NR_028591, intron 1 of 1) | 51834 | NR_028591 | 100043424 | Mm.297222 | NR_028589 | | **Gm14005** | OTTMUSG00000015351 | predicted gene 14005 | ncRNA |
| chr2_128368425_128368989 | chr2 | 128368426 | 128368989 | + | 0.983671 | intron (NR_028589, intron 1 of 2) | intron (NR_028589, intron 1 of 2) | 60644 | NR_028591 | 100043424 | Mm.297222 | NR_028589 | | **Gm14005** | OTTMUSG00000015351 | predicted gene 14005 | ncRNA |
| chr2_128380470_128380998 | chr2 | 128380471 | 128380998 | + | 0.847183 | intron (NR_028591, intron 1 of 1) | intron (NR_028591, intron 1 of 1) | 48617 | NR_028591 | 100043424 | Mm.297222 | NR_028589 | | **Gm14005** | OTTMUSG00000015351 | predicted gene 14005 | ncRNA |
| chr2_128433593_128433956 | chr2 | 128433594 | 128433956 | + | 0.846226 | Intergenic | Intergenic | -4424 | NR_028590 | 100043424 | Mm.297222 | NR_028589 | | **Gm14005** | OTTMUSG00000015351 | predicted gene 14005 | ncRNA |
| chr2_128380612_128380994 | chr2 | 128380613 | 128380994 | + | 0.813604 | intron (NR_028591, intron 1 of 1) | intron (NR_028591, intron 1 of 1) | 48548 | NR_028591 | 100043424 | Mm.297222 | NR_028589 | | **Gm14005** | OTTMUSG00000015351 | predicted gene 14005 | ncRNA |
| chr2_128535870_128536877 | chr2 | 128535871 | 128536877 | + | 0.790695 | Intergenic | Intergenic | -107023 | NR_028590 | 100043424 | Mm.297222 | NR_028589 | | **Gm14005** | OTTMUSG00000015351 | predicted gene 14005 | ncRNA |
| chr2_128430705_128431168 | chr2 | 128430706 | 128431168 | + | 0.787286 | Intergenic | Intergenic | -1586 | NR_028590 | 100043424 | Mm.297222 | NR_028589 | | **Gm14005** | OTTMUSG00000015351 | predicted gene 14005 | ncRNA |
| chr2_128401179_128401762 | chr2 | 128401180 | 128401762 | + | 0.776002 | intron (NR_028591, intron 1 of 1) | MIR\|SINE\|MIR | 27880 | NR_028591 | 100043424 | Mm.297222 | NR_028589 | | **Gm14005** | OTTMUSG00000015351 | predicted gene 14005 | ncRNA |
| chr2_128544903_128545181 | chr2 | 128544904 | 128545181 | + | 0.774455 | Intergenic | Intergenic | -115691 | NR_028590 | 100043424 | Mm.297222 | NR_028589 | | **Gm14005** | OTTMUSG00000015351 | predicted gene 14005 | ncRNA |
| chr2_128377027_128377494 | chr2 | 128377028 | 128377494 | + | 0.762452 | intron (NR_028591, intron 1 of 1) | intron (NR_028591, intron 1 of 1) | 52090 | NR_028591 | 100043424 | Mm.297222 | NR_028589 | | **Gm14005** | OTTMUSG00000015351 | predicted gene 14005 | ncRNA |
| chr2_128372291_128372585 | chr2 | 128372292 | 128372585 | + | 0.639361 | intron (NR_028591, intron 1 of 1) | intron (NR_028591, intron 1 of 1) | 56913 | NR_028591 | 100043424 | Mm.297222 | NR_028589 | | **Gm14005** | OTTMUSG00000015351 | predicted gene 14005 | ncRNA |
| chr2_128387901_128388298 | chr2 | 128387902 | 128388298 | + | 0.617538 | intron (NR_028591, intron 1 of 1) | intron (NR_028591, intron 1 of 1) | 41251 | NR_028591 | 100043424 | Mm.297222 | NR_028589 | | **Gm14005** | OTTMUSG00000015351 | predicted gene 14005 | ncRNA |
| chr2_152349642_152350478 | chr2 | 152349643 | 152350478 | + | 1.00143 | intron (NR_033505, intron 1 of 4) | intron (NR_033505, intron 1 of 4) | 4391 | NR_033505 | 791417 | Mm.451219 | NR_033505 | ENSMUSG00000068062 | **Gm14164** | OTTMUSG00000015846 | predicted gene 14164 | ncRNA |
| chrX_11268040_11268844 | chrX | 11268041 | 11268844 | + | 1.10526 | Intergenic | Intergenic | -30815 | NM_001111037 | 100042922 | Mm.374311 | NM_001111037 | ENSMUSG00000095208 | **Gm14483** | OTTMUSG00000016789 | predicted gene 14483 | protein-coding |
| chrX_11253957_11254420 | chrX | 11253958 | 11254420 | + | 1.02612 | Intergenic | Intergenic | -45068 | NM_001111037 | 100042922 | Mm.374311 | NM_001111037 | ENSMUSG00000095208 | **Gm14483** | OTTMUSG00000016789 | predicted gene 14483 | protein-coding |
| chrX_11251527_11251993 | chrX | 11251528 | 11251993 | + | 0.750102 | Intergenic | LTR16E2\|LTR\|ERVL | -47497 | NM_001111037 | 100042922 | Mm.374311 | NM_001111037 | ENSMUSG00000095208 | **Gm14483** | OTTMUSG00000016789 | predicted gene 14483 | protein-coding |
| chrX_57182536_57182850 | chrX | 57182537 | 57182850 | + | 0.86692 | Intergenic | Intergenic | -17944 | NR_038463 | 100502842 | Mm.164733 | NR_038463 | ENSMUSG00000085808 | **Gm14718** | OTTMUSG00000017790 | predicted gene 14718 | ncRNA |
| chrX_57183410_57183671 | chrX | 57183411 | 57183671 | + | 0.742111 | Intergenic | Intergenic | -17096 | NR_038463 | 100502842 | Mm.164733 | NR_038463 | ENSMUSG00000085808 | **Gm14718** | OTTMUSG00000017790 | predicted gene 14718 | ncRNA |
| chrX_57177166_57177745 | chrX | 57177167 | 57177745 | + | 0.699372 | Intergenic | Intergenic | -23181 | NR_038463 | 100502842 | Mm.164733 | NR_038463 | ENSMUSG00000085808 | **Gm14718** | OTTMUSG00000017790 | predicted gene 14718 | ncRNA |
| chrX_57186319_57186753 | chrX | 57186320 | 57186753 | + | 0.692556 | Intergenic | Intergenic | -14101 | NR_038463 | 100502842 | Mm.164733 | NR_038463 | ENSMUSG00000085808 | **Gm14718** | OTTMUSG00000017790 | predicted gene 14718 | ncRNA |
| chrX_102341713_102342170 | chrX | 102341714 | 102342170 | + | 0.709378 | intron (NM_027382, intron 9 of 10) | intron (NM_027382, intron 9 of 10) | 89291 | NR_040285 | 434825 | Mm.447373 | NR_040285 | ENSMUSG00000085568 | **Gm14858** | OTTMUSG00000018301 | predicted gene 14858 | ncRNA |
| chr7_104296539_104297086 | chr7 | 104296540 | 104297086 | + | 1.00874 | Intergenic | Intergenic | -18218 | NR_040749 | 622070 | Mm.486406 | NR_040749 | ENSMUSG00000087175 | **Gm15133** | OTTMUSG00000019186 | predicted gene 15133 | ncRNA |
| chr8_12821330_12821916 | chr8 | 12821331 | 12821916 | + | 1.01591 | intron (NR_121603, intron 6 of 29) | intron (NR_121603, intron 6 of 29) | 10296 | NR_045775 | 100504249 | Mm.441494 | NR_045775 | ENSMUSG00000084780 | **Gm15350** | OTTMUSG00000020611 | predicted gene 15350 | ncRNA |
| chr8_12821109_12821927 | chr8 | 12821110 | 12821927 | + | 0.987414 | intron (NR_121603, intron 6 of 29) | intron (NR_121603, intron 6 of 29) | 10401 | NR_045775 | 100504249 | Mm.441494 | NR_045775 | ENSMUSG00000084780 | **Gm15350** | OTTMUSG00000020611 | predicted gene 15350 | ncRNA |
| chr6_72659833_72660388 | chr6 | 72659834 | 72660388 | + | 1.09656 | intron (NM_001079822, intron 3 of 12) | intron (NM_001079822, intron 3 of 12) | 30280 | NR_040421 | 100038714 | Mm.321330 | NR_040421 | | **Gm15401** | ENSMUSG00000072999\|OTTMUSG00000021199 | predicted gene 15401 | ncRNA |
| chr7_96390845_96391309 | chr7 | 96390846 | 96391309 | + | 0.885593 | intron (NM_011858, intron 2 of 32) | intron (NM_011858, intron 2 of 32) | -49113 | NR_046043 | 670727 | Mm.421352 | NR_046043 | ENSMUSG00000085751 | **Gm15412** | OTTMUSG00000021245 | predicted gene 15412 | ncRNA |
| chr7_96818265_96819111 | chr7 | 96818266 | 96819111 | + | 1.14991 | intron (NM_011858, intron 17 of 32) | L1MEd\|LINE\|L1 | -17139 | NR_045874 | 791381 | Mm.402112 | NR_045874 | ENSMUSG00000053049 | **Gm15413** | ENSMUSG00000053049\|OTTMUSG00000021246 | predicted gene 15413 | ncRNA |
| chr7_96707079_96707686 | chr7 | 96707080 | 96707686 | + | 1.0205 | intron (NM_011858, intron 8 of 32) | intron (NM_011858, intron 8 of 32) | 94166 | NR_045874 | 791381 | Mm.402112 | NR_045874 | ENSMUSG00000053049 | **Gm15413** | ENSMUSG00000053049\|OTTMUSG00000021246 | predicted gene 15413 | ncRNA |
| chr7_96706593_96707688 | chr7 | 96706594 | 96707688 | + | 0.816359 | intron (NM_011858, intron 8 of 32) | MIRb\|SINE\|MIR | 94408 | NR_045874 | 791381 | Mm.402112 | NR_045874 | ENSMUSG00000053049 | **Gm15413** | ENSMUSG00000053049\|OTTMUSG00000021246 | predicted gene 15413 | ncRNA |
| chr7_96814595_96814999 | chr7 | 96814596 | 96814999 | + | 0.68129 | intron (NM_011858, intron 17 of 32) | intron (NM_011858, intron 17 of 32) | -13248 | NR_045874 | 791381 | Mm.402112 | NR_045874 | ENSMUSG00000053049 | **Gm15413** | ENSMUSG00000053049\|OTTMUSG00000021246 | predicted gene 15413 | ncRNA |
| chr7_96701140_96701506 | chr7 | 96701141 | 96701506 | + | 0.644881 | intron (NM_011858, intron 6 of 32) | intron (NM_011858, intron 6 of 32) | 100226 | NR_045874 | 791381 | Mm.402112 | NR_045874 | ENSMUSG00000053049 | **Gm15413** | ENSMUSG00000053049\|OTTMUSG00000021246 | predicted gene 15413 | ncRNA |
| chr3_89391127_89391408 | chr3 | 89391128 | 89391408 | + | 0.738057 | promoter-TSS (NR_040403) | promoter-TSS (NR_040403) | -596 | NR_040403 | 545539 | Mm.393389 | NR_040403 | ENSMUSG00000074466 | **Gm15417** | ENSMUSG00000074466\|OTTMUSG00000021274 | predicted gene 15417 | ncRNA |
| chr5_22451746_22452090 | chr5 | 22451747 | 22452090 | + | 0.795636 | Intergenic | Intergenic | 77112 | NR_004442 | 100042049 | Mm.21724 | NR_004442 | | **Gm15421** | OTTMUSG00000021442 | ribosomal protein L22 like 1 pseudogene | pseudo |
| chr5_22511275_22511757 | chr5 | 22511276 | 22511757 | + | 0.709466 | intron (NM_011959, intron 12 of 13) | intron (NM_011959, intron 12 of 13) | 17514 | NR_004442 | 100042049 | Mm.21724 | NR_004442 | | **Gm15421** | OTTMUSG00000021442 | ribosomal protein L22 like 1 pseudogene | pseudo |
| chr6_145274084_145274599 | chr6 | 145274085 | 145274599 | + | 1.41741 | Intergenic | Intergenic | 23790 | NR_045598 | 100039495 | Mm.121014 | NR_045598 | ENSMUSG00000086013 | **Gm15706** | OTTMUSG00000025463 | predicted gene 15706 | ncRNA |
| chr6_145272571_145272818 | chr6 | 145272572 | 145272818 | + | 1.12096 | Intergenic | Intergenic | 22143 | NR_045598 | 100039495 | Mm.121014 | NR_045598 | ENSMUSG00000086013 | **Gm15706** | OTTMUSG00000025463 | predicted gene 15706 | ncRNA |
| chr6_145275807_145276167 | chr6 | 145275808 | 145276167 | + | 0.768041 | Intergenic | Intergenic | 25435 | NR_045598 | 100039495 | Mm.121014 | NR_045598 | ENSMUSG00000086013 | **Gm15706** | OTTMUSG00000025463 | predicted gene 15706 | ncRNA |
| chr5_121228920_121229715 | chr5 | 121228921 | 121229715 | + | 1.37124 | intron (NM_181421, intron 1 of 75) | intron (NM_181421, intron 1 of 75) | 9099 | NM_181421 | 269700 | Mm.184589 | NM_181421 | ENSMUSG00000042744 | **Gm15800** | A130010L24\|AU042671\|BC019639\|EG545802\|OTTMUSG00000026283\|mKIAA0614 | predicted gene 15800 | protein-coding |
| chr1_136116936_136117584 | chr1 | 136116937 | 136117584 | + | 1.58971 | intron (NM_014193, intron 39 of 42) | MER5B\|DNA\|hAT-Charlie | 13923 | NR_046168 | 329253 | Mm.487858 | NR_046167 | ENSMUSG00000086264 | **Gm15850** | 6530439I21\|OTTMUSG00000026868 | predicted gene 15850 | ncRNA |
| chr5_149585940_149586794 | chr5 | 149585941 | 149586794 | + | 1.42027 | intron (NM_029440, intron 14 of 22) | MLT1A\|LTR\|ERVL-MaLR | -48337 | NR_045423 | 100415916 | Mm.145528 | NR_045423 | ENSMUSG00000086744 | **Gm15997** | OTTMUSG00000028862 | predicted gene 15997 | ncRNA |
| chr6_8299073_8299508 | chr6 | 8299074 | 8299508 | + | 1.352 | intron (NR_033518, intron 2 of 3) | intron (NR_033518, intron 2 of 3) | 40003 | NR_033518 | 100036521 | Mm.446106 | NR_033518 | ENSMUSG00000089862 | **Gm16039** | OTTMUSG00000029336 | predicted gene 16039 | ncRNA |
| chr6_8288999_8289497 | chr6 | 8289000 | 8289497 | + | 0.697736 | intron (NR_033518, intron 2 of 3) | intron (NR_033518, intron 2 of 3) | 29960 | NR_033518 | 100036521 | Mm.446106 | NR_033518 | ENSMUSG00000089862 | **Gm16039** | OTTMUSG00000029336 | predicted gene 16039 | ncRNA |
| chr9_58027806_58028378 | chr9 | 58027807 | 58028378 | + | 0.908263 | TTS (NM_029212) | TTS (NM_029212) | -4182 | NM_001243265 | 330951 | Mm.406992 | NM_001243265 | | **Gm16130** | E330033L03\|OTTMUSG00000030814 | predicted gene 16130 | protein-coding |
| chr2_71696776_71697115 | chr2 | 71696777 | 71697115 | + | 0.719124 | Intergenic | ID_B1\|SINE\|B4 | -22471 | NR_037979 | 381371 | Mm.330284 | NM_201366 | ENSMUSG00000064263 | **Gm1631** | - | predicted gene 1631 | ncRNA |
| chr2_71643988_71644551 | chr2 | 71643989 | 71644551 | + | 0.643454 | Intergenic | Intergenic | -75147 | NR_037979 | 381371 | Mm.330284 | NM_201366 | ENSMUSG00000064263 | **Gm1631** | - | predicted gene 1631 | ncRNA |
| chr7_110919244_110919666 | chr7 | 110919245 | 110919666 | + | 1.09641 | intron (NM_010826, intron 9 of 19) | intron (NM_010826, intron 9 of 19) | 14470 | NR_045267 | 100504184 | Mm.441327 | NR_045267 | ENSMUSG00000087160 | **Gm16336** | OTTMUSG00000033852 | predicted gene 16336 | ncRNA |

| | | | | | | | | | | | | | | | | | |
|---|---|---|---|---|---|---|---|---|---|---|---|---|---|---|---|---|---|
| chr7_110907871_110908326 | chr7 | 110907872 | 110908326 | + | 0.635125 | intron (NM_010826, intron 9 of 19) | intron (NM_010826, intron 9 of 19) | 3114 | NR_045267 | 100504184 | Mm.441327 | NR_045267 | ENSMUSG00000087160 | **Gm16336** | OTTMUSG00000033852 | predicted gene 16336 | ncRNA |
| chr7_110907854_110908472 | chr7 | 110907855 | 110908472 | + | 0.632787 | intron (NM_010826, intron 9 of 19) | intron (NM_010826, intron 9 of 19) | 3178 | NR_045267 | 100504184 | Mm.441327 | NR_045267 | ENSMUSG00000087160 | **Gm16336** | OTTMUSG00000033852 | predicted gene 16336 | ncRNA |
| chr3_69131070_69131315 | chr3 | 69131071 | 69131315 | + | 0.833518 | promoter-TSS (NM_001243000) | promoter-TSS (NM_001243000) | -40 | NM_001243000 | 381452 | Mm.331067 | NM_001243000 | ENSMUSG00000094763 | **Gm1647** | - | predicted gene 1647 | protein-coding |
| chr2_39123923_39124341 | chr2 | 39123924 | 39124341 | + | 0.783277 | intron (NM_178778, intron 4 of 17) | MRLTR33\|LTR\|ERVK | 38103 | NR_033526 | 100042584 | Mm.451696 | NR_033526 | | **Gm16523** | A130054J05Rik | mitochondrial ribosomal protein L40 pseudogene | pseudo |
| chr2_39123608_39124341 | chr2 | 39123609 | 39124341 | + | 0.781496 | intron (NM_178778, intron 4 of 17) | intron (NM_178778, intron 4 of 17) | 38260 | NR_033526 | 100042584 | Mm.451696 | NR_033526 | | **Gm16523** | A130054J05Rik | mitochondrial ribosomal protein L40 pseudogene | pseudo |
| chr2_39140927_39141792 | chr2 | 39140928 | 39141792 | + | 0.682277 | intron (NM_178778, intron 2 of 17) | intron (NM_178778, intron 2 of 17) | 20875 | NR_033526 | 100042584 | Mm.451696 | NR_033526 | | **Gm16523** | A130054J05Rik | mitochondrial ribosomal protein L40 pseudogene | pseudo |
| chr2_39140958_39141395 | chr2 | 39140959 | 39141395 | + | 0.673853 | intron (NM_178778, intron 2 of 17) | B3A\|SINE\|B2 | 21058 | NR_033526 | 100042584 | Mm.451696 | NR_033526 | | **Gm16523** | A130054J05Rik | mitochondrial ribosomal protein L40 pseudogene | pseudo |
| chr3_149338609_149339106 | chr3 | 149338610 | 149339106 | + | 1.21135 | Intergenic | Intergenic | 64121 | NR_040591 | 381482 | Mm.393866 | NR_040591 | | **Gm1653** | - | predicted gene 1653 | ncRNA |
| chr3_149339372_149339758 | chr3 | 149339373 | 149339758 | + | 1.1743 | Intergenic | Intergenic | 64828 | NR_040591 | 381482 | Mm.393866 | NR_040591 | | **Gm1653** | - | predicted gene 1653 | ncRNA |
| chr12_108557718_108558251 | chr12 | 108557719 | 108558251 | + | 1.62451 | intron (NM_007965, intron 1 of 12) | L1MEg\|LINE\|L1 | -2367 | NR_045753 | 100504467 | Mm.394980 | NR_045751 | ENSMUSG00000097788 | **Gm16596** | - | predicted gene, 16596 | ncRNA |
| chr8_46714349_46714749 | chr8 | 46714350 | 46714749 | + | 1.42532 | Intergenic | Intergenic | 24966 | NR_045750 | 100503498 | Mm.187358 | NR_045750 | ENSMUSG00000097534 | **Gm16675** | - | predicted gene, 16675 | ncRNA |
| chr8_46688316_46688591 | chr8 | 46688317 | 46688591 | + | 0.816727 | Intergenic | Intergenic | 51061 | NR_045750 | 100503498 | Mm.187358 | NR_045750 | ENSMUSG00000097534 | **Gm16675** | - | predicted gene, 16675 | ncRNA |
| chr17_8385463_8385832 | chr17 | 8385464 | 8385832 | + | 0.824828 | intron (NR_045795, intron 2 of 2) | Lx8b\|LINE\|L1 | 3820 | NR_045795 | 100504601 | Mm.365131 | NR_045795 | ENSMUSG00000097796 | **Gm16702** | AU016609 | predicted gene, 16702 | ncRNA |
| chr17_55961300_55961728 | chr17 | 55961301 | 55961728 | + | 0.863606 | intron (NM_028381, intron 2 of 7) | ORR1C2\|LTR\|ERVL-MaLR | -2113 | NR_108021 | 100504333 | Mm.391908 | NR_108021 | ENSMUSG00000097077 | **Gm16712** | - | predicted gene, 16712 | ncRNA |
| chr8_35614033_35614645 | chr8 | 35614034 | 35614645 | + | 1.60657 | intron (NM_001081279, intron 1 of 2) | L1ME1\|LINE\|L1 | -25319 | NR_040376 | 100504714 | Mm.381744 | NR_040376 | ENSMUSG00000097357 | **Gm16793** | - | predicted gene, 16793 | ncRNA |
| chr8_35625839_35626218 | chr8 | 35625840 | 35626218 | + | 1.53716 | intron (NM_001081279, intron 1 of 2) | intron (NM_001081279, intron 1 of 2) | -37009 | NR_040376 | 100504714 | Mm.381744 | NR_040376 | ENSMUSG00000097357 | **Gm16793** | - | predicted gene, 16793 | ncRNA |
| chr8_35632085_35632749 | chr8 | 35632086 | 35632749 | + | 1.34642 | intron (NM_001081279, intron 1 of 2) | intron (NM_001081279, intron 1 of 2) | -43397 | NR_040376 | 100504714 | Mm.381744 | NR_040376 | ENSMUSG00000097357 | **Gm16793** | - | predicted gene, 16793 | ncRNA |
| chr8_35591626_35592182 | chr8 | 35591627 | 35592182 | + | 1.16897 | intron (NM_001081279, intron 1 of 2) | intron (NM_001081279, intron 1 of 2) | -2884 | NR_040376 | 100504714 | Mm.381744 | NR_040376 | ENSMUSG00000097357 | **Gm16793** | - | predicted gene, 16793 | ncRNA |
| chr1_136696272_136696693 | chr1 | 136696273 | 136696693 | + | 0.834257 | intron (NR_037986, intron 1 of 4) | intron (NR_037986, intron 1 of 4) | 272 | NR_037986 | 100503474 | Mm.428276 | NR_037986 | | **Gm16880** | - | predicted gene, 16880 | ncRNA |
| chr1_40109223_40109693 | chr1 | 40109224 | 40109693 | + | 1.0353 | intron (NM_010555, intron 3 of 8) | L1_Mus3\|LINE\|L1 | -23607 | NR_037980 | 100047133 | Mm.447052 | NR_037980 | ENSMUSG00000097899 | **Gm16894** | - | predicted gene, 16894 | ncRNA |
| chr1_194980871_194981248 | chr1 | 194980872 | 194981248 | + | 0.701429 | Intergenic | Intergenic | -4101 | NR_033564 | 320400 | Mm.417608 | NR_033564 | ENSMUSG00000097325 | **Gm16897** | C030002c11rik | predicted gene, 16897 | ncRNA |
| chr13_63274452_63275181 | chr13 | 63274453 | 63275181 | + | 0.776799 | intron (NR_110520, intron 2 of 5) | intron (NR_110520, intron 2 of 5) | 22330 | NR_045794 | 100504114 | Mm.437070 | NR_045794 | ENSMUSG00000097242 | **Gm16907** | - | predicted gene, 16907 | ncRNA |
| chr10_54254812_54255610 | chr10 | 54254813 | 54255610 | + | 1.62117 | Intergenic | MIRb\|SINE\|MIR | 179289 | NR_038016 | 100502920 | Mm.403982 | NR_038016 | ENSMUSG00000097069 | **Gm16998** | - | predicted gene, 16998 | ncRNA |
| chr10_54255906_54256422 | chr10 | 54255907 | 54256422 | + | 1.5041 | Intergenic | Intergenic | 180242 | NR_038016 | 100502920 | Mm.403982 | NR_038016 | ENSMUSG00000097069 | **Gm16998** | - | predicted gene, 16998 | ncRNA |
| chr10_54245037_54245350 | chr10 | 54245038 | 54245350 | + | 0.68785 | Intergenic | Intergenic | 169272 | NR_038016 | 100502920 | Mm.403982 | NR_038016 | ENSMUSG00000097069 | **Gm16998** | - | predicted gene, 16998 | ncRNA |
| chrX_166808088_166808648 | chrX | 166808089 | 166808648 | + | 1.244 | Intergenic | MLT2B3\|LTR\|ERVL | 146343 | NR_046015 | 382255 | Mm.336320 | NR_046015 | ENSMUSG00000086860 | **Gm1720** | - | predicted gene 1720 | ncRNA |
| chrX_166718067_166718626 | chrX | 166718068 | 166718626 | + | 0.931107 | Intergenic | Intergenic | 56322 | NR_046015 | 382255 | Mm.336320 | NR_046015 | ENSMUSG00000086860 | **Gm1720** | - | predicted gene 1720 | ncRNA |
| chr3_79335990_79336510 | chr3 | 79335991 | 79336510 | + | 1.51266 | Intergenic | Intergenic | -9126 | NM_001142953 | 100233207 | Mm.373801 | NM_001142953 | ENSMUSG00000091685 | **Gm17359** | - | predicted gene, 17359 | protein-coding |
| chr3_79451266_79451742 | chr3 | 79451267 | 79451742 | + | 0.851828 | intron (NM_001142953, intron 4 of 4) | intron (NM_001142953, intron 4 of 4) | 106128 | NM_001142953 | 100233207 | Mm.373801 | NM_001142953 | ENSMUSG00000091685 | **Gm17359** | - | predicted gene, 17359 | protein-coding |
| chr3_79280365_79280769 | chr3 | 79280366 | 79280769 | + | 0.826654 | Intergenic | Intergenic | -64809 | NM_001142953 | 100233207 | Mm.373801 | NM_001142953 | ENSMUSG00000091685 | **Gm17359** | - | predicted gene, 17359 | protein-coding |
| chr10_60395607_60396115 | chr10 | 60395608 | 60396115 | + | 1.26102 | intron (NM_023370, intron 28 of 68) | RMER15-int\|LTR\|ERVL | -3865 | NM_001164374 | 100302688 | Mm.444135 | NM_001164374 | ENSMUSG00000090439 | **Gm17455** | - | predicted gene, 17455 | protein-coding |
| chr10_60416847_60417309 | chr10 | 60416848 | 60417309 | + | 1.12732 | intron (NM_023370, intron 22 of 68) | intron (NM_023370, intron 22 of 68) | 17352 | NM_001164374 | 100302688 | Mm.444135 | NM_001164374 | ENSMUSG00000090439 | **Gm17455** | - | predicted gene, 17455 | protein-coding |
| chr10_93343414_93344035 | chr10 | 93343415 | 93344035 | + | 1.54153 | intron (NR_038014, intron 1 of 2) | intron (NR_038014, intron 1 of 2) | 7493 | NR_038014 | 432488 | Mm.32833 | NR_038014 | | **Gm17745** | - | predicted gene, 17745 | ncRNA |
| chr10_93360193_93361029 | chr10 | 93360194 | 93361029 | + | 1.12438 | Intergenic | Intergenic | 24379 | NR_038014 | 432488 | Mm.32833 | NR_038014 | | **Gm17745** | - | predicted gene, 17745 | ncRNA |
| chr10_93345212_93345799 | chr10 | 93345213 | 93345799 | + | 1.01756 | intron (NR_038014, intron 1 of 2) | intron (NR_038014, intron 1 of 2) | 9274 | NR_038014 | 432488 | Mm.32833 | NR_038014 | | **Gm17745** | - | predicted gene, 17745 | ncRNA |
| chr10_93361794_93362151 | chr10 | 93361795 | 93362151 | + | 0.970393 | Intergenic | Plat_L3\|LINE\|CR1 | 25741 | NR_038014 | 432488 | Mm.32833 | NR_038014 | | **Gm17745** | - | predicted gene, 17745 | ncRNA |
| chr10_93353966_93354312 | chr10 | 93353967 | 93354312 | + | 0.904728 | Intergenic | Intergenic | 17907 | NR_038014 | 432488 | Mm.32833 | NR_038014 | | **Gm17745** | - | predicted gene, 17745 | ncRNA |
| chr10_93360696_93361028 | chr10 | 93360697 | 93361028 | + | 0.868342 | Intergenic | MamRep488\|DNA\|hAT-Tip100 | 24630 | NR_038014 | 432488 | Mm.32833 | NR_038014 | | **Gm17745** | - | predicted gene, 17745 | ncRNA |
| chr10_93346228_93346505 | chr10 | 93346229 | 93346505 | + | 0.851708 | intron (NR_038014, intron 1 of 2) | MLT1A\|LTR\|ERVL-MaLR | 10135 | NR_038014 | 432488 | Mm.32833 | NR_038014 | | **Gm17745** | - | predicted gene, 17745 | ncRNA |
| chr12_25438544_25439235 | chr12 | 25438545 | 25439235 | + | 1.34331 | Intergenic | Intergenic | 309716 | NR_045844 | 432641 | Mm.112955 | NR_045844 | | **Gm17746** | - | predicted gene, 17746 | ncRNA |
| chr12_25438400_25439563 | chr12 | 25438401 | 25439563 | + | 1.34145 | Intergenic | Intergenic | 309808 | NR_045844 | 432641 | Mm.112955 | NR_045844 | | **Gm17746** | - | predicted gene, 17746 | ncRNA |
| chr12_25258782_25259615 | chr12 | 25258783 | 25259615 | + | 1.3249 | Intergenic | L1ME3Cz\|LINE\|L1 | 130025 | NR_045844 | 432641 | Mm.112955 | NR_045844 | | **Gm17746** | - | predicted gene, 17746 | ncRNA |
| chr12_25305711_25306678 | chr12 | 25305712 | 25306678 | + | 1.23063 | Intergenic | Intergenic | 177021 | NR_045844 | 432641 | Mm.112955 | NR_045844 | | **Gm17746** | - | predicted gene, 17746 | ncRNA |
| chr12_25190355_25190974 | chr12 | 25190356 | 25190974 | + | 1.04537 | Intergenic | Intergenic | 61491 | NR_045844 | 432641 | Mm.112955 | NR_045844 | | **Gm17746** | - | predicted gene, 17746 | ncRNA |
| chr12_25190412_25190754 | chr12 | 25190413 | 25190754 | + | 1.02232 | Intergenic | Intergenic | 61409 | NR_045844 | 432641 | Mm.112955 | NR_045844 | | **Gm17746** | - | predicted gene, 17746 | ncRNA |
| chr12_25240076_25240501 | chr12 | 25240077 | 25240501 | + | 0.990378 | Intergenic | Intergenic | 111115 | NR_045844 | 432641 | Mm.112955 | NR_045844 | | **Gm17746** | - | predicted gene, 17746 | ncRNA |
| chr12_25436275_25436520 | chr12 | 25436276 | 25436520 | + | 0.976108 | Intergenic | Intergenic | 307224 | NR_045844 | 432641 | Mm.112955 | NR_045844 | | **Gm17746** | - | predicted gene, 17746 | ncRNA |
| chr12_25239779_25240559 | chr12 | 25239780 | 25240559 | + | 0.969437 | Intergenic | Intergenic | 110995 | NR_045844 | 432641 | Mm.112955 | NR_045844 | | **Gm17746** | - | predicted gene, 17746 | ncRNA |
| chr12_25143424_25143915 | chr12 | 25143425 | 25143915 | + | 0.864672 | Intergenic | Intergenic | 14496 | NR_045844 | 432641 | Mm.112955 | NR_045844 | | **Gm17746** | - | predicted gene, 17746 | ncRNA |
| chr12_25246832_25247358 | chr12 | 25246833 | 25247358 | + | 0.7823 | Intergenic | Intergenic | 117921 | NR_045844 | 432641 | Mm.112955 | NR_045844 | | **Gm17746** | - | predicted gene, 17746 | ncRNA |
| chr12_25320332_25320709 | chr12 | 25320333 | 25320709 | + | 0.729992 | Intergenic | Intergenic | 191347 | NR_045844 | 432641 | Mm.112955 | NR_045844 | | **Gm17746** | - | predicted gene, 17746 | ncRNA |
| chr12_25311753_25312110 | chr12 | 25311754 | 25312110 | + | 0.717222 | Intergenic | Intergenic | 182758 | NR_045844 | 432641 | Mm.112955 | NR_045844 | | **Gm17746** | - | predicted gene, 17746 | ncRNA |
| chr12_25320282_25320852 | chr12 | 25320283 | 25320852 | + | 0.700903 | Intergenic | Intergenic | 191393 | NR_045844 | 432641 | Mm.112955 | NR_045844 | | **Gm17746** | - | predicted gene, 17746 | ncRNA |
| chr12_25155321_25155982 | chr12 | 25155322 | 25155982 | + | 0.667836 | Intergenic | Intergenic | 26478 | NR_045844 | 432641 | Mm.112955 | NR_045844 | | **Gm17746** | - | predicted gene, 17746 | ncRNA |
| chr12_25311512_25312189 | chr12 | 25311513 | 25312189 | + | 0.660094 | Intergenic | Intergenic | 182677 | NR_045844 | 432641 | Mm.112955 | NR_045844 | | **Gm17746** | - | predicted gene, 17746 | ncRNA |
| chr10_77430590_77431137 | chr10 | 77430591 | 77431137 | + | 1.03449 | Intergenic | Intergenic | 12488 | NR_027377 | 100045653 | Mm.363175 | NR_027377 | | **Gm17769** | - | predicted gene, 17769 | ncRNA |
| chr17_49269058_49269477 | chr17 | 49269059 | 49269477 | + | 0.689791 | Intergenic | Intergenic | 43692 | NR_033486 | 100415784 | Mm.400421 | NR_033486 | | **Gm17830** | - | predicted gene, 17830 | ncRNA |
| chr15_51176075_51176799 | chr15 | 51176076 | 51176799 | + | 1.18425 | Intergenic | Intergenic | 219173 | NR_045092 | 100502643 | Mm.131987 | NR_045092 | | **Gm19303** | - | predicted gene, 19303 | ncRNA |
| chr15_51175478_51176799 | chr15 | 51175479 | 51176799 | + | 1.0615 | Intergenic | Intergenic | 219471 | NR_045092 | 100502643 | Mm.131987 | NR_045092 | | **Gm19303** | - | predicted gene, 19303 | ncRNA |
| chr15_51206068_51206486 | chr15 | 51206069 | 51206486 | + | 1.05699 | Intergenic | Intergenic | 189333 | NR_045092 | 100502643 | Mm.131987 | NR_045092 | | **Gm19303** | - | predicted gene, 19303 | ncRNA |
| chr15_51175478_51176030 | chr15 | 51175479 | 51176030 | + | 0.618358 | Intergenic | RMER1B\|Other\|Other | 219856 | NR_045092 | 100502643 | Mm.131987 | NR_045092 | | **Gm19303** | - | predicted gene, 19303 | ncRNA |
| chr15_59799269_59799802 | chr15 | 59799270 | 59799802 | + | 1.62578 | Intergenic | Intergenic | -4577 | NR_045076 | 100503021 | Mm.402924 | NR_045076 | | **Gm19510** | - | predicted gene, 19510 | ncRNA |
| chr15_59759526_59760043 | chr15 | 59759527 | 59760043 | + | 1.5742 | Intergenic | Intergenic | 35174 | NR_045076 | 100503021 | Mm.402924 | NR_045076 | | **Gm19510** | - | predicted gene, 19510 | ncRNA |
| chr15_59737244_59738163 | chr15 | 59737245 | 59738163 | + | 1.41129 | Intergenic | MER21C\|LTR\|ERVL | 57255 | NR_045076 | 100503021 | Mm.402924 | NR_045076 | | **Gm19510** | - | predicted gene, 19510 | ncRNA |
| chr15_59822858_59823783 | chr15 | 59822859 | 59823783 | + | 1.27184 | Intergenic | Intergenic | -28362 | NR_045076 | 100503021 | Mm.402924 | NR_045076 | | **Gm19510** | - | predicted gene, 19510 | ncRNA |
| chr15_59739222_59739679 | chr15 | 59739223 | 59739679 | + | 1.1155 | Intergenic | Intergenic | 55508 | NR_045076 | 100503021 | Mm.402924 | NR_045076 | | **Gm19510** | - | predicted gene, 19510 | ncRNA |
| chr15_59728072_59728805 | chr15 | 59728073 | 59728805 | + | 1.05446 | Intergenic | Intergenic | 66520 | NR_045076 | 100503021 | Mm.402924 | NR_045076 | | **Gm19510** | - | predicted gene, 19510 | ncRNA |
| chr15_59802300_59802768 | chr15 | 59802301 | 59802768 | + | 0.993695 | Intergenic | Intergenic | -7575 | NR_045076 | 100503021 | Mm.402924 | NR_045076 | | **Gm19510** | - | predicted gene, 19510 | ncRNA |
| chr15_59795680_59796117 | chr15 | 59795681 | 59796117 | + | 0.758142 | promoter-TSS (NR_045076) | promoter-TSS (NR_045076) | -940 | NR_045076 | 100503021 | Mm.402924 | NR_045076 | | **Gm19510** | - | predicted gene, 19510 | ncRNA |
| chr15_59731106_59731477 | chr15 | 59731107 | 59731477 | + | 0.643068 | Intergenic | Intergenic | 63667 | NR_045076 | 100503021 | Mm.402924 | NR_045076 | | **Gm19510** | - | predicted gene, 19510 | ncRNA |
| chr1_88543065_88544010 | chr1 | 88543066 | 88544010 | + | 0.691389 | Intergenic | Intergenic | -7979 | NR_037971 | 100503197 | Mm.99352 | NR_037971 | | **Gm19589** | - | predicted gene, 19589 | ncRNA |
| chr16_29882209_29882983 | chr16 | 29882210 | 29882983 | + | 0.78467 | Intergenic | Intergenic | 96528 | NR_037689 | 328657 | Mm.425982 | NM_001033403 | | **Gm1968** | - | predicted gene 1968 | ncRNA |

| | | | | | | | | | | | | | | | | | |
|---|---|---|---|---|---|---|---|---|---|---|---|---|---|---|---|---|---|
| chr16_29865134_29865530 | chr16 | 29865135 | 29865530 | + | 0.671463 | Intergenic | Intergenic | 113792 | NR_037689 | 328657 | Mm.425982 | NM_001033403 | | **Gm1968** | - | predicted gene 1968 | ncRNA |
| chr16_29813747_29814498 | chr16 | 29813748 | 29814498 | + | 0.599513 | Intergenic | CpG | 165001 | NR_037689 | 328657 | Mm.425982 | NM_001033403 | | **Gm1968** | - | predicted gene 1968 | ncRNA |
| chr17_52076726_52077277 | chr17 | 52076727 | 52077277 | + | 1.63079 | Intergenic | Intergenic | 194275 | NR_045095 | 100504169 | Mm.417731 | NR_045095 | | **Gm20098** | - | predicted gene, 20098 | ncRNA |
| chr17_52076498_52077275 | chr17 | 52076499 | 52077275 | + | 1.53273 | Intergenic | Intergenic | 194160 | NR_045095 | 100504169 | Mm.417731 | NR_045095 | | **Gm20098** | - | predicted gene, 20098 | ncRNA |
| chr17_52151378_52151608 | chr17 | 52151379 | 52151608 | + | 0.698911 | Intergenic | Intergenic | 268766 | NR_045095 | 100504169 | Mm.417731 | NR_045095 | | **Gm20098** | - | predicted gene, 20098 | ncRNA |
| chr17_52151311_52151625 | chr17 | 52151312 | 52151625 | + | 0.69304 | Intergenic | Intergenic | 268741 | NR_045095 | 100504169 | Mm.417731 | NR_045095 | | **Gm20098** | - | predicted gene, 20098 | ncRNA |
| chr17_52152069_52152455 | chr17 | 52152070 | 52152455 | + | 0.692749 | Intergenic | LTR73\|LTR\|ERV1 | 269535 | NR_045095 | 100504169 | Mm.417731 | NR_045095 | | **Gm20098** | - | predicted gene, 20098 | ncRNA |
| chr17_52125319_52125644 | chr17 | 52125320 | 52125644 | + | 0.647564 | Intergenic | Intergenic | 242755 | NR_045095 | 100504169 | Mm.417731 | NR_045095 | | **Gm20098** | - | predicted gene, 20098 | ncRNA |
| chr17_52125199_52125682 | chr17 | 52125200 | 52125682 | + | 0.645858 | Intergenic | Intergenic | 242714 | NR_045095 | 100504169 | Mm.417731 | NR_045095 | | **Gm20098** | - | predicted gene, 20098 | ncRNA |
| chr10_99447919_99448518 | chr10 | 99447920 | 99448518 | + | 1.66075 | Intergenic | MLTR14\|LTR\|ERV1 | -160952 | NR_038019 | 100504202 | Mm.403801 | NR_038019 | | **Gm20110** | - | predicted gene, 20110 | ncRNA |
| chr10_99555757_99556577 | chr10 | 99555758 | 99556577 | + | 1.58957 | Intergenic | Intergenic | -53004 | NR_038019 | 100504202 | Mm.403801 | NR_038019 | | **Gm20110** | - | predicted gene, 20110 | ncRNA |
| chr10_99561280_99561900 | chr10 | 99561281 | 99561900 | + | 1.29463 | Intergenic | L2\|LINE\|L2 | -47581 | NR_038019 | 100504202 | Mm.403801 | NR_038019 | | **Gm20110** | - | predicted gene, 20110 | ncRNA |
| chr10_99447562_99447852 | chr10 | 99447563 | 99447852 | + | 0.874555 | Intergenic | Intergenic | -161464 | NR_038019 | 100504202 | Mm.403801 | NR_038019 | | **Gm20110** | - | predicted gene, 20110 | ncRNA |
| chr10_99459493_99460422 | chr10 | 99459494 | 99460422 | + | 0.72269 | Intergenic | Intergenic | -149213 | NR_038019 | 100504202 | Mm.403801 | NR_038019 | | **Gm20110** | - | predicted gene, 20110 | ncRNA |
| chr10_16714062_16714478 | chr10 | 16714063 | 16714478 | + | 0.641248 | Intergenic | Intergenic | 381512 | NR_038020 | 100504235 | Mm.84044 | NR_038020 | | **Gm20125** | - | predicted gene, 20125 | ncRNA |
| chr10_19165585_19165879 | chr10 | 19165586 | 19165879 | + | 1.19288 | intron (NR_038021, intron 1 of 3) | intron (NR_038021, intron 1 of 3) | -1506 | NR_038022 | 100504265 | Mm.383417 | NR_038021 | | **Gm20139** | - | predicted gene, 20139 | ncRNA |
| chr12_80937453_80938184 | chr12 | 80937454 | 80938184 | + | 1.12911 | Intergenic | Intergenic | 7580 | NR_045057 | 100504661 | Mm.442322 | NR_045057 | | **Gm20337** | - | predicted gene, 20337 | ncRNA |
| chr15_84743253_84743691 | chr15 | 84743254 | 84743691 | + | 0.661694 | intron (NM_001164627, intron 5 of 12) | intron (NM_001164627, intron 5 of 12) | -23206 | NR_040347 | 328576 | Mm.400211 | NR_040347 | | **Gm20556** | A430075N02 | predicted gene, 20556 | ncRNA |
| chr10_53527389_53527660 | chr10 | 53527390 | 53527660 | + | 1.61204 | Intergenic | Intergenic | -8536 | NR_037973 | 100502868 | Mm.297722 | NR_037973 | | **Gm20597** | B230314j19 | predicted gene, 20597 | ncRNA |
| chr10_53526195_53526768 | chr10 | 53526196 | 53526768 | + | 1.54945 | Intergenic | MERV1_LTR\|LTR\|ERV 1 | -7493 | NR_037973 | 100502868 | Mm.297722 | NR_037973 | | **Gm20597** | B230314j19 | predicted gene, 20597 | ncRNA |
| chr10_53525428_53526778 | chr10 | 53525429 | 53526778 | + | 1.19081 | Intergenic | RMER19C\|LTR\|ERVK | -7114 | NR_037973 | 100502868 | Mm.297722 | NR_037973 | | **Gm20597** | B230314j19 | predicted gene, 20597 | ncRNA |
| chr10_53527865_53528172 | chr10 | 53527866 | 53528172 | + | 1.17223 | Intergenic | Intergenic | -9030 | NR_037973 | 100502868 | Mm.297722 | NR_037973 | | **Gm20597** | B230314j19 | predicted gene, 20597 | ncRNA |
| chr10_53525426_53526108 | chr10 | 53525427 | 53526108 | + | 0.909669 | Intergenic | ORR1C2-int\|LTR\|ERVL-MaLR | -6778 | NR_037973 | 100502868 | Mm.297722 | NR_037973 | | **Gm20597** | B230314j19 | predicted gene, 20597 | ncRNA |
| chr15_63268484_63269040 | chr15 | 63268485 | 63269040 | + | 1.19088 | Intergenic | Intergenic | 210846 | NR_045279 | 432958 | Mm.255105 | NR_045279 | | **Gm20740** | - | predicted gene, 20740 | ncRNA |
| chr15_63268447_63269440 | chr15 | 63268448 | 63269440 | + | 1.18198 | Intergenic | Intergenic | 210664 | NR_045279 | 432958 | Mm.255105 | NR_045279 | | **Gm20740** | - | predicted gene, 20740 | ncRNA |
| chr3_52866121_52866708 | chr3 | 52866122 | 52866708 | + | 0.723404 | Intergenic | Intergenic | 61251 | NR_040555 | 622124 | Mm.389434 | NR_040555 | | **Gm20750** | - | predicted gene, 20750 | ncRNA |
| chr1_106145416_106146121 | chr1 | 106145417 | 106146121 | + | 1.24768 | Intergenic | Intergenic | 25755 | NR_040630 | 626049 | Mm.247410 | NR_040630 | | **Gm20753** | - | predicted gene, 20753 | ncRNA |
| chr1_106145129_106146123 | chr1 | 106145130 | 106146123 | + | 0.98054 | Intergenic | Intergenic | 25898 | NR_040630 | 626049 | Mm.247410 | NR_040630 | | **Gm20753** | - | predicted gene, 20753 | ncRNA |
| chr3_37980493_37980763 | chr3 | 37980494 | 37980763 | + | 0.782377 | intron (NR_040559, intron 2 of 3) | intron (NR_040559, intron 2 of 3) | 81815 | NR_040559 | 626410 | Mm.389417 | NR_040559 | | **Gm20755** | - | predicted gene, 20755 | ncRNA |
| chr3_37974754_37975192 | chr3 | 37974755 | 37975192 | + | 0.750638 | intron (NR_040559, intron 2 of 3) | intron (NR_040559, intron 2 of 3) | 76160 | NR_040559 | 626410 | Mm.389417 | NR_040559 | | **Gm20755** | - | predicted gene, 20755 | ncRNA |
| chr3_37845722_37846101 | chr3 | 37845723 | 37846101 | + | 0.740296 | Intergenic | Intergenic | -52901 | NR_040559 | 626410 | Mm.389417 | NR_040559 | | **Gm20755** | - | predicted gene, 20755 | ncRNA |
| chr3_37845713_37846539 | chr3 | 37845714 | 37846539 | + | 0.720469 | Intergenic | Intergenic | -52687 | NR_040559 | 626410 | Mm.389417 | NR_040559 | | **Gm20755** | - | predicted gene, 20755 | ncRNA |
| chr9_83342437_83343378 | chr9 | 83342438 | 83343378 | + | 1.44697 | Intergenic | Intergenic | -88639 | NR_102437 | 100039186 | Mm.212721 | NR_102437 | | **Gm2087** | - | predicted gene 2087 | ncRNA |
| chr9_83375397_83376077 | chr9 | 83375398 | 83376077 | + | 0.827036 | Intergenic | Intergenic | -55810 | NR_102437 | 100039186 | Mm.212721 | NR_102437 | | **Gm2087** | - | predicted gene 2087 | ncRNA |
| chr9_83341226_83341560 | chr9 | 83341227 | 83341560 | + | 0.630443 | Intergenic | L1_Mur3\|LINE\|L1 | -90154 | NR_102437 | 100039186 | Mm.212721 | NR_102437 | | **Gm2087** | - | predicted gene 2087 | ncRNA |
| chr9_83324332_83324755 | chr9 | 83324333 | 83324755 | + | 0.614947 | Intergenic | Intergenic | -107003 | NR_102437 | 100039186 | Mm.212721 | NR_102437 | | **Gm2087** | - | predicted gene 2087 | ncRNA |
| chr7_80067126_80067478 | chr7 | 80067127 | 80067478 | + | 1.07976 | intron (NR_045695, intron 2 of 2) | MER2\|DNA\|TcMar-Tigger | 7955 | NR_045695 | 100859962 | Mm.392575 | NR_045695 | | **Gm21057** | - | predicted gene, 21057 | ncRNA |
| chr6_83569001_83570329 | chr6 | 83569002 | 83570329 | + | 1.31755 | intron (NR_078345, intron 1 of 7) | intron (NR_078345, intron 1 of 7) | 1626 | NR_078345 | 100861870 | Mm.461230 | NR_078345 | | **Gm21284** | - | predicted gene, 21284 | ncRNA |
| chr18_80616191_80616576 | chr18 | 80616192 | 80616576 | + | 1.85231 | intron (NM_198429, intron 9 of 9) | intron (NM_198429, intron 9 of 9) | -71838 | NR_028424 | 100039348 | Mm.254722 | NR_028424 | | **Gm2176** | A430076E10Rik | glucosamine-6-phosphate deaminase 1 pseudogene | pseudo |
| chr18_80619528_80619833 | chr18 | 80619529 | 80619833 | + | 0.899533 | intron (NM_198429, intron 9 of 9) | intron (NM_198429, intron 9 of 9) | -75135 | NR_028424 | 100039348 | Mm.254722 | NR_028424 | | **Gm2176** | A430076E10Rik | glucosamine-6-phosphate deaminase 1 pseudogene | pseudo |
| chr18_80610119_80610430 | chr18 | 80610120 | 80610430 | + | 0.876751 | intron (NM_198429, intron 9 of 9) | intron (NM_198429, intron 9 of 9) | -65729 | NR_028424 | 100039348 | Mm.254722 | NR_028424 | | **Gm2176** | A430076E10Rik | glucosamine-6-phosphate deaminase 1 pseudogene | pseudo |
| chr9_88721570_88722367 | chr9 | 88721571 | 88722367 | + | 1.38757 | Intergenic | Intergenic | -2171 | NM_001128601 | 100039707 | Mm.347377 | NM_001128601 | ENSMUSG00000079427 | **Gm2382** | - | predicted gene 2382 | protein-coding |
| chr3_52790848_52791489 | chr3 | 52790849 | 52791489 | + | 1.343 | Intergenic | Intergenic | 51445 | NR_038079 | 329639 | Mm.131245 | NR_038079 | | **Gm2447** | 100039832\|A530017F20 | predicted gene 2447 | ncRNA |
| chr3_52828801_52829385 | chr3 | 52828802 | 52829385 | + | 1.05921 | Intergenic | Intergenic | 89369 | NR_038079 | 329639 | Mm.131245 | NR_038079 | | **Gm2447** | 100039832\|A530017F20 | predicted gene 2447 | ncRNA |
| chr3_52828742_52829592 | chr3 | 52828743 | 52829592 | + | 0.970903 | Intergenic | Intergenic | 89443 | NR_038079 | 329639 | Mm.131245 | NR_038079 | | **Gm2447** | 100039832\|A530017F20 | predicted gene 2447 | ncRNA |
| chr8_87472319_87473201 | chr8 | 87472320 | 87473201 | + | 0.725493 | promoter-TSS (NR_033430) | promoter-TSS (NR_033430) | -52 | NR_033430 | 100040294 | Mm.213632 | NR_033430 | ENSMUSG00000097248 | **Gm2694** | C230036H16Rik | predicted gene 2694 | ncRNA |
| chr11_55102209_55102590 | chr11 | 55102210 | 55102590 | + | 1.17093 | intron (NM_010299, intron 1 of 3) | intron (NM_010299, intron 1 of 3) | 4415 | NM_010299 | 14667 | Mm.287807 | NM_010299 | ENSMUSG00000000594 | **Gm2a** | AA408702\|AW215435\|GM2-AP\|SAP-3 | GM2 ganglioside activator protein | protein-coding |
| chr2_19628048_19628659 | chr2 | 19628049 | 19628659 | + | 1.00315 | Intergenic | L1MB7\|LINE\|L1 | 29543 | NR_033642 | 100041244 | Mm.407223 | NR_033642 | | **Gm3230** | - | predicted gene 3230 | ncRNA |
| chr2_19646855_19647343 | chr2 | 19646856 | 19647343 | + | 0.989673 | Intergenic | Intergenic | 10798 | NR_033642 | 100041244 | Mm.407223 | NR_033642 | | **Gm3230** | - | predicted gene 3230 | ncRNA |
| chr2_19630558_19631185 | chr2 | 19630559 | 19631185 | + | 0.957187 | Intergenic | MTEb\|LTR\|ERVL-MaLR | 27025 | NR_033642 | 100041244 | Mm.407223 | NR_033642 | | **Gm3230** | - | predicted gene 3230 | ncRNA |
| chr2_19628901_19629167 | chr2 | 19628902 | 19629167 | + | 0.647506 | Intergenic | Intergenic | 28863 | NR_033642 | 100041244 | Mm.407223 | NR_033642 | | **Gm3230** | - | predicted gene 3230 | ncRNA |
| chr5_64567588_64568221 | chr5 | 64567589 | 64568221 | + | 1.52439 | Intergenic | Intergenic | 42794 | NR_045078 | 100042198 | Mm.25191 | NR_045078 | | **Gm3716** | Gm1683 | predicted gene 3716 | ncRNA |
| chr5_64661912_64662385 | chr5 | 64661913 | 64662385 | + | 1.05914 | Intergenic | Intergenic | -51450 | NR_045078 | 100042198 | Mm.25191 | NR_045078 | | **Gm3716** | Gm1683 | predicted gene 3716 | ncRNA |
| chr5_64624390_64624649 | chr5 | 64624391 | 64624649 | + | 0.833623 | Intergenic | ID_B1\|SINE\|B4 | -13821 | NR_045078 | 100042198 | Mm.25191 | NR_045078 | | **Gm3716** | Gm1683 | predicted gene 3716 | ncRNA |
| chr5_64621373_64621634 | chr5 | 64621374 | 64621634 | + | 0.770098 | Intergenic | Intergenic | -10805 | NR_045078 | 100042198 | Mm.25191 | NR_045078 | | **Gm3716** | Gm1683 | predicted gene 3716 | ncRNA |
| chr5_64650554_64650888 | chr5 | 64650555 | 64650888 | + | 0.676185 | Intergenic | Intergenic | -40022 | NR_045078 | 100042198 | Mm.25191 | NR_045078 | | **Gm3716** | Gm1683 | predicted gene 3716 | ncRNA |
| chr5_64677250_64677790 | chr5 | 64677251 | 64677790 | + | 0.659047 | Intergenic | Intergenic | -66821 | NR_045078 | 100042198 | Mm.25191 | NR_045078 | | **Gm3716** | Gm1683 | predicted gene 3716 | ncRNA |
| chr14_71103943_71104519 | chr14 | 71103944 | 71104519 | + | 1.43226 | Intergenic | Intergenic | -147042 | NR_046075 | 100043132 | Mm.399336 | NR_046075 | | **Gm4251** | - | predicted gene 4251 | ncRNA |
| chr14_71137545_71138439 | chr14 | 71137546 | 71138439 | + | 0.793558 | Intergenic | Intergenic | -113281 | NR_046075 | 100043132 | Mm.399336 | NR_046075 | | **Gm4251** | - | predicted gene 4251 | ncRNA |
| chr14_75819110_75819467 | chr14 | 75819111 | 75819467 | + | 0.732696 | Intergenic | ORR1A4\|LTR\|ERVL-MaLR | 25675 | NR_045294 | 100043189 | Mm.426329 | NR_045294 | | **Gm4285** | - | predicted gene 4285 | ncRNA |
| chr17_32251865_32252332 | chr17 | 32251866 | 32252332 | + | 0.727517 | intron (NM_198094, intron 2 of 12) | intron (NM_198094, intron 2 of 12) | 10179 | NR_102329 | 100043434 | | NR_102329 | | **Gm4432** | - | predicted gene 4432 | pseudo |
| chr10_122193890_122194583 | chr10 | 122193891 | 122194583 | + | 1.33544 | Intergenic | Intergenic | 74540 | NR_027637 | 100043513 | Mm.429141 | NR_027637 | | **Gm4489** | Gm241 | predicted gene 4489 | ncRNA |
| chr10_122198657_122198958 | chr10 | 122198658 | 122198958 | + | 0.741686 | Intergenic | Intergenic | 79111 | NR_027637 | 100043513 | Mm.429141 | NR_027637 | | **Gm4489** | Gm241 | predicted gene 4489 | ncRNA |
| chr16_50204792_50205319 | chr16 | 50204793 | 50205319 | + | 1.28442 | intron (NM_027444, intron 14 of 16) | intron (NM_027444, intron 14 of 16) | -132204 | NR_045935 | 224180 | Mm.146439 | NM_175010 | | **Gm4827** | B130050K08\|EG224180 | predicted gene 4827 | ncRNA |
| chr16_50027116_50027595 | chr16 | 50027117 | 50027595 | + | 0.783333 | intron (NR_045935, intron 2 of 3) | intron (NR_045935, intron 2 of 3) | 45496 | NR_045935 | 224180 | Mm.146439 | NM_175010 | | **Gm4827** | B130050K08\|EG224180 | predicted gene 4827 | ncRNA |
| chr16_50070577_50071113 | chr16 | 50070578 | 50071113 | + | 0.709468 | intron (NR_045935, intron 1 of 3) | intron (NR_045935, intron 1 of 3) | 2007 | NR_045935 | 224180 | Mm.146439 | NM_175010 | | **Gm4827** | B130050K08\|EG224180 | predicted gene 4827 | ncRNA |
| chr17_88195196_88196036 | chr17 | 88195197 | 88196036 | + | 1.48493 | Intergenic | ERVL-B4-int\|LTR\|ERVL | 70104 | NM_001190356 | 225058 | Mm.312053 | NM_001190356 | | **Gm4832** | EG225058 | predicted gene 4832 | protein-coding |
| chr17_88113265_88114093 | chr17 | 88113266 | 88114093 | + | 1.08206 | Intergenic | Intergenic | -11833 | NM_001190356 | 225058 | Mm.312053 | NM_001190356 | | **Gm4832** | EG225058 | predicted gene 4832 | protein-coding |
| chr17_88213506_88213966 | chr17 | 88213507 | 88213966 | + | 0.804825 | Intergenic | Intergenic | 88224 | NM_001190356 | 225058 | Mm.312053 | NM_001190356 | | **Gm4832** | EG225058 | predicted gene 4832 | protein-coding |
| chr17_88112253_88112723 | chr17 | 88112254 | 88112723 | + | 0.74693 | Intergenic | Intergenic | -13024 | NM_001190356 | 225058 | Mm.312053 | NM_001190356 | | **Gm4832** | EG225058 | predicted gene 4832 | protein-coding |
| chr17_88111872_88112750 | chr17 | 88111873 | 88112750 | + | 0.699838 | Intergenic | RMER15\|LTR\|ERVL | -13201 | NM_001190356 | 225058 | Mm.312053 | NM_001190356 | | **Gm4832** | EG225058 | predicted gene 4832 | protein-coding |
| chrX_11665130_11666034 | chrX | 11665131 | 11666034 | + | 0.697541 | Intergenic | Intergenic | 337760 | NM_001114529 | 236663 | Mm.473680 | NM_001114529 | ENSMUSG00000069038 | **Gm4906** | EG236663 | predicted gene 4906 | protein-coding |
| chr1_21309180_21309625 | chr1 | 21309181 | 21309625 | + | 0.931227 | Intergenic | Lx8\|LINE\|L1 | -11091 | NR_002858 | 241041 | Mm.290634 | NR_002858 | ENSMUSG00000025936 | **Gm4956** | EG241041 | predicted gene 4956 | pseudo |
| chr7_73637601_73638098 | chr7 | 73637602 | 73638098 | + | 1.31091 | Intergenic | Intergenic | 48713 | NR_033604 | 244061 | Mm.484605 | NR_033604 | | **Gm4971** | EG244061 | Smad nuclear interacting protein 1 pseudogene | pseudo |
| chr13_41662594_41663262 | chr13 | 41662595 | 41663262 | + | 0.997189 | Intergenic | Intergenic | 12166 | NM_001145878 | 328231 | Mm.227476 | NM_177808 | | **Gm5082** | A030008J09\|EG328231 | predicted gene 5082 | protein-coding |
| chr13_41670923_41671931 | chr13 | 41670924 | 41671931 | + | 0.957081 | Intergenic | ORR1F\|LTR\|ERVL-MaLR | 20665 | NM_001145878 | 328231 | Mm.227476 | NM_177808 | | **Gm5082** | A030008J09\|EG328231 | predicted gene 5082 | protein-coding |
| chr13_41663446_41663826 | chr13 | 41663447 | 41663826 | + | 0.736913 | Intergenic | URR1B\|DNA\|hAT-Charlie | 12874 | NM_001145878 | 328231 | Mm.227476 | NM_177808 | | **Gm5082** | A030008J09\|EG328231 | predicted gene 5082 | protein-coding |

| | | | | | | | | | | | | | | | | | |
|---|---|---|---|---|---|---|---|---|---|---|---|---|---|---|---|---|---|
| chr13_41638109_41638488 | chr13 | 41638110 | 41638488 | + | 0.734953 | 3' UTR (NM_001163572, exon 3 of 3) | 3' UTR (NM_001163572, exon 3 of 3) | -11144 | NM_177808 | 328231 | Mm.227476 | NM_177808 | | **Gm5082** | A030008J09\|EG328231 | predicted gene 5082 | protein-coding |
| chr13_97680364_97681150 | chr13 | 97680365 | 97681150 | + | 1.19874 | Intergenic | Intergenic | 120757 | NR_046157 | 328314 | Mm.138047 | NR_046157 | | **Gm5086** | B230311A21\|EG328314 | predicted gene 5086 | ncRNA |
| chr13_97550851_97551447 | chr13 | 97550852 | 97551447 | + | 0.779723 | Intergenic | Intergenic | -8851 | NR_046157 | 328314 | Mm.138047 | NR_046157 | | **Gm5086** | B230311A21\|EG328314 | predicted gene 5086 | ncRNA |
| chr3_138062685_138063020 | chr3 | 138062686 | 138063020 | + | 0.8627 | intron (NR_037975, intron 1 of 3) | intron (NR_037975, intron 1 of 3) | 4535 | NR_037975 | 329763 | Mm.169710 | NM_177860 | ENSMUSG00000049349 | **Gm5105** | D830013H12\|EG329763 | predicted gene 5105 | ncRNA |
| chr3_138061586_138062024 | chr3 | 138061587 | 138062024 | + | 0.766546 | intron (NR_037975, intron 1 of 3) | intron (NR_037975, intron 1 of 3) | 5583 | NR_037975 | 329763 | Mm.169710 | NM_177860 | ENSMUSG00000049349 | **Gm5105** | D830013H12\|EG329763 | predicted gene 5105 | ncRNA |
| chr3_138061283_138062027 | chr3 | 138061284 | 138062027 | + | 0.737714 | intron (NR_037975, intron 1 of 3) | intron (NR_037975, intron 1 of 3) | 5733 | NR_037975 | 329763 | Mm.169710 | NM_177860 | ENSMUSG00000049349 | **Gm5105** | D830013H12\|EG329763 | predicted gene 5105 | ncRNA |
| chr6_48591201_48591756 | chr6 | 48591202 | 48591756 | + | 0.701845 | TTS (NM_183309) | TTS (NM_183309) | 2034 | NM_183309 | 330305 | Mm.123868 | NM_183309 | ENSMUSG00000052730 | **Gm5111** | A630038D02\|EG330305 | predicted gene 5111 | protein-coding |
| chr6_48591444_48591767 | chr6 | 48591445 | 48591767 | + | 0.643434 | Intergenic | Intergenic | 2161 | NM_183309 | 330305 | Mm.123868 | NM_183309 | ENSMUSG00000052730 | **Gm5111** | A630038D02\|EG330305 | predicted gene 5111 | protein-coding |
| chr9_61014460_61014894 | chr9 | 61014461 | 61014894 | + | 1.05542 | Intergenic | Intergenic | -2392 | NR_040767 | 330956 | Mm.207985 | NR_040767 | | **Gm5122** | A430102L10 | predicted gene 5122 | ncRNA |
| chrX_21349216_21349601 | chrX | 21349217 | 21349601 | + | 0.659532 | Intergenic | B3\|SINE\|B2 | -11456 | NR_045668 | 331392 | Mm.23126 | NM_198633 | ENSMUSG00000051537 | **Gm5124** | EG331392 | nucleolar and coiled-body phosphoprotein 1 pseudogene | pseudo |
| chr5_29739507_29739864 | chr5 | 29739508 | 29739864 | + | 1.24567 | intron (NM_001127367, intron 1 of 7) | intron (NM_001127367, intron 1 of 7) | -3750 | NR_028426 | 332993 | Mm.473746 | NR_028426 | | **Gm5129** | EG332993\|ENSMUSG00000053121 | predicted gene 5129 | ncRNA |
| chr10_75968266_75968605 | chr10 | 75968267 | 75968605 | + | 0.627338 | intron (NM_198635, intron 3 of 13) | RMER1A\|Other\|Other | 13922 | NM_198635 | 333669 | Mm.31077 | NM_198635 | ENSMUSG00000033255 | **Gm5134** | EG333669 | predicted gene 5134 | protein-coding |
| chr3_37754746_37755129 | chr3 | 37754747 | 37755129 | + | 0.769205 | Intergenic | ORR1C1\|LTR\|ERVL-MaLR | -30578 | NM_198657 | 381438 | Mm.490549 | NM_198657 | ENSMUSG00000058174 | **Gm5148** | EG381438 | predicted gene 5148 | protein-coding |
| chr11_89095324_89095801 | chr11 | 89095325 | 89095801 | + | 0.821904 | Intergenic | Intergenic | 21722 | NM_001033266 | 217071 | Mm.342127 | NM_001033266 | ENSMUSG00000072553 | **Gm525** | - | predicted gene 525 | protein-coding |
| chr10_21670062_21670541 | chr10 | 21670063 | 21670541 | + | 0.773308 | Intergenic | Intergenic | -20543 | NR_045843 | 432436 | Mm.244553 | NM_001004162 | ENSMUSG00000056716 | **Gm5420** | EG432436 | predicted gene 5420 | ncRNA |
| chr12_36073223_36073823 | chr12 | 36073224 | 36073823 | + | 0.735767 | Intergenic | Intergenic | -16856 | NR_003649 | 432649 | Mm.347860 | NM_001014396 | ENSMUSG00000059301 | **Gm5434** | EG432649 | ubiquitin-conjugating enzyme E2F (putative) pseudogene | pseudo |
| chr12_117406771_117407363 | chr12 | 117406772 | 117407363 | + | 1.07061 | Intergenic | MER5A\|DNA\|hAT-Charlie | -70055 | NR_044987 | 432713 | Mm.459210 | NR_044987 | | **Gm5441** | - | predicted gene 5441 | ncRNA |
| chr1_13752475_13753052 | chr1 | 13752476 | 13753052 | + | 1.06829 | Intergenic | L1MA4\|LINE\|L1 | 34165 | NR_004447 | 433273 | Mm.304088 | NR_004447 | ENSMUSG00000097711 | **Gm5523** | EG433273 | glyceraldehyde-3-phosphate dehydrogenase pseudogene | pseudo |
| chr2_144213787_144214218 | chr2 | 144213788 | 144214218 | + | 1.11269 | Intergenic | L1MB2\|LINE\|L1 | -24713 | NM_001033778 | 433481 | Mm.330630 | NM_001033778 | | **Gm5535** | EG433481 | predicted gene 5535 | protein-coding |
| chr2_144213865_144214108 | chr2 | 144213866 | 144214108 | + | 1.08859 | Intergenic | L1MB2\|LINE\|L1 | -24697 | NM_001033778 | 433481 | Mm.330630 | NM_001033778 | | **Gm5535** | EG433481 | predicted gene 5535 | protein-coding |
| chr2_144188032_144188656 | chr2 | 144188033 | 144188656 | + | 0.817053 | intron (NM_001033778, intron 1 of 3) | MIR\|SINE\|MIR | 946 | NM_001033778 | 433481 | Mm.330630 | NM_001033778 | | **Gm5535** | EG433481 | predicted gene 5535 | protein-coding |
| chr3_97948453_97949047 | chr3 | 97948454 | 97949047 | + | 1.48227 | intron (NM_001033779, intron 1 of 1) | intron (NM_001033779, intron 1 of 1) | 18577 | NM_001033779 | 433632 | Mm.212421 | NM_001033779 | | **Gm5544** | EG433632 | predicted gene 5544 | protein-coding |
| chr3_97934869_97935278 | chr3 | 97934870 | 97935278 | + | 1.45107 | intron (NM_001033779, intron 1 of 1) | intron (NM_001033779, intron 1 of 1) | 4901 | NM_001033779 | 433632 | Mm.212421 | NM_001033779 | | **Gm5544** | EG433632 | predicted gene 5544 | protein-coding |
| chr3_97930095_97930875 | chr3 | 97930096 | 97930875 | + | 1.33962 | intron (NM_001033779, intron 1 of 1) | intron (NM_001033779, intron 1 of 1) | 312 | NM_001033779 | 433632 | Mm.212421 | NM_001033779 | | **Gm5544** | EG433632 | predicted gene 5544 | protein-coding |
| chr3_97937707_97937997 | chr3 | 97937708 | 97937997 | + | 0.865376 | intron (NM_001033779, intron 1 of 1) | L1MB8\|LINE\|L1 | 7679 | NM_001033779 | 433632 | Mm.212421 | NM_001033779 | | **Gm5544** | EG433632 | predicted gene 5544 | protein-coding |
| chr3_97947508_97947891 | chr3 | 97947509 | 97947891 | + | 0.835698 | intron (NM_001033779, intron 1 of 1) | MIRb\|SINE\|MIR | 17527 | NM_001033779 | 433632 | Mm.212421 | NM_001033779 | | **Gm5544** | EG433632 | predicted gene 5544 | protein-coding |
| chr3_97955487_97955946 | chr3 | 97955488 | 97955946 | + | 0.835512 | intron (NM_001033779, intron 1 of 1) | intron (NM_001033779, intron 1 of 1) | 25544 | NM_001033779 | 433632 | Mm.212421 | NM_001033779 | | **Gm5544** | EG433632 | predicted gene 5544 | protein-coding |
| chr3_97967676_97967953 | chr3 | 97967677 | 97967953 | + | 0.774264 | TTS (NM_001033779) | TTS (NM_001033779) | 37642 | NM_001033779 | 433632 | Mm.212421 | NM_001033779 | | **Gm5544** | EG433632 | predicted gene 5544 | protein-coding |
| chr3_97965944_97966334 | chr3 | 97965945 | 97966334 | + | 0.71806 | intron (NM_001033779, intron 1 of 1) | intron (NM_001033779, intron 1 of 1) | 35966 | NM_001033779 | 433632 | Mm.212421 | NM_001033779 | | **Gm5544** | EG433632 | predicted gene 5544 | protein-coding |
| chr3_105818091_105818872 | chr3 | 105818092 | 105818872 | + | 1.1594 | Intergenic | LTR81B\|LTR\|Gypsy | 2915 | NR_045845 | 433637 | Mm.331285 | NR_045845 | | **Gm5547** | EG433637 | predicted gene 5547 | ncRNA |
| chr1_138741139_138741528 | chr1 | 138741140 | 138741528 | + | 0.727123 | Intergenic | Intergenic | -59259 | NR_040304 | 545363 | Mm.362817 | NR_040304 | | **Gm5833** | - | predicted gene 5833 | ncRNA |
| chr1_138743460_138744141 | chr1 | 138743461 | 138744141 | + | 0.62195 | Intergenic | Intergenic | -61726 | NR_040304 | 545363 | Mm.362817 | NR_040304 | | **Gm5833** | - | predicted gene 5833 | ncRNA |
| chrX_49046226_49046860 | chrX | 49046227 | 49046860 | + | 1.10339 | intron (NM_145951, intron 9 of 15) | intron (NM_145951, intron 9 of 15) | -168830 | NM_001085499 | 209005 | Mm.336054 | NM_001085499 | ENSMUSG00000079606 | **Gm595** | - | predicted gene 595 | protein-coding |
| chrX_49028535_49029027 | chrX | 49028536 | 49029027 | + | 0.727099 | intron (NM_145951, intron 10 of 15) | intron (NM_145951, intron 10 of 15) | -151068 | NM_001085499 | 209005 | Mm.336054 | NM_001085499 | ENSMUSG00000079606 | **Gm595** | - | predicted gene 595 | protein-coding |
| chr16_44210430_44211134 | chr16 | 44210431 | 44211134 | + | 1.15944 | intron (NM_001029889, intron 6 of 7) | intron (NM_001029889, intron 6 of 7) | 37385 | NM_001029889 | 207806 | Mm.310566 | NM_001029889 | ENSMUSG00000068284 | **Gm608** | 5530400K22Rik\|Gm1779\|mKIAA2018 | predicted gene 608 | protein-coding |
| chr16_45506347_45506701 | chr16 | 45506348 | 45506701 | + | 0.661586 | Intergenic | Lx2A1\|LINE\|L1 | -13555 | NM_001005854 | 208166 | Mm.451852 | NM_001005854 | ENSMUSG00000053182 | **Gm609** | Gm1749\|isec1 | predicted gene 609 | protein-coding |
| chr5_74903785_74904249 | chr5 | 74903786 | 74904249 | + | 1.2487 | Intergenic | Intergenic | -45177 | NR_045866 | 619945 | Mm.359372 | NM_001037908 | ENSMUSG00000072874 | **Gm6116** | EG619945 | predicted gene 6116 | ncRNA |
| chr5_74903494_74904283 | chr5 | 74903495 | 74904283 | + | 1.15553 | Intergenic | Intergenic | -45305 | NR_045866 | 619945 | Mm.359372 | NM_001037908 | ENSMUSG00000072874 | **Gm6116** | EG619945 | predicted gene 6116 | ncRNA |
| chr3_143539820_143540293 | chr3 | 143539821 | 143540293 | + | 0.840759 | Intergenic | MTD\|LTR\|ERVL-MaLR | 172086 | NR_040405 | 621809 | Mm.360134 | NR_040405 | | **Gm6260** | EG621809 | predicted gene 6260 | ncRNA |
| chr3_143912394_143913008 | chr3 | 143912395 | 143913008 | + | 0.734843 | Intergenic | Intergenic | -200558 | NR_040405 | 621809 | Mm.360134 | NR_040405 | | **Gm6260** | EG621809 | predicted gene 6260 | ncRNA |
| chrX_36385234_36385665 | chrX | 36385235 | 36385665 | + | 0.926174 | Intergenic | Intergenic | -18027 | NR_044989 | 621901 | Mm.491051 | NR_044989 | ENSMUSG00000079642 | **Gm6268** | EG621901 | predicted gene 6268 | ncRNA |
| chrX_109207857_109208245 | chrX | 109207858 | 109208245 | + | 0.949605 | Intergenic | Intergenic | 11295 | NM_001037917 | 622976 | Mm.482747 | NM_001037917 | ENSMUSG00000048621 | **Gm6377** | EG622976 | predicted gene 6377 | protein-coding |
| chrX_109207551_109208540 | chrX | 109207552 | 109208540 | + | 0.607947 | Intergenic | Intergenic | 11290 | NM_001037917 | 622976 | Mm.482747 | NM_001037917 | ENSMUSG00000048621 | **Gm6377** | EG622976 | predicted gene 6377 | protein-coding |
| chr13_117127067_117127689 | chr13 | 117127068 | 117127689 | + | 0.993805 | Intergenic | Intergenic | -2647 | NR_046023 | 623312 | Mm.131279 | NR_046023 | | **Gm6416** | - | predicted gene 6416 | ncRNA |
| chr14_48235899_48236754 | chr14 | 48235900 | 48236754 | + | 1.59108 | intron (NM_033602, intron 2 of 5) | intron (NM_033602, intron 2 of 5) | 37611 | NR_003630 | 624367 | Mm.445185 | NM_001037921 | | **Gm6498** | EG624367 | glyceraldehyde-3-phosphate dehydrogenase pseudogene | pseudo |
| chr3_84169359_84169986 | chr3 | 84169360 | 84169986 | + | 1.36819 | intron (NM_001271725, intron 10 of 11) | MLT1A1\|LTR\|ERVL-MaLR | -4965 | NR_036654 | 624713 | Mm.442604 | NR_036654 | | **Gm6525** | EG624713 | ribosomal protein L36a pseudogene | pseudo |
| chr19_30924059_30924706 | chr19 | 30924060 | 30924706 | + | 0.938779 | intron (NM_001013833, intron 4 of 17) | intron (NM_001013833, intron 4 of 17) | 6579 | NR_033643 | 625995 | Mm.454508 | NR_033643 | | **Gm6642** | EG625995 | ubiquitin-like 5 pseudogene | pseudo |
| chr19_30924037_30925128 | chr19 | 30924038 | 30925128 | + | 0.832521 | intron (NM_001013833, intron 4 of 17) | intron (NM_001013833, intron 4 of 17) | 6379 | NR_033643 | 625995 | Mm.454508 | NR_033643 | | **Gm6642** | EG625995 | ubiquitin-like 5 pseudogene | pseudo |
| chr6_34274491_34274988 | chr6 | 34274492 | 34274988 | + | 1.20465 | Intergenic | Intergenic | 42703 | NR_037965 | 626009 | Mm.451 | NR_028277 | | **Gm6644** | Akr1b3p\|EG626009 | Akr1b3 pseudogene | pseudo |
| chr6_34272470_34272886 | chr6 | 34272471 | 34272886 | + | 1.13329 | Intergenic | RLTR53_Mm\|LTR\|ERVK | 44765 | NR_037965 | 626009 | Mm.451 | NR_028277 | | **Gm6644** | Akr1b3p\|EG626009 | Akr1b3 pseudogene | pseudo |
| chr6_34276476_34276740 | chr6 | 34276477 | 34276740 | + | 1.03393 | Intergenic | Intergenic | 40835 | NR_037965 | 626009 | Mm.451 | NR_028277 | | **Gm6644** | Akr1b3p\|EG626009 | Akr1b3 pseudogene | pseudo |
| chr9_51285982_51286438 | chr9 | 51285983 | 51286438 | + | 0.904329 | Intergenic | Intergenic | -7656 | NM_001195681 | 100502940 | Mm.235705 | NM_001195681 | ENSMUSG00000079559 | **Gm684** | - | predicted gene 684 | protein-coding |
| chrX_77611948_77612487 | chrX | 77611949 | 77612487 | + | 0.754353 | intron (NM_020601, intron 1 of 15) | intron (NM_020601, intron 1 of 15) | 63234 | NM_001101585 | 628893 | Mm.477437 | NM_001101585 | | **Gm6927** | EG628893 | predicted pseudogene 6927 | protein-coding |
| chrX_77611948_77612348 | chrX | 77611949 | 77612348 | + | 0.745889 | intron (NM_020601, intron 1 of 15) | MTE2a\|LTR\|ERVL-MaLR | 63304 | NM_001101585 | 628893 | Mm.477437 | NM_001101585 | | **Gm6927** | EG628893 | predicted pseudogene 6927 | protein-coding |
| chr16_49982131_49982830 | chr16 | 49982132 | 49982830 | + | 1.62064 | intron (NR_045001, intron 1 of 3) | intron (NR_045001, intron 1 of 3) | 2021 | NR_045001 | 628951 | Mm.428032 | NR_045001 | | **Gm6936** | - | predicted gene 6936 | ncRNA |
| chr16_49969975_49971615 | chr16 | 49969976 | 49971615 | + | 1.18138 | Intergenic | Intergenic | -9665 | NR_045001 | 628951 | Mm.428032 | NR_045001 | | **Gm6936** | - | predicted gene 6936 | ncRNA |
| chr16_49951853_49952210 | chr16 | 49951854 | 49952210 | + | 0.886783 | Intergenic | Intergenic | -28428 | NR_045001 | 628951 | Mm.428032 | NR_045001 | | **Gm6936** | - | predicted gene 6936 | ncRNA |
| chr16_49938668_49939276 | chr16 | 49938669 | 49939276 | + | 0.801599 | Intergenic | Intergenic | -41488 | NR_045001 | 628951 | Mm.428032 | NR_045001 | | **Gm6936** | - | predicted gene 6936 | ncRNA |

| | | | | | | | | | | | | | | | | | |
|---|---|---|---|---|---|---|---|---|---|---|---|---|---|---|---|---|---|
| chr16_49935952_49937049 | chr16 | 49935953 | 49937049 | + | 0.763788 | Intergenic | Intergenic | -43959 | NR_045001 | 628951 | Mm.428032 | NR_045001 | | **Gm6936** | - | predicted gene 6936 | ncRNA |
| chr16_49951837_49952557 | chr16 | 49951838 | 49952557 | + | 0.723376 | Intergenic | Intergenic | -28263 | NR_045001 | 628951 | Mm.428032 | NR_045001 | | **Gm6936** | - | predicted gene 6936 | ncRNA |
| chr16_49943535_49944371 | chr16 | 49943536 | 49944371 | + | 0.699733 | Intergenic | Intergenic | -36507 | NR_045001 | 628951 | Mm.428032 | NR_045001 | | **Gm6936** | - | predicted gene 6936 | ncRNA |
| chr9_52059548_52060591 | chr9 | 52059549 | 52060591 | + | 1.29283 | intron (NM_009041, intron 1 of 13) | intron (NM_009041, intron 1 of 13) | -11365 | NR_023357 | 629557 | Mm.422775 | NR_023357 | | **Gm6981** | EG629557 | glyceraldehyde-3-phosphate dehydrogenase pseudogene | pseudo |
| chr12_40263017_40263336 | chr12 | 40263018 | 40263336 | + | 0.816025 | Intergenic | Intergenic | 39813 | NR_045157 | 629820 | Mm.435540 | NR_045157 | ENSMUSG00000035983 | **Gm7008** | EG629820 | predicted gene 7008 | ncRNA |
| chr12_40223885_40224081 | chr12 | 40223886 | 40224081 | + | 0.6393 | promoter-TSS (NM_013562) | promoter-TSS (NM_013562) | 619 | NR_045157 | 629820 | Mm.435540 | NR_045157 | ENSMUSG00000035983 | **Gm7008** | EG629820 | predicted gene 7008 | ncRNA |
| chr17_45621401_45621729 | chr17 | 45621402 | 45621729 | + | 0.820704 | Intergenic | ORR1A4\|LTR\|ERVL-MaLR | -19463 | NM_001177469 | 653016 | Mm.45676 | NM_001177468 | ENSMUSG00000079471 | **Gm7325** | EG653016\|Esgp | predicted gene 7325 | protein-coding |
| chr17_45616631_45617107 | chr17 | 45616632 | 45617107 | + | 0.783521 | Intergenic | ID_B1\|SINE\|B4 | -14767 | NM_001177469 | 653016 | Mm.45676 | NM_001177468 | ENSMUSG00000079471 | **Gm7325** | EG653016\|Esgp | predicted gene 7325 | protein-coding |
| chr6_142823030_142823713 | chr6 | 142823031 | 142823713 | + | 1.31445 | 3' UTR (NM_011374, exon 5 of 5) | 3' UTR (NM_011374, exon 5 of 5) | 9141 | NR_045707 | 665037 | Mm.401493 | NR_045707 | ENSMUSG00000072683 | **Gm7457** | EG665037\|ENSMUSG00000072683 | predicted gene 7457 | ncRNA |
| chr6_142845726_142846183 | chr6 | 142845727 | 142846183 | + | 1.27328 | intron (NM_011374, intron 4 of 4) | intron (NM_011374, intron 4 of 4) | 31724 | NR_045707 | 665037 | Mm.401493 | NR_045707 | ENSMUSG00000072683 | **Gm7457** | EG665037\|ENSMUSG00000072683 | predicted gene 7457 | ncRNA |
| chr6_142847977_142848783 | chr6 | 142847978 | 142848783 | + | 1.19649 | intron (NM_011374, intron 4 of 4) | intron (NM_011374, intron 4 of 4) | 34149 | NR_045707 | 665037 | Mm.401493 | NR_045707 | ENSMUSG00000072683 | **Gm7457** | EG665037\|ENSMUSG00000072683 | predicted gene 7457 | ncRNA |
| chr6_142834727_142835565 | chr6 | 142834728 | 142835565 | + | 1.10218 | intron (NM_011374, intron 4 of 4) | intron (NM_011374, intron 4 of 4) | 20915 | NR_045707 | 665037 | Mm.401493 | NR_045707 | ENSMUSG00000072683 | **Gm7457** | EG665037\|ENSMUSG00000072683 | predicted gene 7457 | ncRNA |
| chr6_142825347_142825812 | chr6 | 142825348 | 142825812 | + | 0.920788 | 3' UTR (NM_011374, exon 5 of 5) | 3' UTR (NM_011374, exon 5 of 5) | 11349 | NR_045707 | 665037 | Mm.401493 | NR_045707 | ENSMUSG00000072683 | **Gm7457** | EG665037\|ENSMUSG00000072683 | predicted gene 7457 | ncRNA |
| chr6_142881397_142882183 | chr6 | 142881398 | 142882183 | + | 0.694795 | intron (NM_011374, intron 2 of 4) | intron (NM_011374, intron 2 of 4) | 67559 | NR_045707 | 665037 | Mm.401493 | NR_045707 | ENSMUSG00000072683 | **Gm7457** | EG665037\|ENSMUSG00000072683 | predicted gene 7457 | ncRNA |
| chr6_142848005_142848234 | chr6 | 142848006 | 142848234 | + | 0.652464 | intron (NM_011374, intron 4 of 4) | ORR1E\|LTR\|ERVL-MaLR | 33889 | NR_045707 | 665037 | Mm.401493 | NR_045707 | ENSMUSG00000072683 | **Gm7457** | EG665037\|ENSMUSG00000072683 | predicted gene 7457 | ncRNA |
| chr12_54369208_54369942 | chr12 | 54369209 | 54369942 | + | 1.05643 | Intergenic | RSINE1\|SINE\|B4 | 21743 | NR_033689 | 665239 | Mm.26856 | NR_033689 | | **Gm7550** | EG665239\|Gm666 | predicted gene 7550 | ncRNA |
| chr12_54487680_54488167 | chr12 | 54487681 | 54488167 | + | 1.03425 | Intergenic | HAL1b\|LINE\|L1 | -96606 | NR_033689 | 665239 | Mm.26856 | NR_033689 | | **Gm7550** | EG665239\|Gm666 | predicted gene 7550 | ncRNA |
| chr12_54516400_54516979 | chr12 | 54516401 | 54516979 | + | 0.975287 | Intergenic | MLTR25A\|LTR\|ERVK | -125372 | NR_033689 | 665239 | Mm.26856 | NR_033689 | | **Gm7550** | EG665239\|Gm666 | predicted gene 7550 | ncRNA |
| chr12_54312600_54312970 | chr12 | 54312601 | 54312970 | + | 0.856011 | Intergenic | Intergenic | 78533 | NR_033689 | 665239 | Mm.26856 | NR_033689 | | **Gm7550** | EG665239\|Gm666 | predicted gene 7550 | ncRNA |
| chr12_54489521_54490019 | chr12 | 54489522 | 54490019 | + | 0.730979 | Intergenic | Intergenic | -98452 | NR_033689 | 665239 | Mm.26856 | NR_033689 | | **Gm7550** | EG665239\|Gm666 | predicted gene 7550 | ncRNA |
| chr12_54488612_54489053 | chr12 | 54488613 | 54489053 | + | 0.68988 | Intergenic | MTEb\|LTR\|ERVL-MaLR | -97515 | NR_033689 | 665239 | Mm.26856 | NR_033689 | | **Gm7550** | EG665239\|Gm666 | predicted gene 7550 | ncRNA |
| chr6_98597023_98598036 | chr6 | 98597024 | 98598036 | + | 1.10448 | Intergenic | Intergenic | -254776 | NM_001128092 | 330390 | Mm.484161 | NM_001128092 | ENSMUSG00000090667 | **Gm765** | - | predicted gene 765 | protein-coding |
| chr6_98597613_98598020 | chr6 | 98597614 | 98598020 | + | 1.04611 | Intergenic | Intergenic | -255063 | NM_001128092 | 330390 | Mm.484161 | NM_001128092 | ENSMUSG00000090667 | **Gm765** | - | predicted gene 765 | protein-coding |
| chr6_98597028_98597559 | chr6 | 98597029 | 98597559 | + | 0.869028 | Intergenic | Intergenic | -254540 | NM_001128092 | 330390 | Mm.484161 | NM_001128092 | ENSMUSG00000090667 | **Gm765** | - | predicted gene 765 | protein-coding |
| chr6_98651356_98651617 | chr6 | 98651357 | 98651617 | + | 0.729329 | Intergenic | Intergenic | -308733 | NM_001128092 | 330390 | Mm.484161 | NM_001128092 | ENSMUSG00000090667 | **Gm765** | - | predicted gene 765 | protein-coding |
| chr9_21601134_21601445 | chr9 | 21601135 | 21601445 | + | 0.72738 | Intergenic | Intergenic | 7039 | NR_003372 | 666043 | Mm.247282 | NR_003372 | | **Gm7904** | EG666043 | predicted gene 7904 | pseudo |
| chr9_21609863_21610338 | chr9 | 21609864 | 21610338 | + | 0.716854 | Intergenic | Intergenic | -1772 | NR_003372 | 666043 | Mm.247282 | NR_003372 | | **Gm7904** | EG666043 | predicted gene 7904 | pseudo |
| chr12_86142314_86142949 | chr12 | 86142315 | 86142949 | + | 0.873504 | intron (NM_029601, intron 4 of 7) | intron (NM_029601, intron 4 of 7) | -26709 | NR_046081 | 328136 | Mm.312050 | NR_046081 | ENSMUSG00000097543 | **Gm805** | - | predicted gene 805 | ncRNA |
| chr16_58584761_58585117 | chr16 | 58584762 | 58585117 | + | 0.677024 | Intergenic | Intergenic | 32039 | NM_001033404 | 328695 | Mm.293308 | NM_001033404 | ENSMUSG00000075002 | **Gm813** | Gm1781 | predicted gene 813 | protein-coding |
| chr2_160336392_160337096 | chr2 | 160336393 | 160337096 | + | 0.734356 | Intergenic | Intergenic | -9250 | NM_001033411 | 329554 | Mm.291144 | NM_001033411 | | **Gm826** | - | predicted gene 826 | protein-coding |
| chr2_160336058_160337134 | chr2 | 160336059 | 160337134 | + | 0.70889 | Intergenic | Intergenic | -9102 | NM_001033411 | 329554 | Mm.291144 | NM_001033411 | | **Gm826** | - | predicted gene 826 | protein-coding |
| chr6_3343543_3343949 | chr6 | 3343544 | 3343949 | + | 0.735197 | Intergenic | B4A\|SINE\|B4 | 55227 | NR_036696 | 667335 | Mm.380219 | NR_036696 | | **Gm8579** | EG667335 | spermatogenesis associated glutamate (E)-rich protein 4a pseudogene | pseudo |
| chr10_75939833_75940179 | chr10 | 75939834 | 75940179 | + | 1.0196 | exon (NM_001277132, exon 3 of 7) | exon (NM_001277132, exon 3 of 7) | 666 | NM_001277132 | 333670 | Mm.291320 | NM_001037714 | | **Gm867** | Gm736 | predicted gene 867 | protein-coding |
| chr10_75937972_75938334 | chr10 | 75937973 | 75938334 | + | 0.677474 | TTS (NM_175329) | TTS (NM_175329) | 2519 | NM_001277132 | 333670 | Mm.291320 | NM_001037714 | | **Gm867** | Gm736 | predicted gene 867 | protein-coding |
| chr10_26544620_26545231 | chr10 | 26544621 | 26545231 | + | 1.4579 | Intergenic | Intergenic | 129725 | NR_033633 | 667572 | Mm.482728 | NR_033633 | ENSMUSG00000098179 | **Gm8709** | EG667572 | glyceraldehyde-3-phosphate dehydrogenase pseudogene | pseudo |
| chr10_26577054_26577449 | chr10 | 26577055 | 26577449 | + | 0.806081 | Intergenic | Intergenic | 97399 | NR_033633 | 667572 | Mm.482728 | NR_033633 | ENSMUSG00000098179 | **Gm8709** | EG667572 | glyceraldehyde-3-phosphate dehydrogenase pseudogene | pseudo |
| chr10_26544132_26544394 | chr10 | 26544133 | 26544394 | + | 0.753392 | Intergenic | Intergenic | 130388 | NR_033633 | 667572 | Mm.482728 | NR_033633 | ENSMUSG00000098179 | **Gm8709** | EG667572 | glyceraldehyde-3-phosphate dehydrogenase pseudogene | pseudo |
| chrX_166982161_166982927 | chrX | 166982162 | 166982927 | + | 1.74211 | Intergenic | Intergenic | 87826 | NM_001101606 | 667794 | Mm.302232 | NM_001101606 | | **Gm8817** | EG667794 | predicted gene 8817 | protein-coding |
| chrX_167130795_167131679 | chrX | 167130796 | 167131679 | + | 1.37367 | Intergenic | Intergenic | -60867 | NM_001101606 | 667794 | Mm.302232 | NM_001101606 | | **Gm8817** | EG667794 | predicted gene 8817 | protein-coding |
| chr3_95957199_95957795 | chr3 | 95957200 | 95957795 | + | 0.659773 | Intergenic | Intergenic | -28013 | NR_045872 | 668224 | Mm.357766 | NR_045872 | | **Gm9054** | EG668224 | predicted gene 9054 | ncRNA |
| chr14_72886692_72887394 | chr14 | 72886693 | 72887394 | + | 1.1718 | Intergenic | L1MD3\|LINE\|L1 | -138560 | NR_027860 | 668489 | Mm.158324 | NR_027860 | | **Gm9199** | EG668489 | glycine cleavage system protein H (aminomethyl carrier) pseudogene | pseudo |
| chr14_72869194_72869795 | chr14 | 72869195 | 72869795 | + | 1.02536 | Intergenic | Intergenic | -156108 | NR_027860 | 668489 | Mm.158324 | NR_027860 | | **Gm9199** | EG668489 | glycine cleavage system protein H (aminomethyl carrier) pseudogene | pseudo |
| chr19_4701064_4701735 | chr19 | 4701065 | 4701735 | + | 0.711797 | Intergenic | Intergenic | -2732 | NM_001033447 | 381196 | Mm.297789 | NM_001033447 | ENSMUSG00000071691 | **Gm960** | - | predicted gene 960 | protein-coding |
| chr19_4701094_4701567 | chr19 | 4701095 | 4701567 | + | 0.66538 | Intergenic | Intergenic | -2663 | NM_001033447 | 381196 | Mm.297789 | NM_001033447 | ENSMUSG00000071691 | **Gm960** | - | predicted gene 960 | protein-coding |
| chr3_15228588_15229218 | chr3 | 15228589 | 15229218 | + | 1.24438 | Intergenic | Intergenic | 103399 | NM_001076679 | 751864 | Mm.431296 | NM_001076679 | ENSMUSG00000078783 | **Gm9733** | - | predicted gene 9733 | protein-coding |
| chr13_94327989_94328542 | chr13 | 94327990 | 94328542 | + | 1.09079 | Intergenic | Intergenic | 30657 | NR_045619 | 328309 | Mm.174191 | NM_183205 | | **Gm9776** | A530089L17\|ENSMUSG00000042857 | predicted gene 9776 | ncRNA |
| chr10_82632723_82633247 | chr10 | 82632724 | 82633247 | + | 1.2625 | intron (NM_172552, intron 1 of 9) | RSINE1\|SINE\|B4 | 3143 | NR_037190 | 624784 | Mm.284252 | NR_037190 | ENSMUSG00000085666 | **Gm9855** | ENSMUSG00000051559 | thymine DNA glycosylase pseudogene | pseudo |
| chr10_82643525_82644463 | chr10 | 82643526 | 82644463 | + | 1.2171 | intron (NM_172552, intron 4 of 9) | intron (NM_172552, intron 4 of 9) | 14152 | NR_037190 | 624784 | Mm.284252 | NR_037190 | ENSMUSG00000085666 | **Gm9855** | ENSMUSG00000051559 | thymine DNA glycosylase pseudogene | pseudo |
| chr10_82643530_82643940 | chr10 | 82643531 | 82643940 | + | 1.05048 | intron (NM_172552, intron 4 of 9) | intron (NM_172552, intron 4 of 9) | 13893 | NR_037190 | 624784 | Mm.284252 | NR_037190 | ENSMUSG00000085666 | **Gm9855** | ENSMUSG00000051559 | thymine DNA glycosylase pseudogene | pseudo |
| chr10_82643992_82644448 | chr10 | 82643993 | 82644448 | + | 0.942329 | exon (NM_011561, exon 5 of 10) | exon (NM_011561, exon 5 of 10) | 14378 | NR_037190 | 624784 | Mm.284252 | NR_037190 | ENSMUSG00000085666 | **Gm9855** | ENSMUSG00000051559 | thymine DNA glycosylase pseudogene | pseudo |
| chr10_82631049_82631453 | chr10 | 82631050 | 82631453 | + | 0.758175 | intron (NM_172552, intron 1 of 9) | B1F1\|SINE\|Alu | 1409 | NR_037190 | 624784 | Mm.284252 | NR_037190 | ENSMUSG00000085666 | **Gm9855** | ENSMUSG00000051559 | thymine DNA glycosylase pseudogene | pseudo |
| chr16_11868373_11868920 | chr16 | 11868374 | 11868920 | + | 0.705618 | intron (NM_146067, intron 2 of 3) | RLTR17\|LTR\|ERVK | -32717 | NR_033509 | 791359 | Mm.227592 | NR_033509 | ENSMUSG00000055015 | **Gm9961** | ENSMUSG00000055015 | predicted gene 9961 | ncRNA |
| chr17_7398205_7398516 | chr17 | 7398206 | 7398516 | + | 0.854299 | Intergenic | Intergenic | -13056 | NM_001142539 | 667055 | Mm.327371 | NM_001142539 | ENSMUSG00000056133 | **Gm9992** | ENSMUSG00000056133 | predicted gene 9992 | protein-coding |
| chr13_32428089_32428911 | chr13 | 32428090 | 32428911 | + | 1.31837 | Intergenic | Intergenic | -89956 | NM_146041 | 218138 | Mm.247143 | NM_146041 | ENSMUSG00000038372 | **Gmds** | BC031788\|C87208 | GDP-mannose 4, 6-dehydratase | protein-coding |
| chr13_32428334_32428873 | chr13 | 32428335 | 32428873 | + | 1.24674 | Intergenic | Intergenic | -90060 | NM_146041 | 218138 | Mm.247143 | NM_146041 | ENSMUSG00000038372 | **Gmds** | BC031788\|C87208 | GDP-mannose 4, 6-dehydratase | protein-coding |
| chr13_32364410_32365377 | chr13 | 32364411 | 32365377 | + | 1.21693 | Intergenic | ORR1A2-int\|LTR\|ERVL-MaLR | -26350 | NM_146041 | 218138 | Mm.247143 | NM_146041 | ENSMUSG00000038372 | **Gmds** | BC031788\|C87208 | GDP-mannose 4, 6-dehydratase | protein-coding |
| chr13_32364410_32365811 | chr13 | 32364411 | 32365811 | + | 1.16836 | Intergenic | ORR1A2-int\|LTR\|ERVL-MaLR | -26567 | NM_146041 | 218138 | Mm.247143 | NM_146041 | ENSMUSG00000038372 | **Gmds** | BC031788\|C87208 | GDP-mannose 4, 6-dehydratase | protein-coding |
| chr13_32268972_32269241 | chr13 | 32268973 | 32269241 | + | 0.990287 | intron (NM_146041, intron 1 of 10) | intron (NM_146041, intron 1 of 10) | 69437 | NM_146041 | 218138 | Mm.247143 | NM_146041 | ENSMUSG00000038372 | **Gmds** | BC031788\|C87208 | GDP-mannose 4, 6-dehydratase | protein-coding |
| chr13_32268253_32268594 | chr13 | 32268254 | 32268594 | + | 0.926368 | intron (NM_146041, intron 1 of 10) | intron (NM_146041, intron 1 of 10) | 70120 | NM_146041 | 218138 | Mm.247143 | NM_146041 | ENSMUSG00000038372 | **Gmds** | BC031788\|C87208 | GDP-mannose 4, 6-dehydratase | protein-coding |
| chr13_32219944_32220281 | chr13 | 32219945 | 32220281 | + | 0.846985 | intron (NM_146041, intron 4 of 10) | intron (NM_146041, intron 4 of 10) | 118431 | NM_146041 | 218138 | Mm.247143 | NM_146041 | ENSMUSG00000038372 | **Gmds** | BC031788\|C87208 | GDP-mannose 4, 6-dehydratase | protein-coding |
| chr13_32267617_32268006 | chr13 | 32267618 | 32268006 | + | 0.781486 | intron (NM_146041, intron 1 of 10) | intron (NM_146041, intron 1 of 10) | 70732 | NM_146041 | 218138 | Mm.247143 | NM_146041 | ENSMUSG00000038372 | **Gmds** | BC031788\|C87208 | GDP-mannose 4, 6-dehydratase | protein-coding |
| chr13_32242205_32242587 | chr13 | 32242206 | 32242587 | + | 0.725613 | intron (NM_146041, intron 1 of 10) | intron (NM_146041, intron 1 of 10) | 96148 | NM_146041 | 218138 | Mm.247143 | NM_146041 | ENSMUSG00000038372 | **Gmds** | BC031788\|C87208 | GDP-mannose 4, 6-dehydratase | protein-coding |
| chr2_181299265_181299823 | chr2 | 181299266 | 181299823 | + | 1.05482 | Intergenic | RMER1A\|Other\|Other | -11578 | NM_198169 | 229004 | Mm.220423 | NM_198169 | ENSMUSG00000038705 | **Gmeb2** | AI839884 | glucocorticoid modulatory element binding protein 2 | protein-coding |
| chr14_46818683_46819144 | chr14 | 46818684 | 46819144 | + | 1.79047 | intron (NM_022023, intron 1 of 6) | intron (NM_022023, intron 1 of 6) | 3328 | NM_022023 | 63985 | Mm.87312 | NM_022023 | ENSMUSG00000062014 | **Gmfb** | 3110001H22Rik\|3110001O16Rik\|AI851627\|C79176\|D14Ertd630e | glia maturation factor, beta | protein-coding |

| | | | | | | | | | | | | | | | | | |
|---|---|---|---|---|---|---|---|---|---|---|---|---|---|---|---|---|---|
| chr7_28444334_28445051 | chr7 | 28444335 | 28445051 | + | 0.644123 | intron (NM_001290550, intron 3 of 5) | MER45C\|DNA\|hAT-Tip100 | 3756 | NM_001039192 | 63986 | Mm.194536 | NM_022024 | | **Gmfg** | 0610039G16Rik\|2310057N07Rik\|AI324845 | glia maturation factor, gamma | protein-coding |
| chr9_108035169_108035968 | chr9 | 108035170 | 108035968 | + | 0.974173 | intron (NM_013785, intron 3 of 5) | RLTR30E_MM\|LTR\|ERV1 | -13721 | NM_177910 | 331026 | Mm.22554 | NM_177910 | ENSMUSG00000070284 | **Gmppb** | AI317178\|E430010H19 | GDP-mannose pyrophosphorylase B | protein-coding |
| chr13_45685507_45686565 | chr13 | 45685508 | 45686565 | + | 1.7344 | intron (NM_001199304, intron 5 of 6) | intron (NM_001199304, intron 5 of 6) | 178567 | NM_025508 | 66355 | Mm.290802 | NM_025508 | ENSMUSG00000000253 | **Gmpr** | 2310004P21Rik\|AV028449 | guanosine monophosphate reductase | protein-coding |
| chr13_45687649_45688009 | chr13 | 45687650 | 45688009 | + | 1.10902 | intron (NM_001199304, intron 5 of 6) | intron (NM_001199304, intron 5 of 6) | 180360 | NM_025508 | 66355 | Mm.290802 | NM_025508 | ENSMUSG00000000253 | **Gmpr** | 2310004P21Rik\|AV028449 | guanosine monophosphate reductase | protein-coding |
| chr13_45734537_45735054 | chr13 | 45734538 | 45735054 | + | 1.06485 | intron (NM_001199305, intron 5 of 7) | intron (NM_001199305, intron 5 of 7) | 227327 | NM_025508 | 66355 | Mm.290802 | NM_025508 | ENSMUSG00000000253 | **Gmpr** | 2310004P21Rik\|AV028449 | guanosine monophosphate reductase | protein-coding |
| chr13_45685197_45685429 | chr13 | 45685198 | 45685429 | + | 1.04841 | intron (NM_001199304, intron 5 of 6) | intron (NM_001199304, intron 5 of 6) | 177844 | NM_025508 | 66355 | Mm.290802 | NM_025508 | ENSMUSG00000000253 | **Gmpr** | 2310004P21Rik\|AV028449 | guanosine monophosphate reductase | protein-coding |
| chr13_45684873_45685046 | chr13 | 45684874 | 45685046 | + | 0.913081 | intron (NM_001199304, intron 5 of 6) | intron (NM_001199304, intron 5 of 6) | 177491 | NM_025508 | 66355 | Mm.290802 | NM_025508 | ENSMUSG00000000253 | **Gmpr** | 2310004P21Rik\|AV028449 | guanosine monophosphate reductase | protein-coding |
| chr13_45651825_45652225 | chr13 | 45651826 | 45652225 | + | 0.875924 | intron (NM_001199304, intron 5 of 6) | intron (NM_001199304, intron 5 of 6) | 144556 | NM_025508 | 66355 | Mm.290802 | NM_025508 | ENSMUSG00000000253 | **Gmpr** | 2310004P21Rik\|AV028449 | guanosine monophosphate reductase | protein-coding |
| chr13_45694364_45694735 | chr13 | 45694365 | 45694735 | + | 0.848881 | intron (NM_001199304, intron 5 of 6) | URR1B\|DNA\|hAT-Charlie | 187081 | NM_025508 | 66355 | Mm.290802 | NM_025508 | ENSMUSG00000000253 | **Gmpr** | 2310004P21Rik\|AV028449 | guanosine monophosphate reductase | protein-coding |
| chr13_45649134_45649773 | chr13 | 45649135 | 45649773 | + | 0.779868 | intron (NM_001199304, intron 5 of 6) | intron (NM_001199304, intron 5 of 6) | 141985 | NM_025508 | 66355 | Mm.290802 | NM_025508 | ENSMUSG00000000253 | **Gmpr** | 2310004P21Rik\|AV028449 | guanosine monophosphate reductase | protein-coding |
| chr13_45708243_45708915 | chr13 | 45708244 | 45708915 | + | 0.642692 | intron (NM_001199305, intron 5 of 7) | intron (NM_001199305, intron 5 of 7) | 201110 | NM_025508 | 66355 | Mm.290802 | NM_025508 | ENSMUSG00000000253 | **Gmpr** | 2310004P21Rik\|AV028449 | guanosine monophosphate reductase | protein-coding |
| chr3_108124096_108124706 | chr3 | 108124097 | 108124706 | + | 0.834916 | intron (NM_010306, intron 1 of 8) | L2\|LINE\|L2 | 21751 | NM_010306 | 14679 | Mm.271703 | NM_010306 | ENSMUSG00000000001 | **Gnai3** | AI158965\|AW537698\|Galphai3\|Gnai-3 | guanine nucleotide binding protein (G protein), alpha inhibiting 3 | protein-coding |
| chr3_108124093_108125156 | chr3 | 108124094 | 108125156 | + | 0.810886 | intron (NM_010306, intron 1 of 8) | L2\|LINE\|L2 | 21527 | NM_010306 | 14679 | Mm.271703 | NM_010306 | ENSMUSG00000000001 | **Gnai3** | AI158965\|AW537698\|Galphai3\|Gnai-3 | guanine nucleotide binding protein (G protein), alpha inhibiting 3 | protein-coding |
| chr2_174297132_174297633 | chr2 | 174297133 | 174297633 | + | 0.781219 | promoter-TSS (NM_201618) | promoter-TSS (NM_201618) | -476 | NM_001077507 | 14683 | Mm.125770 | NM_010309 | ENSMUSG00000027523 | **Gnas** | 5530400H20Rik\|A930027G11Rik\|C130027O20Rik\|GPSA\|GSP\|Galphas\|Gnas1\|Gnasxl\|Gsa\|Nesp\|Nespl\|Oed-Sml\|Oedsml\|P1\|P2\|P3\|PHP1A\|PHP1B\|POH | GNAS (guanine nucleotide binding protein, alpha stimulating) complex locus | protein-coding |
| chr10_74922764_74923199 | chr10 | 74922765 | 74923199 | + | 0.864758 | Intergenic | Intergenic | -44249 | NM_010311 | 14687 | Mm.32595 | NM_010311 | ENSMUSG00000040009 | **Gnaz** | AI847979\|Gz | guanine nucleotide binding protein, alpha z subunit | protein-coding |
| chr10_74969278_74969552 | chr10 | 74969279 | 74969552 | + | 0.842735 | intron (NM_010311, intron 1 of 3) | intron (NM_010311, intron 1 of 3) | 2184 | NM_010311 | 14687 | Mm.32595 | NM_010311 | ENSMUSG00000040009 | **Gnaz** | AI847979\|Gz | guanine nucleotide binding protein, alpha z subunit | protein-coding |
| chr4_155508620_155508968 | chr4 | 155508621 | 155508968 | + | 0.77039 | intron (NM_001160017, intron 1 of 10) | intron (NM_001160017, intron 1 of 10) | 17433 | NM_001160016 | 14688 | Mm.2344 | NM_008142 | ENSMUSG00000029064 | **Gnb1** | AA409223\|C77571\|Gnb-1 | guanine nucleotide binding protein (G protein), beta 1 | protein-coding |
| chr9_75318261_75318745 | chr9 | 75318262 | 75318745 | + | 1.36941 | intron (NM_138719, intron 1 of 10) | MLT1D\|LTR\|ERVL-MaLR | 4435 | NM_138719 | 14697 | Mm.17604 | NM_010313 | | **Gnb5** | GBS\|Gbeta5\|flr | guanine nucleotide binding protein (G protein), beta 5 | protein-coding |
| chr6_66929424_66930099 | chr6 | 66929425 | 66930099 | + | 0.92197 | intron (NM_001177556, intron 2 of 3) | Lx8\|LINE\|L1 | 32871 | NM_001177558 | 14701 | Mm.234342 | NM_025278 | ENSMUSG00000036402 | **Gng12** | 2010305F15Rik\|AA536815\|AI115529\|AI314170\|AI842738 | guanine nucleotide binding protein (G protein), gamma 12 | protein-coding |
| chr6_66916711_66917900 | chr6 | 66916712 | 66917900 | + | 0.746858 | intron (NM_001177556, intron 2 of 3) | intron (NM_001177556, intron 2 of 3) | 20415 | NM_001177558 | 14701 | Mm.234342 | NM_025278 | ENSMUSG00000036402 | **Gng12** | 2010305F15Rik\|AA536815\|AI115529\|AI314170\|AI842738 | guanine nucleotide binding protein (G protein), gamma 12 | protein-coding |
| chr6_66944239_66945435 | chr6 | 66944240 | 66945435 | + | 0.619349 | intron (NM_001177560, intron 3 of 4) | intron (NM_001177560, intron 3 of 4) | 47946 | NM_001177558 | 14701 | Mm.234342 | NM_025278 | ENSMUSG00000036402 | **Gng12** | 2010305F15Rik\|AA536815\|AI115529\|AI314170\|AI842738 | guanine nucleotide binding protein (G protein), gamma 12 | protein-coding |
| chr14_19970269_19970713 | chr14 | 19970270 | 19970713 | + | 0.832198 | intron (NM_001285908, intron 1 of 3) | L2a\|LINE\|L2 | 6202 | NM_001285911 | 14702 | Mm.41737 | NM_010315 | ENSMUSG00000043004 | **Gng2** | 82 | guanine nucleotide binding protein (G protein), gamma 2 | protein-coding |
| chr14_19969587_19969897 | chr14 | 19969588 | 19969897 | + | 0.713846 | intron (NM_001285908, intron 1 of 3) | intron (NM_001285908, intron 1 of 3) | 6951 | NM_001285911 | 14702 | Mm.41737 | NM_010315 | ENSMUSG00000043004 | **Gng2** | 82 | guanine nucleotide binding protein (G protein), gamma 2 | protein-coding |
| chr14_19964218_19964474 | chr14 | 19964219 | 19964474 | + | 0.635438 | intron (NM_001285908, intron 1 of 3) | intron (NM_001285908, intron 1 of 3) | 12347 | NM_001285911 | 14702 | Mm.41737 | NM_010315 | ENSMUSG00000043004 | **Gng2** | 82 | guanine nucleotide binding protein (G protein), gamma 2 | protein-coding |
| chr13_13785082_13785332 | chr13 | 13785083 | 13785332 | + | 0.730903 | intron (NM_010317, intron 1 of 3) | intron (NM_010317, intron 1 of 3) | 668 | NM_010317 | 14706 | Mm.215394 | NM_010317 | ENSMUSG00000021303 | **Gng4** | - | guanine nucleotide binding protein (G protein), gamma 4 | protein-coding |
| chr13_13783846_13785332 | chr13 | 13783847 | 13785332 | + | 0.670695 | promoter-TSS (NM_010317) | promoter-TSS (NM_010317) | 50 | NM_010317 | 14706 | Mm.215394 | NM_010317 | ENSMUSG00000021303 | **Gng4** | - | guanine nucleotide binding protein (G protein), gamma 4 | protein-coding |
| chr10_80967342_80967925 | chr10 | 80967343 | 80967925 | + | 0.628578 | intron (NM_001038655, intron 2 of 3) | intron (NM_001038655, intron 2 of 3) | 33731 | NM_001038655 | 14708 | Mm.222496 | NM_010319 | ENSMUSG00000048240 | **Gng7** | AI840417 | guanine nucleotide binding protein (G protein), gamma 7 | protein-coding |
| chrX_151007872_151008397 | chrX | 151007873 | 151008397 | + | 0.74813 | intron (NM_198110, intron 2 of 15) | intron (NM_198110, intron 2 of 15) | 9158 | NM_001168600 | 237107 | Mm.45987 | NM_198110 | ENSMUSG00000025266 | **Gnl3l** | BC020354 | guanine nucleotide binding protein-like 3 (nucleolar)-like | protein-coding |
| chrX_150956490_150957278 | chrX | 150956491 | 150957278 | + | 0.747409 | Intergenic | Intergenic | 60409 | NM_001168600 | 237107 | Mm.45987 | NM_198110 | ENSMUSG00000025266 | **Gnl3l** | BC020354 | guanine nucleotide binding protein-like 3 (nucleolar)-like | protein-coding |
| chr14_45385642_45385902 | chr14 | 45385643 | 45385902 | + | 1.03341 | intron (NM_019425, intron 1 of 5) | intron (NM_019425, intron 1 of 5) | 3024 | NM_019425 | 54342 | Mm.312945 | NM_019425 | ENSMUSG00000037722 | **Gnpnat1** | AU017428\|AU040593\|EMeg32\|Gpnat1\|Gsnpat | glucosamine-phosphate N-acetyltransferase 1 | protein-coding |
| chr10_88404112_88404560 | chr10 | 88404113 | 88404560 | + | 1.24001 | intron (NM_001004164, intron 2 of 20) | intron (NM_001004164, intron 2 of 20) | 24924 | NM_001004164 | 432486 | Mm.335190 | NM_001004164 | ENSMUSG00000035311 | **Gnptab** | EG432486\|mKIAA1208 | N-acetylglucosamine-1-phosphate transferase, alpha and beta subunits | protein-coding |
| chr10_88388936_88389198 | chr10 | 88388937 | 88389198 | + | 0.680045 | intron (NM_001004164, intron 1 of 20) | intron (NM_001004164, intron 1 of 20) | 9655 | NM_001004164 | 432486 | Mm.335190 | NM_001004164 | ENSMUSG00000035311 | **Gnptab** | EG432486\|mKIAA1208 | N-acetylglucosamine-1-phosphate transferase, alpha and beta subunits | protein-coding |
| chr9_118512665_118513009 | chr9 | 118512666 | 118513009 | + | 0.70776 | intron (NM_018748, intron 1 of 23) | intron (NM_018748, intron 1 of 23) | 6519 | NM_018748 | 54214 | Mm.10409 | NM_018748 | ENSMUSG00000038708 | **Golga4** | AI225887\|AU019508\|Olp-1 | golgi autoantigen, golgin subfamily a, 4 | protein-coding |
| chr9_118512658_118513275 | chr9 | 118512659 | 118513275 | + | 0.706485 | intron (NM_018748, intron 1 of 23) | intron (NM_018748, intron 1 of 23) | 6649 | NM_018748 | 54214 | Mm.10409 | NM_018748 | ENSMUSG00000038708 | **Golga4** | AI225887\|AU019508\|Olp-1 | golgi autoantigen, golgin subfamily a, 4 | protein-coding |
| chr8_23256020_23256829 | chr8 | 23256021 | 23256829 | + | 0.762958 | intron (NM_001042484, intron 2 of 5) | intron (NM_001042484, intron 2 of 5) | 655 | NM_020585 | 57437 | Mm.196269 | NM_020585 | ENSMUSG00000015341 | **Golga7** | AB041568\|C130038N16Rik\|GCP16\|GOLGA3AP1\|HSPC041\|R75586 | golgi autoantigen, golgin subfamily a, 7 | protein-coding |
| chr3_75939509_75940003 | chr3 | 75939510 | 75940003 | + | 0.785273 | intron (NM_001291069, intron 1 of 15) | intron (NM_001291069, intron 1 of 15) | 17193 | NM_175193 | 73124 | Mm.249232 | NM_175193 | ENSMUSG00000034109 | **Golim4** | 3110027H23Rik\|GIMPC\|GPP130\|Golph4\|P138 | golgi integral membrane protein 4 | protein-coding |
| chr3_75948630_75949136 | chr3 | 75948631 | 75949136 | + | 0.682538 | intron (NM_001291069, intron 1 of 15) | intron (NM_001291069, intron 1 of 15) | 8066 | NM_175193 | 73124 | Mm.249232 | NM_175193 | ENSMUSG00000034109 | **Golim4** | 3110027H23Rik\|GIMPC\|GPP130\|Golph4\|P138 | golgi integral membrane protein 4 | protein-coding |

<table>
<tr><td>chr15_12425112_12425673</td><td>chr15</td><td>12425113</td><td>12425673</td><td>+</td><td>0.862663</td><td>intron (NM_001081064, intron 5 of 23)</td><td>intron (NM_001081064, intron 5 of 23)</td><td>103897</td><td>NM_025673</td><td>66629</td><td>Mm.157655</td><td>NM_025673</td><td>ENSMUSG00000022200</td><td>Golph3</td><td>4733401N08Rik|5730410D03Rik|AW413496</td><td>golgi phosphoprotein 3</td><td>protein-coding</td></tr>
<tr><td>chr6_142389115_142389464</td><td>chr6</td><td>142389116</td><td>142389464</td><td>+</td><td>1.47878</td><td>intron (NM_025872, intron 1 of 4)</td><td>intron (NM_025872, intron 1 of 4)</td><td>2047</td><td>NM_025872</td><td>66964</td><td>Mm.3676</td><td>NM_025872</td><td>ENSMUSG00000030245</td><td>Golt1b</td><td>2310061A22Rik|AA407874|AU024291|CGI-141</td><td>golgi transport 1 homolog B (S. cerevisiae)</td><td>protein-coding</td></tr>
<tr><td>chr1_163374435_163375134</td><td>chr1</td><td>163374436</td><td>163375134</td><td>+</td><td>1.08976</td><td>Intergenic</td><td>Intergenic</td><td>28856</td><td>NM_178883</td><td>98376</td><td>Mm.32901</td><td>NM_178883</td><td>ENSMUSG00000040124</td><td>Gorab</td><td>AI467484|NTKL-BP1|Scyl1bp1</td><td>golgin, RAB6-interacting</td><td>protein-coding</td></tr>
<tr><td>chr1_163381963_163382772</td><td>chr1</td><td>163381964</td><td>163382772</td><td>+</td><td>1.03857</td><td>Intergenic</td><td>Intergenic</td><td>21273</td><td>NM_178883</td><td>98376</td><td>Mm.32901</td><td>NM_178883</td><td>ENSMUSG00000040124</td><td>Gorab</td><td>AI467484|NTKL-BP1|Scyl1bp1</td><td>golgin, RAB6-interacting</td><td>protein-coding</td></tr>
<tr><td>chr1_163383507_163383929</td><td>chr1</td><td>163383508</td><td>163383929</td><td>+</td><td>0.749091</td><td>Intergenic</td><td>Intergenic</td><td>19923</td><td>NM_178883</td><td>98376</td><td>Mm.32901</td><td>NM_178883</td><td>ENSMUSG00000040124</td><td>Gorab</td><td>AI467484|NTKL-BP1|Scyl1bp1</td><td>golgin, RAB6-interacting</td><td>protein-coding</td></tr>
<tr><td>chr11_103685106_103685751</td><td>chr11</td><td>103685107</td><td>103685751</td><td>+</td><td>1.28663</td><td>intron (NM_019650, intron 4 of 5)</td><td>intron (NM_019650, intron 4 of 5)</td><td>12281</td><td>NM_019650</td><td>56494</td><td>Mm.195451</td><td>NM_019650</td><td>ENSMUSG00000020946</td><td>Gosr2</td><td>2310032N09Rik|C76855|Gs27|SNARE|membrin</td><td>golgi SNAP receptor complex member 2</td><td>protein-coding</td></tr>
<tr><td>chr1_187216946_187217502</td><td>chr1</td><td>187216947</td><td>187217502</td><td>+</td><td>0.866135</td><td>intron (NM_026367, intron 1 of 9)</td><td>intron (NM_026367, intron 1 of 9)</td><td>1713</td><td>NM_026367</td><td>67769</td><td>Mm.138321</td><td>NM_026367</td><td>ENSMUSG00000039210</td><td>Gpatch2</td><td>5830433G22Rik|5830436K05Rik|AI427714|AI447508|AW107440|AW491060|Gpatc2</td><td>G patch domain containing 2</td><td>protein-coding</td></tr>
<tr><td>chr1_187216761_187217484</td><td>chr1</td><td>187216762</td><td>187217484</td><td>+</td><td>0.841426</td><td>intron (NM_026367, intron 1 of 9)</td><td>intron (NM_026367, intron 1 of 9)</td><td>1612</td><td>NM_026367</td><td>67769</td><td>Mm.138321</td><td>NM_026367</td><td>ENSMUSG00000039210</td><td>Gpatch2</td><td>5830433G22Rik|5830436K05Rik|AI427714|AI447508|AW107440|AW491060|Gpatc2</td><td>G patch domain containing 2</td><td>protein-coding</td></tr>
<tr><td>chr13_111568693_111569553</td><td>chr13</td><td>111568694</td><td>111569553</td><td>+</td><td>1.15653</td><td>Intergenic</td><td>Intergenic</td><td>-79082</td><td>NM_001122963</td><td>73274</td><td>Mm.57286</td><td>NM_028487</td><td>ENSMUSG00000032745</td><td>Gpbp1</td><td>1700034P14Rik|AU019836|D230035M11Rik|Gpbp|mGPBP</td><td>GC-rich promoter binding protein 1</td><td>protein-coding</td></tr>
<tr><td>chr13_111525418_111526293</td><td>chr13</td><td>111525419</td><td>111526293</td><td>+</td><td>0.985051</td><td>Intergenic</td><td>MER20B|DNA|hAT-Charlie</td><td>-35815</td><td>NM_001122963</td><td>73274</td><td>Mm.57286</td><td>NM_028487</td><td>ENSMUSG00000032745</td><td>Gpbp1</td><td>1700034P14Rik|AU019836|D230035M11Rik|Gpbp|mGPBP</td><td>GC-rich promoter binding protein 1</td><td>protein-coding</td></tr>
<tr><td>chr1_92806539_92807189</td><td>chr1</td><td>92806540</td><td>92807189</td><td>+</td><td>0.625395</td><td>Intergenic</td><td>Intergenic</td><td>-24822</td><td>NM_016696</td><td>14733</td><td>Mm.297976</td><td>NM_016696</td><td>ENSMUSG00000034220</td><td>Gpc1</td><td>AI462976</td><td>glypican 1</td><td>protein-coding</td></tr>
<tr><td>chr14_116975229_116975547</td><td>chr14</td><td>116975230</td><td>116975547</td><td>+</td><td>0.82109</td><td>intron (NM_001079844, intron 1 of 9)</td><td>intron (NM_001079844, intron 1 of 9)</td><td>50091</td><td>NM_011821</td><td>23888</td><td>Mm.440025</td><td>NM_011821</td><td>ENSMUSG00000058571</td><td>Gpc6</td><td>-</td><td>glypican 6</td><td>protein-coding</td></tr>
<tr><td>chr9_114955622_114956463</td><td>chr9</td><td>114955623</td><td>114956463</td><td>+</td><td>1.52417</td><td>Intergenic</td><td>Intergenic</td><td>-22056</td><td>NM_175380</td><td>333433</td><td>Mm.38198</td><td>NM_175380</td><td>ENSMUSG00000050627</td><td>Gpd1l</td><td>2210409H23Rik|D9Ertd660e</td><td>glycerol-3-phosphate dehydrogenase 1-like</td><td>protein-coding</td></tr>
<tr><td>chr2_57268640_57269227</td><td>chr2</td><td>57268641</td><td>57269227</td><td>+</td><td>0.856151</td><td>intron (NM_010274, intron 2 of 16)</td><td>intron (NM_010274, intron 2 of 16)</td><td>30552</td><td>NM_010274</td><td>14571</td><td>Mm.3711</td><td>NM_010274</td><td>ENSMUSG00000026827</td><td>Gpd2</td><td>AA408484|AI448216|AU021455|AW494132|GPDH|Gdm1|Gpd-m|Gpdh-m|TISP38</td><td>glycerol phosphate dehydrogenase 2, mitochondrial</td><td>protein-coding</td></tr>
<tr><td>chr2_57268757_57269161</td><td>chr2</td><td>57268758</td><td>57269161</td><td>+</td><td>0.831321</td><td>intron (NM_010274, intron 2 of 16)</td><td>intron (NM_010274, intron 2 of 16)</td><td>30577</td><td>NM_010274</td><td>14571</td><td>Mm.3711</td><td>NM_010274</td><td>ENSMUSG00000026827</td><td>Gpd2</td><td>AA408484|AI448216|AU021455|AW494132|GPDH|Gdm1|Gpd-m|Gpdh-m|TISP38</td><td>glycerol phosphate dehydrogenase 2, mitochondrial</td><td>protein-coding</td></tr>
<tr><td>chr2_57239607_57239801</td><td>chr2</td><td>57239608</td><td>57239801</td><td>+</td><td>0.825576</td><td>intron (NM_010274, intron 1 of 16)</td><td>intron (NM_010274, intron 1 of 16)</td><td>1322</td><td>NM_010274</td><td>14571</td><td>Mm.3711</td><td>NM_010274</td><td>ENSMUSG00000026827</td><td>Gpd2</td><td>AA408484|AI448216|AU021455|AW494132|GPDH|Gdm1|Gpd-m|Gpdh-m|TISP38</td><td>glycerol phosphate dehydrogenase 2, mitochondrial</td><td>protein-coding</td></tr>
<tr><td>chr2_57230806_57231288</td><td>chr2</td><td>57230807</td><td>57231288</td><td>+</td><td>0.754666</td><td>Intergenic</td><td>(CA)n|Simple_repeat|Simple_repeat</td><td>-6631</td><td>NM_001145820</td><td>14571</td><td>Mm.3711</td><td>NM_010274</td><td>ENSMUSG00000026827</td><td>Gpd2</td><td>AA408484|AI448216|AU021455|AW494132|GPDH|Gdm1|Gpd-m|Gpdh-m|TISP38</td><td>glycerol phosphate dehydrogenase 2, mitochondrial</td><td>protein-coding</td></tr>
<tr><td>chr2_57275946_57276562</td><td>chr2</td><td>57275947</td><td>57276562</td><td>+</td><td>0.745036</td><td>intron (NM_010274, intron 2 of 16)</td><td>intron (NM_010274, intron 2 of 16)</td><td>37872</td><td>NM_010274</td><td>14571</td><td>Mm.3711</td><td>NM_010274</td><td>ENSMUSG00000026827</td><td>Gpd2</td><td>AA408484|AI448216|AU021455|AW494132|GPDH|Gdm1|Gpd-m|Gpdh-m|TISP38</td><td>glycerol phosphate dehydrogenase 2, mitochondrial</td><td>protein-coding</td></tr>
<tr><td>chr2_57356168_57356772</td><td>chr2</td><td>57356169</td><td>57356772</td><td>+</td><td>0.731135</td><td>intron (NM_010274, intron 11 of 16)</td><td>intron (NM_010274, intron 11 of 16)</td><td>118088</td><td>NM_010274</td><td>14571</td><td>Mm.3711</td><td>NM_010274</td><td>ENSMUSG00000026827</td><td>Gpd2</td><td>AA408484|AI448216|AU021455|AW494132|GPDH|Gdm1|Gpd-m|Gpdh-m|TISP38</td><td>glycerol phosphate dehydrogenase 2, mitochondrial</td><td>protein-coding</td></tr>
<tr><td>chr2_57362391_57362837</td><td>chr2</td><td>57362392</td><td>57362837</td><td>+</td><td>0.684502</td><td>intron (NM_010274, intron 13 of 16)</td><td>B3A|SINE|B2</td><td>124232</td><td>NM_010274</td><td>14571</td><td>Mm.3711</td><td>NM_010274</td><td>ENSMUSG00000026827</td><td>Gpd2</td><td>AA408484|AI448216|AU021455|AW494132|GPDH|Gdm1|Gpd-m|Gpdh-m|TISP38</td><td>glycerol phosphate dehydrogenase 2, mitochondrial</td><td>protein-coding</td></tr>
<tr><td>chr2_57251980_57252267</td><td>chr2</td><td>57251981</td><td>57252267</td><td>+</td><td>0.643405</td><td>intron (NM_010274, intron 1 of 16)</td><td>intron (NM_010274, intron 1 of 16)</td><td>13742</td><td>NM_010274</td><td>14571</td><td>Mm.3711</td><td>NM_010274</td><td>ENSMUSG00000026827</td><td>Gpd2</td><td>AA408484|AI448216|AU021455|AW494132|GPDH|Gdm1|Gpd-m|Gpdh-m|TISP38</td><td>glycerol phosphate dehydrogenase 2, mitochondrial</td><td>protein-coding</td></tr>
<tr><td>chr12_78298880_78299485</td><td>chr12</td><td>78298881</td><td>78299485</td><td>+</td><td>1.22496</td><td>intron (NM_172952, intron 1 of 23)</td><td>MERV1_LTR|LTR|ERV1</td><td>72528</td><td>NM_145965</td><td>268566</td><td>Mm.341742</td><td>NM_145965</td><td>ENSMUSG00000047454</td><td>Gphn</td><td>5730552E08Rik|AI662856|BC027112|C230040D23|GPH|GPHRYN|geph</td><td>gephyrin</td><td>protein-coding</td></tr>
<tr><td>chr12_78298875_78299831</td><td>chr12</td><td>78298876</td><td>78299831</td><td>+</td><td>1.09453</td><td>intron (NM_172952, intron 1 of 23)</td><td>MERV1_LTR|LTR|ERV1</td><td>72698</td><td>NM_145965</td><td>268566</td><td>Mm.341742</td><td>NM_145965</td><td>ENSMUSG00000047454</td><td>Gphn</td><td>5730552E08Rik|AI662856|BC027112|C230040D23|GPH|GPHRYN|geph</td><td>gephyrin</td><td>protein-coding</td></tr>
<tr><td>chr12_77887338_77887778</td><td>chr12</td><td>77887339</td><td>77887778</td><td>+</td><td>0.89737</td><td>Intergenic</td><td>Intergenic</td><td>-339097</td><td>NM_172952</td><td>268566</td><td>Mm.341742</td><td>NM_145965</td><td>ENSMUSG00000047454</td><td>Gphn</td><td>5730552E08Rik|AI662856|BC027112|C230040D23|GPH|GPHRYN|geph</td><td>gephyrin</td><td>protein-coding</td></tr>
<tr><td>chr12_78089633_78090162</td><td>chr12</td><td>78089634</td><td>78090162</td><td>+</td><td>0.70917</td><td>Intergenic</td><td>Intergenic</td><td>-136757</td><td>NM_172952</td><td>268566</td><td>Mm.341742</td><td>NM_145965</td><td>ENSMUSG00000047454</td><td>Gphn</td><td>5730552E08Rik|AI662856|BC027112|C230040D23|GPH|GPHRYN|geph</td><td>gephyrin</td><td>protein-coding</td></tr>
<tr><td>chr13_24962144_24962526</td><td>chr13</td><td>24962145</td><td>24962526</td><td>+</td><td>1.26197</td><td>exon (NM_008156, exon 9 of 26)</td><td>exon (NM_008156, exon 9 of 26)</td><td>19183</td><td>NM_008156</td><td>14756</td><td>Mm.291831</td><td>NM_008156</td><td></td><td>Gpld1</td><td>6330541J12Rik|AW546131</td><td>glycosylphosphatidylinositol specific phospholipase D1</td><td>protein-coding</td></tr>
<tr><td>chrX_166226684_166227214</td><td>chrX</td><td>166226685</td><td>166227214</td><td>+</td><td>0.791632</td><td>Intergenic</td><td>Intergenic</td><td>-11994</td><td>NM_001177961</td><td>14758</td><td>Mm.222320</td><td>NM_023122</td><td>ENSMUSG00000031342</td><td>Gpm6b</td><td>AI593561|Gpm6|M6B</td><td>glycoprotein m6b</td><td>protein-coding</td></tr>
<tr><td>chrX_166214461_166215247</td><td>chrX</td><td>166214462</td><td>166215247</td><td>+</td><td>0.699832</td><td>Intergenic</td><td>Intergenic</td><td>-24089</td><td>NM_001177961</td><td>14758</td><td>Mm.222320</td><td>NM_023122</td><td>ENSMUSG00000031342</td><td>Gpm6b</td><td>AI593561|Gpm6|M6B</td><td>glycoprotein m6b</td><td>protein-coding</td></tr>
<tr><td>chrX_166226541_166227272</td><td>chrX</td><td>166226542</td><td>166227272</td><td>+</td><td>0.683758</td><td>Intergenic</td><td>Intergenic</td><td>-12036</td><td>NM_001177961</td><td>14758</td><td>Mm.222320</td><td>NM_023122</td><td>ENSMUSG00000031342</td><td>Gpm6b</td><td>AI593561|Gpm6|M6B</td><td>glycoprotein m6b</td><td>protein-coding</td></tr>
<tr><td>chr2_31180265_31180626</td><td>chr2</td><td>31180266</td><td>31180626</td><td>+</td><td>0.681752</td><td>intron (NM_178760, intron 9 of 17)</td><td>intron (NM_178760, intron 9 of 17)</td><td>28130</td><td>NM_178760</td><td>277463</td><td>Mm.236074</td><td>NM_178760</td><td></td><td>Gpr107</td><td>9930033D15Rik|AI790205|C530034M11|mKIAA1624</td><td>G protein-coupled receptor 107</td><td>protein-coding</td></tr>
<tr><td>chr10_14442224_14442943</td><td>chr10</td><td>14442225</td><td>14442943</td><td>+</td><td>0.935153</td><td>intron (NM_001002268, intron 9 of 24)</td><td>intron (NM_001002268, intron 9 of 24)</td><td>102452</td><td>NM_001002268</td><td>215798</td><td>Mm.440142</td><td>NM_001002268</td><td>ENSMUSG00000039116</td><td>Gpr126</td><td>1190004A11Rik|AI449247|AW045736|DREG|Gm222</td><td>G protein-coupled receptor 126</td><td>protein-coding</td></tr>
<tr><td>chr10_14385619_14386159</td><td>chr10</td><td>14385620</td><td>14386159</td><td>+</td><td>0.748987</td><td>Intergenic</td><td>MLT2B1|LTR|ERVL</td><td>159147</td><td>NM_001002268</td><td>215798</td><td>Mm.440142</td><td>NM_001002268</td><td>ENSMUSG00000039116</td><td>Gpr126</td><td>1190004A11Rik|AI449247|AW045736|DREG|Gm222</td><td>G protein-coupled receptor 126</td><td>protein-coding</td></tr>
<tr><td>chr10_14385613_14385917</td><td>chr10</td><td>14385614</td><td>14385917</td><td>+</td><td>0.738644</td><td>Intergenic</td><td>Intergenic</td><td>159271</td><td>NM_001002268</td><td>215798</td><td>Mm.440142</td><td>NM_001002268</td><td>ENSMUSG00000039116</td><td>Gpr126</td><td>1190004A11Rik|AI449247|AW045736|DREG|Gm222</td><td>G protein-coupled receptor 126</td><td>protein-coding</td></tr>
<tr><td>chr12_112864921_112865215</td><td>chr12</td><td>112864922</td><td>112865215</td><td>+</td><td>0.87519</td><td>Intergenic</td><td>Intergenic</td><td>-4152</td><td>NM_019925</td><td>56696</td><td>Mm.20455</td><td>NM_019925</td><td>ENSMUSG00000021298</td><td>Gpr132</td><td>G2a</td><td>G protein-coupled receptor 132</td><td>protein-coding</td></tr>
<tr><td>chr12_112861844_112862268</td><td>chr12</td><td>112861845</td><td>112862268</td><td>+</td><td>0.814558</td><td>Intergenic</td><td>Intergenic</td><td>-1140</td><td>NM_019925</td><td>56696</td><td>Mm.20455</td><td>NM_019925</td><td>ENSMUSG00000021298</td><td>Gpr132</td><td>G2a</td><td>G protein-coupled receptor 132</td><td>protein-coding</td></tr>
<tr><td>chr12_112867159_112867631</td><td>chr12</td><td>112867160</td><td>112867631</td><td>+</td><td>0.626198</td><td>Intergenic</td><td>Intergenic</td><td>-6479</td><td>NM_019925</td><td>56696</td><td>Mm.20455</td><td>NM_019925</td><td>ENSMUSG00000021298</td><td>Gpr132</td><td>G2a</td><td>G protein-coupled receptor 132</td><td>protein-coding</td></tr>
<tr><td>chr5_129190033_129190528</td><td>chr5</td><td>129190034</td><td>129190528</td><td>+</td><td>0.996835</td><td>intron (NM_001081342, intron 21 of 25)</td><td>intron (NM_001081342, intron 21 of 25)</td><td>93531</td><td>NM_001081342</td><td>243277</td><td>Mm.440501</td><td>NM_177734</td><td>ENSMUSG00000044017</td><td>Gpr133</td><td>E230012M21Rik</td><td>G protein-coupled receptor 133</td><td>protein-coding</td></tr>
<tr><td>chr5_139395908_139396306</td><td>chr5</td><td>139395909</td><td>139396306</td><td>+</td><td>0.766486</td><td>3' UTR (NM_001038703, exon 2 of 2)</td><td>3' UTR (NM_001038703, exon 2 of 2)</td><td>6203</td><td>NM_001038703</td><td>80290</td><td>Mm.270003</td><td>NM_030258</td><td>ENSMUSG00000044197</td><td>Gpr146</td><td>BC003323|PGR8</td><td>G protein-coupled receptor 146</td><td>protein-coding</td></tr>
<tr><td>chr16_58696675_58697175</td><td>chr16</td><td>58696676</td><td>58697175</td><td>+</td><td>2.04257</td><td>Intergenic</td><td>MLT1F|LTR|ERVL-MaLR</td><td>21799</td><td>NM_001162955</td><td>71223</td><td>Mm.359970</td><td>NM_001162955</td><td>ENSMUSG00000047293</td><td>Gpr15</td><td>4933439K08Rik</td><td>G protein-coupled receptor 15</td><td>protein-coding</td></tr>
<tr><td>chr16_58714982_58715434</td><td>chr16</td><td>58714983</td><td>58715434</td><td>+</td><td>1.56885</td><td>Intergenic</td><td>Intergenic</td><td>3516</td><td>NM_001162955</td><td>71223</td><td>Mm.359970</td><td>NM_001162955</td><td>ENSMUSG00000047293</td><td>Gpr15</td><td>4933439K08Rik</td><td>G protein-coupled receptor 15</td><td>protein-coding</td></tr>
<tr><td>chr16_58703317_58703708</td><td>chr16</td><td>58703318</td><td>58703708</td><td>+</td><td>1.54938</td><td>Intergenic</td><td>Intergenic</td><td>15211</td><td>NM_001162955</td><td>71223</td><td>Mm.359970</td><td>NM_001162955</td><td>ENSMUSG00000047293</td><td>Gpr15</td><td>4933439K08Rik</td><td>G protein-coupled receptor 15</td><td>protein-coding</td></tr>
<tr><td>chr16_58703029_58703708</td><td>chr16</td><td>58703030</td><td>58703708</td><td>+</td><td>1.5143</td><td>Intergenic</td><td>Intergenic</td><td>15355</td><td>NM_001162955</td><td>71223</td><td>Mm.359970</td><td>NM_001162955</td><td>ENSMUSG00000047293</td><td>Gpr15</td><td>4933439K08Rik</td><td>G protein-coupled receptor 15</td><td>protein-coding</td></tr>
<tr><td>chr16_58697288_58697569</td><td>chr16</td><td>58697289</td><td>58697569</td><td>+</td><td>1.19041</td><td>Intergenic</td><td>Intergenic</td><td>21295</td><td>NM_001162955</td><td>71223</td><td>Mm.359970</td><td>NM_001162955</td><td>ENSMUSG00000047293</td><td>Gpr15</td><td>4933439K08Rik</td><td>G protein-coupled receptor 15</td><td>protein-coding</td></tr>
<tr><td>chr16_58713353_58713622</td><td>chr16</td><td>58713354</td><td>58713622</td><td>+</td><td>0.882219</td><td>Intergenic</td><td>Intergenic</td><td>5236</td><td>NM_001162955</td><td>71223</td><td>Mm.359970</td><td>NM_001162955</td><td>ENSMUSG00000047293</td><td>Gpr15</td><td>4933439K08Rik</td><td>G protein-coupled receptor 15</td><td>protein-coding</td></tr>
<tr><td>chr16_58703771_58703936</td><td>chr16</td><td>58703772</td><td>58703936</td><td>+</td><td>0.648825</td><td>Intergenic</td><td>Intergenic</td><td>14870</td><td>NM_001162955</td><td>71223</td><td>Mm.359970</td><td>NM_001162955</td><td>ENSMUSG00000047293</td><td>Gpr15</td><td>4933439K08Rik</td><td>G protein-coupled receptor 15</td><td>protein-coding</td></tr>
<tr><td>chr18_42591215_42591455</td><td>chr18</td><td>42591216</td><td>42591455</td><td>+</td><td>0.718042</td><td>Intergenic</td><td>B3|SINE|B2</td><td>-11683</td><td>NM_181543</td><td>240239</td><td>Mm.186779</td><td>NM_181543</td><td>ENSMUSG00000042816</td><td>Gpr151</td><td>C130082O03Rik|GalRL|PGR7|nGPCR-2037</td><td>G protein-coupled receptor 151</td><td>protein-coding</td></tr>
<tr><td>chr18_42591091_42591474</td><td>chr18</td><td>42591092</td><td>42591474</td><td>+</td><td>0.694415</td><td>Intergenic</td><td>B3|SINE|B2</td><td>-11631</td><td>NM_181543</td><td>240239</td><td>Mm.186779</td><td>NM_181543</td><td>ENSMUSG00000042816</td><td>Gpr151</td><td>C130082O03Rik|GalRL|PGR7|nGPCR-2037</td><td>G protein-coupled receptor 151</td><td>protein-coding</td></tr>
<tr><td>chr2_73404491_73404787</td><td>chr2</td><td>73404492</td><td>73404787</td><td>+</td><td>0.893515</td><td>Intergenic</td><td>Intergenic</td><td>-18159</td><td>NM_001276444</td><td>68526</td><td>Mm.451014</td><td>NM_001080707</td><td>ENSMUSG00000041762</td><td>Gpr155</td><td>1110017O10Rik|DEPDC3|F730029F15Rik|PGR22</td><td>G protein-coupled receptor 155</td><td>protein-coding</td></tr>
<tr><td>chr4_150096177_150096712</td><td>chr4</td><td>150096178</td><td>150096712</td><td>+</td><td>0.988008</td><td>intron (NM_177366, intron 1 of 3)</td><td>PB1D7|SINE|Alu</td><td>8942</td><td>NM_177366</td><td>269604</td><td>Mm.219307</td><td>NM_177366</td><td>ENSMUSG00000047875</td><td>Gpr157</td><td>F730108M23Rik</td><td>G protein-coupled receptor 157</td><td>protein-coding</td></tr>
</table>

| | | | | | | | | | | | | | | | | | |
|---|---|---|---|---|---|---|---|---|---|---|---|---|---|---|---|---|---|
| chr3_30888885_30889446 | chr3 | 30888886 | 30889446 | + | 0.755082 | intron (NM_001286995, intron 2 of 2) | RLTR20B5_MM\|LTR\|ERVK | 33216 | NM_001286994 | 71862 | Mm.272019 | NM_027965 | ENSMUSG00000037661 | **Gpr160** | 1700025D19Rik\|GPCR150 | G protein-coupled receptor 160 | protein-coding |
| chr3_30868675_30869059 | chr3 | 30868676 | 30869059 | + | 0.707089 | intron (NM_001286994, intron 1 of 3) | MTEa\|LTR\|ERVL-MaLR | 12917 | NM_001286994 | 71862 | Mm.272019 | NM_027965 | ENSMUSG00000037661 | **Gpr160** | 1700025D19Rik\|GPCR150 | G protein-coupled receptor 160 | protein-coding |
| chr1_165322392_165322708 | chr1 | 165322393 | 165322708 | + | 0.823478 | TTS (NM_001081126) | TTS (NM_001081126) | 26784 | NM_001081126 | 240888 | Mm.329929 | NM_001081126 | | **Gpr161** | Gm208\|Gm208Gpr\|vl | G protein-coupled receptor 161 | protein-coding |
| chr3_59087989_59088849 | chr3 | 59087990 | 59088849 | + | 1.18516 | intron (NM_177855, intron 12 of 43) | intron (NM_177855, intron 12 of 43) | 13402 | NM_173398 | 229323 | Mm.123648 | NM_173398 | ENSMUSG00000050075 | **Gpr171** | A630006E05\|F730001G15Rik\|H963 | G protein-coupled receptor 171 | protein-coding |
| chr3_59113885_59114846 | chr3 | 59113886 | 59114846 | + | 0.789191 | TTS (NM_133200) | TTS (NM_133200) | -12545 | NM_173398 | 229323 | Mm.123648 | NM_173398 | ENSMUSG00000050075 | **Gpr171** | A630006E05\|F730001G15Rik\|H963 | G protein-coupled receptor 171 | protein-coding |
| chr3_59089618_59090046 | chr3 | 59089619 | 59090046 | + | 0.733071 | intron (NM_177855, intron 12 of 43) | MTB_Mm-int\|LTR\|ERVL-MaLR | 11989 | NM_173398 | 229323 | Mm.123648 | NM_173398 | ENSMUSG00000050075 | **Gpr171** | A630006E05\|F730001G15Rik\|H963 | G protein-coupled receptor 171 | protein-coding |
| chr3_59093996_59094419 | chr3 | 59093997 | 59094419 | + | 0.703758 | intron (NM_177855, intron 15 of 43) | intron (NM_177855, intron 15 of 43) | 7613 | NM_173398 | 229323 | Mm.123648 | NM_173398 | ENSMUSG00000050075 | **Gpr171** | A630006E05\|F730001G15Rik\|H963 | G protein-coupled receptor 171 | protein-coding |
| chrX_107251037_107251398 | chrX | 107251038 | 107251398 | + | 1.20053 | Intergenic | Intergenic | -4660 | NM_001177781 | 213439 | Mm.383562 | NM_001033251 | ENSMUSG00000073008 | **Gpr174** | Gm376\|Lypsr3 | G protein-coupled receptor 174 | protein-coding |
| chrX_107294155_107294531 | chrX | 107294156 | 107294531 | + | 0.827004 | 3' UTR (NM_001177782, exon 3 of 3) | 3' UTR (NM_001177782, exon 3 of 3) | 9997 | NM_001177782 | 213439 | Mm.383562 | NM_001033251 | ENSMUSG00000073008 | **Gpr174** | Gm376\|Lypsr3 | G protein-coupled receptor 174 | protein-coding |
| chrX_107294110_107294707 | chrX | 107294111 | 107294707 | + | 0.760131 | 3' UTR (NM_001177782, exon 3 of 3) | 3' UTR (NM_001177782, exon 3 of 3) | 10063 | NM_001177782 | 213439 | Mm.383562 | NM_001033251 | ENSMUSG00000073008 | **Gpr174** | Gm376\|Lypsr3 | G protein-coupled receptor 174 | protein-coding |
| chrX_107260569_107260946 | chrX | 107260570 | 107260946 | + | 0.707527 | intron (NM_001033251, intron 2 of 4) | intron (NM_001033251, intron 2 of 4) | 4730 | NM_001033251 | 213439 | Mm.383562 | NM_001033251 | ENSMUSG00000073008 | **Gpr174** | Gm376\|Lypsr3 | G protein-coupled receptor 174 | protein-coding |
| chrX_107252441_107252637 | chrX | 107252442 | 107252637 | + | 0.605214 | Intergenic | Intergenic | -3339 | NM_001177781 | 213439 | Mm.383562 | NM_001033251 | ENSMUSG00000073008 | **Gpr174** | Gm376\|Lypsr3 | G protein-coupled receptor 174 | protein-coding |
| chr2_118375697_118376224 | chr2 | 118375698 | 118376224 | + | 1.48496 | Intergenic | MLT1I\|LTR\|ERVL-MaLR | -2542 | NM_201367 | 381413 | Mm.70979 | NM_201367 | ENSMUSG00000040133 | **Gpr176** | Gm1012 | G protein-coupled receptor 176 | protein-coding |
| chr14_121905567_121906722 | chr14 | 121905568 | 121906722 | + | 0.764102 | intron (NM_026861, intron 2 of 8) | RSINE1\|SINE\|B4 | 9629 | NM_182806 | 110168 | Mm.37405 | NM_182806 | ENSMUSG00000050350 | **Gpr18** | - | G protein-coupled receptor 18 | protein-coding |
| chr14_121901619_121901789 | chr14 | 121901620 | 121901789 | + | 0.746961 | intron (NM_026861, intron 1 of 8) | intron (NM_026861, intron 1 of 8) | 14070 | NM_182806 | 110168 | Mm.37405 | NM_182806 | ENSMUSG00000050350 | **Gpr18** | - | G protein-coupled receptor 18 | protein-coding |
| chr14_121901600_121901838 | chr14 | 121901601 | 121901838 | + | 0.71046 | intron (NM_026861, intron 1 of 8) | intron (NM_026861, intron 1 of 8) | 14055 | NM_182806 | 110168 | Mm.37405 | NM_182806 | ENSMUSG00000050350 | **Gpr18** | - | G protein-coupled receptor 18 | protein-coding |
| chr14_118161809_118162257 | chr14 | 118161810 | 118162257 | + | 0.965904 | intron (NM_021434, intron 8 of 8) | intron (NM_021434, intron 8 of 8) | 24906 | NM_021434 | 58245 | Mm.275411 | NM_021434 | ENSMUSG00000022131 | **Gpr180** | AB041545\|E130016I23Rik\|Itr | G protein-coupled receptor 180 | protein-coding |
| chr14_118161815_118162766 | chr14 | 118161816 | 118162766 | + | 0.670161 | intron (NM_021434, intron 8 of 8) | intron (NM_021434, intron 8 of 8) | 25164 | NM_021434 | 58245 | Mm.275411 | NM_021434 | ENSMUSG00000022131 | **Gpr180** | AB041545\|E130016I23Rik\|Itr | G protein-coupled receptor 180 | protein-coding |
| chr14_121983343_121983805 | chr14 | 121983344 | 121983805 | + | 1.12369 | intron (NM_026861, intron 6 of 8) | intron (NM_026861, intron 6 of 8) | -18381 | NM_183031 | 321019 | Mm.265618 | NM_183031 | ENSMUSG00000051212 | **Gpr183** | Ebi2 | G protein-coupled receptor 183 | protein-coding |
| chr14_121982971_121983836 | chr14 | 121982972 | 121983836 | + | 0.86119 | intron (NM_026861, intron 6 of 8) | intron (NM_026861, intron 6 of 8) | -18211 | NM_183031 | 321019 | Mm.265618 | NM_183031 | ENSMUSG00000051212 | **Gpr183** | Ebi2 | G protein-coupled receptor 183 | protein-coding |
| chr14_121959617_121959954 | chr14 | 121959618 | 121959954 | + | 0.814779 | intron (NM_183031, intron 1 of 1) | intron (NM_183031, intron 1 of 1) | 5407 | NM_183031 | 321019 | Mm.265618 | NM_183031 | ENSMUSG00000051212 | **Gpr183** | Ebi2 | G protein-coupled receptor 183 | protein-coding |
| chr12_31813802_31814216 | chr12 | 31813803 | 31814216 | + | 0.799682 | intron (NM_001163126, intron 10 of 22) | intron (NM_001163126, intron 10 of 22) | -100083 | NM_175191 | 73010 | Mm.441517 | NM_175191 | ENSMUSG00000044067 | **Gpr22** | 2900068K05Rik\|AW061316 | G protein-coupled receptor 22 | protein-coding |
| chr1_136255120_136255635 | chr1 | 136255121 | 136255635 | + | 1.4446 | Intergenic | Intergenic | 5495 | NM_001101516 | 383563 | Mm.40367 | NM_001101516 | ENSMUSG00000052759 | **Gpr25** | Gm1300 | G protein-coupled receptor 25 | protein-coding |
| chr1_136258584_136258940 | chr1 | 136258585 | 136258940 | + | 1.22729 | TTS (NM_001101516) | TTS (NM_001101516) | 2111 | NM_001101516 | 383563 | Mm.40367 | NM_001101516 | ENSMUSG00000052759 | **Gpr25** | Gm1300 | G protein-coupled receptor 25 | protein-coding |
| chr12_52030784_52031394 | chr12 | 52030785 | 52031394 | + | 1.50264 | Intergenic | Intergenic | -3026 | NM_008159 | 14762 | Mm.12890 | NM_008159 | ENSMUSG00000035148 | **Gpr33** | - | G protein-coupled receptor 33 | protein-coding |
| chrX_13622688_13623362 | chrX | 13622689 | 13623362 | + | 0.646366 | intron (NM_009806, intron 6 of 24) | intron (NM_009806, intron 6 of 24) | -9744 | NM_011823 | 23890 | Mm.487326 | NM_011823 | ENSMUSG00000040229 | **Gpr34** | Lypsr1 | G protein-coupled receptor 34 | protein-coding |
| chr1_85973372_85973996 | chr1 | 85973373 | 85973996 | + | 1.75584 | Intergenic | Intergenic | -12629 | NM_001033290 | 227326 | Mm.244158 | NM_001033290 | ENSMUSG00000049608 | **Gpr55** | Gm218\|Lpir1 | G protein-coupled receptor 55 | protein-coding |
| chr1_85958104_85958749 | chr1 | 85958105 | 85958749 | + | 1.07659 | intron (NM_001033290, intron 1 of 1) | intron (NM_001033290, intron 1 of 1) | 2628 | NM_001033290 | 227326 | Mm.244158 | NM_001033290 | ENSMUSG00000049608 | **Gpr55** | Gm218\|Lpir1 | G protein-coupled receptor 55 | protein-coding |
| chr1_85985772_85986477 | chr1 | 85985773 | 85986477 | + | 1.04123 | Intergenic | Intergenic | -25070 | NM_001033290 | 227326 | Mm.244158 | NM_001033290 | ENSMUSG00000049608 | **Gpr55** | Gm218\|Lpir1 | G protein-coupled receptor 55 | protein-coding |
| chr1_85953276_85953504 | chr1 | 85953277 | 85953504 | + | 0.856703 | intron (NM_001033290, intron 1 of 1) | intron (NM_001033290, intron 1 of 1) | 7665 | NM_001033290 | 227326 | Mm.244158 | NM_001033290 | ENSMUSG00000049608 | **Gpr55** | Gm218\|Lpir1 | G protein-coupled receptor 55 | protein-coding |
| chr1_85957279_85957672 | chr1 | 85957280 | 85957672 | + | 0.816974 | intron (NM_001033290, intron 1 of 1) | intron (NM_001033290, intron 1 of 1) | 3579 | NM_001033290 | 227326 | Mm.244158 | NM_001033290 | ENSMUSG00000049608 | **Gpr55** | Gm218\|Lpir1 | G protein-coupled receptor 55 | protein-coding |
| chr8_94984023_94984599 | chr8 | 94984024 | 94984599 | + | 1.11135 | promoter-TSS (NM_018882) | promoter-TSS (NM_018882) | -88 | NM_018882 | 14766 | Mm.290834 | NM_018882 | ENSMUSG00000031785 | **Gpr56** | Cyt28\|TM7LN4\|TM7XN1 | G protein-coupled receptor 56 | protein-coding |
| chr3_108153971_108154319 | chr3 | 108153972 | 108154319 | + | 0.836693 | intron (NM_175470, intron 1 of 1) | intron (NM_175470, intron 1 of 1) | 737 | NM_175470 | 229714 | Mm.73666 | NM_175470 | ENSMUSG00000046793 | **Gpr61** | - | G protein-coupled receptor 61 | protein-coding |
| chr3_108153957_108154478 | chr3 | 108153958 | 108154478 | + | 0.720296 | intron (NM_175470, intron 1 of 1) | intron (NM_175470, intron 1 of 1) | 664 | NM_175470 | 229714 | Mm.73666 | NM_175470 | ENSMUSG00000046793 | **Gpr61** | - | G protein-coupled receptor 61 | protein-coding |
| chr12_98283107_98283914 | chr12 | 98283108 | 98283914 | + | 1.73603 | Intergenic | Intergenic | 14876 | NM_008152 | 14744 | Mm.378924 | NM_008152 | ENSMUSG00000021886 | **Gpr65** | Dig1\|Gpcr25\|TDAG8 | G-protein coupled receptor 65 | protein-coding |
| chr12_100871601_100872287 | chr12 | 100871602 | 100872287 | + | 1.85359 | 3' UTR (NM_145448, exon 13 of 13) | 3' UTR (NM_145448, exon 13 of 13) | 27580 | NM_001177674 | 238377 | Mm.32160 | NM_175493 | ENSMUSG00000047415 | **Gpr68** | BB131428\|Ogr1 | G protein-coupled receptor 68 | protein-coding |
| chr12_100871372_100872295 | chr12 | 100871373 | 100872295 | + | 1.41742 | 3' UTR (NM_145448, exon 13 of 13) | 3' UTR (NM_145448, exon 13 of 13) | 27690 | NM_001177674 | 238377 | Mm.32160 | NM_175493 | ENSMUSG00000047415 | **Gpr68** | BB131428\|Ogr1 | G protein-coupled receptor 68 | protein-coding |
| chr12_100880858_100881121 | chr12 | 100880859 | 100881121 | + | 0.696282 | intron (NM_001177673, intron 2 of 2) | intron (NM_001177673, intron 2 of 2) | 18534 | NM_001177674 | 238377 | Mm.32160 | NM_175493 | ENSMUSG00000047415 | **Gpr68** | BB131428\|Ogr1 | G protein-coupled receptor 68 | protein-coding |
| chr3_96881471_96882796 | chr3 | 96881472 | 96882796 | + | 0.855027 | intron (NM_026229, intron 8 of 13) | intron (NM_026229, intron 8 of 13) | 23164 | NM_026229 | 67549 | Mm.46722 | NM_026229 | | **Gpr89** | 4933412D19Rik\|Gpr89a\|SH120 | G protein-coupled receptor 89 | protein-coding |
| chr3_96882064_96882618 | chr3 | 96882065 | 96882618 | + | 0.7964 | intron (NM_026229, intron 8 of 13) | MIR3\|SINE\|MIR | 22957 | NM_026229 | 67549 | Mm.46722 | NM_026229 | | **Gpr89** | 4933412D19Rik\|Gpr89a\|SH120 | G protein-coupled receptor 89 | protein-coding |
| chr6_135060769_135061038 | chr6 | 135060770 | 135061038 | + | 0.79812 | Intergenic | RMER15\|LTR\|ERVL | -4758 | NM_181444 | 232431 | Mm.23575 | NM_181444 | ENSMUSG00000046733 | **Gprc5a** | Rai3\|Raig1 | G protein-coupled receptor, family C, group 5, member A | protein-coding |
| chr14_34188664_34188992 | chr14 | 34188665 | 34188992 | + | 0.767257 | Intergenic | Intergenic | 12805 | NM_183209 | 432839 | Mm.294643 | NM_183209 | ENSMUSG00000071531 | **Gprin2** | C130040D06Rik\|C230073P13\|Gm286\|mKIAA0514 | G protein regulated inducer of neurite outgrowth 2 | protein-coding |
| chr6_59418498_59419008 | chr6 | 59418499 | 59419008 | + | 1.43301 | intron (NM_183183, intron 1 of 1) | intron (NM_183183, intron 1 of 1) | 7537 | NM_183183 | 243385 | Mm.138080 | NM_183183 | ENSMUSG00000045441 | **Gprin3** | C030038J10Rik\|C730021L23 | GPRIN family member 3 | protein-coding |
| chr6_59348547_59349254 | chr6 | 59348548 | 59349254 | + | 1.09719 | Intergenic | Intergenic | 77389 | NM_183183 | 243385 | Mm.138080 | NM_183183 | ENSMUSG00000045441 | **Gprin3** | C030038J10Rik\|C730021L23 | GPRIN family member 3 | protein-coding |
| chr6_59450866_59451638 | chr6 | 59450867 | 59451638 | + | 1.07817 | Intergenic | Intergenic | -24962 | NM_183183 | 243385 | Mm.138080 | NM_183183 | ENSMUSG00000045441 | **Gprin3** | C030038J10Rik\|C730021L23 | GPRIN family member 3 | protein-coding |
| chr6_59424687_59424988 | chr6 | 59424688 | 59424988 | + | 0.928859 | intron (NM_183183, intron 1 of 1) | intron (NM_183183, intron 1 of 1) | 1452 | NM_183183 | 243385 | Mm.138080 | NM_183183 | ENSMUSG00000045441 | **Gprin3** | C030038J10Rik\|C730021L23 | GPRIN family member 3 | protein-coding |
| chr6_59421814_59422011 | chr6 | 59421815 | 59422011 | + | 0.867745 | intron (NM_183183, intron 1 of 1) | intron (NM_183183, intron 1 of 1) | 4377 | NM_183183 | 243385 | Mm.138080 | NM_183183 | ENSMUSG00000045441 | **Gprin3** | C030038J10Rik\|C730021L23 | GPRIN family member 3 | protein-coding |
| chr11_69915078_69915348 | chr11 | 69915079 | 69915348 | + | 1.12231 | exon (NM_019726, exon 4 of 10) | exon (NM_019726, exon 4 of 10) | 1021 | NM_019726 | 56310 | Mm.477738 | NM_019726 | ENSMUSG00000023170 | **Gps2** | AI505953 | G protein pathway suppressor 2 | protein-coding |
| chr11_69915692_69916223 | chr11 | 69915693 | 69916223 | + | 0.768139 | TTS (NM_181582) | TTS (NM_181582) | 1766 | NM_019726 | 56310 | Mm.477738 | NM_019726 | ENSMUSG00000023170 | **Gps2** | AI505953 | G protein pathway suppressor 2 | protein-coding |
| chr2_26327649_26328151 | chr2 | 26327650 | 26328151 | + | 0.768873 | intron (NM_001199147, intron 9 of 13) | ORR1C2\|LTR\|ERVL-MaLR | 8130 | NM_001199147 | 67839 | Mm.266611 | NM_153410 | ENSMUSG00000026930 | **Gpsm1** | 1810037C22Rik\|AW107933\|Ags3 | G-protein signalling modulator 1 (AGS3-like, C. elegans) | protein-coding |
| chr13_21249035_21249521 | chr13 | 21249036 | 21249521 | + | 0.894889 | Intergenic | Intergenic | 43408 | NM_010343 | 14780 | Mm.1332 | NM_010343 | ENSMUSG00000004344 | **Gpx5** | AV379049\|Arep | glutathione peroxidase 5 | protein-coding |
| chr13_21240806_21241197 | chr13 | 21240807 | 21241197 | + | 0.773051 | Intergenic | tRNA-Ala-GCA\|tRNA\|tRNA | 51684 | NM_010343 | 14780 | Mm.1332 | NM_010343 | ENSMUSG00000004344 | **Gpx5** | AV379049\|Arep | glutathione peroxidase 5 | protein-coding |

| | | | | | | | | | | | | | | | | | |
|---|---|---|---|---|---|---|---|---|---|---|---|---|---|---|---|---|---|
| chr13_21259698_21260177 | chr13 | 21259699 | 21260177 | + | 0.74587 | Intergenic | Intergenic | 32748 | NM_010343 | 14780 | Mm.1332 | NM_010343 | ENSMUSG00000004344 | **Gpx5** | AV379049\|Arep | glutathione peroxidase 5 | protein-coding |
| chr13_21259700_21260423 | chr13 | 21259701 | 21260423 | + | 0.724127 | Intergenic | Intergenic | 32624 | NM_010343 | 14780 | Mm.1332 | NM_010343 | ENSMUSG00000004344 | **Gpx5** | AV379049\|Arep | glutathione peroxidase 5 | protein-coding |
| chr13_21254025_21254335 | chr13 | 21254026 | 21254335 | + | 0.658871 | Intergenic | Intergenic | 38506 | NM_010343 | 14780 | Mm.1332 | NM_010343 | ENSMUSG00000004344 | **Gpx5** | AV379049\|Arep | glutathione peroxidase 5 | protein-coding |
| chr9_40537006_40537699 | chr9 | 40537007 | 40537699 | + | 1.41128 | Intergenic | Intergenic | -81589 | NM_172768 | 235283 | Mm.138434 | NM_172768 | ENSMUSG00000040111 | **Gramd1b** | 3222402H23\|A930008A22Rik\|AI593249\|mKIAA1201 | GRAM domain containing 1B | protein-coding |
| chr9_40554132_40554452 | chr9 | 40554133 | 40554452 | + | 1.2186 | Intergenic | L1_Mur2\|LINE\|L1 | -98528 | NM_172768 | 235283 | Mm.138434 | NM_172768 | ENSMUSG00000040111 | **Gramd1b** | 3222402H23\|A930008A22Rik\|AI593249\|mKIAA1201 | GRAM domain containing 1B | protein-coding |
| chr9_40434957_40435435 | chr9 | 40434958 | 40435435 | + | 1.01249 | intron (NM_172768, intron 1 of 19) | intron (NM_172768, intron 1 of 19) | 20568 | NM_172768 | 235283 | Mm.138434 | NM_172768 | ENSMUSG00000040111 | **Gramd1b** | 3222402H23\|A930008A22Rik\|AI593249\|mKIAA1201 | GRAM domain containing 1B | protein-coding |
| chr9_40553908_40554465 | chr9 | 40553909 | 40554465 | + | 0.993229 | Intergenic | Intergenic | -98423 | NM_172768 | 235283 | Mm.138434 | NM_172768 | ENSMUSG00000040111 | **Gramd1b** | 3222402H23\|A930008A22Rik\|AI593249\|mKIAA1201 | GRAM domain containing 1B | protein-coding |
| chr9_40429800_40430299 | chr9 | 40429801 | 40430299 | + | 0.944109 | intron (NM_172768, intron 1 of 19) | intron (NM_172768, intron 1 of 19) | 25714 | NM_172768 | 235283 | Mm.138434 | NM_172768 | ENSMUSG00000040111 | **Gramd1b** | 3222402H23\|A930008A22Rik\|AI593249\|mKIAA1201 | GRAM domain containing 1B | protein-coding |
| chr9_40449635_40450618 | chr9 | 40449636 | 40450618 | + | 0.894031 | intron (NM_172768, intron 1 of 19) | intron (NM_172768, intron 1 of 19) | 5637 | NM_172768 | 235283 | Mm.138434 | NM_172768 | ENSMUSG00000040111 | **Gramd1b** | 3222402H23\|A930008A22Rik\|AI593249\|mKIAA1201 | GRAM domain containing 1B | protein-coding |
| chr9_40449887_40450549 | chr9 | 40449888 | 40450549 | + | 0.856418 | intron (NM_172768, intron 1 of 19) | intron (NM_172768, intron 1 of 19) | 5546 | NM_172768 | 235283 | Mm.138434 | NM_172768 | ENSMUSG00000040111 | **Gramd1b** | 3222402H23\|A930008A22Rik\|AI593249\|mKIAA1201 | GRAM domain containing 1B | protein-coding |
| chr9_40448855_40449161 | chr9 | 40448856 | 40449161 | + | 0.761589 | intron (NM_172768, intron 1 of 19) | intron (NM_172768, intron 1 of 19) | 6756 | NM_172768 | 235283 | Mm.138434 | NM_172768 | ENSMUSG00000040111 | **Gramd1b** | 3222402H23\|A930008A22Rik\|AI593249\|mKIAA1201 | GRAM domain containing 1B | protein-coding |
| chr16_44002082_44002429 | chr16 | 44002083 | 44002429 | + | 1.02047 | intron (NM_153528, intron 4 of 12) | intron (NM_153528, intron 4 of 12) | 14180 | NM_153528 | 207798 | Mm.19316 | NM_153528 | ENSMUSG00000036292 | **Gramd1c** | 4921521N14Rik | GRAM domain containing 1C | protein-coding |
| chr18_56429099_56429417 | chr18 | 56429100 | 56429417 | + | 1.05669 | Intergenic | Intergenic | -2874 | NM_026240 | 107022 | Mm.24356 | NM_026240 | ENSMUSG00000001700 | **Gramd3** | 9030613F08Rik\|9130427A09Rik\|AW061306\|NS3TP2 | GRAM domain containing 3 | protein-coding |
| chr18_56028485_56028828 | chr18 | 56028486 | 56028828 | + | 0.762588 | Intergenic | MIR\|SINE\|MIR | -403475 | NM_026240 | 107022 | Mm.24356 | NM_026240 | ENSMUSG00000001700 | **Gramd3** | 9030613F08Rik\|9130427A09Rik\|AW061306\|NS3TP2 | GRAM domain containing 3 | protein-coding |
| chr15_86051088_86051668 | chr15 | 86051089 | 86051668 | + | 0.658683 | Intergenic | MER3\|DNA\|hAT-Charlie | -6317 | NM_172611 | 223752 | Mm.24442 | NM_172611 | ENSMUSG00000035900 | **Gramd4** | 9930016O13 | GRAM domain containing 4 | protein-coding |
| chr11_61644538_61644924 | chr11 | 61644539 | 61644924 | + | 1.592 | Intergenic | L1M4b\|LINE\|L1 | -8590 | NM_027817 | 71520 | Mm.202760 | NM_027817 | ENSMUSG00000004837 | **Grap** | 8430435N19Rik\|BB233594 | GRB2-related adaptor protein | protein-coding |
| chr11_61638007_61638407 | chr11 | 61638008 | 61638407 | + | 0.92323 | Intergenic | MER76\|LTR\|ERVL | -15114 | NM_027817 | 71520 | Mm.202760 | NM_027817 | ENSMUSG00000004837 | **Grap** | 8430435N19Rik\|BB233594 | GRB2-related adaptor protein | protein-coding |
| chr11_61639757_61640008 | chr11 | 61639758 | 61640008 | + | 0.805115 | Intergenic | B4\|SINE\|B4 | -13438 | NM_027817 | 71520 | Mm.202760 | NM_027817 | ENSMUSG00000004837 | **Grap** | 8430435N19Rik\|BB233594 | GRB2-related adaptor protein | protein-coding |
| chr15_80593797_80594218 | chr15 | 80593798 | 80594218 | + | 0.757227 | intron (NM_001289442, intron 1 of 7) | intron (NM_001289442, intron 1 of 7) | 21411 | NM_001289442 | 17444 | Mm.12947 | NM_010815 | ENSMUSG00000042351 | **Grap2** | GRAP-2\|GRB2L\|GRBLG\|GRID\|Gads\|GrbX\|Grf40\|GrpL\|Mona\|P38 | GRB2-related adaptor protein 2 | protein-coding |
| chr15_80607428_80607690 | chr15 | 80607429 | 80607690 | + | 0.70848 | intron (NM_001289442, intron 1 of 7) | intron (NM_001289442, intron 1 of 7) | -15924 | NM_010815 | 17444 | Mm.12947 | NM_010815 | ENSMUSG00000042351 | **Grap2** | GRAP-2\|GRB2L\|GRBLG\|GRID\|Gads\|GrbX\|Grf40\|GrpL\|Mona\|P38 | GRB2-related adaptor protein 2 | protein-coding |
| chr2_65104455_65104959 | chr2 | 65104456 | 65104959 | + | 1.16018 | intron (NM_027225, intron 8 of 12) | intron (NM_027225, intron 8 of 12) | -81941 | NM_016719 | 50915 | Mm.214554 | NM_016719 | ENSMUSG00000026888 | **Grb14** | AI505286 | growth factor receptor bound protein 14 | protein-coding |
| chr11_98472722_98473012 | chr11 | 98472723 | 98473012 | + | 0.781157 | intron (NM_011771, intron 7 of 7) | intron (NM_011771, intron 7 of 7) | 26033 | NM_010346 | 14786 | Mm.276702 | NM_010346 | ENSMUSG00000019312 | **Grb7** | mKIAA4028 | growth factor receptor bound protein 7 | protein-coding |
| chr11_98476828_98478189 | chr11 | 98476829 | 98478189 | + | 0.600313 | intron (NM_011771, intron 6 of 7) | intron (NM_011771, intron 6 of 7) | 30675 | NM_010346 | 14786 | Mm.276702 | NM_010346 | ENSMUSG00000019312 | **Grb7** | mKIAA4028 | growth factor receptor bound protein 7 | protein-coding |
| chr4_44948653_44949087 | chr4 | 44948654 | 44949087 | + | 0.807585 | Intergenic | Intergenic | -32541 | NM_080289 | 76238 | Mm.196574 | NM_080289 | ENSMUSG00000035637 | **Grhpr** | 1110059D05Rik\|6430629L09Rik | glyoxylate reductase/hydroxypyruvate reductase | protein-coding |
| chrX_41398485_41398806 | chrX | 41398486 | 41398806 | + | 0.688705 | Intergenic | Intergenic | -2655 | NM_016886 | 53623 | Mm.327681 | NM_016886 | ENSMUSG00000001986 | **Gria3** | 2900064I19Rik\|GluA3\|GluR-C\|GluR-K3\|Glur-3\|Glur3\|Gluralpha3 | glutamate receptor, ionotropic, AMPA3 (alpha 3) | protein-coding |
| chr10_119803652_119804215 | chr10 | 119803653 | 119804215 | + | 0.677642 | intron (NM_133442, intron 1 of 21) | intron (NM_133442, intron 1 of 21) | -15455 | NM_001277294 | 74053 | Mm.196692 | NM_028736 | ENSMUSG00000034813 | **Grip1** | 4931400F03Rik\|GRIP\|eb | glutamate receptor interacting protein 1 | protein-coding |
| chr19_60950494_60950842 | chr19 | 60950495 | 60950842 | + | 1.1964 | intron (NM_018869, intron 1 of 15) | intron (NM_018869, intron 1 of 15) | 60919 | NM_018869 | 14773 | Mm.279400 | NM_018869 | ENSMUSG00000003228 | **Grk5** | Gprk5 | G protein-coupled receptor kinase 5 | protein-coding |
| chr5_21122948_21123781 | chr5 | 21122949 | 21123781 | + | 0.791997 | Intergenic | Intergenic | -62902 | NM_175437 | 212167 | Mm.121705 | NM_175437 | ENSMUSG00000039934 | **Gsap** | A530088I07Rik\|Pion | gamma-secretase activating protein | protein-coding |
| chr5_21253934_21254179 | chr5 | 21253935 | 21254179 | + | 0.677592 | intron (NM_175437, intron 18 of 30) | intron (NM_175437, intron 18 of 30) | 67790 | NM_175437 | 212167 | Mm.121705 | NM_175437 | ENSMUSG00000039934 | **Gsap** | A530088I07Rik\|Pion | gamma-secretase activating protein | protein-coding |
| chr8_120436432_120437635 | chr8 | 120436433 | 120437635 | + | 0.911602 | Intergenic | Intergenic | -51832 | NM_198671 | 382034 | Mm.334856 | NM_198671 | ENSMUSG00000031822 | **Gse1** | 2210013I18Rik\|mKIAA0182 | genetic suppressor element 1 | protein-coding |
| chr2_35271763_35272172 | chr2 | 35271764 | 35272172 | + | 0.827872 | intron (NM_001206367, intron 2 of 17) | intron (NM_001206367, intron 2 of 17) | 5224 | NM_001206369 | 227753 | Mm.21109 | NM_146120 | ENSMUSG00000026879 | **Gsn** | ADF | gelsolin | protein-coding |
| chr9_78204726_78205201 | chr9 | 78204727 | 78205201 | + | 0.775074 | intron (NM_010357, intron 4 of 6) | intron (NM_010357, intron 4 of 6) | 12998 | NM_010357 | 14860 | Mm.2662 | NM_010357 | ENSMUSG00000032348 | **Gsta4** | GST5.7\|mGsta4 | glutathione S-transferase, alpha 4 | protein-coding |
| chr3_133078158_133078798 | chr3 | 133078159 | 133078798 | + | 1.99689 | intron (NM_026231, intron 3 of 11) | intron (NM_026231, intron 3 of 11) | 13262 | NM_026231 | 67553 | Mm.304899 | NM_026231 | ENSMUSG00000028018 | **Gstcd** | 4933434L15Rik | glutathione S-transferase, C-terminal domain containing | protein-coding |
| chr19_4043957_4044423 | chr19 | 4043958 | 4044423 | + | 0.75765 | Intergenic | RLTR45\|LTR\|ERVK | -1969 | NM_181796 | 14869 | Mm.441542 | NM_181796 | ENSMUSG00000038155 | **Gstp2** | GSTpiA\|Gst-3\|Gst3 | glutathione S-transferase, pi 2 | protein-coding |
| chr19_4044018_4044352 | chr19 | 4044019 | 4044352 | + | 0.717064 | Intergenic | RLTR45\|LTR\|ERVK | -1964 | NM_181796 | 14869 | Mm.441542 | NM_181796 | ENSMUSG00000038155 | **Gstp2** | GSTpiA\|Gst-3\|Gst3 | glutathione S-transferase, pi 2 | protein-coding |
| chr10_75829546_75830270 | chr10 | 75829547 | 75830270 | + | 1.50745 | Intergenic | MTB_Mm-int\|LTR\|ERVL-MaLR | 4973 | NM_010361 | 14872 | Mm.24118 | NM_010361 | ENSMUSG00000033318 | **Gstt2** | AI266894\|Yrs\|mGSTT2 | glutathione S-transferase, theta 2 | protein-coding |
| chr10_75839012_75839800 | chr10 | 75839013 | 75839800 | + | 0.885821 | Intergenic | Arthur1B\|DNA\|hAT-Tip100 | -4525 | NM_010361 | 14872 | Mm.24118 | NM_010361 | ENSMUSG00000033318 | **Gstt2** | AI266894\|Yrs\|mGSTT2 | glutathione S-transferase, theta 2 | protein-coding |
| chr10_75824470_75824969 | chr10 | 75824471 | 75824969 | + | 1.21503 | Intergenic | RLTR48B\|LTR\|ERV1 | -2177 | NM_029472 | 75886 | Mm.268996 | NM_029472 | ENSMUSG00000009093 | **Gstt4** | 4930583C14Rik\|AI507011 | glutathione S-transferase, theta 4 | protein-coding |
| chr10_75824274_75824969 | chr10 | 75824275 | 75824969 | + | 1.11391 | Intergenic | RLTR48B\|LTR\|ERV1 | -2079 | NM_029472 | 75886 | Mm.268996 | NM_029472 | ENSMUSG00000009093 | **Gstt4** | 4930583C14Rik\|AI507011 | glutathione S-transferase, theta 4 | protein-coding |
| chr12_87130067_87130879 | chr12 | 87130068 | 87130879 | + | 1.24903 | intron (NM_153415, intron 6 of 20) | intron (NM_153415, intron 6 of 20) | -16692 | NM_001252555 | 14874 | Mm.29652 | NM_010363 | ENSMUSG00000021033 | **Gstz1** | MAAI | glutathione transferase zeta 1 (maleylacetoacetate isomerase) | protein-coding |
| chr5_75070615_75070838 | chr5 | 75070616 | 75070838 | + | 0.787599 | Intergenic | Intergenic | -4874 | NM_133256 | 14843 | Mm.218752 | NM_133256 | ENSMUSG00000035946 | **Gsx2** | Gsh-2\|Gsh2 | GS homeobox 2 | protein-coding |
| chr2_44886488_44887388 | chr2 | 44886489 | 44887388 | + | 2.0078 | Intergenic | L2b\|LINE\|L2 | -25316 | NM_172662 | 227835 | Mm.314312 | NM_172662 | ENSMUSG00000036890 | **Gtdc1** | E330008O22Rik | glycosyltransferase-like domain containing 1 | protein-coding |
| chr2_44878866_44879922 | chr2 | 44878867 | 44879922 | + | 1.14758 | Intergenic | ORR1B2\|LTR\|ERVL-MaLR | -17772 | NM_172662 | 227835 | Mm.314312 | NM_172662 | ENSMUSG00000036890 | **Gtdc1** | E330008O22Rik | glycosyltransferase-like domain containing 1 | protein-coding |
| chr2_44873734_44874060 | chr2 | 44873735 | 44874060 | + | 0.883388 | Intergenic | L1_Mur3\|LINE\|L1 | -12275 | NM_172662 | 227835 | Mm.314312 | NM_172662 | ENSMUSG00000036890 | **Gtdc1** | E330008O22Rik | glycosyltransferase-like domain containing 1 | protein-coding |
| chr2_44818609_44819091 | chr2 | 44818610 | 44819091 | + | 0.702564 | intron (NM_172662, intron 3 of 10) | intron (NM_172662, intron 3 of 10) | 42772 | NM_172662 | 227835 | Mm.314312 | NM_172662 | ENSMUSG00000036890 | **Gtdc1** | E330008O22Rik | glycosyltransferase-like domain containing 1 | protein-coding |
| chr2_44849725_44850196 | chr2 | 44849726 | 44850196 | + | 0.651106 | intron (NM_172662, intron 2 of 10) | intron (NM_172662, intron 2 of 10) | 11661 | NM_172662 | 227835 | Mm.314312 | NM_172662 | ENSMUSG00000036890 | **Gtdc1** | E330008O22Rik | glycosyltransferase-like domain containing 1 | protein-coding |
| chr2_44864765_44865298 | chr2 | 44864766 | 44865298 | + | 0.619677 | Intergenic | Intergenic | -3410 | NM_172662 | 227835 | Mm.314312 | NM_172662 | ENSMUSG00000036890 | **Gtdc1** | E330008O22Rik | glycosyltransferase-like domain containing 1 | protein-coding |
| chr12_91585117_91585516 | chr12 | 91585118 | 91585516 | + | 1.41485 | intron (NM_031391, intron 2 of 8) | B4A\|SINE\|B4 | 4332 | NM_031391 | 83602 | Mm.275728 | NM_031391 | ENSMUSG00000020962 | **Gtf2a1** | 6330549H03Rik\|AA536742\|AA959775\|AW060250\|TfIIAa/b\|Tfiia1 | general transcription factor II A, 1 | protein-coding |
| chr12_91661411_91661861 | chr12 | 91661412 | 91661861 | + | 1.00284 | intron (NM_175367, intron 4 of 6) | RMER4B\|LTR\|ERVK | -71149 | NM_175335 | 83602 | Mm.275728 | NM_031391 | ENSMUSG00000020962 | **Gtf2a1** | 6330549H03Rik\|AA536742\|AA959775\|AW060250\|TfIIAa/b\|Tfiia1 | general transcription factor II A, 1 | protein-coding |
| chr12_91642591_91643098 | chr12 | 91642592 | 91643098 | + | 0.845097 | intron (NM_175367, intron 5 of 6) | intron (NM_175367, intron 5 of 6) | -52358 | NM_175335 | 83602 | Mm.275728 | NM_031391 | ENSMUSG00000020962 | **Gtf2a1** | 6330549H03Rik\|AA536742\|AA959775\|AW060250\|TfIIAa/b\|Tfiia1 | general transcription factor II A, 1 | protein-coding |
| chr12_91668488_91669182 | chr12 | 91668489 | 91669182 | + | 0.830527 | intron (NM_175367, intron 4 of 6) | intron (NM_175367, intron 4 of 6) | -78348 | NM_175335 | 83602 | Mm.275728 | NM_031391 | ENSMUSG00000020962 | **Gtf2a1** | 6330549H03Rik\|AA536742\|AA959775\|AW060250\|TfIIAa/b\|Tfiia1 | general transcription factor II A, 1 | protein-coding |
| chr12_91663247_91663689 | chr12 | 91663248 | 91663689 | + | 0.782807 | intron (NM_175367, intron 4 of 6) | intron (NM_175367, intron 4 of 6) | -72981 | NM_175335 | 83602 | Mm.275728 | NM_031391 | ENSMUSG00000020962 | **Gtf2a1** | 6330549H03Rik\|AA536742\|AA959775\|AW060250\|TfIIAa/b\|Tfiia1 | general transcription factor II A, 1 | protein-coding |
| chr12_91662359_91662650 | chr12 | 91662360 | 91662650 | + | 0.729778 | intron (NM_175367, intron 4 of 6) | intron (NM_175367, intron 4 of 6) | -72018 | NM_175335 | 83602 | Mm.275728 | NM_031391 | ENSMUSG00000020962 | **Gtf2a1** | 6330549H03Rik\|AA536742\|AA959775\|AW060250\|TfIIAa/b\|Tfiia1 | general transcription factor II A, 1 | protein-coding |
| chr13_100510843_100511832 | chr13 | 100510844 | 100511832 | + | 1.31467 | intron (NM_008756, intron 6 of 8) | intron (NM_008756, intron 6 of 8) | -18729 | NM_022011 | 23894 | Mm.272533 | NM_022011 | ENSMUSG00000021639 | **Gtf2h2** | 44kDa\|BTF2 p44\|BTF2-p44\|Btf2p44\|p44 | general transcription factor II H, polypeptide 2 | protein-coding |
| chr17_35676827_35677172 | chr17 | 35676828 | 35677172 | + | 0.832443 | Intergenic | B1_Mur3\|SINE\|Alu | -3318 | NM_010364 | 14885 | Mm.10182 | NM_010364 | ENSMUSG00000001524 | **Gtf2h4** | AW545633\|BTF2 p52\|TFIIH\|p44\|p52 | general transcription factor II H, polypeptide 4 | protein-coding |
| chr7_125705120_125705471 | chr7 | 125705121 | 125705471 | + | 0.972189 | intron (NM_207239, intron 1 of 36) | intron (NM_207239, intron 1 of 36) | 2392 | NM_207239 | 233863 | Mm.294173 | NM_207239 | ENSMUSG00000032777 | **Gtf3c1** | - | general transcription factor III C 1 | protein-coding |
| chr5_5561097_5561382 | chr5 | 5561098 | 5561382 | + | 0.766969 | Intergenic | B3A\|SINE\|B2 | -1739 | NM_153116 | 207704 | Mm.323984 | NM_153116 | ENSMUSG00000040464 | **Gtpbp10** | 4930545J22Rik\|BC034507 | GTP-binding protein 10 (putative) | protein-coding |

| | | | | | | | | | | | | | | | | | |
|---|---|---|---|---|---|---|---|---|---|---|---|---|---|---|---|---|---|
| chr13_9026794_9027206 | chr13 | 9026795 | 9027206 | + | 0.720873 | Intergenic | Intergenic | -30988 | NM_027000 | 69237 | Mm.41800 | NM_027000 | ENSMUSG00000021149 | **Gtpbp4** | 2610028C09Rik\|Crfg\|Gtpbp3\|NGB\|Nog1 | GTP binding protein 4 | protein-coding |
| chr2_163102929_163103778 | chr2 | 163102930 | 163103778 | + | 1.47927 | Intergenic | Intergenic | -13753 | NM_026630 | 68236 | Mm.33648 | NM_026630 | | **Gtsf1l** | 2410116G06Rik | gametocyte specific factor 1-like | protein-coding |
| chr4_119614067_119614811 | chr4 | 119614068 | 119614811 | + | 0.736498 | intron (NM_172699, intron 6 of 12) | B4\|SINE\|B4 | -23293 | NM_008190 | 14915 | Mm.2614 | NM_008190 | ENSMUSG00000023247 | **Guca2a** | Guca2 | guanylate cyclase activator 2a (guanylin) | protein-coding |
| chr9_3851223_3851859 | chr9 | 3851224 | 3851859 | + | 0.695592 | intron (NM_001033322, intron 6 of 7) | intron (NM_001033322, intron 6 of 7) | 319192 | NM_001033322 | 234889 | Mm.91240 | NM_001033322 | | **Gucy1a2** | 6330407I18Rik\|A230060L24Rik | guanylate cyclase 1, soluble, alpha 2 | protein-coding |
| chr3_82140859_82141735 | chr3 | 82140860 | 82141735 | + | 1.03886 | intron (NM_021896, intron 1 of 8) | intron (NM_021896, intron 1 of 8) | 4580 | NM_021896 | 60596 | Mm.143831 | NM_021896 | ENSMUSG00000033910 | **Gucy1a3** | 1200016O07Rik\|sGC-alpha1 | guanylate cyclase 1, soluble, alpha 3 | protein-coding |
| chr3_82158393_82158952 | chr3 | 82158394 | 82158952 | + | 0.686623 | Intergenic | Intergenic | -12796 | NM_021896 | 60596 | Mm.143831 | NM_021896 | ENSMUSG00000033910 | **Gucy1a3** | 1200016O07Rik\|sGC-alpha1 | guanylate cyclase 1, soluble, alpha 3 | protein-coding |
| chr11_69232832_69233659 | chr11 | 69232833 | 69233659 | + | 0.655579 | intron (NM_008192, intron 3 of 18) | intron (NM_008192, intron 3 of 18) | 3776 | NM_008192 | 14919 | Mm.291816 | NM_008192 | ENSMUSG00000020890 | **Gucy2e** | GC-E\|GC1\|ROS-GC1 | guanylate cyclase 2e | protein-coding |
| chr7_106235669_106237379 | chr7 | 106235670 | 106237379 | + | 0.840028 | Intergenic | Intergenic | -21184 | NM_029000 | 74558 | Mm.458429 | NM_029000 | ENSMUSG00000045868 | **Gvin1** | 9130002C22Rik\|9830104F22Rik\|Iigs1\|VLIG\|VLIG-1 | GTPase, very large interferon inducible 1 | protein-coding |
| chr7_106236131_106237040 | chr7 | 106236132 | 106237040 | + | 0.826943 | Intergenic | RMER5\|LTR\|ERV1 | -21246 | NM_029000 | 74558 | Mm.458429 | NM_029000 | ENSMUSG00000045868 | **Gvin1** | 9130002C22Rik\|9830104F22Rik\|Iigs1\|VLIG\|VLIG-1 | GTPase, very large interferon inducible 1 | protein-coding |
| chr7_106256730_106257072 | chr7 | 106256731 | 106257072 | + | 0.820483 | Intergenic | Intergenic | -41561 | NM_029000 | 74558 | Mm.458429 | NM_029000 | ENSMUSG00000045868 | **Gvin1** | 9130002C22Rik\|9830104F22Rik\|Iigs1\|VLIG\|VLIG-1 | GTPase, very large interferon inducible 1 | protein-coding |
| chr18_32564080_32564776 | chr18 | 32564081 | 32564776 | + | 1.09056 | Intergenic | Intergenic | -4394 | NM_001048207 | 71683 | Mm.292145 | NM_027863 | ENSMUSG00000090523 | **Gypc** | 0610037F22Rik\|Cd236r | glycophorin C | protein-coding |
| chr18_32569431_32570017 | chr18 | 32569432 | 32570017 | + | 0.765373 | Intergenic | Intergenic | -9690 | NM_001048207 | 71683 | Mm.292145 | NM_027863 | ENSMUSG00000090523 | **Gypc** | 0610037F22Rik\|Cd236r | glycophorin C | protein-coding |
| chr6_142455445_142455962 | chr6 | 142455446 | 142455962 | + | 0.683466 | intron (NM_145572, intron 5 of 16) | MTD\|LTR\|ERVL-MaLR | 17405 | NM_145572 | 232493 | Mm.275975 | NM_145572 | ENSMUSG00000030244 | **Gys2** | BC021322\|LGS | glycogen synthase 2 | protein-coding |
| chr2_148692347_148692832 | chr2 | 148692348 | 148692832 | + | 1.38083 | 3' UTR (NM_028986, exon 6 of 6) | 3' UTR (NM_028986, exon 6 of 6) | 11470 | NM_028986 | 74533 | Mm.136913 | NM_028986 | ENSMUSG00000027439 | **Gzf1** | 8430437G08Rik\|Zfp336\|mGZF1 | GDNF-inducible zinc finger protein 1 | protein-coding |
| chr2_148686260_148686818 | chr2 | 148686261 | 148686818 | + | 1.31249 | exon (NM_028986, exon 3 of 6) | exon (NM_028986, exon 3 of 6) | 5419 | NM_028986 | 74533 | Mm.136913 | NM_028986 | ENSMUSG00000027439 | **Gzf1** | 8430437G08Rik\|Zfp336\|mGZF1 | GDNF-inducible zinc finger protein 1 | protein-coding |
| chr2_148685722_148685986 | chr2 | 148685723 | 148685986 | + | 1.22418 | intron (NM_028986, intron 2 of 5) | intron (NM_028986, intron 2 of 5) | 4734 | NM_028986 | 74533 | Mm.136913 | NM_028986 | ENSMUSG00000027439 | **Gzf1** | 8430437G08Rik\|Zfp336\|mGZF1 | GDNF-inducible zinc finger protein 1 | protein-coding |
| chr2_148683113_148683290 | chr2 | 148683114 | 148683290 | + | 1.01162 | intron (NM_028986, intron 1 of 5) | B3A\|SINE\|B2 | 2082 | NM_028986 | 74533 | Mm.136913 | NM_028986 | ENSMUSG00000027439 | **Gzf1** | 8430437G08Rik\|Zfp336\|mGZF1 | GDNF-inducible zinc finger protein 1 | protein-coding |
| chr2_148689452_148690062 | chr2 | 148689453 | 148690062 | + | 0.918649 | intron (NM_028986, intron 4 of 5) | intron (NM_028986, intron 4 of 5) | 8637 | NM_028986 | 74533 | Mm.136913 | NM_028986 | ENSMUSG00000027439 | **Gzf1** | 8430437G08Rik\|Zfp336\|mGZF1 | GDNF-inducible zinc finger protein 1 | protein-coding |
| chr13_113134510_113135277 | chr13 | 113134511 | 113135277 | + | 1.01615 | Intergenic | Intergenic | -33913 | NM_010370 | 14938 | Mm.15510 | NM_010370 | ENSMUSG00000023132 | **Gzma** | AW494114\|Ctla-3\|Ctla3\|Hf\|SE1\|TSP-1\|TSP1 | granzyme A | protein-coding |
| chr13_113134863_113135211 | chr13 | 113134864 | 113135211 | + | 0.951806 | Intergenic | Intergenic | -34056 | NM_010370 | 14938 | Mm.15510 | NM_010370 | ENSMUSG00000023132 | **Gzma** | AW494114\|Ctla-3\|Ctla3\|Hf\|SE1\|TSP-1\|TSP1 | granzyme A | protein-coding |
| chr13_113120223_113121015 | chr13 | 113120224 | 113121015 | + | 0.630647 | Intergenic | Intergenic | -19638 | NM_010370 | 14938 | Mm.15510 | NM_010370 | ENSMUSG00000023132 | **Gzma** | AW494114\|Ctla-3\|Ctla3\|Hf\|SE1\|TSP-1\|TSP1 | granzyme A | protein-coding |
| chr14_56281557_56282007 | chr14 | 56281558 | 56282007 | + | 1.73231 | Intergenic | Intergenic | -19522 | NM_013542 | 14939 | Mm.14874 | NM_013542 | ENSMUSG00000015437 | **Gzmb** | AI553453\|CCP-1/C11\|CCP1\|Ctla-1\|Ctla1\|GZB | granzyme B | protein-coding |
| chr14_56261818_56262843 | chr14 | 56261819 | 56262843 | + | 0.849498 | promoter-TSS (NM_013542) | promoter-TSS (NM_013542) | -71 | NM_013542 | 14939 | Mm.14874 | NM_013542 | ENSMUSG00000015437 | **Gzmb** | AI553453\|CCP-1/C11\|CCP1\|Ctla-1\|Ctla1\|GZB | granzyme B | protein-coding |
| chr14_56275673_56276521 | chr14 | 56275674 | 56276521 | + | 0.838804 | Intergenic | Intergenic | -13837 | NM_013542 | 14939 | Mm.14874 | NM_013542 | ENSMUSG00000015437 | **Gzmb** | AI553453\|CCP-1/C11\|CCP1\|Ctla-1\|Ctla1\|GZB | granzyme B | protein-coding |
| chr14_56261981_56262558 | chr14 | 56261982 | 56262558 | + | 0.759039 | promoter-TSS (NM_013542) | promoter-TSS (NM_013542) | -10 | NM_013542 | 14939 | Mm.14874 | NM_013542 | ENSMUSG00000015437 | **Gzmb** | AI553453\|CCP-1/C11\|CCP1\|Ctla-1\|Ctla1\|GZB | granzyme B | protein-coding |
| chr14_56291333_56291625 | chr14 | 56291334 | 56291625 | + | 0.727035 | Intergenic | Lx6\|LINE\|L1 | -29219 | NM_013542 | 14939 | Mm.14874 | NM_013542 | ENSMUSG00000015437 | **Gzmb** | AI553453\|CCP-1/C11\|CCP1\|Ctla-1\|Ctla1\|GZB | granzyme B | protein-coding |
| chr14_56310251_56310633 | chr14 | 56310252 | 56310633 | + | 0.645787 | Intergenic | Intergenic | -48182 | NM_013542 | 14939 | Mm.14874 | NM_013542 | ENSMUSG00000015437 | **Gzmb** | AI553453\|CCP-1/C11\|CCP1\|Ctla-1\|Ctla1\|GZB | granzyme B | protein-coding |
| chr14_56205640_56206029 | chr14 | 56205641 | 56206029 | + | 0.811376 | intron (NM_010374, intron 4 of 4) | intron (NM_010374, intron 4 of 4) | 5572 | NM_010374 | 14943 | Mm.457976 | NM_010374 | ENSMUSG00000015441 | **Gzmf** | CCP4\|Ctla-7\|Ctla7\|MCSP-3 | granzyme F | protein-coding |
| chr14_56205083_56205394 | chr14 | 56205084 | 56205394 | + | 0.653763 | TTS (NM_010374) | TTS (NM_010374) | 6168 | NM_010374 | 14943 | Mm.457976 | NM_010374 | ENSMUSG00000015441 | **Gzmf** | CCP4\|Ctla-7\|Ctla7\|MCSP-3 | granzyme F | protein-coding |
| chr13_113183263_113183776 | chr13 | 113183264 | 113183776 | + | 0.769816 | Intergenic | Intergenic | -2623 | NM_008196 | 14945 | Mm.56993 | NM_008196 | | **Gzmk** | - | granzyme K | protein-coding |
| chr13_113183263_113183974 | chr13 | 113183264 | 113183974 | + | 0.763821 | Intergenic | Intergenic | -2722 | NM_008196 | 14945 | Mm.56993 | NM_008196 | | **Gzmk** | - | granzyme K | protein-coding |
| chr2_152649912_152650433 | chr2 | 152649913 | 152650433 | + | 0.734467 | Intergenic | Intergenic | -19288 | NM_010376 | 14950 | Mm.277327 | NM_010376 | ENSMUSG00000019188 | **H13** | 1200006O09Rik\|4930443L17Rik\|5031424B04Rik\|AV020344\|H-13\|Hm13\|PSL3\|Spp | histocompatibility 13 | protein-coding |
| chr6_115945647_115946129 | chr6 | 115945648 | 115946129 | + | 1.34507 | intron (NM_138311, intron 1 of 4) | intron (NM_138311, intron 1 of 4) | 949 | NM_138311 | 171506 | Mm.21899 | NM_138311 | ENSMUSG00000042279 | **H1foo** | C86609\|H1fo\|H1oo | H1 histone family, member O, oocyte-specific | protein-coding |
| chr6_115944078_115944428 | chr6 | 115944079 | 115944428 | + | 0.97871 | promoter-TSS (NM_138311) | promoter-TSS (NM_138311) | -686 | NM_138311 | 171506 | Mm.21899 | NM_138311 | ENSMUSG00000042279 | **H1foo** | C86609\|H1fo\|H1oo | H1 histone family, member O, oocyte-specific | protein-coding |
| chr6_115955739_115956130 | chr6 | 115955740 | 115956130 | + | 0.875797 | intron (NM_026376, intron 35 of 35) | intron (NM_026376, intron 35 of 35) | 10996 | NM_138311 | 171506 | Mm.21899 | NM_138311 | ENSMUSG00000042279 | **H1foo** | C86609\|H1fo\|H1oo | H1 histone family, member O, oocyte-specific | protein-coding |
| chr6_115955542_115956255 | chr6 | 115955543 | 115956255 | + | 0.686529 | intron (NM_026376, intron 35 of 35) | intron (NM_026376, intron 35 of 35) | 10960 | NM_138311 | 171506 | Mm.21899 | NM_138311 | ENSMUSG00000042279 | **H1foo** | C86609\|H1fo\|H1oo | H1 histone family, member O, oocyte-specific | protein-coding |
| chr17_34150169_34150456 | chr17 | 34150170 | 34150456 | + | 0.668965 | intron (NM_010388, intron 3 of 5) | intron (NM_010388, intron 3 of 5) | -2878 | NM_010387 | 14999 | Mm.460938 | NM_010387 | ENSMUSG00000079547 | **H2-DMb1** | AI385589\|H-2Mb1\|H2-Mb1 | histocompatibility 2, class II, locus Mb1 | protein-coding |
| chr17_34134381_34134716 | chr17 | 34134382 | 34134716 | + | 0.971676 | intron (NM_010386, intron 1 of 5) | Tigger7\|DNA\|TcMar-Tigger | -10867 | NM_010388 | 15000 | Mm.195060 | NM_010388 | | **H2-DMb2** | H-2Mb2\|H2-Mb2 | histocompatibility 2, class II, locus Mb2 | protein-coding |
| chr17_34068841_34069943 | chr17 | 34068842 | 34069943 | + | 0.834274 | Intergenic | Intergenic | -22987 | NM_008206 | 15001 | Mm.116 | NM_008206 | ENSMUSG00000024334 | **H2-Oa** | H-2Oa | histocompatibility 2, O region alpha locus | protein-coding |
| chr17_35322867_35323225 | chr17 | 35322868 | 35323225 | + | 0.654366 | intron (NM_010390, intron 3 of 7) | intron (NM_010390, intron 3 of 7) | 2488 | NM_010390 | 15006 | Mm.466882 | NM_010390 | ENSMUSG00000079507 | **H2-Q1** | H-2Q1\|Q1\|Q1b\|Q1d\|Q1k\|Qa-1\|Qa1\|Qed-1 | histocompatibility 2, Q region locus 1 | protein-coding |
| chr17_35468938_35469351 | chr17 | 35468939 | 35469351 | + | 0.911175 | promoter-TSS (NM_010391) | promoter-TSS (NM_010391) | -944 | NM_010391 | 15007 | Mm.88795 | NM_010391 | ENSMUSG00000067235 | **H2-Q10** | H-2Q10\|Q10\|Qa10 | histocompatibility 2, Q region locus 10 | protein-coding |
| chr17_36025920_36026371 | chr17 | 36025921 | 36026371 | + | 1.09597 | intron (NR_015618, intron 1 of 1) | MTD\|LTR\|ERVL-MaLR | -5586 | NM_008207 | 15042 | Mm.439654 | NM_008207 | ENSMUSG00000053835 | **H2-T24** | H-2T24 | histocompatibility 2, T region locus 24 | protein-coding |
| chr17_36024926_36025225 | chr17 | 36024927 | 36025225 | + | 0.894939 | intron (NR_015618, intron 1 of 1) | intron (NR_015618, intron 1 of 1) | -4516 | NM_008207 | 15042 | Mm.439654 | NM_008207 | ENSMUSG00000053835 | **H2-T24** | H-2T24 | histocompatibility 2, T region locus 24 | protein-coding |
| chr2_17889281_17890139 | chr2 | 17889282 | 17890139 | + | 0.825087 | Intergenic | Intergenic | 107236 | NM_026627 | 68231 | Mm.23887 | NM_026627 | ENSMUSG00000062651 | **H2afb1** | 1700113O17Rik\|H2A.Bbd\|H2al2\|TSEG-1\|Tseg1 | H2A histone family, member B1 | protein-coding |
| chr13_56115941_56116655 | chr13 | 56115942 | 56116655 | + | 1.08918 | intron (NM_001159514, intron 2 of 8) | intron (NM_001159514, intron 2 of 8) | 19252 | NM_012015 | 26914 | Mm.283802 | NM_012015 | ENSMUSG00000015937 | **H2afy** | H2AF12M\|mH2a1\|macroH2A1 | H2A histone family, member Y | protein-coding |
| chr13_56142446_56143327 | chr13 | 56142447 | 56143327 | + | 0.678248 | Intergenic | Intergenic | -7337 | NM_001159513 | 26914 | Mm.283802 | NM_012015 | ENSMUSG00000015937 | **H2afy** | H2AF12M\|mH2a1\|macroH2A1 | H2A histone family, member Y | protein-coding |
| chr15_62268081_62268634 | chr15 | 62268082 | 62268634 | + | 1.88721 | Intergenic | Intergenic | -48907 | NR_003523 | 67552 | Mm.484027 | NM_026230 | | **H2afy3** | 4933432H23Rik | H2A histone family, member Y3 | pseudo |
| chr15_62239818_62240475 | chr15 | 62239819 | 62240475 | + | 1.70422 | intron (NR_003368, intron 5 of 8) | intron (NR_003368, intron 5 of 8) | -20696 | NR_003523 | 67552 | Mm.484027 | NM_026230 | | **H2afy3** | 4933432H23Rik | H2A histone family, member Y3 | pseudo |
| chr15_62271204_62271894 | chr15 | 62271205 | 62271894 | + | 1.68834 | Intergenic | L1ME2\|LINE\|L1 | -52098 | NR_003523 | 67552 | Mm.484027 | NM_026230 | | **H2afy3** | 4933432H23Rik | H2A histone family, member Y3 | pseudo |
| chr15_62726553_62727066 | chr15 | 62726554 | 62727066 | + | 1.68223 | Intergenic | Intergenic | -507359 | NR_003523 | 67552 | Mm.484027 | NM_026230 | | **H2afy3** | 4933432H23Rik | H2A histone family, member Y3 | pseudo |
| chr15_62189564_62190857 | chr15 | 62189565 | 62190857 | + | 1.68137 | intron (NR_003368, intron 5 of 8) | intron (NR_003368, intron 5 of 8) | 29240 | NR_003523 | 67552 | Mm.484027 | NM_026230 | | **H2afy3** | 4933432H23Rik | H2A histone family, member Y3 | pseudo |
| chr15_62374420_62375085 | chr15 | 62374421 | 62375085 | + | 1.56385 | Intergenic | Intergenic | -155302 | NR_003523 | 67552 | Mm.484027 | NM_026230 | | **H2afy3** | 4933432H23Rik | H2A histone family, member Y3 | pseudo |
| chr15_62568832_62569984 | chr15 | 62568833 | 62569984 | + | 1.52615 | Intergenic | Intergenic | -349957 | NR_003523 | 67552 | Mm.484027 | NM_026230 | | **H2afy3** | 4933432H23Rik | H2A histone family, member Y3 | pseudo |
| chr15_62244553_62245914 | chr15 | 62244554 | 62245914 | + | 1.48278 | intron (NR_003368, intron 8 of 8) | Lx8\|LINE\|L1 | -25783 | NR_003523 | 67552 | Mm.484027 | NM_026230 | | **H2afy3** | 4933432H23Rik | H2A histone family, member Y3 | pseudo |
| chr15_62325346_62326220 | chr15 | 62325347 | 62326220 | + | 1.40684 | Intergenic | Intergenic | -106332 | NR_003523 | 67552 | Mm.484027 | NM_026230 | | **H2afy3** | 4933432H23Rik | H2A histone family, member Y3 | pseudo |
| chr15_62245381_62245677 | chr15 | 62245382 | 62245677 | + | 1.3581 | intron (NR_003368, intron 8 of 8) | intron (NR_003368, intron 8 of 8) | -26078 | NR_003523 | 67552 | Mm.484027 | NM_026230 | | **H2afy3** | 4933432H23Rik | H2A histone family, member Y3 | pseudo |
| chr15_62833013_62833570 | chr15 | 62833014 | 62833570 | + | 1.34431 | Intergenic | Intergenic | -613841 | NR_003523 | 67552 | Mm.484027 | NM_026230 | | **H2afy3** | 4933432H23Rik | H2A histone family, member Y3 | pseudo |

| | | | | | | | | | | | | | | | | | |
|---|---|---|---|---|---|---|---|---|---|---|---|---|---|---|---|---|---|
| chr15_62145027_62145566 | chr15 | 62145028 | 62145566 | + | 1.30876 | intron (NR_003368, intron 2 of 8) | intron (NR_003368, intron 2 of 8) | 74154 | NR_003523 | 67552 | Mm.484027 | NM_026230 | | **H2afy3** | 4933432H23Rik | H2A histone family, member Y3 | pseudo |
| chr15_62832770_62833581 | chr15 | 62832771 | 62833581 | + | 1.30634 | Intergenic | Intergenic | -613725 | NR_003523 | 67552 | Mm.484027 | NM_026230 | | **H2afy3** | 4933432H23Rik | H2A histone family, member Y3 | pseudo |
| chr15_62393005_62393906 | chr15 | 62393006 | 62393906 | + | 1.30412 | Intergenic | ID4_\|SINE\|ID | -174005 | NR_003523 | 67552 | Mm.484027 | NM_026230 | | **H2afy3** | 4933432H23Rik | H2A histone family, member Y3 | pseudo |
| chr15_62787490_62788202 | chr15 | 62787491 | 62788202 | + | 1.30086 | Intergenic | Intergenic | -568395 | NR_003523 | 67552 | Mm.484027 | NM_026230 | | **H2afy3** | 4933432H23Rik | H2A histone family, member Y3 | pseudo |
| chr15_62288241_62288728 | chr15 | 62288242 | 62288728 | + | 1.26512 | Intergenic | L1M4a2\|LINE\|L1 | -69034 | NR_003523 | 67552 | Mm.484027 | NM_026230 | | **H2afy3** | 4933432H23Rik | H2A histone family, member Y3 | pseudo |
| chr15_62821684_62822087 | chr15 | 62821685 | 62822087 | + | 1.21154 | Intergenic | Intergenic | -602435 | NR_003523 | 67552 | Mm.484027 | NM_026230 | | **H2afy3** | 4933432H23Rik | H2A histone family, member Y3 | pseudo |
| chr15_62366266_62366655 | chr15 | 62366267 | 62366655 | + | 1.16045 | Intergenic | MIR\|SINE\|MIR | -147010 | NR_003523 | 67552 | Mm.484027 | NM_026230 | | **H2afy3** | 4933432H23Rik | H2A histone family, member Y3 | pseudo |
| chr15_62366263_62367097 | chr15 | 62366264 | 62367097 | + | 1.15471 | Intergenic | Intergenic | -147229 | NR_003523 | 67552 | Mm.484027 | NM_026230 | | **H2afy3** | 4933432H23Rik | H2A histone family, member Y3 | pseudo |
| chr15_62823798_62824466 | chr15 | 62823799 | 62824466 | + | 1.15053 | Intergenic | Intergenic | -604681 | NR_003523 | 67552 | Mm.484027 | NM_026230 | | **H2afy3** | 4933432H23Rik | H2A histone family, member Y3 | pseudo |
| chr15_62838202_62838683 | chr15 | 62838203 | 62838683 | + | 1.10167 | Intergenic | Intergenic | -618992 | NR_003523 | 67552 | Mm.484027 | NM_026230 | | **H2afy3** | 4933432H23Rik | H2A histone family, member Y3 | pseudo |
| chr15_62734753_62735614 | chr15 | 62734754 | 62735614 | + | 1.04105 | Intergenic | Intergenic | -515733 | NR_003523 | 67552 | Mm.484027 | NM_026230 | | **H2afy3** | 4933432H23Rik | H2A histone family, member Y3 | pseudo |
| chr15_62734748_62735947 | chr15 | 62734749 | 62735947 | + | 1.01428 | Intergenic | Intergenic | -515897 | NR_003523 | 67552 | Mm.484027 | NM_026230 | | **H2afy3** | 4933432H23Rik | H2A histone family, member Y3 | pseudo |
| chr15_62804417_62804838 | chr15 | 62804418 | 62804838 | + | 0.983445 | Intergenic | Intergenic | -585177 | NR_003523 | 67552 | Mm.484027 | NM_026230 | | **H2afy3** | 4933432H23Rik | H2A histone family, member Y3 | pseudo |
| chr15_62273828_62274485 | chr15 | 62273829 | 62274485 | + | 0.968619 | Intergenic | MLT1J1\|LTR\|ERVL-MaLR | -54706 | NR_003523 | 67552 | Mm.484027 | NM_026230 | | **H2afy3** | 4933432H23Rik | H2A histone family, member Y3 | pseudo |
| chr15_62824644_62825825 | chr15 | 62824645 | 62825825 | + | 0.952287 | Intergenic | Intergenic | -605784 | NR_003523 | 67552 | Mm.484027 | NM_026230 | | **H2afy3** | 4933432H23Rik | H2A histone family, member Y3 | pseudo |
| chr15_62261358_62261823 | chr15 | 62261359 | 62261823 | + | 0.931515 | Intergenic | Intergenic | -42140 | NR_003523 | 67552 | Mm.484027 | NM_026230 | | **H2afy3** | 4933432H23Rik | H2A histone family, member Y3 | pseudo |
| chr15_62807394_62807774 | chr15 | 62807395 | 62807774 | + | 0.918938 | Intergenic | Intergenic | -588133 | NR_003523 | 67552 | Mm.484027 | NM_026230 | | **H2afy3** | 4933432H23Rik | H2A histone family, member Y3 | pseudo |
| chr15_62286781_62287693 | chr15 | 62286782 | 62287693 | + | 0.916328 | Intergenic | Intergenic | -67786 | NR_003523 | 67552 | Mm.484027 | NM_026230 | | **H2afy3** | 4933432H23Rik | H2A histone family, member Y3 | pseudo |
| chr15_62458021_62458525 | chr15 | 62458022 | 62458525 | + | 0.897206 | Intergenic | Intergenic | -238822 | NR_003523 | 67552 | Mm.484027 | NM_026230 | | **H2afy3** | 4933432H23Rik | H2A histone family, member Y3 | pseudo |
| chr15_62786365_62786737 | chr15 | 62786366 | 62786737 | + | 0.896638 | Intergenic | Intergenic | -567100 | NR_003523 | 67552 | Mm.484027 | NM_026230 | | **H2afy3** | 4933432H23Rik | H2A histone family, member Y3 | pseudo |
| chr15_62265437_62265876 | chr15 | 62265438 | 62265876 | + | 0.881597 | Intergenic | Intergenic | -46206 | NR_003523 | 67552 | Mm.484027 | NM_026230 | | **H2afy3** | 4933432H23Rik | H2A histone family, member Y3 | pseudo |
| chr15_62839466_62839762 | chr15 | 62839467 | 62839762 | + | 0.838386 | Intergenic | LTR40a\|LTR\|ERVL | -620163 | NR_003523 | 67552 | Mm.484027 | NM_026230 | | **H2afy3** | 4933432H23Rik | H2A histone family, member Y3 | pseudo |
| chr15_62358346_62358897 | chr15 | 62358347 | 62358897 | + | 0.825814 | Intergenic | Intergenic | -139171 | NR_003523 | 67552 | Mm.484027 | NM_026230 | | **H2afy3** | 4933432H23Rik | H2A histone family, member Y3 | pseudo |
| chr15_62835278_62835583 | chr15 | 62835279 | 62835583 | + | 0.806341 | Intergenic | Intergenic | -615980 | NR_003523 | 67552 | Mm.484027 | NM_026230 | | **H2afy3** | 4933432H23Rik | H2A histone family, member Y3 | pseudo |
| chr15_62173321_62173555 | chr15 | 62173322 | 62173555 | + | 0.774564 | intron (NR_003368, intron 4 of 8) | intron (NR_003368, intron 4 of 8) | 46013 | NR_003523 | 67552 | Mm.484027 | NM_026230 | | **H2afy3** | 4933432H23Rik | H2A histone family, member Y3 | pseudo |
| chr15_62438785_62439331 | chr15 | 62438786 | 62439331 | + | 0.764968 | Intergenic | Intergenic | -219607 | NR_003523 | 67552 | Mm.484027 | NM_026230 | | **H2afy3** | 4933432H23Rik | H2A histone family, member Y3 | pseudo |
| chr15_62273901_62274169 | chr15 | 62273902 | 62274169 | + | 0.72274 | Intergenic | Intergenic | -54584 | NR_003523 | 67552 | Mm.484027 | NM_026230 | | **H2afy3** | 4933432H23Rik | H2A histone family, member Y3 | pseudo |
| chr15_62222779_62223165 | chr15 | 62222780 | 62223165 | + | 0.690987 | intron (NR_003368, intron 5 of 8) | MIR\|SINE\|MIR | -3521 | NR_003523 | 67552 | Mm.484027 | NM_026230 | | **H2afy3** | 4933432H23Rik | H2A histone family, member Y3 | pseudo |
| chr15_62483602_62483931 | chr15 | 62483603 | 62483931 | + | 0.664903 | Intergenic | Intergenic | -264316 | NR_003523 | 67552 | Mm.484027 | NM_026230 | | **H2afy3** | 4933432H23Rik | H2A histone family, member Y3 | pseudo |
| chr3_137820541_137821527 | chr3 | 137820542 | 137821527 | + | 0.955883 | Intergenic | Intergenic | -43453 | NM_016750 | 51788 | Mm.117541 | NM_016750 | ENSMUSG00000037894 | **H2afz** | H2A.Z\|H2a.z-1 | H2A histone family, member Z | protein-coding |
| chr10_3238403_3238817 | chr10 | 3238404 | 3238817 | + | 0.760252 | Intergenic | Intergenic | 29161 | NM_001204916 | 670558 | Mm.398582 | NM_001204916 | ENSMUSG00000091618 | **H60c** | 4632413I24Rik\|Gm9491 | histocompatibility 60c | protein-coding |
| chr4_150012041_150012347 | chr4 | 150012042 | 150012347 | + | 0.744156 | Intergenic | Intergenic | -3171 | NM_173371 | 100198 | Mm.22183 | NM_173371 | ENSMUSG00000028980 | **H6pd** | AI785303\|G6pd1\|Gpd-1\|Gpd1 | hexose-6-phosphate dehydrogenase (glucose 1-dehydrogenase) | protein-coding |
| chr17_83816963_83817327 | chr17 | 83816964 | 83817327 | + | 0.913178 | Intergenic | Intergenic | 29645 | NM_025325 | 107766 | Mm.30100 | NM_025325 | ENSMUSG00000000673 | **Haao** | 0610007K21Rik\|0610012J07Rik\|3-HAO\|3-HAOxase\|3HAO | 3-hydroxyanthranilate 3,4-dioxygenase | protein-coding |
| chr17_83850379_83851325 | chr17 | 83850380 | 83851325 | + | 0.772682 | Intergenic | Intergenic | -4062 | NM_025325 | 107766 | Mm.30100 | NM_025325 | ENSMUSG00000000673 | **Haao** | 0610007K21Rik\|0610012J07Rik\|3-HAO\|3-HAOxase\|3HAO | 3-hydroxyanthranilate 3,4-dioxygenase | protein-coding |
| chr17_83854347_83854811 | chr17 | 83854348 | 83854811 | + | 0.620138 | Intergenic | Intergenic | -7789 | NM_025325 | 107766 | Mm.30100 | NM_025325 | ENSMUSG00000000673 | **Haao** | 0610007K21Rik\|0610012J07Rik\|3-HAO\|3-HAOxase\|3HAO | 3-hydroxyanthranilate 3,4-dioxygenase | protein-coding |
| chr3_131250274_131250694 | chr3 | 131250275 | 131250694 | + | 0.716099 | intron (NM_008212, intron 1 of 7) | intron (NM_008212, intron 1 of 7) | 21617 | NM_008212 | 15107 | Mm.260164 | NM_008212 | ENSMUSG00000027984 | **Hadh** | AA409008\|AU019341\|AW742602\|HCDH\|Hadhsc\|Schad | hydroxyacyl-Coenzyme A dehydrogenase | protein-coding |
| chr3_131250248_131250816 | chr3 | 131250249 | 131250816 | + | 0.687241 | intron (NM_008212, intron 1 of 7) | intron (NM_008212, intron 1 of 7) | 21569 | NM_008212 | 15107 | Mm.260164 | NM_008212 | ENSMUSG00000027984 | **Hadh** | AA409008\|AU019341\|AW742602\|HCDH\|Hadhsc\|Schad | hydroxyacyl-Coenzyme A dehydrogenase | protein-coding |
| chr7_30935419_30936024 | chr7 | 30935420 | 30936024 | + | 0.698367 | Intergenic | MTA_Mm\|LTR\|ERVL-MaLR | 8295 | NM_032541 | 84506 | Mm.439939 | NM_032541 | ENSMUSG00000050440 | **Hamp** | Hamp1\|Hepc\|Hepc1 | hepcidin antimicrobial peptide | protein-coding |
| chr11_57911541_57912289 | chr11 | 57911542 | 57912289 | + | 1.01989 | Intergenic | Intergenic | -79768 | NM_008213 | 15110 | Mm.4746 | NM_008213 | ENSMUSG00000037335 | **Hand1** | Ehand1\|Hxt\|Th1\|Thing1\|bHLHa27\|eHAND | heart and neural crest derivatives expressed transcript 1 | protein-coding |
| chr11_57917891_57918300 | chr11 | 57917892 | 57918300 | + | 0.753972 | Intergenic | B3A\|SINE\|B2 | -85949 | NM_008213 | 15110 | Mm.4746 | NM_008213 | ENSMUSG00000037335 | **Hand1** | Ehand1\|Hxt\|Th1\|Thing1\|bHLHa27\|eHAND | heart and neural crest derivatives expressed transcript 1 | protein-coding |
| chr2_134545018_134545587 | chr2 | 134545019 | 134545587 | + | 0.711216 | intron (NM_010403, intron 2 of 7) | intron (NM_010403, intron 2 of 7) | 9049 | NM_010403 | 15112 | Mm.26634 | NM_010403 | ENSMUSG00000027261 | **Hao1** | GOX\|Gox1\|Hao-1 | hydroxyacid oxidase 1, liver | protein-coding |
| chr13_89521441_89522119 | chr13 | 89521442 | 89522119 | + | 1.64791 | Intergenic | L1M4\|LINE\|L1 | -18856 | NM_013500 | 12950 | Mm.266790 | NM_013500 | ENSMUSG00000021613 | **Hapln1** | BB099155\|CLP\|Crtl1\|Crtl1l\|LP\|LP-1 | hyaluronan and proteoglycan link protein 1 | protein-coding |
| chr13_89324096_89324771 | chr13 | 89324097 | 89324771 | + | 1.29928 | Intergenic | Intergenic | -216202 | NM_013500 | 12950 | Mm.266790 | NM_013500 | ENSMUSG00000021613 | **Hapln1** | BB099155\|CLP\|Crtl1\|Crtl1l\|LP\|LP-1 | hyaluronan and proteoglycan link protein 1 | protein-coding |
| chr13_89453539_89454191 | chr13 | 89453540 | 89454191 | + | 1.2895 | Intergenic | Intergenic | -86771 | NM_013500 | 12950 | Mm.266790 | NM_013500 | ENSMUSG00000021613 | **Hapln1** | BB099155\|CLP\|Crtl1\|Crtl1l\|LP\|LP-1 | hyaluronan and proteoglycan link protein 1 | protein-coding |
| chr13_89454674_89455006 | chr13 | 89454675 | 89455006 | + | 0.969509 | Intergenic | B1F\|SINE\|Alu | -85796 | NM_013500 | 12950 | Mm.266790 | NM_013500 | ENSMUSG00000021613 | **Hapln1** | BB099155\|CLP\|Crtl1\|Crtl1l\|LP\|LP-1 | hyaluronan and proteoglycan link protein 1 | protein-coding |
| chr17_17845277_17846059 | chr17 | 17845278 | 17846059 | + | 1.10376 | intron (NM_008215, intron 4 of 4) | intron (NM_008215, intron 4 of 4) | 9520 | NM_008215 | 15116 | Mm.255701 | NM_008215 | ENSMUSG00000003665 | **Has1** | HAS | hyaluronan synthase1 | protein-coding |
| chr8_106802747_106803157 | chr8 | 106802748 | 106803157 | + | 0.889576 | intron (NM_173037, intron 15 of 17) | intron (NM_173037, intron 15 of 17) | -67290 | NM_008217 | 15118 | Mm.56986 | NM_008217 | ENSMUSG00000031910 | **Has3** | - | hyaluronan synthase 3 | protein-coding |
| chr8_106794870_106795200 | chr8 | 106794871 | 106795200 | + | 0.756736 | intron (NM_173037, intron 15 of 17) | MLT1F2\|LTR\|ERVL-MaLR | -75207 | NM_008217 | 15118 | Mm.56986 | NM_008217 | ENSMUSG00000031910 | **Has3** | - | hyaluronan synthase 3 | protein-coding |
| chr8_106802755_106803413 | chr8 | 106802756 | 106803413 | + | 0.708935 | intron (NM_173037, intron 15 of 17) | intron (NM_173037, intron 15 of 17) | -67158 | NM_008217 | 15118 | Mm.56986 | NM_008217 | ENSMUSG00000031910 | **Has3** | - | hyaluronan synthase 3 | protein-coding |
| chr2_120658378_120658949 | chr2 | 120658379 | 120658949 | + | 1.57121 | Intergenic | Intergenic | 49069 | NM_001290807 | 66296 | Mm.221900 | NM_025475 | | **Haus2** | 1700101G24Rik\|2410007P03Rik\|AU020217\|Cep27 | HAUS augmin-like complex, subunit 2 | protein-coding |
| chr11_46474053_46474731 | chr11 | 46474054 | 46474731 | + | 1.56449 | intron (NM_134250, intron 5 of 6) | intron (NM_134250, intron 5 of 6) | 19461 | NM_134250 | 171285 | Mm.72168 | NM_134250 | ENSMUSG00000020399 | **Havcr2** | TIM-3\|Tim3\|Timd3 | hepatitis A virus cellular receptor 2 | protein-coding |
| chr11_46466777_46467364 | chr11 | 46466778 | 46467364 | + | 1.42058 | intron (NM_134250, intron 4 of 6) | intron (NM_134250, intron 4 of 6) | 12140 | NM_134250 | 171285 | Mm.72168 | NM_134250 | ENSMUSG00000020399 | **Havcr2** | TIM-3\|Tim3\|Timd3 | hepatitis A virus cellular receptor 2 | protein-coding |
| chr11_46461413_46462340 | chr11 | 46461414 | 46462340 | + | 1.08654 | intron (NM_134250, intron 3 of 6) | intron (NM_134250, intron 3 of 6) | 6946 | NM_134250 | 171285 | Mm.72168 | NM_134250 | ENSMUSG00000020399 | **Havcr2** | TIM-3\|Tim3\|Timd3 | hepatitis A virus cellular receptor 2 | protein-coding |
| chr11_46461824_46462339 | chr11 | 46461825 | 46462339 | + | 1.01289 | intron (NM_134250, intron 3 of 6) | intron (NM_134250, intron 3 of 6) | 7151 | NM_134250 | 171285 | Mm.72168 | NM_134250 | ENSMUSG00000020399 | **Havcr2** | TIM-3\|Tim3\|Timd3 | hepatitis A virus cellular receptor 2 | protein-coding |
| chr11_46461442_46461786 | chr11 | 46461443 | 46461786 | + | 0.978992 | intron (NM_134250, intron 3 of 6) | intron (NM_134250, intron 3 of 6) | 6683 | NM_134250 | 171285 | Mm.72168 | NM_134250 | ENSMUSG00000020399 | **Havcr2** | TIM-3\|Tim3\|Timd3 | hepatitis A virus cellular receptor 2 | protein-coding |
| chr11_46454581_46455122 | chr11 | 46454582 | 46455122 | + | 0.802096 | promoter-TSS (NM_134250) | promoter-TSS (NM_134250) | -79 | NM_134250 | 171285 | Mm.72168 | NM_134250 | ENSMUSG00000020399 | **Havcr2** | TIM-3\|Tim3\|Timd3 | hepatitis A virus cellular receptor 2 | protein-coding |
| chr18_36513888_36514341 | chr18 | 36513889 | 36514341 | + | 1.01614 | intron (NM_010415, intron 2 of 5) | intron (NM_010415, intron 2 of 5) | 1690 | NM_010415 | 15200 | Mm.289681 | NM_010415 | ENSMUSG00000024486 | **Hbegf** | AW047313\|Dtr\|Dts\|Hegfl | heparin-binding EGF-like growth factor | protein-coding |
| chr10_21277648_21278081 | chr10 | 21277649 | 21278081 | + | 1.08734 | Intergenic | Lx\|LINE\|L1 | -18114 | NM_001042593 | 56422 | Mm.25527 | NM_019702 | ENSMUSG00000019977 | **Hbs1l** | 2810035F15Rik\|AI326327\|eRFS | Hbs1-like (S. cerevisiae) | protein-coding |
| chr10_21318287_21319024 | chr10 | 21318288 | 21319024 | + | 0.93407 | TTS (NR_045352) | TTS (NR_045352) | 22677 | NM_001145209 | 56422 | Mm.25527 | NM_019702 | ENSMUSG00000019977 | **Hbs1l** | 2810035F15Rik\|AI326327\|eRFS | Hbs1-like (S. cerevisiae) | protein-coding |
| chr10_21246059_21246389 | chr10 | 21246060 | 21246389 | + | 0.740514 | Intergenic | Intergenic | -49755 | NM_001042593 | 56422 | Mm.25527 | NM_019702 | ENSMUSG00000019977 | **Hbs1l** | 2810035F15Rik\|AI326327\|eRFS | Hbs1-like (S. cerevisiae) | protein-coding |
| chr10_21244978_21245832 | chr10 | 21244979 | 21245832 | + | 0.650488 | Intergenic | Intergenic | -50574 | NM_001042593 | 56422 | Mm.25527 | NM_019702 | ENSMUSG00000019977 | **Hbs1l** | 2810035F15Rik\|AI326327\|eRFS | Hbs1-like (S. cerevisiae) | protein-coding |
| chr2_35016076_35016701 | chr2 | 35016077 | 35016701 | + | 1.1688 | intron (NM_010406, intron 22 of 40) | RMER15\|LTR\|ERVL | 45052 | NM_010406 | 15139 | Mm.2168 | NM_010406 | ENSMUSG00000026874 | **Hc** | C5\|C5a\|He | hemolytic complement | protein-coding |
| chr2_35023472_35023985 | chr2 | 35023473 | 35023985 | + | 1.072 | intron (NM_010406, intron 20 of 40) | intron (NM_010406, intron 20 of 40) | 37712 | NM_010406 | 15139 | Mm.2168 | NM_010406 | ENSMUSG00000026874 | **Hc** | C5\|C5a\|He | hemolytic complement | protein-coding |
| chrX_73985648_73986179 | chrX | 73985649 | 73986179 | + | 1.09818 | Intergenic | L1MB3\|LINE\|L1 | -19557 | NM_008224 | 15161 | Mm.491126 | NM_008224 | ENSMUSG00000031386 | **Hcfc1** | AW060991\|HCF\|HCF1 | host cell factor C1 | protein-coding |
| chrX_73976259_73976712 | chrX | 73976260 | 73976712 | + | 0.934643 | Intergenic | Intergenic | -10129 | NM_008224 | 15161 | Mm.491126 | NM_008224 | ENSMUSG00000031386 | **Hcfc1** | AW060991\|HCF\|HCF1 | host cell factor C1 | protein-coding |
| chrX_73989073_73989658 | chrX | 73989074 | 73989658 | + | 0.783888 | Intergenic | Intergenic | -23009 | NM_008224 | 15161 | Mm.491126 | NM_008224 | ENSMUSG00000031386 | **Hcfc1** | AW060991\|HCF\|HCF1 | host cell factor C1 | protein-coding |
| chrX_73990853_73991186 | chrX | 73990854 | 73991186 | + | 0.692363 | Intergenic | Intergenic | -24663 | NM_008224 | 15161 | Mm.491126 | NM_008224 | ENSMUSG00000031386 | **Hcfc1** | AW060991\|HCF\|HCF1 | host cell factor C1 | protein-coding |
| chr2_153121853_153122252 | chr2 | 153121854 | 153122252 | + | 0.795125 | intron (NM_001172117, intron 1 of 12) | intron (NM_001172117, intron 1 of 12) | 13585 | NM_010407 | 15162 | Mm.715 | NM_010407 | ENSMUSG00000003283 | **Hck** | AI849071\|Bmk | hemopoietic cell kinase | protein-coding |

| | | | | | | | | | | | | | | | | | |
|---|---|---|---|---|---|---|---|---|---|---|---|---|---|---|---|---|---|
| chr2_153123022_153123302 | chr2 | 153123023 | 153123302 | + | 0.673703 | intron (NM_001172117, intron 1 of 12) | PB1D10\|SINE\|Alu | 14694 | NM_010407 | 15162 | Mm.715 | NM_010407 | ENSMUSG00000003283 | **Hck** | AI849071\|Bmk | hemopoietic cell kinase | protein-coding |
| chr2_153125831_153126292 | chr2 | 153125832 | 153126292 | + | 0.647989 | intron (NM_001172117, intron 2 of 12) | intron (NM_001172117, intron 2 of 12) | 17594 | NM_010407 | 15162 | Mm.715 | NM_010407 | ENSMUSG00000003283 | **Hck** | AI849071\|Bmk | hemopoietic cell kinase | protein-coding |
| chr13_117602261_117602571 | chr13 | 117602262 | 117602571 | + | 0.688426 | promoter-TSS (NM_010408) | promoter-TSS (NM_010408) | 96 | NM_010408 | 15165 | Mm.343429 | NM_010408 | | **Hcn1** | Bcng1\|C630013B14Rik\|HAC2 | hyperpolarization-activated, cyclic nucleotide-gated K+ 1 | protein-coding |
| chr13_117602122_117602812 | chr13 | 117602123 | 117602812 | + | 0.616738 | 5' UTR (NM_010408, exon 1 of 8) | 5' UTR (NM_010408, exon 1 of 8) | 147 | NM_010408 | 15165 | Mm.343429 | NM_010408 | | **Hcn1** | Bcng1\|C630013B14Rik\|HAC2 | hyperpolarization-activated, cyclic nucleotide-gated K+ 1 | protein-coding |
| chr10_36987286_36987625 | chr10 | 36987287 | 36987625 | + | 1.17129 | intron (NM_008229, intron 3 of 13) | intron (NM_008229, intron 3 of 13) | 12912 | NM_008229 | 15182 | Mm.19806 | NM_008229 | ENSMUSG00000019777 | **Hdac2** | D10Wsu179e\|YAF1\|Yy1bp\|mRPD3 | histone deacetylase 2 | protein-coding |
| chr10_36985993_36986381 | chr10 | 36985994 | 36986381 | + | 0.651526 | intron (NM_008229, intron 2 of 13) | intron (NM_008229, intron 2 of 13) | 11643 | NM_008229 | 15182 | Mm.19806 | NM_008229 | ENSMUSG00000019777 | **Hdac2** | D10Wsu179e\|YAF1\|Yy1bp\|mRPD3 | histone deacetylase 2 | protein-coding |
| chr1_92082584_92083576 | chr1 | 92082585 | 92083576 | + | 0.993966 | intron (NM_207225, intron 1 of 25) | intron (NM_207225, intron 1 of 25) | 65313 | NM_207225 | 208727 | Mm.318567 | NM_207225 | ENSMUSG00000026313 | **Hdac4** | 4932408F19Rik\|AI047285 | histone deacetylase 4 | protein-coding |
| chr1_92003814_92004627 | chr1 | 92003815 | 92004627 | + | 0.940456 | intron (NM_207225, intron 4 of 25) | intron (NM_207225, intron 4 of 25) | 144172 | NM_207225 | 208727 | Mm.318567 | NM_207225 | ENSMUSG00000026313 | **Hdac4** | 4932408F19Rik\|AI047285 | histone deacetylase 4 | protein-coding |
| chr1_92015189_92015693 | chr1 | 92015190 | 92015693 | + | 0.872576 | intron (NM_207225, intron 3 of 25) | intron (NM_207225, intron 3 of 25) | 132952 | NM_207225 | 208727 | Mm.318567 | NM_207225 | ENSMUSG00000026313 | **Hdac4** | 4932408F19Rik\|AI047285 | histone deacetylase 4 | protein-coding |
| chr1_92015157_92015997 | chr1 | 92015158 | 92015997 | + | 0.756856 | intron (NM_207225, intron 3 of 25) | intron (NM_207225, intron 3 of 25) | 132816 | NM_207225 | 208727 | Mm.318567 | NM_207225 | ENSMUSG00000026313 | **Hdac4** | 4932408F19Rik\|AI047285 | histone deacetylase 4 | protein-coding |
| chr1_92128379_92128683 | chr1 | 92128380 | 92128683 | + | 0.726082 | intron (NM_207225, intron 1 of 25) | intron (NM_207225, intron 1 of 25) | 19862 | NM_207225 | 208727 | Mm.318567 | NM_207225 | ENSMUSG00000026313 | **Hdac4** | 4932408F19Rik\|AI047285 | histone deacetylase 4 | protein-coding |
| chr1_92118956_92119834 | chr1 | 92118957 | 92119834 | + | 0.636765 | intron (NM_207225, intron 1 of 25) | intron (NM_207225, intron 1 of 25) | 28998 | NM_207225 | 208727 | Mm.318567 | NM_207225 | ENSMUSG00000026313 | **Hdac4** | 4932408F19Rik\|AI047285 | histone deacetylase 4 | protein-coding |
| chrX_102446906_102447450 | chrX | 102446907 | 102447450 | + | 0.752968 | intron (NM_027382, intron 4 of 10) | intron (NM_027382, intron 4 of 10) | 57970 | NM_027382 | 70315 | Mm.328128 | NM_027382 | ENSMUSG00000067567 | **Hdac8** | 2610007D20Rik | histone deacetylase 8 | protein-coding |
| chr12_34489652_34490157 | chr12 | 34489653 | 34490157 | + | 0.794165 | intron (NM_024124, intron 2 of 11) | intron (NM_024124, intron 2 of 11) | 38984 | NM_024124 | 79221 | Mm.310551 | NM_024124 | ENSMUSG00000004698 | **Hdac9** | AV022454\|D030072B18Rik\|HD7B\|HD9\|HDRP\|Hdac7b\|Mitr\|mKIAA0744 | histone deacetylase 9 | protein-coding |
| chr3_87907278_87907804 | chr3 | 87907279 | 87907804 | + | 0.599942 | intron (NM_008231, intron 1 of 5) | intron (NM_008231, intron 1 of 5) | 1220 | NM_008231 | 15191 | Mm.292208 | NM_008231 | ENSMUSG00000004897 | **Hdgf** | AI118077\|D3Ertd299e | hepatoma-derived growth factor | protein-coding |
| chr11_83799934_83800438 | chr11 | 83799935 | 83800438 | + | 0.930598 | Intergenic | Intergenic | 46549 | NM_145432 | 217026 | Mm.117874 | NM_145432 | ENSMUSG00000000976 | **Heatr6** | 2700008B19Rik\|BB135312\|BB249351 | HEAT repeat containing 6 | protein-coding |
| chr11_83789270_83789612 | chr11 | 83789271 | 83789612 | + | 0.76889 | Intergenic | Intergenic | 35804 | NM_145432 | 217026 | Mm.117874 | NM_145432 | ENSMUSG00000000976 | **Heatr6** | 2700008B19Rik\|BB135312\|BB249351 | HEAT repeat containing 6 | protein-coding |
| chr11_83789115_83789774 | chr11 | 83789116 | 83789774 | + | 0.7059 | Intergenic | Intergenic | 35808 | NM_145432 | 217026 | Mm.117874 | NM_145432 | ENSMUSG00000000976 | **Heatr6** | 2700008B19Rik\|BB135312\|BB249351 | HEAT repeat containing 6 | protein-coding |
| chr12_51865098_51865623 | chr12 | 51865099 | 51865623 | + | 1.31275 | Intergenic | Intergenic | -35825 | NM_144788 | 207304 | Mm.249391 | NM_144788 | ENSMUSG00000035247 | **Hectd1** | A630086P08Rik\|AI844876\|b2b327Clo\|opm | HECT domain containing 1 | protein-coding |
| chr12_51864887_51865648 | chr12 | 51864888 | 51865648 | + | 1.29712 | Intergenic | Intergenic | -35732 | NM_144788 | 207304 | Mm.249391 | NM_144788 | ENSMUSG00000035247 | **Hectd1** | A630086P08Rik\|AI844876\|b2b327Clo\|opm | HECT domain containing 1 | protein-coding |
| chr12_51811347_51812237 | chr12 | 51811348 | 51812237 | + | 0.952598 | intron (NM_144788, intron 2 of 42) | ORR1G\|LTR\|ERVL-MaLR | 17744 | NM_144788 | 207304 | Mm.249391 | NM_144788 | ENSMUSG00000035247 | **Hectd1** | A630086P08Rik\|AI844876\|b2b327Clo\|opm | HECT domain containing 1 | protein-coding |
| chr16_33606122_33606585 | chr16 | 33606123 | 33606585 | + | 1.19163 | intron (NM_134251, intron 6 of 12) | intron (NM_134251, intron 6 of 12) | -78112 | NM_175256 | 77446 | Mm.245741 | NM_172934 | ENSMUSG00000075254 | **Heg1** | 4632417D23Rik\|5530401I02Rik\|9530025L16Rik\|BC038614\|Gm629\|OTTMUSG00000027315 | HEG homolog 1 (zebrafish) | protein-coding |
| chr16_33605782_33607456 | chr16 | 33605783 | 33607456 | + | 0.992473 | intron (NM_134251, intron 6 of 12) | intron (NM_134251, intron 6 of 12) | -77847 | NM_175256 | 77446 | Mm.245741 | NM_172934 | ENSMUSG00000075254 | **Heg1** | 4632417D23Rik\|5530401I02Rik\|9530025L16Rik\|BC038614\|Gm629\|OTTMUSG00000027315 | HEG homolog 1 (zebrafish) | protein-coding |
| chr16_33606651_33607429 | chr16 | 33606652 | 33607429 | + | 0.873203 | intron (NM_134251, intron 6 of 12) | intron (NM_134251, intron 6 of 12) | -77426 | NM_175256 | 77446 | Mm.245741 | NM_172934 | ENSMUSG00000075254 | **Heg1** | 4632417D23Rik\|5530401I02Rik\|9530025L16Rik\|BC038614\|Gm629\|OTTMUSG00000027315 | HEG homolog 1 (zebrafish) | protein-coding |
| chr16_33606747_33607150 | chr16 | 33606748 | 33607150 | + | 0.858835 | intron (NM_134251, intron 6 of 12) | intron (NM_134251, intron 6 of 12) | -77517 | NM_175256 | 77446 | Mm.245741 | NM_172934 | ENSMUSG00000075254 | **Heg1** | 4632417D23Rik\|5530401I02Rik\|9530025L16Rik\|BC038614\|Gm629\|OTTMUSG00000027315 | HEG homolog 1 (zebrafish) | protein-coding |
| chr16_33708835_33709178 | chr16 | 33708836 | 33709178 | + | 0.80156 | intron (NM_175256, intron 1 of 14) | Lx4A\|LINE\|L1 | 24541 | NM_175256 | 77446 | Mm.245741 | NM_172934 | ENSMUSG00000075254 | **Heg1** | 4632417D23Rik\|5530401I02Rik\|9530025L16Rik\|BC038614\|Gm629\|OTTMUSG00000027315 | HEG homolog 1 (zebrafish) | protein-coding |
| chr4_46410243_46410820 | chr4 | 46410244 | 46410820 | + | 0.840361 | Intergenic | Intergenic | -6349 | NM_053149 | 93966 | Mm.25793 | NM_053149 | ENSMUSG00000028332 | **Hemgn** | 4921524M03Rik\|AI317176\|EDAG\|Hgn\|mNDR | hemogen | protein-coding |
| chr9_107334120_107334932 | chr9 | 107334121 | 107334932 | + | 0.74217 | intron (NM_133984, intron 4 of 10) | intron (NM_133984, intron 4 of 10) | 3824 | NM_133984 | 69536 | Mm.259467 | NM_133984 | ENSMUSG00000032579 | **Hemk1** | 2310008M14Rik\|AW049265 | HemK methyltransferase family member 1 | protein-coding |
| chr9_107334117_107335399 | chr9 | 107334118 | 107335399 | + | 0.7365 | intron (NM_133984, intron 4 of 10) | intron (NM_133984, intron 4 of 10) | 3592 | NM_133984 | 69536 | Mm.259467 | NM_133984 | ENSMUSG00000032579 | **Hemk1** | 2310008M14Rik\|AW049265 | HemK methyltransferase family member 1 | protein-coding |
| chr9_107335545_107335848 | chr9 | 107335546 | 107335848 | + | 0.735551 | intron (NM_133984, intron 4 of 10) | intron (NM_133984, intron 4 of 10) | 2653 | NM_133984 | 69536 | Mm.259467 | NM_133984 | ENSMUSG00000032579 | **Hemk1** | 2310008M14Rik\|AW049265 | HemK methyltransferase family member 1 | protein-coding |
| chr9_107333679_107334010 | chr9 | 107333680 | 107334010 | + | 0.688294 | intron (NM_133984, intron 4 of 10) | intron (NM_133984, intron 4 of 10) | 4505 | NM_133984 | 69536 | Mm.259467 | NM_133984 | ENSMUSG00000032579 | **Hemk1** | 2310008M14Rik\|AW049265 | HemK methyltransferase family member 1 | protein-coding |
| chr9_107335097_107335393 | chr9 | 107335098 | 107335393 | + | 0.663925 | intron (NM_133984, intron 4 of 10) | B3\|SINE\|B2 | 3105 | NM_133984 | 69536 | Mm.259467 | NM_133984 | ENSMUSG00000032579 | **Hemk1** | 2310008M14Rik\|AW049265 | HemK methyltransferase family member 1 | protein-coding |
| chr9_15103557_15104300 | chr9 | 15103558 | 15104300 | + | 1.62772 | intron (NM_001164797, intron 1 of 19) | intron (NM_001164797, intron 1 of 19) | 8179 | NM_001164797 | 244698 | Mm.325134 | NM_001164797 | ENSMUSG00000031936 | **Hephl1** | Gm509\|Zp\|zyklopen | hephaestin-like 1 | protein-coding |
| chr9_66309499_66309844 | chr9 | 66309500 | 66309844 | + | 0.732996 | Intergenic | Intergenic | -40778 | NM_145617 | 235439 | Mm.244179 | NM_145617 | ENSMUSG00000038664 | **Herc1** | 2810449H11Rik\|B230218H07\|D130015N03Rik\|tbl | hect (homologous to the E6-AP (UBE3A) carboxyl terminus) domain and RCC1 (CHC1)-like domain (RLD) 1 | protein-coding |
| chr9_66306526_66306784 | chr9 | 66306527 | 66306784 | + | 0.673893 | Intergenic | Intergenic | -43795 | NM_145617 | 235439 | Mm.244179 | NM_145617 | ENSMUSG00000038664 | **Herc1** | 2810449H11Rik\|B230218H07\|D130015N03Rik\|tbl | hect (homologous to the E6-AP (UBE3A) carboxyl terminus) domain and RCC1 (CHC1)-like domain (RLD) 1 | protein-coding |
| chr9_66267298_66267667 | chr9 | 66267299 | 66267667 | + | 0.601452 | intron (NM_010019, intron 9 of 11) | intron (NM_010019, intron 9 of 11) | -82967 | NM_145617 | 235439 | Mm.244179 | NM_145617 | ENSMUSG00000038664 | **Herc1** | 2810449H11Rik\|B230218H07\|D130015N03Rik\|tbl | hect (homologous to the E6-AP (UBE3A) carboxyl terminus) domain and RCC1 (CHC1)-like domain (RLD) 1 | protein-coding |
| chr6_58828604_58828885 | chr6 | 58828605 | 58828885 | + | 0.995565 | Intergenic | Intergenic | -4955 | NM_028705 | 73998 | Mm.191892 | NM_028705 | ENSMUSG00000029804 | **Herc3** | 5730409F18Rik\|AI646603\|mKIAA0032 | hect domain and RLD 3 | protein-coding |
| chr6_58841266_58841499 | chr6 | 58841267 | 58841499 | + | 0.778056 | intron (NM_028705, intron 1 of 24) | B2_Mm2\|SINE\|B2 | 7683 | NM_028705 | 73998 | Mm.191892 | NM_028705 | ENSMUSG00000029804 | **Herc3** | 5730409F18Rik\|AI646603\|mKIAA0032 | hect domain and RLD 3 | protein-coding |
| chr10_63240859_63241258 | chr10 | 63240860 | 63241258 | + | 1.20271 | Intergenic | RMER2\|LTR\|ERV1 | -2738 | NM_030114 | 67345 | Mm.234437 | NM_026101 | ENSMUSG00000020064 | **Herc4** | 1700056O17Rik\|4921531D01Rik\|9530080M15Rik\|mKIAA1593 | hect domain and RLD 4 | protein-coding |
| chr9_25161336_25161656 | chr9 | 25161337 | 25161656 | + | 1.17534 | Intergenic | PB1D7\|SINE\|Alu | -9715 | NM_020586 | 80517 | Mm.142843 | NM_020586 | ENSMUSG00000008429 | **Herpud2** | 5031400M07Rik\|AB041580 | HERPUD family member 2 | protein-coding |
| chr9_25068892_25069479 | chr9 | 25068893 | 25069479 | + | 1.0289 | Intergenic | Intergenic | 82595 | NM_020586 | 80517 | Mm.142843 | NM_020586 | ENSMUSG00000008429 | **Herpud2** | 5031400M07Rik\|AB041580 | HERPUD family member 2 | protein-coding |
| chr9_25072694_25073086 | chr9 | 25072695 | 25073086 | + | 0.763555 | Intergenic | Intergenic | 78891 | NM_020586 | 80517 | Mm.142843 | NM_020586 | ENSMUSG00000008429 | **Herpud2** | 5031400M07Rik\|AB041580 | HERPUD family member 2 | protein-coding |
| chr4_123225982_123226362 | chr4 | 123225983 | 123226362 | + | 0.862957 | Intergenic | RMER15\|LTR\|ERVL | -7384 | NM_013905 | 56198 | Mm.103615 | NM_013905 | ENSMUSG00000032744 | **Heyl** | Hey3\|Hrt3\|bHLHb33\|hesr3 | hairy/enhancer-of-split related with YRPW motif-like | protein-coding |
| chr4_123225766_123226371 | chr4 | 123225767 | 123226371 | + | 0.858547 | Intergenic | RMER15\|LTR\|ERVL | -7487 | NM_013905 | 56198 | Mm.103615 | NM_013905 | ENSMUSG00000032744 | **Heyl** | Hey3\|Hrt3\|bHLHb33\|hesr3 | hairy/enhancer-of-split related with YRPW motif-like | protein-coding |
| chr5_106885558_106885968 | chr5 | 106885559 | 106885968 | + | 0.913368 | intron (NM_177873, intron 21 of 39) | intron (NM_177873, intron 21 of 39) | 40127 | NM_177873 | 330149 | Mm.121664 | NM_177873 | ENSMUSG00000043410 | **Hfm1** | A330009G12Rik\|Gm1046\|Mer3\|Sec63d1 | HFM1, ATP-dependent DNA helicase homolog (S. cerevisiae) | protein-coding |
| chr5_106885348_106885975 | chr5 | 106885349 | 106885975 | + | 0.842688 | intron (NM_177873, intron 21 of 39) | intron (NM_177873, intron 21 of 39) | 40228 | NM_177873 | 330149 | Mm.121664 | NM_177873 | ENSMUSG00000043410 | **Hfm1** | A330009G12Rik\|Gm1046\|Mer3\|Sec63d1 | HFM1, ATP-dependent DNA helicase homolog (S. cerevisiae) | protein-coding |

| | | | | | | | | | | | | | | | | | |
|---|---|---|---|---|---|---|---|---|---|---|---|---|---|---|---|---|---|
| chr5_16350965_16351330 | chr5 | 16350966 | 16351330 | + | 0.858837 | intron (NM_009784, intron 24 of 37) | intron (NM_009784, intron 24 of 37) | -202347 | NM_001289460 | 15234 | Mm.267078 | NM_010427 | ENSMUSG00000028864 | **Hgf** | C230052L06Rik\|HGF/SF\|NK1\|NK2\|SF\|SF/HGF | hepatocyte growth factor | protein-coding |
| chr5_16355843_16356300 | chr5 | 16355844 | 16356300 | + | 0.785102 | intron (NM_009784, intron 28 of 37) | intron (NM_009784, intron 28 of 37) | -197423 | NM_001289460 | 15234 | Mm.267078 | NM_010427 | ENSMUSG00000028864 | **Hgf** | C230052L06Rik\|HGF/SF\|NK1\|NK2\|SF\|SF/HGF | hepatocyte growth factor | protein-coding |
| chr5_16350482_16351318 | chr5 | 16350483 | 16351318 | + | 0.60093 | intron (NM_009784, intron 24 of 37) | Lx2\|LINE\|L1 | -202595 | NM_001289460 | 15234 | Mm.267078 | NM_010427 | ENSMUSG00000028864 | **Hgf** | C230052L06Rik\|HGF/SF\|NK1\|NK2\|SF\|SF/HGF | hepatocyte growth factor | protein-coding |
| chr8_25965524_25965883 | chr8 | 25965525 | 25965883 | + | 0.66406 | intron (NM_029884, intron 5 of 17) | intron (NM_029884, intron 5 of 17) | 11040 | NM_029884 | 52120 | Mm.28326 | NM_029884 | ENSMUSG00000037260 | **Hgsnat** | 9430010M12Rik\|AW208455\|D8Ertd354e\|Tmem76 | heparan-alpha-glucosaminide N-acetyltransferase | protein-coding |
| chr1_192742910_192743185 | chr1 | 192742911 | 192743185 | + | 0.921799 | intron (NM_144881, intron 3 of 11) | Lx9\|LINE\|L1 | 28171 | NM_144881 | 226861 | Mm.145857 | NM_144881 | ENSMUSG00000037375 | **Hhat** | 2810432O22Rik\|AI462858\|AP-2CRE\|Skn\|Tg(TFAP2A-cre)1Will | hedgehog acyltransferase | protein-coding |
| chr1_192786080_192786721 | chr1 | 192786081 | 192786721 | + | 0.803301 | Intergenic | MLT1B\|LTR\|ERVL-MaLR | -15182 | NM_144881 | 226861 | Mm.145857 | NM_144881 | ENSMUSG00000037375 | **Hhat** | 2810432O22Rik\|AI462858\|AP-2CRE\|Skn\|Tg(TFAP2A-cre)1Will | hedgehog acyltransferase | protein-coding |
| chr1_192745492_192746524 | chr1 | 192745493 | 192746524 | + | 0.793534 | intron (NM_144881, intron 3 of 11) | intron (NM_144881, intron 3 of 11) | 25211 | NM_144881 | 226861 | Mm.145857 | NM_144881 | ENSMUSG00000037375 | **Hhat** | 2810432O22Rik\|AI462858\|AP-2CRE\|Skn\|Tg(TFAP2A-cre)1Will | hedgehog acyltransferase | protein-coding |
| chr1_192785921_192786811 | chr1 | 192785922 | 192786811 | + | 0.648206 | Intergenic | Intergenic | -15147 | NM_144881 | 226861 | Mm.145857 | NM_144881 | ENSMUSG00000037375 | **Hhat** | 2810432O22Rik\|AI462858\|AP-2CRE\|Skn\|Tg(TFAP2A-cre)1Will | hedgehog acyltransferase | protein-coding |
| chr1_192745497_192745976 | chr1 | 192745498 | 192745976 | + | 0.634285 | intron (NM_144881, intron 3 of 11) | intron (NM_144881, intron 3 of 11) | 25482 | NM_144881 | 226861 | Mm.145857 | NM_144881 | ENSMUSG00000037375 | **Hhat** | 2810432O22Rik\|AI462858\|AP-2CRE\|Skn\|Tg(TFAP2A-cre)1Will | hedgehog acyltransferase | protein-coding |
| chr6_52578063_52578822 | chr6 | 52578064 | 52578822 | + | 1.19697 | intron (NM_145567, intron 4 of 7) | intron (NM_145567, intron 4 of 7) | 61857 | NM_145567 | 58875 | Mm.286458 | NM_145567 | ENSMUSG00000029776 | **Hibadh** | 6430402H10Rik\|AI265272 | 3-hydroxyisobutyrate dehydrogenase | protein-coding |
| chr6_52594701_52595059 | chr6 | 52594702 | 52595059 | + | 1.01025 | intron (NM_145567, intron 4 of 7) | intron (NM_145567, intron 4 of 7) | 45420 | NM_145567 | 58875 | Mm.286458 | NM_145567 | ENSMUSG00000029776 | **Hibadh** | 6430402H10Rik\|AI265272 | 3-hydroxyisobutyrate dehydrogenase | protein-coding |
| chr6_52594690_52595363 | chr6 | 52594691 | 52595363 | + | 0.989201 | intron (NM_145567, intron 4 of 7) | B1F2\|SINE\|Alu | 45273 | NM_145567 | 58875 | Mm.286458 | NM_145567 | ENSMUSG00000029776 | **Hibadh** | 6430402H10Rik\|AI265272 | 3-hydroxyisobutyrate dehydrogenase | protein-coding |
| chr1_52856273_52857023 | chr1 | 52856274 | 52857023 | + | 1.37385 | intron (NM_146108, intron 2 of 13) | RLTR21\|LTR\|ERVK | 11602 | NM_146108 | 227095 | Mm.222063 | NM_146108 | ENSMUSG00000041426 | **Hibch** | 2610509I15Rik\|AI648812 | 3-hydroxyisobutyryl-Coenzyme A hydrolase | protein-coding |
| chr1_52856277_52856683 | chr1 | 52856278 | 52856683 | + | 1.06555 | intron (NM_146108, intron 2 of 13) | RLTR21\|LTR\|ERVK | 11434 | NM_146108 | 227095 | Mm.222063 | NM_146108 | ENSMUSG00000041426 | **Hibch** | 2610509I15Rik\|AI648812 | 3-hydroxyisobutyryl-Coenzyme A hydrolase | protein-coding |
| chr1_52856757_52857079 | chr1 | 52856758 | 52857079 | + | 0.929219 | intron (NM_146108, intron 2 of 13) | intron (NM_146108, intron 2 of 13) | 11872 | NM_146108 | 227095 | Mm.222063 | NM_146108 | ENSMUSG00000041426 | **Hibch** | 2610509I15Rik\|AI648812 | 3-hydroxyisobutyryl-Coenzyme A hydrolase | protein-coding |
| chr1_52863376_52864166 | chr1 | 52863377 | 52864166 | + | 0.6823 | intron (NM_146108, intron 4 of 13) | intron (NM_146108, intron 4 of 13) | 18725 | NM_146108 | 227095 | Mm.222063 | NM_146108 | ENSMUSG00000041426 | **Hibch** | 2610509I15Rik\|AI648812 | 3-hydroxyisobutyryl-Coenzyme A hydrolase | protein-coding |
| chr11_75154004_75154665 | chr11 | 75154005 | 75154665 | + | 1.46636 | intron (NM_001002764, intron 15 of 18) | intron (NM_001002764, intron 15 of 18) | 13817 | NM_001098203 | 15248 | Mm.57250 | NM_010430 | ENSMUSG00000043099 | **Hic1** | AA408311\|HIC-1 | hypermethylated in cancer 1 | protein-coding |
| chr11_75137104_75137468 | chr11 | 75137105 | 75137468 | + | 0.65299 | intron (NM_001002764, intron 13 of 18) | intron (NM_001002764, intron 13 of 18) | 30866 | NM_001098203 | 15248 | Mm.57250 | NM_010430 | ENSMUSG00000043099 | **Hic1** | AA408311\|HIC-1 | hypermethylated in cancer 1 | protein-coding |
| chr12_73887606_73888722 | chr12 | 73887607 | 73888722 | + | 1.78912 | Intergenic | Intergenic | -19703 | NM_010431 | 15251 | Mm.3879 | NM_010431 | ENSMUSG00000021109 | **Hif1a** | AA959795\|HIF1alpha\|MOP1\|bHLHe78 | hypoxia inducible factor 1, alpha subunit | protein-coding |
| chr12_73901195_73901651 | chr12 | 73901196 | 73901651 | + | 1.05906 | Intergenic | L1MCa\|LINE\|L1 | -6444 | NM_010431 | 15251 | Mm.3879 | NM_010431 | ENSMUSG00000021109 | **Hif1a** | AA959795\|HIF1alpha\|MOP1\|bHLHe78 | hypoxia inducible factor 1, alpha subunit | protein-coding |
| chr12_73779262_73779548 | chr12 | 73779263 | 73779548 | + | 1.00764 | Intergenic | Intergenic | -128462 | NM_010431 | 15251 | Mm.3879 | NM_010431 | ENSMUSG00000021109 | **Hif1a** | AA959795\|HIF1alpha\|MOP1\|bHLHe78 | hypoxia inducible factor 1, alpha subunit | protein-coding |
| chr12_73912089_73912282 | chr12 | 73912090 | 73912282 | + | 0.988687 | intron (NM_010431, intron 1 of 14) | L1MB2\|LINE\|L1 | 4319 | NM_010431 | 15251 | Mm.3879 | NM_010431 | ENSMUSG00000021109 | **Hif1a** | AA959795\|HIF1alpha\|MOP1\|bHLHe78 | hypoxia inducible factor 1, alpha subunit | protein-coding |
| chr12_73876344_73876995 | chr12 | 73876345 | 73876995 | + | 0.962746 | Intergenic | Intergenic | -31197 | NM_010431 | 15251 | Mm.3879 | NM_010431 | ENSMUSG00000021109 | **Hif1a** | AA959795\|HIF1alpha\|MOP1\|bHLHe78 | hypoxia inducible factor 1, alpha subunit | protein-coding |
| chr12_73895745_73896123 | chr12 | 73895746 | 73896123 | + | 0.888643 | Intergenic | Intergenic | -11933 | NM_010431 | 15251 | Mm.3879 | NM_010431 | ENSMUSG00000021109 | **Hif1a** | AA959795\|HIF1alpha\|MOP1\|bHLHe78 | hypoxia inducible factor 1, alpha subunit | protein-coding |
| chr12_73880329_73880727 | chr12 | 73880330 | 73880727 | + | 0.877701 | Intergenic | Intergenic | -27339 | NM_010431 | 15251 | Mm.3879 | NM_010431 | ENSMUSG00000021109 | **Hif1a** | AA959795\|HIF1alpha\|MOP1\|bHLHe78 | hypoxia inducible factor 1, alpha subunit | protein-coding |
| chr12_73779211_73779908 | chr12 | 73779212 | 73779908 | + | 0.876561 | Intergenic | Intergenic | -128307 | NM_010431 | 15251 | Mm.3879 | NM_010431 | ENSMUSG00000021109 | **Hif1a** | AA959795\|HIF1alpha\|MOP1\|bHLHe78 | hypoxia inducible factor 1, alpha subunit | protein-coding |
| chr12_73880426_73880722 | chr12 | 73880427 | 73880722 | + | 0.873188 | Intergenic | Intergenic | -27293 | NM_010431 | 15251 | Mm.3879 | NM_010431 | ENSMUSG00000021109 | **Hif1a** | AA959795\|HIF1alpha\|MOP1\|bHLHe78 | hypoxia inducible factor 1, alpha subunit | protein-coding |
| chr6_29247109_29247496 | chr6 | 29247110 | 29247496 | + | 1.1217 | Intergenic | Intergenic | -25185 | NM_001190461 | 69573 | Mm.45161 | NM_023516 | ENSMUSG00000043421 | **Hilpda** | 2310016C08Rik\|AW049004\|Hig2 | hypoxia inducible lipid droplet associated | protein-coding |
| chr6_29257467_29257848 | chr6 | 29257468 | 29257848 | + | 1.09086 | Intergenic | Intergenic | -14830 | NM_001190461 | 69573 | Mm.45161 | NM_023516 | ENSMUSG00000043421 | **Hilpda** | 2310016C08Rik\|AW049004\|Hig2 | hypoxia inducible lipid droplet associated | protein-coding |
| chr6_29259348_29259723 | chr6 | 29259349 | 29259723 | + | 0.736916 | Intergenic | Intergenic | -12952 | NM_001190461 | 69573 | Mm.45161 | NM_023516 | ENSMUSG00000043421 | **Hilpda** | 2310016C08Rik\|AW049004\|Hig2 | hypoxia inducible lipid droplet associated | protein-coding |
| chr6_29270784_29271384 | chr6 | 29270785 | 29271384 | + | 0.645553 | Intergenic | Intergenic | -1404 | NM_001190461 | 69573 | Mm.45161 | NM_023516 | ENSMUSG00000043421 | **Hilpda** | 2310016C08Rik\|AW049004\|Hig2 | hypoxia inducible lipid droplet associated | protein-coding |
| chr10_30636604_30637381 | chr10 | 30636605 | 30637381 | + | 0.981453 | Intergenic | MERV1_LTR\|LTR\|ERV1 | -18627 | NM_025798 | 66847 | Mm.46218 | NM_025798 | | **Hint3** | 0610010I17Rik\|AV009015\|HINT-3\|HINT-4\|HINT4 | histidine triad nucleotide binding protein 3 | protein-coding |
| chr5_135537294_135538367 | chr5 | 135537295 | 135538367 | + | 0.68277 | intron (NM_146001, intron 1 of 30) | MTEa\|LTR\|ERVL-MaLR | 7291 | NM_146001 | 215114 | Mm.280805 | NM_146001 | ENSMUSG00000039959 | **Hip1** | 2610109B09Rik\|A930014B11Rik\|E130315I21Rik\|mKIAA4113 | huntingtin interacting protein 1 | protein-coding |
| chr5_123980517_123981157 | chr5 | 123980518 | 123981157 | + | 0.824285 | intron (NM_145070, intron 1 of 31) | intron (NM_145070, intron 1 of 31) | 7209 | NM_145070 | 29816 | Mm.149954 | NM_145070 | ENSMUSG00000000915 | **Hip1r** | AA410023\|mKIAA0655 | huntingtin interacting protein 1 related | protein-coding |
| chr13_21780168_21780763 | chr13 | 21780169 | 21780763 | + | 1.05021 | exon (NM_020034, exon 1 of 1) | exon (NM_020034, exon 1 of 1) | 159 | NM_020034 | 56702 | Mm.221314 | NM_020034 | ENSMUSG00000058773 | **Hist1h1b** | H1.5\|H1B\|H1f5\|H1s-3 | histone cluster 1, H1b | protein-coding |
| chr13_21768595_21768931 | chr13 | 21768596 | 21768931 | + | 0.784748 | Intergenic | Intergenic | 11862 | NM_020034 | 56702 | Mm.221314 | NM_020034 | ENSMUSG00000058773 | **Hist1h1b** | H1.5\|H1B\|H1f5\|H1s-3 | histone cluster 1, H1b | protein-coding |
| chr13_23554872_23556371 | chr13 | 23554873 | 23556371 | + | 0.605293 | exon (NM_145713, exon 1 of 1) | exon (NM_145713, exon 1 of 1) | 590 | NM_145713 | 14957 | Mm.247046 | NM_145713 | | **Hist1h1d** | H1.3\|H1D\|H1f3\|H1s-4 | histone cluster 1, H1d | protein-coding |
| chr13_23691654_23692367 | chr13 | 23691655 | 23692367 | + | 1.88305 | intron (NM_023422, intron 1 of 1) | intron (NM_023422, intron 1 of 1) | -3800 | NM_010377 | 107970 | Mm.8048 | NM_010377 | ENSMUSG00000036211 | **Hist1h1t** | H1ft\|H1t | histone cluster 1, H1t | protein-coding |
| chr13_23749697_23749919 | chr13 | 23749698 | 23749919 | + | 1.3438 | Intergenic | Intergenic | -1280 | NM_175660 | 319172 | Mm.423402 | NM_175660 | ENSMUSG00000061615 | **Hist1h2ab** | H2A.1\|H2a-53 | histone cluster 1, H2ab | protein-coding |
| chr13_22042053_22042309 | chr13 | 22042054 | 22042309 | + | 0.760315 | promoter-TSS (NM_175656) | promoter-TSS (NM_175656) | 768 | NM_178186 | 319167 | Mm.263165 | NM_178186 | ENSMUSG00000069301 | **Hist1h2ag** | - | histone cluster 1, H2ag | protein-coding |
| chr13_21716683_21717130 | chr13 | 21716684 | 21717130 | + | 0.703103 | promoter-TSS (NM_178199) | promoter-TSS (NM_178199) | 495 | NM_178182 | 319191 | Mm.14767 | NM_178182 | | **Hist1h2ai** | H2a-291A | histone cluster 1, H2ai | protein-coding |
| chr13_23747058_23747500 | chr13 | 23747059 | 23747500 | + | 1.06362 | TTS (NM_175664) | TTS (NM_175664) | 545 | NM_175664 | 319178 | Mm.390567 | NM_175664 | ENSMUSG00000075031 | **Hist1h2bb** | H2b-143 | histone cluster 1, H2bb | protein-coding |
| chr13_23593064_23593674 | chr13 | 23593065 | 23593674 | + | 0.734326 | intron (NM_001177653, intron 1 of 1) | intron (NM_001177653, intron 1 of 1) | -7399 | NM_001290530 | 319179 | Mm.440391 | NM_178194 | ENSMUSG00000047246 | **Hist1h2be** | - | histone cluster 1, H2be | protein-coding |
| chr13_22053820_22054054 | chr13 | 22053821 | 22054054 | + | 1.05338 | Intergenic | Intergenic | 10707 | NM_178198 | 319183 | Mm.371766 | NM_178198 | ENSMUSG00000069300 | **Hist1h2bj** | - | histone cluster 1, H2bj | protein-coding |
| chr13_22036177_22036635 | chr13 | 22036178 | 22036635 | + | 0.821234 | promoter-TSS (NM_175659) | promoter-TSS (NM_175659) | 585 | NM_175665 | 319184 | Mm.358955 | NM_175665 | | **Hist1h2bk** | - | histone cluster 1, H2bk | protein-coding |
| chr13_21715598_21715869 | chr13 | 21715599 | 21715869 | + | 0.778402 | promoter-TSS (NM_178182) | promoter-TSS (NM_178182) | 409 | NM_178199 | 319185 | | NM_178199 | ENSMUSG00000094338 | **Hist1h2bl** | H2b-291a | histone cluster 1, H2bl | protein-coding |
| chr13_21852302_21852983 | chr13 | 21852303 | 21852983 | + | 0.674166 | Intergenic | Intergenic | 18900 | NM_001110555 | 665622 | Mm.471247 | NM_001110555 | | **Hist1h2br** | Gm11277\|OTTMUSG00000000449 | histone cluster 1 H2br | protein-coding |
| chr13_21855102_21855588 | chr13 | 21855103 | 21855588 | + | 0.658029 | Intergenic | tRNA-Ile-ATT\|tRNA\|tRNA | 21602 | NM_001110555 | 665622 | Mm.471247 | NM_001110555 | | **Hist1h2br** | Gm11277\|OTTMUSG00000000449 | histone cluster 1 H2br | protein-coding |
| chr13_21847927_21848188 | chr13 | 21847928 | 21848188 | + | 0.637117 | Intergenic | tRNA-Leu-TTA\|tRNA\|tRNA | 14315 | NM_001110555 | 665622 | Mm.471247 | NM_001110555 | | **Hist1h2br** | Gm11277\|OTTMUSG00000000449 | histone cluster 1 H2br | protein-coding |
| chr13_23752107_23752502 | chr13 | 23752108 | 23752502 | + | 0.917824 | promoter-TSS (NM_178203) | promoter-TSS (NM_178203) | -75 | NM_178203 | 319150 | | NM_178203 | | **Hist1h3b** | H3-53 | histone cluster 1, H3b | protein-coding |
| chr13_23742061_23742286 | chr13 | 23742062 | 23742286 | + | 1.3601 | Intergenic | Intergenic | 3347 | NM_175653 | 319148 | Mm.262517 | NM_175653 | | **Hist1h3c** | H3-143 | histone cluster 1, H3c | protein-coding |
| chr13_23534794_23535055 | chr13 | 23534795 | 23535055 | + | 0.758044 | promoter-TSS (NM_145073) | promoter-TSS (NM_145073) | -493 | NM_145073 | 97908 | Mm.261657 | NM_145073 | ENSMUSG00000062417 | **Hist1h3g** | H3.1-221\|M32460 | histone cluster 1, H3g | protein-coding |
| chr13_23759968_23760399 | chr13 | 23759969 | 23760399 | + | 1.14121 | TTS (NM_178192) | TTS (NM_178192) | 1065 | NM_178192 | 326619 | Mm.228709 | NM_178192 | | **Hist1h4a** | - | histone cluster 1, H4a | protein-coding |
| chr13_21812157_21812544 | chr13 | 21812158 | 21812544 | + | 0.728318 | TTS (NM_175657) | TTS (NM_175657) | 567 | NM_175657 | 319161 | Mm.261664 | NM_175657 | | **Hist1h4n** | - | histone cluster 1, H4n | protein-coding |
| chr13_21812152_21812701 | chr13 | 21812153 | 21812701 | + | 0.684143 | TTS (NM_175657) | TTS (NM_175657) | 643 | NM_175657 | 319161 | Mm.261664 | NM_175657 | | **Hist1h4n** | - | histone cluster 1, H4n | protein-coding |

| | | | | | | | | | | | | | | | | | |
|---|---|---|---|---|---|---|---|---|---|---|---|---|---|---|---|---|---|
| chr3_96246075_96246285 | chr3 | 96246076 | 96246285 | + | 0.681548 | promoter-TSS (NM_178216).2 | promoter-TSS (NM_178216).2 | -505 | NM_178216 | 15077 | Mm.422680 | NM_178216 | ENSMUSG00000093769 | **Hist2h3c1** | BE691662\|H3-615\|H3f2\|Hist2h3ca1 | histone cluster 2, H3c1 | protein-coding |
| chr3_96239541_96239754 | chr3 | 96239542 | 96239754 | + | 0.627084 | promoter-TSS (NM_178216) | promoter-TSS (NM_178216) | -506 | NM_178216 | 15077 | Mm.422680 | NM_178216 | ENSMUSG00000093769 | **Hist2h3c1** | BE691662\|H3-615\|H3f2\|Hist2h3ca1 | histone cluster 2, H3c1 | protein-coding |
| chr3_96237956_96238486 | chr3 | 96237957 | 96238486 | + | 0.955091 | TTS (NM_178216) | TTS (NM_178216) | 906 | NM_054045 | 97114 | Mm.358797 | NM_054045 | ENSMUSG00000081058 | **Hist2h3c2** | H3-614\|H3.2-614\|Hist2h3c2-ps\|Hist2h3ca1\|M32461 | histone cluster 2, H3c2 | protein-coding |
| chr3_96237551_96238517 | chr3 | 96237552 | 96238517 | + | 0.941023 | TTS (NM_178216) | TTS (NM_178216) | 1093 | NM_054045 | 97114 | Mm.358797 | NM_054045 | ENSMUSG00000081058 | **Hist2h3c2** | H3-614\|H3.2-614\|Hist2h3c2-ps\|Hist2h3ca1\|M32461 | histone cluster 2, H3c2 | protein-coding |
| chr6_136804947_136805297 | chr6 | 136804948 | 136805297 | + | 0.837496 | promoter-TSS (NM_175652) | promoter-TSS (NM_175652) | -691 | NM_175652 | 320332 | Mm.246720 | NM_175652 | ENSMUSG00000096010 | **Hist4h4** | B130044J01Rik | histone cluster 4, H4 | protein-coding |
| chr13_42113564_42114153 | chr13 | 42113565 | 42114153 | + | 1.17184 | intron (NM_007772, intron 2 of 8) | intron (NM_007772, intron 2 of 8) | 61838 | NM_007772 | 110521 | Mm.4808 | NM_007772 | ENSMUSG00000021366 | **Hivep1** | Cryabp1 | human immunodeficiency virus type I enhancer binding protein 1 | protein-coding |
| chr13_42110232_42110718 | chr13 | 42110233 | 42110718 | + | 1.14879 | intron (NM_007772, intron 2 of 8) | intron (NM_007772, intron 2 of 8) | 58454 | NM_007772 | 110521 | Mm.4808 | NM_007772 | ENSMUSG00000021366 | **Hivep1** | Cryabp1 | human immunodeficiency virus type I enhancer binding protein 1 | protein-coding |
| chr13_42064275_42064692 | chr13 | 42064276 | 42064692 | + | 1.13325 | intron (NM_007772, intron 2 of 8) | intron (NM_007772, intron 2 of 8) | 12463 | NM_007772 | 110521 | Mm.4808 | NM_007772 | ENSMUSG00000021366 | **Hivep1** | Cryabp1 | human immunodeficiency virus type I enhancer binding protein 1 | protein-coding |
| chr13_42064030_42065111 | chr13 | 42064031 | 42065111 | + | 0.972861 | intron (NM_007772, intron 2 of 8) | intron (NM_007772, intron 2 of 8) | 12550 | NM_007772 | 110521 | Mm.4808 | NM_007772 | ENSMUSG00000021366 | **Hivep1** | Cryabp1 | human immunodeficiency virus type I enhancer binding protein 1 | protein-coding |
| chr13_42064794_42065107 | chr13 | 42064795 | 42065107 | + | 0.954555 | intron (NM_007772, intron 2 of 8) | intron (NM_007772, intron 2 of 8) | 12930 | NM_007772 | 110521 | Mm.4808 | NM_007772 | ENSMUSG00000021366 | **Hivep1** | Cryabp1 | human immunodeficiency virus type I enhancer binding protein 1 | protein-coding |
| chr13_42093654_42094020 | chr13 | 42093655 | 42094020 | + | 0.664507 | intron (NM_007772, intron 2 of 8) | intron (NM_007772, intron 2 of 8) | 41816 | NM_007772 | 110521 | Mm.4808 | NM_007772 | ENSMUSG00000021366 | **Hivep1** | Cryabp1 | human immunodeficiency virus type I enhancer binding protein 1 | protein-coding |
| chr10_14087293_14087725 | chr10 | 14087294 | 14087725 | + | 1.26288 | intron (NM_010437, intron 2 of 8) | intron (NM_010437, intron 2 of 8) | 121130 | NM_010437 | 15273 | Mm.42157 | NM_010437 | ENSMUSG00000015501 | **Hivep2** | MIBP1\|Schnurri-2\|Shn-2 | human immunodeficiency virus type I enhancer binding protein 2 | protein-coding |
| chr10_13997300_13997792 | chr10 | 13997301 | 13997792 | + | 1.17479 | intron (NM_010437, intron 1 of 8) | intron (NM_010437, intron 1 of 8) | 31167 | NM_010437 | 15273 | Mm.42157 | NM_010437 | ENSMUSG00000015501 | **Hivep2** | MIBP1\|Schnurri-2\|Shn-2 | human immunodeficiency virus type I enhancer binding protein 2 | protein-coding |
| chr10_14019308_14019689 | chr10 | 14019309 | 14019689 | + | 1.07876 | intron (NM_010437, intron 1 of 8) | intron (NM_010437, intron 1 of 8) | 53120 | NM_010437 | 15273 | Mm.42157 | NM_010437 | ENSMUSG00000015501 | **Hivep2** | MIBP1\|Schnurri-2\|Shn-2 | human immunodeficiency virus type I enhancer binding protein 2 | protein-coding |
| chr10_14019307_14019892 | chr10 | 14019308 | 14019892 | + | 0.872206 | intron (NM_010437, intron 1 of 8) | intron (NM_010437, intron 1 of 8) | 53221 | NM_010437 | 15273 | Mm.42157 | NM_010437 | ENSMUSG00000015501 | **Hivep2** | MIBP1\|Schnurri-2\|Shn-2 | human immunodeficiency virus type I enhancer binding protein 2 | protein-coding |
| chr10_14033995_14034521 | chr10 | 14033996 | 14034521 | + | 0.83125 | intron (NM_010437, intron 1 of 8) | intron (NM_010437, intron 1 of 8) | 67879 | NM_010437 | 15273 | Mm.42157 | NM_010437 | ENSMUSG00000015501 | **Hivep2** | MIBP1\|Schnurri-2\|Shn-2 | human immunodeficiency virus type I enhancer binding protein 2 | protein-coding |
| chr4_119922918_119923586 | chr4 | 119922919 | 119923586 | + | 1.77161 | intron (NM_010657, intron 1 of 8) | intron (NM_010657, intron 1 of 8) | 108574 | NM_010657 | 16656 | Mm.302758 | NM_010657 | ENSMUSG00000028634 | **Hivep3** | 2900056N03Rik\|A130075N07\|AI848000\|E030045D18Rik\|KBP-1\|KBP1\|Krc\|Rc\|Schnurri-3\|Shn3\|Zas3 | human immunodeficiency virus type I enhancer binding protein 3 | protein-coding |
| chr4_119921055_119921694 | chr4 | 119921056 | 119921694 | + | 1.35938 | intron (NM_010657, intron 1 of 8) | ORR1E\|LTR\|ERVL-MaLR | 106697 | NM_010657 | 16656 | Mm.302758 | NM_010657 | ENSMUSG00000028634 | **Hivep3** | 2900056N03Rik\|A130075N07\|AI848000\|E030045D18Rik\|KBP-1\|KBP1\|Krc\|Rc\|Schnurri-3\|Shn3\|Zas3 | human immunodeficiency virus type I enhancer binding protein 3 | protein-coding |
| chr4_119974053_119974605 | chr4 | 119974054 | 119974605 | + | 1.28612 | intron (NM_010657, intron 1 of 8) | intron (NM_010657, intron 1 of 8) | 159651 | NM_010657 | 16656 | Mm.302758 | NM_010657 | ENSMUSG00000028634 | **Hivep3** | 2900056N03Rik\|A130075N07\|AI848000\|E030045D18Rik\|KBP-1\|KBP1\|Krc\|Rc\|Schnurri-3\|Shn3\|Zas3 | human immunodeficiency virus type I enhancer binding protein 3 | protein-coding |
| chr4_119784860_119785518 | chr4 | 119784861 | 119785518 | + | 1.25189 | Intergenic | ERV3-16A3_I-int\|LTR\|ERVL | -29489 | NM_010657 | 16656 | Mm.302758 | NM_010657 | ENSMUSG00000028634 | **Hivep3** | 2900056N03Rik\|A130075N07\|AI848000\|E030045D18Rik\|KBP-1\|KBP1\|Krc\|Rc\|Schnurri-3\|Shn3\|Zas3 | human immunodeficiency virus type I enhancer binding protein 3 | protein-coding |
| chr4_119974275_119974599 | chr4 | 119974276 | 119974599 | + | 1.2434 | intron (NM_010657, intron 1 of 8) | intron (NM_010657, intron 1 of 8) | 159759 | NM_010657 | 16656 | Mm.302758 | NM_010657 | ENSMUSG00000028634 | **Hivep3** | 2900056N03Rik\|A130075N07\|AI848000\|E030045D18Rik\|KBP-1\|KBP1\|Krc\|Rc\|Schnurri-3\|Shn3\|Zas3 | human immunodeficiency virus type I enhancer binding protein 3 | protein-coding |
| chr4_119786242_119786661 | chr4 | 119786243 | 119786661 | + | 1.1635 | Intergenic | Intergenic | -28226 | NM_010657 | 16656 | Mm.302758 | NM_010657 | ENSMUSG00000028634 | **Hivep3** | 2900056N03Rik\|A130075N07\|AI848000\|E030045D18Rik\|KBP-1\|KBP1\|Krc\|Rc\|Schnurri-3\|Shn3\|Zas3 | human immunodeficiency virus type I enhancer binding protein 3 | protein-coding |
| chr4_119897233_119897932 | chr4 | 119897234 | 119897932 | + | 1.10983 | intron (NM_010657, intron 1 of 8) | intron (NM_010657, intron 1 of 8) | 82905 | NM_010657 | 16656 | Mm.302758 | NM_010657 | ENSMUSG00000028634 | **Hivep3** | 2900056N03Rik\|A130075N07\|AI848000\|E030045D18Rik\|KBP-1\|KBP1\|Krc\|Rc\|Schnurri-3\|Shn3\|Zas3 | human immunodeficiency virus type I enhancer binding protein 3 | protein-coding |
| chr4_119812730_119813923 | chr4 | 119812731 | 119813923 | + | 1.05652 | Intergenic | Intergenic | -1351 | NM_010657 | 16656 | Mm.302758 | NM_010657 | ENSMUSG00000028634 | **Hivep3** | 2900056N03Rik\|A130075N07\|AI848000\|E030045D18Rik\|KBP-1\|KBP1\|Krc\|Rc\|Schnurri-3\|Shn3\|Zas3 | human immunodeficiency virus type I enhancer binding protein 3 | protein-coding |
| chr4_119896667_119897960 | chr4 | 119896668 | 119897960 | + | 1.03062 | intron (NM_010657, intron 1 of 8) | intron (NM_010657, intron 1 of 8) | 82636 | NM_010657 | 16656 | Mm.302758 | NM_010657 | ENSMUSG00000028634 | **Hivep3** | 2900056N03Rik\|A130075N07\|AI848000\|E030045D18Rik\|KBP-1\|KBP1\|Krc\|Rc\|Schnurri-3\|Shn3\|Zas3 | human immunodeficiency virus type I enhancer binding protein 3 | protein-coding |
| chr4_119966306_119967031 | chr4 | 119966307 | 119967031 | + | 0.956078 | intron (NM_010657, intron 1 of 8) | intron (NM_010657, intron 1 of 8) | 151991 | NM_010657 | 16656 | Mm.302758 | NM_010657 | ENSMUSG00000028634 | **Hivep3** | 2900056N03Rik\|A130075N07\|AI848000\|E030045D18Rik\|KBP-1\|KBP1\|Krc\|Rc\|Schnurri-3\|Shn3\|Zas3 | human immunodeficiency virus type I enhancer binding protein 3 | protein-coding |
| chr4_119932558_119933296 | chr4 | 119932559 | 119933296 | + | 0.808096 | intron (NM_010657, intron 1 of 8) | intron (NM_010657, intron 1 of 8) | 118249 | NM_010657 | 16656 | Mm.302758 | NM_010657 | ENSMUSG00000028634 | **Hivep3** | 2900056N03Rik\|A130075N07\|AI848000\|E030045D18Rik\|KBP-1\|KBP1\|Krc\|Rc\|Schnurri-3\|Shn3\|Zas3 | human immunodeficiency virus type I enhancer binding protein 3 | protein-coding |
| chr4_119962779_119963136 | chr4 | 119962780 | 119963136 | + | 0.793705 | intron (NM_010657, intron 1 of 8) | intron (NM_010657, intron 1 of 8) | 148280 | NM_010657 | 16656 | Mm.302758 | NM_010657 | ENSMUSG00000028634 | **Hivep3** | 2900056N03Rik\|A130075N07\|AI848000\|E030045D18Rik\|KBP-1\|KBP1\|Krc\|Rc\|Schnurri-3\|Shn3\|Zas3 | human immunodeficiency virus type I enhancer binding protein 3 | protein-coding |
| chr4_119932616_119933036 | chr4 | 119932617 | 119933036 | + | 0.783978 | intron (NM_010657, intron 1 of 8) | intron (NM_010657, intron 1 of 8) | 118148 | NM_010657 | 16656 | Mm.302758 | NM_010657 | ENSMUSG00000028634 | **Hivep3** | 2900056N03Rik\|A130075N07\|AI848000\|E030045D18Rik\|KBP-1\|KBP1\|Krc\|Rc\|Schnurri-3\|Shn3\|Zas3 | human immunodeficiency virus type I enhancer binding protein 3 | protein-coding |
| chr4_119962119_119962706 | chr4 | 119962120 | 119962706 | + | 0.753739 | intron (NM_010657, intron 1 of 8) | intron (NM_010657, intron 1 of 8) | 147735 | NM_010657 | 16656 | Mm.302758 | NM_010657 | ENSMUSG00000028634 | **Hivep3** | 2900056N03Rik\|A130075N07\|AI848000\|E030045D18Rik\|KBP-1\|KBP1\|Krc\|Rc\|Schnurri-3\|Shn3\|Zas3 | human immunodeficiency virus type I enhancer binding protein 3 | protein-coding |
| chr4_119976474_119977107 | chr4 | 119976475 | 119977107 | + | 0.654169 | intron (NM_010657, intron 1 of 8) | (TC)n\|Simple_repeat\|Simple_repeat | 162113 | NM_010657 | 16656 | Mm.302758 | NM_010657 | ENSMUSG00000028634 | **Hivep3** | 2900056N03Rik\|A130075N07\|AI848000\|E030045D18Rik\|KBP-1\|KBP1\|Krc\|Rc\|Schnurri-3\|Shn3\|Zas3 | human immunodeficiency virus type I enhancer binding protein 3 | protein-coding |
| chr6_82744378_82745150 | chr6 | 82744379 | 82745150 | + | 1.92732 | intron (NM_013820, intron 4 of 17) | intron (NM_013820, intron 4 of 17) | 29690 | NM_013820 | 15277 | Mm.255848 | NM_013820 | ENSMUSG00000000628 | **Hk2** | AI642394\|HKII | hexokinase 2 | protein-coding |
| chr6_82753017_82754024 | chr6 | 82753018 | 82754024 | + | 1.14541 | intron (NM_013820, intron 2 of 17) | intron (NM_013820, intron 2 of 17) | 20933 | NM_013820 | 15277 | Mm.255848 | NM_013820 | ENSMUSG00000000628 | **Hk2** | AI642394\|HKII | hexokinase 2 | protein-coding |
| chr6_82747127_82747428 | chr6 | 82747128 | 82747428 | + | 0.924139 | intron (NM_013820, intron 3 of 17) | intron (NM_013820, intron 3 of 17) | 27176 | NM_013820 | 15277 | Mm.255848 | NM_013820 | ENSMUSG00000000628 | **Hk2** | AI642394\|HKII | hexokinase 2 | protein-coding |
| chr6_82769737_82770036 | chr6 | 82769738 | 82770036 | + | 0.69843 | intron (NM_013820, intron 1 of 17) | intron (NM_013820, intron 1 of 17) | 4567 | NM_013820 | 15277 | Mm.255848 | NM_013820 | ENSMUSG00000000628 | **Hk2** | AI642394\|HKII | hexokinase 2 | protein-coding |
| chr6_82773209_82773904 | chr6 | 82773210 | 82773904 | + | 0.645981 | intron (NM_013820, intron 1 of 17) | CpG | 897 | NM_013820 | 15277 | Mm.255848 | NM_013820 | ENSMUSG00000000628 | **Hk2** | AI642394\|HKII | hexokinase 2 | protein-coding |

| | | | | | | | | | | | | | | | | | |
|---|---|---|---|---|---|---|---|---|---|---|---|---|---|---|---|---|---|
| chr16_94299179_94299610 | chr16 | 94299180 | 94299610 | + | 0.769401 | Intergenic | Intergenic | -11539 | NM_139145 | 110948 | Mm.30921 | NM_139145 | ENSMUSG00000040820 | **Hlcs** | 410I21.SP6\|D16Jhu34 | holocarboxylase synthetase (biotin- [propriony-Coenzyme A-carboxylase (ATP-hydrolysing)] ligase) | protein-coding |
| chr11_90349356_90350670 | chr11 | 90349357 | 90350670 | + | 0.813763 | intron (NM_172563, intron 2 of 3) | intron (NM_172563, intron 2 of 3) | 40904 | NM_172563 | 217082 | Mm.158903 | NM_172563 | ENSMUSG00000003949 | **Hlf** | E230015K02Rik | hepatic leukemia factor | protein-coding |
| chr14_64926948_64927290 | chr14 | 64926949 | 64927290 | + | 0.722553 | intron (NM_177338, intron 1 of 9) | B3\|SINE\|B2 | 22728 | NM_177338 | 219150 | Mm.344074 | NM_177338 | ENSMUSG00000021972 | **Hmbox1** | AI451877\|AI604847\|F830020C16Rik | homeobox containing 1 | protein-coding |
| chr6_87916024_87916543 | chr6 | 87916025 | 87916543 | + | 0.937831 | intron (NM_173737, intron 2 of 6) | intron (NM_173737, intron 2 of 6) | 2308 | NM_173737 | 232210 | Mm.246241 | NM_173737 | ENSMUSG00000030060 | **Hmces** | 8430410A17Rik\|C85376 | 5-hydroxymethylcytosine (hmC) binding, ES cell specific | protein-coding |
| chr1_150774410_150774943 | chr1 | 150774411 | 150774943 | + | 0.790072 | intron (NM_001024720, intron 11 of 106) | intron (NM_001024720, intron 11 of 106) | 218758 | NM_001024720 | 545370 | Mm.379425 | NM_001024720 | ENSMUSG00000066842 | **Hmcn1** | EG545370\|Gm201 | hemicentin 1 | protein-coding |
| chr9_56497274_56497650 | chr9 | 56497275 | 56497650 | + | 0.821589 | TTS (NM_025812) | TTS (NM_025812) | 78616 | NM_025812 | 66867 | Mm.150856 | NM_025812 | ENSMUSG00000032329 | **Hmg20a** | 1200004E06Rik\|5730490E10Rik\|Hmgxb1\|Ibraf | high mobility group 20A | protein-coding |
| chr1_176996797_176997317 | chr1 | 176996798 | 176997317 | + | 1.1558 | intron (NM_029756, intron 16 of 17) | intron (NM_029756, intron 16 of 17) | -160924 | NR_037996 | 15365 | Mm.441435 | NR_037996 | | **Hmga2-ps1** | EG666235\|Gm7996\|Hmgic-ps1\|[y]i-c | high mobility group AT-hook 2, pseudogene 1 | pseudo |
| chr1_176952697_176952997 | chr1 | 176952698 | 176952997 | + | 1.08392 | intron (NM_029756, intron 15 of 17) | intron (NM_029756, intron 15 of 17) | -116714 | NR_037996 | 15365 | Mm.441435 | NR_037996 | | **Hmga2-ps1** | EG666235\|Gm7996\|Hmgic-ps1\|[y]i-c | high mobility group AT-hook 2, pseudogene 1 | pseudo |
| chr1_176995388_176995961 | chr1 | 176995389 | 176995961 | + | 1.01673 | intron (NM_029756, intron 16 of 17) | intron (NM_029756, intron 16 of 17) | -159542 | NR_037996 | 15365 | Mm.441435 | NR_037996 | | **Hmga2-ps1** | EG666235\|Gm7996\|Hmgic-ps1\|[y]i-c | high mobility group AT-hook 2, pseudogene 1 | pseudo |
| chr1_176995607_176995930 | chr1 | 176995608 | 176995930 | + | 1.01377 | intron (NM_029756, intron 16 of 17) | intron (NM_029756, intron 16 of 17) | -159636 | NR_037996 | 15365 | Mm.441435 | NR_037996 | | **Hmga2-ps1** | EG666235\|Gm7996\|Hmgic-ps1\|[y]i-c | high mobility group AT-hook 2, pseudogene 1 | pseudo |
| chr1_176832633_176833052 | chr1 | 176832634 | 176833052 | + | 0.769434 | intron (NM_029756, intron 5 of 17) | intron (NM_029756, intron 5 of 17) | 3290 | NR_037996 | 15365 | Mm.441435 | NR_037996 | | **Hmga2-ps1** | EG666235\|Gm7996\|Hmgic-ps1\|[y]i-c | high mobility group AT-hook 2, pseudogene 1 | pseudo |
| chr1_176939698_176940119 | chr1 | 176939699 | 176940119 | + | 0.66976 | intron (NM_029756, intron 13 of 17) | intron (NM_029756, intron 13 of 17) | -103776 | NR_037996 | 15365 | Mm.441435 | NR_037996 | | **Hmga2-ps1** | EG666235\|Gm7996\|Hmgic-ps1\|[y]i-c | high mobility group AT-hook 2, pseudogene 1 | pseudo |
| chr8_33482435_33482836 | chr8 | 33482436 | 33482836 | + | 0.896781 | Intergenic | Lx3_Mus\|LINE\|L1 | -1675 | NR_033589 | 628431 | Mm.376526 | NR_033589 | | **Hmgb1-rs17** | Hmg1-rs17 | high mobility group box 1, related sequence 17 | pseudo |
| chr8_33435245_33435784 | chr8 | 33435246 | 33435784 | + | 0.792295 | Intergenic | Intergenic | -48796 | NR_033589 | 628431 | Mm.376526 | NR_033589 | | **Hmgb1-rs17** | Hmg1-rs17 | high mobility group box 1, related sequence 17 | pseudo |
| chr8_57577179_57577583 | chr8 | 57577180 | 57577583 | + | 1.29285 | intron (NM_001167981, intron 2 of 11) | intron (NM_001167981, intron 2 of 11) | 65538 | NM_008252 | 97165 | Mm.276881 | NM_008252 | ENSMUSG00000054717 | **Hmgb2** | C80539\|HMG-2\|Hmg2 | high mobility group box 2 | protein-coding |
| chr8_57577736_57578107 | chr8 | 57577737 | 57578107 | + | 1.24199 | intron (NM_001167981, intron 2 of 11) | intron (NM_001167981, intron 2 of 11) | 66079 | NM_008252 | 97165 | Mm.276881 | NM_008252 | ENSMUSG00000054717 | **Hmgb2** | C80539\|HMG-2\|Hmg2 | high mobility group box 2 | protein-coding |
| chr8_57579782_57580182 | chr8 | 57579783 | 57580182 | + | 0.652846 | intron (NM_001167981, intron 2 of 11) | intron (NM_001167981, intron 2 of 11) | 68139 | NM_008252 | 97165 | Mm.276881 | NM_008252 | ENSMUSG00000054717 | **Hmgb2** | C80539\|HMG-2\|Hmg2 | high mobility group box 2 | protein-coding |
| chrX_71524207_71524666 | chrX | 71524208 | 71524666 | + | 0.82855 | Intergenic | L1_Mur3\|LINE\|L1 | -31480 | NM_001293624 | 15354 | Mm.336087 | NM_008253 | ENSMUSG00000015217 | **Hmgb3** | Hmg2a\|Hmg4 | high mobility group box 3 | protein-coding |
| chr16_96126730_96126992 | chr16 | 96126731 | 96126992 | + | 0.77459 | intron (NM_008251, intron 4 of 5) | B3A\|SINE\|B2 | 864 | NM_008251 | 15312 | Mm.2756 | NM_008251 | | **Hmgn1** | HMG-14\|Hmg14 | high mobility group nucleosomal binding domain 1 | protein-coding |
| chr18_61183923_61184317 | chr18 | 61183924 | 61184317 | + | 0.814032 | Intergenic | Intergenic | -7070 | NM_134134 | 106894 | Mm.22574 | NM_134134 | ENSMUSG00000024622 | **Hmgxb3** | 2510002C16Rik\|A630042L21Rik\|AI413166\|mKIAA0194 | HMG box domain containing 3 | protein-coding |
| chr16_4762318_4762860 | chr16 | 4762319 | 4762860 | + | 1.02167 | 5' UTR (NM_010443, exon 2 of 6) | 5' UTR (NM_010443, exon 2 of 6) | 20976 | NM_010443 | 15369 | Mm.272866 | NM_010443 | ENSMUSG00000004070 | **Hmox2** | HO-2\|HO2 | heme oxygenase (decycling) 2 | protein-coding |
| chr16_4741647_4742580 | chr16 | 4741648 | 4742580 | + | 0.917441 | intron (NM_010443, intron 1 of 5) | intron (NM_010443, intron 1 of 5) | 501 | NM_010443 | 15369 | Mm.272866 | NM_010443 | ENSMUSG00000004070 | **Hmox2** | HO-2\|HO2 | heme oxygenase (decycling) 2 | protein-coding |
| chr11_83828490_83829285 | chr11 | 83828491 | 83829285 | + | 1.31956 | Intergenic | Intergenic | -21981 | NM_001291268 | 21410 | Mm.7226 | NM_009330 | ENSMUSG00000020679 | **Hnf1b** | HNF-1-beta\|HNF-1B\|HNF-1Beta\|Hnf1beta\|LFB3\|Tcf-2\|Tcf2\|vHNF1 | HNF1 homeobox B | protein-coding |
| chr14_52094513_52094995 | chr14 | 52094514 | 52094995 | + | 1.15928 | intron (NM_001170982, intron 1 of 7) | URR1A\|DNA\|hAT-Charlie | 9274 | NM_001170982 | 15381 | Mm.427321 | NM_016884 | ENSMUSG00000060373 | **Hnrnpc** | AL022939\|D14Wsu171e\|Hnrpc\|Hnrpc1\|Hnrpc2\|hnRNPC1\|hnRNPC2\|hnrnp-C | heterogeneous nuclear ribonucleoprotein C | protein-coding |
| chr14_52098451_52098732 | chr14 | 52098452 | 52098732 | + | 0.958245 | intron (NM_001170981, intron 1 of 8) | intron (NM_001170981, intron 1 of 8) | 5436 | NM_001170982 | 15381 | Mm.427321 | NM_016884 | ENSMUSG00000060373 | **Hnrnpc** | AL022939\|D14Wsu171e\|Hnrpc\|Hnrpc1\|Hnrpc2\|hnRNPC1\|hnRNPC2\|hnrnp-C | heterogeneous nuclear ribonucleoprotein C | protein-coding |
| chr14_52100668_52100877 | chr14 | 52100669 | 52100877 | + | 0.883444 | intron (NM_001170981, intron 1 of 8) | intron (NM_001170981, intron 1 of 8) | 3255 | NM_001170982 | 15381 | Mm.427321 | NM_016884 | ENSMUSG00000060373 | **Hnrnpc** | AL022939\|D14Wsu171e\|Hnrpc\|Hnrpc1\|Hnrpc2\|hnRNPC1\|hnRNPC2\|hnrnp-C | heterogeneous nuclear ribonucleoprotein C | protein-coding |
| chr5_99973973_99974330 | chr5 | 99973974 | 99974330 | + | 0.79902 | intron (NM_001077267, intron 1 of 6) | intron (NM_001077267, intron 1 of 6) | 4786 | NM_001077266 | 11991 | Mm.150231 | NM_007516 | | **Hnrnpd** | Auf1\|Hnrpd | heterogeneous nuclear ribonucleoprotein D | protein-coding |
| chr11_50369290_50369717 | chr11 | 50369291 | 50369717 | + | 1.00408 | Intergenic | (TC)n\|Simple_repeat\|Simple_repeat | -8215 | NM_021510 | 59013 | Mm.21740 | NM_021510 | ENSMUSG00000007850 | **Hnrnph1** | AI642080\|E430005G16Rik\|Hnrnph\|Hnrph1 | heterogeneous nuclear ribonucleoprotein H1 | protein-coding |
| chr13_58453555_58454138 | chr13 | 58453556 | 58454138 | + | 0.945446 | Intergenic | Intergenic | -50504 | NM_001301341 | 15387 | Mm.142872 | NM_025279 | ENSMUSG00000021546 | **Hnrnpk** | Hnrpk\|KBBP\|NOVA | heterogeneous nuclear ribonucleoprotein K | protein-coding |
| chr13_93331643_93332242 | chr13 | 93331644 | 93332242 | + | 1.20509 | intron (NM_152134, intron 1 of 8) | L1M5\|LINE\|L1 | 23909 | NM_152134 | 26556 | Mm.37533 | NM_011982 | | **Homer1** | PSD-Zip45\|SYN47\|Ves-1\|homer-1\|vesl-1 | homer homolog 1 (Drosophila) | protein-coding |
| chr13_93330726_93331266 | chr13 | 93330727 | 93331266 | + | 1.1772 | intron (NM_152134, intron 1 of 8) | ORR1D1\|LTR\|ERVL-MaLR | 22962 | NM_152134 | 26556 | Mm.37533 | NM_011982 | | **Homer1** | PSD-Zip45\|SYN47\|Ves-1\|homer-1\|vesl-1 | homer homolog 1 (Drosophila) | protein-coding |
| chr11_4506365_4506826 | chr11 | 4506366 | 4506826 | + | 1.61072 | intron (NM_028860, intron 7 of 17) | intron (NM_028860, intron 7 of 17) | -65514 | NM_029458 | 75828 | Mm.159592 | NM_029458 | ENSMUSG00000020419 | **Hormad2** | 4930529M09Rik | HORMA domain containing 2 | protein-coding |
| chr11_4501735_4502269 | chr11 | 4501736 | 4502269 | + | 1.15299 | intron (NM_028860, intron 8 of 17) | intron (NM_028860, intron 8 of 17) | -60920 | NM_029458 | 75828 | Mm.159592 | NM_029458 | ENSMUSG00000020419 | **Hormad2** | 4930529M09Rik | HORMA domain containing 2 | protein-coding |
| chr11_4497454_4498056 | chr11 | 4497455 | 4498056 | + | 1.14671 | intron (NM_028860, intron 10 of 17) | intron (NM_028860, intron 10 of 17) | -56673 | NM_029458 | 75828 | Mm.159592 | NM_029458 | ENSMUSG00000020419 | **Hormad2** | 4930529M09Rik | HORMA domain containing 2 | protein-coding |
| chr11_4438086_4438808 | chr11 | 4438087 | 4438808 | + | 1.10562 | intron (NM_029458, intron 1 of 10) | intron (NM_029458, intron 1 of 10) | 2635 | NM_029458 | 75828 | Mm.159592 | NM_029458 | ENSMUSG00000020419 | **Hormad2** | 4930529M09Rik | HORMA domain containing 2 | protein-coding |
| chr11_4459466_4459802 | chr11 | 4459467 | 4459802 | + | 0.683281 | Intergenic | Intergenic | -18552 | NM_029458 | 75828 | Mm.159592 | NM_029458 | ENSMUSG00000020419 | **Hormad2** | 4930529M09Rik | HORMA domain containing 2 | protein-coding |
| chr11_96280315_96280582 | chr11 | 96280316 | 96280582 | + | 0.780638 | Intergenic | Intergenic | -1456 | NM_010461 | 15416 | Mm.4822 | NM_010461 | ENSMUSG00000056648 | **Hoxb8** | Hox-2.4 | homeobox B8 | protein-coding |
| chr4_138213892_138214252 | chr4 | 138213893 | 138214252 | + | 1.52334 | Intergenic | Intergenic | -2540 | NM_001122896 | 15441 | Mm.348429 | NM_010470 | ENSMUSG00000028759 | **Hp1bp3** | Hp1bp74 | heterochromatin protein 1, binding protein 3 | protein-coding |
| chr4_129113059_129113492 | chr4 | 129113060 | 129113492 | + | 0.599243 | intron (NM_001286081, intron 2 of 3) | intron (NM_001286081, intron 2 of 3) | 6896 | NM_001286083 | 15444 | Mm.384452 | NM_010471 | ENSMUSG00000028785 | **Hpca** | - | hippocalcin | protein-coding |
| chr12_17688987_17689227 | chr12 | 17688988 | 17689227 | + | 1.10159 | Intergenic | MLT1J1\|LTR\|ERVL-MaLR | -1707 | NM_016677 | 53602 | Mm.313619 | NM_016677 | ENSMUSG00000071379 | **Hpcal1** | NVP-3\|Nvp3\|VILIP3\|Vnsl3\|Vsnl3 | hippocalcin-like 1 | protein-coding |
| chr12_17745882_17746367 | chr12 | 17745883 | 17746367 | + | 0.911776 | intron (NM_016677, intron 1 of 3) | intron (NM_016677, intron 1 of 3) | 55311 | NM_016677 | 53602 | Mm.313619 | NM_016677 | ENSMUSG00000071379 | **Hpcal1** | NVP-3\|Nvp3\|VILIP3\|Vnsl3\|Vsnl3 | hippocalcin-like 1 | protein-coding |
| chr12_17688980_17689683 | chr12 | 17688981 | 17689683 | + | 0.835315 | Intergenic | Intergenic | -1482 | NM_016677 | 53602 | Mm.313619 | NM_016677 | ENSMUSG00000071379 | **Hpcal1** | NVP-3\|Nvp3\|VILIP3\|Vnsl3\|Vsnl3 | hippocalcin-like 1 | protein-coding |
| chr12_17737147_17738143 | chr12 | 17737148 | 17738143 | + | 0.764428 | intron (NM_016677, intron 1 of 3) | (TG)n\|Simple_repeat\|Simple_repeat | 46831 | NM_016677 | 53602 | Mm.313619 | NM_016677 | ENSMUSG00000071379 | **Hpcal1** | NVP-3\|Nvp3\|VILIP3\|Vnsl3\|Vsnl3 | hippocalcin-like 1 | protein-coding |
| chr12_17787626_17788228 | chr12 | 17787627 | 17788228 | + | 0.729491 | intron (NM_016677, intron 3 of 3) | MIRc\|SINE\|MIR | 97113 | NM_016677 | 53602 | Mm.313619 | NM_016677 | ENSMUSG00000071379 | **Hpcal1** | NVP-3\|Nvp3\|VILIP3\|Vnsl3\|Vsnl3 | hippocalcin-like 1 | protein-coding |
| chr12_17733163_17733457 | chr12 | 17733164 | 17733457 | + | 0.649761 | intron (NM_016677, intron 1 of 3) | intron (NM_016677, intron 1 of 3) | 42496 | NM_016677 | 53602 | Mm.313619 | NM_016677 | ENSMUSG00000071379 | **Hpcal1** | NVP-3\|Nvp3\|VILIP3\|Vnsl3\|Vsnl3 | hippocalcin-like 1 | protein-coding |
| chr6_65190328_65190712 | chr6 | 65190329 | 65190712 | + | 1.67411 | Intergenic | Intergenic | -45790 | NM_019455 | 54486 | Mm.410688 | NM_019455 | ENSMUSG00000029919 | **Hpgds** | H-PGDS\|Ptgds2 | hematopoietic prostaglandin D synthase | protein-coding |
| chr6_65193052_65193351 | chr6 | 65193053 | 65193351 | + | 0.693987 | Intergenic | MTD\|LTR\|ERVL-MaLR | -48472 | NM_019455 | 54486 | Mm.410688 | NM_019455 | ENSMUSG00000029919 | **Hpgds** | H-PGDS\|Ptgds2 | hematopoietic prostaglandin D synthase | protein-coding |
| chr7_31100809_31101098 | chr7 | 31100810 | 31101098 | + | 0.868419 | intron (NM_008281, intron 9 of 12) | intron (NM_008281, intron 9 of 12) | 10043 | NM_001276269 | 15451 | Mm.19182 | NM_008281 | ENSMUSG00000001249 | **Hpn** | Hlb320 | hepsin | protein-coding |
| chrX_52972099_52972491 | chrX | 52972100 | 52972491 | + | 1.1667 | Intergenic | Intergenic | -15783 | NM_013556 | 15452 | Mm.299381 | NM_013556 | ENSMUSG00000025630 | **Hprt** | C81579\|HPGRT\|Hprt1 | hypoxanthine guanine phosphoribosyl transferase | protein-coding |

| | | | | | | | | | | | | | | | | | |
|---|---|---|---|---|---|---|---|---|---|---|---|---|---|---|---|---|---|
| chrX_52972542_52973063 | chrX | 52972543 | 52973063 | + | 0.706649 | Intergenic | Intergenic | -15275 | NM_013556 | 15452 | Mm.299381 | NM_013556 | ENSMUSG00000025630 | **Hprt** | C81579\|HPGRT\|Hprt1 | hypoxanthine guanine phosphoribosyl transferase | protein-coding |
| chrX_52971024_52971429 | chrX | 52971025 | 52971429 | + | 0.686421 | Intergenic | RMER1A\|Other\|Other | -16851 | NM_013556 | 15452 | Mm.299381 | NM_013556 | ENSMUSG00000025630 | **Hprt** | C81579\|HPGRT\|Hprt1 | hypoxanthine guanine phosphoribosyl transferase | protein-coding |
| chr19_42796234_42796603 | chr19 | 42796235 | 42796603 | + | 1.02089 | intron (NM_001081257, intron 11 of 11) | intron (NM_001081257, intron 11 of 11) | -16443 | NM_019424 | 192236 | Mm.218381 | NM_019424 | ENSMUSG00000025188 | **Hps1** | 6030422N11Rik\|BB405864\|Gm21361\|Hps\|ep | Hermansky-Pudlak syndrome 1 homolog (human) | protein-coding |
| chr19_46002233_46002655 | chr19 | 46002234 | 46002655 | + | 1.14928 | Intergenic | ORR1C2\|LTR\|ERVL-MaLR | -1034 | NM_176785 | 20170 | Mm.101110 | NM_176785 | ENSMUSG00000074811 | **Hps6** | 5330434M19Rik\|BLOC2\|ru | Hermansky-Pudlak syndrome 6 | protein-coding |
| chr19_46006622_46006807 | chr19 | 46006623 | 46006807 | + | 0.744837 | TTS (NM_176785) | TTS (NM_176785) | 3237 | NM_176785 | 20170 | Mm.101110 | NM_176785 | ENSMUSG00000074811 | **Hps6** | 5330434M19Rik\|BLOC2\|ru | Hermansky-Pudlak syndrome 6 | protein-coding |
| chr19_43193599_43194062 | chr19 | 43193600 | 43194062 | + | 0.787838 | intron (NM_001081257, intron 3 of 11) | AmnSINE1\|SINE\|Deu | 194524 | NM_001081257 | 545291 | Mm.442200 | NM_001081257 | ENSMUSG00000074852 | **Hpse2** | Gm968\|Hpa2 | heparanase 2 | protein-coding |
| chr7_45338128_45338605 | chr7 | 45338129 | 45338605 | + | 0.769674 | TTS (NM_029741) | TTS (NM_029741) | 3098 | NM_010473 | 15464 | Mm.39968 | NM_010473 | ENSMUSG00000038239 | **Hrc** | - | histidine rich calcium binding protein | protein-coding |
| chr7_45338116_45338796 | chr7 | 45338117 | 45338796 | + | 0.696206 | TTS (NM_029741) | TTS (NM_029741) | 3187 | NM_010473 | 15464 | Mm.39968 | NM_010473 | ENSMUSG00000038239 | **Hrc** | - | histidine rich calcium binding protein | protein-coding |
| chr6_114347844_114348758 | chr6 | 114347845 | 114348758 | + | 1.39341 | Intergenic | Intergenic | -49635 | NM_008285 | 15465 | Mm.333327 | NM_008285 | ENSMUSG00000053004 | **Hrh1** | Bphs\|H1R\|HH1R\|Hir | histamine receptor H1 | protein-coding |
| chr3_144431964_144432652 | chr3 | 144431965 | 144432652 | + | 1.62372 | 3' UTR (NM_011828, exon 7 of 7) | 3' UTR (NM_011828, exon 7 of 7) | 137908 | NM_011828 | 23908 | Mm.12863 | NM_011828 | ENSMUSG00000040151 | **Hs2st1** | AW214369\|Hs2st\|mKIAA0448 | heparan sulfate 2-O-sulfotransferase 1 | protein-coding |
| chr3_144475810_144476499 | chr3 | 144475811 | 144476499 | + | 1.0802 | intron (NM_011828, intron 1 of 6) | intron (NM_011828, intron 1 of 6) | 94061 | NM_011828 | 23908 | Mm.12863 | NM_011828 | ENSMUSG00000040151 | **Hs2st1** | AW214369\|Hs2st\|mKIAA0448 | heparan sulfate 2-O-sulfotransferase 1 | protein-coding |
| chr3_144448864_144449563 | chr3 | 144448865 | 144449563 | + | 0.902179 | intron (NM_011828, intron 3 of 6) | L2a\|LINE\|L2 | 121002 | NM_011828 | 23908 | Mm.12863 | NM_011828 | ENSMUSG00000040151 | **Hs2st1** | AW214369\|Hs2st\|mKIAA0448 | heparan sulfate 2-O-sulfotransferase 1 | protein-coding |
| chr1_35796893_35797488 | chr1 | 35796894 | 35797488 | + | 0.892378 | Intergenic | Intergenic | -271209 | NM_015818 | 50785 | Mm.213566 | NM_015818 | ENSMUSG00000045216 | **Hs6st1** | 6Ost1 | heparan sulfate 6-O-sulfotransferase 1 | protein-coding |
| chr1_35797103_35797473 | chr1 | 35797104 | 35797473 | + | 0.799791 | Intergenic | L1MDa\|LINE\|L1 | -271112 | NM_015818 | 50785 | Mm.213566 | NM_015818 | ENSMUSG00000045216 | **Hs6st1** | 6Ost1 | heparan sulfate 6-O-sulfotransferase 1 | protein-coding |
| chr1_35913510_35913843 | chr1 | 35913511 | 35913843 | + | 0.650594 | Intergenic | RLTR30D_MM\|LTR\|ERV1 | -154723 | NM_015818 | 50785 | Mm.213566 | NM_015818 | ENSMUSG00000045216 | **Hs6st1** | 6Ost1 | heparan sulfate 6-O-sulfotransferase 1 | protein-coding |
| chr18_80228391_80228876 | chr18 | 80228392 | 80228876 | + | 1.15277 | Intergenic | Intergenic | 18468 | NM_001136181 | 66255 | Mm.272341 | NM_001136181 | ENSMUSG00000078963 | **Hsbp1l1** | 1810005K13Rik | heat shock factor binding protein 1-like 1 | protein-coding |
| chr1_193253831_193254310 | chr1 | 193253832 | 193254310 | + | 0.750974 | intron (NM_008288, intron 1 of 6) | RMER4A\|LTR\|ERVK | 9974 | NM_008288 | 15483 | Mm.28328 | NM_008288 | ENSMUSG00000016194 | **Hsd11b1** | - | hydroxysteroid 11-beta dehydrogenase 1 | protein-coding |
| chr1_193225503_193226154 | chr1 | 193225504 | 193226154 | + | 0.621352 | intron (NM_008288, intron 5 of 6) | intron (NM_008288, intron 5 of 6) | 16531 | NM_001044751 | 15483 | Mm.28328 | NM_008288 | ENSMUSG00000016194 | **Hsd11b1** | - | hydroxysteroid 11-beta dehydrogenase 1 | protein-coding |
| chrX_151975534_151976170 | chrX | 151975535 | 151976170 | + | 0.780999 | Intergenic | Intergenic | -26044 | NM_016763 | 15108 | Mm.6994 | NM_016763 | ENSMUSG00000025260 | **Hsd17b10** | 17bHSD10\|Ads9\|ERAB\|Hadh2 | hydroxysteroid (17-beta) dehydrogenase 10 | protein-coding |
| chr2_94114553_94115011 | chr2 | 94114554 | 94115011 | + | 0.982454 | intron (NM_019657, intron 3 of 10) | Tigger13a\|DNA\|TcMar-Tigger | 43127 | NM_019657 | 56348 | Mm.22505 | NM_019657 | ENSMUSG00000027195 | **Hsd17b12** | 2610510O05Rik\|AI172963\|KIK-I\|Kik1 | hydroxysteroid (17-beta) dehydrogenase 12 | protein-coding |
| chr2_94142379_94142798 | chr2 | 94142380 | 94142798 | + | 0.951188 | intron (NM_019657, intron 1 of 10) | L1M2\|LINE\|L1 | 15320 | NM_019657 | 56348 | Mm.22505 | NM_019657 | ENSMUSG00000027195 | **Hsd17b12** | 2610510O05Rik\|AI172963\|KIK-I\|Kik1 | hydroxysteroid (17-beta) dehydrogenase 12 | protein-coding |
| chr2_94136892_94137234 | chr2 | 94136893 | 94137234 | + | 0.826325 | intron (NM_019657, intron 1 of 10) | intron (NM_019657, intron 1 of 10) | 20846 | NM_019657 | 56348 | Mm.22505 | NM_019657 | ENSMUSG00000027195 | **Hsd17b12** | 2610510O05Rik\|AI172963\|KIK-I\|Kik1 | hydroxysteroid (17-beta) dehydrogenase 12 | protein-coding |
| chr13_64077645_64077929 | chr13 | 64077646 | 64077929 | + | 0.666592 | intron (NM_008291, intron 2 of 10) | B3\|SINE\|B2 | 11414 | NM_008291 | 15487 | Mm.5109 | NM_008291 | ENSMUSG00000033122 | **Hsd17b3** | - | hydroxysteroid (17-beta) dehydrogenase 3 | protein-coding |
| chr18_50096288_50097072 | chr18 | 50096289 | 50097072 | + | 1.33597 | Intergenic | Intergenic | -31521 | NM_008292 | 15488 | Mm.277857 | NM_008292 | ENSMUSG00000024507 | **Hsd17b4** | 17-beta-HSD\|17[b]-HSD\|4\|DBP\|MFE-2\|MFP2\|MPF-2\|Mfp-2\|perMFE-2 | hydroxysteroid (17-beta) dehydrogenase 4 | protein-coding |
| chr18_50095677_50095995 | chr18 | 50095678 | 50095995 | + | 0.785864 | Intergenic | ID_B1\|SINE\|B4 | -32365 | NM_008292 | 15488 | Mm.277857 | NM_008292 | ENSMUSG00000024507 | **Hsd17b4** | 17-beta-HSD\|17[b]-HSD\|4\|DBP\|MFE-2\|MFP2\|MPF-2\|Mfp-2\|perMFE-2 | hydroxysteroid (17-beta) dehydrogenase 4 | protein-coding |
| chr4_59575517_59575998 | chr4 | 59575518 | 59575998 | + | 0.779375 | Intergenic | ORR1E\|LTR\|ERVL-MaLR | -5805 | NM_001301367 | 72479 | Mm.272905 | NM_024255 | ENSMUSG00000028383 | **Hsdl2** | 2610207I16Rik | hydroxysteroid dehydrogenase like 2 | protein-coding |
| chr10_57487228_57487740 | chr10 | 57487229 | 57487740 | + | 1.09445 | intron (NM_008297, intron 1 of 11) | intron (NM_008297, intron 1 of 11) | 1099 | NM_008297 | 15500 | Mm.261395 | NM_008297 | ENSMUSG00000019878 | **Hsf2** | AI661205 | heat shock factor 2 | protein-coding |
| chr16_75784735_75785115 | chr16 | 75784736 | 75785115 | + | 0.777364 | Intergenic | MTC\|LTR\|ERVL-MaLR | -18107 | NM_030201 | 110920 | Mm.26704 | NM_030201 | ENSMUSG00000032932 | **Hspa13** | 1600002I10Rik\|AV006182\|B230217N24Rik\|Stch | heat shock protein 70 family, member 13 | protein-coding |
| chr17_34961113_34961553 | chr17 | 34961114 | 34961553 | + | 0.89236 | Intergenic | Intergenic | -2095 | NM_010478 | 15511 | Mm.372314 | NM_010478 | ENSMUSG00000090877 | **Hspa1b** | Hsp70\|Hsp70-1\|Hsp70.1\|hsp68 | heat shock protein 1B | protein-coding |
| chr3_40761350_40761789 | chr3 | 40761351 | 40761789 | + | 1.82098 | intron (NM_011020, intron 6 of 18) | RMER15\|LTR\|ERVL | 15957 | NM_011020 | 18415 | Mm.39330 | NM_011020 | | **Hspa4l** | 94kDa\|AI461691\|APG-1\|Osp94 | heat shock protein 4 like | protein-coding |
| chr3_40746276_40746570 | chr3 | 40746277 | 40746570 | + | 0.692755 | intron (NM_011020, intron 1 of 18) | intron (NM_011020, intron 1 of 18) | 810 | NM_011020 | 18415 | Mm.39330 | NM_011020 | | **Hspa4l** | 94kDa\|AI461691\|APG-1\|Osp94 | heat shock protein 4 like | protein-coding |
| chr9_40743448_40744135 | chr9 | 40743449 | 40744135 | + | 0.905998 | intron (NM_133733, intron 1 of 6) | intron (NM_133733, intron 1 of 6) | -57481 | NM_031165 | 15481 | Mm.290774 | NM_031165 | ENSMUSG00000015656 | **Hspa8** | 2410008N15Rik\|Hsc70\|Hsc71\|Hsc73\|Hsp73\|Hspa10 | heat shock protein 8 | protein-coding |
| chr9_40806362_40807978 | chr9 | 40806363 | 40807978 | + | 0.753708 | Intergenic | ID_B1\|SINE\|B4 | 5897 | NM_031165 | 15481 | Mm.290774 | NM_031165 | ENSMUSG00000015656 | **Hspa8** | 2410008N15Rik\|Hsc70\|Hsc71\|Hsc73\|Hsp73\|Hspa10 | heat shock protein 8 | protein-coding |
| chr9_40743269_40744678 | chr9 | 40743270 | 40744678 | + | 0.649408 | intron (NM_133733, intron 1 of 6) | intron (NM_133733, intron 1 of 6) | -57299 | NM_031165 | 15481 | Mm.290774 | NM_031165 | ENSMUSG00000015656 | **Hspa8** | 2410008N15Rik\|Hsc70\|Hsc71\|Hsc73\|Hsp73\|Hspa10 | heat shock protein 8 | protein-coding |
| chr4_141425519_141426220 | chr4 | 141425520 | 141426220 | + | 0.871403 | TTS (NM_013868) | TTS (NM_013868) | 5091 | NM_013868 | 29818 | Mm.46181 | NM_013868 | ENSMUSG00000006221 | **Hspb7** | 27kDa\|Hsp25-2\|cvHsp | heat shock protein family, member 7 (cardiovascular) | protein-coding |
| chr4_141425305_141426422 | chr4 | 141425306 | 141426422 | + | 0.832698 | TTS (NM_013868) | TTS (NM_013868) | 5085 | NM_013868 | 29818 | Mm.46181 | NM_013868 | ENSMUSG00000006221 | **Hspb7** | 27kDa\|Hsp25-2\|cvHsp | heat shock protein family, member 7 (cardiovascular) | protein-coding |
| chr4_141419839_141420292 | chr4 | 141419840 | 141420292 | + | 0.67458 | promoter-TSS (NM_013868) | promoter-TSS (NM_013868) | -713 | NM_013868 | 29818 | Mm.46181 | NM_013868 | ENSMUSG00000006221 | **Hspb7** | 27kDa\|Hsp25-2\|cvHsp | heat shock protein family, member 7 (cardiovascular) | protein-coding |
| chr16_35813159_35813594 | chr16 | 35813160 | 35813594 | + | 1.10789 | intron (NM_175111, intron 3 of 7) | intron (NM_175111, intron 3 of 7) | 42991 | NM_175111 | 66667 | Mm.45272 | NM_175111 | ENSMUSG00000022849 | **Hspbap1** | 3830421G21Rik | Hspb associated protein 1 | protein-coding |
| chr13_105377057_105377590 | chr13 | 105377058 | 105377590 | + | 0.704115 | Intergenic | Intergenic | -66369 | NM_008308 | 15550 | Mm.4716 | NM_008308 | ENSMUSG00000021721 | **Htr1a** | Gpcr18 | 5-hydroxytryptamine (serotonin) receptor 1A | protein-coding |
| chr6_83053599_83054019 | chr6 | 83053600 | 83054019 | + | 0.708508 | promoter-TSS (NM_007517) | promoter-TSS (NM_007517) | 762 | NM_019752 | 64704 | Mm.21880 | NM_019752 | ENSMUSG00000068329 | **Htra2** | AI481710\|Omi\|Prss25\|mnd2 | HtrA serine peptidase 2 | protein-coding |
| chr5_35672883_35673438 | chr5 | 35672884 | 35673438 | + | 0.972034 | intron (NM_001042615, intron 1 of 6) | intron (NM_001042615, intron 1 of 6) | 6619 | NM_001042615 | 78558 | Mm.274255 | NM_030127 | ENSMUSG00000029096 | **Htra3** | 2210021K23Rik\|9530081K03Rik\|Prsp\|Tasp | HtrA serine peptidase 3 | protein-coding |
| chr5_34784848_34785523 | chr5 | 34784849 | 34785523 | + | 1.51102 | intron (NM_010414, intron 2 of 66) | MER21C\|LTR\|ERVL | 23446 | NM_010414 | 15194 | Mm.209071 | NM_010414 | ENSMUSG00000029104 | **Htt** | AI256365\|C430023I11Rik\|Hd\|Hdh\|IT15 | huntingtin | protein-coding |
| chr5_34785028_34785522 | chr5 | 34785029 | 34785522 | + | 1.4813 | intron (NM_010414, intron 2 of 66) | MER21C\|LTR\|ERVL | 23535 | NM_010414 | 15194 | Mm.209071 | NM_010414 | ENSMUSG00000029104 | **Htt** | AI256365\|C430023I11Rik\|Hd\|Hdh\|IT15 | huntingtin | protein-coding |
| chr5_34762940_34763207 | chr5 | 34762941 | 34763207 | + | 0.792067 | intron (NM_010414, intron 1 of 66) | intron (NM_010414, intron 1 of 66) | 1334 | NM_010414 | 15194 | Mm.209071 | NM_010414 | ENSMUSG00000029104 | **Htt** | AI256365\|C430023I11Rik\|Hd\|Hdh\|IT15 | huntingtin | protein-coding |
| chr16_90455621_90456092 | chr16 | 90455622 | 90456092 | + | 0.796022 | intron (NM_015755, intron 4 of 10) | MLT1A\|LTR\|ERVL-MaLR | 69460 | NM_015755 | 26559 | Mm.125874 | NM_015755 | ENSMUSG00000053414 | **Hunk** | Bstk1\|Mak-v | hormonally upregulated Neu-associated kinase | protein-coding |
| chr16_90378809_90379240 | chr16 | 90378810 | 90379240 | + | 0.739451 | Intergenic | Intergenic | -7372 | NM_015755 | 26559 | Mm.125874 | NM_015755 | ENSMUSG00000053414 | **Hunk** | Bstk1\|Mak-v | hormonally upregulated Neu-associated kinase | protein-coding |
| chr16_90455614_90456347 | chr16 | 90455615 | 90456347 | + | 0.736419 | intron (NM_015755, intron 4 of 10) | MLT1A\|LTR\|ERVL-MaLR | 69584 | NM_015755 | 26559 | Mm.125874 | NM_015755 | ENSMUSG00000053414 | **Hunk** | Bstk1\|Mak-v | hormonally upregulated Neu-associated kinase | protein-coding |
| chr13_30865520_30865922 | chr13 | 30865521 | 30865922 | + | 1.13145 | intron (NM_025588, intron 21 of 27) | intron (NM_025588, intron 21 of 27) | 82040 | NM_153072 | 210554 | Mm.450571 | NM_153072 | ENSMUSG00000076430 | **Hus1b** | - | Hus1 homolog b (S. pombe) | protein-coding |
| chr13_30879162_30879694 | chr13 | 30879163 | 30879694 | + | 1.12893 | intron (NM_025588, intron 13 of 27) | intron (NM_025588, intron 13 of 27) | 68333 | NM_153072 | 210554 | Mm.450571 | NM_153072 | ENSMUSG00000076430 | **Hus1b** | - | Hus1 homolog b (S. pombe) | protein-coding |
| chr13_30878889_30879727 | chr13 | 30878890 | 30879727 | + | 1.09276 | intron (NM_025588, intron 13 of 27) | intron (NM_025588, intron 13 of 27) | 68453 | NM_153072 | 210554 | Mm.450571 | NM_153072 | ENSMUSG00000076430 | **Hus1b** | - | Hus1 homolog b (S. pombe) | protein-coding |
| chr13_30865321_30865931 | chr13 | 30865322 | 30865931 | + | 0.913935 | intron (NM_025588, intron 21 of 27) | intron (NM_025588, intron 21 of 27) | 82135 | NM_153072 | 210554 | Mm.450571 | NM_153072 | ENSMUSG00000076430 | **Hus1b** | - | Hus1 homolog b (S. pombe) | protein-coding |
| chr13_30921890_30922252 | chr13 | 30921891 | 30922252 | + | 0.802503 | intron (NM_025588, intron 6 of 27) | L1MA4\|LINE\|L1 | 25690 | NM_153072 | 210554 | Mm.450571 | NM_153072 | ENSMUSG00000076430 | **Hus1b** | - | Hus1 homolog b (S. pombe) | protein-coding |
| chrX_151780131_151780896 | chrX | 151780132 | 151780896 | + | 0.938876 | Intergenic | Intergenic | -22768 | NM_021523 | 59026 | Mm.27372 | NM_021523 | ENSMUSG00000025261 | **Huwe1** | 5430439H10Rik\|AU041296\|Arf-bp1\|C430014N20Rik\|C80292\|Gm1718\|Ib772\|LASU1\|Mule\|Ureb1 | HECT, UBA and WWE domain containing 1 | protein-coding |

| | | | | | | | | | | | | | | | | | |
|---|---|---|---|---|---|---|---|---|---|---|---|---|---|---|---|---|---|
| chr5_122230115_122230526 | chr5 | 122230116 | 122230526 | + | 0.733494 | intron (NM_001042489, intron 3 of 7) | L2a\|LINE\|L2 | 20034 | NM_028752 | 74096 | Mm.28804 | NM_028752 | ENSMUSG00000064267 | **Hvcn1** | 0610039P13Rik\|AI450555\|BTS\|HV1\|Vsop.\|mVSOP | hydrogen voltage-gated channel 1 | protein-coding |
| chr9_61215614_61216703 | chr9 | 61215615 | 61216703 | + | 0.896389 | Intergenic | Intergenic | 77644 | NR_045705 | 619328 | Mm.325722 | NR_045705 | | **I730028E13Rik** | - | RIKEN cDNA I730028E13 gene | ncRNA |
| chr12_21335806_21336475 | chr12 | 21335807 | 21336475 | + | 1.09786 | intron (NR_120376, intron 11 of 18) | intron (NR_120376, intron 11 of 18) | 19749 | NM_026347 | 67732 | Mm.335830 | NM_026347 | ENSMUSG00000062054 | **Iah1** | 1500019E20Rik\|4833421E05Rik | isoamyl acetate-hydrolyzing esterase 1 homolog (S. cerevisiae) | protein-coding |
| chr13_49700348_49700767 | chr13 | 49700349 | 49700767 | + | 0.750447 | intron (NM_172015, intron 7 of 33) | intron (NM_172015, intron 7 of 33) | 18428 | NM_172015 | 105148 | Mm.21118 | NM_172015 | ENSMUSG00000037851 | **Iars** | 2510016L12Rik\|AI327140\|AU044614\|E430001P04Rik\|ILRS | isoleucine-tRNA synthetase | protein-coding |
| chr13_49700209_49700951 | chr13 | 49700210 | 49700951 | + | 0.734658 | intron (NM_172015, intron 7 of 33) | intron (NM_172015, intron 7 of 33) | 18450 | NM_172015 | 105148 | Mm.21118 | NM_172015 | ENSMUSG00000037851 | **Iars** | 2510016L12Rik\|AI327140\|AU044614\|E430001P04Rik\|ILRS | isoleucine-tRNA synthetase | protein-coding |
| chr6_8648207_8649043 | chr6 | 8648208 | 8649043 | + | 1.03484 | intron (NM_001252266, intron 10 of 11) | intron (NM_001252266, intron 10 of 11) | 109833 | NM_001252266 | 15893 | Mm.275683 | NM_010492 | ENSMUSG00000062995 | **Ica1** | 69kDa\|ICA69 | islet cell autoantigen 1 | protein-coding |
| chr6_8647771_8648962 | chr6 | 8647772 | 8648962 | + | 0.98136 | intron (NM_001252266, intron 10 of 11) | intron (NM_001252266, intron 10 of 11) | 110091 | NM_001252266 | 15893 | Mm.275683 | NM_010492 | ENSMUSG00000062995 | **Ica1** | 69kDa\|ICA69 | islet cell autoantigen 1 | protein-coding |
| chr6_8738497_8738729 | chr6 | 8738498 | 8738729 | + | 0.806678 | intron (NM_001252266, intron 4 of 11) | B3A\|SINE\|B2 | 19845 | NM_001252266 | 15893 | Mm.275683 | NM_010492 | ENSMUSG00000062995 | **Ica1** | 69kDa\|ICA69 | islet cell autoantigen 1 | protein-coding |
| chr6_8649317_8649613 | chr6 | 8649318 | 8649613 | + | 0.805217 | intron (NM_001252266, intron 10 of 11) | intron (NM_001252266, intron 10 of 11) | 108993 | NM_001252266 | 15893 | Mm.275683 | NM_010492 | ENSMUSG00000062995 | **Ica1** | 69kDa\|ICA69 | islet cell autoantigen 1 | protein-coding |
| chr6_8639491_8639961 | chr6 | 8639492 | 8639961 | + | 0.70826 | intron (NM_001252266, intron 11 of 11) | MTD\|LTR\|ERVL-MaLR | 118732 | NM_001252266 | 15893 | Mm.275683 | NM_010492 | ENSMUSG00000062995 | **Ica1** | 69kDa\|ICA69 | islet cell autoantigen 1 | protein-coding |
| chr1_60016066_60016821 | chr1 | 60016067 | 60016821 | + | 0.935323 | intron (NM_027407, intron 3 of 10) | intron (NM_027407, intron 3 of 10) | 26643 | NM_027407 | 70375 | Mm.17834 | NM_027407 | ENSMUSG00000026018 | **Ica1l** | 1700030B17Rik\|Als2cr15 | islet cell autoantigen 1-like | protein-coding |
| chr1_60007313_60008045 | chr1 | 60007314 | 60008045 | + | 0.66093 | intron (NM_027407, intron 7 of 10) | B3A\|SINE\|B2 | 35408 | NM_027407 | 70375 | Mm.17834 | NM_027407 | ENSMUSG00000026018 | **Ica1l** | 1700030B17Rik\|Als2cr15 | islet cell autoantigen 1-like | protein-coding |
| chr1_60016047_60017500 | chr1 | 60016048 | 60017500 | + | 0.602524 | intron (NM_027407, intron 3 of 10) | B1_Mm\|SINE\|Alu | 26313 | NM_027407 | 70375 | Mm.17834 | NM_027407 | ENSMUSG00000026018 | **Ica1l** | 1700030B17Rik\|Als2cr15 | islet cell autoantigen 1-like | protein-coding |
| chr13_70635817_70636081 | chr13 | 70635818 | 70636081 | + | 0.720901 | intron (NM_144837, intron 1 of 18) | L2\|LINE\|L2 | 1685 | NM_144837 | 218333 | Mm.139738 | NM_144837 | ENSMUSG00000034525 | **Ice1** | C77245\|mKIAA0947 | interactor of little elongation complex ELL subunit 1 | protein-coding |
| chr13_70615081_70615349 | chr13 | 70615082 | 70615349 | + | 0.696805 | intron (NM_144837, intron 8 of 18) | intron (NM_144837, intron 8 of 18) | 22419 | NM_144837 | 218333 | Mm.139738 | NM_144837 | ENSMUSG00000034525 | **Ice1** | C77245\|mKIAA0947 | interactor of little elongation complex ELL subunit 1 | protein-coding |
| chr9_69349532_69349939 | chr9 | 69349533 | 69349939 | + | 1.32975 | intron (NM_001289916, intron 2 of 9) | RSINE1\|SINE\|B4 | -48262 | NM_145618 | 93697 | Mm.35911 | NM_145618 | ENSMUSG00000032235 | **Ice2** | B230343B06Rik\|Narg2 | interactor of little elongation complex ELL subunit 2 | protein-coding |
| chr9_69349021_69349944 | chr9 | 69349022 | 69349944 | + | 1.16967 | intron (NM_001289916, intron 2 of 9) | intron (NM_001289916, intron 2 of 9) | -48515 | NM_145618 | 93697 | Mm.35911 | NM_145618 | ENSMUSG00000032235 | **Ice2** | B230343B06Rik\|Narg2 | interactor of little elongation complex ELL subunit 2 | protein-coding |
| chr9_69345481_69345939 | chr9 | 69345482 | 69345939 | + | 1.09837 | intron (NM_001289916, intron 1 of 9) | intron (NM_001289916, intron 1 of 9) | -52288 | NM_145618 | 93697 | Mm.35911 | NM_145618 | ENSMUSG00000032235 | **Ice2** | B230343B06Rik\|Narg2 | interactor of little elongation complex ELL subunit 2 | protein-coding |
| chr1_61022731_61023342 | chr1 | 61022732 | 61023342 | + | 1.53853 | Intergenic | Intergenic | 45123 | NM_017480 | 54167 | Mm.42044 | NM_017480 | ENSMUSG00000026009 | **Icos** | AILIM\|CCLP\|CRP-1\|H4\|Ly115 | inducible T cell co-stimulator | protein-coding |
| chr1_61050182_61050836 | chr1 | 61050183 | 61050836 | + | 1.46605 | Intergenic | Intergenic | 72595 | NM_017480 | 54167 | Mm.42044 | NM_017480 | ENSMUSG00000026009 | **Icos** | AILIM\|CCLP\|CRP-1\|H4\|Ly115 | inducible T cell co-stimulator | protein-coding |
| chr1_61160444_61160945 | chr1 | 61160445 | 61160945 | + | 1.39691 | Intergenic | Intergenic | 182781 | NM_017480 | 54167 | Mm.42044 | NM_017480 | ENSMUSG00000026009 | **Icos** | AILIM\|CCLP\|CRP-1\|H4\|Ly115 | inducible T cell co-stimulator | protein-coding |
| chr1_61139745_61140267 | chr1 | 61139746 | 61140267 | + | 1.32678 | Intergenic | Intergenic | 162092 | NM_017480 | 54167 | Mm.42044 | NM_017480 | ENSMUSG00000026009 | **Icos** | AILIM\|CCLP\|CRP-1\|H4\|Ly115 | inducible T cell co-stimulator | protein-coding |
| chr1_61137559_61137832 | chr1 | 61137560 | 61137832 | + | 1.10431 | Intergenic | Intergenic | 159782 | NM_017480 | 54167 | Mm.42044 | NM_017480 | ENSMUSG00000026009 | **Icos** | AILIM\|CCLP\|CRP-1\|H4\|Ly115 | inducible T cell co-stimulator | protein-coding |
| chr1_61026721_61027083 | chr1 | 61026722 | 61027083 | + | 1.09955 | Intergenic | Intergenic | 48988 | NM_017480 | 54167 | Mm.42044 | NM_017480 | ENSMUSG00000026009 | **Icos** | AILIM\|CCLP\|CRP-1\|H4\|Ly115 | inducible T cell co-stimulator | protein-coding |
| chr1_60975042_60975490 | chr1 | 60975043 | 60975490 | + | 0.760356 | Intergenic | Intergenic | -2648 | NM_017480 | 54167 | Mm.42044 | NM_017480 | ENSMUSG00000026009 | **Icos** | AILIM\|CCLP\|CRP-1\|H4\|Ly115 | inducible T cell co-stimulator | protein-coding |
| chr1_61029816_61030255 | chr1 | 61029817 | 61030255 | + | 0.73861 | Intergenic | Intergenic | 52122 | NM_017480 | 54167 | Mm.42044 | NM_017480 | ENSMUSG00000026009 | **Icos** | AILIM\|CCLP\|CRP-1\|H4\|Ly115 | inducible T cell co-stimulator | protein-coding |
| chr1_61141011_61141213 | chr1 | 61141012 | 61141213 | + | 0.717665 | Intergenic | Intergenic | 163198 | NM_017480 | 54167 | Mm.42044 | NM_017480 | ENSMUSG00000026009 | **Icos** | AILIM\|CCLP\|CRP-1\|H4\|Ly115 | inducible T cell co-stimulator | protein-coding |
| chr10_78064064_78064533 | chr10 | 78064065 | 78064533 | + | 0.972078 | TTS (NM_001284200) | TTS (NM_001284200) | -5069 | NM_015790 | 50723 | Mm.17819 | NM_015790 | ENSMUSG00000000732 | **Icosl** | AU044799\|B7-H2\|B7RP-1\|B7h\|BG071784\|GI50\|GL50\|GL50-B\|ICOS-L\|Icoslg\|LICOS\|Ly115l\|mKIAA0653 | icos ligand | protein-coding |
| chr4_136123453_136123914 | chr4 | 136123454 | 136123914 | + | 1.27287 | Intergenic | MIR3\|SINE\|MIR | -20138 | NM_008321 | 15903 | Mm.110 | NM_008321 | | **Id3** | Hlh462\|Idb3\|bHLHb25 | inhibitor of DNA binding 3 | protein-coding |
| chr4_136122897_136123943 | chr4 | 136122898 | 136123943 | + | 1.11421 | Intergenic | Intergenic | -20402 | NM_008321 | 15903 | Mm.110 | NM_008321 | | **Id3** | Hlh462\|Idb3\|bHLHb25 | inhibitor of DNA binding 3 | protein-coding |
| chr4_136122955_136123407 | chr4 | 136122956 | 136123407 | + | 0.867784 | Intergenic | Intergenic | -20641 | NM_008321 | 15903 | Mm.110 | NM_008321 | | **Id3** | Hlh462\|Idb3\|bHLHb25 | inhibitor of DNA binding 3 | protein-coding |
| chr4_136106765_136107183 | chr4 | 136106766 | 136107183 | + | 0.786428 | Intergenic | Intergenic | -36848 | NM_008321 | 15903 | Mm.110 | NM_008321 | | **Id3** | Hlh462\|Idb3\|bHLHb25 | inhibitor of DNA binding 3 | protein-coding |
| chr4_136106718_136107435 | chr4 | 136106719 | 136107435 | + | 0.742195 | Intergenic | Intergenic | -36745 | NM_008321 | 15903 | Mm.110 | NM_008321 | | **Id3** | Hlh462\|Idb3\|bHLHb25 | inhibitor of DNA binding 3 | protein-coding |
| chr4_136110537_136110962 | chr4 | 136110538 | 136110962 | + | 0.706877 | Intergenic | Intergenic | -33072 | NM_008321 | 15903 | Mm.110 | NM_008321 | | **Id3** | Hlh462\|Idb3\|bHLHb25 | inhibitor of DNA binding 3 | protein-coding |
| chr4_136110580_136110863 | chr4 | 136110581 | 136110863 | + | 0.681538 | Intergenic | Intergenic | -33100 | NM_008321 | 15903 | Mm.110 | NM_008321 | | **Id3** | Hlh462\|Idb3\|bHLHb25 | inhibitor of DNA binding 3 | protein-coding |
| chr8_84648622_84649006 | chr8 | 84648623 | 84649006 | + | 1.21118 | Intergenic | MIRc\|SINE\|MIR | 14038 | NM_010499 | 15936 | Mm.399 | NM_010499 | ENSMUSG00000053560 | **Ier2** | AI317238\|Ch1\|Pip92 | immediate early response 2 | protein-coding |
| chr17_35819884_35820145 | chr17 | 35819885 | 35820145 | + | 0.870849 | Intergenic | Intergenic | -1698 | NM_133662 | 15937 | Mm.25613 | NM_133662 | ENSMUSG00000003541 | **Ier3** | AI663993\|IEX-1\|cI-3\|gly96 | immediate early response 3 | protein-coding |
| chr1_155108397_155108922 | chr1 | 155108398 | 155108922 | + | 0.988692 | Intergenic | Intergenic | -9024 | NM_010500 | 15939 | Mm.12246 | NM_010500 | ENSMUSG00000056708 | **Ier5** | - | immediate early response 5 | protein-coding |
| chr1_155091265_155091833 | chr1 | 155091266 | 155091833 | + | 0.883149 | Intergenic | Intergenic | 8087 | NM_010500 | 15939 | Mm.12246 | NM_010500 | ENSMUSG00000056708 | **Ier5** | - | immediate early response 5 | protein-coding |
| chr1_155050853_155051433 | chr1 | 155050854 | 155051433 | + | 0.721015 | Intergenic | Intergenic | 48493 | NM_010500 | 15939 | Mm.12246 | NM_010500 | ENSMUSG00000056708 | **Ier5** | - | immediate early response 5 | protein-coding |
| chr4_139578989_139579461 | chr4 | 139578990 | 139579461 | + | 0.618627 | intron (NM_001205173, intron 1 of 8) | intron (NM_001205173, intron 1 of 8) | 4505 | NM_001205173 | 212632 | Mm.30464 | NM_183148 | ENSMUSG00000041025 | **Iffo2** | C79267 | intermediate filament family orphan 2 | protein-coding |
| chr1_173802790_173803313 | chr1 | 173802791 | 173803313 | + | 1.06808 | Intergenic | Intergenic | -36133 | NM_008329 | 15951 | Mm.261270 | NM_008329 | ENSMUSG00000073489 | **Ifi204** | Ifi16\|p204 | interferon activated gene 204 | protein-coding |
| chr12_103445086_103445405 | chr12 | 103445087 | 103445405 | + | 0.862716 | Intergenic | Intergenic | -1566 | NM_001281830 | 76933 | Mm.271275 | NM_029803 | ENSMUSG00000079017 | **Ifi27l2a** | 2310061N23Rik\|Ifi27\|Isg12\|Isg12(b1) | interferon, alpha-inducible protein 27 like 2A | protein-coding |
| chr11_101446306_101446772 | chr11 | 101446307 | 101446772 | + | 1.25887 | TTS (NM_011289) | TTS (NM_011289) | -1873 | NM_027320 | 70110 | Mm.45558 | NM_027320 | ENSMUSG00000010358 | **Ifi35** | 2010008K16Rik\|AW986054\|IFP35\|ifi-35 | interferon-induced protein 35 | protein-coding |
| chr11_49085624_49085917 | chr11 | 49085625 | 49085917 | + | 0.844369 | intron (NM_001271677, intron 1 of 2) | intron (NM_001271677, intron 1 of 2) | -1246 | NM_001271676 | 15953 | Mm.24769 | NM_008330 | ENSMUSG00000078920 | **Ifi47** | 47kDa\|IRG-47\|Ifggc1\|Igrd\|Iigp4\|Iipg4\|Irgd | interferon gamma inducible protein 47 | protein-coding |
| chr11_49098888_49099154 | chr11 | 49098889 | 49099154 | + | 0.82103 | intron (NM_010999, intron 2 of 5) | intron (NM_010999, intron 2 of 5) | 12004 | NM_008330 | 15953 | Mm.24769 | NM_008330 | ENSMUSG00000078920 | **Ifi47** | 47kDa\|IRG-47\|Ifggc1\|Igrd\|Iigp4\|Iipg4\|Irgd | interferon gamma inducible protein 47 | protein-coding |
| chr11_49090210_49090466 | chr11 | 49090211 | 49090466 | + | 0.786223 | intron (NM_010999, intron 2 of 5) | intron (NM_010999, intron 2 of 5) | 3321 | NM_008330 | 15953 | Mm.24769 | NM_008330 | ENSMUSG00000078920 | **Ifi47** | 47kDa\|IRG-47\|Ifggc1\|Igrd\|Iigp4\|Iipg4\|Irgd | interferon gamma inducible protein 47 | protein-coding |
| chr2_62626131_62626630 | chr2 | 62626132 | 62626630 | + | 1.19162 | intron (NM_001164477, intron 2 of 14) | intron (NM_001164477, intron 2 of 14) | 19874 | NM_027835 | 71586 | Mm.136224 | NM_027835 | ENSMUSG00000026896 | **Ifih1** | 9130009C22Rik\|Helicard\|Hlcd\|MDA5\|RLR-2 | interferon induced with helicase C domain 1 | protein-coding |
| chr19_34678211_34678837 | chr19 | 34678212 | 34678837 | + | 1.38756 | intron (NM_172838, intron 3 of 6) | intron (NM_172838, intron 3 of 6) | 37635 | NM_008331 | 15957 | Mm.439751 | NM_008331 | ENSMUSG00000034459 | **Ifit1** | ISG56\|Ifi56\|P56 | interferon-induced protein with tetratricopeptide repeats 1 | protein-coding |
| chr7_142349196_142349735 | chr7 | 142349197 | 142349735 | + | 1.10303 | intron (NM_177265, intron 3 of 3) | ORR1E\|LTR\|ERVL-MaLR | 22793 | NM_177265 | 320802 | Mm.154486 | NM_177265 | ENSMUSG00000045777 | **Ifitm10** | 6330512M04Rik\|AI848576\|DSPA3 | interferon induced transmembrane protein 10 | protein-coding |
| chr7_140998851_140999883 | chr7 | 140998852 | 140999883 | + | 1.03806 | Intergenic | Intergenic | 11377 | NM_025378 | 66141 | Mm.141021 | NM_025378 | ENSMUSG00000025492 | **Ifitm3** | 1110004C05Rik\|Cd225\|Cdw217\|DSPA2b\|Fgls\|IP15\|mil-1 | interferon induced transmembrane protein 3 | protein-coding |
| chr7_141024924_141025468 | chr7 | 141024925 | 141025468 | + | 1.29244 | Intergenic | RMER17C\|LTR\|ERVK | -8304 | NM_001033632 | 213002 | Mm.276440 | NM_001033632 | ENSMUSG00000059108 | **Ifitm6** | A330075D07\|Dspa2f\|Fgls5\|fragilis5 | interferon induced transmembrane protein 6 | protein-coding |
| chr4_88652871_88653178 | chr4 | 88652872 | 88653178 | + | 0.772649 | Intergenic | Intergenic | -8566 | NM_177347 | 230396 | Mm.246592 | NM_177347 | ENSMUSG00000063376 | **Ifna13** | - | interferon alpha 13 | protein-coding |
| chr4_88681226_88681868 | chr4 | 88681227 | 88681868 | + | 1.55503 | Intergenic | Intergenic | 2232 | NM_010503 | 15965 | Mm.14091 | NM_010503 | | **Ifna2** | Ifa2 | interferon alpha 2 | protein-coding |
| chr4_88683666_88684108 | chr4 | 88683667 | 88684108 | + | 0.720824 | promoter-TSS (NM_010503) | promoter-TSS (NM_010503) | -108 | NM_010503 | 15965 | Mm.14091 | NM_010503 | | **Ifna2** | Ifa2 | interferon alpha 2 | protein-coding |
| chr4_88841448_88841764 | chr4 | 88841449 | 88841764 | + | 0.69282 | promoter-TSS (NM_010504) | promoter-TSS (NM_010504) | -210 | NM_010504 | 15967 | Mm.377088 | NM_010504 | ENSMUSG00000070904 | **Ifna4** | Ifa4 | interferon alpha 4 | protein-coding |

| | | | | | | | | | | | | | | | | | |
|---|---|---|---|---|---|---|---|---|---|---|---|---|---|---|---|---|---|
| chr4_88829578_88829988 | chr4 | 88829579 | 88829988 | + | 1.11665 | Intergenic | Intergenic | 2367 | NM_206871 | 15969 | Mm.377090 | NM_206871 | | **Ifna6** | IFNa8/6\|Ifa6\|Ifa8\|Ifna8 | interferon alpha 6 | protein-coding |
| chr16_91500920_91501392 | chr16 | 91500921 | 91501392 | + | 0.643524 | intron (NM_010508, intron 8 of 10) | intron (NM_010508, intron 8 of 10) | 15941 | NM_010508 | 15975 | Mm.502 | NM_010508 | | **Ifnar1** | CD118\|Ifar\|Ifnar\|Ifrc\|Infar | interferon (alpha and beta) receptor 1 | protein-coding |
| chr16_91352759_91353151 | chr16 | 91352760 | 91353151 | + | 0.729969 | Intergenic | Intergenic | -19828 | NM_010509 | 15976 | Mm.6834 | NM_010509 | ENSMUSG00000022971 | **Ifnar2** | AI747302\|Ifnar-2 | interferon (alpha and beta) receptor 2 | protein-coding |
| chr4_88515659_88515939 | chr4 | 88515660 | 88515939 | + | 0.649358 | Intergenic | Intergenic | 6995 | NM_010510 | 15977 | Mm.1245 | NM_010510 | ENSMUSG00000048806 | **Ifnb1** | IFN-beta\|IFNB\|Ifb | interferon beta 1, fibroblast | protein-coding |
| chr4_88875351_88876126 | chr4 | 88875352 | 88876126 | + | 1.22376 | Intergenic | Lx9\|LINE\|L1 | 4462 | NM_177348 | 230405 | Mm.246593 | NM_177348 | ENSMUSG00000045364 | **Ifne** | Ifn-tau-1\|Ifne1\|Ifnt1\|Infe1 | interferon epsilon | protein-coding |
| chr4_88888167_88888579 | chr4 | 88888168 | 88888579 | + | 0.872714 | Intergenic | Intergenic | -8172 | NM_177348 | 230405 | Mm.246593 | NM_177348 | ENSMUSG00000045364 | **Ifne** | Ifn-tau-1\|Ifne1\|Ifnt1\|Infe1 | interferon epsilon | protein-coding |
| chr4_88881137_88881485 | chr4 | 88881138 | 88881485 | + | 0.828835 | Intergenic | ORR1B1\|LTR\|ERVL-MaLR | -1110 | NM_177348 | 230405 | Mm.246593 | NM_177348 | ENSMUSG00000045364 | **Ifne** | Ifn-tau-1\|Ifne1\|Ifnt1\|Infe1 | interferon epsilon | protein-coding |
| chr4_88881168_88881707 | chr4 | 88881169 | 88881707 | + | 0.741025 | Intergenic | ORR1B1\|LTR\|ERVL-MaLR | -1237 | NM_177348 | 230405 | Mm.246593 | NM_177348 | ENSMUSG00000045364 | **Ifne** | Ifn-tau-1\|Ifne1\|Ifnt1\|Infe1 | interferon epsilon | protein-coding |
| chr10_118495166_118495736 | chr10 | 118495167 | 118495736 | + | 1.12942 | Intergenic | Intergenic | 54404 | NM_008337 | 15978 | Mm.240327 | NM_008337 | ENSMUSG00000055170 | **Ifng** | IFN-g\|Ifg | interferon gamma | protein-coding |
| chr10_118442940_118443407 | chr10 | 118442941 | 118443407 | + | 1.12361 | intron (NM_008337, intron 3 of 3) | (TCC)n\|Simple_repeat\|Simple_repeat | 2127 | NM_008337 | 15978 | Mm.240327 | NM_008337 | ENSMUSG00000055170 | **Ifng** | IFN-g\|Ifg | interferon gamma | protein-coding |
| chr10_118386241_118387242 | chr10 | 118386242 | 118387242 | + | 0.938217 | Intergenic | Intergenic | -54305 | NM_008337 | 15978 | Mm.240327 | NM_008337 | ENSMUSG00000055170 | **Ifng** | IFN-g\|Ifg | interferon gamma | protein-coding |
| chr10_118459977_118460654 | chr10 | 118459978 | 118460654 | + | 0.840512 | Intergenic | Intergenic | 19269 | NM_008337 | 15978 | Mm.240327 | NM_008337 | ENSMUSG00000055170 | **Ifng** | IFN-g\|Ifg | interferon gamma | protein-coding |
| chr10_118451675_118452221 | chr10 | 118451676 | 118452221 | + | 0.830105 | Intergenic | L1MC4\|LINE\|L1 | 10901 | NM_008337 | 15978 | Mm.240327 | NM_008337 | ENSMUSG00000055170 | **Ifng** | IFN-g\|Ifg | interferon gamma | protein-coding |
| chr10_118406824_118407421 | chr10 | 118406825 | 118407421 | + | 0.712976 | Intergenic | Intergenic | -33924 | NM_008337 | 15978 | Mm.240327 | NM_008337 | ENSMUSG00000055170 | **Ifng** | IFN-g\|Ifg | interferon gamma | protein-coding |
| chr10_118458575_118459284 | chr10 | 118458576 | 118459284 | + | 0.680839 | Intergenic | Intergenic | 17883 | NM_008337 | 15978 | Mm.240327 | NM_008337 | ENSMUSG00000055170 | **Ifng** | IFN-g\|Ifg | interferon gamma | protein-coding |
| chr10_118458781_118459084 | chr10 | 118458782 | 118459084 | + | 0.654486 | Intergenic | Intergenic | 17886 | NM_008337 | 15978 | Mm.240327 | NM_008337 | ENSMUSG00000055170 | **Ifng** | IFN-g\|Ifg | interferon gamma | protein-coding |
| chr10_19492985_19493394 | chr10 | 19492986 | 19493394 | + | 0.749021 | Intergenic | MTC\|LTR\|ERVL-MaLR | -98768 | NM_010511 | 15979 | Mm.549 | NM_010511 | ENSMUSG00000020009 | **Ifngr1** | CD119\|IFN-gammaR\|Ifgr\|Ifngr\|Nktar | interferon gamma receptor 1 | protein-coding |
| chr16_91555117_91555649 | chr16 | 91555118 | 91555649 | + | 0.988258 | intron (NM_008338, intron 2 of 6) | intron (NM_008338, intron 2 of 6) | 8289 | NM_008338 | 15980 | Mm.249364 | NM_008338 | | **Ifngr2** | Ifgr2\|Ifgt | interferon gamma receptor 2 | protein-coding |
| chr16_91534970_91535158 | chr16 | 91534971 | 91535158 | + | 0.840614 | Intergenic | Intergenic | -12030 | NM_008338 | 15980 | Mm.249364 | NM_008338 | | **Ifngr2** | Ifgr2\|Ifgt | interferon gamma receptor 2 | protein-coding |
| chr4_35060863_35061554 | chr4 | 35060864 | 35061554 | + | 1.87234 | intron (NM_178061, intron 2 of 3) | intron (NM_178061, intron 2 of 3) | -90846 | NM_199157 | 387510 | Mm.386929 | NM_199157 | ENSMUSG00000042993 | **Ifnk** | - | interferon kappa | protein-coding |
| chr2_163020906_163021176 | chr2 | 163020907 | 163021176 | + | 0.718336 | intron (NM_172150, intron 1 of 12) | MER44B\|DNA\|TcMar-Tigger | 3569 | NM_172150 | 245866 | Mm.34034 | NM_172150 | ENSMUSG00000017858 | **Ift52** | BC037708\|NGD5 | intraflagellar transport 52 | protein-coding |
| chr16_49534658_49535099 | chr16 | 49534659 | 49535099 | + | 1.49485 | Intergenic | Intergenic | -164415 | NM_028680 | 73916 | Mm.241276 | NM_028680 | ENSMUSG00000032965 | **Ift57** | 4833420A15Rik\|Esrrbl1\|Hippi\|MHS4R2 | intraflagellar transport 57 | protein-coding |
| chr16_49596511_49597016 | chr16 | 49596512 | 49597016 | + | 1.29549 | Intergenic | Intergenic | -102530 | NM_028680 | 73916 | Mm.241276 | NM_028680 | ENSMUSG00000032965 | **Ift57** | 4833420A15Rik\|Esrrbl1\|Hippi\|MHS4R2 | intraflagellar transport 57 | protein-coding |
| chr16_49775379_49775955 | chr16 | 49775380 | 49775955 | + | 1.27972 | Intergenic | Intergenic | 76373 | NM_028680 | 73916 | Mm.241276 | NM_028680 | ENSMUSG00000032965 | **Ift57** | 4833420A15Rik\|Esrrbl1\|Hippi\|MHS4R2 | intraflagellar transport 57 | protein-coding |
| chr16_49775043_49775956 | chr16 | 49775044 | 49775956 | + | 1.16431 | Intergenic | Intergenic | 76206 | NM_028680 | 73916 | Mm.241276 | NM_028680 | ENSMUSG00000032965 | **Ift57** | 4833420A15Rik\|Esrrbl1\|Hippi\|MHS4R2 | intraflagellar transport 57 | protein-coding |
| chr16_49771620_49771947 | chr16 | 49771621 | 49771947 | + | 0.993139 | Intergenic | Intergenic | 72490 | NM_028680 | 73916 | Mm.241276 | NM_028680 | ENSMUSG00000032965 | **Ift57** | 4833420A15Rik\|Esrrbl1\|Hippi\|MHS4R2 | intraflagellar transport 57 | protein-coding |
| chr9_65084444_65084827 | chr9 | 65084445 | 65084827 | + | 0.690651 | Intergenic | Intergenic | -16850 | NM_001290315 | 56741 | Mm.209041 | NM_020043 | ENSMUSG00000032816 | **Igdcc4** | DDM36\|Nope | immunoglobulin superfamily, DCC subclass, member 4 | protein-coding |
| chr6_49206235_49207065 | chr6 | 49206236 | 49207065 | + | 1.32344 | intron (NM_023670, intron 2 of 14) | intron (NM_023670, intron 2 of 14) | 8304 | NM_023670 | 140488 | Mm.281018 | NM_023670 | ENSMUSG00000029814 | **Igf2bp3** | 2610101N11Rik\|AA522010\|AL022933\|AU045931\|IMP-3\|IMP3\|Koc13\|Neilsen\|mimp3 | insulin-like growth factor 2 mRNA binding protein 3 | protein-coding |
| chr6_49217545_49218331 | chr6 | 49217546 | 49218331 | + | 0.969681 | Intergenic | CpG | -2984 | NM_023670 | 140488 | Mm.281018 | NM_023670 | ENSMUSG00000029814 | **Igf2bp3** | 2610101N11Rik\|AA522010\|AL022933\|AU045931\|IMP-3\|IMP3\|Koc13\|Neilsen\|mimp3 | insulin-like growth factor 2 mRNA binding protein 3 | protein-coding |
| chr6_49215332_49215726 | chr6 | 49215333 | 49215726 | + | 0.626422 | promoter-TSS (NM_023670) | promoter-TSS (NM_023670) | -575 | NM_023670 | 140488 | Mm.281018 | NM_023670 | ENSMUSG00000029814 | **Igf2bp3** | 2610101N11Rik\|AA522010\|AL022933\|AU045931\|IMP-3\|IMP3\|Koc13\|Neilsen\|mimp3 | insulin-like growth factor 2 mRNA binding protein 3 | protein-coding |
| chr17_12758063_12758293 | chr17 | 12758064 | 12758293 | + | 1.25026 | intron (NM_010515, intron 1 of 47) | intron (NM_010515, intron 1 of 47) | 11528 | NM_010515 | 16004 | Mm.26553 | NM_010515 | ENSMUSG00000023830 | **Igf2r** | AI661837\|CD222\|CI-MPR\|M6P/IGF2R\|Mpr300 | insulin-like growth factor 2 receptor | protein-coding |
| chr17_12757925_12758416 | chr17 | 12757926 | 12758416 | + | 1.1283 | intron (NM_010515, intron 1 of 47) | intron (NM_010515, intron 1 of 47) | 11535 | NM_010515 | 16004 | Mm.26553 | NM_010515 | ENSMUSG00000023830 | **Igf2r** | AI661837\|CD222\|CI-MPR\|M6P/IGF2R\|Mpr300 | insulin-like growth factor 2 receptor | protein-coding |
| chr17_12804568_12805480 | chr17 | 12804569 | 12805480 | + | 0.710969 | intron (NR_027784, intron 2 of 2) | intron (NR_027784, intron 2 of 2) | -35318 | NM_010515 | 16004 | Mm.26553 | NM_010515 | ENSMUSG00000023830 | **Igf2r** | AI661837\|CD222\|CI-MPR\|M6P/IGF2R\|Mpr300 | insulin-like growth factor 2 receptor | protein-coding |
| chr5_77367430_77368043 | chr5 | 77367431 | 77368043 | + | 1.65042 | intron (NM_008048, intron 1 of 4) | intron (NM_008048, intron 1 of 4) | 40308 | NM_001159518 | 29817 | Mm.233470 | NM_008048 | ENSMUSG00000036256 | **Igfbp7** | AGM\|Fstl2\|Mac25 | insulin-like growth factor binding protein 7 | protein-coding |
| chr5_77372796_77373260 | chr5 | 77372797 | 77373260 | + | 1.59421 | intron (NM_008048, intron 1 of 4) | intron (NM_008048, intron 1 of 4) | 35017 | NM_001159518 | 29817 | Mm.233470 | NM_008048 | ENSMUSG00000036256 | **Igfbp7** | AGM\|Fstl2\|Mac25 | insulin-like growth factor binding protein 7 | protein-coding |
| chr5_77383909_77384770 | chr5 | 77383910 | 77384770 | + | 1.4517 | intron (NM_008048, intron 1 of 4) | intron (NM_008048, intron 1 of 4) | 23705 | NM_001159518 | 29817 | Mm.233470 | NM_008048 | ENSMUSG00000036256 | **Igfbp7** | AGM\|Fstl2\|Mac25 | insulin-like growth factor binding protein 7 | protein-coding |
| chr5_77374875_77375294 | chr5 | 77374876 | 77375294 | + | 0.938139 | intron (NM_008048, intron 1 of 4) | intron (NM_008048, intron 1 of 4) | 32960 | NM_001159518 | 29817 | Mm.233470 | NM_008048 | ENSMUSG00000036256 | **Igfbp7** | AGM\|Fstl2\|Mac25 | insulin-like growth factor binding protein 7 | protein-coding |
| chr7_30566790_30567073 | chr7 | 30566791 | 30567073 | + | 0.686249 | exon (NM_145580, exon 3 of 5) | exon (NM_145580, exon 3 of 5) | 1505 | NM_145580 | 101883 | Mm.333757 | NM_145580 | ENSMUSG00000036826 | **Igflr1** | F630003I17Rik\|Tmem149 | IGF-like family receptor 1 | protein-coding |
| chr5_88508327_88508995 | chr5 | 88508328 | 88508995 | + | 0.981061 | Intergenic | MTA_Mm\|LTR\|ERVL-MaLR | 19236 | NM_152839 | 16069 | Mm.1192 | NM_152839 | | **Igj** | 9530090F24Rik\|AI323815\|Jch | immunoglobulin joining chain | protein-coding |
| chr5_88508310_88509401 | chr5 | 88508311 | 88509401 | + | 0.772179 | Intergenic | Intergenic | 19041 | NM_152839 | 16069 | Mm.1192 | NM_152839 | | **Igj** | 9530090F24Rik\|AI323815\|Jch | immunoglobulin joining chain | protein-coding |
| chr7_43480850_43481596 | chr7 | 43480851 | 43481596 | + | 1.28911 | intron (NM_001164518, intron 1 of 7) | intron (NM_001164518, intron 1 of 7) | 8800 | NM_001164518 | 210094 | Mm.228087 | NM_001164518 | ENSMUSG00000013367 | **Iglon5** | A230106M20Rik\|Gm1431 | IgLON family member 5 | protein-coding |
| chr18_60356070_60356556 | chr18 | 60356071 | 60356556 | + | 0.828074 | Intergenic | Intergenic | -19716 | NM_021792 | 60440 | Mm.261140 | NM_021792 | ENSMUSG00000054072 | **Iigp1** | 2900074L10Rik\|AI046432\|AW111922\|Ifgga1\|Iigp\|Irga6 | interferon inducible GTPase 1 | protein-coding |
| chr18_60355721_60356556 | chr18 | 60355722 | 60356556 | + | 0.819352 | Intergenic | Intergenic | -19890 | NM_021792 | 60440 | Mm.261140 | NM_021792 | ENSMUSG00000054072 | **Iigp1** | 2900074L10Rik\|AI046432\|AW111922\|Ifgga1\|Iigp\|Irga6 | interferon inducible GTPase 1 | protein-coding |
| chr11_11646389_11646704 | chr11 | 11646390 | 11646704 | + | 1.42583 | Intergenic | Intergenic | -39666 | NM_009578 | 22778 | Mm.103545 | NM_009578 | ENSMUSG00000018654 | **Ikzf1** | 5832432G11Rik\|Ikaros\|LyF-1\|Zfpn1a1\|Znfn1a1\|hlk-1\|mKIAA4227 | IKAROS family zinc finger 1 | protein-coding |
| chr11_11724484_11724920 | chr11 | 11724485 | 11724920 | + | 1.2743 | intron (NM_009578, intron 3 of 6) | PB1D10\|SINE\|Alu | 38489 | NM_001025597 | 22778 | Mm.103545 | NM_009578 | ENSMUSG00000018654 | **Ikzf1** | 5832432G11Rik\|Ikaros\|LyF-1\|Zfpn1a1\|Znfn1a1\|hlk-1\|mKIAA4227 | IKAROS family zinc finger 1 | protein-coding |
| chr11_11721287_11721541 | chr11 | 11721288 | 11721541 | + | 1.12347 | intron (NM_009578, intron 3 of 6) | intron (NM_009578, intron 3 of 6) | 35201 | NM_001025597 | 22778 | Mm.103545 | NM_009578 | ENSMUSG00000018654 | **Ikzf1** | 5832432G11Rik\|Ikaros\|LyF-1\|Zfpn1a1\|Znfn1a1\|hlk-1\|mKIAA4227 | IKAROS family zinc finger 1 | protein-coding |
| chr11_11725263_11725887 | chr11 | 11725264 | 11725887 | + | 1.0544 | intron (NM_009578, intron 3 of 6) | intron (NM_009578, intron 3 of 6) | 39362 | NM_001025597 | 22778 | Mm.103545 | NM_009578 | ENSMUSG00000018654 | **Ikzf1** | 5832432G11Rik\|Ikaros\|LyF-1\|Zfpn1a1\|Znfn1a1\|hlk-1\|mKIAA4227 | IKAROS family zinc finger 1 | protein-coding |
| chr11_11697420_11697830 | chr11 | 11697421 | 11697830 | + | 0.809681 | intron (NM_001025597, intron 1 of 7) | intron (NM_001025597, intron 1 of 7) | 11412 | NM_001025597 | 22778 | Mm.103545 | NM_009578 | ENSMUSG00000018654 | **Ikzf1** | 5832432G11Rik\|Ikaros\|LyF-1\|Zfpn1a1\|Znfn1a1\|hlk-1\|mKIAA4227 | IKAROS family zinc finger 1 | protein-coding |
| chr11_11720942_11721167 | chr11 | 11720943 | 11721167 | + | 0.78199 | intron (NM_009578, intron 3 of 6) | intron (NM_009578, intron 3 of 6) | 34842 | NM_001025597 | 22778 | Mm.103545 | NM_009578 | ENSMUSG00000018654 | **Ikzf1** | 5832432G11Rik\|Ikaros\|LyF-1\|Zfpn1a1\|Znfn1a1\|hlk-1\|mKIAA4227 | IKAROS family zinc finger 1 | protein-coding |
| chr11_11655013_11655420 | chr11 | 11655014 | 11655420 | + | 0.768676 | Intergenic | Intergenic | -30996 | NM_009578 | 22778 | Mm.103545 | NM_009578 | ENSMUSG00000018654 | **Ikzf1** | 5832432G11Rik\|Ikaros\|LyF-1\|Zfpn1a1\|Znfn1a1\|hlk-1\|mKIAA4227 | IKAROS family zinc finger 1 | protein-coding |
| chr11_11683275_11683496 | chr11 | 11683276 | 11683496 | + | 0.723091 | Intergenic | Intergenic | -2827 | NM_009578 | 22778 | Mm.103545 | NM_009578 | ENSMUSG00000018654 | **Ikzf1** | 5832432G11Rik\|Ikaros\|LyF-1\|Zfpn1a1\|Znfn1a1\|hlk-1\|mKIAA4227 | IKAROS family zinc finger 1 | protein-coding |

| | | | | | | | | | | | | | | | | | |
|---|---|---|---|---|---|---|---|---|---|---|---|---|---|---|---|---|---|
| chr11_11656375_11656873 | chr11 | 11656376 | 11656873 | + | 0.704942 | Intergenic | Intergenic | -29589 | NM_009578 | 22778 | Mm.103545 | NM_009578 | ENSMUSG00000018654 | **Ikzf1** | 5832432G11Rik\|Ikaros\|LyF-1\|Zfpn1a1\|Znfn1a1\|hlk-1\|mKIAA4227 | IKAROS family zinc finger 1 | protein-coding |
| chr11_11708210_11708694 | chr11 | 11708211 | 11708694 | + | 0.693847 | intron (NM_009578, intron 3 of 6) | intron (NM_009578, intron 3 of 6) | 22239 | NM_001025597 | 22778 | Mm.103545 | NM_009578 | ENSMUSG00000018654 | **Ikzf1** | 5832432G11Rik\|Ikaros\|LyF-1\|Zfpn1a1\|Znfn1a1\|hlk-1\|mKIAA4227 | IKAROS family zinc finger 1 | protein-coding |
| chr11_11654875_11655420 | chr11 | 11654876 | 11655420 | + | 0.674688 | Intergenic | (TG)n\|Simple_repeat\|Simple_repeat | -31065 | NM_009578 | 22778 | Mm.103545 | NM_009578 | ENSMUSG00000018654 | **Ikzf1** | 5832432G11Rik\|Ikaros\|LyF-1\|Zfpn1a1\|Znfn1a1\|hlk-1\|mKIAA4227 | IKAROS family zinc finger 1 | protein-coding |
| chr11_11716031_11716293 | chr11 | 11716032 | 11716293 | + | 0.651054 | intron (NM_009578, intron 3 of 6) | intron (NM_009578, intron 3 of 6) | 29949 | NM_001025597 | 22778 | Mm.103545 | NM_009578 | ENSMUSG00000018654 | **Ikzf1** | 5832432G11Rik\|Ikaros\|LyF-1\|Zfpn1a1\|Znfn1a1\|hlk-1\|mKIAA4227 | IKAROS family zinc finger 1 | protein-coding |
| chr11_11731358_11732154 | chr11 | 11731359 | 11732154 | + | 0.639832 | intron (NM_009578, intron 3 of 6) | LTR90A\|LTR\|LTR | 45543 | NM_001025597 | 22778 | Mm.103545 | NM_009578 | ENSMUSG00000018654 | **Ikzf1** | 5832432G11Rik\|Ikaros\|LyF-1\|Zfpn1a1\|Znfn1a1\|hlk-1\|mKIAA4227 | IKAROS family zinc finger 1 | protein-coding |
| chr11_11692514_11692921 | chr11 | 11692515 | 11692921 | + | 0.611669 | intron (NM_001025597, intron 1 of 7) | intron (NM_001025597, intron 1 of 7) | 6505 | NM_001025597 | 22778 | Mm.103545 | NM_009578 | ENSMUSG00000018654 | **Ikzf1** | 5832432G11Rik\|Ikaros\|LyF-1\|Zfpn1a1\|Znfn1a1\|hlk-1\|mKIAA4227 | IKAROS family zinc finger 1 | protein-coding |
| chr1_69544090_69544712 | chr1 | 69544091 | 69544712 | + | 1.37314 | intron (NM_011770, intron 6 of 7) | intron (NM_011770, intron 6 of 7) | 141559 | NM_011770 | 22779 | Mm.106343 | NM_011770 | ENSMUSG00000025997 | **Ikzf2** | A730095J18Rik\|Helios\|Zfpn1a2\|Znfn1a2 | IKAROS family zinc finger 2 | protein-coding |
| chr1_69461167_69461612 | chr1 | 69461168 | 69461612 | + | 1.01026 | Intergenic | Intergenic | 224570 | NM_011770 | 22779 | Mm.106343 | NM_011770 | ENSMUSG00000025997 | **Ikzf2** | A730095J18Rik\|Helios\|Zfpn1a2\|Znfn1a2 | IKAROS family zinc finger 2 | protein-coding |
| chr1_69554027_69554791 | chr1 | 69554028 | 69554791 | + | 0.96009 | intron (NM_011770, intron 5 of 7) | L1MEc\|LINE\|L1 | 131551 | NM_011770 | 22779 | Mm.106343 | NM_011770 | ENSMUSG00000025997 | **Ikzf2** | A730095J18Rik\|Helios\|Zfpn1a2\|Znfn1a2 | IKAROS family zinc finger 2 | protein-coding |
| chr1_69498825_69499463 | chr1 | 69498826 | 69499463 | + | 0.780251 | Intergenic | Intergenic | 186816 | NM_011770 | 22779 | Mm.106343 | NM_011770 | ENSMUSG00000025997 | **Ikzf2** | A730095J18Rik\|Helios\|Zfpn1a2\|Znfn1a2 | IKAROS family zinc finger 2 | protein-coding |
| chr1_69548862_69549088 | chr1 | 69548863 | 69549088 | + | 0.766648 | intron (NM_011770, intron 5 of 7) | intron (NM_011770, intron 5 of 7) | 136985 | NM_011770 | 22779 | Mm.106343 | NM_011770 | ENSMUSG00000025997 | **Ikzf2** | A730095J18Rik\|Helios\|Zfpn1a2\|Znfn1a2 | IKAROS family zinc finger 2 | protein-coding |
| chr1_69548535_69549113 | chr1 | 69548536 | 69549113 | + | 0.7525 | intron (NM_011770, intron 5 of 7) | intron (NM_011770, intron 5 of 7) | 137136 | NM_011770 | 22779 | Mm.106343 | NM_011770 | ENSMUSG00000025997 | **Ikzf2** | A730095J18Rik\|Helios\|Zfpn1a2\|Znfn1a2 | IKAROS family zinc finger 2 | protein-coding |
| chr10_128645663_128646039 | chr10 | 128645664 | 128646039 | + | 0.720146 | 5' UTR (NM_011772, exon 1 of 8) | 5' UTR (NM_011772, exon 1 of 8) | 142 | NM_011772 | 22781 | Mm.358648 | NM_011772 | ENSMUSG00000002578 | **Ikzf4** | A630026H08Rik\|Eos\|Zfpn1a4\|Znfn1a4 | IKAROS family zinc finger 4 | protein-coding |
| chr10_128646425_128646740 | chr10 | 128646426 | 128646740 | + | 0.716683 | promoter-TSS (NM_011772) | promoter-TSS (NM_011772) | -590 | NM_011772 | 22781 | Mm.358648 | NM_011772 | ENSMUSG00000002578 | **Ikzf4** | A630026H08Rik\|Eos\|Zfpn1a4\|Znfn1a4 | IKAROS family zinc finger 4 | protein-coding |
| chr10_128645060_128646063 | chr10 | 128645061 | 128646063 | + | 0.643142 | intron (NM_011772, intron 1 of 7) | intron (NM_011772, intron 1 of 7) | 431 | NM_011772 | 22781 | Mm.358648 | NM_011772 | ENSMUSG00000002578 | **Ikzf4** | A630026H08Rik\|Eos\|Zfpn1a4\|Znfn1a4 | IKAROS family zinc finger 4 | protein-coding |
| chr1_131022523_131023260 | chr1 | 131022524 | 131023260 | + | 1.92452 | intron (NM_010548, intron 4 of 4) | intron (NM_010548, intron 4 of 4) | 3047 | NM_010548 | 16153 | Mm.874 | NM_010548 | ENSMUSG00000016529 | **Il10** | CSIF\|Il-10 | interleukin 10 | protein-coding |
| chr1_131038004_131038722 | chr1 | 131038005 | 131038722 | + | 1.58733 | Intergenic | Intergenic | 18518 | NM_010548 | 16153 | Mm.874 | NM_010548 | ENSMUSG00000016529 | **Il10** | CSIF\|Il-10 | interleukin 10 | protein-coding |
| chr1_131010182_131010937 | chr1 | 131010183 | 131010937 | + | 1.4621 | Intergenic | Intergenic | -9285 | NM_010548 | 16153 | Mm.874 | NM_010548 | ENSMUSG00000016529 | **Il10** | CSIF\|Il-10 | interleukin 10 | protein-coding |
| chr1_130996360_130996762 | chr1 | 130996361 | 130996762 | + | 1.19909 | Intergenic | Intergenic | -23284 | NM_010548 | 16153 | Mm.874 | NM_010548 | ENSMUSG00000016529 | **Il10** | CSIF\|Il-10 | interleukin 10 | protein-coding |
| chr1_130993644_130994258 | chr1 | 130993645 | 130994258 | + | 0.998802 | Intergenic | Intergenic | -25894 | NM_010548 | 16153 | Mm.874 | NM_010548 | ENSMUSG00000016529 | **Il10** | CSIF\|Il-10 | interleukin 10 | protein-coding |
| chr1_130993700_130994067 | chr1 | 130993701 | 130994067 | + | 0.959456 | Intergenic | Intergenic | -25961 | NM_010548 | 16153 | Mm.874 | NM_010548 | ENSMUSG00000016529 | **Il10** | CSIF\|Il-10 | interleukin 10 | protein-coding |
| chr1_131049135_131049709 | chr1 | 131049136 | 131049709 | + | 0.953083 | Intergenic | Intergenic | 29577 | NM_010548 | 16153 | Mm.874 | NM_010548 | ENSMUSG00000016529 | **Il10** | CSIF\|Il-10 | interleukin 10 | protein-coding |
| chr1_130999641_131000066 | chr1 | 130999642 | 131000066 | + | 0.928258 | Intergenic | Intergenic | -19991 | NM_010548 | 16153 | Mm.874 | NM_010548 | ENSMUSG00000016529 | **Il10** | CSIF\|Il-10 | interleukin 10 | protein-coding |
| chr1_131000373_131000732 | chr1 | 131000374 | 131000732 | + | 0.831027 | Intergenic | Intergenic | -19292 | NM_010548 | 16153 | Mm.874 | NM_010548 | ENSMUSG00000016529 | **Il10** | CSIF\|Il-10 | interleukin 10 | protein-coding |
| chr1_130999430_130999967 | chr1 | 130999431 | 130999967 | + | 0.819772 | Intergenic | Intergenic | -20146 | NM_010548 | 16153 | Mm.874 | NM_010548 | ENSMUSG00000016529 | **Il10** | CSIF\|Il-10 | interleukin 10 | protein-coding |
| chr1_130999377_131000132 | chr1 | 130999378 | 131000132 | + | 0.798773 | Intergenic | Intergenic | -20090 | NM_010548 | 16153 | Mm.874 | NM_010548 | ENSMUSG00000016529 | **Il10** | CSIF\|Il-10 | interleukin 10 | protein-coding |
| chr1_130993403_130994057 | chr1 | 130993404 | 130994057 | + | 0.792924 | Intergenic | Intergenic | -26115 | NM_010548 | 16153 | Mm.874 | NM_010548 | ENSMUSG00000016529 | **Il10** | CSIF\|Il-10 | interleukin 10 | protein-coding |
| chr1_131028056_131028310 | chr1 | 131028057 | 131028310 | + | 0.730983 | Intergenic | MLT1F1\|LTR\|ERVL-MaLR | 8338 | NM_010548 | 16153 | Mm.874 | NM_010548 | ENSMUSG00000016529 | **Il10** | CSIF\|Il-10 | interleukin 10 | protein-coding |
| chr9_45278167_45278554 | chr9 | 45278168 | 45278554 | + | 0.841052 | Intergenic | Intergenic | -9215 | NM_008348 | 16154 | Mm.379327 | NM_008348 | ENSMUSG00000032089 | **Il10ra** | AW553859\|CDw210\|CDw210a\|Il10r\|mIL-10R | interleukin 10 receptor, alpha | protein-coding |
| chr9_45288029_45288240 | chr9 | 45288030 | 45288240 | + | 0.690031 | Intergenic | L1M5\|LINE\|L1 | -18989 | NM_008348 | 16154 | Mm.379327 | NM_008348 | ENSMUSG00000032089 | **Il10ra** | AW553859\|CDw210\|CDw210a\|Il10r\|mIL-10R | interleukin 10 receptor, alpha | protein-coding |
| chr16_91397970_91398500 | chr16 | 91397971 | 91398500 | + | 1.16271 | intron (NM_010509, intron 7 of 8) | intron (NM_010509, intron 7 of 8) | -8000 | NM_008349 | 16155 | Mm.4154 | NM_008349 | ENSMUSG00000022969 | **Il10rb** | 6620401D04Rik\|AI528744\|CRF2-4\|Crfb4\|D16H21S58\|D21S58h\|IL-10R2\|Il10r2 | interleukin 10 receptor, beta | protein-coding |
| chr16_91413095_91413868 | chr16 | 91413096 | 91413868 | + | 0.882134 | intron (NM_008349, intron 3 of 6) | intron (NM_008349, intron 3 of 6) | 7247 | NM_008349 | 16155 | Mm.4154 | NM_008349 | ENSMUSG00000022969 | **Il10rb** | 6620401D04Rik\|AI528744\|CRF2-4\|Crfb4\|D16H21S58\|D21S58h\|IL-10R2\|Il10r2 | interleukin 10 receptor, beta | protein-coding |
| chr16_91413107_91413660 | chr16 | 91413108 | 91413660 | + | 0.855957 | intron (NM_008349, intron 3 of 6) | intron (NM_008349, intron 3 of 6) | 7149 | NM_008349 | 16155 | Mm.4154 | NM_008349 | ENSMUSG00000022969 | **Il10rb** | 6620401D04Rik\|AI528744\|CRF2-4\|Crfb4\|D16H21S58\|D21S58h\|IL-10R2\|Il10r2 | interleukin 10 receptor, beta | protein-coding |
| chr3_68762937_68763297 | chr3 | 68762938 | 68763297 | + | 1.01781 | Intergenic | Intergenic | 71693 | NM_008351 | 16159 | Mm.103783 | NM_008351 | ENSMUSG00000027776 | **Il12a** | IL-12p35\|Il-12a\|Ll12a\|p35 | interleukin 12a | protein-coding |
| chr3_68761237_68761482 | chr3 | 68761238 | 68761482 | + | 0.909017 | Intergenic | Intergenic | 69936 | NM_008351 | 16159 | Mm.103783 | NM_008351 | ENSMUSG00000027776 | **Il12a** | IL-12p35\|Il-12a\|Ll12a\|p35 | interleukin 12a | protein-coding |
| chr3_68761622_68762427 | chr3 | 68761623 | 68762427 | + | 0.846933 | Intergenic | Intergenic | 70601 | NM_008351 | 16159 | Mm.103783 | NM_008351 | ENSMUSG00000027776 | **Il12a** | IL-12p35\|Il-12a\|Ll12a\|p35 | interleukin 12a | protein-coding |
| chr3_68760865_68761505 | chr3 | 68760866 | 68761505 | + | 0.845712 | Intergenic | Intergenic | 69761 | NM_008351 | 16159 | Mm.103783 | NM_008351 | ENSMUSG00000027776 | **Il12a** | IL-12p35\|Il-12a\|Ll12a\|p35 | interleukin 12a | protein-coding |
| chr3_68761687_68762227 | chr3 | 68761688 | 68762227 | + | 0.825957 | Intergenic | MER92C\|LTR\|ERV1 | 70533 | NM_008351 | 16159 | Mm.103783 | NM_008351 | ENSMUSG00000027776 | **Il12a** | IL-12p35\|Il-12a\|Ll12a\|p35 | interleukin 12a | protein-coding |
| chr3_68772089_68772441 | chr3 | 68772090 | 68772441 | + | 0.803167 | Intergenic | Intergenic | 80841 | NM_008351 | 16159 | Mm.103783 | NM_008351 | ENSMUSG00000027776 | **Il12a** | IL-12p35\|Il-12a\|Ll12a\|p35 | interleukin 12a | protein-coding |
| chr3_68637663_68638392 | chr3 | 68637664 | 68638392 | + | 0.787271 | Intergenic | Intergenic | -52616 | NM_001159424 | 16159 | Mm.103783 | NM_008351 | ENSMUSG00000027776 | **Il12a** | IL-12p35\|Il-12a\|Ll12a\|p35 | interleukin 12a | protein-coding |
| chr3_68637951_68638428 | chr3 | 68637952 | 68638428 | + | 0.731771 | Intergenic | Intergenic | -52454 | NM_001159424 | 16159 | Mm.103783 | NM_008351 | ENSMUSG00000027776 | **Il12a** | IL-12p35\|Il-12a\|Ll12a\|p35 | interleukin 12a | protein-coding |
| chr3_68637663_68639009 | chr3 | 68637664 | 68639009 | + | 0.714907 | Intergenic | Intergenic | -52308 | NM_001159424 | 16159 | Mm.103783 | NM_008351 | ENSMUSG00000027776 | **Il12a** | IL-12p35\|Il-12a\|Ll12a\|p35 | interleukin 12a | protein-coding |
| chr3_68771577_68772443 | chr3 | 68771578 | 68772443 | + | 0.674776 | Intergenic | Intergenic | 80586 | NM_008351 | 16159 | Mm.103783 | NM_008351 | ENSMUSG00000027776 | **Il12a** | IL-12p35\|Il-12a\|Ll12a\|p35 | interleukin 12a | protein-coding |
| chr11_44373812_44374172 | chr11 | 44373813 | 44374172 | + | 0.754991 | Intergenic | Intergenic | -26071 | NM_008352 | 16160 | Mm.239707 | NM_008352 | ENSMUSG00000004296 | **Il12b** | Il-12b\|Il-12p40\|Il12p40\|p40 | interleukin 12b | protein-coding |
| chr11_44256462_44256920 | chr11 | 44256463 | 44256920 | + | 0.624594 | Intergenic | RLTR49\|LTR\|ERVK | -143372 | NM_008352 | 16160 | Mm.239707 | NM_008352 | ENSMUSG00000004296 | **Il12b** | Il-12b\|Il-12p40\|Il12p40\|p40 | interleukin 12b | protein-coding |
| chr6_67395992_67396593 | chr6 | 67395993 | 67396593 | + | 1.70001 | Intergenic | Intergenic | -20156 | NM_008354 | 16162 | Mm.188337 | NM_008354 | ENSMUSG00000018341 | **Il12rb2** | A930027I18Rik\|IL-12RB2\|Ifnm | interleukin 12 receptor, beta 2 | protein-coding |
| chr6_67366950_67367518 | chr6 | 67366951 | 67367518 | + | 1.1758 | intron (NM_008354, intron 1 of 15) | intron (NM_008354, intron 1 of 15) | 8903 | NM_008354 | 16162 | Mm.188337 | NM_008354 | ENSMUSG00000018341 | **Il12rb2** | A930027I18Rik\|IL-12RB2\|Ifnm | interleukin 12 receptor, beta 2 | protein-coding |
| chr6_67366960_67367719 | chr6 | 67366961 | 67367719 | + | 1.15881 | intron (NM_008354, intron 1 of 15) | intron (NM_008354, intron 1 of 15) | 8797 | NM_008354 | 16162 | Mm.188337 | NM_008354 | ENSMUSG00000018341 | **Il12rb2** | A930027I18Rik\|IL-12RB2\|Ifnm | interleukin 12 receptor, beta 2 | protein-coding |
| chr6_67395508_67395839 | chr6 | 67395509 | 67395839 | + | 1.13799 | Intergenic | MLT1B\|LTR\|ERVL-MaLR | -19537 | NM_008354 | 16162 | Mm.188337 | NM_008354 | ENSMUSG00000018341 | **Il12rb2** | A930027I18Rik\|IL-12RB2\|Ifnm | interleukin 12 receptor, beta 2 | protein-coding |
| chr6_67395405_67395880 | chr6 | 67395406 | 67395880 | + | 1.05583 | Intergenic | MLT1B\|LTR\|ERVL-MaLR | -19506 | NM_008354 | 16162 | Mm.188337 | NM_008354 | ENSMUSG00000018341 | **Il12rb2** | A930027I18Rik\|IL-12RB2\|Ifnm | interleukin 12 receptor, beta 2 | protein-coding |
| chr6_67378021_67378407 | chr6 | 67378022 | 67378407 | + | 0.951248 | Intergenic | Intergenic | -2077 | NM_008354 | 16162 | Mm.188337 | NM_008354 | ENSMUSG00000018341 | **Il12rb2** | A930027I18Rik\|IL-12RB2\|Ifnm | interleukin 12 receptor, beta 2 | protein-coding |
| chr6_67378125_67378316 | chr6 | 67378126 | 67378316 | + | 0.944822 | Intergenic | Intergenic | -2084 | NM_008354 | 16162 | Mm.188337 | NM_008354 | ENSMUSG00000018341 | **Il12rb2** | A930027I18Rik\|IL-12RB2\|Ifnm | interleukin 12 receptor, beta 2 | protein-coding |
| chr6_67373725_67373987 | chr6 | 67373726 | 67373987 | + | 0.931938 | intron (NM_008354, intron 1 of 15) | intron (NM_008354, intron 1 of 15) | 2281 | NM_008354 | 16162 | Mm.188337 | NM_008354 | ENSMUSG00000018341 | **Il12rb2** | A930027I18Rik\|IL-12RB2\|Ifnm | interleukin 12 receptor, beta 2 | protein-coding |
| chr6_67373431_67373624 | chr6 | 67373432 | 67373624 | + | 0.783929 | intron (NM_008354, intron 1 of 15) | B3A\|SINE\|B2 | 2609 | NM_008354 | 16162 | Mm.188337 | NM_008354 | ENSMUSG00000018341 | **Il12rb2** | A930027I18Rik\|IL-12RB2\|Ifnm | interleukin 12 receptor, beta 2 | protein-coding |
| chr11_53656658_53657228 | chr11 | 53656659 | 53657228 | + | 1.05472 | intron (NM_009012, intron 21 of 24) | intron (NM_009012, intron 21 of 24) | -22241 | NM_008355 | 16163 | Mm.1284 | NM_008355 | ENSMUSG00000020383 | **Il13** | Il-13 | interleukin 13 | protein-coding |
| chr11_53655045_53655549 | chr11 | 53655046 | 53655549 | + | 0.933967 | intron (NM_009012, intron 22 of 24) | intron (NM_009012, intron 22 of 24) | -20595 | NM_008355 | 16163 | Mm.1284 | NM_008355 | ENSMUSG00000020383 | **Il13** | Il-13 | interleukin 13 | protein-coding |
| chrX_36104665_36105076 | chrX | 36104666 | 36105076 | + | 1.02413 | Intergenic | Intergenic | -7268 | NM_133990 | 16164 | Mm.24208 | NM_133990 | ENSMUSG00000017057 | **Il13ra1** | AI882074\|CD213a1\|IL-13r[a]\|Il13ra\|NR4 | interleukin 13 receptor, alpha 1 | protein-coding |
| chr8_81881314_81882329 | chr8 | 81881315 | 81882329 | + | 1.25008 | intron (NM_001297596, intron 8 of 24) | intron (NM_001297596, intron 8 of 24) | 520764 | NM_001254747 | 16168 | Mm.4392 | NM_008357 | | **Il15** | AI503618\|IL-15 | interleukin 15 | protein-coding |
| chr8_82201654_82202350 | chr8 | 82201655 | 82202350 | + | 0.884644 | Intergenic | Intergenic | 200584 | NM_001254747 | 16168 | Mm.4392 | NM_008357 | | **Il15** | AI503618\|IL-15 | interleukin 15 | protein-coding |

| | | | | | | | | | | | | | | | | |
|---|---|---|---|---|---|---|---|---|---|---|---|---|---|---|---|---|
| chr2_11691122_11691554 | chr2 | 11691123 | 11691554 | + | 1.7016 | 3' UTR (NM_008367, exon 8 of 8) | 3' UTR (NM_008367, exon 8 of 8) | -13955 | NM_001271498 | 16169 | Mm.200196 | NM_008358 | ENSMUSG00000023206 | **Il15ra** | AA690181\|IL-15RA | interleukin 15 receptor, alpha chain | protein-coding |
| chr2_11690246_11690753 | chr2 | 11690247 | 11690753 | + | 1.31663 | 3' UTR (NM_008367, exon 8 of 8) | 3' UTR (NM_008367, exon 8 of 8) | -14793 | NM_001271498 | 16169 | Mm.200196 | NM_008358 | ENSMUSG00000023206 | **Il15ra** | AA690181\|IL-15RA | interleukin 15 receptor, alpha chain | protein-coding |
| chr2_11699978_11700631 | chr2 | 11699979 | 11700631 | + | 0.810183 | Intergenic | ORR1C2\|LTR\|ERVL-MaLR | -4988 | NM_001271498 | 16169 | Mm.200196 | NM_008358 | ENSMUSG00000023206 | **Il15ra** | AA690181\|IL-15RA | interleukin 15 receptor, alpha chain | protein-coding |
| chr1_20740380_20740988 | chr1 | 20740381 | 20740988 | + | 1.12345 | Intergenic | Intergenic | 9779 | NM_010552 | 16171 | Mm.5419 | NM_010552 | ENSMUSG00000025929 | **Il17a** | Ctla-8\|Ctla8\|IL-17\|IL-17A\|Il17 | interleukin 17A | protein-coding |
| chr1_20759740_20760215 | chr1 | 20759741 | 20760215 | + | 1.06088 | Intergenic | Intergenic | 24292 | NM_145856 | 257630 | Mm.222807 | NM_145856 | ENSMUSG00000041872 | **Il17f** | C87042\|IL-17F | interleukin 17F | protein-coding |
| chr1_20759537_20760212 | chr1 | 20759538 | 20760212 | + | 0.87896 | Intergenic | Intergenic | 24395 | NM_145856 | 257630 | Mm.222807 | NM_145856 | ENSMUSG00000041872 | **Il17f** | C87042\|IL-17F | interleukin 17F | protein-coding |
| chr6_120452948_120453634 | chr6 | 120452949 | 120453634 | + | 0.942575 | Intergenic | Intergenic | -9906 | NM_008359 | 16172 | Mm.4481 | NM_008359 | ENSMUSG00000002897 | **Il17ra** | AW538159\|Cdw217\|Il17r\|VDw217 | interleukin 17 receptor A | protein-coding |
| chr6_120470751_120471016 | chr6 | 120470752 | 120471016 | + | 0.632693 | intron (NM_008359, intron 1 of 12) | intron (NM_008359, intron 1 of 12) | 7687 | NM_008359 | 16172 | Mm.4481 | NM_008359 | ENSMUSG00000002897 | **Il17ra** | AW538159\|Cdw217\|Il17r\|VDw217 | interleukin 17 receptor A | protein-coding |
| chr9_50569926_50570526 | chr9 | 50569927 | 50570526 | + | 1.45294 | intron (NM_008360, intron 1 of 5) | intron (NM_008360, intron 1 of 5) | 4858 | NM_008360 | 16173 | Mm.1410 | NM_008360 | ENSMUSG00000039217 | **Il18** | Igif\|Il-18 | interleukin 18 | protein-coding |
| chr7_102003138_102003947 | chr7 | 102003139 | 102003947 | + | 0.73153 | intron (NM_133947, intron 14 of 24) | B3\|SINE\|B2 | 14612 | NM_010531 | 16068 | Mm.422756 | NM_010531 | ENSMUSG00000070427 | **Il18bp** | IL-18BP\|Igifbp\|MC54L | interleukin 18 binding protein | protein-coding |
| chr1_40469651_40469835 | chr1 | 40469652 | 40469835 | + | 0.773151 | intron (NM_008365, intron 1 of 10) | intron (NM_008365, intron 1 of 10) | 3737 | NM_008365 | 16182 | Mm.253664 | NM_008365 | ENSMUSG00000026070 | **Il18r1** | Il18ralpha\|Il1rrp | interleukin 18 receptor 1 | protein-coding |
| chr1_40516075_40516792 | chr1 | 40516076 | 40516792 | + | 1.45434 | 5' UTR (NM_010553, exon 3 of 13) | 5' UTR (NM_010553, exon 3 of 13) | 1072 | NM_010553 | 16174 | Mm.20466 | NM_010553 | ENSMUSG00000026068 | **Il18rap** | AcPL\|IL-18R-beta\|IL-18RAcP\|IL-18Rbeta\|IL-1RAcPL | interleukin 18 receptor accessory protein | protein-coding |
| chr2_129320029_129320616 | chr2 | 129320030 | 129320616 | + | 1.19551 | Intergenic | Intergenic | -10351 | NM_010554 | 16175 | Mm.15534 | NM_010554 | ENSMUSG00000027399 | **Il1a** | Il-1a | interleukin 1 alpha | protein-coding |
| chr2_129385287_129385725 | chr2 | 129385288 | 129385725 | + | 0.996673 | Intergenic | Intergenic | -14386 | NM_008361 | 16176 | Mm.222830 | NM_008361 | ENSMUSG00000027398 | **Il1b** | IL-1beta\|Il-1b | interleukin 1 beta | protein-coding |
| chr2_129385151_129385920 | chr2 | 129385152 | 129385920 | + | 0.901424 | Intergenic | Intergenic | -14416 | NM_008361 | 16176 | Mm.222830 | NM_008361 | ENSMUSG00000027398 | **Il1b** | IL-1beta\|Il-1b | interleukin 1 beta | protein-coding |
| chr1_40235925_40236619 | chr1 | 40235926 | 40236619 | + | 1.47359 | intron (NM_008362, intron 1 of 10) | intron (NM_008362, intron 1 of 10) | 11192 | NM_008362 | 16177 | Mm.896 | NM_008362 | ENSMUSG00000026072 | **Il1r1** | CD121a\|CD121b\|IL-iR\|Il1r-1 | interleukin 1 receptor, type I | protein-coding |
| chr1_40033222_40033744 | chr1 | 40033223 | 40033744 | + | 1.28655 | Intergenic | Intergenic | -51285 | NM_010555 | 16178 | Mm.1349 | NM_010555 | ENSMUSG00000026073 | **Il1r2** | CD121b\|Il1r-2 | interleukin 1 receptor, type II | protein-coding |
| chr1_40055863_40056361 | chr1 | 40055864 | 40056361 | + | 1.26523 | Intergenic | MLT1A1\|LTR\|ERVL-MaLR | -28656 | NM_010555 | 16178 | Mm.1349 | NM_010555 | ENSMUSG00000026073 | **Il1r2** | CD121b\|Il1r-2 | interleukin 1 receptor, type II | protein-coding |
| chr1_40033188_40034091 | chr1 | 40033189 | 40034091 | + | 1.19342 | Intergenic | Intergenic | -51128 | NM_010555 | 16178 | Mm.1349 | NM_010555 | ENSMUSG00000026073 | **Il1r2** | CD121b\|Il1r-2 | interleukin 1 receptor, type II | protein-coding |
| chr1_40053745_40054246 | chr1 | 40053746 | 40054246 | + | 0.879596 | Intergenic | Intergenic | -30772 | NM_010555 | 16178 | Mm.1349 | NM_010555 | ENSMUSG00000026073 | **Il1r2** | CD121b\|Il1r-2 | interleukin 1 receptor, type II | protein-coding |
| chr1_40059129_40059495 | chr1 | 40059130 | 40059495 | + | 0.801254 | Intergenic | MIRc\|SINE\|MIR | -25456 | NM_010555 | 16178 | Mm.1349 | NM_010555 | ENSMUSG00000026073 | **Il1r2** | CD121b\|Il1r-2 | interleukin 1 receptor, type II | protein-coding |
| chr1_40059122_40060039 | chr1 | 40059123 | 40060039 | + | 0.788404 | Intergenic | Intergenic | -25187 | NM_010555 | 16178 | Mm.1349 | NM_010555 | ENSMUSG00000026073 | **Il1r2** | CD121b\|Il1r-2 | interleukin 1 receptor, type II | protein-coding |
| chr1_40022715_40023226 | chr1 | 40022716 | 40023226 | + | 0.725123 | intron (NM_008696, intron 28 of 30) | intron (NM_008696, intron 28 of 30) | -61797 | NM_010555 | 16178 | Mm.1349 | NM_010555 | ENSMUSG00000026073 | **Il1r2** | CD121b\|Il1r-2 | interleukin 1 receptor, type II | protein-coding |
| chr1_40022535_40023164 | chr1 | 40022536 | 40023164 | + | 0.664315 | exon (NM_001252202, exon 27 of 30) | exon (NM_001252202, exon 27 of 30) | -61918 | NM_010555 | 16178 | Mm.1349 | NM_010555 | ENSMUSG00000026073 | **Il1r2** | CD121b\|Il1r-2 | interleukin 1 receptor, type II | protein-coding |
| chr1_40444556_40445258 | chr1 | 40444557 | 40445258 | + | 1.97182 | intron (NM_001294171, intron 6 of 7) | intron (NM_001294171, intron 6 of 7) | 5278 | NM_010743 | 17082 | Mm.289824 | NM_010743 | ENSMUSG00000026069 | **Il1rl1** | DER4\|Fit-1\|Ly84\|ST2L\|St2\|St2-rs1\|T1\|T1/ST2 | interleukin 1 receptor-like 1 | protein-coding |
| chr1_40393098_40393929 | chr1 | 40393099 | 40393929 | + | 1.46671 | Intergenic | Intergenic | -36056 | NM_001294171 | 17082 | Mm.289824 | NM_010743 | ENSMUSG00000026069 | **Il1rl1** | DER4\|Fit-1\|Ly84\|ST2L\|St2\|St2-rs1\|T1\|T1/ST2 | interleukin 1 receptor-like 1 | protein-coding |
| chr1_40412394_40412803 | chr1 | 40412395 | 40412803 | + | 1.38731 | Intergenic | L1_Mus3\|LINE\|L1 | -16971 | NM_001294171 | 17082 | Mm.289824 | NM_010743 | ENSMUSG00000026069 | **Il1rl1** | DER4\|Fit-1\|Ly84\|ST2L\|St2\|St2-rs1\|T1\|T1/ST2 | interleukin 1 receptor-like 1 | protein-coding |
| chr1_40412960_40413544 | chr1 | 40412961 | 40413544 | + | 0.997871 | Intergenic | Intergenic | -16318 | NM_001294171 | 17082 | Mm.289824 | NM_010743 | ENSMUSG00000026069 | **Il1rl1** | DER4\|Fit-1\|Ly84\|ST2L\|St2\|St2-rs1\|T1\|T1/ST2 | interleukin 1 receptor-like 1 | protein-coding |
| chr1_40358686_40359228 | chr1 | 40358687 | 40359228 | + | 1.1629 | intron (NM_133193, intron 9 of 10) | intron (NM_133193, intron 9 of 10) | 34330 | NM_133193 | 107527 | Mm.155583 | NM_133193 | ENSMUSG00000070942 | **Il1rl2** | AI481289\|AW551444\|IL-1Rrp2 | interleukin 1 receptor-like 2 | protein-coding |
| chr1_40315535_40315822 | chr1 | 40315536 | 40315822 | + | 1.14006 | 3' UTR (NM_008362, exon 11 of 11) | 3' UTR (NM_008362, exon 11 of 11) | -8948 | NM_133193 | 107527 | Mm.155583 | NM_133193 | ENSMUSG00000070942 | **Il1rl2** | AI481289\|AW551444\|IL-1Rrp2 | interleukin 1 receptor-like 2 | protein-coding |
| chr1_40315336_40315894 | chr1 | 40315337 | 40315894 | + | 1.12698 | 3' UTR (NM_008362, exon 11 of 11) | 3' UTR (NM_008362, exon 11 of 11) | -9012 | NM_133193 | 107527 | Mm.155583 | NM_133193 | ENSMUSG00000070942 | **Il1rl2** | AI481289\|AW551444\|IL-1Rrp2 | interleukin 1 receptor-like 2 | protein-coding |
| chr2_24341428_24342124 | chr2 | 24341429 | 24342124 | + | 1.01775 | intron (NM_031167, intron 1 of 4) | intron (NM_031167, intron 1 of 4) | -3515 | NM_001159562 | 16181 | Mm.882 | NM_031167 | ENSMUSG00000026981 | **Il1rn** | F630041P17Rik\|IL-1ra | interleukin 1 receptor antagonist | protein-coding |
| chr2_24359334_24359684 | chr2 | 24359335 | 24359684 | + | 0.930247 | Intergenic | MERV1_LTR\|LTR\|ERV1 | 14138 | NM_001039701 | 16181 | Mm.882 | NM_031167 | ENSMUSG00000026981 | **Il1rn** | F630041P17Rik\|IL-1ra | interleukin 1 receptor antagonist | protein-coding |
| chr3_37128262_37128801 | chr3 | 37128263 | 37128801 | + | 1.34329 | Intergenic | Intergenic | -2578 | NM_008366 | 16183 | Mm.14190 | NM_008366 | ENSMUSG00000027720 | **Il2** | Il-2 | interleukin 2 | protein-coding |
| chr3_37174291_37174902 | chr3 | 37174292 | 37174902 | + | 0.755566 | Intergenic | Intergenic | -48643 | NM_008366 | 16183 | Mm.14190 | NM_008366 | ENSMUSG00000027720 | **Il2** | Il-2 | interleukin 2 | protein-coding |
| chr3_37157040_37157404 | chr3 | 37157041 | 37157404 | + | 0.645395 | Intergenic | L4_C_Mam\|LINE\|RTE-X | -31268 | NM_008366 | 16183 | Mm.14190 | NM_008366 | ENSMUSG00000027720 | **Il2** | Il-2 | interleukin 2 | protein-coding |
| chr1_130902459_130902868 | chr1 | 130902460 | 130902868 | + | 0.76788 | Intergenic | Intergenic | 8632 | NM_021380 | 58181 | Mm.103794 | NM_021380 | ENSMUSG00000026416 | **Il20** | Zcyto10 | interleukin 20 | protein-coding |
| chr9_100488906_100489541 | chr9 | 100488907 | 100489541 | + | 1.10304 | Intergenic | Intergenic | -2751 | NM_001033543 | 213208 | Mm.242896 | NM_001033543 | ENSMUSG00000044244 | **Il20rb** | AV228068\|Fndc6\|Gm186\|Il20R2 | interleukin 20 receptor beta | protein-coding |
| chr9_100493510_100493923 | chr9 | 100493511 | 100493923 | + | 0.728077 | Intergenic | Intergenic | -7244 | NM_001033543 | 213208 | Mm.242896 | NM_001033543 | ENSMUSG00000044244 | **Il20rb** | AV228068\|Fndc6\|Gm186\|Il20R2 | interleukin 20 receptor beta | protein-coding |
| chr9_100493679_100493916 | chr9 | 100493680 | 100493916 | + | 0.710413 | Intergenic | Intergenic | -7325 | NM_001033543 | 213208 | Mm.242896 | NM_001033543 | ENSMUSG00000044244 | **Il20rb** | AV228068\|Fndc6\|Gm186\|Il20R2 | interleukin 20 receptor beta | protein-coding |
| chr3_37259349_37260047 | chr3 | 37259350 | 37260047 | + | 1.43741 | Intergenic | Intergenic | -27062 | NM_021782 | 60505 | Mm.157689 | NM_021782 | ENSMUSG00000027718 | **Il21** | IL-21 | interleukin 21 | protein-coding |
| chr3_37231075_37231492 | chr3 | 37231076 | 37231492 | + | 0.854542 | intron (NM_001291041, intron 2 of 3) | intron (NM_001291041, intron 2 of 3) | 1352 | NM_001291041 | 60505 | Mm.157689 | NM_021782 | ENSMUSG00000027718 | **Il21** | IL-21 | interleukin 21 | protein-coding |
| chr7_125607846_125608138 | chr7 | 125607847 | 125608138 | + | 0.660378 | intron (NM_021887, intron 1 of 8) | intron (NM_021887, intron 1 of 8) | 4563 | NM_021887 | 60504 | Mm.155643 | NM_021887 | ENSMUSG00000030745 | **Il21r** | NILR | interleukin 21 receptor | protein-coding |
| chr10_118178672_118179089 | chr10 | 118178673 | 118179089 | + | 0.679284 | Intergenic | Intergenic | -26083 | NM_016971 | 50929 | Mm.103585 | NM_016971 | ENSMUSG00000074695 | **Il22** | IL-22\|IL-22a\|ILTIFa\|Iltif | interleukin 22 | protein-coding |
| chr10_128297813_128298398 | chr10 | 128297814 | 128298398 | + | 1.25567 | promoter-TSS (NM_031252) | promoter-TSS (NM_031252) | -22 | NM_031252 | 83430 | Mm.125482 | NM_031252 | ENSMUSG00000025383 | **Il23a** | IL-23\|p19 | interleukin 23, alpha subunit p19 | protein-coding |
| chr6_67459308_67459804 | chr6 | 67459309 | 67459804 | + | 1.62547 | intron (NM_144548, intron 6 of 10) | RMER15\|LTR\|ERVL | 32299 | NM_144548 | 209590 | Mm.221227 | NM_144548 | ENSMUSG00000049093 | **Il23r** | IL-23R | interleukin 23 receptor | protein-coding |
| chr6_67475437_67476021 | chr6 | 67475438 | 67476021 | + | 1.4756 | intron (NM_144548, intron 4 of 10) | intron (NM_144548, intron 4 of 10) | 16126 | NM_144548 | 209590 | Mm.221227 | NM_144548 | ENSMUSG00000049093 | **Il23r** | IL-23R | interleukin 23 receptor | protein-coding |
| chr6_67483002_67483643 | chr6 | 67483003 | 67483643 | + | 1.27114 | intron (NM_144548, intron 3 of 10) | intron (NM_144548, intron 3 of 10) | 8532 | NM_144548 | 209590 | Mm.221227 | NM_144548 | ENSMUSG00000049093 | **Il23r** | IL-23R | interleukin 23 receptor | protein-coding |
| chr6_67481222_67482213 | chr6 | 67481223 | 67482213 | + | 1.0872 | intron (NM_144548, intron 3 of 10) | B1_Mus1\|SINE\|Alu | 10137 | NM_144548 | 209590 | Mm.221227 | NM_144548 | ENSMUSG00000049093 | **Il23r** | IL-23R | interleukin 23 receptor | protein-coding |
| chr6_67508995_67509337 | chr6 | 67508996 | 67509337 | + | 1.05198 | Intergenic | Intergenic | -17311 | NM_144548 | 209590 | Mm.221227 | NM_144548 | ENSMUSG00000049093 | **Il23r** | IL-23R | interleukin 23 receptor | protein-coding |
| chr6_67481749_67482201 | chr6 | 67481750 | 67482201 | + | 0.871728 | intron (NM_144548, intron 3 of 10) | intron (NM_144548, intron 3 of 10) | 9880 | NM_144548 | 209590 | Mm.221227 | NM_144548 | ENSMUSG00000049093 | **Il23r** | IL-23R | interleukin 23 receptor | protein-coding |
| chr6_67443037_67444086 | chr6 | 67443038 | 67444086 | + | 0.779544 | intron (NM_144548, intron 7 of 10) | intron (NM_144548, intron 7 of 10) | 48293 | NM_144548 | 209590 | Mm.221227 | NM_144548 | ENSMUSG00000049093 | **Il23r** | IL-23R | interleukin 23 receptor | protein-coding |
| chr6_67441375_67441815 | chr6 | 67441376 | 67441815 | + | 0.691908 | intron (NM_144548, intron 7 of 10) | intron (NM_144548, intron 7 of 10) | 50260 | NM_144548 | 209590 | Mm.221227 | NM_144548 | ENSMUSG00000049093 | **Il23r** | IL-23R | interleukin 23 receptor | protein-coding |
| chr1_130879494_130880524 | chr1 | 130879495 | 130880524 | + | 1.68161 | intron (NM_026976, intron 7 of 7) | intron (NM_026976, intron 7 of 7) | 7402 | NM_053095 | 93672 | Mm.196691 | NM_053095 | ENSMUSG00000026420 | **Il24** | FISP\|Mda-7\|Mda7\|St16 | interleukin 24 | protein-coding |
| chr1_130886382_130886961 | chr1 | 130886383 | 130886961 | + | 0.839309 | intron (NM_053095, intron 1 of 5) | intron (NM_053095, intron 1 of 5) | 739 | NM_053095 | 93672 | Mm.196691 | NM_053095 | ENSMUSG00000026420 | **Il24** | FISP\|Mda-7\|Mda7\|St16 | interleukin 24 | protein-coding |
| chr2_11645335_11645759 | chr2 | 11645336 | 11645759 | + | 1.58266 | intron (NM_008367, intron 1 of 7) | intron (NM_008367, intron 1 of 7) | 2755 | NM_008367 | 16184 | Mm.915 | NM_008367 | ENSMUSG00000026770 | **Il2ra** | CD25\|Il2r\|Ly-43 | interleukin 2 receptor, alpha chain | protein-coding |
| chr2_11623113_11623570 | chr2 | 11623114 | 11623570 | + | 1.32098 | Intergenic | Intergenic | -19450 | NM_008367 | 16184 | Mm.915 | NM_008367 | ENSMUSG00000026770 | **Il2ra** | CD25\|Il2r\|Ly-43 | interleukin 2 receptor, alpha chain | protein-coding |
| chr2_11647247_11647520 | chr2 | 11647248 | 11647520 | + | 1.29231 | intron (NM_008367, intron 1 of 7) | intron (NM_008367, intron 1 of 7) | 4592 | NM_008367 | 16184 | Mm.915 | NM_008367 | ENSMUSG00000026770 | **Il2ra** | CD25\|Il2r\|Ly-43 | interleukin 2 receptor, alpha chain | protein-coding |
| chr2_11623790_11624039 | chr2 | 11623791 | 11624039 | + | 1.21234 | Intergenic | Intergenic | -18877 | NM_008367 | 16184 | Mm.915 | NM_008367 | ENSMUSG00000026770 | **Il2ra** | CD25\|Il2r\|Ly-43 | interleukin 2 receptor, alpha chain | protein-coding |
| chr2_11651347_11651910 | chr2 | 11651348 | 11651910 | + | 1.09723 | intron (NM_008367, intron 1 of 7) | intron (NM_008367, intron 1 of 7) | 8837 | NM_008367 | 16184 | Mm.915 | NM_008367 | ENSMUSG00000026770 | **Il2ra** | CD25\|Il2r\|Ly-43 | interleukin 2 receptor, alpha chain | protein-coding |
| chr2_11644982_11645220 | chr2 | 11644983 | 11645220 | + | 1.08393 | intron (NM_008367, intron 1 of 7) | B3\|SINE\|B2 | 2309 | NM_008367 | 16184 | Mm.915 | NM_008367 | ENSMUSG00000026770 | **Il2ra** | CD25\|Il2r\|Ly-43 | interleukin 2 receptor, alpha chain | protein-coding |

| | | | | | | | | | | | | | | | | | |
|---|---|---|---|---|---|---|---|---|---|---|---|---|---|---|---|---|---|
| chr2_11625937_11626286 | chr2 | 11625938 | 11626286 | + | 1.06877 | Intergenic | Intergenic | -16680 | NM_008367 | 16184 | Mm.915 | NM_008367 | ENSMUSG00000026770 | **Il2ra** | CD25\|Il2r\|Ly-43 | interleukin 2 receptor, alpha chain | protein-coding |
| chr2_11630357_11630809 | chr2 | 11630358 | 11630809 | + | 0.931979 | Intergenic | Intergenic | -12209 | NM_008367 | 16184 | Mm.915 | NM_008367 | ENSMUSG00000026770 | **Il2ra** | CD25\|Il2r\|Ly-43 | interleukin 2 receptor, alpha chain | protein-coding |
| chr2_11655005_11655332 | chr2 | 11655006 | 11655332 | + | 0.912108 | intron (NM_008367, intron 1 of 7) | intron (NM_008367, intron 1 of 7) | 12377 | NM_008367 | 16184 | Mm.915 | NM_008367 | ENSMUSG00000026770 | **Il2ra** | CD25\|Il2r\|Ly-43 | interleukin 2 receptor, alpha chain | protein-coding |
| chr2_11655408_11655862 | chr2 | 11655409 | 11655862 | + | 0.896634 | intron (NM_008367, intron 1 of 7) | intron (NM_008367, intron 1 of 7) | 12843 | NM_008367 | 16184 | Mm.915 | NM_008367 | ENSMUSG00000026770 | **Il2ra** | CD25\|Il2r\|Ly-43 | interleukin 2 receptor, alpha chain | protein-coding |
| chr2_11629678_11629970 | chr2 | 11629679 | 11629970 | + | 0.84177 | Intergenic | Intergenic | -12968 | NM_008367 | 16184 | Mm.915 | NM_008367 | ENSMUSG00000026770 | **Il2ra** | CD25\|Il2r\|Ly-43 | interleukin 2 receptor, alpha chain | protein-coding |
| chr2_11662877_11663219 | chr2 | 11662878 | 11663219 | + | 0.836371 | intron (NM_008367, intron 1 of 7) | intron (NM_008367, intron 1 of 7) | 20256 | NM_008367 | 16184 | Mm.915 | NM_008367 | ENSMUSG00000026770 | **Il2ra** | CD25\|Il2r\|Ly-43 | interleukin 2 receptor, alpha chain | protein-coding |
| chr2_11662752_11663722 | chr2 | 11662753 | 11663722 | + | 0.815336 | intron (NM_008367, intron 1 of 7) | intron (NM_008367, intron 1 of 7) | 20445 | NM_008367 | 16184 | Mm.915 | NM_008367 | ENSMUSG00000026770 | **Il2ra** | CD25\|Il2r\|Ly-43 | interleukin 2 receptor, alpha chain | protein-coding |
| chr2_11662730_11663451 | chr2 | 11662731 | 11663451 | + | 0.803009 | intron (NM_008367, intron 1 of 7) | intron (NM_008367, intron 1 of 7) | 20299 | NM_008367 | 16184 | Mm.915 | NM_008367 | ENSMUSG00000026770 | **Il2ra** | CD25\|Il2r\|Ly-43 | interleukin 2 receptor, alpha chain | protein-coding |
| chr2_11655396_11655653 | chr2 | 11655397 | 11655653 | + | 0.749512 | intron (NM_008367, intron 1 of 7) | intron (NM_008367, intron 1 of 7) | 12733 | NM_008367 | 16184 | Mm.915 | NM_008367 | ENSMUSG00000026770 | **Il2ra** | CD25\|Il2r\|Ly-43 | interleukin 2 receptor, alpha chain | protein-coding |
| chr13_112602569_112602968 | chr13 | 112602570 | 112602968 | + | 0.853459 | intron (NM_001145885, intron 18 of 21) | intron (NM_001145885, intron 18 of 21) | -22107 | NM_139299 | 218624 | Mm.380801 | NM_139299 | ENSMUSG00000050377 | **Il31ra** | CRL3\|GLM-R\|GPL\|Glmr\|NR10 | interleukin 31 receptor A | protein-coding |
| chr13_112602520_112603144 | chr13 | 112602521 | 112603144 | + | 0.78932 | intron (NM_001145885, intron 18 of 21) | intron (NM_001145885, intron 18 of 21) | -22170 | NM_139299 | 218624 | Mm.380801 | NM_139299 | ENSMUSG00000050377 | **Il31ra** | CRL3\|GLM-R\|GPL\|Glmr\|NR10 | interleukin 31 receptor A | protein-coding |
| chr19_29887502_29887930 | chr19 | 29887503 | 29887930 | + | 1.6699 | Intergenic | Intergenic | -37398 | NM_133775 | 77125 | Mm.182359 | NM_133775 | ENSMUSG00000024810 | **Il33** | 9230117N10Rik\|Il-33\|Il1f11\|NF-HEV | interleukin 33 | protein-coding |
| chr7_125554485_125554876 | chr7 | 125554486 | 125554876 | + | 0.792302 | intron (NM_001008700, intron 1 of 10) | intron (NM_001008700, intron 1 of 10) | 2399 | NM_001008700 | 16190 | Mm.233802 | NM_010557 | ENSMUSG00000030748 | **Il4ra** | CD124\|Il4r | interleukin 4 receptor, alpha | protein-coding |
| chr7_125554146_125555006 | chr7 | 125554147 | 125555006 | + | 0.731021 | intron (NM_001008700, intron 1 of 10) | intron (NM_001008700, intron 1 of 10) | 2294 | NM_001008700 | 16190 | Mm.233802 | NM_010557 | ENSMUSG00000030748 | **Il4ra** | CD124\|Il4r | interleukin 4 receptor, alpha | protein-coding |
| chr5_29919998_29920768 | chr5 | 29919999 | 29920768 | + | 1.31125 | intron (NR_105781, intron 1 of 2) | intron (NR_105781, intron 1 of 2) | -92778 | NM_031168 | 16193 | Mm.1019 | NM_031168 | ENSMUSG00000025746 | **Il6** | Il-6 | interleukin 6 | protein-coding |
| chr5_29947189_29947747 | chr5 | 29947190 | 29947747 | + | 1.17646 | intron (NR_105781, intron 2 of 2) | intron (NR_105781, intron 2 of 2) | -65693 | NM_031168 | 16193 | Mm.1019 | NM_031168 | ENSMUSG00000025746 | **Il6** | Il-6 | interleukin 6 | protein-coding |
| chr5_29946837_29947742 | chr5 | 29946838 | 29947742 | + | 1.091 | intron (NR_105781, intron 2 of 2) | intron (NR_105781, intron 2 of 2) | -65871 | NM_031168 | 16193 | Mm.1019 | NM_031168 | ENSMUSG00000025746 | **Il6** | Il-6 | interleukin 6 | protein-coding |
| chr3_89879972_89880742 | chr3 | 89879973 | 89880742 | + | 1.4843 | intron (NM_010559, intron 6 of 9) | intron (NM_010559, intron 6 of 9) | 32805 | NM_010559 | 16194 | Mm.2856 | NM_010559 | ENSMUSG00000027947 | **Il6ra** | CD126\|IL-6R\|Il6r | interleukin 6 receptor, alpha | protein-coding |
| chr3_89874031_89874496 | chr3 | 89874032 | 89874496 | + | 0.84214 | intron (NM_010559, intron 9 of 9) | ORR1D1\|LTR\|ERVL-MaLR | 38898 | NM_010559 | 16194 | Mm.2856 | NM_010559 | ENSMUSG00000027947 | **Il6ra** | CD126\|IL-6R\|Il6r | interleukin 6 receptor, alpha | protein-coding |
| chr3_89892494_89893048 | chr3 | 89892495 | 89893048 | + | 0.717085 | intron (NM_010559, intron 1 of 9) | ORR1D1\|LTR\|ERVL-MaLR | 20391 | NM_010559 | 16194 | Mm.2856 | NM_010559 | ENSMUSG00000027947 | **Il6ra** | CD126\|IL-6R\|Il6r | interleukin 6 receptor, alpha | protein-coding |
| chr3_89887615_89888170 | chr3 | 89887616 | 89888170 | + | 0.702688 | intron (NM_010559, intron 3 of 9) | intron (NM_010559, intron 3 of 9) | 25269 | NM_010559 | 16194 | Mm.2856 | NM_010559 | ENSMUSG00000027947 | **Il6ra** | CD126\|IL-6R\|Il6r | interleukin 6 receptor, alpha | protein-coding |
| chr13_112447203_112447491 | chr13 | 112447204 | 112447491 | + | 1.09039 | Intergenic | Intergenic | -16723 | NM_010560 | 16195 | Mm.4364 | NM_010560 | ENSMUSG00000021756 | **Il6st** | 5133400A03Rik\|AA389424\|BB405851\|CD130\|D13Ertd699e\|gp130 | interleukin 6 signal transducer | protein-coding |
| chr13_112447034_112447491 | chr13 | 112447035 | 112447491 | + | 1.01255 | Intergenic | MLT1B\|LTR\|ERVL-MaLR | -16807 | NM_010560 | 16195 | Mm.4364 | NM_010560 | ENSMUSG00000021756 | **Il6st** | 5133400A03Rik\|AA389424\|BB405851\|CD130\|D13Ertd699e\|gp130 | interleukin 6 signal transducer | protein-coding |
| chr13_112425121_112425812 | chr13 | 112425122 | 112425812 | + | 1.00672 | Intergenic | Intergenic | -38603 | NM_010560 | 16195 | Mm.4364 | NM_010560 | ENSMUSG00000021756 | **Il6st** | 5133400A03Rik\|AA389424\|BB405851\|CD130\|D13Ertd699e\|gp130 | interleukin 6 signal transducer | protein-coding |
| chr13_112443153_112443506 | chr13 | 112443154 | 112443506 | + | 1.001 | Intergenic | Intergenic | -20740 | NM_010560 | 16195 | Mm.4364 | NM_010560 | ENSMUSG00000021756 | **Il6st** | 5133400A03Rik\|AA389424\|BB405851\|CD130\|D13Ertd699e\|gp130 | interleukin 6 signal transducer | protein-coding |
| chr13_112396490_112397102 | chr13 | 112396491 | 112397102 | + | 0.856256 | Intergenic | Intergenic | -67274 | NM_010560 | 16195 | Mm.4364 | NM_010560 | ENSMUSG00000021756 | **Il6st** | 5133400A03Rik\|AA389424\|BB405851\|CD130\|D13Ertd699e\|gp130 | interleukin 6 signal transducer | protein-coding |
| chr13_112458587_112459005 | chr13 | 112458588 | 112459005 | + | 0.763545 | Intergenic | Intergenic | -5274 | NM_010560 | 16195 | Mm.4364 | NM_010560 | ENSMUSG00000021756 | **Il6st** | 5133400A03Rik\|AA389424\|BB405851\|CD130\|D13Ertd699e\|gp130 | interleukin 6 signal transducer | protein-coding |
| chr13_112396135_112396436 | chr13 | 112396136 | 112396436 | + | 0.750461 | Intergenic | Intergenic | -67784 | NM_010560 | 16195 | Mm.4364 | NM_010560 | ENSMUSG00000021756 | **Il6st** | 5133400A03Rik\|AA389424\|BB405851\|CD130\|D13Ertd699e\|gp130 | interleukin 6 signal transducer | protein-coding |
| chr13_112443151_112443929 | chr13 | 112443152 | 112443929 | + | 0.745723 | Intergenic | Intergenic | -20530 | NM_010560 | 16195 | Mm.4364 | NM_010560 | ENSMUSG00000021756 | **Il6st** | 5133400A03Rik\|AA389424\|BB405851\|CD130\|D13Ertd699e\|gp130 | interleukin 6 signal transducer | protein-coding |
| chr15_9569416_9569910 | chr15 | 9569417 | 9569910 | + | 2.04149 | Intergenic | Intergenic | -39787 | NM_008372 | 16197 | Mm.389 | NM_008372 | ENSMUSG00000003882 | **Il7r** | CD127\|IL-7Ralpha | interleukin 7 receptor | protein-coding |
| chr15_9570044_9570746 | chr15 | 9570045 | 9570746 | + | 1.53465 | Intergenic | Intergenic | -40519 | NM_008372 | 16197 | Mm.389 | NM_008372 | ENSMUSG00000003882 | **Il7r** | CD127\|IL-7Ralpha | interleukin 7 receptor | protein-coding |
| chr15_9562313_9563342 | chr15 | 9562314 | 9563342 | + | 1.01919 | Intergenic | Intergenic | -32952 | NM_008372 | 16197 | Mm.389 | NM_008372 | ENSMUSG00000003882 | **Il7r** | CD127\|IL-7Ralpha | interleukin 7 receptor | protein-coding |
| chr15_9555933_9556366 | chr15 | 9555934 | 9556366 | + | 1.00045 | Intergenic | Intergenic | -26274 | NM_008372 | 16197 | Mm.389 | NM_008372 | ENSMUSG00000003882 | **Il7r** | CD127\|IL-7Ralpha | interleukin 7 receptor | protein-coding |
| chr15_9555713_9556368 | chr15 | 9555714 | 9556368 | + | 0.928716 | Intergenic | Intergenic | -26165 | NM_008372 | 16197 | Mm.389 | NM_008372 | ENSMUSG00000003882 | **Il7r** | CD127\|IL-7Ralpha | interleukin 7 receptor | protein-coding |
| chr12_41044763_41045127 | chr12 | 41044764 | 41045127 | + | 0.892596 | intron (NM_053122, intron 2 of 6) | intron (NM_053122, intron 2 of 6) | 20855 | NM_053122 | 93757 | Mm.363813 | NM_053122 | ENSMUSG00000056899 | **Immp2l** | AI853880\|IMP2 | IMP2 inner mitochondrial membrane peptidase-like (S. cerevisiae) | protein-coding |
| chr12_41089887_41090456 | chr12 | 41089888 | 41090456 | + | 0.87289 | intron (NM_053122, intron 3 of 6) | MLT1H2\|LTR\|ERVL-MaLR | 66082 | NM_053122 | 93757 | Mm.363813 | NM_053122 | ENSMUSG00000056899 | **Immp2l** | AI853880\|IMP2 | IMP2 inner mitochondrial membrane peptidase-like (S. cerevisiae) | protein-coding |
| chr18_67274249_67274706 | chr18 | 67274250 | 67274706 | + | 1.57328 | Intergenic | Lx8\|LINE\|L1 | -14745 | NM_053261 | 114663 | Mm.34079 | NM_053261 | ENSMUSG00000024525 | **Impa2** | 2210415D20Rik\|AI326924\|AW259601 | inositol (myo)-1(or 4)-monophosphatase 2 | protein-coding |
| chr9_80819061_80819449 | chr9 | 80819062 | 80819449 | + | 0.931973 | Intergenic | Intergenic | -353817 | NM_022016 | 63859 | Mm.422780 | NM_022016 | ENSMUSG00000032343 | **Impg1** | A930015H12Rik\|IMP150\|SPACR | interphotoreceptor matrix proteoglycan 1 | protein-coding |
| chr8_47666261_47666689 | chr8 | 47666262 | 47666689 | + | 0.994941 | TTS (NM_023503) | TTS (NM_023503) | 8684 | NM_023503 | 69260 | Mm.430733 | NM_023503 | ENSMUSG00000063049 | **Ing2** | 2810011M06Rik\|Ing1l\|Ing2b\|P33ING2 | inhibitor of growth family, member 2 | protein-coding |
| chr6_21954928_21955286 | chr6 | 21954929 | 21955286 | + | 0.694756 | intron (NM_023626, intron 4 of 11) | MTD\|LTR\|ERVL-MaLR | 5492 | NM_023626 | 71777 | Mm.39999 | NM_023626 | ENSMUSG00000029670 | **Ing3** | 1300013A07Rik\|P47ING3 | inhibitor of growth family, member 3 | protein-coding |
| chr6_125029801_125030888 | chr6 | 125029802 | 125030888 | + | 0.65327 | intron (NM_001252083, intron 7 of 10) | URR1B\|DNA\|hAT-Charlie | -9503 | NM_133345 | 28019 | Mm.262547 | NM_133345 | ENSMUSG00000030330 | **Ing4** | D6Wsu147e\|D6Xrf92\|p29ING4 | inhibitor of growth family, member 4 | protein-coding |
| chr13_16641041_16642660 | chr13 | 16641042 | 16642660 | + | 1.45616 | Intergenic | Intergenic | 627376 | NM_008380 | 16323 | Mm.8042 | NM_008380 | ENSMUSG00000041324 | **Inhba** | - | inhibin beta-A | protein-coding |
| chr13_16738234_16738728 | chr13 | 16738235 | 16738728 | + | 1.30556 | Intergenic | Intergenic | 724006 | NM_008380 | 16323 | Mm.8042 | NM_008380 | ENSMUSG00000041324 | **Inhba** | - | inhibin beta-A | protein-coding |
| chr13_16738264_16738983 | chr13 | 16738265 | 16738983 | + | 1.18645 | Intergenic | Intergenic | 724149 | NM_008380 | 16323 | Mm.8042 | NM_008380 | ENSMUSG00000041324 | **Inhba** | - | inhibin beta-A | protein-coding |
| chr13_16659218_16660251 | chr13 | 16659219 | 16660251 | + | 1.05655 | Intergenic | Intergenic | 645260 | NM_008380 | 16323 | Mm.8042 | NM_008380 | ENSMUSG00000041324 | **Inhba** | - | inhibin beta-A | protein-coding |
| chr4_59789788_59790215 | chr4 | 59789789 | 59790215 | + | 1.41262 | Intergenic | RMER15\|LTR\|ERVL | -6147 | NM_001013577 | 66209 | Mm.51988 | NM_001013577 | ENSMUSG00000038544 | **Inip** | 1110054O05Rik\|2610312O17Rik\|AA399876\|Ssbip1 | INTS3 and NABP interacting protein | protein-coding |
| chr4_59789617_59790312 | chr4 | 59789618 | 59790312 | + | 1.26411 | Intergenic | RMER15\|LTR\|ERVL | -6110 | NM_001013577 | 66209 | Mm.51988 | NM_001013577 | ENSMUSG00000038544 | **Inip** | 1110054O05Rik\|2610312O17Rik\|AA399876\|Ssbip1 | INTS3 and NABP interacting protein | protein-coding |
| chr18_24116729_24117219 | chr18 | 24116730 | 24117219 | + | 1.5352 | intron (NM_172625, intron 1 of 4) | intron (NM_172625, intron 1 of 4) | 4845 | NM_172625 | 225280 | Mm.258973 | NM_172625 | ENSMUSG00000047989 | **Ino80c** | D030070L09Rik | INO80 complex subunit C | protein-coding |
| chr18_24116557_24117232 | chr18 | 24116558 | 24117232 | + | 1.51913 | intron (NM_172625, intron 1 of 4) | intron (NM_172625, intron 1 of 4) | 4924 | NM_172625 | 225280 | Mm.258973 | NM_172625 | ENSMUSG00000047989 | **Ino80c** | D030070L09Rik | INO80 complex subunit C | protein-coding |
| chr18_24111950_24112328 | chr18 | 24111951 | 24112328 | + | 1.02821 | intron (NM_172625, intron 3 of 4) | intron (NM_172625, intron 3 of 4) | 9680 | NM_172625 | 225280 | Mm.258973 | NM_172625 | ENSMUSG00000047989 | **Ino80c** | D030070L09Rik | INO80 complex subunit C | protein-coding |
| chr18_24088670_24089355 | chr18 | 24088671 | 24089355 | + | 0.987261 | Intergenic | Intergenic | 32806 | NM_172625 | 225280 | Mm.258973 | NM_172625 | ENSMUSG00000047989 | **Ino80c** | D030070L09Rik | INO80 complex subunit C | protein-coding |
| chr18_24101722_24102167 | chr18 | 24101723 | 24102167 | + | 0.937166 | Intergenic | Intergenic | 19874 | NM_172625 | 225280 | Mm.258973 | NM_172625 | ENSMUSG00000047989 | **Ino80c** | D030070L09Rik | INO80 complex subunit C | protein-coding |
| chr18_24104422_24104717 | chr18 | 24104423 | 24104717 | + | 0.640126 | TTS (NM_172625) | TTS (NM_172625) | 17249 | NM_172625 | 225280 | Mm.258973 | NM_172625 | ENSMUSG00000047989 | **Ino80c** | D030070L09Rik | INO80 complex subunit C | protein-coding |
| chr1_63119735_63120065 | chr1 | 63119736 | 63120065 | + | 0.8318 | intron (NR_045914, intron 1 of 3) | intron (NR_045914, intron 1 of 3) | 5160 | NR_045914 | 100380944 | Mm.392955 | NR_045914 | ENSMUSG00000084799 | **Ino80dos** | Gm11602\|OTTMUSG00000002611 | INO80 complex subunit D, opposite strand | ncRNA |
| chr1_63119727_63120277 | chr1 | 63119728 | 63120277 | + | 0.751416 | intron (NR_045914, intron 1 of 3) | intron (NR_045914, intron 1 of 3) | 5262 | NR_045914 | 100380944 | Mm.392955 | NR_045914 | ENSMUSG00000084799 | **Ino80dos** | Gm11602\|OTTMUSG00000002611 | INO80 complex subunit D, opposite strand | ncRNA |
| chr1_52802431_52802881 | chr1 | 52802432 | 52802881 | + | 0.898913 | intron (NM_008384, intron 1 of 5) | intron (NM_008384, intron 1 of 5) | 15032 | NM_008384 | 16329 | Mm.917 | NM_008384 | ENSMUSG00000026102 | **Inpp1** | 2300002C06Rik\|AV137389 | inositol polyphosphate-1-phosphatase | protein-coding |

| | | | | | | | | | | | | | | | | | |
|---|---|---|---|---|---|---|---|---|---|---|---|---|---|---|---|---|---|
| chr1_52803019_52803258 | chr1 | 52803020 | 52803258 | + | 0.706895 | intron (NM_008384, intron 1 of 5) | MLT1B\|LTR\|ERVL-MaLR | 14549 | NM_008384 | 16329 | Mm.917 | NM_008384 | ENSMUSG00000026102 | **Inpp1** | 2300002C06Rik\|AV137389 | inositol polyphosphate-1-phosphatase | protein-coding |
| chr1_37310740_37311428 | chr1 | 37310741 | 37311428 | + | 1.66309 | intron (NM_001290797, intron 1 of 25) | L1MA6\|LINE\|L1 | 11246 | NM_001290797 | 269180 | Mm.5294 | NM_030266 | ENSMUSG00000026113 | **Inpp4a** | 107kDa\|9630012D15\|D130048C09Rik\|R74740 | inositol polyphosphate-4-phosphatase, type I | protein-coding |
| chr1_37310923_37311423 | chr1 | 37310924 | 37311423 | + | 1.62223 | intron (NM_001290797, intron 1 of 25) | intron (NM_001290797, intron 1 of 25) | 11335 | NM_001290797 | 269180 | Mm.5294 | NM_030266 | ENSMUSG00000026113 | **Inpp4a** | 107kDa\|9630012D15\|D130048C09Rik\|R74740 | inositol polyphosphate-4-phosphatase, type I | protein-coding |
| chr1_37303579_37303894 | chr1 | 37303580 | 37303894 | + | 1.26561 | intron (NM_001290797, intron 1 of 25) | L1MC4\|LINE\|L1 | 3899 | NM_001290797 | 269180 | Mm.5294 | NM_030266 | ENSMUSG00000026113 | **Inpp4a** | 107kDa\|9630012D15\|D130048C09Rik\|R74740 | inositol polyphosphate-4-phosphatase, type I | protein-coding |
| chr8_81866965_81867589 | chr8 | 81866966 | 81867589 | + | 1.71546 | intron (NM_001297596, intron 8 of 24) | intron (NM_001297596, intron 8 of 24) | 524715 | NM_001297593 | 234515 | Mm.324468 | NM_001024617 | ENSMUSG00000037940 | **Inpp4b** | E130107I17Rik | inositol polyphosphate-4-phosphatase, type II | protein-coding |
| chr8_81730064_81730810 | chr8 | 81730065 | 81730810 | + | 1.50203 | intron (NM_001024617, intron 3 of 24) | intron (NM_001024617, intron 3 of 24) | 387875 | NM_001297593 | 234515 | Mm.324468 | NM_001024617 | ENSMUSG00000037940 | **Inpp4b** | E130107I17Rik | inositol polyphosphate-4-phosphatase, type II | protein-coding |
| chr8_81729809_81730819 | chr8 | 81729810 | 81730819 | + | 1.30938 | intron (NM_001024617, intron 3 of 24) | intron (NM_001024617, intron 3 of 24) | 387752 | NM_001297593 | 234515 | Mm.324468 | NM_001024617 | ENSMUSG00000037940 | **Inpp4b** | E130107I17Rik | inositol polyphosphate-4-phosphatase, type II | protein-coding |
| chr8_81325856_81326478 | chr8 | 81325857 | 81326478 | + | 1.21471 | Intergenic | ERVL-E-int\|LTR\|ERVL | -16395 | NM_001024617 | 234515 | Mm.324468 | NM_001024617 | ENSMUSG00000037940 | **Inpp4b** | E130107I17Rik | inositol polyphosphate-4-phosphatase, type II | protein-coding |
| chr8_81325972_81326344 | chr8 | 81325973 | 81326344 | + | 1.2003 | Intergenic | ERVL-E-int\|LTR\|ERVL | -16404 | NM_001024617 | 234515 | Mm.324468 | NM_001024617 | ENSMUSG00000037940 | **Inpp4b** | E130107I17Rik | inositol polyphosphate-4-phosphatase, type II | protein-coding |
| chr8_81860593_81860925 | chr8 | 81860594 | 81860925 | + | 1.10487 | intron (NM_001297596, intron 8 of 24) | ORR1E\|LTR\|ERVL-MaLR | 518197 | NM_001297593 | 234515 | Mm.324468 | NM_001024617 | ENSMUSG00000037940 | **Inpp4b** | E130107I17Rik | inositol polyphosphate-4-phosphatase, type II | protein-coding |
| chr8_81859781_81860165 | chr8 | 81859782 | 81860165 | + | 1.07911 | intron (NM_001297596, intron 8 of 24) | intron (NM_001297596, intron 8 of 24) | 517411 | NM_001297593 | 234515 | Mm.324468 | NM_001024617 | ENSMUSG00000037940 | **Inpp4b** | E130107I17Rik | inositol polyphosphate-4-phosphatase, type II | protein-coding |
| chr8_81765384_81765644 | chr8 | 81765385 | 81765644 | + | 1.00049 | intron (NM_001297593, intron 4 of 23) | intron (NM_001297593, intron 4 of 23) | 422952 | NM_001297593 | 234515 | Mm.324468 | NM_001024617 | ENSMUSG00000037940 | **Inpp4b** | E130107I17Rik | inositol polyphosphate-4-phosphatase, type II | protein-coding |
| chr8_81800856_81801206 | chr8 | 81800857 | 81801206 | + | 0.98967 | intron (NM_001024617, intron 7 of 24) | MIR\|SINE\|MIR | 458469 | NM_001297593 | 234515 | Mm.324468 | NM_001024617 | ENSMUSG00000037940 | **Inpp4b** | E130107I17Rik | inositol polyphosphate-4-phosphatase, type II | protein-coding |
| chr8_81803393_81803785 | chr8 | 81803394 | 81803785 | + | 0.961252 | intron (NM_001024617, intron 7 of 24) | intron (NM_001024617, intron 7 of 24) | 461027 | NM_001297593 | 234515 | Mm.324468 | NM_001024617 | ENSMUSG00000037940 | **Inpp4b** | E130107I17Rik | inositol polyphosphate-4-phosphatase, type II | protein-coding |
| chr8_81861332_81861728 | chr8 | 81861333 | 81861728 | + | 0.928919 | intron (NM_001297596, intron 8 of 24) | intron (NM_001297596, intron 8 of 24) | 518968 | NM_001297593 | 234515 | Mm.324468 | NM_001024617 | ENSMUSG00000037940 | **Inpp4b** | E130107I17Rik | inositol polyphosphate-4-phosphatase, type II | protein-coding |
| chr8_81860589_81861102 | chr8 | 81860590 | 81861102 | + | 0.888003 | intron (NM_001297596, intron 8 of 24) | ORR1E\|LTR\|ERVL-MaLR | 518284 | NM_001297593 | 234515 | Mm.324468 | NM_001024617 | ENSMUSG00000037940 | **Inpp4b** | E130107I17Rik | inositol polyphosphate-4-phosphatase, type II | protein-coding |
| chr8_81803393_81803988 | chr8 | 81803394 | 81803988 | + | 0.844269 | intron (NM_001024617, intron 7 of 24) | intron (NM_001024617, intron 7 of 24) | 461129 | NM_001297593 | 234515 | Mm.324468 | NM_001024617 | ENSMUSG00000037940 | **Inpp4b** | E130107I17Rik | inositol polyphosphate-4-phosphatase, type II | protein-coding |
| chr8_81702624_81702880 | chr8 | 81702625 | 81702880 | + | 0.8382 | intron (NM_001297596, intron 2 of 24) | intron (NM_001297596, intron 2 of 24) | 360190 | NM_001297593 | 234515 | Mm.324468 | NM_001024617 | ENSMUSG00000037940 | **Inpp4b** | E130107I17Rik | inositol polyphosphate-4-phosphatase, type II | protein-coding |
| chr8_81563201_81563553 | chr8 | 81563202 | 81563553 | + | 0.828244 | intron (NM_001297596, intron 2 of 24) | RMER6A\|LTR\|ERVK | 220815 | NM_001297593 | 234515 | Mm.324468 | NM_001024617 | ENSMUSG00000037940 | **Inpp4b** | E130107I17Rik | inositol polyphosphate-4-phosphatase, type II | protein-coding |
| chr8_81802338_81802770 | chr8 | 81802339 | 81802770 | + | 0.816832 | intron (NM_001024617, intron 7 of 24) | intron (NM_001024617, intron 7 of 24) | 459992 | NM_001297593 | 234515 | Mm.324468 | NM_001024617 | ENSMUSG00000037940 | **Inpp4b** | E130107I17Rik | inositol polyphosphate-4-phosphatase, type II | protein-coding |
| chr8_81614729_81615428 | chr8 | 81614730 | 81615428 | + | 0.793038 | intron (NM_001297596, intron 2 of 24) | intron (NM_001297596, intron 2 of 24) | 272517 | NM_001297593 | 234515 | Mm.324468 | NM_001024617 | ENSMUSG00000037940 | **Inpp4b** | E130107I17Rik | inositol polyphosphate-4-phosphatase, type II | protein-coding |
| chr8_81514395_81514885 | chr8 | 81514396 | 81514885 | + | 0.728716 | intron (NM_001297596, intron 2 of 24) | intron (NM_001297596, intron 2 of 24) | 172078 | NM_001297593 | 234515 | Mm.324468 | NM_001024617 | ENSMUSG00000037940 | **Inpp4b** | E130107I17Rik | inositol polyphosphate-4-phosphatase, type II | protein-coding |
| chr8_81841566_81842000 | chr8 | 81841567 | 81842000 | + | 0.723932 | intron (NM_001024617, intron 7 of 24) | L2c\|LINE\|L2 | 499221 | NM_001297593 | 234515 | Mm.324468 | NM_001024617 | ENSMUSG00000037940 | **Inpp4b** | E130107I17Rik | inositol polyphosphate-4-phosphatase, type II | protein-coding |
| chr8_81702460_81702882 | chr8 | 81702461 | 81702882 | + | 0.715118 | intron (NM_001297596, intron 2 of 24) | intron (NM_001297596, intron 2 of 24) | 360109 | NM_001297593 | 234515 | Mm.324468 | NM_001024617 | ENSMUSG00000037940 | **Inpp4b** | E130107I17Rik | inositol polyphosphate-4-phosphatase, type II | protein-coding |
| chr8_81823059_81823535 | chr8 | 81823060 | 81823535 | + | 0.653041 | intron (NM_001024617, intron 7 of 24) | intron (NM_001024617, intron 7 of 24) | 480735 | NM_001297593 | 234515 | Mm.324468 | NM_001024617 | ENSMUSG00000037940 | **Inpp4b** | E130107I17Rik | inositol polyphosphate-4-phosphatase, type II | protein-coding |
| chr8_81854505_81854984 | chr8 | 81854506 | 81854984 | + | 0.648827 | intron (NM_001297596, intron 7 of 24) | intron (NM_001297596, intron 7 of 24) | 512183 | NM_001297593 | 234515 | Mm.324468 | NM_001024617 | ENSMUSG00000037940 | **Inpp4b** | E130107I17Rik | inositol polyphosphate-4-phosphatase, type II | protein-coding |
| chr8_81854229_81855003 | chr8 | 81854230 | 81855003 | + | 0.64336 | intron (NM_001297596, intron 7 of 24) | intron (NM_001297596, intron 7 of 24) | 512054 | NM_001297593 | 234515 | Mm.324468 | NM_001024617 | ENSMUSG00000037940 | **Inpp4b** | E130107I17Rik | inositol polyphosphate-4-phosphatase, type II | protein-coding |
| chr7_139476680_139477157 | chr7 | 139476681 | 139477157 | + | 1.03947 | intron (NM_001127363, intron 2 of 15) | intron (NM_001127363, intron 2 of 15) | 87810 | NM_001127363 | 212111 | Mm.28308 | NM_183144 | ENSMUSG00000025477 | **Inpp5a** | - | inositol polyphosphate-5-phosphatase A | protein-coding |
| chr7_139475929_139477155 | chr7 | 139475930 | 139477155 | + | 0.896042 | intron (NM_001127363, intron 2 of 15) | intron (NM_001127363, intron 2 of 15) | 87433 | NM_001127363 | 212111 | Mm.28308 | NM_183144 | ENSMUSG00000025477 | **Inpp5a** | - | inositol polyphosphate-5-phosphatase A | protein-coding |
| chr4_124737874_124739518 | chr4 | 124737875 | 124739518 | + | 0.797875 | Intergenic | Intergenic | -3182 | NM_008385 | 16330 | Mm.296202 | NM_008385 | ENSMUSG00000028894 | **Inpp5b** | 75kDa\|AW260155\|INPP5P | inositol polyphosphate-5-phosphatase B | protein-coding |
| chr7_128622561_128622993 | chr7 | 128622562 | 128622993 | + | 1.29722 | intron (NM_178641, intron 1 of 19) | intron (NM_178641, intron 1 of 19) | 11412 | NM_178641 | 101490 | Mm.490344 | NM_178641 | ENSMUSG00000042105 | **Inpp5f** | 5830435P03Rik\|AI115354\|AW561896\|SAC2\|cl-27\|mKIAA0966 | inositol polyphosphate-5-phosphatase F | protein-coding |
| chr7_128578683_128579019 | chr7 | 128578684 | 128579019 | + | 1.15694 | Intergenic | Intergenic | -32514 | NM_178641 | 101490 | Mm.490344 | NM_178641 | ENSMUSG00000042105 | **Inpp5f** | 5830435P03Rik\|AI115354\|AW561896\|SAC2\|cl-27\|mKIAA0966 | inositol polyphosphate-5-phosphatase F | protein-coding |
| chr7_128578250_128579024 | chr7 | 128578251 | 128579024 | + | 0.874923 | Intergenic | Intergenic | -32728 | NM_178641 | 101490 | Mm.490344 | NM_178641 | ENSMUSG00000042105 | **Inpp5f** | 5830435P03Rik\|AI115354\|AW561896\|SAC2\|cl-27\|mKIAA0966 | inositol polyphosphate-5-phosphatase F | protein-coding |
| chr12_55663331_55663695 | chr12 | 55663332 | 55663695 | + | 0.774429 | intron (NM_001112714, intron 35 of 40) | intron (NM_001112714, intron 35 of 40) | 64596 | NM_020287 | 56856 | Mm.114886 | NM_020287 | ENSMUSG00000045440 | **Insm2** | Mlt1 | insulinoma-associated 2 | protein-coding |
| chr12_55664088_55664432 | chr12 | 55664089 | 55664432 | + | 0.76963 | intron (NM_001112714, intron 35 of 40) | intron (NM_001112714, intron 35 of 40) | 65343 | NM_020287 | 56856 | Mm.114886 | NM_020287 | ENSMUSG00000045440 | **Insm2** | Mlt1 | insulinoma-associated 2 | protein-coding |
| chr12_55663332_55663537 | chr12 | 55663333 | 55663537 | + | 0.697577 | intron (NM_001112714, intron 35 of 40) | intron (NM_001112714, intron 35 of 40) | 64518 | NM_020287 | 56856 | Mm.114886 | NM_020287 | ENSMUSG00000045440 | **Insm2** | Mlt1 | insulinoma-associated 2 | protein-coding |
| chr8_3247026_3247643 | chr8 | 3247027 | 3247643 | + | 0.657352 | intron (NM_010568, intron 2 of 20) | intron (NM_010568, intron 2 of 20) | 32282 | NM_010568 | 16337 | Mm.268003 | NM_010568 | ENSMUSG00000005534 | **Insr** | 4932439J01Rik\|CD220\|D630014A15Rik\|IR\|IR-A\|IR-B | insulin receptor | protein-coding |
| chr7_97492313_97492962 | chr7 | 97492314 | 97492962 | + | 0.860687 | intron (NM_027256, intron 4 of 22) | intron (NM_027256, intron 4 of 22) | 11682 | NM_027256 | 101861 | Mm.27383 | NM_027256 | ENSMUSG00000025133 | **Ints4** | 2610034N24Rik\|AA692684\|AW493223 | integrator complex subunit 4 | protein-coding |

| | | | | | | | | | | | | | | | | | |
|---|---|---|---|---|---|---|---|---|---|---|---|---|---|---|---|---|---|
| chr14_62778144_62778426 | chr14 | 62778145 | 62778426 | + | 0.846733 | intron (NR_045503, intron 2 of 6) | intron (NR_045503, intron 2 of 6) | -17173 | NM_008715 | 18130 | Mm.319684 | NM_008715 | ENSMUSG00000035161 | **Ints6** | 2900075H24Rik\|AI480962\|DICE1\|Ddx26\|HDB\|Notch2l | integrator complex subunit 6 | protein-coding |
| chr14_62775033_62775418 | chr14 | 62775034 | 62775418 | + | 0.82803 | intron (NR_045503, intron 2 of 6) | intron (NR_045503, intron 2 of 6) | -14114 | NM_008715 | 18130 | Mm.319684 | NM_008715 | ENSMUSG00000035161 | **Ints6** | 2900075H24Rik\|AI480962\|DICE1\|Ddx26\|HDB\|Notch2l | integrator complex subunit 6 | protein-coding |
| chr14_62775020_62775691 | chr14 | 62775021 | 62775691 | + | 0.726611 | intron (NR_045503, intron 2 of 6) | MLT1J\|LTR\|ERVL-MaLR | -14244 | NM_008715 | 18130 | Mm.319684 | NM_008715 | ENSMUSG00000035161 | **Ints6** | 2900075H24Rik\|AI480962\|DICE1\|Ddx26\|HDB\|Notch2l | integrator complex subunit 6 | protein-coding |
| chr4_11240268_11240708 | chr4 | 11240269 | 11240708 | + | 1.06179 | intron (NM_178112, intron 7 of 27) | intron (NM_178112, intron 7 of 27) | 13771 | NM_178112 | 72656 | Mm.246713 | NM_178112 | ENSMUSG00000040738 | **Ints8** | 2810013E07Rik\|AV063769\|D130008D20Rik | integrator complex subunit 8 | protein-coding |
| chr14_64984591_64984945 | chr14 | 64984592 | 64984945 | + | 0.814072 | intron (NM_153414, intron 2 of 16) | RSINE1\|SINE\|B4 | 34723 | NM_001253731 | 210925 | Mm.71332 | NM_153414 | ENSMUSG00000021975 | **Ints9** | BC028953\|D14Ertd231e | integrator complex subunit 9 | protein-coding |
| chr10_6953023_6953584 | chr10 | 6953024 | 6953584 | + | 1.84292 | intron (NM_001170801, intron 4 of 6) | intron (NM_001170801, intron 4 of 6) | 27161 | NM_001170800 | 320495 | Mm.331674 | NM_001033391 | ENSMUSG00000064065 | **Ipcef1** | A130090K04Rik | interaction protein for cytohesin exchange factors 1 | protein-coding |
| chr10_6987262_6987846 | chr10 | 6987263 | 6987846 | + | 1.37716 | intron (NM_001039652, intron 4 of 5) | intron (NM_001039652, intron 4 of 5) | -7089 | NM_001170801 | 320495 | Mm.331674 | NM_001033391 | ENSMUSG00000064065 | **Ipcef1** | A130090K04Rik | interaction protein for cytohesin exchange factors 1 | protein-coding |
| chr10_7113793_7114095 | chr10 | 7113794 | 7114095 | + | 1.10828 | Intergenic | Intergenic | -90759 | NM_001033391 | 320495 | Mm.331674 | NM_001033391 | ENSMUSG00000064065 | **Ipcef1** | A130090K04Rik | interaction protein for cytohesin exchange factors 1 | protein-coding |
| chr10_6916109_6917080 | chr10 | 6916110 | 6917080 | + | 1.06759 | TTS (NM_001170802) | TTS (NM_001170802) | 63870 | NM_001170800 | 320495 | Mm.331674 | NM_001033391 | ENSMUSG00000064065 | **Ipcef1** | A130090K04Rik | interaction protein for cytohesin exchange factors 1 | protein-coding |
| chr10_7111208_7111664 | chr10 | 7111209 | 7111664 | + | 0.911496 | Intergenic | Intergenic | -88251 | NM_001033391 | 320495 | Mm.331674 | NM_001033391 | ENSMUSG00000064065 | **Ipcef1** | A130090K04Rik | interaction protein for cytohesin exchange factors 1 | protein-coding |
| chr10_7112881_7113458 | chr10 | 7112882 | 7113458 | + | 0.896716 | Intergenic | Intergenic | -89985 | NM_001033391 | 320495 | Mm.331674 | NM_001033391 | ENSMUSG00000064065 | **Ipcef1** | A130090K04Rik | interaction protein for cytohesin exchange factors 1 | protein-coding |
| chr10_6936535_6937007 | chr10 | 6936536 | 6937007 | + | 0.822394 | intron (NM_001170801, intron 5 of 6) | intron (NM_001170801, intron 5 of 6) | 43694 | NM_001170800 | 320495 | Mm.331674 | NM_001033391 | ENSMUSG00000064065 | **Ipcef1** | A130090K04Rik | interaction protein for cytohesin exchange factors 1 | protein-coding |
| chr13_106691693_106692147 | chr13 | 106691694 | 106692147 | + | 0.778101 | Intergenic | L1M2c\|LINE\|L1 | 244995 | NM_029665 | 76582 | Mm.132208 | NM_029665 | ENSMUSG00000042590 | **Ipo11** | 1700081H05Rik\|2510001A17Rik\|AI314624\|AW555235\|E330021B14Rik\|Ranbp11 | importin 11 | protein-coding |
| chr7_80767009_80767383 | chr7 | 80767010 | 80767383 | + | 0.682593 | intron (NM_016721, intron 3 of 37) | MIR\|SINE\|MIR | 36135 | NM_016721 | 29875 | Mm.207619 | NM_016721 | ENSMUSG00000030536 | **Iqgap1** | AA682088\|D7Ertd237e\|D7Ertd257e\|mKIAA0051 | IQ motif containing GTPase activating protein 1 | protein-coding |
| chr13_96011155_96011673 | chr13 | 96011156 | 96011673 | + | 1.51503 | intron (NM_029210, intron 4 of 12) | intron (NM_029210, intron 4 of 12) | -119492 | NM_027711 | 544963 | Mm.38878 | NM_027711 | ENSMUSG00000021676 | **Iqgap2** | 4933417J23Rik\|A630053O10\|AI788777 | IQ motif containing GTPase activating protein 2 | protein-coding |
| chr13_95845029_95845779 | chr13 | 95845030 | 95845779 | + | 0.842016 | intron (NM_027711, intron 1 of 35) | GA-rich\|Low_complexity\|Low_complexity | 46518 | NM_027711 | 544963 | Mm.38878 | NM_027711 | ENSMUSG00000021676 | **Iqgap2** | 4933417J23Rik\|A630053O10\|AI788777 | IQ motif containing GTPase activating protein 2 | protein-coding |
| chr3_88082640_88083018 | chr3 | 88082641 | 88083018 | + | 0.840797 | intron (NM_001033484, intron 1 of 37) | intron (NM_001033484, intron 1 of 37) | 778 | NM_001033484 | 404710 | Mm.331133 | NM_001033484 | ENSMUSG00000028068 | **Iqgap3** | AI593484\|D030034H08 | IQ motif containing GTPase activating protein 3 | protein-coding |
| chr6_90744072_90744315 | chr6 | 90744073 | 90744315 | + | 0.760084 | intron (NM_001134384, intron 1 of 13) | L1MB8\|LINE\|L1 | -27665 | NM_001134383 | 232227 | Mm.196943 | NM_182784 | ENSMUSG00000034312 | **Iqsec1** | AW561907\|BC026481\|BRAG2\|D6Ertd349e\|cI-43 | IQ motif and Sec7 domain 1 | protein-coding |
| chr9_82877102_82877413 | chr9 | 82877103 | 82877413 | + | 0.866497 | exon (NM_001081216, exon 36 of 40) | exon (NM_001081216, exon 36 of 40) | 47452 | NM_022986 | 65099 | Mm.41415 | NM_022986 | ENSMUSG00000032251 | **Irak1bp1** | 4921528N06Rik\|AI851240\|Aabp3\|Aip70\|Simpl | interleukin-1 receptor-associated kinase 1 binding protein 1 | protein-coding |
| chr6_113643090_113643341 | chr6 | 113643091 | 113643341 | + | 0.958071 | intron (NM_172161, intron 1 of 12) | intron (NM_172161, intron 1 of 12) | 4749 | NM_001113553 | 108960 | Mm.152142 | NM_172161 | ENSMUSG00000060477 | **Irak2** | 6330415L08Rik\|AI649099\|IRAK-2 | interleukin-1 receptor-associated kinase 2 | protein-coding |
| chr10_120204576_120204990 | chr10 | 120204577 | 120204990 | + | 1.1252 | Intergenic | Intergenic | -3246 | NM_028679 | 73914 | Mm.146194 | NM_028679 | ENSMUSG00000020227 | **Irak3** | 4833428C18Rik\|AI563835\|IRAK-M | interleukin-1 receptor-associated kinase 3 | protein-coding |
| chr15_94564821_94565260 | chr15 | 94564822 | 94565260 | + | 0.667641 | intron (NM_029926, intron 10 of 11) | intron (NM_029926, intron 10 of 11) | 21381 | NM_029926 | 266632 | Mm.422858 | NM_029926 | ENSMUSG00000059883 | **Irak4** | 8430405M07Rik\|9330209D03Rik\|IRAK-4\|NY-REN-64 | interleukin-1 receptor-associated kinase 4 | protein-coding |
| chr11_53768775_53769059 | chr11 | 53768776 | 53769059 | + | 0.800749 | Intergenic | Intergenic | -1097 | NM_001159393 | 16362 | Mm.105218 | NM_008390 | ENSMUSG00000018899 | **Irf1** | AU020929\|Irf-1 | interferon regulatory factor 1 | protein-coding |
| chr11_53778266_53779270 | chr11 | 53778267 | 53779270 | + | 0.690018 | TTS (NM_001159393) | TTS (NM_001159393) | 8295 | NM_001159396 | 16362 | Mm.105218 | NM_008390 | ENSMUSG00000018899 | **Irf1** | AU020929\|Irf-1 | interferon regulatory factor 1 | protein-coding |
| chr8_46752321_46752654 | chr8 | 46752322 | 46752654 | + | 0.731693 | intron (NM_008391, intron 1 of 8) | intron (NM_008391, intron 1 of 8) | 12743 | NM_008391 | 16363 | Mm.1149 | NM_008391 | ENSMUSG00000031627 | **Irf2** | 9830146E22Rik\|AI646973\|Irf-2 | interferon regulatory factor 2 | protein-coding |
| chr8_46752970_46753331 | chr8 | 46752971 | 46753331 | + | 0.712284 | intron (NM_008391, intron 1 of 8) | L1MB3\|LINE\|L1 | 13406 | NM_008391 | 16363 | Mm.1149 | NM_008391 | ENSMUSG00000031627 | **Irf2** | 9830146E22Rik\|AI646973\|Irf-2 | interferon regulatory factor 2 | protein-coding |
| chr8_46752970_46753732 | chr8 | 46752971 | 46753732 | + | 0.702373 | intron (NM_008391, intron 1 of 8) | L1MB3\|LINE\|L1 | 13606 | NM_008391 | 16363 | Mm.1149 | NM_008391 | ENSMUSG00000031627 | **Irf2** | 9830146E22Rik\|AI646973\|Irf-2 | interferon regulatory factor 2 | protein-coding |
| chr8_46752291_46752848 | chr8 | 46752292 | 46752848 | + | 0.677925 | intron (NM_008391, intron 1 of 8) | intron (NM_008391, intron 1 of 8) | 12825 | NM_008391 | 16363 | Mm.1149 | NM_008391 | ENSMUSG00000031627 | **Irf2** | 9830146E22Rik\|AI646973\|Irf-2 | interferon regulatory factor 2 | protein-coding |
| chr8_46832483_46832952 | chr8 | 46832484 | 46832952 | + | 0.665048 | intron (NM_008391, intron 6 of 8) | intron (NM_008391, intron 6 of 8) | 92973 | NM_008391 | 16363 | Mm.1149 | NM_008391 | ENSMUSG00000031627 | **Irf2** | 9830146E22Rik\|AI646973\|Irf-2 | interferon regulatory factor 2 | protein-coding |
| chr8_46799187_46799570 | chr8 | 46799188 | 46799570 | + | 0.63933 | intron (NM_008391, intron 2 of 8) | intron (NM_008391, intron 2 of 8) | 59634 | NM_008391 | 16363 | Mm.1149 | NM_008391 | ENSMUSG00000031627 | **Irf2** | 9830146E22Rik\|AI646973\|Irf-2 | interferon regulatory factor 2 | protein-coding |
| chr7_19003189_19003438 | chr7 | 19003190 | 19003438 | + | 0.699489 | promoter-TSS (NM_178757) | promoter-TSS (NM_178757) | -751 | NM_178757 | 272359 | Mm.274237 | NM_178757 | ENSMUSG00000044030 | **Irf2bp1** | 6330414O09Rik\|C530033F24 | interferon regulatory factor 2 binding protein 1 | protein-coding |
| chr8_126550811_126551363 | chr8 | 126550812 | 126551363 | + | 0.930303 | Intergenic | Intergenic | 42349 | NM_001164598 | 270110 | Mm.334918 | NM_001164598 | ENSMUSG00000051495 | **Irf2bp2** | E130305N23Rik\|IRF-2BP2 | interferon regulatory factor 2 binding protein 2 | protein-coding |
| chr8_126539782_126540095 | chr8 | 126539783 | 126540095 | + | 0.672492 | Intergenic | Intergenic | 53497 | NM_001164598 | 270110 | Mm.334918 | NM_001164598 | ENSMUSG00000051495 | **Irf2bp2** | E130305N23Rik\|IRF-2BP2 | interferon regulatory factor 2 binding protein 2 | protein-coding |
| chr12_86859164_86859976 | chr12 | 86859165 | 86859976 | + | 1.20649 | Intergenic | Intergenic | 25244 | NM_145836 | 238330 | Mm.248358 | NM_145836 | ENSMUSG00000034168 | **Irf2bpl** | 6430527G18Rik\|Eap1 | interferon regulatory factor 2 binding protein-like | protein-coding |
| chr13_30831074_30831784 | chr13 | 30831075 | 30831784 | + | 1.67026 | intron (NM_025588, intron 23 of 27) | intron (NM_025588, intron 23 of 27) | 82171 | NM_013674 | 16364 | Mm.4677 | NM_013674 | ENSMUSG00000021356 | **Irf4** | AI385587\|IRF-4\|LSIRF\|Spip | interferon regulatory factor 4 | protein-coding |
| chr13_30793891_30794390 | chr13 | 30793892 | 30794390 | + | 1.41412 | Intergenic | Intergenic | 44883 | NM_013674 | 16364 | Mm.4677 | NM_013674 | ENSMUSG00000021356 | **Irf4** | AI385587\|IRF-4\|LSIRF\|Spip | interferon regulatory factor 4 | protein-coding |
| chr13_30787929_30788259 | chr13 | 30787930 | 30788259 | + | 1.40183 | Intergenic | Intergenic | 38836 | NM_013674 | 16364 | Mm.4677 | NM_013674 | ENSMUSG00000021356 | **Irf4** | AI385587\|IRF-4\|LSIRF\|Spip | interferon regulatory factor 4 | protein-coding |
| chr13_30843031_30843453 | chr13 | 30843032 | 30843453 | + | 1.32744 | intron (NM_025588, intron 23 of 27) | intron (NM_025588, intron 23 of 27) | 93984 | NM_013674 | 16364 | Mm.4677 | NM_013674 | ENSMUSG00000021356 | **Irf4** | AI385587\|IRF-4\|LSIRF\|Spip | interferon regulatory factor 4 | protein-coding |
| chr13_30754077_30754369 | chr13 | 30754078 | 30754369 | + | 0.940688 | intron (NM_013674, intron 5 of 8) | intron (NM_013674, intron 5 of 8) | 4965 | NM_013674 | 16364 | Mm.4677 | NM_013674 | ENSMUSG00000021356 | **Irf4** | AI385587\|IRF-4\|LSIRF\|Spip | interferon regulatory factor 4 | protein-coding |
| chr13_30841446_30841873 | chr13 | 30841447 | 30841873 | + | 0.936744 | intron (NM_025588, intron 23 of 27) | intron (NM_025588, intron 23 of 27) | 92402 | NM_013674 | 16364 | Mm.4677 | NM_013674 | ENSMUSG00000021356 | **Irf4** | AI385587\|IRF-4\|LSIRF\|Spip | interferon regulatory factor 4 | protein-coding |
| chr13_30787620_30787883 | chr13 | 30787621 | 30787883 | + | 0.908464 | Intergenic | Intergenic | 38494 | NM_013674 | 16364 | Mm.4677 | NM_013674 | ENSMUSG00000021356 | **Irf4** | AI385587\|IRF-4\|LSIRF\|Spip | interferon regulatory factor 4 | protein-coding |
| chr13_30814289_30814769 | chr13 | 30814290 | 30814769 | + | 0.898206 | 3' UTR (NM_025588, exon 28 of 28) | 3' UTR (NM_025588, exon 28 of 28) | 65271 | NM_013674 | 16364 | Mm.4677 | NM_013674 | ENSMUSG00000021356 | **Irf4** | AI385587\|IRF-4\|LSIRF\|Spip | interferon regulatory factor 4 | protein-coding |
| chr13_30788494_30788669 | chr13 | 30788495 | 30788669 | + | 0.827292 | Intergenic | MTD\|LTR\|ERVL-MaLR | 39324 | NM_013674 | 16364 | Mm.4677 | NM_013674 | ENSMUSG00000021356 | **Irf4** | AI385587\|IRF-4\|LSIRF\|Spip | interferon regulatory factor 4 | protein-coding |
| chr13_30724895_30725452 | chr13 | 30724896 | 30725452 | + | 0.813324 | Intergenic | Intergenic | -24084 | NM_013674 | 16364 | Mm.4677 | NM_013674 | ENSMUSG00000021356 | **Irf4** | AI385587\|IRF-4\|LSIRF\|Spip | interferon regulatory factor 4 | protein-coding |
| chr13_30841411_30842241 | chr13 | 30841412 | 30842241 | + | 0.80692 | intron (NM_025588, intron 23 of 27) | intron (NM_025588, intron 23 of 27) | 92568 | NM_013674 | 16364 | Mm.4677 | NM_013674 | ENSMUSG00000021356 | **Irf4** | AI385587\|IRF-4\|LSIRF\|Spip | interferon regulatory factor 4 | protein-coding |
| chr13_30726860_30727315 | chr13 | 30726861 | 30727315 | + | 0.782456 | Intergenic | Intergenic | -22170 | NM_013674 | 16364 | Mm.4677 | NM_013674 | ENSMUSG00000021356 | **Irf4** | AI385587\|IRF-4\|LSIRF\|Spip | interferon regulatory factor 4 | protein-coding |
| chr13_30826033_30826379 | chr13 | 30826034 | 30826379 | + | 0.770833 | intron (NM_025588, intron 23 of 27) | intron (NM_025588, intron 23 of 27) | 76948 | NM_013674 | 16364 | Mm.4677 | NM_013674 | ENSMUSG00000021356 | **Irf4** | AI385587\|IRF-4\|LSIRF\|Spip | interferon regulatory factor 4 | protein-coding |
| chr13_30739792_30740078 | chr13 | 30739793 | 30740078 | + | 0.730907 | Intergenic | Intergenic | -9323 | NM_013674 | 16364 | Mm.4677 | NM_013674 | ENSMUSG00000021356 | **Irf4** | AI385587\|IRF-4\|LSIRF\|Spip | interferon regulatory factor 4 | protein-coding |
| chr13_30739670_30740098 | chr13 | 30739671 | 30740098 | + | 0.69564 | Intergenic | Intergenic | -9374 | NM_013674 | 16364 | Mm.4677 | NM_013674 | ENSMUSG00000021356 | **Irf4** | AI385587\|IRF-4\|LSIRF\|Spip | interferon regulatory factor 4 | protein-coding |
| chr13_30765890_30766263 | chr13 | 30765891 | 30766263 | + | 0.616822 | 3' UTR (NM_013674, exon 9 of 9) | 3' UTR (NM_013674, exon 9 of 9) | 16819 | NM_013674 | 16364 | Mm.4677 | NM_013674 | ENSMUSG00000021356 | **Irf4** | AI385587\|IRF-4\|LSIRF\|Spip | interferon regulatory factor 4 | protein-coding |
| chr1_193149472_193149726 | chr1 | 193149473 | 193149726 | + | 0.617164 | Intergenic | Intergenic | -3513 | NM_016851 | 54139 | Mm.273695 | NM_016851 | ENSMUSG00000026638 | **Irf6** | AI876454\|E230028I05Rik\|mirf6 | interferon regulatory factor 6 | protein-coding |
| chr8_120708943_120709550 | chr8 | 120708944 | 120709550 | + | 0.965845 | Intergenic | Intergenic | -27129 | NM_008320 | 15900 | Mm.334861 | NM_008320 | ENSMUSG00000041515 | **Irf8** | AI893568\|ICSBP\|IRF-8\|Icsbp1\|Myls | interferon regulatory factor 8 | protein-coding |
| chr8_120725197_120725583 | chr8 | 120725198 | 120725583 | + | 0.696479 | Intergenic | Intergenic | -10986 | NM_008320 | 15900 | Mm.334861 | NM_008320 | ENSMUSG00000041515 | **Irf8** | AI893568\|ICSBP\|IRF-8\|Icsbp1\|Myls | interferon regulatory factor 8 | protein-coding |

| | | | | | | | | | | | | | | | | | |
|---|---|---|---|---|---|---|---|---|---|---|---|---|---|---|---|---|---|
| chr8_120725063_120725886 | chr8 | 120725064 | 120725886 | + | 0.642089 | Intergenic | Intergenic | -10901 | NM_008320 | 15900 | Mm.334861 | NM_008320 | ENSMUSG00000041515 | **Irf8** | AI893568\|ICSBP\|IRF-8\|Icsbp1\|Myls | interferon regulatory factor 8 | protein-coding |
| chr8_120715954_120716625 | chr8 | 120715955 | 120716625 | + | 0.602384 | Intergenic | Intergenic | -20086 | NM_008320 | 15900 | Mm.334861 | NM_008320 | ENSMUSG00000041515 | **Irf8** | AI893568\|ICSBP\|IRF-8\|Icsbp1\|Myls | interferon regulatory factor 8 | protein-coding |
| chr1_82281757_82281999 | chr1 | 82281758 | 82281999 | + | 0.814597 | intron (NM_010570, intron 1 of 1) | intron (NM_010570, intron 1 of 1) | 9561 | NM_010570 | 16367 | Mm.4952 | NM_010570 | ENSMUSG00000055980 | **Irs1** | G972R\|IRS-1 | insulin receptor substrate 1 | protein-coding |
| chr13_116410714_116411052 | chr13 | 116410715 | 116411052 | + | 0.686045 | Intergenic | ORR1D2\|LTR\|ERVL-MaLR | -101195 | NM_021459 | 16392 | Mm.42242 | NM_021459 | ENSMUSG00000042258 | **Isl1** | - | ISL1 transcription factor, LIM/homeodomain | protein-coding |
| chr2_139670600_139671041 | chr2 | 139670601 | 139671041 | + | 1.62071 | Intergenic | Intergenic | -7357 | NM_001276489 | 319909 | Mm.41113 | NM_001126490 | ENSMUSG00000074766 | **Ism1** | 5430433G21Rik\|Ism | isthmin 1 homolog (zebrafish) | protein-coding |
| chr2_139670412_139671043 | chr2 | 139670413 | 139671043 | + | 1.57673 | Intergenic | Intergenic | -7450 | NM_001276489 | 319909 | Mm.41113 | NM_001126490 | ENSMUSG00000074766 | **Ism1** | 5430433G21Rik\|Ism | isthmin 1 homolog (zebrafish) | protein-coding |
| chr2_139839569_139840256 | chr2 | 139839570 | 139840256 | + | 1.05752 | intron (NR_110351, intron 13 of 13) | intron (NR_110351, intron 13 of 13) | 161735 | NM_001276489 | 319909 | Mm.41113 | NM_001126490 | ENSMUSG00000074766 | **Ism1** | 5430433G21Rik\|Ism | isthmin 1 homolog (zebrafish) | protein-coding |
| chr2_139671854_139672168 | chr2 | 139671855 | 139672168 | + | 0.826291 | Intergenic | Intergenic | -6167 | NM_001276489 | 319909 | Mm.41113 | NM_001126490 | ENSMUSG00000074766 | **Ism1** | 5430433G21Rik\|Ism | isthmin 1 homolog (zebrafish) | protein-coding |
| chr2_139840798_139841867 | chr2 | 139840799 | 139841867 | + | 0.784613 | intron (NR_110351, intron 13 of 13) | intron (NR_110351, intron 13 of 13) | 163155 | NM_001276489 | 319909 | Mm.41113 | NM_001126490 | ENSMUSG00000074766 | **Ism1** | 5430433G21Rik\|Ism | isthmin 1 homolog (zebrafish) | protein-coding |
| chr2_139849677_139850221 | chr2 | 139849678 | 139850221 | + | 0.780532 | intron (NR_110351, intron 13 of 13) | Lx8\|LINE\|L1 | 171771 | NM_001276489 | 319909 | Mm.41113 | NM_001126490 | ENSMUSG00000074766 | **Ism1** | 5430433G21Rik\|Ism | isthmin 1 homolog (zebrafish) | protein-coding |
| chr2_139733942_139734928 | chr2 | 139733943 | 139734928 | + | 0.758956 | intron (NM_001276489, intron 2 of 5) | intron (NM_001276489, intron 2 of 5) | 56257 | NM_001276489 | 319909 | Mm.41113 | NM_001126490 | ENSMUSG00000074766 | **Ism1** | 5430433G21Rik\|Ism | isthmin 1 homolog (zebrafish) | protein-coding |
| chr2_139841450_139841765 | chr2 | 139841451 | 139841765 | + | 0.709918 | intron (NR_110351, intron 13 of 13) | intron (NR_110351, intron 13 of 13) | 163430 | NM_001276489 | 319909 | Mm.41113 | NM_001126490 | ENSMUSG00000074766 | **Ism1** | 5430433G21Rik\|Ism | isthmin 1 homolog (zebrafish) | protein-coding |
| chr2_139840948_139841257 | chr2 | 139840949 | 139841257 | + | 0.707612 | intron (NR_110351, intron 13 of 13) | intron (NR_110351, intron 13 of 13) | 162925 | NM_001276489 | 319909 | Mm.41113 | NM_001126490 | ENSMUSG00000074766 | **Ism1** | 5430433G21Rik\|Ism | isthmin 1 homolog (zebrafish) | protein-coding |
| chr2_139841337_139841850 | chr2 | 139841338 | 139841850 | + | 0.648527 | intron (NR_110351, intron 13 of 13) | intron (NR_110351, intron 13 of 13) | 163416 | NM_001276489 | 319909 | Mm.41113 | NM_001126490 | ENSMUSG00000074766 | **Ism1** | 5430433G21Rik\|Ism | isthmin 1 homolog (zebrafish) | protein-coding |
| chr7_4873385_4873636 | chr7 | 4873386 | 4873636 | + | 1.16574 | Intergenic | Intergenic | -3542 | NM_001101598 | 664994 | Mm.302624 | NM_001101598 | ENSMUSG00000086784 | **Isoc2a** | Isoc2 | isochorismatase domain containing 2a | protein-coding |
| chr7_4861376_4861593 | chr7 | 4861377 | 4861593 | + | 1.10517 | intron (NM_026158, intron 1 of 5) | RLTR14-int\|LTR\|ERV1 | 4694 | NM_026158 | 67441 | Mm.41385 | NM_026158 | ENSMUSG00000052605 | **Isoc2b** | 0610042E07Rik\|AI553388\|Isoc2 | isochorismatase domain containing 2b | protein-coding |
| chr12_36468265_36469148 | chr12 | 36468266 | 36469148 | + | 0.76731 | intron (NM_178629, intron 3 of 9) | HAL1\|LINE\|L1 | 86560 | NM_001289503 | 75847 | Mm.261764 | NM_178629 | ENSMUSG00000043153 | **Ispd** | 4930579E17Rik\|AV040780 | isoprenoid synthase domain containing | protein-coding |
| chr8_109694023_109694339 | chr8 | 109694024 | 109694339 | + | 1.30253 | promoter-TSS (NM_028018) | promoter-TSS (NM_028018) | -887 | NM_028018 | 71955 | Mm.290036 | NM_028018 | ENSMUSG00000031729 | **Ist1** | 2400003C14Rik\|AW536298\|mKIAA0174 | increased sodium tolerance 1 homolog (yeast) | protein-coding |
| chr17_26242838_26243030 | chr17 | 26242839 | 26243030 | + | 0.622629 | intron (NM_001206335, intron 1 of 12) | intron (NM_001206335, intron 1 of 12) | 1308 | NM_207217 | 106581 | Mm.28844 | NM_207217 | ENSMUSG00000024187 | **Itfg3** | - | integrin alpha FG-GAP repeat containing 3 | protein-coding |
| chr9_62732724_62733321 | chr9 | 62732725 | 62733321 | + | 0.655327 | intron (NM_176922, intron 5 of 29) | intron (NM_176922, intron 5 of 29) | 55168 | NM_176922 | 319480 | Mm.34883 | NM_176922 | ENSMUSG00000032243 | **Itga11** | 4732459H24Rik | integrin alpha 11 | protein-coding |
| chr13_114878649_114879478 | chr13 | 114878650 | 114879478 | + | 1.25259 | intron (NM_008396, intron 6 of 29) | intron (NM_008396, intron 6 of 29) | 52977 | NM_008396 | 16398 | Mm.5007 | NM_008396 | ENSMUSG00000015533 | **Itga2** | CD49B\|DX5 | integrin alpha 2 | protein-coding |
| chr13_114935477_114935906 | chr13 | 114935478 | 114935906 | + | 0.964931 | Intergenic | ORR1C2\|LTR\|ERVL-MaLR | -3651 | NM_008396 | 16398 | Mm.5007 | NM_008396 | ENSMUSG00000015533 | **Itga2** | CD49B\|DX5 | integrin alpha 2 | protein-coding |
| chr13_114880675_114881007 | chr13 | 114880676 | 114881007 | + | 0.745095 | intron (NM_008396, intron 5 of 29) | intron (NM_008396, intron 5 of 29) | 51200 | NM_008396 | 16398 | Mm.5007 | NM_008396 | ENSMUSG00000015533 | **Itga2** | CD49B\|DX5 | integrin alpha 2 | protein-coding |
| chr2_79122492_79122920 | chr2 | 79122493 | 79122920 | + | 1.38058 | Intergenic | Intergenic | -132720 | NM_010576 | 16401 | Mm.31903 | NM_010576 | ENSMUSG00000027009 | **Itga4** | CD49D\|Itga4B | integrin alpha 4 | protein-coding |
| chr2_79124255_79124785 | chr2 | 79124256 | 79124785 | + | 0.762309 | Intergenic | Intergenic | -130906 | NM_010576 | 16401 | Mm.31903 | NM_010576 | ENSMUSG00000027009 | **Itga4** | CD49D\|Itga4B | integrin alpha 4 | protein-coding |
| chr2_79124270_79124570 | chr2 | 79124271 | 79124570 | + | 0.709234 | Intergenic | Intergenic | -131006 | NM_010576 | 16401 | Mm.31903 | NM_010576 | ENSMUSG00000027009 | **Itga4** | CD49D\|Itga4B | integrin alpha 4 | protein-coding |
| chr2_79254007_79254217 | chr2 | 79254008 | 79254217 | + | 0.691161 | Intergenic | Intergenic | -1314 | NM_010576 | 16401 | Mm.31903 | NM_010576 | ENSMUSG00000027009 | **Itga4** | CD49D\|Itga4B | integrin alpha 4 | protein-coding |
| chr2_79316269_79316794 | chr2 | 79316270 | 79316794 | + | 0.66014 | intron (NM_010576, intron 21 of 27) | intron (NM_010576, intron 21 of 27) | 61106 | NM_010576 | 16401 | Mm.31903 | NM_010576 | ENSMUSG00000027009 | **Itga4** | CD49D\|Itga4B | integrin alpha 4 | protein-coding |
| chr2_71786157_71786380 | chr2 | 71786158 | 71786380 | + | 0.872732 | promoter-TSS (NM_008397) | promoter-TSS (NM_008397) | -670 | NM_008397 | 16403 | Mm.225096 | NM_008397 | | **Itga6** | 5033401O05Rik\|AI115430\|Cd49f\|VLA-6 | integrin alpha 6 | protein-coding |
| chr11_73092911_73093261 | chr11 | 73092912 | 73093261 | + | 1.10287 | intron (NM_008399, intron 1 of 30) | intron (NM_008399, intron 1 of 30) | 2491 | NM_008399 | 16407 | Mm.96 | NM_008399 | ENSMUSG00000005947 | **Itgae** | A530055J10\|CD103\|aM290\|alpha-E1\|alpha-M290 | integrin alpha E, epithelial-associated | protein-coding |
| chr11_73095587_73095782 | chr11 | 73095588 | 73095782 | + | 0.972461 | intron (NM_008399, intron 1 of 30) | intron (NM_008399, intron 1 of 30) | 5090 | NM_008399 | 16407 | Mm.96 | NM_008399 | ENSMUSG00000005947 | **Itgae** | A530055J10\|CD103\|aM290\|alpha-E1\|alpha-M290 | integrin alpha E, epithelial-associated | protein-coding |
| chr11_73095376_73095889 | chr11 | 73095377 | 73095889 | + | 0.899774 | intron (NM_008399, intron 1 of 30) | intron (NM_008399, intron 1 of 30) | 5038 | NM_008399 | 16407 | Mm.96 | NM_008399 | ENSMUSG00000005947 | **Itgae** | A530055J10\|CD103\|aM290\|alpha-E1\|alpha-M290 | integrin alpha E, epithelial-associated | protein-coding |
| chr11_73098958_73099208 | chr11 | 73098959 | 73099208 | + | 0.884993 | intron (NM_008399, intron 1 of 30) | B1_Mur3\|SINE\|Alu | 8488 | NM_008399 | 16407 | Mm.96 | NM_008399 | ENSMUSG00000005947 | **Itgae** | A530055J10\|CD103\|aM290\|alpha-E1\|alpha-M290 | integrin alpha E, epithelial-associated | protein-coding |
| chr11_73095179_73095905 | chr11 | 73095180 | 73095905 | + | 0.880549 | intron (NM_008399, intron 1 of 30) | intron (NM_008399, intron 1 of 30) | 4947 | NM_008399 | 16407 | Mm.96 | NM_008399 | ENSMUSG00000005947 | **Itgae** | A530055J10\|CD103\|aM290\|alpha-E1\|alpha-M290 | integrin alpha E, epithelial-associated | protein-coding |
| chr11_73109602_73110331 | chr11 | 73109603 | 73110331 | + | 0.777187 | intron (NM_008399, intron 2 of 30) | intron (NM_008399, intron 2 of 30) | 19372 | NM_008399 | 16407 | Mm.96 | NM_008399 | ENSMUSG00000005947 | **Itgae** | A530055J10\|CD103\|aM290\|alpha-E1\|alpha-M290 | integrin alpha E, epithelial-associated | protein-coding |
| chr11_73109296_73110499 | chr11 | 73109297 | 73110499 | + | 0.658609 | intron (NM_008399, intron 2 of 30) | intron (NM_008399, intron 2 of 30) | 19303 | NM_008399 | 16407 | Mm.96 | NM_008399 | ENSMUSG00000005947 | **Itgae** | A530055J10\|CD103\|aM290\|alpha-E1\|alpha-M290 | integrin alpha E, epithelial-associated | protein-coding |
| chr7_127293744_127294098 | chr7 | 127293745 | 127294098 | + | 0.9094 | Intergenic | Intergenic | -2339 | NM_001253873 | 16408 | Mm.1618 | NM_008400 | ENSMUSG00000030830 | **Itgal** | (p180)\|Cd11a\|LFA-1\|LFA-1A\|Ly-15\|Ly-21 | integrin alpha L | protein-coding |
| chr2_83695199_83695938 | chr2 | 83695200 | 83695938 | + | 1.43105 | Intergenic | MLT2B2\|LTR\|ERVL | -28828 | NM_008402 | 16410 | Mm.227 | NM_008402 | ENSMUSG00000027087 | **Itgav** | 1110004F14Rik\|2610028E01Rik\|CD51\|D430040G12Rik | integrin alpha V | protein-coding |
| chr2_83728393_83728702 | chr2 | 83728394 | 83728702 | + | 1.19627 | intron (NM_008402, intron 1 of 29) | intron (NM_008402, intron 1 of 29) | 4151 | NM_008402 | 16410 | Mm.227 | NM_008402 | ENSMUSG00000027087 | **Itgav** | 1110004F14Rik\|2610028E01Rik\|CD51\|D430040G12Rik | integrin alpha V | protein-coding |
| chr2_83700499_83701349 | chr2 | 83700500 | 83701349 | + | 1.13566 | Intergenic | Intergenic | -23473 | NM_008402 | 16410 | Mm.227 | NM_008402 | ENSMUSG00000027087 | **Itgav** | 1110004F14Rik\|2610028E01Rik\|CD51\|D430040G12Rik | integrin alpha V | protein-coding |
| chr8_128535526_128536069 | chr8 | 128535527 | 128536069 | + | 1.90332 | Intergenic | Intergenic | -149856 | NM_010578 | 16412 | Mm.263396 | NM_010578 | ENSMUSG00000025809 | **Itgb1** | 4633401G24Rik\|AA409975\|AA960159\|CD29\|ENSMUSG00000051907\|Fnrb\|Gm9863\|gpIIa | integrin beta 1 (fibronectin receptor beta) | protein-coding |
| chr8_128788088_128788673 | chr8 | 128788089 | 128788673 | + | 1.28255 | Intergenic | L1M3\|LINE\|L1 | 102727 | NM_010578 | 16412 | Mm.263396 | NM_010578 | ENSMUSG00000025809 | **Itgb1** | 4633401G24Rik\|AA409975\|AA960159\|CD29\|ENSMUSG00000051907\|Fnrb\|Gm9863\|gpIIa | integrin beta 1 (fibronectin receptor beta) | protein-coding |
| chr8_128763935_128764628 | chr8 | 128763936 | 128764628 | + | 1.23995 | Intergenic | Lx\|LINE\|L1 | 78628 | NM_010578 | 16412 | Mm.263396 | NM_010578 | ENSMUSG00000025809 | **Itgb1** | 4633401G24Rik\|AA409975\|AA960159\|CD29\|ENSMUSG00000051907\|Fnrb\|Gm9863\|gpIIa | integrin beta 1 (fibronectin receptor beta) | protein-coding |
| chr8_128622712_128623129 | chr8 | 128622713 | 128623129 | + | 1.17307 | Intergenic | Intergenic | -62733 | NM_010578 | 16412 | Mm.263396 | NM_010578 | ENSMUSG00000025809 | **Itgb1** | 4633401G24Rik\|AA409975\|AA960159\|CD29\|ENSMUSG00000051907\|Fnrb\|Gm9863\|gpIIa | integrin beta 1 (fibronectin receptor beta) | protein-coding |
| chr8_128642946_128643464 | chr8 | 128642947 | 128643464 | + | 1.16944 | Intergenic | Intergenic | -42449 | NM_010578 | 16412 | Mm.263396 | NM_010578 | ENSMUSG00000025809 | **Itgb1** | 4633401G24Rik\|AA409975\|AA960159\|CD29\|ENSMUSG00000051907\|Fnrb\|Gm9863\|gpIIa | integrin beta 1 (fibronectin receptor beta) | protein-coding |
| chr8_128647570_128648046 | chr8 | 128647571 | 128648046 | + | 1.08623 | Intergenic | Intergenic | -37846 | NM_010578 | 16412 | Mm.263396 | NM_010578 | ENSMUSG00000025809 | **Itgb1** | 4633401G24Rik\|AA409975\|AA960159\|CD29\|ENSMUSG00000051907\|Fnrb\|Gm9863\|gpIIa | integrin beta 1 (fibronectin receptor beta) | protein-coding |
| chr8_128563105_128563524 | chr8 | 128563106 | 128563524 | + | 1.0652 | Intergenic | Intergenic | -122339 | NM_010578 | 16412 | Mm.263396 | NM_010578 | ENSMUSG00000025809 | **Itgb1** | 4633401G24Rik\|AA409975\|AA960159\|CD29\|ENSMUSG00000051907\|Fnrb\|Gm9863\|gpIIa | integrin beta 1 (fibronectin receptor beta) | protein-coding |
| chr8_128690571_128691017 | chr8 | 128690572 | 128691017 | + | 0.969199 | intron (NM_010578, intron 1 of 15) | intron (NM_010578, intron 1 of 15) | 5140 | NM_010578 | 16412 | Mm.263396 | NM_010578 | ENSMUSG00000025809 | **Itgb1** | 4633401G24Rik\|AA409975\|AA960159\|CD29\|ENSMUSG00000051907\|Fnrb\|Gm9863\|gpIIa | integrin beta 1 (fibronectin receptor beta) | protein-coding |
| chr8_128525886_128526274 | chr8 | 128525887 | 128526274 | + | 0.944223 | Intergenic | HAL1\|LINE\|L1 | -159574 | NM_010578 | 16412 | Mm.263396 | NM_010578 | ENSMUSG00000025809 | **Itgb1** | 4633401G24Rik\|AA409975\|AA960159\|CD29\|ENSMUSG00000051907\|Fnrb\|Gm9863\|gpIIa | integrin beta 1 (fibronectin receptor beta) | protein-coding |

| | | | | | | | | | | | | | | | | | |
|---|---|---|---|---|---|---|---|---|---|---|---|---|---|---|---|---|---|
| chr8_128645500_128645999 | chr8 | 128645501 | 128645999 | + | 0.928894 | Intergenic | Intergenic | -39904 | NM_010578 | 16412 | Mm.263396 | NM_010578 | ENSMUSG00000025809 | **Itgb1** | 4633401G24Rik\|AA409975\|AA960159\|CD29\|ENSMUSG00000051907\|Fnrb\|Gm9863\|gpIIa | integrin beta 1 (fibronectin receptor beta) | protein-coding |
| chr8_128619029_128619362 | chr8 | 128619030 | 128619362 | + | 0.839785 | Intergenic | Intergenic | -66458 | NM_010578 | 16412 | Mm.263396 | NM_010578 | ENSMUSG00000025809 | **Itgb1** | 4633401G24Rik\|AA409975\|AA960159\|CD29\|ENSMUSG00000051907\|Fnrb\|Gm9863\|gpIIa | integrin beta 1 (fibronectin receptor beta) | protein-coding |
| chr8_128694524_128694905 | chr8 | 128694525 | 128694905 | + | 0.778084 | intron (NM_010578, intron 1 of 15) | MTC\|LTR\|ERVL-MaLR | 9061 | NM_010578 | 16412 | Mm.263396 | NM_010578 | ENSMUSG00000025809 | **Itgb1** | 4633401G24Rik\|AA409975\|AA960159\|CD29\|ENSMUSG00000051907\|Fnrb\|Gm9863\|gpIIa | integrin beta 1 (fibronectin receptor beta) | protein-coding |
| chr8_128646823_128647075 | chr8 | 128646824 | 128647075 | + | 0.696264 | Intergenic | Intergenic | -38705 | NM_010578 | 16412 | Mm.263396 | NM_010578 | ENSMUSG00000025809 | **Itgb1** | 4633401G24Rik\|AA409975\|AA960159\|CD29\|ENSMUSG00000051907\|Fnrb\|Gm9863\|gpIIa | integrin beta 1 (fibronectin receptor beta) | protein-coding |
| chr8_128534395_128534675 | chr8 | 128534396 | 128534675 | + | 0.637074 | Intergenic | Intergenic | -151119 | NM_010578 | 16412 | Mm.263396 | NM_010578 | ENSMUSG00000025809 | **Itgb1** | 4633401G24Rik\|AA409975\|AA960159\|CD29\|ENSMUSG00000051907\|Fnrb\|Gm9863\|gpIIa | integrin beta 1 (fibronectin receptor beta) | protein-coding |
| chr4_99813736_99814039 | chr4 | 99813737 | 99814039 | + | 0.697358 | intron (NM_026348, intron 2 of 8) | intron (NM_026348, intron 2 of 8) | 15230 | NM_026348 | 67733 | Mm.257094 | NM_026348 | ENSMUSG00000028549 | **Itgb3bp** | 4930471O16Rik\|AU022583 | integrin beta 3 binding protein (beta3-endonexin) | protein-coding |
| chr2_60699996_60700538 | chr2 | 60699997 | 60700538 | + | 1.48907 | intron (NM_001159564, intron 1 of 17) | intron (NM_001159564, intron 1 of 17) | 22336 | NM_001159564 | 16420 | Mm.98193 | NM_021359 | ENSMUSG00000026971 | **Itgb6** | 2210409C20Rik\|4831415H04Rik | integrin beta 6 | protein-coding |
| chr12_119237728_119238990 | chr12 | 119237729 | 119238990 | + | 0.645302 | promoter-TSS (NM_177290) | promoter-TSS (NM_177290) | -83 | NM_177290 | 320910 | Mm.217000 | NM_177290 | ENSMUSG00000025321 | **Itgb8** | 4832412O06Rik | integrin beta 8 | protein-coding |
| chr2_10121853_10122256 | chr2 | 10121854 | 10122256 | + | 0.776466 | intron (NM_010582, intron 5 of 20) | B3A\|SINE\|B2 | 8628 | NM_010582 | 16425 | Mm.182043 | NM_010582 | ENSMUSG00000037254 | **Itih2** | AI747202\|Intin2\|Itih-2 | inter-alpha trypsin inhibitor, heavy chain 2 | protein-coding |
| chr11_46362556_46363055 | chr11 | 46362557 | 46363055 | + | 0.725799 | intron (NM_001281966, intron 4 of 16) | intron (NM_001281966, intron 4 of 16) | 26709 | NM_001281968 | 16428 | Mm.339927 | NM_010583 | ENSMUSG00000020395 | **Itk** | Emt\|Tcsk\|Tsk | IL2 inducible T cell kinase | protein-coding |
| chr11_46362117_46363073 | chr11 | 46362118 | 46363073 | + | 0.648103 | intron (NM_001281966, intron 4 of 16) | intron (NM_001281966, intron 4 of 16) | 26920 | NM_001281968 | 16428 | Mm.339927 | NM_010583 | ENSMUSG00000020395 | **Itk** | Emt\|Tcsk\|Tsk | IL2 inducible T cell kinase | protein-coding |
| chr14_73379637_73379940 | chr14 | 73379638 | 73379940 | + | 1.44184 | intron (NM_008410, intron 1 of 5) | intron (NM_008410, intron 1 of 5) | 5482 | NM_008410 | 16432 | Mm.4266 | NM_008410 | ENSMUSG00000022108 | **Itm2b** | AI256040\|Bri\|Bri2\|Bricd2b\|D14Sel6\|E25BMM\|imBRI2 | integral membrane protein 2B | protein-coding |
| chr14_73441611_73441907 | chr14 | 73441612 | 73441907 | + | 0.710304 | Intergenic | Intergenic | -56488 | NM_008410 | 16432 | Mm.4266 | NM_008410 | ENSMUSG00000022108 | **Itm2b** | AI256040\|Bri\|Bri2\|Bricd2b\|D14Sel6\|E25BMM\|imBRI2 | integral membrane protein 2B | protein-coding |
| chr12_102695651_102696240 | chr12 | 102695652 | 102696240 | + | 0.617736 | intron (NM_172584, intron 1 of 9) | intron (NM_172584, intron 1 of 9) | 8923 | NM_172584 | 217837 | Mm.482192 | NM_172584 | ENSMUSG00000057963 | **Itpk1** | BC031182 | inositol 1,3,4-triphosphate 5/6 kinase | protein-coding |
| chr1_180312790_180313292 | chr1 | 180312791 | 180313292 | + | 0.926009 | Intergenic | Intergenic | -17435 | NM_001081175 | 320404 | Mm.132535 | NM_001081175 | ENSMUSG00000038855 | **Itpkb** | 1110033J02Rik\|E130307H12Rik | inositol 1,4,5-trisphosphate 3-kinase B | protein-coding |
| chr1_180303465_180304000 | chr1 | 180303466 | 180304000 | + | 0.752881 | Intergenic | Intergenic | -26743 | NM_001081175 | 320404 | Mm.132535 | NM_001081175 | ENSMUSG00000038855 | **Itpkb** | 1110033J02Rik\|E130307H12Rik | inositol 1,4,5-trisphosphate 3-kinase B | protein-coding |
| chr6_108328227_108328625 | chr6 | 108328228 | 108328625 | + | 1.03941 | intron (NM_010585, intron 4 of 61) | intron (NM_010585, intron 4 of 61) | 115330 | NM_010585 | 16438 | Mm.227912 | NM_010585 | ENSMUSG00000030102 | **Itpr1** | D6Pas2\|ENSMUSG00000072853\|Gm10429\|IP3R1\|InsP3R\|Ip3r\|Itpr-1\|P400\|Pcp-1\|Pcp1\|opt | inositol 1,4,5-trisphosphate receptor 1 | protein-coding |
| chr6_146521458_146521886 | chr6 | 146521459 | 146521886 | + | 1.06591 | Intergenic | RMER15\|LTR\|ERVL | -19449 | NM_019923 | 16439 | Mm.7800 | NM_010586 | ENSMUSG00000030287 | **Itpr2** | AI649341\|InsP3R-2\|InsP3R-5\|Ip3r2\|Itpr5\|insP3R2 | inositol 1,4,5-triphosphate receptor 2 | protein-coding |
| chr6_146432363_146433466 | chr6 | 146432364 | 146433466 | + | 1.05536 | intron (NM_019923, intron 2 of 56) | intron (NM_019923, intron 2 of 56) | 69308 | NM_010586 | 16439 | Mm.7800 | NM_010586 | ENSMUSG00000030287 | **Itpr2** | AI649341\|InsP3R-2\|InsP3R-5\|Ip3r2\|Itpr5\|insP3R2 | inositol 1,4,5-triphosphate receptor 2 | protein-coding |
| chr6_146497969_146498299 | chr6 | 146497970 | 146498299 | + | 0.914121 | intron (NM_019923, intron 1 of 56) | intron (NM_019923, intron 1 of 56) | 4089 | NM_010586 | 16439 | Mm.7800 | NM_010586 | ENSMUSG00000030287 | **Itpr2** | AI649341\|InsP3R-2\|InsP3R-5\|Ip3r2\|Itpr5\|insP3R2 | inositol 1,4,5-triphosphate receptor 2 | protein-coding |
| chr6_146453574_146453976 | chr6 | 146453575 | 146453976 | + | 0.769747 | intron (NM_019923, intron 2 of 56) | URR1B\|DNA\|hAT-Charlie | 48448 | NM_010586 | 16439 | Mm.7800 | NM_010586 | ENSMUSG00000030287 | **Itpr2** | AI649341\|InsP3R-2\|InsP3R-5\|Ip3r2\|Itpr5\|insP3R2 | inositol 1,4,5-triphosphate receptor 2 | protein-coding |
| chr6_146497835_146498311 | chr6 | 146497836 | 146498311 | + | 0.703924 | intron (NM_019923, intron 1 of 56) | intron (NM_019923, intron 1 of 56) | 4150 | NM_010586 | 16439 | Mm.7800 | NM_010586 | ENSMUSG00000030287 | **Itpr2** | AI649341\|InsP3R-2\|InsP3R-5\|Ip3r2\|Itpr5\|insP3R2 | inositol 1,4,5-triphosphate receptor 2 | protein-coding |
| chr2_127165740_127166284 | chr2 | 127165741 | 127166284 | + | 1.52215 | Intergenic | Intergenic | -22555 | NM_001163527 | 73338 | Mm.57559 | NM_001163527 | | **Itpripl1** | 1700041B20Rik | inositol 1,4,5-triphosphate receptor interacting protein-like 1 | protein-coding |
| chr2_127139167_127139615 | chr2 | 127139168 | 127139615 | + | 1.15798 | 3' UTR (NM_001163528, exon 2 of 2) | 3' UTR (NM_001163528, exon 2 of 2) | 3199 | NM_001163528 | 73338 | Mm.57559 | NM_001163527 | | **Itpripl1** | 1700041B20Rik | inositol 1,4,5-triphosphate receptor interacting protein-like 1 | protein-coding |
| chr2_127138197_127138418 | chr2 | 127138198 | 127138418 | + | 0.784172 | TTS (NM_001163528) | TTS (NM_001163528) | 4282 | NM_001163528 | 73338 | Mm.57559 | NM_001163527 | | **Itpripl1** | 1700041B20Rik | inositol 1,4,5-triphosphate receptor interacting protein-like 1 | protein-coding |
| chr12_4574101_4574490 | chr12 | 4574102 | 4574490 | + | 1.20981 | Intergenic | MERVL_2A-int\|LTR\|ERVL | -18712 | NM_001198969 | 20403 | Mm.341204 | NM_011365 | ENSMUSG00000020640 | **Itsn2** | AI327390\|Ese2\|Sh3d1B\|Sh3p18\|mKIAA1256 | intersectin 2 | protein-coding |
| chr12_4599619_4600444 | chr12 | 4599620 | 4600444 | + | 1.04787 | intron (NM_011365, intron 1 of 38) | intron (NM_011365, intron 1 of 38) | 7024 | NM_001198968 | 20403 | Mm.341204 | NM_011365 | ENSMUSG00000020640 | **Itsn2** | AI327390\|Ese2\|Sh3d1B\|Sh3p18\|mKIAA1256 | intersectin 2 | protein-coding |
| chr2_118863201_118863610 | chr2 | 118863202 | 118863610 | + | 0.685954 | intron (NM_019826, intron 1 of 11) | MTC-int\|LTR\|ERVL-MaLR | 1406 | NM_019826 | 56357 | Mm.6635 | NM_019826 | ENSMUSG00000027332 | **Ivd** | 1300016K07Rik\|6720455E18Rik\|AI463340 | isovaleryl coenzyme A dehydrogenase | protein-coding |
| chr1_151250699_151251435 | chr1 | 151250700 | 151251435 | + | 1.15863 | Intergenic | MLT1H\|LTR\|ERVL-MaLR | -93431 | NM_001039512 | 117198 | Mm.33764 | NM_028582 | ENSMUSG00000023150 | **Ivns1abp** | 1190004M08Rik\|1700126I16Rik\|AA960440\|HSPC068\|ND1\|NS-1\|NS1-BP\|Nd1-L\|Nd1-S\|mKIAA0850 | influenza virus NS1A binding protein | protein-coding |
| chr1_151248363_151248763 | chr1 | 151248364 | 151248763 | + | 0.860995 | Intergenic | Intergenic | -95935 | NM_001039512 | 117198 | Mm.33764 | NM_028582 | ENSMUSG00000023150 | **Ivns1abp** | 1190004M08Rik\|1700126I16Rik\|AA960440\|HSPC068\|ND1\|NS-1\|NS1-BP\|Nd1-L\|Nd1-S\|mKIAA0850 | influenza virus NS1A binding protein | protein-coding |
| chr3_41606030_41606596 | chr3 | 41606031 | 41606596 | + | 1.21924 | intron (NM_172303, intron 9 of 10) | intron (NM_172303, intron 9 of 10) | 41433 | NM_001130186 | 269424 | Mm.286285 | NM_172303 | ENSMUSG00000025764 | **Jade1** | AU041499\|D530048A03Rik\|Phf17\|mKIAA1807 | jade family PHD finger 1 | protein-coding |
| chr11_51812013_51812721 | chr11 | 51812014 | 51812721 | + | 1.08747 | Intergenic | ORR1F\|LTR\|ERVL-MaLR | 45114 | NM_199299 | 76901 | Mm.259996 | NM_199299 | ENSMUSG00000020387 | **Jade2** | 1200017K05Rik\|AI480685\|Phf15\|mKIAA0239 | jade family PHD finger 2 | protein-coding |
| chr11_51833427_51833988 | chr11 | 51833428 | 51833988 | + | 0.970731 | intron (NM_199299, intron 4 of 11) | MIRb\|SINE\|MIR | 23773 | NM_199299 | 76901 | Mm.259996 | NM_199299 | ENSMUSG00000020387 | **Jade2** | 1200017K05Rik\|AI480685\|Phf15\|mKIAA0239 | jade family PHD finger 2 | protein-coding |
| chr11_51833089_51834003 | chr11 | 51833090 | 51834003 | + | 0.821869 | intron (NM_199299, intron 4 of 11) | intron (NM_199299, intron 4 of 11) | 23935 | NM_199299 | 76901 | Mm.259996 | NM_199299 | ENSMUSG00000020387 | **Jade2** | 1200017K05Rik\|AI480685\|Phf15\|mKIAA0239 | jade family PHD finger 2 | protein-coding |
| chr11_51832414_51832979 | chr11 | 51832415 | 51832979 | + | 0.798249 | intron (NM_199299, intron 4 of 11) | intron (NM_199299, intron 4 of 11) | 24784 | NM_199299 | 76901 | Mm.259996 | NM_199299 | ENSMUSG00000020387 | **Jade2** | 1200017K05Rik\|AI480685\|Phf15\|mKIAA0239 | jade family PHD finger 2 | protein-coding |
| chr11_51880974_51881458 | chr11 | 51880975 | 51881458 | + | 0.695267 | Intergenic | Intergenic | -23735 | NM_199299 | 76901 | Mm.259996 | NM_199299 | ENSMUSG00000020387 | **Jade2** | 1200017K05Rik\|AI480685\|Phf15\|mKIAA0239 | jade family PHD finger 2 | protein-coding |
| chr4_101276509_101277101 | chr4 | 101276510 | 101277101 | + | 1.37081 | Intergenic | Intergenic | -11523 | NM_146145 | 16451 | Mm.289657 | NM_146145 | ENSMUSG00000028530 | **Jak1** | AA960307\|C130039L05Rik | Janus kinase 1 | protein-coding |
| chr4_101272945_101273561 | chr4 | 101272946 | 101273561 | + | 0.793136 | Intergenic | Intergenic | -7971 | NM_146145 | 16451 | Mm.289657 | NM_146145 | ENSMUSG00000028530 | **Jak1** | AA960307\|C130039L05Rik | Janus kinase 1 | protein-coding |
| chr4_101273313_101273552 | chr4 | 101273314 | 101273552 | + | 0.752265 | Intergenic | Intergenic | -8151 | NM_146145 | 16451 | Mm.289657 | NM_146145 | ENSMUSG00000028530 | **Jak1** | AA960307\|C130039L05Rik | Janus kinase 1 | protein-coding |
| chr4_101236757_101237287 | chr4 | 101236758 | 101237287 | + | 0.71286 | intron (NM_146145, intron 1 of 24) | intron (NM_146145, intron 1 of 24) | 28260 | NM_146145 | 16451 | Mm.289657 | NM_146145 | ENSMUSG00000028530 | **Jak1** | AA960307\|C130039L05Rik | Janus kinase 1 | protein-coding |
| chr4_101236797_101237110 | chr4 | 101236798 | 101237110 | + | 0.675069 | intron (NM_146145, intron 1 of 24) | intron (NM_146145, intron 1 of 24) | 28328 | NM_146145 | 16451 | Mm.289657 | NM_146145 | ENSMUSG00000028530 | **Jak1** | AA960307\|C130039L05Rik | Janus kinase 1 | protein-coding |
| chr4_101247720_101248100 | chr4 | 101247721 | 101248100 | + | 0.627106 | intron (NM_146145, intron 1 of 24) | Lx9\|LINE\|L1 | 17372 | NM_146145 | 16451 | Mm.289657 | NM_146145 | ENSMUSG00000028530 | **Jak1** | AA960307\|C130039L05Rik | Janus kinase 1 | protein-coding |
| chr4_101202621_101203077 | chr4 | 101202622 | 101203077 | + | 0.608679 | intron (NM_146145, intron 1 of 24) | intron (NM_146145, intron 1 of 24) | 62433 | NM_146145 | 16451 | Mm.289657 | NM_146145 | ENSMUSG00000028530 | **Jak1** | AA960307\|C130039L05Rik | Janus kinase 1 | protein-coding |
| chr19_29271104_29271591 | chr19 | 29271105 | 29271591 | + | 1.0015 | intron (NM_008413, intron 3 of 24) | intron (NM_008413, intron 3 of 24) | 19545 | NM_008413 | 16452 | Mm.275839 | NM_008413 | ENSMUSG00000024789 | **Jak2** | Fd17 | Janus kinase 2 | protein-coding |
| chr16_84799424_84800397 | chr16 | 84799425 | 84800397 | + | 0.781901 | intron (NM_023844, intron 1 of 9) | ORR1E\|LTR\|ERVL-MaLR | 25788 | NM_023844 | 67374 | Mm.41758 | NM_023844 | ENSMUSG00000053062 | **Jam2** | 1110002N23Rik\|2410030G21Rik\|2410167M24Rik\|JAM-2\|JAM-B\|Jcam2\|VE-JAM | junction adhesion molecule 2 | protein-coding |

| | | | | | | | | | | | | | | | | | |
|---|---|---|---|---|---|---|---|---|---|---|---|---|---|---|---|---|---|
| chr13_44770782_44771559 | chr13 | 44770783 | 44771559 | + | 0.950801 | intron (NM_001205043, intron 1 of 17) | intron (NM_001205043, intron 1 of 17) | 39900 | NM_001205043 | 16468 | Mm.25059 | NM_021878 | ENSMUSG00000038518 | **Jarid2** | Jmj\|jumonji | jumonji, AT rich interactive domain 2 | protein-coding |
| chr13_44770934_44771216 | chr13 | 44770935 | 44771216 | + | 0.882169 | intron (NM_001205043, intron 1 of 17) | intron (NM_001205043, intron 1 of 17) | 39804 | NM_001205043 | 16468 | Mm.25059 | NM_021878 | ENSMUSG00000038518 | **Jarid2** | Jmj\|jumonji | jumonji, AT rich interactive domain 2 | protein-coding |
| chr13_44859233_44859607 | chr13 | 44859234 | 44859607 | + | 0.825566 | intron (NM_001205043, intron 3 of 17) | intron (NM_001205043, intron 3 of 17) | 128149 | NM_001205043 | 16468 | Mm.25059 | NM_021878 | ENSMUSG00000038518 | **Jarid2** | Jmj\|jumonji | jumonji, AT rich interactive domain 2 | protein-coding |
| chr6_52889155_52890235 | chr6 | 52889156 | 52890235 | + | 1.54332 | intron (NM_173406, intron 2 of 4) | intron (NM_173406, intron 2 of 4) | 19925 | NM_001168277 | 231986 | Mm.386865 | NM_173406 | | **Jazf1** | AI591476\|C820002C15\|Jaz1\|Tip27 | JAZF zinc finger 1 | protein-coding |
| chr6_52903175_52903877 | chr6 | 52903176 | 52903877 | + | 1.18719 | intron (NM_173406, intron 1 of 4) | intron (NM_173406, intron 1 of 4) | 6094 | NM_001168277 | 231986 | Mm.386865 | NM_173406 | | **Jazf1** | AI591476\|C820002C15\|Jaz1\|Tip27 | JAZF zinc finger 1 | protein-coding |
| chr6_52825558_52826096 | chr6 | 52825559 | 52826096 | + | 1.04855 | intron (NM_173406, intron 2 of 4) | B3A\|SINE\|B2 | 83793 | NM_001168277 | 231986 | Mm.386865 | NM_173406 | | **Jazf1** | AI591476\|C820002C15\|Jaz1\|Tip27 | JAZF zinc finger 1 | protein-coding |
| chr6_53130723_53131314 | chr6 | 53130724 | 53131314 | + | 0.797311 | Intergenic | Intergenic | -62395 | NM_173406 | 231986 | Mm.386865 | NM_173406 | | **Jazf1** | AI591476\|C820002C15\|Jaz1\|Tip27 | JAZF zinc finger 1 | protein-coding |
| chr6_52965194_52965891 | chr6 | 52965195 | 52965891 | + | 0.636863 | intron (NM_173406, intron 1 of 4) | intron (NM_173406, intron 1 of 4) | -55923 | NM_001168277 | 231986 | Mm.386865 | NM_173406 | | **Jazf1** | AI591476\|C820002C15\|Jaz1\|Tip27 | JAZF zinc finger 1 | protein-coding |
| chr12_85571277_85571792 | chr12 | 85571278 | 85571792 | + | 0.991211 | Intergenic | Intergenic | -27570 | NM_001205052 | 81703 | Mm.103560 | NM_030887 | ENSMUSG00000034271 | **Jdp2** | Jundm2\|Jundp2\|TIF | Jun dimerization protein 2 | protein-coding |
| chr12_85570231_85570639 | chr12 | 85570232 | 85570639 | + | 0.953171 | Intergenic | Intergenic | -28670 | NM_001205052 | 81703 | Mm.103560 | NM_030887 | ENSMUSG00000034271 | **Jdp2** | Jundm2\|Jundp2\|TIF | Jun dimerization protein 2 | protein-coding |
| chr10_67112999_67113276 | chr10 | 67113000 | 67113276 | + | 1.14436 | Intergenic | L1_Mus1\|LINE\|L1 | -14120 | NM_207221 | 108829 | Mm.23846 | NM_207221 | ENSMUSG00000037876 | **Jmjd1c** | 5430433L24Rik\|D630035I23Rik\|Jmjdic\|TRIP8 | jumonji domain containing 1C | protein-coding |
| chr10_67196629_67196987 | chr10 | 67196630 | 67196987 | + | 0.617384 | intron (NM_207221, intron 3 of 25) | intron (NM_207221, intron 3 of 25) | 11058 | NM_001242396 | 108829 | Mm.23846 | NM_207221 | ENSMUSG00000037876 | **Jmjd1c** | 5430433L24Rik\|D630035I23Rik\|Jmjdic\|TRIP8 | jumonji domain containing 1C | protein-coding |
| chr13_93406492_93407154 | chr13 | 93406493 | 93407154 | + | 0.823247 | Intergenic | Lx8b\|LINE\|L1 | 92985 | NM_021310 | 57748 | Mm.100273 | NM_021310 | ENSMUSG00000021690 | **Jmy** | - | junction-mediating and regulatory protein | protein-coding |
| chr10_80814246_80814848 | chr10 | 80814247 | 80814848 | + | 0.67426 | Intergenic | Intergenic | -1049 | NM_028001 | 71912 | Mm.45137 | NM_028001 | ENSMUSG00000020216 | **Jsrp1** | 2300003C06Rik\|2310032K21Rik\|JP-45\|JP45 | junctional sarcoplasmic reticulum protein 1 | protein-coding |
| chr4_95035295_95035976 | chr4 | 95035296 | 95035976 | + | 1.80101 | Intergenic | Intergenic | 16586 | NM_010591 | 16476 | Mm.275071 | NM_010591 | ENSMUSG00000052684 | **Jun** | AP-1\|Junc\|c-jun | jun proto-oncogene | protein-coding |
| chr4_95187944_95188522 | chr4 | 95187945 | 95188522 | + | 1.46269 | Intergenic | Intergenic | -136011 | NM_010591 | 16476 | Mm.275071 | NM_010591 | ENSMUSG00000052684 | **Jun** | AP-1\|Junc\|c-jun | jun proto-oncogene | protein-coding |
| chr4_95105292_95105752 | chr4 | 95105293 | 95105752 | + | 1.26379 | Intergenic | Intergenic | -53300 | NM_010591 | 16476 | Mm.275071 | NM_010591 | ENSMUSG00000052684 | **Jun** | AP-1\|Junc\|c-jun | jun proto-oncogene | protein-coding |
| chr4_95036461_95036917 | chr4 | 95036462 | 95036917 | + | 1.05243 | Intergenic | Intergenic | 15533 | NM_010591 | 16476 | Mm.275071 | NM_010591 | ENSMUSG00000052684 | **Jun** | AP-1\|Junc\|c-jun | jun proto-oncogene | protein-coding |
| chr4_95277760_95278900 | chr4 | 95277761 | 95278900 | + | 1.00226 | Intergenic | Intergenic | -226108 | NM_010591 | 16476 | Mm.275071 | NM_010591 | ENSMUSG00000052684 | **Jun** | AP-1\|Junc\|c-jun | jun proto-oncogene | protein-coding |
| chr4_95100368_95100761 | chr4 | 95100369 | 95100761 | + | 0.995988 | Intergenic | MLT2B4\|LTR\|ERVL | -48343 | NM_010591 | 16476 | Mm.275071 | NM_010591 | ENSMUSG00000052684 | **Jun** | AP-1\|Junc\|c-jun | jun proto-oncogene | protein-coding |
| chr4_95278050_95278661 | chr4 | 95278051 | 95278661 | + | 0.991833 | Intergenic | Intergenic | -226134 | NM_010591 | 16476 | Mm.275071 | NM_010591 | ENSMUSG00000052684 | **Jun** | AP-1\|Junc\|c-jun | jun proto-oncogene | protein-coding |
| chr4_95128011_95128609 | chr4 | 95128012 | 95128609 | + | 0.972964 | Intergenic | Intergenic | -76088 | NM_010591 | 16476 | Mm.275071 | NM_010591 | ENSMUSG00000052684 | **Jun** | AP-1\|Junc\|c-jun | jun proto-oncogene | protein-coding |
| chr4_95159847_95160410 | chr4 | 95159848 | 95160410 | + | 0.885232 | Intergenic | MIR\|SINE\|MIR | -107907 | NM_010591 | 16476 | Mm.275071 | NM_010591 | ENSMUSG00000052684 | **Jun** | AP-1\|Junc\|c-jun | jun proto-oncogene | protein-coding |
| chr4_95147228_95147538 | chr4 | 95147229 | 95147538 | + | 0.675985 | Intergenic | Intergenic | -95161 | NM_010591 | 16476 | Mm.275071 | NM_010591 | ENSMUSG00000052684 | **Jun** | AP-1\|Junc\|c-jun | jun proto-oncogene | protein-coding |
| chr11_100393662_100394175 | chr11 | 100393663 | 100394175 | + | 0.750279 | intron (NM_010593, intron 1 of 13) | intron (NM_010593, intron 1 of 13) | 3871 | NM_010593 | 16480 | Mm.299774 | NM_010593 | ENSMUSG00000001552 | **Jup** | Ctnng\|D930025P04Rik\|PG | junction plakoglobin | protein-coding |
| chr17_33812478_33812866 | chr17 | 33812479 | 33812866 | + | 1.05667 | intron (NM_030697, intron 1 of 10) | ID_B1\|SINE\|B4 | 2149 | NM_030697 | 80880 | Mm.196330 | NM_030697 | ENSMUSG00000042099 | **Kank3** | 0610013D04Rik\|Ankrd47\|D17Ertd288e\|NG28 | KN motif and ankyrin repeat domains 3 | protein-coding |
| chr17_33813176_33813414 | chr17 | 33813177 | 33813414 | + | 0.891149 | intron (NM_030697, intron 1 of 10) | intron (NM_030697, intron 1 of 10) | 2772 | NM_030697 | 80880 | Mm.196330 | NM_030697 | ENSMUSG00000042099 | **Kank3** | 0610013D04Rik\|Ankrd47\|D17Ertd288e\|NG28 | KN motif and ankyrin repeat domains 3 | protein-coding |
| chr11_104412286_104412643 | chr11 | 104412287 | 104412643 | + | 1.30006 | intron (NM_001081045, intron 2 of 13) | B3\|SINE\|B2 | 29826 | NM_001081045 | 76719 | Mm.136418 | NM_001081045 | ENSMUSG00000018412 | **Kansl1** | 1700081L11Rik\|9430041J06Rik\|mKIAA1267 | KAT8 regulatory NSL complex subunit 1 | protein-coding |
| chr11_104364233_104364522 | chr11 | 104364234 | 104364522 | + | 1.29426 | intron (NM_001081045, intron 4 of 13) | intron (NM_001081045, intron 4 of 13) | 77913 | NM_001081045 | 76719 | Mm.136418 | NM_001081045 | ENSMUSG00000018412 | **Kansl1** | 1700081L11Rik\|9430041J06Rik\|mKIAA1267 | KAT8 regulatory NSL complex subunit 1 | protein-coding |
| chr11_104412287_104412863 | chr11 | 104412288 | 104412863 | + | 1.19796 | intron (NM_001081045, intron 2 of 13) | intron (NM_001081045, intron 2 of 13) | 29716 | NM_001081045 | 76719 | Mm.136418 | NM_001081045 | ENSMUSG00000018412 | **Kansl1** | 1700081L11Rik\|9430041J06Rik\|mKIAA1267 | KAT8 regulatory NSL complex subunit 1 | protein-coding |
| chr11_104397127_104397717 | chr11 | 104397128 | 104397717 | + | 0.747993 | intron (NM_001081045, intron 2 of 13) | B2_Mm2\|SINE\|B2 | 44869 | NM_001081045 | 76719 | Mm.136418 | NM_001081045 | ENSMUSG00000018412 | **Kansl1** | 1700081L11Rik\|9430041J06Rik\|mKIAA1267 | KAT8 regulatory NSL complex subunit 1 | protein-coding |
| chr1_66806842_66807150 | chr1 | 66806843 | 66807150 | + | 0.765936 | intron (NM_001122738, intron 1 of 14) | MT2A\|LTR\|ERVL | 10599 | NM_177645 | 68691 | Mm.240912 | NM_177645 | ENSMUSG00000026004 | **Kansl1l** | 1110028C15Rik\|1190030G24\|AI427598\|C430010P07Rik\|mKIAA4189 | KAT8 regulatory NSL complex subunit 1-like | protein-coding |
| chr1_66806703_66807160 | chr1 | 66806704 | 66807160 | + | 0.701496 | intron (NM_001122738, intron 1 of 14) | MT2A\|LTR\|ERVL | 10663 | NM_177645 | 68691 | Mm.240912 | NM_177645 | ENSMUSG00000026004 | **Kansl1l** | 1110028C15Rik\|1190030G24\|AI427598\|C430010P07Rik\|mKIAA4189 | KAT8 regulatory NSL complex subunit 1-like | protein-coding |
| chr17_53577628_53578101 | chr17 | 53577629 | 53578101 | + | 0.872754 | intron (NM_020005, intron 1 of 17) | intron (NM_020005, intron 1 of 17) | -6261 | NM_001190846 | 18519 | Mm.255025 | NM_020005 | ENSMUSG00000000708 | **Kat2b** | A930006P13Rik\|AI461839\|AW536563\|Pcaf\|p/CAF | K(lysine) acetyltransferase 2B | protein-coding |
| chr8_22873245_22873635 | chr8 | 22873246 | 22873635 | + | 0.639163 | intron (NM_001081149, intron 3 of 17) | intron (NM_001081149, intron 3 of 17) | 13901 | NM_001081149 | 244349 | Mm.182776 | NM_001081149 | ENSMUSG00000031540 | **Kat6a** | 1500036M03\|9930021N24Rik\|MOZ\|Myst3\|Zfp220 | K(lysine) acetyltransferase 6A | protein-coding |
| chr7_127915395_127915670 | chr7 | 127915396 | 127915670 | + | 0.788236 | exon (NM_026370, exon 3 of 11) | exon (NM_026370, exon 3 of 11) | 3016 | NM_026370 | 67773 | Mm.425492 | NM_026370 | ENSMUSG00000030801 | **Kat8** | 2010203C02Rik\|5830450F21Rik\|D7Ertd629e\|MOF\|Myst1 | K(lysine) acetyltransferase 8 | protein-coding |
| chr18_77054830_77055280 | chr18 | 77054831 | 77055280 | + | 0.689992 | Intergenic | Intergenic | -7759 | NM_027721 | 71206 | Mm.35803 | NM_027721 | ENSMUSG00000025420 | **Katnal2** | 3110023G01Rik\|4933439B08Rik | katanin p60 subunit A-like 2 | protein-coding |
| chr8_15010113_15010565 | chr8 | 15010114 | 15010565 | + | 0.989322 | promoter-TSS (NM_029116) | promoter-TSS (NM_029116) | -686 | NM_029116 | 74901 | Mm.46675 | NM_029116 | ENSMUSG00000055675 | **Kbtbd11** | 2900016B01Rik\|4930465M17Rik\|mKIAA0711 | kelch repeat and BTB (POZ) domain containing 11 | protein-coding |
| chr8_14965622_14965968 | chr8 | 14965623 | 14965968 | + | 0.952468 | intron (NM_001037736, intron 17 of 27) | intron (NM_001037736, intron 17 of 27) | -45230 | NM_029116 | 74901 | Mm.46675 | NM_029116 | ENSMUSG00000055675 | **Kbtbd11** | 2900016B01Rik\|4930465M17Rik\|mKIAA0711 | kelch repeat and BTB (POZ) domain containing 11 | protein-coding |
| chr6_56803459_56804089 | chr6 | 56803460 | 56804089 | + | 1.34874 | Intergenic | L1MB3\|LINE\|L1 | -5961 | NM_145958 | 210973 | Mm.57295 | NM_145958 | ENSMUSG00000059486 | **Kbtbd2** | BC022962\|Bklhd1\|mKIAA1489 | kelch repeat and BTB (POZ) domain containing 2 | protein-coding |
| chr6_56764877_56765676 | chr6 | 56764878 | 56765676 | + | 0.89706 | Intergenic | Intergenic | 32536 | NM_145958 | 210973 | Mm.57295 | NM_145958 | ENSMUSG00000059486 | **Kbtbd2** | BC022962\|Bklhd1\|mKIAA1489 | kelch repeat and BTB (POZ) domain containing 2 | protein-coding |
| chr6_56784717_56785641 | chr6 | 56784718 | 56785641 | + | 0.859016 | intron (NM_145958, intron 2 of 3) | intron (NM_145958, intron 2 of 3) | 12634 | NM_145958 | 210973 | Mm.57295 | NM_145958 | ENSMUSG00000059486 | **Kbtbd2** | BC022962\|Bklhd1\|mKIAA1489 | kelch repeat and BTB (POZ) domain containing 2 | protein-coding |
| chr6_56764788_56765995 | chr6 | 56764789 | 56765995 | + | 0.789417 | Intergenic | Intergenic | 32421 | NM_145958 | 210973 | Mm.57295 | NM_145958 | ENSMUSG00000059486 | **Kbtbd2** | BC022962\|Bklhd1\|mKIAA1489 | kelch repeat and BTB (POZ) domain containing 2 | protein-coding |
| chr6_72881197_72882038 | chr6 | 72881198 | 72882038 | + | 0.6762 | intron (NM_019715, intron 1 of 6) | intron (NM_019715, intron 1 of 6) | 18361 | NM_019715 | 74287 | Mm.29194 | NM_019715 | ENSMUSG00000055239 | **Kcmf1** | 1700094M07Rik\|Debt91\|Pmcf | potassium channel modulatory factor 1 | protein-coding |
| chr3_107075611_107076038 | chr3 | 107075612 | 107076038 | + | 1.25713 | Intergenic | L3\|LINE\|CR1 | -25742 | NM_008417 | 16490 | Mm.39285 | NM_008417 | | **Kcna2** | Akr6a4\|ENSMUSG00000074335\|Gm10672\|Kca1-2\|Kv1.2\|Mk-2 | potassium voltage-gated channel, shaker-related subfamily, member 2 | protein-coding |
| chr3_107075653_107075925 | chr3 | 107075654 | 107075925 | + | 1.25506 | Intergenic | L3\|LINE\|CR1 | -25778 | NM_008417 | 16490 | Mm.39285 | NM_008417 | | **Kcna2** | Akr6a4\|ENSMUSG00000074335\|Gm10672\|Kca1-2\|Kv1.2\|Mk-2 | potassium voltage-gated channel, shaker-related subfamily, member 2 | protein-coding |
| chr3_107103135_107103526 | chr3 | 107103136 | 107103526 | + | 0.71336 | 5' UTR (NM_008417, exon 2 of 3) | 5' UTR (NM_008417, exon 2 of 3) | 1764 | NM_008417 | 16490 | Mm.39285 | NM_008417 | | **Kcna2** | Akr6a4\|ENSMUSG00000074335\|Gm10672\|Kca1-2\|Kv1.2\|Mk-2 | potassium voltage-gated channel, shaker-related subfamily, member 2 | protein-coding |
| chr3_107076466_107076997 | chr3 | 107076467 | 107076997 | + | 0.655124 | Intergenic | Intergenic | -24835 | NM_008417 | 16490 | Mm.39285 | NM_008417 | | **Kcna2** | Akr6a4\|ENSMUSG00000074335\|Gm10672\|Kca1-2\|Kv1.2\|Mk-2 | potassium voltage-gated channel, shaker-related subfamily, member 2 | protein-coding |
| chr3_107120191_107120540 | chr3 | 107120192 | 107120540 | + | 0.643649 | Intergenic | MIRb\|SINE\|MIR | 18799 | NM_008417 | 16490 | Mm.39285 | NM_008417 | | **Kcna2** | Akr6a4\|ENSMUSG00000074335\|Gm10672\|Kca1-2\|Kv1.2\|Mk-2 | potassium voltage-gated channel, shaker-related subfamily, member 2 | protein-coding |
| chr3_106942900_106943263 | chr3 | 106942901 | 106943263 | + | 1.03135 | Intergenic | Intergenic | -93080 | NM_008418 | 16491 | Mm.30640 | NM_008418 | ENSMUSG00000047959 | **Kcna3** | Kca1-3\|Kv1.3\|Mk-3 | potassium voltage-gated channel, shaker-related subfamily, member 3 | protein-coding |
| chr2_168428438_168428985 | chr2 | 168428439 | 168428985 | + | 0.690826 | Intergenic | Intergenic | -159381 | NM_001081134 | 241794 | Mm.425102 | NM_001081134 | ENSMUSG00000074575 | **Kcng1** | AW536275\|OTTMUSG00000016048 | potassium voltage-gated channel, subfamily G, member 1 | protein-coding |
| chr1_192190413_192191019 | chr1 | 192190414 | 192191019 | + | 0.711925 | promoter-TSS (NM_001038607) | promoter-TSS (NM_001038607) | -61 | NM_001038607 | 16510 | Mm.4489 | NM_010600 | ENSMUSG00000058248 | **Kcnh1** | EAG1\|Kv10.1\|M-eag | potassium voltage-gated channel, subfamily H (eag-related), member 1 | protein-coding |

| | | | | | | | | | | | | | | | | | |
|---|---|---|---|---|---|---|---|---|---|---|---|---|---|---|---|---|---|
| chr12_74894779_74895507 | chr12 | 74894780 | 74895507 | + | 1.11447 | Intergenic | Intergenic | 282189 | NM_172805 | 238271 | Mm.44465 | NM_172805 | ENSMUSG00000034402 | **Kcnh5** | Eag2 | potassium voltage-gated channel, subfamily H (eag-related), member 5 | protein-coding |
| chr17_52497258_52497577 | chr17 | 52497259 | 52497577 | + | 0.663196 | Intergenic | Intergenic | -105291 | NM_001031811 | 211468 | Mm.399068 | NM_001031811 | ENSMUSG00000035580 | **Kcnh8** | C130090D05Rik\|ELK\|ELK1\|ELK3\|Kv12.1 | potassium voltage-gated channel, subfamily H (eag-related), member 8 | protein-coding |
| chr11_33791444_33792448 | chr11 | 33791445 | 33792448 | + | 0.869873 | intron (NM_001190885, intron 1 of 8) | intron (NM_001190885, intron 1 of 8) | 51639 | NM_027398 | 70357 | Mm.252514 | NM_027398 | ENSMUSG00000053519 | **Kcnip1** | KCHIP1\|Kchip1.2 | Kv channel-interacting protein 1 | protein-coding |
| chr2_127488453_127488986 | chr2 | 127488454 | 127488986 | + | 0.975388 | intron (NM_019789, intron 2 of 8) | ORR1F\|LTR\|ERVL-MaLR | -6221 | NM_001111331 | 56461 | Mm.315292 | NM_019789 | ENSMUSG00000079056 | **Kcnip3** | 4933407H12Rik\|AI413860\|Csen\|DREAM\|KChIP3\|R74849 | Kv channel interacting protein 3, calsenilin | protein-coding |
| chr9_32420778_32421093 | chr9 | 32420779 | 32421093 | + | 0.604855 | Intergenic | B3A\|SINE\|B2 | 26951 | NM_001168354 | 56379 | Mm.390168 | NM_019659 | ENSMUSG00000041248 | **Kcnj1** | Kir1.1\|ROMK\|Romk2 | potassium inwardly-rectifying channel, subfamily J, member 1 | protein-coding |
| chr16_95275401_95276142 | chr16 | 95275402 | 95276142 | + | 1.20918 | intron (NM_001271687, intron 3 of 4) | intron (NM_001271687, intron 3 of 4) | 18214 | NM_001271690 | 16516 | Mm.272239 | NM_019664 | ENSMUSG00000062609 | **Kcnj15** | 4930414N08Rik\|AI182284\|AI267127\|IRKK\|Kir4.2 | potassium inwardly-rectifying channel, subfamily J, member 15 | protein-coding |
| chr16_95359984_95361269 | chr16 | 95359985 | 95361269 | + | 0.762576 | 3' UTR (NM_133659, exon 11 of 11) | 3' UTR (NM_133659, exon 11 of 11) | 103069 | NM_001271690 | 16516 | Mm.272239 | NM_019664 | ENSMUSG00000062609 | **Kcnj15** | 4930414N08Rik\|AI182284\|AI267127\|IRKK\|Kir4.2 | potassium inwardly-rectifying channel, subfamily J, member 15 | protein-coding |
| chr16_95264484_95264752 | chr16 | 95264485 | 95264752 | + | 0.669645 | intron (NM_001271687, intron 3 of 4) | intron (NM_001271687, intron 3 of 4) | 7060 | NM_001271690 | 16516 | Mm.272239 | NM_019664 | ENSMUSG00000062609 | **Kcnj15** | 4930414N08Rik\|AI182284\|AI267127\|IRKK\|Kir4.2 | potassium inwardly-rectifying channel, subfamily J, member 15 | protein-coding |
| chr11_111689902_111690236 | chr11 | 111689903 | 111690236 | + | 0.876967 | Intergenic | Intergenic | 623905 | NM_008425 | 16518 | Mm.4951 | NM_008425 | ENSMUSG00000041695 | **Kcnj2** | IRK1\|Kcnf1\|Kir2.1 | potassium inwardly-rectifying channel, subfamily J, member 2 | protein-coding |
| chr6_142578563_142579582 | chr6 | 142578564 | 142579582 | + | 0.936894 | Intergenic | Intergenic | -7717 | NM_008428 | 16523 | Mm.1482 | NM_008428 | ENSMUSG00000030247 | **Kcnj8** | AI448900\|Kir6.1\|gnite\|slmbr\|sltr | potassium inwardly-rectifying channel, subfamily J, member 8 | protein-coding |
| chr12_98572259_98572867 | chr12 | 98572260 | 98572867 | + | 1.69519 | intron (NM_029911, intron 1 of 6) | B1F\|SINE\|Alu | 5377 | NM_029911 | 72258 | Mm.196158 | NM_029911 | ENSMUSG00000033854 | **Kcnk10** | 1700024D23Rik\|3010005K24Rik\|Trek2 | potassium channel, subfamily K, member 10 | protein-coding |
| chr12_98574283_98574848 | chr12 | 98574284 | 98574848 | + | 1.44711 | intron (NM_029911, intron 1 of 6) | CpG | 3374 | NM_029911 | 72258 | Mm.196158 | NM_029911 | ENSMUSG00000033854 | **Kcnk10** | 1700024D23Rik\|3010005K24Rik\|Trek2 | potassium channel, subfamily K, member 10 | protein-coding |
| chr12_98577807_98578346 | chr12 | 98577808 | 98578346 | + | 1.42794 | promoter-TSS (NM_029911) | promoter-TSS (NM_029911) | -137 | NM_029911 | 72258 | Mm.196158 | NM_029911 | ENSMUSG00000033854 | **Kcnk10** | 1700024D23Rik\|3010005K24Rik\|Trek2 | potassium channel, subfamily K, member 10 | protein-coding |
| chr12_98444830_98445215 | chr12 | 98444831 | 98445215 | + | 1.33887 | intron (NM_029911, intron 4 of 6) | intron (NM_029911, intron 4 of 6) | 132917 | NM_029911 | 72258 | Mm.196158 | NM_029911 | ENSMUSG00000033854 | **Kcnk10** | 1700024D23Rik\|3010005K24Rik\|Trek2 | potassium channel, subfamily K, member 10 | protein-coding |
| chr12_98600525_98601393 | chr12 | 98600526 | 98601393 | + | 1.33732 | Intergenic | Intergenic | -23019 | NM_029911 | 72258 | Mm.196158 | NM_029911 | ENSMUSG00000033854 | **Kcnk10** | 1700024D23Rik\|3010005K24Rik\|Trek2 | potassium channel, subfamily K, member 10 | protein-coding |
| chr12_98574277_98575262 | chr12 | 98574278 | 98575262 | + | 1.20335 | intron (NM_029911, intron 1 of 6) | CpG | 3170 | NM_029911 | 72258 | Mm.196158 | NM_029911 | ENSMUSG00000033854 | **Kcnk10** | 1700024D23Rik\|3010005K24Rik\|Trek2 | potassium channel, subfamily K, member 10 | protein-coding |
| chr12_98524872_98525275 | chr12 | 98524873 | 98525275 | + | 1.10228 | intron (NM_029911, intron 1 of 6) | intron (NM_029911, intron 1 of 6) | 52866 | NM_029911 | 72258 | Mm.196158 | NM_029911 | ENSMUSG00000033854 | **Kcnk10** | 1700024D23Rik\|3010005K24Rik\|Trek2 | potassium channel, subfamily K, member 10 | protein-coding |
| chr12_98524687_98525277 | chr12 | 98524688 | 98525277 | + | 0.934401 | intron (NM_029911, intron 1 of 6) | intron (NM_029911, intron 1 of 6) | 52958 | NM_029911 | 72258 | Mm.196158 | NM_029911 | ENSMUSG00000033854 | **Kcnk10** | 1700024D23Rik\|3010005K24Rik\|Trek2 | potassium channel, subfamily K, member 10 | protein-coding |
| chr12_98445604_98445991 | chr12 | 98445605 | 98445991 | + | 0.739587 | intron (NM_029911, intron 4 of 6) | intron (NM_029911, intron 4 of 6) | 132142 | NM_029911 | 72258 | Mm.196158 | NM_029911 | ENSMUSG00000033854 | **Kcnk10** | 1700024D23Rik\|3010005K24Rik\|Trek2 | potassium channel, subfamily K, member 10 | protein-coding |
| chr12_98470037_98470720 | chr12 | 98470038 | 98470720 | + | 0.627252 | intron (NM_029911, intron 4 of 6) | intron (NM_029911, intron 4 of 6) | 107561 | NM_029911 | 72258 | Mm.196158 | NM_029911 | ENSMUSG00000033854 | **Kcnk10** | 1700024D23Rik\|3010005K24Rik\|Trek2 | potassium channel, subfamily K, member 10 | protein-coding |
| chr17_87849008_87849306 | chr17 | 87849009 | 87849306 | + | 1.01015 | Intergenic | PB1D10\|SINE\|Alu | -51163 | NM_199251 | 210741 | Mm.312046 | NM_199251 | ENSMUSG00000050138 | **Kcnk12** | mntk1 | potassium channel, subfamily K, member 12 | protein-coding |
| chr12_99979280_99980206 | chr12 | 99979281 | 99980206 | + | 1.50804 | intron (NM_001164427, intron 2 of 2) | Lx7\|LINE\|L1 | 15244 | NM_001164427 | 217826 | Mm.312335 | NM_146037 | ENSMUSG00000045404 | **Kcnk13** | BB085247\|F730021E22Rik\|Gm1570\|Gm1685 | potassium channel, subfamily K, member 13 | protein-coding |
| chr12_99979518_99980310 | chr12 | 99979519 | 99980310 | + | 1.37387 | intron (NM_001164427, intron 2 of 2) | Lx7\|LINE\|L1 | 15415 | NM_001164427 | 217826 | Mm.312335 | NM_146037 | ENSMUSG00000045404 | **Kcnk13** | BB085247\|F730021E22Rik\|Gm1570\|Gm1685 | potassium channel, subfamily K, member 13 | protein-coding |
| chr1_189499353_189500344 | chr1 | 189499354 | 189500344 | + | 0.830503 | Intergenic | RMER15\|LTR\|ERVL | -97576 | NM_001281848 | 16526 | Mm.33304 | NM_010607 | ENSMUSG00000037624 | **Kcnk2** | A430027H14Rik\|AI848635\|TREK-1 | potassium channel, subfamily K, member 2 | protein-coding |
| chr14_20173756_20174397 | chr14 | 20173757 | 20174397 | + | 1.00093 | intron (NM_021542, intron 1 of 4) | L1MB1\|LINE\|L1 | 7705 | NM_021542 | 16529 | Mm.68998 | NM_021542 | ENSMUSG00000023243 | **Kcnk5** | TASK-2 | potassium channel, subfamily K, member 5 | protein-coding |
| chr14_20173478_20174404 | chr14 | 20173479 | 20174404 | + | 0.998273 | intron (NM_021542, intron 1 of 4) | L1MB1\|LINE\|L1 | 7841 | NM_021542 | 16529 | Mm.68998 | NM_021542 | ENSMUSG00000023243 | **Kcnk5** | TASK-2 | potassium channel, subfamily K, member 5 | protein-coding |
| chr14_20172936_20173418 | chr14 | 20172937 | 20173418 | + | 0.992332 | intron (NM_021542, intron 1 of 4) | intron (NM_021542, intron 1 of 4) | 8605 | NM_021542 | 16529 | Mm.68998 | NM_021542 | ENSMUSG00000023243 | **Kcnk5** | TASK-2 | potassium channel, subfamily K, member 5 | protein-coding |
| chr14_20214137_20214612 | chr14 | 20214138 | 20214612 | + | 0.961237 | Intergenic | L1MB7\|LINE\|L1 | -32593 | NM_021542 | 16529 | Mm.68998 | NM_021542 | ENSMUSG00000023243 | **Kcnk5** | TASK-2 | potassium channel, subfamily K, member 5 | protein-coding |
| chr14_20172949_20173272 | chr14 | 20172950 | 20173272 | + | 0.944654 | intron (NM_021542, intron 1 of 4) | intron (NM_021542, intron 1 of 4) | 8671 | NM_021542 | 16529 | Mm.68998 | NM_021542 | ENSMUSG00000023243 | **Kcnk5** | TASK-2 | potassium channel, subfamily K, member 5 | protein-coding |
| chr14_20213840_20214619 | chr14 | 20213841 | 20214619 | + | 0.897962 | Intergenic | Intergenic | -32448 | NM_021542 | 16529 | Mm.68998 | NM_021542 | ENSMUSG00000023243 | **Kcnk5** | TASK-2 | potassium channel, subfamily K, member 5 | protein-coding |
| chr19_5704168_5704546 | chr19 | 5704169 | 5704546 | + | 0.648403 | promoter-TSS (NM_010609) | promoter-TSS (NM_010609) | -119 | NM_010609 | 16530 | Mm.440033 | NM_010609 | ENSMUSG00000024936 | **Kcnk7** | 2310014G03Rik\|Kcnk6\|Kcnk8\|Knot\|Mlk3 | potassium channel, subfamily K, member 7 | protein-coding |
| chr11_33967530_33968278 | chr11 | 33967531 | 33968278 | + | 0.856061 | intron (NM_031169, intron 3 of 3) | intron (NM_031169, intron 3 of 3) | 4891 | NM_031169 | 16533 | Mm.6206 | NM_031169 | ENSMUSG00000020155 | **Kcnmb1** | BKbeta1 | potassium large conductance calcium-activated channel, subfamily M, beta member 1 | protein-coding |
| chr11_33937481_33937983 | chr11 | 33937482 | 33937983 | + | 0.792881 | intron (NM_001190886, intron 1 of 7) | intron (NM_001190886, intron 1 of 7) | -25281 | NM_031169 | 16533 | Mm.6206 | NM_031169 | ENSMUSG00000020155 | **Kcnmb1** | BKbeta1 | potassium large conductance calcium-activated channel, subfamily M, beta member 1 | protein-coding |
| chr10_116486962_116487717 | chr10 | 116486963 | 116487717 | + | 1.36764 | intron (NR_024327, intron 13 of 13) | B1F1\|SINE\|Alu | -13817 | NM_021452 | 58802 | Mm.440652 | NM_021452 | ENSMUSG00000054934 | **Kcnmb4** | 1700058G18Rik\|2900045G12Rik | potassium large conductance calcium-activated channel, subfamily M, beta member 4 | protein-coding |
| chr10_116470214_116470612 | chr10 | 116470215 | 116470612 | + | 0.958815 | intron (NM_021452, intron 1 of 2) | MER5B\|DNA\|hAT-Charlie | 3110 | NM_021452 | 58802 | Mm.440652 | NM_021452 | ENSMUSG00000054934 | **Kcnmb4** | 1700058G18Rik\|2900045G12Rik | potassium large conductance calcium-activated channel, subfamily M, beta member 4 | protein-coding |
| chr10_116391178_116391543 | chr10 | 116391179 | 116391543 | + | 0.644428 | Intergenic | Intergenic | -26760 | NR_028107 | 67342 | Mm.245800 | NM_026098 | ENSMUSG00000020164 | **Kcnmb4os1** | 1700058G18Rik | potassium large conductance calcium-activated channel, subfamily M, beta member 4, opposite strand 1 | ncRNA |
| chr10_116391029_116391540 | chr10 | 116391030 | 116391540 | + | 0.636108 | Intergenic | Intergenic | -26836 | NR_028107 | 67342 | Mm.245800 | NM_026098 | ENSMUSG00000020164 | **Kcnmb4os1** | 1700058G18Rik | potassium large conductance calcium-activated channel, subfamily M, beta member 4, opposite strand 1 | ncRNA |
| chr3_89491452_89492000 | chr3 | 89491453 | 89492000 | + | 0.813173 | Intergenic | Intergenic | -28438 | NM_080466 | 140493 | Mm.120250 | NM_080466 | | **Kcnn3** | KCa2.3\|SK3\|SKCA3 | potassium intermediate/small conductance calcium-activated channel, subfamily N, member 3 | protein-coding |
| chr7_24365735_24365950 | chr7 | 24365736 | 24365950 | + | 0.838462 | Intergenic | Intergenic | -4420 | NM_001163510 | 16534 | Mm.9911 | NM_008433 | ENSMUSG00000054342 | **Kcnn4** | IK1\|IKCA1\|KCA4\|KCa3.1\|SK4\|SKCas\|mIKCa1 | potassium intermediate/small conductance calcium-activated channel, subfamily N, member 4 | protein-coding |
| chr2_181099206_181099840 | chr2 | 181099207 | 181099840 | + | 1.3042 | intron (NM_001003824, intron 9 of 16) | MLT1C-int\|LTR\|ERVL-MaLR | 35702 | NM_001006679 | 16536 | Mm.40615 | NM_010611 | ENSMUSG00000016346 | **Kcnq2** | HNSPC\|KQT2\|Nmf134 | potassium voltage-gated channel, subfamily Q, member 2 | protein-coding |
| chr2_181100079_181100450 | chr2 | 181100080 | 181100450 | + | 1.04134 | intron (NM_001003824, intron 9 of 16) | intron (NM_001003824, intron 9 of 16) | 34960 | NM_001006679 | 16536 | Mm.40615 | NM_010611 | ENSMUSG00000016346 | **Kcnq2** | HNSPC\|KQT2\|Nmf134 | potassium voltage-gated channel, subfamily Q, member 2 | protein-coding |
| chr15_66313553_66314282 | chr15 | 66313554 | 66314282 | + | 0.911476 | Intergenic | Intergenic | -27694 | NM_152923 | 110862 | Mm.255585 | NM_152923 | | **Kcnq3** | - | potassium voltage-gated channel, subfamily Q, member 3 | protein-coding |
| chr1_21805057_21805606 | chr1 | 21805058 | 21805606 | + | 1.09596 | intron (NM_001160139, intron 1 of 14) | MTE2b\|LTR\|ERVL-MaLR | 156610 | NM_001160139 | 226922 | Mm.336519 | NM_023872 | ENSMUSG00000028033 | **Kcnq5** | 7730402H11\|9230107O05Rik\|AA589396\|D1Mgi1 | potassium voltage-gated channel, subfamily Q, member 5 | protein-coding |
| chr1_21751669_21752198 | chr1 | 21751670 | 21752198 | + | 0.982762 | intron (NM_001160139, intron 1 of 14) | MIRc\|SINE\|MIR | 210008 | NM_001160139 | 226922 | Mm.336519 | NM_023872 | ENSMUSG00000028033 | **Kcnq5** | 7730402H11\|9230107O05Rik\|AA589396\|D1Mgi1 | potassium voltage-gated channel, subfamily Q, member 5 | protein-coding |

| | | | | | | | | | | | | | | | | | |
|---|---|---|---|---|---|---|---|---|---|---|---|---|---|---|---|---|---|
| chr1_21836399_21836765 | chr1 | 21836400 | 21836765 | + | 0.685922 | intron (NM_001160139, intron 1 of 14) | intron (NM_001160139, intron 1 of 14) | 125360 | NM_001160139 | 226922 | Mm.336519 | NM_023872 | ENSMUSG00000028033 | **Kcnq5** | 7730402H11\|9230107O05Rik\|AA589396\|D1Mgi1 | potassium voltage-gated channel, subfamily Q, member 5 | protein-coding |
| chr18_15138405_15138709 | chr18 | 15138406 | 15138709 | + | 0.713694 | intron (NM_134112, intron 1 of 4) | intron (NM_134112, intron 1 of 4) | 12889 | NM_134112 | 106931 | Mm.329299 | NM_134112 | ENSMUSG00000036225 | **Kctd1** | 4933402K10Rik\|AI661543\|AW553000 | potassium channel tetramerisation domain containing 1 | protein-coding |
| chr18_15042690_15043171 | chr18 | 15042691 | 15043171 | + | 0.694944 | intron (NM_134112, intron 1 of 4) | intron (NM_134112, intron 1 of 4) | 20651 | NM_001142731 | 106931 | Mm.329299 | NM_134112 | ENSMUSG00000036225 | **Kctd1** | 4933402K10Rik\|AI661543\|AW553000 | potassium channel tetramerisation domain containing 1 | protein-coding |
| chr1_189006196_189006587 | chr1 | 189006197 | 189006587 | + | 1.29043 | intron (NM_172650, intron 1 of 17) | intron (NM_172650, intron 1 of 17) | 1448 | NM_172650 | 226823 | Mm.209880 | NM_172650 | ENSMUSG00000026608 | **Kctd3** | 4930438A20Rik\|9330185B06\|E330032J19Rik\|NY-REN-45 | potassium channel tetramerisation domain containing 3 | protein-coding |
| chr5_130112704_130112996 | chr5 | 130112705 | 130112996 | + | 0.781429 | intron (NM_013837, intron 2 of 5) | B3\|SINE\|B2 | -32038 | NM_172509 | 212919 | Mm.55812 | NM_172509 | ENSMUSG00000034110 | **Kctd7** | 4932409E18\|9430010P06Rik | potassium channel tetramerisation domain containing 7 | protein-coding |
| chr5_130112476_130113000 | chr5 | 130112477 | 130113000 | + | 0.632536 | intron (NM_013837, intron 2 of 5) | (CA)n\|Simple_repeat\|Simple_repeat | -32150 | NM_172509 | 212919 | Mm.55812 | NM_172509 | ENSMUSG00000034110 | **Kctd7** | 4932409E18\|9430010P06Rik | potassium channel tetramerisation domain containing 7 | protein-coding |
| chr5_69340077_69341069 | chr5 | 69340078 | 69341069 | + | 0.624657 | exon (NM_175519, exon 1 of 2) | exon (NM_175519, exon 1 of 2) | 1136 | NM_175519 | 243043 | Mm.234821 | NM_175519 | ENSMUSG00000037653 | **Kctd8** | A730087N02Rik | potassium channel tetramerisation domain containing 8 | protein-coding |
| chr9_53344500_53345000 | chr9 | 53344501 | 53345000 | + | 0.66128 | intron (NM_176846, intron 3 of 5) | intron (NM_176846, intron 3 of 5) | -39273 | NM_212445 | 68304 | Mm.427519 | NM_212445 | ENSMUSG00000034487 | **Kdelc2** | 2010004J24Rik\|4833410J10Rik\|AW549401 | KDEL (Lys-Asp-Glu-Leu) containing 2 | protein-coding |
| chr5_143425992_143426728 | chr5 | 143425993 | 143426728 | + | 1.13841 | Intergenic | RLTR19\|LTR\|ERVK | 22540 | NM_025841 | 66913 | Mm.181880 | NM_025841 | ENSMUSG00000079111 | **Kdelr2** | 1110007A14Rik | KDEL (Lys-Asp-Glu-Leu) endoplasmic reticulum protein retention receptor 2 | protein-coding |
| chr4_136569897_136570734 | chr4 | 136569898 | 136570734 | + | 1.51165 | intron (NM_133872, intron 7 of 18) | intron (NM_133872, intron 7 of 18) | 32407 | NM_133872 | 99982 | Mm.28540 | NM_133872 | ENSMUSG00000036940 | **Kdm1a** | 1810043O07Rik\|AA408884\|Aof2\|D4Ertd478e\|Kdm1\|Lsd1\|mKIAA0601 | lysine (K)-specific demethylase 1A | protein-coding |
| chr13_47052716_47053283 | chr13 | 47052717 | 47053283 | + | 1.44675 | intron (NM_172262, intron 5 of 20) | L1M5\|LINE\|L1 | 9501 | NM_172262 | 218214 | Mm.31259 | NM_172262 | ENSMUSG00000038080 | **Kdm1b** | 4632428N09Rik\|AI482520\|Aof1 | lysine (K)-specific demethylase 1B | protein-coding |
| chr5_122894659_122895319 | chr5 | 122894660 | 122895319 | + | 1.08261 | intron (NM_001003953, intron 12 of 22) | intron (NM_001003953, intron 12 of 22) | 5361 | NM_013910 | 30841 | Mm.86406 | NM_013910 | ENSMUSG00000029475 | **Kdm2b** | Cxxc2\|E430001G17\|Fbl10\|Fbxl10\|Jhdm1b\|PCCX2\|mKIAA3014 | lysine (K)-specific demethylase 2B | protein-coding |
| chr5_122894324_122895319 | chr5 | 122894325 | 122895319 | + | 1.06103 | intron (NM_001003953, intron 12 of 22) | intron (NM_001003953, intron 12 of 22) | 5528 | NM_013910 | 30841 | Mm.86406 | NM_013910 | ENSMUSG00000029475 | **Kdm2b** | Cxxc2\|E430001G17\|Fbl10\|Fbxl10\|Jhdm1b\|PCCX2\|mKIAA3014 | lysine (K)-specific demethylase 2B | protein-coding |
| chr5_122986167_122986495 | chr5 | 122986168 | 122986495 | + | 0.653297 | intron (NM_001003953, intron 2 of 22) | intron (NM_001003953, intron 2 of 22) | 2202 | NM_001005866 | 30841 | Mm.86406 | NM_013910 | ENSMUSG00000029475 | **Kdm2b** | Cxxc2\|E430001G17\|Fbl10\|Fbxl10\|Jhdm1b\|PCCX2\|mKIAA3014 | lysine (K)-specific demethylase 2B | protein-coding |
| chrX_152218522_152218876 | chrX | 152218523 | 152218876 | + | 0.851101 | intron (NM_001005475, intron 10 of 13) | intron (NM_001005475, intron 10 of 13) | -14528 | NM_013668 | 20591 | Mm.142655 | NM_013668 | ENSMUSG00000025332 | **Kdm5c** | D930009K15Rik\|Jarid1c\|Smcx\|mKIAA0234 | lysine (K)-specific demethylase 5C | protein-coding |
| chr6_39181768_39182197 | chr6 | 39181769 | 39182197 | + | 1.0403 | intron (NM_001033430, intron 1 of 19) | intron (NM_001033430, intron 1 of 19) | 24790 | NM_001033430 | 338523 | Mm.293175 | NM_001033430 | ENSMUSG00000042599 | **Kdm7a** | A630082K20Rik\|BB041802\|ENSMUSG00000073143\|Jhdm1d\|mKIAA1718 | lysine (K)-specific demethylase 7A | protein-coding |
| chr6_39165732_39166196 | chr6 | 39165733 | 39166196 | + | 0.981702 | intron (NM_001033430, intron 8 of 19) | LTR87\|LTR\|ERVL? | 40809 | NM_001033430 | 338523 | Mm.293175 | NM_001033430 | ENSMUSG00000042599 | **Kdm7a** | A630082K20Rik\|BB041802\|ENSMUSG00000073143\|Jhdm1d\|mKIAA1718 | lysine (K)-specific demethylase 7A | protein-coding |
| chr6_39243382_39243703 | chr6 | 39243383 | 39243703 | + | 0.756709 | Intergenic | Intergenic | -36770 | NM_001033430 | 338523 | Mm.293175 | NM_001033430 | ENSMUSG00000042599 | **Kdm7a** | A630082K20Rik\|BB041802\|ENSMUSG00000073143\|Jhdm1d\|mKIAA1718 | lysine (K)-specific demethylase 7A | protein-coding |
| chr7_125425545_125426286 | chr7 | 125425546 | 125426286 | + | 1.43538 | Intergenic | Intergenic | -18704 | NM_029842 | 77035 | Mm.307239 | NM_029842 | ENSMUSG00000030752 | **Kdm8** | 3110005O21Rik\|Jmjd5 | lysine (K)-specific demethylase 8 | protein-coding |
| chr5_75793167_75793538 | chr5 | 75793168 | 75793538 | + | 0.841339 | Intergenic | Intergenic | 185075 | NM_010612 | 16542 | Mm.285 | NM_010612 | ENSMUSG00000062960 | **Kdr** | 6130401C07\|Flk-1\|Flk1\|Krd-1\|Ly73\|VEGFR-2\|VEGFR2\|sVEGFR-2 | kinase insert domain protein receptor | protein-coding |
| chr1_21425661_21426023 | chr1 | 21425662 | 21426023 | + | 1.42433 | intron (NM_001160139, intron 11 of 14) | intron (NM_001160139, intron 11 of 14) | 42286 | NM_001113187 | 98582 | Mm.326336 | NM_001113187 | ENSMUSG00000085079 | **Khdc1b** | AU014713\|AU015226\|AU024076\|AY238603\|C85331\|C86372\|C88118\|Khdc1\|Khdc1c | KH domain containing 1B | protein-coding |
| chr1_21425631_21426221 | chr1 | 21425632 | 21426221 | + | 1.25203 | intron (NM_001160139, intron 11 of 14) | intron (NM_001160139, intron 11 of 14) | 42370 | NM_001113187 | 98582 | Mm.326336 | NM_001113187 | ENSMUSG00000085079 | **Khdc1b** | AU014713\|AU015226\|AU024076\|AY238603\|C85331\|C86372\|C88118\|Khdc1\|Khdc1c | KH domain containing 1B | protein-coding |
| chr1_21603986_21604379 | chr1 | 21603987 | 21604379 | + | 1.0434 | intron (NM_001160139, intron 1 of 14) | L1MA4\|LINE\|L1 | 220627 | NM_001113187 | 98582 | Mm.326336 | NM_001113187 | ENSMUSG00000085079 | **Khdc1b** | AU014713\|AU015226\|AU024076\|AY238603\|C85331\|C86372\|C88118\|Khdc1\|Khdc1c | KH domain containing 1B | protein-coding |
| chr1_21603783_21604444 | chr1 | 21603784 | 21604444 | + | 0.986272 | intron (NM_001160139, intron 1 of 14) | L1MA4\|LINE\|L1 | 220558 | NM_001113187 | 98582 | Mm.326336 | NM_001113187 | ENSMUSG00000085079 | **Khdc1b** | AU014713\|AU015226\|AU024076\|AY238603\|C85331\|C86372\|C88118\|Khdc1\|Khdc1c | KH domain containing 1B | protein-coding |
| chr1_21419232_21419681 | chr1 | 21419233 | 21419681 | + | 0.880092 | intron (NM_001160139, intron 12 of 14) | intron (NM_001160139, intron 12 of 14) | 35901 | NM_001113187 | 98582 | Mm.326336 | NM_001113187 | ENSMUSG00000085079 | **Khdc1b** | AU014713\|AU015226\|AU024076\|AY238603\|C85331\|C86372\|C88118\|Khdc1\|Khdc1c | KH domain containing 1B | protein-coding |
| chr1_21633883_21634426 | chr1 | 21633884 | 21634426 | + | 0.771464 | intron (NM_001160139, intron 1 of 14) | intron (NM_001160139, intron 1 of 14) | 250599 | NM_001113187 | 98582 | Mm.326336 | NM_001113187 | ENSMUSG00000085079 | **Khdc1b** | AU014713\|AU015226\|AU024076\|AY238603\|C85331\|C86372\|C88118\|Khdc1\|Khdc1c | KH domain containing 1B | protein-coding |
| chr4_129746362_129746671 | chr4 | 129746363 | 129746671 | + | 0.64437 | Intergenic | PB1D11\|SINE\|Alu | -4214 | NR_045036 | 20218 | Mm.8256 | NM_011317 | ENSMUSG00000028790 | **Khdrbs1** | Sam68\|p62\|p68 | KH domain containing, RNA binding, signal transduction associated 1 | protein-coding |
| chr13_46945034_46945582 | chr13 | 46945035 | 46945582 | + | 0.64396 | Intergenic | Intergenic | -15590 | NM_010617 | 16553 | Mm.342703 | NM_010617 | ENSMUSG00000021375 | **Kif13a** | 4930505I07Rik\|mKIAA4109 | kinesin family member 13A | protein-coding |
| chr14_64663966_64664278 | chr14 | 64663967 | 64664278 | + | 1.0395 | intron (NM_001081177, intron 1 of 39) | intron (NM_001081177, intron 1 of 39) | 11591 | NM_001081177 | 16554 | Mm.23611 | NM_001081177 | ENSMUSG00000060012 | **Kif13b** | 5330429L19Rik\|6030414C01\|AI429803\|C130021D12Rik\|GAKIN | kinesin family member 13B | protein-coding |
| chr14_64656009_64656301 | chr14 | 64656010 | 64656301 | + | 0.82832 | intron (NM_001081177, intron 1 of 39) | B3\|SINE\|B2 | 3624 | NM_001081177 | 16554 | Mm.23611 | NM_001081177 | ENSMUSG00000060012 | **Kif13b** | 5330429L19Rik\|6030414C01\|AI429803\|C130021D12Rik\|GAKIN | kinesin family member 13B | protein-coding |
| chr14_64678055_64678424 | chr14 | 64678056 | 64678424 | + | 0.801914 | intron (NM_001081177, intron 2 of 39) | MT2B\|LTR\|ERVL | 25709 | NM_001081177 | 16554 | Mm.23611 | NM_001081177 | ENSMUSG00000060012 | **Kif13b** | 5330429L19Rik\|6030414C01\|AI429803\|C130021D12Rik\|GAKIN | kinesin family member 13B | protein-coding |
| chr14_64666627_64667135 | chr14 | 64666628 | 64667135 | + | 0.677487 | intron (NM_001081177, intron 1 of 39) | intron (NM_001081177, intron 1 of 39) | 14350 | NM_001081177 | 16554 | Mm.23611 | NM_001081177 | ENSMUSG00000060012 | **Kif13b** | 5330429L19Rik\|6030414C01\|AI429803\|C130021D12Rik\|GAKIN | kinesin family member 13B | protein-coding |
| chr1_136538156_136538728 | chr1 | 136538157 | 136538728 | + | 0.778537 | Intergenic | Intergenic | 70594 | NM_001287179 | 381293 | Mm.458468 | NM_001081258 | ENSMUSG00000041498 | **Kif14** | D1Ertd367e\|E130203M01 | kinesin family member 14 | protein-coding |
| chr1_136521410_136521868 | chr1 | 136521411 | 136521868 | + | 0.679016 | intron (NM_001287179, intron 26 of 28) | intron (NM_001287179, intron 26 of 28) | 53791 | NM_001287179 | 381293 | Mm.458468 | NM_001081258 | ENSMUSG00000041498 | **Kif14** | D1Ertd367e\|E130203M01 | kinesin family member 14 | protein-coding |
| chr2_142885886_142886487 | chr2 | 142885887 | 142886487 | + | 1.02384 | intron (NM_001081133, intron 1 of 25) | intron (NM_001081133, intron 1 of 25) | 15277 | NM_001081133 | 16558 | Mm.251934 | NM_001081133 | ENSMUSG00000038844 | **Kif16b** | 8430434E15Rik\|AA623607\|AI789011\|C80253\|C80902\|mKIAA1590 | kinesin family member 16B | protein-coding |
| chr2_142757952_142758569 | chr2 | 142757953 | 142758569 | + | 0.704609 | intron (NM_001081133, intron 13 of 25) | intron (NM_001081133, intron 13 of 25) | 143203 | NM_001081133 | 16558 | Mm.251934 | NM_001081133 | ENSMUSG00000038844 | **Kif16b** | 8430434E15Rik\|AA623607\|AI789011\|C80253\|C80902\|mKIAA1590 | kinesin family member 16B | protein-coding |
| chr4_149262282_149262825 | chr4 | 149262283 | 149262825 | + | 1.46145 | intron (NM_001290995, intron 12 of 48) | intron (NM_001290995, intron 12 of 48) | 45144 | NM_207682 | 16561 | Mm.402393 | NM_008441 | ENSMUSG00000063077 | **Kif1b** | A530096N05Rik\|AI448212\|AI506502\|D4Mil1e\|KIF1Bp130\|KIF1Bp204 | kinesin family member 1B | protein-coding |
| chr4_149260927_149261213 | chr4 | 149260928 | 149261213 | + | 0.864254 | intron (NM_001290995, intron 14 of 48) | intron (NM_001290995, intron 14 of 48) | 46628 | NM_207682 | 16561 | Mm.402393 | NM_008441 | ENSMUSG00000063077 | **Kif1b** | A530096N05Rik\|AI448212\|AI506502\|D4Mil1e\|KIF1Bp130\|KIF1Bp204 | kinesin family member 1B | protein-coding |
| chr4_149259445_149260090 | chr4 | 149259446 | 149260090 | + | 0.771658 | intron (NM_001290995, intron 14 of 48) | intron (NM_001290995, intron 14 of 48) | 47930 | NM_207682 | 16561 | Mm.402393 | NM_008441 | ENSMUSG00000063077 | **Kif1b** | A530096N05Rik\|AI448212\|AI506502\|D4Mil1e\|KIF1Bp130\|KIF1Bp204 | kinesin family member 1B | protein-coding |

| | | | | | | | | | | | | | | | | | |
|---|---|---|---|---|---|---|---|---|---|---|---|---|---|---|---|---|---|
| chr1_136136897_136137384 | chr1 | 136136898 | 136137384 | + | 1.0803 | intron (NM_001039472, intron 1 of 34) | intron (NM_001039472, intron 1 of 34) | 5740 | NM_001039472 | 16565 | Mm.276135 | NM_019962 | ENSMUSG00000041642 | **Kif21b** | 2610511N21Rik\|mKIAA0449 | kinesin family member 21B | protein-coding |
| chr1_136136654_136137382 | chr1 | 136136655 | 136137382 | + | 0.970537 | intron (NM_001039472, intron 1 of 34) | intron (NM_001039472, intron 1 of 34) | 5617 | NM_001039472 | 16565 | Mm.276135 | NM_019962 | ENSMUSG00000041642 | **Kif21b** | 2610511N21Rik\|mKIAA0449 | kinesin family member 21B | protein-coding |
| chr1_136161336_136161752 | chr1 | 136161337 | 136161752 | + | 0.644803 | exon (NM_001039472, exon 22 of 35) | exon (NM_001039472, exon 22 of 35) | 30143 | NM_001039472 | 16565 | Mm.276135 | NM_019962 | ENSMUSG00000041642 | **Kif21b** | 2610511N21Rik\|mKIAA0449 | kinesin family member 21B | protein-coding |
| chr9_61963335_61963721 | chr9 | 61963336 | 61963721 | + | 0.873673 | intron (NM_028748, intron 6 of 7) | intron (NM_028748, intron 6 of 7) | -16729 | NM_024245 | 71819 | Mm.259374 | NM_024245 | ENSMUSG00000032254 | **Kif23** | 3110001D19Rik\|C87313\|CHO1\|Knsl5\|MKLP-1\|MKLP1 | kinesin family member 23 | protein-coding |
| chr13_107019335_107019735 | chr13 | 107019336 | 107019735 | + | 1.10252 | intron (NM_008442, intron 1 of 21) | URR1A\|DNA\|hAT-Charlie | 2579 | NM_001145779 | 16563 | Mm.355686 | NM_008442 | ENSMUSG00000021693 | **Kif2a** | C530030B14Rik\|Kif2\|Kns2\|M-kinesin | kinesin family member 2A | protein-coding |
| chr2_153292186_153292962 | chr2 | 153292187 | 153292962 | + | 1.61105 | intron (NM_008444, intron 1 of 8) | intron (NM_008444, intron 1 of 8) | 1158 | NM_008444 | 16569 | Mm.315560 | NM_008444 | ENSMUSG00000027475 | **Kif3b** | AI854312\|AW549267\|mKIAA0359 | kinesin family member 3B | protein-coding |
| chr2_153298547_153299268 | chr2 | 153298548 | 153299268 | + | 0.722627 | intron (NM_008444, intron 1 of 8) | intron (NM_008444, intron 1 of 8) | 7492 | NM_008444 | 16569 | Mm.315560 | NM_008444 | ENSMUSG00000027475 | **Kif3b** | AI854312\|AW549267\|mKIAA0359 | kinesin family member 3B | protein-coding |
| chr12_3338928_3339393 | chr12 | 3338929 | 3339393 | + | 0.903307 | Intergenic | Intergenic | -25971 | NM_008445 | 16570 | Mm.7688 | NM_008445 | ENSMUSG00000020668 | **Kif3c** | mKIAA4058 | kinesin family member 3C | protein-coding |
| chr12_3338666_3339439 | chr12 | 3338667 | 3339439 | + | 0.902022 | Intergenic | Intergenic | -26079 | NM_008445 | 16570 | Mm.7688 | NM_008445 | ENSMUSG00000020668 | **Kif3c** | mKIAA4058 | kinesin family member 3C | protein-coding |
| chr12_3344772_3345626 | chr12 | 3344773 | 3345626 | + | 0.874895 | Intergenic | Lx7\|LINE\|L1 | -19933 | NM_008445 | 16570 | Mm.7688 | NM_008445 | ENSMUSG00000020668 | **Kif3c** | mKIAA4058 | kinesin family member 3C | protein-coding |
| chr12_3359256_3359758 | chr12 | 3359257 | 3359758 | + | 0.822303 | Intergenic | Intergenic | -5625 | NM_008445 | 16570 | Mm.7688 | NM_008445 | ENSMUSG00000020668 | **Kif3c** | mKIAA4058 | kinesin family member 3C | protein-coding |
| chr18_6285898_6286303 | chr18 | 6285899 | 6286303 | + | 1.25434 | Intergenic | Intergenic | -44577 | NM_008448 | 16573 | Mm.223744 | NM_008448 | ENSMUSG00000006740 | **Kif5b** | AL022807\|Khc\|Khcs\|Kns1\|Ukhc | kinesin family member 5B | protein-coding |
| chr18_6289975_6290263 | chr18 | 6289976 | 6290263 | + | 0.869993 | Intergenic | Intergenic | -48595 | NM_008448 | 16573 | Mm.223744 | NM_008448 | ENSMUSG00000006740 | **Kif5b** | AL022807\|Khc\|Khcs\|Kns1\|Ukhc | kinesin family member 5B | protein-coding |
| chr1_163778361_163778589 | chr1 | 163778362 | 163778589 | + | 0.736336 | Intergenic | URR1B\|DNA\|hAT-Charlie | -1204 | NM_010629 | 16579 | Mm.4651 | NM_010629 | ENSMUSG00000026585 | **Kifap3** | KAP3\|SMAP | kinesin-associated protein 3 | protein-coding |
| chr8_95154589_95155188 | chr8 | 95154590 | 95155188 | + | 1.44269 | Intergenic | Intergenic | -12349 | NM_001145831 | 16582 | Mm.378951 | NM_010631 | ENSMUSG00000031788 | **Kifc3** | AI325457\|BB123200 | kinesin family member C3 | protein-coding |
| chr8_95149363_95149857 | chr8 | 95149364 | 95149857 | + | 0.960858 | Intergenic | Intergenic | -7070 | NM_001145831 | 16582 | Mm.378951 | NM_010631 | ENSMUSG00000031788 | **Kifc3** | AI325457\|BB123200 | kinesin family member C3 | protein-coding |
| chr2_146834586_146835178 | chr2 | 146834587 | 146835178 | + | 0.683366 | Intergenic | Intergenic | -21007 | NM_001033298 | 228730 | Mm.292352 | NM_001033298 | ENSMUSG00000074749 | **Kiz** | Gm114\|Ncrna00153\|Plk1s1 | kizuna centrosomal protein | protein-coding |
| chr12_111768011_111768891 | chr12 | 111768012 | 111768891 | + | 1.5233 | intron (NM_001025360, intron 1 of 15) | RLTR48A\|LTR\|ERV1 | 9583 | NM_001025361 | 16593 | Mm.278357 | NM_008450 | ENSMUSG00000021288 | **Klc1** | AI874768\|Kns2 | kinesin light chain 1 | protein-coding |
| chr14_99944234_99945066 | chr14 | 99944235 | 99945066 | + | 0.877769 | intron (NM_010636, intron 4 of 6) | intron (NM_010636, intron 4 of 6) | 205146 | NM_010636 | 16597 | Mm.396499 | NM_010636 | | **Klf12** | 2700063E05Rik\|AI225908\|AI315654\|AP-2rep\|B130052C06Rik\|D530033K05Rik | Kruppel-like factor 12 | protein-coding |
| chr7_63940483_63940652 | chr7 | 63940484 | 63940652 | + | 1.51691 | Intergenic | Intergenic | -1653 | NM_021366 | 50794 | Mm.240473 | NM_021366 | ENSMUSG00000052040 | **Klf13** | 0610043C13Rik\|9430029L20Rik\|Bteb3\|FKLF-2\|FKLF2\|NSLP1\|RFLAT-1\|RFLAT1 | Kruppel-like factor 13 | protein-coding |
| chr7_64005813_64006239 | chr7 | 64005814 | 64006239 | + | 0.874304 | Intergenic | Intergenic | -67111 | NM_021366 | 50794 | Mm.240473 | NM_021366 | ENSMUSG00000052040 | **Klf13** | 0610043C13Rik\|9430029L20Rik\|Bteb3\|FKLF-2\|FKLF2\|NSLP1\|RFLAT-1\|RFLAT1 | Kruppel-like factor 13 | protein-coding |
| chr7_64001513_64002163 | chr7 | 64001514 | 64002163 | + | 0.821533 | Intergenic | Intergenic | -62923 | NM_021366 | 50794 | Mm.240473 | NM_021366 | ENSMUSG00000052040 | **Klf13** | 0610043C13Rik\|9430029L20Rik\|Bteb3\|FKLF-2\|FKLF2\|NSLP1\|RFLAT-1\|RFLAT1 | Kruppel-like factor 13 | protein-coding |
| chr7_63966390_63966739 | chr7 | 63966391 | 63966739 | + | 0.664165 | Intergenic | MER21B\|LTR\|ERVL | -27650 | NM_021366 | 50794 | Mm.240473 | NM_021366 | ENSMUSG00000052040 | **Klf13** | 0610043C13Rik\|9430029L20Rik\|Bteb3\|FKLF-2\|FKLF2\|NSLP1\|RFLAT-1\|RFLAT1 | Kruppel-like factor 13 | protein-coding |
| chr8_72331199_72331477 | chr8 | 72331200 | 72331477 | + | 0.998215 | Intergenic | Intergenic | 12276 | NM_008452 | 16598 | Mm.26938 | NM_008452 | ENSMUSG00000055148 | **Klf2** | Lklf | Kruppel-like factor 2 (lung) | protein-coding |
| chr8_72330653_72331521 | chr8 | 72330654 | 72331521 | + | 0.947559 | Intergenic | Intergenic | 12025 | NM_008452 | 16598 | Mm.26938 | NM_008452 | ENSMUSG00000055148 | **Klf2** | Lklf | Kruppel-like factor 2 (lung) | protein-coding |
| chr5_64765726_64766312 | chr5 | 64765727 | 64766312 | + | 1.7613 | Intergenic | Intergenic | -37504 | NM_008453 | 16599 | Mm.392759 | NM_008453 | ENSMUSG00000029178 | **Klf3** | 9930027G08Rik\|AL024007\|Bklf\|Tef-2 | Kruppel-like factor 3 (basic) | protein-coding |
| chr4_55488549_55488973 | chr4 | 55488550 | 55488973 | + | 1.66814 | Intergenic | LTR16B\|LTR\|ERVL | 43714 | NM_010637 | 16600 | Mm.4325 | NM_010637 | ENSMUSG00000003032 | **Klf4** | EZF\|Gklf\|Zie | Kruppel-like factor 4 (gut) | protein-coding |
| chr4_55489115_55489696 | chr4 | 55489116 | 55489696 | + | 1.2604 | Intergenic | Intergenic | 43069 | NM_010637 | 16600 | Mm.4325 | NM_010637 | ENSMUSG00000003032 | **Klf4** | EZF\|Gklf\|Zie | Kruppel-like factor 4 (gut) | protein-coding |
| chr4_55733253_55733657 | chr4 | 55733254 | 55733657 | + | 0.808424 | Intergenic | Intergenic | -200980 | NM_010637 | 16600 | Mm.4325 | NM_010637 | ENSMUSG00000003032 | **Klf4** | EZF\|Gklf\|Zie | Kruppel-like factor 4 (gut) | protein-coding |
| chr4_55730229_55730649 | chr4 | 55730230 | 55730649 | + | 0.787451 | Intergenic | Intergenic | -197964 | NM_010637 | 16600 | Mm.4325 | NM_010637 | ENSMUSG00000003032 | **Klf4** | EZF\|Gklf\|Zie | Kruppel-like factor 4 (gut) | protein-coding |
| chr4_55732088_55732338 | chr4 | 55732089 | 55732338 | + | 0.645387 | Intergenic | Intergenic | -199738 | NM_010637 | 16600 | Mm.4325 | NM_010637 | ENSMUSG00000003032 | **Klf4** | EZF\|Gklf\|Zie | Kruppel-like factor 4 (gut) | protein-coding |
| chr13_5889754_5890426 | chr13 | 5889755 | 5890426 | + | 1.51116 | Intergenic | Intergenic | 28601 | NM_011803 | 23849 | Mm.275036 | NM_011803 | ENSMUSG00000000078 | **Klf6** | AI448727\|BCD1\|C86813\|CPBP\|Copeb\|FM2\|FM6\|Ierepo1\|Ierepo3\|R75280\|Zf9 | Kruppel-like factor 6 | protein-coding |
| chr13_6159426_6160071 | chr13 | 6159427 | 6160071 | + | 1.44529 | Intergenic | Intergenic | 298260 | NM_011803 | 23849 | Mm.275036 | NM_011803 | ENSMUSG00000000078 | **Klf6** | AI448727\|BCD1\|C86813\|CPBP\|Copeb\|FM2\|FM6\|Ierepo1\|Ierepo3\|R75280\|Zf9 | Kruppel-like factor 6 | protein-coding |
| chr13_5979428_5979852 | chr13 | 5979429 | 5979852 | + | 1.40487 | Intergenic | Intergenic | 118151 | NM_011803 | 23849 | Mm.275036 | NM_011803 | ENSMUSG00000000078 | **Klf6** | AI448727\|BCD1\|C86813\|CPBP\|Copeb\|FM2\|FM6\|Ierepo1\|Ierepo3\|R75280\|Zf9 | Kruppel-like factor 6 | protein-coding |
| chr13_5892730_5893050 | chr13 | 5892731 | 5893050 | + | 1.1212 | Intergenic | Intergenic | 31401 | NM_011803 | 23849 | Mm.275036 | NM_011803 | ENSMUSG00000000078 | **Klf6** | AI448727\|BCD1\|C86813\|CPBP\|Copeb\|FM2\|FM6\|Ierepo1\|Ierepo3\|R75280\|Zf9 | Kruppel-like factor 6 | protein-coding |
| chr13_5979423_5980578 | chr13 | 5979424 | 5980578 | + | 1.11991 | Intergenic | Intergenic | 118512 | NM_011803 | 23849 | Mm.275036 | NM_011803 | ENSMUSG00000000078 | **Klf6** | AI448727\|BCD1\|C86813\|CPBP\|Copeb\|FM2\|FM6\|Ierepo1\|Ierepo3\|R75280\|Zf9 | Kruppel-like factor 6 | protein-coding |
| chr13_6191538_6192389 | chr13 | 6191539 | 6192389 | + | 1.10709 | Intergenic | Intergenic | 330475 | NM_011803 | 23849 | Mm.275036 | NM_011803 | ENSMUSG00000000078 | **Klf6** | AI448727\|BCD1\|C86813\|CPBP\|Copeb\|FM2\|FM6\|Ierepo1\|Ierepo3\|R75280\|Zf9 | Kruppel-like factor 6 | protein-coding |
| chr13_5894947_5895265 | chr13 | 5894948 | 5895265 | + | 1.07052 | Intergenic | Intergenic | 33617 | NM_011803 | 23849 | Mm.275036 | NM_011803 | ENSMUSG00000000078 | **Klf6** | AI448727\|BCD1\|C86813\|CPBP\|Copeb\|FM2\|FM6\|Ierepo1\|Ierepo3\|R75280\|Zf9 | Kruppel-like factor 6 | protein-coding |
| chr13_6111869_6112346 | chr13 | 6111870 | 6112346 | + | 0.996278 | Intergenic | Intergenic | 250619 | NM_011803 | 23849 | Mm.275036 | NM_011803 | ENSMUSG00000000078 | **Klf6** | AI448727\|BCD1\|C86813\|CPBP\|Copeb\|FM2\|FM6\|Ierepo1\|Ierepo3\|R75280\|Zf9 | Kruppel-like factor 6 | protein-coding |
| chr13_6131273_6131633 | chr13 | 6131274 | 6131633 | + | 0.937002 | Intergenic | Intergenic | 269964 | NM_011803 | 23849 | Mm.275036 | NM_011803 | ENSMUSG00000000078 | **Klf6** | AI448727\|BCD1\|C86813\|CPBP\|Copeb\|FM2\|FM6\|Ierepo1\|Ierepo3\|R75280\|Zf9 | Kruppel-like factor 6 | protein-coding |
| chr13_6109004_6109692 | chr13 | 6109005 | 6109692 | + | 0.899703 | Intergenic | MLT1B\|LTR\|ERVL-MaLR | 247859 | NM_011803 | 23849 | Mm.275036 | NM_011803 | ENSMUSG00000000078 | **Klf6** | AI448727\|BCD1\|C86813\|CPBP\|Copeb\|FM2\|FM6\|Ierepo1\|Ierepo3\|R75280\|Zf9 | Kruppel-like factor 6 | protein-coding |
| chr13_6109009_6110174 | chr13 | 6109010 | 6110174 | + | 0.823422 | Intergenic | MLT1B\|LTR\|ERVL-MaLR | 248103 | NM_011803 | 23849 | Mm.275036 | NM_011803 | ENSMUSG00000000078 | **Klf6** | AI448727\|BCD1\|C86813\|CPBP\|Copeb\|FM2\|FM6\|Ierepo1\|Ierepo3\|R75280\|Zf9 | Kruppel-like factor 6 | protein-coding |
| chr13_5886876_5887156 | chr13 | 5886877 | 5887156 | + | 0.789087 | Intergenic | Intergenic | 25527 | NM_011803 | 23849 | Mm.275036 | NM_011803 | ENSMUSG00000000078 | **Klf6** | AI448727\|BCD1\|C86813\|CPBP\|Copeb\|FM2\|FM6\|Ierepo1\|Ierepo3\|R75280\|Zf9 | Kruppel-like factor 6 | protein-coding |
| chr13_6127196_6127855 | chr13 | 6127197 | 6127855 | + | 0.787986 | Intergenic | Intergenic | 266037 | NM_011803 | 23849 | Mm.275036 | NM_011803 | ENSMUSG00000000078 | **Klf6** | AI448727\|BCD1\|C86813\|CPBP\|Copeb\|FM2\|FM6\|Ierepo1\|Ierepo3\|R75280\|Zf9 | Kruppel-like factor 6 | protein-coding |
| chr13_5979933_5980555 | chr13 | 5979934 | 5980555 | + | 0.6858 | Intergenic | Intergenic | 118755 | NM_011803 | 23849 | Mm.275036 | NM_011803 | ENSMUSG00000000078 | **Klf6** | AI448727\|BCD1\|C86813\|CPBP\|Copeb\|FM2\|FM6\|Ierepo1\|Ierepo3\|R75280\|Zf9 | Kruppel-like factor 6 | protein-coding |

| | | | | | | | | | | | | | | | | | |
|---|---|---|---|---|---|---|---|---|---|---|---|---|---|---|---|---|---|
| chr13_6193054_6193307 | chr13 | 6193055 | 6193307 | + | 0.663904 | Intergenic | Intergenic | 331692 | NM_011803 | 23849 | Mm.275036 | NM_011803 | ENSMUSG00000000078 | **Klf6** | AI448727\|BCD1\|C86813\|CPBP\|Copeb\|FM2\|FM6\|Ierepo1\|Ierepo3\|R75280\|Zf9 | Kruppel-like factor 6 | protein-coding |
| chr13_6127194_6128688 | chr13 | 6127195 | 6128688 | + | 0.660651 | Intergenic | Intergenic | 266452 | NM_011803 | 23849 | Mm.275036 | NM_011803 | ENSMUSG00000000078 | **Klf6** | AI448727\|BCD1\|C86813\|CPBP\|Copeb\|FM2\|FM6\|Ierepo1\|Ierepo3\|R75280\|Zf9 | Kruppel-like factor 6 | protein-coding |
| chr1_64191565_64192062 | chr1 | 64191566 | 64192062 | + | 1.10288 | Intergenic | Intergenic | -70425 | NM_033563 | 93691 | Mm.29466 | NM_033563 | ENSMUSG00000025959 | **Klf7** | 9830124P08Rik | Kruppel-like factor 7 (ubiquitous) | protein-coding |
| chr1_64101386_64101825 | chr1 | 64101387 | 64101825 | + | 0.993299 | intron (NM_033563, intron 1 of 3) | intron (NM_033563, intron 1 of 3) | 19783 | NM_033563 | 93691 | Mm.29466 | NM_033563 | ENSMUSG00000025959 | **Klf7** | 9830124P08Rik | Kruppel-like factor 7 (ubiquitous) | protein-coding |
| chr1_64091335_64091871 | chr1 | 64091336 | 64091871 | + | 0.964933 | intron (NM_033563, intron 1 of 3) | intron (NM_033563, intron 1 of 3) | 29786 | NM_033563 | 93691 | Mm.29466 | NM_033563 | ENSMUSG00000025959 | **Klf7** | 9830124P08Rik | Kruppel-like factor 7 (ubiquitous) | protein-coding |
| chr1_64116960_64117213 | chr1 | 64116961 | 64117213 | + | 0.960734 | intron (NM_033563, intron 1 of 3) | intron (NM_033563, intron 1 of 3) | 4302 | NM_033563 | 93691 | Mm.29466 | NM_033563 | ENSMUSG00000025959 | **Klf7** | 9830124P08Rik | Kruppel-like factor 7 (ubiquitous) | protein-coding |
| chr1_64305367_64306013 | chr1 | 64305368 | 64306013 | + | 0.957455 | Intergenic | Intergenic | -184301 | NM_033563 | 93691 | Mm.29466 | NM_033563 | ENSMUSG00000025959 | **Klf7** | 9830124P08Rik | Kruppel-like factor 7 (ubiquitous) | protein-coding |
| chr1_64134264_64134801 | chr1 | 64134265 | 64134801 | + | 0.892528 | Intergenic | Intergenic | -13144 | NM_033563 | 93691 | Mm.29466 | NM_033563 | ENSMUSG00000025959 | **Klf7** | 9830124P08Rik | Kruppel-like factor 7 (ubiquitous) | protein-coding |
| chr1_64090591_64090936 | chr1 | 64090592 | 64090936 | + | 0.82803 | intron (NM_033563, intron 1 of 3) | intron (NM_033563, intron 1 of 3) | 30625 | NM_033563 | 93691 | Mm.29466 | NM_033563 | ENSMUSG00000025959 | **Klf7** | 9830124P08Rik | Kruppel-like factor 7 (ubiquitous) | protein-coding |
| chr1_64105181_64105825 | chr1 | 64105182 | 64105825 | + | 0.784383 | intron (NM_033563, intron 1 of 3) | intron (NM_033563, intron 1 of 3) | 15886 | NM_033563 | 93691 | Mm.29466 | NM_033563 | ENSMUSG00000025959 | **Klf7** | 9830124P08Rik | Kruppel-like factor 7 (ubiquitous) | protein-coding |
| chr1_64301105_64301633 | chr1 | 64301106 | 64301633 | + | 0.658666 | Intergenic | Intergenic | -179980 | NM_033563 | 93691 | Mm.29466 | NM_033563 | ENSMUSG00000025959 | **Klf7** | 9830124P08Rik | Kruppel-like factor 7 (ubiquitous) | protein-coding |
| chr19_23134526_23135320 | chr19 | 23134527 | 23135320 | + | 0.704914 | Intergenic | CpG | -6303 | NM_010638 | 16601 | Mm.291595 | NM_010638 | ENSMUSG00000033863 | **Klf9** | 2310051E17Rik\|AA589643\|BTEB-1\|Bteb1\|ENSMUSG00000055440\|Gm9971 | Kruppel-like factor 9 | protein-coding |
| chr19_23134037_23136236 | chr19 | 23134038 | 23136236 | + | 0.693178 | Intergenic | CpG | -6089 | NM_010638 | 16601 | Mm.291595 | NM_010638 | ENSMUSG00000033863 | **Klf9** | 2310051E17Rik\|AA589643\|BTEB-1\|Bteb1\|ENSMUSG00000055440\|Gm9971 | Kruppel-like factor 9 | protein-coding |
| chr19_23135367_23136186 | chr19 | 23135368 | 23136186 | + | 0.680999 | Intergenic | CpG | -5449 | NM_010638 | 16601 | Mm.291595 | NM_010638 | ENSMUSG00000033863 | **Klf9** | 2310051E17Rik\|AA589643\|BTEB-1\|Bteb1\|ENSMUSG00000055440\|Gm9971 | Kruppel-like factor 9 | protein-coding |
| chr19_23091629_23092023 | chr19 | 23091630 | 23092023 | + | 0.659862 | Intergenic | Intergenic | -49400 | NM_010638 | 16601 | Mm.291595 | NM_010638 | ENSMUSG00000033863 | **Klf9** | 2310051E17Rik\|AA589643\|BTEB-1\|Bteb1\|ENSMUSG00000055440\|Gm9971 | Kruppel-like factor 9 | protein-coding |
| chr6_30427934_30428472 | chr6 | 30427935 | 30428472 | + | 0.792056 | intron (NM_029742, intron 2 of 9) | intron (NM_029742, intron 2 of 9) | 26294 | NM_029742 | 76788 | Mm.277308 | NM_029742 | ENSMUSG00000029775 | **Klhdc10** | 2410127E18Rik\|AI593221\|mKIAA0265 | kelch domain containing 10 | protein-coding |
| chr12_69288659_69289295 | chr12 | 69288660 | 69289295 | + | 1.23927 | Intergenic | Lx9\|LINE\|L1 | -7704 | NM_027117 | 69554 | Mm.234368 | NM_027117 | ENSMUSG00000020978 | **Klhdc2** | 2310022K15Rik\|D12Ertd522e\|HCLP-1\|LCP | kelch domain containing 2 | protein-coding |
| chr12_69290320_69290951 | chr12 | 69290321 | 69290951 | + | 0.719426 | Intergenic | Intergenic | -6045 | NM_027117 | 69554 | Mm.234368 | NM_027117 | ENSMUSG00000020978 | **Klhdc2** | 2310022K15Rik\|D12Ertd522e\|HCLP-1\|LCP | kelch domain containing 2 | protein-coding |
| chr8_121848492_121848967 | chr8 | 121848493 | 121848967 | + | 1.65188 | Intergenic | Intergenic | -19161 | NM_145605 | 234825 | Mm.209329 | NM_145605 | ENSMUSG00000040263 | **Klhdc4** | AA408426\|AV352552\|BC012312\|G430025P05Rik | kelch domain containing 4 | protein-coding |
| chr1_132264581_132265176 | chr1 | 132264582 | 132265176 | + | 1.17208 | Intergenic | Intergenic | -33747 | NM_144810 | 213417 | Mm.39723 | NM_144810 | ENSMUSG00000042115 | **Klhdc8a** | A630065K24Rik | kelch domain containing 8A | protein-coding |
| chrX_94233533_94233759 | chrX | 94233534 | 94233759 | + | 0.798055 | Intergenic | Intergenic | -1284 | NM_001039059 | 236904 | Mm.115533 | NM_153165 | ENSMUSG00000043929 | **Klhl15** | 6330500C13Rik | kelch-like 15 | protein-coding |
| chr8_64841988_64842337 | chr8 | 64841989 | 64842337 | + | 0.992494 | intron (NM_178633, intron 1 of 14) | intron (NM_178633, intron 1 of 14) | 7761 | NM_178633 | 77113 | Mm.350347 | NM_178633 | ENSMUSG00000031605 | **Klhl2** | 6030411N21Rik\|ABP-KELCH\|AU020744\|Mav | kelch-like 2, Mayven | protein-coding |
| chr8_64841716_64842337 | chr8 | 64841717 | 64842337 | + | 0.86799 | intron (NM_178633, intron 1 of 14) | intron (NM_178633, intron 1 of 14) | 7897 | NM_178633 | 77113 | Mm.350347 | NM_178633 | ENSMUSG00000031605 | **Klhl2** | 6030411N21Rik\|ABP-KELCH\|AU020744\|Mav | kelch-like 2, Mayven | protein-coding |
| chr7_75666958_75667597 | chr7 | 75666959 | 75667597 | + | 1.36364 | intron (NM_029332, intron 10 of 36) | intron (NM_029332, intron 10 of 36) | -181060 | NM_001122780 | 207952 | Mm.206223 | NM_029652 | ENSMUSG00000055652 | **Klhl25** | 2810402K13Rik\|BC027373\|Enc-1 | kelch-like 25 | protein-coding |
| chr7_75681129_75681605 | chr7 | 75681130 | 75681605 | + | 1.18013 | intron (NM_029332, intron 12 of 36) | intron (NM_029332, intron 12 of 36) | -166971 | NM_001122780 | 207952 | Mm.206223 | NM_029652 | ENSMUSG00000055652 | **Klhl25** | 2810402K13Rik\|BC027373\|Enc-1 | kelch-like 25 | protein-coding |
| chr7_75744967_75745511 | chr7 | 75744968 | 75745511 | + | 1.06066 | intron (NM_029332, intron 33 of 36) | intron (NM_029332, intron 33 of 36) | -103099 | NM_001122780 | 207952 | Mm.206223 | NM_029652 | ENSMUSG00000055652 | **Klhl25** | 2810402K13Rik\|BC027373\|Enc-1 | kelch-like 25 | protein-coding |
| chr7_75800283_75800885 | chr7 | 75800284 | 75800885 | + | 0.760965 | Intergenic | RLTR41\|LTR\|ERV1 | -47754 | NM_001122780 | 207952 | Mm.206223 | NM_029652 | ENSMUSG00000055652 | **Klhl25** | 2810402K13Rik\|BC027373\|Enc-1 | kelch-like 25 | protein-coding |
| chr7_75684994_75685369 | chr7 | 75684995 | 75685369 | + | 0.715536 | exon (NM_029332, exon 13 of 37) | exon (NM_029332, exon 13 of 37) | -163156 | NM_001122780 | 207952 | Mm.206223 | NM_029652 | ENSMUSG00000055652 | **Klhl25** | 2810402K13Rik\|BC027373\|Enc-1 | kelch-like 25 | protein-coding |
| chr7_75714896_75715537 | chr7 | 75714897 | 75715537 | + | 0.697507 | intron (NM_029332, intron 17 of 36) | intron (NM_029332, intron 17 of 36) | -133121 | NM_001122780 | 207952 | Mm.206223 | NM_029652 | ENSMUSG00000055652 | **Klhl25** | 2810402K13Rik\|BC027373\|Enc-1 | kelch-like 25 | protein-coding |
| chr9_77592403_77592961 | chr9 | 77592404 | 77592961 | + | 0.788894 | Intergenic | Intergenic | -44050 | NM_172925 | 244923 | Mm.138238 | NM_172925 | ENSMUSG00000044938 | **Klhl31** | 9830147P19Rik\|D930047P17Rik\|Kbtbd1 | kelch-like 31 | protein-coding |
| chr6_147117752_147118194 | chr6 | 147117753 | 147118194 | + | 1.27211 | Intergenic | Intergenic | 26898 | NM_001081237 | 232539 | Mm.491206 | NM_001081237 | ENSMUSG00000040102 | **Klhl42** | A330079N23Rik\|AI450355\|C230080I20Rik\|Ctb9\|Klhdc5\|mKIAA1340 | kelch-like 42 | protein-coding |
| chr5_65119170_65119500 | chr5 | 65119171 | 65119500 | + | 0.858992 | Intergenic | Intergenic | -11896 | NM_175174 | 71778 | Mm.10281 | NM_175174 | ENSMUSG00000054920 | **Klhl5** | 1300013C10Rik | kelch-like 5 | protein-coding |
| chr5_65128381_65128836 | chr5 | 65128382 | 65128836 | + | 0.825684 | Intergenic | Intergenic | -2622 | NM_175174 | 71778 | Mm.10281 | NM_175174 | ENSMUSG00000054920 | **Klhl5** | 1300013C10Rik | kelch-like 5 | protein-coding |
| chr5_65128368_65129033 | chr5 | 65128369 | 65129033 | + | 0.722273 | Intergenic | Intergenic | -2530 | NM_175174 | 71778 | Mm.10281 | NM_175174 | ENSMUSG00000054920 | **Klhl5** | 1300013C10Rik | kelch-like 5 | protein-coding |
| chr16_19969493_19969876 | chr16 | 19969494 | 19969876 | + | 1.07388 | intron (NM_183390, intron 1 of 6) | intron (NM_183390, intron 1 of 6) | 13364 | NM_183390 | 239743 | Mm.86699 | NM_183390 | ENSMUSG00000043008 | **Klhl6** | - | kelch-like 6 | protein-coding |
| chr5_103914343_103914836 | chr5 | 103914344 | 103914836 | + | 0.983984 | Intergenic | Intergenic | -3361 | NM_178741 | 246293 | Mm.179871 | NM_178741 | ENSMUSG00000029312 | **Klhl8** | 2310001P09Rik\|D5Ertd431e | kelch-like 8 | protein-coding |
| chr6_130062787_130063246 | chr6 | 130062788 | 130063246 | + | 1.23332 | intron (NM_010649, intron 3 of 6) | intron (NM_010649, intron 3 of 6) | 4241 | NM_053153 | 93970 | Mm.333431 | NM_053153 | | **Klra18** | Ly49r | killer cell lectin-like receptor, subfamily A, member 18 | protein-coding |
| chr6_131261888_131262570 | chr6 | 131261889 | 131262570 | + | 1.15656 | Intergenic | Intergenic | -14867 | NM_008462 | 16633 | Mm.4783 | NM_008462 | ENSMUSG00000030187 | **Klra2** | Klra30\|Ly49b | killer cell lectin-like receptor, subfamily A, member 2 | protein-coding |
| chr6_128826674_128827346 | chr6 | 128826675 | 128827346 | + | 0.743959 | promoter-TSS (NM_030599) | promoter-TSS (NM_030599) | -695 | NM_030599 | 80782 | Mm.6180 | NM_030599 | ENSMUSG00000079298 | **Klrb1b** | Klrb1d\|Ly55b\|Ly55d\|NKR-P1B\|NKR-P1D\|Nkrp1-b\|Nkrp1d | killer cell lectin-like receptor subfamily B member 1B | protein-coding |
| chr6_128783400_128784017 | chr6 | 128783401 | 128784017 | + | 0.778565 | exon (NM_001159904, exon 5 of 6) | exon (NM_001159904, exon 5 of 6) | 4932 | NM_008527 | 17059 | Mm.480764 | NM_008527 | ENSMUSG00000030325 | **Klrb1c** | AI462337\|CD161\|Ly-59\|Ly55c\|Ly59\|NK-RP1\|NK1.1\|NKR-P1.9\|NKRP1\|NKRP140\|Nk-1\|Nk-1.2\|Nk1\|Nk1.2\|Nkrp1c\|ly-55c | killer cell lectin-like receptor subfamily B member 1C | protein-coding |
| chr6_129687097_129687483 | chr6 | 129687098 | 129687483 | + | 1.37048 | Intergenic | Intergenic | -8317 | NM_010652 | 16641 | Mm.56899 | NM_010652 | ENSMUSG00000030167 | **Klrc1** | CD159a\|NKG2A\|NKG2B | killer cell lectin-like receptor subfamily C, member 1 | protein-coding |
| chr6_129565897_129566281 | chr6 | 129565898 | 129566281 | + | 1.16851 | Intergenic | Intergenic | -12197 | NM_153590 | 243655 | Mm.30968 | NM_153590 | ENSMUSG00000050241 | **Klre1** | Klre-1\|NKG2I | killer cell lectin-like receptor family E member 1 | protein-coding |
| chr6_129583026_129583594 | chr6 | 129583027 | 129583594 | + | 1.04897 | intron (NM_153590, intron 5 of 6) | intron (NM_153590, intron 5 of 6) | 5024 | NM_153590 | 243655 | Mm.30968 | NM_153590 | ENSMUSG00000050241 | **Klre1** | Klre-1\|NKG2I | killer cell lectin-like receptor family E member 1 | protein-coding |
| chr6_129583122_129583454 | chr6 | 129583123 | 129583454 | + | 0.97527 | intron (NM_153590, intron 5 of 6) | intron (NM_153590, intron 5 of 6) | 5002 | NM_153590 | 243655 | Mm.30968 | NM_153590 | ENSMUSG00000050241 | **Klre1** | Klre-1\|NKG2I | killer cell lectin-like receptor family E member 1 | protein-coding |
| chr6_129583933_129584577 | chr6 | 129583934 | 129584577 | + | 0.725702 | intron (NM_153590, intron 6 of 6) | intron (NM_153590, intron 6 of 6) | 5969 | NM_153590 | 243655 | Mm.30968 | NM_153590 | ENSMUSG00000050241 | **Klre1** | Klre-1\|NKG2I | killer cell lectin-like receptor family E member 1 | protein-coding |
| chr6_129557641_129557924 | chr6 | 129557642 | 129557924 | + | 0.708523 | Intergenic | RMER17C\|LTR\|ERVK | -20503 | NM_153590 | 243655 | Mm.30968 | NM_153590 | ENSMUSG00000050241 | **Klre1** | Klre-1\|NKG2I | killer cell lectin-like receptor family E member 1 | protein-coding |
| chr6_122258009_122258341 | chr6 | 122258010 | 122258341 | + | 0.700144 | Intergenic | Intergenic | 24658 | NM_016970 | 50928 | Mm.20434 | NM_016970 | ENSMUSG00000030114 | **Klrg1** | 2F1-Ag\|MAFA\|MAFA-L | killer cell lectin-like receptor subfamily G, member 1 | protein-coding |
| chr6_129742257_129742965 | chr6 | 129742258 | 129742965 | + | 1.13984 | Intergenic | Intergenic | -2127 | NM_177155 | 320407 | Mm.123265 | NM_177155 | ENSMUSG00000043932 | **Klri2** | A530090P03Rik | killer cell lectin-like receptor family I member 2 | protein-coding |
| chr6_129743285_129743475 | chr6 | 129743286 | 129743475 | + | 0.891833 | Intergenic | Intergenic | -2896 | NM_177155 | 320407 | Mm.123265 | NM_177155 | ENSMUSG00000043932 | **Klri2** | A530090P03Rik | killer cell lectin-like receptor family I member 2 | protein-coding |
| chr6_129623736_129624133 | chr6 | 129623737 | 129624133 | + | 1.09406 | promoter-TSS (NM_001083322) | promoter-TSS (NM_001083322) | -71 | NM_001286018 | 27007 | Mm.8217 | NM_033078 | ENSMUSG00000030149 | **Klrk1** | D6H12S2489E\|NKG2-D\|Nkg2d | killer cell lectin-like receptor subfamily K, member 1 | protein-coding |
| chr6_129623547_129624133 | chr6 | 129623548 | 129624133 | + | 1.06718 | promoter-TSS (NM_001083322) | promoter-TSS (NM_001083322) | 24 | NM_001083322 | 27007 | Mm.8217 | NM_033078 | ENSMUSG00000030149 | **Klrk1** | D6H12S2489E\|NKG2-D\|Nkg2d | killer cell lectin-like receptor subfamily K, member 1 | protein-coding |
| chr1_175631488_175632300 | chr1 | 175631489 | 175632300 | + | 1.33067 | promoter-TSS (NM_133809) | promoter-TSS (NM_133809) | -299 | NM_133809 | 98256 | Mm.27217 | NM_133809 | ENSMUSG00000039783 | **Kmo** | AI046660 | kynurenine 3-monooxygenase (kynurenine 3-hydroxylase) | protein-coding |
| chr1_175631743_175632257 | chr1 | 175631744 | 175632257 | + | 1.28717 | promoter-TSS (NM_133809) | promoter-TSS (NM_133809) | -193 | NM_133809 | 98256 | Mm.27217 | NM_133809 | ENSMUSG00000039783 | **Kmo** | AI046660 | kynurenine 3-monooxygenase (kynurenine 3-hydroxylase) | protein-coding |

| | | | | | | | | | | | | | | | | | |
|---|---|---|---|---|---|---|---|---|---|---|---|---|---|---|---|---|---|
| chr9_44868732_44869119 | chr9 | 44868733 | 44869119 | + | 0.770923 | intron (NM_001081049, intron 1 of 35) | intron (NM_001081049, intron 1 of 35) | 12348 | NM_001081049 | 214162 | Mm.2389 | NM_001081049 | ENSMUSG00000002028 | **Kmt2a** | 6430520K01\|ALL-1\|All1\|Cxxc7\|HRX\|HTRX1\|Mll\|Mll1\|mKIAA4050 | lysine (K)-specific methyltransferase 2A | protein-coding |
| chr9_44868397_44869116 | chr9 | 44868398 | 44869116 | + | 0.759936 | intron (NM_001081049, intron 1 of 35) | intron (NM_001081049, intron 1 of 35) | 12517 | NM_001081049 | 214162 | Mm.2389 | NM_001081049 | ENSMUSG00000002028 | **Kmt2a** | 6430520K01\|ALL-1\|All1\|Cxxc7\|HRX\|HTRX1\|Mll\|Mll1\|mKIAA4050 | lysine (K)-specific methyltransferase 2A | protein-coding |
| chr9_44854304_44854655 | chr9 | 44854305 | 44854655 | + | 0.745148 | intron (NM_001081049, intron 1 of 35) | B3\|SINE\|B2 | 26794 | NM_001081049 | 214162 | Mm.2389 | NM_001081049 | ENSMUSG00000002028 | **Kmt2a** | 6430520K01\|ALL-1\|All1\|Cxxc7\|HRX\|HTRX1\|Mll\|Mll1\|mKIAA4050 | lysine (K)-specific methyltransferase 2A | protein-coding |
| chr5_23445090_23445793 | chr5 | 23445091 | 23445793 | + | 0.676252 | intron (NM_026984, intron 1 of 25) | L1MD2\|LINE\|L1 | 11013 | NM_026984 | 69188 | Mm.205190 | NM_026984 | ENSMUSG00000029004 | **Kmt2e** | 1810033J14Rik\|9530077A04Rik\|D230038D11Rik\|Mll5\|NKp44L | lysine (K)-specific methyltransferase 2E | protein-coding |
| chr7_139915823_139916360 | chr7 | 139915824 | 139916360 | + | 1.49792 | intron (NM_177261, intron 9 of 30) | intron (NM_177261, intron 9 of 30) | 21396 | NM_177261 | 76484 | Mm.44442 | NM_177261 | ENSMUSG00000066129 | **Kndc1** | 2410012C07Rik\|B830014K08Rik\|VKIND\|very-kind | kinase non-catalytic C-lobe domain (KIND) containing 1 | protein-coding |
| chr7_139914197_139914713 | chr7 | 139914198 | 139914713 | + | 0.809024 | intron (NM_177261, intron 9 of 30) | intron (NM_177261, intron 9 of 30) | 19759 | NM_177261 | 76484 | Mm.44442 | NM_177261 | ENSMUSG00000066129 | **Kndc1** | 2410012C07Rik\|B830014K08Rik\|VKIND\|very-kind | kinase non-catalytic C-lobe domain (KIND) containing 1 | protein-coding |
| chr7_139914771_139915338 | chr7 | 139914772 | 139915338 | + | 0.698719 | intron (NM_177261, intron 9 of 30) | L1MDa\|LINE\|L1 | 20359 | NM_177261 | 76484 | Mm.44442 | NM_177261 | ENSMUSG00000066129 | **Kndc1** | 2410012C07Rik\|B830014K08Rik\|VKIND\|very-kind | kinase non-catalytic C-lobe domain (KIND) containing 1 | protein-coding |
| chr16_35984486_35984870 | chr16 | 35984487 | 35984870 | + | 0.604894 | intron (NM_008465, intron 1 of 13) | intron (NM_008465, intron 1 of 13) | 1315 | NM_008465 | 16646 | Mm.6952 | NM_008465 | ENSMUSG00000022905 | **Kpna1** | AW494490\|IPOA5\|NPI1\|Rch2\|mSRP1 | karyopherin (importin) alpha 1 | protein-coding |
| chr3_69112901_69113262 | chr3 | 69112902 | 69113262 | + | 0.676143 | intron (NM_008467, intron 1 of 16) | intron (NM_008467, intron 1 of 16) | 14010 | NM_008467 | 16649 | Mm.30601 | NM_008467 | | **Kpna4** | 1110058D08Rik\|IPOA3 | karyopherin (importin) alpha 4 | protein-coding |
| chr3_69108906_69109431 | chr3 | 69108907 | 69109431 | + | 0.646079 | intron (NM_008467, intron 1 of 16) | B1F\|SINE\|Alu | 17923 | NM_008467 | 16649 | Mm.30601 | NM_008467 | | **Kpna4** | 1110058D08Rik\|IPOA3 | karyopherin (importin) alpha 4 | protein-coding |
| chr6_48398457_48398789 | chr6 | 48398458 | 48398789 | + | 0.736067 | intron (NM_133922, intron 1 of 17) | ORR1E\|LTR\|ERVL-MaLR | 3037 | NM_133922 | 77827 | Mm.107646 | NM_133922 | ENSMUSG00000042810 | **Krba1** | A930040G15Rik\|AI448780 | KRAB-A domain containing 1 | protein-coding |
| chr6_71273752_71274011 | chr6 | 71273753 | 71274011 | + | 1.07269 | intron (NM_145568, intron 1 of 3) | intron (NM_145568, intron 1 of 3) | 1863 | NM_145568 | 57896 | Mm.250569 | NM_145568 | ENSMUSG00000053012 | **Krcc1** | AA792894 | lysine-rich coiled-coil 1 | protein-coding |
| chr11_5294910_5295272 | chr11 | 5294911 | 5295272 | + | 1.23458 | intron (NM_001080924, intron 2 of 8) | B1F1\|SINE\|Alu | -33481 | NM_032396 | 84035 | Mm.209989 | NM_032396 | ENSMUSG00000020393 | **Kremen1** | AV002070\|Kremen\|Krm1 | kringle containing transmembrane protein 1 | protein-coding |
| chr11_5294878_5295454 | chr11 | 5294879 | 5295454 | + | 1.03933 | intron (NM_001080924, intron 2 of 8) | B1F1\|SINE\|Alu | -33556 | NM_032396 | 84035 | Mm.209989 | NM_032396 | ENSMUSG00000020393 | **Kremen1** | AV002070\|Kremen\|Krm1 | kringle containing transmembrane protein 1 | protein-coding |
| chr15_101459099_101459958 | chr15 | 101459100 | 101459958 | + | 0.705441 | exon (NM_001166157, exon 9 of 9) | exon (NM_001166157, exon 9 of 9) | 4236 | NM_001166157 | 64818 | Mm.347934 | NM_001166157 | ENSMUSG00000067615 | **Krt81** | AA589625\|Krt2-19 | keratin 81 | protein-coding |
| chr5_31256007_31256363 | chr5 | 31256008 | 31256363 | + | 0.742459 | intron (NM_026298, intron 40 of 47) | intron (NM_026298, intron 40 of 47) | 4479 | NM_027221 | 69815 | Mm.389609 | NM_027221 | ENSMUSG00000029149 | **Krtcap3** | 2010001C09Rik\|AW321088\|Kcp3 | keratinocyte associated protein 3 | protein-coding |
| chr7_30794929_30795309 | chr7 | 30794930 | 30795309 | + | 1.07147 | Intergenic | Intergenic | 7214 | NM_001033131 | 64661 | Mm.268157 | NM_001033131 | ENSMUSG00000074199 | **Krtdap** | Kdap\|sk221 | keratinocyte differentiation associated protein | protein-coding |
| chr11_79129431_79130416 | chr11 | 79129432 | 79130416 | + | 1.08828 | intron (NM_013571, intron 1 of 17) | intron (NM_013571, intron 1 of 17) | 16430 | NM_013571 | 16706 | Mm.4745 | NM_013571 | ENSMUSG00000018334 | **Ksr1** | AW492498\|B-KSR1\|D11Bhm183e\|D11Bhm184e\|Ksr\|RSU2\|hb | kinase suppressor of ras 1 | protein-coding |
| chr11_79131692_79132442 | chr11 | 79131693 | 79132442 | + | 0.838873 | intron (NM_013571, intron 1 of 17) | intron (NM_013571, intron 1 of 17) | 14287 | NM_013571 | 16706 | Mm.4745 | NM_013571 | ENSMUSG00000018334 | **Ksr1** | AW492498\|B-KSR1\|D11Bhm183e\|D11Bhm184e\|Ksr\|RSU2\|hb | kinase suppressor of ras 1 | protein-coding |
| chr11_79131890_79132406 | chr11 | 79131891 | 79132406 | + | 0.786047 | intron (NM_013571, intron 1 of 17) | intron (NM_013571, intron 1 of 17) | 14206 | NM_013571 | 16706 | Mm.4745 | NM_013571 | ENSMUSG00000018334 | **Ksr1** | AW492498\|B-KSR1\|D11Bhm183e\|D11Bhm184e\|Ksr\|RSU2\|hb | kinase suppressor of ras 1 | protein-coding |
| chr14_47664454_47664840 | chr14 | 47664455 | 47664840 | + | 1.35985 | intron (NM_001293636, intron 3 of 39) | intron (NM_001293636, intron 3 of 39) | 891 | NM_008477 | 16709 | Mm.3110 | NM_008477 | ENSMUSG00000021843 | **Ktn1** | - | kinectin 1 | protein-coding |
| chr14_47738969_47739497 | chr14 | 47738970 | 47739497 | + | 1.34637 | Intergenic | Lx7\|LINE\|L1 | 75477 | NM_008477 | 16709 | Mm.3110 | NM_008477 | ENSMUSG00000021843 | **Ktn1** | - | kinectin 1 | protein-coding |
| chr14_47654307_47654515 | chr14 | 47654308 | 47654515 | + | 0.730656 | intron (NM_001293636, intron 2 of 39) | intron (NM_001293636, intron 2 of 39) | 5100 | NM_001293635 | 16709 | Mm.3110 | NM_008477 | ENSMUSG00000021843 | **Ktn1** | - | kinectin 1 | protein-coding |
| chr8_70519128_70519386 | chr8 | 70519129 | 70519386 | + | 0.932568 | intron (NM_029366, intron 2 of 4) | B3\|SINE\|B2 | 3923 | NM_029366 | 75620 | Mm.482129 | NM_029366 | ENSMUSG00000055553 | **Kxd1** | 0610030B01Rik\|2810422J05Rik\|C78305 | KxDL motif containing 1 | protein-coding |
| chr10_26500370_26501050 | chr10 | 26500371 | 26501050 | + | 1.57428 | Intergenic | MLT1J2\|LTR\|ERVL-MaLR | -125525 | NM_172787 | 237339 | Mm.338439 | NM_172787 | ENSMUSG00000039089 | **L3mbtl3** | AI481284\|MBT-1 | l(3)mbt-like 3 (Drosophila) | protein-coding |
| chr10_26359730_26360130 | chr10 | 26359731 | 26360130 | + | 1.00144 | intron (NM_172787, intron 1 of 21) | intron (NM_172787, intron 1 of 21) | 15255 | NM_172787 | 237339 | Mm.338439 | NM_172787 | ENSMUSG00000039089 | **L3mbtl3** | AI481284\|MBT-1 | l(3)mbt-like 3 (Drosophila) | protein-coding |
| chr7_89947382_89948174 | chr7 | 89947383 | 89948174 | + | 1.10162 | Intergenic | ORR1E\|LTR\|ERVL-MaLR | -6574 | NM_001291288 | 67669 | Mm.28679 | NM_026304 | | **l7Rn6** | 0610007P06Rik\|1110002N09Rik\|l(7)6Rn | lethal, Chr 7, Rinchik 6 | protein-coding |
| chr7_89954315_89954645 | chr7 | 89954316 | 89954645 | + | 0.625868 | TTS (NM_021876) | TTS (NM_021876) | -13276 | NM_001291288 | 67669 | Mm.28679 | NM_026304 | | **l7Rn6** | 0610007P06Rik\|1110002N09Rik\|l(7)6Rn | lethal, Chr 7, Rinchik 6 | protein-coding |
| chr9_66988721_66989176 | chr9 | 66988722 | 66989176 | + | 1.30479 | Intergenic | Intergenic | -13465 | NM_030717 | 80907 | Mm.157882 | NM_030717 | ENSMUSG00000032370 | **Lactb** | LACT-1\|Lact1\|Mrpl56 | lactamase, beta | protein-coding |
| chr9_66990043_66990531 | chr9 | 66990044 | 66990531 | + | 1.11861 | Intergenic | Intergenic | -14803 | NM_030717 | 80907 | Mm.157882 | NM_030717 | ENSMUSG00000032370 | **Lactb** | LACT-1\|Lact1\|Mrpl56 | lactamase, beta | protein-coding |
| chr9_66990984_66991285 | chr9 | 66990985 | 66991285 | + | 0.853375 | Intergenic | Intergenic | -15651 | NM_030717 | 80907 | Mm.157882 | NM_030717 | ENSMUSG00000032370 | **Lactb** | LACT-1\|Lact1\|Mrpl56 | lactamase, beta | protein-coding |
| chr1_13639201_13640000 | chr1 | 13639202 | 13640000 | + | 1.53149 | intron (NM_145381, intron 3 of 6) | intron (NM_145381, intron 3 of 6) | 20908 | NM_145381 | 212442 | Mm.89572 | NM_145381 | | **Lactb2** | Cgi-83\|E430032H21Rik | lactamase, beta 2 | protein-coding |
| chr1_135815336_135815732 | chr1 | 135815337 | 135815732 | + | 1.30472 | Intergenic | Intergenic | -3064 | NM_133664 | 16763 | Mm.36726 | NM_133664 | ENSMUSG00000041782 | **Lad1** | - | ladinin | protein-coding |
| chr6_124911374_124912311 | chr6 | 124911375 | 124912311 | + | 1.53928 | promoter-TSS (NM_008479) | promoter-TSS (NM_008479) | -138 | NM_008479 | 16768 | Mm.4528 | NM_008479 | ENSMUSG00000030124 | **Lag3** | CD223\|LAG-3\|Ly66 | lymphocyte-activation gene 3 | protein-coding |
| chr6_124901763_124902511 | chr6 | 124901764 | 124902511 | + | 1.33624 | Intergenic | Intergenic | 9568 | NM_008479 | 16768 | Mm.4528 | NM_008479 | ENSMUSG00000030124 | **Lag3** | CD223\|LAG-3\|Ly66 | lymphocyte-activation gene 3 | protein-coding |
| chr6_124907834_124908152 | chr6 | 124907835 | 124908152 | + | 1.04541 | intron (NM_008479, intron 5 of 7) | intron (NM_008479, intron 5 of 7) | 3712 | NM_008479 | 16768 | Mm.4528 | NM_008479 | ENSMUSG00000030124 | **Lag3** | CD223\|LAG-3\|Ly66 | lymphocyte-activation gene 3 | protein-coding |
| chr7_4087353_4087962 | chr7 | 4087354 | 4087962 | + | 0.871537 | intron (NR_033568, intron 6 of 8) | intron (NR_033568, intron 6 of 8) | -24454 | NM_001113474 | 52855 | Mm.290880 | NM_178611 | ENSMUSG00000055541 | **Lair1** | 5133400O11Rik\|BB115266\|D7Bwg0421e\|Lair-1\|mLair-1 | leukocyte-associated Ig-like receptor 1 | protein-coding |
| chr1_153300348_153300924 | chr1 | 153300349 | 153300924 | + | 1.77751 | intron (NM_010683, intron 1 of 27) | intron (NM_010683, intron 1 of 27) | 32150 | NM_010683 | 226519 | Mm.1249 | NM_010683 | ENSMUSG00000026478 | **Lamc1** | Lamb2 | laminin, gamma 1 | protein-coding |
| chr1_153346839_153347459 | chr1 | 153346840 | 153347459 | + | 0.938889 | Intergenic | A-rich\|Low_complexity\|Low_complexity | -14363 | NM_010683 | 226519 | Mm.1249 | NM_010683 | ENSMUSG00000026478 | **Lamc1** | Lamb2 | laminin, gamma 1 | protein-coding |
| chr1_153129754_153130366 | chr1 | 153129755 | 153130366 | + | 1.35652 | exon (NM_008485, exon 20 of 24) | exon (NM_008485, exon 20 of 24) | 56387 | NM_008485 | 16782 | Mm.329272 | NM_008485 | ENSMUSG00000026479 | **Lamc2** | AA589349 | laminin, gamma 2 | protein-coding |
| chr1_153185129_153185672 | chr1 | 153185130 | 153185672 | + | 0.883794 | intron (NM_008485, intron 2 of 23) | intron (NM_008485, intron 2 of 23) | 1046 | NM_008485 | 16782 | Mm.329272 | NM_008485 | ENSMUSG00000026479 | **Lamc2** | AA589349 | laminin, gamma 2 | protein-coding |
| chr1_153092270_153092901 | chr1 | 153092271 | 153092901 | + | 0.829827 | intron (NM_175460, intron 7 of 10) | intron (NM_175460, intron 7 of 10) | 93861 | NM_008485 | 16782 | Mm.329272 | NM_008485 | ENSMUSG00000026479 | **Lamc2** | AA589349 | laminin, gamma 2 | protein-coding |
| chr8_13162179_13162477 | chr8 | 13162180 | 13162477 | + | 0.76286 | intron (NM_010684, intron 1 of 8) | intron (NM_010684, intron 1 of 8) | 3193 | NM_010684 | 16783 | Mm.16716 | NM_010684 | ENSMUSG00000031447 | **Lamp1** | AI196048\|CD107a\|Lamp-1 | lysosomal-associated membrane protein 1 | protein-coding |
| chrX_38431081_38431615 | chrX | 38431082 | 38431615 | + | 0.80991 | intron (NM_001017959, intron 5 of 8) | intron (NM_001017959, intron 5 of 8) | 25112 | NM_010685 | 16784 | Mm.486 | NM_010685 | ENSMUSG00000016534 | **Lamp2** | CD107b\|LGP-B\|Lamp II\|Lamp-2\|Lamp-2a\|Lamp-2b\|Lamp-2c\|Mac3 | lysosomal-associated membrane protein 2 | protein-coding |
| chrX_9178957_9179404 | chrX | 9178958 | 9179404 | + | 0.827861 | Intergenic | UCON29\|DNA\|PiggyBac? | -20792 | NM_173414 | 236285 | Mm.245119 | NM_173414 | ENSMUSG00000047344 | **Lancl3** | 6030463G20Rik | LanC lantibiotic synthetase component C-like 3 (bacterial) | protein-coding |
| chr12_8854285_8854988 | chr12 | 8854286 | 8854988 | + | 0.892393 | Intergenic | Intergenic | -66670 | NM_008640 | 17775 | Mm.30071 | NM_008640 | ENSMUSG00000020585 | **Laptm4a** | AA286466\|LAPTM4\|MTP\|Mtrp\|mKIAA0108 | lysosomal-associated protein transmembrane 4A | protein-coding |
| chr15_34234688_34234941 | chr15 | 34234689 | 34234941 | + | 0.783433 | Intergenic | RLTR1A2_MM\|LTR\|ERV1 | -3211 | NM_033521 | 114128 | Mm.197518 | NM_033521 | ENSMUSG00000022257 | **Laptm4b** | C330023P13Rik\|LAPTM4beta | lysosomal-associated protein transmembrane 4B | protein-coding |
| chr4_130904725_130905746 | chr4 | 130904726 | 130905746 | + | 0.759194 | Intergenic | MIRc\|SINE\|MIR | -8098 | NM_010686 | 16792 | Mm.271868 | NM_010686 | ENSMUSG00000028581 | **Laptm5** | E3 | lysosomal-associated protein transmembrane 5 | protein-coding |

| | | | | | | | | | | | | | | | | | |
|---|---|---|---|---|---|---|---|---|---|---|---|---|---|---|---|---|---|
| chr11_57966608_57967133 | chr11 | 57966609 | 57967133 | + | 1.04658 | Intergenic | Intergenic | -42193 | NM_028451 | 73158 | Mm.248843 | NM_028451 | ENSMUSG00000037331 | **Larp1** | 1810024J12Rik\|3110040D16Rik\|Larp\|mKIAA0731 | La ribonucleoprotein domain family, member 1 | protein-coding |
| chr11_57993085_57993522 | chr11 | 57993086 | 57993522 | + | 0.978681 | Intergenic | ORR1F\|LTR\|ERVL-MaLR | -15760 | NM_028451 | 73158 | Mm.248843 | NM_028451 | ENSMUSG00000037331 | **Larp1** | 1810024J12Rik\|3110040D16Rik\|Larp\|mKIAA0731 | La ribonucleoprotein domain family, member 1 | protein-coding |
| chr11_57964178_57964393 | chr11 | 57964179 | 57964393 | + | 0.683642 | Intergenic | Intergenic | -44778 | NM_028451 | 73158 | Mm.248843 | NM_028451 | ENSMUSG00000037331 | **Larp1** | 1810024J12Rik\|3110040D16Rik\|Larp\|mKIAA0731 | La ribonucleoprotein domain family, member 1 | protein-coding |
| chr11_57952416_57953159 | chr11 | 57952417 | 57953159 | + | 0.635473 | Intergenic | Intergenic | -56276 | NM_028451 | 73158 | Mm.248843 | NM_028451 | ENSMUSG00000037331 | **Larp1** | 1810024J12Rik\|3110040D16Rik\|Larp\|mKIAA0731 | La ribonucleoprotein domain family, member 1 | protein-coding |
| chr3_40980713_40981106 | chr3 | 40980714 | 40981106 | + | 1.41482 | Intergenic | Intergenic | 30279 | NM_001040399 | 214048 | Mm.46782 | NM_027663 | ENSMUSG00000037814 | **Larp1b** | 1700108L22Rik\|4933421B21Rik\|AA690246\|Larp2 | La ribonucleoprotein domain family, member 1B | protein-coding |
| chr13_9099269_9099598 | chr13 | 9099270 | 9099598 | + | 0.910844 | intron (NM_172585, intron 1 of 17) | intron (NM_172585, intron 1 of 17) | 5529 | NM_172585 | 217980 | Mm.234944 | NM_172585 | ENSMUSG00000033499 | **Larp4b** | A630096F19\|AI256361\|D13Wsu64e\|Larp5 | La ribonucleoprotein domain family, member 4B | protein-coding |
| chr13_9099138_9099631 | chr13 | 9099139 | 9099631 | + | 0.779702 | intron (NM_172585, intron 1 of 17) | intron (NM_172585, intron 1 of 17) | 5480 | NM_172585 | 217980 | Mm.234944 | NM_172585 | ENSMUSG00000033499 | **Larp4b** | A630096F19\|AI256361\|D13Wsu64e\|Larp5 | La ribonucleoprotein domain family, member 4B | protein-coding |
| chr13_9139396_9139924 | chr13 | 9139397 | 9139924 | + | 0.687234 | intron (NM_172585, intron 5 of 17) | L1MCa\|LINE\|L1 | 45755 | NM_172585 | 217980 | Mm.234944 | NM_172585 | ENSMUSG00000033499 | **Larp4b** | A630096F19\|AI256361\|D13Wsu64e\|Larp5 | La ribonucleoprotein domain family, member 4B | protein-coding |
| chr13_9139456_9139850 | chr13 | 9139457 | 9139850 | + | 0.678983 | intron (NM_172585, intron 5 of 17) | L1MCa\|LINE\|L1 | 45748 | NM_172585 | 217980 | Mm.234944 | NM_172585 | ENSMUSG00000033499 | **Larp4b** | A630096F19\|AI256361\|D13Wsu64e\|Larp5 | La ribonucleoprotein domain family, member 4B | protein-coding |
| chr9_123333482_123333976 | chr9 | 123333483 | 123333976 | + | 1.10943 | Intergenic | Intergenic | -33211 | NM_153168 | 102436 | Mm.276076 | NM_153168 | ENSMUSG00000035202 | **Lars2** | AI035546\|Kiaa0028\|LEURS | leucyl-tRNA synthetase, mitochondrial | protein-coding |
| chrX_96020243_96020860 | chrX | 96020244 | 96020860 | + | 0.718975 | Intergenic | Intergenic | -63578 | NM_152822 | 76130 | Mm.274318 | NM_152822 | ENSMUSG00000057421 | **Las1l** | 1810030A06Rik\|5830482G23Rik | LAS1-like (S. cerevisiae) | protein-coding |
| chr5_134612185_134612467 | chr5 | 134612186 | 134612467 | + | 0.654392 | intron (NM_022964, intron 1 of 11) | ZP3AR\|Satellite\|Satellite | 2699 | NM_022964 | 56743 | Mm.391375 | NM_020044 | ENSMUSG00000040751 | **Lat2** | AW125574\|LAB\|NTAL\|WSCR5\|Wbscr15\|Wbscr5 | linker for activation of T cells family, member 2 | protein-coding |
| chr17_72955479_72955846 | chr17 | 72955480 | 72955846 | + | 1.03392 | Intergenic | Intergenic | 37358 | NM_029999 | 77889 | Mm.154109 | NM_029999 | ENSMUSG00000024063 | **Lbh** | 1810009F10Rik\|6720416L16Rik | limb-bud and heart | protein-coding |
| chr17_72955343_72955860 | chr17 | 72955344 | 72955860 | + | 0.883771 | Intergenic | Intergenic | 37297 | NM_029999 | 77889 | Mm.154109 | NM_029999 | ENSMUSG00000024063 | **Lbh** | 1810009F10Rik\|6720416L16Rik | limb-bud and heart | protein-coding |
| chr17_72943993_72944338 | chr17 | 72943994 | 72944338 | + | 0.772049 | Intergenic | L1M4\|LINE\|L1 | 25861 | NM_029999 | 77889 | Mm.154109 | NM_029999 | ENSMUSG00000024063 | **Lbh** | 1810009F10Rik\|6720416L16Rik | limb-bud and heart | protein-coding |
| chr17_72955343_72956100 | chr17 | 72955344 | 72956100 | + | 0.714272 | Intergenic | Intergenic | 37417 | NM_029999 | 77889 | Mm.154109 | NM_029999 | ENSMUSG00000024063 | **Lbh** | 1810009F10Rik\|6720416L16Rik | limb-bud and heart | protein-coding |
| chr17_72946678_72946919 | chr17 | 72946679 | 72946919 | + | 0.659958 | Intergenic | Intergenic | 28494 | NM_029999 | 77889 | Mm.154109 | NM_029999 | ENSMUSG00000024063 | **Lbh** | 1810009F10Rik\|6720416L16Rik | limb-bud and heart | protein-coding |
| chr1_181877042_181877534 | chr1 | 181877043 | 181877534 | + | 0.661753 | Intergenic | RMER1B\|Other\|Other | -34887 | NM_133815 | 98386 | Mm.4538 | NM_133815 | ENSMUSG00000004880 | **Lbr** | AI505894\|ic | lamin B receptor | protein-coding |
| chr17_73109308_73109640 | chr17 | 73109309 | 73109640 | + | 1.1564 | intron (NM_001081071, intron 1 of 5) | B3A\|SINE\|B2 | 1489 | NM_001177967 | 225010 | Mm.329194 | NM_001081071 | ENSMUSG00000054469 | **Lclat1** | AI181996\|Agpat8\|Alcat1\|Gm91\|Lycat | lysocardiolipin acyltransferase 1 | protein-coding |
| chr17_73251218_73251525 | chr17 | 73251219 | 73251525 | + | 0.726972 | Intergenic | Intergenic | 143387 | NM_001177967 | 225010 | Mm.329194 | NM_001081071 | ENSMUSG00000054469 | **Lclat1** | AI181996\|Agpat8\|Alcat1\|Gm91\|Lycat | lysocardiolipin acyltransferase 1 | protein-coding |
| chr19_41495480_41495930 | chr19 | 41495481 | 41495930 | + | 1.24859 | Intergenic | Intergenic | -53934 | NM_172154 | 212391 | Mm.422910 | NM_172154 | ENSMUSG00000025019 | **Lcor** | 3110023F06Rik\|A630025C20Rik\|Mlr2\|mKIAA1795 | ligand dependent nuclear receptor corepressor | protein-coding |
| chr14_75176221_75176452 | chr14 | 75176222 | 75176452 | + | 0.860062 | intron (NM_008879, intron 1 of 16) | intron (NM_008879, intron 1 of 16) | 39912 | NM_008879 | 18826 | Mm.153911 | NM_008879 | ENSMUSG00000021998 | **Lcp1** | AW536232\|D14Ertd310e\|LCP-1\|Pls2\|pp65 | lymphocyte cytosolic protein 1 | protein-coding |
| chr14_75182475_75182744 | chr14 | 75182476 | 75182744 | + | 0.708865 | intron (NM_008879, intron 2 of 16) | intron (NM_008879, intron 2 of 16) | 46185 | NM_008879 | 18826 | Mm.153911 | NM_008879 | ENSMUSG00000021998 | **Lcp1** | AW536232\|D14Ertd310e\|LCP-1\|Pls2\|pp65 | lymphocyte cytosolic protein 1 | protein-coding |
| chr11_34111733_34112540 | chr11 | 34111734 | 34112540 | + | 1.80486 | Intergenic | Intergenic | 64936 | NM_010696 | 16822 | Mm.265350 | NM_010696 | ENSMUSG00000002699 | **Lcp2** | AI323664\|BB161688\|SLP-76\|SLP76\|twm | lymphocyte cytosolic protein 2 | protein-coding |
| chr11_34034801_34035099 | chr11 | 34034802 | 34035099 | + | 1.2454 | Intergenic | MIRc\|SINE\|MIR | -12251 | NM_010696 | 16822 | Mm.265350 | NM_010696 | ENSMUSG00000002699 | **Lcp2** | AI323664\|BB161688\|SLP-76\|SLP76\|twm | lymphocyte cytosolic protein 2 | protein-coding |
| chr11_34036341_34036689 | chr11 | 34036342 | 34036689 | + | 1.24109 | Intergenic | Intergenic | -10686 | NM_010696 | 16822 | Mm.265350 | NM_010696 | ENSMUSG00000002699 | **Lcp2** | AI323664\|BB161688\|SLP-76\|SLP76\|twm | lymphocyte cytosolic protein 2 | protein-coding |
| chr11_34101420_34102296 | chr11 | 34101421 | 34102296 | + | 1.047 | Intergenic | Intergenic | 54657 | NM_010696 | 16822 | Mm.265350 | NM_010696 | ENSMUSG00000002699 | **Lcp2** | AI323664\|BB161688\|SLP-76\|SLP76\|twm | lymphocyte cytosolic protein 2 | protein-coding |
| chr11_34074441_34074899 | chr11 | 34074442 | 34074899 | + | 0.909968 | intron (NM_010696, intron 8 of 20) | intron (NM_010696, intron 8 of 20) | 27469 | NM_010696 | 16822 | Mm.265350 | NM_010696 | ENSMUSG00000002699 | **Lcp2** | AI323664\|BB161688\|SLP-76\|SLP76\|twm | lymphocyte cytosolic protein 2 | protein-coding |
| chr11_34022731_34023166 | chr11 | 34022732 | 34023166 | + | 0.904207 | Intergenic | MTD\|LTR\|ERVL-MaLR | -24252 | NM_010696 | 16822 | Mm.265350 | NM_010696 | ENSMUSG00000002699 | **Lcp2** | AI323664\|BB161688\|SLP-76\|SLP76\|twm | lymphocyte cytosolic protein 2 | protein-coding |
| chr11_34086180_34086524 | chr11 | 34086181 | 34086524 | + | 0.827581 | intron (NM_010696, intron 17 of 20) | intron (NM_010696, intron 17 of 20) | 39151 | NM_010696 | 16822 | Mm.265350 | NM_010696 | ENSMUSG00000002699 | **Lcp2** | AI323664\|BB161688\|SLP-76\|SLP76\|twm | lymphocyte cytosolic protein 2 | protein-coding |
| chr11_34033912_34035119 | chr11 | 34033913 | 34035119 | + | 0.719729 | Intergenic | L2\|LINE\|L2 | -12685 | NM_010696 | 16822 | Mm.265350 | NM_010696 | ENSMUSG00000002699 | **Lcp2** | AI323664\|BB161688\|SLP-76\|SLP76\|twm | lymphocyte cytosolic protein 2 | protein-coding |
| chr11_34085829_34086525 | chr11 | 34085830 | 34086525 | + | 0.680244 | intron (NM_010696, intron 17 of 20) | intron (NM_010696, intron 17 of 20) | 38976 | NM_010696 | 16822 | Mm.265350 | NM_010696 | ENSMUSG00000002699 | **Lcp2** | AI323664\|BB161688\|SLP-76\|SLP76\|twm | lymphocyte cytosolic protein 2 | protein-coding |
| chr11_34048437_34049295 | chr11 | 34048438 | 34049295 | + | 0.676817 | intron (NM_010696, intron 1 of 20) | MIRb\|SINE\|MIR | 1665 | NM_010696 | 16822 | Mm.265350 | NM_010696 | ENSMUSG00000002699 | **Lcp2** | AI323664\|BB161688\|SLP-76\|SLP76\|twm | lymphocyte cytosolic protein 2 | protein-coding |
| chr11_34050069_34050395 | chr11 | 34050070 | 34050395 | + | 0.619834 | intron (NM_010696, intron 1 of 20) | intron (NM_010696, intron 1 of 20) | 3031 | NM_010696 | 16822 | Mm.265350 | NM_010696 | ENSMUSG00000002699 | **Lcp2** | AI323664\|BB161688\|SLP-76\|SLP76\|twm | lymphocyte cytosolic protein 2 | protein-coding |
| chr1_128328717_128329121 | chr1 | 128328718 | 128329121 | + | 0.925825 | promoter-TSS (NM_001081078) | promoter-TSS (NM_001081078) | -601 | NM_001081078 | 226413 | Mm.26930 | NM_001081078 | ENSMUSG00000026354 | **Lct** | Gm100\|LAC\|LPH\|Lphl | lactase | protein-coding |
| chr1_128328671_128329221 | chr1 | 128328672 | 128329221 | + | 0.744452 | promoter-TSS (NM_001081078) | promoter-TSS (NM_001081078) | -628 | NM_001081078 | 226413 | Mm.26930 | NM_001081078 | ENSMUSG00000026354 | **Lct** | Gm100\|LAC\|LPH\|Lphl | lactase | protein-coding |
| chr17_5255420_5255872 | chr17 | 5255421 | 5255872 | + | 0.740115 | intron (NM_001085355, intron 5 of 19) | intron (NM_001085355, intron 5 of 19) | 163121 | NM_175349 | 106557 | Mm.432764 | NM_175349 | | **Ldhal6b** | 4933402O15Rik\|AI326310\|LDHL\|Ldhal | lactate dehydrogenase A-like 6B | protein-coding |
| chr2_102117021_102117898 | chr2 | 102117022 | 102117898 | + | 0.723949 | intron (NM_178886, intron 1 of 5) | intron (NM_178886, intron 1 of 5) | 69000 | NM_178886 | 241576 | Mm.489740 | NM_178886 | ENSMUSG00000048058 | **Ldlrad3** | 6430500P08\|AI194318\|Lrad3 | low density lipoprotein receptor class A domain containing 3 | protein-coding |
| chr18_67945470_67946122 | chr18 | 67945471 | 67946122 | + | 1.54149 | intron (NM_172631, intron 1 of 5) | intron (NM_172631, intron 1 of 5) | 12539 | NM_172631 | 52662 | Mm.440183 | NM_172631 | ENSMUSG00000024544 | **Ldlrad4** | 8230401C20Rik\|A430083H02\|A430108L08Rik\|C18orf1\|D18Ertd653e\|D330030L18Rik | low density lipoprotein receptor class A domain containing 4 | protein-coding |
| chr18_67973479_67974015 | chr18 | 67973480 | 67974015 | + | 1.22985 | intron (NM_172631, intron 1 of 5) | intron (NM_172631, intron 1 of 5) | 40490 | NM_172631 | 52662 | Mm.440183 | NM_172631 | ENSMUSG00000024544 | **Ldlrad4** | 8230401C20Rik\|A430083H02\|A430108L08Rik\|C18orf1\|D18Ertd653e\|D330030L18Rik | low density lipoprotein receptor class A domain containing 4 | protein-coding |
| chr3_131169174_131169680 | chr3 | 131169175 | 131169680 | + | 1.32503 | intron (NM_001276402, intron 3 of 9) | intron (NM_001276402, intron 3 of 9) | 56636 | NM_001276403 | 16842 | Mm.255219 | NM_010703 | ENSMUSG00000027985 | **Lef1** | 3000002B05\|AI451430\|Lef-1 | lymphoid enhancer binding factor 1 | protein-coding |
| chr3_131168936_131169728 | chr3 | 131168937 | 131169728 | + | 1.29962 | intron (NM_001276402, intron 3 of 9) | intron (NM_001276402, intron 3 of 9) | 56541 | NM_001276403 | 16842 | Mm.255219 | NM_010703 | ENSMUSG00000027985 | **Lef1** | 3000002B05\|AI451430\|Lef-1 | lymphoid enhancer binding factor 1 | protein-coding |
| chr3_130923413_130924099 | chr3 | 130923414 | 130924099 | + | 0.82217 | Intergenic | Intergenic | -186541 | NM_010703 | 16842 | Mm.255219 | NM_010703 | ENSMUSG00000027985 | **Lef1** | 3000002B05\|AI451430\|Lef-1 | lymphoid enhancer binding factor 1 | protein-coding |
| chr3_131143527_131143869 | chr3 | 131143528 | 131143869 | + | 0.674671 | intron (NM_001276402, intron 3 of 9) | intron (NM_001276402, intron 3 of 9) | 30907 | NM_001276403 | 16842 | Mm.255219 | NM_010703 | ENSMUSG00000027985 | **Lef1** | 3000002B05\|AI451430\|Lef-1 | lymphoid enhancer binding factor 1 | protein-coding |
| chr3_131085161_131085621 | chr3 | 131085162 | 131085621 | + | 0.666609 | Intergenic | Intergenic | -24906 | NM_010703 | 16842 | Mm.255219 | NM_010703 | ENSMUSG00000027985 | **Lef1** | 3000002B05\|AI451430\|Lef-1 | lymphoid enhancer binding factor 1 | protein-coding |
| chr3_131084932_131085622 | chr3 | 131084933 | 131085622 | + | 0.647777 | Intergenic | Intergenic | -25020 | NM_010703 | 16842 | Mm.255219 | NM_010703 | ENSMUSG00000027985 | **Lef1** | 3000002B05\|AI451430\|Lef-1 | lymphoid enhancer binding factor 1 | protein-coding |
| chr3_65793327_65794178 | chr3 | 65793328 | 65794178 | + | 1.0168 | intron (NM_001166659, intron 9 of 12) | ORR1B1-int\|LTR\|ERVL-MaLR | 127525 | NM_001166659 | 624866 | Mm.296317 | NM_001037923 | ENSMUSG00000074579 | **Lekr1** | EG546798\|EG624866\|Gm6534 | leucine, glutamate and lysine rich 1 | protein-coding |
| chr9_75425824_75426269 | chr9 | 75425825 | 75426269 | + | 0.968704 | Intergenic | Intergenic | -15477 | NM_001039522 | 235497 | Mm.41508 | NM_001006122 | ENSMUSG00000042487 | **Leo1** | Gm185 | Leo1, Paf1/RNA polymerase II complex component, homolog (S. cerevisiae) | protein-coding |
| chr16_25970157_25970607 | chr16 | 25970158 | 25970607 | + | 1.16333 | intron (NM_173379, intron 11 of 14) | intron (NM_173379, intron 11 of 14) | 135402 | NM_173379 | 210530 | Mm.326869 | NM_173379 | | **Leprel1** | 4832416N06\|AW553532\|Mlat4\|P3H2 | leprecan-like 1 | protein-coding |
| chr8_34139899_34140213 | chr8 | 34139900 | 34140213 | + | 1.20163 | intron (NM_026609, intron 2 of 3) | intron (NM_026609, intron 2 of 3) | 6683 | NM_026609 | 68192 | Mm.34212 | NM_026609 | ENSMUSG00000031513 | **Leprotl1** | 1110067H13Rik\|1520402O14Rik\|AI854296 | leptin receptor overlapping transcript-like 1 | protein-coding |
| chr14_47357953_47358315 | chr14 | 47357954 | 47358315 | + | 0.823865 | Intergenic | MTD\|LTR\|ERVL-MaLR | -15726 | NM_010705 | 16854 | Mm.248615 | NM_010705 | ENSMUSG00000050335 | **Lgals3** | GBP\|L-34\|Mac-2\|gal3 | lectin, galactose binding, soluble 3 | protein-coding |
| chr14_47369777_47370317 | chr14 | 47369778 | 47370317 | + | 0.806137 | Intergenic | Intergenic | -3813 | NM_010705 | 16854 | Mm.248615 | NM_010705 | ENSMUSG00000050335 | **Lgals3** | GBP\|L-34\|Mac-2\|gal3 | lectin, galactose binding, soluble 3 | protein-coding |
| chr14_47371678_47372024 | chr14 | 47371679 | 47372024 | + | 0.789083 | Intergenic | Intergenic | -2009 | NM_010705 | 16854 | Mm.248615 | NM_010705 | ENSMUSG00000050335 | **Lgals3** | GBP\|L-34\|Mac-2\|gal3 | lectin, galactose binding, soluble 3 | protein-coding |
| chr11_118404496_118404857 | chr11 | 118404497 | 118404857 | + | 1.28689 | Intergenic | Intergenic | -2746 | NM_011150 | 19039 | Mm.3152 | NM_011150 | ENSMUSG00000033880 | **Lgals3bp** | 90K\|CyCAP\|MAC-2BP\|Ppicap\|Tango10b | lectin, galactoside-binding, soluble, 3 binding protein | protein-coding |

| | | | | | | | | | | | | | | | | | |
|---|---|---|---|---|---|---|---|---|---|---|---|---|---|---|---|---|---|
| chr11_118404088_118404848 | chr11 | 118404089 | 118404848 | + | 1.06094 | Intergenic | URR1A\|DNA\|hAT-Charlie | -2537 | NM_011150 | 19039 | Mm.3152 | NM_011150 | ENSMUSG00000033880 | **Lgals3bp** | 90K\|CyCAP\|MAC-2BP\|Ppicap\|Tango10b | lectin, galactoside-binding, soluble, 3 binding protein | protein-coding |
| chr13_94116436_94117238 | chr13 | 94116437 | 94117238 | + | 1.40445 | intron (NM_172589, intron 1 of 3) | intron (NM_172589, intron 1 of 3) | 59041 | NM_172589 | 218454 | Mm.316553 | NM_172589 | ENSMUSG00000045312 | **Lhfpl2** | 6030465B15\|AI447312\|AW050335\|mKIAA0206\|vgim | lipoma HMGIC fusion partner-like 2 | protein-coding |
| chr13_94098186_94099076 | chr13 | 94098187 | 94099076 | + | 1.23282 | intron (NM_172589, intron 1 of 3) | intron (NM_172589, intron 1 of 3) | 40835 | NM_172589 | 218454 | Mm.316553 | NM_172589 | ENSMUSG00000045312 | **Lhfpl2** | 6030465B15\|AI447312\|AW050335\|mKIAA0206\|vgim | lipoma HMGIC fusion partner-like 2 | protein-coding |
| chr13_94153561_94154066 | chr13 | 94153562 | 94154066 | + | 1.11945 | intron (NM_172589, intron 2 of 3) | intron (NM_172589, intron 2 of 3) | 96018 | NM_172589 | 218454 | Mm.316553 | NM_172589 | ENSMUSG00000045312 | **Lhfpl2** | 6030465B15\|AI447312\|AW050335\|mKIAA0206\|vgim | lipoma HMGIC fusion partner-like 2 | protein-coding |
| chr13_94154447_94154796 | chr13 | 94154448 | 94154796 | + | 0.802276 | intron (NM_172589, intron 2 of 3) | intron (NM_172589, intron 2 of 3) | 96826 | NM_172589 | 218454 | Mm.316553 | NM_172589 | ENSMUSG00000045312 | **Lhfpl2** | 6030465B15\|AI447312\|AW050335\|mKIAA0206\|vgim | lipoma HMGIC fusion partner-like 2 | protein-coding |
| chr6_113166619_113167343 | chr6 | 113166620 | 113167343 | + | 1.22303 | Intergenic | Intergenic | 28403 | NM_177763 | 269788 | Mm.37943 | NM_177763 | ENSMUSG00000042873 | **Lhfpl4** | 1190004M23Rik\|AI604880\|B230384L07\|mKIAA4027 | lipoma HMGIC fusion partner-like protein 4 | protein-coding |
| chr7_132672486_132673074 | chr7 | 132672487 | 132673074 | + | 0.959705 | intron (NM_029609, intron 6 of 6) | intron (NM_029609, intron 6 of 6) | 62137 | NM_029609 | 76429 | Mm.276721 | NM_029609 | ENSMUSG00000030946 | **Lhpp** | 2310007H09Rik | phospholysine phosphohistidine inorganic pyrophosphate phosphatase | protein-coding |
| chr11_4339286_4339984 | chr11 | 4339287 | 4339984 | + | 1.17526 | Intergenic | Intergenic | 72839 | NM_001039537 | 16878 | Mm.4964 | NM_008501 | ENSMUSG00000034394 | **Lif** | - | leukemia inhibitory factor | protein-coding |
| chr11_4268457_4268888 | chr11 | 4268458 | 4268888 | + | 1.01473 | intron (NM_001039537, intron 2 of 2) | intron (NM_001039537, intron 2 of 2) | 1877 | NM_001039537 | 16878 | Mm.4964 | NM_008501 | ENSMUSG00000034394 | **Lif** | - | leukemia inhibitory factor | protein-coding |
| chr11_4336319_4337116 | chr11 | 4336320 | 4337116 | + | 0.869331 | Intergenic | Intergenic | 69922 | NM_001039537 | 16878 | Mm.4964 | NM_008501 | ENSMUSG00000034394 | **Lif** | - | leukemia inhibitory factor | protein-coding |
| chr11_4268236_4268956 | chr11 | 4268237 | 4268956 | + | 0.800564 | intron (NM_001039537, intron 2 of 2) | intron (NM_001039537, intron 2 of 2) | 1800 | NM_001039537 | 16878 | Mm.4964 | NM_008501 | ENSMUSG00000034394 | **Lif** | - | leukemia inhibitory factor | protein-coding |
| chr11_4319051_4319315 | chr11 | 4319052 | 4319315 | + | 0.70956 | Intergenic | Intergenic | 52387 | NM_001039537 | 16878 | Mm.4964 | NM_008501 | ENSMUSG00000034394 | **Lif** | - | leukemia inhibitory factor | protein-coding |
| chr11_4262992_4263205 | chr11 | 4262993 | 4263205 | + | 0.705185 | intron (NM_008501, intron 1 of 3) | intron (NM_008501, intron 1 of 3) | -3697 | NM_001039537 | 16878 | Mm.4964 | NM_008501 | ENSMUSG00000034394 | **Lif** | - | leukemia inhibitory factor | protein-coding |
| chr11_4341094_4341336 | chr11 | 4341095 | 4341336 | + | 0.621915 | Intergenic | Lx5b\|LINE\|L1 | 74419 | NM_001039537 | 16878 | Mm.4964 | NM_008501 | ENSMUSG00000034394 | **Lif** | - | leukemia inhibitory factor | protein-coding |
| chr15_7216858_7217457 | chr15 | 7216859 | 7217457 | + | 0.768976 | intron (NM_001289498, intron 20 of 22) | intron (NM_001289498, intron 20 of 22) | 76616 | NM_013584 | 16880 | Mm.149720 | NM_013584 | ENSMUSG00000054263 | **Lifr** | A230075M04Rik\|AW061234\|LIF | leukemia inhibitory factor receptor | protein-coding |
| chr15_7216641_7217510 | chr15 | 7216642 | 7217510 | + | 0.706092 | intron (NM_001289498, intron 20 of 22) | intron (NM_001289498, intron 20 of 22) | 76534 | NM_013584 | 16880 | Mm.149720 | NM_013584 | ENSMUSG00000054263 | **Lifr** | A230075M04Rik\|AW061234\|LIF | leukemia inhibitory factor receptor | protein-coding |
| chr15_7256416_7256972 | chr15 | 7256417 | 7256972 | + | 0.693426 | intron (NM_001289498, intron 6 of 22) | intron (NM_001289498, intron 6 of 22) | 116152 | NM_013584 | 16880 | Mm.149720 | NM_013584 | ENSMUSG00000054263 | **Lifr** | A230075M04Rik\|AW061234\|LIF | leukemia inhibitory factor receptor | protein-coding |
| chr2_181381939_181382387 | chr2 | 181381940 | 181382387 | + | 0.674383 | 5' UTR (NM_023684, exon 1 of 5) | 5' UTR (NM_023684, exon 1 of 5) | 928 | NM_023684 | 72699 | Mm.440138 | NM_023684 | ENSMUSG00000090077 | **Lime1** | 2810038K19Rik\|LIME | Lck interacting transmembrane adaptor 1 | protein-coding |
| chr11_3402942_3403452 | chr11 | 3402943 | 3403452 | + | 0.859347 | intron (NM_010718, intron 1 of 15) | intron (NM_010718, intron 1 of 15) | 6041 | NM_010718 | 16886 | Mm.124176 | NM_010718 | ENSMUSG00000020451 | **Limk2** | Limk-2 | LIM motif-containing protein kinase 2 | protein-coding |
| chr11_3399479_3399796 | chr11 | 3399480 | 3399796 | + | 0.633878 | intron (NM_010718, intron 1 of 15) | intron (NM_010718, intron 1 of 15) | 9600 | NM_010718 | 16886 | Mm.124176 | NM_010718 | ENSMUSG00000020451 | **Limk2** | Limk-2 | LIM motif-containing protein kinase 2 | protein-coding |
| chr10_58333419_58333766 | chr10 | 58333420 | 58333766 | + | 0.838059 | intron (NM_026148, intron 1 of 9) | intron (NM_026148, intron 1 of 9) | 10127 | NM_026148 | 110829 | Mm.57734 | NM_026148 | ENSMUSG00000019920 | **Lims1** | 2310016J22Rik\|4921524A02Rik\|AI507642\|AU021743\|AW551584\|C430041B13Rik\|Lims1l\|PINCH\|PINCH1 | LIM and senescent cell antigen-like domains 1 | protein-coding |
| chr10_58333363_58333901 | chr10 | 58333364 | 58333901 | + | 0.718402 | intron (NM_026148, intron 1 of 9) | intron (NM_026148, intron 1 of 9) | 10166 | NM_026148 | 110829 | Mm.57734 | NM_026148 | ENSMUSG00000019920 | **Lims1** | 2310016J22Rik\|4921524A02Rik\|AI507642\|AU021743\|AW551584\|C430041B13Rik\|Lims1l\|PINCH\|PINCH1 | LIM and senescent cell antigen-like domains 1 | protein-coding |
| chr5_100505838_100506321 | chr5 | 100505839 | 100506321 | + | 1.37367 | Intergenic | MTB\|LTR\|ERVL-MaLR | -5441 | NM_172714 | 231506 | Mm.212568 | NM_172714 | ENSMUSG00000035310 | **Lin54** | 4832414L08\|AI461788\|mKIAA2037 | lin-54 homolog (C. elegans) | protein-coding |
| chr13_97395089_97395686 | chr13 | 97395090 | 97395686 | + | 0.647181 | Intergenic | Intergenic | -39582 | NR_110432 | 100039691 | Mm.219358 | NR_110430 | | **Lincenc1** | ENSMUSG00000078952\|Gm11019\|Gm2373\|linc-Enc1 | long intergenic non-coding RNA, embryonic stem cells expressed 1 | ncRNA |
| chr6_31169339_31169880 | chr6 | 31169340 | 31169880 | + | 1.44433 | intron (NR_110470, intron 2 of 2) | intron (NR_110470, intron 2 of 2) | 48864 | NR_110470 | 232685 | Mm.443019 | NM_144555 | | **Lincpint** | MNCb-1768\|Pint\|linc-Pint | long intergenic non-protein coding RNA, Trp53 induced transcript | ncRNA |
| chr6_31150428_31150900 | chr6 | 31150429 | 31150900 | + | 0.779948 | intron (NR_110471, intron 2 of 3) | B1F\|SINE\|Alu | 67810 | NR_110470 | 232685 | Mm.443019 | NM_144555 | | **Lincpint** | MNCb-1768\|Pint\|linc-Pint | long intergenic non-protein coding RNA, Trp53 induced transcript | ncRNA |
| chr9_70942589_70943112 | chr9 | 70942590 | 70943112 | + | 0.622468 | Intergenic | Intergenic | -8043 | NM_008280 | 15450 | Mm.390187 | NM_008280 | ENSMUSG00000032207 | **Lipc** | AI256194\|HL\|Hpl | lipase, hepatic | protein-coding |
| chr7_25386114_25386796 | chr7 | 25386115 | 25386796 | + | 1.31974 | TTS (NR_073028) | TTS (NR_073028) | 3657 | NM_001039507 | 16890 | Mm.158548 | NM_010719 | ENSMUSG00000003123 | **Lipe** | 4933403G17Rik\|HSL | lipase, hormone sensitive | protein-coding |
| chr7_25385816_25386789 | chr7 | 25385817 | 25386789 | + | 1.25005 | TTS (NR_073028) | TTS (NR_073028) | 3809 | NM_001039507 | 16890 | Mm.158548 | NM_010719 | ENSMUSG00000003123 | **Lipe** | 4933403G17Rik\|HSL | lipase, hormone sensitive | protein-coding |
| chr18_74910957_74911164 | chr18 | 74910958 | 74911164 | + | 1.14397 | Intergenic | L1M4\|LINE\|L1 | 50202 | NM_010720 | 16891 | Mm.299647 | NM_010720 | ENSMUSG00000053846 | **Lipg** | 3110013K01Rik\|EL\|lipase\|mEDL | lipase, endothelial | protein-coding |
| chr18_74910696_74911219 | chr18 | 74910697 | 74911219 | + | 1.1028 | Intergenic | Intergenic | 50305 | NM_010720 | 16891 | Mm.299647 | NM_010720 | ENSMUSG00000053846 | **Lipg** | 3110013K01Rik\|EL\|lipase\|mEDL | lipase, endothelial | protein-coding |
| chr18_74943048_74943905 | chr18 | 74943049 | 74943905 | + | 0.760616 | intron (NM_010720, intron 8 of 9) | intron (NM_010720, intron 8 of 9) | 17786 | NM_010720 | 16891 | Mm.299647 | NM_010720 | ENSMUSG00000053846 | **Lipg** | 3110013K01Rik\|EL\|lipase\|mEDL | lipase, endothelial | protein-coding |
| chr19_34119505_34120536 | chr19 | 34119506 | 34120536 | + | 1.43229 | intron (NM_023903, intron 9 of 9) | intron (NM_023903, intron 9 of 9) | 19078 | NM_023903 | 78753 | Mm.195937 | NM_023903 | ENSMUSG00000056078 | **Lipm** | 4632427C23Rik\|AW227514\|Lipl3 | lipase, family member M | protein-coding |
| chr16_10958534_10959408 | chr16 | 10958535 | 10959408 | + | 0.618561 | TTS (NM_019980) | TTS (NM_019980) | 34150 | NM_019980 | 56722 | Mm.294753 | NM_019980 | ENSMUSG00000022500 | **Litaf** | 3222402J11Rik\|C85531\|N4WBP3\|TBX1 | LPS-induced TN factor | protein-coding |
| chr17_17398701_17399189 | chr17 | 17398702 | 17399189 | + | 0.765339 | Intergenic | Intergenic | -3741 | NM_025681 | 66643 | Mm.268018 | NM_025681 | ENSMUSG00000047786 | **Lix1** | 5730466L18Rik | limb expression 1 homolog (chicken) | protein-coding |
| chr17_17397290_17397596 | chr17 | 17397291 | 17397596 | + | 0.719171 | Intergenic | Intergenic | -5243 | NM_025681 | 66643 | Mm.268018 | NM_025681 | ENSMUSG00000047786 | **Lix1** | 5730466L18Rik | limb expression 1 homolog (chicken) | protein-coding |
| chr17_17405681_17406052 | chr17 | 17405682 | 17406052 | + | 0.694888 | intron (NM_025681, intron 1 of 5) | intron (NM_025681, intron 1 of 5) | 3181 | NM_025681 | 66643 | Mm.268018 | NM_025681 | ENSMUSG00000047786 | **Lix1** | 5730466L18Rik | limb expression 1 homolog (chicken) | protein-coding |
| chr3_88491027_88491526 | chr3 | 88491028 | 88491526 | + | 0.856795 | intron (NM_001002011, intron 1 of 11) | intron (NM_001002011, intron 1 of 11) | 2035 | NM_019390 | 16905 | Mm.243014 | NM_019390 | ENSMUSG00000028063 | **Lmna** | Dhe | lamin A | protein-coding |
| chr18_56685053_56685605 | chr18 | 56685054 | 56685605 | + | 1.02232 | Intergenic | MER34A\|LTR\|ERV1 | -22484 | NM_010721 | 16906 | Mm.4105 | NM_010721 | ENSMUSG00000024590 | **Lmnb1** | - | lamin B1 | protein-coding |
| chr2_103938612_103938959 | chr2 | 103938613 | 103938959 | + | 1.2981 | Intergenic | Intergenic | -19209 | NM_001142337 | 16909 | Mm.29266 | NM_008505 | ENSMUSG00000032698 | **Lmo2** | Rbtn-2\|Rbtn2\|Rhom-2\|Ttg2 | LIM domain only 2 | protein-coding |
| chr2_103938611_103939267 | chr2 | 103938612 | 103939267 | + | 1.22328 | Intergenic | Intergenic | -19056 | NM_001142337 | 16909 | Mm.29266 | NM_008505 | ENSMUSG00000032698 | **Lmo2** | Rbtn-2\|Rbtn2\|Rhom-2\|Ttg2 | LIM domain only 2 | protein-coding |
| chr2_103917241_103917858 | chr2 | 103917242 | 103917858 | + | 0.972969 | Intergenic | L1MC1\|LINE\|L1 | -40445 | NM_001142337 | 16909 | Mm.29266 | NM_008505 | ENSMUSG00000032698 | **Lmo2** | Rbtn-2\|Rbtn2\|Rhom-2\|Ttg2 | LIM domain only 2 | protein-coding |
| chr3_144379867_144380201 | chr3 | 144379868 | 144380201 | + | 0.798625 | Intergenic | Intergenic | -174779 | NM_010723 | 16911 | Mm.29187 | NM_010723 | ENSMUSG00000028266 | **Lmo4** | A730077C12Rik\|Crp3\|Etohi4 | LIM domain only 4 | protein-coding |
| chr14_101879040_101879592 | chr14 | 101879041 | 101879592 | + | 0.867331 | intron (NM_201529, intron 6 of 30) | intron (NM_201529, intron 6 of 30) | 149388 | NM_201529 | 380928 | Mm.218981 | NM_201529 | ENSMUSG00000033060 | **Lmo7** | C78582\|FBXO20\|Gm914\|mKIAA0858 | LIM domain only 7 | protein-coding |
| chr14_101878503_101879593 | chr14 | 101878504 | 101879593 | + | 0.809801 | intron (NM_201529, intron 6 of 30) | intron (NM_201529, intron 6 of 30) | 149120 | NM_201529 | 380928 | Mm.218981 | NM_201529 | ENSMUSG00000033060 | **Lmo7** | C78582\|FBXO20\|Gm914\|mKIAA0858 | LIM domain only 7 | protein-coding |
| chr6_97315958_97316288 | chr6 | 97315959 | 97316288 | + | 1.33802 | intron (NM_145148, intron 12 of 23) | intron (NM_145148, intron 12 of 23) | -63343 | NM_001081157 | 320502 | Mm.68163 | NM_001081157 | ENSMUSG00000044086 | **Lmod3** | 5430424A14Rik | leiomodin 3 (fetal) | protein-coding |
| chr6_97293305_97293959 | chr6 | 97293306 | 97293959 | + | 1.17185 | intron (NM_145148, intron 22 of 23) | MIRb\|SINE\|MIR | -40852 | NM_001081157 | 320502 | Mm.68163 | NM_001081157 | ENSMUSG00000044086 | **Lmod3** | 5430424A14Rik | leiomodin 3 (fetal) | protein-coding |
| chr6_97430258_97430665 | chr6 | 97430259 | 97430665 | + | 0.86354 | intron (NM_145148, intron 2 of 23) | intron (NM_145148, intron 2 of 23) | -177682 | NM_001081157 | 320502 | Mm.68163 | NM_001081157 | ENSMUSG00000044086 | **Lmod3** | 5430424A14Rik | leiomodin 3 (fetal) | protein-coding |
| chr6_97293295_97294829 | chr6 | 97293296 | 97294829 | + | 0.794877 | intron (NM_145148, intron 22 of 23) | intron (NM_145148, intron 22 of 23) | -41282 | NM_001081157 | 320502 | Mm.68163 | NM_001081157 | ENSMUSG00000044086 | **Lmod3** | 5430424A14Rik | leiomodin 3 (fetal) | protein-coding |
| chr6_97430242_97430899 | chr6 | 97430243 | 97430899 | + | 0.769128 | intron (NM_145148, intron 2 of 23) | intron (NM_145148, intron 2 of 23) | -177791 | NM_001081157 | 320502 | Mm.68163 | NM_001081157 | ENSMUSG00000044086 | **Lmod3** | 5430424A14Rik | leiomodin 3 (fetal) | protein-coding |
| chr6_97352488_97353151 | chr6 | 97352489 | 97353151 | + | 0.693128 | intron (NM_145148, intron 8 of 23) | intron (NM_145148, intron 8 of 23) | -100040 | NM_001081157 | 320502 | Mm.68163 | NM_001081157 | ENSMUSG00000044086 | **Lmod3** | 5430424A14Rik | leiomodin 3 (fetal) | protein-coding |
| chr6_97352460_97353394 | chr6 | 97352461 | 97353394 | + | 0.674001 | intron (NM_145148, intron 8 of 23) | intron (NM_145148, intron 8 of 23) | -100147 | NM_001081157 | 320502 | Mm.68163 | NM_001081157 | ENSMUSG00000044086 | **Lmod3** | 5430424A14Rik | leiomodin 3 (fetal) | protein-coding |
| chr17_17592949_17593284 | chr17 | 17592950 | 17593284 | + | 0.833506 | intron (NM_172827, intron 1 of 17) | intron (NM_172827, intron 1 of 17) | 31372 | NM_172827 | 240028 | Mm.328807 | NM_172827 | ENSMUSG00000023845 | **Lnpep** | 2010309L07Rik\|4732490P18Rik\|CAP\|IRAP\|PLAP\|gp160\|vp165 | leucyl/cystinyl aminopeptidase | protein-coding |
| chr5_147055751_147056293 | chr5 | 147055752 | 147056293 | + | 1.68988 | intron (NM_080795, intron 1 of 9) | intron (NM_080795, intron 1 of 9) | 20550 | NM_080795 | 140887 | Mm.272203 | NM_080795 | ENSMUSG00000016520 | **Lnx2** | 9630046H24\|AW209022 | ligand of numb-protein X 2 | protein-coding |

| | | | | | | | | | | | | | | | | | |
|---|---|---|---|---|---|---|---|---|---|---|---|---|---|---|---|---|---|
| chr5_147057599_147057934 | chr5 | 147057600 | 147057934 | + | 1.41903 | intron (NM_080795, intron 1 of 9) | L1M5\|LINE\|L1 | 18805 | NM_080795 | 140887 | Mm.272203 | NM_080795 | ENSMUSG00000016520 | **Lnx2** | 9630046H24\|AW209022 | ligand of numb-protein X 2 | protein-coding |
| chr5_147072025_147072250 | chr5 | 147072026 | 147072250 | + | 1.047 | intron (NM_080795, intron 1 of 9) | intron (NM_080795, intron 1 of 9) | 4434 | NM_080795 | 140887 | Mm.272203 | NM_080795 | ENSMUSG00000016520 | **Lnx2** | 9630046H24\|AW209022 | ligand of numb-protein X 2 | protein-coding |
| chr4_59224876_59225236 | chr4 | 59224877 | 59225236 | + | 0.878789 | Intergenic | Intergenic | -34995 | NR_105031 | 101055769 | Mm.421401 | NR_105031 | | **LOC101055769** | - | uncharacterized LOC101055769 | ncRNA |
| chr4_59338794_59339309 | chr4 | 59338795 | 59339309 | + | 0.856772 | intron (NM_001163288, intron 12 of 16) | intron (NM_001163288, intron 12 of 16) | 79001 | NR_105031 | 101055769 | Mm.421401 | NR_105031 | | **LOC101055769** | - | uncharacterized LOC101055769 | ncRNA |
| chr4_59255959_59256353 | chr4 | 59255960 | 59256353 | + | 0.792762 | Intergenic | Intergenic | -3895 | NR_105031 | 101055769 | Mm.421401 | NR_105031 | | **LOC101055769** | - | uncharacterized LOC101055769 | ncRNA |
| chr4_59224651_59225278 | chr4 | 59224652 | 59225278 | + | 0.739805 | Intergenic | Intergenic | -35086 | NR_105031 | 101055769 | Mm.421401 | NR_105031 | | **LOC101055769** | - | uncharacterized LOC101055769 | ncRNA |
| chr3_132871160_132871879 | chr3 | 132871161 | 132871879 | + | 1.77716 | intron (NR_102297, intron 1 of 3) | intron (NR_102297, intron 1 of 3) | 7076 | NR_102297 | 101243624 | Mm.306256 | NR_102297 | | **LOC101243624** | - | uncharacterized LOC101243624 | ncRNA |
| chr3_132787663_132788117 | chr3 | 132787664 | 132788117 | + | 0.720724 | intron (NM_001163455, intron 23 of 25) | L1MA7\|LINE\|L1 | 90706 | NR_102297 | 101243624 | Mm.306256 | NR_102297 | | **LOC101243624** | - | uncharacterized LOC101243624 | ncRNA |
| chr10_66844291_66845460 | chr10 | 66844292 | 66845460 | + | 1.79381 | Intergenic | (TG)n\|Simple_repeat\|Simple_repeat | -160199 | NR_110511 | 102634101 | Mm.445258 | NR_110511 | | **LOC102634101** | - | uncharacterized LOC102634101 | ncRNA |
| chr10_66955424_66955754 | chr10 | 66955425 | 66955754 | + | 1.2663 | Intergenic | Intergenic | -49486 | NR_110511 | 102634101 | Mm.445258 | NR_110511 | | **LOC102634101** | - | uncharacterized LOC102634101 | ncRNA |
| chr10_66928186_66928612 | chr10 | 66928187 | 66928612 | + | 1.06174 | Intergenic | Intergenic | -76676 | NR_110511 | 102634101 | Mm.445258 | NR_110511 | | **LOC102634101** | - | uncharacterized LOC102634101 | ncRNA |
| chr10_66901132_66901428 | chr10 | 66901133 | 66901428 | + | 0.982514 | Intergenic | Intergenic | -103795 | NR_110511 | 102634101 | Mm.445258 | NR_110511 | | **LOC102634101** | - | uncharacterized LOC102634101 | ncRNA |
| chr10_66870782_66871498 | chr10 | 66870783 | 66871498 | + | 0.913388 | Intergenic | Intergenic | -133935 | NR_110511 | 102634101 | Mm.445258 | NR_110511 | | **LOC102634101** | - | uncharacterized LOC102634101 | ncRNA |
| chr10_67048879_67049592 | chr10 | 67048880 | 67049592 | + | 0.883216 | intron (NM_001204915, intron 2 of 8) | URR1B\|DNA\|hAT-Charlie | 44161 | NR_110511 | 102634101 | Mm.445258 | NR_110511 | | **LOC102634101** | - | uncharacterized LOC102634101 | ncRNA |
| chr10_67048893_67049204 | chr10 | 67048894 | 67049204 | + | 0.8824 | intron (NM_001204915, intron 2 of 8) | intron (NM_001204915, intron 2 of 8) | 43974 | NR_110511 | 102634101 | Mm.445258 | NR_110511 | | **LOC102634101** | - | uncharacterized LOC102634101 | ncRNA |
| chr10_66901123_66901746 | chr10 | 66901124 | 66901746 | + | 0.877342 | Intergenic | Intergenic | -103640 | NR_110511 | 102634101 | Mm.445258 | NR_110511 | | **LOC102634101** | - | uncharacterized LOC102634101 | ncRNA |
| chr10_66953878_66954179 | chr10 | 66953879 | 66954179 | + | 0.768566 | Intergenic | RSINE1\|SINE\|B4 | -51046 | NR_110511 | 102634101 | Mm.445258 | NR_110511 | | **LOC102634101** | - | uncharacterized LOC102634101 | ncRNA |
| chr10_66953850_66954298 | chr10 | 66953851 | 66954298 | + | 0.727079 | Intergenic | Intergenic | -51001 | NR_110511 | 102634101 | Mm.445258 | NR_110511 | | **LOC102634101** | - | uncharacterized LOC102634101 | ncRNA |
| chr10_66852700_66853078 | chr10 | 66852701 | 66853078 | + | 0.720097 | Intergenic | Intergenic | -152186 | NR_110511 | 102634101 | Mm.445258 | NR_110511 | | **LOC102634101** | - | uncharacterized LOC102634101 | ncRNA |
| chr10_67049261_67049570 | chr10 | 67049262 | 67049570 | + | 0.701862 | intron (NM_001204915, intron 2 of 8) | intron (NM_001204915, intron 2 of 8) | 44341 | NR_110511 | 102634101 | Mm.445258 | NR_110511 | | **LOC102634101** | - | uncharacterized LOC102634101 | ncRNA |
| chr10_66869971_66870240 | chr10 | 66869972 | 66870240 | + | 0.681871 | Intergenic | ID_B1\|SINE\|B4 | -134969 | NR_110511 | 102634101 | Mm.445258 | NR_110511 | | **LOC102634101** | - | uncharacterized LOC102634101 | ncRNA |
| chr10_66937822_66938322 | chr10 | 66937823 | 66938322 | + | 0.663874 | Intergenic | Intergenic | -67003 | NR_110511 | 102634101 | Mm.445258 | NR_110511 | | **LOC102634101** | - | uncharacterized LOC102634101 | ncRNA |
| chr10_66953850_66954511 | chr10 | 66953851 | 66954511 | + | 0.65774 | Intergenic | Intergenic | -50894 | NR_110511 | 102634101 | Mm.445258 | NR_110511 | | **LOC102634101** | - | uncharacterized LOC102634101 | ncRNA |
| chr10_66956535_66957119 | chr10 | 66956536 | 66957119 | + | 0.637156 | Intergenic | Lx7\|LINE\|L1 | -48248 | NR_110511 | 102634101 | Mm.445258 | NR_110511 | | **LOC102634101** | - | uncharacterized LOC102634101 | ncRNA |
| chr6_134668967_134669443 | chr6 | 134668968 | 134669443 | + | 0.734614 | intron (NM_001170479, intron 2 of 3) | intron (NM_001170479, intron 2 of 3) | 28265 | NM_026371 | 67774 | Mm.41892 | NM_026371 | ENSMUSG00000042992 | **Loh12cr1** | 5830457J20Rik\|AW124643 | loss of heterozygosity, 12, chromosomal region 1 homolog (human) | protein-coding |
| chr6_134716105_134716552 | chr6 | 134716106 | 134716552 | + | 0.669122 | 3' UTR (NM_001048054, exon 7 of 7) | 3' UTR (NM_001048054, exon 7 of 7) | 75389 | NM_026371 | 67774 | Mm.41892 | NM_026371 | ENSMUSG00000042992 | **Loh12cr1** | 5830457J20Rik\|AW124643 | loss of heterozygosity, 12, chromosomal region 1 homolog (human) | protein-coding |
| chr17_56625505_56625871 | chr17 | 56625506 | 56625871 | + | 1.17658 | intron (NM_028782, intron 1 of 17) | intron (NM_028782, intron 1 of 17) | 1215 | NM_028782 | 74142 | Mm.329136 | NM_028782 | ENSMUSG00000041168 | **Lonp1** | 1200017E13Rik\|LON\|Prss15 | lon peptidase 1, mitochondrial | protein-coding |
| chr8_86641948_86642489 | chr8 | 86641949 | 86642489 | + | 0.857098 | intron (NM_025827, intron 7 of 14) | intron (NM_025827, intron 7 of 14) | -9350 | NM_001168591 | 66887 | Mm.324550 | NM_025827 | ENSMUSG00000047866 | **Lonp2** | 1300002A08Rik\|AU015403\|Lonp | lon peptidase 2, peroxisomal | protein-coding |
| chr8_86641658_86642489 | chr8 | 86641659 | 86642489 | + | 0.819336 | intron (NM_025827, intron 7 of 14) | intron (NM_025827, intron 7 of 14) | -9495 | NM_001168591 | 66887 | Mm.324550 | NM_025827 | ENSMUSG00000047866 | **Lonp2** | 1300002A08Rik\|AU015403\|Lonp | lon peptidase 2, peroxisomal | protein-coding |
| chr8_86630536_86630998 | chr8 | 86630537 | 86630998 | + | 0.733076 | intron (NM_025827, intron 1 of 14) | ORR1D2\|LTR\|ERVL-MaLR | 6724 | NM_025827 | 66887 | Mm.324550 | NM_025827 | ENSMUSG00000047866 | **Lonp2** | 1300002A08Rik\|AU015403\|Lonp | lon peptidase 2, peroxisomal | protein-coding |
| chr8_36190299_36190581 | chr8 | 36190300 | 36190581 | + | 1.00333 | Intergenic | Intergenic | 59076 | NM_001081150 | 244421 | Mm.324032 | NM_001081150 | ENSMUSG00000039633 | **Lonrf1** | - | LON peptidase N-terminal domain and ring finger 1 | protein-coding |
| chr8_36251643_36251941 | chr8 | 36251644 | 36251941 | + | 0.6573 | Intergenic | Intergenic | -2276 | NM_001081150 | 244421 | Mm.324032 | NM_001081150 | ENSMUSG00000039633 | **Lonrf1** | - | LON peptidase N-terminal domain and ring finger 1 | protein-coding |
| chr8_36172862_36173858 | chr8 | 36172863 | 36173858 | + | 0.645207 | Intergenic | Intergenic | 76156 | NM_001081150 | 244421 | Mm.324032 | NM_001081150 | ENSMUSG00000039633 | **Lonrf1** | - | LON peptidase N-terminal domain and ring finger 1 | protein-coding |
| chr1_38777225_38778250 | chr1 | 38777226 | 38778250 | + | 1.15198 | Intergenic | Intergenic | 43477 | NM_001029878 | 381338 | Mm.487137 | NM_028403 | ENSMUSG00000048814 | **Lonrf2** | 2900060P06Rik\|AI851349 | LON peptidase N-terminal domain and ring finger 2 | protein-coding |
| chr1_38777321_38777941 | chr1 | 38777322 | 38777941 | + | 1.1023 | Intergenic | Intergenic | 43584 | NM_001029878 | 381338 | Mm.487137 | NM_028403 | ENSMUSG00000048814 | **Lonrf2** | 2900060P06Rik\|AI851349 | LON peptidase N-terminal domain and ring finger 2 | protein-coding |
| chr1_38771033_38771380 | chr1 | 38771034 | 38771380 | + | 0.818468 | Intergenic | Intergenic | 50008 | NM_001029878 | 381338 | Mm.487137 | NM_028403 | ENSMUSG00000048814 | **Lonrf2** | 2900060P06Rik\|AI851349 | LON peptidase N-terminal domain and ring finger 2 | protein-coding |
| chrX_36359652_36360339 | chrX | 36359653 | 36360339 | + | 0.714095 | intron (NM_028894, intron 10 of 10) | intron (NM_028894, intron 10 of 10) | 31587 | NM_028894 | 74365 | Mm.327654 | NM_028894 | | **Lonrf3** | 4932412G04Rik\|5730439E01Rik\|A830039N02Rik\|AU023707\|Rnf127 | LON peptidase N-terminal domain and ring finger 3 | protein-coding |
| chrX_36358372_36358760 | chrX | 36358373 | 36358760 | + | 0.612085 | intron (NM_028894, intron 9 of 10) | intron (NM_028894, intron 9 of 10) | 30157 | NM_028894 | 74365 | Mm.327654 | NM_028894 | | **Lonrf3** | 4932412G04Rik\|5730439E01Rik\|A830039N02Rik\|AU023707\|Rnf127 | LON peptidase N-terminal domain and ring finger 3 | protein-coding |
| chr18_52512709_52513193 | chr18 | 52512710 | 52513193 | + | 0.872044 | Intergenic | MER2\|DNA\|TcMar-Tigger | 16770 | NM_001286181 | 16948 | Mm.172 | NM_010728 | ENSMUSG00000024529 | **Lox** | AI893619\|TSC-160\|rrg | lysyl oxidase | protein-coding |
| chr4_58471200_58471569 | chr4 | 58471201 | 58471569 | + | 0.72074 | intron (NM_001290486, intron 2 of 2) | intron (NM_001290486, intron 2 of 2) | 28018 | NM_001290486 | 14745 | Mm.4772 | NM_010336 | ENSMUSG00000038668 | **Lpar1** | AI326300\|Edg2\|Gpcr26\|Kdt2\|lpA1\|vzg-1 | lysophosphatidic acid receptor 1 | protein-coding |
| chr4_58470195_58470579 | chr4 | 58470196 | 58470579 | + | 0.683251 | intron (NM_001290486, intron 2 of 2) | intron (NM_001290486, intron 2 of 2) | 29016 | NM_001290486 | 14745 | Mm.4772 | NM_010336 | ENSMUSG00000038668 | **Lpar1** | AI326300\|Edg2\|Gpcr26\|Kdt2\|lpA1\|vzg-1 | lysophosphatidic acid receptor 1 | protein-coding |
| chr3_146251814_146252490 | chr3 | 146251815 | 146252490 | + | 0.73915 | intron (NM_022983, intron 2 of 2) | intron (NM_022983, intron 2 of 2) | 31191 | NM_022983 | 65086 | Mm.155520 | NM_022983 | ENSMUSG00000036832 | **Lpar3** | Edg7\|lpA3 | lysophosphatidic acid receptor 3 | protein-coding |
| chr3_146225439_146225736 | chr3 | 146225440 | 146225736 | + | 0.738812 | intron (NM_022983, intron 1 of 2) | L1MC4\|LINE\|L1 | 4627 | NM_022983 | 65086 | Mm.155520 | NM_022983 | ENSMUSG00000036832 | **Lpar3** | Edg7\|lpA3 | lysophosphatidic acid receptor 3 | protein-coding |
| chr3_146220707_146221234 | chr3 | 146220708 | 146221234 | + | 0.715351 | promoter-TSS (NM_022983) | promoter-TSS (NM_022983) | 10 | NM_022983 | 65086 | Mm.155520 | NM_022983 | ENSMUSG00000036832 | **Lpar3** | Edg7\|lpA3 | lysophosphatidic acid receptor 3 | protein-coding |
| chr3_146211003_146211420 | chr3 | 146211004 | 146211420 | + | 0.675124 | Intergenic | MTA_Mm\|LTR\|ERVL-MaLR | -9749 | NM_022983 | 65086 | Mm.155520 | NM_022983 | ENSMUSG00000036832 | **Lpar3** | Edg7\|lpA3 | lysophosphatidic acid receptor 3 | protein-coding |
| chr14_73274733_73275175 | chr14 | 73274734 | 73275175 | + | 0.835801 | intron (NM_009029, intron 9 of 26) | intron (NM_009029, intron 9 of 26) | 37063 | NM_175116 | 67168 | Mm.390681 | NM_175116 | ENSMUSG00000033446 | **Lpar6** | 2610302I02Rik\|P2ry5\|P2y5 | lysophosphatidic acid receptor 6 | protein-coding |
| chr13_73490937_73491550 | chr13 | 73490938 | 73491550 | + | 0.851207 | intron (NM_145376, intron 3 of 13) | intron (NM_145376, intron 3 of 13) | 23861 | NM_145376 | 210992 | Mm.284649 | NM_145376 | ENSMUSG00000021608 | **Lpcat1** | 2900035H07Rik\|Aytl2\|BB137372\|BC005662\|C87117\|LPCAT\|rd11 | lysophosphatidylcholine acyltransferase 1 | protein-coding |
| chr6_124683595_124684249 | chr6 | 124683596 | 124684249 | + | 1.12274 | intron (NM_145130, intron 1 of 12) | intron (NM_145130, intron 1 of 12) | 20818 | NM_145130 | 14792 | Mm.273915 | NM_145130 | ENSMUSG00000004270 | **Lpcat3** | C3f\|Grcc3f\|Lpcat\|Lpeat\|Lplat5\|Lpsat\|Mboat5\|Moact5\|Oact5\|PTG | lysophosphatidylcholine acyltransferase 3 | protein-coding |
| chr2_112214540_112215131 | chr2 | 112214541 | 112215131 | + | 0.888355 | Intergenic | MIR\|SINE\|MIR | -25005 | NM_207206 | 99010 | Mm.33438 | NM_207206 | ENSMUSG00000027134 | **Lpcat4** | AI505034\|Agpat7\|Aytl3\|LPEAT2 | lysophosphatidylcholine acyltransferase 4 | protein-coding |
| chr1_191662058_191662857 | chr1 | 191662059 | 191662857 | + | 1.14225 | Intergenic | Intergenic | -55566 | NM_172266 | 226856 | Mm.277958 | NM_172266 | ENSMUSG00000026623 | **Lpgat1** | AI649174\|AW112037\|BC013667 | lysophosphatidylglycerol acyltransferase 1 | protein-coding |
| chr1_191711640_191712016 | chr1 | 191711641 | 191712016 | + | 0.629673 | Intergenic | Intergenic | -6196 | NM_172266 | 226856 | Mm.277958 | NM_172266 | ENSMUSG00000026623 | **Lpgat1** | AI649174\|AW112037\|BC013667 | lysophosphatidylglycerol acyltransferase 1 | protein-coding |
| chr3_148899876_148900193 | chr3 | 148899877 | 148900193 | + | 0.870434 | intron (NM_001081298, intron 1 of 23) | intron (NM_001081298, intron 1 of 23) | 54600 | NM_001081298 | 99633 | Mm.9776 | NM_001081298 | ENSMUSG00000028184 | **Lphn2** | AI450192\|Gm619\|Lec1\|Lphh1\|mKIAA0786 | latrophilin 2 | protein-coding |
| chr3_148933075_148933484 | chr3 | 148933076 | 148933484 | + | 0.72088 | intron (NM_001081298, intron 1 of 23) | intron (NM_001081298, intron 1 of 23) | 21355 | NM_001081298 | 99633 | Mm.9776 | NM_001081298 | ENSMUSG00000028184 | **Lphn2** | AI450192\|Gm619\|Lec1\|Lphh1\|mKIAA0786 | latrophilin 2 | protein-coding |

| | | | | | | | | | | | | | | | | | |
|---|---|---|---|---|---|---|---|---|---|---|---|---|---|---|---|---|---|
| chr12_16605679_16606287 | chr12 | 16605680 | 16606287 | + | 1.16822 | Intergenic | CT-rich\|Low_complexity\|Low_complexity | -16213 | NM_172950 | 14245 | Mm.153625 | NM_015763 | ENSMUSG00000020593 | **Lpin1** | 4631420P06\|Kiaa0188\|Lipin1\|fld\|mKIAA0188 | lipin 1 | protein-coding |
| chr12_16531776_16532315 | chr12 | 16531777 | 16532315 | + | 1.16377 | Intergenic | Intergenic | 57724 | NM_015763 | 14245 | Mm.153625 | NM_015763 | ENSMUSG00000020593 | **Lpin1** | 4631420P06\|Kiaa0188\|Lipin1\|fld\|mKIAA0188 | lipin 1 | protein-coding |
| chr12_16532008_16532310 | chr12 | 16532009 | 16532310 | + | 1.1133 | Intergenic | Intergenic | 57611 | NM_015763 | 14245 | Mm.153625 | NM_015763 | ENSMUSG00000020593 | **Lpin1** | 4631420P06\|Kiaa0188\|Lipin1\|fld\|mKIAA0188 | lipin 1 | protein-coding |
| chr12_16605663_16606484 | chr12 | 16605664 | 16606484 | + | 1.11277 | Intergenic | Intergenic | -16304 | NM_172950 | 14245 | Mm.153625 | NM_015763 | ENSMUSG00000020593 | **Lpin1** | 4631420P06\|Kiaa0188\|Lipin1\|fld\|mKIAA0188 | lipin 1 | protein-coding |
| chr12_16533316_16534144 | chr12 | 16533317 | 16534144 | + | 0.830257 | Intergenic | ID_B1\|SINE\|B4 | 56040 | NM_015763 | 14245 | Mm.153625 | NM_015763 | ENSMUSG00000020593 | **Lpin1** | 4631420P06\|Kiaa0188\|Lipin1\|fld\|mKIAA0188 | lipin 1 | protein-coding |
| chr12_16374454_16374980 | chr12 | 16374455 | 16374980 | + | 0.75318 | Intergenic | Intergenic | 215053 | NM_015763 | 14245 | Mm.153625 | NM_015763 | ENSMUSG00000020593 | **Lpin1** | 4631420P06\|Kiaa0188\|Lipin1\|fld\|mKIAA0188 | lipin 1 | protein-coding |
| chr12_16210188_16210516 | chr12 | 16210189 | 16210516 | + | 0.674272 | Intergenic | L1M4\|LINE\|L1 | 379418 | NM_015763 | 14245 | Mm.153625 | NM_015763 | ENSMUSG00000020593 | **Lpin1** | 4631420P06\|Kiaa0188\|Lipin1\|fld\|mKIAA0188 | lipin 1 | protein-coding |
| chr17_71191779_71192362 | chr17 | 71191780 | 71192362 | + | 1.09986 | intron (NM_022882, intron 1 of 19) | intron (NM_022882, intron 1 of 19) | 8093 | NM_022882 | 64898 | Mm.227924 | NM_022882 | ENSMUSG00000024052 | **Lpin2** | 2610511G02Rik\|AI481352\|AW742896 | lipin 2 | protein-coding |
| chr17_71256667_71257509 | chr17 | 71256668 | 71257509 | + | 1.04588 | intron (NM_145158, intron 4 of 7) | intron (NM_145158, intron 4 of 7) | 52434 | NM_001164885 | 64898 | Mm.227924 | NM_022882 | ENSMUSG00000024052 | **Lpin2** | 2610511G02Rik\|AI481352\|AW742896 | lipin 2 | protein-coding |
| chr17_71190576_71190940 | chr17 | 71190577 | 71190940 | + | 1.03823 | intron (NM_022882, intron 1 of 19) | intron (NM_022882, intron 1 of 19) | 6780 | NM_022882 | 64898 | Mm.227924 | NM_022882 | ENSMUSG00000024052 | **Lpin2** | 2610511G02Rik\|AI481352\|AW742896 | lipin 2 | protein-coding |
| chr17_71164950_71165400 | chr17 | 71164951 | 71165400 | + | 0.956526 | Intergenic | Intergenic | -18803 | NM_022882 | 64898 | Mm.227924 | NM_022882 | ENSMUSG00000024052 | **Lpin2** | 2610511G02Rik\|AI481352\|AW742896 | lipin 2 | protein-coding |
| chr17_71187843_71188104 | chr17 | 71187844 | 71188104 | + | 0.851214 | intron (NM_022882, intron 1 of 19) | intron (NM_022882, intron 1 of 19) | 3996 | NM_022882 | 64898 | Mm.227924 | NM_022882 | ENSMUSG00000024052 | **Lpin2** | 2610511G02Rik\|AI481352\|AW742896 | lipin 2 | protein-coding |
| chr16_24859490_24860136 | chr16 | 24859491 | 24860136 | + | 2.20058 | intron (NM_001145952, intron 8 of 11) | intron (NM_001145952, intron 8 of 11) | 137958 | NM_001145954 | 210126 | Mm.209385 | NM_178665 | ENSMUSG00000033306 | **Lpp** | 9430020K16Rik\|AA959454\|AU024130\|B130055L10Rik\|C79715\|D630048H16 | LIM domain containing preferred translocation partner in lipoma | protein-coding |
| chr16_24828378_24828845 | chr16 | 24828379 | 24828845 | + | 1.5692 | intron (NM_001145952, intron 7 of 11) | MIRc\|SINE\|MIR | 106757 | NM_001145954 | 210126 | Mm.209385 | NM_178665 | ENSMUSG00000033306 | **Lpp** | 9430020K16Rik\|AA959454\|AU024130\|B130055L10Rik\|C79715\|D630048H16 | LIM domain containing preferred translocation partner in lipoma | protein-coding |
| chr16_24334450_24334773 | chr16 | 24334451 | 24334773 | + | 1.26896 | Intergenic | MTEa\|LTR\|ERVL-MaLR | -58738 | NM_178665 | 210126 | Mm.209385 | NM_178665 | ENSMUSG00000033306 | **Lpp** | 9430020K16Rik\|AA959454\|AU024130\|B130055L10Rik\|C79715\|D630048H16 | LIM domain containing preferred translocation partner in lipoma | protein-coding |
| chr16_24659110_24659665 | chr16 | 24659111 | 24659665 | + | 1.17644 | intron (NM_001145952, intron 4 of 11) | intron (NM_001145952, intron 4 of 11) | -62467 | NM_001145954 | 210126 | Mm.209385 | NM_178665 | ENSMUSG00000033306 | **Lpp** | 9430020K16Rik\|AA959454\|AU024130\|B130055L10Rik\|C79715\|D630048H16 | LIM domain containing preferred translocation partner in lipoma | protein-coding |
| chr16_24659045_24659862 | chr16 | 24659046 | 24659862 | + | 1.13315 | intron (NM_001145952, intron 4 of 11) | intron (NM_001145952, intron 4 of 11) | -62401 | NM_001145954 | 210126 | Mm.209385 | NM_178665 | ENSMUSG00000033306 | **Lpp** | 9430020K16Rik\|AA959454\|AU024130\|B130055L10Rik\|C79715\|D630048H16 | LIM domain containing preferred translocation partner in lipoma | protein-coding |
| chr16_24659010_24659459 | chr16 | 24659011 | 24659459 | + | 1.07829 | intron (NM_001145952, intron 4 of 11) | intron (NM_001145952, intron 4 of 11) | -62620 | NM_001145954 | 210126 | Mm.209385 | NM_178665 | ENSMUSG00000033306 | **Lpp** | 9430020K16Rik\|AA959454\|AU024130\|B130055L10Rik\|C79715\|D630048H16 | LIM domain containing preferred translocation partner in lipoma | protein-coding |
| chr16_24861341_24861703 | chr16 | 24861342 | 24861703 | + | 1.03379 | intron (NM_001145952, intron 8 of 11) | Tigger7\|DNA\|TcMar-Tigger | 139667 | NM_001145954 | 210126 | Mm.209385 | NM_178665 | ENSMUSG00000033306 | **Lpp** | 9430020K16Rik\|AA959454\|AU024130\|B130055L10Rik\|C79715\|D630048H16 | LIM domain containing preferred translocation partner in lipoma | protein-coding |
| chr16_24637403_24637846 | chr16 | 24637404 | 24637846 | + | 0.990875 | intron (NM_001145952, intron 4 of 11) | intron (NM_001145952, intron 4 of 11) | -84230 | NM_001145954 | 210126 | Mm.209385 | NM_178665 | ENSMUSG00000033306 | **Lpp** | 9430020K16Rik\|AA959454\|AU024130\|B130055L10Rik\|C79715\|D630048H16 | LIM domain containing preferred translocation partner in lipoma | protein-coding |
| chr16_24424744_24425207 | chr16 | 24424745 | 24425207 | + | 0.959507 | intron (NM_178665, intron 1 of 12) | intron (NM_178665, intron 1 of 12) | -23115 | NM_001145952 | 210126 | Mm.209385 | NM_178665 | ENSMUSG00000033306 | **Lpp** | 9430020K16Rik\|AA959454\|AU024130\|B130055L10Rik\|C79715\|D630048H16 | LIM domain containing preferred translocation partner in lipoma | protein-coding |
| chr16_24659507_24659862 | chr16 | 24659508 | 24659862 | + | 0.869993 | intron (NM_001145952, intron 4 of 11) | intron (NM_001145952, intron 4 of 11) | -62170 | NM_001145954 | 210126 | Mm.209385 | NM_178665 | ENSMUSG00000033306 | **Lpp** | 9430020K16Rik\|AA959454\|AU024130\|B130055L10Rik\|C79715\|D630048H16 | LIM domain containing preferred translocation partner in lipoma | protein-coding |
| chr16_24678764_24679335 | chr16 | 24678765 | 24679335 | + | 0.862799 | intron (NM_001145952, intron 5 of 11) | intron (NM_001145952, intron 5 of 11) | -42805 | NM_001145954 | 210126 | Mm.209385 | NM_178665 | ENSMUSG00000033306 | **Lpp** | 9430020K16Rik\|AA959454\|AU024130\|B130055L10Rik\|C79715\|D630048H16 | LIM domain containing preferred translocation partner in lipoma | protein-coding |
| chr16_24427457_24428384 | chr16 | 24427458 | 24428384 | + | 0.84737 | intron (NM_178665, intron 1 of 12) | intron (NM_178665, intron 1 of 12) | -20170 | NM_001145952 | 210126 | Mm.209385 | NM_178665 | ENSMUSG00000033306 | **Lpp** | 9430020K16Rik\|AA959454\|AU024130\|B130055L10Rik\|C79715\|D630048H16 | LIM domain containing preferred translocation partner in lipoma | protein-coding |
| chr16_24757912_24758250 | chr16 | 24757913 | 24758250 | + | 0.704628 | intron (NM_001145954, intron 2 of 7) | intron (NM_001145954, intron 2 of 7) | 36226 | NM_001145954 | 210126 | Mm.209385 | NM_178665 | ENSMUSG00000033306 | **Lpp** | 9430020K16Rik\|AA959454\|AU024130\|B130055L10Rik\|C79715\|D630048H16 | LIM domain containing preferred translocation partner in lipoma | protein-coding |
| chr16_24836478_24837028 | chr16 | 24836479 | 24837028 | + | 0.679609 | intron (NM_001145952, intron 7 of 11) | intron (NM_001145952, intron 7 of 11) | 114898 | NM_001145954 | 210126 | Mm.209385 | NM_178665 | ENSMUSG00000033306 | **Lpp** | 9430020K16Rik\|AA959454\|AU024130\|B130055L10Rik\|C79715\|D630048H16 | LIM domain containing preferred translocation partner in lipoma | protein-coding |
| chr16_24704244_24705073 | chr16 | 24704245 | 24705073 | + | 0.655915 | intron (NM_001145952, intron 6 of 11) | intron (NM_001145952, intron 6 of 11) | -17196 | NM_001145954 | 210126 | Mm.209385 | NM_178665 | ENSMUSG00000033306 | **Lpp** | 9430020K16Rik\|AA959454\|AU024130\|B130055L10Rik\|C79715\|D630048H16 | LIM domain containing preferred translocation partner in lipoma | protein-coding |
| chr16_24474806_24475141 | chr16 | 24474807 | 24475141 | + | 0.646657 | intron (NM_001145952, intron 2 of 11) | intron (NM_001145952, intron 2 of 11) | 26883 | NM_001145952 | 210126 | Mm.209385 | NM_178665 | ENSMUSG00000033306 | **Lpp** | 9430020K16Rik\|AA959454\|AU024130\|B130055L10Rik\|C79715\|D630048H16 | LIM domain containing preferred translocation partner in lipoma | protein-coding |
| chr16_24239200_24239618 | chr16 | 24239201 | 24239618 | + | 0.596185 | Intergenic | Intergenic | -153941 | NM_178665 | 210126 | Mm.209385 | NM_178665 | ENSMUSG00000033306 | **Lpp** | 9430020K16Rik\|AA959454\|AU024130\|B130055L10Rik\|C79715\|D630048H16 | LIM domain containing preferred translocation partner in lipoma | protein-coding |
| chr19_12801075_12801405 | chr19 | 12801076 | 12801405 | + | 1.05702 | intron (NM_134152, intron 1 of 8) | intron (NM_134152, intron 1 of 8) | 2631 | NM_134152 | 107321 | Mm.313136 | NM_134152 | ENSMUSG00000024696 | **Lpxn** | 4933402K05Rik\|A530083L21Rik\|AV278559\|LDPL | leupaxin | protein-coding |
| chr19_12802889_12803147 | chr19 | 12802890 | 12803147 | + | 0.783061 | intron (NM_134152, intron 1 of 8) | intron (NM_134152, intron 1 of 8) | 4409 | NM_134152 | 107321 | Mm.313136 | NM_134152 | ENSMUSG00000024696 | **Lpxn** | 4933402K05Rik\|A530083L21Rik\|AV278559\|LDPL | leupaxin | protein-coding |
| chr3_86213893_86214612 | chr3 | 86213894 | 86214612 | + | 0.741384 | Intergenic | L1_Mus2\|LINE\|L1 | -10437 | NM_001077688 | 80877 | Mm.439825 | NM_030695 | ENSMUSG00000028080 | **Lrba** | C80285\|D3Ertd775e\|Lab300\|Lba | LPS-responsive beige-like anchor | protein-coding |
| chr3_86265546_86266473 | chr3 | 86265547 | 86266473 | + | 0.679738 | intron (NM_001077687, intron 2 of 50) | RMER19B\|LTR\|ERVK | 41320 | NM_001077687 | 80877 | Mm.439825 | NM_030695 | ENSMUSG00000028080 | **Lrba** | C80285\|D3Ertd775e\|Lab300\|Lba | LPS-responsive beige-like anchor | protein-coding |
| chr14_74888657_74889648 | chr14 | 74888658 | 74889648 | + | 1.7221 | intron (NM_001252132, intron 1 of 18) | intron (NM_001252132, intron 1 of 18) | 58724 | NM_001252132 | 380916 | Mm.288868 | NM_001033439 | ENSMUSG00000068015 | **Lrch1** | 4832412D13Rik\|AI552548\|AW215503\|Chdc1\|mKIAA1016 | leucine-rich repeats and calponin homology (CH) domain containing 1 | protein-coding |
| chr14_74880864_74881346 | chr14 | 74880865 | 74881346 | + | 1.09607 | intron (NM_001252132, intron 1 of 18) | L2b\|LINE\|L2 | 66772 | NM_001252132 | 380916 | Mm.288868 | NM_001033439 | ENSMUSG00000068015 | **Lrch1** | 4832412D13Rik\|AI552548\|AW215503\|Chdc1\|mKIAA1016 | leucine-rich repeats and calponin homology (CH) domain containing 1 | protein-coding |
| chr14_74930367_74930716 | chr14 | 74930368 | 74930716 | + | 0.789423 | intron (NM_001252132, intron 1 of 18) | intron (NM_001252132, intron 1 of 18) | 17335 | NM_001252132 | 380916 | Mm.288868 | NM_001033439 | ENSMUSG00000068015 | **Lrch1** | 4832412D13Rik\|AI552548\|AW215503\|Chdc1\|mKIAA1016 | leucine-rich repeats and calponin homology (CH) domain containing 1 | protein-coding |
| chr14_74928882_74929145 | chr14 | 74928883 | 74929145 | + | 0.653079 | intron (NM_001252132, intron 1 of 18) | intron (NM_001252132, intron 1 of 18) | 18863 | NM_001252132 | 380916 | Mm.288868 | NM_001033439 | ENSMUSG00000068015 | **Lrch1** | 4832412D13Rik\|AI552548\|AW215503\|Chdc1\|mKIAA1016 | leucine-rich repeats and calponin homology (CH) domain containing 1 | protein-coding |
| chr16_32934394_32934830 | chr16 | 32934395 | 32934830 | + | 1.43261 | intron (NM_001081255, intron 1 of 20) | intron (NM_001081255, intron 1 of 20) | 20512 | NM_001081255 | 70144 | Mm.491178 | NM_001081255 | ENSMUSG00000022801 | **Lrch3** | 2210409B11Rik\|AW215594\|Gm1742 | leucine-rich repeats and calponin homology (CH) domain containing 3 | protein-coding |
| chr16_32915713_32916002 | chr16 | 32915714 | 32916002 | + | 0.981214 | intron (NM_001081255, intron 1 of 20) | intron (NM_001081255, intron 1 of 20) | 1758 | NM_001081255 | 70144 | Mm.491178 | NM_001081255 | ENSMUSG00000022801 | **Lrch3** | 2210409B11Rik\|AW215594\|Gm1742 | leucine-rich repeats and calponin homology (CH) domain containing 3 | protein-coding |

| | | | | | | | | | | | | | | | | | |
|---|---|---|---|---|---|---|---|---|---|---|---|---|---|---|---|---|---|
| chr7_30347835_30348335 | chr7 | 30347836 | 30348335 | + | 0.661939 | Intergenic | CpG | 14687 | NM_175478 | 233067 | Mm.23157 | NM_175478 | ENSMUSG00000036957 | **Lrfn3** | A530045B06Rik\|SALM4 | leucine rich repeat and fibronectin type III domain containing 3 | protein-coding |
| chr6_33702274_33702885 | chr6 | 33702275 | 33702885 | + | 1.09963 | intron (NM_009148, intron 10 of 17) | intron (NM_009148, intron 10 of 17) | -326868 | NM_028886 | 74354 | Mm.127952 | NM_028886 | ENSMUSG00000056215 | **Lrguk** | 4921528H16Rik | leucine-rich repeats and guanylate kinase domain containing | protein-coding |
| chr6_33727487_33728241 | chr6 | 33727488 | 33728241 | + | 0.642363 | intron (NM_009148, intron 10 of 17) | intron (NM_009148, intron 10 of 17) | -301584 | NM_028886 | 74354 | Mm.127952 | NM_028886 | ENSMUSG00000056215 | **Lrguk** | 4921528H16Rik | leucine-rich repeats and guanylate kinase domain containing | protein-coding |
| chr6_94660204_94660807 | chr6 | 94660205 | 94660807 | + | 1.46363 | intron (NM_008377, intron 2 of 18) | intron (NM_008377, intron 2 of 18) | 39639 | NM_008377 | 16206 | Mm.245210 | NM_008377 | ENSMUSG00000030029 | **Lrig1** | D6Bwg0781e\|Img\|LIG-1 | leucine-rich repeats and immunoglobulin-like domains 1 | protein-coding |
| chr6_94663836_94664281 | chr6 | 94663837 | 94664281 | + | 1.44111 | intron (NM_008377, intron 1 of 18) | intron (NM_008377, intron 1 of 18) | 36086 | NM_008377 | 16206 | Mm.245210 | NM_008377 | ENSMUSG00000030029 | **Lrig1** | D6Bwg0781e\|Img\|LIG-1 | leucine-rich repeats and immunoglobulin-like domains 1 | protein-coding |
| chr6_94670069_94670576 | chr6 | 94670070 | 94670576 | + | 1.15611 | intron (NM_008377, intron 1 of 18) | intron (NM_008377, intron 1 of 18) | 29822 | NM_008377 | 16206 | Mm.245210 | NM_008377 | ENSMUSG00000030029 | **Lrig1** | D6Bwg0781e\|Img\|LIG-1 | leucine-rich repeats and immunoglobulin-like domains 1 | protein-coding |
| chr6_94705841_94706224 | chr6 | 94705842 | 94706224 | + | 0.78213 | Intergenic | Intergenic | -5888 | NM_008377 | 16206 | Mm.245210 | NM_008377 | ENSMUSG00000030029 | **Lrig1** | D6Bwg0781e\|Img\|LIG-1 | leucine-rich repeats and immunoglobulin-like domains 1 | protein-coding |
| chr6_94659975_94661696 | chr6 | 94659976 | 94661696 | + | 0.69596 | intron (NM_008377, intron 2 of 18) | intron (NM_008377, intron 2 of 18) | 39309 | NM_008377 | 16206 | Mm.245210 | NM_008377 | ENSMUSG00000030029 | **Lrig1** | D6Bwg0781e\|Img\|LIG-1 | leucine-rich repeats and immunoglobulin-like domains 1 | protein-coding |
| chr10_125725392_125725629 | chr10 | 125725393 | 125725629 | + | 0.625924 | Intergenic | Intergenic | -240708 | NM_177152 | 320398 | Mm.275586 | NM_177152 | ENSMUSG00000020105 | **Lrig3** | 9030421L11Rik\|9130004I02Rik\|9430095K15Rik\|mKIAA3016 | leucine-rich repeats and immunoglobulin-like domains 3 | protein-coding |
| chr6_145104085_145104878 | chr6 | 145104086 | 145104878 | + | 1.01487 | Intergenic | Intergenic | -11154 | NM_001281980 | 16970 | Mm.843 | NM_008511 | ENSMUSG00000030263 | **Lrmp** | D6Int3\|D6Int4\|D6Int5\|D6Int7\|D6Int8\|Jaw1 | lymphoid-restricted membrane protein | protein-coding |
| chr6_145123616_145124161 | chr6 | 145123617 | 145124161 | + | 0.978325 | intron (NM_008511, intron 1 of 18) | intron (NM_008511, intron 1 of 18) | 2150 | NM_008511 | 16970 | Mm.843 | NM_008511 | ENSMUSG00000030263 | **Lrmp** | D6Int3\|D6Int4\|D6Int5\|D6Int7\|D6Int8\|Jaw1 | lymphoid-restricted membrane protein | protein-coding |
| chr10_7582212_7583097 | chr10 | 7582213 | 7583097 | + | 0.743588 | Intergenic | Intergenic | -7145 | NM_172784 | 237253 | Mm.206759 | NM_172784 | ENSMUSG00000019796 | **Lrp11** | 1700034J19Rik\|6330533B21\|9830160H19Rik | low density lipoprotein receptor-related protein 11 | protein-coding |
| chr10_7582255_7582579 | chr10 | 7582256 | 7582579 | + | 0.703862 | Intergenic | Intergenic | -7383 | NM_172784 | 237253 | Mm.206759 | NM_172784 | ENSMUSG00000019796 | **Lrp11** | 1700034J19Rik\|6330533B21\|9830160H19Rik | low density lipoprotein receptor-related protein 11 | protein-coding |
| chr15_39900778_39901121 | chr15 | 39900779 | 39901121 | + | 0.644149 | intron (NM_172814, intron 2 of 6) | intron (NM_172814, intron 2 of 6) | 42807 | NM_172814 | 239393 | Mm.39874 | NM_172814 | ENSMUSG00000022305 | **Lrp12** | AI848829\|C820005L12Rik\|LRP-12 | low density lipoprotein-related protein 12 | protein-coding |
| chr2_91454481_91454841 | chr2 | 91454482 | 91454841 | + | 0.778189 | Intergenic | Intergenic | -2870 | NM_172668 | 228357 | Mm.275149 | NM_172668 | ENSMUSG00000027253 | **Lrp4** | 6430526J12Rik\|D230026E03\|Megf7\|mdig | low density lipoprotein receptor-related protein 4 | protein-coding |
| chr2_91459726_91460110 | chr2 | 91459727 | 91460110 | + | 0.756025 | intron (NM_001145857, intron 1 of 11) | intron (NM_001145857, intron 1 of 11) | 2387 | NM_001145857 | 228357 | Mm.275149 | NM_172668 | ENSMUSG00000027253 | **Lrp4** | 6430526J12Rik\|D230026E03\|Megf7\|mdig | low density lipoprotein receptor-related protein 4 | protein-coding |
| chr6_134544114_134544464 | chr6 | 134544115 | 134544464 | + | 1.24431 | intron (NM_008514, intron 1 of 22) | intron (NM_008514, intron 1 of 22) | 22624 | NM_008514 | 16974 | Mm.321990 | NM_008514 | ENSMUSG00000030201 | **Lrp6** | C030016K15Rik\|Cd\|Gw\|ska26\|ska<m26Jus>\|skax26 | low density lipoprotein receptor-related protein 6 | protein-coding |
| chr6_134543806_134544471 | chr6 | 134543807 | 134544471 | + | 0.80161 | intron (NM_008514, intron 1 of 22) | intron (NM_008514, intron 1 of 22) | 22774 | NM_008514 | 16974 | Mm.321990 | NM_008514 | ENSMUSG00000030201 | **Lrp6** | C030016K15Rik\|Cd\|Gw\|ska26\|ska<m26Jus>\|skax26 | low density lipoprotein receptor-related protein 6 | protein-coding |
| chr6_134517445_134517815 | chr6 | 134517446 | 134517815 | + | 0.671179 | intron (NM_008514, intron 3 of 22) | L1M4\|LINE\|L1 | 49283 | NM_008514 | 16974 | Mm.321990 | NM_008514 | ENSMUSG00000030201 | **Lrp6** | C030016K15Rik\|Cd\|Gw\|ska26\|ska<m26Jus>\|skax26 | low density lipoprotein receptor-related protein 6 | protein-coding |
| chr4_107757518_107757989 | chr4 | 107757519 | 107757989 | + | 0.800753 | Intergenic | Intergenic | -44505 | NR_033496 | 16975 | Mm.442134 | NM_053073 | | **Lrp8** | 4932703M08Rik\|AA921429\|AI848122\|ApoER2\|Lr8b | low density lipoprotein receptor-related protein 8, apolipoprotein e receptor | protein-coding |
| chr4_107803363_107804249 | chr4 | 107803364 | 107804249 | + | 0.653231 | intron (NM_001080926, intron 2 of 18) | intron (NM_001080926, intron 2 of 18) | 1547 | NM_001080926 | 16975 | Mm.442134 | NM_053073 | | **Lrp8** | 4932703M08Rik\|AA921429\|AI848122\|ApoER2\|Lr8b | low density lipoprotein receptor-related protein 8, apolipoprotein e receptor | protein-coding |
| chr10_117022561_117023025 | chr10 | 117022562 | 117023025 | + | 0.927797 | intron (NM_001007583, intron 9 of 9) | Tigger7\|DNA\|TcMar-Tigger | -22548 | NM_146242 | 237560 | Mm.37320 | NM_146242 | ENSMUSG00000060187 | **Lrrc10** | D330003I11Rik\|Hrlrrp\|Serdin1 | leucine rich repeat containing 10 | protein-coding |
| chr6_124784256_124784796 | chr6 | 124784257 | 124784796 | + | 0.926084 | Intergenic | Intergenic | -4808 | NM_013588 | 16977 | Mm.3207 | NM_013588 | ENSMUSG00000030125 | **Lrrc23** | 4921537K05Rik\|B7\|Lrpb7 | leucine rich repeat containing 23 | protein-coding |
| chr7_98533719_98534141 | chr7 | 98533720 | 98534141 | + | 1.18516 | Intergenic | Intergenic | 39708 | NM_001113379 | 434215 | Mm.323057 | NM_001113379 | ENSMUSG00000090958 | **Lrrc32** | AI426318\|D11S833Eh\|D7H11S833E\|EG434215\|Garp | leucine rich repeat containing 32 | protein-coding |
| chr7_98564595_98565297 | chr7 | 98564596 | 98565297 | + | 0.991189 | Intergenic | Intergenic | 70724 | NM_001113379 | 434215 | Mm.323057 | NM_001113379 | ENSMUSG00000090958 | **Lrrc32** | AI426318\|D11S833Eh\|D7H11S833E\|EG434215\|Garp | leucine rich repeat containing 32 | protein-coding |
| chr7_98490394_98490672 | chr7 | 98490395 | 98490672 | + | 0.909691 | Intergenic | Intergenic | -3689 | NM_001113379 | 434215 | Mm.323057 | NM_001113379 | ENSMUSG00000090958 | **Lrrc32** | AI426318\|D11S833Eh\|D7H11S833E\|EG434215\|Garp | leucine rich repeat containing 32 | protein-coding |
| chr7_98490299_98490672 | chr7 | 98490300 | 98490672 | + | 0.848951 | Intergenic | Intergenic | -3736 | NM_001113379 | 434215 | Mm.323057 | NM_001113379 | ENSMUSG00000090958 | **Lrrc32** | AI426318\|D11S833Eh\|D7H11S833E\|EG434215\|Garp | leucine rich repeat containing 32 | protein-coding |
| chr7_98532998_98533338 | chr7 | 98532999 | 98533338 | + | 0.837208 | Intergenic | Intergenic | 38946 | NM_001113379 | 434215 | Mm.323057 | NM_001113379 | ENSMUSG00000090958 | **Lrrc32** | AI426318\|D11S833Eh\|D7H11S833E\|EG434215\|Garp | leucine rich repeat containing 32 | protein-coding |
| chr7_98554720_98555537 | chr7 | 98554721 | 98555537 | + | 0.834088 | Intergenic | Intergenic | 60907 | NM_001113379 | 434215 | Mm.323057 | NM_001113379 | ENSMUSG00000090958 | **Lrrc32** | AI426318\|D11S833Eh\|D7H11S833E\|EG434215\|Garp | leucine rich repeat containing 32 | protein-coding |
| chr7_98533713_98535013 | chr7 | 98533714 | 98535013 | + | 0.78698 | Intergenic | Intergenic | 40141 | NM_001113379 | 434215 | Mm.323057 | NM_001113379 | ENSMUSG00000090958 | **Lrrc32** | AI426318\|D11S833Eh\|D7H11S833E\|EG434215\|Garp | leucine rich repeat containing 32 | protein-coding |
| chr14_15795298_15796011 | chr14 | 15795299 | 15796011 | + | 0.949772 | Intergenic | Intergenic | -356668 | NM_146052 | 218763 | Mm.138039 | NM_146052 | ENSMUSG00000045201 | **Lrrc3b** | BC019794\|LRP15 | leucine rich repeat containing 3B | protein-coding |
| chr14_15410792_15411703 | chr14 | 15410793 | 15411703 | + | 0.816148 | intron (NM_146052, intron 1 of 1) | intron (NM_146052, intron 1 of 1) | 27739 | NM_146052 | 218763 | Mm.138039 | NM_146052 | ENSMUSG00000045201 | **Lrrc3b** | BC019794\|LRP15 | leucine rich repeat containing 3B | protein-coding |
| chr14_15653259_15653838 | chr14 | 15653260 | 15653838 | + | 0.682453 | Intergenic | Intergenic | -214562 | NM_146052 | 218763 | Mm.138039 | NM_146052 | ENSMUSG00000045201 | **Lrrc3b** | BC019794\|LRP15 | leucine rich repeat containing 3B | protein-coding |
| chr7_101927307_101927732 | chr7 | 101927308 | 101927732 | + | 0.700526 | intron (NM_027053, intron 1 of 5) | B3A\|SINE\|B2 | 6337 | NM_027053 | 69358 | Mm.214841 | NM_027053 | ENSMUSG00000064307 | **Lrrc51** | 1700008D07Rik\|Lrtomt | leucine rich repeat containing 51 | protein-coding |
| chr2_120599274_120600091 | chr2 | 120599275 | 120600091 | + | 1.19397 | 3' UTR (NM_009222, exon 8 of 8) | 3' UTR (NM_009222, exon 8 of 8) | 9632 | NM_025657 | 66606 | Mm.428639 | NM_025657 | ENSMUSG00000027286 | **Lrrc57** | 2810002D13Rik\|AA407405 | leucine rich repeat containing 57 | protein-coding |
| chr16_37875908_37876996 | chr16 | 37875909 | 37876996 | + | 1.2734 | intron (NM_177093, intron 1 of 3) | intron (NM_177093, intron 1 of 3) | 8052 | NM_177093 | 320184 | Mm.390882 | NM_177093 | ENSMUSG00000034158 | **Lrrc58** | 1810012N18Rik\|C330018J07Rik | leucine rich repeat containing 58 | protein-coding |
| chr16_37877076_37877357 | chr16 | 37877077 | 37877357 | + | 0.71631 | intron (NM_177093, intron 2 of 3) | intron (NM_177093, intron 2 of 3) | 8817 | NM_177093 | 320184 | Mm.390882 | NM_177093 | ENSMUSG00000034158 | **Lrrc58** | 1810012N18Rik\|C330018J07Rik | leucine rich repeat containing 58 | protein-coding |
| chr15_66509728_66510070 | chr15 | 66509729 | 66510070 | + | 1.25713 | Intergenic | B4\|SINE\|B4 | -8989 | NM_019457 | 54562 | Mm.244890 | NM_019457 | ENSMUSG00000022375 | **Lrrc6** | LRTP | leucine rich repeat containing 6 (testis) | protein-coding |
| chr15_66522891_66523523 | chr15 | 66522892 | 66523523 | + | 0.747308 | Intergenic | RMER20C_Mm\|LTR\|ERVK | -22297 | NM_019457 | 54562 | Mm.244890 | NM_019457 | ENSMUSG00000022375 | **Lrrc6** | LRTP | leucine rich repeat containing 6 (testis) | protein-coding |
| chr15_66495102_66495658 | chr15 | 66495103 | 66495658 | + | 0.735723 | intron (NM_019457, intron 1 of 11) | intron (NM_019457, intron 1 of 11) | 5530 | NM_019457 | 54562 | Mm.244890 | NM_019457 | ENSMUSG00000022375 | **Lrrc6** | LRTP | leucine rich repeat containing 6 (testis) | protein-coding |
| chr12_36241037_36241755 | chr12 | 36241038 | 36241755 | + | 1.9678 | intron (NM_001177877, intron 3 of 7) | intron (NM_001177877, intron 3 of 7) | 12002 | NM_001177877 | 71156 | Mm.250837 | NM_027699 | ENSMUSG00000020545 | **Lrrc72** | 1700108M19Rik\|4933421E18Rik | leucine rich repeat containing 72 | protein-coding |
| chr12_36267634_36268373 | chr12 | 36267635 | 36268373 | + | 0.796829 | Intergenic | Intergenic | -14606 | NM_001177877 | 71156 | Mm.250837 | NM_027699 | ENSMUSG00000020545 | **Lrrc72** | 1700108M19Rik\|4933421E18Rik | leucine rich repeat containing 72 | protein-coding |
| chr12_86787093_86787613 | chr12 | 86787094 | 86787613 | + | 1.42946 | Intergenic | Intergenic | 52984 | NM_001195767 | 627607 | Mm.341880 | NM_001195767 | ENSMUSG00000059114 | **Lrrc74** | EG627607\|Gm6772 | leucine rich repeat containing 74 | protein-coding |
| chr12_86786836_86787731 | chr12 | 86786837 | 86787731 | + | 1.3199 | Intergenic | Intergenic | 52915 | NM_001195767 | 627607 | Mm.341880 | NM_001195767 | ENSMUSG00000059114 | **Lrrc74** | EG627607\|Gm6772 | leucine rich repeat containing 74 | protein-coding |
| chr5_105444840_105445292 | chr5 | 105444841 | 105445292 | + | 1.22137 | intron (NM_001033550, intron 1 of 4) | intron (NM_001033550, intron 1 of 4) | 29291 | NM_001033550 | 433926 | Mm.439781 | NM_001033550 | ENSMUSG00000070639 | **Lrrc8b** | R75581\|Ta-lrrp\|mKIAA0231 | leucine rich repeat containing 8 family, member B | protein-coding |
| chr5_105444438_105444648 | chr5 | 105444439 | 105444648 | + | 0.86742 | intron (NM_001033550, intron 1 of 4) | intron (NM_001033550, intron 1 of 4) | 28768 | NM_001033550 | 433926 | Mm.439781 | NM_001033550 | ENSMUSG00000070639 | **Lrrc8b** | R75581\|Ta-lrrp\|mKIAA0231 | leucine rich repeat containing 8 family, member B | protein-coding |
| chr5_105500292_105501096 | chr5 | 105500293 | 105501096 | + | 1.0784 | Intergenic | MER21C\|LTR\|ERVL | -18777 | NM_133897 | 100604 | Mm.319847 | NM_133897 | ENSMUSG00000054720 | **Lrrc8c** | AD158\|AI326115\|E430036I04Rik\|fad158 | leucine rich repeat containing 8 family, member C | protein-coding |
| chr5_105500554_105501102 | chr5 | 105500555 | 105501102 | + | 1.04632 | Intergenic | MER21C\|LTR\|ERVL | -18643 | NM_133897 | 100604 | Mm.319847 | NM_133897 | ENSMUSG00000054720 | **Lrrc8c** | AD158\|AI326115\|E430036I04Rik\|fad158 | leucine rich repeat containing 8 family, member C | protein-coding |
| chr5_105510877_105511542 | chr5 | 105510878 | 105511542 | + | 0.97973 | Intergenic | URR1B\|DNA\|hAT-Charlie | -8261 | NM_133897 | 100604 | Mm.319847 | NM_133897 | ENSMUSG00000054720 | **Lrrc8c** | AD158\|AI326115\|E430036I04Rik\|fad158 | leucine rich repeat containing 8 family, member C | protein-coding |
| chr5_105500719_105501084 | chr5 | 105500720 | 105501084 | + | 0.883442 | Intergenic | MER21C\|LTR\|ERVL | -18569 | NM_133897 | 100604 | Mm.319847 | NM_133897 | ENSMUSG00000054720 | **Lrrc8c** | AD158\|AI326115\|E430036I04Rik\|fad158 | leucine rich repeat containing 8 family, member C | protein-coding |
| chr5_105496207_105496391 | chr5 | 105496208 | 105496391 | + | 0.846646 | Intergenic | MTE2a\|LTR\|ERVL-MaLR | -23172 | NM_133897 | 100604 | Mm.319847 | NM_133897 | ENSMUSG00000054720 | **Lrrc8c** | AD158\|AI326115\|E430036I04Rik\|fad158 | leucine rich repeat containing 8 family, member C | protein-coding |

| | | | | | | | | | | | | | | | | | |
|---|---|---|---|---|---|---|---|---|---|---|---|---|---|---|---|---|---|
| chr5_105557742_105558269 | chr5 | 105557743 | 105558269 | + | 0.78556 | intron (NM_133897, intron 2 of 3) | intron (NM_133897, intron 2 of 3) | 38535 | NM_133897 | 100604 | Mm.319847 | NM_133897 | ENSMUSG00000054720 | **Lrrc8c** | AD158\|AI326115\|E430036I04Rik\|fad158 | leucine rich repeat containing 8 family, member C | protein-coding |
| chr5_105501175_105501484 | chr5 | 105501176 | 105501484 | + | 0.766741 | Intergenic | Intergenic | -18141 | NM_133897 | 100604 | Mm.319847 | NM_133897 | ENSMUSG00000054720 | **Lrrc8c** | AD158\|AI326115\|E430036I04Rik\|fad158 | leucine rich repeat containing 8 family, member C | protein-coding |
| chr5_105557742_105558859 | chr5 | 105557743 | 105558859 | + | 0.676787 | intron (NM_133897, intron 2 of 3) | intron (NM_133897, intron 2 of 3) | 38830 | NM_133897 | 100604 | Mm.319847 | NM_133897 | ENSMUSG00000054720 | **Lrrc8c** | AD158\|AI326115\|E430036I04Rik\|fad158 | leucine rich repeat containing 8 family, member C | protein-coding |
| chr5_105499438_105499758 | chr5 | 105499439 | 105499758 | + | 0.622932 | Intergenic | ORR1B2\|LTR\|ERVL-MaLR | -19873 | NM_133897 | 100604 | Mm.319847 | NM_133897 | ENSMUSG00000054720 | **Lrrc8c** | AD158\|AI326115\|E430036I04Rik\|fad158 | leucine rich repeat containing 8 family, member C | protein-coding |
| chr5_105620379_105620920 | chr5 | 105620380 | 105620920 | + | 1.83199 | Intergenic | Intergenic | -79319 | NM_001122768 | 231549 | Mm.23837 | NM_178701 | ENSMUSG00000046079 | **Lrrc8d** | 2810473G09Rik\|4930525N13Rik\|A930019F03\|Lrrc5 | leucine rich repeat containing 8D | protein-coding |
| chr5_105618500_105618944 | chr5 | 105618501 | 105618944 | + | 1.79023 | Intergenic | MLT1G\|LTR\|ERVL-MaLR | -81247 | NM_001122768 | 231549 | Mm.23837 | NM_178701 | ENSMUSG00000046079 | **Lrrc8d** | 2810473G09Rik\|4930525N13Rik\|A930019F03\|Lrrc5 | leucine rich repeat containing 8D | protein-coding |
| chr5_105745778_105746376 | chr5 | 105745779 | 105746376 | + | 0.66958 | intron (NM_178701, intron 2 of 2) | ORR1C2\|LTR\|ERVL-MaLR | 45267 | NM_178701 | 231549 | Mm.23837 | NM_178701 | ENSMUSG00000046079 | **Lrrc8d** | 2810473G09Rik\|4930525N13Rik\|A930019F03\|Lrrc5 | leucine rich repeat containing 8D | protein-coding |
| chr5_105708544_105708985 | chr5 | 105708545 | 105708985 | + | 0.642938 | intron (NM_178701, intron 1 of 2) | ORR1C2\|LTR\|ERVL-MaLR | 7955 | NM_178701 | 231549 | Mm.23837 | NM_178701 | ENSMUSG00000046079 | **Lrrc8d** | 2810473G09Rik\|4930525N13Rik\|A930019F03\|Lrrc5 | leucine rich repeat containing 8D | protein-coding |
| chr1_91069557_91069986 | chr1 | 91069558 | 91069986 | + | 1.08109 | intron (NM_001111311, intron 2 of 7) | intron (NM_001111311, intron 2 of 7) | 16328 | NM_008515 | 16978 | Mm.45039 | NM_008515 | ENSMUSG00000026305 | **Lrrfip1** | AU024550\|Fliiap1 | leucine rich repeat (in FLII) interacting protein 1 | protein-coding |
| chr1_91023923_91024303 | chr1 | 91023924 | 91024303 | + | 0.835462 | intron (NM_001111312, intron 1 of 12) | intron (NM_001111312, intron 1 of 12) | 25386 | NM_001111312 | 16978 | Mm.45039 | NM_008515 | ENSMUSG00000026305 | **Lrrfip1** | AU024550\|Fliiap1 | leucine rich repeat (in FLII) interacting protein 1 | protein-coding |
| chr1_91085911_91086767 | chr1 | 91085912 | 91086767 | + | 0.821559 | intron (NM_008515, intron 7 of 22) | intron (NM_008515, intron 7 of 22) | 32895 | NM_008515 | 16978 | Mm.45039 | NM_008515 | ENSMUSG00000026305 | **Lrrfip1** | AU024550\|Fliiap1 | leucine rich repeat (in FLII) interacting protein 1 | protein-coding |
| chr1_91085916_91086492 | chr1 | 91085917 | 91086492 | + | 0.763821 | intron (NM_008515, intron 7 of 22) | intron (NM_008515, intron 7 of 22) | 32760 | NM_008515 | 16978 | Mm.45039 | NM_008515 | ENSMUSG00000026305 | **Lrrfip1** | AU024550\|Fliiap1 | leucine rich repeat (in FLII) interacting protein 1 | protein-coding |
| chr1_91091537_91091941 | chr1 | 91091538 | 91091941 | + | 0.64695 | intron (NM_008515, intron 9 of 22) | intron (NM_008515, intron 9 of 22) | 38295 | NM_008515 | 16978 | Mm.45039 | NM_008515 | ENSMUSG00000026305 | **Lrrfip1** | AU024550\|Fliiap1 | leucine rich repeat (in FLII) interacting protein 1 | protein-coding |
| chr1_91090510_91090837 | chr1 | 91090511 | 91090837 | + | 0.646507 | intron (NM_008515, intron 8 of 22) | intron (NM_008515, intron 8 of 22) | 37230 | NM_008515 | 16978 | Mm.45039 | NM_008515 | ENSMUSG00000026305 | **Lrrfip1** | AU024550\|Fliiap1 | leucine rich repeat (in FLII) interacting protein 1 | protein-coding |
| chr1_91091531_91092192 | chr1 | 91091532 | 91092192 | + | 0.610763 | intron (NM_008515, intron 9 of 22) | intron (NM_008515, intron 9 of 22) | 38418 | NM_008515 | 16978 | Mm.45039 | NM_008515 | ENSMUSG00000026305 | **Lrrfip1** | AU024550\|Fliiap1 | leucine rich repeat (in FLII) interacting protein 1 | protein-coding |
| chr12_41935384_41935908 | chr12 | 41935385 | 41935908 | + | 1.11015 | intron (NM_053122, intron 6 of 6) | MLT1G3\|LTR\|ERVL-MaLR | -449589 | NM_010733 | 16981 | Mm.404786 | NM_010733 | ENSMUSG00000036295 | **Lrrn3** | NLRR-3 | leucine rich repeat protein 3, neuronal | protein-coding |
| chr12_41949571_41950334 | chr12 | 41949572 | 41950334 | + | 1.10302 | intron (NM_053122, intron 6 of 6) | intron (NM_053122, intron 6 of 6) | -463896 | NM_010733 | 16981 | Mm.404786 | NM_010733 | ENSMUSG00000036295 | **Lrrn3** | NLRR-3 | leucine rich repeat protein 3, neuronal | protein-coding |
| chr12_41606211_41606640 | chr12 | 41606212 | 41606640 | + | 1.00358 | intron (NM_053122, intron 4 of 6) | intron (NM_053122, intron 4 of 6) | -120369 | NM_010733 | 16981 | Mm.404786 | NM_010733 | ENSMUSG00000036295 | **Lrrn3** | NLRR-3 | leucine rich repeat protein 3, neuronal | protein-coding |
| chr12_41500488_41500931 | chr12 | 41500489 | 41500931 | + | 0.870842 | intron (NM_053122, intron 4 of 6) | intron (NM_053122, intron 4 of 6) | -14653 | NM_010733 | 16981 | Mm.404786 | NM_010733 | ENSMUSG00000036295 | **Lrrn3** | NLRR-3 | leucine rich repeat protein 3, neuronal | protein-coding |
| chr12_41942952_41943405 | chr12 | 41942953 | 41943405 | + | 0.713308 | intron (NM_053122, intron 6 of 6) | intron (NM_053122, intron 6 of 6) | -457122 | NM_010733 | 16981 | Mm.404786 | NM_010733 | ENSMUSG00000036295 | **Lrrn3** | NLRR-3 | leucine rich repeat protein 3, neuronal | protein-coding |
| chr6_119369170_119369614 | chr6 | 119369171 | 119369614 | + | 0.715612 | intron (NM_197985, intron 2 of 7) | intron (NM_197985, intron 2 of 7) | -38626 | NM_172492 | 211187 | Mm.121498 | NM_172492 | ENSMUSG00000055003 | **Lrtm2** | A230084J22\|AI841794 | leucine-rich repeats and transmembrane domains 2 | protein-coding |
| chr6_119369735_119370003 | chr6 | 119369736 | 119370003 | + | 0.678757 | intron (NM_197985, intron 2 of 7) | B3\|SINE\|B2 | -39103 | NM_172492 | 211187 | Mm.121498 | NM_172492 | ENSMUSG00000055003 | **Lrtm2** | A230084J22\|AI841794 | leucine-rich repeats and transmembrane domains 2 | protein-coding |
| chr6_119368984_119369599 | chr6 | 119368985 | 119369599 | + | 0.653279 | intron (NM_197985, intron 2 of 7) | B1_Mus2\|SINE\|Alu | -38526 | NM_172492 | 211187 | Mm.121498 | NM_172492 | ENSMUSG00000055003 | **Lrtm2** | A230084J22\|AI841794 | leucine-rich repeats and transmembrane domains 2 | protein-coding |
| chr6_119343614_119343911 | chr6 | 119343615 | 119343911 | + | 0.614781 | intron (NM_001033382, intron 30 of 37) | intron (NM_001033382, intron 30 of 37) | -12997 | NM_172492 | 211187 | Mm.121498 | NM_172492 | ENSMUSG00000055003 | **Lrtm2** | A230084J22\|AI841794 | leucine-rich repeats and transmembrane domains 2 | protein-coding |
| chr11_102172340_102172876 | chr11 | 102172341 | 102172876 | + | 0.655248 | intron (NM_172947, intron 2 of 4) | B1_Mus2\|SINE\|Alu | 12648 | NM_172947 | 268490 | Mm.259079 | NM_172947 | | **Lsm12** | 1110032E16Rik\|1110059P07\|2600001B17Rik | LSM12 homolog (S. cerevisiae) | protein-coding |
| chr11_102172163_102172884 | chr11 | 102172164 | 102172884 | + | 0.643184 | intron (NM_172947, intron 2 of 4) | intron (NM_172947, intron 2 of 4) | 12732 | NM_172947 | 268490 | Mm.259079 | NM_172947 | | **Lsm12** | 1110032E16Rik\|1110059P07\|2600001B17Rik | LSM12 homolog (S. cerevisiae) | protein-coding |
| chr7_34341971_34342478 | chr7 | 34341972 | 34342478 | + | 0.70216 | Intergenic | Intergenic | 47315 | NM_025948 | 67070 | Mm.276503 | NM_025948 | ENSMUSG00000066568 | **Lsm14a** | 2700023B17Rik\|AA407828\|AU017544\|Tral | LSM14 homolog A (SCD6, S. cerevisiae) | protein-coding |
| chr7_142462126_142463361 | chr7 | 142462127 | 142463361 | + | 0.848352 | intron (NM_001271510, intron 1 of 10) | intron (NM_001271510, intron 1 of 10) | 1932 | NM_001136071 | 16985 | Mm.234003 | NM_019391 | ENSMUSG00000018819 | **Lsp1** | Lsp-1\|WP34\|p50\|pp52 | lymphocyte specific 1 | protein-coding |
| chr7_142470729_142471200 | chr7 | 142470730 | 142471200 | + | 0.823187 | promoter-TSS (NM_001271508) | promoter-TSS (NM_001271508) | -878 | NM_001271508 | 16985 | Mm.234003 | NM_019391 | ENSMUSG00000018819 | **Lsp1** | Lsp-1\|WP34\|p50\|pp52 | lymphocyte specific 1 | protein-coding |
| chr7_142470560_142471233 | chr7 | 142470561 | 142471233 | + | 0.75036 | promoter-TSS (NM_001271508) | promoter-TSS (NM_001271508) | -946 | NM_001271508 | 16985 | Mm.234003 | NM_019391 | ENSMUSG00000018819 | **Lsp1** | Lsp-1\|WP34\|p50\|pp52 | lymphocyte specific 1 | protein-coding |
| chr17_35215152_35215354 | chr17 | 35215153 | 35215354 | + | 0.895511 | Intergenic | Intergenic | -9902 | NM_010735 | 16992 | Mm.87787 | NM_010735 | ENSMUSG00000024402 | **Lta** | LT\|LT-[a]\|LT-alpha\|LT[a]\|LTalpha\|Ltx\|TNF-beta\|TNFSF1\|Tnfb\|Tnfsf1b\|hlb382 | lymphotoxin A | protein-coding |
| chr17_35205023_35205683 | chr17 | 35205024 | 35205683 | + | 0.684916 | promoter-TSS (NM_010735) | promoter-TSS (NM_010735) | -2 | NM_010735 | 16992 | Mm.87787 | NM_010735 | ENSMUSG00000024402 | **Lta** | LT\|LT-[a]\|LT-alpha\|LT[a]\|LTalpha\|Ltx\|TNF-beta\|TNFSF1\|Tnfb\|Tnfsf1b\|hlb382 | lymphotoxin A | protein-coding |
| chr17_35219570_35220347 | chr17 | 35219571 | 35220347 | + | 0.642895 | TTS (NM_010909) | TTS (NM_010909) | -14608 | NM_010735 | 16992 | Mm.87787 | NM_010735 | ENSMUSG00000024402 | **Lta** | LT\|LT-[a]\|LT-alpha\|LT[a]\|LTalpha\|Ltx\|TNF-beta\|TNFSF1\|Tnfb\|Tnfsf1b\|hlb382 | lymphotoxin A | protein-coding |
| chr10_93403264_93403732 | chr10 | 93403265 | 93403732 | + | 0.646269 | Intergenic | RLTR31D_MM\|LTR\|ERVK | -49898 | NM_008517 | 16993 | Mm.271071 | NM_008517 | ENSMUSG00000015889 | **Lta4h** | - | leukotriene A4 hydrolase | protein-coding |
| chr17_74869523_74870050 | chr17 | 74869524 | 74870050 | + | 1.64149 | Intergenic | Intergenic | -135742 | NM_019919 | 268977 | Mm.269747 | NM_019919 | ENSMUSG00000001870 | **Ltbp1** | 9430031G15Rik\|9830146M04\|Ltbp-1\|Tgfb\|b2b1000Clo | latent transforming growth factor beta binding protein 1 | protein-coding |
| chr17_74869341_74870371 | chr17 | 74869342 | 74870371 | + | 1.53252 | Intergenic | Intergenic | -135673 | NM_019919 | 268977 | Mm.269747 | NM_019919 | ENSMUSG00000001870 | **Ltbp1** | 9430031G15Rik\|9830146M04\|Ltbp-1\|Tgfb\|b2b1000Clo | latent transforming growth factor beta binding protein 1 | protein-coding |
| chr17_74864769_74865351 | chr17 | 74864770 | 74865351 | + | 1.28171 | Intergenic | Intergenic | -140469 | NM_019919 | 268977 | Mm.269747 | NM_019919 | ENSMUSG00000001870 | **Ltbp1** | 9430031G15Rik\|9830146M04\|Ltbp-1\|Tgfb\|b2b1000Clo | latent transforming growth factor beta binding protein 1 | protein-coding |
| chr17_75001017_75001402 | chr17 | 75001018 | 75001402 | + | 0.967982 | Intergenic | ORR1G\|LTR\|ERVL-MaLR | -4319 | NM_019919 | 268977 | Mm.269747 | NM_019919 | ENSMUSG00000001870 | **Ltbp1** | 9430031G15Rik\|9830146M04\|Ltbp-1\|Tgfb\|b2b1000Clo | latent transforming growth factor beta binding protein 1 | protein-coding |
| chr17_26284898_26285546 | chr17 | 26284899 | 26285546 | + | 1.08095 | 3' UTR (NM_028190, exon 10 of 10) | 3' UTR (NM_028190, exon 10 of 10) | 32312 | NR_037905 | 66978 | Mm.440216 | NM_025881 | ENSMUSG00000024188 | **Luc7l** | 1810045C04Rik\|2410018D03Rik | Luc7 homolog (S. cerevisiae)-like | protein-coding |
| chr15_74932430_74932963 | chr15 | 74932431 | 74932963 | + | 0.969459 | Intergenic | Intergenic | -22354 | NM_001164039 | 17069 | Mm.788 | NM_008529 | ENSMUSG00000022587 | **Ly6e** | 9804\|Ly67\|RIG-E\|Sca-2\|TSA-1\|Tsa1 | lymphocyte antigen 6 complex, locus E | protein-coding |
| chr15_74920593_74921030 | chr15 | 74920594 | 74921030 | + | 0.855917 | Intergenic | MTD\|LTR\|ERVL-MaLR | -34239 | NM_001164039 | 17069 | Mm.788 | NM_008529 | ENSMUSG00000022587 | **Ly6e** | 9804\|Ly67\|RIG-E\|Sca-2\|TSA-1\|Tsa1 | lymphocyte antigen 6 complex, locus E | protein-coding |
| chr15_74919554_74920038 | chr15 | 74919555 | 74920038 | + | 0.671063 | Intergenic | Intergenic | -35255 | NM_001164039 | 17069 | Mm.788 | NM_008529 | ENSMUSG00000022587 | **Ly6e** | 9804\|Ly67\|RIG-E\|Sca-2\|TSA-1\|Tsa1 | lymphocyte antigen 6 complex, locus E | protein-coding |
| chr15_74934789_74935102 | chr15 | 74934790 | 74935102 | + | 0.654009 | Intergenic | MERV1_I-int\|LTR\|ERV1 | -20105 | NM_001164039 | 17069 | Mm.788 | NM_008529 | ENSMUSG00000022587 | **Ly6e** | 9804\|Ly67\|RIG-E\|Sca-2\|TSA-1\|Tsa1 | lymphocyte antigen 6 complex, locus E | protein-coding |
| chr15_75544140_75544526 | chr15 | 75544141 | 75544526 | + | 0.768878 | Intergenic | Intergenic | 22315 | NM_001135689 | 23934 | Mm.22154 | NM_011837 | ENSMUSG00000022577 | **Ly6h** | - | lymphocyte antigen 6 complex, locus H | protein-coding |
| chr2_60370101_60370871 | chr2 | 60370102 | 60370871 | + | 1.69095 | intron (NM_013825, intron 2 of 34) | intron (NM_013825, intron 2 of 34) | 12745 | NM_013825 | 17076 | Mm.2074 | NM_013825 | ENSMUSG00000026980 | **Ly75** | CD205\|DEC-205\|DEC205 | lymphocyte antigen 75 | protein-coding |
| chr2_60369345_60370027 | chr2 | 60369346 | 60370027 | + | 0.997484 | intron (NM_013825, intron 2 of 34) | intron (NM_013825, intron 2 of 34) | 13545 | NM_013825 | 17076 | Mm.2074 | NM_013825 | ENSMUSG00000026980 | **Ly75** | CD205\|DEC-205\|DEC205 | lymphocyte antigen 75 | protein-coding |
| chr2_60369362_60369583 | chr2 | 60369363 | 60369583 | + | 0.759744 | intron (NM_013825, intron 2 of 34) | Lx4A\|LINE\|L1 | 13758 | NM_013825 | 17076 | Mm.2074 | NM_013825 | ENSMUSG00000026980 | **Ly75** | CD205\|DEC-205\|DEC205 | lymphocyte antigen 75 | protein-coding |
| chr13_37290568_37291096 | chr13 | 37290569 | 37291096 | + | 1.35696 | Intergenic | Intergenic | -54513 | NM_010745 | 17084 | Mm.2639 | NM_010745 | ENSMUSG00000021423 | **Ly86** | MD-1\|MD1 | lymphocyte antigen 86 | protein-coding |

| | | | | | | | | | | | | | | | | | |
|---|---|---|---|---|---|---|---|---|---|---|---|---|---|---|---|---|---|
| chr13_37403348_37404071 | chr13 | 37403349 | 37404071 | + | 0.928482 | intron (NM_010745, intron 3 of 4) | L1MA7\|LINE\|L1 | 58365 | NM_010745 | 17084 | Mm.2639 | NM_010745 | ENSMUSG00000021423 | **Ly86** | MD-1\|MD1 | lymphocyte antigen 86 | protein-coding |
| chr13_37302445_37303518 | chr13 | 37302446 | 37303518 | + | 0.905995 | Intergenic | Intergenic | -42363 | NM_010745 | 17084 | Mm.2639 | NM_010745 | ENSMUSG00000021423 | **Ly86** | MD-1\|MD1 | lymphocyte antigen 86 | protein-coding |
| chr13_37241169_37241645 | chr13 | 37241170 | 37241645 | + | 0.633551 | Intergenic | MLT1O\|LTR\|ERVL-MaLR | -103938 | NM_010745 | 17084 | Mm.2639 | NM_010745 | ENSMUSG00000021423 | **Ly86** | MD-1\|MD1 | lymphocyte antigen 86 | protein-coding |
| chr1_171611160_171611441 | chr1 | 171611161 | 171611441 | + | 0.990882 | Intergenic | Lx\|LINE\|L1 | -3891 | NM_008534 | 17085 | Mm.560 | NM_008534 | ENSMUSG00000004707 | **Ly9** | AI893573\|CD229\|Lgp100\|Ly-9\|SLAMF3\|T100 | lymphocyte antigen 9 | protein-coding |
| chr1_171628874_171629524 | chr1 | 171628875 | 171629524 | + | 0.854599 | Intergenic | L2a\|LINE\|L2 | -21789 | NM_008534 | 17085 | Mm.560 | NM_008534 | ENSMUSG00000004707 | **Ly9** | AI893573\|CD229\|Lgp100\|Ly-9\|SLAMF3\|T100 | lymphocyte antigen 9 | protein-coding |
| chr1_171610806_171611460 | chr1 | 171610807 | 171611460 | + | 0.772095 | Intergenic | (TCC)n\|Simple_repeat\|Simple_repeat | -3723 | NM_008534 | 17085 | Mm.560 | NM_008534 | ENSMUSG00000004707 | **Ly9** | AI893573\|CD229\|Lgp100\|Ly-9\|SLAMF3\|T100 | lymphocyte antigen 9 | protein-coding |
| chr1_171628843_171629885 | chr1 | 171628844 | 171629885 | + | 0.766591 | Intergenic | Intergenic | -21954 | NM_008534 | 17085 | Mm.560 | NM_008534 | ENSMUSG00000004707 | **Ly9** | AI893573\|CD229\|Lgp100\|Ly-9\|SLAMF3\|T100 | lymphocyte antigen 9 | protein-coding |
| chr1_16696492_16696992 | chr1 | 16696493 | 16696992 | + | 0.857458 | intron (NM_001159711, intron 2 of 4) | intron (NM_001159711, intron 2 of 4) | 8286 | NM_016923 | 17087 | Mm.116844 | NM_016923 | ENSMUSG00000025779 | **Ly96** | ESOP-1\|MD-2\|MD2 | lymphocyte antigen 96 | protein-coding |
| chr1_16696180_16697034 | chr1 | 16696181 | 16697034 | + | 0.845495 | intron (NM_001159711, intron 2 of 4) | L1_Mus2\|LINE\|L1 | 8151 | NM_016923 | 17087 | Mm.116844 | NM_016923 | ENSMUSG00000025779 | **Ly96** | ESOP-1\|MD-2\|MD2 | lymphocyte antigen 96 | protein-coding |
| chr1_16747068_16747733 | chr1 | 16747069 | 16747733 | + | 0.834454 | Intergenic | Intergenic | 58945 | NM_016923 | 17087 | Mm.116844 | NM_016923 | ENSMUSG00000025779 | **Ly96** | ESOP-1\|MD-2\|MD2 | lymphocyte antigen 96 | protein-coding |
| chr1_16737733_16738325 | chr1 | 16737734 | 16738325 | + | 0.6489 | Intergenic | MERV1_LTR\|LTR\|ERV1 | 49573 | NM_016923 | 17087 | Mm.116844 | NM_016923 | ENSMUSG00000025779 | **Ly96** | ESOP-1\|MD-2\|MD2 | lymphocyte antigen 96 | protein-coding |
| chr1_37931750_37932209 | chr1 | 37931751 | 37932209 | + | 0.694719 | Intergenic | Intergenic | -15487 | NM_001033427 | 332427 | Mm.332981 | NM_001033427 | ENSMUSG00000061584 | **Lyg2** | Gm776 | lysozyme G-like 2 | protein-coding |
| chr2_49744229_49744850 | chr2 | 49744230 | 49744850 | + | 1.43696 | intron (NM_008449, intron 20 of 25) | intron (NM_008449, intron 20 of 25) | -43146 | NM_027990 | 71897 | Mm.41381 | NM_027990 | ENSMUSG00000026765 | **Lypd6b** | 2310010M24Rik\|AW049525 | LY6/PLAUR domain containing 6B | protein-coding |
| chr2_49789414_49789864 | chr2 | 49789415 | 49789864 | + | 0.97061 | intron (NM_027990, intron 1 of 9) | intron (NM_027990, intron 1 of 9) | 1953 | NM_027990 | 71897 | Mm.41381 | NM_027990 | ENSMUSG00000026765 | **Lypd6b** | 2310010M24Rik\|AW049525 | LY6/PLAUR domain containing 6B | protein-coding |
| chr2_49789510_49789847 | chr2 | 49789511 | 49789847 | + | 0.967905 | intron (NM_027990, intron 1 of 9) | intron (NM_027990, intron 1 of 9) | 1993 | NM_027990 | 71897 | Mm.41381 | NM_027990 | ENSMUSG00000026765 | **Lypd6b** | 2310010M24Rik\|AW049525 | LY6/PLAUR domain containing 6B | protein-coding |
| chr2_49840750_49841424 | chr2 | 49840751 | 49841424 | + | 0.900874 | intron (NM_027990, intron 1 of 9) | intron (NM_027990, intron 1 of 9) | 53401 | NM_027990 | 71897 | Mm.41381 | NM_027990 | ENSMUSG00000026765 | **Lypd6b** | 2310010M24Rik\|AW049525 | LY6/PLAUR domain containing 6B | protein-coding |
| chr2_49789155_49789863 | chr2 | 49789156 | 49789863 | + | 0.792854 | intron (NM_027990, intron 1 of 9) | intron (NM_027990, intron 1 of 9) | 1823 | NM_027990 | 71897 | Mm.41381 | NM_027990 | ENSMUSG00000026765 | **Lypd6b** | 2310010M24Rik\|AW049525 | LY6/PLAUR domain containing 6B | protein-coding |
| chr2_49886274_49886570 | chr2 | 49886275 | 49886570 | + | 0.658261 | intron (NM_027990, intron 3 of 9) | intron (NM_027990, intron 3 of 9) | 98736 | NM_027990 | 71897 | Mm.41381 | NM_027990 | ENSMUSG00000026765 | **Lypd6b** | 2310010M24Rik\|AW049525 | LY6/PLAUR domain containing 6B | protein-coding |
| chr7_119906536_119907156 | chr7 | 119906537 | 119907156 | + | 1.44121 | intron (NR_104377, intron 1 of 2) | intron (NR_104377, intron 1 of 2) | 6735 | NR_104377 | 73919 | Mm.46381 | NM_029610 | ENSMUSG00000030922 | **Lyrm1** | 1110065L10Rik\|2310004B22Rik\|4930404J24Rik | LYR motif containing 1 | protein-coding |
| chr7_119920486_119920799 | chr7 | 119920487 | 119920799 | + | 0.723983 | Intergenic | Intergenic | 20532 | NR_104377 | 73919 | Mm.46381 | NM_029610 | ENSMUSG00000030922 | **Lyrm1** | 1110065L10Rik\|2310004B22Rik\|4930404J24Rik | LYR motif containing 1 | protein-coding |
| chr13_36081730_36082094 | chr13 | 36081731 | 36082094 | + | 1.11917 | intron (NM_201358, intron 2 of 2) | intron (NM_201358, intron 2 of 2) | 35445 | NM_201358 | 380840 | Mm.390576 | NM_201358 | ENSMUSG00000046573 | **Lyrm4** | BC034664\|Gm903 | LYR motif containing 4 | protein-coding |
| chr13_36081578_36082111 | chr13 | 36081579 | 36082111 | + | 0.987355 | intron (NM_201358, intron 2 of 2) | intron (NM_201358, intron 2 of 2) | 35512 | NM_201358 | 380840 | Mm.390576 | NM_201358 | ENSMUSG00000046573 | **Lyrm4** | BC034664\|Gm903 | LYR motif containing 4 | protein-coding |
| chr13_36114453_36114723 | chr13 | 36114454 | 36114723 | + | 0.954937 | intron (NM_201358, intron 1 of 2) | ORR1C2\|LTR\|ERVL-MaLR | 2769 | NM_201358 | 380840 | Mm.390576 | NM_201358 | ENSMUSG00000046573 | **Lyrm4** | BC034664\|Gm903 | LYR motif containing 4 | protein-coding |
| chr13_81667616_81668445 | chr13 | 81667617 | 81668445 | + | 1.52332 | intron (NM_030257, intron 2 of 2) | intron (NM_030257, intron 2 of 2) | 10225 | NM_030257 | 80289 | Mm.259969 | NM_030257 | ENSMUSG00000035840 | **Lysmd3** | 1110030H10Rik\|BC003322 | LysM, putative peptidoglycan-binding, domain containing 3 | protein-coding |
| chr7_67112486_67112925 | chr7 | 67112487 | 67112925 | + | 0.873116 | intron (NM_001033877, intron 19 of 22) | intron (NM_001033877, intron 19 of 22) | -109838 | NM_175215 | 75099 | Mm.121691 | NM_175215 | ENSMUSG00000043831 | **Lysmd4** | 4930506D23Rik | LysM, putative peptidoglycan-binding, domain containing 4 | protein-coding |
| chr7_67112474_67113124 | chr7 | 67112475 | 67113124 | + | 0.807895 | intron (NM_001033877, intron 19 of 22) | intron (NM_001033877, intron 19 of 22) | -109745 | NM_175215 | 75099 | Mm.121691 | NM_175215 | ENSMUSG00000043831 | **Lysmd4** | 4930506D23Rik | LysM, putative peptidoglycan-binding, domain containing 4 | protein-coding |
| chr7_67243888_67244269 | chr7 | 67243889 | 67244269 | + | 0.658962 | intron (NM_001291196, intron 9 of 11) | intron (NM_001291196, intron 9 of 11) | 21463 | NM_001191051 | 75099 | Mm.121691 | NM_175215 | ENSMUSG00000043831 | **Lysmd4** | 4930506D23Rik | LysM, putative peptidoglycan-binding, domain containing 4 | protein-coding |
| chr13_13584472_13584711 | chr13 | 13584473 | 13584711 | + | 0.83848 | Intergenic | Intergenic | -5817 | NM_010748 | 17101 | Mm.342337 | NM_010748 | ENSMUSG00000019726 | **Lyst** | D13Sfk13\|beige\|bg | lysosomal trafficking regulator | protein-coding |
| chr13_13583270_13583581 | chr13 | 13583271 | 13583581 | + | 0.807786 | Intergenic | Intergenic | -6983 | NM_010748 | 17101 | Mm.342337 | NM_010748 | ENSMUSG00000019726 | **Lyst** | D13Sfk13\|beige\|bg | lysosomal trafficking regulator | protein-coding |
| chr9_123740670_123741390 | chr9 | 123740671 | 123741390 | + | 1.1485 | Intergenic | Intergenic | -23473 | NM_033322 | 93730 | Mm.378263 | NM_033322 | ENSMUSG00000025245 | **Lztfl1** | 5530402H04Rik\|6130400H19Rik\|AI414725\|AW048545 | leucine zipper transcription factor-like 1 | protein-coding |
| chr9_123732507_123732976 | chr9 | 123732508 | 123732976 | + | 0.776464 | Intergenic | Intergenic | -15185 | NM_033322 | 93730 | Mm.378263 | NM_033322 | ENSMUSG00000025245 | **Lztfl1** | 5530402H04Rik\|6130400H19Rik\|AI414725\|AW048545 | leucine zipper transcription factor-like 1 | protein-coding |
| chr8_69148767_69149716 | chr8 | 69148768 | 69149716 | + | 0.994725 | Intergenic | Intergenic | -8289 | NM_199364 | 211134 | Mm.479807 | NM_199364 | ENSMUSG00000036306 | **Lzts1** | F37\|FEZ1 | leucine zipper, putative tumor suppressor 1 | protein-coding |
| chr8_69123486_69123814 | chr8 | 69123487 | 69123814 | + | 0.634173 | Intergenic | MLTR14\|LTR\|ERV1 | 17303 | NM_199364 | 211134 | Mm.479807 | NM_199364 | ENSMUSG00000036306 | **Lzts1** | F37\|FEZ1 | leucine zipper, putative tumor suppressor 1 | protein-coding |
| chr6_82947986_82948607 | chr6 | 82947987 | 82948607 | + | 1.59912 | intron (NM_033079, intron 1 of 9) | (CATAA)n\|Simple_repeat\|Simple_repeat | 1375 | NM_033079 | 110958 | Mm.100652 | NM_033079 | ENSMUSG00000030041 | **M1ap** | D6Mm5e | meiosis 1 associated protein | protein-coding |
| chr6_82969381_82969977 | chr6 | 82969382 | 82969977 | + | 0.732541 | intron (NM_033079, intron 3 of 9) | L1MCa\|LINE\|L1 | 22757 | NM_033079 | 110958 | Mm.100652 | NM_033079 | ENSMUSG00000030041 | **M1ap** | D6Mm5e | meiosis 1 associated protein | protein-coding |
| chr5_140257316_140257996 | chr5 | 140257317 | 140257996 | + | 1.40428 | intron (NM_010752, intron 10 of 17) | intron (NM_010752, intron 10 of 17) | 63896 | NM_010752 | 17120 | Mm.27250 | NM_010752 | ENSMUSG00000029554 | **Mad1l1** | AI173502\|AW550425\|Mad1 | MAD1 mitotic arrest deficient 1-like 1 | protein-coding |
| chr5_140150821_140151662 | chr5 | 140150822 | 140151662 | + | 0.646107 | intron (NM_010752, intron 11 of 17) | MT2A\|LTR\|ERVL | 170310 | NM_010752 | 17120 | Mm.27250 | NM_010752 | ENSMUSG00000029554 | **Mad1l1** | AI173502\|AW550425\|Mad1 | MAD1 mitotic arrest deficient 1-like 1 | protein-coding |
| chr6_66498243_66498741 | chr6 | 66498244 | 66498741 | + | 1.2471 | Intergenic | Intergenic | -36976 | NM_019499 | 56150 | Mm.489653 | NM_019499 | ENSMUSG00000029910 | **Mad2l1** | AA673185\|MAD2 | MAD2 mitotic arrest deficient-like 1 | protein-coding |
| chr6_66472722_66473032 | chr6 | 66472723 | 66473032 | + | 0.729691 | Intergenic | Intergenic | -62591 | NM_019499 | 56150 | Mm.489653 | NM_019499 | ENSMUSG00000029910 | **Mad2l1** | AA673185\|MAD2 | MAD2 mitotic arrest deficient-like 1 | protein-coding |
| chr6_66472722_66473412 | chr6 | 66472723 | 66473412 | + | 0.685967 | Intergenic | Intergenic | -62401 | NM_019499 | 56150 | Mm.489653 | NM_019499 | ENSMUSG00000029910 | **Mad2l1** | AA673185\|MAD2 | MAD2 mitotic arrest deficient-like 1 | protein-coding |
| chr8_115770860_115771486 | chr8 | 115770861 | 115771486 | + | 1.4355 | Intergenic | Intergenic | -64279 | NM_001025577 | 17132 | Mm.275549 | NM_001025577 | | **Maf** | 2810401A20Rik\|A230108G15Rik\|AW047063\|c-maf | avian musculoaponeurotic fibrosarcoma (v-maf) AS42 oncogene homolog | protein-coding |
| chr8_115183565_115183958 | chr8 | 115183566 | 115183958 | + | 1.02104 | intron (NM_019573, intron 8 of 8) | intron (NM_019573, intron 8 of 8) | 523132 | NM_001025577 | 17132 | Mm.275549 | NM_001025577 | | **Maf** | 2810401A20Rik\|A230108G15Rik\|AW047063\|c-maf | avian musculoaponeurotic fibrosarcoma (v-maf) AS42 oncogene homolog | protein-coding |
| chr8_115625036_115625514 | chr8 | 115625037 | 115625514 | + | 0.998937 | Intergenic | Intergenic | 81619 | NM_001025577 | 17132 | Mm.275549 | NM_001025577 | | **Maf** | 2810401A20Rik\|A230108G15Rik\|AW047063\|c-maf | avian musculoaponeurotic fibrosarcoma (v-maf) AS42 oncogene homolog | protein-coding |
| chr8_115726844_115727200 | chr8 | 115726845 | 115727200 | + | 0.9877 | Intergenic | Intergenic | -20128 | NM_001025577 | 17132 | Mm.275549 | NM_001025577 | | **Maf** | 2810401A20Rik\|A230108G15Rik\|AW047063\|c-maf | avian musculoaponeurotic fibrosarcoma (v-maf) AS42 oncogene homolog | protein-coding |
| chr8_115742577_115743091 | chr8 | 115742578 | 115743091 | + | 0.887797 | Intergenic | Intergenic | -35940 | NM_001025577 | 17132 | Mm.275549 | NM_001025577 | | **Maf** | 2810401A20Rik\|A230108G15Rik\|AW047063\|c-maf | avian musculoaponeurotic fibrosarcoma (v-maf) AS42 oncogene homolog | protein-coding |
| chr8_115496829_115497167 | chr8 | 115496830 | 115497167 | + | 0.79084 | Intergenic | Intergenic | 209896 | NM_001025577 | 17132 | Mm.275549 | NM_001025577 | | **Maf** | 2810401A20Rik\|A230108G15Rik\|AW047063\|c-maf | avian musculoaponeurotic fibrosarcoma (v-maf) AS42 oncogene homolog | protein-coding |
| chr8_115445124_115445606 | chr8 | 115445125 | 115445606 | + | 0.769572 | Intergenic | Intergenic | 261529 | NM_001025577 | 17132 | Mm.275549 | NM_001025577 | | **Maf** | 2810401A20Rik\|A230108G15Rik\|AW047063\|c-maf | avian musculoaponeurotic fibrosarcoma (v-maf) AS42 oncogene homolog | protein-coding |
| chr8_115226878_115227871 | chr8 | 115226879 | 115227871 | + | 0.723797 | intron (NM_019573, intron 8 of 8) | intron (NM_019573, intron 8 of 8) | 479519 | NM_001025577 | 17132 | Mm.275549 | NM_001025577 | | **Maf** | 2810401A20Rik\|A230108G15Rik\|AW047063\|c-maf | avian musculoaponeurotic fibrosarcoma (v-maf) AS42 oncogene homolog | protein-coding |
| chr2_160397436_160398334 | chr2 | 160397437 | 160398334 | + | 0.89717 | Intergenic | MER5A\|DNA\|hAT-Charlie | -30820 | NM_010658 | 16658 | Mm.330745 | NM_010658 | ENSMUSG00000074622 | **Mafb** | Kreisler\|Krml\|Krml1\|kr | v-maf musculoaponeurotic fibrosarcoma oncogene family, protein B (avian) | protein-coding |
| chr2_160467530_160467847 | chr2 | 160467531 | 160467847 | + | 0.778866 | Intergenic | Intergenic | -100624 | NM_010658 | 16658 | Mm.330745 | NM_010658 | ENSMUSG00000074622 | **Mafb** | Kreisler\|Krml\|Krml1\|kr | v-maf musculoaponeurotic fibrosarcoma oncogene family, protein B (avian) | protein-coding |
| chr2_160467530_160468080 | chr2 | 160467531 | 160468080 | + | 0.689442 | Intergenic | RMER17C\|LTR\|ERVK | -100740 | NM_010658 | 16658 | Mm.330745 | NM_010658 | ENSMUSG00000074622 | **Mafb** | Kreisler\|Krml\|Krml1\|kr | v-maf musculoaponeurotic fibrosarcoma oncogene family, protein B (avian) | protein-coding |
| chr15_79346073_79346416 | chr15 | 79346074 | 79346416 | + | 0.894843 | Intergenic | Intergenic | -1433 | NM_010755 | 17133 | Mm.86646 | NM_010755 | ENSMUSG00000042622 | **Maff** | - | v-maf musculoaponeurotic fibrosarcoma oncogene family, protein F (avian) | protein-coding |
| chr5_139794883_139795216 | chr5 | 139794884 | 139795216 | + | 0.684777 | intron (NM_010757, intron 1 of 2) | intron (NM_010757, intron 1 of 2) | 3514 | NM_010757 | 17135 | Mm.157313 | NM_010757 | ENSMUSG00000018143 | **Mafk** | AW061068\|NF-E2\|Nfe2u | v-maf musculoaponeurotic fibrosarcoma oncogene family, protein K (avian) | protein-coding |
| chrX_150817347_150817628 | chrX | 150817348 | 150817628 | + | 0.747191 | Intergenic | Intergenic | -3149 | NM_030700 | 80884 | Mm.22575 | NM_030700 | ENSMUSG00000025268 | **Maged2** | 4833439A22Rik\|Mage-d2\|ORF1 | melanoma antigen, family D, 2 | protein-coding |

| | | | | | | | | | | | | | | | | | |
|---|---|---|---|---|---|---|---|---|---|---|---|---|---|---|---|---|---|
| chr5_20377099_20377956 | chr5 | 20377100 | 20377956 | + | 1.8462 | intron (NM_001170746, intron 7 of 22) | intron (NM_001170746, intron 7 of 22) | 470010 | NM_015823 | 50791 | Mm.332231 | NM_015823 | ENSMUSG00000040003 | **Magi2** | AIP-1\|Acvri1\|Acvrinp1\|Acvrip1\|Magi-2\|S-SCAM\|mKIAA0705 | membrane associated guanylate kinase, WW and PDZ domain containing 2 | protein-coding |
| chr2_127722229_127722918 | chr2 | 127722230 | 127722918 | + | 0.735074 | intron (NM_145532, intron 1 of 3) | intron (NM_145532, intron 1 of 3) | 7323 | NM_145532 | 228576 | Mm.133036 | NM_145532 | ENSMUSG00000027377 | **Mall** | BC012256\|BENE | mal, T cell differentiation protein-like | protein-coding |
| chr6_49130397_49130758 | chr6 | 49130398 | 49130758 | + | 0.95531 | intron (NM_023670, intron 5 of 14) | intron (NM_023670, intron 5 of 14) | 56783 | NM_029353 | 75593 | Mm.439947 | NM_029353 | ENSMUSG00000029815 | **Malsu1** | 2410003K15Rik\|Matsu1 | mitochondrial assembly of ribosomal large subunit 1 | protein-coding |
| chr6_49129813_49130120 | chr6 | 49129814 | 49130120 | + | 0.668585 | intron (NM_023670, intron 5 of 14) | intron (NM_023670, intron 5 of 14) | 56172 | NM_029353 | 75593 | Mm.439947 | NM_029353 | ENSMUSG00000029815 | **Malsu1** | 2410003K15Rik\|Matsu1 | mitochondrial assembly of ribosomal large subunit 1 | protein-coding |
| chr18_65478894_65479516 | chr18 | 65478895 | 65479516 | + | 0.873487 | TTS (NM_172833) | TTS (NM_172833) | 48208 | NM_172833 | 240354 | Mm.132613 | NM_172833 | ENSMUSG00000032688 | **Malt1** | A630046N12\|D430033E09Rik | mucosa associated lymphoid tissue lymphoma translocation gene 1 | protein-coding |
| chr18_65481151_65481749 | chr18 | 65481152 | 65481749 | + | 0.722527 | intron (NR_105742, intron 3 of 6) | intron (NR_105742, intron 3 of 6) | 50453 | NM_172833 | 240354 | Mm.132613 | NM_172833 | ENSMUSG00000032688 | **Malt1** | A630046N12\|D430033E09Rik | mucosa associated lymphoid tissue lymphoma translocation gene 1 | protein-coding |
| chr9_13562660_13563109 | chr9 | 13562661 | 13563109 | + | 1.74865 | Intergenic | Intergenic | -57104 | NM_001013813 | 270118 | Mm.116898 | NM_173776 | ENSMUSG00000031925 | **Maml2** | 5930431H10\|BC032967 | mastermind like 2 (Drosophila) | protein-coding |
| chr9_13563325_13564066 | chr9 | 13563326 | 13564066 | + | 1.72183 | Intergenic | Intergenic | -56293 | NM_001013813 | 270118 | Mm.116898 | NM_173776 | ENSMUSG00000031925 | **Maml2** | 5930431H10\|BC032967 | mastermind like 2 (Drosophila) | protein-coding |
| chr9_13639472_13640230 | chr9 | 13639473 | 13640230 | + | 1.62368 | intron (NM_001013813, intron 1 of 3) | intron (NM_001013813, intron 1 of 3) | 19862 | NM_001013813 | 270118 | Mm.116898 | NM_173776 | ENSMUSG00000031925 | **Maml2** | 5930431H10\|BC032967 | mastermind like 2 (Drosophila) | protein-coding |
| chr9_13651889_13652635 | chr9 | 13651890 | 13652635 | + | 1.38997 | intron (NM_001013813, intron 1 of 3) | intron (NM_001013813, intron 1 of 3) | -10198 | NM_173776 | 270118 | Mm.116898 | NM_173776 | ENSMUSG00000031925 | **Maml2** | 5930431H10\|BC032967 | mastermind like 2 (Drosophila) | protein-coding |
| chr9_13631768_13632248 | chr9 | 13631769 | 13632248 | + | 0.997041 | intron (NM_001013813, intron 1 of 3) | intron (NM_001013813, intron 1 of 3) | 12019 | NM_001013813 | 270118 | Mm.116898 | NM_173776 | ENSMUSG00000031925 | **Maml2** | 5930431H10\|BC032967 | mastermind like 2 (Drosophila) | protein-coding |
| chr9_13645685_13645950 | chr9 | 13645686 | 13645950 | + | 0.768188 | intron (NM_001013813, intron 1 of 3) | intron (NM_001013813, intron 1 of 3) | -16642 | NM_173776 | 270118 | Mm.116898 | NM_173776 | ENSMUSG00000031925 | **Maml2** | 5930431H10\|BC032967 | mastermind like 2 (Drosophila) | protein-coding |
| chr9_13554378_13554793 | chr9 | 13554379 | 13554793 | + | 0.76372 | Intergenic | GA-rich\|Low_complexity\|Low_complexity | -65403 | NM_001013813 | 270118 | Mm.116898 | NM_173776 | ENSMUSG00000031925 | **Maml2** | 5930431H10\|BC032967 | mastermind like 2 (Drosophila) | protein-coding |
| chr3_51934769_51935438 | chr3 | 51934770 | 51935438 | + | 0.896444 | intron (NM_001004176, intron 1 of 4) | intron (NM_001004176, intron 1 of 4) | 169902 | NM_001004176 | 433586 | Mm.234972 | NM_001004176 | ENSMUSG00000061143 | **Maml3** | AV234550\|BC049812\|Mam-2\|mKIAA1816 | mastermind like 3 (Drosophila) | protein-coding |
| chr10_54026546_54027280 | chr10 | 54026547 | 54027280 | + | 1.63967 | intron (NM_008548, intron 3 of 12) | MER44C\|DNA\|TcMar-Tigger | 48883 | NM_008548 | 17155 | Mm.117294 | NM_008548 | ENSMUSG00000003746 | **Man1a** | Man1a1\|PCR1 | mannosidase 1, alpha | protein-coding |
| chr10_53940093_53940638 | chr10 | 53940094 | 53940638 | + | 1.38307 | intron (NM_008548, intron 6 of 12) | intron (NM_008548, intron 6 of 12) | 135430 | NM_008548 | 17155 | Mm.117294 | NM_008548 | ENSMUSG00000003746 | **Man1a** | Man1a1\|PCR1 | mannosidase 1, alpha | protein-coding |
| chr10_53925365_53925830 | chr10 | 53925366 | 53925830 | + | 1.16075 | intron (NM_008548, intron 8 of 12) | intron (NM_008548, intron 8 of 12) | 150198 | NM_008548 | 17155 | Mm.117294 | NM_008548 | ENSMUSG00000003746 | **Man1a** | Man1a1\|PCR1 | mannosidase 1, alpha | protein-coding |
| chr10_54009195_54009992 | chr10 | 54009196 | 54009992 | + | 1.05895 | intron (NM_008548, intron 5 of 12) | intron (NM_008548, intron 5 of 12) | 66202 | NM_008548 | 17155 | Mm.117294 | NM_008548 | ENSMUSG00000003746 | **Man1a** | Man1a1\|PCR1 | mannosidase 1, alpha | protein-coding |
| chr10_54063570_54063860 | chr10 | 54063571 | 54063860 | + | 0.731911 | intron (NM_008548, intron 2 of 12) | MARNA\|DNA\|TcMar-Mariner | 12081 | NM_008548 | 17155 | Mm.117294 | NM_008548 | ENSMUSG00000003746 | **Man1a** | Man1a1\|PCR1 | mannosidase 1, alpha | protein-coding |
| chr10_53918426_53918794 | chr10 | 53918427 | 53918794 | + | 0.687455 | intron (NM_008548, intron 11 of 12) | URR1A\|DNA\|hAT-Charlie | 157186 | NM_008548 | 17155 | Mm.117294 | NM_008548 | ENSMUSG00000003746 | **Man1a** | Man1a1\|PCR1 | mannosidase 1, alpha | protein-coding |
| chr17_64603145_64603590 | chr17 | 64603146 | 64603590 | + | 1.15333 | intron (NM_008549, intron 1 of 21) | intron (NM_008549, intron 1 of 21) | 1719 | NM_008549 | 17158 | Mm.2433 | NM_008549 | ENSMUSG00000024085 | **Man2a1** | Mana-2\|Mana2\|Map-2 | mannosidase 2, alpha 1 | protein-coding |
| chr17_64590453_64590855 | chr17 | 64590454 | 64590855 | + | 0.881637 | Intergenic | Intergenic | -10995 | NM_008549 | 17158 | Mm.2433 | NM_008549 | ENSMUSG00000024085 | **Man2a1** | Mana-2\|Mana2\|Map-2 | mannosidase 2, alpha 1 | protein-coding |
| chr17_64590279_64590856 | chr17 | 64590280 | 64590856 | + | 0.812992 | Intergenic | Intergenic | -11081 | NM_008549 | 17158 | Mm.2433 | NM_008549 | ENSMUSG00000024085 | **Man2a1** | Mana-2\|Mana2\|Map-2 | mannosidase 2, alpha 1 | protein-coding |
| chr17_64620757_64621060 | chr17 | 64620758 | 64621060 | + | 0.799056 | intron (NM_008549, intron 1 of 21) | B3\|SINE\|B2 | 19260 | NM_008549 | 17158 | Mm.2433 | NM_008549 | ENSMUSG00000024085 | **Man2a1** | Mana-2\|Mana2\|Map-2 | mannosidase 2, alpha 1 | protein-coding |
| chr3_135555973_135556613 | chr3 | 135555974 | 135556613 | + | 0.902695 | intron (NM_027288, intron 13 of 16) | intron (NM_027288, intron 13 of 16) | 70682 | NM_027288 | 110173 | Mm.280536 | NM_027288 | ENSMUSG00000028164 | **Manba** | 2410030O07Rik\|B930014J03Rik\|Bmn | mannosidase, beta A, lysosomal | protein-coding |
| chr3_135579268_135579705 | chr3 | 135579269 | 135579705 | + | 0.756868 | Intergenic | (TG)n\|Simple_repeat\|Simple_repeat | 93876 | NM_027288 | 110173 | Mm.280536 | NM_027288 | ENSMUSG00000028164 | **Manba** | 2410030O07Rik\|B930014J03Rik\|Bmn | mannosidase, beta A, lysosomal | protein-coding |
| chr3_135579252_135580290 | chr3 | 135579253 | 135580290 | + | 0.744672 | Intergenic | Intergenic | 94160 | NM_027288 | 110173 | Mm.280536 | NM_027288 | ENSMUSG00000028164 | **Manba** | 2410030O07Rik\|B930014J03Rik\|Bmn | mannosidase, beta A, lysosomal | protein-coding |
| chr3_135527814_135528259 | chr3 | 135527815 | 135528259 | + | 0.718605 | intron (NM_027288, intron 7 of 16) | intron (NM_027288, intron 7 of 16) | 42426 | NM_027288 | 110173 | Mm.280536 | NM_027288 | ENSMUSG00000028164 | **Manba** | 2410030O07Rik\|B930014J03Rik\|Bmn | mannosidase, beta A, lysosomal | protein-coding |
| chr6_134620687_134621842 | chr6 | 134620688 | 134621842 | + | 1.12065 | intron (NM_026345, intron 2 of 3) | B3A\|SINE\|B2 | 11223 | NM_026345 | 67729 | Mm.57648 | NM_026345 | ENSMUSG00000032718 | **Mansc1** | 9130403P13Rik | MANSC domain containing 1 | protein-coding |
| chr8_125671194_125671794 | chr8 | 125671195 | 125671794 | + | 1.33932 | exon (NM_028908, exon 1 of 1) | exon (NM_028908, exon 1 of 1) | 1676 | NM_028908 | 74393 | Mm.41709 | NM_028908 | ENSMUSG00000050930 | **Map10** | 4933403G14Rik\|AI835009\|mKIAA1383 | microtubule-associated protein 10 | protein-coding |
| chr1_66280321_66281236 | chr1 | 66280322 | 66281236 | + | 0.951932 | intron (NM_008632, intron 3 of 13) | intron (NM_008632, intron 3 of 13) | 105450 | NM_001039934 | 17756 | Mm.256966 | NM_008632 | ENSMUSG00000015222 | **Map2** | G1-397-34\|MAP-2\|Mtap-2\|Mtap2\|repro4 | microtubule-associated protein 2 | protein-coding |
| chr11_60937821_60938082 | chr11 | 60937822 | 60938082 | + | 0.734313 | intron (NM_008928, intron 1 of 11) | intron (NM_008928, intron 1 of 11) | 5895 | NM_008928 | 26397 | Mm.18494 | NM_008928 | ENSMUSG00000018932 | **Map2k3** | AW212142\|MEK3\|MKK3\|Prkmk3\|mMKK3b | mitogen-activated protein kinase kinase 3 | protein-coding |
| chr11_60937522_60938109 | chr11 | 60937523 | 60938109 | + | 0.692194 | intron (NM_008928, intron 1 of 11) | intron (NM_008928, intron 1 of 11) | 5759 | NM_008928 | 26397 | Mm.18494 | NM_008928 | ENSMUSG00000018932 | **Map2k3** | AW212142\|MEK3\|MKK3\|Prkmk3\|mMKK3b | mitogen-activated protein kinase kinase 3 | protein-coding |
| chr11_60930756_60930990 | chr11 | 60930757 | 60930990 | + | 0.638676 | intron (NR_027800, intron 1 of 1) | intron (NR_027800, intron 1 of 1) | 994 | NR_027800 | 24082 | Mm.46270 | NM_025293 | ENSMUSG00000042549 | **Map2k3os** | F3-1\|Gm16516\|Gtlf3a | mitogen-activated protein kinase kinase 3, opposite strand | ncRNA |
| chr11_65794253_65795435 | chr11 | 65794254 | 65795435 | + | 0.905297 | Intergenic | L1MB8\|LINE\|L1 | -6547 | NM_009157 | 26398 | Mm.412922 | NM_009157 | ENSMUSG00000033352 | **Map2k4** | JNKK1\|MEK4\|MKK4\|PRKMK4\|Sek1\|Serk1 | mitogen-activated protein kinase kinase 4 | protein-coding |
| chr11_65794601_65795131 | chr11 | 65794602 | 65795131 | + | 0.8656 | Intergenic | L1MB8\|LINE\|L1 | -6569 | NM_009157 | 26398 | Mm.412922 | NM_009157 | ENSMUSG00000033352 | **Map2k4** | JNKK1\|MEK4\|MKK4\|PRKMK4\|Sek1\|Serk1 | mitogen-activated protein kinase kinase 4 | protein-coding |
| chr11_65771436_65771855 | chr11 | 65771437 | 65771855 | + | 0.653447 | intron (NM_009157, intron 1 of 10) | intron (NM_009157, intron 1 of 10) | 16651 | NM_009157 | 26398 | Mm.412922 | NM_009157 | ENSMUSG00000033352 | **Map2k4** | JNKK1\|MEK4\|MKK4\|PRKMK4\|Sek1\|Serk1 | mitogen-activated protein kinase kinase 4 | protein-coding |
| chr13_111826597_111827058 | chr13 | 111826598 | 111827058 | + | 1.21092 | Intergenic | URR1B\|DNA\|hAT-Charlie | -17845 | NM_011945 | 26401 | Mm.15918 | NM_011945 | ENSMUSG00000021754 | **Map3k1** | MAPKKK1\|MEKK1\|Mekk | mitogen-activated protein kinase kinase kinase 1 | protein-coding |
| chr13_111804628_111804997 | chr13 | 111804629 | 111804997 | + | 1.02895 | intron (NM_011945, intron 1 of 19) | intron (NM_011945, intron 1 of 19) | 4170 | NM_011945 | 26401 | Mm.15918 | NM_011945 | ENSMUSG00000021754 | **Map3k1** | MAPKKK1\|MEKK1\|Mekk | mitogen-activated protein kinase kinase kinase 1 | protein-coding |
| chr13_111940322_111940585 | chr13 | 111940323 | 111940585 | + | 0.744101 | Intergenic | Intergenic | -131471 | NM_011945 | 26401 | Mm.15918 | NM_011945 | ENSMUSG00000021754 | **Map3k1** | MAPKKK1\|MEKK1\|Mekk | mitogen-activated protein kinase kinase kinase 1 | protein-coding |
| chr11_103269515_103269753 | chr11 | 103269516 | 103269753 | + | 0.956998 | Intergenic | AmnSINE2\|SINE\|Deu | -2233 | NM_016896 | 53859 | Mm.158981 | NM_016896 | ENSMUSG00000020941 | **Map3k14** | Nik\|aly | mitogen-activated protein kinase kinase kinase 14 | protein-coding |
| chr1_127812927_127813630 | chr1 | 127812928 | 127813630 | + | 0.806063 | Intergenic | MTC\|LTR\|ERVL-MaLR | 27011 | NM_011737 | 22625 | Mm.246566 | NM_011737 | ENSMUSG00000051590 | **Map3k19** | Ysk4 | mitogen-activated protein kinase kinase kinase 19 | protein-coding |
| chr1_127812663_127813648 | chr1 | 127812664 | 127813648 | + | 0.74197 | Intergenic | MTC\|LTR\|ERVL-MaLR | 27134 | NM_011737 | 22625 | Mm.246566 | NM_011737 | ENSMUSG00000051590 | **Map3k19** | Ysk4 | mitogen-activated protein kinase kinase kinase 19 | protein-coding |
| chr18_32192074_32192547 | chr18 | 32192075 | 32192547 | + | 1.296 | intron (NM_011946, intron 2 of 16) | intron (NM_011946, intron 2 of 16) | 29222 | NM_011946 | 26405 | Mm.211762 | NM_011946 | ENSMUSG00000024383 | **Map3k2** | 9630061B06Rik\|AI585793\|Mekk2\|Mekk2b | mitogen-activated protein kinase kinase kinase 2 | protein-coding |
| chr18_32192071_32193068 | chr18 | 32192072 | 32193068 | + | 1.26567 | intron (NM_011946, intron 2 of 16) | intron (NM_011946, intron 2 of 16) | 29481 | NM_011946 | 26405 | Mm.211762 | NM_011946 | ENSMUSG00000024383 | **Map3k2** | 9630061B06Rik\|AI585793\|Mekk2\|Mekk2b | mitogen-activated protein kinase kinase kinase 2 | protein-coding |
| chr18_32185943_32186360 | chr18 | 32185944 | 32186360 | + | 1.10886 | intron (NM_011946, intron 1 of 16) | intron (NM_011946, intron 1 of 16) | 23063 | NM_011946 | 26405 | Mm.211762 | NM_011946 | ENSMUSG00000024383 | **Map3k2** | 9630061B06Rik\|AI585793\|Mekk2\|Mekk2b | mitogen-activated protein kinase kinase kinase 2 | protein-coding |
| chr18_32192647_32193065 | chr18 | 32192648 | 32193065 | + | 0.764267 | intron (NM_011946, intron 2 of 16) | intron (NM_011946, intron 2 of 16) | 29767 | NM_011946 | 26405 | Mm.211762 | NM_011946 | ENSMUSG00000024383 | **Map3k2** | 9630061B06Rik\|AI585793\|Mekk2\|Mekk2b | mitogen-activated protein kinase kinase kinase 2 | protein-coding |
| chr17_12275521_12276259 | chr17 | 12275522 | 12276259 | + | 1.93751 | intron (NM_011948, intron 2 of 26) | L2b\|LINE\|L2 | 42770 | NM_011948 | 26407 | Mm.28587 | NM_011948 | ENSMUSG00000014426 | **Map3k4** | D17Rp17\|D17Rp17e\|MAPKKK4\|MEKK 4\|MTK1\|Mek4b\|Mekk4\|RP17\|Rp17a\|Tas\|mKIAA0213 | mitogen-activated protein kinase kinase kinase 4 | protein-coding |

| | | | | | | | | | | | | | | | | | |
|---|---|---|---|---|---|---|---|---|---|---|---|---|---|---|---|---|---|
| chr10_19999670_20000128 | chr10 | 19999671 | 20000128 | + | 0.766953 | intron (NM_008580, intron 1 of 29) | C-rich\|Low_complexity\|Low_complexity | 65373 | NM_008580 | 26408 | Mm.6595 | NM_008580 | ENSMUSG00000071369 | **Map3k5** | 7420452D20Rik\|ASK\|ASK1\|MAPKKK5\|Mekk5 | mitogen-activated protein kinase kinase kinase 5 | protein-coding |
| chr10_19997481_19997770 | chr10 | 19997482 | 19997770 | + | 0.680939 | intron (NM_008580, intron 1 of 29) | ORR1D1\|LTR\|ERVL-MaLR | 63100 | NM_008580 | 26408 | Mm.6595 | NM_008580 | ENSMUSG00000071369 | **Map3k5** | 7420452D20Rik\|ASK\|ASK1\|MAPKKK5\|Mekk5 | mitogen-activated protein kinase kinase kinase 5 | protein-coding |
| chr10_20000398_20000712 | chr10 | 20000399 | 20000712 | + | 0.612336 | intron (NM_008580, intron 1 of 29) | intron (NM_008580, intron 1 of 29) | 66029 | NM_008580 | 26408 | Mm.6595 | NM_008580 | ENSMUSG00000071369 | **Map3k5** | 7420452D20Rik\|ASK\|ASK1\|MAPKKK5\|Mekk5 | mitogen-activated protein kinase kinase kinase 5 | protein-coding |
| chr4_31941517_31942257 | chr4 | 31941518 | 31942257 | + | 1.52085 | Intergenic | Lx7\|LINE\|L1 | -22220 | NM_009316 | 26409 | Mm.258589 | NM_009316 | ENSMUSG00000028284 | **Map3k7** | B430101B05\|C87327\|Tak1 | mitogen-activated protein kinase kinase kinase 7 | protein-coding |
| chr4_32179631_32180305 | chr4 | 32179632 | 32180305 | + | 1.33666 | Intergenic | Intergenic | 215861 | NM_172688 | 26409 | Mm.258589 | NM_009316 | ENSMUSG00000028284 | **Map3k7** | B430101B05\|C87327\|Tak1 | mitogen-activated protein kinase kinase kinase 7 | protein-coding |
| chr4_32126466_32127360 | chr4 | 32126467 | 32127360 | + | 0.658886 | Intergenic | MTD\|LTR\|ERVL-MaLR | 162806 | NM_172688 | 26409 | Mm.258589 | NM_009316 | ENSMUSG00000028284 | **Map3k7** | B430101B05\|C87327\|Tak1 | mitogen-activated protein kinase kinase kinase 7 | protein-coding |
| chr18_4311243_4311629 | chr18 | 4311244 | 4311629 | + | 1.26112 | Intergenic | Intergenic | 41517 | NM_007746 | 26410 | Mm.3275 | NM_007746 | ENSMUSG00000024235 | **Map3k8** | Cot\|Cot/Tpl2\|Est\|Estf\|Tpl-2\|Tpl2\|c-COT | mitogen-activated protein kinase kinase kinase 8 | protein-coding |
| chr18_4280583_4280963 | chr18 | 4280584 | 4280963 | + | 1.18616 | Intergenic | L1_Mus3\|LINE\|L1 | 72180 | NM_007746 | 26410 | Mm.3275 | NM_007746 | ENSMUSG00000024235 | **Map3k8** | Cot\|Cot/Tpl2\|Est\|Estf\|Tpl-2\|Tpl2\|c-COT | mitogen-activated protein kinase kinase kinase 8 | protein-coding |
| chr18_4349788_4350073 | chr18 | 4349789 | 4350073 | + | 1.01229 | intron (NM_007746, intron 1 of 7) | intron (NM_007746, intron 1 of 7) | 3022 | NM_007746 | 26410 | Mm.3275 | NM_007746 | ENSMUSG00000024235 | **Map3k8** | Cot\|Cot/Tpl2\|Est\|Estf\|Tpl-2\|Tpl2\|c-COT | mitogen-activated protein kinase kinase kinase 8 | protein-coding |
| chr18_4311779_4312328 | chr18 | 4311780 | 4312328 | + | 0.732548 | Intergenic | Intergenic | 40899 | NM_007746 | 26410 | Mm.3275 | NM_007746 | ENSMUSG00000024235 | **Map3k8** | Cot\|Cot/Tpl2\|Est\|Estf\|Tpl-2\|Tpl2\|c-COT | mitogen-activated protein kinase kinase kinase 8 | protein-coding |
| chr18_4279902_4280203 | chr18 | 4279903 | 4280203 | + | 0.686127 | Intergenic | L1_Mus3\|LINE\|L1 | 72900 | NM_007746 | 26410 | Mm.3275 | NM_007746 | ENSMUSG00000024235 | **Map3k8** | Cot\|Cot/Tpl2\|Est\|Estf\|Tpl-2\|Tpl2\|c-COT | mitogen-activated protein kinase kinase kinase 8 | protein-coding |
| chr18_4291588_4291966 | chr18 | 4291589 | 4291966 | + | 0.662447 | Intergenic | Intergenic | 61176 | NM_007746 | 26410 | Mm.3275 | NM_007746 | ENSMUSG00000024235 | **Map3k8** | Cot\|Cot/Tpl2\|Est\|Estf\|Tpl-2\|Tpl2\|c-COT | mitogen-activated protein kinase kinase kinase 8 | protein-coding |
| chr18_4278298_4278812 | chr18 | 4278299 | 4278812 | + | 0.647091 | Intergenic | Intergenic | 74398 | NM_007746 | 26410 | Mm.3275 | NM_007746 | ENSMUSG00000024235 | **Map3k8** | Cot\|Cot/Tpl2\|Est\|Estf\|Tpl-2\|Tpl2\|c-COT | mitogen-activated protein kinase kinase kinase 8 | protein-coding |
| chr7_29026207_29026718 | chr7 | 29026208 | 29026718 | + | 0.889819 | intron (NM_009109, intron 89 of 105) | L2\|LINE\|L2 | 43609 | NM_008279 | 26411 | Mm.148278 | NM_008279 | ENSMUSG00000037337 | **Map4k1** | Hpk1\|mHPK1 | mitogen-activated protein kinase kinase kinase kinase 1 | protein-coding |
| chr17_80809213_80809691 | chr17 | 80809214 | 80809691 | + | 1.11764 | Intergenic | Intergenic | -81427 | NM_001290345 | 225028 | Mm.311948 | NM_001081357 | ENSMUSG00000024242 | **Map4k3** | 4833416M01Rik\|4833416M07Rik\|9530052P13Rik\|Glk\|MAPKKKK3\|MEKKK 3\|MEKKK3\|RAB8IPL1 | mitogen-activated protein kinase kinase kinase kinase 3 | protein-coding |
| chr1_39941279_39941594 | chr1 | 39941280 | 39941594 | + | 0.697671 | intron (NM_008696, intron 2 of 30) | intron (NM_008696, intron 2 of 30) | 40524 | NM_008696 | 26921 | Mm.491139 | NM_008696 | ENSMUSG00000026074 | **Map4k4** | 9430080K19Rik\|AU043147\|AU045934\|AW046177\|HGK\|Nik | mitogen-activated protein kinase kinase kinase kinase 4 | protein-coding |
| chr4_126250407_126250840 | chr4 | 126250408 | 126250840 | + | 0.74406 | intron (NM_001145970, intron 1 of 16) | intron (NM_001145970, intron 1 of 16) | 5695 | NM_144941 | 245877 | Mm.266716 | NM_144941 | ENSMUSG00000028849 | **Map7d1** | AV028413\|BC019977\|Mtap7d1\|Parcc1\|Rprc1\|mKIAA1187 | MAP7 domain containing 1 | protein-coding |
| chrX_159447111_159447518 | chrX | 159447112 | 159447518 | + | 0.635763 | intron (NM_001081124, intron 3 of 16) | Lx6\|LINE\|L1 | 32737 | NM_001081124 | 78283 | Mm.160079 | NM_001081124 | ENSMUSG00000041020 | **Map7d2** | 1600028E09Rik\|2900002G04Rik\|5330432J06Rik\|C030036K04Rik\|Mtap7d2 | MAP7 domain containing 2 | protein-coding |
| chr3_82308636_82309112 | chr3 | 82308637 | 82309112 | + | 1.08115 | Intergenic | Intergenic | -49198 | NM_001081230 | 213582 | Mm.271269 | NM_001081230 | ENSMUSG00000033900 | **Map9** | 5033421J10Rik\|5330427D05Rik\|ASAP\|Mtap9 | microtubule-associated protein 9 | protein-coding |
| chr3_82308481_82309117 | chr3 | 82308482 | 82309117 | + | 0.987466 | Intergenic | Intergenic | -49273 | NM_001081230 | 213582 | Mm.271269 | NM_001081230 | ENSMUSG00000033900 | **Map9** | 5033421J10Rik\|5330427D05Rik\|ASAP\|Mtap9 | microtubule-associated protein 9 | protein-coding |
| chr16_17024434_17024904 | chr16 | 17024435 | 17024904 | + | 1.61539 | intron (NM_001038663, intron 4 of 7) | intron (NM_001038663, intron 4 of 7) | 41287 | NM_001038663 | 26413 | Mm.196581 | NM_011949 | ENSMUSG00000063358 | **Mapk1** | 9030612K14Rik\|AA407128\|AU018647\|C78273\|ERK\|Erk2\|MAPK2\|PRKM2\|Prkm1\|p41mapk\|p42mapk | mitogen-activated protein kinase 1 | protein-coding |
| chr16_17010892_17011532 | chr16 | 17010893 | 17011532 | + | 1.45112 | intron (NM_001038663, intron 1 of 7) | intron (NM_001038663, intron 1 of 7) | 27830 | NM_001038663 | 26413 | Mm.196581 | NM_011949 | ENSMUSG00000063358 | **Mapk1** | 9030612K14Rik\|AA407128\|AU018647\|C78273\|ERK\|Erk2\|MAPK2\|PRKM2\|Prkm1\|p41mapk\|p42mapk | mitogen-activated protein kinase 1 | protein-coding |
| chr7_138841568_138842189 | chr7 | 138841569 | 138842189 | + | 1.224 | intron (NM_001045483, intron 1 of 1) | intron (NM_001045483, intron 1 of 1) | 4388 | NM_001045483 | 69546 | Mm.31598 | NM_027115 | ENSMUSG00000041775 | **Mapk1ip1** | 2310009E07Rik\|AU043776\|MISS | mitogen-activated protein kinase 1 interacting protein 1 | protein-coding |
| chr14_47303348_47303707 | chr14 | 47303349 | 47303707 | + | 0.647584 | intron (NM_178684, intron 1 of 3) | intron (NM_178684, intron 1 of 3) | 5214 | NM_178684 | 218975 | Mm.212297 | NM_178684 | ENSMUSG00000021840 | **Mapk1ip1l** | 9130202B12\|C130032J12Rik | mitogen-activated protein kinase 1 interacting protein 1-like | protein-coding |
| chr14_47327401_47328148 | chr14 | 47327402 | 47328148 | + | 0.614835 | Intergenic | Intergenic | 29461 | NM_178684 | 218975 | Mm.212297 | NM_178684 | ENSMUSG00000021840 | **Mapk1ip1l** | 9130202B12\|C130032J12Rik | mitogen-activated protein kinase 1 interacting protein 1-like | protein-coding |
| chr9_75407868_75408176 | chr9 | 75407869 | 75408176 | + | 0.664071 | intron (NM_015806, intron 1 of 5) | intron (NM_015806, intron 1 of 5) | 1337 | NM_027418 | 50772 | Mm.480076 | NM_015806 | ENSMUSG00000042688 | **Mapk6** | 2610021I23Rik\|D130053K17Rik\|Erk3\|Mapk4\|Mapk63\|Prkm4\|Prkm6 | mitogen-activated protein kinase 6 | protein-coding |
| chr14_33443815_33444333 | chr14 | 33443816 | 33444333 | + | 1.50185 | intron (NM_016700, intron 1 of 11) | intron (NM_016700, intron 1 of 11) | 3084 | NM_016700 | 26419 | Mm.21495 | NM_016700 | ENSMUSG00000021936 | **Mapk8** | AI849689\|JNK\|JNK1\|Prkm8\|SAPK1 | mitogen-activated protein kinase 8 | protein-coding |
| chr2_34508559_34508917 | chr2 | 34508560 | 34508917 | + | 0.696354 | intron (NM_001290625, intron 5 of 11) | intron (NM_001290625, intron 5 of 11) | 101887 | NM_001290626 | 227743 | Mm.270866 | NM_177345 | ENSMUSG00000038696 | **Mapkap1** | AI591529\|D230039K05Rik\|Sin1 | mitogen-activated protein kinase associated protein 1 | protein-coding |
| chr1_131113529_131113888 | chr1 | 131113530 | 131113888 | + | 1.04705 | Intergenic | Intergenic | -16166 | NM_008551 | 17164 | Mm.221235 | NM_008551 | ENSMUSG00000016528 | **Mapkapk2** | AA960234\|MAPKAP-K2\|MK-2\|MK2\|Rps6kc1 | MAP kinase-activated protein kinase 2 | protein-coding |
| chr1_131111630_131111895 | chr1 | 131111631 | 131111895 | + | 0.858001 | Intergenic | (TGGGGG)n\|Simple_repeat\|Simple_repeat | -14220 | NM_008551 | 17164 | Mm.221235 | NM_008551 | ENSMUSG00000016528 | **Mapkapk2** | AA960234\|MAPKAP-K2\|MK-2\|MK2\|Rps6kc1 | MAP kinase-activated protein kinase 2 | protein-coding |
| chr9_107278564_107278931 | chr9 | 107278565 | 107278931 | + | 0.734809 | intron (NM_178907, intron 2 of 10) | intron (NM_178907, intron 2 of 10) | 11129 | NM_178907 | 102626 | Mm.222612 | NM_178907 | ENSMUSG00000032577 | **Mapkapk3** | 3PK\|AI874665\|MAPKAP-K3\|MAPKAP3\|MK-3\|MK3\|MapkKapk3 | mitogen-activated protein kinase-activated protein kinase 3 | protein-coding |
| chr18_23805649_23805981 | chr18 | 23805650 | 23805981 | + | 1.13626 | intron (NM_153058, intron 1 of 6) | intron (NM_153058, intron 1 of 6) | 1845 | NM_153058 | 212307 | Mm.132237 | NM_153058 | ENSMUSG00000024277 | **Mapre2** | AI314113\|C820009F03Rik\|D18Abb1e\|EB1\|EB2\|RP1 | microtubule-associated protein, RP/EB family, member 2 | protein-coding |
| chr18_23844626_23845400 | chr18 | 23844627 | 23845400 | + | 1.06457 | intron (NM_153058, intron 2 of 6) | intron (NM_153058, intron 2 of 6) | 41043 | NM_153058 | 212307 | Mm.132237 | NM_153058 | ENSMUSG00000024277 | **Mapre2** | AI314113\|C820009F03Rik\|D18Abb1e\|EB1\|EB2\|RP1 | microtubule-associated protein, RP/EB family, member 2 | protein-coding |
| chr18_23754220_23754789 | chr18 | 23754221 | 23754789 | + | 1.04785 | intron (NM_001162941, intron 1 of 7) | intron (NM_001162941, intron 1 of 7) | 780 | NM_001162941 | 212307 | Mm.132237 | NM_153058 | ENSMUSG00000024277 | **Mapre2** | AI314113\|C820009F03Rik\|D18Abb1e\|EB1\|EB2\|RP1 | microtubule-associated protein, RP/EB family, member 2 | protein-coding |
| chr18_23762790_23763484 | chr18 | 23762791 | 23763484 | + | 0.77681 | intron (NM_001162941, intron 1 of 7) | intron (NM_001162941, intron 1 of 7) | 9412 | NM_001162941 | 212307 | Mm.132237 | NM_153058 | ENSMUSG00000024277 | **Mapre2** | AI314113\|C820009F03Rik\|D18Abb1e\|EB1\|EB2\|RP1 | microtubule-associated protein, RP/EB family, member 2 | protein-coding |
| chr18_23843371_23843806 | chr18 | 23843372 | 23843806 | + | 0.771136 | intron (NM_153058, intron 2 of 6) | (CCCCCA)n\|Simple_repeat\|Simple_repeat | 39619 | NM_153058 | 212307 | Mm.132237 | NM_153058 | ENSMUSG00000024277 | **Mapre2** | AI314113\|C820009F03Rik\|D18Abb1e\|EB1\|EB2\|RP1 | microtubule-associated protein, RP/EB family, member 2 | protein-coding |
| chr11_105484992_105485419 | chr11 | 105484993 | 105485419 | + | 0.814436 | Intergenic | L1MDa\|LINE\|L1 | -28471 | NM_001039242 | 632687 | Mm.121570 | NM_172568 | ENSMUSG00000078627 | **March10** | 4933417C16Rik\|AV039223\|OTTMUSG00000002847\|Rnf190 | membrane-associated ring finger (C3HC4) 10 | protein-coding |
| chr17_33702209_33702591 | chr17 | 33702210 | 33702591 | + | 1.06322 | intron (NR_045522, intron 3 of 6) | intron (NR_045522, intron 3 of 6) | 16277 | NM_145486 | 224703 | Mm.311439 | NM_145486 | ENSMUSG00000079557 | **March2** | 9530046H09Rik\|MARCH-II | membrane-associated ring finger (C3HC4) 2 | protein-coding |
| chr15_31546293_31546701 | chr15 | 31546294 | 31546701 | + | 1.23866 | Intergenic | Intergenic | -15460 | NM_172606 | 223455 | Mm.272185 | NM_172606 | ENSMUSG00000039100 | **March6** | 3830408G03\|F830029L24Rik\|mKIAA0597 | membrane-associated ring finger (C3HC4) 6 | protein-coding |
| chr15_31542091_31542942 | chr15 | 31542092 | 31542942 | + | 1.07203 | Intergenic | Intergenic | -11480 | NM_172606 | 223455 | Mm.272185 | NM_172606 | ENSMUSG00000039100 | **March6** | 3830408G03\|F830029L24Rik\|mKIAA0597 | membrane-associated ring finger (C3HC4) 6 | protein-coding |
| chr15_31523750_31524354 | chr15 | 31523751 | 31524354 | + | 0.655839 | intron (NM_172606, intron 1 of 25) | MLT1B\|LTR\|ERVL-MaLR | 6985 | NM_172606 | 223455 | Mm.272185 | NM_172606 | ENSMUSG00000039100 | **March6** | 3830408G03\|F830029L24Rik\|mKIAA0597 | membrane-associated ring finger (C3HC4) 6 | protein-coding |
| chr10_37321249_37321514 | chr10 | 37321250 | 37321514 | + | 0.928926 | Intergenic | Intergenic | -182456 | NM_008538 | 17118 | Mm.30059 | NM_008538 | ENSMUSG00000069662 | **Marcks** | Macs\|PKCSL | myristoylated alanine rich protein kinase C substrate | protein-coding |
| chr10_37321177_37321566 | chr10 | 37321178 | 37321566 | + | 0.924002 | Intergenic | Intergenic | -182446 | NM_008538 | 17118 | Mm.30059 | NM_008538 | ENSMUSG00000069662 | **Marcks** | Macs\|PKCSL | myristoylated alanine rich protein kinase C substrate | protein-coding |
| chr10_37145512_37145801 | chr10 | 37145513 | 37145801 | + | 0.70201 | Intergenic | B1F1\|SINE\|Alu | -6731 | NM_008538 | 17118 | Mm.30059 | NM_008538 | ENSMUSG00000069662 | **Marcks** | Macs\|PKCSL | myristoylated alanine rich protein kinase C substrate | protein-coding |
| chr10_37337020_37337698 | chr10 | 37337021 | 37337698 | + | 0.698149 | Intergenic | Intergenic | -198433 | NM_008538 | 17118 | Mm.30059 | NM_008538 | ENSMUSG00000069662 | **Marcks** | Macs\|PKCSL | myristoylated alanine rich protein kinase C substrate | protein-coding |

| | | | | | | | | | | | | | | | | | |
|---|---|---|---|---|---|---|---|---|---|---|---|---|---|---|---|---|---|
| chr10_37320130_37320364 | chr10 | 37320131 | 37320364 | + | 0.645642 | Intergenic | Intergenic | -181321 | NM_008538 | 17118 | Mm.30059 | NM_008538 | ENSMUSG00000069662 | **Marcks** | Macs\|PKCSL | myristoylated alanine rich protein kinase C substrate | protein-coding |
| chr4_129507795_129508383 | chr4 | 129507796 | 129508383 | + | 1.00873 | Intergenic | Intergenic | -5492 | NM_010807 | 17357 | Mm.424974 | NM_010807 | ENSMUSG00000047945 | **Marcksl1** | AL022768\|AW215397\|AW536807\|D4Bc1\|F52\|MacMARCKS\|Macs2\|Macs3\|Mlp\|Mrp | MARCKS-like 1 | protein-coding |
| chr19_7297763_7298096 | chr19 | 7297764 | 7298096 | + | 1.11436 | intron (NM_001080389, intron 1 of 17) | intron (NM_001080389, intron 1 of 17) | -2483 | NM_001080390 | 13728 | Mm.258986 | NM_007928 | ENSMUSG00000024969 | **Mark2** | EMK-1\|Emk\|Par-1\|Par-1b | MAP/microtubule affinity-regulating kinase 2 | protein-coding |
| chr19_7335702_7335991 | chr19 | 7335703 | 7335991 | + | 0.842793 | intron (NM_001080389, intron 1 of 17) | intron (NM_001080389, intron 1 of 17) | 6013 | NM_001080389 | 13728 | Mm.258986 | NM_007928 | ENSMUSG00000024969 | **Mark2** | EMK-1\|Emk\|Par-1\|Par-1b | MAP/microtubule affinity-regulating kinase 2 | protein-coding |
| chr12_111599507_111599958 | chr12 | 111599508 | 111599958 | + | 1.57714 | intron (NM_022801, intron 2 of 15) | intron (NM_022801, intron 2 of 15) | 25223 | NM_022801 | 17169 | Mm.260504 | NM_021516 | ENSMUSG00000007411 | **Mark3** | 1600015G02Rik\|A430080F22Rik\|C-TAK1\|ETK-1\|ETK1\|Emk2\|MPK10 | MAP/microtubule affinity-regulating kinase 3 | protein-coding |
| chr12_111599507_111600183 | chr12 | 111599508 | 111600183 | + | 1.47688 | intron (NM_022801, intron 2 of 15) | intron (NM_022801, intron 2 of 15) | 25335 | NM_022801 | 17169 | Mm.260504 | NM_021516 | ENSMUSG00000007411 | **Mark3** | 1600015G02Rik\|A430080F22Rik\|C-TAK1\|ETK-1\|ETK1\|Emk2\|MPK10 | MAP/microtubule affinity-regulating kinase 3 | protein-coding |
| chr12_111584801_111585537 | chr12 | 111584802 | 111585537 | + | 1.37292 | intron (NM_022801, intron 1 of 15) | B1_Mur2\|SINE\|Alu | 10659 | NM_022801 | 17169 | Mm.260504 | NM_021516 | ENSMUSG00000007411 | **Mark3** | 1600015G02Rik\|A430080F22Rik\|C-TAK1\|ETK-1\|ETK1\|Emk2\|MPK10 | MAP/microtubule affinity-regulating kinase 3 | protein-coding |
| chr12_111584297_111584739 | chr12 | 111584298 | 111584739 | + | 1.15322 | intron (NM_022801, intron 1 of 15) | URR1B\|DNA\|hAT-Charlie | 10008 | NM_022801 | 17169 | Mm.260504 | NM_021516 | ENSMUSG00000007411 | **Mark3** | 1600015G02Rik\|A430080F22Rik\|C-TAK1\|ETK-1\|ETK1\|Emk2\|MPK10 | MAP/microtubule affinity-regulating kinase 3 | protein-coding |
| chr17_12870805_12871315 | chr17 | 12870806 | 12871315 | + | 0.70726 | Intergenic | RSINE1\|SINE\|B4 | -2917 | NM_008552 | 17171 | Mm.425994 | NM_008552 | ENSMUSG00000068037 | **Mas1** | Mas-1\|Mgra | MAS1 oncogene | protein-coding |
| chr16_23492791_23493186 | chr16 | 23492792 | 23493186 | + | 0.954357 | intron (NM_008555, intron 4 of 15) | MIR3\|SINE\|MIR | 27601 | NM_008555 | 17174 | Mm.1213 | NM_008555 | ENSMUSG00000022887 | **Masp1** | AW048060\|CCPII\|Crarf\|Masp1/3 | mannan-binding lectin serine peptidase 1 | protein-coding |
| chr16_23492600_23493199 | chr16 | 23492601 | 23493199 | + | 0.727672 | intron (NM_008555, intron 4 of 15) | intron (NM_008555, intron 4 of 15) | 27690 | NM_008555 | 17174 | Mm.1213 | NM_008555 | ENSMUSG00000022887 | **Masp1** | AW048060\|CCPII\|Crarf\|Masp1/3 | mannan-binding lectin serine peptidase 1 | protein-coding |
| chr4_148604165_148604515 | chr4 | 148604166 | 148604515 | + | 1.14218 | TTS (NM_010767) | TTS (NM_010767) | 1796 | NM_001003893 | 17175 | Mm.378962 | NM_010767 | ENSMUSG00000028979 | **Masp2** | MASP-2\|MAp19 | mannan-binding lectin serine peptidase 2 | protein-coding |
| chr4_116451566_116451960 | chr4 | 116451567 | 116451960 | + | 0.983425 | intron (NM_001042743, intron 1 of 29) | MT2C_Mm\|LTR\|ERVL | 12418 | NM_001042743 | 17776 | Mm.9287 | NM_008641 | ENSMUSG00000003810 | **Mast2** | MAST205\|Mtssk | microtubule associated serine/threonine kinase 2 | protein-coding |
| chr13_103109784_103110250 | chr13 | 103109785 | 103110250 | + | 1.25953 | intron (NM_175171, intron 3 of 28) | intron (NM_175171, intron 3 of 28) | 224475 | NM_175171 | 328329 | Mm.202606 | NM_175171 | ENSMUSG00000034751 | **Mast4** | 4930420O11Rik\|AI642422\|mKIAA0303 | microtubule associated serine/threonine kinase family member 4 | protein-coding |
| chr11_40688648_40688867 | chr11 | 40688649 | 40688867 | + | 1.20389 | intron (NM_001199274, intron 1 of 6) | intron (NM_001199274, intron 1 of 6) | 3924 | NM_134017 | 108645 | Mm.293771 | NM_134017 | ENSMUSG00000042032 | **Mat2b** | 1110064C04Rik\|2410018D16Rik\|AI182287\|AU022853\|MAT-II\|MATIIbeta\|TGR | methionine adenosyltransferase II, beta | protein-coding |
| chr11_40688435_40688879 | chr11 | 40688436 | 40688879 | + | 1.06457 | intron (NM_001199274, intron 1 of 6) | intron (NM_001199274, intron 1 of 6) | 4025 | NM_134017 | 108645 | Mm.293771 | NM_134017 | ENSMUSG00000042032 | **Mat2b** | 1110064C04Rik\|2410018D16Rik\|AI182287\|AU022853\|MAT-II\|MATIIbeta\|TGR | methionine adenosyltransferase II, beta | protein-coding |
| chr12_76950265_76950805 | chr12 | 76950266 | 76950805 | + | 1.28371 | intron (NM_008558, intron 3 of 4) | intron (NM_008558, intron 3 of 4) | 11713 | NM_008558 | 17187 | Mm.268548 | NM_008558 | ENSMUSG00000059436 | **Max** | AA960152\|AI875693\|bHLHd4\|bHLHd5\|bHLHd6\|bHLHd7\|bHLHd8 | Max protein | protein-coding |
| chr15_77058478_77058873 | chr15 | 77058479 | 77058873 | + | 0.613569 | Intergenic | RLTR23\|LTR\|ERV1 | -8008 | NM_013593 | 17189 | Mm.404074 | NM_013593 | ENSMUSG00000018893 | **Mb** | AI325109 | myoglobin | protein-coding |
| chr18_70631849_70632590 | chr18 | 70631850 | 70632590 | + | 1.51222 | Intergenic | Intergenic | 63928 | NM_010773 | 17191 | Mm.322 | NM_010773 | ENSMUSG00000024513 | **Mbd2** | MBD2a | methyl-CpG binding domain protein 2 | protein-coding |
| chr18_70575979_70576536 | chr18 | 70575980 | 70576536 | + | 1.33136 | intron (NM_010773, intron 1 of 6) | intron (NM_010773, intron 1 of 6) | 7966 | NM_010773 | 17191 | Mm.322 | NM_010773 | ENSMUSG00000024513 | **Mbd2** | MBD2a | methyl-CpG binding domain protein 2 | protein-coding |
| chr18_70633449_70633992 | chr18 | 70633450 | 70633992 | + | 1.15378 | Intergenic | MLT1L\|LTR\|ERVL-MaLR | 65429 | NM_010773 | 17191 | Mm.322 | NM_010773 | ENSMUSG00000024513 | **Mbd2** | MBD2a | methyl-CpG binding domain protein 2 | protein-coding |
| chr18_71102222_71103036 | chr18 | 71102223 | 71103036 | + | 1.1119 | Intergenic | MTE2b\|LTR\|ERVL-MaLR | 534337 | NM_010773 | 17191 | Mm.322 | NM_010773 | ENSMUSG00000024513 | **Mbd2** | MBD2a | methyl-CpG binding domain protein 2 | protein-coding |
| chr18_70587644_70588377 | chr18 | 70587645 | 70588377 | + | 0.626785 | intron (NM_010773, intron 2 of 6) | Charlie2b\|DNA\|hAT-Charlie | 19719 | NM_010773 | 17191 | Mm.322 | NM_010773 | ENSMUSG00000024513 | **Mbd2** | MBD2a | methyl-CpG binding domain protein 2 | protein-coding |
| chr2_49204131_49204531 | chr2 | 49204132 | 49204531 | + | 0.730123 | intron (NM_029924, intron 6 of 16) | B1F\|SINE\|Alu | -41569 | NM_001290656 | 109241 | Mm.440436 | NM_029924 | ENSMUSG00000036792 | **Mbd5** | 9430004D19Rik\|AA536666\|AI426407\|C030040A15Rik\|Gm1630\|OTTMUSG00000012483 | methyl-CpG binding domain protein 5 | protein-coding |
| chr12_56325941_56326382 | chr12 | 56325942 | 56326382 | + | 0.69588 | Intergenic | Intergenic | 19732 | NM_145442 | 217588 | Mm.25656 | NM_145442 | ENSMUSG00000021028 | **Mbip** | 4933408E06Rik | MAP3K12 binding inhibitory protein 1 | protein-coding |
| chr13_81741555_81742338 | chr13 | 81741556 | 81742338 | + | 0.648359 | intron (NM_028372, intron 1 of 1) | RMER15\|LTR\|ERVL | 30530 | NM_028372 | 72852 | Mm.31095 | NM_028372 | ENSMUSG00000051098 | **Mblac2** | 2900024O10Rik | metallo-beta-lactamase domain containing 2 | protein-coding |
| chr3_60438707_60439248 | chr3 | 60438708 | 60439248 | + | 1.73761 | Intergenic | B4A\|SINE\|B4 | -33852 | NM_001253713 | 56758 | Mm.255723 | NM_020007 | ENSMUSG00000027763 | **Mbnl1** | Mbnl\|mKIAA0428 | muscleblind-like 1 (Drosophila) | protein-coding |
| chr3_60596968_60597325 | chr3 | 60596969 | 60597325 | + | 1.30972 | intron (NM_001253708, intron 3 of 8) | intron (NM_001253708, intron 3 of 8) | 95895 | NM_001253708 | 56758 | Mm.255723 | NM_020007 | ENSMUSG00000027763 | **Mbnl1** | Mbnl\|mKIAA0428 | muscleblind-like 1 (Drosophila) | protein-coding |
| chr3_60603069_60603542 | chr3 | 60603070 | 60603542 | + | 1.25268 | intron (NM_001253708, intron 3 of 8) | intron (NM_001253708, intron 3 of 8) | 102054 | NM_001253708 | 56758 | Mm.255723 | NM_020007 | ENSMUSG00000027763 | **Mbnl1** | Mbnl\|mKIAA0428 | muscleblind-like 1 (Drosophila) | protein-coding |
| chr3_60567139_60567446 | chr3 | 60567140 | 60567446 | + | 0.897531 | intron (NM_001253708, intron 2 of 8) | intron (NM_001253708, intron 2 of 8) | 66041 | NM_001253708 | 56758 | Mm.255723 | NM_020007 | ENSMUSG00000027763 | **Mbnl1** | Mbnl\|mKIAA0428 | muscleblind-like 1 (Drosophila) | protein-coding |
| chr3_60483240_60483568 | chr3 | 60483241 | 60483568 | + | 0.826435 | intron (NM_001253713, intron 2 of 9) | intron (NM_001253713, intron 2 of 9) | 10574 | NM_001253713 | 56758 | Mm.255723 | NM_020007 | ENSMUSG00000027763 | **Mbnl1** | Mbnl\|mKIAA0428 | muscleblind-like 1 (Drosophila) | protein-coding |
| chr3_60485704_60485931 | chr3 | 60485705 | 60485931 | + | 0.65889 | intron (NM_001253713, intron 2 of 9) | intron (NM_001253713, intron 2 of 9) | 12988 | NM_001253713 | 56758 | Mm.255723 | NM_020007 | ENSMUSG00000027763 | **Mbnl1** | Mbnl\|mKIAA0428 | muscleblind-like 1 (Drosophila) | protein-coding |
| chr14_120295832_120296600 | chr14 | 120295833 | 120296600 | + | 1.30697 | intron (NM_175341, intron 1 of 8) | intron (NM_175341, intron 1 of 8) | 20547 | NM_175341 | 105559 | Mm.238266 | NM_175341 | ENSMUSG00000022139 | **Mbnl2** | 1110002M11Rik\|AI047808\|AI837313\|AI849185\|AL118326\|MBLL\|R75232\|mKIAA4072 | muscleblind-like 2 | protein-coding |
| chr14_120344651_120344937 | chr14 | 120344652 | 120344937 | + | 1.05719 | intron (NM_175341, intron 2 of 8) | ORR1C1\|LTR\|ERVL-MaLR | 69125 | NM_175341 | 105559 | Mm.238266 | NM_175341 | ENSMUSG00000022139 | **Mbnl2** | 1110002M11Rik\|AI047808\|AI837313\|AI849185\|AL118326\|MBLL\|R75232\|mKIAA4072 | muscleblind-like 2 | protein-coding |
| chr14_120344595_120344986 | chr14 | 120344596 | 120344986 | + | 0.973852 | intron (NM_175341, intron 2 of 8) | ORR1C1\|LTR\|ERVL-MaLR | 69122 | NM_175341 | 105559 | Mm.238266 | NM_175341 | ENSMUSG00000022139 | **Mbnl2** | 1110002M11Rik\|AI047808\|AI837313\|AI849185\|AL118326\|MBLL\|R75232\|mKIAA4072 | muscleblind-like 2 | protein-coding |
| chr14_120290800_120291481 | chr14 | 120290801 | 120291481 | + | 0.922598 | intron (NM_175341, intron 1 of 8) | intron (NM_175341, intron 1 of 8) | 15472 | NM_175341 | 105559 | Mm.238266 | NM_175341 | ENSMUSG00000022139 | **Mbnl2** | 1110002M11Rik\|AI047808\|AI837313\|AI849185\|AL118326\|MBLL\|R75232\|mKIAA4072 | muscleblind-like 2 | protein-coding |
| chr14_120325638_120325952 | chr14 | 120325639 | 120325952 | + | 0.754426 | intron (NM_175341, intron 2 of 8) | intron (NM_175341, intron 2 of 8) | 50126 | NM_175341 | 105559 | Mm.238266 | NM_175341 | ENSMUSG00000022139 | **Mbnl2** | 1110002M11Rik\|AI047808\|AI837313\|AI849185\|AL118326\|MBLL\|R75232\|mKIAA4072 | muscleblind-like 2 | protein-coding |
| chr14_120344069_120344992 | chr14 | 120344070 | 120344992 | + | 0.687473 | intron (NM_175341, intron 2 of 8) | intron (NM_175341, intron 2 of 8) | 68862 | NM_175341 | 105559 | Mm.238266 | NM_175341 | ENSMUSG00000022139 | **Mbnl2** | 1110002M11Rik\|AI047808\|AI837313\|AI849185\|AL118326\|MBLL\|R75232\|mKIAA4072 | muscleblind-like 2 | protein-coding |
| chr14_120325217_120326143 | chr14 | 120325218 | 120326143 | + | 0.675071 | intron (NM_175341, intron 2 of 8) | intron (NM_175341, intron 2 of 8) | 50011 | NM_175341 | 105559 | Mm.238266 | NM_175341 | ENSMUSG00000022139 | **Mbnl2** | 1110002M11Rik\|AI047808\|AI837313\|AI849185\|AL118326\|MBLL\|R75232\|mKIAA4072 | muscleblind-like 2 | protein-coding |
| chr13_30109042_30109697 | chr13 | 30109043 | 30109697 | + | 1.10269 | Intergenic | C-rich\|Low_complexity\|Low_complexity | -27120 | NM_153546 | 218121 | Mm.89682 | NM_153546 | ENSMUSG00000038732 | **Mboat1** | 9130215M02Rik\|BC023845\|LPEAT1\|Moact1\|Oact1 | membrane bound O-acyltransferase domain containing 1 | protein-coding |
| chr8_34123316_34123545 | chr8 | 34123317 | 34123545 | + | 0.775611 | intron (NM_001126314, intron 2 of 2) | intron (NM_001126314, intron 2 of 2) | 8401 | NM_001126314 | 234155 | Mm.324008 | NM_001126314 | ENSMUSG00000071113 | **Mboat4** | GOAT\|Gm171 | membrane bound O-acyltransferase domain containing 4 | protein-coding |

| | | | | | | | | | | | | | | | | | |
|---|---|---|---|---|---|---|---|---|---|---|---|---|---|---|---|---|---|
| chr18_82505055_82505255 | chr18 | 82505056 | 82505255 | + | 0.941617 | intron (NM_001025245, intron 2 of 3) | intron (NM_001025245, intron 2 of 3) | 30032 | NM_001025245 | 17196 | Mm.252063 | NM_010777 | ENSMUSG00000041607 | **Mbp** | C76307\|Hmbpr\|R75289\|golli-mbp\|mld\|shi | myelin basic protein | protein-coding |
| chr18_82531878_82532379 | chr18 | 82531879 | 82532379 | + | 0.893424 | intron (NM_001025245, intron 3 of 3) | intron (NM_001025245, intron 3 of 3) | -22334 | NM_001025256 | 17196 | Mm.252063 | NM_010777 | ENSMUSG00000041607 | **Mbp** | C76307\|Hmbpr\|R75289\|golli-mbp\|mld\|shi | myelin basic protein | protein-coding |
| chr18_82508633_82509668 | chr18 | 82508634 | 82509668 | + | 0.87771 | intron (NM_001025245, intron 2 of 3) | intron (NM_001025245, intron 2 of 3) | 34028 | NM_001025245 | 17196 | Mm.252063 | NM_010777 | ENSMUSG00000041607 | **Mbp** | C76307\|Hmbpr\|R75289\|golli-mbp\|mld\|shi | myelin basic protein | protein-coding |
| chr18_82559904_82560325 | chr18 | 82559905 | 82560325 | + | 0.853081 | intron (NM_001025258, intron 1 of 4) | intron (NM_001025258, intron 1 of 4) | 5607 | NM_001025254 | 17196 | Mm.252063 | NM_010777 | ENSMUSG00000041607 | **Mbp** | C76307\|Hmbpr\|R75289\|golli-mbp\|mld\|shi | myelin basic protein | protein-coding |
| chr18_82524635_82524985 | chr18 | 82524636 | 82524985 | + | 0.827245 | intron (NM_001025245, intron 3 of 3) | intron (NM_001025245, intron 3 of 3) | -29653 | NM_001025256 | 17196 | Mm.252063 | NM_010777 | ENSMUSG00000041607 | **Mbp** | C76307\|Hmbpr\|R75289\|golli-mbp\|mld\|shi | myelin basic protein | protein-coding |
| chr18_82559876_82560477 | chr18 | 82559877 | 82560477 | + | 0.807226 | intron (NM_001025258, intron 1 of 4) | intron (NM_001025258, intron 1 of 4) | 5669 | NM_001025254 | 17196 | Mm.252063 | NM_010777 | ENSMUSG00000041607 | **Mbp** | C76307\|Hmbpr\|R75289\|golli-mbp\|mld\|shi | myelin basic protein | protein-coding |
| chr18_82559874_82560748 | chr18 | 82559875 | 82560748 | + | 0.646118 | intron (NM_001025258, intron 1 of 4) | intron (NM_001025258, intron 1 of 4) | 5803 | NM_001025254 | 17196 | Mm.252063 | NM_010777 | ENSMUSG00000041607 | **Mbp** | C76307\|Hmbpr\|R75289\|golli-mbp\|mld\|shi | myelin basic protein | protein-coding |
| chr8_119548588_119549065 | chr8 | 119548589 | 119549065 | + | 0.966175 | intron (NM_001167910, intron 1 of 22) | intron (NM_001167910, intron 1 of 22) | 9934 | NM_001167910 | 56453 | Mm.206934 | NM_019709 | ENSMUSG00000031835 | **Mbtps1** | 0610038M03Rik\|AV003995\|S1P\|SKI-1\|Ski1\|mKIAA0091 | membrane-bound transcription factor peptidase, site 1 | protein-coding |
| chr3_95648079_95648439 | chr3 | 95648080 | 95648439 | + | 1.39733 | Intergenic | Lx8\|LINE\|L1 | -10462 | NM_008562 | 17210 | Mm.1639 | NM_008562 | ENSMUSG00000038612 | **Mcl1** | AW556805\|Mcl-1 | myeloid cell leukemia sequence 1 | protein-coding |
| chr3_95650358_95651289 | chr3 | 95650359 | 95651289 | + | 1.10003 | Intergenic | Intergenic | -7897 | NM_008562 | 17210 | Mm.1639 | NM_008562 | ENSMUSG00000038612 | **Mcl1** | AW556805\|Mcl-1 | myeloid cell leukemia sequence 1 | protein-coding |
| chr3_95663889_95664536 | chr3 | 95663890 | 95664536 | + | 0.755605 | Intergenic | B1F\|SINE\|Alu | 5492 | NM_008562 | 17210 | Mm.1639 | NM_008562 | ENSMUSG00000038612 | **Mcl1** | AW556805\|Mcl-1 | myeloid cell leukemia sequence 1 | protein-coding |
| chr3_95664994_95665333 | chr3 | 95664995 | 95665333 | + | 0.735124 | Intergenic | Intergenic | 6443 | NM_008562 | 17210 | Mm.1639 | NM_008562 | ENSMUSG00000038612 | **Mcl1** | AW556805\|Mcl-1 | myeloid cell leukemia sequence 1 | protein-coding |
| chr1_128357458_128357811 | chr1 | 128357459 | 128357811 | + | 0.617482 | intron (NM_008567, intron 1 of 16) | intron (NM_008567, intron 1 of 16) | 2021 | NM_008567 | 17219 | Mm.4933 | NM_008567 | ENSMUSG00000026355 | **Mcm6** | ASP-I1\|D1Wsu22e\|Mcmd6 | minichromosome maintenance deficient 6 (MIS5 homolog, S. pombe) (S. cerevisiae) | protein-coding |
| chr10_53621359_53622114 | chr10 | 53621360 | 53622114 | + | 1.07856 | intron (NM_027830, intron 4 of 13) | RLTR40\|LTR\|ERVK | 8702 | NM_027830 | 71567 | Mm.278221 | NM_027830 | ENSMUSG00000058298 | **Mcm9** | 9030408O17Rik\|Gm235\|Mcmdc1 | minichromosome maintenance complex component 9 | protein-coding |
| chr3_146172942_146173446 | chr3 | 146172943 | 146173446 | + | 0.974706 | intron (NM_001005846, intron 4 of 13) | RMER15\|LTR\|ERVL | 23020 | NM_001005846 | 68279 | Mm.116862 | NM_026656 | ENSMUSG00000011008 | **Mcoln2** | 3300002C04Rik\|AI549968\|C86638\|TRPML2 | mucolipin 2 | protein-coding |
| chr3_146123113_146123428 | chr3 | 146123114 | 146123428 | + | 0.923537 | intron (NM_134160, intron 1 of 11) | intron (NM_134160, intron 1 of 11) | 1480 | NM_134160 | 171166 | Mm.114683 | NM_134160 | ENSMUSG00000036853 | **Mcoln3** | 6720490O21Rik\|TRPML3\|Va | mucolipin 3 | protein-coding |
| chr3_146122753_146123433 | chr3 | 146122754 | 146123433 | + | 0.749481 | intron (NM_134160, intron 1 of 11) | intron (NM_134160, intron 1 of 11) | 1302 | NM_134160 | 171166 | Mm.114683 | NM_134160 | ENSMUSG00000036853 | **Mcoln3** | 6720490O21Rik\|TRPML3\|Va | mucolipin 3 | protein-coding |
| chr8_18618955_18619497 | chr8 | 18618956 | 18619497 | + | 1.36064 | intron (NM_173189, intron 3 of 13) | B3\|SINE\|B2 | 24053 | NM_173189 | 244329 | Mm.235296 | NM_173189 | ENSMUSG00000039842 | **Mcph1** | 5430437K10Rik\|BRIT1\|D030046N04Rik\|MCT | microcephaly, primary autosomal recessive 1 | protein-coding |
| chr7_72350040_72350438 | chr7 | 72350041 | 72350438 | + | 0.784225 | Intergenic | Intergenic | -43644 | NM_001024703 | 244049 | Mm.217412 | NM_001024703 | ENSMUSG00000032776 | **Mctp2** | Gm489 | multiple C2 domains, transmembrane 2 | protein-coding |
| chr7_72143330_72143702 | chr7 | 72143331 | 72143702 | + | 0.721383 | intron (NM_001024703, intron 17 of 22) | RLTR40\|LTR\|ERVK | 163079 | NM_001024703 | 244049 | Mm.217412 | NM_001024703 | ENSMUSG00000032776 | **Mctp2** | Gm489 | multiple C2 domains, transmembrane 2 | protein-coding |
| chr7_72143319_72143938 | chr7 | 72143320 | 72143938 | + | 0.6693 | intron (NM_001024703, intron 17 of 22) | intron (NM_001024703, intron 17 of 22) | 162966 | NM_001024703 | 244049 | Mm.217412 | NM_001024703 | ENSMUSG00000032776 | **Mctp2** | Gm489 | multiple C2 domains, transmembrane 2 | protein-coding |
| chr10_59580517_59580964 | chr10 | 59580518 | 59580964 | + | 0.973475 | intron (NM_001033259, intron 1 of 7) | L1MD2\|LINE\|L1 | 35951 | NM_001033259 | 215999 | Mm.296470 | NM_001033259 | ENSMUSG00000009647 | **Mcu** | 2010012O16Rik\|AV064928\|C10orf42\|Ccdc109a\|D130073L02Rik\|Gm64 | mitochondrial calcium uniporter | protein-coding |
| chr10_59577811_59578995 | chr10 | 59577812 | 59578995 | + | 0.751956 | intron (NM_001033259, intron 1 of 7) | intron (NM_001033259, intron 1 of 7) | 38289 | NM_001033259 | 215999 | Mm.296470 | NM_001033259 | ENSMUSG00000009647 | **Mcu** | 2010012O16Rik\|AV064928\|C10orf42\|Ccdc109a\|D130073L02Rik\|Gm64 | mitochondrial calcium uniporter | protein-coding |
| chr10_59578300_59578772 | chr10 | 59578301 | 59578772 | + | 0.655217 | intron (NM_001033259, intron 1 of 7) | intron (NM_001033259, intron 1 of 7) | 38156 | NM_001033259 | 215999 | Mm.296470 | NM_001033259 | ENSMUSG00000009647 | **Mcu** | 2010012O16Rik\|AV064928\|C10orf42\|Ccdc109a\|D130073L02Rik\|Gm64 | mitochondrial calcium uniporter | protein-coding |
| chr6_15710746_15711409 | chr6 | 15710747 | 15711409 | + | 1.4672 | Intergenic | Intergenic | -9583 | NM_175088 | 16543 | Mm.1314 | NM_175088 | ENSMUSG00000041390 | **Mdfic** | Kdt1\|Mdfid | MyoD family inhibitor domain containing | protein-coding |
| chr6_15797771_15798252 | chr6 | 15797772 | 15798252 | + | 1.44971 | intron (NM_175088, intron 4 of 4) | intron (NM_175088, intron 4 of 4) | 77351 | NM_175088 | 16543 | Mm.1314 | NM_175088 | ENSMUSG00000041390 | **Mdfic** | Kdt1\|Mdfid | MyoD family inhibitor domain containing | protein-coding |
| chr6_15726471_15726849 | chr6 | 15726472 | 15726849 | + | 1.4362 | intron (NM_175088, intron 2 of 4) | intron (NM_175088, intron 2 of 4) | 5999 | NM_175088 | 16543 | Mm.1314 | NM_175088 | ENSMUSG00000041390 | **Mdfic** | Kdt1\|Mdfid | MyoD family inhibitor domain containing | protein-coding |
| chr6_15751407_15751716 | chr6 | 15751408 | 15751716 | + | 1.32642 | intron (NM_175088, intron 3 of 4) | intron (NM_175088, intron 3 of 4) | 30901 | NM_175088 | 16543 | Mm.1314 | NM_175088 | ENSMUSG00000041390 | **Mdfic** | Kdt1\|Mdfid | MyoD family inhibitor domain containing | protein-coding |
| chr6_15726966_15727253 | chr6 | 15726967 | 15727253 | + | 1.29103 | intron (NM_175088, intron 2 of 4) | intron (NM_175088, intron 2 of 4) | 6449 | NM_175088 | 16543 | Mm.1314 | NM_175088 | ENSMUSG00000041390 | **Mdfic** | Kdt1\|Mdfid | MyoD family inhibitor domain containing | protein-coding |
| chr6_15799262_15799695 | chr6 | 15799263 | 15799695 | + | 1.17893 | intron (NM_175088, intron 4 of 4) | intron (NM_175088, intron 4 of 4) | 78818 | NM_175088 | 16543 | Mm.1314 | NM_175088 | ENSMUSG00000041390 | **Mdfic** | Kdt1\|Mdfid | MyoD family inhibitor domain containing | protein-coding |
| chr6_15683348_15684103 | chr6 | 15683349 | 15684103 | + | 1.16806 | Intergenic | Intergenic | -36935 | NM_175088 | 16543 | Mm.1314 | NM_175088 | ENSMUSG00000041390 | **Mdfic** | Kdt1\|Mdfid | MyoD family inhibitor domain containing | protein-coding |
| chr6_15716794_15717149 | chr6 | 15716795 | 15717149 | + | 1.12733 | Intergenic | Intergenic | -3689 | NM_175088 | 16543 | Mm.1314 | NM_175088 | ENSMUSG00000041390 | **Mdfic** | Kdt1\|Mdfid | MyoD family inhibitor domain containing | protein-coding |
| chr6_15682809_15684093 | chr6 | 15682810 | 15684093 | + | 1.08729 | Intergenic | Intergenic | -37210 | NM_175088 | 16543 | Mm.1314 | NM_175088 | ENSMUSG00000041390 | **Mdfic** | Kdt1\|Mdfid | MyoD family inhibitor domain containing | protein-coding |
| chr6_15728852_15729703 | chr6 | 15728853 | 15729703 | + | 1.08633 | intron (NM_175088, intron 2 of 4) | intron (NM_175088, intron 2 of 4) | 8617 | NM_175088 | 16543 | Mm.1314 | NM_175088 | ENSMUSG00000041390 | **Mdfic** | Kdt1\|Mdfid | MyoD family inhibitor domain containing | protein-coding |
| chr6_15729234_15729644 | chr6 | 15729235 | 15729644 | + | 1.00564 | intron (NM_175088, intron 2 of 4) | intron (NM_175088, intron 2 of 4) | 8778 | NM_175088 | 16543 | Mm.1314 | NM_175088 | ENSMUSG00000041390 | **Mdfic** | Kdt1\|Mdfid | MyoD family inhibitor domain containing | protein-coding |
| chr6_15743313_15743868 | chr6 | 15743314 | 15743868 | + | 0.950143 | intron (NM_175088, intron 3 of 4) | intron (NM_175088, intron 3 of 4) | 22930 | NM_175088 | 16543 | Mm.1314 | NM_175088 | ENSMUSG00000041390 | **Mdfic** | Kdt1\|Mdfid | MyoD family inhibitor domain containing | protein-coding |
| chr6_15780693_15781026 | chr6 | 15780694 | 15781026 | + | 0.902954 | intron (NM_175088, intron 4 of 4) | intron (NM_175088, intron 4 of 4) | 60199 | NM_175088 | 16543 | Mm.1314 | NM_175088 | ENSMUSG00000041390 | **Mdfic** | Kdt1\|Mdfid | MyoD family inhibitor domain containing | protein-coding |
| chr6_15711838_15712143 | chr6 | 15711839 | 15712143 | + | 0.847841 | Intergenic | Intergenic | -8670 | NM_175088 | 16543 | Mm.1314 | NM_175088 | ENSMUSG00000041390 | **Mdfic** | Kdt1\|Mdfid | MyoD family inhibitor domain containing | protein-coding |
| chr6_15793068_15793996 | chr6 | 15793069 | 15793996 | + | 0.843082 | intron (NM_175088, intron 4 of 4) | intron (NM_175088, intron 4 of 4) | 72871 | NM_175088 | 16543 | Mm.1314 | NM_175088 | ENSMUSG00000041390 | **Mdfic** | Kdt1\|Mdfid | MyoD family inhibitor domain containing | protein-coding |
| chr6_15792704_15793990 | chr6 | 15792705 | 15793990 | + | 0.820939 | intron (NM_175088, intron 4 of 4) | intron (NM_175088, intron 4 of 4) | 72686 | NM_175088 | 16543 | Mm.1314 | NM_175088 | ENSMUSG00000041390 | **Mdfic** | Kdt1\|Mdfid | MyoD family inhibitor domain containing | protein-coding |
| chr6_15794047_15794260 | chr6 | 15794048 | 15794260 | + | 0.756917 | intron (NM_175088, intron 4 of 4) | intron (NM_175088, intron 4 of 4) | 73493 | NM_175088 | 16543 | Mm.1314 | NM_175088 | ENSMUSG00000041390 | **Mdfic** | Kdt1\|Mdfid | MyoD family inhibitor domain containing | protein-coding |
| chr6_15794845_15795032 | chr6 | 15794846 | 15795032 | + | 0.752945 | intron (NM_175088, intron 4 of 4) | intron (NM_175088, intron 4 of 4) | 74278 | NM_175088 | 16543 | Mm.1314 | NM_175088 | ENSMUSG00000041390 | **Mdfic** | Kdt1\|Mdfid | MyoD family inhibitor domain containing | protein-coding |
| chr6_15782978_15783453 | chr6 | 15782979 | 15783453 | + | 0.685456 | intron (NM_175088, intron 4 of 4) | intron (NM_175088, intron 4 of 4) | 62555 | NM_175088 | 16543 | Mm.1314 | NM_175088 | ENSMUSG00000041390 | **Mdfic** | Kdt1\|Mdfid | MyoD family inhibitor domain containing | protein-coding |
| chr17_29933959_29934670 | chr17 | 29933960 | 29934670 | + | 0.771324 | Intergenic | Intergenic | -46433 | NM_001081160 | 74762 | Mm.40901 | NM_001081160 | ENSMUSG00000043557 | **Mdga1** | 1200011I03Rik\|GPIM\|Mamdc3 | MAM domain containing glycosylphosphatidylinositol anchor 1 | protein-coding |
| chr10_117721433_117721952 | chr10 | 117721434 | 117721952 | + | 1.71606 | Intergenic | Intergenic | -10935 | NM_001288586 | 17246 | Mm.22670 | NM_010786 | ENSMUSG00000020184 | **Mdm2** | 1700007J15Rik\|AA415488\|Mdm-2 | transformed mouse 3T3 cell double minute 2 | protein-coding |
| chr10_117677499_117677946 | chr10 | 117677500 | 117677946 | + | 0.893765 | intron (NM_027468, intron 7 of 8) | intron (NM_027468, intron 7 of 8) | 33035 | NM_010786 | 17246 | Mm.22670 | NM_010786 | ENSMUSG00000020184 | **Mdm2** | 1700007J15Rik\|AA415488\|Mdm-2 | transformed mouse 3T3 cell double minute 2 | protein-coding |
| chr10_117715141_117715409 | chr10 | 117715142 | 117715409 | + | 0.735983 | Intergenic | RLTR30D_MM\|LTR\|ERV1 | -4517 | NM_001288586 | 17246 | Mm.22670 | NM_010786 | ENSMUSG00000020184 | **Mdm2** | 1700007J15Rik\|AA415488\|Mdm-2 | transformed mouse 3T3 cell double minute 2 | protein-coding |
| chrX_74105825_74106318 | chrX | 74105826 | 74106318 | + | 0.779267 | Intergenic | RMER19B\|LTR\|ERVK | -20382 | NM_001081979 | 17257 | Mm.131408 | NM_010788 | ENSMUSG00000031393 | **Mecp2** | 1500041B07Rik\|D630021H01Rik\|Mbd5\|WBP10 | methyl CpG binding protein 2 | protein-coding |
| chr11_86380350_86380842 | chr11 | 86380351 | 86380842 | + | 0.921261 | Intergenic | MLT1M\|LTR\|ERVL-MaLR | -23071 | NM_001080931 | 327987 | Mm.341886 | NM_001080931 | ENSMUSG00000034297 | **Med13** | 1110067M05Rik\|D030023K18\|Thrap1\|Trap240 | mediator complex subunit 13 | protein-coding |

| | | | | | | | | | | | | | | | | | |
|---|---|---|---|---|---|---|---|---|---|---|---|---|---|---|---|---|---|
| chr5_118494644_118495563 | chr5 | 118494645 | 118495563 | + | 1.19269 | Intergenic | Intergenic | -65615 | NM_172424 | 76199 | Mm.206238 | NM_172424 | ENSMUSG00000018076 | **Med13l** | 2210413I17Rik\|6330591G05Rik\|9030618F05Rik\|Thrap2\|Trap240L\|mKIAA1025 | mediator complex subunit 13-like | protein-coding |
| chr5_118498509_118498839 | chr5 | 118498510 | 118498839 | + | 0.757036 | Intergenic | Intergenic | -62045 | NM_172424 | 76199 | Mm.206238 | NM_172424 | ENSMUSG00000018076 | **Med13l** | 2210413I17Rik\|6330591G05Rik\|9030618F05Rik\|Thrap2\|Trap240L\|mKIAA1025 | mediator complex subunit 13-like | protein-coding |
| chr5_118458791_118459751 | chr5 | 118458792 | 118459751 | + | 0.752782 | Intergenic | RMER17C2\|LTR\|ERVK | -101448 | NM_172424 | 76199 | Mm.206238 | NM_172424 | ENSMUSG00000018076 | **Med13l** | 2210413I17Rik\|6330591G05Rik\|9030618F05Rik\|Thrap2\|Trap240L\|mKIAA1025 | mediator complex subunit 13-like | protein-coding |
| chr16_17694661_17695291 | chr16 | 17694662 | 17695291 | + | 0.949565 | intron (NM_001285886, intron 4 of 16) | intron (NM_001285886, intron 4 of 16) | 27924 | NM_001285886 | 94112 | Mm.208970 | NM_033609 | ENSMUSG00000012114 | **Med15** | A230074L19Rik\|AW536074\|Pcqap\|mPcqap | mediator complex subunit 15 | protein-coding |
| chr2_29383922_29384259 | chr2 | 29383923 | 29384259 | + | 0.740923 | intron (NM_001290489, intron 1 of 6) | intron (NM_001290489, intron 1 of 6) | 5577 | NM_001290489 | 68975 | Mm.330109 | NM_026896 | ENSMUSG00000026799 | **Med27** | 1500015J03Rik\|2310042P07Rik\|AA682045\|Crsp8\|D2Ertd434e | mediator complex subunit 27 | protein-coding |
| chr15_52899401_52900370 | chr15 | 52899402 | 52900370 | + | 0.91758 | Intergenic | Intergenic | 187441 | NM_027212 | 69790 | Mm.195562 | NM_027212 | ENSMUSG00000038622 | **Med30** | 1810038N03Rik\|2510044J04Rik\|TRAP25\|Thrap6 | mediator complex subunit 30 | protein-coding |
| chr15_52591959_52592424 | chr15 | 52591960 | 52592424 | + | 0.835801 | Intergenic | Intergenic | -120253 | NM_027212 | 69790 | Mm.195562 | NM_027212 | ENSMUSG00000038622 | **Med30** | 1810038N03Rik\|2510044J04Rik\|TRAP25\|Thrap6 | mediator complex subunit 30 | protein-coding |
| chr15_52883979_52884266 | chr15 | 52883980 | 52884266 | + | 0.690616 | Intergenic | L1ME3Cz\|LINE\|L1 | 171678 | NM_027212 | 69790 | Mm.195562 | NM_027212 | ENSMUSG00000038622 | **Med30** | 1810038N03Rik\|2510044J04Rik\|TRAP25\|Thrap6 | mediator complex subunit 30 | protein-coding |
| chr15_52565350_52566014 | chr15 | 52565351 | 52566014 | + | 0.679168 | Intergenic | Intergenic | -146763 | NM_027212 | 69790 | Mm.195562 | NM_027212 | ENSMUSG00000038622 | **Med30** | 1810038N03Rik\|2510044J04Rik\|TRAP25\|Thrap6 | mediator complex subunit 30 | protein-coding |
| chr15_52577140_52577699 | chr15 | 52577141 | 52577699 | + | 0.654214 | Intergenic | Intergenic | -135025 | NM_027212 | 69790 | Mm.195562 | NM_027212 | ENSMUSG00000038622 | **Med30** | 1810038N03Rik\|2510044J04Rik\|TRAP25\|Thrap6 | mediator complex subunit 30 | protein-coding |
| chr5_149391872_149392386 | chr5 | 149391873 | 149392386 | + | 0.694911 | Intergenic | Intergenic | -19677 | NM_027519 | 70717 | Mm.252139 | NM_027519 | ENSMUSG00000029659 | **Medag** | 6330406I15Rik\|Awms3\|MEDA-4\|Meda4\|mAWMS3 | mesenteric estrogen dependent adipogenesis | protein-coding |
| chr5_149391625_149392401 | chr5 | 149391626 | 149392401 | + | 0.635229 | Intergenic | Intergenic | -19793 | NM_027519 | 70717 | Mm.252139 | NM_027519 | ENSMUSG00000029659 | **Medag** | 6330406I15Rik\|Awms3\|MEDA-4\|Meda4\|mAWMS3 | mesenteric estrogen dependent adipogenesis | protein-coding |
| chr7_67308096_67308542 | chr7 | 67308097 | 67308542 | + | 1.06417 | intron (NM_001291196, intron 3 of 11) | intron (NM_001291196, intron 3 of 11) | 64539 | NM_001291192 | 17258 | Mm.132788 | NM_001033713 | ENSMUSG00000030557 | **Mef2a** | A430079H05Rik | myocyte enhancer factor 2A | protein-coding |
| chr7_67304801_67305273 | chr7 | 67304802 | 67305273 | + | 0.814757 | intron (NM_001291196, intron 3 of 11) | B3A\|SINE\|B2 | 67821 | NM_001291192 | 17258 | Mm.132788 | NM_001033713 | ENSMUSG00000030557 | **Mef2a** | A430079H05Rik | myocyte enhancer factor 2A | protein-coding |
| chr7_67320378_67320911 | chr7 | 67320379 | 67320911 | + | 0.67856 | intron (NM_001291196, intron 2 of 11) | intron (NM_001291196, intron 2 of 11) | 52213 | NM_001291192 | 17258 | Mm.132788 | NM_001033713 | ENSMUSG00000030557 | **Mef2a** | A430079H05Rik | myocyte enhancer factor 2A | protein-coding |
| chr7_67430070_67430466 | chr7 | 67430071 | 67430466 | + | 0.645377 | Intergenic | MTD\|LTR\|ERVL-MaLR | -57410 | NM_001291195 | 17258 | Mm.132788 | NM_001033713 | ENSMUSG00000030557 | **Mef2a** | A430079H05Rik | myocyte enhancer factor 2A | protein-coding |
| chr13_83597409_83598075 | chr13 | 83597410 | 83598075 | + | 0.78881 | intron (NM_025282, intron 4 of 10) | intron (NM_025282, intron 4 of 10) | 93708 | NM_025282 | 17260 | Mm.24001 | NM_025282 | ENSMUSG00000005583 | **Mef2c** | 5430401D19Rik\|9930028G15Rik\|AV011172\|Mef2 | myocyte enhancer factor 2C | protein-coding |
| chr3_88128166_88128443 | chr3 | 88128167 | 88128443 | + | 0.985505 | Intergenic | MIRb\|SINE\|MIR | -14090 | NM_133665 | 17261 | Mm.28184 | NM_133665 | ENSMUSG00000001419 | **Mef2d** | C80750 | myocyte enhancer factor 2D | protein-coding |
| chr16_3723115_3723685 | chr16 | 3723116 | 3723685 | + | 1.95947 | Intergenic | Intergenic | -5276 | NM_001161791 | 54483 | Mm.143718 | NM_019453 | ENSMUSG00000022534 | **Mefv** | FMF\|TRIM20\|pyrin | Mediterranean fever | protein-coding |
| chr9_81941544_81941827 | chr9 | 81941545 | 81941827 | + | 0.868878 | intron (NM_175213, intron 3 of 4) | intron (NM_175213, intron 3 of 4) | 77942 | NM_175213 | 75033 | Mm.335329 | NM_175213 | ENSMUSG00000043289 | **Mei4** | 4930486G11Rik | meiosis-specific, MEI4 homolog (S. cerevisiae) | protein-coding |
| chr9_82010969_82011385 | chr9 | 82010970 | 82011385 | + | 0.740053 | intron (NM_175213, intron 4 of 4) | intron (NM_175213, intron 4 of 4) | 147433 | NM_175213 | 75033 | Mm.335329 | NM_175213 | ENSMUSG00000043289 | **Mei4** | 4930486G11Rik | meiosis-specific, MEI4 homolog (S. cerevisiae) | protein-coding |
| chr2_115855818_115856216 | chr2 | 115855819 | 115856216 | + | 0.668389 | Intergenic | Intergenic | 208604 | NM_001159569 | 17536 | Mm.247566 | NM_010825 | ENSMUSG00000027210 | **Meis2** | A430109D20Rik\|Mrg1\|Stra10 | Meis homeobox 2 | protein-coding |
| chr4_44272732_44272952 | chr4 | 44272733 | 44272952 | + | 0.736304 | Intergenic | ORR1D1\|LTR\|ERVL-MaLR | -28075 | NM_010790 | 17279 | Mm.268668 | NM_010790 | ENSMUSG00000035683 | **Melk** | AI327312\|MPK38\|mKIAA0175 | maternal embryonic leucine zipper kinase | protein-coding |
| chr2_128693437_128693897 | chr2 | 128693438 | 128693897 | + | 0.740217 | Intergenic | MTC\|LTR\|ERVL-MaLR | -5330 | NM_008587 | 17289 | Mm.239655 | NM_008587 | ENSMUSG00000014361 | **Mertk** | Eyk\|Mer\|Nyk\|nmf12 | c-mer proto-oncogene tyrosine kinase | protein-coding |
| chr6_17478791_17479231 | chr6 | 17478792 | 17479231 | + | 0.627559 | intron (NM_008591, intron 1 of 21) | intron (NM_008591, intron 1 of 21) | 15054 | NM_008591 | 17295 | Mm.86844 | NM_008591 | ENSMUSG00000009376 | **Met** | AI838057\|HGF\|HGFR\|Par4\|c-Met | met proto-oncogene | protein-coding |
| chr10_127042763_127043050 | chr10 | 127042764 | 127043050 | + | 1.34474 | intron (NM_010792, intron 1 of 5) | intron (NM_010792, intron 1 of 5) | 975 | NM_010792 | 17299 | Mm.251593 | NM_010792 | ENSMUSG00000006732 | **Mettl1** | 2810012D02Rik | methyltransferase like 1 | protein-coding |
| chr10_127042765_127043220 | chr10 | 127042766 | 127043220 | + | 1.25611 | intron (NM_010792, intron 1 of 5) | intron (NM_010792, intron 1 of 5) | 1061 | NM_010792 | 17299 | Mm.251593 | NM_010792 | ENSMUSG00000006732 | **Mettl1** | 2810012D02Rik | methyltransferase like 1 | protein-coding |
| chr10_127042761_127043388 | chr10 | 127042762 | 127043388 | + | 1.13785 | exon (NM_010792, exon 2 of 6) | exon (NM_010792, exon 2 of 6) | 1143 | NM_010792 | 17299 | Mm.251593 | NM_010792 | ENSMUSG00000006732 | **Mettl1** | 2810012D02Rik | methyltransferase like 1 | protein-coding |
| chr2_109221519_109221814 | chr2 | 109221520 | 109221814 | + | 0.808187 | intron (NM_029790, intron 2 of 5) | Lx3C\|LINE\|L1 | 56623 | NM_029790 | 76894 | Mm.330451 | NM_029790 | ENSMUSG00000057234 | **Mettl15** | 0610027B03Rik\|AI303697\|AW049952\|Mett5d1 | methyltransferase like 15 | protein-coding |
| chr14_51870761_51871354 | chr14 | 51870762 | 51871354 | + | 1.26436 | Intergenic | Tigger7\|DNA\|TcMar-Tigger | -13784 | NM_001029990 | 52535 | Mm.5750 | NM_001029990 | ENSMUSG00000004561 | **Mettl17** | 2310032K15Rik\|D14Ertd209e\|Mett11d1 | methyltransferase like 17 | protein-coding |
| chr1_163999067_163999368 | chr1 | 163999068 | 163999368 | + | 1.14516 | Intergenic | Intergenic | 4273 | NM_027279 | 69962 | Mm.27284 | NM_027279 | ENSMUSG00000041396 | **Mettl18** | 2810422O20Rik\|C78571 | methyltransferase like 18 | protein-coding |
| chr1_64601307_64601822 | chr1 | 64601308 | 64601822 | + | 1.42205 | 3' UTR (NM_009952, exon 9 of 9) | 3' UTR (NM_009952, exon 9 of 9) | 15603 | NM_025964 | 67099 | Mm.46394 | NM_025964 | ENSMUSG00000025956 | **Mettl21a** | 2310038H17Rik\|AI464204\|Fam119a | methyltransferase like 21A | protein-coding |
| chr1_44032205_44032795 | chr1 | 44032206 | 44032795 | + | 1.08722 | Intergenic | L1MC5a\|LINE\|L1 | -12494 | NM_001013799 | 433294 | Mm.123621 | NM_001013799 | ENSMUSG00000047343 | **Mettl21c** | A530098C11Rik | methyltransferase like 21C | protein-coding |
| chr1_43996966_43997907 | chr1 | 43996967 | 43997907 | + | 1.07335 | intron (NM_009418, intron 27 of 29) | Lx5\|LINE\|L1 | 22569 | NM_001013799 | 433294 | Mm.123621 | NM_001013799 | ENSMUSG00000047343 | **Mettl21c** | A530098C11Rik | methyltransferase like 21C | protein-coding |
| chr1_44047571_44048086 | chr1 | 44047572 | 44048086 | + | 1.06376 | Intergenic | MERV1_LTR\|LTR\|ERV1 | -27823 | NM_001013799 | 433294 | Mm.123621 | NM_001013799 | ENSMUSG00000047343 | **Mettl21c** | A530098C11Rik | methyltransferase like 21C | protein-coding |
| chr1_44047417_44048289 | chr1 | 44047418 | 44048289 | + | 1.01316 | Intergenic | MERV1_LTR\|LTR\|ERV1 | -27847 | NM_001013799 | 433294 | Mm.123621 | NM_001013799 | ENSMUSG00000047343 | **Mettl21c** | A530098C11Rik | methyltransferase like 21C | protein-coding |
| chr1_44049802_44050028 | chr1 | 44049803 | 44050028 | + | 0.725618 | Intergenic | L1_Mus4\|LINE\|L1 | -29909 | NM_001013799 | 433294 | Mm.123621 | NM_001013799 | ENSMUSG00000047343 | **Mettl21c** | A530098C11Rik | methyltransferase like 21C | protein-coding |
| chr16_8471886_8472241 | chr16 | 8471887 | 8472241 | + | 0.882717 | intron (NM_146247, intron 1 of 10) | intron (NM_146247, intron 1 of 10) | 1251 | NM_146247 | 239706 | Mm.214953 | NM_146247 | | **Mettl22** | - | methyltransferase like 22 | protein-coding |
| chr16_8471882_8472747 | chr16 | 8471883 | 8472747 | + | 0.827501 | intron (NM_146247, intron 1 of 10) | intron (NM_146247, intron 1 of 10) | 1502 | NM_146247 | 239706 | Mm.214953 | NM_146247 | | **Mettl22** | - | methyltransferase like 22 | protein-coding |
| chr16_8482475_8482945 | chr16 | 8482476 | 8482945 | + | 0.801452 | intron (NM_146247, intron 5 of 10) | intron (NM_146247, intron 5 of 10) | 11897 | NM_146247 | 239706 | Mm.214953 | NM_146247 | | **Mettl22** | - | methyltransferase like 22 | protein-coding |
| chr10_40688419_40688786 | chr10 | 40688420 | 40688786 | + | 0.838914 | intron (NM_177793, intron 1 of 4) | intron (NM_177793, intron 1 of 4) | 5321 | NM_177793 | 327747 | Mm.80407 | NM_177793 | ENSMUSG00000045555 | **Mettl24** | 9030224M15Rik | methyltransferase like 24 | protein-coding |
| chr10_40688348_40689055 | chr10 | 40688349 | 40689055 | + | 0.802193 | intron (NM_177793, intron 1 of 4) | intron (NM_177793, intron 1 of 4) | 5420 | NM_177793 | 327747 | Mm.80407 | NM_177793 | ENSMUSG00000045555 | **Mettl24** | 9030224M15Rik | methyltransferase like 24 | protein-coding |
| chr10_40712874_40713591 | chr10 | 40712875 | 40713591 | + | 0.732018 | intron (NM_177793, intron 1 of 4) | intron (NM_177793, intron 1 of 4) | 29951 | NM_177793 | 327747 | Mm.80407 | NM_177793 | ENSMUSG00000045555 | **Mettl24** | 9030224M15Rik | methyltransferase like 24 | protein-coding |
| chr10_128972915_128973226 | chr10 | 128972916 | 128973226 | + | 0.856911 | Intergenic | ORR1C1\|LTR\|ERVL-MaLR | -12083 | NM_027853 | 71664 | Mm.337232 | NM_027853 | ENSMUSG00000025347 | **Mettl7b** | 0610006F02Rik\|AI266817 | methyltransferase like 7B | protein-coding |
| chr2_71002487_71002950 | chr2 | 71002488 | 71002950 | + | 0.80933 | intron (NM_001110512, intron 1 of 8) | intron (NM_001110512, intron 1 of 8) | 15655 | NM_001110512 | 228019 | Mm.490343 | NM_145524 | ENSMUSG00000041975 | **Mettl8** | BC004636\|Tip | methyltransferase like 8 | protein-coding |
| chr3_88520088_88520445 | chr3 | 88520089 | 88520445 | + | 1.11323 | Intergenic | Intergenic | -12128 | NM_001029890 | 72640 | Mm.275154 | NM_001029890 | ENSMUSG00000074480 | **Mex3a** | 2700083E18Rik\|Gm411\|Rkhd4 | mex3 homolog A (C. elegans) | protein-coding |
| chr18_73547441_73548114 | chr18 | 73547442 | 73548114 | + | 1.21522 | Intergenic | Intergenic | -24927 | NM_001039214 | 240396 | Mm.24131 | NM_001039214 | ENSMUSG00000037253 | **Mex3c** | A130001D14Rik\|BC035207\|BM-013\|Rkhd2 | mex3 homolog C (C. elegans) | protein-coding |
| chr18_73491752_73492580 | chr18 | 73491753 | 73492580 | + | 1.05729 | Intergenic | MLT1L\|LTR\|ERVL-MaLR | -80539 | NM_001039214 | 240396 | Mm.24131 | NM_001039214 | ENSMUSG00000037253 | **Mex3c** | A130001D14Rik\|BC035207\|BM-013\|Rkhd2 | mex3 homolog C (C. elegans) | protein-coding |
| chr18_73546416_73546718 | chr18 | 73546417 | 73546718 | + | 0.989403 | Intergenic | RMER15\|LTR\|ERVL | -26138 | NM_001039214 | 240396 | Mm.24131 | NM_001039214 | ENSMUSG00000037253 | **Mex3c** | A130001D14Rik\|BC035207\|BM-013\|Rkhd2 | mex3 homolog C (C. elegans) | protein-coding |
| chr18_73491752_73492348 | chr18 | 73491753 | 73492348 | + | 0.984529 | Intergenic | Intergenic | -80655 | NM_001039214 | 240396 | Mm.24131 | NM_001039214 | ENSMUSG00000037253 | **Mex3c** | A130001D14Rik\|BC035207\|BM-013\|Rkhd2 | mex3 homolog C (C. elegans) | protein-coding |
| chr11_57550895_57551485 | chr11 | 57550896 | 57551485 | + | 0.913147 | Intergenic | Intergenic | 32525 | NM_145426 | 216760 | Mm.30501 | NM_145426 | ENSMUSG00000020522 | **Mfap3** | 2610509F16Rik\|2700079M14Rik\|AW536327 | microfibrillar-associated protein 3 | protein-coding |

| | | | | | | | | | | | | | | | | | |
|---|---|---|---|---|---|---|---|---|---|---|---|---|---|---|---|---|---|
| chr11_57550621_57551444 | chr11 | 57550622 | 57551444 | + | 0.904642 | Intergenic | Intergenic | 32368 | NM_145426 | 216760 | Mm.30501 | NM_145426 | ENSMUSG00000020522 | **Mfap3** | 2610509F16Rik\|2700079M14Rik\|AW536327 | microfibrillar-associated protein 3 | protein-coding |
| chr6_122527570_122527846 | chr6 | 122527571 | 122527846 | + | 1.39505 | intron (NM_015776, intron 9 of 9) | intron (NM_015776, intron 9 of 9) | 14032 | NM_015776 | 50530 | Mm.41931 | NM_015776 | ENSMUSG00000030116 | **Mfap5** | MAGP-2 | microfibrillar associated protein 5 | protein-coding |
| chr6_122527540_122528133 | chr6 | 122527541 | 122528133 | + | 1.3462 | intron (NM_015776, intron 9 of 9) | intron (NM_015776, intron 9 of 9) | 14161 | NM_015776 | 50530 | Mm.41931 | NM_015776 | ENSMUSG00000030116 | **Mfap5** | MAGP-2 | microfibrillar associated protein 5 | protein-coding |
| chr6_122532802_122533360 | chr6 | 122532803 | 122533360 | + | 0.840414 | Intergenic | Intergenic | 19405 | NM_015776 | 50530 | Mm.41931 | NM_015776 | ENSMUSG00000030116 | **Mfap5** | MAGP-2 | microfibrillar associated protein 5 | protein-coding |
| chr15_78758020_78758602 | chr15 | 78758021 | 78758602 | + | 1.23761 | intron (NM_008595, intron 6 of 7) | MER2\|DNA\|TcMar-Tigger | 15134 | NM_008595 | 17305 | Mm.149235 | NM_008595 | ENSMUSG00000018169 | **Mfng** | AW546563 | MFNG O-fucosylpeptide 3-beta-N-acetylglucosaminyltransferase | protein-coding |
| chr11_68545766_68546592 | chr11 | 68545767 | 68546592 | + | 1.07768 | intron (NM_001004435, intron 18 of 19) | intron (NM_001004435, intron 18 of 19) | -10007 | NM_146004 | 215723 | Mm.131205 | NM_146004 | ENSMUSG00000048329 | **Mfsd6l** | - | major facilitator superfamily domain containing 6-like | protein-coding |
| chr11_68553674_68554647 | chr11 | 68553675 | 68554647 | + | 0.877557 | Intergenic | Intergenic | -2025 | NM_146004 | 215723 | Mm.131205 | NM_146004 | ENSMUSG00000048329 | **Mfsd6l** | - | major facilitator superfamily domain containing 6-like | protein-coding |
| chr1_191031097_191031397 | chr1 | 191031098 | 191031397 | + | 0.679453 | Intergenic | B3\|SINE\|B2 | -5057 | NM_001081259 | 226844 | Mm.134053 | NM_001081259 | ENSMUSG00000066595 | **Mfsd7b** | 9630055N22Rik\|FLVCR\|Flvcr1 | major facilitator superfamily domain containing 7B | protein-coding |
| chr12_85729408_85730309 | chr12 | 85729409 | 85730309 | + | 0.671561 | Intergenic | Intergenic | -16680 | NM_145447 | 217721 | Mm.301076 | NM_145447 | ENSMUSG00000034258 | **Mfsd7c** | CCT\|Flvcr2 | major facilitator superfamily domain containing 7C | protein-coding |
| chr11_49249237_49249755 | chr11 | 49249238 | 49249755 | + | 0.800427 | intron (NM_001110149, intron 1 of 1) | intron (NM_001110149, intron 1 of 1) | -1056 | NM_001110150 | 17308 | Mm.196933 | NM_010794 | ENSMUSG00000020346 | **Mgat1** | Mgat-1 | mannoside acetylglucosaminyltransferase 1 | protein-coding |
| chr1_127380843_127381585 | chr1 | 127380844 | 127381585 | + | 1.44212 | intron (NM_145128, intron 6 of 17) | intron (NM_145128, intron 6 of 17) | 176228 | NM_145128 | 107895 | Mm.214682 | NM_145128 | ENSMUSG00000036155 | **Mgat5** | 4930471A21Rik\|5330407H02Rik\|AI480971\|GlcNAc-TV | mannoside acetylglucosaminyltransferase 5 | protein-coding |
| chr1_127433039_127433759 | chr1 | 127433040 | 127433759 | + | 1.1403 | intron (NM_145128, intron 12 of 17) | intron (NM_145128, intron 12 of 17) | 228413 | NM_145128 | 107895 | Mm.214682 | NM_145128 | ENSMUSG00000036155 | **Mgat5** | 4930471A21Rik\|5330407H02Rik\|AI480971\|GlcNAc-TV | mannoside acetylglucosaminyltransferase 5 | protein-coding |
| chr1_127433263_127433754 | chr1 | 127433264 | 127433754 | + | 1.13592 | intron (NM_145128, intron 12 of 17) | intron (NM_145128, intron 12 of 17) | 228523 | NM_145128 | 107895 | Mm.214682 | NM_145128 | ENSMUSG00000036155 | **Mgat5** | 4930471A21Rik\|5330407H02Rik\|AI480971\|GlcNAc-TV | mannoside acetylglucosaminyltransferase 5 | protein-coding |
| chr1_127303779_127304122 | chr1 | 127303780 | 127304122 | + | 0.874817 | intron (NM_145128, intron 2 of 17) | MTE2b\|LTR\|ERVL-MaLR | 98965 | NM_145128 | 107895 | Mm.214682 | NM_145128 | ENSMUSG00000036155 | **Mgat5** | 4930471A21Rik\|5330407H02Rik\|AI480971\|GlcNAc-TV | mannoside acetylglucosaminyltransferase 5 | protein-coding |
| chr1_127309070_127309290 | chr1 | 127309071 | 127309290 | + | 0.871468 | intron (NM_145128, intron 3 of 17) | intron (NM_145128, intron 3 of 17) | 104194 | NM_145128 | 107895 | Mm.214682 | NM_145128 | ENSMUSG00000036155 | **Mgat5** | 4930471A21Rik\|5330407H02Rik\|AI480971\|GlcNAc-TV | mannoside acetylglucosaminyltransferase 5 | protein-coding |
| chr1_127311308_127311540 | chr1 | 127311309 | 127311540 | + | 0.732922 | intron (NM_145128, intron 3 of 17) | intron (NM_145128, intron 3 of 17) | 106438 | NM_145128 | 107895 | Mm.214682 | NM_145128 | ENSMUSG00000036155 | **Mgat5** | 4930471A21Rik\|5330407H02Rik\|AI480971\|GlcNAc-TV | mannoside acetylglucosaminyltransferase 5 | protein-coding |
| chr1_127344554_127345486 | chr1 | 127344555 | 127345486 | + | 0.682692 | intron (NM_145128, intron 4 of 17) | intron (NM_145128, intron 4 of 17) | 140034 | NM_145128 | 107895 | Mm.214682 | NM_145128 | ENSMUSG00000036155 | **Mgat5** | 4930471A21Rik\|5330407H02Rik\|AI480971\|GlcNAc-TV | mannoside acetylglucosaminyltransferase 5 | protein-coding |
| chr1_127117901_127118433 | chr1 | 127117902 | 127118433 | + | 0.612713 | Intergenic | Intergenic | -86819 | NM_145128 | 107895 | Mm.214682 | NM_145128 | ENSMUSG00000036155 | **Mgat5** | 4930471A21Rik\|5330407H02Rik\|AI480971\|GlcNAc-TV | mannoside acetylglucosaminyltransferase 5 | protein-coding |
| chr11_70113531_70114916 | chr11 | 70113532 | 70114916 | + | 1.36462 | Intergenic | Intergenic | -16133 | NM_145137 | 216864 | Mm.222465 | NM_145137 | ENSMUSG00000040950 | **Mgl2** | CD301b | macrophage galactose N-acetyl-galactosamine specific lectin 2 | protein-coding |
| chr11_70113587_70114357 | chr11 | 70113588 | 70114357 | + | 1.33176 | Intergenic | PB1D7\|SINE\|Alu | -16385 | NM_145137 | 216864 | Mm.222465 | NM_145137 | ENSMUSG00000040950 | **Mgl2** | CD301b | macrophage galactose N-acetyl-galactosamine specific lectin 2 | protein-coding |
| chr11_70114444_70114868 | chr11 | 70114445 | 70114868 | + | 1.03635 | Intergenic | Intergenic | -15701 | NM_145137 | 216864 | Mm.222465 | NM_145137 | ENSMUSG00000040950 | **Mgl2** | CD301b | macrophage galactose N-acetyl-galactosamine specific lectin 2 | protein-coding |
| chr11_70121315_70121583 | chr11 | 70121316 | 70121583 | + | 0.702449 | Intergenic | L1M2\|LINE\|L1 | -8908 | NM_145137 | 216864 | Mm.222465 | NM_145137 | ENSMUSG00000040950 | **Mgl2** | CD301b | macrophage galactose N-acetyl-galactosamine specific lectin 2 | protein-coding |
| chr6_88756850_88757438 | chr6 | 88756851 | 88757438 | + | 1.12031 | intron (NM_011844, intron 2 of 7) | intron (NM_011844, intron 2 of 7) | 32440 | NM_001166251 | 23945 | Mm.272197 | NM_011844 | ENSMUSG00000033174 | **Mgll** | AA589436\|Magl\|Mgl | monoglyceride lipase | protein-coding |
| chr6_88717687_88718758 | chr6 | 88717688 | 88718758 | + | 1.01283 | Intergenic | Intergenic | -6189 | NM_001166249 | 23945 | Mm.272197 | NM_011844 | ENSMUSG00000033174 | **Mgll** | AA589436\|Magl\|Mgl | monoglyceride lipase | protein-coding |
| chr3_51772175_51772981 | chr3 | 51772176 | 51772981 | + | 1.44239 | intron (NM_001004176, intron 2 of 4) | intron (NM_001004176, intron 2 of 4) | 111385 | NM_174995 | 211666 | Mm.24679 | NM_174995 | ENSMUSG00000074604 | **Mgst2** | GST2\|MGST-II | microsomal glutathione S-transferase 2 | protein-coding |
| chr3_51768205_51768749 | chr3 | 51768206 | 51768749 | + | 0.923666 | intron (NM_001004176, intron 2 of 4) | intron (NM_001004176, intron 2 of 4) | 107284 | NM_174995 | 211666 | Mm.24679 | NM_174995 | ENSMUSG00000074604 | **Mgst2** | GST2\|MGST-II | microsomal glutathione S-transferase 2 | protein-coding |
| chr3_51758208_51758581 | chr3 | 51758209 | 51758581 | + | 0.920675 | intron (NM_001004176, intron 2 of 4) | intron (NM_001004176, intron 2 of 4) | 97202 | NM_174995 | 211666 | Mm.24679 | NM_174995 | ENSMUSG00000074604 | **Mgst2** | GST2\|MGST-II | microsomal glutathione S-transferase 2 | protein-coding |
| chr3_51765229_51765915 | chr3 | 51765230 | 51765915 | + | 0.877596 | intron (NM_001004176, intron 2 of 4) | intron (NM_001004176, intron 2 of 4) | 104379 | NM_174995 | 211666 | Mm.24679 | NM_174995 | ENSMUSG00000074604 | **Mgst2** | GST2\|MGST-II | microsomal glutathione S-transferase 2 | protein-coding |
| chr3_51765028_51765994 | chr3 | 51765029 | 51765994 | + | 0.867394 | intron (NM_001004176, intron 2 of 4) | intron (NM_001004176, intron 2 of 4) | 104318 | NM_174995 | 211666 | Mm.24679 | NM_174995 | ENSMUSG00000074604 | **Mgst2** | GST2\|MGST-II | microsomal glutathione S-transferase 2 | protein-coding |
| chr3_51768215_51768929 | chr3 | 51768216 | 51768929 | + | 0.79782 | intron (NM_001004176, intron 2 of 4) | RSINE1\|SINE\|B4 | 107379 | NM_174995 | 211666 | Mm.24679 | NM_174995 | ENSMUSG00000074604 | **Mgst2** | GST2\|MGST-II | microsomal glutathione S-transferase 2 | protein-coding |
| chr3_51758188_51758819 | chr3 | 51758189 | 51758819 | + | 0.770883 | intron (NM_001004176, intron 2 of 4) | intron (NM_001004176, intron 2 of 4) | 97311 | NM_174995 | 211666 | Mm.24679 | NM_174995 | ENSMUSG00000074604 | **Mgst2** | GST2\|MGST-II | microsomal glutathione S-transferase 2 | protein-coding |
| chr3_51731605_51731960 | chr3 | 51731606 | 51731960 | + | 0.746805 | intron (NM_001004176, intron 2 of 4) | intron (NM_001004176, intron 2 of 4) | 70590 | NM_174995 | 211666 | Mm.24679 | NM_174995 | ENSMUSG00000074604 | **Mgst2** | GST2\|MGST-II | microsomal glutathione S-transferase 2 | protein-coding |
| chr3_51744892_51745628 | chr3 | 51744893 | 51745628 | + | 0.73373 | intron (NM_001004176, intron 2 of 4) | intron (NM_001004176, intron 2 of 4) | 84067 | NM_174995 | 211666 | Mm.24679 | NM_174995 | ENSMUSG00000074604 | **Mgst2** | GST2\|MGST-II | microsomal glutathione S-transferase 2 | protein-coding |
| chr3_51744369_51744794 | chr3 | 51744370 | 51744794 | + | 0.714354 | intron (NM_001004176, intron 2 of 4) | intron (NM_001004176, intron 2 of 4) | 83389 | NM_174995 | 211666 | Mm.24679 | NM_174995 | ENSMUSG00000074604 | **Mgst2** | GST2\|MGST-II | microsomal glutathione S-transferase 2 | protein-coding |
| chr3_51736761_51737378 | chr3 | 51736762 | 51737378 | + | 0.687133 | intron (NM_001004176, intron 2 of 4) | intron (NM_001004176, intron 2 of 4) | 75877 | NM_174995 | 211666 | Mm.24679 | NM_174995 | ENSMUSG00000074604 | **Mgst2** | GST2\|MGST-II | microsomal glutathione S-transferase 2 | protein-coding |
| chr3_51736721_51737214 | chr3 | 51736722 | 51737214 | + | 0.687088 | intron (NM_001004176, intron 2 of 4) | PB1D10\|SINE\|Alu | 75775 | NM_174995 | 211666 | Mm.24679 | NM_174995 | ENSMUSG00000074604 | **Mgst2** | GST2\|MGST-II | microsomal glutathione S-transferase 2 | protein-coding |
| chr3_51788613_51789113 | chr3 | 51788614 | 51789113 | + | 0.668812 | intron (NM_001004176, intron 2 of 4) | intron (NM_001004176, intron 2 of 4) | 127670 | NM_174995 | 211666 | Mm.24679 | NM_174995 | ENSMUSG00000074604 | **Mgst2** | GST2\|MGST-II | microsomal glutathione S-transferase 2 | protein-coding |
| chr1_167392723_167393115 | chr1 | 167392724 | 167393115 | + | 0.711996 | intron (NM_025569, intron 1 of 5) | intron (NM_025569, intron 1 of 5) | 878 | NM_025569 | 66447 | Mm.218286 | NM_025569 | | **Mgst3** | 2010012L10Rik\|2010306B17Rik\|2700004G04Rik\|AA516734\|GST-III | microsomal glutathione S-transferase 3 | protein-coding |
| chr18_10743983_10744361 | chr18 | 10743984 | 10744361 | + | 0.836498 | intron (NM_144860, intron 3 of 20) | intron (NM_144860, intron 3 of 20) | 18547 | NM_144860 | 225164 | Mm.21500 | NM_144860 | ENSMUSG00000024294 | **Mib1** | DIP-1\|E430019M12Rik\|Mib\|mKIAA1323 | mindbomb homolog 1 (Drosophila) | protein-coding |
| chr7_112238452_112239265 | chr7 | 112238453 | 112239265 | + | 0.715729 | intron (NM_177282, intron 2 of 19) | B1F\|SINE\|Alu | 13023 | NM_001193305 | 320878 | Mm.52312 | NM_177282 | ENSMUSG00000038244 | **Mical2** | 5330438E18Rik\|9530064J02\|MICAL-2\|mKIAA0750 | microtubule associated monooxygenase, calponin and LIM domain containing 2 | protein-coding |
| chr7_112238727_112239069 | chr7 | 112238728 | 112239069 | + | 0.694717 | intron (NM_177282, intron 2 of 19) | intron (NM_177282, intron 2 of 19) | 13062 | NM_001193305 | 320878 | Mm.52312 | NM_177282 | ENSMUSG00000038244 | **Mical2** | 5330438E18Rik\|9530064J02\|MICAL-2\|mKIAA0750 | microtubule associated monooxygenase, calponin and LIM domain containing 2 | protein-coding |
| chr13_111740900_111741359 | chr13 | 111740901 | 111741359 | + | 0.658801 | Intergenic | ORR1A4\|LTR\|ERVL-MaLR | 54952 | NM_172593 | 218613 | Mm.31012 | NM_172593 | ENSMUSG00000032727 | **Mier3** | 5730509D14\|AU015422\|D130064H19Rik | mesoderm induction early response 1, family member 3 | protein-coding |
| chr13_111655701_111656277 | chr13 | 111655702 | 111656277 | + | 0.599044 | Intergenic | Intergenic | -30189 | NM_172593 | 218613 | Mm.31012 | NM_172593 | ENSMUSG00000032727 | **Mier3** | 5730509D14\|AU015422\|D130064H19Rik | mesoderm induction early response 1, family member 3 | protein-coding |

| | | | | | | | | | | | | | | | | | |
|---|---|---|---|---|---|---|---|---|---|---|---|---|---|---|---|---|---|
| chr16_59480158_59480731 | chr16 | 59480159 | 59480731 | + | 1.50824 | intron (NM_025910, intron 4 of 9) | intron (NM_025910, intron 4 of 9) | 8670 | NM_025910 | 67014 | Mm.289150 | NM_025910 | ENSMUSG00000022724 | **Mina** | 1810047J07Rik\|2410057H13Rik\|3830408E23Rik\|AI449204 | myc induced nuclear antigen | protein-coding |
| chr16_59473522_59473971 | chr16 | 59473523 | 59473971 | + | 1.50059 | intron (NM_025910, intron 1 of 9) | intron (NM_025910, intron 1 of 9) | 1972 | NM_025910 | 67014 | Mm.289150 | NM_025910 | ENSMUSG00000022724 | **Mina** | 1810047J07Rik\|2410057H13Rik\|3830408E23Rik\|AI449204 | myc induced nuclear antigen | protein-coding |
| chr16_59475890_59476144 | chr16 | 59475891 | 59476144 | + | 1.15711 | intron (NM_025910, intron 2 of 9) | intron (NM_025910, intron 2 of 9) | 4242 | NM_025910 | 67014 | Mm.289150 | NM_025910 | ENSMUSG00000022724 | **Mina** | 1810047J07Rik\|2410057H13Rik\|3830408E23Rik\|AI449204 | myc induced nuclear antigen | protein-coding |
| chr16_59475327_59475487 | chr16 | 59475328 | 59475487 | + | 0.859812 | intron (NM_025910, intron 1 of 9) | intron (NM_025910, intron 1 of 9) | 3632 | NM_025910 | 67014 | Mm.289150 | NM_025910 | ENSMUSG00000022724 | **Mina** | 1810047J07Rik\|2410057H13Rik\|3830408E23Rik\|AI449204 | myc induced nuclear antigen | protein-coding |
| chr16_59483642_59484093 | chr16 | 59483643 | 59484093 | + | 0.682199 | intron (NM_025910, intron 5 of 9) | intron (NM_025910, intron 5 of 9) | 12093 | NM_025910 | 67014 | Mm.289150 | NM_025910 | ENSMUSG00000022724 | **Mina** | 1810047J07Rik\|2410057H13Rik\|3830408E23Rik\|AI449204 | myc induced nuclear antigen | protein-coding |
| chr4_139115336_139115748 | chr4 | 139115337 | 139115748 | + | 0.798816 | intron (NM_001163006, intron 1 of 3) | intron (NM_001163006, intron 1 of 3) | 15495 | NR_033745 | 433771 | Mm.305990 | NM_001163006 | ENSMUSG00000050608 | **Minos1** | 2310028O11Rik | mitochondrial inner membrane organizing system 1 | protein-coding |
| chr4_139115331_139116036 | chr4 | 139115332 | 139116036 | + | 0.727459 | intron (NM_001163006, intron 1 of 3) | intron (NM_001163006, intron 1 of 3) | 15353 | NR_033745 | 433771 | Mm.305990 | NM_001163006 | ENSMUSG00000050608 | **Minos1** | 2310028O11Rik | mitochondrial inner membrane organizing system 1 | protein-coding |
| chr19_32488004_32488209 | chr19 | 32488005 | 32488209 | + | 0.760957 | intron (NM_010799, intron 1 of 4) | RSINE1\|SINE\|B4 | 2338 | NM_010799 | 17330 | Mm.255116 | NM_010799 | ENSMUSG00000024896 | **Minpp1** | AA408516 | multiple inositol polyphosphate histidine phosphatase 1 | protein-coding |
| chr19_29150030_29150733 | chr19 | 29150031 | 29150733 | + | 0.960122 | Intergenic | Intergenic | 15103 | NR_029778 | 724062 | | NR_029778 | | **Mir101b** | Mirn101b\|mir-101b | microRNA 101b | ncRNA |
| chr2_131292022_131292899 | chr2 | 131292023 | 131292899 | + | 1.08731 | intron (NM_153501, intron 5 of 6) | intron (NM_153501, intron 5 of 6) | 4409 | NR_029755 | 723825 | | NR_029755 | | **Mir103-2** | Mirn103-2\|mir-103-2 | microRNA 103-2 | ncRNA |
| chr12_100975684_100976323 | chr12 | 100975685 | 100976323 | + | 1.4695 | intron (NM_026681, intron 4 of 29) | intron (NM_026681, intron 4 of 29) | 45764 | NR_035421 | 100316805 | | NR_035421 | | **Mir1190** | Mirn1190\|mmu-mir-1190 | microRNA 1190 | ncRNA |
| chr12_100993191_100993400 | chr12 | 100993192 | 100993400 | + | 1.07859 | intron (NM_026681, intron 3 of 29) | intron (NM_026681, intron 3 of 29) | 28472 | NR_035421 | 100316805 | | NR_035421 | | **Mir1190** | Mirn1190\|mmu-mir-1190 | microRNA 1190 | ncRNA |
| chr12_100979682_100979951 | chr12 | 100979683 | 100979951 | + | 0.81879 | intron (NM_026681, intron 4 of 29) | intron (NM_026681, intron 4 of 29) | 41951 | NR_035421 | 100316805 | | NR_035421 | | **Mir1190** | Mirn1190\|mmu-mir-1190 | microRNA 1190 | ncRNA |
| chr12_101014264_101014656 | chr12 | 101014265 | 101014656 | + | 0.79086 | intron (NM_026681, intron 3 of 29) | intron (NM_026681, intron 3 of 29) | 7308 | NR_035421 | 100316805 | | NR_035421 | | **Mir1190** | Mirn1190\|mmu-mir-1190 | microRNA 1190 | ncRNA |
| chr12_100992524_100992895 | chr12 | 100992525 | 100992895 | + | 0.790159 | intron (NM_026681, intron 3 of 29) | intron (NM_026681, intron 3 of 29) | 29058 | NR_035421 | 100316805 | | NR_035421 | | **Mir1190** | Mirn1190\|mmu-mir-1190 | microRNA 1190 | ncRNA |
| chr12_101000584_101000865 | chr12 | 101000585 | 101000865 | + | 0.688597 | intron (NM_026681, intron 3 of 29) | intron (NM_026681, intron 3 of 29) | 21043 | NR_035421 | 100316805 | | NR_035421 | | **Mir1190** | Mirn1190\|mmu-mir-1190 | microRNA 1190 | ncRNA |
| chr12_55343194_55344303 | chr12 | 55343195 | 55344303 | + | 1.29501 | intron (NM_025373, intron 4 of 6) | intron (NM_025373, intron 4 of 6) | -26946 | NR_035430 | 100316807 | | NR_035430 | | **Mir1198** | Mirn1198\|mmu-mir-1198 | microRNA 1198 | ncRNA |
| chr18_65169185_65169678 | chr18 | 65169186 | 65169678 | + | 1.1793 | intron (NM_031881, intron 12 of 29) | intron (NM_031881, intron 12 of 29) | -79429 | NR_029600 | 387231 | | NR_029600 | miRBase:MI0000256 | **Mir122a** | Mirn122a\|mmu-mir-122 | microRNA 122a | ncRNA |
| chr18_65260157_65260508 | chr18 | 65260158 | 65260508 | + | 0.698996 | intron (NR_105742, intron 3 of 6) | intron (NR_105742, intron 3 of 6) | 11472 | NR_029600 | 387231 | | NR_029600 | miRBase:MI0000256 | **Mir122a** | Mirn122a\|mmu-mir-122 | microRNA 122a | ncRNA |
| chr9_111918605_111918893 | chr9 | 111918606 | 111918893 | + | 0.645385 | Intergenic | Intergenic | 199962 | NR_029823 | 723815 | | NR_029823 | | **Mir128-2** | Mirn128-2\|Mirn128b\|mir-128b | microRNA 128-2 | ncRNA |
| chr6_28990145_28990668 | chr6 | 28990146 | 28990668 | + | 1.03668 | Intergenic | Intergenic | -32212 | NR_029567 | 387237 | | NR_029567 | miRBase:MI0000222 | **Mir129-1** | Mirn129-1\|Mirn129b\|miR-129b | microRNA 129-1 | ncRNA |
| chr6_28990086_28990918 | chr6 | 28990087 | 28990918 | + | 0.770804 | Intergenic | Intergenic | -32117 | NR_029567 | 387237 | | NR_029567 | miRBase:MI0000222 | **Mir129-1** | Mirn129-1\|Mirn129b\|miR-129b | microRNA 129-1 | ncRNA |
| chr6_29023188_29023847 | chr6 | 29023189 | 29023847 | + | 0.765036 | TTS (NR_029567) | TTS (NR_029567) | 899 | NR_029567 | 387237 | | NR_029567 | miRBase:MI0000222 | **Mir129-1** | Mirn129-1\|Mirn129b\|miR-129b | microRNA 129-1 | ncRNA |
| chr2_29457675_29458053 | chr2 | 29457676 | 29458053 | + | 0.805006 | intron (NM_029615, intron 3 of 4) | B3A\|SINE\|B2 | 17430 | NR_105753 | 102465170 | | NR_105753 | miRBase:MI0021863 | **Mir133c** | mmu-mir-133c | microRNA mir-133c | ncRNA |
| chr2_29439728_29440771 | chr2 | 29439729 | 29440771 | + | 0.747946 | intron (NM_029615, intron 3 of 4) | intron (NM_029615, intron 3 of 4) | 35044 | NR_105753 | 102465170 | | NR_105753 | miRBase:MI0021863 | **Mir133c** | mmu-mir-133c | microRNA mir-133c | ncRNA |
| chr2_29439817_29440303 | chr2 | 29439818 | 29440303 | + | 0.655977 | intron (NM_029615, intron 3 of 4) | intron (NM_029615, intron 3 of 4) | 35234 | NR_105753 | 102465170 | | NR_105753 | miRBase:MI0021863 | **Mir133c** | mmu-mir-133c | microRNA mir-133c | ncRNA |
| chr8_107498752_107499254 | chr8 | 107498753 | 107499254 | + | 0.917804 | intron (NM_025830, intron 6 of 23) | intron (NM_025830, intron 6 of 23) | -52241 | NR_029553 | 387158 | | NR_029553 | | **Mir140** | Mirn140\|mmu-mir-140 | microRNA 140 | ncRNA |
| chr8_107502454_107502777 | chr8 | 107502455 | 107502777 | + | 0.76743 | intron (NM_025830, intron 6 of 23) | intron (NM_025830, intron 6 of 23) | -48628 | NR_029553 | 387158 | | NR_029553 | | **Mir140** | Mirn140\|mmu-mir-140 | microRNA 140 | ncRNA |
| chr8_107498474_107499257 | chr8 | 107498475 | 107499257 | + | 0.720788 | intron (NM_025830, intron 6 of 23) | intron (NM_025830, intron 6 of 23) | -52378 | NR_029553 | 387158 | | NR_029553 | | **Mir140** | Mirn140\|mmu-mir-140 | microRNA 140 | ncRNA |
| chr8_107560375_107561544 | chr8 | 107560376 | 107561544 | + | 0.698934 | Intergenic | Intergenic | 9716 | NR_029553 | 387158 | | NR_029553 | | **Mir140** | Mirn140\|mmu-mir-140 | microRNA 140 | ncRNA |
| chr8_107560790_107561534 | chr8 | 107560791 | 107561534 | + | 0.679646 | Intergenic | C-rich\|Low_complexity\|Low_complexity | 9918 | NR_029553 | 387158 | | NR_029553 | | **Mir140** | Mirn140\|mmu-mir-140 | microRNA 140 | ncRNA |
| chr11_43376891_43377767 | chr11 | 43376892 | 43377767 | + | 1.07018 | Intergenic | Intergenic | -2868 | NR_029558 | 387164 | | NR_029558 | miRBase:MI0000170 | **Mir146** | Mirn146\|miR-146a\|mmu-mir-146 | microRNA 146 | ncRNA |
| chr11_43386956_43387562 | chr11 | 43386957 | 43387562 | + | 1.00636 | Intergenic | Intergenic | -12798 | NR_029558 | 387164 | | NR_029558 | miRBase:MI0000170 | **Mir146** | Mirn146\|miR-146a\|mmu-mir-146 | microRNA 146 | ncRNA |
| chr11_43358302_43359222 | chr11 | 43358303 | 43359222 | + | 0.687394 | Intergenic | Intergenic | 15699 | NR_029558 | 387164 | | NR_029558 | miRBase:MI0000170 | **Mir146** | Mirn146\|miR-146a\|mmu-mir-146 | microRNA 146 | ncRNA |
| chr2_122656154_122656432 | chr2 | 122656155 | 122656432 | + | 0.835282 | Intergenic | Intergenic | 15490 | NR_030547 | 387165 | | NR_030547 | miRBase:MI0005482 | **Mir147** | Mirn147\|mmu-mir-147 | microRNA 147 | ncRNA |
| chr2_122665272_122667024 | chr2 | 122665273 | 122667024 | + | 0.818042 | Intergenic | MER63B\|DNA\|hAT-Blackjack | 25345 | NR_030547 | 387165 | | NR_030547 | miRBase:MI0005482 | **Mir147** | Mirn147\|mmu-mir-147 | microRNA 147 | ncRNA |
| chr6_51287472_51288364 | chr6 | 51287473 | 51288364 | + | 1.26848 | Intergenic | Intergenic | -18008 | NR_029719 | 387166 | | NR_029719 | miRBase:MI0000550 | **Mir148a** | Mirn148\|Mirn148a | microRNA 148a | ncRNA |
| chr6_51199942_51200757 | chr6 | 51199943 | 51200757 | + | 0.966428 | Intergenic | Intergenic | 69560 | NR_029719 | 387166 | | NR_029719 | miRBase:MI0000550 | **Mir148a** | Mirn148\|Mirn148a | microRNA 148a | ncRNA |
| chr6_51178138_51178886 | chr6 | 51178139 | 51178886 | + | 0.7454 | Intergenic | C-rich\|Low_complexity\|Low_complexity | 91398 | NR_029719 | 387166 | | NR_029719 | miRBase:MI0000550 | **Mir148a** | Mirn148\|Mirn148a | microRNA 148a | ncRNA |
| chr11_96859672_96859987 | chr11 | 96859673 | 96859987 | + | 1.06949 | intron (NM_019877, intron 8 of 8) | intron (NM_019877, intron 8 of 8) | 9437 | NR_029562 | 387170 | | NR_029562 | | **Mir152** | Mirn152\|mmu-mir-152 | microRNA 152 | ncRNA |
| chr16_84698822_84699050 | chr16 | 84698823 | 84699050 | + | 1.03872 | Intergenic | Intergenic | -15204 | NR_029565 | 387173 | | NR_029565 | miRBase:MI0000177 | **Mir155** | Mirn155\|mmu-mir-155 | microRNA 155 | ncRNA |
| chr16_84717792_84718243 | chr16 | 84717793 | 84718243 | + | 0.993201 | 3' UTR (NM_017404, exon 10 of 10) | 3' UTR (NM_017404, exon 10 of 10) | 3878 | NR_029565 | 387173 | | NR_029565 | miRBase:MI0000177 | **Mir155** | Mirn155\|mmu-mir-155 | microRNA 155 | ncRNA |
| chr16_84710252_84710606 | chr16 | 84710253 | 84710606 | + | 0.926962 | Intergenic | Intergenic | -3711 | NR_029565 | 387173 | | NR_029565 | miRBase:MI0000177 | **Mir155** | Mirn155\|mmu-mir-155 | microRNA 155 | ncRNA |
| chr16_84712872_84713581 | chr16 | 84712873 | 84713581 | + | 0.740805 | promoter-TSS (NR_029565) | promoter-TSS (NR_029565) | -913 | NR_029565 | 387173 | | NR_029565 | miRBase:MI0000177 | **Mir155** | Mirn155\|mmu-mir-155 | microRNA 155 | ncRNA |
| chr16_84706693_84706909 | chr16 | 84706694 | 84706909 | + | 0.671008 | Intergenic | Intergenic | -7339 | NR_029565 | 387173 | | NR_029565 | miRBase:MI0000177 | **Mir155** | Mirn155\|mmu-mir-155 | microRNA 155 | ncRNA |
| chr16_84713144_84713604 | chr16 | 84713145 | 84713604 | + | 0.663103 | promoter-TSS (NR_029565) | promoter-TSS (NR_029565) | -766 | NR_029565 | 387173 | | NR_029565 | miRBase:MI0000177 | **Mir155** | Mirn155\|mmu-mir-155 | microRNA 155 | ncRNA |
| chr14_114983333_114983810 | chr14 | 114983334 | 114983810 | + | 1.11308 | Intergenic | Lx\|LINE\|L1 | -60099 | NR_029785 | 723905 | | NR_029785 | miRBase:MI0000687 | **Mir17** | Mirn17\|mmu-mir-17 | microRNA 17 | ncRNA |
| chr1_137922650_137923131 | chr1 | 137922651 | 137923131 | + | 0.73499 | Intergenic | Intergenic | -43564 | NR_029795 | 735252 | | NR_029795 | miRBase:MI0000697 | **Mir181a-1** | Mirn181a-1\|Mirn213\|miR-213\|mmu-mir-213 | microRNA 181a-1 | ncRNA |
| chr1_137935692_137936232 | chr1 | 137935693 | 137936232 | + | 0.68293 | Intergenic | Intergenic | -30493 | NR_029795 | 735252 | | NR_029795 | miRBase:MI0000697 | **Mir181a-1** | Mirn181a-1\|Mirn213\|miR-213\|mmu-mir-213 | microRNA 181a-1 | ncRNA |
| chr1_138038517_138039683 | chr1 | 138038518 | 138039683 | + | 1.5324 | Intergenic | Intergenic | 72461 | NR_029820 | 723890 | | NR_029820 | miRBase:MI0000723 | **Mir181b-1** | Mirn181b-1 | microRNA 181b-1 | ncRNA |
| chr1_138039118_138039683 | chr1 | 138039119 | 138039683 | + | 1.50665 | Intergenic | Intergenic | 72762 | NR_029820 | 723890 | | NR_029820 | miRBase:MI0000723 | **Mir181b-1** | Mirn181b-1 | microRNA 181b-1 | ncRNA |
| chr1_138009411_138009904 | chr1 | 138009412 | 138009904 | + | 0.902124 | Intergenic | RMER15\|LTR\|ERVL | 43019 | NR_029820 | 723890 | | NR_029820 | miRBase:MI0000723 | **Mir181b-1** | Mirn181b-1 | microRNA 181b-1 | ncRNA |
| chr6_30162068_30162491 | chr6 | 30162069 | 30162491 | + | 0.749416 | Intergenic | PB1D10\|SINE\|Alu | 3712 | NR_029569 | 387177 | | NR_029569 | miRBase:MI0000224 | **Mir182** | Mirn182\|mmu-mir-182 | microRNA 182 | ncRNA |
| chr6_30161906_30162491 | chr6 | 30161907 | 30162491 | + | 0.724701 | Intergenic | ID_B1\|SINE\|B4 | 3793 | NR_029569 | 387177 | | NR_029569 | miRBase:MI0000224 | **Mir182** | Mirn182\|mmu-mir-182 | microRNA 182 | ncRNA |
| chr12_80386721_80387612 | chr12 | 80386722 | 80387612 | + | 1.40304 | intron (NM_177267, intron 5 of 8) | intron (NM_177267, intron 5 of 8) | 4510 | NR_037207 | 100526530 | | NR_037207 | | **Mir1843a** | Mir1843\|mmu-mir-1843\|mmu-mir-1843a | microRNA 1843a | ncRNA |
| chr12_80387316_80387609 | chr12 | 80387317 | 80387609 | + | 1.31386 | intron (NM_177267, intron 5 of 8) | intron (NM_177267, intron 5 of 8) | 4214 | NR_037207 | 100526530 | | NR_037207 | | **Mir1843a** | Mir1843\|mmu-mir-1843\|mmu-mir-1843a | microRNA 1843a | ncRNA |
| chr12_80386735_80387257 | chr12 | 80386736 | 80387257 | + | 1.01611 | intron (NM_177267, intron 5 of 8) | PB1D7\|SINE\|Alu | 4681 | NR_037207 | 100526530 | | NR_037207 | | **Mir1843a** | Mir1843\|mmu-mir-1843\|mmu-mir-1843a | microRNA 1843a | ncRNA |
| chr12_80360318_80361035 | chr12 | 80360319 | 80361035 | + | 0.887584 | intron (NM_177267, intron 7 of 8) | intron (NM_177267, intron 7 of 8) | 31000 | NR_037207 | 100526530 | | NR_037207 | | **Mir1843a** | Mir1843\|mmu-mir-1843\|mmu-mir-1843a | microRNA 1843a | ncRNA |
| chr18_6389726_6390329 | chr18 | 6389727 | 6390329 | + | 0.748182 | Intergenic | RLTR45\|LTR\|ERVK | 100618 | NR_035446 | 100316773 | | NR_035446 | | **Mir1893** | Mirn1893\|mmu-mir-1893 | microRNA 1893 | ncRNA |
| chr8_128420385_128420821 | chr8 | 128420386 | 128420821 | + | 1.65854 | intron (NM_008737, intron 3 of 16) | UCON12A\|Unknown\|Unknown | 61362 | NR_035436 | 100316680 | | NR_035436 | miRBase:MI0008317 | **Mir1903** | Mirn1903\|mmu-mir-1903 | microRNA 1903 | ncRNA |

| | | | | | | | | | | | | | | | | | |
|---|---|---|---|---|---|---|---|---|---|---|---|---|---|---|---|---|---|
| chr8_128418501_128418918 | chr8 | 128418502 | 128418918 | + | 1.4597 | intron (NM_008737, intron 2 of 16) | intron (NM_008737, intron 2 of 16) | 59469 | NR_035436 | 100316680 | | NR_035436 | miRBase:MI0008317 | **Mir1903** | Mirn1903\|mmu-mir-1903 | microRNA 1903 | ncRNA |
| chr8_128423817_128424114 | chr8 | 128423818 | 128424114 | + | 1.23873 | intron (NM_008737, intron 3 of 16) | intron (NM_008737, intron 3 of 16) | 64725 | NR_035436 | 100316680 | | NR_035436 | miRBase:MI0008317 | **Mir1903** | Mirn1903\|mmu-mir-1903 | microRNA 1903 | ncRNA |
| chr8_128420940_128421331 | chr8 | 128420941 | 128421331 | + | 1.15191 | intron (NM_008737, intron 3 of 16) | intron (NM_008737, intron 3 of 16) | 61895 | NR_035436 | 100316680 | | NR_035436 | miRBase:MI0008317 | **Mir1903** | Mirn1903\|mmu-mir-1903 | microRNA 1903 | ncRNA |
| chr8_128484698_128485228 | chr8 | 128484699 | 128485228 | + | 1.13453 | intron (NM_008737, intron 11 of 16) | intron (NM_008737, intron 11 of 16) | 125722 | NR_035436 | 100316680 | | NR_035436 | miRBase:MI0008317 | **Mir1903** | Mirn1903\|mmu-mir-1903 | microRNA 1903 | ncRNA |
| chr8_128505917_128506307 | chr8 | 128505918 | 128506307 | + | 1.0841 | TTS (NM_008737) | TTS (NM_008737) | 146871 | NR_035436 | 100316680 | | NR_035436 | miRBase:MI0008317 | **Mir1903** | Mirn1903\|mmu-mir-1903 | microRNA 1903 | ncRNA |
| chr8_128504057_128504484 | chr8 | 128504058 | 128504484 | + | 0.933318 | 3' UTR (NM_008737, exon 17 of 17) | 3' UTR (NM_008737, exon 17 of 17) | 145030 | NR_035436 | 100316680 | | NR_035436 | miRBase:MI0008317 | **Mir1903** | Mirn1903\|mmu-mir-1903 | microRNA 1903 | ncRNA |
| chr8_128464079_128464508 | chr8 | 128464080 | 128464508 | + | 0.892962 | intron (NM_008737, intron 8 of 16) | intron (NM_008737, intron 8 of 16) | 105053 | NR_035436 | 100316680 | | NR_035436 | miRBase:MI0008317 | **Mir1903** | Mirn1903\|mmu-mir-1903 | microRNA 1903 | ncRNA |
| chr8_128483420_128483706 | chr8 | 128483421 | 128483706 | + | 0.797086 | intron (NM_008737, intron 11 of 16) | intron (NM_008737, intron 11 of 16) | 124322 | NR_035436 | 100316680 | | NR_035436 | miRBase:MI0008317 | **Mir1903** | Mirn1903\|mmu-mir-1903 | microRNA 1903 | ncRNA |
| chr13_109829488_109829928 | chr13 | 109829489 | 109829928 | + | 0.945686 | intron (NM_011056, intron 7 of 16) | intron (NM_011056, intron 7 of 16) | -74101 | NR_035442 | 100316683 | | NR_035442 | | **Mir1904** | Mirn1904\|mmu-mir-1904 | microRNA 1904 | ncRNA |
| chr13_109789108_109789582 | chr13 | 109789109 | 109789582 | + | 0.63296 | intron (NM_011056, intron 7 of 16) | intron (NM_011056, intron 7 of 16) | -114464 | NR_035442 | 100316683 | | NR_035442 | | **Mir1904** | Mirn1904\|mmu-mir-1904 | microRNA 1904 | ncRNA |
| chr15_50750878_50751698 | chr15 | 50750879 | 50751698 | + | 1.90244 | intron (NM_032000, intron 4 of 5) | intron (NM_032000, intron 4 of 5) | 137826 | NR_035444 | 100316685 | | NR_035444 | miRBase:MI0008325 | **Mir1907** | Mirn1907\|mmu-mir-1907 | microRNA 1907 | ncRNA |
| chr15_50667172_50667988 | chr15 | 50667173 | 50667988 | + | 1.57056 | intron (NM_032000, intron 4 of 5) | intron (NM_032000, intron 4 of 5) | 221534 | NR_035444 | 100316685 | | NR_035444 | miRBase:MI0008325 | **Mir1907** | Mirn1907\|mmu-mir-1907 | microRNA 1907 | ncRNA |
| chr15_50650258_50650581 | chr15 | 50650259 | 50650581 | + | 1.35053 | Intergenic | Intergenic | 238694 | NR_035444 | 100316685 | | NR_035444 | miRBase:MI0008325 | **Mir1907** | Mirn1907\|mmu-mir-1907 | microRNA 1907 | ncRNA |
| chr15_50737012_50737641 | chr15 | 50737013 | 50737641 | + | 1.33646 | intron (NM_032000, intron 4 of 5) | intron (NM_032000, intron 4 of 5) | 151787 | NR_035444 | 100316685 | | NR_035444 | miRBase:MI0008325 | **Mir1907** | Mirn1907\|mmu-mir-1907 | microRNA 1907 | ncRNA |
| chr15_50650055_50650586 | chr15 | 50650056 | 50650586 | + | 1.24657 | Intergenic | Intergenic | 238793 | NR_035444 | 100316685 | | NR_035444 | miRBase:MI0008325 | **Mir1907** | Mirn1907\|mmu-mir-1907 | microRNA 1907 | ncRNA |
| chr15_50664514_50664988 | chr15 | 50664515 | 50664988 | + | 0.940525 | exon (NM_032000, exon 5 of 6) | exon (NM_032000, exon 5 of 6) | 224363 | NR_035444 | 100316685 | | NR_035444 | miRBase:MI0008325 | **Mir1907** | Mirn1907\|mmu-mir-1907 | microRNA 1907 | ncRNA |
| chr15_50749975_50750247 | chr15 | 50749976 | 50750247 | + | 0.702835 | intron (NM_032000, intron 4 of 5) | intron (NM_032000, intron 4 of 5) | 139003 | NR_035444 | 100316685 | | NR_035444 | miRBase:MI0008325 | **Mir1907** | Mirn1907\|mmu-mir-1907 | microRNA 1907 | ncRNA |
| chr15_50153766_50154178 | chr15 | 50153767 | 50154178 | + | 0.702745 | Intergenic | L1MC1\|LINE\|L1 | 735142 | NR_035444 | 100316685 | | NR_035444 | miRBase:MI0008325 | **Mir1907** | Mirn1907\|mmu-mir-1907 | microRNA 1907 | ncRNA |
| chr15_50653958_50654681 | chr15 | 50653959 | 50654681 | + | 0.692517 | TTS (NM_032000) | TTS (NM_032000) | 234794 | NR_035444 | 100316685 | | NR_035444 | miRBase:MI0008325 | **Mir1907** | Mirn1907\|mmu-mir-1907 | microRNA 1907 | ncRNA |
| chr15_50417680_50417983 | chr15 | 50417681 | 50417983 | + | 0.662045 | Intergenic | B4\|SINE\|B4 | 471282 | NR_035444 | 100316685 | | NR_035444 | miRBase:MI0008325 | **Mir1907** | Mirn1907\|mmu-mir-1907 | microRNA 1907 | ncRNA |
| chr15_50653960_50654477 | chr15 | 50653961 | 50654477 | + | 0.643518 | TTS (NM_032000) | TTS (NM_032000) | 234895 | NR_035444 | 100316685 | | NR_035444 | miRBase:MI0008325 | **Mir1907** | Mirn1907\|mmu-mir-1907 | microRNA 1907 | ncRNA |
| chr10_44400590_44400986 | chr10 | 44400591 | 44400986 | + | 0.778774 | Intergenic | Intergenic | 41112 | NR_035450 | 100316688 | | NR_035450 | | **Mir1929** | mmu-mir-1929 | microRNA 1929 | ncRNA |
| chr10_44403257_44403991 | chr10 | 44403258 | 44403991 | + | 0.725812 | Intergenic | Intergenic | 43948 | NR_035450 | 100316688 | | NR_035450 | | **Mir1929** | mmu-mir-1929 | microRNA 1929 | ncRNA |
| chr10_93188383_93188922 | chr10 | 93188384 | 93188922 | + | 0.920067 | intron (NM_146239, intron 1 of 16) | intron (NM_146239, intron 1 of 16) | 25868 | NR_035452 | 100316689 | | NR_035452 | | **Mir1931** | mmu-mir-1931 | microRNA 1931 | ncRNA |
| chr10_93177726_93178228 | chr10 | 93177727 | 93178228 | + | 0.899136 | intron (NM_146239, intron 1 of 16) | MER110A\|LTR\|ERV1 | 15192 | NR_035452 | 100316689 | | NR_035452 | | **Mir1931** | mmu-mir-1931 | microRNA 1931 | ncRNA |
| chr11_21332742_21333468 | chr11 | 21332743 | 21333468 | + | 0.916793 | intron (NM_001290634, intron 4 of 9) | MIRb\|SINE\|MIR | 11570 | NR_035454 | 100316753 | | NR_035454 | | **Mir1933** | mmu-mir-1933 | microRNA 1933 | ncRNA |
| chr16_13428050_13428450 | chr16 | 13428051 | 13428450 | + | 1.38304 | Intergenic | B4A\|SINE\|B4 | -21273 | NR_030549 | 100124432 | | NR_030549 | miRBase:MI0005484 | **Mir193b** | Mir193\|Mirn193b | microRNA 193b | ncRNA |
| chr16_13428050_13428591 | chr16 | 13428051 | 13428591 | + | 1.22191 | Intergenic | B4A\|SINE\|B4 | -21202 | NR_030549 | 100124432 | | NR_030549 | miRBase:MI0005484 | **Mir193b** | Mir193\|Mirn193b | microRNA 193b | ncRNA |
| chr3_96069590_96070451 | chr3 | 96069591 | 96070451 | + | 1.10104 | Intergenic | RLTR13D6\|LTR\|ERVK | -10575 | NR_035496 | 100316714 | | NR_035496 | miRBase:MI0009967 | **Mir1946b** | mmu-mir-1946b | microRNA 1946b | ncRNA |
| chr3_96069592_96070070 | chr3 | 96069593 | 96070070 | + | 1.08609 | Intergenic | RLTR13D6\|LTR\|ERVK | -10765 | NR_035496 | 100316714 | | NR_035496 | miRBase:MI0009967 | **Mir1946b** | mmu-mir-1946b | microRNA 1946b | ncRNA |
| chr2_115610498_115610925 | chr2 | 115610499 | 115610925 | + | 1.17566 | intron (NM_207264, intron 1 of 10) | intron (NM_207264, intron 1 of 10) | -28013 | NR_035476 | 100316703 | | NR_035476 | | **Mir1951** | mmu-mir-1951 | microRNA 1951 | ncRNA |
| chr2_115610274_115610966 | chr2 | 115610275 | 115610966 | + | 1.04682 | intron (NM_207264, intron 1 of 10) | intron (NM_207264, intron 1 of 10) | -28105 | NR_035476 | 100316703 | | NR_035476 | | **Mir1951** | mmu-mir-1951 | microRNA 1951 | ncRNA |
| chr2_151959274_151959819 | chr2 | 151959275 | 151959819 | + | 0.596194 | Intergenic | Intergenic | 8070 | NR_035478 | 100316704 | | NR_035478 | miRBase:MI0009948 | **Mir1953** | mmu-mir-1953 | microRNA 1953 | ncRNA |
| chr2_92197241_92197743 | chr2 | 92197242 | 92197743 | + | 0.934871 | intron (NM_001109691, intron 2 of 18) | B1_Mus1\|SINE\|Alu | 5515 | NR_035480 | 100316756 | | NR_035480 | miRBase:MI0009950 | **Mir1955** | mmu-mir-1955 | microRNA 1955 | ncRNA |
| chr3_138529000_138529215 | chr3 | 138529001 | 138529215 | + | 0.617483 | intron (NM_007917, intron 2 of 7) | intron (NM_007917, intron 2 of 7) | 2687 | NR_035481 | 100316706 | | NR_035481 | miRBase:MI0009952 | **Mir1956** | mmu-mir-1956 | microRNA 1956 | ncRNA |
| chr1_63998850_63999224 | chr1 | 63998851 | 63999224 | + | 1.002 | Intergenic | Intergenic | -14554 | NR_035484 | 100316708 | | NR_035484 | | **Mir1958** | mmu-mir-1958 | microRNA 1958 | ncRNA |
| chr7_135588526_135589253 | chr7 | 135588527 | 135589253 | + | 1.29406 | intron (NM_011212, intron 1 of 20) | intron (NM_011212, intron 1 of 20) | 22728 | NR_035488 | 100316818 | | NR_035488 | | **Mir1962** | mmu-mir-1962 | microRNA 1962 | ncRNA |
| chr8_105625230_105625567 | chr8 | 105625231 | 105625567 | + | 1.05909 | Intergenic | Intergenic | 9933 | NR_035492 | 100316712 | | NR_035492 | | **Mir1966** | mmu-mir-1966 | microRNA 1966 | ncRNA |
| chr13_21914557_21914981 | chr13 | 21914558 | 21914981 | + | 1.1152 | Intergenic | Intergenic | -17720 | NR_035500 | 100316716 | | NR_035500 | miRBase:MI0009990 | **Mir1983** | mmu-mir-1983 | microRNA 1983 | ncRNA |
| chr13_21896831_21897187 | chr13 | 21896832 | 21897187 | + | 0.792186 | promoter-TSS (NR_035500) | promoter-TSS (NR_035500) | 40 | NR_035500 | 100316716 | | NR_035500 | miRBase:MI0009990 | **Mir1983** | mmu-mir-1983 | microRNA 1983 | ncRNA |
| chr13_21919933_21920438 | chr13 | 21919934 | 21920438 | + | 0.757057 | Intergenic | Intergenic | -23137 | NR_035500 | 100316716 | | NR_035500 | miRBase:MI0009990 | **Mir1983** | mmu-mir-1983 | microRNA 1983 | ncRNA |
| chr13_21896847_21897524 | chr13 | 21896848 | 21897524 | + | 0.704934 | promoter-TSS (NR_035500) | promoter-TSS (NR_035500) | -137 | NR_035500 | 100316716 | | NR_035500 | miRBase:MI0009990 | **Mir1983** | mmu-mir-1983 | microRNA 1983 | ncRNA |
| chr13_21911247_21911511 | chr13 | 21911248 | 21911511 | + | 0.67322 | Intergenic | tRNA-Asp-GAY\|tRNA\|tRNA | -14330 | NR_035500 | 100316716 | | NR_035500 | miRBase:MI0009990 | **Mir1983** | mmu-mir-1983 | microRNA 1983 | ncRNA |
| chr13_21917015_21917341 | chr13 | 21917016 | 21917341 | + | 0.635447 | Intergenic | tRNA-Arg-AGA\|tRNA\|tRNA | -20129 | NR_035500 | 100316716 | | NR_035500 | miRBase:MI0009990 | **Mir1983** | mmu-mir-1983 | microRNA 1983 | ncRNA |
| chr13_21918870_21919722 | chr13 | 21918871 | 21919722 | + | 0.612448 | Intergenic | tRNA-Ser-TCY\|tRNA\|tRNA | -22247 | NR_035500 | 100316716 | | NR_035500 | miRBase:MI0009990 | **Mir1983** | mmu-mir-1983 | microRNA 1983 | ncRNA |
| chr12_112125855_112126230 | chr12 | 112125856 | 112126230 | + | 0.626808 | intron (NM_001081169, intron 13 of 14) | (TG)n\|Simple_repeat\|Simple_repeat | -4837 | NR_029590 | 387199 | | NR_029590 | miRBase:MI0000246 | **Mir203** | Mirn203\|mmu-mir-203 | microRNA 203 | ncRNA |
| chr1_193451379_193451858 | chr1 | 193451380 | 193451858 | + | 0.999851 | Intergenic | Intergenic | 55911 | NR_029592 | 387201 | | NR_029592 | miRBase:MI0000248 | **Mir205** | Mirn205\|mmu-mir-205 | microRNA 205 | ncRNA |
| chr1_193556693_193557182 | chr1 | 193556694 | 193557182 | + | 0.635035 | Intergenic | Intergenic | -49408 | NR_029592 | 387201 | | NR_029592 | miRBase:MI0000248 | **Mir205** | Mirn205\|mmu-mir-205 | microRNA 205 | ncRNA |
| chr11_35521216_35521702 | chr11 | 35521217 | 35521702 | + | 0.831432 | intron (NM_011412, intron 5 of 35) | intron (NM_011412, intron 5 of 35) | -95357 | NR_029799 | 723924 | | NR_029799 | | **Mir218-2** | Mirn218-2\|mir-218-2 | microRNA 218-2 | ncRNA |
| chr11_86582666_86582990 | chr11 | 86582667 | 86582990 | + | 1.08403 | Intergenic | URR1B\|DNA\|hAT-Charlie | 1330 | NR_029738 | 387140 | | NR_029738 | miRBase:MI0000569 | **Mir21a** | Mir21\|Mirn21\|mmu-mir-21\|mmu-mir-21a | microRNA 21a | ncRNA |
| chr11_86593009_86593447 | chr11 | 86593010 | 86593447 | + | 0.729116 | intron (NM_029478, intron 10 of 11) | intron (NM_029478, intron 10 of 11) | -9070 | NR_029738 | 387140 | | NR_029738 | miRBase:MI0000569 | **Mir21a** | Mir21\|Mirn21\|mmu-mir-21\|mmu-mir-21a | microRNA 21a | ncRNA |
| chr11_86591273_86591749 | chr11 | 86591274 | 86591749 | + | 0.63061 | intron (NM_029478, intron 10 of 11) | intron (NM_029478, intron 10 of 11) | -7353 | NR_029738 | 387140 | | NR_029738 | miRBase:MI0000569 | **Mir21a** | Mir21\|Mirn21\|mmu-mir-21\|mmu-mir-21a | microRNA 21a | ncRNA |
| chr8_128164750_128165504 | chr8 | 128164751 | 128165504 | + | 1.35199 | Intergenic | Intergenic | 113098 | NR_105822 | 102465212 | | NR_105822 | miRBase:MI0021932 | **Mir21c** | mmu-mir-21c | microRNA mir-21c | ncRNA |
| chr8_128272881_128273513 | chr8 | 128272882 | 128273513 | + | 0.710372 | Intergenic | Intergenic | 5028 | NR_105822 | 102465212 | | NR_105822 | miRBase:MI0021932 | **Mir21c** | mmu-mir-21c | microRNA mir-21c | ncRNA |
| chr11_75464534_75464769 | chr11 | 75464535 | 75464769 | + | 1.25935 | TTS (NR_029739) | TTS (NR_029739) | 936 | NR_029739 | 387141 | | NR_029739 | miRBase:MI0000570 | **Mir22** | Mirn22\|mmu-mir-22 | microRNA 22 | ncRNA |
| chrX_19160259_19160778 | chrX | 19160260 | 19160778 | + | 0.922731 | Intergenic | Intergenic | -13548 | NR_029807 | 723828 | | NR_029807 | miRBase:MI0000710 | **Mir222** | Mirn222\|mmu-mir-222 | microRNA 222 | ncRNA |
| chrX_19282850_19283384 | chrX | 19282851 | 19283384 | + | 0.715849 | Intergenic | Intergenic | -136146 | NR_029807 | 723828 | | NR_029807 | miRBase:MI0000710 | **Mir222** | Mirn222\|mmu-mir-222 | microRNA 222 | ncRNA |
| chrX_19157458_19158188 | chrX | 19157459 | 19158188 | + | 0.669937 | Intergenic | Intergenic | -10852 | NR_029807 | 723828 | | NR_029807 | miRBase:MI0000710 | **Mir222** | Mirn222\|mmu-mir-222 | microRNA 222 | ncRNA |
| chr8_84202582_84202806 | chr8 | 84202583 | 84202806 | + | 0.678444 | Intergenic | Intergenic | -5824 | NR_029740 | 387216 | | NR_029740 | miRBase:MI0000571 | **Mir23a** | Mirn23a\|mmu-mir-23a | microRNA 23a | ncRNA |
| chr8_96552364_96552974 | chr8 | 96552365 | 96552974 | + | 0.811674 | Intergenic | Intergenic | -6696 | NR_039538 | 100628567 | | NR_039538 | | **Mir28c** | mmu-mir-28c | microRNA 28c | ncRNA |
| chr6_31132212_31132762 | chr6 | 31132213 | 31132762 | + | 1.25772 | intron (NR_110471, intron 3 of 3) | intron (NR_110471, intron 3 of 3) | -69394 | NR_029532 | 387223 | | NR_029532 | miRBase:MI0000143 | **Mir29b-1** | Mirn29b\|Mirn29b-1 | microRNA 29b-1 | ncRNA |
| chr6_31088091_31088971 | chr6 | 31088092 | 31088971 | + | 1.13951 | Intergenic | Intergenic | -25438 | NR_029532 | 387223 | | NR_029532 | miRBase:MI0000143 | **Mir29b-1** | Mirn29b\|Mirn29b-1 | microRNA 29b-1 | ncRNA |
| chr6_31080654_31081042 | chr6 | 31080655 | 31081042 | + | 1.13793 | Intergenic | Intergenic | -17755 | NR_029532 | 387223 | | NR_029532 | miRBase:MI0000143 | **Mir29b-1** | Mirn29b\|Mirn29b-1 | microRNA 29b-1 | ncRNA |
| chr6_31117668_31118059 | chr6 | 31117669 | 31118059 | + | 0.726081 | intron (NR_110471, intron 3 of 3) | intron (NR_110471, intron 3 of 3) | -54771 | NR_029532 | 387223 | | NR_029532 | miRBase:MI0000143 | **Mir29b-1** | Mirn29b\|Mirn29b-1 | microRNA 29b-1 | ncRNA |

| | | | | | | | | | | | | | | | | | |
|---|---|---|---|---|---|---|---|---|---|---|---|---|---|---|---|---|---|
| chr11_4138683_4139238 | chr11 | 4138684 | 4139238 | + | 1.0779 | promoter-TSS (NR_037220) | promoter-TSS (NR_037220) | -403 | NR_037220 | 100526466 | | NR_037220 | | **Mir3060** | mmu-mir-3060 | microRNA 3060 | ncRNA |
| chr11_119988929_119989567 | chr11 | 119988930 | 119989567 | + | 0.688155 | intron (NM_001037754, intron 6 of 13) | intron (NM_001037754, intron 6 of 13) | -25519 | NR_037225 | 100526518 | | NR_037225 | | **Mir3065** | mmu-mir-3065 | microRNA 3065 | ncRNA |
| chr12_17373561_17374470 | chr12 | 17373562 | 17374470 | + | 0.873593 | intron (NM_001008421, intron 12 of 20) | intron (NM_001008421, intron 12 of 20) | 18624 | NR_037226 | 100526470 | | NR_037226 | | **Mir3066** | mmu-mir-3066 | microRNA 3066 | ncRNA |
| chr12_87434984_87435351 | chr12 | 87434985 | 87435351 | + | 0.730806 | intron (NM_001102565, intron 3 of 5) | MER77B\|LTR\|ERVL | 2589 | NR_037228 | 100526546 | | NR_037228 | | **Mir3068** | mmu-mir-3068 | microRNA 3068 | ncRNA |
| chr12_87426421_87427204 | chr12 | 87426422 | 87427204 | + | 0.681588 | Intergenic | Intergenic | 10944 | NR_037228 | 100526546 | | NR_037228 | | **Mir3068** | mmu-mir-3068 | microRNA 3068 | ncRNA |
| chr12_105022703_105023184 | chr12 | 105022704 | 105023184 | + | 0.786182 | Intergenic | Intergenic | 8197 | NR_037229 | 100526471 | | NR_037229 | | **Mir3069** | mmu-mir-3069 | microRNA 3069 | ncRNA |
| chr14_25527317_25527653 | chr14 | 25527318 | 25527653 | + | 0.621344 | intron (NM_183208, intron 2 of 23) | intron (NM_183208, intron 2 of 23) | -6954 | NR_037236 | 100526533 | | NR_037236 | miRBase:MI0014038 | **Mir3075** | mmu-mir-3075 | microRNA 3075 | ncRNA |
| chr14_25527283_25527816 | chr14 | 25527284 | 25527816 | + | 0.612342 | intron (NM_183208, intron 2 of 23) | intron (NM_183208, intron 2 of 23) | -6889 | NR_037236 | 100526533 | | NR_037236 | miRBase:MI0014038 | **Mir3075** | mmu-mir-3075 | microRNA 3075 | ncRNA |
| chr14_30584035_30585145 | chr14 | 30584036 | 30585145 | + | 1.36994 | Intergenic | Intergenic | 12441 | NR_037237 | 100526475 | | NR_037237 | | **Mir3076** | mmu-mir-3076 | microRNA 3076 | ncRNA |
| chr14_30584037_30584467 | chr14 | 30584038 | 30584467 | + | 1.31397 | Intergenic | RMER15\|LTR\|ERVL | 12103 | NR_037237 | 100526475 | | NR_037237 | | **Mir3076** | mmu-mir-3076 | microRNA 3076 | ncRNA |
| chr14_30584538_30585111 | chr14 | 30584539 | 30585111 | + | 0.994935 | Intergenic | Intergenic | 12676 | NR_037237 | 100526475 | | NR_037237 | | **Mir3076** | mmu-mir-3076 | microRNA 3076 | ncRNA |
| chr14_64608647_64608950 | chr14 | 64608648 | 64608950 | + | 1.12331 | Intergenic | Intergenic | 17614 | NR_037239 | 100526520 | | NR_037239 | miRBase:MI0014041 | **Mir3078** | mmu-mir-3078 | microRNA 3078 | ncRNA |
| chr14_64608530_64608957 | chr14 | 64608531 | 64608957 | + | 0.836829 | Intergenic | Intergenic | 17559 | NR_037239 | 100526520 | | NR_037239 | miRBase:MI0014041 | **Mir3078** | mmu-mir-3078 | microRNA 3078 | ncRNA |
| chr16_44494516_44494844 | chr16 | 44494517 | 44494844 | + | 0.709215 | intron (NM_172506, intron 8 of 18) | intron (NM_172506, intron 8 of 18) | 63449 | NR_037242 | 100526478 | | NR_037242 | miRBase:MI0014044 | **Mir3081** | mmu-mir-3081 | microRNA 3081 | ncRNA |
| chr2_30723303_30723653 | chr2 | 30723304 | 30723653 | + | 0.775396 | Intergenic | MIR\|SINE\|MIR | 2268 | NR_037272 | 100526499 | | NR_037272 | miRBase:MI0014082 | **Mir3089** | mmu-mir-3089 | microRNA 3089 | ncRNA |
| chr4_58469705_58469977 | chr4 | 58469706 | 58469977 | + | 0.940654 | intron (NM_001290486, intron 2 of 2) | intron (NM_001290486, intron 2 of 2) | -28743 | NR_037278 | 100526502 | | NR_037278 | | **Mir3095** | mmu-mir-3095 | microRNA 3095 | ncRNA |
| chr4_120759734_120760148 | chr4 | 120759735 | 120760148 | + | 0.851087 | intron (NM_001277095, intron 7 of 8) | intron (NM_001277095, intron 7 of 8) | 9681 | NR_029716 | 387227 | | NR_029716 | | **Mir30c-1** | Mirn30c\|Mirn30c-1 | microRNA 30c-1 | ncRNA |
| chr4_88911455_88911748 | chr4 | 88911456 | 88911748 | + | 0.764342 | promoter-TSS (NR_029747) | promoter-TSS (NR_029747) | -940 | NR_029747 | 723895 | | NR_029747 | miRBase:MI0000579 | **Mir31** | Mirn31\|miR-31\|mmu-mir-31 | microRNA 31 | ncRNA |
| chr4_88937782_88938945 | chr4 | 88937783 | 88938945 | + | 0.708148 | Intergenic | L1M4b\|LINE\|L1 | -27702 | NR_029747 | 723895 | | NR_029747 | miRBase:MI0000579 | **Mir31** | Mirn31\|miR-31\|mmu-mir-31 | microRNA 31 | ncRNA |
| chr8_108890850_108891372 | chr8 | 108890851 | 108891372 | + | 0.663186 | intron (NM_007496, intron 4 of 9) | (CAGA)n\|Simple_repeat\|Simple_repeat | -45749 | NR_037295 | 100526511 | | NR_037295 | | **Mir3108** | mmu-mir-3108 | microRNA 3108 | ncRNA |
| chr9_69467184_69467745 | chr9 | 69467185 | 69467745 | + | 1.32633 | intron (NM_007585, intron 3 of 12) | intron (NM_007585, intron 3 of 12) | 10521 | NR_037296 | 100526560 | | NR_037296 | miRBase:MI0014106 | **Mir3109** | mmu-mir-3109 | microRNA 3109 | ncRNA |
| chr9_69457976_69458617 | chr9 | 69457977 | 69458617 | + | 1.30721 | intron (NM_007585, intron 1 of 12) | intron (NM_007585, intron 1 of 12) | 1353 | NR_037296 | 100526560 | | NR_037296 | miRBase:MI0014106 | **Mir3109** | mmu-mir-3109 | microRNA 3109 | ncRNA |
| chr9_69467184_69467959 | chr9 | 69467185 | 69467959 | + | 1.14578 | intron (NM_007585, intron 3 of 12) | intron (NM_007585, intron 3 of 12) | 10628 | NR_037296 | 100526560 | | NR_037296 | miRBase:MI0014106 | **Mir3109** | mmu-mir-3109 | microRNA 3109 | ncRNA |
| chr8_126683899_126684347 | chr8 | 126683900 | 126684347 | + | 1.17592 | Intergenic | Intergenic | 37677 | NR_037297 | 100526512 | | NR_037297 | miRBase:MI0014107 | **Mir3110** | mmu-mir-3110 | microRNA 3110 | ncRNA |
| chr8_126811282_126812033 | chr8 | 126811283 | 126812033 | + | 1.03623 | Intergenic | Intergenic | -89858 | NR_037297 | 100526512 | | NR_037297 | miRBase:MI0014107 | **Mir3110** | mmu-mir-3110 | microRNA 3110 | ncRNA |
| chr8_126767468_126768019 | chr8 | 126767469 | 126768019 | + | 0.95691 | Intergenic | Intergenic | -45944 | NR_037297 | 100526512 | | NR_037297 | miRBase:MI0014107 | **Mir3110** | mmu-mir-3110 | microRNA 3110 | ncRNA |
| chr8_126767250_126768057 | chr8 | 126767251 | 126768057 | + | 0.911885 | Intergenic | Intergenic | -45854 | NR_037297 | 100526512 | | NR_037297 | miRBase:MI0014107 | **Mir3110** | mmu-mir-3110 | microRNA 3110 | ncRNA |
| chr8_126833335_126834281 | chr8 | 126833336 | 126834281 | + | 0.908975 | Intergenic | Intergenic | -112008 | NR_037297 | 100526512 | | NR_037297 | miRBase:MI0014107 | **Mir3110** | mmu-mir-3110 | microRNA 3110 | ncRNA |
| chr8_126826306_126826719 | chr8 | 126826307 | 126826719 | + | 0.710806 | Intergenic | Intergenic | -104713 | NR_037297 | 100526512 | | NR_037297 | miRBase:MI0014107 | **Mir3110** | mmu-mir-3110 | microRNA 3110 | ncRNA |
| chr8_126803993_126804556 | chr8 | 126803994 | 126804556 | + | 0.660372 | Intergenic | Intergenic | -82475 | NR_037297 | 100526512 | | NR_037297 | miRBase:MI0014107 | **Mir3110** | mmu-mir-3110 | microRNA 3110 | ncRNA |
| chr13_107798077_107798490 | chr13 | 107798078 | 107798490 | + | 0.972809 | intron (NM_145456, intron 2 of 14) | intron (NM_145456, intron 2 of 14) | -37197 | NR_029759 | 723929 | | NR_029759 | miRBase:MI0000597 | **Mir325** | Mirn325\|mmu-mir-325 | microRNA 325 | ncRNA |
| chr10_93945954_93946310 | chr10 | 93945955 | 93946310 | + | 0.911384 | Intergenic | Intergenic | 17731 | NR_029764 | 723908 | | NR_029764 | miRBase:MI0000609 | **Mir331** | Mirn331\|mmu-mir-331 | microRNA 331 | ncRNA |
| chr10_93945866_93946699 | chr10 | 93945867 | 93946699 | + | 0.793083 | Intergenic | RLTR19\|LTR\|ERVK | 17580 | NR_029764 | 723908 | | NR_029764 | miRBase:MI0000609 | **Mir331** | Mirn331\|mmu-mir-331 | microRNA 331 | ncRNA |
| chr11_50076792_50077371 | chr11 | 50076793 | 50077371 | + | 1.34378 | intron (NM_001290750, intron 3 of 7) | intron (NM_001290750, intron 3 of 7) | 7380 | NR_029769 | 723845 | | NR_029769 | | **Mir340** | Mirn340\|mmu-mir-340 | microRNA 340 | ncRNA |
| chr7_144417622_144417946 | chr7 | 144417623 | 144417946 | + | 0.819972 | intron (NM_001113373, intron 15 of 15) | intron (NM_001113373, intron 15 of 15) | 25649 | NR_037302 | 100499519 | | NR_037302 | miRBase:MI0014697 | **Mir3470b** | mmu-mir-3470b | microRNA 3470b | ncRNA |
| chr7_144417598_144418089 | chr7 | 144417599 | 144418089 | + | 0.801662 | intron (NM_001113373, intron 15 of 15) | intron (NM_001113373, intron 15 of 15) | 25589 | NR_037302 | 100499519 | | NR_037302 | miRBase:MI0014697 | **Mir3470b** | mmu-mir-3470b | microRNA 3470b | ncRNA |
| chr15_100188691_100189497 | chr15 | 100188692 | 100189497 | + | 1.27896 | intron (NM_172819, intron 18 of 32) | intron (NM_172819, intron 18 of 32) | -1058 | NR_037311 | 100499528 | | NR_037311 | | **Mir3473a** | Mir3473\|mmu-mir-3473\|mmu-mir-3473a | microRNA 3473a | ncRNA |
| chr15_100181186_100181644 | chr15 | 100181187 | 100181644 | + | 0.751203 | exon (NM_001159361, exon 21 of 38) | exon (NM_001159361, exon 21 of 38) | 6621 | NR_037311 | 100499528 | | NR_037311 | | **Mir3473a** | Mir3473\|mmu-mir-3473\|mmu-mir-3473a | microRNA 3473a | ncRNA |
| chr15_100180709_100181074 | chr15 | 100180710 | 100181074 | + | 0.640656 | intron (NM_172819, intron 15 of 32) | RSINE1\|SINE\|B4 | 7144 | NR_037311 | 100499528 | | NR_037311 | | **Mir3473a** | Mir3473\|mmu-mir-3473\|mmu-mir-3473a | microRNA 3473a | ncRNA |
| chr4_150059620_150060723 | chr4 | 150059621 | 150060723 | + | 1.00439 | Intergenic | Intergenic | -8282 | NR_029751 | 723848 | | NR_029751 | miRBase:MI0000584 | **Mir34a** | Mirn34a\|mmu-mir-34a | microRNA 34a | ncRNA |
| chr4_150049673_150049971 | chr4 | 150049674 | 150049971 | + | 0.97285 | Intergenic | Intergenic | -18632 | NR_029751 | 723848 | | NR_029751 | miRBase:MI0000584 | **Mir34a** | Mirn34a\|mmu-mir-34a | microRNA 34a | ncRNA |
| chr4_150049052_150049617 | chr4 | 150049053 | 150049617 | + | 0.965572 | Intergenic | Intergenic | -19119 | NR_029751 | 723848 | | NR_029751 | miRBase:MI0000584 | **Mir34a** | Mirn34a\|mmu-mir-34a | microRNA 34a | ncRNA |
| chr4_150040020_150041178 | chr4 | 150040021 | 150041178 | + | 0.867113 | Intergenic | Intergenic | -27855 | NR_029751 | 723848 | | NR_029751 | miRBase:MI0000584 | **Mir34a** | Mirn34a\|mmu-mir-34a | microRNA 34a | ncRNA |
| chr4_150040053_150040905 | chr4 | 150040054 | 150040905 | + | 0.864555 | Intergenic | RSINE1\|SINE\|B4 | -27975 | NR_029751 | 723848 | | NR_029751 | miRBase:MI0000584 | **Mir34a** | Mirn34a\|mmu-mir-34a | microRNA 34a | ncRNA |
| chr4_150048033_150049973 | chr4 | 150048034 | 150049973 | + | 0.855762 | Intergenic | CpG | -19451 | NR_029751 | 723848 | | NR_029751 | miRBase:MI0000584 | **Mir34a** | Mirn34a\|mmu-mir-34a | microRNA 34a | ncRNA |
| chrX_52739723_52740136 | chrX | 52739724 | 52740136 | + | 1.0861 | Intergenic | Intergenic | 1837 | NR_029853 | 723852 | | NR_029853 | miRBase:MI0000765 | **Mir363** | Mirn363\|mmu-mir-363 | microRNA 363 | ncRNA |
| chrX_52722296_52722935 | chrX | 52722297 | 52722935 | + | 0.80849 | Intergenic | B3A\|SINE\|B2 | 19151 | NR_029853 | 723852 | | NR_029853 | miRBase:MI0000765 | **Mir363** | Mirn363\|mmu-mir-363 | microRNA 363 | ncRNA |
| chrX_52732714_52733210 | chrX | 52732715 | 52733210 | + | 0.724106 | Intergenic | Intergenic | 8805 | NR_029853 | 723852 | | NR_029853 | miRBase:MI0000765 | **Mir363** | Mirn363\|mmu-mir-363 | microRNA 363 | ncRNA |
| chr11_79833729_79834217 | chr11 | 79833730 | 79834217 | + | 1.30981 | Intergenic | Intergenic | 107573 | NR_029959 | 723853 | | NR_029959 | miRBase:MI0001645 | **Mir365-2** | Mirn365-2 | microRNA 365-2 | ncRNA |
| chr11_79793654_79794515 | chr11 | 79793655 | 79794515 | + | 1.23152 | Intergenic | Intergenic | 67685 | NR_029959 | 723853 | | NR_029959 | miRBase:MI0001645 | **Mir365-2** | Mirn365-2 | microRNA 365-2 | ncRNA |
| chr11_79793371_79794547 | chr11 | 79793372 | 79794547 | + | 1.09434 | Intergenic | MIRc\|SINE\|MIR | 67559 | NR_029959 | 723853 | | NR_029959 | miRBase:MI0001645 | **Mir365-2** | Mirn365-2 | microRNA 365-2 | ncRNA |
| chr11_79826954_79827669 | chr11 | 79826955 | 79827669 | + | 0.903259 | Intergenic | Intergenic | 100912 | NR_029959 | 723853 | | NR_029959 | miRBase:MI0001645 | **Mir365-2** | Mirn365-2 | microRNA 365-2 | ncRNA |
| chr11_79761918_79762387 | chr11 | 79761919 | 79762387 | + | 0.695998 | Intergenic | Intergenic | 35753 | NR_029959 | 723853 | | NR_029959 | miRBase:MI0001645 | **Mir365-2** | Mirn365-2 | microRNA 365-2 | ncRNA |
| chr9_71995731_71996135 | chr9 | 71995732 | 71996135 | + | 1.20434 | intron (NM_001253862, intron 5 of 19) | intron (NM_001253862, intron 5 of 19) | 32492 | NR_029879 | 723889 | | NR_029879 | | **Mir378a** | Mir378\|Mirn378\|mmu-mir-378\|mmu-mir-378a | microRNA 378a | ncRNA |
| chr10_97347441_97347768 | chr10 | 97347442 | 97347768 | + | 0.689687 | Intergenic | L1MC1\|LINE\|L1 | -75873 | NR_039547 | 100628616 | | NR_039547 | | **Mir3966** | mmu-mir-3966\|mu-mir-3966 | microRNA 3966 | ncRNA |
| chr2_22679735_22680892 | chr2 | 22679736 | 22680892 | + | 1.83338 | intron (NM_008078, intron 12 of 15) | intron (NM_008078, intron 12 of 15) | 26031 | NR_039548 | 100628608 | | NR_039548 | | **Mir3967** | mmu-mir-3967\|mu-mir-3967 | microRNA 3967 | ncRNA |
| chr2_22687126_22687514 | chr2 | 22687127 | 22687514 | + | 0.762594 | intron (NM_008078, intron 15 of 15) | intron (NM_008078, intron 15 of 15) | 33037 | NR_039548 | 100628608 | | NR_039548 | | **Mir3967** | mmu-mir-3967\|mu-mir-3967 | microRNA 3967 | ncRNA |
| chr11_75564231_75564521 | chr11 | 75564232 | 75564521 | + | 0.830741 | intron (NM_001199284, intron 9 of 13) | intron (NM_001199284, intron 9 of 13) | 12947 | NR_039553 | 100628624 | | NR_039553 | | **Mir3971** | mmu-mir-3971\|mu-mir-3971 | microRNA 3971 | ncRNA |
| chr11_75563775_75564152 | chr11 | 75563776 | 75564152 | + | 0.808191 | intron (NM_001199284, intron 9 of 13) | intron (NM_001199284, intron 9 of 13) | 12535 | NR_039553 | 100628624 | | NR_039553 | | **Mir3971** | mmu-mir-3971\|mu-mir-3971 | microRNA 3971 | ncRNA |
| chr3_142540905_142541335 | chr3 | 142540906 | 142541335 | + | 1.02202 | intron (NM_145545, intron 5 of 10) | URR1B\|DNA\|hAT-Charlie | -1237 | NR_030558 | 100124494 | | NR_030558 | miRBase:MI0005496 | **Mir421** | Mirn421\|mmu-mir-421 | microRNA 421 | ncRNA |
| chrX_77219330_77219845 | chrX | 77219331 | 77219845 | + | 0.641127 | intron (NR_035475, intron 1 of 3) | intron (NR_035475, intron 1 of 3) | -207626 | NR_030558 | 100124494 | | NR_030558 | miRBase:MI0005496 | **Mir421** | Mirn421\|mmu-mir-421 | microRNA 421 | ncRNA |

| | | | | | | | | | | | | | | | | | |
|---|---|---|---|---|---|---|---|---|---|---|---|---|---|---|---|---|---|
| chr2_5395859_5396397 | chr2 | 5395860 | 5396397 | + | 1.05838 | intron (NR_030601, intron 1 of 1) | intron (NR_030601, intron 1 of 1) | 16738 | NR_030601 | 100124465 | | NR_030601 | | **Mir466d** | Mirn466d | microRNA 466d | ncRNA |
| chr2_5401861_5402525 | chr2 | 5401862 | 5402525 | + | 0.942537 | intron (NR_030601, intron 1 of 1) | intron (NR_030601, intron 1 of 1) | 22803 | NR_030601 | 100124465 | | NR_030601 | | **Mir466d** | Mirn466d | microRNA 466d | ncRNA |
| chr2_5296819_5297115 | chr2 | 5296820 | 5297115 | + | 0.841808 | 3' UTR (NM_001290376, exon 11 of 11) | 3' UTR (NM_001290376, exon 11 of 11) | -82423 | NR_030601 | 100124465 | | NR_030601 | | **Mir466d** | Mirn466d | microRNA 466d | ncRNA |
| chr2_5296620_5297120 | chr2 | 5296621 | 5297120 | + | 0.759576 | 3' UTR (NM_001290376, exon 11 of 11) | 3' UTR (NM_001290376, exon 11 of 11) | -82520 | NR_030601 | 100124465 | | NR_030601 | | **Mir466d** | Mirn466d | microRNA 466d | ncRNA |
| chr2_5507923_5508525 | chr2 | 5507924 | 5508525 | + | 0.68226 | intron (NM_177343, intron 2 of 10) | intron (NM_177343, intron 2 of 10) | 128834 | NR_030601 | 100124465 | | NR_030601 | | **Mir466d** | Mirn466d | microRNA 466d | ncRNA |
| chr11_86430265_86430790 | chr11 | 86430266 | 86430790 | + | 1.65239 | intron (NR_030571, intron 1 of 2) | intron (NR_030571, intron 1 of 2) | 14684 | NR_030571 | 100124487 | | NR_030571 | | **Mir467c** | Mirn467c | microRNA 467c | ncRNA |
| chr11_86428936_86429230 | chr11 | 86428937 | 86429230 | + | 1.05386 | intron (NR_030571, intron 1 of 2) | intron (NR_030571, intron 1 of 2) | 13239 | NR_030571 | 100124487 | | NR_030571 | | **Mir467c** | Mirn467c | microRNA 467c | ncRNA |
| chr4_88122115_88122802 | chr4 | 88122116 | 88122802 | + | 1.79711 | TTS (NR_030478) | TTS (NR_030478) | 419 | NR_030478 | 735282 | | NR_030478 | | **Mir491** | Mirn491\|mmu-mir-491 | microRNA 491 | ncRNA |
| chr4_88179347_88179819 | chr4 | 88179348 | 88179819 | + | 1.245 | intron (NM_001081184, intron 8 of 43) | intron (NM_001081184, intron 8 of 43) | 57543 | NR_030478 | 735282 | | NR_030478 | | **Mir491** | Mirn491\|mmu-mir-491 | microRNA 491 | ncRNA |
| chr4_88213871_88214328 | chr4 | 88213872 | 88214328 | + | 1.14363 | intron (NM_001081184, intron 11 of 43) | RMER15\|LTR\|ERVL | 92060 | NR_030478 | 735282 | | NR_030478 | | **Mir491** | Mirn491\|mmu-mir-491 | microRNA 491 | ncRNA |
| chr4_88118046_88118257 | chr4 | 88118047 | 88118257 | + | 1.13714 | intron (NM_001081184, intron 1 of 43) | B3A\|SINE\|B2 | -3888 | NR_030478 | 735282 | | NR_030478 | | **Mir491** | Mirn491\|mmu-mir-491 | microRNA 491 | ncRNA |
| chr4_88120109_88120476 | chr4 | 88120110 | 88120476 | + | 1.00501 | intron (NM_001081184, intron 1 of 43) | intron (NM_001081184, intron 1 of 43) | -1747 | NR_030478 | 735282 | | NR_030478 | | **Mir491** | Mirn491\|mmu-mir-491 | microRNA 491 | ncRNA |
| chr4_88177290_88177811 | chr4 | 88177291 | 88177811 | + | 0.992048 | intron (NM_001081184, intron 7 of 43) | LTRIS_Mm\|LTR\|ERV1 | 55511 | NR_030478 | 735282 | | NR_030478 | | **Mir491** | Mirn491\|mmu-mir-491 | microRNA 491 | ncRNA |
| chr19_16254993_16255776 | chr19 | 16254994 | 16255776 | + | 0.880831 | intron (NM_008139, intron 2 of 6) | intron (NM_008139, intron 2 of 6) | 59619 | NR_105832 | 102465218 | | NR_105832 | miRBase:MI0021942 | **Mir496b** | mmu-mir-496b | microRNA mir-496b | ncRNA |
| chr19_16371762_16372148 | chr19 | 16371763 | 16372148 | + | 0.764698 | intron (NM_008139, intron 5 of 6) | intron (NM_008139, intron 5 of 6) | -56951 | NR_105832 | 102465218 | | NR_105832 | miRBase:MI0021942 | **Mir496b** | mmu-mir-496b | microRNA mir-496b | ncRNA |
| chr19_16329088_16329538 | chr19 | 16329089 | 16329538 | + | 0.657075 | intron (NM_008139, intron 3 of 6) | intron (NM_008139, intron 3 of 6) | -14309 | NR_105832 | 102465218 | | NR_105832 | miRBase:MI0021942 | **Mir496b** | mmu-mir-496b | microRNA mir-496b | ncRNA |
| chr2_155624685_155625131 | chr2 | 155624686 | 155625131 | + | 0.675235 | intron (NM_001085378, intron 21 of 41) | B3A\|SINE\|B2 | 2028 | NR_030757 | 735275 | | NR_030757 | | **Mir499** | Mirn499\|mmu-mir-499 | microRNA 499 | ncRNA |
| chrX_59091687_59092260 | chrX | 59091688 | 59092260 | + | 0.916505 | intron (NM_001290414, intron 5 of 6) | intron (NM_001290414, intron 5 of 6) | 5762 | NR_030574 | 100124476 | | NR_030574 | | **Mir504** | Mirn504\|mmu-mir-504 | microRNA 504 | ncRNA |
| chrX_59090971_59091463 | chrX | 59090972 | 59091463 | + | 0.783172 | intron (NM_001290414, intron 5 of 6) | intron (NM_001290414, intron 5 of 6) | 6519 | NR_030574 | 100124476 | | NR_030574 | | **Mir504** | Mirn504\|mmu-mir-504 | microRNA 504 | ncRNA |
| chrX_60379753_60380264 | chrX | 60379754 | 60380264 | + | 0.780321 | intron (NM_001001798, intron 1 of 28) | B1F\|SINE\|Alu | 14483 | NR_030499 | 751545 | | NR_030499 | | **Mir505** | Mirn505\|mmu-mir-505 | microRNA 505 | ncRNA |
| chr10_7879967_7880637 | chr10 | 7879968 | 7880637 | + | 1.52864 | Intergenic | L1VL2\|LINE\|L1 | 43921 | NR_039563 | 100628581 | | NR_039563 | miRBase:MI0018012 | **Mir5104** | mmu-mir-5104\|mu-mir-5104 | microRNA 5104 | ncRNA |
| chr10_7844739_7845181 | chr10 | 7844740 | 7845181 | + | 0.643262 | intron (NM_172785, intron 2 of 5) | intron (NM_172785, intron 2 of 5) | 8579 | NR_039563 | 100628581 | | NR_039563 | miRBase:MI0018012 | **Mir5104** | mmu-mir-5104\|mu-mir-5104 | microRNA 5104 | ncRNA |
| chr2_14312407_14312964 | chr2 | 14312408 | 14312964 | + | 1.52073 | intron (NM_008625, intron 21 of 29) | intron (NM_008625, intron 21 of 29) | 51683 | NR_030609 | 100124488 | | NR_030609 | | **Mir511** | Mirn511\|mmu-mir-511 | microRNA 511 | ncRNA |
| chr2_14312116_14312969 | chr2 | 14312117 | 14312969 | + | 1.31133 | intron (NM_008625, intron 21 of 29) | intron (NM_008625, intron 21 of 29) | 51540 | NR_030609 | 100124488 | | NR_030609 | | **Mir511** | Mirn511\|mmu-mir-511 | microRNA 511 | ncRNA |
| chr15_80886501_80887069 | chr15 | 80886502 | 80887069 | + | 1.0504 | intron (NM_177124, intron 7 of 21) | intron (NM_177124, intron 7 of 21) | -53254 | NR_039573 | 100628612 | | NR_039573 | miRBase:MI0018022 | **Mir5113** | mmu-mir-5113\|mu-mir-5113 | microRNA 5113 | ncRNA |
| chr4_133399379_133400008 | chr4 | 133399380 | 133400008 | + | 1.09361 | intron (NM_016981, intron 1 of 11) | intron (NM_016981, intron 1 of 11) | 29918 | NR_039582 | 100628620 | | NR_039582 | miRBase:MI0018031 | **Mir5122** | mmu-mir-5122\|mu-mir-5122 | microRNA 5122 | ncRNA |
| chr4_133398654_133399013 | chr4 | 133398655 | 133399013 | + | 0.791891 | intron (NM_016981, intron 1 of 11) | MIR3\|SINE\|MIR | 29058 | NR_039582 | 100628620 | | NR_039582 | miRBase:MI0018031 | **Mir5122** | mmu-mir-5122\|mu-mir-5122 | microRNA 5122 | ncRNA |
| chr4_133378207_133378607 | chr4 | 133378208 | 133378607 | + | 0.623565 | intron (NM_016981, intron 1 of 11) | intron (NM_016981, intron 1 of 11) | 8631 | NR_039582 | 100628620 | | NR_039582 | miRBase:MI0018031 | **Mir5122** | mmu-mir-5122\|mu-mir-5122 | microRNA 5122 | ncRNA |
| chr4_40846582_40846921 | chr4 | 40846583 | 40846921 | + | 0.73507 | intron (NM_022305, intron 1 of 5) | MTD\|LTR\|ERVL-MaLR | 3386 | NR_039584 | 100628614 | | NR_039584 | | **Mir5123** | mmu-mir-5123\|mu-mir-5123 | microRNA 5123 | ncRNA |
| chr9_113984100_113984457 | chr9 | 113984101 | 113984457 | + | 0.765574 | intron (NM_178624, intron 12 of 14) | intron (NM_178624, intron 12 of 14) | 9521 | NR_039590 | 100628597 | | NR_039590 | miRBase:MI0018040 | **Mir5128** | mmu-mir-5128\|mu-mir-5128 | microRNA 5128 | ncRNA |
| chr9_113983985_113984423 | chr9 | 113983986 | 113984423 | + | 0.675996 | intron (NM_178624, intron 12 of 14) | intron (NM_178624, intron 12 of 14) | 9596 | NR_039590 | 100628597 | | NR_039590 | miRBase:MI0018040 | **Mir5128** | mmu-mir-5128\|mu-mir-5128 | microRNA 5128 | ncRNA |
| chr2_44952748_44953383 | chr2 | 44952749 | 44953383 | + | 1.42184 | Intergenic | Intergenic | 70111 | NR_039591 | 100628598 | | NR_039591 | | **Mir5129** | mmu-mir-5129\|mu-mir-5129 | microRNA 5129 | ncRNA |
| chr2_44961106_44961696 | chr2 | 44961107 | 44961696 | + | 1.09323 | Intergenic | Intergenic | 61776 | NR_039591 | 100628598 | | NR_039591 | | **Mir5129** | mmu-mir-5129\|mu-mir-5129 | microRNA 5129 | ncRNA |
| chr2_44961098_44962029 | chr2 | 44961099 | 44962029 | + | 0.935951 | Intergenic | Intergenic | 61613 | NR_039591 | 100628598 | | NR_039591 | | **Mir5129** | mmu-mir-5129\|mu-mir-5129 | microRNA 5129 | ncRNA |
| chr2_45055815_45056520 | chr2 | 45055816 | 45056520 | + | 0.669104 | intron (NM_001289521, intron 1 of 8) | intron (NM_001289521, intron 1 of 8) | -32991 | NR_039591 | 100628598 | | NR_039591 | | **Mir5129** | mmu-mir-5129\|mu-mir-5129 | microRNA 5129 | ncRNA |
| chr2_44956312_44956600 | chr2 | 44956313 | 44956600 | + | 0.650548 | Intergenic | Intergenic | 66721 | NR_039591 | 100628598 | | NR_039591 | | **Mir5129** | mmu-mir-5129\|mu-mir-5129 | microRNA 5129 | ncRNA |
| chr14_102837944_102838481 | chr14 | 102837945 | 102838481 | + | 0.78662 | Intergenic | Intergenic | 144419 | NR_039592 | 100628599 | | NR_039592 | miRBase:MI0018042 | **Mir5130** | mmu-mir-5130\|mu-mir-5130 | microRNA 5130 | ncRNA |
| chr14_102833917_102834290 | chr14 | 102833918 | 102834290 | + | 0.679748 | Intergenic | URR1B\|DNA\|hAT-Charlie | 148528 | NR_039592 | 100628599 | | NR_039592 | miRBase:MI0018042 | **Mir5130** | mmu-mir-5130\|mu-mir-5130 | microRNA 5130 | ncRNA |
| chr14_102674610_102674911 | chr14 | 102674611 | 102674911 | + | 0.648258 | Intergenic | MT2A\|LTR\|ERVL | 307871 | NR_039592 | 100628599 | | NR_039592 | miRBase:MI0018042 | **Mir5130** | mmu-mir-5130\|mu-mir-5130 | microRNA 5130 | ncRNA |
| chr14_45648861_45649531 | chr14 | 45648862 | 45649531 | + | 1.05012 | intron (NM_176845, intron 1 of 12) | intron (NM_176845, intron 1 of 12) | 8857 | NR_039593 | 100628600 | | NR_039593 | | **Mir5131** | mmu-mir-5131\|mu-mir-5131 | microRNA 5131 | ncRNA |
| chr14_45646534_45647127 | chr14 | 45646535 | 45647127 | + | 0.70078 | intron (NM_176845, intron 1 of 12) | intron (NM_176845, intron 1 of 12) | 11222 | NR_039593 | 100628600 | | NR_039593 | | **Mir5131** | mmu-mir-5131\|mu-mir-5131 | microRNA 5131 | ncRNA |
| chr9_62127780_62128578 | chr9 | 62127781 | 62128578 | + | 1.42469 | Intergenic | MLT1F\|LTR\|ERVL-MaLR | -5585 | NR_039595 | 100628602 | | NR_039595 | miRBase:MI0018045 | **Mir5133** | mmu-mir-5133\|mu-mir-5133 | microRNA 5133 | ncRNA |
| chr9_62118836_62119023 | chr9 | 62118837 | 62119023 | + | 1.13392 | Intergenic | Intergenic | 3664 | NR_039595 | 100628602 | | NR_039595 | miRBase:MI0018045 | **Mir5133** | mmu-mir-5133\|mu-mir-5133 | microRNA 5133 | ncRNA |
| chr9_62152115_62152420 | chr9 | 62152116 | 62152420 | + | 0.715075 | Intergenic | Intergenic | -29674 | NR_039595 | 100628602 | | NR_039595 | miRBase:MI0018045 | **Mir5133** | mmu-mir-5133\|mu-mir-5133 | microRNA 5133 | ncRNA |
| chr19_57971068_57971334 | chr19 | 57971069 | 57971334 | + | 0.717348 | intron (NM_181415, intron 27 of 28) | B3A\|SINE\|B2 | -79966 | NR_049196 | 100885835 | | NR_049196 | | **Mir5623** | mmu-mir-5623 | microRNA mir-5623 | ncRNA |
| chr19_57970464_57970972 | chr19 | 57970465 | 57970972 | + | 0.635572 | intron (NM_181415, intron 27 of 28) | intron (NM_181415, intron 27 of 28) | -80449 | NR_049196 | 100885835 | | NR_049196 | | **Mir5623** | mmu-mir-5623 | microRNA mir-5623 | ncRNA |
| chr13_93812968_93813254 | chr13 | 93812969 | 93813254 | + | 0.735679 | intron (NM_009712, intron 4 of 7) | intron (NM_009712, intron 4 of 7) | 22334 | NR_049198 | 100885837 | | NR_049198 | | **Mir5624** | mmu-mir-5624 | microRNA mir-5624 | ncRNA |
| chr5_30643333_30643737 | chr5 | 30643334 | 30643737 | + | 0.693316 | Intergenic | Intergenic | -4401 | NR_049199 | 100885838 | | NR_049199 | miRBase:MI0019194 | **Mir5625** | mmu-mir-5625 | microRNA mir-5625 | ncRNA |
| chr16_43628310_43629050 | chr16 | 43628311 | 43629050 | + | 0.91389 | Intergenic | Intergenic | -11975 | NR_030576 | 100124467 | | NR_030576 | miRBase:MI0005517 | **Mir568** | Mirn568\|mmu-mir-568 | microRNA 568 | ncRNA |
| chr5_64992491_64992978 | chr5 | 64992492 | 64992978 | + | 0.809272 | intron (NM_026667, intron 2 of 13) | intron (NM_026667, intron 2 of 13) | 22417 | NR_030577 | 100124451 | | NR_030577 | miRBase:MI0005518 | **Mir574** | Mirn574\|mmu-mir-574 | microRNA 574 | ncRNA |
| chr9_110272243_110272796 | chr9 | 110272244 | 110272796 | + | 0.763313 | Intergenic | Intergenic | -8767 | NR_105744 | 102465959 | | NR_105744 | miRBase:MI0021583 | **Mir6236** | mmu-mir-6236 | microRNA mir-6236 | ncRNA |

| | | | | | | | | | | | | | | | | | |
|---|---|---|---|---|---|---|---|---|---|---|---|---|---|---|---|---|---|
| chr2_130029730_130030281 | chr2 | 130029731 | 130030281 | + | 1.23149 | exon (NM_009374, exon 7 of 13) | exon (NM_009374, exon 7 of 13) | -13598 | NR_105757 | 102466628 | | NR_105757 | miRBase:MI0021867 | **Mir6339** | mmu-mir-6339 | microRNA mir-6339 | ncRNA |
| chr18_68828978_68829697 | chr18 | 68828979 | 68829697 | + | 1.13476 | Intergenic | Intergenic | 89938 | NR_105774 | 102465183 | | NR_105774 | miRBase:MI0021884 | **Mir6356** | mmu-mir-6356 | microRNA mir-6356 | ncRNA |
| chr18_89075100_89075552 | chr18 | 89075101 | 89075552 | + | 0.635473 | intron (NM_175542, intron 33 of 48) | intron (NM_175542, intron 33 of 48) | 103296 | NR_105777 | 102466633 | | NR_105777 | | **Mir6359** | mmu-mir-6359 | microRNA mir-6359 | ncRNA |
| chr18_23782897_23783756 | chr18 | 23782898 | 23783756 | + | 0.837519 | intron (NM_001162941, intron 2 of 7) | intron (NM_001162941, intron 2 of 7) | -8338 | NR_105778 | 102465185 | | NR_105778 | miRBase:MI0021888 | **Mir6360** | mmu-mir-6360 | microRNA mir-6360 | ncRNA |
| chr18_32902513_32902912 | chr18 | 32902514 | 32902912 | + | 0.755226 | Intergenic | Intergenic | -22040 | NR_105779 | 102465186 | | NR_105779 | miRBase:MI0021889 | **Mir6361** | mmu-mir-6361 | microRNA mir-6361 | ncRNA |
| chr2_166655759_166656044 | chr2 | 166655760 | 166656044 | + | 0.992891 | intron (NM_177782, intron 1 of 39) | intron (NM_177782, intron 1 of 39) | -16816 | NR_105783 | 102465188 | | NR_105783 | | **Mir6364** | mmu-mir-6364 | microRNA mir-6364 | ncRNA |
| chr2_166655687_166656150 | chr2 | 166655688 | 166656150 | + | 0.889324 | intron (NM_177782, intron 1 of 39) | intron (NM_177782, intron 1 of 39) | -16799 | NR_105783 | 102465188 | | NR_105783 | | **Mir6364** | mmu-mir-6364 | microRNA mir-6364 | ncRNA |
| chr2_166647145_166647613 | chr2 | 166647146 | 166647613 | + | 0.743933 | intron (NM_177782, intron 1 of 39) | intron (NM_177782, intron 1 of 39) | -25339 | NR_105783 | 102465188 | | NR_105783 | | **Mir6364** | mmu-mir-6364 | microRNA mir-6364 | ncRNA |
| chr16_13472537_13473219 | chr16 | 13472538 | 13473219 | + | 0.904367 | Intergenic | L2b\|LINE\|L2 | -1698 | NR_105784 | 102466153 | | NR_105784 | miRBase:MI0021894 | **Mir6365** | mmu-mir-6365 | microRNA mir-6365 | ncRNA |
| chr13_28431724_28432527 | chr13 | 28431725 | 28432527 | + | 1.23099 | Intergenic | Intergenic | 278747 | NR_105787 | 102466635 | | NR_105787 | miRBase:MI0021897 | **Mir6368** | mmu-mir-6368 | microRNA mir-6368 | ncRNA |
| chr13_28734905_28735377 | chr13 | 28734906 | 28735377 | + | 1.13783 | intron (NR_045053, intron 3 of 8) | intron (NR_045053, intron 3 of 8) | -24268 | NR_105787 | 102466635 | | NR_105787 | miRBase:MI0021897 | **Mir6368** | mmu-mir-6368 | microRNA mir-6368 | ncRNA |
| chr13_28442122_28442776 | chr13 | 28442123 | 28442776 | + | 0.899162 | Intergenic | ORR1B1\|LTR\|ERVL-MaLR | 268424 | NR_105787 | 102466635 | | NR_105787 | miRBase:MI0021897 | **Mir6368** | mmu-mir-6368 | microRNA mir-6368 | ncRNA |
| chr13_28434284_28434676 | chr13 | 28434285 | 28434676 | + | 0.841969 | Intergenic | L1MA4A\|LINE\|L1 | 276393 | NR_105787 | 102466635 | | NR_105787 | miRBase:MI0021897 | **Mir6368** | mmu-mir-6368 | microRNA mir-6368 | ncRNA |
| chr13_28442057_28442966 | chr13 | 28442058 | 28442966 | + | 0.819627 | Intergenic | ORR1B1\|LTR\|ERVL-MaLR | 268361 | NR_105787 | 102466635 | | NR_105787 | miRBase:MI0021897 | **Mir6368** | mmu-mir-6368 | microRNA mir-6368 | ncRNA |
| chr11_26184807_26185298 | chr11 | 26184808 | 26185298 | + | 1.55427 | intron (NR_027973, intron 1 of 5) | intron (NR_027973, intron 1 of 5) | -17187 | NR_105791 | 102466154 | | NR_105791 | miRBase:MI0021901 | **Mir6372** | mmu-mir-6372 | microRNA mir-6372 | ncRNA |
| chr11_26184804_26185648 | chr11 | 26184805 | 26185648 | + | 1.53539 | intron (NR_027973, intron 1 of 5) | intron (NR_027973, intron 1 of 5) | -17014 | NR_105791 | 102466154 | | NR_105791 | miRBase:MI0021901 | **Mir6372** | mmu-mir-6372 | microRNA mir-6372 | ncRNA |
| chr11_26164616_26165203 | chr11 | 26164617 | 26165203 | + | 1.00567 | intron (NR_027973, intron 1 of 5) | intron (NR_027973, intron 1 of 5) | -37330 | NR_105791 | 102466154 | | NR_105791 | miRBase:MI0021901 | **Mir6372** | mmu-mir-6372 | microRNA mir-6372 | ncRNA |
| chr11_26185356_26185648 | chr11 | 26185357 | 26185648 | + | 0.940117 | intron (NR_027973, intron 1 of 5) | intron (NR_027973, intron 1 of 5) | -16738 | NR_105791 | 102466154 | | NR_105791 | miRBase:MI0021901 | **Mir6372** | mmu-mir-6372 | microRNA mir-6372 | ncRNA |
| chr11_26164601_26165696 | chr11 | 26164602 | 26165696 | + | 0.866854 | intron (NR_027973, intron 1 of 5) | MMAR1\|DNA\|TcMar-Mariner | -37091 | NR_105791 | 102466154 | | NR_105791 | miRBase:MI0021901 | **Mir6372** | mmu-mir-6372 | microRNA mir-6372 | ncRNA |
| chr11_26128453_26128773 | chr11 | 26128454 | 26128773 | + | 0.700491 | intron (NR_027973, intron 1 of 5) | intron (NR_027973, intron 1 of 5) | -73627 | NR_105791 | 102466154 | | NR_105791 | miRBase:MI0021901 | **Mir6372** | mmu-mir-6372 | microRNA mir-6372 | ncRNA |
| chr12_15707731_15708191 | chr12 | 15707732 | 15708191 | + | 1.03266 | Intergenic | Intergenic | 93140 | NR_105809 | 102465204 | | NR_105809 | | **Mir6387** | mmu-mir-6387 | microRNA mir-6387 | ncRNA |
| chr12_15566135_15566421 | chr12 | 15566136 | 15566421 | + | 0.96694 | Intergenic | Intergenic | 234823 | NR_105809 | 102465204 | | NR_105809 | | **Mir6387** | mmu-mir-6387 | microRNA mir-6387 | ncRNA |
| chr12_15592465_15592839 | chr12 | 15592466 | 15592839 | + | 0.718138 | Intergenic | Intergenic | 208449 | NR_105809 | 102465204 | | NR_105809 | | **Mir6387** | mmu-mir-6387 | microRNA mir-6387 | ncRNA |
| chr12_15713640_15714082 | chr12 | 15713641 | 15714082 | + | 0.646616 | Intergenic | Intergenic | 87240 | NR_105809 | 102465204 | | NR_105809 | | **Mir6387** | mmu-mir-6387 | microRNA mir-6387 | ncRNA |
| chr14_105615832_105616204 | chr14 | 105615833 | 105616204 | + | 0.93442 | Intergenic | RMER15\|LTR\|ERVL | 7752 | NR_105814 | 102465207 | | NR_105814 | miRBase:MI0021924 | **Mir6390** | mmu-mir-6390 | microRNA mir-6390 | ncRNA |
| chr14_105612722_105613507 | chr14 | 105612723 | 105613507 | + | 0.866662 | Intergenic | ORR1E\|LTR\|ERVL-MaLR | 10655 | NR_105814 | 102465207 | | NR_105814 | miRBase:MI0021924 | **Mir6390** | mmu-mir-6390 | microRNA mir-6390 | ncRNA |
| chr14_105668915_105669401 | chr14 | 105668916 | 105669401 | + | 0.824906 | Intergenic | Intergenic | -45388 | NR_105814 | 102465207 | | NR_105814 | miRBase:MI0021924 | **Mir6390** | mmu-mir-6390 | microRNA mir-6390 | ncRNA |
| chr14_105612717_105613199 | chr14 | 105612718 | 105613199 | + | 0.779323 | Intergenic | Intergenic | 10812 | NR_105814 | 102465207 | | NR_105814 | miRBase:MI0021924 | **Mir6390** | mmu-mir-6390 | microRNA mir-6390 | ncRNA |
| chr14_105684321_105684709 | chr14 | 105684322 | 105684709 | + | 0.706146 | Intergenic | Intergenic | -60745 | NR_105814 | 102465207 | | NR_105814 | miRBase:MI0021924 | **Mir6390** | mmu-mir-6390 | microRNA mir-6390 | ncRNA |
| chr8_34384268_34384759 | chr8 | 34384269 | 34384759 | + | 1.17322 | Intergenic | Intergenic | -217802 | NR_105820 | 102465211 | | NR_105820 | miRBase:MI0021930 | **Mir6395** | mmu-mir-6395 | microRNA mir-6395 | ncRNA |
| chr8_34191816_34192177 | chr8 | 34191817 | 34192177 | + | 0.889982 | Intergenic | L1M4\|LINE\|L1 | -25285 | NR_105820 | 102465211 | | NR_105820 | miRBase:MI0021930 | **Mir6395** | mmu-mir-6395 | microRNA mir-6395 | ncRNA |
| chr8_34217170_34217416 | chr8 | 34217171 | 34217416 | + | 0.862738 | Intergenic | Intergenic | -50581 | NR_105820 | 102465211 | | NR_105820 | miRBase:MI0021930 | **Mir6395** | mmu-mir-6395 | microRNA mir-6395 | ncRNA |
| chr8_34282299_34282682 | chr8 | 34282300 | 34282682 | + | 0.71123 | Intergenic | RLTR51B_Mm\|LTR\|ERVK? | -115779 | NR_105820 | 102465211 | | NR_105820 | miRBase:MI0021930 | **Mir6395** | mmu-mir-6395 | microRNA mir-6395 | ncRNA |
| chr8_34282163_34282687 | chr8 | 34282164 | 34282687 | + | 0.698525 | Intergenic | RLTR51B_Mm\|LTR\|ERVK? | -115713 | NR_105820 | 102465211 | | NR_105820 | miRBase:MI0021930 | **Mir6395** | mmu-mir-6395 | microRNA mir-6395 | ncRNA |
| chr8_34385313_34385764 | chr8 | 34385314 | 34385764 | + | 0.649286 | Intergenic | Intergenic | -218827 | NR_105820 | 102465211 | | NR_105820 | miRBase:MI0021930 | **Mir6395** | mmu-mir-6395 | microRNA mir-6395 | ncRNA |
| chr4_123611264_123611597 | chr4 | 123611265 | 123611597 | + | 0.631931 | intron (NM_001199136, intron 1 of 97) | intron (NM_001199136, intron 1 of 97) | -4989 | NR_105824 | 102465214 | | NR_105824 | miRBase:MI0021934 | **Mir6398** | mmu-mir-6398 | microRNA mir-6398 | ncRNA |
| chr7_139711777_139712313 | chr7 | 139711778 | 139712313 | + | 0.782598 | Intergenic | Intergenic | -66973 | NR_105827 | 102465215 | | NR_105827 | miRBase:MI0021937 | **Mir6401** | mmu-mir-6401 | microRNA mir-6401 | ncRNA |
| chr11_68213961_68214429 | chr11 | 68213962 | 68214429 | + | 0.632549 | intron (NM_008744, intron 6 of 6) | intron (NM_008744, intron 6 of 6) | -6697 | NR_105833 | 102465219 | | NR_105833 | miRBase:MI0021943 | **Mir6406** | mmu-mir-6406 | microRNA mir-6406 | ncRNA |
| chr10_76016679_76017303 | chr10 | 76016680 | 76017303 | + | 2.11724 | Intergenic | Intergenic | 8065 | NR_105836 | 102466646 | | NR_105836 | miRBase:MI0021946 | **Mir6409** | mmu-mir-6409 | microRNA mir-6409 | ncRNA |
| chr10_76012979_76013502 | chr10 | 76012980 | 76013502 | + | 1.0262 | Intergenic | L1_Mur2\|LINE\|L1 | 4315 | NR_105836 | 102466646 | | NR_105836 | miRBase:MI0021946 | **Mir6409** | mmu-mir-6409 | microRNA mir-6409 | ncRNA |
| chr10_20285632_20285916 | chr10 | 20285633 | 20285916 | + | 0.764529 | Intergenic | Intergenic | 11914 | NR_105840 | 102466647 | | NR_105840 | miRBase:MI0021950 | **Mir6413** | mmu-mir-6413 | microRNA mir-6413 | ncRNA |
| chr5_92467422_92468015 | chr5 | 92467423 | 92468015 | + | 0.856114 | intron (NM_007644, intron 2 of 11) | B1_Mus1\|SINE\|Alu | 9162 | NR_105843 | 102466159 | | NR_105843 | miRBase:MI0021953 | **Mir6415** | mmu-mir-6415 | microRNA mir-6415 | ncRNA |
| chr7_64234351_64234929 | chr7 | 64234352 | 64234929 | + | 0.895736 | intron (NM_001039104, intron 17 of 26) | MTEa\|LTR\|ERVL-MaLR | 15459 | NR_105844 | 102465226 | | NR_105844 | miRBase:MI0021954 | **Mir6416** | mmu-mir-6416 | microRNA mir-6416 | ncRNA |
| chr5_52111102_52111559 | chr5 | 52111103 | 52111559 | + | 1.13874 | Intergenic | (CA)n\|Simple_repeat\|Simple_repeat | 24686 | NR_105845 | 102466648 | | NR_105845 | miRBase:MI0021955 | **Mir6417** | mmu-mir-6417 | microRNA mir-6417 | ncRNA |
| chr5_52111706_52111944 | chr5 | 52111707 | 52111944 | + | 1.04296 | Intergenic | Intergenic | 24192 | NR_105845 | 102466648 | | NR_105845 | miRBase:MI0021955 | **Mir6417** | mmu-mir-6417 | microRNA mir-6417 | ncRNA |
| chr5_52145771_52146049 | chr5 | 52145772 | 52146049 | + | 1.03107 | Intergenic | Intergenic | -9893 | NR_105845 | 102466648 | | NR_105845 | miRBase:MI0021955 | **Mir6417** | mmu-mir-6417 | microRNA mir-6417 | ncRNA |
| chr5_52149215_52149487 | chr5 | 52149216 | 52149487 | + | 0.895486 | TTS (NM_007839) | TTS (NM_007839) | -13334 | NR_105845 | 102466648 | | NR_105845 | miRBase:MI0021955 | **Mir6417** | mmu-mir-6417 | microRNA mir-6417 | ncRNA |
| chr5_52149171_52149592 | chr5 | 52149172 | 52149592 | + | 0.874741 | TTS (NM_007839) | TTS (NM_007839) | -13365 | NR_105845 | 102466648 | | NR_105845 | miRBase:MI0021955 | **Mir6417** | mmu-mir-6417 | microRNA mir-6417 | ncRNA |
| chr5_52149094_52149693 | chr5 | 52149095 | 52149693 | + | 0.867363 | TTS (NM_007839) | TTS (NM_007839) | -13377 | NR_105845 | 102466648 | | NR_105845 | miRBase:MI0021955 | **Mir6417** | mmu-mir-6417 | microRNA mir-6417 | ncRNA |
| chr5_52112226_52112455 | chr5 | 52112227 | 52112455 | + | 0.764732 | Intergenic | Intergenic | 23676 | NR_105845 | 102466648 | | NR_105845 | miRBase:MI0021955 | **Mir6417** | mmu-mir-6417 | microRNA mir-6417 | ncRNA |
| chr5_52112961_52113527 | chr5 | 52112962 | 52113527 | + | 0.735356 | Intergenic | CpG | 22773 | NR_105845 | 102466648 | | NR_105845 | miRBase:MI0021955 | **Mir6417** | mmu-mir-6417 | microRNA mir-6417 | ncRNA |
| chr5_52112226_52112641 | chr5 | 52112227 | 52112641 | + | 0.734184 | Intergenic | Intergenic | 23583 | NR_105845 | 102466648 | | NR_105845 | miRBase:MI0021955 | **Mir6417** | mmu-mir-6417 | microRNA mir-6417 | ncRNA |
| chr5_52112937_52113708 | chr5 | 52112938 | 52113708 | + | 0.691971 | Intergenic | CpG | 22694 | NR_105845 | 102466648 | | NR_105845 | miRBase:MI0021955 | **Mir6417** | mmu-mir-6417 | microRNA mir-6417 | ncRNA |
| chr2_17391437_17391778 | chr2 | 17391438 | 17391778 | + | 0.987736 | intron (NM_028757, intron 4 of 6) | intron (NM_028757, intron 4 of 6) | -171906 | NR_105847 | 102465964 | | NR_105847 | miRBase:MI0021957 | **Mir6419** | mmu-mir-6419 | microRNA mir-6419 | ncRNA |
| chr2_17390867_17391301 | chr2 | 17390868 | 17391301 | + | 0.980252 | intron (NM_028757, intron 4 of 6) | intron (NM_028757, intron 4 of 6) | -172430 | NR_105847 | 102465964 | | NR_105847 | miRBase:MI0021957 | **Mir6419** | mmu-mir-6419 | microRNA mir-6419 | ncRNA |
| chr2_17409265_17409628 | chr2 | 17409266 | 17409628 | + | 0.740662 | intron (NM_028757, intron 4 of 6) | intron (NM_028757, intron 4 of 6) | -154067 | NR_105847 | 102465964 | | NR_105847 | miRBase:MI0021957 | **Mir6419** | mmu-mir-6419 | microRNA mir-6419 | ncRNA |
| chr2_17386414_17386720 | chr2 | 17386415 | 17386720 | + | 0.727376 | intron (NM_028757, intron 4 of 6) | intron (NM_028757, intron 4 of 6) | -176947 | NR_105847 | 102465964 | | NR_105847 | miRBase:MI0021957 | **Mir6419** | mmu-mir-6419 | microRNA mir-6419 | ncRNA |
| chr17_64390864_64391209 | chr17 | 64390865 | 64391209 | + | 0.723966 | Intergenic | B3A\|SINE\|B2 | -49769 | NR_105848 | 102465228 | | NR_105848 | miRBase:MI0021958 | **Mir6420** | mmu-mir-6420 | microRNA mir-6420 | ncRNA |
| chr11_117092668_117093209 | chr11 | 117092669 | 117093209 | + | 0.694742 | Intergenic | Intergenic | 15591 | NR_105853 | 102465258 | | NR_105853 | | **Mir6516** | mmu-mir-6516 | microRNA mir-6516 | ncRNA |
| chr7_25109190_25109772 | chr7 | 25109191 | 25109772 | + | 1.0968 | intron (NM_011138, intron 5 of 13) | intron (NM_011138, intron 5 of 13) | -12306 | NR_105855 | 102465968 | | NR_105855 | | **Mir6537** | mmu-mir-6537 | microRNA mir-6537 | ncRNA |
| chr14_62872005_62872390 | chr14 | 62872006 | 62872390 | + | 1.00485 | intron (NM_175546, intron 1 of 11) | URR1B\|DNA\|hAT-Charlie | 6656 | NR_105860 | 102465311 | | NR_105860 | | **Mir6541** | mmu-mir-6541 | microRNA mir-6541 | ncRNA |
| chr14_62871877_62872490 | chr14 | 62871878 | 62872490 | + | 0.947041 | intron (NM_175546, intron 1 of 11) | URR1B\|DNA\|hAT-Charlie | 6642 | NR_105860 | 102465311 | | NR_105860 | | **Mir6541** | mmu-mir-6541 | microRNA mir-6541 | ncRNA |
| chr14_62874458_62874698 | chr14 | 62874459 | 62874698 | + | 0.677007 | intron (NM_175546, intron 1 of 11) | intron (NM_175546, intron 1 of 11) | 9036 | NR_105860 | 102465311 | | NR_105860 | | **Mir6541** | mmu-mir-6541 | microRNA mir-6541 | ncRNA |
| chr1_171074135_171074666 | chr1 | 171074136 | 171074666 | + | 1.33759 | Intergenic | Intergenic | -9749 | NR_106103 | 102466998 | | NR_106103 | miRBase:MI0024983 | **Mir6546** | mmu-mir-6546 | microRNA mir-6546 | ncRNA |
| chr1_171066675_171066919 | chr1 | 171066676 | 171066919 | + | 1.00115 | Intergenic | Intergenic | -2145 | NR_106103 | 102466998 | | NR_106103 | miRBase:MI0024983 | **Mir6546** | mmu-mir-6546 | microRNA mir-6546 | ncRNA |
| chr1_171081910_171082164 | chr1 | 171081911 | 171082164 | + | 0.925573 | Intergenic | Intergenic | -17385 | NR_106103 | 102466998 | | NR_106103 | miRBase:MI0024983 | **Mir6546** | mmu-mir-6546 | microRNA mir-6546 | ncRNA |
| chr1_171081661_171082186 | chr1 | 171081662 | 171082186 | + | 0.85735 | Intergenic | tRNA-Gly-GGY\|tRNA\|tRNA | -17272 | NR_106103 | 102466998 | | NR_106103 | miRBase:MI0024983 | **Mir6546** | mmu-mir-6546 | microRNA mir-6546 | ncRNA |

| | | | | | | | | | | | | | | | | | |
|---|---|---|---|---|---|---|---|---|---|---|---|---|---|---|---|---|---|
| chr1_171064417_171064770 | chr1 | 171064418 | 171064770 | + | 0.830604 | promoter-TSS (NR_106103) | promoter-TSS (NR_106103) | 58 | NR_106103 | 102466998 | | NR_106103 | miRBase:MI0024983 | **Mir6546** | mmu-mir-6546 | microRNA mir-6546 | ncRNA |
| chr12_91149545_91150078 | chr12 | 91149546 | 91150078 | + | 0.99062 | intron (NM_181815, intron 20 of 25) | intron (NM_181815, intron 20 of 25) | 39583 | NR_030473 | 735273 | | NR_030473 | | **Mir669c** | Mirn669c\|Mirn669c-1 | microRNA 669c | ncRNA |
| chr2_10783090_10783436 | chr2 | 10783091 | 10783436 | + | 1.1711 | Intergenic | Intergenic | 265659 | NR_035417 | 100316668 | | NR_035417 | | **Mir669i** | Mirn669i\|mmu-mir-669i | microRNA 669i | ncRNA |
| chr2_94288148_94288482 | chr2 | 94288149 | 94288482 | + | 1.29577 | Intergenic | Intergenic | -26916 | NR_030431 | 735259 | | NR_030431 | miRBase:MI0004295 | **Mir670** | Mirn670\|mmu-mir-670 | microRNA 670 | ncRNA |
| chr2_94288112_94288646 | chr2 | 94288113 | 94288646 | + | 1.20505 | Intergenic | Intergenic | -26980 | NR_030431 | 735259 | | NR_030431 | miRBase:MI0004295 | **Mir670** | Mirn670\|mmu-mir-670 | microRNA 670 | ncRNA |
| chr16_20192048_20192693 | chr16 | 20192049 | 20192693 | + | 0.820793 | intron (NM_001033237, intron 12 of 29) | RMER15\|LTR\|ERVL | -16137 | NR_030454 | 735274 | | NR_030454 | | **Mir684-1** | Mirn684-1 | microRNA 684-1 | ncRNA |
| chr14_73218247_73218598 | chr14 | 73218248 | 73218598 | + | 1.14798 | intron (NM_009029, intron 17 of 26) | B3A\|SINE\|B2 | -11572 | NR_030459 | 751541 | | NR_030459 | | **Mir687** | Mirn687\|mir-687\|mmu-mir-687 | microRNA 687 | ncRNA |
| chr14_73175573_73176223 | chr14 | 73175574 | 73176223 | + | 0.803552 | intron (NM_001170694, intron 12 of 14) | intron (NM_001170694, intron 12 of 14) | 30953 | NR_030459 | 751541 | | NR_030459 | | **Mir687** | Mirn687\|mir-687\|mmu-mir-687 | microRNA 687 | ncRNA |
| chr15_102671693_102672054 | chr15 | 102671694 | 102672054 | + | 0.646814 | promoter-TSS (NM_026468) | promoter-TSS (NM_026468) | -8 | NR_030460 | 751542 | | NR_030460 | miRBase:MI0004653 | **Mir688** | Mirn688\|mmu-mir-688 | microRNA 688 | ncRNA |
| chr10_60161993_60162899 | chr10 | 60161994 | 60162899 | + | 0.909796 | Intergenic | LTRIS2\|LTR\|ERV1 | 35670 | NR_105871 | 102465546 | | NR_105871 | | **Mir6906** | mmu-mir-6906 | microRNA mir-6906 | ncRNA |
| chr11_74881016_74881363 | chr11 | 74881017 | 74881363 | + | 0.697368 | intron (NM_197943, intron 2 of 22) | RSINE1\|SINE\|B4 | -12996 | NR_105891 | 102465558 | | NR_105891 | | **Mir6926** | mmu-mir-6926 | microRNA mir-6926 | ncRNA |
| chr11_119159517_119160405 | chr11 | 119159518 | 119160405 | + | 0.859443 | intron (NM_172443, intron 3 of 11) | Lx5c\|LINE\|L1 | -5824 | NR_105899 | 102465563 | | NR_105899 | | **Mir6934** | mmu-mir-6934 | microRNA mir-6934 | ncRNA |
| chr11_119142346_119142790 | chr11 | 119142347 | 119142790 | + | 0.722847 | TTS (NM_172443) | TTS (NM_172443) | 11569 | NR_105899 | 102465563 | | NR_105899 | | **Mir6934** | mmu-mir-6934 | microRNA mir-6934 | ncRNA |
| chr12_28678379_28678583 | chr12 | 28678380 | 28678583 | + | 0.881393 | promoter-TSS (NR_105902) | promoter-TSS (NR_105902) | -844 | NR_105902 | 102465980 | | NR_105902 | | **Mir6937** | mmu-mir-6937 | microRNA mir-6937 | ncRNA |
| chr13_21384749_21385485 | chr13 | 21384750 | 21385485 | + | 1.63644 | Intergenic | MERV1_LTR\|LTR\|ERV 1 | 11521 | NR_105907 | 102465568 | | NR_105907 | miRBase:MI0022789 | **Mir6942** | mmu-mir-6942 | microRNA mir-6942 | ncRNA |
| chr14_20686627_20686994 | chr14 | 20686628 | 20686994 | + | 0.80955 | intron (NM_172596, intron 7 of 22) | intron (NM_172596, intron 7 of 22) | -3824 | NR_105911 | 102465571 | | NR_105911 | | **Mir6946** | mmu-mir-6946 | microRNA mir-6946 | ncRNA |
| chr14_20689038_20689394 | chr14 | 20689039 | 20689394 | + | 0.665741 | intron (NM_172596, intron 11 of 22) | intron (NM_172596, intron 11 of 22) | -1419 | NR_105911 | 102465571 | | NR_105911 | | **Mir6946** | mmu-mir-6946 | microRNA mir-6946 | ncRNA |
| chr15_76251683_76252157 | chr15 | 76251684 | 76252157 | + | 0.882382 | Intergenic | Intergenic | 3729 | NR_105918 | 102465575 | | NR_105918 | | **Mir6953** | mmu-mir-6953 | microRNA mir-6953 | ncRNA |
| chr15_89277261_89277919 | chr15 | 89277262 | 89277919 | + | 1.48399 | intron (NM_026813, intron 15 of 24) | intron (NM_026813, intron 15 of 24) | 28142 | NR_105924 | 102465981 | | NR_105924 | | **Mir6959** | mmu-mir-6959 | microRNA mir-6959 | ncRNA |
| chr17_24240056_24240463 | chr17 | 24240057 | 24240463 | + | 0.764874 | TTS (NR_105930) | TTS (NR_105930) | 687 | NR_105930 | 102465582 | | NR_105930 | | **Mir6965** | mmu-mir-6965 | microRNA mir-6965 | ncRNA |
| chr18_37891974_37893051 | chr18 | 37891975 | 37893051 | + | 0.856717 | intron (NM_007858, intron 13 of 26) | intron (NM_007858, intron 13 of 26) | -37853 | NR_105945 | 102465592 | | NR_105945 | | **Mir6979** | mmu-mir-6979 | microRNA mir-6979 | ncRNA |
| chr18_37892264_37893050 | chr18 | 37892265 | 37893050 | + | 0.842806 | intron (NM_007858, intron 13 of 26) | MTC\|LTR\|ERVL-MaLR | -37997 | NR_105945 | 102465592 | | NR_105945 | | **Mir6979** | mmu-mir-6979 | microRNA mir-6979 | ncRNA |
| chr4_124768877_124769760 | chr4 | 124768878 | 124769760 | + | 0.833045 | intron (NM_008385, intron 7 of 23) | MIRc\|SINE\|MIR | 25538 | NR_030480 | 735263 | | NR_030480 | | **Mir698** | Mirn698\|mir-698\|mmu-mir-698 | microRNA 698 | ncRNA |
| chr19_7425405_7425781 | chr19 | 7425406 | 7425781 | + | 1.29105 | TTS (NM_001271487) | TTS (NM_001271487) | 3020 | NR_105957 | 102466781 | | NR_105957 | | **Mir6991** | mmu-mir-6991 | microRNA mir-6991 | ncRNA |
| chr19_7425177_7425781 | chr19 | 7425178 | 7425781 | + | 1.206 | TTS (NM_001271487) | TTS (NM_001271487) | 2906 | NR_105957 | 102466781 | | NR_105957 | | **Mir6991** | mmu-mir-6991 | microRNA mir-6991 | ncRNA |
| chr4_135418856_135419467 | chr4 | 135418857 | 135419467 | + | 1.29142 | intron (NM_022980, intron 3 of 4) | ORR1F\|LTR\|ERVL-MaLR | -2529 | NR_030481 | 735285 | | NR_030481 | | **Mir700** | Mirn700\|mir-700\|mmu-mir-700 | microRNA 700 | ncRNA |
| chr2_93425696_93425976 | chr2 | 93425697 | 93425976 | + | 0.891369 | intron (NM_007656, intron 6 of 9) | intron (NM_007656, intron 6 of 9) | -3829 | NR_105967 | 102465605 | | NR_105967 | | **Mir7001** | mmu-mir-7001 | microRNA mir-7001 | ncRNA |
| chr3_31038658_31039157 | chr3 | 31038659 | 31039157 | + | 1.24341 | promoter-TSS (NR_105975) | promoter-TSS (NR_105975) | -58 | NR_105975 | 102465610 | | NR_105975 | | **Mir7008** | mmu-mir-7008 | microRNA mir-7008 | ncRNA |
| chr3_36467643_36468038 | chr3 | 36467644 | 36468038 | + | 1.51406 | intron (NM_009673, intron 2 of 12) | intron (NM_009673, intron 2 of 12) | 7683 | NR_105976 | 102466786 | | NR_105976 | miRBase:MI0022858 | **Mir7009** | mmu-mir-7009 | microRNA mir-7009 | ncRNA |
| chr5_111124534_111125630 | chr5 | 111124535 | 111125630 | + | 0.830647 | intron (NM_001267622, intron 5 of 22) | intron (NM_001267622, intron 5 of 22) | 120938 | NR_030482 | 735278 | | NR_030482 | | **Mir701** | Mirn701\|mmu-mir-701 | microRNA 701 | ncRNA |
| chr4_120987185_120987700 | chr4 | 120987186 | 120987700 | + | 0.689101 | intron (NM_133716, intron 1 of 9) | intron (NM_133716, intron 1 of 9) | -14121 | NR_105982 | 102465614 | | NR_105982 | | **Mir7015** | mmu-mir-7015 | microRNA mir-7015 | ncRNA |
| chr4_133182436_133183094 | chr4 | 133182437 | 133183094 | + | 0.779198 | intron (NM_153423, intron 2 of 8) | B1F2\|SINE\|Alu | -12890 | NR_105984 | 102466215 | | NR_105984 | | **Mir7017** | mmu-mir-7017 | microRNA mir-7017 | ncRNA |
| chr4_139640739_139641133 | chr4 | 139640740 | 139641133 | + | 1.01786 | intron (NM_175438, intron 9 of 14) | intron (NM_175438, intron 9 of 14) | -3106 | NR_105987 | 102465617 | | NR_105987 | | **Mir7020** | mmu-mir-7020 | microRNA mir-7020 | ncRNA |
| chr4_139639692_139640052 | chr4 | 139639693 | 139640052 | + | 0.710582 | intron (NM_175438, intron 8 of 14) | intron (NM_175438, intron 8 of 14) | -4170 | NR_105987 | 102465617 | | NR_105987 | | **Mir7020** | mmu-mir-7020 | microRNA mir-7020 | ncRNA |
| chr4_143152792_143153602 | chr4 | 143152793 | 143153602 | + | 0.757788 | intron (NM_001256380, intron 6 of 7) | intron (NM_001256380, intron 6 of 7) | -17035 | NR_105988 | 102465618 | | NR_105988 | | **Mir7021** | mmu-mir-7021 | microRNA mir-7021 | ncRNA |
| chr5_75315822_75316347 | chr5 | 75315823 | 75316347 | + | 0.964054 | Intergenic | L1MEd\|LINE\|L1 | 139254 | NR_105992 | 102465620 | | NR_105992 | | **Mir7025** | mmu-mir-7025 | microRNA mir-7025 | ncRNA |
| chr6_94634967_94635793 | chr6 | 94634968 | 94635793 | + | 1.12636 | intron (NM_008377, intron 3 of 18) | intron (NM_008377, intron 3 of 18) | -28965 | NR_106008 | 102466992 | | NR_106008 | | **Mir7041** | mmu-mir-7041 | microRNA mir-7041 | ncRNA |
| chr6_94634421_94635869 | chr6 | 94634422 | 94635869 | + | 0.640668 | intron (NM_008377, intron 3 of 18) | intron (NM_008377, intron 3 of 18) | -28730 | NR_106008 | 102466992 | | NR_106008 | | **Mir7041** | mmu-mir-7041 | microRNA mir-7041 | ncRNA |
| chr7_47103647_47104381 | chr7 | 47103648 | 47104381 | + | 1.24356 | intron (NM_001163565, intron 2 of 13) | MLT1F2\|LTR\|ERVL-MaLR | -20790 | NR_106023 | 102465639 | | NR_106023 | | **Mir7056** | mmu-mir-7056 | microRNA mir-7056 | ncRNA |
| chr7_47098060_47098375 | chr7 | 47098061 | 47098375 | + | 0.829935 | intron (NM_001163565, intron 2 of 13) | intron (NM_001163565, intron 2 of 13) | -14994 | NR_106023 | 102465639 | | NR_106023 | | **Mir7056** | mmu-mir-7056 | microRNA mir-7056 | ncRNA |
| chr7_66304064_66304561 | chr7 | 66304065 | 66304561 | + | 0.783457 | intron (NM_146191, intron 10 of 33) | intron (NM_146191, intron 10 of 33) | 77411 | NR_106024 | 102465640 | | NR_106024 | miRBase:MI0022906 | **Mir7057** | mmu-mir-7057 | microRNA mir-7057 | ncRNA |
| chr7_126348508_126349193 | chr7 | 126348509 | 126349193 | + | 1.31906 | Intergenic | Intergenic | 19194 | NR_106025 | 102466797 | | NR_106025 | | **Mir7058** | mmu-mir-7058 | microRNA mir-7058 | ncRNA |
| chr7_126577958_126578387 | chr7 | 126577959 | 126578387 | + | 0.655413 | intron (NM_009907, intron 8 of 14) | intron (NM_009907, intron 8 of 14) | 1359 | NR_106026 | 102465641 | | NR_106026 | | **Mir7059** | mmu-mir-7059 | microRNA mir-7059 | ncRNA |
| chr6_120010595_120010862 | chr6 | 120010596 | 120010862 | + | 0.741908 | intron (NM_001199083, intron 1 of 27) | RSINE1\|SINE\|B4 | 23582 | NR_030487 | 735290 | | NR_030487 | | **Mir706** | Mirn706\|mir-706\|mmu-mir-706 | microRNA 706 | ncRNA |
| chr8_94493187_94493871 | chr8 | 94493188 | 94493871 | + | 1.16601 | intron (NM_001033207, intron 20 of 48) | intron (NM_001033207, intron 20 of 48) | -11517 | NR_106040 | 102466800 | | NR_106040 | | **Mir7072** | mmu-mir-7072 | microRNA mir-7072 | ncRNA |
| chr9_45262117_45262859 | chr9 | 45262118 | 45262859 | + | 0.954202 | intron (NM_008348, intron 5 of 6) | intron (NM_008348, intron 5 of 6) | 4219 | NR_106054 | 102466803 | | NR_106054 | | **Mir7086** | mmu-mir-7086 | microRNA mir-7086 | ncRNA |
| chr9_45264282_45264712 | chr9 | 45264283 | 45264712 | + | 0.758909 | intron (NM_008348, intron 4 of 6) | intron (NM_008348, intron 4 of 6) | 2210 | NR_106054 | 102466803 | | NR_106054 | | **Mir7086** | mmu-mir-7086 | microRNA mir-7086 | ncRNA |
| chr9_109907925_109908527 | chr9 | 109907926 | 109908527 | + | 0.805211 | Intergenic | Intergenic | 18583 | NR_106057 | 102465660 | | NR_106057 | | **Mir7089** | mmu-mir-7089 | microRNA mir-7089 | ncRNA |
| chr15_27592892_27593306 | chr15 | 27592893 | 27593306 | + | 0.695695 | intron (NM_020332, intron 11 of 11) | intron (NM_020332, intron 11 of 11) | 21605 | NR_106065 | 102466808 | | NR_106065 | | **Mir7117** | mmu-mir-7117 | microRNA mir-7117 | ncRNA |
| chr5_136458419_136459231 | chr5 | 136458420 | 136459231 | + | 0.706602 | intron (NM_001291239, intron 2 of 22) | intron (NM_001291239, intron 2 of 22) | -83022 | NR_030500 | 723975 | | NR_030500 | | **Mir721** | Mirn721\|miRY\|mir-721\|mmu-mir-721 | microRNA 721 | ncRNA |
| chr15_25948819_25949460 | chr15 | 25948820 | 25949460 | + | 0.929849 | TTS (NR_106071) | TTS (NR_106071) | 913 | NR_106071 | 102465698 | | NR_106071 | | **Mir7212** | mmu-mir-7212 | microRNA mir-7212 | ncRNA |
| chr18_68119102_68119668 | chr18 | 68119103 | 68119668 | + | 1.38365 | intron (NM_172631, intron 3 of 5) | intron (NM_172631, intron 3 of 5) | 141588 | NR_106078 | 102465702 | | NR_106078 | miRBase:MI0023714 | **Mir7219** | mmu-mir-7219 | microRNA mir-7219 | ncRNA |
| chr18_68122912_68123517 | chr18 | 68122913 | 68123517 | + | 1.12225 | intron (NM_172631, intron 3 of 5) | intron (NM_172631, intron 3 of 5) | 137758 | NR_106078 | 102465702 | | NR_106078 | miRBase:MI0023714 | **Mir7219** | mmu-mir-7219 | microRNA mir-7219 | ncRNA |

| | | | | | | | | | | | | | | | | | |
|---|---|---|---|---|---|---|---|---|---|---|---|---|---|---|---|---|---|
| chr18_68122995_68123412 | chr18 | 68122996 | 68123412 | + | 1.12073 | intron (NM_172631, intron 3 of 5) | intron (NM_172631, intron 3 of 5) | 137769 | NR_106078 | 102465702 | | NR_106078 | miRBase:MI0023714 | **Mir7219** | mmu-mir-7219 | microRNA mir-7219 | ncRNA |
| chr11_108039027_108039349 | chr11 | 108039028 | 108039349 | + | 0.676236 | intron (NM_011101, intron 5 of 16) | intron (NM_011101, intron 5 of 16) | 42340 | NR_106082 | 102465705 | | NR_106082 | miRBase:MI0023718 | **Mir7223** | mmu-mir-7223 | microRNA mir-7223 | ncRNA |
| chr6_127789907_127790485 | chr6 | 127789908 | 127790485 | + | 0.862978 | Intergenic | Intergenic | -2053 | NR_106092 | 102466996 | | NR_106092 | | **Mir7233** | mmu-mir-7233 | microRNA mir-7233 | ncRNA |
| chr11_97131724_97131954 | chr11 | 97131725 | 97131954 | + | 1.01985 | Intergenic | Intergenic | 14580 | NR_106094 | 102466822 | | NR_106094 | | **Mir7235** | mmu-mir-7235 | microRNA mir-7235 | ncRNA |
| chr11_97136621_97136864 | chr11 | 97136622 | 97136864 | + | 0.665257 | 3' UTR (NM_198100, exon 9 of 9) | 3' UTR (NM_198100, exon 9 of 9) | 9676 | NR_106094 | 102466822 | | NR_106094 | | **Mir7235** | mmu-mir-7235 | microRNA mir-7235 | ncRNA |
| chr11_65733700_65734313 | chr11 | 65733701 | 65734313 | + | 1.1876 | TTS (NR_030417) | TTS (NR_030417) | 825 | NR_030417 | 791070 | | NR_030417 | | **Mir744** | Mirn744\|mir-744\|mmu-mir-744 | microRNA 744 | ncRNA |
| chr11_65735413_65735838 | chr11 | 65735414 | 65735838 | + | 1.07812 | promoter-TSS (NR_030417) | promoter-TSS (NR_030417) | -794 | NR_030417 | 791070 | | NR_030417 | | **Mir744** | Mirn744\|mir-744\|mmu-mir-744 | microRNA 744 | ncRNA |
| chr11_65680284_65680912 | chr11 | 65680285 | 65680912 | + | 0.916149 | Intergenic | L1MB2\|LINE\|L1 | 54234 | NR_030417 | 791070 | | NR_030417 | | **Mir744** | Mirn744\|mir-744\|mmu-mir-744 | microRNA 744 | ncRNA |
| chr11_65734620_65734987 | chr11 | 65734621 | 65734987 | + | 0.780964 | promoter-TSS (NR_030417) | promoter-TSS (NR_030417) | 28 | NR_030417 | 791070 | | NR_030417 | | **Mir744** | Mirn744\|mir-744\|mmu-mir-744 | microRNA 744 | ncRNA |
| chr4_109018272_109018672 | chr4 | 109018273 | 109018672 | + | 0.802586 | TTS (NR_030432) | TTS (NR_030432) | 817 | NR_030432 | 791075 | | NR_030432 | | **Mir761** | Mirn761\|mir-761\|mmu-mir-761 | microRNA 761 | ncRNA |
| chr7_127678826_127679373 | chr7 | 127678827 | 127679373 | + | 1.17265 | Intergenic | Intergenic | -29387 | NR_030428 | 791073 | | NR_030428 | miRBase:MI0004215 | **Mir762** | Mirn762\|mir-762\|mmu-mir-762 | microRNA 762 | ncRNA |
| chr9_15317877_15318680 | chr9 | 15317878 | 15318680 | + | 0.780815 | intron (NM_176976, intron 23 of 26) | intron (NM_176976, intron 23 of 26) | 5668 | NR_106108 | 102465758 | | NR_106108 | | **Mir7650** | mmu-mir-7650 | microRNA mir-7650 | ncRNA |
| chr9_15317588_15318680 | chr9 | 15317589 | 15318680 | + | 0.713954 | intron (NM_176976, intron 23 of 26) | intron (NM_176976, intron 23 of 26) | 5523 | NR_106108 | 102465758 | | NR_106108 | | **Mir7650** | mmu-mir-7650 | microRNA mir-7650 | ncRNA |
| chr11_78179181_78179378 | chr11 | 78179182 | 78179378 | + | 0.730853 | TTS (NR_106113) | TTS (NR_106113) | 453 | NR_106113 | 102465761 | | NR_106113 | | **Mir7653** | mmu-mir-7653 | microRNA mir-7653 | ncRNA |
| chr9_94951816_94952855 | chr9 | 94951817 | 94952855 | + | 1.49535 | intron (NM_177909, intron 8 of 15) | Lx8b\|LINE\|L1 | 200507 | NR_106116 | 102465763 | | NR_106116 | | **Mir7656** | mmu-mir-7656 | microRNA mir-7656 | ncRNA |
| chr9_94803049_94803428 | chr9 | 94803050 | 94803428 | + | 0.715242 | intron (NM_177909, intron 4 of 15) | B3A\|SINE\|B2 | 51410 | NR_106116 | 102465763 | | NR_106116 | | **Mir7656** | mmu-mir-7656 | microRNA mir-7656 | ncRNA |
| chr3_122566034_122566431 | chr3 | 122566035 | 122566431 | + | 0.813841 | intron (NM_153118, intron 5 of 13) | HAL1\|LINE\|L1 | -24552 | NR_106117 | 102465764 | | NR_106117 | | **Mir7657** | mmu-mir-7657 | microRNA mir-7657 | ncRNA |
| chr6_108448012_108448872 | chr6 | 108448013 | 108448872 | + | 1.67289 | intron (NM_010585, intron 44 of 61) | intron (NM_010585, intron 44 of 61) | -41301 | NR_106121 | 102465766 | | NR_106121 | | **Mir7661** | mmu-mir-7661 | microRNA mir-7661 | ncRNA |
| chr6_108544966_108545247 | chr6 | 108544967 | 108545247 | + | 1.37278 | intron (NM_010585, intron 61 of 61) | intron (NM_010585, intron 61 of 61) | 55364 | NR_106121 | 102465766 | | NR_106121 | | **Mir7661** | mmu-mir-7661 | microRNA mir-7661 | ncRNA |
| chr6_108554310_108554706 | chr6 | 108554311 | 108554706 | + | 1.36595 | Intergenic | Intergenic | 64765 | NR_106121 | 102465766 | | NR_106121 | | **Mir7661** | mmu-mir-7661 | microRNA mir-7661 | ncRNA |
| chr6_108481530_108483005 | chr6 | 108481531 | 108483005 | + | 1.30928 | intron (NM_010585, intron 47 of 61) | intron (NM_010585, intron 47 of 61) | -7475 | NR_106121 | 102465766 | | NR_106121 | | **Mir7661** | mmu-mir-7661 | microRNA mir-7661 | ncRNA |
| chr6_108529539_108529841 | chr6 | 108529540 | 108529841 | + | 1.26139 | intron (NM_010585, intron 60 of 61) | intron (NM_010585, intron 60 of 61) | 39947 | NR_106121 | 102465766 | | NR_106121 | | **Mir7661** | mmu-mir-7661 | microRNA mir-7661 | ncRNA |
| chr6_108552653_108553398 | chr6 | 108552654 | 108553398 | + | 1.2089 | Intergenic | Intergenic | 63283 | NR_106121 | 102465766 | | NR_106121 | | **Mir7661** | mmu-mir-7661 | microRNA mir-7661 | ncRNA |
| chr6_108482320_108482994 | chr6 | 108482321 | 108482994 | + | 1.10636 | intron (NM_010585, intron 47 of 61) | intron (NM_010585, intron 47 of 61) | -7086 | NR_106121 | 102465766 | | NR_106121 | | **Mir7661** | mmu-mir-7661 | microRNA mir-7661 | ncRNA |
| chr6_108510274_108510761 | chr6 | 108510275 | 108510761 | + | 1.08603 | intron (NM_010585, intron 54 of 61) | intron (NM_010585, intron 54 of 61) | 20775 | NR_106121 | 102465766 | | NR_106121 | | **Mir7661** | mmu-mir-7661 | microRNA mir-7661 | ncRNA |
| chr6_108542155_108542476 | chr6 | 108542156 | 108542476 | + | 1.01564 | intron (NM_010585, intron 61 of 61) | intron (NM_010585, intron 61 of 61) | 52573 | NR_106121 | 102465766 | | NR_106121 | | **Mir7661** | mmu-mir-7661 | microRNA mir-7661 | ncRNA |
| chr6_108554966_108555182 | chr6 | 108554967 | 108555182 | + | 0.967206 | Intergenic | Intergenic | 65331 | NR_106121 | 102465766 | | NR_106121 | | **Mir7661** | mmu-mir-7661 | microRNA mir-7661 | ncRNA |
| chr6_108545488_108545711 | chr6 | 108545489 | 108545711 | + | 0.90192 | intron (NM_010585, intron 61 of 61) | intron (NM_010585, intron 61 of 61) | 55857 | NR_106121 | 102465766 | | NR_106121 | | **Mir7661** | mmu-mir-7661 | microRNA mir-7661 | ncRNA |
| chr6_108508876_108509212 | chr6 | 108508877 | 108509212 | + | 0.697544 | intron (NM_010585, intron 54 of 61) | intron (NM_010585, intron 54 of 61) | 19301 | NR_106121 | 102465766 | | NR_106121 | | **Mir7661** | mmu-mir-7661 | microRNA mir-7661 | ncRNA |
| chr6_108553471_108554062 | chr6 | 108553472 | 108554062 | + | 0.654772 | Intergenic | Intergenic | 64024 | NR_106121 | 102465766 | | NR_106121 | | **Mir7661** | mmu-mir-7661 | microRNA mir-7661 | ncRNA |
| chr1_53908173_53908733 | chr1 | 53908174 | 53908733 | + | 0.772033 | intron (NM_001001883, intron 13 of 28) | intron (NM_001001883, intron 13 of 28) | 59029 | NR_106149 | 102465783 | | NR_106149 | miRBase:MI0025029 | **Mir7681** | mmu-mir-7681 | microRNA mir-7681 | ncRNA |
| chr19_41596984_41598066 | chr19 | 41596985 | 41598066 | + | 0.67455 | Intergenic | Intergenic | 8874 | NR_106166 | 102465884 | | NR_106166 | miRBase:MI0026018 | **Mir8091** | mmu-mir-8091 | microRNA mir-8091 | ncRNA |
| chr19_41597563_41597806 | chr19 | 41597564 | 41597806 | + | 0.672818 | Intergenic | Intergenic | 9034 | NR_106166 | 102465884 | | NR_106166 | miRBase:MI0026018 | **Mir8091** | mmu-mir-8091 | microRNA mir-8091 | ncRNA |
| chr1_85761693_85762136 | chr1 | 85761694 | 85762136 | + | 1.27198 | Intergenic | Intergenic | 9967 | NR_106172 | 102466882 | | NR_106172 | miRBase:MI0026024 | **Mir8096** | mmu-mir-8096 | microRNA mir-8096 | ncRNA |
| chr12_93462053_93462557 | chr12 | 93462054 | 93462557 | + | 0.875527 | Intergenic | Intergenic | -465033 | NR_106201 | 102466888 | | NR_106201 | miRBase:MI0026053 | **Mir8099-2** | mmu-mir-8099-2 | microRNA mir-8099-2 | ncRNA |
| chr12_92916323_92916694 | chr12 | 92916324 | 92916694 | + | 0.79871 | Intergenic | Intergenic | -1010829 | NR_106201 | 102466888 | | NR_106201 | miRBase:MI0026053 | **Mir8099-2** | mmu-mir-8099-2 | microRNA mir-8099-2 | ncRNA |
| chr11_46126795_46127292 | chr11 | 46126796 | 46127292 | + | 1.25548 | intron (NM_009616, intron 11 of 22) | intron (NM_009616, intron 11 of 22) | 24774 | NR_106177 | 102465996 | | NR_106177 | | **Mir8100** | mmu-mir-8100 | microRNA mir-8100 | ncRNA |
| chr11_46126366_46127360 | chr11 | 46126367 | 46127360 | + | 1.11676 | intron (NM_009616, intron 11 of 22) | L1_Mus4\|LINE\|L1 | 24593 | NR_106177 | 102465996 | | NR_106177 | | **Mir8100** | mmu-mir-8100 | microRNA mir-8100 | ncRNA |
| chr11_46090636_46091240 | chr11 | 46090637 | 46091240 | + | 0.635247 | intron (NM_009616, intron 3 of 22) | intron (NM_009616, intron 3 of 22) | -11332 | NR_106177 | 102465996 | | NR_106177 | | **Mir8100** | mmu-mir-8100 | microRNA mir-8100 | ncRNA |
| chr11_97083595_97084241 | chr11 | 97083596 | 97084241 | + | 0.677461 | Intergenic | Intergenic | 20150 | NR_106181 | 102465893 | | NR_106181 | | **Mir8103** | mmu-mir-8103 | microRNA mir-8103 | ncRNA |
| chr10_122740157_122740401 | chr10 | 122740158 | 122740401 | + | 0.856993 | intron (NM_176919, intron 1 of 8) | MTEa\|LTR\|ERVL-MaLR | 60964 | NR_106182 | 102466884 | | NR_106182 | | **Mir8104** | mmu-mir-8104 | microRNA mir-8104 | ncRNA |
| chr10_122739835_122740034 | chr10 | 122739836 | 122740034 | + | 0.80157 | intron (NM_176919, intron 1 of 8) | MTEa\|LTR\|ERVL-MaLR | 60620 | NR_106182 | 102466884 | | NR_106182 | | **Mir8104** | mmu-mir-8104 | microRNA mir-8104 | ncRNA |
| chr10_122735297_122735833 | chr10 | 122735298 | 122735833 | + | 0.760613 | intron (NM_176919, intron 1 of 8) | intron (NM_176919, intron 1 of 8) | 56250 | NR_106182 | 102466884 | | NR_106182 | | **Mir8104** | mmu-mir-8104 | microRNA mir-8104 | ncRNA |
| chr10_122755077_122755295 | chr10 | 122755078 | 122755295 | + | 0.68102 | intron (NM_176919, intron 1 of 8) | intron (NM_176919, intron 1 of 8) | 75871 | NR_106182 | 102466884 | | NR_106182 | | **Mir8104** | mmu-mir-8104 | microRNA mir-8104 | ncRNA |
| chr10_122829652_122830353 | chr10 | 122829653 | 122830353 | + | 0.652548 | intron (NM_176919, intron 4 of 8) | intron (NM_176919, intron 4 of 8) | 150688 | NR_106182 | 102466884 | | NR_106182 | | **Mir8104** | mmu-mir-8104 | microRNA mir-8104 | ncRNA |
| chr9_122240739_122241392 | chr9 | 122240740 | 122241392 | + | 1.11007 | intron (NM_133979, intron 11 of 12) | intron (NM_133979, intron 11 of 12) | 32962 | NR_106185 | 102465896 | | NR_106185 | | **Mir8106** | mmu-mir-8106 | microRNA mir-8106 | ncRNA |
| chr9_122230793_122231032 | chr9 | 122230794 | 122231032 | + | 0.719531 | intron (NM_133979, intron 11 of 12) | intron (NM_133979, intron 11 of 12) | 43115 | NR_106185 | 102465896 | | NR_106185 | | **Mir8106** | mmu-mir-8106 | microRNA mir-8106 | ncRNA |
| chr8_25149050_25149563 | chr8 | 25149051 | 25149563 | + | 1.32313 | Intergenic | Intergenic | 9926 | NR_106187 | 102467006 | | NR_106187 | miRBase:MI0026039 | **Mir8108** | mmu-mir-8108 | microRNA mir-8108 | ncRNA |
| chr8_25130926_25131347 | chr8 | 25130927 | 25131347 | + | 1.06512 | Intergenic | ORR1F\|LTR\|ERVL-MaLR | 28096 | NR_106187 | 102467006 | | NR_106187 | miRBase:MI0026039 | **Mir8108** | mmu-mir-8108 | microRNA mir-8108 | ncRNA |
| chr11_86652510_86653058 | chr11 | 86652511 | 86653058 | + | 1.06216 | intron (NM_029478, intron 5 of 11) | intron (NM_029478, intron 5 of 11) | -3287 | NR_106195 | 102465902 | | NR_106195 | | **Mir8115** | mmu-mir-8115 | microRNA mir-8115 | ncRNA |
| chr4_33467622_33468058 | chr4 | 33467623 | 33468058 | + | 1.7157 | intron (NM_011884, intron 14 of 15) | intron (NM_011884, intron 14 of 15) | 29759 | NR_106198 | 102465904 | | NR_106198 | | **Mir8118** | mmu-mir-8118 | microRNA mir-8118 | ncRNA |
| chr4_33446239_33446777 | chr4 | 33446240 | 33446777 | + | 1.33975 | intron (NM_011884, intron 14 of 15) | B4A\|SINE\|B4 | 8427 | NR_106198 | 102465904 | | NR_106198 | | **Mir8118** | mmu-mir-8118 | microRNA mir-8118 | ncRNA |
| chr4_33587607_33588654 | chr4 | 33587608 | 33588654 | + | 1.24414 | Intergenic | Intergenic | 150050 | NR_106198 | 102465904 | | NR_106198 | | **Mir8118** | mmu-mir-8118 | microRNA mir-8118 | ncRNA |
| chr4_33588141_33588631 | chr4 | 33588142 | 33588631 | + | 1.14615 | Intergenic | Intergenic | 150305 | NR_106198 | 102465904 | | NR_106198 | | **Mir8118** | mmu-mir-8118 | microRNA mir-8118 | ncRNA |
| chr4_33587602_33588024 | chr4 | 33587603 | 33588024 | + | 1.09721 | Intergenic | Intergenic | 149732 | NR_106198 | 102465904 | | NR_106198 | | **Mir8118** | mmu-mir-8118 | microRNA mir-8118 | ncRNA |
| chr3_65603311_65604034 | chr3 | 65603312 | 65604034 | + | 1.65649 | Intergenic | Intergenic | -55615 | NR_106200 | 102465905 | | NR_106200 | miRBase:MI0026052 | **Mir8120** | mmu-mir-8120 | microRNA mir-8120 | ncRNA |
| chr3_65662080_65662258 | chr3 | 65662081 | 65662258 | + | 1.06584 | Intergenic | B3A\|SINE\|B2 | 2881 | NR_106200 | 102465905 | | NR_106200 | miRBase:MI0026052 | **Mir8120** | mmu-mir-8120 | microRNA mir-8120 | ncRNA |
| chr15_35724392_35725272 | chr15 | 35724393 | 35725272 | + | 1.78593 | intron (NM_177151, intron 34 of 61) | intron (NM_177151, intron 34 of 61) | -63785 | NR_030606 | 100124469 | | NR_030606 | miRBase:MI0005551 | **Mir875** | Mirn875\|mmu-mir-875 | microRNA 875 | ncRNA |
| chr14_115047009_115049073 | chr14 | 115047010 | 115049073 | + | 1.13179 | Intergenic | Intergenic | 3614 | NR_029816 | 751549 | | NR_029816 | miRBase:MI0000719 | **Mir92-1** | Mirn92-1 | microRNA 92-1 | ncRNA |
| chr14_115049130_115049732 | chr14 | 115049131 | 115049732 | + | 0.798315 | Intergenic | Intergenic | 5004 | NR_029816 | 751549 | | NR_029816 | miRBase:MI0000719 | **Mir92-1** | Mirn92-1 | microRNA 92-1 | ncRNA |
| chr14_115049173_115049506 | chr14 | 115049174 | 115049506 | + | 0.72411 | Intergenic | Intergenic | 4913 | NR_029816 | 751549 | | NR_029816 | miRBase:MI0000719 | **Mir92-1** | Mirn92-1 | microRNA 92-1 | ncRNA |
| chrX_151950993_151951612 | chrX | 151950994 | 151951612 | + | 1.20088 | Intergenic | MLT2B3\|LTR\|ERVL | 38089 | NR_029753 | 723947 | | NR_029753 | | **Mir98** | Mirn98\|mmu-mir-98 | microRNA 98 | ncRNA |
| chr10_122982899_122983245 | chr10 | 122982900 | 122983245 | + | 1.16363 | Intergenic | Intergenic | 2652 | NR_029527 | 387251 | | NR_029527 | miRBase:MI0000138 | **Mirlet7i** | Mirnlet7i\|mmu-let-7i | microRNA let7i | ncRNA |
| chr10_122854130_122854614 | chr10 | 122854131 | 122854614 | + | 0.896261 | intron (NM_176919, intron 4 of 8) | URR1A\|DNA\|hAT-Charlie | 131352 | NR_029527 | 387251 | | NR_029527 | miRBase:MI0000138 | **Mirlet7i** | Mirnlet7i\|mmu-let-7i | microRNA let7i | ncRNA |

| | | | | | | | | | | | | | | | | | |
|---|---|---|---|---|---|---|---|---|---|---|---|---|---|---|---|---|---|
| chr10_122853984_122855029 | chr10 | 122853985 | 122855029 | + | 0.733941 | intron (NM_176919, intron 4 of 8) | intron (NM_176919, intron 4 of 8) | 131217 | NR_029527 | 387251 | | NR_029527 | miRBase:MI0000138 | **Mirlet7i** | Mirnlet7i\|mmu-let-7i | microRNA let7i | ncRNA |
| chr10_122940266_122940745 | chr10 | 122940267 | 122940745 | + | 0.706189 | intron (NM_001110218, intron 9 of 9) | intron (NM_001110218, intron 9 of 9) | 45218 | NR_029527 | 387251 | | NR_029527 | miRBase:MI0000138 | **Mirlet7i** | Mirnlet7i\|mmu-let-7i | microRNA let7i | ncRNA |
| chr5_147292575_147292855 | chr5 | 147292576 | 147292855 | + | 0.668412 | Intergenic | Intergenic | -9417 | NR_105854 | 102465309 | | NR_105854 | miRBase:MI0022352 | **Mirlet7k** | mmu-let-7k | microRNA let-7k | ncRNA |
| chr15_81106014_81106339 | chr15 | 81106015 | 81106339 | + | 0.779075 | intron (NM_001082536, intron 2 of 13) | intron (NM_001082536, intron 2 of 13) | -1130 | NM_153049 | 223701 | Mm.439814 | NM_153049 | ENSMUSG00000042292 | **Mkl1** | AMKL\|Bsac\|Mal\|Mrtf-A | MKL (megakaryoblastic leukemia)/myocardin-like 1 | protein-coding |
| chr15_81103864_81104054 | chr15 | 81103865 | 81104054 | + | 0.643714 | intron (NM_001082536, intron 2 of 13) | intron (NM_001082536, intron 2 of 13) | 1088 | NM_153049 | 223701 | Mm.439814 | NM_153049 | ENSMUSG00000042292 | **Mkl1** | AMKL\|Bsac\|Mal\|Mrtf-A | MKL (megakaryoblastic leukemia)/myocardin-like 1 | protein-coding |
| chr6_39447774_39448153 | chr6 | 39447775 | 39448153 | + | 1.14846 | Intergenic | RMER15\|LTR\|ERVL | -27595 | NM_018810 | 54484 | Mm.270484 | NM_018810 | ENSMUSG00000029922 | **Mkrn1** | RFP | makorin, ring finger protein, 1 | protein-coding |
| chr6_115574042_115574497 | chr6 | 115574043 | 115574497 | + | 1.26692 | intron (NM_199033, intron 8 of 11) | intron (NM_199033, intron 8 of 11) | 18306 | NM_001101431 | 70291 | Mm.133694 | NM_001101431 | | **Mkrn2os** | 2510049J12Rik | makorin, ring finger protein 2, opposite strand | protein-coding |
| chr6_115573949_115574566 | chr6 | 115573950 | 115574566 | + | 1.13488 | intron (NM_199033, intron 8 of 11) | intron (NM_199033, intron 8 of 11) | 18318 | NM_001101431 | 70291 | Mm.133694 | NM_001101431 | | **Mkrn2os** | 2510049J12Rik | makorin, ring finger protein 2, opposite strand | protein-coding |
| chr6_115580897_115581424 | chr6 | 115580898 | 115581424 | + | 0.9229 | Intergenic | Intergenic | 11415 | NM_001101431 | 70291 | Mm.133694 | NM_001101431 | | **Mkrn2os** | 2510049J12Rik | makorin, ring finger protein 2, opposite strand | protein-coding |
| chr6_115580887_115581212 | chr6 | 115580888 | 115581212 | + | 0.906385 | Intergenic | Intergenic | 11526 | NM_001101431 | 70291 | Mm.133694 | NM_001101431 | | **Mkrn2os** | 2510049J12Rik | makorin, ring finger protein 2, opposite strand | protein-coding |
| chr6_115577264_115577803 | chr6 | 115577265 | 115577803 | + | 0.794201 | intron (NM_199033, intron 11 of 11) | intron (NM_199033, intron 11 of 11) | 15042 | NM_001101431 | 70291 | Mm.133694 | NM_001101431 | | **Mkrn2os** | 2510049J12Rik | makorin, ring finger protein 2, opposite strand | protein-coding |
| chr3_67373855_67374327 | chr3 | 67373856 | 67374327 | + | 0.684804 | promoter-TSS (NM_010801) | promoter-TSS (NM_010801) | -6 | NM_010801 | 17349 | Mm.10414 | NM_010801 | ENSMUSG00000048416 | **Mlf1** | HLS7 | myeloid leukemia factor 1 | protein-coding |
| chr2_18180373_18180779 | chr2 | 18180374 | 18180779 | + | 1.18864 | intron (NR_045538, intron 13 of 22) | intron (NR_045538, intron 13 of 22) | 115991 | NM_001252561 | 17354 | Mm.209175 | NM_010804 | ENSMUSG00000026743 | **Mllt10** | Af10\|B130021D15Rik\|D630001B22Rik\|mKIAA4140 | myeloid/lymphoid or mixed-lineage leukemia (trithorax homolog, Drosophila); translocated to, 10 | protein-coding |
| chr2_18179122_18179490 | chr2 | 18179123 | 18179490 | + | 1.11001 | intron (NR_045538, intron 13 of 22) | intron (NR_045538, intron 13 of 22) | 114721 | NM_001252561 | 17354 | Mm.209175 | NM_010804 | ENSMUSG00000026743 | **Mllt10** | Af10\|B130021D15Rik\|D630001B22Rik\|mKIAA4140 | myeloid/lymphoid or mixed-lineage leukemia (trithorax homolog, Drosophila); translocated to, 10 | protein-coding |
| chr2_18107507_18107920 | chr2 | 18107508 | 18107920 | + | 0.852529 | intron (NR_045538, intron 4 of 22) | B3\|SINE\|B2 | 43129 | NM_001252561 | 17354 | Mm.209175 | NM_010804 | ENSMUSG00000026743 | **Mllt10** | Af10\|B130021D15Rik\|D630001B22Rik\|mKIAA4140 | myeloid/lymphoid or mixed-lineage leukemia (trithorax homolog, Drosophila); translocated to, 10 | protein-coding |
| chr2_18178616_18179026 | chr2 | 18178617 | 18179026 | + | 0.658022 | intron (NR_045538, intron 13 of 22) | (CCA)n\|Simple_repeat\|Simple_repeat | 114236 | NM_001252561 | 17354 | Mm.209175 | NM_010804 | ENSMUSG00000026743 | **Mllt10** | Af10\|B130021D15Rik\|D630001B22Rik\|mKIAA4140 | myeloid/lymphoid or mixed-lineage leukemia (trithorax homolog, Drosophila); translocated to, 10 | protein-coding |
| chr2_18179632_18180070 | chr2 | 18179633 | 18180070 | + | 0.643587 | intron (NR_045538, intron 13 of 22) | Lx5b\|LINE\|L1 | 115266 | NM_001252561 | 17354 | Mm.209175 | NM_010804 | ENSMUSG00000026743 | **Mllt10** | Af10\|B130021D15Rik\|D630001B22Rik\|mKIAA4140 | myeloid/lymphoid or mixed-lineage leukemia (trithorax homolog, Drosophila); translocated to, 10 | protein-coding |
| chr4_87859047_87859758 | chr4 | 87859048 | 87859758 | + | 1.49178 | intron (NM_027326, intron 4 of 10) | intron (NM_027326, intron 4 of 10) | -53080 | NM_029931 | 70122 | Mm.288898 | NM_027326 | ENSMUSG00000028496 | **Mllt3** | 2210011H10Rik\|2610012I03Rik\|3830408D16Rik\|Af9\|D4Ertd321e | myeloid/lymphoid or mixed-lineage leukemia (trithorax homolog, Drosophila); translocated to, 3 | protein-coding |
| chr4_87987040_87987355 | chr4 | 87987041 | 87987355 | + | 1.44283 | intron (NM_027326, intron 2 of 10) | intron (NM_027326, intron 2 of 10) | 46209 | NM_027326 | 70122 | Mm.288898 | NM_027326 | ENSMUSG00000028496 | **Mllt3** | 2210011H10Rik\|2610012I03Rik\|3830408D16Rik\|Af9\|D4Ertd321e | myeloid/lymphoid or mixed-lineage leukemia (trithorax homolog, Drosophila); translocated to, 3 | protein-coding |
| chr4_87979841_87980081 | chr4 | 87979842 | 87980081 | + | 1.38996 | intron (NM_027326, intron 2 of 10) | intron (NM_027326, intron 2 of 10) | 53446 | NM_027326 | 70122 | Mm.288898 | NM_027326 | ENSMUSG00000028496 | **Mllt3** | 2210011H10Rik\|2610012I03Rik\|3830408D16Rik\|Af9\|D4Ertd321e | myeloid/lymphoid or mixed-lineage leukemia (trithorax homolog, Drosophila); translocated to, 3 | protein-coding |
| chr4_87979104_87980083 | chr4 | 87979105 | 87980083 | + | 1.36437 | intron (NM_027326, intron 2 of 10) | RSINE1\|SINE\|B4 | 53813 | NM_027326 | 70122 | Mm.288898 | NM_027326 | ENSMUSG00000028496 | **Mllt3** | 2210011H10Rik\|2610012I03Rik\|3830408D16Rik\|Af9\|D4Ertd321e | myeloid/lymphoid or mixed-lineage leukemia (trithorax homolog, Drosophila); translocated to, 3 | protein-coding |
| chr4_87976311_87977210 | chr4 | 87976312 | 87977210 | + | 1.32056 | intron (NM_027326, intron 2 of 10) | intron (NM_027326, intron 2 of 10) | 56646 | NM_027326 | 70122 | Mm.288898 | NM_027326 | ENSMUSG00000028496 | **Mllt3** | 2210011H10Rik\|2610012I03Rik\|3830408D16Rik\|Af9\|D4Ertd321e | myeloid/lymphoid or mixed-lineage leukemia (trithorax homolog, Drosophila); translocated to, 3 | protein-coding |
| chr4_87984418_87985031 | chr4 | 87984419 | 87985031 | + | 1.15573 | intron (NM_027326, intron 2 of 10) | intron (NM_027326, intron 2 of 10) | 48682 | NM_027326 | 70122 | Mm.288898 | NM_027326 | ENSMUSG00000028496 | **Mllt3** | 2210011H10Rik\|2610012I03Rik\|3830408D16Rik\|Af9\|D4Ertd321e | myeloid/lymphoid or mixed-lineage leukemia (trithorax homolog, Drosophila); translocated to, 3 | protein-coding |
| chr4_87984675_87985063 | chr4 | 87984676 | 87985063 | + | 1.10374 | intron (NM_027326, intron 2 of 10) | intron (NM_027326, intron 2 of 10) | 48538 | NM_027326 | 70122 | Mm.288898 | NM_027326 | ENSMUSG00000028496 | **Mllt3** | 2210011H10Rik\|2610012I03Rik\|3830408D16Rik\|Af9\|D4Ertd321e | myeloid/lymphoid or mixed-lineage leukemia (trithorax homolog, Drosophila); translocated to, 3 | protein-coding |
| chr4_87979201_87979767 | chr4 | 87979202 | 87979767 | + | 1.05778 | intron (NM_027326, intron 2 of 10) | intron (NM_027326, intron 2 of 10) | 53923 | NM_027326 | 70122 | Mm.288898 | NM_027326 | ENSMUSG00000028496 | **Mllt3** | 2210011H10Rik\|2610012I03Rik\|3830408D16Rik\|Af9\|D4Ertd321e | myeloid/lymphoid or mixed-lineage leukemia (trithorax homolog, Drosophila); translocated to, 3 | protein-coding |
| chr4_87858393_87858755 | chr4 | 87858394 | 87858755 | + | 0.866363 | intron (NM_027326, intron 4 of 10) | intron (NM_027326, intron 4 of 10) | -52251 | NM_029931 | 70122 | Mm.288898 | NM_027326 | ENSMUSG00000028496 | **Mllt3** | 2210011H10Rik\|2610012I03Rik\|3830408D16Rik\|Af9\|D4Ertd321e | myeloid/lymphoid or mixed-lineage leukemia (trithorax homolog, Drosophila); translocated to, 3 | protein-coding |
| chr4_87989918_87990299 | chr4 | 87989919 | 87990299 | + | 0.826155 | intron (NM_027326, intron 2 of 10) | B3A\|SINE\|B2 | 43298 | NM_027326 | 70122 | Mm.288898 | NM_027326 | ENSMUSG00000028496 | **Mllt3** | 2210011H10Rik\|2610012I03Rik\|3830408D16Rik\|Af9\|D4Ertd321e | myeloid/lymphoid or mixed-lineage leukemia (trithorax homolog, Drosophila); translocated to, 3 | protein-coding |
| chr4_88014993_88015760 | chr4 | 88014994 | 88015760 | + | 0.783561 | intron (NM_027326, intron 2 of 10) | intron (NM_027326, intron 2 of 10) | 18030 | NM_027326 | 70122 | Mm.288898 | NM_027326 | ENSMUSG00000028496 | **Mllt3** | 2210011H10Rik\|2610012I03Rik\|3830408D16Rik\|Af9\|D4Ertd321e | myeloid/lymphoid or mixed-lineage leukemia (trithorax homolog, Drosophila); translocated to, 3 | protein-coding |
| chr4_87984147_87984303 | chr4 | 87984148 | 87984303 | + | 0.73167 | intron (NM_027326, intron 2 of 10) | intron (NM_027326, intron 2 of 10) | 49182 | NM_027326 | 70122 | Mm.288898 | NM_027326 | ENSMUSG00000028496 | **Mllt3** | 2210011H10Rik\|2610012I03Rik\|3830408D16Rik\|Af9\|D4Ertd321e | myeloid/lymphoid or mixed-lineage leukemia (trithorax homolog, Drosophila); translocated to, 3 | protein-coding |
| chr4_88010471_88010811 | chr4 | 88010472 | 88010811 | + | 0.709174 | intron (NM_027326, intron 2 of 10) | intron (NM_027326, intron 2 of 10) | 22766 | NM_027326 | 70122 | Mm.288898 | NM_027326 | ENSMUSG00000028496 | **Mllt3** | 2210011H10Rik\|2610012I03Rik\|3830408D16Rik\|Af9\|D4Ertd321e | myeloid/lymphoid or mixed-lineage leukemia (trithorax homolog, Drosophila); translocated to, 3 | protein-coding |
| chr4_88014989_88016013 | chr4 | 88014990 | 88016013 | + | 0.670883 | intron (NM_027326, intron 2 of 10) | intron (NM_027326, intron 2 of 10) | 17906 | NM_027326 | 70122 | Mm.288898 | NM_027326 | ENSMUSG00000028496 | **Mllt3** | 2210011H10Rik\|2610012I03Rik\|3830408D16Rik\|Af9\|D4Ertd321e | myeloid/lymphoid or mixed-lineage leukemia (trithorax homolog, Drosophila); translocated to, 3 | protein-coding |
| chr4_87933959_87934321 | chr4 | 87933960 | 87934321 | + | 0.642301 | intron (NM_027326, intron 2 of 10) | MER5A\|DNA\|hAT-Charlie | 99267 | NM_027326 | 70122 | Mm.288898 | NM_027326 | ENSMUSG00000028496 | **Mllt3** | 2210011H10Rik\|2610012I03Rik\|3830408D16Rik\|Af9\|D4Ertd321e | myeloid/lymphoid or mixed-lineage leukemia (trithorax homolog, Drosophila); translocated to, 3 | protein-coding |
| chr17_13680366_13680706 | chr17 | 13680367 | 13680706 | + | 1.09061 | Intergenic | B4A\|SINE\|B4 | -80005 | NM_010806 | 17356 | Mm.59167 | NM_010806 | ENSMUSG00000068036 | **Mllt4** | 5033403D15Rik\|AF6\|Af-6\|Afadin\|Gm314\|I-afadin\|S-afadin | myeloid/lymphoid or mixed-lineage leukemia (trithorax homolog, Drosophila); translocated to, 4 | protein-coding |
| chr11_97659467_97659808 | chr11 | 97659468 | 97659808 | + | 0.628322 | Intergenic | Intergenic | -3774 | NM_139311 | 246198 | Mm.23685 | NM_139311 | ENSMUSG00000038437 | **Mllt6** | AI315037\|Af17 | myeloid/lymphoid or mixed-lineage leukemia (trithorax homolog, Drosophila); translocated to, 6 | protein-coding |
| chr11_97659445_97659975 | chr11 | 97659446 | 97659975 | + | 0.61622 | Intergenic | Intergenic | -3702 | NM_139311 | 246198 | Mm.23685 | NM_139311 | ENSMUSG00000038437 | **Mllt6** | AI315037\|Af17 | myeloid/lymphoid or mixed-lineage leukemia (trithorax homolog, Drosophila); translocated to, 6 | protein-coding |
| chr8_79273682_79274286 | chr8 | 79273683 | 79274286 | + | 1.3461 | intron (NM_133823, intron 4 of 6) | intron (NM_133823, intron 4 of 6) | 20972 | NM_133823 | 109136 | Mm.26510 | NM_133823 | ENSMUSG00000037022 | **Mmaa** | 2810018E08Rik\|AI840684 | methylmalonic aciduria (cobalamin deficiency) type A | protein-coding |
| chr11_90224852_90225456 | chr11 | 90224853 | 90225456 | + | 0.616823 | Intergenic | Intergenic | -24322 | NM_026178 | 67468 | Mm.277518 | NM_026178 | ENSMUSG00000003948 | **Mmd** | 1200017E07Rik\|1810073C06Rik\|AA690185 | monocyte to macrophage differentiation-associated | protein-coding |
| chr9_7235462_7235847 | chr9 | 7235463 | 7235847 | + | 0.970483 | Intergenic | Intergenic | -36859 | NM_008607 | 17386 | Mm.5022 | NM_008607 | ENSMUSG00000050578 | **Mmp13** | Clg\|MMP-13\|Mmp1 | matrix metallopeptidase 13 | protein-coding |
| chr9_7239928_7240430 | chr9 | 7239929 | 7240430 | + | 0.647462 | Intergenic | Intergenic | -32335 | NM_008607 | 17386 | Mm.5022 | NM_008607 | ENSMUSG00000050578 | **Mmp13** | Clg\|MMP-13\|Mmp1 | matrix metallopeptidase 13 | protein-coding |
| chr10_128779450_128779848 | chr10 | 128779451 | 128779848 | + | 1.24089 | Intergenic | Intergenic | -11261 | NM_001164197 | 58223 | Mm.131266 | NM_021412 | ENSMUSG00000025355 | **Mmp19** | - | matrix metallopeptidase 19 | protein-coding |
| chr10_128780255_128781057 | chr10 | 128780256 | 128781057 | + | 0.862433 | Intergenic | Intergenic | -10254 | NM_001164197 | 58223 | Mm.131266 | NM_021412 | ENSMUSG00000025355 | **Mmp19** | - | matrix metallopeptidase 19 | protein-coding |
| chr9_7553565_7554204 | chr9 | 7553566 | 7554204 | + | 0.630963 | Intergenic | Intergenic | -4544 | NM_008611 | 17394 | Mm.16415 | NM_008611 | ENSMUSG00000005800 | **Mmp8** | BB138268 | matrix metallopeptidase 8 | protein-coding |

| | | | | | | | | | | | | | | | | | |
|---|---|---|---|---|---|---|---|---|---|---|---|---|---|---|---|---|---|
| chr4_24569022_24569514 | chr4 | 24569023 | 24569514 | + | 0.683421 | intron (NM_199467, intron 13 of 23) | intron (NM_199467, intron 13 of 23) | 72806 | NM_199467 | 212377 | Mm.120919 | NM_199467 | ENSMUSG00000045751 | **Mms22l** | F730047E07Rik\|Gm134 | MMS22-like, DNA repair protein | protein-coding |
| chr4_24594632_24594968 | chr4 | 24594633 | 24594968 | + | 0.679992 | intron (NM_199467, intron 20 of 23) | intron (NM_199467, intron 20 of 23) | 98338 | NM_199467 | 212377 | Mm.120919 | NM_199467 | ENSMUSG00000045751 | **Mms22l** | F730047E07Rik\|Gm134 | MMS22-like, DNA repair protein | protein-coding |
| chr14_10397909_10398795 | chr14 | 10397910 | 10398795 | + | 1.42226 | intron (NM_010210, intron 4 of 8) | intron (NM_010210, intron 4 of 8) | 514257 | NR_030680 | 100042415 | Mm.235938 | NR_030680 | | **Mnd1-ps** | Gm3833 | Mnd1 retrotransposed pseudogene | pseudo |
| chr14_10315830_10316584 | chr14 | 10315831 | 10316584 | + | 1.20532 | intron (NM_010210, intron 4 of 8) | intron (NM_010210, intron 4 of 8) | 432112 | NR_030680 | 100042415 | Mm.235938 | NR_030680 | | **Mnd1-ps** | Gm3833 | Mnd1 retrotransposed pseudogene | pseudo |
| chr14_10323022_10323345 | chr14 | 10323023 | 10323345 | + | 1.05473 | intron (NM_010210, intron 4 of 8) | intron (NM_010210, intron 4 of 8) | 439089 | NR_030680 | 100042415 | Mm.235938 | NR_030680 | | **Mnd1-ps** | Gm3833 | Mnd1 retrotransposed pseudogene | pseudo |
| chr14_10322909_10323391 | chr14 | 10322910 | 10323391 | + | 0.979446 | intron (NM_010210, intron 4 of 8) | intron (NM_010210, intron 4 of 8) | 439055 | NR_030680 | 100042415 | Mm.235938 | NR_030680 | | **Mnd1-ps** | Gm3833 | Mnd1 retrotransposed pseudogene | pseudo |
| chr1_173905899_173906455 | chr1 | 173905900 | 173906455 | + | 0.674126 | exon (NM_001033450, exon 3 of 7) | exon (NM_001033450, exon 3 of 7) | 6745 | NM_001033450 | 381308 | Mm.490248 | NM_001033450 | ENSMUSG00000026536 | **Mnda** | A730048F03\|Ifi205b | myeloid cell nuclear differentiation antigen | protein-coding |
| chr9_72418363_72418844 | chr9 | 72418364 | 72418844 | + | 0.983811 | Intergenic | ORR1C2\|LTR\|ERVL-MaLR | -19925 | NM_008613 | 17427 | Mm.387671 | NM_008613 | | **Mns1** | AW546487 | meiosis-specific nuclear structural protein 1 | protein-coding |
| chr5_88717071_88717318 | chr5 | 88717072 | 88717318 | + | 1.24273 | Intergenic | L2\|LINE\|L2 | -3676 | NM_026735 | 68473 | Mm.35764 | NM_026735 | ENSMUSG00000006262 | **Mob1b** | 1110003E08Rik\|AU015450\|B230364F10\|Mobkl1a | MOB kinase activator 1B | protein-coding |
| chr5_88724227_88724425 | chr5 | 88724228 | 88724425 | + | 0.663331 | intron (NM_026735, intron 1 of 5) | intron (NM_026735, intron 1 of 5) | 3455 | NM_026735 | 68473 | Mm.35764 | NM_026735 | ENSMUSG00000006262 | **Mob1b** | 1110003E08Rik\|AU015450\|B230364F10\|Mobkl1a | MOB kinase activator 1B | protein-coding |
| chr10_80695985_80696666 | chr10 | 80695986 | 80696666 | + | 0.868706 | intron (NM_172457, intron 1 of 3) | intron (NM_172457, intron 1 of 3) | 5494 | NM_172457 | 208228 | Mm.49309 | NM_172457 | ENSMUSG00000003348 | **Mob3a** | 5330417K06Rik\|A630029F06\|AV218468\|Mobkl2a | MOB kinase activator 3A | protein-coding |
| chr10_80696758_80696970 | chr10 | 80696759 | 80696970 | + | 0.647858 | intron (NM_172457, intron 1 of 3) | intron (NM_172457, intron 1 of 3) | 4956 | NM_172457 | 208228 | Mm.49309 | NM_172457 | ENSMUSG00000003348 | **Mob3a** | 5330417K06Rik\|A630029F06\|AV218468\|Mobkl2a | MOB kinase activator 3A | protein-coding |
| chr18_24629672_24630310 | chr18 | 24629673 | 24630310 | + | 1.16712 | intron (NM_021448, intron 17 of 21) | ORR1D2\|LTR\|ERVL-MaLR | -23700 | NM_026779 | 68591 | Mm.28252 | NM_026779 | ENSMUSG00000039616 | **Mocos** | 1110018O12Rik\|HMCS\|MCS\|MOS | molybdenum cofactor sulfurase | protein-coding |
| chr7_99248256_99248778 | chr7 | 99248257 | 99248778 | + | 0.87175 | Intergenic | Intergenic | -9906 | NM_177448 | 233549 | Mm.208030 | NM_177448 | ENSMUSG00000052396 | **Mogat2** | DGAT2L5\|MGAT2\|Mgat1l | monoacylglycerol O-acyltransferase 2 | protein-coding |
| chr12_110844867_110845533 | chr12 | 110844868 | 110845533 | + | 0.952043 | Intergenic | Intergenic | -4261 | NM_011973 | 26448 | Mm.140948 | NM_011973 | ENSMUSG00000056458 | **Mok** | RAGE1\|Rage\|Stk30 | MOK protein kinase | protein-coding |
| chr10_123131331_123132114 | chr10 | 123131332 | 123132114 | + | 0.905898 | intron (NM_027604, intron 9 of 21) | Tigger17c\|DNA\|TcMar-Tigger | -55218 | NM_001163025 | 67074 | Mm.169924 | NM_153395 | ENSMUSG00000034602 | **Mon2** | 2610528O22Rik\|AW495628\|Sf21\|mKIAA1040 | MON2 homolog (yeast) | protein-coding |
| chr10_123124377_123124938 | chr10 | 123124378 | 123124938 | + | 0.604621 | intron (NM_027604, intron 17 of 21) | intron (NM_027604, intron 17 of 21) | -48153 | NM_001163025 | 67074 | Mm.169924 | NM_153395 | ENSMUSG00000034602 | **Mon2** | 2610528O22Rik\|AW495628\|Sf21\|mKIAA1040 | MON2 homolog (yeast) | protein-coding |
| chr16_93798514_93799345 | chr16 | 93798515 | 93799345 | + | 1.76109 | exon (NM_027293, exon 28 of 37) | exon (NM_027293, exon 28 of 37) | -33191 | NM_001045529 | 338467 | Mm.287329 | NM_001045529 | ENSMUSG00000039456 | **Morc3** | 1110051N18Rik\|AI452146\|BF318192\|D16Jhu32e\|NXP2\|Zcwcc3 | microrchidia 3 | protein-coding |
| chr16_93778434_93778798 | chr16 | 93778435 | 93778798 | + | 0.98514 | intron (NM_027293, intron 23 of 36) | intron (NM_027293, intron 23 of 36) | -53505 | NM_001045529 | 338467 | Mm.287329 | NM_001045529 | ENSMUSG00000039456 | **Morc3** | 1110051N18Rik\|AI452146\|BF318192\|D16Jhu32e\|NXP2\|Zcwcc3 | microrchidia 3 | protein-coding |
| chr4_155117130_155117931 | chr4 | 155117131 | 155117931 | + | 1.16404 | intron (NM_001081100, intron 10 of 13) | intron (NM_001081100, intron 10 of 13) | 30954 | NM_001081100 | 76866 | Mm.200590 | NM_001081100 | ENSMUSG00000029049 | **Morn1** | 2900057D20Rik\|4930417P05Rik\|AU042805 | MORN repeat containing 1 | protein-coding |
| chr19_42086234_42086498 | chr19 | 42086235 | 42086498 | + | 0.669804 | promoter-TSS (NM_198108) | promoter-TSS (NM_198108) | 4 | NM_198108 | 226123 | Mm.259441 | NM_198108 | ENSMUSG00000049670 | **Morn4** | - | MORN repeat containing 4 | protein-coding |
| chr4_3892937_3893383 | chr4 | 3892938 | 3893383 | + | 1.37797 | Intergenic | Intergenic | -21055 | NM_020021 | 17451 | Mm.459300 | NM_020021 | ENSMUSG00000078365 | **Mos** | c-mos | Moloney sarcoma oncogene | protein-coding |
| chr18_46447401_46448015 | chr18 | 46447402 | 46448015 | + | 1.22545 | Intergenic | Intergenic | 18108 | NR_045438 | 72076 | Mm.82766 | NM_028086 | ENSMUSG00000046487 | **Mospd4** | 2010013H21Rik | motile sperm domain containing 4 | pseudo |
| chr3_104840815_104841604 | chr3 | 104840816 | 104841604 | + | 1.82669 | intron (NM_009797, intron 3 of 9) | L2\|LINE\|L2 | -22647 | NM_001163440 | 17454 | Mm.1597 | NM_008619 | ENSMUSG00000002227 | **Mov10** | C77703\|Mov-10 | Moloney leukemia virus 10 | protein-coding |
| chr3_104819689_104820507 | chr3 | 104819690 | 104820507 | + | 0.709311 | Intergenic | Intergenic | -1535 | NM_001163440 | 17454 | Mm.1597 | NM_008619 | ENSMUSG00000002227 | **Mov10** | C77703\|Mov-10 | Moloney leukemia virus 10 | protein-coding |
| chr19_12472435_12472862 | chr19 | 12472436 | 12472862 | + | 0.891472 | intron (NM_172442, intron 8 of 8) | B4A\|SINE\|B4 | 11870 | NM_010821 | 17476 | Mm.3999 | NM_010821 | ENSMUSG00000046805 | **Mpeg1** | MPS1\|Mpg-1 | macrophage expressed gene 1 | protein-coding |
| chr19_12474162_12474545 | chr19 | 12474163 | 12474545 | + | 0.709203 | intron (NM_172442, intron 7 of 8) | MLT1A1\|LTR\|ERVL-MaLR | 13575 | NM_010821 | 17476 | Mm.3999 | NM_010821 | ENSMUSG00000046805 | **Mpeg1** | MPS1\|Mpg-1 | macrophage expressed gene 1 | protein-coding |
| chr8_117781346_117781801 | chr8 | 117781347 | 117781801 | + | 1.07366 | Intergenic | Intergenic | 20355 | NM_026758 | 68533 | Mm.181836 | NM_026758 | ENSMUSG00000031843 | **Mphosph6** | 1110001M01Rik\|AA536809\|C86426 | M phase phosphoprotein 6 | protein-coding |
| chr8_117772512_117772978 | chr8 | 117772513 | 117772978 | + | 0.808975 | Intergenic | Intergenic | 29184 | NM_026758 | 68533 | Mm.181836 | NM_026758 | ENSMUSG00000031843 | **Mphosph6** | 1110001M01Rik\|AA536809\|C86426 | M phase phosphoprotein 6 | protein-coding |
| chr8_117772163_117773040 | chr8 | 117772164 | 117773040 | + | 0.637152 | Intergenic | Intergenic | 29327 | NM_026758 | 68533 | Mm.181836 | NM_026758 | ENSMUSG00000031843 | **Mphosph6** | 1110001M01Rik\|AA536809\|C86426 | M phase phosphoprotein 6 | protein-coding |
| chr5_124302805_124303241 | chr5 | 124302806 | 124303241 | + | 0.707799 | intron (NR_102690, intron 8 of 22) | URR1B\|DNA\|hAT-Charlie | 24949 | NR_102690 | 269702 | Mm.60224 | NM_001081323 | ENSMUSG00000038126 | **Mphosph9** | 4930548D04Rik\|9630025B04Rik\|AW413446\|AW547060\|B930097C17Rik\|C87456\|MPP-9\|MPP9 | M-phase phosphoprotein 9 | protein-coding |
| chr11_102027671_102028101 | chr11 | 102027672 | 102028101 | + | 0.812169 | promoter-TSS (NM_007863) | promoter-TSS (NM_007863) | -931 | NM_007863 | 13384 | Mm.20449 | NM_007863 | ENSMUSG00000052373 | **Mpp3** | 6430514B01\|Dlgh3 | membrane protein, palmitoylated 3 (MAGUK p55 subfamily member 3) | protein-coding |
| chr1_59177774_59178444 | chr1 | 59177775 | 59178444 | + | 1.38508 | intron (NM_001159948, intron 25 of 33) | intron (NM_001159948, intron 25 of 33) | -14720 | NM_001164682 | 227157 | Mm.474049 | NM_145143 | ENSMUSG00000079550 | **Mpp4** | Dlg6 | membrane protein, palmitoylated 4 (MAGUK p55 subfamily member 4) | protein-coding |
| chr1_59162803_59163516 | chr1 | 59162804 | 59163516 | + | 1.04477 | 5' UTR (NM_001164682, exon 1 of 23) | 5' UTR (NM_001164682, exon 1 of 23) | 229 | NM_001164682 | 227157 | Mm.474049 | NM_145143 | ENSMUSG00000079550 | **Mpp4** | Dlg6 | membrane protein, palmitoylated 4 (MAGUK p55 subfamily member 4) | protein-coding |
| chr1_59180047_59180406 | chr1 | 59180048 | 59180406 | + | 0.751457 | intron (NM_001159948, intron 22 of 33) | intron (NM_001159948, intron 22 of 33) | -16838 | NM_001164682 | 227157 | Mm.474049 | NM_145143 | ENSMUSG00000079550 | **Mpp4** | Dlg6 | membrane protein, palmitoylated 4 (MAGUK p55 subfamily member 4) | protein-coding |
| chr12_78799870_78800532 | chr12 | 78799871 | 78800532 | + | 0.913513 | intron (NM_019579, intron 3 of 14) | MTD-int\|LTR\|ERVL-MaLR | 51254 | NM_019579 | 56217 | Mm.425777 | NM_019579 | ENSMUSG00000021112 | **Mpp5** | 3830420B02Rik\|AI255216\|AI644496\|Pals1 | membrane protein, palmitoylated 5 (MAGUK p55 subfamily member 5) | protein-coding |
| chr6_49987339_49988176 | chr6 | 49987340 | 49988176 | + | 1.14537 | Intergenic | L1MD1\|LINE\|L1 | -122483 | NM_019939 | 56524 | Mm.41288 | NM_008016 | ENSMUSG00000038388 | **Mpp6** | P55t\|Pals2 | membrane protein, palmitoylated 6 (MAGUK p55 subfamily member 6) | protein-coding |
| chr6_50159613_50160284 | chr6 | 50159614 | 50160284 | + | 0.615037 | intron (NM_019939, intron 4 of 12) | intron (NM_019939, intron 4 of 12) | 49708 | NM_001164733 | 56524 | Mm.41288 | NM_008016 | ENSMUSG00000038388 | **Mpp6** | P55t\|Pals2 | membrane protein, palmitoylated 6 (MAGUK p55 subfamily member 6) | protein-coding |
| chr18_7613703_7614088 | chr18 | 7613704 | 7614088 | + | 0.823797 | intron (NM_001081287, intron 1 of 16) | intron (NM_001081287, intron 1 of 16) | 12967 | NM_001081287 | 75739 | Mm.133293 | NM_001081287 | ENSMUSG00000057440 | **Mpp7** | 1110068J02Rik\|2810038M04Rik\|5430426E14Rik\|AI415104\|Gm955 | membrane protein, palmitoylated 7 (MAGUK p55 subfamily member 7) | protein-coding |
| chr1_165603285_165603706 | chr1 | 165603286 | 165603706 | + | 1.03702 | intron (NM_001083897, intron 4 of 5) | intron (NM_001083897, intron 4 of 5) | 31045 | NM_001001880 | 68481 | Mm.46438 | NM_001001880 | ENSMUSG00000026566 | **Mpzl1** | 1110007A10Rik\|PZR | myelin protein zero-like 1 | protein-coding |
| chr1_165603273_165603856 | chr1 | 165603274 | 165603856 | + | 0.917947 | intron (NM_001083897, intron 4 of 5) | intron (NM_001083897, intron 4 of 5) | 30976 | NM_001001880 | 68481 | Mm.46438 | NM_001001880 | ENSMUSG00000026566 | **Mpzl1** | 1110007A10Rik\|PZR | myelin protein zero-like 1 | protein-coding |
| chr9_45058344_45059315 | chr9 | 45058345 | 45059315 | + | 1.04545 | intron (NM_001093749, intron 1 of 5) | intron (NM_001093749, intron 1 of 5) | 3649 | NM_001093749 | 319742 | Mm.49611 | NM_176993 | ENSMUSG00000070305 | **Mpzl3** | 5430427F17Rik\|A530065I17Rik\|rc | myelin protein zero-like 3 | protein-coding |
| chr9_45060206_45060466 | chr9 | 45060207 | 45060466 | + | 0.958639 | intron (NM_001093749, intron 1 of 5) | intron (NM_001093749, intron 1 of 5) | 5155 | NM_001093749 | 319742 | Mm.49611 | NM_176993 | ENSMUSG00000070305 | **Mpzl3** | 5430427F17Rik\|A530065I17Rik\|rc | myelin protein zero-like 3 | protein-coding |
| chr1_155138825_155139268 | chr1 | 155138826 | 155139268 | + | 1.65486 | intron (NM_008209, intron 1 of 5) | intron (NM_008209, intron 1 of 5) | 7733 | NM_008209 | 15064 | Mm.7580 | NM_008209 | ENSMUSG00000026471 | **Mr1** | H2ls | major histocompatibility complex, class I-related | protein-coding |
| chr1_72236863_72237678 | chr1 | 72236864 | 72237678 | + | 0.849982 | Intergenic | Intergenic | -24964 | NM_001005423 | 381269 | Mm.422636 | NM_001005423 | ENSMUSG00000039395 | **Mreg** | Gm974\|Wdt2\|dsu | melanoregulin | protein-coding |
| chr1_72226159_72227243 | chr1 | 72226160 | 72227243 | + | 0.637717 | Intergenic | Intergenic | -14394 | NM_001005423 | 381269 | Mm.422636 | NM_001005423 | ENSMUSG00000039395 | **Mreg** | Gm974\|Wdt2\|dsu | melanoregulin | protein-coding |
| chr7_143786010_143786394 | chr7 | 143786011 | 143786394 | + | 0.849598 | Intergenic | Intergenic | -1702 | NM_175534 | 244238 | Mm.183561 | NM_175534 | ENSMUSG00000048965 | **Mrgpre** | C130069N09Rik\|Ebrt3\|Mgrf\|MrgE | MAS-related GPR, member E | protein-coding |
| chr15_4816650_4817409 | chr15 | 4816651 | 4817409 | + | 0.861631 | Intergenic | Intergenic | -81707 | NM_001166066 | 223825 | Mm.159281 | NM_001166066 | ENSMUSG00000022155 | **Mroh2b** | 4930455B06Rik\|Heatr7b2 | maestro heat-like repeat family member 2B | protein-coding |
| chr15_4816745_4817093 | chr15 | 4816746 | 4817093 | + | 0.796994 | Intergenic | Intergenic | -81818 | NM_001166066 | 223825 | Mm.159281 | NM_001166066 | ENSMUSG00000022155 | **Mroh2b** | 4930455B06Rik\|Heatr7b2 | maestro heat-like repeat family member 2B | protein-coding |
| chr5_96222929_96223350 | chr5 | 96222930 | 96223350 | + | 1.88717 | intron (NM_001039084, intron 2 of 6) | L1M4\|LINE\|L1 | 13025 | NM_053158 | 94061 | Mm.295499 | NM_053158 | ENSMUSG00000029486 | **Mrpl1** | 2410002L03Rik\|5830418D04Rik\|AI462572\|L1mt | mitochondrial ribosomal protein L1 | protein-coding |

| | | | | | | | | | | | | | | | | | |
|---|---|---|---|---|---|---|---|---|---|---|---|---|---|---|---|---|---|
| chr5_96219537_96219995 | chr5 | 96219538 | 96219995 | + | 1.42748 | intron (NM_001039084, intron 2 of 6) | intron (NM_001039084, intron 2 of 6) | 9651 | NM_053158 | 94061 | Mm.295499 | NM_053158 | ENSMUSG00000029486 | **Mrpl1** | 2410002L03Rik\|5830418D04Rik\|AI462572\|L1mt | mitochondrial ribosomal protein L1 | protein-coding |
| chr5_96221302_96221606 | chr5 | 96221303 | 96221606 | + | 0.994013 | intron (NM_001039084, intron 2 of 6) | ORR1E\|LTR\|ERVL-MaLR | 11339 | NM_053158 | 94061 | Mm.295499 | NM_053158 | ENSMUSG00000029486 | **Mrpl1** | 2410002L03Rik\|5830418D04Rik\|AI462572\|L1mt | mitochondrial ribosomal protein L1 | protein-coding |
| chr5_96221812_96222033 | chr5 | 96221813 | 96222033 | + | 0.803407 | intron (NM_001039084, intron 2 of 6) | MLT1H\|LTR\|ERVL-MaLR | 11808 | NM_053158 | 94061 | Mm.295499 | NM_053158 | ENSMUSG00000029486 | **Mrpl1** | 2410002L03Rik\|5830418D04Rik\|AI462572\|L1mt | mitochondrial ribosomal protein L1 | protein-coding |
| chr5_96204626_96205039 | chr5 | 96204627 | 96205039 | + | 0.777925 | Intergenic | Intergenic | -5282 | NM_001039084 | 94061 | Mm.295499 | NM_053158 | ENSMUSG00000029486 | **Mrpl1** | 2410002L03Rik\|5830418D04Rik\|AI462572\|L1mt | mitochondrial ribosomal protein L1 | protein-coding |
| chr5_96232510_96233056 | chr5 | 96232511 | 96233056 | + | 0.666751 | intron (NM_001039084, intron 6 of 6) | intron (NM_001039084, intron 6 of 6) | 22668 | NM_053158 | 94061 | Mm.295499 | NM_053158 | ENSMUSG00000029486 | **Mrpl1** | 2410002L03Rik\|5830418D04Rik\|AI462572\|L1mt | mitochondrial ribosomal protein L1 | protein-coding |
| chr5_96197522_96197880 | chr5 | 96197523 | 96197880 | + | 0.631115 | Intergenic | Intergenic | -12414 | NM_001039084 | 94061 | Mm.295499 | NM_053158 | ENSMUSG00000029486 | **Mrpl1** | 2410002L03Rik\|5830418D04Rik\|AI462572\|L1mt | mitochondrial ribosomal protein L1 | protein-coding |
| chr15_55503233_55503930 | chr15 | 55503234 | 55503930 | + | 0.901217 | intron (NM_181277, intron 45 of 47) | MTE2a\|LTR\|ERVL-MaLR | 53730 | NM_026759 | 68537 | Mm.236795 | NM_026759 | | **Mrpl13** | 1110002D09Rik | mitochondrial ribosomal protein L13 | protein-coding |
| chr15_55502347_55502907 | chr15 | 55502348 | 55502907 | + | 0.69994 | intron (NM_181277, intron 45 of 47) | intron (NM_181277, intron 45 of 47) | 54685 | NM_026759 | 68537 | Mm.236795 | NM_026759 | | **Mrpl13** | 1110002D09Rik | mitochondrial ribosomal protein L13 | protein-coding |
| chr17_45703188_45703744 | chr17 | 45703189 | 45703744 | + | 0.819154 | Intergenic | Intergenic | 17094 | NM_026732 | 68463 | Mm.379158 | NM_026732 | ENSMUSG00000023939 | **Mrpl14** | 1110006I11Rik\|AI846816\|MRP-L32\|Rpml32 | mitochondrial ribosomal protein L14 | protein-coding |
| chr6_71777984_71778368 | chr6 | 71777985 | 71778368 | + | 0.977301 | intron (NM_178608, intron 4 of 6) | intron (NM_178608, intron 4 of 6) | 45608 | NM_025430 | 66223 | Mm.28579 | NM_025430 | ENSMUSG00000052962 | **Mrpl35** | 1110066C01Rik | mitochondrial ribosomal protein L35 | protein-coding |
| chr6_71777766_71778372 | chr6 | 71777767 | 71778372 | + | 0.933371 | intron (NM_178608, intron 4 of 6) | intron (NM_178608, intron 4 of 6) | 45715 | NM_025430 | 66223 | Mm.28579 | NM_025430 | ENSMUSG00000052962 | **Mrpl35** | 1110066C01Rik | mitochondrial ribosomal protein L35 | protein-coding |
| chr6_71799359_71799937 | chr6 | 71799360 | 71799937 | + | 0.770655 | intron (NM_178608, intron 6 of 6) | MLT1F1\|LTR\|ERVL-MaLR | 24136 | NM_025430 | 66223 | Mm.28579 | NM_025430 | ENSMUSG00000052962 | **Mrpl35** | 1110066C01Rik | mitochondrial ribosomal protein L35 | protein-coding |
| chr6_71783226_71783539 | chr6 | 71783227 | 71783539 | + | 0.622896 | intron (NM_178608, intron 5 of 6) | intron (NM_178608, intron 5 of 6) | 40401 | NM_025430 | 66223 | Mm.28579 | NM_025430 | ENSMUSG00000052962 | **Mrpl35** | 1110066C01Rik | mitochondrial ribosomal protein L35 | protein-coding |
| chr11_97321902_97322458 | chr11 | 97321903 | 97322458 | + | 0.925395 | intron (NM_025927, intron 3 of 7) | intron (NM_025927, intron 3 of 7) | 6464 | NM_025927 | 67036 | Mm.5381 | NM_025927 | ENSMUSG00000018882 | **Mrpl45** | 2600005P05Rik\|AU043542\|R75241 | mitochondrial ribosomal protein L45 | protein-coding |
| chr11_97309281_97309721 | chr11 | 97309282 | 97309721 | + | 0.672499 | Intergenic | Intergenic | -6215 | NM_025927 | 67036 | Mm.5381 | NM_025927 | ENSMUSG00000018882 | **Mrpl45** | 2600005P05Rik\|AU043542\|R75241 | mitochondrial ribosomal protein L45 | protein-coding |
| chr7_78757852_78758483 | chr7 | 78757853 | 78758483 | + | 0.856193 | Intergenic | Intergenic | 24921 | NM_023331 | 67308 | Mm.193181 | NM_023331 | ENSMUSG00000030612 | **Mrpl46** | 3110052F15Rik\|C15orf4\|C80107\|LIECG2\|P2ECSL | mitochondrial ribosomal protein L46 | protein-coding |
| chr7_78798113_78798485 | chr7 | 78798114 | 78798485 | + | 0.993665 | Intergenic | Intergenic | 15168 | NM_026498 | 67994 | Mm.297033 | NM_026498 | ENSMUSG00000030611 | **Mrps11** | 0710005I03Rik\|C79873 | mitochondrial ribosomal protein S11 | protein-coding |
| chr1_160179396_160180267 | chr1 | 160179397 | 160180267 | + | 1.04715 | Intergenic | URR1B\|DNA\|hAT-Charlie | -15428 | NM_025474 | 64659 | Mm.29599 | NM_025474 | | **Mrps14** | 1810032L21Rik\|AL022847\|Rpms14 | mitochondrial ribosomal protein S14 | protein-coding |
| chr9_98600164_98600400 | chr9 | 98600165 | 98600400 | + | 0.852428 | intron (NM_025485, intron 1 of 7) | MTC\|LTR\|ERVL-MaLR | 1397 | NM_025485 | 64655 | Mm.17949 | NM_025485 | ENSMUSG00000032459 | **Mrps22** | 3100002P07Rik\|Rpms22 | mitochondrial ribosomal protein S22 | protein-coding |
| chr9_98600150_98600486 | chr9 | 98600151 | 98600486 | + | 0.80625 | intron (NM_025485, intron 1 of 7) | MTC\|LTR\|ERVL-MaLR | 1361 | NM_025485 | 64655 | Mm.17949 | NM_025485 | ENSMUSG00000032459 | **Mrps22** | 3100002P07Rik\|Rpms22 | mitochondrial ribosomal protein S22 | protein-coding |
| chr11_88215168_88215832 | chr11 | 88215169 | 88215832 | + | 0.86891 | Intergenic | Intergenic | 11112 | NM_001291271 | 64656 | Mm.168680 | NM_024174 | ENSMUSG00000023723 | **Mrps23** | D11Bwg1153e\|MRP-S23\|Rpms23\|S23mt | mitochondrial ribosomal protein S23 | protein-coding |
| chr11_88235466_88236114 | chr11 | 88235467 | 88236114 | + | 0.685168 | Intergenic | Intergenic | 31402 | NM_001291271 | 64656 | Mm.168680 | NM_024174 | ENSMUSG00000023723 | **Mrps23** | D11Bwg1153e\|MRP-S23\|Rpms23\|S23mt | mitochondrial ribosomal protein S23 | protein-coding |
| chr3_8845690_8846354 | chr3 | 8845691 | 8846354 | + | 1.96936 | intron (NM_025434, intron 2 of 2) | intron (NM_025434, intron 2 of 2) | 77835 | NM_025434 | 66230 | Mm.38460 | NM_025434 | ENSMUSG00000040269 | **Mrps28** | 1500012D08Rik\|AW061224 | mitochondrial ribosomal protein S28 | protein-coding |
| chr13_100746160_100746391 | chr13 | 100746161 | 100746391 | + | 0.675251 | Intergenic | B3\|SINE\|B2 | -1617 | NM_025369 | 66128 | Mm.306173 | NM_025369 | ENSMUSG00000061474 | **Mrps36** | 1110018B13Rik\|S36mt | mitochondrial ribosomal protein S36 | protein-coding |
| chr19_11527719_11528033 | chr19 | 11527720 | 11528033 | + | 1.05527 | intron (NM_027209, intron 5 of 6) | intron (NM_027209, intron 5 of 6) | -8973 | NM_025658 | 66607 | Mm.441056 | NM_025658 | ENSMUSG00000024678 | **Ms4a4d** | 2210417J23Rik | membrane-spanning 4-domains, subfamily A, member 4D | protein-coding |
| chr19_11527468_11528068 | chr19 | 11527469 | 11528068 | + | 0.888706 | intron (NM_027209, intron 5 of 6) | intron (NM_027209, intron 5 of 6) | -9081 | NM_025658 | 66607 | Mm.441056 | NM_025658 | ENSMUSG00000024678 | **Ms4a4d** | 2210417J23Rik | membrane-spanning 4-domains, subfamily A, member 4D | protein-coding |
| chr19_11559112_11559378 | chr19 | 11559113 | 11559378 | + | 0.700694 | TTS (NM_025658) | TTS (NM_025658) | 22396 | NM_025658 | 66607 | Mm.441056 | NM_025658 | ENSMUSG00000024678 | **Ms4a4d** | 2210417J23Rik | membrane-spanning 4-domains, subfamily A, member 4D | protein-coding |
| chr19_11511010_11511746 | chr19 | 11511011 | 11511746 | + | 2.04589 | intron (NM_001164202, intron 4 of 4) | intron (NM_001164202, intron 4 of 4) | -7181 | NM_027209 | 69774 | Mm.436639 | NM_027209 | ENSMUSG00000024677 | **Ms4a6b** | 1810027D10Rik | membrane-spanning 4-domains, subfamily A, member 6B | protein-coding |
| chr19_11527141_11528071 | chr19 | 11527142 | 11528071 | + | 0.854993 | intron (NM_027209, intron 5 of 6) | intron (NM_027209, intron 5 of 6) | 9047 | NM_027209 | 69774 | Mm.436639 | NM_027209 | ENSMUSG00000024677 | **Ms4a6b** | 1810027D10Rik | membrane-spanning 4-domains, subfamily A, member 6B | protein-coding |
| chr19_11466395_11466864 | chr19 | 11466396 | 11466864 | + | 0.642596 | Intergenic | Intergenic | -2738 | NM_001166376 | 73656 | Mm.440755 | NM_028595 | ENSMUSG00000079419 | **Ms4a6c** | 2200009H22Rik\|2210417N07Rik | membrane-spanning 4-domains, subfamily A, member 6C | protein-coding |
| chr19_11093368_11093831 | chr19 | 11093369 | 11093831 | + | 1.13903 | Intergenic | Intergenic | -12497 | NM_022430 | 64381 | Mm.34460 | NM_022430 | ENSMUSG00000024730 | **Ms4a8a** | 2010004L09Rik\|AI481240\|CD20L5\|Ms4a8 | membrane-spanning 4-domains, subfamily A, member 8A | protein-coding |
| chr1_14747307_14747707 | chr1 | 14747308 | 14747707 | + | 0.823433 | Intergenic | Intergenic | 8459 | NM_010827 | 17681 | Mm.5820 | NM_010827 | ENSMUSG00000025930 | **Msc** | MyoR\|bHLHa22 | musculin | protein-coding |
| chr3_153904605_153904889 | chr3 | 153904606 | 153904889 | + | 0.62881 | 5' UTR (NM_001282054, exon 1 of 21) | 5' UTR (NM_001282054, exon 1 of 21) | 144 | NM_001282054 | 55993 | Mm.272226 | NM_031870 | ENSMUSG00000005493 | **Msh4** | 4930485C04Rik\|AV144863\|mMsh4 | mutS homolog 4 (E. coli) | protein-coding |
| chr17_87979751_87979998 | chr17 | 87979752 | 87979998 | + | 1.12154 | intron (NM_010830, intron 1 of 9) | MTA_Mm\|LTR\|ERVL-MaLR | 4825 | NM_010830 | 17688 | Mm.18210 | NM_010830 | ENSMUSG00000005370 | **Msh6** | AU044881\|AW550279\|GTBP\|Gtmbp | mutS homolog 6 (E. coli) | protein-coding |
| chr5_115431023_115431322 | chr5 | 115431024 | 115431322 | + | 0.774872 | intron (NM_008629, intron 4 of 14) | intron (NM_008629, intron 4 of 14) | 1488 | NM_008629 | 17690 | Mm.5077 | NM_008629 | ENSMUSG00000054256 | **Msi1** | Msi1h\|Musahi1\|m-Msi-1 | musashi RNA-binding protein 1 | protein-coding |
| chr11_88578732_88579108 | chr11 | 88578733 | 88579108 | + | 0.938921 | intron (NM_054043, intron 6 of 13) | intron (NM_054043, intron 6 of 13) | 139309 | NM_054043 | 76626 | Mm.400451 | NM_054043 | ENSMUSG00000069769 | **Msi2** | 1700105C15Rik\|AI451722\|AI563628\|AW489193\|Msi2h | musashi RNA-binding protein 2 | protein-coding |
| chr11_88589260_88590255 | chr11 | 88589261 | 88590255 | + | 0.883695 | intron (NM_054043, intron 6 of 13) | intron (NM_054043, intron 6 of 13) | 128471 | NM_054043 | 76626 | Mm.400451 | NM_054043 | ENSMUSG00000069769 | **Msi2** | 1700105C15Rik\|AI451722\|AI563628\|AW489193\|Msi2h | musashi RNA-binding protein 2 | protein-coding |
| chr11_88552929_88553354 | chr11 | 88552930 | 88553354 | + | 0.880305 | intron (NM_054043, intron 6 of 13) | intron (NM_054043, intron 6 of 13) | 165087 | NM_054043 | 76626 | Mm.400451 | NM_054043 | ENSMUSG00000069769 | **Msi2** | 1700105C15Rik\|AI451722\|AI563628\|AW489193\|Msi2h | musashi RNA-binding protein 2 | protein-coding |
| chr11_88552874_88553773 | chr11 | 88552875 | 88553773 | + | 0.865531 | intron (NM_054043, intron 6 of 13) | intron (NM_054043, intron 6 of 13) | 164905 | NM_054043 | 76626 | Mm.400451 | NM_054043 | ENSMUSG00000069769 | **Msi2** | 1700105C15Rik\|AI451722\|AI563628\|AW489193\|Msi2h | musashi RNA-binding protein 2 | protein-coding |
| chr11_88553053_88553340 | chr11 | 88553054 | 88553340 | + | 0.86464 | intron (NM_054043, intron 6 of 13) | intron (NM_054043, intron 6 of 13) | 165032 | NM_054043 | 76626 | Mm.400451 | NM_054043 | ENSMUSG00000069769 | **Msi2** | 1700105C15Rik\|AI451722\|AI563628\|AW489193\|Msi2h | musashi RNA-binding protein 2 | protein-coding |
| chr11_88552933_88553556 | chr11 | 88552934 | 88553556 | + | 0.845517 | intron (NM_054043, intron 6 of 13) | intron (NM_054043, intron 6 of 13) | 164984 | NM_054043 | 76626 | Mm.400451 | NM_054043 | ENSMUSG00000069769 | **Msi2** | 1700105C15Rik\|AI451722\|AI563628\|AW489193\|Msi2h | musashi RNA-binding protein 2 | protein-coding |
| chr11_88637980_88638900 | chr11 | 88637981 | 88638900 | + | 0.752305 | intron (NM_054043, intron 5 of 13) | intron (NM_054043, intron 5 of 13) | 79789 | NM_054043 | 76626 | Mm.400451 | NM_054043 | ENSMUSG00000069769 | **Msi2** | 1700105C15Rik\|AI451722\|AI563628\|AW489193\|Msi2h | musashi RNA-binding protein 2 | protein-coding |
| chr11_88552417_88552693 | chr11 | 88552418 | 88552693 | + | 0.720209 | intron (NM_054043, intron 6 of 13) | intron (NM_054043, intron 6 of 13) | 165674 | NM_054043 | 76626 | Mm.400451 | NM_054043 | ENSMUSG00000069769 | **Msi2** | 1700105C15Rik\|AI451722\|AI563628\|AW489193\|Msi2h | musashi RNA-binding protein 2 | protein-coding |
| chr11_88690673_88691495 | chr11 | 88690674 | 88691495 | + | 0.688958 | intron (NM_001201341, intron 5 of 7) | intron (NM_001201341, intron 5 of 7) | 27145 | NM_054043 | 76626 | Mm.400451 | NM_054043 | ENSMUSG00000069769 | **Msi2** | 1700105C15Rik\|AI451722\|AI563628\|AW489193\|Msi2h | musashi RNA-binding protein 2 | protein-coding |
| chrX_168658982_168659571 | chrX | 168658983 | 168659571 | + | 0.819057 | intron (NM_010832, intron 9 of 11) | intron (NM_010832, intron 9 of 11) | 14625 | NM_010832 | 17692 | Mm.24248 | NM_010832 | ENSMUSG00000031358 | **Msl3** | AU018931\|Msl31\|Msl3l1 | male-specific lethal 3 homolog (Drosophila) | protein-coding |
| chr8_64725840_64726791 | chr8 | 64725841 | 64726791 | + | 1.33132 | intron (NM_025436, intron 2 of 5) | intron (NM_025436, intron 2 of 5) | 7262 | NM_025436 | 66234 | Mm.30119 | NM_025436 | ENSMUSG00000031604 | **Msmo1** | 1500001G16Rik\|C78600\|DESP4\|ERG25\|Sc4mol | methylsterol monoxygenase 1 | protein-coding |
| chrX_96121888_96122361 | chrX | 96121889 | 96122361 | + | 1.33093 | intron (NM_010833, intron 1 of 12) | intron (NM_010833, intron 1 of 12) | 26080 | NM_010833 | 17698 | Mm.138876 | NM_010833 | ENSMUSG00000031207 | **Msn** | C78546 | moesin | protein-coding |
| chrX_96122595_96122936 | chrX | 96122596 | 96122936 | + | 1.00537 | intron (NM_010833, intron 1 of 12) | intron (NM_010833, intron 1 of 12) | 26721 | NM_010833 | 17698 | Mm.138876 | NM_010833 | ENSMUSG00000031207 | **Msn** | C78546 | moesin | protein-coding |

| | | | | | | | | | | | | | | | | | |
|---|---|---|---|---|---|---|---|---|---|---|---|---|---|---|---|---|---|
| chr14_64368258_64368668 | chr14 | 64368259 | 64368668 | + | 1.10021 | intron (NM_001253715, intron 1 of 4) | intron (NM_001253715, intron 1 of 4) | 48493 | NM_001253716 | 110265 | Mm.26713 | NM_026322 | ENSMUSG00000054733 | **Msra** | 2310045J23Rik\|6530413P12Rik\|MSR-A | methionine sulfoxide reductase A | protein-coding |
| chr1_53049612_53050485 | chr1 | 53049613 | 53050485 | + | 1.03209 | Intergenic | ID_B1\|SINE\|B4 | -11614 | NM_010834 | 17700 | Mm.3514 | NM_010834 | ENSMUSG00000026100 | **Mstn** | Cmpt\|Gdf8 | myostatin | protein-coding |
| chr8_94186541_94186936 | chr8 | 94186542 | 94186936 | + | 1.01598 | Intergenic | Intergenic | 7650 | NM_013602 | 17748 | Mm.192991 | NM_013602 | ENSMUSG00000031765 | **Mt1** | MT-I\|Mt-1 | metallothionein 1 | protein-coding |
| chr8_94178836_94179352 | chr8 | 94178837 | 94179352 | + | 0.856814 | promoter-TSS (NM_013602) | promoter-TSS (NM_013602) | 5 | NM_013602 | 17748 | Mm.192991 | NM_013602 | ENSMUSG00000031765 | **Mt1** | MT-I\|Mt-1 | metallothionein 1 | protein-coding |
| chr8_94172121_94172929 | chr8 | 94172122 | 94172929 | + | 1.1859 | promoter-TSS (NM_008630) | promoter-TSS (NM_008630) | -93 | NM_008630 | 17750 | Mm.147226 | NM_008630 | ENSMUSG00000031762 | **Mt2** | AA409533\|MT-II\|Mt-2 | metallothionein 2 | protein-coding |
| chr8_94172282_94172878 | chr8 | 94172283 | 94172878 | + | 1.16671 | promoter-TSS (NM_008630) | promoter-TSS (NM_008630) | -38 | NM_008630 | 17750 | Mm.147226 | NM_008630 | ENSMUSG00000031762 | **Mt2** | AA409533\|MT-II\|Mt-2 | metallothionein 2 | protein-coding |
| chr8_94134598_94135135 | chr8 | 94134599 | 94135135 | + | 1.29984 | Intergenic | Intergenic | -2337 | NM_008631 | 17752 | Mm.1536 | NM_008631 | ENSMUSG00000031757 | **Mt4** | MT-IV | metallothionein 4 | protein-coding |
| chr17_83775966_83776304 | chr17 | 83775967 | 83776304 | + | 0.714809 | intron (NM_054082, intron 9 of 16) | intron (NM_054082, intron 9 of 16) | 69972 | NM_001171052 | 116871 | Mm.277668 | NM_054082 | ENSMUSG00000055817 | **Mta3** | 1110002J22Rik\|mKIAA1266 | metastasis associated 3 | protein-coding |
| chr4_89177762_89178336 | chr4 | 89177763 | 89178336 | + | 1.29969 | intron (NM_024433, intron 7 of 7) | intron (NM_024433, intron 7 of 7) | 40679 | NM_024433 | 66902 | Mm.28500 | NM_024433 | ENSMUSG00000062937 | **Mtap** | 1300019I21Rik\|MSAP | methylthioadenosine phosphorylase | protein-coding |
| chr4_89163641_89164223 | chr4 | 89163642 | 89164223 | + | 0.810737 | intron (NM_024433, intron 4 of 7) | intron (NM_024433, intron 4 of 7) | 26562 | NM_024433 | 66902 | Mm.28500 | NM_024433 | ENSMUSG00000062937 | **Mtap** | 1300019I21Rik\|MSAP | methylthioadenosine phosphorylase | protein-coding |
| chr4_89178421_89178598 | chr4 | 89178422 | 89178598 | + | 0.718011 | intron (NM_024433, intron 7 of 7) | B4A\|SINE\|B4 | 41140 | NM_024433 | 66902 | Mm.28500 | NM_024433 | ENSMUSG00000062937 | **Mtap** | 1300019I21Rik\|MSAP | methylthioadenosine phosphorylase | protein-coding |
| chr4_89164433_89165164 | chr4 | 89164434 | 89165164 | + | 0.699263 | intron (NM_024433, intron 4 of 7) | intron (NM_024433, intron 4 of 7) | 27429 | NM_024433 | 66902 | Mm.28500 | NM_024433 | ENSMUSG00000062937 | **Mtap** | 1300019I21Rik\|MSAP | methylthioadenosine phosphorylase | protein-coding |
| chrX_56849265_56849651 | chrX | 56849266 | 56849651 | + | 0.725698 | Intergenic | Intergenic | -27133 | NM_177293 | 320923 | Mm.186745 | NM_177293 | ENSMUSG00000067878 | **Mtap7d3** | 4933424A10Rik\|Map7d3 | MAP7 domain containing 3 | protein-coding |
| chr15_34064495_34064829 | chr15 | 34064496 | 34064829 | + | 0.769125 | Intergenic | Intergenic | -18057 | NM_026002 | 67154 | Mm.130883 | NM_026002 | ENSMUSG00000022255 | **Mtdh** | 2610103J23Rik\|3D3\|3D3/Lyric\|AEG-1\|AV353288\|D8Bwg1112e\|Lyric | metadherin | protein-coding |
| chr15_34026560_34027036 | chr15 | 34026561 | 34027036 | + | 0.665053 | Intergenic | RLTR51B_Mm\|LTR\|ERVK? | -55921 | NM_026002 | 67154 | Mm.130883 | NM_026002 | ENSMUSG00000022255 | **Mtdh** | 2610103J23Rik\|3D3\|3D3/Lyric\|AEG-1\|AV353288\|D8Bwg1112e\|Lyric | metadherin | protein-coding |
| chr4_124798443_124798705 | chr4 | 124798444 | 124798705 | + | 0.97097 | intron (NM_008385, intron 22 of 23) | L2\|LINE\|L2 | -3975 | NM_008636 | 17764 | Mm.272397 | NM_008636 | ENSMUSG00000028890 | **Mtf1** | MTF-1\|Thyls | metal response element binding transcription factor 1 | protein-coding |
| chr3_19223191_19223472 | chr3 | 19223192 | 19223472 | + | 0.700439 | 3' UTR (NM_001122759, exon 13 of 13) | 3' UTR (NM_001122759, exon 13 of 13) | 35111 | NR_045575 | 67472 | Mm.45125 | NM_026182 | ENSMUSG00000027601 | **Mtfr1** | 1300002C08Rik\|1700080D04Rik\|2810026O10Rik\|4930579E05Rik\|Chppr\|Fam54a2 | mitochondrial fission regulator 1 | protein-coding |
| chr10_4001521_4001918 | chr10 | 4001522 | 4001918 | + | 0.735783 | intron (NM_001170785, intron 7 of 27) | URR1A\|DNA\|hAT-Charlie | 28645 | NM_172308 | 270685 | Mm.184752 | NM_172308 | ENSMUSG00000040675 | **Mthfd1l** | 2410004L15Rik\|AI647056\|Fthfsdc1 | methylenetetrahydrofolate dehydrogenase (NADP+ dependent) 1-like | protein-coding |
| chr10_3967286_3967570 | chr10 | 3967287 | 3967570 | + | 0.735628 | TTS (NM_001159942) | TTS (NM_001159942) | -5647 | NM_001170786 | 270685 | Mm.184752 | NM_172308 | ENSMUSG00000040675 | **Mthfd1l** | 2410004L15Rik\|AI647056\|Fthfsdc1 | methylenetetrahydrofolate dehydrogenase (NADP+ dependent) 1-like | protein-coding |
| chr6_83311766_83312382 | chr6 | 83311767 | 83312382 | + | 0.881297 | intron (NM_008638, intron 2 of 7) | intron (NM_008638, intron 2 of 7) | 5530 | NM_008638 | 17768 | Mm.443 | NM_008638 | ENSMUSG00000005667 | **Mthfd2** | AW558851\|NMDMC | methylenetetrahydrofolate dehydrogenase (NAD+ dependent), methenyltetrahydrofolate cyclohydrolase | protein-coding |
| chr6_83317964_83318176 | chr6 | 83317965 | 83318176 | + | 0.73825 | promoter-TSS (NM_008638) | promoter-TSS (NM_008638) | -466 | NM_008638 | 17768 | Mm.443 | NM_008638 | ENSMUSG00000005667 | **Mthfd2** | AW558851\|NMDMC | methylenetetrahydrofolate dehydrogenase (NAD+ dependent), methenyltetrahydrofolate cyclohydrolase | protein-coding |
| chr9_89227768_89228500 | chr9 | 89227769 | 89228500 | + | 0.890297 | intron (NM_026829, intron 2 of 2) | intron (NM_026829, intron 2 of 2) | 16944 | NM_026829 | 107885 | Mm.486274 | NM_026829 | ENSMUSG00000066442 | **Mthfs** | 1110034I12Rik\|2310020H23Rik\|AI119695 | 5, 10-methenyltetrahydrofolate synthetase | protein-coding |
| chr9_89208852_89209321 | chr9 | 89208853 | 89209321 | + | 0.810073 | Intergenic | ORR1E\|LTR\|ERVL-MaLR | -2103 | NM_026829 | 107885 | Mm.486274 | NM_026829 | ENSMUSG00000066442 | **Mthfs** | 1110034I12Rik\|2310020H23Rik\|AI119695 | 5, 10-methenyltetrahydrofolate synthetase | protein-coding |
| chr9_89208719_89209374 | chr9 | 89208720 | 89209374 | + | 0.798486 | Intergenic | ORR1E\|LTR\|ERVL-MaLR | -2143 | NM_026829 | 107885 | Mm.486274 | NM_026829 | ENSMUSG00000066442 | **Mthfs** | 1110034I12Rik\|2310020H23Rik\|AI119695 | 5, 10-methenyltetrahydrofolate synthetase | protein-coding |
| chrX_71226512_71227280 | chrX | 71226513 | 71227280 | + | 0.727213 | intron (NM_019926, intron 1 of 14) | intron (NM_019926, intron 1 of 14) | 11814 | NM_019926 | 17772 | Mm.274981 | NM_019926 | ENSMUSG00000031337 | **Mtm1** | AF073996\|Mtm\|mKIAA4176 | X-linked myotubular myopathy gene 1 | protein-coding |
| chr3_96160092_96160759 | chr3 | 96160093 | 96160759 | + | 1.1306 | 3' UTR (NM_001025613, exon 13 of 13) | 3' UTR (NM_001025613, exon 13 of 13) | -1544 | NM_181409 | 194126 | Mm.268536 | NM_181409 | ENSMUSG00000045934 | **Mtmr11** | AI035306\|BC051083\|Cra | myotubularin related protein 11 | protein-coding |
| chr11_4521832_4522614 | chr11 | 4521833 | 4522614 | + | 0.932016 | intron (NM_028860, intron 4 of 17) | intron (NM_028860, intron 4 of 17) | 72592 | NM_028860 | 74302 | Mm.425669 | NM_028860 | ENSMUSG00000034354 | **Mtmr3** | 1700092A20Rik\|AI255150\|AW557713\|FYVE-DSP1\|ZFYVE10\|mKIAA0371 | myotubularin related protein 3 | protein-coding |
| chr18_4366490_4367112 | chr18 | 4366491 | 4367112 | + | 1.48033 | intron (NR_040328, intron 2 of 3) | intron (NR_040328, intron 2 of 3) | -8791 | NM_026157 | 67440 | Mm.49826 | NM_026157 | ENSMUSG00000024234 | **Mtpap** | 0610027A18Rik\|AW551379\|Papd1 | mitochondrial poly(A) polymerase | protein-coding |
| chr18_4366359_4367591 | chr18 | 4366360 | 4367591 | + | 1.29688 | intron (NR_040328, intron 2 of 3) | intron (NR_040328, intron 2 of 3) | -8617 | NM_026157 | 67440 | Mm.49826 | NM_026157 | ENSMUSG00000024234 | **Mtpap** | 0610027A18Rik\|AW551379\|Papd1 | mitochondrial poly(A) polymerase | protein-coding |
| chr18_4367166_4367482 | chr18 | 4367167 | 4367482 | + | 0.638147 | intron (NR_040328, intron 2 of 3) | intron (NR_040328, intron 2 of 3) | -8268 | NM_026157 | 67440 | Mm.49826 | NM_026157 | ENSMUSG00000024234 | **Mtpap** | 0610027A18Rik\|AW551379\|Papd1 | mitochondrial poly(A) polymerase | protein-coding |
| chr15_59105878_59106432 | chr15 | 59105879 | 59106432 | + | 0.972122 | Intergenic | Intergenic | -24129 | NM_144800 | 211401 | Mm.215481 | NM_144800 | ENSMUSG00000022353 | **Mtss1** | 2310003N14Rik\|BC024131\|D130001D01Rik\|Mim\|mKIAA0429 | metastasis suppressor 1 | protein-coding |
| chr15_59105957_59106338 | chr15 | 59105958 | 59106338 | + | 0.970775 | Intergenic | Intergenic | -24122 | NM_144800 | 211401 | Mm.215481 | NM_144800 | ENSMUSG00000022353 | **Mtss1** | 2310003N14Rik\|BC024131\|D130001D01Rik\|Mim\|mKIAA0429 | metastasis suppressor 1 | protein-coding |
| chr15_59105866_59106629 | chr15 | 59105867 | 59106629 | + | 0.947645 | Intergenic | Intergenic | -24222 | NM_144800 | 211401 | Mm.215481 | NM_144800 | ENSMUSG00000022353 | **Mtss1** | 2310003N14Rik\|BC024131\|D130001D01Rik\|Mim\|mKIAA0429 | metastasis suppressor 1 | protein-coding |
| chr15_59075868_59076195 | chr15 | 59075869 | 59076195 | + | 0.79257 | intron (NM_001146180, intron 1 of 13) | intron (NM_001146180, intron 1 of 13) | 5994 | NM_001146180 | 211401 | Mm.215481 | NM_144800 | ENSMUSG00000022353 | **Mtss1** | 2310003N14Rik\|BC024131\|D130001D01Rik\|Mim\|mKIAA0429 | metastasis suppressor 1 | protein-coding |
| chr15_59152730_59153406 | chr15 | 59152731 | 59153406 | + | 0.747002 | Intergenic | MLT1C\|LTR\|ERVL-MaLR | -71042 | NM_144800 | 211401 | Mm.215481 | NM_144800 | ENSMUSG00000022353 | **Mtss1** | 2310003N14Rik\|BC024131\|D130001D01Rik\|Mim\|mKIAA0429 | metastasis suppressor 1 | protein-coding |
| chr15_59145601_59146143 | chr15 | 59145602 | 59146143 | + | 0.65131 | Intergenic | Intergenic | -63846 | NM_144800 | 211401 | Mm.215481 | NM_144800 | ENSMUSG00000022353 | **Mtss1** | 2310003N14Rik\|BC024131\|D130001D01Rik\|Mim\|mKIAA0429 | metastasis suppressor 1 | protein-coding |
| chr2_75113699_75114788 | chr2 | 75113700 | 75114788 | + | 0.670237 | Intergenic | Intergenic | 288432 | NM_016804 | 53375 | Mm.292613 | NM_016804 | | **Mtx2** | 1500012G02Rik | metaxin 2 | protein-coding |
| chr2_75114081_75114782 | chr2 | 75114082 | 75114782 | + | 0.663322 | Intergenic | Intergenic | 288620 | NM_016804 | 53375 | Mm.292613 | NM_016804 | | **Mtx2** | 1500012G02Rik | metaxin 2 | protein-coding |
| chr14_30877691_30878178 | chr14 | 30877692 | 30878178 | + | 1.19973 | TTS (NM_028839) | TTS (NM_028839) | -1322 | NM_181390 | 66175 | Mm.220895 | NM_181390 | ENSMUSG00000042485 | **Mustn1** | 1110028G01Rik\|Mustang | musculoskeletal, embryonic nuclear protein 1 | protein-coding |
| chr14_30877652_30878481 | chr14 | 30877653 | 30878481 | + | 0.980675 | TTS (NM_028839) | TTS (NM_028839) | -1190 | NM_181390 | 66175 | Mm.220895 | NM_181390 | ENSMUSG00000042485 | **Mustn1** | 1110028G01Rik\|Mustang | musculoskeletal, embryonic nuclear protein 1 | protein-coding |
| chr2_34088929_34089195 | chr2 | 34088930 | 34089195 | + | 0.751626 | Intergenic | Intergenic | -201116 | NM_175184 | 72543 | Mm.254914 | NM_175184 | ENSMUSG00000038740 | **Mvb12b** | 2610200O14Rik\|2610528K11Rik\|AI414895\|Fam125b | multivesicular body subunit 12B | protein-coding |
| chr7_127008256_127008622 | chr7 | 127008257 | 127008622 | + | 0.812122 | intron (NM_080638, intron 1 of 14) | intron (NM_080638, intron 1 of 14) | 6155 | NM_080638 | 78388 | Mm.228797 | NM_080638 | ENSMUSG00000030681 | **Mvp** | 2310009M24Rik\|LRP\|VAULT1 | major vault protein | protein-coding |
| chr7_127012407_127012651 | chr7 | 127012408 | 127012651 | + | 0.767806 | intron (NM_080638, intron 1 of 14) | MER5A\|DNA\|hAT-Charlie | 2065 | NM_080638 | 78388 | Mm.228797 | NM_080638 | ENSMUSG00000030681 | **Mvp** | 2310009M24Rik\|LRP\|VAULT1 | major vault protein | protein-coding |
| chr7_127008235_127009004 | chr7 | 127008236 | 127009004 | + | 0.762118 | intron (NM_080638, intron 1 of 14) | intron (NM_080638, intron 1 of 14) | 5974 | NM_080638 | 78388 | Mm.228797 | NM_080638 | ENSMUSG00000030681 | **Mvp** | 2310009M24Rik\|LRP\|VAULT1 | major vault protein | protein-coding |
| chr16_97533876_97534082 | chr16 | 97533877 | 97534082 | + | 0.699618 | Intergenic | ORR1D2\|LTR\|ERVL-MaLR | -2102 | NR_003508 | 17858 | Mm.14157 | NM_013606 | ENSMUSG00000023341 | **Mx2** | AI528743\|Mx-2 | MX dynamin-like GTPase 2 | protein-coding |
| chr13_55339346_55339599 | chr13 | 55339347 | 55339599 | + | 0.894907 | Intergenic | ORR1A4\|LTR\|ERVL-MaLR | -9743 | NM_016662 | 17121 | Mm.20350 | NM_016662 | ENSMUSG00000021485 | **Mxd3** | 4631412E13Rik\|Mad3\|bHLHc13 | Max dimerization protein 3 | protein-coding |
| chr13_55339290_55339869 | chr13 | 55339291 | 55339869 | + | 0.86134 | Intergenic | Intergenic | -9850 | NM_016662 | 17121 | Mm.20350 | NM_016662 | ENSMUSG00000021485 | **Mxd3** | 4631412E13Rik\|Mad3\|bHLHc13 | Max dimerization protein 3 | protein-coding |
| chr19_53318124_53318694 | chr19 | 53318125 | 53318694 | + | 1.15671 | intron (NM_001008542, intron 1 of 5) | intron (NM_001008542, intron 1 of 5) | 7903 | NM_001008542 | 17859 | Mm.2154 | NM_010847 | ENSMUSG00000025025 | **Mxi1** | ENSMUSG00000067085\|Gm10197\|Mad2\|bHLHc11 | Max interacting protein 1 | protein-coding |
| chr10_21226214_21226563 | chr10 | 21226215 | 21226563 | + | 1.22202 | Intergenic | Intergenic | -65405 | NM_010848 | 17863 | Mm.52109 | NM_010848 | ENSMUSG00000019982 | **Myb** | AI550390\|M16449\|c-myb | myeloblastosis oncogene | protein-coding |
| chr10_21226049_21226620 | chr10 | 21226050 | 21226620 | + | 1.19693 | Intergenic | Intergenic | -65351 | NM_010848 | 17863 | Mm.52109 | NM_010848 | ENSMUSG00000019982 | **Myb** | AI550390\|M16449\|c-myb | myeloblastosis oncogene | protein-coding |
| chr10_21204248_21205069 | chr10 | 21204249 | 21205069 | + | 1.09184 | Intergenic | Intergenic | -43675 | NM_010848 | 17863 | Mm.52109 | NM_010848 | ENSMUSG00000019982 | **Myb** | AI550390\|M16449\|c-myb | myeloblastosis oncogene | protein-coding |
| chr10_21204243_21205366 | chr10 | 21204244 | 21205366 | + | 1.0057 | Intergenic | Intergenic | -43821 | NM_010848 | 17863 | Mm.52109 | NM_010848 | ENSMUSG00000019982 | **Myb** | AI550390\|M16449\|c-myb | myeloblastosis oncogene | protein-coding |
| chr10_21187740_21188210 | chr10 | 21187741 | 21188210 | + | 0.825541 | Intergenic | Intergenic | -26991 | NM_010848 | 17863 | Mm.52109 | NM_010848 | ENSMUSG00000019982 | **Myb** | AI550390\|M16449\|c-myb | myeloblastosis oncogene | protein-coding |
| chr10_21178822_21179107 | chr10 | 21178823 | 21179107 | + | 0.737925 | Intergenic | Intergenic | -17981 | NM_010848 | 17863 | Mm.52109 | NM_010848 | ENSMUSG00000019982 | **Myb** | AI550390\|M16449\|c-myb | myeloblastosis oncogene | protein-coding |

| | | | | | | | | | | | | | | | | | |
|---|---|---|---|---|---|---|---|---|---|---|---|---|---|---|---|---|---|
| chr10_21102612_21103059 | chr10 | 21102613 | 21103059 | + | 0.730569 | Intergenic | Intergenic | 58148 | NM_001198914 | 17863 | Mm.52109 | NM_010848 | ENSMUSG00000019982 | **Myb** | AI550390\|M16449\|c-myb | myeloblastosis oncogene | protein-coding |
| chr10_21115296_21115774 | chr10 | 21115297 | 21115774 | + | 0.710607 | Intergenic | RLTR19\|LTR\|ERVK | 45449 | NM_001198914 | 17863 | Mm.52109 | NM_010848 | ENSMUSG00000019982 | **Myb** | AI550390\|M16449\|c-myb | myeloblastosis oncogene | protein-coding |
| chr10_21218313_21218934 | chr10 | 21218314 | 21218934 | + | 0.66299 | Intergenic | Intergenic | -57640 | NM_010848 | 17863 | Mm.52109 | NM_010848 | ENSMUSG00000019982 | **Myb** | AI550390\|M16449\|c-myb | myeloblastosis oncogene | protein-coding |
| chr15_61990926_61991541 | chr15 | 61990927 | 61991541 | + | 0.926076 | TTS (NM_001177352) | TTS (NM_001177352) | 5893 | NM_001177352 | 17869 | Mm.2444 | NM_010849 | ENSMUSG00000022346 | **Myc** | AU016757\|Myc2\|Niard\|Nird\|bHLHe39 | myelocytomatosis oncogene | protein-coding |
| chr15_61998489_61999246 | chr15 | 61998490 | 61999246 | + | 0.896696 | Intergenic | Intergenic | 13527 | NM_001177352 | 17869 | Mm.2444 | NM_010849 | ENSMUSG00000022346 | **Myc** | AU016757\|Myc2\|Niard\|Nird\|bHLHe39 | myelocytomatosis oncogene | protein-coding |
| chr15_61998787_61999152 | chr15 | 61998788 | 61999152 | + | 0.876682 | Intergenic | Intergenic | 13629 | NM_001177352 | 17869 | Mm.2444 | NM_010849 | ENSMUSG00000022346 | **Myc** | AU016757\|Myc2\|Niard\|Nird\|bHLHe39 | myelocytomatosis oncogene | protein-coding |
| chr15_61981536_61981776 | chr15 | 61981537 | 61981776 | + | 0.599442 | Intergenic | Intergenic | -3685 | NM_001177354 | 17869 | Mm.2444 | NM_010849 | ENSMUSG00000022346 | **Myc** | AU016757\|Myc2\|Niard\|Nird\|bHLHe39 | myelocytomatosis oncogene | protein-coding |
| chr14_103375492_103375888 | chr14 | 103375493 | 103375888 | + | 0.606102 | Intergenic | MTC\|LTR\|ERVL-MaLR | -28890 | NM_207215 | 105689 | Mm.6478 | NM_207215 | ENSMUSG00000033004 | **Mycbp2** | AU023734\|AW107953\|AW546647\|C130061D10Rik\|Pam\|Phr1\|R75243 | MYC binding protein 2 | protein-coding |
| chr12_12941700_12942181 | chr12 | 12941701 | 12942181 | + | 0.70617 | promoter-TSS (NM_008709) | promoter-TSS (NM_008709) | -105 | NM_008709 | 18109 | Mm.16469 | NM_008709 | ENSMUSG00000037169 | **Mycn** | N-myc\|Nmyc\|Nmyc-1\|Nmyc1\|bHLHe37\|c-nmyc | v-myc myelocytomatosis viral related oncogene, neuroblastoma derived (avian) | protein-coding |
| chr10_5591464_5592036 | chr10 | 5591465 | 5592036 | + | 0.782329 | Intergenic | Intergenic | -1978 | NM_026793 | 68632 | Mm.338242 | NM_026793 | ENSMUSG00000046916 | **Myct1** | 1110020B04Rik\|AI225941\|AI642973\|Mtmc1 | myc target 1 | protein-coding |
| chr11_68674619_68675729 | chr11 | 68674620 | 68675729 | + | 0.670196 | Intergenic | Intergenic | -16741 | NM_175260 | 77579 | Mm.218233 | NM_175260 | ENSMUSG00000020900 | **Myh10** | 5730504C04Rik\|9330167F11Rik\|Fltn\|Myhn-2\|Myhn2\|NMHC II-B\|NMHC-B\|NMHCII-B\|NMMHC II-b\|NMMHC-B\|NMMHC-IIB\|SMemb\|mKIAA3005 | myosin, heavy polypeptide 10, non-muscle | protein-coding |
| chr16_14303683_14304030 | chr16 | 14303684 | 14304030 | + | 0.711246 | intron (NM_025345, intron 4 of 4) | intron (NM_025345, intron 4 of 4) | -12449 | NM_013607 | 17880 | Mm.250705 | NM_013607 | ENSMUSG00000018830 | **Myh11** | AV071570\|SM1\|SM2\|smMHC | myosin, heavy polypeptide 11, smooth muscle | protein-coding |
| chr16_49197764_49198467 | chr16 | 49197765 | 49198467 | + | 1.22162 | intron (NM_001166210, intron 40 of 40) | intron (NM_001166210, intron 40 of 40) | 140630 | NM_001166210 | 667772 | Mm.301387 | NM_001166210 | ENSMUSG00000092009 | **Myh15** | EG667772 | myosin, heavy chain 15 | protein-coding |
| chr16_49277088_49277391 | chr16 | 49277089 | 49277391 | + | 0.810922 | Intergenic | Intergenic | 219754 | NM_001166210 | 667772 | Mm.301387 | NM_001166210 | ENSMUSG00000092009 | **Myh15** | EG667772 | myosin, heavy chain 15 | protein-coding |
| chr15_77836151_77836452 | chr15 | 77836152 | 77836452 | + | 0.906591 | intron (NM_022410, intron 1 of 40) | intron (NM_022410, intron 1 of 40) | 5873 | NM_022410 | 17886 | Mm.29677 | NM_022410 | ENSMUSG00000022443 | **Myh9** | Fltn\|Myhn-1\|Myhn1\|NMHCIIA\|NMMHC-A\|NMMHC-IIA\|TU72.6 | myosin, heavy polypeptide 9, non-muscle | protein-coding |
| chr15_77811348_77811767 | chr15 | 77811349 | 77811767 | + | 0.869645 | intron (NM_022410, intron 2 of 40) | intron (NM_022410, intron 2 of 40) | 30617 | NM_022410 | 17886 | Mm.29677 | NM_022410 | ENSMUSG00000022443 | **Myh9** | Fltn\|Myhn-1\|Myhn1\|NMHCIIA\|NMMHC-A\|NMMHC-IIA\|TU72.6 | myosin, heavy polypeptide 9, non-muscle | protein-coding |
| chr15_77836015_77836452 | chr15 | 77836016 | 77836452 | + | 0.729065 | intron (NM_022410, intron 1 of 40) | intron (NM_022410, intron 1 of 40) | 5941 | NM_022410 | 17886 | Mm.29677 | NM_022410 | ENSMUSG00000022443 | **Myh9** | Fltn\|Myhn-1\|Myhn1\|NMHCIIA\|NMMHC-A\|NMMHC-IIA\|TU72.6 | myosin, heavy polypeptide 9, non-muscle | protein-coding |
| chr13_45418995_45419527 | chr13 | 45418996 | 45419527 | + | 1.22706 | Intergenic | Intergenic | 29519 | NM_153789 | 218203 | Mm.212855 | NM_153789 | ENSMUSG00000038175 | **Mylip** | 9430057C20Rik\|Mir | myosin regulatory light chain interacting protein | protein-coding |
| chr16_34756140_34756632 | chr16 | 34756141 | 34756632 | + | 0.676363 | Intergenic | Intergenic | -28564 | NM_139300 | 107589 | Mm.33360 | NM_139300 | ENSMUSG00000022836 | **Mylk** | 9530072E15Rik\|A930019C19Rik\|AW489456\|KRP\|MLCK108\|MLCK210\|Mlck\|smMLCK | myosin, light polypeptide kinase | protein-coding |
| chr2_152906265_152906794 | chr2 | 152906266 | 152906794 | + | 0.980961 | Intergenic | Intergenic | -4822 | NM_001081044 | 228785 | Mm.250604 | NM_001081044 | ENSMUSG00000027470 | **Mylk2** | 9830004H17Rik\|BB138278 | myosin, light polypeptide kinase 2, skeletal muscle | protein-coding |
| chr2_152906265_152907225 | chr2 | 152906266 | 152907225 | + | 0.749458 | Intergenic | Intergenic | -4607 | NM_001081044 | 228785 | Mm.250604 | NM_001081044 | ENSMUSG00000027470 | **Mylk2** | 9830004H17Rik\|BB138278 | myosin, light polypeptide kinase 2, skeletal muscle | protein-coding |
| chr2_152903678_152903871 | chr2 | 152903679 | 152903871 | + | 0.712905 | Intergenic | Intergenic | -7577 | NM_001081044 | 228785 | Mm.250604 | NM_001081044 | ENSMUSG00000027470 | **Mylk2** | 9830004H17Rik\|BB138278 | myosin, light polypeptide kinase 2, skeletal muscle | protein-coding |
| chr13_32680029_32680627 | chr13 | 32680030 | 32680627 | + | 0.919386 | Intergenic | Intergenic | 101451 | NM_001166030 | 238564 | Mm.60041 | NM_001166030 | ENSMUSG00000044951 | **Mylk4** | EG238564 | myosin light chain kinase family, member 4 | protein-coding |
| chr13_32692410_32692938 | chr13 | 32692411 | 32692938 | + | 0.740515 | Intergenic | L1M4\|LINE\|L1 | 89105 | NM_001166030 | 238564 | Mm.60041 | NM_001166030 | ENSMUSG00000044951 | **Mylk4** | EG238564 | myosin light chain kinase family, member 4 | protein-coding |
| chr15_25563718_25564085 | chr15 | 25563719 | 25564085 | + | 1.29005 | Intergenic | Intergenic | -58648 | NM_019472 | 17909 | Mm.60590 | NM_019472 | ENSMUSG00000022272 | **Myo10** | AW048724\|D15Ertd600e\|mKIAA0799 | myosin X | protein-coding |
| chr15_25575866_25576342 | chr15 | 25575867 | 25576342 | + | 0.904701 | Intergenic | ORR1D1\|LTR\|ERVL-MaLR | -46446 | NM_019472 | 17909 | Mm.60590 | NM_019472 | ENSMUSG00000022272 | **Myo10** | AW048724\|D15Ertd600e\|mKIAA0799 | myosin X | protein-coding |
| chr15_25561212_25561720 | chr15 | 25561213 | 25561720 | + | 0.885612 | Intergenic | Intergenic | -61084 | NM_019472 | 17909 | Mm.60590 | NM_019472 | ENSMUSG00000022272 | **Myo10** | AW048724\|D15Ertd600e\|mKIAA0799 | myosin X | protein-coding |
| chr15_25560966_25561712 | chr15 | 25560967 | 25561712 | + | 0.865171 | Intergenic | Intergenic | -61211 | NM_019472 | 17909 | Mm.60590 | NM_019472 | ENSMUSG00000022272 | **Myo10** | AW048724\|D15Ertd600e\|mKIAA0799 | myosin X | protein-coding |
| chr1_51820324_51821576 | chr1 | 51820325 | 51821576 | + | 0.775297 | intron (NM_001290982, intron 5 of 28) | intron (NM_001290982, intron 5 of 28) | 95028 | NM_001290982 | 17912 | Mm.3390 | NM_010863 | ENSMUSG00000018417 | **Myo1b** | AA406997\|MIH-L\|MMIa | myosin IB | protein-coding |
| chr11_80711346_80711713 | chr11 | 80711347 | 80711713 | + | 0.923465 | intron (NM_177390, intron 1 of 21) | intron (NM_177390, intron 1 of 21) | 68495 | NM_177390 | 338367 | Mm.151948 | NM_177390 | ENSMUSG00000035441 | **Myo1d** | 9930104H07Rik\|AW544947\|D11Ertd9e\|myosin-1d | myosin ID | protein-coding |
| chr11_80785640_80786292 | chr11 | 80785641 | 80786292 | + | 0.902164 | Intergenic | Intergenic | -5941 | NM_177390 | 338367 | Mm.151948 | NM_177390 | ENSMUSG00000035441 | **Myo1d** | 9930104H07Rik\|AW544947\|D11Ertd9e\|myosin-1d | myosin ID | protein-coding |
| chr9_70241595_70241942 | chr9 | 70241596 | 70241942 | + | 0.752486 | intron (NM_181072, intron 1 of 27) | intron (NM_181072, intron 1 of 27) | 34419 | NM_181072 | 71602 | Mm.249311 | NM_181072 | ENSMUSG00000032220 | **Myo1e** | 2310020N23Rik\|9130023P14Rik\|AA407778\|myosin-1e\|myr 3 | myosin IE | protein-coding |
| chr9_70284585_70284984 | chr9 | 70284586 | 70284984 | + | 0.696171 | intron (NM_181072, intron 1 of 27) | intron (NM_181072, intron 1 of 27) | 77435 | NM_181072 | 71602 | Mm.249311 | NM_181072 | ENSMUSG00000032220 | **Myo1e** | 2310020N23Rik\|9130023P14Rik\|AA407778\|myosin-1e\|myr 3 | myosin IE | protein-coding |
| chr9_70284587_70285253 | chr9 | 70284588 | 70285253 | + | 0.645457 | intron (NM_181072, intron 1 of 27) | intron (NM_181072, intron 1 of 27) | 77570 | NM_181072 | 71602 | Mm.249311 | NM_181072 | ENSMUSG00000032220 | **Myo1e** | 2310020N23Rik\|9130023P14Rik\|AA407778\|myosin-1e\|myr 3 | myosin IE | protein-coding |
| chr17_33552153_33552708 | chr17 | 33552154 | 33552708 | + | 1.57391 | intron (NR_037707, intron 24 of 24) | intron (NR_037707, intron 24 of 24) | -3276 | NM_053214 | 17916 | Mm.42019 | NM_053214 | ENSMUSG00000024300 | **Myo1f** | C330006B10Rik | myosin IF | protein-coding |
| chr11_6522733_6523551 | chr11 | 6522734 | 6523551 | + | 1.20567 | Intergenic | MER58B\|DNA\|hAT-Charlie | -2184 | NM_178440 | 246177 | Mm.239554 | NM_178440 | ENSMUSG00000020437 | **Myo1g** | E430002D17Rik | myosin IG | protein-coding |
| chr2_70118351_70119273 | chr2 | 70118352 | 70119273 | + | 1.48333 | intron (NM_177376, intron 7 of 34) | intron (NM_177376, intron 7 of 34) | 79686 | NM_177376 | 329421 | Mm.99648 | NM_177376 | ENSMUSG00000042064 | **Myo3b** | A430065P19Rik | myosin IIIB | protein-coding |
| chr2_70129526_70130046 | chr2 | 70129527 | 70130046 | + | 1.3502 | intron (NM_177376, intron 8 of 34) | intron (NM_177376, intron 8 of 34) | 90660 | NM_177376 | 329421 | Mm.99648 | NM_177376 | ENSMUSG00000042064 | **Myo3b** | A430065P19Rik | myosin IIIB | protein-coding |
| chr2_70127290_70127679 | chr2 | 70127291 | 70127679 | + | 0.796306 | intron (NM_177376, intron 8 of 34) | intron (NM_177376, intron 8 of 34) | 88359 | NM_177376 | 329421 | Mm.99648 | NM_177376 | ENSMUSG00000042064 | **Myo3b** | A430065P19Rik | myosin IIIB | protein-coding |
| chr9_75198418_75198895 | chr9 | 75198419 | 75198895 | + | 1.11946 | intron (NM_010864, intron 33 of 40) | intron (NM_010864, intron 33 of 40) | -33357 | NM_001081322 | 208943 | Mm.41339 | NM_001081322 | ENSMUSG00000033590 | **Myo5c** | 9130003O20Rik\|BB085560\|C330029I24 | myosin VC | protein-coding |
| chr9_75196431_75196746 | chr9 | 75196432 | 75196746 | + | 1.04295 | intron (NM_010864, intron 32 of 40) | intron (NM_010864, intron 32 of 40) | -35425 | NM_001081322 | 208943 | Mm.41339 | NM_001081322 | ENSMUSG00000033590 | **Myo5c** | 9130003O20Rik\|BB085560\|C330029I24 | myosin VC | protein-coding |
| chr9_75263998_75264344 | chr9 | 75263999 | 75264344 | + | 0.802891 | intron (NM_001081322, intron 12 of 40) | intron (NM_001081322, intron 12 of 40) | 32157 | NM_001081322 | 208943 | Mm.41339 | NM_001081322 | ENSMUSG00000033590 | **Myo5c** | 9130003O20Rik\|BB085560\|C330029I24 | myosin VC | protein-coding |
| chr8_71307270_71307799 | chr8 | 71307271 | 71307799 | + | 0.744796 | intron (NM_015742, intron 2 of 39) | MTD\|LTR\|ERVL-MaLR | 34821 | NM_001142323 | 17925 | Mm.33779 | NM_015742 | ENSMUSG00000004677 | **Myo9b** | - | myosin IXb | protein-coding |
| chr1_162621922_162622567 | chr1 | 162621923 | 162622567 | + | 1.11286 | Intergenic | Intergenic | -16905 | NM_010865 | 17926 | Mm.10694 | NM_010865 | ENSMUSG00000026697 | **Myoc** | AI957332\|GLC1A\|TIGR | myocilin | protein-coding |
| chr1_162621922_162622337 | chr1 | 162621923 | 162622337 | + | 1.00691 | Intergenic | Intergenic | -17020 | NM_010865 | 17926 | Mm.10694 | NM_010865 | ENSMUSG00000026697 | **Myoc** | AI957332\|GLC1A\|TIGR | myocilin | protein-coding |
| chr19_37954804_37955147 | chr19 | 37954805 | 37955147 | + | 0.678201 | intron (NM_001099634, intron 21 of 52) | intron (NM_001099634, intron 21 of 52) | 88601 | NM_001099634 | 226101 | Mm.34674 | NM_177035 | ENSMUSG00000048612 | **Myof** | 2310004N10Rik\|2310051D19Rik\|E030042N20Rik\|Fer1l3 | myoferlin | protein-coding |
| chr17_71089898_71090442 | chr17 | 71089899 | 71090442 | + | 0.910855 | intron (NM_010867, intron 21 of 37) | intron (NM_010867, intron 21 of 37) | 70613 | NM_001083934 | 17929 | Mm.4103 | NM_010867 | ENSMUSG00000024049 | **Myom1** | D430047A17Rik | myomesin 1 | protein-coding |
| chr17_71089249_71090485 | chr17 | 71089250 | 71090485 | + | 0.79791 | exon (NM_001083934, exon 20 of 37) | exon (NM_001083934, exon 20 of 37) | 70310 | NM_001083934 | 17929 | Mm.4103 | NM_010867 | ENSMUSG00000024049 | **Myom1** | D430047A17Rik | myomesin 1 | protein-coding |

| | | | | | | | | | | | | | | | | | |
|---|---|---|---|---|---|---|---|---|---|---|---|---|---|---|---|---|---|
| chr17_71010910_71011247 | chr17 | 71010911 | 71011247 | + | 0.61384 | Intergenic | ORR1A1\|LTR\|ERVL-MaLR | -8478 | NM_010867 | 17929 | Mm.4103 | NM_010867 | ENSMUSG00000024049 | **Myom1** | D430047A17Rik | myomesin 1 | protein-coding |
| chr3_123053592_123053915 | chr3 | 123053593 | 123053915 | + | 0.644277 | Intergenic | Intergenic | -18767 | NM_021503 | 59006 | Mm.141157 | NM_021503 | ENSMUSG00000028116 | **Myoz2** | 1110012I24Rik\|Fatz-2\|Myozl2 | myozenin 2 | protein-coding |
| chr10_63175037_63175712 | chr10 | 63175038 | 63175712 | + | 0.683594 | intron (NM_182992, intron 2 of 19) | intron (NM_182992, intron 2 of 19) | 28577 | NM_182992 | 68802 | Mm.259767 | NM_182992 | ENSMUSG00000020067 | **Mypn** | 1110056A04Rik\|9830112J03\|AI853556\|mKIAA4170 | myopalladin | protein-coding |
| chr19_10261293_10261754 | chr19 | 10261294 | 10261754 | + | 1.02578 | intron (NM_198114, intron 8 of 19) | intron (NM_198114, intron 8 of 19) | -20776 | NM_001033481 | 225908 | Mm.329716 | NM_001033481 | ENSMUSG00000036098 | **Myrf** | 6030439E18\|Gm1804\|Gm98\|Mrf | myelin regulatory factor | protein-coding |
| chr4_95001229_95001814 | chr4 | 95001230 | 95001814 | + | 1.61447 | Intergenic | Intergenic | -22422 | NM_177239 | 320713 | Mm.208868 | NM_177239 | ENSMUSG00000062627 | **Mysm1** | C130067A03Rik\|C530050H10Rik | myb-like, SWIRM and MPN domains 1 | protein-coding |
| chr4_95000145_95000617 | chr4 | 95000146 | 95000617 | + | 1.15879 | Intergenic | Intergenic | -21281 | NM_177239 | 320713 | Mm.208868 | NM_177239 | ENSMUSG00000062627 | **Mysm1** | C130067A03Rik\|C530050H10Rik | myb-like, SWIRM and MPN domains 1 | protein-coding |
| chr8_86819427_86819744 | chr8 | 86819428 | 86819744 | + | 1.10945 | Intergenic | Intergenic | 65672 | NM_030563 | 80750 | Mm.25117 | NM_030563 | ENSMUSG00000031652 | **N4bp1** | AI481586\|C81621 | NEDD4 binding protein 1 | protein-coding |
| chr8_86856283_86856745 | chr8 | 86856284 | 86856745 | + | 1.07474 | intron (NM_030563, intron 2 of 6) | intron (NM_030563, intron 2 of 6) | 28744 | NM_030563 | 80750 | Mm.25117 | NM_030563 | ENSMUSG00000031652 | **N4bp1** | AI481586\|C81621 | NEDD4 binding protein 1 | protein-coding |
| chr8_86870716_86871336 | chr8 | 86870717 | 86871336 | + | 0.959737 | intron (NM_030563, intron 1 of 6) | MTE2a\|LTR\|ERVL-MaLR | 14232 | NM_030563 | 80750 | Mm.25117 | NM_030563 | ENSMUSG00000031652 | **N4bp1** | AI481586\|C81621 | NEDD4 binding protein 1 | protein-coding |
| chr8_86838696_86839593 | chr8 | 86838697 | 86839593 | + | 0.929622 | Intergenic | Intergenic | 46113 | NM_030563 | 80750 | Mm.25117 | NM_030563 | ENSMUSG00000031652 | **N4bp1** | AI481586\|C81621 | NEDD4 binding protein 1 | protein-coding |
| chr8_86853620_86854283 | chr8 | 86853621 | 86854283 | + | 0.799741 | intron (NM_030563, intron 2 of 6) | intron (NM_030563, intron 2 of 6) | 31306 | NM_030563 | 80750 | Mm.25117 | NM_030563 | ENSMUSG00000031652 | **N4bp1** | AI481586\|C81621 | NEDD4 binding protein 1 | protein-coding |
| chr5_150621550_150622162 | chr5 | 150621551 | 150622162 | + | 0.865435 | Intergenic | Intergenic | -27331 | NM_133898 | 100637 | Mm.279932 | NM_133898 | ENSMUSG00000041132 | **N4bp2l1** | 2410024N18Rik\|AI428195\|B230342M21Rik | NEDD4 binding protein 2-like 1 | protein-coding |
| chr5_150660327_150660682 | chr5 | 150660328 | 150660682 | + | 0.964622 | intron (NM_201369, intron 2 of 5) | B3\|SINE\|B2 | 5107 | NM_201369 | 381695 | Mm.246934 | NM_201369 | ENSMUSG00000029655 | **N4bp2l2** | 2700092H06Rik\|zag1 | NEDD4 binding protein 2-like 2 | protein-coding |
| chr3_51425352_51425760 | chr3 | 51425353 | 51425760 | + | 0.988992 | intron (NM_053089, intron 1 of 19) | intron (NM_053089, intron 1 of 19) | 9540 | NM_053089 | 74838 | Mm.275281 | NM_053089 | ENSMUSG00000063273 | **Naa15** | 5730450D16Rik\|6330400I15\|ASTBDN\|Narg1\|Tbdn-1\|mNAT1 | N(alpha)-acetyltransferase 15, NatA auxiliary subunit | protein-coding |
| chr2_145904993_145905695 | chr2 | 145904994 | 145905695 | + | 1.10653 | intron (NM_001141965, intron 1 of 5) | intron (NM_001141965, intron 1 of 5) | 2103 | NM_001141965 | 67877 | Mm.151168 | NM_026425 | ENSMUSG00000002728 | **Naa20** | 1500004D14Rik\|2900026I01Rik\|AU041458\|D2Ertd186e\|Nat5 | N(alpha)-acetyltransferase 20, NatB catalytic subunit | protein-coding |
| chr2_145905248_145905659 | chr2 | 145905249 | 145905659 | + | 1.08978 | intron (NM_001141965, intron 1 of 5) | intron (NM_001141965, intron 1 of 5) | 2213 | NM_001141965 | 67877 | Mm.151168 | NM_026425 | ENSMUSG00000002728 | **Naa20** | 1500004D14Rik\|2900026I01Rik\|AU041458\|D2Ertd186e\|Nat5 | N(alpha)-acetyltransferase 20, NatB catalytic subunit | protein-coding |
| chr2_145900299_145900519 | chr2 | 145900300 | 145900519 | + | 1.04105 | Intergenic | MMERGLN_LTR\|LTR\|ERV1 | -2832 | NM_026425 | 67877 | Mm.151168 | NM_026425 | ENSMUSG00000002728 | **Naa20** | 1500004D14Rik\|2900026I01Rik\|AU041458\|D2Ertd186e\|Nat5 | N(alpha)-acetyltransferase 20, NatB catalytic subunit | protein-coding |
| chr1_52421639_52422171 | chr1 | 52421640 | 52422171 | + | 1.18422 | Intergenic | Intergenic | 78543 | NM_008667 | 17936 | Mm.25903 | NM_008667 | ENSMUSG00000002881 | **Nab1** | - | Ngfi-A binding protein 1 | protein-coding |
| chr1_52454460_52455233 | chr1 | 52454461 | 52455233 | + | 1.13314 | Intergenic | Intergenic | 45601 | NM_008667 | 17936 | Mm.25903 | NM_008667 | ENSMUSG00000002881 | **Nab1** | - | Ngfi-A binding protein 1 | protein-coding |
| chr1_52421481_52422248 | chr1 | 52421482 | 52422248 | + | 1.03833 | Intergenic | Intergenic | 78583 | NM_008667 | 17936 | Mm.25903 | NM_008667 | ENSMUSG00000002881 | **Nab1** | - | Ngfi-A binding protein 1 | protein-coding |
| chr1_52427204_52427642 | chr1 | 52427205 | 52427642 | + | 0.888616 | Intergenic | MTA_Mm\|LTR\|ERVL-MaLR | 73025 | NM_008667 | 17936 | Mm.25903 | NM_008667 | ENSMUSG00000002881 | **Nab1** | - | Ngfi-A binding protein 1 | protein-coding |
| chr1_52470790_52471498 | chr1 | 52470791 | 52471498 | + | 0.764604 | intron (NM_008667, intron 4 of 7) | intron (NM_008667, intron 4 of 7) | 29304 | NM_008667 | 17936 | Mm.25903 | NM_008667 | ENSMUSG00000002881 | **Nab1** | - | Ngfi-A binding protein 1 | protein-coding |
| chr1_52442460_52443263 | chr1 | 52442461 | 52443263 | + | 0.670175 | Intergenic | Intergenic | 57586 | NM_008667 | 17936 | Mm.25903 | NM_008667 | ENSMUSG00000002881 | **Nab1** | - | Ngfi-A binding protein 1 | protein-coding |
| chr1_52419482_52419800 | chr1 | 52419483 | 52419800 | + | 0.653192 | Intergenic | Intergenic | 80807 | NM_008667 | 17936 | Mm.25903 | NM_008667 | ENSMUSG00000002881 | **Nab1** | - | Ngfi-A binding protein 1 | protein-coding |
| chr1_51537647_51538093 | chr1 | 51537648 | 51538093 | + | 1.57853 | Intergenic | MTC\|LTR\|ERVL-MaLR | -59471 | NM_028696 | 109019 | Mm.196290 | NM_028696 | ENSMUSG00000026107 | **Nabp1** | 4930434H03Rik\|4930442A21Rik\|4930488J04Rik\|4933440J18Rik\|5830411E10Rik\|AI852561\|Nbp1\|Obfc2a\|Ssb2 | nucleic acid binding protein 1 | protein-coding |
| chr1_51535165_51535801 | chr1 | 51535166 | 51535801 | + | 1.39928 | Intergenic | (CA)n\|Simple_repeat\|Simple_repeat | -57084 | NM_028696 | 109019 | Mm.196290 | NM_028696 | ENSMUSG00000026107 | **Nabp1** | 4930434H03Rik\|4930442A21Rik\|4930488J04Rik\|4933440J18Rik\|5830411E10Rik\|AI852561\|Nbp1\|Obfc2a\|Ssb2 | nucleic acid binding protein 1 | protein-coding |
| chr1_51538914_51539397 | chr1 | 51538915 | 51539397 | + | 1.28807 | Intergenic | Intergenic | -60757 | NM_028696 | 109019 | Mm.196290 | NM_028696 | ENSMUSG00000026107 | **Nabp1** | 4930434H03Rik\|4930442A21Rik\|4930488J04Rik\|4933440J18Rik\|5830411E10Rik\|AI852561\|Nbp1\|Obfc2a\|Ssb2 | nucleic acid binding protein 1 | protein-coding |
| chr1_51448614_51449275 | chr1 | 51448615 | 51449275 | + | 1.28034 | Intergenic | Intergenic | 29454 | NM_028696 | 109019 | Mm.196290 | NM_028696 | ENSMUSG00000026107 | **Nabp1** | 4930434H03Rik\|4930442A21Rik\|4930488J04Rik\|4933440J18Rik\|5830411E10Rik\|AI852561\|Nbp1\|Obfc2a\|Ssb2 | nucleic acid binding protein 1 | protein-coding |
| chr1_51514289_51515019 | chr1 | 51514290 | 51515019 | + | 1.08831 | Intergenic | Intergenic | -36255 | NM_028696 | 109019 | Mm.196290 | NM_028696 | ENSMUSG00000026107 | **Nabp1** | 4930434H03Rik\|4930442A21Rik\|4930488J04Rik\|4933440J18Rik\|5830411E10Rik\|AI852561\|Nbp1\|Obfc2a\|Ssb2 | nucleic acid binding protein 1 | protein-coding |
| chr1_51512999_51513408 | chr1 | 51513000 | 51513408 | + | 1.03138 | Intergenic | Intergenic | -34805 | NM_028696 | 109019 | Mm.196290 | NM_028696 | ENSMUSG00000026107 | **Nabp1** | 4930434H03Rik\|4930442A21Rik\|4930488J04Rik\|4933440J18Rik\|5830411E10Rik\|AI852561\|Nbp1\|Obfc2a\|Ssb2 | nucleic acid binding protein 1 | protein-coding |
| chr1_51509741_51510079 | chr1 | 51509742 | 51510079 | + | 0.965134 | Intergenic | MLT1K\|LTR\|ERVL-MaLR | -31511 | NM_028696 | 109019 | Mm.196290 | NM_028696 | ENSMUSG00000026107 | **Nabp1** | 4930434H03Rik\|4930442A21Rik\|4930488J04Rik\|4933440J18Rik\|5830411E10Rik\|AI852561\|Nbp1\|Obfc2a\|Ssb2 | nucleic acid binding protein 1 | protein-coding |
| chr1_51536217_51536691 | chr1 | 51536218 | 51536691 | + | 0.875988 | Intergenic | ORR1G\|LTR\|ERVL-MaLR | -58055 | NM_028696 | 109019 | Mm.196290 | NM_028696 | ENSMUSG00000026107 | **Nabp1** | 4930434H03Rik\|4930442A21Rik\|4930488J04Rik\|4933440J18Rik\|5830411E10Rik\|AI852561\|Nbp1\|Obfc2a\|Ssb2 | nucleic acid binding protein 1 | protein-coding |
| chr1_51509841_51510074 | chr1 | 51509842 | 51510074 | + | 0.864228 | Intergenic | MLT1K\|LTR\|ERVL-MaLR | -31559 | NM_028696 | 109019 | Mm.196290 | NM_028696 | ENSMUSG00000026107 | **Nabp1** | 4930434H03Rik\|4930442A21Rik\|4930488J04Rik\|4933440J18Rik\|5830411E10Rik\|AI852561\|Nbp1\|Obfc2a\|Ssb2 | nucleic acid binding protein 1 | protein-coding |
| chr1_51499136_51499457 | chr1 | 51499137 | 51499457 | + | 0.840148 | Intergenic | Intergenic | -20898 | NM_028696 | 109019 | Mm.196290 | NM_028696 | ENSMUSG00000026107 | **Nabp1** | 4930434H03Rik\|4930442A21Rik\|4930488J04Rik\|4933440J18Rik\|5830411E10Rik\|AI852561\|Nbp1\|Obfc2a\|Ssb2 | nucleic acid binding protein 1 | protein-coding |
| chr1_51499012_51499467 | chr1 | 51499013 | 51499467 | + | 0.771403 | Intergenic | Intergenic | -20841 | NM_028696 | 109019 | Mm.196290 | NM_028696 | ENSMUSG00000026107 | **Nabp1** | 4930434H03Rik\|4930442A21Rik\|4930488J04Rik\|4933440J18Rik\|5830411E10Rik\|AI852561\|Nbp1\|Obfc2a\|Ssb2 | nucleic acid binding protein 1 | protein-coding |
| chr10_128041063_128041343 | chr10 | 128041064 | 128041343 | + | 0.700621 | exon (NM_001113199, exon 3 of 9) | exon (NM_001113199, exon 3 of 9) | 5857 | NM_001113199 | 17938 | Mm.3746 | NM_013608 | ENSMUSG00000061315 | **Naca** | AL022831\|AL024382\|Gm1878\|mKIAA0363\|skNAC | nascent polypeptide-associated complex alpha polypeptide | protein-coding |
| chr13_100440778_100441120 | chr13 | 100440779 | 100441120 | + | 0.697881 | intron (NM_008670, intron 4 of 15) | BGLII\|LTR\|ERVK | 11915 | NM_008670 | 17940 | Mm.6898 | NM_008670 | ENSMUSG00000021640 | **Naip1** | AV364616\|Birc1a\|D13Lsd1\|Naip\|Naip-rs1 | NLR family, apoptosis inhibitory protein 1 | protein-coding |
| chr12_32795719_32796098 | chr12 | 32795720 | 32796098 | + | 0.730849 | Intergenic | Intergenic | -24426 | NM_021524 | 59027 | Mm.202727 | NM_021524 | ENSMUSG00000020572 | **Nampt** | 1110035O14Rik\|AI314458\|AI480535\|NAmPRTase\|Pbef\|Pbef1\|Visfatin | nicotinamide phosphoribosyltransferase | protein-coding |
| chr7_18967632_18968127 | chr7 | 18967633 | 18968127 | + | 1.03698 | Intergenic | LTR67B\|LTR\|ERVL | -19644 | NM_194064 | 378430 | Mm.277153 | NM_194064 | ENSMUSG00000051965 | **Nanos2** | nos2 | nanos homolog 2 (Drosophila) | protein-coding |
| chr7_18967521_18968256 | chr7 | 18967522 | 18968256 | + | 0.777924 | Intergenic | LTR67B\|LTR\|ERVL | -19635 | NM_194064 | 378430 | Mm.277153 | NM_194064 | ENSMUSG00000051965 | **Nanos2** | nos2 | nanos homolog 2 (Drosophila) | protein-coding |
| chr7_96920496_96921146 | chr7 | 96920497 | 96921146 | + | 1.34356 | Intergenic | L1MDa\|LINE\|L1 | -30706 | NM_153591 | 244141 | Mm.274864 | NM_153591 | ENSMUSG00000018995 | **Nars2** | AI875199 | asparaginyl-tRNA synthetase 2 (mitochondrial)(putative) | protein-coding |
| chr2_103721526_103722166 | chr2 | 103721527 | 103722166 | + | 0.662989 | exon (NM_153126, exon 29 of 29) | exon (NM_153126, exon 29 of 29) | 39404 | NM_153126 | 98956 | Mm.288005 | NM_153126 | ENSMUSG00000027185 | **Nat10** | AI429152 | N-acetyltransferase 10 | protein-coding |
| chr5_34032112_34032800 | chr5 | 34032113 | 34032800 | + | 0.784276 | intron (NM_181857, intron 19 of 24) | URR1B\|DNA\|hAT-Charlie | 36472 | NM_001001985 | 269642 | Mm.274610 | NM_001001985 | ENSMUSG00000048142 | **Nat8l** | 1110038O08Rik\|Shati | N-acetyltransferase 8-like | protein-coding |

| | | | | | | | | | | | | | | | | | |
|---|---|---|---|---|---|---|---|---|---|---|---|---|---|---|---|---|---|
| chr1_135579644_135580335 | chr1 | 135579645 | 135580335 | + | 0.740025 | intron (NM_173437, intron 1 of 28) | intron (NM_173437, intron 1 of 28) | 5365 | NM_173437 | 215690 | Mm.34977 | NM_173437 | ENSMUSG00000009418 | **Nav1** | 9530089B19\|9930003A20Rik\|C230080M11Rik\|Pomfil3\|Unc53h1\|mKIAA1151\|steerin-1 | neuron navigator 1 | protein-coding |
| chr1_135598399_135598837 | chr1 | 135598400 | 135598837 | + | 0.635981 | Intergenic | Intergenic | -13263 | NM_173437 | 215690 | Mm.34977 | NM_173437 | ENSMUSG00000009418 | **Nav1** | 9530089B19\|9930003A20Rik\|C230080M11Rik\|Pomfil3\|Unc53h1\|mKIAA1151\|steerin-1 | neuron navigator 1 | protein-coding |
| chr7_49245805_49246548 | chr7 | 49245806 | 49246548 | + | 0.712685 | promoter-TSS (NM_175272) | promoter-TSS (NM_175272) | -12 | NM_175272 | 78286 | Mm.333881 | NM_175272 | ENSMUSG00000052512 | **Nav2** | 5330421F07Rik\|AU067780\|BQ960892\|HELAD1\|POMFIL2\|RAINB1\|RAINB2\|Unc53H2\|mKIAA3015 | neuron navigator 2 | protein-coding |
| chr7_49245798_49246790 | chr7 | 49245799 | 49246790 | + | 0.679745 | 5' UTR (NM_175272, exon 1 of 38) | 5' UTR (NM_175272, exon 1 of 38) | 105 | NM_175272 | 78286 | Mm.333881 | NM_175272 | ENSMUSG00000052512 | **Nav2** | 5330421F07Rik\|AU067780\|BQ960892\|HELAD1\|POMFIL2\|RAINB1\|RAINB2\|Unc53H2\|mKIAA3015 | neuron navigator 2 | protein-coding |
| chr12_13462314_13462904 | chr12 | 13462315 | 13462904 | + | 1.10216 | intron (NM_027706, intron 40 of 51) | intron (NM_027706, intron 40 of 51) | 193482 | NM_027706 | 71169 | Mm.177520 | NM_027706 | ENSMUSG00000020576 | **Nbas** | 4933425L03Rik\|Nag | neuroblastoma amplified sequence | protein-coding |
| chr1_60173092_60173640 | chr1 | 60173093 | 60173640 | + | 0.861652 | Intergenic | RLTR11A\|LTR\|ERVK | -7233 | NM_173444 | 269198 | Mm.205224 | NM_173444 | ENSMUSG00000073664 | **Nbeal1** | 2310076G13Rik\|A530050O19Rik\|A530083I02Rik\|AI662228\|ALS2CR17 | neurobeachin like 1 | protein-coding |
| chr15_37607488_37607978 | chr15 | 37607489 | 37607978 | + | 1.29813 | intron (NM_134094, intron 2 of 6) | intron (NM_134094, intron 2 of 6) | -148351 | NM_001170868 | 52589 | Mm.283370 | NM_134094 | ENSMUSG00000051359 | **Ncald** | AI848120\|D030020D09Rik\|D15Ertd412e | neurocalcin delta | protein-coding |
| chr15_37607489_37608211 | chr15 | 37607490 | 37608211 | + | 1.28058 | intron (NM_134094, intron 2 of 6) | intron (NM_134094, intron 2 of 6) | -148468 | NM_001170868 | 52589 | Mm.283370 | NM_134094 | ENSMUSG00000051359 | **Ncald** | AI848120\|D030020D09Rik\|D15Ertd412e | neurocalcin delta | protein-coding |
| chr15_37679394_37680112 | chr15 | 37679395 | 37680112 | + | 0.753392 | intron (NM_134094, intron 1 of 6) | intron (NM_134094, intron 1 of 6) | 112247 | NM_001170866 | 52589 | Mm.283370 | NM_134094 | ENSMUSG00000051359 | **Ncald** | AI848120\|D030020D09Rik\|D15Ertd412e | neurocalcin delta | protein-coding |
| chr15_37656455_37657316 | chr15 | 37656456 | 37657316 | + | 0.714834 | intron (NM_134094, intron 2 of 6) | ORR1D2\|LTR\|ERVL-MaLR | 135114 | NM_001170866 | 52589 | Mm.283370 | NM_134094 | ENSMUSG00000051359 | **Ncald** | AI848120\|D030020D09Rik\|D15Ertd412e | neurocalcin delta | protein-coding |
| chr15_37461354_37461982 | chr15 | 37461355 | 37461982 | + | 0.669135 | intron (NM_134094, intron 4 of 6) | intron (NM_134094, intron 4 of 6) | -2286 | NM_001170868 | 52589 | Mm.283370 | NM_134094 | ENSMUSG00000051359 | **Ncald** | AI848120\|D030020D09Rik\|D15Ertd412e | neurocalcin delta | protein-coding |
| chr16_81365878_81366257 | chr16 | 81365879 | 81366257 | + | 1.17062 | intron (NM_010954, intron 1 of 15) | intron (NM_010954, intron 1 of 15) | 165371 | NM_010954 | 17968 | Mm.433941 | NM_010954 | ENSMUSG00000022762 | **Ncam2** | Ncam-2\|Ocam\|RNCAM | neural cell adhesion molecule 2 | protein-coding |
| chr8_70116971_70117250 | chr8 | 70116972 | 70117250 | + | 0.687084 | intron (NM_007789, intron 1 of 14) | intron (NM_007789, intron 1 of 14) | 3733 | NM_007789 | 13004 | Mm.268079 | NM_007789 | ENSMUSG00000002341 | **Ncan** | C230035B04\|Cspg3\|Cspg3-rs\|Tgfbit | neurocan | protein-coding |
| chr5_134248812_134249460 | chr5 | 134248813 | 134249460 | + | 0.91233 | intron (NM_001080749, intron 19 of 32) | ORR1C2\|LTR\|ERVL-MaLR | -19511 | NR_104383 | 17969 | Mm.425296 | NM_010876 | ENSMUSG00000015950 | **Ncf1** | NCF-47K\|NOXO2\|Ncf-1\|p47-phox\|p47<phox>\|p47phox | neutrophil cytosolic factor 1 | protein-coding |
| chr1_152841050_152841422 | chr1 | 152841051 | 152841422 | + | 0.681634 | intron (NM_001005507, intron 18 of 21) | intron (NM_001005507, intron 18 of 21) | 33346 | NM_010877 | 17970 | Mm.270307 | NM_010877 | ENSMUSG00000026480 | **Ncf2** | NOXA2\|Ncf-2\|p67phox | neutrophil cytosolic factor 2 | protein-coding |
| chr1_43461622_43462118 | chr1 | 43461623 | 43462118 | + | 1.1111 | intron (NM_010879, intron 1 of 4) | MTC\|LTR\|ERVL-MaLR | 16119 | NM_010879 | 17974 | Mm.389903 | NM_010879 | ENSMUSG00000066877 | **Nck2** | 4833426I10Rik\|Grb4\|NCKbeta\|Nck-2 | non-catalytic region of tyrosine kinase adaptor protein 2 | protein-coding |
| chr1_43412864_43413319 | chr1 | 43412865 | 43413319 | + | 1.05543 | Intergenic | MER5A\|DNA\|hAT-Charlie | -32659 | NM_010879 | 17974 | Mm.389903 | NM_010879 | ENSMUSG00000066877 | **Nck2** | 4833426I10Rik\|Grb4\|NCKbeta\|Nck-2 | non-catalytic region of tyrosine kinase adaptor protein 2 | protein-coding |
| chr1_43469029_43469448 | chr1 | 43469030 | 43469448 | + | 0.985092 | intron (NM_010879, intron 1 of 4) | intron (NM_010879, intron 1 of 4) | 23488 | NM_010879 | 17974 | Mm.389903 | NM_010879 | ENSMUSG00000066877 | **Nck2** | 4833426I10Rik\|Grb4\|NCKbeta\|Nck-2 | non-catalytic region of tyrosine kinase adaptor protein 2 | protein-coding |
| chr1_43471321_43471615 | chr1 | 43471322 | 43471615 | + | 0.869771 | intron (NM_010879, intron 1 of 4) | intron (NM_010879, intron 1 of 4) | 25717 | NM_010879 | 17974 | Mm.389903 | NM_010879 | ENSMUSG00000066877 | **Nck2** | 4833426I10Rik\|Grb4\|NCKbeta\|Nck-2 | non-catalytic region of tyrosine kinase adaptor protein 2 | protein-coding |
| chr1_43474093_43475527 | chr1 | 43474094 | 43475527 | + | 0.726459 | intron (NM_010879, intron 1 of 4) | LTR33B\|LTR\|ERVL | 29059 | NM_010879 | 17974 | Mm.389903 | NM_010879 | ENSMUSG00000066877 | **Nck2** | 4833426I10Rik\|Grb4\|NCKbeta\|Nck-2 | non-catalytic region of tyrosine kinase adaptor protein 2 | protein-coding |
| chr1_126747877_126748457 | chr1 | 126747878 | 126748457 | + | 1.35168 | intron (NM_176957, intron 1 of 12) | intron (NM_176957, intron 1 of 12) | 82465 | NM_001081756 | 210356 | Mm.377234 | NM_172484 | ENSMUSG00000049690 | **Nckap5** | 8430408F21\|D130011D22Rik\|E030049G20Rik\|Erih1\|Erih2\|Gm1548\|Nap5 | NCK-associated protein 5 | protein-coding |
| chr1_126817671_126818053 | chr1 | 126817672 | 126818053 | + | 1.01925 | intron (NM_176957, intron 1 of 12) | intron (NM_176957, intron 1 of 12) | 12770 | NM_001081756 | 210356 | Mm.377234 | NM_172484 | ENSMUSG00000049690 | **Nckap5** | 8430408F21\|D130011D22Rik\|E030049G20Rik\|Erih1\|Erih2\|Gm1548\|Nap5 | NCK-associated protein 5 | protein-coding |
| chr1_126817406_126818479 | chr1 | 126817407 | 126818479 | + | 0.924538 | intron (NM_176957, intron 1 of 12) | intron (NM_176957, intron 1 of 12) | 12689 | NM_001081756 | 210356 | Mm.377234 | NM_172484 | ENSMUSG00000049690 | **Nckap5** | 8430408F21\|D130011D22Rik\|E030049G20Rik\|Erih1\|Erih2\|Gm1548\|Nap5 | NCK-associated protein 5 | protein-coding |
| chr12_4433254_4433991 | chr12 | 4433255 | 4433991 | + | 1.68672 | intron (NM_010881, intron 1 of 22) | intron (NM_010881, intron 1 of 22) | 43559 | NM_010881 | 17977 | Mm.301039 | NM_010881 | ENSMUSG00000020647 | **Ncoa1** | KAT13A\|SRC-1\|SRC1\|bHLHe74 | nuclear receptor coactivator 1 | protein-coding |
| chr12_4424709_4425583 | chr12 | 4424710 | 4425583 | + | 0.919781 | intron (NM_010881, intron 1 of 22) | intron (NM_010881, intron 1 of 22) | 52036 | NM_010881 | 17977 | Mm.301039 | NM_010881 | ENSMUSG00000020647 | **Ncoa1** | KAT13A\|SRC-1\|SRC1\|bHLHe74 | nuclear receptor coactivator 1 | protein-coding |
| chr12_4454127_4454674 | chr12 | 4454128 | 4454674 | + | 0.901045 | intron (NM_010881, intron 1 of 22) | ORR1E\|LTR\|ERVL-MaLR | 22781 | NM_010881 | 17977 | Mm.301039 | NM_010881 | ENSMUSG00000020647 | **Ncoa1** | KAT13A\|SRC-1\|SRC1\|bHLHe74 | nuclear receptor coactivator 1 | protein-coding |
| chr12_4424719_4425084 | chr12 | 4424720 | 4425084 | + | 0.856716 | intron (NM_010881, intron 1 of 22) | Tigger7\|DNA\|TcMar-Tigger | 52280 | NM_010881 | 17977 | Mm.301039 | NM_010881 | ENSMUSG00000020647 | **Ncoa1** | KAT13A\|SRC-1\|SRC1\|bHLHe74 | nuclear receptor coactivator 1 | protein-coding |
| chr12_4425178_4425583 | chr12 | 4425179 | 4425583 | + | 0.739169 | intron (NM_010881, intron 1 of 22) | B1_Mur2\|SINE\|Alu | 51801 | NM_010881 | 17977 | Mm.301039 | NM_010881 | ENSMUSG00000020647 | **Ncoa1** | KAT13A\|SRC-1\|SRC1\|bHLHe74 | nuclear receptor coactivator 1 | protein-coding |
| chr12_4403258_4403534 | chr12 | 4403259 | 4403534 | + | 0.718102 | intron (NM_010881, intron 2 of 22) | B3\|SINE\|B2 | 73786 | NM_010881 | 17977 | Mm.301039 | NM_010881 | ENSMUSG00000020647 | **Ncoa1** | KAT13A\|SRC-1\|SRC1\|bHLHe74 | nuclear receptor coactivator 1 | protein-coding |
| chr12_4436159_4436582 | chr12 | 4436160 | 4436582 | + | 0.680837 | intron (NM_010881, intron 1 of 22) | L1M2\|LINE\|L1 | 40811 | NM_010881 | 17977 | Mm.301039 | NM_010881 | ENSMUSG00000020647 | **Ncoa1** | KAT13A\|SRC-1\|SRC1\|bHLHe74 | nuclear receptor coactivator 1 | protein-coding |
| chr1_13383513_13384051 | chr1 | 13383514 | 13384051 | + | 1.2334 | Intergenic | L1M5\|LINE\|L1 | -9699 | NM_008678 | 17978 | Mm.2537 | NM_008678 | ENSMUSG00000005886 | **Ncoa2** | 9530095N19\|D1Ertd433e\|GRIP-1\|Grip1\|KAT13C\|SRC-2\|TIF-2\|TIF2\|bHLHe75 | nuclear receptor coactivator 2 | protein-coding |
| chr1_13465002_13465557 | chr1 | 13465003 | 13465557 | + | 1.01665 | Intergenic | Intergenic | -91197 | NM_008678 | 17978 | Mm.2537 | NM_008678 | ENSMUSG00000005886 | **Ncoa2** | 9530095N19\|D1Ertd433e\|GRIP-1\|Grip1\|KAT13C\|SRC-2\|TIF-2\|TIF2\|bHLHe75 | nuclear receptor coactivator 2 | protein-coding |
| chr1_13304947_13305204 | chr1 | 13304948 | 13305204 | + | 0.919007 | intron (NM_001077695, intron 1 of 21) | intron (NM_001077695, intron 1 of 21) | 69007 | NM_001077695 | 17978 | Mm.2537 | NM_008678 | ENSMUSG00000005886 | **Ncoa2** | 9530095N19\|D1Ertd433e\|GRIP-1\|Grip1\|KAT13C\|SRC-2\|TIF-2\|TIF2\|bHLHe75 | nuclear receptor coactivator 2 | protein-coding |
| chr1_13382563_13382934 | chr1 | 13382564 | 13382934 | + | 0.770318 | Intergenic | L1ME3A\|LINE\|L1 | -8666 | NM_008678 | 17978 | Mm.2537 | NM_008678 | ENSMUSG00000005886 | **Ncoa2** | 9530095N19\|D1Ertd433e\|GRIP-1\|Grip1\|KAT13C\|SRC-2\|TIF-2\|TIF2\|bHLHe75 | nuclear receptor coactivator 2 | protein-coding |
| chr1_13388543_13388972 | chr1 | 13388544 | 13388972 | + | 0.76891 | Intergenic | MLT2D\|LTR\|ERVL | -14675 | NM_008678 | 17978 | Mm.2537 | NM_008678 | ENSMUSG00000005886 | **Ncoa2** | 9530095N19\|D1Ertd433e\|GRIP-1\|Grip1\|KAT13C\|SRC-2\|TIF-2\|TIF2\|bHLHe75 | nuclear receptor coactivator 2 | protein-coding |
| chr2_166020648_166021075 | chr2 | 166020649 | 166021075 | + | 0.950761 | intron (NM_008679, intron 1 of 22) | intron (NM_008679, intron 1 of 22) | 28225 | NM_008679 | 17979 | Mm.469663 | NM_008679 | ENSMUSG00000027678 | **Ncoa3** | 2010305B15Rik\|AW321064\|Actr\|Aib1\|KAT13B\|Rac3\|Src3\|Tram-1\|Tram1\|bHLHe42\|p/Cip\|pCip | nuclear receptor coactivator 3 | protein-coding |
| chr2_166018244_166018574 | chr2 | 166018245 | 166018574 | + | 0.796855 | intron (NM_008679, intron 1 of 22) | intron (NM_008679, intron 1 of 22) | 25772 | NM_008679 | 17979 | Mm.469663 | NM_008679 | ENSMUSG00000027678 | **Ncoa3** | 2010305B15Rik\|AW321064\|Actr\|Aib1\|KAT13B\|Rac3\|Src3\|Tram-1\|Tram1\|bHLHe42\|p/Cip\|pCip | nuclear receptor coactivator 3 | protein-coding |
| chr2_166020155_166020409 | chr2 | 166020156 | 166020409 | + | 0.735262 | intron (NM_008679, intron 1 of 22) | intron (NM_008679, intron 1 of 22) | 27645 | NM_008679 | 17979 | Mm.469663 | NM_008679 | ENSMUSG00000027678 | **Ncoa3** | 2010305B15Rik\|AW321064\|Actr\|Aib1\|KAT13B\|Rac3\|Src3\|Tram-1\|Tram1\|bHLHe42\|p/Cip\|pCip | nuclear receptor coactivator 3 | protein-coding |
| chr10_30712471_30712928 | chr10 | 30712472 | 30712928 | + | 1.64587 | intron (NM_172495, intron 3 of 15) | intron (NM_172495, intron 3 of 15) | -56833 | NM_001111267 | 211329 | Mm.297903 | NM_172495 | ENSMUSG00000039697 | **Ncoa7** | 9030406N13Rik | nuclear receptor coactivator 7 | protein-coding |
| chr10_30709877_30710372 | chr10 | 30709878 | 30710372 | + | 1.56296 | intron (NM_172495, intron 3 of 15) | intron (NM_172495, intron 3 of 15) | -54258 | NM_001111267 | 211329 | Mm.297903 | NM_172495 | ENSMUSG00000039697 | **Ncoa7** | 9030406N13Rik | nuclear receptor coactivator 7 | protein-coding |
| chr10_30662290_30662863 | chr10 | 30662291 | 30662863 | + | 0.943537 | intron (NM_172495, intron 10 of 15) | intron (NM_172495, intron 10 of 15) | -6710 | NM_001111267 | 211329 | Mm.297903 | NM_172495 | ENSMUSG00000039697 | **Ncoa7** | 9030406N13Rik | nuclear receptor coactivator 7 | protein-coding |
| chr10_30638140_30638577 | chr10 | 30638141 | 30638577 | + | 0.900442 | Intergenic | Intergenic | 17508 | NM_001111267 | 211329 | Mm.297903 | NM_172495 | ENSMUSG00000039697 | **Ncoa7** | 9030406N13Rik | nuclear receptor coactivator 7 | protein-coding |
| chr10_30638122_30638772 | chr10 | 30638123 | 30638772 | + | 0.876637 | Intergenic | Intergenic | 17420 | NM_001111267 | 211329 | Mm.297903 | NM_172495 | ENSMUSG00000039697 | **Ncoa7** | 9030406N13Rik | nuclear receptor coactivator 7 | protein-coding |

| | | | | | | | | | | | | | | | | | |
|---|---|---|---|---|---|---|---|---|---|---|---|---|---|---|---|---|---|
| chr10_30662242_30663041 | chr10 | 30662243 | 30663041 | + | 0.861308 | intron (NM_172495, intron 10 of 15) | intron (NM_172495, intron 10 of 15) | -6775 | NM_001111267 | 211329 | Mm.297903 | NM_172495 | ENSMUSG00000039697 | **Ncoa7** | 9030406N13Rik | nuclear receptor coactivator 7 | protein-coding |
| chr10_30729781_30730122 | chr10 | 30729782 | 30730122 | + | 0.726899 | intron (NM_172495, intron 2 of 15) | RMER1B\|Other\|Other | 73155 | NM_172495 | 211329 | Mm.297903 | NM_172495 | ENSMUSG00000039697 | **Ncoa7** | 9030406N13Rik | nuclear receptor coactivator 7 | protein-coding |
| chr5_125189266_125189529 | chr5 | 125189267 | 125189529 | + | 0.69587 | Intergenic | B1_Mur4\|SINE\|Alu | -10184 | NM_011424 | 20602 | Mm.278646 | NM_011424 | ENSMUSG00000029478 | **Ncor2** | N-CoR\|SMRT\|SMRTe | nuclear receptor co-repressor 2 | protein-coding |
| chr2_31303142_31303784 | chr2 | 31303143 | 31303784 | + | 1.97854 | Intergenic | Intergenic | 57540 | NM_019681 | 14299 | Mm.249318 | NM_019681 | ENSMUSG00000062661 | **Ncs1** | 9430075O15Rik\|A730032G13Rik\|AI836659\|Freq\|Mfreq\|NCS-1 | neuronal calcium sensor 1 | protein-coding |
| chr18_38440644_38441068 | chr18 | 38440645 | 38441068 | + | 0.838004 | intron (NM_022996, intron 1 of 7) | intron (NM_022996, intron 1 of 7) | 21881 | NM_022996 | 65113 | Mm.102496 | NM_022996 | ENSMUSG00000024425 | **Ndfip1** | 0610010M22Rik\|N4wbp5 | Nedd4 family interacting protein 1 | protein-coding |
| chr18_38464129_38464434 | chr18 | 38464130 | 38464434 | + | 0.696884 | 3' UTR (NM_022996, exon 8 of 8) | 3' UTR (NM_022996, exon 8 of 8) | 45307 | NM_022996 | 65113 | Mm.102496 | NM_022996 | ENSMUSG00000024425 | **Ndfip1** | 0610010M22Rik\|N4wbp5 | Nedd4 family interacting protein 1 | protein-coding |
| chr18_38439942_38441099 | chr18 | 38439943 | 38441099 | + | 0.695385 | intron (NM_022996, intron 1 of 7) | intron (NM_022996, intron 1 of 7) | 21546 | NM_022996 | 65113 | Mm.102496 | NM_022996 | ENSMUSG00000024425 | **Ndfip1** | 0610010M22Rik\|N4wbp5 | Nedd4 family interacting protein 1 | protein-coding |
| chr14_105287610_105288311 | chr14 | 105287611 | 105288311 | + | 1.40805 | intron (NM_029561, intron 2 of 7) | B1F\|SINE\|Alu | 29288 | NM_001190989 | 76273 | Mm.290669 | NM_029561 | ENSMUSG00000053253 | **Ndfip2** | 0710001O20Rik\|2810436B12Rik\|9130207N19Rik\|N4wbp5a\|mKIAA1165 | Nedd4 family interacting protein 2 | protein-coding |
| chr14_105240121_105240601 | chr14 | 105240122 | 105240601 | + | 1.35814 | Intergenic | Intergenic | -18312 | NM_029561 | 76273 | Mm.290669 | NM_029561 | ENSMUSG00000053253 | **Ndfip2** | 0710001O20Rik\|2810436B12Rik\|9130207N19Rik\|N4wbp5a\|mKIAA1165 | Nedd4 family interacting protein 2 | protein-coding |
| chr14_105288387_105288909 | chr14 | 105288388 | 105288909 | + | 1.27462 | intron (NM_029561, intron 2 of 7) | (CA)n\|Simple_repeat\|Simple_repeat | 29975 | NM_001190989 | 76273 | Mm.290669 | NM_029561 | ENSMUSG00000053253 | **Ndfip2** | 0710001O20Rik\|2810436B12Rik\|9130207N19Rik\|N4wbp5a\|mKIAA1165 | Nedd4 family interacting protein 2 | protein-coding |
| chr14_105288024_105288315 | chr14 | 105288025 | 105288315 | + | 1.20884 | intron (NM_029561, intron 2 of 7) | intron (NM_029561, intron 2 of 7) | 29497 | NM_001190989 | 76273 | Mm.290669 | NM_029561 | ENSMUSG00000053253 | **Ndfip2** | 0710001O20Rik\|2810436B12Rik\|9130207N19Rik\|N4wbp5a\|mKIAA1165 | Nedd4 family interacting protein 2 | protein-coding |
| chr14_105287613_105287974 | chr14 | 105287614 | 105287974 | + | 1.14429 | intron (NM_029561, intron 2 of 7) | intron (NM_029561, intron 2 of 7) | 29121 | NM_001190989 | 76273 | Mm.290669 | NM_029561 | ENSMUSG00000053253 | **Ndfip2** | 0710001O20Rik\|2810436B12Rik\|9130207N19Rik\|N4wbp5a\|mKIAA1165 | Nedd4 family interacting protein 2 | protein-coding |
| chr14_105275259_105275634 | chr14 | 105275260 | 105275634 | + | 0.850472 | intron (NM_029561, intron 1 of 7) | intron (NM_029561, intron 1 of 7) | 16774 | NM_001190989 | 76273 | Mm.290669 | NM_029561 | ENSMUSG00000053253 | **Ndfip2** | 0710001O20Rik\|2810436B12Rik\|9130207N19Rik\|N4wbp5a\|mKIAA1165 | Nedd4 family interacting protein 2 | protein-coding |
| chr15_66997998_66998952 | chr15 | 66997999 | 66998952 | + | 1.98862 | Intergenic | Intergenic | -28834 | NM_008681 | 17988 | Mm.30837 | NM_008681 | ENSMUSG00000005125 | **Ndrg1** | CAP43\|CMT4D\|DRG1\|HMSNL\|NMSL\|Ndr1\|Ndrl\|PROXY1\|RTP\|TDD5 | N-myc downstream regulated gene 1 | protein-coding |
| chr15_66973597_66974190 | chr15 | 66973598 | 66974190 | + | 0.874201 | Intergenic | Intergenic | -4253 | NM_008681 | 17988 | Mm.30837 | NM_008681 | ENSMUSG00000005125 | **Ndrg1** | CAP43\|CMT4D\|DRG1\|HMSNL\|NMSL\|Ndr1\|Ndrl\|PROXY1\|RTP\|TDD5 | N-myc downstream regulated gene 1 | protein-coding |
| chr8_95672687_95673316 | chr8 | 95672688 | 95673316 | + | 0.871677 | Intergenic | MamSINE1\|SINE\|tRNA | -30035 | NM_001195006 | 234593 | Mm.29846 | NM_145602 | ENSMUSG00000036564 | **Ndrg4** | D8Bwg1337e\|Ndr1-rs\|Ndr4\|R74996\|SMAP-8 | N-myc downstream regulated gene 4 | protein-coding |
| chr18_60736617_60736997 | chr18 | 60736618 | 60736997 | + | 0.725384 | Intergenic | Intergenic | -23418 | NM_008306 | 15531 | Mm.181862 | NM_008306 | ENSMUSG00000054008 | **Ndst1** | 1200015G06Rik\|Hsst\|NDST-1\|b2b2230Clo | N-deacetylase/N-sulfotransferase (heparan glucosaminyl) 1 | protein-coding |
| chr10_94242843_94243243 | chr10 | 94242844 | 94243243 | + | 0.761965 | Intergenic | RMER6C\|LTR\|ERVK | 44034 | NM_025551 | 66414 | Mm.27886 | NM_025551 | ENSMUSG00000020022 | **Ndufa12** | 2410011G03Rik\|AW112974 | NADH dehydrogenase (ubiquinone) 1 alpha subcomplex, 12 | protein-coding |
| chr8_69921545_69922117 | chr8 | 69921546 | 69922117 | + | 0.711307 | intron (NM_001286450, intron 3 of 12) | intron (NM_001286450, intron 3 of 12) | -19273 | NM_023312 | 67184 | Mm.21162 | NM_023312 | ENSMUSG00000036199 | **Ndufa13** | 2700054G14Rik\|AU022060\|CDA016\|CGI-39\|GRIM-19\|Grim19 | NADH dehydrogenase (ubiquinone) 1 alpha subcomplex, 13 | protein-coding |
| chr15_82351525_82351832 | chr15 | 82351526 | 82351832 | + | 1.2636 | intron (NM_025987, intron 1 of 2) | intron (NM_025987, intron 1 of 2) | 2612 | NM_025987 | 67130 | Mm.27570 | NM_025987 | ENSMUSG00000022450 | **Ndufa6** | 14kDa\|2700038D15Rik\|B230217P19Rik | NADH dehydrogenase (ubiquinone) 1 alpha subcomplex, 6 (B14) | protein-coding |
| chr7_122086911_122087757 | chr7 | 122086912 | 122087757 | + | 1.12809 | TTS (NM_028177) | TTS (NM_028177) | 14514 | NM_028177 | 70316 | Mm.3014 | NM_028177 | | **Ndufab1** | 2210401F17Rik\|2310039H15Rik\|2610003B19Rik\|8kDa\|9130423F15Rik\|ACP\|CI-SDAP\|SDAP | NADH dehydrogenase (ubiquinone) 1, alpha/beta subcomplex, 1 | protein-coding |
| chr13_108107058_108107408 | chr13 | 108107059 | 108107408 | + | 0.92099 | intron (NM_001127346, intron 1 of 3) | intron (NM_001127346, intron 1 of 3) | 51392 | NM_001127346 | 75597 | Mm.276040 | NM_001127346 | ENSMUSG00000068184 | **Ndufaf2** | 1810058I14Rik\|C86051\|Ndufa12l\|mimitin | NADH dehydrogenase (ubiquinone) 1 alpha subcomplex, assembly factor 2 | protein-coding |
| chr4_11095549_11095828 | chr4 | 11095550 | 11095828 | + | 1.1621 | Intergenic | Intergenic | -19484 | NM_001085493 | 76947 | Mm.275265 | NM_001085493 | ENSMUSG00000050323 | **Ndufaf6** | 2310030N02Rik | NADH dehydrogenase (ubiquinone) complex I, assembly factor 6 | protein-coding |
| chr4_11090573_11091118 | chr4 | 11090574 | 11091118 | + | 1.02691 | Intergenic | Intergenic | -14641 | NM_001085493 | 76947 | Mm.275265 | NM_001085493 | ENSMUSG00000050323 | **Ndufaf6** | 2310030N02Rik | NADH dehydrogenase (ubiquinone) complex I, assembly factor 6 | protein-coding |
| chr6_39626559_39627142 | chr6 | 39626560 | 39627142 | + | 1.71537 | intron (NM_139294, intron 18 of 21) | intron (NM_139294, intron 18 of 21) | 34268 | NM_026612 | 68198 | Mm.29415 | NM_026612 | ENSMUSG00000002416 | **Ndufb2** | 1810011O01Rik\|8kDa\|AGGG\|AI325567\|CI-AGGG | NADH dehydrogenase (ubiquinone) 1 beta subcomplex, 2 | protein-coding |
| chr2_52319180_52319426 | chr2 | 52319181 | 52319426 | + | 1.05963 | intron (NM_010889, intron 10 of 156) | intron (NM_010889, intron 10 of 156) | 19495 | NM_010889 | 17996 | Mm.389247 | NM_010889 | ENSMUSG00000026950 | **Neb** | AI595938 | nebulin | protein-coding |
| chr2_52319162_52319699 | chr2 | 52319163 | 52319699 | + | 1.02609 | intron (NM_010889, intron 10 of 156) | intron (NM_010889, intron 10 of 156) | 19367 | NM_010889 | 17996 | Mm.389247 | NM_010889 | ENSMUSG00000026950 | **Neb** | AI595938 | nebulin | protein-coding |
| chr2_52318959_52319728 | chr2 | 52318960 | 52319728 | + | 0.988228 | intron (NM_010889, intron 10 of 156) | intron (NM_010889, intron 10 of 156) | 19454 | NM_010889 | 17996 | Mm.389247 | NM_010889 | ENSMUSG00000026950 | **Neb** | AI595938 | nebulin | protein-coding |
| chr2_52372095_52372905 | chr2 | 52372096 | 52372905 | + | 0.668456 | Intergenic | Intergenic | -33702 | NM_010889 | 17996 | Mm.389247 | NM_010889 | ENSMUSG00000026950 | **Neb** | AI595938 | nebulin | protein-coding |
| chr2_17844869_17845794 | chr2 | 17844870 | 17845794 | + | 1.9527 | Intergenic | Intergenic | -114264 | NM_028757 | 74103 | Mm.120298 | NM_028757 | ENSMUSG00000053702 | **Nebl** | 1200007O21Rik\|A630080F05Rik\|BB140644\|D830029A09Rik\|Lnebl | nebulette | protein-coding |
| chr2_17741530_17741971 | chr2 | 17741531 | 17741971 | + | 0.950178 | Intergenic | Intergenic | -10683 | NM_028757 | 74103 | Mm.120298 | NM_028757 | ENSMUSG00000053702 | **Nebl** | 1200007O21Rik\|A630080F05Rik\|BB140644\|D830029A09Rik\|Lnebl | nebulette | protein-coding |
| chr2_17845840_17846148 | chr2 | 17845841 | 17846148 | + | 0.94867 | Intergenic | B1_Mur2\|SINE\|Alu | -114926 | NM_028757 | 74103 | Mm.120298 | NM_028757 | ENSMUSG00000053702 | **Nebl** | 1200007O21Rik\|A630080F05Rik\|BB140644\|D830029A09Rik\|Lnebl | nebulette | protein-coding |
| chr2_17741542_17741798 | chr2 | 17741543 | 17741798 | + | 0.883979 | Intergenic | Intergenic | -10602 | NM_028757 | 74103 | Mm.120298 | NM_028757 | ENSMUSG00000053702 | **Nebl** | 1200007O21Rik\|A630080F05Rik\|BB140644\|D830029A09Rik\|Lnebl | nebulette | protein-coding |
| chr2_17730257_17730566 | chr2 | 17730258 | 17730566 | + | 0.841287 | intron (NM_028757, intron 1 of 6) | intron (NM_028757, intron 1 of 6) | 656 | NM_028757 | 74103 | Mm.120298 | NM_028757 | ENSMUSG00000053702 | **Nebl** | 1200007O21Rik\|A630080F05Rik\|BB140644\|D830029A09Rik\|Lnebl | nebulette | protein-coding |
| chr2_17741509_17742738 | chr2 | 17741510 | 17742738 | + | 0.836882 | Intergenic | Intergenic | -11056 | NM_028757 | 74103 | Mm.120298 | NM_028757 | ENSMUSG00000053702 | **Nebl** | 1200007O21Rik\|A630080F05Rik\|BB140644\|D830029A09Rik\|Lnebl | nebulette | protein-coding |
| chr2_17685112_17685509 | chr2 | 17685113 | 17685509 | + | 0.718532 | intron (NM_028757, intron 2 of 6) | intron (NM_028757, intron 2 of 6) | 45757 | NM_028757 | 74103 | Mm.120298 | NM_028757 | ENSMUSG00000053702 | **Nebl** | 1200007O21Rik\|A630080F05Rik\|BB140644\|D830029A09Rik\|Lnebl | nebulette | protein-coding |
| chr10_92719716_92719945 | chr10 | 92719717 | 92719945 | + | 0.862911 | intron (NM_008682, intron 1 of 14) | intron (NM_008682, intron 1 of 14) | 2587 | NM_008682 | 17997 | Mm.2998 | NM_008682 | ENSMUSG00000019988 | **Nedd1** | - | neural precursor cell expressed, developmentally down-regulated gene 1 | protein-coding |
| chr9_72678597_72679145 | chr9 | 72678598 | 72679145 | + | 1.12151 | intron (NM_010890, intron 5 of 28) | intron (NM_010890, intron 5 of 28) | 16524 | NM_010890 | 17999 | Mm.279923 | NM_010890 | ENSMUSG00000032216 | **Nedd4** | AA959633\|AL023035\|AU019897\|E430025J12Rik\|Nedd4-1\|Nedd4a\|mKIAA0093 | neural precursor cell expressed, developmentally down-regulated 4 | protein-coding |
| chr18_65010270_65011068 | chr18 | 65010271 | 65011068 | + | 1.55772 | intron (NM_001114386, intron 2 of 30) | intron (NM_001114386, intron 2 of 30) | -12858 | NM_031881 | 83814 | Mm.98668 | NM_031881 | ENSMUSG00000024589 | **Nedd4l** | 1300012C07Rik\|Nedd4-2\|Nedd4b | neural precursor cell expressed, developmentally down-regulated gene 4-like | protein-coding |
| chr18_65106880_65107483 | chr18 | 65106881 | 65107483 | + | 1.1945 | intron (NM_031881, intron 4 of 29) | intron (NM_031881, intron 4 of 29) | 83655 | NM_031881 | 83814 | Mm.98668 | NM_031881 | ENSMUSG00000024589 | **Nedd4l** | 1300012C07Rik\|Nedd4-2\|Nedd4b | neural precursor cell expressed, developmentally down-regulated gene 4-like | protein-coding |
| chr18_64986599_64987068 | chr18 | 64986600 | 64987068 | + | 0.964093 | intron (NM_001114386, intron 1 of 30) | intron (NM_001114386, intron 1 of 30) | -36693 | NM_031881 | 83814 | Mm.98668 | NM_031881 | ENSMUSG00000024589 | **Nedd4l** | 1300012C07Rik\|Nedd4-2\|Nedd4b | neural precursor cell expressed, developmentally down-regulated gene 4-like | protein-coding |
| chr18_65015899_65016600 | chr18 | 65015900 | 65016600 | + | 0.897789 | intron (NM_001114386, intron 2 of 30) | MLT1A1\|LTR\|ERVL-MaLR | -7277 | NM_031881 | 83814 | Mm.98668 | NM_031881 | ENSMUSG00000024589 | **Nedd4l** | 1300012C07Rik\|Nedd4-2\|Nedd4b | neural precursor cell expressed, developmentally down-regulated gene 4-like | protein-coding |
| chr18_65050775_65051266 | chr18 | 65050776 | 65051266 | + | 0.806813 | intron (NM_031881, intron 1 of 29) | intron (NM_031881, intron 1 of 29) | 27494 | NM_031881 | 83814 | Mm.98668 | NM_031881 | ENSMUSG00000024589 | **Nedd4l** | 1300012C07Rik\|Nedd4-2\|Nedd4b | neural precursor cell expressed, developmentally down-regulated gene 4-like | protein-coding |
| chr18_64962505_64962875 | chr18 | 64962506 | 64962875 | + | 0.722219 | intron (NM_001114386, intron 1 of 30) | intron (NM_001114386, intron 1 of 30) | -60837 | NM_031881 | 83814 | Mm.98668 | NM_031881 | ENSMUSG00000024589 | **Nedd4l** | 1300012C07Rik\|Nedd4-2\|Nedd4b | neural precursor cell expressed, developmentally down-regulated gene 4-like | protein-coding |

| | | | | | | | | | | | | | | | | | |
|---|---|---|---|---|---|---|---|---|---|---|---|---|---|---|---|---|---|
| chr18_64987761_64988011 | chr18 | 64987762 | 64988011 | + | 0.719603 | intron (NM_001114386, intron 1 of 30) | ORR1G\|LTR\|ERVL-MaLR | -35641 | NM_031881 | 83814 | Mm.98668 | NM_031881 | ENSMUSG00000024589 | **Nedd4l** | 1300012C07Rik\|Nedd4-2\|Nedd4b | neural precursor cell expressed, developmentally down-regulated gene 4-like | protein-coding |
| chr18_65099096_65099496 | chr18 | 65099097 | 65099496 | + | 0.716329 | intron (NM_031881, intron 4 of 29) | intron (NM_031881, intron 4 of 29) | 75769 | NM_031881 | 83814 | Mm.98668 | NM_031881 | ENSMUSG00000024589 | **Nedd4l** | 1300012C07Rik\|Nedd4-2\|Nedd4b | neural precursor cell expressed, developmentally down-regulated gene 4-like | protein-coding |
| chr18_65053819_65054088 | chr18 | 65053820 | 65054088 | + | 0.653963 | intron (NM_031881, intron 1 of 29) | intron (NM_031881, intron 1 of 29) | 30427 | NM_031881 | 83814 | Mm.98668 | NM_031881 | ENSMUSG00000024589 | **Nedd4l** | 1300012C07Rik\|Nedd4-2\|Nedd4b | neural precursor cell expressed, developmentally down-regulated gene 4-like | protein-coding |
| chr13_41453314_41453924 | chr13 | 41453315 | 41453924 | + | 1.13489 | intron (NM_001111324, intron 1 of 7) | intron (NM_001111324, intron 1 of 7) | 33741 | NM_001111324 | 18003 | Mm.288980 | NM_017464 | ENSMUSG00000021365 | **Nedd9** | Cas-L\|CasL\|HEF1\|MEF1\|Nedd-9\|p105 | neural precursor cell expressed, developmentally down-regulated gene 9 | protein-coding |
| chr13_41444891_41445956 | chr13 | 41444892 | 41445956 | + | 1.13198 | intron (NM_001111324, intron 1 of 7) | intron (NM_001111324, intron 1 of 7) | 41936 | NM_001111324 | 18003 | Mm.288980 | NM_017464 | ENSMUSG00000021365 | **Nedd9** | Cas-L\|CasL\|HEF1\|MEF1\|Nedd-9\|p105 | neural precursor cell expressed, developmentally down-regulated gene 9 | protein-coding |
| chr13_41469500_41469978 | chr13 | 41469501 | 41469978 | + | 1.13098 | intron (NM_001111324, intron 1 of 7) | intron (NM_001111324, intron 1 of 7) | 17621 | NM_001111324 | 18003 | Mm.288980 | NM_017464 | ENSMUSG00000021365 | **Nedd9** | Cas-L\|CasL\|HEF1\|MEF1\|Nedd-9\|p105 | neural precursor cell expressed, developmentally down-regulated gene 9 | protein-coding |
| chr13_41453034_41453931 | chr13 | 41453035 | 41453931 | + | 1.08477 | intron (NM_001111324, intron 1 of 7) | intron (NM_001111324, intron 1 of 7) | 33877 | NM_001111324 | 18003 | Mm.288980 | NM_017464 | ENSMUSG00000021365 | **Nedd9** | Cas-L\|CasL\|HEF1\|MEF1\|Nedd-9\|p105 | neural precursor cell expressed, developmentally down-regulated gene 9 | protein-coding |
| chr13_41366949_41367808 | chr13 | 41366950 | 41367808 | + | 0.879413 | intron (NM_001111324, intron 2 of 7) | intron (NM_001111324, intron 2 of 7) | -8378 | NM_017464 | 18003 | Mm.288980 | NM_017464 | ENSMUSG00000021365 | **Nedd9** | Cas-L\|CasL\|HEF1\|MEF1\|Nedd-9\|p105 | neural precursor cell expressed, developmentally down-regulated gene 9 | protein-coding |
| chr11_4951367_4951615 | chr11 | 4951368 | 4951615 | + | 0.891289 | intron (NM_001243043, intron 1 of 22) | MTC\|LTR\|ERVL-MaLR | -3427 | NM_010904 | 380684 | Mm.298283 | NM_010904 | ENSMUSG00000020396 | **Nefh** | NF-H\|NF200\|Nfh\|mKIAA0845 | neurofilament, heavy polypeptide | protein-coding |
| chr14_68034273_68034803 | chr14 | 68034274 | 68034803 | + | 1.64652 | Intergenic | Intergenic | -49346 | NM_010910 | 18039 | Mm.1956 | NM_010910 | | **Nefl** | AI847934\|CMT2E\|NF-L\|NF68\|Nfl | neurofilament, light polypeptide | protein-coding |
| chr14_68035262_68035970 | chr14 | 68035263 | 68035970 | + | 1.38076 | Intergenic | Intergenic | -48268 | NM_010910 | 18039 | Mm.1956 | NM_010910 | | **Nefl** | AI847934\|CMT2E\|NF-L\|NF68\|Nfl | neurofilament, light polypeptide | protein-coding |
| chr14_63195043_63195574 | chr14 | 63195044 | 63195574 | + | 1.32086 | Intergenic | Intergenic | -1784 | NM_201610 | 382913 | Mm.239490 | NM_201610 | ENSMUSG00000035121 | **Neil2** | Gm1212\|NEH2 | nei like 2 (E. coli) | protein-coding |
| chr14_14784245_14784592 | chr14 | 14784246 | 14784592 | + | 1.01526 | intron (NM_001033270, intron 21 of 24) | MLT1C\|LTR\|ERVL-MaLR | -36396 | NM_001195229 | 674895 | Mm.317563 | NM_001034865 | ENSMUSG00000042567 | **Nek10** | Gm282 | NIMA (never in mitosis gene a)- related kinase 10 | protein-coding |
| chr9_105368470_105369150 | chr9 | 105368471 | 105369150 | + | 1.25926 | intron (NM_172461, intron 2 of 15) | intron (NM_172461, intron 2 of 15) | 26477 | NM_172461 | 208583 | Mm.347509 | NM_172461 | ENSMUSG00000035032 | **Nek11** | 4932416N14Rik | NIMA (never in mitosis gene a)-related expressed kinase 11 | protein-coding |
| chr9_105387432_105387853 | chr9 | 105387433 | 105387853 | + | 1.06322 | intron (NM_172461, intron 2 of 15) | intron (NM_172461, intron 2 of 15) | 7644 | NM_172461 | 208583 | Mm.347509 | NM_172461 | ENSMUSG00000035032 | **Nek11** | 4932416N14Rik | NIMA (never in mitosis gene a)-related expressed kinase 11 | protein-coding |
| chr9_105384656_105385162 | chr9 | 105384657 | 105385162 | + | 0.799195 | intron (NM_172461, intron 2 of 15) | Lx8\|LINE\|L1 | 10378 | NM_172461 | 208583 | Mm.347509 | NM_172461 | ENSMUSG00000035032 | **Nek11** | 4932416N14Rik | NIMA (never in mitosis gene a)-related expressed kinase 11 | protein-coding |
| chr2_38536249_38536935 | chr2 | 38536250 | 38536935 | + | 1.28497 | intron (NM_001159631, intron 1 of 9) | intron (NM_001159631, intron 1 of 9) | 24716 | NM_001159631 | 59126 | Mm.143818 | NM_021606 | ENSMUSG00000026749 | **Nek6** | 1300007C09Rik | NIMA (never in mitosis gene a)-related expressed kinase 6 | protein-coding |
| chr2_38471713_38472438 | chr2 | 38471714 | 38472438 | + | 1.28139 | Intergenic | Intergenic | -39621 | NM_021606 | 59126 | Mm.143818 | NM_021606 | ENSMUSG00000026749 | **Nek6** | 1300007C09Rik | NIMA (never in mitosis gene a)-related expressed kinase 6 | protein-coding |
| chr2_38471784_38472202 | chr2 | 38471785 | 38472202 | + | 1.21176 | Intergenic | Intergenic | -39704 | NM_021606 | 59126 | Mm.143818 | NM_021606 | ENSMUSG00000026749 | **Nek6** | 1300007C09Rik | NIMA (never in mitosis gene a)-related expressed kinase 6 | protein-coding |
| chr2_38472500_38473293 | chr2 | 38472501 | 38473293 | + | 0.900537 | Intergenic | Intergenic | -38800 | NM_021606 | 59126 | Mm.143818 | NM_021606 | ENSMUSG00000026749 | **Nek6** | 1300007C09Rik | NIMA (never in mitosis gene a)-related expressed kinase 6 | protein-coding |
| chr2_38472854_38473261 | chr2 | 38472855 | 38473261 | + | 0.813913 | Intergenic | Intergenic | -38639 | NM_021606 | 59126 | Mm.143818 | NM_021606 | ENSMUSG00000026749 | **Nek6** | 1300007C09Rik | NIMA (never in mitosis gene a)-related expressed kinase 6 | protein-coding |
| chr2_38510158_38510432 | chr2 | 38510159 | 38510432 | + | 0.686063 | Intergenic | Intergenic | -1402 | NM_021606 | 59126 | Mm.143818 | NM_021606 | ENSMUSG00000026749 | **Nek6** | 1300007C09Rik | NIMA (never in mitosis gene a)-related expressed kinase 6 | protein-coding |
| chr1_138626206_138626795 | chr1 | 138626207 | 138626795 | + | 1.35262 | Intergenic | ORR1F\|LTR\|ERVL-MaLR | -6805 | NM_021605 | 59125 | Mm.143817 | NM_021605 | ENSMUSG00000026393 | **Nek7** | 2810460C19Rik\|AU020186 | NIMA (never in mitosis gene a)-related expressed kinase 7 | protein-coding |
| chr1_138603747_138604458 | chr1 | 138603748 | 138604458 | + | 1.29803 | intron (NM_021605, intron 1 of 9) | MTD\|LTR\|ERVL-MaLR | 15593 | NM_021605 | 59125 | Mm.143817 | NM_021605 | ENSMUSG00000026393 | **Nek7** | 2810460C19Rik\|AU020186 | NIMA (never in mitosis gene a)-related expressed kinase 7 | protein-coding |
| chr1_138595964_138596697 | chr1 | 138595965 | 138596697 | + | 1.18352 | intron (NM_021605, intron 1 of 9) | intron (NM_021605, intron 1 of 9) | 23365 | NM_021605 | 59125 | Mm.143817 | NM_021605 | ENSMUSG00000026393 | **Nek7** | 2810460C19Rik\|AU020186 | NIMA (never in mitosis gene a)-related expressed kinase 7 | protein-coding |
| chr1_138559542_138560106 | chr1 | 138559543 | 138560106 | + | 0.792692 | intron (NM_021605, intron 2 of 9) | intron (NM_021605, intron 2 of 9) | 59872 | NM_021605 | 59125 | Mm.143817 | NM_021605 | ENSMUSG00000026393 | **Nek7** | 2810460C19Rik\|AU020186 | NIMA (never in mitosis gene a)-related expressed kinase 7 | protein-coding |
| chr1_138559312_138560358 | chr1 | 138559313 | 138560358 | + | 0.780071 | intron (NM_021605, intron 2 of 9) | intron (NM_021605, intron 2 of 9) | 59861 | NM_021605 | 59125 | Mm.143817 | NM_021605 | ENSMUSG00000026393 | **Nek7** | 2810460C19Rik\|AU020186 | NIMA (never in mitosis gene a)-related expressed kinase 7 | protein-coding |
| chr1_138590337_138590818 | chr1 | 138590338 | 138590818 | + | 0.730933 | intron (NM_021605, intron 1 of 9) | intron (NM_021605, intron 1 of 9) | 29118 | NM_021605 | 59125 | Mm.143817 | NM_021605 | ENSMUSG00000026393 | **Nek7** | 2810460C19Rik\|AU020186 | NIMA (never in mitosis gene a)-related expressed kinase 7 | protein-coding |
| chr1_138592538_138592836 | chr1 | 138592539 | 138592836 | + | 0.621003 | intron (NM_021605, intron 1 of 9) | Lx7\|LINE\|L1 | 27009 | NM_021605 | 59125 | Mm.143817 | NM_021605 | ENSMUSG00000026393 | **Nek7** | 2810460C19Rik\|AU020186 | NIMA (never in mitosis gene a)-related expressed kinase 7 | protein-coding |
| chr11_78177015_78177715 | chr11 | 78177016 | 78177715 | + | 1.10041 | promoter-TSS (NM_080849) | promoter-TSS (NM_080849) | -699 | NM_080849 | 140859 | Mm.23788 | NM_080849 | ENSMUSG00000017405 | **Nek8** | 4632401F23Rik\|b2b1449Clo\|jck | NIMA (never in mitosis gene a)-related expressed kinase 8 | protein-coding |
| chr1_191275434_191276092 | chr1 | 191275435 | 191276092 | + | 0.722009 | Intergenic | RMER30\|DNA\|hAT-Charlie | 42355 | NM_025424 | 66208 | Mm.46444 | NM_025424 | ENSMUSG00000037499 | **Nenf** | 1110060M21Rik\|SCIRP10\|Spuf | neuron derived neurotrophic factor | protein-coding |
| chr1_191275534_191275912 | chr1 | 191275535 | 191275912 | + | 0.718439 | Intergenic | RMER30\|DNA\|hAT-Charlie | 42395 | NM_025424 | 66208 | Mm.46444 | NM_025424 | ENSMUSG00000037499 | **Nenf** | 1110060M21Rik\|SCIRP10\|Spuf | neuron derived neurotrophic factor | protein-coding |
| chr9_58979036_58979389 | chr9 | 58979037 | 58979389 | + | 0.705824 | intron (NM_001042752, intron 4 of 27) | intron (NM_001042752, intron 4 of 27) | 57228 | NM_008684 | 18007 | Mm.42249 | NM_008684 | ENSMUSG00000032340 | **Neo1** | 2610028H22Rik\|AI327052\|D930014N22Rik\|Igdcc2 | neogenin | protein-coding |
| chr10_52392229_52392871 | chr10 | 52392230 | 52392871 | + | 0.655287 | intron (NM_025684, intron 1 of 2) | intron (NM_025684, intron 1 of 2) | 942 | NM_025684 | 66650 | Mm.90950 | NM_025684 | ENSMUSG00000038624 | **Nepn** | 5730521E12Rik\|Npn\|periolin | nephrocan | protein-coding |
| chr8_85648588_85648944 | chr8 | 85648589 | 85648944 | + | 1.58059 | intron (NM_001081324, intron 6 of 8) | Tigger7\|DNA\|TcMar-Tigger | 42243 | NM_001081324 | 74513 | Mm.126079 | NM_001081324 | ENSMUSG00000036902 | **Neto2** | 5530601C23Rik\|AI854825\|BTCL2 | neuropilin (NRP) and tolloid (TLL)-like 2 | protein-coding |
| chr7_99833668_99834253 | chr7 | 99833669 | 99834253 | + | 1.53573 | Intergenic | Intergenic | -5544 | NM_016720 | 50877 | Mm.103703 | NM_016720 | ENSMUSG00000035239 | **Neu3** | - | neuraminidase 3 | protein-coding |
| chr7_99834366_99834690 | chr7 | 99834367 | 99834690 | + | 1.5101 | Intergenic | L1MA5\|LINE\|L1 | -6111 | NM_016720 | 50877 | Mm.103703 | NM_016720 | ENSMUSG00000035239 | **Neu3** | - | neuraminidase 3 | protein-coding |
| chr1_36266308_36266723 | chr1 | 36266309 | 36266723 | + | 0.924063 | exon (NM_153408, exon 4 of 4) | exon (NM_153408, exon 4 of 4) | 6909 | NM_153408 | 214854 | Mm.389110 | NM_153408 | ENSMUSG00000047180 | **Neurl3** | 2010300P06Rik\|Lincr | neuralized homolog 3 homolog (Drosophila) | protein-coding |
| chr3_127704985_127705758 | chr3 | 127704986 | 127705758 | + | 0.938226 | intron (NM_027808, intron 3 of 13) | intron (NM_027808, intron 3 of 13) | 72227 | NM_009718 | 11924 | Mm.42017 | NM_009718 | ENSMUSG00000027967 | **Neurog2** | Atoh4\|Math4A\|bHLHa8\|ngn-2\|ngn2 | neurogenin 2 | protein-coding |
| chr3_127638088_127638301 | chr3 | 127638089 | 127638301 | + | 0.89665 | Intergenic | Intergenic | 5050 | NM_009718 | 11924 | Mm.42017 | NM_009718 | ENSMUSG00000027967 | **Neurog2** | Atoh4\|Math4A\|bHLHa8\|ngn-2\|ngn2 | neurogenin 2 | protein-coding |
| chr3_152257489_152258053 | chr3 | 152257490 | 152258053 | + | 0.988364 | intron (NM_199465, intron 1 of 12) | intron (NM_199465, intron 1 of 12) | 8549 | NM_199465 | 68810 | Mm.200188 | NM_199465 | ENSMUSG00000039103 | **Nexn** | 1110046H09Rik\|AA553326\|NELIN | nexilin | protein-coding |
| chr11_4826209_4826961 | chr11 | 4826210 | 4826961 | + | 1.55173 | intron (NM_001252251, intron 1 of 16) | intron (NM_001252251, intron 1 of 16) | 6550 | NM_001252253 | 18016 | Mm.297109 | NM_010898 | ENSMUSG00000009073 | **Nf2** | merlin | neurofibromatosis 2 | protein-coding |
| chr11_4804918_4805426 | chr11 | 4804919 | 4805426 | + | 1.27117 | intron (NM_001252251, intron 6 of 16) | intron (NM_001252251, intron 6 of 16) | 27963 | NM_001252253 | 18016 | Mm.297109 | NM_010898 | ENSMUSG00000009073 | **Nf2** | merlin | neurofibromatosis 2 | protein-coding |
| chr11_4803808_4804477 | chr11 | 4803809 | 4804477 | + | 0.759528 | intron (NM_001252251, intron 6 of 16) | intron (NM_001252251, intron 6 of 16) | 28992 | NM_001252253 | 18016 | Mm.297109 | NM_010898 | ENSMUSG00000009073 | **Nf2** | merlin | neurofibromatosis 2 | protein-coding |
| chr11_4803928_4804287 | chr11 | 4803929 | 4804287 | + | 0.710181 | intron (NM_001252251, intron 6 of 16) | ID4\|SINE\|ID | 29027 | NM_001252253 | 18016 | Mm.297109 | NM_010898 | ENSMUSG00000009073 | **Nf2** | merlin | neurofibromatosis 2 | protein-coding |

| | | | | | | | | | | | | | | | | | |
|---|---|---|---|---|---|---|---|---|---|---|---|---|---|---|---|---|---|
| chr8_107290935_107291721 | chr8 | 107290936 | 107291721 | + | 1.8456 | Intergenic | Intergenic | -2142 | NM_018823 | 54446 | Mm.390057 | NM_018823 | ENSMUSG00000003847 | **Nfat5** | AI225870\|B130038B15Rik\|CAG-8\|CAG80\|NFATL1\|OREBP\|TonEBP\|mKIAA0827\|nfatz | nuclear factor of activated T cells 5 | protein-coding |
| chr8_107254805_107255537 | chr8 | 107254806 | 107255537 | + | 1.47418 | Intergenic | MIR\|SINE\|MIR | -38299 | NM_018823 | 54446 | Mm.390057 | NM_018823 | ENSMUSG00000003847 | **Nfat5** | AI225870\|B130038B15Rik\|CAG-8\|CAG80\|NFATL1\|OREBP\|TonEBP\|mKIAA0827\|nfatz | nuclear factor of activated T cells 5 | protein-coding |
| chr8_107254402_107254709 | chr8 | 107254403 | 107254709 | + | 0.603442 | Intergenic | Intergenic | -38914 | NM_018823 | 54446 | Mm.390057 | NM_018823 | ENSMUSG00000003847 | **Nfat5** | AI225870\|B130038B15Rik\|CAG-8\|CAG80\|NFATL1\|OREBP\|TonEBP\|mKIAA0827\|nfatz | nuclear factor of activated T cells 5 | protein-coding |
| chr18_80642475_80642893 | chr18 | 80642476 | 80642893 | + | 1.00311 | intron (NM_198429, intron 8 of 9) | intron (NM_198429, intron 8 of 9) | 65489 | NM_001164111 | 18018 | Mm.329560 | NM_016791 | ENSMUSG00000033016 | **Nfatc1** | 2210017P03Rik\|AI449492\|AV076380\|NF-ATc\|NFAT2\|NFATc\|Nfatcb | nuclear factor of activated T cells, cytoplasmic, calcineurin dependent 1 | protein-coding |
| chr18_80642465_80643093 | chr18 | 80642466 | 80643093 | + | 0.852418 | intron (NM_198429, intron 8 of 9) | intron (NM_198429, intron 8 of 9) | 65394 | NM_001164111 | 18018 | Mm.329560 | NM_016791 | ENSMUSG00000033016 | **Nfatc1** | 2210017P03Rik\|AI449492\|AV076380\|NF-ATc\|NFAT2\|NFATc\|Nfatcb | nuclear factor of activated T cells, cytoplasmic, calcineurin dependent 1 | protein-coding |
| chr18_80802363_80803036 | chr18 | 80802364 | 80803036 | + | 0.671264 | intron (NM_015805, intron 12 of 28) | intron (NM_015805, intron 12 of 28) | -89629 | NM_001164109 | 18018 | Mm.329560 | NM_016791 | ENSMUSG00000033016 | **Nfatc1** | 2210017P03Rik\|AI449492\|AV076380\|NF-ATc\|NFAT2\|NFATc\|Nfatcb | nuclear factor of activated T cells, cytoplasmic, calcineurin dependent 1 | protein-coding |
| chr18_80726931_80727451 | chr18 | 80726932 | 80727451 | + | 0.662902 | Intergenic | Intergenic | -14120 | NM_001164109 | 18018 | Mm.329560 | NM_016791 | ENSMUSG00000033016 | **Nfatc1** | 2210017P03Rik\|AI449492\|AV076380\|NF-ATc\|NFAT2\|NFATc\|Nfatcb | nuclear factor of activated T cells, cytoplasmic, calcineurin dependent 1 | protein-coding |
| chr2_168578228_168578860 | chr2 | 168578229 | 168578860 | + | 1.37497 | intron (NM_001291169, intron 1 of 10) | MIR\|SINE\|MIR | 11821 | NM_001291178 | 18019 | Mm.116802 | NM_010899 | ENSMUSG00000027544 | **Nfatc2** | AI607462\|NF-ATc2\|NF-ATp\|NFAT1\|NFAT1-D\|Nfatp | nuclear factor of activated T cells, cytoplasmic, calcineurin dependent 2 | protein-coding |
| chr2_168579550_168580114 | chr2 | 168579551 | 168580114 | + | 1.26578 | intron (NM_001291169, intron 1 of 10) | intron (NM_001291169, intron 1 of 10) | 10533 | NM_001291178 | 18019 | Mm.116802 | NM_010899 | ENSMUSG00000027544 | **Nfatc2** | AI607462\|NF-ATc2\|NF-ATp\|NFAT1\|NFAT1-D\|Nfatp | nuclear factor of activated T cells, cytoplasmic, calcineurin dependent 2 | protein-coding |
| chr2_168532165_168533012 | chr2 | 168532166 | 168533012 | + | 1.23623 | intron (NM_001291169, intron 5 of 10) | intron (NM_001291169, intron 5 of 10) | 57776 | NM_001291178 | 18019 | Mm.116802 | NM_010899 | ENSMUSG00000027544 | **Nfatc2** | AI607462\|NF-ATc2\|NF-ATp\|NFAT1\|NFAT1-D\|Nfatp | nuclear factor of activated T cells, cytoplasmic, calcineurin dependent 2 | protein-coding |
| chr2_168491761_168492506 | chr2 | 168491762 | 168492506 | + | 1.14008 | intron (NM_001291169, intron 9 of 10) | intron (NM_001291169, intron 9 of 10) | 98231 | NM_001291178 | 18019 | Mm.116802 | NM_010899 | ENSMUSG00000027544 | **Nfatc2** | AI607462\|NF-ATc2\|NF-ATp\|NFAT1\|NFAT1-D\|Nfatp | nuclear factor of activated T cells, cytoplasmic, calcineurin dependent 2 | protein-coding |
| chr2_168594705_168594970 | chr2 | 168594706 | 168594970 | + | 1.07368 | intron (NM_001291169, intron 1 of 10) | intron (NM_001291169, intron 1 of 10) | -4473 | NM_001136073 | 18019 | Mm.116802 | NM_010899 | ENSMUSG00000027544 | **Nfatc2** | AI607462\|NF-ATc2\|NF-ATp\|NFAT1\|NFAT1-D\|Nfatp | nuclear factor of activated T cells, cytoplasmic, calcineurin dependent 2 | protein-coding |
| chr2_168531860_168532903 | chr2 | 168531861 | 168532903 | + | 0.9688 | intron (NM_001291169, intron 5 of 10) | intron (NM_001291169, intron 5 of 10) | 57983 | NM_001291178 | 18019 | Mm.116802 | NM_010899 | ENSMUSG00000027544 | **Nfatc2** | AI607462\|NF-ATc2\|NF-ATp\|NFAT1\|NFAT1-D\|Nfatp | nuclear factor of activated T cells, cytoplasmic, calcineurin dependent 2 | protein-coding |
| chr14_55820996_55821332 | chr14 | 55820997 | 55821332 | + | 0.677705 | Intergenic | Intergenic | -3631 | NM_001168346 | 73181 | Mm.27908 | NM_023699 | ENSMUSG00000023411 | **Nfatc4** | 3110041H08Rik\|AW107667\|AW546455\|Nfat3 | nuclear factor of activated T cells, cytoplasmic, calcineurin dependent 4 | protein-coding |
| chr15_103247328_103249471 | chr15 | 103247329 | 103249471 | + | 1.04127 | 3' UTR (NM_008685, exon 3 of 3) | 3' UTR (NM_008685, exon 3 of 3) | 7016 | NM_008685 | 18022 | Mm.457989 | NM_008685 | ENSMUSG00000058794 | **Nfe2** | 45kDa\|NF-E2\|p45\|p45NFE2\|p45nf-e2 | nuclear factor, erythroid derived 2 | protein-coding |
| chr11_96822621_96823328 | chr11 | 96822622 | 96823328 | + | 1.47291 | intron (NM_008686, intron 2 of 4) | intron (NM_008686, intron 2 of 4) | 1038 | NM_001130453 | 18023 | Mm.6743 | NM_008686 | ENSMUSG00000038615 | **Nfe2l1** | AA408798\|AW212678\|LCR-F1\|Lcrf1\|NRF1\|TCF-11\|TCF11 | nuclear factor, erythroid derived 2,-like 1 | protein-coding |
| chr2_75685418_75685986 | chr2 | 75685419 | 75685986 | + | 0.957717 | intron (NM_010902, intron 1 of 4) | intron (NM_010902, intron 1 of 4) | 18939 | NM_010902 | 18024 | Mm.1025 | NM_010902 | ENSMUSG00000015839 | **Nfe2l2** | AI194320\|Nrf2 | nuclear factor, erythroid derived 2, like 2 | protein-coding |
| chr4_97995674_97996721 | chr4 | 97995675 | 97996721 | + | 1.66827 | intron (NM_001122953, intron 3 of 9) | intron (NM_001122953, intron 3 of 9) | 218572 | NM_010905 | 18027 | Mm.31274 | NM_010905 | ENSMUSG00000028565 | **Nfia** | 1110047K16Rik\|9430022M17Rik\|CTF\|NF1-A\|NF1A | nuclear factor I/A | protein-coding |
| chr4_97860805_97861304 | chr4 | 97860806 | 97861304 | + | 1.43682 | intron (NM_001122953, intron 2 of 9) | intron (NM_001122953, intron 2 of 9) | 83429 | NM_010905 | 18027 | Mm.31274 | NM_010905 | ENSMUSG00000028565 | **Nfia** | 1110047K16Rik\|9430022M17Rik\|CTF\|NF1-A\|NF1A | nuclear factor I/A | protein-coding |
| chr4_97860418_97860719 | chr4 | 97860419 | 97860719 | + | 1.14898 | intron (NM_001122953, intron 2 of 9) | intron (NM_001122953, intron 2 of 9) | 82943 | NM_010905 | 18027 | Mm.31274 | NM_010905 | ENSMUSG00000028565 | **Nfia** | 1110047K16Rik\|9430022M17Rik\|CTF\|NF1-A\|NF1A | nuclear factor I/A | protein-coding |
| chr4_98041497_98042200 | chr4 | 98041498 | 98042200 | + | 0.874296 | intron (NM_010905, intron 7 of 10) | intron (NM_010905, intron 7 of 10) | 264223 | NM_010905 | 18027 | Mm.31274 | NM_010905 | ENSMUSG00000028565 | **Nfia** | 1110047K16Rik\|9430022M17Rik\|CTF\|NF1-A\|NF1A | nuclear factor I/A | protein-coding |
| chr4_97926732_97927512 | chr4 | 97926733 | 97927512 | + | 0.85781 | intron (NM_001122953, intron 2 of 9) | intron (NM_001122953, intron 2 of 9) | 149496 | NM_010905 | 18027 | Mm.31274 | NM_010905 | ENSMUSG00000028565 | **Nfia** | 1110047K16Rik\|9430022M17Rik\|CTF\|NF1-A\|NF1A | nuclear factor I/A | protein-coding |
| chr4_98041861_98042192 | chr4 | 98041862 | 98042192 | + | 0.809147 | intron (NM_010905, intron 7 of 10) | intron (NM_010905, intron 7 of 10) | 264401 | NM_010905 | 18027 | Mm.31274 | NM_010905 | ENSMUSG00000028565 | **Nfia** | 1110047K16Rik\|9430022M17Rik\|CTF\|NF1-A\|NF1A | nuclear factor I/A | protein-coding |
| chr4_97919780_97920076 | chr4 | 97919781 | 97920076 | + | 0.774879 | intron (NM_001122953, intron 2 of 9) | intron (NM_001122953, intron 2 of 9) | 142302 | NM_010905 | 18027 | Mm.31274 | NM_010905 | ENSMUSG00000028565 | **Nfia** | 1110047K16Rik\|9430022M17Rik\|CTF\|NF1-A\|NF1A | nuclear factor I/A | protein-coding |
| chr4_97859079_97859544 | chr4 | 97859080 | 97859544 | + | 0.616043 | intron (NM_001122953, intron 2 of 9) | intron (NM_001122953, intron 2 of 9) | 81686 | NM_010905 | 18027 | Mm.31274 | NM_010905 | ENSMUSG00000028565 | **Nfia** | 1110047K16Rik\|9430022M17Rik\|CTF\|NF1-A\|NF1A | nuclear factor I/A | protein-coding |
| chr13_52976649_52977131 | chr13 | 52976650 | 52977131 | + | 1.06191 | intron (NM_017373, intron 1 of 1) | intron (NM_017373, intron 1 of 1) | 4149 | NM_017373 | 18030 | Mm.136604 | NM_017373 | ENSMUSG00000056749 | **Nfil3** | AV225605\|E4BP4 | nuclear factor, interleukin 3, regulated | protein-coding |
| chr13_52973484_52973748 | chr13 | 52973485 | 52973748 | + | 0.73216 | intron (NM_017373, intron 1 of 1) | intron (NM_017373, intron 1 of 1) | 7423 | NM_017373 | 18030 | Mm.136604 | NM_017373 | ENSMUSG00000056749 | **Nfil3** | AV225605\|E4BP4 | nuclear factor, interleukin 3, regulated | protein-coding |
| chr3_135635592_135636346 | chr3 | 135635593 | 135636346 | + | 1.84111 | intron (NM_008689, intron 6 of 24) | intron (NM_008689, intron 6 of 24) | 55578 | NM_008689 | 18033 | Mm.256765 | NM_008689 | ENSMUSG00000028163 | **Nfkb1** | NF-KB1\|NF-kappaB\|NF-kappaB1\|p105\|p50\|p50/p105 | nuclear factor of kappa light polypeptide gene enhancer in B cells 1, p105 | protein-coding |
| chr3_135720788_135721350 | chr3 | 135720789 | 135721350 | + | 1.70084 | Intergenic | Intergenic | -29522 | NM_008689 | 18033 | Mm.256765 | NM_008689 | ENSMUSG00000028163 | **Nfkb1** | NF-KB1\|NF-kappaB\|NF-kappaB1\|p105\|p50\|p50/p105 | nuclear factor of kappa light polypeptide gene enhancer in B cells 1, p105 | protein-coding |
| chr3_135628682_135629654 | chr3 | 135628683 | 135629654 | + | 1.47907 | intron (NM_008689, intron 6 of 24) | intron (NM_008689, intron 6 of 24) | 62379 | NM_008689 | 18033 | Mm.256765 | NM_008689 | ENSMUSG00000028163 | **Nfkb1** | NF-KB1\|NF-kappaB\|NF-kappaB1\|p105\|p50\|p50/p105 | nuclear factor of kappa light polypeptide gene enhancer in B cells 1, p105 | protein-coding |
| chr3_135640172_135640624 | chr3 | 135640173 | 135640624 | + | 1.35216 | intron (NM_008689, intron 6 of 24) | intron (NM_008689, intron 6 of 24) | 51149 | NM_008689 | 18033 | Mm.256765 | NM_008689 | ENSMUSG00000028163 | **Nfkb1** | NF-KB1\|NF-kappaB\|NF-kappaB1\|p105\|p50\|p50/p105 | nuclear factor of kappa light polypeptide gene enhancer in B cells 1, p105 | protein-coding |
| chr3_135722174_135722599 | chr3 | 135722175 | 135722599 | + | 1.32927 | Intergenic | Intergenic | -30840 | NM_008689 | 18033 | Mm.256765 | NM_008689 | ENSMUSG00000028163 | **Nfkb1** | NF-KB1\|NF-kappaB\|NF-kappaB1\|p105\|p50\|p50/p105 | nuclear factor of kappa light polypeptide gene enhancer in B cells 1, p105 | protein-coding |
| chr3_135699642_135700714 | chr3 | 135699643 | 135700714 | + | 1.29208 | Intergenic | Intergenic | -8631 | NM_008689 | 18033 | Mm.256765 | NM_008689 | ENSMUSG00000028163 | **Nfkb1** | NF-KB1\|NF-kappaB\|NF-kappaB1\|p105\|p50\|p50/p105 | nuclear factor of kappa light polypeptide gene enhancer in B cells 1, p105 | protein-coding |
| chr3_135629968_135630257 | chr3 | 135629969 | 135630257 | + | 1.28976 | intron (NM_008689, intron 6 of 24) | intron (NM_008689, intron 6 of 24) | 61434 | NM_008689 | 18033 | Mm.256765 | NM_008689 | ENSMUSG00000028163 | **Nfkb1** | NF-KB1\|NF-kappaB\|NF-kappaB1\|p105\|p50\|p50/p105 | nuclear factor of kappa light polypeptide gene enhancer in B cells 1, p105 | protein-coding |
| chr3_135670488_135670884 | chr3 | 135670489 | 135670884 | + | 1.17551 | intron (NM_008689, intron 1 of 24) | intron (NM_008689, intron 1 of 24) | 20861 | NM_008689 | 18033 | Mm.256765 | NM_008689 | ENSMUSG00000028163 | **Nfkb1** | NF-KB1\|NF-kappaB\|NF-kappaB1\|p105\|p50\|p50/p105 | nuclear factor of kappa light polypeptide gene enhancer in B cells 1, p105 | protein-coding |
| chr3_135712186_135712975 | chr3 | 135712187 | 135712975 | + | 1.05642 | Intergenic | B3\|SINE\|B2 | -21034 | NM_008689 | 18033 | Mm.256765 | NM_008689 | ENSMUSG00000028163 | **Nfkb1** | NF-KB1\|NF-kappaB\|NF-kappaB1\|p105\|p50\|p50/p105 | nuclear factor of kappa light polypeptide gene enhancer in B cells 1, p105 | protein-coding |
| chr3_135604635_135605148 | chr3 | 135604636 | 135605148 | + | 0.880651 | intron (NM_008689, intron 15 of 24) | intron (NM_008689, intron 15 of 24) | 86655 | NM_008689 | 18033 | Mm.256765 | NM_008689 | ENSMUSG00000028163 | **Nfkb1** | NF-KB1\|NF-kappaB\|NF-kappaB1\|p105\|p50\|p50/p105 | nuclear factor of kappa light polypeptide gene enhancer in B cells 1, p105 | protein-coding |
| chr3_135641122_135641488 | chr3 | 135641123 | 135641488 | + | 0.877096 | intron (NM_008689, intron 6 of 24) | L3\|LINE\|CR1 | 50242 | NM_008689 | 18033 | Mm.256765 | NM_008689 | ENSMUSG00000028163 | **Nfkb1** | NF-KB1\|NF-kappaB\|NF-kappaB1\|p105\|p50\|p50/p105 | nuclear factor of kappa light polypeptide gene enhancer in B cells 1, p105 | protein-coding |
| chr3_135602103_135602634 | chr3 | 135602104 | 135602634 | + | 0.848819 | intron (NM_008689, intron 16 of 24) | intron (NM_008689, intron 16 of 24) | 89178 | NM_008689 | 18033 | Mm.256765 | NM_008689 | ENSMUSG00000028163 | **Nfkb1** | NF-KB1\|NF-kappaB\|NF-kappaB1\|p105\|p50\|p50/p105 | nuclear factor of kappa light polypeptide gene enhancer in B cells 1, p105 | protein-coding |
| chr3_135607039_135607589 | chr3 | 135607040 | 135607589 | + | 0.837757 | intron (NM_008689, intron 13 of 24) | intron (NM_008689, intron 13 of 24) | 84233 | NM_008689 | 18033 | Mm.256765 | NM_008689 | ENSMUSG00000028163 | **Nfkb1** | NF-KB1\|NF-kappaB\|NF-kappaB1\|p105\|p50\|p50/p105 | nuclear factor of kappa light polypeptide gene enhancer in B cells 1, p105 | protein-coding |
| chr3_135636426_135636823 | chr3 | 135636427 | 135636823 | + | 0.832914 | intron (NM_008689, intron 6 of 24) | intron (NM_008689, intron 6 of 24) | 54922 | NM_008689 | 18033 | Mm.256765 | NM_008689 | ENSMUSG00000028163 | **Nfkb1** | NF-KB1\|NF-kappaB\|NF-kappaB1\|p105\|p50\|p50/p105 | nuclear factor of kappa light polypeptide gene enhancer in B cells 1, p105 | protein-coding |
| chr3_135720030_135720276 | chr3 | 135720031 | 135720276 | + | 0.823922 | Intergenic | Intergenic | -28606 | NM_008689 | 18033 | Mm.256765 | NM_008689 | ENSMUSG00000028163 | **Nfkb1** | NF-KB1\|NF-kappaB\|NF-kappaB1\|p105\|p50\|p50/p105 | nuclear factor of kappa light polypeptide gene enhancer in B cells 1, p105 | protein-coding |
| chr3_135673123_135673379 | chr3 | 135673124 | 135673379 | + | 0.796105 | intron (NM_008689, intron 1 of 24) | intron (NM_008689, intron 1 of 24) | 18296 | NM_008689 | 18033 | Mm.256765 | NM_008689 | ENSMUSG00000028163 | **Nfkb1** | NF-KB1\|NF-kappaB\|NF-kappaB1\|p105\|p50\|p50/p105 | nuclear factor of kappa light polypeptide gene enhancer in B cells 1, p105 | protein-coding |

| | | | | | | | | | | | | | | | | | |
|---|---|---|---|---|---|---|---|---|---|---|---|---|---|---|---|---|---|
| chr3_135686398_135686699 | chr3 | 135686399 | 135686699 | + | 0.692943 | intron (NM_008689, intron 1 of 24) | intron (NM_008689, intron 1 of 24) | 4998 | NM_008689 | 18033 | Mm.256765 | NM_008689 | ENSMUSG00000028163 | **Nfkb1** | NF-KB1\|NF-kappaB\|NF-kappaB1\|p105\|p50\|p50/p105 | nuclear factor of kappa light polypeptide gene enhancer in B cells 1, p105 | protein-coding |
| chr3_135621868_135622357 | chr3 | 135621869 | 135622357 | + | 0.647323 | intron (NM_008689, intron 7 of 24) | intron (NM_008689, intron 7 of 24) | 69434 | NM_008689 | 18033 | Mm.256765 | NM_008689 | ENSMUSG00000028163 | **Nfkb1** | NF-KB1\|NF-kappaB\|NF-kappaB1\|p105\|p50\|p50/p105 | nuclear factor of kappa light polypeptide gene enhancer in B cells 1, p105 | protein-coding |
| chr3_135621069_135622368 | chr3 | 135621070 | 135622368 | + | 0.627812 | intron (NM_008689, intron 7 of 24) | intron (NM_008689, intron 7 of 24) | 69828 | NM_008689 | 18033 | Mm.256765 | NM_008689 | ENSMUSG00000028163 | **Nfkb1** | NF-KB1\|NF-kappaB\|NF-kappaB1\|p105\|p50\|p50/p105 | nuclear factor of kappa light polypeptide gene enhancer in B cells 1, p105 | protein-coding |
| chr12_55457622_55458222 | chr12 | 55457623 | 55458222 | + | 0.832781 | intron (NR_035430, intron 1 of 3) | intron (NR_035430, intron 1 of 3) | 34725 | NM_010907 | 18035 | Mm.170515 | NM_010907 | ENSMUSG00000021025 | **Nfkbia** | AI462015\|Nfkbi | nuclear factor of kappa light polypeptide gene enhancer in B cells inhibitor, alpha | protein-coding |
| chr17_35231064_35231503 | chr17 | 35231065 | 35231503 | + | 1.05658 | intron (NM_010909, intron 2 of 3) | intron (NM_010909, intron 2 of 3) | 4531 | NM_010909 | 18038 | Mm.300795 | NM_010909 | ENSMUSG00000042419 | **Nfkbil1** | Def-7\|IKBL | nuclear factor of kappa light polypeptide gene enhancer in B cells inhibitor like 1 | protein-coding |
| chr17_35223498_35223740 | chr17 | 35223499 | 35223740 | + | 0.79494 | intron (NM_010909, intron 2 of 3) | intron (NM_010909, intron 2 of 3) | 12196 | NM_010909 | 18038 | Mm.300795 | NM_010909 | ENSMUSG00000042419 | **Nfkbil1** | Def-7\|IKBL | nuclear factor of kappa light polypeptide gene enhancer in B cells inhibitor like 1 | protein-coding |
| chr17_35225477_35225744 | chr17 | 35225478 | 35225744 | + | 0.786746 | intron (NM_010909, intron 2 of 3) | intron (NM_010909, intron 2 of 3) | 10204 | NM_010909 | 18038 | Mm.300795 | NM_010909 | ENSMUSG00000042419 | **Nfkbil1** | Def-7\|IKBL | nuclear factor of kappa light polypeptide gene enhancer in B cells inhibitor like 1 | protein-coding |
| chr16_55564197_55565330 | chr16 | 55564198 | 55565330 | + | 1.39207 | Intergenic | L1VL2\|LINE\|L1 | 257374 | NM_001159394 | 80859 | Mm.247272 | NM_030612 | ENSMUSG00000035356 | **Nfkbiz** | AA408868\|INAP\|Mail | nuclear factor of kappa light polypeptide gene enhancer in B cells inhibitor, zeta | protein-coding |
| chr16_55747084_55748111 | chr16 | 55747085 | 55748111 | + | 1.32956 | Intergenic | Intergenic | 74540 | NM_001159394 | 80859 | Mm.247272 | NM_030612 | ENSMUSG00000035356 | **Nfkbiz** | AA408868\|INAP\|Mail | nuclear factor of kappa light polypeptide gene enhancer in B cells inhibitor, zeta | protein-coding |
| chr16_55747104_55747695 | chr16 | 55747105 | 55747695 | + | 1.29914 | Intergenic | Intergenic | 74738 | NM_001159394 | 80859 | Mm.247272 | NM_030612 | ENSMUSG00000035356 | **Nfkbiz** | AA408868\|INAP\|Mail | nuclear factor of kappa light polypeptide gene enhancer in B cells inhibitor, zeta | protein-coding |
| chr16_55555182_55555679 | chr16 | 55555183 | 55555679 | + | 1.12713 | Intergenic | Intergenic | 266707 | NM_001159394 | 80859 | Mm.247272 | NM_030612 | ENSMUSG00000035356 | **Nfkbiz** | AA408868\|INAP\|Mail | nuclear factor of kappa light polypeptide gene enhancer in B cells inhibitor, zeta | protein-coding |
| chr16_55555202_55555566 | chr16 | 55555203 | 55555566 | + | 1.1123 | Intergenic | Intergenic | 266754 | NM_001159394 | 80859 | Mm.247272 | NM_030612 | ENSMUSG00000035356 | **Nfkbiz** | AA408868\|INAP\|Mail | nuclear factor of kappa light polypeptide gene enhancer in B cells inhibitor, zeta | protein-coding |
| chr16_55837254_55837468 | chr16 | 55837255 | 55837468 | + | 1.09341 | intron (NM_001159395, intron 1 of 12) | intron (NM_001159395, intron 1 of 12) | 1280 | NM_030612 | 80859 | Mm.247272 | NM_030612 | ENSMUSG00000035356 | **Nfkbiz** | AA408868\|INAP\|Mail | nuclear factor of kappa light polypeptide gene enhancer in B cells inhibitor, zeta | protein-coding |
| chr16_55770190_55770731 | chr16 | 55770191 | 55770731 | + | 1.00391 | Intergenic | Intergenic | 51677 | NM_001159394 | 80859 | Mm.247272 | NM_030612 | ENSMUSG00000035356 | **Nfkbiz** | AA408868\|INAP\|Mail | nuclear factor of kappa light polypeptide gene enhancer in B cells inhibitor, zeta | protein-coding |
| chr16_55835635_55836134 | chr16 | 55835636 | 55836134 | + | 0.947345 | intron (NM_030612, intron 2 of 13) | (TGGGGG)n\|Simple_repeat\|Simple_repeat | 2756 | NM_030612 | 80859 | Mm.247272 | NM_030612 | ENSMUSG00000035356 | **Nfkbiz** | AA408868\|INAP\|Mail | nuclear factor of kappa light polypeptide gene enhancer in B cells inhibitor, zeta | protein-coding |
| chr16_55608476_55609017 | chr16 | 55608477 | 55609017 | + | 0.946706 | Intergenic | B4A\|SINE\|B4 | 213391 | NM_001159394 | 80859 | Mm.247272 | NM_030612 | ENSMUSG00000035356 | **Nfkbiz** | AA408868\|INAP\|Mail | nuclear factor of kappa light polypeptide gene enhancer in B cells inhibitor, zeta | protein-coding |
| chr16_55722853_55723543 | chr16 | 55722854 | 55723543 | + | 0.943479 | Intergenic | Intergenic | 98940 | NM_001159394 | 80859 | Mm.247272 | NM_030612 | ENSMUSG00000035356 | **Nfkbiz** | AA408868\|INAP\|Mail | nuclear factor of kappa light polypeptide gene enhancer in B cells inhibitor, zeta | protein-coding |
| chr16_55690984_55691405 | chr16 | 55690985 | 55691405 | + | 0.879382 | Intergenic | (CAGCC)n\|Simple_repeat\|Simple_repeat | 130943 | NM_001159394 | 80859 | Mm.247272 | NM_030612 | ENSMUSG00000035356 | **Nfkbiz** | AA408868\|INAP\|Mail | nuclear factor of kappa light polypeptide gene enhancer in B cells inhibitor, zeta | protein-coding |
| chr16_55747797_55748111 | chr16 | 55747798 | 55748111 | + | 0.875967 | Intergenic | ORR1A2\|LTR\|ERVL-MaLR | 74184 | NM_001159394 | 80859 | Mm.247272 | NM_030612 | ENSMUSG00000035356 | **Nfkbiz** | AA408868\|INAP\|Mail | nuclear factor of kappa light polypeptide gene enhancer in B cells inhibitor, zeta | protein-coding |
| chr16_55606546_55606840 | chr16 | 55606547 | 55606840 | + | 0.831856 | Intergenic | Intergenic | 215445 | NM_001159394 | 80859 | Mm.247272 | NM_030612 | ENSMUSG00000035356 | **Nfkbiz** | AA408868\|INAP\|Mail | nuclear factor of kappa light polypeptide gene enhancer in B cells inhibitor, zeta | protein-coding |
| chr16_55582865_55583434 | chr16 | 55582866 | 55583434 | + | 0.793833 | Intergenic | Intergenic | 238988 | NM_001159394 | 80859 | Mm.247272 | NM_030612 | ENSMUSG00000035356 | **Nfkbiz** | AA408868\|INAP\|Mail | nuclear factor of kappa light polypeptide gene enhancer in B cells inhibitor, zeta | protein-coding |
| chr16_55723132_55723543 | chr16 | 55723133 | 55723543 | + | 0.778968 | Intergenic | Intergenic | 98800 | NM_001159394 | 80859 | Mm.247272 | NM_030612 | ENSMUSG00000035356 | **Nfkbiz** | AA408868\|INAP\|Mail | nuclear factor of kappa light polypeptide gene enhancer in B cells inhibitor, zeta | protein-coding |
| chr16_55723199_55723507 | chr16 | 55723200 | 55723507 | + | 0.747136 | Intergenic | Intergenic | 98785 | NM_001159394 | 80859 | Mm.247272 | NM_030612 | ENSMUSG00000035356 | **Nfkbiz** | AA408868\|INAP\|Mail | nuclear factor of kappa light polypeptide gene enhancer in B cells inhibitor, zeta | protein-coding |
| chr16_55572653_55573119 | chr16 | 55572654 | 55573119 | + | 0.745878 | Intergenic | Intergenic | 249252 | NM_001159394 | 80859 | Mm.247272 | NM_030612 | ENSMUSG00000035356 | **Nfkbiz** | AA408868\|INAP\|Mail | nuclear factor of kappa light polypeptide gene enhancer in B cells inhibitor, zeta | protein-coding |
| chr16_55582470_55583434 | chr16 | 55582471 | 55583434 | + | 0.712572 | Intergenic | Intergenic | 239186 | NM_001159394 | 80859 | Mm.247272 | NM_030612 | ENSMUSG00000035356 | **Nfkbiz** | AA408868\|INAP\|Mail | nuclear factor of kappa light polypeptide gene enhancer in B cells inhibitor, zeta | protein-coding |
| chr16_55572765_55573101 | chr16 | 55572766 | 55573101 | + | 0.700055 | Intergenic | Intergenic | 249205 | NM_001159394 | 80859 | Mm.247272 | NM_030612 | ENSMUSG00000035356 | **Nfkbiz** | AA408868\|INAP\|Mail | nuclear factor of kappa light polypeptide gene enhancer in B cells inhibitor, zeta | protein-coding |
| chr4_40973848_40974520 | chr4 | 40973849 | 40974520 | + | 1.59739 | intron (NM_001290448, intron 1 of 15) | intron (NM_001290448, intron 1 of 15) | 3241 | NM_023739 | 74164 | Mm.247456 | NM_023739 | ENSMUSG00000028423 | **Nfx1** | 1300017N15Rik\|3000003M19Rik\|NFX.1\|TEG-42\|Tex42 | nuclear transcription factor, X-box binding 1 | protein-coding |
| chr4_40973784_40974691 | chr4 | 40973785 | 40974691 | + | 1.56433 | intron (NM_001290448, intron 1 of 15) | intron (NM_001290448, intron 1 of 15) | 3295 | NM_023739 | 74164 | Mm.247456 | NM_023739 | ENSMUSG00000028423 | **Nfx1** | 1300017N15Rik\|3000003M19Rik\|NFX.1\|TEG-42\|Tex42 | nuclear transcription factor, X-box binding 1 | protein-coding |
| chr10_82780369_82780624 | chr10 | 82780370 | 82780624 | + | 0.810773 | Intergenic | ORR1B1\|LTR\|ERVL-MaLR | -16356 | NM_010914 | 18045 | Mm.245998 | NM_010914 | ENSMUSG00000020248 | **Nfyb** | AA985999\|Cbf-A | nuclear transcription factor-Y beta | protein-coding |
| chr10_82780304_82780740 | chr10 | 82780305 | 82780740 | + | 0.762127 | Intergenic | ORR1B1\|LTR\|ERVL-MaLR | -16381 | NM_010914 | 18045 | Mm.245998 | NM_010914 | ENSMUSG00000020248 | **Nfyb** | AA985999\|Cbf-A | nuclear transcription factor-Y beta | protein-coding |
| chr10_82761157_82761356 | chr10 | 82761158 | 82761356 | + | 0.711954 | intron (NM_010914, intron 2 of 7) | intron (NM_010914, intron 2 of 7) | 2884 | NM_010914 | 18045 | Mm.245998 | NM_010914 | ENSMUSG00000020248 | **Nfyb** | AA985999\|Cbf-A | nuclear transcription factor-Y beta | protein-coding |
| chr1_87550963_87551435 | chr1 | 87550964 | 87551435 | + | 0.857519 | intron (NM_001111314, intron 1 of 14) | intron (NM_001111314, intron 1 of 14) | 22671 | NM_001111314 | 53972 | Mm.435439 | NM_019867 | ENSMUSG00000026259 | **Ngef** | Tims2\|ephexin | neuronal guanine nucleotide exchange factor | protein-coding |
| chr14_16045886_16046771 | chr14 | 16045887 | 16046771 | + | 0.869703 | Intergenic | ORR1C1-int\|LTR\|ERVL-MaLR | -202985 | NM_021504 | 59007 | Mm.258381 | NM_021504 | ENSMUSG00000021785 | **Ngly1** | 1110002C09Rik\|PNGase\|Png1 | N-glycanase 1 | protein-coding |
| chr14_16046486_16046763 | chr14 | 16046487 | 16046763 | + | 0.743677 | Intergenic | ORR1C1-int\|LTR\|ERVL-MaLR | -202689 | NM_021504 | 59007 | Mm.258381 | NM_021504 | ENSMUSG00000021785 | **Ngly1** | 1110002C09Rik\|PNGase\|Png1 | N-glycanase 1 | protein-coding |
| chr14_16107366_16107860 | chr14 | 16107367 | 16107860 | + | 0.698344 | Intergenic | Intergenic | -141701 | NM_021504 | 59007 | Mm.258381 | NM_021504 | ENSMUSG00000021785 | **Ngly1** | 1110002C09Rik\|PNGase\|Png1 | N-glycanase 1 | protein-coding |
| chr14_15936941_15937354 | chr14 | 15936942 | 15937354 | + | 0.653573 | Intergenic | MTC-int\|LTR\|ERVL-MaLR | -312166 | NM_021504 | 59007 | Mm.258381 | NM_021504 | ENSMUSG00000021785 | **Ngly1** | 1110002C09Rik\|PNGase\|Png1 | N-glycanase 1 | protein-coding |
| chr19_56588167_56589036 | chr19 | 56588168 | 56589036 | + | 0.872102 | intron (NM_025811, intron 6 of 10) | MLT1I\|LTR\|ERVL-MaLR | 40341 | NM_025811 | 66866 | Mm.46346 | NM_025811 | ENSMUSG00000025078 | **Nhlrc2** | 1200003G01Rik\|AI835049\|AV002846\|AW496455 | NHL repeat containing 2 | protein-coding |
| chr3_53451716_53452184 | chr3 | 53451717 | 53452184 | + | 0.994203 | TTS (NM_172501) | TTS (NM_172501) | 11308 | NM_172501 | 212114 | Mm.85744 | NM_172501 | | **Nhlrc3** | 4833441N19\|8030451K01Rik\|mKIAA4083 | NHL repeat containing 3 | protein-coding |
| chrX_101882182_101882637 | chrX | 101882183 | 101882637 | + | 1.04353 | intron (NM_001163610, intron 1 of 7) | intron (NM_001163610, intron 1 of 7) | 33025 | NM_001163610 | 100042480 | Mm.232 | NM_001163610 | ENSMUSG00000079481 | **Nhsl2** | 1110062M06Rik\|6430511F03\|AI225852\|AW488865\|EG621083\|ENSMUSG00000073037\|Gm10456 | NHS-like 2 | protein-coding |
| chr13_13493440_13493937 | chr13 | 13493441 | 13493937 | + | 1.52813 | intron (NM_010917, intron 12 of 19) | intron (NM_010917, intron 12 of 19) | 56087 | NM_010917 | 18073 | Mm.4691 | NM_010917 | ENSMUSG00000005397 | **Nid1** | A630025O17\|Nid\|entactin\|entactin-1\|nidogen-1 | nidogen 1 | protein-coding |
| chr6_120190929_120191281 | chr6 | 120190930 | 120191281 | + | 1.60925 | intron (NM_016718, intron 1 of 3) | intron (NM_016718, intron 1 of 3) | 97725 | NM_016718 | 29862 | Mm.103668 | NM_016718 | ENSMUSG00000041377 | **Ninj2** | - | ninjurin 2 | protein-coding |
| chr6_120085725_120086224 | chr6 | 120085726 | 120086224 | + | 1.20033 | Intergenic | Intergenic | -7405 | NM_016718 | 29862 | Mm.103668 | NM_016718 | ENSMUSG00000041377 | **Ninj2** | - | ninjurin 2 | protein-coding |
| chr6_120119901_120120174 | chr6 | 120119902 | 120120174 | + | 1.11519 | intron (NM_016718, intron 1 of 3) | L1M3\|LINE\|L1 | 26658 | NM_016718 | 29862 | Mm.103668 | NM_016718 | ENSMUSG00000041377 | **Ninj2** | - | ninjurin 2 | protein-coding |
| chr6_120179141_120179754 | chr6 | 120179142 | 120179754 | + | 0.989419 | intron (NM_016718, intron 1 of 3) | PB1D10\|SINE\|Alu | 86068 | NM_016718 | 29862 | Mm.103668 | NM_016718 | ENSMUSG00000041377 | **Ninj2** | - | ninjurin 2 | protein-coding |
| chr6_120114671_120114966 | chr6 | 120114672 | 120114966 | + | 0.950494 | intron (NM_016718, intron 1 of 3) | intron (NM_016718, intron 1 of 3) | 21439 | NM_016718 | 29862 | Mm.103668 | NM_016718 | ENSMUSG00000041377 | **Ninj2** | - | ninjurin 2 | protein-coding |
| chr6_120179141_120180069 | chr6 | 120179142 | 120180069 | + | 0.851613 | intron (NM_016718, intron 1 of 3) | intron (NM_016718, intron 1 of 3) | 86225 | NM_016718 | 29862 | Mm.103668 | NM_016718 | ENSMUSG00000041377 | **Ninj2** | - | ninjurin 2 | protein-coding |
| chr6_120122826_120123273 | chr6 | 120122827 | 120123273 | + | 0.807061 | intron (NM_016718, intron 1 of 3) | L1M3\|LINE\|L1 | 29670 | NM_016718 | 29862 | Mm.103668 | NM_016718 | ENSMUSG00000041377 | **Ninj2** | - | ninjurin 2 | protein-coding |
| chr6_120152755_120153009 | chr6 | 120152756 | 120153009 | + | 0.675387 | intron (NM_016718, intron 1 of 3) | intron (NM_016718, intron 1 of 3) | 59502 | NM_016718 | 29862 | Mm.103668 | NM_016718 | ENSMUSG00000041377 | **Ninj2** | - | ninjurin 2 | protein-coding |
| chr7_55938113_55938672 | chr7 | 55938114 | 55938672 | + | 0.904563 | intron (NM_023647, intron 4 of 6) | intron (NM_023647, intron 4 of 6) | 24100 | NM_001256133 | 93790 | Mm.37249 | NM_023647 | ENSMUSG00000030452 | **Nipa2** | 2600017P10Rik\|3830408P04Rik\|AB041581 | non imprinted in Prader-Willi/Angelman syndrome 2 homolog (human) | protein-coding |

| | | | | | | | | | | | | | | | | | |
|---|---|---|---|---|---|---|---|---|---|---|---|---|---|---|---|---|---|
| chr5_72679515_72679851 | chr5 | 72679516 | 72679851 | + | 0.844992 | Intergenic | ORR1B1\|LTR\|ERVL-MaLR | 31887 | NM_001081205 | 70701 | Mm.38884 | NM_001081205 | ENSMUSG00000067219 | **Nipal1** | 3830408G10Rik\|Npal1 | NIPA-like domain containing 1 | protein-coding |
| chr11_46134804_46135365 | chr11 | 46134805 | 46135365 | + | 1.59459 | intron (NM_009616, intron 15 of 22) | intron (NM_009616, intron 15 of 22) | 31274 | NM_172524 | 214112 | Mm.225253 | NM_172524 | ENSMUSG00000020411 | **Nipal4** | 9530066K23Rik | NIPA-like domain containing 4 | protein-coding |
| chr15_8403361_8403720 | chr15 | 8403362 | 8403720 | + | 1.02578 | intron (NM_027707, intron 1 of 46) | intron (NM_027707, intron 1 of 46) | 40922 | NM_027707 | 71175 | Mm.240329 | NM_027707 | ENSMUSG00000022141 | **Nipbl** | Idn3 | Nipped-B homolog (Drosophila) | protein-coding |
| chr15_8403081_8403728 | chr15 | 8403082 | 8403728 | + | 0.912732 | intron (NM_027707, intron 1 of 46) | intron (NM_027707, intron 1 of 46) | 41058 | NM_027707 | 71175 | Mm.240329 | NM_027707 | ENSMUSG00000022141 | **Nipbl** | Idn3 | Nipped-B homolog (Drosophila) | protein-coding |
| chr15_8446483_8446808 | chr15 | 8446484 | 8446808 | + | 0.654328 | Intergenic | Intergenic | -2183 | NM_201232 | 71175 | Mm.240329 | NM_027707 | ENSMUSG00000022141 | **Nipbl** | Idn3 | Nipped-B homolog (Drosophila) | protein-coding |
| chr14_31197865_31198231 | chr14 | 31197866 | 31198231 | + | 0.922689 | intron (NM_022656, intron 3 of 20) | L1MB7\|LINE\|L1 | 8778 | NM_022656 | 64652 | Mm.298728 | NM_022656 | ENSMUSG00000021910 | **Nisch** | 1200007D05Rik\|3202002H23Rik\|AW494485\|I-1 | nischarin | protein-coding |
| chr14_31197865_31198440 | chr14 | 31197866 | 31198440 | + | 0.836671 | intron (NM_022656, intron 3 of 20) | intron (NM_022656, intron 3 of 20) | 8673 | NM_022656 | 64652 | Mm.298728 | NM_022656 | ENSMUSG00000021910 | **Nisch** | 1200007D05Rik\|3202002H23Rik\|AW494485\|I-1 | nischarin | protein-coding |
| chr16_57175528_57176143 | chr16 | 57175529 | 57176143 | + | 0.94654 | intron (NM_026254, intron 15 of 17) | B4\|SINE\|B4 | -8504 | NM_023175 | 52633 | Mm.383203 | NM_023175 | ENSMUSG00000022751 | **Nit2** | 1190017B19Rik\|D16Ertd502e | nitrilase family, member 2 | protein-coding |
| chr16_57175687_57176125 | chr16 | 57175688 | 57176125 | + | 0.861461 | intron (NM_026254, intron 15 of 17) | MTD\|LTR\|ERVL-MaLR | -8574 | NM_023175 | 52633 | Mm.383203 | NM_023175 | ENSMUSG00000022751 | **Nit2** | 1190017B19Rik\|D16Ertd502e | nitrilase family, member 2 | protein-coding |
| chr4_130576328_130576973 | chr4 | 130576329 | 130576973 | + | 0.61411 | Intergenic | Intergenic | -2615 | NM_025998 | 67149 | Mm.423029 | NM_025998 | | **Nkain1** | 2610200G18Rik\|2810426C15Rik | Na+/K+ transporting ATPase interacting 1 | protein-coding |
| chr14_69135758_69136232 | chr14 | 69135759 | 69136232 | + | 0.647389 | Intergenic | Intergenic | -35807 | NM_010920 | 18092 | Mm.390691 | NM_010920 | ENSMUSG00000044186 | **Nkx2-6** | AV044928\|Nkx-2.6\|Nkx2.6\|Tix\|tinman | NK2 homeobox 6 | protein-coding |
| chr14_69184714_69185256 | chr14 | 69184715 | 69185256 | + | 0.956972 | Intergenic | Intergenic | -5707 | NM_010921 | 18095 | Mm.3520 | NM_010921 | ENSMUSG00000022061 | **Nkx3-1** | Bax\|NKX3.1\|NKX3A\|Nkx-3.1\|bagpipe | NK-3 transcription factor, locus 1 (Drosophila) | protein-coding |
| chr14_69184675_69185402 | chr14 | 69184676 | 69185402 | + | 0.849647 | Intergenic | Intergenic | -5653 | NM_010921 | 18095 | Mm.3520 | NM_010921 | ENSMUSG00000022061 | **Nkx3-1** | Bax\|NKX3.1\|NKX3A\|Nkx-3.1\|bagpipe | NK-3 transcription factor, locus 1 (Drosophila) | protein-coding |
| chr14_69186664_69187058 | chr14 | 69186665 | 69187058 | + | 0.784539 | Intergenic | Intergenic | -3831 | NM_010921 | 18095 | Mm.3520 | NM_010921 | ENSMUSG00000022061 | **Nkx3-1** | Bax\|NKX3.1\|NKX3A\|Nkx-3.1\|bagpipe | NK-3 transcription factor, locus 1 (Drosophila) | protein-coding |
| chr5_101665221_101665598 | chr5 | 101665222 | 101665598 | + | 0.856568 | promoter-TSS (NM_144955) | promoter-TSS (NM_144955) | -699 | NM_144955 | 18096 | Mm.193072 | NM_144955 | | **Nkx6-1** | NKX6A\|Nkx6.1 | NK6 homeobox 1 | protein-coding |
| chr11_78649278_78649770 | chr11 | 78649279 | 78649770 | + | 1.08921 | intron (NM_008702, intron 1 of 10) | intron (NM_008702, intron 1 of 10) | 47901 | NM_008702 | 18099 | Mm.9001 | NM_008702 | ENSMUSG00000017376 | **Nlk** | AI194375 | nemo like kinase | protein-coding |
| chr8_94458501_94459226 | chr8 | 94458502 | 94459226 | + | 0.993123 | Intergenic | MLT1A\|LTR\|ERVL-MaLR | -13899 | NM_001033207 | 434341 | Mm.334720 | NM_001033207 | ENSMUSG00000074151 | **Nlrc5** | AI451557\|AK220210\|NOD27 | NLR family, CARD domain containing 5 | protein-coding |
| chr8_94458647_94458990 | chr8 | 94458648 | 94458990 | + | 0.973851 | Intergenic | MLT1A\|LTR\|ERVL-MaLR | -13944 | NM_001033207 | 434341 | Mm.334720 | NM_001033207 | ENSMUSG00000074151 | **Nlrc5** | AI451557\|AK220210\|NOD27 | NLR family, CARD domain containing 5 | protein-coding |
| chr18_34561560_34562039 | chr18 | 34561561 | 34562039 | + | 0.674653 | TTS (NM_080637) | TTS (NM_080637) | 17306 | NM_080637 | 75533 | Mm.275430 | NM_080637 | ENSMUSG00000035984 | **Nme5** | 1700019D05Rik\|Nm23-M5 | NME/NM23 family member 5 | protein-coding |
| chr1_153019265_153019911 | chr1 | 153019266 | 153019911 | + | 0.945777 | intron (NM_175460, intron 1 of 10) | intron (NM_175460, intron 1 of 10) | 64487 | NM_175460 | 226518 | Mm.40548 | NM_175460 | ENSMUSG00000042751 | **Nmnat2** | AI843915\|D030041I09Rik\|PNAT1\|PNAT2 | nicotinamide nucleotide adenylyltransferase 2 | protein-coding |
| chr2_3274054_3274687 | chr2 | 3274055 | 3274687 | + | 0.760812 | Intergenic | MER2\|DNA\|TcMar-Tigger | -9841 | NM_008708 | 18108 | Mm.65021 | NM_008708 | ENSMUSG00000026643 | **Nmt2** | A930001K02Rik\|AI605445\|AU044698\|hNMT-2 | N-myristoyltransferase 2 | protein-coding |
| chr19_38746019_38746613 | chr19 | 38746020 | 38746613 | + | 1.00838 | intron (NM_019588, intron 19 of 31) | (TC)n\|Simple_repeat\|Simple_repeat | 72921 | NM_021315 | 57753 | Mm.86482 | NM_021315 | ENSMUSG00000024999 | **Noc3l** | AF233884\|Fad24 | nucleolar complex associated 3 homolog (S. cerevisiae) | protein-coding |
| chr19_38788410_38789003 | chr19 | 38788411 | 38789003 | + | 0.725168 | 3' UTR (NM_021315, exon 21 of 21) | 3' UTR (NM_021315, exon 21 of 21) | 30530 | NM_021315 | 57753 | Mm.86482 | NM_021315 | ENSMUSG00000024999 | **Noc3l** | AF233884\|Fad24 | nucleolar complex associated 3 homolog (S. cerevisiae) | protein-coding |
| chr6_54956403_54956688 | chr6 | 54956404 | 54956688 | + | 0.876639 | intron (NM_172729, intron 1 of 11) | intron (NM_172729, intron 1 of 11) | 15115 | NM_001171007 | 107607 | Mm.28498 | NM_172729 | ENSMUSG00000038058 | **Nod1** | C230079P11\|Card4\|F830007N14Rik\|Nlrc1 | nucleotide-binding oligomerization domain containing 1 | protein-coding |
| chr6_54922863_54923340 | chr6 | 54922864 | 54923340 | + | 0.855476 | TTS (NM_001171007) | TTS (NM_001171007) | 48559 | NM_001171007 | 107607 | Mm.28498 | NM_172729 | ENSMUSG00000038058 | **Nod1** | C230079P11\|Card4\|F830007N14Rik\|Nlrc1 | nucleotide-binding oligomerization domain containing 1 | protein-coding |
| chr6_54954674_54954973 | chr6 | 54954675 | 54954973 | + | 0.750697 | intron (NM_172729, intron 1 of 11) | intron (NM_172729, intron 1 of 11) | 16837 | NM_001171007 | 107607 | Mm.28498 | NM_172729 | ENSMUSG00000038058 | **Nod1** | C230079P11\|Card4\|F830007N14Rik\|Nlrc1 | nucleotide-binding oligomerization domain containing 1 | protein-coding |
| chr11_107287203_107287898 | chr11 | 107287204 | 107287898 | + | 1.45673 | intron (NM_145823, intron 5 of 9) | intron (NM_145823, intron 5 of 9) | -98170 | NM_001161329 | 68979 | Mm.294617 | NM_133702 | ENSMUSG00000018433 | **Nol11** | 1500002M01Rik\|AU015220\|AU019874 | nucleolar protein 11 | protein-coding |
| chr2_153542642_153543577 | chr2 | 153542643 | 153543577 | + | 1.45819 | Intergenic | L2a\|LINE\|L2 | -13139 | NM_001134300 | 329540 | Mm.50600 | NM_001001986 | ENSMUSG00000061411 | **Nol4l** | 4931410H15\|8430427H17Rik\|AI429157\|Gm1636 | nucleolar protein 4-like | protein-coding |
| chr13_43400884_43401387 | chr13 | 43400885 | 43401387 | + | 1.51768 | intron (NM_023554, intron 5 of 7) | intron (NM_023554, intron 5 of 7) | 2760 | NM_023554 | 70078 | Mm.439927 | NM_023554 | ENSMUSG00000063200 | **Nol7** | 2210008F15Rik\|5730556I21Rik\|NOP27\|RARG-1 | nucleolar protein 7 | protein-coding |
| chr2_112262881_112263122 | chr2 | 112262882 | 112263122 | + | 0.645424 | TTS (NM_025403) | TTS (NM_025403) | 1076 | NM_025403 | 66181 | Mm.272346 | NM_025403 | | **Nop10** | 1110036B12Rik\|NOP10P\|Nola3 | NOP10 ribonucleoprotein | protein-coding |
| chr6_125129622_125130300 | chr6 | 125129623 | 125130300 | + | 0.934638 | exon (NM_145979, exon 39 of 39) | exon (NM_145979, exon 39 of 39) | -1922 | NM_138747 | 110109 | Mm.29203 | NM_138747 | ENSMUSG00000038279 | **Nop2** | 120kDa\|A530002O17\|Nol1 | NOP2 nucleolar protein | protein-coding |
| chr6_125130565_125130779 | chr6 | 125130566 | 125130779 | + | 0.67154 | TTS (NM_145979) | TTS (NM_145979) | -1211 | NM_138747 | 110109 | Mm.29203 | NM_138747 | ENSMUSG00000038279 | **Nop2** | 120kDa\|A530002O17\|Nol1 | NOP2 nucleolar protein | protein-coding |
| chr1_59683955_59684255 | chr1 | 59683956 | 59684255 | + | 0.817253 | promoter-TSS (NM_018868) | promoter-TSS (NM_018868) | -901 | NM_018868 | 55989 | Mm.220367 | NM_018868 | ENSMUSG00000026020 | **Nop58** | MSSP\|Nol5\|SIK\|nop5 | NOP58 ribonucleoprotein | protein-coding |
| chr1_170461894_170462606 | chr1 | 170461895 | 170462606 | + | 0.664098 | intron (NM_001109985, intron 2 of 9) | MTE2a\|LTR\|ERVL-MaLR | 94860 | NM_027528 | 70729 | Mm.487078 | NM_027528 | ENSMUSG00000038473 | **Nos1ap** | 6330408P19Rik\|Capon | nitric oxide synthase 1 (neuronal) adaptor protein | protein-coding |
| chr11_78920384_78921012 | chr11 | 78920385 | 78921012 | + | 1.14985 | promoter-TSS (NM_010927) | promoter-TSS (NM_010927) | -161 | NM_010927 | 18126 | Mm.2893 | NM_010927 | ENSMUSG00000020826 | **Nos2** | NOS-II\|Nos-2\|Nos2a\|i-NOS\|iNOS | nitric oxide synthase 2, inducible | protein-coding |
| chr2_69109756_69110215 | chr2 | 69109757 | 69110215 | + | 0.838614 | 3' UTR (NM_172856, exon 10 of 10) | 3' UTR (NM_172856, exon 10 of 10) | -25814 | NM_181547 | 329416 | Mm.90047 | NM_181547 | ENSMUSG00000034738 | **Nostrin** | Daip2\|mDaIP2 | nitric oxide synthase trafficker | protein-coding |
| chr2_69125358_69125772 | chr2 | 69125359 | 69125772 | + | 0.728164 | Intergenic | Intergenic | -10235 | NM_181547 | 329416 | Mm.90047 | NM_181547 | ENSMUSG00000034738 | **Nostrin** | Daip2\|mDaIP2 | nitric oxide synthase trafficker | protein-coding |
| chr3_98066372_98066884 | chr3 | 98066373 | 98066884 | + | 1.63531 | intron (NM_010928, intron 2 of 33) | B3\|SINE\|B2 | 53090 | NM_010928 | 18129 | Mm.485843 | NM_010928 | ENSMUSG00000027878 | **Notch2** | AI853703\|N2 | notch 2 | protein-coding |
| chr7_18925118_18925685 | chr7 | 18925119 | 18925685 | + | 0.621666 | promoter-TSS (NM_001029877) | promoter-TSS (NM_001029877) | -486 | NM_001029877 | 384569 | Mm.262724 | NM_001029877 | ENSMUSG00000030411 | **Nova2** | Gm1424 | neuro-oncological ventral antigen 2 | protein-coding |
| chr1_39196718_39197321 | chr1 | 39196719 | 39197321 | + | 0.94198 | intron (NM_008719, intron 1 of 20) | intron (NM_008719, intron 1 of 20) | 2748 | NM_008719 | 18143 | Mm.2380 | NM_008719 | ENSMUSG00000026077 | **Npas2** | MOP4\|bHLHe9 | neuronal PAS domain protein 2 | protein-coding |
| chr12_53024590_53024903 | chr12 | 53024591 | 53024903 | + | 0.635289 | intron (NM_198111, intron 8 of 13) | intron (NM_198111, intron 8 of 13) | -223930 | NM_013780 | 27386 | Mm.247044 | NM_013780 | ENSMUSG00000021010 | **Npas3** | 4930423H22Rik\|bHLHe12 | neuronal PAS domain protein 3 | protein-coding |
| chr19_4992969_4993234 | chr19 | 4992970 | 4993234 | + | 0.900494 | Intergenic | Intergenic | -3131 | NM_153553 | 225872 | Mm.287867 | NM_153553 | ENSMUSG00000045903 | **Npas4** | LE-PAS\|Nxf | neuronal PAS domain protein 4 | protein-coding |
| chr19_4992849_4993308 | chr19 | 4992850 | 4993308 | + | 0.822904 | Intergenic | Intergenic | -3108 | NM_153553 | 225872 | Mm.287867 | NM_153553 | ENSMUSG00000045903 | **Npas4** | LE-PAS\|Nxf | neuronal PAS domain protein 4 | protein-coding |
| chr12_84764549_84765112 | chr12 | 84764550 | 84765112 | + | 1.45895 | intron (NM_023409, intron 2 of 3) | intron (NM_023409, intron 2 of 3) | 8281 | NM_023409 | 67963 | Mm.282556 | NM_023409 | ENSMUSG00000021242 | **Npc2** | 2700012J19Rik\|AA408070\|AU045843\|HE1 | Niemann-Pick type C2 | protein-coding |
| chr12_84763930_84764290 | chr12 | 84763931 | 84764290 | + | 1.29662 | intron (NM_023409, intron 2 of 3) | URR1A\|DNA\|hAT-Charlie | 9002 | NM_023409 | 67963 | Mm.282556 | NM_023409 | ENSMUSG00000021242 | **Npc2** | 2700012J19Rik\|AA408070\|AU045843\|HE1 | Niemann-Pick type C2 | protein-coding |
| chr15_102555675_102555953 | chr15 | 102555676 | 102555953 | + | 0.687319 | intron (NM_146065, intron 4 of 10) | URR1A\|DNA\|hAT-Charlie | -30872 | NR_033409 | 54615 | Mm.445915 | NM_018787 | ENSMUSG00000023052 | **Npff** | - | neuropeptide FF-amide peptide precursor | protein-coding |
| chr10_61599492_61599937 | chr10 | 61599493 | 61599937 | + | 1.07569 | intron (NM_001177511, intron 1 of 3) | intron (NM_001177511, intron 1 of 3) | 4231 | NM_001177511 | 237362 | Mm.131297 | NM_001177511 | ENSMUSG00000020090 | **Npffr1** | Gm236\|Gpr147\|NPFF1\|OT7T022 | neuropeptide FF receptor 1 | protein-coding |
| chr1_153585717_153586076 | chr1 | 153585718 | 153586076 | + | 0.919505 | Intergenic | Intergenic | -36183 | NM_028749 | 74091 | Mm.24887 | NM_028749 | ENSMUSG00000042684 | **Npl** | 0610033B02Rik | N-acetylneuraminate pyruvate lyase | protein-coding |
| chr1_153583001_153583730 | chr1 | 153583002 | 153583730 | + | 0.918225 | Intergenic | Intergenic | -33652 | NM_028749 | 74091 | Mm.24887 | NM_028749 | ENSMUSG00000042684 | **Npl** | 0610033B02Rik | N-acetylneuraminate pyruvate lyase | protein-coding |
| chr1_153583959_153584225 | chr1 | 153583960 | 153584225 | + | 0.849036 | Intergenic | Lx\|LINE\|L1 | -34378 | NM_028749 | 74091 | Mm.24887 | NM_028749 | ENSMUSG00000042684 | **Npl** | 0610033B02Rik | N-acetylneuraminate pyruvate lyase | protein-coding |
| chr3_132947442_132947924 | chr3 | 132947443 | 132947924 | + | 1.17684 | intron (NM_001287101, intron 2 of 13) | B3\|SINE\|B2 | 2131 | NM_001287103 | 114249 | Mm.279310 | NM_033525 | ENSMUSG00000040998 | **Npnt** | 1110009H02Rik\|AA682063\|AI314031\|Nctn\|POEM | nephronectin | protein-coding |
| chr1_86671754_86672178 | chr1 | 86671755 | 86672178 | + | 0.785596 | Intergenic | Intergenic | -1393 | NM_010933 | 18159 | Mm.12802 | NM_010933 | ENSMUSG00000026241 | **Nppc** | CNP\|Ibab | natriuretic peptide type C | protein-coding |
| chr11_32271195_32271609 | chr11 | 32271196 | 32271609 | + | 1.15363 | Intergenic | L1M5\|LINE\|L1 | -3695 | NM_001284359 | 17168 | Mm.263161 | NM_181569 | ENSMUSG00000020289 | **Nprl3** | Aag\|CGTHBA\|HS-26\|HS-40\|Mare\|Phg\|Prox1\|m(alpha)RE | nitrogen permease regulator-like 3 | protein-coding |
| chr11_98772638_98773187 | chr11 | 98772639 | 98773187 | + | 0.812735 | intron (NM_145434, intron 1 of 7) | intron (NM_145434, intron 1 of 7) | 2464 | NM_145434 | 217166 | Mm.390397 | NM_145434 | ENSMUSG00000020889 | **Nr1d1** | A530070C09Rik\|R75201 | nuclear receptor subfamily 1, group D, member 1 | protein-coding |
| chr11_98775786_98776083 | chr11 | 98775787 | 98776083 | + | 0.74031 | promoter-TSS (NM_145434) | promoter-TSS (NM_145434) | -558 | NM_145434 | 217166 | Mm.390397 | NM_145434 | ENSMUSG00000020889 | **Nr1d1** | A530070C09Rik\|R75201 | nuclear receptor subfamily 1, group D, member 1 | protein-coding |

| | | | | | | | | | | | | | | | | | |
|---|---|---|---|---|---|---|---|---|---|---|---|---|---|---|---|---|---|
| chr14_18241832_18242154 | chr14 | 18241833 | 18242154 | + | 0.806454 | Intergenic | Intergenic | -2887 | NM_011584 | 353187 | Mm.26587 | NM_011584 | ENSMUSG00000021775 | **Nr1d2** | RVR\|Rev-erb | nuclear receptor subfamily 1, group D, member 2 | protein-coding |
| chr16_38259808_38260324 | chr16 | 38259809 | 38260324 | + | 0.949132 | intron (NM_001098404, intron 2 of 8) | RMER15-int\|LTR\|ERVL | 34758 | NM_010936 | 18171 | Mm.8509 | NM_010936 | ENSMUSG00000022809 | **Nr1i2** | PXR\|PXR.1\|PXR.2\|PXR1\|SXR\|mPXR | nuclear receptor subfamily 1, group I, member 2 | protein-coding |
| chr6_92103724_92104040 | chr6 | 92103725 | 92104040 | + | 0.946054 | intron (NM_011630, intron 1 of 14) | intron (NM_011630, intron 1 of 14) | 12464 | NM_011630 | 22026 | Mm.87062 | NM_011630 | ENSMUSG00000005893 | **Nr2c2** | TAK1\|Tr4\|mKIAA4145 | nuclear receptor subfamily 2, group C, member 2 | protein-coding |
| chr9_59911810_59912292 | chr9 | 59911811 | 59912292 | + | 0.997779 | intron (NM_173018, intron 40 of 43) | MTD\|LTR\|ERVL-MaLR | 38028 | NM_013708 | 23958 | Mm.103641 | NM_013708 | ENSMUSG00000032292 | **Nr2e3** | A930035N01Rik\|PNR\|RNR\|rd7 | nuclear receptor subfamily 2, group E, member 3 | protein-coding |
| chr13_78203846_78204096 | chr13 | 78203847 | 78204096 | + | 0.970513 | intron (NR_045195, intron 1 of 2) | intron (NR_045195, intron 1 of 2) | -4989 | NM_010151 | 13865 | Mm.439653 | NM_010151 | ENSMUSG00000069171 | **Nr2f1** | COUP-TF1\|COUP-TFI\|COUPTFA\|EAR3\|Erbal3\|SVP44\|Tcfcoup1 | nuclear receptor subfamily 2, group F, member 1 | protein-coding |
| chr13_78203822_78204275 | chr13 | 78203823 | 78204275 | + | 0.912881 | intron (NR_045195, intron 1 of 2) | intron (NR_045195, intron 1 of 2) | -5067 | NM_010151 | 13865 | Mm.439653 | NM_010151 | ENSMUSG00000069171 | **Nr2f1** | COUP-TF1\|COUP-TFI\|COUPTFA\|EAR3\|Erbal3\|SVP44\|Tcfcoup1 | nuclear receptor subfamily 2, group F, member 1 | protein-coding |
| chr13_78201582_78201876 | chr13 | 78201583 | 78201876 | + | 0.879009 | intron (NR_045195, intron 1 of 2) | intron (NR_045195, intron 1 of 2) | -2747 | NM_010151 | 13865 | Mm.439653 | NM_010151 | ENSMUSG00000069171 | **Nr2f1** | COUP-TF1\|COUP-TFI\|COUPTFA\|EAR3\|Erbal3\|SVP44\|Tcfcoup1 | nuclear receptor subfamily 2, group F, member 1 | protein-coding |
| chr13_78201564_78202057 | chr13 | 78201565 | 78202057 | + | 0.723723 | intron (NR_045195, intron 1 of 2) | intron (NR_045195, intron 1 of 2) | -2829 | NM_010151 | 13865 | Mm.439653 | NM_010151 | ENSMUSG00000069171 | **Nr2f1** | COUP-TF1\|COUP-TFI\|COUPTFA\|EAR3\|Erbal3\|SVP44\|Tcfcoup1 | nuclear receptor subfamily 2, group F, member 1 | protein-coding |
| chr18_39611614_39612310 | chr18 | 39611615 | 39612310 | + | 1.66825 | Intergenic | L2\|LINE\|L2 | -124717 | NM_008173 | 14815 | Mm.129481 | NM_008173 | ENSMUSG00000024431 | **Nr3c1** | GR\|Grl-1\|Grl1 | nuclear receptor subfamily 3, group C, member 1 | protein-coding |
| chr18_39281523_39282752 | chr18 | 39281524 | 39282752 | + | 1.26331 | intron (NM_175164, intron 17 of 22) | intron (NM_175164, intron 17 of 22) | 205107 | NM_008173 | 14815 | Mm.129481 | NM_008173 | ENSMUSG00000024431 | **Nr3c1** | GR\|Grl-1\|Grl1 | nuclear receptor subfamily 3, group C, member 1 | protein-coding |
| chr18_39319146_39320001 | chr18 | 39319147 | 39320001 | + | 1.01707 | intron (NM_175164, intron 20 of 22) | intron (NM_175164, intron 20 of 22) | 167671 | NM_008173 | 14815 | Mm.129481 | NM_008173 | ENSMUSG00000024431 | **Nr3c1** | GR\|Grl-1\|Grl1 | nuclear receptor subfamily 3, group C, member 1 | protein-coding |
| chr18_39250646_39251308 | chr18 | 39250647 | 39251308 | + | 0.856906 | intron (NM_175164, intron 17 of 22) | intron (NM_175164, intron 17 of 22) | 236268 | NM_008173 | 14815 | Mm.129481 | NM_008173 | ENSMUSG00000024431 | **Nr3c1** | GR\|Grl-1\|Grl1 | nuclear receptor subfamily 3, group C, member 1 | protein-coding |
| chr18_39483616_39484133 | chr18 | 39483617 | 39484133 | + | 0.799239 | intron (NM_008173, intron 1 of 7) | intron (NM_008173, intron 1 of 7) | 3370 | NM_008173 | 14815 | Mm.129481 | NM_008173 | ENSMUSG00000024431 | **Nr3c1** | GR\|Grl-1\|Grl1 | nuclear receptor subfamily 3, group C, member 1 | protein-coding |
| chr18_39308718_39309508 | chr18 | 39308719 | 39309508 | + | 0.75157 | intron (NM_175164, intron 20 of 22) | intron (NM_175164, intron 20 of 22) | 178132 | NM_008173 | 14815 | Mm.129481 | NM_008173 | ENSMUSG00000024431 | **Nr3c1** | GR\|Grl-1\|Grl1 | nuclear receptor subfamily 3, group C, member 1 | protein-coding |
| chr8_77209368_77209790 | chr8 | 77209369 | 77209790 | + | 0.781243 | intron (NM_001083906, intron 6 of 8) | intron (NM_001083906, intron 6 of 8) | 307071 | NM_001083906 | 110784 | Mm.324393 | NM_001083906 | ENSMUSG00000031618 | **Nr3c2** | MR\|Mlr | nuclear receptor subfamily 3, group C, member 2 | protein-coding |
| chr2_57123101_57124478 | chr2 | 57123102 | 57124478 | + | 0.726942 | intron (NM_001139509, intron 1 of 7) | CpG | 213 | NM_001139509 | 18227 | Mm.3507 | NM_013613 | ENSMUSG00000026826 | **Nr4a2** | HZF-3\|NOT\|Nurr1\|RNR-1\|TINOR\|TINUR | nuclear receptor subfamily 4, group A, member 2 | protein-coding |
| chr4_47846818_47847712 | chr4 | 47846819 | 47847712 | + | 1.62176 | Intergenic | Intergenic | -203983 | NM_015743 | 18124 | Mm.247261 | NM_015743 | ENSMUSG00000028341 | **Nr4a3** | AI573420\|CHN\|CSMF\|MINOR\|NOR-1\|Nor1\|TEC | nuclear receptor subfamily 4, group A, member 3 | protein-coding |
| chr4_48002983_48003295 | chr4 | 48002984 | 48003295 | + | 0.979604 | Intergenic | Intergenic | -48109 | NM_015743 | 18124 | Mm.247261 | NM_015743 | ENSMUSG00000028341 | **Nr4a3** | AI573420\|CHN\|CSMF\|MINOR\|NOR-1\|Nor1\|TEC | nuclear receptor subfamily 4, group A, member 3 | protein-coding |
| chr4_47957995_47958304 | chr4 | 47957996 | 47958304 | + | 0.96495 | Intergenic | L1VL2\|LINE\|L1 | -93098 | NM_015743 | 18124 | Mm.247261 | NM_015743 | ENSMUSG00000028341 | **Nr4a3** | AI573420\|CHN\|CSMF\|MINOR\|NOR-1\|Nor1\|TEC | nuclear receptor subfamily 4, group A, member 3 | protein-coding |
| chr4_47957853_47958303 | chr4 | 47957854 | 47958303 | + | 0.784753 | Intergenic | L1VL2\|LINE\|L1 | -93170 | NM_015743 | 18124 | Mm.247261 | NM_015743 | ENSMUSG00000028341 | **Nr4a3** | AI573420\|CHN\|CSMF\|MINOR\|NOR-1\|Nor1\|TEC | nuclear receptor subfamily 4, group A, member 3 | protein-coding |
| chr4_47957853_47958526 | chr4 | 47957854 | 47958526 | + | 0.763887 | Intergenic | L1VL2\|LINE\|L1 | -93058 | NM_015743 | 18124 | Mm.247261 | NM_015743 | ENSMUSG00000028341 | **Nr4a3** | AI573420\|CHN\|CSMF\|MINOR\|NOR-1\|Nor1\|TEC | nuclear receptor subfamily 4, group A, member 3 | protein-coding |
| chr4_47905428_47905942 | chr4 | 47905429 | 47905942 | + | 0.751506 | Intergenic | Intergenic | -145563 | NM_015743 | 18124 | Mm.247261 | NM_015743 | ENSMUSG00000028341 | **Nr4a3** | AI573420\|CHN\|CSMF\|MINOR\|NOR-1\|Nor1\|TEC | nuclear receptor subfamily 4, group A, member 3 | protein-coding |
| chr4_47771333_47771831 | chr4 | 47771334 | 47771831 | + | 0.727473 | Intergenic | Intergenic | -279666 | NM_015743 | 18124 | Mm.247261 | NM_015743 | ENSMUSG00000028341 | **Nr4a3** | AI573420\|CHN\|CSMF\|MINOR\|NOR-1\|Nor1\|TEC | nuclear receptor subfamily 4, group A, member 3 | protein-coding |
| chr4_47998260_47998578 | chr4 | 47998261 | 47998578 | + | 0.687473 | Intergenic | Intergenic | -52829 | NM_015743 | 18124 | Mm.247261 | NM_015743 | ENSMUSG00000028341 | **Nr4a3** | AI573420\|CHN\|CSMF\|MINOR\|NOR-1\|Nor1\|TEC | nuclear receptor subfamily 4, group A, member 3 | protein-coding |
| chr4_47905439_47905773 | chr4 | 47905440 | 47905773 | + | 0.681362 | Intergenic | Intergenic | -145642 | NM_015743 | 18124 | Mm.247261 | NM_015743 | ENSMUSG00000028341 | **Nr4a3** | AI573420\|CHN\|CSMF\|MINOR\|NOR-1\|Nor1\|TEC | nuclear receptor subfamily 4, group A, member 3 | protein-coding |
| chr2_38779836_38780338 | chr2 | 38779837 | 38780338 | + | 0.766558 | intron (NM_010264, intron 3 of 10) | intron (NM_010264, intron 3 of 10) | 4428 | NM_001159548 | 14536 | Mm.439703 | NM_010264 | ENSMUSG00000063972 | **Nr6a1** | 1700113M01Rik\|Gcnf\|NCNF\|RTR | nuclear receptor subfamily 6, group A, member 1 | protein-coding |
| chr6_30081833_30082173 | chr6 | 30081834 | 30082173 | + | 0.970739 | intron (NM_001164229, intron 1 of 12) | MIR\|SINE\|MIR | 5682 | NM_001164229 | 18181 | Mm.259258 | NM_010938 | ENSMUSG00000058440 | **Nrf1** | C87038\|D6Ertd415e | nuclear respiratory factor 1 | protein-coding |
| chr6_30041525_30042388 | chr6 | 30041526 | 30042388 | + | 0.961457 | Intergenic | Intergenic | -6031 | NM_010938 | 18181 | Mm.259258 | NM_010938 | ENSMUSG00000058440 | **Nrf1** | C87038\|D6Ertd415e | nuclear respiratory factor 1 | protein-coding |
| chr6_30041212_30042438 | chr6 | 30041213 | 30042438 | + | 0.918227 | Intergenic | Intergenic | -6163 | NM_010938 | 18181 | Mm.259258 | NM_010938 | ENSMUSG00000058440 | **Nrf1** | C87038\|D6Ertd415e | nuclear respiratory factor 1 | protein-coding |
| chr9_55285247_55285734 | chr9 | 55285248 | 55285734 | + | 1.0283 | Intergenic | ORR1F\|LTR\|ERVL-MaLR | -1866 | NM_032002 | 83961 | Mm.443874 | NM_032002 | ENSMUSG00000032311 | **Nrg4** | AI552600 | neuregulin 4 | protein-coding |
| chr16_76681526_76681749 | chr16 | 76681527 | 76681749 | + | 0.789341 | Intergenic | Intergenic | -308589 | NM_173440 | 268903 | Mm.74711 | NM_173440 | ENSMUSG00000048490 | **Nrip1** | 6030458L20Rik\|8430438I05Rik\|9630050P12\|AA959574\|AW456757\|RIP140 | nuclear receptor interacting protein 1 | protein-coding |
| chr16_76355303_76355686 | chr16 | 76355304 | 76355686 | + | 0.775865 | intron (NM_173440, intron 1 of 3) | intron (NM_173440, intron 1 of 3) | 17554 | NM_173440 | 268903 | Mm.74711 | NM_173440 | ENSMUSG00000048490 | **Nrip1** | 6030458L20Rik\|8430438I05Rik\|9630050P12\|AA959574\|AW456757\|RIP140 | nuclear receptor interacting protein 1 | protein-coding |
| chr16_76575135_76575980 | chr16 | 76575136 | 76575980 | + | 0.752339 | Intergenic | Intergenic | -202509 | NM_173440 | 268903 | Mm.74711 | NM_173440 | ENSMUSG00000048490 | **Nrip1** | 6030458L20Rik\|8430438I05Rik\|9630050P12\|AA959574\|AW456757\|RIP140 | nuclear receptor interacting protein 1 | protein-coding |
| chr16_76334457_76334793 | chr16 | 76334458 | 76334793 | + | 0.714862 | intron (NM_173440, intron 2 of 3) | URR1B\|DNA\|hAT-Charlie | 38424 | NM_173440 | 268903 | Mm.74711 | NM_173440 | ENSMUSG00000048490 | **Nrip1** | 6030458L20Rik\|8430438I05Rik\|9630050P12\|AA959574\|AW456757\|RIP140 | nuclear receptor interacting protein 1 | protein-coding |
| chr13_36534514_36534988 | chr13 | 36534515 | 36534988 | + | 1.38565 | intron (NM_001039189, intron 5 of 5) | intron (NM_001039189, intron 5 of 5) | 199726 | NM_153529 | 68404 | Mm.232930 | NM_153529 | ENSMUSG00000039114 | **Nrn1** | 0710008J23Rik\|Cpg15\|Nrn | neuritin 1 | protein-coding |
| chr13_36534299_36534998 | chr13 | 36534300 | 36534998 | + | 1.24137 | intron (NM_001039189, intron 5 of 5) | ORR1F\|LTR\|ERVL-MaLR | 199828 | NM_153529 | 68404 | Mm.232930 | NM_153529 | ENSMUSG00000039114 | **Nrn1** | 0710008J23Rik\|Cpg15\|Nrn | neuritin 1 | protein-coding |
| chr13_36813806_36814377 | chr13 | 36813807 | 36814377 | + | 1.0869 | Intergenic | Intergenic | -79615 | NM_153529 | 68404 | Mm.232930 | NM_153529 | ENSMUSG00000039114 | **Nrn1** | 0710008J23Rik\|Cpg15\|Nrn | neuritin 1 | protein-coding |
| chr13_36812180_36812472 | chr13 | 36812181 | 36812472 | + | 0.991235 | Intergenic | Intergenic | -77849 | NM_153529 | 68404 | Mm.232930 | NM_153529 | ENSMUSG00000039114 | **Nrn1** | 0710008J23Rik\|Cpg15\|Nrn | neuritin 1 | protein-coding |
| chr13_36807216_36807515 | chr13 | 36807217 | 36807515 | + | 0.915114 | Intergenic | Intergenic | -72889 | NM_153529 | 68404 | Mm.232930 | NM_153529 | ENSMUSG00000039114 | **Nrn1** | 0710008J23Rik\|Cpg15\|Nrn | neuritin 1 | protein-coding |
| chr13_36527860_36528512 | chr13 | 36527861 | 36528512 | + | 0.906591 | intron (NM_001039189, intron 5 of 5) | intron (NM_001039189, intron 5 of 5) | 206291 | NM_153529 | 68404 | Mm.232930 | NM_153529 | ENSMUSG00000039114 | **Nrn1** | 0710008J23Rik\|Cpg15\|Nrn | neuritin 1 | protein-coding |
| chr13_36513068_36513485 | chr13 | 36513069 | 36513485 | + | 0.85997 | intron (NM_001039189, intron 5 of 5) | intron (NM_001039189, intron 5 of 5) | 221200 | NM_153529 | 68404 | Mm.232930 | NM_153529 | ENSMUSG00000039114 | **Nrn1** | 0710008J23Rik\|Cpg15\|Nrn | neuritin 1 | protein-coding |
| chr13_36801157_36801670 | chr13 | 36801158 | 36801670 | + | 0.798569 | Intergenic | Intergenic | -66937 | NM_153529 | 68404 | Mm.232930 | NM_153529 | ENSMUSG00000039114 | **Nrn1** | 0710008J23Rik\|Cpg15\|Nrn | neuritin 1 | protein-coding |
| chr13_36814596_36815205 | chr13 | 36814597 | 36815205 | + | 0.643177 | Intergenic | Intergenic | -80424 | NM_153529 | 68404 | Mm.232930 | NM_153529 | ENSMUSG00000039114 | **Nrn1** | 0710008J23Rik\|Cpg15\|Nrn | neuritin 1 | protein-coding |
| chr2_33818023_33818787 | chr2 | 33818024 | 33818787 | + | 1.13287 | intron (NM_175184, intron 6 of 9) | intron (NM_175184, intron 6 of 9) | 12589 | NR_045729 | 320482 | Mm.442057 | NR_045729 | ENSMUSG00000086484 | **Nron** | A630071L07Rik\|Gm24353 | non-protein coding RNA, repressor of NFAT | ncRNA |
| chr2_33817801_33818832 | chr2 | 33817802 | 33818832 | + | 0.978033 | intron (NM_175184, intron 6 of 9) | intron (NM_175184, intron 6 of 9) | 12501 | NR_045729 | 320482 | Mm.442057 | NR_045729 | ENSMUSG00000086484 | **Nron** | A630071L07Rik\|Gm24353 | non-protein coding RNA, repressor of NFAT | ncRNA |
| chr8_128351020_128351626 | chr8 | 128351021 | 128351626 | + | 1.80127 | Intergenic | B4\|SINE\|B4 | -7750 | NM_008737 | 18186 | Mm.271745 | NM_008737 | ENSMUSG00000025810 | **Nrp1** | C530029I03\|NP-1\|NPN-1\|Npn1\|Nrp | neuropilin 1 | protein-coding |

| | | | | | | | | | | | | | | | | | |
|---|---|---|---|---|---|---|---|---|---|---|---|---|---|---|---|---|---|
| chr8_128352832_128353132 | chr8 | 128352833 | 128353132 | + | 0.950827 | Intergenic | Intergenic | -6091 | NM_008737 | 18186 | Mm.271745 | NM_008737 | ENSMUSG00000025810 | **Nrp1** | C530029I03\|NP-1\|NPN-1\|Npn1\|Nrp | neuropilin 1 | protein-coding |
| chr8_128352712_128353163 | chr8 | 128352713 | 128353163 | + | 0.923473 | Intergenic | Intergenic | -6135 | NM_008737 | 18186 | Mm.271745 | NM_008737 | ENSMUSG00000025810 | **Nrp1** | C530029I03\|NP-1\|NPN-1\|Npn1\|Nrp | neuropilin 1 | protein-coding |
| chr8_128333245_128333857 | chr8 | 128333246 | 128333857 | + | 0.903088 | Intergenic | Intergenic | -25522 | NM_008737 | 18186 | Mm.271745 | NM_008737 | ENSMUSG00000025810 | **Nrp1** | C530029I03\|NP-1\|NPN-1\|Npn1\|Nrp | neuropilin 1 | protein-coding |
| chr8_128333214_128334103 | chr8 | 128333215 | 128334103 | + | 0.834648 | Intergenic | Intergenic | -25414 | NM_008737 | 18186 | Mm.271745 | NM_008737 | ENSMUSG00000025810 | **Nrp1** | C530029I03\|NP-1\|NPN-1\|Npn1\|Nrp | neuropilin 1 | protein-coding |
| chr12_90166092_90166557 | chr12 | 90166093 | 90166557 | + | 0.631916 | intron (NM_172544, intron 15 of 19) | intron (NM_172544, intron 15 of 19) | 353842 | NM_001252074 | 18191 | Mm.425766 | NM_172544 | ENSMUSG00000066392 | **Nrxn3** | - | neurexin III | protein-coding |
| chr4_6443783_6444480 | chr4 | 6443784 | 6444480 | + | 0.902579 | intron (NM_010945, intron 1 of 30) | LTR48\|LTR\|ERV1 | 10139 | NM_010945 | 18201 | Mm.3059 | NM_010945 | ENSMUSG00000028245 | **Nsmaf** | AA959567\|C630007J05\|Fan | neutral sphingomyelinase (N-SMase) activation associated factor | protein-coding |
| chr7_125500659_125501436 | chr7 | 125500660 | 125501436 | + | 0.988085 | Intergenic | Intergenic | -9506 | NM_026330 | 67711 | Mm.4467 | NM_026330 | ENSMUSG00000030750 | **Nsmce1** | 2510027N19Rik | non-SMC element 1 homolog (S. cerevisiae) | protein-coding |
| chr7_125474526_125475133 | chr7 | 125474527 | 125475133 | + | 0.89742 | intron (NM_026330, intron 2 of 7) | intron (NM_026330, intron 2 of 7) | 16712 | NM_026330 | 67711 | Mm.4467 | NM_026330 | ENSMUSG00000030750 | **Nsmce1** | 2510027N19Rik | non-SMC element 1 homolog (S. cerevisiae) | protein-coding |
| chr7_125474581_125474894 | chr7 | 125474582 | 125474894 | + | 0.851275 | intron (NM_026330, intron 2 of 7) | intron (NM_026330, intron 2 of 7) | 16804 | NM_026330 | 67711 | Mm.4467 | NM_026330 | ENSMUSG00000030750 | **Nsmce1** | 2510027N19Rik | non-SMC element 1 homolog (S. cerevisiae) | protein-coding |
| chr15_59410314_59410745 | chr15 | 59410315 | 59410745 | + | 1.43847 | intron (NM_026746, intron 2 of 6) | intron (NM_026746, intron 2 of 6) | 36332 | NM_001164604 | 68501 | Mm.27762 | NM_026746 | ENSMUSG00000059586 | **Nsmce2** | 1110014D18Rik\|AI661537 | non-SMC element 2 homolog (MMS21, S. cerevisiae) | protein-coding |
| chr15_59409996_59410786 | chr15 | 59409997 | 59410786 | + | 1.22084 | intron (NM_026746, intron 2 of 6) | intron (NM_026746, intron 2 of 6) | 36193 | NM_001164604 | 68501 | Mm.27762 | NM_026746 | ENSMUSG00000059586 | **Nsmce2** | 1110014D18Rik\|AI661537 | non-SMC element 2 homolog (MMS21, S. cerevisiae) | protein-coding |
| chr15_59448708_59449051 | chr15 | 59448709 | 59449051 | + | 0.882645 | intron (NM_026746, intron 3 of 6) | intron (NM_026746, intron 3 of 6) | 74682 | NM_001164604 | 68501 | Mm.27762 | NM_026746 | ENSMUSG00000059586 | **Nsmce2** | 1110014D18Rik\|AI661537 | non-SMC element 2 homolog (MMS21, S. cerevisiae) | protein-coding |
| chr15_59486876_59487258 | chr15 | 59486877 | 59487258 | + | 0.830287 | intron (NM_026746, intron 4 of 6) | intron (NM_026746, intron 4 of 6) | 112869 | NM_001164604 | 68501 | Mm.27762 | NM_026746 | ENSMUSG00000059586 | **Nsmce2** | 1110014D18Rik\|AI661537 | non-SMC element 2 homolog (MMS21, S. cerevisiae) | protein-coding |
| chr15_59480379_59480969 | chr15 | 59480380 | 59480969 | + | 0.690015 | intron (NM_026746, intron 4 of 6) | intron (NM_026746, intron 4 of 6) | 106476 | NM_001164604 | 68501 | Mm.27762 | NM_026746 | ENSMUSG00000059586 | **Nsmce2** | 1110014D18Rik\|AI661537 | non-SMC element 2 homolog (MMS21, S. cerevisiae) | protein-coding |
| chr2_25031618_25032107 | chr2 | 25031619 | 25032107 | + | 1.06364 | intron (NM_146251, intron 21 of 34) | RLTR19\|LTR\|ERVK | -22516 | NM_001177654 | 56876 | Mm.42956 | NM_020276 | ENSMUSG00000006476 | **Nsmf** | Jacob\|Nelf | NMDA receptor synaptonuclear signaling and neuronal migration factor | protein-coding |
| chr2_25034920_25035424 | chr2 | 25034921 | 25035424 | + | 1.03631 | intron (NM_146251, intron 21 of 34) | RMER15\|LTR\|ERVL | -19207 | NM_001177654 | 56876 | Mm.42956 | NM_020276 | ENSMUSG00000006476 | **Nsmf** | Jacob\|Nelf | NMDA receptor synaptonuclear signaling and neuronal migration factor | protein-coding |
| chr13_69656149_69656835 | chr13 | 69656150 | 69656835 | + | 0.804602 | Intergenic | MTEa\|LTR\|ERVL-MaLR | 44476 | NM_145354 | 28114 | Mm.260009 | NM_145354 | ENSMUSG00000021595 | **Nsun2** | D13Wsu123e\|Misu | NOL1/NOP2/Sun domain family member 2 | protein-coding |
| chr4_116056316_116056590 | chr4 | 116056317 | 116056590 | + | 0.921831 | Intergenic | MERV1_LTR\|LTR\|ERV1 | -2577 | NM_028142 | 72181 | Mm.282560 | NM_028142 | | **Nsun4** | 2310010O12Rik\|2810405F18Rik\|AI507900\|Shtap | NOL1/NOP2/Sun domain family, member 4 | protein-coding |
| chr2_15050893_15051271 | chr2 | 15050894 | 15051271 | + | 1.3148 | intron (NM_001165941, intron 1 of 11) | intron (NM_001165941, intron 1 of 11) | -1770 | NM_028950 | 74455 | Mm.478923 | NM_028950 | ENSMUSG00000026707 | **Nsun6** | 4933403D21Rik\|4933414E04Rik\|NOPD1 | NOL1/NOP2/Sun domain family member 6 | protein-coding |
| chr2_15035309_15035717 | chr2 | 15035310 | 15035717 | + | 1.18883 | TTS (NM_001165942) | TTS (NM_001165942) | 13799 | NM_001165942 | 74455 | Mm.478923 | NM_028950 | ENSMUSG00000026707 | **Nsun6** | 4933403D21Rik\|4933414E04Rik\|NOPD1 | NOL1/NOP2/Sun domain family member 6 | protein-coding |
| chr2_15035143_15035721 | chr2 | 15035144 | 15035721 | + | 1.17718 | intron (NM_001165941, intron 7 of 11) | intron (NM_001165941, intron 7 of 11) | 13880 | NM_001165942 | 74455 | Mm.478923 | NM_028950 | ENSMUSG00000026707 | **Nsun6** | 4933403D21Rik\|4933414E04Rik\|NOPD1 | NOL1/NOP2/Sun domain family member 6 | protein-coding |
| chr2_15050301_15050786 | chr2 | 15050302 | 15050786 | + | 0.933448 | intron (NM_001165941, intron 1 of 11) | intron (NM_001165941, intron 1 of 11) | -1232 | NM_028950 | 74455 | Mm.478923 | NM_028950 | ENSMUSG00000026707 | **Nsun6** | 4933403D21Rik\|4933414E04Rik\|NOPD1 | NOL1/NOP2/Sun domain family member 6 | protein-coding |
| chr4_123200062_123200395 | chr4 | 123200063 | 123200395 | + | 0.737216 | Intergenic | RMER15\|LTR\|ERVL | -1324 | NM_001085502 | 230718 | Mm.354635 | NM_001085502 | ENSMUSG00000054958 | **Nt5c1a** | Cn1a\|Gm49 | 5'-nucleotidase, cytosolic IA | protein-coding |
| chr19_46985206_46985576 | chr19 | 46985207 | 46985576 | + | 0.677516 | intron (NM_001164363, intron 1 of 17) | intron (NM_001164363, intron 1 of 17) | -15824 | NM_001164365 | 76952 | Mm.40965 | NM_029810 | ENSMUSG00000025041 | **Nt5c2** | 2010002I23Rik\|CnII\|ENSMUSG00000025039\|Gm9751\|Gmp\|Nt5b\|Pnt5\|cN-II | 5'-nucleotidase, cytosolic II | protein-coding |
| chr6_56920612_56921065 | chr6 | 56920613 | 56921065 | + | 0.920758 | intron (NM_026004, intron 1 of 8) | L1MC3\|LINE\|L1 | 3093 | NM_026004 | 107569 | Mm.158936 | NM_026004 | ENSMUSG00000029780 | **Nt5c3** | 1600024P05Rik\|2610206B05Rik\|3110004A18Rik\|AI842776\|Nt5c3a\|Nt5y\|PN-1\|PN-I\|PSN1\|UMPH\|Umph-1\|Umph1\|cN-III\|lupin\|p36 | 5'-nucleotidase, cytosolic III | protein-coding |
| chr6_56934339_56934736 | chr6 | 56934340 | 56934736 | + | 0.65693 | Intergenic | Lx8\|LINE\|L1 | -10606 | NM_026004 | 107569 | Mm.158936 | NM_026004 | ENSMUSG00000029780 | **Nt5c3** | 1600024P05Rik\|2610206B05Rik\|3110004A18Rik\|AI842776\|Nt5c3a\|Nt5y\|PN-1\|PN-I\|PSN1\|UMPH\|Umph-1\|Umph1\|cN-III\|lupin\|p36 | 5'-nucleotidase, cytosolic III | protein-coding |
| chr10_34413551_34413868 | chr10 | 34413552 | 34413868 | + | 0.839197 | intron (NM_176968, intron 1 of 11) | intron (NM_176968, intron 1 of 11) | 4818 | NM_176968 | 319638 | Mm.73467 | NM_176968 | ENSMUSG00000039480 | **Nt5dc1** | 6030401B09Rik\|AW987726\|Nt5c2l1 | 5'-nucleotidase domain containing 1 | protein-coding |
| chr10_86794730_86795623 | chr10 | 86794731 | 86795623 | + | 1.30746 | intron (NM_175331, intron 1 of 13) | intron (NM_175331, intron 1 of 13) | 16172 | NM_175331 | 103466 | Mm.200446 | NM_175331 | ENSMUSG00000054027 | **Nt5dc3** | AU040402\|AW540062\|C630002B14Rik\|TU12B1-TY | 5'-nucleotidase domain containing 3 | protein-coding |
| chr9_88273245_88273692 | chr9 | 88273246 | 88273692 | + | 0.968985 | Intergenic | Intergenic | -54140 | NM_011851 | 23959 | Mm.244235 | NM_011851 | ENSMUSG00000032420 | **Nt5e** | 2210401F01Rik\|AI447961\|CD73\|NT\|Nt5\|eNT | 5' nucleotidase, ecto | protein-coding |
| chr16_13844859_13845418 | chr16 | 13844860 | 13845418 | + | 1.15899 | intron (NM_053181, intron 15 of 22) | intron (NM_053181, intron 15 of 22) | 25862 | NM_010946 | 18203 | Mm.380410 | NM_010946 | ENSMUSG00000022681 | **Ntan1** | - | N-terminal Asn amidase | protein-coding |
| chr8_125727309_125728107 | chr8 | 125727310 | 125728107 | + | 0.98007 | Intergenic | Intergenic | -6495 | NM_025636 | 66566 | Mm.25296 | NM_025636 | ENSMUSG00000031851 | **Ntpcr** | 2310079N02Rik\|AI449709 | nucleoside-triphosphatase, cancer-related | protein-coding |
| chr5_24687790_24688075 | chr5 | 24687791 | 24688075 | + | 0.914988 | intron (NM_016736, intron 1 of 14) | intron (NM_016736, intron 1 of 14) | 2118 | NM_016736 | 53312 | Mm.5856 | NM_016736 | ENSMUSG00000028954 | **Nub1** | 4931404D21Rik\|6330412F12Rik\|BS4\|NY-REN-18 | negative regulator of ubiquitin-like proteins 1 | protein-coding |
| chr7_116493232_116494427 | chr7 | 116493233 | 116494427 | + | 0.999429 | Intergenic | Intergenic | -10539 | NM_001130479 | 53322 | Mm.9901 | NM_016773 | ENSMUSG00000030659 | **Nucb2** | AI607786\|Calnuc\|Nefa\|Nesfatin-1 | nucleobindin 2 | protein-coding |
| chr15_44440791_44441632 | chr15 | 44440792 | 44441632 | + | 0.69696 | Intergenic | Intergenic | -12905 | NM_026149 | 67429 | Mm.490404 | NM_026149 | ENSMUSG00000038736 | **Nudcd1** | 4921532K09Rik\|AA407246\|AW260430\|AW556235\|CML66-L\|Cml66 | NudC domain containing 1 | protein-coding |
| chr16_4913022_4913373 | chr16 | 4913023 | 4913373 | + | 0.678822 | intron (NM_029657, intron 4 of 16) | intron (NM_029657, intron 4 of 16) | -25913 | NM_025839 | 66911 | Mm.428698 | NM_025839 | ENSMUSG00000022516 | **Nudt16l1** | 1110001K21Rik\|5330437I08Rik\|Sdos | nudix (nucleoside diphosphate linked moiety X)-type motif 16-like 1 | protein-coding |
| chr14_70581688_70581944 | chr14 | 70581689 | 70581944 | + | 0.962043 | 3' UTR (NM_153136, exon 3 of 3) | 3' UTR (NM_153136, exon 3 of 3) | 3969 | NM_153136 | 213484 | Mm.34191 | NM_153136 | ENSMUSG00000045211 | **Nudt18** | BC036718 | nudix (nucleoside diphosphate linked moiety X)-type motif 18 | protein-coding |
| chr3_37401532_37401847 | chr3 | 37401533 | 37401847 | + | 0.711207 | intron (NM_008006, intron 2 of 2) | intron (NM_008006, intron 2 of 2) | 17906 | NM_001291044 | 229228 | Mm.436967 | NM_153561 | ENSMUSG00000050174 | **Nudt6** | Asfgf2b | nudix (nucleoside diphosphate linked moiety X)-type motif 6 | protein-coding |
| chr14_76220923_76221305 | chr14 | 76220924 | 76221305 | + | 1.06351 | Intergenic | Intergenic | 110223 | NM_013745 | 27275 | Mm.21138 | NM_013745 | ENSMUSG00000022009 | **Nufip1** | Nufip | nuclear fragile X mental retardation protein interacting protein 1 | protein-coding |
| chr10_117818169_117818531 | chr10 | 117818170 | 117818531 | + | 1.17251 | intron (NM_024457, intron 6 of 7) | B3\|SINE\|B2 | -25645 | NM_134010 | 103468 | Mm.12568 | NM_134010 | ENSMUSG00000052798 | **Nup107** | AW541137\|C76801 | nucleoporin 107 | protein-coding |
| chr10_117818158_117818654 | chr10 | 117818159 | 117818654 | + | 1.05878 | intron (NM_024457, intron 6 of 7) | intron (NM_024457, intron 6 of 7) | -25701 | NM_134010 | 103468 | Mm.12568 | NM_134010 | ENSMUSG00000052798 | **Nup107** | AW541137\|C76801 | nucleoporin 107 | protein-coding |
| chr6_90984128_90984462 | chr6 | 90984129 | 90984462 | + | 1.14561 | Intergenic | Intergenic | 132531 | NM_018815 | 54563 | Mm.28162 | NM_018815 | ENSMUSG00000030091 | **Nup210** | 9830001L10\|AI836801\|Pom210\|gp190\|gp210 | nucleoporin 210 | protein-coding |
| chr6_91066726_91067797 | chr6 | 91066727 | 91067797 | + | 0.985737 | intron (NM_018815, intron 13 of 39) | B3A\|SINE\|B2 | 49564 | NM_018815 | 54563 | Mm.28162 | NM_018815 | ENSMUSG00000030091 | **Nup210** | 9830001L10\|AI836801\|Pom210\|gp190\|gp210 | nucleoporin 210 | protein-coding |
| chr6_91104746_91105311 | chr6 | 91104747 | 91105311 | + | 0.645985 | intron (NM_018815, intron 1 of 39) | intron (NM_018815, intron 1 of 39) | 11797 | NM_018815 | 54563 | Mm.28162 | NM_018815 | ENSMUSG00000030091 | **Nup210** | 9830001L10\|AI836801\|Pom210\|gp190\|gp210 | nucleoporin 210 | protein-coding |
| chr2_80745635_80746150 | chr2 | 80745636 | 80746150 | + | 1.13484 | Intergenic | Intergenic | 106629 | NM_001190179 | 69482 | Mm.29200 | NM_027091 | | **Nup35** | 2310006I24Rik\|35kDa\|5330402E05Rik\|MP44\|NO44 | nucleoporin 35 | protein-coding |
| chr2_80745596_80746300 | chr2 | 80745597 | 80746300 | + | 1.01301 | Intergenic | Intergenic | 106684 | NM_001190179 | 69482 | Mm.29200 | NM_027091 | | **Nup35** | 2310006I24Rik\|35kDa\|5330402E05Rik\|MP44\|NO44 | nucleoporin 35 | protein-coding |
| chr8_94240163_94240632 | chr8 | 94240164 | 94240632 | + | 0.695822 | intron (NM_172410, intron 3 of 22) | intron (NM_172410, intron 3 of 22) | 25797 | NM_172410 | 71805 | Mm.458170 | NM_172410 | ENSMUSG00000032939 | **Nup93** | 2410008G02Rik | nucleoporin 93 | protein-coding |
| chr14_60248009_60248250 | chr14 | 60248010 | 60248250 | + | 1.17193 | intron (NM_170591, intron 1 of 14) | intron (NM_170591, intron 1 of 14) | 3248 | NM_170591 | 71844 | Mm.258051 | NM_170591 | ENSMUSG00000063895 | **Nupl1** | 1700017F11Rik\|mKIAA0410 | nucleoporin like 1 | protein-coding |

| | | | | | | | | | | | | | | | | | |
|---|---|---|---|---|---|---|---|---|---|---|---|---|---|---|---|---|---|
| chr5_24154133_24154703 | chr5 | 24154134 | 24154703 | + | 0.852933 | intron (NM_001161800, intron 8 of 9) | ORR1B1\|LTR\|ERVL-MaLR | -10545 | NM_153092 | 231042 | Mm.45513 | NM_153092 | ENSMUSG00000048439 | **Nupl2** | AV116043\|CG1\|NLP-1\|Nlp1 | nucleoporin like 2 | protein-coding |
| chr7_126619560_126620002 | chr7 | 126619561 | 126620002 | + | 0.617293 | Intergenic | Intergenic | 5689 | NM_019738 | 56312 | Mm.18742 | NM_019738 | | **Nupr1** | 2310032H04Rik\|Com1\|p8 | nuclear protein transcription regulator 1 | protein-coding |
| chr7_126629194_126630258 | chr7 | 126629195 | 126630258 | + | 0.613998 | Intergenic | Intergenic | -4256 | NM_019738 | 56312 | Mm.18742 | NM_019738 | | **Nupr1** | 2310032H04Rik\|Com1\|p8 | nuclear protein transcription regulator 1 | protein-coding |
| chr1_181138585_181139154 | chr1 | 181138586 | 181139154 | + | 1.65417 | intron (NM_026171, intron 4 of 22) | intron (NM_026171, intron 4 of 22) | 5287 | NM_026171 | 67459 | Mm.263464 | NM_026171 | ENSMUSG00000026516 | **Nvl** | 1200009I24Rik | nuclear VCP-like | protein-coding |
| chr8_72682925_72683794 | chr8 | 72682926 | 72683794 | + | 1.10588 | intron (NM_176940, intron 10 of 18) | intron (NM_176940, intron 10 of 18) | 36649 | NM_176940 | 319555 | Mm.212927 | NM_176940 | ENSMUSG00000048148 | **Nwd1** | A230063L24Rik | NACHT and WD repeat domain containing 1 | protein-coding |
| chr8_72682093_72682548 | chr8 | 72682094 | 72682548 | + | 0.769693 | intron (NM_176940, intron 10 of 18) | AmnSINE1\|SINE\|Deu | 35610 | NM_176940 | 319555 | Mm.212927 | NM_176940 | ENSMUSG00000048148 | **Nwd1** | A230063L24Rik | NACHT and WD repeat domain containing 1 | protein-coding |
| chr19_8755499_8756305 | chr19 | 8755500 | 8756305 | + | 0.925961 | TTS (NM_019829) | TTS (NM_019829) | -1201 | NM_016813 | 53319 | Mm.7271 | NM_016813 | ENSMUSG00000010097 | **Nxf1** | Mex67\|Mvb1\|Tap | nuclear RNA export factor 1 | protein-coding |
| chr16_55906809_55907623 | chr16 | 55906810 | 55907623 | + | 0.880448 | intron (NM_028815, intron 10 of 10) | intron (NM_028815, intron 10 of 10) | -11937 | NM_001134457 | 385658 | Mm.310635 | NM_001134457 | ENSMUSG00000075033 | **Nxpe3** | Fam55c\|Gm1752\|Gm611 | neurexophilin and PC-esterase domain family, member 3 | protein-coding |
| chr16_55906800_55907931 | chr16 | 55906801 | 55907931 | + | 0.86356 | intron (NM_028815, intron 10 of 10) | intron (NM_028815, intron 10 of 10) | -12087 | NM_001134457 | 385658 | Mm.310635 | NM_001134457 | ENSMUSG00000075033 | **Nxpe3** | Fam55c\|Gm1752\|Gm611 | neurexophilin and PC-esterase domain family, member 3 | protein-coding |
| chr9_48356941_48357562 | chr9 | 48356942 | 48357562 | + | 1.07369 | Intergenic | Tigger5b\|DNA\|TcMar-Tigger | -4789 | NM_172921 | 244853 | Mm.325522 | NM_172921 | ENSMUSG00000044229 | **Nxpe4** | C130036J11\|D930028F11Rik\|Fam55d | neurexophilin and PC-esterase domain family, member 4 | protein-coding |
| chr5_114911624_114912050 | chr5 | 114911625 | 114912050 | + | 1.2506 | 3' UTR (NM_011854, exon 6 of 6) | 3' UTR (NM_011854, exon 6 of 6) | -11403 | NM_145209 | 231655 | Mm.95479 | NM_145209 | ENSMUSG00000041827 | **Oasl1** | 7530414C13Rik\|oasl9 | 2'-5' oligoadenylate synthetase-like 1 | protein-coding |
| chr19_47526289_47526952 | chr19 | 47526290 | 47526952 | + | 0.616421 | intron (NM_175360, intron 3 of 9) | intron (NM_175360, intron 3 of 9) | 10399 | NM_175360 | 108689 | Mm.218048 | NM_175360 | ENSMUSG00000042694 | **Obfc1** | 0610009H20Rik\|2310057J23Rik\|AAF-44\|AAF44\|AI413458\|RPA-32 | oligonucleotide/oligosaccharide-binding fold containing 1 | protein-coding |
| chr8_71370744_71370994 | chr8 | 71370745 | 71370994 | + | 1.61274 | promoter-TSS (NM_029865) | promoter-TSS (NM_029865) | -429 | NM_029865 | 77090 | Mm.440170 | NM_029865 | ENSMUSG00000002396 | **Ocel1** | 9430098E02Rik\|AI326022\|AI593544 | occludin/ELL domain containing 1 | protein-coding |
| chr5_73344196_73344780 | chr5 | 73344197 | 73344780 | + | 1.35413 | Intergenic | Intergenic | -5849 | NM_026950 | 433904 | Mm.274892 | NM_026950 | ENSMUSG00000029153 | **Ociad2** | 1810027I20Rik | OCIA domain containing 2 | protein-coding |
| chr2_165405315_165405717 | chr2 | 165405316 | 165405717 | + | 1.35077 | 3' UTR (NM_054055, exon 13 of 13) | 3' UTR (NM_054055, exon 13 of 13) | -5122 | NM_029021 | 74614 | Mm.72179 | NM_029021 | | **Ocstamp** | 4833422F24Rik\|OC-STAMP | osteoclast stimulatory transmembrane protein | protein-coding |
| chr15_38197042_38197680 | chr15 | 38197043 | 38197680 | + | 0.713994 | Intergenic | Intergenic | -21842 | NM_008757 | 18285 | Mm.252830 | NM_008757 | ENSMUSG00000061923 | **Odf1** | - | outer dense fiber of sperm tails 1 | protein-coding |
| chr14_8459799_8460225 | chr14 | 8459800 | 8460225 | + | 1.44896 | intron (NM_028934, intron 14 of 15) | intron (NM_028934, intron 14 of 15) | -81249 | NM_146050 | 18300 | Mm.25351 | NM_146050 | ENSMUSG00000021749 | **Oit1** | 2310076N21Rik\|AV067083\|EF-7\|Fam3d | oncoprotein induced transcript 1 | protein-coding |
| chr14_8458587_8459297 | chr14 | 8458588 | 8459297 | + | 0.978231 | intron (NM_028934, intron 14 of 15) | intron (NM_028934, intron 14 of 15) | -80179 | NM_146050 | 18300 | Mm.25351 | NM_146050 | ENSMUSG00000021749 | **Oit1** | 2310076N21Rik\|AV067083\|EF-7\|Fam3d | oncoprotein induced transcript 1 | protein-coding |
| chr14_8501367_8502085 | chr14 | 8501368 | 8502085 | + | 0.694627 | intron (NM_028934, intron 12 of 15) | MLT2B2\|LTR\|ERVL | -122963 | NM_146050 | 18300 | Mm.25351 | NM_146050 | ENSMUSG00000021749 | **Oit1** | 2310076N21Rik\|AV067083\|EF-7\|Fam3d | oncoprotein induced transcript 1 | protein-coding |
| chr14_8460313_8460810 | chr14 | 8460314 | 8460810 | + | 0.631118 | intron (NM_028934, intron 14 of 15) | intron (NM_028934, intron 14 of 15) | -81799 | NM_146050 | 18300 | Mm.25351 | NM_146050 | ENSMUSG00000021749 | **Oit1** | 2310076N21Rik\|AV067083\|EF-7\|Fam3d | oncoprotein induced transcript 1 | protein-coding |
| chr2_38934958_38935336 | chr2 | 38934959 | 38935336 | + | 0.950182 | intron (NM_172854, intron 1 of 7) | intron (NM_172854, intron 1 of 7) | 3167 | NM_172854 | 241327 | Mm.103491 | NM_172854 | | **Olfml2a** | 4932431K08Rik\|mFLJ00237 | olfactomedin-like 2A | protein-coding |
| chr1_170729221_170729978 | chr1 | 170729222 | 170729978 | + | 0.807425 | intron (NM_001081304, intron 15 of 15) | intron (NM_001081304, intron 15 of 15) | 85068 | NM_177068 | 320078 | Mm.287131 | NM_177068 | ENSMUSG00000038463 | **Olfml2b** | 1110018N05Rik\|4832415H08Rik\|AI467542 | olfactomedin-like 2B | protein-coding |
| chr1_170729221_170730348 | chr1 | 170729222 | 170730348 | + | 0.782585 | intron (NM_001081304, intron 15 of 15) | intron (NM_001081304, intron 15 of 15) | 85253 | NM_177068 | 320078 | Mm.287131 | NM_177068 | ENSMUSG00000038463 | **Olfml2b** | 1110018N05Rik\|4832415H08Rik\|AI467542 | olfactomedin-like 2B | protein-coding |
| chr11_49295903_49296198 | chr11 | 49295904 | 49296198 | + | 0.682484 | Intergenic | RMER15\|LTR\|ERVL | 2754 | NM_146470 | 258462 | Mm.377516 | NM_146470 | ENSMUSG00000050959 | **Olfr1392** | MOR256-25 | olfactory receptor 1392 | protein-coding |
| chr11_49266096_49266768 | chr11 | 49266097 | 49266768 | + | 1.04953 | Intergenic | Intergenic | -13718 | NM_146471 | 258463 | Mm.377517 | NM_146471 | ENSMUSG00000059864 | **Olfr1393** | MOR256-24 | olfactory receptor 1393 | protein-coding |
| chr19_12004674_12004972 | chr19 | 12004675 | 12004972 | + | 0.744811 | Intergenic | MER34A1\|LTR\|ERV1 | 31917 | NM_146680 | 258675 | Mm.271517 | NM_146680 | ENSMUSG00000067529 | **Olfr1423** | MOR239-3 | olfactory receptor 1423 | protein-coding |
| chr19_12831404_12831838 | chr19 | 12831405 | 12831838 | + | 0.972459 | intron (NM_134152, intron 7 of 8) | intron (NM_134152, intron 7 of 8) | -30156 | NM_146702 | 258697 | Mm.223132 | NM_146702 | ENSMUSG00000046272 | **Olfr1444** | MOR202-4 | olfactory receptor 1444 | protein-coding |
| chr14_52565527_52565968 | chr14 | 52565528 | 52565968 | + | 1.0796 | Intergenic | Intergenic | -70053 | NM_001170918 | 57269 | Mm.88822 | NM_020512 | ENSMUSG00000059887 | **Olfr1507** | MOR244-1\|Mor28\|Or28 | olfactory receptor 1507 | protein-coding |
| chr14_53039342_53039980 | chr14 | 53039343 | 53039980 | + | 0.911608 | Intergenic | Intergenic | -543966 | NM_001170918 | 57269 | Mm.88822 | NM_020512 | ENSMUSG00000059887 | **Olfr1507** | MOR244-1\|Mor28\|Or28 | olfactory receptor 1507 | protein-coding |
| chr14_53080498_53080929 | chr14 | 53080499 | 53080929 | + | 0.910545 | Intergenic | Intergenic | -585019 | NM_001170918 | 57269 | Mm.88822 | NM_020512 | ENSMUSG00000059887 | **Olfr1507** | MOR244-1\|Mor28\|Or28 | olfactory receptor 1507 | protein-coding |
| chr14_52545314_52545897 | chr14 | 52545315 | 52545897 | + | 0.722314 | Intergenic | Intergenic | -49911 | NM_001170918 | 57269 | Mm.88822 | NM_020512 | ENSMUSG00000059887 | **Olfr1507** | MOR244-1\|Mor28\|Or28 | olfactory receptor 1507 | protein-coding |
| chr16_58826734_58827295 | chr16 | 58826735 | 58827295 | + | 0.677344 | promoter-TSS (NM_147002) | promoter-TSS (NM_147002) | -254 | NM_147002 | 259004 | Mm.223391 | NM_147002 | ENSMUSG00000060663 | **Olfr175-ps1** | MOR184-1\|MOR184-10P\|Olfr174 | olfactory receptor 175, pseudogene 1 | protein-coding |
| chr14_52041280_52041710 | chr14 | 52041281 | 52041710 | + | 0.639064 | Intergenic | RLTR40\|LTR\|ERVK | -5386 | NM_001001808 | 258420 | Mm.377946 | NM_001001808 | ENSMUSG00000044286 | **Olfr221** | MOR205-1\|MOR205-2\|Olfr1514\|Olfr221-ps1 | olfactory receptor 221 | protein-coding |
| chr3_106875592_106875965 | chr3 | 106875593 | 106875965 | + | 1.11501 | Intergenic | Intergenic | -53222 | NM_146489 | 258482 | Mm.377527 | NM_146489 | ENSMUSG00000043529 | **Olfr266** | MOR122-2 | olfactory receptor 266 | protein-coding |
| chr7_84776092_84776447 | chr7 | 84776093 | 84776447 | + | 1.04948 | non-coding (NR_038175, exon 5 of 5) | non-coding (NR_038175, exon 5 of 5) | -77301 | NM_146415 | 258410 | Mm.484702 | NM_146415 | ENSMUSG00000070460 | **Olfr291** | MOR254-2 | olfactory receptor 291 | protein-coding |
| chr8_72037140_72037417 | chr8 | 72037141 | 72037417 | + | 0.658562 | Intergenic | RLTR48A\|LTR\|ERV1 | -20381 | NM_207555 | 404315 | Mm.377936 | NM_207555 | ENSMUSG00000069998 | **Olfr372** | MOR282-1 | olfactory receptor 372 | protein-coding |
| chr11_74420905_74421528 | chr11 | 74420906 | 74421528 | + | 0.686153 | intron (NM_001015046, intron 11 of 24) | intron (NM_001015046, intron 11 of 24) | 56546 | NM_001011851 | 258153 | Mm.376124 | NM_001011851 | ENSMUSG00000058275 | **Olfr412** | MOR127-5P | olfactory receptor 412 | protein-coding |
| chr1_174052880_174053490 | chr1 | 174052881 | 174053490 | + | 1.46723 | Intergenic | Intergenic | 2810 | NM_146716 | 258711 | Mm.377648 | NM_146716 | ENSMUSG00000047048 | **Olfr432** | MOR123-2 | olfactory receptor 432 | protein-coding |
| chr1_174043342_174043763 | chr1 | 174043343 | 174043763 | + | 0.862776 | TTS (NM_146717) | TTS (NM_146717) | 1620 | NM_146717 | 258712 | Mm.223146 | NM_146717 | ENSMUSG00000045381 | **Olfr433** | MOR123-1 | olfactory receptor 433 | protein-coding |
| chr7_99740891_99741531 | chr7 | 99740892 | 99741531 | + | 0.663153 | Intergenic | MTA_Mm\|LTR\|ERVL-MaLR | 6066 | NM_147063 | 259066 | Mm.377818 | NM_147063 | ENSMUSG00000073998 | **Olfr520** | MOR101-1 | olfactory receptor 520 | protein-coding |
| chr7_140329129_140330718 | chr7 | 140329130 | 140330718 | + | 1.26457 | Intergenic | RMER1B\|Other\|Other | -5940 | NM_001011776 | 257939 | Mm.377310 | NM_001011776 | ENSMUSG00000062782 | **Olfr527** | MOR251-5 | olfactory receptor 527 | protein-coding |
| chr7_140539738_140540083 | chr7 | 140539739 | 140540083 | + | 1.18108 | Intergenic | Intergenic | -32593 | NM_146520 | 258513 | Mm.261779 | NM_146520 | ENSMUSG00000052508 | **Olfr536** | MOR252-1\|MOR252-4 | olfactory receptor 536 | protein-coding |
| chr7_140537478_140538030 | chr7 | 140537479 | 140538030 | + | 1.07937 | Intergenic | Intergenic | -30436 | NM_146520 | 258513 | Mm.261779 | NM_146520 | ENSMUSG00000052508 | **Olfr536** | MOR252-1\|MOR252-4 | olfactory receptor 536 | protein-coding |
| chr7_102553291_102553649 | chr7 | 102553292 | 102553649 | + | 1.06387 | promoter-TSS (NM_147101) | promoter-TSS (NM_147101) | -816 | NM_147101 | 259105 | Mm.424092 | NM_147101 | ENSMUSG00000073977 | **Olfr549** | MOR31-3 | olfactory receptor 549 | protein-coding |
| chr7_102553032_102553679 | chr7 | 102553033 | 102553679 | + | 0.932065 | promoter-TSS (NM_147101) | promoter-TSS (NM_147101) | -930 | NM_147101 | 259105 | Mm.424092 | NM_147101 | ENSMUSG00000073977 | **Olfr549** | MOR31-3 | olfactory receptor 549 | protein-coding |
| chr7_102549700_102550110 | chr7 | 102549701 | 102550110 | + | 0.795876 | Intergenic | Intergenic | -4381 | NM_147101 | 259105 | Mm.424092 | NM_147101 | ENSMUSG00000073977 | **Olfr549** | MOR31-3 | olfactory receptor 549 | protein-coding |
| chr7_104105219_104105603 | chr7 | 104105220 | 104105603 | + | 0.810828 | promoter-TSS (NM_147056) | promoter-TSS (NM_147056) | -870 | NM_147056 | 259058 | Mm.377814 | NM_147056 | ENSMUSG00000073931 | **Olfr646** | MOR33-2 | olfactory receptor 646 | protein-coding |
| chr14_50061955_50062977 | chr14 | 50061956 | 50062977 | + | 0.652633 | Intergenic | CpG | 22213 | NM_146316 | 258313 | Mm.463727 | NM_146316 | ENSMUSG00000060523 | **Olfr726** | MOR246-4 | olfactory receptor 726 | protein-coding |
| chr14_50061438_50062970 | chr14 | 50061439 | 50062970 | + | 0.620889 | Intergenic | LSU-rRNA_Hsa\|rRNA\|rRNA | 22475 | NM_146316 | 258313 | Mm.463727 | NM_146316 | ENSMUSG00000060523 | **Olfr726** | MOR246-4 | olfactory receptor 726 | protein-coding |
| chr17_37069871_37070614 | chr17 | 37069872 | 37070614 | + | 0.92513 | intron (NM_019439, intron 19 of 22) | intron (NM_019439, intron 19 of 22) | 15993 | NM_146477 | 258469 | Mm.271392 | NM_146477 | ENSMUSG00000056600 | **Olfr90** | MOR256-21\|bM573K1.2 | olfactory receptor 90 | protein-coding |
| chr10_19406645_19407165 | chr10 | 19406646 | 19407165 | + | 1.09991 | Intergenic | Intergenic | 50366 | NM_053008 | 94222 | Mm.156946 | NM_053008 | ENSMUSG00000045591 | **Olig3** | Bhlhb7\|bHLHe20 | oligodendrocyte transcription factor 3 | protein-coding |
| chr4_103311394_103311625 | chr4 | 103311395 | 103311625 | + | 0.880822 | intron (NR_045887, intron 3 of 3) | Lx8\|LINE\|L1 | -2336 | NM_025909 | 67013 | Mm.30021 | NM_025909 | ENSMUSG00000035069 | **Oma1** | 2010001O09Rik\|MPRP-1\|ZMPOMA1 | OMA1 homolog, zinc metallopeptidase (S. cerevisiae) | protein-coding |
| chr9_74870997_74871287 | chr9 | 74870998 | 74871287 | + | 0.614335 | intron (NM_008262, intron 1 of 1) | intron (NM_008262, intron 1 of 1) | 9221 | NM_008262 | 15379 | Mm.303355 | NM_008262 | | **Onecut1** | D9Ertd423e\|HNF-6\|Hfh12\|Hnf6\|Hnf6b\|OC-1\|Oc1 | one cut domain, family member 1 | protein-coding |
| chr16_29775756_29776352 | chr16 | 29775757 | 29776352 | + | 0.729558 | Intergenic | Intergenic | 196720 | NM_001199177 | 74143 | Mm.274285 | NM_133752 | ENSMUSG00000038084 | **Opa1** | 1200011N24Rik\|AI225888\|AI847218\|lilr3\|mKIAA0567 | optic atrophy 1 | protein-coding |
| chr6_29388585_29389080 | chr6 | 29388586 | 29389080 | + | 0.813867 | Intergenic | tRNA-Pro-CCY\|tRNA\|tRNA | -8320 | NM_007538 | 12057 | Mm.56987 | NM_007538 | ENSMUSG00000058831 | **Opn1sw** | AW551857\|Bcp | opsin 1 (cone pigments), short-wave-sensitive (color blindness, tritan) | protein-coding |
| chr6_29388052_29389113 | chr6 | 29388053 | 29389113 | + | 0.742912 | Intergenic | Intergenic | -8070 | NM_007538 | 12057 | Mm.56987 | NM_007538 | ENSMUSG00000058831 | **Opn1sw** | AW551857\|Bcp | opsin 1 (cone pigments), short-wave-sensitive (color blindness, tritan) | protein-coding |
| chr17_42603405_42603751 | chr17 | 42603406 | 42603751 | + | 0.758928 | intron (NM_181753, intron 2 of 6) | intron (NM_181753, intron 2 of 6) | 7735 | NM_181753 | 353344 | Mm.184937 | NM_181753 | ENSMUSG00000043972 | **Opn5** | Gpr136\|Neuropsin\|PGR12\|TMEM13 | opsin 5 | protein-coding |
| chr17_42613028_42613486 | chr17 | 42613029 | 42613486 | + | 0.703041 | Intergenic | AmnSINE1\|SINE\|Deu | -1944 | NM_181753 | 353344 | Mm.184937 | NM_181753 | ENSMUSG00000043972 | **Opn5** | Gpr136\|Neuropsin\|PGR12\|TMEM13 | opsin 5 | protein-coding |
| chr10_127142183_127142512 | chr10 | 127142184 | 127142512 | + | 0.910023 | Intergenic | MTB_Mm\|LTR\|ERVL-MaLR | -21188 | NM_001171026 | 216440 | Mm.295246 | NM_177614 | ENSMUSG00000040462 | **Os9** | 4632413K17Rik\|AU022351 | amplified in osteosarcoma | protein-coding |

| | | | | | | | | | | | | | | | | | |
|---|---|---|---|---|---|---|---|---|---|---|---|---|---|---|---|---|---|
| chr10_127114303_127114607 | chr10 | 127114304 | 127114607 | + | 0.797484 | intron (NM_177614, intron 5 of 13) | intron (NM_177614, intron 5 of 13) | 6705 | NM_177614 | 216440 | Mm.295246 | NM_177614 | ENSMUSG00000040462 | **Os9** | 4632413K17Rik\|AU022351 | amplified in osteosarcoma | protein-coding |
| chr19_11967039_11967223 | chr19 | 11967040 | 11967223 | + | 0.912662 | intron (NM_001033174, intron 1 of 13) | intron (NM_001033174, intron 1 of 13) | 1287 | NM_001033174 | 76303 | Mm.291279 | NM_001033174 | ENSMUSG00000024687 | **Osbp** | 1110018F06Rik\|AW559088\|mKIAA4220 | oxysterol binding protein | protein-coding |
| chr9_115168526_115168926 | chr9 | 115168527 | 115168926 | + | 0.654017 | intron (NM_148958, intron 3 of 9) | intron (NM_148958, intron 3 of 9) | 101447 | NM_148958 | 74486 | Mm.458474 | NM_148958 | ENSMUSG00000040875 | **Osbpl10** | 4933433D06Rik\|C820004B04Rik\|OPR-10 | oxysterol binding protein-like 10 | protein-coding |
| chr16_33155565_33156657 | chr16 | 33155566 | 33156657 | + | 1.14131 | Intergenic | MTA_Mm\|LTR\|ERVL-MaLR | -28960 | NM_176840 | 106326 | Mm.486405 | NM_176840 | ENSMUSG00000022807 | **Osbpl11** | 9430097N02Rik\|AI132306\|AU019795\|ORP-11 | oxysterol binding protein-like 11 | protein-coding |
| chr18_12936082_12936452 | chr18 | 12936083 | 12936452 | + | 0.678104 | intron (NM_207530, intron 1 of 27) | B3A\|SINE\|B2 | 5574 | NM_207530 | 64291 | Mm.259470 | NM_020573 | ENSMUSG00000044252 | **Osbpl1a** | G430090F17Rik\|Gm753\|ORP-1\|Osbpl1b | oxysterol binding protein-like 1A | protein-coding |
| chr6_50334838_50335283 | chr6 | 50334839 | 50335283 | + | 1.05031 | intron (NM_027881, intron 10 of 21) | intron (NM_027881, intron 10 of 21) | 47776 | NM_027881 | 71720 | Mm.44153 | NM_027881 | ENSMUSG00000029822 | **Osbpl3** | 1200014M06Rik\|6720421I08Rik\|A530055M08\|ORP3\|OSBP3 | oxysterol binding protein-like 3 | protein-coding |
| chr6_50390315_50390693 | chr6 | 50390316 | 50390693 | + | 0.887115 | intron (NM_001163645, intron 2 of 21) | intron (NM_001163645, intron 2 of 21) | -7667 | NM_027881 | 71720 | Mm.44153 | NM_027881 | ENSMUSG00000029822 | **Osbpl3** | 1200014M06Rik\|6720421I08Rik\|A530055M08\|ORP3\|OSBP3 | oxysterol binding protein-like 3 | protein-coding |
| chr6_50391953_50392295 | chr6 | 50391954 | 50392295 | + | 0.86845 | intron (NM_001163645, intron 2 of 21) | intron (NM_001163645, intron 2 of 21) | -9287 | NM_027881 | 71720 | Mm.44153 | NM_027881 | ENSMUSG00000029822 | **Osbpl3** | 1200014M06Rik\|6720421I08Rik\|A530055M08\|ORP3\|OSBP3 | oxysterol binding protein-like 3 | protein-coding |
| chr6_50393896_50394485 | chr6 | 50393897 | 50394485 | + | 0.745281 | intron (NM_001163645, intron 2 of 21) | intron (NM_001163645, intron 2 of 21) | -11354 | NM_027881 | 71720 | Mm.44153 | NM_027881 | ENSMUSG00000029822 | **Osbpl3** | 1200014M06Rik\|6720421I08Rik\|A530055M08\|ORP3\|OSBP3 | oxysterol binding protein-like 3 | protein-coding |
| chr10_111101933_111102286 | chr10 | 111101934 | 111102286 | + | 1.6982 | Intergenic | Intergenic | -62692 | NM_001003717 | 237542 | Mm.220204 | NM_175489 | ENSMUSG00000020189 | **Osbpl8** | AA536976\|AA536995\|C730029P18Rik\|D330025H14Rik\|ORP-8 | oxysterol binding protein-like 8 | protein-coding |
| chr10_111160223_111160504 | chr10 | 111160224 | 111160504 | + | 1.18654 | Intergenic | RLTR28\|LTR\|ERVL | -4438 | NM_001003717 | 237542 | Mm.220204 | NM_175489 | ENSMUSG00000020189 | **Osbpl8** | AA536976\|AA536995\|C730029P18Rik\|D330025H14Rik\|ORP-8 | oxysterol binding protein-like 8 | protein-coding |
| chr10_111213624_111214481 | chr10 | 111213625 | 111214481 | + | 1.09825 | intron (NM_001003717, intron 2 of 22) | (TCCC)n\|Simple_repeat\|Simple_repeat | 49251 | NM_175489 | 237542 | Mm.220204 | NM_175489 | ENSMUSG00000020189 | **Osbpl8** | AA536976\|AA536995\|C730029P18Rik\|D330025H14Rik\|ORP-8 | oxysterol binding protein-like 8 | protein-coding |
| chr10_111103041_111103443 | chr10 | 111103042 | 111103443 | + | 0.979366 | Intergenic | Intergenic | -61560 | NM_001003717 | 237542 | Mm.220204 | NM_175489 | ENSMUSG00000020189 | **Osbpl8** | AA536976\|AA536995\|C730029P18Rik\|D330025H14Rik\|ORP-8 | oxysterol binding protein-like 8 | protein-coding |
| chr10_111128467_111128984 | chr10 | 111128468 | 111128984 | + | 0.804269 | Intergenic | URR1B\|DNA\|hAT-Charlie | -36076 | NM_001003717 | 237542 | Mm.220204 | NM_175489 | ENSMUSG00000020189 | **Osbpl8** | AA536976\|AA536995\|C730029P18Rik\|D330025H14Rik\|ORP-8 | oxysterol binding protein-like 8 | protein-coding |
| chr10_111214111_111214451 | chr10 | 111214112 | 111214451 | + | 0.779511 | intron (NM_001003717, intron 2 of 22) | intron (NM_001003717, intron 2 of 22) | 49479 | NM_175489 | 237542 | Mm.220204 | NM_175489 | ENSMUSG00000020189 | **Osbpl8** | AA536976\|AA536995\|C730029P18Rik\|D330025H14Rik\|ORP-8 | oxysterol binding protein-like 8 | protein-coding |
| chr10_111101933_111102826 | chr10 | 111101934 | 111102826 | + | 0.710386 | Intergenic | Intergenic | -62422 | NM_001003717 | 237542 | Mm.220204 | NM_175489 | ENSMUSG00000020189 | **Osbpl8** | AA536976\|AA536995\|C730029P18Rik\|D330025H14Rik\|ORP-8 | oxysterol binding protein-like 8 | protein-coding |
| chr10_111186507_111186745 | chr10 | 111186508 | 111186745 | + | 0.67754 | intron (NM_175489, intron 1 of 23) | RSINE1\|SINE\|B4 | 21824 | NM_175489 | 237542 | Mm.220204 | NM_175489 | ENSMUSG00000020189 | **Osbpl8** | AA536976\|AA536995\|C730029P18Rik\|D330025H14Rik\|ORP-8 | oxysterol binding protein-like 8 | protein-coding |
| chr4_109104961_109105694 | chr4 | 109104962 | 109105694 | + | 1.88393 | intron (NM_133885, intron 4 of 22) | intron (NM_133885, intron 4 of 22) | 12708 | NM_173350 | 100273 | Mm.366315 | NM_133885 | ENSMUSG00000028559 | **Osbpl9** | 2600011I06Rik\|AU015843\|Orp-9 | oxysterol binding protein-like 9 | protein-coding |
| chr4_109134849_109135609 | chr4 | 109134850 | 109135609 | + | 1.25723 | intron (NM_133885, intron 3 of 22) | URR1B\|DNA\|hAT-Charlie | -17193 | NM_001134791 | 100273 | Mm.366315 | NM_133885 | ENSMUSG00000028559 | **Osbpl9** | 2600011I06Rik\|AU015843\|Orp-9 | oxysterol binding protein-like 9 | protein-coding |
| chr4_109135047_109135584 | chr4 | 109135048 | 109135584 | + | 1.23997 | intron (NM_133885, intron 3 of 22) | URR1B\|DNA\|hAT-Charlie | -17280 | NM_001134791 | 100273 | Mm.366315 | NM_133885 | ENSMUSG00000028559 | **Osbpl9** | 2600011I06Rik\|AU015843\|Orp-9 | oxysterol binding protein-like 9 | protein-coding |
| chr4_109125975_109126507 | chr4 | 109125976 | 109126507 | + | 0.904506 | intron (NM_133885, intron 4 of 22) | Charlie29a\|DNA\|hAT-Charlie | -8205 | NM_001134791 | 100273 | Mm.366315 | NM_133885 | ENSMUSG00000028559 | **Osbpl9** | 2600011I06Rik\|AU015843\|Orp-9 | oxysterol binding protein-like 9 | protein-coding |
| chr4_109126905_109127142 | chr4 | 109126906 | 109127142 | + | 0.833886 | intron (NM_133885, intron 4 of 22) | intron (NM_133885, intron 4 of 22) | -8988 | NM_001134791 | 100273 | Mm.366315 | NM_133885 | ENSMUSG00000028559 | **Osbpl9** | 2600011I06Rik\|AU015843\|Orp-9 | oxysterol binding protein-like 9 | protein-coding |
| chr7_3590590_3590971 | chr7 | 3590591 | 3590971 | + | 1.51593 | Intergenic | Intergenic | 25376 | NM_175632 | 232790 | Mm.358882 | NM_175632 | ENSMUSG00000054594 | **Oscar** | mOSCAR\|mOSCAR-M1\|mOSCAR-M2\|mOSCAR-M3 | osteoclast associated receptor | protein-coding |
| chr7_3594726_3595135 | chr7 | 3594727 | 3595135 | + | 1.24148 | Intergenic | RLTR17\|LTR\|ERVK | 21226 | NM_175632 | 232790 | Mm.358882 | NM_175632 | ENSMUSG00000054594 | **Oscar** | mOSCAR\|mOSCAR-M1\|mOSCAR-M2\|mOSCAR-M3 | osteoclast associated receptor | protein-coding |
| chr8_119436932_119437288 | chr8 | 119436933 | 119437288 | + | 0.696747 | promoter-TSS (NM_027950) | promoter-TSS (NM_027950) | -52 | NM_027950 | 71839 | Mm.272357 | NM_027950 | ENSMUSG00000074063 | **Osgin1** | 1700012B18Rik\|Okl38 | oxidative stress induced growth inhibitor 1 | protein-coding |
| chr4_16071043_16071883 | chr4 | 16071044 | 16071883 | + | 1.38121 | Intergenic | MTE-int\|LTR\|ERVL-MaLR | -57586 | NM_145950 | 209212 | Mm.46455 | NM_145950 | ENSMUSG00000041153 | **Osgin2** | BC010311\|C230027H09Rik | oxidative stress induced growth inhibitor family member 2 | protein-coding |
| chr4_16071238_16071827 | chr4 | 16071239 | 16071827 | + | 1.36528 | Intergenic | Intergenic | -57656 | NM_145950 | 209212 | Mm.46455 | NM_145950 | ENSMUSG00000041153 | **Osgin2** | BC010311\|C230027H09Rik | oxidative stress induced growth inhibitor family member 2 | protein-coding |
| chr4_16015588_16015943 | chr4 | 16015589 | 16015943 | + | 1.17004 | Intergenic | B3A\|SINE\|B2 | -1889 | NM_145950 | 209212 | Mm.46455 | NM_145950 | ENSMUSG00000041153 | **Osgin2** | BC010311\|C230027H09Rik | oxidative stress induced growth inhibitor family member 2 | protein-coding |
| chr4_16078563_16078955 | chr4 | 16078564 | 16078955 | + | 0.975707 | Intergenic | Intergenic | -64882 | NM_145950 | 209212 | Mm.46455 | NM_145950 | ENSMUSG00000041153 | **Osgin2** | BC010311\|C230027H09Rik | oxidative stress induced growth inhibitor family member 2 | protein-coding |
| chr15_35304729_35305097 | chr15 | 35304730 | 35305097 | + | 1.67761 | Intergenic | Intergenic | 8801 | NM_054049 | 107587 | Mm.46336 | NM_054049 | ENSMUSG00000022330 | **Osr2** | 5430409I15Rik\|Osr2A\|Osr2B | odd-skipped related 2 | protein-coding |
| chr15_35304421_35305120 | chr15 | 35304422 | 35305120 | + | 1.46874 | Intergenic | Intergenic | 8659 | NM_054049 | 107587 | Mm.46336 | NM_054049 | ENSMUSG00000022330 | **Osr2** | 5430409I15Rik\|Osr2A\|Osr2B | odd-skipped related 2 | protein-coding |
| chr15_35306441_35306849 | chr15 | 35306442 | 35306849 | + | 1.07848 | TTS (NR_045280) | TTS (NR_045280) | 10533 | NM_054049 | 107587 | Mm.46336 | NM_054049 | ENSMUSG00000022330 | **Osr2** | 5430409I15Rik\|Osr2A\|Osr2B | odd-skipped related 2 | protein-coding |
| chr15_35295742_35296230 | chr15 | 35295743 | 35296230 | + | 0.671219 | promoter-TSS (NM_054049) | promoter-TSS (NM_054049) | -126 | NM_054049 | 107587 | Mm.46336 | NM_054049 | ENSMUSG00000022330 | **Osr2** | 5430409I15Rik\|Osr2A\|Osr2B | odd-skipped related 2 | protein-coding |
| chr3_130694757_130695099 | chr3 | 130694758 | 130695099 | + | 0.798527 | TTS (NM_025509) | TTS (NM_025509) | 14516 | NM_025509 | 66357 | Mm.300697 | NM_025509 | ENSMUSG00000041084 | **Ostc** | 2310008M10Rik\|5730557H03Rik | oligosaccharyltransferase complex subunit | protein-coding |
| chr19_18598290_18598949 | chr19 | 18598291 | 18598949 | + | 1.16706 | intron (NM_017375, intron 2 of 9) | intron (NM_017375, intron 2 of 9) | 33193 | NM_017375 | 20409 | Mm.172222 | NM_017375 | | **Ostf1** | C78236\|SH3P2\|Sh3d3 | osteoclast stimulating factor 1 | protein-coding |
| chr19_18599657_18600075 | chr19 | 18599658 | 18600075 | + | 0.959057 | intron (NM_017375, intron 2 of 9) | intron (NM_017375, intron 2 of 9) | 31947 | NM_017375 | 20409 | Mm.172222 | NM_017375 | | **Ostf1** | C78236\|SH3P2\|Sh3d3 | osteoclast stimulating factor 1 | protein-coding |
| chr19_18599005_18599412 | chr19 | 18599006 | 18599412 | + | 0.791312 | intron (NM_017375, intron 2 of 9) | intron (NM_017375, intron 2 of 9) | 32604 | NM_017375 | 20409 | Mm.172222 | NM_017375 | | **Ostf1** | C78236\|SH3P2\|Sh3d3 | osteoclast stimulating factor 1 | protein-coding |
| chr16_27224005_27224332 | chr16 | 27224006 | 27224332 | + | 0.897775 | Intergenic | Intergenic | -83472 | NM_198112 | 239790 | Mm.390880 | NM_198112 | ENSMUSG00000052276 | **Ostn** | Ostc | osteocrin | protein-coding |
| chr16_27326370_27326820 | chr16 | 27326371 | 27326820 | + | 0.766138 | intron (NM_198112, intron 3 of 4) | RMER15\|LTR\|ERVL | 18954 | NM_198112 | 239790 | Mm.390880 | NM_198112 | ENSMUSG00000052276 | **Ostn** | Ostc | osteocrin | protein-coding |
| chr10_107921272_107921929 | chr10 | 107921273 | 107921929 | + | 1.55565 | Intergenic | URR1A\|DNA\|hAT-Charlie | -9467 | NM_001177567 | 628870 | Mm.336090 | NM_001177567 | ENSMUSG00000091455 | **Otogl** | EG628870\|Gm6924 | otogelin-like | protein-coding |
| chr10_107922107_107922396 | chr10 | 107922108 | 107922396 | + | 1.18416 | Intergenic | Intergenic | -10118 | NM_001177567 | 628870 | Mm.336090 | NM_001177567 | ENSMUSG00000091455 | **Otogl** | EG628870\|Gm6924 | otogelin-like | protein-coding |
| chr10_108033279_108033823 | chr10 | 108033280 | 108033823 | + | 0.794636 | Intergenic | RMER17B\|LTR\|ERVK | -121417 | NM_001177567 | 628870 | Mm.336090 | NM_001177567 | ENSMUSG00000091455 | **Otogl** | EG628870\|Gm6924 | otogelin-like | protein-coding |
| chr10_107951604_107952020 | chr10 | 107951605 | 107952020 | + | 0.765771 | Intergenic | Intergenic | -39678 | NM_001177567 | 628870 | Mm.336090 | NM_001177567 | ENSMUSG00000091455 | **Otogl** | EG628870\|Gm6924 | otogelin-like | protein-coding |
| chr10_107952541_107953208 | chr10 | 107952542 | 107953208 | + | 0.751808 | Intergenic | RCHARR1\|DNA\|hAT-Charlie | -40741 | NM_001177567 | 628870 | Mm.336090 | NM_001177567 | ENSMUSG00000091455 | **Otogl** | EG628870\|Gm6924 | otogelin-like | protein-coding |
| chr2_19669278_19669722 | chr2 | 19669279 | 19669722 | + | 0.675273 | Intergenic | ORR1F\|LTR\|ERVL-MaLR | 11438 | NM_027715 | 71198 | Mm.83981 | NM_027715 | ENSMUSG00000043415 | **Otud1** | 4933428L19Rik | OTU domain containing 1 | protein-coding |
| chr8_79542519_79542899 | chr8 | 79542520 | 79542899 | + | 0.771848 | Intergenic | Intergenic | -96967 | NM_001256033 | 73945 | Mm.34348 | NM_001081164 | ENSMUSG00000036990 | **Otud4** | 4930431L18Rik\|AI449692\|D8Ertd69e\|mKIAA1046 | OTU domain containing 4 | protein-coding |
| chr8_79542434_79543149 | chr8 | 79542435 | 79543149 | + | 0.67829 | Intergenic | Intergenic | -96884 | NM_001256033 | 73945 | Mm.34348 | NM_001081164 | ENSMUSG00000036990 | **Otud4** | 4930431L18Rik\|AI449692\|D8Ertd69e\|mKIAA1046 | OTU domain containing 4 | protein-coding |
| chr3_96105175_96105699 | chr3 | 96105176 | 96105699 | + | 1.10177 | intron (NM_001025614, intron 1 of 11) | intron (NM_001025614, intron 1 of 11) | 910 | NM_001025614 | 229603 | Mm.272336 | NM_001025613 | ENSMUSG00000038495 | **Otud7b** | 2900060B22Rik\|4930463P07Rik\|AI462125\|CEZANNE\|Trabid2\|Za20d1 | OTU domain containing 7B | protein-coding |
| chr15_27625040_27625899 | chr15 | 27625041 | 27625899 | + | 0.660456 | intron (NM_001013792, intron 1 of 6) | intron (NM_001013792, intron 1 of 6) | 5223 | NM_001013792 | 432940 | Mm.309164 | NM_001013792 | ENSMUSG00000046034 | **Otulin** | C79097\|Fam105b\|m3Sapc\|m7-1Sapc | OTU deubiquitinase with linear linkage specificity | protein-coding |
| chr11_22007370_22008683 | chr11 | 22007371 | 22008683 | + | 1.06223 | intron (NM_153078, intron 23 of 24) | intron (NM_153078, intron 23 of 24) | -6376 | NM_011023 | 18423 | Mm.129663 | NM_011023 | ENSMUSG00000005917 | **Otx1** | A730044F23Rik\|jv | orthodenticle homolog 1 | protein-coding |
| chr11_22001423_22002297 | chr11 | 22001424 | 22002297 | + | 0.951799 | promoter-TSS (NM_011023) | promoter-TSS (NM_011023) | -209 | NM_011023 | 18423 | Mm.129663 | NM_011023 | ENSMUSG00000005917 | **Otx1** | A730044F23Rik\|jv | orthodenticle homolog 1 | protein-coding |

| | | | | | | | | | | | | | | | | | |
|---|---|---|---|---|---|---|---|---|---|---|---|---|---|---|---|---|---|
| chr11_21788800_21789295 | chr11 | 21788801 | 21789295 | + | 0.838559 | intron (NM_145425, intron 13 of 17) | RLTR50A\|LTR\|ERVK | 212603 | NM_011023 | 18423 | Mm.129663 | NM_011023 | ENSMUSG00000005917 | **Otx1** | A730044F23Rik\|jv | orthodenticle homolog 1 | protein-coding |
| chr11_22002400_22002963 | chr11 | 22002401 | 22002963 | + | 0.722063 | Intergenic | Intergenic | -1031 | NM_011023 | 18423 | Mm.129663 | NM_011023 | ENSMUSG00000005917 | **Otx1** | A730044F23Rik\|jv | orthodenticle homolog 1 | protein-coding |
| chr2_144342893_144343514 | chr2 | 144342894 | 144343514 | + | 1.03304 | Intergenic | Intergenic | -11124 | NM_026924 | 107586 | Mm.252750 | NM_026924 | ENSMUSG00000037279 | **Ovol2** | 1700108N11Rik\|1810007D21Rik\|M-OVO\|M-OVO-A\|M-OVO-B\|MOVO\|Ovo2\|Zfp339\|movo2 | ovo-like 2 (Drosophila) | protein-coding |
| chr15_4012456_4012916 | chr15 | 4012457 | 4012916 | + | 0.850822 | non-coding (NR_045740, exon 3 of 3) | non-coding (NR_045740, exon 3 of 3) | -13742 | NM_024188 | 67041 | Mm.13445 | NM_024188 | ENSMUSG00000022186 | **Oxct1** | 2610008O03Rik\|Oxct\|Oxct2a\|Scot-s | 3-oxoacid CoA transferase 1 | protein-coding |
| chr14_32086307_32086601 | chr14 | 32086308 | 32086601 | + | 0.860991 | promoter-TSS (NM_001047433) | promoter-TSS (NM_001047433) | 726 | NM_145460 | 218885 | Mm.202257 | NM_145460 | ENSMUSG00000021906 | **Oxnad1** | 2410002F01Rik\|AW492292\|BC019806 | oxidoreductase NAD-binding domain containing 1 | protein-coding |
| chr14_32111998_32112466 | chr14 | 32111999 | 32112466 | + | 0.690493 | Intergenic | RLTR1B\|LTR\|ERV1 | 26504 | NM_145460 | 218885 | Mm.202257 | NM_145460 | ENSMUSG00000021906 | **Oxnad1** | 2410002F01Rik\|AW492292\|BC019806 | oxidoreductase NAD-binding domain containing 1 | protein-coding |
| chr15_41612130_41612658 | chr15 | 41612131 | 41612658 | + | 0.864886 | intron (NM_001130166, intron 2 of 16) | MIR3\|SINE\|MIR | 164912 | NM_001130166 | 170719 | Mm.254267 | NM_130885 | ENSMUSG00000022307 | **Oxr1** | 2210416C20Rik\|C7\|C7B | oxidation resistance 1 | protein-coding |
| chr15_41778262_41778548 | chr15 | 41778263 | 41778548 | + | 0.820819 | intron (NM_001130166, intron 3 of 16) | MER58A\|DNA\|hAT-Charlie | -10629 | NM_001130165 | 170719 | Mm.254267 | NM_130885 | ENSMUSG00000022307 | **Oxr1** | 2210416C20Rik\|C7\|C7B | oxidation resistance 1 | protein-coding |
| chr14_16269791_16270356 | chr14 | 16269792 | 16270356 | + | 1.25309 | intron (NM_021504, intron 3 of 11) | intron (NM_021504, intron 3 of 11) | -20266 | NM_027695 | 71147 | Mm.197960 | NM_027695 | ENSMUSG00000021786 | **Oxsm** | 4933425A18Rik\|C80494 | 3-oxoacyl-ACP synthase, mitochondrial | protein-coding |
| chr14_16251612_16251957 | chr14 | 16251613 | 16251957 | + | 0.726436 | intron (NM_021504, intron 1 of 11) | intron (NM_021504, intron 1 of 11) | -1977 | NM_027695 | 71147 | Mm.197960 | NM_027695 | ENSMUSG00000021786 | **Oxsm** | 4933425A18Rik\|C80494 | 3-oxoacyl-ACP synthase, mitochondrial | protein-coding |
| chr16_17564236_17564744 | chr16 | 17564237 | 17564744 | + | 0.993994 | intron (NM_011028, intron 2 of 11) | RMER15\|LTR\|ERVL | 2605 | NM_001159561 | 18440 | Mm.8032 | NM_011028 | ENSMUSG00000022758 | **P2rx6** | P2rxl1\|P2x6\|P2xm | purinergic receptor P2X, ligand-gated ion channel, 6 | protein-coding |
| chr3_60787824_60788467 | chr3 | 60787825 | 60788467 | + | 1.73018 | Intergenic | Intergenic | -214639 | NM_008772 | 18441 | Mm.281452 | NM_008772 | ENSMUSG00000027765 | **P2ry1** | P2Y1 | purinergic receptor P2Y, G-protein coupled 1 | protein-coding |
| chr3_61053716_61054235 | chr3 | 61053717 | 61054235 | + | 1.15323 | Intergenic | Intergenic | 51191 | NM_001282016 | 18441 | Mm.281452 | NM_008772 | ENSMUSG00000027765 | **P2ry1** | P2Y1 | purinergic receptor P2Y, G-protein coupled 1 | protein-coding |
| chr3_60787551_60787745 | chr3 | 60787552 | 60787745 | + | 0.891989 | Intergenic | URR1B\|DNA\|hAT-Charlie | -215137 | NM_008772 | 18441 | Mm.281452 | NM_008772 | ENSMUSG00000027765 | **P2ry1** | P2Y1 | purinergic receptor P2Y, G-protein coupled 1 | protein-coding |
| chr3_60811924_60812469 | chr3 | 60811925 | 60812469 | + | 0.685646 | Intergenic | Intergenic | -190588 | NM_008772 | 18441 | Mm.281452 | NM_008772 | ENSMUSG00000027765 | **P2ry1** | P2Y1 | purinergic receptor P2Y, G-protein coupled 1 | protein-coding |
| chr3_59295777_59296969 | chr3 | 59295778 | 59296969 | + | 1.10134 | intron (NM_177855, intron 40 of 43) | intron (NM_177855, intron 40 of 43) | -33542 | NM_027571 | 70839 | Mm.422675 | NM_027571 | ENSMUSG00000036353 | **P2ry12** | 2900079B22Rik\|4921504D23Rik\|P2Y12 | purinergic receptor P2Y, G-protein coupled 12 | protein-coding |
| chr3_59277596_59278116 | chr3 | 59277597 | 59278116 | + | 0.673537 | intron (NM_177855, intron 38 of 43) | intron (NM_177855, intron 38 of 43) | -15025 | NM_027571 | 70839 | Mm.422675 | NM_027571 | ENSMUSG00000036353 | **P2ry12** | 2900079B22Rik\|4921504D23Rik\|P2Y12 | purinergic receptor P2Y, G-protein coupled 12 | protein-coding |
| chr3_59219319_59219702 | chr3 | 59219320 | 59219702 | + | 1.54793 | intron (NM_027571, intron 3 of 3) | intron (NM_027571, intron 3 of 3) | -8629 | NM_028808 | 74191 | Mm.90067 | NM_028808 | ENSMUSG00000036362 | **P2ry13** | 2010001L06Rik\|GPCR1\|GPR94\|Gpr86\|P2Y13\|P2yr13\|SP174 | purinergic receptor P2Y, G-protein coupled 13 | protein-coding |
| chr3_59222277_59222646 | chr3 | 59222278 | 59222646 | + | 1.12961 | intron (NM_027571, intron 2 of 3) | intron (NM_027571, intron 2 of 3) | -11580 | NM_028808 | 74191 | Mm.90067 | NM_028808 | ENSMUSG00000036362 | **P2ry13** | 2010001L06Rik\|GPCR1\|GPR94\|Gpr86\|P2Y13\|P2yr13\|SP174 | purinergic receptor P2Y, G-protein coupled 13 | protein-coding |
| chr3_59203742_59204223 | chr3 | 59203743 | 59204223 | + | 0.885945 | intron (NM_177855, intron 15 of 43) | intron (NM_177855, intron 15 of 43) | 6899 | NM_028808 | 74191 | Mm.90067 | NM_028808 | ENSMUSG00000036362 | **P2ry13** | 2010001L06Rik\|GPCR1\|GPR94\|Gpr86\|P2Y13\|P2yr13\|SP174 | purinergic receptor P2Y, G-protein coupled 13 | protein-coding |
| chr3_59203734_59204468 | chr3 | 59203735 | 59204468 | + | 0.761473 | intron (NM_177855, intron 15 of 43) | intron (NM_177855, intron 15 of 43) | 6781 | NM_028808 | 74191 | Mm.90067 | NM_028808 | ENSMUSG00000036362 | **P2ry13** | 2010001L06Rik\|GPCR1\|GPR94\|Gpr86\|P2Y13\|P2yr13\|SP174 | purinergic receptor P2Y, G-protein coupled 13 | protein-coding |
| chr3_59118310_59118574 | chr3 | 59118311 | 59118574 | + | 0.877475 | intron (NM_001287119, intron 1 of 2) | intron (NM_001287119, intron 1 of 2) | 12320 | NM_001008497 | 140795 | Mm.439021 | NM_133200 | ENSMUSG00000036381 | **P2ry14** | A330108O13Rik\|Gpr105\|P2Y14 | purinergic receptor P2Y, G-protein coupled, 14 | protein-coding |
| chr3_59129000_59129245 | chr3 | 59129001 | 59129245 | + | 0.677505 | intron (NM_001287119, intron 1 of 2) | intron (NM_001287119, intron 1 of 2) | 1639 | NM_001008497 | 140795 | Mm.439021 | NM_133200 | ENSMUSG00000036381 | **P2ry14** | A330108O13Rik\|Gpr105\|P2Y14 | purinergic receptor P2Y, G-protein coupled, 14 | protein-coding |
| chr11_54149531_54150475 | chr11 | 54149532 | 54150475 | + | 0.856276 | Intergenic | L2\|LINE\|L2 | 49079 | NM_011031 | 18452 | Mm.3705 | NM_011031 | ENSMUSG00000018906 | **P4ha2** | AA407196\|C76437\|P4hl | procollagen-proline, 2-oxoglutarate 4-dioxygenase (proline 4-hydroxylase), alpha II polypeptide | protein-coding |
| chr11_54127159_54127734 | chr11 | 54127160 | 54127734 | + | 0.779334 | intron (NM_011031, intron 13 of 15) | intron (NM_011031, intron 13 of 15) | 26523 | NM_011031 | 18452 | Mm.3705 | NM_011031 | ENSMUSG00000018906 | **P4ha2** | AA407196\|C76437\|P4hl | procollagen-proline, 2-oxoglutarate 4-dioxygenase (proline 4-hydroxylase), alpha II polypeptide | protein-coding |
| chr15_36604939_36605253 | chr15 | 36604940 | 36605253 | + | 1.4993 | intron (NM_008774, intron 3 of 14) | PB1D10\|SINE\|Alu | 3877 | NM_008774 | 18458 | Mm.371570 | NM_008774 | ENSMUSG00000022283 | **Pabpc1** | PABP\|Pabp1\|Pabpl\|Pabpl1\|ePAB | poly(A) binding protein, cytoplasmic 1 | protein-coding |
| chr15_36606592_36606811 | chr15 | 36606593 | 36606811 | + | 1.03026 | intron (NM_008774, intron 1 of 14) | intron (NM_008774, intron 1 of 14) | 2271 | NM_008774 | 18458 | Mm.371570 | NM_008774 | ENSMUSG00000022283 | **Pabpc1** | PABP\|Pabp1\|Pabpl\|Pabpl1\|ePAB | poly(A) binding protein, cytoplasmic 1 | protein-coding |
| chr18_39653907_39654512 | chr18 | 39653908 | 39654512 | + | 1.04547 | Intergenic | Intergenic | -119287 | NM_011033 | 18459 | Mm.9493 | NM_011033 | ENSMUSG00000051732 | **Pabpc2** | Pabp\|Pabp+\|Pabp2 | poly(A) binding protein, cytoplasmic 2 | protein-coding |
| chr18_39663865_39664760 | chr18 | 39663866 | 39664760 | + | 0.864299 | Intergenic | Intergenic | -109184 | NM_011033 | 18459 | Mm.9493 | NM_011033 | ENSMUSG00000051732 | **Pabpc2** | Pabp\|Pabp+\|Pabp2 | poly(A) binding protein, cytoplasmic 2 | protein-coding |
| chr18_39802119_39802449 | chr18 | 39802120 | 39802449 | + | 0.639551 | Intergenic | Intergenic | 28787 | NM_011033 | 18459 | Mm.9493 | NM_011033 | ENSMUSG00000051732 | **Pabpc2** | Pabp\|Pabp+\|Pabp2 | poly(A) binding protein, cytoplasmic 2 | protein-coding |
| chr18_39877802_39878196 | chr18 | 39877803 | 39878196 | + | 0.638777 | Intergenic | Intergenic | 104502 | NM_011033 | 18459 | Mm.9493 | NM_011033 | ENSMUSG00000051732 | **Pabpc2** | Pabp\|Pabp+\|Pabp2 | poly(A) binding protein, cytoplasmic 2 | protein-coding |
| chr12_113040007_113040244 | chr12 | 113040008 | 113040244 | + | 0.665476 | intron (NM_001291444, intron 1 of 23) | L1M5\|LINE\|L1 | 25618 | NM_001291444 | 217893 | Mm.35754 | NM_198165 | ENSMUSG00000021143 | **Pacs2** | 6720425G15Rik\|AW208651\|AW489976\|E230011J22\|Pacs1l\|mKIAA0602 | phosphofurin acidic cluster sorting protein 2 | protein-coding |
| chr17_27658725_27659146 | chr17 | 27658726 | 27659146 | + | 0.715369 | intron (NM_001286744, intron 1 of 9) | intron (NM_001286744, intron 1 of 9) | 3345 | NM_001286743 | 23969 | Mm.4926 | NM_011861 | ENSMUSG00000040276 | **Pacsin1** | A830061D09Rik\|H74\|mKIAA1379\|syndapin | protein kinase C and casein kinase substrate in neurons 1 | protein-coding |
| chr15_83404033_83404442 | chr15 | 83404034 | 83404442 | + | 0.960514 | intron (NM_001159509, intron 1 of 10) | intron (NM_001159509, intron 1 of 10) | 28612 | NM_011862 | 23970 | Mm.23978 | NM_011862 | ENSMUSG00000016664 | **Pacsin2** | AI197433 | protein kinase C and casein kinase substrate in neurons 2 | protein-coding |
| chr2_91250014_91250507 | chr2 | 91250015 | 91250507 | + | 0.714804 | Intergenic | ID_B1\|SINE\|B4 | -5904 | NM_001289677 | 80708 | Mm.236650 | NM_028733 | ENSMUSG00000027257 | **Pacsin3** | 4921507A02Rik\|6330413E15Rik\|AW413130 | protein kinase C and casein kinase substrate in neurons 3 | protein-coding |
| chr4_140723719_140724329 | chr4 | 140723720 | 140724329 | + | 0.645669 | TTS (NM_173867) | TTS (NM_173867) | 18619 | NM_153106 | 242726 | Mm.271661 | NM_153106 | ENSMUSG00000040935 | **Padi6** | Pad6\|Padi5\|ePAD | peptidyl arginine deiminase, type VI | protein-coding |
| chr3_9766991_9767418 | chr3 | 9766992 | 9767418 | + | 0.884578 | intron (NM_053182, intron 1 of 7) | intron (NM_053182, intron 1 of 7) | 66474 | NM_053182 | 94212 | Mm.23897 | NM_053182 | ENSMUSG00000027508 | **Pag1** | Cbp\|F730007C19Rik\|Pag | phosphoprotein associated with glycosphingolipid microdomains 1 | protein-coding |
| chr3_9765786_9766206 | chr3 | 9765787 | 9766206 | + | 0.884533 | intron (NM_053182, intron 1 of 7) | MTC\|LTR\|ERVL-MaLR | 67683 | NM_053182 | 94212 | Mm.23897 | NM_053182 | ENSMUSG00000027508 | **Pag1** | Cbp\|F730007C19Rik\|Pag | phosphoprotein associated with glycosphingolipid microdomains 1 | protein-coding |
| chr3_9839112_9839549 | chr3 | 9839113 | 9839549 | + | 0.698919 | Intergenic | B1_Mus1\|SINE\|Alu | -5652 | NM_001195031 | 94212 | Mm.23897 | NM_053182 | ENSMUSG00000027508 | **Pag1** | Cbp\|F730007C19Rik\|Pag | phosphoprotein associated with glycosphingolipid microdomains 1 | protein-coding |
| chr3_9756071_9756536 | chr3 | 9756072 | 9756536 | + | 0.649031 | intron (NM_053182, intron 1 of 7) | intron (NM_053182, intron 1 of 7) | 77375 | NM_053182 | 94212 | Mm.23897 | NM_053182 | ENSMUSG00000027508 | **Pag1** | Cbp\|F730007C19Rik\|Pag | phosphoprotein associated with glycosphingolipid microdomains 1 | protein-coding |
| chr5_76948382_76948584 | chr5 | 76948383 | 76948584 | + | 1.03459 | intron (NM_172146, intron 1 of 10) | intron (NM_172146, intron 1 of 10) | -2928 | NM_025939 | 67054 | Mm.182931 | NM_025939 | ENSMUSG00000029247 | **Paics** | 2610511I09Rik\|ADE2H1\|AIRC\|PAIS | phosphoribosylaminoimidazole carboxylase, phosphoribosylaminoribosylaminoimidazole, succinocarboxamide synthetase | protein-coding |
| chr2_118659060_118659477 | chr2 | 118659061 | 118659477 | + | 1.00535 | Intergenic | MIRc\|SINE\|MIR | -4308 | NM_001033254 | 214230 | Mm.241857 | NM_001033254 | ENSMUSG00000074923 | **Pak6** | 4732456M09 | p21 protein (Cdc42/Rac)-activated kinase 6 | protein-coding |
| chr2_118664408_118664746 | chr2 | 118664409 | 118664746 | + | 0.969827 | intron (NM_001033254, intron 1 of 10) | intron (NM_001033254, intron 1 of 10) | 1000 | NM_001033254 | 214230 | Mm.241857 | NM_001033254 | ENSMUSG00000074923 | **Pak6** | 4732456M09 | p21 protein (Cdc42/Rac)-activated kinase 6 | protein-coding |
| chr2_118656330_118656812 | chr2 | 118656331 | 118656812 | + | 0.721339 | Intergenic | Intergenic | -7006 | NM_001033254 | 214230 | Mm.241857 | NM_001033254 | ENSMUSG00000074923 | **Pak6** | 4732456M09 | p21 protein (Cdc42/Rac)-activated kinase 6 | protein-coding |
| chr8_61556890_61557511 | chr8 | 61556891 | 61557511 | + | 1.298 | intron (NM_001293772, intron 10 of 21) | intron (NM_001293772, intron 10 of 21) | 33968 | NM_001293774 | 72333 | Mm.29933 | NM_001081390 | ENSMUSG00000058056 | **Palld** | 2410003B16Rik\|6030492A02 | palladin, cytoskeletal associated protein | protein-coding |
| chr2_131258830_131259147 | chr2 | 131258831 | 131259147 | + | 1.60507 | intron (NR_037281, intron 2 of 2) | intron (NR_037281, intron 2 of 2) | -3511 | NM_153501 | 74450 | Mm.101264 | NM_153501 | ENSMUSG00000037514 | **Pank2** | 4933409I19Rik\|AI642621 | pantothenate kinase 2 | protein-coding |
| chr9_14995449_14995993 | chr9 | 14995450 | 14995993 | + | 0.953309 | Intergenic | Intergenic | 49757 | NM_019482 | 55991 | Mm.142253 | NM_019482 | | **Panx1** | AI847747 | pannexin 1 | protein-coding |
| chr9_15015281_15015537 | chr9 | 15015282 | 15015537 | + | 0.93248 | intron (NM_019482, intron 2 of 4) | intron (NM_019482, intron 2 of 4) | 30069 | NM_019482 | 55991 | Mm.142253 | NM_019482 | | **Panx1** | AI847747 | pannexin 1 | protein-coding |
| chr9_15026428_15026737 | chr9 | 15026429 | 15026737 | + | 0.900181 | intron (NM_019482, intron 1 of 4) | MIRb\|SINE\|MIR | 18895 | NM_019482 | 55991 | Mm.142253 | NM_019482 | | **Panx1** | AI847747 | pannexin 1 | protein-coding |
| chr9_15026356_15026821 | chr9 | 15026357 | 15026821 | + | 0.825348 | intron (NM_019482, intron 1 of 4) | MIRb\|SINE\|MIR | 18889 | NM_019482 | 55991 | Mm.142253 | NM_019482 | | **Panx1** | AI847747 | pannexin 1 | protein-coding |

| | | | | | | | | | | | | | | | | | |
|---|---|---|---|---|---|---|---|---|---|---|---|---|---|---|---|---|---|
| chr8_88177226_88177668 | chr8 | 88177227 | 88177668 | + | 0.950363 | Intergenic | Intergenic | -21766 | NM_001164499 | 214627 | Mm.491211 | NM_001164497 | ENSMUSG00000036779 | **Papd5** | 5730445M16Rik\|5830428A09 | PAP associated domain containing 5 | protein-coding |
| chr8_88202923_88203259 | chr8 | 88202924 | 88203259 | + | 0.886353 | intron (NM_001164498, intron 2 of 12) | RMER15\|LTR\|ERVL | 3878 | NM_001164498 | 214627 | Mm.491211 | NM_001164497 | ENSMUSG00000036779 | **Papd5** | 5730445M16Rik\|5830428A09 | PAP associated domain containing 5 | protein-coding |
| chr13_69485895_69486230 | chr13 | 69485896 | 69486230 | + | 0.799074 | Intergenic | ORR1E\|LTR\|ERVL-MaLR | 47801 | NM_198600 | 210106 | Mm.315959 | NM_198600 | ENSMUSG00000034575 | **Papd7** | LAK-1\|POLK\|Pols\|TRF4\|TRF4-1 | PAP associated domain containing 7 | protein-coding |
| chr13_69485722_69486238 | chr13 | 69485723 | 69486238 | + | 0.619602 | Intergenic | ORR1E\|LTR\|ERVL-MaLR | 47884 | NM_198600 | 210106 | Mm.315959 | NM_198600 | ENSMUSG00000034575 | **Papd7** | LAK-1\|POLK\|Pols\|TRF4\|TRF4-1 | PAP associated domain containing 7 | protein-coding |
| chr12_105762358_105762700 | chr12 | 105762359 | 105762700 | + | 0.774142 | intron (NM_030187, intron 13 of 17) | intron (NM_030187, intron 13 of 17) | -22173 | NM_011112 | 18789 | Mm.255877 | NM_011112 | ENSMUSG00000021111 | **Papola** | Pap\|PapIII\|Plap | poly (A) polymerase alpha | protein-coding |
| chr3_131489661_131491111 | chr3 | 131489662 | 131491111 | + | 0.877277 | Intergenic | CpG | -74382 | NM_001289479 | 23971 | Mm.244912 | NM_011863 | ENSMUSG00000028032 | **Papss1** | AI325286\|Asapk\|SK1 | 3'-phosphoadenosine 5'-phosphosulfate synthase 1 | protein-coding |
| chr19_32596478_32596845 | chr19 | 32596479 | 32596845 | + | 0.811227 | intron (NM_001201470, intron 1 of 11) | intron (NM_001201470, intron 1 of 11) | 947 | NM_001201470 | 23972 | Mm.203916 | NM_011864 | ENSMUSG00000024899 | **Papss2** | 1810018P12Rik\|AI159688\|AtpsU2\|Atpsk2\|Sk2\|bm | 3'-phosphoadenosine 5'-phosphosulfate synthase 2 | protein-coding |
| chr9_62020233_62020912 | chr9 | 62020234 | 62020912 | + | 0.737641 | intron (NM_028748, intron 1 of 7) | intron (NM_028748, intron 1 of 7) | 6217 | NM_028748 | 74090 | Mm.273267 | NM_028748 | ENSMUSG00000032278 | **Paqr5** | 0610010I15Rik\|AV002411\|BB115488\|mPRg | progestin and adipoQ receptor family member V | protein-coding |
| chr1_20912109_20912464 | chr1 | 20912110 | 20912464 | + | 1.24989 | intron (NM_028829, intron 1 of 2) | intron (NM_028829, intron 1 of 2) | 21665 | NM_028829 | 74229 | Mm.40780 | NM_028829 | ENSMUSG00000025931 | **Paqr8** | 1700019B16Rik\|3110001D06Rik | progestin and adipoQ receptor family member VIII | protein-coding |
| chr8_127381277_127381761 | chr8 | 127381278 | 127381761 | + | 1.36767 | intron (NM_001013580, intron 10 of 13) | intron (NM_001013580, intron 10 of 13) | 81867 | NM_001013580 | 93742 | Mm.299254 | NM_033620 | ENSMUSG00000025812 | **Pard3** | AA960621\|AI256638\|ASIP\|D8Ertd580e\|PAR-3\|Par3\|Pard3a | par-3 family cell polarity regulator | protein-coding |
| chr8_127309334_127309802 | chr8 | 127309335 | 127309802 | + | 1.13801 | intron (NM_001013580, intron 2 of 13) | intron (NM_001013580, intron 2 of 13) | 9916 | NM_001013580 | 93742 | Mm.299254 | NM_033620 | ENSMUSG00000025812 | **Pard3** | AA960621\|AI256638\|ASIP\|D8Ertd580e\|PAR-3\|Par3\|Pard3a | par-3 family cell polarity regulator | protein-coding |
| chr8_127655372_127656108 | chr8 | 127655373 | 127656108 | + | 1.04861 | Intergenic | Intergenic | 356088 | NM_001013580 | 93742 | Mm.299254 | NM_033620 | ENSMUSG00000025812 | **Pard3** | AA960621\|AI256638\|ASIP\|D8Ertd580e\|PAR-3\|Par3\|Pard3a | par-3 family cell polarity regulator | protein-coding |
| chr8_127605708_127606446 | chr8 | 127605709 | 127606446 | + | 0.871146 | intron (NM_033620, intron 24 of 24) | MTD\|LTR\|ERVL-MaLR | 306425 | NM_001013580 | 93742 | Mm.299254 | NM_033620 | ENSMUSG00000025812 | **Pard3** | AA960621\|AI256638\|ASIP\|D8Ertd580e\|PAR-3\|Par3\|Pard3a | par-3 family cell polarity regulator | protein-coding |
| chr18_80009243_80009774 | chr18 | 80009244 | 80009774 | + | 0.827555 | Intergenic | MERV1_LTR\|LTR\|ERV1 | -37386 | NM_053117 | 93737 | Mm.24678 | NM_053117 | ENSMUSG00000056214 | **Pard6g** | 2410049N21Rik\|Par6a | par-6 family cell polarity regulator gamma | protein-coding |
| chr5_91517399_91518168 | chr5 | 91517400 | 91518168 | + | 0.629217 | promoter-TSS (NM_145562) | promoter-TSS (NM_145562) | 84 | NM_145562 | 231440 | Mm.5002 | NM_145562 | | **Parm1** | 2210012L08Rik\|9130213B05Rik\|W91666 | prostate androgen-regulated mucin-like protein 1 | protein-coding |
| chr1_180545900_180547137 | chr1 | 180545901 | 180547137 | + | 0.629032 | Intergenic | RMER17C\|LTR\|ERVK | -22456 | NM_007415 | 11545 | Mm.277779 | NM_007415 | ENSMUSG00000026496 | **Parp1** | 5830444G22Rik\|AI893648\|ARTD1\|Adprp\|Adprt1\|C80510\|PARP\|PPOL\|parp-1\|sPARP-1 | poly (ADP-ribose) polymerase family, member 1 | protein-coding |
| chr6_127447766_127448518 | chr6 | 127447767 | 127448518 | + | 0.807724 | Intergenic | Intergenic | -5581 | NM_181402 | 101187 | Mm.110818 | NM_181402 | ENSMUSG00000037997 | **Parp11** | 5330431N24Rik\|AI851877\|ARTD11\|HIN1L | poly (ADP-ribose) polymerase family, member 11 | protein-coding |
| chr6_127447495_127448522 | chr6 | 127447496 | 127448522 | + | 0.799248 | Intergenic | GA-rich\|Low_complexity\|Low_complexity | -5714 | NM_181402 | 101187 | Mm.110818 | NM_181402 | ENSMUSG00000037997 | **Parp11** | 5330431N24Rik\|AI851877\|ARTD11\|HIN1L | poly (ADP-ribose) polymerase family, member 11 | protein-coding |
| chr6_127456827_127457104 | chr6 | 127456828 | 127457104 | + | 0.676545 | intron (NM_181402, intron 1 of 8) | ORR1C2-int\|LTR\|ERVL-MaLR | 3243 | NM_181402 | 101187 | Mm.110818 | NM_181402 | ENSMUSG00000037997 | **Parp11** | 5330431N24Rik\|AI851877\|ARTD11\|HIN1L | poly (ADP-ribose) polymerase family, member 11 | protein-coding |
| chr6_127456743_127457115 | chr6 | 127456744 | 127457115 | + | 0.655585 | intron (NM_181402, intron 1 of 8) | ORR1C2-int\|LTR\|ERVL-MaLR | 3206 | NM_181402 | 101187 | Mm.110818 | NM_181402 | ENSMUSG00000037997 | **Parp11** | 5330431N24Rik\|AI851877\|ARTD11\|HIN1L | poly (ADP-ribose) polymerase family, member 11 | protein-coding |
| chr9_65211544_65211910 | chr9 | 65211545 | 65211910 | + | 0.927378 | Intergenic | B3\|SINE\|B2 | -2963 | NM_177460 | 214424 | Mm.31129 | NM_177460 | ENSMUSG00000032392 | **Parp16** | ARTD15\|BC055447\|C79952\|PARP-16 | poly (ADP-ribose) polymerase family, member 16 | protein-coding |
| chr14_56581005_56581484 | chr14 | 56581006 | 56581484 | + | 0.895513 | intron (NM_001145978, intron 1 of 34) | intron (NM_001145978, intron 1 of 34) | 5626 | NM_001145978 | 328417 | Mm.379357 | NM_001145978 | ENSMUSG00000054509 | **Parp4** | Adprtl1\|C030027K23Rik\|E230037B21Rik\|Gm743\|PARPL\|PH5P\|VAULT3\|VPARP\|p193 | poly (ADP-ribose) polymerase family, member 4 | protein-coding |
| chr13_117018221_117018606 | chr13 | 117018222 | 117018606 | + | 1.30522 | intron (NM_001081009, intron 3 of 26) | RSINE1\|SINE\|B4 | 7102 | NM_001081009 | 52552 | Mm.274466 | NM_027272 | ENSMUSG00000021725 | **Parp8** | 2810430O08Rik\|ARTD16\|D13Ertd275e | poly (ADP-ribose) polymerase family, member 8 | protein-coding |
| chr13_116967521_116967929 | chr13 | 116967522 | 116967929 | + | 0.654075 | intron (NM_001081009, intron 3 of 26) | intron (NM_001081009, intron 3 of 26) | 57791 | NM_001081009 | 52552 | Mm.274466 | NM_027272 | ENSMUSG00000021725 | **Parp8** | 2810430O08Rik\|ARTD16\|D13Ertd275e | poly (ADP-ribose) polymerase family, member 8 | protein-coding |
| chr2_122180960_122181295 | chr2 | 122180961 | 122181295 | + | 1.01546 | intron (NM_026251, intron 1 of 17) | ID4_\|SINE\|ID | 5061 | NM_026251 | 67578 | Mm.150462 | NM_026251 | ENSMUSG00000027233 | **Patl2** | 4930424G05Rik\|Pat1a | protein associated with topoisomerase II homolog 2 (yeast) | protein-coding |
| chr2_122180153_122180397 | chr2 | 122180154 | 122180397 | + | 0.740584 | intron (NM_026251, intron 1 of 17) | B3A\|SINE\|B2 | 5914 | NM_026251 | 67578 | Mm.150462 | NM_026251 | ENSMUSG00000027233 | **Patl2** | 4930424G05Rik\|Pat1a | protein associated with topoisomerase II homolog 2 (yeast) | protein-coding |
| chr2_24440033_24440784 | chr2 | 24440034 | 24440784 | + | 1.3703 | intron (NM_011040, intron 7 of 11) | intron (NM_011040, intron 7 of 11) | 35190 | NM_011040 | 18510 | Mm.2533 | NM_011040 | ENSMUSG00000026976 | **Pax8** | Pax-8 | paired box 8 | protein-coding |
| chr2_34249482_34250295 | chr2 | 34249483 | 34250295 | + | 0.816031 | intron (NM_001290576, intron 2 of 7) | intron (NM_001290576, intron 2 of 7) | 122156 | NM_001290576 | 18516 | Mm.239941 | NM_016768 | ENSMUSG00000038718 | **Pbx3** | - | pre B cell leukemia homeobox 3 | protein-coding |
| chr2_34139184_34140230 | chr2 | 34139185 | 34140230 | + | 0.627402 | Intergenic | Intergenic | 232338 | NM_001290576 | 18516 | Mm.239941 | NM_016768 | ENSMUSG00000038718 | **Pbx3** | - | pre B cell leukemia homeobox 3 | protein-coding |
| chr8_69850646_69851436 | chr8 | 69850647 | 69851436 | + | 0.736769 | intron (NM_001024954, intron 1 of 7) | intron (NM_001024954, intron 1 of 7) | 18337 | NM_001024954 | 80720 | Mm.210823 | NM_030555 | ENSMUSG00000031860 | **Pbx4** | 2410015M21Rik\|AI429113\|AW045266 | pre B cell leukemia homeobox 4 | protein-coding |
| chr14_122713711_122714239 | chr14 | 122713712 | 122714239 | + | 1.47887 | intron (NM_144844, intron 17 of 23) | intron (NM_144844, intron 17 of 23) | 179647 | NM_144844 | 110821 | Mm.23876 | NM_144844 | ENSMUSG00000041650 | **Pcca** | C79630 | propionyl-Coenzyme A carboxylase, alpha polypeptide | protein-coding |
| chr14_122567105_122567823 | chr14 | 122567106 | 122567823 | + | 1.34909 | intron (NM_144844, intron 4 of 23) | intron (NM_144844, intron 4 of 23) | 33136 | NM_144844 | 110821 | Mm.23876 | NM_144844 | ENSMUSG00000041650 | **Pcca** | C79630 | propionyl-Coenzyme A carboxylase, alpha polypeptide | protein-coding |
| chr14_122567110_122567620 | chr14 | 122567111 | 122567620 | + | 1.28763 | intron (NM_144844, intron 4 of 23) | URR1B\|DNA\|hAT-Charlie | 33037 | NM_144844 | 110821 | Mm.23876 | NM_144844 | ENSMUSG00000041650 | **Pcca** | C79630 | propionyl-Coenzyme A carboxylase, alpha polypeptide | protein-coding |
| chr14_122700220_122701072 | chr14 | 122700221 | 122701072 | + | 1.28762 | intron (NM_144844, intron 16 of 23) | intron (NM_144844, intron 16 of 23) | 166318 | NM_144844 | 110821 | Mm.23876 | NM_144844 | ENSMUSG00000041650 | **Pcca** | C79630 | propionyl-Coenzyme A carboxylase, alpha polypeptide | protein-coding |
| chr14_122700227_122700692 | chr14 | 122700228 | 122700692 | + | 1.1938 | intron (NM_144844, intron 16 of 23) | intron (NM_144844, intron 16 of 23) | 166132 | NM_144844 | 110821 | Mm.23876 | NM_144844 | ENSMUSG00000041650 | **Pcca** | C79630 | propionyl-Coenzyme A carboxylase, alpha polypeptide | protein-coding |
| chr14_122710571_122711113 | chr14 | 122710572 | 122711113 | + | 0.831166 | intron (NM_144844, intron 17 of 23) | intron (NM_144844, intron 17 of 23) | 176514 | NM_144844 | 110821 | Mm.23876 | NM_144844 | ENSMUSG00000041650 | **Pcca** | C79630 | propionyl-Coenzyme A carboxylase, alpha polypeptide | protein-coding |
| chr14_122558010_122558722 | chr14 | 122558011 | 122558722 | + | 0.79557 | intron (NM_144844, intron 2 of 23) | intron (NM_144844, intron 2 of 23) | 24038 | NM_144844 | 110821 | Mm.23876 | NM_144844 | ENSMUSG00000041650 | **Pcca** | C79630 | propionyl-Coenzyme A carboxylase, alpha polypeptide | protein-coding |
| chr14_122710795_122711103 | chr14 | 122710796 | 122711103 | + | 0.770293 | intron (NM_144844, intron 17 of 23) | intron (NM_144844, intron 17 of 23) | 176621 | NM_144844 | 110821 | Mm.23876 | NM_144844 | ENSMUSG00000041650 | **Pcca** | C79630 | propionyl-Coenzyme A carboxylase, alpha polypeptide | protein-coding |
| chr14_122642014_122642685 | chr14 | 122642015 | 122642685 | + | 0.730286 | intron (NM_144844, intron 8 of 23) | L1M2\|LINE\|L1 | 108022 | NM_144844 | 110821 | Mm.23876 | NM_144844 | ENSMUSG00000041650 | **Pcca** | C79630 | propionyl-Coenzyme A carboxylase, alpha polypeptide | protein-coding |
| chr9_101042260_101042636 | chr9 | 101042261 | 101042636 | + | 0.704068 | Intergenic | Intergenic | -7573 | NM_025835 | 66904 | Mm.335385 | NM_025835 | ENSMUSG00000032527 | **Pccb** | 1300012P06Rik\|AI314687\|R74805 | propionyl Coenzyme A carboxylase, beta polypeptide | protein-coding |
| chr14_84444221_84444601 | chr14 | 84444222 | 84444601 | + | 0.722183 | 5' UTR (NM_001013753, exon 1 of 4) | 5' UTR (NM_001013753, exon 1 of 4) | 848 | NM_001013753 | 219228 | Mm.153643 | NM_001013753 | ENSMUSG00000035566 | **Pcdh17** | C030033F14Rik\|Gm78 | protocadherin 17 | protein-coding |
| chr5_57973982_57974420 | chr5 | 57973983 | 57974420 | + | 0.898791 | intron (NM_001122758, intron 2 of 2) | intron (NM_001122758, intron 2 of 2) | 256180 | NM_018764 | 54216 | Mm.332387 | NM_018764 | ENSMUSG00000029108 | **Pcdh7** | - | protocadherin 7 | protein-coding |
| chr18_37817849_37818476 | chr18 | 37817850 | 37818476 | + | 1.07056 | intron (NM_033582, intron 1 of 3) | intron (NM_033582, intron 1 of 3) | -1383 | NM_033583 | 93708 | Mm.247203 | NM_033583 | ENSMUSG00000023036 | **Pcdhgc5** | - | protocadherin gamma subfamily C, 5 | protein-coding |
| chr19_36364678_36365183 | chr19 | 36364679 | 36365183 | + | 1.30348 | Intergenic | Intergenic | -14136 | NM_029508 | 76073 | Mm.259180 | NM_029508 | ENSMUSG00000024805 | **Pcgf5** | 0610009F02Rik\|1110054A01Rik\|5830406C17Rik\|5830443C21Rik\|9530023M17Rik\|AI324127 | polycomb group ring finger 5 | protein-coding |

| | | | | | | | | | | | | | | | | | |
|---|---|---|---|---|---|---|---|---|---|---|---|---|---|---|---|---|---|
| chr19_36378463_36378819 | chr19 | 36378464 | 36378819 | + | 1.05117 | promoter-TSS (NM_029508) | promoter-TSS (NM_029508) | -426 | NM_029508 | 76073 | Mm.259180 | NM_029508 | ENSMUSG00000024805 | **Pcgf5** | 0610009F02Rik\|1110054A01Rik\|5830406C17Rik\|5830443C21Rik\|9530023M17Rik\|AI324127 | polycomb group ring finger 5 | protein-coding |
| chr19_36378413_36379181 | chr19 | 36378414 | 36379181 | + | 1.04208 | promoter-TSS (NM_029508) | promoter-TSS (NM_029508) | -270 | NM_029508 | 76073 | Mm.259180 | NM_029508 | ENSMUSG00000024805 | **Pcgf5** | 0610009F02Rik\|1110054A01Rik\|5830406C17Rik\|5830443C21Rik\|9530023M17Rik\|AI324127 | polycomb group ring finger 5 | protein-coding |
| chr2_173150702_173151161 | chr2 | 173150703 | 173151161 | + | 0.629837 | Intergenic | Intergenic | -2141 | NM_011044 | 18534 | Mm.266867 | NM_011044 | ENSMUSG00000027513 | **Pck1** | AI265463\|PEPCK\|Pck-1 | phosphoenolpyruvate carboxykinase 1, cytosolic | protein-coding |
| chr1_7100580_7100872 | chr1 | 7100581 | 7100872 | + | 0.689201 | intron (NM_183028, intron 1 of 5) | intron (NM_183028, intron 1 of 5) | 11806 | NM_183028 | 319263 | Mm.21539 | NM_183028 | ENSMUSG00000051285 | **Pcmtd1** | 8430411F12Rik\|A030012M09Rik | protein-L-isoaspartate (D-aspartate) O-methyltransferase domain containing 1 | protein-coding |
| chr1_7105104_7105612 | chr1 | 7105105 | 7105612 | + | 0.67992 | intron (NM_183028, intron 1 of 5) | intron (NM_183028, intron 1 of 5) | 16438 | NM_183028 | 319263 | Mm.21539 | NM_183028 | ENSMUSG00000051285 | **Pcmtd1** | 8430411F12Rik\|A030012M09Rik | protein-L-isoaspartate (D-aspartate) O-methyltransferase domain containing 1 | protein-coding |
| chr12_81861419_81861661 | chr12 | 81861420 | 81861661 | + | 0.643945 | intron (NM_018814, intron 1 of 35) | intron (NM_018814, intron 1 of 35) | 1510 | NM_018814 | 54604 | Mm.86584 | NM_018814 | ENSMUSG00000021140 | **Pcnx** | 2900024E21Rik\|3526401J03Rik\|AF096286\|AI327143\|AI413187\|Pcnxl1 | pecanex homolog (Drosophila) | protein-coding |
| chr9_95624089_95624510 | chr9 | 95624090 | 95624510 | + | 0.83706 | Intergenic | URR1B\|DNA\|hAT-Charlie | -13328 | NM_029620 | 76477 | Mm.46016 | NM_029620 | | **Pcolce2** | 2400001O18Rik\|Pcpe2 | procollagen C-endopeptidase enhancer 2 | protein-coding |
| chr18_61707081_61708090 | chr18 | 61707082 | 61708090 | + | 0.702543 | promoter-TSS (NM_172832) | promoter-TSS (NM_172832) | 49 | NM_172832 | 240334 | Mm.234457 | NM_172832 | ENSMUSG00000024579 | **Pcyox1l** | BC060677\|C630049M13 | prenylcysteine oxidase 1 like | protein-coding |
| chr1_150251751_150252204 | chr1 | 150251752 | 150252204 | + | 0.640956 | Intergenic | Intergenic | -67439 | NM_024458 | 20028 | Mm.440883 | NM_024458 | ENSMUSG00000006007 | **Pdc** | Rpr-1\|Rpr1 | phosducin | protein-coding |
| chr1_94048034_94048359 | chr1 | 94048035 | 94048359 | + | 0.906087 | intron (NM_008798, intron 1 of 4) | intron (NM_008798, intron 1 of 4) | 4356 | NM_008798 | 18566 | Mm.5024 | NM_008798 | ENSMUSG00000026285 | **Pdcd1** | Ly101\|PD-1\|Pdc1 | programmed cell death 1 | protein-coding |
| chr1_94053227_94054107 | chr1 | 94053228 | 94054107 | + | 0.839168 | Intergenic | Intergenic | -1114 | NM_008798 | 18566 | Mm.5024 | NM_008798 | ENSMUSG00000026285 | **Pdcd1** | Ly101\|PD-1\|Pdc1 | programmed cell death 1 | protein-coding |
| chr1_94055931_94056318 | chr1 | 94055932 | 94056318 | + | 0.741388 | Intergenic | MTC\|LTR\|ERVL-MaLR | -3572 | NM_008798 | 18566 | Mm.5024 | NM_008798 | ENSMUSG00000026285 | **Pdcd1** | Ly101\|PD-1\|Pdc1 | programmed cell death 1 | protein-coding |
| chr1_94051114_94051360 | chr1 | 94051115 | 94051360 | + | 0.698446 | intron (NM_008798, intron 1 of 4) | intron (NM_008798, intron 1 of 4) | 1316 | NM_008798 | 18566 | Mm.5024 | NM_008798 | ENSMUSG00000026285 | **Pdcd1** | Ly101\|PD-1\|Pdc1 | programmed cell death 1 | protein-coding |
| chr3_75491208_75491930 | chr3 | 75491209 | 75491930 | + | 1.10006 | Intergenic | MTEa\|LTR\|ERVL-MaLR | 65283 | NM_019745 | 56426 | Mm.316473 | NM_019745 | ENSMUSG00000027835 | **Pdcd10** | 2410003B13Rik\|Ccm3\|Tfa15\|Tfar15 | programmed cell death 10 | protein-coding |
| chr3_75476759_75477419 | chr3 | 75476760 | 75477419 | + | 1.06269 | Intergenic | Intergenic | 79763 | NM_019745 | 56426 | Mm.316473 | NM_019745 | ENSMUSG00000027835 | **Pdcd10** | 2410003B13Rik\|Ccm3\|Tfa15\|Tfar15 | programmed cell death 10 | protein-coding |
| chr19_53800368_53801324 | chr19 | 53800369 | 53801324 | + | 1.31806 | intron (NM_001170847, intron 1 of 13) | intron (NM_001170847, intron 1 of 13) | -91385 | NM_011050 | 18569 | Mm.1605 | NM_011050 | ENSMUSG00000024975 | **Pdcd4** | D19Ucla1\|Ma3\|Tis | programmed cell death 4 | protein-coding |
| chr19_53791724_53792843 | chr19 | 53791725 | 53792843 | + | 1.05976 | intron (NM_001170847, intron 1 of 13) | intron (NM_001170847, intron 1 of 13) | -99947 | NM_011050 | 18569 | Mm.1605 | NM_011050 | ENSMUSG00000024975 | **Pdcd4** | D19Ucla1\|Ma3\|Tis | programmed cell death 4 | protein-coding |
| chr19_53869288_53870460 | chr19 | 53869289 | 53870460 | + | 0.906693 | Intergenic | Intergenic | -22357 | NM_011050 | 18569 | Mm.1605 | NM_011050 | ENSMUSG00000024975 | **Pdcd4** | D19Ucla1\|Ma3\|Tis | programmed cell death 4 | protein-coding |
| chr17_8504467_8504740 | chr17 | 8504468 | 8504740 | + | 0.688136 | Intergenic | ORR1E\|LTR\|ERVL-MaLR | -22197 | NM_001290707 | 23984 | Mm.87161 | NM_011866 | ENSMUSG00000023868 | **Pde10a** | - | phosphodiesterase 10A | protein-coding |
| chr2_76223712_76224095 | chr2 | 76223713 | 76224095 | + | 1.14945 | intron (NM_001081033, intron 2 of 19) | intron (NM_001081033, intron 2 of 19) | 114749 | NM_001081033 | 241489 | Mm.246613 | NM_001081033 | | **Pde11a** | 6330414F14Rik\|A630086N24Rik\|Gm350\|PDE11A1 | phosphodiesterase 11A | protein-coding |
| chr2_76259301_76259658 | chr2 | 76259302 | 76259658 | + | 0.737671 | intron (NM_001081033, intron 2 of 19) | intron (NM_001081033, intron 2 of 19) | 79173 | NM_001081033 | 241489 | Mm.246613 | NM_001081033 | | **Pde11a** | 6330414F14Rik\|A630086N24Rik\|Gm350\|PDE11A1 | phosphodiesterase 11A | protein-coding |
| chr2_80014133_80014809 | chr2 | 80014134 | 80014809 | + | 0.956775 | intron (NM_001159582, intron 2 of 15) | intron (NM_001159582, intron 2 of 15) | -106072 | NM_001009978 | 18573 | Mm.40678 | NM_016744 | ENSMUSG00000059173 | **Pde1a** | AI987702\|AW125737 | phosphodiesterase 1A, calmodulin-dependent | protein-coding |
| chr2_79966505_79967143 | chr2 | 79966506 | 79967143 | + | 0.9562 | intron (NM_001159582, intron 2 of 15) | intron (NM_001159582, intron 2 of 15) | -58425 | NM_001009978 | 18573 | Mm.40678 | NM_016744 | ENSMUSG00000059173 | **Pde1a** | AI987702\|AW125737 | phosphodiesterase 1A, calmodulin-dependent | protein-coding |
| chr7_114466115_114466806 | chr7 | 114466116 | 114466806 | + | 0.782255 | intron (NM_011055, intron 1 of 15) | L1MA4\|LINE\|L1 | 51207 | NM_011055 | 18576 | Mm.430730 | NM_011055 | | **Pde3b** | 9830102A01Rik\|AI847709 | phosphodiesterase 3B, cGMP-inhibited | protein-coding |
| chr7_114466361_114466784 | chr7 | 114466362 | 114466784 | + | 0.735122 | intron (NM_011055, intron 1 of 15) | intron (NM_011055, intron 1 of 15) | 51319 | NM_011055 | 18576 | Mm.430730 | NM_011055 | | **Pde3b** | 9830102A01Rik\|AI847709 | phosphodiesterase 3B, cGMP-inhibited | protein-coding |
| chr7_114482540_114482749 | chr7 | 114482541 | 114482749 | + | 0.676678 | intron (NM_011055, intron 1 of 15) | ID4_\|SINE\|ID | 67391 | NM_011055 | 18576 | Mm.430730 | NM_011055 | | **Pde3b** | 9830102A01Rik\|AI847709 | phosphodiesterase 3B, cGMP-inhibited | protein-coding |
| chr4_102393312_102394031 | chr4 | 102393313 | 102394031 | + | 2.10064 | intron (NM_001177983, intron 1 of 5) | intron (NM_001177983, intron 1 of 5) | 138733 | NM_019840 | 18578 | Mm.20181 | NM_019840 | ENSMUSG00000028525 | **Pde4b** | Dpde4\|R74983\|dunce | phosphodiesterase 4B, cAMP specific | protein-coding |
| chr4_102393307_102394433 | chr4 | 102393308 | 102394433 | + | 1.82222 | intron (NM_001177983, intron 1 of 5) | MIRc\|SINE\|MIR | 138931 | NM_019840 | 18578 | Mm.20181 | NM_019840 | ENSMUSG00000028525 | **Pde4b** | Dpde4\|R74983\|dunce | phosphodiesterase 4B, cAMP specific | protein-coding |
| chr4_102619036_102619675 | chr4 | 102619037 | 102619675 | + | 1.78809 | Intergenic | Intergenic | 29513 | NM_001177981 | 18578 | Mm.20181 | NM_019840 | ENSMUSG00000028525 | **Pde4b** | Dpde4\|R74983\|dunce | phosphodiesterase 4B, cAMP specific | protein-coding |
| chr4_102523391_102523675 | chr4 | 102523392 | 102523675 | + | 1.40676 | intron (NM_019840, intron 5 of 14) | L1_Mur3\|LINE\|L1 | -46527 | NM_001177980 | 18578 | Mm.20181 | NM_019840 | ENSMUSG00000028525 | **Pde4b** | Dpde4\|R74983\|dunce | phosphodiesterase 4B, cAMP specific | protein-coding |
| chr4_102520870_102521707 | chr4 | 102520871 | 102521707 | + | 1.32867 | intron (NM_019840, intron 5 of 14) | MIRc\|SINE\|MIR | -48771 | NM_001177980 | 18578 | Mm.20181 | NM_019840 | ENSMUSG00000028525 | **Pde4b** | Dpde4\|R74983\|dunce | phosphodiesterase 4B, cAMP specific | protein-coding |
| chr4_102514965_102515270 | chr4 | 102514966 | 102515270 | + | 1.28175 | intron (NM_019840, intron 5 of 14) | intron (NM_019840, intron 5 of 14) | -54942 | NM_001177980 | 18578 | Mm.20181 | NM_019840 | ENSMUSG00000028525 | **Pde4b** | Dpde4\|R74983\|dunce | phosphodiesterase 4B, cAMP specific | protein-coding |
| chr4_102498940_102499330 | chr4 | 102498941 | 102499330 | + | 0.974993 | intron (NM_001177983, intron 4 of 5) | intron (NM_001177983, intron 4 of 5) | -70925 | NM_001177980 | 18578 | Mm.20181 | NM_019840 | ENSMUSG00000028525 | **Pde4b** | Dpde4\|R74983\|dunce | phosphodiesterase 4B, cAMP specific | protein-coding |
| chr4_102563158_102563759 | chr4 | 102563159 | 102563759 | + | 0.870807 | intron (NM_019840, intron 5 of 14) | MIRc\|SINE\|MIR | -6601 | NM_001177980 | 18578 | Mm.20181 | NM_019840 | ENSMUSG00000028525 | **Pde4b** | Dpde4\|R74983\|dunce | phosphodiesterase 4B, cAMP specific | protein-coding |
| chr4_102587705_102588103 | chr4 | 102587706 | 102588103 | + | 0.62912 | intron (NM_001177982, intron 1 of 9) | intron (NM_001177982, intron 1 of 9) | 346 | NM_001177982 | 18578 | Mm.20181 | NM_019840 | ENSMUSG00000028525 | **Pde4b** | Dpde4\|R74983\|dunce | phosphodiesterase 4B, cAMP specific | protein-coding |
| chr13_108603494_108603927 | chr13 | 108603495 | 108603927 | + | 1.57873 | Intergenic | MLT1N2\|LTR\|ERVL-MaLR | -50466 | NM_011056 | 238871 | Mm.434429 | NM_011056 | | **Pde4d** | 9630011N22Rik\|Dpde3 | phosphodiesterase 4D, cAMP specific | protein-coding |
| chr13_108819869_108820325 | chr13 | 108819870 | 108820325 | + | 0.780277 | intron (NM_011056, intron 1 of 16) | intron (NM_011056, intron 1 of 16) | 165920 | NM_011056 | 238871 | Mm.434429 | NM_011056 | | **Pde4d** | 9630011N22Rik\|Dpde3 | phosphodiesterase 4D, cAMP specific | protein-coding |
| chr3_122718602_122718983 | chr3 | 122718603 | 122718983 | + | 1.34916 | Intergenic | MTC\|LTR\|ERVL-MaLR | -10365 | NM_153422 | 242202 | Mm.134911 | NM_153422 | ENSMUSG00000053965 | **Pde5a** | Cgbpde\|Cn5n\|Pde5\|Pde5a1 | phosphodiesterase 5A, cGMP-specific | protein-coding |
| chr3_122721357_122721980 | chr3 | 122721358 | 122721980 | + | 0.811172 | Intergenic | L1_Rod\|LINE\|L1 | -7489 | NM_153422 | 242202 | Mm.134911 | NM_153422 | ENSMUSG00000053965 | **Pde5a** | Cgbpde\|Cn5n\|Pde5\|Pde5a1 | phosphodiesterase 5A, cGMP-specific | protein-coding |
| chr1_86580368_86580850 | chr1 | 86580369 | 86580850 | + | 1.07411 | intron (NM_008801, intron 1 of 4) | intron (NM_008801, intron 1 of 4) | 1892 | NM_008801 | 18582 | Mm.12925 | NM_008801 | ENSMUSG00000026239 | **Pde6d** | AI841218\|PrBP/delta | phosphodiesterase 6D, cGMP-specific, rod, delta | protein-coding |
| chr6_136965883_136966276 | chr6 | 136965884 | 136966276 | + | 1.36576 | Intergenic | Intergenic | 11557 | NM_023898 | 78600 | Mm.290622 | NM_023898 | | **Pde6h** | A930033D18Rik\|PDEgamma | phosphodiesterase 6H, cGMP-specific, cone, gamma | protein-coding |
| chr3_19308697_19309064 | chr3 | 19308698 | 19309064 | + | 1.17752 | intron (NM_001122759, intron 1 of 12) | intron (NM_001122759, intron 1 of 12) | 2441 | NM_001122759 | 18583 | Mm.355614 | NM_008802 | ENSMUSG00000069094 | **Pde7a** | AU015378\|AW047537 | phosphodiesterase 7A | protein-coding |
| chr3_19246948_19247639 | chr3 | 19246949 | 19247639 | + | 0.880463 | intron (NM_008802, intron 3 of 12) | B1_Mur3\|SINE\|Alu | 17714 | NM_008802 | 18583 | Mm.355614 | NM_008802 | ENSMUSG00000069094 | **Pde7a** | AU015378\|AW047537 | phosphodiesterase 7A | protein-coding |
| chr3_19372813_19373737 | chr3 | 19372814 | 19373737 | + | 0.650866 | Intergenic | Intergenic | -61953 | NM_001122759 | 18583 | Mm.355614 | NM_008802 | ENSMUSG00000069094 | **Pde7a** | AU015378\|AW047537 | phosphodiesterase 7A | protein-coding |
| chr7_81184229_81184450 | chr7 | 81184230 | 81184450 | + | 0.735758 | Intergenic | L3\|LINE\|CR1 | -29464 | NM_008803 | 18584 | Mm.371577 | NM_008803 | | **Pde8a** | AI551852\|Pde8 | phosphodiesterase 8A | protein-coding |
| chr15_80024304_80024911 | chr15 | 80024305 | 80024911 | + | 0.735605 | Intergenic | Intergenic | -9800 | NM_011057 | 18591 | Mm.144089 | NM_011057 | ENSMUSG00000000489 | **Pdgfb** | PDGF-B\|Sis | platelet derived growth factor, B polypeptide | protein-coding |
| chr2_102998857_102999669 | chr2 | 102998858 | 102999669 | + | 0.90666 | Intergenic | Intergenic | 74250 | NM_175094 | 27402 | Mm.315011 | NM_175094 | ENSMUSG00000010914 | **Pdhx** | AI481367\|E3bp\|Pdx1 | pyruvate dehydrogenase complex, component X | protein-coding |
| chr2_102999094_102999636 | chr2 | 102999095 | 102999636 | + | 0.887768 | Intergenic | Intergenic | 74148 | NM_175094 | 27402 | Mm.315011 | NM_175094 | ENSMUSG00000010914 | **Pdhx** | AI481367\|E3bp\|Pdx1 | pyruvate dehydrogenase complex, component X | protein-coding |
| chr2_103052381_103052938 | chr2 | 103052382 | 103052938 | + | 0.651133 | intron (NM_175094, intron 2 of 10) | intron (NM_175094, intron 2 of 10) | 20853 | NM_175094 | 27402 | Mm.315011 | NM_175094 | ENSMUSG00000010914 | **Pdhx** | AI481367\|E3bp\|Pdx1 | pyruvate dehydrogenase complex, component X | protein-coding |

| | | | | | | | | | | | | | | | | | |
|---|---|---|---|---|---|---|---|---|---|---|---|---|---|---|---|---|---|
| chr2_121418557_121418770 | chr2 | 121418558 | 121418770 | + | 0.790781 | intron (NM_007952, intron 1 of 12) | intron (NM_007952, intron 1 of 12) | 4762 | NM_007952 | 14827 | Mm.263177 | NM_007952 | ENSMUSG00000027248 | **Pdia3** | 58kDa\|ERp57\|ERp60\|ERp61\|Erp\|Grp58\|PDI\|PDI-Q2\|PI-PLC\|PLC[a]\|Plca | protein disulfide isomerase associated 3 | protein-coding |
| chr2_121423714_121424215 | chr2 | 121423715 | 121424215 | + | 0.683076 | intron (NM_007952, intron 2 of 12) | MTA_Mm\|LTR\|ERVL-MaLR | 10063 | NM_007952 | 14827 | Mm.263177 | NM_007952 | ENSMUSG00000027248 | **Pdia3** | 58kDa\|ERp57\|ERp60\|ERp61\|Erp\|Grp58\|PDI\|PDI-Q2\|PI-PLC\|PLC[a]\|Plca | protein disulfide isomerase associated 3 | protein-coding |
| chr16_35484468_35485412 | chr16 | 35484469 | 35485412 | + | 0.722236 | intron (NM_028295, intron 1 of 16) | intron (NM_028295, intron 1 of 16) | 5933 | NM_028295 | 72599 | Mm.71015 | NM_028295 | ENSMUSG00000022844 | **Pdia5** | 2700053F16Rik\|AU015525\|Pdir | protein disulfide isomerase associated 5 | protein-coding |
| chr6_5442697_5443225 | chr6 | 5442698 | 5443225 | + | 0.724569 | Intergenic | Intergenic | 53317 | NM_013743 | 27273 | Mm.235547 | NM_013743 | ENSMUSG00000019577 | **Pdk4** | AV005916 | pyruvate dehydrogenase kinase, isoenzyme 4 | protein-coding |
| chr19_40259591_40259870 | chr19 | 40259592 | 40259870 | + | 0.978558 | intron (NM_016861, intron 1 of 6) | intron (NM_016861, intron 1 of 6) | 11885 | NM_016861 | 54132 | Mm.5567 | NM_016861 | | **Pdlim1** | Clim1\|mClim1 | PDZ and LIM domain 1 (elfin) | protein-coding |
| chr19_40268668_40269108 | chr19 | 40268669 | 40269108 | + | 0.73315 | intron (NM_016861, intron 1 of 6) | intron (NM_016861, intron 1 of 6) | 2728 | NM_016861 | 54132 | Mm.5567 | NM_016861 | | **Pdlim1** | Clim1\|mClim1 | PDZ and LIM domain 1 (elfin) | protein-coding |
| chr3_142345548_142346035 | chr3 | 142345549 | 142346035 | + | 0.969638 | intron (NM_019808, intron 3 of 12) | intron (NM_019808, intron 3 of 12) | 46058 | NM_022554 | 56376 | Mm.117709 | NM_019808 | ENSMUSG00000028273 | **Pdlim5** | 1110001A05Rik\|AI987914\|C87059\|Enh\|Enh1\|Enh2\|Enh3\|LIM | PDZ and LIM domain 5 | protein-coding |
| chr3_142345751_142346048 | chr3 | 142345752 | 142346048 | + | 0.742642 | intron (NM_019808, intron 3 of 12) | intron (NM_019808, intron 3 of 12) | 45950 | NM_022554 | 56376 | Mm.117709 | NM_019808 | ENSMUSG00000028273 | **Pdlim5** | 1110001A05Rik\|AI987914\|C87059\|Enh\|Enh1\|Enh2\|Enh3\|LIM | PDZ and LIM domain 5 | protein-coding |
| chr3_142345263_142346054 | chr3 | 142345264 | 142346054 | + | 0.735146 | intron (NM_019808, intron 3 of 12) | intron (NM_019808, intron 3 of 12) | 46191 | NM_022554 | 56376 | Mm.117709 | NM_019808 | ENSMUSG00000028273 | **Pdlim5** | 1110001A05Rik\|AI987914\|C87059\|Enh\|Enh1\|Enh2\|Enh3\|LIM | PDZ and LIM domain 5 | protein-coding |
| chr3_142358919_142359360 | chr3 | 142358920 | 142359360 | + | 0.603376 | intron (NM_019808, intron 2 of 12) | intron (NM_019808, intron 2 of 12) | 32710 | NM_022554 | 56376 | Mm.117709 | NM_019808 | ENSMUSG00000028273 | **Pdlim5** | 1110001A05Rik\|AI987914\|C87059\|Enh\|Enh1\|Enh2\|Enh3\|LIM | PDZ and LIM domain 5 | protein-coding |
| chr4_11887676_11888292 | chr4 | 11887677 | 11888292 | + | 0.800315 | Intergenic | Intergenic | 77729 | NM_001098231 | 381511 | Mm.331489 | NM_001033453 | ENSMUSG00000049225 | **Pdp1** | Gm1024\|Ppm2c | pyruvate dehyrogenase phosphatase catalytic subunit 1 | protein-coding |
| chr17_24127604_24127958 | chr17 | 24127605 | 24127958 | + | 0.713732 | intron (NM_001286662, intron 1 of 13) | intron (NM_001286662, intron 1 of 13) | 13813 | NM_011062 | 18607 | Mm.10504 | NM_011062 | ENSMUSG00000024122 | **Pdpk1** | Pdk1 | 3-phosphoinositide dependent protein kinase 1 | protein-coding |
| chr5_65705312_65705775 | chr5 | 65705313 | 65705775 | + | 0.710504 | Intergenic | Intergenic | -7688 | NM_001081321 | 71521 | Mm.278974 | NM_027818 | ENSMUSG00000029202 | **Pds5a** | 9030416H16Rik\|E230024D05Rik\|mKIAA0648 | PDS5, regulator of cohesion maintenance, homolog A (S. cerevisiae) | protein-coding |
| chr5_65705351_65705643 | chr5 | 65705352 | 65705643 | + | 0.664097 | Intergenic | Intergenic | -7641 | NM_001081321 | 71521 | Mm.278974 | NM_027818 | ENSMUSG00000029202 | **Pds5a** | 9030416H16Rik\|E230024D05Rik\|mKIAA0648 | PDS5, regulator of cohesion maintenance, homolog A (S. cerevisiae) | protein-coding |
| chr2_22857518_22858078 | chr2 | 22857519 | 22858078 | + | 0.954297 | intron (NM_019456, intron 9 of 13) | intron (NM_019456, intron 9 of 13) | -37724 | NM_019501 | 56075 | Mm.249752 | NM_019501 | ENSMUSG00000026784 | **Pdss1** | 2610203G20Rik\|2700031G06Rik\|TPT\|TPT 1\|Tprt\|mDLP1\|mSPS1 | prenyl (solanesyl) diphosphate synthase, subunit 1 | protein-coding |
| chr10_43217237_43217834 | chr10 | 43217238 | 43217834 | + | 0.700667 | Intergenic | Intergenic | -3950 | NM_027772 | 71365 | Mm.490430 | NM_027772 | ENSMUSG00000038240 | **Pdss2** | 5430420P03Rik\|Gm60\|Plmp\|kd\|mDLP1 | prenyl (solanesyl) diphosphate synthase, subunit 2 | protein-coding |
| chr15_12588495_12589287 | chr15 | 12588496 | 12589287 | + | 1.23049 | intron (NM_001081064, intron 1 of 23) | intron (NM_001081064, intron 1 of 23) | 3665 | NM_001081064 | 68070 | Mm.423205 | NM_001081064 | ENSMUSG00000022197 | **Pdzd2** | 4930537L06Rik\|A930022H17Rik\|Gm82\|Pdzk3\|mKIAA0300 | PDZ domain containing 2 | protein-coding |
| chr15_12557979_12558564 | chr15 | 12557980 | 12558564 | + | 1.11706 | intron (NM_001081064, intron 1 of 23) | intron (NM_001081064, intron 1 of 23) | 34284 | NM_001081064 | 68070 | Mm.423205 | NM_001081064 | ENSMUSG00000022197 | **Pdzd2** | 4930537L06Rik\|A930022H17Rik\|Gm82\|Pdzk3\|mKIAA0300 | PDZ domain containing 2 | protein-coding |
| chr15_12616310_12616853 | chr15 | 12616311 | 12616853 | + | 0.715357 | Intergenic | Intergenic | -24026 | NM_001081064 | 68070 | Mm.423205 | NM_001081064 | ENSMUSG00000022197 | **Pdzd2** | 4930537L06Rik\|A930022H17Rik\|Gm82\|Pdzk3\|mKIAA0300 | PDZ domain containing 2 | protein-coding |
| chr15_12586014_12586448 | chr15 | 12586015 | 12586448 | + | 0.6572 | intron (NM_001081064, intron 1 of 23) | intron (NM_001081064, intron 1 of 23) | 6325 | NM_001081064 | 68070 | Mm.423205 | NM_001081064 | ENSMUSG00000022197 | **Pdzd2** | 4930537L06Rik\|A930022H17Rik\|Gm82\|Pdzk3\|mKIAA0300 | PDZ domain containing 2 | protein-coding |
| chr4_115088588_115089353 | chr4 | 115088589 | 115089353 | + | 1.17933 | exon (NM_001164557, exon 1 of 4) | exon (NM_001164557, exon 1 of 4) | 263 | NM_001164557 | 67182 | Mm.30181 | NM_026018 | ENSMUSG00000028716 | **Pdzk1ip1** | 0610007F13Rik\|2700030M23Rik\|AU046042\|Map17 | PDZK1 interacting protein 1 | protein-coding |
| chr9_56378552_56379141 | chr9 | 56378553 | 56379141 | + | 1.05446 | intron (NM_172924, intron 1 of 6) | L1_Mur3\|LINE\|L1 | 39203 | NM_172924 | 244895 | Mm.490331 | NM_172924 | ENSMUSG00000074305 | **Peak1** | 1110049L02Rik\|9530046P14\|AA690169\|BE132841\|C230081A13Rik\|SGK269\|mKIAA2002 | pseudopodium-enriched atypical kinase 1 | protein-coding |
| chr9_56370693_56371221 | chr9 | 56370694 | 56371221 | + | 1.00405 | intron (NM_172924, intron 2 of 6) | intron (NM_172924, intron 2 of 6) | 47093 | NM_172924 | 244895 | Mm.490331 | NM_172924 | ENSMUSG00000074305 | **Peak1** | 1110049L02Rik\|9530046P14\|AA690169\|BE132841\|C230081A13Rik\|SGK269\|mKIAA2002 | pseudopodium-enriched atypical kinase 1 | protein-coding |
| chr11_106666282_106667130 | chr11 | 106666283 | 106667130 | + | 0.953505 | intron (NM_001032378, intron 13 of 13) | intron (NM_001032378, intron 13 of 13) | 48575 | NM_001032378 | 18613 | Mm.343951 | NM_008816 | ENSMUSG00000020717 | **Pecam1** | C85791\|Cd31\|PECAM-1\|Pecam | platelet/endothelial cell adhesion molecule 1 | protein-coding |
| chr11_21108078_21108631 | chr11 | 21108079 | 21108631 | + | 1.00094 | intron (NM_023324, intron 1 of 6) | Lx8\|LINE\|L1 | 17031 | NM_023324 | 67245 | Mm.28957 | NM_023324 | ENSMUSG00000020134 | **Peli1** | 2810468L03Rik\|A930031K15Rik\|AA409794\|AI586297\|D11Ertd676e | pellino 1 | protein-coding |
| chr11_21102294_21103008 | chr11 | 21102295 | 21103008 | + | 0.905165 | intron (NM_023324, intron 1 of 6) | intron (NM_023324, intron 1 of 6) | 11327 | NM_023324 | 67245 | Mm.28957 | NM_023324 | ENSMUSG00000020134 | **Peli1** | 2810468L03Rik\|A930031K15Rik\|AA409794\|AI586297\|D11Ertd676e | pellino 1 | protein-coding |
| chr11_21082745_21083024 | chr11 | 21082746 | 21083024 | + | 0.776518 | Intergenic | Intergenic | -8439 | NM_023324 | 67245 | Mm.28957 | NM_023324 | ENSMUSG00000020134 | **Peli1** | 2810468L03Rik\|A930031K15Rik\|AA409794\|AI586297\|D11Ertd676e | pellino 1 | protein-coding |
| chr11_21082714_21083122 | chr11 | 21082715 | 21083122 | + | 0.720083 | Intergenic | Intergenic | -8406 | NM_023324 | 67245 | Mm.28957 | NM_023324 | ENSMUSG00000020134 | **Peli1** | 2810468L03Rik\|A930031K15Rik\|AA409794\|AI586297\|D11Ertd676e | pellino 1 | protein-coding |
| chr11_21110728_21111090 | chr11 | 21110729 | 21111090 | + | 0.654269 | intron (NM_023324, intron 1 of 6) | intron (NM_023324, intron 1 of 6) | 19585 | NM_023324 | 67245 | Mm.28957 | NM_023324 | ENSMUSG00000020134 | **Peli1** | 2810468L03Rik\|A930031K15Rik\|AA409794\|AI586297\|D11Ertd676e | pellino 1 | protein-coding |
| chr11_70385453_70385891 | chr11 | 70385454 | 70385891 | + | 0.686474 | Intergenic | Intergenic | 24359 | NM_029231 | 75273 | Mm.340601 | NM_029231 | ENSMUSG00000018921 | **Pelp1** | 4930563C04Rik\|MNAR | proline, glutamic acid and leucine rich protein 1 | protein-coding |
| chr11_69102403_69102650 | chr11 | 69102404 | 69102650 | + | 0.840559 | intron (NM_011065, intron 8 of 22) | intron (NM_011065, intron 8 of 22) | 2369 | NM_001159367 | 18626 | Mm.7373 | NM_011065 | ENSMUSG00000020893 | **Per1** | Per\|m-rigui\|mPer1 | period circadian clock 1 | protein-coding |
| chr10_18860636_18861415 | chr10 | 18860637 | 18861415 | + | 1.75353 | Intergenic | Intergenic | 15955 | NM_022032 | 64058 | Mm.28209 | NM_022032 | ENSMUSG00000019851 | **Perp** | 1110017A08Rik\|KCP1\|KRTCAP1\|PIGPC1\|THW | PERP, TP53 apoptosis effector | protein-coding |
| chr10_18868186_18868547 | chr10 | 18868187 | 18868547 | + | 0.882632 | Intergenic | Tigger1\|DNA\|TcMar-Tigger | 23296 | NM_022032 | 64058 | Mm.28209 | NM_022032 | ENSMUSG00000019851 | **Perp** | 1110017A08Rik\|KCP1\|KRTCAP1\|PIGPC1\|THW | PERP, TP53 apoptosis effector | protein-coding |
| chr3_85629881_85630291 | chr3 | 85629882 | 85630291 | + | 0.790714 | intron (NM_144896, intron 9 of 12) | intron (NM_144896, intron 9 of 12) | 55957 | NM_144896 | 229487 | Mm.52275 | NM_144896 | ENSMUSG00000028085 | **Pet112** | 9430026F02Rik\|Pet112l | PET112 homolog (S. cerevisiae) | protein-coding |
| chr4_149062977_149063764 | chr4 | 149062978 | 149063764 | + | 1.05904 | intron (NM_019781, intron 2 of 8) | intron (NM_019781, intron 2 of 8) | 36441 | NM_019781 | 56273 | Mm.184172 | NM_019781 | ENSMUSG00000028975 | **Pex14** | Pex14p\|R75137 | peroxisomal biogenesis factor 14 | protein-coding |
| chr10_13675070_13675690 | chr10 | 13675071 | 13675690 | + | 1.0767 | intron (NM_025446, intron 4 of 5) | intron (NM_025446, intron 4 of 5) | -122238 | NM_001164195 | 56535 | Mm.282711 | NM_019961 | ENSMUSG00000019809 | **Pex3** | 1700014F15Rik\|2810027F19Rik\|2900010N04Rik | peroxisomal biogenesis factor 3 | protein-coding |
| chr18_36415146_36415774 | chr18 | 36415147 | 36415774 | + | 1.36195 | intron (NM_026027, intron 3 of 3) | intron (NM_026027, intron 3 of 3) | 39035 | NM_026027 | 67199 | Mm.30184 | NM_026027 | ENSMUSG00000024346 | **Pfdn1** | 2700086I23Rik\|AA408327\|AU044714 | prefoldin 1 | protein-coding |
| chr18_36415225_36415420 | chr18 | 36415226 | 36415420 | + | 1.35571 | intron (NM_026027, intron 3 of 3) | B3A\|SINE\|B2 | 39172 | NM_026027 | 67199 | Mm.30184 | NM_026027 | ENSMUSG00000024346 | **Pfdn1** | 2700086I23Rik\|AA408327\|AU044714 | prefoldin 1 | protein-coding |
| chr18_36415475_36415766 | chr18 | 36415476 | 36415766 | + | 1.04592 | intron (NM_026027, intron 3 of 3) | intron (NM_026027, intron 3 of 3) | 38874 | NM_026027 | 67199 | Mm.30184 | NM_026027 | ENSMUSG00000024346 | **Pfdn1** | 2700086I23Rik\|AA408327\|AU044714 | prefoldin 1 | protein-coding |
| chr2_11530261_11530478 | chr2 | 11530262 | 11530478 | + | 1.23389 | intron (NM_001177758, intron 1 of 14) | intron (NM_001177758, intron 1 of 14) | 23559 | NM_001177758 | 170768 | Mm.19669 | NM_133232 | ENSMUSG00000026773 | **Pfkfb3** | E330010H22Rik\|iPFK-2\|uPFK-2 | 6-phosphofructo-2-kinase/fructose-2,6-biphosphatase 3 | protein-coding |
| chr2_11530079_11530540 | chr2 | 11530080 | 11530540 | + | 1.16582 | intron (NM_001177758, intron 1 of 14) | intron (NM_001177758, intron 1 of 14) | 23619 | NM_001177758 | 170768 | Mm.19669 | NM_133232 | ENSMUSG00000026773 | **Pfkfb3** | E330010H22Rik\|iPFK-2\|uPFK-2 | 6-phosphofructo-2-kinase/fructose-2,6-biphosphatase 3 | protein-coding |
| chr2_11530034_11530717 | chr2 | 11530035 | 11530717 | + | 1.11934 | intron (NM_001177758, intron 1 of 14) | intron (NM_001177758, intron 1 of 14) | 23553 | NM_001177758 | 170768 | Mm.19669 | NM_133232 | ENSMUSG00000026773 | **Pfkfb3** | E330010H22Rik\|iPFK-2\|uPFK-2 | 6-phosphofructo-2-kinase/fructose-2,6-biphosphatase 3 | protein-coding |

| | | | | | | | | | | | | | | | | | |
|---|---|---|---|---|---|---|---|---|---|---|---|---|---|---|---|---|---|
| chr2_11540381_11540988 | chr2 | 11540382 | 11540988 | + | 1.04999 | intron (NM_001177758, intron 1 of 14) | intron (NM_001177758, intron 1 of 14) | 13244 | NM_001177758 | 170768 | Mm.19669 | NM_133232 | ENSMUSG00000026773 | **Pfkfb3** | E330010H22Rik\|iPFK-2\|uPFK-2 | 6-phosphofructo-2-kinase/fructose-2,6-biphosphatase 3 | protein-coding |
| chr15_98066424_98067122 | chr15 | 98066425 | 98067122 | + | 1.4974 | exon (NM_144851, exon 7 of 18) | exon (NM_144851, exon 7 of 18) | -25816 | NM_001163488 | 18642 | Mm.272582 | NM_021514 | ENSMUSG00000033065 | **Pfkm** | AI131669\|ATP-PFK\|PFK-A\|PFK-M\|Pfk-4\|Pfk4\|Pfka\|Pfkx | phosphofructokinase, muscle | protein-coding |
| chr13_6618526_6619103 | chr13 | 6618527 | 6619103 | + | 1.60025 | intron (NM_001291071, intron 7 of 21) | intron (NM_001291071, intron 7 of 21) | 29956 | NM_019703 | 56421 | Mm.273874 | NM_019703 | ENSMUSG00000021196 | **Pfkp** | 1200015H23Rik\|9330125N24Rik\|ATP-PFK\|PFK-C\|PFK-P | phosphofructokinase, platelet | protein-coding |
| chr13_6652528_6652937 | chr13 | 6652529 | 6652937 | + | 0.984232 | Intergenic | L1MA9\|LINE\|L1 | -3962 | NM_001291071 | 56421 | Mm.273874 | NM_019703 | ENSMUSG00000021196 | **Pfkp** | 1200015H23Rik\|9330125N24Rik\|ATP-PFK\|PFK-C\|PFK-P | phosphofructokinase, platelet | protein-coding |
| chr13_6636640_6636854 | chr13 | 6636641 | 6636854 | + | 0.962537 | intron (NM_001291071, intron 1 of 21) | intron (NM_001291071, intron 1 of 21) | 12024 | NM_019703 | 56421 | Mm.273874 | NM_019703 | ENSMUSG00000021196 | **Pfkp** | 1200015H23Rik\|9330125N24Rik\|ATP-PFK\|PFK-C\|PFK-P | phosphofructokinase, platelet | protein-coding |
| chr13_6634743_6635007 | chr13 | 6634744 | 6635007 | + | 0.88182 | intron (NM_001291071, intron 2 of 21) | B4A\|SINE\|B4 | 13896 | NM_019703 | 56421 | Mm.273874 | NM_019703 | ENSMUSG00000021196 | **Pfkp** | 1200015H23Rik\|9330125N24Rik\|ATP-PFK\|PFK-C\|PFK-P | phosphofructokinase, platelet | protein-coding |
| chr13_6626438_6627179 | chr13 | 6626439 | 6627179 | + | 0.727645 | intron (NM_001291071, intron 2 of 21) | intron (NM_001291071, intron 2 of 21) | 21962 | NM_019703 | 56421 | Mm.273874 | NM_019703 | ENSMUSG00000021196 | **Pfkp** | 1200015H23Rik\|9330125N24Rik\|ATP-PFK\|PFK-C\|PFK-P | phosphofructokinase, platelet | protein-coding |
| chr13_6635230_6635485 | chr13 | 6635231 | 6635485 | + | 0.653352 | intron (NM_001291071, intron 2 of 21) | intron (NM_001291071, intron 2 of 21) | 13413 | NM_019703 | 56421 | Mm.273874 | NM_019703 | ENSMUSG00000021196 | **Pfkp** | 1200015H23Rik\|9330125N24Rik\|ATP-PFK\|PFK-C\|PFK-P | phosphofructokinase, platelet | protein-coding |
| chr11_70651456_70651663 | chr11 | 70651457 | 70651663 | + | 1.05987 | TTS (NM_027445) | TTS (NM_027445) | 3090 | NM_011072 | 18643 | Mm.2647 | NM_011072 | ENSMUSG00000018293 | **Pfn1** | Pfn | profilin 1 | protein-coding |
| chr1_54520094_54520575 | chr1 | 54520095 | 54520575 | + | 1.22289 | intron (NM_001163314, intron 13 of 26) | intron (NM_001163314, intron 13 of 26) | 37349 | NM_001163314 | 241062 | Mm.103539 | NM_175508 | ENSMUSG00000073678 | **Pgap1** | 5033403E17Rik\|9030223K07Rik\|A530084K22\|D230012E17Rik\|oto | post-GPI attachment to proteins 1 | protein-coding |
| chr1_54606039_54606502 | chr1 | 54606040 | 54606502 | + | 1.11552 | Intergenic | MERV1_I-int\|LTR\|ERV1 | -48587 | NM_001163314 | 241062 | Mm.103539 | NM_175508 | ENSMUSG00000073678 | **Pgap1** | 5033403E17Rik\|9030223K07Rik\|A530084K22\|D230012E17Rik\|oto | post-GPI attachment to proteins 1 | protein-coding |
| chr1_54585939_54586535 | chr1 | 54585940 | 54586535 | + | 0.821614 | Intergenic | Intergenic | -28553 | NM_001163314 | 241062 | Mm.103539 | NM_175508 | ENSMUSG00000073678 | **Pgap1** | 5033403E17Rik\|9030223K07Rik\|A530084K22\|D230012E17Rik\|oto | post-GPI attachment to proteins 1 | protein-coding |
| chr7_102217808_102218134 | chr7 | 102217809 | 102218134 | + | 1.04905 | intron (NM_001291358, intron 1 of 6) | B3\|SINE\|B2 | -5057 | NR_111938 | 233575 | Mm.302013 | NM_145583 | ENSMUSG00000030990 | **Pgap2** | 1810006G21Rik\|C76649\|Frag1 | post-GPI attachment to proteins 2 | protein-coding |
| chr4_149216822_149217140 | chr4 | 149216823 | 149217140 | + | 0.79663 | intron (NM_001290995, intron 27 of 48) | intron (NM_001290995, intron 27 of 48) | -50274 | NM_001081274 | 110208 | Mm.252080 | NM_025801 | ENSMUSG00000028961 | **Pgd** | 0610042A05Rik\|AU019875\|C78335 | phosphogluconate dehydrogenase | protein-coding |
| chr18_46285781_46286822 | chr18 | 46285782 | 46286822 | + | 0.916182 | intron (NM_001160399, intron 7 of 9) | intron (NM_001160399, intron 7 of 9) | -5452 | NM_172627 | 225467 | Mm.262096 | NM_172627 | ENSMUSG00000024477 | **Pggt1b** | 2010207C17Rik\|2610100E13\|AI451237\|AI551093\|BGG1\|GGT1 | protein geranylgeranyltransferase type I, beta subunit | protein-coding |
| chrX_106165058_106165428 | chrX | 106165059 | 106165428 | + | 0.87951 | Intergenic | B3\|SINE\|B2 | -21857 | NM_008828 | 18655 | Mm.316355 | NM_008828 | ENSMUSG00000062070 | **Pgk1** | Pgk-1 | phosphoglycerate kinase 1 | protein-coding |
| chr5_64085373_64085852 | chr5 | 64085374 | 64085852 | + | 0.750897 | Intergenic | Intergenic | -7337 | NM_025700 | 66681 | Mm.2325 | NM_025700 | ENSMUSG00000029171 | **Pgm1** | 3230402E02Rik\|Pgm-1\|Pgm2 | phosphoglucomutase 1 | protein-coding |
| chr4_99944608_99945550 | chr4 | 99944609 | 99945550 | + | 1.11087 | intron (NM_028132, intron 1 of 10) | intron (NM_028132, intron 1 of 10) | 15628 | NM_028132 | 72157 | Mm.217764 | NM_028132 | ENSMUSG00000025791 | **Pgm2** | 2610020G18Rik\|AA407108\|AI098105\|Pgm-2\|Pgm1 | phosphoglucomutase 2 | protein-coding |
| chr4_99985706_99986582 | chr4 | 99985707 | 99986582 | + | 0.99304 | intron (NM_028132, intron 10 of 10) | B2_Mm2\|SINE\|B2 | 56693 | NM_028132 | 72157 | Mm.217764 | NM_028132 | ENSMUSG00000025791 | **Pgm2** | 2610020G18Rik\|AA407108\|AI098105\|Pgm-2\|Pgm1 | phosphoglucomutase 2 | protein-coding |
| chr4_99942035_99942598 | chr4 | 99942036 | 99942598 | + | 0.871823 | intron (NM_028132, intron 1 of 10) | intron (NM_028132, intron 1 of 10) | 12866 | NM_028132 | 72157 | Mm.217764 | NM_028132 | ENSMUSG00000025791 | **Pgm2** | 2610020G18Rik\|AA407108\|AI098105\|Pgm-2\|Pgm1 | phosphoglucomutase 2 | protein-coding |
| chrX_36609015_36609658 | chrX | 36609016 | 36609658 | + | 1.255 | 3' UTR (NM_001081956, exon 7 of 7) | 3' UTR (NM_001081956, exon 7 of 7) | 11112 | NM_016783 | 53328 | Mm.9052 | NM_016783 | ENSMUSG00000006373 | **Pgrmc1** | AA415812\|HPR6.6\|PPMR\|Vema | progesterone receptor membrane component 1 | protein-coding |
| chr3_41228625_41228830 | chr3 | 41228626 | 41228830 | + | 1.20355 | Intergenic | Intergenic | -145682 | NM_027558 | 70804 | Mm.40321 | NM_027558 | ENSMUSG00000049940 | **Pgrmc2** | 4631434O19Rik\|5730409G06Rik\|DG6\|PMBP | progesterone receptor membrane component 2 | protein-coding |
| chr3_41228509_41228950 | chr3 | 41228510 | 41228950 | + | 1.16486 | Intergenic | Intergenic | -145684 | NM_027558 | 70804 | Mm.40321 | NM_027558 | ENSMUSG00000049940 | **Pgrmc2** | 4631434O19Rik\|5730409G06Rik\|DG6\|PMBP | progesterone receptor membrane component 2 | protein-coding |
| chr11_117988056_117988297 | chr11 | 117988057 | 117988297 | + | 0.769043 | intron (NM_133757, intron 1 of 9) | intron (NM_133757, intron 1 of 9) | 1320 | NM_133757 | 74451 | Mm.28864 | NM_133757 | ENSMUSG00000017715 | **Pgs1** | 2610019F11Rik\|4933424M23Rik\|SAF | phosphatidylglycerophosphate synthase 1 | protein-coding |
| chr11_117989814_117990449 | chr11 | 117989815 | 117990449 | + | 0.733114 | intron (NM_133757, intron 1 of 9) | intron (NM_133757, intron 1 of 9) | 3275 | NM_133757 | 74451 | Mm.28864 | NM_133757 | ENSMUSG00000017715 | **Pgs1** | 2610019F11Rik\|4933424M23Rik\|SAF | phosphatidylglycerophosphate synthase 1 | protein-coding |
| chr11_117989998_117990325 | chr11 | 117989999 | 117990325 | + | 0.714167 | intron (NM_133757, intron 1 of 9) | intron (NM_133757, intron 1 of 9) | 3305 | NM_133757 | 74451 | Mm.28864 | NM_133757 | ENSMUSG00000017715 | **Pgs1** | 2610019F11Rik\|4933424M23Rik\|SAF | phosphatidylglycerophosphate synthase 1 | protein-coding |
| chr13_43085240_43086071 | chr13 | 43085241 | 43086071 | + | 0.875015 | intron (NM_001005740, intron 6 of 12) | intron (NM_001005740, intron 6 of 12) | -37183 | NM_001005748 | 218194 | Mm.160124 | NM_198419 | ENSMUSG00000054728 | **Phactr1** | 9630030F18Rik\|Rpel1 | phosphatase and actin regulator 1 | protein-coding |
| chr10_13438032_13438852 | chr10 | 13438033 | 13438852 | + | 1.33755 | intron (NM_001195065, intron 1 of 12) | intron (NM_001195065, intron 1 of 12) | 35954 | NM_001195065 | 215789 | Mm.283359 | NM_001033257 | ENSMUSG00000062866 | **Phactr2** | AV158170\|BC012871 | phosphatase and actin regulator 2 | protein-coding |
| chr10_13424241_13424852 | chr10 | 13424242 | 13424852 | + | 1.29489 | intron (NM_001195065, intron 1 of 12) | intron (NM_001195065, intron 1 of 12) | -35577 | NM_001195066 | 215789 | Mm.283359 | NM_001033257 | ENSMUSG00000062866 | **Phactr2** | AV158170\|BC012871 | phosphatase and actin regulator 2 | protein-coding |
| chr10_13466897_13467383 | chr10 | 13466898 | 13467383 | + | 0.943115 | intron (NM_001195065, intron 1 of 12) | intron (NM_001195065, intron 1 of 12) | 7256 | NM_001195065 | 215789 | Mm.283359 | NM_001033257 | ENSMUSG00000062866 | **Phactr2** | AV158170\|BC012871 | phosphatase and actin regulator 2 | protein-coding |
| chr10_13468773_13469272 | chr10 | 13468774 | 13469272 | + | 0.854204 | intron (NM_001195065, intron 1 of 12) | intron (NM_001195065, intron 1 of 12) | 5373 | NM_001195065 | 215789 | Mm.283359 | NM_001033257 | ENSMUSG00000062866 | **Phactr2** | AV158170\|BC012871 | phosphatase and actin regulator 2 | protein-coding |
| chr4_132409676_132410219 | chr4 | 132409677 | 132410219 | + | 0.824136 | intron (NM_175306, intron 2 of 13) | B1_Mur2\|SINE\|Alu | 12498 | NM_001161797 | 100169 | Mm.158361 | NM_175306 | ENSMUSG00000066043 | **Phactr4** | 3110001B12Rik\|AI527228\|AW495572\|C330013F19Rik\|N28169\|mKIAA4120 | phosphatase and actin regulator 4 | protein-coding |
| chr11_95668322_95668608 | chr11 | 95668323 | 95668608 | + | 0.863317 | intron (NM_008831, intron 2 of 6) | intron (NM_008831, intron 2 of 6) | 1508 | NM_008831 | 18673 | Mm.263862 | NM_008831 | ENSMUSG00000038845 | **Phb** | Bap32 | prohibitin | protein-coding |
| chr11_95668248_95668634 | chr11 | 95668249 | 95668634 | + | 0.77644 | intron (NM_008831, intron 2 of 6) | intron (NM_008831, intron 2 of 6) | 1484 | NM_008831 | 18673 | Mm.263862 | NM_008831 | ENSMUSG00000038845 | **Phb** | Bap32 | prohibitin | protein-coding |
| chr4_128733562_128734010 | chr4 | 128733563 | 128734010 | + | 0.882923 | intron (NM_001195083, intron 1 of 6) | intron (NM_001195083, intron 1 of 6) | 6215 | NM_001195083 | 54383 | Mm.473738 | NM_018774 | ENSMUSG00000028796 | **Phc2** | A3galt2\|AA415044\|D130050K19Rik\|D4Ertd810e\|Edr2\|Mph2\|p36 | polyhomeotic-like 2 (Drosophila) | protein-coding |
| chr4_128733621_128733845 | chr4 | 128733622 | 128733845 | + | 0.79378 | intron (NM_001195083, intron 1 of 6) | intron (NM_001195083, intron 1 of 6) | 6162 | NM_001195083 | 54383 | Mm.473738 | NM_018774 | ENSMUSG00000028796 | **Phc2** | A3galt2\|AA415044\|D130050K19Rik\|D4Ertd810e\|Edr2\|Mph2\|p36 | polyhomeotic-like 2 (Drosophila) | protein-coding |
| chr4_128713078_128713346 | chr4 | 128713079 | 128713346 | + | 0.680204 | intron (NM_001195130, intron 7 of 14) | intron (NM_001195130, intron 7 of 14) | -14359 | NM_001195083 | 54383 | Mm.473738 | NM_018774 | ENSMUSG00000028796 | **Phc2** | A3galt2\|AA415044\|D130050K19Rik\|D4Ertd810e\|Edr2\|Mph2\|p36 | polyhomeotic-like 2 (Drosophila) | protein-coding |
| chr3_30949448_30949816 | chr3 | 30949449 | 30949816 | + | 1.38621 | intron (NM_153421, intron 5 of 14) | intron (NM_153421, intron 5 of 14) | 19783 | NM_001165954 | 241915 | Mm.233173 | NM_153421 | ENSMUSG00000037652 | **Phc3** | E030046K01Rik\|Edr3\|Hph3 | polyhomeotic-like 3 (Drosophila) | protein-coding |
| chr3_30952305_30953034 | chr3 | 30952306 | 30953034 | + | 1.3747 | intron (NM_153421, intron 4 of 14) | intron (NM_153421, intron 4 of 14) | 16745 | NM_001165954 | 241915 | Mm.233173 | NM_153421 | ENSMUSG00000037652 | **Phc3** | E030046K01Rik\|Edr3\|Hph3 | polyhomeotic-like 3 (Drosophila) | protein-coding |
| chr3_30951715_30952004 | chr3 | 30951716 | 30952004 | + | 1.06972 | intron (NM_153421, intron 4 of 14) | ID_B1\|SINE\|B4 | 17555 | NM_001165954 | 241915 | Mm.233173 | NM_153421 | ENSMUSG00000037652 | **Phc3** | E030046K01Rik\|Edr3\|Hph3 | polyhomeotic-like 3 (Drosophila) | protein-coding |
| chr3_30971606_30971986 | chr3 | 30971607 | 30971986 | + | 0.991181 | Intergenic | MTC-int\|LTR\|ERVL-MaLR | -2381 | NM_001165956 | 241915 | Mm.233173 | NM_153421 | ENSMUSG00000037652 | **Phc3** | E030046K01Rik\|Edr3\|Hph3 | polyhomeotic-like 3 (Drosophila) | protein-coding |
| chr3_30929809_30930266 | chr3 | 30929810 | 30930266 | + | 0.885284 | exon (NM_001165955, exon 9 of 14) | exon (NM_001165955, exon 9 of 14) | 39377 | NM_001165954 | 241915 | Mm.233173 | NM_153421 | ENSMUSG00000037652 | **Phc3** | E030046K01Rik\|Edr3\|Hph3 | polyhomeotic-like 3 (Drosophila) | protein-coding |

| | | | | | | | | | | | | | | | | | |
|---|---|---|---|---|---|---|---|---|---|---|---|---|---|---|---|---|---|
| chr3_30946695_30946915 | chr3 | 30946696 | 30946915 | + | 0.769128 | intron (NM_001165955, intron 5 of 13) | B1F\|SINE\|Alu | 22610 | NM_001165954 | 241915 | Mm.233173 | NM_153421 | ENSMUSG00000037652 | **Phc3** | E030046K01Rik\|Edr3\|Hph3 | polyhomeotic-like 3 (Drosophila) | protein-coding |
| chr17_14960031_14960341 | chr17 | 14960032 | 14960341 | + | 0.633769 | intron (NM_024250, intron 1 of 11) | intron (NM_024250, intron 1 of 11) | 1074 | NM_024250 | 72057 | Mm.440085 | NM_024250 | ENSMUSG00000023883 | **Phf10** | 1810055P05Rik\|AV024533 | PHD finger protein 10 | protein-coding |
| chr13_48879953_48880204 | chr13 | 48879954 | 48880204 | + | 0.755014 | 3' UTR (NM_001033268, exon 18 of 18) | 3' UTR (NM_001033268, exon 18 of 18) | -9194 | NM_011078 | 18676 | Mm.486213 | NM_011078 | ENSMUSG00000038025 | **Phf2** | GRC5 | PHD finger protein 2 | protein-coding |
| chr2_156208885_156209843 | chr2 | 156208886 | 156209843 | + | 0.819793 | intron (NM_172674, intron 1 of 17) | MIRb\|SINE\|MIR | 12717 | NM_172674 | 228829 | Mm.427078 | NM_172674 | ENSMUSG00000038116 | **Phf20** | 6820402O20Rik | PHD finger protein 20 | protein-coding |
| chr2_156208886_156209502 | chr2 | 156208887 | 156209502 | + | 0.814466 | intron (NM_172674, intron 1 of 17) | intron (NM_172674, intron 1 of 17) | 12547 | NM_172674 | 228829 | Mm.427078 | NM_172674 | ENSMUSG00000038116 | **Phf20** | 6820402O20Rik | PHD finger protein 20 | protein-coding |
| chr2_156222045_156222517 | chr2 | 156222046 | 156222517 | + | 0.757605 | intron (NM_172674, intron 1 of 17) | B3\|SINE\|B2 | 25634 | NM_172674 | 228829 | Mm.427078 | NM_172674 | ENSMUSG00000038116 | **Phf20** | 6820402O20Rik | PHD finger protein 20 | protein-coding |
| chr2_92241233_92241679 | chr2 | 92241234 | 92241679 | + | 0.98951 | intron (NM_001109691, intron 7 of 18) | intron (NM_001109691, intron 7 of 18) | 19894 | NM_001109690 | 192285 | Mm.330408 | NM_138755 | ENSMUSG00000058318 | **Phf21a** | 80kDa\|Bhc\|Bhc80\|D030065N23Rik\|PFTF1 | PHD finger protein 21A | protein-coding |
| chr2_92240876_92241690 | chr2 | 92240877 | 92241690 | + | 0.827926 | intron (NM_001109691, intron 7 of 18) | intron (NM_001109691, intron 7 of 18) | 19721 | NM_001109690 | 192285 | Mm.330408 | NM_138755 | ENSMUSG00000058318 | **Phf21a** | 80kDa\|Bhc\|Bhc80\|D030065N23Rik\|PFTF1 | PHD finger protein 21A | protein-coding |
| chr2_92266241_92266596 | chr2 | 92266242 | 92266596 | + | 0.705265 | intron (NM_001109691, intron 7 of 18) | intron (NM_001109691, intron 7 of 18) | 44857 | NM_001109690 | 192285 | Mm.330408 | NM_138755 | ENSMUSG00000058318 | **Phf21a** | 80kDa\|Bhc\|Bhc80\|D030065N23Rik\|PFTF1 | PHD finger protein 21A | protein-coding |
| chr1_30792925_30793378 | chr1 | 30792926 | 30793378 | + | 0.796747 | Intergenic | Intergenic | 70104 | NM_001081080 | 213109 | Mm.194486 | NM_025871 | ENSMUSG00000048874 | **Phf3** | 2310061N19Rik\|mKIAA0244 | PHD finger protein 3 | protein-coding |
| chr1_30812107_30812509 | chr1 | 30812108 | 30812509 | + | 0.683259 | intron (NM_001081080, intron 10 of 14) | intron (NM_001081080, intron 10 of 14) | 50948 | NM_001081080 | 213109 | Mm.194486 | NM_025871 | ENSMUSG00000048874 | **Phf3** | 2310061N19Rik\|mKIAA0244 | PHD finger protein 3 | protein-coding |
| chrX_52928165_52928546 | chrX | 52928166 | 52928546 | + | 1.30614 | intron (NM_027642, intron 3 of 10) | B2_Mm2\|SINE\|B2 | 16142 | NM_001290546 | 70998 | Mm.26870 | NM_027642 | ENSMUSG00000025626 | **Phf6** | 2700007B13Rik\|4931428F02Rik\|mKIAA1823 | PHD finger protein 6 | protein-coding |
| chrX_52928165_52928743 | chrX | 52928166 | 52928743 | + | 1.18442 | intron (NM_027642, intron 3 of 10) | PB1D7\|SINE\|Alu | 16240 | NM_001290546 | 70998 | Mm.26870 | NM_027642 | ENSMUSG00000025626 | **Phf6** | 2700007B13Rik\|4931428F02Rik\|mKIAA1823 | PHD finger protein 6 | protein-coding |
| chrX_52929816_52930120 | chrX | 52929817 | 52930120 | + | 0.895268 | intron (NM_027642, intron 3 of 10) | intron (NM_027642, intron 3 of 10) | 17754 | NM_001290546 | 70998 | Mm.26870 | NM_027642 | ENSMUSG00000025626 | **Phf6** | 2700007B13Rik\|4931428F02Rik\|mKIAA1823 | PHD finger protein 6 | protein-coding |
| chr3_98356701_98357532 | chr3 | 98356702 | 98357532 | + | 0.728414 | Intergenic | RMER15-int\|LTR\|ERVL | -17148 | NM_016966 | 236539 | Mm.16898 | NM_016966 | ENSMUSG00000053398 | **Phgdh** | 3-PGDH\|3PGDH\|4930479N23\|A10\|PGAD\|PGD\|PGDH\|SERA | 3-phosphoglycerate dehydrogenase | protein-coding |
| chr9_82967004_82967553 | chr9 | 82967005 | 82967553 | + | 0.727671 | intron (NM_001081216, intron 4 of 39) | intron (NM_001081216, intron 4 of 39) | 8210 | NM_001081216 | 83946 | Mm.221688 | NM_031879 | ENSMUSG00000032253 | **Phip** | 2810004D21Rik\|4632404O06Rik\|Ndrp\|Wdr11 | pleckstrin homology domain interacting protein | protein-coding |
| chr9_82907244_82907824 | chr9 | 82907245 | 82907824 | + | 0.71919 | intron (NM_001081216, intron 19 of 39) | intron (NM_001081216, intron 19 of 39) | 67955 | NM_001081216 | 83946 | Mm.221688 | NM_031879 | ENSMUSG00000032253 | **Phip** | 2810004D21Rik\|4632404O06Rik\|Ndrp\|Wdr11 | pleckstrin homology domain interacting protein | protein-coding |
| chrX_160603894_160604662 | chrX | 160603895 | 160604662 | + | 1.85652 | Intergenic | Intergenic | 101844 | NM_001177878 | 110094 | Mm.350712 | NM_172783 | ENSMUSG00000031295 | **Phka2** | 6330505C01Rik\|D330034O08\|Phk | phosphorylase kinase alpha 2 | protein-coding |
| chrX_160494422_160494744 | chrX | 160494423 | 160494744 | + | 0.847003 | intron (NM_001290445, intron 26 of 26) | intron (NM_001290445, intron 26 of 26) | -7583 | NM_001177879 | 110094 | Mm.350712 | NM_172783 | ENSMUSG00000031295 | **Phka2** | 6330505C01Rik\|D330034O08\|Phk | phosphorylase kinase alpha 2 | protein-coding |
| chrX_160496252_160496779 | chrX | 160496253 | 160496779 | + | 0.83846 | 3' UTR (NM_178712, exon 28 of 28) | 3' UTR (NM_178712, exon 28 of 28) | -5650 | NM_001177879 | 110094 | Mm.350712 | NM_172783 | ENSMUSG00000031295 | **Phka2** | 6330505C01Rik\|D330034O08\|Phk | phosphorylase kinase alpha 2 | protein-coding |
| chr8_86079173_86079854 | chr8 | 86079174 | 86079854 | + | 0.65918 | Intergenic | MTD\|LTR\|ERVL-MaLR | 238512 | NM_199446 | 102093 | Mm.237296 | NM_199446 | ENSMUSG00000036879 | **Phkb** | AI463271 | phosphorylase kinase beta | protein-coding |
| chr8_85860613_85860991 | chr8 | 85860614 | 85860991 | + | 0.635912 | intron (NM_199446, intron 2 of 31) | intron (NM_199446, intron 2 of 31) | 19800 | NM_199446 | 102093 | Mm.237296 | NM_199446 | ENSMUSG00000036879 | **Phkb** | AI463271 | phosphorylase kinase beta | protein-coding |
| chr10_111543863_111544341 | chr10 | 111543864 | 111544341 | + | 1.13364 | Intergenic | Intergenic | 37816 | NM_009344 | 21664 | Mm.3117 | NM_009344 | ENSMUSG00000020205 | **Phlda1** | DT1P1B11\|TDAG51\|Tdag | pleckstrin homology-like domain, family A, member 1 | protein-coding |
| chr10_111516751_111517290 | chr10 | 111516752 | 111517290 | + | 1.05876 | Intergenic | Intergenic | 10735 | NM_009344 | 21664 | Mm.3117 | NM_009344 | ENSMUSG00000020205 | **Phlda1** | DT1P1B11\|TDAG51\|Tdag | pleckstrin homology-like domain, family A, member 1 | protein-coding |
| chr10_111509338_111509557 | chr10 | 111509339 | 111509557 | + | 0.872603 | TTS (NM_009344) | TTS (NM_009344) | 3162 | NM_009344 | 21664 | Mm.3117 | NM_009344 | ENSMUSG00000020205 | **Phlda1** | DT1P1B11\|TDAG51\|Tdag | pleckstrin homology-like domain, family A, member 1 | protein-coding |
| chr10_111516222_111517311 | chr10 | 111516223 | 111517311 | + | 0.633142 | Intergenic | (TG)n\|Simple_repeat\|Simple_repeat | 10481 | NM_009344 | 21664 | Mm.3117 | NM_009344 | ENSMUSG00000020205 | **Phlda1** | DT1P1B11\|TDAG51\|Tdag | pleckstrin homology-like domain, family A, member 1 | protein-coding |
| chr7_143523489_143524199 | chr7 | 143523490 | 143524199 | + | 0.724846 | intron (NM_001285490, intron 12 of 14) | intron (NM_001285490, intron 12 of 14) | -21320 | NM_009434 | 22113 | Mm.334344 | NM_009434 | ENSMUSG00000010760 | **Phlda2** | Ipl\|Tssc3 | pleckstrin homology-like domain, family A, member 2 | protein-coding |
| chr1_106351542_106351926 | chr1 | 106351543 | 106351926 | + | 1.38066 | intron (NM_133821, intron 10 of 16) | intron (NM_133821, intron 10 of 16) | 179865 | NM_133821 | 98432 | Mm.24115 | NM_133821 | ENSMUSG00000044340 | **Phlpp1** | AI836256\|Phlpp\|Plekhe1\|SCOP\|mKIAA0606 | PH domain and leucine rich repeat protein phosphatase 1 | protein-coding |
| chr1_106320615_106320937 | chr1 | 106320616 | 106320937 | + | 1.37664 | intron (NM_133821, intron 4 of 16) | intron (NM_133821, intron 4 of 16) | 148907 | NM_133821 | 98432 | Mm.24115 | NM_133821 | ENSMUSG00000044340 | **Phlpp1** | AI836256\|Phlpp\|Plekhe1\|SCOP\|mKIAA0606 | PH domain and leucine rich repeat protein phosphatase 1 | protein-coding |
| chr1_106352761_106353116 | chr1 | 106352762 | 106353116 | + | 1.3477 | intron (NM_133821, intron 10 of 16) | intron (NM_133821, intron 10 of 16) | 181070 | NM_133821 | 98432 | Mm.24115 | NM_133821 | ENSMUSG00000044340 | **Phlpp1** | AI836256\|Phlpp\|Plekhe1\|SCOP\|mKIAA0606 | PH domain and leucine rich repeat protein phosphatase 1 | protein-coding |
| chr1_106352767_106353464 | chr1 | 106352768 | 106353464 | + | 1.33467 | intron (NM_133821, intron 10 of 16) | intron (NM_133821, intron 10 of 16) | 181247 | NM_133821 | 98432 | Mm.24115 | NM_133821 | ENSMUSG00000044340 | **Phlpp1** | AI836256\|Phlpp\|Plekhe1\|SCOP\|mKIAA0606 | PH domain and leucine rich repeat protein phosphatase 1 | protein-coding |
| chr1_106345274_106345650 | chr1 | 106345275 | 106345650 | + | 1.27021 | intron (NM_133821, intron 8 of 16) | intron (NM_133821, intron 8 of 16) | 173593 | NM_133821 | 98432 | Mm.24115 | NM_133821 | ENSMUSG00000044340 | **Phlpp1** | AI836256\|Phlpp\|Plekhe1\|SCOP\|mKIAA0606 | PH domain and leucine rich repeat protein phosphatase 1 | protein-coding |
| chr1_106345102_106345651 | chr1 | 106345103 | 106345651 | + | 1.24867 | intron (NM_133821, intron 8 of 16) | intron (NM_133821, intron 8 of 16) | 173508 | NM_133821 | 98432 | Mm.24115 | NM_133821 | ENSMUSG00000044340 | **Phlpp1** | AI836256\|Phlpp\|Plekhe1\|SCOP\|mKIAA0606 | PH domain and leucine rich repeat protein phosphatase 1 | protein-coding |
| chr1_106325275_106325635 | chr1 | 106325276 | 106325635 | + | 0.996756 | intron (NM_133821, intron 4 of 16) | intron (NM_133821, intron 4 of 16) | 153586 | NM_133821 | 98432 | Mm.24115 | NM_133821 | ENSMUSG00000044340 | **Phlpp1** | AI836256\|Phlpp\|Plekhe1\|SCOP\|mKIAA0606 | PH domain and leucine rich repeat protein phosphatase 1 | protein-coding |
| chr1_106307350_106307955 | chr1 | 106307351 | 106307955 | + | 0.98677 | intron (NM_133821, intron 3 of 16) | intron (NM_133821, intron 3 of 16) | 135784 | NM_133821 | 98432 | Mm.24115 | NM_133821 | ENSMUSG00000044340 | **Phlpp1** | AI836256\|Phlpp\|Plekhe1\|SCOP\|mKIAA0606 | PH domain and leucine rich repeat protein phosphatase 1 | protein-coding |
| chr1_106373230_106373593 | chr1 | 106373231 | 106373593 | + | 0.974854 | intron (NM_133821, intron 12 of 16) | intron (NM_133821, intron 12 of 16) | 201543 | NM_133821 | 98432 | Mm.24115 | NM_133821 | ENSMUSG00000044340 | **Phlpp1** | AI836256\|Phlpp\|Plekhe1\|SCOP\|mKIAA0606 | PH domain and leucine rich repeat protein phosphatase 1 | protein-coding |
| chr1_106325089_106325673 | chr1 | 106325090 | 106325673 | + | 0.942052 | intron (NM_133821, intron 4 of 16) | intron (NM_133821, intron 4 of 16) | 153512 | NM_133821 | 98432 | Mm.24115 | NM_133821 | ENSMUSG00000044340 | **Phlpp1** | AI836256\|Phlpp\|Plekhe1\|SCOP\|mKIAA0606 | PH domain and leucine rich repeat protein phosphatase 1 | protein-coding |
| chr1_106321334_106321923 | chr1 | 106321335 | 106321923 | + | 0.742119 | intron (NM_133821, intron 4 of 16) | intron (NM_133821, intron 4 of 16) | 149760 | NM_133821 | 98432 | Mm.24115 | NM_133821 | ENSMUSG00000044340 | **Phlpp1** | AI836256\|Phlpp\|Plekhe1\|SCOP\|mKIAA0606 | PH domain and leucine rich repeat protein phosphatase 1 | protein-coding |
| chr1_106321313_106322148 | chr1 | 106321314 | 106322148 | + | 0.723143 | intron (NM_133821, intron 4 of 16) | intron (NM_133821, intron 4 of 16) | 149862 | NM_133821 | 98432 | Mm.24115 | NM_133821 | ENSMUSG00000044340 | **Phlpp1** | AI836256\|Phlpp\|Plekhe1\|SCOP\|mKIAA0606 | PH domain and leucine rich repeat protein phosphatase 1 | protein-coding |
| chr1_106374000_106374754 | chr1 | 106374001 | 106374754 | + | 0.687555 | intron (NM_133821, intron 12 of 16) | Lx7\|LINE\|L1 | 202508 | NM_133821 | 98432 | Mm.24115 | NM_133821 | ENSMUSG00000044340 | **Phlpp1** | AI836256\|Phlpp\|Plekhe1\|SCOP\|mKIAA0606 | PH domain and leucine rich repeat protein phosphatase 1 | protein-coding |
| chr1_106400702_106401592 | chr1 | 106400703 | 106401592 | + | 0.666641 | Intergenic | Intergenic | 229278 | NM_133821 | 98432 | Mm.24115 | NM_133821 | ENSMUSG00000044340 | **Phlpp1** | AI836256\|Phlpp\|Plekhe1\|SCOP\|mKIAA0606 | PH domain and leucine rich repeat protein phosphatase 1 | protein-coding |
| chr1_106269340_106269684 | chr1 | 106269341 | 106269684 | + | 0.637337 | intron (NM_133821, intron 1 of 16) | intron (NM_133821, intron 1 of 16) | 97643 | NM_133821 | 98432 | Mm.24115 | NM_133821 | ENSMUSG00000044340 | **Phlpp1** | AI836256\|Phlpp\|Plekhe1\|SCOP\|mKIAA0606 | PH domain and leucine rich repeat protein phosphatase 1 | protein-coding |
| chr3_103991284_103991719 | chr3 | 103991285 | 103991719 | + | 0.860154 | intron (NM_001163469, intron 9 of 17) | intron (NM_001163469, intron 9 of 17) | -23186 | NR_030676 | 100125931 | Mm.400392 | NR_030676 | ENSMUSG00000089998 | **Phtf1os** | A130049A11Rik | putative homeodomain transcription factor 1, opposite strand | ncRNA |
| chr3_103990597_103991174 | chr3 | 103990598 | 103991174 | + | 0.859632 | intron (NM_001163469, intron 7 of 17) | intron (NM_001163469, intron 7 of 17) | -22570 | NR_030676 | 100125931 | Mm.400392 | NR_030676 | ENSMUSG00000089998 | **Phtf1os** | A130049A11Rik | putative homeodomain transcription factor 1, opposite strand | ncRNA |
| chr5_20680879_20681559 | chr5 | 20680880 | 20681559 | + | 1.04386 | intron (NM_001170746, intron 22 of 22) | intron (NM_001170746, intron 22 of 22) | 200905 | NM_172992 | 68770 | Mm.86410 | NM_172992 | ENSMUSG00000039987 | **Phtf2** | 1110054G21Rik\|9530062N20Rik\|AI447096\|C77923 | putative homeodomain transcription factor 2 | protein-coding |

| | | | | | | | | | | | | | | | | | |
|---|---|---|---|---|---|---|---|---|---|---|---|---|---|---|---|---|---|
| chr5_20778193_20778891 | chr5 | 20778194 | 20778891 | + | 0.959323 | intron (NM_172992, intron 10 of 17) | intron (NM_172992, intron 10 of 17) | 103582 | NM_172992 | 68770 | Mm.86410 | NM_172992 | ENSMUSG00000039987 | **Phtf2** | 1110054G21Rik\|9530062N20Rik\|AI447096\|C77923 | putative homeodomain transcription factor 2 | protein-coding |
| chr5_20754139_20754523 | chr5 | 20754140 | 20754523 | + | 0.937313 | Intergenic | Intergenic | 127793 | NM_172992 | 68770 | Mm.86410 | NM_172992 | ENSMUSG00000039987 | **Phtf2** | 1110054G21Rik\|9530062N20Rik\|AI447096\|C77923 | putative homeodomain transcription factor 2 | protein-coding |
| chr5_20845214_20845501 | chr5 | 20845215 | 20845501 | + | 0.809691 | intron (NM_172992, intron 2 of 17) | B4\|SINE\|B4 | 36766 | NM_172992 | 68770 | Mm.86410 | NM_172992 | ENSMUSG00000039987 | **Phtf2** | 1110054G21Rik\|9530062N20Rik\|AI447096\|C77923 | putative homeodomain transcription factor 2 | protein-coding |
| chr5_20845121_20845521 | chr5 | 20845122 | 20845521 | + | 0.782647 | intron (NM_172992, intron 2 of 17) | B4\|SINE\|B4 | 36803 | NM_172992 | 68770 | Mm.86410 | NM_172992 | ENSMUSG00000039987 | **Phtf2** | 1110054G21Rik\|9530062N20Rik\|AI447096\|C77923 | putative homeodomain transcription factor 2 | protein-coding |
| chr9_13509358_13510238 | chr9 | 13509359 | 13510238 | + | 0.772185 | Intergenic | Intergenic | 78437 | NR_028271 | 18689 | Mm.41972 | NM_008835 | | **Phxr4** | Sp5 | per-hexamer repeat gene 4 | ncRNA |
| chr9_13473216_13473540 | chr9 | 13473217 | 13473540 | + | 0.719478 | Intergenic | Intergenic | 42017 | NR_028271 | 18689 | Mm.41972 | NM_008835 | | **Phxr4** | Sp5 | per-hexamer repeat gene 4 | ncRNA |
| chr9_13445215_13445494 | chr9 | 13445216 | 13445494 | + | 0.622708 | Intergenic | Intergenic | 13994 | NR_028271 | 18689 | Mm.41972 | NM_008835 | | **Phxr4** | Sp5 | per-hexamer repeat gene 4 | ncRNA |
| chr11_51594726_51595919 | chr11 | 51594727 | 51595919 | + | 0.767646 | intron (NM_028398, intron 8 of 12) | intron (NM_028398, intron 8 of 12) | 10566 | NM_028398 | 72947 | Mm.256875 | NM_028398 | | **Phykpl** | 2900006B13Rik\|Agxt2l2 | 5-phosphohydroxy-L-lysine phospholyase | protein-coding |
| chr16_17414955_17415282 | chr16 | 17414956 | 17415282 | + | 0.826518 | intron (NM_023348, intron 1 of 4) | RMER3D-int\|LTR\|ERVK | -8805 | NM_001001983 | 224020 | Mm.5718 | NM_001001983 | ENSMUSG00000041720 | **Pi4ka** | Pik4ca | phosphatidylinositol 4-kinase, catalytic, alpha polypeptide | protein-coding |
| chr9_62932602_62932949 | chr9 | 62932603 | 62932949 | + | 1.04895 | intron (NM_019663, intron 2 of 13) | intron (NM_019663, intron 2 of 13) | 48103 | NM_019663 | 56469 | Mm.431253 | NM_019663 | ENSMUSG00000032405 | **Pias1** | 2900068C24Rik\|Ddxbp1\|GBP | protein inhibitor of activated STAT 1 | protein-coding |
| chr9_62932439_62932968 | chr9 | 62932440 | 62932968 | + | 0.773469 | intron (NM_019663, intron 2 of 13) | intron (NM_019663, intron 2 of 13) | 48175 | NM_019663 | 56469 | Mm.431253 | NM_019663 | ENSMUSG00000032405 | **Pias1** | 2900068C24Rik\|Ddxbp1\|GBP | protein inhibitor of activated STAT 1 | protein-coding |
| chr9_62941246_62941631 | chr9 | 62941247 | 62941631 | + | 0.668344 | intron (NM_019663, intron 2 of 13) | ORR1D1\|LTR\|ERVL-MaLR | 39440 | NM_019663 | 56469 | Mm.431253 | NM_019663 | ENSMUSG00000032405 | **Pias1** | 2900068C24Rik\|Ddxbp1\|GBP | protein inhibitor of activated STAT 1 | protein-coding |
| chr7_90146250_90146698 | chr7 | 90146251 | 90146698 | + | 0.817877 | intron (NM_001252524, intron 1 of 18) | intron (NM_001252524, intron 1 of 18) | 16242 | NM_001252520 | 233489 | Mm.235175 | NM_146194 | ENSMUSG00000039361 | **Picalm** | CALM\|CLTH\|PAP180\|fit-1\|fit1\|mKIAA4114 | phosphatidylinositol binding clathrin assembly protein | protein-coding |
| chr7_90145981_90146698 | chr7 | 90145982 | 90146698 | + | 0.812729 | intron (NM_001252524, intron 1 of 18) | intron (NM_001252524, intron 1 of 18) | 16108 | NM_001252520 | 233489 | Mm.235175 | NM_146194 | ENSMUSG00000039361 | **Picalm** | CALM\|CLTH\|PAP180\|fit-1\|fit1\|mKIAA4114 | phosphatidylinositol binding clathrin assembly protein | protein-coding |
| chr8_122550217_122550536 | chr8 | 122550218 | 122550536 | + | 0.736704 | intron (NM_001037298, intron 1 of 50) | intron (NM_001037298, intron 1 of 50) | 952 | NM_001037298 | 234839 | Mm.37324 | NM_001037298 | ENSMUSG00000014444 | **Piezo1** | 9630020g22\|Fam38a\|mKIAA0233 | piezo-type mechanosensitive ion channel component 1 | protein-coding |
| chr8_122548804_122549471 | chr8 | 122548805 | 122549471 | + | 0.59691 | intron (NM_001037298, intron 1 of 50) | intron (NM_001037298, intron 1 of 50) | 2191 | NM_001037298 | 234839 | Mm.37324 | NM_001037298 | ENSMUSG00000014444 | **Piezo1** | 9630020g22\|Fam38a\|mKIAA0233 | piezo-type mechanosensitive ion channel component 1 | protein-coding |
| chr3_106030174_106030875 | chr3 | 106030175 | 106030875 | + | 1.08904 | Intergenic | Intergenic | -15879 | NM_001200028 | 100503311 | Mm.386825 | NM_029604 | ENSMUSG00000010136 | **Pifo** | 1700027A23Rik | primary cilia formation | protein-coding |
| chr12_79090454_79091259 | chr12 | 79090455 | 79091259 | + | 1.41366 | intron (NR_110486, intron 2 of 2) | intron (NR_110486, intron 2 of 2) | -1187 | NM_029988 | 110417 | Mm.268317 | NM_029988 | ENSMUSG00000021120 | **Pigh** | 2210416H01Rik\|A930028P05Rik | phosphatidylinositol glycan anchor biosynthesis, class H | protein-coding |
| chr3_152813833_152814286 | chr3 | 152813834 | 152814286 | + | 1.09644 | Intergenic | RMER15\|LTR\|ERVL | 99960 | NM_025662 | 329777 | Mm.331447 | NM_025662 | ENSMUSG00000039047 | **Pigk** | 3000001O05Rik | phosphatidylinositol glycan anchor biosynthesis, class K | protein-coding |
| chr3_152758449_152758950 | chr3 | 152758450 | 152758950 | + | 1.01788 | intron (NM_178016, intron 10 of 11) | intron (NM_178016, intron 10 of 11) | 44600 | NM_025662 | 329777 | Mm.331447 | NM_025662 | ENSMUSG00000039047 | **Pigk** | 3000001O05Rik | phosphatidylinositol glycan anchor biosynthesis, class K | protein-coding |
| chr3_152762308_152762663 | chr3 | 152762309 | 152762663 | + | 0.879076 | intron (NM_178016, intron 10 of 11) | intron (NM_178016, intron 10 of 11) | 48386 | NM_025662 | 329777 | Mm.331447 | NM_025662 | ENSMUSG00000039047 | **Pigk** | 3000001O05Rik | phosphatidylinositol glycan anchor biosynthesis, class K | protein-coding |
| chr1_172390867_172391256 | chr1 | 172390868 | 172391256 | + | 0.805287 | Intergenic | ORR1B1\|LTR\|ERVL-MaLR | 14531 | NM_026234 | 67556 | Mm.26612 | NM_026234 | ENSMUSG00000050229 | **Pigm** | 4933437L05Rik\|AI644422\|AV302080\|C920011G20Rik | phosphatidylinositol glycan anchor biosynthesis, class M | protein-coding |
| chr1_105564726_105565437 | chr1 | 105564727 | 105565437 | + | 1.4032 | intron (NM_013784, intron 23 of 30) | intron (NM_013784, intron 23 of 30) | 98594 | NM_013784 | 27392 | Mm.268911 | NM_013784 | ENSMUSG00000056536 | **Pign** | Gm20308\|PIG-N | phosphatidylinositol glycan anchor biosynthesis, class N | protein-coding |
| chr1_105523698_105524179 | chr1 | 105523699 | 105524179 | + | 1.1214 | intron (NM_013784, intron 30 of 30) | intron (NM_013784, intron 30 of 30) | 139737 | NM_013784 | 27392 | Mm.268911 | NM_013784 | ENSMUSG00000056536 | **Pign** | Gm20308\|PIG-N | phosphatidylinositol glycan anchor biosynthesis, class N | protein-coding |
| chr1_105519092_105519606 | chr1 | 105519093 | 105519606 | + | 1.10125 | 3' UTR (NM_013784, exon 31 of 31) | 3' UTR (NM_013784, exon 31 of 31) | 144327 | NM_013784 | 27392 | Mm.268911 | NM_013784 | ENSMUSG00000056536 | **Pign** | Gm20308\|PIG-N | phosphatidylinositol glycan anchor biosynthesis, class N | protein-coding |
| chr1_105527408_105527986 | chr1 | 105527409 | 105527986 | + | 0.982289 | intron (NM_013784, intron 30 of 30) | L1_Mur1\|LINE\|L1 | 135979 | NM_013784 | 27392 | Mm.268911 | NM_013784 | ENSMUSG00000056536 | **Pign** | Gm20308\|PIG-N | phosphatidylinositol glycan anchor biosynthesis, class N | protein-coding |
| chr1_105652560_105652807 | chr1 | 105652561 | 105652807 | + | 0.747904 | intron (NM_013784, intron 7 of 30) | intron (NM_013784, intron 7 of 30) | 10992 | NM_013784 | 27392 | Mm.268911 | NM_013784 | ENSMUSG00000056536 | **Pign** | Gm20308\|PIG-N | phosphatidylinositol glycan anchor biosynthesis, class N | protein-coding |
| chr16_94362195_94362518 | chr16 | 94362196 | 94362518 | + | 0.671022 | intron (NM_001159620, intron 3 of 3) | intron (NM_001159620, intron 3 of 3) | 7973 | NM_001159619 | 56176 | Mm.474449 | NM_019543 | ENSMUSG00000022940 | **Pigp** | Dcrc\|Dscr5 | phosphatidylinositol glycan anchor biosynthesis, class P | protein-coding |
| chr4_133688345_133688803 | chr4 | 133688346 | 133688803 | + | 0.856301 | intron (NM_001080819, intron 12 of 19) | intron (NM_001080819, intron 12 of 19) | -15927 | NM_001145956 | 230801 | Mm.217004 | NM_178698 | | **Pigv** | B330013B03\|D430024F16Rik | phosphatidylinositol glycan anchor biosynthesis, class V | protein-coding |
| chr4_133680183_133680547 | chr4 | 133680184 | 133680547 | + | 0.678103 | exon (NM_001080819, exon 20 of 20) | exon (NM_001080819, exon 20 of 20) | -7718 | NM_001145956 | 230801 | Mm.217004 | NM_178698 | | **Pigv** | B330013B03\|D430024F16Rik | phosphatidylinositol glycan anchor biosynthesis, class V | protein-coding |
| chr19_41417730_41418242 | chr19 | 41417731 | 41418242 | + | 0.715052 | Intergenic | MLT1J\|LTR\|ERVL-MaLR | -32916 | NM_031376 | 83490 | Mm.222266 | NM_031376 | ENSMUSG00000025017 | **Pik3ap1** | 1810044J04Rik\|BCAP | phosphoinositide-3-kinase adaptor protein 1 | protein-coding |
| chr7_116456987_116457353 | chr7 | 116456988 | 116457353 | + | 0.713608 | Intergenic | L4_A_Mam\|LINE\|RTE-X | -13712 | NM_011083 | 18704 | Mm.3810 | NM_011083 | ENSMUSG00000030660 | **Pik3c2a** | Cpk-m\|PI3KC2 | phosphatidylinositol 3-kinase, C2 domain containing, alpha polypeptide | protein-coding |
| chr18_30312333_30312840 | chr18 | 30312334 | 30312840 | + | 0.815844 | exon (NM_181414, exon 15 of 25) | exon (NM_181414, exon 15 of 25) | 39691 | NM_181414 | 225326 | Mm.194127 | NM_181414 | ENSMUSG00000033628 | **Pik3c3** | 5330434F23Rik\|Vps34 | phosphoinositide-3-kinase, class 3 | protein-coding |
| chr3_32417570_32418271 | chr3 | 32417571 | 32418271 | + | 1.52609 | Intergenic | RMER15\|LTR\|ERVL | -18230 | NM_008839 | 18706 | Mm.260521 | NM_008839 | ENSMUSG00000027665 | **Pik3ca** | 6330412C24Rik\|caPI3K\|p110\|p110alpha | phosphatidylinositol 3-kinase, catalytic, alpha polypeptide | protein-coding |
| chr3_32421771_32422013 | chr3 | 32421772 | 32422013 | + | 1.46744 | Intergenic | Intergenic | -14259 | NM_008839 | 18706 | Mm.260521 | NM_008839 | ENSMUSG00000027665 | **Pik3ca** | 6330412C24Rik\|caPI3K\|p110\|p110alpha | phosphatidylinositol 3-kinase, catalytic, alpha polypeptide | protein-coding |
| chr3_32427468_32428249 | chr3 | 32427469 | 32428249 | + | 0.976388 | Intergenic | ORR1E\|LTR\|ERVL-MaLR | -8292 | NM_008839 | 18706 | Mm.260521 | NM_008839 | ENSMUSG00000027665 | **Pik3ca** | 6330412C24Rik\|caPI3K\|p110\|p110alpha | phosphatidylinositol 3-kinase, catalytic, alpha polypeptide | protein-coding |
| chr3_32437285_32438184 | chr3 | 32437286 | 32438184 | + | 0.737487 | intron (NM_008839, intron 2 of 19) | intron (NM_008839, intron 2 of 19) | 1584 | NM_008839 | 18706 | Mm.260521 | NM_008839 | ENSMUSG00000027665 | **Pik3ca** | 6330412C24Rik\|caPI3K\|p110\|p110alpha | phosphatidylinositol 3-kinase, catalytic, alpha polypeptide | protein-coding |
| chr3_32412036_32412565 | chr3 | 32412037 | 32412565 | + | 0.679418 | Intergenic | Intergenic | -23850 | NM_008839 | 18706 | Mm.260521 | NM_008839 | ENSMUSG00000027665 | **Pik3ca** | 6330412C24Rik\|caPI3K\|p110\|p110alpha | phosphatidylinositol 3-kinase, catalytic, alpha polypeptide | protein-coding |
| chr3_32419863_32420299 | chr3 | 32419864 | 32420299 | + | 0.619452 | Intergenic | Intergenic | -16070 | NM_008839 | 18706 | Mm.260521 | NM_008839 | ENSMUSG00000027665 | **Pik3ca** | 6330412C24Rik\|caPI3K\|p110\|p110alpha | phosphatidylinositol 3-kinase, catalytic, alpha polypeptide | protein-coding |
| chr4_149661979_149662281 | chr4 | 149661980 | 149662281 | + | 0.983146 | intron (NM_001029837, intron 3 of 23) | intron (NM_001029837, intron 3 of 23) | 1186 | NM_001164052 | 18707 | Mm.229108 | NM_008840 | ENSMUSG00000039936 | **Pik3cd** | 2410099E07Rik\|2610208K16Rik\|AW545373\|p110delta | phosphatidylinositol 3-kinase catalytic delta polypeptide | protein-coding |
| chr4_149689841_149690192 | chr4 | 149689842 | 149690192 | + | 0.933381 | intron (NM_001029837, intron 1 of 23) | intron (NM_001029837, intron 1 of 23) | 8691 | NM_001164050 | 18707 | Mm.229108 | NM_008840 | ENSMUSG00000039936 | **Pik3cd** | 2410099E07Rik\|2610208K16Rik\|AW545373\|p110delta | phosphatidylinositol 3-kinase catalytic delta polypeptide | protein-coding |
| chr4_149696158_149696909 | chr4 | 149696159 | 149696909 | + | 0.893284 | intron (NM_001029837, intron 1 of 23) | intron (NM_001029837, intron 1 of 23) | 2174 | NM_001164050 | 18707 | Mm.229108 | NM_008840 | ENSMUSG00000039936 | **Pik3cd** | 2410099E07Rik\|2610208K16Rik\|AW545373\|p110delta | phosphatidylinositol 3-kinase catalytic delta polypeptide | protein-coding |
| chr4_149662366_149662573 | chr4 | 149662367 | 149662573 | + | 0.686599 | intron (NM_001029837, intron 3 of 23) | B4\|SINE\|B4 | 846 | NM_001164052 | 18707 | Mm.229108 | NM_008840 | ENSMUSG00000039936 | **Pik3cd** | 2410099E07Rik\|2610208K16Rik\|AW545373\|p110delta | phosphatidylinositol 3-kinase catalytic delta polypeptide | protein-coding |
| chr12_32201838_32202270 | chr12 | 32201839 | 32202270 | + | 1.09872 | intron (NM_020272, intron 2 of 10) | intron (NM_020272, intron 2 of 10) | 6284 | NM_001146201 | 30955 | Mm.101369 | NM_020272 | ENSMUSG00000020573 | **Pik3cg** | 5830428L06Rik\|PI3Kgamma\|p110gamma | phosphoinositide-3-kinase, catalytic, gamma polypeptide | protein-coding |
| chr12_32203732_32204081 | chr12 | 32203733 | 32204081 | + | 0.778412 | intron (NM_020272, intron 2 of 10) | intron (NM_020272, intron 2 of 10) | 4431 | NM_001146201 | 30955 | Mm.101369 | NM_020272 | ENSMUSG00000020573 | **Pik3cg** | 5830428L06Rik\|PI3Kgamma\|p110gamma | phosphoinositide-3-kinase, catalytic, gamma polypeptide | protein-coding |
| chr13_101549642_101550092 | chr13 | 101549643 | 101550092 | + | 1.07309 | Intergenic | Intergenic | 142763 | NM_001024955 | 18708 | Mm.259333 | NM_001024955 | ENSMUSG00000041417 | **Pik3r1** | PI3K\|p50alpha\|p55alpha\|p85alpha | phosphatidylinositol 3-kinase, regulatory subunit, polypeptide 1 (p85 alpha) | protein-coding |
| chr13_101549644_101549957 | chr13 | 101549645 | 101549957 | + | 1.05412 | Intergenic | Intergenic | 142829 | NM_001024955 | 18708 | Mm.259333 | NM_001024955 | ENSMUSG00000041417 | **Pik3r1** | PI3K\|p50alpha\|p55alpha\|p85alpha | phosphatidylinositol 3-kinase, regulatory subunit, polypeptide 1 (p85 alpha) | protein-coding |

| | | | | | | | | | | | | | | | | | |
|---|---|---|---|---|---|---|---|---|---|---|---|---|---|---|---|---|---|
| chr13_102093699_102094457 | chr13 | 102093700 | 102094457 | + | 1.047 | Intergenic | Intergenic | -325861 | NM_001077495 | 18708 | Mm.259333 | NM_001024955 | ENSMUSG00000041417 | **Pik3r1** | PI3K\|p50alpha\|p55alpha\|p85alpha | phosphatidylinositol 3-kinase, regulatory subunit, polypeptide 1 (p85 alpha) | protein-coding |
| chr13_102093927_102094444 | chr13 | 102093928 | 102094444 | + | 1.01042 | Intergenic | Intergenic | -325969 | NM_001077495 | 18708 | Mm.259333 | NM_001024955 | ENSMUSG00000041417 | **Pik3r1** | PI3K\|p50alpha\|p55alpha\|p85alpha | phosphatidylinositol 3-kinase, regulatory subunit, polypeptide 1 (p85 alpha) | protein-coding |
| chr13_102070677_102071181 | chr13 | 102070678 | 102071181 | + | 0.900634 | Intergenic | Intergenic | -302712 | NM_001077495 | 18708 | Mm.259333 | NM_001024955 | ENSMUSG00000041417 | **Pik3r1** | PI3K\|p50alpha\|p55alpha\|p85alpha | phosphatidylinositol 3-kinase, regulatory subunit, polypeptide 1 (p85 alpha) | protein-coding |
| chr13_101691615_101691884 | chr13 | 101691616 | 101691884 | + | 0.778003 | intron (NM_001077495, intron 7 of 15) | intron (NM_001077495, intron 7 of 15) | 880 | NM_001024955 | 18708 | Mm.259333 | NM_001024955 | ENSMUSG00000041417 | **Pik3r1** | PI3K\|p50alpha\|p55alpha\|p85alpha | phosphatidylinositol 3-kinase, regulatory subunit, polypeptide 1 (p85 alpha) | protein-coding |
| chr13_101547789_101548372 | chr13 | 101547790 | 101548372 | + | 0.762447 | Intergenic | LTR33B\|LTR\|ERVL | 144549 | NM_001024955 | 18708 | Mm.259333 | NM_001024955 | ENSMUSG00000041417 | **Pik3r1** | PI3K\|p50alpha\|p55alpha\|p85alpha | phosphatidylinositol 3-kinase, regulatory subunit, polypeptide 1 (p85 alpha) | protein-coding |
| chr13_101569014_101569774 | chr13 | 101569015 | 101569774 | + | 0.761474 | Intergenic | MER45R\|DNA\|hAT-Tip100 | 123236 | NM_001024955 | 18708 | Mm.259333 | NM_001024955 | ENSMUSG00000041417 | **Pik3r1** | PI3K\|p50alpha\|p55alpha\|p85alpha | phosphatidylinositol 3-kinase, regulatory subunit, polypeptide 1 (p85 alpha) | protein-coding |
| chr13_101996083_101996630 | chr13 | 101996084 | 101996630 | + | 0.731168 | Intergenic | Intergenic | -228140 | NM_001077495 | 18708 | Mm.259333 | NM_001024955 | ENSMUSG00000041417 | **Pik3r1** | PI3K\|p50alpha\|p55alpha\|p85alpha | phosphatidylinositol 3-kinase, regulatory subunit, polypeptide 1 (p85 alpha) | protein-coding |
| chr4_116300833_116301565 | chr4 | 116300834 | 116301565 | + | 0.72023 | 3' UTR (NM_181585, exon 10 of 10) | 3' UTR (NM_181585, exon 10 of 10) | 79285 | NM_181585 | 18710 | Mm.253819 | NM_181585 | ENSMUSG00000028698 | **Pik3r3** | AA414954\|p55pik | phosphatidylinositol 3 kinase, regulatory subunit, polypeptide 3 (p55) | protein-coding |
| chr11_68458048_68458493 | chr11 | 68458049 | 68458493 | + | 1.38496 | intron (NM_177320, intron 1 of 18) | intron (NM_177320, intron 1 of 18) | 26146 | NM_177320 | 320207 | Mm.244960 | NM_177320 | ENSMUSG00000020901 | **Pik3r5** | AV230647\|F730038I15Rik | phosphoinositide-3-kinase, regulatory subunit 5, p101 | protein-coding |
| chr11_68443583_68443937 | chr11 | 68443584 | 68443937 | + | 1.07222 | intron (NM_177320, intron 1 of 18) | intron (NM_177320, intron 1 of 18) | 11635 | NM_177320 | 320207 | Mm.244960 | NM_177320 | ENSMUSG00000020901 | **Pik3r5** | AV230647\|F730038I15Rik | phosphoinositide-3-kinase, regulatory subunit 5, p101 | protein-coding |
| chr11_68445658_68446170 | chr11 | 68445659 | 68446170 | + | 0.881148 | intron (NM_177320, intron 1 of 18) | intron (NM_177320, intron 1 of 18) | 13789 | NM_177320 | 320207 | Mm.244960 | NM_177320 | ENSMUSG00000020901 | **Pik3r5** | AV230647\|F730038I15Rik | phosphoinositide-3-kinase, regulatory subunit 5, p101 | protein-coding |
| chr11_68446556_68446820 | chr11 | 68446557 | 68446820 | + | 0.850503 | intron (NM_177320, intron 1 of 18) | L1M5\|LINE\|L1 | 14563 | NM_177320 | 320207 | Mm.244960 | NM_177320 | ENSMUSG00000020901 | **Pik3r5** | AV230647\|F730038I15Rik | phosphoinositide-3-kinase, regulatory subunit 5, p101 | protein-coding |
| chr11_68502467_68502816 | chr11 | 68502468 | 68502816 | + | 0.785026 | promoter-TSS (NM_001081566) | promoter-TSS (NM_001081566) | -377 | NM_001004435 | 104709 | Mm.234573 | NM_001004435 | ENSMUSG00000046207 | **Pik3r6** | BB220380\|p84\|p84 PIKAP\|p87(PIKAP)\|p87PIKAP | phosphoinositide-3-kinase, regulatory subunit 6 | protein-coding |
| chr5_137816915_137817825 | chr5 | 137816916 | 137817825 | + | 0.88888 | intron (NM_001005426, intron 12 of 16) | intron (NM_001005426, intron 12 of 16) | 18908 | NM_153510 | 231805 | Mm.379385 | NM_153510 | ENSMUSG00000046245 | **Pilra** | AV021745\|FDF03 | paired immunoglobin-like type 2 receptor alpha | protein-coding |
| chr5_137817342_137817800 | chr5 | 137817343 | 137817800 | + | 0.766471 | intron (NM_001005426, intron 13 of 16) | intron (NM_001005426, intron 13 of 16) | 18707 | NM_153510 | 231805 | Mm.379385 | NM_153510 | ENSMUSG00000046245 | **Pilra** | AV021745\|FDF03 | paired immunoglobin-like type 2 receptor alpha | protein-coding |
| chr5_137856868_137857329 | chr5 | 137856869 | 137857329 | + | 0.938355 | intron (NM_133209, intron 2 of 4) | intron (NM_133209, intron 2 of 4) | 950 | NM_133209 | 170741 | Mm.347593 | NM_133209 | ENSMUSG00000066684 | **Pilrb1** | FDFACT\|Fdact\|Pilrb | paired immunoglobin-like type 2 receptor beta 1 | protein-coding |
| chr17_29499607_29500166 | chr17 | 29499608 | 29500166 | + | 1.24161 | Intergenic | Intergenic | 8842 | NM_008842 | 18712 | Mm.405293 | NM_008842 | | **Pim1** | Pim-1 | proviral integration site 1 | protein-coding |
| chr17_29499595_29500621 | chr17 | 29499596 | 29500621 | + | 1.07821 | Intergenic | Intergenic | 9063 | NM_008842 | 18712 | Mm.405293 | NM_008842 | | **Pim1** | Pim-1 | proviral integration site 1 | protein-coding |
| chr17_29500634_29501279 | chr17 | 29500635 | 29501279 | + | 0.838835 | Intergenic | Intergenic | 9912 | NM_008842 | 18712 | Mm.405293 | NM_008842 | | **Pim1** | Pim-1 | proviral integration site 1 | protein-coding |
| chrX_7863119_7863736 | chrX | 7863120 | 7863736 | + | 1.10169 | intron (NM_001290536, intron 2 of 8) | intron (NM_001290536, intron 2 of 8) | -14878 | NM_138606 | 18715 | Mm.347478 | NM_138606 | ENSMUSG00000031155 | **Pim2** | DXCch3\|Pim-2 | proviral integration site 2 | protein-coding |
| chrX_7867276_7867549 | chrX | 7867277 | 7867549 | + | 0.931894 | intron (NM_001290536, intron 2 of 8) | intron (NM_001290536, intron 2 of 8) | -10893 | NM_138606 | 18715 | Mm.347478 | NM_138606 | ENSMUSG00000031155 | **Pim2** | DXCch3\|Pim-2 | proviral integration site 2 | protein-coding |
| chr15_88843391_88844634 | chr15 | 88843392 | 88844634 | + | 1.09007 | Intergenic | Intergenic | -18181 | NM_145478 | 223775 | Mm.400129 | NM_145478 | ENSMUSG00000035828 | **Pim3** | BC026639\|Kid1 | proviral integration site 3 | protein-coding |
| chrX_102121406_102121760 | chrX | 102121407 | 102121760 | + | 0.951772 | intron (NM_027181, intron 1 of 3) | intron (NM_027181, intron 1 of 3) | 2118 | NM_027181 | 69713 | Mm.371635 | NM_027181 | ENSMUSG00000079480 | **Pin4** | 2410002I22Rik\|EPVH\|Par14 | protein (peptidyl-prolyl cis/trans isomerase) NIMA-interacting, 4 (parvulin) | protein-coding |
| chr2_18918758_18919113 | chr2 | 18918759 | 18919113 | + | 1.15632 | intron (NM_008845, intron 1 of 9) | intron (NM_008845, intron 1 of 9) | 79185 | NM_008845 | 18718 | Mm.313977 | NM_008845 | ENSMUSG00000026737 | **Pip4k2a** | AW742916\|Pip5k2a | phosphatidylinositol-5-phosphate 4-kinase, type II, alpha | protein-coding |
| chr2_18911346_18911875 | chr2 | 18911347 | 18911875 | + | 0.76702 | intron (NM_008845, intron 1 of 9) | intron (NM_008845, intron 1 of 9) | 86510 | NM_008845 | 18718 | Mm.313977 | NM_008845 | ENSMUSG00000026737 | **Pip4k2a** | AW742916\|Pip5k2a | phosphatidylinositol-5-phosphate 4-kinase, type II, alpha | protein-coding |
| chr4_135987997_135988509 | chr4 | 135987998 | 135988509 | + | 1.85046 | Intergenic | MIRb\|SINE\|MIR | -1009 | NM_025411 | 66193 | Mm.29131 | NM_025411 | ENSMUSG00000028669 | **Pithd1** | 1110049F12Rik\|AA408179\|Trp26 | PITH (C-terminal proteasome-interacting domain of thioredoxin-like) domain containing 1 | protein-coding |
| chr5_111371895_111372620 | chr5 | 111371896 | 111372620 | + | 1.23655 | intron (NM_019640, intron 8 of 11) | PB1D9\|SINE\|Alu | 41495 | NM_019640 | 56305 | Mm.200516 | NM_019640 | ENSMUSG00000050017 | **Pitpnb** | AI256223\|AU040890 | phosphatidylinositol transfer protein, beta | protein-coding |
| chr5_111338134_111338620 | chr5 | 111338135 | 111338620 | + | 0.986573 | exon (NM_019640, exon 3 of 12) | exon (NM_019640, exon 3 of 12) | 7614 | NM_019640 | 56305 | Mm.200516 | NM_019640 | ENSMUSG00000050017 | **Pitpnb** | AI256223\|AU040890 | phosphatidylinositol transfer protein, beta | protein-coding |
| chr5_111338076_111338804 | chr5 | 111338077 | 111338804 | + | 0.904246 | intron (NM_019640, intron 3 of 11) | intron (NM_019640, intron 3 of 11) | 7677 | NM_019640 | 56305 | Mm.200516 | NM_019640 | ENSMUSG00000050017 | **Pitpnb** | AI256223\|AU040890 | phosphatidylinositol transfer protein, beta | protein-coding |
| chr5_124190819_124191072 | chr5 | 124190820 | 124191072 | + | 0.902805 | intron (NM_011256, intron 2 of 24) | intron (NM_011256, intron 2 of 24) | -3764 | NM_001289472 | 19679 | Mm.44261 | NM_011256 | ENSMUSG00000029406 | **Pitpnm2** | NIR3\|RDGBA2\|Rdgb2\|mKIAA1457 | phosphatidylinositol transfer protein, membrane-associated 2 | protein-coding |
| chr5_124192920_124193233 | chr5 | 124192921 | 124193233 | + | 0.668487 | intron (NM_011256, intron 2 of 24) | intron (NM_011256, intron 2 of 24) | -5895 | NM_001289472 | 19679 | Mm.44261 | NM_011256 | ENSMUSG00000029406 | **Pitpnm2** | NIR3\|RDGBA2\|Rdgb2\|mKIAA1457 | phosphatidylinositol transfer protein, membrane-associated 2 | protein-coding |
| chr5_124221398_124221752 | chr5 | 124221399 | 124221752 | + | 0.630806 | intron (NM_011256, intron 1 of 24) | intron (NM_011256, intron 1 of 24) | -8150 | NR_045369 | 75927 | Mm.410803 | NR_045369 | ENSMUSG00000090220 | **Pitpnm2os1** | 4930568K20Rik | phosphatidylinositol transfer protein, membrane-associated 2, opposite strand 1 | ncRNA |
| chr13_6287729_6288341 | chr13 | 6287730 | 6288341 | + | 0.989624 | Intergenic | Intergenic | -260122 | NM_145131 | 69617 | Mm.41933 | NM_145131 | ENSMUSG00000021193 | **Pitrm1** | 2310012C15Rik\|AA410010\|MP-1\|MP1\|Ntup1\|PreP\|mKIAA1104 | pitrilysin metallepetidase 1 | protein-coding |
| chr13_6212013_6212896 | chr13 | 6212014 | 6212896 | + | 0.919431 | Intergenic | MLT1D\|LTR\|ERVL-MaLR | -335702 | NM_145131 | 69617 | Mm.41933 | NM_145131 | ENSMUSG00000021193 | **Pitrm1** | 2310012C15Rik\|AA410010\|MP-1\|MP1\|Ntup1\|PreP\|mKIAA1104 | pitrilysin metallepetidase 1 | protein-coding |
| chr13_6287730_6288050 | chr13 | 6287731 | 6288050 | + | 0.864688 | Intergenic | Intergenic | -260267 | NM_145131 | 69617 | Mm.41933 | NM_145131 | ENSMUSG00000021193 | **Pitrm1** | 2310012C15Rik\|AA410010\|MP-1\|MP1\|Ntup1\|PreP\|mKIAA1104 | pitrilysin metallepetidase 1 | protein-coding |
| chr13_6595397_6595699 | chr13 | 6595398 | 6595699 | + | 0.74795 | intron (NM_001291071, intron 15 of 21) | MER20\|DNA\|hAT-Charlie | 47391 | NM_145131 | 69617 | Mm.41933 | NM_145131 | ENSMUSG00000021193 | **Pitrm1** | 2310012C15Rik\|AA410010\|MP-1\|MP1\|Ntup1\|PreP\|mKIAA1104 | pitrilysin metallepetidase 1 | protein-coding |
| chr13_6456582_6457209 | chr13 | 6456583 | 6457209 | + | 0.667617 | Intergenic | Intergenic | -91261 | NM_145131 | 69617 | Mm.41933 | NM_145131 | ENSMUSG00000021193 | **Pitrm1** | 2310012C15Rik\|AA410010\|MP-1\|MP1\|Ntup1\|PreP\|mKIAA1104 | pitrilysin metallepetidase 1 | protein-coding |
| chr3_129204138_129205213 | chr3 | 129204139 | 129205213 | + | 0.849443 | intron (NM_001286942, intron 3 of 4) | intron (NM_001286942, intron 3 of 4) | 4798 | NM_001042504 | 18741 | Mm.246804 | NM_011098 | ENSMUSG00000028023 | **Pitx2** | 9430085M16Rik\|Brx1\|Brx1b\|Munc30\|Otlx2\|Ptx2\|Rieg | paired-like homeodomain transcription factor 2 | protein-coding |
| chr3_129204579_129205213 | chr3 | 129204580 | 129205213 | + | 0.804834 | intron (NM_001286942, intron 3 of 4) | intron (NM_001286942, intron 3 of 4) | 5018 | NM_001042504 | 18741 | Mm.246804 | NM_011098 | ENSMUSG00000028023 | **Pitx2** | 9430085M16Rik\|Brx1\|Brx1b\|Munc30\|Otlx2\|Ptx2\|Rieg | paired-like homeodomain transcription factor 2 | protein-coding |
| chr9_14737543_14738291 | chr9 | 14737544 | 14738291 | + | 0.894663 | intron (NM_177905, intron 2 of 18) | B1_Mus1\|SINE\|Alu | 2816 | NM_177905 | 330890 | Mm.334969 | NM_177905 | ENSMUSG00000036912 | **Piwil4** | 9230101H05Rik\|Miwi2 | piwi-like RNA-mediated gene silencing 4 | protein-coding |
| chr17_64334005_64334274 | chr17 | 64334006 | 64334274 | + | 1.39809 | Intergenic | Intergenic | -2257 | NM_001025309 | 224938 | Mm.489681 | NM_144859 | ENSMUSG00000024083 | **Pja2** | AI447901\|AL022700\|Neurodap1\|mKIAA0438 | praja 2, RING-H2 motif containing | protein-coding |
| chr17_64341677_64342094 | chr17 | 64341678 | 64342094 | + | 0.938237 | Intergenic | Intergenic | -10003 | NM_001025309 | 224938 | Mm.489681 | NM_144859 | ENSMUSG00000024083 | **Pja2** | AI447901\|AL022700\|Neurodap1\|mKIAA0438 | praja 2, RING-H2 motif containing | protein-coding |
| chr8_117074142_117075122 | chr8 | 117074143 | 117075122 | + | 1.15586 | intron (NM_029686, intron 4 of 42) | intron (NM_029686, intron 4 of 42) | 7817 | NM_029686 | 76645 | Mm.159753 | NM_029686 | ENSMUSG00000034416 | **Pkd1l2** | 1700126L06Rik | polycystic kidney disease 1 like 2 | protein-coding |
| chr15_44457318_44457955 | chr15 | 44457319 | 44457955 | + | 0.663828 | promoter-TSS (NM_138674) | promoter-TSS (NM_138674) | 84 | NM_138674 | 192190 | Mm.211494 | NM_138674 | ENSMUSG00000038725 | **Pkhd1l1** | AB055648\|PKHDL1 | polycystic kidney and hepatic disease 1-like 1 | protein-coding |
| chr10_57666532_57667326 | chr10 | 57666533 | 57667326 | + | 1.00733 | intron (NM_008863, intron 2 of 5) | ERVL-B4-int\|LTR\|ERVL | 15948 | NM_001190402 | 18768 | Mm.262135 | NM_008863 | ENSMUSG00000019876 | **Pkib** | Pkibeta\|Prkacn2 | protein kinase inhibitor beta, cAMP dependent, testis specific | protein-coding |
| chr10_57666587_57666898 | chr10 | 57666588 | 57666898 | + | 0.862947 | intron (NM_008863, intron 2 of 5) | ERVL-B4-int\|LTR\|ERVL | 15762 | NM_001190402 | 18768 | Mm.262135 | NM_008863 | ENSMUSG00000019876 | **Pkib** | Pkibeta\|Prkacn2 | protein kinase inhibitor beta, cAMP dependent, testis specific | protein-coding |
| chr10_57666967_57667292 | chr10 | 57666968 | 57667292 | + | 0.857948 | intron (NM_008863, intron 2 of 5) | intron (NM_008863, intron 2 of 5) | 16149 | NM_001190402 | 18768 | Mm.262135 | NM_008863 | ENSMUSG00000019876 | **Pkib** | Pkibeta\|Prkacn2 | protein kinase inhibitor beta, cAMP dependent, testis specific | protein-coding |
| chr8_83686225_83686701 | chr8 | 83686226 | 83686701 | + | 1.17678 | intron (NM_177262, intron 4 of 21) | B3A\|SINE\|B2 | 7598 | NM_001199593 | 320795 | Mm.213000 | NM_177262 | ENSMUSG00000057672 | **Pkn1** | DBK\|F730027O18Rik\|PAK1\|PRK1\|Pkn\|Prkcl1\|Stk3 | protein kinase N1 | protein-coding |

| | | | | | | | | | | | | | | | | | |
|---|---|---|---|---|---|---|---|---|---|---|---|---|---|---|---|---|---|
| chr3_143016638_143017260 | chr3 | 143016639 | 143017260 | + | 1.17829 | Intergenic | Intergenic | -134945 | NM_178654 | 109333 | Mm.244236 | NM_178654 | ENSMUSG00000004591 | **Pkn2** | 6030436C20Rik\|AI507382\|PRK2\|Prkcl2\|Stk7 | protein kinase N2 | protein-coding |
| chr3_143058806_143059494 | chr3 | 143058807 | 143059494 | + | 0.804191 | Intergenic | L1M2\|LINE\|L1 | -177146 | NM_178654 | 109333 | Mm.244236 | NM_178654 | ENSMUSG00000004591 | **Pkn2** | 6030436C20Rik\|AI507382\|PRK2\|Prkcl2\|Stk7 | protein kinase N2 | protein-coding |
| chr3_143014974_143015326 | chr3 | 143014975 | 143015326 | + | 0.789657 | Intergenic | Intergenic | -133146 | NM_178654 | 109333 | Mm.244236 | NM_178654 | ENSMUSG00000004591 | **Pkn2** | 6030436C20Rik\|AI507382\|PRK2\|Prkcl2\|Stk7 | protein kinase N2 | protein-coding |
| chr3_143058866_143059238 | chr3 | 143058867 | 143059238 | + | 0.7697 | Intergenic | Tigger5\|DNA\|TcMar-Tigger | -177048 | NM_178654 | 109333 | Mm.244236 | NM_178654 | ENSMUSG00000004591 | **Pkn2** | 6030436C20Rik\|AI507382\|PRK2\|Prkcl2\|Stk7 | protein kinase N2 | protein-coding |
| chr3_142826685_142827312 | chr3 | 142826686 | 142827312 | + | 0.649747 | intron (NM_178654, intron 6 of 21) | MER77B\|LTR\|ERVL | 55005 | NM_178654 | 109333 | Mm.244236 | NM_178654 | ENSMUSG00000004591 | **Pkn2** | 6030436C20Rik\|AI507382\|PRK2\|Prkcl2\|Stk7 | protein kinase N2 | protein-coding |
| chr1_135929635_135930294 | chr1 | 135929636 | 135930294 | + | 0.783148 | Intergenic | Intergenic | -10941 | NM_019645 | 18772 | Mm.4494 | NM_019645 | ENSMUSG00000026413 | **Pkp1** | AI528551 | plakophilin 1 | protein-coding |
| chr1_135929637_135930519 | chr1 | 135929638 | 135930519 | + | 0.680664 | Intergenic | Intergenic | -11054 | NM_019645 | 18772 | Mm.4494 | NM_019645 | ENSMUSG00000026413 | **Pkp1** | AI528551 | plakophilin 1 | protein-coding |
| chr16_16216994_16217294 | chr16 | 16216995 | 16217294 | + | 1.40465 | intron (NM_026163, intron 1 of 12) | intron (NM_026163, intron 1 of 12) | 3799 | NM_026163 | 67451 | Mm.2252 | NM_026163 | ENSMUSG00000041957 | **Pkp2** | 1200008D14Rik\|1200012P04Rik\|AA516617\|Pkp2l | plakophilin 2 | protein-coding |
| chr2_59171560_59172058 | chr2 | 59171561 | 59172058 | + | 0.924716 | intron (NM_026361, intron 1 of 21) | intron (NM_026361, intron 1 of 21) | 10959 | NM_175464 | 227937 | Mm.260938 | NM_026361 | ENSMUSG00000026991 | **Pkp4** | 5031422I09Rik\|9430019K17Rik\|Armrp\|p0071 | plakophilin 4 | protein-coding |
| chr2_59171568_59171921 | chr2 | 59171569 | 59171921 | + | 0.895657 | intron (NM_026361, intron 1 of 21) | intron (NM_026361, intron 1 of 21) | 10895 | NM_175464 | 227937 | Mm.260938 | NM_026361 | ENSMUSG00000026991 | **Pkp4** | 5031422I09Rik\|9430019K17Rik\|Armrp\|p0071 | plakophilin 4 | protein-coding |
| chr2_59171380_59171950 | chr2 | 59171381 | 59171950 | + | 0.721969 | intron (NM_026361, intron 1 of 21) | intron (NM_026361, intron 1 of 21) | 10815 | NM_175464 | 227937 | Mm.260938 | NM_026361 | ENSMUSG00000026991 | **Pkp4** | 5031422I09Rik\|9430019K17Rik\|Armrp\|p0071 | plakophilin 4 | protein-coding |
| chr2_59171246_59172068 | chr2 | 59171247 | 59172068 | + | 0.703233 | intron (NM_026361, intron 1 of 21) | intron (NM_026361, intron 1 of 21) | 10807 | NM_175464 | 227937 | Mm.260938 | NM_026361 | ENSMUSG00000026991 | **Pkp4** | 5031422I09Rik\|9430019K17Rik\|Armrp\|p0071 | plakophilin 4 | protein-coding |
| chr16_38406718_38407151 | chr16 | 38406719 | 38407151 | + | 1.35643 | intron (NM_134102, intron 7 of 10) | intron (NM_134102, intron 7 of 10) | 26210 | NM_134102 | 85031 | Mm.279805 | NM_134102 | ENSMUSG00000002847 | **Pla1a** | AA986889\|Ps-pla1\|Pspla1 | phospholipase A1 member A | protein-coding |
| chr16_38407605_38407932 | chr16 | 38407606 | 38407932 | + | 1.21218 | intron (NM_134102, intron 6 of 10) | intron (NM_134102, intron 6 of 10) | 25376 | NM_134102 | 85031 | Mm.279805 | NM_134102 | ENSMUSG00000002847 | **Pla1a** | AA986889\|Ps-pla1\|Pspla1 | phospholipase A1 member A | protein-coding |
| chr16_38425639_38426029 | chr16 | 38425640 | 38426029 | + | 0.956243 | intron (NM_134102, intron 1 of 10) | intron (NM_134102, intron 1 of 10) | 7311 | NM_134102 | 85031 | Mm.279805 | NM_134102 | ENSMUSG00000002847 | **Pla1a** | AA986889\|Ps-pla1\|Pspla1 | phospholipase A1 member A | protein-coding |
| chr19_7558762_7559112 | chr19 | 7558763 | 7559112 | + | 0.649212 | intron (NM_139269, intron 2 of 4) | MER97c\|DNA\|hAT-Tip100 | 1478 | NM_139269 | 225845 | Mm.274810 | NM_139269 | ENSMUSG00000060675 | **Pla2g16** | C78643\|HRSL3\|Hrasls3\|Hrev107\|MLP-3 | phospholipase A2, group XVI | protein-coding |
| chrX_53162612_53162985 | chrX | 53162613 | 53162985 | + | 0.752038 | Intergenic | Intergenic | -48394 | NM_019538 | 56096 | Mm.274842 | NM_019538 | ENSMUSG00000061082 | **Plac1** | DXWsu72e\|Epcs26\|Oosp2l | placental specific protein 1 | protein-coding |
| chrX_53168610_53169017 | chrX | 53168611 | 53169017 | + | 0.672468 | Intergenic | Intergenic | -54409 | NM_019538 | 56096 | Mm.274842 | NM_019538 | ENSMUSG00000061082 | **Plac1** | DXWsu72e\|Epcs26\|Oosp2l | placental specific protein 1 | protein-coding |
| chr5_100622300_100622791 | chr5 | 100622301 | 100622791 | + | 0.743756 | Intergenic | Intergenic | -50341 | NM_139198 | 231507 | Mm.34609 | NM_139198 | ENSMUSG00000029322 | **Plac8** | C15\|D5Wsu111e | placenta-specific 8 | protein-coding |
| chr5_100597748_100598314 | chr5 | 100597749 | 100598314 | + | 0.667352 | Intergenic | Intergenic | -25826 | NM_139198 | 231507 | Mm.34609 | NM_139198 | ENSMUSG00000029322 | **Plac8** | C15\|D5Wsu111e | placenta-specific 8 | protein-coding |
| chr10_13059911_13060597 | chr10 | 13059912 | 13060597 | + | 0.976551 | Intergenic | Intergenic | -30534 | NM_009538 | 22634 | Mm.287857 | NM_009538 | ENSMUSG00000019817 | **Plagl1** | Lot1\|Zac1 | pleiomorphic adenoma gene-like 1 | protein-coding |
| chr2_153237890_153238343 | chr2 | 153237891 | 153238343 | + | 2.31157 | intron (NM_018807, intron 1 of 2) | intron (NM_018807, intron 1 of 2) | 3241 | NM_018807 | 54711 | Mm.103199 | NM_018807 | ENSMUSG00000051413 | **Plagl2** | AU018672\|AW552839\|mKIAA0198 | pleiomorphic adenoma gene-like 2 | protein-coding |
| chr2_153234754_153235124 | chr2 | 153234755 | 153235124 | + | 1.19662 | intron (NM_018807, intron 2 of 2) | Charlie15a\|DNA\|hAT-Charlie | 6419 | NM_018807 | 54711 | Mm.103199 | NM_018807 | ENSMUSG00000051413 | **Plagl2** | AU018672\|AW552839\|mKIAA0198 | pleiomorphic adenoma gene-like 2 | protein-coding |
| chr2_153238555_153238766 | chr2 | 153238556 | 153238766 | + | 0.856184 | intron (NM_018807, intron 1 of 2) | MIRc\|SINE\|MIR | 2697 | NM_018807 | 54711 | Mm.103199 | NM_018807 | ENSMUSG00000051413 | **Plagl2** | AU018672\|AW552839\|mKIAA0198 | pleiomorphic adenoma gene-like 2 | protein-coding |
| chr2_153239077_153239252 | chr2 | 153239078 | 153239252 | + | 0.797736 | intron (NM_018807, intron 1 of 2) | intron (NM_018807, intron 1 of 2) | 2193 | NM_018807 | 54711 | Mm.103199 | NM_018807 | ENSMUSG00000051413 | **Plagl2** | AU018672\|AW552839\|mKIAA0198 | pleiomorphic adenoma gene-like 2 | protein-coding |
| chr5_32191897_32192580 | chr5 | 32191898 | 32192580 | + | 1.61776 | Intergenic | Intergenic | -40469 | NM_030072 | 665270 | Mm.160067 | NM_030072 | ENSMUSG00000029134 | **Plb1** | 4632413E21Rik\|4930433E17Rik\|4930539A06Rik\|BC033606 | phospholipase B1 | protein-coding |
| chr5_120502032_120502481 | chr5 | 120502033 | 120502481 | + | 0.990902 | intron (NM_023625, intron 1 of 11) | intron (NM_023625, intron 1 of 11) | 1366 | NM_023625 | 71772 | Mm.100065 | NM_023625 | ENSMUSG00000029598 | **Plbd2** | 1300012G16Rik\|AU019810\|P76 | phospholipase B domain containing 2 | protein-coding |
| chr2_135737397_135737912 | chr2 | 135737398 | 135737912 | + | 0.84502 | Intergenic | MLT1H2\|LTR\|ERVL-MaLR | -4175 | NM_013829 | 18798 | Mm.38009 | NM_013829 | ENSMUSG00000039943 | **Plcb4** | A930039J07Rik\|AI854601\|C230058B11Rik | phospholipase C, beta 4 | protein-coding |
| chr1_74550909_74551526 | chr1 | 74550910 | 74551526 | + | 0.998457 | intron (NM_001081456, intron 5 of 16) | intron (NM_001081456, intron 5 of 16) | 6858 | NM_148937 | 18802 | Mm.290731 | NM_148937 | ENSMUSG00000026173 | **Plcd4** | 4921507K24Rik | phospholipase C, delta 4 | protein-coding |
| chr2_160723646_160724354 | chr2 | 160723647 | 160724354 | + | 1.38063 | Intergenic | Intergenic | -7310 | NM_021280 | 18803 | Mm.44463 | NM_021280 | ENSMUSG00000016933 | **Plcg1** | AI894140\|Cded\|Plc-1\|Plc-gamma1\|Plcg-1 | phospholipase C, gamma 1 | protein-coding |
| chr1_55488255_55488658 | chr1 | 55488256 | 55488658 | + | 1.85412 | intron (NM_001114663, intron 1 of 5) | intron (NM_001114663, intron 1 of 5) | 82511 | NM_001114663 | 227120 | Mm.194183 | NM_001114663 | | **Plcl1** | C230017K02Rik\|PLC-L\|PLC-L1\|PLCE\|PLCL\|PLDL1\|PRIP\|PRIP-1 | phospholipase C-like 1 | protein-coding |
| chr1_55410693_55411111 | chr1 | 55410694 | 55411111 | + | 1.40071 | intron (NM_001114663, intron 1 of 5) | intron (NM_001114663, intron 1 of 5) | 4956 | NM_001114663 | 227120 | Mm.194183 | NM_001114663 | | **Plcl1** | C230017K02Rik\|PLC-L\|PLC-L1\|PLCE\|PLCL\|PLDL1\|PRIP\|PRIP-1 | phospholipase C-like 1 | protein-coding |
| chr1_55430393_55431039 | chr1 | 55430394 | 55431039 | + | 1.37425 | intron (NM_001114663, intron 1 of 5) | intron (NM_001114663, intron 1 of 5) | 24770 | NM_001114663 | 227120 | Mm.194183 | NM_001114663 | | **Plcl1** | C230017K02Rik\|PLC-L\|PLC-L1\|PLCE\|PLCL\|PLDL1\|PRIP\|PRIP-1 | phospholipase C-like 1 | protein-coding |
| chr1_55442012_55442379 | chr1 | 55442013 | 55442379 | + | 1.07713 | intron (NM_001114663, intron 1 of 5) | B3A\|SINE\|B2 | 36250 | NM_001114663 | 227120 | Mm.194183 | NM_001114663 | | **Plcl1** | C230017K02Rik\|PLC-L\|PLC-L1\|PLCE\|PLCL\|PLDL1\|PRIP\|PRIP-1 | phospholipase C-like 1 | protein-coding |
| chr1_55489436_55489800 | chr1 | 55489437 | 55489800 | + | 0.966847 | intron (NM_001114663, intron 1 of 5) | intron (NM_001114663, intron 1 of 5) | 83672 | NM_001114663 | 227120 | Mm.194183 | NM_001114663 | | **Plcl1** | C230017K02Rik\|PLC-L\|PLC-L1\|PLCE\|PLCL\|PLDL1\|PRIP\|PRIP-1 | phospholipase C-like 1 | protein-coding |
| chr1_55420301_55420901 | chr1 | 55420302 | 55420901 | + | 0.861062 | intron (NM_001114663, intron 1 of 5) | intron (NM_001114663, intron 1 of 5) | 14655 | NM_001114663 | 227120 | Mm.194183 | NM_001114663 | | **Plcl1** | C230017K02Rik\|PLC-L\|PLC-L1\|PLCE\|PLCL\|PLDL1\|PRIP\|PRIP-1 | phospholipase C-like 1 | protein-coding |
| chr1_55528401_55529192 | chr1 | 55528402 | 55529192 | + | 0.834946 | intron (NM_001114663, intron 1 of 5) | intron (NM_001114663, intron 1 of 5) | 122851 | NM_001114663 | 227120 | Mm.194183 | NM_001114663 | | **Plcl1** | C230017K02Rik\|PLC-L\|PLC-L1\|PLCE\|PLCL\|PLDL1\|PRIP\|PRIP-1 | phospholipase C-like 1 | protein-coding |
| chr1_55528819_55529144 | chr1 | 55528820 | 55529144 | + | 0.759975 | intron (NM_001114663, intron 1 of 5) | intron (NM_001114663, intron 1 of 5) | 123036 | NM_001114663 | 227120 | Mm.194183 | NM_001114663 | | **Plcl1** | C230017K02Rik\|PLC-L\|PLC-L1\|PLCE\|PLCL\|PLDL1\|PRIP\|PRIP-1 | phospholipase C-like 1 | protein-coding |
| chr1_55447766_55448673 | chr1 | 55447767 | 55448673 | + | 0.72438 | intron (NM_001114663, intron 1 of 5) | intron (NM_001114663, intron 1 of 5) | 42274 | NM_001114663 | 227120 | Mm.194183 | NM_001114663 | | **Plcl1** | C230017K02Rik\|PLC-L\|PLC-L1\|PLCE\|PLCL\|PLDL1\|PRIP\|PRIP-1 | phospholipase C-like 1 | protein-coding |
| chr1_55446927_55447259 | chr1 | 55446928 | 55447259 | + | 0.683892 | intron (NM_001114663, intron 1 of 5) | intron (NM_001114663, intron 1 of 5) | 41147 | NM_001114663 | 227120 | Mm.194183 | NM_001114663 | | **Plcl1** | C230017K02Rik\|PLC-L\|PLC-L1\|PLCE\|PLCL\|PLDL1\|PRIP\|PRIP-1 | phospholipase C-like 1 | protein-coding |
| chr17_50570597_50571063 | chr17 | 50570598 | 50571063 | + | 1.00894 | intron (NM_013880, intron 1 of 5) | URR1B\|DNA\|hAT-Charlie | 61283 | NM_013880 | 224860 | Mm.217362 | NM_013880 | ENSMUSG00000038910 | **Plcl2** | PLC-L2\|PRIP-2\|Plce2\|mKIAA1092 | phospholipase C-like 2 | protein-coding |
| chr17_50570592_50571527 | chr17 | 50570593 | 50571527 | + | 0.952205 | intron (NM_013880, intron 1 of 5) | MTD\|LTR\|ERVL-MaLR | 61513 | NM_013880 | 224860 | Mm.217362 | NM_013880 | ENSMUSG00000038910 | **Plcl2** | PLC-L2\|PRIP-2\|Plce2\|mKIAA1092 | phospholipase C-like 2 | protein-coding |
| chr17_50551887_50552489 | chr17 | 50551888 | 50552489 | + | 0.884889 | intron (NM_013880, intron 1 of 5) | intron (NM_013880, intron 1 of 5) | 42641 | NM_013880 | 224860 | Mm.217362 | NM_013880 | ENSMUSG00000038910 | **Plcl2** | PLC-L2\|PRIP-2\|Plce2\|mKIAA1092 | phospholipase C-like 2 | protein-coding |
| chr17_50504134_50504757 | chr17 | 50504135 | 50504757 | + | 0.859448 | Intergenic | MTD\|LTR\|ERVL-MaLR | -5101 | NM_013880 | 224860 | Mm.217362 | NM_013880 | ENSMUSG00000038910 | **Plcl2** | PLC-L2\|PRIP-2\|Plce2\|mKIAA1092 | phospholipase C-like 2 | protein-coding |
| chr17_50478391_50478829 | chr17 | 50478392 | 50478829 | + | 0.793338 | Intergenic | Intergenic | -30937 | NM_013880 | 224860 | Mm.217362 | NM_013880 | ENSMUSG00000038910 | **Plcl2** | PLC-L2\|PRIP-2\|Plce2\|mKIAA1092 | phospholipase C-like 2 | protein-coding |
| chr17_50582693_50583407 | chr17 | 50582694 | 50583407 | + | 0.781929 | intron (NM_013880, intron 1 of 5) | intron (NM_013880, intron 1 of 5) | 73503 | NM_013880 | 224860 | Mm.217362 | NM_013880 | ENSMUSG00000038910 | **Plcl2** | PLC-L2\|PRIP-2\|Plce2\|mKIAA1092 | phospholipase C-like 2 | protein-coding |
| chr17_50454042_50454403 | chr17 | 50454043 | 50454403 | + | 0.708829 | Intergenic | Intergenic | -55324 | NM_013880 | 224860 | Mm.217362 | NM_013880 | ENSMUSG00000038910 | **Plcl2** | PLC-L2\|PRIP-2\|Plce2\|mKIAA1092 | phospholipase C-like 2 | protein-coding |
| chr15_76162392_76162866 | chr15 | 76162393 | 76162866 | + | 1.00155 | Intergenic | Intergenic | 33081 | NM_201394 | 18810 | Mm.234912 | NM_011117 | ENSMUSG00000022565 | **Plec** | AA591047\|AU042537\|EBS1\|PCN\|PLTN\|Plec1 | plectin | protein-coding |
| chr15_76203898_76204569 | chr15 | 76203899 | 76204569 | + | 0.633575 | intron (NM_011117, intron 1 of 31) | intron (NM_011117, intron 1 of 31) | 2087 | NM_201389 | 18810 | Mm.234912 | NM_011117 | ENSMUSG00000022565 | **Plec** | AA591047\|AU042537\|EBS1\|PCN\|PLTN\|Plec1 | plectin | protein-coding |
| chr11_17008313_17009169 | chr11 | 17008314 | 17009169 | + | 0.742317 | promoter-TSS (NM_019549) | promoter-TSS (NM_019549) | -23 | NM_019549 | 56193 | Mm.98232 | NM_019549 | ENSMUSG00000020120 | **Plek** | 2010300B13Rik | pleckstrin | protein-coding |
| chr12_78940042_78940479 | chr12 | 78940043 | 78940479 | + | 1.5523 | Intergenic | Intergenic | -33323 | NM_013738 | 27260 | Mm.103380 | NM_013738 | ENSMUSG00000021118 | **Plek2** | - | pleckstrin 2 | protein-coding |

| | | | | | | | | | | | | | | | | | |
|---|---|---|---|---|---|---|---|---|---|---|---|---|---|---|---|---|---|
| chr12_78939706_78940518 | chr12 | 78939707 | 78940518 | + | 1.43904 | Intergenic | MER21C\|LTR\|ERVL | -33174 | NM_013738 | 27260 | Mm.103380 | NM_013738 | ENSMUSG00000021118 | **Plek2** | - | pleckstrin 2 | protein-coding |
| chr7_130886259_130886753 | chr7 | 130886260 | 130886753 | + | 0.870306 | intron (NM_133942, intron 3 of 9) | intron (NM_133942, intron 3 of 9) | 20596 | NM_133942 | 101476 | Mm.323554 | NM_133942 | ENSMUSG00000040268 | **Plekha1** | AA960558\|C920009D07Rik\|TAPP1 | pleckstrin homology domain containing, family A (phosphoinositide binding specific) member 1 | protein-coding |
| chr7_130886103_130887028 | chr7 | 130886104 | 130887028 | + | 0.772907 | intron (NM_133942, intron 3 of 9) | intron (NM_133942, intron 3 of 9) | 20656 | NM_133942 | 101476 | Mm.323554 | NM_133942 | ENSMUSG00000040268 | **Plekha1** | AA960558\|C920009D07Rik\|TAPP1 | pleckstrin homology domain containing, family A (phosphoinositide binding specific) member 1 | protein-coding |
| chr7_130827695_130828974 | chr7 | 130827696 | 130828974 | + | 0.740843 | Intergenic | Intergenic | -37575 | NM_133942 | 101476 | Mm.323554 | NM_133942 | ENSMUSG00000040268 | **Plekha1** | AA960558\|C920009D07Rik\|TAPP1 | pleckstrin homology domain containing, family A (phosphoinositide binding specific) member 1 | protein-coding |
| chr8_25122423_25122822 | chr8 | 25122424 | 25122822 | + | 0.763754 | Intergenic | Intergenic | -20812 | NM_031257 | 83436 | Mm.261122 | NM_031257 | ENSMUSG00000031557 | **Plekha2** | 6430512N22Rik\|AI851530\|AU041791\|TAPP2\|W91773 | pleckstrin homology domain-containing, family A (phosphoinositide binding specific) member 2 | protein-coding |
| chr8_25122396_25123030 | chr8 | 25122397 | 25123030 | + | 0.696922 | Intergenic | MMERGLN_LTR\|LTR\|ERV1 | -20902 | NM_031257 | 83436 | Mm.261122 | NM_031257 | ENSMUSG00000031557 | **Plekha2** | 6430512N22Rik\|AI851530\|AU041791\|TAPP2\|W91773 | pleckstrin homology domain-containing, family A (phosphoinositide binding specific) member 2 | protein-coding |
| chr2_76676877_76677113 | chr2 | 76676878 | 76677113 | + | 1.00429 | intron (NM_031256, intron 1 of 7) | intron (NM_031256, intron 1 of 7) | 1680 | NM_031256 | 83435 | Mm.41636 | NM_031256 | ENSMUSG00000002733 | **Plekha3** | FAPP1 | pleckstrin homology domain-containing, family A (phosphoinositide binding specific) member 3 | protein-coding |
| chr6_140466899_140467299 | chr6 | 140466900 | 140467299 | + | 1.86984 | intron (NM_144920, intron 3 of 30) | intron (NM_144920, intron 3 of 30) | 43000 | NM_144920 | 109135 | Mm.247670 | NM_144920 | ENSMUSG00000030231 | **Plekha5** | 2810431N21Rik\|AI428202\|AK129423\|AMH-Cre\|Ayu21-9\|Gt(pU21)9Imeg\|Pepp2\|Tg(AMH-cre)1Flor | pleckstrin homology domain containing, family A member 5 | protein-coding |
| chr1_34852442_34852805 | chr1 | 34852443 | 34852805 | + | 1.03455 | intron (NM_175421, intron 1 of 6) | intron (NM_175421, intron 1 of 6) | 2665 | NM_145516 | 226971 | Mm.292751 | NM_145516 | ENSMUSG00000026123 | **Plekhb2** | 2310009M15Rik\|AU018239\|Evt2\|Phdc\|evt-2 | pleckstrin homology domain containing, family B (evectins) member 2 | protein-coding |
| chr1_34852329_34852836 | chr1 | 34852330 | 34852836 | + | 0.944523 | intron (NM_175421, intron 1 of 6) | intron (NM_175421, intron 1 of 6) | 2624 | NM_145516 | 226971 | Mm.292751 | NM_145516 | ENSMUSG00000026123 | **Plekhb2** | 2310009M15Rik\|AU018239\|Evt2\|Phdc\|evt-2 | pleckstrin homology domain containing, family B (evectins) member 2 | protein-coding |
| chr12_80674947_80675383 | chr12 | 80674948 | 80675383 | + | 0.74954 | intron (NM_026244, intron 4 of 6) | MTC\|LTR\|ERVL-MaLR | 17301 | NR_037995 | 68354 | Mm.46194 | NR_037995 | | **Plekhd1os** | 0610009B14Rik | pleckstrin homology domain containing, family D (with coiled-coil domains) member 1, opposite strand | ncRNA |
| chr6_125433664_125434169 | chr6 | 125433665 | 125434169 | + | 1.12739 | Intergenic | Intergenic | -53413 | NM_198604 | 213522 | Mm.333396 | NM_198604 | ENSMUSG00000038167 | **Plekhg6** | BC026778\|Gm155 | pleckstrin homology domain containing, family G (with RhoGef domain) member 6 | protein-coding |
| chr6_125425385_125425696 | chr6 | 125425386 | 125425696 | + | 0.647996 | Intergenic | Intergenic | -45037 | NM_198604 | 213522 | Mm.333396 | NM_198604 | ENSMUSG00000038167 | **Plekhg6** | BC026778\|Gm155 | pleckstrin homology domain containing, family G (with RhoGef domain) member 6 | protein-coding |
| chr17_84535288_84536057 | chr17 | 84535289 | 84536057 | + | 0.983829 | intron (NM_177606, intron 2 of 29) | intron (NM_177606, intron 2 of 29) | 23778 | NM_177606 | 213556 | Mm.215159 | NM_177606 | ENSMUSG00000040852 | **Plekhh2** | AI256725\|E030001K05\|mKIAA2028 | pleckstrin homology domain containing, family H (with MyTH4 domain) member 2 | protein-coding |
| chr1_64875495_64876187 | chr1 | 64875496 | 64876187 | + | 0.694124 | intron (NM_001039493, intron 5 of 7) | L2a\|LINE\|L2 | 80983 | NM_001039493 | 241075 | Mm.239534 | NM_175509 | ENSMUSG00000051344 | **Plekhm3** | 9430067K14Rik\|A230102O09Rik\|AI449570\|DAPR\|Plekhm1l | pleckstrin homology domain containing, family M, member 3 | protein-coding |
| chr3_96010050_96010655 | chr3 | 96010051 | 96010655 | + | 0.846507 | intron (NM_013841, intron 14 of 14) | intron (NM_013841, intron 14 of 14) | -14514 | NM_023320 | 67220 | Mm.458147 | NM_023320 | ENSMUSG00000015745 | **Plekho1** | 2810052M02Rik\|CKIP-1\|Ckip1\|JZA-20\|Jza2 | pleckstrin homology domain containing, family O member 1 | protein-coding |
| chr19_56438162_56438724 | chr19 | 56438163 | 56438724 | + | 0.834993 | intron (NM_007611, intron 6 of 6) | intron (NM_007611, intron 6 of 6) | -23194 | NM_001164263 | 226245 | Mm.156742 | NM_172641 | ENSMUSG00000035818 | **Plekhs1** | 9930023K05Rik | pleckstrin homology domain containing, family S member 1 | protein-coding |
| chr9_50476635_50476999 | chr9 | 50476636 | 50476999 | + | 1.12118 | Intergenic | L2\|LINE\|L2 | -17708 | NM_029639 | 76509 | Mm.29959 | NM_029639 | ENSMUSG00000032068 | **Plet1** | 0610037B23Rik\|1600029D21Rik\|AA407923\|AU043048 | placenta expressed transcript 1 | protein-coding |
| chr9_50324303_50325174 | chr9 | 50324304 | 50325174 | + | 0.864958 | Intergenic | Intergenic | -169786 | NM_029639 | 76509 | Mm.29959 | NM_029639 | ENSMUSG00000032068 | **Plet1** | 0610037B23Rik\|1600029D21Rik\|AA407923\|AU043048 | placenta expressed transcript 1 | protein-coding |
| chr9_50324309_50324718 | chr9 | 50324310 | 50324718 | + | 0.806773 | Intergenic | Intergenic | -170011 | NM_029639 | 76509 | Mm.29959 | NM_029639 | ENSMUSG00000032068 | **Plet1** | 0610037B23Rik\|1600029D21Rik\|AA407923\|AU043048 | placenta expressed transcript 1 | protein-coding |
| chr9_50324768_50325157 | chr9 | 50324769 | 50325157 | + | 0.75886 | Intergenic | B3\|SINE\|B2 | -169562 | NM_029639 | 76509 | Mm.29959 | NM_029639 | ENSMUSG00000032068 | **Plet1** | 0610037B23Rik\|1600029D21Rik\|AA407923\|AU043048 | placenta expressed transcript 1 | protein-coding |
| chr17_12363982_12364703 | chr17 | 12363983 | 12364703 | + | 0.720876 | Intergenic | Tigger5\|DNA\|TcMar-Tigger | -14266 | NM_008877 | 18815 | Mm.971 | NM_008877 | ENSMUSG00000059481 | **Plg** | AI649309\|Pg | plasminogen | protein-coding |
| chr17_12363612_12364715 | chr17 | 12363613 | 12364715 | + | 0.692247 | Intergenic | Tigger5\|DNA\|TcMar-Tigger | -14445 | NM_008877 | 18815 | Mm.971 | NM_008877 | ENSMUSG00000059481 | **Plg** | AI649309\|Pg | plasminogen | protein-coding |
| chr4_86671854_86672411 | chr4 | 86671855 | 86672411 | + | 1.17657 | Intergenic | Intergenic | -2074 | NM_007408 | 11520 | Mm.381 | NM_007408 | ENSMUSG00000028494 | **Plin2** | AA407157\|ADPH\|Adfp\|Adrp | perilipin 2 | protein-coding |
| chr8_94703366_94703757 | chr8 | 94703367 | 94703757 | + | 0.799871 | Intergenic | (TGG)n\|Simple_repeat\|Simple_repeat | -7320 | NM_026385 | 67801 | Mm.279977 | NM_026385 | ENSMUSG00000031775 | **Pllp** | 0610010I06Rik\|AV001002\|Plapi\|Tm4sf11 | plasma membrane proteolipid | protein-coding |
| chr10_53355368_53355897 | chr10 | 53355369 | 53355897 | + | 1.49189 | intron (NM_001204983, intron 2 of 12) | ORR1D2\|LTR\|ERVL-MaLR | 17947 | NM_001141927 | 18821 | Mm.34145 | NM_023129 | ENSMUSG00000038583 | **Pln** | Plb | phospholamban | protein-coding |
| chr10_53355420_53355718 | chr10 | 53355421 | 53355718 | + | 1.47839 | intron (NM_001204983, intron 2 of 12) | ORR1D2\|LTR\|ERVL-MaLR | 17883 | NM_001141927 | 18821 | Mm.34145 | NM_023129 | ENSMUSG00000038583 | **Pln** | Plb | phospholamban | protein-coding |
| chrX_7673494_7674268 | chrX | 7673495 | 7674268 | + | 0.69915 | non-coding (NR_037581, exon 6 of 6) | non-coding (NR_037581, exon 6 of 6) | -2603 | NM_019755 | 18824 | Mm.18565 | NM_019755 | | **Plp2** | mIMA4 | proteolipid protein 2 | protein-coding |
| chrX_7671230_7671538 | chrX | 7671231 | 7671538 | + | 0.680481 | promoter-TSS (NM_019755) | promoter-TSS (NM_019755) | -106 | NM_019755 | 18824 | Mm.18565 | NM_019755 | | **Plp2** | mIMA4 | proteolipid protein 2 | protein-coding |
| chr9_95835601_95836294 | chr9 | 95835602 | 95836294 | + | 1.23039 | intron (NM_001033210, intron 1 of 15) | intron (NM_001033210, intron 1 of 15) | 9331 | NM_001033210 | 102502 | Mm.11869 | NM_001033210 | ENSMUSG00000049493 | **Pls1** | AI427122 | plastin 1 (I-isoform) | protein-coding |
| chr9_95832720_95833340 | chr9 | 95832721 | 95833340 | + | 1.00386 | intron (NM_001033210, intron 1 of 15) | MER44D\|DNA\|TcMar-Tigger | 12249 | NM_001033210 | 102502 | Mm.11869 | NM_001033210 | ENSMUSG00000049493 | **Pls1** | AI427122 | plastin 1 (I-isoform) | protein-coding |
| chr9_95827638_95827979 | chr9 | 95827639 | 95827979 | + | 0.790103 | intron (NM_001033210, intron 1 of 15) | intron (NM_001033210, intron 1 of 15) | 17470 | NM_001033210 | 102502 | Mm.11869 | NM_001033210 | ENSMUSG00000049493 | **Pls1** | AI427122 | plastin 1 (I-isoform) | protein-coding |
| chr9_92249344_92249862 | chr9 | 92249345 | 92249862 | + | 1.48088 | promoter-TSS (NM_011636) | promoter-TSS (NM_011636) | -591 | NM_011636 | 22038 | Mm.421956 | NM_011636 | ENSMUSG00000032369 | **Plscr1** | MmTRA1a\|MmTRA1b\|Nor1\|Tra1\|Tra1a\|Tra1b\|Tras1\|Tras2 | phospholipid scramblase 1 | protein-coding |
| chr9_92228974_92229787 | chr9 | 92228975 | 92229787 | + | 1.16808 | Intergenic | Intergenic | -20813 | NM_011636 | 22038 | Mm.421956 | NM_011636 | ENSMUSG00000032369 | **Plscr1** | MmTRA1a\|MmTRA1b\|Nor1\|Tra1\|Tra1a\|Tra1b\|Tras1\|Tras2 | phospholipid scramblase 1 | protein-coding |
| chr9_92245849_92246299 | chr9 | 92245850 | 92246299 | + | 1.09602 | Intergenic | RMER6A\|LTR\|ERVK | -4120 | NM_011636 | 22038 | Mm.421956 | NM_011636 | ENSMUSG00000032369 | **Plscr1** | MmTRA1a\|MmTRA1b\|Nor1\|Tra1\|Tra1a\|Tra1b\|Tras1\|Tras2 | phospholipid scramblase 1 | protein-coding |
| chr9_92249705_92250807 | chr9 | 92249706 | 92250807 | + | 0.900302 | promoter-TSS (NM_011636) | promoter-TSS (NM_011636) | 62 | NM_011636 | 22038 | Mm.421956 | NM_011636 | ENSMUSG00000032369 | **Plscr1** | MmTRA1a\|MmTRA1b\|Nor1\|Tra1\|Tra1a\|Tra1b\|Tras1\|Tras2 | phospholipid scramblase 1 | protein-coding |
| chr9_92226131_92226503 | chr9 | 92226132 | 92226503 | + | 0.740346 | Intergenic | Intergenic | -23877 | NM_011636 | 22038 | Mm.421956 | NM_011636 | ENSMUSG00000032369 | **Plscr1** | MmTRA1a\|MmTRA1b\|Nor1\|Tra1\|Tra1a\|Tra1b\|Tras1\|Tras2 | phospholipid scramblase 1 | protein-coding |
| chr9_92274150_92274817 | chr9 | 92274151 | 92274817 | + | 0.859798 | Intergenic | Intergenic | -1118 | NM_008880 | 18828 | Mm.10306 | NM_008880 | ENSMUSG00000032372 | **Plscr2** | - | phospholipid scramblase 2 | protein-coding |
| chr9_92384043_92385040 | chr9 | 92384044 | 92385040 | + | 1.51667 | Intergenic | Tigger1\|DNA\|TcMar-Tigger | -72836 | NM_178711 | 235527 | Mm.55289 | NM_178711 | | **Plscr4** | AV245873 | phospholipid scramblase 4 | protein-coding |
| chr9_92217559_92218217 | chr9 | 92217560 | 92218217 | + | 0.746258 | Intergenic | Intergenic | 24952 | NM_001195693 | 100504689 | Mm.464262 | NM_001195693 | ENSMUSG00000095654 | **Plscr5** | 331000 | phospholipid scramblase family, member 5 | protein-coding |
| chr2_164854664_164854876 | chr2 | 164854665 | 164854876 | + | 0.684111 | intron (NM_011125, intron 4 of 15) | Lx8\|LINE\|L1 | 2938 | NM_011125 | 18830 | Mm.6105 | NM_011125 | ENSMUSG00000017754 | **Pltp** | Bpife\|OD107 | phospholipid transfer protein | protein-coding |
| chr2_16352857_16353352 | chr2 | 16352858 | 16353352 | + | 1.66423 | Intergenic | Intergenic | -3199 | NM_026162 | 67448 | Mm.313938 | NM_026162 | ENSMUSG00000026748 | **Plxdc2** | 1200007L24Rik\|5430431D22Rik\|AI646728\|AU022916\|AV231634\|Tem7r | plexin domain containing 2 | protein-coding |
| chr2_16570416_16571343 | chr2 | 16570417 | 16571343 | + | 0.993335 | intron (NM_026162, intron 4 of 13) | intron (NM_026162, intron 4 of 13) | 214576 | NM_026162 | 67448 | Mm.313938 | NM_026162 | ENSMUSG00000026748 | **Plxdc2** | 1200007L24Rik\|5430431D22Rik\|AI646728\|AU022916\|AV231634\|Tem7r | plexin domain containing 2 | protein-coding |

| | | | | | | | | | | | | | | | | | |
|---|---|---|---|---|---|---|---|---|---|---|---|---|---|---|---|---|---|
| chr2_16565858_16566151 | chr2 | 16565859 | 16566151 | + | 0.811622 | intron (NM_026162, intron 4 of 13) | intron (NM_026162, intron 4 of 13) | 209701 | NM_026162 | 67448 | Mm.313938 | NM_026162 | ENSMUSG00000026748 | **Plxdc2** | 1200007L24Rik\|5430431D22Rik\|AI646728\|AU022916\|AV231634\|Tem7r | plexin domain containing 2 | protein-coding |
| chr2_16570381_16571645 | chr2 | 16570382 | 16571645 | + | 0.807964 | intron (NM_026162, intron 4 of 13) | intron (NM_026162, intron 4 of 13) | 214709 | NM_026162 | 67448 | Mm.313938 | NM_026162 | ENSMUSG00000026748 | **Plxdc2** | 1200007L24Rik\|5430431D22Rik\|AI646728\|AU022916\|AV231634\|Tem7r | plexin domain containing 2 | protein-coding |
| chrX_74324872_74325311 | chrX | 74324873 | 74325311 | + | 0.724637 | Intergenic | RMER1A\|Other\|Other | -3974 | NM_008883 | 18846 | Mm.1685 | NM_008883 | ENSMUSG00000031398 | **Plxna3** | PlexA3\|Plxa3\|Plxn3\|Plxn4\|SEX\|mKIAA4078 | plexin A3 | protein-coding |
| chr6_32796113_32796478 | chr6 | 32796114 | 32796478 | + | 0.760535 | intron (NM_025336, intron 7 of 7) | intron (NM_025336, intron 7 of 7) | -208104 | NM_175750 | 243743 | Mm.444029 | NM_175750 | ENSMUSG00000029765 | **Plxna4** | 9330117B14\|Plxa4\|mKIAA1550 | plexin A4 | protein-coding |
| chr10_94860714_94861426 | chr10 | 94860715 | 94861426 | + | 1.78225 | intron (NM_018797, intron 9 of 30) | intron (NM_018797, intron 9 of 30) | 83508 | NM_018797 | 54712 | Mm.256712 | NM_018797 | ENSMUSG00000074785 | **Plxnc1** | 2510048K12Rik\|AW742158\|CD232\|Vespr | plexin C1 | protein-coding |
| chr10_94864284_94864977 | chr10 | 94864285 | 94864977 | + | 1.41957 | intron (NM_018797, intron 9 of 30) | intron (NM_018797, intron 9 of 30) | 79947 | NM_018797 | 54712 | Mm.256712 | NM_018797 | ENSMUSG00000074785 | **Plxnc1** | 2510048K12Rik\|AW742158\|CD232\|Vespr | plexin C1 | protein-coding |
| chr10_94849393_94849751 | chr10 | 94849394 | 94849751 | + | 1.21261 | intron (NM_018797, intron 11 of 30) | intron (NM_018797, intron 11 of 30) | 95006 | NM_018797 | 54712 | Mm.256712 | NM_018797 | ENSMUSG00000074785 | **Plxnc1** | 2510048K12Rik\|AW742158\|CD232\|Vespr | plexin C1 | protein-coding |
| chr10_95119339_95119686 | chr10 | 95119340 | 95119686 | + | 0.883386 | Intergenic | Intergenic | -174935 | NM_018797 | 54712 | Mm.256712 | NM_018797 | ENSMUSG00000074785 | **Plxnc1** | 2510048K12Rik\|AW742158\|CD232\|Vespr | plexin C1 | protein-coding |
| chr10_94919558_94919920 | chr10 | 94919559 | 94919920 | + | 0.859222 | intron (NM_018797, intron 2 of 30) | intron (NM_018797, intron 2 of 30) | 24839 | NM_018797 | 54712 | Mm.256712 | NM_018797 | ENSMUSG00000074785 | **Plxnc1** | 2510048K12Rik\|AW742158\|CD232\|Vespr | plexin C1 | protein-coding |
| chr6_116089341_116089840 | chr6 | 116089342 | 116089840 | + | 1.37763 | intron (NM_177412, intron 3 of 5) | URR1A\|DNA\|hAT-Charlie | -94586 | NM_026376 | 67784 | Mm.3085 | NM_026376 | ENSMUSG00000030123 | **Plxnd1** | 6230425C21Rik\|b2b1863Clo\|b2b553Clo | plexin D1 | protein-coding |
| chr6_115989021_115990050 | chr6 | 115989022 | 115990050 | + | 0.644262 | intron (NM_026376, intron 1 of 35) | intron (NM_026376, intron 1 of 35) | 5469 | NM_026376 | 67784 | Mm.3085 | NM_026376 | ENSMUSG00000030123 | **Plxnd1** | 6230425C21Rik\|b2b1863Clo\|b2b553Clo | plexin D1 | protein-coding |
| chr18_66453180_66453750 | chr18 | 66453181 | 66453750 | + | 0.842189 | Intergenic | PB1D7\|SINE\|Alu | -5139 | NM_021451 | 58801 | Mm.271878 | NM_021451 | ENSMUSG00000024521 | **Pmaip1** | Noxa | phorbol-12-myristate-13-acetate-induced protein 1 | protein-coding |
| chr18_66453189_66453583 | chr18 | 66453190 | 66453583 | + | 0.836239 | Intergenic | PB1D7\|SINE\|Alu | -5218 | NM_021451 | 58801 | Mm.271878 | NM_021451 | ENSMUSG00000024521 | **Pmaip1** | Noxa | phorbol-12-myristate-13-acetate-induced protein 1 | protein-coding |
| chr18_66428624_66429003 | chr18 | 66428625 | 66429003 | + | 0.819457 | Intergenic | B3\|SINE\|B2 | -29790 | NM_021451 | 58801 | Mm.271878 | NM_021451 | ENSMUSG00000024521 | **Pmaip1** | Noxa | phorbol-12-myristate-13-acetate-induced protein 1 | protein-coding |
| chr18_66428562_66429485 | chr18 | 66428563 | 66429485 | + | 0.774034 | Intergenic | B3\|SINE\|B2 | -29580 | NM_021451 | 58801 | Mm.271878 | NM_021451 | ENSMUSG00000024521 | **Pmaip1** | Noxa | phorbol-12-myristate-13-acetate-induced protein 1 | protein-coding |
| chr10_88054100_88054885 | chr10 | 88054101 | 88054885 | + | 0.923768 | Intergenic | Intergenic | -36579 | NM_029971 | 110312 | Mm.391436 | NM_029971 | ENSMUSG00000035383 | **Pmch** | A230109K23Rik | pro-melanin-concentrating hormone | protein-coding |
| chr2_173385642_173386306 | chr2 | 173385643 | 173386306 | + | 1.34989 | Intergenic | Intergenic | -109441 | NM_022995 | 65112 | Mm.73682 | NM_022995 | ENSMUSG00000038400 | **Pmepa1** | 2210418I02Rik\|AW455466\|Erg1.2\|N4wbp4\|Stag1\|Tmepai | prostate transmembrane protein, androgen induced 1 | protein-coding |
| chr2_173273932_173274293 | chr2 | 173273933 | 173274293 | + | 1.20564 | intron (NM_022995, intron 1 of 3) | intron (NM_022995, intron 1 of 3) | 2420 | NM_022995 | 65112 | Mm.73682 | NM_022995 | ENSMUSG00000038400 | **Pmepa1** | 2210418I02Rik\|AW455466\|Erg1.2\|N4wbp4\|Stag1\|Tmepai | prostate transmembrane protein, androgen induced 1 | protein-coding |
| chr2_173385299_173385579 | chr2 | 173385300 | 173385579 | + | 0.911378 | Intergenic | Intergenic | -108906 | NM_022995 | 65112 | Mm.73682 | NM_022995 | ENSMUSG00000038400 | **Pmepa1** | 2210418I02Rik\|AW455466\|Erg1.2\|N4wbp4\|Stag1\|Tmepai | prostate transmembrane protein, androgen induced 1 | protein-coding |
| chr2_173385160_173385576 | chr2 | 173385161 | 173385576 | + | 0.897323 | Intergenic | Intergenic | -108835 | NM_022995 | 65112 | Mm.73682 | NM_022995 | ENSMUSG00000038400 | **Pmepa1** | 2210418I02Rik\|AW455466\|Erg1.2\|N4wbp4\|Stag1\|Tmepai | prostate transmembrane protein, androgen induced 1 | protein-coding |
| chr9_58250301_58250672 | chr9 | 58250302 | 58250672 | + | 1.28235 | promoter-TSS (NM_008884) | promoter-TSS (NM_008884) | -701 | NM_008884 | 18854 | Mm.392123 | NM_008884 | ENSMUSG00000036986 | **Pml** | 1200009E24Rik\|AI661194\|Trim19 | promyelocytic leukemia | protein-coding |
| chr3_89451914_89453013 | chr3 | 89451915 | 89453013 | + | 1.77614 | Intergenic | Intergenic | -6654 | NM_026784 | 68603 | Mm.34242 | NM_026784 | ENSMUSG00000027952 | **Pmvk** | 1110011E12Rik\|2900002L22Rik\|AV001327\|PMK\|PMKA\|PMKASE | phosphomevalonate kinase | protein-coding |
| chr12_59077996_59078454 | chr12 | 59077997 | 59078454 | + | 0.844872 | Intergenic | Intergenic | 11306 | NM_008891 | 18949 | Mm.22347 | NM_008891 | ENSMUSG00000020994 | **Pnn** | AU045199\|D12Ertd512e | pinin | protein-coding |
| chr11_17187675_17187992 | chr11 | 17187676 | 17187992 | + | 0.830482 | intron (NM_024459, intron 2 of 5) | intron (NM_024459, intron 2 of 5) | 23755 | NM_025443 | 66249 | Mm.27831 | NM_025443 | ENSMUSG00000020116 | **Pno1** | 1810003N24Rik\|AA407421\|AU015468\|AU019495\|AU042323\|Emi3\|Imi3 | partner of NOB1 homolog (S. cerevisiae) | protein-coding |
| chr14_65393543_65393861 | chr14 | 65393544 | 65393861 | + | 0.843202 | intron (NM_199029, intron 7 of 9) | intron (NM_199029, intron 7 of 9) | 31770 | NM_001205075 | 18155 | Mm.16347 | NM_010932 | ENSMUSG00000045731 | **Pnoc** | N/OFQ\|N23K\|Npnc1\|OFQ/N | prepronociceptin | protein-coding |
| chr14_65393438_65393872 | chr14 | 65393439 | 65393872 | + | 0.833089 | intron (NM_199029, intron 7 of 9) | intron (NM_199029, intron 7 of 9) | 31817 | NM_001205075 | 18155 | Mm.16347 | NM_010932 | ENSMUSG00000045731 | **Pnoc** | N/OFQ\|N23K\|Npnc1\|OFQ/N | prepronociceptin | protein-coding |
| chr4_33253186_33253412 | chr4 | 33253187 | 33253412 | + | 0.901007 | Intergenic | Intergenic | -4512 | NM_001033225 | 108767 | Mm.27769 | NM_001033225 | ENSMUSG00000040128 | **Pnrc1** | 5730463C12Rik\|B4-2\|PRR2\|Prol2 | proline-rich nuclear receptor coactivator 1 | protein-coding |
| chr10_99081550_99082247 | chr10 | 99081551 | 99082247 | + | 0.860849 | Intergenic | Intergenic | -25272 | NM_027740 | 382406 | Mm.440656 | NM_027740 | ENSMUSG00000019952 | **Poc1b** | 4933430F16Rik\|Wdr51b | POC1 centriolar protein homolog B (Chlamydomonas) | protein-coding |
| chr4_108058117_108058723 | chr4 | 108058118 | 108058723 | + | 1.23961 | intron (NM_011327, intron 13 of 15) | MTEb\|LTR\|ERVL-MaLR | -26330 | NM_172874 | 242608 | Mm.74710 | NM_172874 | ENSMUSG00000028600 | **Podn** | 9430070G18\|Pcan\|SLRR5A | podocan | protein-coding |
| chr6_31494909_31496553 | chr6 | 31494910 | 31496553 | + | 1.3046 | intron (NM_013791, intron 15 of 17) | intron (NM_013791, intron 15 of 17) | 68206 | NM_013723 | 27205 | Mm.89918 | NM_013723 | ENSMUSG00000025608 | **Podxl** | AW121214\|Ly102\|PC\|PCLP-1\|Pclp1 | podocalyxin-like | protein-coding |
| chr6_31489725_31490375 | chr6 | 31489726 | 31490375 | + | 1.14645 | intron (NM_013791, intron 13 of 17) | intron (NM_013791, intron 13 of 17) | 73887 | NM_013723 | 27205 | Mm.89918 | NM_013723 | ENSMUSG00000025608 | **Podxl** | AW121214\|Ly102\|PC\|PCLP-1\|Pclp1 | podocalyxin-like | protein-coding |
| chr6_31640615_31641132 | chr6 | 31640616 | 31641132 | + | 0.729483 | Intergenic | Intergenic | -76937 | NM_013723 | 27205 | Mm.89918 | NM_013723 | ENSMUSG00000025608 | **Podxl** | AW121214\|Ly102\|PC\|PCLP-1\|Pclp1 | podocalyxin-like | protein-coding |
| chr10_77273306_77273831 | chr10 | 77273307 | 77273831 | + | 0.835817 | Intergenic | Intergenic | 14269 | NM_030262 | 80294 | Mm.203556 | NM_030262 | ENSMUSG00000020260 | **Pofut2** | 2310011G23Rik\|AI256847\|BC003494\|C21orf80\|FUT13 | protein O-fucosyltransferase 2 | protein-coding |
| chr10_77273065_77273842 | chr10 | 77273066 | 77273842 | + | 0.758637 | Intergenic | Intergenic | 14154 | NM_030262 | 80294 | Mm.203556 | NM_030262 | ENSMUSG00000020260 | **Pofut2** | 2310011G23Rik\|AI256847\|BC003494\|C21orf80\|FUT13 | protein O-fucosyltransferase 2 | protein-coding |
| chr10_77273023_77274163 | chr10 | 77273024 | 77274163 | + | 0.75484 | Intergenic | Intergenic | 14293 | NM_030262 | 80294 | Mm.203556 | NM_030262 | ENSMUSG00000020260 | **Pofut2** | 2310011G23Rik\|AI256847\|BC003494\|C21orf80\|FUT13 | protein O-fucosyltransferase 2 | protein-coding |
| chr11_5882678_5882958 | chr11 | 5882679 | 5882958 | + | 0.86221 | Intergenic | Intergenic | -4562 | NM_008894 | 18972 | Mm.35788 | NM_008894 | ENSMUSG00000020471 | **Pold2** | 50kDa\|p50\|po1D2 | polymerase (DNA directed), delta 2, regulatory subunit | protein-coding |
| chr11_5882515_5883007 | chr11 | 5882516 | 5883007 | + | 0.773808 | Intergenic | Intergenic | -4505 | NM_008894 | 18972 | Mm.35788 | NM_008894 | ENSMUSG00000020471 | **Pold2** | 50kDa\|p50\|po1D2 | polymerase (DNA directed), delta 2, regulatory subunit | protein-coding |
| chr7_100100030_100100356 | chr7 | 100100031 | 100100356 | + | 0.768999 | intron (NM_133692, intron 6 of 11) | PB1D10\|SINE\|Alu | 21307 | NM_133692 | 67967 | Mm.37562 | NM_133692 | ENSMUSG00000030726 | **Pold3** | 2410142G14Rik\|C85233\|P66\|P68 | polymerase (DNA-directed), delta 3, accessory subunit | protein-coding |
| chr19_4230499_4230791 | chr19 | 4230500 | 4230791 | + | 0.83685 | Intergenic | Intergenic | -1254 | NM_027196 | 69745 | Mm.32518 | NM_027196 | ENSMUSG00000024854 | **Pold4** | 2410012M21Rik\|AI463381\|AW060307\|Polds\|p12 | polymerase (DNA-directed), delta 4 | protein-coding |
| chr5_110300387_110301036 | chr5 | 110300388 | 110301036 | + | 1.2359 | intron (NM_011132, intron 20 of 48) | intron (NM_011132, intron 20 of 48) | 14393 | NM_011132 | 18973 | Mm.35061 | NM_011132 | ENSMUSG00000007080 | **Pole** | - | polymerase (DNA directed), epsilon | protein-coding |
| chr5_110300656_110301039 | chr5 | 110300657 | 110301039 | + | 1.13911 | intron (NM_011132, intron 20 of 48) | intron (NM_011132, intron 20 of 48) | 14529 | NM_011132 | 18973 | Mm.35061 | NM_011132 | ENSMUSG00000007080 | **Pole** | - | polymerase (DNA directed), epsilon | protein-coding |
| chr5_110300656_110301201 | chr5 | 110300657 | 110301201 | + | 1.01087 | intron (NM_011132, intron 20 of 48) | intron (NM_011132, intron 20 of 48) | 14610 | NM_011132 | 18973 | Mm.35061 | NM_011132 | ENSMUSG00000007080 | **Pole** | - | polymerase (DNA directed), epsilon | protein-coding |
| chr13_96533784_96534196 | chr13 | 96533785 | 96534196 | + | 1.02467 | intron (NM_012048, intron 1 of 14) | intron (NM_012048, intron 1 of 14) | 8495 | NM_012048 | 27015 | Mm.89926 | NM_012048 | ENSMUSG00000021668 | **Polk** | Dinb1 | polymerase (DNA directed), kappa | protein-coding |
| chr11_5835092_5835417 | chr11 | 5835093 | 5835417 | + | 1.5328 | intron (NM_017401, intron 4 of 10) | intron (NM_017401, intron 4 of 10) | 2761 | NM_017401 | 54125 | Mm.260194 | NM_017401 | ENSMUSG00000020474 | **Polm** | AI462166\|B230309I03Rik\|Tdt-N | polymerase (DNA directed), mu | protein-coding |
| chr5_147086531_147087030 | chr5 | 147086532 | 147087030 | + | 1.00283 | intron (NM_181730, intron 1 of 2) | MTC\|LTR\|ERVL-MaLR | 9435 | NM_009087 | 20018 | Mm.317557 | NM_009087 | ENSMUSG00000029642 | **Polr1d** | 1110003G10Rik\|AC19\|RPA16\|Rpo1-3 | polymerase (RNA) I polypeptide D | protein-coding |
| chr9_71483928_71484288 | chr9 | 71483929 | 71484288 | + | 0.83527 | intron (NM_178602, intron 1 of 3) | B1_Mus2\|SINE\|Alu | 850 | NM_001164793 | 28015 | Mm.21094 | NM_178602 | ENSMUSG00000032199 | **Polr2m** | AA407243\|AA408799\|AI303746\|AW553238\|D9Wsu138e\|Grinl1a\|mKIAA4185 | polymerase (RNA) II (DNA directed) polypeptide M | protein-coding |
| chr13_81684134_81684722 | chr13 | 81684135 | 81684722 | + | 1.19314 | intron (NM_001081176, intron 5 of 7) | ORR1B1-int\|LTR\|ERVL-MaLR | 26585 | NM_001081176 | 67486 | Mm.279781 | NM_001081176 | ENSMUSG00000035834 | **Polr3g** | 2310047G20Rik\|AV275904\|RPC32\|RPC7 | polymerase (RNA) III (DNA directed) polypeptide G | protein-coding |
| chr13_81684903_81685329 | chr13 | 81684904 | 81685329 | + | 1.15633 | intron (NM_001081176, intron 5 of 7) | B3A\|SINE\|B2 | 25897 | NM_001081176 | 67486 | Mm.279781 | NM_001081176 | ENSMUSG00000035834 | **Polr3g** | 2310047G20Rik\|AV275904\|RPC32\|RPC7 | polymerase (RNA) III (DNA directed) polypeptide G | protein-coding |
| chr2_181878570_181879519 | chr2 | 181878571 | 181879519 | + | 1.09071 | Intergenic | Intergenic | 14685 | NM_025901 | 67005 | Mm.271986 | NM_025901 | ENSMUSG00000038628 | **Polr3k** | 12.3kDa\|1500004O14Rik\|AI196849\|AU014893\|C11\|RPC10\|RPC11 | polymerase (RNA) III (DNA directed) polypeptide K | protein-coding |

| | | | | | | | | | | | | | | | | | |
|---|---|---|---|---|---|---|---|---|---|---|---|---|---|---|---|---|---|
| chr13_21984358_21984993 | chr13 | 21984359 | 21984993 | + | 1.2136 | 3' UTR (NM_001162928, exon 1 of 1) | 3' UTR (NM_001162928, exon 1 of 1) | 3482 | NM_001162929 | 195236 | Mm.204664 | NM_001162928 | ENSMUSG00000016982 | **Pom121l2** | Gm24 | POM121 membrane glycoprotein-like 2 (rat) | protein-coding |
| chr13_21984087_21985184 | chr13 | 21984088 | 21985184 | + | 1.11536 | 3' UTR (NM_001162928, exon 1 of 1) | 3' UTR (NM_001162928, exon 1 of 1) | 3442 | NM_001162929 | 195236 | Mm.204664 | NM_001162928 | ENSMUSG00000016982 | **Pom121l2** | Gm24 | POM121 membrane glycoprotein-like 2 (rat) | protein-coding |
| chr13_21993862_21994611 | chr13 | 21993863 | 21994611 | + | 0.863192 | Intergenic | tRNA-Ser-AGY\|tRNA\|tRNA | 13043 | NM_001162929 | 195236 | Mm.204664 | NM_001162928 | ENSMUSG00000016982 | **Pom121l2** | Gm24 | POM121 membrane glycoprotein-like 2 (rat) | protein-coding |
| chr13_21985418_21985729 | chr13 | 21985419 | 21985729 | + | 0.77896 | 3' UTR (NM_001162928, exon 1 of 1) | 3' UTR (NM_001162928, exon 1 of 1) | 4380 | NM_001162929 | 195236 | Mm.204664 | NM_001162928 | ENSMUSG00000016982 | **Pom121l2** | Gm24 | POM121 membrane glycoprotein-like 2 (rat) | protein-coding |
| chr6_5295113_5295389 | chr6 | 5295114 | 5295389 | + | 0.903419 | intron (NM_183308, intron 1 of 8) | intron (NM_183308, intron 1 of 8) | 3122 | NM_183308 | 330260 | Mm.126984 | NM_183308 | | **Pon2** | 6330405I24Rik\|AI481612 | paraoxonase 2 | protein-coding |
| chr16_38396043_38396685 | chr16 | 38396044 | 38396685 | + | 1.61434 | 3' UTR (NM_134102, exon 11 of 11) | 3' UTR (NM_134102, exon 11 of 11) | 34154 | NM_022318 | 64082 | Mm.390856 | NM_022318 | ENSMUSG00000022803 | **Popdc2** | AV006127\|Pop2 | popeye domain containing 2 | protein-coding |
| chr16_38379000_38379466 | chr16 | 38379001 | 38379466 | + | 1.4912 | Intergenic | Intergenic | 17023 | NM_022318 | 64082 | Mm.390856 | NM_022318 | ENSMUSG00000022803 | **Popdc2** | AV006127\|Pop2 | popeye domain containing 2 | protein-coding |
| chr16_38378606_38379466 | chr16 | 38378607 | 38379466 | + | 1.23774 | TTS (NM_001081984) | TTS (NM_001081984) | 16826 | NM_022318 | 64082 | Mm.390856 | NM_022318 | ENSMUSG00000022803 | **Popdc2** | AV006127\|Pop2 | popeye domain containing 2 | protein-coding |
| chr16_38374818_38375328 | chr16 | 38374819 | 38375328 | + | 1.1528 | intron (NM_001081984, intron 3 of 3) | intron (NM_001081984, intron 3 of 3) | 12863 | NM_022318 | 64082 | Mm.390856 | NM_022318 | ENSMUSG00000022803 | **Popdc2** | AV006127\|Pop2 | popeye domain containing 2 | protein-coding |
| chr16_38358354_38358918 | chr16 | 38358355 | 38358918 | + | 1.1351 | Intergenic | Intergenic | -3537 | NM_001081984 | 64082 | Mm.390856 | NM_022318 | ENSMUSG00000022803 | **Popdc2** | AV006127\|Pop2 | popeye domain containing 2 | protein-coding |
| chr16_38395221_38395602 | chr16 | 38395222 | 38395602 | + | 1.10842 | TTS (NM_134102) | TTS (NM_134102) | 33202 | NM_022318 | 64082 | Mm.390856 | NM_022318 | ENSMUSG00000022803 | **Popdc2** | AV006127\|Pop2 | popeye domain containing 2 | protein-coding |
| chr16_38371905_38372263 | chr16 | 38371906 | 38372263 | + | 1.04762 | intron (NM_022318, intron 2 of 3) | intron (NM_022318, intron 2 of 3) | 9874 | NM_022318 | 64082 | Mm.390856 | NM_022318 | ENSMUSG00000022803 | **Popdc2** | AV006127\|Pop2 | popeye domain containing 2 | protein-coding |
| chr16_38358066_38358916 | chr16 | 38358067 | 38358916 | + | 0.984326 | Intergenic | Intergenic | -3682 | NM_001081984 | 64082 | Mm.390856 | NM_022318 | ENSMUSG00000022803 | **Popdc2** | AV006127\|Pop2 | popeye domain containing 2 | protein-coding |
| chr16_38385322_38386328 | chr16 | 38385323 | 38386328 | + | 0.901149 | Intergenic | Intergenic | 23615 | NM_022318 | 64082 | Mm.390856 | NM_022318 | ENSMUSG00000022803 | **Popdc2** | AV006127\|Pop2 | popeye domain containing 2 | protein-coding |
| chr16_38384887_38386328 | chr16 | 38384888 | 38386328 | + | 0.839602 | Intergenic | Intergenic | 23398 | NM_022318 | 64082 | Mm.390856 | NM_022318 | ENSMUSG00000022803 | **Popdc2** | AV006127\|Pop2 | popeye domain containing 2 | protein-coding |
| chr16_38374372_38374700 | chr16 | 38374373 | 38374700 | + | 0.832265 | intron (NM_001081984, intron 3 of 3) | intron (NM_001081984, intron 3 of 3) | 12326 | NM_022318 | 64082 | Mm.390856 | NM_022318 | ENSMUSG00000022803 | **Popdc2** | AV006127\|Pop2 | popeye domain containing 2 | protein-coding |
| chr16_38378606_38378946 | chr16 | 38378607 | 38378946 | + | 0.734367 | TTS (NM_001081984) | TTS (NM_001081984) | 16566 | NM_022318 | 64082 | Mm.390856 | NM_022318 | ENSMUSG00000022803 | **Popdc2** | AV006127\|Pop2 | popeye domain containing 2 | protein-coding |
| chr16_38369003_38369397 | chr16 | 38369004 | 38369397 | + | 0.69609 | intron (NM_022318, intron 1 of 3) | intron (NM_022318, intron 1 of 3) | 6990 | NM_022318 | 64082 | Mm.390856 | NM_022318 | ENSMUSG00000022803 | **Popdc2** | AV006127\|Pop2 | popeye domain containing 2 | protein-coding |
| chr16_65351083_65351431 | chr16 | 65351084 | 65351431 | + | 0.879876 | Intergenic | Intergenic | -169372 | NM_008849 | 18736 | Mm.355440 | NM_008849 | ENSMUSG00000004842 | **Pou1f1** | GHF-1\|Hmp1\|Pit-1\|Pit1\|Pit1-rs1\|dw\|dwarf | POU domain, class 1, transcription factor 1 | protein-coding |
| chr16_65349902_65350292 | chr16 | 65349903 | 65350292 | + | 0.809803 | Intergenic | ORR1B1\|LTR\|ERVL-MaLR | -170532 | NM_008849 | 18736 | Mm.355440 | NM_008849 | ENSMUSG00000004842 | **Pou1f1** | GHF-1\|Hmp1\|Pit-1\|Pit1\|Pit1-rs1\|dw\|dwarf | POU domain, class 1, transcription factor 1 | protein-coding |
| chr9_51177319_51177836 | chr9 | 51177320 | 51177836 | + | 1.34813 | Intergenic | Intergenic | -36112 | NM_011136 | 18985 | Mm.897 | NM_011136 | ENSMUSG00000032053 | **Pou2af1** | BOB.1\|Bob-1\|Bob1\|OBF-1\|OBF.1\|OCA-B\|OCAB | POU domain, class 2, associating factor 1 | protein-coding |
| chr7_25130572_25130986 | chr7 | 25130573 | 25130986 | + | 1.46036 | intron (NM_011138, intron 1 of 13) | intron (NM_011138, intron 1 of 13) | 1681 | NM_011138 | 18987 | Mm.76683 | NM_011138 | | **Pou2f2** | Oct-2\|Oct2a\|Oct2b\|Otf-2\|Otf2 | POU domain, class 2, transcription factor 2 | protein-coding |
| chr7_25121023_25122040 | chr7 | 25121024 | 25122040 | + | 1.02125 | intron (NM_011138, intron 1 of 13) | intron (NM_011138, intron 1 of 13) | 10928 | NM_011138 | 18987 | Mm.76683 | NM_011138 | | **Pou2f2** | Oct-2\|Oct2a\|Oct2b\|Otf-2\|Otf2 | POU domain, class 2, transcription factor 2 | protein-coding |
| chr13_77908618_77909190 | chr13 | 77908619 | 77909190 | + | 1.37516 | intron (NR_028109, intron 7 of 10) | intron (NR_028109, intron 7 of 10) | -115998 | NM_029315 | 75507 | Mm.443255 | NM_029315 | ENSMUSG00000093668 | **Pou5f2** | 1700013G10Rik\|SPRM-1\|Sprm1 | POU domain class 5, transcription factor 2 | protein-coding |
| chr3_133319656_133320166 | chr3 | 133319657 | 133320166 | + | 1.14636 | intron (NM_146141, intron 1 of 11) | intron (NM_146141, intron 1 of 11) | 9801 | NM_146141 | 74776 | Mm.210305 | NM_146141 | ENSMUSG00000028013 | **Ppa2** | 1110013G13Rik\|Sid6306 | pyrophosphatase (inorganic) 2 | protein-coding |
| chr3_133333582_133334105 | chr3 | 133333583 | 133334105 | + | 0.889645 | intron (NM_146141, intron 5 of 11) | intron (NM_146141, intron 5 of 11) | 23734 | NM_146141 | 74776 | Mm.210305 | NM_146141 | ENSMUSG00000028013 | **Ppa2** | 1110013G13Rik\|Sid6306 | pyrophosphatase (inorganic) 2 | protein-coding |
| chr3_133338807_133339266 | chr3 | 133338808 | 133339266 | + | 0.712178 | intron (NM_146141, intron 5 of 11) | intron (NM_146141, intron 5 of 11) | 28927 | NM_146141 | 74776 | Mm.210305 | NM_146141 | ENSMUSG00000028013 | **Ppa2** | 1110013G13Rik\|Sid6306 | pyrophosphatase (inorganic) 2 | protein-coding |
| chr6_115315771_115316148 | chr6 | 115315772 | 115316148 | + | 1.36276 | Intergenic | Tigger7\|DNA\|TcMar-Tigger | -45261 | NM_001127330 | 19016 | Mm.3020 | NM_011146 | ENSMUSG00000000440 | **Pparg** | Nr1c3\|PPAR-gamma\|PPAR-gamma2\|PPARgamma\|PPARgamma2 | peroxisome proliferator activated receptor gamma | protein-coding |
| chr6_115315702_115316388 | chr6 | 115315703 | 115316388 | + | 1.29129 | Intergenic | Tigger7\|DNA\|TcMar-Tigger | -45176 | NM_001127330 | 19016 | Mm.3020 | NM_011146 | ENSMUSG00000000440 | **Pparg** | Nr1c3\|PPAR-gamma\|PPAR-gamma2\|PPARgamma\|PPARgamma2 | peroxisome proliferator activated receptor gamma | protein-coding |
| chr6_115479984_115480833 | chr6 | 115479985 | 115480833 | + | 0.777315 | intron (NM_011146, intron 6 of 6) | intron (NM_011146, intron 6 of 6) | 58343 | NM_011146 | 19016 | Mm.3020 | NM_011146 | ENSMUSG00000000440 | **Pparg** | Nr1c3\|PPAR-gamma\|PPAR-gamma2\|PPARgamma\|PPARgamma2 | peroxisome proliferator activated receptor gamma | protein-coding |
| chr18_61385121_61385777 | chr18 | 61385122 | 61385777 | + | 0.686425 | intron (NM_133249, intron 1 of 11) | intron (NM_133249, intron 1 of 11) | 14982 | NM_133249 | 170826 | Mm.415302 | NM_133249 | ENSMUSG00000033871 | **Ppargc1b** | 4631412G21Rik\|Perc | peroxisome proliferative activated receptor, gamma, coactivator 1 beta | protein-coding |
| chr9_57427345_57427758 | chr9 | 57427346 | 57427758 | + | 1.10892 | intron (NM_176831, intron 3 of 6) | intron (NM_176831, intron 3 of 6) | 12562 | NM_176831 | 66812 | Mm.250508 | NM_176831 | ENSMUSG00000063849 | **Ppcdc** | 1810057I13Rik\|8430432M10Rik | phosphopantothenoylcysteine decarboxylase | protein-coding |
| chr5_92262572_92263230 | chr5 | 92262573 | 92263230 | + | 0.733015 | intron (NM_025972, intron 7 of 10) | intron (NM_025972, intron 7 of 10) | -9742 | NM_011148 | 19023 | Mm.87155 | NM_011148 | ENSMUSG00000029410 | **Ppef2** | - | protein phosphatase, EF hand calcium-binding domain 2 | protein-coding |
| chr6_146893909_146894434 | chr6 | 146893910 | 146894434 | + | 1.55692 | intron (NM_001170433, intron 1 of 27) | intron (NM_001170433, intron 1 of 27) | 5678 | NM_026221 | 67533 | Mm.103382 | NM_026221 | ENSMUSG00000016487 | **Ppfibp1** | 4632409B19Rik\|AW214454\|AW261454 | PTPRF interacting protein, binding protein 1 (liprin beta 1) | protein-coding |
| chr6_146909666_146910440 | chr6 | 146909667 | 146910440 | + | 0.965734 | intron (NM_001170433, intron 1 of 27) | intron (NM_001170433, intron 1 of 27) | 21559 | NM_026221 | 67533 | Mm.103382 | NM_026221 | ENSMUSG00000016487 | **Ppfibp1** | 4632409B19Rik\|AW214454\|AW261454 | PTPRF interacting protein, binding protein 1 (liprin beta 1) | protein-coding |
| chr6_146909658_146910035 | chr6 | 146909659 | 146910035 | + | 0.891466 | intron (NM_001170433, intron 1 of 27) | intron (NM_001170433, intron 1 of 27) | 21353 | NM_026221 | 67533 | Mm.103382 | NM_026221 | ENSMUSG00000016487 | **Ppfibp1** | 4632409B19Rik\|AW214454\|AW261454 | PTPRF interacting protein, binding protein 1 (liprin beta 1) | protein-coding |
| chr7_107648196_107649446 | chr7 | 107648197 | 107649446 | + | 1.22639 | intron (NM_008905, intron 2 of 24) | intron (NM_008905, intron 2 of 24) | 41977 | NM_001163557 | 19024 | Mm.2817 | NM_008905 | ENSMUSG00000036528 | **Ppfibp2** | Cclp1 | PTPRF interacting protein, binding protein 2 (liprin beta 2) | protein-coding |
| chr7_107609395_107610299 | chr7 | 107609396 | 107610299 | + | 0.987902 | intron (NM_001163557, intron 1 of 25) | intron (NM_001163557, intron 1 of 25) | 3003 | NM_001163557 | 19024 | Mm.2817 | NM_008905 | ENSMUSG00000036528 | **Ppfibp2** | Cclp1 | PTPRF interacting protein, binding protein 2 (liprin beta 2) | protein-coding |
| chr4_123146765_123148201 | chr4 | 123146766 | 123148201 | + | 0.868479 | Intergenic | L2b\|LINE\|L2 | -7542 | NM_019489 | 56031 | Mm.126873 | NM_019489 | ENSMUSG00000028651 | **Ppie** | 2010010D16Rik\|Cyp33 | peptidylprolyl isomerase E (cyclophilin E) | protein-coding |
| chr1_97746271_97746824 | chr1 | 97746272 | 97746824 | + | 1.46867 | intron (NM_173760, intron 10 of 27) | intron (NM_173760, intron 10 of 27) | 23544 | NM_173760 | 227399 | Mm.220817 | NM_173760 | ENSMUSG00000040648 | **Ppip5k2** | AW555814\|D330021B20\|Hisppd1\|Vip2\|mKIAA0433 | diphosphoinositol pentakisphosphate kinase 2 | protein-coding |
| chr1_97745749_97746036 | chr1 | 97745750 | 97746036 | + | 0.863237 | intron (NM_173760, intron 11 of 27) | intron (NM_173760, intron 11 of 27) | 24199 | NM_173760 | 227399 | Mm.220817 | NM_173760 | ENSMUSG00000040648 | **Ppip5k2** | AW555814\|D330021B20\|Hisppd1\|Vip2\|mKIAA0433 | diphosphoinositol pentakisphosphate kinase 2 | protein-coding |
| chr1_97757989_97758342 | chr1 | 97757990 | 97758342 | + | 0.624011 | intron (NM_173760, intron 3 of 27) | intron (NM_173760, intron 3 of 27) | 11926 | NM_173760 | 227399 | Mm.220817 | NM_173760 | ENSMUSG00000040648 | **Ppip5k2** | AW555814\|D330021B20\|Hisppd1\|Vip2\|mKIAA0433 | diphosphoinositol pentakisphosphate kinase 2 | protein-coding |
| chr12_72748357_72749196 | chr12 | 72748358 | 72749196 | + | 0.999019 | Intergenic | RSINE1\|SINE\|B4 | -12434 | NM_008910 | 19042 | Mm.261045 | NM_008910 | ENSMUSG00000021096 | **Ppm1a** | 2310003C21Rik\|2900017D14Rik\|AI427932\|AU017636\|MMPa-2\|MPPa-1 | protein phosphatase 1A, magnesium dependent, alpha isoform | protein-coding |
| chr6_57529062_57529659 | chr6 | 57529063 | 57529659 | + | 0.977062 | intron (NM_175523, intron 1 of 6) | URR1B\|DNA\|hAT-Charlie | 6065 | NM_175523 | 243382 | Mm.396893 | NM_175523 | ENSMUSG00000037826 | **Ppm1k** | 2900063A19Rik\|A930026L03Rik\|PP2Cm | protein phosphatase 1K (PP2C domain containing) | protein-coding |
| chr3_69440605_69441213 | chr3 | 69440606 | 69441213 | + | 1.45626 | intron (NM_178726, intron 1 of 3) | intron (NM_178726, intron 1 of 3) | 123991 | NM_178726 | 242083 | Mm.40577 | NM_178726 | | **Ppm1l** | 3222401G13\|5930404J21Rik\|AI481720\|AW045850\|PP2C\|PP2Cepsilon\|Pp2ce\|Ppp2ce\|mKIAA4175 | protein phosphatase 1 (formerly 2C)-like | protein-coding |
| chr3_69269526_69269963 | chr3 | 69269527 | 69269963 | + | 0.769681 | Intergenic | B3\|SINE\|B2 | -47173 | NM_178726 | 242083 | Mm.40577 | NM_178726 | | **Ppm1l** | 3222401G13\|5930404J21Rik\|AI481720\|AW045850\|PP2C\|PP2Cepsilon\|Pp2ce\|Ppp2ce\|mKIAA4175 | protein phosphatase 1 (formerly 2C)-like | protein-coding |
| chr5_32447201_32447758 | chr5 | 32447202 | 32447758 | + | 1.38613 | Intergenic | L1ME3\|LINE\|L1 | -11490 | NM_172707 | 19046 | Mm.241931 | NM_172707 | | **Ppp1cb** | 1200010B19 | protein phosphatase 1, catalytic subunit, beta isoform | protein-coding |
| chr5_32447012_32447768 | chr5 | 32447013 | 32447768 | + | 1.20233 | Intergenic | L1ME3\|LINE\|L1 | -11580 | NM_172707 | 19046 | Mm.241931 | NM_172707 | | **Ppp1cb** | 1200010B19 | protein phosphatase 1, catalytic subunit, beta isoform | protein-coding |

| | | | | | | | | | | | | | | | | | |
|---|---|---|---|---|---|---|---|---|---|---|---|---|---|---|---|---|---|
| chr5_32430016_32431284 | chr5 | 32430017 | 32431284 | + | 1.0736 | Intergenic | Intergenic | -28320 | NM_172707 | 19046 | Mm.241931 | NM_172707 | | **Ppp1cb** | 1200010B19 | protein phosphatase 1, catalytic subunit, beta isoform | protein-coding |
| chr5_32427501_32427861 | chr5 | 32427502 | 32427861 | + | 1.03662 | Intergenic | Intergenic | -31289 | NM_172707 | 19046 | Mm.241931 | NM_172707 | | **Ppp1cb** | 1200010B19 | protein phosphatase 1, catalytic subunit, beta isoform | protein-coding |
| chr10_108173136_108173612 | chr10 | 108173137 | 108173612 | + | 1.25081 | intron (NM_027892, intron 1 of 25) | B4A\|SINE\|B4 | 10974 | NM_027892 | 17931 | Mm.422959 | NM_027892 | ENSMUSG00000019907 | **Ppp1r12a** | 1200015F06Rik\|5730577I22Rik\|AA792106\|AV099298\|D10Ertd625e\|Mypt1 | protein phosphatase 1, regulatory (inhibitor) subunit 12A | protein-coding |
| chr10_108138348_108138690 | chr10 | 108138349 | 108138690 | + | 1.01803 | Intergenic | MTEa\|LTR\|ERVL-MaLR | -23881 | NM_027892 | 17931 | Mm.422959 | NM_027892 | ENSMUSG00000019907 | **Ppp1r12a** | 1200015F06Rik\|5730577I22Rik\|AA792106\|AV099298\|D10Ertd625e\|Mypt1 | protein phosphatase 1, regulatory (inhibitor) subunit 12A | protein-coding |
| chr10_108051586_108051956 | chr10 | 108051587 | 108051956 | + | 1.00733 | Intergenic | L1MB8\|LINE\|L1 | -110629 | NM_027892 | 17931 | Mm.422959 | NM_027892 | ENSMUSG00000019907 | **Ppp1r12a** | 1200015F06Rik\|5730577I22Rik\|AA792106\|AV099298\|D10Ertd625e\|Mypt1 | protein phosphatase 1, regulatory (inhibitor) subunit 12A | protein-coding |
| chr10_108170839_108171139 | chr10 | 108170840 | 108171139 | + | 0.998519 | intron (NM_027892, intron 1 of 25) | intron (NM_027892, intron 1 of 25) | 8589 | NM_027892 | 17931 | Mm.422959 | NM_027892 | ENSMUSG00000019907 | **Ppp1r12a** | 1200015F06Rik\|5730577I22Rik\|AA792106\|AV099298\|D10Ertd625e\|Mypt1 | protein phosphatase 1, regulatory (inhibitor) subunit 12A | protein-coding |
| chr10_108108610_108109366 | chr10 | 108108611 | 108109366 | + | 0.824223 | Intergenic | MTA_Mm\|LTR\|ERVL-MaLR | -53412 | NM_027892 | 17931 | Mm.422959 | NM_027892 | ENSMUSG00000019907 | **Ppp1r12a** | 1200015F06Rik\|5730577I22Rik\|AA792106\|AV099298\|D10Ertd625e\|Mypt1 | protein phosphatase 1, regulatory (inhibitor) subunit 12A | protein-coding |
| chr10_108190062_108190358 | chr10 | 108190063 | 108190358 | + | 0.806386 | intron (NM_027892, intron 1 of 25) | intron (NM_027892, intron 1 of 25) | 27810 | NM_027892 | 17931 | Mm.422959 | NM_027892 | ENSMUSG00000019907 | **Ppp1r12a** | 1200015F06Rik\|5730577I22Rik\|AA792106\|AV099298\|D10Ertd625e\|Mypt1 | protein phosphatase 1, regulatory (inhibitor) subunit 12A | protein-coding |
| chr10_108108878_108109341 | chr10 | 108108879 | 108109341 | + | 0.788095 | Intergenic | MTA_Mm\|LTR\|ERVL-MaLR | -53290 | NM_027892 | 17931 | Mm.422959 | NM_027892 | ENSMUSG00000019907 | **Ppp1r12a** | 1200015F06Rik\|5730577I22Rik\|AA792106\|AV099298\|D10Ertd625e\|Mypt1 | protein phosphatase 1, regulatory (inhibitor) subunit 12A | protein-coding |
| chr10_108043739_108044351 | chr10 | 108043740 | 108044351 | + | 0.764815 | Intergenic | MTD\|LTR\|ERVL-MaLR | -118355 | NM_027892 | 17931 | Mm.422959 | NM_027892 | ENSMUSG00000019907 | **Ppp1r12a** | 1200015F06Rik\|5730577I22Rik\|AA792106\|AV099298\|D10Ertd625e\|Mypt1 | protein phosphatase 1, regulatory (inhibitor) subunit 12A | protein-coding |
| chr12_111909188_111909526 | chr12 | 111909189 | 111909526 | + | 0.726425 | Intergenic | MIR\|SINE\|MIR | -1302 | NM_011625 | 21981 | Mm.313076 | NM_011625 | ENSMUSG00000021285 | **Ppp1r13b** | AI449786\|ASPP1\|AW545810\|Tp53bp2\|Trp53bp2\|p85 | protein phosphatase 1, regulatory (inhibitor) subunit 13B | protein-coding |
| chr2_158674470_158674968 | chr2 | 158674471 | 158674968 | + | 0.948831 | intron (NM_001159662, intron 1 of 10) | B4\|SINE\|B4 | 7584 | NM_001159662 | 228852 | Mm.150540 | NM_153089 | ENSMUSG00000037754 | **Ppp1r16b** | ANKRD4\|C130078N17Rik\|TIMAP\|Wdt4 | protein phosphatase 1, regulatory (inhibitor) subunit 16B | protein-coding |
| chr2_158700908_158701303 | chr2 | 158700909 | 158701303 | + | 0.687117 | intron (NM_153089, intron 2 of 10) | L1MB7\|LINE\|L1 | 33971 | NM_001159662 | 228852 | Mm.150540 | NM_153089 | ENSMUSG00000037754 | **Ppp1r16b** | ANKRD4\|C130078N17Rik\|TIMAP\|Wdt4 | protein phosphatase 1, regulatory (inhibitor) subunit 16B | protein-coding |
| chr2_79698006_79698873 | chr2 | 79698007 | 79698873 | + | 1.70106 | Intergenic | Intergenic | -9340 | NM_172420 | 75276 | Mm.257373 | NM_172420 | ENSMUSG00000034683 | **Ppp1r1c** | 4930565M23Rik\|4930579P15Rik | protein phosphatase 1, regulatory (inhibitor) subunit 1C | protein-coding |
| chr17_88508793_88509636 | chr17 | 88508794 | 88509636 | + | 0.816366 | Intergenic | Intergenic | -20909 | NM_028658 | 73825 | Mm.271817 | NM_028658 | ENSMUSG00000034709 | **Ppp1r21** | 1110018J12Rik\|AI426045\|AW550781\|Ccdc128\|Klraq1 | protein phosphatase 1, regulatory subunit 21 | protein-coding |
| chr2_28445052_28445487 | chr2 | 28445053 | 28445487 | + | 0.751054 | Intergenic | ORR1D1\|LTR\|ERVL-MaLR | -2671 | NM_001005420 | 241289 | Mm.330095 | NM_001005420 | ENSMUSG00000035829 | **Ppp1r26** | Gm347\|Kiaa0649\|mKIAA0649 | protein phosphatase 1, regulatory subunit 26 | protein-coding |
| chr19_10497459_10498252 | chr19 | 10497460 | 10498252 | + | 0.764474 | Intergenic | MIRb\|SINE\|MIR | -14959 | NM_133689 | 67752 | Mm.25939 | NM_133689 | ENSMUSG00000035179 | **Ppp1r32** | 4930579J09Rik\|AU015816\|IIIG9\|IIIG9L\|IIIG9S | protein phosphatase 1, regulatory subunit 32 | protein-coding |
| chr5_137773241_137773913 | chr5 | 137773242 | 137773913 | + | 1.43615 | Intergenic | Intergenic | -5341 | NM_027242 | 69871 | Mm.276058 | NM_027242 | ENSMUSG00000029725 | **Ppp1r35** | 2010007H12Rik\|2010011D20Rik\|AL024004 | protein phosphatase 1, regulatory subunit 35 | protein-coding |
| chr8_35425915_35426386 | chr8 | 35425916 | 35426386 | + | 1.0761 | Intergenic | Intergenic | 50410 | NM_177741 | 244416 | Mm.247126 | NM_177741 | ENSMUSG00000046794 | **Ppp1r3b** | 6430576E21\|AW821953\|GL\|PTG | protein phosphatase 1, regulatory (inhibitor) subunit 3B | protein-coding |
| chr13_36028815_36029394 | chr13 | 36028816 | 36029394 | + | 0.772487 | intron (NM_201358, intron 2 of 2) | ORR1F\|LTR\|ERVL-MaLR | 61199 | NM_029628 | 76487 | Mm.44745 | NM_029628 | ENSMUSG00000050423 | **Ppp1r3g** | 1600032L17Rik | protein phosphatase 1, regulatory (inhibitor) subunit 3G | protein-coding |
| chr11_52079955_52080452 | chr11 | 52079956 | 52080452 | + | 1.39011 | intron (NM_001166654, intron 13 of 13) | intron (NM_001166654, intron 13 of 13) | -18620 | NM_019411 | 19052 | Mm.260288 | NM_019411 | | **Ppp2ca** | PP2A\|R75353 | protein phosphatase 2 (formerly 2A), catalytic subunit, alpha isoform | protein-coding |
| chr8_33606228_33606630 | chr8 | 33606229 | 33606630 | + | 0.792866 | intron (NM_017374, intron 1 of 6) | RMER6B\|LTR\|ERVK | 6808 | NM_017374 | 19053 | Mm.29301 | NM_017374 | ENSMUSG00000009630 | **Ppp2cb** | AI115466\|D8Ertd766e\|PP2Ac | protein phosphatase 2 (formerly 2A), catalytic subunit, beta isoform | protein-coding |
| chr9_50886672_50887696 | chr9 | 50886673 | 50887696 | + | 0.905361 | intron (NM_001286553, intron 15 of 16) | intron (NM_001286553, intron 15 of 16) | 30260 | NM_001286553 | 73699 | Mm.7726 | NM_028614 | ENSMUSG00000032058 | **Ppp2r1b** | 2410091N08Rik\|AI790395 | protein phosphatase 2, regulatory subunit A, beta | protein-coding |
| chr9_50835784_50837126 | chr9 | 50835785 | 50837126 | + | 0.901028 | intron (NR_045601, intron 13 of 13) | intron (NR_045601, intron 13 of 13) | -20469 | NM_001034085 | 73699 | Mm.7726 | NM_028614 | ENSMUSG00000032058 | **Ppp2r1b** | 2410091N08Rik\|AI790395 | protein phosphatase 2, regulatory subunit A, beta | protein-coding |
| chr9_50823482_50824130 | chr9 | 50823483 | 50824130 | + | 0.809287 | intron (NR_045601, intron 13 of 13) | MLT2B2\|LTR\|ERVL | -33118 | NM_001034085 | 73699 | Mm.7726 | NM_028614 | ENSMUSG00000032058 | **Ppp2r1b** | 2410091N08Rik\|AI790395 | protein phosphatase 2, regulatory subunit A, beta | protein-coding |
| chr14_67045174_67045816 | chr14 | 67045175 | 67045816 | + | 0.771343 | intron (NM_028032, intron 2 of 9) | L1MC3\|LINE\|L1 | 26976 | NM_028032 | 71978 | Mm.273997 | NM_028032 | ENSMUSG00000022052 | **Ppp2r2a** | 2410004D02Rik\|mKIAA1541 | protein phosphatase 2, regulatory subunit B, alpha | protein-coding |
| chr14_67047777_67048166 | chr14 | 67047778 | 67048166 | + | 0.680902 | intron (NM_028032, intron 2 of 9) | intron (NM_028032, intron 2 of 9) | 24499 | NM_028032 | 71978 | Mm.273997 | NM_028032 | ENSMUSG00000022052 | **Ppp2r2a** | 2410004D02Rik\|mKIAA1541 | protein phosphatase 2, regulatory subunit B, alpha | protein-coding |
| chr9_101232046_101232558 | chr9 | 101232047 | 101232558 | + | 0.869817 | intron (NM_001161362, intron 1 of 13) | RMER15\|LTR\|ERVL | 19530 | NM_001161362 | 235542 | Mm.335386 | NM_172144 | ENSMUSG00000043154 | **Ppp2r3a** | 3222402P14Rik\|A730042E07 | protein phosphatase 2, regulatory subunit B'', alpha | protein-coding |
| chr1_191368237_191368608 | chr1 | 191368238 | 191368608 | + | 1.49367 | intron (NM_144880, intron 3 of 12) | RSINE1\|SINE\|B4 | 28618 | NM_144880 | 226849 | Mm.205569 | NM_144880 | ENSMUSG00000026626 | **Ppp2r5a** | PR61alpha | protein phosphatase 2, regulatory subunit B', alpha | protein-coding |
| chr1_191384548_191385431 | chr1 | 191384549 | 191385431 | + | 1.00829 | intron (NM_144880, intron 1 of 12) | intron (NM_144880, intron 1 of 12) | 12051 | NM_144880 | 226849 | Mm.205569 | NM_144880 | ENSMUSG00000026626 | **Ppp2r5a** | PR61alpha | protein phosphatase 2, regulatory subunit B', alpha | protein-coding |
| chr1_191414366_191414739 | chr1 | 191414367 | 191414739 | + | 0.88997 | Intergenic | Charlie5\|DNA\|hAT-Charlie | -17512 | NM_144880 | 226849 | Mm.205569 | NM_144880 | ENSMUSG00000026626 | **Ppp2r5a** | PR61alpha | protein phosphatase 2, regulatory subunit B', alpha | protein-coding |
| chr1_191431952_191433120 | chr1 | 191431953 | 191433120 | + | 0.803185 | Intergenic | Intergenic | -35495 | NM_144880 | 226849 | Mm.205569 | NM_144880 | ENSMUSG00000026626 | **Ppp2r5a** | PR61alpha | protein phosphatase 2, regulatory subunit B', alpha | protein-coding |
| chr1_191432273_191433120 | chr1 | 191432274 | 191433120 | + | 0.789753 | Intergenic | Intergenic | -35656 | NM_144880 | 226849 | Mm.205569 | NM_144880 | ENSMUSG00000026626 | **Ppp2r5a** | PR61alpha | protein phosphatase 2, regulatory subunit B', alpha | protein-coding |
| chr12_110381048_110381882 | chr12 | 110381049 | 110381882 | + | 0.800286 | Intergenic | Intergenic | -65714 | NM_001135001 | 26931 | Mm.240396 | NM_012023 | ENSMUSG00000017843 | **Ppp2r5c** | 2610043M05Rik\|2700063L20Rik\|AI060890\|AW545884\|C85228\|D12Bwg0916e\|mKIAA0044 | protein phosphatase 2, regulatory subunit B', gamma | protein-coding |
| chr12_75544210_75544610 | chr12 | 75544211 | 75544610 | + | 0.886291 | intron (NM_012024, intron 2 of 13) | intron (NM_012024, intron 2 of 13) | 51790 | NM_012024 | 26932 | Mm.259626 | NM_012024 | ENSMUSG00000021051 | **Ppp2r5e** | 4633401M22Rik\|AI449017\|B56beta | protein phosphatase 2, regulatory subunit B', epsilon | protein-coding |
| chr12_75599563_75599729 | chr12 | 75599564 | 75599729 | + | 0.876497 | Intergenic | B4A\|SINE\|B4 | -3446 | NM_012024 | 26932 | Mm.259626 | NM_012024 | ENSMUSG00000021051 | **Ppp2r5e** | 4633401M22Rik\|AI449017\|B56beta | protein phosphatase 2, regulatory subunit B', epsilon | protein-coding |
| chr3_136901866_136902417 | chr3 | 136901867 | 136902417 | + | 1.17032 | intron (NM_008913, intron 8 of 13) | intron (NM_008913, intron 8 of 13) | 232076 | NM_001293622 | 19055 | Mm.331389 | NM_008913 | ENSMUSG00000028161 | **Ppp3ca** | 2900074D19Rik\|CN\|Caln\|Calna\|CnA | protein phosphatase 3, catalytic subunit, alpha isoform | protein-coding |
| chr3_136940783_136941843 | chr3 | 136940784 | 136941843 | + | 1.11029 | Intergenic | Intergenic | 271247 | NM_001293622 | 19055 | Mm.331389 | NM_008913 | ENSMUSG00000028161 | **Ppp3ca** | 2900074D19Rik\|CN\|Caln\|Calna\|CnA | protein phosphatase 3, catalytic subunit, alpha isoform | protein-coding |
| chr3_136941008_136941329 | chr3 | 136941009 | 136941329 | + | 1.1051 | Intergenic | Intergenic | 271103 | NM_001293622 | 19055 | Mm.331389 | NM_008913 | ENSMUSG00000028161 | **Ppp3ca** | 2900074D19Rik\|CN\|Caln\|Calna\|CnA | protein phosphatase 3, catalytic subunit, alpha isoform | protein-coding |
| chr3_136951726_136952722 | chr3 | 136951727 | 136952722 | + | 0.922937 | Intergenic | Intergenic | 282158 | NM_001293622 | 19055 | Mm.331389 | NM_008913 | ENSMUSG00000028161 | **Ppp3ca** | 2900074D19Rik\|CN\|Caln\|Calna\|CnA | protein phosphatase 3, catalytic subunit, alpha isoform | protein-coding |
| chr14_70281832_70282054 | chr14 | 70281833 | 70282054 | + | 1.06491 | intron (NM_008915, intron 1 of 13) | intron (NM_008915, intron 1 of 13) | 7506 | NM_008915 | 19057 | Mm.439683 | NM_008915 | ENSMUSG00000022092 | **Ppp3cc** | Calnc | protein phosphatase 3, catalytic subunit, gamma isoform | protein-coding |
| chr14_70263093_70263795 | chr14 | 70263094 | 70263795 | + | 0.948808 | intron (NM_008915, intron 3 of 13) | intron (NM_008915, intron 3 of 13) | 26005 | NM_008915 | 19057 | Mm.439683 | NM_008915 | ENSMUSG00000022092 | **Ppp3cc** | Calnc | protein phosphatase 3, catalytic subunit, gamma isoform | protein-coding |
| chr11_17176493_17176989 | chr11 | 17176494 | 17176989 | + | 1.21884 | intron (NM_024459, intron 1 of 5) | PB1D9\|SINE\|Alu | 17443 | NM_024459 | 19058 | Mm.41840 | NM_024459 | ENSMUSG00000033953 | **Ppp3r1** | CaNB1\|Cnb1\|MCIP1 | protein phosphatase 3, regulatory subunit B, alpha isoform (calcineurin B, type I) | protein-coding |
| chr17_65818990_65819548 | chr17 | 65818991 | 65819548 | + | 1.12613 | intron (NR_104465, intron 10 of 18) | intron (NR_104465, intron 10 of 18) | 35914 | NR_104465 | 70351 | Mm.48686 | NM_146081 | ENSMUSG00000061950 | **Ppp4r1** | 3110001J10Rik\|Pp4r1 | protein phosphatase 4, regulatory subunit 1 | protein-coding |
| chr6_100807161_100807845 | chr6 | 100807162 | 100807845 | + | 1.26722 | Intergenic | Intergenic | -26135 | NM_182939 | 232314 | Mm.27223 | NM_182939 | ENSMUSG00000052144 | **Ppp4r2** | BE691708\|C230060M08Rik | protein phosphatase 4, regulatory subunit 2 | protein-coding |

| | | | | | | | | | | | | | | | | | |
|---|---|---|---|---|---|---|---|---|---|---|---|---|---|---|---|---|---|
| chr6_100809682_100810487 | chr6 | 100809683 | 100810487 | + | 1.09375 | Intergenic | Intergenic | -23553 | NM_182939 | 232314 | Mm.27223 | NM_182939 | ENSMUSG00000052144 | **Ppp4r2** | BE691708\|C230060M08Rik | protein phosphatase 4, regulatory subunit 2 | protein-coding |
| chr6_100879343_100879985 | chr6 | 100879344 | 100879985 | + | 0.887671 | Intergenic | Intergenic | 46026 | NM_182939 | 232314 | Mm.27223 | NM_182939 | ENSMUSG00000052144 | **Ppp4r2** | BE691708\|C230060M08Rik | protein phosphatase 4, regulatory subunit 2 | protein-coding |
| chr6_100842772_100843067 | chr6 | 100842773 | 100843067 | + | 0.789854 | intron (NM_182939, intron 2 of 8) | intron (NM_182939, intron 2 of 8) | 9282 | NM_182939 | 232314 | Mm.27223 | NM_182939 | ENSMUSG00000052144 | **Ppp4r2** | BE691708\|C230060M08Rik | protein phosphatase 4, regulatory subunit 2 | protein-coding |
| chr6_100800371_100801037 | chr6 | 100800372 | 100801037 | + | 0.672755 | intron (NM_198612, intron 6 of 6) | intron (NM_198612, intron 6 of 6) | -32934 | NM_182939 | 232314 | Mm.27223 | NM_182939 | ENSMUSG00000052144 | **Ppp4r2** | BE691708\|C230060M08Rik | protein phosphatase 4, regulatory subunit 2 | protein-coding |
| chr6_100824249_100825053 | chr6 | 100824250 | 100825053 | + | 0.61564 | Intergenic | Intergenic | -8987 | NM_182939 | 232314 | Mm.27223 | NM_182939 | ENSMUSG00000052144 | **Ppp4r2** | BE691708\|C230060M08Rik | protein phosphatase 4, regulatory subunit 2 | protein-coding |
| chr2_39295466_39296013 | chr2 | 39295467 | 39296013 | + | 1.17902 | Intergenic | RLTR48A\|LTR\|ERV1 | -69402 | NM_024209 | 67857 | Mm.389242 | NM_024209 | | **Ppp6c** | 2310003C10Rik | protein phosphatase 6, catalytic subunit | protein-coding |
| chr2_39377177_39377644 | chr2 | 39377178 | 39377644 | + | 1.02263 | Intergenic | RLTR19\|LTR\|ERVK | -151073 | NM_024209 | 67857 | Mm.389242 | NM_024209 | | **Ppp6c** | 2310003C10Rik | protein phosphatase 6, catalytic subunit | protein-coding |
| chr2_39347359_39347885 | chr2 | 39347360 | 39347885 | + | 0.765746 | Intergenic | MLT2B1\|LTR\|ERVL | -121284 | NM_024209 | 67857 | Mm.389242 | NM_024209 | | **Ppp6c** | 2310003C10Rik | protein phosphatase 6, catalytic subunit | protein-coding |
| chr2_39299369_39299648 | chr2 | 39299370 | 39299648 | + | 0.687696 | Intergenic | Intergenic | -73171 | NM_024209 | 67857 | Mm.389242 | NM_024209 | | **Ppp6c** | 2310003C10Rik | protein phosphatase 6, catalytic subunit | protein-coding |
| chr4_122847977_122848707 | chr4 | 122847978 | 122848707 | + | 1.27031 | intron (NM_008917, intron 4 of 8) | intron (NM_008917, intron 4 of 8) | 12115 | NM_008917 | 19063 | Mm.277719 | NM_008917 | ENSMUSG00000028657 | **Ppt1** | 9530043G02Rik\|AA960502\|C77813\|CLN1\|D4Ertd184e\|INCL\|PPT | palmitoyl-protein thioesterase 1 | protein-coding |
| chr4_122853477_122854142 | chr4 | 122853478 | 122854142 | + | 1.13814 | intron (NM_008917, intron 6 of 8) | intron (NM_008917, intron 6 of 8) | 17583 | NM_008917 | 19063 | Mm.277719 | NM_008917 | ENSMUSG00000028657 | **Ppt1** | 9530043G02Rik\|AA960502\|C77813\|CLN1\|D4Ertd184e\|INCL\|PPT | palmitoyl-protein thioesterase 1 | protein-coding |
| chr4_122810746_122811577 | chr4 | 122810747 | 122811577 | + | 0.993527 | Intergenic | L1MB3\|LINE\|L1 | -25065 | NM_008917 | 19063 | Mm.277719 | NM_008917 | ENSMUSG00000028657 | **Ppt1** | 9530043G02Rik\|AA960502\|C77813\|CLN1\|D4Ertd184e\|INCL\|PPT | palmitoyl-protein thioesterase 1 | protein-coding |
| chr4_122857975_122858474 | chr4 | 122857976 | 122858474 | + | 0.748669 | TTS (NM_001301067) | TTS (NM_001301067) | 21998 | NM_008917 | 19063 | Mm.277719 | NM_008917 | ENSMUSG00000028657 | **Ppt1** | 9530043G02Rik\|AA960502\|C77813\|CLN1\|D4Ertd184e\|INCL\|PPT | palmitoyl-protein thioesterase 1 | protein-coding |
| chr4_122838413_122838855 | chr4 | 122838414 | 122838855 | + | 0.6404 | intron (NM_008917, intron 1 of 8) | L2b\|LINE\|L2 | 2407 | NM_008917 | 19063 | Mm.277719 | NM_008917 | ENSMUSG00000028657 | **Ppt1** | 9530043G02Rik\|AA960502\|C77813\|CLN1\|D4Ertd184e\|INCL\|PPT | palmitoyl-protein thioesterase 1 | protein-coding |
| chr12_17005281_17005711 | chr12 | 17005282 | 17005711 | + | 0.762831 | 3' UTR (NM_001201333, exon 4 of 4) | 3' UTR (NM_001201333, exon 4 of 4) | -5378 | NM_001161111 | 217430 | Mm.379451 | NM_172574 | ENSMUSG00000045679 | **Pqlc3** | C78076\|E030024M05Rik | PQ loop repeat containing | protein-coding |
| chr7_80299548_80299973 | chr7 | 80299549 | 80299973 | + | 0.878253 | intron (NM_145150, intron 1 of 13) | B1_Mur4\|SINE\|Alu | 5310 | NM_001285998 | 233406 | Mm.227274 | NM_145150 | ENSMUSG00000038943 | **Prc1** | D7Ertd348e | protein regulator of cytokinesis 1 | protein-coding |
| chr7_80298827_80299046 | chr7 | 80298828 | 80299046 | + | 0.732514 | intron (NM_145150, intron 1 of 13) | B2_Mm1t\|SINE\|B2 | 4486 | NM_001285998 | 233406 | Mm.227274 | NM_145150 | ENSMUSG00000038943 | **Prc1** | D7Ertd348e | protein regulator of cytokinesis 1 | protein-coding |
| chr10_44432211_44432647 | chr10 | 44432212 | 44432647 | + | 1.3396 | Intergenic | Intergenic | 26258 | NM_007548 | 12142 | Mm.4800 | NM_007548 | ENSMUSG00000038151 | **Prdm1** | Blimp-1\|Blimp1\|PRDI-BF1\|ZNFPR1A1\|b2b1765Clo | PR domain containing 1, with ZNF domain | protein-coding |
| chr10_44444428_44444775 | chr10 | 44444429 | 44444775 | + | 1.20273 | intron (NM_007548, intron 4 of 6) | intron (NM_007548, intron 4 of 6) | 14085 | NM_007548 | 12142 | Mm.4800 | NM_007548 | ENSMUSG00000038151 | **Prdm1** | Blimp-1\|Blimp1\|PRDI-BF1\|ZNFPR1A1\|b2b1765Clo | PR domain containing 1, with ZNF domain | protein-coding |
| chr10_44532465_44532887 | chr10 | 44532466 | 44532887 | + | 0.960231 | Intergenic | Intergenic | -73989 | NM_007548 | 12142 | Mm.4800 | NM_007548 | ENSMUSG00000038151 | **Prdm1** | Blimp-1\|Blimp1\|PRDI-BF1\|ZNFPR1A1\|b2b1765Clo | PR domain containing 1, with ZNF domain | protein-coding |
| chr10_44564936_44565466 | chr10 | 44564937 | 44565466 | + | 0.869986 | Intergenic | Intergenic | -106514 | NM_007548 | 12142 | Mm.4800 | NM_007548 | ENSMUSG00000038151 | **Prdm1** | Blimp-1\|Blimp1\|PRDI-BF1\|ZNFPR1A1\|b2b1765Clo | PR domain containing 1, with ZNF domain | protein-coding |
| chr10_44538761_44539703 | chr10 | 44538762 | 44539703 | + | 0.797317 | Intergenic | Intergenic | -80545 | NM_007548 | 12142 | Mm.4800 | NM_007548 | ENSMUSG00000038151 | **Prdm1** | Blimp-1\|Blimp1\|PRDI-BF1\|ZNFPR1A1\|b2b1765Clo | PR domain containing 1, with ZNF domain | protein-coding |
| chr10_44439640_44439936 | chr10 | 44439641 | 44439936 | + | 0.79001 | exon (NM_007548, exon 7 of 7) | exon (NM_007548, exon 7 of 7) | 18899 | NM_007548 | 12142 | Mm.4800 | NM_007548 | ENSMUSG00000038151 | **Prdm1** | Blimp-1\|Blimp1\|PRDI-BF1\|ZNFPR1A1\|b2b1765Clo | PR domain containing 1, with ZNF domain | protein-coding |
| chr10_44615599_44615993 | chr10 | 44615600 | 44615993 | + | 0.758263 | Intergenic | Intergenic | -157109 | NM_007548 | 12142 | Mm.4800 | NM_007548 | ENSMUSG00000038151 | **Prdm1** | Blimp-1\|Blimp1\|PRDI-BF1\|ZNFPR1A1\|b2b1765Clo | PR domain containing 1, with ZNF domain | protein-coding |
| chr10_44708772_44709276 | chr10 | 44708773 | 44709276 | + | 0.735979 | Intergenic | Intergenic | -250337 | NM_007548 | 12142 | Mm.4800 | NM_007548 | ENSMUSG00000038151 | **Prdm1** | Blimp-1\|Blimp1\|PRDI-BF1\|ZNFPR1A1\|b2b1765Clo | PR domain containing 1, with ZNF domain | protein-coding |
| chr10_44708319_44709488 | chr10 | 44708320 | 44709488 | + | 0.735951 | Intergenic | Intergenic | -250217 | NM_007548 | 12142 | Mm.4800 | NM_007548 | ENSMUSG00000038151 | **Prdm1** | Blimp-1\|Blimp1\|PRDI-BF1\|ZNFPR1A1\|b2b1765Clo | PR domain containing 1, with ZNF domain | protein-coding |
| chr1_13219938_13220300 | chr1 | 13219939 | 13220300 | + | 1.03793 | intron (NM_001077695, intron 4 of 21) | intron (NM_001077695, intron 4 of 21) | -92956 | NM_001081209 | 383491 | Mm.309380 | NM_001081209 | ENSMUSG00000042414 | **Prdm14** | - | PR domain containing 14 | protein-coding |
| chr1_13155349_13156050 | chr1 | 13155350 | 13156050 | + | 0.973909 | intron (NM_001077695, intron 15 of 21) | intron (NM_001077695, intron 15 of 21) | -28537 | NM_001081209 | 383491 | Mm.309380 | NM_001081209 | ENSMUSG00000042414 | **Prdm14** | - | PR domain containing 14 | protein-coding |
| chr1_13231274_13231819 | chr1 | 13231275 | 13231819 | + | 0.626414 | intron (NM_001077695, intron 2 of 21) | intron (NM_001077695, intron 2 of 21) | -104384 | NM_001081209 | 383491 | Mm.309380 | NM_001081209 | ENSMUSG00000042414 | **Prdm14** | - | PR domain containing 14 | protein-coding |
| chr4_143210081_143210339 | chr4 | 143210082 | 143210339 | + | 0.994918 | intron (NM_001256380, intron 1 of 7) | intron (NM_001256380, intron 1 of 7) | 2499 | NM_001081355 | 110593 | Mm.332020 | NM_001081355 | ENSMUSG00000057637 | **Prdm2** | 4833427P12Rik\|E330024L24Rik\|Gm1033\|Gm19732\|KMT8\|Riz\|Riz1\|Znfpr1c1 | PR domain containing 2, with ZNF domain | protein-coding |
| chr4_116676082_116676451 | chr4 | 116676083 | 116676451 | + | 0.689051 | Intergenic | Intergenic | -9332 | NM_011034 | 18477 | Mm.30929 | NM_011034 | | **Prdx1** | MSP23\|NkefA\|OSF-3\|OSF3\|PAG\|Paga\|PrdxI\|PrxI\|TDX2\|TPxA\|Tdpx2\|prx1 | peroxiredoxin 1 | protein-coding |
| chr1_161252196_161252522 | chr1 | 161252197 | 161252522 | + | 1.05688 | Intergenic | Intergenic | -1193 | NM_007453 | 11758 | Mm.186185 | NM_007453 | ENSMUSG00000026701 | **Prdx6** | 1-cysPrx\|9430088D19Rik\|AA690119\|Aop2\|Aop2-rs3\|Brp-12\|CC26\|CP-3\|GPx\|Ltw-4\|Ltw4\|Lvtw-4\|NSGP\|NSGPx\|ORF06\|Prdx5\|Prdx6-rs3\|aiPLA2\|mKIAA0106 | peroxiredoxin 6 | protein-coding |
| chr1_161252045_161252592 | chr1 | 161252046 | 161252592 | + | 0.645323 | Intergenic | Intergenic | -1153 | NM_007453 | 11758 | Mm.186185 | NM_007453 | ENSMUSG00000026701 | **Prdx6** | 1-cysPrx\|9430088D19Rik\|AA690119\|Aop2\|Aop2-rs3\|Brp-12\|CC26\|CP-3\|GPx\|Ltw-4\|Ltw4\|Lvtw-4\|NSGP\|NSGPx\|ORF06\|Prdx5\|Prdx6-rs3\|aiPLA2\|mKIAA0106 | peroxiredoxin 6 | protein-coding |
| chr18_41947080_41947319 | chr18 | 41947081 | 41947319 | + | 1.41365 | intron (NM_029942, intron 1 of 6) | Lx8b\|LINE\|L1 | 3994 | NM_029942 | 77619 | Mm.30649 | NM_029942 | ENSMUSG00000056671 | **Prelid2** | 1700003A01Rik\|C330008K14Rik | PRELI domain containing 2 | protein-coding |
| chr18_41947059_41947394 | chr18 | 41947060 | 41947394 | + | 1.39162 | intron (NM_029942, intron 1 of 6) | Lx8b\|LINE\|L1 | 3967 | NM_029942 | 77619 | Mm.30649 | NM_029942 | ENSMUSG00000056671 | **Prelid2** | 1700003A01Rik\|C330008K14Rik | PRELI domain containing 2 | protein-coding |
| chr18_41922103_41922483 | chr18 | 41922104 | 41922483 | + | 1.17047 | intron (NM_029942, intron 4 of 6) | intron (NM_029942, intron 4 of 6) | 28901 | NM_029942 | 77619 | Mm.30649 | NM_029942 | ENSMUSG00000056671 | **Prelid2** | 1700003A01Rik\|C330008K14Rik | PRELI domain containing 2 | protein-coding |
| chr18_41922037_41922730 | chr18 | 41922038 | 41922730 | + | 1.02245 | intron (NM_029942, intron 4 of 6) | intron (NM_029942, intron 4 of 6) | 28810 | NM_029942 | 77619 | Mm.30649 | NM_029942 | ENSMUSG00000056671 | **Prelid2** | 1700003A01Rik\|C330008K14Rik | PRELI domain containing 2 | protein-coding |
| chr18_41887994_41888873 | chr18 | 41887995 | 41888873 | + | 0.730224 | intron (NM_029942, intron 5 of 6) | intron (NM_029942, intron 5 of 6) | 62760 | NM_029942 | 77619 | Mm.30649 | NM_029942 | ENSMUSG00000056671 | **Prelid2** | 1700003A01Rik\|C330008K14Rik | PRELI domain containing 2 | protein-coding |
| chr10_44821608_44822220 | chr10 | 44821609 | 44822220 | + | 1.04836 | Intergenic | Intergenic | -245300 | NM_011156 | 19072 | Mm.37294 | NM_011156 | ENSMUSG00000019849 | **Prep** | AI047692\|AI450383\|D10Wsu136e\|PEP\|Pop | prolyl endopeptidase | protein-coding |
| chr10_44819395_44819657 | chr10 | 44819396 | 44819657 | + | 0.830158 | Intergenic | Intergenic | -247688 | NM_011156 | 19072 | Mm.37294 | NM_011156 | ENSMUSG00000019849 | **Prep** | AI047692\|AI450383\|D10Wsu136e\|PEP\|Pop | prolyl endopeptidase | protein-coding |
| chr10_45117951_45118480 | chr10 | 45117952 | 45118480 | + | 0.769944 | intron (NM_011156, intron 8 of 14) | intron (NM_011156, intron 8 of 14) | 51002 | NM_011156 | 19072 | Mm.37294 | NM_011156 | ENSMUSG00000019849 | **Prep** | AI047692\|AI450383\|D10Wsu136e\|PEP\|Pop | prolyl endopeptidase | protein-coding |
| chr10_45001420_45001734 | chr10 | 45001421 | 45001734 | + | 0.742524 | Intergenic | Intergenic | -65637 | NM_011156 | 19072 | Mm.37294 | NM_011156 | ENSMUSG00000019849 | **Prep** | AI047692\|AI450383\|D10Wsu136e\|PEP\|Pop | prolyl endopeptidase | protein-coding |
| chr10_45020267_45020781 | chr10 | 45020268 | 45020781 | + | 0.637388 | Intergenic | Intergenic | -46690 | NM_011156 | 19072 | Mm.37294 | NM_011156 | ENSMUSG00000019849 | **Prep** | AI047692\|AI450383\|D10Wsu136e\|PEP\|Pop | prolyl endopeptidase | protein-coding |
| chr2_166715839_166716079 | chr2 | 166715840 | 166716079 | + | 0.722876 | intron (NR_105783, intron 2 of 2) | intron (NR_105783, intron 2 of 2) | -2127 | NM_177782 | 277360 | Mm.34420 | NM_177782 | ENSMUSG00000039621 | **Prex1** | G630042G04\|P-REX1\|Setd6 | phosphatidylinositol-3,4,5-trisphosphate-dependent Rac exchange factor 1 | protein-coding |
| chr15_93513904_93514412 | chr15 | 93513905 | 93514412 | + | 1.21147 | intron (NM_001033217, intron 1 of 7) | intron (NM_001033217, intron 1 of 7) | 81733 | NM_001033217 | 106042 | Mm.150314 | NM_001033217 | ENSMUSG00000036158 | **Prickle1** | 1110058P22Rik\|AW215793\|Prickle\|b2b019Clo\|mpk1 | prickle homolog 1 (Drosophila) | protein-coding |
| chr15_93715201_93715970 | chr15 | 93715202 | 93715970 | + | 0.858773 | Intergenic | Intergenic | -119695 | NM_001033217 | 106042 | Mm.150314 | NM_001033217 | ENSMUSG00000036158 | **Prickle1** | 1110058P22Rik\|AW215793\|Prickle\|b2b019Clo\|mpk1 | prickle homolog 1 (Drosophila) | protein-coding |
| chr15_93715498_93715970 | chr15 | 93715499 | 93715970 | + | 0.799158 | Intergenic | MLT1G3\|LTR\|ERVL-MaLR | -119843 | NM_001033217 | 106042 | Mm.150314 | NM_001033217 | ENSMUSG00000036158 | **Prickle1** | 1110058P22Rik\|AW215793\|Prickle\|b2b019Clo\|mpk1 | prickle homolog 1 (Drosophila) | protein-coding |

| | | | | | | | | | | | | | | | | | |
|---|---|---|---|---|---|---|---|---|---|---|---|---|---|---|---|---|---|
| chr1_33452742_33453118 | chr1 | 33452743 | 33453118 | + | 0.964298 | TTS (NM_008922) | TTS (NM_008922) | 216864 | NM_008922 | 19076 | Mm.27705 | NM_008922 | ENSMUSG00000026134 | **Prim2** | AI323589 | DNA primase, p58 subunit | protein-coding |
| chr1_33612064_33612519 | chr1 | 33612065 | 33612519 | + | 0.959261 | intron (NM_008922, intron 7 of 13) | intron (NM_008922, intron 7 of 13) | 57502 | NM_008922 | 19076 | Mm.27705 | NM_008922 | ENSMUSG00000026134 | **Prim2** | AI323589 | DNA primase, p58 subunit | protein-coding |
| chr1_33584236_33584528 | chr1 | 33584237 | 33584528 | + | 0.7559 | intron (NM_008922, intron 7 of 13) | intron (NM_008922, intron 7 of 13) | 85412 | NM_008922 | 19076 | Mm.27705 | NM_008922 | ENSMUSG00000026134 | **Prim2** | AI323589 | DNA primase, p58 subunit | protein-coding |
| chr1_33452733_33453276 | chr1 | 33452734 | 33453276 | + | 0.701561 | TTS (NM_008922) | TTS (NM_008922) | 216789 | NM_008922 | 19076 | Mm.27705 | NM_008922 | ENSMUSG00000026134 | **Prim2** | AI323589 | DNA primase, p58 subunit | protein-coding |
| chr1_33584230_33584684 | chr1 | 33584231 | 33584684 | + | 0.615365 | intron (NM_008922, intron 7 of 13) | intron (NM_008922, intron 7 of 13) | 85337 | NM_008922 | 19076 | Mm.27705 | NM_008922 | ENSMUSG00000026134 | **Prim2** | AI323589 | DNA primase, p58 subunit | protein-coding |
| chr12_103099192_103099555 | chr12 | 103099193 | 103099555 | + | 0.961794 | intron (NM_001081017, intron 26 of 48) | intron (NM_001081017, intron 26 of 48) | 142772 | NM_133364 | 170952 | Mm.217019 | NM_133364 | ENSMUSG00000041669 | **Prima1** | B230212M13Rik\|BB120497\|PRiMA | proline rich membrane anchor 1 | protein-coding |
| chr8_46600961_46601700 | chr8 | 46600962 | 46601700 | + | 0.841135 | intron (NM_001001184, intron 5 of 14) | intron (NM_001001184, intron 5 of 14) | 15869 | NM_001001184 | 408022 | Mm.217385 | NM_001001184 | ENSMUSG00000038225 | **Primpol** | BC065112\|Ccdc111 | primase and polymerase (DNA-directed) | protein-coding |
| chr8_46590449_46591038 | chr8 | 46590450 | 46591038 | + | 0.837412 | intron (NM_001001184, intron 8 of 14) | URR1A\|DNA\|hAT-Charlie | 26456 | NM_001001184 | 408022 | Mm.217385 | NM_001001184 | ENSMUSG00000038225 | **Primpol** | BC065112\|Ccdc111 | primase and polymerase (DNA-directed) | protein-coding |
| chr15_5145642_5145885 | chr15 | 5145643 | 5145885 | + | 0.662848 | intron (NM_001013367, intron 1 of 8) | intron (NM_001013367, intron 1 of 8) | 1903 | NM_001013367 | 105787 | Mm.207004 | NM_001013367 | ENSMUSG00000050697 | **Prkaa1** | AI194361\|AI450832\|AL024255\|AMPKalpha1\|C130083N04Rik | protein kinase, AMP-activated, alpha 1 catalytic subunit | protein-coding |
| chr3_146757825_146758317 | chr3 | 146757826 | 146758317 | + | 1.30602 | intron (NM_011100, intron 1 of 9) | intron (NM_011100, intron 1 of 9) | 12167 | NM_001164198 | 18749 | Mm.16766 | NM_011100 | ENSMUSG00000005034 | **Prkacb** | Pkacb | protein kinase, cAMP dependent, catalytic, beta | protein-coding |
| chr3_146784281_146784802 | chr3 | 146784282 | 146784802 | + | 0.737691 | intron (NM_011100, intron 1 of 9) | intron (NM_011100, intron 1 of 9) | -3126 | NM_001164200 | 18749 | Mm.16766 | NM_011100 | ENSMUSG00000005034 | **Prkacb** | Pkacb | protein kinase, cAMP dependent, catalytic, beta | protein-coding |
| chr5_24970882_24971288 | chr5 | 24970883 | 24971288 | + | 1.39727 | intron (NM_145401, intron 3 of 15) | intron (NM_145401, intron 3 of 15) | -62576 | NM_001170556 | 108099 | Mm.33649 | NM_145401 | ENSMUSG00000028944 | **Prkag2** | 2410051C13Rik\|AAKG\|AAKG2\|AI854673\|H91620p\|WPWS | protein kinase, AMP-activated, gamma 2 non-catalytic subunit | protein-coding |
| chr5_24969433_24969931 | chr5 | 24969434 | 24969931 | + | 0.718308 | intron (NM_145401, intron 3 of 15) | intron (NM_145401, intron 3 of 15) | -61173 | NM_001170556 | 108099 | Mm.33649 | NM_145401 | ENSMUSG00000028944 | **Prkag2** | 2410051C13Rik\|AAKG\|AAKG2\|AI854673\|H91620p\|WPWS | protein kinase, AMP-activated, gamma 2 non-catalytic subunit | protein-coding |
| chr12_32014967_32015807 | chr12 | 32014968 | 32015807 | + | 1.0939 | intron (NM_011158, intron 2 of 10) | intron (NM_011158, intron 2 of 10) | 45892 | NM_011158 | 19088 | Mm.25594 | NM_011158 | ENSMUSG00000002997 | **Prkar2b** | AI451071\|AW061005\|PKARIIbeta\|Pkarb2\|RII(beta) | protein kinase, cAMP dependent regulatory, type II beta | protein-coding |
| chr11_108261436_108262210 | chr11 | 108261437 | 108262210 | + | 1.50316 | intron (NM_011101, intron 2 of 16) | L1MB2\|LINE\|L1 | 82065 | NM_011101 | 18750 | Mm.222178 | NM_011101 | ENSMUSG00000050965 | **Prkca** | AI875142\|Pkca | protein kinase C, alpha | protein-coding |
| chr11_108170123_108170689 | chr11 | 108170124 | 108170689 | + | 1.43077 | intron (NM_011101, intron 3 of 16) | intron (NM_011101, intron 3 of 16) | 173482 | NM_011101 | 18750 | Mm.222178 | NM_011101 | ENSMUSG00000050965 | **Prkca** | AI875142\|Pkca | protein kinase C, alpha | protein-coding |
| chr11_108282743_108283382 | chr11 | 108282744 | 108283382 | + | 1.42793 | intron (NM_011101, intron 2 of 16) | MIR\|SINE\|MIR | 60825 | NM_011101 | 18750 | Mm.222178 | NM_011101 | ENSMUSG00000050965 | **Prkca** | AI875142\|Pkca | protein kinase C, alpha | protein-coding |
| chr11_108199620_108199967 | chr11 | 108199621 | 108199967 | + | 1.41678 | intron (NM_011101, intron 2 of 16) | intron (NM_011101, intron 2 of 16) | 144094 | NM_011101 | 18750 | Mm.222178 | NM_011101 | ENSMUSG00000050965 | **Prkca** | AI875142\|Pkca | protein kinase C, alpha | protein-coding |
| chr11_108203931_108204583 | chr11 | 108203932 | 108204583 | + | 1.2146 | intron (NM_011101, intron 2 of 16) | intron (NM_011101, intron 2 of 16) | 139631 | NM_011101 | 18750 | Mm.222178 | NM_011101 | ENSMUSG00000050965 | **Prkca** | AI875142\|Pkca | protein kinase C, alpha | protein-coding |
| chr11_108202699_108202998 | chr11 | 108202700 | 108202998 | + | 1.10694 | intron (NM_011101, intron 2 of 16) | MLT1C\|LTR\|ERVL-MaLR | 141039 | NM_011101 | 18750 | Mm.222178 | NM_011101 | ENSMUSG00000050965 | **Prkca** | AI875142\|Pkca | protein kinase C, alpha | protein-coding |
| chr11_108215512_108215936 | chr11 | 108215513 | 108215936 | + | 0.998781 | intron (NM_011101, intron 2 of 16) | intron (NM_011101, intron 2 of 16) | 128164 | NM_011101 | 18750 | Mm.222178 | NM_011101 | ENSMUSG00000050965 | **Prkca** | AI875142\|Pkca | protein kinase C, alpha | protein-coding |
| chr11_108176747_108177059 | chr11 | 108176748 | 108177059 | + | 0.979177 | intron (NM_011101, intron 3 of 16) | intron (NM_011101, intron 3 of 16) | 166985 | NM_011101 | 18750 | Mm.222178 | NM_011101 | ENSMUSG00000050965 | **Prkca** | AI875142\|Pkca | protein kinase C, alpha | protein-coding |
| chr11_108277838_108278113 | chr11 | 108277839 | 108278113 | + | 0.978299 | intron (NM_011101, intron 2 of 16) | intron (NM_011101, intron 2 of 16) | 65912 | NM_011101 | 18750 | Mm.222178 | NM_011101 | ENSMUSG00000050965 | **Prkca** | AI875142\|Pkca | protein kinase C, alpha | protein-coding |
| chr11_108176722_108177352 | chr11 | 108176723 | 108177352 | + | 0.845926 | intron (NM_011101, intron 3 of 16) | intron (NM_011101, intron 3 of 16) | 166851 | NM_011101 | 18750 | Mm.222178 | NM_011101 | ENSMUSG00000050965 | **Prkca** | AI875142\|Pkca | protein kinase C, alpha | protein-coding |
| chr11_108171883_108172245 | chr11 | 108171884 | 108172245 | + | 0.742 | intron (NM_011101, intron 3 of 16) | intron (NM_011101, intron 3 of 16) | 171824 | NM_011101 | 18750 | Mm.222178 | NM_011101 | ENSMUSG00000050965 | **Prkca** | AI875142\|Pkca | protein kinase C, alpha | protein-coding |
| chr11_108246263_108246819 | chr11 | 108246264 | 108246819 | + | 0.740976 | intron (NM_011101, intron 2 of 16) | RMER4B\|LTR\|ERVK | 97347 | NM_011101 | 18750 | Mm.222178 | NM_011101 | ENSMUSG00000050965 | **Prkca** | AI875142\|Pkca | protein kinase C, alpha | protein-coding |
| chr11_108236338_108237356 | chr11 | 108236339 | 108237356 | + | 0.657426 | intron (NM_011101, intron 2 of 16) | intron (NM_011101, intron 2 of 16) | 107041 | NM_011101 | 18750 | Mm.222178 | NM_011101 | ENSMUSG00000050965 | **Prkca** | AI875142\|Pkca | protein kinase C, alpha | protein-coding |
| chr11_108336217_108336796 | chr11 | 108336218 | 108336796 | + | 0.633156 | intron (NM_011101, intron 2 of 16) | intron (NM_011101, intron 2 of 16) | 7381 | NM_011101 | 18750 | Mm.222178 | NM_011101 | ENSMUSG00000050965 | **Prkca** | AI875142\|Pkca | protein kinase C, alpha | protein-coding |
| chr7_122397506_122398146 | chr7 | 122397507 | 122398146 | + | 1.70223 | intron (NM_008855, intron 2 of 16) | intron (NM_008855, intron 2 of 16) | 108701 | NM_008855 | 18751 | Mm.207496 | NM_008855 | ENSMUSG00000052889 | **Prkcb** | A130082F03Rik\|PKC-Beta\|Pkcb\|Prkcb1\|Prkcb2 | protein kinase C, beta | protein-coding |
| chr7_122371935_122372724 | chr7 | 122371936 | 122372724 | + | 1.40093 | intron (NM_008855, intron 2 of 16) | intron (NM_008855, intron 2 of 16) | 83205 | NM_008855 | 18751 | Mm.207496 | NM_008855 | ENSMUSG00000052889 | **Prkcb** | A130082F03Rik\|PKC-Beta\|Pkcb\|Prkcb1\|Prkcb2 | protein kinase C, beta | protein-coding |
| chr7_122395218_122395482 | chr7 | 122395219 | 122395482 | + | 1.11895 | intron (NM_008855, intron 2 of 16) | MIR\|SINE\|MIR | 106225 | NM_008855 | 18751 | Mm.207496 | NM_008855 | ENSMUSG00000052889 | **Prkcb** | A130082F03Rik\|PKC-Beta\|Pkcb\|Prkcb1\|Prkcb2 | protein kinase C, beta | protein-coding |
| chr7_122296683_122297234 | chr7 | 122296684 | 122297234 | + | 0.915667 | intron (NM_008855, intron 2 of 16) | intron (NM_008855, intron 2 of 16) | 7834 | NM_008855 | 18751 | Mm.207496 | NM_008855 | ENSMUSG00000052889 | **Prkcb** | A130082F03Rik\|PKC-Beta\|Pkcb\|Prkcb1\|Prkcb2 | protein kinase C, beta | protein-coding |
| chr7_122296357_122297312 | chr7 | 122296358 | 122297312 | + | 0.868492 | intron (NM_008855, intron 2 of 16) | MER113\|DNA\|hAT-Charlie | 7710 | NM_008855 | 18751 | Mm.207496 | NM_008855 | ENSMUSG00000052889 | **Prkcb** | A130082F03Rik\|PKC-Beta\|Pkcb\|Prkcb1\|Prkcb2 | protein kinase C, beta | protein-coding |
| chr7_122415395_122415815 | chr7 | 122415396 | 122415815 | + | 0.768271 | intron (NM_008855, intron 2 of 16) | MIR\|SINE\|MIR | 126480 | NM_008855 | 18751 | Mm.207496 | NM_008855 | ENSMUSG00000052889 | **Prkcb** | A130082F03Rik\|PKC-Beta\|Pkcb\|Prkcb1\|Prkcb2 | protein kinase C, beta | protein-coding |
| chr7_122389722_122390029 | chr7 | 122389723 | 122390029 | + | 0.753867 | intron (NM_008855, intron 2 of 16) | intron (NM_008855, intron 2 of 16) | 100751 | NM_008855 | 18751 | Mm.207496 | NM_008855 | ENSMUSG00000052889 | **Prkcb** | A130082F03Rik\|PKC-Beta\|Pkcb\|Prkcb1\|Prkcb2 | protein kinase C, beta | protein-coding |
| chr7_122386853_122387135 | chr7 | 122386854 | 122387135 | + | 0.692849 | intron (NM_008855, intron 2 of 16) | intron (NM_008855, intron 2 of 16) | 97869 | NM_008855 | 18751 | Mm.207496 | NM_008855 | ENSMUSG00000052889 | **Prkcb** | A130082F03Rik\|PKC-Beta\|Pkcb\|Prkcb1\|Prkcb2 | protein kinase C, beta | protein-coding |
| chr7_122444948_122445416 | chr7 | 122444949 | 122445416 | + | 0.640939 | intron (NM_008855, intron 3 of 16) | intron (NM_008855, intron 3 of 16) | 156057 | NM_008855 | 18751 | Mm.207496 | NM_008855 | ENSMUSG00000052889 | **Prkcb** | A130082F03Rik\|PKC-Beta\|Pkcb\|Prkcb1\|Prkcb2 | protein kinase C, beta | protein-coding |
| chr7_122388910_122389199 | chr7 | 122388911 | 122389199 | + | 0.633776 | intron (NM_008855, intron 2 of 16) | intron (NM_008855, intron 2 of 16) | 99930 | NM_008855 | 18751 | Mm.207496 | NM_008855 | ENSMUSG00000052889 | **Prkcb** | A130082F03Rik\|PKC-Beta\|Pkcb\|Prkcb1\|Prkcb2 | protein kinase C, beta | protein-coding |
| chr7_105486974_105487496 | chr7 | 105486975 | 105487496 | + | 1.7012 | Intergenic | Intergenic | -5038 | NM_028444 | 109042 | Mm.3124 | NM_028444 | ENSMUSG00000037060 | **Prkcdbp** | 3110015B12Rik\|6330514M23Rik\|SRBC | protein kinase C, delta binding protein | protein-coding |
| chr7_105481510_105482374 | chr7 | 105481511 | 105482374 | + | 0.613462 | exon (NM_028444, exon 1 of 2) | exon (NM_028444, exon 1 of 2) | 255 | NM_028444 | 109042 | Mm.3124 | NM_028444 | ENSMUSG00000037060 | **Prkcdbp** | 3110015B12Rik\|6330514M23Rik\|SRBC | protein kinase C, delta binding protein | protein-coding |
| chr12_73726101_73726637 | chr12 | 73726102 | 73726637 | + | 1.42482 | intron (NM_008856, intron 10 of 13) | intron (NM_008856, intron 10 of 13) | 141328 | NM_008856 | 18755 | Mm.341677 | NM_008856 | ENSMUSG00000021108 | **Prkch** | Pkch | protein kinase C, eta | protein-coding |
| chr12_73725784_73726788 | chr12 | 73725785 | 73726788 | + | 1.08241 | intron (NM_008856, intron 10 of 13) | L3\|LINE\|CR1 | 141245 | NM_008856 | 18755 | Mm.341677 | NM_008856 | ENSMUSG00000021108 | **Prkch** | Pkch | protein kinase C, eta | protein-coding |
| chr12_73741999_73742301 | chr12 | 73742000 | 73742301 | + | 0.927078 | intron (NM_008856, intron 10 of 13) | B3A\|SINE\|B2 | 157109 | NM_008856 | 18755 | Mm.341677 | NM_008856 | ENSMUSG00000021108 | **Prkch** | Pkch | protein kinase C, eta | protein-coding |
| chr12_73641176_73641734 | chr12 | 73641177 | 73641734 | + | 0.918181 | intron (NM_008856, intron 1 of 13) | RSINE1\|SINE\|B4 | 56414 | NM_008856 | 18755 | Mm.341677 | NM_008856 | ENSMUSG00000021108 | **Prkch** | Pkch | protein kinase C, eta | protein-coding |
| chr12_73745992_73746398 | chr12 | 73745993 | 73746398 | + | 0.909604 | intron (NM_008856, intron 10 of 13) | L1MA6\|LINE\|L1 | 161154 | NM_008856 | 18755 | Mm.341677 | NM_008856 | ENSMUSG00000021108 | **Prkch** | Pkch | protein kinase C, eta | protein-coding |
| chr12_73632664_73633216 | chr12 | 73632665 | 73633216 | + | 0.829022 | intron (NM_008856, intron 1 of 13) | intron (NM_008856, intron 1 of 13) | 47899 | NM_008856 | 18755 | Mm.341677 | NM_008856 | ENSMUSG00000021108 | **Prkch** | Pkch | protein kinase C, eta | protein-coding |
| chr12_73684100_73684346 | chr12 | 73684101 | 73684346 | + | 0.771565 | intron (NM_008856, intron 2 of 13) | intron (NM_008856, intron 2 of 13) | 99182 | NM_008856 | 18755 | Mm.341677 | NM_008856 | ENSMUSG00000021108 | **Prkch** | Pkch | protein kinase C, eta | protein-coding |
| chr12_73570561_73570846 | chr12 | 73570562 | 73570846 | + | 0.753232 | Intergenic | MTC\|LTR\|ERVL-MaLR | -14337 | NM_008856 | 18755 | Mm.341677 | NM_008856 | ENSMUSG00000021108 | **Prkch** | Pkch | protein kinase C, eta | protein-coding |
| chr12_73728796_73729086 | chr12 | 73728797 | 73729086 | + | 0.664712 | intron (NM_008856, intron 10 of 13) | intron (NM_008856, intron 10 of 13) | 143900 | NM_008856 | 18755 | Mm.341677 | NM_008856 | ENSMUSG00000021108 | **Prkch** | Pkch | protein kinase C, eta | protein-coding |
| chr12_73673198_73673618 | chr12 | 73673199 | 73673618 | + | 0.64668 | intron (NM_008856, intron 2 of 13) | URR1A\|DNA\|hAT-Charlie | 88367 | NM_008856 | 18755 | Mm.341677 | NM_008856 | ENSMUSG00000021108 | **Prkch** | Pkch | protein kinase C, eta | protein-coding |

| | | | | | | | | | | | | | | | | | |
|---|---|---|---|---|---|---|---|---|---|---|---|---|---|---|---|---|---|
| chr2_11146705_11147010 | chr2 | 11146706 | 11147010 | + | 1.52281 | Intergenic | Charlie6\|DNA\|hAT-Charlie | -25524 | NM_008859 | 18761 | Mm.329993 | NM_008859 | ENSMUSG00000026778 | **Prkcq** | A130035A12Rik\|AW494342\|PKC-0\|PKC-theta\|PKCtheta\|Pkcq | protein kinase C, theta | protein-coding |
| chr2_10978739_10979323 | chr2 | 10978740 | 10979323 | + | 1.4919 | Intergenic | Intergenic | -193351 | NM_008859 | 18761 | Mm.329993 | NM_008859 | ENSMUSG00000026778 | **Prkcq** | A130035A12Rik\|AW494342\|PKC-0\|PKC-theta\|PKCtheta\|Pkcq | protein kinase C, theta | protein-coding |
| chr2_11142541_11143314 | chr2 | 11142542 | 11143314 | + | 1.2567 | Intergenic | Intergenic | -29454 | NM_008859 | 18761 | Mm.329993 | NM_008859 | ENSMUSG00000026778 | **Prkcq** | A130035A12Rik\|AW494342\|PKC-0\|PKC-theta\|PKCtheta\|Pkcq | protein kinase C, theta | protein-coding |
| chr2_11145655_11146003 | chr2 | 11145656 | 11146003 | + | 0.884181 | Intergenic | MTEa\|LTR\|ERVL-MaLR | -26553 | NM_008859 | 18761 | Mm.329993 | NM_008859 | ENSMUSG00000026778 | **Prkcq** | A130035A12Rik\|AW494342\|PKC-0\|PKC-theta\|PKCtheta\|Pkcq | protein kinase C, theta | protein-coding |
| chr2_11145655_11146224 | chr2 | 11145656 | 11146224 | + | 0.81289 | Intergenic | Intergenic | -26442 | NM_008859 | 18761 | Mm.329993 | NM_008859 | ENSMUSG00000026778 | **Prkcq** | A130035A12Rik\|AW494342\|PKC-0\|PKC-theta\|PKCtheta\|Pkcq | protein kinase C, theta | protein-coding |
| chr2_11141901_11142296 | chr2 | 11141902 | 11142296 | + | 0.765732 | Intergenic | MER110A\|LTR\|ERV1 | -30283 | NM_008859 | 18761 | Mm.329993 | NM_008859 | ENSMUSG00000026778 | **Prkcq** | A130035A12Rik\|AW494342\|PKC-0\|PKC-theta\|PKCtheta\|Pkcq | protein kinase C, theta | protein-coding |
| chr2_11013355_11013734 | chr2 | 11013356 | 11013734 | + | 0.757414 | Intergenic | Intergenic | -158837 | NM_008859 | 18761 | Mm.329993 | NM_008859 | ENSMUSG00000026778 | **Prkcq** | A130035A12Rik\|AW494342\|PKC-0\|PKC-theta\|PKCtheta\|Pkcq | protein kinase C, theta | protein-coding |
| chr2_11173085_11173457 | chr2 | 11173086 | 11173457 | + | 0.752729 | intron (NM_008859, intron 1 of 17) | intron (NM_008859, intron 1 of 17) | 889 | NM_008859 | 18761 | Mm.329993 | NM_008859 | ENSMUSG00000026778 | **Prkcq** | A130035A12Rik\|AW494342\|PKC-0\|PKC-theta\|PKCtheta\|Pkcq | protein kinase C, theta | protein-coding |
| chr2_11115822_11116109 | chr2 | 11115823 | 11116109 | + | 0.646787 | Intergenic | Intergenic | -56416 | NM_008859 | 18761 | Mm.329993 | NM_008859 | ENSMUSG00000026778 | **Prkcq** | A130035A12Rik\|AW494342\|PKC-0\|PKC-theta\|PKCtheta\|Pkcq | protein kinase C, theta | protein-coding |
| chr2_11139799_11140105 | chr2 | 11139800 | 11140105 | + | 0.639744 | Intergenic | Intergenic | -32430 | NM_008859 | 18761 | Mm.329993 | NM_008859 | ENSMUSG00000026778 | **Prkcq** | A130035A12Rik\|AW494342\|PKC-0\|PKC-theta\|PKCtheta\|Pkcq | protein kinase C, theta | protein-coding |
| chr7_98698667_98698965 | chr7 | 98698668 | 98698965 | + | 1.02813 | Intergenic | Intergenic | -4547 | NM_028410 | 72981 | Mm.4428 | NM_028410 | ENSMUSG00000030753 | **Prkrir** | 2900052B10Rik\|Dap4\|Rpkrir | protein-kinase, interferon-inducible double stranded RNA dependent inhibitor, repressor of (P58 repressor) | protein-coding |
| chr19_60469513_60469975 | chr19 | 60469514 | 60469975 | + | 1.21501 | Intergenic | Intergenic | -1440 | NM_201615 | 226278 | Mm.377241 | NM_201615 | ENSMUSG00000045052 | **Prlhr** | Gm339\|Gpr10\|Gr3\|Prrpr | prolactin releasing hormone receptor | protein-coding |
| chr15_10073448_10073835 | chr15 | 10073449 | 10073835 | + | 1.01093 | Intergenic | Intergenic | -103596 | NM_011169 | 19116 | Mm.10516 | NM_011169 | ENSMUSG00000005268 | **Prlr** | AI987712\|Pr-1\|Pr-3\|Prlr-rs1 | prolactin receptor | protein-coding |
| chr15_10072914_10073200 | chr15 | 10072915 | 10073200 | + | 0.714016 | Intergenic | Intergenic | -104181 | NM_011169 | 19116 | Mm.10516 | NM_011169 | ENSMUSG00000005268 | **Prlr** | AI987712\|Pr-1\|Pr-3\|Prlr-rs1 | prolactin receptor | protein-coding |
| chr16_10812140_10812484 | chr16 | 10812141 | 10812484 | + | 1.02878 | Intergenic | L1MB3\|LINE\|L1 | -15489 | NM_013637 | 19118 | Mm.42733 | NM_013637 | ENSMUSG00000022501 | **Prm1** | Prm-1 | protamine 1 | protein-coding |
| chr2_131935750_131936126 | chr2 | 131935751 | 131936126 | + | 0.62132 | intron (NM_001278256, intron 3 of 3) | intron (NM_001278256, intron 3 of 3) | -14923 | NM_001278257 | 26434 | Mm.180750 | NM_023043 | ENSMUSG00000027338 | **Prnd** | AI450264\|Dpl\|PrPLP\|doppel | prion protein dublet | protein-coding |
| chr2_131885847_131886121 | chr2 | 131885848 | 131886121 | + | 1.32935 | Intergenic | Intergenic | -23944 | NM_011170 | 19122 | Mm.648 | NM_011170 | ENSMUSG00000079037 | **Prnp** | AA960666\|AI325101\|CD230\|PrP\|PrP<C>\|PrPC\|PrPSc\|Prn-i\|Prn-p\|Sinc\|prP27-30\|prP33-35C | prion protein | protein-coding |
| chr2_131885784_131886349 | chr2 | 131885785 | 131886349 | + | 1.2292 | Intergenic | Intergenic | -23861 | NM_011170 | 19122 | Mm.648 | NM_011170 | ENSMUSG00000079037 | **Prnp** | AA960666\|AI325101\|CD230\|PrP\|PrP<C>\|PrPC\|PrPSc\|Prn-i\|Prn-p\|Sinc\|prP27-30\|prP33-35C | prion protein | protein-coding |
| chr2_155761734_155762415 | chr2 | 155761735 | 155762415 | + | 1.02535 | Intergenic | Intergenic | 10858 | NM_011171 | 19124 | Mm.3243 | NM_011171 | ENSMUSG00000027611 | **Procr** | AI325044\|Ccca\|Ccd41\|Epcr | protein C receptor, endothelial | protein-coding |
| chr2_132379548_132380358 | chr2 | 132379549 | 132380358 | + | 1.63942 | intron (NM_144944, intron 2 of 2) | intron (NM_144944, intron 2 of 2) | 5494 | NM_144944 | 246313 | Mm.283777 | NM_144944 | ENSMUSG00000050558 | **Prokr2** | B830005M06Rik\|EG-VEGFR2\|Gpcr73l1\|Gpr73l1\|PKR2 | prokineticin receptor 2 | protein-coding |
| chr2_132379562_132380087 | chr2 | 132379563 | 132380087 | + | 1.63847 | intron (NM_144944, intron 2 of 2) | intron (NM_144944, intron 2 of 2) | 5622 | NM_144944 | 246313 | Mm.283777 | NM_144944 | ENSMUSG00000050558 | **Prokr2** | B830005M06Rik\|EG-VEGFR2\|Gpcr73l1\|Gpr73l1\|PKR2 | prokineticin receptor 2 | protein-coding |
| chr2_6129612_6130439 | chr2 | 6129613 | 6130439 | + | 0.62181 | 5' UTR (NM_001159657, exon 1 of 4) | 5' UTR (NM_001159657, exon 1 of 4) | 159 | NM_144883 | 227545 | Mm.133542 | NM_144883 | | **Proser2** | 5430407P10Rik\|AU067770 | proline and serine rich 2 | protein-coding |
| chr2_4649637_4650105 | chr2 | 4649638 | 4650105 | + | 0.854259 | intron (NM_026045, intron 1 of 9) | intron (NM_026045, intron 1 of 9) | 2215 | NM_026045 | 67229 | Mm.38529 | NM_026045 | ENSMUSG00000039449 | **Prpf18** | 2810441A10Rik | PRP18 pre-mRNA processing factor 18 homolog (yeast) | protein-coding |
| chr3_108916661_108917362 | chr3 | 108916662 | 108917362 | + | 0.750554 | Intergenic | Intergenic | -5308 | NM_025845 | 66921 | Mm.51049 | NM_025845 | ENSMUSG00000027881 | **Prpf38b** | 1110021E09Rik\|AU018955 | PRP38 pre-mRNA processing factor 38 (yeast) domain containing B | protein-coding |
| chr12_65022692_65023215 | chr12 | 65022693 | 65023215 | + | 1.64184 | TTS (NM_177805) | TTS (NM_177805) | -13380 | NM_177806 | 328110 | Mm.283339 | NM_177806 | ENSMUSG00000035597 | **Prpf39** | Srcs1 | PRP39 pre-mRNA processing factor 39 homolog (yeast) | protein-coding |
| chr12_65002885_65003364 | chr12 | 65002886 | 65003364 | + | 1.1635 | intron (NM_177805, intron 11 of 19) | intron (NM_177805, intron 11 of 19) | -33209 | NM_177806 | 328110 | Mm.283339 | NM_177806 | ENSMUSG00000035597 | **Prpf39** | Srcs1 | PRP39 pre-mRNA processing factor 39 homolog (yeast) | protein-coding |
| chr12_65002876_65003630 | chr12 | 65002877 | 65003630 | + | 1.07194 | intron (NM_177805, intron 11 of 19) | intron (NM_177805, intron 11 of 19) | -33081 | NM_177806 | 328110 | Mm.283339 | NM_177806 | ENSMUSG00000035597 | **Prpf39** | Srcs1 | PRP39 pre-mRNA processing factor 39 homolog (yeast) | protein-coding |
| chr17_46916157_46916760 | chr17 | 46916158 | 46916760 | + | 0.806374 | intron (NM_008938, intron 1 of 2) | intron (NM_008938, intron 1 of 2) | 5974 | NM_008938 | 19133 | Mm.291370 | NM_008938 | ENSMUSG00000023978 | **Prph2** | AOFMD\|AVMD\|Nmf193\|PRPH\|RP7\|Rd-2\|Rd2\|Rds\|Tspan22\|rds | peripherin 2 | protein-coding |
| chr17_46932965_46933491 | chr17 | 46932966 | 46933491 | + | 0.655327 | intron (NM_001177374, intron 45 of 46) | RMER6A\|LTR\|ERVK | 22743 | NM_008938 | 19133 | Mm.291370 | NM_008938 | ENSMUSG00000023978 | **Prph2** | AOFMD\|AVMD\|Nmf193\|PRPH\|RP7\|Rd-2\|Rd2\|Rds\|Tspan22\|rds | peripherin 2 | protein-coding |
| chr12_35005653_35006243 | chr12 | 35005654 | 35006243 | + | 1.30977 | Intergenic | ORR1B2-int\|LTR\|ERVL-MaLR | 21187 | NM_029294 | 75456 | Mm.79179 | NM_029294 | ENSMUSG00000092305 | **Prps1l1** | 1700011K15Rik | phosphoribosyl pyrophosphate synthetase 1-like 1 | protein-coding |
| chr12_35005653_35006657 | chr12 | 35005654 | 35006657 | + | 1.03696 | Intergenic | ORR1B2-int\|LTR\|ERVL-MaLR | 21394 | NM_029294 | 75456 | Mm.79179 | NM_029294 | ENSMUSG00000092305 | **Prps1l1** | 1700011K15Rik | phosphoribosyl pyrophosphate synthetase 1-like 1 | protein-coding |
| chr12_34991947_34992350 | chr12 | 34991948 | 34992350 | + | 0.877353 | Intergenic | Intergenic | 7388 | NM_029294 | 75456 | Mm.79179 | NM_029294 | ENSMUSG00000092305 | **Prps1l1** | 1700011K15Rik | phosphoribosyl pyrophosphate synthetase 1-like 1 | protein-coding |
| chrX_167597645_167598128 | chrX | 167597646 | 167598128 | + | 0.695117 | intron (NM_001033330, intron 7 of 19) | RMER17D\|LTR\|ERVK | -215188 | NM_026662 | 110639 | Mm.272955 | NM_026662 | ENSMUSG00000025742 | **Prps2** | 2610101M19Rik\|AA589463\|AI464149\|Prps-2 | phosphoribosyl pyrophosphate synthetase 2 | protein-coding |
| chr2_101754416_101754762 | chr2 | 101754417 | 101754762 | + | 1.60827 | intron (NM_001083810, intron 4 of 8) | intron (NM_001083810, intron 4 of 8) | 43118 | NM_175181 | 72446 | Mm.24138 | NM_175181 | ENSMUSG00000032841 | **Prr5l** | 2600010E01Rik\|4833411O04Rik\|Protor-2\|Protor2 | proline rich 5 like | protein-coding |
| chr2_101810680_101811272 | chr2 | 101810681 | 101811272 | + | 1.55181 | intron (NM_001110849, intron 2 of 9) | intron (NM_001110849, intron 2 of 9) | -13269 | NM_001083810 | 72446 | Mm.24138 | NM_175181 | ENSMUSG00000032841 | **Prr5l** | 2600010E01Rik\|4833411O04Rik\|Protor-2\|Protor2 | proline rich 5 like | protein-coding |
| chr2_101815257_101815793 | chr2 | 101815258 | 101815793 | + | 1.44147 | intron (NM_001110849, intron 2 of 9) | intron (NM_001110849, intron 2 of 9) | -17818 | NM_001083810 | 72446 | Mm.24138 | NM_175181 | ENSMUSG00000032841 | **Prr5l** | 2600010E01Rik\|4833411O04Rik\|Protor-2\|Protor2 | proline rich 5 like | protein-coding |
| chr2_101796382_101797799 | chr2 | 101796383 | 101797799 | + | 1.41665 | intron (NM_001110849, intron 2 of 9) | CpG | 616 | NM_175181 | 72446 | Mm.24138 | NM_175181 | ENSMUSG00000032841 | **Prr5l** | 2600010E01Rik\|4833411O04Rik\|Protor-2\|Protor2 | proline rich 5 like | protein-coding |
| chr2_101819291_101819822 | chr2 | 101819292 | 101819822 | + | 1.39818 | intron (NM_001110849, intron 2 of 9) | intron (NM_001110849, intron 2 of 9) | 19423 | NM_001110849 | 72446 | Mm.24138 | NM_175181 | ENSMUSG00000032841 | **Prr5l** | 2600010E01Rik\|4833411O04Rik\|Protor-2\|Protor2 | proline rich 5 like | protein-coding |
| chr2_101796641_101797048 | chr2 | 101796642 | 101797048 | + | 1.08919 | intron (NM_001110849, intron 2 of 9) | CpG | 862 | NM_175181 | 72446 | Mm.24138 | NM_175181 | ENSMUSG00000032841 | **Prr5l** | 2600010E01Rik\|4833411O04Rik\|Protor-2\|Protor2 | proline rich 5 like | protein-coding |
| chr2_101830217_101830537 | chr2 | 101830218 | 101830537 | + | 0.723299 | intron (NM_001110849, intron 2 of 9) | intron (NM_001110849, intron 2 of 9) | 8603 | NM_001110849 | 72446 | Mm.24138 | NM_175181 | ENSMUSG00000032841 | **Prr5l** | 2600010E01Rik\|4833411O04Rik\|Protor-2\|Protor2 | proline rich 5 like | protein-coding |
| chr2_101755943_101756758 | chr2 | 101755944 | 101756758 | + | 0.662388 | intron (NM_001083810, intron 4 of 8) | intron (NM_001083810, intron 4 of 8) | 41356 | NM_175181 | 72446 | Mm.24138 | NM_175181 | ENSMUSG00000032841 | **Prr5l** | 2600010E01Rik\|4833411O04Rik\|Protor-2\|Protor2 | proline rich 5 like | protein-coding |
| chr2_101811862_101812287 | chr2 | 101811863 | 101812287 | + | 0.662325 | intron (NM_001110849, intron 2 of 9) | intron (NM_001110849, intron 2 of 9) | -14368 | NM_001083810 | 72446 | Mm.24138 | NM_175181 | ENSMUSG00000032841 | **Prr5l** | 2600010E01Rik\|4833411O04Rik\|Protor-2\|Protor2 | proline rich 5 like | protein-coding |
| chr2_32180046_32180604 | chr2 | 32180047 | 32180604 | + | 1.46854 | intron (NM_001159634, intron 1 of 31) | intron (NM_001159634, intron 1 of 31) | 29177 | NM_001159634 | 227723 | Mm.25504 | NM_172661 | ENSMUSG00000039262 | **Prrc2b** | 5830434P21Rik\|AI173903\|Bat2l\|Bat2l1\|D430039P21\|mKIAA0515 | proline-rich coiled-coil 2B | protein-coding |
| chr2_32146633_32147107 | chr2 | 32146634 | 32147107 | + | 0.891437 | Intergenic | MER2\|DNA\|TcMar-Tigger | -4278 | NM_172661 | 227723 | Mm.25504 | NM_172661 | ENSMUSG00000039262 | **Prrc2b** | 5830434P21Rik\|AI173903\|Bat2l\|Bat2l1\|D430039P21\|mKIAA0515 | proline-rich coiled-coil 2B | protein-coding |

| | | | | | | | | | | | | | | | | | |
|---|---|---|---|---|---|---|---|---|---|---|---|---|---|---|---|---|---|
| chr1_162773543_162774058 | chr1 | 162773544 | 162774058 | + | 1.1775 | Intergenic | Intergenic | -33245 | NM_001081290 | 226562 | Mm.245446 | NM_001081290 | ENSMUSG00000040225 | **Prrc2c** | 1810043M20Rik\|9630039I18Rik\|A630006J20\|Bat2d\|Bat2d1\|Bat2l2\|E130112L15Rik\|Prrc3\|mKIAA1096 | proline-rich coiled-coil 2C | protein-coding |
| chr2_104855753_104856157 | chr2 | 104855754 | 104856157 | + | 1.64779 | Intergenic | Intergenic | -6105 | NM_178695 | 228413 | Mm.259609 | NM_178695 | ENSMUSG00000027171 | **Prrg4** | 9930111I18Rik\|TMG4 | proline rich Gla (G-carboxyglutamic acid) 4 (transmembrane) | protein-coding |
| chr2_104864416_104864929 | chr2 | 104864417 | 104864929 | + | 1.52571 | Intergenic | Intergenic | -14823 | NM_178695 | 228413 | Mm.259609 | NM_178695 | ENSMUSG00000027171 | **Prrg4** | 9930111I18Rik\|TMG4 | proline rich Gla (G-carboxyglutamic acid) 4 (transmembrane) | protein-coding |
| chr2_104848570_104849357 | chr2 | 104848571 | 104849357 | + | 0.933997 | intron (NM_178695, intron 2 of 5) | intron (NM_178695, intron 2 of 5) | 886 | NM_178695 | 228413 | Mm.259609 | NM_178695 | ENSMUSG00000027171 | **Prrg4** | 9930111I18Rik\|TMG4 | proline rich Gla (G-carboxyglutamic acid) 4 (transmembrane) | protein-coding |
| chr2_104848540_104850296 | chr2 | 104848541 | 104850296 | + | 0.844791 | intron (NM_178695, intron 1 of 5) | CpG | 432 | NM_178695 | 228413 | Mm.259609 | NM_178695 | ENSMUSG00000027171 | **Prrg4** | 9930111I18Rik\|TMG4 | proline rich Gla (G-carboxyglutamic acid) 4 (transmembrane) | protein-coding |
| chr2_104856232_104856517 | chr2 | 104856233 | 104856517 | + | 0.793158 | Intergenic | B3\|SINE\|B2 | -6525 | NM_178695 | 228413 | Mm.259609 | NM_178695 | ENSMUSG00000027171 | **Prrg4** | 9930111I18Rik\|TMG4 | proline rich Gla (G-carboxyglutamic acid) 4 (transmembrane) | protein-coding |
| chr2_104859169_104859425 | chr2 | 104859170 | 104859425 | + | 0.774446 | Intergenic | Intergenic | -9447 | NM_178695 | 228413 | Mm.259609 | NM_178695 | ENSMUSG00000027171 | **Prrg4** | 9930111I18Rik\|TMG4 | proline rich Gla (G-carboxyglutamic acid) 4 (transmembrane) | protein-coding |
| chr13_22020818_22021296 | chr13 | 22020819 | 22021296 | + | 0.752732 | Intergenic | Intergenic | -11316 | NM_019429 | 54373 | Mm.32043 | NM_019429 | ENSMUSG00000006179 | **Prss16** | AI448615\|TSSP | protease, serine 16 (thymus) | protein-coding |
| chr6_41522824_41523424 | chr6 | 41522825 | 41523424 | + | 1.10121 | intron (NM_009430, intron 2 of 4) | intron (NM_009430, intron 2 of 4) | 1348 | NM_009430 | 22072 | Mm.276926 | NM_009430 | ENSMUSG00000057163 | **Prss2** | TRY8\|TRYP\|Ta\|Tesp4\|Try2 | protease, serine 2 | protein-coding |
| chr6_41522860_41523219 | chr6 | 41522861 | 41523219 | + | 0.9763 | intron (NM_009430, intron 2 of 4) | intron (NM_009430, intron 2 of 4) | 1264 | NM_009430 | 22072 | Mm.276926 | NM_009430 | ENSMUSG00000057163 | **Prss2** | TRY8\|TRYP\|Ta\|Tesp4\|Try2 | protease, serine 2 | protein-coding |
| chr6_41549260_41549495 | chr6 | 41549261 | 41549495 | + | 0.885034 | Intergenic | Intergenic | 27602 | NM_009430 | 22072 | Mm.276926 | NM_009430 | ENSMUSG00000057163 | **Prss2** | TRY8\|TRYP\|Ta\|Tesp4\|Try2 | protease, serine 2 | protein-coding |
| chr6_41550160_41550468 | chr6 | 41550161 | 41550468 | + | 0.835347 | Intergenic | Intergenic | 28538 | NM_009430 | 22072 | Mm.276926 | NM_009430 | ENSMUSG00000057163 | **Prss2** | TRY8\|TRYP\|Ta\|Tesp4\|Try2 | protease, serine 2 | protein-coding |
| chr6_41548327_41548598 | chr6 | 41548328 | 41548598 | + | 0.780962 | Intergenic | Intergenic | 26687 | NM_009430 | 22072 | Mm.276926 | NM_009430 | ENSMUSG00000057163 | **Prss2** | TRY8\|TRYP\|Ta\|Tesp4\|Try2 | protease, serine 2 | protein-coding |
| chr6_41549839_41550038 | chr6 | 41549840 | 41550038 | + | 0.755984 | Intergenic | Intergenic | 28163 | NM_009430 | 22072 | Mm.276926 | NM_009430 | ENSMUSG00000057163 | **Prss2** | TRY8\|TRYP\|Ta\|Tesp4\|Try2 | protease, serine 2 | protein-coding |
| chr7_89560949_89561469 | chr7 | 89560950 | 89561469 | + | 0.954017 | Intergenic | Intergenic | -43623 | NM_029614 | 76453 | Mm.250438 | NM_029614 | ENSMUSG00000039405 | **Prss23** | 2310046G15Rik\|4930479H08Rik\|AI790574\|Spuve | protease, serine 23 | protein-coding |
| chr17_23968742_23969251 | chr17 | 23968743 | 23969251 | + | 1.55426 | Intergenic | Intergenic | 6233 | NM_013921 | 30943 | Mm.142568 | NM_013921 | ENSMUSG00000024124 | **Prss30** | 2010015P21Rik\|Disp\|Disp1\|Tmprss8 | protease, serine 30 | protein-coding |
| chr17_23968654_23969556 | chr17 | 23968655 | 23969556 | + | 1.25931 | Intergenic | Intergenic | 6125 | NM_013921 | 30943 | Mm.142568 | NM_013921 | ENSMUSG00000024124 | **Prss30** | 2010015P21Rik\|Disp\|Disp1\|Tmprss8 | protease, serine 30 | protein-coding |
| chr17_23983021_23983320 | chr17 | 23983022 | 23983320 | + | 0.86898 | Intergenic | Intergenic | -7941 | NM_013921 | 30943 | Mm.142568 | NM_013921 | ENSMUSG00000024124 | **Prss30** | 2010015P21Rik\|Disp\|Disp1\|Tmprss8 | protease, serine 30 | protein-coding |
| chr1_34564251_34565096 | chr1 | 34564252 | 34565096 | + | 0.713595 | Intergenic | Intergenic | -3731 | NM_009356 | 21756 | Mm.14418 | NM_009356 | ENSMUSG00000037529 | **Prss40** | Tesp2 | protease, serine 40 | protein-coding |
| chr9_110820142_110820621 | chr9 | 110820143 | 110820621 | + | 1.00607 | Intergenic | Intergenic | -6308 | NM_199471 | 272643 | Mm.458525 | NM_199471 | ENSMUSG00000058398 | **Prss43** | Gm661\|Tessp3 | protease, serine 43 | protein-coding |
| chr9_110819950_110820615 | chr9 | 110819951 | 110820615 | + | 0.942004 | Intergenic | Intergenic | 6289 | NM_148940 | 73336 | Mm.200587 | NM_148940 | ENSMUSG00000032493 | **Prss44** | 1700036D21Rik\|79H19A\|AU043574\|AU043597\|TESSP-4\|Tessp4 | protease, serine 44 | protein-coding |
| chr9_110849198_110849612 | chr9 | 110849199 | 110849612 | + | 0.775876 | intron (NM_183103, intron 1 of 4) | intron (NM_183103, intron 1 of 4) | 4899 | NM_183103 | 74306 | Mm.87419 | NM_183103 | ENSMUSG00000049719 | **Prss46** | 1700112C13Rik\|79H19B\|Tessp6 | protease, serine 46 | protein-coding |
| chr1_87180837_87181335 | chr1 | 87180838 | 87181335 | + | 1.17958 | Intergenic | Intergenic | -2407 | NM_027084 | 69453 | Mm.84618 | NM_027084 | ENSMUSG00000036480 | **Prss56** | 1700027L20Rik | protease, serine 56 | protein-coding |
| chr1_87180830_87181632 | chr1 | 87180831 | 87181632 | + | 1.1609 | Intergenic | Intergenic | -2262 | NM_027084 | 69453 | Mm.84618 | NM_027084 | ENSMUSG00000036480 | **Prss56** | 1700027L20Rik | protease, serine 56 | protein-coding |
| chr9_72803017_72803602 | chr9 | 72803018 | 72803602 | + | 1.04839 | Intergenic | B4A\|SINE\|B4 | -3964 | NM_175485 | 235472 | Mm.31696 | NM_175485 | ENSMUSG00000036030 | **Prtg** | A230098A12Rik\|Igdcc5 | protogenin homolog (Gallus gallus) | protein-coding |
| chr9_72803283_72803592 | chr9 | 72803284 | 72803592 | + | 1.02664 | Intergenic | Intergenic | -3836 | NM_175485 | 235472 | Mm.31696 | NM_175485 | ENSMUSG00000036030 | **Prtg** | A230098A12Rik\|Igdcc5 | protogenin homolog (Gallus gallus) | protein-coding |
| chr9_72746121_72746654 | chr9 | 72746122 | 72746654 | + | 0.700611 | intron (NM_010890, intron 27 of 28) | intron (NM_010890, intron 27 of 28) | -60886 | NM_175485 | 235472 | Mm.31696 | NM_175485 | ENSMUSG00000036030 | **Prtg** | A230098A12Rik\|Igdcc5 | protogenin homolog (Gallus gallus) | protein-coding |
| chr19_17024625_17025165 | chr19 | 17024626 | 17025165 | + | 1.46829 | intron (NM_181348, intron 6 of 19) | intron (NM_181348, intron 6 of 19) | 68777 | NM_181348 | 353211 | Mm.257354 | NM_181348 | ENSMUSG00000039126 | **Prune2** | 6330414G02Rik\|A230083H22Rik\|A330102H22Rik\|Bmcc1\|mKIAA0367 | prune homolog 2 (Drosophila) | protein-coding |
| chr10_60296936_60297443 | chr10 | 60296937 | 60297443 | + | 1.25079 | intron (NM_011179, intron 8 of 14) | intron (NM_011179, intron 8 of 14) | 12729 | NM_001146124 | 19156 | Mm.277498 | NM_011179 | ENSMUSG00000004207 | **Psap** | AI037048\|SGP-1 | prosaposin | protein-coding |
| chr10_60283449_60283679 | chr10 | 60283450 | 60283679 | + | 0.988049 | promoter-TSS (NM_001146124) | promoter-TSS (NM_001146124) | -897 | NM_001146124 | 19156 | Mm.277498 | NM_011179 | ENSMUSG00000004207 | **Psap** | AI037048\|SGP-1 | prosaposin | protein-coding |
| chr18_36025499_36025909 | chr18 | 36025500 | 36025909 | + | 0.877646 | intron (NM_001167891, intron 5 of 9) | L2\|LINE\|L2 | 60874 | NR_110349 | 74002 | Mm.196944 | NM_028707 | ENSMUSG00000024347 | **Psd2** | 6330404E20Rik\|AW125584\|EFA6C | pleckstrin and Sec7 domain containing 2 | protein-coding |
| chr2_24365251_24365804 | chr2 | 24365252 | 24365804 | + | 1.10756 | Intergenic | ORR1F\|LTR\|ERVL-MaLR | -19869 | NM_177611 | 215632 | Mm.490434 | NM_177611 | ENSMUSG00000026979 | **Psd4** | BC046518\|EFA6B | pleckstrin and Sec7 domain containing 4 | protein-coding |
| chr2_24392045_24392729 | chr2 | 24392046 | 24392729 | + | 0.771675 | intron (NM_177611, intron 1 of 16) | intron (NM_177611, intron 1 of 16) | 6990 | NM_177611 | 215632 | Mm.490434 | NM_177611 | ENSMUSG00000026979 | **Psd4** | BC046518\|EFA6B | pleckstrin and Sec7 domain containing 4 | protein-coding |
| chr1_180290261_180290894 | chr1 | 180290262 | 180290894 | + | 0.861524 | Intergenic | Intergenic | -34278 | NM_001128605 | 19165 | Mm.330850 | NM_011183 | ENSMUSG00000010609 | **Psen2** | ALG-3\|Ad4h\|Alg3\|PS-2\|PS2\|Psnl2\|STM2 | presenilin 2 | protein-coding |
| chr12_55392618_55393369 | chr12 | 55392619 | 55393369 | + | 1.44279 | intron (NR_035430, intron 1 of 3) | intron (NR_035430, intron 1 of 3) | -5831 | NM_011968 | 26443 | Mm.30210 | NM_011968 | ENSMUSG00000021024 | **Psma6** | IOTA | proteasome (prosome, macropain) subunit, alpha type 6 | protein-coding |
| chr3_94888638_94889092 | chr3 | 94888639 | 94889092 | + | 0.672607 | Intergenic | B3\|SINE\|B2 | -1907 | NM_008945 | 19172 | Mm.368 | NM_008945 | ENSMUSG00000005779 | **Psmb4** | Pros-27 | proteasome (prosome, macropain) subunit, beta type 4 | protein-coding |
| chr2_38610000_38610742 | chr2 | 38610001 | 38610742 | + | 0.953256 | intron (NM_011187, intron 6 of 7) | intron (NM_011187, intron 6 of 7) | 33535 | NM_011187 | 19177 | Mm.389251 | NM_011187 | ENSMUSG00000026750 | **Psmb7** | AU020723\|MC14 | proteasome (prosome, macropain) subunit, beta type 7 | protein-coding |
| chr2_38616058_38616436 | chr2 | 38616059 | 38616436 | + | 0.742178 | intron (NM_011187, intron 5 of 7) | intron (NM_011187, intron 5 of 7) | 27659 | NM_011187 | 19177 | Mm.389251 | NM_011187 | ENSMUSG00000026750 | **Psmb7** | AU020723\|MC14 | proteasome (prosome, macropain) subunit, beta type 7 | protein-coding |
| chr17_34179857_34180317 | chr17 | 34179858 | 34180317 | + | 1.09146 | Intergenic | Intergenic | 7243 | NM_013585 | 16912 | Mm.390983 | NM_013585 | | **Psmb9** | Lmp-2\|Lmp2 | proteasome (prosome, macropain) subunit, beta type 9 (large multifunctional peptidase 2) | protein-coding |
| chr2_91052806_91053074 | chr2 | 91052807 | 91053074 | + | 0.688314 | Intergenic | B2_Mm1t\|SINE\|B2 | -1076 | NM_008948 | 19182 | Mm.289832 | NM_008948 | ENSMUSG00000002102 | **Psmc3** | TBP-1 | proteasome (prosome, macropain) 26S subunit, ATPase 3 | protein-coding |
| chr14_45325450_45325767 | chr14 | 45325451 | 45325767 | + | 1.04075 | Intergenic | MLT1F1\|LTR\|ERVL-MaLR | -4215 | NM_025959 | 67089 | Mm.18472 | NM_025959 | ENSMUSG00000021832 | **Psmc6** | 2300001E01Rik\|AI451058 | proteasome (prosome, macropain) 26S subunit, ATPase, 6 | protein-coding |
| chrX_140978959_140979489 | chrX | 140978960 | 140979489 | + | 0.795983 | Intergenic | Intergenic | -22513 | NM_001164177 | 53380 | Mm.17640 | NM_016883 | ENSMUSG00000031429 | **Psmd10** | AW554874 | proteasome (prosome, macropain) 26S subunit, non-ATPase, 10 | protein-coding |
| chr11_80417103_80417581 | chr11 | 80417104 | 80417581 | + | 1.15295 | Intergenic | RMER6C\|LTR\|ERVK | -11273 | NM_178616 | 69077 | Mm.260539 | NM_178616 | ENSMUSG00000017428 | **Psmd11** | 1700089D09Rik\|1810019E17Rik\|2610024G20Rik\|2810055C24Rik\|C78232\|P44.5\|S9 | proteasome (prosome, macropain) 26S subunit, non-ATPase, 11 | protein-coding |
| chr11_80420128_80420629 | chr11 | 80420129 | 80420629 | + | 0.957685 | Intergenic | Intergenic | -8236 | NM_178616 | 69077 | Mm.260539 | NM_178616 | ENSMUSG00000017428 | **Psmd11** | 1700089D09Rik\|1810019E17Rik\|2610024G20Rik\|2810055C24Rik\|C78232\|P44.5\|S9 | proteasome (prosome, macropain) 26S subunit, non-ATPase, 11 | protein-coding |
| chr11_80412069_80412465 | chr11 | 80412070 | 80412465 | + | 0.641509 | Intergenic | Intergenic | -16348 | NM_178616 | 69077 | Mm.260539 | NM_178616 | ENSMUSG00000017428 | **Psmd11** | 1700089D09Rik\|1810019E17Rik\|2610024G20Rik\|2810055C24Rik\|C78232\|P44.5\|S9 | proteasome (prosome, macropain) 26S subunit, non-ATPase, 11 | protein-coding |
| chr3_95043384_95043604 | chr3 | 95043385 | 95043604 | + | 0.669861 | promoter-TSS (NM_001282017) | promoter-TSS (NM_001282017) | -880 | NM_008951 | 19185 | Mm.2261 | NM_008951 | ENSMUSG00000005625 | **Psmd4** | Af1\|Mcb1\|angiocidin | proteasome (prosome, macropain) 26S subunit, non-ATPase, 4 | protein-coding |
| chr14_14091310_14091730 | chr14 | 14091311 | 14091730 | + | 1.00544 | intron (NM_139227, intron 7 of 11) | intron (NM_139227, intron 7 of 11) | 29384 | NM_025550 | 66413 | Mm.27591 | NM_025550 | ENSMUSG00000021737 | **Psmd6** | 2400006A19Rik | proteasome (prosome, macropain) 26S subunit, non-ATPase, 6 | protein-coding |
| chr14_14087417_14087696 | chr14 | 14087418 | 14087696 | + | 0.794045 | intron (NM_139227, intron 5 of 11) | intron (NM_139227, intron 5 of 11) | 33347 | NM_025550 | 66413 | Mm.27591 | NM_025550 | ENSMUSG00000021737 | **Psmd6** | 2400006A19Rik | proteasome (prosome, macropain) 26S subunit, non-ATPase, 6 | protein-coding |
| chr8_107589219_107589505 | chr8 | 107589220 | 107589505 | + | 1.26959 | promoter-TSS (NM_010817) | promoter-TSS (NM_010817) | -880 | NM_010817 | 17463 | Mm.18347 | NM_010817 | ENSMUSG00000039067 | **Psmd7** | AW107203\|Mov-34\|Mov34 | proteasome (prosome, macropain) 26S subunit, non-ATPase, 7 | protein-coding |
| chr9_56101065_56101399 | chr9 | 56101066 | 56101399 | + | 1.1052 | intron (NM_011193, intron 1 of 14) | intron (NM_011193, intron 1 of 14) | 11256 | NM_011193 | 19200 | Mm.2534 | NM_011193 | ENSMUSG00000032322 | **Pstpip1** | CD2BP1\|def-2 | proline-serine-threonine phosphatase-interacting protein 1 | protein-coding |
| chr9_56104090_56104889 | chr9 | 56104091 | 56104889 | + | 0.812797 | intron (NM_011193, intron 1 of 14) | intron (NM_011193, intron 1 of 14) | 14514 | NM_011193 | 19200 | Mm.2534 | NM_011193 | ENSMUSG00000032322 | **Pstpip1** | CD2BP1\|def-2 | proline-serine-threonine phosphatase-interacting protein 1 | protein-coding |
| chr18_77787545_77788192 | chr18 | 77787546 | 77788192 | + | 0.853018 | Intergenic | Intergenic | -6681 | NM_013831 | 19201 | Mm.57174 | NM_013831 | | **Pstpip2** | MAYP\|cmo | proline-serine-threonine phosphatase-interacting protein 2 | protein-coding |
| chr4_132588497_132588875 | chr4 | 132588498 | 132588875 | + | 0.94493 | Intergenic | Intergenic | 24619 | NM_001081211 | 19204 | Mm.89389 | NM_001081211 | | **Ptafr** | PAFR | platelet-activating factor receptor | protein-coding |
| chr4_132588426_132589070 | chr4 | 132588427 | 132589070 | + | 0.814719 | Intergenic | Intergenic | 24681 | NM_001081211 | 19204 | Mm.89389 | NM_001081211 | | **Ptafr** | PAFR | platelet-activating factor receptor | protein-coding |

| | | | | | | | | | | | | | | | | | |
|---|---|---|---|---|---|---|---|---|---|---|---|---|---|---|---|---|---|
| chr4_59505451_59505886 | chr4 | 59505452 | 59505886 | + | 1.1784 | intron (NM_144904, intron 5 of 14) | ID_B1\|SINE\|B4 | 43695 | NM_178164 | 230257 | Mm.331640 | NM_144904 | ENSMUSG00000028382 | **Ptbp3** | 5830471K22Rik\|AA407443\|AI462022\|AW107884\|C86549\|Rod1 | polypyrimidine tract binding protein 3 | protein-coding |
| chr4_59496176_59497111 | chr4 | 59496177 | 59497111 | + | 0.958315 | intron (NM_144904, intron 6 of 14) | intron (NM_144904, intron 6 of 14) | 52720 | NM_178164 | 230257 | Mm.331640 | NM_144904 | ENSMUSG00000028382 | **Ptbp3** | 5830471K22Rik\|AA407443\|AI462022\|AW107884\|C86549\|Rod1 | polypyrimidine tract binding protein 3 | protein-coding |
| chr4_59552016_59552303 | chr4 | 59552017 | 59552303 | + | 0.795201 | Intergenic | B4A\|SINE\|B4 | -2796 | NM_144904 | 230257 | Mm.331640 | NM_144904 | ENSMUSG00000028382 | **Ptbp3** | 5830471K22Rik\|AA407443\|AI462022\|AW107884\|C86549\|Rod1 | polypyrimidine tract binding protein 3 | protein-coding |
| chr13_63689775_63690384 | chr13 | 63689776 | 63690384 | + | 1.34425 | Intergenic | Intergenic | -124560 | NM_008957 | 19206 | Mm.228798 | NM_008957 | ENSMUSG00000021466 | **Ptch1** | A230106A15Rik\|Ptc\|Ptc1\|Ptch\|mes | patched homolog 1 | protein-coding |
| chr13_63557920_63558276 | chr13 | 63557921 | 63558276 | + | 1.18019 | intron (NM_008957, intron 2 of 22) | intron (NM_008957, intron 2 of 22) | 7422 | NM_008957 | 19206 | Mm.228798 | NM_008957 | ENSMUSG00000021466 | **Ptch1** | A230106A15Rik\|Ptc\|Ptc1\|Ptch\|mes | patched homolog 1 | protein-coding |
| chr13_63541195_63542088 | chr13 | 63541196 | 63542088 | + | 1.14088 | intron (NM_008957, intron 8 of 22) | intron (NM_008957, intron 8 of 22) | 23878 | NM_008957 | 19206 | Mm.228798 | NM_008957 | ENSMUSG00000021466 | **Ptch1** | A230106A15Rik\|Ptc\|Ptc1\|Ptch\|mes | patched homolog 1 | protein-coding |
| chr13_63548297_63548875 | chr13 | 63548298 | 63548875 | + | 1.07102 | intron (NM_008957, intron 2 of 22) | intron (NM_008957, intron 2 of 22) | 16934 | NM_008957 | 19206 | Mm.228798 | NM_008957 | ENSMUSG00000021466 | **Ptch1** | A230106A15Rik\|Ptc\|Ptc1\|Ptch\|mes | patched homolog 1 | protein-coding |
| chr13_63604593_63605286 | chr13 | 63604594 | 63605286 | + | 0.997054 | Intergenic | ORR1C2\|LTR\|ERVL-MaLR | -39420 | NM_008957 | 19206 | Mm.228798 | NM_008957 | ENSMUSG00000021466 | **Ptch1** | A230106A15Rik\|Ptc\|Ptc1\|Ptch\|mes | patched homolog 1 | protein-coding |
| chr13_63662383_63662986 | chr13 | 63662384 | 63662986 | + | 0.96113 | Intergenic | ORR1F\|LTR\|ERVL-MaLR | -97165 | NM_008957 | 19206 | Mm.228798 | NM_008957 | ENSMUSG00000021466 | **Ptch1** | A230106A15Rik\|Ptc\|Ptc1\|Ptch\|mes | patched homolog 1 | protein-coding |
| chr13_63557757_63558293 | chr13 | 63557758 | 63558293 | + | 0.928907 | intron (NM_008957, intron 2 of 22) | intron (NM_008957, intron 2 of 22) | 7495 | NM_008957 | 19206 | Mm.228798 | NM_008957 | ENSMUSG00000021466 | **Ptch1** | A230106A15Rik\|Ptc\|Ptc1\|Ptch\|mes | patched homolog 1 | protein-coding |
| chr13_63502204_63502729 | chr13 | 63502205 | 63502729 | + | 0.684263 | Intergenic | Intergenic | 63053 | NM_008957 | 19206 | Mm.228798 | NM_008957 | ENSMUSG00000021466 | **Ptch1** | A230106A15Rik\|Ptc\|Ptc1\|Ptch\|mes | patched homolog 1 | protein-coding |
| chr13_63553442_63553959 | chr13 | 63553443 | 63553959 | + | 0.66681 | intron (NM_008957, intron 2 of 22) | intron (NM_008957, intron 2 of 22) | 11819 | NM_008957 | 19206 | Mm.228798 | NM_008957 | ENSMUSG00000021466 | **Ptch1** | A230106A15Rik\|Ptc\|Ptc1\|Ptch\|mes | patched homolog 1 | protein-coding |
| chr4_148287861_148288495 | chr4 | 148287862 | 148288495 | + | 0.734032 | promoter-TSS (NM_001083342) | promoter-TSS (NM_001083342) | -213 | NM_001083342 | 242748 | Mm.33873 | NM_001083342 | | **Ptchd2** | G630052C06Rik\|RNDEu-2 | patched domain containing 2 | protein-coding |
| chr4_148287640_148288564 | chr4 | 148287641 | 148288564 | + | 0.63351 | promoter-TSS (NM_001083342) | promoter-TSS (NM_001083342) | -137 | NM_001083342 | 242748 | Mm.33873 | NM_001083342 | | **Ptchd2** | G630052C06Rik\|RNDEu-2 | patched domain containing 2 | protein-coding |
| chr11_121776464_121777408 | chr11 | 121776465 | 121777408 | + | 1.0635 | Intergenic | Intergenic | -53282 | NM_029049 | 74675 | Mm.61213 | NM_029049 | ENSMUSG00000039198 | **Ptchd3** | 4930451E13Rik\|4933440L20Rik\|RNDEu-3 | patched domain containing 3 | protein-coding |
| chr11_121977315_121977861 | chr11 | 121977316 | 121977861 | + | 1.05426 | Intergenic | RLTR30B_MM\|LTR\|ERV1 | 147370 | NM_029049 | 74675 | Mm.61213 | NM_029049 | ENSMUSG00000039198 | **Ptchd3** | 4930451E13Rik\|4933440L20Rik\|RNDEu-3 | patched domain containing 3 | protein-coding |
| chr11_121768842_121769170 | chr11 | 121768843 | 121769170 | + | 1.04903 | Intergenic | Intergenic | -61212 | NM_029049 | 74675 | Mm.61213 | NM_029049 | ENSMUSG00000039198 | **Ptchd3** | 4930451E13Rik\|4933440L20Rik\|RNDEu-3 | patched domain containing 3 | protein-coding |
| chr11_121776986_121777408 | chr11 | 121776987 | 121777408 | + | 1.01065 | Intergenic | Intergenic | -53021 | NM_029049 | 74675 | Mm.61213 | NM_029049 | ENSMUSG00000039198 | **Ptchd3** | 4930451E13Rik\|4933440L20Rik\|RNDEu-3 | patched domain containing 3 | protein-coding |
| chr11_121768366_121769206 | chr11 | 121768367 | 121769206 | + | 0.983165 | Intergenic | L1M2\|LINE\|L1 | -61432 | NM_029049 | 74675 | Mm.61213 | NM_029049 | ENSMUSG00000039198 | **Ptchd3** | 4930451E13Rik\|4933440L20Rik\|RNDEu-3 | patched domain containing 3 | protein-coding |
| chr11_121977277_121978267 | chr11 | 121977278 | 121978267 | + | 0.871967 | Intergenic | RLTR30B_MM\|LTR\|ERV1 | 147554 | NM_029049 | 74675 | Mm.61213 | NM_029049 | ENSMUSG00000039198 | **Ptchd3** | 4930451E13Rik\|4933440L20Rik\|RNDEu-3 | patched domain containing 3 | protein-coding |
| chr11_121777542_121777997 | chr11 | 121777543 | 121777997 | + | 0.783989 | Intergenic | Intergenic | -52448 | NM_029049 | 74675 | Mm.61213 | NM_029049 | ENSMUSG00000039198 | **Ptchd3** | 4930451E13Rik\|4933440L20Rik\|RNDEu-3 | patched domain containing 3 | protein-coding |
| chr19_32988611_32989139 | chr19 | 32988612 | 32989139 | + | 1.67565 | Intergenic | Intergenic | 231298 | NM_008960 | 19211 | Mm.245395 | NM_008960 | ENSMUSG00000013663 | **Pten** | 2310035O07Rik\|A130070J02Rik\|AI463227\|B430203M17Rik\|MMAC1\|TEP1 | phosphatase and tensin homolog | protein-coding |
| chr19_32956929_32957480 | chr19 | 32956930 | 32957480 | + | 1.02754 | Intergenic | Intergenic | 199628 | NM_008960 | 19211 | Mm.245395 | NM_008960 | ENSMUSG00000013663 | **Pten** | 2310035O07Rik\|A130070J02Rik\|AI463227\|B430203M17Rik\|MMAC1\|TEP1 | phosphatase and tensin homolog | protein-coding |
| chr19_32904260_32904948 | chr19 | 32904261 | 32904948 | + | 0.99065 | Intergenic | Intergenic | 147027 | NM_008960 | 19211 | Mm.245395 | NM_008960 | ENSMUSG00000013663 | **Pten** | 2310035O07Rik\|A130070J02Rik\|AI463227\|B430203M17Rik\|MMAC1\|TEP1 | phosphatase and tensin homolog | protein-coding |
| chr19_32952457_32952771 | chr19 | 32952458 | 32952771 | + | 0.978694 | Intergenic | ORR1B2\|LTR\|ERVL-MaLR | 195037 | NM_008960 | 19211 | Mm.245395 | NM_008960 | ENSMUSG00000013663 | **Pten** | 2310035O07Rik\|A130070J02Rik\|AI463227\|B430203M17Rik\|MMAC1\|TEP1 | phosphatase and tensin homolog | protein-coding |
| chr19_32760660_32760959 | chr19 | 32760661 | 32760959 | + | 0.858277 | intron (NM_008960, intron 1 of 8) | intron (NM_008960, intron 1 of 8) | 3233 | NM_008960 | 19211 | Mm.245395 | NM_008960 | ENSMUSG00000013663 | **Pten** | 2310035O07Rik\|A130070J02Rik\|AI463227\|B430203M17Rik\|MMAC1\|TEP1 | phosphatase and tensin homolog | protein-coding |
| chr19_33036700_33037113 | chr19 | 33036701 | 33037113 | + | 0.693893 | Intergenic | Intergenic | 279330 | NM_008960 | 19211 | Mm.245395 | NM_008960 | ENSMUSG00000013663 | **Pten** | 2310035O07Rik\|A130070J02Rik\|AI463227\|B430203M17Rik\|MMAC1\|TEP1 | phosphatase and tensin homolog | protein-coding |
| chr8_83665929_83666454 | chr8 | 83665930 | 83666454 | + | 1.05411 | promoter-TSS (NM_013641) | promoter-TSS (NM_013641) | -448 | NM_013641 | 19216 | Mm.213000 | NM_013641 | | **Ptger1** | 42kDa\|EP1\|Ptgerep1 | prostaglandin E receptor 1 (subtype EP1) | protein-coding |
| chr14_45102009_45102649 | chr14 | 45102010 | 45102649 | + | 1.83856 | Intergenic | RMER1C\|Other\|Other | 114218 | NM_008964 | 19217 | Mm.4630 | NM_008964 | ENSMUSG00000037759 | **Ptger2** | EP2\|Ptgerep2 | prostaglandin E receptor 2 (subtype EP2) | protein-coding |
| chr14_45005341_45006279 | chr14 | 45005342 | 45006279 | + | 1.77549 | Intergenic | Intergenic | 17699 | NM_008964 | 19217 | Mm.4630 | NM_008964 | ENSMUSG00000037759 | **Ptger2** | EP2\|Ptgerep2 | prostaglandin E receptor 2 (subtype EP2) | protein-coding |
| chr14_45049483_45050663 | chr14 | 45049484 | 45050663 | + | 1.59558 | Intergenic | Intergenic | 61962 | NM_008964 | 19217 | Mm.4630 | NM_008964 | ENSMUSG00000037759 | **Ptger2** | EP2\|Ptgerep2 | prostaglandin E receptor 2 (subtype EP2) | protein-coding |
| chr14_45034294_45034671 | chr14 | 45034295 | 45034671 | + | 1.20166 | Intergenic | Lx4A\|LINE\|L1 | 46372 | NM_008964 | 19217 | Mm.4630 | NM_008964 | ENSMUSG00000037759 | **Ptger2** | EP2\|Ptgerep2 | prostaglandin E receptor 2 (subtype EP2) | protein-coding |
| chr14_45009406_45009679 | chr14 | 45009407 | 45009679 | + | 1.18619 | Intergenic | Intergenic | 21432 | NM_008964 | 19217 | Mm.4630 | NM_008964 | ENSMUSG00000037759 | **Ptger2** | EP2\|Ptgerep2 | prostaglandin E receptor 2 (subtype EP2) | protein-coding |
| chr14_45040866_45041617 | chr14 | 45040867 | 45041617 | + | 0.994863 | Intergenic | Intergenic | 53131 | NM_008964 | 19217 | Mm.4630 | NM_008964 | ENSMUSG00000037759 | **Ptger2** | EP2\|Ptgerep2 | prostaglandin E receptor 2 (subtype EP2) | protein-coding |
| chr14_45038054_45038437 | chr14 | 45038055 | 45038437 | + | 0.919832 | Intergenic | L1MB5\|LINE\|L1 | 50135 | NM_008964 | 19217 | Mm.4630 | NM_008964 | ENSMUSG00000037759 | **Ptger2** | EP2\|Ptgerep2 | prostaglandin E receptor 2 (subtype EP2) | protein-coding |
| chr14_45094605_45095167 | chr14 | 45094606 | 45095167 | + | 0.787492 | Intergenic | Lx4A\|LINE\|L1 | 106775 | NM_008964 | 19217 | Mm.4630 | NM_008964 | ENSMUSG00000037759 | **Ptger2** | EP2\|Ptgerep2 | prostaglandin E receptor 2 (subtype EP2) | protein-coding |
| chr3_157635653_157636275 | chr3 | 157635654 | 157636275 | + | 0.760214 | intron (NM_011196, intron 2 of 2) | intron (NM_011196, intron 2 of 2) | 69072 | NM_011196 | 19218 | Mm.30424 | NM_011196 | ENSMUSG00000040016 | **Ptger3** | EP3\|Pgerep3\|Ptgerep3 | prostaglandin E receptor 3 (subtype EP3) | protein-coding |
| chr15_5238585_5238981 | chr15 | 5238586 | 5238981 | + | 0.845067 | intron (NM_001136079, intron 2 of 2) | intron (NM_001136079, intron 2 of 2) | 5404 | NM_001136079 | 19219 | Mm.18509 | NM_008965 | ENSMUSG00000039942 | **Ptger4** | EP4\|Ptgerep4 | prostaglandin E receptor 4 (subtype EP4) | protein-coding |
| chr15_5252341_5252946 | chr15 | 5252342 | 5252946 | + | 0.701713 | Intergenic | Intergenic | -8457 | NM_008965 | 19219 | Mm.18509 | NM_008965 | ENSMUSG00000039942 | **Ptger4** | EP4\|Ptgerep4 | prostaglandin E receptor 4 (subtype EP4) | protein-coding |
| chr3_101115890_101116343 | chr3 | 101115891 | 101116343 | + | 0.802657 | Intergenic | Intergenic | -5951 | NM_011197 | 19221 | Mm.24807 | NM_011197 | ENSMUSG00000027864 | **Ptgfrn** | 4833445A08Rik\|AU042434\|CD9P-1\|FPRP\|Trim45 | prostaglandin F2 receptor negative regulator | protein-coding |
| chr15_73459548_73460025 | chr15 | 73459549 | 73460025 | + | 1.2534 | Intergenic | Intergenic | -36596 | NM_001130409 | 14083 | Mm.254494 | NM_007982 | ENSMUSG00000022607 | **Ptk2** | FAK\|FRNK\|Fadk\|mKIAA4203\|p125FAK | PTK2 protein tyrosine kinase 2 | protein-coding |
| chr15_73318482_73319121 | chr15 | 73318483 | 73319121 | + | 0.767997 | intron (NM_007982, intron 7 of 31) | intron (NM_007982, intron 7 of 31) | 104389 | NM_007982 | 14083 | Mm.254494 | NM_007982 | ENSMUSG00000022607 | **Ptk2** | FAK\|FRNK\|Fadk\|mKIAA4203\|p125FAK | PTK2 protein tyrosine kinase 2 | protein-coding |
| chr15_73461896_73462175 | chr15 | 73461897 | 73462175 | + | 0.654848 | Intergenic | MTE2a\|LTR\|ERVL-MaLR | -38845 | NM_001130409 | 14083 | Mm.254494 | NM_007982 | ENSMUSG00000022607 | **Ptk2** | FAK\|FRNK\|Fadk\|mKIAA4203\|p125FAK | PTK2 protein tyrosine kinase 2 | protein-coding |
| chr14_66268922_66269386 | chr14 | 66268923 | 66269386 | + | 1.36297 | intron (NM_001162365, intron 1 of 30) | L1M4\|LINE\|L1 | 11898 | NM_172498 | 19229 | Mm.21613 | NM_172498 | ENSMUSG00000059456 | **Ptk2b** | CADTK\|CAKB\|CAKbeta\|E430023O05Rik\|FADK2\|FAK2\|PYK2\|Raftk | PTK2 protein tyrosine kinase 2 beta | protein-coding |
| chr14_66220980_66221394 | chr14 | 66220981 | 66221394 | + | 1.1854 | intron (NM_001162365, intron 1 of 30) | intron (NM_001162365, intron 1 of 30) | -7635 | NM_001162366 | 19229 | Mm.21613 | NM_172498 | ENSMUSG00000059456 | **Ptk2b** | CADTK\|CAKB\|CAKbeta\|E430023O05Rik\|FADK2\|FAK2\|PYK2\|Raftk | PTK2 protein tyrosine kinase 2 beta | protein-coding |
| chr14_66259051_66259503 | chr14 | 66259052 | 66259503 | + | 1.00992 | intron (NM_001162365, intron 1 of 30) | intron (NM_001162365, intron 1 of 30) | 21775 | NM_172498 | 19229 | Mm.21613 | NM_172498 | ENSMUSG00000059456 | **Ptk2b** | CADTK\|CAKB\|CAKbeta\|E430023O05Rik\|FADK2\|FAK2\|PYK2\|Raftk | PTK2 protein tyrosine kinase 2 beta | protein-coding |
| chr14_66265879_66266284 | chr14 | 66265880 | 66266284 | + | 0.948741 | intron (NM_001162365, intron 1 of 30) | intron (NM_001162365, intron 1 of 30) | 14970 | NM_172498 | 19229 | Mm.21613 | NM_172498 | ENSMUSG00000059456 | **Ptk2b** | CADTK\|CAKB\|CAKbeta\|E430023O05Rik\|FADK2\|FAK2\|PYK2\|Raftk | PTK2 protein tyrosine kinase 2 beta | protein-coding |
| chr14_66230835_66231287 | chr14 | 66230836 | 66231287 | + | 0.885662 | intron (NM_001162365, intron 1 of 30) | intron (NM_001162365, intron 1 of 30) | -17509 | NM_001162366 | 19229 | Mm.21613 | NM_172498 | ENSMUSG00000059456 | **Ptk2b** | CADTK\|CAKB\|CAKbeta\|E430023O05Rik\|FADK2\|FAK2\|PYK2\|Raftk | PTK2 protein tyrosine kinase 2 beta | protein-coding |
| chr14_66265879_66266554 | chr14 | 66265880 | 66266554 | + | 0.818921 | intron (NM_001162365, intron 1 of 30) | intron (NM_001162365, intron 1 of 30) | 14835 | NM_172498 | 19229 | Mm.21613 | NM_172498 | ENSMUSG00000059456 | **Ptk2b** | CADTK\|CAKB\|CAKbeta\|E430023O05Rik\|FADK2\|FAK2\|PYK2\|Raftk | PTK2 protein tyrosine kinase 2 beta | protein-coding |

| | | | | | | | | | | | | | | | | | |
|---|---|---|---|---|---|---|---|---|---|---|---|---|---|---|---|---|---|
| chr14_66249052_66249717 | chr14 | 66249053 | 66249717 | + | 0.746765 | intron (NM_001162365, intron 1 of 30) | intron (NM_001162365, intron 1 of 30) | 31667 | NM_172498 | 19229 | Mm.21613 | NM_172498 | ENSMUSG00000059456 | **Ptk2b** | CADTK\|CAKB\|CAKbeta\|E430023O05Rik\|FADK2\|FAK2\|PYK2\|Raftk | PTK2 protein tyrosine kinase 2 beta | protein-coding |
| chr14_66228912_66229352 | chr14 | 66228913 | 66229352 | + | 0.682596 | intron (NM_001162365, intron 1 of 30) | intron (NM_001162365, intron 1 of 30) | -15580 | NM_001162366 | 19229 | Mm.21613 | NM_172498 | ENSMUSG00000059456 | **Ptk2b** | CADTK\|CAKB\|CAKbeta\|E430023O05Rik\|FADK2\|FAK2\|PYK2\|Raftk | PTK2 protein tyrosine kinase 2 beta | protein-coding |
| chr14_66248941_66250057 | chr14 | 66248942 | 66250057 | + | 0.654075 | intron (NM_001162365, intron 1 of 30) | intron (NM_001162365, intron 1 of 30) | 31553 | NM_172498 | 19229 | Mm.21613 | NM_172498 | ENSMUSG00000059456 | **Ptk2b** | CADTK\|CAKB\|CAKbeta\|E430023O05Rik\|FADK2\|FAK2\|PYK2\|Raftk | PTK2 protein tyrosine kinase 2 beta | protein-coding |
| chr1_86531006_86531311 | chr1 | 86531007 | 86531311 | + | 0.957393 | TTS (NM_008972) | TTS (NM_008972) | 4423 | NM_008972 | 19231 | Mm.19187 | NM_008972 | ENSMUSG00000026238 | **Ptma** | Thym | prothymosin alpha | protein-coding |
| chr6_37055086_37055539 | chr6 | 37055087 | 37055539 | + | 0.941878 | intron (NM_001081206, intron 10 of 33) | intron (NM_001081206, intron 10 of 33) | -243952 | NM_008973 | 19242 | Mm.279690 | NM_008973 | ENSMUSG00000029838 | **Ptn** | HARP\|HB-GAM\|HBBN\|HBGF-8\|HBNF\|OSF\|Osf-1\|Osf1 | pleiotrophin | protein-coding |
| chr6_37055081_37055863 | chr6 | 37055082 | 37055863 | + | 0.765338 | intron (NM_001081206, intron 10 of 33) | intron (NM_001081206, intron 10 of 33) | -244111 | NM_008973 | 19242 | Mm.279690 | NM_008973 | ENSMUSG00000029838 | **Ptn** | HARP\|HB-GAM\|HBBN\|HBGF-8\|HBNF\|OSF\|Osf-1\|Osf1 | pleiotrophin | protein-coding |
| chr4_129823273_129823693 | chr4 | 129823274 | 129823693 | + | 0.972333 | intron (NM_001164745, intron 1 of 5) | intron (NM_001164745, intron 1 of 5) | 2775 | NM_001164745 | 19244 | Mm.193688 | NM_008974 | ENSMUSG00000028788 | **Ptp4a2** | Prl-2 | protein tyrosine phosphatase 4a2 | protein-coding |
| chr4_129823280_129823471 | chr4 | 129823281 | 129823471 | + | 0.925941 | intron (NM_001164745, intron 1 of 5) | intron (NM_001164745, intron 1 of 5) | 2668 | NM_001164745 | 19244 | Mm.193688 | NM_008974 | ENSMUSG00000028788 | **Ptp4a2** | Prl-2 | protein tyrosine phosphatase 4a2 | protein-coding |
| chr4_129809334_129809629 | chr4 | 129809335 | 129809629 | + | 0.780311 | Intergenic | RSINE1\|SINE\|B4 | -10997 | NM_008974 | 19244 | Mm.193688 | NM_008974 | ENSMUSG00000028788 | **Ptp4a2** | Prl-2 | protein tyrosine phosphatase 4a2 | protein-coding |
| chr4_129827592_129828367 | chr4 | 129827593 | 129828367 | + | 0.709895 | intron (NM_001164745, intron 1 of 5) | intron (NM_001164745, intron 1 of 5) | 7272 | NM_001164745 | 19244 | Mm.193688 | NM_008974 | ENSMUSG00000028788 | **Ptp4a2** | Prl-2 | protein tyrosine phosphatase 4a2 | protein-coding |
| chr4_129818600_129818891 | chr4 | 129818601 | 129818891 | + | 0.686919 | Intergenic | Intergenic | -1733 | NM_008974 | 19244 | Mm.193688 | NM_008974 | ENSMUSG00000028788 | **Ptp4a2** | Prl-2 | protein tyrosine phosphatase 4a2 | protein-coding |
| chr9_65003841_65004573 | chr9 | 65003842 | 65004573 | + | 1.5084 | intron (NM_021345, intron 5 of 10) | intron (NM_021345, intron 5 of 10) | 17510 | NM_021345 | 57874 | Mm.308180 | NM_021345 | ENSMUSG00000033629 | **Ptplad1** | 4930523M17Rik\|AW742319\|B-ind1\|HACD3\|Hspc121 | protein tyrosine phosphatase-like A domain containing 1 | protein-coding |
| chr2_167909640_167910693 | chr2 | 167909641 | 167910693 | + | 1.30342 | Intergenic | Intergenic | -22160 | NM_011201 | 19246 | Mm.277916 | NM_011201 | ENSMUSG00000027540 | **Ptpn1** | PTP-1B\|PTP-HA2\|PTP1B | protein tyrosine phosphatase, non-receptor type 1 | protein-coding |
| chr2_167958794_167959921 | chr2 | 167958795 | 167959921 | + | 0.934377 | intron (NM_011201, intron 1 of 8) | intron (NM_011201, intron 1 of 8) | 27031 | NM_011201 | 19246 | Mm.277916 | NM_011201 | ENSMUSG00000027540 | **Ptpn1** | PTP-1B\|PTP-HA2\|PTP1B | protein tyrosine phosphatase, non-receptor type 1 | protein-coding |
| chr2_167942105_167942622 | chr2 | 167942106 | 167942622 | + | 0.880073 | intron (NM_011201, intron 1 of 8) | intron (NM_011201, intron 1 of 8) | 10037 | NM_011201 | 19246 | Mm.277916 | NM_011201 | ENSMUSG00000027540 | **Ptpn1** | PTP-1B\|PTP-HA2\|PTP1B | protein tyrosine phosphatase, non-receptor type 1 | protein-coding |
| chr2_167959653_167959852 | chr2 | 167959654 | 167959852 | + | 0.785798 | intron (NM_011201, intron 1 of 8) | intron (NM_011201, intron 1 of 8) | 27426 | NM_011201 | 19246 | Mm.277916 | NM_011201 | ENSMUSG00000027540 | **Ptpn1** | PTP-1B\|PTP-HA2\|PTP1B | protein tyrosine phosphatase, non-receptor type 1 | protein-coding |
| chr5_121196803_121197085 | chr5 | 121196804 | 121197085 | + | 0.614095 | Intergenic | Intergenic | -5547 | NM_011202 | 19247 | Mm.8681 | NM_011202 | ENSMUSG00000043733 | **Ptpn11** | 2700084A17Rik\|AW536184\|PTP1D\|PTP2C\|SAP-2\|SH-PTP2\|SH-PTP3\|SHP-2\|Shp2\|Syp | protein tyrosine phosphatase, non-receptor type 11 | protein-coding |
| chr5_21019580_21020094 | chr5 | 21019581 | 21020094 | + | 0.623181 | intron (NM_011203, intron 3 of 17) | intron (NM_011203, intron 3 of 17) | 35960 | NM_011203 | 19248 | Mm.319117 | NM_011203 | ENSMUSG00000028771 | **Ptpn12** | P19-PTP\|PTP-P19\|PTP-PEST\|PTPG1 | protein tyrosine phosphatase, non-receptor type 12 | protein-coding |
| chr5_103442129_103442590 | chr5 | 103442130 | 103442590 | + | 1.65742 | intron (NM_011204, intron 1 of 47) | intron (NM_011204, intron 1 of 47) | 17168 | NM_011204 | 19249 | Mm.3414 | NM_011204 | ENSMUSG00000034573 | **Ptpn13** | AI324989\|PTP-BL\|PTPL1\|Ptpri\|RIP | protein tyrosine phosphatase, non-receptor type 13 | protein-coding |
| chr5_103518654_103519624 | chr5 | 103518655 | 103519624 | + | 1.369 | intron (NM_011204, intron 9 of 47) | intron (NM_011204, intron 9 of 47) | 93947 | NM_011204 | 19249 | Mm.3414 | NM_011204 | ENSMUSG00000034573 | **Ptpn13** | AI324989\|PTP-BL\|PTPL1\|Ptpri\|RIP | protein tyrosine phosphatase, non-receptor type 13 | protein-coding |
| chr5_103365320_103365532 | chr5 | 103365321 | 103365532 | + | 1.04674 | Intergenic | Intergenic | -59766 | NM_011204 | 19249 | Mm.3414 | NM_011204 | ENSMUSG00000034573 | **Ptpn13** | AI324989\|PTP-BL\|PTPL1\|Ptpri\|RIP | protein tyrosine phosphatase, non-receptor type 13 | protein-coding |
| chr5_103379218_103379502 | chr5 | 103379219 | 103379502 | + | 0.82548 | Intergenic | Intergenic | -45832 | NM_011204 | 19249 | Mm.3414 | NM_011204 | ENSMUSG00000034573 | **Ptpn13** | AI324989\|PTP-BL\|PTPL1\|Ptpri\|RIP | protein tyrosine phosphatase, non-receptor type 13 | protein-coding |
| chr5_103463670_103464280 | chr5 | 103463671 | 103464280 | + | 0.773351 | intron (NM_011204, intron 2 of 47) | intron (NM_011204, intron 2 of 47) | 38783 | NM_011204 | 19249 | Mm.3414 | NM_011204 | ENSMUSG00000034573 | **Ptpn13** | AI324989\|PTP-BL\|PTPL1\|Ptpri\|RIP | protein tyrosine phosphatase, non-receptor type 13 | protein-coding |
| chr5_103463691_103464025 | chr5 | 103463692 | 103464025 | + | 0.742978 | intron (NM_011204, intron 2 of 47) | intron (NM_011204, intron 2 of 47) | 38666 | NM_011204 | 19249 | Mm.3414 | NM_011204 | ENSMUSG00000034573 | **Ptpn13** | AI324989\|PTP-BL\|PTPL1\|Ptpri\|RIP | protein tyrosine phosphatase, non-receptor type 13 | protein-coding |
| chr5_103365306_103365692 | chr5 | 103365307 | 103365692 | + | 0.729092 | Intergenic | Intergenic | -59693 | NM_011204 | 19249 | Mm.3414 | NM_011204 | ENSMUSG00000034573 | **Ptpn13** | AI324989\|PTP-BL\|PTPL1\|Ptpri\|RIP | protein tyrosine phosphatase, non-receptor type 13 | protein-coding |
| chr5_103439195_103439494 | chr5 | 103439196 | 103439494 | + | 0.669489 | intron (NM_011204, intron 1 of 47) | intron (NM_011204, intron 1 of 47) | 14153 | NM_011204 | 19249 | Mm.3414 | NM_011204 | ENSMUSG00000034573 | **Ptpn13** | AI324989\|PTP-BL\|PTPL1\|Ptpri\|RIP | protein tyrosine phosphatase, non-receptor type 13 | protein-coding |
| chr1_189819139_189819615 | chr1 | 189819140 | 189819615 | + | 0.877987 | intron (NM_008976, intron 3 of 18) | intron (NM_008976, intron 3 of 18) | 91109 | NM_008976 | 19250 | Mm.4498 | NM_008976 | ENSMUSG00000026604 | **Ptpn14** | C130080N23Rik\|OTTMUSG00000022087\|PTP36 | protein tyrosine phosphatase, non-receptor type 14 | protein-coding |
| chr1_189816355_189817150 | chr1 | 189816356 | 189817150 | + | 0.779321 | intron (NM_008976, intron 3 of 18) | intron (NM_008976, intron 3 of 18) | 88485 | NM_008976 | 19250 | Mm.4498 | NM_008976 | ENSMUSG00000026604 | **Ptpn14** | C130080N23Rik\|OTTMUSG00000022087\|PTP36 | protein tyrosine phosphatase, non-receptor type 14 | protein-coding |
| chr1_189819145_189819855 | chr1 | 189819146 | 189819855 | + | 0.735137 | intron (NM_008976, intron 3 of 18) | intron (NM_008976, intron 3 of 18) | 91232 | NM_008976 | 19250 | Mm.4498 | NM_008976 | ENSMUSG00000026604 | **Ptpn14** | C130080N23Rik\|OTTMUSG00000022087\|PTP36 | protein tyrosine phosphatase, non-receptor type 14 | protein-coding |
| chr1_189824466_189824819 | chr1 | 189824467 | 189824819 | + | 0.708723 | intron (NM_008976, intron 4 of 18) | intron (NM_008976, intron 4 of 18) | 96375 | NM_008976 | 19250 | Mm.4498 | NM_008976 | ENSMUSG00000026604 | **Ptpn14** | C130080N23Rik\|OTTMUSG00000022087\|PTP36 | protein tyrosine phosphatase, non-receptor type 14 | protein-coding |
| chr18_67693261_67694301 | chr18 | 67693262 | 67694301 | + | 0.969355 | intron (NM_001127177, intron 3 of 8) | intron (NM_001127177, intron 3 of 8) | 30840 | NM_001127177 | 19255 | Mm.260433 | NM_008977 | ENSMUSG00000024539 | **Ptpn2** | AI325124\|Ptpt\|TC-PTP | protein tyrosine phosphatase, non-receptor type 2 | protein-coding |
| chr18_67706001_67706463 | chr18 | 67706002 | 67706463 | + | 0.867342 | intron (NM_001127177, intron 2 of 8) | intron (NM_001127177, intron 2 of 8) | 18389 | NM_001127177 | 19255 | Mm.260433 | NM_008977 | ENSMUSG00000024539 | **Ptpn2** | AI325124\|Ptpt\|TC-PTP | protein tyrosine phosphatase, non-receptor type 2 | protein-coding |
| chr18_67699072_67699456 | chr18 | 67699073 | 67699456 | + | 0.817552 | intron (NM_001127177, intron 2 of 8) | intron (NM_001127177, intron 2 of 8) | 25357 | NM_001127177 | 19255 | Mm.260433 | NM_008977 | ENSMUSG00000024539 | **Ptpn2** | AI325124\|Ptpt\|TC-PTP | protein tyrosine phosphatase, non-receptor type 2 | protein-coding |
| chr18_67688439_67688787 | chr18 | 67688440 | 67688787 | + | 0.788695 | intron (NM_001127177, intron 4 of 8) | intron (NM_001127177, intron 4 of 8) | 36008 | NM_001127177 | 19255 | Mm.260433 | NM_008977 | ENSMUSG00000024539 | **Ptpn2** | AI325124\|Ptpt\|TC-PTP | protein tyrosine phosphatase, non-receptor type 2 | protein-coding |
| chr18_67699203_67699428 | chr18 | 67699204 | 67699428 | + | 0.738003 | intron (NM_001127177, intron 2 of 8) | intron (NM_001127177, intron 2 of 8) | 25305 | NM_001127177 | 19255 | Mm.260433 | NM_008977 | ENSMUSG00000024539 | **Ptpn2** | AI325124\|Ptpt\|TC-PTP | protein tyrosine phosphatase, non-receptor type 2 | protein-coding |
| chr18_67699555_67699959 | chr18 | 67699556 | 67699959 | + | 0.687627 | intron (NM_001127177, intron 2 of 8) | intron (NM_001127177, intron 2 of 8) | 24864 | NM_001127177 | 19255 | Mm.260433 | NM_008977 | ENSMUSG00000024539 | **Ptpn2** | AI325124\|Ptpt\|TC-PTP | protein tyrosine phosphatase, non-receptor type 2 | protein-coding |
| chr3_103850657_103851260 | chr3 | 103850658 | 103851260 | + | 1.15032 | intron (NM_001142960, intron 3 of 3) | intron (NM_001142960, intron 3 of 3) | -8836 | NR_104070 | 19260 | Mm.395 | NM_008979 | ENSMUSG00000027843 | **Ptpn22** | 70zpep\|PEP\|Ptpn8 | protein tyrosine phosphatase, non-receptor type 22 (lymphoid) | protein-coding |
| chr3_103870358_103870666 | chr3 | 103870359 | 103870666 | + | 1.0034 | intron (NM_008979, intron 1 of 20) | ORR1F\|LTR\|ERVL-MaLR | 10235 | NM_008979 | 19260 | Mm.395 | NM_008979 | ENSMUSG00000027843 | **Ptpn22** | 70zpep\|PEP\|Ptpn8 | protein tyrosine phosphatase, non-receptor type 22 (lymphoid) | protein-coding |
| chr4_57287817_57288255 | chr4 | 57287818 | 57288255 | + | 0.739453 | intron (NM_011207, intron 1 of 25) | intron (NM_011207, intron 1 of 25) | 13801 | NM_011207 | 545622 | Mm.246552 | NM_011207 | ENSMUSG00000038764 | **Ptpn3** | 9530011I20Rik\|PTP-H1\|PTPCL | protein tyrosine phosphatase, non-receptor type 3 | protein-coding |
| chr4_57300723_57301042 | chr4 | 57300724 | 57301042 | + | 0.711751 | intron (NM_011207, intron 1 of 25) | intron (NM_011207, intron 1 of 25) | 954 | NM_011207 | 545622 | Mm.246552 | NM_011207 | ENSMUSG00000038764 | **Ptpn3** | 9530011I20Rik\|PTP-H1\|PTPCL | protein tyrosine phosphatase, non-receptor type 3 | protein-coding |
| chr4_57306118_57306613 | chr4 | 57306119 | 57306613 | + | 0.657016 | Intergenic | L2a\|LINE\|L2 | -4529 | NM_011207 | 545622 | Mm.246552 | NM_011207 | ENSMUSG00000038764 | **Ptpn3** | 9530011I20Rik\|PTP-H1\|PTPCL | protein tyrosine phosphatase, non-receptor type 3 | protein-coding |
| chr1_119835771_119836029 | chr1 | 119835772 | 119836029 | + | 1.04383 | intron (NM_019933, intron 1 of 26) | intron (NM_019933, intron 1 of 26) | 1171 | NM_019933 | 19258 | Mm.458796 | NM_019933 | | **Ptpn4** | PTPMEG\|Ptn4\|TEP\|TEP/mPTPMEG\|hPTP-MEG | protein tyrosine phosphatase, non-receptor type 4 | protein-coding |
| chr1_119806044_119806331 | chr1 | 119806045 | 119806331 | + | 0.6708 | intron (NM_019933, intron 1 of 26) | RSINE1\|SINE\|B4 | 30883 | NM_019933 | 19258 | Mm.458796 | NM_019933 | | **Ptpn4** | PTPMEG\|Ptn4\|TEP\|TEP/mPTPMEG\|hPTP-MEG | protein tyrosine phosphatase, non-receptor type 4 | protein-coding |
| chr7_47113374_47114026 | chr7 | 47113375 | 47114026 | + | 1.79075 | intron (NM_013643, intron 1 of 13) | intron (NM_013643, intron 1 of 13) | 18898 | NM_001163565 | 19259 | Mm.4654 | NM_013643 | ENSMUSG00000030854 | **Ptpn5** | Step | protein tyrosine phosphatase, non-receptor type 5 | protein-coding |

| | | | | | | | | | | | | | | | | | |
|---|---|---|---|---|---|---|---|---|---|---|---|---|---|---|---|---|---|
| chr7_47110574_47111062 | chr7 | 47110575 | 47111062 | + | 1.36429 | intron (NM_001163565, intron 2 of 13) | intron (NM_001163565, intron 2 of 13) | 21780 | NM_001163565 | 19259 | Mm.4654 | NM_013643 | ENSMUSG00000030854 | **Ptpn5** | Step | protein tyrosine phosphatase, non-receptor type 5 | protein-coding |
| chr7_47110452_47111452 | chr7 | 47110453 | 47111452 | + | 1.32365 | intron (NM_001163565, intron 2 of 13) | L2b\|LINE\|L2 | 21646 | NM_001163565 | 19259 | Mm.4654 | NM_013643 | ENSMUSG00000030854 | **Ptpn5** | Step | protein tyrosine phosphatase, non-receptor type 5 | protein-coding |
| chr7_47111113_47111448 | chr7 | 47111114 | 47111448 | + | 1.27406 | intron (NM_001163565, intron 2 of 13) | L3\|LINE\|CR1 | 21317 | NM_001163565 | 19259 | Mm.4654 | NM_013643 | ENSMUSG00000030854 | **Ptpn5** | Step | protein tyrosine phosphatase, non-receptor type 5 | protein-coding |
| chr1_135133659_135133960 | chr1 | 135133660 | 135133960 | + | 0.780814 | exon (NM_177081, exon 2 of 10) | exon (NM_177081, exon 2 of 10) | 1085 | NM_177081 | 320139 | Mm.258388 | NM_177081 | ENSMUSG00000031506 | **Ptpn7** | BPTP-4\|C920001D21Rik\|HEPTP\|LC-PTP\|LPTP | protein tyrosine phosphatase, non-receptor type 7 | protein-coding |
| chr9_57031254_57032164 | chr9 | 57031255 | 57032164 | + | 1.42933 | intron (NM_019651, intron 5 of 12) | intron (NM_019651, intron 5 of 12) | 36741 | NM_019651 | 56294 | Mm.325643 | NM_019651 | ENSMUSG00000032290 | **Ptpn9** | MEG2 | protein tyrosine phosphatase, non-receptor type 9 | protein-coding |
| chr1_138076459_138076971 | chr1 | 138076460 | 138076971 | + | 1.14626 | intron (NM_011210, intron 23 of 29) | intron (NM_011210, intron 23 of 29) | 98411 | NM_001111316 | 19264 | Mm.391573 | NM_011210 | ENSMUSG00000026395 | **Ptprc** | B220\|CD45R\|Cd45\|L-CA\|Ly-5\|Lyt-4\|T200\|loc | protein tyrosine phosphatase, receptor type, C | protein-coding |
| chr1_138195194_138195517 | chr1 | 138195195 | 138195517 | + | 1.05531 | Intergenic | Intergenic | -20051 | NM_001268286 | 19264 | Mm.391573 | NM_011210 | ENSMUSG00000026395 | **Ptprc** | B220\|CD45R\|Cd45\|L-CA\|Ly-5\|Lyt-4\|T200\|loc | protein tyrosine phosphatase, receptor type, C | protein-coding |
| chr1_138142225_138142649 | chr1 | 138142226 | 138142649 | + | 0.974759 | intron (NM_011210, intron 2 of 29) | intron (NM_011210, intron 2 of 29) | 32689 | NM_001111316 | 19264 | Mm.391573 | NM_011210 | ENSMUSG00000026395 | **Ptprc** | B220\|CD45R\|Cd45\|L-CA\|Ly-5\|Lyt-4\|T200\|loc | protein tyrosine phosphatase, receptor type, C | protein-coding |
| chr1_138176597_138176915 | chr1 | 138176598 | 138176915 | + | 0.842346 | Intergenic | Lx9\|LINE\|L1 | -1451 | NM_001268286 | 19264 | Mm.391573 | NM_011210 | ENSMUSG00000026395 | **Ptprc** | B220\|CD45R\|Cd45\|L-CA\|Ly-5\|Lyt-4\|T200\|loc | protein tyrosine phosphatase, receptor type, C | protein-coding |
| chr1_138144390_138144786 | chr1 | 138144391 | 138144786 | + | 0.807583 | intron (NM_011210, intron 2 of 29) | intron (NM_011210, intron 2 of 29) | 30538 | NM_001111316 | 19264 | Mm.391573 | NM_011210 | ENSMUSG00000026395 | **Ptprc** | B220\|CD45R\|Cd45\|L-CA\|Ly-5\|Lyt-4\|T200\|loc | protein tyrosine phosphatase, receptor type, C | protein-coding |
| chr1_138129536_138129915 | chr1 | 138129537 | 138129915 | + | 0.732192 | intron (NM_011210, intron 2 of 29) | intron (NM_011210, intron 2 of 29) | 45400 | NM_001111316 | 19264 | Mm.391573 | NM_011210 | ENSMUSG00000026395 | **Ptprc** | B220\|CD45R\|Cd45\|L-CA\|Ly-5\|Lyt-4\|T200\|loc | protein tyrosine phosphatase, receptor type, C | protein-coding |
| chr1_138190250_138190447 | chr1 | 138190251 | 138190447 | + | 0.723029 | Intergenic | Intergenic | -15044 | NM_001268286 | 19264 | Mm.391573 | NM_011210 | ENSMUSG00000026395 | **Ptprc** | B220\|CD45R\|Cd45\|L-CA\|Ly-5\|Lyt-4\|T200\|loc | protein tyrosine phosphatase, receptor type, C | protein-coding |
| chr1_138166132_138166793 | chr1 | 138166133 | 138166793 | + | 0.654751 | intron (NM_011210, intron 2 of 29) | intron (NM_011210, intron 2 of 29) | 8663 | NM_001111316 | 19264 | Mm.391573 | NM_011210 | ENSMUSG00000026395 | **Ptprc** | B220\|CD45R\|Cd45\|L-CA\|Ly-5\|Lyt-4\|T200\|loc | protein tyrosine phosphatase, receptor type, C | protein-coding |
| chr1_138190830_138191118 | chr1 | 138190831 | 138191118 | + | 0.632051 | Intergenic | Intergenic | -15669 | NM_001268286 | 19264 | Mm.391573 | NM_011210 | ENSMUSG00000026395 | **Ptprc** | B220\|CD45R\|Cd45\|L-CA\|Ly-5\|Lyt-4\|T200\|loc | protein tyrosine phosphatase, receptor type, C | protein-coding |
| chr2_90542571_90543540 | chr2 | 90542572 | 90543540 | + | 1.59597 | intron (NM_008982, intron 1 of 23) | L2c\|LINE\|L2 | 37591 | NM_008982 | 19271 | Mm.330393 | NM_008982 | ENSMUSG00000025314 | **Ptprj** | AI450271\|BET\|Byp\|CD148\|DEP-1\|PTPbeta2\|Ptpb2\|RPTPJ\|Scc-1\|Scc1 | protein tyrosine phosphatase, receptor type, J | protein-coding |
| chr2_90563107_90563852 | chr2 | 90563108 | 90563852 | + | 1.40867 | intron (NM_008982, intron 1 of 23) | intron (NM_008982, intron 1 of 23) | 17167 | NM_008982 | 19271 | Mm.330393 | NM_008982 | ENSMUSG00000025314 | **Ptprj** | AI450271\|BET\|Byp\|CD148\|DEP-1\|PTPbeta2\|Ptpb2\|RPTPJ\|Scc-1\|Scc1 | protein tyrosine phosphatase, receptor type, J | protein-coding |
| chr2_90490708_90491102 | chr2 | 90490709 | 90491102 | + | 0.940903 | intron (NM_008982, intron 1 of 23) | intron (NM_008982, intron 1 of 23) | -11731 | NM_001135657 | 19271 | Mm.330393 | NM_008982 | ENSMUSG00000025314 | **Ptprj** | AI450271\|BET\|Byp\|CD148\|DEP-1\|PTPbeta2\|Ptpb2\|RPTPJ\|Scc-1\|Scc1 | protein tyrosine phosphatase, receptor type, J | protein-coding |
| chr2_90490348_90490599 | chr2 | 90490349 | 90490599 | + | 0.905186 | intron (NM_008982, intron 1 of 23) | intron (NM_008982, intron 1 of 23) | -11300 | NM_001135657 | 19271 | Mm.330393 | NM_008982 | ENSMUSG00000025314 | **Ptprj** | AI450271\|BET\|Byp\|CD148\|DEP-1\|PTPbeta2\|Ptpb2\|RPTPJ\|Scc-1\|Scc1 | protein tyrosine phosphatase, receptor type, J | protein-coding |
| chr2_90582421_90582626 | chr2 | 90582422 | 90582626 | + | 0.609043 | Intergenic | Intergenic | -1877 | NM_008982 | 19271 | Mm.330393 | NM_008982 | ENSMUSG00000025314 | **Ptprj** | AI450271\|BET\|Byp\|CD148\|DEP-1\|PTPbeta2\|Ptpb2\|RPTPJ\|Scc-1\|Scc1 | protein tyrosine phosphatase, receptor type, J | protein-coding |
| chr10_28074398_28075500 | chr10 | 28074399 | 28075500 | + | 0.65899 | 5' UTR (NM_008983, exon 1 of 32) | 5' UTR (NM_008983, exon 1 of 32) | 129 | NM_008983 | 19272 | Mm.332303 | NM_008983 | ENSMUSG00000019889 | **Ptprk** | AI853699\|PTPk\|RPTPkappa | protein tyrosine phosphatase, receptor type, K | protein-coding |
| chr12_116622653_116623283 | chr12 | 116622654 | 116623283 | + | 1.03511 | intron (NM_011215, intron 2 of 22) | intron (NM_011215, intron 2 of 22) | 137248 | NM_011215 | 19276 | Mm.206054 | NM_011215 | ENSMUSG00000056553 | **Ptprn2** | 4930425H11Rik\|IA2beta\|PTP-NP\|Phol\|mKIAA0387\|phogrin | protein tyrosine phosphatase, receptor type, N polypeptide 2 | protein-coding |
| chr12_116585015_116585485 | chr12 | 116585016 | 116585485 | + | 0.934551 | intron (NM_011215, intron 2 of 22) | intron (NM_011215, intron 2 of 22) | 99530 | NM_011215 | 19276 | Mm.206054 | NM_011215 | ENSMUSG00000056553 | **Ptprn2** | 4930425H11Rik\|IA2beta\|PTP-NP\|Phol\|mKIAA0387\|phogrin | protein tyrosine phosphatase, receptor type, N polypeptide 2 | protein-coding |
| chr12_116629672_116630152 | chr12 | 116629673 | 116630152 | + | 0.889162 | intron (NM_011215, intron 2 of 22) | intron (NM_011215, intron 2 of 22) | 144192 | NM_011215 | 19276 | Mm.206054 | NM_011215 | ENSMUSG00000056553 | **Ptprn2** | 4930425H11Rik\|IA2beta\|PTP-NP\|Phol\|mKIAA0387\|phogrin | protein tyrosine phosphatase, receptor type, N polypeptide 2 | protein-coding |
| chr12_116485534_116486091 | chr12 | 116485535 | 116486091 | + | 0.778126 | promoter-TSS (NM_011215) | promoter-TSS (NM_011215) | 93 | NM_011215 | 19276 | Mm.206054 | NM_011215 | ENSMUSG00000056553 | **Ptprn2** | 4930425H11Rik\|IA2beta\|PTP-NP\|Phol\|mKIAA0387\|phogrin | protein tyrosine phosphatase, receptor type, N polypeptide 2 | protein-coding |
| chr12_116553397_116553739 | chr12 | 116553398 | 116553739 | + | 0.749285 | intron (NM_011215, intron 1 of 22) | intron (NM_011215, intron 1 of 22) | 67848 | NM_011215 | 19276 | Mm.206054 | NM_011215 | ENSMUSG00000056553 | **Ptprn2** | 4930425H11Rik\|IA2beta\|PTP-NP\|Phol\|mKIAA0387\|phogrin | protein tyrosine phosphatase, receptor type, N polypeptide 2 | protein-coding |
| chr12_116539344_116539797 | chr12 | 116539345 | 116539797 | + | 0.731752 | intron (NM_011215, intron 1 of 22) | ORR1B2\|LTR\|ERVL-MaLR | 53851 | NM_011215 | 19276 | Mm.206054 | NM_011215 | ENSMUSG00000056553 | **Ptprn2** | 4930425H11Rik\|IA2beta\|PTP-NP\|Phol\|mKIAA0387\|phogrin | protein tyrosine phosphatase, receptor type, N polypeptide 2 | protein-coding |
| chr12_116485521_116486425 | chr12 | 116485522 | 116486425 | + | 0.623622 | exon (NM_011215, exon 1 of 23) | exon (NM_011215, exon 1 of 23) | 253 | NM_011215 | 19276 | Mm.206054 | NM_011215 | ENSMUSG00000056553 | **Ptprn2** | 4930425H11Rik\|IA2beta\|PTP-NP\|Phol\|mKIAA0387\|phogrin | protein tyrosine phosphatase, receptor type, N polypeptide 2 | protein-coding |
| chr6_137316145_137316779 | chr6 | 137316146 | 137316779 | + | 0.753258 | intron (NM_001164401, intron 1 of 25) | intron (NM_001164401, intron 1 of 25) | 64163 | NM_011216 | 19277 | Mm.186361 | NM_011216 | ENSMUSG00000030223 | **Ptpro** | D28\|GLEPP1\|PTPBK\|PTPROt\|PTPU2\|PTPoc\|PTPphi\|Ptpn15\|R-PTP-O | protein tyrosine phosphatase, receptor type, O | protein-coding |
| chr6_137316382_137316752 | chr6 | 137316383 | 137316752 | + | 0.71549 | intron (NM_001164401, intron 1 of 25) | intron (NM_001164401, intron 1 of 25) | 64268 | NM_011216 | 19277 | Mm.186361 | NM_011216 | ENSMUSG00000030223 | **Ptpro** | D28\|GLEPP1\|PTPBK\|PTPROt\|PTPU2\|PTPoc\|PTPphi\|Ptpn15\|R-PTP-O | protein tyrosine phosphatase, receptor type, O | protein-coding |
| chr17_56478193_56478796 | chr17 | 56478194 | 56478796 | + | 1.26756 | Intergenic | Intergenic | -2015 | NM_011218 | 19280 | Mm.258771 | NM_011218 | ENSMUSG00000013236 | **Ptprs** | AL022616\|PTP\|PTP-NU3\|PTPNU-3\|PTPsigma\|Ptpt9\|R-PTP-S\|RPTPsigma | protein tyrosine phosphatase, receptor type, S | protein-coding |
| chr17_56472106_56472415 | chr17 | 56472107 | 56472415 | + | 0.9657 | intron (NM_001252455, intron 1 of 27) | intron (NM_001252455, intron 1 of 27) | 4219 | NM_001252455 | 19280 | Mm.258771 | NM_011218 | ENSMUSG00000013236 | **Ptprs** | AL022616\|PTP\|PTP-NU3\|PTPNU-3\|PTPsigma\|Ptpt9\|R-PTP-S\|RPTPsigma | protein tyrosine phosphatase, receptor type, S | protein-coding |
| chr17_56472162_56472503 | chr17 | 56472163 | 56472503 | + | 0.922895 | intron (NM_001252455, intron 1 of 27) | intron (NM_001252455, intron 1 of 27) | 4147 | NM_001252455 | 19280 | Mm.258771 | NM_011218 | ENSMUSG00000013236 | **Ptprs** | AL022616\|PTP\|PTP-NU3\|PTPNU-3\|PTPsigma\|Ptpt9\|R-PTP-S\|RPTPsigma | protein tyrosine phosphatase, receptor type, S | protein-coding |
| chr17_56472190_56472409 | chr17 | 56472191 | 56472409 | + | 0.858448 | intron (NM_001252455, intron 1 of 27) | intron (NM_001252455, intron 1 of 27) | 4180 | NM_001252455 | 19280 | Mm.258771 | NM_011218 | ENSMUSG00000013236 | **Ptprs** | AL022616\|PTP\|PTP-NU3\|PTPNU-3\|PTPsigma\|Ptpt9\|R-PTP-S\|RPTPsigma | protein tyrosine phosphatase, receptor type, S | protein-coding |
| chr11_86691447_86691944 | chr11 | 86691448 | 86691944 | + | 0.765936 | 3' UTR (NM_175004, exon 3 of 3) | 3' UTR (NM_175004, exon 3 of 3) | 7615 | NM_001098810 | 217057 | Mm.24868 | NM_175004 | ENSMUSG00000072582 | **Ptrh2** | A230072I16Rik\|Bit1\|CGI-147 | peptidyl-tRNA hydrolase 2 | protein-coding |
| chr9_50537240_50537609 | chr9 | 50537241 | 50537609 | + | 1.13171 | intron (NM_133217, intron 8 of 11) | L1M4\|LINE\|L1 | -8784 | NM_011220 | 19286 | Mm.35856 | NM_011220 | ENSMUSG00000032067 | **Pts** | PTPS | 6-pyruvoyl-tetrahydropterin synthase | protein-coding |
| chr9_50520502_50521103 | chr9 | 50520503 | 50521103 | + | 0.680867 | TTS (NM_011220) | TTS (NM_011220) | 7838 | NM_011220 | 19286 | Mm.35856 | NM_011220 | ENSMUSG00000032067 | **Pts** | PTPS | 6-pyruvoyl-tetrahydropterin synthase | protein-coding |
| chr9_50525617_50525961 | chr9 | 50525618 | 50525961 | + | 0.670711 | intron (NM_011220, intron 2 of 5) | intron (NM_011220, intron 2 of 5) | 2852 | NM_011220 | 19286 | Mm.35856 | NM_011220 | ENSMUSG00000032067 | **Pts** | PTPS | 6-pyruvoyl-tetrahydropterin synthase | protein-coding |
| chr12_8704420_8704983 | chr12 | 8704421 | 8704983 | + | 0.977716 | intron (NR_027670, intron 4 of 21) | intron (NR_027670, intron 4 of 21) | 30021 | NM_030723 | 80913 | Mm.341243 | NM_030723 | ENSMUSG00000020594 | **Pum2** | 5730503J23Rik\|Pumm2\|mKIAA0235 | pumilio RNA-binding family member 2 | protein-coding |
| chr12_8712611_8713009 | chr12 | 8712612 | 8713009 | + | 0.787538 | intron (NR_027670, intron 7 of 21) | intron (NR_027670, intron 7 of 21) | 38129 | NM_030723 | 80913 | Mm.341243 | NM_030723 | ENSMUSG00000020594 | **Pum2** | 5730503J23Rik\|Pumm2\|mKIAA0235 | pumilio RNA-binding family member 2 | protein-coding |
| chr12_8703992_8704290 | chr12 | 8703993 | 8704290 | + | 0.732004 | intron (NR_027670, intron 4 of 21) | intron (NR_027670, intron 4 of 21) | 29460 | NM_030723 | 80913 | Mm.341243 | NM_030723 | ENSMUSG00000020594 | **Pum2** | 5730503J23Rik\|Pumm2\|mKIAA0235 | pumilio RNA-binding family member 2 | protein-coding |
| chr12_8713109_8713509 | chr12 | 8713110 | 8713509 | + | 0.696586 | intron (NR_027670, intron 7 of 21) | intron (NR_027670, intron 7 of 21) | 38628 | NM_030723 | 80913 | Mm.341243 | NM_030723 | ENSMUSG00000020594 | **Pum2** | 5730503J23Rik\|Pumm2\|mKIAA0235 | pumilio RNA-binding family member 2 | protein-coding |
| chr11_23687932_23688304 | chr11 | 23687933 | 23688304 | + | 0.710414 | intron (NM_001033654, intron 4 of 17) | PB1D11\|SINE\|Alu | 22142 | NM_028304 | 74467 | Mm.29536 | NM_028304 | ENSMUSG00000020280 | **Pus10** | 2810013G11Rik\|4933435A13Rik\|AU014648\|C77560\|Ccdc139 | pseudouridylate synthase 10 | protein-coding |
| chr15_94475965_94476375 | chr15 | 94475966 | 94476375 | + | 1.18063 | Intergenic | Intergenic | 67337 | NM_172437 | 78895 | Mm.34344 | NM_172437 | ENSMUSG00000033356 | **Pus7l** | 3000003F02Rik | pseudouridylate synthase 7 homolog (S. cerevisiae)-like | protein-coding |

| | | | | | | | | | | | | | | | | | |
|---|---|---|---|---|---|---|---|---|---|---|---|---|---|---|---|---|---|
| chr9_43548201_43548627 | chr9 | 43548202 | 43548627 | + | 0.749455 | Intergenic | Intergenic | -196162 | NM_021424 | 58235 | Mm.335096 | NM_021424 | ENSMUSG00000032012 | **Pvrl1** | AI835281\|AW549174\|Cd111\|HIgR\|HveC\|PRR\|PRR1\|nectin-1 | poliovirus receptor-related 1 | protein-coding |
| chr15_62095184_62095576 | chr15 | 62095185 | 62095576 | + | 0.944952 | intron (NR_003368, intron 1 of 8) | intron (NR_003368, intron 1 of 8) | 57393 | NR_003368 | 19296 | Mm.4608 | NM_011222 | ENSMUSG00000097039 | **Pvt1** | Gt(pU21)84Imeg\|Mis-1\|Mlvi-1\|Pvt-1 | plasmacytoma variant translocation 1 | ncRNA |
| chr15_62079178_62079919 | chr15 | 62079179 | 62079919 | + | 0.868829 | intron (NR_003368, intron 1 of 8) | intron (NR_003368, intron 1 of 8) | 41562 | NR_003368 | 19296 | Mm.4608 | NM_011222 | ENSMUSG00000097039 | **Pvt1** | Gt(pU21)84Imeg\|Mis-1\|Mlvi-1\|Pvt-1 | plasmacytoma variant translocation 1 | ncRNA |
| chr15_62043611_62044321 | chr15 | 62043612 | 62044321 | + | 0.824101 | intron (NR_003368, intron 1 of 8) | intron (NR_003368, intron 1 of 8) | 5979 | NR_003368 | 19296 | Mm.4608 | NM_011222 | ENSMUSG00000097039 | **Pvt1** | Gt(pU21)84Imeg\|Mis-1\|Mlvi-1\|Pvt-1 | plasmacytoma variant translocation 1 | ncRNA |
| chr5_110283973_110284244 | chr5 | 110283974 | 110284244 | + | 0.990593 | intron (NM_008993, intron 1 of 4) | intron (NM_008993, intron 1 of 4) | 2059 | NM_008993 | 19301 | Mm.21853 | NM_008993 | ENSMUSG00000029499 | **Pxmp2** | 22kDa\|PMP22 | peroxisomal membrane protein 2 | protein-coding |
| chr5_115530391_115530932 | chr5 | 115530392 | 115530932 | + | 0.702378 | intron (NM_011223, intron 1 of 10) | intron (NM_011223, intron 1 of 10) | 23986 | NM_133915 | 19303 | Mm.18714 | NM_011223 | ENSMUSG00000029528 | **Pxn** | AW108311\|AW123232\|Pax | paxillin | protein-coding |
| chr2_150840065_150840744 | chr2 | 150840066 | 150840744 | + | 0.942486 | intron (NM_024465, intron 6 of 12) | intron (NM_024465, intron 6 of 12) | 53609 | NM_153781 | 110078 | Mm.222584 | NM_153781 | ENSMUSG00000033059 | **Pygb** | - | brain glycogen phosphorylase | protein-coding |
| chr1_173624060_173624650 | chr1 | 173624061 | 173624650 | + | 1.19307 | Intergenic | Lx2\|LINE\|L1 | -6504 | NM_175026 | 236312 | Mm.447 | NM_175026 | ENSMUSG00000043263 | **Pyhin1** | 4930422C14\|AI447904\|Ifi209\|Ifix | pyrin and HIN domain family, member 1 | protein-coding |
| chr1_173646306_173646933 | chr1 | 173646307 | 173646933 | + | 0.954337 | 3' UTR (NM_175026, exon 7 of 7) | 3' UTR (NM_175026, exon 7 of 7) | 15761 | NM_175026 | 236312 | Mm.447 | NM_175026 | ENSMUSG00000043263 | **Pyhin1** | 4930422C14\|AI447904\|Ifi209\|Ifix | pyrin and HIN domain family, member 1 | protein-coding |
| chr1_173645402_173645760 | chr1 | 173645403 | 173645760 | + | 0.712018 | intron (NM_175026, intron 6 of 6) | intron (NM_175026, intron 6 of 6) | 14722 | NM_175026 | 236312 | Mm.447 | NM_175026 | ENSMUSG00000043263 | **Pyhin1** | 4930422C14\|AI447904\|Ifi209\|Ifix | pyrin and HIN domain family, member 1 | protein-coding |
| chr5_45295036_45295778 | chr5 | 45295037 | 45295778 | + | 0.67024 | Intergenic | Intergenic | 154822 | NM_024236 | 110391 | Mm.30204 | NM_024236 | ENSMUSG00000015806 | **Qdpr** | 2610008L04Rik\|D5Ertd371e\|Dhpr\|PKU2 | quinoid dihydropteridine reductase | protein-coding |
| chr7_127112163_127112449 | chr7 | 127112164 | 127112449 | + | 0.825482 | intron (NM_133686, intron 1 of 3) | intron (NM_133686, intron 1 of 3) | 9723 | NM_133686 | 67375 | Mm.26928 | NM_133686 | | **Qprt** | 2410027J01Rik\|AI647766\|QPRTase | quinolinate phosphoribosyltransferase | protein-coding |
| chr9_108521702_108522165 | chr9 | 108521703 | 108522165 | + | 1.23656 | intron (NM_001114119, intron 2 of 10) | intron (NM_001114119, intron 2 of 10) | 4847 | NM_175143 | 69232 | Mm.335449 | NM_175143 | ENSMUSG00000006673 | **Qrich1** | 2610028H07Rik | glutamine-rich 1 | protein-coding |
| chr2_104813184_104813536 | chr2 | 104813185 | 104813536 | + | 1.25823 | intron (NM_001123327, intron 1 of 11) | intron (NM_001123327, intron 1 of 11) | 3336 | NM_001123327 | 99003 | Mm.274314 | NM_001123327 | ENSMUSG00000074994 | **Qser1** | 4732486I23Rik\|AI502974 | glutamine and serine rich 1 | protein-coding |
| chr2_104817547_104817814 | chr2 | 104817548 | 104817814 | + | 1.12138 | promoter-TSS (NM_001123327) | promoter-TSS (NM_001123327) | -985 | NM_001123327 | 99003 | Mm.274314 | NM_001123327 | ENSMUSG00000074994 | **Qser1** | 4732486I23Rik\|AI502974 | glutamine and serine rich 1 | protein-coding |
| chr1_155836085_155836411 | chr1 | 155836086 | 155836411 | + | 0.624684 | Intergenic | L1_Rod\|LINE\|L1 | -23349 | NM_001024945 | 104009 | Mm.27035 | NM_023268 | ENSMUSG00000033684 | **Qsox1** | 1300003H02Rik\|QSOX\|Qscn6\|SOx | quiescin Q6 sulfhydryl oxidase 1 | protein-coding |
| chr2_26231276_26231989 | chr2 | 26231277 | 26231989 | + | 0.887599 | intron (NM_153559, intron 1 of 12) | intron (NM_153559, intron 1 of 12) | 5787 | NM_153559 | 227638 | Mm.116769 | NM_153559 | ENSMUSG00000036327 | **Qsox2** | BC030934\|Qscn6l1\|SOXN | quiescin Q6 sulfhydryl oxidase 2 | protein-coding |
| chr1_128124374_128124855 | chr1 | 128124375 | 128124855 | + | 1.51005 | intron (NM_181750, intron 1 of 26) | intron (NM_181750, intron 1 of 26) | 21309 | NM_181750 | 226412 | Mm.221041 | NM_181750 | ENSMUSG00000056211 | **R3hdm1** | R3hdm | R3H domain containing 1 | protein-coding |
| chr12_3282964_3283339 | chr12 | 3282965 | 3283339 | + | 1.30508 | intron (NM_016676, intron 1 of 5) | Charlie1a\|DNA\|hAT-Charlie | 26817 | NM_016676 | 19325 | Mm.378993 | NM_016676 | ENSMUSG00000020671 | **Rab10** | AW107754 | RAB10, member RAS oncogene family | protein-coding |
| chr12_3283960_3284260 | chr12 | 3283961 | 3284260 | + | 0.758512 | intron (NM_016676, intron 1 of 5) | B1F2\|SINE\|Alu | 25859 | NM_016676 | 19325 | Mm.378993 | NM_016676 | ENSMUSG00000020671 | **Rab10** | AW107754 | RAB10, member RAS oncogene family | protein-coding |
| chr12_3283923_3284467 | chr12 | 3283924 | 3284467 | + | 0.722655 | intron (NM_016676, intron 1 of 5) | B1F2\|SINE\|Alu | 25774 | NM_016676 | 19325 | Mm.378993 | NM_016676 | ENSMUSG00000020671 | **Rab10** | AW107754 | RAB10, member RAS oncogene family | protein-coding |
| chr12_3283921_3284755 | chr12 | 3283922 | 3284755 | + | 0.651972 | intron (NM_016676, intron 1 of 5) | ID4_\|SINE\|ID | 25631 | NM_016676 | 19325 | Mm.378993 | NM_016676 | ENSMUSG00000020671 | **Rab10** | AW107754 | RAB10, member RAS oncogene family | protein-coding |
| chr9_64564118_64564897 | chr9 | 64564119 | 64564897 | + | 1.5739 | intron (NM_001134399, intron 5 of 24) | intron (NM_001134399, intron 5 of 24) | 173248 | NM_017382 | 53869 | Mm.1387 | NM_017382 | ENSMUSG00000004771 | **Rab11a** | - | RAB11A, member RAS oncogene family | protein-coding |
| chr9_64568258_64568582 | chr9 | 64568259 | 64568582 | + | 1.11973 | intron (NM_001134399, intron 5 of 24) | intron (NM_001134399, intron 5 of 24) | 169336 | NM_017382 | 53869 | Mm.1387 | NM_017382 | ENSMUSG00000004771 | **Rab11a** | - | RAB11A, member RAS oncogene family | protein-coding |
| chr9_64716748_64717184 | chr9 | 64716749 | 64717184 | + | 0.792921 | exon (NM_017382, exon 5 of 5) | exon (NM_017382, exon 5 of 5) | 20790 | NM_017382 | 53869 | Mm.1387 | NM_017382 | ENSMUSG00000004771 | **Rab11a** | - | RAB11A, member RAS oncogene family | protein-coding |
| chr8_27168190_27168564 | chr8 | 27168191 | 27168564 | + | 1.5904 | intron (NM_029423, intron 1 of 4) | intron (NM_029423, intron 1 of 4) | 6269 | NM_029423 | 75767 | Mm.205335 | NM_029423 | ENSMUSG00000031488 | **Rab11fip1** | 2010200K21Rik\|4833414G05Rik\|Rcp | RAB11 family interacting protein 1 (class I) | protein-coding |
| chr8_27172535_27172777 | chr8 | 27172536 | 27172777 | + | 1.06149 | intron (NM_029423, intron 1 of 4) | intron (NM_029423, intron 1 of 4) | 1990 | NM_029423 | 75767 | Mm.205335 | NM_029423 | ENSMUSG00000031488 | **Rab11fip1** | 2010200K21Rik\|4833414G05Rik\|Rcp | RAB11 family interacting protein 1 (class I) | protein-coding |
| chr8_27172031_27172822 | chr8 | 27172032 | 27172822 | + | 1.05388 | intron (NM_029423, intron 1 of 4) | intron (NM_029423, intron 1 of 4) | 2219 | NM_029423 | 75767 | Mm.205335 | NM_029423 | ENSMUSG00000031488 | **Rab11fip1** | 2010200K21Rik\|4833414G05Rik\|Rcp | RAB11 family interacting protein 1 (class I) | protein-coding |
| chr8_27172044_27172479 | chr8 | 27172045 | 27172479 | + | 0.966781 | intron (NM_029423, intron 1 of 4) | intron (NM_029423, intron 1 of 4) | 2384 | NM_029423 | 75767 | Mm.205335 | NM_029423 | ENSMUSG00000031488 | **Rab11fip1** | 2010200K21Rik\|4833414G05Rik\|Rcp | RAB11 family interacting protein 1 (class I) | protein-coding |
| chr19_59905636_59906074 | chr19 | 59905637 | 59906074 | + | 1.0407 | 3' UTR (NM_001164367, exon 5 of 5) | 3' UTR (NM_001164367, exon 5 of 5) | 37509 | NM_001033172 | 74998 | Mm.24167 | NM_001033172 | ENSMUSG00000040022 | **Rab11fip2** | 4930470G04Rik\|A830046J09Rik\|AW558126\|Nrip11 | RAB11 family interacting protein 2 (class I) | protein-coding |
| chr11_79676476_79676847 | chr11 | 79676477 | 79676847 | + | 0.950024 | intron (NM_175543, intron 3 of 14) | intron (NM_175543, intron 3 of 14) | -1576 | NR_045898 | 72218 | Mm.443271 | NR_045898 | ENSMUSG00000086569 | **Rab11fip4os2** | 1700020F10Rik\|4930542H20Rik\|Rab11fib4os2 | RAB11 family interacting protein 4 (class II), opposite strand 2 | ncRNA |
| chr11_79676510_79677086 | chr11 | 79676511 | 79677086 | + | 0.876491 | intron (NM_175543, intron 3 of 14) | intron (NM_175543, intron 3 of 14) | -1712 | NR_045898 | 72218 | Mm.443271 | NR_045898 | ENSMUSG00000086569 | **Rab11fip4os2** | 1700020F10Rik\|4930542H20Rik\|Rab11fib4os2 | RAB11 family interacting protein 4 (class II), opposite strand 2 | ncRNA |
| chr11_79678284_79679184 | chr11 | 79678285 | 79679184 | + | 0.759989 | intron (NM_175543, intron 3 of 14) | intron (NM_175543, intron 3 of 14) | -3648 | NR_045898 | 72218 | Mm.443271 | NR_045898 | ENSMUSG00000086569 | **Rab11fip4os2** | 1700020F10Rik\|4930542H20Rik\|Rab11fib4os2 | RAB11 family interacting protein 4 (class II), opposite strand 2 | ncRNA |
| chr11_79654295_79654731 | chr11 | 79654296 | 79654731 | + | 0.717669 | intron (NM_175543, intron 3 of 14) | intron (NM_175543, intron 3 of 14) | 20573 | NR_045898 | 72218 | Mm.443271 | NR_045898 | ENSMUSG00000086569 | **Rab11fip4os2** | 1700020F10Rik\|4930542H20Rik\|Rab11fib4os2 | RAB11 family interacting protein 4 (class II), opposite strand 2 | ncRNA |
| chr2_35203839_35204172 | chr2 | 35203840 | 35204172 | + | 1.13649 | Intergenic | ORR1C2\|LTR\|ERVL-MaLR | -2886 | NM_026697 | 68365 | Mm.198264 | NM_026697 | ENSMUSG00000026878 | **Rab14** | 0610030G24Rik\|2810475J17Rik\|A830021G03Rik\|AI314285\|AI649155\|D030017L14Rik | RAB14, member RAS oncogene family | protein-coding |
| chr8_11452347_11453363 | chr8 | 11452348 | 11453363 | + | 0.917272 | Intergenic | Intergenic | 25644 | NM_011227 | 19332 | Mm.37632 | NM_011227 | | **Rab20** | AA536966\|D8Ertd350e | RAB20, member RAS oncogene family | protein-coding |
| chr8_11476589_11477255 | chr8 | 11476590 | 11477255 | + | 0.664453 | intron (NM_011227, intron 1 of 1) | intron (NM_011227, intron 1 of 1) | 1577 | NM_011227 | 19332 | Mm.37632 | NM_011227 | | **Rab20** | AA536966\|D8Ertd350e | RAB20, member RAS oncogene family | protein-coding |
| chr10_115332115_115333418 | chr10 | 115332116 | 115333418 | + | 1.3158 | Intergenic | ID_B1\|SINE\|B4 | -17176 | NM_024454 | 216344 | Mm.21936 | NM_024454 | ENSMUSG00000020132 | **Rab21** | 9630024B22 | RAB21, member RAS oncogene family | protein-coding |
| chr10_115332132_115332824 | chr10 | 115332133 | 115332824 | + | 1.27039 | Intergenic | Intergenic | -16887 | NM_024454 | 216344 | Mm.21936 | NM_024454 | ENSMUSG00000020132 | **Rab21** | 9630024B22 | RAB21, member RAS oncogene family | protein-coding |
| chr10_115332907_115333408 | chr10 | 115332908 | 115333408 | + | 1.11158 | Intergenic | L1MCa\|LINE\|L1 | -17567 | NM_024454 | 216344 | Mm.21936 | NM_024454 | ENSMUSG00000020132 | **Rab21** | 9630024B22 | RAB21, member RAS oncogene family | protein-coding |
| chr10_115328455_115328887 | chr10 | 115328456 | 115328887 | + | 0.72495 | Intergenic | MER70B\|LTR\|ERVL | -13080 | NM_024454 | 216344 | Mm.21936 | NM_024454 | ENSMUSG00000020132 | **Rab21** | 9630024B22 | RAB21, member RAS oncogene family | protein-coding |
| chr2_173662604_173662912 | chr2 | 173662605 | 173662912 | + | 1.78556 | intron (NM_024436, intron 2 of 6) | intron (NM_024436, intron 2 of 6) | 2913 | NM_024436 | 19334 | Mm.275304 | NM_024436 | | **Rab22a** | 3732413A17Rik\|AI662177\|AW319644\|AW550514\|E130120E14Rik\|Rab22 | RAB22A, member RAS oncogene family | protein-coding |
| chr2_173680012_173680626 | chr2 | 173680013 | 173680626 | + | 1.07147 | intron (NM_024436, intron 2 of 6) | intron (NM_024436, intron 2 of 6) | 20474 | NM_024436 | 19334 | Mm.275304 | NM_024436 | | **Rab22a** | 3732413A17Rik\|AI662177\|AW319644\|AW550514\|E130120E14Rik\|Rab22 | RAB22A, member RAS oncogene family | protein-coding |
| chr1_33710995_33711630 | chr1 | 33710996 | 33711630 | + | 1.25333 | Intergenic | Intergenic | -8583 | NM_008999 | 19335 | Mm.86744 | NM_008999 | ENSMUSG00000004768 | **Rab23** | AW545388\|opb\|opb2 | RAB23, member RAS oncogene family | protein-coding |
| chr1_33710469_33711652 | chr1 | 33710470 | 33711652 | + | 1.24317 | Intergenic | Intergenic | -8835 | NM_008999 | 19335 | Mm.86744 | NM_008999 | ENSMUSG00000004768 | **Rab23** | AW545388\|opb\|opb2 | RAB23, member RAS oncogene family | protein-coding |
| chr1_33710485_33710923 | chr1 | 33710486 | 33710923 | + | 0.910373 | Intergenic | Intergenic | -9192 | NM_008999 | 19335 | Mm.86744 | NM_008999 | ENSMUSG00000004768 | **Rab23** | AW545388\|opb\|opb2 | RAB23, member RAS oncogene family | protein-coding |
| chr9_73060291_73060752 | chr9 | 73060292 | 73060752 | + | 1.21629 | intron (NM_023635, intron 1 of 5) | HAL1\|LINE\|L1 | -3626 | NM_001301232 | 11891 | Mm.34867 | NM_023635 | ENSMUSG00000032202 | **Rab27a** | 2210402C08Rik\|2410003M20Rik\|4933437C11Rik\|ash | RAB27A, member RAS oncogene family | protein-coding |
| chr9_73053388_73053842 | chr9 | 73053389 | 73053842 | + | 1.12804 | intron (NM_023635, intron 1 of 5) | intron (NM_023635, intron 1 of 5) | 8805 | NM_001301230 | 11891 | Mm.34867 | NM_023635 | ENSMUSG00000032202 | **Rab27a** | 2210402C08Rik\|2410003M20Rik\|4933437C11Rik\|ash | RAB27A, member RAS oncogene family | protein-coding |
| chr9_73079793_73080423 | chr9 | 73079794 | 73080423 | + | 1.10476 | intron (NM_001301230, intron 3 of 6) | intron (NM_001301230, intron 3 of 6) | 15960 | NM_001301232 | 11891 | Mm.34867 | NM_023635 | ENSMUSG00000032202 | **Rab27a** | 2210402C08Rik\|2410003M20Rik\|4933437C11Rik\|ash | RAB27A, member RAS oncogene family | protein-coding |
| chr9_73047961_73048258 | chr9 | 73047962 | 73048258 | + | 1.03787 | intron (NM_023635, intron 1 of 5) | intron (NM_023635, intron 1 of 5) | 3300 | NM_001301230 | 11891 | Mm.34867 | NM_023635 | ENSMUSG00000032202 | **Rab27a** | 2210402C08Rik\|2410003M20Rik\|4933437C11Rik\|ash | RAB27A, member RAS oncogene family | protein-coding |

| | | | | | | | | | | | | | | | | | |
|---|---|---|---|---|---|---|---|---|---|---|---|---|---|---|---|---|---|
| chr9_73071051_73071635 | chr9 | 73071052 | 73071635 | + | 0.649389 | intron (NM_001301232, intron 1 of 5) | intron (NM_001301232, intron 1 of 5) | 7195 | NM_001301232 | 11891 | Mm.34867 | NM_023635 | ENSMUSG00000032202 | **Rab27a** | 2210402C08Rik\|2410003M20Rik\|4933437C11Rik\|ash | RAB27A, member RAS oncogene family | protein-coding |
| chr7_92758046_92758523 | chr7 | 92758047 | 92758523 | + | 0.663317 | intron (NM_029494, intron 1 of 4) | MIRb\|SINE\|MIR | 16571 | NM_029494 | 75985 | Mm.389966 | NM_029494 | ENSMUSG00000030643 | **Rab30** | 5033421K01Rik\|AI323892\|Rsb30 | RAB30, member RAS oncogene family | protein-coding |
| chr17_65741555_65741849 | chr17 | 65741556 | 65741849 | + | 0.876549 | intron (NM_133685, intron 1 of 6) | MER21B\|LTR\|ERVL | 31050 | NM_133685 | 106572 | Mm.29274 | NM_133685 | ENSMUSG00000056515 | **Rab31** | 1700093E07Rik\|AI415285\|Rab22B | RAB31, member RAS oncogene family | protein-coding |
| chr17_65740924_65741857 | chr17 | 65740925 | 65741857 | + | 0.865478 | intron (NM_133685, intron 1 of 6) | RMER6A\|LTR\|ERVK | 31361 | NM_133685 | 106572 | Mm.29274 | NM_133685 | ENSMUSG00000056515 | **Rab31** | 1700093E07Rik\|AI415285\|Rab22B | RAB31, member RAS oncogene family | protein-coding |
| chr3_51475783_51476165 | chr3 | 51475784 | 51476165 | + | 0.695166 | TTS (NM_053089) | TTS (NM_053089) | -7992 | NM_016858 | 19338 | Mm.1664 | NM_016858 | ENSMUSG00000027739 | **Rab33b** | - | RAB33B, member RAS oncogene family | protein-coding |
| chr11_115104897_115105553 | chr11 | 115104898 | 115105553 | + | 0.774239 | intron (NM_001163753, intron 1 of 8) | intron (NM_001163753, intron 1 of 8) | 13794 | NM_001163753 | 58222 | Mm.143789 | NM_021411 | ENSMUSG00000020732 | **Rab37** | B230331O03Rik\|B230354I04Rik | RAB37, member RAS oncogene family | protein-coding |
| chr7_88459636_88460258 | chr7 | 88459637 | 88460258 | + | 1.27734 | intron (NM_028238, intron 2 of 2) | MIRb\|SINE\|MIR | 29674 | NM_028238 | 72433 | Mm.276669 | NM_028238 | ENSMUSG00000030559 | **Rab38** | 2310011F14Rik\|AU043391\|cht | RAB38, member RAS oncogene family | protein-coding |
| chr9_53695712_53696357 | chr9 | 53695713 | 53696357 | + | 0.815263 | intron (NM_175562, intron 1 of 1) | PB1D7\|SINE\|Alu | 10197 | NM_175562 | 270160 | Mm.290285 | NM_175562 | ENSMUSG00000055069 | **Rab39** | C230094F14Rik\|Rab39a | RAB39, member RAS oncogene family | protein-coding |
| chr1_185205122_185205403 | chr1 | 185205123 | 185205403 | + | 0.939732 | intron (NM_001163754, intron 1 of 34) | intron (NM_001163754, intron 1 of 34) | 1095 | NM_001163754 | 98732 | Mm.275841 | NM_172649 | ENSMUSG00000039318 | **Rab3gap2** | 1110059F07Rik\|2010002H18Rik\|5830469C09\|AI851069\|AW743433\|RAB3-GAP150\|mKIAA0839 | RAB3 GTPase activating protein subunit 2 | protein-coding |
| chr10_116928885_116929170 | chr10 | 116928886 | 116929170 | + | 0.71032 | intron (NM_001003950, intron 3 of 9) | intron (NM_001003950, intron 3 of 9) | 21352 | NM_001003950 | 216363 | Mm.336394 | NM_001003950 | ENSMUSG00000064181 | **Rab3ip** | B230311A06\|Gtpat12\|Rabin3 | RAB3A interacting protein | protein-coding |
| chr4_132299936_132300239 | chr4 | 132299937 | 132300239 | + | 0.878666 | Intergenic | Intergenic | 3268 | NM_001081651 | 242681 | Mm.331927 | NM_001081651 | ENSMUSG00000089687 | **Rab42** | 9530096D07Rik\|Rab42-ps | RAB42, member RAS oncogene family | protein-coding |
| chr4_132299834_132300285 | chr4 | 132299835 | 132300285 | + | 0.825033 | Intergenic | Intergenic | 3296 | NM_001081651 | 242681 | Mm.331927 | NM_001081651 | ENSMUSG00000089687 | **Rab42** | 9530096D07Rik\|Rab42-ps | RAB42, member RAS oncogene family | protein-coding |
| chr17_29169055_29169297 | chr17 | 29169056 | 29169297 | + | 1.01893 | intron (NM_153166, intron 13 of 20) | intron (NM_153166, intron 13 of 20) | 34120 | NM_001002786 | 442827 | Mm.440341 | NM_001002786 | ENSMUSG00000064147 | **Rab44** | 9830134C10Rik | RAB44, member RAS oncogene family | protein-coding |
| chr6_88056852_88057379 | chr6 | 88056853 | 88057379 | + | 0.726357 | Intergenic | Intergenic | -11846 | NM_009005 | 19349 | Mm.333233 | NM_009005 | ENSMUSG00000079477 | **Rab7** | Rab7a | RAB7, member RAS oncogene family | protein-coding |
| chr9_66918649_66918925 | chr9 | 66918650 | 66918925 | + | 0.945949 | intron (NM_173413, intron 1 of 7) | intron (NM_173413, intron 1 of 7) | 918 | NM_173413 | 235442 | Mm.260376 | NM_173413 | ENSMUSG00000036943 | **Rab8b** | 5930437D16\|D330025I23Rik | RAB8B, member RAS oncogene family | protein-coding |
| chr9_66917224_66917684 | chr9 | 66917225 | 66917684 | + | 0.941248 | intron (NM_173413, intron 1 of 7) | intron (NM_173413, intron 1 of 7) | 2251 | NM_173413 | 235442 | Mm.260376 | NM_173413 | ENSMUSG00000036943 | **Rab8b** | 5930437D16\|D330025I23Rik | RAB8B, member RAS oncogene family | protein-coding |
| chr11_70866550_70866906 | chr11 | 70866551 | 70866906 | + | 0.668578 | intron (NM_001291143, intron 2 of 18) | Tigger7\|DNA\|TcMar-Tigger | 21967 | NM_001291142 | 54189 | Mm.7087 | NM_019400 | ENSMUSG00000020817 | **Rabep1** | Rab5ep\|rabaptin-5\|rabaptin-5alpha | rabaptin, RAB GTPase binding effector protein 1 | protein-coding |
| chr1_160765564_160766288 | chr1 | 160765565 | 160766288 | + | 1.31874 | intron (NM_013862, intron 1 of 20) | intron (NM_013862, intron 1 of 20) | 27012 | NM_013862 | 29809 | Mm.25833 | NM_013862 | ENSMUSG00000026721 | **Rabgap1l** | 5830411O09Rik\|8430421H08Rik\|9630005B12Rik\|AW049894\|HHL\|Hh1\|mKIAA0471 | RAB GTPase activating protein 1-like | protein-coding |
| chr1_160799924_160800395 | chr1 | 160799925 | 160800395 | + | 1.10323 | Intergenic | L1MA4A\|LINE\|L1 | -7222 | NM_013862 | 29809 | Mm.25833 | NM_013862 | ENSMUSG00000026721 | **Rabgap1l** | 5830411O09Rik\|8430421H08Rik\|9630005B12Rik\|AW049894\|HHL\|Hh1\|mKIAA0471 | RAB GTPase activating protein 1-like | protein-coding |
| chr1_160791323_160791650 | chr1 | 160791324 | 160791650 | + | 1.09828 | intron (NM_013862, intron 1 of 20) | intron (NM_013862, intron 1 of 20) | 1451 | NM_013862 | 29809 | Mm.25833 | NM_013862 | ENSMUSG00000026721 | **Rabgap1l** | 5830411O09Rik\|8430421H08Rik\|9630005B12Rik\|AW049894\|HHL\|Hh1\|mKIAA0471 | RAB GTPase activating protein 1-like | protein-coding |
| chr1_160805257_160805675 | chr1 | 160805258 | 160805675 | + | 1.06156 | Intergenic | Intergenic | -12528 | NM_013862 | 29809 | Mm.25833 | NM_013862 | ENSMUSG00000026721 | **Rabgap1l** | 5830411O09Rik\|8430421H08Rik\|9630005B12Rik\|AW049894\|HHL\|Hh1\|mKIAA0471 | RAB GTPase activating protein 1-like | protein-coding |
| chr1_160793831_160794090 | chr1 | 160793832 | 160794090 | + | 0.94417 | Intergenic | Intergenic | -1023 | NM_013862 | 29809 | Mm.25833 | NM_013862 | ENSMUSG00000026721 | **Rabgap1l** | 5830411O09Rik\|8430421H08Rik\|9630005B12Rik\|AW049894\|HHL\|Hh1\|mKIAA0471 | RAB GTPase activating protein 1-like | protein-coding |
| chr1_160286420_160286803 | chr1 | 160286421 | 160286803 | + | 0.861012 | intron (NM_013862, intron 19 of 20) | intron (NM_013862, intron 19 of 20) | 64959 | NM_001038621 | 29809 | Mm.25833 | NM_013862 | ENSMUSG00000026721 | **Rabgap1l** | 5830411O09Rik\|8430421H08Rik\|9630005B12Rik\|AW049894\|HHL\|Hh1\|mKIAA0471 | RAB GTPase activating protein 1-like | protein-coding |
| chr1_160805971_160806467 | chr1 | 160805972 | 160806467 | + | 0.832178 | Intergenic | B4\|SINE\|B4 | -13281 | NM_013862 | 29809 | Mm.25833 | NM_013862 | ENSMUSG00000026721 | **Rabgap1l** | 5830411O09Rik\|8430421H08Rik\|9630005B12Rik\|AW049894\|HHL\|Hh1\|mKIAA0471 | RAB GTPase activating protein 1-like | protein-coding |
| chr1_160794463_160794625 | chr1 | 160794464 | 160794625 | + | 0.786363 | Intergenic | Intergenic | -1606 | NM_013862 | 29809 | Mm.25833 | NM_013862 | ENSMUSG00000026721 | **Rabgap1l** | 5830411O09Rik\|8430421H08Rik\|9630005B12Rik\|AW049894\|HHL\|Hh1\|mKIAA0471 | RAB GTPase activating protein 1-like | protein-coding |
| chr1_160286409_160286998 | chr1 | 160286410 | 160286998 | + | 0.782192 | intron (NM_013862, intron 19 of 20) | intron (NM_013862, intron 19 of 20) | 64867 | NM_001038621 | 29809 | Mm.25833 | NM_013862 | ENSMUSG00000026721 | **Rabgap1l** | 5830411O09Rik\|8430421H08Rik\|9630005B12Rik\|AW049894\|HHL\|Hh1\|mKIAA0471 | RAB GTPase activating protein 1-like | protein-coding |
| chr1_160416699_160417113 | chr1 | 160416700 | 160417113 | + | 0.767269 | intron (NM_013862, intron 17 of 20) | RSINE1\|SINE\|B4 | -65335 | NM_001038621 | 29809 | Mm.25833 | NM_013862 | ENSMUSG00000026721 | **Rabgap1l** | 5830411O09Rik\|8430421H08Rik\|9630005B12Rik\|AW049894\|HHL\|Hh1\|mKIAA0471 | RAB GTPase activating protein 1-like | protein-coding |
| chr5_143518140_143518624 | chr5 | 143518141 | 143518624 | + | 1.11036 | intron (NM_009007, intron 1 of 5) | intron (NM_009007, intron 1 of 5) | 9611 | NM_009007 | 19353 | Mm.292510 | NM_009007 | ENSMUSG00000001847 | **Rac1** | AL023026\|D5Ertd559e | RAS-related C3 botulinum substrate 1 | protein-coding |
| chr5_143517946_143518760 | chr5 | 143517947 | 143518760 | + | 1.08195 | intron (NM_009007, intron 1 of 5) | intron (NM_009007, intron 1 of 5) | 9640 | NM_009007 | 19353 | Mm.292510 | NM_009007 | ENSMUSG00000001847 | **Rac1** | AL023026\|D5Ertd559e | RAS-related C3 botulinum substrate 1 | protein-coding |
| chr5_143511922_143512678 | chr5 | 143511923 | 143512678 | + | 0.809028 | intron (NM_009007, intron 3 of 5) | intron (NM_009007, intron 3 of 5) | 15693 | NM_009007 | 19353 | Mm.292510 | NM_009007 | ENSMUSG00000001847 | **Rac1** | AL023026\|D5Ertd559e | RAS-related C3 botulinum substrate 1 | protein-coding |
| chr5_143499343_143500203 | chr5 | 143499344 | 143500203 | + | 0.714396 | intron (NM_144915, intron 11 of 14) | intron (NM_144915, intron 11 of 14) | 28220 | NM_009007 | 19353 | Mm.292510 | NM_009007 | ENSMUSG00000001847 | **Rac1** | AL023026\|D5Ertd559e | RAS-related C3 botulinum substrate 1 | protein-coding |
| chr15_78577446_78577803 | chr15 | 78577447 | 78577803 | + | 0.792909 | Intergenic | Intergenic | -4842 | NM_009008 | 19354 | Mm.1972 | NM_009008 | ENSMUSG00000033220 | **Rac2** | AI323801\|AI452260 | RAS-related C3 botulinum substrate 2 | protein-coding |
| chr4_55353147_55353313 | chr4 | 55353148 | 55353313 | + | 0.672136 | intron (NM_009011, intron 1 of 9) | intron (NM_009011, intron 1 of 9) | 3188 | NM_009011 | 19359 | Mm.196846 | NM_009011 | ENSMUSG00000028426 | **Rad23b** | 0610007D13Rik\|AV001138\|HR23B\|mHR23B\|p58 | RAD23b homolog (S. cerevisiae) | protein-coding |
| chr12_79436906_79437342 | chr12 | 79436907 | 79437342 | + | 1.22971 | intron (NM_009014, intron 7 of 10) | intron (NM_009014, intron 7 of 10) | 139773 | NM_001252562 | 19363 | Mm.341756 | NM_009014 | ENSMUSG00000059060 | **Rad51b** | AI553500\|R51H2\|Rad51l1\|mREC2 | RAD51 homolog B | protein-coding |
| chr12_79650713_79651046 | chr12 | 79650714 | 79651046 | + | 1.18631 | intron (NM_009014, intron 7 of 10) | Lx8\|LINE\|L1 | 353529 | NM_001252562 | 19363 | Mm.341756 | NM_009014 | ENSMUSG00000059060 | **Rad51b** | AI553500\|R51H2\|Rad51l1\|mREC2 | RAD51 homolog B | protein-coding |
| chr12_79436893_79437475 | chr12 | 79436894 | 79437475 | + | 1.16221 | intron (NM_009014, intron 7 of 10) | intron (NM_009014, intron 7 of 10) | 139833 | NM_001252562 | 19363 | Mm.341756 | NM_009014 | ENSMUSG00000059060 | **Rad51b** | AI553500\|R51H2\|Rad51l1\|mREC2 | RAD51 homolog B | protein-coding |
| chr12_79600109_79600753 | chr12 | 79600110 | 79600753 | + | 1.0733 | intron (NM_009014, intron 7 of 10) | intron (NM_009014, intron 7 of 10) | 303080 | NM_001252562 | 19363 | Mm.341756 | NM_009014 | ENSMUSG00000059060 | **Rad51b** | AI553500\|R51H2\|Rad51l1\|mREC2 | RAD51 homolog B | protein-coding |
| chr12_79550184_79550654 | chr12 | 79550185 | 79550654 | + | 1.07291 | intron (NM_009014, intron 7 of 10) | intron (NM_009014, intron 7 of 10) | 253068 | NM_001252562 | 19363 | Mm.341756 | NM_009014 | ENSMUSG00000059060 | **Rad51b** | AI553500\|R51H2\|Rad51l1\|mREC2 | RAD51 homolog B | protein-coding |
| chr12_79550030_79550855 | chr12 | 79550031 | 79550855 | + | 0.775284 | intron (NM_009014, intron 7 of 10) | intron (NM_009014, intron 7 of 10) | 253092 | NM_001252562 | 19363 | Mm.341756 | NM_009014 | ENSMUSG00000059060 | **Rad51b** | AI553500\|R51H2\|Rad51l1\|mREC2 | RAD51 homolog B | protein-coding |
| chr12_79439828_79440310 | chr12 | 79439829 | 79440310 | + | 0.709854 | intron (NM_009014, intron 7 of 10) | intron (NM_009014, intron 7 of 10) | 142718 | NM_001252562 | 19363 | Mm.341756 | NM_009014 | ENSMUSG00000059060 | **Rad51b** | AI553500\|R51H2\|Rad51l1\|mREC2 | RAD51 homolog B | protein-coding |
| chr12_79492157_79492558 | chr12 | 79492158 | 79492558 | + | 0.703931 | intron (NM_009014, intron 7 of 10) | MIR3\|SINE\|MIR | 195007 | NM_001252562 | 19363 | Mm.341756 | NM_009014 | ENSMUSG00000059060 | **Rad51b** | AI553500\|R51H2\|Rad51l1\|mREC2 | RAD51 homolog B | protein-coding |
| chr12_79393499_79394056 | chr12 | 79393500 | 79394056 | + | 0.654534 | intron (NM_009014, intron 7 of 10) | intron (NM_009014, intron 7 of 10) | 96427 | NM_001252562 | 19363 | Mm.341756 | NM_009014 | ENSMUSG00000059060 | **Rad51b** | AI553500\|R51H2\|Rad51l1\|mREC2 | RAD51 homolog B | protein-coding |
| chr6_119924224_119924705 | chr6 | 119924225 | 119924705 | + | 1.41975 | 3' UTR (NM_001185021, exon 26 of 26) | 3' UTR (NM_001185021, exon 26 of 26) | 21767 | NM_001166381 | 19365 | Mm.149 | NM_011236 | ENSMUSG00000030166 | **Rad52** | Rad52yh | RAD52 homolog (S. cerevisiae) | protein-coding |
| chr6_119908479_119908827 | chr6 | 119908480 | 119908827 | + | 1.32518 | intron (NM_001166382, intron 1 of 11) | MTE2a\|LTR\|ERVL-MaLR | 5955 | NM_001166381 | 19365 | Mm.149 | NM_011236 | ENSMUSG00000030166 | **Rad52** | Rad52yh | RAD52 homolog (S. cerevisiae) | protein-coding |

| | | | | | | | | | | | | | | | | | |
|---|---|---|---|---|---|---|---|---|---|---|---|---|---|---|---|---|---|
| chr6_119908105_119908396 | chr6 | 119908106 | 119908396 | + | 1.17214 | intron (NM_001166382, intron 1 of 11) | B3\|SINE\|B2 | 5553 | NM_001166381 | 19365 | Mm.149 | NM_011236 | ENSMUSG00000030166 | **Rad52** | Rad52yh | RAD52 homolog (S. cerevisiae) | protein-coding |
| chr6_119908133_119908336 | chr6 | 119908134 | 119908336 | + | 1.16687 | intron (NM_001166382, intron 1 of 11) | B3\|SINE\|B2 | 5537 | NM_001166381 | 19365 | Mm.149 | NM_011236 | ENSMUSG00000030166 | **Rad52** | Rad52yh | RAD52 homolog (S. cerevisiae) | protein-coding |
| chr4_116122657_116123184 | chr4 | 116122658 | 116123184 | + | 0.917142 | exon (NM_001122958, exon 3 of 19) | exon (NM_001122958, exon 3 of 19) | 768 | NM_009015 | 19366 | Mm.3655 | NM_009015 | ENSMUSG00000028702 | **Rad54l** | RAD54 | RAD54 like (S. cerevisiae) | protein-coding |
| chr10_22201442_22201781 | chr10 | 22201443 | 22201781 | + | 0.777943 | intron (NM_009018, intron 5 of 5) | intron (NM_009018, intron 5 of 5) | 27710 | NM_009018 | 19370 | Mm.439724 | NM_009018 | | **Raet1c** | RAE-1gamma | retinoic acid early transcript gamma | protein-coding |
| chr6_115677643_115678353 | chr6 | 115677644 | 115678353 | + | 1.82632 | Intergenic | L1MB5\|LINE\|L1 | -1363 | NM_029780 | 110157 | Mm.184163 | NM_029780 | ENSMUSG00000000441 | **Raf1** | 6430402F14Rik\|AA990557\|BB129353\|Craf1\|D830050J10Rik\|Raf-1\|c-Raf\|v-Raf | v-raf-leukemia viral oncogene 1 | protein-coding |
| chr6_115677389_115678368 | chr6 | 115677390 | 115678368 | + | 1.67397 | Intergenic | L1MB5\|LINE\|L1 | -1244 | NM_029780 | 110157 | Mm.184163 | NM_029780 | ENSMUSG00000000441 | **Raf1** | 6430402F14Rik\|AA990557\|BB129353\|Craf1\|D830050J10Rik\|Raf-1\|c-Raf\|v-Raf | v-raf-leukemia viral oncogene 1 | protein-coding |
| chr6_115678086_115678358 | chr6 | 115678087 | 115678358 | + | 1.56147 | Intergenic | B3\|SINE\|B2 | -1587 | NM_029780 | 110157 | Mm.184163 | NM_029780 | ENSMUSG00000000441 | **Raf1** | 6430402F14Rik\|AA990557\|BB129353\|Craf1\|D830050J10Rik\|Raf-1\|c-Raf\|v-Raf | v-raf-leukemia viral oncogene 1 | protein-coding |
| chr6_115677390_115678019 | chr6 | 115677391 | 115678019 | + | 1.44768 | Intergenic | L1MB5\|LINE\|L1 | -1070 | NM_029780 | 110157 | Mm.184163 | NM_029780 | ENSMUSG00000000441 | **Raf1** | 6430402F14Rik\|AA990557\|BB129353\|Craf1\|D830050J10Rik\|Raf-1\|c-Raf\|v-Raf | v-raf-leukemia viral oncogene 1 | protein-coding |
| chr6_115693070_115693825 | chr6 | 115693071 | 115693825 | + | 0.831235 | Intergenic | Intergenic | -16813 | NM_029780 | 110157 | Mm.184163 | NM_029780 | ENSMUSG00000000441 | **Raf1** | 6430402F14Rik\|AA990557\|BB129353\|Craf1\|D830050J10Rik\|Raf-1\|c-Raf\|v-Raf | v-raf-leukemia viral oncogene 1 | protein-coding |
| chr6_115653991_115654272 | chr6 | 115653992 | 115654272 | + | 0.808822 | intron (NM_029780, intron 1 of 16) | Tigger7\|DNA\|TcMar-Tigger | 22503 | NM_029780 | 110157 | Mm.184163 | NM_029780 | ENSMUSG00000000441 | **Raf1** | 6430402F14Rik\|AA990557\|BB129353\|Craf1\|D830050J10Rik\|Raf-1\|c-Raf\|v-Raf | v-raf-leukemia viral oncogene 1 | protein-coding |
| chr6_115684387_115685311 | chr6 | 115684388 | 115685311 | + | 0.667157 | Intergenic | MT2B2\|LTR\|ERVL | -8214 | NM_029780 | 110157 | Mm.184163 | NM_029780 | ENSMUSG00000000441 | **Raf1** | 6430402F14Rik\|AA990557\|BB129353\|Craf1\|D830050J10Rik\|Raf-1\|c-Raf\|v-Raf | v-raf-leukemia viral oncogene 1 | protein-coding |
| chr6_115684799_115685544 | chr6 | 115684800 | 115685544 | + | 0.611012 | Intergenic | RLTR17\|LTR\|ERVK | -8537 | NM_029780 | 110157 | Mm.184163 | NM_029780 | ENSMUSG00000000441 | **Raf1** | 6430402F14Rik\|AA990557\|BB129353\|Craf1\|D830050J10Rik\|Raf-1\|c-Raf\|v-Raf | v-raf-leukemia viral oncogene 1 | protein-coding |
| chr2_101623065_101623549 | chr2 | 101623066 | 101623549 | + | 0.674757 | intron (NM_026592, intron 1 of 6) | Tigger7\|DNA\|TcMar-Tigger | -1441 | NM_009020 | 19374 | Mm.4988 | NM_009020 | ENSMUSG00000032864 | **Rag2** | Rag-2 | recombination activating gene 2 | protein-coding |
| chr1_119470530_119470999 | chr1 | 119470531 | 119470999 | + | 0.825343 | 3' UTR (NM_022327, exon 5 of 5) | 3' UTR (NM_022327, exon 5 of 5) | 34017 | NM_022327 | 64143 | Mm.27832 | NM_022327 | ENSMUSG00000004451 | **Ralb** | 5730472O18Rik | v-ral simian leukemia viral oncogene homolog B (ras related) | protein-coding |
| chr2_146416124_146416916 | chr2 | 146416125 | 146416916 | + | 0.953118 | intron (NM_001033348, intron 16 of 40) | intron (NM_001033348, intron 16 of 40) | 95484 | NM_001033348 | 241694 | Mm.222310 | NM_001033348 | ENSMUSG00000037110 | **Ralgapa2** | A230067G21Rik\|AS250\|BC053994\|RGC2\|pp250 | Ral GTPase activating protein, alpha subunit 2 (catalytic) | protein-coding |
| chr2_146516990_146517552 | chr2 | 146516991 | 146517552 | + | 0.902583 | Intergenic | Intergenic | -5267 | NM_001033348 | 241694 | Mm.222310 | NM_001033348 | ENSMUSG00000037110 | **Ralgapa2** | A230067G21Rik\|AS250\|BC053994\|RGC2\|pp250 | Ral GTPase activating protein, alpha subunit 2 (catalytic) | protein-coding |
| chr2_146457371_146458267 | chr2 | 146457372 | 146458267 | + | 0.78545 | intron (NM_001033348, intron 4 of 40) | intron (NM_001033348, intron 4 of 40) | 54185 | NM_001033348 | 241694 | Mm.222310 | NM_001033348 | ENSMUSG00000037110 | **Ralgapa2** | A230067G21Rik\|AS250\|BC053994\|RGC2\|pp250 | Ral GTPase activating protein, alpha subunit 2 (catalytic) | protein-coding |
| chr2_146476050_146476630 | chr2 | 146476051 | 146476630 | + | 0.62075 | intron (NM_001033348, intron 3 of 40) | intron (NM_001033348, intron 3 of 40) | 35664 | NM_001033348 | 241694 | Mm.222310 | NM_001033348 | ENSMUSG00000037110 | **Ralgapa2** | A230067G21Rik\|AS250\|BC053994\|RGC2\|pp250 | Ral GTPase activating protein, alpha subunit 2 (catalytic) | protein-coding |
| chr2_158450698_158451179 | chr2 | 158450699 | 158451179 | + | 0.975443 | intron (NM_001291137, intron 17 of 30) | L1MB5\|LINE\|L1 | 41086 | NM_001291138 | 228850 | Mm.398185 | NM_177227 | ENSMUSG00000027652 | **Ralgapb** | 9330195P16\|AI507211\|B230339M05Rik\|mKIAA1219\|p170 | Ral GTPase activating protein, beta subunit (non-catalytic) | protein-coding |
| chr2_28523770_28524111 | chr2 | 28523771 | 28524111 | + | 0.998769 | intron (NM_009058, intron 1 of 17) | intron (NM_009058, intron 1 of 17) | -9523 | NM_001145835 | 19730 | Mm.5236 | NM_009058 | ENSMUSG00000026821 | **Ralgds** | Gnds\|Rgds\|mKIAA1308 | ral guanine nucleotide dissociation stimulator | protein-coding |
| chr2_33346064_33346348 | chr2 | 33346065 | 33346348 | + | 0.719869 | intron (NM_001290570, intron 1 of 16) | intron (NM_001290570, intron 1 of 16) | 25288 | NM_001290570 | 241308 | Mm.274249 | NM_175211 | ENSMUSG00000038831 | **Ralgps1** | 5830418G11Rik\|AI853783\|AI854138\|RALGEF2\|RALGPS1A\|RalGEF2\|mKIAA0351 | Ral GEF with PH domain and SH3 binding motif 1 | protein-coding |
| chr1_157019804_157020319 | chr1 | 157019805 | 157020319 | + | 0.946144 | Intergenic | Intergenic | -80436 | NM_023884 | 78255 | Mm.28376 | NM_023884 | ENSMUSG00000026594 | **Ralgps2** | 1810020P17Rik\|2210408F11Rik\|4921528G01Rik\|9130014M22Rik\|AU043409\|AW046161 | Ral GEF with PH domain and SH3 binding motif 2 | protein-coding |
| chr1_156910958_156911354 | chr1 | 156910959 | 156911354 | + | 0.752584 | intron (NR_027640, intron 1 of 19) | intron (NR_027640, intron 1 of 19) | 28231 | NM_001159966 | 78255 | Mm.28376 | NM_023884 | ENSMUSG00000026594 | **Ralgps2** | 1810020P17Rik\|2210408F11Rik\|4921528G01Rik\|9130014M22Rik\|AU043409\|AW046161 | Ral GEF with PH domain and SH3 binding motif 2 | protein-coding |
| chr1_156995093_156995655 | chr1 | 156995094 | 156995655 | + | 0.676033 | Intergenic | Intergenic | -55748 | NM_023884 | 78255 | Mm.28376 | NM_023884 | ENSMUSG00000026594 | **Ralgps2** | 1810020P17Rik\|2210408F11Rik\|4921528G01Rik\|9130014M22Rik\|AU043409\|AW046161 | Ral GEF with PH domain and SH3 binding motif 2 | protein-coding |
| chr2_154757676_154758050 | chr2 | 154757677 | 154758050 | + | 1.00991 | Intergenic | Intergenic | -33247 | NM_001139512 | 19383 | Mm.221440 | NM_023130 | ENSMUSG00000027593 | **Raly** | AI663842\|Merc | hnRNP-associated with lethal yellow | protein-coding |
| chr1_91187997_91188389 | chr1 | 91187998 | 91188389 | + | 0.796111 | intron (NM_001168392, intron 1 of 3) | intron (NM_001168392, intron 1 of 3) | 8371 | NM_001168392 | 51801 | Mm.3272 | NM_016894 | ENSMUSG00000034353 | **Ramp1** | 9130218E19Rik | receptor (calcitonin) activity modifying protein 1 | protein-coding |
| chr5_129025008_129025320 | chr5 | 129025009 | 129025320 | + | 0.716329 | TTS (NM_009391) | TTS (NM_009391) | 5008 | NM_009391 | 19384 | Mm.297440 | NM_009391 | ENSMUSG00000029430 | **Ran** | - | RAN, member RAS oncogene family | protein-coding |
| chr10_58457368_58457831 | chr10 | 58457369 | 58457831 | + | 0.677697 | exon (NM_011240, exon 7 of 29) | exon (NM_011240, exon 7 of 29) | 10748 | NM_011240 | 19386 | Mm.431695 | NM_011240 | ENSMUSG00000003226 | **Ranbp2** | A430087B05Rik\|AI256741\|NUP358 | RAN binding protein 2 | protein-coding |
| chr17_56667579_56668127 | chr17 | 56667580 | 56668127 | + | 0.734907 | Intergenic | ORR1C1\|LTR\|ERVL-MaLR | -5372 | NM_001252466 | 71810 | Mm.29608 | NM_027933 | ENSMUSG00000002372 | **Ranbp3** | 2610024N24Rik\|AA408221 | RAN binding protein 3 | protein-coding |
| chr19_29849543_29850642 | chr19 | 29849544 | 29850642 | + | 1.06913 | Intergenic | Intergenic | -37119 | NM_177721 | 240614 | Mm.125503 | NM_177721 | ENSMUSG00000074909 | **Ranbp6** | C630001B19 | RAN binding protein 6 | protein-coding |
| chr19_29850184_29850617 | chr19 | 29850185 | 29850617 | + | 1.01872 | Intergenic | Intergenic | -37427 | NM_177721 | 240614 | Mm.125503 | NM_177721 | ENSMUSG00000074909 | **Ranbp6** | C630001B19 | RAN binding protein 6 | protein-coding |
| chr19_29849818_29850617 | chr19 | 29849819 | 29850617 | + | 1.00446 | Intergenic | Intergenic | -37244 | NM_177721 | 240614 | Mm.125503 | NM_177721 | ENSMUSG00000074909 | **Ranbp6** | C630001B19 | RAN binding protein 6 | protein-coding |
| chr13_43461169_43461613 | chr13 | 43461170 | 43461613 | + | 1.37017 | intron (NM_019930, intron 2 of 13) | intron (NM_019930, intron 2 of 13) | 19582 | NM_019930 | 56705 | Mm.148781 | NM_019930 | ENSMUSG00000038546 | **Ranbp9** | IBAP-1\|Ibap1\|RanBPM | RAN binding protein 9 | protein-coding |
| chr13_43463564_43463865 | chr13 | 43463565 | 43463865 | + | 0.813517 | intron (NM_019930, intron 1 of 13) | intron (NM_019930, intron 1 of 13) | 17258 | NM_019930 | 56705 | Mm.148781 | NM_019930 | ENSMUSG00000038546 | **Ranbp9** | IBAP-1\|Ibap1\|RanBPM | RAN binding protein 9 | protein-coding |
| chr3_105806645_105807253 | chr3 | 105806646 | 105807253 | + | 1.32232 | Intergenic | Intergenic | -5568 | NM_145541 | 109905 | Mm.101922 | NM_145541 | ENSMUSG00000068798 | **Rap1a** | AI848598\|G-22K\|Krev-1\|Rap1 | RAS-related protein-1a | protein-coding |
| chr3_105775289_105776054 | chr3 | 105775290 | 105776054 | + | 0.857982 | intron (NM_145541, intron 1 of 7) | MTA_Mm\|LTR\|ERVL-MaLR | 25709 | NM_145541 | 109905 | Mm.101922 | NM_145541 | ENSMUSG00000068798 | **Rap1a** | AI848598\|G-22K\|Krev-1\|Rap1 | RAS-related protein-1a | protein-coding |
| chr10_117867304_117867700 | chr10 | 117867305 | 117867700 | + | 1.27534 | Intergenic | Intergenic | -21528 | NM_024457 | 215449 | Mm.233009 | NM_024457 | ENSMUSG00000052681 | **Rap1b** | 2810443E11Rik | RAS related protein 1b | protein-coding |
| chr10_117872479_117873787 | chr10 | 117872480 | 117873787 | + | 1.11394 | Intergenic | Intergenic | -27159 | NM_024457 | 215449 | Mm.233009 | NM_024457 | ENSMUSG00000052681 | **Rap1b** | 2810443E11Rik | RAS related protein 1b | protein-coding |
| chr10_117840451_117840754 | chr10 | 117840452 | 117840754 | + | 1.01579 | intron (NM_024457, intron 1 of 7) | intron (NM_024457, intron 1 of 7) | 5371 | NM_024457 | 215449 | Mm.233009 | NM_024457 | ENSMUSG00000052681 | **Rap1b** | 2810443E11Rik | RAS related protein 1b | protein-coding |
| chr10_117838942_117839295 | chr10 | 117838943 | 117839295 | + | 0.720075 | intron (NM_024457, intron 1 of 7) | B3\|SINE\|B2 | 6855 | NM_024457 | 215449 | Mm.233009 | NM_024457 | ENSMUSG00000052681 | **Rap1b** | 2810443E11Rik | RAS related protein 1b | protein-coding |
| chr4_137648115_137648702 | chr4 | 137648116 | 137648702 | + | 0.838742 | intron (NM_130879, intron 24 of 27) | intron (NM_130879, intron 24 of 27) | -16317 | NM_029563 | 110351 | Mm.180763 | NM_029563 | ENSMUSG00000041351 | **Rap1gap** | 1300019I11Rik\|2310004O14Rik\|AI427470\|ARPP-90\|Rap1GAP1\|Rap1ga1 | Rap1 GTPase-activating protein | protein-coding |
| chr11_74523600_74523901 | chr11 | 74523601 | 74523901 | + | 0.694334 | intron (NM_001015046, intron 2 of 24) | intron (NM_001015046, intron 2 of 24) | 66407 | NM_001015046 | 380711 | Mm.425690 | NM_001015046 | ENSMUSG00000038807 | **Rap1gap2** | AU067654\|Garnl4\|Gm1561\|mKIAA1039 | RAP1 GTPase activating protein 2 | protein-coding |
| chr3_139000129_139000942 | chr3 | 139000130 | 139000942 | + | 0.879895 | intron (NM_001286759, intron 4 of 14) | L1_Mur3\|LINE\|L1 | 74665 | NM_145544 | 229877 | Mm.28893 | NM_145544 | ENSMUSG00000028149 | **Rap1gds1** | BC011279\|GDS1 | RAP1, GTP-GDP dissociation stimulator 1 | protein-coding |

<table>
<tr><td></td><td></td><td></td><td></td><td></td><td></td><td></td><td></td><td></td><td></td><td></td><td></td><td></td><td></td><td></td><td></td><td></td><td></td></tr>
<tr><td>chr3_139073786_139074107</td><td>chr3</td><td>139073787</td><td>139074107</td><td>+</td><td>0.790673</td><td>intron (NM_001286759, intron 1 of 14)</td><td>intron (NM_001286759, intron 1 of 14)</td><td>1254</td><td>NM_145544</td><td>229877</td><td>Mm.28893</td><td>NM_145544</td><td>ENSMUSG00000028149</td><td>Rap1gds1</td><td>BC011279|GDS1</td><td>RAP1, GTP-GDP dissociation stimulator 1</td><td>protein-coding</td></tr>
<tr><td>chr14_120535853_120536317</td><td>chr14</td><td>120535854</td><td>120536317</td><td>+</td><td>0.962387</td><td>Intergenic</td><td>Intergenic</td><td>57624</td><td>NM_029519</td><td>76108</td><td>Mm.261448</td><td>NM_029519</td><td>ENSMUSG00000051615</td><td>Rap2a</td><td>5830461H18Rik</td><td>RAS related protein 2a</td><td>protein-coding</td></tr>
<tr><td>chr14_120520706_120521521</td><td>chr14</td><td>120520707</td><td>120521521</td><td>+</td><td>0.766154</td><td>Intergenic</td><td>Intergenic</td><td>42653</td><td>NM_029519</td><td>76108</td><td>Mm.261448</td><td>NM_029519</td><td>ENSMUSG00000051615</td><td>Rap2a</td><td>5830461H18Rik</td><td>RAS related protein 2a</td><td>protein-coding</td></tr>
<tr><td>chr14_120400211_120400498</td><td>chr14</td><td>120400212</td><td>120400498</td><td>+</td><td>0.717323</td><td>intron (NM_175341, intron 7 of 8)</td><td>intron (NM_175341, intron 7 of 8)</td><td>-78106</td><td>NM_029519</td><td>76108</td><td>Mm.261448</td><td>NM_029519</td><td>ENSMUSG00000051615</td><td>Rap2a</td><td>5830461H18Rik</td><td>RAS related protein 2a</td><td>protein-coding</td></tr>
<tr><td>chr3_61368566_61369139</td><td>chr3</td><td>61368567</td><td>61369139</td><td>+</td><td>1.41487</td><td>TTS (NM_028712)</td><td>TTS (NM_028712)</td><td>4346</td><td>NM_028712</td><td>74012</td><td>Mm.422674</td><td>NM_028712</td><td>ENSMUSG00000036894</td><td>Rap2b</td><td>4021402C18Rik|AA408554</td><td>RAP2B, member of RAS oncogene family</td><td>protein-coding</td></tr>
<tr><td>chr3_61370072_61370418</td><td>chr3</td><td>61370073</td><td>61370418</td><td>+</td><td>1.1523</td><td>Intergenic</td><td>Intergenic</td><td>5738</td><td>NM_028712</td><td>74012</td><td>Mm.422674</td><td>NM_028712</td><td>ENSMUSG00000036894</td><td>Rap2b</td><td>4021402C18Rik|AA408554</td><td>RAP2B, member of RAS oncogene family</td><td>protein-coding</td></tr>
<tr><td>chr3_61509746_61510127</td><td>chr3</td><td>61509747</td><td>61510127</td><td>+</td><td>0.975746</td><td>Intergenic</td><td>Lx5c|LINE|L1</td><td>145430</td><td>NM_028712</td><td>74012</td><td>Mm.422674</td><td>NM_028712</td><td>ENSMUSG00000036894</td><td>Rap2b</td><td>4021402C18Rik|AA408554</td><td>RAP2B, member of RAS oncogene family</td><td>protein-coding</td></tr>
<tr><td>chr3_61509616_61510165</td><td>chr3</td><td>61509617</td><td>61510165</td><td>+</td><td>0.89745</td><td>Intergenic</td><td>Lx5c|LINE|L1</td><td>145384</td><td>NM_028712</td><td>74012</td><td>Mm.422674</td><td>NM_028712</td><td>ENSMUSG00000036894</td><td>Rap2b</td><td>4021402C18Rik|AA408554</td><td>RAP2B, member of RAS oncogene family</td><td>protein-coding</td></tr>
<tr><td>chr3_61506443_61506943</td><td>chr3</td><td>61506444</td><td>61506943</td><td>+</td><td>0.871066</td><td>Intergenic</td><td>Intergenic</td><td>142186</td><td>NM_028712</td><td>74012</td><td>Mm.422674</td><td>NM_028712</td><td>ENSMUSG00000036894</td><td>Rap2b</td><td>4021402C18Rik|AA408554</td><td>RAP2B, member of RAS oncogene family</td><td>protein-coding</td></tr>
<tr><td>chr3_61343827_61344461</td><td>chr3</td><td>61343828</td><td>61344461</td><td>+</td><td>0.624465</td><td>Intergenic</td><td>Intergenic</td><td>-20363</td><td>NM_028712</td><td>74012</td><td>Mm.422674</td><td>NM_028712</td><td>ENSMUSG00000036894</td><td>Rap2b</td><td>4021402C18Rik|AA408554</td><td>RAP2B, member of RAS oncogene family</td><td>protein-coding</td></tr>
<tr><td>chr2_29629258_29629797</td><td>chr2</td><td>29629259</td><td>29629797</td><td>+</td><td>1.04482</td><td>intron (NM_001039086, intron 1 of 25)</td><td>intron (NM_001039086, intron 1 of 25)</td><td>9808</td><td>NM_001039086</td><td>107746</td><td>Mm.298274</td><td>NM_054050</td><td>ENSMUSG00000039844</td><td>Rapgef1</td><td>4932418O06Rik|C3G|Grf2</td><td>Rap guanine nucleotide exchange factor (GEF) 1</td><td>protein-coding</td></tr>
<tr><td>chr2_29628747_29629798</td><td>chr2</td><td>29628748</td><td>29629798</td><td>+</td><td>0.788875</td><td>intron (NM_001039086, intron 1 of 25)</td><td>intron (NM_001039086, intron 1 of 25)</td><td>9553</td><td>NM_001039086</td><td>107746</td><td>Mm.298274</td><td>NM_054050</td><td>ENSMUSG00000039844</td><td>Rapgef1</td><td>4932418O06Rik|C3G|Grf2</td><td>Rap guanine nucleotide exchange factor (GEF) 1</td><td>protein-coding</td></tr>
<tr><td>chr3_78889650_78890272</td><td>chr3</td><td>78889651</td><td>78890272</td><td>+</td><td>1.5878</td><td>Intergenic</td><td>Intergenic</td><td>255914</td><td>NM_001099624</td><td>76089</td><td>Mm.31220</td><td>NM_001099624</td><td>ENSMUSG00000062232</td><td>Rapgef2</td><td>5830453M24Rik|CNRasGEF|Pdzgef1|RA-GEF-1|mKIAA0313|nRapGEP</td><td>Rap guanine nucleotide exchange factor (GEF) 2</td><td>protein-coding</td></tr>
<tr><td>chr3_79113882_79114368</td><td>chr3</td><td>79113883</td><td>79114368</td><td>+</td><td>0.762707</td><td>intron (NM_001099624, intron 1 of 23)</td><td>intron (NM_001099624, intron 1 of 23)</td><td>31750</td><td>NM_001099624</td><td>76089</td><td>Mm.31220</td><td>NM_001099624</td><td>ENSMUSG00000062232</td><td>Rapgef2</td><td>5830453M24Rik|CNRasGEF|Pdzgef1|RA-GEF-1|mKIAA0313|nRapGEP</td><td>Rap guanine nucleotide exchange factor (GEF) 2</td><td>protein-coding</td></tr>
<tr><td>chr3_79113923_79114175</td><td>chr3</td><td>79113924</td><td>79114175</td><td>+</td><td>0.669481</td><td>intron (NM_001099624, intron 1 of 23)</td><td>intron (NM_001099624, intron 1 of 23)</td><td>31826</td><td>NM_001099624</td><td>76089</td><td>Mm.31220</td><td>NM_001099624</td><td>ENSMUSG00000062232</td><td>Rapgef2</td><td>5830453M24Rik|CNRasGEF|Pdzgef1|RA-GEF-1|mKIAA0313|nRapGEP</td><td>Rap guanine nucleotide exchange factor (GEF) 2</td><td>protein-coding</td></tr>
<tr><td>chr3_78947103_78947663</td><td>chr3</td><td>78947104</td><td>78947663</td><td>+</td><td>0.65387</td><td>Intergenic</td><td>Intergenic</td><td>198492</td><td>NM_001099624</td><td>76089</td><td>Mm.31220</td><td>NM_001099624</td><td>ENSMUSG00000062232</td><td>Rapgef2</td><td>5830453M24Rik|CNRasGEF|Pdzgef1|RA-GEF-1|mKIAA0313|nRapGEP</td><td>Rap guanine nucleotide exchange factor (GEF) 2</td><td>protein-coding</td></tr>
<tr><td>chr2_71964165_71965039</td><td>chr2</td><td>71964166</td><td>71965039</td><td>+</td><td>1.2178</td><td>Intergenic</td><td>L1M5|LINE|L1</td><td>-16613</td><td>NM_001204167</td><td>56508</td><td>Mm.196153</td><td>NM_019688</td><td>ENSMUSG00000049044</td><td>Rapgef4</td><td>1300003D15Rik|5730402K07Rik|6330581N18Rik|EPAC 2|Epac-2|Epac2|mKIAA4040</td><td>Rap guanine nucleotide exchange factor (GEF) 4</td><td>protein-coding</td></tr>
<tr><td>chr2_71974205_71974870</td><td>chr2</td><td>71974206</td><td>71974870</td><td>+</td><td>0.935892</td><td>Intergenic</td><td>Intergenic</td><td>-6677</td><td>NM_001204167</td><td>56508</td><td>Mm.196153</td><td>NM_019688</td><td>ENSMUSG00000049044</td><td>Rapgef4</td><td>1300003D15Rik|5730402K07Rik|6330581N18Rik|EPAC 2|Epac-2|Epac2|mKIAA4040</td><td>Rap guanine nucleotide exchange factor (GEF) 4</td><td>protein-coding</td></tr>
<tr><td>chr2_72078077_72078452</td><td>chr2</td><td>72078078</td><td>72078452</td><td>+</td><td>0.810349</td><td>intron (NM_001204166, intron 1 of 27)</td><td>intron (NM_001204166, intron 1 of 27)</td><td>23648</td><td>NM_001204166</td><td>56508</td><td>Mm.196153</td><td>NM_019688</td><td>ENSMUSG00000049044</td><td>Rapgef4</td><td>1300003D15Rik|5730402K07Rik|6330581N18Rik|EPAC 2|Epac-2|Epac2|mKIAA4040</td><td>Rap guanine nucleotide exchange factor (GEF) 4</td><td>protein-coding</td></tr>
<tr><td>chr14_16519625_16519898</td><td>chr14</td><td>16519626</td><td>16519898</td><td>+</td><td>0.707774</td><td>intron (NM_001289761, intron 3 of 8)</td><td>intron (NM_001289761, intron 3 of 8)</td><td>55714</td><td>NM_011243</td><td>218772</td><td>Mm.259318</td><td>NM_011243</td><td>ENSMUSG00000017491</td><td>Rarb</td><td>A830025K23|Hap|Nr1b2</td><td>retinoic acid receptor, beta</td><td>protein-coding</td></tr>
<tr><td>chr13_85459242_85459791</td><td>chr13</td><td>85459243</td><td>85459791</td><td>+</td><td>1.35487</td><td>Intergenic</td><td>MERV1_LTR|LTR|ERV1</td><td>-170031</td><td>NM_145452</td><td>218397</td><td>Mm.259653</td><td>NM_145452</td><td>ENSMUSG00000021549</td><td>Rasa1</td><td>Gap|RasGAP|Rasa</td><td>RAS p21 protein activator 1</td><td>protein-coding</td></tr>
<tr><td>chr13_85428139_85428628</td><td>chr13</td><td>85428140</td><td>85428628</td><td>+</td><td>1.28818</td><td>Intergenic</td><td>Intergenic</td><td>-138898</td><td>NM_145452</td><td>218397</td><td>Mm.259653</td><td>NM_145452</td><td>ENSMUSG00000021549</td><td>Rasa1</td><td>Gap|RasGAP|Rasa</td><td>RAS p21 protein activator 1</td><td>protein-coding</td></tr>
<tr><td>chr13_85458661_85458917</td><td>chr13</td><td>85458662</td><td>85458917</td><td>+</td><td>1.21406</td><td>Intergenic</td><td>MTC|LTR|ERVL-MaLR</td><td>-169303</td><td>NM_145452</td><td>218397</td><td>Mm.259653</td><td>NM_145452</td><td>ENSMUSG00000021549</td><td>Rasa1</td><td>Gap|RasGAP|Rasa</td><td>RAS p21 protein activator 1</td><td>protein-coding</td></tr>
<tr><td>chr13_85458615_85459103</td><td>chr13</td><td>85458616</td><td>85459103</td><td>+</td><td>1.13771</td><td>Intergenic</td><td>MTC|LTR|ERVL-MaLR</td><td>-169373</td><td>NM_145452</td><td>218397</td><td>Mm.259653</td><td>NM_145452</td><td>ENSMUSG00000021549</td><td>Rasa1</td><td>Gap|RasGAP|Rasa</td><td>RAS p21 protein activator 1</td><td>protein-coding</td></tr>
<tr><td>chr13_85485175_85485433</td><td>chr13</td><td>85485176</td><td>85485433</td><td>+</td><td>0.710878</td><td>Intergenic</td><td>Intergenic</td><td>-195818</td><td>NM_145452</td><td>218397</td><td>Mm.259653</td><td>NM_145452</td><td>ENSMUSG00000021549</td><td>Rasa1</td><td>Gap|RasGAP|Rasa</td><td>RAS p21 protein activator 1</td><td>protein-coding</td></tr>
<tr><td>chr13_85359153_85359900</td><td>chr13</td><td>85359154</td><td>85359900</td><td>+</td><td>0.704469</td><td>Intergenic</td><td>Intergenic</td><td>-70041</td><td>NM_145452</td><td>218397</td><td>Mm.259653</td><td>NM_145452</td><td>ENSMUSG00000021549</td><td>Rasa1</td><td>Gap|RasGAP|Rasa</td><td>RAS p21 protein activator 1</td><td>protein-coding</td></tr>
<tr><td>chr13_85398753_85399214</td><td>chr13</td><td>85398754</td><td>85399214</td><td>+</td><td>0.626872</td><td>Intergenic</td><td>Intergenic</td><td>-109498</td><td>NM_145452</td><td>218397</td><td>Mm.259653</td><td>NM_145452</td><td>ENSMUSG00000021549</td><td>Rasa1</td><td>Gap|RasGAP|Rasa</td><td>RAS p21 protein activator 1</td><td>protein-coding</td></tr>
<tr><td>chr13_85282107_85282542</td><td>chr13</td><td>85282108</td><td>85282542</td><td>+</td><td>0.60791</td><td>intron (NM_145452, intron 1 of 24)</td><td>Lx8|LINE|L1</td><td>7161</td><td>NM_145452</td><td>218397</td><td>Mm.259653</td><td>NM_145452</td><td>ENSMUSG00000021549</td><td>Rasa1</td><td>Gap|RasGAP|Rasa</td><td>RAS p21 protein activator 1</td><td>protein-coding</td></tr>
<tr><td>chr9_96649870_96650486</td><td>chr9</td><td>96649871</td><td>96650486</td><td>+</td><td>0.743531</td><td>Intergenic</td><td>Intergenic</td><td>-18675</td><td>NM_053268</td><td>114713</td><td>Mm.124502</td><td>NM_053268</td><td>ENSMUSG00000032413</td><td>Rasa2</td><td>5430433H21Rik|AA517451|AU023900|GAP1m</td><td>RAS p21 protein activator 2</td><td>protein-coding</td></tr>
<tr><td>chr8_13687034_13687555</td><td>chr8</td><td>13687035</td><td>13687555</td><td>+</td><td>1.11422</td><td>Intergenic</td><td>Intergenic</td><td>-9708</td><td>NM_009025</td><td>19414</td><td>Mm.18517</td><td>NM_009025</td><td>ENSMUSG00000031453</td><td>Rasa3</td><td>AI326412|C86362|E130011G04|GAPIII|scat</td><td>RAS p21 protein activator 3</td><td>protein-coding</td></tr>
<tr><td>chr8_13625495_13625835</td><td>chr8</td><td>13625496</td><td>13625835</td><td>+</td><td>1.04207</td><td>intron (NM_009025, intron 2 of 23)</td><td>intron (NM_009025, intron 2 of 23)</td><td>51922</td><td>NM_009025</td><td>19414</td><td>Mm.18517</td><td>NM_009025</td><td>ENSMUSG00000031453</td><td>Rasa3</td><td>AI326412|C86362|E130011G04|GAPIII|scat</td><td>RAS p21 protein activator 3</td><td>protein-coding</td></tr>
<tr><td>chr8_13663034_13663346</td><td>chr8</td><td>13663035</td><td>13663346</td><td>+</td><td>0.642841</td><td>intron (NM_009025, intron 1 of 23)</td><td>intron (NM_009025, intron 1 of 23)</td><td>14397</td><td>NM_009025</td><td>19414</td><td>Mm.18517</td><td>NM_009025</td><td>ENSMUSG00000031453</td><td>Rasa3</td><td>AI326412|C86362|E130011G04|GAPIII|scat</td><td>RAS p21 protein activator 3</td><td>protein-coding</td></tr>
<tr><td>chr8_75212694_75213629</td><td>chr8</td><td>75212695</td><td>75213629</td><td>+</td><td>0.692</td><td>promoter-TSS (NM_029182)</td><td>promoter-TSS (NM_029182)</td><td>-782</td><td>NM_029182</td><td>75141</td><td>Mm.179267</td><td>NM_029182</td><td>ENSMUSG00000034472</td><td>Rasd2</td><td>4930526B11Rik|AU045414|Rhes|TEM-2|TEM2</td><td>RASD family, member 2</td><td>protein-coding</td></tr>
<tr><td>chr5_99214292_99214958</td><td>chr5</td><td>99214293</td><td>99214958</td><td>+</td><td>1.01574</td><td>Intergenic</td><td>Intergenic</td><td>38302</td><td>NM_181318</td><td>320292</td><td>Mm.259074</td><td>NM_145839</td><td>ENSMUSG00000029333</td><td>Rasgef1b</td><td>4732452O09Rik|GPIgamma4|Gpig4</td><td>RasGEF domain family, member 1B</td><td>protein-coding</td></tr>
<tr><td>chr5_99203477_99203856</td><td>chr5</td><td>99203478</td><td>99203856</td><td>+</td><td>0.828088</td><td>Intergenic</td><td>Intergenic</td><td>49260</td><td>NM_181318</td><td>320292</td><td>Mm.259074</td><td>NM_145839</td><td>ENSMUSG00000029333</td><td>Rasgef1b</td><td>4732452O09Rik|GPIgamma4|Gpig4</td><td>RasGEF domain family, member 1B</td><td>protein-coding</td></tr>
<tr><td>chr5_99365196_99365924</td><td>chr5</td><td>99365197</td><td>99365924</td><td>+</td><td>0.709936</td><td>intron (NR_030692, intron 3 of 3)</td><td>intron (NR_030692, intron 3 of 3)</td><td>-112633</td><td>NM_145839</td><td>320292</td><td>Mm.259074</td><td>NM_145839</td><td>ENSMUSG00000029333</td><td>Rasgef1b</td><td>4732452O09Rik|GPIgamma4|Gpig4</td><td>RasGEF domain family, member 1B</td><td>protein-coding</td></tr>
<tr><td>chr5_99329075_99329379</td><td>chr5</td><td>99329076</td><td>99329379</td><td>+</td><td>0.707063</td><td>intron (NR_030692, intron 3 of 3)</td><td>intron (NR_030692, intron 3 of 3)</td><td>-76300</td><td>NM_145839</td><td>320292</td><td>Mm.259074</td><td>NM_145839</td><td>ENSMUSG00000029333</td><td>Rasgef1b</td><td>4732452O09Rik|GPIgamma4|Gpig4</td><td>RasGEF domain family, member 1B</td><td>protein-coding</td></tr>
<tr><td>chr5_99460926_99461837</td><td>chr5</td><td>99460927</td><td>99461837</td><td>+</td><td>0.647961</td><td>intron (NR_030692, intron 2 of 3)</td><td>intron (NR_030692, intron 2 of 3)</td><td>-208455</td><td>NM_145839</td><td>320292</td><td>Mm.259074</td><td>NM_145839</td><td>ENSMUSG00000029333</td><td>Rasgef1b</td><td>4732452O09Rik|GPIgamma4|Gpig4</td><td>RasGEF domain family, member 1B</td><td>protein-coding</td></tr>
<tr><td>chr9_89874292_89874850</td><td>chr9</td><td>89874293</td><td>89874850</td><td>+</td><td>1.38019</td><td>Intergenic</td><td>Intergenic</td><td>-35204</td><td>NM_011245</td><td>19417</td><td>Mm.44561</td><td>NM_011245</td><td>ENSMUSG00000032356</td><td>Rasgrf1</td><td>AI844718|CDC25|CDC25Mm|Gnrp|Grf1|Grfbeta|P190-A|Ras-GRF1|p190|p190RhoGEF</td><td>RAS protein-specific guanine nucleotide-releasing factor 1</td><td>protein-coding</td></tr>
<tr><td>chr13_91933643_91934086</td><td>chr13</td><td>91933644</td><td>91934086</td><td>+</td><td>0.946556</td><td>intron (NM_009027, intron 5 of 14)</td><td>Lx8|LINE|L1</td><td>54177</td><td>NM_009027</td><td>19418</td><td>Mm.248630</td><td>NM_009027</td><td></td><td>Rasgrf2</td><td>6330417G04Rik|AW048350|Grf2|Ras-GRF2</td><td>RAS protein-specific guanine nucleotide-releasing factor 2</td><td>protein-coding</td></tr>
<tr><td>chr13_91933497_91934334</td><td>chr13</td><td>91933498</td><td>91934334</td><td>+</td><td>0.786123</td><td>intron (NM_009027, intron 5 of 14)</td><td>Lx8|LINE|L1</td><td>54126</td><td>NM_009027</td><td>19418</td><td>Mm.248630</td><td>NM_009027</td><td></td><td>Rasgrf2</td><td>6330417G04Rik|AW048350|Grf2|Ras-GRF2</td><td>RAS protein-specific guanine nucleotide-releasing factor 2</td><td>protein-coding</td></tr>
<tr><td>chr13_91995720_91996026</td><td>chr13</td><td>91995721</td><td>91996026</td><td>+</td><td>0.659144</td><td>Intergenic</td><td>Intergenic</td><td>-7831</td><td>NM_009027</td><td>19418</td><td>Mm.248630</td><td>NM_009027</td><td></td><td>Rasgrf2</td><td>6330417G04Rik|AW048350|Grf2|Ras-GRF2</td><td>RAS protein-specific guanine nucleotide-releasing factor 2</td><td>protein-coding</td></tr>
<tr><td>chr2_117458406_117459123</td><td>chr2</td><td>117458407</td><td>117459123</td><td>+</td><td>1.61352</td><td>Intergenic</td><td>Intergenic</td><td>-115888</td><td>NM_011246</td><td>19419</td><td>Mm.42150</td><td>NM_011246</td><td>ENSMUSG00000027347</td><td>Rasgrp1</td><td>Rasgrp</td><td>RAS guanyl releasing protein 1</td><td>protein-coding</td></tr>
<tr><td>chr2_117451730_117452199</td><td>chr2</td><td>117451731</td><td>117452199</td><td>+</td><td>1.08524</td><td>Intergenic</td><td>Intergenic</td><td>-109088</td><td>NM_011246</td><td>19419</td><td>Mm.42150</td><td>NM_011246</td><td>ENSMUSG00000027347</td><td>Rasgrp1</td><td>Rasgrp</td><td>RAS guanyl releasing protein 1</td><td>protein-coding</td></tr>
<tr><td>chr2_117354778_117355210</td><td>chr2</td><td>117354779</td><td>117355210</td><td>+</td><td>0.845178</td><td>Intergenic</td><td>Intergenic</td><td>-12117</td><td>NM_011246</td><td>19419</td><td>Mm.42150</td><td>NM_011246</td><td>ENSMUSG00000027347</td><td>Rasgrp1</td><td>Rasgrp</td><td>RAS guanyl releasing protein 1</td><td>protein-coding</td></tr>
<tr><td>chr2_117459773_117460197</td><td>chr2</td><td>117459774</td><td>117460197</td><td>+</td><td>0.830783</td><td>Intergenic</td><td>Intergenic</td><td>-117108</td><td>NM_011246</td><td>19419</td><td>Mm.42150</td><td>NM_011246</td><td>ENSMUSG00000027347</td><td>Rasgrp1</td><td>Rasgrp</td><td>RAS guanyl releasing protein 1</td><td>protein-coding</td></tr>
<tr><td>chr2_117432663_117433278</td><td>chr2</td><td>117432664</td><td>117433278</td><td>+</td><td>0.726913</td><td>Intergenic</td><td>Intergenic</td><td>-90094</td><td>NM_011246</td><td>19419</td><td>Mm.42150</td><td>NM_011246</td><td>ENSMUSG00000027347</td><td>Rasgrp1</td><td>Rasgrp</td><td>RAS guanyl releasing protein 1</td><td>protein-coding</td></tr>
<tr><td>chr2_117431704_117433336</td><td>chr2</td><td>117431705</td><td>117433336</td><td>+</td><td>0.613166</td><td>Intergenic</td><td>Intergenic</td><td>-89643</td><td>NM_011246</td><td>19419</td><td>Mm.42150</td><td>NM_011246</td><td>ENSMUSG00000027347</td><td>Rasgrp1</td><td>Rasgrp</td><td>RAS guanyl releasing protein 1</td><td>protein-coding</td></tr>
<tr><td>chr2_117461438_117461916</td><td>chr2</td><td>117461439</td><td>117461916</td><td>+</td><td>0.607906</td><td>Intergenic</td><td>Intergenic</td><td>-118800</td><td>NM_011246</td><td>19419</td><td>Mm.42150</td><td>NM_011246</td><td>ENSMUSG00000027347</td><td>Rasgrp1</td><td>Rasgrp</td><td>RAS guanyl releasing protein 1</td><td>protein-coding</td></tr>
<tr><td>chr17_75504841_75505550</td><td>chr17</td><td>75504842</td><td>75505550</td><td>+</td><td>0.809941</td><td>intron (NM_207246, intron 10 of 17)</td><td>intron (NM_207246, intron 10 of 17)</td><td>39658</td><td>NM_207246</td><td>240168</td><td>Mm.329203</td><td>NM_207246</td><td>ENSMUSG00000071042</td><td>Rasgrp3</td><td>BC066069|Gm327</td><td>RAS, guanyl releasing protein 3</td><td>protein-coding</td></tr>
<tr><td>chr5_74348206_74349034</td><td>chr5</td><td>74348207</td><td>74349034</td><td>+</td><td>1.50916</td><td>intron (NM_001286437, intron 5 of 7)</td><td>intron (NM_001286437, intron 5 of 7)</td><td>153294</td><td>NM_026878</td><td>68939</td><td>Mm.293316</td><td>NM_026878</td><td>ENSMUSG00000049907</td><td>Rasl11b</td><td>1190017B18Rik</td><td>RAS-like, family 11, member B</td><td>protein-coding</td></tr>
<tr><td>chr5_74261591_74262006</td><td>chr5</td><td>74261592</td><td>74262006</td><td>+</td><td>0.796435</td><td>intron (NM_001286437, intron 5 of 7)</td><td>intron (NM_001286437, intron 5 of 7)</td><td>66473</td><td>NM_026878</td><td>68939</td><td>Mm.293316</td><td>NM_026878</td><td>ENSMUSG00000049907</td><td>Rasl11b</td><td>1190017B18Rik</td><td>RAS-like, family 11, member B</td><td>protein-coding</td></tr>
<tr><td>chr5_74349758_74350008</td><td>chr5</td><td>74349759</td><td>74350008</td><td>+</td><td>0.739529</td><td>intron (NM_001286437, intron 5 of 7)</td><td>intron (NM_001286437, intron 5 of 7)</td><td>154557</td><td>NM_026878</td><td>68939</td><td>Mm.293316</td><td>NM_026878</td><td>ENSMUSG00000049907</td><td>Rasl11b</td><td>1190017B18Rik</td><td>RAS-like, family 11, member B</td><td>protein-coding</td></tr>
<tr><td>chr10_121517450_121517789</td><td>chr10</td><td>121517451</td><td>121517789</td><td>+</td><td>0.928589</td><td>Intergenic</td><td>ORR1B1|LTR|ERVL-MaLR</td><td>-41370</td><td>NM_138956</td><td>192678</td><td>Mm.41265</td><td>NM_138956</td><td>ENSMUSG00000025795</td><td>Rassf3</td><td>AW212023|AW322379</td><td>Ras association (RalGDS/AF-6) domain family member 3</td><td>protein-coding</td></tr>
</table>

| | | | | | | | | | | | | | | | | | |
|---|---|---|---|---|---|---|---|---|---|---|---|---|---|---|---|---|---|
| chr10_121520837_121521527 | chr10 | 121520838 | 121521527 | + | 0.856382 | Intergenic | Intergenic | -44932 | NM_138956 | 192678 | Mm.41265 | NM_138956 | ENSMUSG00000025795 | **Rassf3** | AW212023\|AW322379 | Ras association (RalGDS/AF-6) domain family member 3 | protein-coding |
| chr10_121443756_121444396 | chr10 | 121443757 | 121444396 | + | 0.614148 | intron (NM_138956, intron 1 of 4) | intron (NM_138956, intron 1 of 4) | 32174 | NM_138956 | 192678 | Mm.41265 | NM_138956 | ENSMUSG00000025795 | **Rassf3** | AW212023\|AW322379 | Ras association (RalGDS/AF-6) domain family member 3 | protein-coding |
| chr4_101141857_101142174 | chr4 | 101141858 | 101142174 | + | 0.747216 | intron (NM_183024, intron 11 of 11) | intron (NM_183024, intron 11 of 11) | 72978 | NM_183024 | 242570 | Mm.185636 | NM_183024 | ENSMUSG00000035275 | **Raver2** | A430091O22Rik | ribonucleoprotein, PTB-binding 2 | protein-coding |
| chr4_129324807_129325189 | chr4 | 129324808 | 129325189 | + | 0.832166 | intron (NM_009030, intron 3 of 11) | B3\|SINE\|B2 | 10372 | NM_009030 | 19646 | Mm.12145 | NM_009030 | ENSMUSG00000057236 | **Rbbp4** | mRbAp48 | retinoblastoma binding protein 4 | protein-coding |
| chr7_123025189_123025493 | chr7 | 123025190 | 123025493 | + | 0.910279 | Intergenic | Intergenic | 54777 | NM_175023 | 19647 | Mm.4480 | NM_011247 | ENSMUSG00000030779 | **Rbbp6** | 4933422O15Rik\|AI316869\|BB233631\|C030034J04Rik\|C77662\|P2P-R\|PACT | retinoblastoma binding protein 6 | protein-coding |
| chr7_123025162_123025777 | chr7 | 123025163 | 123025777 | + | 0.708267 | Intergenic | ID_B1\|SINE\|B4 | 54906 | NM_175023 | 19647 | Mm.4480 | NM_011247 | ENSMUSG00000030779 | **Rbbp6** | 4933422O15Rik\|AI316869\|BB233631\|C030034J04Rik\|C77662\|P2P-R\|PACT | retinoblastoma binding protein 6 | protein-coding |
| chr18_11659775_11659937 | chr18 | 11659776 | 11659937 | + | 0.828865 | intron (NR_045526, intron 1 of 17) | B1F\|SINE\|Alu | 2507 | NM_001081223 | 225182 | Mm.154275 | NM_175458 | ENSMUSG00000041238 | **Rbbp8** | 9930104E21Rik\|CtIP\|RBBP-8\|RIM\|SAE2 | retinoblastoma binding protein 8 | protein-coding |
| chr2_180307746_180308070 | chr2 | 180307747 | 180308070 | + | 0.839949 | Intergenic | Intergenic | -18029 | NM_173031 | 271887 | Mm.246640 | NM_173031 | ENSMUSG00000038980 | **Rbbp8nl** | 4732416F18 | RBBP8 N-terminal like | protein-coding |
| chr2_180307745_180308469 | chr2 | 180307746 | 180308469 | + | 0.750487 | Intergenic | Intergenic | -18228 | NM_173031 | 271887 | Mm.246640 | NM_173031 | ENSMUSG00000038980 | **Rbbp8nl** | 4732416F18 | RBBP8 N-terminal like | protein-coding |
| chr8_91073567_91074265 | chr8 | 91073568 | 91074265 | + | 0.964106 | intron (NM_001282001, intron 1 of 21) | MLT1G3\|LTR\|ERVL-MaLR | 3859 | NM_001282000 | 19651 | Mm.235580 | NM_011250 | ENSMUSG00000031666 | **Rbl2** | PRB2\|RBR-2\|Rb2\|p130 | retinoblastoma-like 2 | protein-coding |
| chr2_156089650_156090264 | chr2 | 156089651 | 156090264 | + | 0.662875 | intron (NM_170590, intron 1 of 15) | intron (NM_170590, intron 1 of 15) | 22008 | NM_170598 | 75710 | Mm.27660 | NM_029397 | | **Rbm12** | 5730420G12Rik\|9430070C08Rik\|AI852903\|SWAN\|mKIAA0765 | RNA binding motif protein 12 | protein-coding |
| chr19_4803826_4804625 | chr19 | 4803827 | 4804625 | + | 0.782274 | intron (NM_019869, intron 1 of 2) | intron (NM_019869, intron 1 of 2) | 7408 | NM_001290127 | 102902673 | Mm.426069 | NM_001290127 | | **Rbm14-rbm4** | - | Rbm14-Rbm4 readthrough | protein-coding |
| chr2_11620263_11620658 | chr2 | 11620264 | 11620658 | + | 1.25808 | Intergenic | Intergenic | -17262 | NM_152824 | 76938 | Mm.182769 | NM_152824 | ENSMUSG00000037197 | **Rbm17** | 2700027J02Rik | RNA binding motif protein 17 | protein-coding |
| chr2_11618662_11618950 | chr2 | 11618663 | 11618950 | + | 1.00572 | Intergenic | Intergenic | -15607 | NM_152824 | 76938 | Mm.182769 | NM_152824 | ENSMUSG00000037197 | **Rbm17** | 2700027J02Rik | RNA binding motif protein 17 | protein-coding |
| chr2_11616617_11617394 | chr2 | 11616618 | 11617394 | + | 0.782362 | Intergenic | Intergenic | -13807 | NM_152824 | 76938 | Mm.182769 | NM_152824 | ENSMUSG00000037197 | **Rbm17** | 2700027J02Rik | RNA binding motif protein 17 | protein-coding |
| chr19_53740738_53741337 | chr19 | 53740739 | 53741337 | + | 0.931452 | intron (NM_001170847, intron 1 of 13) | intron (NM_001170847, intron 1 of 13) | 63732 | NM_001170847 | 73713 | Mm.313695 | NM_001170847 | ENSMUSG00000043639 | **Rbm20** | 1110018J23Rik\|2010003H22Rik\|AI646761 | RNA binding motif protein 20 | protein-coding |
| chr14_105179879_105180114 | chr14 | 105179880 | 105180114 | + | 0.763058 | Intergenic | Intergenic | -2670 | NM_134077 | 74213 | Mm.474531 | NM_134077 | ENSMUSG00000022119 | **Rbm26** | 1700009P03Rik\|AI447318\|C230097K14Rik\|C86225\|MLZ-393\|Pro1777\|Se70-2 | RNA binding motif protein 26 | protein-coding |
| chr18_42307857_42308455 | chr18 | 42307858 | 42308455 | + | 1.57429 | intron (NM_172626, intron 8 of 19) | intron (NM_172626, intron 8 of 19) | 32803 | NM_172626 | 225432 | Mm.329400 | NM_172626 | ENSMUSG00000024491 | **Rbm27** | A730010I06\|AI043120\|Psc1\|mKIAA1311 | RNA binding motif protein 27 | protein-coding |
| chr2_173047951_173048646 | chr2 | 173047952 | 173048646 | + | 0.666054 | Intergenic | Intergenic | 26397 | NM_019547 | 56190 | Mm.485782 | NM_019547 | ENSMUSG00000027510 | **Rbm38** | Rnpc1\|Seb4\|Seb4l | RNA binding motif protein 38 | protein-coding |
| chr2_173047973_173048394 | chr2 | 173047974 | 173048394 | + | 0.60591 | Intergenic | Intergenic | 26282 | NM_019547 | 56190 | Mm.485782 | NM_019547 | ENSMUSG00000027510 | **Rbm38** | Rnpc1\|Seb4\|Seb4l | RNA binding motif protein 38 | protein-coding |
| chr19_4797788_4798253 | chr19 | 4797789 | 4798253 | + | 1.26356 | intron (NM_001290127, intron 1 of 3) | intron (NM_001290127, intron 1 of 3) | -4012 | NM_001290123 | 19653 | Mm.426069 | NM_009032 | ENSMUSG00000094936 | **Rbm4** | 4921506I22Rik\|Lark1\|Mlark\|Rbm4a\|lark | RNA binding motif protein 4 | protein-coding |
| chr1_91153309_91153714 | chr1 | 91153310 | 91153714 | + | 0.710672 | exon (NM_001033408, exon 3 of 16) | exon (NM_001033408, exon 3 of 16) | 8409 | NM_001033408 | 329207 | Mm.295940 | NM_001033408 | ENSMUSG00000070732 | **Rbm44** | Gm817 | RNA binding motif protein 44 | protein-coding |
| chr5_66168691_66169571 | chr5 | 66168692 | 66169571 | + | 1.31469 | intron (NR_040695, intron 2 of 2) | RSINE1\|SINE\|B4 | -17177 | NM_001127382 | 245945 | Mm.36863 | NM_139065 | ENSMUSG00000070780 | **Rbm47** | 9530077J19Rik | RNA binding motif protein 47 | protein-coding |
| chr5_66169026_66169548 | chr5 | 66169027 | 66169548 | + | 1.26743 | intron (NR_040695, intron 2 of 2) | RSINE1\|SINE\|B4 | -17333 | NM_001127382 | 245945 | Mm.36863 | NM_139065 | ENSMUSG00000070780 | **Rbm47** | 9530077J19Rik | RNA binding motif protein 47 | protein-coding |
| chr5_66168656_66169836 | chr5 | 66168657 | 66169836 | + | 1.1327 | intron (NR_040695, intron 2 of 2) | RSINE1\|SINE\|B4 | -17292 | NM_001127382 | 245945 | Mm.36863 | NM_139065 | ENSMUSG00000070780 | **Rbm47** | 9530077J19Rik | RNA binding motif protein 47 | protein-coding |
| chr5_66137222_66137698 | chr5 | 66137223 | 66137698 | + | 0.923126 | intron (NM_001127382, intron 2 of 7) | intron (NM_001127382, intron 2 of 7) | 13518 | NM_001291226 | 245945 | Mm.36863 | NM_139065 | ENSMUSG00000070780 | **Rbm47** | 9530077J19Rik | RNA binding motif protein 47 | protein-coding |
| chr2_60867109_60867821 | chr2 | 60867110 | 60867821 | + | 0.771848 | intron (NM_020296, intron 1 of 13) | intron (NM_020296, intron 1 of 13) | 13973 | NM_001141932 | 56878 | Mm.259667 | NM_020296 | ENSMUSG00000026970 | **Rbms1** | 2600014B10Rik\|AI255215\|MSSP-1\|MSSP-2\|MSSP-3\|YC1 | RNA binding motif, single stranded interacting protein 1 | protein-coding |
| chr2_60878066_60878560 | chr2 | 60878067 | 60878560 | + | 0.655597 | intron (NM_020296, intron 1 of 13) | intron (NM_020296, intron 1 of 13) | 3125 | NM_001141932 | 56878 | Mm.259667 | NM_020296 | ENSMUSG00000026970 | **Rbms1** | 2600014B10Rik\|AI255215\|MSSP-1\|MSSP-2\|MSSP-3\|YC1 | RNA binding motif, single stranded interacting protein 1 | protein-coding |
| chr2_60877786_60878633 | chr2 | 60877787 | 60878633 | + | 0.613316 | intron (NM_020296, intron 1 of 13) | intron (NM_020296, intron 1 of 13) | 3228 | NM_001141932 | 56878 | Mm.259667 | NM_020296 | ENSMUSG00000026970 | **Rbms1** | 2600014B10Rik\|AI255215\|MSSP-1\|MSSP-2\|MSSP-3\|YC1 | RNA binding motif, single stranded interacting protein 1 | protein-coding |
| chrX_48778050_48778982 | chrX | 48778051 | 48778982 | + | 0.661694 | Intergenic | Lx6\|LINE\|L1 | 83512 | NM_173376 | 209003 | Mm.32561 | NM_173376 | ENSMUSG00000031107 | **Rbmx2** | 2700089A17Rik\|2810411A03Rik\|CGI-79 | RNA binding motif protein, X-linked 2 | protein-coding |
| chr5_53543491_53543852 | chr5 | 53543492 | 53543852 | + | 1.26277 | Intergenic | Intergenic | -12107 | NM_001080928 | 19664 | Mm.209292 | NM_009035 | ENSMUSG00000039191 | **Rbpj** | AI843960\|CBF1\|Igkjrb\|Igkrsbp\|RBP-J\|RBP-J kappa\|RBPjk\|Rbpsuh | recombination signal binding protein for immunoglobulin kappa J region | protein-coding |
| chr5_53413599_53414119 | chr5 | 53413600 | 53414119 | + | 1.25877 | Intergenic | Intergenic | -141920 | NM_001080928 | 19664 | Mm.209292 | NM_009035 | ENSMUSG00000039191 | **Rbpj** | AI843960\|CBF1\|Igkjrb\|Igkrsbp\|RBP-J\|RBP-J kappa\|RBPjk\|Rbpsuh | recombination signal binding protein for immunoglobulin kappa J region | protein-coding |
| chr5_53543095_53543840 | chr5 | 53543096 | 53543840 | + | 1.10711 | Intergenic | Intergenic | -12311 | NM_001080928 | 19664 | Mm.209292 | NM_009035 | ENSMUSG00000039191 | **Rbpj** | AI843960\|CBF1\|Igkjrb\|Igkrsbp\|RBP-J\|RBP-J kappa\|RBPjk\|Rbpsuh | recombination signal binding protein for immunoglobulin kappa J region | protein-coding |
| chr5_53553401_53553967 | chr5 | 53553402 | 53553967 | + | 1.06761 | Intergenic | Intergenic | -2095 | NM_001080928 | 19664 | Mm.209292 | NM_009035 | ENSMUSG00000039191 | **Rbpj** | AI843960\|CBF1\|Igkjrb\|Igkrsbp\|RBP-J\|RBP-J kappa\|RBPjk\|Rbpsuh | recombination signal binding protein for immunoglobulin kappa J region | protein-coding |
| chr5_53441093_53441664 | chr5 | 53441094 | 53441664 | + | 1.02088 | Intergenic | Intergenic | -114400 | NM_001080928 | 19664 | Mm.209292 | NM_009035 | ENSMUSG00000039191 | **Rbpj** | AI843960\|CBF1\|Igkjrb\|Igkrsbp\|RBP-J\|RBP-J kappa\|RBPjk\|Rbpsuh | recombination signal binding protein for immunoglobulin kappa J region | protein-coding |
| chr5_53490480_53491398 | chr5 | 53490481 | 53491398 | + | 0.895654 | Intergenic | Intergenic | -64840 | NM_001080928 | 19664 | Mm.209292 | NM_009035 | ENSMUSG00000039191 | **Rbpj** | AI843960\|CBF1\|Igkjrb\|Igkrsbp\|RBP-J\|RBP-J kappa\|RBPjk\|Rbpsuh | recombination signal binding protein for immunoglobulin kappa J region | protein-coding |
| chr5_53539980_53540638 | chr5 | 53539981 | 53540638 | + | 0.881305 | Intergenic | Intergenic | -15470 | NM_001080928 | 19664 | Mm.209292 | NM_009035 | ENSMUSG00000039191 | **Rbpj** | AI843960\|CBF1\|Igkjrb\|Igkrsbp\|RBP-J\|RBP-J kappa\|RBPjk\|Rbpsuh | recombination signal binding protein for immunoglobulin kappa J region | protein-coding |
| chr5_53421306_53421791 | chr5 | 53421307 | 53421791 | + | 0.819698 | Intergenic | Intergenic | -134230 | NM_001080928 | 19664 | Mm.209292 | NM_009035 | ENSMUSG00000039191 | **Rbpj** | AI843960\|CBF1\|Igkjrb\|Igkrsbp\|RBP-J\|RBP-J kappa\|RBPjk\|Rbpsuh | recombination signal binding protein for immunoglobulin kappa J region | protein-coding |
| chr5_53572505_53572867 | chr5 | 53572506 | 53572867 | + | 0.791896 | intron (NM_001080928, intron 1 of 10) | L1MB7\|LINE\|L1 | 16907 | NM_001080928 | 19664 | Mm.209292 | NM_009035 | ENSMUSG00000039191 | **Rbpj** | AI843960\|CBF1\|Igkjrb\|Igkrsbp\|RBP-J\|RBP-J kappa\|RBPjk\|Rbpsuh | recombination signal binding protein for immunoglobulin kappa J region | protein-coding |
| chr5_53420767_53421791 | chr5 | 53420768 | 53421791 | + | 0.739038 | Intergenic | Intergenic | -134500 | NM_001080928 | 19664 | Mm.209292 | NM_009035 | ENSMUSG00000039191 | **Rbpj** | AI843960\|CBF1\|Igkjrb\|Igkrsbp\|RBP-J\|RBP-J kappa\|RBPjk\|Rbpsuh | recombination signal binding protein for immunoglobulin kappa J region | protein-coding |
| chr5_53566480_53567282 | chr5 | 53566481 | 53567282 | + | 0.680814 | intron (NM_001080928, intron 1 of 10) | intron (NM_001080928, intron 1 of 10) | 11102 | NM_001080928 | 19664 | Mm.209292 | NM_009035 | ENSMUSG00000039191 | **Rbpj** | AI843960\|CBF1\|Igkjrb\|Igkrsbp\|RBP-J\|RBP-J kappa\|RBPjk\|Rbpsuh | recombination signal binding protein for immunoglobulin kappa J region | protein-coding |
| chr5_53566155_53567272 | chr5 | 53566156 | 53567272 | + | 0.674001 | intron (NM_001080928, intron 1 of 10) | intron (NM_001080928, intron 1 of 10) | 10935 | NM_001080928 | 19664 | Mm.209292 | NM_009035 | ENSMUSG00000039191 | **Rbpj** | AI843960\|CBF1\|Igkjrb\|Igkrsbp\|RBP-J\|RBP-J kappa\|RBPjk\|Rbpsuh | recombination signal binding protein for immunoglobulin kappa J region | protein-coding |
| chr8_33968233_33968779 | chr8 | 33968234 | 33968779 | + | 1.18322 | Intergenic | L1MB4\|LINE\|L1 | -38643 | NM_001042675 | 19663 | Mm.323997 | NM_019733 | ENSMUSG00000031586 | **Rbpms** | 2010300K22Rik\|2700019M19Rik\|AU017537\|RBP-MS\|hermes | RNA binding protein gene with multiple splicing | protein-coding |
| chr9_65622747_65623448 | chr9 | 65622748 | 65623448 | + | 0.798654 | Intergenic | Intergenic | -7484 | NM_028030 | 71973 | Mm.29148 | NM_028030 | ENSMUSG00000032387 | **Rbpms2** | 2400008B06Rik\|AI316523 | RNA binding protein with multiple splicing 2 | protein-coding |
| chr15_81437827_81438223 | chr15 | 81437828 | 81438223 | + | 0.609751 | intron (NM_177310, intron 5 of 8) | intron (NM_177310, intron 5 of 8) | -28291 | NM_019712 | 56438 | Mm.29405 | NM_019712 | ENSMUSG00000022400 | **Rbx1** | 1500002P15Rik\|AA517855\|ROC1 | ring-box 1 | protein-coding |
| chr16_92443497_92444206 | chr16 | 92443498 | 92444206 | + | 0.792398 | intron (NM_001081549, intron 1 of 3) | intron (NM_001081549, intron 1 of 3) | 22317 | NM_001081549 | 54720 | Mm.265744 | NM_019466 | ENSMUSG00000022951 | **Rcan1** | Adapt78\|CALP1L\|CSP1\|DSC1\|Dscr1\|MCIP1\|RCN1 | regulator of calcineurin 1 | protein-coding |
| chr4_135439988_135440515 | chr4 | 135439989 | 135440515 | + | 0.778391 | Intergenic | ORR1C1\|LTR\|ERVL-MaLR | -6447 | NM_022980 | 53902 | Mm.331970 | NM_022980 | ENSMUSG00000059713 | **Rcan3** | AU041093\|Csp3\|Dscr1l2 | regulator of calcineurin 3 | protein-coding |
| chr14_73169896_73170577 | chr14 | 73169897 | 73170577 | + | 1.01111 | intron (NM_001170694, intron 10 of 14) | intron (NM_001170694, intron 10 of 14) | 27452 | NR_033185 | 105670 | Mm.280068 | NM_134083 | ENSMUSG00000022106 | **Rcbtb2** | 2610028E02Rik\|2810420M18Rik\|AI448345\|AW240694\|Chc1l\|RC/BTB2 | regulator of chromosome condensation (RCC1) and BTB (POZ) domain containing protein 2 | protein-coding |

| | | | | | | | | | | | | | | | | | |
|---|---|---|---|---|---|---|---|---|---|---|---|---|---|---|---|---|---|
| chr14_73146636_73147075 | chr14 | 73146637 | 73147075 | + | 0.686076 | intron (NR_033185, intron 1 of 15) | RMER6A\|LTR\|ERVK | 4071 | NR_033185 | 105670 | Mm.280068 | NM_134083 | ENSMUSG00000022106 | **Rcbtb2** | 2610028E02Rik\|2810420M18Rik\|AI448345\|AW240694\|Chc1l\|RC/BTB2 | regulator of chromosome condensation (RCC1) and BTB (POZ) domain containing protein 2 | protein-coding |
| chr5_91973992_91974423 | chr5 | 91973993 | 91974423 | + | 1.3579 | Intergenic | Intergenic | -11140 | NM_026557 | 68098 | Mm.159453 | NM_026557 | ENSMUSG00000029397 | **Rchy1** | ARNIP\|CHIMP\|Pirh2\|Zfp363\|Znf363 | ring finger and CHY zinc finger domain containing 1 | protein-coding |
| chr5_91973992_91974579 | chr5 | 91973993 | 91974579 | + | 1.32864 | Intergenic | Intergenic | -11218 | NM_026557 | 68098 | Mm.159453 | NM_026557 | ENSMUSG00000029397 | **Rchy1** | ARNIP\|CHIMP\|Pirh2\|Zfp363\|Znf363 | ring finger and CHY zinc finger domain containing 1 | protein-coding |
| chr5_91961398_91961710 | chr5 | 91961399 | 91961710 | + | 0.764807 | intron (NR_073442, intron 1 of 9) | intron (NR_073442, intron 1 of 9) | 1002 | NR_073442 | 68098 | Mm.159453 | NM_026557 | ENSMUSG00000029397 | **Rchy1** | ARNIP\|CHIMP\|Pirh2\|Zfp363\|Znf363 | ring finger and CHY zinc finger domain containing 1 | protein-coding |
| chr5_91960010_91960267 | chr5 | 91960011 | 91960267 | + | 0.665272 | intron (NM_026557, intron 1 of 8) | Charlie1a\|DNA\|hAT-Charlie | 2417 | NR_073442 | 68098 | Mm.159453 | NM_026557 | ENSMUSG00000029397 | **Rchy1** | ARNIP\|CHIMP\|Pirh2\|Zfp363\|Znf363 | ring finger and CHY zinc finger domain containing 1 | protein-coding |
| chr12_111048232_111048731 | chr12 | 111048233 | 111048731 | + | 0.746685 | intron (NM_198023, intron 2 of 11) | B1_Mus1\|SINE\|Alu | 8684 | NM_198023 | 217864 | Mm.23808 | NM_198023 | ENSMUSG00000037896 | **Rcor1** | 5730409O11\|6720480E22Rik\|AU042633\|D12Wsu95e\|Rocr1\|mKIAA0071 | REST corepressor 1 | protein-coding |
| chr12_111048280_111048531 | chr12 | 111048281 | 111048531 | + | 0.725709 | intron (NM_198023, intron 2 of 11) | intron (NM_198023, intron 2 of 11) | 8608 | NM_198023 | 217864 | Mm.23808 | NM_198023 | ENSMUSG00000037896 | **Rcor1** | 5730409O11\|6720480E22Rik\|AU042633\|D12Wsu95e\|Rocr1\|mKIAA0071 | REST corepressor 1 | protein-coding |
| chr1_165722298_165723092 | chr1 | 165722299 | 165723092 | + | 1.07765 | Intergenic | Intergenic | -14601 | NM_001038846 | 226594 | Mm.265552 | NM_178593 | ENSMUSG00000040723 | **Rcsd1** | A430105K13Rik\|BC025872 | RCSD domain containing 1 | protein-coding |
| chr1_165689437_165690169 | chr1 | 165689438 | 165690169 | + | 0.842692 | intron (NM_001038846, intron 1 of 5) | intron (NM_001038846, intron 1 of 5) | 18291 | NM_178593 | 226594 | Mm.265552 | NM_178593 | ENSMUSG00000040723 | **Rcsd1** | A430105K13Rik\|BC025872 | RCSD domain containing 1 | protein-coding |
| chr1_191960858_191961292 | chr1 | 191960859 | 191961292 | + | 1.07888 | Intergenic | Intergenic | -16298 | NM_001177900 | 74023 | Mm.274393 | NM_023727 | ENSMUSG00000049353 | **Rd3** | 3322402L07Rik\|rd-3\|rd3 | retinal degeneration 3 | protein-coding |
| chr1_16227061_16227458 | chr1 | 16227062 | 16227458 | + | 0.740638 | Intergenic | Intergenic | 121378 | NM_133832 | 98711 | Mm.274376 | NM_133832 | | **Rdh10** | 3110069K09Rik\|4921506A21Rik\|AI875664\|AW549993\|D1Ertd762e\|m366Asp | retinol dehydrogenase 10 (all-trans) | protein-coding |
| chr9_20835534_20835951 | chr9 | 20835535 | 20835951 | + | 0.850835 | Intergenic | RLTR30B_MM\|LTR\|ERV1 | 17239 | NM_001030290 | 235033 | Mm.297661 | NM_001030290 | ENSMUSG00000053773 | **Rdh8** | Gm182\|prRDH | retinol dehydrogenase 8 | protein-coding |
| chr4_43897267_43897558 | chr4 | 43897268 | 43897558 | + | 0.955044 | intron (NM_016678, intron 4 of 20) | intron (NM_016678, intron 4 of 20) | 21883 | NM_016678 | 53614 | Mm.331573 | NM_016678 | ENSMUSG00000028476 | **Reck** | St15\|mRECK | reversion-inducing-cysteine-rich protein with kazal motifs | protein-coding |
| chr10_67100180_67100656 | chr10 | 67100181 | 67100656 | + | 1.34051 | Intergenic | Intergenic | -3430 | NM_178606 | 28193 | Mm.41272 | NM_178606 | ENSMUSG00000019873 | **Reep3** | D10Ucla1 | receptor accessory protein 3 | protein-coding |
| chr10_67102796_67103105 | chr10 | 67102797 | 67103105 | + | 0.796391 | Intergenic | L1MC1\|LINE\|L1 | -5963 | NM_178606 | 28193 | Mm.41272 | NM_178606 | ENSMUSG00000019873 | **Reep3** | D10Ucla1 | receptor accessory protein 3 | protein-coding |
| chr11_23754314_23754930 | chr11 | 23754315 | 23754930 | + | 1.07694 | intron (NM_009044, intron 3 of 9) | intron (NM_009044, intron 3 of 9) | 16348 | NM_009044 | 19696 | Mm.4869 | NM_009044 | ENSMUSG00000020275 | **Rel** | c-Rel | reticuloendotheliosis oncogene | protein-coding |
| chr11_23754317_23754672 | chr11 | 23754318 | 23754672 | + | 1.0162 | intron (NM_009044, intron 3 of 9) | intron (NM_009044, intron 3 of 9) | 16475 | NM_009044 | 19696 | Mm.4869 | NM_009044 | ENSMUSG00000020275 | **Rel** | c-Rel | reticuloendotheliosis oncogene | protein-coding |
| chr11_23820576_23820889 | chr11 | 23820577 | 23820889 | + | 0.796011 | Intergenic | Intergenic | -49763 | NM_009044 | 19696 | Mm.4869 | NM_009044 | ENSMUSG00000020275 | **Rel** | c-Rel | reticuloendotheliosis oncogene | protein-coding |
| chr11_23772878_23773127 | chr11 | 23772879 | 23773127 | + | 0.782674 | Intergenic | Intergenic | -2033 | NM_009044 | 19696 | Mm.4869 | NM_009044 | ENSMUSG00000020275 | **Rel** | c-Rel | reticuloendotheliosis oncogene | protein-coding |
| chr11_23734886_23735555 | chr11 | 23734887 | 23735555 | + | 0.706737 | Intergenic | Intergenic | 35749 | NM_009044 | 19696 | Mm.4869 | NM_009044 | ENSMUSG00000020275 | **Rel** | c-Rel | reticuloendotheliosis oncogene | protein-coding |
| chr11_23823000_23823633 | chr11 | 23823001 | 23823633 | + | 0.63932 | Intergenic | Intergenic | -52347 | NM_009044 | 19696 | Mm.4869 | NM_009044 | ENSMUSG00000020275 | **Rel** | c-Rel | reticuloendotheliosis oncogene | protein-coding |
| chr11_23734142_23735569 | chr11 | 23734143 | 23735569 | + | 0.625126 | Intergenic | Intergenic | 36114 | NM_009044 | 19696 | Mm.4869 | NM_009044 | ENSMUSG00000020275 | **Rel** | c-Rel | reticuloendotheliosis oncogene | protein-coding |
| chr11_23749883_23750276 | chr11 | 23749884 | 23750276 | + | 0.601689 | intron (NM_009044, intron 4 of 9) | B4\|SINE\|B4 | 20890 | NM_009044 | 19696 | Mm.4869 | NM_009044 | ENSMUSG00000020275 | **Rel** | c-Rel | reticuloendotheliosis oncogene | protein-coding |
| chrX_162541712_162542391 | chrX | 162541713 | 162542391 | + | 1.01555 | intron (NM_178256, intron 5 of 17) | intron (NM_178256, intron 5 of 17) | 101606 | NM_001290633 | 194590 | Mm.442441 | NM_178256 | ENSMUSG00000040855 | **Reps2** | POB1 | RALBP1 associated Eps domain containing protein 2 | protein-coding |
| chr4_150351211_150352052 | chr4 | 150351212 | 150352052 | + | 0.803741 | intron (NM_001085492, intron 1 of 22) | intron (NM_001085492, intron 1 of 22) | 69716 | NM_001085492 | 68703 | Mm.291274 | NM_001085492 | ENSMUSG00000039852 | **Rere** | 1110033A15Rik\|AI414665\|ARG\|ARP\|ATN1L\|AW742570\|Atr2\|DNB1\|eye<m03Jus>\|eyes3\|mKIAA0458 | arginine glutamic acid dipeptide (RE) repeats | protein-coding |
| chr4_150351689_150352052 | chr4 | 150351690 | 150352052 | + | 0.797608 | intron (NM_001085492, intron 1 of 22) | intron (NM_001085492, intron 1 of 22) | 69955 | NM_001085492 | 68703 | Mm.291274 | NM_001085492 | ENSMUSG00000039852 | **Rere** | 1110033A15Rik\|AI414665\|ARG\|ARP\|ATN1L\|AW742570\|Atr2\|DNB1\|eye<m03Jus>\|eyes3\|mKIAA0458 | arginine glutamic acid dipeptide (RE) repeats | protein-coding |
| chr4_150450016_150450789 | chr4 | 150450017 | 150450789 | + | 0.788269 | intron (NM_001085492, intron 4 of 22) | intron (NM_001085492, intron 4 of 22) | 168487 | NM_001085492 | 68703 | Mm.291274 | NM_001085492 | ENSMUSG00000039852 | **Rere** | 1110033A15Rik\|AI414665\|ARG\|ARP\|ATN1L\|AW742570\|Atr2\|DNB1\|eye<m03Jus>\|eyes3\|mKIAA0458 | arginine glutamic acid dipeptide (RE) repeats | protein-coding |
| chr4_150379850_150380105 | chr4 | 150379851 | 150380105 | + | 0.619525 | intron (NM_001085492, intron 1 of 22) | intron (NM_001085492, intron 1 of 22) | 98062 | NM_001085492 | 68703 | Mm.291274 | NM_001085492 | ENSMUSG00000039852 | **Rere** | 1110033A15Rik\|AI414665\|ARG\|ARP\|ATN1L\|AW742570\|Atr2\|DNB1\|eye<m03Jus>\|eyes3\|mKIAA0458 | arginine glutamic acid dipeptide (RE) repeats | protein-coding |
| chr5_77252126_77252915 | chr5 | 77252127 | 77252915 | + | 1.73926 | Intergenic | B2_Mm2\|SINE\|B2 | -12973 | NM_011263 | 19712 | Mm.28840 | NM_011263 | ENSMUSG00000029249 | **Rest** | 2610008J04Rik\|AA407358\|NRSF | RE1-silencing transcription factor | protein-coding |
| chr5_77253234_77253514 | chr5 | 77253235 | 77253514 | + | 0.759738 | Intergenic | B3\|SINE\|B2 | -12120 | NM_011263 | 19712 | Mm.28840 | NM_011263 | ENSMUSG00000029249 | **Rest** | 2610008J04Rik\|AA407358\|NRSF | RE1-silencing transcription factor | protein-coding |
| chr6_118196746_118197639 | chr6 | 118196747 | 118197639 | + | 0.630119 | 5' UTR (NM_001080780, exon 1 of 19) | 5' UTR (NM_001080780, exon 1 of 19) | 551 | NM_009050 | 19713 | Mm.57199 | NM_009050 | ENSMUSG00000030110 | **Ret** | PTC\|RET51\|RET9\|c-Ret | ret proto-oncogene | protein-coding |
| chr16_48798270_48798645 | chr16 | 48798271 | 48798645 | + | 1.14307 | Intergenic | Intergenic | -18398 | NM_023881 | 57263 | Mm.21123 | NM_023881 | ENSMUSG00000022650 | **Retnlb** | 9030012B21Rik\|Fizz2\|RELMbeta\|Relmb\|Xcp3 | resistin like beta | protein-coding |
| chr10_39742097_39742374 | chr10 | 39742098 | 39742374 | + | 0.979363 | intron (NM_011264, intron 1 of 31) | intron (NM_011264, intron 1 of 31) | 10076 | NM_011264 | 19714 | Mm.439723 | NM_011264 | ENSMUSG00000019841 | **Rev3l** | Rev\|Rev3\|Sez4 | REV3-like, catalytic subunit of DNA polymerase zeta RAD54 like (S. cerevisiae) | protein-coding |
| chr10_39815131_39815522 | chr10 | 39815132 | 39815522 | + | 0.647406 | intron (NM_011264, intron 10 of 31) | intron (NM_011264, intron 10 of 31) | 83167 | NM_011264 | 19714 | Mm.439723 | NM_011264 | ENSMUSG00000019841 | **Rev3l** | Rev\|Rev3\|Sez4 | REV3-like, catalytic subunit of DNA polymerase zeta RAD54 like (S. cerevisiae) | protein-coding |
| chr16_23129973_23130357 | chr16 | 23129974 | 23130357 | + | 0.836552 | Intergenic | Intergenic | -2435 | NM_145480 | 106344 | Mm.386835 | NM_145480 | ENSMUSG00000022881 | **Rfc4** | A1\|AI894123\|AU040575\|RFC37 | replication factor C (activator 1) 4 | protein-coding |
| chr19_17385471_17385816 | chr19 | 17385472 | 17385816 | + | 1.13408 | Intergenic | Intergenic | -8399 | NM_019437 | 54391 | Mm.7013 | NM_019437 | ENSMUSG00000024712 | **Rfk** | 0610038L10Rik\|AF031381\|KOI-4 | riboflavin kinase | protein-coding |
| chr19_17385086_17385404 | chr19 | 17385087 | 17385404 | + | 0.864742 | Intergenic | Intergenic | -8798 | NM_019437 | 54391 | Mm.7013 | NM_019437 | ENSMUSG00000024712 | **Rfk** | 0610038L10Rik\|AF031381\|KOI-4 | riboflavin kinase | protein-coding |
| chr17_50086032_50086577 | chr17 | 50086033 | 50086577 | + | 1.80069 | intron (NM_181397, intron 4 of 9) | RLTR20A\|LTR\|ERVK | 104192 | NM_181397 | 76438 | Mm.41854 | NM_181397 | | **Rftn1** | 2310015N21Rik\|AI853526 | raftlin lipid raft linker 1 | protein-coding |
| chr17_50085706_50085962 | chr17 | 50085707 | 50085962 | + | 1.24632 | intron (NM_181397, intron 4 of 9) | intron (NM_181397, intron 4 of 9) | 104663 | NM_181397 | 76438 | Mm.41854 | NM_181397 | | **Rftn1** | 2310015N21Rik\|AI853526 | raftlin lipid raft linker 1 | protein-coding |
| chr17_50083936_50084341 | chr17 | 50083937 | 50084341 | + | 0.941367 | intron (NM_181397, intron 4 of 9) | intron (NM_181397, intron 4 of 9) | 106358 | NM_181397 | 76438 | Mm.41854 | NM_181397 | | **Rftn1** | 2310015N21Rik\|AI853526 | raftlin lipid raft linker 1 | protein-coding |
| chr17_50085090_50085390 | chr17 | 50085091 | 50085390 | + | 0.867528 | intron (NM_181397, intron 4 of 9) | intron (NM_181397, intron 4 of 9) | 105257 | NM_181397 | 76438 | Mm.41854 | NM_181397 | | **Rftn1** | 2310015N21Rik\|AI853526 | raftlin lipid raft linker 1 | protein-coding |
| chr17_49992545_49992975 | chr17 | 49992546 | 49992975 | + | 0.744572 | TTS (NM_181397) | TTS (NM_181397) | 197737 | NM_181397 | 76438 | Mm.41854 | NM_181397 | | **Rftn1** | 2310015N21Rik\|AI853526 | raftlin lipid raft linker 1 | protein-coding |
| chr17_49992531_49993447 | chr17 | 49992532 | 49993447 | + | 0.718731 | TTS (NM_181397) | TTS (NM_181397) | 197508 | NM_181397 | 76438 | Mm.41854 | NM_181397 | | **Rftn1** | 2310015N21Rik\|AI853526 | raftlin lipid raft linker 1 | protein-coding |
| chr17_50181799_50182165 | chr17 | 50181800 | 50182165 | + | 0.675261 | intron (NM_181397, intron 1 of 9) | intron (NM_181397, intron 1 of 9) | 8515 | NM_181397 | 76438 | Mm.41854 | NM_181397 | | **Rftn1** | 2310015N21Rik\|AI853526 | raftlin lipid raft linker 1 | protein-coding |
| chr17_50097605_50098530 | chr17 | 50097606 | 50098530 | + | 0.622539 | intron (NM_181397, intron 3 of 9) | intron (NM_181397, intron 3 of 9) | 92429 | NM_181397 | 76438 | Mm.41854 | NM_181397 | | **Rftn1** | 2310015N21Rik\|AI853526 | raftlin lipid raft linker 1 | protein-coding |
| chr17_50097790_50098343 | chr17 | 50097791 | 50098343 | + | 0.60314 | intron (NM_181397, intron 3 of 9) | intron (NM_181397, intron 3 of 9) | 92430 | NM_181397 | 76438 | Mm.41854 | NM_181397 | | **Rftn1** | 2310015N21Rik\|AI853526 | raftlin lipid raft linker 1 | protein-coding |
| chr17_56819240_56819507 | chr17 | 56819241 | 56819507 | + | 0.766474 | intron (NM_027787, intron 1 of 17) | intron (NM_027787, intron 1 of 17) | 11634 | NM_009056 | 19725 | Mm.102 | NM_009056 | ENSMUSG00000024206 | **Rfx2** | 5430432H19Rik | regulatory factor X, 2 (influences HLA class II expression) | protein-coding |
| chr17_56819108_56819513 | chr17 | 56819109 | 56819513 | + | 0.722161 | intron (NM_027787, intron 1 of 17) | intron (NM_027787, intron 1 of 17) | 11697 | NM_009056 | 19725 | Mm.102 | NM_009056 | ENSMUSG00000024206 | **Rfx2** | 5430432H19Rik | regulatory factor X, 2 (influences HLA class II expression) | protein-coding |
| chr19_27872249_27872625 | chr19 | 27872250 | 27872625 | + | 0.718439 | intron (NM_001166414, intron 3 of 17) | L1_Mus3\|LINE\|L1 | 110511 | NM_001166414 | 19726 | Mm.313321 | NM_011265 | ENSMUSG00000040929 | **Rfx3** | C230093O12Rik\|MRFX3 | regulatory factor X, 3 (influences HLA class II expression) | protein-coding |
| chr19_27906928_27907323 | chr19 | 27906929 | 27907323 | + | 0.685606 | intron (NM_011265, intron 1 of 16) | intron (NM_011265, intron 1 of 16) | 75822 | NM_001166414 | 19726 | Mm.313321 | NM_011265 | ENSMUSG00000040929 | **Rfx3** | C230093O12Rik\|MRFX3 | regulatory factor X, 3 (influences HLA class II expression) | protein-coding |
| chr19_27976326_27976662 | chr19 | 27976327 | 27976662 | + | 0.658829 | intron (NM_001166414, intron 1 of 17) | ORR1D2\|LTR\|ERVL-MaLR | 6454 | NM_001166414 | 19726 | Mm.313321 | NM_011265 | ENSMUSG00000040929 | **Rfx3** | C230093O12Rik\|MRFX3 | regulatory factor X, 3 (influences HLA class II expression) | protein-coding |
| chr10_84816016_84816617 | chr10 | 84816017 | 84816617 | + | 0.995314 | intron (NM_001024918, intron 4 of 17) | MER3\|DNA\|hAT-Charlie | -21831 | NM_027689 | 71137 | Mm.79070 | NM_027689 | ENSMUSG00000020037 | **Rfx4** | 4933412G19Rik\|NYD-sp10 | regulatory factor X, 4 (influences HLA class II expression) | protein-coding |

| | | | | | | | | | | | | | | | | | |
|---|---|---|---|---|---|---|---|---|---|---|---|---|---|---|---|---|---|
| chr10_84816101_84816433 | chr10 | 84816102 | 84816433 | + | 0.980454 | intron (NM_001024918, intron 4 of 17) | MER3\|DNA\|hAT-Charlie | -21881 | NM_027689 | 71137 | Mm.79070 | NM_027689 | ENSMUSG00000020037 | **Rfx4** | 4933412G19Rik\|NYD-sp10 | regulatory factor X, 4 (influences HLA class II expression) | protein-coding |
| chr1_39743702_39744283 | chr1 | 39743703 | 39744283 | + | 1.14281 | Intergenic | Intergenic | -23004 | NM_001145660 | 619289 | Mm.334107 | NM_001145660 | ENSMUSG00000057173 | **Rfx8** | 4933400N17Rik | regulatory factor X 8 | protein-coding |
| chr1_39714744_39715071 | chr1 | 39714745 | 39715071 | + | 0.828556 | intron (NM_001145660, intron 2 of 14) | intron (NM_001145660, intron 2 of 14) | 6081 | NM_001145660 | 619289 | Mm.334107 | NM_001145660 | ENSMUSG00000057173 | **Rfx8** | 4933400N17Rik | regulatory factor X 8 | protein-coding |
| chr14_79290553_79291084 | chr14 | 79290554 | 79291084 | + | 0.782828 | intron (NM_025427, intron 3 of 4) | intron (NM_025427, intron 3 of 4) | 10816 | NM_025427 | 66214 | Mm.29811 | NM_025427 | ENSMUSG00000022018 | **Rgcc** | 1190002H23Rik\|Rgc-32\|Rgc32 | regulator of cell cycle | protein-coding |
| chr1_144316257_144316834 | chr1 | 144316258 | 144316834 | + | 1.99064 | Intergenic | Intergenic | -67442 | NM_015811 | 50778 | Mm.103701 | NM_015811 | ENSMUSG00000026358 | **Rgs1** | BL34 | regulator of G-protein signaling 1 | protein-coding |
| chr1_144238954_144239638 | chr1 | 144238955 | 144239638 | + | 1.88396 | Intergenic | Lx8b\|LINE\|L1 | 9808 | NM_015811 | 50778 | Mm.103701 | NM_015811 | ENSMUSG00000026358 | **Rgs1** | BL34 | regulator of G-protein signaling 1 | protein-coding |
| chr1_144241397_144241928 | chr1 | 144241398 | 144241928 | + | 1.61973 | Intergenic | Intergenic | 7441 | NM_015811 | 50778 | Mm.103701 | NM_015811 | ENSMUSG00000026358 | **Rgs1** | BL34 | regulator of G-protein signaling 1 | protein-coding |
| chr1_144263980_144264515 | chr1 | 144263981 | 144264515 | + | 1.46052 | Intergenic | L2a\|LINE\|L2 | -15144 | NM_015811 | 50778 | Mm.103701 | NM_015811 | ENSMUSG00000026358 | **Rgs1** | BL34 | regulator of G-protein signaling 1 | protein-coding |
| chr1_144380728_144381983 | chr1 | 144380729 | 144381983 | + | 1.45906 | Intergenic | (CA)n\|Simple_repeat\|Simple_repeat | -132252 | NM_015811 | 50778 | Mm.103701 | NM_015811 | ENSMUSG00000026358 | **Rgs1** | BL34 | regulator of G-protein signaling 1 | protein-coding |
| chr1_144215236_144215762 | chr1 | 144215237 | 144215762 | + | 1.27699 | Intergenic | Intergenic | 33605 | NM_015811 | 50778 | Mm.103701 | NM_015811 | ENSMUSG00000026358 | **Rgs1** | BL34 | regulator of G-protein signaling 1 | protein-coding |
| chr1_144215219_144216023 | chr1 | 144215220 | 144216023 | + | 1.18642 | Intergenic | Intergenic | 33483 | NM_015811 | 50778 | Mm.103701 | NM_015811 | ENSMUSG00000026358 | **Rgs1** | BL34 | regulator of G-protein signaling 1 | protein-coding |
| chr1_144216215_144216457 | chr1 | 144216216 | 144216457 | + | 1.06056 | Intergenic | L1_Mus3\|LINE\|L1 | 32768 | NM_015811 | 50778 | Mm.103701 | NM_015811 | ENSMUSG00000026358 | **Rgs1** | BL34 | regulator of G-protein signaling 1 | protein-coding |
| chr1_144216083_144216462 | chr1 | 144216084 | 144216462 | + | 1.00753 | Intergenic | Intergenic | 32831 | NM_015811 | 50778 | Mm.103701 | NM_015811 | ENSMUSG00000026358 | **Rgs1** | BL34 | regulator of G-protein signaling 1 | protein-coding |
| chr1_144307451_144307698 | chr1 | 144307452 | 144307698 | + | 0.8465 | Intergenic | Intergenic | -58471 | NM_015811 | 50778 | Mm.103701 | NM_015811 | ENSMUSG00000026358 | **Rgs1** | BL34 | regulator of G-protein signaling 1 | protein-coding |
| chr1_144293286_144293623 | chr1 | 144293287 | 144293623 | + | 0.752764 | Intergenic | L1_Mus3\|LINE\|L1 | -44351 | NM_015811 | 50778 | Mm.103701 | NM_015811 | ENSMUSG00000026358 | **Rgs1** | BL34 | regulator of G-protein signaling 1 | protein-coding |
| chr1_144292844_144293707 | chr1 | 144292845 | 144293707 | + | 0.739757 | Intergenic | L1_Mus3\|LINE\|L1 | -44172 | NM_015811 | 50778 | Mm.103701 | NM_015811 | ENSMUSG00000026358 | **Rgs1** | BL34 | regulator of G-protein signaling 1 | protein-coding |
| chr1_144306322_144306631 | chr1 | 144306323 | 144306631 | + | 0.719804 | Intergenic | Intergenic | -57373 | NM_015811 | 50778 | Mm.103701 | NM_015811 | ENSMUSG00000026358 | **Rgs1** | BL34 | regulator of G-protein signaling 1 | protein-coding |
| chr1_144373680_144374463 | chr1 | 144373681 | 144374463 | + | 0.673449 | Intergenic | Lx4A\|LINE\|L1 | -124968 | NM_015811 | 50778 | Mm.103701 | NM_015811 | ENSMUSG00000026358 | **Rgs1** | BL34 | regulator of G-protein signaling 1 | protein-coding |
| chr7_128410539_128411054 | chr7 | 128410540 | 128411054 | + | 0.796217 | intron (NM_026418, intron 1 of 4) | intron (NM_026418, intron 1 of 4) | 7375 | NM_026418 | 67865 | Mm.18635 | NM_026418 | ENSMUSG00000030844 | **Rgs10** | 2310010N19Rik | regulator of G-protein signalling 10 | protein-coding |
| chr1_144206226_144206867 | chr1 | 144206227 | 144206867 | + | 1.96265 | Intergenic | Intergenic | -29175 | NM_153171 | 246709 | Mm.38271 | NM_153171 | ENSMUSG00000051079 | **Rgs13** | - | regulator of G-protein signaling 13 | protein-coding |
| chr1_144204757_144205093 | chr1 | 144204758 | 144205093 | + | 1.23955 | Intergenic | Intergenic | -27553 | NM_153171 | 246709 | Mm.38271 | NM_153171 | ENSMUSG00000051079 | **Rgs13** | - | regulator of G-protein signaling 13 | protein-coding |
| chr1_144202907_144203545 | chr1 | 144202908 | 144203545 | + | 0.998462 | Intergenic | Intergenic | -25854 | NM_153171 | 246709 | Mm.38271 | NM_153171 | ENSMUSG00000051079 | **Rgs13** | - | regulator of G-protein signaling 13 | protein-coding |
| chr1_144202952_144203342 | chr1 | 144202953 | 144203342 | + | 0.918617 | Intergenic | Intergenic | -25775 | NM_153171 | 246709 | Mm.38271 | NM_153171 | ENSMUSG00000051079 | **Rgs13** | - | regulator of G-protein signaling 13 | protein-coding |
| chr1_144168937_144169310 | chr1 | 144168938 | 144169310 | + | 0.848762 | intron (NM_153171, intron 3 of 5) | intron (NM_153171, intron 3 of 5) | 8248 | NM_153171 | 246709 | Mm.38271 | NM_153171 | ENSMUSG00000051079 | **Rgs13** | - | regulator of G-protein signaling 13 | protein-coding |
| chr1_144142586_144142969 | chr1 | 144142587 | 144142969 | + | 0.719989 | intron (NM_153171, intron 4 of 5) | intron (NM_153171, intron 4 of 5) | 34594 | NM_153171 | 246709 | Mm.38271 | NM_153171 | ENSMUSG00000051079 | **Rgs13** | - | regulator of G-protein signaling 13 | protein-coding |
| chr1_153740043_153740525 | chr1 | 153740044 | 153740525 | + | 0.847754 | promoter-TSS (NM_011267) | promoter-TSS (NM_011267) | -69 | NM_011267 | 19734 | Mm.181709 | NM_011267 | ENSMUSG00000026475 | **Rgs16** | Rgs14\|Rgsr | regulator of G-protein signaling 16 | protein-coding |
| chr1_153739822_153740544 | chr1 | 153739823 | 153740544 | + | 0.802564 | promoter-TSS (NM_011267) | promoter-TSS (NM_011267) | -170 | NM_011267 | 19734 | Mm.181709 | NM_011267 | ENSMUSG00000026475 | **Rgs16** | Rgs14\|Rgsr | regulator of G-protein signaling 16 | protein-coding |
| chr10_6035864_6036173 | chr10 | 6035865 | 6036173 | + | 0.647687 | Intergenic | Intergenic | -113619 | NM_019958 | 56533 | Mm.44606 | NM_019958 | ENSMUSG00000019775 | **Rgs17** | 6430507P11Rik\|Rgsz2 | regulator of G-protein signaling 17 | protein-coding |
| chr1_144688781_144689438 | chr1 | 144688782 | 144689438 | + | 0.982492 | Intergenic | Intergenic | 86311 | NM_022881 | 64214 | Mm.253927 | NM_022881 | ENSMUSG00000026357 | **Rgs18** | - | regulator of G-protein signaling 18 | protein-coding |
| chr1_144075968_144076474 | chr1 | 144075969 | 144076474 | + | 1.49211 | Intergenic | Intergenic | -72072 | NM_009061 | 19735 | Mm.28262 | NM_009061 | ENSMUSG00000026360 | **Rgs2** | GOS8 | regulator of G-protein signaling 2 | protein-coding |
| chr1_144064670_144065055 | chr1 | 144064671 | 144065055 | + | 0.714045 | Intergenic | Charlie13a\|DNA\|hAT-Charlie | -60714 | NM_009061 | 19735 | Mm.28262 | NM_009061 | ENSMUSG00000026360 | **Rgs2** | GOS8 | regulator of G-protein signaling 2 | protein-coding |
| chr1_5031605_5031897 | chr1 | 5031606 | 5031897 | + | 0.721359 | intron (NM_001177795, intron 1 of 5) | intron (NM_001177795, intron 1 of 5) | -12212 | NM_021374 | 58175 | Mm.103771 | NM_021374 | ENSMUSG00000002459 | **Rgs20** | 2900073E09Rik\|Rgsz1 | regulator of G-protein signaling 20 | protein-coding |
| chr1_144605520_144606522 | chr1 | 144605521 | 144606522 | + | 1.43512 | Intergenic | Intergenic | -38354 | NR_110891 | 624910 | Mm.389199 | NM_001290269 | ENSMUSG00000098509 | **Rgs21** | EG624910 | regulator of G-protein signalling 21 | protein-coding |
| chr1_144502251_144502549 | chr1 | 144502252 | 144502549 | + | 0.872097 | Intergenic | Intergenic | 65267 | NM_001290269 | 624910 | Mm.389199 | NM_001290269 | ENSMUSG00000098509 | **Rgs21** | EG624910 | regulator of G-protein signalling 21 | protein-coding |
| chr1_144502084_144502549 | chr1 | 144502085 | 144502549 | + | 0.77813 | Intergenic | LTR40b\|LTR\|ERVL | 65350 | NM_001290269 | 624910 | Mm.389199 | NM_001290269 | ENSMUSG00000098509 | **Rgs21** | EG624910 | regulator of G-protein signalling 21 | protein-coding |
| chr4_62679210_62679559 | chr4 | 62679211 | 62679559 | + | 1.04778 | intron (NM_134257, intron 10 of 15) | L1ME3\|LINE\|L1 | 59713 | NM_134257 | 50780 | Mm.286753 | NM_019492 | ENSMUSG00000059810 | **Rgs3** | 4930506N09Rik\|C2PA-RGS3\|C2pa\|PDZ-RGS3\|RGS3S | regulator of G-protein signaling 3 | protein-coding |
| chr4_62543669_62544418 | chr4 | 62543670 | 62544418 | + | 0.978996 | intron (NM_177607, intron 12 of 14) | PB1D10\|SINE\|Alu | -15803 | NM_019492 | 50780 | Mm.286753 | NM_019492 | ENSMUSG00000059810 | **Rgs3** | 4930506N09Rik\|C2PA-RGS3\|C2pa\|PDZ-RGS3\|RGS3S | regulator of G-protein signaling 3 | protein-coding |
| chr4_62731300_62731902 | chr4 | 62731301 | 62731902 | + | 0.879624 | Intergenic | Intergenic | 111929 | NM_134257 | 50780 | Mm.286753 | NM_019492 | ENSMUSG00000059810 | **Rgs3** | 4930506N09Rik\|C2PA-RGS3\|C2pa\|PDZ-RGS3\|RGS3S | regulator of G-protein signaling 3 | protein-coding |
| chr4_62709498_62709999 | chr4 | 62709499 | 62709999 | + | 0.858438 | Intergenic | L1MD2\|LINE\|L1 | 90077 | NM_134257 | 50780 | Mm.286753 | NM_019492 | ENSMUSG00000059810 | **Rgs3** | 4930506N09Rik\|C2PA-RGS3\|C2pa\|PDZ-RGS3\|RGS3S | regulator of G-protein signaling 3 | protein-coding |
| chr4_62553267_62553464 | chr4 | 62553268 | 62553464 | + | 0.845282 | Intergenic | PB1D10\|SINE\|Alu | -6481 | NM_019492 | 50780 | Mm.286753 | NM_019492 | ENSMUSG00000059810 | **Rgs3** | 4930506N09Rik\|C2PA-RGS3\|C2pa\|PDZ-RGS3\|RGS3S | regulator of G-protein signaling 3 | protein-coding |
| chr4_62731091_62731902 | chr4 | 62731092 | 62731902 | + | 0.813346 | Intergenic | Intergenic | 111825 | NM_134257 | 50780 | Mm.286753 | NM_019492 | ENSMUSG00000059810 | **Rgs3** | 4930506N09Rik\|C2PA-RGS3\|C2pa\|PDZ-RGS3\|RGS3S | regulator of G-protein signaling 3 | protein-coding |
| chr4_62653579_62654140 | chr4 | 62653580 | 62654140 | + | 0.75866 | intron (NM_134257, intron 10 of 15) | MIR\|SINE\|MIR | 34188 | NM_134257 | 50780 | Mm.286753 | NM_019492 | ENSMUSG00000059810 | **Rgs3** | 4930506N09Rik\|C2PA-RGS3\|C2pa\|PDZ-RGS3\|RGS3S | regulator of G-protein signaling 3 | protein-coding |
| chr4_62691564_62691978 | chr4 | 62691565 | 62691978 | + | 0.739992 | intron (NM_134257, intron 11 of 15) | MIR3\|SINE\|MIR | 72099 | NM_134257 | 50780 | Mm.286753 | NM_019492 | ENSMUSG00000059810 | **Rgs3** | 4930506N09Rik\|C2PA-RGS3\|C2pa\|PDZ-RGS3\|RGS3S | regulator of G-protein signaling 3 | protein-coding |
| chr4_62638247_62638599 | chr4 | 62638248 | 62638599 | + | 0.635276 | intron (NM_019492, intron 16 of 19) | intron (NM_019492, intron 16 of 19) | 18751 | NM_134257 | 50780 | Mm.286753 | NM_019492 | ENSMUSG00000059810 | **Rgs3** | 4930506N09Rik\|C2PA-RGS3\|C2pa\|PDZ-RGS3\|RGS3S | regulator of G-protein signaling 3 | protein-coding |
| chr1_153617121_153617895 | chr1 | 153617122 | 153617895 | + | 1.17363 | Intergenic | Intergenic | -35529 | NM_026380 | 67792 | Mm.379143 | NM_026380 | ENSMUSG00000042671 | **Rgs8** | 6530413N01Rik | regulator of G-protein signaling 8 | protein-coding |
| chr1_153617266_153617713 | chr1 | 153617267 | 153617713 | + | 1.12984 | Intergenic | Intergenic | -35547 | NM_026380 | 67792 | Mm.379143 | NM_026380 | ENSMUSG00000042671 | **Rgs8** | 6530413N01Rik | regulator of G-protein signaling 8 | protein-coding |
| chr11_109271398_109271878 | chr11 | 109271399 | 109271878 | + | 0.82655 | intron (NM_001165934, intron 7 of 16) | MTEa\|LTR\|ERVL-MaLR | 26543 | NM_001165934 | 19739 | Mm.38548 | NM_011268 | ENSMUSG00000020599 | **Rgs9** | RGS9-1\|Rgs9-2 | regulator of G-protein signaling 9 | protein-coding |
| chr11_109269211_109270004 | chr11 | 109269212 | 109270004 | + | 0.714952 | intron (NM_001165934, intron 7 of 16) | intron (NM_001165934, intron 7 of 16) | 28573 | NM_001165934 | 19739 | Mm.38548 | NM_011268 | ENSMUSG00000020599 | **Rgs9** | RGS9-1\|Rgs9-2 | regulator of G-protein signaling 9 | protein-coding |
| chr1_82372559_82372890 | chr1 | 82372560 | 82372890 | + | 0.812123 | intron (NM_001122685, intron 5 of 6) | MTEa\|LTR\|ERVL-MaLR | 33676 | NM_001122685 | 76867 | Mm.226994 | NM_029777 | ENSMUSG00000026142 | **Rhbdd1** | 4930418P06Rik\|RRP4\|mRHBDD1 | rhomboid domain containing 1 | protein-coding |
| chr9_108318861_108319421 | chr9 | 108318862 | 108319421 | + | 0.783747 | intron (NM_016802, intron 2 of 5) | intron (NM_016802, intron 2 of 5) | 12936 | NM_016802 | 11848 | Mm.757 | NM_016802 | ENSMUSG00000007815 | **Rhoa** | Arha\|Arha1\|Arha2 | ras homolog gene family, member A | protein-coding |
| chr10_69185926_69186211 | chr10 | 69185927 | 69186211 | + | 0.838605 | Intergenic | Intergenic | -22483 | NM_001252636 | 69288 | Mm.26659 | NM_001081347 | ENSMUSG00000019944 | **Rhobtb1** | 1700008H16Rik\|3110048G13Rik\|AI173445\|AI573858\|AV350930 | Rho-related BTB domain containing 1 | protein-coding |
| chr5_65914329_65914652 | chr5 | 65914330 | 65914652 | + | 1.0821 | Intergenic | Intergenic | 50922 | NM_001081105 | 74734 | Mm.20323 | NM_001081105 | ENSMUSG00000029204 | **Rhoh** | 5830400A04Rik\|AU019774\|Arhh | ras homolog gene family, member H | protein-coding |
| chr12_75292897_75293359 | chr12 | 75292898 | 75293359 | + | 1.3858 | Intergenic | Intergenic | -15185 | NM_023275 | 80837 | Mm.27467 | NM_023275 | ENSMUSG00000046768 | **Rhoj** | 1110005O19Rik\|AW210585\|Arhj\|TC10L\|TCL | ras homolog gene family, member J | protein-coding |
| chr17_86961288_86961678 | chr17 | 86961289 | 86961678 | + | 1.20248 | Intergenic | Intergenic | -1628 | NM_145491 | 104215 | Mm.826 | NM_145491 | ENSMUSG00000024143 | **Rhoq** | Arhq\|TC10A\|Tc10 | ras homolog gene family, member Q | protein-coding |
| chr10_84926965_84927222 | chr10 | 84926966 | 84927222 | + | 1.00614 | intron (NM_001013441, intron 2 of 8) | intron (NM_001013441, intron 2 of 8) | 9481 | NM_183172 | 237422 | Mm.18936 | NM_183172 | | **Ric8b** | BC051080\|Ric-8\|Ric-8b | resistance to inhibitors of cholinesterase 8 homolog B (C. elegans) | protein-coding |
| chr10_84895796_84896127 | chr10 | 84895797 | 84896127 | + | 0.9921 | exon (NM_027689, exon 13 of 14) | exon (NM_027689, exon 13 of 14) | -21651 | NM_001013441 | 237422 | Mm.18936 | NM_183172 | | **Ric8b** | BC051080\|Ric-8\|Ric-8b | resistance to inhibitors of cholinesterase 8 homolog B (C. elegans) | protein-coding |
| chr10_84926738_84927227 | chr10 | 84926739 | 84927227 | + | 0.94697 | intron (NM_001013441, intron 2 of 8) | intron (NM_001013441, intron 2 of 8) | 9370 | NM_183172 | 237422 | Mm.18936 | NM_183172 | | **Ric8b** | BC051080\|Ric-8\|Ric-8b | resistance to inhibitors of cholinesterase 8 homolog B (C. elegans) | protein-coding |
| chr10_84945232_84945737 | chr10 | 84945233 | 84945737 | + | 0.900225 | intron (NM_001013441, intron 2 of 8) | intron (NM_001013441, intron 2 of 8) | 27872 | NM_183172 | 237422 | Mm.18936 | NM_183172 | | **Ric8b** | BC051080\|Ric-8\|Ric-8b | resistance to inhibitors of cholinesterase 8 homolog B (C. elegans) | protein-coding |
| chr10_84928360_84928890 | chr10 | 84928361 | 84928890 | + | 0.790246 | intron (NM_001013441, intron 2 of 8) | URR1B\|DNA\|hAT-Charlie | 11012 | NM_183172 | 237422 | Mm.18936 | NM_183172 | | **Ric8b** | BC051080\|Ric-8\|Ric-8b | resistance to inhibitors of cholinesterase 8 homolog B (C. elegans) | protein-coding |
| chr5_124478566_124478768 | chr5 | 124478567 | 124478768 | + | 0.727635 | promoter-TSS (NM_030259) | promoter-TSS (NM_030259) | -432 | NM_030259 | 80291 | Mm.425280 | NM_030259 | ENSMUSG00000029401 | **Rilpl2** | - | Rab interacting lysosomal protein-like 2 | protein-coding |

| | | | | | | | | | | | | | | | | | |
|---|---|---|---|---|---|---|---|---|---|---|---|---|---|---|---|---|---|
| chr15_39354534_39355081 | chr15 | 39354535 | 39355081 | + | 1.04534 | intron (NM_053271, intron 5 of 26) | URR1B\|DNA\|hAT-Charlie | 156522 | NM_053271 | 116838 | Mm.309296 | NM_053271 | | **Rims2** | 2810036I15Rik\|AW048769\|RIM2\|Rab3ip2\|Rim2(+40A)\|Rim2(+44A)\|Rim2(+4A)\|Serg2\|Syt3-rs\|mKIAA0751 | regulating synaptic membrane exocytosis 2 | protein-coding |
| chr19_5050004_5050695 | chr19 | 5050005 | 5050695 | + | 0.833368 | promoter-TSS (NM_145495) | promoter-TSS (NM_145495) | -458 | NM_145495 | 225870 | Mm.271922 | NM_145495 | ENSMUSG00000024883 | **Rin1** | - | Ras and Rab interactor 1 | protein-coding |
| chr12_102309894_102310137 | chr12 | 102309895 | 102310137 | + | 0.702081 | intron (NM_001161365, intron 1 of 9) | L1ME2z\|LINE\|L1 | 26375 | NM_001161365 | 217835 | Mm.24145 | NM_177620 | ENSMUSG00000044456 | **Rin3** | 6430500K07\|C86933 | Ras and Rab interactor 3 | protein-coding |
| chr18_12131278_12131667 | chr18 | 12131279 | 12131667 | + | 0.876666 | intron (NM_024182, intron 1 of 12) | intron (NM_024182, intron 1 of 12) | 2623 | NM_024182 | 66878 | Mm.28551 | NM_024182 | ENSMUSG00000024404 | **Riok3** | 1200013N13Rik\|D18Ertd331e\|E130306C24Rik\|Sudd | RIO kinase 3 | protein-coding |
| chr4_16101566_16101942 | chr4 | 16101567 | 16101942 | + | 1.09122 | Intergenic | Intergenic | 61744 | NM_138952 | 192656 | Mm.112765 | NM_138952 | ENSMUSG00000041135 | **Ripk2** | 2210420D18Rik\|CARD3\|CARDIAK\|CCK\|D4Bwg0615e\|RICK\|RIP2 | receptor (TNFRSF)-interacting serine-threonine kinase 2 | protein-coding |
| chr4_16101576_16102577 | chr4 | 16101577 | 16102577 | + | 1.03388 | Intergenic | Intergenic | 61421 | NM_138952 | 192656 | Mm.112765 | NM_138952 | ENSMUSG00000041135 | **Ripk2** | 2210420D18Rik\|CARD3\|CARDIAK\|CCK\|D4Bwg0615e\|RICK\|RIP2 | receptor (TNFRSF)-interacting serine-threonine kinase 2 | protein-coding |
| chr4_16101566_16102907 | chr4 | 16101567 | 16102907 | + | 0.99619 | Intergenic | Intergenic | 61261 | NM_138952 | 192656 | Mm.112765 | NM_138952 | ENSMUSG00000041135 | **Ripk2** | 2210420D18Rik\|CARD3\|CARDIAK\|CCK\|D4Bwg0615e\|RICK\|RIP2 | receptor (TNFRSF)-interacting serine-threonine kinase 2 | protein-coding |
| chr4_16112451_16112904 | chr4 | 16112452 | 16112904 | + | 0.939314 | Intergenic | Intergenic | 50820 | NM_138952 | 192656 | Mm.112765 | NM_138952 | ENSMUSG00000041135 | **Ripk2** | 2210420D18Rik\|CARD3\|CARDIAK\|CCK\|D4Bwg0615e\|RICK\|RIP2 | receptor (TNFRSF)-interacting serine-threonine kinase 2 | protein-coding |
| chr4_16101990_16102547 | chr4 | 16101991 | 16102547 | + | 0.770372 | Intergenic | Intergenic | 61229 | NM_138952 | 192656 | Mm.112765 | NM_138952 | ENSMUSG00000041135 | **Ripk2** | 2210420D18Rik\|CARD3\|CARDIAK\|CCK\|D4Bwg0615e\|RICK\|RIP2 | receptor (TNFRSF)-interacting serine-threonine kinase 2 | protein-coding |
| chr7_79387184_79387614 | chr7 | 79387185 | 79387614 | + | 1.30769 | promoter-TSS (NM_001173483) | promoter-TSS (NM_001173483) | -372 | NM_001173483 | 19771 | Mm.41653 | NM_020599 | ENSMUSG00000039194 | **Rlbp1** | 3110056M11Rik\|CRALBP | retinaldehyde binding protein 1 | protein-coding |
| chrX_104033824_104034211 | chrX | 104033825 | 104034211 | + | 1.53084 | Intergenic | Intergenic | -52734 | NM_011276 | 19820 | Mm.427762 | NM_011276 | ENSMUSG00000056537 | **Rlim** | AL022832\|AW743871\|Ha1r\|Rlim1\|Rnf12 | ring finger protein, LIM domain interacting | protein-coding |
| chrX_104033820_104034426 | chrX | 104033821 | 104034426 | + | 1.46574 | Intergenic | Intergenic | -52839 | NM_011276 | 19820 | Mm.427762 | NM_011276 | ENSMUSG00000056537 | **Rlim** | AL022832\|AW743871\|Ha1r\|Rlim1\|Rnf12 | ring finger protein, LIM domain interacting | protein-coding |
| chr19_29342364_29342915 | chr19 | 29342365 | 29342915 | + | 1.15347 | Intergenic | LTR53\|LTR\|ERVL | -7970 | NM_011272 | 19773 | Mm.5033 | NM_011272 | ENSMUSG00000039097 | **Rln1** | Rln\|rlx | relaxin 1 | protein-coding |
| chr19_29341062_29341678 | chr19 | 29341063 | 29341678 | + | 0.871091 | Intergenic | Intergenic | -6700 | NM_011272 | 19773 | Mm.5033 | NM_011272 | ENSMUSG00000039097 | **Rln1** | Rln\|rlx | relaxin 1 | protein-coding |
| chr17_79599225_79599636 | chr17 | 79599226 | 79599636 | + | 1.10596 | Intergenic | MTC\|LTR\|ERVL-MaLR | -15469 | NM_201361 | 381110 | Mm.293063 | NM_201361 | ENSMUSG00000036368 | **Rmdn2** | AW061290\|Fam82a1\|RMD-2\|mRMD-2 | regulator of microtubule dynamics 2 | protein-coding |
| chr17_79584399_79584752 | chr17 | 79584400 | 79584752 | + | 1.06468 | Intergenic | Intergenic | -30324 | NM_201361 | 381110 | Mm.293063 | NM_201361 | ENSMUSG00000036368 | **Rmdn2** | AW061290\|Fam82a1\|RMD-2\|mRMD-2 | regulator of microtubule dynamics 2 | protein-coding |
| chr17_79584383_79584885 | chr17 | 79584384 | 79584885 | + | 1.05062 | Intergenic | Intergenic | -30266 | NM_201361 | 381110 | Mm.293063 | NM_201361 | ENSMUSG00000036368 | **Rmdn2** | AW061290\|Fam82a1\|RMD-2\|mRMD-2 | regulator of microtubule dynamics 2 | protein-coding |
| chr16_10829067_10829665 | chr16 | 10829068 | 10829665 | + | 0.90165 | Intergenic | RSINE1\|SINE\|B4 | -5693 | NM_001162932 | 223970 | Mm.325777 | NM_001033278 | ENSMUSG00000037991 | **Rmi2** | A630055G03Rik\|Gm537\|Gm929 | RMI2, RecQ mediated genome instability 2, homolog (S. cerevisiae) | protein-coding |
| chr16_10842342_10842704 | chr16 | 10842343 | 10842704 | + | 0.711636 | 3' UTR (NM_001162932, exon 2 of 2) | 3' UTR (NM_001162932, exon 2 of 2) | 7464 | NM_001162932 | 223970 | Mm.325777 | NM_001033278 | ENSMUSG00000037991 | **Rmi2** | A630055G03Rik\|Gm537\|Gm929 | RMI2, RecQ mediated genome instability 2, homolog (S. cerevisiae) | protein-coding |
| chr6_47635648_47636223 | chr6 | 47635649 | 47636223 | + | 1.4144 | Intergenic | Intergenic | 19158 | NR_002841 | 19799 | | NR_002841 | | **Rn4.5s** | - | 4.5S RNA | rRNA |
| chr6_47633533_47633782 | chr6 | 47633534 | 47633782 | + | 0.6232 | Intergenic | B3\|SINE\|B2 | 21436 | NR_002841 | 19799 | | NR_002841 | | **Rn4.5s** | - | 4.5S RNA | rRNA |
| chr14_62353401_62354272 | chr14 | 62353402 | 62354272 | + | 1.48385 | intron (NM_026001, intron 5 of 10) | intron (NM_026001, intron 5 of 10) | 21732 | NM_026001 | 67153 | Mm.26040 | NM_026001 | ENSMUSG00000021932 | **Rnaseh2b** | 1110019N06Rik\|2610207P08Rik\|Deu8 | ribonuclease H2, subunit B | protein-coding |
| chr14_62389485_62389911 | chr14 | 62389486 | 62389911 | + | 1.08495 | Intergenic | Intergenic | 57593 | NM_026001 | 67153 | Mm.26040 | NM_026001 | ENSMUSG00000021932 | **Rnaseh2b** | 1110019N06Rik\|2610207P08Rik\|Deu8 | ribonuclease H2, subunit B | protein-coding |
| chr14_62389399_62390211 | chr14 | 62389400 | 62390211 | + | 0.666396 | Intergenic | Intergenic | 57700 | NM_026001 | 67153 | Mm.26040 | NM_026001 | ENSMUSG00000021932 | **Rnaseh2b** | 1110019N06Rik\|2610207P08Rik\|Deu8 | ribonuclease H2, subunit B | protein-coding |
| chr4_109484625_109485004 | chr4 | 109484626 | 109485004 | + | 1.03011 | Intergenic | Intergenic | -8310 | NM_013876 | 29864 | Mm.392580 | NM_013876 | ENSMUSG00000028557 | **Rnf11** | - | ring finger protein 11 | protein-coding |
| chr4_109469713_109470123 | chr4 | 109469714 | 109470123 | + | 0.937483 | intron (NM_013876, intron 1 of 2) | intron (NM_013876, intron 1 of 2) | 6587 | NM_013876 | 29864 | Mm.392580 | NM_013876 | ENSMUSG00000028557 | **Rnf11** | - | ring finger protein 11 | protein-coding |
| chr4_109470899_109471348 | chr4 | 109470900 | 109471348 | + | 0.659994 | intron (NM_013876, intron 1 of 2) | intron (NM_013876, intron 1 of 2) | 5381 | NM_013876 | 29864 | Mm.392580 | NM_013876 | ENSMUSG00000028557 | **Rnf11** | - | ring finger protein 11 | protein-coding |
| chr2_167485664_167486087 | chr2 | 167485665 | 167486087 | + | 0.924995 | intron (NM_170756, intron 1 of 2) | intron (NM_170756, intron 1 of 2) | -6769 | NM_030743 | 81018 | Mm.22225 | NM_030743 | ENSMUSG00000006418 | **Rnf114** | 1110008J21Rik\|AI225886\|AW549494\|Zfp228\|Zfp313\|Znf228 | ring finger protein 114 | protein-coding |
| chr2_167485636_167486281 | chr2 | 167485637 | 167486281 | + | 0.771568 | intron (NM_170756, intron 1 of 2) | intron (NM_170756, intron 1 of 2) | -6686 | NM_030743 | 81018 | Mm.22225 | NM_030743 | ENSMUSG00000006418 | **Rnf114** | 1110008J21Rik\|AI225886\|AW549494\|Zfp228\|Zfp313\|Znf228 | ring finger protein 114 | protein-coding |
| chr3_96759568_96760134 | chr3 | 96759569 | 96760134 | + | 1.28644 | intron (NM_026406, intron 3 of 8) | intron (NM_026406, intron 3 of 8) | 32240 | NM_026406 | 67845 | Mm.386792 | NM_026406 | ENSMUSG00000028098 | **Rnf115** | 2610028E05Rik\|AU042696\|Zfp364 | ring finger protein 115 | protein-coding |
| chr3_96765418_96765802 | chr3 | 96765419 | 96765802 | + | 0.746195 | intron (NM_026406, intron 3 of 8) | MTE2b\|LTR\|ERVL-MaLR | 37999 | NM_026406 | 67845 | Mm.386792 | NM_026406 | ENSMUSG00000028098 | **Rnf115** | 2610028E05Rik\|AU042696\|Zfp364 | ring finger protein 115 | protein-coding |
| chr3_96729906_96730164 | chr3 | 96729907 | 96730164 | + | 0.641872 | intron (NM_026406, intron 1 of 8) | intron (NM_026406, intron 1 of 8) | 2424 | NM_026406 | 67845 | Mm.386792 | NM_026406 | ENSMUSG00000028098 | **Rnf115** | 2610028E05Rik\|AU042696\|Zfp364 | ring finger protein 115 | protein-coding |
| chr7_102063778_102064131 | chr7 | 102063779 | 102064131 | + | 1.01733 | intron (NM_029211, intron 1 of 8) | intron (NM_029211, intron 1 of 8) | 1177 | NM_029211 | 75212 | Mm.101141 | NM_029211 | ENSMUSG00000070426 | **Rnf121** | 4930544L10Rik | ring finger protein 121 | protein-coding |
| chr11_49977974_49978721 | chr11 | 49977975 | 49978721 | + | 0.730051 | intron (NM_029004, intron 12 of 13) | intron (NM_029004, intron 12 of 13) | -46983 | NM_021540 | 59044 | Mm.359004 | NM_021540 | ENSMUSG00000020376 | **Rnf130** | G1RZFP\|G1rp\|GOLIATH\|GP | ring finger protein 130 | protein-coding |
| chr18_20982419_20982853 | chr18 | 20982420 | 20982853 | + | 0.692994 | intron (NM_026301, intron 5 of 5) | intron (NM_026301, intron 5 of 5) | -18705 | NM_019706 | 56515 | Mm.253542 | NM_019706 | ENSMUSG00000024317 | **Rnf138** | 2410015A17Rik\|2810480D20Rik\|STRIN\|Trif\|Trif-d | ring finger protein 138 | protein-coding |
| chr13_47133036_47133904 | chr13 | 47133037 | 47133904 | + | 1.06129 | intron (NM_146042, intron 1 of 7) | B1_Mur1\|SINE\|Alu | 10750 | NM_146042 | 218215 | Mm.287609 | NM_146042 | ENSMUSG00000038068 | **Rnf144b** | BC025007\|E130105P19Rik\|Ibrdc2\|Pir2 | ring finger protein 144B | protein-coding |
| chr13_47272877_47273247 | chr13 | 47272878 | 47273247 | + | 0.988527 | Intergenic | Intergenic | 79059 | NM_001170643 | 218215 | Mm.287609 | NM_146042 | ENSMUSG00000038068 | **Rnf144b** | BC025007\|E130105P19Rik\|Ibrdc2\|Pir2 | ring finger protein 144B | protein-coding |
| chr13_47272830_47273712 | chr13 | 47272831 | 47273712 | + | 0.797125 | Intergenic | Intergenic | 79268 | NM_001170643 | 218215 | Mm.287609 | NM_146042 | ENSMUSG00000038068 | **Rnf144b** | BC025007\|E130105P19Rik\|Ibrdc2\|Pir2 | ring finger protein 144B | protein-coding |
| chr13_47142812_47143289 | chr13 | 47142813 | 47143289 | + | 0.670706 | intron (NM_146042, intron 1 of 7) | intron (NM_146042, intron 1 of 7) | 20331 | NM_146042 | 218215 | Mm.287609 | NM_146042 | ENSMUSG00000038068 | **Rnf144b** | BC025007\|E130105P19Rik\|Ibrdc2\|Pir2 | ring finger protein 144B | protein-coding |
| chr11_44504737_44505391 | chr11 | 44504738 | 44505391 | + | 0.810645 | Intergenic | Intergenic | -13900 | NM_028862 | 74315 | Mm.156771 | NM_028862 | ENSMUSG00000019189 | **Rnf145** | 3732413I11Rik\|TMRF1 | ring finger protein 145 | protein-coding |
| chr1_39578540_39578879 | chr1 | 39578541 | 39578879 | + | 0.795404 | Intergenic | Intergenic | -1363 | NM_001033135 | 67702 | Mm.28614 | NM_001033135 | ENSMUSG00000048234 | **Rnf149** | 1600023E10Rik\|AV037261\|C76763\|Gm15832\|Greul4\|OTTMUSG00000026591 | ring finger protein 149 | protein-coding |
| chr1_105434839_105435420 | chr1 | 105434840 | 105435420 | + | 0.869716 | Intergenic | Intergenic | -78420 | NM_001160368 | 320311 | Mm.389162 | NM_178779 | ENSMUSG00000047496 | **Rnf152** | A930029B02Rik | ring finger protein 152 | protein-coding |
| chr1_105443548_105444143 | chr1 | 105443549 | 105444143 | + | 0.835058 | Intergenic | MLT1A\|LTR\|ERVL-MaLR | -87136 | NM_001160368 | 320311 | Mm.389162 | NM_178779 | ENSMUSG00000047496 | **Rnf152** | A930029B02Rik | ring finger protein 152 | protein-coding |
| chr11_116379445_116379886 | chr11 | 116379446 | 116379886 | + | 0.979408 | intron (NM_027258, intron 2 of 18) | intron (NM_027258, intron 2 of 18) | 33366 | NM_027258 | 217340 | Mm.426344 | NM_027258 | ENSMUSG00000052949 | **Rnf157** | 2610036E23Rik\|A130073L17Rik\|mKIAA1917 | ring finger protein 157 | protein-coding |
| chr11_116406124_116407109 | chr11 | 116406125 | 116407109 | + | 0.812184 | intron (NM_027258, intron 1 of 18) | intron (NM_027258, intron 1 of 18) | 6415 | NM_027258 | 217340 | Mm.426344 | NM_027258 | ENSMUSG00000052949 | **Rnf157** | 2610036E23Rik\|A130073L17Rik\|mKIAA1917 | ring finger protein 157 | protein-coding |
| chr11_70648702_70648935 | chr11 | 70648703 | 70648935 | + | 0.77122 | intron (NM_027445, intron 2 of 8) | intron (NM_027445, intron 2 of 8) | 1230 | NM_027445 | 70510 | Mm.261818 | NM_027445 | ENSMUSG00000040746 | **Rnf167** | 0610010G05Rik\|5730408C10Rik\|AU019305\|AV328608 | ring finger protein 167 | protein-coding |
| chr11_70648665_70648976 | chr11 | 70648666 | 70648976 | + | 0.752885 | intron (NM_027445, intron 2 of 8) | intron (NM_027445, intron 2 of 8) | 1232 | NM_027445 | 70510 | Mm.261818 | NM_027445 | ENSMUSG00000040746 | **Rnf167** | 0610010G05Rik\|5730408C10Rik\|AU019305\|AV328608 | ring finger protein 167 | protein-coding |
| chr8_26137933_26138706 | chr8 | 26137934 | 26138706 | + | 1.12362 | intron (NM_029965, intron 6 of 7) | intron (NM_029965, intron 6 of 7) | 18940 | NM_029965 | 77733 | Mm.291906 | NM_029965 | ENSMUSG00000013878 | **Rnf170** | 6720407G21Rik\|AI481227 | ring finger protein 170 | protein-coding |
| chr15_36364864_36365203 | chr15 | 36364865 | 36365203 | + | 1.11596 | Intergenic | Intergenic | -81887 | NM_013923 | 30945 | Mm.5181 | NM_013923 | ENSMUSG00000022280 | **Rnf19a** | AA032313\|Dorfin\|Rnf19\|UIP117\|Ube7ip2\|XYbp | ring finger protein 19A | protein-coding |
| chr15_36360393_36360639 | chr15 | 36360394 | 36360639 | + | 1.0396 | Intergenic | Intergenic | -77369 | NM_013923 | 30945 | Mm.5181 | NM_013923 | ENSMUSG00000022280 | **Rnf19a** | AA032313\|Dorfin\|Rnf19\|UIP117\|Ube7ip2\|XYbp | ring finger protein 19A | protein-coding |
| chr15_36363213_36363460 | chr15 | 36363214 | 36363460 | + | 0.718769 | Intergenic | Intergenic | -80190 | NM_013923 | 30945 | Mm.5181 | NM_013923 | ENSMUSG00000022280 | **Rnf19a** | AA032313\|Dorfin\|Rnf19\|UIP117\|Ube7ip2\|XYbp | ring finger protein 19A | protein-coding |
| chr15_36363014_36363460 | chr15 | 36363015 | 36363460 | + | 0.691279 | Intergenic | Intergenic | -80090 | NM_013923 | 30945 | Mm.5181 | NM_013923 | ENSMUSG00000022280 | **Rnf19a** | AA032313\|Dorfin\|Rnf19\|UIP117\|Ube7ip2\|XYbp | ring finger protein 19A | protein-coding |

| | | | | | | | | | | | | | | | | | |
|---|---|---|---|---|---|---|---|---|---|---|---|---|---|---|---|---|---|
| chr4_129067589_129068444 | chr4 | 129067590 | 129068444 | + | 0.93875 | intron (NM_029219, intron 1 of 8) | intron (NM_029219, intron 1 of 8) | 9746 | NM_029219 | 75234 | Mm.259672 | NM_029219 | ENSMUSG00000028793 | **Rnf19b** | 4930534K13Rik\|4930555L03Rik\|Ibrdc3 | ring finger protein 19B | protein-coding |
| chr4_129067618_129067994 | chr4 | 129067619 | 129067994 | + | 0.844521 | intron (NM_029219, intron 1 of 8) | B1_Mus2\|SINE\|Alu | 9535 | NM_029219 | 75234 | Mm.259672 | NM_029219 | ENSMUSG00000028793 | **Rnf19b** | 4930534K13Rik\|4930555L03Rik\|Ibrdc3 | ring finger protein 19B | protein-coding |
| chr4_129068053_129068435 | chr4 | 129068054 | 129068435 | + | 0.75392 | intron (NM_029219, intron 1 of 8) | intron (NM_029219, intron 1 of 8) | 9973 | NM_029219 | 75234 | Mm.259672 | NM_029219 | ENSMUSG00000028793 | **Rnf19b** | 4930534K13Rik\|4930555L03Rik\|Ibrdc3 | ring finger protein 19B | protein-coding |
| chr4_129070748_129071163 | chr4 | 129070749 | 129071163 | + | 0.723949 | intron (NM_029219, intron 1 of 8) | B3\|SINE\|B2 | 12685 | NM_029219 | 75234 | Mm.259672 | NM_029219 | ENSMUSG00000028793 | **Rnf19b** | 4930534K13Rik\|4930555L03Rik\|Ibrdc3 | ring finger protein 19B | protein-coding |
| chr4_129033482_129034060 | chr4 | 129033483 | 129034060 | + | 0.610468 | Intergenic | Intergenic | -24500 | NM_029219 | 75234 | Mm.259672 | NM_029219 | ENSMUSG00000028793 | **Rnf19b** | 4930534K13Rik\|4930555L03Rik\|Ibrdc3 | ring finger protein 19B | protein-coding |
| chr4_129032649_129033014 | chr4 | 129032650 | 129033014 | + | 0.591371 | Intergenic | Intergenic | -25439 | NM_029219 | 75234 | Mm.259672 | NM_029219 | ENSMUSG00000028793 | **Rnf19b** | 4930534K13Rik\|4930555L03Rik\|Ibrdc3 | ring finger protein 19B | protein-coding |
| chr14_104516951_104517858 | chr14 | 104516952 | 104517858 | + | 1.01588 | intron (NM_026047, intron 1 of 5) | L1MA4\|LINE\|L1 | 5261 | NM_026047 | 72486 | Mm.291811 | NM_026047 | ENSMUSG00000022120 | **Rnf219** | 2610206B13Rik\|2810449K13Rik\|AI451544 | ring finger protein 219 | protein-coding |
| chr14_104517509_104517842 | chr14 | 104517510 | 104517842 | + | 0.893424 | intron (NM_026047, intron 1 of 5) | L1MA4\|LINE\|L1 | 4990 | NM_026047 | 72486 | Mm.291811 | NM_026047 | ENSMUSG00000022120 | **Rnf219** | 2610206B13Rik\|2810449K13Rik\|AI451544 | ring finger protein 219 | protein-coding |
| chr14_104516961_104517328 | chr14 | 104516962 | 104517328 | + | 0.751651 | intron (NM_026047, intron 1 of 5) | ORR1C2\|LTR\|ERVL-MaLR | 5521 | NM_026047 | 72486 | Mm.291811 | NM_026047 | ENSMUSG00000022120 | **Rnf219** | 2610206B13Rik\|2810449K13Rik\|AI451544 | ring finger protein 219 | protein-coding |
| chr4_117447640_117448322 | chr4 | 117447641 | 117448322 | + | 1.17393 | intron (NM_025739, intron 2 of 14) | B1_Mus1\|SINE\|Alu | 48934 | NM_025739 | 66743 | Mm.318331 | NM_025739 | ENSMUSG00000028677 | **Rnf220** | 4732477A13\|4931406I20Rik\|5730503K05Rik\|AV302541 | ring finger protein 220 | protein-coding |
| chr4_117476772_117477112 | chr4 | 117476773 | 117477112 | + | 0.890718 | intron (NM_025739, intron 2 of 14) | intron (NM_025739, intron 2 of 14) | 19973 | NM_025739 | 66743 | Mm.318331 | NM_025739 | ENSMUSG00000028677 | **Rnf220** | 4732477A13\|4931406I20Rik\|5730503K05Rik\|AV302541 | ring finger protein 220 | protein-coding |
| chr11_68889485_68890086 | chr11 | 68889486 | 68890086 | + | 1.15465 | intron (NM_177060, intron 1 of 2) | intron (NM_177060, intron 1 of 2) | 1233 | NM_177060 | 320040 | Mm.305696 | NM_177060 | | **Rnf222** | 9930039A11Rik | ring finger protein 222 | protein-coding |
| chr2_131386147_131386809 | chr2 | 131386148 | 131386809 | + | 1.09007 | intron (NR_037281, intron 2 of 2) | intron (NR_037281, intron 2 of 2) | -33586 | NM_178607 | 51902 | Mm.477523 | NM_178607 | ENSMUSG00000048911 | **Rnf24** | 2810473M14Rik\|4930505A13Rik\|AI317164\|C86507\|D2Ertd504e | ring finger protein 24 | protein-coding |
| chr10_128433867_128434132 | chr10 | 128433868 | 128434132 | + | 0.903633 | intron (NR_104290, intron 4 of 7) | intron (NR_104290, intron 4 of 7) | 22384 | NM_026259 | 67588 | Mm.294120 | NM_026259 | ENSMUSG00000025373 | **Rnf41** | 2210404G21Rik\|4930511A05Rik\|4933415P08Rik\|D10Ertd722e\|FLRF\|Nrdp1 | ring finger protein 41 | protein-coding |
| chr19_33159952_33160539 | chr19 | 33159953 | 33160539 | + | 1.29169 | intron (NM_001167818, intron 6 of 6) | intron (NM_001167818, intron 6 of 6) | 232049 | NM_001167818 | 67795 | Mm.204591 | NM_001146342 | ENSMUSG00000071573 | **Rnls** | 6530404N21Rik\|AI452315\|AW060440\|C10orf59 | renalase, FAD-dependent amine oxidase | protein-coding |
| chr19_33339852_33340495 | chr19 | 33339853 | 33340495 | + | 1.15605 | intron (NM_001167818, intron 4 of 6) | intron (NM_001167818, intron 4 of 6) | 52121 | NM_001167818 | 67795 | Mm.204591 | NM_001146342 | ENSMUSG00000071573 | **Rnls** | 6530404N21Rik\|AI452315\|AW060440\|C10orf59 | renalase, FAD-dependent amine oxidase | protein-coding |
| chr19_33157091_33157441 | chr19 | 33157092 | 33157441 | + | 0.846118 | intron (NM_001167818, intron 6 of 6) | L1MB8\|LINE\|L1 | 235029 | NM_001167818 | 67795 | Mm.204591 | NM_001146342 | ENSMUSG00000071573 | **Rnls** | 6530404N21Rik\|AI452315\|AW060440\|C10orf59 | renalase, FAD-dependent amine oxidase | protein-coding |
| chr19_33159292_33159538 | chr19 | 33159293 | 33159538 | + | 0.713868 | intron (NM_001167818, intron 6 of 6) | intron (NM_001167818, intron 6 of 6) | 232880 | NM_001167818 | 67795 | Mm.204591 | NM_001146342 | ENSMUSG00000071573 | **Rnls** | 6530404N21Rik\|AI452315\|AW060440\|C10orf59 | renalase, FAD-dependent amine oxidase | protein-coding |
| chr19_33223297_33223751 | chr19 | 33223298 | 33223751 | + | 0.688689 | intron (NM_001167818, intron 4 of 6) | intron (NM_001167818, intron 4 of 6) | 168771 | NM_001167818 | 67795 | Mm.204591 | NM_001146342 | ENSMUSG00000071573 | **Rnls** | 6530404N21Rik\|AI452315\|AW060440\|C10orf59 | renalase, FAD-dependent amine oxidase | protein-coding |
| chr15_83155470_83155864 | chr15 | 83155471 | 83155864 | + | 1.14326 | intron (NM_029787, intron 8 of 8) | intron (NM_029787, intron 8 of 8) | 6022 | NR_004432 | 104307 | Mm.471434 | NR_004432 | ENSMUSG00000065176 | **Rnu12** | - | RNA U12, small nuclear | snRNA |
| chr15_83155436_83156027 | chr15 | 83155437 | 83156027 | + | 0.933555 | intron (NM_029787, intron 8 of 8) | intron (NM_029787, intron 8 of 8) | 6087 | NR_004432 | 104307 | Mm.471434 | NR_004432 | ENSMUSG00000065176 | **Rnu12** | - | RNA U12, small nuclear | snRNA |
| chr19_14451813_14452209 | chr19 | 14451814 | 14452209 | + | 0.74316 | intron (NM_011600, intron 18 of 19) | intron (NM_011600, intron 18 of 19) | -13505 | NR_003027 | 19862 | | NR_003027 | ENSMUSG00000095132 | **Rnu6** | - | U6 small nuclear RNA | snRNA |
| chr19_14437212_14437697 | chr19 | 14437213 | 14437697 | + | 0.708512 | Intergenic | Intergenic | 1051 | NR_003027 | 19862 | | NR_003027 | ENSMUSG00000095132 | **Rnu6** | - | U6 small nuclear RNA | snRNA |
| chr19_14451550_14452194 | chr19 | 14451551 | 14452194 | + | 0.688219 | intron (NM_011600, intron 18 of 19) | intron (NM_011600, intron 18 of 19) | -13366 | NR_003027 | 19862 | | NR_003027 | ENSMUSG00000095132 | **Rnu6** | - | U6 small nuclear RNA | snRNA |
| chr18_10178181_10178453 | chr18 | 10178182 | 10178453 | + | 1.12871 | intron (NM_009071, intron 1 of 32) | B2_Mm2\|SINE\|B2 | 3475 | NM_009071 | 19877 | Mm.6710 | NM_009071 | ENSMUSG00000024290 | **Rock1** | 1110055K06Rik\|Rock-I | Rho-associated coiled-coil containing protein kinase 1 | protein-coding |
| chr12_16940740_16941312 | chr12 | 16940741 | 16941312 | + | 1.34602 | intron (NM_009072, intron 4 of 32) | intron (NM_009072, intron 4 of 32) | 46048 | NM_009072 | 19878 | Mm.276024 | NM_009072 | ENSMUSG00000020580 | **Rock2** | B230113H15Rik\|ROKalpha\|Rho-kinase\|Rock-II\|Rock2m\|mKIAA0619 | Rho-associated coiled-coil containing protein kinase 2 | protein-coding |
| chr12_16940721_16941539 | chr12 | 16940722 | 16941539 | + | 1.23218 | intron (NM_009072, intron 4 of 32) | intron (NM_009072, intron 4 of 32) | 46152 | NM_009072 | 19878 | Mm.276024 | NM_009072 | ENSMUSG00000020580 | **Rock2** | B230113H15Rik\|ROKalpha\|Rho-kinase\|Rock-II\|Rock2m\|mKIAA0619 | Rho-associated coiled-coil containing protein kinase 2 | protein-coding |
| chr12_16940946_16941509 | chr12 | 16940947 | 16941509 | + | 1.19819 | intron (NM_009072, intron 4 of 32) | B1_Mus1\|SINE\|Alu | 46250 | NM_009072 | 19878 | Mm.276024 | NM_009072 | ENSMUSG00000020580 | **Rock2** | B230113H15Rik\|ROKalpha\|Rho-kinase\|Rock-II\|Rock2m\|mKIAA0619 | Rho-associated coiled-coil containing protein kinase 2 | protein-coding |
| chr12_16920390_16920897 | chr12 | 16920391 | 16920897 | + | 0.834496 | intron (NM_009072, intron 1 of 32) | L2a\|LINE\|L2 | 25666 | NM_009072 | 19878 | Mm.276024 | NM_009072 | ENSMUSG00000020580 | **Rock2** | B230113H15Rik\|ROKalpha\|Rho-kinase\|Rock-II\|Rock2m\|mKIAA0619 | Rho-associated coiled-coil containing protein kinase 2 | protein-coding |
| chr12_16930667_16931027 | chr12 | 16930668 | 16931027 | + | 0.786261 | intron (NM_009072, intron 3 of 32) | PB1D9\|SINE\|Alu | 35869 | NM_009072 | 19878 | Mm.276024 | NM_009072 | ENSMUSG00000020580 | **Rock2** | B230113H15Rik\|ROKalpha\|Rho-kinase\|Rock-II\|Rock2m\|mKIAA0619 | Rho-associated coiled-coil containing protein kinase 2 | protein-coding |
| chr12_16942010_16942403 | chr12 | 16942011 | 16942403 | + | 0.732286 | intron (NM_009072, intron 4 of 32) | intron (NM_009072, intron 4 of 32) | 47229 | NM_009072 | 19878 | Mm.276024 | NM_009072 | ENSMUSG00000020580 | **Rock2** | B230113H15Rik\|ROKalpha\|Rho-kinase\|Rock-II\|Rock2m\|mKIAA0619 | Rho-associated coiled-coil containing protein kinase 2 | protein-coding |
| chr16_34665613_34665849 | chr16 | 34665614 | 34665849 | + | 0.664776 | intron (NM_030744, intron 1 of 5) | intron (NM_030744, intron 1 of 5) | 14520 | NM_030744 | 76378 | Mm.23515 | NM_030744 | ENSMUSG00000022832 | **Ropn1** | 1700008N21Rik\|ODF6\|RHPNAP1\|Rppn\|ropporin | ropporin, rhophilin associated protein 1 | protein-coding |
| chr9_69300310_69300626 | chr9 | 69300311 | 69300626 | + | 1.06517 | intron (NM_001289916, intron 1 of 9) | intron (NM_001289916, intron 1 of 9) | 10627 | NM_001289916 | 19883 | Mm.378450 | NM_013646 | ENSMUSG00000032238 | **Rora** | 9530021D13Rik\|Nr1f1\|ROR1\|ROR2\|ROR3\|nmf267\|sg\|staggerer\|tmgc26 | RAR-related orphan receptor alpha | protein-coding |
| chr9_69155788_69157174 | chr9 | 69155789 | 69157174 | + | 0.929485 | intron (NM_013646, intron 1 of 10) | intron (NM_013646, intron 1 of 10) | -133360 | NM_001289916 | 19883 | Mm.378450 | NM_013646 | ENSMUSG00000032238 | **Rora** | 9530021D13Rik\|Nr1f1\|ROR1\|ROR2\|ROR3\|nmf267\|sg\|staggerer\|tmgc26 | RAR-related orphan receptor alpha | protein-coding |
| chr9_69319298_69320160 | chr9 | 69319299 | 69320160 | + | 0.771168 | intron (NM_001289916, intron 1 of 9) | MIR\|SINE\|MIR | 29888 | NM_001289916 | 19883 | Mm.378450 | NM_013646 | ENSMUSG00000032238 | **Rora** | 9530021D13Rik\|Nr1f1\|ROR1\|ROR2\|ROR3\|nmf267\|sg\|staggerer\|tmgc26 | RAR-related orphan receptor alpha | protein-coding |
| chr9_69304403_69304968 | chr9 | 69304404 | 69304968 | + | 0.639018 | intron (NM_001289916, intron 1 of 9) | Charlie1a\|DNA\|hAT-Charlie | 14845 | NM_001289916 | 19883 | Mm.378450 | NM_013646 | ENSMUSG00000032238 | **Rora** | 9530021D13Rik\|Nr1f1\|ROR1\|ROR2\|ROR3\|nmf267\|sg\|staggerer\|tmgc26 | RAR-related orphan receptor alpha | protein-coding |
| chr3_94379562_94380136 | chr3 | 94379563 | 94380136 | + | 1.15313 | intron (NR_121656, intron 1 of 9) | intron (NR_121656, intron 1 of 9) | 2417 | NR_121656 | 19885 | Mm.4372 | NM_011281 | ENSMUSG00000028150 | **Rorc** | Nr1f3\|RORgamma\|TOR\|Thor | RAR-related orphan receptor gamma | protein-coding |
| chr3_94379545_94380532 | chr3 | 94379546 | 94380532 | + | 1.09375 | intron (NR_121656, intron 1 of 9) | intron (NR_121656, intron 1 of 9) | 2607 | NR_121656 | 19885 | Mm.4372 | NM_011281 | ENSMUSG00000028150 | **Rorc** | Nr1f3\|RORgamma\|TOR\|Thor | RAR-related orphan receptor gamma | protein-coding |
| chr3_94375337_94375610 | chr3 | 94375338 | 94375610 | + | 0.882829 | intron (NM_011281, intron 2 of 10) | intron (NM_011281, intron 2 of 10) | -1958 | NM_001293734 | 19885 | Mm.4372 | NM_011281 | ENSMUSG00000028150 | **Rorc** | Nr1f3\|RORgamma\|TOR\|Thor | RAR-related orphan receptor gamma | protein-coding |
| chr15_97499150_97499691 | chr15 | 97499151 | 97499691 | + | 1.27558 | Intergenic | URR1B\|DNA\|hAT-Charlie | 206401 | NM_028003 | 71919 | Mm.12255 | NM_028003 | ENSMUSG00000022466 | **Rpap3** | 2310042P20Rik\|D15Ertd682e | RNA polymerase II associated protein 3 | protein-coding |
| chr1_66648305_66648726 | chr1 | 66648306 | 66648726 | + | 1.07001 | intron (NM_175510, intron 45 of 64) | intron (NM_175510, intron 45 of 64) | -52377 | NM_025683 | 66646 | Mm.240912 | NM_025683 | ENSMUSG00000026005 | **Rpe** | 2810429B02Rik\|5730518J08Rik | ribulose-5-phosphate-3-epimerase | protein-coding |
| chr1_66649601_66649940 | chr1 | 66649602 | 66649940 | + | 0.930346 | intron (NM_175510, intron 47 of 64) | intron (NM_175510, intron 47 of 64) | -51122 | NM_025683 | 66646 | Mm.240912 | NM_025683 | ENSMUSG00000026005 | **Rpe** | 2810429B02Rik\|5730518J08Rik | ribulose-5-phosphate-3-epimerase | protein-coding |
| chr1_66650036_66650374 | chr1 | 66650037 | 66650374 | + | 0.887497 | intron (NM_175510, intron 47 of 64) | intron (NM_175510, intron 47 of 64) | -50688 | NM_025683 | 66646 | Mm.240912 | NM_025683 | ENSMUSG00000026005 | **Rpe** | 2810429B02Rik\|5730518J08Rik | ribulose-5-phosphate-3-epimerase | protein-coding |
| chr6_70725590_70726163 | chr6 | 70725591 | 70726163 | + | 1.03907 | Intergenic | Intergenic | 66298 | NM_009075 | 19895 | Mm.17905 | NM_009075 | ENSMUSG00000053604 | **Rpia** | RPI | ribose 5-phosphate isomerase A | protein-coding |

| | | | | | | | | | | | | | | | | | |
|---|---|---|---|---|---|---|---|---|---|---|---|---|---|---|---|---|---|
| chr6_70743667_70744393 | chr6 | 70743668 | 70744393 | + | 0.806602 | Intergenic | Intergenic | 48145 | NM_009075 | 19895 | Mm.17905 | NM_009075 | ENSMUSG00000053604 | **Rpia** | RPI | ribose 5-phosphate isomerase A | protein-coding |
| chr17_28332003_28332750 | chr17 | 28332004 | 28332750 | + | 0.728724 | 3' UTR (NM_001098226, exon 13 of 13) | 3' UTR (NM_001098226, exon 13 of 13) | 3906 | NM_011287 | 19896 | Mm.336955 | NM_011287 | ENSMUSG00000037805 | **Rpl10a** | CsA-19\|Nedd6 | ribosomal protein L10A | protein-coding |
| chr17_28331932_28333409 | chr17 | 28331933 | 28333409 | + | 0.701453 | 3' UTR (NM_001098226, exon 13 of 13) | 3' UTR (NM_001098226, exon 13 of 13) | 4200 | NM_011287 | 19896 | Mm.336955 | NM_011287 | ENSMUSG00000037805 | **Rpl10a** | CsA-19\|Nedd6 | ribosomal protein L10A | protein-coding |
| chr5_146818718_146819442 | chr5 | 146818719 | 146819442 | + | 1.23682 | Intergenic | RSINE1\|SINE\|B4 | -13810 | NM_019647 | 19933 | Mm.316592 | NM_019647 | ENSMUSG00000041453 | **Rpl21** | 8430440E03Rik\|L21 | ribosomal protein L21 | protein-coding |
| chr16_55954240_55954618 | chr16 | 55954241 | 55954618 | + | 0.84837 | Intergenic | LTRIS_Mus\|LTR\|ERV1 | -11846 | NM_024218 | 68193 | Mm.29221 | NM_024218 | ENSMUSG00000098274 | **Rpl24** | 0610008L05Rik\|Bst | ribosomal protein L24 | protein-coding |
| chr16_55963141_55963378 | chr16 | 55963142 | 55963378 | + | 0.81688 | Intergenic | RSINE1\|SINE\|B4 | -3015 | NM_024218 | 68193 | Mm.29221 | NM_024218 | ENSMUSG00000098274 | **Rpl24** | 0610008L05Rik\|Bst | ribosomal protein L24 | protein-coding |
| chr1_39281896_39282232 | chr1 | 39281897 | 39282232 | + | 1.40853 | intron (NM_008719, intron 2 of 20) | intron (NM_008719, intron 2 of 20) | -85787 | NM_053257 | 114641 | Mm.379302 | NM_053257 | ENSMUSG00000073702 | **Rpl31** | - | ribosomal protein L31 | protein-coding |
| chr1_39281845_39282414 | chr1 | 39281846 | 39282414 | + | 1.39876 | intron (NM_008719, intron 2 of 20) | intron (NM_008719, intron 2 of 20) | -85721 | NM_053257 | 114641 | Mm.379302 | NM_053257 | ENSMUSG00000073702 | **Rpl31** | - | ribosomal protein L31 | protein-coding |
| chr1_39281613_39282456 | chr1 | 39281614 | 39282456 | + | 1.369 | intron (NM_008719, intron 2 of 20) | intron (NM_008719, intron 2 of 20) | -85816 | NM_053257 | 114641 | Mm.379302 | NM_053257 | ENSMUSG00000073702 | **Rpl31** | - | ribosomal protein L31 | protein-coding |
| chr3_130894043_130894678 | chr3 | 130894044 | 130894678 | + | 0.884045 | Intergenic | Intergenic | -163963 | NM_001005859 | 68436 | Mm.140568 | NM_026724 | ENSMUSG00000062006 | **Rpl34** | 1100001I22Rik | ribosomal protein L34 | protein-coding |
| chr3_130841725_130842123 | chr3 | 130841726 | 130842123 | + | 0.652878 | Intergenic | Intergenic | -111526 | NM_001005859 | 68436 | Mm.140568 | NM_026724 | ENSMUSG00000062006 | **Rpl34** | 1100001I22Rik | ribosomal protein L34 | protein-coding |
| chr17_56614518_56615056 | chr17 | 56614519 | 56615056 | + | 0.740492 | TTS (NM_018730) | TTS (NM_018730) | 1392 | NM_018730 | 54217 | Mm.11376 | NM_018730 | ENSMUSG00000057863 | **Rpl36** | - | ribosomal protein L36 | protein-coding |
| chr17_56615180_56616162 | chr17 | 56615181 | 56616162 | + | 0.681258 | intron (NM_028782, intron 13 of 17) | intron (NM_028782, intron 13 of 17) | 2276 | NM_018730 | 54217 | Mm.11376 | NM_018730 | ENSMUSG00000057863 | **Rpl36** | - | ribosomal protein L36 | protein-coding |
| chr1_72724413_72724791 | chr1 | 72724414 | 72724791 | + | 0.626929 | Intergenic | Intergenic | 13342 | NM_009084 | 19981 | Mm.21529 | NM_009084 | ENSMUSG00000046330 | **Rpl37a** | - | ribosomal protein L37a | protein-coding |
| chr10_128556333_128556779 | chr10 | 128556334 | 128556779 | + | 0.738841 | Intergenic | B2_Mm2\|SINE\|B2 | -7388 | NM_018860 | 67945 | Mm.290786 | NM_018860 | ENSMUSG00000093674 | **Rpl41** | 1810055P16Rik\|2210411K19Rik | ribosomal protein L41 | protein-coding |
| chr9_61827603_61828131 | chr9 | 61827604 | 61828131 | + | 1.39604 | Intergenic | Intergenic | 86643 | NM_018853 | 56040 | Mm.3158 | NM_018853 | ENSMUSG00000007892 | **Rplp1** | 2410042H16Rik\|Arpp1\|C430017H15Rik | ribosomal protein, large, P1 | protein-coding |
| chr9_61778604_61779866 | chr9 | 61778605 | 61779866 | + | 0.764113 | Intergenic | Intergenic | 135275 | NM_018853 | 56040 | Mm.3158 | NM_018853 | ENSMUSG00000007892 | **Rplp1** | 2410042H16Rik\|Arpp1\|C430017H15Rik | ribosomal protein, large, P1 | protein-coding |
| chr9_61664021_61664570 | chr9 | 61664022 | 61664570 | + | 0.660454 | Intergenic | Intergenic | 250214 | NM_018853 | 56040 | Mm.3158 | NM_018853 | ENSMUSG00000007892 | **Rplp1** | 2410042H16Rik\|Arpp1\|C430017H15Rik | ribosomal protein, large, P1 | protein-coding |
| chr9_61892895_61893369 | chr9 | 61892896 | 61893369 | + | 0.651205 | Intergenic | L1M5\|LINE\|L1 | 21378 | NM_018853 | 56040 | Mm.3158 | NM_018853 | ENSMUSG00000007892 | **Rplp1** | 2410042H16Rik\|Arpp1\|C430017H15Rik | ribosomal protein, large, P1 | protein-coding |
| chr7_116328870_116329351 | chr7 | 116328871 | 116329351 | + | 0.961049 | Intergenic | Intergenic | 5079 | NM_026533 | 68052 | Mm.14798 | NM_026533 | ENSMUSG00000090862 | **Rps13** | 2700063M04Rik | ribosomal protein S13 | protein-coding |
| chr7_116335228_116335430 | chr7 | 116335229 | 116335430 | + | 0.681883 | Intergenic | Intergenic | -1139 | NM_026533 | 68052 | Mm.14798 | NM_026533 | ENSMUSG00000090862 | **Rps13** | 2700063M04Rik | ribosomal protein S13 | protein-coding |
| chr7_117981995_117982559 | chr7 | 117981996 | 117982559 | + | 1.06651 | Intergenic | Intergenic | 133870 | NM_170669 | 267019 | Mm.288212 | NM_170669 | ENSMUSG00000008683 | **Rps15a** | A630031B11Rik | ribosomal protein S15A | protein-coding |
| chr7_118102624_118104309 | chr7 | 118102625 | 118104309 | + | 0.734903 | TTS (NM_170669) | TTS (NM_170669) | 12680 | NM_170669 | 267019 | Mm.288212 | NM_170669 | ENSMUSG00000008683 | **Rps15a** | A630031B11Rik | ribosomal protein S15A | protein-coding |
| chr7_118103678_118104287 | chr7 | 118103679 | 118104287 | + | 0.70707 | TTS (NM_170669) | TTS (NM_170669) | 12164 | NM_170669 | 267019 | Mm.288212 | NM_170669 | ENSMUSG00000008683 | **Rps15a** | A630031B11Rik | ribosomal protein S15A | protein-coding |
| chr7_118102624_118103532 | chr7 | 118102625 | 118103532 | + | 0.699962 | Intergenic | RMER1B\|Other\|Other | 13069 | NM_170669 | 267019 | Mm.288212 | NM_170669 | ENSMUSG00000008683 | **Rps15a** | A630031B11Rik | ribosomal protein S15A | protein-coding |
| chr11_6130579_6131262 | chr11 | 6130580 | 6131262 | + | 1.18647 | intron (NM_173748, intron 3 of 5) | intron (NM_173748, intron 3 of 5) | 42410 | NR_029471 | 100042173 | Mm.441799 | NR_029471 | | **Rps15a-ps6** | Gm11968\|OTTMUSG00000005107 | ribosomal protein S15A, pseudogene 6 | pseudo |
| chr11_6130590_6130923 | chr11 | 6130591 | 6130923 | + | 1.01344 | intron (NM_173748, intron 3 of 5) | intron (NM_173748, intron 3 of 5) | 42574 | NR_029471 | 100042173 | Mm.441799 | NR_029471 | | **Rps15a-ps6** | Gm11968\|OTTMUSG00000005107 | ribosomal protein S15A, pseudogene 6 | pseudo |
| chr11_6130981_6131260 | chr11 | 6130982 | 6131260 | + | 0.753257 | intron (NM_173748, intron 3 of 5) | L1M4\|LINE\|L1 | 42210 | NR_029471 | 100042173 | Mm.441799 | NR_029471 | | **Rps15a-ps6** | Gm11968\|OTTMUSG00000005107 | ribosomal protein S15A, pseudogene 6 | pseudo |
| chr7_81319703_81320194 | chr7 | 81319704 | 81320194 | + | 0.975815 | intron (NM_008803, intron 16 of 21) | intron (NM_008803, intron 16 of 21) | 25285 | NM_009092 | 20068 | Mm.42767 | NM_009092 | ENSMUSG00000061787 | **Rps17** | - | ribosomal protein S17 | protein-coding |
| chr4_3824587_3825191 | chr4 | 3824588 | 3825191 | + | 1.25126 | Intergenic | Intergenic | 10711 | NM_026147 | 67427 | Mm.21938 | NM_026147 | | **Rps20** | 4632426K06Rik\|Dsk4 | ribosomal protein S20 | protein-coding |
| chr2_180248945_180249422 | chr2 | 180248946 | 180249422 | + | 0.648597 | Intergenic | Intergenic | -8195 | NM_025587 | 66481 | Mm.289669 | NM_025587 | ENSMUSG00000039001 | **Rps21** | 1810049N11Rik\|2410030A14Rik | ribosomal protein S21 | protein-coding |
| chr13_90957505_90957946 | chr13 | 90957506 | 90957946 | + | 0.977405 | intron (NM_025770, intron 5 of 7) | intron (NM_025770, intron 5 of 7) | 34604 | NM_024175 | 66475 | Mm.306302 | NM_024175 | | **Rps23** | 2410044J15Rik | ribosomal protein S23 | protein-coding |
| chr13_90958307_90958611 | chr13 | 90958308 | 90958611 | + | 0.857645 | intron (NM_025770, intron 5 of 7) | intron (NM_025770, intron 5 of 7) | 35337 | NM_024175 | 66475 | Mm.306302 | NM_024175 | | **Rps23** | 2410044J15Rik | ribosomal protein S23 | protein-coding |
| chr13_90927064_90927594 | chr13 | 90927065 | 90927594 | + | 0.793169 | Intergenic | B3\|SINE\|B2 | 4207 | NM_024175 | 66475 | Mm.306302 | NM_024175 | | **Rps23** | 2410044J15Rik | ribosomal protein S23 | protein-coding |
| chr13_90927055_90928012 | chr13 | 90927056 | 90928012 | + | 0.696023 | Intergenic | MTC\|LTR\|ERVL-MaLR | 4412 | NM_024175 | 66475 | Mm.306302 | NM_024175 | | **Rps23** | 2410044J15Rik | ribosomal protein S23 | protein-coding |
| chr12_69101853_69102374 | chr12 | 69101854 | 69102374 | + | 0.720785 | Intergenic | Intergenic | 57072 | NM_009093 | 20090 | Mm.379006 | NM_009093 | ENSMUSG00000034892 | **Rps29** | - | ribosomal protein S29 | protein-coding |
| chr12_69101816_69102669 | chr12 | 69101817 | 69102669 | + | 0.643866 | Intergenic | Intergenic | 56943 | NM_009093 | 20090 | Mm.379006 | NM_009093 | ENSMUSG00000034892 | **Rps29** | - | ribosomal protein S29 | protein-coding |
| chr4_133839320_133839569 | chr4 | 133839321 | 133839569 | + | 1.52915 | Intergenic | Intergenic | 35286 | NM_001285506 | 20111 | Mm.301827 | NM_009097 | ENSMUSG00000003644 | **Rps6ka1** | Mapkapk-1a\|Rsk\|Rsk-1\|Rsk1\|S6K-alpha-1\|p90-Rsk1\|p90Rsk1\|p90S6K\|p90rsk | ribosomal protein S6 kinase polypeptide 1 | protein-coding |
| chr4_133842916_133843765 | chr4 | 133842917 | 133843765 | + | 1.34228 | Intergenic | Intergenic | 31390 | NM_001285506 | 20111 | Mm.301827 | NM_009097 | ENSMUSG00000003644 | **Rps6ka1** | Mapkapk-1a\|Rsk\|Rsk-1\|Rsk1\|S6K-alpha-1\|p90-Rsk1\|p90Rsk1\|p90S6K\|p90rsk | ribosomal protein S6 kinase polypeptide 1 | protein-coding |
| chr4_133897155_133897764 | chr4 | 133897156 | 133897764 | + | 1.04644 | Intergenic | Intergenic | -9663 | NM_009097 | 20111 | Mm.301827 | NM_009097 | ENSMUSG00000003644 | **Rps6ka1** | Mapkapk-1a\|Rsk\|Rsk-1\|Rsk1\|S6K-alpha-1\|p90-Rsk1\|p90Rsk1\|p90S6K\|p90rsk | ribosomal protein S6 kinase polypeptide 1 | protein-coding |
| chr4_133825598_133826065 | chr4 | 133825599 | 133826065 | + | 0.951457 | Intergenic | Intergenic | 48899 | NM_001285506 | 20111 | Mm.301827 | NM_009097 | ENSMUSG00000003644 | **Rps6ka1** | Mapkapk-1a\|Rsk\|Rsk-1\|Rsk1\|S6K-alpha-1\|p90-Rsk1\|p90Rsk1\|p90S6K\|p90rsk | ribosomal protein S6 kinase polypeptide 1 | protein-coding |
| chr17_7163911_7164451 | chr17 | 7163912 | 7164451 | + | 1.00943 | Intergenic | Intergenic | -5934 | NM_011299 | 20112 | Mm.268383 | NM_011299 | ENSMUSG00000023809 | **Rps6ka2** | 90kDa\|D17Wsu134e\|Rps6ka-rs1\|Rsk3\|p90rsk\|pp90rsk | ribosomal protein S6 kinase, polypeptide 2 | protein-coding |
| chrX_159085981_159086638 | chrX | 159085982 | 159086638 | + | 1.10157 | Intergenic | MLT1K\|LTR\|ERVL-MaLR | -169472 | NM_148945 | 110651 | Mm.328476 | NM_148945 | ENSMUSG00000031309 | **Rps6ka3** | C130006E23\|MAPKAPK-1b\|MPK-9\|Rsk2\|S6K-alpha3\|p90RSK3\|pp90RSK2 | ribosomal protein S6 kinase polypeptide 3 | protein-coding |
| chrX_159085985_159086298 | chrX | 159085986 | 159086298 | + | 1.02078 | Intergenic | MLT1K\|LTR\|ERVL-MaLR | -169640 | NM_148945 | 110651 | Mm.328476 | NM_148945 | ENSMUSG00000031309 | **Rps6ka3** | C130006E23\|MAPKAPK-1b\|MPK-9\|Rsk2\|S6K-alpha3\|p90RSK3\|pp90RSK2 | ribosomal protein S6 kinase polypeptide 3 | protein-coding |
| chrX_158940635_158941374 | chrX | 158940636 | 158941374 | + | 0.775985 | Intergenic | Intergenic | -314777 | NM_148945 | 110651 | Mm.328476 | NM_148945 | ENSMUSG00000031309 | **Rps6ka3** | C130006E23\|MAPKAPK-1b\|MPK-9\|Rsk2\|S6K-alpha3\|p90RSK3\|pp90RSK2 | ribosomal protein S6 kinase polypeptide 3 | protein-coding |
| chrX_158969316_158969695 | chrX | 158969317 | 158969695 | + | 0.762649 | Intergenic | Intergenic | -286276 | NM_148945 | 110651 | Mm.328476 | NM_148945 | ENSMUSG00000031309 | **Rps6ka3** | C130006E23\|MAPKAPK-1b\|MPK-9\|Rsk2\|S6K-alpha3\|p90RSK3\|pp90RSK2 | ribosomal protein S6 kinase polypeptide 3 | protein-coding |
| chrX_159189232_159189661 | chrX | 159189233 | 159189661 | + | 0.713868 | Intergenic | ID4\|SINE\|ID | -66335 | NM_148945 | 110651 | Mm.328476 | NM_148945 | ENSMUSG00000031309 | **Rps6ka3** | C130006E23\|MAPKAPK-1b\|MPK-9\|Rsk2\|S6K-alpha3\|p90RSK3\|pp90RSK2 | ribosomal protein S6 kinase polypeptide 3 | protein-coding |
| chr12_100747082_100747552 | chr12 | 100747083 | 100747552 | + | 0.937459 | Intergenic | Intergenic | -22289 | NM_153587 | 73086 | Mm.220417 | NM_153587 | ENSMUSG00000021180 | **Rps6ka5** | 3110005L17Rik\|6330404E13Rik\|AI854034\|MSK1\|MSPK1\|RLPK | ribosomal protein S6 kinase, polypeptide 5 | protein-coding |
| chr12_100668935_100669534 | chr12 | 100668936 | 100669534 | + | 0.744019 | intron (NM_153587, intron 2 of 17) | intron (NM_153587, intron 2 of 17) | 55793 | NM_153587 | 73086 | Mm.220417 | NM_153587 | ENSMUSG00000021180 | **Rps6ka5** | 3110005L17Rik\|6330404E13Rik\|AI854034\|MSK1\|MSPK1\|RLPK | ribosomal protein S6 kinase, polypeptide 5 | protein-coding |
| chr12_100668933_100669780 | chr12 | 100668934 | 100669780 | + | 0.72004 | intron (NM_153587, intron 2 of 17) | intron (NM_153587, intron 2 of 17) | 55671 | NM_153587 | 73086 | Mm.220417 | NM_153587 | ENSMUSG00000021180 | **Rps6ka5** | 3110005L17Rik\|6330404E13Rik\|AI854034\|MSK1\|MSPK1\|RLPK | ribosomal protein S6 kinase, polypeptide 5 | protein-coding |
| chr12_100704918_100705377 | chr12 | 100704919 | 100705377 | + | 0.651952 | intron (NM_153587, intron 1 of 17) | LTR16E2\|LTR\|ERVL | 19880 | NM_153587 | 73086 | Mm.220417 | NM_153587 | ENSMUSG00000021180 | **Rps6ka5** | 3110005L17Rik\|6330404E13Rik\|AI854034\|MSK1\|MSPK1\|RLPK | ribosomal protein S6 kinase, polypeptide 5 | protein-coding |
| chr1_190899820_190900304 | chr1 | 190899821 | 190900304 | + | 1.54363 | intron (NM_178775, intron 2 of 14) | B3A\|SINE\|B2 | 11708 | NM_178775 | 320119 | Mm.220912 | NM_178775 | ENSMUSG00000089872 | **Rps6kc1** | AA682037\|B130003F20Rik\|C80612\|Rpk118 | ribosomal protein S6 kinase polypeptide 1 | protein-coding |
| chr1_190914156_190914401 | chr1 | 190914157 | 190914401 | + | 1.05128 | Intergenic | B3\|SINE\|B2 | -2509 | NM_178775 | 320119 | Mm.220912 | NM_178775 | ENSMUSG00000089872 | **Rps6kc1** | AA682037\|B130003F20Rik\|C80612\|Rpk118 | ribosomal protein S6 kinase polypeptide 1 | protein-coding |
| chr7_3707353_3707847 | chr7 | 3707354 | 3707847 | + | 0.783226 | TTS (NM_029767) | TTS (NM_029767) | 3559 | NM_029767 | 76846 | Mm.13944 | NM_029767 | ENSMUSG00000006333 | **Rps9** | 3010033P07Rik\|AL022771\|AL022885 | ribosomal protein S9 | protein-coding |
| chr7_3709081_3709290 | chr7 | 3709082 | 3709290 | + | 0.735466 | Intergenic | PB1D11\|SINE\|Alu | 5145 | NM_029767 | 76846 | Mm.13944 | NM_029767 | ENSMUSG00000006333 | **Rps9** | 3010033P07Rik\|AL022771\|AL022885 | ribosomal protein S9 | protein-coding |

| | | | | | | | | | | | | | | | | | |
|---|---|---|---|---|---|---|---|---|---|---|---|---|---|---|---|---|---|
| chr7_3708177_3708663 | chr7 | 3708178 | 3708663 | + | 0.734282 | Intergenic | Intergenic | 4379 | NM_029767 | 76846 | Mm.13944 | NM_029767 | ENSMUSG00000006333 | **Rps9** | 3010033P07Rik\|AL022771\|AL022885 | ribosomal protein S9 | protein-coding |
| chr7_3709048_3709414 | chr7 | 3709049 | 3709414 | + | 0.671246 | Intergenic | PB1D11\|SINE\|Alu | 5190 | NM_029767 | 76846 | Mm.13944 | NM_029767 | ENSMUSG00000006333 | **Rps9** | 3010033P07Rik\|AL022771\|AL022885 | ribosomal protein S9 | protein-coding |
| chr7_3707087_3707860 | chr7 | 3707088 | 3707860 | + | 0.667416 | TTS (NM_029767) | TTS (NM_029767) | 3433 | NM_029767 | 76846 | Mm.13944 | NM_029767 | ENSMUSG00000006333 | **Rps9** | 3010033P07Rik\|AL022771\|AL022885 | ribosomal protein S9 | protein-coding |
| chr11_119788945_119789225 | chr11 | 119788946 | 119789225 | + | 0.650145 | intron (NM_028898, intron 7 of 33) | intron (NM_028898, intron 7 of 33) | 45280 | NR_045313 | 319454 | Mm.442144 | NR_045312 | ENSMUSG00000085081 | **Rptoros** | A430071A18Rik | regulatory associated protein of MTOR, complex 1, opposite strand | ncRNA |
| chr7_114100179_114100701 | chr7 | 114100180 | 114100701 | + | 0.85806 | intron (NM_025846, intron 1 of 5) | intron (NM_025846, intron 1 of 5) | 17341 | NM_025846 | 66922 | Mm.276572 | NM_025846 | ENSMUSG00000055723 | **Rras2** | 2610016H24Rik\|C86394\|TC21 | related RAS viral (r-ras) oncogene homolog 2 | protein-coding |
| chr7_114139103_114140368 | chr7 | 114139104 | 114140368 | + | 0.848339 | Intergenic | Intergenic | -21955 | NM_025846 | 66922 | Mm.276572 | NM_025846 | ENSMUSG00000055723 | **Rras2** | 2610016H24Rik\|C86394\|TC21 | related RAS viral (r-ras) oncogene homolog 2 | protein-coding |
| chr7_114139877_114140368 | chr7 | 114139878 | 114140368 | + | 0.798553 | Intergenic | B3A\|SINE\|B2 | -22342 | NM_025846 | 66922 | Mm.276572 | NM_025846 | ENSMUSG00000055723 | **Rras2** | 2610016H24Rik\|C86394\|TC21 | related RAS viral (r-ras) oncogene homolog 2 | protein-coding |
| chr7_114175684_114175946 | chr7 | 114175685 | 114175946 | + | 0.772114 | Intergenic | Intergenic | -58034 | NM_025846 | 66922 | Mm.276572 | NM_025846 | ENSMUSG00000055723 | **Rras2** | 2610016H24Rik\|C86394\|TC21 | related RAS viral (r-ras) oncogene homolog 2 | protein-coding |
| chr7_114172688_114173037 | chr7 | 114172689 | 114173037 | + | 0.695967 | Intergenic | MLT1J\|LTR\|ERVL-MaLR | -55082 | NM_025846 | 66922 | Mm.276572 | NM_025846 | ENSMUSG00000055723 | **Rras2** | 2610016H24Rik\|C86394\|TC21 | related RAS viral (r-ras) oncogene homolog 2 | protein-coding |
| chr7_114159078_114159302 | chr7 | 114159079 | 114159302 | + | 0.685338 | Intergenic | RMER4B\|LTR\|ERVK | -41409 | NM_025846 | 66922 | Mm.276572 | NM_025846 | ENSMUSG00000055723 | **Rras2** | 2610016H24Rik\|C86394\|TC21 | related RAS viral (r-ras) oncogene homolog 2 | protein-coding |
| chr7_114171284_114172183 | chr7 | 114171285 | 114172183 | + | 0.67228 | Intergenic | Intergenic | -53953 | NM_025846 | 66922 | Mm.276572 | NM_025846 | ENSMUSG00000055723 | **Rras2** | 2610016H24Rik\|C86394\|TC21 | related RAS viral (r-ras) oncogene homolog 2 | protein-coding |
| chr13_37622233_37623085 | chr13 | 37622234 | 37623085 | + | 1.41208 | Intergenic | Intergenic | -203379 | NM_026830 | 68750 | Mm.491109 | NM_026830 | ENSMUSG00000039087 | **Rreb1** | 1110037N09Rik\|AA414966\|B930013M22Rik | ras responsive element binding protein 1 | protein-coding |
| chr13_37792814_37793713 | chr13 | 37792815 | 37793713 | + | 1.24583 | Intergenic | Intergenic | -32774 | NM_026830 | 68750 | Mm.491109 | NM_026830 | ENSMUSG00000039087 | **Rreb1** | 1110037N09Rik\|AA414966\|B930013M22Rik | ras responsive element binding protein 1 | protein-coding |
| chr13_37691758_37692427 | chr13 | 37691759 | 37692427 | + | 0.997652 | Intergenic | Intergenic | -133945 | NM_026830 | 68750 | Mm.491109 | NM_026830 | ENSMUSG00000039087 | **Rreb1** | 1110037N09Rik\|AA414966\|B930013M22Rik | ras responsive element binding protein 1 | protein-coding |
| chr13_37759743_37760392 | chr13 | 37759744 | 37760392 | + | 0.861948 | Intergenic | Intergenic | -65970 | NM_026830 | 68750 | Mm.491109 | NM_026830 | ENSMUSG00000039087 | **Rreb1** | 1110037N09Rik\|AA414966\|B930013M22Rik | ras responsive element binding protein 1 | protein-coding |
| chr13_37940772_37941475 | chr13 | 37940773 | 37941475 | + | 0.849025 | intron (NR_033218, intron 10 of 11) | (C)n\|Simple_repeat\|Simple_repeat | 52253 | NM_001177869 | 68750 | Mm.491109 | NM_026830 | ENSMUSG00000039087 | **Rreb1** | 1110037N09Rik\|AA414966\|B930013M22Rik | ras responsive element binding protein 1 | protein-coding |
| chr13_37801137_37801454 | chr13 | 37801138 | 37801454 | + | 0.77272 | Intergenic | Intergenic | -24742 | NM_026830 | 68750 | Mm.491109 | NM_026830 | ENSMUSG00000039087 | **Rreb1** | 1110037N09Rik\|AA414966\|B930013M22Rik | ras responsive element binding protein 1 | protein-coding |
| chr13_37680823_37681528 | chr13 | 37680824 | 37681528 | + | 0.769267 | Intergenic | Intergenic | -144862 | NM_026830 | 68750 | Mm.491109 | NM_026830 | ENSMUSG00000039087 | **Rreb1** | 1110037N09Rik\|AA414966\|B930013M22Rik | ras responsive element binding protein 1 | protein-coding |
| chr13_37852907_37853159 | chr13 | 37852908 | 37853159 | + | 0.707305 | intron (NM_001039188, intron 1 of 9) | intron (NM_001039188, intron 1 of 9) | 25640 | NM_001039188 | 68750 | Mm.491109 | NM_026830 | ENSMUSG00000039087 | **Rreb1** | 1110037N09Rik\|AA414966\|B930013M22Rik | ras responsive element binding protein 1 | protein-coding |
| chr13_37633058_37633416 | chr13 | 37633059 | 37633416 | + | 0.693973 | Intergenic | Intergenic | -192801 | NM_026830 | 68750 | Mm.491109 | NM_026830 | ENSMUSG00000039087 | **Rreb1** | 1110037N09Rik\|AA414966\|B930013M22Rik | ras responsive element binding protein 1 | protein-coding |
| chr13_37783606_37784300 | chr13 | 37783607 | 37784300 | + | 0.678721 | Intergenic | Intergenic | -42085 | NM_026830 | 68750 | Mm.491109 | NM_026830 | ENSMUSG00000039087 | **Rreb1** | 1110037N09Rik\|AA414966\|B930013M22Rik | ras responsive element binding protein 1 | protein-coding |
| chr13_37671172_37671485 | chr13 | 37671173 | 37671485 | + | 0.660324 | Intergenic | ORR1C2\|LTR\|ERVL-MaLR | -154709 | NM_026830 | 68750 | Mm.491109 | NM_026830 | ENSMUSG00000039087 | **Rreb1** | 1110037N09Rik\|AA414966\|B930013M22Rik | ras responsive element binding protein 1 | protein-coding |
| chr7_102402062_102402555 | chr7 | 102402063 | 102402555 | + | 0.931477 | intron (NM_009287, intron 3 of 11) | Charlie29a\|DNA\|hAT-Charlie | -39386 | NM_009103 | 20133 | Mm.197486 | NM_009103 | ENSMUSG00000030978 | **Rrm1** | RnrM1 | ribonucleotide reductase M1 | protein-coding |
| chr12_24706794_24707139 | chr12 | 24706795 | 24707139 | + | 0.835759 | Intergenic | Intergenic | -1287 | NM_009104 | 20135 | Mm.99 | NM_009104 | ENSMUSG00000020649 | **Rrm2** | AA407299\|R2 | ribonucleotide reductase M2 | protein-coding |
| chr12_24707570_24707819 | chr12 | 24707571 | 24707819 | + | 0.651978 | promoter-TSS (NM_009104) | promoter-TSS (NM_009104) | -559 | NM_009104 | 20135 | Mm.99 | NM_009104 | ENSMUSG00000020649 | **Rrm2** | AA407299\|R2 | ribonucleotide reductase M2 | protein-coding |
| chr15_37899671_37900307 | chr15 | 37899672 | 37900307 | + | 0.775288 | Intergenic | Intergenic | 61066 | NM_199476 | 382985 | Mm.24738 | NM_199476 | ENSMUSG00000022292 | **Rrm2b** | p53R2 | ribonucleotide reductase M2 B (TP53 inducible) | protein-coding |
| chr16_13772074_13772434 | chr16 | 13772075 | 13772434 | + | 0.909728 | Intergenic | Intergenic | -8445 | NM_001039521 | 106298 | Mm.490362 | NM_001039521 | ENSMUSG00000022682 | **Rrn3** | AL023001\|E130302O19Rik\|R75565\|TIF-1A\|Tif1a | RRN3 RNA polymerase I transcription factor homolog (yeast) | protein-coding |
| chr19_41894931_41895349 | chr19 | 41894932 | 41895349 | + | 1.43993 | intron (NM_199447, intron 2 of 33) | intron (NM_199447, intron 2 of 33) | 1013 | NM_199447 | 107094 | Mm.276044 | NM_199447 | ENSMUSG00000035049 | **Rrp12** | AA408556\|AA536972\|mKIAA0690 | ribosomal RNA processing 12 homolog (S. cerevisiae) | protein-coding |
| chr3_103908827_103909148 | chr3 | 103908828 | 103909148 | + | 1.04301 | intron (NM_008979, intron 19 of 20) | intron (NM_008979, intron 19 of 20) | -5132 | NM_172684 | 229675 | Mm.29366 | NM_172684 | ENSMUSG00000044098 | **Rsbn1** | C230004D03Rik\|E330004N01\|Rosbin\|Rsbn\|Rsbp\|mKIAA3002 | rosbin, round spermatid basic protein 1 | protein-coding |
| chr3_103907333_103908145 | chr3 | 103907334 | 103908145 | + | 1.03539 | intron (NM_008979, intron 19 of 20) | intron (NM_008979, intron 19 of 20) | -6381 | NM_172684 | 229675 | Mm.29366 | NM_172684 | ENSMUSG00000044098 | **Rsbn1** | C230004D03Rik\|E330004N01\|Rosbin\|Rsbn\|Rsbp\|mKIAA3002 | rosbin, round spermatid basic protein 1 | protein-coding |
| chr9_73150362_73150926 | chr9 | 73150363 | 73150926 | + | 1.11351 | Intergenic | Intergenic | 37175 | NM_198609 | 225215 | Mm.379366 | NM_198609 | ENSMUSG00000032215 | **Rsl24d1** | 2410159K22Rik | ribosomal L24 domain containing 1 | protein-coding |
| chr17_7939081_7939360 | chr17 | 7939082 | 7939360 | + | 0.901372 | Intergenic | Intergenic | -6393 | NM_025789 | 66832 | Mm.29830 | NM_025789 | ENSMUSG00000073471 | **Rsph3a** | 1700012G05Rik\|4930524H12Rik\|R74860\|Rshl2\|Rshl2a | radial spoke 3A homolog (Chlamydomonas) | protein-coding |
| chr3_67059677_67060121 | chr3 | 67059678 | 67060121 | + | 1.17783 | intron (NM_025822, intron 3 of 9) | L1ME3B\|LINE\|L1 | 74227 | NM_025822 | 66880 | Mm.316418 | NM_025822 | ENSMUSG00000034544 | **Rsrc1** | 1200013F24Rik\|SRrp53 | arginine/serine-rich coiled-coil 1 | protein-coding |
| chr3_67163677_67164075 | chr3 | 67163678 | 67164075 | + | 0.860577 | intron (NM_025822, intron 4 of 9) | intron (NM_025822, intron 4 of 9) | 178204 | NM_025822 | 66880 | Mm.316418 | NM_025822 | ENSMUSG00000034544 | **Rsrc1** | 1200013F24Rik\|SRrp53 | arginine/serine-rich coiled-coil 1 | protein-coding |
| chr3_67087200_67087625 | chr3 | 67087201 | 67087625 | + | 0.827181 | intron (NM_025822, intron 4 of 9) | LTR37A\|LTR\|ERV1 | 101741 | NM_025822 | 66880 | Mm.316418 | NM_025822 | ENSMUSG00000034544 | **Rsrc1** | 1200013F24Rik\|SRrp53 | arginine/serine-rich coiled-coil 1 | protein-coding |
| chr3_67086649_67087628 | chr3 | 67086650 | 67087628 | + | 0.817569 | intron (NM_025822, intron 4 of 9) | B2_Mm2\|SINE\|B2 | 101467 | NM_025822 | 66880 | Mm.316418 | NM_025822 | ENSMUSG00000034544 | **Rsrc1** | 1200013F24Rik\|SRrp53 | arginine/serine-rich coiled-coil 1 | protein-coding |
| chr3_67108396_67109047 | chr3 | 67108397 | 67109047 | + | 0.752924 | intron (NM_025822, intron 4 of 9) | intron (NM_025822, intron 4 of 9) | 123050 | NM_025822 | 66880 | Mm.316418 | NM_025822 | ENSMUSG00000034544 | **Rsrc1** | 1200013F24Rik\|SRrp53 | arginine/serine-rich coiled-coil 1 | protein-coding |
| chr3_116474620_116474956 | chr3 | 116474621 | 116474956 | + | 1.46375 | Intergenic | Charlie29a\|DNA\|hAT-Charlie | 33387 | NM_025517 | 66368 | Mm.251228 | NM_025517 | ENSMUSG00000000339 | **Rtca** | 2310009A18Rik\|AI450277\|RPC\|Rtcd1 | RNA 3'-terminal phosphate cyclase | protein-coding |
| chr3_116466243_116466820 | chr3 | 116466244 | 116466820 | + | 0.660581 | Intergenic | ORR1B1\|LTR\|ERVL-MaLR | 41643 | NM_025517 | 66368 | Mm.251228 | NM_025517 | ENSMUSG00000000339 | **Rtca** | 2310009A18Rik\|AI450277\|RPC\|Rtcd1 | RNA 3'-terminal phosphate cyclase | protein-coding |
| chr2_172435568_172436185 | chr2 | 172435569 | 172436185 | + | 0.96204 | Intergenic | Intergenic | -4701 | NM_025542 | 66404 | Mm.25157 | NM_025542 | ENSMUSG00000027502 | **Rtfdc1** | 1700067C04Rik\|2410001C21Rik\|5730427M17Rik\|AA589417 | replication termination factor 2 domain containing 1 | protein-coding |
| chr10_68039598_68040274 | chr10 | 68039599 | 68040274 | + | 1.5951 | intron (NM_001081346, intron 11 of 12) | intron (NM_001081346, intron 11 of 12) | 60338 | NM_001081346 | 170799 | Mm.209844 | NM_133244 | ENSMUSG00000037846 | **Rtkn2** | B130039D23Rik\|Mbf\|Plekhk1 | rhotekin 2 | protein-coding |
| chr10_68035763_68036256 | chr10 | 68035764 | 68036256 | + | 1.113 | intron (NM_001081346, intron 11 of 12) | intron (NM_001081346, intron 11 of 12) | 56412 | NM_001081346 | 170799 | Mm.209844 | NM_133244 | ENSMUSG00000037846 | **Rtkn2** | B130039D23Rik\|Mbf\|Plekhk1 | rhotekin 2 | protein-coding |
| chr10_68066028_68066562 | chr10 | 68066029 | 68066562 | + | 0.962061 | Intergenic | Intergenic | 86697 | NM_001081346 | 170799 | Mm.209844 | NM_133244 | ENSMUSG00000037846 | **Rtkn2** | B130039D23Rik\|Mbf\|Plekhk1 | rhotekin 2 | protein-coding |
| chr11_29705231_29705659 | chr11 | 29705232 | 29705659 | + | 0.643988 | intron (NM_194051, intron 1 of 8) | intron (NM_194051, intron 1 of 8) | 11671 | NM_194051 | 68585 | Mm.192580 | NM_024226 | ENSMUSG00000020458 | **Rtn4** | 1110020G17Rik\|AA407876\|AA409940\|AA960376\|ASY\|C130026I10Rik\|NOGO\|NSP-CL\|NgA\|Nogo-A\|Nogo-B\|Nogo-C\|mKIAA0886\|mKIAA4153 | reticulon 4 | protein-coding |
| chr11_75199051_75199809 | chr11 | 75199052 | 75199809 | + | 1.05387 | intron (NM_177708, intron 1 of 1) | intron (NM_177708, intron 1 of 1) | 5437 | NM_177708 | 237847 | Mm.82661 | NM_177708 | ENSMUSG00000045287 | **Rtn4rl1** | Ngr3\|Ngrh2\|Ngrl2 | reticulon 4 receptor-like 1 | protein-coding |
| chr11_75199938_75200269 | chr11 | 75199939 | 75200269 | + | 0.991682 | intron (NM_177708, intron 1 of 1) | intron (NM_177708, intron 1 of 1) | 6111 | NM_177708 | 237847 | Mm.82661 | NM_177708 | ENSMUSG00000045287 | **Rtn4rl1** | Ngr3\|Ngrh2\|Ngrl2 | reticulon 4 receptor-like 1 | protein-coding |
| chr11_75221108_75221537 | chr11 | 75221109 | 75221537 | + | 0.975609 | intron (NM_177708, intron 1 of 1) | (CAGA)n\|Simple_repeat\|Simple_repeat | 27330 | NM_177708 | 237847 | Mm.82661 | NM_177708 | ENSMUSG00000045287 | **Rtn4rl1** | Ngr3\|Ngrh2\|Ngrl2 | reticulon 4 receptor-like 1 | protein-coding |
| chr11_75199949_75200552 | chr11 | 75199950 | 75200552 | + | 0.847068 | intron (NM_177708, intron 1 of 1) | intron (NM_177708, intron 1 of 1) | 6258 | NM_177708 | 237847 | Mm.82661 | NM_177708 | ENSMUSG00000045287 | **Rtn4rl1** | Ngr3\|Ngrh2\|Ngrl2 | reticulon 4 receptor-like 1 | protein-coding |
| chr11_75254609_75255040 | chr11 | 75254610 | 75255040 | + | 0.674938 | intron (NM_177708, intron 1 of 1) | intron (NM_177708, intron 1 of 1) | 60832 | NM_177708 | 237847 | Mm.82661 | NM_177708 | ENSMUSG00000045287 | **Rtn4rl1** | Ngr3\|Ngrh2\|Ngrl2 | reticulon 4 receptor-like 1 | protein-coding |
| chr11_50439743_50440319 | chr11 | 50439744 | 50440319 | + | 1.48274 | Intergenic | Intergenic | -8920 | NM_172557 | 216724 | Mm.390400 | NM_172557 | ENSMUSG00000020375 | **Rufy1** | 3000002E04Rik\|Rabip4\|ZFYVE12 | RUN and FYVE domain containing 1 | protein-coding |
| chr5_88599424_88599988 | chr5 | 88599425 | 88599988 | + | 0.921298 | intron (NM_001289777, intron 3 of 14) | L1M4\|LINE\|L1 | 16193 | NM_027530 | 52822 | Mm.270469 | NM_027530 | ENSMUSG00000029291 | **Rufy3** | 2810428M05Rik\|6330416M07Rik\|AW455998\|AW538594\|D5Bwg0860e\|Ripx\|Rpipx | RUN and FYVE domain containing 3 | protein-coding |
| chr5_8542384_8543068 | chr5 | 8542385 | 8543068 | + | 1.24162 | intron (NM_198620, intron 6 of 9) | intron (NM_198620, intron 6 of 9) | 80226 | NM_198620 | 242819 | Mm.332192 | NM_198620 | ENSMUSG00000040570 | **Rundc3b** | Gm440 | RUN domain containing 3B | protein-coding |

| | | | | | | | | | | | | | | | | | |
|---|---|---|---|---|---|---|---|---|---|---|---|---|---|---|---|---|---|
| chr16_92864220_92864762 | chr16 | 92864221 | 92864762 | + | 1.43535 | Intergenic | Intergenic | -38417 | NM_001111022 | 12394 | Mm.4081 | NM_009821 | ENSMUSG00000022952 | **Runx1** | AML1\|CBF-alpha-2\|Cbfa2\|Pebp2a2\|Pebpa2b | runt related transcription factor 1 | protein-coding |
| chr16_92748195_92748670 | chr16 | 92748196 | 92748670 | + | 1.2322 | intron (NM_001111022, intron 2 of 6) | intron (NM_001111022, intron 2 of 6) | -51104 | NM_009821 | 12394 | Mm.4081 | NM_009821 | ENSMUSG00000022952 | **Runx1** | AML1\|CBF-alpha-2\|Cbfa2\|Pebp2a2\|Pebpa2b | runt related transcription factor 1 | protein-coding |
| chr16_92823664_92823926 | chr16 | 92823665 | 92823926 | + | 1.05553 | intron (NM_001111022, intron 2 of 6) | intron (NM_001111022, intron 2 of 6) | 2279 | NM_001111021 | 12394 | Mm.4081 | NM_009821 | ENSMUSG00000022952 | **Runx1** | AML1\|CBF-alpha-2\|Cbfa2\|Pebp2a2\|Pebpa2b | runt related transcription factor 1 | protein-coding |
| chr16_92743549_92744252 | chr16 | 92743550 | 92744252 | + | 0.847394 | intron (NM_001111022, intron 2 of 6) | intron (NM_001111022, intron 2 of 6) | -46572 | NM_009821 | 12394 | Mm.4081 | NM_009821 | ENSMUSG00000022952 | **Runx1** | AML1\|CBF-alpha-2\|Cbfa2\|Pebp2a2\|Pebpa2b | runt related transcription factor 1 | protein-coding |
| chr16_92997682_92998081 | chr16 | 92997683 | 92998081 | + | 0.811885 | Intergenic | Intergenic | -171808 | NM_001111022 | 12394 | Mm.4081 | NM_009821 | ENSMUSG00000022952 | **Runx1** | AML1\|CBF-alpha-2\|Cbfa2\|Pebp2a2\|Pebpa2b | runt related transcription factor 1 | protein-coding |
| chr16_92896831_92897506 | chr16 | 92896832 | 92897506 | + | 0.687204 | Intergenic | Intergenic | -71095 | NM_001111022 | 12394 | Mm.4081 | NM_009821 | ENSMUSG00000022952 | **Runx1** | AML1\|CBF-alpha-2\|Cbfa2\|Pebp2a2\|Pebpa2b | runt related transcription factor 1 | protein-coding |
| chr16_92620579_92621543 | chr16 | 92620580 | 92621543 | + | 0.686253 | intron (NM_001111022, intron 5 of 6) | intron (NM_001111022, intron 5 of 6) | 76268 | NM_001111023 | 12394 | Mm.4081 | NM_009821 | ENSMUSG00000022952 | **Runx1** | AML1\|CBF-alpha-2\|Cbfa2\|Pebp2a2\|Pebpa2b | runt related transcription factor 1 | protein-coding |
| chr16_92997522_92998093 | chr16 | 92997523 | 92998093 | + | 0.646115 | Intergenic | Intergenic | -171734 | NM_001111022 | 12394 | Mm.4081 | NM_009821 | ENSMUSG00000022952 | **Runx1** | AML1\|CBF-alpha-2\|Cbfa2\|Pebp2a2\|Pebpa2b | runt related transcription factor 1 | protein-coding |
| chr17_44536419_44536963 | chr17 | 44536420 | 44536963 | + | 1.62201 | intron (NR_073425, intron 5 of 5) | intron (NR_073425, intron 5 of 5) | 199485 | NR_073425 | 12393 | Mm.391013 | NM_009820 | ENSMUSG00000039153 | **Runx2** | AML3\|Cbf\|Cbfa-1\|Cbfa1\|LS3\|Osf2\|PEBP2aA\|Pebp2a1\|Pebpa2a | runt related transcription factor 2 | protein-coding |
| chr17_44536565_44536917 | chr17 | 44536566 | 44536917 | + | 1.59219 | intron (NR_073425, intron 5 of 5) | intron (NR_073425, intron 5 of 5) | 199435 | NR_073425 | 12393 | Mm.391013 | NM_009820 | ENSMUSG00000039153 | **Runx2** | AML3\|Cbf\|Cbfa-1\|Cbfa1\|LS3\|Osf2\|PEBP2aA\|Pebp2a1\|Pebpa2a | runt related transcription factor 2 | protein-coding |
| chr17_44714149_44714520 | chr17 | 44714150 | 44714520 | + | 1.2936 | intron (NR_073392, intron 4 of 7) | intron (NR_073392, intron 4 of 7) | 21841 | NR_073425 | 12393 | Mm.391013 | NM_009820 | ENSMUSG00000039153 | **Runx2** | AML3\|Cbf\|Cbfa-1\|Cbfa1\|LS3\|Osf2\|PEBP2aA\|Pebp2a1\|Pebpa2a | runt related transcription factor 2 | protein-coding |
| chr17_44727667_44728213 | chr17 | 44727668 | 44728213 | + | 1.2774 | intron (NR_073392, intron 2 of 7) | intron (NR_073392, intron 2 of 7) | 8236 | NR_073425 | 12393 | Mm.391013 | NM_009820 | ENSMUSG00000039153 | **Runx2** | AML3\|Cbf\|Cbfa-1\|Cbfa1\|LS3\|Osf2\|PEBP2aA\|Pebp2a1\|Pebpa2a | runt related transcription factor 2 | protein-coding |
| chr17_44713298_44713887 | chr17 | 44713299 | 44713887 | + | 1.12387 | intron (NR_073392, intron 4 of 7) | intron (NR_073392, intron 4 of 7) | 22583 | NR_073425 | 12393 | Mm.391013 | NM_009820 | ENSMUSG00000039153 | **Runx2** | AML3\|Cbf\|Cbfa-1\|Cbfa1\|LS3\|Osf2\|PEBP2aA\|Pebp2a1\|Pebpa2a | runt related transcription factor 2 | protein-coding |
| chr17_44674570_44675435 | chr17 | 44674571 | 44675435 | + | 1.05914 | intron (NR_073392, intron 4 of 7) | intron (NR_073392, intron 4 of 7) | 61173 | NR_073425 | 12393 | Mm.391013 | NM_009820 | ENSMUSG00000039153 | **Runx2** | AML3\|Cbf\|Cbfa-1\|Cbfa1\|LS3\|Osf2\|PEBP2aA\|Pebp2a1\|Pebpa2a | runt related transcription factor 2 | protein-coding |
| chr17_44728523_44728808 | chr17 | 44728524 | 44728808 | + | 0.976885 | intron (NR_073392, intron 2 of 7) | intron (NR_073392, intron 2 of 7) | 7510 | NR_073425 | 12393 | Mm.391013 | NM_009820 | ENSMUSG00000039153 | **Runx2** | AML3\|Cbf\|Cbfa-1\|Cbfa1\|LS3\|Osf2\|PEBP2aA\|Pebp2a1\|Pebpa2a | runt related transcription factor 2 | protein-coding |
| chr17_44728339_44728829 | chr17 | 44728340 | 44728829 | + | 0.938812 | intron (NR_073392, intron 2 of 7) | intron (NR_073392, intron 2 of 7) | 7592 | NR_073425 | 12393 | Mm.391013 | NM_009820 | ENSMUSG00000039153 | **Runx2** | AML3\|Cbf\|Cbfa-1\|Cbfa1\|LS3\|Osf2\|PEBP2aA\|Pebp2a1\|Pebpa2a | runt related transcription factor 2 | protein-coding |
| chr17_44532794_44533195 | chr17 | 44532795 | 44533195 | + | 0.809506 | intron (NR_073425, intron 5 of 5) | intron (NR_073425, intron 5 of 5) | 203181 | NR_073425 | 12393 | Mm.391013 | NM_009820 | ENSMUSG00000039153 | **Runx2** | AML3\|Cbf\|Cbfa-1\|Cbfa1\|LS3\|Osf2\|PEBP2aA\|Pebp2a1\|Pebpa2a | runt related transcription factor 2 | protein-coding |
| chr17_44718598_44718893 | chr17 | 44718599 | 44718893 | + | 0.722801 | intron (NR_073425, intron 2 of 5) | intron (NR_073425, intron 2 of 5) | 17430 | NR_073425 | 12393 | Mm.391013 | NM_009820 | ENSMUSG00000039153 | **Runx2** | AML3\|Cbf\|Cbfa-1\|Cbfa1\|LS3\|Osf2\|PEBP2aA\|Pebp2a1\|Pebpa2a | runt related transcription factor 2 | protein-coding |
| chr17_44555914_44556277 | chr17 | 44555915 | 44556277 | + | 0.685276 | intron (NR_073425, intron 5 of 5) | intron (NR_073425, intron 5 of 5) | 180080 | NR_073425 | 12393 | Mm.391013 | NM_009820 | ENSMUSG00000039153 | **Runx2** | AML3\|Cbf\|Cbfa-1\|Cbfa1\|LS3\|Osf2\|PEBP2aA\|Pebp2a1\|Pebpa2a | runt related transcription factor 2 | protein-coding |
| chr17_44718491_44718898 | chr17 | 44718492 | 44718898 | + | 0.640578 | intron (NR_073425, intron 2 of 5) | intron (NR_073425, intron 2 of 5) | 17481 | NR_073425 | 12393 | Mm.391013 | NM_009820 | ENSMUSG00000039153 | **Runx2** | AML3\|Cbf\|Cbfa-1\|Cbfa1\|LS3\|Osf2\|PEBP2aA\|Pebp2a1\|Pebpa2a | runt related transcription factor 2 | protein-coding |
| chr4_135118559_135118881 | chr4 | 135118560 | 135118881 | + | 1.47607 | Intergenic | Intergenic | -1925 | NM_019732 | 12399 | Mm.378894 | NM_019732 | ENSMUSG00000070691 | **Runx3** | AML2\|Cbfa3\|Pebp2a3 | runt related transcription factor 3 | protein-coding |
| chr4_135118555_135119173 | chr4 | 135118556 | 135119173 | + | 1.31874 | Intergenic | Intergenic | -1781 | NM_019732 | 12399 | Mm.378894 | NM_019732 | ENSMUSG00000070691 | **Runx3** | AML2\|Cbfa3\|Pebp2a3 | runt related transcription factor 3 | protein-coding |
| chr4_135073381_135073937 | chr4 | 135073382 | 135073937 | + | 0.959868 | Intergenic | Intergenic | -46986 | NM_019732 | 12399 | Mm.378894 | NM_019732 | ENSMUSG00000070691 | **Runx3** | AML2\|Cbfa3\|Pebp2a3 | runt related transcription factor 3 | protein-coding |
| chr4_135118927_135119161 | chr4 | 135118928 | 135119161 | + | 0.935117 | Intergenic | Intergenic | -1601 | NM_019732 | 12399 | Mm.378894 | NM_019732 | ENSMUSG00000070691 | **Runx3** | AML2\|Cbfa3\|Pebp2a3 | runt related transcription factor 3 | protein-coding |
| chr4_135086047_135086476 | chr4 | 135086048 | 135086476 | + | 0.930484 | Intergenic | Intergenic | -34383 | NM_019732 | 12399 | Mm.378894 | NM_019732 | ENSMUSG00000070691 | **Runx3** | AML2\|Cbfa3\|Pebp2a3 | runt related transcription factor 3 | protein-coding |
| chr4_135114366_135114765 | chr4 | 135114367 | 135114765 | + | 0.720357 | Intergenic | Intergenic | -6079 | NM_019732 | 12399 | Mm.378894 | NM_019732 | ENSMUSG00000070691 | **Runx3** | AML2\|Cbfa3\|Pebp2a3 | runt related transcription factor 3 | protein-coding |
| chr4_135158598_135159405 | chr4 | 135158599 | 135159405 | + | 0.643492 | intron (NM_019732, intron 3 of 5) | intron (NM_019732, intron 3 of 5) | 38357 | NM_019732 | 12399 | Mm.378894 | NM_019732 | ENSMUSG00000070691 | **Runx3** | AML2\|Cbfa3\|Pebp2a3 | runt related transcription factor 3 | protein-coding |
| chr3_89046853_89047221 | chr3 | 89046854 | 89047221 | + | 1.14778 | intron (NM_138679, intron 11 of 26) | intron (NM_138679, intron 11 of 26) | 42994 | NM_001083808 | 72296 | Mm.27687 | NM_028188 | ENSMUSG00000041263 | **Rusc1** | 2210403N08Rik\|AA408288\|NESCA | RUN and SH3 domain containing 1 | protein-coding |
| chr9_86624773_86625217 | chr9 | 86624774 | 86625217 | + | 0.981259 | intron (NM_001198933, intron 6 of 13) | MIRb\|SINE\|MIR | 52942 | NM_027100 | 69519 | Mm.272517 | NM_027100 | ENSMUSG00000032417 | **Rwdd2a** | 1700030C20Rik\|AI848608\|Rwdd2 | RWD domain containing 2A | protein-coding |
| chr9_86624044_86624552 | chr9 | 86624045 | 86624552 | + | 0.650211 | intron (NM_001198933, intron 6 of 13) | intron (NM_001198933, intron 6 of 13) | 52245 | NM_027100 | 69519 | Mm.272517 | NM_027100 | ENSMUSG00000032417 | **Rwdd2a** | 1700030C20Rik\|AI848608\|Rwdd2 | RWD domain containing 2A | protein-coding |
| chr8_47566537_47566922 | chr8 | 47566538 | 47566922 | + | 0.778986 | Intergenic | ORR1B1\|LTR\|ERVL-MaLR | 33085 | NM_203507 | 192174 | Mm.273081 | NM_203507 | ENSMUSG00000031568 | **Rwdd4a** | BC016198\|Gm1942\|Rwdd4 | RWD domain containing 4A | protein-coding |
| chr2_27728015_27729348 | chr2 | 27728016 | 27729348 | + | 0.772571 | intron (NM_001290482, intron 1 of 9) | intron (NM_001290482, intron 1 of 9) | 18099 | NM_001290482 | 20181 | Mm.24624 | NM_011305 | ENSMUSG00000015846 | **Rxra** | 9530071D11Rik\|Nr2b1\|RXRalpha1 | retinoid X receptor alpha | protein-coding |
| chr2_27712388_27713022 | chr2 | 27712389 | 27713022 | + | 0.735484 | intron (NM_001290482, intron 1 of 9) | intron (NM_001290482, intron 1 of 9) | 2122 | NM_001290482 | 20181 | Mm.24624 | NM_011305 | ENSMUSG00000015846 | **Rxra** | 9530071D11Rik\|Nr2b1\|RXRalpha1 | retinoid X receptor alpha | protein-coding |
| chr2_27728495_27729276 | chr2 | 27728496 | 27729276 | + | 0.653741 | intron (NM_001290482, intron 1 of 9) | intron (NM_001290482, intron 1 of 9) | 18303 | NM_001290482 | 20181 | Mm.24624 | NM_011305 | ENSMUSG00000015846 | **Rxra** | 9530071D11Rik\|Nr2b1\|RXRalpha1 | retinoid X receptor alpha | protein-coding |
| chr6_100280443_100280887 | chr6 | 100280444 | 100280887 | + | 1.39196 | intron (NM_019743, intron 2 of 4) | intron (NM_019743, intron 2 of 4) | 6693 | NM_019743 | 56353 | Mm.321633 | NM_019743 | ENSMUSG00000072872 | **Rybp** | 2410018J24Rik\|DEDAF\|YEAF1 | RING1 and YY1 binding protein | protein-coding |
| chr6_100248351_100248898 | chr6 | 100248352 | 100248898 | + | 1.05345 | intron (NM_019743, intron 2 of 4) | intron (NM_019743, intron 2 of 4) | 38733 | NM_019743 | 56353 | Mm.321633 | NM_019743 | ENSMUSG00000072872 | **Rybp** | 2410018J24Rik\|DEDAF\|YEAF1 | RING1 and YY1 binding protein | protein-coding |
| chr6_100314061_100314648 | chr6 | 100314062 | 100314648 | + | 0.977224 | Intergenic | Intergenic | -26997 | NM_019743 | 56353 | Mm.321633 | NM_019743 | ENSMUSG00000072872 | **Rybp** | 2410018J24Rik\|DEDAF\|YEAF1 | RING1 and YY1 binding protein | protein-coding |
| chr6_100363625_100364103 | chr6 | 100363626 | 100364103 | + | 0.944106 | Intergenic | Intergenic | -76506 | NM_019743 | 56353 | Mm.321633 | NM_019743 | ENSMUSG00000072872 | **Rybp** | 2410018J24Rik\|DEDAF\|YEAF1 | RING1 and YY1 binding protein | protein-coding |
| chr6_100275497_100276009 | chr6 | 100275498 | 100276009 | + | 0.851842 | intron (NM_019743, intron 2 of 4) | intron (NM_019743, intron 2 of 4) | 11605 | NM_019743 | 56353 | Mm.321633 | NM_019743 | ENSMUSG00000072872 | **Rybp** | 2410018J24Rik\|DEDAF\|YEAF1 | RING1 and YY1 binding protein | protein-coding |
| chr6_100152910_100153507 | chr6 | 100152911 | 100153507 | + | 0.745257 | Intergenic | Intergenic | 134149 | NM_019743 | 56353 | Mm.321633 | NM_019743 | ENSMUSG00000072872 | **Rybp** | 2410018J24Rik\|DEDAF\|YEAF1 | RING1 and YY1 binding protein | protein-coding |
| chr6_100259596_100259962 | chr6 | 100259597 | 100259962 | + | 0.739082 | intron (NM_019743, intron 2 of 4) | L1MB7\|LINE\|L1 | 27579 | NM_019743 | 56353 | Mm.321633 | NM_019743 | ENSMUSG00000072872 | **Rybp** | 2410018J24Rik\|DEDAF\|YEAF1 | RING1 and YY1 binding protein | protein-coding |
| chr6_100275474_100276217 | chr6 | 100275475 | 100276217 | + | 0.727268 | intron (NM_019743, intron 2 of 4) | intron (NM_019743, intron 2 of 4) | 11512 | NM_019743 | 56353 | Mm.321633 | NM_019743 | ENSMUSG00000072872 | **Rybp** | 2410018J24Rik\|DEDAF\|YEAF1 | RING1 and YY1 binding protein | protein-coding |
| chr6_100084151_100084517 | chr6 | 100084152 | 100084517 | + | 0.723464 | Intergenic | Intergenic | 203024 | NM_019743 | 56353 | Mm.321633 | NM_019743 | ENSMUSG00000072872 | **Rybp** | 2410018J24Rik\|DEDAF\|YEAF1 | RING1 and YY1 binding protein | protein-coding |
| chr6_100072550_100073206 | chr6 | 100072551 | 100073206 | + | 0.67654 | Intergenic | Intergenic | 214480 | NM_019743 | 56353 | Mm.321633 | NM_019743 | ENSMUSG00000072872 | **Rybp** | 2410018J24Rik\|DEDAF\|YEAF1 | RING1 and YY1 binding protein | protein-coding |
| chr3_93571871_93572807 | chr3 | 93571872 | 93572807 | + | 1.84975 | Intergenic | Intergenic | 17222 | NM_009112 | 20194 | Mm.1 | NM_009112 | ENSMUSG00000041959 | **S100a10** | 42C\|AA409961\|AL024248\|CAL12\|CLP11\|Cal1l\|p10\|p11 | S100 calcium binding protein A10 (calpactin) | protein-coding |
| chr3_93552076_93552263 | chr3 | 93552077 | 93552263 | + | 1.012 | Intergenic | MLT1J\|LTR\|ERVL-MaLR | -2947 | NM_009112 | 20194 | Mm.1 | NM_009112 | ENSMUSG00000041959 | **S100a10** | 42C\|AA409961\|AL024248\|CAL12\|CLP11\|Cal1l\|p10\|p11 | S100 calcium binding protein A10 (calpactin) | protein-coding |

| | | | | | | | | | | | | | | | | | |
|---|---|---|---|---|---|---|---|---|---|---|---|---|---|---|---|---|---|
| chr3_93598043_93598325 | chr3 | 93598044 | 93598325 | + | 0.923974 | Intergenic | Intergenic | 43067 | NM_009112 | 20194 | Mm.1 | NM_009112 | ENSMUSG00000041959 | **S100a10** | 42C\|AA409961\|AL024248\|CAL12\|CLP11\|Cal1l\|p10\|p11 | S100 calcium binding protein A10 (calpactin) | protein-coding |
| chr3_93597141_93597803 | chr3 | 93597142 | 93597803 | + | 0.916864 | Intergenic | Intergenic | 42355 | NM_009112 | 20194 | Mm.1 | NM_009112 | ENSMUSG00000041959 | **S100a10** | 42C\|AA409961\|AL024248\|CAL12\|CLP11\|Cal1l\|p10\|p11 | S100 calcium binding protein A10 (calpactin) | protein-coding |
| chr3_93597167_93597500 | chr3 | 93597168 | 93597500 | + | 0.885031 | Intergenic | Intergenic | 42217 | NM_009112 | 20194 | Mm.1 | NM_009112 | ENSMUSG00000041959 | **S100a10** | 42C\|AA409961\|AL024248\|CAL12\|CLP11\|Cal1l\|p10\|p11 | S100 calcium binding protein A10 (calpactin) | protein-coding |
| chr3_93562356_93562716 | chr3 | 93562357 | 93562716 | + | 0.71709 | intron (NM_009112, intron 2 of 2) | intron (NM_009112, intron 2 of 2) | 7419 | NM_009112 | 20194 | Mm.1 | NM_009112 | ENSMUSG00000041959 | **S100a10** | 42C\|AA409961\|AL024248\|CAL12\|CLP11\|Cal1l\|p10\|p11 | S100 calcium binding protein A10 (calpactin) | protein-coding |
| chr3_93562329_93562919 | chr3 | 93562330 | 93562919 | + | 0.713526 | intron (NM_009112, intron 2 of 2) | intron (NM_009112, intron 2 of 2) | 7507 | NM_009112 | 20194 | Mm.1 | NM_009112 | ENSMUSG00000041959 | **S100a10** | 42C\|AA409961\|AL024248\|CAL12\|CLP11\|Cal1l\|p10\|p11 | S100 calcium binding protein A10 (calpactin) | protein-coding |
| chr3_93504910_93505105 | chr3 | 93504911 | 93505105 | + | 0.677097 | Intergenic | Looper\|DNA\|PiggyBac | -15488 | NM_016740 | 20195 | Mm.280038 | NM_016740 | ENSMUSG00000027907 | **S100a11** | EMAPI\|Emap1\|S100a14\|S100c\|cal | S100 calcium binding protein A11 (calgizzarin) | protein-coding |
| chr3_93496000_93496352 | chr3 | 93496001 | 93496352 | + | 0.628698 | Intergenic | B3\|SINE\|B2 | -24320 | NM_016740 | 20195 | Mm.280038 | NM_016740 | ENSMUSG00000027907 | **S100a11** | EMAPI\|Emap1\|S100a14\|S100c\|cal | S100 calcium binding protein A11 (calgizzarin) | protein-coding |
| chr3_90587739_90588136 | chr3 | 90587740 | 90588136 | + | 0.902524 | Intergenic | Intergenic | -2309 | NM_001195760 | 628324 | Mm.33816 | NM_001195760 | ENSMUSG00000094018 | **S100a2** | CaN19\|EG628324\|S100L | S100 calcium binding protein A2 | protein-coding |
| chr3_90585591_90585959 | chr3 | 90585592 | 90585959 | + | 0.722453 | Intergenic | Intergenic | -4472 | NM_001195760 | 628324 | Mm.33816 | NM_001195760 | ENSMUSG00000094018 | **S100a2** | CaN19\|EG628324\|S100L | S100 calcium binding protein A2 | protein-coding |
| chr3_90604345_90605024 | chr3 | 90604346 | 90605024 | + | 0.754493 | intron (NM_011311, intron 1 of 2) | intron (NM_011311, intron 1 of 2) | 915 | NM_011311 | 20198 | Mm.3925 | NM_011311 | ENSMUSG00000001020 | **S100a4** | 18A2\|42a\|Capl\|FSp1\|Mts1\|PeL98\|metastasin\|pk9a | S100 calcium binding protein A4 | protein-coding |
| chr3_90629316_90629640 | chr3 | 90629317 | 90629640 | + | 1.03283 | Intergenic | L2b\|LINE\|L2 | 16584 | NM_011313 | 20200 | Mm.100144 | NM_011313 | ENSMUSG00000001025 | **S100a6** | 2A9\|5B10\|CALCYCLIN\|Cacy\|PRA | S100 calcium binding protein A6 (calcyclin) | protein-coding |
| chr3_90629140_90629910 | chr3 | 90629141 | 90629910 | + | 0.979094 | Intergenic | Intergenic | 16631 | NM_011313 | 20200 | Mm.100144 | NM_011313 | ENSMUSG00000001025 | **S100a6** | 2A9\|5B10\|CALCYCLIN\|Cacy\|PRA | S100 calcium binding protein A6 (calcyclin) | protein-coding |
| chr3_90638918_90639248 | chr3 | 90638919 | 90639248 | + | 0.711087 | Intergenic | Intergenic | -15219 | NM_199422 | 381493 | Mm.291525 | NM_199422 | ENSMUSG00000063767 | **S100a7a** | AY465109\|Gm1020\|S100A7L1\|S100A7f\|S100a15\|S100a15a\|S100a17l1 | S100 calcium binding protein A7A | protein-coding |
| chr3_115730586_115731228 | chr3 | 115730587 | 115731228 | + | 1.24142 | Intergenic | L1M2\|LINE\|L1 | -15852 | NM_007901 | 13609 | Mm.982 | NM_007901 | ENSMUSG00000045092 | **S1pr1** | AI849002\|Edg1\|Lpb1\|S1p\|S1p1 | sphingosine-1-phosphate receptor 1 | protein-coding |
| chr3_115728305_115728570 | chr3 | 115728306 | 115728570 | + | 0.887725 | Intergenic | HAL1\|LINE\|L1 | -13383 | NM_007901 | 13609 | Mm.982 | NM_007901 | ENSMUSG00000045092 | **S1pr1** | AI849002\|Edg1\|Lpb1\|S1p\|S1p1 | sphingosine-1-phosphate receptor 1 | protein-coding |
| chr3_115761296_115761712 | chr3 | 115761297 | 115761712 | + | 0.878529 | Intergenic | Intergenic | -46449 | NM_007901 | 13609 | Mm.982 | NM_007901 | ENSMUSG00000045092 | **S1pr1** | AI849002\|Edg1\|Lpb1\|S1p\|S1p1 | sphingosine-1-phosphate receptor 1 | protein-coding |
| chr9_123505091_123505473 | chr9 | 123505092 | 123505473 | + | 0.972702 | intron (NM_013860, intron 2 of 7) | intron (NM_013860, intron 2 of 7) | -24600 | NM_030692 | 83493 | Mm.273671 | NM_030692 | ENSMUSG00000025240 | **Sacm1l** | SAC1\|Sac1p\|mKIAA0851 | SAC1 (suppressor of actin mutations 1, homolog)-like (S. cerevisiae) | protein-coding |
| chr9_123511052_123511259 | chr9 | 123511053 | 123511259 | + | 0.922834 | intron (NM_013860, intron 3 of 7) | intron (NM_013860, intron 3 of 7) | -18726 | NM_030692 | 83493 | Mm.273671 | NM_030692 | ENSMUSG00000025240 | **Sacm1l** | SAC1\|Sac1p\|mKIAA0851 | SAC1 (suppressor of actin mutations 1, homolog)-like (S. cerevisiae) | protein-coding |
| chr9_123535779_123536194 | chr9 | 123535780 | 123536194 | + | 0.864373 | intron (NM_030692, intron 1 of 19) | intron (NM_030692, intron 1 of 19) | 6105 | NM_030692 | 83493 | Mm.273671 | NM_030692 | ENSMUSG00000025240 | **Sacm1l** | SAC1\|Sac1p\|mKIAA0851 | SAC1 (suppressor of actin mutations 1, homolog)-like (S. cerevisiae) | protein-coding |
| chr9_123510981_123511308 | chr9 | 123510982 | 123511308 | + | 0.861691 | intron (NM_013860, intron 3 of 7) | intron (NM_013860, intron 3 of 7) | -18737 | NM_030692 | 83493 | Mm.273671 | NM_030692 | ENSMUSG00000025240 | **Sacm1l** | SAC1\|Sac1p\|mKIAA0851 | SAC1 (suppressor of actin mutations 1, homolog)-like (S. cerevisiae) | protein-coding |
| chr1_87811103_87811876 | chr1 | 87811104 | 87811876 | + | 0.676127 | intron (NM_009118, intron 3 of 15) | L1MD1\|LINE\|L1 | 7810 | NM_009118 | 20215 | Mm.1276 | NM_009118 | ENSMUSG00000056055 | **Sag** | A930001K18Rik\|Arr1\|Irbp\|arrestin | S-antigen, retina and pineal gland (arrestin) | protein-coding |
| chr18_81319377_81320150 | chr18 | 81319378 | 81320150 | + | 0.895245 | Intergenic | Intergenic | -333186 | NM_178280 | 20689 | Mm.215917 | NM_178280 | ENSMUSG00000024565 | **Sall3** | B130022O04Rik\|Msal\|Msal-1\|Sal\|Salt\|Spalt | sal-like 3 (Drosophila) | protein-coding |
| chr15_54071790_54072135 | chr15 | 54071791 | 54072135 | + | 0.896906 | Intergenic | MER97c\|DNA\|hAT-Tip100 | -169527 | NM_177225 | 320679 | Mm.442455 | NM_177225 | ENSMUSG00000058656 | **Samd12** | A830094I09Rik | sterile alpha motif domain containing 12 | protein-coding |
| chr10_26264029_26264559 | chr10 | 26264030 | 26264559 | + | 1.06192 | intron (NM_001115154, intron 8 of 10) | intron (NM_001115154, intron 8 of 10) | 34587 | NM_001115154 | 268288 | Mm.213322 | NM_001013766 | ENSMUSG00000051354 | **Samd3** | Gm623 | sterile alpha motif domain containing 3 | protein-coding |
| chr14_46891317_46891694 | chr14 | 46891318 | 46891694 | + | 0.928504 | intron (NM_028966, intron 2 of 11) | intron (NM_028966, intron 2 of 11) | 8541 | NM_001037221 | 74480 | Mm.214454 | NM_028966 | ENSMUSG00000021838 | **Samd4** | 1700024G08Rik\|1700111L17Rik\|4933436G17Rik\|Samd4a\|Smaug\|Smaug1 | sterile alpha motif domain containing 4 | protein-coding |
| chr7_28422877_28423760 | chr7 | 28422878 | 28423760 | + | 1.28975 | intron (NM_175021, intron 2 of 12) | intron (NM_175021, intron 2 of 12) | 12872 | NM_175021 | 233033 | Mm.54120 | NM_175021 | ENSMUSG00000037513 | **Samd4b** | BC042901 | sterile alpha motif domain containing 4B | protein-coding |
| chr6_3395707_3396126 | chr6 | 3395708 | 3396126 | + | 1.36951 | intron (NM_010156, intron 1 of 1) | B4A\|SINE\|B4 | 3654 | NM_010156 | 209086 | Mm.196013 | NM_010156 | ENSMUSG00000047735 | **Samd9l** | AA175286\|ESTM25\|mKIAA2005 | sterile alpha motif domain containing 9-like | protein-coding |
| chr6_3395702_3396551 | chr6 | 3395703 | 3396551 | + | 0.908786 | intron (NM_010156, intron 1 of 1) | RMER30\|DNA\|hAT-Charlie | 3444 | NM_010156 | 209086 | Mm.196013 | NM_010156 | ENSMUSG00000047735 | **Samd9l** | AA175286\|ESTM25\|mKIAA2005 | sterile alpha motif domain containing 9-like | protein-coding |
| chr16_75837718_75838372 | chr16 | 75837719 | 75838372 | + | 1.38689 | Intergenic | L1MB7\|LINE\|L1 | 71221 | NM_023380 | 67742 | Mm.131406 | NM_023380 | ENSMUSG00000022876 | **Samsn1** | 4930571B16Rik\|930571B16Rik\|Hacs1\|SLy2 | SAM domain, SH3 domain and nuclear localization signals, 1 | protein-coding |
| chr16_75894177_75894883 | chr16 | 75894178 | 75894883 | + | 1.01448 | intron (NM_023380, intron 1 of 7) | intron (NM_023380, intron 1 of 7) | 14736 | NM_023380 | 67742 | Mm.131406 | NM_023380 | ENSMUSG00000022876 | **Samsn1** | 4930571B16Rik\|930571B16Rik\|Hacs1\|SLy2 | SAM domain, SH3 domain and nuclear localization signals, 1 | protein-coding |
| chr18_31692833_31693263 | chr18 | 31692834 | 31693263 | + | 0.943769 | intron (NM_172965, intron 15 of 20) | Tigger6b\|DNA\|TcMar-Tigger | 58665 | NM_172965 | 269003 | Mm.194538 | NM_172965 | ENSMUSG00000024260 | **Sap130** | 2610304F09Rik\|6720406D06 | Sin3A associated protein | protein-coding |
| chr18_31692554_31693266 | chr18 | 31692555 | 31693266 | + | 0.828435 | intron (NM_172965, intron 15 of 20) | Tigger6b\|DNA\|TcMar-Tigger | 58527 | NM_172965 | 269003 | Mm.194538 | NM_172965 | ENSMUSG00000024260 | **Sap130** | 2610304F09Rik\|6720406D06 | Sin3A associated protein | protein-coding |
| chr8_57476857_57477300 | chr8 | 57476858 | 57477300 | + | 0.978229 | intron (NM_009136, intron 2 of 2) | intron (NM_009136, intron 2 of 2) | 10781 | NM_021788 | 60406 | Mm.118 | NM_021788 | ENSMUSG00000031609 | **Sap30** | 30kDa | sin3 associated polypeptide | protein-coding |
| chr8_57476291_57476658 | chr8 | 57476292 | 57476658 | + | 0.728325 | intron (NM_009136, intron 2 of 2) | intron (NM_009136, intron 2 of 2) | 11385 | NM_021788 | 60406 | Mm.118 | NM_021788 | ENSMUSG00000031609 | **Sap30** | 30kDa | sin3 associated polypeptide | protein-coding |
| chr11_51760052_51760757 | chr11 | 51760053 | 51760757 | + | 1.89194 | Intergenic | URR1B\|DNA\|hAT-Charlie | -3258 | NM_025535 | 66397 | Mm.196592 | NM_025535 | ENSMUSG00000020386 | **Sar1b** | 2310075M17Rik\|2900019I22Rik\|CMRD\|Sara1b\|Sara2\|Sarb | SAR1 gene homolog B (S. cerevisiae) | protein-coding |
| chr11_51761312_51761556 | chr11 | 51761313 | 51761556 | + | 1.06424 | Intergenic | Intergenic | -2229 | NM_025535 | 66397 | Mm.196592 | NM_025535 | ENSMUSG00000020386 | **Sar1b** | 2310075M17Rik\|2900019I22Rik\|CMRD\|Sara1b\|Sara2\|Sarb | SAR1 gene homolog B (S. cerevisiae) | protein-coding |
| chr2_27322123_27322700 | chr2 | 27322124 | 27322700 | + | 0.774039 | intron (NM_009500, intron 3 of 27) | MTEb\|LTR\|ERVL-MaLR | -75109 | NM_138665 | 192166 | Mm.278467 | NM_138665 | ENSMUSG00000009614 | **Sardh** | - | sarcosine dehydrogenase | protein-coding |
| chrX_155218382_155218621 | chrX | 155218383 | 155218621 | + | 1.02811 | Intergenic | Intergenic | -2053 | NM_001291865 | 20229 | Mm.2734 | NM_009121 | ENSMUSG00000025283 | **Sat1** | AA617398\|SSAT\|Sat | spermidine/spermine N1-acetyl transferase 1 | protein-coding |
| chr14_20074783_20075339 | chr14 | 20074784 | 20075339 | + | 1.18289 | TTS (NM_026209) | TTS (NM_026209) | 8111 | NM_026209 | 67509 | Mm.292423 | NM_026209 | ENSMUSG00000045107 | **Saysd1** | 1810063B07Rik\|4930488P03Rik | SAYSVFN motif domain containing 1 | protein-coding |
| chr14_20098008_20098465 | chr14 | 20098009 | 20098465 | + | 0.733595 | Intergenic | Intergenic | -15065 | NM_026209 | 67509 | Mm.292423 | NM_026209 | ENSMUSG00000045107 | **Saysd1** | 1810063B07Rik\|4930488P03Rik | SAYSVFN motif domain containing 1 | protein-coding |
| chr7_110619121_110619839 | chr7 | 110619122 | 110619839 | + | 0.755741 | Intergenic | Intergenic | -4560 | NM_177324 | 319934 | Mm.102970 | NM_177324 | ENSMUSG00000038371 | **Sbf2** | 4833411B01Rik\|AA414977\|AI317167\|B430219L04Rik\|Mtmr13\|mMTMH1 | SET binding factor 2 | protein-coding |
| chr7_126282598_126282913 | chr7 | 126282599 | 126282913 | + | 0.745436 | intron (NM_145587, intron 1 of 3) | intron (NM_145587, intron 1 of 3) | 10137 | NM_145587 | 104175 | Mm.29660 | NM_145587 | ENSMUSG00000042978 | **Sbk1** | Sbk | SH3-binding kinase 1 | protein-coding |
| chr7_126289415_126289731 | chr7 | 126289416 | 126289731 | + | 0.735181 | intron (NM_145587, intron 1 of 3) | intron (NM_145587, intron 1 of 3) | 16954 | NM_145587 | 104175 | Mm.29660 | NM_145587 | ENSMUSG00000042978 | **Sbk1** | Sbk | SH3-binding kinase 1 | protein-coding |
| chr5_124411552_124412079 | chr5 | 124411553 | 124412079 | + | 1.26135 | intron (NM_001081203, intron 3 of 31) | intron (NM_001081203, intron 3 of 31) | 14098 | NM_001081203 | 243272 | Mm.23879 | NM_001081203 | ENSMUSG00000038095 | **Sbno1** | 9330180L10Rik\|AI849128\|AW557836\|BC021875\|Sbno\|mSno1\|sno | sno, strawberry notch homolog 1 (Drosophila) | protein-coding |
| chr5_124412297_124412647 | chr5 | 124412298 | 124412647 | + | 0.778616 | intron (NM_001081203, intron 3 of 31) | intron (NM_001081203, intron 3 of 31) | 13442 | NM_001081203 | 243272 | Mm.23879 | NM_001081203 | ENSMUSG00000038095 | **Sbno1** | 9330180L10Rik\|AI849128\|AW557836\|BC021875\|Sbno\|mSno1\|sno | sno, strawberry notch homolog 1 (Drosophila) | protein-coding |
| chr9_42313844_42314270 | chr9 | 42313845 | 42314270 | + | 0.862673 | Intergenic | Intergenic | -49757 | NM_172769 | 235293 | Mm.32700 | NM_172769 | ENSMUSG00000032018 | **Sc5d** | A830037K02\|A830073K23Rik\|Sc5dl | sterol-C5-desaturase (fungal ERG3, delta-5-desaturase) homolog (S. cerevisae) | protein-coding |
| chr9_42252921_42253269 | chr9 | 42252922 | 42253269 | + | 0.762942 | Intergenic | Intergenic | 11205 | NM_172769 | 235293 | Mm.32700 | NM_172769 | ENSMUSG00000032018 | **Sc5d** | A830037K02\|A830073K23Rik\|Sc5dl | sterol-C5-desaturase (fungal ERG3, delta-5-desaturase) homolog (S. cerevisae) | protein-coding |
| chr9_42252921_42253452 | chr9 | 42252922 | 42253452 | + | 0.650947 | TTS (NM_172769) | TTS (NM_172769) | 11113 | NM_172769 | 235293 | Mm.32700 | NM_172769 | ENSMUSG00000032018 | **Sc5d** | A830037K02\|A830073K23Rik\|Sc5dl | sterol-C5-desaturase (fungal ERG3, delta-5-desaturase) homolog (S. cerevisae) | protein-coding |
| chr15_96520396_96520803 | chr15 | 96520397 | 96520803 | + | 0.98618 | Intergenic | Intergenic | -59757 | NM_028148 | 72193 | Mm.324474 | NM_028148 | ENSMUSG00000033228 | **Scaf11** | 1110061H03Rik\|2610510E10Rik\|AI462454\|CASP11\|SIP1\|SRRP129\|Sfrs2ip\|Srsf2ip\|mKIAA3013 | SR-related CTD-associated factor 11 | protein-coding |
| chr16_90331008_90331495 | chr16 | 90331009 | 90331495 | + | 0.642213 | Intergenic | MER3\|DNA\|hAT-Charlie | -46827 | NM_178923 | 224432 | Mm.439811 | NM_178923 | ENSMUSG00000022983 | **Scaf4** | AA517739\|Sfrs15\|Sra4\|Srsf15 | SR-related CTD-associated factor 4 | protein-coding |
| chr17_3104912_3105410 | chr17 | 3104913 | 3105410 | + | 1.2181 | Intergenic | Lx8\|LINE\|L1 | -9811 | NM_134123 | 106583 | Mm.124373 | NM_134123 | ENSMUSG00000046201 | **Scaf8** | A630086M08Rik\|A930036P18Rik\|AA408306\|AI447644\|AI448652\|AU015035\|Rbm16\|mKIAA1116 | SR-related CTD-associated factor 8 | protein-coding |

| | | | | | | | | | | | | | | | | | |
|---|---|---|---|---|---|---|---|---|---|---|---|---|---|---|---|---|---|
| chr17_3102896_3103173 | chr17 | 3102897 | 3103173 | + | 0.964983 | Intergenic | Intergenic | -11937 | NM_134123 | 106583 | Mm.124373 | NM_134123 | ENSMUSG00000046201 | **Scaf8** | A630086M08Rik\|A930036P18Rik\|AA408306\|AI447644\|AI448652\|AU015035\|Rbm16\|mKIAA1116 | SR-related CTD-associated factor 8 | protein-coding |
| chr17_3102747_3103181 | chr17 | 3102748 | 3103181 | + | 0.773894 | Intergenic | L1MB4\|LINE\|L1 | -12008 | NM_134123 | 106583 | Mm.124373 | NM_134123 | ENSMUSG00000046201 | **Scaf8** | A630086M08Rik\|A930036P18Rik\|AA408306\|AI447644\|AI448652\|AU015035\|Rbm16\|mKIAA1116 | SR-related CTD-associated factor 8 | protein-coding |
| chr17_3204004_3204340 | chr17 | 3204005 | 3204340 | + | 0.7694 | Intergenic | Intergenic | 89200 | NM_134123 | 106583 | Mm.124373 | NM_134123 | ENSMUSG00000046201 | **Scaf8** | A630086M08Rik\|A930036P18Rik\|AA408306\|AI447644\|AI448652\|AU015035\|Rbm16\|mKIAA1116 | SR-related CTD-associated factor 8 | protein-coding |
| chr17_3203492_3204382 | chr17 | 3203493 | 3204382 | + | 0.741409 | Intergenic | Intergenic | 88965 | NM_134123 | 106583 | Mm.124373 | NM_134123 | ENSMUSG00000046201 | **Scaf8** | A630086M08Rik\|A930036P18Rik\|AA408306\|AI447644\|AI448652\|AU015035\|Rbm16\|mKIAA1116 | SR-related CTD-associated factor 8 | protein-coding |
| chr9_55754456_55755054 | chr9 | 55754457 | 55755054 | + | 0.965467 | intron (NM_001081341, intron 23 of 32) | intron (NM_001081341, intron 23 of 32) | 183360 | NM_001081341 | 244891 | Mm.86588 | NM_175536 | ENSMUSG00000034007 | **Scaper** | C430017I08\|D530014O03Rik\|Zfp291 | S phase cyclin A-associated protein in the ER | protein-coding |
| chr9_55786740_55787064 | chr9 | 55786741 | 55787064 | + | 0.801424 | intron (NM_001081341, intron 21 of 32) | L1VL4\|LINE\|L1 | 151213 | NM_001081341 | 244891 | Mm.86588 | NM_175536 | ENSMUSG00000034007 | **Scaper** | C430017I08\|D530014O03Rik\|Zfp291 | S phase cyclin A-associated protein in the ER | protein-coding |
| chr14_65900694_65901050 | chr14 | 65900695 | 65901050 | + | 0.865903 | Intergenic | Intergenic | 52872 | NM_172604 | 219151 | Mm.344095 | NM_172604 | ENSMUSG00000034463 | **Scara3** | APC7\|C130058N24Rik\|CSR\|CSR1\|MSLR1\|MSRL1 | scavenger receptor class A, member 3 | protein-coding |
| chr14_65900206_65901050 | chr14 | 65900207 | 65901050 | + | 0.852364 | Intergenic | Intergenic | 53116 | NM_172604 | 219151 | Mm.344095 | NM_172604 | ENSMUSG00000034463 | **Scara3** | APC7\|C130058N24Rik\|CSR\|CSR1\|MSLR1\|MSRL1 | scavenger receptor class A, member 3 | protein-coding |
| chr5_92489017_92489429 | chr5 | 92489018 | 92489429 | + | 0.67021 | intron (NM_007644, intron 1 of 11) | intron (NM_007644, intron 1 of 11) | 16385 | NM_007644 | 12492 | Mm.297964 | NM_007644 | ENSMUSG00000029426 | **Scarb2** | 9330185J12Rik\|Cd36l2\|LGP85\|LIMP-2\|MLGP85 | scavenger receptor class B, member 2 | protein-coding |
| chr19_44393822_44394448 | chr19 | 44393823 | 44394448 | + | 0.938868 | TTS (NM_009127) | TTS (NM_009127) | 13574 | NM_009127 | 20249 | Mm.267377 | NM_009127 | ENSMUSG00000037071 | **Scd1** | AA589638\|AI265570\|Scd\|Scd-1\|ab | stearoyl-Coenzyme A desaturase 1 | protein-coding |
| chr14_103532344_103532844 | chr14 | 103532345 | 103532844 | + | 0.843711 | intron (NM_022886, intron 3 of 30) | intron (NM_022886, intron 3 of 30) | 19253 | NM_022886 | 64929 | Mm.244003 | NM_022886 | ENSMUSG00000022123 | **Scel** | 9230114I02Rik | sciellin | protein-coding |
| chr14_103474704_103475006 | chr14 | 103474705 | 103475006 | + | 0.740115 | Intergenic | Intergenic | -38486 | NM_022886 | 64929 | Mm.244003 | NM_022886 | ENSMUSG00000022123 | **Scel** | 9230114I02Rik | sciellin | protein-coding |
| chr14_103472005_103472507 | chr14 | 103472006 | 103472507 | + | 0.723051 | Intergenic | Intergenic | -41085 | NM_022886 | 64929 | Mm.244003 | NM_022886 | ENSMUSG00000022123 | **Scel** | 9230114I02Rik | sciellin | protein-coding |
| chr14_103512954_103513676 | chr14 | 103512955 | 103513676 | + | 0.655414 | promoter-TSS (NM_022886) | promoter-TSS (NM_022886) | -26 | NM_022886 | 64929 | Mm.244003 | NM_022886 | ENSMUSG00000022123 | **Scel** | 9230114I02Rik | sciellin | protein-coding |
| chr14_103531804_103532276 | chr14 | 103531805 | 103532276 | + | 0.638209 | intron (NM_022886, intron 3 of 30) | intron (NM_022886, intron 3 of 30) | 18699 | NM_022886 | 64929 | Mm.244003 | NM_022886 | ENSMUSG00000022123 | **Scel** | 9230114I02Rik | sciellin | protein-coding |
| chr5_74523462_74523800 | chr5 | 74523463 | 74523800 | + | 0.815526 | intron (NM_001286437, intron 1 of 7) | intron (NM_001286437, intron 1 of 7) | 8111 | NM_178672 | 212986 | Mm.134147 | NM_178672 | ENSMUSG00000062110 | **Scfd2** | E430013M20Rik\|STXBP1L1 | Sec1 family domain containing 2 | protein-coding |
| chr19_9067187_9067707 | chr19 | 9067188 | 9067707 | + | 0.662433 | intron (NM_001286518, intron 6 of 6) | intron (NM_001286518, intron 6 of 6) | 20509 | NM_011681 | 22287 | Mm.2258 | NM_011681 | ENSMUSG00000024653 | **Scgb1a1** | CC10\|CC16\|CCSP\|PCB-BP\|UG\|UGB\|Utg | secretoglobin, family 1A, member 1 (uteroglobin) | protein-coding |
| chr3_68608784_68609458 | chr3 | 68608785 | 68609458 | + | 1.20562 | intron (NM_001282044, intron 3 of 7) | intron (NM_001282044, intron 3 of 7) | 36753 | NM_013928 | 30953 | Mm.331064 | NM_013928 | ENSMUSG00000027777 | **Schip1** | Nf2ip\|Schip-1 | schwannomin interacting protein 1 | protein-coding |
| chr11_70790960_70791206 | chr11 | 70790961 | 70791206 | + | 0.616705 | 3' UTR (NM_001045526, exon 5 of 5) | 3' UTR (NM_001045526, exon 5 of 5) | 21478 | NM_001045526 | 327957 | Mm.164325 | NM_001045526 | ENSMUSG00000057135 | **Scimp** | A430084P05Rik | SLP adaptor and CSK interacting membrane protein | protein-coding |
| chr12_40154156_40154562 | chr12 | 40154157 | 40154562 | + | 0.893474 | Intergenic | MTD\|LTR\|ERVL-MaLR | -20131 | NM_001146196 | 20259 | Mm.2416 | NM_009132 | ENSMUSG00000002565 | **Scin** | AW545522\|adseverin | scinderin | protein-coding |
| chr10_42977612_42978275 | chr10 | 42977613 | 42978275 | + | 0.627318 | Intergenic | ORR1C1\|LTR\|ERVL-MaLR | 117432 | NM_172938 | 268297 | Mm.98731 | NM_172938 | ENSMUSG00000044770 | **Scml4** | 9330161D17\|mFLJ00197 | sex comb on midleg-like 4 (Drosophila) | protein-coding |
| chr15_100844274_100844753 | chr15 | 100844275 | 100844753 | + | 0.892423 | Intergenic | Intergenic | -26131 | NM_001077499 | 20273 | Mm.385012 | NM_011323 | ENSMUSG00000023033 | **Scn8a** | AI853486\|C630029C19Rik\|NaCh6\|Nav1.6\|dmu\|med\|mnd-2\|mnd2\|nmf2\|nmf335\|nmf58\|nur14\|seal | sodium channel, voltage-gated, type VIII, alpha | protein-coding |
| chr11_67054820_67055543 | chr11 | 67054821 | 67055543 | + | 1.2747 | intron (NM_001040026, intron 2 of 5) | MTE2a\|LTR\|ERVL-MaLR | 2512 | NM_001040026 | 52892 | Mm.129731 | NM_001040026 | | **Sco1** | 2610001C07Rik\|D11Bwg1310e | SCO cytochrome oxidase deficient homolog 1 (yeast) | protein-coding |
| chr11_88952199_88952478 | chr11 | 88952200 | 88952478 | + | 0.828581 | intron (NM_029023, intron 2 of 12) | intron (NM_029023, intron 2 of 12) | 3103 | NM_029023 | 74617 | Mm.34126 | NM_029023 | ENSMUSG00000000278 | **Scpep1** | 2410018F01Rik\|4833411K15Rik\|AI957256\|Risc | serine carboxypeptidase 1 | protein-coding |
| chr8_57430938_57431206 | chr8 | 57430939 | 57431206 | + | 0.723336 | Intergenic | Intergenic | -24851 | NM_009136 | 20284 | Mm.12886 | NM_009136 | ENSMUSG00000031610 | **Scrg1** | AW124307 | scrapie responsive gene 1 | protein-coding |
| chr10_89677788_89678527 | chr10 | 89677789 | 89678527 | + | 1.56165 | intron (NM_198021, intron 2 of 17) | intron (NM_198021, intron 2 of 17) | 8127 | NM_198021 | 213326 | Mm.27651 | NM_198021 | ENSMUSG00000069539 | **Scyl2** | AU015688\|AW536445\|BC030932\|CVAK104\|D10Ertd802e | SCY1-like 2 (S. cerevisiae) | protein-coding |
| chr10_89680529_89680863 | chr10 | 89680530 | 89680863 | + | 1.50915 | intron (NM_198021, intron 1 of 17) | intron (NM_198021, intron 1 of 17) | 5589 | NM_198021 | 213326 | Mm.27651 | NM_198021 | ENSMUSG00000069539 | **Scyl2** | AU015688\|AW536445\|BC030932\|CVAK104\|D10Ertd802e | SCY1-like 2 (S. cerevisiae) | protein-coding |
| chr12_8829223_8830068 | chr12 | 8829224 | 8830068 | + | 1.02109 | Intergenic | L1MB2\|LINE\|L1 | 58250 | NM_011519 | 20969 | Mm.2580 | NM_011519 | ENSMUSG00000020592 | **Sdc1** | AA408134\|AA409076\|CD138\|Sstn\|Synd\|Synd1\|syn-1 | syndecan 1 | protein-coding |
| chr2_151569499_151569974 | chr2 | 151569500 | 151569974 | + | 1.62291 | Intergenic | Intergenic | -2886 | NM_145535 | 228765 | Mm.32068 | NM_145535 | ENSMUSG00000027456 | **Sdcbp2** | - | syndecan binding protein (syntenin) 2 | protein-coding |
| chr2_151564983_151565241 | chr2 | 151564984 | 151565241 | + | 0.92767 | Intergenic | Intergenic | -7511 | NM_145535 | 228765 | Mm.32068 | NM_145535 | ENSMUSG00000027456 | **Sdcbp2** | - | syndecan binding protein (syntenin) 2 | protein-coding |
| chr2_151564896_151565248 | chr2 | 151564897 | 151565248 | + | 0.85347 | Intergenic | Intergenic | -7551 | NM_145535 | 228765 | Mm.32068 | NM_145535 | ENSMUSG00000027456 | **Sdcbp2** | - | syndecan binding protein (syntenin) 2 | protein-coding |
| chr2_26387817_26388100 | chr2 | 26387818 | 26388100 | + | 0.648454 | intron (NM_001085408, intron 1 of 7) | intron (NM_001085408, intron 1 of 7) | 1357 | NM_001085407 | 68112 | Mm.253771 | NM_026563 | ENSMUSG00000026927 | **Sdccag3** | C330016H24Rik\|C630038K21Rik | serologically defined colon cancer antigen 3 | protein-coding |
| chr7_30322823_30323336 | chr7 | 30322824 | 30323336 | + | 0.633916 | promoter-TSS (NM_001033140) | promoter-TSS (NM_001033140) | -705 | NM_001033140 | 68332 | Mm.6844 | NM_001033140 | ENSMUSG00000074211 | **Sdhaf1** | 0610010E21Rik\|AI430885\|AW490662\|Lyrm8 | succinate dehydrogenase complex assembly factor 1 | protein-coding |
| chr9_50590835_50591522 | chr9 | 50590836 | 50591522 | + | 1.4556 | Intergenic | Intergenic | 12670 | NM_025848 | 66925 | Mm.10406 | NM_025848 | ENSMUSG00000000171 | **Sdhd** | 3110001M13Rik\|AVLL5809\|C78570\|PRO19626 | succinate dehydrogenase complex, subunit D, integral membrane protein | protein-coding |
| chr11_114069921_114070436 | chr11 | 114069922 | 114070436 | + | 0.727493 | Intergenic | Intergenic | -4228 | NM_172800 | 237979 | Mm.130504 | NM_172800 | ENSMUSG00000041592 | **Sdk2** | 4632412F08Rik\|5330435L01Rik\|Sdk-2\|mKIAA1514 | sidekick homolog 2 (chicken) | protein-coding |
| chr18_65808401_65808681 | chr18 | 65808402 | 65808681 | + | 0.769191 | intron (NM_025468, intron 1 of 5) | intron (NM_025468, intron 1 of 5) | 7963 | NM_025468 | 66286 | Mm.27800 | NM_025468 | ENSMUSG00000024516 | **Sec11c** | 1810029G24Rik\|Sec11l3 | SEC11 homolog C (S. cerevisiae) | protein-coding |
| chr18_65808778_65809272 | chr18 | 65808779 | 65809272 | + | 0.741852 | intron (NM_025468, intron 1 of 5) | intron (NM_025468, intron 1 of 5) | 8447 | NM_025468 | 66286 | Mm.27800 | NM_025468 | ENSMUSG00000024516 | **Sec11c** | 1810029G24Rik\|Sec11l3 | SEC11 homolog C (S. cerevisiae) | protein-coding |
| chr18_65801754_65802050 | chr18 | 65801755 | 65802050 | + | 0.712422 | intron (NM_025468, intron 1 of 5) | intron (NM_025468, intron 1 of 5) | 1324 | NM_025468 | 66286 | Mm.27800 | NM_025468 | ENSMUSG00000024516 | **Sec11c** | 1810029G24Rik\|Sec11l3 | SEC11 homolog C (S. cerevisiae) | protein-coding |
| chr18_65803142_65803532 | chr18 | 65803143 | 65803532 | + | 0.640311 | intron (NM_025468, intron 1 of 5) | intron (NM_025468, intron 1 of 5) | 2759 | NM_025468 | 66286 | Mm.27800 | NM_025468 | ENSMUSG00000024516 | **Sec11c** | 1810029G24Rik\|Sec11l3 | SEC11 homolog C (S. cerevisiae) | protein-coding |
| chr16_5165451_5166059 | chr16 | 5165452 | 5166059 | + | 0.626752 | intron (NM_001127725, intron 3 of 15) | intron (NM_001127725, intron 3 of 15) | 18646 | NM_001127725 | 665119 | Mm.102072 | NM_001127725 | ENSMUSG00000091712 | **Sec14l5** | - | SEC14-like 5 (S. cerevisiae) | protein-coding |
| chr9_121680137_121680332 | chr9 | 121680138 | 121680332 | + | 0.635265 | 3' UTR (NM_001164562, exon 7 of 7) | 3' UTR (NM_001164562, exon 7 of 7) | 24794 | NM_001164562 | 215474 | Mm.31421 | NM_178677 | ENSMUSG00000061536 | **Sec22c** | 4932412K21\|5930407I15Rik\|C530046H07\|Sec22l3 | SEC22 vesicle trafficking protein homolog C (S. cerevisiae) | protein-coding |
| chr7_128744046_128744203 | chr7 | 128744047 | 128744203 | + | 0.749025 | promoter-TSS (NM_001029982) | promoter-TSS (NM_001029982) | -745 | NM_001029982 | 207352 | Mm.297105 | NM_001029982 | ENSMUSG00000055319 | **Sec23ip** | D7Ertd373e\|p125 | Sec23 interacting protein | protein-coding |
| chr11_51746957_51747404 | chr11 | 51746958 | 51747404 | + | 1.1693 | intron (NM_001290785, intron 1 of 22) | ORR1C2\|LTR\|ERVL-MaLR | 9653 | NM_001290785 | 77371 | Mm.33878 | NM_175255 | ENSMUSG00000036391 | **Sec24a** | - | Sec24 related gene family, member A (S. cerevisiae) | protein-coding |
| chr11_51746936_51747564 | chr11 | 51746937 | 51747564 | + | 1.06811 | intron (NM_001290785, intron 1 of 22) | ORR1C2\|LTR\|ERVL-MaLR | 9584 | NM_001290785 | 77371 | Mm.33878 | NM_175255 | ENSMUSG00000036391 | **Sec24a** | - | Sec24 related gene family, member A (S. cerevisiae) | protein-coding |
| chr3_130075836_130076081 | chr3 | 130075837 | 130076081 | + | 0.766726 | Intergenic | Intergenic | -15052 | NM_207209 | 99683 | Mm.192303 | NM_207209 | ENSMUSG00000001052 | **Sec24b** | AI605202\|SEC24 | Sec24 related gene family, member B (S. cerevisiae) | protein-coding |

| | | | | | | | | | | | | | | | | | |
|---|---|---|---|---|---|---|---|---|---|---|---|---|---|---|---|---|---|
| chr5_100341047_100341487 | chr5 | 100341048 | 100341487 | + | 0.7408 | Intergenic | Intergenic | 74967 | NM_026969 | 69162 | Mm.18634 | NM_026969 | ENSMUSG00000035325 | **Sec31a** | 1810024J13Rik\|ABP125\|ABP130\|AU042160\|HSPC275\|Sec31l1 | Sec31 homolog A (S. cerevisiae) | protein-coding |
| chr5_100320490_100320706 | chr5 | 100320491 | 100320706 | + | 0.649326 | Intergenic | B2_Mm2\|SINE\|B2 | 95636 | NM_026969 | 69162 | Mm.18634 | NM_026969 | ENSMUSG00000035325 | **Sec31a** | 1810024J13Rik\|ABP125\|ABP130\|AU042160\|HSPC275\|Sec31l1 | Sec31 homolog A (S. cerevisiae) | protein-coding |
| chr4_47603770_47604428 | chr4 | 47603771 | 47604428 | + | 0.758725 | Intergenic | Intergenic | 129438 | NM_024171 | 66212 | Mm.20188 | NM_024171 | ENSMUSG00000053317 | **Sec61b** | 1190006C12Rik\|AI326121\|AW122942 | Sec61 beta subunit | protein-coding |
| chr13_51712023_51712436 | chr13 | 51712024 | 51712436 | + | 1.08799 | intron (NM_013660, intron 9 of 15) | intron (NM_013660, intron 9 of 15) | 60516 | NM_029279 | 75420 | Mm.275981 | NM_029279 | ENSMUSG00000035139 | **Secisbp2** | 2210413N07Rik\|2810012K13Rik\|SBP2 | SECIS binding protein 2 | protein-coding |
| chr13_51716332_51717299 | chr13 | 51716333 | 51717299 | + | 0.901407 | intron (NM_013660, intron 6 of 15) | intron (NM_013660, intron 6 of 15) | 65102 | NM_029279 | 75420 | Mm.275981 | NM_029279 | ENSMUSG00000035139 | **Secisbp2** | 2210413N07Rik\|2810012K13Rik\|SBP2 | SECIS binding protein 2 | protein-coding |
| chr13_51716332_51716999 | chr13 | 51716333 | 51716999 | + | 0.88882 | intron (NM_013660, intron 6 of 15) | intron (NM_013660, intron 6 of 15) | 64952 | NM_029279 | 75420 | Mm.275981 | NM_029279 | ENSMUSG00000035139 | **Secisbp2** | 2210413N07Rik\|2810012K13Rik\|SBP2 | SECIS binding protein 2 | protein-coding |
| chr12_91845371_91845681 | chr12 | 91845372 | 91845681 | + | 1.23052 | intron (NM_001039089, intron 1 of 20) | intron (NM_001039089, intron 1 of 20) | 3631 | NM_001039089 | 20338 | Mm.250605 | NM_011344 | ENSMUSG00000020964 | **Sel1l** | AW493766\|Sel1h\|mKIAA4137 | sel-1 suppressor of lin-12-like (C. elegans) | protein-coding |
| chr12_92486774_92487354 | chr12 | 92486775 | 92487354 | + | 0.910594 | Intergenic | Intergenic | -637907 | NM_011344 | 20338 | Mm.250605 | NM_011344 | ENSMUSG00000020964 | **Sel1l** | AW493766\|Sel1h\|mKIAA4137 | sel-1 suppressor of lin-12-like (C. elegans) | protein-coding |
| chr12_92204804_92205172 | chr12 | 92204805 | 92205172 | + | 0.746469 | Intergenic | L1_Mur2\|LINE\|L1 | -355831 | NM_011344 | 20338 | Mm.250605 | NM_011344 | ENSMUSG00000020964 | **Sel1l** | AW493766\|Sel1h\|mKIAA4137 | sel-1 suppressor of lin-12-like (C. elegans) | protein-coding |
| chr1_164030852_164031342 | chr1 | 164030853 | 164031342 | + | 1.07127 | Intergenic | Intergenic | -17137 | NM_011345 | 20339 | Mm.5245 | NM_011345 | ENSMUSG00000026582 | **Sele** | CD62E\|E-selectin\|Elam | selectin, endothelial cell | protein-coding |
| chr3_144589558_144590045 | chr3 | 144589559 | 144590045 | + | 1.10092 | intron (NM_053102, intron 2 of 4) | RMER1A\|Other\|Other | 19375 | NM_053102 | 93684 | Mm.29812 | NM_053102 | ENSMUSG00000037072 | **Selenof** | 9430015P09Rik | selenoprotein | protein-coding |
| chr3_144589159_144590186 | chr3 | 144589160 | 144590186 | + | 0.875154 | intron (NM_053102, intron 2 of 4) | RMER1A\|Other\|Other | 19246 | NM_053102 | 93684 | Mm.29812 | NM_053102 | ENSMUSG00000037072 | **Selenof** | 9430015P09Rik | selenoprotein | protein-coding |
| chr14_29946712_29947345 | chr14 | 29946713 | 29947345 | + | 0.909059 | Intergenic | Tigger7\|DNA\|TcMar-Tigger | -21351 | NM_019979 | 80795 | Mm.104491 | NM_019979 | ENSMUSG00000042682 | **Selk** | 1110001C03Rik\|AA673253\|HSPC030\|Hsp30 | selenoprotein K | protein-coding |
| chr1_164073074_164073471 | chr1 | 164073075 | 164073471 | + | 1.02758 | intron (NM_011346, intron 7 of 8) | intron (NM_011346, intron 7 of 8) | 11197 | NM_001164059 | 20343 | Mm.1461 | NM_011346 | ENSMUSG00000026581 | **Sell** | AI528707\|CD62L\|L-selectin\|LECAM-1\|Lnhr\|Ly-22\|Ly-m22\|Lyam-1\|Lyam1 | selectin, lymphocyte | protein-coding |
| chr1_164081840_164082575 | chr1 | 164081841 | 164082575 | + | 0.850796 | Intergenic | Intergenic | 20132 | NM_001164059 | 20343 | Mm.1461 | NM_011346 | ENSMUSG00000026581 | **Sell** | AI528707\|CD62L\|L-selectin\|LECAM-1\|Lnhr\|Ly-22\|Ly-m22\|Lyam-1\|Lyam1 | selectin, lymphocyte | protein-coding |
| chr1_164075449_164075961 | chr1 | 164075450 | 164075961 | + | 0.67893 | intron (NM_011346, intron 7 of 8) | intron (NM_011346, intron 7 of 8) | 13629 | NM_001164059 | 20343 | Mm.1461 | NM_011346 | ENSMUSG00000026581 | **Sell** | AI528707\|CD62L\|L-selectin\|LECAM-1\|Lnhr\|Ly-22\|Ly-m22\|Lyam-1\|Lyam1 | selectin, lymphocyte | protein-coding |
| chr15_89090622_89090965 | chr15 | 89090623 | 89090965 | + | 0.80106 | intron (NM_027905, intron 1 of 8) | intron (NM_027905, intron 1 of 8) | 1687 | NM_027905 | 223776 | Mm.44763 | NM_027905 | ENSMUSG00000035757 | **Selo** | 1300018J18Rik | selenoprotein O | protein-coding |
| chr9_107600320_107600676 | chr9 | 107600321 | 107600676 | + | 0.683675 | exon (NM_001042779, exon 15 of 20) | exon (NM_001042779, exon 15 of 20) | 4904 | NM_009153 | 20347 | Mm.4083 | NM_009153 | ENSMUSG00000057969 | **Sema3b** | SemA\|Semaa\|sema5\|semaV | sema domain, immunoglobulin domain (Ig), short basic domain, secreted, (semaphorin) 3B | protein-coding |
| chr5_17556044_17556521 | chr5 | 17556045 | 17556521 | + | 0.6347 | Intergenic | Intergenic | -18533 | NM_013657 | 20348 | Mm.5071 | NM_013657 | ENSMUSG00000028780 | **Sema3c** | 1110036B02Rik\|SemE\|Semae | sema domain, immunoglobulin domain (Ig), short basic domain, secreted, (semaphorin) 3C | protein-coding |
| chr3_88457198_88457603 | chr3 | 88457199 | 88457603 | + | 1.32247 | intron (NM_001163489, intron 1 of 14) | intron (NM_001163489, intron 1 of 14) | 1483 | NM_001163491 | 20351 | Mm.439752 | NM_013658 | ENSMUSG00000028064 | **Sema4a** | AI132332\|SemB\|Semab | sema domain, immunoglobulin domain (Ig), transmembrane domain (TM) and short cytoplasmic domain, (semaphorin) 4A | protein-coding |
| chr3_88457193_88457807 | chr3 | 88457194 | 88457807 | + | 1.24555 | intron (NM_001163489, intron 1 of 14) | intron (NM_001163489, intron 1 of 14) | 1384 | NM_001163491 | 20351 | Mm.439752 | NM_013658 | ENSMUSG00000028064 | **Sema4a** | AI132332\|SemB\|Semab | sema domain, immunoglobulin domain (Ig), transmembrane domain (TM) and short cytoplasmic domain, (semaphorin) 4A | protein-coding |
| chr7_80166686_80167153 | chr7 | 80166687 | 80167153 | + | 1.76524 | Intergenic | Intergenic | -19921 | NM_013659 | 20352 | Mm.275909 | NM_013659 | ENSMUSG00000030539 | **Sema4b** | SemC\|Semac\|mKIAA1745 | sema domain, immunoglobulin domain (Ig), transmembrane domain (TM) and short cytoplasmic domain, (semaphorin) 4B | protein-coding |
| chr7_80206275_80206547 | chr7 | 80206276 | 80206547 | + | 0.801337 | intron (NM_013659, intron 2 of 14) | intron (NM_013659, intron 2 of 14) | 19570 | NM_013659 | 20352 | Mm.275909 | NM_013659 | ENSMUSG00000030539 | **Sema4b** | SemC\|Semac\|mKIAA1745 | sema domain, immunoglobulin domain (Ig), transmembrane domain (TM) and short cytoplasmic domain, (semaphorin) 4B | protein-coding |
| chr1_36556043_36556401 | chr1 | 36556044 | 36556401 | + | 1.03918 | intron (NM_001126047, intron 1 of 14) | intron (NM_001126047, intron 1 of 14) | 2159 | NM_001126047 | 20353 | Mm.29558 | NM_001126047 | ENSMUSG00000026121 | **Sema4c** | AI426163\|Semacl1\|Semaf\|Semai\|mKIAA1739 | sema domain, immunoglobulin domain (Ig), transmembrane domain (TM) and short cytoplasmic domain, (semaphorin) 4C | protein-coding |
| chr1_36555838_36556420 | chr1 | 36555839 | 36556420 | + | 0.905757 | intron (NM_001126047, intron 1 of 14) | intron (NM_001126047, intron 1 of 14) | 2252 | NM_001126047 | 20353 | Mm.29558 | NM_001126047 | ENSMUSG00000026121 | **Sema4c** | AI426163\|Semacl1\|Semaf\|Semai\|mKIAA1739 | sema domain, immunoglobulin domain (Ig), transmembrane domain (TM) and short cytoplasmic domain, (semaphorin) 4C | protein-coding |
| chr1_36555778_36556782 | chr1 | 36555779 | 36556782 | + | 0.685934 | intron (NM_001126047, intron 1 of 14) | intron (NM_001126047, intron 1 of 14) | 2101 | NM_001126047 | 20353 | Mm.29558 | NM_001126047 | ENSMUSG00000026121 | **Sema4c** | AI426163\|Semacl1\|Semaf\|Semai\|mKIAA1739 | sema domain, immunoglobulin domain (Ig), transmembrane domain (TM) and short cytoplasmic domain, (semaphorin) 4C | protein-coding |
| chr13_51802977_51803256 | chr13 | 51802978 | 51803256 | + | 1.15187 | Intergenic | Intergenic | -9370 | NM_013660 | 20354 | Mm.33903 | NM_013660 | ENSMUSG00000021451 | **Sema4d** | CD100\|Semacl2\|Semaj\|Semcl2\|coll-4 | sema domain, immunoglobulin domain (Ig), transmembrane domain (TM) and short cytoplasmic domain, (semaphorin) 4D | protein-coding |
| chr13_51814634_51815091 | chr13 | 51814635 | 51815091 | + | 1.08018 | Intergenic | Intergenic | -21116 | NM_013660 | 20354 | Mm.33903 | NM_013660 | ENSMUSG00000021451 | **Sema4d** | CD100\|Semacl2\|Semaj\|Semcl2\|coll-4 | sema domain, immunoglobulin domain (Ig), transmembrane domain (TM) and short cytoplasmic domain, (semaphorin) 4D | protein-coding |
| chr2_124732438_124733206 | chr2 | 124732439 | 124733206 | + | 0.91645 | Intergenic | Intergenic | 79241 | NM_199239 | 214968 | Mm.330536 | NM_172537 | ENSMUSG00000027200 | **Sema6d** | 1110067B02Rik\|AA409156\|D330011G23\|mKIAA1479 | sema domain, transmembrane domain (TM), and cytoplasmic domain, (semaphorin) 6D | protein-coding |
| chr2_124704484_124705327 | chr2 | 124704485 | 124705327 | + | 0.708909 | Intergenic | Intergenic | 51325 | NM_199239 | 214968 | Mm.330536 | NM_172537 | ENSMUSG00000027200 | **Sema6d** | 1110067B02Rik\|AA409156\|D330011G23\|mKIAA1479 | sema domain, transmembrane domain (TM), and cytoplasmic domain, (semaphorin) 6D | protein-coding |
| chr9_57973711_57974102 | chr9 | 57973712 | 57974102 | + | 1.2679 | Intergenic | Intergenic | 33772 | NM_011352 | 20361 | Mm.335187 | NM_011352 | ENSMUSG00000038264 | **Sema7a** | 2900057C09Rik\|CDw108\|H-Sema-L\|M-Sema-L\|Semal | sema domain, immunoglobulin domain (Ig), and GPI membrane anchor, (semaphorin) 7A | protein-coding |
| chr9_57968734_57969211 | chr9 | 57968735 | 57969211 | + | 0.722702 | Intergenic | Intergenic | 28838 | NM_011352 | 20361 | Mm.335187 | NM_011352 | ENSMUSG00000038264 | **Sema7a** | 2900057C09Rik\|CDw108\|H-Sema-L\|M-Sema-L\|Semal | sema domain, immunoglobulin domain (Ig), and GPI membrane anchor, (semaphorin) 7A | protein-coding |
| chr16_32009861_32010324 | chr16 | 32009862 | 32010324 | + | 1.12094 | Intergenic | RMER15\|LTR\|ERVL | -6806 | NM_177103 | 320213 | Mm.152890 | NM_023801 | | **Senp5** | 6230429P13Rik\|A730063F07Rik\|AI851888\|BB189556\|SMT3IP3 | SUMO/sentrin specific peptidase 5 | protein-coding |
| chr9_80057549_80058357 | chr9 | 80057550 | 80058357 | + | 1.32388 | Intergenic | MLT1D\|LTR\|ERVL-MaLR | -8950 | NM_146003 | 215351 | Mm.28232 | NM_146003 | ENSMUSG00000034252 | **Senp6** | 2810017C20Rik\|E130319N12Rik\|Susp1\|mKIAA0797 | SUMO/sentrin specific peptidase 6 | protein-coding |
| chr4_134559042_134559594 | chr4 | 134559043 | 134559594 | + | 0.661061 | Intergenic | Intergenic | -7152 | NM_029100 | 74777 | Mm.29773 | NM_029100 | ENSMUSG00000050989 | **Sepn1** | 1110019I12Rik\|AI414492 | selenoprotein N, 1 | protein-coding |
| chr7_127216450_127216761 | chr7 | 127216451 | 127216761 | + | 0.930944 | intron (NM_017461, intron 6 of 10) | intron (NM_017461, intron 6 of 10) | 1839 | NM_017461 | 54204 | Mm.18 | NM_017461 | ENSMUSG00000000486 | **Sept1** | Diff6\|Pnutl3 | septin 1 | protein-coding |
| chr15_82282453_82283062 | chr15 | 82282454 | 82283062 | + | 0.874778 | intron (NM_011889, intron 2 of 9) | intron (NM_011889, intron 2 of 9) | 7823 | NM_011889 | 24050 | Mm.309707 | NM_011889 | ENSMUSG00000022456 | **Sept3** | 3110018K01Rik\|AV154067\|B530002E20Rik\|Sep3 | septin 3 | protein-coding |
| chr16_18686886_18687355 | chr16 | 18686887 | 18687355 | + | 0.868588 | Intergenic | Intergenic | -57183 | NM_213614 | 18951 | Mm.20365 | NM_213614 | ENSMUSG00000072214 | **Sept5** | Cdcrel-1\|Cdcrel1\|Pnutl1 | septin 5 | protein-coding |
| chr16_18686886_18687737 | chr16 | 18686887 | 18687737 | + | 0.826926 | Intergenic | Intergenic | -57374 | NM_213614 | 18951 | Mm.20365 | NM_213614 | ENSMUSG00000072214 | **Sept5** | Cdcrel-1\|Cdcrel1\|Pnutl1 | septin 5 | protein-coding |
| chr9_25230241_25230938 | chr9 | 25230242 | 25230938 | + | 1.56444 | Intergenic | Intergenic | -21849 | NM_009859 | 235072 | Mm.270259 | NM_009859 | ENSMUSG00000001833 | **Sept7** | Cdc10\|E430034N22 | septin 7 | protein-coding |
| chr9_25260990_25261339 | chr9 | 25260991 | 25261339 | + | 0.970063 | intron (NM_001205367, intron 1 of 12) | intron (NM_001205367, intron 1 of 12) | 8726 | NM_001205367 | 235072 | Mm.270259 | NM_009859 | ENSMUSG00000001833 | **Sept7** | Cdc10\|E430034N22 | septin 7 | protein-coding |
| chr9_25260988_25261616 | chr9 | 25260989 | 25261616 | + | 0.889886 | intron (NM_001205367, intron 1 of 12) | intron (NM_001205367, intron 1 of 12) | 8863 | NM_001205367 | 235072 | Mm.270259 | NM_009859 | ENSMUSG00000001833 | **Sept7** | Cdc10\|E430034N22 | septin 7 | protein-coding |
| chr9_25243897_25244271 | chr9 | 25243898 | 25244271 | + | 0.797291 | Intergenic | Intergenic | -8355 | NM_009859 | 235072 | Mm.270259 | NM_009859 | ENSMUSG00000001833 | **Sept7** | Cdc10\|E430034N22 | septin 7 | protein-coding |
| chr9_25318174_25318533 | chr9 | 25318175 | 25318533 | + | 0.723492 | Intergenic | Intergenic | 65915 | NM_001205367 | 235072 | Mm.270259 | NM_009859 | ENSMUSG00000001833 | **Sept7** | Cdc10\|E430034N22 | septin 7 | protein-coding |
| chr9_25317962_25318551 | chr9 | 25317963 | 25318551 | + | 0.708723 | Intergenic | Intergenic | 65818 | NM_001205367 | 235072 | Mm.270259 | NM_009859 | ENSMUSG00000001833 | **Sept7** | Cdc10\|E430034N22 | septin 7 | protein-coding |
| chr11_117239673_117239945 | chr11 | 117239674 | 117239945 | + | 1.04129 | intron (NM_001113486, intron 2 of 11) | L1M5\|LINE\|L1 | -26437 | NM_001113487 | 53860 | Mm.38450 | NM_017380 | ENSMUSG00000059248 | **Sept9** | MSF1\|Msf\|PNUTL4\|Sint1 | septin 9 | protein-coding |
| chr11_117311041_117311414 | chr11 | 117311042 | 117311414 | + | 0.866474 | intron (NM_001113487, intron 2 of 10) | intron (NM_001113487, intron 2 of 10) | -20487 | NM_017380 | 53860 | Mm.38450 | NM_017380 | ENSMUSG00000059248 | **Sept9** | MSF1\|Msf\|PNUTL4\|Sint1 | septin 9 | protein-coding |

| | | | | | | | | | | | | | | | | | |
|---|---|---|---|---|---|---|---|---|---|---|---|---|---|---|---|---|---|
| chr11_117345811_117346142 | chr11 | 117345812 | 117346142 | + | 0.842614 | intron (NM_001113488, intron 1 of 9) | intron (NM_001113488, intron 1 of 9) | 13664 | NM_001113488 | 53860 | Mm.38450 | NM_017380 | ENSMUSG00000059248 | **Sept9** | MSF1\|Msf\|PNUTL4\|Sint1 | septin 9 | protein-coding |
| chr11_117341775_117342221 | chr11 | 117341776 | 117342221 | + | 0.741158 | intron (NM_001113488, intron 1 of 9) | intron (NM_001113488, intron 1 of 9) | 9685 | NM_001113488 | 53860 | Mm.38450 | NM_017380 | ENSMUSG00000059248 | **Sept9** | MSF1\|Msf\|PNUTL4\|Sint1 | septin 9 | protein-coding |
| chr11_117250165_117250620 | chr11 | 117250166 | 117250620 | + | 0.740886 | intron (NM_001113486, intron 2 of 11) | ID_B1\|SINE\|B4 | -15853 | NM_001113487 | 53860 | Mm.38450 | NM_017380 | ENSMUSG00000059248 | **Sept9** | MSF1\|Msf\|PNUTL4\|Sint1 | septin 9 | protein-coding |
| chr11_117220747_117221142 | chr11 | 117220748 | 117221142 | + | 0.717267 | intron (NM_001113486, intron 2 of 11) | intron (NM_001113486, intron 2 of 11) | 21284 | NM_001113486 | 53860 | Mm.38450 | NM_017380 | ENSMUSG00000059248 | **Sept9** | MSF1\|Msf\|PNUTL4\|Sint1 | septin 9 | protein-coding |
| chr7_15921719_15922092 | chr7 | 15921720 | 15922092 | + | 0.704596 | intron (NM_009156, intron 1 of 5) | intron (NM_009156, intron 1 of 5) | 465 | NM_009156 | 20364 | Mm.42829 | NM_009156 | ENSMUSG00000041571 | **Sepw1** | selW | selenoprotein W, muscle 1 | protein-coding |
| chr6_67186911_67187417 | chr6 | 67186912 | 67187417 | + | 1.22735 | Intergenic | LTR67B\|LTR\|ERVL | -79815 | NM_001113564 | 66870 | Mm.240490 | NM_025814 | ENSMUSG00000036371 | **Serbp1** | 1200009K13Rik\|9330147J08Rik\|AL022786\|Pairbp1 | serpine1 mRNA binding protein 1 | protein-coding |
| chr6_67291673_67292760 | chr6 | 67291674 | 67292760 | + | 0.958339 | exon (NM_008354, exon 16 of 16) | exon (NM_008354, exon 16 of 16) | 25238 | NM_025814 | 66870 | Mm.240490 | NM_025814 | ENSMUSG00000036371 | **Serbp1** | 1200009K13Rik\|9330147J08Rik\|AL022786\|Pairbp1 | serpine1 mRNA binding protein 1 | protein-coding |
| chr6_67291954_67292684 | chr6 | 67291955 | 67292684 | + | 0.95823 | exon (NM_008354, exon 16 of 16) | exon (NM_008354, exon 16 of 16) | 25340 | NM_025814 | 66870 | Mm.240490 | NM_025814 | ENSMUSG00000036371 | **Serbp1** | 1200009K13Rik\|9330147J08Rik\|AL022786\|Pairbp1 | serpine1 mRNA binding protein 1 | protein-coding |
| chr10_57530569_57530850 | chr10 | 57530570 | 57530850 | + | 0.827161 | intron (NM_019760, intron 1 of 8) | intron (NM_019760, intron 1 of 8) | 1819 | NM_019760 | 56442 | Mm.29344 | NM_019760 | ENSMUSG00000019877 | **Serinc1** | 1500011D18Rik\|AI315070\|AIGP-2\|TMS-2\|Tde1l\|Tde2\|Tms2\|mKIAA1253 | serine incorporator 1 | protein-coding |
| chr13_92613185_92613471 | chr13 | 92613186 | 92613471 | + | 1.29681 | intron (NM_172588, intron 1 of 11) | LTR72_RN\|LTR\|ERV1 | 2190 | NM_172588 | 218442 | Mm.297970 | NM_172588 | ENSMUSG00000021703 | **Serinc5** | A130038L21Rik\|AIGP3\|C86123\|TPO1 | serine incorporator 5 | protein-coding |
| chr13_92612550_92613002 | chr13 | 92612551 | 92613002 | + | 1.28545 | intron (NM_172588, intron 1 of 11) | intron (NM_172588, intron 1 of 11) | 1638 | NM_172588 | 218442 | Mm.297970 | NM_172588 | ENSMUSG00000021703 | **Serinc5** | A130038L21Rik\|AIGP3\|C86123\|TPO1 | serine incorporator 5 | protein-coding |
| chr13_92629513_92630077 | chr13 | 92629514 | 92630077 | + | 1.26165 | intron (NM_172588, intron 1 of 11) | intron (NM_172588, intron 1 of 11) | 18657 | NM_172588 | 218442 | Mm.297970 | NM_172588 | ENSMUSG00000021703 | **Serinc5** | A130038L21Rik\|AIGP3\|C86123\|TPO1 | serine incorporator 5 | protein-coding |
| chr13_92629322_92630110 | chr13 | 92629323 | 92630110 | + | 1.19215 | intron (NM_172588, intron 1 of 11) | intron (NM_172588, intron 1 of 11) | 18578 | NM_172588 | 218442 | Mm.297970 | NM_172588 | ENSMUSG00000021703 | **Serinc5** | A130038L21Rik\|AIGP3\|C86123\|TPO1 | serine incorporator 5 | protein-coding |
| chr13_92639002_92639475 | chr13 | 92639003 | 92639475 | + | 0.828555 | intron (NM_172588, intron 1 of 11) | intron (NM_172588, intron 1 of 11) | 28101 | NM_172588 | 218442 | Mm.297970 | NM_172588 | ENSMUSG00000021703 | **Serinc5** | A130038L21Rik\|AIGP3\|C86123\|TPO1 | serine incorporator 5 | protein-coding |
| chr13_32831878_32832281 | chr13 | 32831879 | 32832281 | + | 0.628556 | Intergenic | MLTR11B\|LTR\|ERVK | 19105 | NM_025429 | 66222 | Mm.20144 | NM_025429 | ENSMUSG00000044734 | **Serpinb1a** | 1190005M04Rik\|AI325983\|EI\|EIA\|ELANH2\|LEI\|M/NEI\|MNEI\|PI2 | serine (or cysteine) peptidase inhibitor, clade B, member 1a | protein-coding |
| chr13_33077021_33077716 | chr13 | 33077022 | 33077716 | + | 1.71352 | Intergenic | MLT1A1\|LTR\|ERVL-MaLR | -6734 | NM_173052 | 282663 | Mm.314490 | NM_173052 | ENSMUSG00000051029 | **Serpinb1b** | 6330533H24Rik\|EIB\|Serpin1b1\|ovalbumin | serine (or cysteine) peptidase inhibitor, clade B, member 1b | protein-coding |
| chr13_33998524_33998758 | chr13 | 33998525 | 33998758 | + | 1.06076 | intron (NM_001243192, intron 1 of 7) | ORR1B2\|LTR\|ERVL-MaLR | 4153 | NM_001243192 | 20719 | Mm.252210 | NM_009254 | ENSMUSG00000060147 | **Serpinb6a** | 4930482L21Rik\|D330015H01Rik\|PI-6\|Serpinb6\|Spi3\|ovalbumin | serine (or cysteine) peptidase inhibitor, clade B, member 6a | protein-coding |
| chr13_32981762_32982445 | chr13 | 32981763 | 32982445 | + | 1.85936 | Intergenic | Intergenic | 16591 | NM_011454 | 20708 | Mm.36526 | NM_011454 | ENSMUSG00000042842 | **Serpinb6b** | NK13\|Spi12\|ovalbumin | serine (or cysteine) peptidase inhibitor, clade B, member 6b | protein-coding |
| chr13_32982656_32983164 | chr13 | 32982657 | 32983164 | + | 1.60475 | Intergenic | Intergenic | 17397 | NM_011454 | 20708 | Mm.36526 | NM_011454 | ENSMUSG00000042842 | **Serpinb6b** | NK13\|Spi12\|ovalbumin | serine (or cysteine) peptidase inhibitor, clade B, member 6b | protein-coding |
| chr13_32982656_32983377 | chr13 | 32982657 | 32983377 | + | 1.58004 | Intergenic | Intergenic | 17504 | NM_011454 | 20708 | Mm.36526 | NM_011454 | ENSMUSG00000042842 | **Serpinb6b** | NK13\|Spi12\|ovalbumin | serine (or cysteine) peptidase inhibitor, clade B, member 6b | protein-coding |
| chr13_32980811_32981062 | chr13 | 32980812 | 32981062 | + | 0.782545 | Intergenic | MER67B\|LTR\|ERV1 | 15424 | NM_011454 | 20708 | Mm.36526 | NM_011454 | ENSMUSG00000042842 | **Serpinb6b** | NK13\|Spi12\|ovalbumin | serine (or cysteine) peptidase inhibitor, clade B, member 6b | protein-coding |
| chr13_32992212_32992465 | chr13 | 32992213 | 32992465 | + | 1.34563 | Intergenic | MLTR73\|LTR\|ERV1 | -12202 | NM_009256 | 20723 | Mm.272569 | NM_009256 | ENSMUSG00000045827 | **Serpinb9** | BB283241\|CAP-3\|CAP3\|PI-9\|PI9\|Spi6\|ovalbumin | serine (or cysteine) peptidase inhibitor, clade B, member 9 | protein-coding |
| chr13_33001837_33002089 | chr13 | 33001838 | 33002089 | + | 0.965845 | Intergenic | L1MB7\|LINE\|L1 | -2578 | NM_009256 | 20723 | Mm.272569 | NM_009256 | ENSMUSG00000045827 | **Serpinb9** | BB283241\|CAP-3\|CAP3\|PI-9\|PI9\|Spi6\|ovalbumin | serine (or cysteine) peptidase inhibitor, clade B, member 9 | protein-coding |
| chr1_160984289_160985227 | chr1 | 160984290 | 160985227 | + | 0.966979 | intron (NM_080844, intron 1 of 7) | MTA_Mm\|LTR\|ERVL-MaLR | 6152 | NM_080844 | 11905 | Mm.260770 | NM_080844 | ENSMUSG00000026715 | **Serpinc1** | AI114908\|ATIII\|At-3\|At3 | serine (or cysteine) peptidase inhibitor, clade C (antithrombin), member 1 | protein-coding |
| chr5_137065468_137065816 | chr5 | 137065469 | 137065816 | + | 1.00022 | intron (NM_008871, intron 5 of 8) | intron (NM_008871, intron 5 of 8) | 6630 | NM_008871 | 18787 | Mm.250422 | NM_008871 | ENSMUSG00000037411 | **Serpine1** | PAI-1\|PAI1\|Planh1 | serine (or cysteine) peptidase inhibitor, clade E, member 1 | protein-coding |
| chr5_137061845_137062136 | chr5 | 137061846 | 137062136 | + | 0.764999 | 3' UTR (NM_008871, exon 9 of 9) | 3' UTR (NM_008871, exon 9 of 9) | 10281 | NM_008871 | 18787 | Mm.250422 | NM_008871 | ENSMUSG00000037411 | **Serpine1** | PAI-1\|PAI1\|Planh1 | serine (or cysteine) peptidase inhibitor, clade E, member 1 | protein-coding |
| chr5_137068737_137069006 | chr5 | 137068738 | 137069006 | + | 0.733206 | intron (NM_008871, intron 3 of 8) | intron (NM_008871, intron 3 of 8) | 3400 | NM_008871 | 18787 | Mm.250422 | NM_008871 | ENSMUSG00000037411 | **Serpine1** | PAI-1\|PAI1\|Planh1 | serine (or cysteine) peptidase inhibitor, clade E, member 1 | protein-coding |
| chr5_137072049_137072539 | chr5 | 137072050 | 137072539 | + | 0.670974 | promoter-TSS (NM_008871) | promoter-TSS (NM_008871) | -22 | NM_008871 | 18787 | Mm.250422 | NM_008871 | ENSMUSG00000037411 | **Serpine1** | PAI-1\|PAI1\|Planh1 | serine (or cysteine) peptidase inhibitor, clade E, member 1 | protein-coding |
| chr1_79859269_79860145 | chr1 | 79859270 | 79860145 | + | 1.25946 | Intergenic | ID4\|SINE\|ID | -1042 | NM_009255 | 20720 | Mm.3093 | NM_009255 | ENSMUSG00000026249 | **Serpine2** | B230326M24Rik\|PAI-1\|PI-7\|PI7\|PN-1\|Spi4 | serine (or cysteine) peptidase inhibitor, clade E, member 2 | protein-coding |
| chr1_79858885_79860149 | chr1 | 79858886 | 79860149 | + | 1.0267 | promoter-TSS (NM_009255) | promoter-TSS (NM_009255) | -852 | NM_009255 | 20720 | Mm.3093 | NM_009255 | ENSMUSG00000026249 | **Serpine2** | B230326M24Rik\|PAI-1\|PI-7\|PI7\|PN-1\|Spi4 | serine (or cysteine) peptidase inhibitor, clade E, member 2 | protein-coding |
| chr1_79858104_79860156 | chr1 | 79858105 | 79860156 | + | 0.688946 | promoter-TSS (NM_009255) | promoter-TSS (NM_009255) | -465 | NM_009255 | 20720 | Mm.3093 | NM_009255 | ENSMUSG00000026249 | **Serpine2** | B230326M24Rik\|PAI-1\|PI-7\|PI7\|PN-1\|Spi4 | serine (or cysteine) peptidase inhibitor, clade E, member 2 | protein-coding |
| chr11_75410067_75410777 | chr11 | 75410068 | 75410777 | + | 0.746729 | intron (NM_011340, intron 7 of 7) | intron (NM_011340, intron 7 of 7) | 12201 | NM_011340 | 20317 | Mm.2044 | NM_011340 | ENSMUSG00000000753 | **Serpinf1** | AI195227\|EPC-1\|Pedf\|Pedfl\|Sdf3 | serine (or cysteine) peptidase inhibitor, clade F, member 1 | protein-coding |
| chr7_99311399_99311820 | chr7 | 99311400 | 99311820 | + | 0.973122 | intron (NM_001048167, intron 1 of 2) | intron (NM_001048167, intron 1 of 2) | 41629 | NM_009825 | 12406 | Mm.22708 | NM_009825 | ENSMUSG00000070436 | **Serpinh1** | BERF-1\|Cbp1\|Cbp2\|Hsp47\|J6\|Serpinh2\|gp46 | serine (or cysteine) peptidase inhibitor, clade H, member 1 | protein-coding |
| chr11_20564535_20565544 | chr11 | 20564536 | 20565544 | + | 1.66678 | intron (NM_021372, intron 1 of 1) | intron (NM_021372, intron 1 of 1) | 21787 | NM_021372 | 58172 | Mm.339676 | NM_021372 | ENSMUSG00000049800 | **Sertad2** | AB041541\|AU021878\|Kiaa0127\|SEI-2\|Sei2\|TRIP-Br2\|mKIAA0127 | SERTA domain containing 2 | protein-coding |
| chr11_20635304_20635494 | chr11 | 20635305 | 20635494 | + | 0.691399 | intron (NM_001038625, intron 1 of 1) | intron (NM_001038625, intron 1 of 1) | 3422 | NM_001038625 | 58172 | Mm.339676 | NM_021372 | ENSMUSG00000049800 | **Sertad2** | AB041541\|AU021878\|Kiaa0127\|SEI-2\|Sei2\|TRIP-Br2\|mKIAA0127 | SERTA domain containing 2 | protein-coding |
| chr11_20608414_20608846 | chr11 | 20608415 | 20608846 | + | 0.66719 | intron (NM_021372, intron 1 of 1) | intron (NM_021372, intron 1 of 1) | -23347 | NM_001038625 | 58172 | Mm.339676 | NM_021372 | ENSMUSG00000049800 | **Sertad2** | AB041541\|AU021878\|Kiaa0127\|SEI-2\|Sei2\|TRIP-Br2\|mKIAA0127 | SERTA domain containing 2 | protein-coding |
| chr10_41822421_41823176 | chr10 | 41822422 | 41823176 | + | 0.90253 | intron (NM_001162908, intron 1 of 9) | intron (NM_001162908, intron 1 of 9) | 12225 | NM_001162908 | 140742 | Mm.139418 | NM_001013370 | ENSMUSG00000038332 | **Sesn1** | 1110002G11Rik\|AU044290\|Pa26\|Sest1 | sestrin 1 | protein-coding |
| chr9_14240965_14241378 | chr9 | 14240966 | 14241378 | + | 1.48407 | Intergenic | Intergenic | -35129 | NM_030261 | 75747 | Mm.325126 | NM_030261 | ENSMUSG00000032009 | **Sesn3** | 5630400E15Rik\|BC003348\|SEST3 | sestrin 3 | protein-coding |
| chr9_14240796_14241377 | chr9 | 14240797 | 14241377 | + | 1.45887 | Intergenic | Intergenic | -35214 | NM_030261 | 75747 | Mm.325126 | NM_030261 | ENSMUSG00000032009 | **Sesn3** | 5630400E15Rik\|BC003348\|SEST3 | sestrin 3 | protein-coding |
| chr9_14247912_14248558 | chr9 | 14247913 | 14248558 | + | 1.35561 | Intergenic | Intergenic | -28066 | NM_030261 | 75747 | Mm.325126 | NM_030261 | ENSMUSG00000032009 | **Sesn3** | 5630400E15Rik\|BC003348\|SEST3 | sestrin 3 | protein-coding |
| chr9_14248137_14248559 | chr9 | 14248138 | 14248559 | + | 1.30823 | Intergenic | Intergenic | -27953 | NM_030261 | 75747 | Mm.325126 | NM_030261 | ENSMUSG00000032009 | **Sesn3** | 5630400E15Rik\|BC003348\|SEST3 | sestrin 3 | protein-coding |
| chr9_14185543_14186122 | chr9 | 14185544 | 14186122 | + | 1.12054 | Intergenic | Intergenic | -90468 | NM_030261 | 75747 | Mm.325126 | NM_030261 | ENSMUSG00000032009 | **Sesn3** | 5630400E15Rik\|BC003348\|SEST3 | sestrin 3 | protein-coding |
| chr9_14140243_14141324 | chr9 | 14140244 | 14141324 | + | 1.04998 | Intergenic | Intergenic | -135517 | NM_030261 | 75747 | Mm.325126 | NM_030261 | ENSMUSG00000032009 | **Sesn3** | 5630400E15Rik\|BC003348\|SEST3 | sestrin 3 | protein-coding |
| chr2_77268600_77268886 | chr2 | 77268601 | 77268886 | + | 1.47982 | intron (NM_175465, intron 1 of 17) | intron (NM_175465, intron 1 of 17) | 11849 | NM_175465 | 228071 | Mm.27120 | NM_175465 | ENSMUSG00000042272 | **Sestd1** | 1500031J16Rik\|A930010O20\|Solo/Trio8 | SEC14 and spectrin domains 1 | protein-coding |
| chr2_77271797_77272292 | chr2 | 77271798 | 77272292 | + | 1.33851 | intron (NM_175465, intron 1 of 17) | intron (NM_175465, intron 1 of 17) | 8547 | NM_175465 | 228071 | Mm.27120 | NM_175465 | ENSMUSG00000042272 | **Sestd1** | 1500031J16Rik\|A930010O20\|Solo/Trio8 | SEC14 and spectrin domains 1 | protein-coding |
| chr2_77288005_77288535 | chr2 | 77288006 | 77288535 | + | 1.31728 | Intergenic | Intergenic | -7678 | NM_175465 | 228071 | Mm.27120 | NM_175465 | ENSMUSG00000042272 | **Sestd1** | 1500031J16Rik\|A930010O20\|Solo/Trio8 | SEC14 and spectrin domains 1 | protein-coding |

| | | | | | | | | | | | | | | | | | |
|---|---|---|---|---|---|---|---|---|---|---|---|---|---|---|---|---|---|
| chr2_77271797_77272159 | chr2 | 77271798 | 77272159 | + | 1.2903 | intron (NM_175465, intron 1 of 17) | L1MD3\|LINE\|L1 | 8614 | NM_175465 | 228071 | Mm.27120 | NM_175465 | ENSMUSG00000042272 | **Sestd1** | 1500031J16Rik\|A930010O20\|Solo/Trio8 | SEC14 and spectrin domains 1 | protein-coding |
| chr2_77267789_77268531 | chr2 | 77267790 | 77268531 | + | 1.06808 | intron (NM_175465, intron 1 of 17) | intron (NM_175465, intron 1 of 17) | 12432 | NM_175465 | 228071 | Mm.27120 | NM_175465 | ENSMUSG00000042272 | **Sestd1** | 1500031J16Rik\|A930010O20\|Solo/Trio8 | SEC14 and spectrin domains 1 | protein-coding |
| chr2_77268290_77268531 | chr2 | 77268291 | 77268531 | + | 0.815648 | intron (NM_175465, intron 1 of 17) | MLT1A0\|LTR\|ERVL-MaLR | 12181 | NM_175465 | 228071 | Mm.27120 | NM_175465 | ENSMUSG00000042272 | **Sestd1** | 1500031J16Rik\|A930010O20\|Solo/Trio8 | SEC14 and spectrin domains 1 | protein-coding |
| chr2_30057218_30057510 | chr2 | 30057219 | 30057510 | + | 0.66833 | Intergenic | Intergenic | -4632 | NM_023871 | 56086 | Mm.335942 | NM_023871 | ENSMUSG00000054766 | **Set** | 2610030F17Rik\|5730420M11Rik\|AA407739\|I-2PP2A\|StF-IT-1\|TAF-I | SET nuclear oncogene | protein-coding |
| chr18_79185975_79186472 | chr18 | 79185976 | 79186472 | + | 1.24171 | Intergenic | Intergenic | -76833 | NM_053099 | 240427 | Mm.312871 | NM_053099 | ENSMUSG00000024548 | **Setbp1** | C130092E12\|Seb\|mKIAA0437 | SET binding protein 1 | protein-coding |
| chr18_79262939_79263463 | chr18 | 79262940 | 79263463 | + | 1.16807 | Intergenic | ORR1D2\|LTR\|ERVL-MaLR | -153810 | NM_053099 | 240427 | Mm.312871 | NM_053099 | ENSMUSG00000024548 | **Setbp1** | C130092E12\|Seb\|mKIAA0437 | SET binding protein 1 | protein-coding |
| chr18_79466857_79467423 | chr18 | 79466858 | 79467423 | + | 1.1522 | Intergenic | RMER15\|LTR\|ERVL | -357749 | NM_053099 | 240427 | Mm.312871 | NM_053099 | ENSMUSG00000024548 | **Setbp1** | C130092E12\|Seb\|mKIAA0437 | SET binding protein 1 | protein-coding |
| chr18_79300593_79301128 | chr18 | 79300594 | 79301128 | + | 1.08436 | Intergenic | Intergenic | -191470 | NM_053099 | 240427 | Mm.312871 | NM_053099 | ENSMUSG00000024548 | **Setbp1** | C130092E12\|Seb\|mKIAA0437 | SET binding protein 1 | protein-coding |
| chr18_79238464_79239195 | chr18 | 79238465 | 79239195 | + | 1.06611 | Intergenic | Intergenic | -129439 | NM_053099 | 240427 | Mm.312871 | NM_053099 | ENSMUSG00000024548 | **Setbp1** | C130092E12\|Seb\|mKIAA0437 | SET binding protein 1 | protein-coding |
| chr18_79184610_79184978 | chr18 | 79184611 | 79184978 | + | 0.999605 | Intergenic | Intergenic | -75403 | NM_053099 | 240427 | Mm.312871 | NM_053099 | ENSMUSG00000024548 | **Setbp1** | C130092E12\|Seb\|mKIAA0437 | SET binding protein 1 | protein-coding |
| chr18_79434525_79435241 | chr18 | 79434526 | 79435241 | + | 0.93482 | Intergenic | Intergenic | -325492 | NM_053099 | 240427 | Mm.312871 | NM_053099 | ENSMUSG00000024548 | **Setbp1** | C130092E12\|Seb\|mKIAA0437 | SET binding protein 1 | protein-coding |
| chr18_79294828_79295211 | chr18 | 79294829 | 79295211 | + | 0.884829 | Intergenic | Intergenic | -185629 | NM_053099 | 240427 | Mm.312871 | NM_053099 | ENSMUSG00000024548 | **Setbp1** | C130092E12\|Seb\|mKIAA0437 | SET binding protein 1 | protein-coding |
| chr18_79352840_79353196 | chr18 | 79352841 | 79353196 | + | 0.84127 | Intergenic | MIR3\|SINE\|MIR | -243627 | NM_053099 | 240427 | Mm.312871 | NM_053099 | ENSMUSG00000024548 | **Setbp1** | C130092E12\|Seb\|mKIAA0437 | SET binding protein 1 | protein-coding |
| chr9_110530789_110531079 | chr9 | 110530790 | 110531079 | + | 0.619684 | Intergenic | PB1D10\|SINE\|Alu | -1663 | NM_001081340 | 235626 | Mm.288949 | NM_001081340 | ENSMUSG00000044791 | **Setd2** | 4921524K10Rik\|BC031601\|KMT3A | SET domain containing 2 | protein-coding |
| chr12_108152441_108153090 | chr12 | 108152442 | 108153090 | + | 0.882324 | intron (NM_028262, intron 6 of 12) | intron (NM_028262, intron 6 of 12) | 26518 | NM_028262 | 52690 | Mm.159185 | NM_028262 | ENSMUSG00000056770 | **Setd3** | 2610102I01Rik\|2610305M23Rik\|6030490B17Rik\|D12Ertd771e | SET domain containing 3 | protein-coding |
| chr12_108176529_108176693 | chr12 | 108176530 | 108176693 | + | 0.698837 | intron (NM_028262, intron 1 of 12) | intron (NM_028262, intron 1 of 12) | 2673 | NM_028262 | 52690 | Mm.159185 | NM_028262 | ENSMUSG00000056770 | **Setd3** | 2610102I01Rik\|2610305M23Rik\|6030490B17Rik\|D12Ertd771e | SET domain containing 3 | protein-coding |
| chr12_108130943_108131251 | chr12 | 108130944 | 108131251 | + | 0.633665 | intron (NM_028262, intron 6 of 12) | intron (NM_028262, intron 6 of 12) | 48187 | NM_028262 | 52690 | Mm.159185 | NM_028262 | ENSMUSG00000056770 | **Setd3** | 2610102I01Rik\|2610305M23Rik\|6030490B17Rik\|D12Ertd771e | SET domain containing 3 | protein-coding |
| chr3_95360224_95360856 | chr3 | 95360225 | 95360856 | + | 0.90629 | Intergenic | RLTR48A\|LTR\|ERV1 | -3338 | NM_001163642 | 84505 | Mm.490259 | NM_018877 | ENSMUSG00000015697 | **Setdb1** | AU022152\|ESET\|KMT1E\|mKIAA0067 | SET domain, bifurcated 1 | protein-coding |
| chr11_4164739_4165029 | chr11 | 4164740 | 4165029 | + | 0.880676 | intron (NM_026175, intron 2 of 15) | intron (NM_026175, intron 2 of 15) | 4530 | NM_026175 | 67465 | Mm.156914 | NM_026175 | ENSMUSG00000002129 | **Sf3a1** | 1200014H24Rik\|5930416L09Rik\|AI159724 | splicing factor 3a, subunit 1 | protein-coding |
| chr11_4164730_4165249 | chr11 | 4164731 | 4165249 | + | 0.869839 | intron (NM_026175, intron 2 of 15) | intron (NM_026175, intron 2 of 15) | 4636 | NM_026175 | 67465 | Mm.156914 | NM_026175 | ENSMUSG00000002129 | **Sf3a1** | 1200014H24Rik\|5930416L09Rik\|AI159724 | splicing factor 3a, subunit 1 | protein-coding |
| chr4_124709034_124709436 | chr4 | 124709035 | 124709436 | + | 0.632122 | TTS (NM_010213) | TTS (NM_010213) | -5626 | NM_029157 | 75062 | Mm.25779 | NM_029157 | ENSMUSG00000028902 | **Sf3a3** | 4930512K19Rik\|60kDa | splicing factor 3a, subunit 3 | protein-coding |
| chr10_13035243_13035600 | chr10 | 13035244 | 13035600 | + | 0.769946 | Intergenic | Intergenic | 26972 | NM_175102 | 66125 | Mm.29743 | NM_175102 | ENSMUSG00000078348 | **Sf3b5** | 10kDa\|1110005L13Rik\|AU043053\|Sf3b10 | splicing factor 3b, subunit 5 | protein-coding |
| chr10_12993358_12994113 | chr10 | 12993359 | 12994113 | + | 0.657647 | Intergenic | L1_Rod\|LINE\|L1 | -14714 | NM_175102 | 66125 | Mm.29743 | NM_175102 | ENSMUSG00000078348 | **Sf3b5** | 10kDa\|1110005L13Rik\|AU043053\|Sf3b10 | splicing factor 3b, subunit 5 | protein-coding |
| chr2_10314211_10314587 | chr2 | 10314212 | 10314587 | + | 1.48478 | Intergenic | Intergenic | -56052 | NM_001198808 | 353282 | Mm.329991 | NM_177386 | ENSMUSG00000061186 | **Sfmbt2** | D2Wsu23e\|D330030P06Rik | Scm-like with four mbt domains 2 | protein-coding |
| chr2_10351136_10351616 | chr2 | 10351137 | 10351616 | + | 1.47548 | Intergenic | Intergenic | -19075 | NM_001198808 | 353282 | Mm.329991 | NM_177386 | ENSMUSG00000061186 | **Sfmbt2** | D2Wsu23e\|D330030P06Rik | Scm-like with four mbt domains 2 | protein-coding |
| chr2_10323091_10323813 | chr2 | 10323092 | 10323813 | + | 1.16474 | Intergenic | Intergenic | -46999 | NM_001198808 | 353282 | Mm.329991 | NM_177386 | ENSMUSG00000061186 | **Sfmbt2** | D2Wsu23e\|D330030P06Rik | Scm-like with four mbt domains 2 | protein-coding |
| chr2_10346876_10347491 | chr2 | 10346877 | 10347491 | + | 0.877162 | Intergenic | MTEb\|LTR\|ERVL-MaLR | -23267 | NM_001198808 | 353282 | Mm.329991 | NM_177386 | ENSMUSG00000061186 | **Sfmbt2** | D2Wsu23e\|D330030P06Rik | Scm-like with four mbt domains 2 | protein-coding |
| chr2_10314173_10315549 | chr2 | 10314174 | 10315549 | + | 0.775055 | Intergenic | Intergenic | -55590 | NM_001198808 | 353282 | Mm.329991 | NM_177386 | ENSMUSG00000061186 | **Sfmbt2** | D2Wsu23e\|D330030P06Rik | Scm-like with four mbt domains 2 | protein-coding |
| chr4_133614277_133614847 | chr4 | 133614278 | 133614847 | + | 0.85186 | intron (NM_001017968, intron 3 of 7) | intron (NM_001017968, intron 3 of 7) | -12394 | NM_018754 | 55948 | Mm.44482 | NM_018754 | ENSMUSG00000047281 | **Sfn** | Er\|Mme1\|Ywhas | stratifin | protein-coding |
| chr19_47754735_47755389 | chr19 | 47754736 | 47755389 | + | 1.33678 | intron (NM_027559, intron 28 of 37) | MTE2a\|LTR\|ERVL-MaLR | 23306 | NM_026377 | 67788 | Mm.181610 | NM_026377 | ENSMUSG00000025066 | **Sfr1** | 6330577E15Rik\|Meir5 | SWI5 dependent recombination repair 1 | protein-coding |
| chr3_83795644_83796135 | chr3 | 83795645 | 83796135 | + | 1.18955 | Intergenic | Intergenic | 29569 | NM_009144 | 20319 | Mm.19155 | NM_009144 | ENSMUSG00000027996 | **Sfrp2** | AI851596\|Sdf5 | secreted frizzled-related protein 2 | protein-coding |
| chr3_83781816_83782446 | chr3 | 83781817 | 83782446 | + | 1.04916 | Intergenic | Intergenic | 15810 | NM_009144 | 20319 | Mm.19155 | NM_009144 | ENSMUSG00000027996 | **Sfrp2** | AI851596\|Sdf5 | secreted frizzled-related protein 2 | protein-coding |
| chr3_83746199_83746622 | chr3 | 83746200 | 83746622 | + | 1.02305 | Intergenic | ORR1D2\|LTR\|ERVL-MaLR | -19910 | NM_009144 | 20319 | Mm.19155 | NM_009144 | ENSMUSG00000027996 | **Sfrp2** | AI851596\|Sdf5 | secreted frizzled-related protein 2 | protein-coding |
| chr3_83746021_83746716 | chr3 | 83746022 | 83746716 | + | 0.960658 | Intergenic | ORR1D2\|LTR\|ERVL-MaLR | -19952 | NM_009144 | 20319 | Mm.19155 | NM_009144 | ENSMUSG00000027996 | **Sfrp2** | AI851596\|Sdf5 | secreted frizzled-related protein 2 | protein-coding |
| chr5_129456275_129457044 | chr5 | 129456276 | 129457044 | + | 1.13671 | Intergenic | Intergenic | -44571 | NM_172276 | 231769 | Mm.288714 | NM_172276 | ENSMUSG00000029439 | **Sfswap** | 1190005N23Rik\|6330437E22Rik\|AI197402\|AW212079\|SWAP\|Sfrs8\|Srsf8 | splicing factor, suppressor of white-apricot homolog (Drosophila) | protein-coding |
| chr17_8305642_8306275 | chr17 | 8305643 | 8306275 | + | 0.878607 | intron (NR_073374, intron 1 of 4) | intron (NR_073374, intron 1 of 4) | -5144 | NM_134114 | 106489 | Mm.311080 | NM_134114 | ENSMUSG00000073468 | **Sft2d1** | 5630401J11Rik\|AA409698 | SFT2 domain containing 1 | protein-coding |
| chr1_165195549_165196010 | chr1 | 165195550 | 165196010 | + | 1.37915 | Intergenic | ORR1B1\|LTR\|ERVL-MaLR | -1347 | NM_145512 | 108735 | Mm.288369 | NM_145512 | ENSMUSG00000040848 | **Sft2d2** | 2010005O13Rik\|AI506168\|AW561909\|cI-45 | SFT2 domain containing 2 | protein-coding |
| chr13_54101565_54102515 | chr13 | 54101566 | 54102515 | + | 0.978926 | intron (NM_027324, intron 8 of 10) | intron (NM_027324, intron 8 of 10) | 30195 | NM_027324 | 14057 | Mm.134191 | NM_027324 | ENSMUSG00000021474 | **Sfxn1** | 2810002O05Rik\|A930015P12Rik\|f | sideroflexin 1 | protein-coding |
| chr13_54101612_54102049 | chr13 | 54101613 | 54102049 | + | 0.792899 | intron (NM_027324, intron 8 of 10) | intron (NM_027324, intron 8 of 10) | 29986 | NM_027324 | 14057 | Mm.134191 | NM_027324 | ENSMUSG00000021474 | **Sfxn1** | 2810002O05Rik\|A930015P12Rik\|f | sideroflexin 1 | protein-coding |
| chr6_85331404_85331681 | chr6 | 85331405 | 85331681 | + | 0.725093 | intron (NM_178639, intron 1 of 13) | intron (NM_178639, intron 1 of 13) | 1879 | NM_178639 | 94282 | Mm.121485 | NM_178639 | ENSMUSG00000033720 | **Sfxn5** | C230001H08Rik | sideroflexin 5 | protein-coding |
| chr6_4708901_4709449 | chr6 | 4708902 | 4709449 | + | 1.82044 | intron (NM_001130188, intron 5 of 11) | intron (NM_001130188, intron 5 of 11) | 38029 | NM_001130188 | 20392 | Mm.8739 | NM_011360 | ENSMUSG00000004631 | **Sgce** | e-SG | sarcoglycan, epsilon | protein-coding |
| chr6_4708893_4709764 | chr6 | 4708894 | 4709764 | + | 1.80735 | intron (NM_001130188, intron 5 of 11) | intron (NM_001130188, intron 5 of 11) | 37875 | NM_001130188 | 20392 | Mm.8739 | NM_011360 | ENSMUSG00000004631 | **Sgce** | e-SG | sarcoglycan, epsilon | protein-coding |
| chr4_102682103_102682613 | chr4 | 102682104 | 102682613 | + | 0.952652 | Intergenic | Intergenic | -77777 | NM_001285860 | 73094 | Mm.54201 | NM_144906 | ENSMUSG00000028524 | **Sgip1** | 3110007P09Rik\|SGIP1alpha\|gag | SH3-domain GRB2-like (endophilin) interacting protein 1 | protein-coding |
| chr2_162969125_162969486 | chr2 | 162969126 | 162969486 | + | 0.765289 | intron (NM_001081338, intron 20 of 22) | intron (NM_001081338, intron 20 of 22) | -18175 | NM_001291154 | 27219 | Mm.26462 | NM_013731 | ENSMUSG00000017868 | **Sgk2** | AI098171\|AW146006\|Sgkl | serum/glucocorticoid regulated kinase 2 | protein-coding |
| chr1_9833403_9833771 | chr1 | 9833404 | 9833771 | + | 0.683652 | intron (NM_133220, intron 2 of 17) | intron (NM_133220, intron 2 of 17) | -14694 | NM_001037759 | 170755 | Mm.336410 | NM_133220 | ENSMUSG00000025915 | **Sgk3** | 2510015P22Rik\|A330005P07Rik\|Cisk\|fy\|fz | serum/glucocorticoid regulated kinase 3 | protein-coding |
| chr1_9833287_9833884 | chr1 | 9833288 | 9833884 | + | 0.659545 | intron (NM_133220, intron 2 of 17) | intron (NM_133220, intron 2 of 17) | -14695 | NM_001037759 | 170755 | Mm.336410 | NM_133220 | ENSMUSG00000025915 | **Sgk3** | 2510015P22Rik\|A330005P07Rik\|Cisk\|fy\|fz | serum/glucocorticoid regulated kinase 3 | protein-coding |
| chr19_32263905_32264268 | chr19 | 32263906 | 32264268 | + | 1.07544 | intron (NM_001168525, intron 2 of 9) | intron (NM_001168525, intron 2 of 9) | -53074 | NM_001168526 | 208449 | Mm.329810 | NM_144792 | ENSMUSG00000040451 | **Sgms1** | 9530058O11Rik\|AI841905\|C80702\|Mob\|Sms1\|Sor1\|Tmem23 | sphingomyelin synthase 1 | protein-coding |
| chr19_32198619_32198907 | chr19 | 32198620 | 32198907 | + | 1.05814 | intron (NM_144792, intron 5 of 9) | intron (NM_144792, intron 5 of 9) | 12250 | NM_001168526 | 208449 | Mm.329810 | NM_144792 | ENSMUSG00000040451 | **Sgms1** | 9530058O11Rik\|AI841905\|C80702\|Mob\|Sms1\|Sor1\|Tmem23 | sphingomyelin synthase 1 | protein-coding |
| chr19_32274156_32274551 | chr19 | 32274157 | 32274551 | + | 0.978946 | intron (NM_001168525, intron 2 of 9) | intron (NM_001168525, intron 2 of 9) | -63341 | NM_001168526 | 208449 | Mm.329810 | NM_144792 | ENSMUSG00000040451 | **Sgms1** | 9530058O11Rik\|AI841905\|C80702\|Mob\|Sms1\|Sor1\|Tmem23 | sphingomyelin synthase 1 | protein-coding |
| chr19_32308345_32308817 | chr19 | 32308346 | 32308817 | + | 0.780128 | intron (NM_144792, intron 2 of 9) | intron (NM_144792, intron 2 of 9) | 40735 | NM_001168525 | 208449 | Mm.329810 | NM_144792 | ENSMUSG00000040451 | **Sgms1** | 9530058O11Rik\|AI841905\|C80702\|Mob\|Sms1\|Sor1\|Tmem23 | sphingomyelin synthase 1 | protein-coding |
| chr19_32262804_32263155 | chr19 | 32262805 | 32263155 | + | 0.59233 | intron (NM_001168525, intron 2 of 9) | RLTR21\|LTR\|ERVK | -51967 | NM_001168526 | 208449 | Mm.329810 | NM_144792 | ENSMUSG00000040451 | **Sgms1** | 9530058O11Rik\|AI841905\|C80702\|Mob\|Sms1\|Sor1\|Tmem23 | sphingomyelin synthase 1 | protein-coding |
| chr3_131364602_131364987 | chr3 | 131364603 | 131364987 | + | 1.21177 | Intergenic | Intergenic | -19872 | NM_028943 | 74442 | Mm.273360 | NM_028943 | ENSMUSG00000050931 | **Sgms2** | 4933405A16Rik\|5133401H06Rik\|AI854299 | sphingomyelin synthase 2 | protein-coding |
| chr3_131364422_131365073 | chr3 | 131364423 | 131365073 | + | 0.972439 | Intergenic | Intergenic | -19825 | NM_028943 | 74442 | Mm.273360 | NM_028943 | ENSMUSG00000050931 | **Sgms2** | 4933405A16Rik\|5133401H06Rik\|AI854299 | sphingomyelin synthase 2 | protein-coding |
| chrX_42365629_42366371 | chrX | 42365630 | 42366371 | + | 1.32155 | Intergenic | Intergenic | -136608 | NM_011364 | 20400 | Mm.441197 | NM_011364 | ENSMUSG00000005696 | **Sh2d1a** | Gm686\|SAP | SH2 domain protein 1A | protein-coding |
| chrX_42403189_42403840 | chrX | 42403190 | 42403840 | + | 1.20316 | Intergenic | Intergenic | -99093 | NM_011364 | 20400 | Mm.441197 | NM_011364 | ENSMUSG00000005696 | **Sh2d1a** | Gm686\|SAP | SH2 domain protein 1A | protein-coding |
| chrX_42408066_42408599 | chrX | 42408067 | 42408599 | + | 1.07438 | Intergenic | Intergenic | -94275 | NM_011364 | 20400 | Mm.441197 | NM_011364 | ENSMUSG00000005696 | **Sh2d1a** | Gm686\|SAP | SH2 domain protein 1A | protein-coding |

| | | | | | | | | | | | | | | | | | |
|---|---|---|---|---|---|---|---|---|---|---|---|---|---|---|---|---|---|
| chrX_42480233_42480729 | chrX | 42480234 | 42480729 | + | 0.969541 | Intergenic | Intergenic | -22127 | NM_011364 | 20400 | Mm.441197 | NM_011364 | ENSMUSG00000005696 | **Sh2d1a** | Gm686\|SAP | SH2 domain protein 1A | protein-coding |
| chrX_42481898_42482715 | chrX | 42481899 | 42482715 | + | 0.720782 | Intergenic | Lx8\|LINE\|L1 | -20301 | NM_011364 | 20400 | Mm.441197 | NM_011364 | ENSMUSG00000005696 | **Sh2d1a** | Gm686\|SAP | SH2 domain protein 1A | protein-coding |
| chrX_42481905_42482481 | chrX | 42481906 | 42482481 | + | 0.685546 | Intergenic | Lx8\|LINE\|L1 | -20415 | NM_011364 | 20400 | Mm.441197 | NM_011364 | ENSMUSG00000005696 | **Sh2d1a** | Gm686\|SAP | SH2 domain protein 1A | protein-coding |
| chr8_68157116_68157643 | chr8 | 68157117 | 68157643 | + | 1.91754 | Intergenic | Intergenic | -119148 | NM_028182 | 72281 | Mm.40974 | NM_028182 | ENSMUSG00000053886 | **Sh2d4a** | 2210402M20Rik\|SH2A | SH2 domain containing 4A | protein-coding |
| chr8_68248522_68249199 | chr8 | 68248523 | 68249199 | + | 1.12198 | Intergenic | URR1A\|DNA\|hAT-Charlie | -27667 | NM_028182 | 72281 | Mm.40974 | NM_028182 | ENSMUSG00000053886 | **Sh2d4a** | 2210402M20Rik\|SH2A | SH2 domain containing 4A | protein-coding |
| chr8_68212105_68212624 | chr8 | 68212106 | 68212624 | + | 0.969171 | Intergenic | Intergenic | -64163 | NM_028182 | 72281 | Mm.40974 | NM_028182 | ENSMUSG00000053886 | **Sh2d4a** | 2210402M20Rik\|SH2A | SH2 domain containing 4A | protein-coding |
| chr8_68126042_68126479 | chr8 | 68126043 | 68126479 | + | 0.967301 | Intergenic | Intergenic | -150267 | NM_028182 | 72281 | Mm.40974 | NM_028182 | ENSMUSG00000053886 | **Sh2d4a** | 2210402M20Rik\|SH2A | SH2 domain containing 4A | protein-coding |
| chr8_68246977_68247418 | chr8 | 68246978 | 68247418 | + | 0.818749 | Intergenic | Intergenic | -29330 | NM_028182 | 72281 | Mm.40974 | NM_028182 | ENSMUSG00000053886 | **Sh2d4a** | 2210402M20Rik\|SH2A | SH2 domain containing 4A | protein-coding |
| chr8_68206788_68207535 | chr8 | 68206789 | 68207535 | + | 0.649996 | Intergenic | Intergenic | -69366 | NM_028182 | 72281 | Mm.40974 | NM_028182 | ENSMUSG00000053886 | **Sh2d4a** | 2210402M20Rik\|SH2A | SH2 domain containing 4A | protein-coding |
| chr4_138247211_138247693 | chr4 | 138247212 | 138247693 | + | 0.989705 | Intergenic | Intergenic | -2959 | NM_001099631 | 230863 | Mm.482408 | NM_001099631 | ENSMUSG00000045349 | **Sh2d5** | BC036961 | SH2 domain containing 5 | protein-coding |
| chrX_109119719_109120219 | chrX | 109119720 | 109120219 | + | 1.15876 | intron (NM_019989, intron 1 of 3) | intron (NM_019989, intron 1 of 3) | 24562 | NM_019989 | 56726 | Mm.260760 | NM_019989 | ENSMUSG00000031246 | **Sh3bgrl** | 1190008F14Rik | SH3-binding domain glutamic acid-rich protein like | protein-coding |
| chr9_83563821_83564555 | chr9 | 83563822 | 83564555 | + | 1.06312 | intron (NM_172507, intron 1 of 3) | intron (NM_172507, intron 1 of 3) | 15850 | NM_172507 | 212531 | Mm.491246 | NM_172507 | ENSMUSG00000032261 | **Sh3bgrl2** | A930014C21Rik\|AA408038\|B930089I21 | SH3 domain binding glutamic acid-rich protein like 2 | protein-coding |
| chr1_89141557_89141969 | chr1 | 89141558 | 89141969 | + | 1.27085 | intron (NM_133816, intron 3 of 5) | intron (NM_133816, intron 3 of 5) | 71301 | NM_133816 | 98402 | Mm.170983 | NM_133816 | | **Sh3bp4** | AI594717\|AW227605\|BOG25 | SH3-domain binding protein 4 | protein-coding |
| chr1_89141082_89141979 | chr1 | 89141083 | 89141979 | + | 1.08189 | intron (NM_133816, intron 3 of 5) | intron (NM_133816, intron 3 of 5) | 71069 | NM_133816 | 98402 | Mm.170983 | NM_133816 | | **Sh3bp4** | AI594717\|AW227605\|BOG25 | SH3-domain binding protein 4 | protein-coding |
| chr1_89093659_89094137 | chr1 | 89093660 | 89094137 | + | 0.70094 | intron (NM_133816, intron 1 of 5) | intron (NM_133816, intron 1 of 5) | 23436 | NM_133816 | 98402 | Mm.170983 | NM_133816 | | **Sh3bp4** | AI594717\|AW227605\|BOG25 | SH3-domain binding protein 4 | protein-coding |
| chr14_31417176_31417545 | chr14 | 31417177 | 31417545 | + | 0.842876 | intron (NM_011894, intron 2 of 8) | intron (NM_011894, intron 2 of 8) | 18672 | NM_011894 | 24056 | Mm.383198 | NM_011894 | ENSMUSG00000021892 | **Sh3bp5** | AI606498\|SH3BP-5\|Sab | SH3-domain binding protein 5 (BTK-associated) | protein-coding |
| chr3_86108586_86108948 | chr3 | 86108587 | 86108948 | + | 0.697866 | intron (NM_001082414, intron 7 of 14) | RMER6A\|LTR\|ERVK | 24333 | NM_001082414 | 27059 | Mm.2454 | NM_012059 | ENSMUSG00000028082 | **Sh3d19** | AW011754\|Kryn | SH3 domain protein D19 | protein-coding |
| chr7_82275838_82276207 | chr7 | 82275839 | 82276207 | + | 1.24674 | intron (NM_001277955, intron 5 of 8) | intron (NM_001277955, intron 5 of 8) | 16115 | NM_001277954 | 20408 | Mm.736 | NM_017400 | | **Sh3gl3** | EEN-B2\|SH3P13\|Sh3d2c\|Sh3d2c2 | SH3-domain GRB2-like 3 | protein-coding |
| chr7_82204402_82205092 | chr7 | 82204403 | 82205092 | + | 0.663944 | Intergenic | Intergenic | -55161 | NM_001277955 | 20408 | Mm.736 | NM_017400 | | **Sh3gl3** | EEN-B2\|SH3P13\|Sh3d2c\|Sh3d2c2 | SH3-domain GRB2-like 3 | protein-coding |
| chr3_144669380_144669890 | chr3 | 144669381 | 144669890 | + | 0.781096 | Intergenic | Intergenic | 50700 | NM_001282042 | 54673 | Mm.271775 | NM_019464 | | **Sh3glb1** | AA409932\|AI314629\|AU015566\|Bif-1\|mKIAA0491 | SH3-domain GRB2-like B1 (endophilin) | protein-coding |
| chr3_144669121_144669890 | chr3 | 144669122 | 144669890 | + | 0.654068 | Intergenic | Intergenic | 50829 | NM_001282042 | 54673 | Mm.271775 | NM_019464 | | **Sh3glb1** | AA409932\|AI314629\|AU015566\|Bif-1\|mKIAA0491 | SH3-domain GRB2-like B1 (endophilin) | protein-coding |
| chrX_159850718_159851105 | chrX | 159850719 | 159851105 | + | 1.09871 | intron (NM_001135728, intron 1 of 12) | intron (NM_001135728, intron 1 of 12) | 10444 | NM_001135728 | 58194 | Mm.286495 | NM_021389 | ENSMUSG00000040990 | **Sh3kbp1** | 1200007H22Rik\|1700125L08Rik\|5830464D22Rik\|AI447724\|IN85\|Ruk\|Seta | SH3-domain kinase binding protein 1 | protein-coding |
| chrX_159910641_159911225 | chrX | 159910642 | 159911225 | + | 1.00814 | intron (NM_021389, intron 8 of 16) | intron (NM_021389, intron 8 of 16) | -31760 | NM_001290661 | 58194 | Mm.286495 | NM_021389 | ENSMUSG00000040990 | **Sh3kbp1** | 1200007H22Rik\|1700125L08Rik\|5830464D22Rik\|AI447724\|IN85\|Ruk\|Seta | SH3-domain kinase binding protein 1 | protein-coding |
| chrX_159911727_159912122 | chrX | 159911728 | 159912122 | + | 0.829365 | intron (NM_021389, intron 8 of 16) | intron (NM_021389, intron 8 of 16) | -30768 | NM_001290661 | 58194 | Mm.286495 | NM_021389 | ENSMUSG00000040990 | **Sh3kbp1** | 1200007H22Rik\|1700125L08Rik\|5830464D22Rik\|AI447724\|IN85\|Ruk\|Seta | SH3-domain kinase binding protein 1 | protein-coding |
| chrX_159950650_159951219 | chrX | 159950651 | 159951219 | + | 0.757178 | intron (NR_110969, intron 13 of 16) | intron (NR_110969, intron 13 of 16) | 8242 | NM_001290664 | 58194 | Mm.286495 | NM_021389 | ENSMUSG00000040990 | **Sh3kbp1** | 1200007H22Rik\|1700125L08Rik\|5830464D22Rik\|AI447724\|IN85\|Ruk\|Seta | SH3-domain kinase binding protein 1 | protein-coding |
| chr8_61227217_61227478 | chr8 | 61227218 | 61227478 | + | 0.804341 | intron (NM_021506, intron 2 of 11) | B3\|SINE\|B2 | 3177 | NM_021506 | 59009 | Mm.27949 | NM_021506 | ENSMUSG00000031642 | **Sh3rf1** | 2200003J05Rik\|Posh\|R75531\|Sh3md2 | SH3 domain containing ring finger 1 | protein-coding |
| chr18_62037615_62038460 | chr18 | 62037616 | 62038460 | + | 0.828647 | Intergenic | Intergenic | 84963 | NM_172628 | 225608 | Mm.262320 | NM_172628 | ENSMUSG00000045629 | **Sh3tc2** | D430044G18Rik | SH3 domain and tetratricopeptide repeats 2 | protein-coding |
| chr18_62037625_62037896 | chr18 | 62037626 | 62037896 | + | 0.822357 | Intergenic | Intergenic | 84686 | NM_172628 | 225608 | Mm.262320 | NM_172628 | ENSMUSG00000045629 | **Sh3tc2** | D430044G18Rik | SH3 domain and tetratricopeptide repeats 2 | protein-coding |
| chr2_125735175_125736080 | chr2 | 125735176 | 125736080 | + | 1.07443 | Intergenic | Tigger1\|DNA\|TcMar-Tigger | -11480 | NM_199022 | 271849 | Mm.425101 | NM_199022 | ENSMUSG00000035109 | **Shc4** | 6230417E10Rik\|9930029B02Rik\|Gm685\|RaLP | SHC (Src homology 2 domain containing) family, member 4 | protein-coding |
| chr3_89861055_89861561 | chr3 | 89861056 | 89861561 | + | 0.845622 | Intergenic | Intergenic | 29938 | NM_172530 | 214547 | Mm.225 | NM_172530 | ENSMUSG00000046280 | **She** | 9430022A14 | src homology 2 domain-containing transforming protein E | protein-coding |
| chr5_67643544_67644210 | chr5 | 67643545 | 67644210 | + | 1.55723 | intron (NM_001284345, intron 33 of 36) | intron (NM_001284345, intron 33 of 36) | 35994 | NM_001033415 | 330096 | Mm.442747 | NM_001033415 | ENSMUSG00000050010 | **Shisa3** | D830007B15Rik | shisa homolog 3 (Xenopus laevis) | protein-coding |
| chr9_109050118_109050358 | chr9 | 109050119 | 109050358 | + | 0.854426 | intron (NM_025858, intron 2 of 5) | RLTR16\|LTR\|ERVK | -4628 | NM_026381 | 66940 | Mm.196533 | NM_025858 | ENSMUSG00000025647 | **Shisa5** | 2310008D10Rik\|6430628I05Rik\|Scotin\|mShisa5 | shisa homolog 5 (Xenopus laevis) | protein-coding |
| chr9_109049927_109050366 | chr9 | 109049928 | 109050366 | + | 0.655298 | intron (NM_025858, intron 2 of 5) | RLTR16\|LTR\|ERVK | -4719 | NM_026381 | 66940 | Mm.196533 | NM_025858 | ENSMUSG00000025647 | **Shisa5** | 2310008D10Rik\|6430628I05Rik\|Scotin\|mShisa5 | shisa homolog 5 (Xenopus laevis) | protein-coding |
| chr3_66959969_66960378 | chr3 | 66959970 | 66960378 | + | 0.66445 | Intergenic | Intergenic | 21597 | NM_013665 | 20429 | Mm.39093 | NM_013665 | | **Shox2** | 6330543G17Rik\|OG12\|Og12x\|Prx3\|SHOT | short stature homeobox 2 | protein-coding |
| chr5_92970475_92970842 | chr5 | 92970476 | 92970842 | + | 1.19714 | Intergenic | MER47A\|DNA\|TcMar-Tigger | 72666 | NM_001077596 | 27428 | Mm.46014 | NM_015756 | ENSMUSG00000029381 | **Shroom3** | AL022960\|D5Ertd287e\|Shrm\|Shrm3 | shroom family member 3 | protein-coding |
| chr5_92897644_92898466 | chr5 | 92897645 | 92898466 | + | 1.11171 | promoter-TSS (NM_001077596) | promoter-TSS (NM_001077596) | 62 | NM_001077596 | 27428 | Mm.46014 | NM_015756 | ENSMUSG00000029381 | **Shroom3** | AL022960\|D5Ertd287e\|Shrm\|Shrm3 | shroom family member 3 | protein-coding |
| chr5_92964452_92965073 | chr5 | 92964453 | 92965073 | + | 0.8945 | 3' UTR (NM_015756, exon 11 of 11) | 3' UTR (NM_015756, exon 11 of 11) | 66770 | NM_001077596 | 27428 | Mm.46014 | NM_015756 | ENSMUSG00000029381 | **Shroom3** | AL022960\|D5Ertd287e\|Shrm\|Shrm3 | shroom family member 3 | protein-coding |
| chr5_92967148_92967553 | chr5 | 92967149 | 92967553 | + | 0.787871 | Intergenic | Intergenic | 69358 | NM_001077596 | 27428 | Mm.46014 | NM_015756 | ENSMUSG00000029381 | **Shroom3** | AL022960\|D5Ertd287e\|Shrm\|Shrm3 | shroom family member 3 | protein-coding |
| chr5_92964125_92965020 | chr5 | 92964126 | 92965020 | + | 0.759595 | exon (NM_015756, exon 11 of 11) | exon (NM_015756, exon 11 of 11) | 66580 | NM_001077596 | 27428 | Mm.46014 | NM_015756 | ENSMUSG00000029381 | **Shroom3** | AL022960\|D5Ertd287e\|Shrm\|Shrm3 | shroom family member 3 | protein-coding |
| chr14_75521785_75522393 | chr14 | 75521786 | 75522393 | + | 0.953725 | intron (NM_001128093, intron 1 of 1) | intron (NM_001128093, intron 1 of 1) | 66107 | NM_001128093 | 380918 | Mm.484360 | NM_001128093 | ENSMUSG00000091722 | **Siah3** | Gm912 | seven in absentia homolog 3 (Drosophila) | protein-coding |
| chr2_131096474_131096803 | chr2 | 131096475 | 131096803 | + | 0.764793 | Intergenic | MIR\|SINE\|MIR | -9874 | NM_011426 | 20612 | Mm.1374 | NM_011426 | ENSMUSG00000027322 | **Siglec1** | Cd169\|Siglec-1\|Sn | sialic acid binding Ig-like lectin 1, sialoadhesin | protein-coding |
| chr2_131096223_131096808 | chr2 | 131096224 | 131096808 | + | 0.755463 | Intergenic | Intergenic | -9751 | NM_011426 | 20612 | Mm.1374 | NM_011426 | ENSMUSG00000027322 | **Siglec1** | Cd169\|Siglec-1\|Sn | sialic acid binding Ig-like lectin 1, sialoadhesin | protein-coding |
| chr7_43399435_43400245 | chr7 | 43399436 | 43400245 | + | 1.36258 | Intergenic | L2\|LINE\|L2 | -8440 | NM_172900 | 243958 | Mm.24444 | NM_172900 | ENSMUSG00000030468 | **Siglecg** | 9830164H23\|A630096C01Rik\|Siglec-G\|Siglec10\|mSiglec-G | sialic acid binding Ig-like lectin G | protein-coding |
| chr4_41738003_41738230 | chr4 | 41738004 | 41738230 | + | 0.715034 | TTS (NM_001286538) | TTS (NM_001286538) | 3242 | NM_001286540 | 18391 | Mm.425181 | NM_011014 | ENSMUSG00000036078 | **Sigmar1** | AL024364\|Oprs1\|sigma1R | sigma non-opioid intracellular receptor 1 | protein-coding |
| chr17_31875325_31875861 | chr17 | 31875326 | 31875861 | + | 1.04964 | Intergenic | Intergenic | -19801 | NM_010831 | 17691 | Mm.290941 | NM_010831 | ENSMUSG00000024042 | **Sik1** | Msk\|Sik\|Snf1lk | salt inducible kinase 1 | protein-coding |
| chr17_31831045_31831648 | chr17 | 31831046 | 31831648 | + | 0.916868 | Intergenic | Intergenic | 24445 | NM_010831 | 17691 | Mm.290941 | NM_010831 | ENSMUSG00000024042 | **Sik1** | Msk\|Sik\|Snf1lk | salt inducible kinase 1 | protein-coding |
| chr17_31831222_31831624 | chr17 | 31831223 | 31831624 | + | 0.91466 | Intergenic | Intergenic | 24369 | NM_010831 | 17691 | Mm.290941 | NM_010831 | ENSMUSG00000024042 | **Sik1** | Msk\|Sik\|Snf1lk | salt inducible kinase 1 | protein-coding |
| chr17_31772167_31772775 | chr17 | 31772168 | 31772775 | + | 0.72418 | Intergenic | Intergenic | 83321 | NM_010831 | 17691 | Mm.290941 | NM_010831 | ENSMUSG00000024042 | **Sik1** | Msk\|Sik\|Snf1lk | salt inducible kinase 1 | protein-coding |
| chr17_31914373_31914911 | chr17 | 31914374 | 31914911 | + | 0.719177 | Intergenic | MTC\|LTR\|ERVL-MaLR | -58850 | NM_010831 | 17691 | Mm.290941 | NM_010831 | ENSMUSG00000024042 | **Sik1** | Msk\|Sik\|Snf1lk | salt inducible kinase 1 | protein-coding |
| chr9_51032867_51033568 | chr9 | 51032868 | 51033568 | + | 1.24324 | Intergenic | Intergenic | -24145 | NM_178710 | 235344 | Mm.104932 | NM_178710 | ENSMUSG00000037112 | **Sik2** | G630080D20Rik\|Snf1lk2 | salt inducible kinase 2 | protein-coding |
| chr9_51015700_51016068 | chr9 | 51015701 | 51016068 | + | 0.875186 | Intergenic | RSINE1\|SINE\|B4 | -6811 | NM_178710 | 235344 | Mm.104932 | NM_178710 | ENSMUSG00000037112 | **Sik2** | G630080D20Rik\|Snf1lk2 | salt inducible kinase 2 | protein-coding |
| chr9_46083326_46083995 | chr9 | 46083327 | 46083995 | + | 1.99452 | intron (NM_027498, intron 1 of 24) | intron (NM_027498, intron 1 of 24) | 70841 | NM_027498 | 70661 | Mm.219459 | NM_027498 | ENSMUSG00000034135 | **Sik3** | 5730525O22Rik\|9030204A07Rik\|AI447252\|AI846254\|AI849445\|Qsk\|SIK-3\|mKIAA0999 | SIK family kinase 3 | protein-coding |
| chr9_46084415_46084703 | chr9 | 46084416 | 46084703 | + | 1.06485 | intron (NM_027498, intron 1 of 24) | intron (NM_027498, intron 1 of 24) | 71739 | NM_027498 | 70661 | Mm.219459 | NM_027498 | ENSMUSG00000034135 | **Sik3** | 5730525O22Rik\|9030204A07Rik\|AI447252\|AI846254\|AI849445\|Qsk\|SIK-3\|mKIAA0999 | SIK family kinase 3 | protein-coding |
| chr9_46067623_46068157 | chr9 | 46067624 | 46068157 | + | 0.983482 | intron (NM_027498, intron 1 of 24) | intron (NM_027498, intron 1 of 24) | 55070 | NM_027498 | 70661 | Mm.219459 | NM_027498 | ENSMUSG00000034135 | **Sik3** | 5730525O22Rik\|9030204A07Rik\|AI447252\|AI846254\|AI849445\|Qsk\|SIK-3\|mKIAA0999 | SIK family kinase 3 | protein-coding |
| chr9_46116845_46117198 | chr9 | 46116846 | 46117198 | + | 0.936539 | intron (NM_027498, intron 1 of 24) | intron (NM_027498, intron 1 of 24) | 104202 | NM_027498 | 70661 | Mm.219459 | NM_027498 | ENSMUSG00000034135 | **Sik3** | 5730525O22Rik\|9030204A07Rik\|AI447252\|AI846254\|AI849445\|Qsk\|SIK-3\|mKIAA0999 | SIK family kinase 3 | protein-coding |

| | | | | | | | | | | | | | | | | | |
|---|---|---|---|---|---|---|---|---|---|---|---|---|---|---|---|---|---|
| chr9_46057090_46057772 | chr9 | 46057091 | 46057772 | + | 0.886853 | intron (NM_027498, intron 1 of 24) | intron (NM_027498, intron 1 of 24) | 44611 | NM_027498 | 70661 | Mm.219459 | NM_027498 | ENSMUSG00000034135 | **Sik3** | 5730525O22Rik\|9030204A07Rik\|AI447252\|AI846254\|AI849445\|Qsk\|SIK-3\|mKIAA0999 | SIK family kinase 3 | protein-coding |
| chr9_46073229_46073938 | chr9 | 46073230 | 46073938 | + | 0.86128 | intron (NM_027498, intron 1 of 24) | intron (NM_027498, intron 1 of 24) | 60764 | NM_027498 | 70661 | Mm.219459 | NM_027498 | ENSMUSG00000034135 | **Sik3** | 5730525O22Rik\|9030204A07Rik\|AI447252\|AI846254\|AI849445\|Qsk\|SIK-3\|mKIAA0999 | SIK family kinase 3 | protein-coding |
| chr9_46110263_46111045 | chr9 | 46110264 | 46111045 | + | 0.824201 | intron (NM_027498, intron 1 of 24) | PB1D7\|SINE\|Alu | 97834 | NM_027498 | 70661 | Mm.219459 | NM_027498 | ENSMUSG00000034135 | **Sik3** | 5730525O22Rik\|9030204A07Rik\|AI447252\|AI846254\|AI849445\|Qsk\|SIK-3\|mKIAA0999 | SIK family kinase 3 | protein-coding |
| chr9_46046453_46047467 | chr9 | 46046454 | 46047467 | + | 0.821318 | intron (NM_027498, intron 1 of 24) | intron (NM_027498, intron 1 of 24) | 34140 | NM_027498 | 70661 | Mm.219459 | NM_027498 | ENSMUSG00000034135 | **Sik3** | 5730525O22Rik\|9030204A07Rik\|AI447252\|AI846254\|AI849445\|Qsk\|SIK-3\|mKIAA0999 | SIK family kinase 3 | protein-coding |
| chr9_46116309_46116597 | chr9 | 46116310 | 46116597 | + | 0.669377 | intron (NM_027498, intron 1 of 24) | intron (NM_027498, intron 1 of 24) | 103633 | NM_027498 | 70661 | Mm.219459 | NM_027498 | ENSMUSG00000034135 | **Sik3** | 5730525O22Rik\|9030204A07Rik\|AI447252\|AI846254\|AI849445\|Qsk\|SIK-3\|mKIAA0999 | SIK family kinase 3 | protein-coding |
| chr8_72721248_72721646 | chr8 | 72721249 | 72721646 | + | 0.673642 | Intergenic | Intergenic | -1823 | NM_001113248 | 20467 | Mm.2137 | NM_009188 | ENSMUSG00000031622 | **Sin3b** | 2810430C10Rik | transcriptional regulator, SIN3B (yeast) | protein-coding |
| chr12_82285866_82286337 | chr12 | 82285867 | 82286337 | + | 1.10971 | intron (NM_001167983, intron 3 of 23) | intron (NM_001167983, intron 3 of 23) | -25247 | NM_172579 | 217692 | Mm.261333 | NM_172579 | ENSMUSG00000042700 | **Sipa1l1** | 4931426N11Rik\|AW213287\|Spar\|mKIAA0440 | signal-induced proliferation-associated 1 like 1 | protein-coding |
| chr12_82387650_82388074 | chr12 | 82387651 | 82388074 | + | 0.946462 | intron (NM_172579, intron 7 of 20) | intron (NM_172579, intron 7 of 20) | 76513 | NM_172579 | 217692 | Mm.261333 | NM_172579 | ENSMUSG00000042700 | **Sipa1l1** | 4931426N11Rik\|AW213287\|Spar\|mKIAA0440 | signal-induced proliferation-associated 1 like 1 | protein-coding |
| chr12_82323135_82323431 | chr12 | 82323136 | 82323431 | + | 0.641215 | intron (NM_172579, intron 1 of 20) | intron (NM_172579, intron 1 of 20) | 11934 | NM_172579 | 217692 | Mm.261333 | NM_172579 | ENSMUSG00000042700 | **Sipa1l1** | 4931426N11Rik\|AW213287\|Spar\|mKIAA0440 | signal-induced proliferation-associated 1 like 1 | protein-coding |
| chr8_125333469_125333971 | chr8 | 125333470 | 125333971 | + | 1.2001 | Intergenic | Intergenic | 158990 | NM_001081337 | 244668 | Mm.271668 | NM_001081337 | ENSMUSG00000001995 | **Sipa1l2** | BC058408\|mKIAA1389 | signal-induced proliferation-associated 1 like 2 | protein-coding |
| chr8_125333007_125333964 | chr8 | 125333008 | 125333964 | + | 1.00571 | Intergenic | Intergenic | 159224 | NM_001081337 | 244668 | Mm.271668 | NM_001081337 | ENSMUSG00000001995 | **Sipa1l2** | BC058408\|mKIAA1389 | signal-induced proliferation-associated 1 like 2 | protein-coding |
| chr7_29443946_29444436 | chr7 | 29443947 | 29444436 | + | 0.74679 | intron (NM_001081028, intron 2 of 22) | intron (NM_001081028, intron 2 of 22) | 61269 | NM_001081028 | 74206 | Mm.447042 | NM_001081028 | ENSMUSG00000030583 | **Sipa1l3** | 2610511M17Rik | signal-induced proliferation-associated 1 like 3 | protein-coding |
| chr10_63326935_63327464 | chr10 | 63326936 | 63327464 | + | 0.664038 | intron (NM_019812, intron 4 of 8) | L1MC4a\|LINE\|L1 | 11832 | NM_019812 | 93759 | Mm.351459 | NM_019812 | ENSMUSG00000020063 | **Sirt1** | AA673258\|SIR2L1\|Sir2\|Sir2a\|Sir2alpha | sirtuin 1 | protein-coding |
| chr5_115491838_115492310 | chr5 | 115491839 | 115492310 | + | 0.681487 | Intergenic | CpG | -7590 | NM_133760 | 75387 | Mm.332616 | NM_133760 | ENSMUSG00000029524 | **Sirt4** | 4930596O17Rik | sirtuin 4 | protein-coding |
| chr17_85664904_85665277 | chr17 | 85664905 | 85665277 | + | 0.765452 | Intergenic | Intergenic | 23163 | NM_011380 | 20472 | Mm.5039 | NM_011380 | ENSMUSG00000024134 | **Six2** | - | sine oculis-related homeobox 2 | protein-coding |
| chr17_85664682_85665290 | chr17 | 85664683 | 85665290 | + | 0.638694 | Intergenic | (CCCCCA)n\|Simple_repeat\|Simple_repeat | 23268 | NM_011380 | 20472 | Mm.5039 | NM_011380 | ENSMUSG00000024134 | **Six2** | - | sine oculis-related homeobox 2 | protein-coding |
| chr17_85645742_85646076 | chr17 | 85645743 | 85646076 | + | 0.752416 | Intergenic | Intergenic | 25075 | NM_011381 | 20473 | Mm.370208 | NM_011381 | | **Six3** | E130112M24Rik\|Six3a\|Six3alpha\|Six3b\|Six3beta | sine oculis-related homeobox 3 | protein-coding |
| chr17_85645737_85646187 | chr17 | 85645738 | 85646187 | + | 0.674257 | Intergenic | Intergenic | 25128 | NM_011381 | 20473 | Mm.370208 | NM_011381 | | **Six3** | E130112M24Rik\|Six3a\|Six3alpha\|Six3b\|Six3beta | sine oculis-related homeobox 3 | protein-coding |
| chr3_31090044_31090231 | chr3 | 31090045 | 31090231 | + | 0.75829 | Intergenic | Intergenic | -4920 | NM_011386 | 20482 | Mm.15406 | NM_011386 | ENSMUSG00000027660 | **Skil** | Skir\|SnoN\|sno | SKI-like | protein-coding |
| chr3_31106596_31106890 | chr3 | 31106597 | 31106890 | + | 0.67919 | intron (NM_011386, intron 2 of 6) | intron (NM_011386, intron 2 of 6) | 11685 | NM_001039090 | 20482 | Mm.15406 | NM_011386 | ENSMUSG00000027660 | **Skil** | Skir\|SnoN\|sno | SKI-like | protein-coding |
| chr15_9189754_9190361 | chr15 | 9189755 | 9190361 | + | 1.07266 | intron (NM_177178, intron 16 of 17) | RMER4B\|LTR\|ERVK | -34633 | NM_001285980 | 27401 | Mm.35584 | NM_013787 | ENSMUSG00000054115 | **Skp2** | FBXL1\|FWD1\|p45 | S-phase kinase-associated protein 2 (p45) | protein-coding |
| chr15_66795265_66795994 | chr15 | 66795266 | 66795994 | + | 1.46976 | intron (NM_001029841, intron 2 of 7) | intron (NM_001029841, intron 2 of 7) | 36199 | NM_001029841 | 20491 | Mm.7601 | NM_009192 | ENSMUSG00000022372 | **Sla** | Slap\|Slap-1 | src-like adaptor | protein-coding |
| chr15_66795273_66795773 | chr15 | 66795274 | 66795773 | + | 1.44068 | intron (NM_001029841, intron 2 of 7) | intron (NM_001029841, intron 2 of 7) | 36306 | NM_001029841 | 20491 | Mm.7601 | NM_009192 | ENSMUSG00000022372 | **Sla** | Slap\|Slap-1 | src-like adaptor | protein-coding |
| chr15_66845837_66846335 | chr15 | 66845838 | 66846335 | + | 1.42476 | intron (NM_009375, intron 45 of 47) | intron (NM_009375, intron 45 of 47) | -14257 | NM_009192 | 20491 | Mm.7601 | NM_009192 | ENSMUSG00000022372 | **Sla** | Slap\|Slap-1 | src-like adaptor | protein-coding |
| chr15_66851468_66852154 | chr15 | 66851469 | 66852154 | + | 1.26028 | Intergenic | Intergenic | -19982 | NM_009192 | 20491 | Mm.7601 | NM_009192 | ENSMUSG00000022372 | **Sla** | Slap\|Slap-1 | src-like adaptor | protein-coding |
| chr15_66798461_66798878 | chr15 | 66798462 | 66798878 | + | 1.16363 | intron (NM_001029841, intron 2 of 7) | intron (NM_001029841, intron 2 of 7) | 33159 | NM_001029841 | 20491 | Mm.7601 | NM_009192 | ENSMUSG00000022372 | **Sla** | Slap\|Slap-1 | src-like adaptor | protein-coding |
| chr15_66859188_66859655 | chr15 | 66859189 | 66859655 | + | 0.971382 | Intergenic | Intergenic | -27593 | NM_009192 | 20491 | Mm.7601 | NM_009192 | ENSMUSG00000022372 | **Sla** | Slap\|Slap-1 | src-like adaptor | protein-coding |
| chr15_66778518_66779357 | chr15 | 66778519 | 66779357 | + | 0.945784 | intron (NM_009375, intron 41 of 47) | ORR1B2\|LTR\|ERVL-MaLR | 52891 | NM_001029841 | 20491 | Mm.7601 | NM_009192 | ENSMUSG00000022372 | **Sla** | Slap\|Slap-1 | src-like adaptor | protein-coding |
| chr15_66807309_66807565 | chr15 | 66807310 | 66807565 | + | 0.934703 | intron (NM_001029841, intron 1 of 7) | intron (NM_001029841, intron 1 of 7) | 24392 | NM_001029841 | 20491 | Mm.7601 | NM_009192 | ENSMUSG00000022372 | **Sla** | Slap\|Slap-1 | src-like adaptor | protein-coding |
| chr15_66818768_66819201 | chr15 | 66818769 | 66819201 | + | 0.928982 | intron (NM_009192, intron 1 of 8) | URR1B\|DNA\|hAT-Charlie | 12844 | NM_001029841 | 20491 | Mm.7601 | NM_009192 | ENSMUSG00000022372 | **Sla** | Slap\|Slap-1 | src-like adaptor | protein-coding |
| chr15_66818553_66819201 | chr15 | 66818554 | 66819201 | + | 0.772801 | intron (NM_009192, intron 1 of 8) | intron (NM_009192, intron 1 of 8) | 12952 | NM_001029841 | 20491 | Mm.7601 | NM_009192 | ENSMUSG00000022372 | **Sla** | Slap\|Slap-1 | src-like adaptor | protein-coding |
| chr15_66799699_66800256 | chr15 | 66799700 | 66800256 | + | 0.75827 | intron (NM_001029841, intron 2 of 7) | intron (NM_001029841, intron 2 of 7) | 31851 | NM_001029841 | 20491 | Mm.7601 | NM_009192 | ENSMUSG00000022372 | **Sla** | Slap\|Slap-1 | src-like adaptor | protein-coding |
| chr15_66861392_66861593 | chr15 | 66861393 | 66861593 | + | 0.724782 | Intergenic | Intergenic | -29664 | NM_009192 | 20491 | Mm.7601 | NM_009192 | ENSMUSG00000022372 | **Sla** | Slap\|Slap-1 | src-like adaptor | protein-coding |
| chr15_66815533_66815813 | chr15 | 66815534 | 66815813 | + | 0.721722 | intron (NM_009192, intron 1 of 8) | intron (NM_009192, intron 1 of 8) | 16156 | NM_001029841 | 20491 | Mm.7601 | NM_009192 | ENSMUSG00000022372 | **Sla** | Slap\|Slap-1 | src-like adaptor | protein-coding |
| chr15_66799730_66800053 | chr15 | 66799731 | 66800053 | + | 0.715505 | intron (NM_001029841, intron 2 of 7) | intron (NM_001029841, intron 2 of 7) | 31937 | NM_001029841 | 20491 | Mm.7601 | NM_009192 | ENSMUSG00000022372 | **Sla** | Slap\|Slap-1 | src-like adaptor | protein-coding |
| chr15_66806353_66806672 | chr15 | 66806354 | 66806672 | + | 0.704401 | intron (NM_001029841, intron 1 of 7) | intron (NM_001029841, intron 1 of 7) | 25316 | NM_001029841 | 20491 | Mm.7601 | NM_009192 | ENSMUSG00000022372 | **Sla** | Slap\|Slap-1 | src-like adaptor | protein-coding |
| chr14_103666379_103667058 | chr14 | 103666380 | 103667058 | + | 1.62551 | intron (NM_198014, intron 2 of 6) | intron (NM_198014, intron 2 of 6) | 16476 | NM_198014 | 105439 | Mm.27548 | NM_198014 | ENSMUSG00000055717 | **Slain1** | 9630044O09Rik\|AA675320\|AW742596 | SLAIN motif family, member 1 | protein-coding |
| chr14_103667113_103667381 | chr14 | 103667114 | 103667381 | + | 1.28076 | intron (NM_198014, intron 2 of 6) | intron (NM_198014, intron 2 of 6) | 17004 | NM_198014 | 105439 | Mm.27548 | NM_198014 | ENSMUSG00000055717 | **Slain1** | 9630044O09Rik\|AA675320\|AW742596 | SLAIN motif family, member 1 | protein-coding |
| chr14_103619447_103620199 | chr14 | 103619448 | 103620199 | + | 1.24592 | Intergenic | LTR84a\|LTR\|ERVL | -30420 | NM_198014 | 105439 | Mm.27548 | NM_198014 | ENSMUSG00000055717 | **Slain1** | 9630044O09Rik\|AA675320\|AW742596 | SLAIN motif family, member 1 | protein-coding |
| chr14_103619747_103620183 | chr14 | 103619748 | 103620183 | + | 1.18721 | Intergenic | Intergenic | -30278 | NM_198014 | 105439 | Mm.27548 | NM_198014 | ENSMUSG00000055717 | **Slain1** | 9630044O09Rik\|AA675320\|AW742596 | SLAIN motif family, member 1 | protein-coding |
| chr14_103661582_103661963 | chr14 | 103661583 | 103661963 | + | 1.18231 | intron (NM_198014, intron 2 of 6) | intron (NM_198014, intron 2 of 6) | 11530 | NM_198014 | 105439 | Mm.27548 | NM_198014 | ENSMUSG00000055717 | **Slain1** | 9630044O09Rik\|AA675320\|AW742596 | SLAIN motif family, member 1 | protein-coding |
| chr14_103621262_103621771 | chr14 | 103621263 | 103621771 | + | 0.982359 | Intergenic | Intergenic | -28726 | NM_198014 | 105439 | Mm.27548 | NM_198014 | ENSMUSG00000055717 | **Slain1** | 9630044O09Rik\|AA675320\|AW742596 | SLAIN motif family, member 1 | protein-coding |
| chr14_103614340_103614839 | chr14 | 103614341 | 103614839 | + | 0.81279 | Intergenic | B4A\|SINE\|B4 | -35653 | NM_198014 | 105439 | Mm.27548 | NM_198014 | ENSMUSG00000055717 | **Slain1** | 9630044O09Rik\|AA675320\|AW742596 | SLAIN motif family, member 1 | protein-coding |
| chr14_103657931_103658199 | chr14 | 103657932 | 103658199 | + | 0.800964 | intron (NM_198014, intron 2 of 6) | intron (NM_198014, intron 2 of 6) | 7822 | NM_198014 | 105439 | Mm.27548 | NM_198014 | ENSMUSG00000055717 | **Slain1** | 9630044O09Rik\|AA675320\|AW742596 | SLAIN motif family, member 1 | protein-coding |
| chr14_103606941_103607231 | chr14 | 103606942 | 103607231 | + | 0.688523 | intron (NM_022886, intron 28 of 30) | intron (NM_022886, intron 28 of 30) | -43157 | NM_198014 | 105439 | Mm.27548 | NM_198014 | ENSMUSG00000055717 | **Slain1** | 9630044O09Rik\|AA675320\|AW742596 | SLAIN motif family, member 1 | protein-coding |
| chr5_72940784_72941433 | chr5 | 72940785 | 72941433 | + | 0.800476 | intron (NM_153567, intron 1 of 8) | intron (NM_153567, intron 1 of 8) | 26747 | NM_153567 | 75991 | Mm.121285 | NM_153567 | ENSMUSG00000036087 | **Slain2** | 5033405K12Rik\|8030444K12Rik\|AI596370 | SLAIN motif family, member 2 | protein-coding |
| chr5_72947984_72948495 | chr5 | 72947985 | 72948495 | + | 0.733351 | intron (NM_153567, intron 2 of 8) | intron (NM_153567, intron 2 of 8) | 33878 | NM_153567 | 75991 | Mm.121285 | NM_153567 | ENSMUSG00000036087 | **Slain2** | 5033405K12Rik\|8030444K12Rik\|AI596370 | SLAIN motif family, member 2 | protein-coding |
| chr5_72939200_72939479 | chr5 | 72939201 | 72939479 | + | 0.709473 | intron (NM_153567, intron 1 of 8) | B3\|SINE\|B2 | 24978 | NM_153567 | 75991 | Mm.121285 | NM_153567 | ENSMUSG00000036087 | **Slain2** | 5033405K12Rik\|8030444K12Rik\|AI596370 | SLAIN motif family, member 2 | protein-coding |

| | | | | | | | | | | | | | | | | | |
|---|---|---|---|---|---|---|---|---|---|---|---|---|---|---|---|---|---|
| chr5_72937774_72938067 | chr5 | 72937775 | 72938067 | + | 0.634682 | intron (NM_153567, intron 1 of 8) | intron (NM_153567, intron 1 of 8) | 23559 | NM_153567 | 75991 | Mm.121285 | NM_153567 | ENSMUSG00000036087 | **Slain2** | 5033405K12Rik\|8030444K12Rik\|AI596370 | SLAIN motif family, member 2 | protein-coding |
| chr1_171770865_171771395 | chr1 | 171770866 | 171771395 | + | 0.926744 | intron (NM_013730, intron 1 of 6) | intron (NM_013730, intron 1 of 6) | 3998 | NM_013730 | 27218 | Mm.103648 | NM_013730 | ENSMUSG00000015316 | **Slamf1** | 4933415F16\|AA177906\|CD150\|CDw150\|ESTM51\|IPO-3\|Slam | signaling lymphocytic activation molecule family member 1 | protein-coding |
| chr1_171761612_171762397 | chr1 | 171761613 | 171762397 | + | 0.713643 | Intergenic | Intergenic | -5127 | NM_013730 | 27218 | Mm.103648 | NM_013730 | ENSMUSG00000015316 | **Slamf1** | 4933415F16\|AA177906\|CD150\|CDw150\|ESTM51\|IPO-3\|Slam | signaling lymphocytic activation molecule family member 1 | protein-coding |
| chr1_171761624_171762851 | chr1 | 171761625 | 171762851 | + | 0.677064 | Intergenic | Intergenic | -4894 | NM_013730 | 27218 | Mm.103648 | NM_013730 | ENSMUSG00000015316 | **Slamf1** | 4933415F16\|AA177906\|CD150\|CDw150\|ESTM51\|IPO-3\|Slam | signaling lymphocytic activation molecule family member 1 | protein-coding |
| chr1_171924994_171925258 | chr1 | 171924995 | 171925258 | + | 0.925482 | intron (NM_030710, intron 1 of 7) | Lx8\|LINE\|L1 | 7589 | NM_030710 | 30925 | Mm.486404 | NM_030710 | ENSMUSG00000015314 | **Slamf6** | KAL1\|KAL1b\|Ly108\|NTB-A\|NTBA\|SF2000 | SLAM family member 6 | protein-coding |
| chr1_171642505_171642766 | chr1 | 171642506 | 171642766 | + | 0.657471 | intron (NM_144539, intron 2 of 7) | intron (NM_144539, intron 2 of 7) | 10401 | NM_144539 | 75345 | Mm.164642 | NM_144539 | | **Slamf7** | 19A\|19A24\|4930560D03Rik\|CRACC\|CS1 | SLAM family member 7 | protein-coding |
| chr16_33600889_33601797 | chr16 | 33600890 | 33601797 | + | 1.04039 | intron (NM_134251, intron 5 of 12) | intron (NM_134251, intron 5 of 12) | 82748 | NM_001083902 | 171286 | Mm.35253 | NM_134251 | ENSMUSG00000035506 | **Slc12a8** | CCC9\|E330020C02Rik | solute carrier family 12 (potassium/chloride transporters), member 8 | protein-coding |
| chr6_24311434_24311997 | chr6 | 24311435 | 24311997 | + | 1.02579 | Intergenic | Intergenic | -143624 | NM_019481 | 55961 | Mm.19299 | NM_019481 | ENSMUSG00000029700 | **Slc13a1** | AI314066\|AI987826\|NaSi-1\|Nas1 | solute carrier family 13 (sodium/sulfate symporters), member 1 | protein-coding |
| chr6_24166746_24167300 | chr6 | 24166747 | 24167300 | + | 1.01002 | intron (NM_019481, intron 1 of 14) | MIR\|SINE\|MIR | 1069 | NM_019481 | 55961 | Mm.19299 | NM_019481 | ENSMUSG00000029700 | **Slc13a1** | AI314066\|AI987826\|NaSi-1\|Nas1 | solute carrier family 13 (sodium/sulfate symporters), member 1 | protein-coding |
| chr6_24166759_24167553 | chr6 | 24166760 | 24167553 | + | 0.991137 | intron (NM_019481, intron 1 of 14) | intron (NM_019481, intron 1 of 14) | 936 | NM_019481 | 55961 | Mm.19299 | NM_019481 | ENSMUSG00000029700 | **Slc13a1** | AI314066\|AI987826\|NaSi-1\|Nas1 | solute carrier family 13 (sodium/sulfate symporters), member 1 | protein-coding |
| chr18_78480423_78481133 | chr18 | 78480424 | 78481133 | + | 1.11831 | intron (NM_030683, intron 2 of 11) | intron (NM_030683, intron 2 of 11) | 116172 | NM_207651 | 27411 | Mm.44158 | NM_030683 | ENSMUSG00000024552 | **Slc14a2** | UT-A1\|UT-A3 | solute carrier family 14 (urea transporter), member 2 | protein-coding |
| chr18_78480794_78481128 | chr18 | 78480795 | 78481128 | + | 0.816185 | intron (NM_030683, intron 2 of 11) | Lx8\|LINE\|L1 | 115989 | NM_207651 | 27411 | Mm.44158 | NM_030683 | ENSMUSG00000024552 | **Slc14a2** | UT-A1\|UT-A3 | solute carrier family 14 (urea transporter), member 2 | protein-coding |
| chr18_78238474_78238885 | chr18 | 78238475 | 78238885 | + | 0.71374 | intron (NM_030683, intron 3 of 11) | intron (NM_030683, intron 3 of 11) | -32240 | NM_001110274 | 27411 | Mm.44158 | NM_030683 | ENSMUSG00000024552 | **Slc14a2** | UT-A1\|UT-A3 | solute carrier family 14 (urea transporter), member 2 | protein-coding |
| chr10_40152317_40152988 | chr10 | 40152318 | 40152988 | + | 1.71818 | Intergenic | Intergenic | -10399 | NM_001114332 | 72472 | Mm.186778 | NM_028247 | ENSMUSG00000019838 | **Slc16a10** | 2610103N14Rik\|9830169E08\|Mct10\|PRO0813\|TAT1 | solute carrier family 16 (monocarboxylic acid transporters), member 10 | protein-coding |
| chr10_40128323_40128819 | chr10 | 40128324 | 40128819 | + | 1.19943 | intron (NM_028247, intron 1 of 2) | B4A\|SINE\|B4 | 13683 | NM_028247 | 72472 | Mm.186778 | NM_028247 | ENSMUSG00000019838 | **Slc16a10** | 2610103N14Rik\|9830169E08\|Mct10\|PRO0813\|TAT1 | solute carrier family 16 (monocarboxylic acid transporters), member 10 | protein-coding |
| chr10_40128335_40129066 | chr10 | 40128336 | 40129066 | + | 1.17509 | intron (NM_028247, intron 1 of 2) | intron (NM_028247, intron 1 of 2) | 13553 | NM_028247 | 72472 | Mm.186778 | NM_028247 | ENSMUSG00000019838 | **Slc16a10** | 2610103N14Rik\|9830169E08\|Mct10\|PRO0813\|TAT1 | solute carrier family 16 (monocarboxylic acid transporters), member 10 | protein-coding |
| chr10_40098406_40099016 | chr10 | 40098407 | 40099016 | + | 0.765117 | intron (NM_028247, intron 1 of 2) | L1M4\|LINE\|L1 | 43543 | NM_028247 | 72472 | Mm.186778 | NM_028247 | ENSMUSG00000019838 | **Slc16a10** | 2610103N14Rik\|9830169E08\|Mct10\|PRO0813\|TAT1 | solute carrier family 16 (monocarboxylic acid transporters), member 10 | protein-coding |
| chr10_40098456_40098739 | chr10 | 40098457 | 40098739 | + | 0.73767 | intron (NM_028247, intron 1 of 2) | RSINE1\|SINE\|B4 | 43656 | NM_028247 | 72472 | Mm.186778 | NM_028247 | ENSMUSG00000019838 | **Slc16a10** | 2610103N14Rik\|9830169E08\|Mct10\|PRO0813\|TAT1 | solute carrier family 16 (monocarboxylic acid transporters), member 10 | protein-coding |
| chr10_40098420_40098802 | chr10 | 40098421 | 40098802 | + | 0.66744 | intron (NM_028247, intron 1 of 2) | RSINE1\|SINE\|B4 | 43643 | NM_028247 | 72472 | Mm.186778 | NM_028247 | ENSMUSG00000019838 | **Slc16a10** | 2610103N14Rik\|9830169E08\|Mct10\|PRO0813\|TAT1 | solute carrier family 16 (monocarboxylic acid transporters), member 10 | protein-coding |
| chr11_70198288_70198872 | chr11 | 70198289 | 70198872 | + | 0.774175 | Intergenic | RLTR19B\|LTR\|ERVK | -15330 | NM_153081 | 216867 | Mm.289238 | NM_153081 | ENSMUSG00000040938 | **Slc16a11** | AB041591\|MNCb-2717 | solute carrier family 16 (monocarboxylic acid transporters), member 11 | protein-coding |
| chr11_70191879_70192447 | chr11 | 70191880 | 70192447 | + | 0.648198 | Intergenic | B1_Mur1\|SINE\|Alu | -21747 | NM_153081 | 216867 | Mm.289238 | NM_153081 | ENSMUSG00000040938 | **Slc16a11** | AB041591\|MNCb-2717 | solute carrier family 16 (monocarboxylic acid transporters), member 11 | protein-coding |
| chr11_70192140_70192444 | chr11 | 70192141 | 70192444 | + | 0.616608 | Intergenic | Intergenic | -21618 | NM_153081 | 216867 | Mm.289238 | NM_153081 | ENSMUSG00000040938 | **Slc16a11** | AB041591\|MNCb-2717 | solute carrier family 16 (monocarboxylic acid transporters), member 11 | protein-coding |
| chr19_34773656_34774903 | chr19 | 34773657 | 34774903 | + | 1.30617 | Intergenic | Intergenic | -27169 | NM_172838 | 240638 | Mm.74636 | NM_172838 | ENSMUSG00000009378 | **Slc16a12** | 3230401A21\|AW210596 | solute carrier family 16 (monocarboxylic acid transporters), member 12 | protein-coding |
| chr11_109479668_109480654 | chr11 | 109479669 | 109480654 | + | 0.743766 | intron (NM_028710, intron 1 of 13) | L2a\|LINE\|L2 | -6565 | NM_001029842 | 104681 | Mm.265874 | NM_134038 | ENSMUSG00000041920 | **Slc16a6** | AW743111\|ESTM12\|MCT 6\|MCT 7\|MCT6 | solute carrier family 16 (monocarboxylic acid transporters), member 6 | protein-coding |
| chr7_51723930_51724382 | chr7 | 51723931 | 51724382 | + | 1.19732 | Intergenic | MER44D\|DNA\|TcMar-Tigger | 102326 | NM_080853 | 140919 | Mm.256618 | NM_080853 | ENSMUSG00000030500 | **Slc17a6** | 2900073D12Rik\|DNPI\|VGLUT2 | solute carrier family 17 (sodium-dependent inorganic phosphate cotransporter), member 6 | protein-coding |
| chr10_89589625_89590506 | chr10 | 89589626 | 89590506 | + | 1.34906 | intron (NM_182959, intron 8 of 11) | intron (NM_182959, intron 8 of 11) | 31183 | NM_182959 | 216227 | Mm.233921 | NM_182959 | ENSMUSG00000019935 | **Slc17a8** | BC042593\|Vglut3 | solute carrier family 17 (sodium-dependent inorganic phosphate cotransporter), member 8 | protein-coding |
| chr10_89599116_89599993 | chr10 | 89599117 | 89599993 | + | 1.27063 | intron (NM_182959, intron 3 of 11) | intron (NM_182959, intron 3 of 11) | 21694 | NM_182959 | 216227 | Mm.233921 | NM_182959 | ENSMUSG00000019935 | **Slc17a8** | BC042593\|Vglut3 | solute carrier family 17 (sodium-dependent inorganic phosphate cotransporter), member 8 | protein-coding |
| chr19_59289742_59290180 | chr19 | 59289743 | 59290180 | + | 0.739521 | intron (NM_172523, intron 15 of 15) | intron (NM_172523, intron 15 of 15) | 29083 | NM_172523 | 214084 | Mm.268797 | NM_172523 | ENSMUSG00000025094 | **Slc18a2** | 1110037L13Rik\|9330105E13\|Vmat2 | solute carrier family 18 (vesicular monoamine), member 2 | protein-coding |
| chr1_83077045_83077754 | chr1 | 83077046 | 83077754 | + | 1.43492 | Intergenic | Intergenic | -38952 | NM_030556 | 80721 | Mm.261542 | NM_030556 | ENSMUSG00000038496 | **Slc19a3** | A230084E24Rik\|AI788884\|ThTr2 | solute carrier family 19, member 3 | protein-coding |
| chr1_83006076_83006749 | chr1 | 83006077 | 83006749 | + | 0.988182 | Intergenic | Intergenic | 32035 | NM_030556 | 80721 | Mm.261542 | NM_030556 | ENSMUSG00000038496 | **Slc19a3** | A230084E24Rik\|AI788884\|ThTr2 | solute carrier family 19, member 3 | protein-coding |
| chr1_83006151_83006526 | chr1 | 83006152 | 83006526 | + | 0.905877 | Intergenic | Intergenic | 32109 | NM_030556 | 80721 | Mm.261542 | NM_030556 | ENSMUSG00000038496 | **Slc19a3** | A230084E24Rik\|AI788884\|ThTr2 | solute carrier family 19, member 3 | protein-coding |
| chr1_83076197_83076629 | chr1 | 83076198 | 83076629 | + | 0.683586 | Intergenic | Intergenic | -37965 | NM_030556 | 80721 | Mm.261542 | NM_030556 | ENSMUSG00000038496 | **Slc19a3** | A230084E24Rik\|AI788884\|ThTr2 | solute carrier family 19, member 3 | protein-coding |
| chr19_28869010_28869516 | chr19 | 28869011 | 28869516 | + | 0.701237 | intron (NM_009199, intron 1 of 11) | intron (NM_009199, intron 1 of 11) | 34097 | NM_009199 | 20510 | Mm.246670 | NM_009199 | ENSMUSG00000024935 | **Slc1a1** | D130048G10Rik\|EAAC1\|EAAC2\|EAAT3\|MEAAC1 | solute carrier family 1 (neuronal/epithelial high affinity glutamate transporter, system Xag), member 1 | protein-coding |
| chr2_102701143_102701870 | chr2 | 102701144 | 102701870 | + | 1.81255 | intron (NM_001077514, intron 1 of 10) | intron (NM_001077514, intron 1 of 10) | -4931 | NM_011393 | 20511 | Mm.267547 | NM_011393 | ENSMUSG00000005089 | **Slc1a2** | 1700091C19Rik\|2900019G14Rik\|AI159670\|Eaat2\|GLT-1\|GLT1\|MGLT1 | solute carrier family 1 (glial high affinity glutamate transporter), member 2 | protein-coding |
| chr2_102732972_102733702 | chr2 | 102732973 | 102733702 | + | 1.5843 | intron (NM_001077515, intron 1 of 10) | intron (NM_001077515, intron 1 of 10) | 26899 | NM_001077515 | 20511 | Mm.267547 | NM_011393 | ENSMUSG00000005089 | **Slc1a2** | 1700091C19Rik\|2900019G14Rik\|AI159670\|Eaat2\|GLT-1\|GLT1\|MGLT1 | solute carrier family 1 (glial high affinity glutamate transporter), member 2 | protein-coding |
| chr2_102738749_102739262 | chr2 | 102738750 | 102739262 | + | 1.44381 | intron (NM_001077515, intron 3 of 10) | intron (NM_001077515, intron 3 of 10) | 32568 | NM_001077515 | 20511 | Mm.267547 | NM_011393 | ENSMUSG00000005089 | **Slc1a2** | 1700091C19Rik\|2900019G14Rik\|AI159670\|Eaat2\|GLT-1\|GLT1\|MGLT1 | solute carrier family 1 (glial high affinity glutamate transporter), member 2 | protein-coding |
| chr2_102735261_102735523 | chr2 | 102735262 | 102735523 | + | 1.01696 | intron (NM_001077515, intron 1 of 10) | intron (NM_001077515, intron 1 of 10) | 28954 | NM_001077515 | 20511 | Mm.267547 | NM_011393 | ENSMUSG00000005089 | **Slc1a2** | 1700091C19Rik\|2900019G14Rik\|AI159670\|Eaat2\|GLT-1\|GLT1\|MGLT1 | solute carrier family 1 (glial high affinity glutamate transporter), member 2 | protein-coding |
| chr2_102739400_102739692 | chr2 | 102739401 | 102739692 | + | 0.985832 | intron (NM_001077515, intron 3 of 10) | intron (NM_001077515, intron 3 of 10) | 33108 | NM_001077515 | 20511 | Mm.267547 | NM_011393 | ENSMUSG00000005089 | **Slc1a2** | 1700091C19Rik\|2900019G14Rik\|AI159670\|Eaat2\|GLT-1\|GLT1\|MGLT1 | solute carrier family 1 (glial high affinity glutamate transporter), member 2 | protein-coding |
| chr2_102694192_102694603 | chr2 | 102694193 | 102694603 | + | 0.951613 | intron (NM_001077514, intron 1 of 10) | intron (NM_001077514, intron 1 of 10) | -12040 | NM_011393 | 20511 | Mm.267547 | NM_011393 | ENSMUSG00000005089 | **Slc1a2** | 1700091C19Rik\|2900019G14Rik\|AI159670\|Eaat2\|GLT-1\|GLT1\|MGLT1 | solute carrier family 1 (glial high affinity glutamate transporter), member 2 | protein-coding |
| chr2_102692260_102692759 | chr2 | 102692261 | 102692759 | + | 0.943945 | intron (NM_001077514, intron 1 of 10) | intron (NM_001077514, intron 1 of 10) | -13928 | NM_011393 | 20511 | Mm.267547 | NM_011393 | ENSMUSG00000005089 | **Slc1a2** | 1700091C19Rik\|2900019G14Rik\|AI159670\|Eaat2\|GLT-1\|GLT1\|MGLT1 | solute carrier family 1 (glial high affinity glutamate transporter), member 2 | protein-coding |
| chr2_102736555_102736948 | chr2 | 102736556 | 102736948 | + | 0.844891 | intron (NM_001077515, intron 2 of 10) | intron (NM_001077515, intron 2 of 10) | 30314 | NM_001077515 | 20511 | Mm.267547 | NM_011393 | ENSMUSG00000005089 | **Slc1a2** | 1700091C19Rik\|2900019G14Rik\|AI159670\|Eaat2\|GLT-1\|GLT1\|MGLT1 | solute carrier family 1 (glial high affinity glutamate transporter), member 2 | protein-coding |
| chr2_102712953_102713947 | chr2 | 102712954 | 102713947 | + | 0.822008 | intron (NM_001077515, intron 1 of 10) | intron (NM_001077515, intron 1 of 10) | 7012 | NM_001077515 | 20511 | Mm.267547 | NM_011393 | ENSMUSG00000005089 | **Slc1a2** | 1700091C19Rik\|2900019G14Rik\|AI159670\|Eaat2\|GLT-1\|GLT1\|MGLT1 | solute carrier family 1 (glial high affinity glutamate transporter), member 2 | protein-coding |
| chr2_102693510_102693930 | chr2 | 102693511 | 102693930 | + | 0.663341 | intron (NM_001077514, intron 1 of 10) | intron (NM_001077514, intron 1 of 10) | -12718 | NM_011393 | 20511 | Mm.267547 | NM_011393 | ENSMUSG00000005089 | **Slc1a2** | 1700091C19Rik\|2900019G14Rik\|AI159670\|Eaat2\|GLT-1\|GLT1\|MGLT1 | solute carrier family 1 (glial high affinity glutamate transporter), member 2 | protein-coding |
| chr7_16784521_16784999 | chr7 | 16784522 | 16784999 | + | 0.894348 | intron (NM_009201, intron 1 of 8) | intron (NM_009201, intron 1 of 8) | 3414 | NM_009201 | 20514 | Mm.1056 | NM_009201 | ENSMUSG00000001918 | **Slc1a5** | AAAT\|ASCT2\|ATBO\|M7V1\|M7VS1\|R16\|RDRC\|Slc1a7 | solute carrier family 1 (neutral amino acid transporter), member 5 | protein-coding |
| chr9_119265462_119266009 | chr9 | 119265463 | 119266009 | + | 1.17601 | intron (NR_045693, intron 8 of 18) | ID4_\|SINE\|ID | -34044 | NR_033303 | 109280 | Mm.261557 | NM_175420 | ENSMUSG00000092212 | **Slc22a13b-ps** | 9330176C04Rik\|Slc22a13b | solute carrier family 22 (organic cation transporter), member 13b, pseudogene | pseudo |
| chr9_119271362_119271951 | chr9 | 119271363 | 119271951 | + | 1.04723 | intron (NR_045693, intron 6 of 18) | MTE2a\|LTR\|ERVL-MaLR | -39965 | NR_033303 | 109280 | Mm.261557 | NM_175420 | ENSMUSG00000092212 | **Slc22a13b-ps** | 9330176C04Rik\|Slc22a13b | solute carrier family 22 (organic cation transporter), member 13b, pseudogene | pseudo |
| chr9_119266072_119266484 | chr9 | 119266073 | 119266484 | + | 0.771185 | intron (NR_045693, intron 8 of 18) | B1_Mus2\|SINE\|Alu | -34586 | NR_033303 | 109280 | Mm.261557 | NM_175420 | ENSMUSG00000092212 | **Slc22a13b-ps** | 9330176C04Rik\|Slc22a13b | solute carrier family 22 (organic cation transporter), member 13b, pseudogene | pseudo |

| | | | | | | | | | | | | | | | | | |
|---|---|---|---|---|---|---|---|---|---|---|---|---|---|---|---|---|---|
| chr3_101948342_101949049 | chr3 | 101948343 | 101949049 | + | 1.06804 | Intergenic | Intergenic | -24243 | NM_001039371 | 242126 | Mm.90204 | NM_001039371 | | **Slc22a15** | 2610034P21Rik\|4930477M19\|A530052I06Rik\|BB219042 | solute carrier family 22 (organic anion/cation transporter), member 15 | protein-coding |
| chr3_101917704_101918091 | chr3 | 101917705 | 101918091 | + | 0.908768 | intron (NM_001039371, intron 1 of 11) | intron (NM_001039371, intron 1 of 11) | 6555 | NM_001039371 | 242126 | Mm.90204 | NM_001039371 | | **Slc22a15** | 2610034P21Rik\|4930477M19\|A530052I06Rik\|BB219042 | solute carrier family 22 (organic anion/cation transporter), member 15 | protein-coding |
| chr17_12505988_12506306 | chr17 | 12505989 | 12506306 | + | 0.71138 | intron (NM_011395, intron 1 of 10) | intron (NM_011395, intron 1 of 10) | 1557 | NM_011395 | 20519 | Mm.99252 | NM_011395 | ENSMUSG00000023828 | **Slc22a3** | EMT\|Oct3\|Orct3\|Slca22a3 | solute carrier family 22 (organic cation transporter), member 3 | protein-coding |
| chr12_102108835_102109292 | chr12 | 102108836 | 102109292 | + | 1.04826 | Intergenic | MTD\|LTR\|ERVL-MaLR | -20355 | NM_172152 | 238384 | Mm.312448 | NM_172152 | ENSMUSG00000041771 | **Slc24a4** | A930002M03Rik\|AW492432\|NCKX4 | solute carrier family 24 (sodium/potassium/calcium exchanger), member 4 | protein-coding |
| chr12_102127905_102128513 | chr12 | 102127906 | 102128513 | + | 0.91282 | Intergenic | Intergenic | -1210 | NM_172152 | 238384 | Mm.312448 | NM_172152 | ENSMUSG00000041771 | **Slc24a4** | A930002M03Rik\|AW492432\|NCKX4 | solute carrier family 24 (sodium/potassium/calcium exchanger), member 4 | protein-coding |
| chr2_71327146_71327531 | chr2 | 71327147 | 71327531 | + | 1.02449 | intron (NM_172436, intron 4 of 17) | intron (NM_172436, intron 4 of 17) | 40215 | NM_172436 | 78830 | Mm.30928 | NM_172436 | | **Slc25a12** | 2610002D09Rik\|AI839531\|B230107K20Rik\|BB129864 | solute carrier family 25 (mitochondrial carrier, Aralar), member 12 | protein-coding |
| chr2_71327114_71327687 | chr2 | 71327115 | 71327687 | + | 0.996678 | intron (NM_172436, intron 4 of 17) | intron (NM_172436, intron 4 of 17) | 40153 | NM_172436 | 78830 | Mm.30928 | NM_172436 | | **Slc25a12** | 2610002D09Rik\|AI839531\|B230107K20Rik\|BB129864 | solute carrier family 25 (mitochondrial carrier, Aralar), member 12 | protein-coding |
| chr2_71365892_71366137 | chr2 | 71365893 | 71366137 | + | 0.729103 | intron (NM_172436, intron 2 of 17) | intron (NM_172436, intron 2 of 17) | 1539 | NM_172436 | 78830 | Mm.30928 | NM_172436 | | **Slc25a12** | 2610002D09Rik\|AI839531\|B230107K20Rik\|BB129864 | solute carrier family 25 (mitochondrial carrier, Aralar), member 12 | protein-coding |
| chr6_6187214_6187694 | chr6 | 6187215 | 6187694 | + | 1.48485 | intron (NM_001177572, intron 2 of 17) | intron (NM_001177572, intron 2 of 17) | 29719 | NM_015829 | 50799 | Mm.24513 | NM_015829 | ENSMUSG00000015112 | **Slc25a13** | AI785475\|Ctrn | solute carrier family 25 (mitochondrial carrier, adenine nucleotide translocator), member 13 | protein-coding |
| chr6_6230309_6230673 | chr6 | 6230310 | 6230673 | + | 0.810175 | Intergenic | Intergenic | -13318 | NM_001177572 | 50799 | Mm.24513 | NM_015829 | ENSMUSG00000015112 | **Slc25a13** | AI785475\|Ctrn | solute carrier family 25 (mitochondrial carrier, adenine nucleotide translocator), member 13 | protein-coding |
| chr6_6230993_6231453 | chr6 | 6230994 | 6231453 | + | 0.760366 | Intergenic | Intergenic | -14050 | NM_001177572 | 50799 | Mm.24513 | NM_015829 | ENSMUSG00000015112 | **Slc25a13** | AI785475\|Ctrn | solute carrier family 25 (mitochondrial carrier, adenine nucleotide translocator), member 13 | protein-coding |
| chr8_22396091_22396498 | chr8 | 22396092 | 22396498 | + | 0.761115 | intron (NM_181325, intron 1 of 6) | intron (NM_181325, intron 1 of 6) | 2326 | NM_181325 | 18408 | Mm.200907 | NM_181325 | ENSMUSG00000031482 | **Slc25a15** | D630044L02Rik\|Ornt1 | solute carrier family 25 (mitochondrial carrier ornithine transporter), member 15 | protein-coding |
| chr15_81353043_81353779 | chr15 | 81353044 | 81353779 | + | 0.992533 | intron (NM_011399, intron 1 of 8) | L1M4\|LINE\|L1 | 7354 | NM_011399 | 20524 | Mm.222536 | NM_011399 | ENSMUSG00000022404 | **Slc25a17** | 34kDa\|47kDa\|PMP34\|Pmp47 | solute carrier family 25 (mitochondrial carrier, peroxisomal membrane protein), member 17 | protein-coding |
| chr15_81353270_81353726 | chr15 | 81353271 | 81353726 | + | 0.923901 | intron (NM_011399, intron 1 of 8) | L1M4\|LINE\|L1 | 7267 | NM_011399 | 20524 | Mm.222536 | NM_011399 | ENSMUSG00000022404 | **Slc25a17** | 34kDa\|47kDa\|PMP34\|Pmp47 | solute carrier family 25 (mitochondrial carrier, peroxisomal membrane protein), member 17 | protein-coding |
| chr11_115660355_115660818 | chr11 | 115660356 | 115660818 | + | 0.771634 | intron (NM_008163, intron 2 of 5) | intron (NM_008163, intron 2 of 5) | -32292 | NM_001252395 | 67283 | Mm.383426 | NM_026071 | ENSMUSG00000020744 | **Slc25a19** | 2900089E13Rik\|DNC\|MUP1\|TPC | solute carrier family 25 (mitochondrial thiamine pyrophosphate carrier), member 19 | protein-coding |
| chr11_115631050_115631556 | chr11 | 115631051 | 115631556 | + | 0.745928 | Intergenic | Intergenic | -3008 | NM_001252395 | 67283 | Mm.383426 | NM_026071 | ENSMUSG00000020744 | **Slc25a19** | 2900089E13Rik\|DNC\|MUP1\|TPC | solute carrier family 25 (mitochondrial thiamine pyrophosphate carrier), member 19 | protein-coding |
| chr12_57213372_57213725 | chr12 | 57213373 | 57213725 | + | 0.708917 | Intergenic | L1MB5\|LINE\|L1 | -16077 | NM_172577 | 217593 | Mm.32835 | NM_172577 | ENSMUSG00000035472 | **Slc25a21** | 9930033G19Rik\|A630030I10\|BB158148 | solute carrier family 25 (mitochondrial oxodicarboxylate carrier), member 21 | protein-coding |
| chr3_109165134_109165781 | chr3 | 109165135 | 109165781 | + | 1.38292 | intron (NM_172685, intron 9 of 9) | RMER15\|LTR\|ERVL | 42309 | NM_172685 | 229731 | Mm.33574 | NM_172685 | ENSMUSG00000040322 | **Slc25a24** | 2610016M12Rik | solute carrier family 25 (mitochondrial carrier, phosphate carrier), member 24 | protein-coding |
| chr3_109195881_109196801 | chr3 | 109195882 | 109196801 | + | 1.0696 | Intergenic | Intergenic | 73192 | NM_172685 | 229731 | Mm.33574 | NM_172685 | ENSMUSG00000040322 | **Slc25a24** | 2610016M12Rik | solute carrier family 25 (mitochondrial carrier, phosphate carrier), member 24 | protein-coding |
| chr3_109173085_109173411 | chr3 | 109173086 | 109173411 | + | 0.727044 | Intergenic | Intergenic | 50099 | NM_172685 | 229731 | Mm.33574 | NM_172685 | ENSMUSG00000040322 | **Slc25a24** | 2610016M12Rik | solute carrier family 25 (mitochondrial carrier, phosphate carrier), member 24 | protein-coding |
| chr3_109172954_109173411 | chr3 | 109172955 | 109173411 | + | 0.691855 | Intergenic | Intergenic | 50034 | NM_172685 | 229731 | Mm.33574 | NM_172685 | ENSMUSG00000040322 | **Slc25a24** | 2610016M12Rik | solute carrier family 25 (mitochondrial carrier, phosphate carrier), member 24 | protein-coding |
| chr2_32428281_32428689 | chr2 | 32428282 | 32428689 | + | 0.794529 | intron (NM_146118, intron 1 of 10) | intron (NM_146118, intron 1 of 10) | 2666 | NM_001164358 | 227731 | Mm.37395 | NM_146118 | ENSMUSG00000026819 | **Slc25a25** | 1110030N17Rik\|MCSC\|mKIAA1896 | solute carrier family 25 (mitochondrial carrier, phosphate carrier), member 25 | protein-coding |
| chr6_94552623_94553126 | chr6 | 94552624 | 94553126 | + | 1.10081 | intron (NM_026255, intron 5 of 9) | MTB\|LTR\|ERVL-MaLR | 52561 | NM_026255 | 67582 | Mm.280725 | NM_026255 | ENSMUSG00000045100 | **Slc25a26** | 4930433D19Rik\|4933433F13Rik\|AW557176\|D6Bwg0781e\|Slc25a6 | solute carrier family 25 (mitochondrial carrier, phosphate carrier), member 26 | protein-coding |
| chr6_94508798_94509263 | chr6 | 94508799 | 94509263 | + | 0.709803 | intron (NM_026255, intron 2 of 9) | intron (NM_026255, intron 2 of 9) | 8717 | NM_026255 | 67582 | Mm.280725 | NM_026255 | ENSMUSG00000045100 | **Slc25a26** | 4930433D19Rik\|4933433F13Rik\|AW557176\|D6Bwg0781e\|Slc25a6 | solute carrier family 25 (mitochondrial carrier, phosphate carrier), member 26 | protein-coding |
| chr6_94508793_94509471 | chr6 | 94508794 | 94509471 | + | 0.682932 | intron (NM_026255, intron 2 of 9) | intron (NM_026255, intron 2 of 9) | 8818 | NM_026255 | 67582 | Mm.280725 | NM_026255 | ENSMUSG00000045100 | **Slc25a26** | 4930433D19Rik\|4933433F13Rik\|AW557176\|D6Bwg0781e\|Slc25a6 | solute carrier family 25 (mitochondrial carrier, phosphate carrier), member 26 | protein-coding |
| chr10_91138286_91138712 | chr10 | 91138287 | 91138712 | + | 0.647215 | Intergenic | ORR1B2\|LTR\|ERVL-MaLR | -14536 | NM_133668 | 18674 | Mm.298 | NM_133668 | ENSMUSG00000061904 | **Slc25a3** | 5730556H19Rik\|PTP\|Phc | solute carrier family 25 (mitochondrial carrier, phosphate carrier), member 3 | protein-coding |
| chr5_8457838_8458272 | chr5 | 8457839 | 8458272 | + | 1.35308 | Intergenic | L1_Mur1\|LINE\|L1 | 35217 | NM_001289595 | 319653 | Mm.146262 | NM_178766 | ENSMUSG00000054099 | **Slc25a40** | B230315F11Rik | solute carrier family 25, member 40 | protein-coding |
| chr18_61212469_61212718 | chr18 | 61212470 | 61212718 | + | 0.935468 | promoter-TSS (NM_007885) | promoter-TSS (NM_007885) | -998 | NM_007885 | 13521 | Mm.24803 | NM_007885 | | **Slc26a2** | Dtd\|ST-OB | solute carrier family 26 (sulfate transporter), member 2 | protein-coding |
| chr18_61208242_61208555 | chr18 | 61208243 | 61208555 | + | 0.842855 | intron (NM_007885, intron 1 of 3) | Lx9\|LINE\|L1 | 3197 | NM_007885 | 13521 | Mm.24803 | NM_007885 | | **Slc26a2** | Dtd\|ST-OB | solute carrier family 26 (sulfate transporter), member 2 | protein-coding |
| chr12_31415850_31416341 | chr12 | 31415851 | 31416341 | + | 1.61528 | Intergenic | MLT2C1\|LTR\|ERVL | -22123 | NM_021353 | 13487 | Mm.490454 | NM_021353 | ENSMUSG00000001225 | **Slc26a3** | 9030623B18Rik\|9130013M11Rik\|AV376035\|Dra | solute carrier family 26, member 3 | protein-coding |
| chr12_31580644_31581041 | chr12 | 31580645 | 31581041 | + | 0.868596 | Intergenic | URR1B\|DNA\|hAT-Charlie | -20874 | NM_011867 | 23985 | Mm.100187 | NM_011867 | ENSMUSG00000020651 | **Slc26a4** | Pds\|pendrin | solute carrier family 26, member 4 | protein-coding |
| chr12_31580637_31581272 | chr12 | 31580638 | 31581272 | + | 0.836326 | Intergenic | URR1B\|DNA\|hAT-Charlie | -20986 | NM_011867 | 23985 | Mm.100187 | NM_011867 | ENSMUSG00000020651 | **Slc26a4** | Pds\|pendrin | solute carrier family 26, member 4 | protein-coding |
| chr12_31563995_31564282 | chr12 | 31563996 | 31564282 | + | 0.686921 | Intergenic | Intergenic | -4170 | NM_011867 | 23985 | Mm.100187 | NM_011867 | ENSMUSG00000020651 | **Slc26a4** | Pds\|pendrin | solute carrier family 26, member 4 | protein-coding |
| chr1_131746931_131747633 | chr1 | 131746932 | 131747633 | + | 1.17533 | intron (NM_177243, intron 1 of 20) | intron (NM_177243, intron 1 of 20) | 3260 | NM_177243 | 320718 | Mm.482704 | NM_177243 | ENSMUSG00000042268 | **Slc26a9** | E030002L01Rik | solute carrier family 26, member 9 | protein-coding |
| chr1_131760070_131760558 | chr1 | 131760071 | 131760558 | + | 0.967869 | intron (NM_177243, intron 12 of 20) | intron (NM_177243, intron 12 of 20) | 16292 | NM_177243 | 320718 | Mm.482704 | NM_177243 | ENSMUSG00000042268 | **Slc26a9** | E030002L01Rik | solute carrier family 26, member 9 | protein-coding |
| chr10_60746534_60747115 | chr10 | 60746535 | 60747115 | + | 1.59405 | intron (NM_023596, intron 2 of 5) | MIR3\|SINE\|MIR | 5957 | NM_023596 | 71279 | Mm.284462 | NM_023596 | ENSMUSG00000020100 | **Slc29a3** | 4933435C21Rik\|AW987637\|Ent3 | solute carrier family 29 (nucleoside transporters), member 3 | protein-coding |
| chr10_60744588_60744931 | chr10 | 60744589 | 60744931 | + | 1.35084 | intron (NM_023596, intron 2 of 5) | intron (NM_023596, intron 2 of 5) | 8022 | NM_023596 | 71279 | Mm.284462 | NM_023596 | ENSMUSG00000020100 | **Slc29a3** | 4933435C21Rik\|AW987637\|Ent3 | solute carrier family 29 (nucleoside transporters), member 3 | protein-coding |
| chr10_60749060_60749334 | chr10 | 60749061 | 60749334 | + | 1.30995 | intron (NM_023596, intron 2 of 5) | intron (NM_023596, intron 2 of 5) | 3585 | NM_023596 | 71279 | Mm.284462 | NM_023596 | ENSMUSG00000020100 | **Slc29a3** | 4933435C21Rik\|AW987637\|Ent3 | solute carrier family 29 (nucleoside transporters), member 3 | protein-coding |
| chr4_119126007_119126295 | chr4 | 119126008 | 119126295 | + | 0.632334 | intron (NM_011400, intron 2 of 9) | intron (NM_011400, intron 2 of 9) | 17406 | NM_011400 | 20525 | Mm.21002 | NM_011400 | ENSMUSG00000028645 | **Slc2a1** | Glut-1\|Glut1 | solute carrier family 2 (facilitated glucose transporter), member 1 | protein-coding |
| chr15_91619394_91619886 | chr15 | 91619395 | 91619886 | + | 1.355 | Intergenic | L1M2\|LINE\|L1 | -46379 | NM_001033633 | 239606 | Mm.360596 | NM_001033633 | ENSMUSG00000036298 | **Slc2a13** | 6530403A04\|A630029G22Rik\|AI505012\|Gm308 | solute carrier family 2 (facilitated glucose transporter), member 13 | protein-coding |
| chr6_122758579_122759131 | chr6 | 122758580 | 122759131 | + | 0.717194 | Intergenic | Intergenic | -16110 | NM_011401 | 20527 | Mm.395108 | NM_011401 | ENSMUSG00000003153 | **Slc2a3** | AA408729\|AL023014\|AL024341\|AU040424\|C78366\|Glut-3\|Glut3 | solute carrier family 2 (facilitated glucose transporter), member 3 | protein-coding |
| chr4_150138032_150138492 | chr4 | 150138033 | 150138492 | + | 0.959702 | intron (NM_019741, intron 7 of 13) | intron (NM_019741, intron 7 of 13) | -10710 | NM_001085529 | 435818 | Mm.359648 | NM_001085529 | ENSMUSG00000062064 | **Slc2a7** | OTTMUSG00000010396 | solute carrier family 2 (facilitated glucose transporter), member 7 | protein-coding |
| chr2_32983657_32984134 | chr2 | 32983658 | 32984134 | + | 1.13573 | Intergenic | Intergenic | -1840 | NM_019488 | 56017 | Mm.305754 | NM_019488 | ENSMUSG00000026791 | **Slc2a8** | D2Ertd44e\|GLUT8\|GlutX1 | solute carrier family 2, (facilitated glucose transporter), member 8 | protein-coding |
| chr2_33012254_33013083 | chr2 | 33012255 | 33013083 | + | 0.718528 | intron (NM_178888, intron 17 of 28) | intron (NM_178888, intron 17 of 28) | -30613 | NM_019488 | 56017 | Mm.305754 | NM_019488 | ENSMUSG00000026791 | **Slc2a8** | D2Ertd44e\|GLUT8\|GlutX1 | solute carrier family 2, (facilitated glucose transporter), member 8 | protein-coding |
| chr2_122719170_122719672 | chr2 | 122719171 | 122719672 | + | 1.48369 | Intergenic | L1M4\|LINE\|L1 | -16758 | NM_001290993 | 22785 | Mm.27801 | NM_011774 | ENSMUSG00000005802 | **Slc30a4** | Znt4\|lm\|znT-4 | solute carrier family 30 (zinc transporter), member 4 | protein-coding |
| chr2_122716383_122716766 | chr2 | 122716384 | 122716766 | + | 0.878539 | Intergenic | ORR1E\|LTR\|ERVL-MaLR | -13912 | NM_001290993 | 22785 | Mm.27801 | NM_011774 | ENSMUSG00000005802 | **Slc30a4** | Znt4\|lm\|znT-4 | solute carrier family 30 (zinc transporter), member 4 | protein-coding |
| chr3_115952281_115952710 | chr3 | 115952282 | 115952710 | + | 1.1233 | intron (NM_023214, intron 10 of 10) | RSINE1\|SINE\|B4 | 54910 | NM_023214 | 66500 | Mm.34550 | NM_023214 | ENSMUSG00000054414 | **Slc30a7** | 1810059J10Rik\|2610034N15Rik\|4833428C12Rik\|AI467242\|ZnT-7\|ZnT7\|Zntl2 | solute carrier family 30 (zinc transporter), member 7 | protein-coding |
| chr3_115953114_115953694 | chr3 | 115953115 | 115953694 | + | 0.961065 | intron (NM_023214, intron 10 of 10) | intron (NM_023214, intron 10 of 10) | 54002 | NM_023214 | 66500 | Mm.34550 | NM_023214 | ENSMUSG00000054414 | **Slc30a7** | 1810059J10Rik\|2610034N15Rik\|4833428C12Rik\|AI467242\|ZnT-7\|ZnT7\|Zntl2 | solute carrier family 30 (zinc transporter), member 7 | protein-coding |

| | | | | | | | | | | | | | | | | | |
|---|---|---|---|---|---|---|---|---|---|---|---|---|---|---|---|---|---|
| chr3_115961718_115961983 | chr3 | 115961719 | 115961983 | + | 0.653889 | intron (NM_023214, intron 8 of 10) | intron (NM_023214, intron 8 of 10) | 45555 | NM_023214 | 66500 | Mm.34550 | NM_023214 | ENSMUSG00000054414 | **Slc30a7** | 1810059J10Rik\|2610034N15Rik\|4833428C12Rik\|AI467242\|ZnT-7\|ZnT7\|Zntl2 | solute carrier family 30 (zinc transporter), member 7 | protein-coding |
| chr15_52341058_52341639 | chr15 | 52341059 | 52341639 | + | 0.952753 | Intergenic | Intergenic | 45796 | NM_172816 | 239436 | Mm.208831 | NM_172816 | ENSMUSG00000022315 | **Slc30a8** | C820002P14Rik\|ZnT-8\|ZnT8 | solute carrier family 30 (zinc transporter), member 8 | protein-coding |
| chr15_52341018_52341897 | chr15 | 52341019 | 52341897 | + | 0.822406 | Intergenic | Intergenic | 45905 | NM_172816 | 239436 | Mm.208831 | NM_172816 | ENSMUSG00000022315 | **Slc30a8** | C820002P14Rik\|ZnT-8\|ZnT8 | solute carrier family 30 (zinc transporter), member 8 | protein-coding |
| chr16_45141775_45142489 | chr16 | 45141776 | 45142489 | + | 1.03959 | 3' UTR (NM_028756, exon 7 of 7) | 3' UTR (NM_028756, exon 7 of 7) | 16541 | NM_028756 | 74102 | Mm.264984 | NM_028756 | ENSMUSG00000022664 | **Slc35a5** | 1010001J06Rik\|AU021179\|BB097433\|D16Ertd450e\|D730043G07Rik | solute carrier family 35, member A5 | protein-coding |
| chr17_45566253_45566608 | chr17 | 45566254 | 45566608 | + | 0.779692 | exon (NM_028662, exon 3 of 3) | exon (NM_028662, exon 3 of 3) | 2279 | NM_028662 | 73836 | Mm.289716 | NM_028662 | ENSMUSG00000037089 | **Slc35b2** | 1110003M08Rik\|AA407792\|AI842275\|PAPST1\|Slc35b1 | solute carrier family 35, member B2 | protein-coding |
| chr17_45566219_45567584 | chr17 | 45566220 | 45567584 | + | 0.736245 | TTS (NM_008302) | TTS (NM_008302) | 2750 | NM_028662 | 73836 | Mm.289716 | NM_028662 | ENSMUSG00000037089 | **Slc35b2** | 1110003M08Rik\|AA407792\|AI842275\|PAPST1\|Slc35b1 | solute carrier family 35, member B2 | protein-coding |
| chr13_38950387_38950969 | chr13 | 38950388 | 38950969 | + | 1.57282 | intron (NM_134060, intron 3 of 9) | L1M2\|LINE\|L1 | 9859 | NM_001170430 | 108652 | Mm.255506 | NM_134060 | ENSMUSG00000021432 | **Slc35b3** | 4921526O06Rik\|AI428480\|CGI-19\|PABST2\|PAPST2 | solute carrier family 35, member B3 | protein-coding |
| chr6_34242184_34243111 | chr6 | 34242185 | 34243111 | + | 0.818668 | Intergenic | Intergenic | -65594 | NM_021435 | 58246 | Mm.245527 | NM_021435 | | **Slc35b4** | 4930474D06Rik\|AB041549\|AW494345 | solute carrier family 35, member B4 | protein-coding |
| chr6_34225082_34225625 | chr6 | 34225083 | 34225625 | + | 0.801579 | Intergenic | Intergenic | -48300 | NM_021435 | 58246 | Mm.245527 | NM_021435 | | **Slc35b4** | 4930474D06Rik\|AB041549\|AW494345 | solute carrier family 35, member B4 | protein-coding |
| chr1_125503817_125504370 | chr1 | 125503818 | 125504370 | + | 1.11738 | Intergenic | Intergenic | -56922 | NM_028787 | 74150 | Mm.28654 | NM_028787 | ENSMUSG00000026342 | **Slc35f5** | 1300003P13Rik\|AI646727 | solute carrier family 35, member F5 | protein-coding |
| chr1_125503322_125504419 | chr1 | 125503323 | 125504419 | + | 1.05297 | Intergenic | B4\|SINE\|B4 | -57145 | NM_028787 | 74150 | Mm.28654 | NM_028787 | ENSMUSG00000026342 | **Slc35f5** | 1300003P13Rik\|AI646727 | solute carrier family 35, member F5 | protein-coding |
| chr11_55134491_55134741 | chr11 | 55134492 | 55134741 | + | 1.03633 | intron (NM_172258, intron 7 of 10) | intron (NM_172258, intron 7 of 10) | 17090 | NM_172258 | 215332 | Mm.87054 | NM_172258 | ENSMUSG00000049491 | **Slc36a3** | PAT3\|TRAMD2\|tramdorin2 | solute carrier family 36 (proton/amino acid symporter), member 3 | protein-coding |
| chr11_55131839_55132286 | chr11 | 55131840 | 55132286 | + | 0.649866 | intron (NM_172258, intron 7 of 10) | L1MB4\|LINE\|L1 | 19643 | NM_172258 | 215332 | Mm.87054 | NM_172258 | ENSMUSG00000049491 | **Slc36a3** | PAT3\|TRAMD2\|tramdorin2 | solute carrier family 36 (proton/amino acid symporter), member 3 | protein-coding |
| chr17_31297361_31297703 | chr17 | 31297362 | 31297703 | + | 0.796345 | intron (NM_001242427, intron 1 of 19) | intron (NM_001242427, intron 1 of 19) | 1308 | NM_001242427 | 224674 | Mm.311395 | NM_153062 | ENSMUSG00000024036 | **Slc37a1** | G3PP | solute carrier family 37 (glycerol-3-phosphate transporter), member 1 | protein-coding |
| chr17_31309495_31309896 | chr17 | 31309496 | 31309896 | + | 0.729152 | intron (NM_153062, intron 3 of 19) | intron (NM_153062, intron 3 of 19) | 13472 | NM_001242427 | 224674 | Mm.311395 | NM_153062 | ENSMUSG00000024036 | **Slc37a1** | G3PP | solute carrier family 37 (glycerol-3-phosphate transporter), member 1 | protein-coding |
| chr9_37235724_37236466 | chr9 | 37235725 | 37236466 | + | 0.741842 | intron (NM_001145960, intron 10 of 18) | intron (NM_001145960, intron 10 of 18) | 19643 | NM_020258 | 56857 | Mm.440037 | NM_020258 | ENSMUSG00000032122 | **Slc37a2** | G3PP\|Slc37a1\|cl-2\|ci2 | solute carrier family 37 (glycerol-3-phosphate transporter), member 2 | protein-coding |
| chr15_96708006_96708318 | chr15 | 96708007 | 96708318 | + | 0.787516 | Intergenic | Intergenic | -8464 | NM_175121 | 67760 | Mm.46754 | NM_175121 | ENSMUSG00000022462 | **Slc38a2** | 5033402L14Rik\|AI316867\|ATA2\|SAT2\|mKIAA1382 | solute carrier family 38, member 2 | protein-coding |
| chr12_73324068_73324660 | chr12 | 73324069 | 73324660 | + | 1.2169 | intron (NM_001037717, intron 5 of 15) | intron (NM_001037717, intron 5 of 15) | 37516 | NM_001037717 | 625098 | Mm.32075 | NM_001037717 | ENSMUSG00000044712 | **Slc38a6** | AW322671\|EG625098 | solute carrier family 38, member 6 | protein-coding |
| chr12_73324013_73325019 | chr12 | 73324014 | 73325019 | + | 1.05706 | intron (NM_001037717, intron 5 of 15) | intron (NM_001037717, intron 5 of 15) | 37668 | NM_001037717 | 625098 | Mm.32075 | NM_001037717 | ENSMUSG00000044712 | **Slc38a6** | AW322671\|EG625098 | solute carrier family 38, member 6 | protein-coding |
| chr12_73327350_73327679 | chr12 | 73327351 | 73327679 | + | 0.776888 | intron (NM_001037717, intron 5 of 15) | intron (NM_001037717, intron 5 of 15) | 40667 | NM_001037717 | 625098 | Mm.32075 | NM_001037717 | ENSMUSG00000044712 | **Slc38a6** | AW322671\|EG625098 | solute carrier family 38, member 6 | protein-coding |
| chr1_46912273_46913206 | chr1 | 46912274 | 46913206 | + | 1.251 | Intergenic | (TCCC)n\|Simple_repeat\|Simple_repeat | -59231 | NM_172653 | 227059 | Mm.233889 | NM_172653 | ENSMUSG00000025986 | **Slc39a10** | 2900042E17Rik\|5430433I10\|mKIAA1265 | solute carrier family 39 (zinc transporter), member 10 | protein-coding |
| chr1_46864072_46864769 | chr1 | 46864073 | 46864769 | + | 0.826152 | Intergenic | Intergenic | -10912 | NM_172653 | 227059 | Mm.233889 | NM_172653 | ENSMUSG00000025986 | **Slc39a10** | 2900042E17Rik\|5430433I10\|mKIAA1265 | solute carrier family 39 (zinc transporter), member 10 | protein-coding |
| chr1_46917128_46917622 | chr1 | 46917129 | 46917622 | + | 0.786579 | Intergenic | ORR1D2\|LTR\|ERVL-MaLR | -63866 | NM_172653 | 227059 | Mm.233889 | NM_172653 | ENSMUSG00000025986 | **Slc39a10** | 2900042E17Rik\|5430433I10\|mKIAA1265 | solute carrier family 39 (zinc transporter), member 10 | protein-coding |
| chr1_46916033_46916257 | chr1 | 46916034 | 46916257 | + | 0.670756 | Intergenic | ORR1D2-int\|LTR\|ERVL-MaLR | -62636 | NM_172653 | 227059 | Mm.233889 | NM_172653 | ENSMUSG00000025986 | **Slc39a10** | 2900042E17Rik\|5430433I10\|mKIAA1265 | solute carrier family 39 (zinc transporter), member 10 | protein-coding |
| chr1_46843458_46843795 | chr1 | 46843459 | 46843795 | + | 0.617095 | intron (NM_172653, intron 1 of 9) | intron (NM_172653, intron 1 of 9) | 9882 | NM_172653 | 227059 | Mm.233889 | NM_172653 | ENSMUSG00000025986 | **Slc39a10** | 2900042E17Rik\|5430433I10\|mKIAA1265 | solute carrier family 39 (zinc transporter), member 10 | protein-coding |
| chr11_113474114_113474415 | chr11 | 113474115 | 113474415 | + | 0.689953 | intron (NM_027216, intron 4 of 9) | intron (NM_027216, intron 4 of 9) | 91550 | NM_001166503 | 69806 | Mm.341021 | NM_027216 | ENSMUSG00000041654 | **Slc39a11** | 1810074D23Rik | solute carrier family 39 (metal ion transporter), member 11 | protein-coding |
| chr11_113487321_113487957 | chr11 | 113487322 | 113487957 | + | 0.661551 | intron (NM_027216, intron 4 of 9) | intron (NM_027216, intron 4 of 9) | 78176 | NM_001166503 | 69806 | Mm.341021 | NM_027216 | ENSMUSG00000041654 | **Slc39a11** | 1810074D23Rik | solute carrier family 39 (metal ion transporter), member 11 | protein-coding |
| chr14_70370746_70371298 | chr14 | 70370747 | 70371298 | + | 1.66571 | Intergenic | ORR1B1\|LTR\|ERVL-MaLR | -19598 | NM_001135151 | 213053 | Mm.270647 | NM_144808 | ENSMUSG00000022094 | **Slc39a14** | FAD-123\|ZIP-14\|Zip14\|fad123 | solute carrier family 39 (zinc transporter), member 14 | protein-coding |
| chr14_70370720_70371476 | chr14 | 70370721 | 70371476 | + | 1.40204 | Intergenic | ORR1B1\|LTR\|ERVL-MaLR | -19674 | NM_001135151 | 213053 | Mm.270647 | NM_144808 | ENSMUSG00000022094 | **Slc39a14** | FAD-123\|ZIP-14\|Zip14\|fad123 | solute carrier family 39 (zinc transporter), member 14 | protein-coding |
| chr19_8701343_8701802 | chr19 | 8701344 | 8701802 | + | 1.14256 | Intergenic | RSINE1\|SINE\|B4 | 12324 | NM_008577 | 17254 | Mm.4114 | NM_008577 | ENSMUSG00000010095 | **Slc3a2** | 4F2\|4F2HC\|AI314110\|Cd98\|Ly-10\|Ly-m10\|Ly10\|Mdu1\|Mgp-2hc\|NACAE | solute carrier family 3 (activators of dibasic and neutral amino acid transport), member 2 | protein-coding |
| chr19_8706126_8706603 | chr19 | 8706127 | 8706603 | + | 0.627333 | TTS (NM_001161413) | TTS (NM_001161413) | 7532 | NM_008577 | 17254 | Mm.4114 | NM_008577 | ENSMUSG00000010095 | **Slc3a2** | 4F2\|4F2HC\|AI314110\|Cd98\|Ly-10\|Ly-m10\|Ly10\|Mdu1\|Mgp-2hc\|NACAE | solute carrier family 3 (activators of dibasic and neutral amino acid transport), member 2 | protein-coding |
| chr1_131822875_131823156 | chr1 | 131822876 | 131823156 | + | 1.23263 | Intergenic | Intergenic | -4996 | NM_173865 | 98396 | Mm.389181 | NM_173865 | ENSMUSG00000013275 | **Slc41a1** | AI573938\|B230315F01Rik | solute carrier family 41, member 1 | protein-coding |
| chr1_131820723_131821151 | chr1 | 131820724 | 131821151 | + | 0.786921 | Intergenic | Intergenic | -7075 | NM_173865 | 98396 | Mm.389181 | NM_173865 | ENSMUSG00000013275 | **Slc41a1** | AI573938\|B230315F01Rik | solute carrier family 41, member 1 | protein-coding |
| chr1_131823621_131823891 | chr1 | 131823622 | 131823891 | + | 0.665526 | Intergenic | Intergenic | -4256 | NM_173865 | 98396 | Mm.389181 | NM_173865 | ENSMUSG00000013275 | **Slc41a1** | AI573938\|B230315F01Rik | solute carrier family 41, member 1 | protein-coding |
| chr10_83266135_83266669 | chr10 | 83266136 | 83266669 | + | 1.54277 | intron (NM_177388, intron 8 of 10) | intron (NM_177388, intron 8 of 10) | 71415 | NM_177388 | 338365 | Mm.67073 | NM_177388 | ENSMUSG00000034591 | **Slc41a2** | A230035L05Rik\|SLC41A1-L1 | solute carrier family 41, member 2 | protein-coding |
| chr10_83265572_83266083 | chr10 | 83265573 | 83266083 | + | 1.23621 | intron (NM_177388, intron 8 of 10) | MTC\|LTR\|ERVL-MaLR | 71989 | NM_177388 | 338365 | Mm.67073 | NM_177388 | ENSMUSG00000034591 | **Slc41a2** | A230035L05Rik\|SLC41A1-L1 | solute carrier family 41, member 2 | protein-coding |
| chr6_90608253_90608485 | chr6 | 90608254 | 90608485 | + | 0.656558 | intron (NM_027868, intron 1 of 10) | PB1D11\|SINE\|Alu | 3489 | NM_027868 | 71699 | Mm.272633 | NM_027868 | ENSMUSG00000030089 | **Slc41a3** | 1010001P06Rik\|AI480742\|SLC41A1-L2 | solute carrier family 41, member 3 | protein-coding |
| chr2_84938298_84938976 | chr2 | 84938299 | 84938976 | + | 1.34933 | intron (NM_021398, intron 3 of 12) | intron (NM_021398, intron 3 of 12) | 1991 | NM_021398 | 58207 | Mm.290729 | NM_021398 | ENSMUSG00000027074 | **Slc43a3** | Eeg1\|SEEEG-1 | solute carrier family 43, member 3 | protein-coding |
| chr2_84939750_84940027 | chr2 | 84939751 | 84940027 | + | 0.811566 | intron (NM_021398, intron 3 of 12) | ORR1D2\|LTR\|ERVL-MaLR | 3243 | NM_021398 | 58207 | Mm.290729 | NM_021398 | ENSMUSG00000027074 | **Slc43a3** | Eeg1\|SEEEG-1 | solute carrier family 43, member 3 | protein-coding |
| chr4_53446115_53446578 | chr4 | 53446116 | 53446578 | + | 1.17539 | intron (NM_001159633, intron 1 of 12) | intron (NM_001159633, intron 1 of 12) | 5934 | NM_133891 | 100434 | Mm.482207 | NM_133891 | ENSMUSG00000028412 | **Slc44a1** | 2210409B22Rik\|4833416H08Rik\|AW547365\|CHTL1\|CTL1\|Cdw92 | solute carrier family 44, member 1 | protein-coding |
| chr9_21324287_21324532 | chr9 | 21324288 | 21324532 | + | 0.700605 | intron (NM_001199186, intron 1 of 21) | intron (NM_001199186, intron 1 of 21) | 3691 | NM_001199186 | 68682 | Mm.148425 | NM_152808 | ENSMUSG00000057193 | **Slc44a2** | 1110028E10Rik\|CTL2 | solute carrier family 44, member 2 | protein-coding |
| chr3_121584520_121585296 | chr3 | 121584521 | 121585296 | + | 1.29284 | Intergenic | Intergenic | -52564 | NM_145394 | 213603 | Mm.33062 | NM_145394 | ENSMUSG00000039865 | **Slc44a3** | BC010552 | solute carrier family 44, member 3 | protein-coding |
| chr3_121490842_121491540 | chr3 | 121490843 | 121491540 | + | 1.18852 | intron (NM_145394, intron 10 of 14) | intron (NM_145394, intron 10 of 14) | 41153 | NM_145394 | 213603 | Mm.33062 | NM_145394 | ENSMUSG00000039865 | **Slc44a3** | BC010552 | solute carrier family 44, member 3 | protein-coding |
| chr3_153971906_153972205 | chr3 | 153971907 | 153972205 | + | 1.21774 | Intergenic | Intergenic | -1380 | NM_001081263 | 242259 | Mm.239962 | NM_001081263 | ENSMUSG00000028360 | **Slc44a5** | AA117069\|Gm422 | solute carrier family 44, member 5 | protein-coding |
| chr4_150692431_150693329 | chr4 | 150692432 | 150693329 | + | 0.916542 | Intergenic | Intergenic | -40706 | NM_173774 | 242773 | Mm.29280 | NM_173774 | | **Slc45a1** | C230078B22\|Dnb5 | solute carrier family 45, member 1 | protein-coding |
| chr4_59894993_59895546 | chr4 | 59894994 | 59895546 | + | 1.26337 | intron (NM_172468, intron 8 of 8) | intron (NM_172468, intron 8 of 8) | 19786 | NM_021053 | 30936 | Mm.284433 | NM_021053 | | **Slc46a2** | 5430429N04Rik\|Ly110\|TSO-1C12\|Tscot | solute carrier family 46, member 2 | protein-coding |
| chr2_62134670_62135291 | chr2 | 62134671 | 62135291 | + | 0.94431 | intron (NM_001242378, intron 1 of 25) | intron (NM_001242378, intron 1 of 25) | 88466 | NM_033552 | 94229 | Mm.314497 | NM_033552 | ENSMUSG00000026904 | **Slc4a10** | NCBE\|mKIAA4136 | solute carrier family 4, sodium bicarbonate cotransporter-like, member 10 | protein-coding |
| chr2_130688960_130689741 | chr2 | 130688961 | 130689741 | + | 0.979811 | intron (NM_001081162, intron 8 of 20) | ID4\|SINE\|ID | 8168 | NM_001081162 | 269356 | Mm.441391 | NM_001081162 | ENSMUSG00000074796 | **Slc4a11** | AI503023\|BTR1\|NaBC1 | solute carrier family 4, sodium bicarbonate transporter-like, member 11 | protein-coding |

| | | | | | | | | | | | | | | | | | |
|---|---|---|---|---|---|---|---|---|---|---|---|---|---|---|---|---|---|
| chr5_89147157_89147819 | chr5 | 89147158 | 89147819 | + | 1.46713 | intron (NM_018760, intron 12 of 25) | intron (NM_018760, intron 12 of 25) | 260228 | NM_018760 | 54403 | Mm.41044 | NM_018760 | ENSMUSG00000060961 | **Slc4a4** | AI835705\|NBC\|NBC1 | solute carrier family 4 (anion exchanger), member 4 | protein-coding |
| chr5_89146857_89147826 | chr5 | 89146858 | 89147826 | + | 1.3644 | intron (NM_018760, intron 12 of 25) | intron (NM_018760, intron 12 of 25) | 260082 | NM_018760 | 54403 | Mm.41044 | NM_018760 | ENSMUSG00000060961 | **Slc4a4** | AI835705\|NBC\|NBC1 | solute carrier family 4 (anion exchanger), member 4 | protein-coding |
| chr14_14607098_14607598 | chr14 | 14607099 | 14607598 | + | 0.879224 | Intergenic | Intergenic | -95677 | NM_001033270 | 218756 | Mm.258893 | NM_001033270 | ENSMUSG00000021733 | **Slc4a7** | E430014N10Rik\|NBC3\|NBCn1\|NBCn1-G | solute carrier family 4, sodium bicarbonate cotransporter, member 7 | protein-coding |
| chr14_14632915_14633705 | chr14 | 14632916 | 14633705 | + | 0.645879 | Intergenic | Intergenic | -69715 | NM_001033270 | 218756 | Mm.258893 | NM_001033270 | ENSMUSG00000021733 | **Slc4a7** | E430014N10Rik\|NBC3\|NBCn1\|NBCn1-G | solute carrier family 4, sodium bicarbonate cotransporter, member 7 | protein-coding |
| chr15_100766148_100766477 | chr15 | 100766149 | 100766477 | + | 0.893313 | intron (NM_021530, intron 1 of 24) | B3\|SINE\|B2 | 4566 | NM_021530 | 59033 | Mm.209856 | NM_021530 | ENSMUSG00000023032 | **Slc4a8** | AW493845\|C230026C11\|NDCBE\|k NBC-3 | solute carrier family 4 (anion exchanger), member 8 | protein-coding |
| chr5_33121853_33122416 | chr5 | 33121854 | 33122416 | + | 0.679484 | intron (NM_019810, intron 2 of 14) | intron (NM_019810, intron 2 of 14) | 17916 | NM_019810 | 20537 | Mm.25237 | NM_019810 | ENSMUSG00000011034 | **Slc5a1** | Sglt1 | solute carrier family 5 (sodium/glucose cotransporter), member 1 | protein-coding |
| chr7_123189819_123190141 | chr7 | 123189820 | 123190141 | + | 0.974536 | exon (NM_144925, exon 21 of 24) | exon (NM_144925, exon 21 of 24) | -24886 | NM_146198 | 233836 | Mm.211838 | NM_146198 | ENSMUSG00000030769 | **Slc5a11** | 2010013B02Rik\|Kst1\|RKST2\|SMIT 2\|Slc5a10 | solute carrier family 5 (sodium/glucose cotransporter), member 11 | protein-coding |
| chr7_123188445_123189111 | chr7 | 123188446 | 123189111 | + | 0.639688 | intron (NM_144925, intron 19 of 23) | intron (NM_144925, intron 19 of 23) | -26088 | NM_146198 | 233836 | Mm.211838 | NM_146198 | ENSMUSG00000030769 | **Slc5a11** | 2010013B02Rik\|Kst1\|RKST2\|SMIT 2\|Slc5a10 | solute carrier family 5 (sodium/glucose cotransporter), member 11 | protein-coding |
| chr7_123191397_123192105 | chr7 | 123191398 | 123192105 | + | 0.616913 | intron (NM_144925, intron 22 of 23) | intron (NM_144925, intron 22 of 23) | -23115 | NM_146198 | 233836 | Mm.211838 | NM_146198 | ENSMUSG00000030769 | **Slc5a11** | 2010013B02Rik\|Kst1\|RKST2\|SMIT 2\|Slc5a10 | solute carrier family 5 (sodium/glucose cotransporter), member 11 | protein-coding |
| chr13_73685448_73686179 | chr13 | 73685449 | 73686179 | + | 0.92914 | exon (NM_028878, exon 8 of 12) | exon (NM_028878, exon 8 of 12) | -7791 | NM_001168643 | 22598 | Mm.30370 | NM_011730 | ENSMUSG00000021612 | **Slc6a18** | B0AT3\|D630001K16Rik\|Xt2\|Xtrp2 | solute carrier family 6 (neurotransmitter transporter), member 18 | protein-coding |
| chr13_73688770_73689289 | chr13 | 73688771 | 73689289 | + | 0.76131 | intron (NM_028878, intron 6 of 11) | intron (NM_028878, intron 6 of 11) | -10796 | NM_027780 | 71390 | Mm.105251 | NM_027780 | | **Slc6a19os** | 5430425J12Rik | solute carrier family 6 (neurotransmitter transporter), member 19, opposite strand | protein-coding |
| chr13_73689069_73689249 | chr13 | 73689070 | 73689249 | + | 0.718545 | intron (NM_028878, intron 5 of 11) | intron (NM_028878, intron 5 of 11) | -10667 | NM_027780 | 71390 | Mm.105251 | NM_027780 | | **Slc6a19os** | 5430425J12Rik | solute carrier family 6 (neurotransmitter transporter), member 19, opposite strand | protein-coding |
| chr9_123598844_123599485 | chr9 | 123598845 | 123599485 | + | 0.911861 | intron (NM_011731, intron 7 of 10) | intron (NM_011731, intron 7 of 10) | 33400 | NM_011731 | 22599 | Mm.41963 | NM_011731 | ENSMUSG00000025243 | **Slc6a20b** | AA986306\|Sit1\|Slc6a20\|XT3\|Xtrp3 | solute carrier family 6 (neurotransmitter transporter), member 20B | protein-coding |
| chr6_91712405_91713341 | chr6 | 91712406 | 91713341 | + | 1.05572 | intron (NM_009320, intron 2 of 14) | intron (NM_009320, intron 2 of 14) | 28806 | NM_009320 | 21366 | Mm.395650 | NM_009320 | ENSMUSG00000030096 | **Slc6a6** | AA589629\|C80501\|Taut | solute carrier family 6 (neurotransmitter transporter, taurine), member 6 | protein-coding |
| chr6_91712083_91713369 | chr6 | 91712084 | 91713369 | + | 0.92957 | intron (NM_009320, intron 2 of 14) | MIR3\|SINE\|MIR | 28659 | NM_009320 | 21366 | Mm.395650 | NM_009320 | ENSMUSG00000030096 | **Slc6a6** | AA589629\|C80501\|Taut | solute carrier family 6 (neurotransmitter transporter, taurine), member 6 | protein-coding |
| chr6_91705359_91706634 | chr6 | 91705360 | 91706634 | + | 0.703217 | intron (NM_009320, intron 2 of 14) | intron (NM_009320, intron 2 of 14) | 21930 | NM_009320 | 21366 | Mm.395650 | NM_009320 | ENSMUSG00000030096 | **Slc6a6** | AA589629\|C80501\|Taut | solute carrier family 6 (neurotransmitter transporter, taurine), member 6 | protein-coding |
| chr5_148397542_148397718 | chr5 | 148397543 | 148397718 | + | 0.934894 | intron (NM_007513, intron 1 of 12) | intron (NM_007513, intron 1 of 12) | 2274 | NM_007513 | 11987 | Mm.275489 | NM_007513 | ENSMUSG00000041313 | **Slc7a1** | 4831426K01Rik\|AI447493\|Atrc-1\|Atrc1\|Cat1\|Rec-1\|Rev-1\|mCAT-1 | solute carrier family 7 (cationic amino acid transporter, y+ system), member 1 | protein-coding |
| chr5_148377457_148378250 | chr5 | 148377458 | 148378250 | + | 0.873349 | intron (NM_007513, intron 1 of 12) | intron (NM_007513, intron 1 of 12) | 22050 | NM_007513 | 11987 | Mm.275489 | NM_007513 | ENSMUSG00000041313 | **Slc7a1** | 4831426K01Rik\|AI447493\|Atrc-1\|Atrc1\|Cat1\|Rec-1\|Rev-1\|mCAT-1 | solute carrier family 7 (cationic amino acid transporter, y+ system), member 1 | protein-coding |
| chr5_148377457_148377983 | chr5 | 148377458 | 148377983 | + | 0.820008 | intron (NM_007513, intron 1 of 12) | intron (NM_007513, intron 1 of 12) | 22184 | NM_007513 | 11987 | Mm.275489 | NM_007513 | ENSMUSG00000041313 | **Slc7a1** | 4831426K01Rik\|AI447493\|Atrc-1\|Atrc1\|Cat1\|Rec-1\|Rev-1\|mCAT-1 | solute carrier family 7 (cationic amino acid transporter, y+ system), member 1 | protein-coding |
| chr7_35176568_35177276 | chr7 | 35176569 | 35177276 | + | 0.994791 | Intergenic | Intergenic | -9463 | NM_017394 | 53896 | Mm.35567 | NM_017394 | ENSMUSG00000030495 | **Slc7a10** | AL024237\|Asc-1\|D7Bwg0847e | solute carrier family 7 (cationic amino acid transporter, y+ system), member 10 | protein-coding |
| chr4_19850566_19851041 | chr4 | 19850567 | 19851041 | + | 1.3747 | Intergenic | Intergenic | 32077 | NM_028746 | 74087 | Mm.173833 | NM_028746 | ENSMUSG00000041052 | **Slc7a13** | 0610009O04Rik\|AGT-1\|AGT1\|XAT2 | solute carrier family 7, (cationic amino acid transporter, y+ system) member 13 | protein-coding |
| chr4_19848415_19848830 | chr4 | 19848416 | 19848830 | + | 1.00312 | Intergenic | ORR1C1\|LTR\|ERVL-MaLR | 29896 | NM_028746 | 74087 | Mm.173833 | NM_028746 | ENSMUSG00000041052 | **Slc7a13** | 0610009O04Rik\|AGT-1\|AGT1\|XAT2 | solute carrier family 7, (cationic amino acid transporter, y+ system) member 13 | protein-coding |
| chr4_19851470_19851774 | chr4 | 19851471 | 19851774 | + | 0.729427 | Intergenic | MTC\|LTR\|ERVL-MaLR | 32895 | NM_028746 | 74087 | Mm.173833 | NM_028746 | ENSMUSG00000041052 | **Slc7a13** | 0610009O04Rik\|AGT-1\|AGT1\|XAT2 | solute carrier family 7, (cationic amino acid transporter, y+ system) member 13 | protein-coding |
| chr12_8571268_8572124 | chr12 | 8571269 | 8572124 | + | 1.33384 | intron (NM_177802, intron 1 of 6) | L1M4\|LINE\|L1 | 27370 | NM_177802 | 328059 | Mm.46326 | NM_177802 | ENSMUSG00000020600 | **Slc7a15** | Arpat | solute carrier family 7 (cationic amino acid transporter, y+ system), member 15 | protein-coding |
| chr8_121898119_121899072 | chr8 | 121898120 | 121899072 | + | 1.49341 | intron (NM_011404, intron 1 of 9) | intron (NM_011404, intron 1 of 9) | 9090 | NM_011404 | 20539 | Mm.27943 | NM_011404 | ENSMUSG00000040010 | **Slc7a5** | 4F2LC\|D0H16S474E\|LAT1\|TA1 | solute carrier family 7 (cationic amino acid transporter, y+ system), member 5 | protein-coding |
| chr8_121878061_121878309 | chr8 | 121878062 | 121878309 | + | 0.860394 | Intergenic | Intergenic | 29501 | NM_011404 | 20539 | Mm.27943 | NM_011404 | ENSMUSG00000040010 | **Slc7a5** | 4F2LC\|D0H16S474E\|LAT1\|TA1 | solute carrier family 7 (cationic amino acid transporter, y+ system), member 5 | protein-coding |
| chr11_115172844_115173382 | chr11 | 115172845 | 115173382 | + | 0.657227 | intron (NM_012030, intron 1 of 5) | URR1B\|DNA\|hAT-Charlie | 9772 | NM_012030 | 26941 | Mm.27842 | NM_012030 | ENSMUSG00000020733 | **Slc9a3r1** | EBP-50\|NHE-RF\|NHERF-1\|NHERF1 | solute carrier family 9 (sodium/hydrogen exchanger), member 3 regulator 1 | protein-coding |
| chr1_40586772_40587293 | chr1 | 40586773 | 40587293 | + | 0.916182 | intron (NM_177084, intron 2 of 11) | intron (NM_177084, intron 2 of 11) | 6806 | NM_177084 | 110895 | Mm.108602 | NM_177084 | ENSMUSG00000026065 | **Slc9a4** | AW990558\|D730009J23Rik\|NHE4 | solute carrier family 9 (sodium/hydrogen exchanger), member 4 | protein-coding |
| chr1_40585502_40586129 | chr1 | 40585503 | 40586129 | + | 0.667543 | intron (NM_177084, intron 2 of 11) | ORR1D1\|LTR\|ERVL-MaLR | 5589 | NM_177084 | 110895 | Mm.108602 | NM_177084 | ENSMUSG00000026065 | **Slc9a4** | AW990558\|D730009J23Rik\|NHE4 | solute carrier family 9 (sodium/hydrogen exchanger), member 4 | protein-coding |
| chrX_20221336_20221868 | chrX | 20221337 | 20221868 | + | 0.714593 | intron (NM_177353, intron 1 of 16) | intron (NM_177353, intron 1 of 16) | 70162 | NM_177353 | 236727 | Mm.132983 | NM_177353 | ENSMUSG00000037341 | **Slc9a7** | A530087D17Rik\|NHE7 | solute carrier family 9 (sodium/hydrogen exchanger), member 7 | protein-coding |
| chr3_135298401_135299136 | chr3 | 135298402 | 135299136 | + | 1.36942 | intron (NM_027208, intron 7 of 9) | intron (NM_027208, intron 7 of 9) | -8931 | NM_178877 | 97086 | Mm.441764 | NM_178877 | ENSMUSG00000037994 | **Slc9b2** | C80638\|NHA2\|NHE10\|Nhedc2\|nha-oc\|nhaoc | solute carrier family 9, subfamily B (NHA2, cation proton antiporter 2), member 2 | protein-coding |
| chr3_135325029_135325457 | chr3 | 135325030 | 135325457 | + | 0.824836 | intron (NM_178877, intron 6 of 12) | MIR\|SINE\|MIR | 17543 | NM_178877 | 97086 | Mm.441764 | NM_178877 | ENSMUSG00000037994 | **Slc9b2** | C80638\|NHA2\|NHE10\|Nhedc2\|nha-oc\|nhaoc | solute carrier family 9, subfamily B (NHA2, cation proton antiporter 2), member 2 | protein-coding |
| chr9_103040343_103040699 | chr9 | 103040344 | 103040699 | + | 0.634019 | intron (NM_033314, intron 1 of 13) | B4\|SINE\|B4 | 32032 | NM_033314 | 24059 | Mm.207106 | NM_033314 | ENSMUSG00000032548 | **Slco2a1** | 2310021C19Rik\|Pgt\|Slc21a2\|mPgt | solute carrier organic anion transporter family, member 2a1 | protein-coding |
| chr7_74646985_74647994 | chr7 | 74646986 | 74647994 | + | 1.32312 | Intergenic | Intergenic | -92710 | NM_001038643 | 108116 | Mm.425467 | NM_023908 | ENSMUSG00000025790 | **Slco3a1** | 5830414C08Rik\|Anr1\|MJAM\|OATP-D\|R75096\|Slc21a11 | solute carrier organic anion transporter family, member 3a1 | protein-coding |
| chr7_74527415_74528088 | chr7 | 74527416 | 74528088 | + | 1.28747 | intron (NM_001038643, intron 1 of 10) | intron (NM_001038643, intron 1 of 10) | 27028 | NM_023908 | 108116 | Mm.425467 | NM_023908 | ENSMUSG00000025790 | **Slco3a1** | 5830414C08Rik\|Anr1\|MJAM\|OATP-D\|R75096\|Slc21a11 | solute carrier organic anion transporter family, member 3a1 | protein-coding |
| chr7_74531529_74532008 | chr7 | 74531530 | 74532008 | + | 1.05178 | intron (NM_001038643, intron 1 of 10) | intron (NM_001038643, intron 1 of 10) | 23011 | NM_023908 | 108116 | Mm.425467 | NM_023908 | ENSMUSG00000025790 | **Slco3a1** | 5830414C08Rik\|Anr1\|MJAM\|OATP-D\|R75096\|Slc21a11 | solute carrier organic anion transporter family, member 3a1 | protein-coding |
| chr7_74696149_74696677 | chr7 | 74696150 | 74696677 | + | 0.870188 | Intergenic | ORR1B2\|LTR\|ERVL-MaLR | -141633 | NM_001038643 | 108116 | Mm.425467 | NM_023908 | ENSMUSG00000025790 | **Slco3a1** | 5830414C08Rik\|Anr1\|MJAM\|OATP-D\|R75096\|Slc21a11 | solute carrier organic anion transporter family, member 3a1 | protein-coding |
| chr7_74696284_74696667 | chr7 | 74696285 | 74696667 | + | 0.840514 | Intergenic | ORR1B2\|LTR\|ERVL-MaLR | -141696 | NM_001038643 | 108116 | Mm.425467 | NM_023908 | ENSMUSG00000025790 | **Slco3a1** | 5830414C08Rik\|Anr1\|MJAM\|OATP-D\|R75096\|Slc21a11 | solute carrier organic anion transporter family, member 3a1 | protein-coding |
| chr7_74323164_74323552 | chr7 | 74323165 | 74323552 | + | 0.757707 | intron (NM_001038643, intron 5 of 10) | intron (NM_001038643, intron 5 of 10) | 231422 | NM_023908 | 108116 | Mm.425467 | NM_023908 | ENSMUSG00000025790 | **Slco3a1** | 5830414C08Rik\|Anr1\|MJAM\|OATP-D\|R75096\|Slc21a11 | solute carrier organic anion transporter family, member 3a1 | protein-coding |
| chr7_74338186_74338516 | chr7 | 74338187 | 74338516 | + | 0.744584 | intron (NM_001038643, intron 4 of 10) | intron (NM_001038643, intron 4 of 10) | 216429 | NM_023908 | 108116 | Mm.425467 | NM_023908 | ENSMUSG00000025790 | **Slco3a1** | 5830414C08Rik\|Anr1\|MJAM\|OATP-D\|R75096\|Slc21a11 | solute carrier organic anion transporter family, member 3a1 | protein-coding |
| chr7_74341118_74341810 | chr7 | 74341119 | 74341810 | + | 0.673182 | intron (NM_001038643, intron 4 of 10) | intron (NM_001038643, intron 4 of 10) | 213316 | NM_023908 | 108116 | Mm.425467 | NM_023908 | ENSMUSG00000025790 | **Slco3a1** | 5830414C08Rik\|Anr1\|MJAM\|OATP-D\|R75096\|Slc21a11 | solute carrier organic anion transporter family, member 3a1 | protein-coding |
| chr2_180453651_180453905 | chr2 | 180453652 | 180453905 | + | 1.21945 | Intergenic | Intergenic | -7200 | NM_148933 | 108115 | Mm.133687 | NM_148933 | ENSMUSG00000038963 | **Slco4a1** | OATP-E\|Slc21a12 | solute carrier organic anion transporter family, member 4a1 | protein-coding |
| chr2_180438185_180438610 | chr2 | 180438186 | 180438610 | + | 1.0271 | Intergenic | Intergenic | -22580 | NM_148933 | 108115 | Mm.133687 | NM_148933 | ENSMUSG00000038963 | **Slco4a1** | OATP-E\|Slc21a12 | solute carrier organic anion transporter family, member 4a1 | protein-coding |
| chr2_180444367_180444746 | chr2 | 180444368 | 180444746 | + | 0.919146 | Intergenic | Intergenic | -16421 | NM_148933 | 108115 | Mm.133687 | NM_148933 | ENSMUSG00000038963 | **Slco4a1** | OATP-E\|Slc21a12 | solute carrier organic anion transporter family, member 4a1 | protein-coding |
| chr2_180435588_180435951 | chr2 | 180435589 | 180435951 | + | 0.725052 | Intergenic | Intergenic | -25208 | NM_148933 | 108115 | Mm.133687 | NM_148933 | ENSMUSG00000038963 | **Slco4a1** | OATP-E\|Slc21a12 | solute carrier organic anion transporter family, member 4a1 | protein-coding |
| chr2_180436978_180438609 | chr2 | 180436979 | 180438609 | + | 0.707636 | Intergenic | Intergenic | -23184 | NM_148933 | 108115 | Mm.133687 | NM_148933 | ENSMUSG00000038963 | **Slco4a1** | OATP-E\|Slc21a12 | solute carrier organic anion transporter family, member 4a1 | protein-coding |
| chr2_180443384_180444799 | chr2 | 180443385 | 180444799 | + | 0.686446 | Intergenic | Intergenic | -16886 | NM_148933 | 108115 | Mm.133687 | NM_148933 | ENSMUSG00000038963 | **Slco4a1** | OATP-E\|Slc21a12 | solute carrier organic anion transporter family, member 4a1 | protein-coding |
| chr1_12991456_12992912 | chr1 | 12991457 | 12992912 | + | 0.851633 | Intergenic | CpG | -1049 | NM_172841 | 240726 | Mm.24481 | NM_172841 | ENSMUSG00000025938 | **Slco5a1** | 4732454B15\|A630033C23Rik\|AA4 07151 | solute carrier organic anion transporter family, member 5A1 | protein-coding |

| | | | | | | | | | | | | | | | | | |
|---|---|---|---|---|---|---|---|---|---|---|---|---|---|---|---|---|---|
| chr1_13052049_13052672 | chr1 | 13052050 | 13052672 | + | 0.67023 | Intergenic | Intergenic | -61226 | NM_172841 | 240726 | Mm.24481 | NM_172841 | ENSMUSG00000025938 | **Slco5a1** | 4732454B15\|A630033C23Rik\|AA407151 | solute carrier organic anion transporter family, member 5A1 | protein-coding |
| chr11_83043838_83044290 | chr11 | 83043839 | 83044290 | + | 1.65838 | Intergenic | RLTR14_RN\|LTR\|ERV1 | -3531 | NR_073524 | 237887 | Mm.341645 | NM_181542 | ENSMUSG00000072621 | **Slfn10-ps** | Slfn10\|mFLJ00257 | schlafen 10, pseudogene | pseudo |
| chr11_83043838_83044704 | chr11 | 83043839 | 83044704 | + | 1.59375 | Intergenic | Intergenic | -3738 | NR_073524 | 237887 | Mm.341645 | NM_181542 | ENSMUSG00000072621 | **Slfn10-ps** | Slfn10\|mFLJ00257 | schlafen 10, pseudogene | pseudo |
| chr11_83044391_83044691 | chr11 | 83044392 | 83044691 | + | 1.08533 | Intergenic | L1_Mus3\|LINE\|L1 | -4008 | NR_073524 | 237887 | Mm.341645 | NM_181542 | ENSMUSG00000072621 | **Slfn10-ps** | Slfn10\|mFLJ00257 | schlafen 10, pseudogene | pseudo |
| chr11_83050114_83050393 | chr11 | 83050115 | 83050393 | + | 0.752213 | Intergenic | L1VL2\|LINE\|L1 | -9721 | NR_073524 | 237887 | Mm.341645 | NM_181542 | ENSMUSG00000072621 | **Slfn10-ps** | Slfn10\|mFLJ00257 | schlafen 10, pseudogene | pseudo |
| chr11_83063913_83064502 | chr11 | 83063914 | 83064502 | + | 0.922316 | promoter-TSS (NM_011408) | promoter-TSS (NM_011408) | -904 | NM_011408 | 20556 | Mm.278689 | NM_011408 | ENSMUSG00000072620 | **Slfn2** | Shlf2 | schlafen 2 | protein-coding |
| chr11_83060439_83061114 | chr11 | 83060440 | 83061114 | + | 0.850302 | Intergenic | Intergenic | -4335 | NM_011408 | 20556 | Mm.278689 | NM_011408 | ENSMUSG00000072620 | **Slfn2** | Shlf2 | schlafen 2 | protein-coding |
| chr11_35178170_35178949 | chr11 | 35178171 | 35178949 | + | 0.772397 | intron (NM_011412, intron 3 of 35) | intron (NM_011412, intron 3 of 35) | 57104 | NM_011412 | 20564 | Mm.478604 | NM_011412 | ENSMUSG00000056427 | **Slit3** | Slil2\|Slit1 | slit homolog 3 (Drosophila) | protein-coding |
| chr14_111674551_111675014 | chr14 | 111674552 | 111675014 | + | 0.951375 | promoter-TSS (NM_198865) | promoter-TSS (NM_198865) | -332 | NM_198865 | 75409 | Mm.235097 | NM_029273 | ENSMUSG00000033214 | **Slitrk5** | 2610019D03Rik | SLIT and NTRK-like family, member 5 | protein-coding |
| chr14_111674692_111674960 | chr14 | 111674693 | 111674960 | + | 0.929442 | promoter-TSS (NM_198865) | promoter-TSS (NM_198865) | -289 | NM_198865 | 75409 | Mm.235097 | NM_029273 | ENSMUSG00000033214 | **Slitrk5** | 2610019D03Rik | SLIT and NTRK-like family, member 5 | protein-coding |
| chr2_174514923_174515319 | chr2 | 174514924 | 174515319 | + | 1.56729 | Intergenic | Intergenic | -42180 | NM_025531 | 66390 | Mm.182294 | NM_025531 | ENSMUSG00000016257 | **Slmo2** | 2310042G06Rik\|AA407013 | slowmo homolog 2 (Drosophila) | protein-coding |
| chr2_174517171_174517746 | chr2 | 174517172 | 174517746 | + | 0.986082 | Intergenic | Intergenic | -44518 | NM_025531 | 66390 | Mm.182294 | NM_025531 | ENSMUSG00000016257 | **Slmo2** | 2310042G06Rik\|AA407013 | slowmo homolog 2 (Drosophila) | protein-coding |
| chr2_174517826_174518043 | chr2 | 174517827 | 174518043 | + | 0.677462 | Intergenic | Intergenic | -44994 | NM_025531 | 66390 | Mm.182294 | NM_025531 | ENSMUSG00000016257 | **Slmo2** | 2310042G06Rik\|AA407013 | slowmo homolog 2 (Drosophila) | protein-coding |
| chr9_70581002_70581335 | chr9 | 70581003 | 70581335 | + | 0.661916 | intron (NM_025690, intron 13 of 20) | B3A\|SINE\|B2 | 38299 | NM_025690 | 66660 | Mm.22379 | NM_025690 | ENSMUSG00000032212 | **Sltm** | 5730455C01Rik\|5730555F13Rik\|9130215G10Rik\|Met | SAFB-like, transcription modulator | protein-coding |
| chr8_79452062_79452443 | chr8 | 79452063 | 79452443 | + | 1.04565 | Intergenic | ORR1D1\|LTR\|ERVL-MaLR | -52785 | NM_008539 | 17125 | Mm.223717 | NM_008539 | ENSMUSG00000031681 | **Smad1** | AI528653\|Madh1\|Madr1\|Mlp1\|MusMLP | SMAD family member 1 | protein-coding |
| chr8_79450215_79450521 | chr8 | 79450216 | 79450521 | + | 0.664179 | Intergenic | Intergenic | -50900 | NM_008539 | 17125 | Mm.223717 | NM_008539 | ENSMUSG00000031681 | **Smad1** | AI528653\|Madh1\|Madr1\|Mlp1\|MusMLP | SMAD family member 1 | protein-coding |
| chr9_63718150_63718554 | chr9 | 63718151 | 63718554 | + | 0.854735 | intron (NM_016769, intron 1 of 8) | intron (NM_016769, intron 1 of 8) | 39642 | NM_016769 | 17127 | Mm.7320 | NM_016769 | ENSMUSG00000032402 | **Smad3** | AU022421\|Madh3 | SMAD family member 3 | protein-coding |
| chr9_63718840_63719074 | chr9 | 63718841 | 63719074 | + | 0.821718 | intron (NM_016769, intron 1 of 8) | intron (NM_016769, intron 1 of 8) | 39037 | NM_016769 | 17127 | Mm.7320 | NM_016769 | ENSMUSG00000032402 | **Smad3** | AU022421\|Madh3 | SMAD family member 3 | protein-coding |
| chr18_73680578_73680957 | chr18 | 73680579 | 73680957 | + | 0.82348 | intron (NM_008540, intron 1 of 11) | ORR1D1\|LTR\|ERVL-MaLR | 22973 | NM_008540 | 17128 | Mm.100399 | NM_008540 | ENSMUSG00000024515 | **Smad4** | AW743858\|D18Wsu70e\|DPC4\|Madh4 | SMAD family member 4 | protein-coding |
| chr13_56740012_56740781 | chr13 | 56740013 | 56740781 | + | 0.900196 | 3' UTR (NM_001164042, exon 7 of 7) | 3' UTR (NM_001164042, exon 7 of 7) | 36729 | NM_001164042 | 17129 | Mm.272920 | NM_008541 | ENSMUSG00000021540 | **Smad5** | 1110051M15Rik\|AI451355\|Dwf-C\|Madh5\|MusMLP | SMAD family member 5 | protein-coding |
| chr18_75371340_75371636 | chr18 | 75371341 | 75371636 | + | 0.614848 | intron (NM_001042660, intron 2 of 3) | intron (NM_001042660, intron 2 of 3) | 4123 | NM_001042660 | 17131 | Mm.34407 | NM_008543 | ENSMUSG00000025880 | **Smad7** | Madh7 | SMAD family member 7 | protein-coding |
| chr19_26414584_26415010 | chr19 | 26414585 | 26415010 | + | 0.644177 | Intergenic | Intergenic | -190363 | NM_011416 | 67155 | Mm.491146 | NM_011416 | ENSMUSG00000024921 | **Smarca2** | 2610209L14Rik\|SNF2alpha\|Snf2l2\|brahma\|brm | SWI/SNF related, matrix associated, actin dependent regulator of chromatin, subfamily a, member 2 | protein-coding |
| chr8_80742866_80743243 | chr8 | 80742867 | 80743243 | + | 1.66418 | Intergenic | URR1B\|DNA\|hAT-Charlie | -3596 | NM_053124 | 93762 | Mm.246803 | NM_053124 | | **Smarca5** | 4933427E24Rik\|D030040M08Rik\|D330027N15Rik\|MommeD4\|Snf2h | SWI/SNF related, matrix associated, actin dependent regulator of chromatin, subfamily a, member 5 | protein-coding |
| chr8_80732212_80732700 | chr8 | 80732213 | 80732700 | + | 0.807678 | intron (NM_053124, intron 3 of 23) | Lx5c\|LINE\|L1 | 7003 | NM_053124 | 93762 | Mm.246803 | NM_053124 | | **Smarca5** | 4933427E24Rik\|D030040M08Rik\|D330027N15Rik\|MommeD4\|Snf2h | SWI/SNF related, matrix associated, actin dependent regulator of chromatin, subfamily a, member 5 | protein-coding |
| chr9_110142211_110142428 | chr9 | 110142212 | 110142428 | + | 0.66995 | intron (NM_009211, intron 3 of 27) | intron (NM_009211, intron 3 of 27) | 10296 | NM_009211 | 20588 | Mm.85410 | NM_009211 | ENSMUSG00000032481 | **Smarcc1** | AI115498\|BAF155\|Rsc8\|SRG3 | SWI/SNF related, matrix associated, actin dependent regulator of chromatin, subfamily c, member 1 | protein-coding |
| chr10_128463886_128464173 | chr10 | 128463887 | 128464173 | + | 0.702314 | intron (NM_001114097, intron 5 of 27) | intron (NM_001114097, intron 5 of 27) | 4794 | NM_001114097 | 68094 | Mm.417338 | NM_198160 | ENSMUSG00000025369 | **Smarcc2** | 5930405J04Rik | SWI/SNF related, matrix associated, actin dependent regulator of chromatin, subfamily c, member 2 | protein-coding |
| chr19_53565795_53566594 | chr19 | 53565796 | 53566594 | + | 0.820896 | Intergenic | Intergenic | -34201 | NM_007790 | 13006 | Mm.14910 | NM_007790 | ENSMUSG00000024974 | **Smc3** | Bamacan\|Cspg6\|HCAP\|Mmip1\|SMC-3\|SmcD | structural maintenance of chromosomes 3 | protein-coding |
| chr19_23350759_23351227 | chr19 | 23350760 | 23351227 | + | 1.09696 | intron (NM_174857, intron 8 of 13) | URR1A\|DNA\|hAT-Charlie | -77096 | NM_001252684 | 226026 | Mm.23267 | NM_153808 | ENSMUSG00000024943 | **Smc5** | MLZ-453\|SMC-5\|Smc5l1\|mKIAA0594\|mSMC5 | structural maintenance of chromosomes 5 | protein-coding |
| chr19_23327050_23327342 | chr19 | 23327051 | 23327342 | + | 0.695301 | intron (NM_174857, intron 11 of 13) | intron (NM_174857, intron 11 of 13) | -53299 | NM_001252684 | 226026 | Mm.23267 | NM_153808 | ENSMUSG00000024943 | **Smc5** | MLZ-453\|SMC-5\|Smc5l1\|mKIAA0594\|mSMC5 | structural maintenance of chromosomes 5 | protein-coding |
| chr11_29166554_29166840 | chr11 | 29166555 | 29166840 | + | 0.656642 | Intergenic | Intergenic | -6210 | NM_134034 | 104570 | Mm.276386 | NM_134034 | ENSMUSG00000020463 | **Smek2** | AW011752\|AW557776\|mKIAA1387 | SMEK homolog 2, suppressor of mek1 (Dictyostelium) | protein-coding |
| chr3_88337337_88337681 | chr3 | 88337338 | 88337681 | + | 0.831044 | intron (NM_178246, intron 1 of 21) | intron (NM_178246, intron 1 of 21) | 1249 | NM_178246 | 229512 | Mm.439830 | NM_173188 | ENSMUSG00000001415 | **Smg5** | BC024683\|mKIAA1089 | Smg-5 homolog, nonsense mediated mRNA decay factor (C. elegans) | protein-coding |
| chr3_88337430_88337662 | chr3 | 88337431 | 88337662 | + | 0.810861 | intron (NM_178246, intron 1 of 21) | intron (NM_178246, intron 1 of 21) | 1286 | NM_178246 | 229512 | Mm.439830 | NM_173188 | ENSMUSG00000001415 | **Smg5** | BC024683\|mKIAA1089 | Smg-5 homolog, nonsense mediated mRNA decay factor (C. elegans) | protein-coding |
| chr11_74973179_74973735 | chr11 | 74973180 | 74973735 | + | 1.37719 | intron (NM_001002764, intron 8 of 18) | intron (NM_001002764, intron 8 of 18) | 47585 | NM_001002764 | 103677 | Mm.288460 | NM_001002764 | ENSMUSG00000038290 | **Smg6** | AI317223\|AU041178 | Smg-6 homolog, nonsense mediated mRNA decay factor (C. elegans) | protein-coding |
| chr11_74973112_74974061 | chr11 | 74973113 | 74974061 | + | 1.32516 | intron (NM_001002764, intron 8 of 18) | intron (NM_001002764, intron 8 of 18) | 47715 | NM_001002764 | 103677 | Mm.288460 | NM_001002764 | ENSMUSG00000038290 | **Smg6** | AI317223\|AU041178 | Smg-6 homolog, nonsense mediated mRNA decay factor (C. elegans) | protein-coding |
| chr11_74947288_74947645 | chr11 | 74947289 | 74947645 | + | 0.722006 | intron (NM_001002764, intron 8 of 18) | B2_Mm1t\|SINE\|B2 | 21595 | NM_001002764 | 103677 | Mm.288460 | NM_001002764 | ENSMUSG00000038290 | **Smg6** | AI317223\|AU041178 | Smg-6 homolog, nonsense mediated mRNA decay factor (C. elegans) | protein-coding |
| chr4_154024606_154025060 | chr4 | 154024607 | 154025060 | + | 0.689699 | intron (NM_001163721, intron 2 of 3) | intron (NM_001163721, intron 2 of 3) | 1211 | NM_001163722 | 68859 | Mm.477488 | NM_001033144 | ENSMUSG00000078350 | **Smim1** | 0610011H04Rik\|1190007F08Rik\|2700009I11Rik | small integral membrane protein 1 | protein-coding |
| chr13_41271816_41272221 | chr13 | 41271817 | 41272221 | + | 0.892657 | intron (NM_001135577, intron 1 of 1) | intron (NM_001135577, intron 1 of 1) | 22175 | NM_001135577 | 108934 | Mm.38578 | NM_001135577 | ENSMUSG00000091264 | **Smim13** | 2900036K02Rik | small integral membrane protein 13 | protein-coding |
| chr5_65483657_65484401 | chr5 | 65483658 | 65484401 | + | 1.07155 | intron (NM_133697, intron 1 of 4) | intron (NM_133697, intron 1 of 4) | 8806 | NM_133697 | 68552 | Mm.10709 | NM_133697 | ENSMUSG00000037822 | **Smim14** | 1110003E01Rik\|1700127H04Rik\|5430439C17Rik\|MAd4 | small integral membrane protein 14 | protein-coding |
| chr5_65483933_65484304 | chr5 | 65483934 | 65484304 | + | 1.04611 | intron (NM_133697, intron 1 of 4) | MLT-int\|LTR\|ERVL-MaLR | 8716 | NM_133697 | 68552 | Mm.10709 | NM_133697 | ENSMUSG00000037822 | **Smim14** | 1110003E01Rik\|1700127H04Rik\|5430439C17Rik\|MAd4 | small integral membrane protein 14 | protein-coding |
| chr13_108092156_108092683 | chr13 | 108092157 | 108092683 | + | 0.906487 | intron (NM_001127346, intron 1 of 3) | intron (NM_001127346, intron 1 of 3) | 47946 | NM_001048250 | 75616 | Mm.491164 | NM_001048250 | | **Smim15** | 2810008M24Rik | small integral membrane protein 15 | protein-coding |
| chr13_107967986_107968346 | chr13 | 107967987 | 107968346 | + | 0.619391 | Intergenic | ORR1A2\|LTR\|ERVL-MaLR | -76308 | NM_001048250 | 75616 | Mm.491164 | NM_001048250 | | **Smim15** | 2810008M24Rik | small integral membrane protein 15 | protein-coding |
| chr5_53402066_53403166 | chr5 | 53402067 | 53403166 | + | 1.84285 | Intergenic | Intergenic | 135510 | NM_001145433 | 66278 | Mm.27900 | NM_001145433 | | **Smim20** | 1110067B18Rik\|1810013D10Rik\|N28078 | small integral membrane protein 20 | protein-coding |
| chr5_53394134_53395000 | chr5 | 53394135 | 53395000 | + | 1.32822 | Intergenic | Intergenic | 127461 | NM_001145433 | 66278 | Mm.27900 | NM_001145433 | | **Smim20** | 1110067B18Rik\|1810013D10Rik\|N28078 | small integral membrane protein 20 | protein-coding |
| chr5_53393539_53395008 | chr5 | 53393540 | 53395008 | + | 1.2653 | Intergenic | Intergenic | 127168 | NM_001145433 | 66278 | Mm.27900 | NM_001145433 | | **Smim20** | 1110067B18Rik\|1810013D10Rik\|N28078 | small integral membrane protein 20 | protein-coding |
| chr5_53409502_53410133 | chr5 | 53409503 | 53410133 | + | 1.24141 | Intergenic | Intergenic | 142712 | NM_001145433 | 66278 | Mm.27900 | NM_001145433 | | **Smim20** | 1110067B18Rik\|1810013D10Rik\|N28078 | small integral membrane protein 20 | protein-coding |
| chr5_53400927_53401382 | chr5 | 53400928 | 53401382 | + | 1.12 | Intergenic | Intergenic | 134049 | NM_001145433 | 66278 | Mm.27900 | NM_001145433 | | **Smim20** | 1110067B18Rik\|1810013D10Rik\|N28078 | small integral membrane protein 20 | protein-coding |
| chr5_53401130_53401366 | chr5 | 53401131 | 53401366 | + | 1.00618 | Intergenic | Intergenic | 134142 | NM_001145433 | 66278 | Mm.27900 | NM_001145433 | | **Smim20** | 1110067B18Rik\|1810013D10Rik\|N28078 | small integral membrane protein 20 | protein-coding |
| chr5_53410850_53411356 | chr5 | 53410851 | 53411356 | + | 0.801448 | Intergenic | MLT1D\|LTR\|ERVL-MaLR | 143997 | NM_001145433 | 66278 | Mm.27900 | NM_001145433 | | **Smim20** | 1110067B18Rik\|1810013D10Rik\|N28078 | small integral membrane protein 20 | protein-coding |

| | | | | | | | | | | | | | | | | | |
|---|---|---|---|---|---|---|---|---|---|---|---|---|---|---|---|---|---|
| chr11_32762479_32763300 | chr11 | 32762480 | 32763300 | + | 1.04646 | Intergenic | MIRb\|SINE\|MIR | 61704 | NM_027050 | 69351 | Mm.65488 | NM_027050 | ENSMUSG00000020270 | **Smim23** | 1700008A04Rik | small integral membrane protein 23 | protein-coding |
| chr18_60502317_60502918 | chr18 | 60502318 | 60502918 | + | 1.10724 | promoter-TSS (NM_134133) | promoter-TSS (NM_134133) | -635 | NM_134133 | 106878 | Mm.273197 | NM_134133 | ENSMUSG00000038059 | **Smim3** | 2010002N04Rik\|AI195350\|AW561905\|Nid67\|cI-41 | small integral membrane protein 3 | protein-coding |
| chr18_60499794_60500741 | chr18 | 60499795 | 60500741 | + | 1.0714 | intron (NM_134133, intron 1 of 1) | intron (NM_134133, intron 1 of 1) | 1715 | NM_134133 | 106878 | Mm.273197 | NM_134133 | ENSMUSG00000038059 | **Smim3** | 2010002N04Rik\|AI195350\|AW561905\|Nid67\|cI-41 | small integral membrane protein 3 | protein-coding |
| chr18_60480855_60481592 | chr18 | 60480856 | 60481592 | + | 0.948184 | intron (NM_134133, intron 1 of 1) | intron (NM_134133, intron 1 of 1) | 20759 | NM_134133 | 106878 | Mm.273197 | NM_134133 | ENSMUSG00000038059 | **Smim3** | 2010002N04Rik\|AI195350\|AW561905\|Nid67\|cI-41 | small integral membrane protein 3 | protein-coding |
| chr14_31088564_31088876 | chr14 | 31088565 | 31088876 | + | 0.739336 | intron (NM_001081251, intron 24 of 32) | L2a\|LINE\|L2 | 40210 | NR_024069 | 66487 | Mm.379116 | NR_024069 | ENSMUSG00000058351 | **Smim4** | 2010107H07Rik\|Snhg8 | small itegral membrane protein 4 | ncRNA |
| chr11_115869309_115869837 | chr11 | 115869310 | 115869837 | + | 0.716717 | Intergenic | Intergenic | -30566 | NM_183259 | 66528 | Mm.275740 | NM_183259 | ENSMUSG00000048442 | **Smim5** | 2210020M01Rik | small integral membrane protein 5 | protein-coding |
| chr4_34819882_34820324 | chr4 | 34819883 | 34820324 | + | 0.95562 | intron (NM_013889, intron 5 of 7) | intron (NM_013889, intron 5 of 7) | -41766 | NM_025471 | 66291 | Mm.28071 | NM_025471 | ENSMUSG00000028295 | **Smim8** | 1810030N24Rik\|2810406B13Rik | small integral membrane protein 8 | protein-coding |
| chr19_53408385_53408751 | chr19 | 53408386 | 53408751 | + | 1.45366 | Intergenic | L2a\|LINE\|L2 | -17995 | NM_172429 | 76479 | Mm.313687 | NM_172429 | ENSMUSG00000025024 | **Smndc1** | 2410004J23Rik\|4933440I19Rik | survival motor neuron domain containing 1 | protein-coding |
| chr19_53408215_53408793 | chr19 | 53408216 | 53408793 | + | 1.35825 | Intergenic | L2a\|LINE\|L2 | -17931 | NM_172429 | 76479 | Mm.313687 | NM_172429 | ENSMUSG00000025024 | **Smndc1** | 2410004J23Rik\|4933440I19Rik | survival motor neuron domain containing 1 | protein-coding |
| chr19_53412109_53412439 | chr19 | 53412110 | 53412439 | + | 0.776845 | Intergenic | Intergenic | -21701 | NM_172429 | 76479 | Mm.313687 | NM_172429 | ENSMUSG00000025024 | **Smndc1** | 2410004J23Rik\|4933440I19Rik | survival motor neuron domain containing 1 | protein-coding |
| chr19_53404290_53404581 | chr19 | 53404291 | 53404581 | + | 0.678264 | Intergenic | Intergenic | -13863 | NM_172429 | 76479 | Mm.313687 | NM_172429 | ENSMUSG00000025024 | **Smndc1** | 2410004J23Rik\|4933440I19Rik | survival motor neuron domain containing 1 | protein-coding |
| chr2_131423596_131424049 | chr2 | 131423597 | 131424049 | + | 1.33721 | intron (NR_037281, intron 1 of 2) | ORR1F\|LTR\|ERVL-MaLR | -68039 | NM_001177834 | 228608 | Mm.136586 | NM_145533 | ENSMUSG00000027333 | **Smox** | B130066H01Rik\|PAO\|PAOh1\|SMO | spermine oxidase | protein-coding |
| chr2_131427022_131427294 | chr2 | 131427023 | 131427294 | + | 1.01364 | intron (NR_037281, intron 1 of 2) | RMER1B\|Other\|Other | -64704 | NM_001177834 | 228608 | Mm.136586 | NM_145533 | ENSMUSG00000027333 | **Smox** | B130066H01Rik\|PAO\|PAOh1\|SMO | spermine oxidase | protein-coding |
| chr2_131427012_131427386 | chr2 | 131427013 | 131427386 | + | 0.804769 | intron (NR_037281, intron 1 of 2) | RMER1B\|Other\|Other | -64663 | NM_001177834 | 228608 | Mm.136586 | NM_145533 | ENSMUSG00000027333 | **Smox** | B130066H01Rik\|PAO\|PAOh1\|SMO | spermine oxidase | protein-coding |
| chr7_105519390_105519767 | chr7 | 105519391 | 105519767 | + | 1.48294 | Intergenic | Intergenic | -34781 | NM_011421 | 20597 | Mm.4628 | NM_011421 | ENSMUSG00000037049 | **Smpd1** | A-SMase\|ASM\|Zn-SMase\|aSMase | sphingomyelin phosphodiesterase 1, acid lysosomal | protein-coding |
| chr7_105519265_105519737 | chr7 | 105519266 | 105519737 | + | 1.42861 | Intergenic | Chap1_Mam\|DNA\|hAT-Charlie | -34859 | NM_011421 | 20597 | Mm.4628 | NM_011421 | ENSMUSG00000037049 | **Smpd1** | A-SMase\|ASM\|Zn-SMase\|aSMase | sphingomyelin phosphodiesterase 1, acid lysosomal | protein-coding |
| chr7_105526290_105526740 | chr7 | 105526291 | 105526740 | + | 0.967219 | Intergenic | Intergenic | -27845 | NM_011421 | 20597 | Mm.4628 | NM_011421 | ENSMUSG00000037049 | **Smpd1** | A-SMase\|ASM\|Zn-SMase\|aSMase | sphingomyelin phosphodiesterase 1, acid lysosomal | protein-coding |
| chr10_57819031_57819587 | chr10 | 57819032 | 57819587 | + | 1.33864 | Intergenic | L1M3\|LINE\|L1 | 24765 | NM_020561 | 57319 | Mm.2379 | NM_020561 | ENSMUSG00000019872 | **Smpdl3a** | 0610010C24Rik\|AI529588\|ASM3A\|ASML3\|ASML3A | sphingomyelin phosphodiesterase, acid-like 3A | protein-coding |
| chr10_57818360_57818878 | chr10 | 57818361 | 57818878 | + | 0.870427 | Intergenic | B1F\|SINE\|Alu | 24075 | NM_020561 | 57319 | Mm.2379 | NM_020561 | ENSMUSG00000019872 | **Smpdl3a** | 0610010C24Rik\|AI529588\|ASM3A\|ASML3\|ASML3A | sphingomyelin phosphodiesterase, acid-like 3A | protein-coding |
| chr10_57818368_57818689 | chr10 | 57818369 | 57818689 | + | 0.843024 | Intergenic | Intergenic | 23985 | NM_020561 | 57319 | Mm.2379 | NM_020561 | ENSMUSG00000019872 | **Smpdl3a** | 0610010C24Rik\|AI529588\|ASM3A\|ASML3\|ASML3A | sphingomyelin phosphodiesterase, acid-like 3A | protein-coding |
| chr4_132757681_132758231 | chr4 | 132757682 | 132758231 | + | 0.925838 | promoter-TSS (NM_133888) | promoter-TSS (NM_133888) | -785 | NM_133888 | 100340 | Mm.287187 | NM_133888 | ENSMUSG00000028885 | **Smpdl3b** | 1110054A24Rik\|AU045240\|Asml3b | sphingomyelin phosphodiesterase, acid-like 3B | protein-coding |
| chr4_132757501_132758248 | chr4 | 132757502 | 132758248 | + | 0.897144 | promoter-TSS (NM_133888) | promoter-TSS (NM_133888) | -704 | NM_133888 | 100340 | Mm.287187 | NM_133888 | ENSMUSG00000028885 | **Smpdl3b** | 1110054A24Rik\|AU045240\|Asml3b | sphingomyelin phosphodiesterase, acid-like 3B | protein-coding |
| chrX_157731945_157732312 | chrX | 157731946 | 157732312 | + | 0.651912 | intron (NM_001252591, intron 4 of 4) | intron (NM_001252591, intron 4 of 4) | 33156 | NR_103722 | 66106 | Mm.140340 | NM_025357 | ENSMUSG00000041476 | **Smpx** | 1010001C09Rik\|Csl | small muscle protein, X-linked | protein-coding |
| chrX_157476157_157476743 | chrX | 157476158 | 157476743 | + | 1.2227 | intron (NM_009214, intron 1 of 10) | L1MB7\|LINE\|L1 | 15596 | NM_009214 | 20603 | Mm.18652 | NM_009214 | ENSMUSG00000071708 | **Sms** | AI427066\|Gy\|SPMSY\|SpmST\|gyro | spermine synthase | protein-coding |
| chrX_157468029_157468572 | chrX | 157468030 | 157468572 | + | 0.998475 | intron (NM_009214, intron 2 of 10) | intron (NM_009214, intron 2 of 10) | 23745 | NM_009214 | 20603 | Mm.18652 | NM_009214 | ENSMUSG00000071708 | **Sms** | AI427066\|Gy\|SPMSY\|SpmST\|gyro | spermine synthase | protein-coding |
| chrX_157468029_157469008 | chrX | 157468030 | 157469008 | + | 0.727932 | intron (NM_009214, intron 2 of 10) | intron (NM_009214, intron 2 of 10) | 23527 | NM_009214 | 20603 | Mm.18652 | NM_009214 | ENSMUSG00000071708 | **Sms** | AI427066\|Gy\|SPMSY\|SpmST\|gyro | spermine synthase | protein-coding |
| chr4_40777886_40778186 | chr4 | 40777887 | 40778186 | + | 1.24366 | Intergenic | RMER17C\|LTR\|ERVK | -20151 | NM_021535 | 74255 | Mm.289929 | NM_021535 | | **Smu1** | 2600001O03Rik\|2610203K23Rik\|AB044414\|AI845086\|AW556129\|Bwd | smu-1 suppressor of mec-8 and unc-52 homolog (C. elegans) | protein-coding |
| chr5_144935992_144936685 | chr5 | 144935993 | 144936685 | + | 0.862384 | intron (NM_029438, intron 1 of 17) | MER94\|DNA\|hAT-Blackjack | 29491 | NM_029438 | 75788 | Mm.27735 | NM_029438 | ENSMUSG00000038780 | **Smurf1** | 4930431E10Rik\|mKIAA1625 | SMAD specific E3 ubiquitin protein ligase 1 | protein-coding |
| chr5_144935892_144936933 | chr5 | 144935893 | 144936933 | + | 0.805964 | intron (NM_029438, intron 1 of 17) | intron (NM_029438, intron 1 of 17) | 29417 | NM_029438 | 75788 | Mm.27735 | NM_029438 | ENSMUSG00000038780 | **Smurf1** | 4930431E10Rik\|mKIAA1625 | SMAD specific E3 ubiquitin protein ligase 1 | protein-coding |
| chr5_144935990_144936315 | chr5 | 144935991 | 144936315 | + | 0.727723 | intron (NM_029438, intron 1 of 17) | intron (NM_029438, intron 1 of 17) | 29677 | NM_029438 | 75788 | Mm.27735 | NM_029438 | ENSMUSG00000038780 | **Smurf1** | 4930431E10Rik\|mKIAA1625 | SMAD specific E3 ubiquitin protein ligase 1 | protein-coding |
| chr6_71233978_71234359 | chr6 | 71233979 | 71234359 | + | 0.768527 | intron (NM_009762, intron 4 of 8) | intron (NM_009762, intron 4 of 8) | 28112 | NM_001160127 | 12180 | Mm.234274 | NM_009762 | ENSMUSG00000055027 | **Smyd1** | 4632404M21Rik\|Bop\|C78565\|Zmynd18 | SET and MYND domain containing 1 | protein-coding |
| chr1_189868033_189868454 | chr1 | 189868034 | 189868454 | + | 1.07427 | intron (NM_008976, intron 18 of 18) | intron (NM_008976, intron 18 of 18) | 54044 | NM_026796 | 226830 | Mm.156895 | NM_026796 | ENSMUSG00000026603 | **Smyd2** | 1110020E07Rik\|4930402C15\|KMT3C\|Zmynd14 | SET and MYND domain containing 2 | protein-coding |
| chr1_179492591_179493773 | chr1 | 179492592 | 179493773 | + | 1.52814 | intron (NM_027188, intron 1 of 11) | intron (NM_027188, intron 1 of 11) | 24821 | NM_027188 | 69726 | Mm.222338 | NM_027188 | | **Smyd3** | 2410008A19Rik\|Zmynd1 | SET and MYND domain containing 3 | protein-coding |
| chr1_179511728_179512435 | chr1 | 179511729 | 179512435 | + | 0.920375 | intron (NM_027188, intron 1 of 11) | intron (NM_027188, intron 1 of 11) | 5921 | NM_027188 | 69726 | Mm.222338 | NM_027188 | | **Smyd3** | 2410008A19Rik\|Zmynd1 | SET and MYND domain containing 3 | protein-coding |
| chr1_179130032_179130384 | chr1 | 179130033 | 179130384 | + | 0.668452 | intron (NM_027188, intron 5 of 11) | intron (NM_027188, intron 5 of 11) | 387795 | NM_027188 | 69726 | Mm.222338 | NM_027188 | | **Smyd3** | 2410008A19Rik\|Zmynd1 | SET and MYND domain containing 3 | protein-coding |
| chr6_85428920_85429390 | chr6 | 85428921 | 85429390 | + | 1.22258 | TTS (NM_001007472) | TTS (NM_001007472) | -2821 | NM_144918 | 232187 | Mm.219946 | NM_144918 | ENSMUSG00000033706 | **Smyd5** | AW536703\|NN8-4AG\|Rai15\|Rrg1 | SET and MYND domain containing 5 | protein-coding |
| chr8_122461527_122461975 | chr8 | 122461528 | 122461975 | + | 1.04801 | Intergenic | Intergenic | -1059 | NM_013914 | 30927 | Mm.103673 | NM_013914 | ENSMUSG00000006587 | **Snai3** | AI643946\|Smuc\|Zfp293 | snail family zinc finger 3 | protein-coding |
| chr8_122460454_122461982 | chr8 | 122460455 | 122461982 | + | 0.637243 | promoter-TSS (NM_013914) | promoter-TSS (NM_013914) | -526 | NM_013914 | 30927 | Mm.103673 | NM_013914 | ENSMUSG00000006587 | **Snai3** | AI643946\|Smuc\|Zfp293 | snail family zinc finger 3 | protein-coding |
| chr12_73956056_73956411 | chr12 | 73956057 | 73956411 | + | 0.852623 | Intergenic | Intergenic | -8296 | NM_178392 | 75627 | Mm.25937 | NM_178392 | ENSMUSG00000021113 | **Snapc1** | 2700033G17Rik\|AU015787 | small nuclear RNA activating complex, polypeptide 1 | protein-coding |
| chr12_73974518_73975128 | chr12 | 73974519 | 73975128 | + | 0.742609 | intron (NM_178392, intron 7 of 9) | intron (NM_178392, intron 7 of 9) | 10293 | NM_178392 | 75627 | Mm.25937 | NM_178392 | ENSMUSG00000021113 | **Snapc1** | 2700033G17Rik\|AU015787 | small nuclear RNA activating complex, polypeptide 1 | protein-coding |
| chr6_28583236_28583829 | chr6 | 28583237 | 28583829 | + | 1.46865 | intron (NM_019776, intron 10 of 23) | intron (NM_019776, intron 10 of 23) | 103185 | NM_019776 | 56463 | Mm.439987 | NM_019776 | ENSMUSG00000001424 | **Snd1** | AL033314\|Tudor-SN | staphylococcal nuclease and tudor domain containing 1 | protein-coding |
| chr6_28559413_28559789 | chr6 | 28559414 | 28559789 | + | 1.1091 | intron (NM_019776, intron 10 of 23) | intron (NM_019776, intron 10 of 23) | 79253 | NM_019776 | 56463 | Mm.439987 | NM_019776 | ENSMUSG00000001424 | **Snd1** | AL033314\|Tudor-SN | staphylococcal nuclease and tudor domain containing 1 | protein-coding |
| chr6_28552824_28553398 | chr6 | 28552825 | 28553398 | + | 1.02891 | intron (NM_019776, intron 10 of 23) | intron (NM_019776, intron 10 of 23) | 72763 | NM_019776 | 56463 | Mm.439987 | NM_019776 | ENSMUSG00000001424 | **Snd1** | AL033314\|Tudor-SN | staphylococcal nuclease and tudor domain containing 1 | protein-coding |
| chr6_28609399_28609768 | chr6 | 28609400 | 28609768 | + | 0.9448 | intron (NM_019776, intron 10 of 23) | intron (NM_019776, intron 10 of 23) | 129236 | NM_019776 | 56463 | Mm.439987 | NM_019776 | ENSMUSG00000001424 | **Snd1** | AL033314\|Tudor-SN | staphylococcal nuclease and tudor domain containing 1 | protein-coding |
| chr6_28469699_28470023 | chr6 | 28469700 | 28470023 | + | 0.892378 | Intergenic | Intergenic | -10487 | NM_019776 | 56463 | Mm.439987 | NM_019776 | ENSMUSG00000001424 | **Snd1** | AL033314\|Tudor-SN | staphylococcal nuclease and tudor domain containing 1 | protein-coding |
| chr6_28548193_28548683 | chr6 | 28548194 | 28548683 | + | 0.797853 | intron (NM_019776, intron 10 of 23) | intron (NM_019776, intron 10 of 23) | 68090 | NM_019776 | 56463 | Mm.439987 | NM_019776 | ENSMUSG00000001424 | **Snd1** | AL033314\|Tudor-SN | staphylococcal nuclease and tudor domain containing 1 | protein-coding |
| chr6_28483928_28484370 | chr6 | 28483929 | 28484370 | + | 0.654162 | intron (NM_019776, intron 1 of 23) | intron (NM_019776, intron 1 of 23) | 3801 | NM_019776 | 56463 | Mm.439987 | NM_019776 | ENSMUSG00000001424 | **Snd1** | AL033314\|Tudor-SN | staphylococcal nuclease and tudor domain containing 1 | protein-coding |
| chr4_132354606_132354894 | chr4 | 132354607 | 132354894 | + | 0.717861 | Intergenic | Intergenic | -1064 | NR_003270 | 399101 | Mm.486919 | NR_002904 | ENSMUSG00000085241 | **Snhg3** | Rnu17d\|U17HG | small nucleolar RNA host gene 3 | ncRNA |
| chr9_88516407_88516931 | chr9 | 88516408 | 88516931 | + | 0.885357 | Intergenic | Intergenic | 6228 | NR_040721 | 72655 | Mm.21798 | NR_040721 | ENSMUSG00000097195 | **Snhg5** | 2810026P18Rik\|4933428I09Rik\|mU50HG-b | small nucleolar RNA host gene 5 | ncRNA |
| chr14_75851884_75852449 | chr14 | 75851885 | 75852449 | + | 1.30464 | Intergenic | ORR1F\|LTR\|ERVL-MaLR | 4244 | NR_028481 | 100303751 | Mm.484707 | NR_028481 | | **Snora31** | MBI-161 | small nucleolar RNA, H/ACA box 31 | snoRNA |
| chr14_75850292_75850583 | chr14 | 75850293 | 75850583 | + | 0.68712 | Intergenic | MTD\|LTR\|ERVL-MaLR | 2515 | NR_028481 | 100303751 | Mm.484707 | NR_028481 | | **Snora31** | MBI-161 | small nucleolar RNA, H/ACA box 31 | snoRNA |
| chr10_23783310_23783732 | chr10 | 23783311 | 23783732 | + | 0.877347 | Intergenic | B4A\|SINE\|B4 | 1930 | NR_037680 | 100529074 | Mm.371846 | NR_037680 | | **Snora33** | ACA33\|MBI-42 | small nucleolar RNA, H/ACA box 33 | snoRNA |
| chr10_23782953_23783738 | chr10 | 23782954 | 23783738 | + | 0.716132 | Intergenic | Intergenic | 2105 | NR_037680 | 100529074 | Mm.371846 | NR_037680 | | **Snora33** | ACA33\|MBI-42 | small nucleolar RNA, H/ACA box 33 | snoRNA |
| chr15_98512714_98513286 | chr15 | 98512715 | 98513286 | + | 1.03018 | Intergenic | Intergenic | 6834 | NR_034051 | 100217417 | Mm.486440 | NR_034051 | | **Snora34** | MBI-89 | small nucleolar RNA, H/ACA box 34 | snoRNA |
| chr18_35555959_35556134 | chr18 | 35555960 | 35556134 | + | 0.851631 | promoter-TSS (NR_002905) | promoter-TSS (NR_002905) | -983 | NR_002905 | 436583 | Mm.482118 | NR_002905 | ENSMUSG00000065649 | **Snora74a** | EG436583\|RNU19\|U19 | small nucleolar RNA, H/ACA box 74A | snoRNA |
| chr1_86342417_86342844 | chr1 | 86342418 | 86342844 | + | 0.684074 | Intergenic | ID4_\|SINE\|ID | 8654 | NR_028478 | 100303740 | Mm.154378 | NR_028478 | | **Snora75** | MBI-55 | small nucleolar RNA, H/ACA box 75 | snoRNA |

| | | | | | | | | | | | | | | | | | |
|---|---|---|---|---|---|---|---|---|---|---|---|---|---|---|---|---|---|
| chr11_106545801_106546592 | chr11 | 106545802 | 106546592 | + | 1.2192 | intron (NM_198292, intron 4 of 11) | intron (NM_198292, intron 4 of 11) | 45206 | NR_030703 | 100216537 | | NR_030703 | ENSMUSG00000065126 | **Snord104** | - | small nucleolar RNA, C/D box 104 | snoRNA |
| chr19_7456820_7457428 | chr19 | 7456821 | 7457428 | + | 1.63725 | exon (NM_001271486, exon 3 of 9) | exon (NM_001271486, exon 3 of 9) | -21394 | NR_028566 | 100216530 | Mm.453052 | NR_028566 | ENSMUSG00000064899 | **Snord118** | MBI-34 | small nucleolar RNA, C/D box 118 | snoRNA |
| chr19_7457610_7457899 | chr19 | 7457611 | 7457899 | + | 1.12538 | exon (NM_001003934, exon 3 of 9) | exon (NM_001003934, exon 3 of 9) | -20763 | NR_028566 | 100216530 | Mm.453052 | NR_028566 | ENSMUSG00000064899 | **Snord118** | MBI-34 | small nucleolar RNA, C/D box 118 | snoRNA |
| chr19_8727593_8728287 | chr19 | 8727594 | 8728287 | + | 0.875457 | Intergenic | B1_Mus1\|SINE\|Alu | 2074 | NR_004445 | 100127111 | Mm.292489 | NR_004445 | ENSMUSG00000065087 | **Snord22** | Rnu22\|U22 | small nucleolar RNA, C/D box 22 | snoRNA |
| chr19_8726841_8727224 | chr19 | 8726842 | 8727224 | + | 0.665265 | TTS (NR_002896) | TTS (NR_002896) | 1167 | NR_004445 | 100127111 | Mm.292489 | NR_004445 | ENSMUSG00000065087 | **Snord22** | Rnu22\|U22 | small nucleolar RNA, C/D box 22 | snoRNA |
| chr1_161039737_161040843 | chr1 | 161039738 | 161040843 | + | 1.06221 | TTS (NM_172644) | TTS (NM_172644) | 2198 | NR_028543 | 100217446 | | NR_028543 | ENSMUSG00000064968 | **Snord47** | MBII-96 | small nucleolar RNA, C/D box 47 | snoRNA |
| chr1_161039539_161041251 | chr1 | 161039540 | 161041251 | + | 1.02361 | TTS (NM_172644) | TTS (NM_172644) | 2303 | NR_028543 | 100217446 | | NR_028543 | ENSMUSG00000064968 | **Snord47** | MBII-96 | small nucleolar RNA, C/D box 47 | snoRNA |
| chr2_130279350_130280306 | chr2 | 130279351 | 130280306 | + | 1.01647 | TTS (NM_024193) | TTS (NM_024193) | 1802 | NR_028528 | 100217428 | | NR_028528 | | **Snord57** | MBII-434 | small nucleolar RNA, C/D box 57 | snoRNA |
| chr14_52059370_52059925 | chr14 | 52059371 | 52059925 | + | 0.716696 | Intergenic | Intergenic | -9667 | NR_028552 | 100217457 | | NR_028552 | ENSMUSG00000064871 | **Snord58b** | MBII-199 | small nucleolar RNA, C/D box 58B | snoRNA |
| chr11_62610476_62610722 | chr11 | 62610477 | 62610722 | + | 0.806105 | intron (NM_198861, intron 2 of 3) | MIRb\|SINE\|MIR | 6069 | NR_028541 | 100217444 | | NR_028541 | | **Snord65** | MBII-135\|snoRNA135 | small nucleolar RNA, C/D box 65 | snoRNA |
| chr15_5127478_5128042 | chr15 | 5127479 | 5128042 | + | 0.758326 | Intergenic | ORR1C2\|LTR\|ERVL-MaLR | 9339 | NR_028091 | 100302529 | | NR_028091 | | **Snord72** | MBII-240 | small nucleolar RNA, C/D box 72 | snoRNA |
| chr15_80060964_80061594 | chr15 | 80060965 | 80061594 | + | 1.59925 | Intergenic | Intergenic | 17300 | NR_028282 | 100302601 | | NR_028282 | | **Snord83b** | MBII-10 | small nucleolar RNA, C/D box 83B | snoRNA |
| chr15_80062798_80063206 | chr15 | 80062799 | 80063206 | + | 0.706599 | Intergenic | MIRb\|SINE\|MIR | 15577 | NR_028282 | 100302601 | | NR_028282 | | **Snord83b** | MBII-10 | small nucleolar RNA, C/D box 83B | snoRNA |
| chr15_80075382_80077320 | chr15 | 80075383 | 80077320 | + | 0.673885 | Intergenic | Intergenic | 2228 | NR_028282 | 100302601 | | NR_028282 | | **Snord83b** | MBII-10 | small nucleolar RNA, C/D box 83B | snoRNA |
| chr5_23893960_23894783 | chr5 | 23893961 | 23894783 | + | 1.5456 | intron (NR_037304, intron 2 of 4) | intron (NR_037304, intron 2 of 4) | 42139 | NR_028536 | 100217436 | | NR_028536 | | **Snord93** | Gm22397\|MBII-336 | small nucleolar RNA, C/D box 93 | snoRNA |
| chr5_23871002_23871716 | chr5 | 23871003 | 23871716 | + | 0.814179 | intron (NR_037304, intron 3 of 4) | MERV1_LTR\|LTR\|ERV 1 | 19126 | NR_028536 | 100217436 | | NR_028536 | | **Snord93** | Gm22397\|MBII-336 | small nucleolar RNA, C/D box 93 | snoRNA |
| chr5_23874418_23874926 | chr5 | 23874419 | 23874926 | + | 0.695787 | intron (NR_037304, intron 3 of 4) | intron (NR_037304, intron 3 of 4) | 22439 | NR_028536 | 100217436 | | NR_028536 | | **Snord93** | Gm22397\|MBII-336 | small nucleolar RNA, C/D box 93 | snoRNA |
| chr10_75527215_75527680 | chr10 | 75527216 | 75527680 | + | 0.816997 | intron (NM_026095, intron 2 of 3) | intron (NM_026095, intron 2 of 3) | 9406 | NM_026095 | 67332 | Mm.45151 | NM_026095 | ENSMUSG00000020180 | **Snrpd3** | 1700043E15Rik\|2310009E13Rik\|AW046420\|SMD3 | small nuclear ribonucleoprotein D3 | protein-coding |
| chr15_55862441_55862796 | chr15 | 55862442 | 55862796 | + | 1.49471 | intron (NM_016667, intron 1 of 6) | MTEb\|LTR\|ERVL-MaLR | 44330 | NM_016667 | 20649 | Mm.309425 | NM_016667 | ENSMUSG00000060429 | **Sntb1** | - | syntrophin, basic 1 | protein-coding |
| chr15_55806136_55807025 | chr15 | 55806137 | 55807025 | + | 1.34051 | intron (NM_016667, intron 1 of 6) | intron (NM_016667, intron 1 of 6) | 100368 | NM_016667 | 20649 | Mm.309425 | NM_016667 | ENSMUSG00000060429 | **Sntb1** | - | syntrophin, basic 1 | protein-coding |
| chr15_55794376_55795238 | chr15 | 55794377 | 55795238 | + | 1.13496 | intron (NM_016667, intron 1 of 6) | intron (NM_016667, intron 1 of 6) | 112142 | NM_016667 | 20649 | Mm.309425 | NM_016667 | ENSMUSG00000060429 | **Sntb1** | - | syntrophin, basic 1 | protein-coding |
| chr15_55794397_55794833 | chr15 | 55794398 | 55794833 | + | 1.01935 | intron (NM_016667, intron 1 of 6) | intron (NM_016667, intron 1 of 6) | 112334 | NM_016667 | 20649 | Mm.309425 | NM_016667 | ENSMUSG00000060429 | **Sntb1** | - | syntrophin, basic 1 | protein-coding |
| chr15_55866156_55866563 | chr15 | 55866157 | 55866563 | + | 0.89991 | intron (NM_016667, intron 1 of 6) | intron (NM_016667, intron 1 of 6) | 40589 | NM_016667 | 20649 | Mm.309425 | NM_016667 | ENSMUSG00000060429 | **Sntb1** | - | syntrophin, basic 1 | protein-coding |
| chr15_55863267_55863575 | chr15 | 55863268 | 55863575 | + | 0.819901 | intron (NM_016667, intron 1 of 6) | intron (NM_016667, intron 1 of 6) | 43528 | NM_016667 | 20649 | Mm.309425 | NM_016667 | ENSMUSG00000060429 | **Sntb1** | - | syntrophin, basic 1 | protein-coding |
| chr15_55794890_55795219 | chr15 | 55794891 | 55795219 | + | 0.790005 | intron (NM_016667, intron 1 of 6) | intron (NM_016667, intron 1 of 6) | 111894 | NM_016667 | 20649 | Mm.309425 | NM_016667 | ENSMUSG00000060429 | **Sntb1** | - | syntrophin, basic 1 | protein-coding |
| chr15_55852716_55853209 | chr15 | 55852717 | 55853209 | + | 0.731971 | intron (NM_016667, intron 1 of 6) | intron (NM_016667, intron 1 of 6) | 53986 | NM_016667 | 20649 | Mm.309425 | NM_016667 | ENSMUSG00000060429 | **Sntb1** | - | syntrophin, basic 1 | protein-coding |
| chr8_106937188_106937509 | chr8 | 106937189 | 106937509 | + | 0.811038 | intron (NM_009229, intron 1 of 6) | intron (NM_009229, intron 1 of 6) | 1599 | NM_009229 | 20650 | Mm.30228 | NM_009229 | ENSMUSG00000041308 | **Sntb2** | Snt2 | syntrophin, basic 2 | protein-coding |
| chr12_87470152_87470736 | chr12 | 87470153 | 87470736 | + | 0.777769 | intron (NM_025507, intron 1 of 13) | MTB_Mm\|LTR\|ERVL-MaLR | 1855 | NM_025507 | 66354 | Mm.271174 | NM_025507 | ENSMUSG00000021039 | **Snw1** | 2310008B08Rik\|AW048543\|NCoA-62\|SKIP\|Skiip | SNW domain containing 1 | protein-coding |
| chr11_96679111_96679639 | chr11 | 96679112 | 96679639 | + | 0.780736 | intron (NM_001177898, intron 4 of 10) | intron (NM_001177898, intron 4 of 10) | 98180 | NM_028965 | 74479 | Mm.270781 | NM_028965 | ENSMUSG00000020876 | **Snx11** | 4933439F10Rik\|A930041K09Rik\|AI117694 | sorting nexin 11 | protein-coding |
| chr12_35199298_35199904 | chr12 | 35199299 | 35199904 | + | 1.12406 | Intergenic | Intergenic | 152412 | NM_001014973 | 217463 | Mm.213991 | NM_001014973 | ENSMUSG00000020590 | **Snx13** | Rgs-px1\|mKIAA0713 | sorting nexin 13 | protein-coding |
| chr12_35051023_35051321 | chr12 | 35051024 | 35051321 | + | 0.718476 | intron (NM_001014973, intron 1 of 25) | intron (NM_001014973, intron 1 of 25) | 3983 | NM_001014973 | 217463 | Mm.213991 | NM_001014973 | ENSMUSG00000020590 | **Snx13** | Rgs-px1\|mKIAA0713 | sorting nexin 13 | protein-coding |
| chr9_88424955_88425500 | chr9 | 88424956 | 88425500 | + | 1.11153 | intron (NM_172926, intron 1 of 27) | intron (NM_172926, intron 1 of 27) | 13723 | NM_172926 | 244962 | Mm.439873 | NM_172926 | ENSMUSG00000032422 | **Snx14** | B830022K16\|C330035N22Rik\|YR-14 | sorting nexin 14 | protein-coding |
| chr3_10446369_10447023 | chr3 | 10446370 | 10447023 | + | 1.28574 | Intergenic | Intergenic | -6566 | NM_001127191 | 74718 | Mm.440548 | NM_029068 | ENSMUSG00000027534 | **Snx16** | 4930522N22Rik\|AI117577\|AI747675 | sorting nexin 16 | protein-coding |
| chr13_113630156_113630826 | chr13 | 113630157 | 113630826 | + | 0.9306 | Intergenic | Intergenic | -11927 | NM_130796 | 170625 | Mm.33721 | NM_130796 | ENSMUSG00000042364 | **Snx18** | Snag1 | sorting nexin 18 | protein-coding |
| chr8_46142138_46143020 | chr8 | 46142139 | 46143020 | + | 0.736353 | Intergenic | Intergenic | -18433 | NM_207213 | 102141 | Mm.267258 | NM_207213 | ENSMUSG00000038291 | **Snx25** | AI661919\|Gm1699\|Gm173\|Sbbi31 | sorting nexin 25 | protein-coding |
| chr3_94563964_94564340 | chr3 | 94563965 | 94564340 | + | 1.5938 | intron (NM_029721, intron 1 of 11) | intron (NM_029721, intron 1 of 11) | 18564 | NM_001082484 | 76742 | Mm.28160 | NM_029721 | ENSMUSG00000028136 | **Snx27** | 5730552M22Rik\|ESTM45\|ESTM47\|R75405 | sorting nexin family member 27 | protein-coding |
| chr3_94565619_94565966 | chr3 | 94565620 | 94565966 | + | 1.36622 | intron (NM_029721, intron 1 of 11) | B3A\|SINE\|B2 | 16923 | NM_001082484 | 76742 | Mm.28160 | NM_029721 | ENSMUSG00000028136 | **Snx27** | 5730552M22Rik\|ESTM45\|ESTM47\|R75405 | sorting nexin family member 27 | protein-coding |
| chr16_11475265_11476360 | chr16 | 11475266 | 11476360 | + | 1.4218 | intron (NM_001290148, intron 6 of 13) | intron (NM_001290148, intron 6 of 13) | 70165 | NM_001290148 | 74478 | Mm.325831 | NM_028964 | ENSMUSG00000071669 | **Snx29** | 4933437K13Rik\|ENSMUSG00000079738\|Gm11170\|Gm1737\|Gm930\|Rundc2a | sorting nexin 29 | protein-coding |
| chr19_5512088_5512558 | chr19 | 5512089 | 5512558 | + | 0.66399 | Intergenic | Intergenic | -1834 | NM_001024560 | 225861 | Mm.20071 | NM_001024560 | ENSMUSG00000056185 | **Snx32** | B930037P14Rik\|Snx6b | sorting nexin 32 | protein-coding |
| chr5_140379127_140379925 | chr5 | 140379128 | 140379925 | + | 1.09563 | intron (NM_172277, intron 1 of 10) | intron (NM_172277, intron 1 of 10) | 9721 | NM_172277 | 231834 | Mm.27098 | NM_172277 | ENSMUSG00000029560 | **Snx8** | B130023O14Rik | sorting nexin 8 | protein-coding |
| chr17_5827848_5828823 | chr17 | 5827849 | 5828823 | + | 1.26972 | Intergenic | Intergenic | -13044 | NM_025664 | 66616 | Mm.89515 | NM_025664 | ENSMUSG00000002365 | **Snx9** | 2700073N08Rik\|SDP1\|SH3PX1 | sorting nexin 9 | protein-coding |
| chr1_156454414_156454741 | chr1 | 156454415 | 156454741 | + | 0.725035 | intron (NM_009230, intron 3 of 15) | intron (NM_009230, intron 3 of 15) | 19750 | NM_009230 | 20652 | Mm.28099 | NM_009230 | ENSMUSG00000026600 | **Soat1** | 8430426K15Rik\|ACAT-1\|AW550831\|Acact\|ald\|hid | sterol O-acyltransferase 1 | protein-coding |
| chr15_102150043_102151299 | chr15 | 102150044 | 102151299 | + | 1.2323 | promoter-TSS (NM_146064) | promoter-TSS (NM_146064) | 96 | NM_146064 | 223920 | Mm.358862 | NM_146064 | ENSMUSG00000023045 | **Soat2** | ACAT2\|D15Wsu97e | sterol O-acyltransferase 2 | protein-coding |
| chr15_102150225_102150843 | chr15 | 102150226 | 102150843 | + | 1.0639 | promoter-TSS (NM_146064) | promoter-TSS (NM_146064) | -41 | NM_146064 | 223920 | Mm.358862 | NM_146064 | ENSMUSG00000023045 | **Soat2** | ACAT2\|D15Wsu97e | sterol O-acyltransferase 2 | protein-coding |
| chr16_10754349_10755082 | chr16 | 10754350 | 10755082 | + | 1.33836 | Intergenic | Intergenic | 30820 | NM_009896 | 12703 | Mm.130 | NM_009896 | ENSMUSG00000038037 | **Socs1** | Cish1\|Cish7\|JAB\|SOCS-1\|SSI-1 | suppressor of cytokine signaling 1 | protein-coding |
| chr16_10668373_10668808 | chr16 | 10668374 | 10668808 | + | 1.10919 | intron (NM_177562, intron 18 of 22) | intron (NM_177562, intron 18 of 22) | 116945 | NM_009896 | 12703 | Mm.130 | NM_009896 | ENSMUSG00000038037 | **Socs1** | Cish1\|Cish7\|JAB\|SOCS-1\|SSI-1 | suppressor of cytokine signaling 1 | protein-coding |
| chr16_10782571_10782793 | chr16 | 10782572 | 10782793 | + | 0.931665 | Intergenic | Intergenic | 2854 | NM_009896 | 12703 | Mm.130 | NM_009896 | ENSMUSG00000038037 | **Socs1** | Cish1\|Cish7\|JAB\|SOCS-1\|SSI-1 | suppressor of cytokine signaling 1 | protein-coding |
| chr16_10758847_10759311 | chr16 | 10758848 | 10759311 | + | 0.774224 | Intergenic | L2b\|LINE\|L2 | 26457 | NM_009896 | 12703 | Mm.130 | NM_009896 | ENSMUSG00000038037 | **Socs1** | Cish1\|Cish7\|JAB\|SOCS-1\|SSI-1 | suppressor of cytokine signaling 1 | protein-coding |
| chr16_10702369_10702960 | chr16 | 10702370 | 10702960 | + | 0.773608 | intron (NM_177562, intron 21 of 22) | intron (NM_177562, intron 21 of 22) | 82871 | NM_009896 | 12703 | Mm.130 | NM_009896 | ENSMUSG00000038037 | **Socs1** | Cish1\|Cish7\|JAB\|SOCS-1\|SSI-1 | suppressor of cytokine signaling 1 | protein-coding |
| chr16_10758720_10759586 | chr16 | 10758721 | 10759586 | + | 0.660026 | Intergenic | L2b\|LINE\|L2 | 26383 | NM_009896 | 12703 | Mm.130 | NM_009896 | ENSMUSG00000038037 | **Socs1** | Cish1\|Cish7\|JAB\|SOCS-1\|SSI-1 | suppressor of cytokine signaling 1 | protein-coding |
| chr16_10676561_10677052 | chr16 | 10676562 | 10677052 | + | 0.625486 | intron (NM_177562, intron 18 of 22) | intron (NM_177562, intron 18 of 22) | 108729 | NM_009896 | 12703 | Mm.130 | NM_009896 | ENSMUSG00000038037 | **Socs1** | Cish1\|Cish7\|JAB\|SOCS-1\|SSI-1 | suppressor of cytokine signaling 1 | protein-coding |
| chr10_95416435_95417660 | chr10 | 95416436 | 95417660 | + | 0.89094 | promoter-TSS (NM_001168655) | promoter-TSS (NM_001168655) | -191 | NM_007706 | 216233 | Mm.474283 | NM_007706 | ENSMUSG00000020027 | **Socs2** | 8030460M17\|AI527257\|AW108012\|CIS2\|Cish2\|D130043N08Rik\|JAB\|SOCS-2\|SSI-2\|hg | suppressor of cytokine signaling 2 | protein-coding |
| chr17_87118663_87119584 | chr17 | 87118664 | 87119584 | + | 1.15023 | intron (NM_019654, intron 1 of 2) | URR1B\|DNA\|hAT-Charlie | 11445 | NM_019654 | 56468 | Mm.126885 | NM_019654 | ENSMUSG00000037104 | **Socs5** | 1810018L08Rik\|Cish5\|mKIAA0671 | suppressor of cytokine signaling 5 | protein-coding |
| chr18_88845609_88845932 | chr18 | 88845610 | 88845932 | + | 0.700394 | Intergenic | Intergenic | 48436 | NM_018821 | 54607 | Mm.91920 | NM_018821 | ENSMUSG00000056153 | **Socs6** | 1500012M23Rik\|5830401B18Rik\|AI447482\|Cis4\|Cish4\|HSPC060\|SOCS-4\|SOCS-6\|SSI4\|STAI4\|STATI4\|Socs4 | suppressor of cytokine signaling 6 | protein-coding |
| chr11_97401041_97401601 | chr11 | 97401042 | 97401601 | + | 0.722333 | Intergenic | Intergenic | 38770 | NM_138657 | 192157 | Mm.34339 | NM_138657 | ENSMUSG00000038485 | **Socs7** | 2310063P06Rik\|C85125\|Nap4 | suppressor of cytokine signaling 7 | protein-coding |
| chr11_97401009_97402056 | chr11 | 97401010 | 97402056 | + | 0.708473 | Intergenic | Intergenic | 38982 | NM_138657 | 192157 | Mm.34339 | NM_138657 | ENSMUSG00000038485 | **Socs7** | 2310063P06Rik\|C85125\|Nap4 | suppressor of cytokine signaling 7 | protein-coding |

| | | | | | | | | | | | | | | | | | |
|---|---|---|---|---|---|---|---|---|---|---|---|---|---|---|---|---|---|
| chr14_70187209_70187564 | chr14 | 70187210 | 70187564 | + | 0.625606 | intron (NM_011366, intron 16 of 20) | ORR1C2\|LTR\|ERVL-MaLR | 4859 | NM_001271409 | 20410 | Mm.5068 | NM_011366 | ENSMUSG00000022091 | **Sorbs3** | SCAM-1\|SH3P3\|Sh3d4\|vinexin-g | sorbin and SH3 domain containing 3 | protein-coding |
| chr9_41967483_41968182 | chr9 | 41967484 | 41968182 | + | 0.979573 | TTS (NM_011436) | TTS (NM_011436) | 156456 | NM_011436 | 20660 | Mm.121920 | NM_011436 | | **Sorl1** | 2900010L19Rik\|AI596264\|AW261561\|LR11\|SorLA\|gp250\|mSorLA | sortilin-related receptor, LDLR class A repeats-containing | protein-coding |
| chr9_42022215_42022713 | chr9 | 42022216 | 42022713 | + | 0.929947 | intron (NM_011436, intron 22 of 47) | intron (NM_011436, intron 22 of 47) | 101825 | NM_011436 | 20660 | Mm.121920 | NM_011436 | | **Sorl1** | 2900010L19Rik\|AI596264\|AW261561\|LR11\|SorLA\|gp250\|mSorLA | sortilin-related receptor, LDLR class A repeats-containing | protein-coding |
| chr9_42022197_42022915 | chr9 | 42022198 | 42022915 | + | 0.838705 | intron (NM_011436, intron 22 of 47) | intron (NM_011436, intron 22 of 47) | 101733 | NM_011436 | 20660 | Mm.121920 | NM_011436 | | **Sorl1** | 2900010L19Rik\|AI596264\|AW261561\|LR11\|SorLA\|gp250\|mSorLA | sortilin-related receptor, LDLR class A repeats-containing | protein-coding |
| chr9_42031301_42031545 | chr9 | 42031302 | 42031545 | + | 0.783203 | intron (NM_011436, intron 20 of 47) | intron (NM_011436, intron 20 of 47) | 92866 | NM_011436 | 20660 | Mm.121920 | NM_011436 | | **Sorl1** | 2900010L19Rik\|AI596264\|AW261561\|LR11\|SorLA\|gp250\|mSorLA | sortilin-related receptor, LDLR class A repeats-containing | protein-coding |
| chr9_41985548_41985957 | chr9 | 41985549 | 41985957 | + | 0.680538 | intron (NM_011436, intron 38 of 47) | intron (NM_011436, intron 38 of 47) | 138536 | NM_011436 | 20660 | Mm.121920 | NM_011436 | | **Sorl1** | 2900010L19Rik\|AI596264\|AW261561\|LR11\|SorLA\|gp250\|mSorLA | sortilin-related receptor, LDLR class A repeats-containing | protein-coding |
| chr10_59236476_59236881 | chr10 | 59236477 | 59236881 | + | 1.13842 | Intergenic | Intergenic | 14757 | NM_172939 | 268301 | Mm.3036 | NM_172939 | ENSMUSG00000098188 | **Sowahc** | 4921515A04Rik\|9430099J10Rik\|Ankrd57\|C820004L04Rik | sosondowah ankyrin repeat domain family member C | protein-coding |
| chr10_59236289_59236875 | chr10 | 59236290 | 59236875 | + | 1.02816 | Intergenic | Intergenic | 14660 | NM_172939 | 268301 | Mm.3036 | NM_172939 | ENSMUSG00000098188 | **Sowahc** | 4921515A04Rik\|9430099J10Rik\|Ankrd57\|C820004L04Rik | sosondowah ankyrin repeat domain family member C | protein-coding |
| chr6_144020004_144020449 | chr6 | 144020005 | 144020449 | + | 1.13851 | intron (NM_001113559, intron 6 of 13) | intron (NM_001113559, intron 6 of 13) | 189341 | NM_001113559 | 20678 | Mm.1752 | NM_011444 | ENSMUSG00000041540 | **Sox5** | A730017D01Rik\|AI528773 | SRY (sex determining region Y)-box 5 | protein-coding |
| chr6_144048148_144048522 | chr6 | 144048149 | 144048522 | + | 0.618798 | intron (NM_011444, intron 3 of 14) | ID_B1\|SINE\|B4 | 161233 | NM_001113559 | 20678 | Mm.1752 | NM_011444 | ENSMUSG00000041540 | **Sox5** | A730017D01Rik\|AI528773 | SRY (sex determining region Y)-box 5 | protein-coding |
| chr6_144688940_144689571 | chr6 | 144688941 | 144689571 | + | 1.94639 | intron (NR_040519, intron 1 of 2) | intron (NR_040519, intron 1 of 2) | 16388 | NR_040519 | 66760 | Mm.233631 | NR_040519 | ENSMUSG00000085785 | **Sox5os3** | 4933425H06Rik | SRY (sex determining region Y)-box 5, opposite strand 3 | ncRNA |
| chr6_144672419_144673003 | chr6 | 144672420 | 144673003 | + | 0.966407 | promoter-TSS (NR_040519) | promoter-TSS (NR_040519) | -157 | NR_040519 | 66760 | Mm.233631 | NR_040519 | ENSMUSG00000085785 | **Sox5os3** | 4933425H06Rik | SRY (sex determining region Y)-box 5, opposite strand 3 | ncRNA |
| chr6_144773059_144773379 | chr6 | 144773060 | 144773379 | + | 0.947236 | Intergenic | ID_B1\|SINE\|B4 | 100351 | NR_040519 | 66760 | Mm.233631 | NR_040519 | ENSMUSG00000085785 | **Sox5os3** | 4933425H06Rik | SRY (sex determining region Y)-box 5, opposite strand 3 | ncRNA |
| chr6_144693580_144693839 | chr6 | 144693581 | 144693839 | + | 0.866382 | non-coding (NR_040519, exon 3 of 3) | non-coding (NR_040519, exon 3 of 3) | 20842 | NR_040519 | 66760 | Mm.233631 | NR_040519 | ENSMUSG00000085785 | **Sox5os3** | 4933425H06Rik | SRY (sex determining region Y)-box 5, opposite strand 3 | ncRNA |
| chr6_144772590_144773400 | chr6 | 144772591 | 144773400 | + | 0.842412 | Intergenic | Intergenic | 100127 | NR_040519 | 66760 | Mm.233631 | NR_040519 | ENSMUSG00000085785 | **Sox5os3** | 4933425H06Rik | SRY (sex determining region Y)-box 5, opposite strand 3 | ncRNA |
| chr6_144804336_144804859 | chr6 | 144804337 | 144804859 | + | 0.839813 | Intergenic | Intergenic | 131730 | NR_040519 | 66760 | Mm.233631 | NR_040519 | ENSMUSG00000085785 | **Sox5os3** | 4933425H06Rik | SRY (sex determining region Y)-box 5, opposite strand 3 | ncRNA |
| chr6_144804158_144804840 | chr6 | 144804159 | 144804840 | + | 0.77997 | Intergenic | Intergenic | 131631 | NR_040519 | 66760 | Mm.233631 | NR_040519 | ENSMUSG00000085785 | **Sox5os3** | 4933425H06Rik | SRY (sex determining region Y)-box 5, opposite strand 3 | ncRNA |
| chr6_144640027_144641269 | chr6 | 144640028 | 144641269 | + | 0.769105 | Intergenic | Intergenic | -32220 | NR_040519 | 66760 | Mm.233631 | NR_040519 | ENSMUSG00000085785 | **Sox5os3** | 4933425H06Rik | SRY (sex determining region Y)-box 5, opposite strand 3 | ncRNA |
| chr7_115494418_115495443 | chr7 | 115494419 | 115495443 | + | 2.07431 | intron (NM_001277328, intron 11 of 13) | intron (NM_001277328, intron 11 of 13) | 306678 | NM_001277328 | 20679 | Mm.323365 | NM_011445 | ENSMUSG00000051910 | **Sox6** | AI987981\|SOX-LZ | SRY (sex determining region Y)-box 6 | protein-coding |
| chr7_115494398_115495876 | chr7 | 115494399 | 115495876 | + | 2.04216 | intron (NM_001277328, intron 11 of 13) | intron (NM_001277328, intron 11 of 13) | 306472 | NM_001277328 | 20679 | Mm.323365 | NM_011445 | ENSMUSG00000051910 | **Sox6** | AI987981\|SOX-LZ | SRY (sex determining region Y)-box 6 | protein-coding |
| chr7_115474050_115474576 | chr7 | 115474051 | 115474576 | + | 1.49895 | 3' UTR (NM_011445, exon 17 of 17) | 3' UTR (NM_011445, exon 17 of 17) | 327296 | NM_001277328 | 20679 | Mm.323365 | NM_011445 | ENSMUSG00000051910 | **Sox6** | AI987981\|SOX-LZ | SRY (sex determining region Y)-box 6 | protein-coding |
| chr7_115448403_115449037 | chr7 | 115448404 | 115449037 | + | 1.23728 | Intergenic | Intergenic | 352889 | NM_001277328 | 20679 | Mm.323365 | NM_011445 | ENSMUSG00000051910 | **Sox6** | AI987981\|SOX-LZ | SRY (sex determining region Y)-box 6 | protein-coding |
| chr7_115448647_115449033 | chr7 | 115448648 | 115449033 | + | 1.21501 | Intergenic | Intergenic | 352769 | NM_001277328 | 20679 | Mm.323365 | NM_011445 | ENSMUSG00000051910 | **Sox6** | AI987981\|SOX-LZ | SRY (sex determining region Y)-box 6 | protein-coding |
| chr7_115495586_115495876 | chr7 | 115495587 | 115495876 | + | 1.14384 | intron (NM_001277328, intron 11 of 13) | intron (NM_001277328, intron 11 of 13) | 305878 | NM_001277328 | 20679 | Mm.323365 | NM_011445 | ENSMUSG00000051910 | **Sox6** | AI987981\|SOX-LZ | SRY (sex determining region Y)-box 6 | protein-coding |
| chr7_115476992_115477384 | chr7 | 115476993 | 115477384 | + | 0.698523 | exon (NM_011445, exon 17 of 17) | exon (NM_011445, exon 17 of 17) | 324421 | NM_001277328 | 20679 | Mm.323365 | NM_011445 | ENSMUSG00000051910 | **Sox6** | AI987981\|SOX-LZ | SRY (sex determining region Y)-box 6 | protein-coding |
| chr7_115494006_115494205 | chr7 | 115494007 | 115494205 | + | 0.690536 | intron (NM_001277328, intron 11 of 13) | intron (NM_001277328, intron 11 of 13) | 307503 | NM_001277328 | 20679 | Mm.323365 | NM_011445 | ENSMUSG00000051910 | **Sox6** | AI987981\|SOX-LZ | SRY (sex determining region Y)-box 6 | protein-coding |
| chr1_85644202_85644484 | chr1 | 85644203 | 85644484 | + | 0.677649 | intron (NM_001013817, intron 16 of 16) | B4A\|SINE\|B4 | -5707 | NM_013673 | 20684 | Mm.290906 | NM_013673 | ENSMUSG00000026222 | **Sp100** | A430075G10Rik | nuclear antigen Sp100 | protein-coding |
| chr2_72961744_72962120 | chr2 | 72961745 | 72962120 | + | 0.910283 | intron (NM_001098425, intron 2 of 4) | intron (NM_001098425, intron 2 of 4) | 17349 | NM_001098425 | 20687 | Mm.124328 | NM_011450 | ENSMUSG00000027109 | **Sp3** | D130027J01Rik | trans-acting transcription factor 3 | protein-coding |
| chr12_118358906_118359949 | chr12 | 118358907 | 118359949 | + | 1.22281 | Intergenic | Intergenic | -57988 | NM_001166385 | 20688 | Mm.34699 | NM_009239 | ENSMUSG00000025323 | **Sp4** | 5730497N03Rik\|HF-1b\|HF1-b | trans-acting transcription factor 4 | protein-coding |
| chr12_118372000_118372240 | chr12 | 118372001 | 118372240 | + | 0.688272 | Intergenic | Intergenic | -70680 | NM_001166385 | 20688 | Mm.34699 | NM_009239 | ENSMUSG00000025323 | **Sp4** | 5730497N03Rik\|HF-1b\|HF1-b | trans-acting transcription factor 4 | protein-coding |
| chr12_118855014_118855499 | chr12 | 118855015 | 118855499 | + | 0.881241 | Intergenic | Intergenic | 8928 | NM_177082 | 320145 | Mm.51682 | NM_177082 | ENSMUSG00000048562 | **Sp8** | D930049B17Rik\|mBtd | trans-acting transcription factor 8 | protein-coding |
| chr12_118849641_118850248 | chr12 | 118849642 | 118850248 | + | 0.692859 | 3' UTR (NM_177082, exon 3 of 3) | 3' UTR (NM_177082, exon 3 of 3) | 3616 | NM_177082 | 320145 | Mm.51682 | NM_177082 | ENSMUSG00000048562 | **Sp8** | D930049B17Rik\|mBtd | trans-acting transcription factor 8 | protein-coding |
| chr2_73247122_73247609 | chr2 | 73247123 | 73247609 | + | 0.809839 | Intergenic | ORR1D2\|LTR\|ERVL-MaLR | -24560 | NM_001005343 | 381373 | Mm.330293 | NM_001005343 | | **Sp9** | - | trans-acting transcription factor 9 | protein-coding |
| chr11_94098863_94099363 | chr11 | 94098864 | 94099363 | + | 1.30813 | exon (NM_001199203, exon 22 of 30) | exon (NM_001199203, exon 22 of 30) | 54908 | NM_001025429 | 70834 | Mm.260737 | NM_027569 | ENSMUSG00000020859 | **Spag9** | 3110018C07Rik\|4733401I23Rik\|4831406C20Rik\|AW552012\|JLP\|JSAP2\|JSAP2a\|Jip4\|Mapk8ip4\|syd1 | sperm associated antigen 9 | protein-coding |
| chr17_74354765_74355408 | chr17 | 74354766 | 74355408 | + | 0.759225 | intron (NM_016962, intron 2 of 16) | intron (NM_016962, intron 2 of 16) | 16100 | NM_016962 | 50850 | Mm.19804 | NM_016962 | ENSMUSG00000024068 | **Spast** | Spg4\|mKIAA1083 | spastin | protein-coding |
| chr17_74354245_74355419 | chr17 | 74354246 | 74355419 | + | 0.6979 | intron (NM_016962, intron 2 of 16) | intron (NM_016962, intron 2 of 16) | 15845 | NM_016962 | 50850 | Mm.19804 | NM_016962 | ENSMUSG00000024068 | **Spast** | Spg4\|mKIAA1083 | spastin | protein-coding |
| chr14_60524851_60525206 | chr14 | 60524852 | 60525206 | + | 0.76856 | Intergenic | Intergenic | -109700 | NM_001033272 | 219140 | Mm.149776 | NM_001033272 | ENSMUSG00000021990 | **Spata13** | C79050\|ESTM11\|X83327\|mFLJ00298 | spermatogenesis associated 13 | protein-coding |
| chr2_167496364_167496721 | chr2 | 167496365 | 167496721 | + | 1.39788 | intron (NM_030743, intron 1 of 7) | intron (NM_030743, intron 1 of 7) | -3669 | NM_170756 | 263876 | Mm.34342 | NM_170756 | ENSMUSG00000047030 | **Spata2** | AI504642\|mKIAA0757 | spermatogenesis associated 2 | protein-coding |
| chr1_86006060_86006760 | chr1 | 86006061 | 86006760 | + | 1.01842 | Intergenic | Intergenic | -15532 | NM_001122732 | 70060 | Mm.196313 | NM_027029 | ENSMUSG00000026226 | **Spata3** | 1700011N12Rik\|1700029H01Rik\|4930424D10Rik\|Mtsarg1\|Tsarg1 | spermatogenesis associated 3 | protein-coding |
| chr1_86000924_86001151 | chr1 | 86000925 | 86001151 | + | 0.847572 | Intergenic | L1MC5a\|LINE\|L1 | -20904 | NM_001122732 | 70060 | Mm.196313 | NM_027029 | ENSMUSG00000026226 | **Spata3** | 1700011N12Rik\|1700029H01Rik\|4930424D10Rik\|Mtsarg1\|Tsarg1 | spermatogenesis associated 3 | protein-coding |
| chr13_59713683_59714206 | chr13 | 59713684 | 59714206 | + | 1.06955 | intron (NM_001167593, intron 2 of 3) | intron (NM_001167593, intron 2 of 3) | 1661 | NM_001167593 | 238662 | Mm.295939 | NM_001167593 | ENSMUSG00000091311 | **Spata31d1b** | EG238662\|Fam75d1b\|Gm4934 | spermatogenesis associated 31 subfamily D, member 1B | protein-coding |
| chr3_37456014_37456627 | chr3 | 37456015 | 37456627 | + | 1.38565 | intron (NM_001163511, intron 11 of 15) | intron (NM_001163511, intron 11 of 15) | 36041 | NM_021343 | 57815 | Mm.172679 | NM_021343 | ENSMUSG00000027722 | **Spata5** | 2510048F20Rik\|C78064\|Spaf | spermatogenesis associated 5 | protein-coding |
| chr3_37479812_37480356 | chr3 | 37479813 | 37480356 | + | 1.03459 | intron (NM_001163511, intron 14 of 15) | intron (NM_001163511, intron 14 of 15) | 59804 | NM_021343 | 57815 | Mm.172679 | NM_021343 | ENSMUSG00000027722 | **Spata5** | 2510048F20Rik\|C78064\|Spaf | spermatogenesis associated 5 | protein-coding |
| chr3_37434223_37434884 | chr3 | 37434224 | 37434884 | + | 0.78672 | intron (NM_001163511, intron 7 of 15) | MTE2a\|LTR\|ERVL-MaLR | 14274 | NM_021343 | 57815 | Mm.172679 | NM_021343 | ENSMUSG00000027722 | **Spata5** | 2510048F20Rik\|C78064\|Spaf | spermatogenesis associated 5 | protein-coding |
| chr12_98605155_98605593 | chr12 | 98605156 | 98605593 | + | 1.35364 | Intergenic | Intergenic | -22768 | NM_001289573 | 104871 | Mm.210053 | NM_178914 | ENSMUSG00000021007 | **Spata7** | AI661438\|B230306G18Rik\|HSD3 | spermatogenesis associated 7 | protein-coding |
| chr8_54534545_54534841 | chr8 | 54534546 | 54534841 | + | 1.23874 | Intergenic | ORR1B2\|LTR\|ERVL-MaLR | -4695 | NM_029701 | 76687 | Mm.86437 | NM_029701 | ENSMUSG00000054408 | **Spcs3** | 1810011E08Rik | signal peptidase complex subunit 3 homolog (S. cerevisiae) | protein-coding |
| chr11_34879623_34880201 | chr11 | 34879624 | 34880201 | + | 0.630683 | Intergenic | Intergenic | -46271 | NM_027411 | 70385 | Mm.194368 | NM_027411 | ENSMUSG00000069910 | **Spdl1** | 1700018I02Rik\|2600001J17Rik\|2810049B11Rik\|AA409762\|Ccdc99 | spindle apparatus coiled-coil protein 1 | protein-coding |

| | | | | | | | | | | | | | | | | | |
|---|---|---|---|---|---|---|---|---|---|---|---|---|---|---|---|---|---|
| chr11_62148181_62149063 | chr11 | 62148182 | 62149063 | + | 1.2585 | intron (NM_001281818, intron 9 of 14) | intron (NM_001281818, intron 9 of 14) | 71598 | NM_001029936 | 432572 | Mm.458025 | NM_001029936 | ENSMUSG00000042331 | **Specc1** | 2810012G08Rik\|B230396K10Rik\|Cytsb | sperm antigen with calponin homology and coiled-coil domains 1 | protein-coding |
| chr11_62007438_62007787 | chr11 | 62007439 | 62007787 | + | 1.0422 | intron (NM_001281818, intron 1 of 14) | intron (NM_001281818, intron 1 of 14) | 50850 | NM_001281818 | 432572 | Mm.458025 | NM_001029936 | ENSMUSG00000042331 | **Specc1** | 2810012G08Rik\|B230396K10Rik\|Cytsb | sperm antigen with calponin homology and coiled-coil domains 1 | protein-coding |
| chr11_62005219_62006208 | chr11 | 62005220 | 62006208 | + | 0.812613 | intron (NM_001281818, intron 1 of 14) | intron (NM_001281818, intron 1 of 14) | 48951 | NM_001281818 | 432572 | Mm.458025 | NM_001029936 | ENSMUSG00000042331 | **Specc1** | 2810012G08Rik\|B230396K10Rik\|Cytsb | sperm antigen with calponin homology and coiled-coil domains 1 | protein-coding |
| chr11_62090336_62090659 | chr11 | 62090337 | 62090659 | + | 0.731212 | intron (NM_001029936, intron 1 of 12) | intron (NM_001029936, intron 1 of 12) | 13474 | NM_001029936 | 432572 | Mm.458025 | NM_001029936 | ENSMUSG00000042331 | **Specc1** | 2810012G08Rik\|B230396K10Rik\|Cytsb | sperm antigen with calponin homology and coiled-coil domains 1 | protein-coding |
| chr11_62092465_62093021 | chr11 | 62092466 | 62093021 | + | 0.639956 | intron (NM_001029936, intron 1 of 12) | intron (NM_001029936, intron 1 of 12) | 15719 | NM_001029936 | 432572 | Mm.458025 | NM_001029936 | ENSMUSG00000042331 | **Specc1** | 2810012G08Rik\|B230396K10Rik\|Cytsb | sperm antigen with calponin homology and coiled-coil domains 1 | protein-coding |
| chr10_75222203_75223080 | chr10 | 75222204 | 75223080 | + | 1.00314 | intron (NM_001145826, intron 2 of 18) | intron (NM_001145826, intron 2 of 18) | 10252 | NM_001145826 | 74392 | Mm.27716 | NM_153406 | ENSMUSG00000033444 | **Specc1l** | 4930470P14Rik\|4932439K10Rik\|9530057A13Rik\|AI317249\|Cytsa\|mKIAA0376 | sperm antigen with calponin homology and coiled-coil domains 1-like | protein-coding |
| chr10_75170282_75170935 | chr10 | 75170283 | 75170935 | + | 0.725669 | intron (NM_001081412, intron 16 of 22) | intron (NM_001081412, intron 16 of 22) | -41464 | NM_153406 | 74392 | Mm.27716 | NM_153406 | ENSMUSG00000033444 | **Specc1l** | 4930470P14Rik\|4932439K10Rik\|9530057A13Rik\|AI317249\|Cytsa\|mKIAA0376 | sperm antigen with calponin homology and coiled-coil domains 1-like | protein-coding |
| chr15_9869964_9870711 | chr15 | 9869965 | 9870711 | + | 1.82507 | Intergenic | Intergenic | -121532 | NM_177123 | 320277 | Mm.127822 | NM_177123 | ENSMUSG00000072663 | **Spef2** | C230086A09Rik | sperm flagellar 2 | protein-coding |
| chr15_9713487_9714051 | chr15 | 9713488 | 9714051 | + | 0.834458 | intron (NM_177123, intron 7 of 17) | intron (NM_177123, intron 7 of 17) | 35037 | NM_177123 | 320277 | Mm.127822 | NM_177123 | ENSMUSG00000072663 | **Spef2** | C230086A09Rik | sperm flagellar 2 | protein-coding |
| chr15_9667564_9668119 | chr15 | 9667565 | 9668119 | + | 0.667244 | intron (NM_177123, intron 16 of 17) | intron (NM_177123, intron 16 of 17) | 80964 | NM_177123 | 320277 | Mm.127822 | NM_177123 | ENSMUSG00000072663 | **Spef2** | C230086A09Rik | sperm flagellar 2 | protein-coding |
| chr13_51107397_51107820 | chr13 | 51107398 | 51107820 | + | 0.91782 | intron (NM_001283029, intron 1 of 5) | intron (NM_001283029, intron 1 of 5) | 6062 | NM_001283029 | 20729 | Mm.188432 | NM_011462 | ENSMUSG00000021395 | **Spin1** | Spin | spindlin 1 | protein-coding |
| chrX_95077000_95077570 | chrX | 95077001 | 95077570 | + | 1.48968 | intron (NM_001290385, intron 6 of 9) | intron (NM_001290385, intron 6 of 9) | -50603 | NM_178753 | 270624 | Mm.115205 | NM_178753 | ENSMUSG00000071722 | **Spin4** | 9630042H07Rik | spindlin family, member 4 | protein-coding |
| chrX_95074518_95075046 | chrX | 95074519 | 95075046 | + | 1.34252 | intron (NM_001290385, intron 6 of 9) | intron (NM_001290385, intron 6 of 9) | -48100 | NM_178753 | 270624 | Mm.115205 | NM_178753 | ENSMUSG00000071722 | **Spin4** | 9630042H07Rik | spindlin family, member 4 | protein-coding |
| chrX_95087717_95088345 | chrX | 95087718 | 95088345 | + | 0.699096 | intron (NM_001290385, intron 4 of 9) | Lx9\|LINE\|L1 | -61349 | NM_178753 | 270624 | Mm.115205 | NM_178753 | ENSMUSG00000071722 | **Spin4** | 9630042H07Rik | spindlin family, member 4 | protein-coding |
| chr18_62717652_62718024 | chr18 | 62717653 | 62718024 | + | 0.866825 | intron (NM_001168423, intron 1 of 4) | ORR1F\|LTR\|ERVL-MaLR | 23549 | NM_001168423 | 100038417 | Mm.387557 | NM_001168423 | ENSMUSG00000073551 | **Spink13** | ENSMUSG00000073551\|Gm10534\|Spink5l3 | serine peptidase inhibitor, Kazal type 13 | protein-coding |
| chr5_77217693_77218276 | chr5 | 77217694 | 77218276 | + | 1.6334 | Intergenic | MLTR11B\|LTR\|ERVK | -6514 | NM_001289768 | 69982 | Mm.46106 | NM_183284 | ENSMUSG00000053030 | **Spink2** | 1700007F22Rik\|AV038945\|HUSI-II | serine peptidase inhibitor, Kazal type 2 | protein-coding |
| chr5_77174733_77175147 | chr5 | 77174734 | 77175147 | + | 1.10701 | Intergenic | Intergenic | 36265 | NM_001289767 | 69982 | Mm.46106 | NM_183284 | ENSMUSG00000053030 | **Spink2** | 1700007F22Rik\|AV038945\|HUSI-II | serine peptidase inhibitor, Kazal type 2 | protein-coding |
| chr18_67536603_67536946 | chr18 | 67536604 | 67536946 | + | 0.635179 | intron (NM_176832, intron 2 of 13) | intron (NM_176832, intron 2 of 13) | 12398 | NM_176832 | 68166 | Mm.208723 | NM_176832 | ENSMUSG00000024533 | **Spire1** | 6030430B19Rik\|AI415299\|AU022898\|AW550622\|Spir-1 | spire homolog 1 (Drosophila) | protein-coding |
| chr7_127153401_127154146 | chr7 | 127153402 | 127154146 | + | 1.64438 | Intergenic | RLTR42-int\|LTR\|ERVK | -15951 | NM_001037810 | 20737 | Mm.283714 | NM_009259 | ENSMUSG00000051457 | **Spn** | A630014B01Rik\|Cd43\|Galgp\|Ly-48\|Ly48 | sialophorin | protein-coding |
| chr7_127163485_127163959 | chr7 | 127163486 | 127163959 | + | 0.955985 | Intergenic | MERV1_LTR\|LTR\|ERV1 | -25899 | NM_001037810 | 20737 | Mm.283714 | NM_009259 | ENSMUSG00000051457 | **Spn** | A630014B01Rik\|Cd43\|Galgp\|Ly-48\|Ly48 | sialophorin | protein-coding |
| chr11_72491113_72491460 | chr11 | 72491114 | 72491460 | + | 0.689796 | Intergenic | Intergenic | -1367 | NM_153060 | 216892 | Mm.212812 | NM_153060 | ENSMUSG00000040447 | **Spns2** | - | spinster homolog 2 | protein-coding |
| chr2_172971813_172972344 | chr2 | 172971814 | 172972344 | + | 1.96974 | Intergenic | Intergenic | -7763 | NM_012046 | 26972 | Mm.23495 | NM_012046 | ENSMUSG00000005883 | **Spo11** | AI449549 | SPO11 meiotic protein covalently bound to DSB homolog (S. cerevisiae) | protein-coding |
| chr5_115045136_115045670 | chr5 | 115045137 | 115045670 | + | 0.945345 | intron (NM_029012, intron 1 of 10) | intron (NM_029012, intron 1 of 10) | 33879 | NM_029012 | 74585 | Mm.355894 | NM_029012 | ENSMUSG00000029550 | **Sppl3** | 4833416I09Rik\|Psl4\|Usmg3 | signal peptide peptidase 3 | protein-coding |
| chr5_115043941_115044321 | chr5 | 115043942 | 115044321 | + | 0.867849 | intron (NM_029012, intron 1 of 10) | intron (NM_029012, intron 1 of 10) | 32607 | NM_029012 | 74585 | Mm.355894 | NM_029012 | ENSMUSG00000029550 | **Sppl3** | 4833416I09Rik\|Psl4\|Usmg3 | signal peptide peptidase 3 | protein-coding |
| chr5_115002702_115003769 | chr5 | 115002703 | 115003769 | + | 0.679067 | Intergenic | Intergenic | -8288 | NM_029012 | 74585 | Mm.355894 | NM_029012 | ENSMUSG00000029550 | **Sppl3** | 4833416I09Rik\|Psl4\|Usmg3 | signal peptide peptidase 3 | protein-coding |
| chr2_117023250_117023852 | chr2 | 117023251 | 117023852 | + | 1.04668 | Intergenic | Intergenic | -97520 | NM_001277256 | 114715 | Mm.245890 | NM_033524 | ENSMUSG00000027351 | **Spred1** | 5730461F13Rik\|AL024345\|AW047647 | sprouty protein with EVH-1 domain 1, related sequence | protein-coding |
| chr2_117105215_117105832 | chr2 | 117105216 | 117105832 | + | 0.806464 | Intergenic | MIR\|SINE\|MIR | -15547 | NM_001277256 | 114715 | Mm.245890 | NM_033524 | ENSMUSG00000027351 | **Spred1** | 5730461F13Rik\|AL024345\|AW047647 | sprouty protein with EVH-1 domain 1, related sequence | protein-coding |
| chr2_117021820_117022237 | chr2 | 117021821 | 117022237 | + | 0.762149 | Intergenic | Intergenic | -99042 | NM_001277256 | 114715 | Mm.245890 | NM_033524 | ENSMUSG00000027351 | **Spred1** | 5730461F13Rik\|AL024345\|AW047647 | sprouty protein with EVH-1 domain 1, related sequence | protein-coding |
| chr2_117151219_117151479 | chr2 | 117151220 | 117151479 | + | 0.674789 | intron (NM_033524, intron 1 of 6) | B3A\|SINE\|B2 | 30278 | NM_033524 | 114715 | Mm.245890 | NM_033524 | ENSMUSG00000027351 | **Spred1** | 5730461F13Rik\|AL024345\|AW047647 | sprouty protein with EVH-1 domain 1, related sequence | protein-coding |
| chr2_117018361_117018813 | chr2 | 117018362 | 117018813 | + | 0.674393 | Intergenic | Intergenic | -102484 | NM_001277256 | 114715 | Mm.245890 | NM_033524 | ENSMUSG00000027351 | **Spred1** | 5730461F13Rik\|AL024345\|AW047647 | sprouty protein with EVH-1 domain 1, related sequence | protein-coding |
| chr2_117121514_117122234 | chr2 | 117121515 | 117122234 | + | 0.60317 | intron (NM_033524, intron 1 of 6) | CpG | 803 | NM_033524 | 114715 | Mm.245890 | NM_033524 | ENSMUSG00000027351 | **Spred1** | 5730461F13Rik\|AL024345\|AW047647 | sprouty protein with EVH-1 domain 1, related sequence | protein-coding |
| chr11_19922249_19922602 | chr11 | 19922250 | 19922602 | + | 1.90182 | Intergenic | Intergenic | -2016 | NM_033523 | 114716 | Mm.266627 | NM_033523 | ENSMUSG00000045671 | **Spred2** | C79158 | sprouty-related, EVH1 domain containing 2 | protein-coding |
| chr11_19950898_19951331 | chr11 | 19950899 | 19951331 | + | 0.972235 | intron (NM_033523, intron 1 of 5) | intron (NM_033523, intron 1 of 5) | 26673 | NM_033523 | 114716 | Mm.266627 | NM_033523 | ENSMUSG00000045671 | **Spred2** | C79158 | sprouty-related, EVH1 domain containing 2 | protein-coding |
| chr11_19961211_19961536 | chr11 | 19961212 | 19961536 | + | 0.828554 | intron (NM_033523, intron 1 of 5) | intron (NM_033523, intron 1 of 5) | 36932 | NM_033523 | 114716 | Mm.266627 | NM_033523 | ENSMUSG00000045671 | **Spred2** | C79158 | sprouty-related, EVH1 domain containing 2 | protein-coding |
| chr11_19956476_19956998 | chr11 | 19956477 | 19956998 | + | 0.635191 | intron (NM_033523, intron 1 of 5) | B3A\|SINE\|B2 | 32295 | NM_033523 | 114716 | Mm.266627 | NM_033523 | ENSMUSG00000045671 | **Spred2** | C79158 | sprouty-related, EVH1 domain containing 2 | protein-coding |
| chr3_37680193_37681375 | chr3 | 37680194 | 37681375 | + | 1.55958 | Intergenic | B4\|SINE\|B4 | 40834 | NM_011896 | 24063 | Mm.330986 | NM_011896 | ENSMUSG00000037211 | **Spry1** | sprouty1 | sprouty homolog 1 (Drosophila) | protein-coding |
| chr3_37649886_37650385 | chr3 | 37649887 | 37650385 | + | 1.47072 | Intergenic | ORR1D2\|LTR\|ERVL-MaLR | 10186 | NM_011896 | 24063 | Mm.330986 | NM_011896 | ENSMUSG00000037211 | **Spry1** | sprouty1 | sprouty homolog 1 (Drosophila) | protein-coding |
| chr3_37645107_37645476 | chr3 | 37645108 | 37645476 | + | 0.955213 | TTS (NM_011896) | TTS (NM_011896) | 5342 | NM_011896 | 24063 | Mm.330986 | NM_011896 | ENSMUSG00000037211 | **Spry1** | sprouty1 | sprouty homolog 1 (Drosophila) | protein-coding |
| chr3_37674784_37675811 | chr3 | 37674785 | 37675811 | + | 0.852384 | Intergenic | Intergenic | 35348 | NM_011896 | 24063 | Mm.330986 | NM_011896 | ENSMUSG00000037211 | **Spry1** | sprouty1 | sprouty homolog 1 (Drosophila) | protein-coding |
| chr3_37654010_37654835 | chr3 | 37654011 | 37654835 | + | 0.795142 | Intergenic | Intergenic | 14473 | NM_011896 | 24063 | Mm.330986 | NM_011896 | ENSMUSG00000037211 | **Spry1** | sprouty1 | sprouty homolog 1 (Drosophila) | protein-coding |
| chr3_37672343_37672663 | chr3 | 37672344 | 37672663 | + | 0.649446 | Intergenic | Intergenic | 32553 | NM_011896 | 24063 | Mm.330986 | NM_011896 | ENSMUSG00000037211 | **Spry1** | sprouty1 | sprouty homolog 1 (Drosophila) | protein-coding |
| chr14_105922851_105923485 | chr14 | 105922852 | 105923485 | + | 1.18957 | Intergenic | Intergenic | -26349 | NM_011897 | 24064 | Mm.89982 | NM_011897 | ENSMUSG00000022114 | **Spry2** | sprouty2 | sprouty homolog 2 (Drosophila) | protein-coding |
| chr14_105805307_105805763 | chr14 | 105805308 | 105805763 | + | 0.852761 | Intergenic | Intergenic | 91284 | NM_011897 | 24064 | Mm.89982 | NM_011897 | ENSMUSG00000022114 | **Spry2** | sprouty2 | sprouty homolog 2 (Drosophila) | protein-coding |
| chr14_105922516_105922787 | chr14 | 105922517 | 105922787 | + | 0.852143 | Intergenic | Intergenic | -25833 | NM_011897 | 24064 | Mm.89982 | NM_011897 | ENSMUSG00000022114 | **Spry2** | sprouty2 | sprouty homolog 2 (Drosophila) | protein-coding |
| chr4_149914442_149915049 | chr4 | 149914443 | 149915049 | + | 1.08771 | intron (NM_029035, intron 1 of 2) | intron (NM_029035, intron 1 of 2) | 40260 | NM_029035 | 74646 | Mm.30 | NM_029035 | ENSMUSG00000039911 | **Spsb1** | 1110014L01Rik\|4930422J18Rik\|AI596360\|AI854583\|SSB-1\|SSB1 | splA/ryanodine receptor domain and SOCS box containing 1 | protein-coding |
| chr4_149917955_149918272 | chr4 | 149917956 | 149918272 | + | 0.908519 | intron (NM_029035, intron 1 of 2) | B4A\|SINE\|B4 | 36892 | NM_029035 | 74646 | Mm.30 | NM_029035 | ENSMUSG00000039911 | **Spsb1** | 1110014L01Rik\|4930422J18Rik\|AI596360\|AI854583\|SSB-1\|SSB1 | splA/ryanodine receptor domain and SOCS box containing 1 | protein-coding |
| chr4_149941825_149942101 | chr4 | 149941826 | 149942101 | + | 0.806554 | intron (NM_029035, intron 1 of 2) | intron (NM_029035, intron 1 of 2) | 13043 | NM_029035 | 74646 | Mm.30 | NM_029035 | ENSMUSG00000039911 | **Spsb1** | 1110014L01Rik\|4930422J18Rik\|AI596360\|AI854583\|SSB-1\|SSB1 | splA/ryanodine receptor domain and SOCS box containing 1 | protein-coding |
| chr4_149896102_149896538 | chr4 | 149896103 | 149896538 | + | 0.751636 | TTS (NM_029035) | TTS (NM_029035) | 58686 | NM_029035 | 74646 | Mm.30 | NM_029035 | ENSMUSG00000039911 | **Spsb1** | 1110014L01Rik\|4930422J18Rik\|AI596360\|AI854583\|SSB-1\|SSB1 | splA/ryanodine receptor domain and SOCS box containing 1 | protein-coding |
| chr4_149914066_149915024 | chr4 | 149914067 | 149915024 | + | 0.744589 | intron (NM_029035, intron 1 of 2) | intron (NM_029035, intron 1 of 2) | 40461 | NM_029035 | 74646 | Mm.30 | NM_029035 | ENSMUSG00000039911 | **Spsb1** | 1110014L01Rik\|4930422J18Rik\|AI596360\|AI854583\|SSB-1\|SSB1 | splA/ryanodine receptor domain and SOCS box containing 1 | protein-coding |
| chr1_174180849_174181948 | chr1 | 174180850 | 174181948 | + | 0.739168 | intron (NM_011465, intron 6 of 51) | intron (NM_011465, intron 6 of 51) | 8623 | NM_011465 | 20739 | Mm.200611 | NM_011465 | ENSMUSG00000026532 | **Spta1** | AF093576\|AI451697\|Spna-1\|Spna1\|ha\|ihj\|nmf4\|sph | spectrin alpha, erythrocytic 1 | protein-coding |
| chr13_53374068_53374625 | chr13 | 53374069 | 53374625 | + | 1.23408 | intron (NM_009269, intron 1 of 14) | intron (NM_009269, intron 1 of 14) | 3014 | NM_009269 | 268656 | Mm.240336 | NM_009269 | ENSMUSG00000021468 | **Sptlc1** | AW552086\|C77762\|E030036H05\|Lcb1 | serine palmitoyltransferase, long chain base subunit 1 | protein-coding |

| | | | | | | | | | | | | | | | | | |
|---|---|---|---|---|---|---|---|---|---|---|---|---|---|---|---|---|---|
| chr7_47022280_47022954 | chr7 | 47022281 | 47022954 | + | 0.760678 | Intergenic | URR1B\|DNA\|hAT-Charlie | -14203 | NM_175318 | 101685 | Mm.155687 | NM_175318 | ENSMUSG00000049516 | **Spty2d1** | 5830435K17Rik\|AI852426\|P16H6 | SPT2, Suppressor of Ty, domain containing 1 (S. cerevisiae) | protein-coding |
| chr15_59338476_59338902 | chr15 | 59338477 | 59338902 | + | 1.42241 | intron (NM_153548, intron 25 of 28) | intron (NM_153548, intron 25 of 28) | 23597 | NM_009270 | 20775 | Mm.296169 | NM_009270 | ENSMUSG00000022351 | **Sqle** | AI323792 | squalene epoxidase | protein-coding |
| chr11_50213024_50213360 | chr11 | 50213025 | 50213360 | + | 0.780415 | Intergenic | Intergenic | -2372 | NM_011018 | 18412 | Mm.40828 | NM_011018 | ENSMUSG00000015837 | **Sqstm1** | A170\|OSF-6\|Osi\|STAP\|STONE14\|p62 | sequestosome 1 | protein-coding |
| chr11_50213024_50213731 | chr11 | 50213025 | 50213731 | + | 0.777636 | Intergenic | Intergenic | -2558 | NM_011018 | 18412 | Mm.40828 | NM_011018 | ENSMUSG00000015837 | **Sqstm1** | A170\|OSF-6\|Osi\|STAP\|STONE14\|p62 | sequestosome 1 | protein-coding |
| chr17_86134521_86134958 | chr17 | 86134522 | 86134958 | + | 0.967269 | intron (NM_030133, intron 4 of 20) | intron (NM_030133, intron 4 of 20) | 10435 | NM_030133 | 78586 | Mm.146236 | NM_030133 | ENSMUSG00000024135 | **Srbd1** | AI461933\|C85414\|D530025C17Rik | S1 RNA binding domain 1 | protein-coding |
| chr17_86052338_86053066 | chr17 | 86052339 | 86053066 | + | 0.941358 | intron (NM_030133, intron 15 of 20) | Lx7\|LINE\|L1 | 92473 | NM_030133 | 78586 | Mm.146236 | NM_030133 | ENSMUSG00000024135 | **Srbd1** | AI461933\|C85414\|D530025C17Rik | S1 RNA binding domain 1 | protein-coding |
| chr2_157400539_157401093 | chr2 | 157400540 | 157401093 | + | 1.0146 | Intergenic | Intergenic | -23477 | NM_009271 | 20779 | Mm.22845 | NM_009271 | ENSMUSG00000027646 | **Src** | AW259666\|pp60c-src | Rous sarcoma oncogene | protein-coding |
| chr2_157400265_157401115 | chr2 | 157400266 | 157401115 | + | 0.800349 | Intergenic | Intergenic | -23603 | NM_009271 | 20779 | Mm.22845 | NM_009271 | ENSMUSG00000027646 | **Src** | AW259666\|pp60c-src | Rous sarcoma oncogene | protein-coding |
| chr13_104825527_104825871 | chr13 | 104825528 | 104825871 | + | 1.28531 | intron (NM_026075, intron 1 of 4) | intron (NM_026075, intron 1 of 4) | 8460 | NM_026075 | 67288 | Mm.125465 | NM_026075 | | **Srek1ip1** | 3110031B13Rik\|AI462702\|AV348305\|AW121048\|Sfrs12ip1\|Srsf12ip1 | splicing regulatory glutamine/lysine-rich protein 1interacting protein 1 | protein-coding |
| chr13_104829481_104830100 | chr13 | 104829482 | 104830100 | + | 0.683038 | intron (NM_026075, intron 2 of 4) | MTE2b\|LTR\|ERVL-MaLR | 12552 | NM_026075 | 67288 | Mm.125465 | NM_026075 | | **Srek1ip1** | 3110031B13Rik\|AI462702\|AV348305\|AW121048\|Sfrs12ip1\|Srsf12ip1 | splicing regulatory glutamine/lysine-rich protein 1interacting protein 1 | protein-coding |
| chr1_131473721_131474256 | chr1 | 131473722 | 131474256 | + | 0.926893 | intron (NM_001081011, intron 2 of 22) | intron (NM_001081011, intron 2 of 22) | 53372 | NM_001081011 | 14270 | Mm.276259 | NM_019520 | ENSMUSG00000026425 | **Srgap2** | 9930124L22Rik\|AI448945\|FBP2\|Fnbp2\|srGAP3 | SLIT-ROBO Rho GTPase activating protein 2 | protein-coding |
| chr6_112856622_112856976 | chr6 | 112856623 | 112856976 | + | 0.729108 | intron (NM_080448, intron 1 of 21) | intron (NM_080448, intron 1 of 21) | 90467 | NM_080448 | 259302 | Mm.236401 | NM_080448 | ENSMUSG00000030257 | **Srgap3** | AI452337\|Arhgap14\|D130026O08Rik\|Gbi\|MEGAP\|WRP\|mKIAA0411 | SLIT-ROBO Rho GTPase activating protein 3 | protein-coding |
| chr6_112861877_112862279 | chr6 | 112861878 | 112862279 | + | 0.714133 | intron (NM_080448, intron 1 of 21) | intron (NM_080448, intron 1 of 21) | 85188 | NM_080448 | 259302 | Mm.236401 | NM_080448 | ENSMUSG00000030257 | **Srgap3** | AI452337\|Arhgap14\|D130026O08Rik\|Gbi\|MEGAP\|WRP\|mKIAA0411 | SLIT-ROBO Rho GTPase activating protein 3 | protein-coding |
| chr6_112861398_112862458 | chr6 | 112861399 | 112862458 | + | 0.669621 | intron (NM_080448, intron 1 of 21) | intron (NM_080448, intron 1 of 21) | 85338 | NM_080448 | 259302 | Mm.236401 | NM_080448 | ENSMUSG00000030257 | **Srgap3** | AI452337\|Arhgap14\|D130026O08Rik\|Gbi\|MEGAP\|WRP\|mKIAA0411 | SLIT-ROBO Rho GTPase activating protein 3 | protein-coding |
| chr10_62505473_62505792 | chr10 | 62505474 | 62505792 | + | 0.751395 | intron (NM_011157, intron 1 of 2) | intron (NM_011157, intron 1 of 2) | 2122 | NM_011157 | 19073 | Mm.338790 | NM_011157 | ENSMUSG00000020077 | **Srgn** | Prg\|Prg1\|Sgc | serglycin | protein-coding |
| chr10_62509290_62509505 | chr10 | 62509291 | 62509505 | + | 0.66923 | Intergenic | MTE2a\|LTR\|ERVL-MaLR | -1643 | NM_011157 | 19073 | Mm.338790 | NM_011157 | ENSMUSG00000020077 | **Srgn** | Prg\|Prg1\|Sgc | serglycin | protein-coding |
| chr2_118448279_118448654 | chr2 | 118448280 | 118448654 | + | 1.36292 | exon (NM_001177806, exon 20 of 38) | exon (NM_001177806, exon 20 of 38) | 31229 | NM_009273 | 20813 | Mm.363404 | NM_009273 | ENSMUSG00000009549 | **Srp14** | 14kDa\|AW536328 | signal recognition particle 14 | protein-coding |
| chr2_118469406_118470073 | chr2 | 118469407 | 118470073 | + | 1.35574 | intron (NM_013719, intron 33 of 38) | intron (NM_013719, intron 33 of 38) | 9956 | NM_009273 | 20813 | Mm.363404 | NM_009273 | ENSMUSG00000009549 | **Srp14** | 14kDa\|AW536328 | signal recognition particle 14 | protein-coding |
| chr2_118454494_118455192 | chr2 | 118454495 | 118455192 | + | 0.694823 | exon (NM_001177806, exon 23 of 38) | exon (NM_001177806, exon 23 of 38) | 24853 | NM_009273 | 20813 | Mm.363404 | NM_009273 | ENSMUSG00000009549 | **Srp14** | 14kDa\|AW536328 | signal recognition particle 14 | protein-coding |
| chr5_76975722_76976099 | chr5 | 76975723 | 76976099 | + | 1.03914 | intron (NM_025691, intron 1 of 18) | intron (NM_025691, intron 1 of 18) | 1210 | NM_025691 | 66661 | Mm.296976 | NM_025691 | ENSMUSG00000036323 | **Srp72** | 5730576P14Rik\|72kDa\|AI132477\|BC019196\|C77589 | signal recognition particle 72 | protein-coding |
| chr1_182101524_182101921 | chr1 | 182101525 | 182101921 | + | 0.734653 | Intergenic | Intergenic | -23014 | NM_012058 | 27058 | Mm.303071 | NM_012058 | ENSMUSG00000026511 | **Srp9** | 9kDa | signal recognition particle 9 | protein-coding |
| chr17_28618792_28619106 | chr17 | 28618793 | 28619106 | + | 0.884484 | intron (NM_016795, intron 2 of 15) | intron (NM_016795, intron 2 of 15) | 3505 | NM_016795 | 20815 | Mm.15252 | NM_016795 | | **Srpk1** | AU017960 | serine/arginine-rich protein specific kinase 1 | protein-coding |
| chr5_23626099_23626530 | chr5 | 23626100 | 23626530 | + | 0.703197 | Intergenic | Intergenic | -9744 | NM_009274 | 20817 | Mm.288728 | NM_009274 | ENSMUSG00000062604 | **Srpk2** | AW226533\|AW492537\|AW547358\|Wbp6\|mSRPK2 | serine/arginine-rich protein specific kinase 2 | protein-coding |
| chrX_133920251_133920793 | chrX | 133920252 | 133920793 | + | 1.89833 | intron (NM_001083895, intron 3 of 10) | L3\|LINE\|CR1 | 12097 | NM_001083895 | 68792 | Mm.263553 | NM_026838 | ENSMUSG00000031253 | **Srpx2** | 1110039C07Rik\|SRP\|SRPUL | sushi-repeat-containing protein, X-linked 2 | protein-coding |
| chr11_74916464_74917161 | chr11 | 74916465 | 74917161 | + | 1.2749 | intron (NM_001163311, intron 1 of 7) | B3\|SINE\|B2 | 8985 | NM_013761 | 27364 | Mm.131443 | NM_013761 | ENSMUSG00000001323 | **Srr** | Srs | serine racemase | protein-coding |
| chr11_88039279_88039615 | chr11 | 88039280 | 88039615 | + | 0.758044 | Intergenic | B3\|SINE\|B2 | -7926 | NM_001078167 | 110809 | Mm.391719 | NM_173374 | ENSMUSG00000018379 | **Srsf1** | 1110054N12Rik\|5730507C05Rik\|6330415C05Rik\|AI482334\|AW491331\|Asf\|Sf2\|Sfrs1 | serine/arginine-rich splicing factor 1 | protein-coding |
| chr11_88045937_88046157 | chr11 | 88045938 | 88046157 | + | 0.682169 | Intergenic | RSINE1\|SINE\|B4 | -1326 | NM_001078167 | 110809 | Mm.391719 | NM_173374 | ENSMUSG00000018379 | **Srsf1** | 1110054N12Rik\|5730507C05Rik\|6330415C05Rik\|AI482334\|AW491331\|Asf\|Sf2\|Sfrs1 | serine/arginine-rich splicing factor 1 | protein-coding |
| chr4_135860295_135860546 | chr4 | 135860296 | 135860546 | + | 1.06161 | intron (NM_010178, intron 2 of 5) | intron (NM_010178, intron 2 of 5) | 4350 | NM_010178 | 14105 | Mm.10229 | NM_010178 | ENSMUSG00000028676 | **Srsf10** | FUSIP2\|Fusip1\|NSSR1\|NSSR2\|Nssr\|SRrp40\|Sfrs13a\|Srsf13a\|TASR\|TASR1\|TASR2 | serine/arginine-rich splicing factor 10 | protein-coding |
| chr17_29049908_29050197 | chr17 | 29049909 | 29050197 | + | 0.733294 | Intergenic | Intergenic | 17393 | NR_036613 | 20383 | Mm.6787 | NM_013663 | ENSMUSG00000071172 | **Srsf3** | AL024116\|Sfrs3\|X16 | serine/arginine-rich splicing factor 3 | protein-coding |
| chr17_80167703_80168281 | chr17 | 80167704 | 80168281 | + | 1.24424 | intron (NM_176963, intron 4 of 6) | intron (NM_176963, intron 4 of 6) | 39313 | NM_001195486 | 225027 | Mm.292016 | NM_146083 | ENSMUSG00000024097 | **Srsf7** | 35kDa\|9430065L19Rik\|9G8\|NX-96\|Sfrs7 | serine/arginine-rich splicing factor 7 | protein-coding |
| chr17_80169031_80169234 | chr17 | 80169032 | 80169234 | + | 0.69448 | intron (NM_176963, intron 4 of 6) | ID_B1\|SINE\|B4 | 38172 | NM_001195486 | 225027 | Mm.292016 | NM_146083 | ENSMUSG00000024097 | **Srsf7** | 35kDa\|9430065L19Rik\|9G8\|NX-96\|Sfrs7 | serine/arginine-rich splicing factor 7 | protein-coding |
| chr2_152105224_152106072 | chr2 | 152105225 | 152106072 | + | 0.902854 | 5' UTR (NM_029688, exon 1 of 2) | 5' UTR (NM_029688, exon 1 of 2) | 124 | NM_029688 | 76650 | Mm.218639 | NM_029688 | ENSMUSG00000032802 | **Srxn1** | 1700127B04Rik\|AI854065\|AW488194\|Npn3\|Srx\|Srx1\|TX01 | sulfiredoxin 1 homolog (S. cerevisiae) | protein-coding |
| chr2_152101826_152102272 | chr2 | 152101827 | 152102272 | + | 0.86404 | Intergenic | Intergenic | -3475 | NM_029688 | 76650 | Mm.218639 | NM_029688 | ENSMUSG00000032802 | **Srxn1** | 1700127B04Rik\|AI854065\|AW488194\|Npn3\|Srx\|Srx1\|TX01 | sulfiredoxin 1 homolog (S. cerevisiae) | protein-coding |
| chr2_152101584_152102306 | chr2 | 152101585 | 152102306 | + | 0.779177 | Intergenic | Intergenic | -3579 | NM_029688 | 76650 | Mm.218639 | NM_029688 | ENSMUSG00000032802 | **Srxn1** | 1700127B04Rik\|AI854065\|AW488194\|Npn3\|Srx\|Srx1\|TX01 | sulfiredoxin 1 homolog (S. cerevisiae) | protein-coding |
| chr18_14613283_14614083 | chr18 | 14613284 | 14614083 | + | 1.81674 | Intergenic | Intergenic | 69231 | NM_001161369 | 268996 | Mm.31761 | NM_009280 | ENSMUSG00000037013 | **Ss18** | D130059H17\|Ssxt\|Syt | synovial sarcoma translocation, Chromosome 18 | protein-coding |
| chr18_14511474_14511748 | chr18 | 14511475 | 14511748 | + | 1.01792 | Intergenic | L1M4b\|LINE\|L1 | 171303 | NM_001161369 | 268996 | Mm.31761 | NM_009280 | ENSMUSG00000037013 | **Ss18** | D130059H17\|Ssxt\|Syt | synovial sarcoma translocation, Chromosome 18 | protein-coding |
| chr18_14654748_14655287 | chr18 | 14654749 | 14655287 | + | 0.840131 | intron (NM_001161369, intron 4 of 9) | intron (NM_001161369, intron 4 of 9) | 27896 | NM_001161369 | 268996 | Mm.31761 | NM_009280 | ENSMUSG00000037013 | **Ss18** | D130059H17\|Ssxt\|Syt | synovial sarcoma translocation, Chromosome 18 | protein-coding |
| chr13_91582297_91582897 | chr13 | 91582298 | 91582897 | + | 0.907632 | intron (NM_024186, intron 4 of 15) | L1VL2\|LINE\|L1 | 121500 | NM_024186 | 66970 | Mm.343095 | NM_024186 | ENSMUSG00000003992 | **Ssbp2** | 1500004K09Rik\|2310079I02Rik\|9330163K02Rik\|A830008M03Rik\|AU067692\|AW558684\|Hspc116\|SSDP2 | single-stranded DNA binding protein 2 | protein-coding |
| chr13_91582019_91582894 | chr13 | 91582020 | 91582894 | + | 0.887324 | intron (NM_024186, intron 4 of 15) | L1VL2\|LINE\|L1 | 121360 | NM_024186 | 66970 | Mm.343095 | NM_024186 | ENSMUSG00000003992 | **Ssbp2** | 1500004K09Rik\|2310079I02Rik\|9330163K02Rik\|A830008M03Rik\|AU067692\|AW558684\|Hspc116\|SSDP2 | single-stranded DNA binding protein 2 | protein-coding |
| chr13_91564653_91565144 | chr13 | 91564654 | 91565144 | + | 0.744187 | intron (NM_024186, intron 4 of 15) | intron (NM_024186, intron 4 of 15) | 103802 | NM_024186 | 66970 | Mm.343095 | NM_024186 | ENSMUSG00000003992 | **Ssbp2** | 1500004K09Rik\|2310079I02Rik\|9330163K02Rik\|A830008M03Rik\|AU067692\|AW558684\|Hspc116\|SSDP2 | single-stranded DNA binding protein 2 | protein-coding |
| chr13_91579720_91580263 | chr13 | 91579721 | 91580263 | + | 0.680306 | intron (NM_024186, intron 4 of 15) | intron (NM_024186, intron 4 of 15) | 118895 | NM_024186 | 66970 | Mm.343095 | NM_024186 | ENSMUSG00000003992 | **Ssbp2** | 1500004K09Rik\|2310079I02Rik\|9330163K02Rik\|A830008M03Rik\|AU067692\|AW558684\|Hspc116\|SSDP2 | single-stranded DNA binding protein 2 | protein-coding |

| | | | | | | | | | | | | | | | | | |
|---|---|---|---|---|---|---|---|---|---|---|---|---|---|---|---|---|---|
| chr4_106983666_106984052 | chr4 | 106983667 | 106984052 | + | 0.672412 | intron (NM_198438, intron 4 of 16) | intron (NM_198438, intron 4 of 16) | 72389 | NM_023672 | 72475 | Mm.195635 | NM_023672 | ENSMUSG00000061887 | **Ssbp3** | 2610021L12Rik\|2610200M23Rik\|5730488C10Rik\|AI854733\|AW551939\|LAST\|SSDP1 | single-stranded DNA binding protein 3 | protein-coding |
| chr7_4924816_4925288 | chr7 | 4924817 | 4925288 | + | 0.80484 | promoter-TSS (NM_173008) | promoter-TSS (NM_173008) | -792 | NM_173008 | 269855 | Mm.44190 | NM_173008 | ENSMUSG00000035279 | **Ssc5d** | A430110N23Rik\|S5D-SRCRB | scavenger receptor cysteine rich family, 5 domains | protein-coding |
| chr7_4924816_4925029 | chr7 | 4924817 | 4925029 | + | 0.75964 | promoter-TSS (NM_173008) | promoter-TSS (NM_173008) | -921 | NM_173008 | 269855 | Mm.44190 | NM_173008 | ENSMUSG00000035279 | **Ssc5d** | A430110N23Rik\|S5D-SRCRB | scavenger receptor cysteine rich family, 5 domains | protein-coding |
| chr2_79555412_79555865 | chr2 | 79555413 | 79555865 | + | 1.9421 | Intergenic | Intergenic | -79786 | NM_080558 | 70599 | Mm.272881 | NM_080558 | ENSMUSG00000027007 | **Ssfa2** | 5730488C15Rik\|CS-1\|CS1\|KRAP\|SPAG13\|mKIAA1927 | sperm specific antigen 2 | protein-coding |
| chr2_79555403_79556164 | chr2 | 79555404 | 79556164 | + | 1.92536 | Intergenic | Intergenic | -79641 | NM_080558 | 70599 | Mm.272881 | NM_080558 | ENSMUSG00000027007 | **Ssfa2** | 5730488C15Rik\|CS-1\|CS1\|KRAP\|SPAG13\|mKIAA1927 | sperm specific antigen 2 | protein-coding |
| chr2_79622485_79622863 | chr2 | 79622486 | 79622863 | + | 0.658472 | Intergenic | L2\|LINE\|L2 | -12751 | NM_080558 | 70599 | Mm.272881 | NM_080558 | ENSMUSG00000027007 | **Ssfa2** | 5730488C15Rik\|CS-1\|CS1\|KRAP\|SPAG13\|mKIAA1927 | sperm specific antigen 2 | protein-coding |
| chr11_77394782_77395768 | chr11 | 77394783 | 77395768 | + | 0.685619 | intron (NM_177710, intron 3 of 14) | intron (NM_177710, intron 3 of 14) | 46995 | NM_001291190 | 237860 | Mm.440381 | NM_177710 | ENSMUSG00000037926 | **Ssh2** | SSH-2\|SSH-2L\|mSSH-2L | slingshot homolog 2 (Drosophila) | protein-coding |
| chr11_77240563_77241467 | chr11 | 77240564 | 77241467 | + | 0.653625 | intron (NM_177710, intron 1 of 14) | RMER19A\|LTR\|ERVK | 24590 | NM_177710 | 237860 | Mm.440381 | NM_177710 | ENSMUSG00000037926 | **Ssh2** | SSH-2\|SSH-2L\|mSSH-2L | slingshot homolog 2 (Drosophila) | protein-coding |
| chr6_145981079_145981580 | chr6 | 145981080 | 145981580 | + | 1.60636 | Intergenic | MER34A1\|LTR\|ERV1 | 47183 | NM_010656 | 16651 | Mm.49689 | NM_010656 | ENSMUSG00000030255 | **Sspn** | Krag | sarcospan | protein-coding |
| chr6_145980833_145981585 | chr6 | 145980834 | 145981585 | + | 1.313 | Intergenic | Intergenic | 47062 | NM_010656 | 16651 | Mm.49689 | NM_010656 | ENSMUSG00000030255 | **Sspn** | Krag | sarcospan | protein-coding |
| chr6_146185487_146186018 | chr6 | 146185488 | 146186018 | + | 1.07162 | intron (NM_019923, intron 48 of 56) | intron (NM_019923, intron 48 of 56) | 251606 | NM_010656 | 16651 | Mm.49689 | NM_010656 | ENSMUSG00000030255 | **Sspn** | Krag | sarcospan | protein-coding |
| chr6_146000780_146001369 | chr6 | 146000781 | 146001369 | + | 0.608153 | Intergenic | Intergenic | 66928 | NM_010656 | 16651 | Mm.49689 | NM_010656 | ENSMUSG00000030255 | **Sspn** | Krag | sarcospan | protein-coding |
| chr3_65434898_65435685 | chr3 | 65434899 | 65435685 | + | 0.822323 | Intergenic | Intergenic | -42739 | NM_026155 | 67437 | Mm.232728 | NM_026155 | ENSMUSG00000027828 | **Ssr3** | 0610038P07Rik\|AL022999\|AU022074\|AW553833\|TRAPG | signal sequence receptor, gamma | protein-coding |
| chr1_6453931_6454720 | chr1 | 6453932 | 6454720 | + | 0.897336 | Intergenic | Intergenic | -32905 | NM_001244693 | 240690 | Mm.234612 | NM_173868 | ENSMUSG00000033740 | **St18** | AV348974\|Myt3\|NZF-3\|Nzf3\|mKIAA0535 | suppression of tumorigenicity 18 | protein-coding |
| chr15_67138581_67139603 | chr15 | 67138582 | 67139603 | + | 1.13581 | intron (NM_009177, intron 2 of 8) | intron (NM_009177, intron 2 of 8) | 37790 | NM_009177 | 20442 | Mm.248334 | NM_009177 | ENSMUSG00000013846 | **St3gal1** | 5330418N22Rik\|AI467004\|ST3GalI\|Siat4\|Siat4a\|St3gal-1 | ST3 beta-galactoside alpha-2,3-sialyltransferase 1 | protein-coding |
| chr15_67129841_67130387 | chr15 | 67129842 | 67130387 | + | 1.11114 | intron (NM_009177, intron 2 of 8) | MIR\|SINE\|MIR | 46768 | NM_009177 | 20442 | Mm.248334 | NM_009177 | ENSMUSG00000013846 | **St3gal1** | 5330418N22Rik\|AI467004\|ST3GalI\|Siat4\|Siat4a\|St3gal-1 | ST3 beta-galactoside alpha-2,3-sialyltransferase 1 | protein-coding |
| chr15_67128882_67129373 | chr15 | 67128883 | 67129373 | + | 0.91883 | intron (NM_009177, intron 2 of 8) | intron (NM_009177, intron 2 of 8) | 47754 | NM_009177 | 20442 | Mm.248334 | NM_009177 | ENSMUSG00000013846 | **St3gal1** | 5330418N22Rik\|AI467004\|ST3GalI\|Siat4\|Siat4a\|St3gal-1 | ST3 beta-galactoside alpha-2,3-sialyltransferase 1 | protein-coding |
| chr15_67196870_67197423 | chr15 | 67196871 | 67197423 | + | 0.876729 | Intergenic | LTRIS_Mus\|LTR\|ERV1 | -20265 | NM_009177 | 20442 | Mm.248334 | NM_009177 | ENSMUSG00000013846 | **St3gal1** | 5330418N22Rik\|AI467004\|ST3GalI\|Siat4\|Siat4a\|St3gal-1 | ST3 beta-galactoside alpha-2,3-sialyltransferase 1 | protein-coding |
| chr15_67116587_67117427 | chr15 | 67116588 | 67117427 | + | 0.797749 | intron (NM_009177, intron 3 of 8) | intron (NM_009177, intron 3 of 8) | 59875 | NM_009177 | 20442 | Mm.248334 | NM_009177 | ENSMUSG00000013846 | **St3gal1** | 5330418N22Rik\|AI467004\|ST3GalI\|Siat4\|Siat4a\|St3gal-1 | ST3 beta-galactoside alpha-2,3-sialyltransferase 1 | protein-coding |
| chr15_67197080_67197403 | chr15 | 67197081 | 67197403 | + | 0.750428 | Intergenic | LTRIS_Mus\|LTR\|ERV1 | -20360 | NM_009177 | 20442 | Mm.248334 | NM_009177 | ENSMUSG00000013846 | **St3gal1** | 5330418N22Rik\|AI467004\|ST3GalI\|Siat4\|Siat4a\|St3gal-1 | ST3 beta-galactoside alpha-2,3-sialyltransferase 1 | protein-coding |
| chr15_67138583_67138839 | chr15 | 67138584 | 67138839 | + | 0.737548 | intron (NM_009177, intron 2 of 8) | intron (NM_009177, intron 2 of 8) | 38171 | NM_009177 | 20442 | Mm.248334 | NM_009177 | ENSMUSG00000013846 | **St3gal1** | 5330418N22Rik\|AI467004\|ST3GalI\|Siat4\|Siat4a\|St3gal-1 | ST3 beta-galactoside alpha-2,3-sialyltransferase 1 | protein-coding |
| chr15_67161581_67162010 | chr15 | 67161582 | 67162010 | + | 0.724067 | intron (NM_009177, intron 1 of 8) | intron (NM_009177, intron 1 of 8) | 15086 | NM_009177 | 20442 | Mm.248334 | NM_009177 | ENSMUSG00000013846 | **St3gal1** | 5330418N22Rik\|AI467004\|ST3GalI\|Siat4\|Siat4a\|St3gal-1 | ST3 beta-galactoside alpha-2,3-sialyltransferase 1 | protein-coding |
| chr8_110954717_110955159 | chr8 | 110954718 | 110955159 | + | 1.22154 | intron (NM_009179, intron 1 of 6) | intron (NM_009179, intron 1 of 6) | 35073 | NM_009179 | 20444 | Mm.200388 | NM_009179 | ENSMUSG00000031749 | **St3gal2** | AI429591\|AW822065\|ST3GalII\|Siat5 | ST3 beta-galactoside alpha-2,3-sialyltransferase 2 | protein-coding |
| chr4_118070472_118071130 | chr4 | 118070473 | 118071130 | + | 0.74092 | intron (NM_009176, intron 2 of 11) | MER44B\|DNA\|TcMar-Tigger | 64145 | NM_001285520 | 20441 | Mm.251002 | NM_009176 | ENSMUSG00000028538 | **St3gal3** | ST3GalIII\|ST3N\|Siat3\|Siat6 | ST3 beta-galactoside alpha-2,3-sialyltransferase 3 | protein-coding |
| chr9_35123844_35124781 | chr9 | 35123845 | 35124781 | + | 0.71346 | TTS (NM_027030) | TTS (NM_027030) | -7503 | NM_009178 | 20443 | Mm.275973 | NM_009178 | ENSMUSG00000032038 | **St3gal4** | Siat4c | ST3 beta-galactoside alpha-2,3-sialyltransferase 4 | protein-coding |
| chr9_35123509_35124818 | chr9 | 35123510 | 35124818 | + | 0.679355 | TTS (NM_027030) | TTS (NM_027030) | -7354 | NM_009178 | 20443 | Mm.275973 | NM_009178 | ENSMUSG00000032038 | **St3gal4** | Siat4c | ST3 beta-galactoside alpha-2,3-sialyltransferase 4 | protein-coding |
| chr16_58475667_58476258 | chr16 | 58475668 | 58476258 | + | 1.88228 | intron (NM_018784, intron 6 of 10) | intron (NM_018784, intron 6 of 10) | 47349 | NM_018784 | 54613 | Mm.212742 | NM_018784 | ENSMUSG00000022747 | **St3gal6** | 1700023B24Rik\|AI930218\|AW552396\|Siat10\|St3galVI | ST3 beta-galactoside alpha-2,3-sialyltransferase 6 | protein-coding |
| chr16_58479957_58480468 | chr16 | 58479958 | 58480468 | + | 1.15735 | intron (NM_018784, intron 6 of 10) | intron (NM_018784, intron 6 of 10) | 43099 | NM_018784 | 54613 | Mm.212742 | NM_018784 | ENSMUSG00000022747 | **St3gal6** | 1700023B24Rik\|AI930218\|AW552396\|Siat10\|St3galVI | ST3 beta-galactoside alpha-2,3-sialyltransferase 6 | protein-coding |
| chr7_109587727_109588481 | chr7 | 109587728 | 109588481 | + | 0.916439 | intron (NM_001001326, intron 1 of 19) | intron (NM_001001326, intron 1 of 19) | 28464 | NR_037617 | 76954 | Mm.252009 | NM_029811 | ENSMUSG00000031024 | **St5** | 2010004M01Rik\|2610305K15Rik | suppression of tumorigenicity 5 | protein-coding |
| chr16_23272170_23272774 | chr16 | 23272171 | 23272774 | + | 1.09792 | intron (NM_001252505, intron 2 of 7) | intron (NM_001252505, intron 2 of 7) | -17998 | NM_001252506 | 20440 | Mm.149029 | NM_145933 | ENSMUSG00000022885 | **St6gal1** | AW742324\|Siat1\|St6Gal-I\|St6gal\|St6galI | beta galactoside alpha 2,6 sialyltransferase 1 | protein-coding |
| chr16_23274973_23275237 | chr16 | 23274974 | 23275237 | + | 0.90255 | intron (NM_001252505, intron 2 of 7) | ORR1D1\|LTR\|ERVL-MaLR | -15365 | NM_001252506 | 20440 | Mm.149029 | NM_145933 | ENSMUSG00000022885 | **St6gal1** | AW742324\|Siat1\|St6Gal-I\|St6gal\|St6galI | beta galactoside alpha 2,6 sialyltransferase 1 | protein-coding |
| chr16_23301811_23302129 | chr16 | 23301812 | 23302129 | + | 0.869336 | intron (NM_001252506, intron 1 of 6) | L1M3\|LINE\|L1 | 11500 | NM_001252506 | 20440 | Mm.149029 | NM_145933 | ENSMUSG00000022885 | **St6gal1** | AW742324\|Siat1\|St6Gal-I\|St6gal\|St6galI | beta galactoside alpha 2,6 sialyltransferase 1 | protein-coding |
| chr16_23294786_23295109 | chr16 | 23294787 | 23295109 | + | 0.771313 | intron (NM_001252506, intron 1 of 6) | intron (NM_001252506, intron 1 of 6) | 4478 | NM_001252506 | 20440 | Mm.149029 | NM_145933 | ENSMUSG00000022885 | **St6gal1** | AW742324\|Siat1\|St6Gal-I\|St6gal\|St6galI | beta galactoside alpha 2,6 sialyltransferase 1 | protein-coding |
| chr16_23351546_23351946 | chr16 | 23351547 | 23351946 | + | 0.65239 | intron (NM_001252505, intron 5 of 7) | intron (NM_001252505, intron 5 of 7) | 61276 | NM_001252506 | 20440 | Mm.149029 | NM_145933 | ENSMUSG00000022885 | **St6gal1** | AW742324\|Siat1\|St6Gal-I\|St6gal\|St6galI | beta galactoside alpha 2,6 sialyltransferase 1 | protein-coding |
| chr16_23274077_23274467 | chr16 | 23274078 | 23274467 | + | 0.629158 | intron (NM_001252505, intron 2 of 7) | intron (NM_001252505, intron 2 of 7) | -16198 | NM_001252506 | 20440 | Mm.149029 | NM_145933 | ENSMUSG00000022885 | **St6gal1** | AW742324\|Siat1\|St6Gal-I\|St6gal\|St6galI | beta galactoside alpha 2,6 sialyltransferase 1 | protein-coding |
| chr2_32585306_32585696 | chr2 | 32585307 | 32585696 | + | 1.40873 | Intergenic | Intergenic | -1577 | NM_001276425 | 20448 | Mm.27446 | NM_011373 | ENSMUSG00000079442 | **St6galnac4** | SIAT7-D\|ST6GalNAcIV\|Siat7d | ST6 (alpha-N-acetyl-neuraminyl-2,3-beta-galactosyl-1,3)-N-acetylgalactosaminide alpha-2,6-sialyltransferase 4 | protein-coding |
| chr3_152907732_152908117 | chr3 | 152907733 | 152908117 | + | 1.28725 | intron (NM_012028, intron 2 of 4) | intron (NM_012028, intron 2 of 4) | 74282 | NM_012028 | 26938 | Mm.40915 | NM_012028 | | **St6galnac5** | AI851940\|Siat7e | ST6 (alpha-N-acetyl-neuraminyl-2,3-beta-galactosyl-1,3)-N-acetylgalactosaminide alpha-2,6-sialyltransferase 5 | protein-coding |
| chr3_152906504_152906744 | chr3 | 152906505 | 152906744 | + | 0.76738 | intron (NM_012028, intron 2 of 4) | intron (NM_012028, intron 2 of 4) | 75583 | NM_012028 | 26938 | Mm.40915 | NM_012028 | | **St6galnac5** | AI851940\|Siat7e | ST6 (alpha-N-acetyl-neuraminyl-2,3-beta-galactosyl-1,3)-N-acetylgalactosaminide alpha-2,6-sialyltransferase 5 | protein-coding |
| chr3_152886436_152887181 | chr3 | 152886437 | 152887181 | + | 0.699847 | intron (NM_012028, intron 2 of 4) | intron (NM_012028, intron 2 of 4) | 95398 | NM_012028 | 26938 | Mm.40915 | NM_012028 | | **St6galnac5** | AI851940\|Siat7e | ST6 (alpha-N-acetyl-neuraminyl-2,3-beta-galactosyl-1,3)-N-acetylgalactosaminide alpha-2,6-sialyltransferase 5 | protein-coding |
| chr6_17731673_17732076 | chr6 | 17731674 | 17732076 | + | 1.22461 | intron (NM_001289626, intron 1 of 14) | intron (NM_001289626, intron 1 of 14) | -11717 | NM_001289629 | 64213 | Mm.12051 | NM_022332 | ENSMUSG00000029534 | **St7** | 9430001H04Rik\|Fam4a2\|HELG\|RAY1\|SEN4\|TSG7 | suppression of tumorigenicity 7 | protein-coding |
| chr1_95624166_95624786 | chr1 | 95624167 | 95624786 | + | 1.35503 | intron (NM_009183, intron 4 of 4) | intron (NM_009183, intron 4 of 4) | 43118 | NM_009183 | 20452 | Mm.306228 | NM_009183 | ENSMUSG00000040710 | **St8sia4** | PST\|PST-1\|SIAT8-D\|ST8SiaIV\|Siat8d | ST8 alpha-N-acetyl-neuraminide alpha-2,8-sialyltransferase 4 | protein-coding |
| chr1_95673412_95673909 | chr1 | 95673413 | 95673909 | + | 0.721237 | Intergenic | Intergenic | -6067 | NM_001159745 | 20452 | Mm.306228 | NM_009183 | ENSMUSG00000040710 | **St8sia4** | PST\|PST-1\|SIAT8-D\|ST8SiaIV\|Siat8d | ST8 alpha-N-acetyl-neuraminide alpha-2,8-sialyltransferase 4 | protein-coding |
| chr14_31160475_31160969 | chr14 | 31160476 | 31160969 | + | 0.921946 | intron (NM_138672, intron 11 of 68) | intron (NM_138672, intron 11 of 68) | 7919 | NM_138672 | 192187 | Mm.220821 | NM_138672 | ENSMUSG00000042286 | **Stab1** | FEEL-1\|FELE-1\|MFEEL-1\|MS-1\|STAB-1\|mKIAA0246 | stabilin 1 | protein-coding |
| chr11_98057982_98058346 | chr11 | 98057983 | 98058346 | + | 0.956971 | Intergenic | MTD\|LTR\|ERVL-MaLR | -4702 | NM_146028 | 217154 | Mm.181008 | NM_146028 | ENSMUSG00000017400 | **Stac2** | 24b2\|24b2/STAC2\|AW240854 | SH3 and cysteine rich domain 2 | protein-coding |
| chr11_98057421_98057860 | chr11 | 98057422 | 98057860 | + | 0.754846 | Intergenic | Intergenic | -4179 | NM_146028 | 217154 | Mm.181008 | NM_146028 | ENSMUSG00000017400 | **Stac2** | 24b2\|24b2/STAC2\|AW240854 | SH3 and cysteine rich domain 2 | protein-coding |

| | | | | | | | | | | | | | | | | | |
|---|---|---|---|---|---|---|---|---|---|---|---|---|---|---|---|---|---|
| chr9_100724942_100725234 | chr9 | 100724943 | 100725234 | + | 0.657442 | intron (NM_009282, intron 3 of 33) | Lx2B2\|LINE\|L1 | 81465 | NM_009282 | 20842 | Mm.42135 | NM_009282 | ENSMUSG00000037286 | **Stag1** | AU045003\|SA-1\|Scc3 | stromal antigen 1 | protein-coding |
| chr9_100724294_100725290 | chr9 | 100724295 | 100725290 | + | 0.630529 | intron (NM_009282, intron 3 of 33) | Lx2B2\|LINE\|L1 | 81169 | NM_009282 | 20842 | Mm.42135 | NM_009282 | ENSMUSG00000037286 | **Stag1** | AU045003\|SA-1\|Scc3 | stromal antigen 1 | protein-coding |
| chr9_100724675_100725237 | chr9 | 100724676 | 100725237 | + | 0.629464 | intron (NM_009282, intron 3 of 33) | Lx2B2\|LINE\|L1 | 81333 | NM_009282 | 20842 | Mm.42135 | NM_009282 | ENSMUSG00000037286 | **Stag1** | AU045003\|SA-1\|Scc3 | stromal antigen 1 | protein-coding |
| chrX_42342144_42342698 | chrX | 42342145 | 42342698 | + | 1.75653 | Intergenic | Intergenic | 140566 | NM_001290713 | 20843 | Mm.290422 | NM_021465 | ENSMUSG00000025862 | **Stag2** | 9230105L23Rik\|B230112I07Rik\|SA-2\|SAP2 | stromal antigen 2 | protein-coding |
| chrX_42336341_42337176 | chrX | 42336342 | 42337176 | + | 1.24276 | Intergenic | B3\|SINE\|B2 | 134904 | NM_001290713 | 20843 | Mm.290422 | NM_021465 | ENSMUSG00000025862 | **Stag2** | 9230105L23Rik\|B230112I07Rik\|SA-2\|SAP2 | stromal antigen 2 | protein-coding |
| chrX_42344298_42344597 | chrX | 42344299 | 42344597 | + | 1.14485 | Intergenic | Intergenic | 142593 | NM_001290713 | 20843 | Mm.290422 | NM_021465 | ENSMUSG00000025862 | **Stag2** | 9230105L23Rik\|B230112I07Rik\|SA-2\|SAP2 | stromal antigen 2 | protein-coding |
| chrX_42188904_42189763 | chrX | 42188905 | 42189763 | + | 0.687519 | intron (NM_021465, intron 1 of 32) | L1MA5A\|LINE\|L1 | -12521 | NM_001290713 | 20843 | Mm.290422 | NM_021465 | ENSMUSG00000025862 | **Stag2** | 9230105L23Rik\|B230112I07Rik\|SA-2\|SAP2 | stromal antigen 2 | protein-coding |
| chr6_83582057_83582811 | chr6 | 83582058 | 83582811 | + | 1.24144 | intron (NR_078345, intron 5 of 7) | Lx8\|LINE\|L1 | -9930 | NM_024239 | 70527 | Mm.32801 | NM_024239 | ENSMUSG00000006906 | **Stambp** | 5330424L14Rik\|5730422L11Rik\|AW107289\|Amsh\|mKIAA4198 | STAM binding protein | protein-coding |
| chr6_83597992_83598427 | chr6 | 83597993 | 83598427 | + | 1.06905 | Intergenic | Intergenic | -25706 | NM_024239 | 70527 | Mm.32801 | NM_024239 | ENSMUSG00000006906 | **Stambp** | 5330424L14Rik\|5730422L11Rik\|AW107289\|Amsh\|mKIAA4198 | STAM binding protein | protein-coding |
| chr19_34179151_34179600 | chr19 | 34179152 | 34179600 | + | 1.14857 | Intergenic | Intergenic | -12894 | NM_029682 | 76630 | Mm.130952 | NM_029682 | ENSMUSG00000024776 | **Stambpl1** | 1700095N21Rik\|8230401J17Rik\|ALMalpha\|AMSH-FP | STAM binding protein like 1 | protein-coding |
| chr19_34216914_34217445 | chr19 | 34216915 | 34217445 | + | 0.604576 | intron (NM_029682, intron 1 of 10) | MLT1N2\|LTR\|ERVL-MaLR | 24910 | NM_029682 | 76630 | Mm.130952 | NM_029682 | ENSMUSG00000024776 | **Stambpl1** | 1700095N21Rik\|8230401J17Rik\|ALMalpha\|AMSH-FP | STAM binding protein like 1 | protein-coding |
| chr5_86078630_86079327 | chr5 | 86078631 | 86079327 | + | 1.00631 | intron (NM_019992, intron 2 of 8) | intron (NM_019992, intron 2 of 8) | 7151 | NM_019992 | 56792 | Mm.131237 | NM_019992 | ENSMUSG00000029254 | **Stap1** | AI586015\|Brdg1\|STAP-1 | signal transducing adaptor family member 1 | protein-coding |
| chr5_86096650_86097062 | chr5 | 86096651 | 86097062 | + | 0.766395 | intron (NM_019992, intron 8 of 8) | intron (NM_019992, intron 8 of 8) | 25028 | NM_019992 | 56792 | Mm.131237 | NM_019992 | ENSMUSG00000029254 | **Stap1** | AI586015\|Brdg1\|STAP-1 | signal transducing adaptor family member 1 | protein-coding |
| chr5_151252127_151252644 | chr5 | 151252128 | 151252644 | + | 0.996674 | Intergenic | Intergenic | -62193 | NM_146258 | 243362 | Mm.209272 | NM_146258 | ENSMUSG00000016128 | **Stard13** | DLC2\|GT650 | StAR-related lipid transfer (START) domain containing 13 | protein-coding |
| chr5_151251666_151252049 | chr5 | 151251667 | 151252049 | + | 0.921543 | Intergenic | Intergenic | -61665 | NM_146258 | 243362 | Mm.209272 | NM_146258 | ENSMUSG00000016128 | **Stard13** | DLC2\|GT650 | StAR-related lipid transfer (START) domain containing 13 | protein-coding |
| chr5_151210648_151210907 | chr5 | 151210649 | 151210907 | + | 0.733165 | Intergenic | Intergenic | -20585 | NM_146258 | 243362 | Mm.209272 | NM_146258 | ENSMUSG00000016128 | **Stard13** | DLC2\|GT650 | StAR-related lipid transfer (START) domain containing 13 | protein-coding |
| chr5_151210950_151211383 | chr5 | 151210951 | 151211383 | + | 0.700267 | Intergenic | Intergenic | -20974 | NM_146258 | 243362 | Mm.209272 | NM_146258 | ENSMUSG00000016128 | **Stard13** | DLC2\|GT650 | StAR-related lipid transfer (START) domain containing 13 | protein-coding |
| chr5_151210976_151211289 | chr5 | 151210977 | 151211289 | + | 0.690351 | Intergenic | Intergenic | -20940 | NM_146258 | 243362 | Mm.209272 | NM_146258 | ENSMUSG00000016128 | **Stard13** | DLC2\|GT650 | StAR-related lipid transfer (START) domain containing 13 | protein-coding |
| chr18_33088329_33088919 | chr18 | 33088330 | 33088919 | + | 1.10163 | intron (NM_009793, intron 2 of 10) | L1MA9\|LINE\|L1 | 125192 | NM_133774 | 170459 | Mm.127058 | NM_133774 | | **Stard4** | 4632419C16Rik\|9030213J02Rik\|AA517649\|AW324468 | StAR-related lipid transfer (START) domain containing 4 | protein-coding |
| chr18_33084153_33084409 | chr18 | 33084154 | 33084409 | + | 0.769539 | intron (NM_009793, intron 2 of 10) | intron (NM_009793, intron 2 of 10) | 129535 | NM_133774 | 170459 | Mm.127058 | NM_133774 | | **Stard4** | 4632419C16Rik\|9030213J02Rik\|AA517649\|AW324468 | StAR-related lipid transfer (START) domain containing 4 | protein-coding |
| chr18_33084632_33085229 | chr18 | 33084633 | 33085229 | + | 0.726421 | intron (NM_009793, intron 2 of 10) | MLT1H2-int\|LTR\|ERVL-MaLR | 128885 | NM_133774 | 170459 | Mm.127058 | NM_133774 | | **Stard4** | 4632419C16Rik\|9030213J02Rik\|AA517649\|AW324468 | StAR-related lipid transfer (START) domain containing 4 | protein-coding |
| chr18_33084869_33085137 | chr18 | 33084870 | 33085137 | + | 0.722121 | intron (NM_009793, intron 2 of 10) | MLT1H2-int\|LTR\|ERVL-MaLR | 128813 | NM_133774 | 170459 | Mm.127058 | NM_133774 | | **Stard4** | 4632419C16Rik\|9030213J02Rik\|AA517649\|AW324468 | StAR-related lipid transfer (START) domain containing 4 | protein-coding |
| chr18_33192778_33193404 | chr18 | 33192779 | 33193404 | + | 0.694551 | 3' UTR (NM_009793, exon 11 of 11) | 3' UTR (NM_009793, exon 11 of 11) | 20725 | NM_133774 | 170459 | Mm.127058 | NM_133774 | | **Stard4** | 4632419C16Rik\|9030213J02Rik\|AA517649\|AW324468 | StAR-related lipid transfer (START) domain containing 4 | protein-coding |
| chr7_83639058_83639791 | chr7 | 83639059 | 83639791 | + | 0.727874 | intron (NM_023377, intron 5 of 5) | intron (NM_023377, intron 5 of 5) | 7408 | NM_023377 | 170460 | Mm.357953 | NM_023377 | ENSMUSG00000046027 | **Stard5** | 2310058G22Rik\|AI607880\|D7Ertd152e | StAR-related lipid transfer (START) domain containing 5 | protein-coding |
| chr18_70459708_70460062 | chr18 | 70459709 | 70460062 | + | 1.28791 | intron (NM_172967, intron 6 of 8) | L1_Rod\|LINE\|L1 | -12569 | NM_029019 | 170461 | Mm.83623 | NM_029019 | ENSMUSG00000079608 | **Stard6** | 1700011K09Rik\|4833424I06Rik\|4933429L05Rik\|AI644424 | StAR-related lipid transfer (START) domain containing 6 | protein-coding |
| chr18_70391178_70391907 | chr18 | 70391179 | 70391907 | + | 1.24975 | Intergenic | Intergenic | -80911 | NM_029019 | 170461 | Mm.83623 | NM_029019 | ENSMUSG00000079608 | **Stard6** | 1700011K09Rik\|4833424I06Rik\|4933429L05Rik\|AI644424 | StAR-related lipid transfer (START) domain containing 6 | protein-coding |
| chr18_70462650_70463111 | chr18 | 70462651 | 70463111 | + | 0.894203 | intron (NM_172967, intron 6 of 8) | intron (NM_172967, intron 6 of 8) | -9573 | NM_029019 | 170461 | Mm.83623 | NM_029019 | ENSMUSG00000079608 | **Stard6** | 1700011K09Rik\|4833424I06Rik\|4933429L05Rik\|AI644424 | StAR-related lipid transfer (START) domain containing 6 | protein-coding |
| chr18_70462660_70462979 | chr18 | 70462661 | 70462979 | + | 0.892398 | intron (NM_172967, intron 6 of 8) | intron (NM_172967, intron 6 of 8) | -9634 | NM_029019 | 170461 | Mm.83623 | NM_029019 | ENSMUSG00000079608 | **Stard6** | 1700011K09Rik\|4833424I06Rik\|4933429L05Rik\|AI644424 | StAR-related lipid transfer (START) domain containing 6 | protein-coding |
| chr18_70462660_70463366 | chr18 | 70462661 | 70463366 | + | 0.769686 | intron (NM_172967, intron 6 of 8) | intron (NM_172967, intron 6 of 8) | -9441 | NM_029019 | 170461 | Mm.83623 | NM_029019 | ENSMUSG00000079608 | **Stard6** | 1700011K09Rik\|4833424I06Rik\|4933429L05Rik\|AI644424 | StAR-related lipid transfer (START) domain containing 6 | protein-coding |
| chr2_127300710_127301379 | chr2 | 127300711 | 127301379 | + | 0.677998 | Intergenic | Intergenic | 30816 | NM_139308 | 99138 | Mm.41271 | NM_139308 | ENSMUSG00000027367 | **Stard7** | AI852671\|AL022671\|AW544915 | START domain containing 7 | protein-coding |
| chr1_52066257_52067048 | chr1 | 52066258 | 52067048 | + | 1.76029 | intron (NM_011487, intron 4 of 23) | intron (NM_011487, intron 4 of 23) | -52785 | NM_009283 | 20846 | Mm.277406 | NM_009283 | ENSMUSG00000026104 | **Stat1** | 2010005J02Rik\|AA408197 | signal transducer and activator of transcription 1 | protein-coding |
| chr1_52133078_52133828 | chr1 | 52133079 | 52133828 | + | 1.10172 | intron (NM_009283, intron 6 of 24) | intron (NM_009283, intron 6 of 24) | 13831 | NM_001205313 | 20846 | Mm.277406 | NM_009283 | ENSMUSG00000026104 | **Stat1** | 2010005J02Rik\|AA408197 | signal transducer and activator of transcription 1 | protein-coding |
| chr1_52143605_52144068 | chr1 | 52143606 | 52144068 | + | 0.761387 | intron (NM_009283, intron 12 of 24) | intron (NM_009283, intron 12 of 24) | 24215 | NM_001205313 | 20846 | Mm.277406 | NM_009283 | ENSMUSG00000026104 | **Stat1** | 2010005J02Rik\|AA408197 | signal transducer and activator of transcription 1 | protein-coding |
| chr1_52144839_52145190 | chr1 | 52144840 | 52145190 | + | 0.669977 | intron (NM_009283, intron 14 of 24) | intron (NM_009283, intron 14 of 24) | 25393 | NM_001205313 | 20846 | Mm.277406 | NM_009283 | ENSMUSG00000026104 | **Stat1** | 2010005J02Rik\|AA408197 | signal transducer and activator of transcription 1 | protein-coding |
| chr1_52078609_52079161 | chr1 | 52078610 | 52079161 | + | 0.626865 | intron (NM_011487, intron 9 of 23) | intron (NM_011487, intron 9 of 23) | -40553 | NM_009283 | 20846 | Mm.277406 | NM_009283 | ENSMUSG00000026104 | **Stat1** | 2010005J02Rik\|AA408197 | signal transducer and activator of transcription 1 | protein-coding |
| chr11_100921272_100921849 | chr11 | 100921273 | 100921849 | + | 1.6064 | intron (NM_213659, intron 1 of 23) | intron (NM_213659, intron 1 of 23) | 17950 | NM_213660 | 20848 | Mm.249934 | NM_011486 | ENSMUSG00000004040 | **Stat3** | 1110034C02Rik\|AW109958\|Aprf | signal transducer and activator of transcription 3 | protein-coding |
| chr1_52039178_52040069 | chr1 | 52039179 | 52040069 | + | 1.71446 | intron (NM_011487, intron 3 of 23) | intron (NM_011487, intron 3 of 23) | 31380 | NM_011487 | 20849 | Mm.1550 | NM_011487 | ENSMUSG00000062939 | **Stat4** | - | signal transducer and activator of transcription 4 | protein-coding |
| chr1_52052961_52053346 | chr1 | 52052962 | 52053346 | + | 1.39422 | intron (NM_011487, intron 3 of 23) | intron (NM_011487, intron 3 of 23) | 44910 | NM_011487 | 20849 | Mm.1550 | NM_011487 | ENSMUSG00000062939 | **Stat4** | - | signal transducer and activator of transcription 4 | protein-coding |
| chr1_52040715_52041005 | chr1 | 52040716 | 52041005 | + | 1.12836 | intron (NM_011487, intron 3 of 23) | intron (NM_011487, intron 3 of 23) | 32616 | NM_011487 | 20849 | Mm.1550 | NM_011487 | ENSMUSG00000062939 | **Stat4** | - | signal transducer and activator of transcription 4 | protein-coding |
| chr1_52040153_52040609 | chr1 | 52040154 | 52040609 | + | 1.10515 | intron (NM_011487, intron 3 of 23) | intron (NM_011487, intron 3 of 23) | 32137 | NM_011487 | 20849 | Mm.1550 | NM_011487 | ENSMUSG00000062939 | **Stat4** | - | signal transducer and activator of transcription 4 | protein-coding |
| chr1_52027615_52028145 | chr1 | 52027616 | 52028145 | + | 0.924504 | intron (NM_011487, intron 3 of 23) | intron (NM_011487, intron 3 of 23) | 19636 | NM_011487 | 20849 | Mm.1550 | NM_011487 | ENSMUSG00000062939 | **Stat4** | - | signal transducer and activator of transcription 4 | protein-coding |
| chr1_52055802_52056435 | chr1 | 52055803 | 52056435 | + | 0.919773 | intron (NM_011487, intron 3 of 23) | intron (NM_011487, intron 3 of 23) | 47875 | NM_011487 | 20849 | Mm.1550 | NM_011487 | ENSMUSG00000062939 | **Stat4** | - | signal transducer and activator of transcription 4 | protein-coding |
| chr1_51970117_51970550 | chr1 | 51970118 | 51970550 | + | 0.869368 | Intergenic | Intergenic | -37910 | NM_011487 | 20849 | Mm.1550 | NM_011487 | ENSMUSG00000062939 | **Stat4** | - | signal transducer and activator of transcription 4 | protein-coding |
| chr11_100861787_100862247 | chr11 | 100861788 | 100862247 | + | 0.6199 | intron (NM_001164062, intron 3 of 17) | intron (NM_001164062, intron 3 of 17) | 1533 | NM_001164062 | 20850 | Mm.277403 | NM_011488 | ENSMUSG00000004043 | **Stat5a** | AA959963\|STAT5 | signal transducer and activator of transcription 5A | protein-coding |
| chr11_100852080_100852617 | chr11 | 100852081 | 100852617 | + | 0.797094 | Intergenic | Intergenic | -1764 | NM_001113563 | 20851 | Mm.34064 | NM_011489 | ENSMUSG00000020919 | **Stat5b** | - | signal transducer and activator of transcription 5B | protein-coding |
| chr11_100844184_100844418 | chr11 | 100844185 | 100844418 | + | 0.730717 | intron (NM_001113563, intron 1 of 18) | intron (NM_001113563, intron 1 of 18) | 6284 | NM_001113563 | 20851 | Mm.34064 | NM_011489 | ENSMUSG00000020919 | **Stat5b** | - | signal transducer and activator of transcription 5B | protein-coding |
| chr1_120236412_120236776 | chr1 | 120236413 | 120236776 | + | 0.689217 | intron (NM_001085409, intron 4 of 5) | intron (NM_001085409, intron 4 of 5) | 28686 | NM_133186 | 68428 | Mm.181033 | NM_133186 | ENSMUSG00000026389 | **Steap3** | 1010001D01Rik\|Tsap6\|pHyde | STEAP family member 3 | protein-coding |
| chr16_36505842_36506279 | chr16 | 36505843 | 36506279 | + | 1.05588 | intron (NM_013803, intron 4 of 6) | intron (NM_013803, intron 4 of 6) | -50669 | NM_025288 | 20863 | Mm.136573 | NM_025288 | ENSMUSG00000054905 | **Stfa3** | Stf3 | stefin A3 | protein-coding |
| chr16_36505214_36505519 | chr16 | 36505215 | 36505519 | + | 1.05103 | intron (NM_013803, intron 4 of 6) | L1M2\|LINE\|L1 | -49975 | NM_025288 | 20863 | Mm.136573 | NM_025288 | ENSMUSG00000054905 | **Stfa3** | Stf3 | stefin A3 | protein-coding |
| chr7_102347325_102347898 | chr7 | 102347326 | 102347898 | + | 0.857219 | intron (NM_009287, intron 1 of 11) | intron (NM_009287, intron 1 of 11) | 79788 | NM_009287 | 20866 | Mm.645 | NM_009287 | ENSMUSG00000030987 | **Stim1** | SIM | stromal interaction molecule 1 | protein-coding |

| | | | | | | | | | | | | | | | | | |
|---|---|---|---|---|---|---|---|---|---|---|---|---|---|---|---|---|---|
| chr5_54105279_54106033 | chr5 | 54105280 | 54106033 | + | 1.4174 | intron (NM_001081103, intron 7 of 11) | intron (NM_001081103, intron 7 of 11) | 107133 | NM_001081103 | 116873 | Mm.265985 | NM_054083 | ENSMUSG00000039156 | **Stim2** | - | stromal interaction molecule 2 | protein-coding |
| chr5_54024655_54025270 | chr5 | 54024656 | 54025270 | + | 1.33666 | intron (NM_001081103, intron 1 of 11) | intron (NM_001081103, intron 1 of 11) | 26440 | NM_001081103 | 116873 | Mm.265985 | NM_054083 | ENSMUSG00000039156 | **Stim2** | - | stromal interaction molecule 2 | protein-coding |
| chr5_54013290_54013892 | chr5 | 54013291 | 54013892 | + | 0.9252 | intron (NM_001081103, intron 1 of 11) | intron (NM_001081103, intron 1 of 11) | 15068 | NM_001081103 | 116873 | Mm.265985 | NM_054083 | ENSMUSG00000039156 | **Stim2** | - | stromal interaction molecule 2 | protein-coding |
| chr5_54029800_54030163 | chr5 | 54029801 | 54030163 | + | 0.924112 | intron (NM_001081103, intron 1 of 11) | ERVL-B4-int\|LTR\|ERVL | 31459 | NM_001081103 | 116873 | Mm.265985 | NM_054083 | ENSMUSG00000039156 | **Stim2** | - | stromal interaction molecule 2 | protein-coding |
| chr5_54061403_54061869 | chr5 | 54061404 | 54061869 | + | 0.857517 | intron (NM_001081103, intron 2 of 11) | RLTR15\|LTR\|ERVK | 63113 | NM_001081103 | 116873 | Mm.265985 | NM_054083 | ENSMUSG00000039156 | **Stim2** | - | stromal interaction molecule 2 | protein-coding |
| chr5_54012486_54012901 | chr5 | 54012487 | 54012901 | + | 0.793765 | intron (NM_001081103, intron 1 of 11) | intron (NM_001081103, intron 1 of 11) | 14171 | NM_001081103 | 116873 | Mm.265985 | NM_054083 | ENSMUSG00000039156 | **Stim2** | - | stromal interaction molecule 2 | protein-coding |
| chr5_54108276_54108508 | chr5 | 54108277 | 54108508 | + | 0.776185 | intron (NM_001081103, intron 7 of 11) | intron (NM_001081103, intron 7 of 11) | 109869 | NM_001081103 | 116873 | Mm.265985 | NM_054083 | ENSMUSG00000039156 | **Stim2** | - | stromal interaction molecule 2 | protein-coding |
| chr5_53971065_53971574 | chr5 | 53971066 | 53971574 | + | 0.768358 | Intergenic | Intergenic | -27203 | NM_001081103 | 116873 | Mm.265985 | NM_054083 | ENSMUSG00000039156 | **Stim2** | - | stromal interaction molecule 2 | protein-coding |
| chr5_54046487_54046835 | chr5 | 54046488 | 54046835 | + | 0.668661 | intron (NM_001081103, intron 1 of 11) | intron (NM_001081103, intron 1 of 11) | 48138 | NM_001081103 | 116873 | Mm.265985 | NM_054083 | ENSMUSG00000039156 | **Stim2** | - | stromal interaction molecule 2 | protein-coding |
| chr5_54097949_54098705 | chr5 | 54097950 | 54098705 | + | 0.650679 | intron (NM_001081103, intron 3 of 11) | intron (NM_001081103, intron 3 of 11) | 99804 | NM_001081103 | 116873 | Mm.265985 | NM_054083 | ENSMUSG00000039156 | **Stim2** | - | stromal interaction molecule 2 | protein-coding |
| chr5_54059986_54060426 | chr5 | 54059987 | 54060426 | + | 0.648596 | intron (NM_001081103, intron 2 of 11) | MER5A\|DNA\|hAT-Charlie | 61683 | NM_001081103 | 116873 | Mm.265985 | NM_054083 | ENSMUSG00000039156 | **Stim2** | - | stromal interaction molecule 2 | protein-coding |
| chr5_54110861_54111173 | chr5 | 54110862 | 54111173 | + | 0.636995 | exon (NM_001081103, exon 10 of 12) | exon (NM_001081103, exon 10 of 12) | 112494 | NM_001081103 | 116873 | Mm.265985 | NM_054083 | ENSMUSG00000039156 | **Stim2** | - | stromal interaction molecule 2 | protein-coding |
| chr11_32558008_32558683 | chr11 | 32558009 | 32558683 | + | 1.15024 | intron (NM_009288, intron 2 of 18) | intron (NM_009288, intron 2 of 18) | 25080 | NM_009288 | 20868 | Mm.8235 | NM_009288 | ENSMUSG00000020272 | **Stk10** | Gek1\|Lok\|mKIAA4026 | serine/threonine kinase 10 | protein-coding |
| chr14_121344935_121345493 | chr14 | 121344936 | 121345493 | + | 1.31935 | intron (NM_145465, intron 1 of 10) | intron (NM_145465, intron 1 of 10) | 34016 | NM_145465 | 223255 | Mm.390756 | NM_145465 | ENSMUSG00000063410 | **Stk24** | 1810013H02Rik\|C76483\|MST-3\|Mst3\|STE20 | serine/threonine kinase 24 | protein-coding |
| chr14_121327028_121327272 | chr14 | 121327029 | 121327272 | + | 0.674921 | intron (NM_145465, intron 2 of 10) | intron (NM_145465, intron 2 of 10) | 52080 | NM_145465 | 223255 | Mm.390756 | NM_145465 | ENSMUSG00000063410 | **Stk24** | 1810013H02Rik\|C76483\|MST-3\|Mst3\|STE20 | serine/threonine kinase 24 | protein-coding |
| chr15_35175811_35176223 | chr15 | 35175812 | 35176223 | + | 0.954505 | Intergenic | Lx\|LINE\|L1 | -20211 | NM_019635 | 56274 | Mm.262330 | NM_019635 | ENSMUSG00000022329 | **Stk3** | 0610042I06Rik\|MST\|Mst2\|Mst3\|Ste20\|mess1 | serine/threonine kinase 3 | protein-coding |
| chr15_35157362_35157641 | chr15 | 35157363 | 35157641 | + | 0.803201 | Intergenic | Intergenic | -1696 | NM_019635 | 56274 | Mm.262330 | NM_019635 | ENSMUSG00000022329 | **Stk3** | 0610042I06Rik\|MST\|Mst2\|Mst3\|Ste20\|mess1 | serine/threonine kinase 3 | protein-coding |
| chr7_139151100_139151644 | chr7 | 139151101 | 139151644 | + | 0.997369 | intron (NM_001162540, intron 1 of 11) | intron (NM_001162540, intron 1 of 11) | 37122 | NM_021302 | 57740 | Mm.65336 | NM_021302 | ENSMUSG00000015981 | **Stk32c** | PKE\|Pkek\|YANK3 | serine/threonine kinase 32C | protein-coding |
| chr7_139151094_139151865 | chr7 | 139151095 | 139151865 | + | 0.845881 | intron (NM_001162540, intron 1 of 11) | intron (NM_001162540, intron 1 of 11) | 37014 | NM_021302 | 57740 | Mm.65336 | NM_021302 | ENSMUSG00000015981 | **Stk32c** | PKE\|Pkek\|YANK3 | serine/threonine kinase 32C | protein-coding |
| chr7_109456525_109457260 | chr7 | 109456526 | 109457260 | + | 1.38807 | intron (NM_181853, intron 16 of 19) | ORR1F\|LTR\|ERVL-MaLR | -17840 | NM_054103 | 117229 | Mm.389950 | NM_054103 | ENSMUSG00000031027 | **Stk33** | 4921505G21Rik\|AW061264 | serine/threonine kinase 33 | protein-coding |
| chr7_109417923_109418584 | chr7 | 109417924 | 109418584 | + | 0.794036 | intron (NM_054103, intron 1 of 13) | Lx4B\|LINE\|L1 | 20799 | NM_054103 | 117229 | Mm.389950 | NM_054103 | ENSMUSG00000031027 | **Stk33** | 4921505G21Rik\|AW061264 | serine/threonine kinase 33 | protein-coding |
| chr6_146748321_146748666 | chr6 | 146748322 | 146748666 | + | 1.39431 | intron (NM_172734, intron 1 of 13) | intron (NM_172734, intron 1 of 13) | 23564 | NM_172734 | 232533 | Mm.322121 | NM_172734 | ENSMUSG00000001630 | **Stk38l** | 4930473A22Rik\|B230328I19\|Ndr2\|Ndr54 | serine/threonine kinase 38 like | protein-coding |
| chr6_146733965_146734720 | chr6 | 146733966 | 146734720 | + | 1.10398 | intron (NM_172734, intron 1 of 13) | intron (NM_172734, intron 1 of 13) | 9413 | NM_172734 | 232533 | Mm.322121 | NM_172734 | ENSMUSG00000001630 | **Stk38l** | 4930473A22Rik\|B230328I19\|Ndr2\|Ndr54 | serine/threonine kinase 38 like | protein-coding |
| chr6_146747666_146747879 | chr6 | 146747667 | 146747879 | + | 0.810924 | intron (NM_172734, intron 1 of 13) | intron (NM_172734, intron 1 of 13) | 22843 | NM_172734 | 232533 | Mm.322121 | NM_172734 | ENSMUSG00000001630 | **Stk38l** | 4930473A22Rik\|B230328I19\|Ndr2\|Ndr54 | serine/threonine kinase 38 like | protein-coding |
| chr6_146747585_146747921 | chr6 | 146747586 | 146747921 | + | 0.804903 | intron (NM_172734, intron 1 of 13) | intron (NM_172734, intron 1 of 13) | 22823 | NM_172734 | 232533 | Mm.322121 | NM_172734 | ENSMUSG00000001630 | **Stk38l** | 4930473A22Rik\|B230328I19\|Ndr2\|Ndr54 | serine/threonine kinase 38 like | protein-coding |
| chr6_146746691_146747328 | chr6 | 146746692 | 146747328 | + | 0.726445 | intron (NM_172734, intron 1 of 13) | intron (NM_172734, intron 1 of 13) | 22080 | NM_172734 | 232533 | Mm.322121 | NM_172734 | ENSMUSG00000001630 | **Stk38l** | 4930473A22Rik\|B230328I19\|Ndr2\|Ndr54 | serine/threonine kinase 38 like | protein-coding |
| chr2_68434507_68435104 | chr2 | 68434508 | 68435104 | + | 1.22244 | intron (NM_016866, intron 1 of 17) | intron (NM_016866, intron 1 of 17) | 37175 | NM_016866 | 53416 | Mm.198414 | NM_016866 | ENSMUSG00000027030 | **Stk39** | AW227544\|AW556857\|DCHT\|RF005\|Rnl5\|SPAK | serine/threonine kinase 39 | protein-coding |
| chr2_68303577_68303977 | chr2 | 68303578 | 68303977 | + | 1.16787 | intron (NM_016866, intron 14 of 17) | RLTR24\|LTR\|ERV1 | 168204 | NM_016866 | 53416 | Mm.198414 | NM_016866 | ENSMUSG00000027030 | **Stk39** | AW227544\|AW556857\|DCHT\|RF005\|Rnl5\|SPAK | serine/threonine kinase 39 | protein-coding |
| chr2_68398305_68398584 | chr2 | 68398306 | 68398584 | + | 1.11393 | exon (NM_016866, exon 3 of 18) | exon (NM_016866, exon 3 of 18) | 73536 | NM_016866 | 53416 | Mm.198414 | NM_016866 | ENSMUSG00000027030 | **Stk39** | AW227544\|AW556857\|DCHT\|RF005\|Rnl5\|SPAK | serine/threonine kinase 39 | protein-coding |
| chr2_68336201_68337213 | chr2 | 68336202 | 68337213 | + | 1.02745 | intron (NM_016866, intron 10 of 17) | intron (NM_016866, intron 10 of 17) | 135274 | NM_016866 | 53416 | Mm.198414 | NM_016866 | ENSMUSG00000027030 | **Stk39** | AW227544\|AW556857\|DCHT\|RF005\|Rnl5\|SPAK | serine/threonine kinase 39 | protein-coding |
| chr2_68436357_68436820 | chr2 | 68436358 | 68436820 | + | 0.937602 | intron (NM_016866, intron 1 of 17) | intron (NM_016866, intron 1 of 17) | 35392 | NM_016866 | 53416 | Mm.198414 | NM_016866 | ENSMUSG00000027030 | **Stk39** | AW227544\|AW556857\|DCHT\|RF005\|Rnl5\|SPAK | serine/threonine kinase 39 | protein-coding |
| chr2_68336226_68336648 | chr2 | 68336227 | 68336648 | + | 0.923891 | intron (NM_016866, intron 10 of 17) | intron (NM_016866, intron 10 of 17) | 135544 | NM_016866 | 53416 | Mm.198414 | NM_016866 | ENSMUSG00000027030 | **Stk39** | AW227544\|AW556857\|DCHT\|RF005\|Rnl5\|SPAK | serine/threonine kinase 39 | protein-coding |
| chr2_68289094_68289600 | chr2 | 68289095 | 68289600 | + | 0.821112 | intron (NM_016866, intron 14 of 17) | intron (NM_016866, intron 14 of 17) | 182634 | NM_016866 | 53416 | Mm.198414 | NM_016866 | ENSMUSG00000027030 | **Stk39** | AW227544\|AW556857\|DCHT\|RF005\|Rnl5\|SPAK | serine/threonine kinase 39 | protein-coding |
| chr2_68301853_68302311 | chr2 | 68301854 | 68302311 | + | 0.729245 | intron (NM_016866, intron 14 of 17) | intron (NM_016866, intron 14 of 17) | 169899 | NM_016866 | 53416 | Mm.198414 | NM_016866 | ENSMUSG00000027030 | **Stk39** | AW227544\|AW556857\|DCHT\|RF005\|Rnl5\|SPAK | serine/threonine kinase 39 | protein-coding |
| chr2_68398085_68398840 | chr2 | 68398086 | 68398840 | + | 0.71314 | exon (NM_016866, exon 3 of 18) | exon (NM_016866, exon 3 of 18) | 73518 | NM_016866 | 53416 | Mm.198414 | NM_016866 | ENSMUSG00000027030 | **Stk39** | AW227544\|AW556857\|DCHT\|RF005\|Rnl5\|SPAK | serine/threonine kinase 39 | protein-coding |
| chr2_68301866_68302155 | chr2 | 68301867 | 68302155 | + | 0.705619 | intron (NM_016866, intron 14 of 17) | intron (NM_016866, intron 14 of 17) | 169970 | NM_016866 | 53416 | Mm.198414 | NM_016866 | ENSMUSG00000027030 | **Stk39** | AW227544\|AW556857\|DCHT\|RF005\|Rnl5\|SPAK | serine/threonine kinase 39 | protein-coding |
| chr2_68425971_68426327 | chr2 | 68425972 | 68426327 | + | 0.655701 | intron (NM_016866, intron 1 of 17) | intron (NM_016866, intron 1 of 17) | 45832 | NM_016866 | 53416 | Mm.198414 | NM_016866 | ENSMUSG00000027030 | **Stk39** | AW227544\|AW556857\|DCHT\|RF005\|Rnl5\|SPAK | serine/threonine kinase 39 | protein-coding |
| chr4_126102525_126102780 | chr4 | 126102526 | 126102780 | + | 0.766851 | Intergenic | Intergenic | -1304 | NM_028800 | 74178 | Mm.440269 | NM_028800 | ENSMUSG00000042608 | **Stk40** | 2310004N11Rik\|AI428792 | serine/threonine kinase 40 | protein-coding |
| chr9_115236292_115237114 | chr9 | 115236293 | 115237114 | + | 0.791612 | Intergenic | Intergenic | 45367 | NR_029430 | 111186 | Mm.484753 | NR_029430 | | **Stmn1-rs1** | Lag-rs1\|Lap18-rs1 | stathmin 1, related sequence 1 | pseudo |
| chr2_181303232_181303585 | chr2 | 181303233 | 181303585 | + | 1.03298 | Intergenic | MLT1F2\|LTR\|ERVL-MaLR | 11091 | NM_009133 | 20262 | Mm.2319 | NM_009133 | ENSMUSG00000027581 | **Stmn3** | 9330161A03Rik\|AI604839\|R75323\|Sclip | stathmin-like 3 | protein-coding |
| chr2_181303225_181303982 | chr2 | 181303226 | 181303982 | + | 0.973886 | Intergenic | Intergenic | 10896 | NM_009133 | 20262 | Mm.2319 | NM_009133 | ENSMUSG00000027581 | **Stmn3** | 9330161A03Rik\|AI604839\|R75323\|Sclip | stathmin-like 3 | protein-coding |
| chr14_66340778_66341499 | chr14 | 66340779 | 66341499 | + | 0.687904 | Intergenic | Intergenic | -3236 | NM_019675 | 56471 | Mm.35474 | NM_019675 | ENSMUSG00000022044 | **Stmn4** | Lagl\|R75359\|RB3 | stathmin-like 4 | protein-coding |
| chr13_46212976_46213427 | chr13 | 46212977 | 46213427 | + | 0.794436 | Intergenic | ORR1E\|LTR\|ERVL-MaLR | -60519 | NM_001005422 | 380842 | Mm.188108 | NM_001005422 | ENSMUSG00000063529 | **Stmnd1** | Gm1574 | stathmin domain containing 1 | protein-coding |
| chr9_58274622_58274905 | chr9 | 58274623 | 58274905 | + | 0.826145 | Intergenic | Intergenic | 21600 | NM_026942 | 69106 | Mm.216502 | NM_026942 | ENSMUSG00000032333 | **Stoml1** | 1810015E19Rik\|AI847431\|SLP1\|STORP\|UNC24\|WPB72 | stomatin-like 1 | protein-coding |
| chr9_58265404_58265968 | chr9 | 58265405 | 58265968 | + | 0.703714 | Intergenic | Intergenic | 12522 | NM_026942 | 69106 | Mm.216502 | NM_026942 | ENSMUSG00000032333 | **Stoml1** | 1810015E19Rik\|AI847431\|SLP1\|STORP\|UNC24\|WPB72 | stomatin-like 1 | protein-coding |
| chr9_58255759_58256400 | chr9 | 58255760 | 58256400 | + | 0.677323 | intron (NR_028146, intron 3 of 7) | intron (NR_028146, intron 3 of 7) | 2916 | NM_026942 | 69106 | Mm.216502 | NM_026942 | ENSMUSG00000032333 | **Stoml1** | 1810015E19Rik\|AI847431\|SLP1\|STORP\|UNC24\|WPB72 | stomatin-like 1 | protein-coding |

| | | | | | | | | | | | | | | | | | |
|---|---|---|---|---|---|---|---|---|---|---|---|---|---|---|---|---|---|
| chr9_58275089_58275558 | chr9 | 58275090 | 58275558 | + | 0.627271 | Intergenic | Intergenic | 22160 | NM_026942 | 69106 | Mm.216502 | NM_026942 | ENSMUSG00000032333 | **Stoml1** | 1810015E19Rik\|AI847431\|SLP1\|STORP\|UNC24\|WPB72 | stomatin-like 1 | protein-coding |
| chr9_58262986_58263223 | chr9 | 58262987 | 58263223 | + | 0.59961 | TTS (NM_026942) | TTS (NM_026942) | 9941 | NM_026942 | 69106 | Mm.216502 | NM_026942 | ENSMUSG00000032333 | **Stoml1** | 1810015E19Rik\|AI847431\|SLP1\|STORP\|UNC24\|WPB72 | stomatin-like 1 | protein-coding |
| chr17_88598816_88599549 | chr17 | 88598817 | 88599549 | + | 0.859551 | Intergenic | Intergenic | -27372 | NM_029858 | 77057 | Mm.482203 | NM_029858 | ENSMUSG00000033855 | **Ston1** | 4921524J06Rik\|C86231\|Salf\|Sblf | stonin 1 | protein-coding |
| chr12_91803197_91804002 | chr12 | 91803198 | 91804002 | + | 1.75761 | Intergenic | Intergenic | -17164 | NM_175367 | 108800 | Mm.23125 | NM_175367 | | **Ston2** | 4933401N24Rik\|AI425999\|AW544599 | stonin 2 | protein-coding |
| chr12_91802771_91804019 | chr12 | 91802772 | 91804019 | + | 1.6894 | Intergenic | Intergenic | -16959 | NM_175367 | 108800 | Mm.23125 | NM_175367 | | **Ston2** | 4933401N24Rik\|AI425999\|AW544599 | stonin 2 | protein-coding |
| chr12_91774636_91775008 | chr12 | 91774637 | 91775008 | + | 1.04446 | intron (NM_175367, intron 1 of 6) | intron (NM_175367, intron 1 of 6) | 11614 | NM_175367 | 108800 | Mm.23125 | NM_175367 | | **Ston2** | 4933401N24Rik\|AI425999\|AW544599 | stonin 2 | protein-coding |
| chr12_91802805_91803121 | chr12 | 91802806 | 91803121 | + | 0.905493 | Intergenic | Intergenic | -16527 | NM_175367 | 108800 | Mm.23125 | NM_175367 | | **Ston2** | 4933401N24Rik\|AI425999\|AW544599 | stonin 2 | protein-coding |
| chr8_47217969_47218423 | chr8 | 47217970 | 47218423 | + | 0.792353 | intron (NM_175162, intron 1 of 4) | intron (NM_175162, intron 1 of 4) | 71206 | NM_001286376 | 71069 | Mm.23973 | NM_175162 | ENSMUSG00000038143 | **Stox2** | 4933409N07Rik\|AI449080\|AI835114\|mKIAA1392 | storkhead box 2 | protein-coding |
| chr6_34923776_34924126 | chr6 | 34923777 | 34924126 | + | 1.19144 | intron (NM_009292, intron 1 of 8) | intron (NM_009292, intron 1 of 8) | 2991 | NM_009292 | 20899 | Mm.5171 | NM_009292 | ENSMUSG00000029848 | **Stra8** | - | stimulated by retinoic acid gene 8 | protein-coding |
| chr17_78665005_78665678 | chr17 | 78665006 | 78665678 | + | 1.16838 | intron (NM_011500, intron 10 of 17) | intron (NM_011500, intron 10 of 17) | 71218 | NM_011500 | 268980 | Mm.311915 | NM_011500 | ENSMUSG00000024077 | **Strn** | AU022939\|D130055P19 | striatin, calmodulin binding protein | protein-coding |
| chr7_16815006_16815572 | chr7 | 16815007 | 16815572 | + | 0.733645 | promoter-TSS (NM_133789) | promoter-TSS (NM_133789) | -600 | NM_133789 | 97387 | Mm.21612 | NM_133789 | ENSMUSG00000030374 | **Strn4** | C80611\|ZIN\|zinedin | striatin, calmodulin binding protein 4 | protein-coding |
| chr9_115299053_115299431 | chr9 | 115299054 | 115299431 | + | 0.824742 | intron (NM_024222, intron 1 of 15) | intron (NM_024222, intron 1 of 15) | 11179 | NM_024222 | 68292 | Mm.296158 | NM_024222 | ENSMUSG00000032437 | **Stt3b** | 1300006C19Rik\|Simp | STT3, subunit of the oligosaccharyltransferase complex, homolog B (S. cerevisiae) | protein-coding |
| chr9_115300349_115300709 | chr9 | 115300350 | 115300709 | + | 0.823527 | intron (NM_024222, intron 1 of 15) | URR1B\|DNA\|hAT-Charlie | 9892 | NM_024222 | 68292 | Mm.296158 | NM_024222 | ENSMUSG00000032437 | **Stt3b** | 1300006C19Rik\|Simp | STT3, subunit of the oligosaccharyltransferase complex, homolog B (S. cerevisiae) | protein-coding |
| chr10_12977512_12978335 | chr10 | 12977513 | 12978335 | + | 1.17489 | Intergenic | Intergenic | -13665 | NM_001163591 | 74732 | Mm.248648 | NM_029075 | ENSMUSG00000039232 | **Stx11** | 5830405C08Rik | syntaxin 11 | protein-coding |
| chr10_12965718_12966056 | chr10 | 12965719 | 12966056 | + | 1.16804 | Intergenic | Intergenic | -1628 | NM_001163591 | 74732 | Mm.248648 | NM_029075 | ENSMUSG00000039232 | **Stx11** | 5830405C08Rik | syntaxin 11 | protein-coding |
| chr7_127815886_127816718 | chr7 | 127815887 | 127816718 | + | 0.726036 | intron (NM_024414, intron 1 of 9) | intron (NM_024414, intron 1 of 9) | 8229 | NM_024414 | 56216 | Mm.439759 | NM_024414 | | **Stx1b** | Stx1b2\|Stx1bl\|Syn1b | syntaxin 1B | protein-coding |
| chr5_129011450_129011830 | chr5 | 129011451 | 129011830 | + | 1.02354 | Intergenic | Intergenic | -3068 | NM_001286034 | 13852 | Mm.260716 | NM_007941 | ENSMUSG00000029428 | **Stx2** | AW538950\|Epim\|G1-536-1\|Syn-2\|repro34 | syntaxin 2 | protein-coding |
| chr19_11812986_11813562 | chr19 | 11812987 | 11813562 | + | 1.09117 | intron (NM_001286543, intron 1 of 11) | intron (NM_001286543, intron 1 of 11) | 6129 | NM_001286543 | 20908 | Mm.272264 | NM_011502 | ENSMUSG00000041488 | **Stx3** | Syn-3 | syntaxin 3 | protein-coding |
| chr1_155201058_155201865 | chr1 | 155201059 | 155201865 | + | 1.29729 | intron (NM_021433, intron 7 of 7) | intron (NM_021433, intron 7 of 7) | 42759 | NM_021433 | 58244 | Mm.66264 | NM_021433 | ENSMUSG00000026470 | **Stx6** | 2310039E05Rik\|2410005I16Rik\|AA437559\|AI415714\|AI845861 | syntaxin 6 | protein-coding |
| chr3_108850434_108850913 | chr3 | 108850435 | 108850913 | + | 1.28948 | Intergenic | Intergenic | -10172 | NM_011504 | 20912 | Mm.316894 | NM_011504 | ENSMUSG00000027882 | **Stxbp3a** | Munc-18c\|Stxbp3\|Sxtbp3\|Unc18-3 | syntaxin binding protein 3A | protein-coding |
| chr3_108855881_108856241 | chr3 | 108855882 | 108856241 | + | 0.869142 | 3' UTR (NM_001165965, exon 12 of 12) | 3' UTR (NM_001165965, exon 12 of 12) | -15559 | NM_011504 | 20912 | Mm.316894 | NM_011504 | ENSMUSG00000027882 | **Stxbp3a** | Munc-18c\|Stxbp3\|Sxtbp3\|Unc18-3 | syntaxin binding protein 3A | protein-coding |
| chr3_108794927_108795144 | chr3 | 108794928 | 108795144 | + | 0.766136 | intron (NM_011504, intron 17 of 18) | intron (NM_011504, intron 17 of 18) | 45466 | NM_011504 | 20912 | Mm.316894 | NM_011504 | ENSMUSG00000027882 | **Stxbp3a** | Munc-18c\|Stxbp3\|Sxtbp3\|Unc18-3 | syntaxin binding protein 3A | protein-coding |
| chr15_12034335_12034675 | chr15 | 12034336 | 12034675 | + | 1.01835 | Intergenic | Intergenic | -38498 | NM_011294 | 20024 | Mm.41746 | NM_011294 | ENSMUSG00000022205 | **Sub1** | AI842364\|P15\|P9\|Pc4\|Rpo2tc1 | SUB1 homolog (S. cerevisiae) | protein-coding |
| chr15_12030135_12030679 | chr15 | 12030136 | 12030679 | + | 0.887502 | Intergenic | (CCA)n\|Simple_repeat\|Simple_repeat | -34400 | NM_011294 | 20024 | Mm.41746 | NM_011294 | ENSMUSG00000022205 | **Sub1** | AI842364\|P15\|P9\|Pc4\|Rpo2tc1 | SUB1 homolog (S. cerevisiae) | protein-coding |
| chr15_12013956_12014382 | chr15 | 12013957 | 12014382 | + | 0.815209 | Intergenic | RMER4A\|LTR\|ERVK | -18162 | NM_011294 | 20024 | Mm.41746 | NM_011294 | ENSMUSG00000022205 | **Sub1** | AI842364\|P15\|P9\|Pc4\|Rpo2tc1 | SUB1 homolog (S. cerevisiae) | protein-coding |
| chr15_12039103_12039517 | chr15 | 12039104 | 12039517 | + | 0.73299 | Intergenic | Intergenic | -43303 | NM_011294 | 20024 | Mm.41746 | NM_011294 | ENSMUSG00000022205 | **Sub1** | AI842364\|P15\|P9\|Pc4\|Rpo2tc1 | SUB1 homolog (S. cerevisiae) | protein-coding |
| chr15_12026094_12026348 | chr15 | 12026095 | 12026348 | + | 0.611155 | Intergenic | L1MB5\|LINE\|L1 | -30214 | NM_011294 | 20024 | Mm.41746 | NM_011294 | ENSMUSG00000022205 | **Sub1** | AI842364\|P15\|P9\|Pc4\|Rpo2tc1 | SUB1 homolog (S. cerevisiae) | protein-coding |
| chr14_73578186_73578709 | chr14 | 73578187 | 73578709 | + | 1.38085 | intron (NM_011506, intron 4 of 10) | intron (NM_011506, intron 4 of 10) | 25662 | NM_011506 | 20916 | Mm.38951 | NM_011506 | ENSMUSG00000022110 | **Sucla2** | 4930547K18Rik | succinate-Coenzyme A ligase, ADP-forming, beta subunit | protein-coding |
| chr14_73577603_73577891 | chr14 | 73577604 | 73577891 | + | 0.710134 | intron (NM_011506, intron 4 of 10) | ORR1B1\|LTR\|ERVL-MaLR | 24961 | NM_011506 | 20916 | Mm.38951 | NM_011506 | ENSMUSG00000022110 | **Sucla2** | 4930547K18Rik | succinate-Coenzyme A ligase, ADP-forming, beta subunit | protein-coding |
| chr6_73249724_73250024 | chr6 | 73249725 | 73250024 | + | 0.737615 | intron (NM_019879, intron 1 of 8) | intron (NM_019879, intron 1 of 8) | 1369 | NM_019879 | 56451 | Mm.29845 | NM_019879 | ENSMUSG00000052738 | **Suclg1** | 1500000I01Rik\|Sucla1 | succinate-CoA ligase, GDP-forming, alpha subunit | protein-coding |
| chr6_95613293_95613677 | chr6 | 95613294 | 95613677 | + | 0.873447 | intron (NM_011507, intron 2 of 10) | intron (NM_011507, intron 2 of 10) | 105361 | NM_011507 | 20917 | Mm.371585 | NM_011507 | ENSMUSG00000061838 | **Suclg2** | AF171077\|AW556404\|D6Wsu120e\|SCS-betaG | succinate-Coenzyme A ligase, GDP-forming, beta subunit | protein-coding |
| chr6_95621718_95622259 | chr6 | 95621719 | 95622259 | + | 0.863976 | intron (NM_011507, intron 2 of 10) | intron (NM_011507, intron 2 of 10) | 96857 | NM_011507 | 20917 | Mm.371585 | NM_011507 | ENSMUSG00000061838 | **Suclg2** | AF171077\|AW556404\|D6Wsu120e\|SCS-betaG | succinate-Coenzyme A ligase, GDP-forming, beta subunit | protein-coding |
| chr3_60123851_60124851 | chr3 | 60123852 | 60124851 | + | 1.19616 | Intergenic | Intergenic | 42482 | NM_032400 | 84112 | Mm.125110 | NM_032400 | ENSMUSG00000027762 | **Sucnr1** | Gpr91 | succinate receptor 1 | protein-coding |
| chr3_60128159_60128563 | chr3 | 60128160 | 60128563 | + | 1.00362 | Intergenic | Intergenic | 46492 | NM_032400 | 84112 | Mm.125110 | NM_032400 | ENSMUSG00000027762 | **Sucnr1** | Gpr91 | succinate receptor 1 | protein-coding |
| chr3_60119833_60120182 | chr3 | 60119834 | 60120182 | + | 0.724763 | Intergenic | Lx3C\|LINE\|L1 | 38139 | NM_032400 | 84112 | Mm.125110 | NM_032400 | ENSMUSG00000027762 | **Sucnr1** | Gpr91 | succinate receptor 1 | protein-coding |
| chr6_108186144_108186506 | chr6 | 108186145 | 108186506 | + | 0.72939 | promoter-TSS (NM_145937) | promoter-TSS (NM_145937) | -742 | NM_145937 | 58911 | Mm.439876 | NM_145937 | ENSMUSG00000030101 | **Sumf1** | AA543204\|AI463102\|AI851573\|FGE | sulfatase modifying factor 1 | protein-coding |
| chr1_59673629_59674209 | chr1 | 59673630 | 59674209 | + | 1.82491 | Intergenic | ORR1F\|LTR\|ERVL-MaLR | -3085 | NM_009460 | 22218 | Mm.362118 | NM_009460 | ENSMUSG00000026021 | **Sumo1** | GMP1\|PIC1\|SENTRIN\|SMT3\|SMT3H3\|SMTP3\|SUMO-1\|Smt3C\|Ubl1 | SMT3 suppressor of mif two 3 homolog 1 (yeast) | protein-coding |
| chr5_139215046_139215743 | chr5 | 139215047 | 139215743 | + | 1.30827 | intron (NM_024451, intron 2 of 22) | intron (NM_024451, intron 2 of 22) | 237 | NM_001256116 | 77053 | Mm.210845 | NM_024451 | ENSMUSG00000036817 | **Sun1** | 4632417G13Rik\|5730434D03Rik\|Unc84a\|mKIAA0810 | Sad1 and UNC84 domain containing 1 | protein-coding |
| chr11_9048355_9049297 | chr11 | 9048356 | 9049297 | + | 0.670103 | 5' UTR (NM_001037928, exon 1 of 7) | 5' UTR (NM_001037928, exon 1 of 7) | 165 | NM_001290520 | 194974 | Mm.79210 | NM_177576 | ENSMUSG00000040985 | **Sun3** | Sunc1 | Sad1 and UNC84 domain containing 3 | protein-coding |
| chr10_128665373_128665736 | chr10 | 128665374 | 128665736 | + | 1.00257 | Intergenic | B3\|SINE\|B2 | 8363 | NM_173733 | 211389 | Mm.23352 | NM_173733 | ENSMUSG00000049858 | **Suox** | SO | sulfite oxidase | protein-coding |
| chr17_44766273_44766819 | chr17 | 44766274 | 44766819 | + | 0.861635 | intron (NM_001146038, intron 2 of 8) | intron (NM_001146038, intron 2 of 8) | -10625 | NM_178652 | 109115 | Mm.431327 | NM_178652 | ENSMUSG00000038954 | **Supt3** | 2310066G22Rik\|AI315192\|SPT3\|SPT3L\|Supt3h | suppressor of Ty 3 | protein-coding |
| chr17_44767380_44767743 | chr17 | 44767381 | 44767743 | + | 0.838192 | intron (NM_001146038, intron 2 of 8) | intron (NM_001146038, intron 2 of 8) | -9609 | NM_178652 | 109115 | Mm.431327 | NM_178652 | ENSMUSG00000038954 | **Supt3** | 2310066G22Rik\|AI315192\|SPT3\|SPT3L\|Supt3h | suppressor of Ty 3 | protein-coding |
| chr11_87724786_87725248 | chr11 | 87724787 | 87725248 | + | 1.27659 | intron (NM_172448, intron 3 of 9) | intron (NM_172448, intron 3 of 9) | -12548 | NM_009296 | 20922 | Mm.622 | NM_009296 | ENSMUSG00000020485 | **Supt4a** | AL022777\|Supt4h\|Supt4h1\|Supt4h1a | suppressor of Ty 4A | protein-coding |
| chr4_59452328_59452966 | chr4 | 59452329 | 59452966 | + | 1.88175 | Intergenic | RLTR40\|LTR\|ERVK | -14014 | NM_001163288 | 634731 | Mm.483589 | NM_001163288 | ENSMUSG00000038578 | **Susd1** | A530080P10\|Gm12528\|OTTMUSG00000007332 | sushi domain containing 1 | protein-coding |
| chr4_59366140_59366733 | chr4 | 59366141 | 59366733 | + | 0.99749 | intron (NM_001163288, intron 9 of 16) | intron (NM_001163288, intron 9 of 16) | 72196 | NM_001163288 | 634731 | Mm.483589 | NM_001163288 | ENSMUSG00000038578 | **Susd1** | A530080P10\|Gm12528\|OTTMUSG00000007332 | sushi domain containing 1 | protein-coding |
| chr4_59454115_59454417 | chr4 | 59454116 | 59454417 | + | 0.985494 | Intergenic | Intergenic | -15633 | NM_001163288 | 634731 | Mm.483589 | NM_001163288 | ENSMUSG00000038578 | **Susd1** | A530080P10\|Gm12528\|OTTMUSG00000007332 | sushi domain containing 1 | protein-coding |
| chr4_59455044_59455303 | chr4 | 59455045 | 59455303 | + | 0.893497 | Intergenic | Intergenic | -16541 | NM_001163288 | 634731 | Mm.483589 | NM_001163288 | ENSMUSG00000038578 | **Susd1** | A530080P10\|Gm12528\|OTTMUSG00000007332 | sushi domain containing 1 | protein-coding |
| chr4_59458831_59459052 | chr4 | 59458832 | 59459052 | + | 0.687796 | Intergenic | RSINE1\|SINE\|B4 | -20309 | NM_001163288 | 634731 | Mm.483589 | NM_001163288 | ENSMUSG00000038578 | **Susd1** | A530080P10\|Gm12528\|OTTMUSG00000007332 | sushi domain containing 1 | protein-coding |
| chr13_49250963_49251485 | chr13 | 49250964 | 49251485 | + | 0.893578 | Intergenic | Intergenic | -2518 | NM_028340 | 66329 | Mm.34308 | NM_025491 | ENSMUSG00000021384 | **Susd3** | 1700017I11Rik\|2810440J20Rik | sushi domain containing 3 | protein-coding |
| chr13_49272280_49273180 | chr13 | 49272281 | 49273180 | + | 0.855041 | intron (NM_015759, intron 14 of 17) | intron (NM_015759, intron 14 of 17) | -24024 | NM_028340 | 66329 | Mm.34308 | NM_025491 | ENSMUSG00000021384 | **Susd3** | 1700017I11Rik\|2810440J20Rik | sushi domain containing 3 | protein-coding |
| chr13_49241182_49241661 | chr13 | 49241183 | 49241661 | + | 0.854862 | intron (NM_028340, intron 1 of 4) | intron (NM_028340, intron 1 of 4) | 6741 | NM_025491 | 66329 | Mm.34308 | NM_025491 | ENSMUSG00000021384 | **Susd3** | 1700017I11Rik\|2810440J20Rik | sushi domain containing 3 | protein-coding |
| chr13_49241247_49241557 | chr13 | 49241248 | 49241557 | + | 0.853614 | intron (NM_028340, intron 1 of 4) | B3A\|SINE\|B2 | 6761 | NM_025491 | 66329 | Mm.34308 | NM_025491 | ENSMUSG00000021384 | **Susd3** | 1700017I11Rik\|2810440J20Rik | sushi domain containing 3 | protein-coding |
| chr13_49272259_49272761 | chr13 | 49272260 | 49272761 | + | 0.840586 | intron (NM_015759, intron 14 of 17) | intron (NM_015759, intron 14 of 17) | -23804 | NM_028340 | 66329 | Mm.34308 | NM_025491 | ENSMUSG00000021384 | **Susd3** | 1700017I11Rik\|2810440J20Rik | sushi domain containing 3 | protein-coding |
| chr13_49270562_49270845 | chr13 | 49270563 | 49270845 | + | 0.778404 | intron (NM_015759, intron 14 of 17) | intron (NM_015759, intron 14 of 17) | -21998 | NM_028340 | 66329 | Mm.34308 | NM_025491 | ENSMUSG00000021384 | **Susd3** | 1700017I11Rik\|2810440J20Rik | sushi domain containing 3 | protein-coding |

| | | | | | | | | | | | | | | | | |
|---|---|---|---|---|---|---|---|---|---|---|---|---|---|---|---|---|
| chr13_49272851_49273174 | chr13 | 49272852 | 49273174 | + | 0.640381 | intron (NM_015759, intron 14 of 17) | intron (NM_015759, intron 14 of 17) | -24307 | NM_028340 | 66329 | Mm.34308 | NM_025491 | ENSMUSG00000021384 | **Susd3** | 1700017I11Rik\|2810440J20Rik | sushi domain containing 3 | protein-coding |
| chr13_96015556_96015980 | chr13 | 96015557 | 96015980 | + | 1.40857 | intron (NM_029210, intron 4 of 12) | intron (NM_029210, intron 4 of 12) | 116809 | NM_029210 | 75209 | Mm.40436 | NM_029210 | ENSMUSG00000051111 | **Sv2c** | 4930527L09Rik\|AI841971 | synaptic vesicle glycoprotein 2c | protein-coding |
| chr18_5020281_5020855 | chr18 | 5020282 | 5020855 | + | 0.952647 | Intergenic | Intergenic | -26021 | NM_153153 | 225115 | Mm.136791 | NM_153153 | ENSMUSG00000024236 | **Svil** | AU024053\|B430302E16Rik | supervillin | protein-coding |
| chr18_5020093_5020861 | chr18 | 5020094 | 5020861 | + | 0.895787 | Intergenic | Intergenic | -26112 | NM_153153 | 225115 | Mm.136791 | NM_153153 | ENSMUSG00000024236 | **Svil** | AU024053\|B430302E16Rik | supervillin | protein-coding |
| chr5_114061659_114062003 | chr5 | 114061660 | 114062003 | + | 0.765765 | intron (NM_026805, intron 4 of 15) | intron (NM_026805, intron 4 of 15) | 29549 | NM_026805 | 68666 | Mm.63584 | NM_026805 | ENSMUSG00000042078 | **Svop** | 1110030H18Rik\|AI415691\|msvop | SV2 related protein | protein-coding |
| chr5_114061515_114062023 | chr5 | 114061516 | 114062023 | + | 0.635751 | intron (NM_026805, intron 4 of 15) | intron (NM_026805, intron 4 of 15) | 29611 | NM_026805 | 68666 | Mm.63584 | NM_026805 | ENSMUSG00000042078 | **Svop** | 1110030H18Rik\|AI415691\|msvop | SV2 related protein | protein-coding |
| chr7_110222904_110223215 | chr7 | 110222905 | 110223215 | + | 0.6842 | intron (NM_009302, intron 1 of 11) | L1MD2\|LINE\|L1 | 1357 | NM_009302 | 20947 | Mm.334144 | NM_009302 | ENSMUSG00000031015 | **Swap70** | 70kDa\|AV235546 | SWA-70 protein | protein-coding |
| chr8_113655998_113656717 | chr8 | 113655999 | 113656717 | + | 1.50727 | TTS (NM_001048145) | TTS (NM_001048145) | 13145 | NM_001048145 | 668110 | Mm.390078 | NM_029139 | ENSMUSG00000033409 | **Syce1l** | 4930481F22Rik\|MMRP\|PSESS\|mmrp2 | synaptonemal complex central element protein 1 like | protein-coding |
| chr8_113655979_113656949 | chr8 | 113655980 | 113656949 | + | 1.47217 | TTS (NM_001048145) | TTS (NM_001048145) | 13251 | NM_001048145 | 668110 | Mm.390078 | NM_029139 | ENSMUSG00000033409 | **Syce1l** | 4930481F22Rik\|MMRP\|PSESS\|mmrp2 | synaptonemal complex central element protein 1 like | protein-coding |
| chr10_88420063_88421457 | chr10 | 88420064 | 88421457 | + | 1.04284 | intron (NM_001004164, intron 7 of 20) | intron (NM_001004164, intron 7 of 20) | -38827 | NM_011517 | 20962 | Mm.297977 | NM_011517 | ENSMUSG00000020059 | **Sycp3** | Cor1\|Scp3 | synaptonemal complex protein 3 | protein-coding |
| chr10_88420070_88421057 | chr10 | 88420071 | 88421057 | + | 1.02448 | intron (NM_001004164, intron 7 of 20) | CT-rich\|Low_complexity\|Low_complexity | -39023 | NM_011517 | 20962 | Mm.297977 | NM_011517 | ENSMUSG00000020059 | **Sycp3** | Cor1\|Scp3 | synaptonemal complex protein 3 | protein-coding |
| chr10_88421113_88421414 | chr10 | 88421114 | 88421414 | + | 0.866528 | intron (NM_001004164, intron 7 of 20) | intron (NM_001004164, intron 7 of 20) | -38323 | NM_011517 | 20962 | Mm.297977 | NM_011517 | ENSMUSG00000020059 | **Sycp3** | Cor1\|Scp3 | synaptonemal complex protein 3 | protein-coding |
| chr4_135019091_135019546 | chr4 | 135019092 | 135019546 | + | 0.631497 | Intergenic | Intergenic | 88339 | NM_026780 | 68592 | Mm.29989 | NM_026780 | ENSMUSG00000028821 | **Syf2** | 1110018L13Rik\|Cbpin\|D4Bwg1551e\|Gcipip\|Ntc31\|mp29\|p29 | SYF2 homolog, RNA splicing factor (S. cerevisiae) | protein-coding |
| chr6_115141321_115141754 | chr6 | 115141322 | 115141754 | + | 1.05378 | intron (NM_001111015, intron 1 of 12) | B3A\|SINE\|B2 | 6636 | NM_001111015 | 20965 | Mm.441431 | NM_013681 | ENSMUSG00000009394 | **Syn2** | 2900074L19Rik\|AI836018\|AI841723 | synapsin II | protein-coding |
| chr6_115123210_115123618 | chr6 | 115123211 | 115123618 | + | 0.987372 | Intergenic | Intergenic | -11488 | NM_013681 | 20965 | Mm.441431 | NM_013681 | ENSMUSG00000009394 | **Syn2** | 2900074L19Rik\|AI836018\|AI841723 | synapsin II | protein-coding |
| chr6_115123187_115123744 | chr6 | 115123188 | 115123744 | + | 0.969264 | Intergenic | Intergenic | -11436 | NM_013681 | 20965 | Mm.441431 | NM_013681 | ENSMUSG00000009394 | **Syn2** | 2900074L19Rik\|AI836018\|AI841723 | synapsin II | protein-coding |
| chr10_86498337_86499096 | chr10 | 86498338 | 86499096 | + | 0.923982 | intron (NM_013722, intron 1 of 13) | intron (NM_013722, intron 1 of 13) | 179 | NM_001164495 | 27204 | Mm.394931 | NM_013722 | ENSMUSG00000059602 | **Syn3** | - | synapsin III | protein-coding |
| chr10_86498381_86499356 | chr10 | 86498382 | 86499356 | + | 0.908653 | promoter-TSS (NM_001164495) | promoter-TSS (NM_001164495) | 27 | NM_001164495 | 27204 | Mm.394931 | NM_013722 | ENSMUSG00000059602 | **Syn3** | - | synapsin III | protein-coding |
| chr10_4988631_4989414 | chr10 | 4988632 | 4989414 | + | 1.10984 | intron (NM_007956, intron 7 of 8) | L1VL2\|LINE\|L1 | 80040 | NM_022027 | 64009 | Mm.331626 | NM_022027 | ENSMUSG00000019769 | **Syne1** | 8B\|A330049M09Rik\|BE692247\|C130039F11Rik\|CPG2\|Myne1\|mKIAA1756 | spectrin repeat containing, nuclear envelope 1 | protein-coding |
| chr10_4976962_4977885 | chr10 | 4976963 | 4977885 | + | 0.93422 | intron (NM_007956, intron 7 of 8) | intron (NM_007956, intron 7 of 8) | 91639 | NM_022027 | 64009 | Mm.331626 | NM_022027 | ENSMUSG00000019769 | **Syne1** | 8B\|A330049M09Rik\|BE692247\|C130039F11Rik\|CPG2\|Myne1\|mKIAA1756 | spectrin repeat containing, nuclear envelope 1 | protein-coding |
| chr10_4971583_4972460 | chr10 | 4971584 | 4972460 | + | 0.933913 | intron (NM_007956, intron 7 of 8) | intron (NM_007956, intron 7 of 8) | 97041 | NM_022027 | 64009 | Mm.331626 | NM_022027 | ENSMUSG00000019769 | **Syne1** | 8B\|A330049M09Rik\|BE692247\|C130039F11Rik\|CPG2\|Myne1\|mKIAA1756 | spectrin repeat containing, nuclear envelope 1 | protein-coding |
| chr10_4989074_4989373 | chr10 | 4989075 | 4989373 | + | 0.771785 | intron (NM_007956, intron 7 of 8) | L1VL2\|LINE\|L1 | 79839 | NM_022027 | 64009 | Mm.331626 | NM_022027 | ENSMUSG00000019769 | **Syne1** | 8B\|A330049M09Rik\|BE692247\|C130039F11Rik\|CPG2\|Myne1\|mKIAA1756 | spectrin repeat containing, nuclear envelope 1 | protein-coding |
| chr12_75834400_75834666 | chr12 | 75834401 | 75834666 | + | 0.635214 | intron (NM_001005510, intron 1 of 115) | intron (NM_001005510, intron 1 of 115) | 16215 | NM_001005510 | 319565 | Mm.26652 | NM_001005510 | ENSMUSG00000063450 | **Syne2** | 6820443O06Rik\|AW546258\|D12Ertd777e\|NUA\|Nesp2g\|mKIAA1011 | spectrin repeat containing, nuclear envelope 2 | protein-coding |
| chr12_105019262_105019794 | chr12 | 105019263 | 105019794 | + | 0.953208 | Intergenic | Intergenic | -9719 | NM_172500 | 212073 | Mm.212688 | NM_172500 | ENSMUSG00000054150 | **Syne3** | nesprin-3 | spectrin repeat containing, nuclear envelope family member 3 | protein-coding |
| chr12_105019262_105020277 | chr12 | 105019263 | 105020277 | + | 0.748215 | Intergenic | Intergenic | -9961 | NM_172500 | 212073 | Mm.212688 | NM_172500 | ENSMUSG00000054150 | **Syne3** | nesprin-3 | spectrin repeat containing, nuclear envelope family member 3 | protein-coding |
| chr12_104983398_104983721 | chr12 | 104983399 | 104983721 | + | 0.729907 | intron (NM_172500, intron 1 of 16) | intron (NM_172500, intron 1 of 16) | 15117 | NM_001042699 | 212073 | Mm.212688 | NM_172500 | ENSMUSG00000054150 | **Syne3** | nesprin-3 | spectrin repeat containing, nuclear envelope family member 3 | protein-coding |
| chr12_105000501_105001273 | chr12 | 105000502 | 105001273 | + | 0.647152 | intron (NM_172500, intron 1 of 16) | intron (NM_172500, intron 1 of 16) | -2210 | NM_001042699 | 212073 | Mm.212688 | NM_172500 | ENSMUSG00000054150 | **Syne3** | nesprin-3 | spectrin repeat containing, nuclear envelope family member 3 | protein-coding |
| chr11_117808153_117808346 | chr11 | 117808154 | 117808346 | + | 0.683737 | TTS (NM_029964) | TTS (NM_029964) | -1417 | NM_009304 | 20973 | Mm.468228 | NM_009304 | ENSMUSG00000048277 | **Syngr2** | Clast2\|cellugyrin | synaptogyrin 2 | protein-coding |
| chr16_90997554_90998105 | chr16 | 90997555 | 90998105 | + | 1.04596 | intron (NM_001164483, intron 2 of 31) | intron (NM_001164483, intron 2 of 31) | 13265 | NM_001045515 | 104015 | Mm.187079 | NM_001045515 | ENSMUSG00000022973 | **Synj1** | A930006D20Rik\|AA675315\|mKIAA0910 | synaptojanin 1 | protein-coding |
| chr17_5993156_5993688 | chr17 | 5993157 | 5993688 | + | 1.87867 | intron (NM_001113351, intron 3 of 24) | intron (NM_001113351, intron 3 of 24) | -14158 | NM_001290698 | 20975 | Mm.236068 | NM_011523 | ENSMUSG00000023805 | **Synj2** | AI481647\|SJ2\|mKIAA0348 | synaptojanin 2 | protein-coding |
| chr17_5994580_5994902 | chr17 | 5994581 | 5994902 | + | 1.04137 | intron (NM_001113351, intron 3 of 24) | intron (NM_001113351, intron 3 of 24) | -12839 | NM_001290698 | 20975 | Mm.236068 | NM_011523 | ENSMUSG00000023805 | **Synj2** | AI481647\|SJ2\|mKIAA0348 | synaptojanin 2 | protein-coding |
| chr17_5977006_5977352 | chr17 | 5977007 | 5977352 | + | 0.659196 | intron (NM_001113351, intron 1 of 24) | intron (NM_001113351, intron 1 of 24) | 1593 | NM_011523 | 20975 | Mm.236068 | NM_011523 | ENSMUSG00000023805 | **Synj2** | AI481647\|SJ2\|mKIAA0348 | synaptojanin 2 | protein-coding |
| chr17_6008065_6008354 | chr17 | 6008066 | 6008354 | + | 0.645791 | intron (NM_001290698, intron 2 of 19) | intron (NM_001290698, intron 2 of 19) | 630 | NM_001290698 | 20975 | Mm.236068 | NM_011523 | ENSMUSG00000023805 | **Synj2** | AI481647\|SJ2\|mKIAA0348 | synaptojanin 2 | protein-coding |
| chr18_60625390_60625751 | chr18 | 60625391 | 60625751 | + | 1.15826 | Intergenic | Intergenic | -1266 | NM_177340 | 104027 | Mm.252321 | NM_177340 | ENSMUSG00000043079 | **Synpo** | 9030217H17Rik\|9130229N11\|9330140I15Rik\|AW046661 | synaptopodin | protein-coding |
| chr18_60623669_60625759 | chr18 | 60623670 | 60625759 | + | 0.865926 | promoter-TSS (NM_177340) | promoter-TSS (NM_177340) | -409 | NM_177340 | 104027 | Mm.252321 | NM_177340 | ENSMUSG00000043079 | **Synpo** | 9030217H17Rik\|9130229N11\|9330140I15Rik\|AW046661 | synaptopodin | protein-coding |
| chr18_60623776_60625178 | chr18 | 60623777 | 60625178 | + | 0.792189 | promoter-TSS (NM_177340) | promoter-TSS (NM_177340) | -172 | NM_177340 | 104027 | Mm.252321 | NM_177340 | ENSMUSG00000043079 | **Synpo** | 9030217H17Rik\|9130229N11\|9330140I15Rik\|AW046661 | synaptopodin | protein-coding |
| chr11_84015967_84016514 | chr11 | 84015968 | 84016514 | + | 1.11724 | intron (NM_194341, intron 13 of 19) | intron (NM_194341, intron 13 of 19) | 51810 | NM_194341 | 217030 | Mm.82680 | NM_194341 | ENSMUSG00000034940 | **Synrg** | Ap1gbp1\|SYNG | synergin, gamma | protein-coding |
| chr2_164456692_164457235 | chr2 | 164456693 | 164457235 | + | 0.67523 | Intergenic | Intergenic | -4007 | NM_025575 | 66460 | Mm.44218 | NM_025575 | ENSMUSG00000045503 | **Sys1** | 2610042O14Rik\|AI266964\|C87389 | SYS1 Golgi-localized integral membrane protein homolog (S. cerevisiae) | protein-coding |
| chr2_92945171_92945683 | chr2 | 92945172 | 92945683 | + | 0.97045 | intron (NM_030725, intron 3 of 5) | ORR1F\|LTR\|ERVL-MaLR | 30326 | NM_030725 | 80976 | Mm.33490 | NM_030725 | ENSMUSG00000027220 | **Syt13** | 5730409J20Rik\|AI549909\|mKIAA1427 | synaptotagmin XIII | protein-coding |
| chr1_193018417_193018841 | chr1 | 193018418 | 193018841 | + | 0.734718 | intron (NM_181546, intron 2 of 7) | MusHAL1\|LINE\|L1 | 17146 | NM_001301370 | 329324 | Mm.76710 | NM_181546 | ENSMUSG00000016200 | **Syt14** | B230320I09Rik\|sytXIV | synaptotagmin XIV | protein-coding |
| chr12_74093084_74093563 | chr12 | 74093085 | 74093563 | + | 0.711059 | intron (NM_172804, intron 1 of 6) | intron (NM_172804, intron 1 of 6) | 95563 | NM_172804 | 238266 | Mm.311393 | NM_172804 | ENSMUSG00000044912 | **Syt16** | 6430703N11\|Strep14\|Syt14l\|syt14r | synaptotagmin XVI | protein-coding |
| chr7_44401734_44402052 | chr7 | 44401735 | 44402052 | + | 0.793355 | Intergenic | L1M3\|LINE\|L1 | 17767 | NM_016663 | 20981 | Mm.4824 | NM_016663 | ENSMUSG00000030731 | **Syt3** | AI385753\|sytIII | synaptotagmin III | protein-coding |
| chr19_10356787_10357212 | chr19 | 10356788 | 10357212 | + | 0.935977 | Intergenic | Intergenic | -32090 | NM_018801 | 54525 | Mm.182654 | NM_018801 | ENSMUSG00000024743 | **Syt7** | AI851541\|B230112P13Rik | synaptotagmin VII | protein-coding |
| chr19_10356787_10357433 | chr19 | 10356788 | 10357433 | + | 0.87412 | Intergenic | Intergenic | -31980 | NM_018801 | 54525 | Mm.182654 | NM_018801 | ENSMUSG00000024743 | **Syt7** | AI851541\|B230112P13Rik | synaptotagmin VII | protein-coding |
| chr4_133264605_133264956 | chr4 | 133264606 | 133264956 | + | 1.1155 | Intergenic | B3\|SINE\|B2 | -1694 | NM_031393 | 269589 | Mm.25660 | NM_031393 | ENSMUSG00000028860 | **Sytl1** | 9030417L18\|PSGL-1\|Slp1 | synaptotagmin-like 1 | protein-coding |
| chr4_141142591_141142840 | chr4 | 141142592 | 141142840 | + | 0.806213 | Intergenic | Intergenic | -2920 | NR_102343 | 213491 | Mm.332006 | NM_174996 | ENSMUSG00000040842 | **Szrd1** | 1110022I03\|D4Ertd22e | SUZ RNA binding domain containing 1 | protein-coding |
| chr10_7992552_7993099 | chr10 | 7992553 | 7993099 | + | 1.78823 | Intergenic | Intergenic | -36703 | NM_138667 | 68652 | Mm.193041 | NM_138667 | ENSMUSG00000015755 | **Tab2** | 1110030N06Rik\|A530078N03Rik\|Map3k7ip2\|mKIAA0733 | TGF-beta activated kinase 1/MAP3K7 binding protein 2 | protein-coding |
| chr10_8129933_8131400 | chr10 | 8129934 | 8131400 | + | 1.36135 | Intergenic | Intergenic | -174544 | NM_138667 | 68652 | Mm.193041 | NM_138667 | ENSMUSG00000015755 | **Tab2** | 1110030N06Rik\|A530078N03Rik\|Map3k7ip2\|mKIAA0733 | TGF-beta activated kinase 1/MAP3K7 binding protein 2 | protein-coding |

| | | | | | | | | | | | | | | | | | |
|---|---|---|---|---|---|---|---|---|---|---|---|---|---|---|---|---|---|
| chr10_8130474_8131393 | chr10 | 8130475 | 8131393 | + | 1.28256 | Intergenic | Intergenic | -174811 | NM_138667 | 68652 | Mm.193041 | NM_138667 | ENSMUSG00000015755 | **Tab2** | 1110030N06Rik\|A530078N03Rik\|Map3k7ip2\|mKIAA0733 | TGF-beta activated kinase 1/MAP3K7 binding protein 2 | protein-coding |
| chr10_8129968_8130425 | chr10 | 8129969 | 8130425 | + | 1.18373 | Intergenic | Intergenic | -174074 | NM_138667 | 68652 | Mm.193041 | NM_138667 | ENSMUSG00000015755 | **Tab2** | 1110030N06Rik\|A530078N03Rik\|Map3k7ip2\|mKIAA0733 | TGF-beta activated kinase 1/MAP3K7 binding protein 2 | protein-coding |
| chr10_7966138_7966542 | chr10 | 7966139 | 7966542 | + | 1.09767 | Intergenic | Intergenic | -10217 | NM_138667 | 68652 | Mm.193041 | NM_138667 | ENSMUSG00000015755 | **Tab2** | 1110030N06Rik\|A530078N03Rik\|Map3k7ip2\|mKIAA0733 | TGF-beta activated kinase 1/MAP3K7 binding protein 2 | protein-coding |
| chr10_7953715_7954279 | chr10 | 7953716 | 7954279 | + | 0.95828 | intron (NM_138667, intron 1 of 6) | intron (NM_138667, intron 1 of 6) | 2126 | NM_138667 | 68652 | Mm.193041 | NM_138667 | ENSMUSG00000015755 | **Tab2** | 1110030N06Rik\|A530078N03Rik\|Map3k7ip2\|mKIAA0733 | TGF-beta activated kinase 1/MAP3K7 binding protein 2 | protein-coding |
| chr10_8007290_8008275 | chr10 | 8007291 | 8008275 | + | 0.948054 | Intergenic | Intergenic | -51660 | NM_138667 | 68652 | Mm.193041 | NM_138667 | ENSMUSG00000015755 | **Tab2** | 1110030N06Rik\|A530078N03Rik\|Map3k7ip2\|mKIAA0733 | TGF-beta activated kinase 1/MAP3K7 binding protein 2 | protein-coding |
| chr10_7923403_7923835 | chr10 | 7923404 | 7923835 | + | 0.908228 | intron (NM_138667, intron 2 of 6) | MTD\|LTR\|ERVL-MaLR | 32504 | NM_138667 | 68652 | Mm.193041 | NM_138667 | ENSMUSG00000015755 | **Tab2** | 1110030N06Rik\|A530078N03Rik\|Map3k7ip2\|mKIAA0733 | TGF-beta activated kinase 1/MAP3K7 binding protein 2 | protein-coding |
| chr10_7907078_7907603 | chr10 | 7907079 | 7907603 | + | 0.804842 | 3' UTR (NM_138667, exon 7 of 7) | 3' UTR (NM_138667, exon 7 of 7) | 48782 | NM_138667 | 68652 | Mm.193041 | NM_138667 | ENSMUSG00000015755 | **Tab2** | 1110030N06Rik\|A530078N03Rik\|Map3k7ip2\|mKIAA0733 | TGF-beta activated kinase 1/MAP3K7 binding protein 2 | protein-coding |
| chr10_7991662_7991876 | chr10 | 7991663 | 7991876 | + | 0.709107 | Intergenic | Intergenic | -35646 | NM_138667 | 68652 | Mm.193041 | NM_138667 | ENSMUSG00000015755 | **Tab2** | 1110030N06Rik\|A530078N03Rik\|Map3k7ip2\|mKIAA0733 | TGF-beta activated kinase 1/MAP3K7 binding protein 2 | protein-coding |
| chr2_9992251_9993183 | chr2 | 9992252 | 9993183 | + | 0.959185 | intron (NM_027748, intron 2 of 6) | (CTCA)n\|Simple_repeat\|Simple_repeat | 55892 | NM_027748 | 209361 | Mm.86343 | NM_027748 | ENSMUSG00000025782 | **Taf3** | 140kDa\|4933439M23Rik\|AW539625\|TAF140\|TAFII-140\|TAFII140\|mTAFII140 | TAF3 RNA polymerase II, TATA box binding protein (TBP)-associated factor | protein-coding |
| chr18_14847535_14847962 | chr18 | 14847536 | 14847962 | + | 0.882686 | intron (NM_001100449, intron 13 of 14) | intron (NM_001100449, intron 13 of 14) | 64504 | NM_001100449 | 72504 | Mm.312233 | NM_001100449 | ENSMUSG00000054321 | **Taf4b** | 105kDa\|2610524B04Rik\|4932409F03Rik\|AW987595\|TAFII105\|Taf2c2 | TAF4B RNA polymerase II, TATA box binding protein (TBP)-associated factor | protein-coding |
| chr17_7888904_7889524 | chr17 | 7888905 | 7889524 | + | 1.61914 | Intergenic | Intergenic | -36786 | NM_145968 | 72536 | Mm.119680 | NM_145968 | ENSMUSG00000033450 | **Tagap** | 2610315E15Rik | T cell activation Rho GTPase activating protein | protein-coding |
| chr17_7875968_7876538 | chr17 | 7875969 | 7876538 | + | 1.32277 | Intergenic | LTR72_RN\|LTR\|ERV1 | -49747 | NM_145968 | 72536 | Mm.119680 | NM_145968 | ENSMUSG00000033450 | **Tagap** | 2610315E15Rik | T cell activation Rho GTPase activating protein | protein-coding |
| chr17_7888357_7888670 | chr17 | 7888358 | 7888670 | + | 1.18438 | Intergenic | Intergenic | -37486 | NM_145968 | 72536 | Mm.119680 | NM_145968 | ENSMUSG00000033450 | **Tagap** | 2610315E15Rik | T cell activation Rho GTPase activating protein | protein-coding |
| chr17_7882982_7883459 | chr17 | 7882983 | 7883459 | + | 1.03368 | Intergenic | Intergenic | -42779 | NM_145968 | 72536 | Mm.119680 | NM_145968 | ENSMUSG00000033450 | **Tagap** | 2610315E15Rik | T cell activation Rho GTPase activating protein | protein-coding |
| chr17_7904893_7905335 | chr17 | 7904894 | 7905335 | + | 1.01813 | Intergenic | Intergenic | -20886 | NM_145968 | 72536 | Mm.119680 | NM_145968 | ENSMUSG00000033450 | **Tagap** | 2610315E15Rik | T cell activation Rho GTPase activating protein | protein-coding |
| chr17_7871030_7871457 | chr17 | 7871031 | 7871457 | + | 0.867625 | Intergenic | Intergenic | -54756 | NM_145968 | 72536 | Mm.119680 | NM_145968 | ENSMUSG00000033450 | **Tagap** | 2610315E15Rik | T cell activation Rho GTPase activating protein | protein-coding |
| chr17_7896127_7896766 | chr17 | 7896128 | 7896766 | + | 0.857567 | Intergenic | Intergenic | -29553 | NM_145968 | 72536 | Mm.119680 | NM_145968 | ENSMUSG00000033450 | **Tagap** | 2610315E15Rik | T cell activation Rho GTPase activating protein | protein-coding |
| chr11_105625824_105626264 | chr11 | 105625825 | 105626264 | + | 0.809399 | intron (NM_181071, intron 2 of 25) | intron (NM_181071, intron 2 of 25) | 36058 | NM_181071 | 77097 | Mm.22501 | NM_175492 | ENSMUSG00000053580 | **Tanc2** | 3526402J09Rik\|5730590C14Rik | tetratricopeptide repeat, ankyrin repeat and coiled-coil containing 2 | protein-coding |
| chr11_105530860_105531206 | chr11 | 105530861 | 105531206 | + | 0.665085 | Intergenic | Intergenic | -58953 | NM_181071 | 77097 | Mm.22501 | NM_175492 | ENSMUSG00000053580 | **Tanc2** | 3526402J09Rik\|5730590C14Rik | tetratricopeptide repeat, ankyrin repeat and coiled-coil containing 2 | protein-coding |
| chr8_106670566_106671210 | chr8 | 106670567 | 106671210 | + | 0.697761 | TTS (NM_009864) | TTS (NM_009864) | -12180 | NM_173037 | 272538 | Mm.87151 | NM_173037 | ENSMUSG00000041949 | **Tango6** | AW413431\|E330010G16\|Tmco7 | transport and golgi organization 6 | protein-coding |
| chr2_61309007_61309780 | chr2 | 61309008 | 61309780 | + | 1.65183 | Intergenic | Intergenic | -269192 | NM_011529 | 21353 | Mm.244393 | NM_011529 | ENSMUSG00000064289 | **Tank** | C86182\|E430026L09Rik\|I-TRAF | TRAF family member-associated Nf-kappa B activator | protein-coding |
| chr2_61406218_61406919 | chr2 | 61406219 | 61406919 | + | 1.29309 | Intergenic | URR1A\|DNA\|hAT-Charlie | -172017 | NM_011529 | 21353 | Mm.244393 | NM_011529 | ENSMUSG00000064289 | **Tank** | C86182\|E430026L09Rik\|I-TRAF | TRAF family member-associated Nf-kappa B activator | protein-coding |
| chr2_61595952_61596227 | chr2 | 61595953 | 61596227 | + | 0.928315 | intron (NM_001164071, intron 2 of 8) | ORR1D1\|LTR\|ERVL-MaLR | 2993 | NM_001164071 | 21353 | Mm.244393 | NM_011529 | ENSMUSG00000064289 | **Tank** | C86182\|E430026L09Rik\|I-TRAF | TRAF family member-associated Nf-kappa B activator | protein-coding |
| chr2_61489769_61490129 | chr2 | 61489770 | 61490129 | + | 0.86123 | Intergenic | Intergenic | -88637 | NM_011529 | 21353 | Mm.244393 | NM_011529 | ENSMUSG00000064289 | **Tank** | C86182\|E430026L09Rik\|I-TRAF | TRAF family member-associated Nf-kappa B activator | protein-coding |
| chr2_61496658_61497454 | chr2 | 61496659 | 61497454 | + | 0.803447 | Intergenic | Intergenic | -81530 | NM_011529 | 21353 | Mm.244393 | NM_011529 | ENSMUSG00000064289 | **Tank** | C86182\|E430026L09Rik\|I-TRAF | TRAF family member-associated Nf-kappa B activator | protein-coding |
| chr2_61605442_61606177 | chr2 | 61605443 | 61606177 | + | 0.713283 | intron (NM_001164071, intron 2 of 8) | intron (NM_001164071, intron 2 of 8) | 12713 | NM_001164071 | 21353 | Mm.244393 | NM_011529 | ENSMUSG00000064289 | **Tank** | C86182\|E430026L09Rik\|I-TRAF | TRAF family member-associated Nf-kappa B activator | protein-coding |
| chr2_61490491_61491062 | chr2 | 61490492 | 61491062 | + | 0.621383 | Intergenic | Intergenic | -87809 | NM_011529 | 21353 | Mm.244393 | NM_011529 | ENSMUSG00000064289 | **Tank** | C86182\|E430026L09Rik\|I-TRAF | TRAF family member-associated Nf-kappa B activator | protein-coding |
| chr5_117158449_117159213 | chr5 | 117158450 | 117159213 | + | 1.29282 | intron (NM_001081308, intron 1 of 19) | intron (NM_001081308, intron 1 of 19) | 25243 | NM_183306 | 330177 | Mm.24343 | NM_183306 | ENSMUSG00000061288 | **Taok3** | 2900006A08Rik\|A130052D22\|A430105I05Rik | TAO kinase 3 | protein-coding |
| chr17_34218287_34219430 | chr17 | 34218288 | 34219430 | + | 0.989113 | Intergenic | Intergenic | 14380 | NM_011530 | 21355 | Mm.14814 | NM_011530 | | **Tap2** | ABC18\|AI462429\|APT2\|Abcb3\|Ham-2\|Ham2\|MTP2\|PSF2\|RING11\|Tap-2\|Y1\|jas | transporter 2, ATP-binding cassette, sub-family B (MDR/TAP) | protein-coding |
| chr17_34203127_34203850 | chr17 | 34203128 | 34203850 | + | 0.947626 | promoter-TSS (NM_011530) | promoter-TSS (NM_011530) | -990 | NM_011530 | 21355 | Mm.14814 | NM_011530 | | **Tap2** | ABC18\|AI462429\|APT2\|Abcb3\|Ham-2\|Ham2\|MTP2\|PSF2\|RING11\|Tap-2\|Y1\|jas | transporter 2, ATP-binding cassette, sub-family B (MDR/TAP) | protein-coding |
| chr17_34203374_34203832 | chr17 | 34203375 | 34203832 | + | 0.933935 | promoter-TSS (NM_011530) | promoter-TSS (NM_011530) | -876 | NM_011530 | 21355 | Mm.14814 | NM_011530 | | **Tap2** | ABC18\|AI462429\|APT2\|Abcb3\|Ham-2\|Ham2\|MTP2\|PSF2\|RING11\|Tap-2\|Y1\|jas | transporter 2, ATP-binding cassette, sub-family B (MDR/TAP) | protein-coding |
| chr17_33924109_33924475 | chr17 | 33924110 | 33924475 | + | 1.12676 | intron (NM_001025313, intron 3 of 7) | RMER4A\|LTR\|ERVK | 4814 | NM_009318 | 21356 | Mm.154457 | NM_009318 | ENSMUSG00000024308 | **Tapbp** | D17Wsu91e\|TPN | TAP binding protein | protein-coding |
| chr5_44254264_44254901 | chr5 | 44254265 | 44254901 | + | 2.14955 | Intergenic | L2a\|LINE\|L2 | -27977 | NM_173764 | 231225 | Mm.30688 | NM_173764 | ENSMUSG00000046985 | **Tapt1** | 4932414K18Rik | transmembrane anterior posterior transformation 1 | protein-coding |
| chr5_44255832_44256176 | chr5 | 44255833 | 44256176 | + | 1.62818 | Intergenic | Intergenic | -29398 | NM_173764 | 231225 | Mm.30688 | NM_173764 | ENSMUSG00000046985 | **Tapt1** | 4932414K18Rik | transmembrane anterior posterior transformation 1 | protein-coding |
| chr5_44253510_44253836 | chr5 | 44253511 | 44253836 | + | 1.25078 | Intergenic | ORR1B1\|LTR\|ERVL-MaLR | -27067 | NM_173764 | 231225 | Mm.30688 | NM_173764 | ENSMUSG00000046985 | **Tapt1** | 4932414K18Rik | transmembrane anterior posterior transformation 1 | protein-coding |
| chr15_102520555_102520822 | chr15 | 102520556 | 102520822 | + | 1.22803 | intron (NM_009319, intron 3 of 8) | intron (NM_009319, intron 3 of 8) | 2497 | NM_009319 | 21357 | Mm.291485 | NM_009319 | ENSMUSG00000023051 | **Tarbp2** | Prbp\|TRBP | TAR (HIV) RNA binding protein 2 | protein-coding |
| chr15_11524370_11524967 | chr15 | 11524371 | 11524967 | + | 0.846036 | Intergenic | Intergenic | -125011 | NM_033074 | 110960 | Mm.286061 | NM_033074 | ENSMUSG00000022241 | **Tars** | D15Wsu59e\|ThrRS | threonyl-tRNA synthetase | protein-coding |
| chr15_11524375_11524764 | chr15 | 11524376 | 11524764 | + | 0.727021 | Intergenic | Intergenic | -124912 | NM_033074 | 110960 | Mm.286061 | NM_033074 | ENSMUSG00000022241 | **Tars** | D15Wsu59e\|ThrRS | threonyl-tRNA synthetase | protein-coding |
| chr15_11519055_11519404 | chr15 | 11519056 | 11519404 | + | 0.649227 | Intergenic | Intergenic | -119572 | NM_033074 | 110960 | Mm.286061 | NM_033074 | ENSMUSG00000022241 | **Tars** | D15Wsu59e\|ThrRS | threonyl-tRNA synthetase | protein-coding |
| chr3_95760594_95761064 | chr3 | 95760595 | 95761064 | + | 1.06293 | 3' UTR (NM_001081293, exon 11 of 11) | 3' UTR (NM_001081293, exon 11 of 11) | -5852 | NM_001163618 | 71807 | Mm.271754 | NM_027931 | ENSMUSG00000028107 | **Tars2** | 2610024N01Rik\|AI429208\|Tarsl1 | threonyl-tRNA synthetase 2, mitochondrial (putative) | protein-coding |
| chr4_152038709_152039048 | chr4 | 152038710 | 152039048 | + | 0.748749 | promoter-TSS (NM_031867) | promoter-TSS (NM_031867) | -389 | NM_031867 | 110326 | Mm.29232 | NM_031867 | ENSMUSG00000028950 | **Tas1r1** | Gpr70\|T1r1\|TR1 | taste receptor, type 1, member 1 | protein-coding |
| chr6_42419455_42419782 | chr6 | 42419456 | 42419782 | + | 0.825154 | Intergenic | Intergenic | 14090 | NM_199159 | 387512 | Mm.377919 | NM_199159 | ENSMUSG00000056203 | **Tas2r135** | Tas2r35\|mt2r38 | taste receptor, type 2, member 135 | protein-coding |
| chr6_42221189_42221648 | chr6 | 42221190 | 42221648 | + | 0.963141 | Intergenic | Intergenic | 6091 | NM_001001453 | 387515 | Mm.377922 | NM_001001453 | ENSMUSG00000051917 | **Tas2r144** | Tas2r40\|Tas2r44\|mt2r33 | taste receptor, type 2, member 144 | protein-coding |
| chr6_42221179_42221819 | chr6 | 42221180 | 42221819 | + | 0.94328 | Intergenic | Intergenic | 6171 | NM_001001453 | 387515 | Mm.377922 | NM_001001453 | ENSMUSG00000051917 | **Tas2r144** | Tas2r40\|Tas2r44\|mt2r33 | taste receptor, type 2, member 144 | protein-coding |
| chr2_140100809_140101714 | chr2 | 140100810 | 140101714 | + | 2.05863 | Intergenic | Intergenic | -34457 | NR_110351 | 75812 | Mm.293449 | NM_175225 | ENSMUSG00000039033 | **Tasp1** | 4930485D02Rik\|AW986064 | taspase, threonine aspartase 1 | protein-coding |
| chr2_139935166_139935499 | chr2 | 139935167 | 139935499 | + | 0.791187 | intron (NR_110351, intron 10 of 13) | intron (NR_110351, intron 10 of 13) | 131472 | NM_001159640 | 75812 | Mm.293449 | NM_175225 | ENSMUSG00000039033 | **Tasp1** | 4930485D02Rik\|AW986064 | taspase, threonine aspartase 1 | protein-coding |
| chr2_139935204_139935439 | chr2 | 139935205 | 139935439 | + | 0.767801 | intron (NR_110351, intron 10 of 13) | intron (NR_110351, intron 10 of 13) | 131483 | NM_001159640 | 75812 | Mm.293449 | NM_175225 | ENSMUSG00000039033 | **Tasp1** | 4930485D02Rik\|AW986064 | taspase, threonine aspartase 1 | protein-coding |
| chr6_113675249_113675688 | chr6 | 113675250 | 113675688 | + | 0.852388 | intron (NM_172161, intron 6 of 12) | intron (NM_172161, intron 6 of 12) | -22030 | NM_001033463 | 381801 | Mm.333322 | NM_001033463 | ENSMUSG00000056952 | **Tatdn2** | AI646012\|mKIAA0218 | TatD DNase domain containing 2 | protein-coding |

| | | | | | | | | | | | | | | | | | |
|---|---|---|---|---|---|---|---|---|---|---|---|---|---|---|---|---|---|
| chr6_113674986_113675724 | chr6 | 113674987 | 113675724 | + | 0.816483 | intron (NM_172161, intron 6 of 12) | intron (NM_172161, intron 6 of 12) | -22144 | NM_001033463 | 381801 | Mm.333322 | NM_001033463 | ENSMUSG00000056952 | **Tatdn2** | AI646012\|mKIAA0218 | TatD DNase domain containing 2 | protein-coding |
| chr6_52703094_52703716 | chr6 | 52703095 | 52703716 | + | 1.27565 | Intergenic | LTR33\|LTR\|ERVL | -10324 | NM_025816 | 52440 | Mm.389757 | NM_025816 | ENSMUSG00000004535 | **Tax1bp1** | 1200003J11Rik\|1700069J21Rik\|AA930106\|D6Ertd404e\|D6Ertd772e\|T6bp\|TXBP151 | Tax1 (human T cell leukemia virus type I) binding protein 1 | protein-coding |
| chr6_52703251_52703677 | chr6 | 52703252 | 52703677 | + | 1.26991 | Intergenic | Intergenic | -10265 | NM_025816 | 52440 | Mm.389757 | NM_025816 | ENSMUSG00000004535 | **Tax1bp1** | 1200003J11Rik\|1700069J21Rik\|AA930106\|D6Ertd404e\|D6Ertd772e\|T6bp\|TXBP151 | Tax1 (human T cell leukemia virus type I) binding protein 1 | protein-coding |
| chr6_52680487_52681071 | chr6 | 52680488 | 52681071 | + | 0.809823 | Intergenic | MLTR25A\|LTR\|ERVK | -32950 | NM_025816 | 52440 | Mm.389757 | NM_025816 | ENSMUSG00000004535 | **Tax1bp1** | 1200003J11Rik\|1700069J21Rik\|AA930106\|D6Ertd404e\|D6Ertd772e\|T6bp\|TXBP151 | Tax1 (human T cell leukemia virus type I) binding protein 1 | protein-coding |
| chr6_52801790_52802789 | chr6 | 52801791 | 52802789 | + | 0.737332 | intron (NM_173406, intron 3 of 4) | intron (NM_173406, intron 3 of 4) | 88561 | NM_025816 | 52440 | Mm.389757 | NM_025816 | ENSMUSG00000004535 | **Tax1bp1** | 1200003J11Rik\|1700069J21Rik\|AA930106\|D6Ertd404e\|D6Ertd772e\|T6bp\|TXBP151 | Tax1 (human T cell leukemia virus type I) binding protein 1 | protein-coding |
| chr10_61175785_61176173 | chr10 | 61175786 | 61176173 | + | 0.692431 | intron (NM_001017419, intron 3 of 9) | intron (NM_001017419, intron 3 of 9) | 717 | NM_023064 | 65971 | Mm.30227 | NM_023064 | ENSMUSG00000020096 | **Tbata** | 1700021K02Rik\|AI428928\|Spatial\|Titest | thymus, brain and testes associated | protein-coding |
| chr5_64238993_64239904 | chr5 | 64238994 | 64239904 | + | 1.4913 | intron (NM_001289514, intron 1 of 18) | intron (NM_001289514, intron 1 of 18) | 9092 | NM_001289514 | 57915 | Mm.286353 | NM_019636 | ENSMUSG00000029174 | **Tbc1d1** | 1110062G02Rik\|AI385682\|AW555803\|Tbc1\|mKIAA1108 | TBC1 domain family, member 1 | protein-coding |
| chr5_64224700_64225133 | chr5 | 64224701 | 64225133 | + | 0.864991 | intron (NM_019636, intron 2 of 19) | intron (NM_019636, intron 2 of 19) | -5440 | NM_001289514 | 57915 | Mm.286353 | NM_019636 | ENSMUSG00000029174 | **Tbc1d1** | 1110062G02Rik\|AI385682\|AW555803\|Tbc1\|mKIAA1108 | TBC1 domain family, member 1 | protein-coding |
| chr5_64222090_64222496 | chr5 | 64222091 | 64222496 | + | 0.761735 | intron (NM_019636, intron 2 of 19) | intron (NM_019636, intron 2 of 19) | -8064 | NM_001289514 | 57915 | Mm.286353 | NM_019636 | ENSMUSG00000029174 | **Tbc1d1** | 1110062G02Rik\|AI385682\|AW555803\|Tbc1\|mKIAA1108 | TBC1 domain family, member 1 | protein-coding |
| chr11_4195404_4196068 | chr11 | 4195405 | 4196068 | + | 1.09614 | intron (NM_134023, intron 1 of 8) | intron (NM_134023, intron 1 of 8) | 8903 | NM_134023 | 103724 | Mm.28140 | NM_134023 | ENSMUSG00000034412 | **Tbc1d10a** | AI447804\|EPI64\|Tbc1d10\|mFLJ00288 | TBC1 domain family, member 10a | protein-coding |
| chr5_36561504_36561901 | chr5 | 36561505 | 36561901 | + | 0.88665 | intron (NM_001113362, intron 3 of 14) | MER3\|DNA\|hAT-Charlie | 20694 | NM_133910 | 100855 | Mm.486203 | NM_133910 | ENSMUSG00000029192 | **Tbc1d14** | 2810413P16Rik\|AU043625\|C86258\|D5Ertd110e\|mKIAA1322 | TBC1 domain family, member 14 | protein-coding |
| chr5_36561506_36561765 | chr5 | 36561507 | 36561765 | + | 0.878299 | intron (NM_001113362, intron 3 of 14) | MER3\|DNA\|hAT-Charlie | 20761 | NM_133910 | 100855 | Mm.486203 | NM_133910 | ENSMUSG00000029192 | **Tbc1d14** | 2810413P16Rik\|AU043625\|C86258\|D5Ertd110e\|mKIAA1322 | TBC1 domain family, member 14 | protein-coding |
| chr10_115245491_115245901 | chr10 | 115245492 | 115245901 | + | 0.645906 | intron (NM_025706, intron 1 of 16) | intron (NM_025706, intron 1 of 16) | 5797 | NM_025706 | 66687 | Mm.22252 | NM_025706 | ENSMUSG00000020130 | **Tbc1d15** | 4432405K22Rik\|AU022358\|AU023730\|AW557774\|Ly6dl\|Rab7-GAP | TBC1 domain family, member 15 | protein-coding |
| chr2_152309652_152310154 | chr2 | 152309653 | 152310154 | + | 0.643104 | intron (NM_024196, intron 5 of 7) | (TTTTTG)n\|Simple_repeat\|Simple_repeat | 16031 | NM_024196 | 67231 | Mm.288733 | NM_024196 | | **Tbc1d20** | 1110028I04Rik\|2810442O16Rik\|AI414693 | TBC1 domain family, member 20 | protein-coding |
| chr15_86253519_86253833 | chr15 | 86253520 | 86253833 | + | 0.656384 | intron (NM_145476, intron 4 of 12) | intron (NM_145476, intron 4 of 12) | 39217 | NM_145476 | 223754 | Mm.28904 | NM_145476 | ENSMUSG00000051864 | **Tbc1d22a** | BC023106\|D15Ertd781e | TBC1 domain family, member 22a | protein-coding |
| chr16_57215568_57216421 | chr16 | 57215569 | 57216421 | + | 0.685161 | intron (NM_026254, intron 2 of 17) | intron (NM_026254, intron 2 of 17) | 15471 | NM_026254 | 67581 | Mm.257819 | NM_026254 | ENSMUSG00000022749 | **Tbc1d23** | 4930451A13Rik\|AU015720\|AU043671\|AU043778\|D030022P07Rik\|W51689 | TBC1 domain family, member 23 | protein-coding |
| chr9_90253618_90253878 | chr9 | 90253619 | 90253878 | + | 0.784982 | intron (NM_194334, intron 1 of 12) | intron (NM_194334, intron 1 of 12) | 17021 | NM_194334 | 67016 | Mm.335347 | NM_194334 | ENSMUSG00000037410 | **Tbc1d2b** | 1810061M12Rik\|AI480956\|AV021536\|AV023399\|AW491493\|mKIAA1055 | TBC1 domain family, member 2B | protein-coding |
| chr10_121188562_121189118 | chr10 | 121188563 | 121189118 | + | 0.98094 | Intergenic | MLT1B\|LTR\|ERVL-MaLR | 122349 | NM_029057 | 74694 | Mm.233833 | NM_029057 | ENSMUSG00000052302 | **Tbc1d30** | 4930505D03Rik | TBC1 domain family, member 30 | protein-coding |
| chr10_121265051_121265466 | chr10 | 121265052 | 121265466 | + | 0.874109 | 3' UTR (NM_029057, exon 12 of 12) | 3' UTR (NM_029057, exon 12 of 12) | 45930 | NM_029057 | 74694 | Mm.233833 | NM_029057 | ENSMUSG00000052302 | **Tbc1d30** | 4930505D03Rik | TBC1 domain family, member 30 | protein-coding |
| chr10_121197064_121197638 | chr10 | 121197065 | 121197638 | + | 0.823177 | Intergenic | MLT1J2\|LTR\|ERVL-MaLR | 113838 | NM_029057 | 74694 | Mm.233833 | NM_029057 | ENSMUSG00000052302 | **Tbc1d30** | 4930505D03Rik | TBC1 domain family, member 30 | protein-coding |
| chr10_121293804_121294345 | chr10 | 121293805 | 121294345 | + | 0.754624 | intron (NM_029057, intron 7 of 11) | RLTR16C_MM\|LTR\|ERVK | 17114 | NM_029057 | 74694 | Mm.233833 | NM_029057 | ENSMUSG00000052302 | **Tbc1d30** | 4930505D03Rik | TBC1 domain family, member 30 | protein-coding |
| chr10_121300642_121301072 | chr10 | 121300643 | 121301072 | + | 0.739868 | intron (NM_029057, intron 5 of 11) | intron (NM_029057, intron 5 of 11) | 10332 | NM_029057 | 74694 | Mm.233833 | NM_029057 | ENSMUSG00000052302 | **Tbc1d30** | 4930505D03Rik | TBC1 domain family, member 30 | protein-coding |
| chr10_121300386_121301101 | chr10 | 121300387 | 121301101 | + | 0.701134 | intron (NM_029057, intron 5 of 11) | ORR1C2\|LTR\|ERVL-MaLR | 10445 | NM_029057 | 74694 | Mm.233833 | NM_029057 | ENSMUSG00000052302 | **Tbc1d30** | 4930505D03Rik | TBC1 domain family, member 30 | protein-coding |
| chr15_57906509_57907117 | chr15 | 57906510 | 57907117 | + | 1.45939 | Intergenic | Intergenic | -5386 | NM_001081396 | 210544 | Mm.390835 | NM_199300 | ENSMUSG00000022364 | **Tbc1d31** | 4B3\|D330013L20Rik\|Gm85\|Wdr67 | TBC1 domain family, member 31 | protein-coding |
| chr14_101480532_101480942 | chr14 | 101480533 | 101480942 | + | 1.07474 | intron (NM_001081278, intron 8 of 18) | URR1A\|DNA\|hAT-Charlie | 128454 | NM_001081278 | 210789 | Mm.320639 | NM_173380 | ENSMUSG00000033083 | **Tbc1d4** | 5930406J04Rik\|A930035N22\|AV295684\|As160 | TBC1 domain family, member 4 | protein-coding |
| chr14_101466396_101467021 | chr14 | 101466397 | 101467021 | + | 0.80564 | intron (NM_001081278, intron 10 of 18) | intron (NM_001081278, intron 10 of 18) | 142482 | NM_001081278 | 210789 | Mm.320639 | NM_173380 | ENSMUSG00000033083 | **Tbc1d4** | 5930406J04Rik\|A930035N22\|AV295684\|As160 | TBC1 domain family, member 4 | protein-coding |
| chr13_43202360_43202796 | chr13 | 43202361 | 43202796 | + | 0.974365 | intron (NM_001033399, intron 1 of 1) | intron (NM_001033399, intron 1 of 1) | -31077 | NM_001252640 | 67046 | Mm.269213 | NM_025935 | ENSMUSG00000021368 | **Tbc1d7** | 2610009C09Rik | TBC1 domain family, member 7 | protein-coding |
| chr13_43218286_43218788 | chr13 | 43218287 | 43218788 | + | 0.86039 | intron (NM_001033399, intron 1 of 1) | MLT1D\|LTR\|ERVL-MaLR | -47036 | NM_001252640 | 67046 | Mm.269213 | NM_025935 | ENSMUSG00000021368 | **Tbc1d7** | 2610009C09Rik | TBC1 domain family, member 7 | protein-coding |
| chr8_83086602_83087048 | chr8 | 83086603 | 83087048 | + | 0.742768 | 3' UTR (NM_177378, exon 7 of 7) | 3' UTR (NM_177378, exon 7 of 7) | -78527 | NM_001111304 | 71310 | Mm.24031 | NM_027758 | ENSMUSG00000031709 | **Tbc1d9** | 4933431N12Rik\|AI847101\|AW490653\|C76116 | TBC1 domain family, member 9 | protein-coding |
| chr8_83131424_83132138 | chr8 | 83131425 | 83132138 | + | 0.691222 | Intergenic | Intergenic | -33571 | NM_001111304 | 71310 | Mm.24031 | NM_027758 | ENSMUSG00000031709 | **Tbc1d9** | 4933431N12Rik\|AI847101\|AW490653\|C76116 | TBC1 domain family, member 9 | protein-coding |
| chr13_94806331_94806726 | chr13 | 94806332 | 94806726 | + | 1.18023 | intron (NM_009321, intron 1 of 3) | intron (NM_009321, intron 1 of 3) | 17586 | NM_009321 | 21371 | Mm.379025 | NM_009321 | ENSMUSG00000042043 | **Tbca** | Tbca13 | tubulin cofactor A | protein-coding |
| chr13_94794646_94795223 | chr13 | 94794647 | 94795223 | + | 0.853459 | intron (NM_009321, intron 1 of 3) | intron (NM_009321, intron 1 of 3) | 5992 | NM_009321 | 21371 | Mm.379025 | NM_009321 | ENSMUSG00000042043 | **Tbca** | Tbca13 | tubulin cofactor A | protein-coding |
| chr13_94808170_94808451 | chr13 | 94808171 | 94808451 | + | 0.852777 | intron (NM_009321, intron 1 of 3) | intron (NM_009321, intron 1 of 3) | 19368 | NM_009321 | 21371 | Mm.379025 | NM_009321 | ENSMUSG00000042043 | **Tbca** | Tbca13 | tubulin cofactor A | protein-coding |
| chr17_46866788_46867315 | chr17 | 46866789 | 46867315 | + | 1.49714 | Intergenic | Intergenic | -23569 | NM_178385 | 72726 | Mm.33780 | NM_178385 | ENSMUSG00000036430 | **Tbcc** | 2810055C19Rik | tubulin-specific chaperone C | protein-coding |
| chr17_46879762_46880375 | chr17 | 46879763 | 46880375 | + | 0.901391 | Intergenic | ORR1F\|LTR\|ERVL-MaLR | -10552 | NM_178385 | 72726 | Mm.33780 | NM_178385 | ENSMUSG00000036430 | **Tbcc** | 2810055C19Rik | tubulin-specific chaperone C | protein-coding |
| chr16_22839046_22839840 | chr16 | 22839047 | 22839840 | + | 1.145 | intron (NM_001081368, intron 2 of 7) | intron (NM_001081368, intron 2 of 7) | 18126 | NM_001081368 | 70573 | Mm.68291 | NM_001081368 | ENSMUSG00000004462 | **Tbccd1** | 5730478M09Rik\|AU015083\|BB165076 | TBCC domain containing 1 | protein-coding |
| chr13_14040295_14040728 | chr13 | 14040296 | 14040728 | + | 1.03671 | promoter-TSS (NM_178337) | promoter-TSS (NM_178337) | -912 | NM_178337 | 70430 | Mm.260209 | NM_178337 | ENSMUSG00000039233 | **Tbce** | 2610206D02Rik\|C530005D02Rik\|pmn | tubulin-specific chaperone E | protein-coding |
| chr13_14034494_14034855 | chr13 | 14034495 | 14034855 | + | 0.782476 | intron (NM_178337, intron 1 of 16) | intron (NM_178337, intron 1 of 16) | 4925 | NM_178337 | 70430 | Mm.260209 | NM_178337 | ENSMUSG00000039233 | **Tbce** | 2610206D02Rik\|C530005D02Rik\|pmn | tubulin-specific chaperone E | protein-coding |
| chr9_42437785_42438326 | chr9 | 42437786 | 42438326 | + | 0.90765 | intron (NM_173038, intron 6 of 7) | intron (NM_173038, intron 6 of 7) | 34170 | NM_173038 | 272589 | Mm.34483 | NM_173038 | ENSMUSG00000037287 | **Tbcel** | D330014K23\|E130107N23Rik\|Lrrc35 | tubulin folding cofactor E-like | protein-coding |
| chr10_121580363_121580724 | chr10 | 121580364 | 121580724 | + | 1.08297 | intron (NM_019786, intron 2 of 20) | intron (NM_019786, intron 2 of 20) | 6250 | NM_019786 | 56480 | Mm.34580 | NM_019786 | ENSMUSG00000020115 | **Tbk1** | 1200008B05Rik\|AI462036\|AW048562 | TANK-binding kinase 1 | protein-coding |
| chr10_121580242_121580901 | chr10 | 121580243 | 121580901 | + | 0.960616 | intron (NM_019786, intron 2 of 20) | intron (NM_019786, intron 2 of 20) | 6222 | NM_019786 | 56480 | Mm.34580 | NM_019786 | ENSMUSG00000020115 | **Tbk1** | 1200008B05Rik\|AI462036\|AW048562 | TANK-binding kinase 1 | protein-coding |
| chr11_97158993_97159635 | chr11 | 97158994 | 97159635 | + | 1.17964 | TTS (NM_008379) | TTS (NM_008379) | -9602 | NM_198100 | 73174 | Mm.27882 | NM_198100 | ENSMUSG00000038517 | **Tbkbp1** | 3110043L15Rik\|ProSAPiP2\|SINTBAD | TBK1 binding protein 1 | protein-coding |
| chr11_97158981_97159468 | chr11 | 97158982 | 97159468 | + | 1.11871 | TTS (NM_008379) | TTS (NM_008379) | -9513 | NM_198100 | 73174 | Mm.27882 | NM_198100 | ENSMUSG00000038517 | **Tbkbp1** | 3110043L15Rik\|ProSAPiP2\|SINTBAD | TBK1 binding protein 1 | protein-coding |
| chr11_97152795_97153221 | chr11 | 97152796 | 97153221 | + | 0.657394 | Intergenic | Intergenic | -3296 | NM_198100 | 73174 | Mm.27882 | NM_198100 | ENSMUSG00000038517 | **Tbkbp1** | 3110043L15Rik\|ProSAPiP2\|SINTBAD | TBK1 binding protein 1 | protein-coding |

<table>
<tr><td>chr3_21965407_21965959</td><td>chr3</td><td>21965408</td><td>21965959</td><td>+</td><td>1.2481</td><td>Intergenic</td><td>Intergenic</td><td>-110969</td><td>NM_030732</td><td>81004</td><td>Mm.202966</td><td>NM_030732</td><td>ENSMUSG00000027630</td><td>Tbl1xr1</td><td>8030499H02Rik|A630076E03Rik|AW539987|C21|C230089I12Rik|DC42|Ira1|TBLR1</td><td>transducin (beta)-like 1X-linked receptor 1</td><td>protein-coding</td></tr>
<tr><td>chr3_21965590_21966020</td><td>chr3</td><td>21965591</td><td>21966020</td><td>+</td><td>1.19453</td><td>Intergenic</td><td>Intergenic</td><td>-110847</td><td>NM_030732</td><td>81004</td><td>Mm.202966</td><td>NM_030732</td><td>ENSMUSG00000027630</td><td>Tbl1xr1</td><td>8030499H02Rik|A630076E03Rik|AW539987|C21|C230089I12Rik|DC42|Ira1|TBLR1</td><td>transducin (beta)-like 1X-linked receptor 1</td><td>protein-coding</td></tr>
<tr><td>chr3_21562764_21563252</td><td>chr3</td><td>21562765</td><td>21563252</td><td>+</td><td>1.17438</td><td>Intergenic</td><td>MLTR14|LTR|ERV1</td><td>-513644</td><td>NM_030732</td><td>81004</td><td>Mm.202966</td><td>NM_030732</td><td>ENSMUSG00000027630</td><td>Tbl1xr1</td><td>8030499H02Rik|A630076E03Rik|AW539987|C21|C230089I12Rik|DC42|Ira1|TBLR1</td><td>transducin (beta)-like 1X-linked receptor 1</td><td>protein-coding</td></tr>
<tr><td>chr3_21969702_21970143</td><td>chr3</td><td>21969703</td><td>21970143</td><td>+</td><td>1.0324</td><td>Intergenic</td><td>Intergenic</td><td>-106729</td><td>NM_030732</td><td>81004</td><td>Mm.202966</td><td>NM_030732</td><td>ENSMUSG00000027630</td><td>Tbl1xr1</td><td>8030499H02Rik|A630076E03Rik|AW539987|C21|C230089I12Rik|DC42|Ira1|TBLR1</td><td>transducin (beta)-like 1X-linked receptor 1</td><td>protein-coding</td></tr>
<tr><td>chr3_21950750_21951268</td><td>chr3</td><td>21950751</td><td>21951268</td><td>+</td><td>0.94205</td><td>Intergenic</td><td>Intergenic</td><td>-125643</td><td>NM_030732</td><td>81004</td><td>Mm.202966</td><td>NM_030732</td><td>ENSMUSG00000027630</td><td>Tbl1xr1</td><td>8030499H02Rik|A630076E03Rik|AW539987|C21|C230089I12Rik|DC42|Ira1|TBLR1</td><td>transducin (beta)-like 1X-linked receptor 1</td><td>protein-coding</td></tr>
<tr><td>chr3_21949535_21949743</td><td>chr3</td><td>21949536</td><td>21949743</td><td>+</td><td>0.935175</td><td>Intergenic</td><td>Intergenic</td><td>-127013</td><td>NM_030732</td><td>81004</td><td>Mm.202966</td><td>NM_030732</td><td>ENSMUSG00000027630</td><td>Tbl1xr1</td><td>8030499H02Rik|A630076E03Rik|AW539987|C21|C230089I12Rik|DC42|Ira1|TBLR1</td><td>transducin (beta)-like 1X-linked receptor 1</td><td>protein-coding</td></tr>
<tr><td>chr3_21961734_21962835</td><td>chr3</td><td>21961735</td><td>21962835</td><td>+</td><td>0.932634</td><td>Intergenic</td><td>Intergenic</td><td>-114367</td><td>NM_030732</td><td>81004</td><td>Mm.202966</td><td>NM_030732</td><td>ENSMUSG00000027630</td><td>Tbl1xr1</td><td>8030499H02Rik|A630076E03Rik|AW539987|C21|C230089I12Rik|DC42|Ira1|TBLR1</td><td>transducin (beta)-like 1X-linked receptor 1</td><td>protein-coding</td></tr>
<tr><td>chr3_22058498_22059016</td><td>chr3</td><td>22058499</td><td>22059016</td><td>+</td><td>0.921972</td><td>Intergenic</td><td>Intergenic</td><td>-17895</td><td>NM_030732</td><td>81004</td><td>Mm.202966</td><td>NM_030732</td><td>ENSMUSG00000027630</td><td>Tbl1xr1</td><td>8030499H02Rik|A630076E03Rik|AW539987|C21|C230089I12Rik|DC42|Ira1|TBLR1</td><td>transducin (beta)-like 1X-linked receptor 1</td><td>protein-coding</td></tr>
<tr><td>chr3_22039437_22040130</td><td>chr3</td><td>22039438</td><td>22040130</td><td>+</td><td>0.918101</td><td>Intergenic</td><td>RLTR17|LTR|ERVK</td><td>-36868</td><td>NM_030732</td><td>81004</td><td>Mm.202966</td><td>NM_030732</td><td>ENSMUSG00000027630</td><td>Tbl1xr1</td><td>8030499H02Rik|A630076E03Rik|AW539987|C21|C230089I12Rik|DC42|Ira1|TBLR1</td><td>transducin (beta)-like 1X-linked receptor 1</td><td>protein-coding</td></tr>
<tr><td>chr3_22117559_22118131</td><td>chr3</td><td>22117560</td><td>22118131</td><td>+</td><td>0.87456</td><td>intron (NM_030732, intron 1 of 15)</td><td>RMER6A|LTR|ERVK</td><td>41193</td><td>NM_030732</td><td>81004</td><td>Mm.202966</td><td>NM_030732</td><td>ENSMUSG00000027630</td><td>Tbl1xr1</td><td>8030499H02Rik|A630076E03Rik|AW539987|C21|C230089I12Rik|DC42|Ira1|TBLR1</td><td>transducin (beta)-like 1X-linked receptor 1</td><td>protein-coding</td></tr>
<tr><td>chr3_21972977_21973521</td><td>chr3</td><td>21972978</td><td>21973521</td><td>+</td><td>0.863765</td><td>Intergenic</td><td>Intergenic</td><td>-103403</td><td>NM_030732</td><td>81004</td><td>Mm.202966</td><td>NM_030732</td><td>ENSMUSG00000027630</td><td>Tbl1xr1</td><td>8030499H02Rik|A630076E03Rik|AW539987|C21|C230089I12Rik|DC42|Ira1|TBLR1</td><td>transducin (beta)-like 1X-linked receptor 1</td><td>protein-coding</td></tr>
<tr><td>chr3_21722631_21722982</td><td>chr3</td><td>21722632</td><td>21722982</td><td>+</td><td>0.771714</td><td>Intergenic</td><td>Intergenic</td><td>-353845</td><td>NM_030732</td><td>81004</td><td>Mm.202966</td><td>NM_030732</td><td>ENSMUSG00000027630</td><td>Tbl1xr1</td><td>8030499H02Rik|A630076E03Rik|AW539987|C21|C230089I12Rik|DC42|Ira1|TBLR1</td><td>transducin (beta)-like 1X-linked receptor 1</td><td>protein-coding</td></tr>
<tr><td>chr3_21927665_21928225</td><td>chr3</td><td>21927666</td><td>21928225</td><td>+</td><td>0.690051</td><td>Intergenic</td><td>RMER15|LTR|ERVL</td><td>-148707</td><td>NM_030732</td><td>81004</td><td>Mm.202966</td><td>NM_030732</td><td>ENSMUSG00000027630</td><td>Tbl1xr1</td><td>8030499H02Rik|A630076E03Rik|AW539987|C21|C230089I12Rik|DC42|Ira1|TBLR1</td><td>transducin (beta)-like 1X-linked receptor 1</td><td>protein-coding</td></tr>
<tr><td>chr3_21917203_21918207</td><td>chr3</td><td>21917204</td><td>21918207</td><td>+</td><td>0.63543</td><td>Intergenic</td><td>Intergenic</td><td>-158947</td><td>NM_030732</td><td>81004</td><td>Mm.202966</td><td>NM_030732</td><td>ENSMUSG00000027630</td><td>Tbl1xr1</td><td>8030499H02Rik|A630076E03Rik|AW539987|C21|C230089I12Rik|DC42|Ira1|TBLR1</td><td>transducin (beta)-like 1X-linked receptor 1</td><td>protein-coding</td></tr>
<tr><td>chr11_97129041_97129356</td><td>chr11</td><td>97129042</td><td>97129356</td><td>+</td><td>1.30802</td><td>Intergenic</td><td>Intergenic</td><td>-13868</td><td>NM_019507</td><td>57765</td><td>Mm.94519</td><td>NM_019507</td><td>ENSMUSG00000001444</td><td>Tbx21</td><td>TBT1|Tbet|Tblym</td><td>T-box 21</td><td>protein-coding</td></tr>
<tr><td>chr11_97112652_97113222</td><td>chr11</td><td>97112653</td><td>97113222</td><td>+</td><td>0.870986</td><td>intron (NM_019507, intron 1 of 5)</td><td>intron (NM_019507, intron 1 of 5)</td><td>2394</td><td>NM_019507</td><td>57765</td><td>Mm.94519</td><td>NM_019507</td><td>ENSMUSG00000001444</td><td>Tbx21</td><td>TBT1|Tbet|Tblym</td><td>T-box 21</td><td>protein-coding</td></tr>
<tr><td>chr11_97128673_97128954</td><td>chr11</td><td>97128674</td><td>97128954</td><td>+</td><td>0.783846</td><td>Intergenic</td><td>Intergenic</td><td>-13483</td><td>NM_019507</td><td>57765</td><td>Mm.94519</td><td>NM_019507</td><td>ENSMUSG00000001444</td><td>Tbx21</td><td>TBT1|Tbet|Tblym</td><td>T-box 21</td><td>protein-coding</td></tr>
<tr><td>chr11_97119404_97119830</td><td>chr11</td><td>97119405</td><td>97119830</td><td>+</td><td>0.68692</td><td>Intergenic</td><td>Intergenic</td><td>-4286</td><td>NM_019507</td><td>57765</td><td>Mm.94519</td><td>NM_019507</td><td>ENSMUSG00000001444</td><td>Tbx21</td><td>TBT1|Tbet|Tblym</td><td>T-box 21</td><td>protein-coding</td></tr>
<tr><td>chr11_97109711_97110119</td><td>chr11</td><td>97109712</td><td>97110119</td><td>+</td><td>0.659999</td><td>intron (NM_019507, intron 1 of 5)</td><td>intron (NM_019507, intron 1 of 5)</td><td>5416</td><td>NM_019507</td><td>57765</td><td>Mm.94519</td><td>NM_019507</td><td>ENSMUSG00000001444</td><td>Tbx21</td><td>TBT1|Tbet|Tblym</td><td>T-box 21</td><td>protein-coding</td></tr>
<tr><td>chr5_119670154_119671079</td><td>chr5</td><td>119670155</td><td>119671079</td><td>+</td><td>0.709058</td><td>promoter-TSS (NM_011535)</td><td>promoter-TSS (NM_011535)</td><td>-245</td><td>NM_198052</td><td>21386</td><td>Mm.219139</td><td>NM_011535</td><td>ENSMUSG00000018604</td><td>Tbx3</td><td>D5Ertd189e</td><td>T-box 3</td><td>protein-coding</td></tr>
<tr><td>chr11_85929479_85930127</td><td>chr11</td><td>85929480</td><td>85930127</td><td>+</td><td>1.09503</td><td>Intergenic</td><td>Intergenic</td><td>33038</td><td>NM_172798</td><td>21387</td><td>Mm.275336</td><td>NM_011536</td><td>ENSMUSG00000000094</td><td>Tbx4</td><td>3930401C23</td><td>T-box 4</td><td>protein-coding</td></tr>
<tr><td>chr6_38954420_38955061</td><td>chr6</td><td>38954421</td><td>38955061</td><td>+</td><td>0.736737</td><td>intron (NM_011539, intron 3 of 12)</td><td>(CACCAT)n|Simple_repeat|Simple_repeat</td><td>35755</td><td>NM_011539</td><td>21391</td><td>Mm.4054</td><td>NM_011539</td><td>ENSMUSG00000029925</td><td>Tbxas1</td><td>CYP5|CYP5A1|THAS|TS|TXAS|TXS</td><td>thromboxane A synthase 1, platelet</td><td>protein-coding</td></tr>
<tr><td>chr4_136030947_136031915</td><td>chr4</td><td>136030948</td><td>136031915</td><td>+</td><td>1.53219</td><td>Intergenic</td><td>Intergenic</td><td>-9782</td><td>NM_013736</td><td>27224</td><td>Mm.27663</td><td>NM_013736</td><td>ENSMUSG00000028668</td><td>Tceb3</td><td>110kDa|AA408125|Tceb3a</td><td>transcription elongation factor B (SIII), polypeptide 3</td><td>protein-coding</td></tr>
<tr><td>chr18_42529659_42530056</td><td>chr18</td><td>42529660</td><td>42530056</td><td>+</td><td>0.831506</td><td>intron (NR_110344, intron 4 of 22)</td><td>intron (NR_110344, intron 4 of 22)</td><td>18371</td><td>NR_110344</td><td>56070</td><td>Mm.270511</td><td>NM_001039474</td><td>ENSMUSG00000024498</td><td>Tcerg1</td><td>2410022J09Rik|2900090C16Rik|AI428505|CA150b|FBP23|FBP28|Taf2s|ca150|p144</td><td>transcription elongation regulator 1 (CA150)</td><td>protein-coding</td></tr>
<tr><td>chr9_72076159_72076568</td><td>chr9</td><td>72076160</td><td>72076568</td><td>+</td><td>0.925585</td><td>intron (NM_001253862, intron 3 of 19)</td><td>intron (NM_001253862, intron 3 of 19)</td><td>34837</td><td>NM_001253862</td><td>21406</td><td>Mm.171615</td><td>NM_011544</td><td>ENSMUSG00000032228</td><td>Tcf12</td><td>A130037E08Rik|ALF1|HEB|HEBAlt|HTF-4|HTF4|ME1|REB|bHLHb20</td><td>transcription factor 12</td><td>protein-coding</td></tr>
<tr><td>chr11_52260906_52261298</td><td>chr11</td><td>52260907</td><td>52261298</td><td>+</td><td>0.777421</td><td>intron (NM_009331, intron 3 of 9)</td><td>intron (NM_009331, intron 3 of 9)</td><td>21469</td><td>NM_009331</td><td>21414</td><td>Mm.31630</td><td>NM_009331</td><td>ENSMUSG00000000782</td><td>Tcf7</td><td>AI465550|TCF-1|Tcf1</td><td>transcription factor 7, T cell specific</td><td>protein-coding</td></tr>
<tr><td>chr19_55750391_55750962</td><td>chr19</td><td>55750392</td><td>55750962</td><td>+</td><td>1.37358</td><td>intron (NM_009333, intron 3 of 12)</td><td>intron (NM_009333, intron 3 of 12)</td><td>8867</td><td>NM_001142923</td><td>21416</td><td>Mm.139815</td><td>NM_009333</td><td>ENSMUSG00000024985</td><td>Tcf7l2</td><td>TCF4B|TCF4E|Tcf-4|Tcf4|mTcf-4B|mTcf-4E</td><td>transcription factor 7 like 2, T cell specific, HMG box</td><td>protein-coding</td></tr>
<tr><td>chr19_55768290_55768648</td><td>chr19</td><td>55768291</td><td>55768648</td><td>+</td><td>1.20051</td><td>intron (NM_001142920, intron 4 of 11)</td><td>intron (NM_001142920, intron 4 of 11)</td><td>26659</td><td>NM_001142923</td><td>21416</td><td>Mm.139815</td><td>NM_009333</td><td>ENSMUSG00000024985</td><td>Tcf7l2</td><td>TCF4B|TCF4E|Tcf-4|Tcf4|mTcf-4B|mTcf-4E</td><td>transcription factor 7 like 2, T cell specific, HMG box</td><td>protein-coding</td></tr>
<tr><td>chr19_55791569_55792245</td><td>chr19</td><td>55791570</td><td>55792245</td><td>+</td><td>0.662791</td><td>intron (NM_001142920, intron 4 of 11)</td><td>intron (NM_001142920, intron 4 of 11)</td><td>50097</td><td>NM_001142923</td><td>21416</td><td>Mm.139815</td><td>NM_009333</td><td>ENSMUSG00000024985</td><td>Tcf7l2</td><td>TCF4B|TCF4E|Tcf-4|Tcf4|mTcf-4B|mTcf-4E</td><td>transcription factor 7 like 2, T cell specific, HMG box</td><td>protein-coding</td></tr>
<tr><td>chr18_60849359_60849687</td><td>chr18</td><td>60849360</td><td>60849687</td><td>+</td><td>0.611378</td><td>promoter-TSS (NM_011552)</td><td>promoter-TSS (NM_011552)</td><td>-559</td><td>NM_001198984</td><td>21453</td><td>Mm.486251</td><td>NM_011552</td><td>ENSMUSG00000024613</td><td>Tcof1</td><td>AA408847|AW209012|treacle</td><td>Treacher Collins Franceschetti syndrome 1, homolog</td><td>protein-coding</td></tr>
<tr><td>chr2_104687158_104687688</td><td>chr2</td><td>104687159</td><td>104687688</td><td>+</td><td>0.873703</td><td>intron (NM_177190, intron 7 of 9)</td><td>intron (NM_177190, intron 7 of 9)</td><td>24739</td><td>NM_177190</td><td>320554</td><td>Mm.249065</td><td>NM_177190</td><td>ENSMUSG00000027175</td><td>Tcp11l1</td><td>C130096D04Rik</td><td>t-complex 11 like 1</td><td>protein-coding</td></tr>
<tr><td>chr4_102886883_102887488</td><td>chr4</td><td>102886884</td><td>102887488</td><td>+</td><td>1.14663</td><td>intron (NM_001285862, intron 6 of 19)</td><td>MIR|SINE|MIR</td><td>-99193</td><td>NM_001163767</td><td>67344</td><td>Mm.111346</td><td>NM_026100</td><td>ENSMUSG00000028523</td><td>Tctex1d1</td><td>1700055O19Rik</td><td>Tctex1 domain containing 1</td><td>protein-coding</td></tr>
<tr><td>chr4_102886204_102886784</td><td>chr4</td><td>102886205</td><td>102886784</td><td>+</td><td>0.868766</td><td>intron (NM_001285862, intron 6 of 19)</td><td>intron (NM_001285862, intron 6 of 19)</td><td>-99885</td><td>NM_001163767</td><td>67344</td><td>Mm.111346</td><td>NM_026100</td><td>ENSMUSG00000028523</td><td>Tctex1d1</td><td>1700055O19Rik</td><td>Tctex1 domain containing 1</td><td>protein-coding</td></tr>
<tr><td>chr14_87739504_87739822</td><td>chr14</td><td>87739505</td><td>87739822</td><td>+</td><td>0.757833</td><td>Intergenic</td><td>ORR1D1|LTR|ERVL-MaLR</td><td>323080</td><td>NM_001253755</td><td>219249</td><td>Mm.35168</td><td>NM_172605</td><td>ENSMUSG00000022019</td><td>Tdrd3</td><td>4732418C03Rik|6720468N18</td><td>tudor domain containing 3</td><td>protein-coding</td></tr>
<tr><td>chr8_14013679_14013965</td><td>chr8</td><td>14013680</td><td>14013965</td><td>+</td><td>0.800287</td><td>Intergenic</td><td>Intergenic</td><td>-39045</td><td>NM_173744</td><td>72148</td><td>Mm.5727</td><td>NM_173744</td><td>ENSMUSG00000050052</td><td>Tdrp</td><td>2610019F03Rik</td><td>testis development related protein</td><td>protein-coding</td></tr>
<tr><td>chr8_14001442_14002099</td><td>chr8</td><td>14001443</td><td>14002099</td><td>+</td><td>0.783433</td><td>Intergenic</td><td>MERV1_LTR|LTR|ERV1</td><td>-26994</td><td>NM_173744</td><td>72148</td><td>Mm.5727</td><td>NM_173744</td><td>ENSMUSG00000050052</td><td>Tdrp</td><td>2610019F03Rik</td><td>testis development related protein</td><td>protein-coding</td></tr>
<tr><td>chr5_72844169_72844786</td><td>chr5</td><td>72844170</td><td>72844786</td><td>+</td><td>0.832346</td><td>intron (NM_013689, intron 1 of 17)</td><td>L1MB5|LINE|L1</td><td>23970</td><td>NM_001113461</td><td>21682</td><td>Mm.319581</td><td>NM_013689</td><td>ENSMUSG00000029217</td><td>Tec</td><td>-</td><td>tec protein tyrosine kinase</td><td>protein-coding</td></tr>
<tr><td>chr8_83599098_83599401</td><td>chr8</td><td>83599099</td><td>83599401</td><td>+</td><td>0.984266</td><td>Intergenic</td><td>ORR1B1|LTR|ERVL-MaLR</td><td>-4759</td><td>NM_134118</td><td>106529</td><td>Mm.352239</td><td>NM_027179</td><td>ENSMUSG00000031708</td><td>Tecr</td><td>2410016D23Rik|A230102P12Rik|AI173355|D17Ertd178e|Gpsn2|SC2</td><td>trans-2,3-enoyl-CoA reductase</td><td>protein-coding</td></tr>
<tr><td>chr11_80145670_80146301</td><td>chr11</td><td>80145671</td><td>80146301</td><td>+</td><td>0.864823</td><td>Intergenic</td><td>Intergenic</td><td>-3833</td><td>NM_183275</td><td>68550</td><td>Mm.18466</td><td>NM_183275</td><td>ENSMUSG00000046909</td><td>Tefm</td><td>1110002N22Rik</td><td>transcription elongation factor, mitochondrial</td><td>protein-coding</td></tr>
<tr><td>chr3_96401632_96402549</td><td>chr3</td><td>96401633</td><td>96402549</td><td>+</td><td>0.963582</td><td>Intergenic</td><td>CpG</td><td>12742</td><td>NR_001579</td><td>21748</td><td></td><td>NR_001579</td><td>ENSMUSG00000064796</td><td>Terc</td><td>mTER|mTR</td><td>telomerase RNA component</td><td>ncRNA</td></tr>
<tr><td>chr3_96409246_96410272</td><td>chr3</td><td>96409247</td><td>96410272</td><td>+</td><td>0.782891</td><td>Intergenic</td><td>Intergenic</td><td>5074</td><td>NR_001579</td><td>21748</td><td></td><td>NR_001579</td><td>ENSMUSG00000064796</td><td>Terc</td><td>mTER|mTR</td><td>telomerase RNA component</td><td>ncRNA</td></tr>
<tr><td>chr3_96427820_96428523</td><td>chr3</td><td>96427821</td><td>96428523</td><td>+</td><td>0.771002</td><td>Intergenic</td><td>B1_Mur4|SINE|Alu</td><td>-13339</td><td>NR_001579</td><td>21748</td><td></td><td>NR_001579</td><td>ENSMUSG00000064796</td><td>Terc</td><td>mTER|mTR</td><td>telomerase RNA component</td><td>ncRNA</td></tr>
<tr><td>chr8_107113932_107114318</td><td>chr8</td><td>107113933</td><td>107114318</td><td>+</td><td>0.807833</td><td>Intergenic</td><td>B3A|SINE|B2</td><td>-17580</td><td>NR_104410</td><td>21750</td><td>Mm.6402</td><td>NM_009353</td><td>ENSMUSG00000031921</td><td>Terf2</td><td>TRF2</td><td>telomeric repeat binding factor 2</td><td>protein-coding</td></tr>
<tr><td>chr6_17060021_17060607</td><td>chr6</td><td>17060022</td><td>17060607</td><td>+</td><td>2.01711</td><td>Intergenic</td><td>Intergenic</td><td>-4835</td><td>NM_207176</td><td>21753</td><td>Mm.436548</td><td>NM_011570</td><td>ENSMUSG00000029552</td><td>Tes</td><td>D6Ertd352e|TESS|Tes1|Tes2|testin2</td><td>testis derived transcript</td><td>protein-coding</td></tr>
<tr><td>chr6_17059067_17059382</td><td>chr6</td><td>17059068</td><td>17059382</td><td>+</td><td>0.883681</td><td>Intergenic</td><td>HAL1M8|LINE|L1</td><td>-5924</td><td>NM_207176</td><td>21753</td><td>Mm.436548</td><td>NM_011570</td><td>ENSMUSG00000029552</td><td>Tes</td><td>D6Ertd352e|TESS|Tes1|Tes2|testin2</td><td>testis derived transcript</td><td>protein-coding</td></tr>
</table>

| | | | | | | | | | | | | | | | | | |
|---|---|---|---|---|---|---|---|---|---|---|---|---|---|---|---|---|---|
| chr6_17052844_17053518 | chr6 | 17052845 | 17053518 | + | 0.800711 | Intergenic | MLT1G3-int\|LTR\|ERVL-MaLR | -11968 | NM_207176 | 21753 | Mm.436548 | NM_011570 | ENSMUSG00000029552 | **Tes** | D6Ertd352e\|TESS\|Tes1\|Tes2\|testin2 | testis derived transcript | protein-coding |
| chr6_17053048_17053513 | chr6 | 17053049 | 17053513 | + | 0.791666 | Intergenic | MLT1G3-int\|LTR\|ERVL-MaLR | -11868 | NM_207176 | 21753 | Mm.436548 | NM_011570 | ENSMUSG00000029552 | **Tes** | D6Ertd352e\|TESS\|Tes1\|Tes2\|testin2 | testis derived transcript | protein-coding |
| chr5_118079385_118079693 | chr5 | 118079386 | 118079693 | + | 0.829419 | intron (NM_172721, intron 7 of 10) | intron (NM_172721, intron 7 of 10) | 51715 | NM_021344 | 57816 | Mm.273285 | NM_021344 | ENSMUSG00000029359 | **Tesc** | 1010001A17Rik\|2410011K10Rik\|TE-1 | tescalcin | protein-coding |
| chr5_118079390_118080282 | chr5 | 118079391 | 118080282 | + | 0.640945 | intron (NM_172721, intron 7 of 10) | intron (NM_172721, intron 7 of 10) | 52012 | NM_021344 | 57816 | Mm.273285 | NM_021344 | ENSMUSG00000029359 | **Tesc** | 1010001A17Rik\|2410011K10Rik\|TE-1 | tescalcin | protein-coding |
| chr10_130340124_130340438 | chr10 | 130340125 | 130340438 | + | 0.827048 | intron (NM_183264, intron 1 of 8) | Lx\|LINE\|L1 | 17429 | NM_183264 | 67596 | Mm.131264 | NM_183264 | ENSMUSG00000034833 | **Tespa1** | 5830405N20Rik\|A430001F24Rik\|AI450549 | thymocyte expressed, positive selection associated 1 | protein-coding |
| chr10_130347326_130347721 | chr10 | 130347327 | 130347721 | + | 0.734873 | intron (NM_183264, intron 2 of 8) | intron (NM_183264, intron 2 of 8) | 24672 | NM_183264 | 67596 | Mm.131264 | NM_183264 | ENSMUSG00000034833 | **Tespa1** | 5830405N20Rik\|A430001F24Rik\|AI450549 | thymocyte expressed, positive selection associated 1 | protein-coding |
| chr3_133436795_133437437 | chr3 | 133436796 | 133437437 | + | 1.6956 | Intergenic | Intergenic | 107274 | NM_001040400 | 214133 | Mm.347816 | NM_145989 | ENSMUSG00000040943 | **Tet2** | Ayu17-449\|E130014J05Rik\|mKIAA1546 | tet methylcytosine dioxygenase 2 | protein-coding |
| chr3_133626354_133627164 | chr3 | 133626355 | 133627164 | + | 1.59424 | Intergenic | Intergenic | -82369 | NM_001040400 | 214133 | Mm.347816 | NM_145989 | ENSMUSG00000040943 | **Tet2** | Ayu17-449\|E130014J05Rik\|mKIAA1546 | tet methylcytosine dioxygenase 2 | protein-coding |
| chr3_133515078_133515596 | chr3 | 133515079 | 133515596 | + | 1.5579 | intron (NM_001040400, intron 1 of 10) | intron (NM_001040400, intron 1 of 10) | 29053 | NM_001040400 | 214133 | Mm.347816 | NM_145989 | ENSMUSG00000040943 | **Tet2** | Ayu17-449\|E130014J05Rik\|mKIAA1546 | tet methylcytosine dioxygenase 2 | protein-coding |
| chr3_133626299_133627460 | chr3 | 133626300 | 133627460 | + | 1.49003 | Intergenic | Intergenic | -82490 | NM_001040400 | 214133 | Mm.347816 | NM_145989 | ENSMUSG00000040943 | **Tet2** | Ayu17-449\|E130014J05Rik\|mKIAA1546 | tet methylcytosine dioxygenase 2 | protein-coding |
| chr3_133758024_133758911 | chr3 | 133758025 | 133758911 | + | 1.48723 | Intergenic | Intergenic | -214078 | NM_001040400 | 214133 | Mm.347816 | NM_145989 | ENSMUSG00000040943 | **Tet2** | Ayu17-449\|E130014J05Rik\|mKIAA1546 | tet methylcytosine dioxygenase 2 | protein-coding |
| chr3_133582057_133582527 | chr3 | 133582058 | 133582527 | + | 1.19304 | Intergenic | Intergenic | -37902 | NM_001040400 | 214133 | Mm.347816 | NM_145989 | ENSMUSG00000040943 | **Tet2** | Ayu17-449\|E130014J05Rik\|mKIAA1546 | tet methylcytosine dioxygenase 2 | protein-coding |
| chr3_133542115_133542449 | chr3 | 133542116 | 133542449 | + | 1.16669 | intron (NM_001040400, intron 1 of 10) | intron (NM_001040400, intron 1 of 10) | 2108 | NM_001040400 | 214133 | Mm.347816 | NM_145989 | ENSMUSG00000040943 | **Tet2** | Ayu17-449\|E130014J05Rik\|mKIAA1546 | tet methylcytosine dioxygenase 2 | protein-coding |
| chr3_133581552_133581902 | chr3 | 133581553 | 133581902 | + | 1.02035 | Intergenic | Intergenic | -37337 | NM_001040400 | 214133 | Mm.347816 | NM_145989 | ENSMUSG00000040943 | **Tet2** | Ayu17-449\|E130014J05Rik\|mKIAA1546 | tet methylcytosine dioxygenase 2 | protein-coding |
| chr3_133582749_133583422 | chr3 | 133582750 | 133583422 | + | 1.02012 | Intergenic | Intergenic | -38696 | NM_001040400 | 214133 | Mm.347816 | NM_145989 | ENSMUSG00000040943 | **Tet2** | Ayu17-449\|E130014J05Rik\|mKIAA1546 | tet methylcytosine dioxygenase 2 | protein-coding |
| chr3_133506084_133506835 | chr3 | 133506085 | 133506835 | + | 0.948429 | intron (NM_001040400, intron 2 of 10) | MER21B\|LTR\|ERVL | 37930 | NM_001040400 | 214133 | Mm.347816 | NM_145989 | ENSMUSG00000040943 | **Tet2** | Ayu17-449\|E130014J05Rik\|mKIAA1546 | tet methylcytosine dioxygenase 2 | protein-coding |
| chr3_133527589_133528125 | chr3 | 133527590 | 133528125 | + | 0.792078 | intron (NM_001040400, intron 1 of 10) | B1_Mur2\|SINE\|Alu | 16533 | NM_001040400 | 214133 | Mm.347816 | NM_145989 | ENSMUSG00000040943 | **Tet2** | Ayu17-449\|E130014J05Rik\|mKIAA1546 | tet methylcytosine dioxygenase 2 | protein-coding |
| chr3_133711330_133711985 | chr3 | 133711331 | 133711985 | + | 0.699971 | Intergenic | Intergenic | -167268 | NM_001040400 | 214133 | Mm.347816 | NM_145989 | ENSMUSG00000040943 | **Tet2** | Ayu17-449\|E130014J05Rik\|mKIAA1546 | tet methylcytosine dioxygenase 2 | protein-coding |
| chr3_133623506_133623770 | chr3 | 133623507 | 133623770 | + | 0.667881 | Intergenic | Intergenic | -79248 | NM_001040400 | 214133 | Mm.347816 | NM_145989 | ENSMUSG00000040943 | **Tet2** | Ayu17-449\|E130014J05Rik\|mKIAA1546 | tet methylcytosine dioxygenase 2 | protein-coding |
| chr3_133467177_133467743 | chr3 | 133467178 | 133467743 | + | 0.667853 | exon (NM_001040400, exon 11 of 11) | exon (NM_001040400, exon 11 of 11) | 76930 | NM_001040400 | 214133 | Mm.347816 | NM_145989 | ENSMUSG00000040943 | **Tet2** | Ayu17-449\|E130014J05Rik\|mKIAA1546 | tet methylcytosine dioxygenase 2 | protein-coding |
| chr3_133625036_133625296 | chr3 | 133625037 | 133625296 | + | 0.652897 | Intergenic | Intergenic | -80776 | NM_001040400 | 214133 | Mm.347816 | NM_145989 | ENSMUSG00000040943 | **Tet2** | Ayu17-449\|E130014J05Rik\|mKIAA1546 | tet methylcytosine dioxygenase 2 | protein-coding |
| chr6_83423629_83424450 | chr6 | 83423630 | 83424450 | + | 1.1625 | intron (NM_183138, intron 2 of 10) | intron (NM_183138, intron 2 of 10) | 17638 | NM_183138 | 194388 | Mm.211030 | NM_183138 | ENSMUSG00000034832 | **Tet3** | B430006D22Rik\|BC037432\|D230004J03Rik | tet methylcytosine dioxygenase 3 | protein-coding |
| chr6_83440003_83440759 | chr6 | 83440004 | 83440759 | + | 0.835802 | intron (NM_183138, intron 2 of 10) | intron (NM_183138, intron 2 of 10) | 1297 | NM_183138 | 194388 | Mm.211030 | NM_183138 | ENSMUSG00000034832 | **Tet3** | B430006D22Rik\|BC037432\|D230004J03Rik | tet methylcytosine dioxygenase 3 | protein-coding |
| chr6_83439998_83440415 | chr6 | 83439999 | 83440415 | + | 0.75911 | intron (NM_183138, intron 2 of 10) | intron (NM_183138, intron 2 of 10) | 1471 | NM_183138 | 194388 | Mm.211030 | NM_183138 | ENSMUSG00000034832 | **Tet3** | B430006D22Rik\|BC037432\|D230004J03Rik | tet methylcytosine dioxygenase 3 | protein-coding |
| chr6_83440466_83440746 | chr6 | 83440467 | 83440746 | + | 0.756614 | intron (NM_183138, intron 2 of 10) | intron (NM_183138, intron 2 of 10) | 1072 | NM_183138 | 194388 | Mm.211030 | NM_183138 | ENSMUSG00000034832 | **Tet3** | B430006D22Rik\|BC037432\|D230004J03Rik | tet methylcytosine dioxygenase 3 | protein-coding |
| chr6_83426278_83426636 | chr6 | 83426279 | 83426636 | + | 0.627807 | intron (NM_183138, intron 2 of 10) | intron (NM_183138, intron 2 of 10) | 15221 | NM_183138 | 194388 | Mm.211030 | NM_183138 | ENSMUSG00000034832 | **Tet3** | B430006D22Rik\|BC037432\|D230004J03Rik | tet methylcytosine dioxygenase 3 | protein-coding |
| chr11_87483173_87483709 | chr11 | 87483174 | 87483709 | + | 0.851939 | intron (NM_031386, intron 3 of 30) | intron (NM_031386, intron 3 of 30) | 55850 | NM_001199293 | 83560 | Mm.440758 | NM_031386 | ENSMUSG00000010342 | **Tex14** | C85585 | testis expressed gene 14 | protein-coding |
| chr11_87459770_87460086 | chr11 | 87459771 | 87460086 | + | 0.848663 | intron (NM_031386, intron 2 of 30) | LTRIS_Mus\|LTR\|ERV1 | 32337 | NM_001199293 | 83560 | Mm.440758 | NM_031386 | ENSMUSG00000010342 | **Tex14** | C85585 | testis expressed gene 14 | protein-coding |
| chr11_87483235_87483517 | chr11 | 87483236 | 87483517 | + | 0.835599 | intron (NM_031386, intron 3 of 30) | PB1D10\|SINE\|Alu | 55785 | NM_001199293 | 83560 | Mm.440758 | NM_031386 | ENSMUSG00000010342 | **Tex14** | C85585 | testis expressed gene 14 | protein-coding |
| chr11_87483176_87483559 | chr11 | 87483177 | 87483559 | + | 0.776174 | intron (NM_031386, intron 3 of 30) | PB1D10\|SINE\|Alu | 55777 | NM_001199293 | 83560 | Mm.440758 | NM_031386 | ENSMUSG00000010342 | **Tex14** | C85585 | testis expressed gene 14 | protein-coding |
| chr11_87453584_87454321 | chr11 | 87453585 | 87454321 | + | 0.760178 | intron (NM_031386, intron 2 of 30) | intron (NM_031386, intron 2 of 30) | 26362 | NM_001199293 | 83560 | Mm.440758 | NM_031386 | ENSMUSG00000010342 | **Tex14** | C85585 | testis expressed gene 14 | protein-coding |
| chr8_33548378_33548873 | chr8 | 33548379 | 33548873 | + | 1.16866 | Intergenic | Intergenic | -21918 | NM_031374 | 104271 | Mm.280624 | NM_031374 | ENSMUSG00000009628 | **Tex15** | 2210014E14Rik\|AL022622\|AU022940 | testis expressed gene 15 | protein-coding |
| chr8_33529694_33530120 | chr8 | 33529695 | 33530120 | + | 0.989477 | Intergenic | RSINE1\|SINE\|B4 | -40637 | NM_031374 | 104271 | Mm.280624 | NM_031374 | ENSMUSG00000009628 | **Tex15** | 2210014E14Rik\|AL022622\|AU022940 | testis expressed gene 15 | protein-coding |
| chr8_33549154_33549470 | chr8 | 33549155 | 33549470 | + | 0.9235 | Intergenic | Intergenic | -21232 | NM_031374 | 104271 | Mm.280624 | NM_031374 | ENSMUSG00000009628 | **Tex15** | 2210014E14Rik\|AL022622\|AU022940 | testis expressed gene 15 | protein-coding |
| chr8_33547186_33547814 | chr8 | 33547187 | 33547814 | + | 0.654594 | Intergenic | RMER1A\|Other\|Other | -23044 | NM_031374 | 104271 | Mm.280624 | NM_031374 | ENSMUSG00000009628 | **Tex15** | 2210014E14Rik\|AL022622\|AU022940 | testis expressed gene 15 | protein-coding |
| chr5_149427323_149427726 | chr5 | 149427324 | 149427726 | + | 0.792675 | intron (NM_027519, intron 4 of 4) | intron (NM_027519, intron 4 of 4) | -12181 | NM_029464 | 75860 | Mm.159605 | NM_029464 | ENSMUSG00000029660 | **Tex26** | 4930588N13Rik | testis expressed 26 | protein-coding |
| chr1_157248070_157248445 | chr1 | 157248071 | 157248445 | + | 1.27715 | intron (NM_177644, intron 3 of 17) | intron (NM_177644, intron 3 of 17) | -139613 | NM_028540 | 73435 | Mm.116705 | NM_028540 | ENSMUSG00000026592 | **Tex35** | 1700057K13Rik | testis expressed 35 | protein-coding |
| chr1_157232375_157232742 | chr1 | 157232376 | 157232742 | + | 0.796678 | intron (NM_177644, intron 3 of 17) | intron (NM_177644, intron 3 of 17) | -123914 | NM_028540 | 73435 | Mm.116705 | NM_028540 | ENSMUSG00000026592 | **Tex35** | 1700057K13Rik | testis expressed 35 | protein-coding |
| chr1_157090846_157091290 | chr1 | 157090847 | 157091290 | + | 0.734244 | Intergenic | Intergenic | 17577 | NM_028540 | 73435 | Mm.116705 | NM_028540 | ENSMUSG00000026592 | **Tex35** | 1700057K13Rik | testis expressed 35 | protein-coding |
| chr18_56600948_56601465 | chr18 | 56600949 | 56601465 | + | 1.31806 | Intergenic | RMER15\|LTR\|ERVL | 12859 | NM_026099 | 67343 | Mm.87057 | NM_026099 | ENSMUSG00000032900 | **Tex43** | 1700065I17Rik\|Tseg7 | testis expressed 43 | protein-coding |
| chr8_13326568_13327025 | chr8 | 13326569 | 13327025 | + | 0.877254 | Intergenic | RSINE1\|SINE\|B4 | -11954 | NM_001291765 | 21781 | Mm.925 | NM_009361 | ENSMUSG00000038482 | **Tfdp1** | Dp1\|Drtf1 | transcription factor Dp 1 | protein-coding |
| chr9_96214216_96214615 | chr9 | 96214217 | 96214615 | + | 1.04748 | intron (NM_178667, intron 2 of 13) | Lx8b\|LINE\|L1 | 18141 | NM_001184706 | 211586 | Mm.17977 | NM_178667 | ENSMUSG00000032411 | **Tfdp2** | 1110029I05Rik\|A330080J22Rik\|DP-3\|DP3 | transcription factor Dp 2 | protein-coding |
| chr17_31165110_31165595 | chr17 | 31165111 | 31165595 | + | 1.03924 | promoter-TSS (NM_009362) | promoter-TSS (NM_009362) | -300 | NM_009362 | 21784 | Mm.2854 | NM_009362 | ENSMUSG00000024032 | **Tff1** | Bcei\|PS2 | trefoil factor 1 | protein-coding |
| chr17_31162536_31163067 | chr17 | 31162537 | 31163067 | + | 0.91798 | exon (NM_009362, exon 2 of 3) | exon (NM_009362, exon 2 of 3) | 2251 | NM_009362 | 21784 | Mm.2854 | NM_009362 | ENSMUSG00000024032 | **Tff1** | Bcei\|PS2 | trefoil factor 1 | protein-coding |
| chr15_66644936_66645490 | chr15 | 66644937 | 66645490 | + | 1.47978 | TTS (NM_001081409) | TTS (NM_001081409) | -25557 | NM_009375 | 21819 | Mm.441333 | NM_009375 | ENSMUSG00000053469 | **Tg** | Tgn\|cog | thyroglobulin | protein-coding |
| chr14_118109716_118110290 | chr14 | 118109717 | 118110290 | + | 1.86397 | Intergenic | Intergenic | 22762 | NM_029578 | 76355 | Mm.251537 | NM_029578 | ENSMUSG00000022130 | **Tgds** | 2610017J16Rik\|2610025M23Rik\|AI648925 | TDP-glucose 4,6-dehydratase | protein-coding |
| chr1_186676007_186676591 | chr1 | 186676008 | 186676591 | + | 0.93214 | intron (NM_009367, intron 1 of 6) | intron (NM_009367, intron 1 of 6) | 29693 | NM_009367 | 21808 | Mm.18213 | NM_009367 | ENSMUSG00000039239 | **Tgfb2** | BB105277\|Tgf-beta2\|Tgfb-2 | transforming growth factor, beta 2 | protein-coding |
| chr1_186706530_186707000 | chr1 | 186706531 | 186707000 | + | 0.810135 | promoter-TSS (NM_009367) | promoter-TSS (NM_009367) | -773 | NM_009367 | 21808 | Mm.18213 | NM_009367 | ENSMUSG00000039239 | **Tgfb2** | BB105277\|Tgf-beta2\|Tgfb-2 | transforming growth factor, beta 2 | protein-coding |
| chr12_86060930_86061281 | chr12 | 86060931 | 86061281 | + | 0.710294 | intron (NM_009368, intron 5 of 6) | MTE2a\|LTR\|ERVL-MaLR | 17935 | NM_009368 | 21809 | Mm.3992 | NM_009368 | ENSMUSG00000021253 | **Tgfb3** | Tgfb-3 | transforming growth factor, beta 3 | protein-coding |
| chr4_47322308_47322827 | chr4 | 47322309 | 47322827 | + | 0.955233 | Intergenic | MTE2a\|LTR\|ERVL-MaLR | -30737 | NM_009370 | 21812 | Mm.197552 | NM_009370 | ENSMUSG00000007613 | **Tgfbr1** | ALK5\|AU017191\|Alk-5\|TbetaR-I\|TbetaRI | transforming growth factor, beta receptor I | protein-coding |

| | | | | | | | | | | | | | | | | | |
|---|---|---|---|---|---|---|---|---|---|---|---|---|---|---|---|---|---|
| chr4_47313925_47314530 | chr4 | 47313926 | 47314530 | + | 0.901378 | Intergenic | Intergenic | -39077 | NM_009370 | 21812 | Mm.197552 | NM_009370 | ENSMUSG00000007613 | **Tgfbr1** | ALK5\|AU017191\|Alk-5\|TbetaR-I\|TbetaRI | transforming growth factor, beta receptor I | protein-coding |
| chr4_47313904_47314819 | chr4 | 47313905 | 47314819 | + | 0.749347 | Intergenic | Intergenic | -38943 | NM_009370 | 21812 | Mm.197552 | NM_009370 | ENSMUSG00000007613 | **Tgfbr1** | ALK5\|AU017191\|Alk-5\|TbetaR-I\|TbetaRI | transforming growth factor, beta receptor I | protein-coding |
| chr9_116136841_116137199 | chr9 | 116136842 | 116137199 | + | 1.47674 | intron (NM_029575, intron 1 of 6) | intron (NM_029575, intron 1 of 6) | 38343 | NM_029575 | 21813 | Mm.172346 | NM_009371 | ENSMUSG00000032440 | **Tgfbr2** | 1110020H15Rik\|AU042018\|DNIIR\|RIIDN\|TBR-II\|TbetaR-II\|TbetaRII | transforming growth factor, beta receptor II | protein-coding |
| chr9_116332472_116333113 | chr9 | 116332473 | 116333113 | + | 1.35109 | Intergenic | Intergenic | -157430 | NM_009371 | 21813 | Mm.172346 | NM_009371 | ENSMUSG00000032440 | **Tgfbr2** | 1110020H15Rik\|AU042018\|DNIIR\|RIIDN\|TBR-II\|TbetaR-II\|TbetaRII | transforming growth factor, beta receptor II | protein-coding |
| chr9_116296352_116296630 | chr9 | 116296353 | 116296630 | + | 0.859837 | Intergenic | Intergenic | -121128 | NM_009371 | 21813 | Mm.172346 | NM_009371 | ENSMUSG00000032440 | **Tgfbr2** | 1110020H15Rik\|AU042018\|DNIIR\|RIIDN\|TBR-II\|TbetaR-II\|TbetaRII | transforming growth factor, beta receptor II | protein-coding |
| chr9_116243414_116244452 | chr9 | 116243415 | 116244452 | + | 0.717272 | Intergenic | Intergenic | -68570 | NM_009371 | 21813 | Mm.172346 | NM_009371 | ENSMUSG00000032440 | **Tgfbr2** | 1110020H15Rik\|AU042018\|DNIIR\|RIIDN\|TBR-II\|TbetaR-II\|TbetaRII | transforming growth factor, beta receptor II | protein-coding |
| chr9_116125219_116125572 | chr9 | 116125220 | 116125572 | + | 0.695445 | intron (NM_029575, intron 3 of 6) | intron (NM_029575, intron 3 of 6) | 49967 | NM_029575 | 21813 | Mm.172346 | NM_009371 | ENSMUSG00000032440 | **Tgfbr2** | 1110020H15Rik\|AU042018\|DNIIR\|RIIDN\|TBR-II\|TbetaR-II\|TbetaRII | transforming growth factor, beta receptor II | protein-coding |
| chr9_116193258_116193459 | chr9 | 116193259 | 116193459 | + | 0.679856 | Intergenic | LTR16A1\|LTR\|ERVL | -17996 | NM_009371 | 21813 | Mm.172346 | NM_009371 | ENSMUSG00000032440 | **Tgfbr2** | 1110020H15Rik\|AU042018\|DNIIR\|RIIDN\|TBR-II\|TbetaR-II\|TbetaRII | transforming growth factor, beta receptor II | protein-coding |
| chr9_116214747_116215023 | chr9 | 116214748 | 116215023 | + | 0.654515 | Intergenic | RLTR30D_RN\|LTR\|ERV1 | -39522 | NM_009371 | 21813 | Mm.172346 | NM_009371 | ENSMUSG00000032440 | **Tgfbr2** | 1110020H15Rik\|AU042018\|DNIIR\|RIIDN\|TBR-II\|TbetaR-II\|TbetaRII | transforming growth factor, beta receptor II | protein-coding |
| chr9_116152774_116153257 | chr9 | 116152775 | 116153257 | + | 0.629031 | intron (NM_029575, intron 1 of 6) | intron (NM_029575, intron 1 of 6) | 22347 | NM_029575 | 21813 | Mm.172346 | NM_009371 | ENSMUSG00000032440 | **Tgfbr2** | 1110020H15Rik\|AU042018\|DNIIR\|RIIDN\|TBR-II\|TbetaR-II\|TbetaRII | transforming growth factor, beta receptor II | protein-coding |
| chr5_107283392_107284466 | chr5 | 107283393 | 107284466 | + | 1.58713 | intron (NM_011578, intron 1 of 16) | intron (NM_011578, intron 1 of 16) | 5666 | NM_011578 | 21814 | Mm.200775 | NM_011578 | ENSMUSG00000029287 | **Tgfbr3** | 1110036H20Rik\|AU015626\|AW215636\|TBRIII | transforming growth factor, beta receptor III | protein-coding |
| chr5_107260687_107261090 | chr5 | 107260688 | 107261090 | + | 0.955296 | intron (NM_011578, intron 2 of 16) | MIR3\|SINE\|MIR | 28706 | NM_011578 | 21814 | Mm.200775 | NM_011578 | ENSMUSG00000029287 | **Tgfbr3** | 1110036H20Rik\|AU015626\|AW215636\|TBRIII | transforming growth factor, beta receptor III | protein-coding |
| chr5_107282370_107282885 | chr5 | 107282371 | 107282885 | + | 0.914205 | intron (NM_011578, intron 1 of 16) | intron (NM_011578, intron 1 of 16) | 6967 | NM_011578 | 21814 | Mm.200775 | NM_011578 | ENSMUSG00000029287 | **Tgfbr3** | 1110036H20Rik\|AU015626\|AW215636\|TBRIII | transforming growth factor, beta receptor III | protein-coding |
| chr17_70728684_70729405 | chr17 | 70728685 | 70729405 | + | 1.15187 | intron (NM_001128181, intron 5 of 9) | intron (NM_001128181, intron 5 of 9) | 117732 | NM_001164077 | 21815 | Mm.101034 | NM_009372 | ENSMUSG00000047407 | **Tgif1** | AA959811\|AI462167\|Tgif | TGFB-induced factor homeobox 1 | protein-coding |
| chr17_70885268_70885517 | chr17 | 70885269 | 70885517 | + | 0.929571 | Intergenic | Intergenic | -31861 | NM_001164074 | 21815 | Mm.101034 | NM_009372 | ENSMUSG00000047407 | **Tgif1** | AA959811\|AI462167\|Tgif | TGFB-induced factor homeobox 1 | protein-coding |
| chr17_70813469_70813837 | chr17 | 70813470 | 70813837 | + | 0.898481 | intron (NM_027712, intron 9 of 9) | intron (NM_027712, intron 9 of 9) | 33124 | NM_001164077 | 21815 | Mm.101034 | NM_009372 | ENSMUSG00000047407 | **Tgif1** | AA959811\|AI462167\|Tgif | TGFB-induced factor homeobox 1 | protein-coding |
| chr17_70836865_70837158 | chr17 | 70836866 | 70837158 | + | 0.83468 | Intergenic | Tigger7\|DNA\|TcMar-Tigger | 9765 | NM_001164077 | 21815 | Mm.101034 | NM_009372 | ENSMUSG00000047407 | **Tgif1** | AA959811\|AI462167\|Tgif | TGFB-induced factor homeobox 1 | protein-coding |
| chr17_70746383_70747200 | chr17 | 70746384 | 70747200 | + | 0.763271 | intron (NM_001128181, intron 5 of 9) | intron (NM_001128181, intron 5 of 9) | 99985 | NM_001164077 | 21815 | Mm.101034 | NM_009372 | ENSMUSG00000047407 | **Tgif1** | AA959811\|AI462167\|Tgif | TGFB-induced factor homeobox 1 | protein-coding |
| chr17_70886016_70886251 | chr17 | 70886017 | 70886251 | + | 0.666306 | Intergenic | Intergenic | -32602 | NM_001164074 | 21815 | Mm.101034 | NM_009372 | ENSMUSG00000047407 | **Tgif1** | AA959811\|AI462167\|Tgif | TGFB-induced factor homeobox 1 | protein-coding |
| chr2_158161540_158162069 | chr2 | 158161541 | 158162069 | + | 0.662925 | Intergenic | Intergenic | -15413 | NM_009373 | 21817 | Mm.330731 | NM_009373 | ENSMUSG00000037820 | **Tgm2** | G[a]h\|TG2\|TGase2\|tTG\|tTGas | transglutaminase 2, C polypeptide | protein-coding |
| chr17_84422098_84422371 | chr17 | 84422099 | 84422371 | + | 1.12398 | intron (NM_183021, intron 21 of 37) | intron (NM_183021, intron 21 of 37) | 43973 | NM_183021 | 240174 | Mm.246767 | NM_177826 | ENSMUSG00000024251 | **Thada** | 6530405K19\|A930021O22\|AU019899\|BC052885\|GITA | thyroid adenoma associated | protein-coding |
| chr17_84434633_84435465 | chr17 | 84434634 | 84435465 | + | 1.07048 | intron (NM_183021, intron 16 of 37) | intron (NM_183021, intron 16 of 37) | 31159 | NM_183021 | 240174 | Mm.246767 | NM_177826 | ENSMUSG00000024251 | **Thada** | 6530405K19\|A930021O22\|AU019899\|BC052885\|GITA | thyroid adenoma associated | protein-coding |
| chr17_84405655_84406065 | chr17 | 84405656 | 84406065 | + | 1.06767 | intron (NM_183021, intron 22 of 37) | intron (NM_183021, intron 22 of 37) | 60348 | NM_183021 | 240174 | Mm.246767 | NM_177826 | ENSMUSG00000024251 | **Thada** | 6530405K19\|A930021O22\|AU019899\|BC052885\|GITA | thyroid adenoma associated | protein-coding |
| chr17_84421492_84421704 | chr17 | 84421493 | 84421704 | + | 1.03742 | intron (NM_183021, intron 21 of 37) | intron (NM_183021, intron 21 of 37) | 44610 | NM_183021 | 240174 | Mm.246767 | NM_177826 | ENSMUSG00000024251 | **Thada** | 6530405K19\|A930021O22\|AU019899\|BC052885\|GITA | thyroid adenoma associated | protein-coding |
| chr2_118111174_118111518 | chr2 | 118111175 | 118111518 | + | 0.599734 | promoter-TSS (NM_011580) | promoter-TSS (NM_011580) | -576 | NM_011580 | 21825 | Mm.4159 | NM_011580 | ENSMUSG00000040152 | **Thbs1** | TSP-1\|TSP1\|Thbs-1\|tbsp1 | thrombospondin 1 | protein-coding |
| chr10_28595673_28595973 | chr10 | 28595674 | 28595973 | + | 1.50754 | exon (NM_008983, exon 32 of 32) | exon (NM_008983, exon 32 of 32) | -72572 | NM_178666 | 210757 | Mm.123021 | NM_178666 | ENSMUSG00000049109 | **Themis** | A430108D01\|BB160712\|E430004N04Rik\|Gasp\|Tsepa\|thylex | thymocyte selection associated | protein-coding |
| chr10_28592717_28593163 | chr10 | 28592718 | 28593163 | + | 1.27423 | intron (NM_008983, intron 31 of 31) | intron (NM_008983, intron 31 of 31) | -75455 | NM_178666 | 210757 | Mm.123021 | NM_178666 | ENSMUSG00000049109 | **Themis** | A430108D01\|BB160712\|E430004N04Rik\|Gasp\|Tsepa\|thylex | thymocyte selection associated | protein-coding |
| chr10_28375054_28375735 | chr10 | 28375055 | 28375735 | + | 1.20121 | intron (NM_008983, intron 6 of 31) | intron (NM_008983, intron 6 of 31) | -293000 | NM_178666 | 210757 | Mm.123021 | NM_178666 | ENSMUSG00000049109 | **Themis** | A430108D01\|BB160712\|E430004N04Rik\|Gasp\|Tsepa\|thylex | thymocyte selection associated | protein-coding |
| chr10_28776541_28777131 | chr10 | 28776542 | 28777131 | + | 0.978548 | intron (NM_178666, intron 3 of 5) | intron (NM_178666, intron 3 of 5) | 108441 | NM_178666 | 210757 | Mm.123021 | NM_178666 | ENSMUSG00000049109 | **Themis** | A430108D01\|BB160712\|E430004N04Rik\|Gasp\|Tsepa\|thylex | thymocyte selection associated | protein-coding |
| chr10_28776478_28777518 | chr10 | 28776479 | 28777518 | + | 0.970758 | intron (NM_178666, intron 3 of 5) | L1MCa\|LINE\|L1 | 108603 | NM_178666 | 210757 | Mm.123021 | NM_178666 | ENSMUSG00000049109 | **Themis** | A430108D01\|BB160712\|E430004N04Rik\|Gasp\|Tsepa\|thylex | thymocyte selection associated | protein-coding |
| chr10_28682499_28682736 | chr10 | 28682500 | 28682736 | + | 0.849357 | intron (NM_178666, intron 1 of 5) | intron (NM_178666, intron 1 of 5) | 14223 | NM_178666 | 210757 | Mm.123021 | NM_178666 | ENSMUSG00000049109 | **Themis** | A430108D01\|BB160712\|E430004N04Rik\|Gasp\|Tsepa\|thylex | thymocyte selection associated | protein-coding |
| chr10_28564237_28564779 | chr10 | 28564238 | 28564779 | + | 0.831601 | intron (NM_008983, intron 16 of 31) | intron (NM_008983, intron 16 of 31) | -103887 | NM_178666 | 210757 | Mm.123021 | NM_178666 | ENSMUSG00000049109 | **Themis** | A430108D01\|BB160712\|E430004N04Rik\|Gasp\|Tsepa\|thylex | thymocyte selection associated | protein-coding |
| chr10_28375813_28376171 | chr10 | 28375814 | 28376171 | + | 0.795628 | intron (NM_008983, intron 6 of 31) | L1Md_F2\|LINE\|L1 | -292403 | NM_178666 | 210757 | Mm.123021 | NM_178666 | ENSMUSG00000049109 | **Themis** | A430108D01\|BB160712\|E430004N04Rik\|Gasp\|Tsepa\|thylex | thymocyte selection associated | protein-coding |
| chr10_28682445_28682834 | chr10 | 28682446 | 28682834 | + | 0.767775 | intron (NM_178666, intron 1 of 5) | intron (NM_178666, intron 1 of 5) | 14245 | NM_178666 | 210757 | Mm.123021 | NM_178666 | ENSMUSG00000049109 | **Themis** | A430108D01\|BB160712\|E430004N04Rik\|Gasp\|Tsepa\|thylex | thymocyte selection associated | protein-coding |
| chr10_28694603_28695393 | chr10 | 28694604 | 28695393 | + | 0.72451 | intron (NM_178666, intron 1 of 5) | B3\|SINE\|B2 | 26603 | NM_178666 | 210757 | Mm.123021 | NM_178666 | ENSMUSG00000049109 | **Themis** | A430108D01\|BB160712\|E430004N04Rik\|Gasp\|Tsepa\|thylex | thymocyte selection associated | protein-coding |
| chr10_28600708_28601227 | chr10 | 28600709 | 28601227 | + | 0.721597 | Intergenic | Intergenic | -67427 | NM_178666 | 210757 | Mm.123021 | NM_178666 | ENSMUSG00000049109 | **Themis** | A430108D01\|BB160712\|E430004N04Rik\|Gasp\|Tsepa\|thylex | thymocyte selection associated | protein-coding |
| chr10_28694840_28695417 | chr10 | 28694841 | 28695417 | + | 0.706796 | intron (NM_178666, intron 1 of 5) | L1MDa\|LINE\|L1 | 26734 | NM_178666 | 210757 | Mm.123021 | NM_178666 | ENSMUSG00000049109 | **Themis** | A430108D01\|BB160712\|E430004N04Rik\|Gasp\|Tsepa\|thylex | thymocyte selection associated | protein-coding |
| chr10_28747876_28748401 | chr10 | 28747877 | 28748401 | + | 0.695284 | intron (NM_178666, intron 2 of 5) | MTA_Mm\|LTR\|ERVL-MaLR | 79744 | NM_178666 | 210757 | Mm.123021 | NM_178666 | ENSMUSG00000049109 | **Themis** | A430108D01\|BB160712\|E430004N04Rik\|Gasp\|Tsepa\|thylex | thymocyte selection associated | protein-coding |
| chr4_132799824_132800342 | chr4 | 132799825 | 132800342 | + | 1.92702 | Intergenic | Intergenic | -3719 | NM_001033308 | 230787 | Mm.290912 | NM_001033308 | ENSMUSG00000037731 | **Themis2** | ICB-1 | thymocyte selection associated family member 2 | protein-coding |
| chr11_45956437_45956800 | chr11 | 45956438 | 45956800 | + | 1.23532 | Intergenic | L1MC5a\|LINE\|L1 | -1116 | NM_001080969 | 66628 | Mm.269083 | NM_001080969 | ENSMUSG00000011254 | **Thg1l** | - | tRNA-histidine guanylyltransferase 1-like (S. cerevisiae) | protein-coding |
| chrX_41803663_41804119 | chrX | 41803664 | 41804119 | + | 0.628245 | intron (NM_001033422, intron 36 of 38) | L1MB5\|LINE\|L1 | 108010 | NM_001033422 | 331401 | Mm.259498 | NM_001033422 | ENSMUSG00000037475 | **Thoc2** | 6330441O12Rik\|D130005M13Rik\|Gm1139\|Gm1793\|Tho2 | THO complex 2 | protein-coding |
| chr14_13895214_13895975 | chr14 | 13895215 | 13895975 | + | 0.889426 | Intergenic | Intergenic | 65718 | NM_001285780 | 66231 | Mm.295875 | NM_025435 | ENSMUSG00000053453 | **Thoc7** | Nif3l1bp1 | THO complex 7 homolog (Drosophila) | protein-coding |
| chr14_18083928_18084640 | chr14 | 18083929 | 18084640 | + | 1.05537 | Intergenic | Intergenic | 102640 | NM_009380 | 21834 | Mm.32563 | NM_009380 | ENSMUSG00000021779 | **Thrb** | Nr1a2\|T3R[b]\|T3Rbeta\|Thrb1\|Thrb2\|c-erbAbeta | thyroid hormone receptor beta | protein-coding |
| chr14_18083938_18084991 | chr14 | 18083939 | 18084991 | + | 0.96255 | Intergenic | Intergenic | 102821 | NM_009380 | 21834 | Mm.32563 | NM_009380 | ENSMUSG00000021779 | **Thrb** | Nr1a2\|T3R[b]\|T3Rbeta\|Thrb1\|Thrb2\|c-erbAbeta | thyroid hormone receptor beta | protein-coding |
| chr14_17409496_17409984 | chr14 | 17409497 | 17409984 | + | 0.811095 | Intergenic | Intergenic | -251220 | NM_001113417 | 21834 | Mm.32563 | NM_009380 | ENSMUSG00000021779 | **Thrb** | Nr1a2\|T3R[b]\|T3Rbeta\|Thrb1\|Thrb2\|c-erbAbeta | thyroid hormone receptor beta | protein-coding |
| chr14_18100186_18100526 | chr14 | 18100187 | 18100526 | + | 0.6972 | Intergenic | Intergenic | 118712 | NM_009380 | 21834 | Mm.32563 | NM_009380 | ENSMUSG00000021779 | **Thrb** | Nr1a2\|T3R[b]\|T3Rbeta\|Thrb1\|Thrb2\|c-erbAbeta | thyroid hormone receptor beta | protein-coding |
| chr14_18100010_18100553 | chr14 | 18100011 | 18100553 | + | 0.685162 | Intergenic | Intergenic | 118638 | NM_009380 | 21834 | Mm.32563 | NM_009380 | ENSMUSG00000021779 | **Thrb** | Nr1a2\|T3R[b]\|T3Rbeta\|Thrb1\|Thrb2\|c-erbAbeta | thyroid hormone receptor beta | protein-coding |
| chr14_17981255_17982532 | chr14 | 17981256 | 17982532 | + | 0.623531 | exon (NM_009380, exon 1 of 7) | exon (NM_009380, exon 1 of 7) | 250 | NM_009380 | 21834 | Mm.32563 | NM_009380 | ENSMUSG00000021779 | **Thrb** | Nr1a2\|T3R[b]\|T3Rbeta\|Thrb1\|Thrb2\|c-erbAbeta | thyroid hormone receptor beta | protein-coding |
| chr6_113047397_113047727 | chr6 | 113047398 | 113047727 | + | 0.970473 | exon (NM_008188, exon 2 of 10) | exon (NM_008188, exon 2 of 10) | 1235 | NM_008188 | 14911 | Mm.781 | NM_008188 | ENSMUSG00000030264 | **Thumpd3** | AW556087\|Gt(ROSA)26asSor\|Gtrosa26as | THUMP domain containing 3 | protein-coding |
| chr9_44009276_44009828 | chr9 | 44009277 | 44009828 | + | 1.92563 | Intergenic | MLT1A1\|LTR\|ERVL-MaLR | -33832 | NM_009382 | 21838 | Mm.3951 | NM_009382 | ENSMUSG00000032011 | **Thy1** | CD90\|T25\|Thy-1\|Thy-1.2\|Thy1.1\|Thy1.2 | thymus cell antigen 1, theta | protein-coding |
| chr9_44039788_44040290 | chr9 | 44039789 | 44040290 | + | 1.24487 | Intergenic | Intergenic | -3345 | NM_009382 | 21838 | Mm.3951 | NM_009382 | ENSMUSG00000032011 | **Thy1** | CD90\|T25\|Thy-1\|Thy-1.2\|Thy1.1\|Thy1.2 | thymus cell antigen 1, theta | protein-coding |

| | | | | | | | | | | | | | | | | |
|---|---|---|---|---|---|---|---|---|---|---|---|---|---|---|---|---|
| chr9_44005372_44005765 | chr9 | 44005373 | 44005765 | + | 1.1258 | Intergenic | RMER15-int\|LTR\|ERVL | -37815 | NM_009382 | 21838 | Mm.3951 | NM_009382 | ENSMUSG00000032011 | **Thy1** | CD90\|T25\|Thy-1\|Thy-1.2\|Thy1.1\|Thy1.2 | thymus cell antigen 1, theta | protein-coding |
| chr9_44032032_44032372 | chr9 | 44032033 | 44032372 | + | 0.889147 | Intergenic | Intergenic | -11182 | NM_009382 | 21838 | Mm.3951 | NM_009382 | ENSMUSG00000032011 | **Thy1** | CD90\|T25\|Thy-1\|Thy-1.2\|Thy1.1\|Thy1.2 | thymus cell antigen 1, theta | protein-coding |
| chr9_44032038_44032250 | chr9 | 44032039 | 44032250 | + | 0.860129 | Intergenic | Intergenic | -11240 | NM_009382 | 21838 | Mm.3951 | NM_009382 | ENSMUSG00000032011 | **Thy1** | CD90\|T25\|Thy-1\|Thy-1.2\|Thy1.1\|Thy1.2 | thymus cell antigen 1, theta | protein-coding |
| chr9_44010935_44011538 | chr9 | 44010936 | 44011538 | + | 0.698662 | Intergenic | Lx5c\|LINE\|L1 | -32147 | NM_009382 | 21838 | Mm.3951 | NM_009382 | ENSMUSG00000032011 | **Thy1** | CD90\|T25\|Thy-1\|Thy-1.2\|Thy1.1\|Thy1.2 | thymus cell antigen 1, theta | protein-coding |
| chr16_89848319_89849554 | chr16 | 89848320 | 89849554 | + | 1.39601 | intron (NM_001145886, intron 12 of 27) | intron (NM_001145886, intron 12 of 27) | -30585 | NM_001145887 | 21844 | Mm.310902 | NM_009384 | ENSMUSG00000002489 | **Tiam1** | AI847750\|D16Ium10\|D16Ium10e | T cell lymphoma invasion and metastasis 1 | protein-coding |
| chr16_89835639_89835900 | chr16 | 89835640 | 89835900 | + | 0.76345 | intron (NM_001145886, intron 14 of 27) | intron (NM_001145886, intron 14 of 27) | -17418 | NM_001145887 | 21844 | Mm.310902 | NM_009384 | ENSMUSG00000002489 | **Tiam1** | AI847750\|D16Ium10\|D16Ium10e | T cell lymphoma invasion and metastasis 1 | protein-coding |
| chr16_89974361_89974751 | chr16 | 89974362 | 89974751 | + | 0.698533 | intron (NM_001145886, intron 1 of 27) | intron (NM_001145886, intron 1 of 27) | 143 | NM_001145886 | 21844 | Mm.310902 | NM_009384 | ENSMUSG00000002489 | **Tiam1** | AI847750\|D16Ium10\|D16Ium10e | T cell lymphoma invasion and metastasis 1 | protein-coding |
| chr6_59315764_59316286 | chr6 | 59315765 | 59316286 | + | 1.59833 | Intergenic | Intergenic | 107155 | NM_001081145 | 68140 | Mm.25843 | NM_001081145 | ENSMUSG00000049232 | **Tigd2** | 3632410O17Rik | tigger transposable element derived 2 | protein-coding |
| chr6_59312928_59313891 | chr6 | 59312929 | 59313891 | + | 1.47038 | Intergenic | RLTR14_RN\|LTR\|ERV1 | 104540 | NM_001081145 | 68140 | Mm.25843 | NM_001081145 | ENSMUSG00000049232 | **Tigd2** | 3632410O17Rik | tigger transposable element derived 2 | protein-coding |
| chr6_59314916_59315499 | chr6 | 59314917 | 59315499 | + | 1.43874 | Intergenic | Intergenic | 106338 | NM_001081145 | 68140 | Mm.25843 | NM_001081145 | ENSMUSG00000049232 | **Tigd2** | 3632410O17Rik | tigger transposable element derived 2 | protein-coding |
| chr6_59244830_59245324 | chr6 | 59244831 | 59245324 | + | 1.42667 | Intergenic | Arthur1B\|DNA\|hAT-Tip100 | 36207 | NM_001081145 | 68140 | Mm.25843 | NM_001081145 | ENSMUSG00000049232 | **Tigd2** | 3632410O17Rik | tigger transposable element derived 2 | protein-coding |
| chr16_43697796_43698059 | chr16 | 43697797 | 43698059 | + | 1.12464 | Intergenic | B3A\|SINE\|B2 | -33744 | NM_001146325 | 100043314 | Mm.390896 | NM_001146325 | ENSMUSG00000071552 | **Tigit** | ENSMUSG00000071552\|Vstm3 | T cell immunoreceptor with Ig and ITIM domains | protein-coding |
| chr16_43695822_43696193 | chr16 | 43695823 | 43696193 | + | 0.976385 | Intergenic | Intergenic | -31824 | NM_001146325 | 100043314 | Mm.390896 | NM_001146325 | ENSMUSG00000071552 | **Tigit** | ENSMUSG00000071552\|Vstm3 | T cell immunoreceptor with Ig and ITIM domains | protein-coding |
| chr16_43701854_43702519 | chr16 | 43701855 | 43702519 | + | 0.930596 | Intergenic | Intergenic | -38003 | NM_001146325 | 100043314 | Mm.390896 | NM_001146325 | ENSMUSG00000071552 | **Tigit** | ENSMUSG00000071552\|Vstm3 | T cell immunoreceptor with Ig and ITIM domains | protein-coding |
| chr16_43661535_43661962 | chr16 | 43661536 | 43661962 | + | 0.830039 | intron (NM_001146325, intron 2 of 3) | intron (NM_001146325, intron 2 of 3) | 2435 | NM_001146325 | 100043314 | Mm.390896 | NM_001146325 | ENSMUSG00000071552 | **Tigit** | ENSMUSG00000071552\|Vstm3 | T cell immunoreceptor with Ig and ITIM domains | protein-coding |
| chr16_43653817_43654127 | chr16 | 43653818 | 43654127 | + | 0.796309 | intron (NM_001146325, intron 3 of 3) | intron (NM_001146325, intron 3 of 3) | 10212 | NM_001146325 | 100043314 | Mm.390896 | NM_001146325 | ENSMUSG00000071552 | **Tigit** | ENSMUSG00000071552\|Vstm3 | T cell immunoreceptor with Ig and ITIM domains | protein-coding |
| chr11_46667920_46668598 | chr11 | 46667921 | 46668598 | + | 1.63394 | TTS (NM_001161356) | TTS (NM_001161356) | 22098 | NM_001161356 | 171284 | Mm.234654 | NM_134249 | ENSMUSG00000040413 | **Timd2** | AA816106\|AU019358\|AU019662\|C78111\|G9D3\|TIM-2\|Tim2 | T cell immunoglobulin and mucin domain containing 2 | protein-coding |
| chr11_46682844_46683285 | chr11 | 46682845 | 46683285 | + | 1.17726 | intron (NM_134249, intron 3 of 8) | intron (NM_134249, intron 3 of 8) | 7292 | NM_001161356 | 171284 | Mm.234654 | NM_134249 | ENSMUSG00000040413 | **Timd2** | AA816106\|AU019358\|AU019662\|C78111\|G9D3\|TIM-2\|Tim2 | T cell immunoglobulin and mucin domain containing 2 | protein-coding |
| chr11_46687864_46688308 | chr11 | 46687865 | 46688308 | + | 0.913429 | intron (NM_134249, intron 2 of 8) | ID_B1\|SINE\|B4 | 2271 | NM_001161356 | 171284 | Mm.234654 | NM_134249 | ENSMUSG00000040413 | **Timd2** | AA816106\|AU019358\|AU019662\|C78111\|G9D3\|TIM-2\|Tim2 | T cell immunoglobulin and mucin domain containing 2 | protein-coding |
| chr11_76434593_76435792 | chr11 | 76434594 | 76435792 | + | 0.709598 | intron (NM_198894, intron 16 of 22) | intron (NM_198894, intron 16 of 22) | 28050 | NM_019818 | 56322 | Mm.18803 | NM_019818 | ENSMUSG00000020843 | **Timm22** | Tim22 | translocase of inner mitochondrial membrane 22 | protein-coding |
| chr8_4274054_4274594 | chr8 | 4274055 | 4274594 | + | 1.26139 | intron (NM_011592, intron 1 of 12) | intron (NM_011592, intron 1 of 12) | 1581 | NM_011592 | 21856 | Mm.195249 | NM_011592 | ENSMUSG00000002949 | **Timm44** | 0710005E20Rik\|D8Ertd118e\|Mimt44\|Tim44 | translocase of inner mitochondrial membrane 44 | protein-coding |
| chr14_122047016_122047409 | chr14 | 122047017 | 122047409 | + | 0.919523 | Intergenic | Intergenic | 12539 | NM_001037744 | 223262 | Mm.297637 | NM_001037744 | ENSMUSG00000071229 | **Timm8a2** | - | translocase of inner mitochondrial membrane 8A2 | protein-coding |
| chr14_122041979_122042714 | chr14 | 122041980 | 122042714 | + | 0.694362 | Intergenic | Intergenic | 7673 | NM_001037744 | 223262 | Mm.297637 | NM_001037744 | ENSMUSG00000071229 | **Timm8a2** | - | translocase of inner mitochondrial membrane 8A2 | protein-coding |
| chr16_38503250_38503953 | chr16 | 38503251 | 38503953 | + | 1.22189 | intron (NM_024273, intron 5 of 6) | RMER1A\|Other\|Other | 19061 | NM_024273 | 76916 | Mm.132231 | NM_024273 | | **Timmdc1** | 2810021C21Rik\|4930455C21Rik\|AV135763 | translocase of inner mitochondrial membrane domain containing 1 | protein-coding |
| chr10_86302704_86303188 | chr10 | 86302705 | 86303188 | + | 0.789655 | intron (NM_011595, intron 1 of 4) | intron (NM_011595, intron 1 of 4) | 2534 | NM_011595 | 21859 | Mm.4871 | NM_011595 | ENSMUSG00000020044 | **Timp3** | Timp-3 | tissue inhibitor of metalloproteinase 3 | protein-coding |
| chr10_86302726_86303035 | chr10 | 86302727 | 86303035 | + | 0.716568 | intron (NM_011595, intron 1 of 4) | intron (NM_011595, intron 1 of 4) | 2469 | NM_011595 | 21859 | Mm.4871 | NM_011595 | ENSMUSG00000020044 | **Timp3** | Timp-3 | tissue inhibitor of metalloproteinase 3 | protein-coding |
| chr6_115283796_115284686 | chr6 | 115283797 | 115284686 | + | 1.45159 | Intergenic | Intergenic | -32392 | NM_080639 | 110595 | Mm.255607 | NM_080639 | ENSMUSG00000030317 | **Timp4** | TIMP-4 | tissue inhibitor of metalloproteinase 4 | protein-coding |
| chr6_115288292_115288589 | chr6 | 115288293 | 115288589 | + | 0.676541 | Intergenic | MIRb\|SINE\|MIR | -36592 | NM_080639 | 110595 | Mm.255607 | NM_080639 | ENSMUSG00000030317 | **Timp4** | TIMP-4 | tissue inhibitor of metalloproteinase 4 | protein-coding |
| chr3_65515703_65516219 | chr3 | 65515704 | 65516219 | + | 1.16326 | Intergenic | Intergenic | -12486 | NM_178892 | 99929 | Mm.246398 | NM_178892 | ENSMUSG00000034640 | **Tiparp** | ARTD14\|AW558171\|PARP7 | TCDD-inducible poly(ADP-ribose) polymerase | protein-coding |
| chr1_165250504_165251216 | chr1 | 165250505 | 165251216 | + | 1.21758 | Intergenic | Intergenic | -13902 | NM_145513 | 226591 | Mm.21520 | NM_145513 | ENSMUSG00000040843 | **Tiprl** | 1810011K17Rik | TIP41, TOR signalling pathway regulator-like (S. cerevisiae) | protein-coding |
| chr14_30556492_30557080 | chr14 | 30556493 | 30557080 | + | 0.92841 | intron (NM_009388, intron 1 of 13) | intron (NM_009388, intron 1 of 13) | 7655 | NM_009388 | 21881 | Mm.290692 | NM_009388 | ENSMUSG00000021957 | **Tkt** | p68 | transketolase | protein-coding |
| chr4_72184416_72184976 | chr4 | 72184417 | 72184976 | + | 1.38127 | intron (NM_001285530, intron 4 of 17) | intron (NM_001285530, intron 4 of 17) | 16197 | NM_011599 | 21885 | Mm.278444 | NM_011599 | ENSMUSG00000008305 | **Tle1** | C230057C06Rik\|Estm14\|Grg-1\|Grg1\|Tle4l | transducin-like enhancer of split 1, homolog of Drosophila E(spl) | protein-coding |
| chr4_72184339_72185174 | chr4 | 72184340 | 72185174 | + | 1.2457 | intron (NM_001285530, intron 4 of 17) | intron (NM_001285530, intron 4 of 17) | 16136 | NM_011599 | 21885 | Mm.278444 | NM_011599 | ENSMUSG00000008305 | **Tle1** | C230057C06Rik\|Estm14\|Grg-1\|Grg1\|Tle4l | transducin-like enhancer of split 1, homolog of Drosophila E(spl) | protein-coding |
| chr9_61367096_61367311 | chr9 | 61367097 | 61367311 | + | 0.834681 | Intergenic | Intergenic | -5162 | NM_001083928 | 21887 | Mm.24255 | NM_009389 | ENSMUSG00000032280 | **Tle3** | 2610103N05Rik\|ESG\|Grg3a\|Grg3b\|mKIAA1547 | transducin-like enhancer of split 3, homolog of Drosophila E(spl) | protein-coding |
| chr19_14620264_14620943 | chr19 | 14620265 | 14620943 | + | 1.42817 | Intergenic | MLT1H\|LTR\|ERVL-MaLR | -22621 | NM_011600 | 21888 | Mm.103638 | NM_011600 | ENSMUSG00000024642 | **Tle4** | 5730411M05Rik\|AA792082\|Bce-1\|Bce1\|ESTM13\|ESTM14\|Grg4\|X83332\|X83333 | transducin-like enhancer of split 4, homolog of Drosophila E(spl) | protein-coding |
| chr2_70816969_70817279 | chr2 | 70816970 | 70817279 | + | 0.931868 | intron (NM_172664, intron 1 of 19) | intron (NM_172664, intron 1 of 19) | 8356 | NM_172664 | 228012 | Mm.136511 | NM_172664 | ENSMUSG00000041997 | **Tlk1** | 4930545J15Rik | tousled-like kinase 1 | protein-coding |
| chr2_70844455_70845683 | chr2 | 70844456 | 70845683 | + | 0.778215 | Intergenic | Intergenic | -19589 | NM_172664 | 228012 | Mm.136511 | NM_172664 | ENSMUSG00000041997 | **Tlk1** | 4930545J15Rik | tousled-like kinase 1 | protein-coding |
| chr2_70844779_70845669 | chr2 | 70844780 | 70845669 | + | 0.751064 | Intergenic | Intergenic | -19744 | NM_172664 | 228012 | Mm.136511 | NM_172664 | ENSMUSG00000041997 | **Tlk1** | 4930545J15Rik | tousled-like kinase 1 | protein-coding |
| chr2_70919806_70920380 | chr2 | 70919807 | 70920380 | + | 0.696751 | Intergenic | ORR1E\|LTR\|ERVL-MaLR | -94613 | NM_172664 | 228012 | Mm.136511 | NM_172664 | ENSMUSG00000041997 | **Tlk1** | 4930545J15Rik | tousled-like kinase 1 | protein-coding |
| chr2_70816685_70817285 | chr2 | 70816686 | 70817285 | + | 0.665163 | intron (NM_172664, intron 1 of 19) | intron (NM_172664, intron 1 of 19) | 8495 | NM_172664 | 228012 | Mm.136511 | NM_172664 | ENSMUSG00000041997 | **Tlk1** | 4930545J15Rik | tousled-like kinase 1 | protein-coding |
| chr5_64886079_64886446 | chr5 | 64886080 | 64886446 | + | 0.841591 | Intergenic | Intergenic | 46436 | NM_030682 | 21897 | Mm.273024 | NM_030682 | ENSMUSG00000044827 | **Tlr1** | - | toll-like receptor 1 | protein-coding |
| chrX_106135476_106136475 | chrX | 106135477 | 106136475 | + | 0.867428 | Intergenic | Intergenic | -7299 | NM_205820 | 279572 | Mm.336203 | NM_205820 | ENSMUSG00000033777 | **Tlr13** | AI666735\|Gm713 | toll-like receptor 13 | protein-coding |
| chrX_106164024_106164364 | chrX | 106164025 | 106164364 | + | 0.841531 | Intergenic | Intergenic | 20919 | NM_205820 | 279572 | Mm.336203 | NM_205820 | ENSMUSG00000033777 | **Tlr13** | AI666735\|Gm713 | toll-like receptor 13 | protein-coding |
| chrX_106155938_106156352 | chrX | 106155939 | 106156352 | + | 0.820583 | intron (NM_205820, intron 2 of 2) | intron (NM_205820, intron 2 of 2) | 12870 | NM_205820 | 279572 | Mm.336203 | NM_205820 | ENSMUSG00000033777 | **Tlr13** | AI666735\|Gm713 | toll-like receptor 13 | protein-coding |
| chrX_106135956_106136374 | chrX | 106135957 | 106136374 | + | 0.805925 | Intergenic | Intergenic | -7110 | NM_205820 | 279572 | Mm.336203 | NM_205820 | ENSMUSG00000033777 | **Tlr13** | AI666735\|Gm713 | toll-like receptor 13 | protein-coding |
| chr1_182989875_182990749 | chr1 | 182989876 | 182990749 | + | 0.853488 | Intergenic | Intergenic | 35524 | NM_016928 | 53791 | Mm.116894 | NM_016928 | ENSMUSG00000079164 | **Tlr5** | - | toll-like receptor 5 | protein-coding |
| chr7_65714309_65714792 | chr7 | 65714310 | 65714792 | + | 1.03507 | Intergenic | MER2B\|DNA\|TcMar-Tigger | 21134 | NM_026795 | 68634 | Mm.476828 | NM_026795 | ENSMUSG00000078681 | **Tm2d3** | 1110025I09Rik\|5930422O05Rik\|Blp2 | TM2 domain containing 3 | protein-coding |
| chr16_32401585_32401919 | chr16 | 32401586 | 32401919 | + | 1.23309 | intron (NM_001160402, intron 1 of 4) | intron (NM_001160402, intron 1 of 4) | 1246 | NM_001160402 | 277203 | Mm.484240 | NM_001160402 | ENSMUSG00000079625 | **Tm4sf19** | EG277203 | transmembrane 4 L six family member 19 | protein-coding |
| chr16_32398128_32398600 | chr16 | 32398129 | 32398600 | + | 1.22677 | Intergenic | Intergenic | -2142 | NM_001160402 | 277203 | Mm.484240 | NM_001160402 | ENSMUSG00000079625 | **Tm4sf19** | EG277203 | transmembrane 4 L six family member 19 | protein-coding |
| chr16_32395522_32396173 | chr16 | 32395523 | 32396173 | + | 0.971279 | Intergenic | Intergenic | -4658 | NM_001160402 | 277203 | Mm.484240 | NM_001160402 | ENSMUSG00000079625 | **Tm4sf19** | EG277203 | transmembrane 4 L six family member 19 | protein-coding |
| chr16_32401377_32401924 | chr16 | 32401378 | 32401924 | + | 0.962532 | intron (NM_001160402, intron 1 of 4) | intron (NM_001160402, intron 1 of 4) | 1145 | NM_001160402 | 277203 | Mm.484240 | NM_001160402 | ENSMUSG00000079625 | **Tm4sf19** | EG277203 | transmembrane 4 L six family member 19 | protein-coding |
| chr16_32395284_32396244 | chr16 | 32395285 | 32396244 | + | 0.842587 | Intergenic | Intergenic | -4742 | NM_001160402 | 277203 | Mm.484240 | NM_001160402 | ENSMUSG00000079625 | **Tm4sf19** | EG277203 | transmembrane 4 L six family member 19 | protein-coding |
| chr16_32387437_32387854 | chr16 | 32387438 | 32387854 | + | 0.689282 | 3' UTR (NM_177633, exon 11 of 11) | 3' UTR (NM_177633, exon 11 of 11) | -12860 | NM_001160402 | 277203 | Mm.484240 | NM_001160402 | ENSMUSG00000079625 | **Tm4sf19** | EG277203 | transmembrane 4 L six family member 19 | protein-coding |

| | | | | | | | | | | | | | | | | | |
|---|---|---|---|---|---|---|---|---|---|---|---|---|---|---|---|---|---|
| chr16_32387564_32387819 | chr16 | 32387565 | 32387819 | + | 0.62389 | 3' UTR (NM_177633, exon 11 of 11) | 3' UTR (NM_177633, exon 11 of 11) | -12814 | NM_001160402 | 277203 | Mm.484240 | NM_001160402 | ENSMUSG00000079625 | **Tm4sf19** | EG277203 | transmembrane 4 L six family member 19 | protein-coding |
| chr1_82795288_82796664 | chr1 | 82795289 | 82796664 | + | 0.720406 | Intergenic | Intergenic | -27520 | NM_025453 | 66261 | Mm.46325 | NM_025453 | ENSMUSG00000026149 | **Tm4sf20** | 1810018L02Rik | transmembrane 4 L six family member 20 | protein-coding |
| chr6_146612756_146613129 | chr6 | 146612757 | 146613129 | + | 1.15759 | intron (NM_026281, intron 7 of 11) | intron (NM_026281, intron 7 of 11) | 21649 | NM_026281 | 67623 | Mm.278037 | NM_026281 | ENSMUSG00000040234 | **Tm7sf3** | 2010003B14Rik\|NARP1 | transmembrane 7 superfamily member 3 | protein-coding |
| chr19_41323130_41323687 | chr19 | 41323131 | 41323687 | + | 1.47966 | intron (NM_031376, intron 7 of 16) | intron (NM_031376, intron 7 of 16) | -59405 | NM_133352 | 107358 | Mm.246440 | NM_133352 | ENSMUSG00000025016 | **Tm9sf3** | 1810073M23Rik\|2810031D16Rik\|AI115521\|AI413748\|AW146116\|AW549777\|Smbp\|mKIAA4036 | transmembrane 9 superfamily member 3 | protein-coding |
| chr19_41321159_41321486 | chr19 | 41321160 | 41321486 | + | 1.05716 | intron (NM_031376, intron 8 of 16) | intron (NM_031376, intron 8 of 16) | -57319 | NM_133352 | 107358 | Mm.246440 | NM_133352 | ENSMUSG00000025016 | **Tm9sf3** | 1810073M23Rik\|2810031D16Rik\|AI115521\|AI413748\|AW146116\|AW549777\|Smbp\|mKIAA4036 | transmembrane 9 superfamily member 3 | protein-coding |
| chr2_153162885_153163386 | chr2 | 153162886 | 153163386 | + | 1.43177 | intron (NM_133847, intron 1 of 17) | intron (NM_133847, intron 1 of 17) | 1835 | NM_133847 | 99237 | Mm.330674 | NM_133847 | ENSMUSG00000068040 | **Tm9sf4** | AA986553\|AU045326\|B930079E06\|mKIAA0255 | transmembrane 9 superfamily protein member 4 | protein-coding |
| chr2_153162309_153162577 | chr2 | 153162310 | 153162577 | + | 0.785516 | intron (NM_133847, intron 1 of 17) | intron (NM_133847, intron 1 of 17) | 1142 | NM_133847 | 99237 | Mm.330674 | NM_133847 | ENSMUSG00000068040 | **Tm9sf4** | AA986553\|AU045326\|B930079E06\|mKIAA0255 | transmembrane 9 superfamily protein member 4 | protein-coding |
| chr2_153175848_153176260 | chr2 | 153175849 | 153176260 | + | 0.703897 | intron (NM_133847, intron 1 of 17) | intron (NM_133847, intron 1 of 17) | 14753 | NM_133847 | 99237 | Mm.330674 | NM_133847 | ENSMUSG00000068040 | **Tm9sf4** | AA986553\|AU045326\|B930079E06\|mKIAA0255 | transmembrane 9 superfamily protein member 4 | protein-coding |
| chr2_153164489_153164723 | chr2 | 153164490 | 153164723 | + | 0.646649 | intron (NM_133847, intron 1 of 17) | intron (NM_133847, intron 1 of 17) | 3305 | NM_133847 | 99237 | Mm.330674 | NM_133847 | ENSMUSG00000068040 | **Tm9sf4** | AA986553\|AU045326\|B930079E06\|mKIAA0255 | transmembrane 9 superfamily protein member 4 | protein-coding |
| chr1_74295180_74296048 | chr1 | 74295181 | 74296048 | + | 1.17201 | intron (NM_027154, intron 1 of 11) | intron (NM_027154, intron 1 of 11) | 8722 | NM_027154 | 69660 | Mm.261182 | NM_027154 | ENSMUSG00000006301 | **Tmbim1** | 2310061B02Rik\|AA960455\|AU024746\|C78899\|RECS1\|Tmbib1\|mKIAA4161 | transmembrane BAX inhibitor motif containing 1 | protein-coding |
| chr15_99400651_99401013 | chr15 | 99400652 | 99401013 | + | 1.09183 | intron (NM_001171034, intron 2 of 10) | B3\|SINE\|B2 | 7613 | NM_001171035 | 110213 | Mm.16769 | NM_026669 | ENSMUSG00000023010 | **Tmbim6** | 5031406P05Rik\|Bi1\|Tegt | transmembrane BAX inhibitor motif containing 6 | protein-coding |
| chr15_99394636_99394868 | chr15 | 99394637 | 99394868 | + | 0.926065 | intron (NM_001171036, intron 2 of 10) | intron (NM_001171036, intron 2 of 10) | 1533 | NM_001171035 | 110213 | Mm.16769 | NM_026669 | ENSMUSG00000023010 | **Tmbim6** | 5031406P05Rik\|Bi1\|Tegt | transmembrane BAX inhibitor motif containing 6 | protein-coding |
| chr7_83438507_83438907 | chr7 | 83438508 | 83438907 | + | 0.738218 | Intergenic | Intergenic | -146224 | NM_177695 | 233424 | Mm.486619 | NM_177695 | | **Tmc3** | 9030203A06 | transmembrane channel-like gene family 3 | protein-coding |
| chr6_116127927_116128418 | chr6 | 116127928 | 116128418 | + | 1.34194 | intron (NM_177412, intron 3 of 5) | intron (NM_177412, intron 3 of 5) | 65201 | NM_177412 | 330401 | Mm.425352 | NM_177412 | ENSMUSG00000030126 | **Tmcc1** | 3632431M01Rik\|mKIAA0779 | transmembrane and coiled coil domains 1 | protein-coding |
| chr1_132383796_132384414 | chr1 | 132383797 | 132384414 | + | 0.823089 | intron (NM_178874, intron 1 of 4) | intron (NM_178874, intron 1 of 4) | 7176 | NM_178874 | 68875 | Mm.273785 | NM_178874 | ENSMUSG00000042066 | **Tmcc2** | 1110063G11Rik | transmembrane and coiled-coil domains 2 | protein-coding |
| chr10_94554134_94554474 | chr10 | 94554135 | 94554474 | + | 0.91878 | intron (NM_172051, intron 1 of 3) | intron (NM_172051, intron 1 of 3) | -20953 | NM_001168684 | 319880 | Mm.23047 | NM_172051 | ENSMUSG00000020023 | **Tmcc3** | AW488095\|C630016B22Rik\|C88213\|Tmcc1 | transmembrane and coiled coil domains 3 | protein-coding |
| chr10_94525859_94526143 | chr10 | 94525860 | 94526143 | + | 0.766206 | intron (NM_172051, intron 1 of 3) | Tigger5\|DNA\|TcMar-Tigger | 11144 | NM_172051 | 319880 | Mm.23047 | NM_172051 | ENSMUSG00000020023 | **Tmcc3** | AW488095\|C630016B22Rik\|C88213\|Tmcc1 | transmembrane and coiled coil domains 3 | protein-coding |
| chr12_85362444_85362902 | chr12 | 85362445 | 85362902 | + | 0.756348 | intron (NM_026775, intron 1 of 4) | L1MC3\|LINE\|L1 | 12044 | NM_026775 | 68581 | Mm.379159 | NM_026775 | ENSMUSG00000021248 | **Tmed10** | 1110014C03Rik\|Tmp21\|p23\|p24delta1 | transmembrane emp24-like trafficking protein 10 (yeast) | protein-coding |
| chr6_13075728_13076171 | chr6 | 13075729 | 13076171 | + | 0.710977 | intron (NM_027992, intron 4 of 8) | ORR1B2\|LTR\|ERVL-MaLR | 6191 | NM_027992 | 71900 | Mm.27742 | NM_027992 | ENSMUSG00000029571 | **Tmem106b** | 2310036D22Rik\|5830455K21Rik\|6430519M21Rik\|AI428776\|AI661344 | transmembrane protein 106B | protein-coding |
| chr6_13061186_13062618 | chr6 | 13061187 | 13062618 | + | 0.683569 | Intergenic | Intergenic | -7857 | NM_027992 | 71900 | Mm.27742 | NM_027992 | ENSMUSG00000029571 | **Tmem106b** | 2310036D22Rik\|5830455K21Rik\|6430519M21Rik\|AI428776\|AI661344 | transmembrane protein 106B | protein-coding |
| chr14_30829035_30829393 | chr14 | 30829036 | 30829393 | + | 1.08466 | intron (NM_028839, intron 1 of 7) | intron (NM_028839, intron 1 of 7) | 3620 | NM_028839 | 69179 | Mm.143025 | NM_028839 | ENSMUSG00000006526 | **Tmem110** | 1810038N08Rik\|2310014H19Rik\|AW554125 | transmembrane protein 110 | protein-coding |
| chr14_30828726_30828938 | chr14 | 30828727 | 30828938 | + | 0.836767 | intron (NM_028839, intron 1 of 7) | intron (NM_028839, intron 1 of 7) | 3238 | NM_028839 | 69179 | Mm.143025 | NM_028839 | ENSMUSG00000006526 | **Tmem110** | 1810038N08Rik\|2310014H19Rik\|AW554125 | transmembrane protein 110 | protein-coding |
| chr9_107519296_107519835 | chr9 | 107519297 | 107519835 | + | 0.728304 | intron (NM_020263, intron 23 of 38) | intron (NM_020263, intron 23 of 38) | -14379 | NM_019704 | 56395 | Mm.260153 | NM_019704 | ENSMUSG00000010045 | **Tmem115** | C78915\|Pl6\|Pp6 | transmembrane protein 115 | protein-coding |
| chr5_121469428_121469803 | chr5 | 121469429 | 121469803 | + | 0.741423 | intron (NM_001161626, intron 4 of 9) | MIRc\|SINE\|MIR | 6435 | NM_001161627 | 77462 | Mm.358781 | NM_029912 | ENSMUSG00000029452 | **Tmem116** | 4930406A18Rik\|4930513P12Rik\|C030022K24Rik | transmembrane protein 116 | protein-coding |
| chr5_113814016_113815123 | chr5 | 113814017 | 113815123 | + | 0.757052 | Intergenic | L1M5\|LINE\|L1 | -14218 | NM_146162 | 231633 | Mm.41681 | NM_146162 | ENSMUSG00000054675 | **Tmem119** | AW208946\|BC025600\|obif | transmembrane protein 119 | protein-coding |
| chr12_113193240_113193813 | chr12 | 113193241 | 113193813 | + | 0.727062 | Intergenic | Intergenic | 7624 | NM_153776 | 69195 | Mm.313244 | NM_153776 | ENSMUSG00000049036 | **Tmem121** | 2410008J05Rik\|Hole | transmembrane protein 121 | protein-coding |
| chr12_113306049_113306327 | chr12 | 113306050 | 113306327 | + | 0.652868 | Intergenic | Intergenic | 120285 | NM_153776 | 69195 | Mm.313244 | NM_153776 | ENSMUSG00000049036 | **Tmem121** | 2410008J05Rik\|Hole | transmembrane protein 121 | protein-coding |
| chr7_90464639_90465648 | chr7 | 90464640 | 90465648 | + | 1.69282 | Intergenic | Intergenic | -7936 | NM_025460 | 66271 | Mm.28963 | NM_025460 | ENSMUSG00000030615 | **Tmem126a** | 1810020E01Rik | transmembrane protein 126A | protein-coding |
| chr7_90539461_90539785 | chr7 | 90539462 | 90539785 | + | 0.750204 | Intergenic | Intergenic | -63628 | NM_026734 | 68472 | Mm.289791 | NM_026734 | | **Tmem126b** | 1110001A23Rik | transmembrane protein 126B | protein-coding |
| chr7_90541170_90541504 | chr7 | 90541171 | 90541504 | + | 0.73977 | Intergenic | ORR1A4\|LTR\|ERVL-MaLR | -65342 | NM_026734 | 68472 | Mm.289791 | NM_026734 | | **Tmem126b** | 1110001A23Rik | transmembrane protein 126B | protein-coding |
| chr1_36965821_36966216 | chr1 | 36965822 | 36966216 | + | 0.965886 | Intergenic | Intergenic | -26492 | NM_018872 | 56030 | Mm.186943 | NM_018872 | ENSMUSG00000026116 | **Tmem131** | 2610524E03Rik\|CC28\|D1Bwg0491e\|Neg\|RW1\|YR-23\|mKIAA0257 | transmembrane protein 131 | protein-coding |
| chr1_37002605_37002981 | chr1 | 37002606 | 37002981 | + | 0.922832 | Intergenic | Intergenic | -63266 | NM_018872 | 56030 | Mm.186943 | NM_018872 | ENSMUSG00000026116 | **Tmem131** | 2610524E03Rik\|CC28\|D1Bwg0491e\|Neg\|RW1\|YR-23\|mKIAA0257 | transmembrane protein 131 | protein-coding |
| chr1_36886420_36887010 | chr1 | 36886421 | 36887010 | + | 0.858828 | intron (NM_018872, intron 2 of 40) | intron (NM_018872, intron 2 of 40) | 52812 | NM_018872 | 56030 | Mm.186943 | NM_018872 | ENSMUSG00000026116 | **Tmem131** | 2610524E03Rik\|CC28\|D1Bwg0491e\|Neg\|RW1\|YR-23\|mKIAA0257 | transmembrane protein 131 | protein-coding |
| chr1_36892379_36892960 | chr1 | 36892380 | 36892960 | + | 0.822445 | intron (NM_018872, intron 1 of 40) | MTEa\|LTR\|ERVL-MaLR | 46857 | NM_018872 | 56030 | Mm.186943 | NM_018872 | ENSMUSG00000026116 | **Tmem131** | 2610524E03Rik\|CC28\|D1Bwg0491e\|Neg\|RW1\|YR-23\|mKIAA0257 | transmembrane protein 131 | protein-coding |
| chr1_36879179_36879457 | chr1 | 36879180 | 36879457 | + | 0.780678 | intron (NM_018872, intron 3 of 40) | intron (NM_018872, intron 3 of 40) | 60209 | NM_018872 | 56030 | Mm.186943 | NM_018872 | ENSMUSG00000026116 | **Tmem131** | 2610524E03Rik\|CC28\|D1Bwg0491e\|Neg\|RW1\|YR-23\|mKIAA0257 | transmembrane protein 131 | protein-coding |
| chr1_36962301_36963071 | chr1 | 36962302 | 36963071 | + | 0.73107 | Intergenic | PB1\|SINE\|Alu | -23159 | NM_018872 | 56030 | Mm.186943 | NM_018872 | ENSMUSG00000026116 | **Tmem131** | 2610524E03Rik\|CC28\|D1Bwg0491e\|Neg\|RW1\|YR-23\|mKIAA0257 | transmembrane protein 131 | protein-coding |
| chr1_36923118_36923724 | chr1 | 36923119 | 36923724 | + | 0.693802 | intron (NM_018872, intron 1 of 40) | intron (NM_018872, intron 1 of 40) | 16106 | NM_018872 | 56030 | Mm.186943 | NM_018872 | ENSMUSG00000026116 | **Tmem131** | 2610524E03Rik\|CC28\|D1Bwg0491e\|Neg\|RW1\|YR-23\|mKIAA0257 | transmembrane protein 131 | protein-coding |
| chr7_89268246_89268884 | chr7 | 89268247 | 89268884 | + | 1.47619 | intron (NM_028343, intron 4 of 14) | intron (NM_028343, intron 4 of 14) | 70222 | NM_028343 | 72759 | Mm.208477 | NM_028343 | ENSMUSG00000039428 | **Tmem135** | 2810439K08Rik\|AW319712\|PMP52 | transmembrane protein 135 | protein-coding |
| chr7_89235281_89235675 | chr7 | 89235282 | 89235675 | + | 1.05823 | intron (NM_028343, intron 5 of 14) | intron (NM_028343, intron 5 of 14) | 103309 | NM_028343 | 72759 | Mm.208477 | NM_028343 | ENSMUSG00000039428 | **Tmem135** | 2810439K08Rik\|AW319712\|PMP52 | transmembrane protein 135 | protein-coding |
| chr7_89146594_89147355 | chr7 | 89146595 | 89147355 | + | 1.00811 | intron (NM_028343, intron 12 of 14) | intron (NM_028343, intron 12 of 14) | 191812 | NM_028343 | 72759 | Mm.208477 | NM_028343 | ENSMUSG00000039428 | **Tmem135** | 2810439K08Rik\|AW319712\|PMP52 | transmembrane protein 135 | protein-coding |
| chr7_89234813_89235067 | chr7 | 89234814 | 89235067 | + | 0.781143 | intron (NM_028343, intron 5 of 14) | intron (NM_028343, intron 5 of 14) | 103847 | NM_028343 | 72759 | Mm.208477 | NM_028343 | ENSMUSG00000039428 | **Tmem135** | 2810439K08Rik\|AW319712\|PMP52 | transmembrane protein 135 | protein-coding |
| chr7_89312845_89313274 | chr7 | 89312846 | 89313274 | + | 0.650632 | intron (NM_028343, intron 1 of 14) | intron (NM_028343, intron 1 of 14) | 25727 | NM_028343 | 72759 | Mm.208477 | NM_028343 | ENSMUSG00000039428 | **Tmem135** | 2810439K08Rik\|AW319712\|PMP52 | transmembrane protein 135 | protein-coding |
| chr7_89335313_89335674 | chr7 | 89335314 | 89335674 | + | 0.630507 | intron (NM_028343, intron 1 of 14) | intron (NM_028343, intron 1 of 14) | 3293 | NM_028343 | 72759 | Mm.208477 | NM_028343 | ENSMUSG00000039428 | **Tmem135** | 2810439K08Rik\|AW319712\|PMP52 | transmembrane protein 135 | protein-coding |
| chr7_45901605_45901880 | chr7 | 45901606 | 45901880 | + | 0.644114 | intron (NM_144801, intron 3 of 7) | intron (NM_144801, intron 3 of 7) | 4674 | NM_144801 | 70209 | Mm.163766 | NM_144801 | ENSMUSG00000002781 | **Tmem143** | 2310076O21Rik\|AI481604 | transmembrane protein 143 | protein-coding |
| chr3_79853921_79854268 | chr3 | 79853922 | 79854268 | + | 1.02422 | Intergenic | Intergenic | -11433 | NM_027495 | 70652 | Mm.46635 | NM_027495 | ENSMUSG00000027956 | **Tmem144** | 1110057I03Rik\|5730537D05Rik | transmembrane protein 144 | protein-coding |
| chr3_84660380_84661287 | chr3 | 84660381 | 84661287 | + | 0.881495 | Intergenic | Intergenic | -5358 | NM_177260 | 320782 | Mm.107370 | NM_177260 | ENSMUSG00000056498 | **Tmem154** | 9930117H01Rik | transmembrane protein 154 | protein-coding |
| chr3_84722666_84723059 | chr3 | 84722667 | 84723059 | + | 0.629902 | Intergenic | Intergenic | 56671 | NM_177260 | 320782 | Mm.107370 | NM_177260 | ENSMUSG00000056498 | **Tmem154** | 9930117H01Rik | transmembrane protein 154 | protein-coding |
| chr3_84685957_84686278 | chr3 | 84685958 | 84686278 | + | 0.620478 | intron (NM_177260, intron 3 of 6) | intron (NM_177260, intron 3 of 6) | 19926 | NM_177260 | 320782 | Mm.107370 | NM_177260 | ENSMUSG00000056498 | **Tmem154** | 9930117H01Rik | transmembrane protein 154 | protein-coding |
| chr9_123240441_123240753 | chr9 | 123240442 | 123240753 | + | 0.833444 | Intergenic | B3A\|SINE\|B2 | 20192 | NM_001002267 | 72309 | Mm.8569 | NM_001002267 | ENSMUSG00000054871 | **Tmem158** | 2310037P21Rik\|Ris1 | transmembrane protein 158 | protein-coding |
| chr8_70174061_70174460 | chr8 | 70174062 | 70174460 | + | 0.873426 | intron (NM_001293797, intron 1 of 10) | intron (NM_001293797, intron 1 of 10) | 1853 | NR_122028 | 234371 | Mm.23488 | NM_145597 | ENSMUSG00000002342 | **Tmem161a** | AI428876\|BB161850\|BC021367 | transmembrane protein 161A | protein-coding |
| chr13_84293961_84294395 | chr13 | 84293962 | 84294395 | + | 0.956073 | intron (NM_175187, intron 11 of 11) | intron (NM_175187, intron 11 of 11) | 71882 | NM_175187 | 72745 | Mm.295063 | NM_175187 | ENSMUSG00000035762 | **Tmem161b** | 2810446P07Rik\|AI843389 | transmembrane protein 161B | protein-coding |
| chr1_127637894_127638338 | chr1 | 127637895 | 127638338 | + | 1.27017 | intron (NM_028135, intron 2 of 7) | intron (NM_028135, intron 2 of 7) | 39905 | NM_028135 | 72160 | Mm.281287 | NM_028135 | ENSMUSG00000026347 | **Tmem163** | 2610024A01Rik\|SV31 | transmembrane protein 163 | protein-coding |

| | | | | | | | | | | | | | | | | |
|---|---|---|---|---|---|---|---|---|---|---|---|---|---|---|---|---|
| chr1_127627381_127627775 | chr1 | 127627382 | 127627775 | + | 1.09243 | intron (NM_028135, intron 2 of 7) | intron (NM_028135, intron 2 of 7) | 50443 | NM_028135 | 72160 | Mm.281287 | NM_028135 | ENSMUSG00000026347 | **Tmem163** | 2610024A01Rik\|SV31 | transmembrane protein 163 | protein-coding |
| chr1_127646212_127646664 | chr1 | 127646213 | 127646664 | + | 0.617035 | intron (NM_028135, intron 2 of 7) | intron (NM_028135, intron 2 of 7) | 31583 | NM_028135 | 72160 | Mm.281287 | NM_028135 | ENSMUSG00000026347 | **Tmem163** | 2610024A01Rik\|SV31 | transmembrane protein 163 | protein-coding |
| chrX_142737722_142738173 | chrX | 142737723 | 142738173 | + | 0.82673 | intron (NM_001199360, intron 2 of 6) | intron (NM_001199360, intron 2 of 6) | 56222 | NM_001199360 | 209497 | Mm.249648 | NM_177592 | ENSMUSG00000047045 | **Tmem164** | AI316850\|AI852450\|AW547186\|F730011B02 | transmembrane protein 164 | protein-coding |
| chrX_142694865_142695374 | chrX | 142694866 | 142695374 | + | 0.767302 | intron (NM_001199360, intron 2 of 6) | intron (NM_001199360, intron 2 of 6) | 13394 | NM_001199360 | 209497 | Mm.249648 | NM_177592 | ENSMUSG00000047045 | **Tmem164** | AI316850\|AI852450\|AW547186\|F730011B02 | transmembrane protein 164 | protein-coding |
| chrX_142735733_142736048 | chrX | 142735734 | 142736048 | + | 0.758364 | intron (NM_001199360, intron 2 of 6) | intron (NM_001199360, intron 2 of 6) | 54165 | NM_001199360 | 209497 | Mm.249648 | NM_177592 | ENSMUSG00000047045 | **Tmem164** | AI316850\|AI852450\|AW547186\|F730011B02 | transmembrane protein 164 | protein-coding |
| chrX_142694842_142695589 | chrX | 142694843 | 142695589 | + | 0.688935 | intron (NM_001199360, intron 2 of 6) | intron (NM_001199360, intron 2 of 6) | 13490 | NM_001199360 | 209497 | Mm.249648 | NM_177592 | ENSMUSG00000047045 | **Tmem164** | AI316850\|AI852450\|AW547186\|F730011B02 | transmembrane protein 164 | protein-coding |
| chr5_76243699_76244335 | chr5 | 76243700 | 76244335 | + | 1.3824 | intron (NM_007715, intron 11 of 22) | intron (NM_007715, intron 11 of 22) | 60137 | NM_011626 | 21982 | Mm.790 | NM_011626 | ENSMUSG00000029234 | **Tmem165** | AV026557\|Tpardl\|Tparl\|pFT27 | transmembrane protein 165 | protein-coding |
| chr6_13610906_13611279 | chr6 | 13610907 | 13611279 | + | 0.937529 | Intergenic | MLTR11A\|LTR\|ERVK | -3030 | NM_028990 | 101118 | Mm.254845 | NM_028990 | ENSMUSG00000029569 | **Tmem168** | 5730526F17Rik\|8430437G11Rik\|AI462344\|AI504145 | transmembrane protein 168 | protein-coding |
| chr11_22626281_22626801 | chr11 | 22626282 | 22626801 | + | 1.29505 | Intergenic | Intergenic | 114258 | NM_153596 | 103765 | Mm.221274 | NM_153596 | ENSMUSG00000049904 | **Tmem17** | AI503894\|AW049890 | transmembrane protein 17 | protein-coding |
| chr11_22669179_22669650 | chr11 | 22669180 | 22669650 | + | 1.14641 | Intergenic | Intergenic | 157132 | NM_153596 | 103765 | Mm.221274 | NM_153596 | ENSMUSG00000049904 | **Tmem17** | AI503894\|AW049890 | transmembrane protein 17 | protein-coding |
| chr11_22643981_22644549 | chr11 | 22643982 | 22644549 | + | 1.05877 | Intergenic | Intergenic | 131982 | NM_153596 | 103765 | Mm.221274 | NM_153596 | ENSMUSG00000049904 | **Tmem17** | AI503894\|AW049890 | transmembrane protein 17 | protein-coding |
| chr11_22675754_22676287 | chr11 | 22675755 | 22676287 | + | 0.845054 | Intergenic | MLT1F1\|LTR\|ERVL-MaLR | 163738 | NM_153596 | 103765 | Mm.221274 | NM_153596 | ENSMUSG00000049904 | **Tmem17** | AI503894\|AW049890 | transmembrane protein 17 | protein-coding |
| chr11_22632237_22632753 | chr11 | 22632238 | 22632753 | + | 0.833455 | Intergenic | Intergenic | 120212 | NM_153596 | 103765 | Mm.221274 | NM_153596 | ENSMUSG00000049904 | **Tmem17** | AI503894\|AW049890 | transmembrane protein 17 | protein-coding |
| chr18_35745268_35745612 | chr18 | 35745269 | 35745612 | + | 0.77106 | Intergenic | RLTR11D\|LTR\|ERVK | -4886 | NM_001289591 | 72512 | Mm.45995 | NM_028261 | ENSMUSG00000024349 | **Tmem173** | 2610307O08Rik\|ERIS\|MPYS\|Mita\|STING | transmembrane protein 173 | protein-coding |
| chr18_35744221_35744726 | chr18 | 35744222 | 35744726 | + | 0.723803 | Intergenic | Intergenic | -3920 | NM_001289591 | 72512 | Mm.45995 | NM_028261 | ENSMUSG00000024349 | **Tmem173** | 2610307O08Rik\|ERIS\|MPYS\|Mita\|STING | transmembrane protein 173 | protein-coding |
| chr12_30598944_30599833 | chr12 | 30598945 | 30599833 | + | 1.08727 | Intergenic | Intergenic | 14946 | NM_172049 | 211986 | Mm.34035 | NM_172049 | ENSMUSG00000043061 | **Tmem18** | BC024093 | transmembrane protein 18 | protein-coding |
| chr12_30557419_30557838 | chr12 | 30557420 | 30557838 | + | 1.07557 | Intergenic | L1_Mur2\|LINE\|L1 | -26814 | NM_172049 | 211986 | Mm.34035 | NM_172049 | ENSMUSG00000043061 | **Tmem18** | BC024093 | transmembrane protein 18 | protein-coding |
| chr12_30598918_30600273 | chr12 | 30598919 | 30600273 | + | 0.907368 | Intergenic | Intergenic | 15153 | NM_172049 | 211986 | Mm.34035 | NM_172049 | ENSMUSG00000043061 | **Tmem18** | BC024093 | transmembrane protein 18 | protein-coding |
| chr1_40847550_40847959 | chr1 | 40847551 | 40847959 | + | 1.88183 | intron (NM_001081198, intron 4 of 4) | intron (NM_001081198, intron 4 of 4) | 42154 | NM_001081198 | 381339 | Mm.334678 | NM_001081198 | | **Tmem182** | 2310079P10Rik\|AI414494 | transmembrane protein 182 | protein-coding |
| chr1_40845273_40845670 | chr1 | 40845274 | 40845670 | + | 0.92217 | intron (NM_001081198, intron 4 of 4) | intron (NM_001081198, intron 4 of 4) | 39871 | NM_001081198 | 381339 | Mm.334678 | NM_001081198 | | **Tmem182** | 2310079P10Rik\|AI414494 | transmembrane protein 182 | protein-coding |
| chr8_64934805_64935629 | chr8 | 64934806 | 64935629 | + | 1.37268 | Intergenic | Intergenic | -11968 | NM_001163747 | 73067 | Mm.294445 | NM_028427 | ENSMUSG00000025521 | **Tmem192** | 3110005G23Rik | transmembrane protein 192 | protein-coding |
| chr8_64934553_64935711 | chr8 | 64934554 | 64935711 | + | 1.32284 | Intergenic | Intergenic | -12053 | NM_001163747 | 73067 | Mm.294445 | NM_028427 | ENSMUSG00000025521 | **Tmem192** | 3110005G23Rik | transmembrane protein 192 | protein-coding |
| chr1_52571292_52571907 | chr1 | 52571293 | 52571907 | + | 0.737032 | Intergenic | MTE2b\|LTR\|ERVL-MaLR | -59105 | NR_024590 | 227094 | Mm.83541 | NM_172654 | ENSMUSG00000043015 | **Tmem194b** | 5330401P04Rik\|9830163O08 | transmembrane protein 194B | protein-coding |
| chr1_52571475_52571855 | chr1 | 52571476 | 52571855 | + | 0.701371 | Intergenic | MTE2b\|LTR\|ERVL-MaLR | -59040 | NR_024590 | 227094 | Mm.83541 | NM_172654 | ENSMUSG00000043015 | **Tmem194b** | 5330401P04Rik\|9830163O08 | transmembrane protein 194B | protein-coding |
| chr19_21913766_21914158 | chr19 | 21913767 | 21914158 | + | 1.24035 | Intergenic | Intergenic | 135622 | NM_031997 | 83921 | Mm.329776 | NM_031997 | ENSMUSG00000024754 | **Tmem2** | - | transmembrane protein 2 | protein-coding |
| chr19_21863538_21863954 | chr19 | 21863539 | 21863954 | + | 1.08798 | Intergenic | Intergenic | 85406 | NM_031997 | 83921 | Mm.329776 | NM_031997 | ENSMUSG00000024754 | **Tmem2** | - | transmembrane protein 2 | protein-coding |
| chr19_21866274_21866558 | chr19 | 21866275 | 21866558 | + | 1.02717 | Intergenic | Intergenic | 88076 | NM_031997 | 83921 | Mm.329776 | NM_031997 | ENSMUSG00000024754 | **Tmem2** | - | transmembrane protein 2 | protein-coding |
| chr19_21947844_21948558 | chr19 | 21947845 | 21948558 | + | 0.990369 | Intergenic | Intergenic | 169861 | NM_031997 | 83921 | Mm.329776 | NM_031997 | ENSMUSG00000024754 | **Tmem2** | - | transmembrane protein 2 | protein-coding |
| chr19_21946521_21946926 | chr19 | 21946522 | 21946926 | + | 0.629734 | Intergenic | Intergenic | 168384 | NM_031997 | 83921 | Mm.329776 | NM_031997 | ENSMUSG00000024754 | **Tmem2** | - | transmembrane protein 2 | protein-coding |
| chr4_131923703_131924106 | chr4 | 131923704 | 131924106 | + | 0.700866 | TTS (NM_001201367) | TTS (NM_001201367) | 2134 | NM_001201367 | 623230 | Mm.421179 | NM_001201367 | ENSMUSG00000070720 | **Tmem200b** | EG623230 | transmembrane protein 200B | protein-coding |
| chr9_21923920_21924399 | chr9 | 21923921 | 21924399 | + | 0.832097 | intron (NR_045606, intron 3 of 4) | intron (NR_045606, intron 3 of 4) | 3128 | NM_001253869 | 235043 | Mm.253329 | NM_178577 | ENSMUSG00000040883 | **Tmem205** | - | transmembrane protein 205 | protein-coding |
| chr6_30507449_30507987 | chr6 | 30507450 | 30507987 | + | 2.09726 | intron (NM_178625, intron 3 of 14) | B3\|SINE\|B2 | 1988 | NM_178625 | 72649 | Mm.45418 | NM_178625 | ENSMUSG00000029782 | **Tmem209** | 2700094F01Rik\|AI428435 | transmembrane protein 209 | protein-coding |
| chr2_132230421_132230704 | chr2 | 132230422 | 132230704 | + | 0.807653 | Intergenic | PB1D7\|SINE\|Alu | 17225 | NM_027478 | 70612 | Mm.30116 | NM_027478 | ENSMUSG00000027341 | **Tmem230** | 5730494N06Rik\|AA407821 | transmembrane protein 230 | protein-coding |
| chr17_65556876_65557716 | chr17 | 65556877 | 65557716 | + | 1.23605 | Intergenic | Intergenic | -16514 | NM_001289486 | 381107 | Mm.141275 | NM_001008973 | ENSMUSG00000045036 | **Tmem232** | E130009J12Rik\|Gm940\|Tes13-L\|Tes13-S | transmembrane protein 232 | protein-coding |
| chr2_14129753_14130022 | chr2 | 14129754 | 14130022 | + | 0.675688 | intron (NM_011484, intron 7 of 13) | MLT1A0\|LTR\|ERVL-MaLR | -44636 | NM_001081310 | 625286 | Mm.158781 | NM_001081310 | ENSMUSG00000061531 | **Tmem236** | 2010003H20Rik\|Fam23a\|Gm613 | transmembrane protein 236 | protein-coding |
| chr5_9059084_9059641 | chr5 | 9059085 | 9059641 | + | 1.30979 | Intergenic | Intergenic | -41374 | NM_001081029 | 652925 | Mm.439954 | NM_001081029 | ENSMUSG00000079659 | **Tmem243** | 4930420K17Rik\|E030031F02Rik | transmembrane protein 243, mitochondrial | protein-coding |
| chr5_9069804_9070176 | chr5 | 9069805 | 9070176 | + | 0.826865 | Intergenic | Intergenic | -30747 | NM_001081029 | 652925 | Mm.439954 | NM_001081029 | ENSMUSG00000079659 | **Tmem243** | 4930420K17Rik\|E030031F02Rik | transmembrane protein 243, mitochondrial | protein-coding |
| chr5_9085150_9086023 | chr5 | 9085151 | 9086023 | + | 0.759262 | Intergenic | Intergenic | -15150 | NM_001081029 | 652925 | Mm.439954 | NM_001081029 | ENSMUSG00000079659 | **Tmem243** | 4930420K17Rik\|E030031F02Rik | transmembrane protein 243, mitochondrial | protein-coding |
| chr5_9068772_9069286 | chr5 | 9068773 | 9069286 | + | 0.646393 | Intergenic | Intergenic | -31708 | NM_001081029 | 652925 | Mm.439954 | NM_001081029 | ENSMUSG00000079659 | **Tmem243** | 4930420K17Rik\|E030031F02Rik | transmembrane protein 243, mitochondrial | protein-coding |
| chr4_56944363_56944726 | chr4 | 56944364 | 56944726 | + | 1.49354 | intron (NM_175518, intron 1 of 17) | intron (NM_175518, intron 1 of 17) | 2884 | NM_175518 | 242474 | Mm.296632 | NM_175518 | | **Tmem245** | A630051L19Rik\|AI957324\|D730040F13Rik | transmembrane protein 245 | protein-coding |
| chr17_86850692_86851282 | chr17 | 86850693 | 86851282 | + | 1.53798 | Intergenic | Intergenic | -66361 | NM_001277980 | 78469 | Mm.67234 | NM_030104 | ENSMUSG00000037689 | **Tmem247** | 1700090G07Rik | transmembrane protein 247 | protein-coding |
| chr17_86852603_86853052 | chr17 | 86852604 | 86853052 | + | 1.00328 | Intergenic | MER21B\|LTR\|ERVL | -64520 | NM_001277980 | 78469 | Mm.67234 | NM_030104 | ENSMUSG00000037689 | **Tmem247** | 1700090G07Rik | transmembrane protein 247 | protein-coding |
| chr17_86872583_86873059 | chr17 | 86872584 | 86873059 | + | 0.643938 | Intergenic | L1M2\|LINE\|L1 | -44527 | NM_001277980 | 78469 | Mm.67234 | NM_030104 | ENSMUSG00000037689 | **Tmem247** | 1700090G07Rik | transmembrane protein 247 | protein-coding |
| chr17_86872528_86873345 | chr17 | 86872529 | 86873345 | + | 0.637656 | Intergenic | L1M2\|LINE\|L1 | -44411 | NM_001277980 | 78469 | Mm.67234 | NM_030104 | ENSMUSG00000037689 | **Tmem247** | 1700090G07Rik | transmembrane protein 247 | protein-coding |
| chr17_86872643_86872989 | chr17 | 86872644 | 86872989 | + | 0.62856 | Intergenic | L1M2\|LINE\|L1 | -44532 | NM_001277980 | 78469 | Mm.67234 | NM_030104 | ENSMUSG00000037689 | **Tmem247** | 1700090G07Rik | transmembrane protein 247 | protein-coding |
| chr19_24720462_24720904 | chr19 | 24720463 | 24720904 | + | 1.00554 | intron (NM_175013, intron 9 of 10) | intron (NM_175013, intron 9 of 10) | 46675 | NM_183160 | 226040 | Mm.391170 | NM_183160 | ENSMUSG00000048572 | **Tmem252** | E030010A14Rik | transmembrane protein 252 | protein-coding |
| chrX_38235901_38236490 | chrX | 38235902 | 38236490 | + | 1.17838 | intron (NM_172930, intron 2 of 9) | intron (NM_172930, intron 2 of 9) | 16285 | NM_001289727 | 245386 | Mm.72979 | NM_172930 | ENSMUSG00000036502 | **Tmem255a** | 4933417N17\|6430550H21Rik\|Fam70a | transmembrane protein 255A | protein-coding |
| chrX_38246914_38247520 | chrX | 38246915 | 38247520 | + | 0.967207 | intron (NM_172930, intron 1 of 9) | intron (NM_172930, intron 1 of 9) | 5264 | NM_001289727 | 245386 | Mm.72979 | NM_172930 | ENSMUSG00000036502 | **Tmem255a** | 4933417N17\|6430550H21Rik\|Fam70a | transmembrane protein 255A | protein-coding |
| chrX_38259567_38259889 | chrX | 38259568 | 38259889 | + | 0.71213 | Intergenic | Intergenic | -7247 | NM_001289728 | 245386 | Mm.72979 | NM_172930 | ENSMUSG00000036502 | **Tmem255a** | 4933417N17\|6430550H21Rik\|Fam70a | transmembrane protein 255A | protein-coding |
| chrX_134301271_134301937 | chrX | 134301272 | 134301937 | + | 1.49804 | intron (NM_026239, intron 1 of 1) | intron (NM_026239, intron 1 of 1) | 6379 | NM_026239 | 67564 | Mm.41489 | NM_026239 | ENSMUSG00000033578 | **Tmem35** | 9030603L14Rik\|AI841526\|PMP52 | transmembrane protein 35 | protein-coding |
| chrX_134300913_134301947 | chrX | 134300914 | 134301947 | + | 1.48599 | intron (NM_026239, intron 1 of 1) | intron (NM_026239, intron 1 of 1) | 6205 | NM_026239 | 67564 | Mm.41489 | NM_026239 | ENSMUSG00000033578 | **Tmem35** | 9030603L14Rik\|AI841526\|PMP52 | transmembrane protein 35 | protein-coding |
| chr4_53832329_53832692 | chr4 | 53832330 | 53832692 | + | 0.59085 | intron (NM_028053, intron 1 of 5) | intron (NM_028053, intron 1 of 5) | 6466 | NM_028053 | 52076 | Mm.27184 | NM_028053 | ENSMUSG00000028420 | **Tmem38b** | 1600017F22Rik\|AA437809\|AV051057\|D4Ertd89e\|TRIC-B\|mg33b | transmembrane protein 38B | protein-coding |
| chr9_123024264_123024574 | chr9 | 123024265 | 123024574 | + | 0.859283 | TTS (NM_001164823) | TTS (NM_001164823) | 3093 | NM_001164823 | 66079 | Mm.482265 | NM_025339 | ENSMUSG00000066233 | **Tmem42** | 0610027O18Rik\|4933429E06Rik\|AV003444\|D9Ertd662e | transmembrane protein 42 | protein-coding |
| chr6_91488496_91489274 | chr6 | 91488497 | 91489274 | + | 1.03772 | TTS (NM_009531) | TTS (NM_009531) | 15134 | NM_028766 | 74122 | Mm.38801 | NM_028766 | ENSMUSG00000030095 | **Tmem43** | 1200015A22Rik\|LUMA | transmembrane protein 43 | protein-coding |
| chr6_91474178_91474669 | chr6 | 91474179 | 91474669 | + | 0.926123 | intron (NM_028766, intron 1 of 11) | intron (NM_028766, intron 1 of 11) | 673 | NM_028766 | 74122 | Mm.38801 | NM_028766 | ENSMUSG00000030095 | **Tmem43** | 1200015A22Rik\|LUMA | transmembrane protein 43 | protein-coding |
| chr16_30504451_30505136 | chr16 | 30504452 | 30505136 | + | 0.655775 | Intergenic | Intergenic | 45784 | NM_172614 | 224090 | Mm.241459 | NM_172614 | | **Tmem44** | 1700007N03Rik\|9330161C17Rik\|B230220N21 | transmembrane protein 44 | protein-coding |
| chr6_129514298_129514945 | chr6 | 129514299 | 129514945 | + | 0.760555 | intron (NM_001081186, intron 2 of 4) | intron (NM_001081186, intron 2 of 4) | 2065 | NM_001081186 | 330428 | Mm.44237 | NM_001081186 | ENSMUSG00000030160 | **Tmem52b** | D630042F21Rik | transmembrane protein 52B | protein-coding |
| chr4_134841910_134842355 | chr4 | 134841911 | 134842355 | + | 0.721033 | intron (NM_025382, intron 1 of 10) | B3\|SINE\|B2 | 11212 | NM_025382 | 66146 | Mm.99793 | NM_025382 | ENSMUSG00000028826 | **Tmem57** | 1110007C24Rik\|9230118A01Rik\|AI317300\|AI606104\|C61 | transmembrane protein 57 | protein-coding |
| chr2_120981021_120981489 | chr2 | 120981022 | 120981489 | + | 1.27137 | intron (NM_175285, intron 4 of 13) | intron (NM_175285, intron 4 of 13) | 4193 | NM_175285 | 96957 | Mm.315212 | NM_175285 | ENSMUSG00000054484 | **Tmem62** | B830009D23Rik\|C77355 | transmembrane protein 62 | protein-coding |
| chr2_120981188_120981489 | chr2 | 120981189 | 120981489 | + | 1.1802 | intron (NM_175285, intron 4 of 13) | intron (NM_175285, intron 4 of 13) | 4277 | NM_175285 | 96957 | Mm.315212 | NM_175285 | ENSMUSG00000054484 | **Tmem62** | B830009D23Rik\|C77355 | transmembrane protein 62 | protein-coding |
| chr15_58749019_58749575 | chr15 | 58749020 | 58749575 | + | 1.53829 | Intergenic | Intergenic | 74130 | NM_175212 | 74868 | Mm.293754 | NM_175212 | ENSMUSG00000062373 | **Tmem65** | 2610029O13Rik\|4930438D12Rik | transmembrane protein 65 | protein-coding |

| | | | | | | | | | | | | | | | | | |
|---|---|---|---|---|---|---|---|---|---|---|---|---|---|---|---|---|---|
| chr15_58669390_58670644 | chr15 | 58669391 | 58670644 | + | 0.776017 | Intergenic | Intergenic | 153410 | NM_175212 | 74868 | Mm.293754 | NM_175212 | ENSMUSG00000062373 | **Tmem65** | 2610029O13Rik\|4930438D12Rik | transmembrane protein 65 | protein-coding |
| chr15_58817999_58818739 | chr15 | 58818000 | 58818739 | + | 0.726843 | intron (NM_175212, intron 1 of 6) | MLT1D\|LTR\|ERVL-MaLR | 5058 | NM_175212 | 74868 | Mm.293754 | NM_175212 | ENSMUSG00000062373 | **Tmem65** | 2610029O13Rik\|4930438D12Rik | transmembrane protein 65 | protein-coding |
| chr4_12036407_12036985 | chr4 | 12036408 | 12036985 | + | 0.653955 | Intergenic | Intergenic | 51261 | NM_177861 | 329795 | Mm.475125 | NM_177861 | ENSMUSG00000049488 | **Tmem67** | 5330408M12Rik\|B230117O07\|b2b1163.1Clo\|b2b1291.1Clo | transmembrane protein 67 | protein-coding |
| chr3_88334982_88335250 | chr3 | 88334983 | 88335250 | + | 0.877468 | promoter-TSS (NM_024246) | promoter-TSS (NM_024246) | -683 | NM_024246 | 71913 | Mm.34256 | NM_024246 | ENSMUSG00000001420 | **Tmem79** | 2310042N02Rik\|2310074C17Rik | transmembrane protein 79 | protein-coding |
| chr7_47031199_47031929 | chr7 | 47031200 | 47031929 | + | 1.00416 | Intergenic | URR1B\|DNA\|hAT-Charlie | -19076 | NM_026436 | 67893 | Mm.27338 | NM_026436 | ENSMUSG00000010307 | **Tmem86a** | 1810054O13Rik\|AI414959\|AW742698 | transmembrane protein 86A | protein-coding |
| chr2_128832768_128833416 | chr2 | 128832769 | 128833416 | + | 1.16791 | intron (NM_028248, intron 8 of 18) | intron (NM_028248, intron 8 of 18) | 14789 | NM_028248 | 72477 | Mm.259688 | NM_028248 | ENSMUSG00000014353 | **Tmem87b** | 2610301K12Rik\|2810431I02Rik\|AU014804 | transmembrane protein 87B | protein-coding |
| chr1_136010230_136010761 | chr1 | 136010231 | 136010761 | + | 0.782247 | intron (NM_001160145, intron 1 of 5) | intron (NM_001160145, intron 1 of 5) | 2277 | NM_001160145 | 66241 | Mm.41773 | NM_025439 | ENSMUSG00000026411 | **Tmem9** | 1500015G18Rik\|AW545782 | transmembrane protein 9 | protein-coding |
| chr10_118506110_118506568 | chr10 | 118506111 | 118506568 | + | 0.685422 | intron (NR_104123, intron 4 of 4) | intron (NR_104123, intron 4 of 4) | 50186 | NR_104123 | 103214 | Mm.30529 | NR_104123 | | **Tmevpg1** | AI447422 | Theiler's murine encephalomyelitis virus persistence candidate gene 1 | ncRNA |
| chr10_91177128_91177779 | chr10 | 91177129 | 91177779 | + | 1.26192 | Intergenic | ORR1F\|LTR\|ERVL-MaLR | -5835 | NM_001080130 | 21917 | Mm.159684 | NM_011605 | ENSMUSG00000019961 | **Tmpo** | 5630400D24Rik\|AI195756\|AI606875\|AW214352\|AW547477\|LAP2\|TP | thymopoietin | protein-coding |
| chr9_114446319_114446645 | chr9 | 114446320 | 114446645 | + | 0.779585 | intron (NR_108101, intron 9 of 14) | intron (NR_108101, intron 9 of 14) | 45387 | NM_001200002 | 100504715 | Mm.290516 | NM_001200002 | ENSMUSG00000079260 | **Tmppe** | - | transmembrane protein with metallophosphoesterase domain | protein-coding |
| chr9_114442872_114443150 | chr9 | 114442873 | 114443150 | + | 0.700894 | non-coding (NR_108102, exon 9 of 15) | non-coding (NR_108102, exon 9 of 15) | 41916 | NM_001200002 | 100504715 | Mm.290516 | NM_001200002 | ENSMUSG00000079260 | **Tmppe** | - | transmembrane protein with metallophosphoesterase domain | protein-coding |
| chr9_114446194_114446655 | chr9 | 114446195 | 114446655 | + | 0.636585 | intron (NR_108101, intron 9 of 14) | intron (NR_108101, intron 9 of 14) | 45330 | NM_001200002 | 100504715 | Mm.290516 | NM_001200002 | ENSMUSG00000079260 | **Tmppe** | - | transmembrane protein with metallophosphoesterase domain | protein-coding |
| chr9_114447730_114448084 | chr9 | 114447731 | 114448084 | + | 0.617568 | intron (NR_108101, intron 9 of 14) | intron (NR_108101, intron 9 of 14) | 46812 | NM_001200002 | 100504715 | Mm.290516 | NM_001200002 | ENSMUSG00000079260 | **Tmppe** | - | transmembrane protein with metallophosphoesterase domain | protein-coding |
| chr9_45297896_45298120 | chr9 | 45297897 | 45298120 | + | 0.691686 | Intergenic | MER20\|DNA\|hAT-Charlie | -21092 | NM_001013373 | 214531 | Mm.394987 | NM_001013373 | ENSMUSG00000037129 | **Tmprss13** | BC010843\|Tpmrss13 | transmembrane protease, serine 13 | protein-coding |
| chr9_45208279_45209039 | chr9 | 45208280 | 45209039 | + | 0.837113 | intron (NM_001198971, intron 1 of 4) | (TC)n\|Simple_repeat\|Simple_repeat | -4584 | NM_145403 | 214523 | Mm.436756 | NM_145403 | ENSMUSG00000032091 | **Tmprss4** | mCAP2 | transmembrane protease, serine 4 | protein-coding |
| chr9_45208579_45209026 | chr9 | 45208580 | 45209026 | + | 0.806592 | intron (NM_001198971, intron 1 of 4) | MTC\|LTR\|ERVL-MaLR | -4728 | NM_145403 | 214523 | Mm.436756 | NM_145403 | ENSMUSG00000032091 | **Tmprss4** | mCAP2 | transmembrane protease, serine 4 | protein-coding |
| chr15_78476217_78476622 | chr15 | 78476218 | 78476622 | + | 1.1169 | Intergenic | Intergenic | -7786 | NM_027902 | 71753 | Mm.34056 | NM_027902 | ENSMUSG00000016942 | **Tmprss6** | 1300008A22Rik | transmembrane serine protease 6 | protein-coding |
| chr15_78450040_78450743 | chr15 | 78450041 | 78450743 | + | 0.906179 | intron (NM_027902, intron 10 of 17) | MLT1A1\|LTR\|ERVL-MaLR | 18242 | NM_027902 | 71753 | Mm.34056 | NM_027902 | ENSMUSG00000016942 | **Tmprss6** | 1300008A22Rik | transmembrane serine protease 6 | protein-coding |
| chr15_78449189_78450820 | chr15 | 78449190 | 78450820 | + | 0.773467 | intron (NM_027902, intron 10 of 17) | intron (NM_027902, intron 10 of 17) | 18629 | NM_027902 | 71753 | Mm.34056 | NM_027902 | ENSMUSG00000016942 | **Tmprss6** | 1300008A22Rik | transmembrane serine protease 6 | protein-coding |
| chr15_78450113_78450359 | chr15 | 78450114 | 78450359 | + | 0.665565 | intron (NM_027902, intron 10 of 17) | intron (NM_027902, intron 10 of 17) | 18398 | NM_027902 | 71753 | Mm.34056 | NM_027902 | ENSMUSG00000016942 | **Tmprss6** | 1300008A22Rik | transmembrane serine protease 6 | protein-coding |
| chrX_167175330_167175718 | chrX | 167175331 | 167175718 | + | 0.712911 | Intergenic | Intergenic | 33694 | NM_021278 | 19241 | Mm.142729 | NM_021278 | ENSMUSG00000049775 | **Tmsb4x** | Ptmb4\|Tb4 | thymosin, beta 4, X chromosome | protein-coding |
| chrX_167173014_167173243 | chrX | 167173015 | 167173243 | + | 0.701605 | Intergenic | Intergenic | 36089 | NM_021278 | 19241 | Mm.142729 | NM_021278 | ENSMUSG00000049775 | **Tmsb4x** | Ptmb4\|Tb4 | thymosin, beta 4, X chromosome | protein-coding |
| chrX_167172968_167173287 | chrX | 167172969 | 167173287 | + | 0.627523 | Intergenic | Intergenic | 36090 | NM_021278 | 19241 | Mm.142729 | NM_021278 | ENSMUSG00000049775 | **Tmsb4x** | Ptmb4\|Tb4 | thymosin, beta 4, X chromosome | protein-coding |
| chr6_148436295_148436886 | chr6 | 148436296 | 148436886 | + | 0.765315 | intron (NM_198967, intron 1 of 19) | intron (NM_198967, intron 1 of 19) | 7761 | NM_198967 | 387314 | Mm.137858 | NM_198967 | ENSMUSG00000030306 | **Tmtc1** | Arg99\|BC023818 | transmembrane and tetratricopeptide repeat containing 1 | protein-coding |
| chr6_148435912_148436912 | chr6 | 148435913 | 148436912 | + | 0.697443 | intron (NM_198967, intron 1 of 19) | ORR1F\|LTR\|ERVL-MaLR | 7940 | NM_198967 | 387314 | Mm.137858 | NM_198967 | ENSMUSG00000030306 | **Tmtc1** | Arg99\|BC023818 | transmembrane and tetratricopeptide repeat containing 1 | protein-coding |
| chr6_148436173_148436734 | chr6 | 148436174 | 148436734 | + | 0.672374 | intron (NM_198967, intron 1 of 19) | ORR1F\|LTR\|ERVL-MaLR | 7898 | NM_198967 | 387314 | Mm.137858 | NM_198967 | ENSMUSG00000030306 | **Tmtc1** | Arg99\|BC023818 | transmembrane and tetratricopeptide repeat containing 1 | protein-coding |
| chr10_105572809_105573236 | chr10 | 105572810 | 105573236 | + | 0.861337 | intron (NM_177368, intron 1 of 11) | intron (NM_177368, intron 1 of 11) | 1456 | NM_177368 | 278279 | Mm.25225 | NM_177368 | ENSMUSG00000036019 | **Tmtc2** | 8430438D04Rik\|D330034A10Rik | transmembrane and tetratricopeptide repeat containing 2 | protein-coding |
| chr10_105281052_105281338 | chr10 | 105281053 | 105281338 | + | 0.784632 | intron (NM_177368, intron 8 of 11) | RMER15\|LTR\|ERVL | 293284 | NM_177368 | 278279 | Mm.25225 | NM_177368 | ENSMUSG00000036019 | **Tmtc2** | 8430438D04Rik\|D330034A10Rik | transmembrane and tetratricopeptide repeat containing 2 | protein-coding |
| chr10_105280955_105281343 | chr10 | 105280956 | 105281343 | + | 0.673422 | intron (NM_177368, intron 8 of 11) | intron (NM_177368, intron 8 of 11) | 293330 | NM_177368 | 278279 | Mm.25225 | NM_177368 | ENSMUSG00000036019 | **Tmtc2** | 8430438D04Rik\|D330034A10Rik | transmembrane and tetratricopeptide repeat containing 2 | protein-coding |
| chr14_123004743_123005704 | chr14 | 123004744 | 123005704 | + | 0.784636 | Intergenic | Intergenic | -21963 | NM_028651 | 70551 | Mm.271988 | NM_028651 | ENSMUSG00000041594 | **Tmtc4** | 4930403J22Rik\|5730419O14Rik | transmembrane and tetratricopeptide repeat containing 4 | protein-coding |
| chr14_122992907_122993317 | chr14 | 122992908 | 122993317 | + | 0.69564 | Intergenic | RMER1A\|Other\|Other | -9851 | NM_028651 | 70551 | Mm.271988 | NM_028651 | ENSMUSG00000041594 | **Tmtc4** | 4930403J22Rik\|5730419O14Rik | transmembrane and tetratricopeptide repeat containing 4 | protein-coding |
| chr12_70469963_70471119 | chr12 | 70469964 | 70471119 | + | 0.772943 | intron (NR_037248, intron 2 of 4) | intron (NR_037248, intron 2 of 4) | 17387 | NM_028339 | 72736 | Mm.204670 | NM_028339 | ENSMUSG00000021072 | **Tmx1** | 2810425A04Rik\|AI465319\|Txndc1 | thioredoxin-related transmembrane protein 1 | protein-coding |
| chr18_90585472_90586562 | chr18 | 90585473 | 90586562 | + | 1.45592 | Intergenic | MTD-int\|LTR\|ERVL-MaLR | 75863 | NM_198295 | 67988 | Mm.268041 | NM_198295 | ENSMUSG00000024614 | **Tmx3** | 6430411B10Rik\|A730024F05Rik\|AV259382\|Txndc10\|mKIAA1830 | thioredoxin-related transmembrane protein 3 | protein-coding |
| chr18_90586112_90586548 | chr18 | 90586113 | 90586548 | + | 1.38857 | Intergenic | MTD-int\|LTR\|ERVL-MaLR | 76176 | NM_198295 | 67988 | Mm.268041 | NM_198295 | ENSMUSG00000024614 | **Tmx3** | 6430411B10Rik\|A730024F05Rik\|AV259382\|Txndc10\|mKIAA1830 | thioredoxin-related transmembrane protein 3 | protein-coding |
| chr18_90586154_90586438 | chr18 | 90586155 | 90586438 | + | 1.35427 | Intergenic | MTD-int\|LTR\|ERVL-MaLR | 76142 | NM_198295 | 67988 | Mm.268041 | NM_198295 | ENSMUSG00000024614 | **Tmx3** | 6430411B10Rik\|A730024F05Rik\|AV259382\|Txndc10\|mKIAA1830 | thioredoxin-related transmembrane protein 3 | protein-coding |
| chr18_90585470_90586061 | chr18 | 90585471 | 90586061 | + | 1.29746 | Intergenic | Intergenic | 75612 | NM_198295 | 67988 | Mm.268041 | NM_198295 | ENSMUSG00000024614 | **Tmx3** | 6430411B10Rik\|A730024F05Rik\|AV259382\|Txndc10\|mKIAA1830 | thioredoxin-related transmembrane protein 3 | protein-coding |
| chr10_18987793_18988238 | chr10 | 18987794 | 18988238 | + | 1.32411 | Intergenic | Intergenic | 23947 | NM_001166402 | 21929 | Mm.116683 | NM_009397 | ENSMUSG00000019850 | **Tnfaip3** | A20\|Tnfip3 | tumor necrosis factor, alpha-induced protein 3 | protein-coding |
| chr10_18987177_18987748 | chr10 | 18987178 | 18987748 | + | 1.10905 | Intergenic | Intergenic | 24500 | NM_001166402 | 21929 | Mm.116683 | NM_009397 | ENSMUSG00000019850 | **Tnfaip3** | A20\|Tnfip3 | tumor necrosis factor, alpha-induced protein 3 | protein-coding |
| chr10_18992952_18993402 | chr10 | 18992953 | 18993402 | + | 0.830153 | Intergenic | Intergenic | 18786 | NM_001166402 | 21929 | Mm.116683 | NM_009397 | ENSMUSG00000019850 | **Tnfaip3** | A20\|Tnfip3 | tumor necrosis factor, alpha-induced protein 3 | protein-coding |
| chr10_19009196_19009761 | chr10 | 19009197 | 19009761 | + | 0.74934 | intron (NM_009397, intron 2 of 8) | intron (NM_009397, intron 2 of 8) | 2484 | NM_001166402 | 21929 | Mm.116683 | NM_009397 | ENSMUSG00000019850 | **Tnfaip3** | A20\|Tnfip3 | tumor necrosis factor, alpha-induced protein 3 | protein-coding |
| chr10_18949196_18949665 | chr10 | 18949197 | 18949665 | + | 0.724786 | Intergenic | Intergenic | 62532 | NM_001166402 | 21929 | Mm.116683 | NM_009397 | ENSMUSG00000019850 | **Tnfaip3** | A20\|Tnfip3 | tumor necrosis factor, alpha-induced protein 3 | protein-coding |
| chr10_19047324_19047808 | chr10 | 19047325 | 19047808 | + | 0.658608 | Intergenic | Intergenic | -32156 | NM_009397 | 21929 | Mm.116683 | NM_009397 | ENSMUSG00000019850 | **Tnfaip3** | A20\|Tnfip3 | tumor necrosis factor, alpha-induced protein 3 | protein-coding |
| chr2_52037816_52038278 | chr2 | 52037817 | 52038278 | + | 0.757354 | promoter-TSS (NM_009398) | promoter-TSS (NM_009398) | -66 | NM_009398 | 21930 | Mm.3509 | NM_009398 | ENSMUSG00000053475 | **Tnfaip6** | TSG-6\|Tnfip6\|Tsg6 | tumor necrosis factor alpha induced protein 6 | protein-coding |
| chr18_50065805_50066645 | chr18 | 50065806 | 50066645 | + | 1.32175 | intron (NM_134131, intron 1 of 1) | intron (NM_134131, intron 1 of 1) | 12943 | NM_134131 | 106869 | Mm.27740 | NM_134131 | ENSMUSG00000062210 | **Tnfaip8** | AA987150\|E130304C20Rik\|ENSMUSG00000073567\|Gg2-1\|Gm10539\|Nded\|Ssc-2 | tumor necrosis factor, alpha-induced protein 8 | protein-coding |
| chr18_49986640_49987406 | chr18 | 49986641 | 49987406 | + | 1.14503 | intron (NM_001177760, intron 1 of 2) | intron (NM_001177760, intron 1 of 2) | 7596 | NM_001177759 | 106869 | Mm.27740 | NM_134131 | ENSMUSG00000062210 | **Tnfaip8** | AA987150\|E130304C20Rik\|ENSMUSG00000073567\|Gg2-1\|Gm10539\|Nded\|Ssc-2 | tumor necrosis factor, alpha-induced protein 8 | protein-coding |
| chr18_50077991_50078787 | chr18 | 50077992 | 50078787 | + | 1.07812 | intron (NM_134131, intron 1 of 1) | intron (NM_134131, intron 1 of 1) | 25107 | NM_134131 | 106869 | Mm.27740 | NM_134131 | ENSMUSG00000062210 | **Tnfaip8** | AA987150\|E130304C20Rik\|ENSMUSG00000073567\|Gg2-1\|Gm10539\|Nded\|Ssc-2 | tumor necrosis factor, alpha-induced protein 8 | protein-coding |
| chr18_49981904_49982132 | chr18 | 49981905 | 49982132 | + | 0.655046 | intron (NM_001177760, intron 1 of 2) | RLTR12E\|LTR\|ERVK | 2591 | NM_001177759 | 106869 | Mm.27740 | NM_134131 | ENSMUSG00000062210 | **Tnfaip8** | AA987150\|E130304C20Rik\|ENSMUSG00000073567\|Gg2-1\|Gm10539\|Nded\|Ssc-2 | tumor necrosis factor, alpha-induced protein 8 | protein-coding |
| chr1_105880638_105881321 | chr1 | 105880639 | 105881321 | + | 0.789302 | Intergenic | Intergenic | 100257 | NM_009399 | 21934 | Mm.6251 | NM_009399 | ENSMUSG00000026321 | **Tnfrsf11a** | Ly109\|ODFR\|OFE\|Rank\|TRANCE-R\|mRANK | tumor necrosis factor receptor superfamily, member 11a, NFKB activator | protein-coding |
| chr15_54113997_54114915 | chr15 | 54113998 | 54114915 | + | 1.35796 | Intergenic | Intergenic | 164028 | NM_008764 | 18383 | Mm.15383 | NM_008764 | ENSMUSG00000063727 | **Tnfrsf11b** | OCIF\|Opg\|TR1 | tumor necrosis factor receptor superfamily, member 11b (osteoprotegerin) | protein-coding |
| chr17_23675768_23676014 | chr17 | 23675769 | 23676014 | + | 0.68342 | TTS (NM_181821) | TTS (NM_181821) | 1558 | NM_001161746 | 27279 | Mm.28518 | NM_013749 | ENSMUSG00000023905 | **Tnfrsf12a** | AI255180\|C87282\|Fn14\|HPIP\|TWEAK-R\|TweakR | tumor necrosis factor receptor superfamily, member 12a | protein-coding |
| chr11_61119727_61120614 | chr11 | 61119728 | 61120614 | + | 1.00139 | Intergenic | Intergenic | -20664 | NM_021349 | 57916 | Mm.265915 | NM_021349 | ENSMUSG00000010142 | **Tnfrsf13b** | 1200009E08Rik\|Taci | tumor necrosis factor receptor superfamily, member 13b | protein-coding |

| | | | | | | | | | | | | | | | | | |
|---|---|---|---|---|---|---|---|---|---|---|---|---|---|---|---|---|---|
| chr11_61122502_61123369 | chr11 | 61122503 | 61123369 | + | 0.682594 | Intergenic | Intergenic | -17899 | NM_021349 | 57916 | Mm.265915 | NM_021349 | ENSMUSG00000010142 | **Tnfrsf13b** | 1200009E08Rik\|Taci | tumor necrosis factor receptor superfamily, member 13b | protein-coding |
| chr16_11285721_11286384 | chr16 | 11285722 | 11286384 | + | 1.00451 | Intergenic | Tigger7\|DNA\|TcMar-Tigger | -27756 | NM_011608 | 21935 | Mm.12935 | NM_011608 | ENSMUSG00000022496 | **Tnfrsf17** | BCM\|BCMA\|Tnfrsf13\|Tnfrsf13a | tumor necrosis factor receptor superfamily, member 17 | protein-coding |
| chr16_11285404_11286377 | chr16 | 11285405 | 11286377 | + | 0.847635 | Intergenic | Tigger7\|DNA\|TcMar-Tigger | -27918 | NM_011608 | 21935 | Mm.12935 | NM_011608 | ENSMUSG00000022496 | **Tnfrsf17** | BCM\|BCMA\|Tnfrsf13\|Tnfrsf13a | tumor necrosis factor receptor superfamily, member 17 | protein-coding |
| chr14_60937465_60938107 | chr14 | 60937466 | 60938107 | + | 0.739012 | Intergenic | Intergenic | 100201 | NM_001164155 | 29820 | Mm.281356 | NM_013869 | ENSMUSG00000060548 | **Tnfrsf19** | AL023044\|AW123854\|TAJ\|TAJ-ALPHA\|TRADE\|Troy | tumor necrosis factor receptor superfamily, member 19 | protein-coding |
| chr4_145274851_145275351 | chr4 | 145274852 | 145275351 | + | 0.663777 | intron (NM_009401, intron 11 of 12) | intron (NM_009401, intron 11 of 12) | -28231 | NM_011610 | 21938 | Mm.235328 | NM_011610 | | **Tnfrsf1b** | CD120b\|TNF-R-II\|TNF-R2\|TNF-R75\|TNF-alphaR2\|TNFBR\|TNFR80\|TNFRII\|TNFalpha-R2\|Tnfr-1\|Tnfr2\|p75 | tumor necrosis factor receptor superfamily, member 1b | protein-coding |
| chr17_42989990_42990530 | chr17 | 42989991 | 42990530 | + | 1.07016 | Intergenic | Intergenic | -26295 | NM_178589 | 94185 | Mm.200792 | NM_178589 | ENSMUSG00000023915 | **Tnfrsf21** | AA959878\|DR6\|R74815\|TR7 | tumor necrosis factor receptor superfamily, member 21 | protein-coding |
| chr7_143631522_143631854 | chr7 | 143631523 | 143631854 | + | 1.17865 | Intergenic | RMER17C-int\|LTR\|ERVK | -3843 | NM_175649 | 244237 | Mm.247498 | NM_175649 | ENSMUSG00000045362 | **Tnfrsf26** | Tnfrh3\|Tnfrsf24 | tumor necrosis factor receptor superfamily, member 26 | protein-coding |
| chr4_156008038_156008885 | chr4 | 156008039 | 156008885 | + | 1.25253 | intron (NM_011341, intron 4 of 6) | intron (NM_011341, intron 4 of 6) | -5233 | NM_011659 | 22163 | Mm.13885 | NM_011659 | ENSMUSG00000029075 | **Tnfrsf4** | ACT35\|CD134\|Ly-70\|Ox40\|TXGP1L\|Txgp1 | tumor necrosis factor receptor superfamily, member 4 | protein-coding |
| chr4_156010289_156010720 | chr4 | 156010290 | 156010720 | + | 0.778404 | 3' UTR (NM_011341, exon 7 of 7) | 3' UTR (NM_011341, exon 7 of 7) | -3190 | NM_011659 | 22163 | Mm.13885 | NM_011659 | ENSMUSG00000029075 | **Tnfrsf4** | ACT35\|CD134\|Ly-70\|Ox40\|TXGP1L\|Txgp1 | tumor necrosis factor receptor superfamily, member 4 | protein-coding |
| chr4_145341659_145342575 | chr4 | 145341660 | 145342575 | + | 1.23749 | Intergenic | Intergenic | -26970 | NM_009401 | 21941 | Mm.12810 | NM_009401 | ENSMUSG00000028602 | **Tnfrsf8** | Cd30\|D1S166E\|Ki\|Ki-1 | tumor necrosis factor receptor superfamily, member 8 | protein-coding |
| chr4_145315111_145315380 | chr4 | 145315112 | 145315380 | + | 1.04982 | promoter-TSS (NM_009401) | promoter-TSS (NM_009401) | -99 | NM_009401 | 21941 | Mm.12810 | NM_009401 | ENSMUSG00000028602 | **Tnfrsf8** | Cd30\|D1S166E\|Ki\|Ki-1 | tumor necrosis factor receptor superfamily, member 8 | protein-coding |
| chr4_145314985_145315478 | chr4 | 145314986 | 145315478 | + | 1.02988 | promoter-TSS (NM_009401) | promoter-TSS (NM_009401) | -85 | NM_009401 | 21941 | Mm.12810 | NM_009401 | ENSMUSG00000028602 | **Tnfrsf8** | Cd30\|D1S166E\|Ki\|Ki-1 | tumor necrosis factor receptor superfamily, member 8 | protein-coding |
| chr4_145323107_145323740 | chr4 | 145323108 | 145323740 | + | 0.958626 | Intergenic | Intergenic | -8277 | NM_009401 | 21941 | Mm.12810 | NM_009401 | ENSMUSG00000028602 | **Tnfrsf8** | Cd30\|D1S166E\|Ki\|Ki-1 | tumor necrosis factor receptor superfamily, member 8 | protein-coding |
| chr4_145335825_145336357 | chr4 | 145335826 | 145336357 | + | 0.900069 | Intergenic | Intergenic | -20944 | NM_009401 | 21941 | Mm.12810 | NM_009401 | ENSMUSG00000028602 | **Tnfrsf8** | Cd30\|D1S166E\|Ki\|Ki-1 | tumor necrosis factor receptor superfamily, member 8 | protein-coding |
| chr4_145319135_145319781 | chr4 | 145319136 | 145319781 | + | 0.856778 | Intergenic | Intergenic | -4311 | NM_009401 | 21941 | Mm.12810 | NM_009401 | ENSMUSG00000028602 | **Tnfrsf8** | Cd30\|D1S166E\|Ki\|Ki-1 | tumor necrosis factor receptor superfamily, member 8 | protein-coding |
| chr4_145323107_145323979 | chr4 | 145323108 | 145323979 | + | 0.846849 | Intergenic | PB1D10\|SINE\|Alu | -8396 | NM_009401 | 21941 | Mm.12810 | NM_009401 | ENSMUSG00000028602 | **Tnfrsf8** | Cd30\|D1S166E\|Ki\|Ki-1 | tumor necrosis factor receptor superfamily, member 8 | protein-coding |
| chr4_145310931_145311206 | chr4 | 145310932 | 145311206 | + | 0.790044 | intron (NM_009401, intron 1 of 12) | intron (NM_009401, intron 1 of 12) | 4078 | NM_009401 | 21941 | Mm.12810 | NM_009401 | ENSMUSG00000028602 | **Tnfrsf8** | Cd30\|D1S166E\|Ki\|Ki-1 | tumor necrosis factor receptor superfamily, member 8 | protein-coding |
| chr4_145338506_145338753 | chr4 | 145338507 | 145338753 | + | 0.664874 | Intergenic | Intergenic | -23483 | NM_009401 | 21941 | Mm.12810 | NM_009401 | ENSMUSG00000028602 | **Tnfrsf8** | Cd30\|D1S166E\|Ki\|Ki-1 | tumor necrosis factor receptor superfamily, member 8 | protein-coding |
| chr4_150917359_150917691 | chr4 | 150917360 | 150917691 | + | 1.01849 | Intergenic | Intergenic | -2630 | NM_011612 | 21942 | Mm.244187 | NM_011612 | ENSMUSG00000028965 | **Tnfrsf9** | 4-1BB\|A930040I11Rik\|AA408498\|AI325004\|CDw137\|Cd137\|ILA\|Ly63 | tumor necrosis factor receptor superfamily, member 9 | protein-coding |
| chr4_150915286_150915863 | chr4 | 150915287 | 150915863 | + | 0.990637 | Intergenic | Intergenic | -4580 | NM_011612 | 21942 | Mm.244187 | NM_011612 | ENSMUSG00000028965 | **Tnfrsf9** | 4-1BB\|A930040I11Rik\|AA408498\|AI325004\|CDw137\|Cd137\|ILA\|Ly63 | tumor necrosis factor receptor superfamily, member 9 | protein-coding |
| chr4_150927810_150928163 | chr4 | 150927811 | 150928163 | + | 0.682883 | promoter-TSS (NM_001077509) | promoter-TSS (NM_001077509) | -12 | NM_001077509 | 21942 | Mm.244187 | NM_011612 | ENSMUSG00000028965 | **Tnfrsf9** | 4-1BB\|A930040I11Rik\|AA408498\|AI325004\|CDw137\|Cd137\|ILA\|Ly63 | tumor necrosis factor receptor superfamily, member 9 | protein-coding |
| chr3_27330724_27331563 | chr3 | 27330725 | 27331563 | + | 0.853456 | intron (NM_009425, intron 3 of 4) | L1MA4\|LINE\|L1 | 14067 | NM_009425 | 22035 | Mm.1062 | NM_009425 | ENSMUSG00000039304 | **Tnfsf10** | A330042I21Rik\|AI448571\|APO-2L\|Ly81\|TL2\|Trail | tumor necrosis factor (ligand) superfamily, member 10 | protein-coding |
| chr3_27330731_27331301 | chr3 | 27330732 | 27331301 | + | 0.836996 | intron (NM_009425, intron 3 of 4) | L1MA4\|LINE\|L1 | 13939 | NM_009425 | 22035 | Mm.1062 | NM_009425 | ENSMUSG00000039304 | **Tnfsf10** | A330042I21Rik\|AI448571\|APO-2L\|Ly81\|TL2\|Trail | tumor necrosis factor (ligand) superfamily, member 10 | protein-coding |
| chr3_27310985_27311770 | chr3 | 27310986 | 27311770 | + | 0.678016 | Intergenic | Intergenic | -5699 | NM_009425 | 22035 | Mm.1062 | NM_009425 | ENSMUSG00000039304 | **Tnfsf10** | A330042I21Rik\|AI448571\|APO-2L\|Ly81\|TL2\|Trail | tumor necrosis factor (ligand) superfamily, member 10 | protein-coding |
| chr14_78420146_78420605 | chr14 | 78420147 | 78420605 | + | 1.89987 | Intergenic | Intergenic | -112333 | NM_011613 | 21943 | Mm.249221 | NM_011613 | ENSMUSG00000022015 | **Tnfsf11** | Ly109l\|ODF\|OPG\|OPGL\|RANKL\|Trance | tumor necrosis factor (ligand) superfamily, member 11 | protein-coding |
| chr14_78416647_78416961 | chr14 | 78416648 | 78416961 | + | 0.651826 | Intergenic | Intergenic | -108761 | NM_011613 | 21943 | Mm.249221 | NM_011613 | ENSMUSG00000022015 | **Tnfsf11** | Ly109l\|ODF\|OPG\|OPGL\|RANKL\|Trance | tumor necrosis factor (ligand) superfamily, member 11 | protein-coding |
| chr8_10006183_10006783 | chr8 | 10006184 | 10006783 | + | 0.647697 | promoter-TSS (NM_033622) | promoter-TSS (NM_033622) | -150 | NM_033622 | 24099 | Mm.28835 | NM_033622 | | **Tnfsf13b** | BAFF\|BLyS\|D8Ertd387e\|TALL-1\|TALL1\|THANK\|TNFSF20\|zTNF4 | tumor necrosis factor (ligand) superfamily, member 13b | protein-coding |
| chr8_10006324_10006938 | chr8 | 10006325 | 10006938 | + | 0.62145 | promoter-TSS (NM_033622) | promoter-TSS (NM_033622) | -2 | NM_033622 | 24099 | Mm.28835 | NM_033622 | | **Tnfsf13b** | BAFF\|BLyS\|D8Ertd387e\|TALL-1\|TALL1\|THANK\|TNFSF20\|zTNF4 | tumor necrosis factor (ligand) superfamily, member 13b | protein-coding |
| chr17_57198162_57198641 | chr17 | 57198163 | 57198641 | + | 1.15679 | Intergenic | RMER15\|LTR\|ERVL | -4221 | NM_019418 | 50930 | Mm.483369 | NM_019418 | ENSMUSG00000005824 | **Tnfsf14** | HVEM-L\|HVEML\|LIGHT\|LTg\|Ly113 | tumor necrosis factor (ligand) superfamily, member 14 | protein-coding |
| chr17_57201606_57202198 | chr17 | 57201607 | 57202198 | + | 1.15542 | Intergenic | Lx8b\|LINE\|L1 | -7721 | NM_019418 | 50930 | Mm.483369 | NM_019418 | ENSMUSG00000005824 | **Tnfsf14** | HVEM-L\|HVEML\|LIGHT\|LTg\|Ly113 | tumor necrosis factor (ligand) superfamily, member 14 | protein-coding |
| chr17_57182810_57183369 | chr17 | 57182811 | 57183369 | + | 1.12935 | Intergenic | Intergenic | 11091 | NM_019418 | 50930 | Mm.483369 | NM_019418 | ENSMUSG00000005824 | **Tnfsf14** | HVEM-L\|HVEML\|LIGHT\|LTg\|Ly113 | tumor necrosis factor (ligand) superfamily, member 14 | protein-coding |
| chr17_57198331_57198639 | chr17 | 57198332 | 57198639 | + | 1.09152 | Intergenic | RMER15\|LTR\|ERVL | -4304 | NM_019418 | 50930 | Mm.483369 | NM_019418 | ENSMUSG00000005824 | **Tnfsf14** | HVEM-L\|HVEML\|LIGHT\|LTg\|Ly113 | tumor necrosis factor (ligand) superfamily, member 14 | protein-coding |
| chr4_63716087_63716580 | chr4 | 63716088 | 63716580 | + | 1.29339 | Intergenic | Intergenic | 28779 | NM_177371 | 326623 | Mm.208152 | NM_177371 | | **Tnfsf15** | Tl1\|Tl1a\|Vegi | tumor necrosis factor (ligand) superfamily, member 15 | protein-coding |
| chr4_63715849_63716584 | chr4 | 63715850 | 63716584 | + | 1.25502 | Intergenic | Intergenic | 28896 | NM_177371 | 326623 | Mm.208152 | NM_177371 | | **Tnfsf15** | Tl1\|Tl1a\|Vegi | tumor necrosis factor (ligand) superfamily, member 15 | protein-coding |
| chr4_63772496_63772868 | chr4 | 63772497 | 63772868 | + | 1.10735 | Intergenic | MIRb\|SINE\|MIR | -27569 | NM_177371 | 326623 | Mm.208152 | NM_177371 | | **Tnfsf15** | Tl1\|Tl1a\|Vegi | tumor necrosis factor (ligand) superfamily, member 15 | protein-coding |
| chr4_63772359_63772868 | chr4 | 63772360 | 63772868 | + | 1.04974 | Intergenic | MIRb\|SINE\|MIR | -27501 | NM_177371 | 326623 | Mm.208152 | NM_177371 | | **Tnfsf15** | Tl1\|Tl1a\|Vegi | tumor necrosis factor (ligand) superfamily, member 15 | protein-coding |
| chr4_63717118_63717804 | chr4 | 63717119 | 63717804 | + | 1.0086 | Intergenic | Intergenic | 27652 | NM_177371 | 326623 | Mm.208152 | NM_177371 | | **Tnfsf15** | Tl1\|Tl1a\|Vegi | tumor necrosis factor (ligand) superfamily, member 15 | protein-coding |
| chr1_161487074_161487464 | chr1 | 161487075 | 161487464 | + | 0.615947 | Intergenic | Intergenic | -7388 | NM_183391 | 240873 | Mm.276823 | NM_183391 | ENSMUSG00000066755 | **Tnfsf18** | Gitrl | tumor necrosis factor (ligand) superfamily, member 18 | protein-coding |
| chr1_161375713_161376697 | chr1 | 161375714 | 161376697 | + | 1.52198 | Intergenic | Intergenic | -19233 | NM_009452 | 22164 | Mm.4994 | NM_009452 | ENSMUSG00000026700 | **Tnfsf4** | Ath-1\|Ath1\|CD134L\|OX-40L\|Ox40l\|TXGP1\|Txgp1l\|gp34 | tumor necrosis factor (ligand) superfamily, member 4 | protein-coding |
| chr1_161339948_161340686 | chr1 | 161339949 | 161340686 | + | 1.40356 | Intergenic | Intergenic | -55121 | NM_009452 | 22164 | Mm.4994 | NM_009452 | ENSMUSG00000026700 | **Tnfsf4** | Ath-1\|Ath1\|CD134L\|OX-40L\|Ox40l\|TXGP1\|Txgp1l\|gp34 | tumor necrosis factor (ligand) superfamily, member 4 | protein-coding |
| chr1_161436393_161437130 | chr1 | 161436394 | 161437130 | + | 1.16988 | Intergenic | Intergenic | 41324 | NM_009452 | 22164 | Mm.4994 | NM_009452 | ENSMUSG00000026700 | **Tnfsf4** | Ath-1\|Ath1\|CD134L\|OX-40L\|Ox40l\|TXGP1\|Txgp1l\|gp34 | tumor necrosis factor (ligand) superfamily, member 4 | protein-coding |
| chr1_161378727_161379388 | chr1 | 161378728 | 161379388 | + | 1.11747 | Intergenic | Intergenic | -16380 | NM_009452 | 22164 | Mm.4994 | NM_009452 | ENSMUSG00000026700 | **Tnfsf4** | Ath-1\|Ath1\|CD134L\|OX-40L\|Ox40l\|TXGP1\|Txgp1l\|gp34 | tumor necrosis factor (ligand) superfamily, member 4 | protein-coding |
| chr1_161327324_161327861 | chr1 | 161327325 | 161327861 | + | 0.917967 | Intergenic | Intergenic | -67845 | NM_009452 | 22164 | Mm.4994 | NM_009452 | ENSMUSG00000026700 | **Tnfsf4** | Ath-1\|Ath1\|CD134L\|OX-40L\|Ox40l\|TXGP1\|Txgp1l\|gp34 | tumor necrosis factor (ligand) superfamily, member 4 | protein-coding |
| chr1_161375291_161375591 | chr1 | 161375292 | 161375591 | + | 0.912756 | Intergenic | ORR1F\|LTR\|ERVL-MaLR | -19997 | NM_009452 | 22164 | Mm.4994 | NM_009452 | ENSMUSG00000026700 | **Tnfsf4** | Ath-1\|Ath1\|CD134L\|OX-40L\|Ox40l\|TXGP1\|Txgp1l\|gp34 | tumor necrosis factor (ligand) superfamily, member 4 | protein-coding |
| chr1_161406374_161406791 | chr1 | 161406375 | 161406791 | + | 0.647249 | intron (NM_009452, intron 1 of 2) | intron (NM_009452, intron 1 of 2) | 11145 | NM_009452 | 22164 | Mm.4994 | NM_009452 | ENSMUSG00000026700 | **Tnfsf4** | Ath-1\|Ath1\|CD134L\|OX-40L\|Ox40l\|TXGP1\|Txgp1l\|gp34 | tumor necrosis factor (ligand) superfamily, member 4 | protein-coding |
| chr4_63874413_63874993 | chr4 | 63874414 | 63874993 | + | 1.67372 | Intergenic | Intergenic | -13419 | NM_009403 | 21949 | Mm.4664 | NM_009403 | | **Tnfsf8** | CD153\|CD30LG\|Cd30l | tumor necrosis factor (ligand) superfamily, member 8 | protein-coding |
| chr4_63935314_63935907 | chr4 | 63935315 | 63935907 | + | 1.55363 | Intergenic | Intergenic | -74327 | NM_009403 | 21949 | Mm.4664 | NM_009403 | | **Tnfsf8** | CD153\|CD30LG\|Cd30l | tumor necrosis factor (ligand) superfamily, member 8 | protein-coding |
| chr4_63875524_63875789 | chr4 | 63875525 | 63875789 | + | 1.52614 | Intergenic | Intergenic | -14373 | NM_009403 | 21949 | Mm.4664 | NM_009403 | | **Tnfsf8** | CD153\|CD30LG\|Cd30l | tumor necrosis factor (ligand) superfamily, member 8 | protein-coding |

| | | | | | | | | | | | | | | | | | |
|---|---|---|---|---|---|---|---|---|---|---|---|---|---|---|---|---|---|
| chr4_63928181_63928729 | chr4 | 63928182 | 63928729 | + | 1.22035 | Intergenic | Intergenic | -67171 | NM_009403 | 21949 | Mm.4664 | NM_009403 | | **Tnfsf8** | CD153\|CD30LG\|Cd30l | tumor necrosis factor (ligand) superfamily, member 8 | protein-coding |
| chr4_63867911_63868203 | chr4 | 63867912 | 63868203 | + | 1.17971 | Intergenic | Intergenic | -6773 | NM_009403 | 21949 | Mm.4664 | NM_009403 | | **Tnfsf8** | CD153\|CD30LG\|Cd30l | tumor necrosis factor (ligand) superfamily, member 8 | protein-coding |
| chr4_63898321_63898652 | chr4 | 63898322 | 63898652 | + | 1.03994 | Intergenic | Intergenic | -37203 | NM_009403 | 21949 | Mm.4664 | NM_009403 | | **Tnfsf8** | CD153\|CD30LG\|Cd30l | tumor necrosis factor (ligand) superfamily, member 8 | protein-coding |
| chr4_63867799_63868215 | chr4 | 63867800 | 63868215 | + | 1.0375 | Intergenic | Intergenic | -6723 | NM_009403 | 21949 | Mm.4664 | NM_009403 | | **Tnfsf8** | CD153\|CD30LG\|Cd30l | tumor necrosis factor (ligand) superfamily, member 8 | protein-coding |
| chr4_63898850_63899909 | chr4 | 63898851 | 63899909 | + | 0.979453 | Intergenic | Intergenic | -38096 | NM_009403 | 21949 | Mm.4664 | NM_009403 | | **Tnfsf8** | CD153\|CD30LG\|Cd30l | tumor necrosis factor (ligand) superfamily, member 8 | protein-coding |
| chr4_63898841_63899567 | chr4 | 63898842 | 63899567 | + | 0.926888 | Intergenic | Intergenic | -37920 | NM_009403 | 21949 | Mm.4664 | NM_009403 | | **Tnfsf8** | CD153\|CD30LG\|Cd30l | tumor necrosis factor (ligand) superfamily, member 8 | protein-coding |
| chr4_63841786_63842938 | chr4 | 63841787 | 63842938 | + | 0.821336 | intron (NM_009403, intron 2 of 3) | intron (NM_009403, intron 2 of 3) | 18922 | NM_009403 | 21949 | Mm.4664 | NM_009403 | | **Tnfsf8** | CD153\|CD30LG\|Cd30l | tumor necrosis factor (ligand) superfamily, member 8 | protein-coding |
| chr4_63868287_63868587 | chr4 | 63868288 | 63868587 | + | 0.780181 | Intergenic | B3\|SINE\|B2 | -7153 | NM_009403 | 21949 | Mm.4664 | NM_009403 | | **Tnfsf8** | CD153\|CD30LG\|Cd30l | tumor necrosis factor (ligand) superfamily, member 8 | protein-coding |
| chr4_63924677_63925204 | chr4 | 63924678 | 63925204 | + | 0.760959 | Intergenic | Intergenic | -63657 | NM_009403 | 21949 | Mm.4664 | NM_009403 | | **Tnfsf8** | CD153\|CD30LG\|Cd30l | tumor necrosis factor (ligand) superfamily, member 8 | protein-coding |
| chr4_63924851_63925182 | chr4 | 63924852 | 63925182 | + | 0.759317 | Intergenic | Intergenic | -63733 | NM_009403 | 21949 | Mm.4664 | NM_009403 | | **Tnfsf8** | CD153\|CD30LG\|Cd30l | tumor necrosis factor (ligand) superfamily, member 8 | protein-coding |
| chr4_63862695_63863445 | chr4 | 63862696 | 63863445 | + | 0.751696 | Intergenic | Intergenic | -1786 | NM_009403 | 21949 | Mm.4664 | NM_009403 | | **Tnfsf8** | CD153\|CD30LG\|Cd30l | tumor necrosis factor (ligand) superfamily, member 8 | protein-coding |
| chr4_63862907_63863427 | chr4 | 63862908 | 63863427 | + | 0.728777 | Intergenic | Intergenic | -1883 | NM_009403 | 21949 | Mm.4664 | NM_009403 | | **Tnfsf8** | CD153\|CD30LG\|Cd30l | tumor necrosis factor (ligand) superfamily, member 8 | protein-coding |
| chr4_63853774_63854236 | chr4 | 63853775 | 63854236 | + | 0.727842 | intron (NM_009403, intron 1 of 3) | intron (NM_009403, intron 1 of 3) | 7279 | NM_009403 | 21949 | Mm.4664 | NM_009403 | | **Tnfsf8** | CD153\|CD30LG\|Cd30l | tumor necrosis factor (ligand) superfamily, member 8 | protein-coding |
| chr4_63840457_63841017 | chr4 | 63840458 | 63841017 | + | 0.690822 | intron (NM_009403, intron 2 of 3) | intron (NM_009403, intron 2 of 3) | 20547 | NM_009403 | 21949 | Mm.4664 | NM_009403 | | **Tnfsf8** | CD153\|CD30LG\|Cd30l | tumor necrosis factor (ligand) superfamily, member 8 | protein-coding |
| chr4_63871301_63871624 | chr4 | 63871302 | 63871624 | + | 0.687317 | Intergenic | Intergenic | -10179 | NM_009403 | 21949 | Mm.4664 | NM_009403 | | **Tnfsf8** | CD153\|CD30LG\|Cd30l | tumor necrosis factor (ligand) superfamily, member 8 | protein-coding |
| chr4_63840481_63840817 | chr4 | 63840482 | 63840817 | + | 0.638968 | intron (NM_009403, intron 2 of 3) | intron (NM_009403, intron 2 of 3) | 20635 | NM_009403 | 21949 | Mm.4664 | NM_009403 | | **Tnfsf8** | CD153\|CD30LG\|Cd30l | tumor necrosis factor (ligand) superfamily, member 8 | protein-coding |
| chr3_28286835_28287208 | chr3 | 28286836 | 28287208 | + | 1.06879 | intron (NM_001163007, intron 1 of 31) | intron (NM_001163007, intron 1 of 31) | 23808 | NM_001163009 | 665113 | Mm.126193 | NM_026910 | ENSMUSG00000027692 | **Tnik** | 1500031A17Rik\|4831440I19Rik\|AI451411\|C530008O15Rik\|C630040K21Rik | TRAF2 and NCK interacting kinase | protein-coding |
| chr3_28437446_28437834 | chr3 | 28437447 | 28437834 | + | 1.03001 | intron (NM_001163007, intron 2 of 31) | intron (NM_001163007, intron 2 of 31) | 174426 | NM_001163009 | 665113 | Mm.126193 | NM_026910 | ENSMUSG00000027692 | **Tnik** | 1500031A17Rik\|4831440I19Rik\|AI451411\|C530008O15Rik\|C630040K21Rik | TRAF2 and NCK interacting kinase | protein-coding |
| chr3_28211324_28211730 | chr3 | 28211325 | 28211730 | + | 0.674217 | Intergenic | Intergenic | -51687 | NM_001163007 | 665113 | Mm.126193 | NM_026910 | ENSMUSG00000027692 | **Tnik** | 1500031A17Rik\|4831440I19Rik\|AI451411\|C530008O15Rik\|C630040K21Rik | TRAF2 and NCK interacting kinase | protein-coding |
| chr11_54952355_54952956 | chr11 | 54952356 | 54952956 | + | 0.933443 | intron (NM_001271456, intron 1 of 18) | L2a\|LINE\|L2 | 10284 | NM_021327 | 57783 | Mm.259671 | NM_021327 | ENSMUSG00000020400 | **Tnip1** | ABIN\|ABIN-1\|ABIN1\|AU018810\|Naf1\|Nef\|VAN | TNFAIP3 interacting protein 1 | protein-coding |
| chr11_54952715_54952947 | chr11 | 54952716 | 54952947 | + | 0.888561 | intron (NM_001271456, intron 1 of 18) | intron (NM_001271456, intron 1 of 18) | 10109 | NM_021327 | 57783 | Mm.259671 | NM_021327 | ENSMUSG00000020400 | **Tnip1** | ABIN\|ABIN-1\|ABIN1\|AU018810\|Naf1\|Nef\|VAN | TNFAIP3 interacting protein 1 | protein-coding |
| chr11_54977961_54978339 | chr11 | 54977962 | 54978339 | + | 0.849974 | TTS (NM_001110211) | TTS (NM_001110211) | -15210 | NM_001271455 | 57783 | Mm.259671 | NM_021327 | ENSMUSG00000020400 | **Tnip1** | ABIN\|ABIN-1\|ABIN1\|AU018810\|Naf1\|Nef\|VAN | TNFAIP3 interacting protein 1 | protein-coding |
| chr11_54952415_54952653 | chr11 | 54952416 | 54952653 | + | 0.843728 | intron (NM_001271456, intron 1 of 18) | L2a\|LINE\|L2 | 10406 | NM_021327 | 57783 | Mm.259671 | NM_021327 | ENSMUSG00000020400 | **Tnip1** | ABIN\|ABIN-1\|ABIN1\|AU018810\|Naf1\|Nef\|VAN | TNFAIP3 interacting protein 1 | protein-coding |
| chr11_54935983_54936780 | chr11 | 54935984 | 54936780 | + | 0.674532 | intron (NM_021327, intron 5 of 17) | intron (NM_021327, intron 5 of 17) | 26558 | NM_021327 | 57783 | Mm.259671 | NM_021327 | ENSMUSG00000020400 | **Tnip1** | ABIN\|ABIN-1\|ABIN1\|AU018810\|Naf1\|Nef\|VAN | TNFAIP3 interacting protein 1 | protein-coding |
| chr5_34501659_34502028 | chr5 | 34501660 | 34502028 | + | 0.713463 | intron (NM_139064, intron 2 of 5) | intron (NM_139064, intron 2 of 5) | 12135 | NM_139064 | 231130 | Mm.28615 | NM_139064 | ENSMUSG00000059866 | **Tnip2** | 1810020H16Rik\|ABIN-2\|AI428870 | TNFAIP3 interacting protein 2 | protein-coding |
| chr6_65516729_65517191 | chr6 | 65516730 | 65517191 | + | 1.61699 | Intergenic | Intergenic | -73438 | NM_001001495 | 414084 | Mm.117558 | NM_001001495 | ENSMUSG00000044162 | **Tnip3** | 9030611K07Rik\|ABIN-3\|LIND | TNFAIP3 interacting protein 3 | protein-coding |
| chr6_65603637_65604054 | chr6 | 65603638 | 65604054 | + | 1.26173 | intron (NM_001001495, intron 5 of 7) | intron (NM_001001495, intron 5 of 7) | 13448 | NM_001001495 | 414084 | Mm.117558 | NM_001001495 | ENSMUSG00000044162 | **Tnip3** | 9030611K07Rik\|ABIN-3\|LIND | TNFAIP3 interacting protein 3 | protein-coding |
| chr6_65563293_65563806 | chr6 | 65563294 | 65563806 | + | 1.09597 | Intergenic | Intergenic | -26848 | NM_001001495 | 414084 | Mm.117558 | NM_001001495 | ENSMUSG00000044162 | **Tnip3** | 9030611K07Rik\|ABIN-3\|LIND | TNFAIP3 interacting protein 3 | protein-coding |
| chr6_65513871_65514480 | chr6 | 65513872 | 65514480 | + | 1.09016 | Intergenic | RMER15\|LTR\|ERVL | -76222 | NM_001001495 | 414084 | Mm.117558 | NM_001001495 | ENSMUSG00000044162 | **Tnip3** | 9030611K07Rik\|ABIN-3\|LIND | TNFAIP3 interacting protein 3 | protein-coding |
| chr6_65530116_65530504 | chr6 | 65530117 | 65530504 | + | 1.02602 | Intergenic | Intergenic | -60088 | NM_001001495 | 414084 | Mm.117558 | NM_001001495 | ENSMUSG00000044162 | **Tnip3** | 9030611K07Rik\|ABIN-3\|LIND | TNFAIP3 interacting protein 3 | protein-coding |
| chr6_65590085_65590517 | chr6 | 65590086 | 65590517 | + | 0.799524 | promoter-TSS (NM_001001495) | promoter-TSS (NM_001001495) | -97 | NM_001001495 | 414084 | Mm.117558 | NM_001001495 | ENSMUSG00000044162 | **Tnip3** | 9030611K07Rik\|ABIN-3\|LIND | TNFAIP3 interacting protein 3 | protein-coding |
| chr6_65524902_65525697 | chr6 | 65524903 | 65525697 | + | 0.711963 | Intergenic | Intergenic | -65098 | NM_001001495 | 414084 | Mm.117558 | NM_001001495 | ENSMUSG00000044162 | **Tnip3** | 9030611K07Rik\|ABIN-3\|LIND | TNFAIP3 interacting protein 3 | protein-coding |
| chr6_65517988_65518310 | chr6 | 65517989 | 65518310 | + | 0.664559 | Intergenic | Intergenic | -72249 | NM_001001495 | 414084 | Mm.117558 | NM_001001495 | ENSMUSG00000044162 | **Tnip3** | 9030611K07Rik\|ABIN-3\|LIND | TNFAIP3 interacting protein 3 | protein-coding |
| chr16_32672169_32672818 | chr16 | 32672170 | 32672818 | + | 1.06298 | promoter-TSS (NM_001289443) | promoter-TSS (NM_001289443) | -660 | NM_001289443 | 51789 | Mm.251115 | NM_016788 | ENSMUSG00000022791 | **Tnk2** | Ack\|Ack-1\|Ack1\|Cdgip\|Pyk1 | tyrosine kinase, non-receptor, 2 | protein-coding |
| chr13_98901659_98902122 | chr13 | 98901660 | 98902122 | + | 0.871672 | intron (NM_178716, intron 1 of 23) | intron (NM_178716, intron 1 of 23) | -10849 | NM_001048267 | 238799 | Mm.173286 | NM_178716 | ENSMUSG00000009470 | **Tnpo1** | AU021749\|D13Ertd688e\|IPO2\|Kpnb2\|MIP\|MIP1\|TRN | transportin 1 | protein-coding |
| chr13_98923972_98924361 | chr13 | 98923973 | 98924361 | + | 0.85939 | intron (NM_178716, intron 1 of 23) | intron (NM_178716, intron 1 of 23) | 2217 | NM_178716 | 238799 | Mm.173286 | NM_178716 | ENSMUSG00000009470 | **Tnpo1** | AU021749\|D13Ertd688e\|IPO2\|Kpnb2\|MIP\|MIP1\|TRN | transportin 1 | protein-coding |
| chr13_98923508_98923742 | chr13 | 98923509 | 98923742 | + | 0.774053 | intron (NM_178716, intron 1 of 23) | intron (NM_178716, intron 1 of 23) | 2759 | NM_178716 | 238799 | Mm.173286 | NM_178716 | ENSMUSG00000009470 | **Tnpo1** | AU021749\|D13Ertd688e\|IPO2\|Kpnb2\|MIP\|MIP1\|TRN | transportin 1 | protein-coding |
| chr1_159674020_159674689 | chr1 | 159674021 | 159674689 | + | 1.66814 | intron (NM_022312, intron 1 of 22) | intron (NM_022312, intron 1 of 22) | 150586 | NM_022312 | 21960 | Mm.44701 | NM_022312 | ENSMUSG00000015829 | **Tnr** | J1-tenascin\|TN-R\|janusin\|restrictin | tenascin R | protein-coding |
| chr1_159666361_159667092 | chr1 | 159666362 | 159667092 | + | 1.01511 | intron (NM_022312, intron 1 of 22) | MIR3\|SINE\|MIR | 142958 | NM_022312 | 21960 | Mm.44701 | NM_022312 | ENSMUSG00000015829 | **Tnr** | J1-tenascin\|TN-R\|janusin\|restrictin | tenascin R | protein-coding |
| chr1_159666385_159666895 | chr1 | 159666386 | 159666895 | + | 0.973361 | intron (NM_022312, intron 1 of 22) | intron (NM_022312, intron 1 of 22) | 142871 | NM_022312 | 21960 | Mm.44701 | NM_022312 | ENSMUSG00000015829 | **Tnr** | J1-tenascin\|TN-R\|janusin\|restrictin | tenascin R | protein-coding |
| chr1_159673602_159673902 | chr1 | 159673603 | 159673902 | + | 0.971405 | intron (NM_022312, intron 1 of 22) | intron (NM_022312, intron 1 of 22) | 149983 | NM_022312 | 21960 | Mm.44701 | NM_022312 | ENSMUSG00000015829 | **Tnr** | J1-tenascin\|TN-R\|janusin\|restrictin | tenascin R | protein-coding |
| chr7_123108001_123109182 | chr7 | 123108002 | 123109182 | + | 1.3954 | Intergenic | MIR\|SINE\|MIR | -15293 | NM_144925 | 233833 | Mm.102305 | NM_144925 | ENSMUSG00000052707 | **Tnrc6a** | 2010321I05Rik\|3110054G10Rik\|AW557223\|CAGH26\|D130023A07Rik\|GW182\|Tnrc6 | trinucleotide repeat containing 6a | protein-coding |
| chr7_123120548_123121098 | chr7 | 123120549 | 123121098 | + | 1.05221 | Intergenic | Intergenic | -3062 | NM_144925 | 233833 | Mm.102305 | NM_144925 | ENSMUSG00000052707 | **Tnrc6a** | 2010321I05Rik\|3110054G10Rik\|AW557223\|CAGH26\|D130023A07Rik\|GW182\|Tnrc6 | trinucleotide repeat containing 6a | protein-coding |
| chr7_123120561_123120882 | chr7 | 123120562 | 123120882 | + | 0.94753 | Intergenic | L1ME2\|LINE\|L1 | -3163 | NM_144925 | 233833 | Mm.102305 | NM_144925 | ENSMUSG00000052707 | **Tnrc6a** | 2010321I05Rik\|3110054G10Rik\|AW557223\|CAGH26\|D130023A07Rik\|GW182\|Tnrc6 | trinucleotide repeat containing 6a | protein-coding |
| chr15_80744092_80744284 | chr15 | 80744093 | 80744284 | + | 0.698327 | intron (NM_144812, intron 1 of 24) | ID_B1\|SINE\|B4 | 32875 | NM_144812 | 213988 | Mm.131328 | NM_144812 | ENSMUSG00000047888 | **Tnrc6b** | 2700090M07Rik\|A730065C02Rik\|AI848765\|D230019K20Rik | trinucleotide repeat containing 6b | protein-coding |
| chr11_8688720_8689700 | chr11 | 8688721 | 8689700 | + | 0.819765 | Intergenic | Intergenic | -24675 | NM_001083587 | 319939 | Mm.337820 | NM_001083587 | ENSMUSG00000020422 | **Tns3** | BC023928\|F830010I22Rik\|TEM6\|Tens1 | tensin 3 | protein-coding |
| chr11_8685663_8686362 | chr11 | 8685664 | 8686362 | + | 0.64959 | Intergenic | RMER15\|LTR\|ERVL | -21478 | NM_001083587 | 319939 | Mm.337820 | NM_001083587 | ENSMUSG00000020422 | **Tns3** | BC023928\|F830010I22Rik\|TEM6\|Tens1 | tensin 3 | protein-coding |
| chr11_99098767_99099342 | chr11 | 99098768 | 99099342 | + | 1.52981 | Intergenic | Intergenic | -9749 | NM_172564 | 217169 | Mm.262067 | NM_172564 | ENSMUSG00000017607 | **Tns4** | 9930017A07Rik\|AA589547\|AU016405\|Cten | tensin 4 | protein-coding |
| chr17_34681026_34681544 | chr17 | 34681027 | 34681544 | + | 0.650988 | intron (NM_031176, intron 5 of 32) | intron (NM_031176, intron 5 of 32) | 10750 | NM_031176 | 81877 | Mm.290527 | NM_031176 | ENSMUSG00000033327 | **Tnxb** | Tn-mhc\|Tnx | tenascin XB | protein-coding |

| | | | | | | | | | | | | | | | | | |
|---|---|---|---|---|---|---|---|---|---|---|---|---|---|---|---|---|---|
| chr11_94141498_94141965 | chr11 | 94141499 | 94141965 | + | 0.857809 | Intergenic | L2a\|LINE\|L2 | -69722 | NM_009427 | 22057 | Mm.259673 | NM_009427 | ENSMUSG00000037573 | **Tob1** | Tob\|Trob | transducer of ErbB-2.1 | protein-coding |
| chr11_94160372_94160969 | chr11 | 94160373 | 94160969 | + | 0.650073 | Intergenic | Intergenic | -50783 | NM_009427 | 22057 | Mm.259673 | NM_009427 | ENSMUSG00000037573 | **Tob1** | Tob\|Trob | transducer of ErbB-2.1 | protein-coding |
| chr11_90675540_90676031 | chr11 | 90675541 | 90676031 | + | 0.750773 | intron (NM_028011, intron 3 of 15) | intron (NM_028011, intron 3 of 15) | 11815 | NM_028011 | 71943 | Mm.167868 | NM_028011 | ENSMUSG00000020541 | **Tom1l1** | 2310045L10Rik\|C80573\|Srcasm | target of myb1-like 1 (chicken) | protein-coding |
| chr11_60290814_60291456 | chr11 | 60290815 | 60291456 | + | 0.780023 | intron (NM_001039092, intron 1 of 13) | intron (NM_001039092, intron 1 of 13) | 61770 | NM_153080 | 216810 | Mm.218875 | NM_153080 | ENSMUSG00000000538 | **Tom1l2** | 2900016I08Rik\|A730055F12Rik\|AU042072\|Srebf1 | target of myb1-like 2 (chicken) | protein-coding |
| chr8_126881649_126882179 | chr8 | 126881650 | 126882179 | + | 1.66192 | Intergenic | Intergenic | 64007 | NM_024214 | 67952 | Mm.6932 | NM_024214 | ENSMUSG00000093904 | **Tomm20** | 1810060K07Rik\|BB284719\|Gm19268\|MAS20\|MOM19\|TOM20\|mKIAA0016 | translocase of outer mitochondrial membrane 20 homolog (yeast) | protein-coding |
| chr8_126919168_126919689 | chr8 | 126919169 | 126919689 | + | 1.65672 | Intergenic | Intergenic | 26492 | NM_024214 | 67952 | Mm.6932 | NM_024214 | ENSMUSG00000093904 | **Tomm20** | 1810060K07Rik\|BB284719\|Gm19268\|MAS20\|MOM19\|TOM20\|mKIAA0016 | translocase of outer mitochondrial membrane 20 homolog (yeast) | protein-coding |
| chr8_126854397_126854745 | chr8 | 126854398 | 126854745 | + | 1.44089 | Intergenic | Intergenic | 91350 | NM_024214 | 67952 | Mm.6932 | NM_024214 | ENSMUSG00000093904 | **Tomm20** | 1810060K07Rik\|BB284719\|Gm19268\|MAS20\|MOM19\|TOM20\|mKIAA0016 | translocase of outer mitochondrial membrane 20 homolog (yeast) | protein-coding |
| chr8_126894947_126895792 | chr8 | 126894948 | 126895792 | + | 1.38568 | Intergenic | Intergenic | 50551 | NM_024214 | 67952 | Mm.6932 | NM_024214 | ENSMUSG00000093904 | **Tomm20** | 1810060K07Rik\|BB284719\|Gm19268\|MAS20\|MOM19\|TOM20\|mKIAA0016 | translocase of outer mitochondrial membrane 20 homolog (yeast) | protein-coding |
| chr8_126896380_126896783 | chr8 | 126896381 | 126896783 | + | 1.29143 | Intergenic | ORR1E\|LTR\|ERVL-MaLR | 49339 | NM_024214 | 67952 | Mm.6932 | NM_024214 | ENSMUSG00000093904 | **Tomm20** | 1810060K07Rik\|BB284719\|Gm19268\|MAS20\|MOM19\|TOM20\|mKIAA0016 | translocase of outer mitochondrial membrane 20 homolog (yeast) | protein-coding |
| chr8_126875500_126876216 | chr8 | 126875501 | 126876216 | + | 1.05944 | Intergenic | Intergenic | 70063 | NM_024214 | 67952 | Mm.6932 | NM_024214 | ENSMUSG00000093904 | **Tomm20** | 1810060K07Rik\|BB284719\|Gm19268\|MAS20\|MOM19\|TOM20\|mKIAA0016 | translocase of outer mitochondrial membrane 20 homolog (yeast) | protein-coding |
| chr8_126833742_126834276 | chr8 | 126833743 | 126834276 | + | 0.979194 | Intergenic | Intergenic | 111912 | NM_024214 | 67952 | Mm.6932 | NM_024214 | ENSMUSG00000093904 | **Tomm20** | 1810060K07Rik\|BB284719\|Gm19268\|MAS20\|MOM19\|TOM20\|mKIAA0016 | translocase of outer mitochondrial membrane 20 homolog (yeast) | protein-coding |
| chr8_126852207_126852539 | chr8 | 126852208 | 126852539 | + | 0.923111 | Intergenic | Intergenic | 93548 | NM_024214 | 67952 | Mm.6932 | NM_024214 | ENSMUSG00000093904 | **Tomm20** | 1810060K07Rik\|BB284719\|Gm19268\|MAS20\|MOM19\|TOM20\|mKIAA0016 | translocase of outer mitochondrial membrane 20 homolog (yeast) | protein-coding |
| chr8_126834800_126835059 | chr8 | 126834801 | 126835059 | + | 0.838145 | Intergenic | Intergenic | 110991 | NM_024214 | 67952 | Mm.6932 | NM_024214 | ENSMUSG00000093904 | **Tomm20** | 1810060K07Rik\|BB284719\|Gm19268\|MAS20\|MOM19\|TOM20\|mKIAA0016 | translocase of outer mitochondrial membrane 20 homolog (yeast) | protein-coding |
| chr8_126854808_126855022 | chr8 | 126854809 | 126855022 | + | 0.829252 | Intergenic | Intergenic | 91006 | NM_024214 | 67952 | Mm.6932 | NM_024214 | ENSMUSG00000093904 | **Tomm20** | 1810060K07Rik\|BB284719\|Gm19268\|MAS20\|MOM19\|TOM20\|mKIAA0016 | translocase of outer mitochondrial membrane 20 homolog (yeast) | protein-coding |
| chr8_126849599_126849839 | chr8 | 126849600 | 126849839 | + | 0.802811 | Intergenic | AmnSINE1\|SINE\|Deu | 96202 | NM_024214 | 67952 | Mm.6932 | NM_024214 | ENSMUSG00000093904 | **Tomm20** | 1810060K07Rik\|BB284719\|Gm19268\|MAS20\|MOM19\|TOM20\|mKIAA0016 | translocase of outer mitochondrial membrane 20 homolog (yeast) | protein-coding |
| chr8_126848462_126848957 | chr8 | 126848463 | 126848957 | + | 0.628782 | Intergenic | Intergenic | 97211 | NM_024214 | 67952 | Mm.6932 | NM_024214 | ENSMUSG00000093904 | **Tomm20** | 1810060K07Rik\|BB284719\|Gm19268\|MAS20\|MOM19\|TOM20\|mKIAA0016 | translocase of outer mitochondrial membrane 20 homolog (yeast) | protein-coding |
| chr4_45106338_45106653 | chr4 | 45106339 | 45106653 | + | 0.878678 | intron (NM_001099675, intron 1 of 1) | intron (NM_001099675, intron 1 of 1) | 1617 | NM_001134646 | 68512 | Mm.282081 | NM_001099675 | ENSMUSG00000078713 | **Tomm5** | 1110019J04Rik\|Tom5 | translocase of outer mitochondrial membrane 5 homolog (yeast) | protein-coding |
| chr16_57130747_57131252 | chr16 | 57130748 | 57131252 | + | 0.677915 | intron (NM_138599, intron 1 of 11) | B4\|SINE\|B4 | 9286 | NM_138599 | 28185 | Mm.213292 | NM_138599 | ENSMUSG00000022752 | **Tomm70a** | 2610044B22Rik\|D16Ium22\|D16Ium22e\|D16Wsu109e\|Tom70\|mKIAA0719 | translocase of outer mitochondrial membrane 70 homolog A (yeast) | protein-coding |
| chr2_160620335_160620905 | chr2 | 160620336 | 160620905 | + | 1.81707 | Intergenic | Intergenic | -25277 | NM_009408 | 21969 | Mm.217233 | NM_009408 | ENSMUSG00000070544 | **Top1** | AI467334\|D130064I21Rik\|Top-1 | topoisomerase (DNA) I | protein-coding |
| chr2_160620182_160621414 | chr2 | 160620183 | 160621414 | + | 1.47284 | Intergenic | Intergenic | -25099 | NM_009408 | 21969 | Mm.217233 | NM_009408 | ENSMUSG00000070544 | **Top1** | AI467334\|D130064I21Rik\|Top-1 | topoisomerase (DNA) I | protein-coding |
| chr14_16447564_16447958 | chr14 | 16447565 | 16447958 | + | 0.723024 | intron (NM_001289761, intron 4 of 8) | intron (NM_001289761, intron 4 of 8) | 82555 | NM_009409 | 21974 | Mm.130362 | NM_009409 | ENSMUSG00000017485 | **Top2b** | D230016L12Rik\|Top-2 | topoisomerase (DNA) II beta | protein-coding |
| chr14_16433253_16433622 | chr14 | 16433254 | 16433622 | + | 0.721996 | intron (NM_001289761, intron 8 of 8) | intron (NM_001289761, intron 8 of 8) | 68232 | NM_009409 | 21974 | Mm.130362 | NM_009409 | ENSMUSG00000017485 | **Top2b** | D230016L12Rik\|Top-2 | topoisomerase (DNA) II beta | protein-coding |
| chr14_16432658_16433025 | chr14 | 16432659 | 16433025 | + | 0.641144 | intron (NM_001289761, intron 8 of 8) | intron (NM_001289761, intron 8 of 8) | 67636 | NM_009409 | 21974 | Mm.130362 | NM_009409 | ENSMUSG00000017485 | **Top2b** | D230016L12Rik\|Top-2 | topoisomerase (DNA) II beta | protein-coding |
| chr1_156008843_156009407 | chr1 | 156008844 | 156009407 | + | 1.32888 | intron (NM_144791, intron 6 of 6) | PB1D9\|SINE\|Alu | 25692 | NM_001160019 | 208263 | Mm.211654 | NM_144791 | ENSMUSG00000026466 | **Tor1aip1** | LAP1\|Lap1B | torsin A interacting protein 1 | protein-coding |
| chr1_156010955_156011339 | chr1 | 156010956 | 156011339 | + | 1.06037 | intron (NM_144791, intron 6 of 6) | Charlie1a\|DNA\|hAT-Charlie | 23670 | NM_001160019 | 208263 | Mm.211654 | NM_144791 | ENSMUSG00000026466 | **Tor1aip1** | LAP1\|Lap1B | torsin A interacting protein 1 | protein-coding |
| chr1_156066017_156066662 | chr1 | 156066018 | 156066662 | + | 1.18224 | 3' UTR (NM_172843, exon 7 of 7) | 3' UTR (NM_172843, exon 7 of 7) | 25479 | NM_022329 | 240832 | Mm.253335 | NM_022329 | ENSMUSG00000050565 | **Tor1aip2** | 1110020D10Rik\|15kDa\|A130072J07\|AA103493\|AW060462\|AW610675\|C77739\|Ifrg15\|Lull1 | torsin A interacting protein 2 | protein-coding |
| chr1_156107376_156107763 | chr1 | 156107377 | 156107763 | + | 0.706748 | intron (NM_177838, intron 1 of 4) | B4A\|SINE\|B4 | 66709 | NM_022329 | 240832 | Mm.253335 | NM_022329 | ENSMUSG00000050565 | **Tor1aip2** | 1110020D10Rik\|15kDa\|A130072J07\|AA103493\|AW060462\|AW610675\|C77739\|Ifrg15\|Lull1 | torsin A interacting protein 2 | protein-coding |
| chr1_156667502_156667946 | chr1 | 156667503 | 156667946 | + | 0.842746 | intron (NM_023141, intron 3 of 5) | intron (NM_023141, intron 3 of 5) | 6615 | NM_023141 | 30935 | Mm.206737 | NM_023141 | ENSMUSG00000060519 | **Tor3a** | Adir | torsin family 3, member A | protein-coding |
| chr1_156667182_156667972 | chr1 | 156667183 | 156667972 | + | 0.840916 | intron (NM_023141, intron 3 of 5) | intron (NM_023141, intron 3 of 5) | 6762 | NM_023141 | 30935 | Mm.206737 | NM_023141 | ENSMUSG00000060519 | **Tor3a** | Adir | torsin family 3, member A | protein-coding |
| chr4_6940812_6941451 | chr4 | 6940813 | 6941451 | + | 0.954314 | intron (NM_145711, intron 1 of 8) | intron (NM_145711, intron 1 of 8) | 49591 | NM_145711 | 252838 | Mm.435828 | NM_145711 | ENSMUSG00000041272 | **Tox** | 1700007F02Rik | thymocyte selection-associated high mobility group box | protein-coding |
| chr4_6959963_6960339 | chr4 | 6959964 | 6960339 | + | 0.942656 | intron (NM_145711, intron 1 of 8) | intron (NM_145711, intron 1 of 8) | 30572 | NM_145711 | 252838 | Mm.435828 | NM_145711 | ENSMUSG00000041272 | **Tox** | 1700007F02Rik | thymocyte selection-associated high mobility group box | protein-coding |
| chr4_6883248_6883580 | chr4 | 6883249 | 6883580 | + | 0.876011 | intron (NM_145711, intron 1 of 8) | intron (NM_145711, intron 1 of 8) | 107309 | NM_145711 | 252838 | Mm.435828 | NM_145711 | ENSMUSG00000041272 | **Tox** | 1700007F02Rik | thymocyte selection-associated high mobility group box | protein-coding |
| chr4_7218176_7218804 | chr4 | 7218177 | 7218804 | + | 0.858688 | Intergenic | Intergenic | -227767 | NM_145711 | 252838 | Mm.435828 | NM_145711 | ENSMUSG00000041272 | **Tox** | 1700007F02Rik | thymocyte selection-associated high mobility group box | protein-coding |
| chr4_6914937_6915290 | chr4 | 6914938 | 6915290 | + | 0.803685 | intron (NM_145711, intron 1 of 8) | intron (NM_145711, intron 1 of 8) | 75609 | NM_145711 | 252838 | Mm.435828 | NM_145711 | ENSMUSG00000041272 | **Tox** | 1700007F02Rik | thymocyte selection-associated high mobility group box | protein-coding |
| chr4_7217934_7219019 | chr4 | 7217935 | 7219019 | + | 0.797714 | Intergenic | Intergenic | -227754 | NM_145711 | 252838 | Mm.435828 | NM_145711 | ENSMUSG00000041272 | **Tox** | 1700007F02Rik | thymocyte selection-associated high mobility group box | protein-coding |
| chr4_6957951_6958931 | chr4 | 6957952 | 6958931 | + | 0.761233 | intron (NM_145711, intron 1 of 8) | intron (NM_145711, intron 1 of 8) | 32282 | NM_145711 | 252838 | Mm.435828 | NM_145711 | ENSMUSG00000041272 | **Tox** | 1700007F02Rik | thymocyte selection-associated high mobility group box | protein-coding |
| chr4_6895877_6896335 | chr4 | 6895878 | 6896335 | + | 0.669659 | intron (NM_145711, intron 1 of 8) | intron (NM_145711, intron 1 of 8) | 94617 | NM_145711 | 252838 | Mm.435828 | NM_145711 | ENSMUSG00000041272 | **Tox** | 1700007F02Rik | thymocyte selection-associated high mobility group box | protein-coding |
| chr4_7112181_7112927 | chr4 | 7112182 | 7112927 | + | 0.66594 | Intergenic | Intergenic | -121831 | NM_145711 | 252838 | Mm.435828 | NM_145711 | ENSMUSG00000041272 | **Tox** | 1700007F02Rik | thymocyte selection-associated high mobility group box | protein-coding |
| chr2_163176332_163176887 | chr2 | 163176333 | 163176887 | + | 0.875997 | Intergenic | MTD-int\|LTR\|ERVL-MaLR | -48844 | NM_001098799 | 269389 | Mm.207709 | NM_001098799 | ENSMUSG00000074607 | **Tox2** | AI851523\|AV026525\|BI987407\|Gcx1\|RxHMG1 | TOX high mobility group box family member 2 | protein-coding |
| chr2_163198687_163198967 | chr2 | 163198688 | 163198967 | + | 0.732252 | Intergenic | L2b\|LINE\|L2 | -26627 | NM_001098799 | 269389 | Mm.207709 | NM_001098799 | ENSMUSG00000074607 | **Tox2** | AI851523\|AV026525\|BI987407\|Gcx1\|RxHMG1 | TOX high mobility group box family member 2 | protein-coding |
| chr5_120583509_120583846 | chr5 | 120583510 | 120583846 | + | 0.800166 | intron (NM_145853, intron 2 of 27) | intron (NM_145853, intron 2 of 27) | 4935 | NM_145853 | 252972 | Mm.114054 | NM_145853 | ENSMUSG00000032741 | **Tpcn1** | 5730403B01Rik\|Tpc1\|mKIAA1169 | two pore channel 1 | protein-coding |
| chr3_8962593_8963149 | chr3 | 8962594 | 8963149 | + | 0.790439 | intron (NM_009412, intron 1 of 5) | intron (NM_009412, intron 1 of 5) | 1183 | NM_001025262 | 21985 | Mm.371590 | NM_009412 | ENSMUSG00000027506 | **Tpd52** | AI043038\|mD52 | tumor protein D52 | protein-coding |
| chr2_181498623_181498907 | chr2 | 181498624 | 181498907 | + | 1.00507 | intron (NR_111899, intron 1 of 5) | RMER15\|LTR\|ERVL | 1623 | NM_001291204 | 66314 | Mm.136648 | NM_025482 | | **Tpd52l2** | 2810411G23Rik\|AU016537\|C86069\|D54 | tumor protein D52-like 2 | protein-coding |

| | | | | | | | | | | | | | | | | | |
|---|---|---|---|---|---|---|---|---|---|---|---|---|---|---|---|---|---|
| chr6_43645105_43645574 | chr6 | 43645106 | 43645574 | + | 0.697485 | intron (NM_013861, intron 2 of 8) | intron (NM_013861, intron 2 of 8) | 20835 | NM_013861 | 29807 | Mm.320979 | NM_013861 | ENSMUSG00000029735 | **Tpk1** | - | thiamine pyrophosphokinase | protein-coding |
| chr4_43522193_43522621 | chr4 | 43522194 | 43522621 | + | 0.615156 | intron (NM_001277876, intron 2 of 8) | intron (NM_001277876, intron 2 of 8) | 1147 | NM_009416 | 22004 | Mm.646 | NM_009416 | | **Tpm2** | Tpm-2\|Trop-2 | tropomyosin 2, beta | protein-coding |
| chr1_43969319_43969698 | chr1 | 43969320 | 43969698 | + | 0.752672 | intron (NM_009418, intron 11 of 29) | intron (NM_009418, intron 11 of 29) | 35502 | NM_009418 | 22019 | Mm.401675 | NM_009418 | ENSMUSG00000041763 | **Tpp2** | TPP-2\|TppII | tripeptidyl peptidase II | protein-coding |
| chr16_25249954_25250811 | chr16 | 25249955 | 25250811 | + | 1.38819 | Intergenic | MTEb\|LTR\|ERVL-MaLR | -36434 | NM_175165 | 71338 | Mm.126851 | NM_175165 | ENSMUSG00000048399 | **Tprg** | 5430420C16Rik\|Tprg1 | transformation related protein 63 regulated | protein-coding |
| chr5_130097354_130098624 | chr5 | 130097355 | 130098624 | + | 1.09322 | intron (NM_001130476, intron 1 of 5) | intron (NM_001130476, intron 1 of 5) | 18619 | NM_001130476 | 22021 | Mm.332720 | NM_013837 | ENSMUSG00000034118 | **Tpst1** | R75054\|Tango13a | protein-tyrosine sulfotransferase 1 | protein-coding |
| chr5_130089208_130090090 | chr5 | 130089209 | 130090090 | + | 1.05619 | intron (NM_001130476, intron 1 of 5) | ID_B1\|SINE\|B4 | 10279 | NM_001130476 | 22021 | Mm.332720 | NM_013837 | ENSMUSG00000034118 | **Tpst1** | R75054\|Tango13a | protein-tyrosine sulfotransferase 1 | protein-coding |
| chr5_130097354_130098184 | chr5 | 130097355 | 130098184 | + | 1.05353 | intron (NM_001130476, intron 1 of 5) | L1M4\|LINE\|L1 | 18399 | NM_001130476 | 22021 | Mm.332720 | NM_013837 | ENSMUSG00000034118 | **Tpst1** | R75054\|Tango13a | protein-tyrosine sulfotransferase 1 | protein-coding |
| chr5_112295158_112295479 | chr5 | 112295159 | 112295479 | + | 0.769112 | intron (NM_009419, intron 1 of 6) | intron (NM_009419, intron 1 of 6) | 18612 | NM_009419 | 22022 | Mm.250631 | NM_009419 | ENSMUSG00000029344 | **Tpst2** | AI448750\|D5Ucla3\|Tango13b\|grm\|grt | protein-tyrosine sulfotransferase 2 | protein-coding |
| chr16_22241261_22242473 | chr16 | 22241262 | 22242473 | + | 0.70692 | Intergenic | Intergenic | 24062 | NM_009186 | 20462 | Mm.210352 | NM_009186 | | **Tra2b** | 5730405G21Rik\|D16Ertd266e\|SIG-41\|Sfrs10\|Silg41\|TRA2beta | transformer 2 beta homolog (Drosophila) | protein-coding |
| chr2_34954370_34954876 | chr2 | 34954371 | 34954876 | + | 0.785722 | intron (NM_009421, intron 6 of 10) | intron (NM_009421, intron 6 of 10) | 7149 | NM_009421 | 22029 | Mm.239514 | NM_009421 | ENSMUSG00000026875 | **Traf1** | 4732496E14Rik | TNF receptor-associated factor 1 | protein-coding |
| chr2_25537403_25537676 | chr2 | 25537404 | 25537676 | + | 0.830349 | intron (NM_009422, intron 2 of 10) | intron (NM_009422, intron 2 of 10) | 9400 | NM_001290413 | 22030 | Mm.3399 | NM_009422 | ENSMUSG00000026942 | **Traf2** | AI325259 | TNF receptor-associated factor 2 | protein-coding |
| chr12_111181839_111182368 | chr12 | 111181840 | 111182368 | + | 0.656836 | intron (NM_001286122, intron 1 of 9) | intron (NM_001286122, intron 1 of 9) | 15734 | NM_011632 | 22031 | Mm.27431 | NM_011632 | ENSMUSG00000021277 | **Traf3** | AI528849\|CAP-1\|CD40bp\|CRAF1\|LAP1\|T-BAM\|amn | TNF receptor-associated factor 3 | protein-coding |
| chr10_39505509_39506113 | chr10 | 39505510 | 39506113 | + | 1.02122 | intron (NM_008054, intron 3 of 13) | intron (NM_008054, intron 3 of 13) | -107123 | NM_134000 | 103213 | Mm.436686 | NM_134000 | ENSMUSG00000019842 | **Traf3ip2** | AI429613\|Act1\|CIKS | TRAF3 interacting protein 2 | protein-coding |
| chr10_39505497_39506394 | chr10 | 39505498 | 39506394 | + | 0.912276 | intron (NM_008054, intron 3 of 13) | intron (NM_008054, intron 3 of 13) | -106988 | NM_134000 | 103213 | Mm.436686 | NM_134000 | ENSMUSG00000019842 | **Traf3ip2** | AI429613\|Act1\|CIKS | TRAF3 interacting protein 2 | protein-coding |
| chr10_39506473_39506850 | chr10 | 39506474 | 39506850 | + | 0.68574 | intron (NM_008054, intron 3 of 13) | MLT1E3\|LTR\|ERVL-MaLR | -106272 | NM_134000 | 103213 | Mm.436686 | NM_134000 | ENSMUSG00000019842 | **Traf3ip2** | AI429613\|Act1\|CIKS | TRAF3 interacting protein 2 | protein-coding |
| chr10_39614813_39614991 | chr10 | 39614814 | 39614991 | + | 0.67819 | intron (NM_134000, intron 1 of 8) | intron (NM_134000, intron 1 of 8) | 1968 | NM_134000 | 103213 | Mm.436686 | NM_134000 | ENSMUSG00000019842 | **Traf3ip2** | AI429613\|Act1\|CIKS | TRAF3 interacting protein 2 | protein-coding |
| chr1_193213714_193214274 | chr1 | 193213715 | 193214274 | + | 0.879613 | Intergenic | Intergenic | -12448 | NM_153137 | 215243 | Mm.261259 | NM_153137 | ENSMUSG00000037318 | **Traf3ip3** | 6030423D04Rik\|AI115021\|T3jam | TRAF3 interacting protein 3 | protein-coding |
| chr1_192090926_192091517 | chr1 | 192090927 | 192091517 | + | 0.794303 | intron (NM_011633, intron 1 of 9) | intron (NM_011633, intron 1 of 9) | 1377 | NM_011633 | 22033 | Mm.380491 | NM_011633 | ENSMUSG00000026637 | **Traf5** | - | TNF receptor-associated factor 5 | protein-coding |
| chr1_192061637_192062188 | chr1 | 192061638 | 192062188 | + | 0.663762 | intron (NM_011633, intron 4 of 9) | intron (NM_011633, intron 4 of 9) | 30686 | NM_011633 | 22033 | Mm.380491 | NM_011633 | ENSMUSG00000026637 | **Traf5** | - | TNF receptor-associated factor 5 | protein-coding |
| chr2_101736451_101736735 | chr2 | 101736452 | 101736735 | + | 1.31574 | intron (NM_001083810, intron 6 of 8) | intron (NM_001083810, intron 6 of 8) | 58153 | NM_009424 | 22034 | Mm.292729 | NM_009424 | ENSMUSG00000027164 | **Traf6** | 2310003F17Rik\|AI851288\|C630032O20Rik | TNF receptor-associated factor 6 | protein-coding |
| chr2_101736439_101736875 | chr2 | 101736440 | 101736875 | + | 1.25314 | intron (NM_001083810, intron 6 of 8) | intron (NM_001083810, intron 6 of 8) | 58217 | NM_009424 | 22034 | Mm.292729 | NM_009424 | ENSMUSG00000027164 | **Traf6** | 2310003F17Rik\|AI851288\|C630032O20Rik | TNF receptor-associated factor 6 | protein-coding |
| chr2_101737654_101738037 | chr2 | 101737655 | 101738037 | + | 1.04293 | intron (NM_001083810, intron 6 of 8) | intron (NM_001083810, intron 6 of 8) | 59406 | NM_009424 | 22034 | Mm.292729 | NM_009424 | ENSMUSG00000027164 | **Traf6** | 2310003F17Rik\|AI851288\|C630032O20Rik | TNF receptor-associated factor 6 | protein-coding |
| chr2_101679156_101679732 | chr2 | 101679157 | 101679732 | + | 0.983908 | intron (NM_009424, intron 1 of 7) | intron (NM_009424, intron 1 of 7) | 1004 | NM_009424 | 22034 | Mm.292729 | NM_009424 | ENSMUSG00000027164 | **Traf6** | 2310003F17Rik\|AI851288\|C630032O20Rik | TNF receptor-associated factor 6 | protein-coding |
| chr5_121383963_121384175 | chr5 | 121383964 | 121384175 | + | 0.816912 | exon (NM_001163470, exon 2 of 12) | exon (NM_001163470, exon 2 of 12) | 1546 | NM_001163470 | 231712 | Mm.20488 | NM_172275 | ENSMUSG00000042726 | **Trafd1** | 1110008K06Rik\|Fln29 | TRAF type zinc finger domain containing 1 | protein-coding |
| chr1_13615901_13616406 | chr1 | 13615902 | 13616406 | + | 1.11534 | Intergenic | Tigger7\|DNA\|TcMar-Tigger | -26290 | NM_028173 | 72265 | Mm.28765 | NM_028173 | ENSMUSG00000025935 | **Tram1** | 1810049E02Rik\|TRAMP | translocating chain-associating membrane protein 1 | protein-coding |
| chr1_13572411_13572963 | chr1 | 13572412 | 13572963 | + | 0.848681 | intron (NM_028173, intron 6 of 10) | intron (NM_028173, intron 6 of 10) | 17177 | NM_028173 | 72265 | Mm.28765 | NM_028173 | ENSMUSG00000025935 | **Tram1** | 1810049E02Rik\|TRAMP | translocating chain-associating membrane protein 1 | protein-coding |
| chr1_13521431_13522125 | chr1 | 13521432 | 13522125 | + | 0.688406 | Intergenic | Intergenic | 68086 | NM_028173 | 72265 | Mm.28765 | NM_028173 | ENSMUSG00000025935 | **Tram1** | 1810049E02Rik\|TRAMP | translocating chain-associating membrane protein 1 | protein-coding |
| chr1_21064099_21064919 | chr1 | 21064100 | 21064919 | + | 1.16668 | intron (NM_177409, intron 1 of 10) | intron (NM_177409, intron 1 of 10) | 14716 | NM_177409 | 170829 | Mm.185680 | NM_177409 | | **Tram2** | C330003D03Rik | translocating chain-associating membrane protein 2 | protein-coding |
| chr9_111314247_111314592 | chr9 | 111314248 | 111314592 | + | 0.846379 | intron (NM_001164659, intron 1 of 21) | B3\|SINE\|B2 | 2681 | NM_001164659 | 320429 | Mm.270754 | NM_001164659 | ENSMUSG00000062296 | **Trank1** | A230061D21Rik\|C030048J01Rik\|Gm187\|Lba1 | tetratricopeptide repeat and ankyrin repeat containing 1 | protein-coding |
| chr10_78241290_78241523 | chr10 | 78241291 | 78241523 | + | 0.873678 | intron (NM_001081055, intron 1 of 22) | intron (NM_001081055, intron 1 of 22) | 3235 | NM_001081055 | 216131 | Mm.27539 | NM_001081055 | ENSMUSG00000000374 | **Trappc10** | B230307C21Rik\|Gm870\|Tmem1 | trafficking protein particle complex 10 | protein-coding |
| chr10_78227210_78227535 | chr10 | 78227211 | 78227535 | + | 0.691425 | intron (NM_001081055, intron 3 of 22) | intron (NM_001081055, intron 3 of 22) | 17269 | NM_001081055 | 216131 | Mm.27539 | NM_001081055 | ENSMUSG00000000374 | **Trappc10** | B230307C21Rik\|Gm870\|Tmem1 | trafficking protein particle complex 10 | protein-coding |
| chr4_126260826_126261094 | chr4 | 126260827 | 126261094 | + | 0.734184 | Intergenic | Intergenic | -1445 | NM_013718 | 27096 | Mm.8392 | NM_013718 | ENSMUSG00000028847 | **Trappc3** | 1110058K12Rik\|AI835942\|AU019197\|Bet3 | trafficking protein particle complex 3 | protein-coding |
| chr12_59039685_59041185 | chr12 | 59039686 | 59041185 | + | 0.746505 | Intergenic | U2\|snRNA\|snRNA | 21026 | NM_030057 | 78232 | Mm.290070 | NM_030057 | ENSMUSG00000020993 | **Trappc6b** | 5830498C14Rik\|C79212 | trafficking protein particle complex 6B | protein-coding |
| chr15_73006259_73006892 | chr15 | 73006260 | 73006892 | + | 1.28355 | intron (NM_001164641, intron 8 of 17) | intron (NM_001164641, intron 8 of 17) | 49236 | NM_029640 | 76510 | Mm.158902 | NM_029640 | ENSMUSG00000047921 | **Trappc9** | 1810044A24Rik\|2900005P22Rik\|4632408O18Rik\|Ibp\|Nibp\|TRS130\|mKIAA1882 | trafficking protein particle complex 9 | protein-coding |
| chr15_73006236_73007086 | chr15 | 73006237 | 73007086 | + | 1.1789 | intron (NM_001164641, intron 8 of 17) | intron (NM_001164641, intron 8 of 17) | 49151 | NM_029640 | 76510 | Mm.158902 | NM_029640 | ENSMUSG00000047921 | **Trappc9** | 1810044A24Rik\|2900005P22Rik\|4632408O18Rik\|Ibp\|Nibp\|TRS130\|mKIAA1882 | trafficking protein particle complex 9 | protein-coding |
| chr16_48677949_48678495 | chr16 | 48677950 | 48678495 | + | 1.42719 | Intergenic | Intergenic | 93734 | NM_198297 | 77647 | Mm.167298 | NM_198297 | ENSMUSG00000030775 | **Trat1** | C030046M14Rik\|Tcrim\|Trim | T cell receptor associated transmembrane adaptor 1 | protein-coding |
| chr16_48738406_48739098 | chr16 | 48738407 | 48739098 | + | 0.861245 | intron (NM_198297, intron 5 of 5) | intron (NM_198297, intron 5 of 5) | 33204 | NM_198297 | 77647 | Mm.167298 | NM_198297 | ENSMUSG00000030775 | **Trat1** | C030046M14Rik\|Tcrim\|Trim | T cell receptor associated transmembrane adaptor 1 | protein-coding |
| chr16_48739948_48740646 | chr16 | 48739949 | 48740646 | + | 0.853515 | intron (NM_198297, intron 4 of 5) | intron (NM_198297, intron 4 of 5) | 31659 | NM_198297 | 77647 | Mm.167298 | NM_198297 | ENSMUSG00000030775 | **Trat1** | C030046M14Rik\|Tcrim\|Trim | T cell receptor associated transmembrane adaptor 1 | protein-coding |
| chr16_48759024_48760163 | chr16 | 48759025 | 48760163 | + | 0.845277 | intron (NM_198297, intron 2 of 5) | intron (NM_198297, intron 2 of 5) | 12362 | NM_198297 | 77647 | Mm.167298 | NM_198297 | ENSMUSG00000030775 | **Trat1** | C030046M14Rik\|Tcrim\|Trim | T cell receptor associated transmembrane adaptor 1 | protein-coding |
| chr16_48716368_48716818 | chr16 | 48716369 | 48716818 | + | 0.845061 | Intergenic | ORR1F-int\|LTR\|ERVL-MaLR | 55363 | NM_198297 | 77647 | Mm.167298 | NM_198297 | ENSMUSG00000030775 | **Trat1** | C030046M14Rik\|Tcrim\|Trim | T cell receptor associated transmembrane adaptor 1 | protein-coding |
| chr16_48716140_48716841 | chr16 | 48716141 | 48716841 | + | 0.80631 | Intergenic | ORR1F-int\|LTR\|ERVL-MaLR | 55465 | NM_198297 | 77647 | Mm.167298 | NM_198297 | ENSMUSG00000030775 | **Trat1** | C030046M14Rik\|Tcrim\|Trim | T cell receptor associated transmembrane adaptor 1 | protein-coding |
| chr16_48783301_48784028 | chr16 | 48783302 | 48784028 | + | 0.792953 | Intergenic | RMER6D\|LTR\|ERVK | -11709 | NM_198297 | 77647 | Mm.167298 | NM_198297 | ENSMUSG00000030775 | **Trat1** | C030046M14Rik\|Tcrim\|Trim | T cell receptor associated transmembrane adaptor 1 | protein-coding |
| chr16_48784678_48784958 | chr16 | 48784679 | 48784958 | + | 0.792643 | Intergenic | L1MA4\|LINE\|L1 | -12862 | NM_198297 | 77647 | Mm.167298 | NM_198297 | ENSMUSG00000030775 | **Trat1** | C030046M14Rik\|Tcrim\|Trim | T cell receptor associated transmembrane adaptor 1 | protein-coding |
| chr16_48759736_48760152 | chr16 | 48759737 | 48760152 | + | 0.77094 | intron (NM_198297, intron 2 of 5) | intron (NM_198297, intron 2 of 5) | 12012 | NM_198297 | 77647 | Mm.167298 | NM_198297 | ENSMUSG00000030775 | **Trat1** | C030046M14Rik\|Tcrim\|Trim | T cell receptor associated transmembrane adaptor 1 | protein-coding |
| chr16_48783311_48783749 | chr16 | 48783312 | 48783749 | + | 0.741972 | Intergenic | RMER6D\|LTR\|ERVK | -11574 | NM_198297 | 77647 | Mm.167298 | NM_198297 | ENSMUSG00000030775 | **Trat1** | C030046M14Rik\|Tcrim\|Trim | T cell receptor associated transmembrane adaptor 1 | protein-coding |

| | | | | | | | | | | | | | | | | | |
|---|---|---|---|---|---|---|---|---|---|---|---|---|---|---|---|---|---|
| chr16_48759063_48759375 | chr16 | 48759064 | 48759375 | + | 0.719609 | intron (NM_198297, intron 2 of 5) | intron (NM_198297, intron 2 of 5) | 12737 | NM_198297 | 77647 | Mm.167298 | NM_198297 | ENSMUSG00000030775 | **Trat1** | C030046M14Rik\|Tcrim\|Trim | T cell receptor associated transmembrane adaptor 1 | protein-coding |
| chr17_48332393_48333407 | chr17 | 48332394 | 48333407 | + | 0.963345 | Intergenic | RLTR19-int\|LTR\|ERVK | -13501 | NM_031254 | 83433 | Mm.261623 | NM_031254 | ENSMUSG00000023992 | **Trem2** | TREM-2\|Trem2a\|Trem2b\|Trem2c | triggering receptor expressed on myeloid cells 2 | protein-coding |
| chr17_48332403_48332976 | chr17 | 48332404 | 48332976 | + | 0.90015 | Intergenic | RLTR19-int\|LTR\|ERVK | -13711 | NM_031254 | 83433 | Mm.261623 | NM_031254 | ENSMUSG00000023992 | **Trem2** | TREM-2\|Trem2a\|Trem2b\|Trem2c | triggering receptor expressed on myeloid cells 2 | protein-coding |
| chr17_48333108_48333405 | chr17 | 48333109 | 48333405 | + | 0.725948 | Intergenic | RLTR19-int\|LTR\|ERVK | -13144 | NM_031254 | 83433 | Mm.261623 | NM_031254 | ENSMUSG00000023992 | **Trem2** | TREM-2\|Trem2a\|Trem2b\|Trem2c | triggering receptor expressed on myeloid cells 2 | protein-coding |
| chr17_48298673_48299023 | chr17 | 48298674 | 48299023 | + | 0.864214 | Intergenic | RMER5\|LTR\|ERV1 | -1190 | NM_001033405 | 328833 | Mm.329104 | NM_001033405 | ENSMUSG00000071068 | **Treml2** | AW049306\|Gm750\|Tlt2 | triggering receptor expressed on myeloid cells-like 2 | protein-coding |
| chr17_48293412_48294396 | chr17 | 48293413 | 48294396 | + | 0.619191 | Intergenic | B3\|SINE\|B2 | -6134 | NM_001033405 | 328833 | Mm.329104 | NM_001033405 | ENSMUSG00000071068 | **Treml2** | AW049306\|Gm750\|Tlt2 | triggering receptor expressed on myeloid cells-like 2 | protein-coding |
| chr17_47129681_47130448 | chr17 | 47129682 | 47130448 | + | 1.35579 | Intergenic | Intergenic | -10877 | NM_001097623 | 224829 | Mm.260989 | NM_172622 | ENSMUSG00000064043 | **Trerf1** | 9430096I18Rik\|AI429294\|B830015H24\|RAPA\|Trep-132\|Trep132 | transcriptional regulating factor 1 | protein-coding |
| chr17_47234355_47234730 | chr17 | 47234356 | 47234730 | + | 1.30467 | intron (NM_001097623, intron 4 of 19) | intron (NM_001097623, intron 4 of 19) | 4313 | NM_172622 | 224829 | Mm.260989 | NM_172622 | ENSMUSG00000064043 | **Trerf1** | 9430096I18Rik\|AI429294\|B830015H24\|RAPA\|Trep-132\|Trep132 | transcriptional regulating factor 1 | protein-coding |
| chr17_47186166_47186767 | chr17 | 47186167 | 47186767 | + | 1.05464 | intron (NM_001097623, intron 3 of 19) | intron (NM_001097623, intron 3 of 19) | -43763 | NM_172622 | 224829 | Mm.260989 | NM_172622 | ENSMUSG00000064043 | **Trerf1** | 9430096I18Rik\|AI429294\|B830015H24\|RAPA\|Trep-132\|Trep132 | transcriptional regulating factor 1 | protein-coding |
| chr17_47234333_47235135 | chr17 | 47234334 | 47235135 | + | 1.00669 | intron (NM_001097623, intron 4 of 19) | intron (NM_001097623, intron 4 of 19) | 4504 | NM_172622 | 224829 | Mm.260989 | NM_172622 | ENSMUSG00000064043 | **Trerf1** | 9430096I18Rik\|AI429294\|B830015H24\|RAPA\|Trep-132\|Trep132 | transcriptional regulating factor 1 | protein-coding |
| chr17_47294011_47294346 | chr17 | 47294012 | 47294346 | + | 0.949232 | intron (NM_001097623, intron 5 of 19) | intron (NM_001097623, intron 5 of 19) | 63949 | NM_172622 | 224829 | Mm.260989 | NM_172622 | ENSMUSG00000064043 | **Trerf1** | 9430096I18Rik\|AI429294\|B830015H24\|RAPA\|Trep-132\|Trep132 | transcriptional regulating factor 1 | protein-coding |
| chr17_47234190_47235282 | chr17 | 47234191 | 47235282 | + | 0.868068 | intron (NM_001097623, intron 4 of 19) | intron (NM_001097623, intron 4 of 19) | 4506 | NM_172622 | 224829 | Mm.260989 | NM_172622 | ENSMUSG00000064043 | **Trerf1** | 9430096I18Rik\|AI429294\|B830015H24\|RAPA\|Trep-132\|Trep132 | transcriptional regulating factor 1 | protein-coding |
| chr17_47294008_47294632 | chr17 | 47294009 | 47294632 | + | 0.833118 | intron (NM_001097623, intron 5 of 19) | intron (NM_001097623, intron 5 of 19) | 64090 | NM_172622 | 224829 | Mm.260989 | NM_172622 | ENSMUSG00000064043 | **Trerf1** | 9430096I18Rik\|AI429294\|B830015H24\|RAPA\|Trep-132\|Trep132 | transcriptional regulating factor 1 | protein-coding |
| chr17_47171165_47171633 | chr17 | 47171166 | 47171633 | + | 0.815857 | intron (NM_001097623, intron 3 of 19) | intron (NM_001097623, intron 3 of 19) | 30457 | NM_001097623 | 224829 | Mm.260989 | NM_172622 | ENSMUSG00000064043 | **Trerf1** | 9430096I18Rik\|AI429294\|B830015H24\|RAPA\|Trep-132\|Trep132 | transcriptional regulating factor 1 | protein-coding |
| chr17_47225235_47225726 | chr17 | 47225236 | 47225726 | + | 0.795959 | intron (NM_001097623, intron 3 of 19) | intron (NM_001097623, intron 3 of 19) | -4749 | NM_172622 | 224829 | Mm.260989 | NM_172622 | ENSMUSG00000064043 | **Trerf1** | 9430096I18Rik\|AI429294\|B830015H24\|RAPA\|Trep-132\|Trep132 | transcriptional regulating factor 1 | protein-coding |
| chr17_47177513_47178239 | chr17 | 47177514 | 47178239 | + | 0.681998 | intron (NM_001097623, intron 3 of 19) | intron (NM_001097623, intron 3 of 19) | 36934 | NM_001097623 | 224829 | Mm.260989 | NM_172622 | ENSMUSG00000064043 | **Trerf1** | 9430096I18Rik\|AI429294\|B830015H24\|RAPA\|Trep-132\|Trep132 | transcriptional regulating factor 1 | protein-coding |
| chr17_47216532_47216904 | chr17 | 47216533 | 47216904 | + | 0.659684 | intron (NM_001097623, intron 3 of 19) | intron (NM_001097623, intron 3 of 19) | -13512 | NM_172622 | 224829 | Mm.260989 | NM_172622 | ENSMUSG00000064043 | **Trerf1** | 9430096I18Rik\|AI429294\|B830015H24\|RAPA\|Trep-132\|Trep132 | transcriptional regulating factor 1 | protein-coding |
| chr17_47189407_47190174 | chr17 | 47189408 | 47190174 | + | 0.624848 | intron (NM_001097623, intron 3 of 19) | intron (NM_001097623, intron 3 of 19) | -40439 | NM_172622 | 224829 | Mm.260989 | NM_172622 | ENSMUSG00000064043 | **Trerf1** | 9430096I18Rik\|AI429294\|B830015H24\|RAPA\|Trep-132\|Trep132 | transcriptional regulating factor 1 | protein-coding |
| chr8_122359260_122359798 | chr8 | 122359261 | 122359798 | + | 1.20333 | intron (NM_133202, intron 1 of 2) | B1F\|SINE\|Alu | 1217 | NM_133202 | 170732 | Mm.154808 | NM_133202 | ENSMUSG00000039079 | **Trhr2** | TRH-R2 | thyrotropin releasing hormone receptor 2 | protein-coding |
| chr8_122355669_122356482 | chr8 | 122355670 | 122356482 | + | 0.983083 | TTS (NM_133202) | TTS (NM_133202) | 4670 | NM_133202 | 170732 | Mm.154808 | NM_133202 | ENSMUSG00000039079 | **Trhr2** | TRH-R2 | thyrotropin releasing hormone receptor 2 | protein-coding |
| chr15_59585133_59585433 | chr15 | 59585134 | 59585433 | + | 1.39257 | intron (NM_026746, intron 4 of 6) | intron (NM_026746, intron 4 of 6) | -63371 | NM_144549 | 211770 | Mm.40298 | NM_144549 | ENSMUSG00000032501 | **Trib1** | A530090O15Rik\|TRB-1\|Trb1 | tribbles homolog 1 (Drosophila) | protein-coding |
| chr15_59540830_59541434 | chr15 | 59540831 | 59541434 | + | 1.17777 | intron (NM_026746, intron 4 of 6) | intron (NM_026746, intron 4 of 6) | -107522 | NM_144549 | 211770 | Mm.40298 | NM_144549 | ENSMUSG00000032501 | **Trib1** | A530090O15Rik\|TRB-1\|Trb1 | tribbles homolog 1 (Drosophila) | protein-coding |
| chr15_59585978_59586606 | chr15 | 59585979 | 59586606 | + | 1.13293 | intron (NM_026746, intron 4 of 6) | intron (NM_026746, intron 4 of 6) | -62362 | NM_144549 | 211770 | Mm.40298 | NM_144549 | ENSMUSG00000032501 | **Trib1** | A530090O15Rik\|TRB-1\|Trb1 | tribbles homolog 1 (Drosophila) | protein-coding |
| chr15_59719360_59719702 | chr15 | 59719361 | 59719702 | + | 1.04192 | Intergenic | Intergenic | 70877 | NM_144549 | 211770 | Mm.40298 | NM_144549 | ENSMUSG00000032501 | **Trib1** | A530090O15Rik\|TRB-1\|Trb1 | tribbles homolog 1 (Drosophila) | protein-coding |
| chr15_59707457_59708263 | chr15 | 59707458 | 59708263 | + | 1.00556 | Intergenic | Intergenic | 59206 | NM_144549 | 211770 | Mm.40298 | NM_144549 | ENSMUSG00000032501 | **Trib1** | A530090O15Rik\|TRB-1\|Trb1 | tribbles homolog 1 (Drosophila) | protein-coding |
| chr15_59661815_59662270 | chr15 | 59661816 | 59662270 | + | 0.964636 | Intergenic | URR1B\|DNA\|hAT-Charlie | 13389 | NM_144549 | 211770 | Mm.40298 | NM_144549 | ENSMUSG00000032501 | **Trib1** | A530090O15Rik\|TRB-1\|Trb1 | tribbles homolog 1 (Drosophila) | protein-coding |
| chr15_59593196_59593429 | chr15 | 59593197 | 59593429 | + | 0.736194 | intron (NM_026746, intron 6 of 6) | intron (NM_026746, intron 6 of 6) | -55341 | NM_144549 | 211770 | Mm.40298 | NM_144549 | ENSMUSG00000032501 | **Trib1** | A530090O15Rik\|TRB-1\|Trb1 | tribbles homolog 1 (Drosophila) | protein-coding |
| chr15_59669748_59670145 | chr15 | 59669749 | 59670145 | + | 0.717586 | Intergenic | Lx6\|LINE\|L1 | 21293 | NM_144549 | 211770 | Mm.40298 | NM_144549 | ENSMUSG00000032501 | **Trib1** | A530090O15Rik\|TRB-1\|Trb1 | tribbles homolog 1 (Drosophila) | protein-coding |
| chr15_59524079_59524401 | chr15 | 59524080 | 59524401 | + | 0.645768 | intron (NM_026746, intron 4 of 6) | intron (NM_026746, intron 4 of 6) | -124414 | NM_144549 | 211770 | Mm.40298 | NM_144549 | ENSMUSG00000032501 | **Trib1** | A530090O15Rik\|TRB-1\|Trb1 | tribbles homolog 1 (Drosophila) | protein-coding |
| chr12_15932049_15932439 | chr12 | 15932050 | 15932439 | + | 1.42736 | Intergenic | Intergenic | -115459 | NM_144551 | 217410 | Mm.491136 | NM_144551 | ENSMUSG00000020601 | **Trib2** | AW319517\|TRB-2\|TRB2 | tribbles homolog 2 (Drosophila) | protein-coding |
| chr12_15821096_15821379 | chr12 | 15821097 | 15821379 | + | 1.35408 | Intergenic | Intergenic | -4453 | NM_144551 | 217410 | Mm.491136 | NM_144551 | ENSMUSG00000020601 | **Trib2** | AW319517\|TRB-2\|TRB2 | tribbles homolog 2 (Drosophila) | protein-coding |
| chr12_16012055_16012417 | chr12 | 16012056 | 16012417 | + | 1.19062 | Intergenic | Intergenic | -195451 | NM_144551 | 217410 | Mm.491136 | NM_144551 | ENSMUSG00000020601 | **Trib2** | AW319517\|TRB-2\|TRB2 | tribbles homolog 2 (Drosophila) | protein-coding |
| chr12_15832441_15833179 | chr12 | 15832442 | 15833179 | + | 0.981402 | Intergenic | Intergenic | -16025 | NM_144551 | 217410 | Mm.491136 | NM_144551 | ENSMUSG00000020601 | **Trib2** | AW319517\|TRB-2\|TRB2 | tribbles homolog 2 (Drosophila) | protein-coding |
| chr12_15913053_15913355 | chr12 | 15913054 | 15913355 | + | 0.824707 | Intergenic | B4A\|SINE\|B4 | -96419 | NM_144551 | 217410 | Mm.491136 | NM_144551 | ENSMUSG00000020601 | **Trib2** | AW319517\|TRB-2\|TRB2 | tribbles homolog 2 (Drosophila) | protein-coding |
| chr12_16006657_16007026 | chr12 | 16006658 | 16007026 | + | 0.75865 | Intergenic | Intergenic | -190057 | NM_144551 | 217410 | Mm.491136 | NM_144551 | ENSMUSG00000020601 | **Trib2** | AW319517\|TRB-2\|TRB2 | tribbles homolog 2 (Drosophila) | protein-coding |
| chr12_15970254_15970808 | chr12 | 15970255 | 15970808 | + | 0.70813 | Intergenic | Intergenic | -153746 | NM_144551 | 217410 | Mm.491136 | NM_144551 | ENSMUSG00000020601 | **Trib2** | AW319517\|TRB-2\|TRB2 | tribbles homolog 2 (Drosophila) | protein-coding |
| chr12_15896483_15897361 | chr12 | 15896484 | 15897361 | + | 0.68436 | Intergenic | Intergenic | -80137 | NM_144551 | 217410 | Mm.491136 | NM_144551 | ENSMUSG00000020601 | **Trib2** | AW319517\|TRB-2\|TRB2 | tribbles homolog 2 (Drosophila) | protein-coding |
| chr12_16006337_16007188 | chr12 | 16006338 | 16007188 | + | 0.648328 | Intergenic | Intergenic | -189978 | NM_144551 | 217410 | Mm.491136 | NM_144551 | ENSMUSG00000020601 | **Trib2** | AW319517\|TRB-2\|TRB2 | tribbles homolog 2 (Drosophila) | protein-coding |
| chr3_84312247_84312881 | chr3 | 84312248 | 84312881 | + | 1.03636 | Intergenic | Intergenic | -7789 | NM_001271726 | 80890 | Mm.44876 | NM_030706 | ENSMUSG00000027993 | **Trim2** | mKIAA0517\|narf | tripartite motif-containing 2 | protein-coding |
| chr3_84253120_84253603 | chr3 | 84253121 | 84253603 | + | 0.997467 | intron (NM_001271726, intron 1 of 11) | intron (NM_001271726, intron 1 of 11) | 6417 | NM_001271727 | 80890 | Mm.44876 | NM_030706 | ENSMUSG00000027993 | **Trim2** | mKIAA0517\|narf | tripartite motif-containing 2 | protein-coding |
| chr3_84305340_84305704 | chr3 | 84305341 | 84305704 | + | 0.845261 | promoter-TSS (NM_001271726) | promoter-TSS (NM_001271726) | -747 | NM_001271726 | 80890 | Mm.44876 | NM_030706 | ENSMUSG00000027993 | **Trim2** | mKIAA0517\|narf | tripartite motif-containing 2 | protein-coding |
| chr6_37705592_37706154 | chr6 | 37705593 | 37706154 | + | 1.02215 | Intergenic | Intergenic | -164938 | NM_001272064 | 21848 | Mm.41063 | NM_145076 | | **Trim24** | A130082H20Rik\|AI447469\|D430004I05Rik\|TIF1\|TIF1-alpha\|TIF1alpha\|Tif1a | tripartite motif-containing 24 | protein-coding |
| chr6_37842849_37844553 | chr6 | 37842850 | 37844553 | + | 0.835432 | Intergenic | Intergenic | -27110 | NM_001272064 | 21848 | Mm.41063 | NM_145076 | | **Trim24** | A130082H20Rik\|AI447469\|D430004I05Rik\|TIF1\|TIF1-alpha\|TIF1alpha\|Tif1a | tripartite motif-containing 24 | protein-coding |
| chr6_37817742_37818092 | chr6 | 37817743 | 37818092 | + | 0.731817 | Intergenic | ORR1D1\|LTR\|ERVL-MaLR | -52894 | NM_001272064 | 21848 | Mm.41063 | NM_145076 | | **Trim24** | A130082H20Rik\|AI447469\|D430004I05Rik\|TIF1\|TIF1-alpha\|TIF1alpha\|Tif1a | tripartite motif-containing 24 | protein-coding |
| chr13_21172549_21172996 | chr13 | 21172550 | 21172996 | + | 0.825847 | Intergenic | Intergenic | -7158 | NM_009054 | 19720 | Mm.314056 | NM_009054 | ENSMUSG00000021326 | **Trim27** | AW538890\|Gm19403\|Rfp | tripartite motif-containing 27 | protein-coding |
| chr7_104462883_104463201 | chr7 | 104462884 | 104463201 | + | 0.740826 | intron (NM_009099, intron 1 of 7) | Lx4A\|LINE\|L1 | 2151 | NM_009099 | 20128 | Mm.295578 | NM_009099 | ENSMUSG00000030921 | **Trim30a** | Rpt-1\|Rpt1\|Trim30 | tripartite motif-containing 30A | protein-coding |
| chr3_103375585_103376086 | chr3 | 103375586 | 103376086 | + | 1.03462 | intron (NR_106164, intron 1 of 4) | intron (NR_106164, intron 1 of 4) | 96543 | NM_001079830 | 94093 | Mm.195036 | NM_053170 | ENSMUSG00000033014 | **Trim33** | 8030451N04Rik\|AI413936\|Ecto\|Tif1g | tripartite motif-containing 33 | protein-coding |
| chr7_104240690_104241165 | chr7 | 104240691 | 104241165 | + | 0.79508 | Intergenic | L1MCa\|LINE\|L1 | -3529 | NM_030684 | 94094 | Mm.326945 | NM_030684 | ENSMUSG00000056144 | **Trim34a** | Trim34\|Trim34-1 | tripartite motif-containing 34A | protein-coding |
| chr18_46181686_46182664 | chr18 | 46181687 | 46182664 | + | 0.92598 | intron (NM_001170855, intron 4 of 9) | intron (NM_001170855, intron 4 of 9) | 16643 | NM_001170855 | 28105 | Mm.277382 | NM_178872 | ENSMUSG00000033949 | **Trim36** | D18Wsu100e\|haprin | tripartite motif-containing 36 | protein-coding |
| chr17_36274749_36275239 | chr17 | 36274750 | 36275239 | + | 0.783964 | Intergenic | Intergenic | -2990 | NM_024468 | 79263 | Mm.40624 | NM_024468 | ENSMUSG00000045409 | **Trim39** | 1100001D15Rik\|E130103K13Rik\|RBCC-B30.2\|Rnf23\|mKIAA4179\|tfp | tripartite motif-containing 39 | protein-coding |

| | | | | | | | | | | | | | | | | | |
|---|---|---|---|---|---|---|---|---|---|---|---|---|---|---|---|---|---|
| chr11_48820446_48820666 | chr11 | 48820447 | 48820666 | + | 0.931635 | Intergenic | RMER6A\|LTR\|ERVK | -3165 | NM_145377 | 211007 | Mm.794 | NM_145377 | ENSMUSG00000040365 | **Trim41** | AW552703\|BC020156\|R75223\|RINCK | tripartite motif-containing 41 | protein-coding |
| chr2_102378944_102379303 | chr2 | 102378945 | 102379303 | + | 1.25401 | intron (NM_020267, intron 2 of 4) | intron (NM_020267, intron 2 of 4) | 21776 | NM_020267 | 80985 | Mm.315002 | NM_020267 | ENSMUSG00000027189 | **Trim44** | Dipb\|Mc7 | tripartite motif-containing 44 | protein-coding |
| chr2_102312947_102313274 | chr2 | 102312948 | 102313274 | + | 0.801353 | intron (NM_020267, intron 4 of 4) | intron (NM_020267, intron 4 of 4) | 87789 | NM_020267 | 80985 | Mm.315002 | NM_020267 | ENSMUSG00000027189 | **Trim44** | Dipb\|Mc7 | tripartite motif-containing 44 | protein-coding |
| chr2_102312871_102313430 | chr2 | 102312872 | 102313430 | + | 0.778686 | intron (NM_020267, intron 4 of 4) | MIR\|SINE\|MIR | 87749 | NM_020267 | 80985 | Mm.315002 | NM_020267 | ENSMUSG00000027189 | **Trim44** | Dipb\|Mc7 | tripartite motif-containing 44 | protein-coding |
| chr3_19620143_19620775 | chr3 | 19620144 | 19620775 | + | 1.09863 | Intergenic | MT2A\|LTR\|ERVL | -24001 | NM_001081281 | 381485 | Mm.35419 | NM_001081281 | ENSMUSG00000060913 | **Trim55** | D830041C10Rik\|Murf2\|Rnf29 | tripartite motif-containing 55 | protein-coding |
| chr3_19619514_19619978 | chr3 | 19619515 | 19619978 | + | 0.752135 | Intergenic | Intergenic | -24714 | NM_001081281 | 381485 | Mm.35419 | NM_001081281 | ENSMUSG00000060913 | **Trim55** | D830041C10Rik\|Murf2\|Rnf29 | tripartite motif-containing 55 | protein-coding |
| chr4_128848947_128849330 | chr4 | 128848948 | 128849330 | + | 0.79788 | Intergenic | L2\|LINE\|L2 | -35001 | NM_178110 | 67525 | Mm.30653 | NM_178110 | ENSMUSG00000041000 | **Trim62** | 6330414G21Rik\|AI450348 | tripartite motif-containing 62 | protein-coding |
| chr7_109484074_109484747 | chr7 | 109484075 | 109484747 | + | 0.924714 | intron (NM_181853, intron 3 of 19) | intron (NM_181853, intron 3 of 19) | 9327 | NM_001170913 | 330627 | Mm.183528 | NM_181853 | ENSMUSG00000031026 | **Trim66** | Kiaa0298-hp\|Tif1d | tripartite motif-containing 66 | protein-coding |
| chr7_109470658_109471065 | chr7 | 109470659 | 109471065 | + | 0.888519 | intron (NM_181853, intron 10 of 19) | intron (NM_181853, intron 10 of 19) | 22876 | NM_001170913 | 330627 | Mm.183528 | NM_181853 | ENSMUSG00000031026 | **Trim66** | Kiaa0298-hp\|Tif1d | tripartite motif-containing 66 | protein-coding |
| chr7_109484277_109484523 | chr7 | 109484278 | 109484523 | + | 0.837453 | intron (NM_181853, intron 3 of 19) | intron (NM_181853, intron 3 of 19) | 9338 | NM_001170913 | 330627 | Mm.183528 | NM_181853 | ENSMUSG00000031026 | **Trim66** | Kiaa0298-hp\|Tif1d | tripartite motif-containing 66 | protein-coding |
| chr7_109484268_109484719 | chr7 | 109484269 | 109484719 | + | 0.82508 | intron (NM_181853, intron 3 of 19) | intron (NM_181853, intron 3 of 19) | 9244 | NM_001170913 | 330627 | Mm.183528 | NM_181853 | ENSMUSG00000031026 | **Trim66** | Kiaa0298-hp\|Tif1d | tripartite motif-containing 66 | protein-coding |
| chr7_109513495_109513772 | chr7 | 109513496 | 109513772 | + | 0.712569 | Intergenic | Intergenic | -5500 | NM_181853 | 330627 | Mm.183528 | NM_181853 | ENSMUSG00000031026 | **Trim66** | Kiaa0298-hp\|Tif1d | tripartite motif-containing 66 | protein-coding |
| chr8_124794255_124794905 | chr8 | 124794256 | 124794905 | + | 0.909997 | exon (NM_198632, exon 1 of 10) | exon (NM_198632, exon 1 of 10) | 1561 | NM_198632 | 330863 | Mm.422769 | NM_198632 | ENSMUSG00000036913 | **Trim67** | D130049O21Rik\|TNL | tripartite motif-containing 67 | protein-coding |
| chr8_124792734_124793688 | chr8 | 124792735 | 124793688 | + | 0.778414 | 5' UTR (NM_198632, exon 1 of 10) | 5' UTR (NM_198632, exon 1 of 10) | 192 | NM_198632 | 330863 | Mm.422769 | NM_198632 | ENSMUSG00000036913 | **Trim67** | D130049O21Rik\|TNL | tripartite motif-containing 67 | protein-coding |
| chr2_122161537_122161900 | chr2 | 122161538 | 122161900 | + | 1.8872 | intron (NM_080510, intron 1 of 6) | intron (NM_080510, intron 1 of 6) | 1019 | NM_080510 | 70928 | Mm.379188 | NM_080510 | ENSMUSG00000033368 | **Trim69** | 4921519C19Rik\|Rnf36\|Trif\|Trimless | tripartite motif-containing 69 | protein-coding |
| chr19_46522452_46523054 | chr19 | 46522453 | 46523054 | + | 0.790038 | Intergenic | Intergenic | 21105 | NM_053100 | 93679 | Mm.392177 | NM_053100 | ENSMUSG00000025034 | **Trim8** | AA408830\|BB006755\|GERP\|Rnf27 | tripartite motif-containing 8 | protein-coding |
| chr19_46522481_46522877 | chr19 | 46522482 | 46522877 | + | 0.73085 | Intergenic | L2a\|LINE\|L2 | 21031 | NM_053100 | 93679 | Mm.392177 | NM_053100 | ENSMUSG00000025034 | **Trim8** | AA408830\|BB006755\|GERP\|Rnf27 | tripartite motif-containing 8 | protein-coding |
| chr15_78962930_78963278 | chr15 | 78962931 | 78963278 | + | 0.662214 | intron (NM_001039156, intron 7 of 24) | intron (NM_001039156, intron 7 of 24) | 15380 | NM_001039155 | 110253 | Mm.123714 | NM_138579 | ENSMUSG00000033088 | **Triobp** | AI428493\|EST478828\|Tara\|mKIAA1662 | TRIO and F-actin binding protein | protein-coding |
| chr1_84791366_84791978 | chr1 | 84791367 | 84791978 | + | 1.43071 | intron (NM_133975, intron 3 of 40) | intron (NM_133975, intron 3 of 40) | 47632 | NM_133975 | 14897 | Mm.209265 | NM_133975 | ENSMUSG00000026219 | **Trip12** | 1110036I07Rik\|6720416K24Rik\|AA410158\|Gtl6\|TRIP-12 | thyroid hormone receptor interactor 12 | protein-coding |
| chr4_123042937_123043211 | chr4 | 123042938 | 123043211 | + | 0.670429 | intron (NM_025873, intron 2 of 10) | intron (NM_025873, intron 2 of 10) | 26477 | NM_025873 | 66966 | Mm.235030 | NM_025873 | ENSMUSG00000028653 | **Trit1** | 2310075G14Rik\|AI314189\|AI314635\|AV099619\|IPT\|MOD5 | tRNA isopentenyltransferase 1 | protein-coding |
| chr10_30535294_30536071 | chr10 | 30535295 | 30536071 | + | 1.92464 | intron (NM_028604, intron 12 of 12) | intron (NM_028604, intron 12 of 12) | 65066 | NM_028604 | 73681 | Mm.294503 | NM_028604 | ENSMUSG00000019792 | **Trmt11** | 2410075D05Rik\|3110045I18Rik\|AW213713 | tRNA methyltransferase 11 | protein-coding |
| chr10_30529204_30530255 | chr10 | 30529205 | 30530255 | + | 1.31297 | Intergenic | LTR40b\|LTR\|ERVL | 71019 | NM_028604 | 73681 | Mm.294503 | NM_028604 | ENSMUSG00000019792 | **Trmt11** | 2410075D05Rik\|3110045I18Rik\|AW213713 | tRNA methyltransferase 11 | protein-coding |
| chr10_30549999_30550456 | chr10 | 30550000 | 30550456 | + | 0.882012 | intron (NM_028604, intron 11 of 12) | L1MB7\|LINE\|L1 | 50521 | NM_028604 | 73681 | Mm.294503 | NM_028604 | ENSMUSG00000019792 | **Trmt11** | 2410075D05Rik\|3110045I18Rik\|AW213713 | tRNA methyltransferase 11 | protein-coding |
| chr15_58854251_58854846 | chr15 | 58854252 | 58854846 | + | 0.922563 | Intergenic | L1M5\|LINE\|L1 | -18100 | NM_026642 | 68260 | Mm.278805 | NM_026642 | ENSMUSG00000037085 | **Trmt12** | 4632406N01Rik\|AI449275\|Tyw2 | tRNA methyltranferase 12 | protein-coding |
| chr12_73210488_73210842 | chr12 | 73210489 | 73210842 | + | 0.722183 | intron (NM_008612, intron 6 of 7) | intron (NM_008612, intron 6 of 7) | 76046 | NM_029580 | 76357 | Mm.311344 | NM_029580 | ENSMUSG00000034442 | **Trmt5** | 2610027O18Rik\|mKIAA1393 | TRM5 tRNA methyltransferase 5 | protein-coding |
| chr12_111686297_111687006 | chr12 | 111686298 | 111687006 | + | 0.723986 | Intergenic | Intergenic | 8547 | NM_177374 | 328162 | Mm.425747 | NM_177374 | ENSMUSG00000060950 | **Trmt61a** | 6720458F09Rik\|AI606093\|Gcd14\|Trm61 | tRNA methyltransferase 61A | protein-coding |
| chr4_133501458_133501926 | chr4 | 133501459 | 133501926 | + | 1.10005 | Intergenic | Intergenic | -3142 | NM_001081156 | 69539 | Mm.151594 | NM_001081156 | ENSMUSG00000056596 | **Trnp1** | 2300002D11Rik\|Trnp | TMF1-regulated nuclear protein 1 | protein-coding |
| chr4_133501273_133501926 | chr4 | 133501274 | 133501926 | + | 0.959503 | Intergenic | L2b\|LINE\|L2 | -3050 | NM_001081156 | 69539 | Mm.151594 | NM_001081156 | ENSMUSG00000056596 | **Trnp1** | 2300002D11Rik\|Trnp | TMF1-regulated nuclear protein 1 | protein-coding |
| chr17_29078869_29079315 | chr17 | 29078870 | 29079315 | + | 1.09848 | promoter-TSS (NR_036469) | promoter-TSS (NR_036469) | 34 | NR_036469 | 100504267 | Mm.438550 | NR_036469 | ENSMUSG00000085912 | **Trp53cor1** | Gm16197\|OTTMUSG00000031656\|linc-p21\|lincRNA-p21 | tumor protein p53 pathway corepressor 1 | ncRNA |
| chr17_29078875_29079484 | chr17 | 29078876 | 29079484 | + | 1.00586 | promoter-TSS (NR_036469) | promoter-TSS (NR_036469) | -54 | NR_036469 | 100504267 | Mm.438550 | NR_036469 | ENSMUSG00000085912 | **Trp53cor1** | Gm16197\|OTTMUSG00000031656\|linc-p21\|lincRNA-p21 | tumor protein p53 pathway corepressor 1 | ncRNA |
| chr17_29080761_29081097 | chr17 | 29080762 | 29081097 | + | 0.907112 | Intergenic | Intergenic | -1803 | NR_036469 | 100504267 | Mm.438550 | NR_036469 | ENSMUSG00000085912 | **Trp53cor1** | Gm16197\|OTTMUSG00000031656\|linc-p21\|lincRNA-p21 | tumor protein p53 pathway corepressor 1 | ncRNA |
| chr17_29065702_29066411 | chr17 | 29065703 | 29066411 | + | 0.735616 | intron (NR_036469, intron 1 of 1) | B3A\|SINE\|B2 | 13069 | NR_036469 | 100504267 | Mm.438550 | NR_036469 | ENSMUSG00000085912 | **Trp53cor1** | Gm16197\|OTTMUSG00000031656\|linc-p21\|lincRNA-p21 | tumor protein p53 pathway corepressor 1 | ncRNA |
| chr17_29056310_29056689 | chr17 | 29056311 | 29056689 | + | 0.732449 | TTS (NR_036469) | TTS (NR_036469) | 22626 | NR_036469 | 100504267 | Mm.438550 | NR_036469 | ENSMUSG00000085912 | **Trp53cor1** | Gm16197\|OTTMUSG00000031656\|linc-p21\|lincRNA-p21 | tumor protein p53 pathway corepressor 1 | ncRNA |
| chr2_93213832_93214647 | chr2 | 93213833 | 93214647 | + | 0.61167 | intron (NM_183180, intron 5 of 9) | intron (NM_183180, intron 5 of 9) | 26656 | NM_001025246 | 277414 | Mm.41033 | NM_001025246 | ENSMUSG00000068735 | **Trp53i11** | Tp53i11 | transformation related protein 53 inducible protein 11 | protein-coding |
| chr2_166757861_166758504 | chr2 | 166757862 | 166758504 | + | 1.06521 | Intergenic | Intergenic | -35584 | NM_023815 | 76367 | Mm.330796 | NM_023815 | ENSMUSG00000042854 | **Trp53rk** | 2810408M09Rik\|4933401B08Rik\|5630401H01Rik\|Nori-2\|PRPK\|Tp53rk\|Trp53tk\|mNori-2p | transformation related protein 53 regulating kinase | protein-coding |
| chr16_25484991_25485660 | chr16 | 25484992 | 25485660 | + | 1.11769 | Intergenic | Intergenic | -198439 | NM_001127259 | 22061 | Mm.20894 | NM_011641 | ENSMUSG00000022510 | **Trp63** | AI462811\|Ket\|P51/P63\|P63\|P73l\|Tp63\|Trp53rp1 | transformation related protein 63 | protein-coding |
| chr16_25587349_25588199 | chr16 | 25587350 | 25588199 | + | 1.10153 | Intergenic | Intergenic | -95991 | NM_001127259 | 22061 | Mm.20894 | NM_011641 | ENSMUSG00000022510 | **Trp63** | AI462811\|Ket\|P51/P63\|P63\|P73l\|Tp63\|Trp53rp1 | transformation related protein 63 | protein-coding |
| chr4_154137146_154137466 | chr4 | 154137147 | 154137466 | + | 0.762682 | intron (NM_011642, intron 1 of 13) | intron (NM_011642, intron 1 of 13) | 2902 | NM_011642 | 22062 | Mm.78015 | NM_011642 | ENSMUSG00000029026 | **Trp73** | Tp73\|p73 | transformation related protein 73 | protein-coding |
| chr1_14924681_14925265 | chr1 | 14924682 | 14925265 | + | 0.98003 | Intergenic | ORR1E\|LTR\|ERVL-MaLR | -6111 | NM_177781 | 277328 | Mm.186329 | NM_177781 | ENSMUSG00000032769 | **Trpa1** | Anktm1\|TRPA1b | transient receptor potential cation channel, subfamily A, member 1 | protein-coding |
| chr10_77939571_77939989 | chr10 | 77939572 | 77939989 | + | 1.16578 | intron (NM_138301, intron 11 of 33) | RMER15\|LTR\|ERVL | 30092 | NM_138301 | 28240 | Mm.276762 | NM_138301 | ENSMUSG00000009292 | **Trpm2** | 9830168K16Rik\|C79133\|LTRPC2\|TRPC7\|Trp7\|Trrp7 | transient receptor potential cation channel, subfamily M, member 2 | protein-coding |
| chr10_77939041_77939990 | chr10 | 77939042 | 77939990 | + | 1.00041 | intron (NM_138301, intron 11 of 33) | RMER15\|LTR\|ERVL | 30356 | NM_138301 | 28240 | Mm.276762 | NM_138301 | ENSMUSG00000009292 | **Trpm2** | 9830168K16Rik\|C79133\|LTRPC2\|TRPC7\|Trp7\|Trrp7 | transient receptor potential cation channel, subfamily M, member 2 | protein-coding |
| chr10_77958107_77958965 | chr10 | 77958108 | 77958965 | + | 0.795648 | intron (NM_138301, intron 5 of 33) | ORR1B1\|LTR\|ERVL-MaLR | 11336 | NM_138301 | 28240 | Mm.276762 | NM_138301 | ENSMUSG00000009292 | **Trpm2** | 9830168K16Rik\|C79133\|LTRPC2\|TRPC7\|Trp7\|Trrp7 | transient receptor potential cation channel, subfamily M, member 2 | protein-coding |
| chr19_18746471_18746939 | chr19 | 18746472 | 18746939 | + | 1.50643 | Intergenic | Intergenic | -3278 | NM_153417 | 225997 | Mm.215171 | NM_153417 | ENSMUSG00000024727 | **Trpm6** | CHAK2\|TRPM6/CHAK2 | transient receptor potential cation channel, subfamily M, member 6 | protein-coding |
| chr19_18747288_18747786 | chr19 | 18747289 | 18747786 | + | 1.28412 | Intergenic | L2b\|LINE\|L2 | -2446 | NM_153417 | 225997 | Mm.215171 | NM_153417 | ENSMUSG00000024727 | **Trpm6** | CHAK2\|TRPM6/CHAK2 | transient receptor potential cation channel, subfamily M, member 6 | protein-coding |
| chr19_18766384_18766930 | chr19 | 18766385 | 18766930 | + | 1.06364 | intron (NM_153417, intron 1 of 38) | intron (NM_153417, intron 1 of 38) | 16674 | NM_153417 | 225997 | Mm.215171 | NM_153417 | ENSMUSG00000024727 | **Trpm6** | CHAK2\|TRPM6/CHAK2 | transient receptor potential cation channel, subfamily M, member 6 | protein-coding |
| chr2_126878570_126879066 | chr2 | 126878571 | 126879066 | + | 1.4335 | Intergenic | Intergenic | -2557 | NM_021450 | 58800 | Mm.244705 | NM_021450 | ENSMUSG00000027365 | **Trpm7** | 2310022G15Rik\|4833414K03Rik\|5033407O22Rik\|CHAK\|CHAK1\|LTrpC-7\|Ltpr7\|Ltrpc7\|TRPPLIK | transient receptor potential cation channel, subfamily M, member 7 | protein-coding |
| chr2_126847759_126848078 | chr2 | 126847760 | 126848078 | + | 0.82088 | intron (NM_001164325, intron 6 of 38) | intron (NM_001164325, intron 6 of 38) | 28342 | NM_001164325 | 58800 | Mm.244705 | NM_021450 | ENSMUSG00000027365 | **Trpm7** | 2310022G15Rik\|4833414K03Rik\|5033407O22Rik\|CHAK\|CHAK1\|LTrpC-7\|Ltpr7\|Ltrpc7\|TRPPLIK | transient receptor potential cation channel, subfamily M, member 7 | protein-coding |
| chr1_88345396_88346013 | chr1 | 88345397 | 88346013 | + | 0.729571 | intron (NM_134252, intron 12 of 26) | intron (NM_134252, intron 12 of 26) | 38993 | NM_134252 | 171382 | Mm.218753 | NM_134252 | ENSMUSG00000036251 | **Trpm8** | CMR1\|LTRPC6\|LTrpC-6\|TRPP8\|Trp-p8 | transient receptor potential cation channel, subfamily M, member 8 | protein-coding |
| chr15_51122499_51122861 | chr15 | 51122500 | 51122861 | + | 0.775749 | Intergenic | B3\|SINE\|B2 | -232639 | NM_032000 | 83925 | Mm.30466 | NM_032000 | ENSMUSG00000038679 | **Trps1** | AI115454\|AI447310\|D15Ertd586e | trichorhinophalangeal syndrome I (human) | protein-coding |
| chr15_50986970_50987426 | chr15 | 50986971 | 50987426 | + | 0.74907 | Intergenic | RMER6C\|LTR\|ERVK | -97157 | NM_032000 | 83925 | Mm.30466 | NM_032000 | ENSMUSG00000038679 | **Trps1** | AI115454\|AI447310\|D15Ertd586e | trichorhinophalangeal syndrome I (human) | protein-coding |
| chr6_41179683_41179976 | chr6 | 41179684 | 41179976 | + | 0.924243 | Intergenic | Intergenic | -122442 | NM_011646 | 22074 | Mm.383263 | NM_011646 | ENSMUSG00000054106 | **Try4** | 0910001B19Rik\|Td | trypsin 4 | protein-coding |
| chr6_41266470_41267300 | chr6 | 41266471 | 41267300 | + | 0.813696 | Intergenic | Intergenic | -35387 | NM_011646 | 22074 | Mm.383263 | NM_011646 | ENSMUSG00000054106 | **Try4** | 0910001B19Rik\|Td | trypsin 4 | protein-coding |
| chr6_41266798_41267295 | chr6 | 41266799 | 41267295 | + | 0.757154 | Intergenic | Intergenic | -35225 | NM_011646 | 22074 | Mm.383263 | NM_011646 | ENSMUSG00000054106 | **Try4** | 0910001B19Rik\|Td | trypsin 4 | protein-coding |
| chr6_41299365_41300081 | chr6 | 41299366 | 41300081 | + | 0.684186 | Intergenic | RMER1A\|Other\|Other | -2549 | NM_011646 | 22074 | Mm.383263 | NM_011646 | ENSMUSG00000054106 | **Try4** | 0910001B19Rik\|Td | trypsin 4 | protein-coding |
| chr17_24610679_24611058 | chr17 | 24610680 | 24611058 | + | 0.656317 | intron (NM_001039363, intron 20 of 39) | ORR1B2\|LTR\|ERVL-MaLR | 21758 | NM_011647 | 22084 | Mm.30435 | NM_011647 | ENSMUSG00000002496 | **Tsc2** | Nafld\|Tcs2 | tuberous sclerosis 2 | protein-coding |

| | | | | | | | | | | | | | | | | | |
|---|---|---|---|---|---|---|---|---|---|---|---|---|---|---|---|---|---|
| chr17_24610679_24611243 | chr17 | 24610680 | 24611243 | + | 0.639263 | intron (NM_001039363, intron 20 of 39) | ORR1B2\|LTR\|ERVL-MaLR | 21666 | NM_011647 | 22084 | Mm.30435 | NM_011647 | ENSMUSG00000002496 | **Tsc2** | Nafld\|Tcs2 | tuberous sclerosis 2 | protein-coding |
| chr14_76330013_76330419 | chr14 | 76330014 | 76330419 | + | 0.617184 | Intergenic | Intergenic | -85605 | NM_207652 | 21807 | Mm.153272 | NM_009366 | ENSMUSG00000022010 | **Tsc22d1** | AA589566\|AW105905\|Egr5\|TSC-22\|Tgfb1i4\|Tsc\|Tsc22 | TSC22 domain family, member 1 | protein-coding |
| chr3_58449509_58449858 | chr3 | 58449510 | 58449858 | + | 0.839233 | intron (NM_001081229, intron 2 of 3) | intron (NM_001081229, intron 2 of 3) | 33995 | NM_001081229 | 72033 | Mm.218409 | NM_028067 | ENSMUSG00000027806 | **Tsc22d2** | 1810043J12Rik\|5530402M19Rik | TSC22 domain family, member 2 | protein-coding |
| chrX_140552458_140552897 | chrX | 140552459 | 140552897 | + | 0.867663 | intron (NM_001077364, intron 2 of 3) | intron (NM_001077364, intron 2 of 3) | -9491 | NM_010286 | 14605 | Mm.22216 | NM_010286 | ENSMUSG00000031431 | **Tsc22d3** | DIP\|Dsip1\|Gilz\|TSC-22R\|Tilz3 | TSC22 domain family, member 3 | protein-coding |
| chr6_115501461_115501908 | chr6 | 115501462 | 115501908 | + | 0.679693 | Intergenic | L2a\|LINE\|L2 | -43019 | NM_199033 | 381802 | Mm.291208 | NM_199033 | ENSMUSG00000042389 | **Tsen2** | AU067695 | tRNA splicing endonuclease 2 homolog (S. cerevisiae) | protein-coding |
| chr7_46916472_46916679 | chr7 | 46916473 | 46916679 | + | 0.710374 | intron (NM_021884, intron 1 of 9) | MER44B\|DNA\|TcMar-Tigger | 3354 | NM_021884 | 22088 | Mm.241334 | NM_021884 | ENSMUSG00000014402 | **Tsg101** | AI255943\|CC2 | tumor susceptibility gene 101 | protein-coding |
| chr12_91424627_91424986 | chr12 | 91424628 | 91424986 | + | 0.687381 | intron (NM_011648, intron 1 of 9) | B3\|SINE\|B2 | 23814 | NM_011648 | 22095 | Mm.173847 | NM_011648 | ENSMUSG00000020963 | **Tshr** | AI481368\|hypothroid\|hyt\|pet | thyroid stimulating hormone receptor | protein-coding |
| chr2_169388659_169389301 | chr2 | 169388660 | 169389301 | + | 0.858091 | Intergenic | Intergenic | -244666 | NM_080455 | 228911 | Mm.315789 | NM_080455 | ENSMUSG00000047907 | **Tshz2** | 2900073F20Rik\|B830045G17\|Mtsh2\|Sdccag33l\|Tsh2\|Zfp218\|mKIAA4248\|teashirt2 | teashirt zinc finger family member 2 | protein-coding |
| chr2_169756271_169756742 | chr2 | 169756272 | 169756742 | + | 0.712937 | intron (NM_080455, intron 1 of 1) | intron (NM_080455, intron 1 of 1) | 122861 | NM_080455 | 228911 | Mm.315789 | NM_080455 | ENSMUSG00000047907 | **Tshz2** | 2900073F20Rik\|B830045G17\|Mtsh2\|Sdccag33l\|Tsh2\|Zfp218\|mKIAA4248\|teashirt2 | teashirt zinc finger family member 2 | protein-coding |
| chr2_169396021_169396712 | chr2 | 169396022 | 169396712 | + | 0.651728 | Intergenic | Intergenic | -237279 | NM_080455 | 228911 | Mm.315789 | NM_080455 | ENSMUSG00000047907 | **Tshz2** | 2900073F20Rik\|B830045G17\|Mtsh2\|Sdccag33l\|Tsh2\|Zfp218\|mKIAA4248\|teashirt2 | teashirt zinc finger family member 2 | protein-coding |
| chr1_118311885_118312416 | chr1 | 118311886 | 118312416 | + | 0.649194 | Intergenic | Intergenic | -1019 | NM_011650 | 22099 | Mm.426637 | NM_011650 | ENSMUSG00000026374 | **Tsn** | 2610034C24Rik\|AU040286\|C3PO\|TB-RBP | translin | protein-coding |
| chr12_36026169_36026501 | chr12 | 36026170 | 36026501 | + | 0.603098 | intron (NM_025359, intron 1 of 5) | intron (NM_025359, intron 1 of 5) | 16143 | NM_025359 | 66109 | Mm.254663 | NM_025359 | ENSMUSG00000020577 | **Tspan13** | 1100001I23Rik\|NET-6\|Tm4sf13 | tetraspanin 13 | protein-coding |
| chr14_40949760_40950028 | chr14 | 40949761 | 40950028 | + | 0.868318 | intron (NM_145928, intron 1 of 8) | L1MC4\|LINE\|L1 | 16913 | NM_145928 | 52588 | Mm.28876 | NM_145928 | ENSMUSG00000037824 | **Tspan14** | AA958793\|D14Ertd226e\|Tm4sf14 | tetraspanin 14 | protein-coding |
| chr14_40949714_40950569 | chr14 | 40949715 | 40950569 | + | 0.758635 | intron (NM_145928, intron 1 of 8) | intron (NM_145928, intron 1 of 8) | 16665 | NM_145928 | 52588 | Mm.28876 | NM_145928 | ENSMUSG00000037824 | **Tspan14** | AA958793\|D14Ertd226e\|Tm4sf14 | tetraspanin 14 | protein-coding |
| chr14_40938365_40938993 | chr14 | 40938366 | 40938993 | + | 0.671042 | intron (NM_145928, intron 1 of 8) | intron (NM_145928, intron 1 of 8) | 28128 | NM_145928 | 52588 | Mm.28876 | NM_145928 | ENSMUSG00000037824 | **Tspan14** | AA958793\|D14Ertd226e\|Tm4sf14 | tetraspanin 14 | protein-coding |
| chr3_102733051_102733687 | chr3 | 102733052 | 102733687 | + | 1.33401 | intron (NR_040588, intron 1 of 3) | intron (NR_040588, intron 1 of 3) | -1401 | NM_027533 | 70747 | Mm.27469 | NM_027533 | ENSMUSG00000027858 | **Tspan2** | 6330415F13Rik\|B230119D02Rik\|tspan-2 | tetraspanin 2 | protein-coding |
| chr3_102724815_102725557 | chr3 | 102724816 | 102725557 | + | 1.07165 | intron (NR_040588, intron 3 of 3) | intron (NR_040588, intron 3 of 3) | -9584 | NM_027533 | 70747 | Mm.27469 | NM_027533 | ENSMUSG00000027858 | **Tspan2** | 6330415F13Rik\|B230119D02Rik\|tspan-2 | tetraspanin 2 | protein-coding |
| chr3_102659414_102659681 | chr3 | 102659415 | 102659681 | + | 0.886813 | Intergenic | Intergenic | -75222 | NM_027533 | 70747 | Mm.27469 | NM_027533 | ENSMUSG00000027858 | **Tspan2** | 6330415F13Rik\|B230119D02Rik\|tspan-2 | tetraspanin 2 | protein-coding |
| chr3_102719717_102720039 | chr3 | 102719718 | 102720039 | + | 0.690258 | TTS (NR_040588) | TTS (NR_040588) | -14892 | NM_027533 | 70747 | Mm.27469 | NM_027533 | ENSMUSG00000027858 | **Tspan2** | 6330415F13Rik\|B230119D02Rik\|tspan-2 | tetraspanin 2 | protein-coding |
| chr3_102659314_102659774 | chr3 | 102659315 | 102659774 | + | 0.656007 | Intergenic | Intergenic | -75226 | NM_027533 | 70747 | Mm.27469 | NM_027533 | ENSMUSG00000027858 | **Tspan2** | 6330415F13Rik\|B230119D02Rik\|tspan-2 | tetraspanin 2 | protein-coding |
| chr3_102659352_102660047 | chr3 | 102659353 | 102660047 | + | 0.649092 | Intergenic | Intergenic | -75070 | NM_027533 | 70747 | Mm.27469 | NM_027533 | ENSMUSG00000027858 | **Tspan2** | 6330415F13Rik\|B230119D02Rik\|tspan-2 | tetraspanin 2 | protein-coding |
| chr9_56265567_56266116 | chr9 | 56265568 | 56266116 | + | 1.4045 | intron (NM_172924, intron 3 of 6) | intron (NM_172924, intron 3 of 6) | -104772 | NM_019793 | 56434 | Mm.28484 | NM_019793 | ENSMUSG00000032324 | **Tspan3** | 1700055K04Rik\|TM4-A\|Tm4sf8 | tetraspanin 3 | protein-coding |
| chr9_56196864_56197269 | chr9 | 56196865 | 56197269 | + | 1.25977 | Intergenic | MTC\|LTR\|ERVL-MaLR | -35997 | NM_019793 | 56434 | Mm.28484 | NM_019793 | ENSMUSG00000032324 | **Tspan3** | 1700055K04Rik\|TM4-A\|Tm4sf8 | tetraspanin 3 | protein-coding |
| chr9_56174719_56175202 | chr9 | 56174720 | 56175202 | + | 0.905599 | Intergenic | RSINE1\|SINE\|B4 | -13891 | NM_019793 | 56434 | Mm.28484 | NM_019793 | ENSMUSG00000032324 | **Tspan3** | 1700055K04Rik\|TM4-A\|Tm4sf8 | tetraspanin 3 | protein-coding |
| chr9_56133919_56134315 | chr9 | 56133920 | 56134315 | + | 0.788873 | Intergenic | L1MC4\|LINE\|L1 | 26953 | NM_019793 | 56434 | Mm.28484 | NM_019793 | ENSMUSG00000032324 | **Tspan3** | 1700055K04Rik\|TM4-A\|Tm4sf8 | tetraspanin 3 | protein-coding |
| chr9_56144428_56145073 | chr9 | 56144429 | 56145073 | + | 0.783151 | intron (NM_019793, intron 6 of 6) | ID2\|SINE\|ID | 16319 | NM_019793 | 56434 | Mm.28484 | NM_019793 | ENSMUSG00000032324 | **Tspan3** | 1700055K04Rik\|TM4-A\|Tm4sf8 | tetraspanin 3 | protein-coding |
| chr9_56260911_56261221 | chr9 | 56260912 | 56261221 | + | 0.727321 | intron (NM_172924, intron 3 of 6) | intron (NM_172924, intron 3 of 6) | -99996 | NM_019793 | 56434 | Mm.28484 | NM_019793 | ENSMUSG00000032324 | **Tspan3** | 1700055K04Rik\|TM4-A\|Tm4sf8 | tetraspanin 3 | protein-coding |
| chr9_56273364_56274053 | chr9 | 56273365 | 56274053 | + | 0.721277 | intron (NM_172924, intron 3 of 6) | intron (NM_172924, intron 3 of 6) | -112639 | NM_019793 | 56434 | Mm.28484 | NM_019793 | ENSMUSG00000032324 | **Tspan3** | 1700055K04Rik\|TM4-A\|Tm4sf8 | tetraspanin 3 | protein-coding |
| chr3_138864046_138864500 | chr3 | 138864047 | 138864500 | + | 1.07734 | intron (NR_121195, intron 1 of 7) | intron (NR_121195, intron 1 of 7) | 122065 | NM_019571 | 56224 | Mm.31927 | NM_019571 | ENSMUSG00000028152 | **Tspan5** | 2810455A09Rik\|4930505M03Rik\|NET-4\|Tm4sf9 | tetraspanin 5 | protein-coding |
| chr3_138841063_138841750 | chr3 | 138841064 | 138841750 | + | 0.981319 | intron (NR_121195, intron 1 of 7) | intron (NR_121195, intron 1 of 7) | 99199 | NM_019571 | 56224 | Mm.31927 | NM_019571 | ENSMUSG00000028152 | **Tspan5** | 2810455A09Rik\|4930505M03Rik\|NET-4\|Tm4sf9 | tetraspanin 5 | protein-coding |
| chr3_138893343_138893891 | chr3 | 138893344 | 138893891 | + | 0.941234 | intron (NR_121195, intron 3 of 7) | intron (NR_121195, intron 3 of 7) | 151409 | NM_019571 | 56224 | Mm.31927 | NM_019571 | ENSMUSG00000028152 | **Tspan5** | 2810455A09Rik\|4930505M03Rik\|NET-4\|Tm4sf9 | tetraspanin 5 | protein-coding |
| chr3_138893216_138893988 | chr3 | 138893217 | 138893988 | + | 0.929047 | intron (NR_121195, intron 3 of 7) | intron (NR_121195, intron 3 of 7) | 151394 | NM_019571 | 56224 | Mm.31927 | NM_019571 | ENSMUSG00000028152 | **Tspan5** | 2810455A09Rik\|4930505M03Rik\|NET-4\|Tm4sf9 | tetraspanin 5 | protein-coding |
| chr3_138746541_138746850 | chr3 | 138746542 | 138746850 | + | 0.905784 | intron (NR_121195, intron 1 of 7) | MTC\|LTR\|ERVL-MaLR | 4488 | NM_019571 | 56224 | Mm.31927 | NM_019571 | ENSMUSG00000028152 | **Tspan5** | 2810455A09Rik\|4930505M03Rik\|NET-4\|Tm4sf9 | tetraspanin 5 | protein-coding |
| chr3_138740450_138740625 | chr3 | 138740451 | 138740625 | + | 0.771667 | Intergenic | L1MC1\|LINE\|L1 | -1670 | NR_121195 | 56224 | Mm.31927 | NM_019571 | ENSMUSG00000028152 | **Tspan5** | 2810455A09Rik\|4930505M03Rik\|NET-4\|Tm4sf9 | tetraspanin 5 | protein-coding |
| chr3_138751987_138752431 | chr3 | 138751988 | 138752431 | + | 0.652006 | intron (NR_121195, intron 1 of 7) | ORR1F\|LTR\|ERVL-MaLR | 10001 | NM_019571 | 56224 | Mm.31927 | NM_019571 | ENSMUSG00000028152 | **Tspan5** | 2810455A09Rik\|4930505M03Rik\|NET-4\|Tm4sf9 | tetraspanin 5 | protein-coding |
| chr10_34307086_34307606 | chr10 | 34307087 | 34307606 | + | 0.876288 | intron (NM_176968, intron 11 of 11) | intron (NM_176968, intron 11 of 11) | 9925 | NM_030203 | 72480 | Mm.21485 | NM_030203 | ENSMUSG00000039485 | **Tspyl4** | 2610102M01Rik\|B230210I21Rik\|D10Bwg0791e | TSPY-like 4 | protein-coding |
| chr10_34306942_34307700 | chr10 | 34306943 | 34307700 | + | 0.712284 | intron (NM_176968, intron 11 of 11) | intron (NM_176968, intron 11 of 11) | 9900 | NM_030203 | 72480 | Mm.21485 | NM_030203 | ENSMUSG00000039485 | **Tspyl4** | 2610102M01Rik\|B230210I21Rik\|D10Bwg0791e | TSPY-like 4 | protein-coding |
| chr4_46095474_46095904 | chr4 | 46095475 | 46095904 | + | 0.771479 | intron (NM_021883, intron 7 of 9) | intron (NM_021883, intron 7 of 9) | 42786 | NM_173033 | 272027 | Mm.331588 | NM_173033 | ENSMUSG00000035495 | **Tstd2** | 3010020C06 | thiosulfate sulfurtransferase (rhodanese)-like domain containing 2 | protein-coding |
| chr4_46099767_46100303 | chr4 | 46099768 | 46100303 | + | 0.759583 | intron (NM_021883, intron 8 of 9) | intron (NM_021883, intron 8 of 9) | 38440 | NM_173033 | 272027 | Mm.331588 | NM_173033 | ENSMUSG00000035495 | **Tstd2** | 3010020C06 | thiosulfate sulfurtransferase (rhodanese)-like domain containing 2 | protein-coding |
| chr2_120823846_120824240 | chr2 | 120823847 | 120824240 | + | 0.684136 | intron (NM_001024857, intron 1 of 14) | MTC\|LTR\|ERVL-MaLR | 26375 | NM_001024857 | 140810 | Mm.275698 | NM_080788 | ENSMUSG00000090100 | **Ttbk2** | 2610507N02Rik\|AI326283\|B930008N24Rik\|TTK\|Ttbk\|Ttbk1\|mKIAA0847 | tau tubulin kinase 2 | protein-coding |
| chr11_43716997_43717915 | chr11 | 43716998 | 43717915 | + | 0.799787 | intron (NM_001164231, intron 3 of 3) | intron (NM_001164231, intron 3 of 3) | 30517 | NM_133795 | 66827 | Mm.271974 | NM_133795 | ENSMUSG00000041278 | **Ttc1** | 4833412C19Rik\|C79328\|TPR1 | tetratricopeptide repeat domain 1 | protein-coding |
| chr11_43728690_43729401 | chr11 | 43728691 | 43729401 | + | 0.771981 | TTS (NM_133795) | TTS (NM_133795) | 18927 | NM_133795 | 66827 | Mm.271974 | NM_133795 | ENSMUSG00000041278 | **Ttc1** | 4833412C19Rik\|C79328\|TPR1 | tetratricopeptide repeat domain 1 | protein-coding |
| chr2_32772950_32773274 | chr2 | 32772951 | 32773274 | + | 0.857443 | intron (NM_001290564, intron 5 of 14) | RSINE1\|SINE\|B4 | 2227 | NM_001290564 | 338348 | Mm.475130 | NM_177384 | ENSMUSG00000039021 | **Ttc16** | 1200002K10Rik | tetratricopeptide repeat domain 16 | protein-coding |
| chr2_32772945_32773401 | chr2 | 32772946 | 32773401 | + | 0.784821 | intron (NM_001290564, intron 5 of 14) | RSINE1\|SINE\|B4 | 2166 | NM_001290564 | 338348 | Mm.475130 | NM_177384 | ENSMUSG00000039021 | **Ttc16** | 1200002K10Rik | tetratricopeptide repeat domain 16 | protein-coding |
| chr2_94396415_94396822 | chr2 | 94396416 | 94396822 | + | 0.769776 | intron (NM_183106, intron 1 of 24) | intron (NM_183106, intron 1 of 24) | 10070 | NM_183106 | 74569 | Mm.248504 | NM_183106 | ENSMUSG00000027194 | **Ttc17** | 9130020K17Rik\|D2Bwg1005e | tetratricopeptide repeat domain 17 | protein-coding |
| chr6_38374294_38374943 | chr6 | 38374295 | 38374943 | + | 0.655376 | Intergenic | Intergenic | -6905 | NM_153600 | 264134 | Mm.253446 | NM_153600 | ENSMUSG00000056832 | **Ttc26** | 9330141E21Rik\|9430097H08Rik | tetratricopeptide repeat domain 26 | protein-coding |
| chr17_74667636_74668378 | chr17 | 74667637 | 74668378 | + | 1.50484 | intron (NM_007566, intron 62 of 72) | Lx5\|LINE\|L1 | -49743 | NM_152817 | 74196 | Mm.209503 | NM_152817 | ENSMUSG00000024078 | **Ttc27** | 2610511O17Rik | tetratricopeptide repeat domain 27 | protein-coding |
| chr17_74841706_74842109 | chr17 | 74841707 | 74842109 | + | 0.681435 | intron (NM_152817, intron 16 of 19) | MIR\|SINE\|MIR | 124158 | NM_152817 | 74196 | Mm.209503 | NM_152817 | ENSMUSG00000024078 | **Ttc27** | 2610511O17Rik | tetratricopeptide repeat domain 27 | protein-coding |
| chr8_78113083_78113480 | chr8 | 78113084 | 78113480 | + | 0.746051 | Intergenic | Intergenic | -100060 | NM_183096 | 73301 | Mm.61243 | NM_183096 | ENSMUSG00000037101 | **Ttc29** | 1700031F13Rik | tetratricopeptide repeat domain 29 | protein-coding |

| | | | | | | | | | | | | | | | | | |
|---|---|---|---|---|---|---|---|---|---|---|---|---|---|---|---|---|---|
| chr2_76031623_76032321 | chr2 | 76031624 | 76032321 | + | 1.14101 | intron (NM_001081033, intron 15 of 19) | intron (NM_001081033, intron 15 of 19) | -50005 | NM_030188 | 78802 | Mm.282339 | NM_030188 | ENSMUSG00000075271 | **Ttc30a1** | 4930506L13Rik | tetratricopeptide repeat domain 30A1 | protein-coding |
| chr2_75891872_75893005 | chr2 | 75891873 | 75893005 | + | 0.796022 | intron (NM_172666, intron 13 of 19) | intron (NM_172666, intron 13 of 19) | 46023 | NM_028235 | 72421 | Mm.275109 | NM_028235 | ENSMUSG00000075273 | **Ttc30b** | 2510042P03Rik\|Tcc30b | tetratricopeptide repeat domain 30B | protein-coding |
| chr2_75891891_75892725 | chr2 | 75891892 | 75892725 | + | 0.784716 | intron (NM_172666, intron 13 of 19) | intron (NM_172666, intron 13 of 19) | 46154 | NM_028235 | 72421 | Mm.275109 | NM_028235 | ENSMUSG00000075273 | **Ttc30b** | 2510042P03Rik\|Tcc30b | tetratricopeptide repeat domain 30B | protein-coding |
| chr2_75891922_75892383 | chr2 | 75891923 | 75892383 | + | 0.731298 | intron (NM_172666, intron 13 of 19) | intron (NM_172666, intron 13 of 19) | 46309 | NM_028235 | 72421 | Mm.275109 | NM_028235 | ENSMUSG00000075273 | **Ttc30b** | 2510042P03Rik\|Tcc30b | tetratricopeptide repeat domain 30B | protein-coding |
| chr2_75904863_75905174 | chr2 | 75904864 | 75905174 | + | 0.688114 | intron (NM_172666, intron 17 of 19) | RSINE1\|SINE\|B4 | 33443 | NM_028235 | 72421 | Mm.275109 | NM_028235 | ENSMUSG00000075273 | **Ttc30b** | 2510042P03Rik\|Tcc30b | tetratricopeptide repeat domain 30B | protein-coding |
| chr4_83223209_83223565 | chr4 | 83223210 | 83223565 | + | 1.85786 | 3' UTR (NM_027238, exon 20 of 20) | 3' UTR (NM_027238, exon 20 of 20) | 100802 | NM_027238 | 69863 | Mm.5540 | NM_025782 | ENSMUSG00000038172 | **Ttc39b** | 1810054D07Rik\|9130422G05Rik\|AI450603\|AV103077\|AW215585 | tetratricopeptide repeat domain 39B | protein-coding |
| chr4_83319701_83320112 | chr4 | 83319702 | 83320112 | + | 1.60204 | intron (NM_027238, intron 1 of 19) | intron (NM_027238, intron 1 of 19) | 4282 | NM_027238 | 69863 | Mm.5540 | NM_025782 | ENSMUSG00000038172 | **Ttc39b** | 1810054D07Rik\|9130422G05Rik\|AI450603\|AV103077\|AW215585 | tetratricopeptide repeat domain 39B | protein-coding |
| chr4_83321346_83321715 | chr4 | 83321347 | 83321715 | + | 1.19449 | intron (NM_027238, intron 1 of 19) | intron (NM_027238, intron 1 of 19) | 2658 | NM_027238 | 69863 | Mm.5540 | NM_025782 | ENSMUSG00000038172 | **Ttc39b** | 1810054D07Rik\|9130422G05Rik\|AI450603\|AV103077\|AW215585 | tetratricopeptide repeat domain 39B | protein-coding |
| chr4_83227489_83227937 | chr4 | 83227490 | 83227937 | + | 1.0908 | intron (NM_027238, intron 19 of 19) | intron (NM_027238, intron 19 of 19) | 96476 | NM_027238 | 69863 | Mm.5540 | NM_025782 | ENSMUSG00000038172 | **Ttc39b** | 1810054D07Rik\|9130422G05Rik\|AI450603\|AV103077\|AW215585 | tetratricopeptide repeat domain 39B | protein-coding |
| chr4_83327428_83327774 | chr4 | 83327429 | 83327774 | + | 0.971723 | Intergenic | Intergenic | -3412 | NM_027238 | 69863 | Mm.5540 | NM_025782 | ENSMUSG00000038172 | **Ttc39b** | 1810054D07Rik\|9130422G05Rik\|AI450603\|AV103077\|AW215585 | tetratricopeptide repeat domain 39B | protein-coding |
| chr4_83328045_83328291 | chr4 | 83328046 | 83328291 | + | 0.619642 | Intergenic | Intergenic | -3979 | NM_027238 | 69863 | Mm.5540 | NM_025782 | ENSMUSG00000038172 | **Ttc39b** | 1810054D07Rik\|9130422G05Rik\|AI450603\|AV103077\|AW215585 | tetratricopeptide repeat domain 39B | protein-coding |
| chr18_12584755_12585587 | chr18 | 12584756 | 12585587 | + | 1.56515 | Intergenic | Intergenic | -58362 | NM_028341 | 72747 | Mm.52526 | NM_028341 | ENSMUSG00000024424 | **Ttc39c** | 1700008N02Rik\|2810439F02Rik | tetratricopeptide repeat domain 39C | protein-coding |
| chr18_12640452_12641310 | chr18 | 12640453 | 12641310 | + | 1.38639 | Intergenic | Intergenic | -2652 | NM_028341 | 72747 | Mm.52526 | NM_028341 | ENSMUSG00000024424 | **Ttc39c** | 1700008N02Rik\|2810439F02Rik | tetratricopeptide repeat domain 39C | protein-coding |
| chr18_12611095_12611819 | chr18 | 12611096 | 12611819 | + | 0.996169 | Intergenic | MLTR73\|LTR\|ERV1 | -32076 | NM_028341 | 72747 | Mm.52526 | NM_028341 | ENSMUSG00000024424 | **Ttc39c** | 1700008N02Rik\|2810439F02Rik | tetratricopeptide repeat domain 39C | protein-coding |
| chr17_87302354_87302667 | chr17 | 87302355 | 87302667 | + | 0.611363 | intron (NM_028639, intron 3 of 19) | intron (NM_028639, intron 3 of 19) | 19625 | NM_028639 | 225049 | Mm.77396 | NM_028639 | ENSMUSG00000036918 | **Ttc7** | 1110035E02Rik\|1700007L07Rik\|Ttc7a\|fsn\|hea | tetratricopeptide repeat domain 7 | protein-coding |
| chr12_100420437_100421270 | chr12 | 100420438 | 100421270 | + | 0.727509 | intron (NM_001033213, intron 5 of 19) | intron (NM_001033213, intron 5 of 19) | 99968 | NM_001033213 | 104718 | Mm.295441 | NM_001033213 | ENSMUSG00000033530 | **Ttc7b** | AA408451\|Ttc7l1 | tetratricopeptide repeat domain 7B | protein-coding |
| chr7_4098854_4099206 | chr7 | 4098855 | 4099206 | + | 0.801481 | intron (NR_033568, intron 5 of 8) | ORR1B1\|LTR\|ERVL-MaLR | -20503 | NM_001109765 | 57776 | Mm.29729 | NM_021324 | ENSMUSG00000030428 | **Ttyh1** | 4930459B04Rik\|6330408P11Rik\|tty | tweety homolog 1 (Drosophila) | protein-coding |
| chr15_99030709_99030951 | chr15 | 99030710 | 99030951 | + | 0.702738 | intron (NM_009448, intron 1 of 3) | intron (NM_009448, intron 1 of 3) | 939 | NM_009448 | 22146 | Mm.88212 | NM_009448 | ENSMUSG00000043091 | **Tuba1c** | M[a]6\|Tuba6 | tubulin, alpha 1C | protein-coding |
| chr6_145677417_145677971 | chr6 | 145677418 | 145677971 | + | 0.77671 | Intergenic | Intergenic | 61731 | NM_009449 | 22147 | Mm.270295 | NM_009449 | ENSMUSG00000067338 | **Tuba3b** | M[a]7\|Tuba7 | tubulin, alpha 3B | protein-coding |
| chr6_145610081_145610625 | chr6 | 145610082 | 145610625 | + | 0.676143 | Intergenic | Intergenic | -5610 | NM_009449 | 22147 | Mm.270295 | NM_009449 | ENSMUSG00000067338 | **Tuba3b** | M[a]7\|Tuba7 | tubulin, alpha 3B | protein-coding |
| chr13_34066336_34066679 | chr13 | 34066337 | 34066679 | + | 0.691235 | intron (NM_026512, intron 6 of 6) | intron (NM_026512, intron 6 of 6) | 11500 | NM_009450 | 22151 | Mm.422827 | NM_009450 | ENSMUSG00000058672 | **Tubb2a** | M(beta)2\|Tubb2 | tubulin, beta 2A class IIA | protein-coding |
| chr18_67391069_67391628 | chr18 | 67391070 | 67391628 | + | 0.658822 | exon (NM_026473, exon 2 of 4) | exon (NM_026473, exon 2 of 4) | 618 | NM_026473 | 67951 | Mm.181860 | NM_026473 | ENSMUSG00000001473 | **Tubb6** | 2310057H16Rik\|BB220206 | tubulin, beta 6 class V | protein-coding |
| chr11_101142039_101142343 | chr11 | 101142040 | 101142343 | + | 0.844226 | Intergenic | Intergenic | -13693 | NM_134028 | 103768 | Mm.142348 | NM_134028 | ENSMUSG00000045007 | **Tubg2** | AI504772\|Tubgl | tubulin, gamma 2 | protein-coding |
| chr2_121220078_121220556 | chr2 | 121220079 | 121220556 | + | 0.911529 | intron (NM_013735, intron 16 of 27) | intron (NM_013735, intron 16 of 27) | 49149 | NM_001290824 | 51885 | Mm.383499 | NM_153387 | ENSMUSG00000027263 | **Tubgcp4** | 4932441P04Rik\|76p\|D2Ertd435e\|GCP-4 | tubulin, gamma complex associated protein 4 | protein-coding |
| chr7_55799967_55800515 | chr7 | 55799968 | 55800515 | + | 2.00278 | intron (NM_146190, intron 5 of 22) | intron (NM_146190, intron 5 of 22) | 6093 | NM_146190 | 233276 | Mm.284306 | NM_146190 | ENSMUSG00000033790 | **Tubgcp5** | B130010C12Rik\|GCP5\|mKIAA1899 | tubulin, gamma complex associated protein 5 | protein-coding |
| chr3_94648800_94649658 | chr3 | 94648801 | 94649658 | + | 0.752695 | intron (NM_011656, intron 1 of 12) | intron (NM_011656, intron 1 of 12) | 9643 | NM_001293728 | 22156 | Mm.10214 | NM_011656 | ENSMUSG00000005968 | **Tuft1** | - | tuftelin 1 | protein-coding |
| chr3_94649189_94649583 | chr3 | 94649190 | 94649583 | + | 0.671167 | intron (NM_011656, intron 1 of 12) | intron (NM_011656, intron 1 of 12) | 9486 | NM_001293728 | 22156 | Mm.10214 | NM_011656 | ENSMUSG00000005968 | **Tuft1** | - | tuftelin 1 | protein-coding |
| chr6_128345518_128346051 | chr6 | 128345519 | 128346051 | + | 0.731512 | intron (NM_011657, intron 1 of 10) | intron (NM_011657, intron 1 of 10) | 10066 | NM_011657 | 22158 | Mm.291783 | NM_011657 | ENSMUSG00000001521 | **Tulp3** | 2310022L06Rik\|AI316887 | tubby-like protein 3 | protein-coding |
| chr17_6155903_6156143 | chr17 | 6155904 | 6156143 | + | 0.84258 | intron (NM_054040, intron 1 of 13) | intron (NM_054040, intron 1 of 13) | 17875 | NM_054040 | 68842 | Mm.449307 | NM_054040 | ENSMUSG00000034377 | **Tulp4** | 1110057P05Rik\|2210038L17Rik\|Tusp | tubby like protein 4 | protein-coding |
| chr17_6145920_6146522 | chr17 | 6145921 | 6146522 | + | 0.683516 | intron (NM_054040, intron 1 of 13) | MTD\|LTR\|ERVL-MaLR | 8073 | NM_054040 | 68842 | Mm.449307 | NM_054040 | ENSMUSG00000034377 | **Tulp4** | 1110057P05Rik\|2210038L17Rik\|Tusp | tubby like protein 4 | protein-coding |
| chr17_6145720_6146518 | chr17 | 6145721 | 6146518 | + | 0.682417 | intron (NM_054040, intron 1 of 13) | MTD\|LTR\|ERVL-MaLR | 7971 | NM_054040 | 68842 | Mm.449307 | NM_054040 | ENSMUSG00000034377 | **Tulp4** | 1110057P05Rik\|2210038L17Rik\|Tusp | tubby like protein 4 | protein-coding |
| chr17_6154405_6154777 | chr17 | 6154406 | 6154777 | + | 0.610961 | intron (NM_054040, intron 1 of 13) | intron (NM_054040, intron 1 of 13) | 16443 | NM_054040 | 68842 | Mm.449307 | NM_054040 | ENSMUSG00000034377 | **Tulp4** | 1110057P05Rik\|2210038L17Rik\|Tusp | tubby like protein 4 | protein-coding |
| chr8_39011053_39011411 | chr8 | 39011054 | 39011411 | + | 0.69419 | intron (NM_030254, intron 1 of 7) | intron (NM_030254, intron 1 of 7) | 5365 | NM_030254 | 80286 | Mm.272722 | NM_030254 | | **Tusc3** | AU022242\|BC003311\|N33 | tumor suppressor candidate 3 | protein-coding |
| chr16_10448819_10449391 | chr16 | 10448820 | 10449391 | + | 0.857783 | Intergenic | Intergenic | -1755 | NM_001013778 | 383103 | Mm.325732 | NM_001013778 | ENSMUSG00000050908 | **Tvp23a** | Fam18a | trans-golgi network vesicle protein 23A | protein-coding |
| chr16_10448818_10449615 | chr16 | 10448819 | 10449615 | + | 0.682236 | Intergenic | Intergenic | -1867 | NM_001013778 | 383103 | Mm.325732 | NM_001013778 | ENSMUSG00000050908 | **Tvp23a** | Fam18a | trans-golgi network vesicle protein 23A | protein-coding |
| chr11_62876191_62876836 | chr11 | 62876192 | 62876836 | + | 1.59365 | intron (NM_001033669, intron 12 of 12) | intron (NM_001033669, intron 12 of 12) | -2976 | NM_026210 | 67510 | Mm.9806 | NM_026210 | ENSMUSG00000014177 | **Tvp23b** | 1810036I24Rik\|Fam18b\|Fam18b1 | trans-golgi network vesicle protein 23B | protein-coding |
| chr15_94574653_94575279 | chr15 | 94574654 | 94575279 | + | 1.1318 | Intergenic | RMER15\|LTR\|ERVL | 14858 | NM_008971 | 19230 | Mm.490250 | NM_008971 | ENSMUSG00000022451 | **Twf1** | A6\|Ptk9\|twinfilin | twinfilin, actin-binding protein, homolog 1 (Drosophila) | protein-coding |
| chr15_94573528_94573898 | chr15 | 94573529 | 94573898 | + | 1.07749 | Intergenic | Lx8\|LINE\|L1 | 16111 | NM_008971 | 19230 | Mm.490250 | NM_008971 | ENSMUSG00000022451 | **Twf1** | A6\|Ptk9\|twinfilin | twinfilin, actin-binding protein, homolog 1 (Drosophila) | protein-coding |
| chr12_34075492_34076460 | chr12 | 34075493 | 34076460 | + | 1.24565 | intron (NM_001271386, intron 27 of 27) | intron (NM_001271386, intron 27 of 27) | 118305 | NM_011658 | 22160 | Mm.3280 | NM_011658 | ENSMUSG00000035799 | **Twist1** | AA960487\|M-Twist\|Pde\|Ska10\|Ska<m10Jus>\|Twist\|bHLHa38\|pdt | twist basic helix-loop-helix transcription factor 1 | protein-coding |
| chr12_34075884_34076460 | chr12 | 34075885 | 34076460 | + | 1.13517 | intron (NM_001271386, intron 27 of 27) | intron (NM_001271386, intron 27 of 27) | 118501 | NM_011658 | 22160 | Mm.3280 | NM_011658 | ENSMUSG00000035799 | **Twist1** | AA960487\|M-Twist\|Pde\|Ska10\|Ska<m10Jus>\|Twist\|bHLHa38\|pdt | twist basic helix-loop-helix transcription factor 1 | protein-coding |
| chr12_34040871_34041362 | chr12 | 34040872 | 34041362 | + | 1.12764 | Intergenic | L1M2\|LINE\|L1 | 83446 | NM_011658 | 22160 | Mm.3280 | NM_011658 | ENSMUSG00000035799 | **Twist1** | AA960487\|M-Twist\|Pde\|Ska10\|Ska<m10Jus>\|Twist\|bHLHa38\|pdt | twist basic helix-loop-helix transcription factor 1 | protein-coding |
| chr1_91692699_91693190 | chr1 | 91692700 | 91693190 | + | 0.796479 | Intergenic | Intergenic | -108532 | NM_007855 | 13345 | Mm.9474 | NM_007855 | ENSMUSG00000007805 | **Twist2** | Dermo1\|bHLHa39 | twist basic helix-loop-helix transcription factor 2 | protein-coding |
| chr17_65939102_65940564 | chr17 | 65939103 | 65940564 | + | 0.885524 | intron (NM_023053, intron 2 of 4) | intron (NM_023053, intron 2 of 4) | 11354 | NM_023053 | 65960 | Mm.10153 | NM_023053 | ENSMUSG00000024098 | **Twsg1** | 1810013J15Rik\|9030422N06Rik\|AW552143\|D17Ertd403e\|Tsg\|Twg | twisted gastrulation homolog 1 (Drosophila) | protein-coding |
| chr5_72725461_72725813 | chr5 | 72725462 | 72725813 | + | 1.12972 | intron (NM_001289494, intron 4 of 13) | intron (NM_001289494, intron 4 of 13) | 10965 | NM_013698 | 22165 | Mm.3264 | NM_013698 | ENSMUSG00000054892 | **Txk** | A130089B16Rik\|Btkl\|PTK-RL-18\|PTK4\|Rlk | TXK tyrosine kinase | protein-coding |
| chr5_72723466_72724123 | chr5 | 72723467 | 72724123 | + | 0.990208 | intron (NM_001289495, intron 6 of 14) | intron (NM_001289495, intron 6 of 14) | 12807 | NM_013698 | 22165 | Mm.3264 | NM_013698 | ENSMUSG00000054892 | **Txk** | A130089B16Rik\|Btkl\|PTK-RL-18\|PTK4\|Rlk | TXK tyrosine kinase | protein-coding |
| chr5_72723466_72723920 | chr5 | 72723467 | 72723920 | + | 0.946306 | intron (NM_001289495, intron 6 of 14) | intron (NM_001289495, intron 6 of 14) | 12909 | NM_013698 | 22165 | Mm.3264 | NM_013698 | ENSMUSG00000054892 | **Txk** | A130089B16Rik\|Btkl\|PTK-RL-18\|PTK4\|Rlk | TXK tyrosine kinase | protein-coding |
| chr15_77938410_77938949 | chr15 | 77938411 | 77938949 | + | 1.47721 | Intergenic | Intergenic | -9686 | NM_019913 | 56551 | Mm.291917 | NM_019913 | ENSMUSG00000005354 | **Txn2** | 2510006J11Rik\|AI788873\|Trx2 | thioredoxin 2 | protein-coding |
| chr14_45211468_45212102 | chr14 | 45211469 | 45212102 | + | 0.678882 | intron (NM_172597, intron 3 of 20) | intron (NM_172597, intron 3 of 20) | 7609 | NM_172597 | 70561 | Mm.28129 | NM_172597 | ENSMUSG00000021830 | **Txndc16** | 5730420B22Rik\|C77647 | thioredoxin domain containing 16 | protein-coding |
| chr13_38547663_38548088 | chr13 | 38547664 | 38548088 | + | 1.27055 | Intergenic | Intergenic | -19052 | NM_001289598 | 105245 | Mm.28622 | NM_145367 | ENSMUSG00000038991 | **Txndc5** | AL022641\|ERp46\|PC-TRP | thioredoxin domain containing 5 | protein-coding |
| chr13_38551994_38552419 | chr13 | 38551995 | 38552419 | + | 0.653592 | Intergenic | Lx7\|LINE\|L1 | -23383 | NM_001289598 | 105245 | Mm.28622 | NM_145367 | ENSMUSG00000038991 | **Txndc5** | AL022641\|ERp46\|PC-TRP | thioredoxin domain containing 5 | protein-coding |
| chr1_37982146_37982685 | chr1 | 37982147 | 37982685 | + | 1.01316 | Intergenic | L1M4b\|LINE\|L1 | 14792 | NM_172054 | 98258 | Mm.28438 | NM_172054 | ENSMUSG00000058407 | **Txndc9** | AI098020\|APACD | thioredoxin domain containing 9 | protein-coding |
| chr8_109570754_109571127 | chr8 | 109570755 | 109571127 | + | 0.711148 | intron (NM_175646, intron 2 of 3) | ORR1D1\|LTR\|ERVL-MaLR | 4955 | NM_175646 | 234723 | Mm.37667 | NM_175646 | ENSMUSG00000031723 | **Txnl4b** | AI595343\|D530025J19Rik\|DLP\|Dim2 | thioredoxin-like 4B | protein-coding |
| chr17_31667254_31668294 | chr17 | 31667255 | 31668294 | + | 1.11143 | Intergenic | Intergenic | -9020 | NM_001163769 | 108121 | Mm.379289 | NM_024187 | ENSMUSG00000061613 | **U2af1** | 2010107D16Rik\|35kDa | U2 small nuclear ribonucleoprotein auxiliary factor (U2AF) 1 | protein-coding |

| | | | | | | | | | | | | | | | | | |
|---|---|---|---|---|---|---|---|---|---|---|---|---|---|---|---|---|---|
| chr17_31657517_31657745 | chr17 | 31657518 | 31657745 | + | 0.873463 | intron (NM_024187, intron 1 of 7) | B3A\|SINE\|B2 | 1123 | NM_024187 | 108121 | Mm.379289 | NM_024187 | ENSMUSG00000061613 | **U2af1** | 2010107D16Rik\|35kDa | U2 small nuclear ribonucleoprotein auxiliary factor (U2AF) 1 | protein-coding |
| chr1_170174183_170174448 | chr1 | 170174184 | 170174448 | + | 0.660288 | intron (NM_133806, intron 1 of 10) | intron (NM_133806, intron 1 of 10) | 630 | NM_133806 | 107652 | Mm.27969 | NM_133806 | ENSMUSG00000026670 | **Uap1** | AA420407\|AA437972\|AGX-1\|AGX-2\|AGX1\|AgX\|ESTM38\|SPAG2 | UDP-N-acetylglucosamine pyrophosphorylase 1 | protein-coding |
| chr6_97204564_97204816 | chr6 | 97204565 | 97204816 | + | 0.891502 | intron (NM_001111106, intron 1 of 16) | intron (NM_001111106, intron 1 of 16) | 952 | NM_001111106 | 22200 | Mm.277626 | NM_011666 | ENSMUSG00000030061 | **Uba3** | A830034N06Rik\|AI256736\|AI848246\|AW546539\|Ube1c | ubiquitin-like modifier activating enzyme 3 | protein-coding |
| chr6_97204391_97204840 | chr6 | 97204392 | 97204840 | + | 0.74975 | intron (NM_001111106, intron 1 of 16) | intron (NM_001111106, intron 1 of 16) | 1026 | NM_001111106 | 22200 | Mm.277626 | NM_011666 | ENSMUSG00000030061 | **Uba3** | A830034N06Rik\|AI256736\|AI848246\|AW546539\|Ube1c | ubiquitin-like modifier activating enzyme 3 | protein-coding |
| chr14_121895240_121895808 | chr14 | 121895241 | 121895808 | + | 0.672826 | intron (NM_026861, intron 1 of 8) | intron (NM_026861, intron 1 of 8) | 16918 | NM_026861 | 68889 | Mm.441151 | NM_026861 | ENSMUSG00000041765 | **Ubac2** | 1190008A14Rik\|AW554959\|Phgdhl1 | ubiquitin associated domain containing 2 | protein-coding |
| chr16_4878642_4878914 | chr16 | 4878643 | 4878914 | + | 0.648307 | intron (NM_145359, intron 1 of 2) | intron (NM_145359, intron 1 of 2) | 1073 | NM_145359 | 207740 | Mm.268895 | NM_145359 | | **Ubald1** | 1500031H01Rik\|BC013706\|Fam100a | UBA-like domain containing 1 | protein-coding |
| chr4_41255584_41256250 | chr4 | 41255585 | 41256250 | + | 1.16308 | intron (NM_026872, intron 1 of 28) | intron (NM_026872, intron 1 of 28) | 19218 | NM_026872 | 68926 | Mm.41864 | NM_026872 | ENSMUSG00000028433 | **Ubap2** | 1190005K07Rik\|AA408600\|AU045235\|UBAP-2\|mKIAA1491 | ubiquitin-associated protein 2 | protein-coding |
| chr4_41240578_41241457 | chr4 | 41240579 | 41241457 | + | 0.930774 | intron (NM_026872, intron 2 of 28) | intron (NM_026872, intron 2 of 28) | 34117 | NM_026872 | 68926 | Mm.41864 | NM_026872 | ENSMUSG00000028433 | **Ubap2** | 1190005K07Rik\|AA408600\|AU045235\|UBAP-2\|mKIAA1491 | ubiquitin-associated protein 2 | protein-coding |
| chr4_41240902_41241404 | chr4 | 41240903 | 41241404 | + | 0.92756 | intron (NM_026872, intron 2 of 28) | intron (NM_026872, intron 2 of 28) | 33982 | NM_026872 | 68926 | Mm.41864 | NM_026872 | ENSMUSG00000028433 | **Ubap2** | 1190005K07Rik\|AA408600\|AU045235\|UBAP-2\|mKIAA1491 | ubiquitin-associated protein 2 | protein-coding |
| chr4_41240947_41241298 | chr4 | 41240948 | 41241298 | + | 0.841474 | intron (NM_026872, intron 2 of 28) | intron (NM_026872, intron 2 of 28) | 34012 | NM_026872 | 68926 | Mm.41864 | NM_026872 | ENSMUSG00000028433 | **Ubap2** | 1190005K07Rik\|AA408600\|AU045235\|UBAP-2\|mKIAA1491 | ubiquitin-associated protein 2 | protein-coding |
| chr17_31222930_31223468 | chr17 | 31222931 | 31223468 | + | 0.731643 | intron (NM_177823, intron 6 of 13) | intron (NM_177823, intron 6 of 13) | 15133 | NM_177823 | 328795 | Mm.387930 | NM_177823 | ENSMUSG00000042345 | **Ubash3a** | 5830413C03Rik\|C330001M22\|Sts-2\|TULA\|TULA-1 | ubiquitin associated and SH3 domain containing, A | protein-coding |
| chr9_41179045_41179448 | chr9 | 41179046 | 41179448 | + | 1.46858 | Intergenic | Intergenic | -21753 | NM_176860 | 72828 | Mm.133615 | NM_176860 | ENSMUSG00000032020 | **Ubash3b** | 2810457I06Rik\|BB125008\|TULA-2\|p70 | ubiquitin associated and SH3 domain containing, B | protein-coding |
| chr9_41169519_41169989 | chr9 | 41169520 | 41169989 | + | 1.34593 | Intergenic | Intergenic | -12260 | NM_176860 | 72828 | Mm.133615 | NM_176860 | ENSMUSG00000032020 | **Ubash3b** | 2810457I06Rik\|BB125008\|TULA-2\|p70 | ubiquitin associated and SH3 domain containing, B | protein-coding |
| chr9_41085278_41085482 | chr9 | 41085279 | 41085482 | + | 0.734551 | intron (NM_176860, intron 1 of 13) | intron (NM_176860, intron 1 of 13) | 72114 | NM_176860 | 72828 | Mm.133615 | NM_176860 | ENSMUSG00000032020 | **Ubash3b** | 2810457I06Rik\|BB125008\|TULA-2\|p70 | ubiquitin associated and SH3 domain containing, B | protein-coding |
| chr9_41112455_41113019 | chr9 | 41112456 | 41113019 | + | 0.692943 | intron (NM_176860, intron 1 of 13) | intron (NM_176860, intron 1 of 13) | 44757 | NM_176860 | 72828 | Mm.133615 | NM_176860 | ENSMUSG00000032020 | **Ubash3b** | 2810457I06Rik\|BB125008\|TULA-2\|p70 | ubiquitin associated and SH3 domain containing, B | protein-coding |
| chr9_41170103_41170478 | chr9 | 41170104 | 41170478 | + | 0.68856 | Intergenic | Intergenic | -12797 | NM_176860 | 72828 | Mm.133615 | NM_176860 | ENSMUSG00000032020 | **Ubash3b** | 2810457I06Rik\|BB125008\|TULA-2\|p70 | ubiquitin associated and SH3 domain containing, B | protein-coding |
| chr5_125375616_125376277 | chr5 | 125375617 | 125376277 | + | 1.12072 | Intergenic | B3\|SINE\|B2 | 14070 | NM_019639 | 22190 | Mm.331 | NM_019639 | ENSMUSG00000008348 | **Ubc** | 2700054O04Rik\|AI194771\|Rps27a\|TI-225\|Uba52\|Ubb | ubiquitin C | protein-coding |
| chr5_125383200_125383640 | chr5 | 125383201 | 125383640 | + | 0.988427 | Intergenic | Intergenic | 6597 | NM_019639 | 22190 | Mm.331 | NM_019639 | ENSMUSG00000008348 | **Ubc** | 2700054O04Rik\|AI194771\|Rps27a\|TI-225\|Uba52\|Ubb | ubiquitin C | protein-coding |
| chr5_125377088_125377578 | chr5 | 125377089 | 125377578 | + | 0.745701 | Intergenic | Intergenic | 12684 | NM_019639 | 22190 | Mm.331 | NM_019639 | ENSMUSG00000008348 | **Ubc** | 2700054O04Rik\|AI194771\|Rps27a\|TI-225\|Uba52\|Ubb | ubiquitin C | protein-coding |
| chr5_125382627_125383745 | chr5 | 125382628 | 125383745 | + | 0.673922 | Intergenic | Intergenic | 6831 | NM_019639 | 22190 | Mm.331 | NM_019639 | ENSMUSG00000008348 | **Ubc** | 2700054O04Rik\|AI194771\|Rps27a\|TI-225\|Uba52\|Ubb | ubiquitin C | protein-coding |
| chr17_37185363_37185921 | chr17 | 37185364 | 37185921 | + | 1.28258 | Intergenic | Intergenic | -8250 | NM_023137 | 24108 | Mm.140210 | NM_023137 | ENSMUSG00000035186 | **Ubd** | FAT10 | ubiquitin D | protein-coding |
| chr9_86357665_86358337 | chr9 | 86357666 | 86358337 | + | 0.767487 | intron (NM_027394, intron 9 of 9) | intron (NM_027394, intron 9 of 9) | 106915 | NM_027394 | 70348 | Mm.53469 | NM_027394 | ENSMUSG00000032415 | **Ube2cbp** | 2610018I03Rik\|Ube3d | ubiquitin-conjugating enzyme E2C binding protein | protein-coding |
| chr10_71266025_71266517 | chr10 | 71266026 | 71266517 | + | 0.645361 | intron (NM_145420, intron 1 of 6) | intron (NM_145420, intron 1 of 6) | 18991 | NM_145420 | 216080 | Mm.270530 | NM_145420 | ENSMUSG00000019927 | **Ube2d1** | UBCH5 | ubiquitin-conjugating enzyme E2D 1 | protein-coding |
| chr18_35797484_35798063 | chr18 | 35797485 | 35798063 | + | 0.966995 | intron (NM_019912, intron 2 of 6) | RSINE1\|SINE\|B4 | 26215 | NM_019912 | 56550 | Mm.180052 | NM_019912 | ENSMUSG00000091896 | **Ube2d2a** | 1500034D03Rik\|Ubc2e\|Ube2d2\|ubc4 | ubiquitin-conjugating enzyme E2D 2A | protein-coding |
| chr5_107778812_107779222 | chr5 | 107778813 | 107779222 | + | 0.926399 | intron (NM_007964, intron 16 of 17) | intron (NM_007964, intron 16 of 17) | -51145 | NM_001276397 | 73318 | Mm.441500 | NM_028502 | | **Ube2d2b** | 1700013N18Rik | ubiquitin-conjugating enzyme E2D 2B | protein-coding |
| chr5_107802623_107802985 | chr5 | 107802624 | 107802985 | + | 0.62834 | intron (NM_007964, intron 11 of 17) | ORR1C2\|LTR\|ERVL-MaLR | -27358 | NM_001276397 | 73318 | Mm.441500 | NM_028502 | | **Ube2d2b** | 1700013N18Rik | ubiquitin-conjugating enzyme E2D 2B | protein-coding |
| chr14_18529843_18530366 | chr14 | 18529844 | 18530366 | + | 0.942449 | Intergenic | Intergenic | -198261 | NM_009455 | 22194 | Mm.4429 | NM_009455 | ENSMUSG00000021774 | **Ube2e1** | UbcM3\|Ubce5\|ubcM2 | ubiquitin-conjugating enzyme E2E 1 | protein-coding |
| chr14_18529791_18530625 | chr14 | 18529792 | 18530625 | + | 0.876577 | Intergenic | Intergenic | -198364 | NM_009455 | 22194 | Mm.4429 | NM_009455 | ENSMUSG00000021774 | **Ube2e1** | UbcM3\|Ubce5\|ubcM2 | ubiquitin-conjugating enzyme E2E 1 | protein-coding |
| chr14_18390944_18391585 | chr14 | 18390945 | 18391585 | + | 0.794517 | Intergenic | Intergenic | -59421 | NM_009455 | 22194 | Mm.4429 | NM_009455 | ENSMUSG00000021774 | **Ube2e1** | UbcM3\|Ubce5\|ubcM2 | ubiquitin-conjugating enzyme E2E 1 | protein-coding |
| chr14_18335684_18336007 | chr14 | 18335685 | 18336007 | + | 0.721721 | Intergenic | Intergenic | -4002 | NM_009455 | 22194 | Mm.4429 | NM_009455 | ENSMUSG00000021774 | **Ube2e1** | UbcM3\|Ubce5\|ubcM2 | ubiquitin-conjugating enzyme E2E 1 | protein-coding |
| chr14_18939807_18940533 | chr14 | 18939808 | 18940533 | + | 0.889516 | Intergenic | Intergenic | -46543 | NM_144839 | 218793 | Mm.23551 | NM_144839 | ENSMUSG00000058317 | **Ube2e2** | BC016265 | ubiquitin-conjugating enzyme E2E 2 | protein-coding |
| chr14_18820359_18821177 | chr14 | 18820360 | 18821177 | + | 0.631073 | intron (NM_144839, intron 3 of 5) | RMER15\|LTR\|ERVL | 72859 | NM_144839 | 218793 | Mm.23551 | NM_144839 | ENSMUSG00000058317 | **Ube2e2** | BC016265 | ubiquitin-conjugating enzyme E2E 2 | protein-coding |
| chr14_18820600_18821186 | chr14 | 18820601 | 18821186 | + | 0.614111 | intron (NM_144839, intron 3 of 5) | RMER15\|LTR\|ERVL | 72734 | NM_144839 | 218793 | Mm.23551 | NM_144839 | ENSMUSG00000058317 | **Ube2e2** | BC016265 | ubiquitin-conjugating enzyme E2E 2 | protein-coding |
| chr2_78961577_78962185 | chr2 | 78961578 | 78962185 | + | 1.49377 | Intergenic | Intergenic | 92834 | NM_009454 | 22193 | Mm.1485 | NM_009454 | | **Ube2e3** | Ubce4\|ubcM2 | ubiquitin-conjugating enzyme E2E 3 | protein-coding |
| chr2_79009754_79010225 | chr2 | 79009755 | 79010225 | + | 1.35492 | Intergenic | Intergenic | 140943 | NM_009454 | 22193 | Mm.1485 | NM_009454 | | **Ube2e3** | Ubce4\|ubcM2 | ubiquitin-conjugating enzyme E2E 3 | protein-coding |
| chr2_78980731_78981453 | chr2 | 78980732 | 78981453 | + | 1.28335 | Intergenic | L2\|LINE\|L2 | 112045 | NM_009454 | 22193 | Mm.1485 | NM_009454 | | **Ube2e3** | Ubce4\|ubcM2 | ubiquitin-conjugating enzyme E2E 3 | protein-coding |
| chr2_79052251_79052594 | chr2 | 79052252 | 79052594 | + | 0.985198 | Intergenic | Intergenic | 183376 | NM_009454 | 22193 | Mm.1485 | NM_009454 | | **Ube2e3** | Ubce4\|ubcM2 | ubiquitin-conjugating enzyme E2E 3 | protein-coding |
| chr2_78979922_78980357 | chr2 | 78979923 | 78980357 | + | 0.918194 | Intergenic | Intergenic | 111093 | NM_009454 | 22193 | Mm.1485 | NM_009454 | | **Ube2e3** | Ubce4\|ubcM2 | ubiquitin-conjugating enzyme E2E 3 | protein-coding |
| chr1_91259794_91260178 | chr1 | 91259795 | 91260178 | + | 0.780273 | intron (NM_026454, intron 2 of 9) | URR1A\|DNA\|hAT-Charlie | 9667 | NM_026454 | 67921 | Mm.337238 | NM_026454 | ENSMUSG00000034343 | **Ube2f** | 2510010F15Rik\|AI851109 | ubiquitin-conjugating enzyme E2F (putative) | protein-coding |
| chr11_72594946_72595309 | chr11 | 72594947 | 72595309 | + | 1.04674 | Intergenic | Intergenic | -12133 | NM_025985 | 67128 | Mm.458052 | NM_025985 | ENSMUSG00000020794 | **Ube2g1** | 2700059C12Rik\|AI256795\|AU014992\|AW552068\|D130023C12Rik | ubiquitin-conjugating enzyme E2G 1 | protein-coding |
| chr11_72594946_72595616 | chr11 | 72594947 | 72595616 | + | 1.03629 | Intergenic | T-rich\|Low_complexity\|Low_complexity | -11980 | NM_025985 | 67128 | Mm.458052 | NM_025985 | ENSMUSG00000020794 | **Ube2g1** | 2700059C12Rik\|AI256795\|AU014992\|AW552068\|D130023C12Rik | ubiquitin-conjugating enzyme E2G 1 | protein-coding |
| chr11_72597870_72598221 | chr11 | 72597871 | 72598221 | + | 0.793406 | Intergenic | Intergenic | -9215 | NM_025985 | 67128 | Mm.458052 | NM_025985 | ENSMUSG00000020794 | **Ube2g1** | 2700059C12Rik\|AI256795\|AU014992\|AW552068\|D130023C12Rik | ubiquitin-conjugating enzyme E2G 1 | protein-coding |
| chr11_72597177_72597560 | chr11 | 72597178 | 72597560 | + | 0.719468 | Intergenic | Intergenic | -9892 | NM_025985 | 67128 | Mm.458052 | NM_025985 | ENSMUSG00000020794 | **Ube2g1** | 2700059C12Rik\|AI256795\|AU014992\|AW552068\|D130023C12Rik | ubiquitin-conjugating enzyme E2G 1 | protein-coding |
| chr10_77623184_77623490 | chr10 | 77623185 | 77623490 | + | 0.633153 | intron (NM_019803, intron 1 of 5) | MLT1E2\|LTR\|ERVL-MaLR | 1016 | NM_019803 | 22213 | Mm.458125 | NM_019803 | ENSMUSG00000009293 | **Ube2g2** | 1110003O05Rik\|D10Xrf369\|UBC7\|Ubc7p | ubiquitin-conjugating enzyme E2G 2 | protein-coding |
| chr6_30282174_30282694 | chr6 | 30282175 | 30282694 | + | 1.03269 | intron (NM_001169576, intron 1 of 4) | intron (NM_001169576, intron 1 of 4) | 22105 | NM_009459 | 22214 | Mm.332967 | NM_009459 | ENSMUSG00000039159 | **Ube2h** | 1500009C23Rik\|AI181839\|AI326965\|AI462483\|AW227540\|E2-20K\|Gid3\|N28148\|Ubc8 | ubiquitin-conjugating enzyme E2H | protein-coding |
| chr6_30282378_30282678 | chr6 | 30282379 | 30282678 | + | 0.936979 | intron (NM_001169576, intron 1 of 4) | intron (NM_001169576, intron 1 of 4) | 22011 | NM_009459 | 22214 | Mm.332967 | NM_009459 | ENSMUSG00000039159 | **Ube2h** | 1500009C23Rik\|AI181839\|AI326965\|AI462483\|AW227540\|E2-20K\|Gid3\|N28148\|Ubc8 | ubiquitin-conjugating enzyme E2H | protein-coding |
| chr6_30300854_30301320 | chr6 | 30300855 | 30301320 | + | 0.776855 | intron (NM_001169576, intron 1 of 4) | intron (NM_001169576, intron 1 of 4) | 3452 | NM_009459 | 22214 | Mm.332967 | NM_009459 | ENSMUSG00000039159 | **Ube2h** | 1500009C23Rik\|AI181839\|AI326965\|AI462483\|AW227540\|E2-20K\|Gid3\|N28148\|Ubc8 | ubiquitin-conjugating enzyme E2H | protein-coding |
| chr16_17177581_17178130 | chr16 | 17177582 | 17178130 | + | 1.17732 | intron (NM_009456, intron 1 of 3) | intron (NM_009456, intron 1 of 3) | 23636 | NM_009456 | 22195 | Mm.3074 | NM_009456 | ENSMUSG00000038965 | **Ube2l3** | C79827\|UbcM4\|Ubce7 | ubiquitin-conjugating enzyme E2L 3 | protein-coding |
| chr4_41187122_41187626 | chr4 | 41187123 | 41187626 | + | 1.20802 | intron (NM_026275, intron 3 of 4) | Lx5c\|LINE\|L1 | 51353 | NM_026275 | 67615 | Mm.389540 | NM_026275 | ENSMUSG00000036241 | **Ube2r2** | 1200003M11Rik\|Cdc34b\|Ubc3b | ubiquitin-conjugating enzyme E2R 2 | protein-coding |
| chr4_41186760_41187639 | chr4 | 41186761 | 41187639 | + | 0.912217 | intron (NM_026275, intron 3 of 4) | Lx5c\|LINE\|L1 | 51179 | NM_026275 | 67615 | Mm.389540 | NM_026275 | ENSMUSG00000036241 | **Ube2r2** | 1200003M11Rik\|Cdc34b\|Ubc3b | ubiquitin-conjugating enzyme E2R 2 | protein-coding |
| chr11_96058226_96058961 | chr11 | 96058227 | 96058961 | + | 1.2477 | intron (NM_172300, intron 3 of 6) | intron (NM_172300, intron 3 of 6) | 6770 | NM_172300 | 268470 | Mm.38802 | NM_172300 | ENSMUSG00000014349 | **Ube2z** | AW049199\|C030047H17Rik\|D11Moh35 | ubiquitin-conjugating enzyme E2Z (putative) | protein-coding |
| chr5_29587889_29588319 | chr5 | 29587890 | 29588319 | + | 1.06474 | intron (NM_133907, intron 2 of 22) | intron (NM_133907, intron 2 of 22) | 18862 | NM_133907 | 100763 | Mm.137746 | NM_133907 | ENSMUSG00000039000 | **Ube3c** | AI853514\|mKIAA0010 | ubiquitin protein ligase E3C | protein-coding |
| chr5_29581837_29582566 | chr5 | 29581838 | 29582566 | + | 0.938024 | intron (NM_133907, intron 1 of 22) | B1_Mur1\|SINE\|Alu | 12960 | NM_133907 | 100763 | Mm.137746 | NM_133907 | ENSMUSG00000039000 | **Ube3c** | AI853514\|mKIAA0010 | ubiquitin protein ligase E3C | protein-coding |

| | | | | | | | | | | | | | | | | | |
|---|---|---|---|---|---|---|---|---|---|---|---|---|---|---|---|---|---|
| chr4_149422206_149422449 | chr4 | 149422207 | 149422449 | + | 0.853243 | intron (NM_022022, intron 1 of 26) | intron (NM_022022, intron 1 of 26) | 4303 | NM_022022 | 63958 | Mm.415260 | NM_022022 | ENSMUSG00000028960 | **Ube4b** | 4930551I19Rik\|4933406G05Rik\|AU014668\|D4Bwg0973e\|UFD2\|UFD2a\|Ufd2p\|mKIAA0684 | ubiquitination factor E4B | protein-coding |
| chr4_149401645_149401901 | chr4 | 149401646 | 149401901 | + | 0.852936 | intron (NM_022022, intron 1 of 26) | B3A\|SINE\|B2 | 24858 | NM_022022 | 63958 | Mm.415260 | NM_022022 | ENSMUSG00000028960 | **Ube4b** | 4930551I19Rik\|4933406G05Rik\|AU014668\|D4Bwg0973e\|UFD2\|UFD2a\|Ufd2p\|mKIAA0684 | ubiquitination factor E4B | protein-coding |
| chr4_149421979_149422450 | chr4 | 149421980 | 149422450 | + | 0.765045 | intron (NM_022022, intron 1 of 26) | intron (NM_022022, intron 1 of 26) | 4416 | NM_022022 | 63958 | Mm.415260 | NM_022022 | ENSMUSG00000028960 | **Ube4b** | 4930551I19Rik\|4933406G05Rik\|AU014668\|D4Bwg0973e\|UFD2\|UFD2a\|Ufd2p\|mKIAA0684 | ubiquitination factor E4B | protein-coding |
| chr4_149403055_149403403 | chr4 | 149403056 | 149403403 | + | 0.69845 | intron (NM_022022, intron 1 of 26) | intron (NM_022022, intron 1 of 26) | 23402 | NM_022022 | 63958 | Mm.415260 | NM_022022 | ENSMUSG00000028960 | **Ube4b** | 4930551I19Rik\|4933406G05Rik\|AU014668\|D4Bwg0973e\|UFD2\|UFD2a\|Ufd2p\|mKIAA0684 | ubiquitination factor E4B | protein-coding |
| chr4_148438205_148438886 | chr4 | 148438206 | 148438886 | + | 0.962465 | intron (NM_027873, intron 1 of 1) | intron (NM_027873, intron 1 of 1) | 6205 | NM_027873 | 71707 | Mm.292503 | NM_027873 | ENSMUSG00000047719 | **Ubiad1** | 1200002M06Rik\|AI426463\|AW320947\|Tere1 | UbiA prenyltransferase domain containing 1 | protein-coding |
| chr4_148437851_148438914 | chr4 | 148437852 | 148438914 | + | 0.808866 | intron (NM_027873, intron 1 of 1) | intron (NM_027873, intron 1 of 1) | 6368 | NM_027873 | 71707 | Mm.292503 | NM_027873 | ENSMUSG00000047719 | **Ubiad1** | 1200002M06Rik\|AI426463\|AW320947\|Tere1 | UbiA prenyltransferase domain containing 1 | protein-coding |
| chr13_58189421_58189824 | chr13 | 58189422 | 58189824 | + | 0.963798 | intron (NM_152234, intron 6 of 9) | L1_Mur3\|LINE\|L1 | 26030 | NM_152234 | 56085 | Mm.182053 | NM_026842 | ENSMUSG00000005312 | **Ubqln1** | 1110046H03Rik\|1810030E05Rik\|AU019746\|C77538\|D13Ertd372e\|Da41\|Dsk2\|Plic-1\|Plic1\|Xdrp1 | ubiquilin 1 | protein-coding |
| chrX_153433577_153434000 | chrX | 153433578 | 153434000 | + | 0.622651 | Intergenic | Intergenic | -64443 | NM_018798 | 54609 | Mm.430772 | NM_018798 | ENSMUSG00000050148 | **Ubqln2** | Chap1\|Dsk2\|HRIHFB2157\|Plic-2\|Plic2 | ubiquilin 2 | protein-coding |
| chr2_120933049_120933716 | chr2 | 120933050 | 120933716 | + | 1.21868 | intron (NM_009461, intron 14 of 46) | B4A\|SINE\|B4 | 37332 | NM_009461 | 22222 | Mm.389330 | NM_009461 | ENSMUSG00000027272 | **Ubr1** | AI504731 | ubiquitin protein ligase E3 component n-recognin 1 | protein-coding |
| chr17_47047672_47048108 | chr17 | 47047673 | 47048108 | + | 0.716567 | Intergenic | Intergenic | -37358 | NM_001177374 | 224826 | Mm.28234 | NM_146078 | ENSMUSG00000023977 | **Ubr2** | 9930021A08Rik\|AI462103\|AW540746\|E130209G04Rik\|mKIAA0349 | ubiquitin protein ligase E3 component n-recognin 2 | protein-coding |
| chr15_38122722_38123124 | chr15 | 38122723 | 38123124 | + | 1.21915 | Intergenic | MTD\|LTR\|ERVL-MaLR | -44069 | NM_001081359 | 70790 | Mm.476840 | NM_027553 | ENSMUSG00000037487 | **Ubr5** | Edd\|Edd1 | ubiquitin protein ligase E3 component n-recognin 5 | protein-coding |
| chr15_38120318_38120733 | chr15 | 38120319 | 38120733 | + | 1.02975 | Intergenic | B4\|SINE\|B4 | -41672 | NM_001081359 | 70790 | Mm.476840 | NM_027553 | ENSMUSG00000037487 | **Ubr5** | Edd\|Edd1 | ubiquitin protein ligase E3 component n-recognin 5 | protein-coding |
| chr15_38129395_38129771 | chr15 | 38129396 | 38129771 | + | 0.778552 | Intergenic | L1MD1\|LINE\|L1 | -50729 | NM_001081359 | 70790 | Mm.476840 | NM_027553 | ENSMUSG00000037487 | **Ubr5** | Edd\|Edd1 | ubiquitin protein ligase E3 component n-recognin 5 | protein-coding |
| chr15_38124511_38124895 | chr15 | 38124512 | 38124895 | + | 0.663108 | Intergenic | Intergenic | -45849 | NM_001081359 | 70790 | Mm.476840 | NM_027553 | ENSMUSG00000037487 | **Ubr5** | Edd\|Edd1 | ubiquitin protein ligase E3 component n-recognin 5 | protein-coding |
| chr11_32442411_32442981 | chr11 | 32442412 | 32442981 | + | 0.7895 | Intergenic | L1MB8\|LINE\|L1 | -12676 | NM_173784 | 327900 | Mm.260956 | NM_173784 | | **Ubtd2** | 4930571L24\|9630054F20Rik\|AI645720 | ubiquitin domain containing 2 | protein-coding |
| chr4_6228968_6229416 | chr4 | 6228969 | 6229416 | + | 0.687248 | Intergenic | Intergenic | 38087 | NM_026534 | 68053 | Mm.404043 | NM_026534 | | **Ubxn2b** | 3110003A22Rik\|6430407D20Rik\|AI451665\|p37 | UBX domain protein 2B | protein-coding |
| chr8_33636497_33637120 | chr8 | 33636498 | 33637120 | + | 0.958377 | intron (NM_178648, intron 2 of 7) | AT_rich\|Low_complexity\|Low_complexity | 5167 | NM_178648 | 108159 | Mm.29301 | NM_178648 | ENSMUSG00000052906 | **Ubxn8** | D0H8S2298E\|Rep-8\|Rep8h\|Ubxd6 | UBX domain protein 8 | protein-coding |
| chr14_101661046_101662129 | chr14 | 101661047 | 101662129 | + | 0.620221 | intron (NM_016723, intron 2 of 8) | intron (NM_016723, intron 2 of 8) | 7621 | NM_016723 | 50933 | Mm.275970 | NM_016723 | ENSMUSG00000022111 | **Uchl3** | - | ubiquitin carboxyl-terminal esterase L3 (ubiquitin thiolesterase) | protein-coding |
| chr1_143818940_143819385 | chr1 | 143818941 | 143819385 | + | 0.604056 | Intergenic | Intergenic | 41885 | NM_019562 | 56207 | Mm.261004 | NM_019562 | ENSMUSG00000018189 | **Uchl5** | 5830413B11Rik\|Uch37 | ubiquitin carboxyl-terminal esterase L5 | protein-coding |
| chr1_167255539_167255881 | chr1 | 167255540 | 167255881 | + | 0.769601 | intron (NM_030724, intron 1 of 6) | intron (NM_030724, intron 1 of 6) | 29417 | NM_030724 | 80914 | Mm.280895 | NM_030724 | ENSMUSG00000026558 | **Uck2** | AA407809\|AI481316\|AU018180\|AU020720\|TSA903\|UK\|UMK\|Umpk | uridine-cytidine kinase 2 | protein-coding |
| chr4_59214630_59215301 | chr4 | 59214631 | 59215301 | + | 0.709915 | intron (NM_011673, intron 5 of 8) | intron (NM_011673, intron 5 of 8) | 25416 | NM_011673 | 22234 | Mm.198803 | NM_011673 | ENSMUSG00000028381 | **Ugcg** | AU043821\|C80537\|Epcs21\|GlcT-1\|Ugcgl | UDP-glucose ceramide glucosyltransferase | protein-coding |
| chr1_36235073_36235714 | chr1 | 36235074 | 36235714 | + | 1.27335 | intron (NM_198899, intron 1 of 40) | MLT1B\|LTR\|ERVL-MaLR | 8908 | NM_198899 | 320011 | Mm.261022 | NM_198899 | ENSMUSG00000037470 | **Uggt1** | 0910001L17Rik\|A930007H10Rik\|AA589501\|AI414429\|AI448372\|C820010P03Rik\|GT\|UGT1\|Ugcgl1 | UDP-glucose glycoprotein glucosyltransferase 1 | protein-coding |
| chr1_88055447_88055966 | chr1 | 88055448 | 88055966 | + | 1.08057 | exon (NM_201641, exon 1 of 5) | exon (NM_201641, exon 1 of 5) | 296 | NM_201641 | 394430 | Mm.300095 | NM_201641 | | **Ugt1a10** | A13\|Ugt1a13 | UDP glycosyltransferase 1 family, polypeptide A10 | protein-coding |
| chr1_88054122_88054611 | chr1 | 88054123 | 88054611 | + | 1.04783 | Intergenic | Intergenic | -1044 | NM_201641 | 394430 | Mm.300095 | NM_201641 | | **Ugt1a10** | A13\|Ugt1a13 | UDP glycosyltransferase 1 family, polypeptide A10 | protein-coding |
| chr17_56302916_56303756 | chr17 | 56302917 | 56303756 | + | 0.855194 | promoter-TSS (NM_010931) | promoter-TSS (NM_010931) | -1 | NM_010931 | 18140 | Mm.42196 | NM_010931 | ENSMUSG00000001228 | **Uhrf1** | AL022808\|ICBP90\|Np95\|RNF106 | ubiquitin-like, containing PHD and RING finger domains, 1 | protein-coding |
| chr10_7483677_7484039 | chr10 | 7483678 | 7484039 | + | 1.26725 | Intergenic | Intergenic | -10381 | NM_029975 | 77777 | Mm.442442 | NM_029975 | ENSMUSG00000079685 | **Ulbp1** | A430108B07Rik\|MULT1 | UL16 binding protein 1 | protein-coding |
| chr10_7484341_7484648 | chr10 | 7484342 | 7484648 | + | 1.07716 | Intergenic | Intergenic | -11018 | NM_029975 | 77777 | Mm.442442 | NM_029975 | ENSMUSG00000079685 | **Ulbp1** | A430108B07Rik\|MULT1 | UL16 binding protein 1 | protein-coding |
| chr10_7483033_7484039 | chr10 | 7483034 | 7484039 | + | 1.03519 | Intergenic | Intergenic | -10059 | NM_029975 | 77777 | Mm.442442 | NM_029975 | ENSMUSG00000079685 | **Ulbp1** | A430108B07Rik\|MULT1 | UL16 binding protein 1 | protein-coding |
| chr10_7388248_7388565 | chr10 | 7388249 | 7388565 | + | 0.99293 | Intergenic | (CAGA)n\|Simple_repeat\|Simple_repeat | 85070 | NM_029975 | 77777 | Mm.442442 | NM_029975 | ENSMUSG00000079685 | **Ulbp1** | A430108B07Rik\|MULT1 | UL16 binding protein 1 | protein-coding |
| chr10_7491813_7492331 | chr10 | 7491814 | 7492331 | + | 0.862942 | Intergenic | Intergenic | -18595 | NM_029975 | 77777 | Mm.442442 | NM_029975 | ENSMUSG00000079685 | **Ulbp1** | A430108B07Rik\|MULT1 | UL16 binding protein 1 | protein-coding |
| chr10_7491654_7492724 | chr10 | 7491655 | 7492724 | + | 0.746874 | Intergenic | Intergenic | -18712 | NM_029975 | 77777 | Mm.442442 | NM_029975 | ENSMUSG00000079685 | **Ulbp1** | A430108B07Rik\|MULT1 | UL16 binding protein 1 | protein-coding |
| chr11_82920441_82921144 | chr11 | 82920442 | 82921144 | + | 1.04193 | intron (NM_178680, intron 8 of 19) | MER21C\|LTR\|ERVL | 9540 | NM_178680 | 217012 | Mm.295205 | NM_178680 | ENSMUSG00000018845 | **Unc45b** | AA445617\|Cmya4\|D230041A13Rik\|Unc45 | unc-45 homolog B (C. elegans) | protein-coding |
| chr10_60808485_60808983 | chr10 | 60808486 | 60808983 | + | 0.685389 | intron (NM_029770, intron 1 of 16) | intron (NM_029770, intron 1 of 16) | 22847 | NM_029770 | 107449 | Mm.290433 | NM_029770 | ENSMUSG00000020099 | **Unc5b** | 6330415E02Rik\|A630020F16\|D10Bwg0792e\|Unc5h2 | unc-5 homolog B (C. elegans) | protein-coding |
| chr12_102947291_102947606 | chr12 | 102947292 | 102947606 | + | 0.708178 | Intergenic | Intergenic | -1410 | NM_001081017 | 217843 | Mm.387073 | NM_177621 | ENSMUSG00000021198 | **Unc79** | 9030205A07Rik\|Mlca3 | unc-79 homolog (C. elegans) | protein-coding |
| chr12_102947188_102947621 | chr12 | 102947189 | 102947621 | + | 0.61192 | Intergenic | (TTTTG)n\|Simple_repeat\|Simple_repeat | -1454 | NM_001081017 | 217843 | Mm.387073 | NM_177621 | ENSMUSG00000021198 | **Unc79** | 9030205A07Rik\|Mlca3 | unc-79 homolog (C. elegans) | protein-coding |
| chr2_6014378_6014877 | chr2 | 6014379 | 6014877 | + | 0.7851 | intron (NM_001081132, intron 10 of 21) | MTA_Mm\|LTR\|ERVL-MaLR | 63159 | NM_001081132 | 326622 | Mm.31102 | NM_001081132 | ENSMUSG00000043241 | **Upf2** | - | UPF2 regulator of nonsense transcripts homolog (yeast) | protein-coding |
| chrX_37117371_37117766 | chrX | 37117372 | 37117766 | + | 0.739038 | Intergenic | RSINE1\|SINE\|B4 | -7247 | NM_026573 | 68134 | Mm.271160 | NM_026573 | ENSMUSG00000036572 | **Upf3b** | 5730594O13Rik\|AI317193\|AW541158\|RENT3B\|UPF3X | UPF3 regulator of nonsense transcripts homolog B (yeast) | protein-coding |
| chr2_58589382_58590332 | chr2 | 58589383 | 58590332 | + | 1.31277 | intron (NM_001289660, intron 1 of 7) | intron (NM_001289660, intron 1 of 7) | 22523 | NM_001289660 | 76654 | Mm.200370 | NM_029692 | ENSMUSG00000026839 | **Upp2** | 1700124F02Rik\|AI266885\|UDRPASE2\|UP2\|UPASE2 | uridine phosphorylase 2 | protein-coding |
| chr2_58580019_58580857 | chr2 | 58580020 | 58580857 | + | 0.825896 | intron (NM_001289660, intron 1 of 7) | intron (NM_001289660, intron 1 of 7) | 13104 | NM_001289660 | 76654 | Mm.200370 | NM_029692 | ENSMUSG00000026839 | **Upp2** | 1700124F02Rik\|AI266885\|UDRPASE2\|UP2\|UPASE2 | uridine phosphorylase 2 | protein-coding |
| chrX_104488046_104488412 | chrX | 104488047 | 104488412 | + | 1.06856 | intron (NM_001081189, intron 1 of 6) | intron (NM_001081189, intron 1 of 6) | 5447 | NM_001081189 | 331487 | Mm.438275 | NM_001081189 | | **Uprt** | Gm774 | uracil phosphoribosyltransferase (FUR1) homolog (S. cerevisiae) | protein-coding |
| chrX_104488041_104488628 | chrX | 104488042 | 104488628 | + | 1.05258 | intron (NM_001081189, intron 1 of 6) | MER5A\|DNA\|hAT-Charlie | 5553 | NM_001081189 | 331487 | Mm.438275 | NM_001081189 | | **Uprt** | Gm774 | uracil phosphoribosyltransferase (FUR1) homolog (S. cerevisiae) | protein-coding |
| chr17_27131182_27131781 | chr17 | 27131183 | 27131781 | + | 0.922957 | intron (NM_026063, intron 1 of 3) | intron (NM_026063, intron 1 of 3) | 2409 | NM_026063 | 67267 | Mm.272389 | NM_026063 | ENSMUSG00000024208 | **Uqcc2** | 2900010M23Rik\|Cbp6\|M19\|Mnf1 | ubiquinol-cytochrome c reductase complex assembly factor 2 | protein-coding |
| chr17_27131146_27131977 | chr17 | 27131147 | 27131977 | + | 0.79006 | intron (NM_026063, intron 1 of 3) | intron (NM_026063, intron 1 of 3) | 2329 | NM_026063 | 67267 | Mm.272389 | NM_026063 | ENSMUSG00000024208 | **Uqcc2** | 2900010M23Rik\|Cbp6\|M19\|Mnf1 | ubiquinol-cytochrome c reductase complex assembly factor 2 | protein-coding |
| chr13_30564845_30565460 | chr13 | 30564846 | 30565460 | + | 1.45082 | Intergenic | MIR3\|SINE\|MIR | -19837 | NM_025710 | 66694 | Mm.181933 | NM_025710 | ENSMUSG00000038462 | **Uqcrfs1** | 4430402G14Rik\|AI875505 | ubiquinol-cytochrome c reductase, Rieske iron-sulfur polypeptide 1 | protein-coding |
| chr13_30567219_30567738 | chr13 | 30567220 | 30567738 | + | 1.15386 | Intergenic | Intergenic | -22163 | NM_025710 | 66694 | Mm.181933 | NM_025710 | ENSMUSG00000038462 | **Uqcrfs1** | 4430402G14Rik\|AI875505 | ubiquinol-cytochrome c reductase, Rieske iron-sulfur polypeptide 1 | protein-coding |
| chr13_30573047_30573495 | chr13 | 30573048 | 30573495 | + | 1.01129 | Intergenic | Intergenic | -27955 | NM_025710 | 66694 | Mm.181933 | NM_025710 | ENSMUSG00000038462 | **Uqcrfs1** | 4430402G14Rik\|AI875505 | ubiquinol-cytochrome c reductase, Rieske iron-sulfur polypeptide 1 | protein-coding |
| chr13_30553990_30554240 | chr13 | 30553991 | 30554240 | + | 0.870823 | Intergenic | L1M2\|LINE\|L1 | -8799 | NM_025710 | 66694 | Mm.181933 | NM_025710 | ENSMUSG00000038462 | **Uqcrfs1** | 4430402G14Rik\|AI875505 | ubiquinol-cytochrome c reductase, Rieske iron-sulfur polypeptide 1 | protein-coding |
| chr13_30551815_30552159 | chr13 | 30551816 | 30552159 | + | 0.843679 | Intergenic | L1MB3\|LINE\|L1 | -6671 | NM_025710 | 66694 | Mm.181933 | NM_025710 | ENSMUSG00000038462 | **Uqcrfs1** | 4430402G14Rik\|AI875505 | ubiquinol-cytochrome c reductase, Rieske iron-sulfur polypeptide 1 | protein-coding |

| | | | | | | | | | | | | | | | | | |
|---|---|---|---|---|---|---|---|---|---|---|---|---|---|---|---|---|---|
| chr8_124085467_124086006 | chr8 | 124085468 | 124086006 | + | 1.25529 | Intergenic | Intergenic | 62264 | NM_001029876 | 382038 | Mm.334891 | NM_001029876 | ENSMUSG00000031976 | **Urb2** | C77575\|mKIAA0133 | URB2 ribosome biogenesis 2 homolog (S. cerevisiae) | protein-coding |
| chr11_5737203_5737490 | chr11 | 5737204 | 5737490 | + | 1.00838 | intron (NM_001077661, intron 1 of 5) | MER44B\|DNA\|TcMar-Tigger | 3829 | NM_178623 | 72046 | Mm.271657 | NM_178623 | ENSMUSG00000049680 | **Urgcp** | 2010005J08Rik\|9130001I21Rik\|AW060359\|Urg4\|mKIAA1507 | upregulator of cell proliferation | protein-coding |
| chr7_38032317_38032907 | chr7 | 38032318 | 38032907 | + | 0.629999 | Intergenic | Intergenic | -13060 | NM_011274 | 19777 | Mm.387220 | NM_011274 | ENSMUSG00000030421 | **Uri1** | C80913\|NNX3\|Rmp | URI1, prefoldin-like chaperone | protein-coding |
| chr8_71324697_71325222 | chr8 | 71324698 | 71325222 | + | 0.610528 | intron (NM_015742, intron 7 of 39) | intron (NM_015742, intron 7 of 39) | -41888 | NM_001145780 | 67023 | Mm.41890 | NM_025917 | ENSMUSG00000002395 | **Use1** | 2010315L10Rik\|5730403H22Rik\|AV002165\|D12\|Ed2\|Q-snare\|p31 | unconventional SNARE in the ER 1 homolog (S. cerevisiae) | protein-coding |
| chr8_119948294_119948973 | chr8 | 119948295 | 119948973 | + | 0.668383 | intron (NM_009462, intron 7 of 13) | intron (NM_009462, intron 7 of 13) | 37782 | NM_009462 | 22224 | Mm.256910 | NM_009462 | ENSMUSG00000031826 | **Usp10** | 2610014N07Rik\|UBPO\|Uchrp\|mKIAA0190 | ubiquitin specific peptidase 10 | protein-coding |
| chrX_20771743_20772447 | chrX | 20771744 | 20772447 | + | 1.04706 | Intergenic | Lx8\|LINE\|L1 | 68186 | NM_145628 | 236733 | Mm.34489 | NM_145628 | ENSMUSG00000031066 | **Usp11** | 6230415D12Rik\|mKIAA4085 | ubiquitin specific peptidase 11 | protein-coding |
| chr2_31026646_31027168 | chr2 | 31026647 | 31027168 | + | 1.05102 | 3' UTR (NM_001177650, exon 14 of 14) | 3' UTR (NM_001177650, exon 14 of 14) | 44628 | NM_028846 | 74270 | Mm.346654 | NM_028846 | ENSMUSG00000026854 | **Usp20** | 1700055M05Rik\|AI467231\|Vdu2 | ubiquitin specific peptidase 20 | protein-coding |
| chr2_31036532_31037285 | chr2 | 31036533 | 31037285 | + | 0.943252 | intron (NM_001038700, intron 11 of 14) | intron (NM_001038700, intron 11 of 14) | 54630 | NM_028846 | 74270 | Mm.346654 | NM_028846 | ENSMUSG00000026854 | **Usp20** | 1700055M05Rik\|AI467231\|Vdu2 | ubiquitin specific peptidase 20 | protein-coding |
| chr4_106279161_106279823 | chr4 | 106279162 | 106279823 | + | 1.43906 | Intergenic | Intergenic | -36721 | NM_183225 | 329908 | Mm.234544 | NM_183225 | ENSMUSG00000028514 | **Usp24** | 2700066K03Rik\|2810030C21Rik\|AI414051\|B130021E18\|C79851 | ubiquitin specific peptidase 24 | protein-coding |
| chr4_106281150_106282003 | chr4 | 106281151 | 106282003 | + | 1.36992 | Intergenic | Intergenic | -34636 | NM_183225 | 329908 | Mm.234544 | NM_183225 | ENSMUSG00000028514 | **Usp24** | 2700066K03Rik\|2810030C21Rik\|AI414051\|B130021E18\|C79851 | ubiquitin specific peptidase 24 | protein-coding |
| chr16_77037872_77038668 | chr16 | 77037873 | 77038668 | + | 1.5361 | intron (NM_013918, intron 3 of 23) | intron (NM_013918, intron 3 of 23) | 24201 | NM_013918 | 30940 | Mm.40986 | NM_013918 | ENSMUSG00000022867 | **Usp25** | - | ubiquitin specific peptidase 25 | protein-coding |
| chr16_77015573_77015763 | chr16 | 77015574 | 77015763 | + | 0.674203 | intron (NM_013918, intron 1 of 23) | intron (NM_013918, intron 1 of 23) | 1599 | NM_013918 | 30940 | Mm.40986 | NM_013918 | ENSMUSG00000022867 | **Usp25** | - | ubiquitin specific peptidase 25 | protein-coding |
| chr9_48986715_48987042 | chr9 | 48986716 | 48987042 | + | 0.927023 | intron (NM_175482, intron 1 of 23) | intron (NM_175482, intron 1 of 23) | 1494 | NM_175482 | 235323 | Mm.21630 | NM_175482 | ENSMUSG00000032267 | **Usp28** | 9830148O20Rik\|AU022237\|mKIAA1515 | ubiquitin specific peptidase 28 | protein-coding |
| chr9_66571966_66572274 | chr9 | 66571967 | 66572274 | + | 0.870162 | intron (NM_144937, intron 1 of 14) | intron (NM_144937, intron 1 of 14) | 20860 | NM_144937 | 235441 | Mm.38976 | NM_144937 | ENSMUSG00000032376 | **Usp3** | AA409661\|BC017156 | ubiquitin specific peptidase 3 | protein-coding |
| chr9_66571945_66572621 | chr9 | 66571946 | 66572621 | + | 0.740888 | intron (NM_144937, intron 1 of 14) | intron (NM_144937, intron 1 of 14) | 20697 | NM_144937 | 235441 | Mm.38976 | NM_144937 | ENSMUSG00000032376 | **Usp3** | AA409661\|BC017156 | ubiquitin specific peptidase 3 | protein-coding |
| chr7_97210033_97210640 | chr7 | 97210034 | 97210640 | + | 1.4191 | intron (NM_001162477, intron 1 of 9) | intron (NM_001162477, intron 1 of 9) | 115627 | NM_001177412 | 244144 | Mm.297541 | NM_001177412 | ENSMUSG00000035713 | **Usp35** | Gm1088\|Gm493 | ubiquitin specific peptidase 35 | protein-coding |
| chr7_97211010_97211513 | chr7 | 97211011 | 97211513 | + | 0.880026 | intron (NM_001162477, intron 1 of 9) | intron (NM_001162477, intron 1 of 9) | 114702 | NM_001177412 | 244144 | Mm.297541 | NM_001177412 | ENSMUSG00000035713 | **Usp35** | Gm1088\|Gm493 | ubiquitin specific peptidase 35 | protein-coding |
| chr8_81004817_81005618 | chr8 | 81004818 | 81005618 | + | 1.16724 | intron (NM_027554, intron 3 of 9) | intron (NM_027554, intron 3 of 9) | 9688 | NM_027554 | 74841 | Mm.246018 | NM_027554 | ENSMUSG00000038250 | **Usp38** | 4631402N15Rik\|4833420O05Rik\|AA536967\|AU044701\|AW544820\|mKIAA1891 | ubiquitin specific peptidase 38 | protein-coding |
| chr8_81069299_81069750 | chr8 | 81069300 | 81069750 | + | 0.611426 | Intergenic | Intergenic | -54619 | NM_027554 | 74841 | Mm.246018 | NM_027554 | ENSMUSG00000038250 | **Usp38** | 4631402N15Rik\|4833420O05Rik\|AA536967\|AU044701\|AW544820\|mKIAA1891 | ubiquitin specific peptidase 38 | protein-coding |
| chr6_72345687_72346008 | chr6 | 72345688 | 72346008 | + | 0.945698 | promoter-TSS (NM_138592) | promoter-TSS (NM_138592) | -673 | NM_138592 | 28035 | Mm.281900 | NM_138592 | ENSMUSG00000056305 | **Usp39** | AA408960\|AI894154\|CGI-21\|D6Wsu157e\|SAD1 | ubiquitin specific peptidase 39 | protein-coding |
| chr10_93829841_93830273 | chr10 | 93829842 | 93830273 | + | 0.854188 | Intergenic | Intergenic | -1498 | NM_183199 | 327799 | Mm.491598 | NM_183199 | ENSMUSG00000020020 | **Usp44** | E430004F17Rik | ubiquitin specific peptidase 44 | protein-coding |
| chr7_112053489_112054342 | chr7 | 112053490 | 112054342 | + | 0.607087 | intron (NM_133758, intron 2 of 27) | intron (NM_133758, intron 2 of 27) | 30410 | NM_177249 | 74996 | Mm.16974 | NM_133758 | ENSMUSG00000059263 | **Usp47** | 4930502N04Rik\|A630020C16Rik | ubiquitin specific peptidase 47 | protein-coding |
| chr2_6593895_6594630 | chr2 | 6593896 | 6594630 | + | 1.78882 | intron (NM_001110232, intron 10 of 15) | intron (NM_001110232, intron 10 of 15) | 241527 | NM_181399 | 98910 | Mm.40673 | NM_181399 | ENSMUSG00000039046 | **Usp6nl** | AI316785\|RNTRE\|TRE2NL\|mKIAA0019 | USP6 N-terminal like | protein-coding |
| chr2_6594777_6595047 | chr2 | 6594778 | 6595047 | + | 1.6277 | intron (NM_001110232, intron 10 of 15) | intron (NM_001110232, intron 10 of 15) | 242176 | NM_181399 | 98910 | Mm.40673 | NM_181399 | ENSMUSG00000039046 | **Usp6nl** | AI316785\|RNTRE\|TRE2NL\|mKIAA0019 | USP6 N-terminal like | protein-coding |
| chr2_6557656_6558201 | chr2 | 6557657 | 6558201 | + | 1.50532 | intron (NM_001110231, intron 13 of 15) | intron (NM_001110231, intron 13 of 15) | 205193 | NM_181399 | 98910 | Mm.40673 | NM_181399 | ENSMUSG00000039046 | **Usp6nl** | AI316785\|RNTRE\|TRE2NL\|mKIAA0019 | USP6 N-terminal like | protein-coding |
| chr2_6592033_6592599 | chr2 | 6592034 | 6592599 | + | 1.44954 | intron (NM_001110232, intron 10 of 15) | intron (NM_001110232, intron 10 of 15) | 239580 | NM_181399 | 98910 | Mm.40673 | NM_181399 | ENSMUSG00000039046 | **Usp6nl** | AI316785\|RNTRE\|TRE2NL\|mKIAA0019 | USP6 N-terminal like | protein-coding |
| chr2_6320190_6320568 | chr2 | 6320191 | 6320568 | + | 1.29192 | non-coding (NR_040333, exon 7 of 7) | non-coding (NR_040333, exon 7 of 7) | -2378 | NM_001080548 | 98910 | Mm.40673 | NM_181399 | ENSMUSG00000039046 | **Usp6nl** | AI316785\|RNTRE\|TRE2NL\|mKIAA0019 | USP6 N-terminal like | protein-coding |
| chr2_6333554_6333968 | chr2 | 6333555 | 6333968 | + | 1.27769 | intron (NM_001080548, intron 1 of 14) | intron (NM_001080548, intron 1 of 14) | 11004 | NM_001080548 | 98910 | Mm.40673 | NM_181399 | ENSMUSG00000039046 | **Usp6nl** | AI316785\|RNTRE\|TRE2NL\|mKIAA0019 | USP6 N-terminal like | protein-coding |
| chr2_6560894_6561146 | chr2 | 6560895 | 6561146 | + | 1.27285 | intron (NM_001110232, intron 11 of 15) | intron (NM_001110232, intron 11 of 15) | 208284 | NM_181399 | 98910 | Mm.40673 | NM_181399 | ENSMUSG00000039046 | **Usp6nl** | AI316785\|RNTRE\|TRE2NL\|mKIAA0019 | USP6 N-terminal like | protein-coding |
| chr2_6598479_6598840 | chr2 | 6598480 | 6598840 | + | 1.26318 | intron (NM_001110232, intron 10 of 15) | intron (NM_001110232, intron 10 of 15) | 245924 | NM_181399 | 98910 | Mm.40673 | NM_181399 | ENSMUSG00000039046 | **Usp6nl** | AI316785\|RNTRE\|TRE2NL\|mKIAA0019 | USP6 N-terminal like | protein-coding |
| chr2_6598910_6599136 | chr2 | 6598911 | 6599136 | + | 1.1578 | intron (NM_001110232, intron 10 of 15) | intron (NM_001110232, intron 10 of 15) | 246287 | NM_181399 | 98910 | Mm.40673 | NM_181399 | ENSMUSG00000039046 | **Usp6nl** | AI316785\|RNTRE\|TRE2NL\|mKIAA0019 | USP6 N-terminal like | protein-coding |
| chr2_6366449_6367294 | chr2 | 6366450 | 6367294 | + | 1.14527 | intron (NM_181399, intron 1 of 13) | intron (NM_181399, intron 1 of 13) | 14136 | NM_181399 | 98910 | Mm.40673 | NM_181399 | ENSMUSG00000039046 | **Usp6nl** | AI316785\|RNTRE\|TRE2NL\|mKIAA0019 | USP6 N-terminal like | protein-coding |
| chr2_6366464_6367002 | chr2 | 6366465 | 6367002 | + | 1.06849 | intron (NM_181399, intron 1 of 13) | intron (NM_181399, intron 1 of 13) | 13997 | NM_181399 | 98910 | Mm.40673 | NM_181399 | ENSMUSG00000039046 | **Usp6nl** | AI316785\|RNTRE\|TRE2NL\|mKIAA0019 | USP6 N-terminal like | protein-coding |
| chr2_6599254_6599504 | chr2 | 6599255 | 6599504 | + | 0.99461 | intron (NM_001110232, intron 10 of 15) | intron (NM_001110232, intron 10 of 15) | 246643 | NM_181399 | 98910 | Mm.40673 | NM_181399 | ENSMUSG00000039046 | **Usp6nl** | AI316785\|RNTRE\|TRE2NL\|mKIAA0019 | USP6 N-terminal like | protein-coding |
| chr2_6545457_6545935 | chr2 | 6545458 | 6545935 | + | 0.955748 | 3' UTR (NM_001110231, exon 16 of 16) | 3' UTR (NM_001110231, exon 16 of 16) | 192960 | NM_181399 | 98910 | Mm.40673 | NM_181399 | ENSMUSG00000039046 | **Usp6nl** | AI316785\|RNTRE\|TRE2NL\|mKIAA0019 | USP6 N-terminal like | protein-coding |
| chr2_6592765_6593031 | chr2 | 6592766 | 6593031 | + | 0.946772 | intron (NM_001110232, intron 10 of 15) | intron (NM_001110232, intron 10 of 15) | 240162 | NM_181399 | 98910 | Mm.40673 | NM_181399 | ENSMUSG00000039046 | **Usp6nl** | AI316785\|RNTRE\|TRE2NL\|mKIAA0019 | USP6 N-terminal like | protein-coding |
| chr2_6379740_6380619 | chr2 | 6379741 | 6380619 | + | 0.938456 | intron (NM_181399, intron 1 of 13) | intron (NM_181399, intron 1 of 13) | 27444 | NM_181399 | 98910 | Mm.40673 | NM_181399 | ENSMUSG00000039046 | **Usp6nl** | AI316785\|RNTRE\|TRE2NL\|mKIAA0019 | USP6 N-terminal like | protein-coding |
| chr2_6378699_6379031 | chr2 | 6378700 | 6379031 | + | 0.822062 | intron (NM_181399, intron 1 of 13) | intron (NM_181399, intron 1 of 13) | 26129 | NM_181399 | 98910 | Mm.40673 | NM_181399 | ENSMUSG00000039046 | **Usp6nl** | AI316785\|RNTRE\|TRE2NL\|mKIAA0019 | USP6 N-terminal like | protein-coding |
| chr2_6378699_6379153 | chr2 | 6378700 | 6379153 | + | 0.813038 | intron (NM_181399, intron 1 of 13) | intron (NM_181399, intron 1 of 13) | 26190 | NM_181399 | 98910 | Mm.40673 | NM_181399 | ENSMUSG00000039046 | **Usp6nl** | AI316785\|RNTRE\|TRE2NL\|mKIAA0019 | USP6 N-terminal like | protein-coding |
| chr2_6559462_6559907 | chr2 | 6559463 | 6559907 | + | 0.794166 | intron (NM_001110231, intron 13 of 15) | intron (NM_001110231, intron 13 of 15) | 206949 | NM_181399 | 98910 | Mm.40673 | NM_181399 | ENSMUSG00000039046 | **Usp6nl** | AI316785\|RNTRE\|TRE2NL\|mKIAA0019 | USP6 N-terminal like | protein-coding |
| chr2_6565277_6565833 | chr2 | 6565278 | 6565833 | + | 0.718313 | intron (NM_001110232, intron 11 of 15) | intron (NM_001110232, intron 11 of 15) | 212819 | NM_181399 | 98910 | Mm.40673 | NM_181399 | ENSMUSG00000039046 | **Usp6nl** | AI316785\|RNTRE\|TRE2NL\|mKIAA0019 | USP6 N-terminal like | protein-coding |

| | | | | | | | | | | | | | | | | | |
|---|---|---|---|---|---|---|---|---|---|---|---|---|---|---|---|---|---|
| chr2_6602245_6602500 | chr2 | 6602246 | 6602500 | + | 0.714385 | intron (NM_001110232, intron 9 of 15) | Lx9\|LINE\|L1 | 249637 | NM_181399 | 98910 | Mm.40673 | NM_181399 | ENSMUSG00000039046 | **Usp6nl** | AI316785\|RNTRE\|TRE2NL\|mKIAA0019 | USP6 N-terminal like | protein-coding |
| chr2_6565290_6565567 | chr2 | 6565291 | 6565567 | + | 0.66176 | intron (NM_001110232, intron 11 of 15) | Tigger5\|DNA\|TcMar-Tigger | 212693 | NM_181399 | 98910 | Mm.40673 | NM_181399 | ENSMUSG00000039046 | **Usp6nl** | AI316785\|RNTRE\|TRE2NL\|mKIAA0019 | USP6 N-terminal like | protein-coding |
| chr16_8728169_8728838 | chr16 | 8728170 | 8728838 | + | 0.680625 | intron (NM_001003918, intron 1 of 30) | intron (NM_001003918, intron 1 of 30) | 9838 | NM_001003918 | 252870 | Mm.295330 | NM_001003918 | ENSMUSG00000022710 | **Usp7** | 2210010O09Rik\|AA409944\|AA617399\|AU019296\|AW548146\|C80752\|Hausp | ubiquitin specific peptidase 7 | protein-coding |
| chr16_8731418_8732076 | chr16 | 8731419 | 8732076 | + | 0.679558 | intron (NM_001003918, intron 1 of 30) | intron (NM_001003918, intron 1 of 30) | 6595 | NM_001003918 | 252870 | Mm.295330 | NM_001003918 | ENSMUSG00000022710 | **Usp7** | 2210010O09Rik\|AA409944\|AA617399\|AU019296\|AW548146\|C80752\|Hausp | ubiquitin specific peptidase 7 | protein-coding |
| chr5_149202854_149203325 | chr5 | 149202855 | 149203325 | + | 1.15199 | intron (NM_001013378, intron 6 of 9) | intron (NM_001013378, intron 6 of 9) | 18530 | NM_001115149 | 231915 | Mm.29402 | NM_001013378 | ENSMUSG00000041264 | **Uspl1** | E430026A01Rik | ubiquitin specific peptidase like 1 | protein-coding |
| chr5_149202597_149203490 | chr5 | 149202598 | 149203490 | + | 0.808065 | intron (NM_001013378, intron 6 of 9) | intron (NM_001013378, intron 6 of 9) | 18484 | NM_001115149 | 231915 | Mm.29402 | NM_001013378 | ENSMUSG00000041264 | **Uspl1** | E430026A01Rik | ubiquitin specific peptidase like 1 | protein-coding |
| chr10_8363247_8363596 | chr10 | 8363248 | 8363596 | + | 0.806934 | intron (NM_177387, intron 2 of 7) | intron (NM_177387, intron 2 of 7) | 155403 | NM_177387 | 338362 | Mm.41163 | NM_177387 | ENSMUSG00000047712 | **Ust** | D930010O20Rik\|UA2OST | uronyl-2-sulfotransferase | protein-coding |
| chr10_8415055_8415802 | chr10 | 8415056 | 8415802 | + | 0.695023 | intron (NM_177387, intron 1 of 7) | intron (NM_177387, intron 1 of 7) | 103396 | NM_177387 | 338362 | Mm.41163 | NM_177387 | ENSMUSG00000047712 | **Ust** | D930010O20Rik\|UA2OST | uronyl-2-sulfotransferase | protein-coding |
| chr1_78682022_78682728 | chr1 | 78682023 | 78682728 | + | 1.01582 | intron (NM_028817, intron 3 of 15) | intron (NM_028817, intron 3 of 15) | 23935 | NM_001136226 | 195434 | Mm.276016 | NM_001001981 | ENSMUSG00000079470 | **Utp14b** | 4932411L21Rik\|jsd | UTP14, U3 small nucleolar ribonucleoprotein, homolog B (yeast) | protein-coding |
| chr10_12821172_12821788 | chr10 | 12821173 | 12821788 | + | 1.28347 | intron (NM_011682, intron 1 of 73) | intron (NM_011682, intron 1 of 73) | 40255 | NM_011682 | 22288 | Mm.331784 | NM_011682 | ENSMUSG00000019820 | **Utrn** | AA589569\|DRP\|Dmdl | utrophin | protein-coding |
| chr10_12798214_12798733 | chr10 | 12798215 | 12798733 | + | 1.04422 | intron (NM_011682, intron 1 of 73) | intron (NM_011682, intron 1 of 73) | 63261 | NM_011682 | 22288 | Mm.331784 | NM_011682 | ENSMUSG00000019820 | **Utrn** | AA589569\|DRP\|Dmdl | utrophin | protein-coding |
| chr10_12754309_12754879 | chr10 | 12754310 | 12754879 | + | 0.860928 | intron (NM_011682, intron 2 of 73) | intron (NM_011682, intron 2 of 73) | 107141 | NM_011682 | 22288 | Mm.331784 | NM_011682 | ENSMUSG00000019820 | **Utrn** | AA589569\|DRP\|Dmdl | utrophin | protein-coding |
| chr10_12704239_12704726 | chr10 | 12704240 | 12704726 | + | 0.819853 | intron (NM_011682, intron 22 of 73) | intron (NM_011682, intron 22 of 73) | 157252 | NM_011682 | 22288 | Mm.331784 | NM_011682 | ENSMUSG00000019820 | **Utrn** | AA589569\|DRP\|Dmdl | utrophin | protein-coding |
| chr10_12608250_12608559 | chr10 | 12608251 | 12608559 | + | 0.776472 | intron (NM_011682, intron 50 of 73) | ORR1B1\|LTR\|ERVL-MaLR | 253330 | NM_011682 | 22288 | Mm.331784 | NM_011682 | ENSMUSG00000019820 | **Utrn** | AA589569\|DRP\|Dmdl | utrophin | protein-coding |
| chr10_12609969_12610244 | chr10 | 12609970 | 12610244 | + | 0.773126 | intron (NM_011682, intron 50 of 73) | intron (NM_011682, intron 50 of 73) | 251628 | NM_011682 | 22288 | Mm.331784 | NM_011682 | ENSMUSG00000019820 | **Utrn** | AA589569\|DRP\|Dmdl | utrophin | protein-coding |
| chr10_12754117_12755264 | chr10 | 12754118 | 12755264 | + | 0.771662 | intron (NM_011682, intron 2 of 73) | intron (NM_011682, intron 2 of 73) | 107044 | NM_011682 | 22288 | Mm.331784 | NM_011682 | ENSMUSG00000019820 | **Utrn** | AA589569\|DRP\|Dmdl | utrophin | protein-coding |
| chr10_12601203_12601601 | chr10 | 12601204 | 12601601 | + | 0.742122 | intron (NM_011682, intron 50 of 73) | B3A\|SINE\|B2 | 260333 | NM_011682 | 22288 | Mm.331784 | NM_011682 | ENSMUSG00000019820 | **Utrn** | AA589569\|DRP\|Dmdl | utrophin | protein-coding |
| chr10_12494006_12494827 | chr10 | 12494007 | 12494827 | + | 0.721569 | intron (NM_011682, intron 53 of 73) | intron (NM_011682, intron 53 of 73) | 367318 | NM_011682 | 22288 | Mm.331784 | NM_011682 | ENSMUSG00000019820 | **Utrn** | AA589569\|DRP\|Dmdl | utrophin | protein-coding |
| chr10_12809663_12809917 | chr10 | 12809664 | 12809917 | + | 0.700511 | intron (NM_011682, intron 1 of 73) | intron (NM_011682, intron 1 of 73) | 51945 | NM_011682 | 22288 | Mm.331784 | NM_011682 | ENSMUSG00000019820 | **Utrn** | AA589569\|DRP\|Dmdl | utrophin | protein-coding |
| chr10_12888398_12888707 | chr10 | 12888399 | 12888707 | + | 0.700317 | Intergenic | Intergenic | -26818 | NM_011682 | 22288 | Mm.331784 | NM_011682 | ENSMUSG00000019820 | **Utrn** | AA589569\|DRP\|Dmdl | utrophin | protein-coding |
| chr10_12812673_12813048 | chr10 | 12812674 | 12813048 | + | 0.683514 | intron (NM_011682, intron 1 of 73) | PB1D10\|SINE\|Alu | 48874 | NM_011682 | 22288 | Mm.331784 | NM_011682 | ENSMUSG00000019820 | **Utrn** | AA589569\|DRP\|Dmdl | utrophin | protein-coding |
| chr10_12716087_12716574 | chr10 | 12716088 | 12716574 | + | 0.675365 | intron (NM_011682, intron 18 of 73) | intron (NM_011682, intron 18 of 73) | 145404 | NM_011682 | 22288 | Mm.331784 | NM_011682 | ENSMUSG00000019820 | **Utrn** | AA589569\|DRP\|Dmdl | utrophin | protein-coding |
| chr10_12888350_12888808 | chr10 | 12888351 | 12888808 | + | 0.665942 | Intergenic | Intergenic | -26844 | NM_011682 | 22288 | Mm.331784 | NM_011682 | ENSMUSG00000019820 | **Utrn** | AA589569\|DRP\|Dmdl | utrophin | protein-coding |
| chr4_151006065_151006340 | chr4 | 151006066 | 151006340 | + | 0.879226 | intron (NM_001289877, intron 20 of 20) | MTD\|LTR\|ERVL-MaLR | 9106 | NM_011910 | 24111 | Mm.89984 | NM_011910 | ENSMUSG00000028963 | **Uts2** | UII\|Ucn2\|prepro-UII | urotensin 2 | protein-coding |
| chr4_151005708_151006499 | chr4 | 151005709 | 151006499 | + | 0.791112 | intron (NM_001289877, intron 20 of 20) | MTD\|LTR\|ERVL-MaLR | 9007 | NM_011910 | 24111 | Mm.89984 | NM_011910 | ENSMUSG00000028963 | **Uts2** | UII\|Ucn2\|prepro-UII | urotensin 2 | protein-coding |
| chr16_27349956_27350579 | chr16 | 27349957 | 27350579 | + | 1.47634 | intron (NM_198112, intron 4 of 4) | intron (NM_198112, intron 4 of 4) | 19971 | NM_198166 | 224065 | Mm.326918 | NM_198166 | ENSMUSG00000056423 | **Uts2b** | Gm538\|U2B\|Urp\|Uts2d | urotensin 2B | protein-coding |
| chr16_27350214_27350548 | chr16 | 27350215 | 27350548 | + | 1.44838 | intron (NM_198112, intron 4 of 4) | intron (NM_198112, intron 4 of 4) | 19858 | NM_198166 | 224065 | Mm.326918 | NM_198166 | ENSMUSG00000056423 | **Uts2b** | Gm538\|U2B\|Urp\|Uts2d | urotensin 2B | protein-coding |
| chr16_27355174_27355768 | chr16 | 27355175 | 27355768 | + | 1.4182 | intron (NM_198166, intron 4 of 4) | intron (NM_198166, intron 4 of 4) | 14768 | NM_198166 | 224065 | Mm.326918 | NM_198166 | ENSMUSG00000056423 | **Uts2b** | Gm538\|U2B\|Urp\|Uts2d | urotensin 2B | protein-coding |
| chr16_27349180_27349694 | chr16 | 27349181 | 27349694 | + | 1.0674 | intron (NM_198112, intron 4 of 4) | intron (NM_198112, intron 4 of 4) | 20802 | NM_198166 | 224065 | Mm.326918 | NM_198166 | ENSMUSG00000056423 | **Uts2b** | Gm538\|U2B\|Urp\|Uts2d | urotensin 2B | protein-coding |
| chr16_27349231_27349871 | chr16 | 27349232 | 27349871 | + | 1.0504 | intron (NM_198112, intron 4 of 4) | intron (NM_198112, intron 4 of 4) | 20688 | NM_198166 | 224065 | Mm.326918 | NM_198166 | ENSMUSG00000056423 | **Uts2b** | Gm538\|U2B\|Urp\|Uts2d | urotensin 2B | protein-coding |
| chr16_27354545_27354907 | chr16 | 27354546 | 27354907 | + | 0.898302 | intron (NM_198166, intron 4 of 4) | intron (NM_198166, intron 4 of 4) | 15513 | NM_198166 | 224065 | Mm.326918 | NM_198166 | ENSMUSG00000056423 | **Uts2b** | Gm538\|U2B\|Urp\|Uts2d | urotensin 2B | protein-coding |
| chr16_27354313_27354909 | chr16 | 27354314 | 27354909 | + | 0.873977 | intron (NM_198166, intron 4 of 4) | intron (NM_198166, intron 4 of 4) | 15628 | NM_198166 | 224065 | Mm.326918 | NM_198166 | ENSMUSG00000056423 | **Uts2b** | Gm538\|U2B\|Urp\|Uts2d | urotensin 2B | protein-coding |
| chr1_43818036_43818678 | chr1 | 43818037 | 43818678 | + | 0.74885 | intron (NM_026430, intron 1 of 14) | intron (NM_026430, intron 1 of 14) | 9351 | NM_026430 | 67883 | Mm.387202 | NM_026430 | | **Uxs1** | 1600025I13Rik\|AI451869\|AI649125\|AW550562 | UDP-glucuronate decarboxylase 1 | protein-coding |
| chr1_43818388_43818673 | chr1 | 43818389 | 43818673 | + | 0.739663 | intron (NM_026430, intron 1 of 14) | intron (NM_026430, intron 1 of 14) | 9177 | NM_026430 | 67883 | Mm.387202 | NM_026430 | | **Uxs1** | 1600025I13Rik\|AI451869\|AI649125\|AW550562 | UDP-glucuronate decarboxylase 1 | protein-coding |
| chr1_43819053_43819719 | chr1 | 43819054 | 43819719 | + | 0.657478 | intron (NM_026430, intron 1 of 14) | MLT1A0\|LTR\|ERVL-MaLR | 8322 | NM_026430 | 67883 | Mm.387202 | NM_026430 | | **Uxs1** | 1600025I13Rik\|AI451869\|AI649125\|AW550562 | UDP-glucuronate decarboxylase 1 | protein-coding |
| chr4_151108161_151108858 | chr4 | 151108162 | 151108858 | + | 0.668633 | intron (NM_001081557, intron 11 of 23) | CpG | -50557 | NM_009498 | 22319 | Mm.273930 | NM_009498 | ENSMUSG00000028955 | **Vamp3** | D130027G05Rik\|VAMP-3\|ceb | vesicle-associated membrane protein 3 | protein-coding |
| chr1_171981949_171982572 | chr1 | 171981950 | 171982572 | + | 1.78358 | Intergenic | Intergenic | 45007 | NM_033509 | 93840 | Mm.36148 | NM_033509 | ENSMUSG00000026556 | **Vangl2** | C530001F03Rik\|Lp\|Lpp1\|Ltap\|Vang1l2\|loop-tail\|ska17\|ska<m17Jus>\|stbm\|strabismus | vang-like 2 (van gogh, Drosophila) | protein-coding |
| chr1_171982776_171983360 | chr1 | 171982777 | 171983360 | + | 0.830843 | Intergenic | Intergenic | 44200 | NM_033509 | 93840 | Mm.36148 | NM_033509 | ENSMUSG00000026556 | **Vangl2** | C530001F03Rik\|Lp\|Lpp1\|Ltap\|Vang1l2\|loop-tail\|ska17\|ska<m17Jus>\|stbm\|strabismus | vang-like 2 (van gogh, Drosophila) | protein-coding |
| chr2_173730257_173730878 | chr2 | 173730258 | 173730878 | + | 1.08831 | Intergenic | RMER5\|LTR\|ERV1 | -7003 | NM_019806 | 56491 | Mm.260456 | NM_019806 | ENSMUSG00000054455 | **Vapb** | AI225786\|AI840687\|AI848259\|D2Abb2e\|R75548\|VAP33b\|Vamp33b | vesicle-associated membrane protein, associated protein B and C | protein-coding |
| chr2_173730039_173730916 | chr2 | 173730040 | 173730916 | + | 0.948251 | Intergenic | RMER5\|LTR\|ERV1 | -7093 | NM_019806 | 56491 | Mm.260456 | NM_019806 | ENSMUSG00000054455 | **Vapb** | AI225786\|AI840687\|AI848259\|D2Abb2e\|R75548\|VAP33b\|Vamp33b | vesicle-associated membrane protein, associated protein B and C | protein-coding |
| chr2_173762836_173763372 | chr2 | 173762837 | 173763372 | + | 0.905196 | intron (NM_019806, intron 2 of 5) | intron (NM_019806, intron 2 of 5) | 25533 | NM_019806 | 56491 | Mm.260456 | NM_019806 | ENSMUSG00000054455 | **Vapb** | AI225786\|AI840687\|AI848259\|D2Abb2e\|R75548\|VAP33b\|Vamp33b | vesicle-associated membrane protein, associated protein B and C | protein-coding |
| chr2_173762639_173763475 | chr2 | 173762640 | 173763475 | + | 0.789526 | intron (NM_019806, intron 2 of 5) | intron (NM_019806, intron 2 of 5) | 25486 | NM_019806 | 56491 | Mm.260456 | NM_019806 | ENSMUSG00000054455 | **Vapb** | AI225786\|AI840687\|AI848259\|D2Abb2e\|R75548\|VAP33b\|Vamp33b | vesicle-associated membrane protein, associated protein B and C | protein-coding |
| chr17_35002344_35002897 | chr17 | 35002345 | 35002897 | + | 0.7121 | intron (NM_011690, intron 2 of 29) | intron (NM_011690, intron 2 of 29) | 1714 | NM_011690 | 22321 | Mm.28420 | NM_011690 | ENSMUSG00000007029 | **Vars** | Bat6\|D17H6S56E\|G7a\|Vars2 | valyl-tRNA synthetase | protein-coding |

| | | | | | | | | | | | | | | | | | |
|---|---|---|---|---|---|---|---|---|---|---|---|---|---|---|---|---|---|
| chr12_86694424_86694813 | chr12 | 86694425 | 86694813 | + | 0.890196 | 3' UTR (NM_177354, exon 7 of 7) | 3' UTR (NM_177354, exon 7 of 7) | 15919 | NM_177354 | 238328 | Mm.216101 | NM_177354 | | **Vash1** | AI834978\|D930046M13Rik\|G630009D10Rik | vasohibin 1 | protein-coding |
| chr12_86694292_86694801 | chr12 | 86694293 | 86694801 | + | 0.83181 | 3' UTR (NM_177354, exon 7 of 7) | 3' UTR (NM_177354, exon 7 of 7) | 15847 | NM_177354 | 238328 | Mm.216101 | NM_177354 | | **Vash1** | AI834978\|D930046M13Rik\|G630009D10Rik | vasohibin 1 | protein-coding |
| chr12_86693432_86693735 | chr12 | 86693433 | 86693735 | + | 0.740402 | 3' UTR (NM_177354, exon 7 of 7) | 3' UTR (NM_177354, exon 7 of 7) | 14884 | NM_177354 | 238328 | Mm.216101 | NM_177354 | | **Vash1** | AI834978\|D930046M13Rik\|G630009D10Rik | vasohibin 1 | protein-coding |
| chr7_19267121_19267405 | chr7 | 19267122 | 19267405 | + | 0.667595 | intron (NM_001282021, intron 1 of 12) | PB1D10\|SINE\|Alu | 4591 | NM_001282021 | 22323 | Mm.9684 | NM_009499 | ENSMUSG00000030403 | **Vasp** | - | vasodilator-stimulated phosphoprotein | protein-coding |
| chr17_57284995_57285738 | chr17 | 57284996 | 57285738 | + | 0.922116 | intron (NM_011691, intron 1 of 26) | intron (NM_011691, intron 1 of 26) | 6267 | NM_001163815 | 22324 | Mm.248172 | NM_011691 | | **Vav1** | Vav\|vav-T | vav 1 oncogene | protein-coding |
| chr2_27404482_27405812 | chr2 | 27404483 | 27405812 | + | 0.829319 | intron (NM_009500, intron 1 of 27) | intron (NM_009500, intron 1 of 27) | 21678 | NM_009500 | 22325 | Mm.179011 | NM_009500 | ENSMUSG00000009621 | **Vav2** | 2810040F13Rik\|AI847175 | vav 2 oncogene | protein-coding |
| chr2_27406932_27407277 | chr2 | 27406933 | 27407277 | + | 0.665558 | intron (NM_009500, intron 1 of 27) | intron (NM_009500, intron 1 of 27) | 19720 | NM_009500 | 22325 | Mm.179011 | NM_009500 | ENSMUSG00000009621 | **Vav2** | 2810040F13Rik\|AI847175 | vav 2 oncogene | protein-coding |
| chr3_109273043_109273794 | chr3 | 109273044 | 109273794 | + | 1.17783 | Intergenic | Intergenic | -67264 | NM_020505 | 57257 | Mm.282257 | NM_020505 | ENSMUSG00000033721 | **Vav3** | A530094I06Rik\|AA986410\|Idd18.1 | vav 3 oncogene | protein-coding |
| chr3_109273039_109273374 | chr3 | 109273040 | 109273374 | + | 1.02095 | Intergenic | Intergenic | -67476 | NM_020505 | 57257 | Mm.282257 | NM_020505 | ENSMUSG00000033721 | **Vav3** | A530094I06Rik\|AA986410\|Idd18.1 | vav 3 oncogene | protein-coding |
| chr3_109434880_109435698 | chr3 | 109434881 | 109435698 | + | 0.843829 | intron (NM_020505, intron 2 of 26) | intron (NM_020505, intron 2 of 26) | 94606 | NM_020505 | 57257 | Mm.282257 | NM_020505 | ENSMUSG00000033721 | **Vav3** | A530094I06Rik\|AA986410\|Idd18.1 | vav 3 oncogene | protein-coding |
| chr3_109420815_109421437 | chr3 | 109420816 | 109421437 | + | 0.768088 | intron (NM_020505, intron 1 of 26) | intron (NM_020505, intron 1 of 26) | 80443 | NM_020505 | 57257 | Mm.282257 | NM_020505 | ENSMUSG00000033721 | **Vav3** | A530094I06Rik\|AA986410\|Idd18.1 | vav 3 oncogene | protein-coding |
| chr3_109747651_109748268 | chr3 | 109747652 | 109748268 | + | 0.723867 | Intergenic | ORR1F\|LTR\|ERVL-MaLR | 173953 | NM_146139 | 57257 | Mm.282257 | NM_020505 | ENSMUSG00000033721 | **Vav3** | A530094I06Rik\|AA986410\|Idd18.1 | vav 3 oncogene | protein-coding |
| chr3_109311347_109311781 | chr3 | 109311348 | 109311781 | + | 0.716995 | Intergenic | Intergenic | -29119 | NM_020505 | 57257 | Mm.282257 | NM_020505 | ENSMUSG00000033721 | **Vav3** | A530094I06Rik\|AA986410\|Idd18.1 | vav 3 oncogene | protein-coding |
| chr3_109273423_109273792 | chr3 | 109273424 | 109273792 | + | 0.69073 | Intergenic | Intergenic | -67075 | NM_020505 | 57257 | Mm.282257 | NM_020505 | ENSMUSG00000033721 | **Vav3** | A530094I06Rik\|AA986410\|Idd18.1 | vav 3 oncogene | protein-coding |
| chr3_109443871_109444439 | chr3 | 109443872 | 109444439 | + | 0.675444 | intron (NM_020505, intron 2 of 26) | intron (NM_020505, intron 2 of 26) | 103472 | NM_020505 | 57257 | Mm.282257 | NM_020505 | ENSMUSG00000033721 | **Vav3** | A530094I06Rik\|AA986410\|Idd18.1 | vav 3 oncogene | protein-coding |
| chr3_109326088_109326363 | chr3 | 109326089 | 109326363 | + | 0.668922 | Intergenic | Intergenic | -14457 | NM_020505 | 57257 | Mm.282257 | NM_020505 | ENSMUSG00000033721 | **Vav3** | A530094I06Rik\|AA986410\|Idd18.1 | vav 3 oncogene | protein-coding |
| chr3_109384084_109384789 | chr3 | 109384085 | 109384789 | + | 0.658069 | intron (NM_020505, intron 1 of 26) | intron (NM_020505, intron 1 of 26) | 43754 | NM_020505 | 57257 | Mm.282257 | NM_020505 | ENSMUSG00000033721 | **Vav3** | A530094I06Rik\|AA986410\|Idd18.1 | vav 3 oncogene | protein-coding |
| chr3_109420249_109421465 | chr3 | 109420250 | 109421465 | + | 0.636431 | intron (NM_020505, intron 1 of 26) | intron (NM_020505, intron 1 of 26) | 80174 | NM_020505 | 57257 | Mm.282257 | NM_020505 | ENSMUSG00000033721 | **Vav3** | A530094I06Rik\|AA986410\|Idd18.1 | vav 3 oncogene | protein-coding |
| chr3_109384249_109384647 | chr3 | 109384250 | 109384647 | + | 0.63152 | intron (NM_020505, intron 1 of 26) | intron (NM_020505, intron 1 of 26) | 43765 | NM_020505 | 57257 | Mm.282257 | NM_020505 | ENSMUSG00000033721 | **Vav3** | A530094I06Rik\|AA986410\|Idd18.1 | vav 3 oncogene | protein-coding |
| chr3_116129198_116129664 | chr3 | 116129199 | 116129664 | + | 0.714285 | 5' UTR (NM_011693, exon 1 of 9) | 5' UTR (NM_011693, exon 1 of 9) | 257 | NM_011693 | 22329 | Mm.76649 | NM_011693 | ENSMUSG00000027962 | **Vcam1** | CD106\|Vcam-1 | vascular cell adhesion molecule 1 | protein-coding |
| chr13_89888978_89889526 | chr13 | 89888979 | 89889526 | + | 1.38004 | intron (NM_028012, intron 7 of 7) | intron (NM_028012, intron 7 of 7) | -146740 | NM_001134475 | 13003 | Mm.158700 | NM_019389 | ENSMUSG00000021614 | **Vcan** | 5430420N07Rik\|9430051N09\|Cspg2\|DPEAAE\|NG2\|PG-M\|PG-M(V0)\|PG-M(V1)\|hdf | versican | protein-coding |
| chr14_20967223_20967645 | chr14 | 20967224 | 20967645 | + | 1.42799 | intron (NM_009502, intron 1 of 20) | B1F1\|SINE\|Alu | 38001 | NM_009502 | 22330 | Mm.279361 | NM_009502 | ENSMUSG00000021823 | **Vcl** | 9430097D22\|AA571387\|AI462105\|AW545629 | vinculin | protein-coding |
| chr14_20977196_20977978 | chr14 | 20977197 | 20977978 | + | 1.36381 | intron (NM_009502, intron 2 of 20) | intron (NM_009502, intron 2 of 20) | 48154 | NM_009502 | 22330 | Mm.279361 | NM_009502 | ENSMUSG00000021823 | **Vcl** | 9430097D22\|AA571387\|AI462105\|AW545629 | vinculin | protein-coding |
| chr14_20903011_20903519 | chr14 | 20903012 | 20903519 | + | 0.7261 | Intergenic | MLT1F\|LTR\|ERVL-MaLR | -26168 | NM_009502 | 22330 | Mm.279361 | NM_009502 | ENSMUSG00000021823 | **Vcl** | 9430097D22\|AA571387\|AI462105\|AW545629 | vinculin | protein-coding |
| chr14_20973804_20974249 | chr14 | 20973805 | 20974249 | + | 0.68738 | intron (NM_009502, intron 2 of 20) | intron (NM_009502, intron 2 of 20) | 44594 | NM_009502 | 22330 | Mm.279361 | NM_009502 | ENSMUSG00000021823 | **Vcl** | 9430097D22\|AA571387\|AI462105\|AW545629 | vinculin | protein-coding |
| chr12_69537399_69538519 | chr12 | 69537400 | 69538519 | + | 0.77763 | Intergenic | Intergenic | 45069 | NM_001033236 | 207965 | Mm.102487 | NM_001033236 | ENSMUSG00000049882 | **Vcpkmt** | Gm71\|Mettl21d\|Mettl221d\|VCP-KMT | valosin containing protein lysine (K) methyltransferase | protein-coding |
| chr12_69595951_69596568 | chr12 | 69595952 | 69596568 | + | 0.777554 | intron (NM_001135559, intron 18 of 22) | intron (NM_001135559, intron 18 of 22) | -13232 | NM_001033236 | 207965 | Mm.102487 | NM_001033236 | ENSMUSG00000049882 | **Vcpkmt** | Gm71\|Mettl21d\|Mettl221d\|VCP-KMT | valosin containing protein lysine (K) methyltransferase | protein-coding |
| chr11_52324319_52324870 | chr11 | 52324320 | 52324870 | + | 0.951721 | Intergenic | Intergenic | -36520 | NM_011694 | 22333 | Mm.3555 | NM_011694 | ENSMUSG00000020402 | **Vdac1** | AL033343\|Vdac5 | voltage-dependent anion channel 1 | protein-coding |
| chr11_52324315_52325242 | chr11 | 52324316 | 52325242 | + | 0.869597 | Intergenic | Intergenic | -36336 | NM_011694 | 22333 | Mm.3555 | NM_011694 | ENSMUSG00000020402 | **Vdac1** | AL033343\|Vdac5 | voltage-dependent anion channel 1 | protein-coding |
| chr11_52333739_52334034 | chr11 | 52333740 | 52334034 | + | 0.773436 | Intergenic | Intergenic | -27228 | NM_011694 | 22333 | Mm.3555 | NM_011694 | ENSMUSG00000020402 | **Vdac1** | AL033343\|Vdac5 | voltage-dependent anion channel 1 | protein-coding |
| chr11_52333702_52334186 | chr11 | 52333703 | 52334186 | + | 0.652105 | Intergenic | Intergenic | -27171 | NM_011694 | 22333 | Mm.3555 | NM_011694 | ENSMUSG00000020402 | **Vdac1** | AL033343\|Vdac5 | voltage-dependent anion channel 1 | protein-coding |
| chr14_21829952_21830191 | chr14 | 21829953 | 21830191 | + | 0.947486 | Intergenic | Intergenic | -1489 | NM_011695 | 22334 | Mm.262327 | NM_011695 | ENSMUSG00000021771 | **Vdac2** | Vdac6\|mVDAC2\|mVDAC6 | voltage-dependent anion channel 2 | protein-coding |
| chr14_21806069_21806709 | chr14 | 21806070 | 21806709 | + | 0.855758 | Intergenic | Intergenic | -25172 | NM_011695 | 22334 | Mm.262327 | NM_011695 | ENSMUSG00000021771 | **Vdac2** | Vdac6\|mVDAC2\|mVDAC6 | voltage-dependent anion channel 2 | protein-coding |
| chr14_21806064_21807032 | chr14 | 21806065 | 21807032 | + | 0.67924 | Intergenic | Intergenic | -25013 | NM_011695 | 22334 | Mm.262327 | NM_011695 | ENSMUSG00000021771 | **Vdac2** | Vdac6\|mVDAC2\|mVDAC6 | voltage-dependent anion channel 2 | protein-coding |
| chr8_22535996_22536435 | chr8 | 22535997 | 22536435 | + | 0.709294 | intron (NM_011394, intron 2 of 10) | intron (NM_011394, intron 2 of 10) | 57597 | NM_001198998 | 22335 | Mm.227704 | NM_011696 | ENSMUSG00000008892 | **Vdac3** | - | voltage-dependent anion channel 3 | protein-coding |
| chr10_94021084_94021418 | chr10 | 94021085 | 94021418 | + | 1.49577 | intron (NM_172538, intron 1 of 11) | intron (NM_172538, intron 1 of 11) | 14497 | NM_172538 | 215008 | Mm.256845 | NM_172538 | ENSMUSG00000036099 | **Vezt** | 6330418D12\|AI854408\|vezatin | vezatin, adherens junctions transmembrane protein | protein-coding |
| chr10_94021609_94022013 | chr10 | 94021610 | 94022013 | + | 1.35383 | intron (NM_172538, intron 1 of 11) | intron (NM_172538, intron 1 of 11) | 13937 | NM_172538 | 215008 | Mm.256845 | NM_172538 | ENSMUSG00000036099 | **Vezt** | 6330418D12\|AI854408\|vezatin | vezatin, adherens junctions transmembrane protein | protein-coding |
| chr10_94032995_94033220 | chr10 | 94032996 | 94033220 | + | 0.754704 | intron (NM_172538, intron 1 of 11) | L1ME3A\|LINE\|L1 | 2640 | NM_172538 | 215008 | Mm.256845 | NM_172538 | ENSMUSG00000036099 | **Vezt** | 6330418D12\|AI854408\|vezatin | vezatin, adherens junctions transmembrane protein | protein-coding |
| chrX_57106908_57107356 | chrX | 57106909 | 57107356 | + | 1.21006 | Intergenic | Intergenic | 19026 | NM_133251 | 170828 | Mm.190616 | NM_133251 | ENSMUSG00000031131 | **Vgll1** | Tondu | vestigial like 1 homolog (Drosophila) | protein-coding |
| chrX_57143876_57144539 | chrX | 57143877 | 57144539 | + | 1.15955 | Intergenic | Intergenic | 56102 | NM_133251 | 170828 | Mm.190616 | NM_133251 | ENSMUSG00000031131 | **Vgll1** | Tondu | vestigial like 1 homolog (Drosophila) | protein-coding |
| chrX_57106731_57107418 | chrX | 57106732 | 57107418 | + | 1.09365 | Intergenic | Intergenic | 18969 | NM_133251 | 170828 | Mm.190616 | NM_133251 | ENSMUSG00000031131 | **Vgll1** | Tondu | vestigial like 1 homolog (Drosophila) | protein-coding |
| chrX_57137689_57138163 | chrX | 57137690 | 57138163 | + | 1.0612 | Intergenic | Intergenic | 49820 | NM_133251 | 170828 | Mm.190616 | NM_133251 | ENSMUSG00000031131 | **Vgll1** | Tondu | vestigial like 1 homolog (Drosophila) | protein-coding |
| chrX_57137394_57138182 | chrX | 57137395 | 57138182 | + | 0.953334 | Intergenic | Intergenic | 49682 | NM_133251 | 170828 | Mm.190616 | NM_133251 | ENSMUSG00000031131 | **Vgll1** | Tondu | vestigial like 1 homolog (Drosophila) | protein-coding |
| chrX_57141522_57141881 | chrX | 57141523 | 57141881 | + | 0.859014 | Intergenic | Intergenic | 53596 | NM_133251 | 170828 | Mm.190616 | NM_133251 | ENSMUSG00000031131 | **Vgll1** | Tondu | vestigial like 1 homolog (Drosophila) | protein-coding |
| chr6_114901427_114901786 | chr6 | 114901428 | 114901786 | + | 0.866901 | intron (NM_177683, intron 1 of 4) | intron (NM_177683, intron 1 of 4) | 20145 | NM_177683 | 232334 | Mm.268357 | NM_177683 | | **Vgll4** | BC048841\|VGL-4 | vestigial like 4 (Drosophila) | protein-coding |
| chr6_114861076_114861545 | chr6 | 114861077 | 114861545 | + | 0.813899 | TTS (NM_177683) | TTS (NM_177683) | 60441 | NM_177683 | 232334 | Mm.268357 | NM_177683 | | **Vgll4** | BC048841\|VGL-4 | vestigial like 4 (Drosophila) | protein-coding |
| chr6_114931209_114931608 | chr6 | 114931210 | 114931608 | + | 0.806368 | Intergenic | Intergenic | -9657 | NM_177683 | 232334 | Mm.268357 | NM_177683 | | **Vgll4** | BC048841\|VGL-4 | vestigial like 4 (Drosophila) | protein-coding |
| chr6_114842605_114843277 | chr6 | 114842606 | 114843277 | + | 0.770776 | intron (NM_001253717, intron 19 of 19) | intron (NM_001253717, intron 19 of 19) | 78811 | NM_177683 | 232334 | Mm.268357 | NM_177683 | | **Vgll4** | BC048841\|VGL-4 | vestigial like 4 (Drosophila) | protein-coding |
| chr1_74438060_74438516 | chr1 | 74438061 | 74438516 | + | 1.14986 | 3' UTR (NM_176972, exon 26 of 26) | 3' UTR (NM_176972, exon 26 of 26) | 28904 | NM_009509 | 22349 | Mm.471601 | NM_009509 | ENSMUSG00000026175 | **Vil1** | Vil | villin 1 | protein-coding |
| chr1_74437840_74438521 | chr1 | 74437841 | 74438521 | + | 1.10787 | 3' UTR (NM_176972, exon 26 of 26) | 3' UTR (NM_176972, exon 26 of 26) | 28797 | NM_009509 | 22349 | Mm.471601 | NM_009509 | ENSMUSG00000026175 | **Vil1** | Vil | villin 1 | protein-coding |
| chr9_119055410_119055741 | chr9 | 119055411 | 119055741 | + | 0.656395 | intron (NM_001164567, intron 1 of 19) | MLTR14\|LTR\|ERV1 | 2798 | NM_001164567 | 22351 | Mm.83817 | NM_011700 | ENSMUSG00000038775 | **Vill** | Villp | villin-like | protein-coding |
| chr2_13559793_13560062 | chr2 | 13559794 | 13560062 | + | 1.39056 | Intergenic | Intergenic | -14383 | NM_011701 | 22352 | Mm.268000 | NM_011701 | ENSMUSG00000026728 | **Vim** | - | vimentin | protein-coding |
| chr2_13559647_13560113 | chr2 | 13559648 | 13560113 | + | 1.37568 | Intergenic | Intergenic | -14431 | NM_011701 | 22352 | Mm.268000 | NM_011701 | ENSMUSG00000026728 | **Vim** | - | vimentin | protein-coding |
| chr12_116132959_116133314 | chr12 | 116132960 | 116133314 | + | 0.657828 | intron (NM_009511, intron 6 of 12) | intron (NM_009511, intron 6 of 12) | 55411 | NM_009511 | 22355 | Mm.395245 | NM_009511 | ENSMUSG00000011171 | **Vipr2** | VPAC2\|VPAC2R\|Vip2 | vasoactive intestinal peptide receptor 2 | protein-coding |
| chr19_27106770_27107419 | chr19 | 27106771 | 27107419 | + | 0.813326 | Intergenic | ORR1D2-int\|LTR\|ERVL-MaLR | -109925 | NM_013703 | 22359 | Mm.4141 | NM_013703 | ENSMUSG00000024924 | **Vldlr** | AA408956\|AI451093\|AW047288 | very low density lipoprotein receptor | protein-coding |
| chrX_71853748_71854174 | chrX | 71853749 | 71854174 | + | 1.4683 | Intergenic | Intergenic | 37203 | NM_001290780 | 67048 | Mm.41102 | NM_001081356 | ENSMUSG00000073131 | **Vma21** | 2610030H06Rik\|AI840175 | VMA21 vacuolar H+-ATPase homolog (S. cerevisiae) | protein-coding |
| chrX_71832166_71832743 | chrX | 71832167 | 71832743 | + | 0.867713 | Intergenic | Intergenic | 15697 | NM_001290780 | 67048 | Mm.41102 | NM_001081356 | ENSMUSG00000073131 | **Vma21** | 2610030H06Rik\|AI840175 | VMA21 vacuolar H+-ATPase homolog (S. cerevisiae) | protein-coding |

| | | | | | | | | | | | | | | | | | |
|---|---|---|---|---|---|---|---|---|---|---|---|---|---|---|---|---|---|
| chr17_19764437_19764772 | chr17 | 19764438 | 19764772 | + | 0.944516 | Intergenic | ORR1A2\|LTR\|ERVL-MaLR | -8758 | NM_001104565 | 627636 | Mm.469855 | NM_001104565 | ENSMUSG00000091771 | **Vmn2r103** | EG627636\|Vmn103 | vomeronasal 2, receptor 103 | protein-coding |
| chr10_130370700_130371774 | chr10 | 130370701 | 130371774 | + | 1.49371 | Intergenic | LTRIS2\|LTR\|ERV1 | 23004 | NM_001081448 | 625068 | Mm.359168 | NM_001081448 | ENSMUSG00000070601 | **Vmn2r84** | EG625068 | vomeronasal 2, receptor 84 | protein-coding |
| chr11_70512549_70513358 | chr11 | 70512550 | 70513358 | + | 1.12597 | TTS (NM_001013607) | TTS (NM_001013607) | 1662 | NM_001013607 | 327956 | Mm.244514 | NM_001013607 | ENSMUSG00000020830 | **Vmo1** | Gm741 | vitelline membrane outer layer 1 homolog (chicken) | protein-coding |
| chr19_16789970_16790434 | chr19 | 16789971 | 16790434 | + | 1.38221 | Intergenic | Intergenic | -9269 | NM_173028 | 271564 | Mm.211963 | NM_173028 | ENSMUSG00000046230 | **Vps13a** | 4930425F11\|4930516E05Rik\|4930543C13Rik\|9930023P20\|Chac\|Chorein\|D330038K10Rik\|mKIAA0986 | vacuolar protein sorting 13A (yeast) | protein-coding |
| chr19_16684759_16685475 | chr19 | 16684760 | 16685475 | + | 0.790052 | intron (NM_173028, intron 39 of 71) | intron (NM_173028, intron 39 of 71) | 95816 | NM_173028 | 271564 | Mm.211963 | NM_173028 | ENSMUSG00000046230 | **Vps13a** | 4930425F11\|4930516E05Rik\|4930543C13Rik\|9930023P20\|Chac\|Chorein\|D330038K10Rik\|mKIAA0986 | vacuolar protein sorting 13A (yeast) | protein-coding |
| chr19_16684861_16685222 | chr19 | 16684862 | 16685222 | + | 0.778782 | intron (NM_173028, intron 39 of 71) | intron (NM_173028, intron 39 of 71) | 95891 | NM_173028 | 271564 | Mm.211963 | NM_173028 | ENSMUSG00000046230 | **Vps13a** | 4930425F11\|4930516E05Rik\|4930543C13Rik\|9930023P20\|Chac\|Chorein\|D330038K10Rik\|mKIAA0986 | vacuolar protein sorting 13A (yeast) | protein-coding |
| chr15_35427799_35428171 | chr15 | 35427800 | 35428171 | + | 0.75135 | intron (NM_177151, intron 6 of 61) | Lx4A\|LINE\|L1 | 56439 | NM_177151 | 666173 | Mm.309250 | NM_177151 | ENSMUSG00000037646 | **Vps13b** | 1810042B05Rik\|2310042E16Rik\|4732488H20\|C330002D13Rik\|C87206\|Coh1\|D230005K13\|mKIAA0532\|mKIAA5032 | vacuolar protein sorting 13B (yeast) | protein-coding |
| chr15_35467009_35467510 | chr15 | 35467010 | 35467510 | + | 0.722037 | intron (NM_177151, intron 16 of 61) | intron (NM_177151, intron 16 of 61) | 95714 | NM_177151 | 666173 | Mm.309250 | NM_177151 | ENSMUSG00000037646 | **Vps13b** | 1810042B05Rik\|2310042E16Rik\|4732488H20\|C330002D13Rik\|C87206\|Coh1\|D230005K13\|mKIAA0532\|mKIAA5032 | vacuolar protein sorting 13B (yeast) | protein-coding |
| chr4_145219705_145220379 | chr4 | 145219706 | 145220379 | + | 0.946016 | intron (NM_011610, intron 8 of 9) | intron (NM_011610, intron 8 of 9) | -25037 | NM_001276465 | 230895 | Mm.240310 | NM_001033190 | ENSMUSG00000020220 | **Vps13d** | - | vacuolar protein sorting 13 D (yeast) | protein-coding |
| chr4_145209378_145209677 | chr4 | 145209379 | 145209677 | + | 0.943318 | Intergenic | B3\|SINE\|B2 | -14523 | NM_001276465 | 230895 | Mm.240310 | NM_001033190 | ENSMUSG00000020220 | **Vps13d** | - | vacuolar protein sorting 13 D (yeast) | protein-coding |
| chr4_145220586_145220925 | chr4 | 145220587 | 145220925 | + | 0.811012 | intron (NM_011610, intron 8 of 9) | intron (NM_011610, intron 8 of 9) | -25751 | NM_001276465 | 230895 | Mm.240310 | NM_001033190 | ENSMUSG00000020220 | **Vps13d** | - | vacuolar protein sorting 13 D (yeast) | protein-coding |
| chr4_145209149_145209682 | chr4 | 145209150 | 145209682 | + | 0.797723 | Intergenic | B3\|SINE\|B2 | -14411 | NM_001276465 | 230895 | Mm.240310 | NM_001033190 | ENSMUSG00000020220 | **Vps13d** | - | vacuolar protein sorting 13 D (yeast) | protein-coding |
| chr4_145201435_145201847 | chr4 | 145201436 | 145201847 | + | 0.694471 | Intergenic | ORR1D2\|LTR\|ERVL-MaLR | -6636 | NM_001276465 | 230895 | Mm.240310 | NM_001033190 | ENSMUSG00000020220 | **Vps13d** | - | vacuolar protein sorting 13 D (yeast) | protein-coding |
| chr4_145201556_145201763 | chr4 | 145201557 | 145201763 | + | 0.692411 | Intergenic | ORR1D2\|LTR\|ERVL-MaLR | -6655 | NM_001276465 | 230895 | Mm.240310 | NM_001033190 | ENSMUSG00000020220 | **Vps13d** | - | vacuolar protein sorting 13 D (yeast) | protein-coding |
| chr10_62490963_62491174 | chr10 | 62490964 | 62491174 | + | 0.711821 | Intergenic | Intergenic | -4264 | NM_133672 | 30930 | Mm.260703 | NM_133672 | ENSMUSG00000020078 | **Vps26a** | AA407240\|HB58\|Vps26 | vacuolar protein sorting 26 homolog A (yeast) | protein-coding |
| chr10_62477160_62477630 | chr10 | 62477161 | 62477630 | + | 0.619471 | intron (NM_133672, intron 2 of 8) | L1M5\|LINE\|L1 | 9203 | NM_001113355 | 30930 | Mm.260703 | NM_133672 | ENSMUSG00000020078 | **Vps26a** | AA407240\|HB58\|Vps26 | vacuolar protein sorting 26 homolog A (yeast) | protein-coding |
| chr3_96027959_96028232 | chr3 | 96027960 | 96028232 | + | 0.603747 | intron (NM_013841, intron 13 of 14) | intron (NM_013841, intron 13 of 14) | 30359 | NM_013841 | 22365 | Mm.263185 | NM_013841 | ENSMUSG00000015747 | **Vps45** | AI462172\|AW554165\|mVps45 | vacuolar protein sorting 45 (yeast) | protein-coding |
| chr11_76151033_76151432 | chr11 | 76151034 | 76151432 | + | 0.654115 | intron (NM_026664, intron 4 of 21) | intron (NM_026664, intron 4 of 21) | 28397 | NM_026664 | 68299 | Mm.46397 | NM_026664 | ENSMUSG00000017288 | **Vps53** | 2010002A08Rik\|2310040I21Rik\|3100002B05Rik\|Hccs1 | vacuolar protein sorting 53 (yeast) | protein-coding |
| chr11_21169397_21169969 | chr11 | 21169398 | 21169969 | + | 1.12438 | Intergenic | MERV1_LTR\|LTR\|ERV1 | -69349 | NM_139061 | 245944 | Mm.170103 | NM_139061 | ENSMUSG00000020128 | **Vps54** | Hcc8\|Vps54l\|mSLP8\|wr | vacuolar protein sorting 54 (yeast) | protein-coding |
| chr11_21221335_21221820 | chr11 | 21221336 | 21221820 | + | 0.996901 | Intergenic | Intergenic | -17454 | NM_139061 | 245944 | Mm.170103 | NM_139061 | ENSMUSG00000020128 | **Vps54** | Hcc8\|Vps54l\|mSLP8\|wr | vacuolar protein sorting 54 (yeast) | protein-coding |
| chr11_21224249_21224768 | chr11 | 21224250 | 21224768 | + | 0.659425 | Intergenic | Intergenic | -14523 | NM_139061 | 245944 | Mm.170103 | NM_139061 | ENSMUSG00000020128 | **Vps54** | Hcc8\|Vps54l\|mSLP8\|wr | vacuolar protein sorting 54 (yeast) | protein-coding |
| chr12_106030247_106030827 | chr12 | 106030248 | 106030827 | + | 1.00803 | intron (NM_001029843, intron 1 of 13) | intron (NM_001029843, intron 1 of 13) | 20274 | NM_001029843 | 22367 | Mm.2981 | NM_011705 | ENSMUSG00000021115 | **Vrk1** | 51PK | vaccinia related kinase 1 | protein-coding |
| chr12_106033016_106033366 | chr12 | 106033017 | 106033366 | + | 0.715419 | intron (NM_001029843, intron 1 of 13) | MER44B\|DNA\|TcMar-Tigger | 22928 | NM_001029843 | 22367 | Mm.2981 | NM_011705 | ENSMUSG00000021115 | **Vrk1** | 51PK | vaccinia related kinase 1 | protein-coding |
| chr11_26657092_26658134 | chr11 | 26657093 | 26658134 | + | 1.54501 | Intergenic | Intergenic | -63693 | NM_001252447 | 69922 | Mm.285351 | NM_027260 | ENSMUSG00000064090 | **Vrk2** | 2810003O05Rik\|AI447698 | vaccinia related kinase 2 | protein-coding |
| chr11_26601569_26602154 | chr11 | 26601570 | 26602154 | + | 1.53729 | Intergenic | MER34A\|LTR\|ERV1 | -7942 | NM_001252447 | 69922 | Mm.285351 | NM_027260 | ENSMUSG00000064090 | **Vrk2** | 2810003O05Rik\|AI447698 | vaccinia related kinase 2 | protein-coding |
| chr11_26640203_26640872 | chr11 | 26640204 | 26640872 | + | 0.754032 | Intergenic | Intergenic | -46618 | NM_001252447 | 69922 | Mm.285351 | NM_027260 | ENSMUSG00000064090 | **Vrk2** | 2810003O05Rik\|AI447698 | vaccinia related kinase 2 | protein-coding |
| chr11_26586441_26587476 | chr11 | 26586442 | 26587476 | + | 0.617771 | intron (NM_027260, intron 2 of 12) | MER130\|Unknown\|Unknown | 6961 | NM_027260 | 69922 | Mm.285351 | NM_027260 | ENSMUSG00000064090 | **Vrk2** | 2810003O05Rik\|AI447698 | vaccinia related kinase 2 | protein-coding |
| chr9_15225222_15225634 | chr9 | 15225223 | 15225634 | + | 0.749138 | Intergenic | LTRIS2\|LTR\|ERV1 | -13617 | NM_026955 | 69137 | Mm.41396 | NM_026955 | ENSMUSG00000031937 | **Vstm5** | 2200002K05Rik | V-set and transmembrane domain containing 5 | protein-coding |
| chr9_15241555_15241888 | chr9 | 15241556 | 15241888 | + | 0.71296 | intron (NM_026955, intron 1 of 3) | MLT1A0-int\|LTR\|ERVL-MaLR | 2677 | NM_026955 | 69137 | Mm.41396 | NM_026955 | ENSMUSG00000031937 | **Vstm5** | 2200002K05Rik | V-set and transmembrane domain containing 5 | protein-coding |
| chr19_55382742_55383266 | chr19 | 55382743 | 55383266 | + | 0.781092 | intron (NM_016862, intron 4 of 7) | intron (NM_016862, intron 4 of 7) | 66947 | NM_001293686 | 53611 | Mm.258637 | NM_016862 | ENSMUSG00000024983 | **Vti1a** | 1110014F16Rik\|1110018K19Rik\|4921537J05Rik\|MVti1\|MVti1a\|Vti1\|Vti1-rp2 | vesicle transport through interaction with t-SNAREs 1A | protein-coding |
| chr19_55382741_55383595 | chr19 | 55382742 | 55383595 | + | 0.719736 | intron (NM_016862, intron 4 of 7) | intron (NM_016862, intron 4 of 7) | 67111 | NM_001293686 | 53611 | Mm.258637 | NM_016862 | ENSMUSG00000024983 | **Vti1a** | 1110014F16Rik\|1110018K19Rik\|4921537J05Rik\|MVti1\|MVti1a\|Vti1\|Vti1-rp2 | vesicle transport through interaction with t-SNAREs 1A | protein-coding |
| chr4_155775648_155776089 | chr4 | 155775649 | 155776089 | + | 0.75007 | Intergenic | Intergenic | -1308 | NM_147776 | 246228 | Mm.26515 | NM_147776 | ENSMUSG00000042116 | **Vwa1** | 4932416A11Rik\|WARP | von Willebrand factor A domain containing 1 | protein-coding |
| chr7_120722673_120723227 | chr7 | 120722674 | 120723227 | + | 0.685874 | intron (NM_001164580, intron 1 of 2) | intron (NM_001164580, intron 1 of 2) | -16607 | NM_177697 | 233813 | Mm.389939 | NM_177697 | ENSMUSG00000030889 | **Vwa3a** | E030013G06Rik | von Willebrand factor A domain containing 3A | protein-coding |
| chr9_38728238_38728717 | chr9 | 38728239 | 38728717 | + | 0.914293 | intron (NM_172767, intron 10 of 19) | intron (NM_172767, intron 10 of 19) | 10210 | NM_172767 | 67776 | Mm.290950 | NM_172767 | ENSMUSG00000023186 | **Vwa5a** | 5830475I06Rik\|AW552491\|BCSC-1\|E130119J22\|Loh11cr2a | von Willebrand factor A domain containing 5A | protein-coding |
| chr9_38725140_38725574 | chr9 | 38725141 | 38725574 | + | 0.63324 | intron (NM_172767, intron 7 of 19) | intron (NM_172767, intron 7 of 19) | 7089 | NM_172767 | 67776 | Mm.290950 | NM_172767 | ENSMUSG00000023186 | **Vwa5a** | 5830475I06Rik\|AW552491\|BCSC-1\|E130119J22\|Loh11cr2a | von Willebrand factor A domain containing 5A | protein-coding |
| chr4_138549774_138550253 | chr4 | 138549775 | 138550253 | + | 0.632105 | Intergenic | Intergenic | 73978 | NM_029401 | 75718 | Mm.251530 | NM_029401 | | **Vwa5b1** | 4931403E03Rik | von Willebrand factor A domain containing 5B1 | protein-coding |
| chr14_78951488_78951869 | chr14 | 78951489 | 78951869 | + | 0.653006 | intron (NM_027906, intron 12 of 44) | MTD\|LTR\|ERVL-MaLR | 102501 | NM_173758 | 219189 | Mm.159651 | NM_027906 | ENSMUSG00000058997 | **Vwa8** | 1300010F03Rik\|4932416F07Rik\|AI957255\|Kiaa0564\|mKIAA0564 | von Willebrand factor A domain containing 8 | protein-coding |
| chr14_78951477_78952280 | chr14 | 78951478 | 78952280 | + | 0.636218 | intron (NM_027906, intron 12 of 44) | MTA_Mm\|LTR\|ERVL-MaLR | 102701 | NM_173758 | 219189 | Mm.159651 | NM_027906 | ENSMUSG00000058997 | **Vwa8** | 1300010F03Rik\|4932416F07Rik\|AI957255\|Kiaa0564\|mKIAA0564 | von Willebrand factor A domain containing 8 | protein-coding |
| chr11_10159542_10160083 | chr11 | 10159543 | 10160083 | + | 1.09878 | Intergenic | MTA_Mm\|LTR\|ERVL-MaLR | -954203 | NM_177033 | 319922 | Mm.337851 | NM_177033 | ENSMUSG00000050830 | **Vwc2** | A930041G11Rik\|G11\|PSST739\|UNQ739\|cradin | von Willebrand factor C domain containing 2 | protein-coding |
| chr11_10194654_10195142 | chr11 | 10194655 | 10195142 | + | 1.07835 | Intergenic | MER74A\|LTR\|ERVL | -919118 | NM_177033 | 319922 | Mm.337851 | NM_177033 | ENSMUSG00000050830 | **Vwc2** | A930041G11Rik\|G11\|PSST739\|UNQ739\|cradin | von Willebrand factor C domain containing 2 | protein-coding |
| chr11_10160141_10160406 | chr11 | 10160142 | 10160406 | + | 0.98717 | Intergenic | Lx2B\|LINE\|L1 | -953742 | NM_177033 | 319922 | Mm.337851 | NM_177033 | ENSMUSG00000050830 | **Vwc2** | A930041G11Rik\|G11\|PSST739\|UNQ739\|cradin | von Willebrand factor C domain containing 2 | protein-coding |
| chr6_125561531_125562059 | chr6 | 125561532 | 125562059 | + | 1.09478 | intron (NM_011708, intron 3 of 51) | intron (NM_011708, intron 3 of 51) | 8847 | NM_011708 | 22371 | Mm.22339 | NM_011708 | ENSMUSG00000001930 | **Vwf** | 6820430P06Rik\|AI551257\|B130011O06Rik\|C630030D09\|F8VWF\|VWD | Von Willebrand factor homolog | protein-coding |
| chr6_125540702_125541190 | chr6 | 125540703 | 125541190 | + | 0.939995 | Intergenic | Intergenic | -12002 | NM_011708 | 22371 | Mm.22339 | NM_011708 | ENSMUSG00000001930 | **Vwf** | 6820430P06Rik\|AI551257\|B130011O06Rik\|C630030D09\|F8VWF\|VWD | Von Willebrand factor homolog | protein-coding |
| chr18_7905350_7905879 | chr18 | 7905351 | 7905879 | + | 1.48674 | intron (NM_153085, intron 5 of 13) | intron (NM_153085, intron 5 of 13) | 36418 | NM_153085 | 225131 | Mm.204524 | NM_153085 | ENSMUSG00000024283 | **Wac** | 1110067P07Rik\|A230035H12Rik\|AI256735\|AI256776\|Wwp4 | WW domain containing adaptor with coiled-coil | protein-coding |
| chr18_7905345_7906088 | chr18 | 7905346 | 7906088 | + | 1.4303 | intron (NM_153085, intron 5 of 13) | intron (NM_153085, intron 5 of 13) | 36520 | NM_153085 | 225131 | Mm.204524 | NM_153085 | ENSMUSG00000024283 | **Wac** | 1110067P07Rik\|A230035H12Rik\|AI256735\|AI256776\|Wwp4 | WW domain containing adaptor with coiled-coil | protein-coding |
| chr18_7888596_7888948 | chr18 | 7888597 | 7888948 | + | 0.688612 | intron (NM_153085, intron 3 of 13) | L1ME1\|LINE\|L1 | 19575 | NM_153085 | 225131 | Mm.204524 | NM_153085 | ENSMUSG00000024283 | **Wac** | 1110067P07Rik\|A230035H12Rik\|AI256735\|AI256776\|Wwp4 | WW domain containing adaptor with coiled-coil | protein-coding |
| chr10_40969363_40970033 | chr10 | 40969364 | 40970033 | + | 0.787274 | Intergenic | Intergenic | 86164 | NM_031877 | 83767 | Mm.41353 | NM_031877 | ENSMUSG00000019831 | **Wasf1** | AI195380\|AI838537\|Scar\|WAVE\|WAVE-1 | WAS protein family, member 1 | protein-coding |
| chr17_66106554_66106871 | chr17 | 66106555 | 66106871 | + | 0.952941 | Intergenic | RMER15\|LTR\|ERVL | -4833 | NM_026833 | 68767 | Mm.296545 | NM_026833 | ENSMUSG00000024101 | **Wash** | 1110049F14Rik\|ORF19\|Wash1 | WAS protein family homolog | protein-coding |

| | | | | | | | | | | | | | | | | | |
|---|---|---|---|---|---|---|---|---|---|---|---|---|---|---|---|---|---|
| chr19_46629662_46629996 | chr19 | 46629663 | 46629996 | + | 0.686643 | intron (NM_001177813, intron 1 of 3) | intron (NM_001177813, intron 1 of 3) | 6428 | NM_001177813 | 226178 | Mm.329895 | NM_146099 | ENSMUSG00000047731 | **Wbp1l** | D19Wsu162e\|Opal1 | WW domain binding protein 1 like | protein-coding |
| chr19_46629657_46630198 | chr19 | 46629658 | 46630198 | + | 0.620146 | intron (NM_001177813, intron 1 of 3) | intron (NM_001177813, intron 1 of 3) | 6527 | NM_001177813 | 226178 | Mm.329895 | NM_146099 | ENSMUSG00000047731 | **Wbp1l** | D19Wsu162e\|Opal1 | WW domain binding protein 1 like | protein-coding |
| chrX_136243127_136243489 | chrX | 136243128 | 136243489 | + | 0.693534 | Intergenic | Intergenic | -1772 | NM_011712 | 22381 | Mm.512 | NM_011712 | ENSMUSG00000042712 | **Wbp5** | AL024233 | WW domain binding protein 5 | protein-coding |
| chr5_134878380_134879223 | chr5 | 134878381 | 134879223 | + | 0.830117 | Intergenic | B4\|SINE\|B4 | 27931 | NM_029681 | 76629 | Mm.159750 | NM_029681 | ENSMUSG00000040576 | **Wbscr28** | 1700111I05Rik | Williams-Beuren syndrome chromosome region 28 (human) | protein-coding |
| chr5_134877913_134879286 | chr5 | 134877914 | 134879286 | + | 0.760183 | Intergenic | Intergenic | 28133 | NM_029681 | 76629 | Mm.159750 | NM_029681 | ENSMUSG00000040576 | **Wbscr28** | 1700111I05Rik | Williams-Beuren syndrome chromosome region 28 (human) | protein-coding |
| chr1_79766303_79766552 | chr1 | 79766304 | 79766552 | + | 0.741329 | Intergenic | MER57E2\|LTR\|ERV1 | -4659 | NM_027057 | 69368 | Mm.293273 | NM_027057 | ENSMUSG00000073643 | **Wdfy1** | 1700013B03Rik\|1700120F24Rik\|FENS-1\|Jr1\|WDF1\|ZFYVE17\|mKIAA1435 | WD repeat and FYVE domain containing 1 | protein-coding |
| chr1_79736542_79736816 | chr1 | 79736543 | 79736816 | + | 0.694457 | intron (NM_027057, intron 2 of 4) | intron (NM_027057, intron 2 of 4) | 25090 | NM_001111279 | 69368 | Mm.293273 | NM_027057 | ENSMUSG00000073643 | **Wdfy1** | 1700013B03Rik\|1700120F24Rik\|FENS-1\|Jr1\|WDF1\|ZFYVE17\|mKIAA1435 | WD repeat and FYVE domain containing 1 | protein-coding |
| chr1_79766126_79766543 | chr1 | 79766127 | 79766543 | + | 0.673992 | Intergenic | Intergenic | -4566 | NM_027057 | 69368 | Mm.293273 | NM_027057 | ENSMUSG00000073643 | **Wdfy1** | 1700013B03Rik\|1700120F24Rik\|FENS-1\|Jr1\|WDF1\|ZFYVE17\|mKIAA1435 | WD repeat and FYVE domain containing 1 | protein-coding |
| chr14_62812789_62813102 | chr14 | 62812790 | 62813102 | + | 0.925441 | intron (NR_045503, intron 3 of 6) | intron (NR_045503, intron 3 of 6) | -24744 | NM_175546 | 268752 | Mm.1960 | NM_175546 | ENSMUSG00000014547 | **Wdfy2** | 5830485M08\|B130024L21Rik\|ProF\|ZFYVE22 | WD repeat and FYVE domain containing 2 | protein-coding |
| chr11_21688365_21688804 | chr11 | 21688366 | 21688804 | + | 0.769534 | intron (NM_145425, intron 6 of 17) | intron (NM_145425, intron 6 of 17) | 116304 | NM_145425 | 216560 | Mm.138090 | NM_145425 | ENSMUSG00000020319 | **Wdpcp** | AV249152 | WD repeat containing planar cell polarity effector | protein-coding |
| chr5_38566427_38566870 | chr5 | 38566428 | 38566870 | + | 1.6626 | Intergenic | MTD\|LTR\|ERVL-MaLR | -5054 | NM_011715 | 22388 | Mm.2654 | NM_011715 | ENSMUSG00000005103 | **Wdr1** | Aip1\|D5Wsu185e\|rede | WD repeat domain 1 | protein-coding |
| chr5_38566914_38567327 | chr5 | 38566915 | 38567327 | + | 1.36417 | Intergenic | RMER19B\|LTR\|ERVK | -5526 | NM_011715 | 22388 | Mm.2654 | NM_011715 | ENSMUSG00000005103 | **Wdr1** | Aip1\|D5Wsu185e\|rede | WD repeat domain 1 | protein-coding |
| chr5_38559867_38560119 | chr5 | 38559868 | 38560119 | + | 1.07907 | intron (NM_011715, intron 2 of 14) | intron (NM_011715, intron 2 of 14) | 1602 | NM_011715 | 22388 | Mm.2654 | NM_011715 | ENSMUSG00000005103 | **Wdr1** | Aip1\|D5Wsu185e\|rede | WD repeat domain 1 | protein-coding |
| chr5_38553900_38554441 | chr5 | 38553901 | 38554441 | + | 0.654919 | intron (NM_011715, intron 2 of 14) | intron (NM_011715, intron 2 of 14) | 7424 | NM_011715 | 22388 | Mm.2654 | NM_011715 | ENSMUSG00000005103 | **Wdr1** | Aip1\|D5Wsu185e\|rede | WD repeat domain 1 | protein-coding |
| chr7_129764388_129764889 | chr7 | 129764389 | 129764889 | + | 1.08712 | Intergenic | Intergenic | 172776 | NM_172255 | 207425 | Mm.229323 | NM_172255 | ENSMUSG00000042055 | **Wdr11** | 2900055P10Rik\|AW489876\|Brwd2\|mKIAA1351 | WD repeat domain 11 | protein-coding |
| chr1_181247167_181247731 | chr1 | 181247168 | 181247731 | + | 0.726345 | Intergenic | Intergenic | -35471 | NM_145514 | 226757 | Mm.289082 | NM_145514 | ENSMUSG00000038733 | **Wdr26** | 1600024A01Rik\|AA693241\|AI447817\|AU044014\|C77982\|Gid7 | WD repeat domain 26 | protein-coding |
| chr1_181203023_181203474 | chr1 | 181203024 | 181203474 | + | 0.648612 | intron (NM_145514, intron 3 of 13) | intron (NM_145514, intron 3 of 13) | 8729 | NM_145514 | 226757 | Mm.289082 | NM_145514 | ENSMUSG00000038733 | **Wdr26** | 1600024A01Rik\|AA693241\|AI447817\|AU044014\|C77982\|Gid7 | WD repeat domain 26 | protein-coding |
| chr4_62473416_62473665 | chr4 | 62473417 | 62473665 | + | 0.738057 | Intergenic | Intergenic | -2646 | NM_001290521 | 71354 | Mm.155573 | NM_023597 | ENSMUSG00000028391 | **Wdr31** | 5430402I10Rik\|spWD | WD repeat domain 31 | protein-coding |
| chr18_31840482_31840949 | chr18 | 31840483 | 31840949 | + | 1.03393 | intron (NM_028866, intron 7 of 21) | ORR1D1-int\|LTR\|ERVL-MaLR | 36659 | NM_001170966 | 74320 | Mm.277705 | NM_028866 | ENSMUSG00000024400 | **Wdr33** | 1110001N06Rik\|2310011G05Rik\|2810021O11Rik\|8430413N20Rik\|WDC146 | WD repeat domain 33 | protein-coding |
| chr13_8645684_8646362 | chr13 | 8645685 | 8646362 | + | 1.36438 | intron (NM_001289530, intron 3 of 9) | L1_Mus3\|LINE\|L1 | 225021 | NM_001039389 | 207615 | Mm.284654 | NM_172445 | ENSMUSG00000021147 | **Wdr37** | - | WD repeat domain 37 | protein-coding |
| chr13_8542717_8543312 | chr13 | 8542718 | 8543312 | + | 0.626358 | intron (NM_001289530, intron 1 of 9) | intron (NM_001289530, intron 1 of 9) | 328029 | NM_001039389 | 207615 | Mm.284654 | NM_172445 | ENSMUSG00000021147 | **Wdr37** | - | WD repeat domain 37 | protein-coding |
| chr2_38980489_38980856 | chr2 | 38980490 | 38980856 | + | 1.09707 | Intergenic | MamRep1879\|DNA\|hAT-Tip100? | -17636 | NM_029687 | 76646 | Mm.358777 | NM_029687 | ENSMUSG00000035295 | **Wdr38** | 1700123D08Rik | WD repeat domain 38 | protein-coding |
| chr2_38974150_38974627 | chr2 | 38974151 | 38974627 | + | 0.866791 | Intergenic | Intergenic | -23920 | NM_029687 | 76646 | Mm.358777 | NM_029687 | ENSMUSG00000035295 | **Wdr38** | 1700123D08Rik | WD repeat domain 38 | protein-coding |
| chr17_31482618_31483020 | chr17 | 31482619 | 31483020 | + | 0.657508 | Intergenic | MTE2b\|LTR\|ERVL-MaLR | 29668 | NM_021322 | 57773 | Mm.143771 | NM_021322 | ENSMUSG00000024037 | **Wdr4** | AI415180\|AI448349\|D530049K22Rik | WD repeat domain 4 | protein-coding |
| chr8_111520628_111520835 | chr8 | 111520629 | 111520835 | + | 0.715786 | intron (NM_001170743, intron 1 of 26) | LTRIS_Mus\|LTR\|ERV1 | 1375 | NM_001170742 | 319481 | Mm.247294 | NM_176923 | ENSMUSG00000031959 | **Wdr59** | 5430401O09Rik\|Cdw12 | WD repeat domain 59 | protein-coding |
| chr12_116266658_116267104 | chr12 | 116266659 | 116267104 | + | 0.96076 | Intergenic | Intergenic | -3856 | NM_146039 | 217935 | Mm.31382 | NM_146039 | ENSMUSG00000042050 | **Wdr60** | D430033N04Rik | WD repeat domain 60 | protein-coding |
| chr12_116170791_116171369 | chr12 | 116170792 | 116171369 | + | 0.737383 | Intergenic | Intergenic | 91945 | NM_146039 | 217935 | Mm.31382 | NM_146039 | ENSMUSG00000042050 | **Wdr60** | D430033N04Rik | WD repeat domain 60 | protein-coding |
| chr12_116269479_116269848 | chr12 | 116269480 | 116269848 | + | 0.695436 | Intergenic | MTE2b\|LTR\|ERVL-MaLR | -6639 | NM_146039 | 217935 | Mm.31382 | NM_146039 | ENSMUSG00000042050 | **Wdr60** | D430033N04Rik | WD repeat domain 60 | protein-coding |
| chr18_63896211_63896745 | chr18 | 63896212 | 63896745 | + | 0.717531 | intron (NM_001014981, intron 21 of 27) | intron (NM_001014981, intron 21 of 27) | 187783 | NM_001014981 | 104082 | Mm.30850 | NM_001014981 | ENSMUSG00000040560 | **Wdr7** | AI462727\|TRAG\|mKIAA0541 | WD repeat domain 7 | protein-coding |
| chr2_121570009_121570453 | chr2 | 121570010 | 121570453 | + | 1.33389 | intron (NM_172673, intron 6 of 14) | B3\|SINE\|B2 | 63508 | NM_001290986 | 241627 | Mm.293284 | NM_030234 | ENSMUSG00000027242 | **Wdr76** | 5830411K18Rik\|6030457D09\|AA517836 | WD repeat domain 76 | protein-coding |
| chr3_105958284_105958518 | chr3 | 105958285 | 105958518 | + | 0.707461 | intron (NM_009725, intron 4 of 8) | intron (NM_009725, intron 4 of 8) | -1097 | NM_027432 | 70465 | Mm.5110 | NM_027432 | ENSMUSG00000000561 | **Wdr77** | 2610003I18Rik\|2610312E17Rik\|C79984\|p44/MEP50 | WD repeat domain 77 | protein-coding |
| chr4_103098473_103098912 | chr4 | 103098474 | 103098912 | + | 1.08277 | intron (NM_146254, intron 2 of 16) | L1_Rod\|LINE\|L1 | 15606 | NM_146254 | 242584 | Mm.374877 | NM_146254 | ENSMUSG00000035126 | **Wdr78** | BC028975 | WD repeat domain 78 | protein-coding |
| chr11_17216493_17216809 | chr11 | 17216494 | 17216809 | + | 0.827803 | intron (NM_178909, intron 1 of 7) | B4A\|SINE\|B4 | 4758 | NM_178909 | 103784 | Mm.298132 | NM_178909 | | **Wdr92** | AI553587\|HZGJ | WD repeat domain 92 | protein-coding |
| chr2_59969053_59969480 | chr2 | 59969054 | 59969480 | + | 1.20657 | exon (NM_001001182, exon 7 of 36) | exon (NM_001001182, exon 7 of 36) | -86661 | NM_028118 | 72137 | Mm.288734 | NM_028118 | ENSMUSG00000026988 | **Wdsub1** | 1700048E19Rik\|2610014F08Rik | WD repeat, SAM and U-box domain containing 1 | protein-coding |
| chr2_59937687_59938033 | chr2 | 59937688 | 59938033 | + | 1.20009 | intron (NM_001001182, intron 15 of 35) | intron (NM_001001182, intron 15 of 35) | -55254 | NM_028118 | 72137 | Mm.288734 | NM_028118 | ENSMUSG00000026988 | **Wdsub1** | 1700048E19Rik\|2610014F08Rik | WD repeat, SAM and U-box domain containing 1 | protein-coding |
| chr2_59933044_59933430 | chr2 | 59933045 | 59933430 | + | 1.16983 | intron (NM_001001182, intron 18 of 35) | intron (NM_001001182, intron 18 of 35) | -50631 | NM_028118 | 72137 | Mm.288734 | NM_028118 | ENSMUSG00000026988 | **Wdsub1** | 1700048E19Rik\|2610014F08Rik | WD repeat, SAM and U-box domain containing 1 | protein-coding |
| chr7_110108952_110109605 | chr7 | 110108953 | 110109605 | + | 1.34293 | Intergenic | Intergenic | -12780 | NM_009516 | 22390 | Mm.287173 | NM_009516 | | **Wee1** | Wee1A | WEE 1 homolog 1 (S. pombe) | protein-coding |
| chr7_110120356_110120621 | chr7 | 110120357 | 110120621 | + | 1.00971 | Intergenic | Intergenic | -1570 | NM_009516 | 22390 | Mm.287173 | NM_009516 | | **Wee1** | Wee1A | WEE 1 homolog 1 (S. pombe) | protein-coding |
| chr11_83737125_83737863 | chr11 | 83737126 | 83737863 | + | 0.896589 | Intergenic | L1MA4\|LINE\|L1 | -9446 | NM_183249 | 66107 | Mm.902 | NM_183249 | ENSMUSG00000051748 | **Wfdc21** | 1100001G20Rik\|Wdnm1-like | WAP four-disulfide core domain 21 | protein-coding |
| chr4_63468886_63469468 | chr4 | 63468887 | 63469468 | + | 0.728802 | intron (NM_028640, intron 2 of 11) | intron (NM_028640, intron 2 of 11) | 26814 | NM_001008793 | 73750 | Mm.300397 | NM_028640 | ENSMUSG00000039137 | **Whrn** | 1110035G07Rik\|AW122018\|AW742671\|C430046P22Rik\|Dfnb31\|wi | whirlin | protein-coding |
| chr5_33822469_33822722 | chr5 | 33822470 | 33822722 | + | 0.980763 | Intergenic | Intergenic | -20516 | NM_175231 | 107823 | Mm.19892 | NM_175231 | ENSMUSG00000057406 | **Whsc1** | 5830445G22Rik\|9430010A17Rik\|AW555663\|C130020C13Rik\|D030027O06Rik\|D930023B08Rik\|MMSET\|NSD2\|Whsc1l\|mKIAA1090 | Wolf-Hirschhorn syndrome candidate 1 (human) | protein-coding |
| chr2_73546776_73547606 | chr2 | 73546777 | 73547606 | + | 0.993063 | Intergenic | Intergenic | -17704 | NM_153138 | 215280 | Mm.223504 | NM_153138 | ENSMUSG00000075284 | **Wipf1** | AI115543\|D2Ertd120e\|WIP\|Waspip | WAS/WASL interacting protein family, member 1 | protein-coding |
| chr2_73487732_73487936 | chr2 | 73487733 | 73487936 | + | 0.984571 | intron (NM_153138, intron 1 of 7) | intron (NM_153138, intron 1 of 7) | -2081 | NM_001289723 | 215280 | Mm.223504 | NM_153138 | ENSMUSG00000075284 | **Wipf1** | AI115543\|D2Ertd120e\|WIP\|Waspip | WAS/WASL interacting protein family, member 1 | protein-coding |
| chr2_73546769_73548088 | chr2 | 73546770 | 73548088 | + | 0.956924 | Intergenic | URR1B\|DNA\|hAT-Charlie | -17942 | NM_153138 | 215280 | Mm.223504 | NM_153138 | ENSMUSG00000075284 | **Wipf1** | AI115543\|D2Ertd120e\|WIP\|Waspip | WAS/WASL interacting protein family, member 1 | protein-coding |
| chr2_73459364_73460556 | chr2 | 73459365 | 73460556 | + | 0.823196 | intron (NM_153138, intron 1 of 7) | intron (NM_153138, intron 1 of 7) | -6071 | NM_001289722 | 215280 | Mm.223504 | NM_153138 | ENSMUSG00000075284 | **Wipf1** | AI115543\|D2Ertd120e\|WIP\|Waspip | WAS/WASL interacting protein family, member 1 | protein-coding |
| chr2_73460210_73460544 | chr2 | 73460211 | 73460544 | + | 0.80498 | intron (NM_153138, intron 1 of 7) | intron (NM_153138, intron 1 of 7) | -6488 | NM_001289722 | 215280 | Mm.223504 | NM_153138 | ENSMUSG00000075284 | **Wipf1** | AI115543\|D2Ertd120e\|WIP\|Waspip | WAS/WASL interacting protein family, member 1 | protein-coding |
| chr2_73459548_73460152 | chr2 | 73459549 | 73460152 | + | 0.759792 | intron (NM_153138, intron 1 of 7) | intron (NM_153138, intron 1 of 7) | -5961 | NM_001289722 | 215280 | Mm.223504 | NM_153138 | ENSMUSG00000075284 | **Wipf1** | AI115543\|D2Ertd120e\|WIP\|Waspip | WAS/WASL interacting protein family, member 1 | protein-coding |

| | | | | | | | | | | | | | | | | | |
|---|---|---|---|---|---|---|---|---|---|---|---|---|---|---|---|---|---|
| chr15_66862543_66862820 | chr15 | 66862544 | 66862820 | + | 1.41622 | Intergenic | Intergenic | -28711 | NM_018865 | 22402 | Mm.10222 | NM_018865 | ENSMUSG00000005124 | **Wisp1** | AW146261\|CCN4\|Elm1 | WNT1 inducible signaling pathway protein 1 | protein-coding |
| chr15_66861773_66862161 | chr15 | 66861774 | 66862161 | + | 1.33403 | Intergenic | Intergenic | -29426 | NM_018865 | 22402 | Mm.10222 | NM_018865 | ENSMUSG00000005124 | **Wisp1** | AW146261\|CCN4\|Elm1 | WNT1 inducible signaling pathway protein 1 | protein-coding |
| chr15_66868525_66868980 | chr15 | 66868526 | 66868980 | + | 1.25712 | Intergenic | Intergenic | -22640 | NM_018865 | 22402 | Mm.10222 | NM_018865 | ENSMUSG00000005124 | **Wisp1** | AW146261\|CCN4\|Elm1 | WNT1 inducible signaling pathway protein 1 | protein-coding |
| chr15_66877729_66878269 | chr15 | 66877730 | 66878269 | + | 0.802361 | Intergenic | Intergenic | -13394 | NM_018865 | 22402 | Mm.10222 | NM_018865 | ENSMUSG00000005124 | **Wisp1** | AW146261\|CCN4\|Elm1 | WNT1 inducible signaling pathway protein 1 | protein-coding |
| chr15_66890894_66891405 | chr15 | 66890895 | 66891405 | + | 0.791273 | promoter-TSS (NM_018865) | promoter-TSS (NM_018865) | -243 | NM_018865 | 22402 | Mm.10222 | NM_018865 | ENSMUSG00000005124 | **Wisp1** | AW146261\|CCN4\|Elm1 | WNT1 inducible signaling pathway protein 1 | protein-coding |
| chr2_163831122_163831885 | chr2 | 163831123 | 163831885 | + | 1.08705 | intron (NM_016873, intron 4 of 4) | intron (NM_016873, intron 4 of 4) | 10670 | NM_016873 | 22403 | Mm.13828 | NM_016873 | ENSMUSG00000027656 | **Wisp2** | Ccn5\|Crgr4\|Ctgfl\|Rcop1 | WNT1 inducible signaling pathway protein 2 | protein-coding |
| chr3_159831378_159831901 | chr3 | 159831379 | 159831901 | + | 1.15595 | Intergenic | Intergenic | -8055 | NR_037590 | 68151 | Mm.6766 | NM_026582 | | **Wls** | 5031439A09Rik\|AI173978\|AI987742\|Gpr177 | wntless homolog (Drosophila) | protein-coding |
| chr1_74806434_74807503 | chr1 | 74806435 | 74807503 | + | 0.783462 | Intergenic | Intergenic | 14950 | NM_009518 | 22409 | Mm.5130 | NM_009518 | | **Wnt10a** | - | wingless-type MMTV integration site family, member 10A | protein-coding |
| chr1_74806543_74806901 | chr1 | 74806544 | 74806901 | + | 0.674197 | Intergenic | Intergenic | 14703 | NM_009518 | 22409 | Mm.5130 | NM_009518 | | **Wnt10a** | - | wingless-type MMTV integration site family, member 10A | protein-coding |
| chr15_98777059_98777495 | chr15 | 98777060 | 98777495 | + | 0.667051 | intron (NM_011718, intron 1 of 4) | intron (NM_011718, intron 1 of 4) | 873 | NM_011718 | 22410 | Mm.4709 | NM_011718 | ENSMUSG00000022996 | **Wnt10b** | Wnt12 | wingless-type MMTV integration site family, member 10B | protein-coding |
| chr7_98878508_98878861 | chr7 | 98878509 | 98878861 | + | 0.77801 | Intergenic | Intergenic | 39840 | NM_001285795 | 22411 | Mm.22182 | NM_009519 | ENSMUSG00000015957 | **Wnt11** | - | wingless-type MMTV integration site family, member 11 | protein-coding |
| chr7_98878357_98878895 | chr7 | 98878358 | 98878895 | + | 0.692436 | Intergenic | BGLII_Mus\|LTR\|ERVK | 39781 | NM_001285795 | 22411 | Mm.22182 | NM_009519 | ENSMUSG00000015957 | **Wnt11** | - | wingless-type MMTV integration site family, member 11 | protein-coding |
| chr7_98797276_98797993 | chr7 | 98797277 | 98797993 | + | 0.682609 | Intergenic | Intergenic | -37477 | NM_001285794 | 22411 | Mm.22182 | NM_009519 | ENSMUSG00000015957 | **Wnt11** | - | wingless-type MMTV integration site family, member 11 | protein-coding |
| chr6_22138882_22139200 | chr6 | 22138883 | 22139200 | + | 0.984747 | intron (NM_001081351, intron 13 of 21) | intron (NM_001081351, intron 13 of 21) | -149186 | NM_053116 | 93735 | Mm.137403 | NM_053116 | ENSMUSG00000029671 | **Wnt16** | E130309I19Rik | wingless-type MMTV integration site family, member 16 | protein-coding |
| chr3_104937804_104938553 | chr3 | 104937805 | 104938553 | + | 0.990092 | Intergenic | Intergenic | 23530 | NM_009520 | 22414 | Mm.10740 | NM_009520 | ENSMUSG00000027840 | **Wnt2b** | Wnt13 | wingless-type MMTV integration site family, member 2B | protein-coding |
| chr3_104937884_104938269 | chr3 | 104937885 | 104938269 | + | 0.928601 | Intergenic | Intergenic | 23632 | NM_009520 | 22414 | Mm.10740 | NM_009520 | ENSMUSG00000027840 | **Wnt2b** | Wnt13 | wingless-type MMTV integration site family, member 2B | protein-coding |
| chr4_137312609_137312998 | chr4 | 137312610 | 137312998 | + | 0.82438 | Intergenic | Intergenic | 35169 | NM_009523 | 22417 | Mm.20355 | NM_009523 | ENSMUSG00000036856 | **Wnt4** | Wnt-4 | wingless-type MMTV integration site family, member 4 | protein-coding |
| chr4_137311666_137311955 | chr4 | 137311667 | 137311955 | + | 0.790352 | Intergenic | URR1B\|DNA\|hAT-Charlie | 34176 | NM_009523 | 22417 | Mm.20355 | NM_009523 | ENSMUSG00000036856 | **Wnt4** | Wnt-4 | wingless-type MMTV integration site family, member 4 | protein-coding |
| chr8_115045521_115046287 | chr8 | 115045522 | 115046287 | + | 1.05035 | intron (NM_019573, intron 8 of 8) | intron (NM_019573, intron 8 of 8) | 606252 | NM_019573 | 80707 | Mm.440420 | NM_019573 | ENSMUSG00000004637 | **Wwox** | 5330426P09Rik\|9030416C10Rik\|WOX1 | WW domain-containing oxidoreductase | protein-coding |
| chr8_114977773_114978214 | chr8 | 114977774 | 114978214 | + | 0.776275 | intron (NM_019573, intron 8 of 8) | intron (NM_019573, intron 8 of 8) | 538342 | NM_019573 | 80707 | Mm.440420 | NM_019573 | ENSMUSG00000004637 | **Wwox** | 5330426P09Rik\|9030416C10Rik\|WOX1 | WW domain-containing oxidoreductase | protein-coding |
| chr4_19685976_19686497 | chr4 | 19685977 | 19686497 | + | 1.47382 | intron (NR_074089, intron 2 of 23) | intron (NR_074089, intron 2 of 23) | 22767 | NM_177327 | 107568 | Mm.78312 | NM_177327 | ENSMUSG00000041058 | **Wwp1** | 8030445B08Rik\|AIP5\|SDRP1\|Tiul1 | WW domain containing E3 ubiquitin protein ligase 1 | protein-coding |
| chr4_19681198_19681660 | chr4 | 19681199 | 19681660 | + | 1.21022 | intron (NR_074089, intron 2 of 23) | intron (NR_074089, intron 2 of 23) | 27575 | NM_177327 | 107568 | Mm.78312 | NM_177327 | ENSMUSG00000041058 | **Wwp1** | 8030445B08Rik\|AIP5\|SDRP1\|Tiul1 | WW domain containing E3 ubiquitin protein ligase 1 | protein-coding |
| chr4_19682541_19682820 | chr4 | 19682542 | 19682820 | + | 0.815701 | intron (NR_074089, intron 2 of 23) | intron (NR_074089, intron 2 of 23) | 26323 | NM_177327 | 107568 | Mm.78312 | NM_177327 | ENSMUSG00000041058 | **Wwp1** | 8030445B08Rik\|AIP5\|SDRP1\|Tiul1 | WW domain containing E3 ubiquitin protein ligase 1 | protein-coding |
| chr4_19699858_19700219 | chr4 | 19699859 | 19700219 | + | 0.730393 | intron (NR_074089, intron 1 of 23) | intron (NR_074089, intron 1 of 23) | 8965 | NM_177327 | 107568 | Mm.78312 | NM_177327 | ENSMUSG00000041058 | **Wwp1** | 8030445B08Rik\|AIP5\|SDRP1\|Tiul1 | WW domain containing E3 ubiquitin protein ligase 1 | protein-coding |
| chr8_107444160_107444755 | chr8 | 107444161 | 107444755 | + | 1.25996 | intron (NM_025830, intron 1 of 23) | intron (NM_025830, intron 1 of 23) | 8060 | NM_025830 | 66894 | Mm.390058 | NM_025830 | ENSMUSG00000031930 | **Wwp2** | 1300010O06Rik\|AA690238\|AIP2\|AW554328 | WW domain containing E3 ubiquitin protein ligase 2 | protein-coding |
| chr11_5509314_5510121 | chr11 | 5509315 | 5510121 | + | 0.763559 | Intergenic | Intergenic | -10923 | NM_013842 | 22433 | Mm.469937 | NM_013842 | ENSMUSG00000020484 | **Xbp1** | D11Ertd39e\|TREB-5\|TREB5\|XBP-1 | X-box binding protein 1 | protein-coding |
| chr1_164945426_164946098 | chr1 | 164945427 | 164946098 | + | 1.07836 | Intergenic | Intergenic | -10252 | NM_008510 | 16963 | Mm.190 | NM_008510 | ENSMUSG00000026573 | **Xcl1** | AI661682\|ATAC\|LTN\|Lptn\|SCM-1\|SCM-1a\|Scyc1 | chemokine (C motif) ligand 1 | protein-coding |
| chr1_164903929_164904422 | chr1 | 164903930 | 164904422 | + | 0.916569 | Intergenic | Intergenic | 31334 | NM_008510 | 16963 | Mm.190 | NM_008510 | ENSMUSG00000026573 | **Xcl1** | AI661682\|ATAC\|LTN\|Lptn\|SCM-1\|SCM-1a\|Scyc1 | chemokine (C motif) ligand 1 | protein-coding |
| chr1_164975732_164976308 | chr1 | 164975733 | 164976308 | + | 0.853366 | Intergenic | Intergenic | -40510 | NM_008510 | 16963 | Mm.190 | NM_008510 | ENSMUSG00000026573 | **Xcl1** | AI661682\|ATAC\|LTN\|Lptn\|SCM-1\|SCM-1a\|Scyc1 | chemokine (C motif) ligand 1 | protein-coding |
| chr1_164903310_164903835 | chr1 | 164903311 | 164903835 | + | 0.836684 | Intergenic | Intergenic | 31937 | NM_008510 | 16963 | Mm.190 | NM_008510 | ENSMUSG00000026573 | **Xcl1** | AI661682\|ATAC\|LTN\|Lptn\|SCM-1\|SCM-1a\|Scyc1 | chemokine (C motif) ligand 1 | protein-coding |
| chr1_164975664_164976147 | chr1 | 164975665 | 164976147 | + | 0.824219 | Intergenic | Intergenic | -40396 | NM_008510 | 16963 | Mm.190 | NM_008510 | ENSMUSG00000026573 | **Xcl1** | AI661682\|ATAC\|LTN\|Lptn\|SCM-1\|SCM-1a\|Scyc1 | chemokine (C motif) ligand 1 | protein-coding |
| chr1_164910725_164911218 | chr1 | 164910726 | 164911218 | + | 0.798599 | Intergenic | Intergenic | 24538 | NM_008510 | 16963 | Mm.190 | NM_008510 | ENSMUSG00000026573 | **Xcl1** | AI661682\|ATAC\|LTN\|Lptn\|SCM-1\|SCM-1a\|Scyc1 | chemokine (C motif) ligand 1 | protein-coding |
| chr17_73918849_73919421 | chr17 | 73918850 | 73919421 | + | 1.62989 | intron (NM_011723, intron 13 of 35) | intron (NM_011723, intron 13 of 35) | 31061 | NM_011723 | 22436 | Mm.11223 | NM_011723 | ENSMUSG00000024066 | **Xdh** | XO\|Xor\|Xox-1\|Xox1 | xanthine dehydrogenase | protein-coding |
| chr9_120012698_120013212 | chr9 | 120012699 | 120013212 | + | 1.90602 | TTS (NM_011724) | TTS (NM_011724) | 10643 | NM_011724 | 22437 | Mm.10117 | NM_011724 | ENSMUSG00000079243 | **Xirp1** | AI415219\|Cmya1 | xin actin-binding repeat containing 1 | protein-coding |
| chr9_120009326_120009567 | chr9 | 120009327 | 120009567 | + | 1.06381 | Intergenic | RMER15\|LTR\|ERVL | 14151 | NM_011724 | 22437 | Mm.10117 | NM_011724 | ENSMUSG00000079243 | **Xirp1** | AI415219\|Cmya1 | xin actin-binding repeat containing 1 | protein-coding |
| chr7_126222781_126223098 | chr7 | 126222782 | 126223098 | + | 0.926184 | Intergenic | Intergenic | -22532 | NM_028816 | 74204 | Mm.235663 | NM_028816 | ENSMUSG00000000131 | **Xpo6** | 2610005L19Rik\|AI854665\|AL022631\|C230091E20Rik\|R75304\|Ranbp20\|mKIAA0370 | exportin 6 | protein-coding |
| chr1_155393367_155393826 | chr1 | 155393368 | 155393826 | + | 1.29529 | intron (NM_011273, intron 1 of 14) | B3A\|SINE\|B2 | 23847 | NM_011273 | 19775 | Mm.266215 | NM_011273 | ENSMUSG00000026469 | **Xpr1** | Rmc-1\|Rmc1\|Sxv\|Syg1\|XR | xenotropic and polytropic retrovirus receptor 1 | protein-coding |
| chr1_155472448_155473162 | chr1 | 155472449 | 155473162 | + | 1.00866 | Intergenic | Intergenic | -55361 | NM_011273 | 19775 | Mm.266215 | NM_011273 | ENSMUSG00000026469 | **Xpr1** | Rmc-1\|Rmc1\|Sxv\|Syg1\|XR | xenotropic and polytropic retrovirus receptor 1 | protein-coding |
| chr1_155472470_155472937 | chr1 | 155472471 | 155472937 | + | 0.96794 | Intergenic | Intergenic | -55260 | NM_011273 | 19775 | Mm.266215 | NM_011273 | ENSMUSG00000026469 | **Xpr1** | Rmc-1\|Rmc1\|Sxv\|Syg1\|XR | xenotropic and polytropic retrovirus receptor 1 | protein-coding |
| chr9_96023820_96024498 | chr9 | 96023821 | 96024498 | + | 1.27244 | exon (NM_011916, exon 31 of 42) | exon (NM_011916, exon 31 of 42) | 69399 | NM_011916 | 24127 | Mm.291179 | NM_011916 | ENSMUSG00000032410 | **Xrn1** | Dhm2\|exo\|mXrn1 | 5'-3' exoribonuclease 1 | protein-coding |
| chr7_117393828_117394636 | chr7 | 117393829 | 117394636 | + | 0.813841 | intron (NM_175645, intron 1 of 11) | intron (NM_175645, intron 1 of 11) | 13253 | NM_175645 | 233781 | Mm.461427 | NM_175645 | ENSMUSG00000030657 | **Xylt1** | 8030490L12 | xylosyltransferase 1 | protein-coding |
| chr7_117394066_117394505 | chr7 | 117394067 | 117394505 | + | 0.770878 | intron (NM_175645, intron 1 of 11) | intron (NM_175645, intron 1 of 11) | 13307 | NM_175645 | 233781 | Mm.461427 | NM_175645 | ENSMUSG00000030657 | **Xylt1** | 8030490L12 | xylosyltransferase 1 | protein-coding |
| chr7_117470139_117470810 | chr7 | 117470140 | 117470810 | + | 0.723523 | intron (NM_175645, intron 1 of 11) | intron (NM_175645, intron 1 of 11) | 89496 | NM_175645 | 233781 | Mm.461427 | NM_175645 | ENSMUSG00000030657 | **Xylt1** | 8030490L12 | xylosyltransferase 1 | protein-coding |
| chr7_117465993_117466836 | chr7 | 117465994 | 117466836 | + | 0.709549 | intron (NM_175645, intron 1 of 11) | B4A\|SINE\|B4 | 85436 | NM_175645 | 233781 | Mm.461427 | NM_175645 | ENSMUSG00000030657 | **Xylt1** | 8030490L12 | xylosyltransferase 1 | protein-coding |
| chr7_117470139_117470497 | chr7 | 117470140 | 117470497 | + | 0.656127 | intron (NM_175645, intron 1 of 11) | intron (NM_175645, intron 1 of 11) | 89339 | NM_175645 | 233781 | Mm.461427 | NM_175645 | ENSMUSG00000030657 | **Xylt1** | 8030490L12 | xylosyltransferase 1 | protein-coding |
| chr9_8003316_8003878 | chr9 | 8003317 | 8003878 | + | 0.615651 | intron (NM_009534, intron 1 of 7) | CpG | 999 | NM_001171147 | 22601 | Mm.221992 | NM_009534 | ENSMUSG00000053110 | **Yap1** | AI325207\|Yap\|Yap65\|Yki\|Yorkie | yes-associated protein 1 | protein-coding |
| chr6_131360725_131361093 | chr6 | 131360726 | 131361093 | + | 0.964551 | Intergenic | ORR1B1\|LTR\|ERVL-MaLR | 27541 | NM_139117 | 56449 | Mm.458000 | NM_011733 | ENSMUSG00000030189 | **Ybx3** | Csda\|Dpba\|MSY3\|MSY4\|Yb2\|dbpA\|oxyR | Y box protein 3 | protein-coding |
| chr6_131411191_131411459 | chr6 | 131411192 | 131411459 | + | 0.64978 | Intergenic | Intergenic | -22875 | NM_011733 | 56449 | Mm.458000 | NM_011733 | ENSMUSG00000030189 | **Ybx3** | Csda\|Dpba\|MSY3\|MSY4\|Yb2\|dbpA\|oxyR | Y box protein 3 | protein-coding |
| chr16_17148123_17148441 | chr16 | 17148124 | 17148441 | + | 0.665311 | exon (NM_026940, exon 5 of 5) | exon (NM_026940, exon 5 of 5) | 1315 | NM_026940 | 69101 | Mm.101656 | NM_026940 | ENSMUSG00000041774 | **Ydjc** | 1810015A11Rik\|4930521M19Rik | YdjC homolog (bacterial) | protein-coding |
| chr16_17157712_17158395 | chr16 | 17157713 | 17158395 | + | 0.653769 | intron (NM_009456, intron 3 of 3) | intron (NM_009456, intron 3 of 3) | 11087 | NM_026940 | 69101 | Mm.101656 | NM_026940 | ENSMUSG00000041774 | **Ydjc** | 1810015A11Rik\|4930521M19Rik | YdjC homolog (bacterial) | protein-coding |
| chr11_6001912_6002299 | chr11 | 6001913 | 6002299 | + | 0.634692 | intron (NM_007595, intron 3 of 20) | intron (NM_007595, intron 3 of 20) | 46348 | NM_019661 | 56418 | Mm.294821 | NM_019661 | ENSMUSG00000002741 | **Ykt6** | 0610042I15Rik\|1810013M05Rik\|AW105923 | YKT6 homolog (S. Cerevisiae) | protein-coding |
| chr12_85050938_85051481 | chr12 | 85050939 | 85051481 | + | 1.02276 | intron (NM_178363, intron 16 of 20) | intron (NM_178363, intron 16 of 20) | 54889 | NM_178363 | 56531 | Mm.153183 | NM_178363 | ENSMUSG00000021244 | **Ylpm1** | A930013E17Rik\|ZAP\|Zap3 | YLP motif containing 1 | protein-coding |
| chr12_85051186_85051474 | chr12 | 85051187 | 85051474 | + | 0.89666 | intron (NM_178363, intron 16 of 20) | intron (NM_178363, intron 16 of 20) | 55009 | NM_178363 | 56531 | Mm.153183 | NM_178363 | ENSMUSG00000021244 | **Ylpm1** | A930013E17Rik\|ZAP\|Zap3 | YLP motif containing 1 | protein-coding |
| chr12_85050918_85051746 | chr12 | 85050919 | 85051746 | + | 0.80028 | intron (NM_178363, intron 16 of 20) | intron (NM_178363, intron 16 of 20) | 55011 | NM_178363 | 56531 | Mm.153183 | NM_178363 | ENSMUSG00000021244 | **Ylpm1** | A930013E17Rik\|ZAP\|Zap3 | YLP motif containing 1 | protein-coding |

| | | | | | | | | | | | | | | | | | |
|---|---|---|---|---|---|---|---|---|---|---|---|---|---|---|---|---|---|
| chr16_17031426_17031834 | chr16 | 17031427 | 17031834 | + | 0.696152 | intron (NM_001038663, intron 6 of 7) | intron (NM_001038663, intron 6 of 7) | -38509 | NM_023249 | 106369 | Mm.237941 | NM_023249 | ENSMUSG00000022773 | **Ypel1** | 0610009L05Rik\|1700016N17Rik\|1700019O22Rik\|4921520K19Rik\|4930511F14Rik\|AV118478\|Dgl1\|Ppil2\|mdgl-1 | yippee-like 1 (Drosophila) | protein-coding |
| chr11_86914353_86914726 | chr11 | 86914354 | 86914726 | + | 0.995873 | intron (NR_106195, intron 4 of 8) | intron (NR_106195, intron 4 of 8) | 79222 | NM_001005341 | 77864 | Mm.191894 | NM_001005341 | ENSMUSG00000018427 | **Ypel2** | 6430570G24\|E130113K08Rik | yippee-like 2 (Drosophila) | protein-coding |
| chr11_86994877_86995170 | chr11 | 86994878 | 86995170 | + | 0.789148 | intron (NR_106195, intron 4 of 8) | intron (NR_106195, intron 4 of 8) | -1262 | NM_001005341 | 77864 | Mm.191894 | NM_001005341 | ENSMUSG00000018427 | **Ypel2** | 6430570G24\|E130113K08Rik | yippee-like 2 (Drosophila) | protein-coding |
| chr17_72821837_72822633 | chr17 | 72821838 | 72822633 | + | 1.69733 | Intergenic | L1MD\|LINE\|L1 | -14469 | NM_027166 | 383295 | Mm.277533 | NM_027166 | ENSMUSG00000039770 | **Ypel5** | 2310076K21Rik\|CGI-127 | yippee-like 5 (Drosophila) | protein-coding |
| chr5_33038674_33039171 | chr5 | 33038675 | 33039171 | + | 0.979205 | Intergenic | B1_Mus2\|SINE\|Alu | 20107 | NM_011738 | 22629 | Mm.332314 | NM_011738 | ENSMUSG00000018965 | **Ywhah** | - | tyrosine 3-monooxygenase/tryptophan 5-monooxygenase activation protein, eta polypeptide | protein-coding |
| chr12_21450309_21450636 | chr12 | 21450310 | 21450636 | + | 0.908238 | Intergenic | Intergenic | -33037 | NM_011739 | 22630 | Mm.289630 | NM_011739 | ENSMUSG00000076432 | **Ywhaq** | 2700028P07Rik\|AA409740\|AU021156\|R74690 | tyrosine 3-monooxygenase/tryptophan 5-monooxygenase activation protein, theta polypeptide | protein-coding |
| chr12_21428949_21429534 | chr12 | 21428950 | 21429534 | + | 0.800331 | Intergenic | MERV1_LTR\|LTR\|ERV1 | -11806 | NM_011739 | 22630 | Mm.289630 | NM_011739 | ENSMUSG00000076432 | **Ywhaq** | 2700028P07Rik\|AA409740\|AU021156\|R74690 | tyrosine 3-monooxygenase/tryptophan 5-monooxygenase activation protein, theta polypeptide | protein-coding |
| chr12_21463308_21463834 | chr12 | 21463309 | 21463834 | + | 0.759926 | Intergenic | Intergenic | -46135 | NM_011739 | 22630 | Mm.289630 | NM_011739 | ENSMUSG00000076432 | **Ywhaq** | 2700028P07Rik\|AA409740\|AU021156\|R74690 | tyrosine 3-monooxygenase/tryptophan 5-monooxygenase activation protein, theta polypeptide | protein-coding |
| chr12_21451964_21452248 | chr12 | 21451965 | 21452248 | + | 0.624499 | Intergenic | Intergenic | -34670 | NM_011739 | 22630 | Mm.289630 | NM_011739 | ENSMUSG00000076432 | **Ywhaq** | 2700028P07Rik\|AA409740\|AU021156\|R74690 | tyrosine 3-monooxygenase/tryptophan 5-monooxygenase activation protein, theta polypeptide | protein-coding |
| chr15_36884680_36885310 | chr15 | 36884681 | 36885310 | + | 1.17872 | Intergenic | Intergenic | -88066 | NM_001253806 | 22631 | Mm.3360 | NM_011740 | ENSMUSG00000022285 | **Ywhaz** | 1110013I11Rik\|14-3-3zeta\|AI596267\|AL022924\|AU020854 | tyrosine 3-monooxygenase/tryptophan 5-monooxygenase activation protein, zeta polypeptide | protein-coding |
| chr15_36758904_36759672 | chr15 | 36758905 | 36759672 | + | 0.88098 | Intergenic | Intergenic | 31644 | NM_001253807 | 22631 | Mm.3360 | NM_011740 | ENSMUSG00000022285 | **Ywhaz** | 1110013I11Rik\|14-3-3zeta\|AI596267\|AL022924\|AU020854 | tyrosine 3-monooxygenase/tryptophan 5-monooxygenase activation protein, zeta polypeptide | protein-coding |
| chr15_36729216_36729476 | chr15 | 36729217 | 36729476 | + | 0.85347 | Intergenic | RLTR16B_MM\|LTR\|ERVK | 61586 | NM_001253807 | 22631 | Mm.3360 | NM_011740 | ENSMUSG00000022285 | **Ywhaz** | 1110013I11Rik\|14-3-3zeta\|AI596267\|AL022924\|AU020854 | tyrosine 3-monooxygenase/tryptophan 5-monooxygenase activation protein, zeta polypeptide | protein-coding |
| chr2_72187424_72187822 | chr2 | 72187425 | 72187822 | + | 1.21575 | intron (NM_019688, intron 9 of 29) | intron (NM_019688, intron 9 of 29) | -98014 | NM_023057 | 65964 | Mm.304143 | NM_023057 | ENSMUSG00000004085 | **Zak** | AV006891\|B230120H23Rik\|MLTK\|MLTKalpha\|MLTKbeta | sterile alpha motif and leucine zipper containing kinase AZK | protein-coding |
| chr2_72253181_72253496 | chr2 | 72253182 | 72253496 | + | 0.737377 | intron (NM_019688, intron 29 of 29) | intron (NM_019688, intron 29 of 29) | -32298 | NM_023057 | 65964 | Mm.304143 | NM_023057 | ENSMUSG00000004085 | **Zak** | AV006891\|B230120H23Rik\|MLTK\|MLTKalpha\|MLTKbeta | sterile alpha motif and leucine zipper containing kinase AZK | protein-coding |
| chr2_72373572_72374087 | chr2 | 72373573 | 72374087 | + | 0.604068 | intron (NM_001164791, intron 7 of 11) | intron (NM_001164791, intron 7 of 11) | 88129 | NM_178084 | 65964 | Mm.304143 | NM_023057 | ENSMUSG00000004085 | **Zak** | AV006891\|B230120H23Rik\|MLTK\|MLTKalpha\|MLTKbeta | sterile alpha motif and leucine zipper containing kinase AZK | protein-coding |
| chr1_36768547_36769273 | chr1 | 36768548 | 36769273 | + | 1.37071 | intron (NM_001289765, intron 1 of 12) | intron (NM_001289765, intron 1 of 12) | 1976 | NM_001289765 | 22637 | Mm.8038 | NM_009539 | ENSMUSG00000026117 | **Zap70** | Srk\|ZAP-70\|mrtle\|mur | zeta-chain (TCR) associated protein kinase | protein-coding |
| chr1_36772264_36772868 | chr1 | 36772265 | 36772868 | + | 0.984975 | intron (NM_009539, intron 3 of 12) | intron (NM_009539, intron 3 of 12) | 5632 | NM_001289765 | 22637 | Mm.8038 | NM_009539 | ENSMUSG00000026117 | **Zap70** | Srk\|ZAP-70\|mrtle\|mur | zeta-chain (TCR) associated protein kinase | protein-coding |
| chr12_76375568_76375915 | chr12 | 76375569 | 76375915 | + | 1.32767 | intron (NM_178744, intron 1 of 1) | intron (NM_178744, intron 1 of 1) | 5476 | NM_178744 | 268564 | Mm.77065 | NM_178744 | | **Zbtb1** | 9430077A12\|AI256369\|Bsg2\|C430003J21Rik | zinc finger and BTB domain containing 1 | protein-coding |
| chr12_76374016_76374439 | chr12 | 76374017 | 76374439 | + | 1.15575 | intron (NM_178744, intron 1 of 1) | intron (NM_178744, intron 1 of 1) | 3962 | NM_178744 | 268564 | Mm.77065 | NM_178744 | | **Zbtb1** | 9430077A12\|AI256369\|Bsg2\|C430003J21Rik | zinc finger and BTB domain containing 1 | protein-coding |
| chr12_76380757_76381089 | chr12 | 76380758 | 76381089 | + | 0.774702 | intron (NM_178744, intron 1 of 1) | intron (NM_178744, intron 1 of 1) | 10657 | NM_178744 | 268564 | Mm.77065 | NM_178744 | | **Zbtb1** | 9430077A12\|AI256369\|Bsg2\|C430003J21Rik | zinc finger and BTB domain containing 1 | protein-coding |
| chr3_9245971_9246461 | chr3 | 9245972 | 9246461 | + | 1.1456 | Intergenic | Intergenic | -4351 | NM_177660 | 229055 | Mm.103262 | NM_177660 | ENSMUSG00000069114 | **Zbtb10** | 4832414A18\|AU021107\|RINZF | zinc finger and BTB domain containing 10 | protein-coding |
| chr9_48783051_48783808 | chr9 | 48783052 | 48783808 | + | 0.878317 | intron (NM_001033324, intron 2 of 6) | intron (NM_001033324, intron 2 of 6) | 52515 | NM_001033324 | 235320 | Mm.457803 | NM_001033324 | ENSMUSG00000066687 | **Zbtb16** | AI467657\|PLZF\|Zfp145\|lu | zinc finger and BTB domain containing 16 | protein-coding |
| chr1_177382591_177383177 | chr1 | 177382592 | 177383177 | + | 1.08043 | Intergenic | Intergenic | -61777 | NM_001012330 | 30928 | Mm.330700 | NM_013915 | ENSMUSG00000063659 | **Zbtb18** | RP58\|Zfp238\|Znf238 | zinc finger and BTB domain containing 18 | protein-coding |
| chr1_177456666_177457381 | chr1 | 177456667 | 177457381 | + | 1.06312 | Intergenic | Intergenic | 11203 | NM_013915 | 30928 | Mm.330700 | NM_013915 | ENSMUSG00000063659 | **Zbtb18** | RP58\|Zfp238\|Znf238 | zinc finger and BTB domain containing 18 | protein-coding |
| chr10_4384892_4385141 | chr10 | 4384893 | 4385141 | + | 0.885454 | intron (NM_001033466, intron 1 of 2) | ORR1C2\|LTR\|ERVL-MaLR | 3091 | NM_001033466 | 381990 | Mm.262466 | NM_001033466 | ENSMUSG00000075327 | **Zbtb2** | Gm1103 | zinc finger and BTB domain containing 2 | protein-coding |
| chr10_4389029_4389378 | chr10 | 4389030 | 4389378 | + | 0.759156 | Intergenic | Intergenic | -1096 | NM_001033466 | 381990 | Mm.262466 | NM_001033466 | ENSMUSG00000075327 | **Zbtb2** | Gm1103 | zinc finger and BTB domain containing 2 | protein-coding |
| chr16_43253011_43253439 | chr16 | 43253012 | 43253439 | + | 1.50349 | intron (NM_181058, intron 1 of 6) | intron (NM_181058, intron 1 of 6) | 5941 | NM_019778 | 56490 | Mm.136238 | NM_019778 | ENSMUSG00000022708 | **Zbtb20** | 1300017A20Rik\|7330412A13Rik\|A930017C21Rik\|D16Wsu73e\|DPZF\|HOF\|ODA-8S\|Oda8\|Zfp288 | zinc finger and BTB domain containing 20 | protein-coding |
| chr16_42900367_42900742 | chr16 | 42900368 | 42900742 | + | 1.37658 | intron (NR_040402, intron 1 of 3) | intron (NR_040402, intron 1 of 3) | -7090 | NM_001285805 | 56490 | Mm.136238 | NM_019778 | ENSMUSG00000022708 | **Zbtb20** | 1300017A20Rik\|7330412A13Rik\|A930017C21Rik\|D16Wsu73e\|DPZF\|HOF\|ODA-8S\|Oda8\|Zfp288 | zinc finger and BTB domain containing 20 | protein-coding |
| chr16_43190168_43190798 | chr16 | 43190169 | 43190798 | + | 1.10815 | intron (NM_001285805, intron 2 of 10) | intron (NM_001285805, intron 2 of 10) | -56801 | NM_181058 | 56490 | Mm.136238 | NM_019778 | ENSMUSG00000022708 | **Zbtb20** | 1300017A20Rik\|7330412A13Rik\|A930017C21Rik\|D16Wsu73e\|DPZF\|HOF\|ODA-8S\|Oda8\|Zfp288 | zinc finger and BTB domain containing 20 | protein-coding |
| chr16_43131099_43131498 | chr16 | 43131100 | 43131498 | + | 1.09444 | intron (NM_001285805, intron 2 of 10) | intron (NM_001285805, intron 2 of 10) | -115985 | NM_181058 | 56490 | Mm.136238 | NM_019778 | ENSMUSG00000022708 | **Zbtb20** | 1300017A20Rik\|7330412A13Rik\|A930017C21Rik\|D16Wsu73e\|DPZF\|HOF\|ODA-8S\|Oda8\|Zfp288 | zinc finger and BTB domain containing 20 | protein-coding |
| chr16_43303259_43304126 | chr16 | 43303260 | 43304126 | + | 0.934001 | intron (NM_181058, intron 2 of 6) | intron (NM_181058, intron 2 of 6) | 56409 | NM_019778 | 56490 | Mm.136238 | NM_019778 | ENSMUSG00000022708 | **Zbtb20** | 1300017A20Rik\|7330412A13Rik\|A930017C21Rik\|D16Wsu73e\|DPZF\|HOF\|ODA-8S\|Oda8\|Zfp288 | zinc finger and BTB domain containing 20 | protein-coding |
| chr16_97961375_97961871 | chr16 | 97961376 | 97961871 | + | 0.775995 | intron (NM_175428, intron 1 of 2) | intron (NM_175428, intron 1 of 2) | 998 | NM_001081685 | 114565 | Mm.474489 | NM_175428 | ENSMUSG00000046962 | **Zbtb21** | 5430437K12Rik\|B430213I24Rik\|Zfp295\|Znf295\|mKIAA1227 | zinc finger and BTB domain containing 21 | protein-coding |
| chr7_30592867_30593878 | chr7 | 30592868 | 30593878 | + | 1.55796 | promoter-TSS (NM_021397) | promoter-TSS (NM_021397) | -431 | NM_021397 | 58206 | Mm.116789 | NM_021397 | ENSMUSG00000006310 | **Zbtb32** | 4930524C15Rik\|FAXF\|FAZF\|PLZP\|Rog\|Tzfp | zinc finger and BTB domain containing 32 | protein-coding |
| chr7_30594524_30594932 | chr7 | 30594525 | 30594932 | + | 1.54025 | Intergenic | Intergenic | -1786 | NM_021397 | 58206 | Mm.116789 | NM_021397 | ENSMUSG00000006310 | **Zbtb32** | 4930524C15Rik\|FAXF\|FAZF\|PLZP\|Rog\|Tzfp | zinc finger and BTB domain containing 32 | protein-coding |
| chr7_30594015_30594932 | chr7 | 30594016 | 30594932 | + | 1.47757 | Intergenic | Intergenic | -1532 | NM_021397 | 58206 | Mm.116789 | NM_021397 | ENSMUSG00000006310 | **Zbtb32** | 4930524C15Rik\|FAXF\|FAZF\|PLZP\|Rog\|Tzfp | zinc finger and BTB domain containing 32 | protein-coding |
| chr7_30594188_30594473 | chr7 | 30594189 | 30594473 | + | 1.28383 | Intergenic | Intergenic | -1389 | NM_021397 | 58206 | Mm.116789 | NM_021397 | ENSMUSG00000006310 | **Zbtb32** | 4930524C15Rik\|FAXF\|FAZF\|PLZP\|Rog\|Tzfp | zinc finger and BTB domain containing 32 | protein-coding |
| chrX_38185109_38185307 | chrX | 38185110 | 38185307 | + | 0.720385 | Intergenic | RLTR30E_MM\|LTR\|ERV1 | -4585 | NM_020256 | 56805 | Mm.391248 | NM_020256 | ENSMUSG00000048047 | **Zbtb33** | AW260253\|E130014G12Rik\|Kaiso\|Znf-kaiso | zinc finger and BTB domain containing 33 | protein-coding |
| chr9_96714157_96714518 | chr9 | 96714158 | 96714518 | + | 0.983183 | intron (NM_175537, intron 2 of 2) | intron (NM_175537, intron 2 of 2) | 17337 | NM_175537 | 245007 | Mm.485954 | NM_175537 | ENSMUSG00000040433 | **Zbtb38** | A930014K01Rik\|CIBZ | zinc finger and BTB domain containing 38 | protein-coding |
| chr9_96713974_96714578 | chr9 | 96713975 | 96714578 | + | 0.952178 | intron (NM_175537, intron 2 of 2) | intron (NM_175537, intron 2 of 2) | 17399 | NM_175537 | 245007 | Mm.485954 | NM_175537 | ENSMUSG00000040433 | **Zbtb38** | A930014K01Rik\|CIBZ | zinc finger and BTB domain containing 38 | protein-coding |
| chr1_139407758_139408130 | chr1 | 139407759 | 139408130 | + | 1.10832 | Intergenic | B3A\|SINE\|B2 | -14439 | NM_172643 | 226470 | Mm.119711 | NM_172643 | ENSMUSG00000033964 | **Zbtb41** | 8430415N23Rik\|9430031N01\|9830132G07Rik\|AI316857 | zinc finger and BTB domain containing 41 homolog | protein-coding |
| chr1_139407760_139408673 | chr1 | 139407761 | 139408673 | + | 1.04386 | Intergenic | MTC\|LTR\|ERVL-MaLR | -14166 | NM_172643 | 226470 | Mm.119711 | NM_172643 | ENSMUSG00000033964 | **Zbtb41** | 8430415N23Rik\|9430031N01\|9830132G07Rik\|AI316857 | zinc finger and BTB domain containing 41 homolog | protein-coding |
| chr1_139413081_139413842 | chr1 | 139413082 | 139413842 | + | 0.893809 | Intergenic | Intergenic | -8921 | NM_172643 | 226470 | Mm.119711 | NM_172643 | ENSMUSG00000033964 | **Zbtb41** | 8430415N23Rik\|9430031N01\|9830132G07Rik\|AI316857 | zinc finger and BTB domain containing 41 homolog | protein-coding |
| chr10_81146124_81146871 | chr10 | 81146125 | 81146871 | + | 1.11428 | intron (NM_010731, intron 2 of 2) | intron (NM_010731, intron 2 of 2) | 10227 | NM_010731 | 16969 | Mm.20920 | NM_010731 | ENSMUSG00000035011 | **Zbtb7a** | 9030619K07Rik\|9130006G12Rik\|AI452336\|FBI-1\|Lrf\|Pokemon\|Zbtb7 | zinc finger and BTB domain containing 7a | protein-coding |

| | | | | | | | | | | | | | | | | | |
|---|---|---|---|---|---|---|---|---|---|---|---|---|---|---|---|---|---|
| chr10_81145778_81146917 | chr10 | 81145779 | 81146917 | + | 0.937637 | intron (NM_010731, intron 2 of 2) | intron (NM_010731, intron 2 of 2) | 10077 | NM_010731 | 16969 | Mm.20920 | NM_010731 | ENSMUSG00000035011 | **Zbtb7a** | 9030619K07Rik\|9130006G12Rik\|AI452336\|FBI-1\|Lrf\|Pokemon\|Zbtb7 | zinc finger and BTB domain containing 7a | protein-coding |
| chr9_52210349_52210681 | chr9 | 52210350 | 52210681 | + | 0.776579 | Intergenic | L1M4\|LINE\|L1 | -42404 | NM_001162921 | 244871 | Mm.295686 | NM_001162921 | ENSMUSG00000035164 | **Zc3h12c** | A230108E06\|C230027N18Rik\|mKIAA1726 | zinc finger CCCH type containing 12C | protein-coding |
| chr9_52348770_52349595 | chr9 | 52348771 | 52349595 | + | 0.64208 | Intergenic | Intergenic | -181072 | NM_001162921 | 244871 | Mm.295686 | NM_001162921 | ENSMUSG00000035164 | **Zc3h12c** | A230108E06\|C230027N18Rik\|mKIAA1726 | zinc finger CCCH type containing 12C | protein-coding |
| chr10_7818722_7819071 | chr10 | 7818723 | 7819071 | + | 0.903776 | intron (NM_026141, intron 12 of 12) | B4\|SINE\|B4 | -13573 | NM_172785 | 237256 | Mm.140055 | NM_172785 | ENSMUSG00000039981 | **Zc3h12d** | D730019B10Rik | zinc finger CCCH type containing 12D | protein-coding |
| chr10_7816464_7817273 | chr10 | 7816465 | 7817273 | + | 0.715232 | intron (NM_026141, intron 12 of 12) | intron (NM_026141, intron 12 of 12) | -15601 | NM_172785 | 237256 | Mm.140055 | NM_172785 | ENSMUSG00000039981 | **Zc3h12d** | D730019B10Rik | zinc finger CCCH type containing 12D | protein-coding |
| chr15_81760272_81760778 | chr15 | 81760273 | 81760778 | + | 0.707238 | intron (NM_001081016, intron 1 of 22) | intron (NM_001081016, intron 1 of 22) | 15677 | NM_001081016 | 20286 | Mm.432075 | NM_001081016 | ENSMUSG00000022390 | **Zc3h7b** | Gm627\|Scrg3\|Zc2h7b\|mKIAA1031 | zinc finger CCCH type containing 7B | protein-coding |
| chr6_30353665_30354114 | chr6 | 30353666 | 30354114 | + | 0.876282 | Intergenic | Intergenic | 37120 | NM_172735 | 232679 | Mm.29780 | NM_172735 | ENSMUSG00000039130 | **Zc3hc1** | 1110054L24Rik\|AU018540\|AU019789\|HSPC216\|Nipa | zinc finger, C3HC type 1 | protein-coding |
| chr1_106043435_106043991 | chr1 | 106043436 | 106043991 | + | 0.910585 | Intergenic | Intergenic | 53307 | NM_001122675 | 227449 | Mm.210188 | NM_001122675 | ENSMUSG00000038866 | **Zcchc2** | 9930114B20Rik\|AW212015 | zinc finger, CCHC domain containing 2 | protein-coding |
| chr1_106045398_106045960 | chr1 | 106045399 | 106045960 | + | 0.856314 | Intergenic | Intergenic | 55273 | NM_001122675 | 227449 | Mm.210188 | NM_001122675 | ENSMUSG00000038866 | **Zcchc2** | 9930114B20Rik\|AW212015 | zinc finger, CCHC domain containing 2 | protein-coding |
| chr1_106043242_106044028 | chr1 | 106043243 | 106044028 | + | 0.791597 | Intergenic | Intergenic | 53229 | NM_001122675 | 227449 | Mm.210188 | NM_001122675 | ENSMUSG00000038866 | **Zcchc2** | 9930114B20Rik\|AW212015 | zinc finger, CCHC domain containing 2 | protein-coding |
| chr5_52786520_52786906 | chr5 | 52786521 | 52786906 | + | 0.889068 | intron (NM_030185, intron 3 of 11) | intron (NM_030185, intron 3 of 11) | 3646 | NM_030185 | 78796 | Mm.266886 | NM_030185 | ENSMUSG00000029179 | **Zcchc4** | 4930449I23Rik | zinc finger, CCHC domain containing 4 | protein-coding |
| chr4_44788638_44789400 | chr4 | 44788639 | 44789400 | + | 1.33466 | intron (NM_138590, intron 2 of 8) | L1M3\|LINE\|L1 | 32460 | NM_138590 | 319885 | Mm.331576 | NM_138590 | ENSMUSG00000035649 | **Zcchc7** | 4930572I07Rik\|D4Wsu132e | zinc finger, CCHC domain containing 7 | protein-coding |
| chr4_44785301_44785592 | chr4 | 44785302 | 44785592 | + | 0.812991 | intron (NM_138590, intron 2 of 8) | RSINE1\|SINE\|B4 | 28888 | NM_138590 | 319885 | Mm.331576 | NM_138590 | ENSMUSG00000035649 | **Zcchc7** | 4930572I07Rik\|D4Wsu132e | zinc finger, CCHC domain containing 7 | protein-coding |
| chr4_44849303_44849885 | chr4 | 44849304 | 44849885 | + | 0.628762 | intron (NM_138590, intron 2 of 8) | intron (NM_138590, intron 2 of 8) | 93035 | NM_138590 | 319885 | Mm.331576 | NM_138590 | ENSMUSG00000035649 | **Zcchc7** | 4930572I07Rik\|D4Wsu132e | zinc finger, CCHC domain containing 7 | protein-coding |
| chr15_93374550_93375082 | chr15 | 93374551 | 93375082 | + | 0.687558 | Intergenic | Intergenic | 23474 | NM_026025 | 67197 | Mm.293181 | NM_026025 | ENSMUSG00000022635 | **Zcrb1** | 2700088M22Rik\|Madp-1 | zinc finger CCHC-type and RNA binding motif 1 | protein-coding |
| chr7_48760538_48761246 | chr7 | 48760539 | 48761246 | + | 1.12837 | Intergenic | Intergenic | -28111 | NM_028031 | 243983 | Mm.279116 | NM_028031 | ENSMUSG00000030471 | **Zdhhc13** | 2410004E01Rik\|C530010M18\|HIP3RP\|Hip14l\|kojak\|skc4 | zinc finger, DHHC domain containing 13 | protein-coding |
| chrX_104652725_104653048 | chrX | 104652726 | 104653048 | + | 0.825045 | intron (NM_175358, intron 1 of 11) | Lx2\|LINE\|L1 | 18177 | NM_175358 | 108672 | Mm.30574 | NM_175358 | ENSMUSG00000033906 | **Zdhhc15** | 6030457O13Rik | zinc finger, DHHC domain containing 15 | protein-coding |
| chr10_111047909_111048439 | chr10 | 111047910 | 111048439 | + | 0.734378 | Intergenic | Intergenic | -38108 | NM_172554 | 320150 | Mm.339281 | NM_172554 | ENSMUSG00000035798 | **Zdhhc17** | A230053P19Rik\|BB187739\|D130071N24Rik\|Hip14 | zinc finger, DHHC domain containing 17 | protein-coding |
| chr10_111031400_111032031 | chr10 | 111031401 | 111032031 | + | 0.669107 | Intergenic | Intergenic | -21650 | NM_172554 | 320150 | Mm.339281 | NM_172554 | ENSMUSG00000035798 | **Zdhhc17** | A230053P19Rik\|BB187739\|D130071N24Rik\|Hip14 | zinc finger, DHHC domain containing 17 | protein-coding |
| chr8_40394651_40395494 | chr8 | 40394652 | 40395494 | + | 1.25849 | Intergenic | Lx4A\|LINE\|L1 | -28742 | NM_178395 | 70546 | Mm.34326 | NM_178395 | | **Zdhhc2** | 5730415P04Rik\|6430583A19Rik | zinc finger, DHHC domain containing 2 | protein-coding |
| chr8_40422285_40422621 | chr8 | 40422286 | 40422621 | + | 0.745983 | Intergenic | L1_Mus3\|LINE\|L1 | -1362 | NM_178395 | 70546 | Mm.34326 | NM_178395 | | **Zdhhc2** | 5730415P04Rik\|6430583A19Rik | zinc finger, DHHC domain containing 2 | protein-coding |
| chr4_82848627_82849553 | chr4 | 82848628 | 82849553 | + | 1.38947 | intron (NM_026647, intron 2 of 8) | intron (NM_026647, intron 2 of 8) | 10571 | NM_026647 | 68268 | Mm.32047 | NM_026647 | ENSMUSG00000028403 | **Zdhhc21** | 9130404H11Rik\|AL024349\|D130004H04Rik\|dep | zinc finger, DHHC domain containing 21 | protein-coding |
| chr4_82848625_82849223 | chr4 | 82848626 | 82849223 | + | 1.35383 | intron (NM_026647, intron 2 of 8) | L2c\|LINE\|L2 | 10737 | NM_026647 | 68268 | Mm.32047 | NM_026647 | ENSMUSG00000028403 | **Zdhhc21** | 9130404H11Rik\|AL024349\|D130004H04Rik\|dep | zinc finger, DHHC domain containing 21 | protein-coding |
| chr4_82833231_82834069 | chr4 | 82833232 | 82834069 | + | 1.19455 | intron (NM_026647, intron 6 of 8) | B1F\|SINE\|Alu | 26011 | NM_026647 | 68268 | Mm.32047 | NM_026647 | ENSMUSG00000028403 | **Zdhhc21** | 9130404H11Rik\|AL024349\|D130004H04Rik\|dep | zinc finger, DHHC domain containing 21 | protein-coding |
| chr12_86985335_86985825 | chr12 | 86985336 | 86985825 | + | 0.736869 | intron (NM_001080943, intron 1 of 1) | intron (NM_001080943, intron 1 of 1) | 3096 | NM_001080943 | 238331 | Mm.185890 | NM_001080943 | ENSMUSG00000048483 | **Zdhhc22** | Gm262 | zinc finger, DHHC-type containing 22 | protein-coding |
| chr18_5464524_5465119 | chr18 | 5464525 | 5465119 | + | 0.973993 | Intergenic | Intergenic | -127038 | NM_011546 | 21417 | Mm.3929 | NM_011546 | ENSMUSG00000024238 | **Zeb1** | 3110032K11Rik\|AREB6\|BZP\|MEB1\|Nil2\|TCF-8\|Tcf18\|Tcf8\|Tw\|ZEB\|Zfhep\|Zfhx1a\|Zfx1a\|Zfx1ha\|[delta]EF1 | zinc finger E-box binding homeobox 1 | protein-coding |
| chr18_5464458_5465272 | chr18 | 5464459 | 5465272 | + | 0.828643 | Intergenic | L1MEc\|LINE\|L1 | -126995 | NM_011546 | 21417 | Mm.3929 | NM_011546 | ENSMUSG00000024238 | **Zeb1** | 3110032K11Rik\|AREB6\|BZP\|MEB1\|Nil2\|TCF-8\|Tcf18\|Tcf8\|Tw\|ZEB\|Zfhep\|Zfhx1a\|Zfx1a\|Zfx1ha\|[delta]EF1 | zinc finger E-box binding homeobox 1 | protein-coding |
| chr2_45172497_45173422 | chr2 | 45172498 | 45173422 | + | 1.24706 | Intergenic | (TG)n\|Simple_repeat\|Simple_repeat | -59681 | NM_015753 | 24136 | Mm.440702 | NM_015753 | ENSMUSG00000026872 | **Zeb2** | 9130203F04Rik\|D130016B08Rik\|SIP1\|Zfhx1b\|Zfx1b\|Zfxh1b | zinc finger E-box binding homeobox 2 | protein-coding |
| chr2_45197157_45197689 | chr2 | 45197158 | 45197689 | + | 0.944109 | Intergenic | Intergenic | -84144 | NM_015753 | 24136 | Mm.440702 | NM_015753 | ENSMUSG00000026872 | **Zeb2** | 9130203F04Rik\|D130016B08Rik\|SIP1\|Zfhx1b\|Zfx1b\|Zfxh1b | zinc finger E-box binding homeobox 2 | protein-coding |
| chr2_45196964_45197741 | chr2 | 45196965 | 45197741 | + | 0.873277 | Intergenic | Intergenic | -84074 | NM_015753 | 24136 | Mm.440702 | NM_015753 | ENSMUSG00000026872 | **Zeb2** | 9130203F04Rik\|D130016B08Rik\|SIP1\|Zfhx1b\|Zfx1b\|Zfxh1b | zinc finger E-box binding homeobox 2 | protein-coding |
| chr2_45068111_45068687 | chr2 | 45068112 | 45068687 | + | 0.751716 | intron (NM_001289521, intron 1 of 8) | intron (NM_001289521, intron 1 of 8) | 41878 | NM_001289521 | 24136 | Mm.440702 | NM_015753 | ENSMUSG00000026872 | **Zeb2** | 9130203F04Rik\|D130016B08Rik\|SIP1\|Zfhx1b\|Zfx1b\|Zfxh1b | zinc finger E-box binding homeobox 2 | protein-coding |
| chr2_45102981_45103562 | chr2 | 45102982 | 45103562 | + | 0.729363 | intron (NM_001289521, intron 1 of 8) | (TC)n\|Simple_repeat\|Simple_repeat | 7005 | NM_001289521 | 24136 | Mm.440702 | NM_015753 | ENSMUSG00000026872 | **Zeb2** | 9130203F04Rik\|D130016B08Rik\|SIP1\|Zfhx1b\|Zfx1b\|Zfxh1b | zinc finger E-box binding homeobox 2 | protein-coding |
| chr17_30041435_30041869 | chr17 | 30041436 | 30041869 | + | 0.954221 | intron (NM_148926, intron 1 of 4) | intron (NM_148926, intron 1 of 4) | 36565 | NM_148926 | 21769 | Mm.220972 | NM_148926 | ENSMUSG00000044477 | **Zfand3** | AW539211\|TEG-27\|Tex27 | zinc finger, AN1-type domain 3 | protein-coding |
| chr17_30052320_30052740 | chr17 | 30052321 | 30052740 | + | 0.727236 | intron (NM_148926, intron 1 of 4) | intron (NM_148926, intron 1 of 4) | 47443 | NM_148926 | 21769 | Mm.220972 | NM_148926 | ENSMUSG00000044477 | **Zfand3** | AW539211\|TEG-27\|Tex27 | zinc finger, AN1-type domain 3 | protein-coding |
| chr6_116240330_116240893 | chr6 | 116240331 | 116240893 | + | 1.38748 | intron (NM_026585, intron 19 of 30) | MTD\|LTR\|ERVL-MaLR | -23607 | NM_001290338 | 67492 | Mm.272772 | NM_001081317 | ENSMUSG00000042213 | **Zfand4** | Anubl1 | zinc finger, AN1-type domain 4 | protein-coding |
| chr7_84650443_84650955 | chr7 | 84650444 | 84650955 | + | 0.796246 | intron (NM_022985, intron 2 of 7) | intron (NM_022985, intron 2 of 7) | 28652 | NM_022985 | 65098 | Mm.209349 | NM_022985 | ENSMUSG00000030629 | **Zfand6** | 3110005P07Rik\|Awp1\|Za20d3 | zinc finger, AN1-type domain 6 | protein-coding |
| chr7_84681013_84681286 | chr7 | 84681014 | 84681286 | + | 0.787118 | Intergenic | Intergenic | -1799 | NM_022985 | 65098 | Mm.209349 | NM_022985 | ENSMUSG00000030629 | **Zfand6** | 3110005P07Rik\|Awp1\|Za20d3 | zinc finger, AN1-type domain 6 | protein-coding |
| chr15_67758573_67758988 | chr15 | 67758574 | 67758988 | + | 0.754683 | Intergenic | Intergenic | 500075 | NM_001145888 | 380993 | Mm.309526 | NM_198644 | ENSMUSG00000022335 | **Zfat** | Gm922\|Zfat1\|Zfp406 | zinc finger and AT hook domain containing | protein-coding |
| chr2_120561582_120561929 | chr2 | 120561583 | 120561929 | + | 1.13186 | intron (NM_011743, intron 1 of 21) | intron (NM_011743, intron 1 of 21) | 2075 | NM_011743 | 20402 | Mm.485295 | NM_011743 | ENSMUSG00000027288 | **Zfp106** | Cd-1\|D2Dcr28\|H3a\|Sh3bp3\|Znf106\|sirm | zinc finger protein 106 | protein-coding |
| chr17_55875746_55876039 | chr17 | 55875747 | 55876039 | + | 0.742497 | intron (NM_144546, intron 1 of 3) | L1VL2\|LINE\|L1 | 3060 | NM_144546 | 104349 | Mm.460659 | NM_144546 | | **Zfp119a** | AL024077\|Mzf13\|Zfp119 | zinc finger protein 119a | protein-coding |
| chr17_55948652_55948870 | chr17 | 55948653 | 55948870 | + | 0.67991 | Intergenic | Intergenic | -3502 | NM_146249 | 240120 | Mm.441018 | NM_146249 | | **Zfp119b** | - | zinc finger protein 119b | protein-coding |
| chr17_23584732_23585129 | chr17 | 23584733 | 23585129 | + | 0.939313 | intron (NM_011747, intron 2 of 5) | MIRc\|SINE\|MIR | 14556 | NM_011747 | 22654 | Mm.40326 | NM_011747 | ENSMUSG00000062012 | **Zfp13** | 4933429B21\|AI835008\|Krox-8\|Rhit\|Zfp-13 | zinc finger protein 13 | protein-coding |
| chr17_23584674_23585221 | chr17 | 23584675 | 23585221 | + | 0.916988 | intron (NM_011747, intron 2 of 5) | MIRc\|SINE\|MIR | 14539 | NM_011747 | 22654 | Mm.40326 | NM_011747 | ENSMUSG00000062012 | **Zfp13** | 4933429B21\|AI835008\|Krox-8\|Rhit\|Zfp-13 | zinc finger protein 13 | protein-coding |
| chr13_21924594_21925128 | chr13 | 21924595 | 21925128 | + | 0.993047 | Intergenic | Intergenic | -20233 | NM_183014 | 193452 | Mm.187446 | NM_183014 | ENSMUSG00000006720 | **Zfp184** | 4930500C15Rik | zinc finger protein 184 (Kruppel-like) | protein-coding |
| chr18_24052106_24052516 | chr18 | 24052107 | 24052516 | + | 0.665713 | Intergenic | ORR1E\|LTR\|ERVL-MaLR | -31540 | NM_021559 | 59057 | Mm.417427 | NM_021559 | | **Zfp191** | 3526401F17Rik\|5033419P20Rik\|AI480505\|AI642085\|AL024422\|KOX17\|ZF-12\|Znf191\|Znf24\|Znp191 | zinc finger protein 191 | protein-coding |
| chr17_23538610_23539038 | chr17 | 23538611 | 23539038 | + | 1.14811 | Intergenic | tRNA-Pro-CCA\|tRNA\|tRNA | 25402 | NM_001033496 | 449521 | Mm.268582 | NM_001033496 | ENSMUSG00000071256 | **Zfp213** | BC038178\|D17Ertd197e | zinc finger protein 213 | protein-coding |
| chr17_23538357_23539060 | chr17 | 23538358 | 23539060 | + | 1.11674 | Intergenic | Intergenic | 25517 | NM_001033496 | 449521 | Mm.268582 | NM_001033496 | ENSMUSG00000071256 | **Zfp213** | BC038178\|D17Ertd197e | zinc finger protein 213 | protein-coding |
| chr17_23534911_23535392 | chr17 | 23534912 | 23535392 | + | 0.857532 | Intergenic | Intergenic | 29074 | NM_001033496 | 449521 | Mm.268582 | NM_001033496 | ENSMUSG00000071256 | **Zfp213** | BC038178\|D17Ertd197e | zinc finger protein 213 | protein-coding |
| chr17_23533935_23534561 | chr17 | 23533936 | 23534561 | + | 0.764645 | Intergenic | Intergenic | 29978 | NM_001033496 | 449521 | Mm.268582 | NM_001033496 | ENSMUSG00000071256 | **Zfp213** | BC038178\|D17Ertd197e | zinc finger protein 213 | protein-coding |
| chr17_23546099_23546629 | chr17 | 23546100 | 23546629 | + | 0.758645 | Intergenic | Intergenic | 17862 | NM_001033496 | 449521 | Mm.268582 | NM_001033496 | ENSMUSG00000071256 | **Zfp213** | BC038178\|D17Ertd197e | zinc finger protein 213 | protein-coding |
| chr17_23548652_23549258 | chr17 | 23548653 | 23549258 | + | 0.688313 | Intergenic | Intergenic | 15271 | NM_001033496 | 449521 | Mm.268582 | NM_001033496 | ENSMUSG00000071256 | **Zfp213** | BC038178\|D17Ertd197e | zinc finger protein 213 | protein-coding |
| chr17_23548657_23549064 | chr17 | 23548658 | 23549064 | + | 0.652489 | Intergenic | Intergenic | 15365 | NM_001033496 | 449521 | Mm.268582 | NM_001033496 | ENSMUSG00000071256 | **Zfp213** | BC038178\|D17Ertd197e | zinc finger protein 213 | protein-coding |

| | | | | | | | | | | | | | | | | | |
|---|---|---|---|---|---|---|---|---|---|---|---|---|---|---|---|---|---|
| chr2_170129320_170129587 | chr2 | 170129321 | 170129587 | + | 1.20061 | intron (NM_001159683, intron 2 of 6) | intron (NM_001159683, intron 2 of 6) | 1766 | NM_001033299 | 228913 | Mm.22858 | NM_001033299 | ENSMUSG00000052056 | **Zfp217** | 4933431C08Rik\|AW987152\|Gm562\|ZABC1\|Znf217 | zinc finger protein 217 | protein-coding |
| chr2_170158404_170158612 | chr2 | 170158405 | 170158612 | + | 0.671103 | Intergenic | Intergenic | -15833 | NM_001159683 | 228913 | Mm.22858 | NM_001033299 | ENSMUSG00000052056 | **Zfp217** | 4933431C08Rik\|AW987152\|Gm562\|ZABC1\|Znf217 | zinc finger protein 217 | protein-coding |
| chr2_170158319_170158675 | chr2 | 170158320 | 170158675 | + | 0.653846 | Intergenic | Intergenic | -15822 | NM_001159683 | 228913 | Mm.22858 | NM_001033299 | ENSMUSG00000052056 | **Zfp217** | 4933431C08Rik\|AW987152\|Gm562\|ZABC1\|Znf217 | zinc finger protein 217 | protein-coding |
| chr2_170247710_170248025 | chr2 | 170247711 | 170248025 | + | 0.631688 | Intergenic | Intergenic | -105193 | NM_001159683 | 228913 | Mm.22858 | NM_001033299 | ENSMUSG00000052056 | **Zfp217** | 4933431C08Rik\|AW987152\|Gm562\|ZABC1\|Znf217 | zinc finger protein 217 | protein-coding |
| chr2_170247693_170248226 | chr2 | 170247694 | 170248226 | + | 0.63113 | Intergenic | Intergenic | -105285 | NM_001159683 | 228913 | Mm.22858 | NM_001033299 | ENSMUSG00000052056 | **Zfp217** | 4933431C08Rik\|AW987152\|Gm562\|ZABC1\|Znf217 | zinc finger protein 217 | protein-coding |
| chr12_40391287_40391630 | chr12 | 40391288 | 40391630 | + | 0.805074 | intron (NM_178845, intron 1 of 11) | intron (NM_178845, intron 1 of 11) | 54331 | NM_172575 | 246196 | Mm.26852 | NM_172575 | ENSMUSG00000055917 | **Zfp277** | 2410017E24Rik\|NIRF4 | zinc finger protein 277 | protein-coding |
| chr12_40383754_40384083 | chr12 | 40383755 | 40384083 | + | 0.77807 | intron (NM_178845, intron 1 of 11) | LTR16B\|LTR\|ERVL | 61871 | NM_172575 | 246196 | Mm.26852 | NM_172575 | ENSMUSG00000055917 | **Zfp277** | 2410017E24Rik\|NIRF4 | zinc finger protein 277 | protein-coding |
| chr10_76034743_76035094 | chr10 | 76034744 | 76035094 | + | 1.13653 | intron (NM_177475, intron 1 of 1) | L1MD\|LINE\|L1 | 2307 | NM_177475 | 64453 | Mm.80609 | NM_177475 | ENSMUSG00000049764 | **Zfp280b** | D10Jhu82e\|Suhw2 | zinc finger protein 280B | protein-coding |
| chr10_76071524_76072035 | chr10 | 76071525 | 76072035 | + | 0.987073 | intron (NM_023219, intron 10 of 14) | intron (NM_023219, intron 10 of 14) | 39168 | NM_177475 | 64453 | Mm.80609 | NM_177475 | ENSMUSG00000049764 | **Zfp280b** | D10Jhu82e\|Suhw2 | zinc finger protein 280B | protein-coding |
| chr10_76035379_76035600 | chr10 | 76035380 | 76035600 | + | 0.78688 | intron (NM_177475, intron 1 of 1) | L1_Rod\|LINE\|L1 | 2878 | NM_177475 | 64453 | Mm.80609 | NM_177475 | ENSMUSG00000049764 | **Zfp280b** | D10Jhu82e\|Suhw2 | zinc finger protein 280B | protein-coding |
| chr1_136601502_136601750 | chr1 | 136601503 | 136601750 | + | 0.669216 | Intergenic | Intergenic | -23275 | NM_177643 | 226442 | Mm.62521 | NM_177643 | ENSMUSG00000041483 | **Zfp281** | Znf281 | zinc finger protein 281 | protein-coding |
| chr11_62732682_62733004 | chr11 | 62732683 | 62733004 | + | 0.991062 | Intergenic | B3A\|SINE\|B2 | -3750 | NM_133208 | 170740 | Mm.339982 | NM_133208 | | **Zfp287** | B230333C16Rik\|SKAT-2\|Skat2\|Znf287\|mszf16\|mszf74 | zinc finger protein 287 | protein-coding |
| chr17_46376671_46377190 | chr17 | 46376672 | 46377190 | + | 0.748971 | Intergenic | RLTR19D\|LTR\|ERVK | -6835 | NM_207671 | 57908 | Mm.439916 | NM_021346 | ENSMUSG00000015597 | **Zfp318** | 2610034E08Rik\|D530032D06Rik\|TZF\|Znf318 | zinc finger protein 318 | protein-coding |
| chr5_105798482_105799186 | chr5 | 105798483 | 105799186 | + | 1.45428 | intron (NM_178701, intron 2 of 2) | (GA)n\|Simple_repeat\|Simple_repeat | -77734 | NR_030777 | 54367 | Mm.248876 | NM_018759 | ENSMUSG00000029290 | **Zfp326** | 5730470H14Rik\|ZAN75\|Znf326 | zinc finger protein 326 | protein-coding |
| chr5_105798145_105799191 | chr5 | 105798146 | 105799191 | + | 1.38817 | intron (NM_178701, intron 2 of 2) | intron (NM_178701, intron 2 of 2) | -77900 | NR_030777 | 54367 | Mm.248876 | NM_018759 | ENSMUSG00000029290 | **Zfp326** | 5730470H14Rik\|ZAN75\|Znf326 | zinc finger protein 326 | protein-coding |
| chr5_105839792_105840393 | chr5 | 105839793 | 105840393 | + | 0.691572 | Intergenic | ORR1D2\|LTR\|ERVL-MaLR | -36475 | NR_030777 | 54367 | Mm.248876 | NM_018759 | ENSMUSG00000029290 | **Zfp326** | 5730470H14Rik\|ZAN75\|Znf326 | zinc finger protein 326 | protein-coding |
| chr5_105877778_105878106 | chr5 | 105877779 | 105878106 | + | 0.651084 | intron (NR_030777, intron 1 of 9) | intron (NR_030777, intron 1 of 9) | 1374 | NM_018759 | 54367 | Mm.248876 | NM_018759 | ENSMUSG00000029290 | **Zfp326** | 5730470H14Rik\|ZAN75\|Znf326 | zinc finger protein 326 | protein-coding |
| chr8_82782304_82782865 | chr8 | 82782305 | 82782865 | + | 1.08295 | Intergenic | ORR1C1\|LTR\|ERVL-MaLR | -8459 | NM_145600 | 30932 | Mm.269248 | NM_145600 | ENSMUSG00000031711 | **Zfp330** | BC008086\|Noa36\|Znf330 | zinc finger protein 330 | protein-coding |
| chr8_82782306_82783098 | chr8 | 82782307 | 82783098 | + | 0.741819 | Intergenic | B3\|SINE\|B2 | -8576 | NM_145600 | 30932 | Mm.269248 | NM_145600 | ENSMUSG00000031711 | **Zfp330** | BC008086\|Noa36\|Znf330 | zinc finger protein 330 | protein-coding |
| chr4_128786861_128787701 | chr4 | 128786862 | 128787701 | + | 0.864267 | intron (NM_001081098, intron 4 of 8) | intron (NM_001081098, intron 4 of 8) | 18831 | NM_001081098 | 230761 | Mm.22616 | NM_001081098 | ENSMUSG00000028799 | **Zfp362** | - | zinc finger protein 362 | protein-coding |
| chr10_67840667_67841401 | chr10 | 67840668 | 67841401 | + | 1.68864 | Intergenic | Intergenic | 71628 | NM_178679 | 216049 | Mm.268346 | NM_178679 | ENSMUSG00000037855 | **Zfp365** | AI839779\|AV340892\|DBZ\|Su48\|Znf365\|talanin | zinc finger protein 365 | protein-coding |
| chr10_67890772_67891629 | chr10 | 67890773 | 67891629 | + | 1.20124 | intron (NM_178679, intron 3 of 4) | intron (NM_178679, intron 3 of 4) | 21461 | NM_178679 | 216049 | Mm.268346 | NM_178679 | ENSMUSG00000037855 | **Zfp365** | AI839779\|AV340892\|DBZ\|Su48\|Znf365\|talanin | zinc finger protein 365 | protein-coding |
| chr10_67898985_67899650 | chr10 | 67898986 | 67899650 | + | 1.16187 | intron (NM_178679, intron 2 of 4) | intron (NM_178679, intron 2 of 4) | 13344 | NM_178679 | 216049 | Mm.268346 | NM_178679 | ENSMUSG00000037855 | **Zfp365** | AI839779\|AV340892\|DBZ\|Su48\|Znf365\|talanin | zinc finger protein 365 | protein-coding |
| chr10_67881685_67882210 | chr10 | 67881686 | 67882210 | + | 1.04622 | Intergenic | Intergenic | 30714 | NM_178679 | 216049 | Mm.268346 | NM_178679 | ENSMUSG00000037855 | **Zfp365** | AI839779\|AV340892\|DBZ\|Su48\|Znf365\|talanin | zinc finger protein 365 | protein-coding |
| chr10_67899353_67899651 | chr10 | 67899354 | 67899651 | + | 1.01823 | intron (NM_178679, intron 2 of 4) | intron (NM_178679, intron 2 of 4) | 13160 | NM_178679 | 216049 | Mm.268346 | NM_178679 | ENSMUSG00000037855 | **Zfp365** | AI839779\|AV340892\|DBZ\|Su48\|Znf365\|talanin | zinc finger protein 365 | protein-coding |
| chr13_99158768_99159269 | chr13 | 99158769 | 99159269 | + | 1.03568 | Intergenic | Intergenic | -25804 | NM_001004149 | 238803 | Mm.343207 | NM_001004149 | ENSMUSG00000050919 | **Zfp366** | DC-SCRIPT\|Znf366 | zinc finger protein 366 | protein-coding |
| chr13_99158552_99159286 | chr13 | 99158553 | 99159286 | + | 0.93094 | Intergenic | Intergenic | -25904 | NM_001004149 | 238803 | Mm.343207 | NM_001004149 | ENSMUSG00000050919 | **Zfp366** | DC-SCRIPT\|Znf366 | zinc finger protein 366 | protein-coding |
| chr13_99152291_99152942 | chr13 | 99152292 | 99152942 | + | 0.65718 | Intergenic | Intergenic | -32206 | NM_001004149 | 238803 | Mm.343207 | NM_001004149 | ENSMUSG00000050919 | **Zfp366** | DC-SCRIPT\|Znf366 | zinc finger protein 366 | protein-coding |
| chr12_79959718_79960476 | chr12 | 79959719 | 79960476 | + | 1.32132 | Intergenic | Intergenic | 152916 | NM_007564 | 12192 | Mm.235132 | NM_007564 | ENSMUSG00000021127 | **Zfp36l1** | AW742437\|AW743212\|Berg36\|Brf1\|D530020L18Rik\|ERF1\|TIS11b\|cMG1 | zinc finger protein 36, C3H type-like 1 | protein-coding |
| chr12_79832607_79833027 | chr12 | 79832608 | 79833027 | + | 0.969479 | Intergenic | L1MB4\|LINE\|L1 | 280196 | NM_007564 | 12192 | Mm.235132 | NM_007564 | ENSMUSG00000021127 | **Zfp36l1** | AW742437\|AW743212\|Berg36\|Brf1\|D530020L18Rik\|ERF1\|TIS11b\|cMG1 | zinc finger protein 36, C3H type-like 1 | protein-coding |
| chr12_80064931_80065459 | chr12 | 80064932 | 80065459 | + | 0.890076 | Intergenic | Intergenic | 47818 | NM_007564 | 12192 | Mm.235132 | NM_007564 | ENSMUSG00000021127 | **Zfp36l1** | AW742437\|AW743212\|Berg36\|Brf1\|D530020L18Rik\|ERF1\|TIS11b\|cMG1 | zinc finger protein 36, C3H type-like 1 | protein-coding |
| chr12_79832601_79833218 | chr12 | 79832602 | 79833218 | + | 0.825717 | Intergenic | Lx8\|LINE\|L1 | 280103 | NM_007564 | 12192 | Mm.235132 | NM_007564 | ENSMUSG00000021127 | **Zfp36l1** | AW742437\|AW743212\|Berg36\|Brf1\|D530020L18Rik\|ERF1\|TIS11b\|cMG1 | zinc finger protein 36, C3H type-like 1 | protein-coding |
| chr12_80069075_80069570 | chr12 | 80069076 | 80069570 | + | 0.818018 | Intergenic | Intergenic | 43690 | NM_007564 | 12192 | Mm.235132 | NM_007564 | ENSMUSG00000021127 | **Zfp36l1** | AW742437\|AW743212\|Berg36\|Brf1\|D530020L18Rik\|ERF1\|TIS11b\|cMG1 | zinc finger protein 36, C3H type-like 1 | protein-coding |
| chr12_80069116_80069461 | chr12 | 80069117 | 80069461 | + | 0.801962 | Intergenic | Intergenic | 43724 | NM_007564 | 12192 | Mm.235132 | NM_007564 | ENSMUSG00000021127 | **Zfp36l1** | AW742437\|AW743212\|Berg36\|Brf1\|D530020L18Rik\|ERF1\|TIS11b\|cMG1 | zinc finger protein 36, C3H type-like 1 | protein-coding |
| chr12_79841120_79841677 | chr12 | 79841121 | 79841677 | + | 0.770556 | Intergenic | Intergenic | 271614 | NM_007564 | 12192 | Mm.235132 | NM_007564 | ENSMUSG00000021127 | **Zfp36l1** | AW742437\|AW743212\|Berg36\|Brf1\|D530020L18Rik\|ERF1\|TIS11b\|cMG1 | zinc finger protein 36, C3H type-like 1 | protein-coding |
| chr17_84291180_84291680 | chr17 | 84291181 | 84291680 | + | 1.02142 | intron (NM_183021, intron 29 of 37) | L1M4\|LINE\|L1 | -103483 | NM_001001806 | 12193 | Mm.259321 | NM_001001806 | ENSMUSG00000045817 | **Zfp36l2** | Brf2\|ERF2\|Tis11d | zinc finger protein 36, C3H type-like 2 | protein-coding |
| chr17_84320375_84321180 | chr17 | 84320376 | 84321180 | + | 0.877139 | intron (NM_183021, intron 28 of 37) | intron (NM_183021, intron 28 of 37) | -132831 | NM_001001806 | 12193 | Mm.259321 | NM_001001806 | ENSMUSG00000045817 | **Zfp36l2** | Brf2\|ERF2\|Tis11d | zinc finger protein 36, C3H type-like 2 | protein-coding |
| chr17_84148667_84149271 | chr17 | 84148668 | 84149271 | + | 0.710182 | Intergenic | Intergenic | 38978 | NM_001001806 | 12193 | Mm.259321 | NM_001001806 | ENSMUSG00000045817 | **Zfp36l2** | Brf2\|ERF2\|Tis11d | zinc finger protein 36, C3H type-like 2 | protein-coding |
| chr17_84148858_84149221 | chr17 | 84148859 | 84149221 | + | 0.685318 | Intergenic | Intergenic | 38907 | NM_001001806 | 12193 | Mm.259321 | NM_001001806 | ENSMUSG00000045817 | **Zfp36l2** | Brf2\|ERF2\|Tis11d | zinc finger protein 36, C3H type-like 2 | protein-coding |
| chr17_84215196_84215661 | chr17 | 84215197 | 84215661 | + | 0.621013 | intron (NM_183021, intron 36 of 37) | intron (NM_183021, intron 36 of 37) | -27482 | NM_001001806 | 12193 | Mm.259321 | NM_001001806 | ENSMUSG00000045817 | **Zfp36l2** | Brf2\|ERF2\|Tis11d | zinc finger protein 36, C3H type-like 2 | protein-coding |
| chr14_65374559_65375185 | chr14 | 65374560 | 65375185 | + | 1.07108 | intron (NM_199029, intron 1 of 9) | intron (NM_199029, intron 1 of 9) | 16196 | NM_199029 | 380912 | Mm.270087 | NM_199029 | ENSMUSG00000034522 | **Zfp395** | BC053701\|Gm910 | zinc finger protein 395 | protein-coding |
| chr14_65330508_65331008 | chr14 | 65330509 | 65331008 | + | 0.905623 | Intergenic | L1MC4\|LINE\|L1 | -27918 | NM_199029 | 380912 | Mm.270087 | NM_199029 | ENSMUSG00000034522 | **Zfp395** | BC053701\|Gm910 | zinc finger protein 395 | protein-coding |
| chr18_23912914_23913901 | chr18 | 23912915 | 23913901 | + | 1.02791 | Intergenic | B4\|SINE\|B4 | -41280 | NM_027007 | 69256 | Mm.218343 | NM_027007 | ENSMUSG00000024276 | **Zfp397** | 2810411K16Rik\|6720480F11Rik\|AI461664 | zinc finger protein 397 | protein-coding |
| chr18_23921138_23921922 | chr18 | 23921139 | 23921922 | + | 0.857262 | Intergenic | Intergenic | -33158 | NM_027007 | 69256 | Mm.218343 | NM_027007 | ENSMUSG00000024276 | **Zfp397** | 2810411K16Rik\|6720480F11Rik\|AI461664 | zinc finger protein 397 | protein-coding |
| chr18_84414238_84414577 | chr18 | 84414239 | 84414577 | + | 1.30843 | intron (NM_001033341, intron 4 of 8) | intron (NM_001033341, intron 4 of 8) | 175096 | NM_001033341 | 240476 | Mm.329575 | NM_001033341 | ENSMUSG00000048410 | **Zfp407** | 6430585N13Rik\|Gm333\|Gm334\|Gm948\|ZNF407 | zinc finger protein 407 | protein-coding |
| chr18_84413675_84413983 | chr18 | 84413676 | 84413983 | + | 1.1101 | intron (NM_001033341, intron 4 of 8) | intron (NM_001033341, intron 4 of 8) | 175675 | NM_001033341 | 240476 | Mm.329575 | NM_001033341 | ENSMUSG00000048410 | **Zfp407** | 6430585N13Rik\|Gm333\|Gm334\|Gm948\|ZNF407 | zinc finger protein 407 | protein-coding |
| chr18_84520725_84521499 | chr18 | 84520726 | 84521499 | + | 0.918846 | intron (NM_001033341, intron 3 of 8) | intron (NM_001033341, intron 3 of 8) | 68392 | NM_001033341 | 240476 | Mm.329575 | NM_001033341 | ENSMUSG00000048410 | **Zfp407** | 6430585N13Rik\|Gm333\|Gm334\|Gm948\|ZNF407 | zinc finger protein 407 | protein-coding |
| chr18_84393679_84394060 | chr18 | 84393680 | 84394060 | + | 0.726489 | intron (NM_001033341, intron 4 of 8) | intron (NM_001033341, intron 4 of 8) | 195634 | NM_001033341 | 240476 | Mm.329575 | NM_001033341 | ENSMUSG00000048410 | **Zfp407** | 6430585N13Rik\|Gm333\|Gm334\|Gm948\|ZNF407 | zinc finger protein 407 | protein-coding |

| | | | | | | | | | | | | | | | | | |
|---|---|---|---|---|---|---|---|---|---|---|---|---|---|---|---|---|---|
| chr18_84418921_84419380 | chr18 | 84418922 | 84419380 | + | 0.722932 | intron (NM_001033341, intron 4 of 8) | intron (NM_001033341, intron 4 of 8) | 170353 | NM_001033341 | 240476 | Mm.329575 | NM_001033341 | ENSMUSG00000048410 | **Zfp407** | 6430585N13Rik\|Gm333\|Gm334\|Gm948\|ZNF407 | zinc finger protein 407 | protein-coding |
| chr9_20500095_20500702 | chr9 | 20500096 | 20500702 | + | 1.15184 | exon (NM_001082485, exon 9 of 9) | exon (NM_001082485, exon 9 of 9) | -7653 | NM_001110309 | 235028 | Mm.297660 | NM_146221 | ENSMUSG00000059475 | **Zfp426** | 2900057C04Rik\|KRAB1\|Zfo61\|Zfp68-rs1\|Znf426\|mKIAA4237 | zinc finger protein 426 | protein-coding |
| chr18_5395928_5396360 | chr18 | 5395929 | 5396360 | + | 0.739798 | Intergenic | Intergenic | -61705 | NM_178722 | 240186 | Mm.118257 | NM_178722 | ENSMUSG00000050945 | **Zfp438** | 9430031G09\|9430091M14Rik\|B830013J05Rik | zinc finger protein 438 | protein-coding |
| chr18_5397296_5397768 | chr18 | 5397297 | 5397768 | + | 0.659897 | Intergenic | Intergenic | -63093 | NM_178722 | 240186 | Mm.118257 | NM_178722 | ENSMUSG00000050945 | **Zfp438** | 9430031G09\|9430091M14Rik\|B830013J05Rik | zinc finger protein 438 | protein-coding |
| chr5_38637902_38638750 | chr5 | 38637903 | 38638750 | + | 1.31379 | Intergenic | Intergenic | 46500 | NM_001081144 | 100515 | Mm.38277 | NM_001081144 | ENSMUSG00000046572 | **Zfp518b** | 6820424L24Rik\|AA410078\|AI661722\|Znf518b | zinc finger protein 518B | protein-coding |
| chr18_13760369_13760737 | chr18 | 13760370 | 13760737 | + | 0.591699 | intron (NM_145492, intron 5 of 7) | B3A\|SINE\|B2 | 212180 | NM_145492 | 225207 | Mm.40325 | NM_145492 | ENSMUSG00000024420 | **Zfp521** | B930086A16Rik\|Evi3\|Znf521 | zinc finger protein 521 | protein-coding |
| chr7_16055581_16055933 | chr7 | 16055582 | 16055933 | + | 1.22854 | Intergenic | L1MD\|LINE\|L1 | -16185 | NM_001099277 | 666528 | Mm.290718 | NM_001099277 | ENSMUSG00000078796 | **Zfp541** | EG666528\|Znf541 | zinc finger protein 541 | protein-coding |
| chr7_30086337_30086792 | chr7 | 30086338 | 30086792 | + | 1.00393 | intron (NM_152814, intron 2 of 3) | RLTR31D_MM\|LTR\|ERVK | 3945 | NM_152814 | 72556 | Mm.441512 | NM_152814 | ENSMUSG00000078768 | **Zfp566** | 2700043M03Rik\|mszf4 | zinc finger protein 566 | protein-coding |
| chr15_59256023_59256332 | chr15 | 59256024 | 59256332 | + | 0.726385 | Intergenic | Charlie1a\|DNA\|hAT-Charlie | -50770 | NR_045613 | 100416830 | Mm.490788 | NR_045613 | ENSMUSG00000098528 | **Zfp572** | 4930472F24Rik | zinc finger protein 572 | pseudo |
| chr13_67509031_67509335 | chr13 | 67509032 | 67509335 | + | 0.830416 | Intergenic | Intergenic | -8661 | NM_001007575 | 238693 | Mm.478914 | NM_001007575 | ENSMUSG00000071291 | **Zfp58** | A530094I17Rik\|Mfg-1\|Mfg1\|Rslcan5\|Zfp817 | zinc finger protein 58 | protein-coding |
| chr13_67509020_67509795 | chr13 | 67509021 | 67509795 | + | 0.681568 | Intergenic | Intergenic | -8886 | NM_001007575 | 238693 | Mm.478914 | NM_001007575 | ENSMUSG00000071291 | **Zfp58** | A530094I17Rik\|Mfg-1\|Mfg1\|Rslcan5\|Zfp817 | zinc finger protein 58 | protein-coding |
| chr18_55409197_55409983 | chr18 | 55409198 | 55409983 | + | 1.71981 | Intergenic | Intergenic | -419410 | NM_175751 | 269023 | Mm.192984 | NM_175751 | ENSMUSG00000052713 | **Zfp608** | 4932417D18Rik\|D430007A19Rik\|Znf608 | zinc finger protein 608 | protein-coding |
| chr18_54921847_54922388 | chr18 | 54921848 | 54922388 | + | 1.7159 | intron (NM_175751, intron 3 of 9) | intron (NM_175751, intron 3 of 9) | 68062 | NM_175751 | 269023 | Mm.192984 | NM_175751 | ENSMUSG00000052713 | **Zfp608** | 4932417D18Rik\|D430007A19Rik\|Znf608 | zinc finger protein 608 | protein-coding |
| chr18_54970607_54971190 | chr18 | 54970608 | 54971190 | + | 1.45179 | intron (NM_175751, intron 2 of 9) | intron (NM_175751, intron 2 of 9) | 19281 | NM_175751 | 269023 | Mm.192984 | NM_175751 | ENSMUSG00000052713 | **Zfp608** | 4932417D18Rik\|D430007A19Rik\|Znf608 | zinc finger protein 608 | protein-coding |
| chr18_55055612_55056092 | chr18 | 55055613 | 55056092 | + | 1.44782 | Intergenic | ORR1B2\|LTR\|ERVL-MaLR | -65672 | NM_175751 | 269023 | Mm.192984 | NM_175751 | ENSMUSG00000052713 | **Zfp608** | 4932417D18Rik\|D430007A19Rik\|Znf608 | zinc finger protein 608 | protein-coding |
| chr18_54921082_54921677 | chr18 | 54921083 | 54921677 | + | 1.44098 | intron (NM_175751, intron 3 of 9) | intron (NM_175751, intron 3 of 9) | 68800 | NM_175751 | 269023 | Mm.192984 | NM_175751 | ENSMUSG00000052713 | **Zfp608** | 4932417D18Rik\|D430007A19Rik\|Znf608 | zinc finger protein 608 | protein-coding |
| chr18_54936714_54937242 | chr18 | 54936715 | 54937242 | + | 1.40042 | intron (NM_175751, intron 3 of 9) | ORR1E\|LTR\|ERVL-MaLR | 53202 | NM_175751 | 269023 | Mm.192984 | NM_175751 | ENSMUSG00000052713 | **Zfp608** | 4932417D18Rik\|D430007A19Rik\|Znf608 | zinc finger protein 608 | protein-coding |
| chr18_54921279_54921764 | chr18 | 54921280 | 54921764 | + | 1.3754 | intron (NM_175751, intron 3 of 9) | intron (NM_175751, intron 3 of 9) | 68658 | NM_175751 | 269023 | Mm.192984 | NM_175751 | ENSMUSG00000052713 | **Zfp608** | 4932417D18Rik\|D430007A19Rik\|Znf608 | zinc finger protein 608 | protein-coding |
| chr18_54972950_54973252 | chr18 | 54972951 | 54973252 | + | 1.08116 | intron (NM_175751, intron 2 of 9) | intron (NM_175751, intron 2 of 9) | 17079 | NM_175751 | 269023 | Mm.192984 | NM_175751 | ENSMUSG00000052713 | **Zfp608** | 4932417D18Rik\|D430007A19Rik\|Znf608 | zinc finger protein 608 | protein-coding |
| chr18_55175971_55176542 | chr18 | 55175972 | 55176542 | + | 1.07166 | Intergenic | MTEb\|LTR\|ERVL-MaLR | -186077 | NM_175751 | 269023 | Mm.192984 | NM_175751 | ENSMUSG00000052713 | **Zfp608** | 4932417D18Rik\|D430007A19Rik\|Znf608 | zinc finger protein 608 | protein-coding |
| chr18_54992418_54992837 | chr18 | 54992419 | 54992837 | + | 0.958336 | Intergenic | Intergenic | -2448 | NM_175751 | 269023 | Mm.192984 | NM_175751 | ENSMUSG00000052713 | **Zfp608** | 4932417D18Rik\|D430007A19Rik\|Znf608 | zinc finger protein 608 | protein-coding |
| chr18_54936375_54936670 | chr18 | 54936376 | 54936670 | + | 0.860308 | intron (NM_175751, intron 3 of 9) | Lx8\|LINE\|L1 | 53657 | NM_175751 | 269023 | Mm.192984 | NM_175751 | ENSMUSG00000052713 | **Zfp608** | 4932417D18Rik\|D430007A19Rik\|Znf608 | zinc finger protein 608 | protein-coding |
| chr18_54860512_54861263 | chr18 | 54860513 | 54861263 | + | 0.774064 | Intergenic | Intergenic | 129292 | NM_175751 | 269023 | Mm.192984 | NM_175751 | ENSMUSG00000052713 | **Zfp608** | 4932417D18Rik\|D430007A19Rik\|Znf608 | zinc finger protein 608 | protein-coding |
| chr18_54822619_54823145 | chr18 | 54822620 | 54823145 | + | 0.699393 | Intergenic | Intergenic | 167298 | NM_175751 | 269023 | Mm.192984 | NM_175751 | ENSMUSG00000052713 | **Zfp608** | 4932417D18Rik\|D430007A19Rik\|Znf608 | zinc finger protein 608 | protein-coding |
| chr9_65811676_65812803 | chr9 | 65811677 | 65812803 | + | 1.4505 | intron (NM_172536, intron 1 of 9) | B3\|SINE\|B2 | 15324 | NM_172536 | 214812 | Mm.139231 | NM_172536 | ENSMUSG00000040524 | **Zfp609** | 9830165N24\|BC016271\|Znf609\|mKIAA0295 | zinc finger protein 609 | protein-coding |
| chr9_65831225_65831684 | chr9 | 65831226 | 65831684 | + | 1.23339 | 3' UTR (NM_001170907, exon 12 of 12) | 3' UTR (NM_001170907, exon 12 of 12) | -3891 | NM_172536 | 214812 | Mm.139231 | NM_172536 | ENSMUSG00000040524 | **Zfp609** | 9830165N24\|BC016271\|Znf609\|mKIAA0295 | zinc finger protein 609 | protein-coding |
| chr4_62877221_62877768 | chr4 | 62877222 | 62877768 | + | 0.812963 | Intergenic | Intergenic | -88079 | NM_028326 | 72701 | Mm.26479 | NM_028326 | ENSMUSG00000028358 | **Zfp618** | 2810031P15Rik\|2810040O04Rik\|AI415580\|AI505139\|D430033D05Rik\|Nedd10\|Znf618\|bM257P7.1 | zinc finger protein 618 | protein-coding |
| chr4_63018277_63019051 | chr4 | 63018278 | 63019051 | + | 0.706176 | intron (NM_028326, intron 1 of 14) | B4A\|SINE\|B4 | 53090 | NM_028326 | 72701 | Mm.26479 | NM_028326 | ENSMUSG00000028358 | **Zfp618** | 2810031P15Rik\|2810040O04Rik\|AI415580\|AI505139\|D430033D05Rik\|Nedd10\|Znf618\|bM257P7.1 | zinc finger protein 618 | protein-coding |
| chr4_62812612_62813684 | chr4 | 62812613 | 62813684 | + | 0.68806 | Intergenic | RMER1B\|Other\|Other | -152426 | NM_028326 | 72701 | Mm.26479 | NM_028326 | ENSMUSG00000028358 | **Zfp618** | 2810031P15Rik\|2810040O04Rik\|AI415580\|AI505139\|D430033D05Rik\|Nedd10\|Znf618\|bM257P7.1 | zinc finger protein 618 | protein-coding |
| chr11_49203862_49204312 | chr11 | 49203863 | 49204312 | + | 0.672058 | intron (NM_001024846, intron 1 of 4) | intron (NM_001024846, intron 1 of 4) | 587 | NM_009562 | 22720 | Mm.16650 | NM_009562 | ENSMUSG00000046311 | **Zfp62** | - | zinc finger protein 62 | protein-coding |
| chr15_75944133_75944611 | chr15 | 75944134 | 75944611 | + | 0.731118 | intron (NM_030199, intron 1 of 1) | L2\|LINE\|L2 | 3420 | NM_030199 | 78834 | Mm.273264 | NM_030199 | ENSMUSG00000050846 | **Zfp623** | 2610029D06Rik\|AI847036 | zinc finger protein 623 | protein-coding |
| chr11_95748121_95748792 | chr11 | 95748122 | 95748792 | + | 1.12465 | promoter-TSS (NM_201609) | promoter-TSS (NM_201609) | -610 | NM_201609 | 268469 | Mm.389038 | NM_201609 | ENSMUSG00000075595 | **Zfp652** | 9530033F24Rik\|ZNF652 | zinc finger protein 652 | protein-coding |
| chr7_43554432_43555113 | chr7 | 43554433 | 43555113 | + | 1.44779 | Intergenic | Intergenic | -7597 | NM_001008549 | 210104 | Mm.333821 | NM_001008549 | ENSMUSG00000056592 | **Zfp658** | - | zinc finger protein 658 | protein-coding |
| chr7_6285459_6285678 | chr7 | 6285460 | 6285678 | + | 0.677713 | Intergenic | Intergenic | -1011 | NM_001024928 | 384763 | Mm.276279 | NM_001024928 | ENSMUSG00000054893 | **Zfp667** | A830025F02Rik\|Mip1\|ZNF667 | zinc finger protein 667 | protein-coding |
| chr8_26943186_26943574 | chr8 | 26943187 | 26943574 | + | 1.15724 | Intergenic | Intergenic | -33956 | NM_001101502 | 353310 | Mm.212469 | NM_001101502 | ENSMUSG00000085795 | **Zfp703** | 1110032O19Rik\|AI430822\|AL022941\|Csmn1\|End2\|Zeppo1\|Znf703\|Zpo1 | zinc finger protein 703 | protein-coding |
| chr8_26943174_26943721 | chr8 | 26943175 | 26943721 | + | 1.08533 | Intergenic | ORR1F\|LTR\|ERVL-MaLR | -33888 | NM_001101502 | 353310 | Mm.212469 | NM_001101502 | ENSMUSG00000085795 | **Zfp703** | 1110032O19Rik\|AI430822\|AL022941\|Csmn1\|End2\|Zeppo1\|Znf703\|Zpo1 | zinc finger protein 703 | protein-coding |
| chr8_26952769_26953333 | chr8 | 26952770 | 26953333 | + | 0.82648 | Intergenic | Intergenic | -24285 | NM_001101502 | 353310 | Mm.212469 | NM_001101502 | ENSMUSG00000085795 | **Zfp703** | 1110032O19Rik\|AI430822\|AL022941\|Csmn1\|End2\|Zeppo1\|Znf703\|Zpo1 | zinc finger protein 703 | protein-coding |
| chr8_26963066_26963839 | chr8 | 26963067 | 26963839 | + | 0.763124 | Intergenic | Intergenic | -13883 | NM_001101502 | 353310 | Mm.212469 | NM_001101502 | ENSMUSG00000085795 | **Zfp703** | 1110032O19Rik\|AI430822\|AL022941\|Csmn1\|End2\|Zeppo1\|Znf703\|Zpo1 | zinc finger protein 703 | protein-coding |
| chr3_9581140_9581641 | chr3 | 9581141 | 9581641 | + | 0.835311 | intron (NM_133218, intron 1 of 8) | intron (NM_133218, intron 1 of 8) | 28694 | NM_133218 | 170753 | Mm.212908 | NM_133218 | ENSMUSG00000040209 | **Zfp704** | C030026M03\|C030034L19Rik\|Gig1\|Znf704 | zinc finger protein 704 | protein-coding |
| chr3_9566702_9567365 | chr3 | 9566703 | 9567365 | + | 0.759498 | intron (NM_133218, intron 1 of 8) | intron (NM_133218, intron 1 of 8) | 43051 | NM_133218 | 170753 | Mm.212908 | NM_133218 | ENSMUSG00000040209 | **Zfp704** | C030026M03\|C030034L19Rik\|Gig1\|Znf704 | zinc finger protein 704 | protein-coding |
| chr3_9566906_9567260 | chr3 | 9566907 | 9567260 | + | 0.71737 | intron (NM_133218, intron 1 of 8) | intron (NM_133218, intron 1 of 8) | 43002 | NM_133218 | 170753 | Mm.212908 | NM_133218 | ENSMUSG00000040209 | **Zfp704** | C030026M03\|C030034L19Rik\|Gig1\|Znf704 | zinc finger protein 704 | protein-coding |
| chr13_67080521_67080778 | chr13 | 67080522 | 67080778 | + | 0.733838 | intron (NM_001012449, intron 1 of 2) | intron (NM_001012449, intron 1 of 2) | 895 | NM_001012448 | 432769 | Mm.458849 | NM_001012325 | ENSMUSG00000058883 | **Zfp708** | Rslcan11 | zinc finger protein 708 | protein-coding |
| chr6_48096626_48097106 | chr6 | 48096627 | 48097106 | + | 1.23851 | Intergenic | Mam_R4\|LINE\|Dong-R4 | -10273 | NM_001163475 | 69228 | Mm.207555 | NM_001163475 | ENSMUSG00000057691 | **Zfp746** | 2810407L07Rik\|AI317225\|Znf746 | zinc finger protein 746 | protein-coding |
| chr11_121554699_121555429 | chr11 | 121554700 | 121555429 | + | 1.53066 | intron (NM_029878, intron 15 of 38) | L1MD\|LINE\|L1 | -35722 | NM_178763 | 319530 | Mm.11395 | NM_178763 | ENSMUSG00000039238 | **Zfp750** | A030007D23Rik\|Znf750 | zinc finger protein 750 | protein-coding |
| chr6_28292526_28293152 | chr6 | 28292527 | 28293152 | + | 1.07758 | Intergenic | Intergenic | -31238 | NM_001081678 | 627049 | Mm.84441 | NM_001081678 | ENSMUSG00000039841 | **Zfp800** | AA407452\|Znf800 | zinc finger protein 800 | protein-coding |
| chr6_28292299_28293150 | chr6 | 28292300 | 28293150 | + | 0.866482 | Intergenic | Intergenic | -31124 | NM_001081678 | 627049 | Mm.84441 | NM_001081678 | ENSMUSG00000039841 | **Zfp800** | AA407452\|Znf800 | zinc finger protein 800 | protein-coding |
| chr6_28252840_28253375 | chr6 | 28252841 | 28253375 | + | 0.788912 | intron (NM_001081678, intron 3 of 5) | intron (NM_001081678, intron 3 of 5) | 8493 | NM_001081678 | 627049 | Mm.84441 | NM_001081678 | ENSMUSG00000039841 | **Zfp800** | AA407452\|Znf800 | zinc finger protein 800 | protein-coding |

| | | | | | | | | | | | | | | | | | |
|---|---|---|---|---|---|---|---|---|---|---|---|---|---|---|---|---|---|
| chr8_79044636_79045264 | chr8 | 79044637 | 79045264 | + | 1.04769 | intron (NM_001294279, intron 1 of 14) | MTD\|LTR\|ERVL-MaLR | 16513 | NM_178267 | 622675 | Mm.48724 | NM_178267 | ENSMUSG00000071064 | **Zfp827** | 2810449M09Rik\|D630040G17Rik\|Znf827 | zinc finger protein 827 | protein-coding |
| chr8_79027330_79027560 | chr8 | 79027331 | 79027560 | + | 0.736281 | promoter-TSS (NM_178267) | promoter-TSS (NM_178267) | -992 | NM_001294279 | 622675 | Mm.48724 | NM_178267 | ENSMUSG00000071064 | **Zfp827** | 2810449M09Rik\|D630040G17Rik\|Znf827 | zinc finger protein 827 | protein-coding |
| chr8_79041366_79041772 | chr8 | 79041367 | 79041772 | + | 0.674434 | intron (NM_001294279, intron 1 of 14) | intron (NM_001294279, intron 1 of 14) | 13132 | NM_178267 | 622675 | Mm.48724 | NM_178267 | ENSMUSG00000071064 | **Zfp827** | 2810449M09Rik\|D630040G17Rik\|Znf827 | zinc finger protein 827 | protein-coding |
| chr2_174569887_174570281 | chr2 | 174569888 | 174570281 | + | 0.620564 | Intergenic | Intergenic | -73450 | NM_001099328 | 100043757 | Mm.48711 | NM_001099328 | ENSMUSG00000050600 | **Zfp831** | ENSMUSG00000050600\|OTTMUSG00000017459 | zinc finger protein 831 | protein-coding |
| chr6_48508058_48508461 | chr6 | 48508059 | 48508461 | + | 1.09771 | intron (NR_027808, intron 1 of 6) | intron (NR_027808, intron 1 of 6) | 3921 | NR_027808 | 58894 | Mm.31983 | NR_015597 | | **Zfp862-ps** | 4732460K03Rik\|AI448302\|Zfp862\|mKIAA0543 | zinc finger protein 862, pseudogene | pseudo |
| chr8_69202079_69202394 | chr8 | 69202080 | 69202394 | + | 0.799304 | Intergenic | B3A\|SINE\|B2 | -6809 | NM_001013379 | 234358 | Mm.334577 | NM_001013379 | | **Zfp930** | - | zinc finger protein 930 | protein-coding |
| chr17_22358255_22358824 | chr17 | 22358256 | 22358824 | + | 0.771922 | intron (NM_176962, intron 3 of 5) | Lx8\|LINE\|L1 | 2860 | NM_176962 | 319615 | Mm.442162 | NM_176962 | ENSMUSG00000033972 | **Zfp944** | 6330416L07Rik | zinc finger protein 944 | protein-coding |
| chr17_22358462_22358819 | chr17 | 22358463 | 22358819 | + | 0.771896 | intron (NM_176962, intron 3 of 5) | Lx8\|LINE\|L1 | 2759 | NM_176962 | 319615 | Mm.442162 | NM_176962 | ENSMUSG00000033972 | **Zfp944** | 6330416L07Rik | zinc finger protein 944 | protein-coding |
| chr5_104918448_104918949 | chr5 | 104918449 | 104918949 | + | 1.13659 | Intergenic | Intergenic | -58631 | NM_001039231 | 626391 | Mm.385405 | NM_001039231 | | **Zfp951** | C230055K05Rik | zinc finger protein 951 | protein-coding |
| chr6_47978860_47979525 | chr6 | 47978861 | 47979525 | + | 0.73056 | Intergenic | Intergenic | 25803 | NM_178898 | 101197 | Mm.241063 | NM_178898 | ENSMUSG00000045466 | **Zfp956** | AI894139 | zinc finger protein 956 | protein-coding |
| chr14_79141547_79142667 | chr14 | 79141548 | 79142667 | + | 1.59154 | intron (NM_027906, intron 37 of 44) | intron (NM_027906, intron 37 of 44) | 105260 | NM_001033215 | 105590 | Mm.486536 | NM_001033215 | ENSMUSG00000071262 | **Zfp957** | AU017455 | zinc finger protein 957 | protein-coding |
| chr14_79248648_79249355 | chr14 | 79248649 | 79249355 | + | 1.22369 | Intergenic | Intergenic | -1635 | NM_001033215 | 105590 | Mm.486536 | NM_001033215 | ENSMUSG00000071262 | **Zfp957** | AU017455 | zinc finger protein 957 | protein-coding |
| chr14_79114868_79115329 | chr14 | 79114869 | 79115329 | + | 1.21704 | intron (NM_027906, intron 37 of 44) | intron (NM_027906, intron 37 of 44) | 132268 | NM_001033215 | 105590 | Mm.486536 | NM_001033215 | ENSMUSG00000071262 | **Zfp957** | AU017455 | zinc finger protein 957 | protein-coding |
| chr14_79181757_79182118 | chr14 | 79181758 | 79182118 | + | 1.05669 | intron (NM_027906, intron 41 of 44) | intron (NM_027906, intron 41 of 44) | 65429 | NM_001033215 | 105590 | Mm.486536 | NM_001033215 | ENSMUSG00000071262 | **Zfp957** | AU017455 | zinc finger protein 957 | protein-coding |
| chr14_79089458_79089912 | chr14 | 79089459 | 79089912 | + | 0.892439 | intron (NM_027906, intron 29 of 44) | intron (NM_027906, intron 29 of 44) | 157682 | NM_001033215 | 105590 | Mm.486536 | NM_001033215 | ENSMUSG00000071262 | **Zfp957** | AU017455 | zinc finger protein 957 | protein-coding |
| chr8_4689673_4690143 | chr8 | 4689674 | 4690143 | + | 0.940672 | Intergenic | Intergenic | 76738 | NM_145591 | 233987 | Mm.102799 | NM_145591 | | **Zfp958** | - | zinc finger protein 958 | protein-coding |
| chr8_122285750_122286173 | chr8 | 122285751 | 122286173 | + | 1.15518 | intron (NM_009569, intron 1 of 9) | intron (NM_009569, intron 1 of 9) | 3821 | NM_009569 | 22761 | Mm.3105 | NM_009569 | ENSMUSG00000049577 | **Zfpm1** | FOG\|Fog1 | zinc finger protein, multitype 1 | protein-coding |
| chr8_122174040_122174877 | chr8 | 122174041 | 122174877 | + | 0.883676 | Intergenic | Intergenic | -107682 | NM_009569 | 22761 | Mm.3105 | NM_009569 | ENSMUSG00000049577 | **Zfpm1** | FOG\|Fog1 | zinc finger protein, multitype 1 | protein-coding |
| chr15_12103668_12104383 | chr15 | 12103669 | 12104383 | + | 1.03097 | Intergenic | Intergenic | -13825 | NM_011767 | 22763 | Mm.273496 | NM_011767 | ENSMUSG00000022201 | **Zfr** | C920030H05Rik | zinc finger RNA binding protein | protein-coding |
| chr15_12115861_12116265 | chr15 | 12115862 | 12116265 | + | 0.720264 | Intergenic | Intergenic | -1788 | NM_011767 | 22763 | Mm.273496 | NM_011767 | ENSMUSG00000022201 | **Zfr** | C920030H05Rik | zinc finger RNA binding protein | protein-coding |
| chr15_12116034_12116265 | chr15 | 12116035 | 12116265 | + | 0.603573 | Intergenic | Intergenic | -1701 | NM_011767 | 22763 | Mm.273496 | NM_011767 | ENSMUSG00000022201 | **Zfr** | C920030H05Rik | zinc finger RNA binding protein | protein-coding |
| chrX_94106060_94106624 | chrX | 94106061 | 94106624 | + | 1.17272 | intron (NM_011768, intron 1 of 7) | intron (NM_011768, intron 1 of 7) | 17065 | NM_011768 | 22764 | Mm.919 | NM_011768 | ENSMUSG00000079509 | **Zfx** | Zfx5\|Zfx5,6\|Zfx6 | zinc finger protein X-linked | protein-coding |
| chrX_94105812_94106633 | chrX | 94105813 | 94106633 | + | 0.871361 | intron (NM_011768, intron 1 of 7) | intron (NM_011768, intron 1 of 7) | 17184 | NM_011768 | 22764 | Mm.919 | NM_011768 | ENSMUSG00000079509 | **Zfx** | Zfx5\|Zfx5,6\|Zfx6 | zinc finger protein X-linked | protein-coding |
| chr9_21062986_21063346 | chr9 | 21062987 | 21063346 | + | 0.984517 | intron (NM_001103168, intron 3 of 4) | intron (NM_001103168, intron 3 of 4) | 3927 | NM_001103168 | 100009600 | Mm.483362 | NM_001103168 | ENSMUSG00000079681 | **Zglp1** | Glp1 | zinc finger, GATA-like protein 1 | protein-coding |
| chr9_21062670_21063403 | chr9 | 21062671 | 21063403 | + | 0.951751 | intron (NM_001103168, intron 3 of 4) | intron (NM_001103168, intron 3 of 4) | 4056 | NM_001103168 | 100009600 | Mm.483362 | NM_001103168 | ENSMUSG00000079681 | **Zglp1** | Glp1 | zinc finger, GATA-like protein 1 | protein-coding |
| chr9_91361747_91362638 | chr9 | 91361748 | 91362638 | + | 0.689372 | intron (NM_009573, intron 2 of 2) | intron (NM_009573, intron 2 of 2) | 3606 | NM_009573 | 22771 | Mm.335350 | NM_009573 | ENSMUSG00000032368 | **Zic1** | ZIC\|ZNF201 | zinc finger protein of the cerebellum 1 | protein-coding |
| chr13_21417367_21418016 | chr13 | 21417368 | 21418016 | + | 0.86139 | Intergenic | Intergenic | -14937 | NM_023685 | 72739 | Mm.296071 | NM_023685 | ENSMUSG00000021327 | **Zkscan3** | 2810435N07Rik\|AA408594\|AI132359\|Skz1\|Zf47\|Zfp-47\|Zfp306\|Zfp307\|mszf35 | zinc finger with KRAB and SCAN domains 3 | protein-coding |
| chr11_65818114_65818535 | chr11 | 65818115 | 65818535 | + | 1.2244 | intron (NM_026107, intron 5 of 7) | intron (NM_026107, intron 5 of 7) | 11081 | NM_026107 | 52712 | Mm.340052 | NM_026107 | ENSMUSG00000018347 | **Zkscan6** | 1700128E15Rik\|D11Ertd714e\|KOX11\|Zfp535\|Znf18 | zinc finger with KRAB and SCAN domains 6 | protein-coding |
| chr11_65818100_65818817 | chr11 | 65818101 | 65818817 | + | 1.19555 | intron (NM_026107, intron 5 of 7) | B1F1\|SINE\|Alu | 11215 | NM_026107 | 52712 | Mm.340052 | NM_026107 | ENSMUSG00000018347 | **Zkscan6** | 1700128E15Rik\|D11Ertd714e\|KOX11\|Zfp535\|Znf18 | zinc finger with KRAB and SCAN domains 6 | protein-coding |
| chr11_65844525_65845343 | chr11 | 65844526 | 65845343 | + | 1.02797 | intron (NM_001099633, intron 66 of 68) | intron (NM_001099633, intron 66 of 68) | 37690 | NM_026107 | 52712 | Mm.340052 | NM_026107 | ENSMUSG00000018347 | **Zkscan6** | 1700128E15Rik\|D11Ertd714e\|KOX11\|Zfp535\|Znf18 | zinc finger with KRAB and SCAN domains 6 | protein-coding |
| chr11_65844870_65845342 | chr11 | 65844871 | 65845342 | + | 0.978127 | intron (NM_001099633, intron 66 of 68) | intron (NM_001099633, intron 66 of 68) | 37862 | NM_026107 | 52712 | Mm.340052 | NM_026107 | ENSMUSG00000018347 | **Zkscan6** | 1700128E15Rik\|D11Ertd714e\|KOX11\|Zfp535\|Znf18 | zinc finger with KRAB and SCAN domains 6 | protein-coding |
| chr11_65852024_65852446 | chr11 | 65852025 | 65852446 | + | 0.670108 | intron (NM_001099633, intron 63 of 68) | intron (NM_001099633, intron 63 of 68) | 44991 | NM_026107 | 52712 | Mm.340052 | NM_026107 | ENSMUSG00000018347 | **Zkscan6** | 1700128E15Rik\|D11Ertd714e\|KOX11\|Zfp535\|Znf18 | zinc finger with KRAB and SCAN domains 6 | protein-coding |
| chr14_25464594_25464944 | chr14 | 25464595 | 25464944 | + | 1.07705 | intron (NM_183208, intron 1 of 23) | intron (NM_183208, intron 1 of 23) | 5584 | NM_183208 | 328365 | Mm.227484 | NM_183208 | ENSMUSG00000007817 | **Zmiz1** | BC065120\|E330020C23\|ENSMUSG00000072684\|Gm10397\|I920194n01\|Rai17\|Zimp10 | zinc finger, MIZ-type containing 1 | protein-coding |
| chr14_25464589_25465086 | chr14 | 25464590 | 25465086 | + | 0.982678 | intron (NM_183208, intron 1 of 23) | intron (NM_183208, intron 1 of 23) | 5653 | NM_183208 | 328365 | Mm.227484 | NM_183208 | ENSMUSG00000007817 | **Zmiz1** | BC065120\|E330020C23\|ENSMUSG00000072684\|Gm10397\|I920194n01\|Rai17\|Zimp10 | zinc finger, MIZ-type containing 1 | protein-coding |
| chr14_25371525_25372300 | chr14 | 25371526 | 25372300 | + | 0.760723 | Intergenic | Intergenic | -87272 | NM_183208 | 328365 | Mm.227484 | NM_183208 | ENSMUSG00000007817 | **Zmiz1** | BC065120\|E330020C23\|ENSMUSG00000072684\|Gm10397\|I920194n01\|Rai17\|Zimp10 | zinc finger, MIZ-type containing 1 | protein-coding |
| chr4_127041068_127041590 | chr4 | 127041069 | 127041590 | + | 1.25255 | Intergenic | B2_Mm2\|SINE\|B2 | 19803 | NM_026670 | 68310 | Mm.273806 | NM_026670 | ENSMUSG00000043872 | **Zmym1** | 5830412B09Rik\|AW209050 | zinc finger, MYM domain containing 1 | protein-coding |
| chr14_56890222_56890540 | chr14 | 56890223 | 56890540 | + | 0.754237 | intron (NM_029498, intron 1 of 23) | intron (NM_029498, intron 1 of 23) | 2586 | NM_029498 | 76007 | Mm.31417 | NM_029498 | ENSMUSG00000021945 | **Zmym2** | 5830413P05Rik\|FIM\|MYM\|RAMP\|SCLL\|Zfp198\|Znf198 | zinc finger, MYM-type 2 | protein-coding |
| chr13_9767258_9767835 | chr13 | 9767259 | 9767835 | + | 1.15695 | Intergenic | Intergenic | -2233 | NM_144516 | 66505 | Mm.397307 | NM_144516 | ENSMUSG00000021156 | **Zmynd11** | 2210402G22Rik\|BS69 | zinc finger, MYND domain containing 11 | protein-coding |
| chr13_9749301_9750232 | chr13 | 9749302 | 9750232 | + | 1.07085 | intron (NM_144516, intron 1 of 13) | intron (NM_144516, intron 1 of 13) | 15547 | NM_001199141 | 66505 | Mm.397307 | NM_144516 | ENSMUSG00000021156 | **Zmynd11** | 2210402G22Rik\|BS69 | zinc finger, MYND domain containing 11 | protein-coding |
| chr13_9589006_9589722 | chr13 | 9589007 | 9589722 | + | 0.888012 | intron (NM_001081426, intron 17 of 36) | intron (NM_001081426, intron 17 of 36) | 175950 | NM_001199141 | 66505 | Mm.397307 | NM_144516 | ENSMUSG00000021156 | **Zmynd11** | 2210402G22Rik\|BS69 | zinc finger, MYND domain containing 11 | protein-coding |
| chr13_9749448_9749700 | chr13 | 9749449 | 9749700 | + | 0.857706 | intron (NM_144516, intron 1 of 13) | intron (NM_144516, intron 1 of 13) | 15740 | NM_001199141 | 66505 | Mm.397307 | NM_144516 | ENSMUSG00000021156 | **Zmynd11** | 2210402G22Rik\|BS69 | zinc finger, MYND domain containing 11 | protein-coding |
| chr8_111534167_111534443 | chr8 | 111534168 | 111534443 | + | 1.55131 | Intergenic | B3\|SINE\|B2 | -2335 | NM_001168622 | 170737 | Mm.271578 | NM_133206 | ENSMUSG00000033545 | **Znrf1** | B830022L21Rik\|Rnf42\|Zrfp1\|nin283 | zinc and ring finger 1 | protein-coding |
| chr8_111544174_111544751 | chr8 | 111544175 | 111544751 | + | 0.866795 | intron (NM_001168623, intron 1 of 4) | intron (NM_001168623, intron 1 of 4) | 6120 | NM_001168623 | 170737 | Mm.271578 | NM_133206 | ENSMUSG00000033545 | **Znrf1** | B830022L21Rik\|Rnf42\|Zrfp1\|nin283 | zinc and ring finger 1 | protein-coding |
| chr8_111555338_111555671 | chr8 | 111555339 | 111555671 | + | 0.682235 | intron (NM_001168623, intron 1 of 4) | intron (NM_001168623, intron 1 of 4) | 17162 | NM_001168623 | 170737 | Mm.271578 | NM_133206 | ENSMUSG00000033545 | **Znrf1** | B830022L21Rik\|Rnf42\|Zrfp1\|nin283 | zinc and ring finger 1 | protein-coding |
| chr6_54862322_54862615 | chr6 | 54862323 | 54862615 | + | 1.30938 | intron (NM_199143, intron 1 of 4) | intron (NM_199143, intron 1 of 4) | 45553 | NM_199143 | 387524 | Mm.286149 | NM_199143 | | **Znrf2** | 1190002C14Rik\|AU044205\|D6Ertd365e | zinc and ring finger 2 | protein-coding |
| chr6_54862115_54862622 | chr6 | 54862116 | 54862622 | + | 1.23995 | intron (NM_199143, intron 1 of 4) | intron (NM_199143, intron 1 of 4) | 45453 | NM_199143 | 387524 | Mm.286149 | NM_199143 | | **Znrf2** | 1190002C14Rik\|AU044205\|D6Ertd365e | zinc and ring finger 2 | protein-coding |
| chr6_54792678_54793067 | chr6 | 54792679 | 54793067 | + | 0.870647 | Intergenic | Intergenic | -24043 | NM_199143 | 387524 | Mm.286149 | NM_199143 | | **Znrf2** | 1190002C14Rik\|AU044205\|D6Ertd365e | zinc and ring finger 2 | protein-coding |
| chr6_54769634_54770007 | chr6 | 54769635 | 54770007 | + | 0.685501 | Intergenic | Intergenic | -47095 | NM_199143 | 387524 | Mm.286149 | NM_199143 | | **Znrf2** | 1190002C14Rik\|AU044205\|D6Ertd365e | zinc and ring finger 2 | protein-coding |
| chr11_5339807_5340218 | chr11 | 5339808 | 5340218 | + | 1.1022 | intron (NM_001080924, intron 1 of 8) | intron (NM_001080924, intron 1 of 8) | 41727 | NM_001290501 | 407821 | Mm.216313 | NM_001080924 | ENSMUSG00000041961 | **Znrf3** | Gm1167 | zinc and ring finger 3 | protein-coding |

| | | | | | | | | | | | | | | | | | |
|---|---|---|---|---|---|---|---|---|---|---|---|---|---|---|---|---|---|
| chr11_5434815_5435170 | chr11 | 5434816 | 5435170 | + | 0.880265 | intron (NM_001080924, intron 1 of 8) | intron (NM_001080924, intron 1 of 8) | 9854 | NM_001080924 | 407821 | Mm.216313 | NM_001080924 | ENSMUSG00000041961 | **Znrf3** | Gm1167 | zinc and ring finger 3 | protein-coding |
| chr11_5333619_5334371 | chr11 | 5333620 | 5334371 | + | 0.724744 | intron (NM_001080924, intron 2 of 8) | intron (NM_001080924, intron 2 of 8) | 47745 | NM_001290501 | 407821 | Mm.216313 | NM_001080924 | ENSMUSG00000041961 | **Znrf3** | Gm1167 | zinc and ring finger 3 | protein-coding |
| chr11_5436703_5437020 | chr11 | 5436704 | 5437020 | + | 0.723801 | intron (NM_001080924, intron 1 of 8) | B4\|SINE\|B4 | 7985 | NM_001080924 | 407821 | Mm.216313 | NM_001080924 | ENSMUSG00000041961 | **Znrf3** | Gm1167 | zinc and ring finger 3 | protein-coding |
| chr11_5336339_5337127 | chr11 | 5336340 | 5337127 | + | 0.708843 | intron (NM_001080924, intron 2 of 8) | intron (NM_001080924, intron 2 of 8) | 45007 | NM_001290501 | 407821 | Mm.216313 | NM_001080924 | ENSMUSG00000041961 | **Znrf3** | Gm1167 | zinc and ring finger 3 | protein-coding |
| chr11_5398396_5398785 | chr11 | 5398397 | 5398785 | + | 0.672851 | intron (NM_001080924, intron 1 of 8) | intron (NM_001080924, intron 1 of 8) | -16851 | NM_001290501 | 407821 | Mm.216313 | NM_001080924 | ENSMUSG00000041961 | **Znrf3** | Gm1167 | zinc and ring finger 3 | protein-coding |
| chr11_5434793_5435374 | chr11 | 5434794 | 5435374 | + | 0.666514 | intron (NM_001080924, intron 1 of 8) | B1F\|SINE\|Alu | 9763 | NM_001080924 | 407821 | Mm.216313 | NM_001080924 | ENSMUSG00000041961 | **Znrf3** | Gm1167 | zinc and ring finger 3 | protein-coding |
| chr11_5416583_5416811 | chr11 | 5416584 | 5416811 | + | 0.665656 | intron (NM_001080924, intron 1 of 8) | intron (NM_001080924, intron 1 of 8) | 28150 | NM_001080924 | 407821 | Mm.216313 | NM_001080924 | ENSMUSG00000041961 | **Znrf3** | Gm1167 | zinc and ring finger 3 | protein-coding |
| chr11_5396075_5396474 | chr11 | 5396076 | 5396474 | + | 0.632278 | intron (NM_001080924, intron 1 of 8) | intron (NM_001080924, intron 1 of 8) | -14535 | NM_001290501 | 407821 | Mm.216313 | NM_001080924 | ENSMUSG00000041961 | **Znrf3** | Gm1167 | zinc and ring finger 3 | protein-coding |
| chr11_98563047_98563489 | chr11 | 98563048 | 98563489 | + | 0.98625 | Intergenic | Intergenic | 12171 | NM_001166494 | 69376 | Mm.55996 | NM_027061 | ENSMUSG00000017195 | **Zpbp2** | 1700017D11Rik\|2610022C02Rik\|ZPBPL | zona pellucida binding protein 2 | protein-coding |
| chr16_55525993_55526856 | chr16 | 55525994 | 55526856 | + | 1.39532 | Intergenic | Intergenic | -243188 | NM_178720 | 239852 | Mm.245045 | NM_178720 | ENSMUSG00000064310 | **Zpld1** | 9430016A21Rik | zona pellucida like domain containing 1 | protein-coding |
| chr16_55526277_55526860 | chr16 | 55526278 | 55526860 | + | 1.35684 | Intergenic | Intergenic | -243332 | NM_178720 | 239852 | Mm.245045 | NM_178720 | ENSMUSG00000064310 | **Zpld1** | 9430016A21Rik | zona pellucida like domain containing 1 | protein-coding |
| chr16_55547079_55547424 | chr16 | 55547080 | 55547424 | + | 1.26401 | Intergenic | Tigger7\|DNA\|TcMar-Tigger | -264015 | NM_178720 | 239852 | Mm.245045 | NM_178720 | ENSMUSG00000064310 | **Zpld1** | 9430016A21Rik | zona pellucida like domain containing 1 | protein-coding |
| chr16_55484358_55484741 | chr16 | 55484359 | 55484741 | + | 1.02392 | Intergenic | ORR1E\|LTR\|ERVL-MaLR | -201313 | NM_178720 | 239852 | Mm.245045 | NM_178720 | ENSMUSG00000064310 | **Zpld1** | 9430016A21Rik | zona pellucida like domain containing 1 | protein-coding |
| chr16_55550795_55551289 | chr16 | 55550796 | 55551289 | + | 0.982068 | Intergenic | Intergenic | -267805 | NM_178720 | 239852 | Mm.245045 | NM_178720 | ENSMUSG00000064310 | **Zpld1** | 9430016A21Rik | zona pellucida like domain containing 1 | protein-coding |
| chr7_80841010_80841403 | chr7 | 80841011 | 80841403 | + | 1.11388 | Intergenic | Intergenic | -20901 | NM_009553 | 22691 | Mm.358645 | NM_009553 | ENSMUSG00000038797 | **Zscan2** | Zfp-29\|Zfp29 | zinc finger and SCAN domain containing 2 | protein-coding |
| chr7_80839814_80840098 | chr7 | 80839815 | 80840098 | + | 0.975293 | Intergenic | Intergenic | -22152 | NM_009553 | 22691 | Mm.358645 | NM_009553 | ENSMUSG00000038797 | **Zscan2** | Zfp-29\|Zfp29 | zinc finger and SCAN domain containing 2 | protein-coding |
| chr7_80837312_80837501 | chr7 | 80837313 | 80837501 | + | 0.708292 | Intergenic | Intergenic | -24701 | NM_009553 | 22691 | Mm.358645 | NM_009553 | ENSMUSG00000038797 | **Zscan2** | Zfp-29\|Zfp29 | zinc finger and SCAN domain containing 2 | protein-coding |
| chr13_107856855_107857565 | chr13 | 107856856 | 107857565 | + | 1.45791 | intron (NM_145456, intron 2 of 14) | intron (NM_145456, intron 2 of 14) | 32854 | NM_145456 | 67263 | Mm.490418 | NM_145456 | ENSMUSG00000032846 | **Zswim6** | 2900036G02Rik\|mKIAA1577 | zinc finger SWIM-type containing 6 | protein-coding |
| chr13_107871660_107872058 | chr13 | 107871661 | 107872058 | + | 1.44485 | intron (NM_145456, intron 2 of 14) | intron (NM_145456, intron 2 of 14) | 18205 | NM_145456 | 67263 | Mm.490418 | NM_145456 | ENSMUSG00000032846 | **Zswim6** | 2900036G02Rik\|mKIAA1577 | zinc finger SWIM-type containing 6 | protein-coding |
| chr6_42348199_42348459 | chr6 | 42348200 | 42348459 | + | 1.24149 | Intergenic | Intergenic | -1499 | NM_011777 | 22793 | Mm.282303 | NM_011777 | ENSMUSG00000029860 | **Zyx** | 9530098H06Rik\|R75157 | zyxin | protein-coding |
| chr11_72842745_72843437 | chr11 | 72842746 | 72843437 | + | 1.46593 | intron (NM_001045536, intron 9 of 54) | intron (NM_001045536, intron 9 of 54) | 46865 | NM_001045536 | 195018 | Mm.272030 | NM_001045536 | ENSMUSG00000055670 | **Zzef1** | 8430405D05Rik\|BC051585\|C130099L13Rik\|C85424\|mKIAA0399 | zinc finger, ZZ-type with EF hand domain 1 | protein-coding |
| chr11_72847897_72848557 | chr11 | 72847898 | 72848557 | + | 1.2098 | intron (NM_001045536, intron 11 of 54) | intron (NM_001045536, intron 11 of 54) | 52001 | NM_001045536 | 195018 | Mm.272030 | NM_001045536 | ENSMUSG00000055670 | **Zzef1** | 8430405D05Rik\|BC051585\|C130099L13Rik\|C85424\|mKIAA0399 | zinc finger, ZZ-type with EF hand domain 1 | protein-coding |
| chr11_72810384_72810764 | chr11 | 72810385 | 72810764 | + | 0.664375 | intron (NM_001045536, intron 1 of 54) | B3\|SINE\|B2 | 14348 | NM_001045536 | 195018 | Mm.272030 | NM_001045536 | ENSMUSG00000055670 | **Zzef1** | 8430405D05Rik\|BC051585\|C130099L13Rik\|C85424\|mKIAA0399 | zinc finger, ZZ-type with EF hand domain 1 | protein-coding |
| chr3_152444106_152444513 | chr3 | 152444107 | 152444513 | + | 1.27108 | intron (NM_001287139, intron 4 of 13) | intron (NM_001287139, intron 4 of 13) | 47626 | NM_001287139 | 108946 | Mm.379292 | NM_198416 | ENSMUSG00000039068 | **Zzz3** | 3110065C23Rik\|6430567E01Rik\|AA408566\|AI481057\|AW556059\|R75514 | zinc finger, ZZ domain containing 3 | protein-coding |

**Table S1C. HOMER Opening peaks in both Cytokine-primed and Ag-stimulated cells**

Data is sorted alphabetically by the gene name; Only peaks above the peak score value of 0.5 are shown.

| PeakID (cmd=annotatePeaks.pl ../averagescore_opening_peaks_BallCA.bed mm10 -go go_opening_peaks_BallCvA) | Chr | Start | End | Strand | Peak Score | Annotation | Detailed Annotation | Distance to TSS | Nearest PromoterID | Entrez ID | Nearest Unigene | Nearest Refseq | Nearest Ensembl | Gene Name | Gene Alias | Gene Description | Gene Type |
|---|---|---|---|---|---|---|---|---|---|---|---|---|---|---|---|---|---|
| chr2_46371045_46371404 | chr2 | 46371046 | 46371404 | + | 1.02429 | Intergenic | Intergenic | 674620 | NR_040497 | 69405 | Mm.396184 | NR_040497 | ENSMUSG00000086202 | **1700019E08Rik** | - | RIKEN cDNA 1700019E08 gene | ncRNA |
| chr2_46370821_46371458 | chr2 | 46370822 | 46371458 | + | 1.00873 | Intergenic | Intergenic | 674535 | NR_040497 | 69405 | Mm.396184 | NR_040497 | ENSMUSG00000086202 | **1700019E08Rik** | - | RIKEN cDNA 1700019E08 gene | ncRNA |
| chr12_12547387_12548252 | chr12 | 12547388 | 12548252 | + | 0.822163 | Intergenic | Intergenic | -155345 | NR_045829 | 70858 | Mm.410838 | NR_045829 | | **4921511I17Rik** | - | RIKEN cDNA 4921511I17 gene | ncRNA |
| chr9_118360838_118361565 | chr9 | 118360839 | 118361565 | + | 0.682352 | Intergenic | Intergenic | -68637 | NR_040716 | 71196 | Mm.73218 | NR_040716 | | **4933432G23Rik** | - | RIKEN cDNA 4933432G23 gene | ncRNA |
| chr3_115857750_115858282 | chr3 | 115857751 | 115858282 | + | 0.859895 | Intergenic | Intergenic | 30043 | NR_027894 | 68161 | Mm.392043 | NR_015487 | | **A930005H10Rik** | - | RIKEN cDNA A930005H10 gene | ncRNA |
| chr3_115857603_115858461 | chr3 | 115857604 | 115858461 | + | 0.81318 | Intergenic | Intergenic | 30027 | NR_027894 | 68161 | Mm.392043 | NR_015487 | | **A930005H10Rik** | - | RIKEN cDNA A930005H10 gene | ncRNA |
| chrX_142907468_142908077 | chrX | 142907469 | 142908077 | + | 0.99965 | intron (NM_019496, intron 2 of 5) | intron (NM_019496, intron 2 of 5) | 58955 | NM_019496 | 56068 | Mm.143724 | NM_019496 | | **Ammecr1** | 6230420G18Rik | Alport syndrome, mental retardation, midface hypoplasia and elliptocytosis chromosomal region gene 1 | protein-coding |
| chr7_58475100_58475677 | chr7 | 58475101 | 58475677 | + | 0.913423 | Intergenic | Intergenic | -182813 | NM_009728 | 11982 | Mm.135129 | NM_009728 | ENSMUSG00000025324 | **Atp10a** | Atp10c\|pfatp | ATPase, class V, type 10A | protein-coding |
| chr16_45038590_45039050 | chr16 | 45038591 | 45039050 | + | 1.10968 | Intergenic | Intergenic | -55233 | NM_026439 | 67896 | Mm.181074 | NM_026439 | ENSMUSG00000022665 | **Ccdc80** | 2610001E17Rik\|DRO1\|Ssg1\|Urb | coiled-coil domain containing 80 | protein-coding |
| chr16_45038390_45039051 | chr16 | 45038391 | 45039051 | + | 1.06367 | Intergenic | MLT2B2\|LTR\|ERVL | -55332 | NM_026439 | 67896 | Mm.181074 | NM_026439 | ENSMUSG00000022665 | **Ccdc80** | 2610001E17Rik\|DRO1\|Ssg1\|Urb | coiled-coil domain containing 80 | protein-coding |
| chr9_114487351_114488780 | chr9 | 114487352 | 114488780 | + | 1.13284 | Intergenic | LTR16A\|LTR\|ERVL | 8478 | NM_009916 | 12773 | Mm.1337 | NM_009916 | ENSMUSG00000047898 | **Ccr4** | C-C CKR-4\|CHEMR1\|Cmkbr4\|LESTR\|Sdf1r | chemokine (C-C motif) receptor 4 | protein-coding |
| chr9_114488332_114488757 | chr9 | 114488333 | 114488757 | + | 1.05555 | Intergenic | Intergenic | 7999 | NM_009916 | 12773 | Mm.1337 | NM_009916 | ENSMUSG00000047898 | **Ccr4** | C-C CKR-4\|CHEMR1\|Cmkbr4\|LESTR\|Sdf1r | chemokine (C-C motif) receptor 4 | protein-coding |
| chr9_114488171_114488772 | chr9 | 114488172 | 114488772 | + | 1.04079 | Intergenic | Intergenic | 8072 | NM_009916 | 12773 | Mm.1337 | NM_009916 | ENSMUSG00000047898 | **Ccr4** | C-C CKR-4\|CHEMR1\|Cmkbr4\|LESTR\|Sdf1r | chemokine (C-C motif) receptor 4 | protein-coding |
| chr11_99180054_99180723 | chr11 | 99180055 | 99180723 | + | 1.00928 | Intergenic | Intergenic | -25312 | NM_007719 | 12775 | Mm.2932 | NM_007719 | ENSMUSG00000037944 | **Ccr7** | CD197\|Cdw197\|Cmkbr7\|EBI1\|Ebi1h | chemokine (C-C motif) receptor 7 | protein-coding |
| chr4_89388116_89388489 | chr4 | 89388117 | 89388489 | + | 1.05541 | Intergenic | Intergenic | -77271 | NM_007670 | 12579 | Mm.423094 | NM_007670 | ENSMUSG00000073802 | **Cdkn2b** | AV083695\|INK4b\|MTS2\|p15\|p15(INK4b)\|p15INK4b | cyclin-dependent kinase inhibitor 2B (p15, inhibits CDK4) | protein-coding |
| chr17_78271345_78271924 | chr17 | 78271346 | 78271924 | + | 1.31401 | intron (NM_015800, intron 2 of 16) | intron (NM_015800, intron 2 of 16) | 71387 | NM_015800 | 50766 | Mm.311912 | NM_015800 | ENSMUSG00000024074 | **Crim1** | AU015004 | cysteine rich transmembrane BMP regulator 1 (chordin like) | protein-coding |
| chr18_84872385_84872663 | chr18 | 84872386 | 84872663 | + | 0.819584 | intron (NM_025797, intron 2 of 4) | intron (NM_025797, intron 2 of 4) | 21110 | NM_025797 | 109672 | Mm.31018 | NM_025797 | | **Cyb5a** | 0610009N12Rik\|Cyb5 | cytochrome b5 type A (microsomal) | protein-coding |
| chr2_62331849_62332356 | chr2 | 62331850 | 62332356 | + | 1.05245 | 3' UTR (NM_010074, exon 26 of 26) | 3' UTR (NM_010074, exon 26 of 26) | 80128 | NM_010074 | 13482 | Mm.1151 | NM_010074 | ENSMUSG00000035000 | **Dpp4** | Cd26\|Dpp-4\|THAM | dipeptidylpeptidase 4 | protein-coding |
| chr7_90067888_90068928 | chr7 | 90067889 | 90068928 | + | 0.741812 | Intergenic | Intergenic | 39249 | NR_110364 | 319711 | Mm.31753 | NR_015614 | | **E230029C05Rik** | AI452102\|B230337J15 | RIKEN cDNA E230029C05 gene | ncRNA |
| chr7_90067893_90068425 | chr7 | 90067894 | 90068425 | + | 0.723427 | Intergenic | Intergenic | 39000 | NR_110364 | 319711 | Mm.31753 | NR_015614 | | **E230029C05Rik** | AI452102\|B230337J15 | RIKEN cDNA E230029C05 gene | ncRNA |
| chr7_90067998_90068378 | chr7 | 90067999 | 90068378 | + | 0.706886 | Intergenic | Intergenic | 39029 | NR_110364 | 319711 | Mm.31753 | NR_015614 | | **E230029C05Rik** | AI452102\|B230337J15 | RIKEN cDNA E230029C05 gene | ncRNA |
| chr9_77955595_77955870 | chr9 | 77955596 | 77955870 | + | 0.68443 | intron (NM_134255, intron 1 of 7) | intron (NM_134255, intron 1 of 7) | 38368 | NM_134255 | 68801 | Mm.430736 | NM_134255 | ENSMUSG00000032349 | **Elovl5** | 1110059L23Rik\|AI747313\|AU043003\|HELO1 | ELOVL family member 5, elongation of long chain fatty acids (yeast) | protein-coding |
| chr5_3477687_3478305 | chr5 | 3477688 | 3478305 | + | 1.1884 | intron (NM_009873, intron 4 of 6) | intron (NM_009873, intron 4 of 6) | -65837 | NM_001042501 | 68152 | Mm.240265 | NM_026583 | | **Fam133b** | 2900022K02Rik\|5830415L20Rik | family with sequence similarity 133, member B | protein-coding |
| chr5_3477606_3478593 | chr5 | 3477607 | 3478593 | + | 1.13954 | intron (NM_009873, intron 4 of 6) | intron (NM_009873, intron 4 of 6) | -65733 | NM_001042501 | 68152 | Mm.240265 | NM_026583 | | **Fam133b** | 2900022K02Rik\|5830415L20Rik | family with sequence similarity 133, member B | protein-coding |
| chr5_3460776_3461381 | chr5 | 3460777 | 3461381 | + | 1.01678 | intron (NM_009873, intron 3 of 6) | intron (NM_009873, intron 3 of 6) | -82754 | NM_001042501 | 68152 | Mm.240265 | NM_026583 | | **Fam133b** | 2900022K02Rik\|5830415L20Rik | family with sequence similarity 133, member B | protein-coding |
| chr5_3460214_3461370 | chr5 | 3460215 | 3461370 | + | 0.997305 | intron (NM_009873, intron 3 of 6) | intron (NM_009873, intron 3 of 6) | -83041 | NM_001042501 | 68152 | Mm.240265 | NM_026583 | | **Fam133b** | 2900022K02Rik\|5830415L20Rik | family with sequence similarity 133, member B | protein-coding |
| chr11_109758486_109759304 | chr11 | 109758487 | 109759304 | + | 0.725995 | Intergenic | Lx9\|LINE\|L1 | -36639 | NM_153782 | 208659 | Mm.208662 | NM_153782 | ENSMUSG00000020614 | **Fam20a** | AI606893 | family with sequence similarity 20, member A | protein-coding |
| chr9_85107251_85107927 | chr9 | 85107252 | 85107927 | + | 0.825215 | Intergenic | Intergenic | 219535 | NM_001160378 | 212943 | Mm.74610 | NM_001160378 | ENSMUSG00000032265 | **Fam46a** | D930050G01Rik | family with sequence similarity 46, member A | protein-coding |
| chr9_85107295_85107691 | chr9 | 85107296 | 85107691 | + | 0.768774 | Intergenic | Intergenic | 219631 | NM_001160378 | 212943 | Mm.74610 | NM_001160378 | ENSMUSG00000032265 | **Fam46a** | D930050G01Rik | family with sequence similarity 46, member A | protein-coding |
| chr17_28488204_28488676 | chr17 | 28488205 | 28488676 | + | 0.723907 | Intergenic | Intergenic | -2291 | NM_010220 | 14229 | Mm.276405 | NM_010220 | ENSMUSG00000024222 | **Fkbp5** | D17Ertd592e\|Dit1\|FKBP-5\|FKBP51 | FK506 binding protein 5 | protein-coding |
| chr12_99510119_99510476 | chr12 | 99510120 | 99510476 | + | 1.1985 | Intergenic | B3\|SINE\|B2 | -60224 | NM_183186 | 71375 | Mm.341972 | NM_183186 | ENSMUSG00000033713 | **Foxn3** | 5430426H20Rik\|AA589593\|AW556347\|Ches1\|Ches1l\|HTLFL1 | forkhead box N3 | protein-coding |
| chr12_99509971_99510622 | chr12 | 99509972 | 99510622 | + | 1.18551 | Intergenic | B3\|SINE\|B2 | -60223 | NM_183186 | 71375 | Mm.341972 | NM_183186 | ENSMUSG00000033713 | **Foxn3** | 5430426H20Rik\|AA589593\|AW556347\|Ches1\|Ches1l\|HTLFL1 | forkhead box N3 | protein-coding |
| chr3_79335990_79336510 | chr3 | 79335991 | 79336510 | + | 1.14104 | Intergenic | Intergenic | -9126 | NM_001142953 | 100233207 | Mm.373801 | NM_001142953 | ENSMUSG00000091685 | **Gm17359** | - | predicted gene, 17359 | protein-coding |
| chr14_56275673_56276521 | chr14 | 56275674 | 56276521 | + | 0.717193 | Intergenic | Intergenic | -13837 | NM_013542 | 14939 | Mm.14874 | NM_013542 | ENSMUSG00000015437 | **Gzmb** | AI553453\|CCP-1/C11\|CCP1\|Ctla-1\|Ctla1\|GZB | granzyme B | protein-coding |
| chr13_23593064_23593674 | chr13 | 23593065 | 23593674 | + | 0.670901 | intron (NM_001177653, intron 1 of 1) | intron (NM_001177653, intron 1 of 1) | -7399 | NM_001290530 | 319179 | Mm.440391 | NM_178194 | ENSMUSG00000047246 | **Hist1h2be** | - | histone cluster 1, H2be | protein-coding |
| chr6_48398457_48398789 | chr6 | 48398458 | 48398789 | + | 0.670235 | intron (NM_133922, intron 1 of 17) | ORR1E\|LTR\|ERVL-MaLR | 3037 | NM_133922 | 77827 | Mm.107646 | NM_133922 | ENSMUSG00000042810 | **Krba1** | A930040G15Rik\|AI448780 | KRAB-A domain containing 1 | protein-coding |
| chr8_34139899_34140213 | chr8 | 34139900 | 34140213 | + | 0.946487 | intron (NM_026609, intron 2 of 3) | intron (NM_026609, intron 2 of 3) | 6683 | NM_026609 | 68192 | Mm.34212 | NM_026609 | ENSMUSG00000031513 | **Leprotl1** | 1110067H13Rik\|1520402O14Rik\|AI854296 | leptin receptor overlapping transcript-like 1 | protein-coding |
| chr6_124683595_124684249 | chr6 | 124683596 | 124684249 | + | 0.921817 | intron (NM_145130, intron 1 of 12) | intron (NM_145130, intron 1 of 12) | 20818 | NM_145130 | 14792 | Mm.273915 | NM_145130 | ENSMUSG00000004270 | **Lpcat3** | C3f\|Grcc3f\|Lpcat\|Lpeat\|Lplat5\|Lpsat\|Mboat5\|Moact5\|Oact5\|PTG | lysophosphatidylcholine acyltransferase 3 | protein-coding |
| chr6_119343614_119343911 | chr6 | 119343615 | 119343911 | + | 0.610866 | intron (NM_001033382, intron 30 of 37) | intron (NM_001033382, intron 30 of 37) | -12997 | NM_172492 | 211187 | Mm.121498 | NM_172492 | ENSMUSG00000055003 | **Lrtm2** | A230084J22\|AI841794 | leucine-rich repeats and transmembrane domains 2 | protein-coding |
| chr18_4278298_4278812 | chr18 | 4278299 | 4278812 | + | 0.7489 | Intergenic | Intergenic | 74398 | NM_007746 | 26410 | Mm.3275 | NM_007746 | ENSMUSG00000024235 | **Map3k8** | Cot\|Cot/Tpl2\|Est\|Estf\|Tpl-2\|Tpl2\|c-COT | mitogen-activated protein kinase kinase kinase 8 | protein-coding |
| chr12_105022703_105023184 | chr12 | 105022704 | 105023184 | + | 0.696501 | Intergenic | Intergenic | 8197 | NR_037229 | 100526471 | | NR_037229 | | **Mir3069** | mmu-mir-3069 | microRNA 3069 | ncRNA |
| chr8_108890850_108891372 | chr8 | 108890851 | 108891372 | + | 0.64507 | intron (NM_007496, intron 4 of 9) | (CAGA)n\|Simple_repeat\|Simple_repeat | -45749 | NR_037295 | 100526511 | | NR_037295 | | **Mir3108** | mmu-mir-3108 | microRNA 3108 | ncRNA |
| chr1_51514289_51515019 | chr1 | 51514290 | 51515019 | + | 0.94139 | Intergenic | Intergenic | -36255 | NM_028696 | 109019 | Mm.196290 | NM_028696 | ENSMUSG00000026107 | **Nabp1** | 4930434H03Rik\|4930442A21Rik\|4930488J04Rik\|4933440J18Rik\|5830411E10Rik\|AI852561\|Nbp1\|Obfc2a\|Ssb2 | nucleic acid binding protein 1 | protein-coding |
| chr2_52319180_52319426 | chr2 | 52319181 | 52319426 | + | 0.870781 | intron (NM_010889, intron 10 of 156) | intron (NM_010889, intron 10 of 156) | 19495 | NM_010889 | 17996 | Mm.389247 | NM_010889 | ENSMUSG00000026950 | **Neb** | AI595938 | nebulin | protein-coding |
| chr2_52319162_52319699 | chr2 | 52319163 | 52319699 | + | 0.842868 | intron (NM_010889, intron 10 of 156) | intron (NM_010889, intron 10 of 156) | 19367 | NM_010889 | 17996 | Mm.389247 | NM_010889 | ENSMUSG00000026950 | **Neb** | AI595938 | nebulin | protein-coding |
| chr2_52318959_52319728 | chr2 | 52318960 | 52319728 | + | 0.810805 | intron (NM_010889, intron 10 of 156) | intron (NM_010889, intron 10 of 156) | 19454 | NM_010889 | 17996 | Mm.389247 | NM_010889 | ENSMUSG00000026950 | **Neb** | AI595938 | nebulin | protein-coding |
| chr16_55582865_55583434 | chr16 | 55582866 | 55583434 | + | 0.781145 | Intergenic | Intergenic | 238988 | NM_001159394 | 80859 | Mm.247272 | NM_030612 | ENSMUSG00000035356 | **Nfkbiz** | AA408868\|INAP\|Mail | nuclear factor of kappa light polypeptide gene enhancer in B cells inhibitor, zeta | protein-coding |
| chr16_55582470_55583434 | chr16 | 55582471 | 55583434 | + | 0.712203 | Intergenic | Intergenic | 239186 | NM_001159394 | 80859 | Mm.247272 | NM_030612 | ENSMUSG00000035356 | **Nfkbiz** | AA408868\|INAP\|Mail | nuclear factor of kappa light polypeptide gene enhancer in B cells inhibitor, zeta | protein-coding |
| chr6_120122826_120123273 | chr6 | 120122827 | 120123273 | + | 0.766281 | intron (NM_016718, intron 1 of 3) | L1M3\|LINE\|L1 | 29670 | NM_016718 | 29862 | Mm.103668 | NM_016718 | ENSMUSG00000041377 | **Ninj2** | - | ninjurin 2 | protein-coding |
| chr19_53791724_53792843 | chr19 | 53791725 | 53792843 | + | 0.832579 | intron (NM_001170847, intron 1 of 13) | intron (NM_001170847, intron 1 of 13) | -99947 | NM_011050 | 18569 | Mm.1605 | NM_011050 | ENSMUSG00000024975 | **Pdcd4** | D19Ucla1\|Ma3\|Tis | programmed cell death 4 | protein-coding |
| chr17_29500634_29501279 | chr17 | 29500635 | 29501279 | + | 0.76991 | Intergenic | Intergenic | 9912 | NM_008842 | 18712 | Mm.405293 | NM_008842 | | **Pim1** | Pim-1 | proviral integration site 1 | protein-coding |
| chr2_59171560_59172058 | chr2 | 59171561 | 59172058 | + | 0.794447 | intron (NM_026361, intron 1 of 21) | intron (NM_026361, intron 1 of 21) | 10959 | NM_175464 | 227937 | Mm.260938 | NM_026361 | ENSMUSG00000026991 | **Pkp4** | 5031422I09Rik\|9430019K17Rik\|Armrp\|p0071 | plakophilin 4 | protein-coding |
| chr2_59171568_59171921 | chr2 | 59171569 | 59171921 | + | 0.776069 | intron (NM_026361, intron 1 of 21) | intron (NM_026361, intron 1 of 21) | 10895 | NM_175464 | 227937 | Mm.260938 | NM_026361 | ENSMUSG00000026991 | **Pkp4** | 5031422I09Rik\|9430019K17Rik\|Armrp\|p0071 | plakophilin 4 | protein-coding |
| chr2_59171380_59171950 | chr2 | 59171381 | 59171950 | + | 0.673682 | intron (NM_026361, intron 1 of 21) | intron (NM_026361, intron 1 of 21) | 10815 | NM_175464 | 227937 | Mm.260938 | NM_026361 | ENSMUSG00000026991 | **Pkp4** | 5031422I09Rik\|9430019K17Rik\|Armrp\|p0071 | plakophilin 4 | protein-coding |

| | | | | | | | | | | | | | | | | | |
|---|---|---|---|---|---|---|---|---|---|---|---|---|---|---|---|---|---|
| chr2_59171246_59172068 | chr2 | 59171247 | 59172068 | + | 0.668804 | intron (NM_026361, intron 1 of 21) | intron (NM_026361, intron 1 of 21) | 10807 | NM_175464 | 227937 | Mm.260938 | NM_026361 | ENSMUSG00000026991 | **Pkp4** | 5031422I09Rik\|9430019K17Rik\|Armrp\|p0071 | plakophilin 4 | protein-coding |
| chr2_11173085_11173457 | chr2 | 11173086 | 11173457 | + | 0.705917 | intron (NM_008859, intron 1 of 17) | intron (NM_008859, intron 1 of 17) | 889 | NM_008859 | 18761 | Mm.329993 | NM_008859 | ENSMUSG00000026778 | **Prkcq** | A130035A12Rik\|AW494342\|PKC-0\|PKC-theta\|PKCtheta\|Pkcq | protein kinase C, theta | protein-coding |
| chr17_50097790_50098343 | chr17 | 50097791 | 50098343 | + | 0.595274 | intron (NM_181397, intron 3 of 9) | intron (NM_181397, intron 3 of 9) | 92430 | NM_181397 | 76438 | Mm.41854 | NM_181397 | | **Rftn1** | 2310015N21Rik\|AI853526 | raftlin lipid raft linker 1 | protein-coding |
| chr7_114171284_114172183 | chr7 | 114171285 | 114172183 | + | 0.655987 | Intergenic | Intergenic | -53953 | NM_025846 | 66922 | Mm.276572 | NM_025846 | ENSMUSG00000055723 | **Rras2** | 2610016H24Rik\|C86394\|TC21 | related RAS viral (r-ras) oncogene homolog 2 | protein-coding |
| chr18_79434525_79435241 | chr18 | 79434526 | 79435241 | + | 0.816524 | Intergenic | Intergenic | -325492 | NM_053099 | 240427 | Mm.312871 | NM_053099 | ENSMUSG00000024548 | **Setbp1** | C130092E12\|Seb\|mKIAA0437 | SET binding protein 1 | protein-coding |
| chr1_46864072_46864769 | chr1 | 46864073 | 46864769 | + | 0.783076 | Intergenic | Intergenic | -10912 | NM_172653 | 227059 | Mm.233889 | NM_172653 | ENSMUSG00000025986 | **Slc39a10** | 2900042E17Rik\|5430433I10\|mKIAA1265 | solute carrier family 39 (zinc transporter), member 10 | protein-coding |
| chr11_62148181_62149063 | chr11 | 62148182 | 62149063 | + | 1.20623 | intron (NM_001281818, intron 9 of 14) | intron (NM_001281818, intron 9 of 14) | 71598 | NM_001029936 | 432572 | Mm.458025 | NM_001029936 | ENSMUSG00000042331 | **Specc1** | 2810012G08Rik\|B230396K10Rik\|Cytsb | sperm antigen with calponin homology and coiled-coil domains 1 | protein-coding |
| chr12_105019262_105019794 | chr12 | 105019263 | 105019794 | + | 0.804234 | Intergenic | Intergenic | -9719 | NM_172500 | 212073 | Mm.212688 | NM_172500 | ENSMUSG00000054150 | **Syne3** | nesprin-3 | spectrin repeat containing, nuclear envelope family member 3 | protein-coding |
| chr6_52801790_52802789 | chr6 | 52801791 | 52802789 | + | 0.677644 | intron (NM_173406, intron 3 of 4) | intron (NM_173406, intron 3 of 4) | 88561 | NM_025816 | 52440 | Mm.389757 | NM_025816 | ENSMUSG00000004535 | **Tax1bp1** | 1200003J11Rik\|1700069J21Rik\|AA930106\|D6Ertd404e\|D6Ertd772e\|T6bp\|TXBP151 | Tax1 (human T cell leukemia virus type I) binding protein 1 | protein-coding |
| chr3_21965407_21965959 | chr3 | 21965408 | 21965959 | + | 1.0854 | Intergenic | Intergenic | -110969 | NM_030732 | 81004 | Mm.202966 | NM_030732 | ENSMUSG00000027630 | **Tbl1xr1** | 8030499H02Rik\|A630076E03Rik\|AW539987\|C21\|C230089I12Rik\|DC42\|Ira1\|TBLR1 | transducin (beta)-like 1X-linked receptor 1 | protein-coding |
| chr3_21965590_21966020 | chr3 | 21965591 | 21966020 | + | 1.03397 | Intergenic | Intergenic | -110847 | NM_030732 | 81004 | Mm.202966 | NM_030732 | ENSMUSG00000027630 | **Tbl1xr1** | 8030499H02Rik\|A630076E03Rik\|AW539987\|C21\|C230089I12Rik\|DC42\|Ira1\|TBLR1 | transducin (beta)-like 1X-linked receptor 1 | protein-coding |
| chr6_59312928_59313891 | chr6 | 59312929 | 59313891 | + | 1.03185 | Intergenic | RLTR14_RN\|LTR\|ERV1 | 104540 | NM_001081145 | 68140 | Mm.25843 | NM_001081145 | ENSMUSG00000049232 | **Tigd2** | 3632410O17Rik | tigger transposable element derived 2 | protein-coding |
| chr1_36892379_36892960 | chr1 | 36892380 | 36892960 | + | 0.731518 | intron (NM_018872, intron 1 of 40) | MTEa\|LTR\|ERVL-MaLR | 46857 | NM_018872 | 56030 | Mm.186943 | NM_018872 | ENSMUSG00000026116 | **Tmem131** | 2610524E03Rik\|CC28\|D1Bwg0491e\|Neg\|RW1\|YR-23\|mKIAA0257 | transmembrane protein 131 | protein-coding |
| chr4_63841786_63842938 | chr4 | 63841787 | 63842938 | + | 0.710108 | intron (NM_009403, intron 2 of 3) | intron (NM_009403, intron 2 of 3) | 18922 | NM_009403 | 21949 | Mm.4664 | NM_009403 | | **Tnfsf8** | CD153\|CD30LG\|Cd30l | tumor necrosis factor (ligand) superfamily, member 8 | protein-coding |
| chr9_65831225_65831684 | chr9 | 65831226 | 65831684 | + | 0.986403 | 3' UTR (NM_001170907, exon 12 of 12) | 3' UTR (NM_001170907, exon 12 of 12) | -3891 | NM_172536 | 214812 | Mm.139231 | NM_172536 | ENSMUSG00000040524 | **Zfp609** | 9830165N24\|BC016271\|Znf609\|mKIAA0295 | zinc finger protein 609 | protein-coding |
| chr11_95748121_95748792 | chr11 | 95748122 | 95748792 | + | 0.896636 | promoter-TSS (NM_201609) | promoter-TSS (NM_201609) | -610 | NM_201609 | 268469 | Mm.389038 | NM_201609 | ENSMUSG00000075595 | **Zfp652** | 9530033F24Rik\|ZNF652 | zinc finger protein 652 | protein-coding |

**Table S2A_HOMER Closing peaks in Cytokine-primed cells**
Data is sorted alphabetically by the gene name; Only peaks above the peak score value of 0.5 are shown.

| PeakID (cmd=annotatePeaks.pl ../closing_peaks_BA.bed mm10 -go go_closing_peaks_BA) | Chr | Start | End | Strand | Peak Score | Annotation | Detailed Annotation | Distance to TSS | Nearest PromoterID | Entrez ID | Nearest Unigene | Nearest Refseq | Nearest Ensembl | Gene Name | Gene Alias | Gene Description | Gene Type |
|---|---|---|---|---|---|---|---|---|---|---|---|---|---|---|---|---|---|
| chr2_132691701_132692100 | chr2 | 132691702 | 132692100 | + | -0.939148 | intron (NM_028637, intron 1 of 2) | intron (NM_028637, intron 1 of 2) | 1618 | NM_028637 | 73747 | Mm.284114 | NM_028637 | ENSMUSG00000044991 | **1110034G24Rik** | - | RIKEN cDNA 1110034G24 gene | protein-coding |
| chr16_95901395_95901682 | chr16 | 95901396 | 95901682 | + | -0.612542 | intron (NR_040484, intron 1 of 3) | intron (NR_040484, intron 1 of 3) | -2826 | NR_040485 | 69776 | Mm.120470 | NR_040484 | ENSMUSG00000086899 | **1600002D24Rik** | - | RIKEN cDNA 1600002D24 gene | ncRNA |
| chr16_95860698_95861353 | chr16 | 95860699 | 95861353 | + | -0.624739 | intron (NR_040484, intron 1 of 3) | intron (NR_040484, intron 1 of 3) | 37687 | NR_040485 | 69776 | Mm.120470 | NR_040484 | ENSMUSG00000086899 | **1600002D24Rik** | - | RIKEN cDNA 1600002D24 gene | ncRNA |
| chr16_95860787_95861174 | chr16 | 95860788 | 95861174 | + | -0.645614 | intron (NR_040484, intron 1 of 3) | intron (NR_040484, intron 1 of 3) | 37732 | NR_040485 | 69776 | Mm.120470 | NR_040484 | ENSMUSG00000086899 | **1600002D24Rik** | - | RIKEN cDNA 1600002D24 gene | ncRNA |
| chr16_95899720_95900588 | chr16 | 95899721 | 95900588 | + | -0.700049 | intron (NR_040484, intron 1 of 3) | intron (NR_040484, intron 1 of 3) | -1441 | NR_040485 | 69776 | Mm.120470 | NR_040484 | ENSMUSG00000086899 | **1600002D24Rik** | - | RIKEN cDNA 1600002D24 gene | ncRNA |
| chr16_95900040_95900625 | chr16 | 95900041 | 95900625 | + | -0.755249 | intron (NR_040484, intron 1 of 3) | intron (NR_040484, intron 1 of 3) | -1620 | NR_040485 | 69776 | Mm.120470 | NR_040484 | ENSMUSG00000086899 | **1600002D24Rik** | - | RIKEN cDNA 1600002D24 gene | ncRNA |
| chr1_183987633_183988190 | chr1 | 183987634 | 183988190 | + | -0.864347 | Intergenic | Intergenic | 46086 | NM_028516 | 73363 | Mm.50115 | NM_028516 | ENSMUSG00000044854 | **1700056E22Rik** | - | RIKEN cDNA 1700056E22 gene | protein-coding |
| chr7_73583327_73583806 | chr7 | 73583328 | 73583806 | + | -0.769368 | Intergenic | Intergenic | -25172 | NR_037569 | 69170 | Mm.34539 | NR_037569 | | **1810026B05Rik** | - | RIKEN cDNA 1810026B05 gene | ncRNA |
| chr18_75304375_75305489 | chr18 | 75304376 | 75305489 | + | -0.589506 | Intergenic | Intergenic | -7603 | NR_045393 | 100503117 | Mm.250517 | NR_045393 | | **2010010A06Rik** | 5033420E22Rik | RIKEN cDNA 2010010A06 gene | ncRNA |
| chr18_75323821_75324179 | chr18 | 75323822 | 75324179 | + | -0.607654 | Intergenic | Intergenic | -26671 | NR_045393 | 100503117 | Mm.250517 | NR_045393 | | **2010010A06Rik** | 5033420E22Rik | RIKEN cDNA 2010010A06 gene | ncRNA |
| chr18_75304315_75305067 | chr18 | 75304316 | 75305067 | + | -0.671616 | Intergenic | Intergenic | -7362 | NR_045393 | 100503117 | Mm.250517 | NR_045393 | | **2010010A06Rik** | 5033420E22Rik | RIKEN cDNA 2010010A06 gene | ncRNA |
| chr6_31221602_31221903 | chr6 | 31221603 | 31221903 | + | -0.778298 | intron (NR_040260, intron 1 of 5) | intron (NR_040260, intron 1 of 5) | 1402 | NR_040261 | 73652 | Mm.441297 | NR_040257 | ENSMUSG00000087380 | **2210408F21Rik** | - | RIKEN cDNA 2210408F21 gene | ncRNA |
| chr6_125265106_125265641 | chr6 | 125265107 | 125265641 | + | -0.587566 | promoter-TSS (NR_015527) | promoter-TSS (NR_015527) | -151 | NR_015527 | 73870 | Mm.440681 | NR_015527 | | **4930417O13Rik** | - | RIKEN cDNA 4930417O13 gene | ncRNA |
| chr6_125265122_125265859 | chr6 | 125265123 | 125265859 | + | -0.591097 | promoter-TSS (NR_015527) | promoter-TSS (NR_015527) | -34 | NR_015527 | 73870 | Mm.440681 | NR_015527 | | **4930417O13Rik** | - | RIKEN cDNA 4930417O13 gene | ncRNA |
| chr19_5405985_5406340 | chr19 | 5405986 | 5406340 | + | -0.625645 | promoter-TSS (NR_015545) | promoter-TSS (NR_015545) | -652 | NR_015545 | 74931 | Mm.388511 | NR_015545 | ENSMUSG00000086938 | **4930481A15Rik** | - | RIKEN cDNA 4930481A15 gene | ncRNA |
| chr1_160042289_160042752 | chr1 | 160042290 | 160042752 | + | -0.656981 | Intergenic | Intergenic | -1859 | NR_028111 | 67647 | Mm.171518 | NM_001024470 | ENSMUSG00000090394 | **4930523C07Rik** | - | RIKEN cDNA 4930523C07 gene | protein-coding |
| chr9_124425516_124425974 | chr9 | 124425517 | 124425974 | + | -0.716907 | TTS (NR_015516) | TTS (NR_015516) | 1958 | NR_015516 | 75135 | Mm.476814 | NR_015516 | | **4930526I15Rik** | 1600027J15Rik | RIKEN cDNA 4930526I15 gene | ncRNA |
| chr10_117918508_117919211 | chr10 | 117918509 | 117919211 | + | -0.781604 | Intergenic | Intergenic | -6599 | NR_045342 | 71067 | Mm.410793 | NR_045342 | | **4933411E08Rik** | - | RIKEN cDNA 4933411E08 gene | ncRNA |
| chr10_117906187_117906940 | chr10 | 117906188 | 117906940 | + | -0.88986 | Intergenic | Intergenic | -18895 | NR_045342 | 71067 | Mm.410793 | NR_045342 | | **4933411E08Rik** | - | RIKEN cDNA 4933411E08 gene | ncRNA |
| chr1_36722477_36723686 | chr1 | 36722478 | 36723686 | + | -0.61017 | intron (NR_040290, intron 2 of 8) | B2_Mm2\|SINE\|B2 | 8544 | NR_040290 | 71166 | Mm.157049 | NR_040290 | | **4933424G06Rik** | 1700104I15Rik | RIKEN cDNA 4933424G06 gene | ncRNA |
| chr1_36723118_36723653 | chr1 | 36723119 | 36723653 | + | -0.617532 | non-coding (NR_040290, exon 3 of 9) | non-coding (NR_040290, exon 3 of 9) | 8848 | NR_040290 | 71166 | Mm.157049 | NR_040290 | | **4933424G06Rik** | 1700104I15Rik | RIKEN cDNA 4933424G06 gene | ncRNA |
| chr9_118390703_118391123 | chr9 | 118390704 | 118391123 | + | -0.609477 | Intergenic | Intergenic | -38926 | NR_040716 | 71196 | Mm.73218 | NR_040716 | | **4933432G23Rik** | - | RIKEN cDNA 4933432G23 gene | ncRNA |
| chr17_84006786_84007167 | chr17 | 84006787 | 84007167 | + | -0.586951 | Intergenic | L1MA10\|LINE\|L1 | -71683 | NR_045458 | 74473 | Mm.399575 | NR_045458 | | **4933433H22Rik** | - | RIKEN cDNA 4933433H22 gene | ncRNA |
| chr2_59338374_59338812 | chr2 | 59338375 | 59338812 | + | -0.703029 | intron (NM_026361, intron 13 of 21) | intron (NM_026361, intron 13 of 21) | 48089 | NR_040510 | 76676 | Mm.110038 | NR_040510 | ENSMUSG00000087455 | **5330411J11Rik** | - | RIKEN cDNA 5330411J11 gene | ncRNA |
| chr19_53441362_53441961 | chr19 | 53441363 | 53441961 | + | -0.642268 | non-coding (NR_028427, exon 10 of 10) | non-coding (NR_028427, exon 10 of 10) | 23134 | NR_028427 | 381232 | Mm.248624 | NR_028427 | ENSMUSG00000097636 | **5830416P10Rik** | 4833407H14Rik\|5430437C04Rik\|ENSMUSG00000074788\|Gm10758 | RIKEN cDNA 5830416P10 gene | ncRNA |
| chr19_53443084_53443703 | chr19 | 53443085 | 53443703 | + | -0.764273 | non-coding (NR_028427, exon 10 of 10) | non-coding (NR_028427, exon 10 of 10) | 21402 | NR_028427 | 381232 | Mm.248624 | NR_028427 | ENSMUSG00000097636 | **5830416P10Rik** | 4833407H14Rik\|5430437C04Rik\|ENSMUSG00000074788\|Gm10758 | RIKEN cDNA 5830416P10 gene | ncRNA |
| chr15_81302159_81302724 | chr15 | 81302160 | 81302724 | + | -0.685521 | Intergenic | Intergenic | -50580 | NR_077230 | 78102 | Mm.296832 | NR_077229 | | **8430426J06Rik** | - | RIKEN cDNA 8430426J06 gene | ncRNA |
| chr15_81302366_81302722 | chr15 | 81302367 | 81302722 | + | -0.740845 | Intergenic | ID_B1\|SINE\|B4 | -50682 | NR_077230 | 78102 | Mm.296832 | NR_077229 | | **8430426J06Rik** | - | RIKEN cDNA 8430426J06 gene | ncRNA |
| chr13_24921296_24921856 | chr13 | 24921297 | 24921856 | + | -0.586372 | intron (NM_172532, intron 4 of 9) | intron (NM_172532, intron 4 of 9) | -15825 | NR_102323 | 493800 | Mm.458692 | NR_102323 | | **9330162012Rik** | - | cDNA RIKEN 9330162012 gene | ncRNA |
| chr2_75458078_75458547 | chr2 | 75458079 | 75458547 | + | -0.671187 | Intergenic | Intergenic | -14134 | NR_040635 | 77269 | Mm.389278 | NR_040635 | | **9430019J16Rik** | - | RIKEN cDNA 9430019J16 gene | ncRNA |
| chr19_10740510_10741377 | chr19 | 10740511 | 10741377 | + | -0.591136 | promoter-TSS (NR_027805) | promoter-TSS (NR_027805) | -3 | NR_027805 | 403202 | Mm.34482 | NM_207529 | ENSMUSG00000067577 | **A430093F15Rik** | Gm674 | RIKEN cDNA A430093F15 gene | ncRNA |
| chr19_10740509_10741085 | chr19 | 10740510 | 10741085 | + | -0.619013 | promoter-TSS (NR_027805) | promoter-TSS (NR_027805) | -150 | NR_027805 | 403202 | Mm.34482 | NM_207529 | ENSMUSG00000067577 | **A430093F15Rik** | Gm674 | RIKEN cDNA A430093F15 gene | ncRNA |
| chr14_53878677_53878921 | chr14 | 53878678 | 53878921 | + | -0.715199 | Intergenic | RMER15-int\|LTR\|ERVL | 212859 | NM_001205242 | 100568452 | Mm.213248 | NM_001205242 | | **A430107P09Rik** | Va10 | RIKEN cDNA A430107P09 gene | protein-coding |
| chr2_167777342_167778653 | chr2 | 167777343 | 167778653 | + | -0.586074 | Intergenic | Intergenic | 86821 | NR_015500 | 329562 | Mm.123101 | NM_177851 | ENSMUSG00000006462 | **A530013C23Rik** | - | RIKEN cDNA A530013C23 gene | ncRNA |
| chr2_167777689_167778551 | chr2 | 167777690 | 167778551 | + | -0.618057 | Intergenic | Intergenic | 86943 | NR_015500 | 329562 | Mm.123101 | NM_177851 | ENSMUSG00000006462 | **A530013C23Rik** | - | RIKEN cDNA A530013C23 gene | ncRNA |
| chr2_167777349_167778277 | chr2 | 167777350 | 167778277 | + | -0.618259 | Intergenic | Intergenic | 86636 | NR_015500 | 329562 | Mm.123101 | NM_177851 | ENSMUSG00000006462 | **A530013C23Rik** | - | RIKEN cDNA A530013C23 gene | ncRNA |
| chr2_167777703_167778191 | chr2 | 167777704 | 167778191 | + | -0.642047 | Intergenic | Intergenic | 86770 | NR_015500 | 329562 | Mm.123101 | NM_177851 | ENSMUSG00000006462 | **A530013C23Rik** | - | RIKEN cDNA A530013C23 gene | ncRNA |
| chr5_122990470_122990823 | chr5 | 122990471 | 122990823 | + | -0.912242 | intron (NR_045820, intron 1 of 2) | intron (NR_045820, intron 1 of 2) | 1293 | NR_045820 | 109202 | Mm.30769 | NM_175405 | ENSMUSG00000056735 | **A930024E05Rik** | - | RIKEN cDNA A930024E05 gene | ncRNA |
| chr5_122989013_122989663 | chr5 | 122989014 | 122989663 | + | -0.917634 | promoter-TSS (NR_045820) | promoter-TSS (NR_045820) | -16 | NR_045820 | 109202 | Mm.30769 | NM_175405 | ENSMUSG00000056735 | **A930024E05Rik** | - | RIKEN cDNA A930024E05 gene | ncRNA |
| chr5_122990341_122990929 | chr5 | 122990342 | 122990929 | + | -0.954641 | intron (NR_045820, intron 1 of 2) | intron (NR_045820, intron 1 of 2) | 1281 | NR_045820 | 109202 | Mm.30769 | NM_175405 | ENSMUSG00000056735 | **A930024E05Rik** | - | RIKEN cDNA A930024E05 gene | ncRNA |
| chr5_8838752_8839476 | chr5 | 8838753 | 8839476 | + | -0.648135 | intron (NM_011075, intron 21 of 27) | intron (NM_011075, intron 21 of 27) | 40967 | NM_011075 | 18669 | Mm.146649 | NM_011075 | ENSMUSG00000028970 | **Abcb1b** | Abcb1\|Mdr1\|Mdr1b\|Pgy-1\|Pgy1\|mdr | ATP-binding cassette, sub-family B (MDR/TAP), member 1B | protein-coding |
| chr5_124074886_124075396 | chr5 | 124074887 | 124075396 | + | -0.914221 | intron (NM_019875, intron 9 of 11) | intron (NM_019875, intron 9 of 11) | 20657 | NM_019875 | 56325 | Mm.254839 | NM_019875 | ENSMUSG00000029408 | **Abcb9** | TAPL\|mKIAA1520 | ATP-binding cassette, sub-family B (MDR/TAP), member 9 | protein-coding |
| chr7_79300507_79301293 | chr7 | 79300508 | 79301293 | + | -0.677344 | intron (NM_018811, intron 3 of 10) | intron (NM_018811, intron 3 of 10) | 27634 | NM_018811 | 54608 | Mm.365490 | NM_018811 | | **Abhd2** | 2210009N18Rik\|LABH2\|Labh-2 | abhydrolase domain containing 2 | protein-coding |
| chr16_23170786_23171252 | chr16 | 23170787 | 23171252 | + | -0.586603 | Intergenic | RMER19C\|LTR\|ERVK | 24483 | NM_009605 | 11450 | Mm.3969 | NM_009605 | ENSMUSG00000022878 | **Adipoq** | 30kDa\|APN\|Acdc\|Acrp30\|GBP28\|adipo\|apM1 | adiponectin, C1Q and collagen domain containing | protein-coding |
| chr6_140819347_140819929 | chr6 | 140819348 | 140819929 | + | -0.604921 | Intergenic | ORR1D2\|LTR\|ERVL-MaLR | 195340 | NM_009637 | 11569 | Mm.86453 | NM_009637 | ENSMUSG00000030232 | **Aebp2** | AU023766\|B230313N05Rik | AE binding protein 2 | protein-coding |
| chr6_140816877_140817496 | chr6 | 140816878 | 140817496 | + | -0.611217 | Intergenic | Intergenic | 192889 | NM_009637 | 11569 | Mm.86453 | NM_009637 | ENSMUSG00000030232 | **Aebp2** | AU023766\|B230313N05Rik | AE binding protein 2 | protein-coding |
| chr11_53351474_53352040 | chr11 | 53351475 | 53352040 | + | -0.600972 | intron (NM_033565, intron 1 of 20) | intron (NM_033565, intron 1 of 20) | 990 | NM_033565 | 93736 | Mm.395281 | NM_033565 | ENSMUSG00000049470 | **Aff4** | AF5Q31\|Alf4\|Laf4l\|MCEF | AF4/FMR2 family, member 4 | protein-coding |
| chr11_53351715_53352000 | chr11 | 53351716 | 53352000 | + | -0.608271 | intron (NM_033565, intron 1 of 20) | intron (NM_033565, intron 1 of 20) | 1091 | NM_033565 | 93736 | Mm.395281 | NM_033565 | ENSMUSG00000049470 | **Aff4** | AF5Q31\|Alf4\|Laf4l\|MCEF | AF4/FMR2 family, member 4 | protein-coding |
| chr15_73137469_73138256 | chr15 | 73137470 | 73138256 | + | -0.625396 | intron (NM_153178, intron 3 of 18) | intron (NM_153178, intron 3 of 18) | 47084 | NM_153178 | 239528 | Mm.23095 | NM_153178 | | **Ago2** | 1110029L17Rik\|2310051F07Rik\|AI225898\|AL022874\|AW546247\|ENSMUSG00000072493\|Eif2c2\|Gerp95\|Gm10365\|mKIAA4215 | argonaute RISC catalytic subunit 2 | protein-coding |
| chr15_73137732_73138206 | chr15 | 73137733 | 73138206 | + | -0.795699 | intron (NM_153178, intron 3 of 18) | intron (NM_153178, intron 3 of 18) | 46978 | NM_153178 | 239528 | Mm.23095 | NM_153178 | | **Ago2** | 1110029L17Rik\|2310051F07Rik\|AI225898\|AL022874\|AW546247\|ENSMUSG00000072493\|Eif2c2\|Gerp95\|Gm10365\|mKIAA4215 | argonaute RISC catalytic subunit 2 | protein-coding |
| chr8_123000248_123000708 | chr8 | 123000249 | 123000708 | + | -0.587959 | intron (NR_037865, intron 1 of 13) | intron (NR_037865, intron 1 of 13) | 41806 | NR_037865 | 77087 | Mm.31567 | NM_001081379 | ENSMUSG00000035569 | **Ankrd11** | 2410104C19Rik\|3010027A04Rik\|6330578C09Rik\|9530048I21Rik\|AA930108\|Gm176\|Yod | ankyrin repeat domain 11 | protein-coding |
| chr2_43727557_43728279 | chr2 | 43727558 | 43728279 | + | -0.656733 | Intergenic | Intergenic | -20906 | NM_153820 | 76117 | Mm.217350 | NM_153820 | ENSMUSG00000049744 | **Arhgap15** | 5830480G12Rik | Rho GTPase activating protein 15 | protein-coding |
| chr6_87502863_87503288 | chr6 | 87502864 | 87503288 | + | -0.691669 | intron (NM_001037727, intron 1 of 10) | intron (NM_001037727, intron 1 of 10) | -6749 | NM_175476 | 232201 | Mm.119564 | NM_175476 | | **Arhgap25** | A130039I20Rik | Rho GTPase activating protein 25 | protein-coding |
| chr7_101537627_101538556 | chr7 | 101537628 | 101538556 | + | -0.744896 | Intergenic | Intergenic | 17710 | NM_007490 | 11871 | Mm.439746 | NM_007490 | ENSMUSG00000092517 | **Art2a-ps** | ARTC2\|Art2a\|Ly92a\|Rt-6\|Rt6\|Rt6-1 | ADP-ribosyltransferase 2a, pseudogene | protein-coding |

| | | | | | | | | | | | | | | | | | |
|---|---|---|---|---|---|---|---|---|---|---|---|---|---|---|---|---|---|
| chr7_101537952_101538573 | chr7 | 101537953 | 101538573 | + | -0.775676 | Intergenic | Intergenic | 17539 | NM_007490 | 11871 | Mm.439746 | NM_007490 | ENSMUSG00000092517 | **Art2a-ps** | ARTC2\|Art2a\|Ly92a\|Rt-6\|Rt6\|Rt6-1 | ADP-ribosyltransferase 2a, pseudogene | protein-coding |
| chr7_101538256_101538486 | chr7 | 101538257 | 101538486 | + | -0.800638 | Intergenic | Intergenic | 17431 | NM_007490 | 11871 | Mm.439746 | NM_007490 | ENSMUSG00000092517 | **Art2a-ps** | ARTC2\|Art2a\|Ly92a\|Rt-6\|Rt6\|Rt6-1 | ADP-ribosyltransferase 2a, pseudogene | protein-coding |
| chr10_81194063_81194682 | chr10 | 81194064 | 81194682 | + | -0.67509 | promoter-TSS (NR_015477) | promoter-TSS (NR_015477) | -318 | NR_015477 | 69678 | Mm.81562 | NR_015477 | | **Atcayos** | 2310050B05Rik | ataxia, cerebellar, Cayman type homolog (human), opposite strand | ncRNA |
| chr1_133762852_133763673 | chr1 | 133762853 | 133763673 | + | -0.612846 | Intergenic | Intergenic | -9516 | NM_213616 | 381290 | Mm.188617 | NM_213616 | ENSMUSG00000026463 | **Atp2b4** | Pmca4 | ATPase, Ca++ transporting, plasma membrane 4 | protein-coding |
| chr1_133762997_133763604 | chr1 | 133762998 | 133763604 | + | -0.613347 | Intergenic | Intergenic | -9554 | NM_213616 | 381290 | Mm.188617 | NM_213616 | ENSMUSG00000026463 | **Atp2b4** | Pmca4 | ATPase, Ca++ transporting, plasma membrane 4 | protein-coding |
| chr12_33302725_33303374 | chr12 | 33302726 | 33303374 | + | -0.75515 | intron (NM_001033436, intron 1 of 7) | intron (NM_001033436, intron 1 of 7) | 535 | NM_001033436 | 380753 | Mm.41540 | NM_028139 | ENSMUSG00000020564 | **Atxn7l1** | 2810423G08Rik\|Atxn7l4 | ataxin 7-like 1 | protein-coding |
| chr4_32343297_32344526 | chr4 | 32343298 | 32344526 | + | -0.589575 | Intergenic | Intergenic | -73523 | NM_001109661 | 12014 | Mm.491223 | NM_007521 | ENSMUSG00000040270 | **Bach2** | E030004N02Rik | BTB and CNC homology 2 | protein-coding |
| chr4_32351934_32352244 | chr4 | 32351935 | 32352244 | + | -0.600273 | Intergenic | MER119\|DNA\|hAT-Charlie | -65346 | NM_001109661 | 12014 | Mm.491223 | NM_007521 | ENSMUSG00000040270 | **Bach2** | E030004N02Rik | BTB and CNC homology 2 | protein-coding |
| chr4_32363583_32364191 | chr4 | 32363584 | 32364191 | + | -0.653421 | Intergenic | Intergenic | -53548 | NM_001109661 | 12014 | Mm.491223 | NM_007521 | ENSMUSG00000040270 | **Bach2** | E030004N02Rik | BTB and CNC homology 2 | protein-coding |
| chr4_32343650_32344700 | chr4 | 32343651 | 32344700 | + | -0.669454 | Intergenic | Intergenic | -73260 | NM_001109661 | 12014 | Mm.491223 | NM_007521 | ENSMUSG00000040270 | **Bach2** | E030004N02Rik | BTB and CNC homology 2 | protein-coding |
| chr4_32363349_32364219 | chr4 | 32363350 | 32364219 | + | -0.676601 | Intergenic | Intergenic | -53651 | NM_001109661 | 12014 | Mm.491223 | NM_007521 | ENSMUSG00000040270 | **Bach2** | E030004N02Rik | BTB and CNC homology 2 | protein-coding |
| chr19_4060521_4061410 | chr19 | 4060522 | 4061410 | + | -0.630734 | Intergenic | (TCTA)n\|Simple_repeat\|Simple_repeat | -1672 | NM_144869 | 225884 | Mm.440201 | NM_144869 | ENSMUSG00000058216 | **BC021614** | - | cDNA sequence BC021614 | protein-coding |
| chr3_145925618_145925830 | chr3 | 145925619 | 145925830 | + | -0.669864 | intron (NM_009740, intron 1 of 2) | intron (NM_009740, intron 1 of 2) | 1462 | NM_009740 | 12042 | Mm.239141 | NM_009740 | ENSMUSG00000028191 | **Bcl10** | AI132454\|BCL-10\|C81403\|CARMEN\|CIPER\|CLAP\|ME10\|cE10 | B cell leukemia/lymphoma 10 | protein-coding |
| chr5_67394006_67394358 | chr5 | 67394007 | 67394358 | + | -0.642428 | 3' UTR (NM_001164806, exon 5 of 5) | 3' UTR (NM_001164806, exon 5 of 5) | 33617 | NM_001164806 | 666938 | Mm.101339 | NM_001164806 | ENSMUSG00000092060 | **Bend4** | D330027G24Rik\|EG666938 | BEN domain containing 4 | protein-coding |
| chr5_67393860_67394502 | chr5 | 67393861 | 67394502 | + | -0.663434 | 3' UTR (NM_001164806, exon 5 of 5) | 3' UTR (NM_001164806, exon 5 of 5) | 33618 | NM_001164806 | 666938 | Mm.101339 | NM_001164806 | ENSMUSG00000092060 | **Bend4** | D330027G24Rik\|EG666938 | BEN domain containing 4 | protein-coding |
| chr5_67394730_67395861 | chr5 | 67394731 | 67395861 | + | -0.675601 | 3' UTR (NM_001164806, exon 5 of 5) | 3' UTR (NM_001164806, exon 5 of 5) | 32503 | NM_001164806 | 666938 | Mm.101339 | NM_001164806 | ENSMUSG00000092060 | **Bend4** | D330027G24Rik\|EG666938 | BEN domain containing 4 | protein-coding |
| chr5_67395197_67395896 | chr5 | 67395198 | 67395896 | + | -0.755933 | 3' UTR (NM_001164806, exon 5 of 5) | 3' UTR (NM_001164806, exon 5 of 5) | 32252 | NM_001164806 | 666938 | Mm.101339 | NM_001164806 | ENSMUSG00000092060 | **Bend4** | D330027G24Rik\|EG666938 | BEN domain containing 4 | protein-coding |
| chr10_96690648_96691340 | chr10 | 96690649 | 96691340 | + | -0.586489 | Intergenic | Intergenic | 73993 | NM_007569 | 12226 | Mm.23811 | NM_007569 | ENSMUSG00000036478 | **Btg1** | AI426953\|AW546738 | B cell translocation gene 1, anti-proliferative | protein-coding |
| chr1_134058883_134059281 | chr1 | 134058884 | 134059281 | + | -0.841073 | Intergenic | Intergenic | 20073 | NM_007570 | 12227 | Mm.392646 | NM_007570 | ENSMUSG00000020423 | **Btg2** | AA959598\|APRO1\|Pc3\|TIS21 | B cell translocation gene 2, anti-proliferative | protein-coding |
| chr5_64518862_64519395 | chr5 | 64518863 | 64519395 | + | -0.608723 | Intergenic | Intergenic | 42121 | NR_045413 | 77382 | Mm.366742 | NR_045411 | | **C030018K13Rik** | - | RIKEN cDNA C030018K13 gene | ncRNA |
| chr5_64498340_64498840 | chr5 | 64498341 | 64498840 | + | -0.653264 | Intergenic | Intergenic | 21582 | NR_045413 | 77382 | Mm.366742 | NR_045411 | | **C030018K13Rik** | - | RIKEN cDNA C030018K13 gene | ncRNA |
| chr15_100604623_100605434 | chr15 | 100604624 | 100605434 | + | -0.877565 | Intergenic | Intergenic | 10081 | NR_045701 | 78445 | Mm.44383 | NR_045701 | ENSMUSG00000097093 | **C330013E15Rik** | - | RIKEN cDNA C330013E15 gene | ncRNA |
| chr9_96772857_96773556 | chr9 | 96772858 | 96773556 | + | -0.662578 | intron (NR_102293, intron 4 of 4) | intron (NR_102293, intron 4 of 4) | 7645 | NR_102296 | 319707 | Mm.392191 | NR_102293 | | **C430002N11Rik** | - | RIKEN cDNA C430002N11 gene | ncRNA |
| chr9_96773111_96773563 | chr9 | 96773112 | 96773563 | + | -0.676304 | non-coding (NR_102296, exon 5 of 5) | non-coding (NR_102296, exon 5 of 5) | 7775 | NR_102296 | 319707 | Mm.392191 | NR_102293 | | **C430002N11Rik** | - | RIKEN cDNA C430002N11 gene | ncRNA |
| chr10_22299428_22299683 | chr10 | 22299429 | 22299683 | + | -0.668397 | intron (NM_009018, intron 5 of 5) | intron (NM_009018, intron 5 of 5) | -7164 | NR_015465 | 606736 | Mm.387042 | NR_015465 | ENSMUSG00000097352 | **C920009B18Rik** | - | RIKEN cDNA C920009B18 gene | ncRNA |
| chr10_22299319_22299716 | chr10 | 22299320 | 22299716 | + | -0.7071 | intron (NM_009018, intron 5 of 5) | intron (NM_009018, intron 5 of 5) | -7202 | NR_015465 | 606736 | Mm.387042 | NR_015465 | ENSMUSG00000097352 | **C920009B18Rik** | - | RIKEN cDNA C920009B18 gene | ncRNA |
| chr18_32970174_32970666 | chr18 | 32970175 | 32970666 | + | -0.822927 | intron (NM_009793, intron 1 of 10) | intron (NM_009793, intron 1 of 10) | 31379 | NM_009793 | 12326 | Mm.222329 | NM_009793 | ENSMUSG00000038128 | **Camk4** | A430110E23Rik\|AI666733\|CaMKIV\|CaMKIV/Gr\|D18Bwg0362e | calcium/calmodulin-dependent protein kinase IV | protein-coding |
| chr18_32970174_32970909 | chr18 | 32970175 | 32970909 | + | -0.849383 | intron (NM_009793, intron 1 of 10) | intron (NM_009793, intron 1 of 10) | 31501 | NM_009793 | 12326 | Mm.222329 | NM_009793 | ENSMUSG00000038128 | **Camk4** | A430110E23Rik\|AI666733\|CaMKIV\|CaMKIV/Gr\|D18Bwg0362e | calcium/calmodulin-dependent protein kinase IV | protein-coding |
| chr18_32958142_32958609 | chr18 | 32958143 | 32958609 | + | -0.980416 | intron (NM_009793, intron 1 of 10) | L2\|LINE\|L2 | 19335 | NM_009793 | 12326 | Mm.222329 | NM_009793 | ENSMUSG00000038128 | **Camk4** | A430110E23Rik\|AI666733\|CaMKIV\|CaMKIV/Gr\|D18Bwg0362e | calcium/calmodulin-dependent protein kinase IV | protein-coding |
| chr16_52147278_52147856 | chr16 | 52147279 | 52147856 | + | -0.588092 | intron (NM_001033238, intron 9 of 17) | intron (NM_001033238, intron 9 of 17) | 116018 | NM_001033238 | 208650 | Mm.328206 | NM_001033238 | ENSMUSG00000022637 | **Cblb** | AI429560\|AI851073\|Cbl-b | Casitas B-lineage lymphoma b | protein-coding |
| chr16_52147074_52147844 | chr16 | 52147075 | 52147844 | + | -0.591675 | intron (NM_001033238, intron 9 of 17) | intron (NM_001033238, intron 9 of 17) | 115910 | NM_001033238 | 208650 | Mm.328206 | NM_001033238 | ENSMUSG00000022637 | **Cblb** | AI429560\|AI851073\|Cbl-b | Casitas B-lineage lymphoma b | protein-coding |
| chr16_52060897_52061678 | chr16 | 52060898 | 52061678 | + | -0.708816 | intron (NM_001033238, intron 3 of 17) | intron (NM_001033238, intron 3 of 17) | 29739 | NM_001033238 | 208650 | Mm.328206 | NM_001033238 | ENSMUSG00000022637 | **Cblb** | AI429560\|AI851073\|Cbl-b | Casitas B-lineage lymphoma b | protein-coding |
| chr16_52142791_52143602 | chr16 | 52142792 | 52143602 | + | -0.863634 | intron (NM_001033238, intron 9 of 17) | intron (NM_001033238, intron 9 of 17) | 111648 | NM_001033238 | 208650 | Mm.328206 | NM_001033238 | ENSMUSG00000022637 | **Cblb** | AI429560\|AI851073\|Cbl-b | Casitas B-lineage lymphoma b | protein-coding |
| chr16_52143084_52143560 | chr16 | 52143085 | 52143560 | + | -0.86694 | intron (NM_001033238, intron 9 of 17) | intron (NM_001033238, intron 9 of 17) | 111773 | NM_001033238 | 208650 | Mm.328206 | NM_001033238 | ENSMUSG00000022637 | **Cblb** | AI429560\|AI851073\|Cbl-b | Casitas B-lineage lymphoma b | protein-coding |
| chr9_114451546_114452014 | chr9 | 114451547 | 114452014 | + | -0.674066 | intron (NR_108101, intron 10 of 14) | intron (NR_108101, intron 10 of 14) | 44764 | NM_009916 | 12773 | Mm.1337 | NM_009916 | ENSMUSG00000047898 | **Ccr4** | C-C CKR-4\|CHEMR1\|Cmkbr4\|LESTR\|Sdf1r | chemokine (C-C motif) receptor 4 | protein-coding |
| chr11_99168068_99168618 | chr11 | 99168069 | 99168618 | + | -1.27289 | Intergenic | Intergenic | -13266 | NM_007719 | 12775 | Mm.2932 | NM_007719 | ENSMUSG00000037944 | **Ccr7** | CD197\|Cdw197\|Cmkbr7\|EBI1\|Ebi1h | chemokine (C-C motif) receptor 7 | protein-coding |
| chr3_101010471_101011126 | chr3 | 101010472 | 101011126 | + | -0.600818 | intron (NM_001099332, intron 6 of 8) | intron (NM_001099332, intron 6 of 8) | 11377 | NM_001099332 | 630146 | Mm.190915 | NM_001099332 | ENSMUSG00000086564 | **Cd101** | Gm1016\|Gm734\|Igsf2 | CD101 antigen | protein-coding |
| chr1_165837439_165838249 | chr1 | 165837440 | 165838249 | + | -0.592383 | intron (NM_031162, intron 1 of 7) | intron (NM_031162, intron 1 of 7) | 49163 | NM_001113392 | 12503 | Mm.217308 | NM_031162 | ENSMUSG00000005763 | **Cd247** | 4930549J05Rik\|A430104F18Rik\|AW552088\|Cd3\|Cd3-eta\|Cd3-zeta\|Cd3h\|Cd3z\|Cd3zeta\|T3z\|Tcrk\|Tcrz | CD247 antigen | protein-coding |
| chr1_60776919_60777374 | chr1 | 60776920 | 60777374 | + | -0.678554 | Intergenic | Intergenic | 30759 | NM_007642 | 12487 | Mm.255003 | NM_007642 | ENSMUSG00000026012 | **Cd28** | - | CD28 antigen | protein-coding |
| chr1_60776944_60777573 | chr1 | 60776945 | 60777573 | + | -0.687261 | Intergenic | (GGGA)n\|Simple_repeat\|Simple_repeat | 30871 | NM_007642 | 12487 | Mm.255003 | NM_007642 | ENSMUSG00000026012 | **Cd28** | - | CD28 antigen | protein-coding |
| chr1_60755819_60756231 | chr1 | 60755820 | 60756231 | + | -0.766756 | intron (NM_007642, intron 1 of 3) | intron (NM_007642, intron 1 of 3) | 9637 | NM_007642 | 12487 | Mm.255003 | NM_007642 | ENSMUSG00000026012 | **Cd28** | - | CD28 antigen | protein-coding |
| chr16_49792406_49793000 | chr16 | 49792407 | 49793000 | + | -0.796609 | Intergenic | Intergenic | -62951 | NM_010581 | 16423 | Mm.31752 | NM_010581 | ENSMUSG00000055447 | **Cd47** | 9130415E20Rik\|AA407862\|AI848868\|AW108519\|B430305P08Rik\|IAP\|Itgp | CD47 antigen (Rh-related antigen, integrin-associated signal transducer) | protein-coding |
| chr8_83747715_83748376 | chr8 | 83747716 | 83748376 | + | -0.723554 | Intergenic | Intergenic | -6735 | NM_001163031 | 26364 | Mm.334648 | NM_011925 | ENSMUSG00000002885 | **Cd97** | AA409984\|TM7LN1 | CD97 antigen | protein-coding |
| chr2_6705477_6706564 | chr2 | 6705478 | 6706564 | + | -0.612025 | intron (NM_001110232, intron 4 of 15) | intron (NM_001110232, intron 4 of 15) | 178975 | NM_010160 | 14007 | Mm.398543 | NM_010160 | ENSMUSG00000002107 | **Celf2** | B230218O03\|B230345P09Rik\|C88023\|CELF-2\|CUG-BP2\|Cugbp2\|D230046B21Rik\|Etr-3\|Napor\|Napor-2\|mETR-3 | CUGBP, Elav-like family member 2 | protein-coding |
| chr2_6705962_6706564 | chr2 | 6705963 | 6706564 | + | -0.67703 | intron (NM_001110232, intron 4 of 15) | intron (NM_001110232, intron 4 of 15) | 178733 | NM_010160 | 14007 | Mm.398543 | NM_010160 | ENSMUSG00000002107 | **Celf2** | B230218O03\|B230345P09Rik\|C88023\|CELF-2\|CUG-BP2\|Cugbp2\|D230046B21Rik\|Etr-3\|Napor\|Napor-2\|mETR-3 | CUGBP, Elav-like family member 2 | protein-coding |
| chr1_161071325_161071880 | chr1 | 161071326 | 161071880 | + | -0.6637 | promoter-TSS (NM_172644) | promoter-TSS (NM_172644) | 711 | NM_001159930 | 70454 | Mm.248212 | NM_027429 | ENSMUSG00000026708 | **Cenpl** | 2610300B10Rik\|AW121806\|AW550697 | centromere protein L | protein-coding |

| | | | | | | | | | | | | | | | | | |
|---|---|---|---|---|---|---|---|---|---|---|---|---|---|---|---|---|---|
| chr1_161071310_161071716 | chr1 | 161071311 | 161071716 | + | -0.680545 | promoter-TSS (NM_172644) | promoter-TSS (NM_172644) | 621 | NM_001159930 | 70454 | Mm.248212 | NM_027429 | ENSMUSG00000026708 | **Cenpl** | 2610300B10Rik\|AW121806\|AW550697 | centromere protein L | protein-coding |
| chr2_68959564_68959979 | chr2 | 68959565 | 68959979 | + | -0.63436 | intron (NM_172856, intron 3 of 9) | intron (NM_172856, intron 3 of 9) | 98215 | NM_172856 | 241447 | Mm.222222 | NM_172856 | ENSMUSG00000027035 | **Cers6** | 4732462C07Rik\|AW544719\|Lass6\|T1L | ceramide synthase 6 | protein-coding |
| chr7_73523287_73523501 | chr7 | 73523288 | 73523501 | + | -0.853891 | intron (NM_001081345, intron 2 of 38) | intron (NM_001081345, intron 2 of 38) | 18352 | NM_001081345 | 244059 | Mm.34539 | NM_001081345 | ENSMUSG00000078671 | **Chd2** | 2810013C04Rik\|2810040A01Rik\|5630401D06Rik\|AI851092\|BC029703\|CHD-2 | chromodomain helicase DNA binding protein 2 | protein-coding |
| chr7_73523174_73523601 | chr7 | 73523175 | 73523601 | + | -0.92602 | intron (NM_001081345, intron 2 of 38) | intron (NM_001081345, intron 2 of 38) | 18358 | NM_001081345 | 244059 | Mm.34539 | NM_001081345 | ENSMUSG00000078671 | **Chd2** | 2810013C04Rik\|2810040A01Rik\|5630401D06Rik\|AI851092\|BC029703\|CHD-2 | chromodomain helicase DNA binding protein 2 | protein-coding |
| chr4_8812369_8812802 | chr4 | 8812370 | 8812802 | + | -0.636749 | intron (NM_001277149, intron 7 of 37) | intron (NM_001277149, intron 7 of 37) | 122180 | NM_001277149 | 320790 | Mm.138792 | NM_001033395 | ENSMUSG00000041235 | **Chd7** | A730019I05Rik\|Cycn\|Cyn\|Dz\|Edy\|Flo\|GENA 47\|GENA 60\|Gena 52\|Lda\|Mt\|Obt\|Todo\|WBE1\|Whi\|metis | chromodomain helicase DNA binding protein 7 | protein-coding |
| chr5_65955295_65955786 | chr5 | 65955296 | 65955786 | + | -0.654553 | Intergenic | Intergenic | -11583 | NM_001081104 | 231252 | Mm.486671 | NM_001081104 | ENSMUSG00000029205 | **Chrna9** | 2410015I05Rik\|Acra9\|EG666827\|Gm8311 | cholinergic receptor, nicotinic, alpha polypeptide 9 | protein-coding |
| chr9_95314858_95315516 | chr9 | 95314859 | 95315516 | + | -0.669768 | Intergenic | Intergenic | 92083 | NM_018763 | 54371 | Mm.212446 | NM_018763 | ENSMUSG00000033350 | **Chst2** | AI428561\|AW121776\|C130041E03Rik\|Chts2\|GST-2\|GlcNAc6ST\|Gn6st | carbohydrate sulfotransferase 2 | protein-coding |
| chr9_95113255_95113827 | chr9 | 95113256 | 95113827 | + | -0.759051 | intron (NM_177909, intron 12 of 15) | intron (NM_177909, intron 12 of 15) | 293729 | NM_018763 | 54371 | Mm.212446 | NM_018763 | ENSMUSG00000033350 | **Chst2** | AI428561\|AW121776\|C130041E03Rik\|Chts2\|GST-2\|GlcNAc6ST\|Gn6st | carbohydrate sulfotransferase 2 | protein-coding |
| chr4_135203188_135203802 | chr4 | 135203189 | 135203802 | + | -0.771963 | Intergenic | Intergenic | 69265 | NM_013885 | 29876 | Mm.257765 | NM_013885 | ENSMUSG00000037242 | **Clic4** | D0Jmb3\|TU-74\|mc3s5\|mtCLIC | chloride intracellular channel 4 (mitochondrial) | protein-coding |
| chr13_24486959_24488025 | chr13 | 24486960 | 24488025 | + | -0.593281 | Intergenic | Intergenic | 72370 | NM_001284520 | 12763 | Mm.8396 | NM_007717 | ENSMUSG00000016756 | **Cmah** | - | cytidine monophospho-N-acetylneuraminic acid hydroxylase | protein-coding |
| chr13_24415259_24416182 | chr13 | 24415260 | 24416182 | + | -0.632391 | intron (NM_001284520, intron 1 of 14) | intron (NM_001284520, intron 1 of 14) | 599 | NM_001284520 | 12763 | Mm.8396 | NM_007717 | ENSMUSG00000016756 | **Cmah** | - | cytidine monophospho-N-acetylneuraminic acid hydroxylase | protein-coding |
| chr6_142757611_142757914 | chr6 | 142757612 | 142757914 | + | -0.631294 | intron (NM_009908, intron 1 of 7) | intron (NM_009908, intron 1 of 7) | 1077 | NM_009908 | 12764 | Mm.3820 | NM_009908 | ENSMUSG00000030282 | **Cmas** | AW208911\|CMPNeu5Ac\|D6Bwg0250e | cytidine monophospho-N-acetylneuraminic acid synthetase | protein-coding |
| chr1_179549409_179549748 | chr1 | 179549410 | 179549748 | + | -0.616864 | intron (NM_146105, intron 1 of 10) | intron (NM_146105, intron 1 of 10) | 3050 | NM_146105 | 226744 | Mm.87662 | NM_146105 | ENSMUSG00000038949 | **Cnst** | 9630058J23Rik | consortin, connexin sorting protein | protein-coding |
| chr6_106792891_106793895 | chr6 | 106792892 | 106793895 | + | -0.883561 | intron (NM_175357, intron 4 of 10) | intron (NM_175357, intron 4 of 10) | 6694 | NM_021449 | 58799 | Mm.290085 | NM_021449 | | **Crbn** | 2610203G15Rik\|2900045O07Rik\|AF229032\|AW108261\|piL | cereblon | protein-coding |
| chr6_106792919_106793563 | chr6 | 106792920 | 106793563 | + | -1.04726 | intron (NM_175357, intron 4 of 10) | intron (NM_175357, intron 4 of 10) | 6846 | NM_021449 | 58799 | Mm.290085 | NM_021449 | | **Crbn** | 2610203G15Rik\|2900045O07Rik\|AF229032\|AW108261\|piL | cereblon | protein-coding |
| chr18_3281717_3281910 | chr18 | 3281718 | 3281910 | + | -0.713143 | promoter-TSS (NM_001110855) | promoter-TSS (NM_001110855) | -67 | NM_001110855 | 12916 | Mm.5244 | NM_013498 | ENSMUSG00000063889 | **Crem** | ICER\|ICERI | cAMP responsive element modulator | protein-coding |
| chr18_3281669_3282064 | chr18 | 3281670 | 3282064 | + | -0.724842 | promoter-TSS (NM_001110855) | promoter-TSS (NM_001110855) | -120 | NM_001110855 | 12916 | Mm.5244 | NM_013498 | ENSMUSG00000063889 | **Crem** | ICER\|ICERI | cAMP responsive element modulator | protein-coding |
| chr17_31693304_31694399 | chr17 | 31693305 | 31694399 | + | -0.936866 | Intergenic | Intergenic | 15921 | NM_001278570 | 12954 | Mm.1228 | NM_013501 | ENSMUSG00000024041 | **Cryaa** | Acry-1\|Crya-1\|Crya1\|DAcry-1\|lop18 | crystallin, alpha A | protein-coding |
| chr17_31692974_31693974 | chr17 | 31692975 | 31693974 | + | -0.991822 | Intergenic | Intergenic | 15543 | NM_001278570 | 12954 | Mm.1228 | NM_013501 | ENSMUSG00000024041 | **Cryaa** | Acry-1\|Crya-1\|Crya1\|DAcry-1\|lop18 | crystallin, alpha A | protein-coding |
| chr17_31693314_31693912 | chr17 | 31693315 | 31693912 | + | -1.02062 | Intergenic | Intergenic | 15682 | NM_001278570 | 12954 | Mm.1228 | NM_013501 | ENSMUSG00000024041 | **Cryaa** | Acry-1\|Crya-1\|Crya1\|DAcry-1\|lop18 | crystallin, alpha A | protein-coding |
| chr2_30625214_30626629 | chr2 | 30625215 | 30626629 | + | -0.675188 | Intergenic | MTE2a\|LTR\|ERVL-MaLR | 30878 | NM_030137 | 78617 | Mm.40413 | NM_030137 | ENSMUSG00000047363 | **Cstad** | 9530039B02Rik | CSA-conditional, T cell activation-dependent protein | protein-coding |
| chr2_30625873_30626611 | chr2 | 30625874 | 30626611 | + | -0.719401 | Intergenic | ORR1D2-int\|LTR\|ERVL-MaLR | 31198 | NM_030137 | 78617 | Mm.40413 | NM_030137 | ENSMUSG00000047363 | **Cstad** | 9530039B02Rik | CSA-conditional, T cell activation-dependent protein | protein-coding |
| chr11_105901427_105901989 | chr11 | 105901428 | 105901989 | + | -0.611432 | intron (NM_181071, intron 17 of 25) | intron (NM_181071, intron 17 of 25) | 42439 | NM_007805 | 13056 | Mm.149403 | NM_007805 | ENSMUSG00000019590 | **Cyb561** | - | cytochrome b-561 | protein-coding |
| chr11_46308254_46308622 | chr11 | 46308255 | 46308622 | + | -0.591325 | intron (NM_001252459, intron 1 of 30) | intron (NM_001252459, intron 1 of 30) | 3920 | NM_001252460 | 76884 | Mm.154358 | NM_133769 | ENSMUSG00000020340 | **Cyfip2** | 1500004I01Rik\|6430511D02Rik\|AA930218\|AU022376\|Pir121\|mKIAA1168 | cytoplasmic FMR1 interacting protein 2 | protein-coding |
| chr7_114494055_114494463 | chr7 | 114494056 | 114494463 | + | -0.628812 | intron (NM_011055, intron 2 of 15) | intron (NM_011055, intron 2 of 15) | 68713 | NR_110407 | 244209 | Mm.108037 | NM_177382 | ENSMUSG00000030670 | **Cyp2r1** | - | cytochrome P450, family 2, subfamily r, polypeptide 1 | protein-coding |
| chr15_78594782_78595208 | chr15 | 78594783 | 78595208 | + | -0.78476 | Intergenic | MTE2b\|LTR\|ERVL-MaLR | -2052 | NM_028195 | 72318 | Mm.32911 | NM_028195 | ENSMUSG00000018008 | **Cyth4** | 2510004M07Rik\|5830469K17Rik\|AI467541\|Pscd4\|mFLJ00017 | cytohesin 4 | protein-coding |
| chr2_58161031_58161256 | chr2 | 58161032 | 58161256 | + | -0.692544 | Intergenic | Intergenic | -1022 | NM_139200 | 227929 | Mm.273905 | NM_139200 | ENSMUSG00000026832 | **Cytip** | A130053M09Rik\|AI462064\|C80816\|Cbp\|Cybr\|Pscdbp | cytohesin 1 interacting protein | protein-coding |
| chr14_54231123_54231478 | chr14 | 54231124 | 54231478 | + | -0.7377 | Intergenic | RSINE1\|SINE\|B4 | 22628 | NM_001113358 | 13135 | Mm.319038 | NM_010015 | ENSMUSG00000022174 | **Dad1** | AI323713 | defender against cell death 1 | protein-coding |
| chr11_106026770_106028187 | chr11 | 106026771 | 106028187 | + | -0.683957 | intron (NM_001037712, intron 11 of 12) | intron (NM_001037712, intron 11 of 12) | -9393 | NM_027946 | 71833 | Mm.307455 | NM_027946 | ENSMUSG00000049354 | **Dcaf7** | 1700012F10Rik\|2610037L01Rik\|C86529\|HAN11\|Wdr68 | DDB1 and CUL4 associated factor 7 | protein-coding |
| chr11_106033501_106033721 | chr11 | 106033502 | 106033721 | + | -0.775444 | intron (NM_001037712, intron 12 of 12) | intron (NM_001037712, intron 12 of 12) | -3261 | NM_027946 | 71833 | Mm.307455 | NM_027946 | ENSMUSG00000049354 | **Dcaf7** | 1700012F10Rik\|2610037L01Rik\|C86529\|HAN11\|Wdr68 | DDB1 and CUL4 associated factor 7 | protein-coding |
| chr11_106033340_106033755 | chr11 | 106033341 | 106033755 | + | -0.842685 | intron (NM_001037712, intron 12 of 12) | IAPLTR1_Mm\|LTR\|ERVK | -3324 | NM_027946 | 71833 | Mm.307455 | NM_027946 | ENSMUSG00000049354 | **Dcaf7** | 1700012F10Rik\|2610037L01Rik\|C86529\|HAN11\|Wdr68 | DDB1 and CUL4 associated factor 7 | protein-coding |
| chr9_44579320_44579786 | chr9 | 44579321 | 44579786 | + | -0.810822 | Intergenic | Intergenic | -25339 | NM_001110826 | 13209 | Mm.267061 | NM_007841 | ENSMUSG00000032097 | **Ddx6** | 1110001P04Rik\|E230023J21Rik\|HLR2\|mRCK/P54\|p54\|rck | DEAD (Asp-Glu-Ala-Asp) box polypeptide 6 | protein-coding |
| chr11_86822354_86823068 | chr11 | 86822355 | 86823068 | + | -0.770838 | intron (NR_106195, intron 2 of 8) | RMER19B\|LTR\|ERVK | -15051 | NM_026191 | 67487 | Mm.260627 | NM_026191 | ENSMUSG00000018425 | **Dhx40** | 2410016C14Rik\|ARG147\|DDX40\|PAD | DEAH (Asp-Glu-Ala-His) box polypeptide 40 | protein-coding |
| chr11_86810271_86811754 | chr11 | 86810272 | 86811754 | + | -0.879455 | intron (NR_106195, intron 2 of 8) | intron (NR_106195, intron 2 of 8) | -3353 | NM_026191 | 67487 | Mm.260627 | NM_026191 | ENSMUSG00000018425 | **Dhx40** | 2410016C14Rik\|ARG147\|DDX40\|PAD | DEAH (Asp-Glu-Ala-His) box polypeptide 40 | protein-coding |
| chr15_100041368_100041561 | chr15 | 100041369 | 100041561 | + | -0.585377 | intron (NM_001159361, intron 1 of 37) | intron (NM_001159361, intron 1 of 37) | 2801 | NM_001159361 | 239667 | Mm.243658 | NM_172819 | ENSMUSG00000023026 | **Dip2b** | 4932422C22\|AI317237\|AI854602\|mKIAA1463 | DIP2 disco-interacting protein 2 homolog B (Drosophila) | protein-coding |
| chr11_118145697_118146415 | chr11 | 118145698 | 118146415 | + | -0.628699 | Intergenic | Intergenic | -16837 | NM_001167746 | 69926 | Mm.473893 | NM_001167746 | ENSMUSG00000033987 | **Dnah17** | 2810003K23Rik\|Dnahc17\|Dnahcl1\|Gm1178\|mKIAA3028 | dynein, axonemal, heavy chain 17 | protein-coding |
| chr4_101495790_101496157 | chr4 | 101495791 | 101496157 | + | -0.775299 | promoter-TSS (NM_198412) | promoter-TSS (NM_198412) | -674 | NM_198412 | 72685 | Mm.76494 | NM_198412 | ENSMUSG00000028528 | **Dnajc6** | 2810027M23Rik\|mKIAA0473 | DnaJ (Hsp40) homolog, subfamily C, member 6 | protein-coding |
| chr1_80755697_80756095 | chr1 | 80755698 | 80756095 | + | -1.14115 | intron (NM_175291, intron 1 of 55) | intron (NM_175291, intron 1 of 55) | 2657 | NM_175291 | 210293 | Mm.133473 | NM_175291 | ENSMUSG00000038608 | **Dock10** | 9330153B10Rik\|A630054M16Rik\|Jr4\|Jr5\|R75174\|ZIZ3\|Zizimin3\|mKIAA0694 | dedicator of cytokinesis 10 | protein-coding |
| chr1_80755731_80756009 | chr1 | 80755732 | 80756009 | + | -1.17058 | intron (NM_175291, intron 1 of 55) | intron (NM_175291, intron 1 of 55) | 2683 | NM_175291 | 210293 | Mm.133473 | NM_175291 | ENSMUSG00000038608 | **Dock10** | 9330153B10Rik\|A630054M16Rik\|Jr4\|Jr5\|R75174\|ZIZ3\|Zizimin3\|mKIAA0694 | dedicator of cytokinesis 10 | protein-coding |
| chr10_80751491_80751840 | chr10 | 80751492 | 80751840 | + | -0.591195 | Intergenic | MLT2D\|LTR\|ERVL | -3540 | NM_199322 | 208266 | Mm.152371 | NM_199322 | ENSMUSG00000061589 | **Dot1l** | A630076O07\|AW907654\|Dot1\|KMT4\|mDot1 | DOT1-like, histone H3 methyltransferase (S. cerevisiae) | protein-coding |
| chr17_26502785_26503609 | chr17 | 26502786 | 26503609 | + | -0.585539 | Intergenic | Intergenic | 5275 | NM_013642 | 19252 | Mm.239041 | NM_013642 | ENSMUSG00000024190 | **Dusp1** | 3CH134\|MKP1\|Ptpn16\|U19515\|erp\|mkp-1 | dual specificity phosphatase 1 | protein-coding |
| chr1_184066590_184067145 | chr1 | 184066591 | 184067145 | + | -0.602289 | intron (NM_022019, intron 2 of 3) | intron (NM_022019, intron 2 of 3) | 32407 | NM_022019 | 63953 | Mm.404024 | NM_022019 | ENSMUSG00000039384 | **Dusp10** | 2610306G15Rik\|AI158871\|MKP-5\|MKP5 | dual specificity phosphatase 10 | protein-coding |
| chr1_184065970_184066448 | chr1 | 184065971 | 184066448 | + | -0.800524 | intron (NM_022019, intron 2 of 3) | intron (NM_022019, intron 2 of 3) | 31748 | NM_022019 | 63953 | Mm.404024 | NM_022019 | ENSMUSG00000039384 | **Dusp10** | 2610306G15Rik\|AI158871\|MKP-5\|MKP5 | dual specificity phosphatase 10 | protein-coding |
| chr2_127312825_127313145 | chr2 | 127312826 | 127313145 | + | -0.699871 | Intergenic | L1MB7\|LINE\|L1 | -23174 | NM_010090 | 13537 | Mm.4729 | NM_010090 | ENSMUSG00000027368 | **Dusp2** | PAC1 | dual specificity phosphatase 2 | protein-coding |
| chr10_118714040_118714879 | chr10 | 118714041 | 118714879 | + | -0.595205 | Intergenic | Intergenic | 154443 | NM_001014390 | 69181 | Mm.45565 | NM_001014390 | ENSMUSG00000028630 | **Dyrk2** | 1810038L18Rik | dual-specificity tyrosine-(Y)-phosphorylation regulated kinase 2 | protein-coding |
| chr10_18197009_18197833 | chr10 | 18197010 | 18197833 | + | -0.591057 | intron (NM_001195036, intron 3 of 20) | intron (NM_001195036, intron 3 of 20) | 13469 | NM_001195036 | 100045792 | Mm.380617 | NM_001195036 | ENSMUSG00000071392 | **Ect2l** | C330021H03Rik\|ENSMUSG00000071392\|Gm10331 | epithelial cell transforming sequence 2 oncogene-like | protein-coding |

| | | | | | | | | | | | | | | | | | |
|---|---|---|---|---|---|---|---|---|---|---|---|---|---|---|---|---|---|
| chr10_18197009_18197618 | chr10 | 18197010 | 18197618 | + | -0.611839 | intron (NM_001195036, intron 3 of 20) | intron (NM_001195036, intron 3 of 20) | 13576 | NM_001195036 | 100045792 | Mm.380617 | NM_001195036 | ENSMUSG00000071392 | **Ect2l** | C330021H03Rik\|ENSMUSG00000071392\|Gm10331 | epithelial cell transforming sequence 2 oncogene-like | protein-coding |
| chr6_99643430_99644106 | chr6 | 99643431 | 99644106 | + | -0.740275 | intron (NM_025829, intron 2 of 6) | intron (NM_025829, intron 2 of 6) | 23003 | NM_025829 | 66892 | Mm.32889 | NM_025829 | ENSMUSG00000093661 | **Eif4e3** | 1300018P11Rik\|AI451927\|eIF4E-3 | eukaryotic translation initiation factor 4E member 3 | protein-coding |
| chr14_79530052_79530466 | chr14 | 79530053 | 79530466 | + | -0.587802 | intron (NM_001286411, intron 1 of 8) | ID4_\|SINE\|ID | 14585 | NM_001286411 | 13709 | Mm.20938 | NM_007920 | ENSMUSG00000036461 | **Elf1** | Elf-1\|Sts1\|mElf-1\|p70 | E74-like factor 1 | protein-coding |
| chr14_79530048_79530760 | chr14 | 79530049 | 79530760 | + | -0.621828 | intron (NM_001286411, intron 1 of 8) | MIR3\|SINE\|MIR | 14730 | NM_001286411 | 13709 | Mm.20938 | NM_007920 | ENSMUSG00000036461 | **Elf1** | Elf-1\|Sts1\|mElf-1\|p70 | E74-like factor 1 | protein-coding |
| chr14_79526136_79526408 | chr14 | 79526137 | 79526408 | + | -0.654636 | intron (NM_001286411, intron 1 of 8) | intron (NM_001286411, intron 1 of 8) | 10598 | NM_001286411 | 13709 | Mm.20938 | NM_007920 | ENSMUSG00000036461 | **Elf1** | Elf-1\|Sts1\|mElf-1\|p70 | E74-like factor 1 | protein-coding |
| chr12_84221032_84221484 | chr12 | 84221033 | 84221484 | + | -0.837653 | Intergenic | MLT1E1A\|LTR\|ERVL-MaLR | -2377 | NM_001163501 | 238317 | Mm.31256 | NM_001163501 | ENSMUSG00000042507 | **Elmsan1** | 9430029N19Rik\|C130039O16Rik\|Gm260 | ELM2 and Myb/SANT-like domain containing 1 | protein-coding |
| chr15_80562444_80562786 | chr15 | 80562445 | 80562786 | + | -0.594237 | Intergenic | Intergenic | -2145 | NM_001163189 | 383075 | Mm.297626 | NM_001163189 | ENSMUSG00000050439 | **Enthd1** | Gm1242\|Gm86 | ENTH domain containing 1 | protein-coding |
| chr15_80559545_80560047 | chr15 | 80559546 | 80560047 | + | -0.942777 | intron (NM_001163189, intron 1 of 5) | intron (NM_001163189, intron 1 of 5) | 674 | NM_001163189 | 383075 | Mm.297626 | NM_001163189 | ENSMUSG00000050439 | **Enthd1** | Gm1242\|Gm86 | ENTH domain containing 1 | protein-coding |
| chr15_80559640_80559964 | chr15 | 80559641 | 80559964 | + | -0.956397 | intron (NM_001163189, intron 1 of 5) | intron (NM_001163189, intron 1 of 5) | 668 | NM_001163189 | 383075 | Mm.297626 | NM_001163189 | ENSMUSG00000050439 | **Enthd1** | Gm1242\|Gm86 | ENTH domain containing 1 | protein-coding |
| chr4_132007857_132009093 | chr4 | 132007858 | 132009093 | + | -0.762303 | intron (NM_183428, intron 1 of 20) | intron (NM_183428, intron 1 of 20) | 40603 | NM_001128606 | 269587 | Mm.30038 | NM_183428 | ENSMUSG00000028906 | **Epb4.1** | 4.1R\|AI415518\|D4Ertd442e\|Elp-1\|Elp1\|Epb41\|mKIAA4056 | erythrocyte protein band 4.1 | protein-coding |
| chr4_132008054_132008655 | chr4 | 132008055 | 132008655 | + | -0.808211 | intron (NM_183428, intron 1 of 20) | intron (NM_183428, intron 1 of 20) | 40723 | NM_001128606 | 269587 | Mm.30038 | NM_183428 | ENSMUSG00000028906 | **Epb4.1** | 4.1R\|AI415518\|D4Ertd442e\|Elp-1\|Elp1\|Epb41\|mKIAA4056 | erythrocyte protein band 4.1 | protein-coding |
| chr16_13158975_13159867 | chr16 | 13158976 | 13159867 | + | -0.596928 | Intergenic | Intergenic | 49685 | NM_015769 | 50505 | Mm.287837 | NM_015769 | | **Ercc4** | AI606920\|Xpf | excision repair cross-complementing rodent repair deficiency, complementation group 4 | protein-coding |
| chr16_13158926_13159472 | chr16 | 13158927 | 13159472 | + | -0.686146 | Intergenic | Intergenic | 49463 | NM_015769 | 50505 | Mm.287837 | NM_015769 | | **Ercc4** | AI606920\|Xpf | excision repair cross-complementing rodent repair deficiency, complementation group 4 | protein-coding |
| chrY_90740034_90740593 | chrY | 90740035 | 90740593 | + | -0.7855 | Intergenic | Intergenic | -45128 | NM_133362 | 170942 | Mm.391385 | NM_133362 | ENSMUSG00000096768 | **Erdr1** | edr | erythroid differentiation regulator 1 | protein-coding |
| chr9_32731843_32732301 | chr9 | 32731844 | 32732301 | + | -0.585562 | intron (NM_001038642, intron 3 of 6) | intron (NM_001038642, intron 3 of 6) | 36030 | NM_001038642 | 23871 | Mm.292415 | NM_011808 | ENSMUSG00000032035 | **Ets1** | AI196000\|AI448617\|D230050P06\|Ets-1\|Tpl1 | E26 avian leukemia oncogene 1, 5' domain | protein-coding |
| chr9_32906299_32906819 | chr9 | 32906300 | 32906819 | + | -0.681131 | Intergenic | Intergenic | 210517 | NM_001038642 | 23871 | Mm.292415 | NM_011808 | ENSMUSG00000032035 | **Ets1** | AI196000\|AI448617\|D230050P06\|Ets-1\|Tpl1 | E26 avian leukemia oncogene 1, 5' domain | protein-coding |
| chr9_32743621_32744194 | chr9 | 32743622 | 32744194 | + | -0.691444 | intron (NM_011808, intron 6 of 7) | intron (NM_011808, intron 6 of 7) | 47866 | NM_001038642 | 23871 | Mm.292415 | NM_011808 | ENSMUSG00000032035 | **Ets1** | AI196000\|AI448617\|D230050P06\|Ets-1\|Tpl1 | E26 avian leukemia oncogene 1, 5' domain | protein-coding |
| chr9_32743680_32744030 | chr9 | 32743681 | 32744030 | + | -0.714436 | intron (NM_011808, intron 6 of 7) | intron (NM_011808, intron 6 of 7) | 47813 | NM_001038642 | 23871 | Mm.292415 | NM_011808 | ENSMUSG00000032035 | **Ets1** | AI196000\|AI448617\|D230050P06\|Ets-1\|Tpl1 | E26 avian leukemia oncogene 1, 5' domain | protein-coding |
| chr16_95728819_95729211 | chr16 | 95728820 | 95729211 | + | -0.594187 | Intergenic | Intergenic | 26608 | NM_011809 | 23872 | Mm.290207 | NM_011809 | ENSMUSG00000022895 | **Ets2** | AU022856\|Ets-2 | E26 avian leukemia oncogene 2, 3' domain | protein-coding |
| chr16_95728547_95729395 | chr16 | 95728548 | 95729395 | + | -0.609146 | Intergenic | Intergenic | 26564 | NM_011809 | 23872 | Mm.290207 | NM_011809 | ENSMUSG00000022895 | **Ets2** | AU022856\|Ets-2 | E26 avian leukemia oncogene 2, 3' domain | protein-coding |
| chr15_53156861_53157781 | chr15 | 53156862 | 53157781 | + | -0.593814 | intron (NM_010162, intron 1 of 10) | intron (NM_010162, intron 1 of 10) | 188862 | NM_010162 | 14042 | Mm.309395 | NM_010162 | | **Ext1** | AA409028 | exostoses (multiple) 1 | protein-coding |
| chr2_91638837_91639308 | chr2 | 91638838 | 91639308 | + | -0.634592 | Intergenic | Intergenic | -2659 | NM_010168 | 14061 | Mm.89048 | NM_010168 | ENSMUSG00000027249 | **F2** | Cf-2\|Cf2\|FII\|thrombin | coagulation factor II | protein-coding |
| chr2_3726423_3727253 | chr2 | 3726424 | 3727253 | + | -0.777223 | intron (NM_025626, intron 1 of 3) | intron (NM_025626, intron 1 of 3) | 13380 | NM_025626 | 66540 | Mm.277864 | NM_025626 | ENSMUSG00000026655 | **Fam107b** | 3110001A13Rik\|AA589595\|AA959793\|AI132312 | family with sequence similarity 107, member B | protein-coding |
| chr7_79392856_79393189 | chr7 | 79392857 | 79393189 | + | -0.658401 | intron (NM_145946, intron 1 of 37) | ID_B1\|SINE\|B4 | 685 | NM_145946 | 208836 | Mm.349493 | NM_145946 | ENSMUSG00000039187 | **Fanci** | - | Fanconi anemia, complementation group I | protein-coding |
| chr1_161704549_161705621 | chr1 | 161704550 | 161705621 | + | -0.611753 | Intergenic | MIR3\|SINE\|MIR | 83410 | NM_001205243 | 14103 | Mm.3355 | NM_010177 | ENSMUSG00000000817 | **Fasl** | APT1LG1\|CD178\|CD95-L\|CD95L\|Fas-L\|Faslg\|Tnfsf6\|gld | Fas ligand (TNF superfamily, member 6) | protein-coding |
| chr1_161704513_161705291 | chr1 | 161704514 | 161705291 | + | -0.633532 | Intergenic | Intergenic | 83593 | NM_001205243 | 14103 | Mm.3355 | NM_010177 | ENSMUSG00000000817 | **Fasl** | APT1LG1\|CD178\|CD95-L\|CD95L\|Fas-L\|Faslg\|Tnfsf6\|gld | Fas ligand (TNF superfamily, member 6) | protein-coding |
| chr1_161765913_161766325 | chr1 | 161765914 | 161766325 | + | -0.719196 | Intergenic | Intergenic | 22376 | NM_001205243 | 14103 | Mm.3355 | NM_010177 | ENSMUSG00000000817 | **Fasl** | APT1LG1\|CD178\|CD95-L\|CD95L\|Fas-L\|Faslg\|Tnfsf6\|gld | Fas ligand (TNF superfamily, member 6) | protein-coding |
| chr1_161767101_161767600 | chr1 | 161767102 | 161767600 | + | -0.753763 | Intergenic | Intergenic | 21144 | NM_001205243 | 14103 | Mm.3355 | NM_010177 | ENSMUSG00000000817 | **Fasl** | APT1LG1\|CD178\|CD95-L\|CD95L\|Fas-L\|Faslg\|Tnfsf6\|gld | Fas ligand (TNF superfamily, member 6) | protein-coding |
| chr9_20645202_20645493 | chr9 | 20645203 | 20645493 | + | -0.624531 | promoter-TSS (NM_001286530) | promoter-TSS (NM_001286530) | -581 | NM_001286530 | 30843 | Mm.24608 | NM_013911 | ENSMUSG00000066892 | **Fbxl12** | 3110048D16Rik\|Fbl12 | F-box and leucine-rich repeat protein 12 | protein-coding |
| chr9_32507068_32507844 | chr9 | 32507069 | 32507844 | + | -0.620958 | intron (NM_008026, intron 1 of 8) | intron (NM_008026, intron 1 of 8) | 33996 | NM_008026 | 14247 | Mm.258908 | NM_008026 | ENSMUSG00000016087 | **Fli1** | EWSR2\|Fli-1\|SIC-1\|Sic1 | Friend leukemia integration 1 | protein-coding |
| chr5_32132864_32133883 | chr5 | 32132865 | 32133883 | + | -0.712501 | Intergenic | CpG | -3098 | NM_008037 | 14284 | Mm.24684 | NM_008037 | ENSMUSG00000029135 | **Fosl2** | Fra-2 | fos-like antigen 2 | protein-coding |
| chr5_32133508_32133880 | chr5 | 32133509 | 32133880 | + | -0.882388 | Intergenic | Intergenic | -2778 | NM_008037 | 14284 | Mm.24684 | NM_008037 | ENSMUSG00000029135 | **Fosl2** | Fra-2 | fos-like antigen 2 | protein-coding |
| chr6_99350333_99350990 | chr6 | 99350334 | 99350990 | + | -0.601138 | intron (NM_053202, intron 1 of 18) | intron (NM_053202, intron 1 of 18) | -84147 | NM_001197321 | 108655 | Mm.234965 | NM_053202 | ENSMUSG00000030067 | **Foxp1** | 3110052D19Rik\|4932443N09Rik\|AI461938\|AW494214 | forkhead box P1 | protein-coding |
| chr6_99095208_99095908 | chr6 | 99095209 | 99095908 | + | -0.663077 | intron (NM_053202, intron 4 of 18) | intron (NM_053202, intron 4 of 18) | -67260 | NM_001197322 | 108655 | Mm.234965 | NM_053202 | ENSMUSG00000030067 | **Foxp1** | 3110052D19Rik\|4932443N09Rik\|AI461938\|AW494214 | forkhead box P1 | protein-coding |
| chr6_99095573_99095908 | chr6 | 99095574 | 99095908 | + | -0.663872 | intron (NM_053202, intron 4 of 18) | intron (NM_053202, intron 4 of 18) | -67443 | NM_001197322 | 108655 | Mm.234965 | NM_053202 | ENSMUSG00000030067 | **Foxp1** | 3110052D19Rik\|4932443N09Rik\|AI461938\|AW494214 | forkhead box P1 | protein-coding |
| chr6_99171421_99172601 | chr6 | 99171422 | 99172601 | + | -0.665152 | intron (NM_053202, intron 3 of 18) | intron (NM_053202, intron 3 of 18) | 94504 | NM_001197321 | 108655 | Mm.234965 | NM_053202 | ENSMUSG00000030067 | **Foxp1** | 3110052D19Rik\|4932443N09Rik\|AI461938\|AW494214 | forkhead box P1 | protein-coding |
| chr6_99023634_99023960 | chr6 | 99023635 | 99023960 | + | -0.666641 | intron (NM_053202, intron 5 of 18) | intron (NM_053202, intron 5 of 18) | 4501 | NM_001197322 | 108655 | Mm.234965 | NM_053202 | ENSMUSG00000030067 | **Foxp1** | 3110052D19Rik\|4932443N09Rik\|AI461938\|AW494214 | forkhead box P1 | protein-coding |
| chr5_73265322_73265878 | chr5 | 73265323 | 73265878 | + | -0.774644 | Intergenic | L2b\|LINE\|L2 | -8982 | NM_028194 | 72313 | Mm.260594 | NM_028194 | ENSMUSG00000070733 | **Fryl** | 2010313D22Rik\|2310004H21Rik\|2510002A14Rik\|9030227G01Rik\|mKIAA0826 | furry homolog-like (Drosophila) | protein-coding |
| chr15_6592556_6593127 | chr15 | 6592557 | 6593127 | + | -0.591045 | intron (NM_011815, intron 1 of 17) | intron (NM_011815, intron 1 of 17) | 12995 | NM_011815 | 23880 | Mm.170905 | NM_011815 | | **Fyb** | ADAP\|B630013F22Rik\|FYB-120/130\|Slap130\|p120/130\|p120/p130 | FYN binding protein | protein-coding |
| chrX_170017203_170017868 | chrX | 170017204 | 170017868 | + | -0.65842 | Intergenic | Intergenic | -38619 | NR_029457 | 654820 | Mm.34441 | NM_001039559 | ENSMUSG00000072844 | **G530011O06Rik** | - | RIKEN cDNA G530011O06 gene | ncRNA |
| chrX_169991082_169991551 | chrX | 169991083 | 169991551 | + | -0.903685 | TTS (NM_001290512) | TTS (NM_001290512) | -12400 | NR_029457 | 654820 | Mm.34441 | NM_001039559 | ENSMUSG00000072844 | **G530011O06Rik** | - | RIKEN cDNA G530011O06 gene | ncRNA |
| chrX_169990905_169991756 | chrX | 169990906 | 169991756 | + | -0.906835 | TTS (NM_001290512) | TTS (NM_001290512) | -12414 | NR_029457 | 654820 | Mm.34441 | NM_001039559 | ENSMUSG00000072844 | **G530011O06Rik** | - | RIKEN cDNA G530011O06 gene | ncRNA |
| chrX_169996857_169998570 | chrX | 169996858 | 169998570 | + | -0.956519 | Intergenic | CpG | -18797 | NR_029457 | 654820 | Mm.34441 | NM_001039559 | ENSMUSG00000072844 | **G530011O06Rik** | - | RIKEN cDNA G530011O06 gene | ncRNA |
| chrX_169989845_169990751 | chrX | 169989846 | 169990751 | + | -1.11526 | 3' UTR (NM_001290512, exon 10 of 10) | 3' UTR (NM_001290512, exon 10 of 10) | -11381 | NR_029457 | 654820 | Mm.34441 | NM_001039559 | ENSMUSG00000072844 | **G530011O06Rik** | - | RIKEN cDNA G530011O06 gene | ncRNA |
| chrX_169990112_169990772 | chrX | 169990113 | 169990772 | + | -1.14841 | 3' UTR (NM_001290512, exon 10 of 10) | 3' UTR (NM_001290512, exon 10 of 10) | -11525 | NR_029457 | 654820 | Mm.34441 | NM_001039559 | ENSMUSG00000072844 | **G530011O06Rik** | - | RIKEN cDNA G530011O06 gene | ncRNA |
| chr13_3526226_3526919 | chr13 | 3526227 | 3526919 | + | -0.593823 | Intergenic | Intergenic | -11502 | NM_008112 | 14569 | Mm.153226 | NM_008112 | ENSMUSG00000021218 | **Gdi2** | GDI-B\|GDIB\|Gdi3 | guanosine diphosphate (GDP) dissociation inhibitor 2 | protein-coding |
| chr2_35507133_35507605 | chr2 | 35507134 | 35507605 | + | -0.60583 | Intergenic | MIRb\|SINE\|MIR | -45944 | NM_001145821 | 14594 | Mm.281124 | NM_010283 | ENSMUSG00000035778 | **Ggta1** | AW108479\|GALT\|Gal\|Ggta\|Ggta-1\|alpha3GalT | glycoprotein galactosyltransferase alpha 1, 3 | protein-coding |
| chr18_5593691_5594020 | chr18 | 5593692 | 5594020 | + | -0.669132 | intron (NM_011546, intron 1 of 7) | intron (NM_011546, intron 1 of 7) | -1419 | NR_033552 | 791318 | Mm.84339 | NR_033552 | ENSMUSG00000063087 | **Gm10125** | ENSMUSG00000063087\|Zeb1os | predicted gene 10125 | ncRNA |
| chr18_5593653_5594312 | chr18 | 5593654 | 5594312 | + | -0.835285 | intron (NM_011546, intron 1 of 7) | intron (NM_011546, intron 1 of 7) | -1546 | NR_033552 | 791318 | Mm.84339 | NR_033552 | ENSMUSG00000063087 | **Gm10125** | ENSMUSG00000063087\|Zeb1os | predicted gene 10125 | ncRNA |

| | | | | | | | | | | | | | | | | | |
|---|---|---|---|---|---|---|---|---|---|---|---|---|---|---|---|---|---|
| chr11_6679691_6680777 | chr11 | 6679692 | 6680777 | + | -0.807904 | Intergenic | RSINE1\|SINE\|B4 | -19998 | NR_028586 | 100041311 | Mm.388018 | NR_028586 | ENSMUSG00000086373 | **Gm11978** | OTTMUSG00000005125\|OTTMUSG00000005127 | predicted gene 11978 | pseudo |
| chr11_6694948_6695746 | chr11 | 6694949 | 6695746 | + | -1.1789 | Intergenic | Tigger7\|DNA\|TcMar-Tigger | 24690 | NR_046025 | 626419 | Mm.390334 | NR_046025 | ENSMUSG00000086808 | **Gm11981** | OTTMUSG00000005129 | predicted gene 11981 | ncRNA |
| chr2_128294653_128295473 | chr2 | 128294654 | 128295473 | + | -0.626176 | Intergenic | Intergenic | 134288 | NR_028591 | 100043424 | Mm.297222 | NR_028589 | | **Gm14005** | OTTMUSG00000015351 | predicted gene 14005 | ncRNA |
| chr6_145306665_145307079 | chr6 | 145306666 | 145307079 | + | -0.710653 | Intergenic | Intergenic | 56320 | NR_045598 | 100039495 | Mm.121014 | NR_045598 | ENSMUSG00000086013 | **Gm15706** | OTTMUSG00000025463 | predicted gene 15706 | ncRNA |
| chr8_35678774_35679367 | chr8 | 35678775 | 35679367 | + | -0.685664 | 3' UTR (NM_001081279, exon 3 of 3) | 3' UTR (NM_001081279, exon 3 of 3) | -90051 | NR_040376 | 100504714 | Mm.381744 | NR_040376 | ENSMUSG00000097357 | **Gm16793** | - | predicted gene, 16793 | ncRNA |
| chr19_47512763_47513353 | chr19 | 47512764 | 47513353 | + | -0.788906 | promoter-TSS (NR_045322) | promoter-TSS (NR_045322) | 76 | NR_045322 | 100503104 | Mm.442735 | NR_045322 | ENSMUSG00000097990 | **Gm19557** | - | predicted gene, 19557 | ncRNA |
| chr17_51903825_51904308 | chr17 | 51903826 | 51904308 | + | -0.588733 | intron (NR_045095, intron 1 of 2) | intron (NR_045095, intron 1 of 2) | 21340 | NR_045095 | 100504169 | Mm.417731 | NR_045095 | | **Gm20098** | - | predicted gene, 20098 | ncRNA |
| chr17_52162267_52163411 | chr17 | 52162268 | 52163411 | + | -0.802091 | Intergenic | Intergenic | 280112 | NR_045095 | 100504169 | Mm.417731 | NR_045095 | | **Gm20098** | - | predicted gene, 20098 | ncRNA |
| chr17_52177915_52178461 | chr17 | 52177916 | 52178461 | + | -0.853897 | Intergenic | Intergenic | 295461 | NR_045095 | 100504169 | Mm.417731 | NR_045095 | | **Gm20098** | - | predicted gene, 20098 | ncRNA |
| chr17_52168131_52169101 | chr17 | 52168132 | 52169101 | + | -0.912526 | Intergenic | Intergenic | 285889 | NR_045095 | 100504169 | Mm.417731 | NR_045095 | | **Gm20098** | - | predicted gene, 20098 | ncRNA |
| chr10_19104540_19105371 | chr10 | 19104541 | 19105371 | + | -0.599941 | Intergenic | Intergenic | -36793 | NR_038021 | 100504265 | Mm.383417 | NR_038021 | | **Gm20139** | - | predicted gene, 20139 | ncRNA |
| chr10_19171826_19172093 | chr10 | 19171827 | 19172093 | + | -0.684093 | intron (NR_038022, intron 1 of 1) | intron (NR_038022, intron 1 of 1) | 4722 | NR_038022 | 100504265 | Mm.383417 | NR_038021 | | **Gm20139** | - | predicted gene, 20139 | ncRNA |
| chr10_19171830_19172216 | chr10 | 19171831 | 19172216 | + | -0.757275 | intron (NR_038022, intron 1 of 1) | intron (NR_038022, intron 1 of 1) | 4785 | NR_038022 | 100504265 | Mm.383417 | NR_038021 | | **Gm20139** | - | predicted gene, 20139 | ncRNA |
| chr10_19171773_19172315 | chr10 | 19171774 | 19172315 | + | -0.777152 | intron (NR_038022, intron 1 of 1) | intron (NR_038022, intron 1 of 1) | 4806 | NR_038022 | 100504265 | Mm.383417 | NR_038021 | | **Gm20139** | - | predicted gene, 20139 | ncRNA |
| chr10_19185775_19186634 | chr10 | 19185776 | 19186634 | + | -0.777511 | intron (NR_038022, intron 1 of 1) | intron (NR_038022, intron 1 of 1) | 18967 | NR_038022 | 100504265 | Mm.383417 | NR_038021 | | **Gm20139** | - | predicted gene, 20139 | ncRNA |
| chr10_19122123_19122601 | chr10 | 19122124 | 19122601 | + | -0.862036 | Intergenic | Intergenic | -19387 | NR_038021 | 100504265 | Mm.383417 | NR_038021 | | **Gm20139** | - | predicted gene, 20139 | ncRNA |
| chr10_19116814_19117545 | chr10 | 19116815 | 19117545 | + | -0.95548 | Intergenic | Intergenic | -24569 | NR_038021 | 100504265 | Mm.383417 | NR_038021 | | **Gm20139** | - | predicted gene, 20139 | ncRNA |
| chr3_52927334_52927956 | chr3 | 52927335 | 52927956 | + | -0.655338 | promoter-TSS (NR_040555) | promoter-TSS (NR_040555) | 21 | NR_040555 | 622124 | Mm.389434 | NR_040555 | | **Gm20750** | - | predicted gene, 20750 | ncRNA |
| chr3_52954575_52955768 | chr3 | 52954576 | 52955768 | + | -0.77178 | Intergenic | Intergenic | -27506 | NR_040555 | 622124 | Mm.389434 | NR_040555 | | **Gm20750** | - | predicted gene, 20750 | ncRNA |
| chr3_52954509_52955041 | chr3 | 52954510 | 52955041 | + | -0.853593 | Intergenic | Intergenic | -27109 | NR_040555 | 622124 | Mm.389434 | NR_040555 | | **Gm20750** | - | predicted gene, 20750 | ncRNA |
| chr18_80608440_80608714 | chr18 | 80608441 | 80608714 | + | -0.664229 | intron (NM_198429, intron 9 of 9) | intron (NM_198429, intron 9 of 9) | -64031 | NR_028424 | 100039348 | Mm.254722 | NR_028424 | | **Gm2176** | A430076E10Rik | glucosamine-6-phosphate deaminase 1 pseudogene | pseudo |
| chr18_80608384_80608782 | chr18 | 80608385 | 80608782 | + | -0.668397 | intron (NM_198429, intron 9 of 9) | intron (NM_198429, intron 9 of 9) | -64037 | NR_028424 | 100039348 | Mm.254722 | NR_028424 | | **Gm2176** | A430076E10Rik | glucosamine-6-phosphate deaminase 1 pseudogene | pseudo |
| chr5_64684153_64684889 | chr5 | 64684154 | 64684889 | + | -0.710593 | Intergenic | Intergenic | -73822 | NR_045078 | 100042198 | Mm.25191 | NR_045078 | | **Gm3716** | Gm1683 | predicted gene 3716 | ncRNA |
| chr1_180403251_180404193 | chr1 | 180403252 | 180404193 | + | -0.673665 | intron (NM_001081175, intron 2 of 7) | intron (NM_001081175, intron 2 of 7) | -73173 | NR_003623 | 277333 | Mm.389223 | NM_199472 | | **Gm5069** | EG277333 | glyceraldehyde-3-phosphate dehydrogenase pseudogene | pseudo |
| chr1_180359353_180359735 | chr1 | 180359354 | 180359735 | + | -0.710447 | intron (NM_001081175, intron 2 of 7) | MLT1L\|LTR\|ERVL-MaLR | -28995 | NR_003623 | 277333 | Mm.389223 | NM_199472 | | **Gm5069** | EG277333 | glyceraldehyde-3-phosphate dehydrogenase pseudogene | pseudo |
| chr1_180401758_180402093 | chr1 | 180401759 | 180402093 | + | -0.778458 | intron (NM_001081175, intron 2 of 7) | intron (NM_001081175, intron 2 of 7) | -71377 | NR_003623 | 277333 | Mm.389223 | NM_199472 | | **Gm5069** | EG277333 | glyceraldehyde-3-phosphate dehydrogenase pseudogene | pseudo |
| chr9_107675658_107676704 | chr9 | 107675659 | 107676704 | + | -0.779163 | intron (NM_008140, intron 7 of 8) | CpG | 3411 | NM_008140 | 14685 | Mm.284853 | NM_008140 | ENSMUSG00000034837 | **Gnat1** | Gnat-1\|Ird1\|Ird2\|Tralpha\|irdc\|irdr\|transducin | guanine nucleotide binding protein, alpha transducing 1 | protein-coding |
| chr1_74262410_74262906 | chr1 | 74262411 | 74262906 | + | -0.861964 | intron (NM_029711, intron 7 of 9) | ID2\|SINE\|ID | -15942 | NM_174985 | 227289 | Mm.246587 | NM_174985 | ENSMUSG00000064272 | **Gpbar1** | BG37\|GPCR\|GPR131\|M-BAR\|TGR5 | G protein-coupled bile acid receptor 1 | protein-coding |
| chr1_74262508_74262868 | chr1 | 74262509 | 74262868 | + | -0.885216 | intron (NM_029711, intron 7 of 9) | intron (NM_029711, intron 7 of 9) | -15912 | NM_174985 | 227289 | Mm.246587 | NM_174985 | ENSMUSG00000064272 | **Gpbar1** | BG37\|GPCR\|GPR131\|M-BAR\|TGR5 | G protein-coupled bile acid receptor 1 | protein-coding |
| chr12_112854986_112855390 | chr12 | 112854987 | 112855390 | + | -0.609736 | intron (NM_019925, intron 2 of 2) | intron (NM_019925, intron 2 of 2) | 5728 | NM_019925 | 56696 | Mm.20455 | NM_019925 | ENSMUSG00000021298 | **Gpr132** | G2a | G protein-coupled receptor 132 | protein-coding |
| chr12_112860791_112861427 | chr12 | 112860792 | 112861427 | + | -0.84521 | promoter-TSS (NM_019925) | promoter-TSS (NM_019925) | -193 | NM_019925 | 56696 | Mm.20455 | NM_019925 | ENSMUSG00000021298 | **Gpr132** | G2a | G protein-coupled receptor 132 | protein-coding |
| chr12_112860822_112861177 | chr12 | 112860823 | 112861177 | + | -0.857465 | promoter-TSS (NM_019925) | promoter-TSS (NM_019925) | -84 | NM_019925 | 56696 | Mm.20455 | NM_019925 | ENSMUSG00000021298 | **Gpr132** | G2a | G protein-coupled receptor 132 | protein-coding |
| chr14_121930700_121931031 | chr14 | 121930701 | 121931031 | + | -0.592395 | intron (NM_026861, intron 4 of 8) | L1ME3A\|LINE\|L1 | -15092 | NM_182806 | 110168 | Mm.37405 | NM_182806 | ENSMUSG00000050350 | **Gpr18** | - | G protein-coupled receptor 18 | protein-coding |
| chr14_121930687_121931346 | chr14 | 121930688 | 121931346 | + | -0.618788 | intron (NM_026861, intron 4 of 8) | intron (NM_026861, intron 4 of 8) | -15243 | NM_182806 | 110168 | Mm.37405 | NM_182806 | ENSMUSG00000050350 | **Gpr18** | - | G protein-coupled receptor 18 | protein-coding |
| chr14_121938658_121939268 | chr14 | 121938659 | 121939268 | + | -0.650911 | intron (NM_026861, intron 4 of 8) | intron (NM_026861, intron 4 of 8) | -23189 | NM_182806 | 110168 | Mm.37405 | NM_182806 | ENSMUSG00000050350 | **Gpr18** | - | G protein-coupled receptor 18 | protein-coding |
| chr14_121938253_121939239 | chr14 | 121938254 | 121939239 | + | -0.651617 | intron (NM_026861, intron 4 of 8) | intron (NM_026861, intron 4 of 8) | -22972 | NM_182806 | 110168 | Mm.37405 | NM_182806 | ENSMUSG00000050350 | **Gpr18** | - | G protein-coupled receptor 18 | protein-coding |
| chr14_121929282_121929879 | chr14 | 121929283 | 121929879 | + | -0.812046 | intron (NM_026861, intron 4 of 8) | intron (NM_026861, intron 4 of 8) | -13807 | NM_182806 | 110168 | Mm.37405 | NM_182806 | ENSMUSG00000050350 | **Gpr18** | - | G protein-coupled receptor 18 | protein-coding |
| chr18_56468961_56469469 | chr18 | 56468962 | 56469469 | + | -0.595081 | intron (NM_026240, intron 1 of 13) | intron (NM_026240, intron 1 of 13) | 37083 | NM_026240 | 107022 | Mm.24356 | NM_026240 | ENSMUSG00000001700 | **Gramd3** | 9030613F08Rik\|9130427A09Rik\|AW061306\|NS3TP2 | GRAM domain containing 3 | protein-coding |
| chr5_134481933_134482727 | chr5 | 134481934 | 134482727 | + | -0.60432 | Intergenic | Intergenic | -25614 | NM_001081464 | 57080 | Mm.332735 | NM_020331 | ENSMUSG00000023079 | **Gtf2ird1** | 1700012P16Rik\|BEN\|Cream1\|ESTM9\|GTF3\|Gtf2il\|MusTRD1\|Tg(Alb1-Myc)166.8Sst\|WBSCR11\|X83320 | general transcription factor II I repeat domain-containing 1 | protein-coding |
| chr6_136743701_136744184 | chr6 | 136743702 | 136744184 | + | -0.83254 | exon (NM_001127318, exon 10 of 27) | exon (NM_001127318, exon 10 of 27) | 37799 | NM_145067 | 14917 | Mm.222729 | NM_145067 | ENSMUSG00000042638 | **Gucy2c** | AI893437\|GC-C\|Gcc | guanylate cyclase 2c | protein-coding |
| chr6_136742308_136742630 | chr6 | 136742309 | 136742630 | + | -0.902219 | intron (NM_145067, intron 10 of 25) | intron (NM_145067, intron 10 of 25) | 39273 | NM_145067 | 14917 | Mm.222729 | NM_145067 | ENSMUSG00000042638 | **Gucy2c** | AI893437\|GC-C\|Gcc | guanylate cyclase 2c | protein-coding |
| chr6_136742252_136742694 | chr6 | 136742253 | 136742694 | + | -0.939701 | intron (NM_145067, intron 10 of 25) | intron (NM_145067, intron 10 of 25) | 39269 | NM_145067 | 14917 | Mm.222729 | NM_145067 | ENSMUSG00000042638 | **Gucy2c** | AI893437\|GC-C\|Gcc | guanylate cyclase 2c | protein-coding |
| chr17_34293420_34293946 | chr17 | 34293421 | 34293946 | + | -0.645198 | Intergenic | Intergenic | -5912 | NM_010378 | 14960 | Mm.235338 | NM_010378 | ENSMUSG00000036594 | **H2-Aa** | Aalpha\|H-2Aa\|H2Aa\|I-Aalpha\|IAalpha\|Ia-1\|Ia1 | histocompatibility 2, class II antigen A, alpha | protein-coding |
| chr17_34347593_34348276 | chr17 | 34347594 | 34348276 | + | -0.663857 | Intergenic | Intergenic | -3290 | NM_010381 | 100504404 | Mm.15680 | NM_010381 | | **H2-Ea-ps** | AI323765\|E-alpha-f\|H-2Ea\|H2-Ea\|I-Ealpha\|Ia-3\|Ia3 | histocompatibility 2, class II antigen E alpha, pseudogene | protein-coding |
| chr17_34239429_34240003 | chr17 | 34239430 | 34240003 | + | -0.665143 | intron (NM_010389, intron 1 of 5) | intron (NM_010389, intron 1 of 5) | 811 | NM_010389 | 15002 | Mm.327358 | NM_010389 | ENSMUSG00000041538 | **H2-Ob** | A-beta-2\|A-beta2\|H-2Ab\|H-2I\|H-2Ob\|H2-Ab\|H2-Ab2\|H2-IAb2\|Ob | histocompatibility 2, O region beta locus | protein-coding |
| chr10_21248615_21249072 | chr10 | 21248616 | 21249072 | + | -0.632525 | Intergenic | Intergenic | -47135 | NM_001042593 | 56422 | Mm.25527 | NM_019702 | ENSMUSG00000019977 | **Hbs1l** | 2810035F15Rik\|AI326327\|eRFS | Hbs1-like (S. cerevisiae) | protein-coding |
| chr10_21254618_21255194 | chr10 | 21254619 | 21255194 | + | -0.784426 | Intergenic | Intergenic | -41073 | NM_001042593 | 56422 | Mm.25527 | NM_019702 | ENSMUSG00000019977 | **Hbs1l** | 2810035F15Rik\|AI326327\|eRFS | Hbs1-like (S. cerevisiae) | protein-coding |
| chr12_51865759_51866206 | chr12 | 51865760 | 51866206 | + | -0.591093 | Intergenic | Intergenic | -36447 | NM_144788 | 207304 | Mm.249391 | NM_144788 | ENSMUSG00000035247 | **Hectd1** | A630086P08Rik\|AI844876\|b2b327Clo\|opm | HECT domain containing 1 | protein-coding |
| chr12_51866253_51866791 | chr12 | 51866254 | 51866791 | + | -0.595306 | Intergenic | Intergenic | -36986 | NM_144788 | 207304 | Mm.249391 | NM_144788 | ENSMUSG00000035247 | **Hectd1** | A630086P08Rik\|AI844876\|b2b327Clo\|opm | HECT domain containing 1 | protein-coding |
| chr12_51865759_51866801 | chr12 | 51865760 | 51866801 | + | -0.633146 | Intergenic | Intergenic | -36744 | NM_144788 | 207304 | Mm.249391 | NM_144788 | ENSMUSG00000035247 | **Hectd1** | A630086P08Rik\|AI844876\|b2b327Clo\|opm | HECT domain containing 1 | protein-coding |
| chr3_8766590_8767289 | chr3 | 8766591 | 8767289 | + | -0.693148 | Intergenic | Intergenic | -99902 | NM_010423 | 15213 | Mm.29581 | NM_010423 | ENSMUSG00000040289 | **Hey1** | AI316788\|AI414254\|CHF2\|HRT1\|Herp2\|Hesr1\|bHLHb31\|hesr-1 | hairy/enhancer-of-split related with YRPW motif 1 | protein-coding |
| chr13_23551220_23551881 | chr13 | 23551221 | 23551881 | + | -0.642747 | promoter-TSS (NM_175655) | promoter-TSS (NM_175655) | 92 | NM_175655 | 319157 | Mm.442307 | NM_175655 | | **Hist1h4f** | - | histone cluster 1, H4f | protein-coding |
| chr10_13993235_13993904 | chr10 | 13993236 | 13993904 | + | -0.70078 | intron (NM_010437, intron 1 of 8) | intron (NM_010437, intron 1 of 8) | 27191 | NM_010437 | 15273 | Mm.42157 | NM_010437 | ENSMUSG00000015501 | **Hivep2** | MIBP1\|Schnurri-2\|Shn-2 | human immunodeficiency virus type I enhancer binding protein 2 | protein-coding |
| chr16_91532982_91534276 | chr16 | 91532983 | 91534276 | + | -0.600613 | Intergenic | Intergenic | -13465 | NM_008338 | 15980 | Mm.249364 | NM_008338 | | **Ifngr2** | Ifgr2\|Ifgt | interferon gamma receptor 2 | protein-coding |

| | | | | | | | | | | | | | | | | | |
|---|---|---|---|---|---|---|---|---|---|---|---|---|---|---|---|---|---|
| chr16_91538144_91538390 | chr16 | 91538145 | 91538390 | + | -0.764928 | Intergenic | Intergenic | -8827 | NM_008338 | 15980 | Mm.249364 | NM_008338 | | **Ifngr2** | Ifgr2\|Ifgt | interferon gamma receptor 2 | protein-coding |
| chr2_11718225_11718632 | chr2 | 11718226 | 11718632 | + | -0.955423 | exon (NM_008358, exon 2 of 7) | exon (NM_008358, exon 2 of 7) | 12717 | NM_001271500 | 16169 | Mm.200196 | NM_008358 | ENSMUSG00000023206 | **Il15ra** | AA690181\|IL-15RA | interleukin 15 receptor, alpha chain | protein-coding |
| chr7_125580991_125581804 | chr7 | 125580992 | 125581804 | + | -0.618884 | Intergenic | CpG | -22031 | NM_021887 | 60504 | Mm.155643 | NM_021887 | ENSMUSG00000030745 | **Il21r** | NILR | interleukin 21 receptor | protein-coding |
| chr13_112427556_112427943 | chr13 | 112427557 | 112427943 | + | -0.585427 | Intergenic | Intergenic | -36320 | NM_010560 | 16195 | Mm.4364 | NM_010560 | ENSMUSG00000021756 | **Il6st** | 5133400A03Rik\|AA389424\|BB405851\|CD130\|D13Ertd699e\|gp130 | interleukin 6 signal transducer | protein-coding |
| chr13_112427470_112428001 | chr13 | 112427471 | 112428001 | + | -0.597674 | Intergenic | Intergenic | -36334 | NM_010560 | 16195 | Mm.4364 | NM_010560 | ENSMUSG00000021756 | **Il6st** | 5133400A03Rik\|AA389424\|BB405851\|CD130\|D13Ertd699e\|gp130 | interleukin 6 signal transducer | protein-coding |
| chr8_81558420_81559100 | chr8 | 81558421 | 81559100 | + | -0.990312 | intron (NM_001297596, intron 2 of 24) | RMER19A\|LTR\|ERVK | 216198 | NM_001297593 | 234515 | Mm.324468 | NM_001024617 | ENSMUSG00000037940 | **Inpp4b** | E130107I17Rik | inositol polyphosphate-4-phosphatase, type II | protein-coding |
| chr1_191579503_191579925 | chr1 | 191579504 | 191579925 | + | -0.61809 | intron (NM_178632, intron 1 of 19) | intron (NM_178632, intron 1 of 19) | 3980 | NM_178632 | 77065 | Mm.338175 | NM_178632 | ENSMUSG00000037461 | **Ints7** | 5930412E23Rik | integrator complex subunit 7 | protein-coding |
| chr9_108003652_108003926 | chr9 | 108003653 | 108003926 | + | -0.688755 | intron (NM_013785, intron 1 of 5) | intron (NM_013785, intron 1 of 5) | 1141 | NM_013785 | 27399 | Mm.276155 | NM_013785 | ENSMUSG00000032594 | **Ip6k1** | 1200016D08Rik\|AU022776\|Ihpk1\|InsP6\|InsP6k1\|Itpk6 | inositol hexaphosphate kinase 1 | protein-coding |
| chr8_109662987_109663545 | chr8 | 109662988 | 109663545 | + | -0.614162 | intron (NM_001039700, intron 27 of 33) | intron (NM_001039700, intron 27 of 33) | 30028 | NM_028018 | 71955 | Mm.290036 | NM_028018 | ENSMUSG00000031729 | **Ist1** | 2400003C14Rik\|AW536298\|mKIAA0174 | increased sodium tolerance 1 homolog (yeast) | protein-coding |
| chr8_109663284_109663532 | chr8 | 109663285 | 109663532 | + | -0.681842 | intron (NM_001039700, intron 27 of 33) | intron (NM_001039700, intron 27 of 33) | 29886 | NM_028018 | 71955 | Mm.290036 | NM_028018 | ENSMUSG00000031729 | **Ist1** | 2400003C14Rik\|AW536298\|mKIAA0174 | increased sodium tolerance 1 homolog (yeast) | protein-coding |
| chr8_109663239_109663572 | chr8 | 109663240 | 109663572 | + | -0.690867 | intron (NM_001039700, intron 27 of 33) | intron (NM_001039700, intron 27 of 33) | 29888 | NM_028018 | 71955 | Mm.290036 | NM_028018 | ENSMUSG00000031729 | **Ist1** | 2400003C14Rik\|AW536298\|mKIAA0174 | increased sodium tolerance 1 homolog (yeast) | protein-coding |
| chr11_46372236_46372775 | chr11 | 46372237 | 46372775 | + | -0.620823 | intron (NM_001281966, intron 1 of 16) | Lx8b\|LINE\|L1 | 17009 | NM_001281968 | 16428 | Mm.339927 | NM_010583 | ENSMUSG00000020395 | **Itk** | Emt\|Tcsk\|Tsk | IL2 inducible T cell kinase | protein-coding |
| chr11_46372236_46373030 | chr11 | 46372237 | 46373030 | + | -0.642167 | intron (NM_001281966, intron 1 of 16) | Lx8b\|LINE\|L1 | 16882 | NM_001281968 | 16428 | Mm.339927 | NM_010583 | ENSMUSG00000020395 | **Itk** | Emt\|Tcsk\|Tsk | IL2 inducible T cell kinase | protein-coding |
| chr4_101185286_101185604 | chr4 | 101185287 | 101185604 | + | -0.613153 | intron (NM_146145, intron 4 of 24) | intron (NM_146145, intron 4 of 24) | 79837 | NM_146145 | 16451 | Mm.289657 | NM_146145 | ENSMUSG00000028530 | **Jak1** | AA960307\|C130039L05Rik | Janus kinase 1 | protein-coding |
| chr13_44799702_44800666 | chr13 | 44799703 | 44800666 | + | -0.716237 | intron (NM_001205043, intron 1 of 17) | intron (NM_001205043, intron 1 of 17) | 68913 | NM_001205043 | 16468 | Mm.25059 | NM_021878 | ENSMUSG00000038518 | **Jarid2** | Jmj\|jumonji | jumonji, AT rich interactive domain 2 | protein-coding |
| chr17_52506633_52507595 | chr17 | 52506634 | 52507595 | + | -0.596847 | Intergenic | Intergenic | -95595 | NM_001031811 | 211468 | Mm.399068 | NM_001031811 | ENSMUSG00000035580 | **Kcnh8** | C130090D05Rik\|ELK\|ELK1\|ELK3\|Kv12.1 | potassium voltage-gated channel, subfamily H (eag-related), member 8 | protein-coding |
| chr17_52506784_52507483 | chr17 | 52506785 | 52507483 | + | -0.644788 | Intergenic | Intergenic | -95575 | NM_001031811 | 211468 | Mm.399068 | NM_001031811 | ENSMUSG00000035580 | **Kcnh8** | C130090D05Rik\|ELK\|ELK1\|ELK3\|Kv12.1 | potassium voltage-gated channel, subfamily H (eag-related), member 8 | protein-coding |
| chr17_52506636_52507189 | chr17 | 52506637 | 52507189 | + | -0.68202 | Intergenic | MIR\|SINE\|MIR | -95796 | NM_001031811 | 211468 | Mm.399068 | NM_001031811 | ENSMUSG00000035580 | **Kcnh8** | C130090D05Rik\|ELK\|ELK1\|ELK3\|Kv12.1 | potassium voltage-gated channel, subfamily H (eag-related), member 8 | protein-coding |
| chr17_52506779_52507244 | chr17 | 52506780 | 52507244 | + | -0.682031 | Intergenic | MIR\|SINE\|MIR | -95697 | NM_001031811 | 211468 | Mm.399068 | NM_001031811 | ENSMUSG00000035580 | **Kcnh8** | C130090D05Rik\|ELK\|ELK1\|ELK3\|Kv12.1 | potassium voltage-gated channel, subfamily H (eag-related), member 8 | protein-coding |
| chr5_122949024_122949658 | chr5 | 122949025 | 122949658 | + | -0.605275 | intron (NM_001003953, intron 6 of 22) | intron (NM_001003953, intron 6 of 22) | 39192 | NM_001005866 | 30841 | Mm.86406 | NM_013910 | ENSMUSG00000029475 | **Kdm2b** | Cxxc2\|E430001G17\|Fbl10\|Fbxl10\|Jhdm1b\|PCCX2\|mKIAA3014 | lysine (K)-specific demethylase 2B | protein-coding |
| chr8_72295545_72296217 | chr8 | 72295546 | 72296217 | + | -0.63972 | Intergenic | Intergenic | -23181 | NM_008452 | 16598 | Mm.26938 | NM_008452 | ENSMUSG00000055148 | **Klf2** | Lklf | Kruppel-like factor 2 (lung) | protein-coding |
| chr5_64806144_64806522 | chr5 | 64806145 | 64806522 | + | -0.898129 | intron (NM_008453, intron 1 of 5) | intron (NM_008453, intron 1 of 5) | 2810 | NM_008453 | 16599 | Mm.392759 | NM_008453 | ENSMUSG00000029178 | **Klf3** | 9930027G08Rik\|AL024007\|Bklf\|Tef-2 | Kruppel-like factor 3 (basic) | protein-coding |
| chr5_64806013_64806530 | chr5 | 64806014 | 64806530 | + | -0.918655 | intron (NM_008453, intron 1 of 5) | intron (NM_008453, intron 1 of 5) | 2749 | NM_008453 | 16599 | Mm.392759 | NM_008453 | ENSMUSG00000029178 | **Klf3** | 9930027G08Rik\|AL024007\|Bklf\|Tef-2 | Kruppel-like factor 3 (basic) | protein-coding |
| chr13_5864196_5864577 | chr13 | 5864197 | 5864577 | + | -0.599174 | intron (NM_011803, intron 1 of 3) | intron (NM_011803, intron 1 of 3) | 2898 | NM_011803 | 23849 | Mm.275036 | NM_011803 | ENSMUSG00000000078 | **Klf6** | AI448727\|BCD1\|C86813\|CPBP\|Copeb\|FM2\|FM6\|Ierepo1\|Ierepo3\|R75280\|Zf9 | Kruppel-like factor 6 | protein-coding |
| chr12_69255967_69256325 | chr12 | 69255968 | 69256325 | + | -0.607038 | intron (NM_178253, intron 5 of 12) | intron (NM_178253, intron 5 of 12) | 14314 | NM_178253 | 271005 | Mm.180969 | NM_178253 | | **Klhdc1** | - | kelch domain containing 1 | protein-coding |
| chr3_131170152_131170622 | chr3 | 131170153 | 131170622 | + | -0.705317 | intron (NM_001276402, intron 3 of 9) | RLTR51B_Mm\|LTR\|ERVK? | 57596 | NM_001276403 | 16842 | Mm.255219 | NM_010703 | ENSMUSG00000027985 | **Lef1** | 3000002B05\|AI451430\|Lef-1 | lymphoid enhancer binding factor 1 | protein-coding |
| chr6_31201168_31201921 | chr6 | 31201169 | 31201921 | + | -0.603343 | intron (NR_110469, intron 1 of 3) | Lx8\|LINE\|L1 | 16929 | NR_110470 | 232685 | Mm.443019 | NM_144555 | | **Lincpint** | MNCb-1768\|Pint\|linc-Pint | long intergenic non-protein coding RNA, Trp53 induced transcript | ncRNA |
| chr7_45778961_45779867 | chr7 | 45778962 | 45779867 | + | -0.788727 | Intergenic | L1MA8\|LINE\|L1 | -4533 | NM_001005511 | 381983 | Mm.44928 | NM_001005511 | ENSMUSG00000062044 | **Lmtk3** | BC059845\|aatyk3 | lemur tyrosine kinase 3 | protein-coding |
| chr5_147063215_147063791 | chr5 | 147063216 | 147063791 | + | -0.617529 | intron (NM_080795, intron 1 of 9) | intron (NM_080795, intron 1 of 9) | 13069 | NM_080795 | 140887 | Mm.272203 | NM_080795 | ENSMUSG00000016520 | **Lnx2** | 9630046H24\|AW209022 | ligand of numb-protein X 2 | protein-coding |
| chr8_36194064_36194571 | chr8 | 36194065 | 36194571 | + | -0.636637 | Intergenic | Intergenic | 55198 | NM_001081150 | 244421 | Mm.324032 | NM_001081150 | ENSMUSG00000039633 | **Lonrf1** | - | LON peptidase N-terminal domain and ring finger 1 | protein-coding |
| chr5_105717851_105718475 | chr5 | 105717852 | 105718475 | + | -0.76636 | intron (NM_178701, intron 1 of 2) | intron (NM_178701, intron 1 of 2) | 17353 | NM_178701 | 231549 | Mm.23837 | NM_178701 | ENSMUSG00000046079 | **Lrrc8d** | 2810473G09Rik\|4930525N13Rik\|A930019F03\|Lrrc5 | leucine rich repeat containing 8D | protein-coding |
| chr5_105717796_105719115 | chr5 | 105717797 | 105719115 | + | -0.892767 | intron (NM_178701, intron 1 of 2) | intron (NM_178701, intron 1 of 2) | 17646 | NM_178701 | 231549 | Mm.23837 | NM_178701 | ENSMUSG00000046079 | **Lrrc8d** | 2810473G09Rik\|4930525N13Rik\|A930019F03\|Lrrc5 | leucine rich repeat containing 8D | protein-coding |
| chr5_105718092_105718770 | chr5 | 105718093 | 105718770 | + | -0.959372 | intron (NM_178701, intron 1 of 2) | intron (NM_178701, intron 1 of 2) | 17621 | NM_178701 | 231549 | Mm.23837 | NM_178701 | ENSMUSG00000046079 | **Lrrc8d** | 2810473G09Rik\|4930525N13Rik\|A930019F03\|Lrrc5 | leucine rich repeat containing 8D | protein-coding |
| chr9_13536663_13537278 | chr9 | 13536664 | 13537278 | + | -0.685664 | Intergenic | Intergenic | -83018 | NM_001013813 | 270118 | Mm.116898 | NM_173776 | ENSMUSG00000031925 | **Maml2** | 5930431H10\|BC032967 | mastermind like 2 (Drosophila) | protein-coding |
| chr9_13536663_13537067 | chr9 | 13536664 | 13537067 | + | -0.693213 | Intergenic | Intergenic | -83124 | NM_001013813 | 270118 | Mm.116898 | NM_173776 | ENSMUSG00000031925 | **Maml2** | 5930431H10\|BC032967 | mastermind like 2 (Drosophila) | protein-coding |
| chr13_111807667_111807884 | chr13 | 111807668 | 111807884 | + | -0.659404 | intron (NM_011945, intron 1 of 19) | CpG | 1207 | NM_011945 | 26401 | Mm.15918 | NM_011945 | ENSMUSG00000021754 | **Map3k1** | MAPKKK1\|MEKK1\|Mekk | mitogen-activated protein kinase kinase kinase 1 | protein-coding |
| chr13_111807162_111807954 | chr13 | 111807163 | 111807954 | + | -0.702269 | intron (NM_011945, intron 1 of 19) | intron (NM_011945, intron 1 of 19) | 1425 | NM_011945 | 26401 | Mm.15918 | NM_011945 | ENSMUSG00000021754 | **Map3k1** | MAPKKK1\|MEKK1\|Mekk | mitogen-activated protein kinase kinase kinase 1 | protein-coding |
| chr13_111807453_111807951 | chr13 | 111807454 | 111807951 | + | -0.76461 | intron (NM_011945, intron 1 of 19) | intron (NM_011945, intron 1 of 19) | 1281 | NM_011945 | 26401 | Mm.15918 | NM_011945 | ENSMUSG00000021754 | **Map3k1** | MAPKKK1\|MEKK1\|Mekk | mitogen-activated protein kinase kinase kinase 1 | protein-coding |
| chr17_28697294_28697955 | chr17 | 28697295 | 28697955 | + | -0.599182 | intron (NM_001168514, intron 2 of 12) | intron (NM_001168514, intron 2 of 12) | 5057 | NM_001168513 | 26416 | Mm.311337 | NM_011951 | ENSMUSG00000053436 | **Mapk14** | CSBP2\|Crk1\|Csbp1\|Mxi2\|PRKM14\|PRKM15\|p38\|p38-alpha\|p38MAPK\|p38a\|p38alpha | mitogen-activated protein kinase 14 | protein-coding |
| chr11_49837689_49838208 | chr11 | 49837690 | 49838208 | + | -0.619643 | 3' UTR (NM_013529, exon 19 of 19) | 3' UTR (NM_013529, exon 19 of 19) | -8802 | NM_001163671 | 26420 | Mm.68933 | NM_016961 | ENSMUSG00000020366 | **Mapk9** | AI851083\|JNK2\|Prkm9\|p54aSAPK | mitogen-activated protein kinase 9 | protein-coding |
| chr18_70571230_70571498 | chr18 | 70571231 | 70571498 | + | -0.624582 | intron (NM_010773, intron 1 of 6) | RSINE1\|SINE\|B4 | 3072 | NM_010773 | 17191 | Mm.322 | NM_010773 | ENSMUSG00000024513 | **Mbd2** | MBD2a | methyl-CpG binding domain protein 2 | protein-coding |
| chr3_60501635_60502133 | chr3 | 60501636 | 60502133 | + | -0.631744 | intron (NM_001253709, intron 1 of 8) | intron (NM_001253709, intron 1 of 8) | 632 | NM_001253708 | 56758 | Mm.255723 | NM_020007 | ENSMUSG00000027763 | **Mbnl1** | Mbnl\|mKIAA0428 | muscleblind-like 1 (Drosophila) | protein-coding |
| chr2_144275660_144276139 | chr2 | 144275661 | 144276139 | + | -0.59569 | intron (NM_001289631, intron 2 of 4) | B1_Mur2\|SINE\|Alu | 4996 | NM_001289630 | 74528 | Mm.386817 | NM_028984 | ENSMUSG00000027424 | **Mgme1** | 8430406I07Rik\|AI426476 | mitochondrial genome maintainance exonuclease 1 | protein-coding |
| chr4_101328960_101329600 | chr4 | 101328961 | 101329600 | + | -0.673132 | Intergenic | MTD\|LTR\|ERVL-MaLR | 17747 | NR_029537 | 387143 | | NR_029537 | miRBase:MI0000148 | **Mir101a** | Mirn101\|Mirn101a\|miR-101 | microRNA 101a | ncRNA |
| chr18_56489940_56490223 | chr18 | 56489941 | 56490223 | + | -0.622848 | intron (NM_026240, intron 10 of 13) | ORR1B1\|LTR\|ERVL-MaLR | -48057 | NR_106160 | 102465790 | | NR_106160 | miRBase:MI0025040 | **Mir1258** | mmu-mir-1258 | microRNA mir-1258 | ncRNA |
| chr18_56489797_56490223 | chr18 | 56489798 | 56490223 | + | -0.644004 | intron (NM_026240, intron 10 of 13) | ORR1B1\|LTR\|ERVL-MaLR | -48129 | NR_106160 | 102465790 | | NR_106160 | miRBase:MI0025040 | **Mir1258** | mmu-mir-1258 | microRNA mir-1258 | ncRNA |
| chr18_56504305_56505231 | chr18 | 56504306 | 56505231 | + | -0.722922 | TTS (NM_026240) | TTS (NM_026240) | -33371 | NR_106160 | 102465790 | | NR_106160 | miRBase:MI0025040 | **Mir1258** | mmu-mir-1258 | microRNA mir-1258 | ncRNA |
| chr18_56504283_56504776 | chr18 | 56504284 | 56504776 | + | -0.820091 | TTS (NM_026240) | TTS (NM_026240) | -33609 | NR_106160 | 102465790 | | NR_106160 | miRBase:MI0025040 | **Mir1258** | mmu-mir-1258 | microRNA mir-1258 | ncRNA |

| | | | | | | | | | | | | | | | | | |
|---|---|---|---|---|---|---|---|---|---|---|---|---|---|---|---|---|---|
| chr7_101466101_101467148 | chr7 | 101466102 | 101467148 | + | -0.707226 | intron (NM_001243757, intron 2 of 30) | intron (NM_001243757, intron 2 of 30) | -8751 | NR_029791 | 387157 | | NR_029791 | | **Mir139** | Mirn139\|mmu-mir-139 | microRNA 139 | ncRNA |
| chr18_11931345_11932050 | chr18 | 11931346 | 11932050 | + | -0.732811 | intron (NM_022021, intron 6 of 8) | intron (NM_022021, intron 6 of 8) | -91260 | NR_035447 | 100316686 | | NR_035447 | miRBase:MI0008328 | **Mir1901** | Mirn1901\|mmu-mir-1901 | microRNA 1901 | ncRNA |
| chr15_100181744_100182377 | chr15 | 100181745 | 100182377 | + | -0.725889 | intron (NM_172819, intron 16 of 32) | intron (NM_172819, intron 16 of 32) | 5975 | NR_037311 | 100499528 | | NR_037311 | | **Mir3473a** | Mir3473\|mmu-mir-3473\|mmu-mir-3473a | microRNA 3473a | ncRNA |
| chr15_100182653_100183245 | chr15 | 100182654 | 100183245 | + | -0.775622 | intron (NM_172819, intron 16 of 32) | intron (NM_172819, intron 16 of 32) | 5087 | NR_037311 | 100499528 | | NR_037311 | | **Mir3473a** | Mir3473\|mmu-mir-3473\|mmu-mir-3473a | microRNA 3473a | ncRNA |
| chr15_80930716_80931040 | chr15 | 80930717 | 80931040 | + | -0.602933 | 3' UTR (NM_144812, exon 25 of 25) | 3' UTR (NM_144812, exon 25 of 25) | -9161 | NR_039573 | 100628612 | | NR_039573 | miRBase:MI0018022 | **Mir5113** | mmu-mir-5113\|mu-mir-5113 | microRNA 5113 | ncRNA |
| chr2_166672555_166673015 | chr2 | 166672556 | 166673015 | + | -0.665809 | promoter-TSS (NR_105783) | promoter-TSS (NR_105783) | 67 | NR_105783 | 102465188 | | NR_105783 | | **Mir6364** | mmu-mir-6364 | microRNA mir-6364 | ncRNA |
| chr5_124002395_124002879 | chr5 | 124002396 | 124002879 | + | -0.638019 | 3' UTR (NM_145070, exon 32 of 32) | 3' UTR (NM_145070, exon 32 of 32) | 5785 | NR_105999 | 102465625 | | NR_105999 | | **Mir7032** | mmu-mir-7032 | microRNA mir-7032 | ncRNA |
| chr5_124003124_124003804 | chr5 | 124003125 | 124003804 | + | -0.979271 | TTS (NM_145070) | TTS (NM_145070) | 6612 | NR_105999 | 102465625 | | NR_105999 | | **Mir7032** | mmu-mir-7032 | microRNA mir-7032 | ncRNA |
| chr5_136459874_136460385 | chr5 | 136459875 | 136460385 | + | -0.653042 | intron (NM_001291239, intron 2 of 22) | intron (NM_001291239, intron 2 of 22) | -84327 | NR_030500 | 723975 | | NR_030500 | | **Mir721** | Mirn721\|miRY\|mir-721\|mmu-mir-721 | microRNA 721 | ncRNA |
| chr1_53852284_53852501 | chr1 | 53852285 | 53852501 | + | -0.776795 | intron (NM_001001883, intron 20 of 28) | intron (NM_001001883, intron 20 of 28) | 2969 | NR_106149 | 102465783 | | NR_106149 | miRBase:MI0025029 | **Mir7681** | mmu-mir-7681 | microRNA mir-7681 | ncRNA |
| chr1_53852283_53852630 | chr1 | 53852284 | 53852630 | + | -0.813485 | intron (NM_001001883, intron 20 of 28) | intron (NM_001001883, intron 20 of 28) | 3033 | NR_106149 | 102465783 | | NR_106149 | miRBase:MI0025029 | **Mir7681** | mmu-mir-7681 | microRNA mir-7681 | ncRNA |
| chr1_53850849_53851293 | chr1 | 53850850 | 53851293 | + | -0.899729 | intron (NM_001001883, intron 21 of 28) | intron (NM_001001883, intron 21 of 28) | 1647 | NR_106149 | 102465783 | | NR_106149 | miRBase:MI0025029 | **Mir7681** | mmu-mir-7681 | microRNA mir-7681 | ncRNA |
| chr10_80678002_80678547 | chr10 | 80678003 | 80678547 | + | -0.648004 | Intergenic | Intergenic | -6299 | NM_021462 | 17347 | Mm.42126 | NM_021462 | ENSMUSG00000020190 | **Mknk2** | 2010016G11Rik\|Gprk7\|Mnk2 | MAP kinase-interacting serine/threonine kinase 2 | protein-coding |
| chr1_78538462_78538696 | chr1 | 78538463 | 78538696 | + | -0.68357 | TTS (NM_026713) | TTS (NM_026713) | 27519 | NM_026713 | 68393 | Mm.41325 | NM_026713 | ENSMUSG00000012187 | **Mogat1** | 0610030A14Rik\|1110064N14Rik\|Dgat2l\|Dgat2l1\|MGAT1\|mDC2 | monoacylglycerol O-acyltransferase 1 | protein-coding |
| chr1_78538279_78538786 | chr1 | 78538280 | 78538786 | + | -0.716835 | TTS (NM_026713) | TTS (NM_026713) | 27473 | NM_026713 | 68393 | Mm.41325 | NM_026713 | ENSMUSG00000012187 | **Mogat1** | 0610030A14Rik\|1110064N14Rik\|Dgat2l\|Dgat2l1\|MGAT1\|mDC2 | monoacylglycerol O-acyltransferase 1 | protein-coding |
| chr10_120895714_120896348 | chr10 | 120895715 | 120896348 | + | -0.705093 | intron (NM_177092, intron 1 of 6) | intron (NM_177092, intron 1 of 6) | 2940 | NM_177092 | 320183 | Mm.169755 | NM_177092 | ENSMUSG00000051236 | **Msrb3** | D430026P16Rik\|I-8-23\|MsrB2 | methionine sulfoxide reductase B3 | protein-coding |
| chr15_12229607_12229878 | chr15 | 12229608 | 12229878 | + | -0.612175 | intron (NM_172958, intron 1 of 15) | B4A\|SINE\|B4 | 24649 | NM_172958 | 268783 | Mm.54460 | NM_172958 | ENSMUSG00000039458 | **Mtmr12** | 3Pap\|4932703C11\|C730015A02Rik\|Pip3ap\|mKIAA1682 | myotubularin related protein 12 | protein-coding |
| chr15_12229597_12230145 | chr15 | 12229598 | 12230145 | + | -0.666342 | intron (NM_172958, intron 1 of 15) | ID\|SINE\|ID | 24777 | NM_172958 | 268783 | Mm.54460 | NM_172958 | ENSMUSG00000039458 | **Mtmr12** | 3Pap\|4932703C11\|C730015A02Rik\|Pip3ap\|mKIAA1682 | myotubularin related protein 12 | protein-coding |
| chr1_51929491_51930119 | chr1 | 51929492 | 51930119 | + | -0.745108 | Intergenic | Intergenic | -13827 | NM_001161817 | 17912 | Mm.3390 | NM_010863 | ENSMUSG00000018417 | **Myo1b** | AA406997\|MIH-L\|MMIa | myosin IB | protein-coding |
| chr5_134269556_134270311 | chr5 | 134269557 | 134270311 | + | -0.604742 | intron (NM_001080749, intron 10 of 32) | intron (NM_001080749, intron 10 of 32) | -40309 | NR_104383 | 17969 | Mm.425296 | NM_010876 | ENSMUSG00000015950 | **Ncf1** | NCF-47K\|NOXO2\|Ncf-1\|p47-phox\|p47<phox>\|p47phox | neutrophil cytosolic factor 1 | protein-coding |
| chr8_85623259_85623571 | chr8 | 85623260 | 85623571 | + | -0.643346 | Intergenic | Intergenic | 67594 | NM_001081324 | 74513 | Mm.126079 | NM_001081324 | ENSMUSG00000036902 | **Neto2** | 5530601C23Rik\|AI854825\|BTCL2 | neuropilin (NRP) and tolloid (TLL)-like 2 | protein-coding |
| chr8_85623174_85623788 | chr8 | 85623175 | 85623788 | + | -0.711063 | Intergenic | Intergenic | 67528 | NM_001081324 | 74513 | Mm.126079 | NM_001081324 | ENSMUSG00000036902 | **Neto2** | 5530601C23Rik\|AI854825\|BTCL2 | neuropilin (NRP) and tolloid (TLL)-like 2 | protein-coding |
| chr17_26370397_26371291 | chr17 | 26370398 | 26371291 | + | -0.721553 | Intergenic | Intergenic | -44121 | NM_001081656 | 240055 | Mm.133211 | NM_001081656 | ENSMUSG00000034413 | **Neurl1b** | C230078M08Rik\|EG240055 | neuralized homolog 1b (Drosophila) | protein-coding |
| chr18_80706073_80706363 | chr18 | 80706074 | 80706363 | + | -0.627279 | intron (NM_198429, intron 1 of 9) | intron (NM_198429, intron 1 of 9) | 1955 | NM_001164111 | 18018 | Mm.329560 | NM_016791 | ENSMUSG00000033016 | **Nfatc1** | 2210017P03Rik\|AI449492\|AV076380\|NF-ATc\|NFAT2\|NFATc\|Nfatcb | nuclear factor of activated T cells, cytoplasmic, calcineurin dependent 1 | protein-coding |
| chr6_120082215_120082857 | chr6 | 120082216 | 120082857 | + | -0.591238 | Intergenic | Intergenic | -10844 | NM_016718 | 29862 | Mm.103668 | NM_016718 | ENSMUSG00000041377 | **Ninj2** | - | ninjurin 2 | protein-coding |
| chr6_120082552_120082837 | chr6 | 120082553 | 120082837 | + | -0.65482 | Intergenic | Intergenic | -10685 | NM_016718 | 29862 | Mm.103668 | NM_016718 | ENSMUSG00000041377 | **Ninj2** | - | ninjurin 2 | protein-coding |
| chr15_8569688_8570104 | chr15 | 8569689 | 8570104 | + | -0.749503 | Intergenic | Intergenic | -125433 | NM_201232 | 71175 | Mm.240329 | NM_027707 | ENSMUSG00000022141 | **Nipbl** | Idn3 | Nipped-B homolog (Drosophila) | protein-coding |
| chr16_76345951_76346481 | chr16 | 76345952 | 76346481 | + | -0.608376 | intron (NM_173440, intron 2 of 3) | intron (NM_173440, intron 2 of 3) | 26833 | NM_173440 | 268903 | Mm.74711 | NM_173440 | ENSMUSG00000048490 | **Nrip1** | 6030458L20Rik\|8430438I05Rik\|9630050P12\|AA959574\|AW456757\|RIP140 | nuclear receptor interacting protein 1 | protein-coding |
| chr7_125507192_125508055 | chr7 | 125507193 | 125508055 | + | -0.599507 | Intergenic | Intergenic | -16082 | NM_026330 | 67711 | Mm.4467 | NM_026330 | ENSMUSG00000030750 | **Nsmce1** | 2510027N19Rik | non-SMC element 1 homolog (S. cerevisiae) | protein-coding |
| chr10_127135239_127135502 | chr10 | 127135240 | 127135502 | + | -0.639827 | Intergenic | B4A\|SINE\|B4 | -14211 | NM_001171026 | 216440 | Mm.295246 | NM_177614 | ENSMUSG00000040462 | **Os9** | 4632413K17Rik\|AU022351 | amplified in osteosarcoma | protein-coding |
| chr10_127135106_127135507 | chr10 | 127135107 | 127135507 | + | -0.698912 | Intergenic | B4A\|SINE\|B4 | -14147 | NM_001171026 | 216440 | Mm.295246 | NM_177614 | ENSMUSG00000040462 | **Os9** | 4632413K17Rik\|AU022351 | amplified in osteosarcoma | protein-coding |
| chr15_27605243_27605725 | chr15 | 27605244 | 27605725 | + | -0.624043 | TTS (NM_001013792) | TTS (NM_001013792) | 25209 | NM_001013792 | 432940 | Mm.309164 | NM_001013792 | ENSMUSG00000046034 | **Otulin** | C79097\|Fam105b\|m3Sapc\|m7-1Sapc | OTU deubiquitinase with linear linkage specificity | protein-coding |
| chr5_122718879_122719648 | chr5 | 122718880 | 122719648 | + | -0.6393 | intron (NM_011026, intron 5 of 11) | intron (NM_011026, intron 5 of 11) | 11707 | NM_011026 | 18438 | Mm.290884 | NM_011026 | ENSMUSG00000029470 | **P2rx4** | AI504491\|AW555605\|D5Ertd444e\|P2X4 | purinergic receptor P2X, ligand-gated ion channel 4 | protein-coding |
| chr5_122719103_122719577 | chr5 | 122719104 | 122719577 | + | -0.642566 | intron (NM_011026, intron 5 of 11) | intron (NM_011026, intron 5 of 11) | 11783 | NM_011026 | 18438 | Mm.290884 | NM_011026 | ENSMUSG00000029470 | **P2rx4** | AI504491\|AW555605\|D5Ertd444e\|P2X4 | purinergic receptor P2X, ligand-gated ion channel 4 | protein-coding |
| chr9_21164901_21165542 | chr9 | 21164902 | 21165542 | + | -0.65503 | promoter-TSS (NM_183408) | promoter-TSS (NM_183408) | -492 | NM_183408 | 18577 | Mm.191749 | NM_019798 | ENSMUSG00000032177 | **Pde4a** | D9Ertd60e\|Dpde2 | phosphodiesterase 4A, cAMP specific | protein-coding |
| chr9_21164912_21165362 | chr9 | 21164913 | 21165362 | + | -0.661003 | promoter-TSS (NM_183408) | promoter-TSS (NM_183408) | -577 | NM_183408 | 18577 | Mm.191749 | NM_019798 | ENSMUSG00000032177 | **Pde4a** | D9Ertd60e\|Dpde2 | phosphodiesterase 4A, cAMP specific | protein-coding |
| chr11_69097039_69097338 | chr11 | 69097040 | 69097338 | + | -0.810334 | Intergenic | Intergenic | -1767 | NM_011065 | 18626 | Mm.7373 | NM_011065 | ENSMUSG00000020893 | **Per1** | Per\|m-rigui\|mPer1 | period circadian clock 1 | protein-coding |
| chr2_18984007_18984373 | chr2 | 18984008 | 18984373 | + | -0.754149 | intron (NM_008845, intron 1 of 9) | intron (NM_008845, intron 1 of 9) | 13931 | NM_008845 | 18718 | Mm.313977 | NM_008845 | ENSMUSG00000026737 | **Pip4k2a** | AW742916\|Pip5k2a | phosphatidylinositol-5-phosphate 4-kinase, type II, alpha | protein-coding |
| chr16_46001586_46002133 | chr16 | 46001587 | 46002133 | + | -0.960635 | intron (NM_001134480, intron 1 of 3) | intron (NM_001134480, intron 1 of 3) | 8553 | NM_001134480 | 433022 | Mm.380993 | NM_001134480 | ENSMUSG00000087141 | **Plcxd2** | AW494639\|EG433022 | phosphatidylinositol-specific phospholipase C, X domain containing 2 | protein-coding |
| chr11_106778955_106780258 | chr11 | 106778956 | 106780258 | + | -0.602651 | promoter-TSS (NM_015810) | promoter-TSS (NM_015810) | -70 | NM_015810 | 50776 | Mm.859 | NM_015810 | ENSMUSG00000020718 | **Polg2** | - | polymerase (DNA directed), gamma 2, accessory subunit | protein-coding |
| chr11_106779956_106780218 | chr11 | 106779957 | 106780218 | + | -0.684412 | promoter-TSS (NM_015810) | promoter-TSS (NM_015810) | -550 | NM_015810 | 50776 | Mm.859 | NM_015810 | ENSMUSG00000020718 | **Polg2** | - | polymerase (DNA directed), gamma 2, accessory subunit | protein-coding |
| chr5_122163142_122163640 | chr5 | 122163143 | 122163640 | + | -0.96456 | intron (NM_013636, intron 1 of 6) | intron (NM_013636, intron 1 of 6) | 5112 | NM_013636 | 19047 | Mm.280784 | NM_013636 | ENSMUSG00000004455 | **Ppp1cc** | PP-1G\|PP1\|dis2m1 | protein phosphatase 1, catalytic subunit, gamma isoform | protein-coding |
| chr1_191425839_191426311 | chr1 | 191425840 | 191426311 | + | -0.81229 | Intergenic | Intergenic | -29034 | NM_144880 | 226849 | Mm.205569 | NM_144880 | ENSMUSG00000026626 | **Ppp2r5a** | PR61alpha | protein phosphatase 2, regulatory subunit B', alpha | protein-coding |
| chr1_191425930_191426225 | chr1 | 191425931 | 191426225 | + | -0.859041 | Intergenic | Intergenic | -29037 | NM_144880 | 226849 | Mm.205569 | NM_144880 | ENSMUSG00000026626 | **Ppp2r5a** | PR61alpha | protein phosphatase 2, regulatory subunit B', alpha | protein-coding |
| chr15_93712466_93713017 | chr15 | 93712467 | 93713017 | + | -0.701573 | Intergenic | MIR3\|SINE\|MIR | -116851 | NM_001033217 | 106042 | Mm.150314 | NM_001033217 | ENSMUSG00000036158 | **Prickle1** | 1110058P22Rik\|AW215793\|Prickle\|b2b019Clo\|mpk1 | prickle homolog 1 (Drosophila) | protein-coding |
| chr11_108273579_108274081 | chr11 | 108273580 | 108274081 | + | -0.623429 | intron (NM_011101, intron 2 of 16) | intron (NM_011101, intron 2 of 16) | 70058 | NM_011101 | 18750 | Mm.222178 | NM_011101 | ENSMUSG00000050965 | **Prkca** | AI875142\|Pkca | protein kinase C, alpha | protein-coding |
| chr12_73603612_73604189 | chr12 | 73603613 | 73604189 | + | -0.764329 | intron (NM_008856, intron 1 of 13) | intron (NM_008856, intron 1 of 13) | 18860 | NM_008856 | 18755 | Mm.341677 | NM_008856 | ENSMUSG00000021108 | **Prkch** | Pkch | protein kinase C, eta | protein-coding |
| chr13_28417358_28418588 | chr13 | 28417359 | 28418588 | + | -0.591309 | Intergenic | (TC)n\|Simple_repeat\|Simple_repeat | 275489 | NM_023746 | 28078 | Mm.155058 | NM_023746 | ENSMUSG00000017064 | **Prl5a1** | 1600013P04Rik\|D13Wsu14e\|PLP-L\|Prlpl | prolactin family 5, subfamily a, member 1 | protein-coding |
| chr13_28417363_28418009 | chr13 | 28417364 | 28418009 | + | -0.742444 | Intergenic | Intergenic | 275202 | NM_023746 | 28078 | Mm.155058 | NM_023746 | ENSMUSG00000017064 | **Prl5a1** | 1600013P04Rik\|D13Wsu14e\|PLP-L\|Prlpl | prolactin family 5, subfamily a, member 1 | protein-coding |
| chr2_32167829_32168216 | chr2 | 32167830 | 32168216 | + | -0.686414 | intron (NM_001159634, intron 1 of 31) | intron (NM_001159634, intron 1 of 31) | 16875 | NM_001159634 | 227723 | Mm.25504 | NM_172661 | ENSMUSG00000039262 | **Prrc2b** | 5830434P21Rik\|AI173903\|Bat2l\|Bat2l1\|D430039P21\|mKIAA0515 | proline-rich coiled-coil 2B | protein-coding |
| chr1_34484376_34484821 | chr1 | 34484377 | 34484821 | + | -0.639954 | Intergenic | MTE2b\|LTR\|ERVL-MaLR | -13831 | NM_009355 | 21755 | Mm.10725 | NM_009355 | ENSMUSG00000026125 | **Prss39** | Tesp1 | protease, serine 39 | protein-coding |
| chr4_129793603_129794339 | chr4 | 129793604 | 129794339 | + | -0.861608 | Intergenic | Intergenic | -26508 | NM_008974 | 19244 | Mm.193688 | NM_008974 | ENSMUSG00000028788 | **Ptp4a2** | Prl-2 | protein tyrosine phosphatase 4a2 | protein-coding |
| chr4_129793881_129794276 | chr4 | 129793882 | 129794276 | + | -0.862479 | Intergenic | Intergenic | -26400 | NM_008974 | 19244 | Mm.193688 | NM_008974 | ENSMUSG00000028788 | **Ptp4a2** | Prl-2 | protein tyrosine phosphatase 4a2 | protein-coding |

| | | | | | | | | | | | | | | | | | |
|---|---|---|---|---|---|---|---|---|---|---|---|---|---|---|---|---|---|
| chr11_98959433_98960017 | chr11 | 98959434 | 98960017 | + | -0.92456 | promoter-TSS (NM_001176528) | promoter-TSS (NM_001176528) | -746 | NM_001176528 | 19401 | Mm.439744 | NM_009024 | ENSMUSG00000037992 | **Rara** | Nr1b1\|RAR\|RARalpha1 | retinoic acid receptor, alpha | protein-coding |
| chr11_98959532_98959813 | chr11 | 98959533 | 98959813 | + | -0.926385 | promoter-TSS (NM_001176528) | promoter-TSS (NM_001176528) | -798 | NM_001176528 | 19401 | Mm.439744 | NM_009024 | ENSMUSG00000037992 | **Rara** | Nr1b1\|RAR\|RARalpha1 | retinoic acid receptor, alpha | protein-coding |
| chr19_6400327_6400732 | chr19 | 6400328 | 6400732 | + | -0.853454 | promoter-TSS (NM_011242) | promoter-TSS (NM_011242) | -53 | NM_011242 | 19395 | Mm.77017 | NM_011242 | ENSMUSG00000032946 | **Rasgrp2** | CDC25L\|Caldaggef1 | RAS, guanyl releasing protein 2 | protein-coding |
| chr19_6400392_6400679 | chr19 | 6400393 | 6400679 | + | -0.857973 | promoter-TSS (NM_011242) | promoter-TSS (NM_011242) | -47 | NM_011242 | 19395 | Mm.77017 | NM_011242 | ENSMUSG00000032946 | **Rasgrp2** | CDC25L\|Caldaggef1 | RAS, guanyl releasing protein 2 | protein-coding |
| chr7_128373861_128374645 | chr7 | 128373862 | 128374645 | + | -0.605676 | intron (NM_026418, intron 4 of 4) | RSINE1\|SINE\|B4 | 43919 | NM_026418 | 67865 | Mm.18635 | NM_026418 | ENSMUSG00000030844 | **Rgs10** | 2310010N19Rik | regulator of G-protein signalling 10 | protein-coding |
| chr7_128376099_128376449 | chr7 | 128376100 | 128376449 | + | -0.741231 | intron (NM_026418, intron 4 of 4) | intron (NM_026418, intron 4 of 4) | 41898 | NM_026418 | 67865 | Mm.18635 | NM_026418 | ENSMUSG00000030844 | **Rgs10** | 2310010N19Rik | regulator of G-protein signalling 10 | protein-coding |
| chr7_128374131_128374534 | chr7 | 128374132 | 128374534 | + | -0.742853 | intron (NM_026418, intron 4 of 4) | RSINE1\|SINE\|B4 | 43839 | NM_026418 | 67865 | Mm.18635 | NM_026418 | ENSMUSG00000030844 | **Rgs10** | 2310010N19Rik | regulator of G-protein signalling 10 | protein-coding |
| chr7_128376524_128376832 | chr7 | 128376525 | 128376832 | + | -0.754328 | intron (NM_026418, intron 4 of 4) | RSINE1\|SINE\|B4 | 41494 | NM_026418 | 67865 | Mm.18635 | NM_026418 | ENSMUSG00000030844 | **Rgs10** | 2310010N19Rik | regulator of G-protein signalling 10 | protein-coding |
| chr7_128376082_128376853 | chr7 | 128376083 | 128376853 | + | -0.874099 | intron (NM_026418, intron 4 of 4) | intron (NM_026418, intron 4 of 4) | 41704 | NM_026418 | 67865 | Mm.18635 | NM_026418 | ENSMUSG00000030844 | **Rgs10** | 2310010N19Rik | regulator of G-protein signalling 10 | protein-coding |
| chr13_38037844_38038300 | chr13 | 38037845 | 38038300 | + | -0.913851 | intron (NM_024242, intron 1 of 16) | intron (NM_024242, intron 1 of 16) | 1083 | NM_024242 | 71340 | Mm.297130 | NM_024242 | ENSMUSG00000021428 | **Riok1** | 3110046C13Rik\|5430416A05Rik\|Ad034 | RIO kinase 1 (yeast) | protein-coding |
| chr18_20962647_20962856 | chr18 | 20962648 | 20962856 | + | -0.725153 | intron (NM_026301, intron 1 of 5) | intron (NM_026301, intron 1 of 5) | 18127 | NM_026301 | 67664 | Mm.45980 | NM_026301 | ENSMUSG00000033107 | **Rnf125** | 4930553F04Rik | ring finger protein 125 | protein-coding |
| chr18_20962629_20963056 | chr18 | 20962630 | 20963056 | + | -0.834728 | intron (NM_026301, intron 1 of 5) | intron (NM_026301, intron 1 of 5) | 18218 | NM_026301 | 67664 | Mm.45980 | NM_026301 | ENSMUSG00000033107 | **Rnf125** | 4930553F04Rik | ring finger protein 125 | protein-coding |
| chr6_70813932_70814833 | chr6 | 70813933 | 70814833 | + | -0.706599 | Intergenic | B4A\|SINE\|B4 | -22208 | NM_009075 | 19895 | Mm.17905 | NM_009075 | ENSMUSG00000053604 | **Rpia** | RPI | ribose 5-phosphate isomerase A | protein-coding |
| chr6_70813932_70814413 | chr6 | 70813933 | 70814413 | + | -0.76051 | Intergenic | Intergenic | -21998 | NM_009075 | 19895 | Mm.17905 | NM_009075 | ENSMUSG00000053604 | **Rpia** | RPI | ribose 5-phosphate isomerase A | protein-coding |
| chr7_114069019_114069665 | chr7 | 114069020 | 114069665 | + | -0.816693 | intron (NM_025846, intron 1 of 5) | intron (NM_025846, intron 1 of 5) | 48439 | NM_025846 | 66922 | Mm.276572 | NM_025846 | ENSMUSG00000055723 | **Rras2** | 2610016H24Rik\|C86394\|TC21 | related RAS viral (r-ras) oncogene homolog 2 | protein-coding |
| chr7_114093540_114094123 | chr7 | 114093541 | 114094123 | + | -0.95309 | intron (NM_025846, intron 1 of 5) | intron (NM_025846, intron 1 of 5) | 23949 | NM_025846 | 66922 | Mm.276572 | NM_025846 | ENSMUSG00000055723 | **Rras2** | 2610016H24Rik\|C86394\|TC21 | related RAS viral (r-ras) oncogene homolog 2 | protein-coding |
| chr7_114093619_114093886 | chr7 | 114093620 | 114093886 | + | -1.04692 | intron (NM_025846, intron 1 of 5) | (CA)n\|Simple_repeat\|Simple_repeat | 24028 | NM_025846 | 66922 | Mm.276572 | NM_025846 | ENSMUSG00000055723 | **Rras2** | 2610016H24Rik\|C86394\|TC21 | related RAS viral (r-ras) oncogene homolog 2 | protein-coding |
| chr10_68081671_68082192 | chr10 | 68081672 | 68082192 | + | -0.594332 | Intergenic | Intergenic | 102334 | NM_001081346 | 170799 | Mm.209844 | NM_133244 | ENSMUSG00000037846 | **Rtkn2** | B130039D23Rik\|Mbf\|Plekhk1 | rhotekin 2 | protein-coding |
| chr3_115793316_115794039 | chr3 | 115793317 | 115794039 | + | -0.650075 | Intergenic | Intergenic | -78623 | NM_007901 | 13609 | Mm.982 | NM_007901 | ENSMUSG00000045092 | **S1pr1** | AI849002\|Edg1\|Lpb1\|S1p\|S1p1 | sphingosine-1-phosphate receptor 1 | protein-coding |
| chr19_44282417_44283497 | chr19 | 44282418 | 44283497 | + | -0.667031 | Intergenic | ID4\|SINE\|ID | -10719 | NM_009128 | 20250 | Mm.487021 | NM_009128 | ENSMUSG00000025203 | **Scd2** | Scd-2\|swty | stearoyl-Coenzyme A desaturase 2 | protein-coding |
| chr19_44283080_44283398 | chr19 | 44283081 | 44283398 | + | -0.800072 | Intergenic | Intergenic | -10437 | NM_009128 | 20250 | Mm.487021 | NM_009128 | ENSMUSG00000025203 | **Scd2** | Scd-2\|swty | stearoyl-Coenzyme A desaturase 2 | protein-coding |
| chr10_42838768_42839441 | chr10 | 42838769 | 42839441 | + | -0.613083 | Intergenic | L1MB3\|LINE\|L1 | -21407 | NM_172938 | 268297 | Mm.98731 | NM_172938 | ENSMUSG00000044770 | **Scml4** | 9330161D17\|mFLJ00197 | sex comb on midleg-like 4 (Drosophila) | protein-coding |
| chr3_58552051_58552413 | chr3 | 58552052 | 58552413 | + | -0.70092 | intron (NM_001005509, intron 10 of 13) | intron (NM_001005509, intron 10 of 13) | -24426 | NM_001040396 | 69227 | Mm.270950 | NM_001040396 | ENSMUSG00000075700 | **Selt** | 2810407C02Rik\|5730408P04Rik | selenoprotein T | protein-coding |
| chr19_32182104_32182466 | chr19 | 32182105 | 32182466 | + | -0.867028 | intron (NM_144792, intron 5 of 9) | intron (NM_144792, intron 5 of 9) | 28728 | NM_001168526 | 208449 | Mm.329810 | NM_144792 | ENSMUSG00000040451 | **Sgms1** | 9530058O11Rik\|AI841905\|C80702\|Mob\|Sms1\|Sor1\|Tmem23 | sphingomyelin synthase 1 | protein-coding |
| chr7_126470094_126470563 | chr7 | 126470095 | 126470563 | + | -0.633367 | intron (NR_110346, intron 4 of 9) | intron (NR_110346, intron 4 of 9) | 2765 | NR_110346 | 20399 | Mm.8538 | NM_011363 | ENSMUSG00000030733 | **Sh2b1** | AI425885\|C530001K22Rik\|Irip\|Psm\|SH2-B\|SH2-Bb\|Sh2bpsm1\|mKIAA1299 | SH2B adaptor protein 1 | protein-coding |
| chr5_115488758_115489306 | chr5 | 115488759 | 115489306 | + | -0.831129 | Intergenic | Intergenic | -4548 | NM_133760 | 75387 | Mm.332616 | NM_133760 | ENSMUSG00000029524 | **Sirt4** | 4930596O17Rik | sirtuin 4 | protein-coding |
| chr4_155168612_155169528 | chr4 | 155168613 | 155169528 | + | -0.622023 | intron (NM_011385, intron 1 of 6) | intron (NM_011385, intron 1 of 6) | 53465 | NM_011385 | 20481 | Mm.28520 | NM_011385 | ENSMUSG00000029050 | **Ski** | 2310012I02Rik\|2610001A11Rik\|AA062172\|AA589460\|BC004088 | ski sarcoma viral oncogene homolog (avian) | protein-coding |
| chr4_155169046_155169577 | chr4 | 155169047 | 155169577 | + | -0.732629 | intron (NM_011385, intron 1 of 6) | intron (NM_011385, intron 1 of 6) | 53223 | NM_011385 | 20481 | Mm.28520 | NM_011385 | ENSMUSG00000029050 | **Ski** | 2310012I02Rik\|2610001A11Rik\|AA062172\|AA589460\|BC004088 | ski sarcoma viral oncogene homolog (avian) | protein-coding |
| chr4_155169048_155169438 | chr4 | 155169049 | 155169438 | + | -0.739567 | intron (NM_011385, intron 1 of 6) | intron (NM_011385, intron 1 of 6) | 53292 | NM_011385 | 20481 | Mm.28520 | NM_011385 | ENSMUSG00000029050 | **Ski** | 2310012I02Rik\|2610001A11Rik\|AA062172\|AA589460\|BC004088 | ski sarcoma viral oncogene homolog (avian) | protein-coding |
| chr15_66803794_66804260 | chr15 | 66803795 | 66804260 | + | -0.676625 | intron (NM_001029841, intron 1 of 7) | intron (NM_001029841, intron 1 of 7) | 27802 | NM_001029841 | 20491 | Mm.7601 | NM_009192 | ENSMUSG00000022372 | **Sla** | Slap\|Slap-1 | src-like adaptor | protein-coding |
| chr6_34176096_34176379 | chr6 | 34176097 | 34176379 | + | -0.601726 | intron (NM_021435, intron 1 of 9) | intron (NM_021435, intron 1 of 9) | 816 | NM_021435 | 58246 | Mm.245527 | NM_021435 | | **Slc35b4** | 4930474D06Rik\|AB041549\|AW494345 | solute carrier family 35, member B4 | protein-coding |
| chr15_96638108_96638665 | chr15 | 96638109 | 96638665 | + | -0.604225 | intron (NM_001166458, intron 1 of 15) | intron (NM_001166458, intron 1 of 15) | 3575 | NM_134086 | 105727 | Mm.103568 | NM_134086 | ENSMUSG00000023169 | **Slc38a1** | AA408026\|AA409865\|AL022800\|AU015942\|NAT2 | solute carrier family 38, member 1 | protein-coding |
| chr19_8712799_8713197 | chr19 | 8712800 | 8713197 | + | -1.08662 | intron (NM_008577, intron 1 of 8) | intron (NM_008577, intron 1 of 8) | 899 | NM_008577 | 17254 | Mm.4114 | NM_008577 | ENSMUSG00000010095 | **Slc3a2** | 4F2\|4F2HC\|AI314110\|Cd98\|Ly-10\|Ly-m10\|Ly10\|Mdu1\|Mgp-2hc\|NACAE | solute carrier family 3 (activators of dibasic and neutral amino acid transport), member 2 | protein-coding |
| chrX_101147485_101147962 | chrX | 101147486 | 101147962 | + | -0.84308 | intron (NM_001110311, intron 4 of 5) | PB1\|SINE\|Alu | -62372 | NM_007515 | 11989 | Mm.5140 | NM_007515 | ENSMUSG00000031297 | **Slc7a3** | Atrc3\|CAT3\|SLC7A1\|SLC7A2 | solute carrier family 7 (cationic amino acid transporter, y+ system), member 3 | protein-coding |
| chr11_99203842_99204380 | chr11 | 99203843 | 99204380 | + | -0.891868 | Intergenic | Intergenic | 26906 | NM_020618 | 57376 | Mm.379086 | NM_020618 | ENSMUSG00000037935 | **Smarce1** | 2810417B20Rik\|5830412H02Rik\|9030408N19Rik\|Baf27 | SWI/SNF related, matrix associated, actin dependent regulator of chromatin, subfamily e, member 1 | protein-coding |
| chr11_99203585_99204395 | chr11 | 99203586 | 99204395 | + | -0.904371 | Intergenic | Intergenic | 27027 | NM_020618 | 57376 | Mm.379086 | NM_020618 | ENSMUSG00000037935 | **Smarce1** | 2810417B20Rik\|5830412H02Rik\|9030408N19Rik\|Baf27 | SWI/SNF related, matrix associated, actin dependent regulator of chromatin, subfamily e, member 1 | protein-coding |
| chr19_53420953_53421631 | chr19 | 53420954 | 53421631 | + | -0.655632 | Intergenic | Intergenic | -30719 | NM_172429 | 76479 | Mm.313687 | NM_172429 | ENSMUSG00000025024 | **Smndc1** | 2410004J23Rik\|4933440I19Rik | survival motor neuron domain containing 1 | protein-coding |
| chr19_53423285_53424050 | chr19 | 53423286 | 53424050 | + | -0.673111 | Intergenic | Intergenic | -33095 | NM_172429 | 76479 | Mm.313687 | NM_172429 | ENSMUSG00000025024 | **Smndc1** | 2410004J23Rik\|4933440I19Rik | survival motor neuron domain containing 1 | protein-coding |
| chr19_53423785_53423998 | chr19 | 53423786 | 53423998 | + | -0.730596 | Intergenic | Intergenic | -33319 | NM_172429 | 76479 | Mm.313687 | NM_172429 | ENSMUSG00000025024 | **Smndc1** | 2410004J23Rik\|4933440I19Rik | survival motor neuron domain containing 1 | protein-coding |
| chr2_131509652_131510440 | chr2 | 131509653 | 131510440 | + | -0.633135 | intron (NM_001177840, intron 1 of 3) | intron (NM_001177840, intron 1 of 3) | 18184 | NM_001177838 | 228608 | Mm.136586 | NM_145533 | ENSMUSG00000027333 | **Smox** | B130066H01Rik\|PAO\|PAOh1\|SMO | spermine oxidase | protein-coding |
| chr11_106535914_106536654 | chr11 | 106535915 | 106536654 | + | -0.7666 | intron (NM_198292, intron 5 of 11) | intron (NM_198292, intron 5 of 11) | 35293 | NR_030703 | 100216537 | | NR_030703 | ENSMUSG00000065126 | **Snord104** | - | small nucleolar RNA, C/D box 104 | snoRNA |
| chr13_113574184_113574790 | chr13 | 113574185 | 113574790 | + | -0.747291 | Intergenic | Intergenic | 44077 | NM_130796 | 170625 | Mm.33721 | NM_130796 | ENSMUSG00000042364 | **Snx18** | Snag1 | sorting nexin 18 | protein-coding |
| chr16_90221557_90222012 | chr16 | 90221558 | 90222012 | + | -0.760998 | intron (NM_011434, intron 1 of 4) | intron (NM_011434, intron 1 of 4) | 1043 | NM_011434 | 20655 | Mm.276325 | NM_011434 | ENSMUSG00000022982 | **Sod1** | B430204E11Rik\|Cu/Zn-SOD\|CuZnSOD\|Ipo-1\|Ipo1\|SODC\|Sod-1 | superoxide dismutase 1, soluble | protein-coding |
| chr7_45742219_45742834 | chr7 | 45742220 | 45742834 | + | -0.716893 | intron (NM_017465, intron 1 of 6) | intron (NM_017465, intron 1 of 6) | -16711 | NM_027055 | 69363 | Mm.45607 | NM_027055 | ENSMUSG00000070563 | **Spaca4** | 1700008E09Rik\|AV043694\|Samp14 | sperm acrosome associated 4 | protein-coding |
| chr11_93995051_93995518 | chr11 | 93995052 | 93995518 | + | -0.794039 | promoter-TSS (NM_001199203) | promoter-TSS (NM_001199203) | -806 | NM_001199203 | 70834 | Mm.260737 | NM_027569 | ENSMUSG00000020859 | **Spag9** | 3110018C07Rik\|4733401I23Rik\|4831406C20Rik\|AW552012\|JLP\|JSAP2\|JSAP2a\|Jip4\|Mapk8ip4\|syd1 | sperm associated antigen 9 | protein-coding |
| chr7_47022280_47022954 | chr7 | 47022281 | 47022954 | + | -0.65418 | Intergenic | URR1B\|DNA\|hAT-Charlie | -14203 | NM_175318 | 101685 | Mm.155687 | NM_175318 | ENSMUSG00000049516 | **Spty2d1** | 5830435K17Rik\|AI852426\|P16H6 | SPT2, Suppressor of Ty, domain containing 1 (S. cerevisiae) | protein-coding |
| chr18_14656710_14657755 | chr18 | 14656711 | 14657755 | + | -0.714973 | intron (NM_001161369, intron 3 of 9) | intron (NM_001161369, intron 3 of 9) | 25681 | NM_001161369 | 268996 | Mm.31761 | NM_009280 | ENSMUSG00000037013 | **Ss18** | D130059H17\|Ssxt\|Syt | synovial sarcoma translocation, Chromosome 18 | protein-coding |
| chr18_14656705_14657281 | chr18 | 14656706 | 14657281 | + | -0.826731 | intron (NM_001161369, intron 3 of 9) | intron (NM_001161369, intron 3 of 9) | 25921 | NM_001161369 | 268996 | Mm.31761 | NM_009280 | ENSMUSG00000037013 | **Ss18** | D130059H17\|Ssxt\|Syt | synovial sarcoma translocation, Chromosome 18 | protein-coding |
| chr6_145992531_145993564 | chr6 | 145992532 | 145993564 | + | -0.708956 | Intergenic | Intergenic | 58901 | NM_010656 | 16651 | Mm.49689 | NM_010656 | ENSMUSG00000030255 | **Sspn** | Krag | sarcospan | protein-coding |
| chr6_145992806_145993553 | chr6 | 145992807 | 145993553 | + | -0.780612 | Intergenic | Intergenic | 59033 | NM_010656 | 16651 | Mm.49689 | NM_010656 | ENSMUSG00000030255 | **Sspn** | Krag | sarcospan | protein-coding |
| chr6_142915336_142915820 | chr6 | 142915337 | 142915820 | + | -0.599052 | intron (NM_011374, intron 1 of 4) | intron (NM_011374, intron 1 of 4) | 48874 | NM_011374 | 20449 | Mm.260838 | NM_011374 | ENSMUSG00000030283 | **St8sia1** | 9330109E03Rik\|GD3S\|Sia-T\|Siat8\|Siat8a | ST8 alpha-N-acetyl-neuraminide alpha-2,8-sialyltransferase 1 | protein-coding |

| | | | | | | | | | | | | | | | | | |
|---|---|---|---|---|---|---|---|---|---|---|---|---|---|---|---|---|---|
| chr6_142917112_142918188 | chr6 | 142917113 | 142918188 | + | -0.743827 | intron (NM_011374, intron 1 of 4) | L1MA5\|LINE\|L1 | 46802 | NM_011374 | 20449 | Mm.260838 | NM_011374 | ENSMUSG00000030283 | **St8sia1** | 9330109E03Rik\|GD3S\|Sia-T\|Siat8\|Siat8a | ST8 alpha-N-acetyl-neuraminide alpha-2,8-sialyltransferase 1 | protein-coding |
| chr1_52008149_52009255 | chr1 | 52008150 | 52009255 | + | -0.597726 | 5' UTR (NM_011487, exon 1 of 24) | 5' UTR (NM_011487, exon 1 of 24) | 458 | NM_011487 | 20849 | Mm.1550 | NM_011487 | ENSMUSG00000062939 | **Stat4** | - | signal transducer and activator of transcription 4 | protein-coding |
| chr1_52008168_52008819 | chr1 | 52008169 | 52008819 | + | -0.677219 | 5' UTR (NM_011487, exon 1 of 24) | 5' UTR (NM_011487, exon 1 of 24) | 250 | NM_011487 | 20849 | Mm.1550 | NM_011487 | ENSMUSG00000062939 | **Stat4** | - | signal transducer and activator of transcription 4 | protein-coding |
| chr1_52008481_52008758 | chr1 | 52008482 | 52008758 | + | -0.757996 | 5' UTR (NM_011487, exon 1 of 24) | 5' UTR (NM_011487, exon 1 of 24) | 376 | NM_011487 | 20849 | Mm.1550 | NM_011487 | ENSMUSG00000062939 | **Stat4** | - | signal transducer and activator of transcription 4 | protein-coding |
| chr11_100848081_100848593 | chr11 | 100848082 | 100848593 | + | -0.758353 | intron (NM_001113563, intron 1 of 18) | intron (NM_001113563, intron 1 of 18) | 2248 | NM_001113563 | 20851 | Mm.34064 | NM_011489 | ENSMUSG00000020919 | **Stat5b** | - | signal transducer and activator of transcription 5B | protein-coding |
| chr11_100845784_100846384 | chr11 | 100845785 | 100846384 | + | -1.04672 | intron (NM_001113563, intron 1 of 18) | intron (NM_001113563, intron 1 of 18) | 4501 | NM_001113563 | 20851 | Mm.34064 | NM_011489 | ENSMUSG00000020919 | **Stat5b** | - | signal transducer and activator of transcription 5B | protein-coding |
| chr11_32535640_32536064 | chr11 | 32535641 | 32536064 | + | -0.844081 | intron (NM_009288, intron 1 of 18) | intron (NM_009288, intron 1 of 18) | 2586 | NM_009288 | 20868 | Mm.8235 | NM_009288 | ENSMUSG00000020272 | **Stk10** | Gek1\|Lok\|mKIAA4026 | serine/threonine kinase 10 | protein-coding |
| chr5_44223227_44223602 | chr5 | 44223228 | 44223602 | + | -0.617034 | intron (NM_173764, intron 1 of 13) | intron (NM_173764, intron 1 of 13) | 3191 | NM_173764 | 231225 | Mm.30688 | NM_173764 | ENSMUSG00000046985 | **Tapt1** | 4932414K18Rik | transmembrane anterior posterior transformation 1 | protein-coding |
| chr3_95758482_95759067 | chr3 | 95758483 | 95759067 | + | -0.668247 | Intergenic | Intergenic | -3798 | NM_001163618 | 71807 | Mm.271754 | NM_027931 | ENSMUSG00000028107 | **Tars2** | 2610024N01Rik\|AI429208\|Tarsl1 | threonyl-tRNA synthetase 2, mitochondrial (putative) | protein-coding |
| chr9_42463712_42463977 | chr9 | 42463713 | 42463977 | + | -0.674349 | intron (NM_173038, intron 1 of 7) | intron (NM_173038, intron 1 of 7) | 8381 | NM_173038 | 272589 | Mm.34483 | NM_173038 | ENSMUSG00000037287 | **Tbcel** | D330014K23\|E130107N23Rik\|Lrrc35 | tubulin folding cofactor E-like | protein-coding |
| chr4_102967676_102968473 | chr4 | 102967677 | 102968473 | + | -0.883561 | non-coding (NR_040444, exon 4 of 4) | non-coding (NR_040444, exon 4 of 4) | -18304 | NM_001163767 | 67344 | Mm.111346 | NM_026100 | ENSMUSG00000028523 | **Tctex1d1** | 1700055O19Rik | Tctex1 domain containing 1 | protein-coding |
| chr8_13964406_13965163 | chr8 | 13964407 | 13965163 | + | -0.625197 | intron (NM_173744, intron 1 of 2) | intron (NM_173744, intron 1 of 2) | 9992 | NM_173744 | 72148 | Mm.5727 | NM_173744 | ENSMUSG00000050052 | **Tdrp** | 2610019F03Rik | testis development related protein | protein-coding |
| chr9_116160974_116161287 | chr9 | 116160975 | 116161287 | + | -0.601636 | intron (NM_009371, intron 1 of 7) | intron (NM_009371, intron 1 of 7) | 14232 | NM_029575 | 21813 | Mm.172346 | NM_009371 | ENSMUSG00000032440 | **Tgfbr2** | 1110020H15Rik\|AU042018\|DNIIR\|RIIDN\|TBR-II\|TbetaR-II\|TbetaRII | transforming growth factor, beta receptor II | protein-coding |
| chr9_116160841_116161289 | chr9 | 116160842 | 116161289 | + | -0.614947 | intron (NM_009371, intron 1 of 7) | intron (NM_009371, intron 1 of 7) | 14298 | NM_029575 | 21813 | Mm.172346 | NM_009371 | ENSMUSG00000032440 | **Tgfbr2** | 1110020H15Rik\|AU042018\|DNIIR\|RIIDN\|TBR-II\|TbetaR-II\|TbetaRII | transforming growth factor, beta receptor II | protein-coding |
| chr9_116163871_116164319 | chr9 | 116163872 | 116164319 | + | -0.632895 | intron (NM_009371, intron 1 of 7) | intron (NM_009371, intron 1 of 7) | 11268 | NM_029575 | 21813 | Mm.172346 | NM_009371 | ENSMUSG00000032440 | **Tgfbr2** | 1110020H15Rik\|AU042018\|DNIIR\|RIIDN\|TBR-II\|TbetaR-II\|TbetaRII | transforming growth factor, beta receptor II | protein-coding |
| chr9_116101767_116102237 | chr9 | 116101768 | 116102237 | + | -0.664754 | intron (NM_029575, intron 5 of 6) | intron (NM_029575, intron 5 of 6) | 73361 | NM_029575 | 21813 | Mm.172346 | NM_009371 | ENSMUSG00000032440 | **Tgfbr2** | 1110020H15Rik\|AU042018\|DNIIR\|RIIDN\|TBR-II\|TbetaR-II\|TbetaRII | transforming growth factor, beta receptor II | protein-coding |
| chr9_116091050_116091449 | chr9 | 116091051 | 116091449 | + | -0.666641 | intron (NM_029575, intron 6 of 6) | intron (NM_029575, intron 6 of 6) | 84113 | NM_029575 | 21813 | Mm.172346 | NM_009371 | ENSMUSG00000032440 | **Tgfbr2** | 1110020H15Rik\|AU042018\|DNIIR\|RIIDN\|TBR-II\|TbetaR-II\|TbetaRII | transforming growth factor, beta receptor II | protein-coding |
| chr9_116090899_116091769 | chr9 | 116090900 | 116091769 | + | -0.689612 | intron (NM_029575, intron 6 of 6) | intron (NM_029575, intron 6 of 6) | 84029 | NM_029575 | 21813 | Mm.172346 | NM_009371 | ENSMUSG00000032440 | **Tgfbr2** | 1110020H15Rik\|AU042018\|DNIIR\|RIIDN\|TBR-II\|TbetaR-II\|TbetaRII | transforming growth factor, beta receptor II | protein-coding |
| chr9_116187401_116187876 | chr9 | 116187402 | 116187876 | + | -0.855934 | Intergenic | Intergenic | -12276 | NM_009371 | 21813 | Mm.172346 | NM_009371 | ENSMUSG00000032440 | **Tgfbr2** | 1110020H15Rik\|AU042018\|DNIIR\|RIIDN\|TBR-II\|TbetaR-II\|TbetaRII | transforming growth factor, beta receptor II | protein-coding |
| chr9_116084183_116084953 | chr9 | 116084184 | 116084953 | + | -0.948067 | Intergenic | Intergenic | 90795 | NM_029575 | 21813 | Mm.172346 | NM_009371 | ENSMUSG00000032440 | **Tgfbr2** | 1110020H15Rik\|AU042018\|DNIIR\|RIIDN\|TBR-II\|TbetaR-II\|TbetaRII | transforming growth factor, beta receptor II | protein-coding |
| chr9_116084411_116085026 | chr9 | 116084412 | 116085026 | + | -0.96118 | Intergenic | Intergenic | 90644 | NM_029575 | 21813 | Mm.172346 | NM_009371 | ENSMUSG00000032440 | **Tgfbr2** | 1110020H15Rik\|AU042018\|DNIIR\|RIIDN\|TBR-II\|TbetaR-II\|TbetaRII | transforming growth factor, beta receptor II | protein-coding |
| chr17_84352152_84352509 | chr17 | 84352153 | 84352509 | + | -0.598221 | intron (NM_183021, intron 26 of 37) | RSINE1\|SINE\|B4 | 113877 | NM_183021 | 240174 | Mm.246767 | NM_177826 | ENSMUSG00000024251 | **Thada** | 6530405K19\|A930021O22\|AU019899\|BC052885\|GITA | thyroid adenoma associated | protein-coding |
| chr17_84419716_84420868 | chr17 | 84419717 | 84420868 | + | -1.02857 | intron (NM_183021, intron 21 of 37) | intron (NM_183021, intron 21 of 37) | 45916 | NM_183021 | 240174 | Mm.246767 | NM_177826 | ENSMUSG00000024251 | **Thada** | 6530405K19\|A930021O22\|AU019899\|BC052885\|GITA | thyroid adenoma associated | protein-coding |
| chr17_84419716_84420245 | chr17 | 84419717 | 84420245 | + | -1.16168 | intron (NM_183021, intron 21 of 37) | intron (NM_183021, intron 21 of 37) | 46227 | NM_183021 | 240174 | Mm.246767 | NM_177826 | ENSMUSG00000024251 | **Thada** | 6530405K19\|A930021O22\|AU019899\|BC052885\|GITA | thyroid adenoma associated | protein-coding |
| chr16_43663800_43664646 | chr16 | 43663801 | 43664646 | + | -0.725792 | promoter-TSS (NM_001146325) | promoter-TSS (NM_001146325) | -39 | NM_001146325 | 100043314 | Mm.390896 | NM_001146325 | ENSMUSG00000071552 | **Tigit** | ENSMUSG00000071552\|Vstm3 | T cell immunoreceptor with Ig and ITIM domains | protein-coding |
| chr14_122062825_122063070 | chr14 | 122062826 | 122063070 | + | -0.662361 | Intergenic | MTD\|LTR\|ERVL-MaLR | 28274 | NM_001037744 | 223262 | Mm.297637 | NM_001037744 | ENSMUSG00000071229 | **Timm8a2** | - | translocase of inner mitochondrial membrane 8A2 | protein-coding |
| chr14_122062567_122063111 | chr14 | 122062568 | 122063111 | + | -0.679745 | Intergenic | B4A\|SINE\|B4 | 28165 | NM_001037744 | 223262 | Mm.297637 | NM_001037744 | ENSMUSG00000071229 | **Timm8a2** | - | translocase of inner mitochondrial membrane 8A2 | protein-coding |
| chr11_115187728_115188235 | chr11 | 115187729 | 115188235 | + | -0.719985 | promoter-TSS (NM_025400) | promoter-TSS (NM_025400) | 495 | NM_001033393 | 320534 | Mm.260587 | NM_001033393 | ENSMUSG00000045980 | **Tmem104** | C630005D06Rik\|mFLJ00021 | transmembrane protein 104 | protein-coding |
| chr9_7764650_7764936 | chr9 | 7764651 | 7764936 | + | -0.743014 | intron (NM_133739, intron 1 of 4) | intron (NM_133739, intron 1 of 4) | 716 | NM_133739 | 71929 | Mm.283293 | NM_133739 | ENSMUSG00000050912 | **Tmem123** | 2310075C12Rik | transmembrane protein 123 | protein-coding |
| chr1_52653261_52653799 | chr1 | 52653262 | 52653799 | + | -0.671568 | Intergenic | Intergenic | 22825 | NM_001142647 | 227094 | Mm.83541 | NM_172654 | ENSMUSG00000043015 | **Tmem194b** | 5330401P04Rik\|9830163O08 | transmembrane protein 194B | protein-coding |
| chr4_133298519_133298893 | chr4 | 133298520 | 133298893 | + | -0.606235 | intron (NM_199306, intron 12 of 15) | intron (NM_199306, intron 12 of 15) | -20916 | NR_110978 | 52174 | Mm.244748 | NM_025667 | ENSMUSG00000028857 | **Tmem222** | 5730406H10Rik\|AA409341\|D4Ertd196e | transmembrane protein 222 | protein-coding |
| chr4_15273203_15273560 | chr4 | 15273204 | 15273560 | + | -0.784087 | intron (NM_181401, intron 1 of 2) | intron (NM_181401, intron 1 of 2) | 7562 | NM_181401 | 100201 | Mm.394111 | NM_181401 | ENSMUSG00000043252 | **Tmem64** | 9630015D15Rik\|AI790744\|AV300874 | transmembrane protein 64 | protein-coding |
| chr4_15273187_15273723 | chr4 | 15273188 | 15273723 | + | -0.792385 | intron (NM_181401, intron 1 of 2) | intron (NM_181401, intron 1 of 2) | 7635 | NM_181401 | 100201 | Mm.394111 | NM_181401 | ENSMUSG00000043252 | **Tmem64** | 9630015D15Rik\|AI790744\|AV300874 | transmembrane protein 64 | protein-coding |
| chr10_118633378_118634010 | chr10 | 118633379 | 118634010 | + | -0.883987 | Intergenic | Intergenic | -77169 | NR_104123 | 103214 | Mm.30529 | NR_104123 | | **Tmevpg1** | AI447422 | Theiler's murine encephalomyelitis virus persistence candidate gene 1 | ncRNA |
| chr10_118599120_118599607 | chr10 | 118599121 | 118599607 | + | -0.919615 | Intergenic | Intergenic | -42839 | NR_104123 | 103214 | Mm.30529 | NR_104123 | | **Tmevpg1** | AI447422 | Theiler's murine encephalomyelitis virus persistence candidate gene 1 | ncRNA |
| chr10_18999278_18999761 | chr10 | 18999279 | 18999761 | + | -0.653017 | Intergenic | Intergenic | 12443 | NM_001166402 | 21929 | Mm.116683 | NM_009397 | ENSMUSG00000019850 | **Tnfaip3** | A20\|Tnfip3 | tumor necrosis factor, alpha-induced protein 3 | protein-coding |
| chr10_19061928_19063110 | chr10 | 19061929 | 19063110 | + | -0.794629 | Intergenic | Intergenic | -47109 | NM_009397 | 21929 | Mm.116683 | NM_009397 | ENSMUSG00000019850 | **Tnfaip3** | A20\|Tnfip3 | tumor necrosis factor, alpha-induced protein 3 | protein-coding |
| chr6_125351328_125351741 | chr6 | 125351329 | 125351741 | + | -0.887942 | intron (NM_011609, intron 1 of 9) | intron (NM_011609, intron 1 of 9) | 1812 | NM_011609 | 21937 | Mm.1258 | NM_011609 | ENSMUSG00000030341 | **Tnfrsf1a** | CD120a\|FPF\|TNF-R\|TNF-R-I\|TNF-R1\|TNF-R55\|TNF-alphaR1\|TNFAR\|TNFR60\|TNFRI\|TNFRp55\|TNFalpha-R1\|Tnfr-2\|Tnfr1\|p55\|p55-R | tumor necrosis factor receptor superfamily, member 1a | protein-coding |
| chr4_145241058_145241326 | chr4 | 145241059 | 145241326 | + | -0.604891 | intron (NM_011610, intron 1 of 9) | intron (NM_011610, intron 1 of 9) | 5678 | NM_011610 | 21938 | Mm.235328 | NM_011610 | | **Tnfrsf1b** | CD120b\|TNF-R-II\|TNF-R2\|TNF-R75\|TNF-alphaR2\|TNFBR\|TNFR80\|TNFRII\|TNFalpha-R2\|Tnfr-1\|Tnfr2\|p75 | tumor necrosis factor receptor superfamily, member 1b | protein-coding |
| chr4_145240725_145240978 | chr4 | 145240726 | 145240978 | + | -0.67329 | intron (NM_011610, intron 1 of 9) | intron (NM_011610, intron 1 of 9) | 6018 | NM_011610 | 21938 | Mm.235328 | NM_011610 | | **Tnfrsf1b** | CD120b\|TNF-R-II\|TNF-R2\|TNF-R75\|TNF-alphaR2\|TNFBR\|TNFR80\|TNFRII\|TNFalpha-R2\|Tnfr-1\|Tnfr2\|p75 | tumor necrosis factor receptor superfamily, member 1b | protein-coding |
| chr4_145240652_145241342 | chr4 | 145240653 | 145241342 | + | -0.691832 | intron (NM_011610, intron 1 of 9) | intron (NM_011610, intron 1 of 9) | 5873 | NM_011610 | 21938 | Mm.235328 | NM_011610 | | **Tnfrsf1b** | CD120b\|TNF-R-II\|TNF-R2\|TNF-R75\|TNF-alphaR2\|TNFBR\|TNFR80\|TNFRII\|TNFalpha-R2\|Tnfr-1\|Tnfr2\|p75 | tumor necrosis factor receptor superfamily, member 1b | protein-coding |
| chr14_78356724_78357235 | chr14 | 78356725 | 78357235 | + | -0.588229 | Intergenic | L4_A_Mam\|LINE\|RTE-X | -48937 | NM_011613 | 21943 | Mm.249221 | NM_011613 | ENSMUSG00000022015 | **Tnfsf11** | Ly109l\|ODF\|OPG\|OPGL\|RANKL\|Trance | tumor necrosis factor (ligand) superfamily, member 11 | protein-coding |
| chr15_80743579_80743988 | chr15 | 80743580 | 80743988 | + | -0.60577 | intron (NM_144812, intron 1 of 24) | intron (NM_144812, intron 1 of 24) | 32471 | NM_144812 | 213988 | Mm.131328 | NM_144812 | ENSMUSG00000047888 | **Tnrc6b** | 2700090M07Rik\|A730065C02Rik\|AI848765\|D230019K20Rik | trinucleotide repeat containing 6b | protein-coding |
| chr15_59652839_59653665 | chr15 | 59652840 | 59653665 | + | -1.01069 | intron (NM_144549, intron 2 of 2) | intron (NM_144549, intron 2 of 2) | 4598 | NM_144549 | 211770 | Mm.40298 | NM_144549 | ENSMUSG00000032501 | **Trib1** | A530090O15Rik\|TRB-1\|Trb1 | tribbles homolog 1 (Drosophila) | protein-coding |
| chr11_89024212_89024566 | chr11 | 89024213 | 89024566 | + | -0.591564 | Intergenic | Intergenic | 24986 | NM_009546 | 217069 | Mm.248445 | NM_009546 | ENSMUSG00000000275 | **Trim25** | AA960166\|AL022677\|EFP\|Zfp147 | tripartite motif-containing 25 | protein-coding |
| chr6_41196244_41197189 | chr6 | 41196245 | 41197189 | + | -0.616144 | Intergenic | L1ME3D\|LINE\|L1 | -105555 | NM_011646 | 22074 | Mm.383263 | NM_011646 | ENSMUSG00000054106 | **Try4** | 0910001B19Rik\|Td | trypsin 4 | protein-coding |
| chr8_109566105_109566382 | chr8 | 109566106 | 109566382 | + | -0.626327 | promoter-TSS (NM_178380) | promoter-TSS (NM_178380) | 258 | NM_175646 | 234723 | Mm.37667 | NM_175646 | ENSMUSG00000031723 | **Txnl4b** | AI595343\|D530025J19Rik\|DLP\|Dim2 | thioredoxin-like 4B | protein-coding |

| | | | | | | | | | | | | | | | | | |
|---|---|---|---|---|---|---|---|---|---|---|---|---|---|---|---|---|---|
| chr14_121879453_121879826 | chr14 | 121879454 | 121879826 | + | -0.782935 | intron (NM_026861, intron 1 of 8) | intron (NM_026861, intron 1 of 8) | 1034 | NM_026861 | 68889 | Mm.441151 | NM_026861 | ENSMUSG00000041765 | **Ubac2** | 1190008A14Rik\|AW554959\|Phgdhl1 | ubiquitin associated domain containing 2 | protein-coding |
| chr14_121879351_121879995 | chr14 | 121879352 | 121879995 | + | -0.785221 | intron (NM_026861, intron 1 of 8) | intron (NM_026861, intron 1 of 8) | 1067 | NM_026861 | 68889 | Mm.441151 | NM_026861 | ENSMUSG00000041765 | **Ubac2** | 1190008A14Rik\|AW554959\|Phgdhl1 | ubiquitin associated domain containing 2 | protein-coding |
| chr16_4878977_4879679 | chr16 | 4878978 | 4879679 | + | -0.591045 | intron (NM_145359, intron 1 of 2) | CpG | 523 | NM_145359 | 207740 | Mm.268895 | NM_145359 | | **Ubald1** | 1500031H01Rik\|BC013706\|Fam100a | UBA-like domain containing 1 | protein-coding |
| chr16_4878642_4879475 | chr16 | 4878643 | 4879475 | + | -0.605764 | intron (NM_145359, intron 1 of 2) | intron (NM_145359, intron 1 of 2) | 792 | NM_145359 | 207740 | Mm.268895 | NM_145359 | | **Ubald1** | 1500031H01Rik\|BC013706\|Fam100a | UBA-like domain containing 1 | protein-coding |
| chr16_4878960_4879500 | chr16 | 4878961 | 4879500 | + | -0.663867 | intron (NM_145359, intron 1 of 2) | intron (NM_145359, intron 1 of 2) | 621 | NM_145359 | 207740 | Mm.268895 | NM_145359 | | **Ubald1** | 1500031H01Rik\|BC013706\|Fam100a | UBA-like domain containing 1 | protein-coding |
| chr5_148554695_148555114 | chr5 | 148554696 | 148555114 | + | -0.619114 | Intergenic | Intergenic | -2117 | NM_011908 | 24109 | Mm.21846 | NM_011908 | ENSMUSG00000001687 | **Ubl3** | AW108023\|HCG | ubiquitin-like 3 | protein-coding |
| chr17_46966852_46967427 | chr17 | 46966853 | 46967427 | + | -0.629933 | intron (NM_001177374, intron 19 of 46) | intron (NM_001177374, intron 19 of 46) | 43392 | NM_146078 | 224826 | Mm.28234 | NM_146078 | ENSMUSG00000023977 | **Ubr2** | 9930021A08Rik\|AI462103\|AW540746\|E130209G04Rik\|mKIAA0349 | ubiquitin protein ligase E3 component n-recognin 2 | protein-coding |
| chr2_6478111_6479186 | chr2 | 6478112 | 6479186 | + | -0.826871 | Intergenic | Intergenic | 125913 | NM_181399 | 98910 | Mm.40673 | NM_181399 | ENSMUSG00000039046 | **Usp6nl** | AI316785\|RNTRE\|TRE2NL\|mKIAA0019 | USP6 N-terminal like | protein-coding |
| chr2_6478111_6478914 | chr2 | 6478112 | 6478914 | + | -0.841864 | Intergenic | Intergenic | 125777 | NM_181399 | 98910 | Mm.40673 | NM_181399 | ENSMUSG00000039046 | **Usp6nl** | AI316785\|RNTRE\|TRE2NL\|mKIAA0019 | USP6 N-terminal like | protein-coding |
| chr1_162571181_162571726 | chr1 | 162571182 | 162571726 | + | -0.789618 | intron (NM_016796, intron 1 of 7) | CpG | 626 | NM_016796 | 53330 | Mm.10699 | NM_016796 | ENSMUSG00000026696 | **Vamp4** | D1Ertd147e | vesicle-associated membrane protein 4 | protein-coding |
| chr1_162571357_162571682 | chr1 | 162571358 | 162571682 | + | -0.8575 | intron (NM_016796, intron 1 of 7) | CpG | 692 | NM_016796 | 53330 | Mm.10699 | NM_016796 | ENSMUSG00000026696 | **Vamp4** | D1Ertd147e | vesicle-associated membrane protein 4 | protein-coding |
| chr6_114904798_114905179 | chr6 | 114904799 | 114905179 | + | -0.748953 | intron (NM_177683, intron 1 of 4) | intron (NM_177683, intron 1 of 4) | 16763 | NM_177683 | 232334 | Mm.268357 | NM_177683 | | **Vgll4** | BC048841\|VGL-4 | vestigial like 4 (Drosophila) | protein-coding |
| chr5_124035460_124035764 | chr5 | 124035461 | 124035764 | + | -0.626327 | Intergenic | Intergenic | -3352 | NM_177876 | 330192 | Mm.4079 | NM_177876 | ENSMUSG00000066278 | **Vps37b** | 2300007F24Rik\|AI415429\|BC026744 | vacuolar protein sorting 37B (yeast) | protein-coding |
| chr4_133162175_133162677 | chr4 | 133162176 | 133162677 | + | -0.753638 | intron (NM_153423, intron 1 of 8) | intron (NM_153423, intron 1 of 8) | 31793 | NM_153423 | 242687 | Mm.23566 | NM_153423 | ENSMUSG00000028868 | **Wasf2** | AW742646\|D4Ertd13e\|WAVE2 | WAS protein family, member 2 | protein-coding |
| chr1_181227800_181228230 | chr1 | 181227801 | 181228230 | + | -0.598192 | Intergenic | Intergenic | -16037 | NM_145514 | 226757 | Mm.289082 | NM_145514 | ENSMUSG00000038733 | **Wdr26** | 1600024A01Rik\|AA693241\|AI447817\|AU044014\|C77982\|Gid7 | WD repeat domain 26 | protein-coding |
| chr15_58165135_58165664 | chr15 | 58165136 | 58165664 | + | -0.753443 | Intergenic | Intergenic | 23964 | NM_029734 | 76773 | Mm.26771 | NM_029734 | ENSMUSG00000022359 | **Wdyhv1** | 2410187C16Rik\|AU014961\|AW550036\|Ntaq1 | WDYHV motif containing 1 | protein-coding |
| chr8_114803112_114803622 | chr8 | 114803113 | 114803622 | + | -0.607505 | intron (NM_019573, intron 8 of 8) | intron (NM_019573, intron 8 of 8) | 363715 | NM_019573 | 80707 | Mm.440420 | NM_019573 | ENSMUSG00000004637 | **Wwox** | 5330426P09Rik\|9030416C10Rik\|WOX1 | WW domain-containing oxidoreductase | protein-coding |
| chr8_114801755_114802112 | chr8 | 114801756 | 114802112 | + | -0.63393 | intron (NM_019573, intron 8 of 8) | intron (NM_019573, intron 8 of 8) | 362282 | NM_019573 | 80707 | Mm.440420 | NM_019573 | ENSMUSG00000004637 | **Wwox** | 5330426P09Rik\|9030416C10Rik\|WOX1 | WW domain-containing oxidoreductase | protein-coding |
| chr8_114973000_114973511 | chr8 | 114973001 | 114973511 | + | -0.919955 | intron (NM_019573, intron 8 of 8) | intron (NM_019573, intron 8 of 8) | 533604 | NM_019573 | 80707 | Mm.440420 | NM_019573 | ENSMUSG00000004637 | **Wwox** | 5330426P09Rik\|9030416C10Rik\|WOX1 | WW domain-containing oxidoreductase | protein-coding |
| chr11_5521514_5522334 | chr11 | 5521515 | 5522334 | + | -0.607773 | exon (NM_013842, exon 2 of 5) | exon (NM_013842, exon 2 of 5) | 1283 | NM_001271730 | 22433 | Mm.469937 | NM_013842 | ENSMUSG00000020484 | **Xbp1** | D11Ertd39e\|TREB-5\|TREB5\|XBP-1 | X-box binding protein 1 | protein-coding |
| chr9_123869249_123869740 | chr9 | 123869250 | 123869740 | + | -0.605424 | Intergenic | Intergenic | -7466 | NM_011798 | 23832 | Mm.390241 | NM_011798 | ENSMUSG00000060509 | **Xcr1** | Ccxcr1\|Gpr5\|mXcr1 | chemokine (C motif) receptor 1 | protein-coding |
| chr17_73893262_73893776 | chr17 | 73893263 | 73893776 | + | -1.01248 | intron (NM_011723, intron 32 of 35) | intron (NM_011723, intron 32 of 35) | 56677 | NM_011723 | 22436 | Mm.11223 | NM_011723 | ENSMUSG00000024066 | **Xdh** | XO\|Xor\|Xox-1\|Xox1 | xanthine dehydrogenase | protein-coding |
| chrX_48096979_48097439 | chrX | 48096980 | 48097439 | + | -0.648166 | Intergenic | Intergenic | -11516 | NM_001289729 | 170745 | Mm.129279 | NM_133213 | ENSMUSG00000037005 | **Xpnpep2** | 9030008G12Rik\|mAPP | X-prolyl aminopeptidase (aminopeptidase P) 2, membrane-bound | protein-coding |
| chrX_48098458_48098833 | chrX | 48098459 | 48098833 | + | -0.657551 | Intergenic | Intergenic | -10079 | NM_001289729 | 170745 | Mm.129279 | NM_133213 | ENSMUSG00000037005 | **Xpnpep2** | 9030008G12Rik\|mAPP | X-prolyl aminopeptidase (aminopeptidase P) 2, membrane-bound | protein-coding |
| chrX_48098405_48099066 | chrX | 48098406 | 48099066 | + | -0.698402 | Intergenic | Intergenic | -9989 | NM_001289729 | 170745 | Mm.129279 | NM_133213 | ENSMUSG00000037005 | **Xpnpep2** | 9030008G12Rik\|mAPP | X-prolyl aminopeptidase (aminopeptidase P) 2, membrane-bound | protein-coding |
| chr7_117428686_117429333 | chr7 | 117428687 | 117429333 | + | -0.937848 | intron (NM_175645, intron 1 of 11) | intron (NM_175645, intron 1 of 11) | 48031 | NM_175645 | 233781 | Mm.461427 | NM_175645 | ENSMUSG00000030657 | **Xylt1** | 8030490L12 | xylosyltransferase 1 | protein-coding |
| chr2_84733625_84734215 | chr2 | 84733626 | 84734215 | + | -0.604492 | promoter-TSS (NM_001005342) | promoter-TSS (NM_001005342) | -284 | NM_001005342 | 241525 | Mm.486398 | NM_001005342 | | **Ypel4** | - | yippee-like 4 (Drosophila) | protein-coding |
| chr9_96696211_96696876 | chr9 | 96696212 | 96696876 | + | -0.783977 | intron (NM_175537, intron 2 of 2) | intron (NM_175537, intron 2 of 2) | 35131 | NM_175537 | 245007 | Mm.485954 | NM_175537 | ENSMUSG00000040433 | **Zbtb38** | A930014K01Rik\|CIBZ | zinc finger and BTB domain containing 38 | protein-coding |
| chr9_96696012_96696960 | chr9 | 96696013 | 96696960 | + | -0.814557 | intron (NM_175537, intron 2 of 2) | intron (NM_175537, intron 2 of 2) | 35189 | NM_175537 | 245007 | Mm.485954 | NM_175537 | ENSMUSG00000040433 | **Zbtb38** | A930014K01Rik\|CIBZ | zinc finger and BTB domain containing 38 | protein-coding |
| chr5_139483151_139483450 | chr5 | 139483152 | 139483450 | + | -0.649041 | intron (NM_001159908, intron 1 of 6) | intron (NM_001159908, intron 1 of 6) | 1190 | NM_133349 | 100494 | Mm.24521 | NM_133349 | ENSMUSG00000053581 | **Zfand2a** | AA407930\|AU016206\|Airap | zinc finger, AN1-type domain 2A | protein-coding |
| chr13_48515791_48516429 | chr13 | 48515792 | 48516429 | + | -0.586896 | Intergenic | Intergenic | -2674 | NM_001164576 | 67911 | Mm.251375 | NM_026450 | ENSMUSG00000050954 | **Zfp169** | 1700025J14Rik\|4930429A13Rik\|BE688704 | zinc finger protein 169 | protein-coding |
| chr11_70762685_70763066 | chr11 | 70762686 | 70763066 | + | -0.61987 | Intergenic | ORR1D-int\|LTR\|ERVL-MaLR | -1571 | NM_177565 | 193043 | Mm.40966 | NM_177565 | ENSMUSG00000043602 | **Zfp3** | AI854635\|B230309J24\|Fnp-1\|Zfp-3 | zinc finger protein 3 | protein-coding |
| chr2_154615715_154616074 | chr2 | 154615716 | 154616074 | + | -0.796319 | intron (NM_199304, intron 1 of 14) | intron (NM_199304, intron 1 of 14) | 2527 | NM_199304 | 228807 | Mm.103555 | NM_199304 | ENSMUSG00000059842 | **Zfp341** | Znf341 | zinc finger protein 341 | protein-coding |
| chr17_21473491_21474369 | chr17 | 21473492 | 21474369 | + | -0.592892 | Intergenic | MMERGLN-int\|LTR\|ERV1 | -15058 | NM_013843 | 24132 | Mm.42140 | NM_013843 | | **Zfp53** | D030067O06Rik\|KRAZ1\|Zfp-53\|Zfp118\|zfas8 | zinc finger protein 53 | protein-coding |
| chr17_23617035_23617592 | chr17 | 23617036 | 23617592 | + | -0.746569 | Intergenic | MLTR11B\|LTR\|ERVK | 16488 | NM_001289482 | 332221 | Mm.270315 | NM_001033425 | ENSMUSG00000023902 | **Zscan10** | BC068284\|Zfp206\|Zkscan10\|Znf206 | zinc finger and SCAN domain containing 10 | protein-coding |
| chr17_23620278_23620796 | chr17 | 23620279 | 23620796 | + | -0.837726 | Intergenic | Intergenic | 19711 | NM_001289482 | 332221 | Mm.270315 | NM_001033425 | ENSMUSG00000023902 | **Zscan10** | BC068284\|Zfp206\|Zkscan10\|Znf206 | zinc finger and SCAN domain containing 10 | protein-coding |

**Table S2B_HOMER Closing peaks in Ag-stimulated cells**
Data is sorted alphabetically by the gene name; Only peaks above the peak score value of 0.5 are shown.

| PeakID (cmd=annotatePeaks.pl ../averagescore_closing_peaks_a [illegible] | Chr | Start | End | Strand | Peak Score | Annotation | Detailed Annotation | Distance to TSS | Nearest PromoterID | Entrez ID | Nearest Unigene | Nearest Refseq | Nearest Ensembl | Gene Name | Gene Alias | Gene Description | Gene Type |
|---|---|---|---|---|---|---|---|---|---|---|---|---|---|---|---|---|---|
| chr6_108572725_108573053 | chr6 | 108572726 | 108573053 | + | -0.644557 | Intergenic | ID_B1\|SINE\|B4 | 50278 | NR_040757 | 75394 | Mm.395061 | NR_040757 | ENSMUSG00000087341 | **0610040F04Rik** | - | RIKEN cDNA 0610040F04 gene | ncRNA |
| chr6_108572794_108573038 | chr6 | 108572795 | 108573038 | + | -0.688422 | Intergenic | B1F\|SINE\|Alu | 50251 | NR_040757 | 75394 | Mm.395061 | NR_040757 | ENSMUSG00000087341 | **0610040F04Rik** | - | RIKEN cDNA 0610040F04 gene | ncRNA |
| chr2_156863815_156864115 | chr2 | 156863816 | 156864115 | + | -0.647432 | intron (NM_026124, intron 1 of 3) | intron (NM_026124, intron 1 of 3) | 843 | NM_026124 | 67388 | Mm.29264 | NM_026124 | ENSMUSG00000027637 | **1110008F13Rik** | AI316789 | RIKEN cDNA 1110008F13 gene | protein-coding |
| chr3_68784068_68784464 | chr3 | 68784069 | 68784464 | + | -0.636287 | Intergenic | (TG)n\|Simple_repeat\|Simple_repeat | -85320 | NM_001167996 | 68725 | Mm.45741 | NM_001167996 | ENSMUSG00000046999 | **1110032F04Rik** | AI115547 | RIKEN cDNA 1110032F04 gene | protein-coding |
| chr2_132658334_132659328 | chr2 | 132658335 | 132659328 | + | -0.760789 | Intergenic | MT2A\|LTR\|ERVL | -31452 | NM_028637 | 73747 | Mm.284114 | NM_028637 | ENSMUSG00000044991 | **1110034G24Rik** | - | RIKEN cDNA 1110034G24 gene | protein-coding |
| chr2_132658397_132658871 | chr2 | 132658398 | 132658871 | + | -0.878571 | Intergenic | Intergenic | -31649 | NM_028637 | 73747 | Mm.284114 | NM_028637 | ENSMUSG00000044991 | **1110034G24Rik** | - | RIKEN cDNA 1110034G24 gene | protein-coding |
| chr2_132691701_132692100 | chr2 | 132691702 | 132692100 | + | -0.911321 | intron (NM_028637, intron 1 of 2) | intron (NM_028637, intron 1 of 2) | 1618 | NM_028637 | 73747 | Mm.284114 | NM_028637 | ENSMUSG00000044991 | **1110034G24Rik** | - | RIKEN cDNA 1110034G24 gene | protein-coding |
| chr4_11488886_11489117 | chr4 | 11488887 | 11489117 | + | -0.636125 | intron (NM_001081183, intron 1 of 24) | RSINE1\|SINE\|B4 | 3044 | NM_001081183 | 66185 | Mm.331487 | NM_001081183 | ENSMUSG00000040720 | **1110037F02Rik** | 4930422M05Rik\|Kiaa1429\|mKIAA1429 | RIKEN cDNA 1110037F02 gene | protein-coding |
| chr4_11492597_11493366 | chr4 | 11492598 | 11493366 | + | -0.988686 | intron (NM_001081183, intron 1 of 24) | intron (NM_001081183, intron 1 of 24) | 7024 | NM_001081183 | 66185 | Mm.331487 | NM_001081183 | ENSMUSG00000040720 | **1110037F02Rik** | 4930422M05Rik\|Kiaa1429\|mKIAA1429 | RIKEN cDNA 1110037F02 gene | protein-coding |
| chr4_11493054_11493355 | chr4 | 11493055 | 11493355 | + | -1.10082 | intron (NM_001081183, intron 1 of 24) | ID_B1\|SINE\|B4 | 7247 | NM_001081183 | 66185 | Mm.331487 | NM_001081183 | ENSMUSG00000040720 | **1110037F02Rik** | 4930422M05Rik\|Kiaa1429\|mKIAA1429 | RIKEN cDNA 1110037F02 gene | protein-coding |
| chr19_21630649_21631116 | chr19 | 21630650 | 21631116 | + | -0.708568 | intron (NM_025423, intron 1 of 3) | intron (NM_025423, intron 1 of 3) | 21908 | NM_025423 | 66206 | Mm.272430 | NM_025423 | ENSMUSG00000035171 | **1110059E24Rik** | - | RIKEN cDNA 1110059E24 gene | protein-coding |
| chr19_21646117_21646692 | chr19 | 21646118 | 21646692 | + | -0.7255 | intron (NM_025423, intron 1 of 3) | L2\|LINE\|L2 | 6386 | NM_025423 | 66206 | Mm.272430 | NM_025423 | ENSMUSG00000035171 | **1110059E24Rik** | - | RIKEN cDNA 1110059E24 gene | protein-coding |
| chr19_21646127_21646507 | chr19 | 21646128 | 21646507 | + | -0.783106 | intron (NM_025423, intron 1 of 3) | MTD\|LTR\|ERVL-MaLR | 6474 | NM_025423 | 66206 | Mm.272430 | NM_025423 | ENSMUSG00000035171 | **1110059E24Rik** | - | RIKEN cDNA 1110059E24 gene | protein-coding |
| chr6_134932565_134932839 | chr6 | 134932566 | 134932839 | + | -0.762448 | intron (NR_037956, intron 1 of 1) | ID_B1\|SINE\|B4 | 3610 | NR_037955 | 381822 | Mm.5935 | NR_037955 | ENSMUSG00000098318 | **1190002F15Rik** | AI120476 | RIKEN cDNA 1190002F15 gene | ncRNA |
| chr9_94440782_94441821 | chr9 | 94440783 | 94441821 | + | -1.01507 | Intergenic | LTR16B2\|LTR\|ERVL | 96779 | NM_001033145 | 68861 | Mm.258746 | NM_001033145 | ENSMUSG00000045414 | **1190002N15Rik** | GoPro49\|HASF | RIKEN cDNA 1190002N15 gene | protein-coding |
| chr9_94441428_94441798 | chr9 | 94441429 | 94441798 | + | -1.04619 | Intergenic | Intergenic | 96468 | NM_001033145 | 68861 | Mm.258746 | NM_001033145 | ENSMUSG00000045414 | **1190002N15Rik** | GoPro49\|HASF | RIKEN cDNA 1190002N15 gene | protein-coding |
| chr9_94441296_94441844 | chr9 | 94441297 | 94441844 | + | -1.05084 | Intergenic | Intergenic | 96511 | NM_001033145 | 68861 | Mm.258746 | NM_001033145 | ENSMUSG00000045414 | **1190002N15Rik** | GoPro49\|HASF | RIKEN cDNA 1190002N15 gene | protein-coding |
| chr16_21835404_21835959 | chr16 | 21835405 | 21835959 | + | -0.847283 | Intergenic | ID_B1\|SINE\|B4 | 41335 | NR_037957 | 100043489 | Mm.326535 | NR_037957 | ENSMUSG00000097290 | **1300002E11Rik** | 1500026P21Rik | RIKEN cDNA 1300002E11 gene | ncRNA |
| chr16_95899720_95900588 | chr16 | 95899721 | 95900588 | + | -0.627482 | intron (NR_040484, intron 1 of 3) | intron (NR_040484, intron 1 of 3) | -1441 | NR_040485 | 69776 | Mm.120470 | NR_040484 | ENSMUSG00000086899 | **1600002D24Rik** | - | RIKEN cDNA 1600002D24 gene | ncRNA |
| chr16_95815231_95816406 | chr16 | 95815232 | 95816406 | + | -0.747297 | Intergenic | Intergenic | 82894 | NR_040485 | 69776 | Mm.120470 | NR_040484 | ENSMUSG00000086899 | **1600002D24Rik** | - | RIKEN cDNA 1600002D24 gene | ncRNA |
| chr16_95815780_95816438 | chr16 | 95815781 | 95816438 | + | -0.881282 | Intergenic | Intergenic | 82604 | NR_040485 | 69776 | Mm.120470 | NR_040484 | ENSMUSG00000086899 | **1600002D24Rik** | - | RIKEN cDNA 1600002D24 gene | ncRNA |
| chr16_95821025_95821543 | chr16 | 95821026 | 95821543 | + | -0.89766 | Intergenic | Intergenic | 77429 | NR_040485 | 69776 | Mm.120470 | NR_040484 | ENSMUSG00000086899 | **1600002D24Rik** | - | RIKEN cDNA 1600002D24 gene | ncRNA |
| chr16_95900040_95900625 | chr16 | 95900041 | 95900625 | + | -0.973217 | intron (NR_040484, intron 1 of 3) | intron (NR_040484, intron 1 of 3) | -1620 | NR_040485 | 69776 | Mm.120470 | NR_040484 | ENSMUSG00000086899 | **1600002D24Rik** | - | RIKEN cDNA 1600002D24 gene | ncRNA |
| chr16_95901395_95901682 | chr16 | 95901396 | 95901682 | + | -1.01738 | intron (NR_040484, intron 1 of 3) | intron (NR_040484, intron 1 of 3) | -2826 | NR_040485 | 69776 | Mm.120470 | NR_040484 | ENSMUSG00000086899 | **1600002D24Rik** | - | RIKEN cDNA 1600002D24 gene | ncRNA |
| chr16_95860698_95861353 | chr16 | 95860699 | 95861353 | + | -1.06914 | intron (NR_040484, intron 1 of 3) | intron (NR_040484, intron 1 of 3) | 37687 | NR_040485 | 69776 | Mm.120470 | NR_040484 | ENSMUSG00000086899 | **1600002D24Rik** | - | RIKEN cDNA 1600002D24 gene | ncRNA |
| chr16_95860787_95861174 | chr16 | 95860788 | 95861174 | + | -1.07979 | intron (NR_040484, intron 1 of 3) | intron (NR_040484, intron 1 of 3) | 37732 | NR_040485 | 69776 | Mm.120470 | NR_040484 | ENSMUSG00000086899 | **1600002D24Rik** | - | RIKEN cDNA 1600002D24 gene | ncRNA |
| chr19_53263758_53264546 | chr19 | 53263759 | 53264546 | + | -0.663395 | Intergenic | Intergenic | -8947 | NR_036590 | 69319 | Mm.57452 | NR_036590 | | **1700001K23Rik** | - | RIKEN cDNA 1700001K23 gene | ncRNA |
| chr2_30769292_30769666 | chr2 | 30769293 | 30769666 | + | -0.652292 | Intergenic | ID_B1\|SINE\|B4 | 32258 | NM_198000 | 73598 | Mm.34166 | NM_198000 | ENSMUSG00000044320 | **1700001O22Rik** | 1700113K14Rik\|4930549G18Rik | RIKEN cDNA 1700001O22 gene | protein-coding |
| chr13_107645976_107646578 | chr13 | 107645977 | 107646578 | + | -0.658475 | Intergenic | Intergenic | -41120 | NR_045900 | 72257 | Mm.234155 | NR_045900 | | **1700006H21Rik** | - | RIKEN cDNA 1700006H21 gene | ncRNA |
| chr16_33337774_33338045 | chr16 | 33337775 | 33338045 | + | -0.629857 | Intergenic | RSINE1\|SINE\|B4 | 42826 | NR_045709 | 100502742 | Mm.217880 | NR_045709 | ENSMUSG00000097318 | **1700007L15Rik** | - | RIKEN cDNA 1700007L15 gene | ncRNA |
| chr16_66276683_66277130 | chr16 | 66276684 | 66277130 | + | -1.17026 | Intergenic | Intergenic | -380211 | NR_040512 | 75501 | Mm.394300 | NR_040512 | ENSMUSG00000092599 | **1700010K23Rik** | - | RIKEN cDNA 1700010K23 gene | ncRNA |
| chr14_49221596_49222098 | chr14 | 49221597 | 49222098 | + | -0.654069 | Intergenic | RLTR33\|LTR\|ERVK | 23581 | NM_025956 | 67082 | Mm.87439 | NM_025956 | ENSMUSG00000021850 | **1700011H14Rik** | - | RIKEN cDNA 1700011H14 gene | protein-coding |
| chr14_49221760_49222111 | chr14 | 49221761 | 49222111 | + | -0.703473 | Intergenic | RLTR33\|LTR\|ERVK | 23492 | NM_025956 | 67082 | Mm.87439 | NM_025956 | ENSMUSG00000021850 | **1700011H14Rik** | - | RIKEN cDNA 1700011H14 gene | protein-coding |
| chr14_49221758_49222332 | chr14 | 49221759 | 49222332 | + | -0.7227 | Intergenic | RLTR33\|LTR\|ERVK | 23383 | NM_025956 | 67082 | Mm.87439 | NM_025956 | ENSMUSG00000021850 | **1700011H14Rik** | - | RIKEN cDNA 1700011H14 gene | protein-coding |
| chr15_67289716_67290327 | chr15 | 67289717 | 67290327 | + | -0.84473 | intron (NR_045140, intron 5 of 8) | intron (NR_045140, intron 5 of 8) | 63253 | NR_045140 | 69334 | Mm.63770 | NR_045140 | | **1700012I11Rik** | - | RIKEN cDNA 1700012I11 gene | ncRNA |
| chr6_124375368_124376043 | chr6 | 124375369 | 124376043 | + | -0.993324 | Intergenic | Intergenic | -18620 | NM_001177502 | 76921 | Mm.425335 | NM_001177502 | ENSMUSG00000079346 | **1700013D24Rik** | - | RIKEN cDNA 1700013D24 gene | protein-coding |
| chr13_5756858_5757933 | chr13 | 5756859 | 5757933 | + | -0.68661 | Intergenic | Intergenic | 100696 | NR_045891 | 71867 | Mm.179181 | NR_045891 | | **1700016G22Rik** | - | RIKEN cDNA 1700016G22 gene | ncRNA |
| chr13_5806183_5806554 | chr13 | 5806184 | 5806554 | + | -0.880736 | Intergenic | B1F2\|SINE\|Alu | 51723 | NR_045891 | 71867 | Mm.179181 | NR_045891 | | **1700016G22Rik** | - | RIKEN cDNA 1700016G22 gene | ncRNA |
| chr11_75992575_75992856 | chr11 | 75992576 | 75992856 | + | -0.777239 | Intergenic | Intergenic | -7196 | NM_198637 | 74230 | Mm.271247 | NM_198637 | ENSMUSG00000053783 | **1700016K19Rik** | - | RIKEN cDNA 1700016K19 gene | protein-coding |
| chr11_75989517_75989896 | chr11 | 75989518 | 75989896 | + | -0.824028 | Intergenic | PB1D10\|SINE\|Alu | -10205 | NM_198637 | 74230 | Mm.271247 | NM_198637 | ENSMUSG00000053783 | **1700016K19Rik** | - | RIKEN cDNA 1700016K19 gene | protein-coding |
| chr11_75989674_75989926 | chr11 | 75989675 | 75989926 | + | -0.884587 | Intergenic | ID_B1\|SINE\|B4 | -10112 | NM_198637 | 74230 | Mm.271247 | NM_198637 | ENSMUSG00000053783 | **1700016K19Rik** | - | RIKEN cDNA 1700016K19 gene | protein-coding |
| chr11_75976740_75977394 | chr11 | 75976741 | 75977394 | + | -1.18196 | Intergenic | Intergenic | -22845 | NM_198637 | 74230 | Mm.271247 | NM_198637 | ENSMUSG00000053783 | **1700016K19Rik** | - | RIKEN cDNA 1700016K19 gene | protein-coding |
| chr11_75980416_75980840 | chr11 | 75980417 | 75980840 | + | -1.30423 | Intergenic | Intergenic | -19284 | NM_198637 | 74230 | Mm.271247 | NM_198637 | ENSMUSG00000053783 | **1700016K19Rik** | - | RIKEN cDNA 1700016K19 gene | protein-coding |
| chr1_80410245_80410571 | chr1 | 80410246 | 80410571 | + | -0.801453 | Intergenic | B4A\|SINE\|B4 | -35524 | NR_040460 | 72208 | Mm.396181 | NR_040460 | | **1700016L21Rik** | Jr3 | RIKEN cDNA 1700016L21 gene | ncRNA |
| chr1_80409777_80409987 | chr1 | 80409778 | 80409987 | + | -0.806387 | Intergenic | Intergenic | -36050 | NR_040460 | 72208 | Mm.396181 | NR_040460 | | **1700016L21Rik** | Jr3 | RIKEN cDNA 1700016L21 gene | ncRNA |
| chr1_80440617_80441502 | chr1 | 80440618 | 80441502 | + | -0.950836 | Intergenic | MTE-int\|LTR\|ERVL-MaLR | -4872 | NR_040460 | 72208 | Mm.396181 | NR_040460 | | **1700016L21Rik** | Jr3 | RIKEN cDNA 1700016L21 gene | ncRNA |
| chr1_80440934_80441513 | chr1 | 80440935 | 80441513 | + | -0.994201 | Intergenic | Intergenic | -4708 | NR_040460 | 72208 | Mm.396181 | NR_040460 | | **1700016L21Rik** | Jr3 | RIKEN cDNA 1700016L21 gene | ncRNA |
| chr1_80440807_80441283 | chr1 | 80440808 | 80441283 | + | -1.08424 | Intergenic | MTE-int\|LTR\|ERVL-MaLR | -4887 | NR_040460 | 72208 | Mm.396181 | NR_040460 | | **1700016L21Rik** | Jr3 | RIKEN cDNA 1700016L21 gene | ncRNA |
| chr9_57278293_57278723 | chr9 | 57278294 | 57278723 | + | -0.723883 | Intergenic | Intergenic | -15909 | NM_028820 | 74211 | Mm.428905 | NM_028820 | ENSMUSG00000032300 | **1700017B05Rik** | AA682102\|D9Ertd278e | RIKEN cDNA 1700017B05 gene | protein-coding |
| chr9_57278043_57278777 | chr9 | 57278044 | 57278777 | + | -0.740717 | Intergenic | Intergenic | -15811 | NM_028820 | 74211 | Mm.428905 | NM_028820 | ENSMUSG00000032300 | **1700017B05Rik** | AA682102\|D9Ertd278e | RIKEN cDNA 1700017B05 gene | protein-coding |
| chr10_100770604_100770867 | chr10 | 100770605 | 100770867 | + | -0.630946 | Intergenic | Intergenic | 178350 | NM_001081246 | 66605 | Mm.72646 | NM_001081246 | ENSMUSG00000056912 | **1700017N19Rik** | - | RIKEN cDNA 1700017N19 gene | protein-coding |
| chr10_100701700_100702336 | chr10 | 100701701 | 100702336 | + | -0.637865 | Intergenic | Intergenic | 109632 | NM_001081246 | 66605 | Mm.72646 | NM_001081246 | ENSMUSG00000056912 | **1700017N19Rik** | - | RIKEN cDNA 1700017N19 gene | protein-coding |
| chr10_100770254_100770932 | chr10 | 100770255 | 100770932 | + | -0.884938 | Intergenic | B3\|SINE\|B2 | 178207 | NM_001081246 | 66605 | Mm.72646 | NM_001081246 | ENSMUSG00000056912 | **1700017N19Rik** | - | RIKEN cDNA 1700017N19 gene | protein-coding |
| chr10_100716017_100716798 | chr10 | 100716018 | 100716798 | + | -0.938254 | Intergenic | Lx8\|LINE\|L1 | 124022 | NM_001081246 | 66605 | Mm.72646 | NM_001081246 | ENSMUSG00000056912 | **1700017N19Rik** | - | RIKEN cDNA 1700017N19 gene | protein-coding |
| chr10_110890636_110891243 | chr10 | 110890637 | 110891243 | + | -1.24206 | intron (NR_045979, intron 2 of 2) | intron (NR_045979, intron 2 of 2) | 5567 | NR_045979 | 75630 | Mm.392786 | NR_045979 | | **1700020G17Rik** | 4930480D05Rik | RIKEN cDNA 1700020G17 gene | ncRNA |
| chr10_110890791_110891274 | chr10 | 110890792 | 110891274 | + | -1.3384 | intron (NR_045979, intron 2 of 2) | intron (NR_045979, intron 2 of 2) | 5474 | NR_045979 | 75630 | Mm.392786 | NR_045979 | | **1700020G17Rik** | 4930480D05Rik | RIKEN cDNA 1700020G17 gene | ncRNA |
| chr2_173436613_173437167 | chr2 | 173436614 | 173437167 | + | -0.948685 | Intergenic | L1M4\|LINE\|L1 | -85702 | NM_028158 | 72221 | Mm.46132 | NM_028158 | ENSMUSG00000027518 | **1700021F07Rik** | - | RIKEN cDNA 1700021F07 gene | protein-coding |
| chr2_173455138_173455576 | chr2 | 173455139 | 173455576 | + | -0.951991 | Intergenic | Intergenic | -67235 | NM_028158 | 72221 | Mm.46132 | NM_028158 | ENSMUSG00000027518 | **1700021F07Rik** | - | RIKEN cDNA 1700021F07 gene | protein-coding |
| chr2_173534503_173535038 | chr2 | 173534504 | 173535038 | + | -1.13978 | Intergenic | Intergenic | 12179 | NM_028158 | 72221 | Mm.46132 | NM_028158 | ENSMUSG00000027518 | **1700021F07Rik** | - | RIKEN cDNA 1700021F07 gene | protein-coding |
| chr1_152038062_152038615 | chr1 | 152038063 | 152038615 | + | -0.861407 | intron (NM_197990, intron 1 of 5) | intron (NM_197990, intron 1 of 5) | 51981 | NM_197990 | 69399 | Mm.278031 | NM_197990 | ENSMUSG00000032666 | **1700025G04Rik** | 2610510E02Rik\|2900042O04Rik\|ENSMUSG00000073540 | RIKEN cDNA 1700025G04 gene | protein-coding |
| chr5_112082975_112083611 | chr5 | 112082976 | 112083611 | + | -0.959871 | Intergenic | Intergenic | -123470 | NR_045378 | 75569 | Mm.243721 | NR_045377 | | **1700028D13Rik** | - | RIKEN cDNA 1700028D13 gene | ncRNA |
| chr5_107545835_107546320 | chr5 | 107545836 | 107546320 | + | -1.0984 | intron (NM_182745, intron 4 of 5) | intron (NM_182745, intron 4 of 5) | 11367 | NM_175241 | 76421 | Mm.67037 | NM_175241 | ENSMUSG00000089798 | **1700028K03Rik** | - | RIKEN cDNA 1700028K03 gene | protein-coding |
| chr13_96995218_96995650 | chr13 | 96995219 | 96995650 | + | -0.846182 | Intergenic | Intergenic | 38928 | NM_025585 | 66479 | Mm.390629 | NM_025585 | ENSMUSG00000052075 | **1700029F12Rik** | - | RIKEN cDNA 1700029F12 gene | protein-coding |
| chr8_13600087_13600732 | chr8 | 13600088 | 13600732 | + | -0.651216 | intron (NM_009025, intron 4 of 23) | intron (NM_009025, intron 4 of 23) | -37949 | NM_001080781 | 66501 | Mm.46136 | NM_025601 | ENSMUSG00000031452 | **1700029H14Rik** | - | RIKEN cDNA 1700029H14 gene | protein-coding |
| chr8_13619437_13619717 | chr8 | 13619438 | 13619717 | + | -0.865999 | intron (NM_009025, intron 2 of 23) | intron (NM_009025, intron 2 of 23) | -57116 | NM_001080781 | 66501 | Mm.46136 | NM_025601 | ENSMUSG00000031452 | **1700029H14Rik** | - | RIKEN cDNA 1700029H14 gene | protein-coding |
| chr8_13601236_13601553 | chr8 | 13601237 | 13601553 | + | -0.948104 | intron (NM_009025, intron 4 of 23) | L1MC\|LINE\|L1 | -38934 | NM_001080781 | 66501 | Mm.46136 | NM_025601 | ENSMUSG00000031452 | **1700029H14Rik** | - | RIKEN cDNA 1700029H14 gene | protein-coding |
| chr4_135618164_135618519 | chr4 | 135618165 | 135618519 | + | -0.753082 | Intergenic | Intergenic | -8313 | NR_015613 | 73937 | Mm.292326 | NM_028688 | ENSMUSG00000086788 | **1700029M20Rik** | 4930406E24Rik | RIKEN cDNA 1700029M20 gene | ncRNA |
| chr10_68552626_68553025 | chr10 | 68552627 | 68553025 | + | -0.766916 | Intergenic | RLTR24\|LTR\|ERV1 | -10951 | NM_028491 | 73287 | Mm.141830 | NM_028491 | ENSMUSG00000019945 | **1700040L02Rik** | - | RIKEN cDNA 1700040L02 gene | protein-coding |

| | | | | | | | | | | | | | | | | | |
|---|---|---|---|---|---|---|---|---|---|---|---|---|---|---|---|---|---|
| chr10_68552485_68552943 | chr10 | 68552486 | 68552943 | + | -0.786996 | Intergenic | ORR1D1\|LTR\|ERVL-MaLR | -10839 | NM_028491 | 73287 | Mm.141830 | NM_028491 | ENSMUSG00000019945 | **1700040L02Rik** | - | RIKEN cDNA 1700040L02 gene | protein-coding |
| chr16_76765686_76765975 | chr16 | 76765687 | 76765975 | + | -0.787199 | Intergenic | Intergenic | 245050 | NR_040573 | 76601 | Mm.396217 | NR_040573 | | **1700041M19Rik** | - | RIKEN cDNA 1700041M19 gene | ncRNA |
| chr16_76765747_76765930 | chr16 | 76765748 | 76765930 | + | -0.813675 | Intergenic | Intergenic | 245042 | NR_040573 | 76601 | Mm.396217 | NR_040573 | | **1700041M19Rik** | - | RIKEN cDNA 1700041M19 gene | ncRNA |
| chr4_57350536_57350871 | chr4 | 57350537 | 57350871 | + | -1.16033 | Intergenic | ORR1A3-int\|LTR\|ERVL-MaLR | 13588 | NR_038179 | 73349 | Mm.393774 | NR_038178 | | **1700042G15Rik** | - | RIKEN cDNA 1700042G15 gene | ncRNA |
| chr6_100580844_100581729 | chr6 | 100580845 | 100581729 | + | -0.650561 | intron (NM_181590, intron 10 of 10) | intron (NM_181590, intron 10 of 10) | 53887 | NR_040525 | 73395 | Mm.69413 | NR_040525 | | **1700049E22Rik** | - | RIKEN cDNA 1700049E22 gene | ncRNA |
| chr6_100580839_100581388 | chr6 | 100580840 | 100581388 | + | -0.679672 | intron (NM_181590, intron 10 of 10) | intron (NM_181590, intron 10 of 10) | 53714 | NR_040525 | 73395 | Mm.69413 | NR_040525 | | **1700049E22Rik** | - | RIKEN cDNA 1700049E22 gene | ncRNA |
| chr1_184005351_184006085 | chr1 | 184005352 | 184006085 | + | -0.636777 | Intergenic | Intergenic | 28280 | NM_028516 | 73363 | Mm.50115 | NM_028516 | ENSMUSG00000044854 | **1700056E22Rik** | - | RIKEN cDNA 1700056E22 gene | protein-coding |
| chr1_184013134_184013801 | chr1 | 184013135 | 184013801 | + | -0.688513 | Intergenic | Intergenic | 20530 | NM_028516 | 73363 | Mm.50115 | NM_028516 | ENSMUSG00000044854 | **1700056E22Rik** | - | RIKEN cDNA 1700056E22 gene | protein-coding |
| chr1_183989754_183989990 | chr1 | 183989755 | 183989990 | + | -0.721289 | Intergenic | ORR1F\|LTR\|ERVL-MaLR | 44126 | NM_028516 | 73363 | Mm.50115 | NM_028516 | ENSMUSG00000044854 | **1700056E22Rik** | - | RIKEN cDNA 1700056E22 gene | protein-coding |
| chr1_183898854_183899144 | chr1 | 183898855 | 183899144 | + | -0.762261 | Intergenic | Intergenic | 134999 | NM_028516 | 73363 | Mm.50115 | NM_028516 | ENSMUSG00000044854 | **1700056E22Rik** | - | RIKEN cDNA 1700056E22 gene | protein-coding |
| chr1_184013148_184013493 | chr1 | 184013149 | 184013493 | + | -0.767987 | Intergenic | MIRb\|SINE\|MIR | 20677 | NM_028516 | 73363 | Mm.50115 | NM_028516 | ENSMUSG00000044854 | **1700056E22Rik** | - | RIKEN cDNA 1700056E22 gene | protein-coding |
| chr1_184013023_184013505 | chr1 | 184013024 | 184013505 | + | -0.779683 | Intergenic | PB1D10\|SINE\|Alu | 20734 | NM_028516 | 73363 | Mm.50115 | NM_028516 | ENSMUSG00000044854 | **1700056E22Rik** | - | RIKEN cDNA 1700056E22 gene | protein-coding |
| chr1_183987633_183988190 | chr1 | 183987634 | 183988190 | + | -0.876153 | Intergenic | Intergenic | 46086 | NM_028516 | 73363 | Mm.50115 | NM_028516 | ENSMUSG00000044854 | **1700056E22Rik** | - | RIKEN cDNA 1700056E22 gene | protein-coding |
| chr1_184025049_184025505 | chr1 | 184025050 | 184025505 | + | -1.03146 | Intergenic | Intergenic | 8721 | NM_028516 | 73363 | Mm.50115 | NM_028516 | ENSMUSG00000044854 | **1700056E22Rik** | - | RIKEN cDNA 1700056E22 gene | protein-coding |
| chr1_183854006_183854511 | chr1 | 183854007 | 183854511 | + | -1.11884 | Intergenic | Intergenic | 179739 | NM_028516 | 73363 | Mm.50115 | NM_028516 | ENSMUSG00000044854 | **1700056E22Rik** | - | RIKEN cDNA 1700056E22 gene | protein-coding |
| chr1_184016499_184016768 | chr1 | 184016500 | 184016768 | + | -1.17801 | Intergenic | Intergenic | 17364 | NM_028516 | 73363 | Mm.50115 | NM_028516 | ENSMUSG00000044854 | **1700056E22Rik** | - | RIKEN cDNA 1700056E22 gene | protein-coding |
| chr1_184016414_184016819 | chr1 | 184016415 | 184016819 | + | -1.17908 | Intergenic | Intergenic | 17381 | NM_028516 | 73363 | Mm.50115 | NM_028516 | ENSMUSG00000044854 | **1700056E22Rik** | - | RIKEN cDNA 1700056E22 gene | protein-coding |
| chr1_184024577_184025005 | chr1 | 184024578 | 184025005 | + | -1.36 | Intergenic | ORR1G\|LTR\|ERVL-MaLR | 9207 | NM_028516 | 73363 | Mm.50115 | NM_028516 | ENSMUSG00000044854 | **1700056E22Rik** | - | RIKEN cDNA 1700056E22 gene | protein-coding |
| chr2_9519863_9520472 | chr2 | 9519864 | 9520472 | + | -0.948817 | Intergenic | Intergenic | 330653 | NR_038180 | 73438 | Mm.23558 | NR_038180 | ENSMUSG00000085580 | **1700061F12Rik** | - | RIKEN cDNA 1700061F12 gene | ncRNA |
| chr10_119031319_119031701 | chr10 | 119031320 | 119031701 | + | -0.709106 | Intergenic | Intergenic | -61395 | NR_045348 | 73400 | Mm.416864 | NR_045348 | | **1700064J06Rik** | - | RIKEN cDNA 1700064J06 gene | ncRNA |
| chr10_119010825_119011349 | chr10 | 119010826 | 119011349 | + | -0.926887 | Intergenic | Intergenic | -81818 | NR_045348 | 73400 | Mm.416864 | NR_045348 | | **1700064J06Rik** | - | RIKEN cDNA 1700064J06 gene | ncRNA |
| chr10_119011035_119011347 | chr10 | 119011036 | 119011347 | + | -0.983001 | Intergenic | Intergenic | -81714 | NR_045348 | 73400 | Mm.416864 | NR_045348 | | **1700064J06Rik** | - | RIKEN cDNA 1700064J06 gene | ncRNA |
| chr12_99559499_99559845 | chr12 | 99559500 | 99559845 | + | -0.827102 | Intergenic | Intergenic | 68302 | NR_045288 | 73457 | Mm.383439 | NR_045288 | | **1700064M15Rik** | - | RIKEN cDNA 1700064M15 gene | ncRNA |
| chr6_145339848_145340364 | chr6 | 145339849 | 145340364 | + | -0.783378 | Intergenic | Intergenic | -47519 | NR_003625 | 381827 | Mm.179075 | NM_001001987 | | **1700073E17Rik** | - | ribosomal protein L7 pseudogene | pseudo |
| chr6_145339902_145340168 | chr6 | 145339903 | 145340168 | + | -0.850124 | Intergenic | Intergenic | -47590 | NR_003625 | 381827 | Mm.179075 | NM_001001987 | | **1700073E17Rik** | - | ribosomal protein L7 pseudogene | pseudo |
| chr14_76664834_76665167 | chr14 | 76664835 | 76665167 | + | -0.801974 | Intergenic | Intergenic | -12567 | NR_015485 | 73272 | Mm.116804 | NR_015485 | | **1700108F19Rik** | 1700020I02Rik\|BB065030 | RIKEN cDNA 1700108F19 gene | ncRNA |
| chr14_76664522_76665161 | chr14 | 76664523 | 76665161 | + | -0.86171 | Intergenic | Intergenic | -12726 | NR_015485 | 73272 | Mm.116804 | NR_015485 | | **1700108F19Rik** | 1700020I02Rik\|BB065030 | RIKEN cDNA 1700108F19 gene | ncRNA |
| chr4_10753066_10753631 | chr4 | 10753067 | 10753631 | + | -0.888654 | intron (NR_045185, intron 2 of 4) | intron (NR_045185, intron 2 of 4) | 44453 | NR_045185 | 73624 | Mm.67392 | NR_045185 | ENSMUSG00000086080 | **1700123O12Rik** | - | RIKEN cDNA 1700123O12 gene | ncRNA |
| chr4_10753122_10753808 | chr4 | 10753123 | 10753808 | + | -0.889453 | intron (NR_045185, intron 2 of 4) | intron (NR_045185, intron 2 of 4) | 44337 | NR_045185 | 73624 | Mm.67392 | NR_045185 | ENSMUSG00000086080 | **1700123O12Rik** | - | RIKEN cDNA 1700123O12 gene | ncRNA |
| chr4_10804759_10805059 | chr4 | 10804760 | 10805059 | + | -0.980089 | Intergenic | Intergenic | -7107 | NR_045185 | 73624 | Mm.67392 | NR_045185 | ENSMUSG00000086080 | **1700123O12Rik** | - | RIKEN cDNA 1700123O12 gene | ncRNA |
| chr4_10804658_10805068 | chr4 | 10804659 | 10805068 | + | -1.00968 | Intergenic | Intergenic | -7061 | NR_045185 | 73624 | Mm.67392 | NR_045185 | ENSMUSG00000086080 | **1700123O12Rik** | - | RIKEN cDNA 1700123O12 gene | ncRNA |
| chr4_10769595_10770630 | chr4 | 10769596 | 10770630 | + | -1.16699 | intron (NR_045185, intron 2 of 4) | intron (NR_045185, intron 2 of 4) | 27689 | NR_045185 | 73624 | Mm.67392 | NR_045185 | ENSMUSG00000086080 | **1700123O12Rik** | - | RIKEN cDNA 1700123O12 gene | ncRNA |
| chr14_54682512_54682711 | chr14 | 54682513 | 54682711 | + | -0.826341 | intron (NM_001085473, intron 2 of 18) | B3\|SINE\|B2 | -3559 | NM_021437 | 58248 | Mm.197509 | NM_021437 | ENSMUSG00000040822 | **1700123O20Rik** | - | RIKEN cDNA 1700123O20 gene | protein-coding |
| chr8_68336552_68336888 | chr8 | 68336553 | 68336888 | + | -0.626779 | intron (NM_028182, intron 7 of 8) | intron (NM_028182, intron 7 of 8) | -31739 | NR_038189 | 73609 | Mm.443247 | NR_038181 | ENSMUSG00000085197 | **1700125H03Rik** | 4930510D22Rik | RIKEN cDNA 1700125H03 gene | ncRNA |
| chr1_98129888_98130423 | chr1 | 98129889 | 98130423 | + | -0.889527 | Intergenic | URR1A\|DNA\|hAT-Charlie | -1311 | NR_040439 | 69030 | Mm.402692 | NR_040439 | | **1810006J02Rik** | - | RIKEN cDNA 1810006J02 gene | ncRNA |
| chr7_73583897_73584344 | chr7 | 73583898 | 73584344 | + | -0.746883 | Intergenic | Intergenic | -25726 | NR_037569 | 69170 | Mm.34539 | NR_037569 | | **1810026B05Rik** | - | RIKEN cDNA 1810026B05 gene | ncRNA |
| chr7_73600058_73600299 | chr7 | 73600059 | 73600299 | + | -0.838704 | Intergenic | RLTR50B\|LTR\|ERVK | -41784 | NR_037569 | 69170 | Mm.34539 | NR_037569 | | **1810026B05Rik** | - | RIKEN cDNA 1810026B05 gene | ncRNA |
| chr19_3730953_3731702 | chr19 | 3730954 | 3731702 | + | -0.720319 | Intergenic | Intergenic | 22995 | NM_028077 | 72056 | Mm.23780 | NM_028077 | ENSMUSG00000035372 | **1810055G02Rik** | AI256204 | RIKEN cDNA 1810055G02 gene | protein-coding |
| chr6_35257798_35258132 | chr6 | 35257799 | 35258132 | + | -0.949504 | intron (NR_027875, intron 1 of 2) | L1MB8\|LINE\|L1 | 5233 | NR_027875 | 67705 | Mm.228046 | NR_015608 | ENSMUSG00000073155 | **1810058I24Rik** | AI414869\|AI848234 | RIKEN cDNA 1810058I24 gene | ncRNA |
| chr18_75330967_75331316 | chr18 | 75330968 | 75331316 | + | -0.672831 | Intergenic | Intergenic | -33813 | NR_045393 | 100503117 | Mm.250517 | NR_045393 | | **2010010A06Rik** | 5033420E22Rik | RIKEN cDNA 2010010A06 gene | ncRNA |
| chr18_75323821_75324179 | chr18 | 75323822 | 75324179 | + | -1.04185 | Intergenic | Intergenic | -26671 | NR_045393 | 100503117 | Mm.250517 | NR_045393 | | **2010010A06Rik** | 5033420E22Rik | RIKEN cDNA 2010010A06 gene | ncRNA |
| chr18_75304375_75305489 | chr18 | 75304376 | 75305489 | + | -1.15738 | Intergenic | Intergenic | -7603 | NR_045393 | 100503117 | Mm.250517 | NR_045393 | | **2010010A06Rik** | 5033420E22Rik | RIKEN cDNA 2010010A06 gene | ncRNA |
| chr18_75322124_75322710 | chr18 | 75322125 | 75322710 | + | -1.19242 | Intergenic | Intergenic | -25088 | NR_045393 | 100503117 | Mm.250517 | NR_045393 | | **2010010A06Rik** | 5033420E22Rik | RIKEN cDNA 2010010A06 gene | ncRNA |
| chr18_75304315_75305067 | chr18 | 75304316 | 75305067 | + | -1.24685 | Intergenic | Intergenic | -7362 | NR_045393 | 100503117 | Mm.250517 | NR_045393 | | **2010010A06Rik** | 5033420E22Rik | RIKEN cDNA 2010010A06 gene | ncRNA |
| chr4_155414680_155415013 | chr4 | 155414681 | 155415013 | + | -0.806081 | non-coding (NR_102287, exon 2 of 4) | non-coding (NR_102287, exon 2 of 4) | 5657 | NM_001166029 | 544678 | Mm.472798 | NM_177674 | ENSMUSG00000042233 | **2010015L04Rik** | A530045M11 | RIKEN cDNA 2010015L04 gene | protein-coding |
| chr17_86322955_86323794 | chr17 | 86322956 | 86323794 | + | -0.629792 | intron (NM_011104, intron 1 of 14) | intron (NM_011104, intron 1 of 14) | -36197 | NR_045435 | 72104 | Mm.443401 | NR_045435 | | **2010106C02Rik** | - | RIKEN cDNA 2010106C02 gene | ncRNA |
| chr17_86336591_86337770 | chr17 | 86336592 | 86337770 | + | -0.898823 | intron (NM_011104, intron 1 of 14) | intron (NM_011104, intron 1 of 14) | -50003 | NR_045435 | 72104 | Mm.443401 | NR_045435 | | **2010106C02Rik** | - | RIKEN cDNA 2010106C02 gene | ncRNA |
| chr17_86434810_86435221 | chr17 | 86434811 | 86435221 | + | -0.963213 | intron (NM_011104, intron 2 of 14) | intron (NM_011104, intron 2 of 14) | -147838 | NR_045435 | 72104 | Mm.443401 | NR_045435 | | **2010106C02Rik** | - | RIKEN cDNA 2010106C02 gene | ncRNA |
| chr13_63261079_63262001 | chr13 | 63261080 | 63262001 | + | -1.0194 | intron (NR_110520, intron 2 of 5) | intron (NR_110520, intron 2 of 5) | 21511 | NR_110521 | 72061 | Mm.188105 | NM_028079 | ENSMUSG00000021458 | **2010111I01Rik** | 2300006M17Rik\|AP-O\|Aopep | RIKEN cDNA 2010111I01 gene | protein-coding |
| chr13_63261032_63261733 | chr13 | 63261033 | 63261733 | + | -1.01963 | intron (NR_110520, intron 2 of 5) | intron (NR_110520, intron 2 of 5) | 21354 | NR_110521 | 72061 | Mm.188105 | NM_028079 | ENSMUSG00000021458 | **2010111I01Rik** | 2300006M17Rik\|AP-O\|Aopep | RIKEN cDNA 2010111I01 gene | protein-coding |
| chr9_124257523_124257722 | chr9 | 124257524 | 124257722 | + | -0.707674 | Intergenic | Intergenic | 55071 | NM_001243118 | 630836 | Mm.458553 | NM_001243117 | | **2010315B03Rik** | BC029716 | RIKEN cDNA 2010315B03 gene | protein-coding |
| chr7_29251205_29251476 | chr7 | 29251206 | 29251476 | + | -0.770591 | Intergenic | Intergenic | -2874 | NM_028179 | 72275 | Mm.441142 | NM_028179 | | **2200002D01Rik** | 2010005J02Rik\|H2RSP\|IMUP\|IMUP-1\|IMUP-2 | RIKEN cDNA 2200002D01 gene | protein-coding |
| chr6_31240079_31240537 | chr6 | 31240080 | 31240537 | + | -0.727228 | intron (NR_040260, intron 1 of 5) | intron (NR_040260, intron 1 of 5) | 19957 | NR_040261 | 73652 | Mm.441297 | NR_040257 | ENSMUSG00000087380 | **2210408F21Rik** | - | RIKEN cDNA 2210408F21 gene | ncRNA |
| chr6_31221602_31221903 | chr6 | 31221603 | 31221903 | + | -1.10124 | intron (NR_040260, intron 1 of 5) | intron (NR_040260, intron 1 of 5) | 1402 | NR_040261 | 73652 | Mm.441297 | NR_040257 | ENSMUSG00000087380 | **2210408F21Rik** | - | RIKEN cDNA 2210408F21 gene | ncRNA |
| chr5_148708969_148709435 | chr5 | 148708970 | 148709435 | + | -0.741498 | Intergenic | Intergenic | -32638 | NR_028285 | 70138 | Mm.417834 | NR_028285 | ENSMUSG00000086494 | **2210417A02Rik** | AV081004 | RIKEN cDNA 2210417A02 gene | ncRNA |
| chr6_129210242_129210789 | chr6 | 129210243 | 129210789 | + | -0.770686 | Intergenic | Intergenic | 27529 | NR_040265 | 76432 | Mm.261738 | NR_040265 | ENSMUSG00000097354 | **2310001H17Rik** | AI646108 | RIKEN cDNA 2310001H17 gene | ncRNA |
| chr8_26716903_26717271 | chr8 | 26716904 | 26717271 | + | -0.873641 | Intergenic | ORR1F\|LTR\|ERVL-MaLR | 107336 | NR_045904 | 72316 | Mm.490534 | NR_045904 | | **2310008N11Rik** | - | RIKEN cDNA 2310008N11 gene | ncRNA |
| chr10_70180774_70181372 | chr10 | 70180775 | 70181372 | + | -1.03517 | intron (NM_001111121, intron 6 of 8) | PB1D10\|SINE\|Alu | -23594 | NR_038041 | 69563 | Mm.64672 | NR_038041 | ENSMUSG00000019933 | **2310015B20Rik** | - | RIKEN cDNA 2310015B20 gene | ncRNA |
| chr10_70180709_70181184 | chr10 | 70180710 | 70181184 | + | -1.10376 | intron (NM_001111121, intron 6 of 8) | intron (NM_001111121, intron 6 of 8) | -23720 | NR_038041 | 69563 | Mm.64672 | NR_038041 | ENSMUSG00000019933 | **2310015B20Rik** | - | RIKEN cDNA 2310015B20 gene | ncRNA |
| chr11_115751794_115752028 | chr11 | 115751795 | 115752028 | + | -0.708732 | Intergenic | ID_B1\|SINE\|B4 | -13522 | NM_028014 | 71947 | Mm.23168 | NM_028014 | ENSMUSG00000020747 | **2310067B10Rik** | Kiaa0195\|mKIAA0195 | RIKEN cDNA 2310067B10 gene | protein-coding |
| chr11_62598660_62598927 | chr11 | 62598661 | 62598927 | + | -0.635356 | intron (NM_011706, intron 15 of 15) | intron (NM_011706, intron 15 of 15) | -4083 | NR_030738 | 69221 | Mm.270874 | NR_030738 | ENSMUSG00000086841 | **2410006H16Rik** | - | RIKEN cDNA 2410006H16 gene | ncRNA |
| chr3_75713644_75714294 | chr3 | 75713645 | 75714294 | + | -0.850161 | Intergenic | Intergenic | -58238 | NR_040539 | 69193 | Mm.374088 | NR_040539 | | **2410007B07Rik** | - | RIKEN cDNA 2410007B07 gene | ncRNA |
| chr3_75713998_75714228 | chr3 | 75713999 | 75714228 | + | -0.99861 | Intergenic | Intergenic | -58382 | NR_040539 | 69193 | Mm.374088 | NR_040539 | | **2410007B07Rik** | - | RIKEN cDNA 2410007B07 gene | ncRNA |
| chr3_75682681_75683221 | chr3 | 75682682 | 75683221 | + | -1.20033 | Intergenic | MTC\|LTR\|ERVL-MaLR | -27220 | NR_040539 | 69193 | Mm.374088 | NR_040539 | | **2410007B07Rik** | - | RIKEN cDNA 2410007B07 gene | ncRNA |
| chr3_75682914_75683186 | chr3 | 75682915 | 75683186 | + | -1.20036 | Intergenic | MTC\|LTR\|ERVL-MaLR | -27319 | NR_040539 | 69193 | Mm.374088 | NR_040539 | | **2410007B07Rik** | - | RIKEN cDNA 2410007B07 gene | ncRNA |
| chr15_8168445_8169769 | chr15 | 8168446 | 8169769 | + | -0.602087 | promoter-TSS (NM_001162906) | promoter-TSS (NM_001162906) | 1 | NM_001162906 | 73692 | Mm.164436 | NM_001162906 | ENSMUSG00000039801 | **2410089E03Rik** | 4732468D17Rik\|b2b012Clo | RIKEN cDNA 2410089E03 gene | protein-coding |
| chr17_36972788_36973162 | chr17 | 36972789 | 36973162 | + | -0.649766 | Intergenic | Intergenic | 8262 | NM_029747 | 76797 | Mm.159787 | NM_029747 | ENSMUSG00000064308 | **2410137M14Rik** | - | RIKEN cDNA 2410137M14 gene | protein-coding |
| chr2_153338296_153338564 | chr2 | 153338297 | 153338564 | + | -0.723332 | Intergenic | Intergenic | 7380 | NR_040318 | 72326 | Mm.400102 | NR_040318 | ENSMUSG00000073236 | **2500004C02Rik** | - | RIKEN cDNA 2500004C02 gene | ncRNA |
| chr16_21689191_21689544 | chr16 | 21689192 | 21689544 | + | -0.70037 | intron (NM_001001881, intron 1 of 1) | ORR1D2\|LTR\|ERVL-MaLR | 5297 | NM_001001881 | 72190 | Mm.131606 | NM_001001881 | ENSMUSG00000043391 | **2510009E07Rik** | AI194952 | RIKEN cDNA 2510009E07 gene | protein-coding |

| | | | | | | | | | | | | | | | | | |
|---|---|---|---|---|---|---|---|---|---|---|---|---|---|---|---|---|---|
| chrX_50835662_50836670 | chrX | 50835663 | 50836670 | + | -0.78179 | Intergenic | Intergenic | -5270 | NM_133729 | 70415 | Mm.391246 | NM_133729 | | **2610018G03Rik** | Mst4 | RIKEN cDNA 2610018G03 gene | protein-coding |
| chrX_50836109_50836678 | chrX | 50836110 | 50836678 | + | -0.79026 | Intergenic | Intergenic | -5042 | NM_133729 | 70415 | Mm.391246 | NM_133729 | | **2610018G03Rik** | Mst4 | RIKEN cDNA 2610018G03 gene | protein-coding |
| chr7_119835838_119836837 | chr7 | 119835839 | 119836837 | + | -0.642621 | intron (NM_001004187, intron 12 of 12) | Lx5b\|LINE\|L1 | 42151 | NM_001004187 | 434234 | Mm.38237 | NM_028129 | ENSMUSG00000030924 | **2610020H08Rik** | - | RIKEN cDNA 2610020H08 gene | protein-coding |
| chr11_113173964_113174882 | chr11 | 113173965 | 113174882 | + | -0.818459 | intron (NR_015556, intron 1 of 2) | intron (NR_015556, intron 1 of 2) | 27415 | NR_015556 | 72386 | Mm.266961 | NR_015556 | ENSMUSG00000087259 | **2610035D17Rik** | - | RIKEN cDNA 2610035D17 gene | ncRNA |
| chr11_113190569_113190936 | chr11 | 113190570 | 113190936 | + | -0.821165 | intron (NR_015556, intron 1 of 2) | intron (NR_015556, intron 1 of 2) | 11085 | NR_015556 | 72386 | Mm.266961 | NR_015556 | ENSMUSG00000087259 | **2610035D17Rik** | - | RIKEN cDNA 2610035D17 gene | ncRNA |
| chr11_113153349_113153802 | chr11 | 113153350 | 113153802 | + | -0.841604 | intron (NR_015556, intron 1 of 2) | intron (NR_015556, intron 1 of 2) | 48262 | NR_015556 | 72386 | Mm.266961 | NR_015556 | ENSMUSG00000087259 | **2610035D17Rik** | - | RIKEN cDNA 2610035D17 gene | ncRNA |
| chr11_113173905_113174486 | chr11 | 113173906 | 113174486 | + | -0.900738 | intron (NR_015556, intron 1 of 2) | intron (NR_015556, intron 1 of 2) | 27642 | NR_015556 | 72386 | Mm.266961 | NR_015556 | ENSMUSG00000087259 | **2610035D17Rik** | - | RIKEN cDNA 2610035D17 gene | ncRNA |
| chr19_32406612_32407122 | chr19 | 32406613 | 32407122 | + | -0.668811 | Intergenic | ORR1E\|LTR\|ERVL-MaLR | 17651 | NR_033198 | 67188 | Mm.391160 | NR_033198 | ENSMUSG00000097787 | **2700046G09Rik** | - | RIKEN cDNA 2700046G09 gene | ncRNA |
| chr19_32406788_32407111 | chr19 | 32406789 | 32407111 | + | -0.698677 | Intergenic | ORR1E\|LTR\|ERVL-MaLR | 17734 | NR_033198 | 67188 | Mm.391160 | NR_033198 | ENSMUSG00000097787 | **2700046G09Rik** | - | RIKEN cDNA 2700046G09 gene | ncRNA |
| chr19_32409527_32410030 | chr19 | 32409528 | 32410030 | + | -1.36998 | Intergenic | RMER12B\|LTR\|ERVK | 20563 | NR_033198 | 67188 | Mm.391160 | NR_033198 | ENSMUSG00000097787 | **2700046G09Rik** | - | RIKEN cDNA 2700046G09 gene | ncRNA |
| chr12_71192732_71193067 | chr12 | 71192733 | 71193067 | + | -1.40658 | intron (NM_029818, intron 25 of 27) | intron (NM_029818, intron 25 of 27) | 56052 | NM_001163378 | 76967 | Mm.87290 | NM_029818 | ENSMUSG00000034601 | **2700049A03Rik** | Talpid3\|mKIAA0586 | RIKEN cDNA 2700049A03 gene | protein-coding |
| chr12_71192669_71193298 | chr12 | 71192670 | 71193298 | + | -1.40697 | intron (NM_029818, intron 25 of 27) | intron (NM_029818, intron 25 of 27) | 56136 | NM_001163378 | 76967 | Mm.87290 | NM_029818 | ENSMUSG00000034601 | **2700049A03Rik** | Talpid3\|mKIAA0586 | RIKEN cDNA 2700049A03 gene | protein-coding |
| chr11_64123259_64123463 | chr11 | 64123260 | 64123463 | + | -0.707402 | Intergenic | Intergenic | 43877 | NR_033780 | 66456 | Mm.124828 | NR_033780 | ENSMUSG00000087497 | **2810001G20Rik** | AW045514 | RIKEN cDNA 2810001G20 gene | ncRNA |
| chr11_64116449_64117426 | chr11 | 64116450 | 64117426 | + | -0.806582 | Intergenic | Intergenic | 37454 | NR_033780 | 66456 | Mm.124828 | NR_033780 | ENSMUSG00000087497 | **2810001G20Rik** | AW045514 | RIKEN cDNA 2810001G20 gene | ncRNA |
| chr11_64085878_64086261 | chr11 | 64085879 | 64086261 | + | -0.852731 | Intergenic | ORR1E\|LTR\|ERVL-MaLR | 6586 | NR_033780 | 66456 | Mm.124828 | NR_033780 | ENSMUSG00000087497 | **2810001G20Rik** | AW045514 | RIKEN cDNA 2810001G20 gene | ncRNA |
| chr1_64688093_64688401 | chr1 | 64688094 | 64688401 | + | -0.895345 | intron (NR_038009, intron 1 of 1) | MTC\|LTR\|ERVL-MaLR | 2412 | NR_038009 | 69941 | Mm.69044 | NR_038009 | ENSMUSG00000087213 | **2810408I11Rik** | - | RIKEN cDNA 2810408I11 gene | ncRNA |
| chr4_35226389_35226620 | chr4 | 35226390 | 35226620 | + | -0.977212 | promoter-TSS (NM_001081343) | promoter-TSS (NM_001081343) | -625 | NM_001081343 | 73205 | Mm.331544 | NM_001081343 | ENSMUSG00000028300 | **3110043O21Rik** | AI840585 | RIKEN cDNA 3110043O21 gene | protein-coding |
| chr17_21617219_21617960 | chr17 | 21617220 | 21617960 | + | -0.694029 | Intergenic | Intergenic | -33021 | NM_001166497 | 73229 | Mm.296733 | NM_001166497 | | **3110052M02Rik** | mszf57\|mszf71 | RIKEN cDNA 3110052M02 gene | protein-coding |
| chr12_71006728_71007252 | chr12 | 71006729 | 71007252 | + | -1.31072 | intron (NR_045055, intron 3 of 3) | Lx8\|LINE\|L1 | 8833 | NR_045056 | 73204 | Mm.363770 | NR_045055 | ENSMUSG00000085622 | **3110056K07Rik** | 4921536K06Rik | RIKEN cDNA 3110056K07 gene | ncRNA |
| chr12_88408038_88408849 | chr12 | 88408039 | 88408849 | + | -0.651971 | intron (NM_001277296, intron 4 of 10) | intron (NM_001277296, intron 4 of 10) | -31121 | NR_045888 | 71818 | Mm.490789 | NR_045888 | | **3200001D21Rik** | - | RIKEN cDNA 3200001D21 gene | ncRNA |
| chr13_107059367_107059859 | chr13 | 107059368 | 107059859 | + | -0.829648 | Intergenic | Intergenic | 37043 | NR_015471 | 100040322 | Mm.133867 | NR_015471 | | **3830408C21Rik** | - | RIKEN cDNA 3830408C21 gene | ncRNA |
| chr10_60370494_60371275 | chr10 | 60370495 | 60371275 | + | -0.985717 | 3' UTR (NM_028732, exon 7 of 7) | 3' UTR (NM_028732, exon 7 of 7) | 24034 | NM_028732 | 74048 | Mm.273584 | NM_028732 | ENSMUSG00000020101 | **4632428N05Rik** | Dies1\|PD-1H\|VISTA | RIKEN cDNA 4632428N05 gene | protein-coding |
| chr11_113251067_113251745 | chr11 | 113251068 | 113251745 | + | -0.666603 | intron (NM_027216, intron 7 of 9) | intron (NM_027216, intron 7 of 9) | -41737 | NR_040276 | 319871 | Mm.383743 | NR_040276 | ENSMUSG00000085421 | **4732490B19Rik** | - | RIKEN cDNA 4732490B19 gene | ncRNA |
| chr11_113207956_113208321 | chr11 | 113207957 | 113208321 | + | -0.70484 | intron (NR_040276, intron 2 of 3) | intron (NR_040276, intron 2 of 3) | 1530 | NR_040276 | 319871 | Mm.383743 | NR_040276 | ENSMUSG00000085421 | **4732490B19Rik** | - | RIKEN cDNA 4732490B19 gene | ncRNA |
| chr11_113339802_113340239 | chr11 | 113339803 | 113340239 | + | -0.719724 | intron (NM_027216, intron 6 of 9) | B4A\|SINE\|B4 | -130352 | NR_040276 | 319871 | Mm.383743 | NR_040276 | ENSMUSG00000085421 | **4732490B19Rik** | - | RIKEN cDNA 4732490B19 gene | ncRNA |
| chr11_113207815_113208317 | chr11 | 113207816 | 113208317 | + | -0.733526 | intron (NR_040276, intron 2 of 3) | intron (NR_040276, intron 2 of 3) | 1603 | NR_040276 | 319871 | Mm.383743 | NR_040276 | ENSMUSG00000085421 | **4732490B19Rik** | - | RIKEN cDNA 4732490B19 gene | ncRNA |
| chr11_113251161_113251562 | chr11 | 113251162 | 113251562 | + | -0.754252 | intron (NM_027216, intron 7 of 9) | intron (NM_027216, intron 7 of 9) | -41693 | NR_040276 | 319871 | Mm.383743 | NR_040276 | ENSMUSG00000085421 | **4732490B19Rik** | - | RIKEN cDNA 4732490B19 gene | ncRNA |
| chr11_113267581_113268495 | chr11 | 113267582 | 113268495 | + | -0.883873 | intron (NM_027216, intron 7 of 9) | L1MB3\|LINE\|L1 | -58369 | NR_040276 | 319871 | Mm.383743 | NR_040276 | ENSMUSG00000085421 | **4732490B19Rik** | - | RIKEN cDNA 4732490B19 gene | ncRNA |
| chr13_119463262_119463544 | chr13 | 119463263 | 119463544 | + | -0.730196 | intron (NM_001113550, intron 1 of 12) | intron (NM_001113550, intron 1 of 12) | 644 | NM_026127 | 67392 | Mm.390641 | NM_026127 | ENSMUSG00000062822 | **4833420G17Rik** | 2210017J06Rik\|5730533D17Rik\|AW107518\|C85193 | RIKEN cDNA 4833420G17 gene | protein-coding |
| chr17_35741921_35742267 | chr17 | 35741922 | 35742267 | + | -0.67693 | Intergenic | B4A\|SINE\|B4 | -30356 | NR_045459 | 74601 | Mm.410840 | NR_045459 | ENSMUSG00000092171 | **4833427F10Rik** | - | RIKEN cDNA 4833427F10 gene | ncRNA |
| chr17_35766414_35766809 | chr17 | 35766415 | 35766809 | + | -0.67805 | Intergenic | Intergenic | -5838 | NR_045459 | 74601 | Mm.410840 | NR_045459 | ENSMUSG00000092171 | **4833427F10Rik** | - | RIKEN cDNA 4833427F10 gene | ncRNA |
| chr17_35752184_35752590 | chr17 | 35752185 | 35752590 | + | -0.74065 | Intergenic | MER91A\|DNA\|hAT-Tip100 | -20063 | NR_045459 | 74601 | Mm.410840 | NR_045459 | ENSMUSG00000092171 | **4833427F10Rik** | - | RIKEN cDNA 4833427F10 gene | ncRNA |
| chr3_79252830_79253407 | chr3 | 79252831 | 79253407 | + | -0.815819 | intron (NR_045502, intron 2 of 2) | intron (NR_045502, intron 2 of 2) | 40200 | NR_045502 | 70924 | Mm.444622 | NR_045502 | | **4921511C10Rik** | - | RIKEN cDNA 4921511C10 gene | ncRNA |
| chr12_12372894_12373398 | chr12 | 12372895 | 12373398 | + | -0.65283 | 3' UTR (NM_029758, exon 13 of 13) | 3' UTR (NM_029758, exon 13 of 13) | 19329 | NR_045829 | 70858 | Mm.410838 | NR_045829 | | **4921511I17Rik** | - | RIKEN cDNA 4921511I17 gene | ncRNA |
| chr10_120625278_120625899 | chr10 | 120625279 | 120625899 | + | -1.23944 | Intergenic | Intergenic | -140143 | NR_038000 | 70874 | Mm.427286 | NR_038000 | ENSMUSG00000044544 | **4921513I03Rik** | - | RIKEN cDNA 4921513I03 gene | ncRNA |
| chr13_59695776_59696139 | chr13 | 59695777 | 59696139 | + | -0.738357 | Intergenic | Intergenic | -1857 | NM_183290 | 70900 | Mm.73124 | NM_183290 | ENSMUSG00000049902 | **4921517D22Rik** | - | RIKEN cDNA 4921517D22 gene | protein-coding |
| chr13_52144087_52144606 | chr13 | 52144088 | 52144606 | + | -0.990786 | Intergenic | ORR1A2-int\|LTR\|ERVL-MaLR | 47606 | NR_045061 | 74050 | Mm.84860 | NR_045061 | | **4921525O09Rik** | - | RIKEN cDNA 4921525O09 gene | ncRNA |
| chr13_52144137_52144387 | chr13 | 52144138 | 52144387 | + | -1.01607 | Intergenic | ORR1A2-int\|LTR\|ERVL-MaLR | 47521 | NR_045061 | 74050 | Mm.84860 | NR_045061 | | **4921525O09Rik** | - | RIKEN cDNA 4921525O09 gene | ncRNA |
| chr13_52100040_52100555 | chr13 | 52100041 | 52100555 | + | -1.04179 | intron (NR_045061, intron 3 of 7) | intron (NR_045061, intron 3 of 7) | 3557 | NR_045061 | 74050 | Mm.84860 | NR_045061 | | **4921525O09Rik** | - | RIKEN cDNA 4921525O09 gene | ncRNA |
| chr13_52026331_52026787 | chr13 | 52026332 | 52026787 | + | -1.1512 | Intergenic | Intergenic | -70182 | NR_045061 | 74050 | Mm.84860 | NR_045061 | | **4921525O09Rik** | - | RIKEN cDNA 4921525O09 gene | ncRNA |
| chr13_101538129_101538472 | chr13 | 101538130 | 101538472 | + | -0.724726 | Intergenic | ORR1E\|LTR\|ERVL-MaLR | -140721 | NR_046187 | 78136 | Mm.399065 | NR_046187 | | **4922502H24Rik** | - | RIKEN cDNA 4922502H24 gene | ncRNA |
| chr2_35422398_35422913 | chr2 | 35422399 | 35422913 | + | -0.711781 | intron (NM_010283, intron 3 of 8) | intron (NM_010283, intron 3 of 8) | -25482 | NM_001272040 | 74854 | Mm.159338 | NM_001080709 | ENSMUSG00000079421 | **4930402F06Rik** | - | RIKEN cDNA 4930402F06 gene | protein-coding |
| chr2_35423730_35424332 | chr2 | 35423731 | 35424332 | + | -1.03589 | intron (NM_010283, intron 2 of 8) | ORR1A1-int\|LTR\|ERVL-MaLR | -26857 | NM_001272040 | 74854 | Mm.159338 | NM_001080709 | ENSMUSG00000079421 | **4930402F06Rik** | - | RIKEN cDNA 4930402F06 gene | protein-coding |
| chr7_122932049_122932672 | chr7 | 122932050 | 122932672 | + | -1.23641 | Intergenic | MT2B1\|LTR\|ERVL | 38098 | NR_045614 | 73951 | Mm.389936 | NR_045614 | | **4930413G21Rik** | - | RIKEN cDNA 4930413G21 gene | ncRNA |
| chr9_115617793_115618333 | chr9 | 115617794 | 115618333 | + | -0.686055 | Intergenic | Intergenic | 281902 | NR_040730 | 73856 | Mm.399156 | NR_040730 | | **4930428G15Rik** | - | RIKEN cDNA 4930428G15 gene | ncRNA |
| chr9_115407980_115408517 | chr9 | 115407981 | 115408517 | + | -0.741902 | Intergenic | Intergenic | 72088 | NR_040730 | 73856 | Mm.399156 | NR_040730 | | **4930428G15Rik** | - | RIKEN cDNA 4930428G15 gene | ncRNA |
| chr9_115481122_115481867 | chr9 | 115481123 | 115481867 | + | -0.747892 | Intergenic | Intergenic | 145334 | NR_040730 | 73856 | Mm.399156 | NR_040730 | | **4930428G15Rik** | - | RIKEN cDNA 4930428G15 gene | ncRNA |
| chr9_115473904_115474637 | chr9 | 115473905 | 115474637 | + | -0.786717 | Intergenic | MLT1A1-int\|LTR\|ERVL-MaLR | 138110 | NR_040730 | 73856 | Mm.399156 | NR_040730 | | **4930428G15Rik** | - | RIKEN cDNA 4930428G15 gene | ncRNA |
| chr9_115415048_115415395 | chr9 | 115415049 | 115415395 | + | -0.791032 | Intergenic | L1MCb\|LINE\|L1 | 79061 | NR_040730 | 73856 | Mm.399156 | NR_040730 | | **4930428G15Rik** | - | RIKEN cDNA 4930428G15 gene | ncRNA |
| chr9_115480916_115481569 | chr9 | 115480917 | 115481569 | + | -0.830478 | Intergenic | Intergenic | 145082 | NR_040730 | 73856 | Mm.399156 | NR_040730 | | **4930428G15Rik** | - | RIKEN cDNA 4930428G15 gene | ncRNA |
| chr9_115526145_115526849 | chr9 | 115526146 | 115526849 | + | -0.866497 | Intergenic | Intergenic | 190336 | NR_040730 | 73856 | Mm.399156 | NR_040730 | | **4930428G15Rik** | - | RIKEN cDNA 4930428G15 gene | ncRNA |
| chr9_115481204_115481589 | chr9 | 115481205 | 115481589 | + | -0.878854 | Intergenic | Intergenic | 145236 | NR_040730 | 73856 | Mm.399156 | NR_040730 | | **4930428G15Rik** | - | RIKEN cDNA 4930428G15 gene | ncRNA |
| chr9_115471584_115471953 | chr9 | 115471585 | 115471953 | + | -0.976803 | Intergenic | Intergenic | 135608 | NR_040730 | 73856 | Mm.399156 | NR_040730 | | **4930428G15Rik** | - | RIKEN cDNA 4930428G15 gene | ncRNA |
| chr9_115468618_115469153 | chr9 | 115468619 | 115469153 | + | -1.02682 | Intergenic | Intergenic | 132725 | NR_040730 | 73856 | Mm.399156 | NR_040730 | | **4930428G15Rik** | - | RIKEN cDNA 4930428G15 gene | ncRNA |
| chr9_115434723_115435092 | chr9 | 115434724 | 115435092 | + | -1.10749 | Intergenic | Intergenic | 98747 | NR_040730 | 73856 | Mm.399156 | NR_040730 | | **4930428G15Rik** | - | RIKEN cDNA 4930428G15 gene | ncRNA |
| chr9_115434565_115435080 | chr9 | 115434566 | 115435080 | + | -1.11185 | Intergenic | Intergenic | 98662 | NR_040730 | 73856 | Mm.399156 | NR_040730 | | **4930428G15Rik** | - | RIKEN cDNA 4930428G15 gene | ncRNA |
| chr9_115468617_115468973 | chr9 | 115468618 | 115468973 | + | -1.12614 | Intergenic | Intergenic | 132634 | NR_040730 | 73856 | Mm.399156 | NR_040730 | | **4930428G15Rik** | - | RIKEN cDNA 4930428G15 gene | ncRNA |
| chr9_115560962_115561629 | chr9 | 115560963 | 115561629 | + | -1.1507 | Intergenic | Intergenic | 225135 | NR_040730 | 73856 | Mm.399156 | NR_040730 | | **4930428G15Rik** | - | RIKEN cDNA 4930428G15 gene | ncRNA |
| chr9_115330732_115331332 | chr9 | 115330733 | 115331332 | + | -1.28962 | Intergenic | Intergenic | -5129 | NR_040730 | 73856 | Mm.399156 | NR_040730 | | **4930428G15Rik** | - | RIKEN cDNA 4930428G15 gene | ncRNA |
| chr9_79795114_79795447 | chr9 | 79795115 | 79795447 | + | -0.931069 | intron (NR_040734, intron 1 of 2) | intron (NR_040734, intron 1 of 2) | 1640 | NR_040734 | 74633 | Mm.262175 | NR_040734 | ENSMUSG00000086918 | **4930429F24Rik** | - | RIKEN cDNA 4930429F24 gene | ncRNA |
| chr15_30662070_30662425 | chr15 | 30662071 | 30662425 | + | -1.15991 | intron (NM_008729, intron 7 of 22) | intron (NM_008729, intron 7 of 22) | -376878 | NR_045186 | 73784 | Mm.66947 | NR_045186 | | **4930430F21Rik** | - | RIKEN cDNA 4930430F21 gene | ncRNA |
| chr9_57343810_57344061 | chr9 | 57343811 | 57344061 | + | -0.984073 | Intergenic | Intergenic | -47134 | NR_040729 | 73772 | Mm.444684 | NR_040729 | | **4930430J02Rik** | - | RIKEN cDNA 4930430J02 gene | ncRNA |
| chr9_57407046_57407492 | chr9 | 57407047 | 57407492 | + | -1.05538 | Intergenic | Intergenic | 16199 | NR_040729 | 73772 | Mm.444684 | NR_040729 | | **4930430J02Rik** | - | RIKEN cDNA 4930430J02 gene | ncRNA |
| chr9_57404012_57404413 | chr9 | 57404013 | 57404413 | + | -1.19724 | Intergenic | RLTR14\|LTR\|ERV1 | 13143 | NR_040729 | 73772 | Mm.444684 | NR_040729 | | **4930430J02Rik** | - | RIKEN cDNA 4930430J02 gene | ncRNA |
| chr9_57403851_57404439 | chr9 | 57403852 | 57404439 | + | -1.21911 | Intergenic | Intergenic | 13075 | NR_040729 | 73772 | Mm.444684 | NR_040729 | | **4930430J02Rik** | - | RIKEN cDNA 4930430J02 gene | ncRNA |

| | | | | | | | | | | | | | | | | | |
|---|---|---|---|---|---|---|---|---|---|---|---|---|---|---|---|---|---|
| chr18_82846885_82847470 | chr18 | 82846886 | 82847470 | + | -1.04982 | Intergenic | Intergenic | -8719 | NR_125720 | 73985 | Mm.399211 | NR_125720 | | **4930445N18Rik** | 3010027C24Rik | RIKEN cDNA 4930445N18 gene | ncRNA |
| chr12_72843985_72844648 | chr12 | 72843986 | 72844648 | + | -0.881184 | Intergenic | Intergenic | 73448 | NM_029444 | 75801 | Mm.242892 | NM_029444 | ENSMUSG00000021098 | **4930447C04Rik** | 4921504I02Rik\|A930035O15Rik\|Six6OS\|Six6as\|Six6os1 | RIKEN cDNA 4930447C04 gene | protein-coding |
| chr13_34114486_34115084 | chr13 | 34114487 | 34115084 | + | -1.13194 | Intergenic | MTD\|LTR\|ERVL-MaLR | -1513 | NR_046184 | 432745 | Mm.159104 | NR_046184 | | **4930447K03Rik** | - | RIKEN cDNA 4930447K03 gene | ncRNA |
| chr16_64758550_64759310 | chr16 | 64758551 | 64759310 | + | -0.834333 | Intergenic | Intergenic | 12009 | NM_026273 | 67609 | Mm.212157 | NM_026273 | ENSMUSG00000059920 | **4930453N24Rik** | AI649104 | RIKEN cDNA 4930453N24 gene | protein-coding |
| chr10_21327577_21328054 | chr10 | 21327578 | 21328054 | + | -0.907545 | intron (NR_045352, intron 1 of 3) | intron (NR_045352, intron 1 of 3) | 14239 | NR_045352 | 73990 | Mm.399240 | NR_045352 | | **4930455C13Rik** | - | RIKEN cDNA 4930455C13 gene | ncRNA |
| chr10_96214757_96215355 | chr10 | 96214758 | 96215355 | + | -0.664213 | Intergenic | Intergenic | -11151 | NR_110448 | 74881 | Mm.410819 | NR_110448 | | **4930459C07Rik** | - | RIKEN cDNA 4930459C07 gene | ncRNA |
| chr10_96180741_96181334 | chr10 | 96180742 | 96181334 | + | -0.74569 | Intergenic | MIRb\|SINE\|MIR | -45169 | NR_110448 | 74881 | Mm.410819 | NR_110448 | | **4930459C07Rik** | - | RIKEN cDNA 4930459C07 gene | ncRNA |
| chr10_96136499_96136982 | chr10 | 96136500 | 96136982 | + | -0.81064 | Intergenic | Intergenic | -89466 | NR_110448 | 74881 | Mm.410819 | NR_110448 | | **4930459C07Rik** | - | RIKEN cDNA 4930459C07 gene | ncRNA |
| chr10_96240583_96240967 | chr10 | 96240584 | 96240967 | + | -0.960642 | intron (NR_110448, intron 1 of 2) | intron (NR_110448, intron 1 of 2) | 14568 | NR_110448 | 74881 | Mm.410819 | NR_110448 | | **4930459C07Rik** | - | RIKEN cDNA 4930459C07 gene | ncRNA |
| chr10_96240656_96240908 | chr10 | 96240657 | 96240908 | + | -1.00304 | intron (NR_110448, intron 1 of 2) | intron (NR_110448, intron 1 of 2) | 14575 | NR_110448 | 74881 | Mm.410819 | NR_110448 | | **4930459C07Rik** | - | RIKEN cDNA 4930459C07 gene | ncRNA |
| chr10_96204298_96204619 | chr10 | 96204299 | 96204619 | + | -1.07308 | Intergenic | RLTR18-int\|LTR\|ERVK | -21748 | NR_110448 | 74881 | Mm.410819 | NR_110448 | | **4930459C07Rik** | - | RIKEN cDNA 4930459C07 gene | ncRNA |
| chr10_96200716_96201846 | chr10 | 96200717 | 96201846 | + | -1.17265 | Intergenic | Intergenic | -24926 | NR_110448 | 74881 | Mm.410819 | NR_110448 | | **4930459C07Rik** | - | RIKEN cDNA 4930459C07 gene | ncRNA |
| chr10_96200990_96201824 | chr10 | 96200991 | 96201824 | + | -1.21045 | Intergenic | Intergenic | -24800 | NR_110448 | 74881 | Mm.410819 | NR_110448 | | **4930459C07Rik** | - | RIKEN cDNA 4930459C07 gene | ncRNA |
| chr12_107361954_107362710 | chr12 | 107361955 | 107362710 | + | -0.82694 | Intergenic | Intergenic | 373156 | NR_045973 | 74968 | Mm.359821 | NR_045973 | ENSMUSG00000085641 | **4930465M20Rik** | 4930431H15Rik | RIKEN cDNA 4930465M20 gene | ncRNA |
| chr12_107566361_107566897 | chr12 | 107566362 | 107566897 | + | -0.94 | Intergenic | Intergenic | 168859 | NR_045973 | 74968 | Mm.359821 | NR_045973 | ENSMUSG00000085641 | **4930465M20Rik** | 4930431H15Rik | RIKEN cDNA 4930465M20 gene | ncRNA |
| chr12_107748899_107749171 | chr12 | 107748900 | 107749171 | + | -0.997757 | Intergenic | Intergenic | -13547 | NR_045973 | 74968 | Mm.359821 | NR_045973 | ENSMUSG00000085641 | **4930465M20Rik** | 4930431H15Rik | RIKEN cDNA 4930465M20 gene | ncRNA |
| chr12_107779295_107780123 | chr12 | 107779296 | 107780123 | + | -1.01639 | Intergenic | Intergenic | -44221 | NR_045973 | 74968 | Mm.359821 | NR_045973 | ENSMUSG00000085641 | **4930465M20Rik** | 4930431H15Rik | RIKEN cDNA 4930465M20 gene | ncRNA |
| chr12_107729222_107729671 | chr12 | 107729223 | 107729671 | + | -1.01877 | intron (NR_045973, intron 2 of 2) | intron (NR_045973, intron 2 of 2) | 6041 | NR_045973 | 74968 | Mm.359821 | NR_045973 | ENSMUSG00000085641 | **4930465M20Rik** | 4930431H15Rik | RIKEN cDNA 4930465M20 gene | ncRNA |
| chr12_107729248_107729858 | chr12 | 107729249 | 107729858 | + | -1.01911 | non-coding (NR_045973, exon 2 of 3) | non-coding (NR_045973, exon 2 of 3) | 5935 | NR_045973 | 74968 | Mm.359821 | NR_045973 | ENSMUSG00000085641 | **4930465M20Rik** | 4930431H15Rik | RIKEN cDNA 4930465M20 gene | ncRNA |
| chr12_107779324_107779648 | chr12 | 107779325 | 107779648 | + | -1.11692 | Intergenic | Intergenic | -43998 | NR_045973 | 74968 | Mm.359821 | NR_045973 | ENSMUSG00000085641 | **4930465M20Rik** | 4930431H15Rik | RIKEN cDNA 4930465M20 gene | ncRNA |
| chr12_107779318_107779543 | chr12 | 107779319 | 107779543 | + | -1.11918 | Intergenic | Intergenic | -43943 | NR_045973 | 74968 | Mm.359821 | NR_045973 | ENSMUSG00000085641 | **4930465M20Rik** | 4930431H15Rik | RIKEN cDNA 4930465M20 gene | ncRNA |
| chr16_78557641_78557969 | chr16 | 78557642 | 78557969 | + | -0.900771 | intron (NM_001252438, intron 2 of 4) | intron (NM_001252438, intron 2 of 4) | -2511 | NR_040569 | 75818 | Mm.410817 | NR_040569 | | **4930478L05Rik** | - | RIKEN cDNA 4930478L05 gene | ncRNA |
| chr12_26185118_26185432 | chr12 | 26185119 | 26185432 | + | -0.695254 | Intergenic | Intergenic | -25992 | NR_046278 | 74930 | Mm.398932 | NR_046278 | | **4930480M12Rik** | - | RIKEN cDNA 4930480M12 gene | ncRNA |
| chr12_26187426_26187890 | chr12 | 26187427 | 26187890 | + | -0.707773 | Intergenic | Intergenic | -23609 | NR_046278 | 74930 | Mm.398932 | NR_046278 | | **4930480M12Rik** | - | RIKEN cDNA 4930480M12 gene | ncRNA |
| chr12_25882513_25882789 | chr12 | 25882514 | 25882789 | + | -0.837179 | Intergenic | ORR1D2\|LTR\|ERVL-MaLR | -328616 | NR_046278 | 74930 | Mm.398932 | NR_046278 | | **4930480M12Rik** | - | RIKEN cDNA 4930480M12 gene | ncRNA |
| chr12_26008031_26008339 | chr12 | 26008032 | 26008339 | + | -0.862441 | Intergenic | Intergenic | -203082 | NR_046278 | 74930 | Mm.398932 | NR_046278 | | **4930480M12Rik** | - | RIKEN cDNA 4930480M12 gene | ncRNA |
| chr19_5405985_5406340 | chr19 | 5405986 | 5406340 | + | -0.76116 | promoter-TSS (NR_015545) | promoter-TSS (NR_015545) | -652 | NR_015545 | 74931 | Mm.388511 | NR_015545 | ENSMUSG00000086938 | **4930481A15Rik** | - | RIKEN cDNA 4930481A15 gene | ncRNA |
| chr13_60855069_60855326 | chr13 | 60855070 | 60855326 | + | -0.733929 | intron (NM_178098, intron 1 of 7) | intron (NM_178098, intron 1 of 7) | 9218 | NM_178098 | 214639 | Mm.19839 | NM_178098 | ENSMUSG00000050345 | **4930486L24Rik** | AW545031 | RIKEN cDNA 4930486L24 gene | protein-coding |
| chr13_60842729_60843218 | chr13 | 60842730 | 60843218 | + | -0.920608 | exon (NM_178098, exon 8 of 8) | exon (NM_178098, exon 8 of 8) | 21442 | NM_178098 | 214639 | Mm.19839 | NM_178098 | ENSMUSG00000050345 | **4930486L24Rik** | AW545031 | RIKEN cDNA 4930486L24 gene | protein-coding |
| chr13_60848742_60849123 | chr13 | 60848743 | 60849123 | + | -1.02113 | intron (NM_178098, intron 5 of 7) | intron (NM_178098, intron 5 of 7) | 15483 | NM_178098 | 214639 | Mm.19839 | NM_178098 | ENSMUSG00000050345 | **4930486L24Rik** | AW545031 | RIKEN cDNA 4930486L24 gene | protein-coding |
| chr13_60839325_60839907 | chr13 | 60839326 | 60839907 | + | -1.235 | Intergenic | Intergenic | 24800 | NM_178098 | 214639 | Mm.19839 | NM_178098 | ENSMUSG00000050345 | **4930486L24Rik** | AW545031 | RIKEN cDNA 4930486L24 gene | protein-coding |
| chr1_62875548_62876224 | chr1 | 62875549 | 62876224 | + | -0.636295 | Intergenic | Intergenic | -81154 | NR_040601 | 74992 | Mm.401979 | NR_040601 | | **4930487H11Rik** | - | RIKEN cDNA 4930487H11 gene | ncRNA |
| chr1_62876783_62877042 | chr1 | 62876784 | 62877042 | + | -0.743344 | Intergenic | Intergenic | -82181 | NR_040601 | 74992 | Mm.401979 | NR_040601 | | **4930487H11Rik** | - | RIKEN cDNA 4930487H11 gene | ncRNA |
| chr1_62829678_62830336 | chr1 | 62829679 | 62830336 | + | -0.852861 | Intergenic | Intergenic | -35275 | NR_040601 | 74992 | Mm.401979 | NR_040601 | | **4930487H11Rik** | - | RIKEN cDNA 4930487H11 gene | ncRNA |
| chr4_140868441_140868817 | chr4 | 140868442 | 140868817 | + | -0.704554 | Intergenic | C-rich\|Low_complexity\|Low_complexity | -4064 | NR_040654 | 75128 | Mm.67085 | NR_040654 | ENSMUSG00000087045 | **4930515B02Rik** | - | RIKEN cDNA 4930515B02 gene | ncRNA |
| chr1_159928739_159929130 | chr1 | 159928740 | 159929130 | + | -0.766358 | Intergenic | Intergenic | -115445 | NR_028111 | 67647 | Mm.171518 | NM_001024470 | ENSMUSG00000090394 | **4930523C07Rik** | - | RIKEN cDNA 4930523C07 gene | protein-coding |
| chr1_160053497_160053983 | chr1 | 160053498 | 160053983 | + | -1.03033 | intron (NR_028111, intron 1 of 2) | intron (NR_028111, intron 1 of 2) | 9360 | NM_001162896 | 67647 | Mm.171518 | NM_001024470 | ENSMUSG00000090394 | **4930523C07Rik** | - | RIKEN cDNA 4930523C07 gene | protein-coding |
| chr11_31963918_31964440 | chr11 | 31963919 | 31964440 | + | -0.987113 | Intergenic | RMER19B\|LTR\|ERVK | -1454 | NM_026262 | 67592 | Mm.78654 | NM_026262 | ENSMUSG00000020299 | **4930524B15Rik** | - | RIKEN cDNA 4930524B15 gene | protein-coding |
| chr19_10647891_10648572 | chr19 | 10647892 | 10648572 | + | -0.880469 | promoter-TSS (NR_045316) | promoter-TSS (NR_045316) | -179 | NR_045316 | 75096 | Mm.399287 | NR_045316 | | **4930524O05Rik** | - | RIKEN cDNA 4930524O05 gene | ncRNA |
| chr9_124412950_124413262 | chr9 | 124412951 | 124413262 | + | -0.767157 | Intergenic | Intergenic | -10145 | NR_027950 | 75135 | Mm.476814 | NR_015516 | | **4930526I15Rik** | 1600027J15Rik | RIKEN cDNA 4930526I15 gene | ncRNA |
| chr14_86681925_86682315 | chr14 | 86681926 | 86682315 | + | -1.12381 | intron (NM_019670, intron 27 of 27) | intron (NM_019670, intron 27 of 27) | 436150 | NR_040457 | 75155 | Mm.399239 | NR_040457 | | **4930529K09Rik** | 4930507G21Rik | RIKEN cDNA 4930529K09 gene | ncRNA |
| chr14_86681934_86682499 | chr14 | 86681935 | 86682499 | + | -1.12916 | intron (NM_019670, intron 27 of 27) | intron (NM_019670, intron 27 of 27) | 436247 | NR_040457 | 75155 | Mm.399239 | NR_040457 | | **4930529K09Rik** | 4930507G21Rik | RIKEN cDNA 4930529K09 gene | ncRNA |
| chr9_15582039_15582362 | chr9 | 15582040 | 15582362 | + | -0.825217 | intron (NM_181816, intron 9 of 11) | intron (NM_181816, intron 9 of 11) | -37656 | NR_040746 | 78025 | Mm.55983 | NR_040746 | | **4930540M03Rik** | - | RIKEN cDNA 4930540M03 gene | ncRNA |
| chr9_15587229_15587868 | chr9 | 15587230 | 15587868 | + | -0.889451 | intron (NM_181816, intron 9 of 11) | RMER15\|LTR\|ERVL | -32308 | NR_040746 | 78025 | Mm.55983 | NR_040746 | | **4930540M03Rik** | - | RIKEN cDNA 4930540M03 gene | ncRNA |
| chr9_15580224_15580521 | chr9 | 15580225 | 15580521 | + | -0.965467 | intron (NM_181816, intron 9 of 11) | intron (NM_181816, intron 9 of 11) | -39484 | NR_040746 | 78025 | Mm.55983 | NR_040746 | | **4930540M03Rik** | - | RIKEN cDNA 4930540M03 gene | ncRNA |
| chr6_66154748_66155293 | chr6 | 66154749 | 66155293 | + | -0.836182 | Intergenic | Intergenic | 202450 | NM_001161773 | 67653 | Mm.389756 | NM_001161773 | ENSMUSG00000036463 | **4930544G11Rik** | - | RIKEN cDNA 4930544G11 gene | protein-coding |
| chr6_65998329_65998721 | chr6 | 65998330 | 65998721 | + | -0.906378 | Intergenic | Intergenic | 45954 | NM_001161773 | 67653 | Mm.389756 | NM_001161773 | ENSMUSG00000036463 | **4930544G11Rik** | - | RIKEN cDNA 4930544G11 gene | protein-coding |
| chr18_68847250_68848123 | chr18 | 68847251 | 68848123 | + | -0.664328 | Intergenic | B1F1\|SINE\|Alu | 103841 | NR_038051 | 78931 | Mm.160155 | NR_038051 | | **4930546C10Rik** | - | RIKEN cDNA 4930546C10 gene | ncRNA |
| chr18_68847260_68847810 | chr18 | 68847261 | 68847810 | + | -0.685321 | Intergenic | B3A\|SINE\|B2 | 103993 | NR_038051 | 78931 | Mm.160155 | NR_038051 | | **4930546C10Rik** | - | RIKEN cDNA 4930546C10 gene | ncRNA |
| chr18_68968851_68969903 | chr18 | 68968852 | 68969903 | + | -0.959961 | Intergenic | Intergenic | -17849 | NR_038051 | 78931 | Mm.160155 | NR_038051 | | **4930546C10Rik** | - | RIKEN cDNA 4930546C10 gene | ncRNA |
| chr18_68951788_68952143 | chr18 | 68951789 | 68952143 | + | -1.07644 | promoter-TSS (NR_038051) | promoter-TSS (NR_038051) | -438 | NR_038051 | 78931 | Mm.160155 | NR_038051 | | **4930546C10Rik** | - | RIKEN cDNA 4930546C10 gene | ncRNA |
| chr18_68951553_68952237 | chr18 | 68951554 | 68952237 | + | -1.08461 | promoter-TSS (NR_038051) | promoter-TSS (NR_038051) | -367 | NR_038051 | 78931 | Mm.160155 | NR_038051 | | **4930546C10Rik** | - | RIKEN cDNA 4930546C10 gene | ncRNA |
| chr19_55544446_55544839 | chr19 | 55544447 | 55544839 | + | -0.741535 | intron (NM_001293686, intron 7 of 7) | L1MB8\|LINE\|L1 | 157561 | NR_045318 | 77594 | Mm.329924 | NR_045318 | ENSMUSG00000097632 | **4930552P12Rik** | - | RIKEN cDNA 4930552P12 gene | ncRNA |
| chr10_96955029_96955338 | chr10 | 96955030 | 96955338 | + | -0.933423 | Intergenic | Intergenic | -80584 | NR_045358 | 75301 | Mm.108650 | NR_045358 | | **4930556N09Rik** | - | RIKEN cDNA 4930556N09 gene | ncRNA |
| chr10_96838318_96838877 | chr10 | 96838319 | 96838877 | + | -0.988574 | Intergenic | Intergenic | -197170 | NR_045358 | 75301 | Mm.108650 | NR_045358 | | **4930556N09Rik** | - | RIKEN cDNA 4930556N09 gene | ncRNA |
| chr10_96838269_96838580 | chr10 | 96838270 | 96838580 | + | -0.996302 | Intergenic | Intergenic | -197343 | NR_045358 | 75301 | Mm.108650 | NR_045358 | | **4930556N09Rik** | - | RIKEN cDNA 4930556N09 gene | ncRNA |
| chr3_105132754_105133076 | chr3 | 105132755 | 105133076 | + | -0.622642 | Intergenic | Intergenic | 66071 | NM_029228 | 75269 | Mm.307994 | NM_029228 | ENSMUSG00000051788 | **4930564D02Rik** | - | RIKEN cDNA 4930564D02 gene | protein-coding |
| chr16_76202191_76202537 | chr16 | 76202192 | 76202537 | + | -0.613526 | Intergenic | Intergenic | 79751 | NR_040575 | 78215 | Mm.243885 | NR_040575 | | **4930578N18Rik** | - | RIKEN cDNA 4930578N18 gene | ncRNA |
| chr16_76246172_76246664 | chr16 | 76246173 | 76246664 | + | -0.655697 | Intergenic | MARNA\|DNA\|TcMar-Mariner | 123805 | NR_040575 | 78215 | Mm.243885 | NR_040575 | | **4930578N18Rik** | - | RIKEN cDNA 4930578N18 gene | ncRNA |
| chr7_118254048_118254353 | chr7 | 118254049 | 118254353 | + | -0.624425 | Intergenic | ID_B1\|SINE\|B4 | 10294 | NR_027879 | 67745 | Mm.483404 | NM_183267 | ENSMUSG00000055159 | **4930583K01Rik** | - | RIKEN cDNA 4930583K01 gene | ncRNA |
| chr1_37495856_37496292 | chr1 | 37495857 | 37496292 | + | -0.748066 | intron (NM_001290801, intron 2 of 14) | intron (NM_001290801, intron 2 of 14) | -7469 | NR_024017 | 77633 | Mm.483177 | NR_024017 | | **4930594C11Rik** | - | G1 to S phase transition pseudogene | pseudo |
| chr1_37496553_37497043 | chr1 | 37496554 | 37497043 | + | -0.748692 | intron (NM_001290801, intron 2 of 14) | intron (NM_001290801, intron 2 of 14) | -8193 | NR_024017 | 77633 | Mm.483177 | NR_024017 | | **4930594C11Rik** | - | G1 to S phase transition pseudogene | pseudo |
| chr1_37496035_37496309 | chr1 | 37496036 | 37496309 | + | -0.791969 | intron (NM_001290801, intron 2 of 14) | intron (NM_001290801, intron 2 of 14) | -7567 | NR_024017 | 77633 | Mm.483177 | NR_024017 | | **4930594C11Rik** | - | G1 to S phase transition pseudogene | pseudo |
| chr1_37473164_37473625 | chr1 | 37473165 | 37473625 | + | -0.863442 | intron (NM_001290801, intron 4 of 14) | intron (NM_001290801, intron 4 of 14) | 15210 | NR_024017 | 77633 | Mm.483177 | NR_024017 | | **4930594C11Rik** | - | G1 to S phase transition pseudogene | pseudo |

| | | | | | | | | | | | | | | | | | |
|---|---|---|---|---|---|---|---|---|---|---|---|---|---|---|---|---|---|
| chr1_37481272_37481710 | chr1 | 37481273 | 37481710 | + | -1.10907 | intron (NM_001290801, intron 4 of 14) | intron (NM_001290801, intron 4 of 14) | 7114 | NR_024017 | 77633 | Mm.483177 | NR_024017 | | **4930594C11Rik** | - | G1 to S phase transition pseudogene | pseudo |
| chr1_139036365_139037445 | chr1 | 139036366 | 139037445 | + | -0.608984 | intron (NM_001166501, intron 3 of 21) | intron (NM_001166501, intron 3 of 21) | 59248 | NR_108103 | 78229 | Mm.437838 | NR_108103 | | **4930596I21Rik** | - | RIKEN cDNA 4930596I21 gene | ncRNA |
| chr6_39142238_39142524 | chr6 | 39142239 | 39142524 | + | -0.821312 | 3' UTR (NM_001033430, exon 20 of 20) | 3' UTR (NM_001033430, exon 20 of 20) | 23908 | NR_045813 | 75379 | Mm.348014 | NR_045813 | ENSMUSG00000073144 | **4930599N23Rik** | - | RIKEN cDNA 4930599N23 gene | ncRNA |
| chr3_65537358_65537683 | chr3 | 65537359 | 65537683 | + | -0.789967 | intron (NM_178892, intron 2 of 5) | intron (NM_178892, intron 2 of 5) | -8107 | NR_027955 | 71004 | Mm.475465 | NM_001033527 | ENSMUSG00000074580 | **4931440P22Rik** | - | RIKEN cDNA 4931440P22 gene | ncRNA |
| chr2_129815998_129816463 | chr2 | 129815999 | 129816463 | + | -0.858514 | intron (NM_183262, intron 3 of 3) | intron (NM_183262, intron 3 of 3) | -14772 | NR_029452 | 654409 | Mm.305555 | NR_029452 | | **4932416H05Rik** | - | RIKEN cDNA 4932416H05 gene | ncRNA |
| chr3_36853724_36854605 | chr3 | 36853725 | 36854605 | + | -1.17776 | Intergenic | Intergenic | -8941 | NM_172679 | 229227 | Mm.207907 | NM_172679 | ENSMUSG00000037270 | **4932438A13Rik** | 4732443H21\|B830039D19Rik\|D630029K19Rik\|FSA\|Kiaa1109\|Tweek | RIKEN cDNA 4932438A13 gene | protein-coding |
| chr11_54162896_54163484 | chr11 | 54162897 | 54163484 | + | -0.851038 | Intergenic | Intergenic | 50098 | NR_045506 | 71056 | Mm.56332 | NR_045506 | ENSMUSG00000086209 | **4933405E24Rik** | - | RIKEN cDNA 4933405E24 gene | ncRNA |
| chr8_105727213_105728232 | chr8 | 105727214 | 105728232 | + | -0.675709 | intron (NM_027469, intron 2 of 2) | intron (NM_027469, intron 2 of 2) | 19420 | NM_027655 | 71046 | Mm.280888 | NM_027655 | ENSMUSG00000013158 | **4933405L10Rik** | - | RIKEN cDNA 4933405L10 gene | protein-coding |
| chr8_105727267_105728029 | chr8 | 105727268 | 105728029 | + | -0.686609 | intron (NM_027469, intron 2 of 2) | intron (NM_027469, intron 2 of 2) | 19345 | NM_027655 | 71046 | Mm.280888 | NM_027655 | ENSMUSG00000013158 | **4933405L10Rik** | - | RIKEN cDNA 4933405L10 gene | protein-coding |
| chr7_114347092_114347351 | chr7 | 114347093 | 114347351 | + | -0.776253 | intron (NR_029437, intron 1 of 3) | intron (NR_029437, intron 1 of 3) | 67939 | NR_029437 | 71045 | Mm.235682 | NR_029437 | ENSMUSG00000087475 | **4933406I18Rik** | 4933403D24Rik | RIKEN cDNA 4933406I18 gene | ncRNA |
| chr7_82773574_82774422 | chr7 | 82773575 | 82774422 | + | -0.951279 | intron (NM_175317, intron 19 of 19) | intron (NM_175317, intron 19 of 19) | -18354 | NR_046004 | 77800 | Mm.444811 | NR_046004 | ENSMUSG00000086690 | **4933406J10Rik** | A930009F24Rik | RIKEN cDNA 4933406J10 gene | ncRNA |
| chr7_82773834_82774390 | chr7 | 82773835 | 82774390 | + | -0.956555 | intron (NM_175317, intron 19 of 19) | intron (NM_175317, intron 19 of 19) | -18468 | NR_046004 | 77800 | Mm.444811 | NR_046004 | ENSMUSG00000086690 | **4933406J10Rik** | A930009F24Rik | RIKEN cDNA 4933406J10 gene | ncRNA |
| chr12_107159977_107160656 | chr12 | 107159978 | 107160656 | + | -0.651372 | Intergenic | Intergenic | -443993 | NR_015512 | 71033 | Mm.159404 | NR_015512 | ENSMUSG00000085820 | **4933406K04Rik** | 4930426M03Rik\|4930485I22Rik\|4933427G14Rik | RIKEN cDNA 4933406K04 gene | ncRNA |
| chr12_107170986_107171277 | chr12 | 107170987 | 107171277 | + | -0.662942 | Intergenic | Lx8\|LINE\|L1 | -454808 | NR_015512 | 71033 | Mm.159404 | NR_015512 | ENSMUSG00000085820 | **4933406K04Rik** | 4930426M03Rik\|4930485I22Rik\|4933427G14Rik | RIKEN cDNA 4933406K04 gene | ncRNA |
| chr12_106658075_106658733 | chr12 | 106658076 | 106658733 | + | -0.761506 | Intergenic | Intergenic | 57920 | NR_015512 | 71033 | Mm.159404 | NR_015512 | ENSMUSG00000085820 | **4933406K04Rik** | 4930426M03Rik\|4930485I22Rik\|4933427G14Rik | RIKEN cDNA 4933406K04 gene | ncRNA |
| chr12_107016850_107017149 | chr12 | 107016851 | 107017149 | + | -0.893201 | Intergenic | Intergenic | -300676 | NR_015512 | 71033 | Mm.159404 | NR_015512 | ENSMUSG00000085820 | **4933406K04Rik** | 4930426M03Rik\|4930485I22Rik\|4933427G14Rik | RIKEN cDNA 4933406K04 gene | ncRNA |
| chr12_107016714_107017227 | chr12 | 107016715 | 107017227 | + | -0.895295 | Intergenic | Intergenic | -300647 | NR_015512 | 71033 | Mm.159404 | NR_015512 | ENSMUSG00000085820 | **4933406K04Rik** | 4930426M03Rik\|4930485I22Rik\|4933427G14Rik | RIKEN cDNA 4933406K04 gene | ncRNA |
| chr12_107050989_107051710 | chr12 | 107050990 | 107051710 | + | -0.915054 | Intergenic | Intergenic | -335026 | NR_015512 | 71033 | Mm.159404 | NR_015512 | ENSMUSG00000085820 | **4933406K04Rik** | 4930426M03Rik\|4930485I22Rik\|4933427G14Rik | RIKEN cDNA 4933406K04 gene | ncRNA |
| chr12_107140862_107141579 | chr12 | 107140863 | 107141579 | + | -1.0363 | Intergenic | Intergenic | -424897 | NR_015512 | 71033 | Mm.159404 | NR_015512 | ENSMUSG00000085820 | **4933406K04Rik** | 4930426M03Rik\|4930485I22Rik\|4933427G14Rik | RIKEN cDNA 4933406K04 gene | ncRNA |
| chr10_117947682_117948269 | chr10 | 117947683 | 117948269 | + | -0.724333 | non-coding (NR_045342, exon 4 of 4) | non-coding (NR_045342, exon 4 of 4) | 22517 | NR_045342 | 71067 | Mm.410793 | NR_045342 | | **4933411E08Rik** | - | RIKEN cDNA 4933411E08 gene | ncRNA |
| chr10_117947895_117948243 | chr10 | 117947896 | 117948243 | + | -0.758896 | non-coding (NR_045342, exon 4 of 4) | non-coding (NR_045342, exon 4 of 4) | 22610 | NR_045342 | 71067 | Mm.410793 | NR_045342 | | **4933411E08Rik** | - | RIKEN cDNA 4933411E08 gene | ncRNA |
| chr10_117943768_117944106 | chr10 | 117943769 | 117944106 | + | -0.760445 | intron (NR_045342, intron 3 of 3) | intron (NR_045342, intron 3 of 3) | 18478 | NR_045342 | 71067 | Mm.410793 | NR_045342 | | **4933411E08Rik** | - | RIKEN cDNA 4933411E08 gene | ncRNA |
| chr10_117982600_117982866 | chr10 | 117982601 | 117982866 | + | -0.849291 | Intergenic | Intergenic | 57274 | NR_045342 | 71067 | Mm.410793 | NR_045342 | | **4933411E08Rik** | - | RIKEN cDNA 4933411E08 gene | ncRNA |
| chr10_117982466_117982856 | chr10 | 117982467 | 117982856 | + | -0.893034 | Intergenic | Intergenic | 57202 | NR_045342 | 71067 | Mm.410793 | NR_045342 | | **4933411E08Rik** | - | RIKEN cDNA 4933411E08 gene | ncRNA |
| chr10_117992370_117992720 | chr10 | 117992371 | 117992720 | + | -1.09611 | Intergenic | Intergenic | 67086 | NR_045342 | 71067 | Mm.410793 | NR_045342 | | **4933411E08Rik** | - | RIKEN cDNA 4933411E08 gene | ncRNA |
| chr15_59948277_59948562 | chr15 | 59948278 | 59948562 | + | -0.668429 | Intergenic | Intergenic | 68198 | NM_027668 | 71088 | Mm.425950 | NM_027668 | ENSMUSG00000071749 | **4933412E24Rik** | - | RIKEN cDNA 4933412E24 gene | protein-coding |
| chr15_59950934_59951210 | chr15 | 59950935 | 59951210 | + | -0.686279 | Intergenic | CT-rich\|Low_complexity\|Low_complexity | 65546 | NM_027668 | 71088 | Mm.425950 | NM_027668 | ENSMUSG00000071749 | **4933412E24Rik** | - | RIKEN cDNA 4933412E24 gene | protein-coding |
| chr15_60087384_60087769 | chr15 | 60087385 | 60087769 | + | -0.821128 | Intergenic | B4A\|SINE\|B4 | -70959 | NM_027668 | 71088 | Mm.425950 | NM_027668 | ENSMUSG00000071749 | **4933412E24Rik** | - | RIKEN cDNA 4933412E24 gene | protein-coding |
| chr15_59937365_59937764 | chr15 | 59937366 | 59937764 | + | -0.864359 | Intergenic | Intergenic | 79053 | NM_027668 | 71088 | Mm.425950 | NM_027668 | ENSMUSG00000071749 | **4933412E24Rik** | - | RIKEN cDNA 4933412E24 gene | protein-coding |
| chr15_60169720_60170047 | chr15 | 60169721 | 60170047 | + | -0.87471 | Intergenic | L1VL2\|LINE\|L1 | -153266 | NM_027668 | 71088 | Mm.425950 | NM_027668 | ENSMUSG00000071749 | **4933412E24Rik** | - | RIKEN cDNA 4933412E24 gene | protein-coding |
| chr15_59936605_59936966 | chr15 | 59936606 | 59936966 | + | -0.875624 | Intergenic | Intergenic | 79832 | NM_027668 | 71088 | Mm.425950 | NM_027668 | ENSMUSG00000071749 | **4933412E24Rik** | - | RIKEN cDNA 4933412E24 gene | protein-coding |
| chr15_59978937_59979296 | chr15 | 59978938 | 59979296 | + | -0.907467 | Intergenic | Intergenic | 37501 | NM_027668 | 71088 | Mm.425950 | NM_027668 | ENSMUSG00000071749 | **4933412E24Rik** | - | RIKEN cDNA 4933412E24 gene | protein-coding |
| chr15_60136701_60137048 | chr15 | 60136702 | 60137048 | + | -0.990986 | Intergenic | Intergenic | -120257 | NM_027668 | 71088 | Mm.425950 | NM_027668 | ENSMUSG00000071749 | **4933412E24Rik** | - | RIKEN cDNA 4933412E24 gene | protein-coding |
| chr15_60119918_60120512 | chr15 | 60119919 | 60120512 | + | -1.0281 | Intergenic | Intergenic | -103597 | NM_027668 | 71088 | Mm.425950 | NM_027668 | ENSMUSG00000071749 | **4933412E24Rik** | - | RIKEN cDNA 4933412E24 gene | protein-coding |
| chr15_60124508_60124867 | chr15 | 60124509 | 60124867 | + | -1.13871 | Intergenic | MER92A\|LTR\|ERV1 | -108070 | NM_027668 | 71088 | Mm.425950 | NM_027668 | ENSMUSG00000071749 | **4933412E24Rik** | - | RIKEN cDNA 4933412E24 gene | protein-coding |
| chr15_60124283_60124807 | chr15 | 60124284 | 60124807 | + | -1.17083 | Intergenic | (GA)n\|Simple_repeat\|Simple_repeat | -107927 | NM_027668 | 71088 | Mm.425950 | NM_027668 | ENSMUSG00000071749 | **4933412E24Rik** | - | RIKEN cDNA 4933412E24 gene | protein-coding |
| chr15_60111770_60112436 | chr15 | 60111771 | 60112436 | + | -1.30445 | Intergenic | Intergenic | -95485 | NM_027668 | 71088 | Mm.425950 | NM_027668 | ENSMUSG00000071749 | **4933412E24Rik** | - | RIKEN cDNA 4933412E24 gene | protein-coding |
| chr1_36720170_36720545 | chr1 | 36720171 | 36720545 | + | -0.707271 | intron (NR_040290, intron 2 of 8) | intron (NR_040290, intron 2 of 8) | 5820 | NR_040290 | 71166 | Mm.157049 | NR_040290 | | **4933424G06Rik** | 1700104I15Rik | RIKEN cDNA 4933424G06 gene | ncRNA |
| chr1_36720212_36720482 | chr1 | 36720213 | 36720482 | + | -0.712538 | intron (NR_040290, intron 2 of 8) | intron (NR_040290, intron 2 of 8) | 5809 | NR_040290 | 71166 | Mm.157049 | NR_040290 | | **4933424G06Rik** | 1700104I15Rik | RIKEN cDNA 4933424G06 gene | ncRNA |
| chr1_36723118_36723653 | chr1 | 36723119 | 36723653 | + | -0.794203 | non-coding (NR_040290, exon 3 of 9) | non-coding (NR_040290, exon 3 of 9) | 8848 | NR_040290 | 71166 | Mm.157049 | NR_040290 | | **4933424G06Rik** | 1700104I15Rik | RIKEN cDNA 4933424G06 gene | ncRNA |
| chr7_82988874_82989167 | chr7 | 82988875 | 82989167 | + | -0.802072 | Intergenic | Intergenic | -17630 | NR_045855 | 71285 | Mm.76818 | NR_045855 | | **4933430H16Rik** | - | RIKEN cDNA 4933430H16 gene | ncRNA |
| chr9_118390703_118391123 | chr9 | 118390704 | 118391123 | + | -1.4761 | Intergenic | Intergenic | -38926 | NR_040716 | 71196 | Mm.73218 | NR_040716 | | **4933432G23Rik** | - | RIKEN cDNA 4933432G23 gene | ncRNA |
| chr14_103017056_103017412 | chr14 | 103017057 | 103017412 | + | -0.827354 | intron (NR_045657, intron 2 of 2) | intron (NR_045657, intron 2 of 2) | 16569 | NR_045657 | 71264 | Mm.399572 | NR_045657 | | **4933432I03Rik** | - | RIKEN cDNA 4933432I03 gene | ncRNA |
| chr17_84035991_84036353 | chr17 | 84035992 | 84036353 | + | -0.681007 | Intergenic | Intergenic | -42488 | NR_045458 | 74473 | Mm.399575 | NR_045458 | | **4933433H22Rik** | - | RIKEN cDNA 4933433H22 gene | ncRNA |
| chr17_84126242_84126449 | chr17 | 84126243 | 84126449 | + | -0.699071 | Intergenic | Intergenic | 47686 | NR_045458 | 74473 | Mm.399575 | NR_045458 | | **4933433H22Rik** | - | RIKEN cDNA 4933433H22 gene | ncRNA |
| chr17_84032354_84032983 | chr17 | 84032355 | 84032983 | + | -0.717626 | Intergenic | Intergenic | -45991 | NR_045458 | 74473 | Mm.399575 | NR_045458 | | **4933433H22Rik** | - | RIKEN cDNA 4933433H22 gene | ncRNA |
| chr17_84032433_84032790 | chr17 | 84032434 | 84032790 | + | -0.762418 | Intergenic | Intergenic | -46048 | NR_045458 | 74473 | Mm.399575 | NR_045458 | | **4933433H22Rik** | - | RIKEN cDNA 4933433H22 gene | ncRNA |
| chr17_84006786_84007167 | chr17 | 84006787 | 84007167 | + | -0.898787 | Intergenic | L1MA10\|LINE\|L1 | -71683 | NR_045458 | 74473 | Mm.399575 | NR_045458 | | **4933433H22Rik** | - | RIKEN cDNA 4933433H22 gene | ncRNA |
| chr17_84128259_84128716 | chr17 | 84128260 | 84128716 | + | -0.953011 | Intergenic | BGLII_B\|LTR\|ERVK | 49828 | NR_045458 | 74473 | Mm.399575 | NR_045458 | | **4933433H22Rik** | - | RIKEN cDNA 4933433H22 gene | ncRNA |
| chr14_75034074_75034733 | chr14 | 75034075 | 75034733 | + | -0.843655 | intron (NM_198642, intron 2 of 13) | intron (NM_198642, intron 2 of 13) | 18377 | NM_198642 | 271221 | Mm.320280 | NM_198642 | ENSMUSG00000034959 | **5031414D18Rik** | Gm744\|Gm911\|Kiaa0226l | RIKEN cDNA 5031414D18 gene | protein-coding |
| chr2_166491899_166492365 | chr2 | 166491900 | 166492365 | + | -0.671623 | Intergenic | ORR1E\|LTR\|ERVL-MaLR | 44681 | NR_015558 | 319684 | Mm.399746 | NR_015558 | | **5031425F14Rik** | - | RIKEN cDNA 5031425F14 gene | ncRNA |
| chr2_166467183_166467538 | chr2 | 166467184 | 166467538 | + | -0.862549 | Intergenic | Intergenic | 19910 | NR_015558 | 319684 | Mm.399746 | NR_015558 | | **5031425F14Rik** | - | RIKEN cDNA 5031425F14 gene | ncRNA |
| chr2_59353116_59353509 | chr2 | 59353117 | 59353509 | + | -1.00414 | intron (NM_026361, intron 21 of 21) | intron (NM_026361, intron 21 of 21) | 33369 | NR_040510 | 76676 | Mm.110038 | NR_040510 | ENSMUSG00000087455 | **5330411J11Rik** | - | RIKEN cDNA 5330411J11 gene | ncRNA |
| chr2_59353146_59353433 | chr2 | 59353147 | 59353433 | + | -1.00666 | intron (NM_026361, intron 21 of 21) | intron (NM_026361, intron 21 of 21) | 33392 | NR_040510 | 76676 | Mm.110038 | NR_040510 | ENSMUSG00000087455 | **5330411J11Rik** | - | RIKEN cDNA 5330411J11 gene | ncRNA |
| chr2_59338374_59338812 | chr2 | 59338375 | 59338812 | + | -1.07769 | intron (NM_026361, intron 13 of 21) | intron (NM_026361, intron 13 of 21) | 48089 | NR_040510 | 76676 | Mm.110038 | NR_040510 | ENSMUSG00000087455 | **5330411J11Rik** | - | RIKEN cDNA 5330411J11 gene | ncRNA |
| chr8_25274072_25275007 | chr8 | 25274073 | 25275007 | + | -0.732696 | Intergenic | Intergenic | 31730 | NR_040352 | 74489 | Mm.256468 | NR_040352 | | **5430421F17Rik** | - | RIKEN cDNA 5430421F17 gene | ncRNA |
| chr8_25353821_25354477 | chr8 | 25353822 | 25354477 | + | -0.749895 | Intergenic | Intergenic | -47879 | NR_040352 | 74489 | Mm.256468 | NR_040352 | | **5430421F17Rik** | - | RIKEN cDNA 5430421F17 gene | ncRNA |
| chr3_38282928_38283201 | chr3 | 38282929 | 38283201 | + | -0.712659 | Intergenic | Intergenic | -74946 | NR_040541 | 71342 | Mm.410782 | NR_040541 | | **5430434I15Rik** | - | RIKEN cDNA 5430434I15 gene | ncRNA |
| chr3_38283655_38284216 | chr3 | 38283656 | 38284216 | + | -0.78346 | Intergenic | MIR\|SINE\|MIR | -75817 | NR_040541 | 71342 | Mm.410782 | NR_040541 | | **5430434I15Rik** | - | RIKEN cDNA 5430434I15 gene | ncRNA |
| chr3_38282771_38283319 | chr3 | 38282772 | 38283319 | + | -0.80264 | Intergenic | Intergenic | -74926 | NR_040541 | 71342 | Mm.410782 | NR_040541 | | **5430434I15Rik** | - | RIKEN cDNA 5430434I15 gene | ncRNA |
| chr3_38283657_38283970 | chr3 | 38283658 | 38283970 | + | -0.894449 | Intergenic | Intergenic | -75695 | NR_040541 | 71342 | Mm.410782 | NR_040541 | | **5430434I15Rik** | - | RIKEN cDNA 5430434I15 gene | ncRNA |
| chr3_38280085_38280581 | chr3 | 38280086 | 38280581 | + | -1.13263 | Intergenic | Intergenic | -72214 | NR_040541 | 71342 | Mm.410782 | NR_040541 | | **5430434I15Rik** | - | RIKEN cDNA 5430434I15 gene | ncRNA |

| | | | | | | | | | | | | | | | | | |
|---|---|---|---|---|---|---|---|---|---|---|---|---|---|---|---|---|---|
| chr15_5740691_5740970 | chr15 | 5740692 | 5740970 | + | -1.0586 | Intergenic | MIR\|SINE\|MIR | 244513 | NR_045274 | 71432 | Mm.291087 | NR_045274 | | **5430437J10Rik** | - | RIKEN cDNA 5430437J10 gene | ncRNA |
| chr9_77466867_77467226 | chr9 | 77466868 | 77467226 | + | -0.778959 | intron (NM_172528, intron 4 of 13) | intron (NM_172528, intron 4 of 13) | -67631 | NR_040378 | 70487 | Mm.58958 | NR_040378 | ENSMUSG00000099170 | **5730403I07Rik** | - | RIKEN cDNA 5730403I07 gene | ncRNA |
| chr9_77466968_77467237 | chr9 | 77466969 | 77467237 | + | -0.802256 | intron (NM_172528, intron 4 of 13) | intron (NM_172528, intron 4 of 13) | -67687 | NR_040378 | 70487 | Mm.58958 | NR_040378 | ENSMUSG00000099170 | **5730403I07Rik** | - | RIKEN cDNA 5730403I07 gene | ncRNA |
| chr9_77380441_77380744 | chr9 | 77380442 | 77380744 | + | -0.982133 | intron (NR_040378, intron 3 of 3) | intron (NR_040378, intron 3 of 3) | 18823 | NR_040378 | 70487 | Mm.58958 | NR_040378 | ENSMUSG00000099170 | **5730403I07Rik** | - | RIKEN cDNA 5730403I07 gene | ncRNA |
| chr4_44672647_44672888 | chr4 | 44672648 | 44672888 | + | -0.654712 | intron (NM_008782, intron 5 of 9) | intron (NM_008782, intron 5 of 9) | -30032 | NR_073462 | 70602 | Mm.491142 | NR_073462 | | **5730488B01Rik** | - | RIKEN cDNA 5730488B01 gene | ncRNA |
| chr3_127868100_127868783 | chr3 | 127868101 | 127868783 | + | -1.35363 | Intergenic | MER63A\|DNA\|hAT-Blackjack | 27881 | NM_027482 | 70617 | Mm.317049 | NM_027482 | ENSMUSG00000050549 | **5730508B09Rik** | 2700063A17Rik\|C80506 | RIKEN cDNA 5730508B09 gene | protein-coding |
| chr19_53464594_53465275 | chr19 | 53464595 | 53465275 | + | -0.778291 | promoter-TSS (NR_028427) | promoter-TSS (NR_028427) | -139 | NR_028427 | 381232 | Mm.248624 | NR_028427 | ENSMUSG00000097636 | **5830416P10Rik** | 4833407H14Rik\|5430437C04Rik\|ENSMUSG00000074788\|Gm10758 | RIKEN cDNA 5830416P10 gene | ncRNA |
| chr9_120679786_120680172 | chr9 | 120679787 | 120680172 | + | -0.835174 | Intergenic | RLTR9A\|LTR\|ERVK | 102648 | NR_073359 | 76100 | Mm.48212 | NR_073359 | ENSMUSG00000052658 | **5830454E08Rik** | - | RIKEN cDNA 5830454E08 gene | ncRNA |
| chr9_120679448_120680182 | chr9 | 120679449 | 120680182 | + | -0.891235 | Intergenic | RLTR9A\|LTR\|ERVK | 102484 | NR_073359 | 76100 | Mm.48212 | NR_073359 | ENSMUSG00000052658 | **5830454E08Rik** | - | RIKEN cDNA 5830454E08 gene | ncRNA |
| chr5_148976797_148977319 | chr5 | 148976798 | 148977319 | + | -0.844735 | Intergenic | Intergenic | 18156 | NR_102383 | 319982 | Mm.33303 | NR_102383 | ENSMUSG00000041298 | **5930430L01Rik** | - | RIKEN cDNA 5930430L01 gene | ncRNA |
| chr15_12800321_12801212 | chr15 | 12800322 | 12801212 | + | -0.654753 | Intergenic | ID_B1\|SINE\|B4 | 23890 | NM_029998 | 77877 | Mm.207311 | NM_029998 | ENSMUSG00000022195 | **6030458C11Rik** | - | RIKEN cDNA 6030458C11 gene | protein-coding |
| chr11_117794056_117794534 | chr11 | 117794057 | 117794534 | + | -1.05512 | Intergenic | Intergenic | -3365 | NM_029964 | 77727 | Mm.159981 | NM_029964 | ENSMUSG00000025573 | **6030468B19Rik** | - | RIKEN cDNA 6030468B19 gene | protein-coding |
| chr6_135217999_135219011 | chr6 | 135218000 | 135219011 | + | -0.755457 | intron (NM_028982, intron 4 of 12) | intron (NM_028982, intron 4 of 12) | 20518 | NM_028982 | 74525 | Mm.34182 | NM_028982 | ENSMUSG00000030207 | **8430419L09Rik** | Kiaa1467\|mKIAA1467 | RIKEN cDNA 8430419L09 gene | protein-coding |
| chr6_135218232_135218883 | chr6 | 135218233 | 135218883 | + | -0.837423 | intron (NM_028982, intron 4 of 12) | intron (NM_028982, intron 4 of 12) | 20571 | NM_028982 | 74525 | Mm.34182 | NM_028982 | ENSMUSG00000030207 | **8430419L09Rik** | Kiaa1467\|mKIAA1467 | RIKEN cDNA 8430419L09 gene | protein-coding |
| chr19_31210431_31211053 | chr19 | 31210432 | 31211053 | + | -0.759642 | intron (NM_001013833, intron 3 of 17) | intron (NM_001013833, intron 3 of 17) | 5654 | NR_002849 | 78103 | Mm.445198 | NR_002849 | | **8430431K14Rik** | - | RIKEN cDNA 8430431K14 gene | ncRNA |
| chr19_31210397_31210863 | chr19 | 31210398 | 31210863 | + | -0.794199 | intron (NM_001013833, intron 3 of 17) | intron (NM_001013833, intron 3 of 17) | 5766 | NR_002849 | 78103 | Mm.445198 | NR_002849 | | **8430431K14Rik** | - | RIKEN cDNA 8430431K14 gene | ncRNA |
| chr19_31337289_31337831 | chr19 | 31337290 | 31337831 | + | -1.02773 | intron (NM_001013833, intron 2 of 17) | intron (NM_001013833, intron 2 of 17) | -121164 | NR_002849 | 78103 | Mm.445198 | NR_002849 | | **8430431K14Rik** | - | RIKEN cDNA 8430431K14 gene | ncRNA |
| chr4_7363615_7364079 | chr4 | 7363616 | 7364079 | + | -0.68983 | Intergenic | Intergenic | -196841 | NR_040645 | 71510 | Mm.410775 | NR_040645 | ENSMUSG00000086070 | **8430436N08Rik** | - | RIKEN cDNA 8430436N08 gene | ncRNA |
| chr4_7364129_7364787 | chr4 | 7364130 | 7364787 | + | -0.770095 | Intergenic | Intergenic | -196230 | NR_040645 | 71510 | Mm.410775 | NR_040645 | ENSMUSG00000086070 | **8430436N08Rik** | - | RIKEN cDNA 8430436N08 gene | ncRNA |
| chr4_7454988_7455611 | chr4 | 7454989 | 7455611 | + | -0.900676 | Intergenic | Intergenic | -105388 | NR_040645 | 71510 | Mm.410775 | NR_040645 | ENSMUSG00000086070 | **8430436N08Rik** | - | RIKEN cDNA 8430436N08 gene | ncRNA |
| chr12_35690952_35691341 | chr12 | 35690953 | 35691341 | + | -0.71613 | intron (NR_045050, intron 3 of 4) | intron (NR_045050, intron 3 of 4) | 3971 | NR_045050 | 71581 | Mm.100110 | NR_045050 | | **9130015A21Rik** | - | RIKEN cDNA 9130015A21 gene | ncRNA |
| chr13_24921296_24921856 | chr13 | 24921297 | 24921856 | + | -1.65797 | intron (NM_172532, intron 4 of 9) | intron (NM_172532, intron 4 of 9) | -15825 | NR_102323 | 493800 | Mm.458692 | NR_102323 | | **9330162012Rik** | - | cDNA RIKEN 9330162012 gene | ncRNA |
| chr8_94422275_94422758 | chr8 | 94422276 | 94422758 | + | -0.914195 | Intergenic | Intergenic | 12586 | NR_015514 | 320377 | Mm.486022 | NR_015514 | ENSMUSG00000097194 | **9330175E14Rik** | A630085K21 | RIKEN cDNA 9330175E14 gene | ncRNA |
| chr2_158334214_158334709 | chr2 | 158334215 | 158334709 | + | -0.620002 | Intergenic | LTR16C\|LTR\|ERVL | 27047 | NR_015463 | 68108 | Mm.119198 | NR_015463 | | **9430008C03Rik** | - | RIKEN cDNA 9430008C03 gene | ncRNA |
| chr15_93963096_93963793 | chr15 | 93963097 | 93963793 | + | -0.820729 | Intergenic | Lx7\|LINE\|L1 | -36334 | NR_045737 | 77248 | Mm.410768 | NR_045737 | | **9430014N10Rik** | - | RIKEN cDNA 9430014N10 gene | ncRNA |
| chr15_93963132_93963543 | chr15 | 93963133 | 93963543 | + | -0.835064 | Intergenic | Lx7\|LINE\|L1 | -36227 | NR_045737 | 77248 | Mm.410768 | NR_045737 | | **9430014N10Rik** | - | RIKEN cDNA 9430014N10 gene | ncRNA |
| chr15_93974246_93974652 | chr15 | 93974247 | 93974652 | + | -0.958948 | Intergenic | Intergenic | -47338 | NR_045737 | 77248 | Mm.410768 | NR_045737 | | **9430014N10Rik** | - | RIKEN cDNA 9430014N10 gene | ncRNA |
| chr15_93974355_93974642 | chr15 | 93974356 | 93974642 | + | -0.975443 | Intergenic | Intergenic | -47388 | NR_045737 | 77248 | Mm.410768 | NR_045737 | | **9430014N10Rik** | - | RIKEN cDNA 9430014N10 gene | ncRNA |
| chr2_75458078_75458547 | chr2 | 75458079 | 75458547 | + | -1.44845 | Intergenic | Intergenic | -14134 | NR_040635 | 77269 | Mm.389278 | NR_040635 | | **9430019J16Rik** | - | RIKEN cDNA 9430019J16 gene | ncRNA |
| chr5_28402413_28403291 | chr5 | 28402414 | 28403291 | + | -0.958752 | intron (NM_028234, intron 15 of 18) | RSINE1\|SINE\|B4 | -64132 | NR_015562 | 654796 | Mm.57202 | NR_015562 | ENSMUSG00000097726 | **9530036O11Rik** | - | RIKEN cDNA 9530036O11Rik | ncRNA |
| chr5_28402730_28403280 | chr5 | 28402731 | 28403280 | + | -0.96654 | intron (NM_028234, intron 15 of 18) | intron (NM_028234, intron 15 of 18) | -63979 | NR_015562 | 654796 | Mm.57202 | NR_015562 | ENSMUSG00000097726 | **9530036O11Rik** | - | RIKEN cDNA 9530036O11Rik | ncRNA |
| chr8_11001210_11001654 | chr8 | 11001211 | 11001654 | + | -0.683213 | intron (NM_001081212, intron 1 of 1) | intron (NM_001081212, intron 1 of 1) | -6418 | NR_046017 | 619321 | Mm.403007 | NR_046017 | ENSMUSG00000096938 | **9530052E02Rik** | - | RIKEN cDNA 9530052E02 gene | ncRNA |
| chr3_127924554_127924798 | chr3 | 127924555 | 127924798 | + | -0.845828 | intron (NR_040552, intron 1 of 4) | MLT1A0\|LTR\|ERVL-MaLR | 8505 | NR_040552 | 329753 | Mm.119720 | NR_040552 | | **9830132P13Rik** | 9830132P13 | RIKEN cDNA 9830132P13 gene | ncRNA |
| chr3_127924490_127924917 | chr3 | 127924491 | 127924917 | + | -0.864069 | intron (NR_040552, intron 1 of 4) | MLT1A0\|LTR\|ERVL-MaLR | 8533 | NR_040552 | 329753 | Mm.119720 | NR_040552 | | **9830132P13Rik** | 9830132P13 | RIKEN cDNA 9830132P13 gene | ncRNA |
| chr2_155033421_155033658 | chr2 | 155033422 | 155033658 | + | -0.607249 | intron (NM_015770, intron 1 of 3) | intron (NM_015770, intron 1 of 3) | 19970 | NM_015770 | 50518 | Mm.315593 | NM_015770 | ENSMUSG00000027596 | **a** | A<y>\|ASIP\|ASP\|As | nonagouti | protein-coding |
| chr2_155011191_155011646 | chr2 | 155011192 | 155011646 | + | -0.818832 | Intergenic | MTD\|LTR\|ERVL-MaLR | -2151 | NM_015770 | 50518 | Mm.315593 | NM_015770 | ENSMUSG00000027596 | **a** | A<y>\|ASIP\|ASP\|As | nonagouti | protein-coding |
| chr2_154991281_154991723 | chr2 | 154991282 | 154991723 | + | -1.00416 | Intergenic | Intergenic | -22068 | NM_015770 | 50518 | Mm.315593 | NM_015770 | ENSMUSG00000027596 | **a** | A<y>\|ASIP\|ASP\|As | nonagouti | protein-coding |
| chr2_154991437_154991725 | chr2 | 154991438 | 154991725 | + | -1.06327 | Intergenic | Intergenic | -21989 | NM_015770 | 50518 | Mm.315593 | NM_015770 | ENSMUSG00000027596 | **a** | A<y>\|ASIP\|ASP\|As | nonagouti | protein-coding |
| chr10_122566666_122567237 | chr10 | 122566667 | 122567237 | + | -0.677876 | non-coding (NR_040616, exon 3 of 3) | non-coding (NR_040616, exon 3 of 3) | 2939 | NR_040616 | 319272 | Mm.411763 | NR_040616 | | **A130077B15Rik** | - | RIKEN cDNA A130077B15 gene | ncRNA |
| chr10_122599937_122600297 | chr10 | 122599938 | 122600297 | + | -0.890786 | Intergenic | Intergenic | -30226 | NR_040616 | 319272 | Mm.411763 | NR_040616 | | **A130077B15Rik** | - | RIKEN cDNA A130077B15 gene | ncRNA |
| chr10_122604850_122605434 | chr10 | 122604851 | 122605434 | + | -1.2286 | Intergenic | RMER5\|LTR\|ERV1 | -35251 | NR_040616 | 319272 | Mm.411763 | NR_040616 | | **A130077B15Rik** | - | RIKEN cDNA A130077B15 gene | ncRNA |
| chr10_122599311_122600237 | chr10 | 122599312 | 122600237 | + | -1.31886 | Intergenic | Intergenic | -29883 | NR_040616 | 319272 | Mm.411763 | NR_040616 | | **A130077B15Rik** | - | RIKEN cDNA A130077B15 gene | ncRNA |
| chr10_122599570_122599811 | chr10 | 122599571 | 122599811 | + | -1.45073 | Intergenic | Intergenic | -29800 | NR_040616 | 319272 | Mm.411763 | NR_040616 | | **A130077B15Rik** | - | RIKEN cDNA A130077B15 gene | ncRNA |
| chr16_24947557_24947861 | chr16 | 24947558 | 24947861 | + | -0.766882 | intron (NM_001145952, intron 9 of 11) | L3\|LINE\|CR1 | -111930 | NR_040374 | 319487 | Mm.137341 | NR_040374 | ENSMUSG00000096960 | **A230028O05Rik** | - | RIKEN cDNA A230028O05 gene | ncRNA |
| chr16_24947543_24947956 | chr16 | 24947544 | 24947956 | + | -0.822383 | intron (NM_001145952, intron 9 of 11) | L3\|LINE\|CR1 | -111889 | NR_040374 | 319487 | Mm.137341 | NR_040374 | ENSMUSG00000096960 | **A230028O05Rik** | - | RIKEN cDNA A230028O05 gene | ncRNA |
| chr16_24909663_24909858 | chr16 | 24909664 | 24909858 | + | -0.917951 | intron (NM_001145952, intron 9 of 11) | intron (NM_001145952, intron 9 of 11) | -149878 | NR_040374 | 319487 | Mm.137341 | NR_040374 | ENSMUSG00000096960 | **A230028O05Rik** | - | RIKEN cDNA A230028O05 gene | ncRNA |
| chr13_44210214_44210542 | chr13 | 44210215 | 44210542 | + | -0.635181 | intron (NR_045088, intron 2 of 3) | intron (NR_045088, intron 2 of 3) | 6109 | NR_045088 | 100503592 | Mm.184925 | NR_045088 | | **A330076C08Rik** | - | RIKEN cDNA A330076C08 gene | ncRNA |
| chr18_46092488_46093239 | chr18 | 46092489 | 46093239 | + | -1.11905 | Intergenic | Intergenic | -47604 | NR_040342 | 107003 | Mm.395148 | NR_040342 | | **A330093E20Rik** | AU043608 | RIKEN cDNA A330093E20 gene | ncRNA |
| chr6_8485016_8486188 | chr6 | 8485017 | 8486188 | + | -0.648514 | Intergenic | Intergenic | 23559 | NR_040452 | 320312 | Mm.374890 | NR_040452 | ENSMUSG00000087305 | **A430035B10Rik** | 2810488G03Rik | RIKEN cDNA A430035B10 gene | ncRNA |
| chr6_8504745_8505236 | chr6 | 8504746 | 8505236 | + | -0.935704 | Intergenic | Intergenic | 4170 | NR_040452 | 320312 | Mm.374890 | NR_040452 | ENSMUSG00000087305 | **A430035B10Rik** | 2810488G03Rik | RIKEN cDNA A430035B10 gene | ncRNA |
| chr15_80697520_80698146 | chr15 | 80697521 | 80698146 | + | -1.10451 | promoter-TSS (NR_045309) | promoter-TSS (NR_045309) | -223 | NR_045310 | 106059 | Mm.442457 | NR_045309 | | **A430088P11Rik** | - | RIKEN cDNA A430088P11 gene | ncRNA |
| chr19_10751604_10751964 | chr19 | 10751605 | 10751964 | + | -0.784127 | intron (NR_027805, intron 1 of 3) | intron (NR_027805, intron 1 of 3) | 10837 | NR_027805 | 403202 | Mm.34482 | NM_207529 | ENSMUSG00000067577 | **A430093F15Rik** | Gm674 | RIKEN cDNA A430093F15 gene | ncRNA |
| chr14_53134635_53135356 | chr14 | 53134636 | 53135356 | + | -0.665861 | Intergenic | (TG)n\|Simple_repeat\|Simple_repeat | -530944 | NM_001205242 | 100568452 | Mm.213248 | NM_001205242 | | **A430107P09Rik** | Va10 | RIKEN cDNA A430107P09 gene | protein-coding |
| chr14_53545320_53545815 | chr14 | 53545321 | 53545815 | + | -0.705428 | Intergenic | Intergenic | -120372 | NM_001205242 | 100568452 | Mm.213248 | NM_001205242 | | **A430107P09Rik** | Va10 | RIKEN cDNA A430107P09 gene | protein-coding |
| chr14_53640497_53640716 | chr14 | 53640498 | 53640716 | + | -0.776552 | Intergenic | Intergenic | -25333 | NM_001205242 | 100568452 | Mm.213248 | NM_001205242 | | **A430107P09Rik** | Va10 | RIKEN cDNA A430107P09 gene | protein-coding |
| chr14_53762862_53763255 | chr14 | 53762863 | 53763255 | + | -0.823073 | Intergenic | Intergenic | 97119 | NM_001205242 | 100568452 | Mm.213248 | NM_001205242 | | **A430107P09Rik** | Va10 | RIKEN cDNA A430107P09 gene | protein-coding |
| chr14_53553766_53554124 | chr14 | 53553767 | 53554124 | + | -0.873083 | Intergenic | Intergenic | -111995 | NM_001205242 | 100568452 | Mm.213248 | NM_001205242 | | **A430107P09Rik** | Va10 | RIKEN cDNA A430107P09 gene | protein-coding |
| chr14_53369878_53370163 | chr14 | 53369879 | 53370163 | + | -0.951641 | Intergenic | Intergenic | -295919 | NM_001205242 | 100568452 | Mm.213248 | NM_001205242 | | **A430107P09Rik** | Va10 | RIKEN cDNA A430107P09 gene | protein-coding |
| chr14_53802503_53802866 | chr14 | 53802504 | 53802866 | + | -0.979905 | Intergenic | ERVL-E-int\|LTR\|ERVL | 136745 | NM_001205242 | 100568452 | Mm.213248 | NM_001205242 | | **A430107P09Rik** | Va10 | RIKEN cDNA A430107P09 gene | protein-coding |
| chr14_53369735_53370145 | chr14 | 53369736 | 53370145 | + | -1.02372 | Intergenic | Intergenic | -296000 | NM_001205242 | 100568452 | Mm.213248 | NM_001205242 | | **A430107P09Rik** | Va10 | RIKEN cDNA A430107P09 gene | protein-coding |
| chr14_53364011_53364345 | chr14 | 53364012 | 53364345 | + | -1.05714 | Intergenic | Intergenic | -301762 | NM_001205242 | 100568452 | Mm.213248 | NM_001205242 | | **A430107P09Rik** | Va10 | RIKEN cDNA A430107P09 gene | protein-coding |
| chr14_53878367_53878586 | chr14 | 53878368 | 53878586 | + | -1.26 | Intergenic | RMER15-int\|LTR\|ERVL | 212537 | NM_001205242 | 100568452 | Mm.213248 | NM_001205242 | | **A430107P09Rik** | Va10 | RIKEN cDNA A430107P09 gene | protein-coding |
| chr14_53878677_53878921 | chr14 | 53878678 | 53878921 | + | -1.31233 | Intergenic | RMER15-int\|LTR\|ERVL | 212859 | NM_001205242 | 100568452 | Mm.213248 | NM_001205242 | | **A430107P09Rik** | Va10 | RIKEN cDNA A430107P09 gene | protein-coding |

| | | | | | | | | | | | | | | | | | |
|---|---|---|---|---|---|---|---|---|---|---|---|---|---|---|---|---|---|
| chr14_53878313_53879006 | chr14 | 53878314 | 53879006 | + | -1.39773 | Intergenic | RMER15-int\|LTR\|ERVL | 212720 | NM_001205242 | 100568452 | Mm.213248 | NM_001205242 | | **A430107P09Rik** | Va10 | RIKEN cDNA A430107P09 gene | protein-coding |
| chr2_167777342_167778653 | chr2 | 167777343 | 167778653 | + | -0.786147 | Intergenic | Intergenic | 86821 | NR_015500 | 329562 | Mm.123101 | NM_177851 | ENSMUSG00000006462 | **A530013C23Rik** | - | RIKEN cDNA A530013C23 gene | ncRNA |
| chr2_167777349_167778277 | chr2 | 167777350 | 167778277 | + | -0.790092 | Intergenic | Intergenic | 86636 | NR_015500 | 329562 | Mm.123101 | NM_177851 | ENSMUSG00000006462 | **A530013C23Rik** | - | RIKEN cDNA A530013C23 gene | ncRNA |
| chr2_167777703_167778191 | chr2 | 167777704 | 167778191 | + | -0.860497 | Intergenic | Intergenic | 86770 | NR_015500 | 329562 | Mm.123101 | NM_177851 | ENSMUSG00000006462 | **A530013C23Rik** | - | RIKEN cDNA A530013C23 gene | ncRNA |
| chr2_167777689_167778551 | chr2 | 167777690 | 167778551 | + | -0.869272 | Intergenic | Intergenic | 86943 | NR_015500 | 329562 | Mm.123101 | NM_177851 | ENSMUSG00000006462 | **A530013C23Rik** | - | RIKEN cDNA A530013C23 gene | ncRNA |
| chr13_15845666_15846521 | chr13 | 15845667 | 15846521 | + | -0.752758 | Intergenic | Intergenic | 38851 | NR_046131 | 328190 | Mm.446957 | NR_046131 | | **A530046M15Rik** | A530046M15 | RIKEN cDNA A530046M15 gene | ncRNA |
| chr1_85720413_85721246 | chr1 | 85720414 | 85721246 | + | -0.931607 | intron (NM_177055, intron 4 of 5) | intron (NM_177055, intron 4 of 5) | 15776 | NM_177055 | 319997 | Mm.116691 | NM_177055 | ENSMUSG00000052760 | **A630001G21Rik** | - | RIKEN cDNA A630001G21 gene | protein-coding |
| chr5_110514717_110514981 | chr5 | 110514718 | 110514981 | + | -0.639422 | promoter-TSS (NR_102290) | promoter-TSS (NR_102290) | -74 | NR_102290 | 231603 | Mm.22836 | NM_173766 | ENSMUSG00000048215 | **A630023P12Rik** | - | RIKEN cDNA A630023P12 gene | ncRNA |
| chr2_170052808_170053085 | chr2 | 170052809 | 170053085 | + | -0.654145 | Intergenic | Charlie4a\|DNA\|hAT-Charlie | 17724 | NR_033634 | 100043910 | Mm.468546 | NR_033632 | ENSMUSG00000097065 | **A630075F10Rik** | 4831414D02\|Gm4722 | RIKEN cDNA A630075F10 gene | ncRNA |
| chr2_170024765_170025399 | chr2 | 170024766 | 170025399 | + | -0.710568 | Intergenic | ORR1D2\|LTR\|ERVL-MaLR | 45589 | NR_033634 | 100043910 | Mm.468546 | NR_033632 | ENSMUSG00000097065 | **A630075F10Rik** | 4831414D02\|Gm4722 | RIKEN cDNA A630075F10 gene | ncRNA |
| chr2_169958527_169959221 | chr2 | 169958528 | 169959221 | + | -1.02907 | Intergenic | ORR1E\|LTR\|ERVL-MaLR | 111797 | NR_033634 | 100043910 | Mm.468546 | NR_033632 | ENSMUSG00000097065 | **A630075F10Rik** | 4831414D02\|Gm4722 | RIKEN cDNA A630075F10 gene | ncRNA |
| chr2_170050443_170051271 | chr2 | 170050444 | 170051271 | + | -1.31608 | Intergenic | Intergenic | 19814 | NR_033634 | 100043910 | Mm.468546 | NR_033632 | ENSMUSG00000097065 | **A630075F10Rik** | 4831414D02\|Gm4722 | RIKEN cDNA A630075F10 gene | ncRNA |
| chr2_170050692_170051022 | chr2 | 170050693 | 170051022 | + | -1.35033 | Intergenic | Intergenic | 19814 | NR_033634 | 100043910 | Mm.468546 | NR_033632 | ENSMUSG00000097065 | **A630075F10Rik** | 4831414D02\|Gm4722 | RIKEN cDNA A630075F10 gene | ncRNA |
| chr2_129243823_129244266 | chr2 | 129243824 | 129244266 | + | -0.884854 | Intergenic | Intergenic | 25076 | NR_045838 | 329513 | Mm.473529 | NM_177847 | ENSMUSG00000056738 | **A730036I17Rik** | - | RIKEN cDNA A730036I17 gene | ncRNA |
| chr9_62260756_62261303 | chr9 | 62260757 | 62261303 | + | -0.860491 | Intergenic | Intergenic | -16930 | NR_040769 | 330958 | Mm.411888 | NR_040769 | | **A730043L09Rik** | A730043L09 | RIKEN cDNA A730043L09 gene | ncRNA |
| chr9_62260977_62261349 | chr9 | 62260978 | 62261349 | + | -0.880072 | Intergenic | Intergenic | -17063 | NR_040769 | 330958 | Mm.411888 | NR_040769 | | **A730043L09Rik** | A730043L09 | RIKEN cDNA A730043L09 gene | ncRNA |
| chr16_24538988_24539471 | chr16 | 24538989 | 24539471 | + | -0.603837 | intron (NM_001145952, intron 2 of 11) | intron (NM_001145952, intron 2 of 11) | 9916 | NR_038091 | 624582 | Mm.439775 | NR_038091 | | **A730098P11Rik** | - | mortality factor 4 like 1 pseudogene | pseudo |
| chr13_91377426_91378196 | chr13 | 91377427 | 91378196 | + | -0.889791 | intron (NR_045161, intron 1 of 2) | intron (NR_045161, intron 1 of 2) | 10274 | NR_045161 | 320297 | Mm.375866 | NR_045161 | ENSMUSG00000097182 | **A830009L08Rik** | - | RIKEN cDNA A830009L08 gene | ncRNA |
| chr13_91377405_91378000 | chr13 | 91377406 | 91378000 | + | -0.924681 | intron (NR_045161, intron 1 of 2) | intron (NR_045161, intron 1 of 2) | 10382 | NR_045161 | 320297 | Mm.375866 | NR_045161 | ENSMUSG00000097182 | **A830009L08Rik** | - | RIKEN cDNA A830009L08 gene | ncRNA |
| chr13_91377426_91377868 | chr13 | 91377427 | 91377868 | + | -0.95872 | intron (NR_045161, intron 1 of 2) | intron (NR_045161, intron 1 of 2) | 10438 | NR_045161 | 320297 | Mm.375866 | NR_045161 | ENSMUSG00000097182 | **A830009L08Rik** | - | RIKEN cDNA A830009L08 gene | ncRNA |
| chr3_115820157_115820392 | chr3 | 115820158 | 115820392 | + | -0.74895 | Intergenic | Intergenic | 67784 | NR_027894 | 68161 | Mm.392043 | NR_015487 | | **A930005H10Rik** | - | RIKEN cDNA A930005H10 gene | ncRNA |
| chr3_115804548_115804967 | chr3 | 115804549 | 115804967 | + | -0.947978 | Intergenic | Intergenic | 83301 | NR_027894 | 68161 | Mm.392043 | NR_015487 | | **A930005H10Rik** | - | RIKEN cDNA A930005H10 gene | ncRNA |
| chr3_115815165_115816215 | chr3 | 115815166 | 115816215 | + | -1.02586 | Intergenic | Intergenic | 72369 | NR_027894 | 68161 | Mm.392043 | NR_015487 | | **A930005H10Rik** | - | RIKEN cDNA A930005H10 gene | ncRNA |
| chr3_115804690_115804947 | chr3 | 115804691 | 115804947 | + | -1.24992 | Intergenic | Intergenic | 83240 | NR_027894 | 68161 | Mm.392043 | NR_015487 | | **A930005H10Rik** | - | RIKEN cDNA A930005H10 gene | ncRNA |
| chr3_115801382_115801977 | chr3 | 115801383 | 115801977 | + | -1.31314 | Intergenic | Intergenic | 86379 | NR_027894 | 68161 | Mm.392043 | NR_015487 | | **A930005H10Rik** | - | RIKEN cDNA A930005H10 gene | ncRNA |
| chr3_115801641_115801818 | chr3 | 115801642 | 115801818 | + | -1.38881 | Intergenic | Intergenic | 86329 | NR_027894 | 68161 | Mm.392043 | NR_015487 | | **A930005H10Rik** | - | RIKEN cDNA A930005H10 gene | ncRNA |
| chr3_115801617_115801883 | chr3 | 115801618 | 115801883 | + | -1.43365 | Intergenic | Intergenic | 86309 | NR_027894 | 68161 | Mm.392043 | NR_015487 | | **A930005H10Rik** | - | RIKEN cDNA A930005H10 gene | ncRNA |
| chr16_91445613_91445982 | chr16 | 91445614 | 91445982 | + | -0.616676 | Intergenic | Intergenic | -19306 | NR_077219 | 100503120 | Mm.387130 | NR_077219 | ENSMUSG00000084866 | **A930006K02Rik** | - | RIKEN cDNA A930006K02 gene | ncRNA |
| chr16_91458090_91458767 | chr16 | 91458091 | 91458767 | + | -0.624646 | Intergenic | Intergenic | -6675 | NR_077219 | 100503120 | Mm.387130 | NR_077219 | ENSMUSG00000084866 | **A930006K02Rik** | - | RIKEN cDNA A930006K02 gene | ncRNA |
| chr16_91447938_91448558 | chr16 | 91447939 | 91448558 | + | -0.74089 | Intergenic | Intergenic | -16856 | NR_077219 | 100503120 | Mm.387130 | NR_077219 | ENSMUSG00000084866 | **A930006K02Rik** | - | RIKEN cDNA A930006K02 gene | ncRNA |
| chr19_29475388_29475705 | chr19 | 29475389 | 29475705 | + | -0.758619 | Intergenic | Intergenic | 46440 | NR_015567 | 77779 | Mm.458738 | NR_015567 | ENSMUSG00000097855 | **A930007I19Rik** | - | RIKEN cDNA A930007I19 gene | ncRNA |
| chr19_29475273_29475719 | chr19 | 29475274 | 29475719 | + | -0.7658 | Intergenic | Intergenic | 46491 | NR_015567 | 77779 | Mm.458738 | NR_015567 | ENSMUSG00000097855 | **A930007I19Rik** | - | RIKEN cDNA A930007I19 gene | ncRNA |
| chr19_29466858_29467527 | chr19 | 29466859 | 29467527 | + | -1.0073 | intron (NM_021396, intron 5 of 5) | MLTR18C_MM\|LTR\|ERVK | 54794 | NR_015567 | 77779 | Mm.458738 | NR_015567 | ENSMUSG00000097855 | **A930007I19Rik** | - | RIKEN cDNA A930007I19 gene | ncRNA |
| chr19_29466894_29467234 | chr19 | 29466895 | 29467234 | + | -1.05404 | intron (NM_021396, intron 5 of 5) | intron (NM_021396, intron 5 of 5) | 54923 | NR_015567 | 77779 | Mm.458738 | NR_015567 | ENSMUSG00000097855 | **A930007I19Rik** | - | RIKEN cDNA A930007I19 gene | ncRNA |
| chr17_35999368_35999737 | chr17 | 35999369 | 35999737 | + | -0.895968 | intron (NR_015618, intron 1 of 1) | intron (NR_015618, intron 1 of 1) | 5049 | NR_015618 | 77810 | Mm.483143 | NR_015618 | | **A930015D03Rik** | - | RIKEN cDNA A930015D03 gene | ncRNA |
| chr17_35999405_35999652 | chr17 | 35999406 | 35999652 | + | -0.903815 | intron (NR_015618, intron 1 of 1) | intron (NR_015618, intron 1 of 1) | 5025 | NR_015618 | 77810 | Mm.483143 | NR_015618 | | **A930015D03Rik** | - | RIKEN cDNA A930015D03 gene | ncRNA |
| chr5_122989013_122989663 | chr5 | 122989014 | 122989663 | + | -0.75846 | promoter-TSS (NR_045820) | promoter-TSS (NR_045820) | -16 | NR_045820 | 109202 | Mm.30769 | NM_175405 | ENSMUSG00000056735 | **A930024E05Rik** | - | RIKEN cDNA A930024E05 gene | ncRNA |
| chr5_122990341_122990929 | chr5 | 122990342 | 122990929 | + | -0.908069 | intron (NR_045820, intron 1 of 2) | intron (NR_045820, intron 1 of 2) | 1281 | NR_045820 | 109202 | Mm.30769 | NM_175405 | ENSMUSG00000056735 | **A930024E05Rik** | - | RIKEN cDNA A930024E05 gene | ncRNA |
| chr5_122990470_122990823 | chr5 | 122990471 | 122990823 | + | -0.911391 | intron (NR_045820, intron 1 of 2) | intron (NR_045820, intron 1 of 2) | 1293 | NR_045820 | 109202 | Mm.30769 | NM_175405 | ENSMUSG00000056735 | **A930024E05Rik** | - | RIKEN cDNA A930024E05 gene | ncRNA |
| chr1_130743785_130744250 | chr1 | 130743786 | 130744250 | + | -0.650105 | 3' UTR (NM_177604, exon 4 of 4) | 3' UTR (NM_177604, exon 4 of 4) | 12042 | NM_177604 | 212439 | Mm.268547 | NM_177604 | ENSMUSG00000042510 | **AA986860** | E030025M07\|Sarg | expressed sequence AA986860 | protein-coding |
| chr6_86909949_86910324 | chr6 | 86909950 | 86910324 | + | -0.732042 | intron (NM_177762, intron 1 of 19) | intron (NM_177762, intron 1 of 19) | 60620 | NM_001040106 | 269774 | Mm.221038 | NM_177762 | ENSMUSG00000057230 | **Aak1** | 5530400K14Rik\|9630042K20\|AU067724\|AU067726\|BC028270\|C79663\|D6Ertd245e\|R75501\|mKIAA1048 | AP2 associated kinase 1 | protein-coding |
| chr6_86850654_86850992 | chr6 | 86850655 | 86850992 | + | -0.867522 | intron (NM_177762, intron 1 of 19) | intron (NM_177762, intron 1 of 19) | 1306 | NM_001040106 | 269774 | Mm.221038 | NM_177762 | ENSMUSG00000057230 | **Aak1** | 5530400K14Rik\|9630042K20\|AU067724\|AU067726\|BC028270\|C79663\|D6Ertd245e\|R75501\|mKIAA1048 | AP2 associated kinase 1 | protein-coding |
| chr5_8669631_8670462 | chr5 | 8669632 | 8670462 | + | -0.743112 | intron (NM_011076, intron 3 of 27) | intron (NM_011076, intron 3 of 27) | 9955 | NM_011076 | 18671 | Mm.207354 | NM_011076 | ENSMUSG00000040584 | **Abcb1a** | Abcb4\|Evi32\|Mdr1a\|Mdr3\|P-gp\|Pgp\|Pgy-3\|Pgy3\|mdr-3 | ATP-binding cassette, sub-family B (MDR/TAP), member 1A | protein-coding |
| chr5_8652036_8652482 | chr5 | 8652037 | 8652482 | + | -0.801509 | Intergenic | MIR\|SINE\|MIR | -7833 | NM_011076 | 18671 | Mm.207354 | NM_011076 | ENSMUSG00000040584 | **Abcb1a** | Abcb4\|Evi32\|Mdr1a\|Mdr3\|P-gp\|Pgp\|Pgy-3\|Pgy3\|mdr-3 | ATP-binding cassette, sub-family B (MDR/TAP), member 1A | protein-coding |
| chr5_8675611_8676224 | chr5 | 8675612 | 8676224 | + | -0.832633 | intron (NM_011076, intron 4 of 27) | intron (NM_011076, intron 4 of 27) | 15826 | NM_011076 | 18671 | Mm.207354 | NM_011076 | ENSMUSG00000040584 | **Abcb1a** | Abcb4\|Evi32\|Mdr1a\|Mdr3\|P-gp\|Pgp\|Pgy-3\|Pgy3\|mdr-3 | ATP-binding cassette, sub-family B (MDR/TAP), member 1A | protein-coding |
| chr5_8675331_8675561 | chr5 | 8675332 | 8675561 | + | -0.839186 | intron (NM_011076, intron 4 of 27) | intron (NM_011076, intron 4 of 27) | 15354 | NM_011076 | 18671 | Mm.207354 | NM_011076 | ENSMUSG00000040584 | **Abcb1a** | Abcb4\|Evi32\|Mdr1a\|Mdr3\|P-gp\|Pgp\|Pgy-3\|Pgy3\|mdr-3 | ATP-binding cassette, sub-family B (MDR/TAP), member 1A | protein-coding |
| chr5_8675325_8676229 | chr5 | 8675326 | 8676229 | + | -0.858594 | intron (NM_011076, intron 4 of 27) | Tigger7\|DNA\|TcMar-Tigger | 15685 | NM_011076 | 18671 | Mm.207354 | NM_011076 | ENSMUSG00000040584 | **Abcb1a** | Abcb4\|Evi32\|Mdr1a\|Mdr3\|P-gp\|Pgp\|Pgy-3\|Pgy3\|mdr-3 | ATP-binding cassette, sub-family B (MDR/TAP), member 1A | protein-coding |
| chr5_8669454_8670072 | chr5 | 8669455 | 8670072 | + | -1.04059 | intron (NM_011076, intron 3 of 27) | L2a\|LINE\|L2 | 9671 | NM_011076 | 18671 | Mm.207354 | NM_011076 | ENSMUSG00000040584 | **Abcb1a** | Abcb4\|Evi32\|Mdr1a\|Mdr3\|P-gp\|Pgp\|Pgy-3\|Pgy3\|mdr-3 | ATP-binding cassette, sub-family B (MDR/TAP), member 1A | protein-coding |
| chr5_8839711_8840017 | chr5 | 8839712 | 8840017 | + | -0.765175 | intron (NM_011075, intron 21 of 27) | intron (NM_011075, intron 21 of 27) | 41717 | NM_011075 | 18669 | Mm.146649 | NM_011075 | ENSMUSG00000028970 | **Abcb1b** | Abcb1\|Mdr1\|Mdr1b\|Pgy-1\|Pgy1\|mdr | ATP-binding cassette, sub-family B (MDR/TAP), member 1B | protein-coding |
| chr5_8838752_8839476 | chr5 | 8838753 | 8839476 | + | -0.823301 | intron (NM_011075, intron 21 of 27) | intron (NM_011075, intron 21 of 27) | 40967 | NM_011075 | 18669 | Mm.146649 | NM_011075 | ENSMUSG00000028970 | **Abcb1b** | Abcb1\|Mdr1\|Mdr1b\|Pgy-1\|Pgy1\|mdr | ATP-binding cassette, sub-family B (MDR/TAP), member 1B | protein-coding |
| chr5_8863656_8863955 | chr5 | 8863657 | 8863955 | + | -0.841848 | intron (NM_011075, intron 26 of 27) | intron (NM_011075, intron 26 of 27) | -29915 | NM_008830 | 18670 | Mm.297825 | NM_008830 | ENSMUSG00000042476 | **Abcb4** | Mdr2\|Pgy-2\|Pgy2\|mdr-2 | ATP-binding cassette, sub-family B (MDR/TAP), member 4 | protein-coding |
| chr5_124079006_124079417 | chr5 | 124079007 | 124079417 | + | -0.741846 | intron (NM_019875, intron 6 of 11) | intron (NM_019875, intron 6 of 11) | 16586 | NM_019875 | 56325 | Mm.254839 | NM_019875 | ENSMUSG00000029408 | **Abcb9** | TAPL\|mKIAA1520 | ATP-binding cassette, sub-family B (MDR/TAP), member 9 | protein-coding |
| chr17_31039496_31040053 | chr17 | 31039497 | 31040053 | + | -0.795834 | Intergenic | Intergenic | -17919 | NM_009593 | 11307 | Mm.15691 | NM_009593 | | **Abcg1** | AW413978\|Abc8\|White | ATP-binding cassette, sub-family G (WHITE), member 1 | protein-coding |
| chr17_31060724_31061483 | chr17 | 31060725 | 31061483 | + | -0.83684 | intron (NM_009593, intron 1 of 14) | intron (NM_009593, intron 1 of 14) | 3410 | NM_009593 | 11307 | Mm.15691 | NM_009593 | | **Abcg1** | AW413978\|Abc8\|White | ATP-binding cassette, sub-family G (WHITE), member 1 | protein-coding |
| chr17_31060741_31061211 | chr17 | 31060742 | 31061211 | + | -0.855483 | intron (NM_009593, intron 1 of 14) | intron (NM_009593, intron 1 of 14) | 3282 | NM_009593 | 11307 | Mm.15691 | NM_009593 | | **Abcg1** | AW413978\|Abc8\|White | ATP-binding cassette, sub-family G (WHITE), member 1 | protein-coding |
| chr17_31058696_31059173 | chr17 | 31058697 | 31059173 | + | -0.961508 | intron (NM_009593, intron 1 of 14) | intron (NM_009593, intron 1 of 14) | 1241 | NM_009593 | 11307 | Mm.15691 | NM_009593 | | **Abcg1** | AW413978\|Abc8\|White | ATP-binding cassette, sub-family G (WHITE), member 1 | protein-coding |
| chr17_31077377_31078198 | chr17 | 31077378 | 31078198 | + | -1.2183 | intron (NM_009593, intron 2 of 14) | intron (NM_009593, intron 2 of 14) | 20094 | NM_009593 | 11307 | Mm.15691 | NM_009593 | | **Abcg1** | AW413978\|Abc8\|White | ATP-binding cassette, sub-family G (WHITE), member 1 | protein-coding |
| chr16_45786543_45787137 | chr16 | 45786544 | 45787137 | + | -0.683812 | intron (NM_153412, intron 6 of 17) | intron (NM_153412, intron 6 of 17) | -43885 | NM_172511 | 213012 | Mm.247453 | NM_172511 | ENSMUSG00000033157 | **Abhd10** | - | abhydrolase domain containing 10 | protein-coding |
| chr16_45763740_45764104 | chr16 | 45763741 | 45764104 | + | -0.769802 | intron (NM_153412, intron 12 of 17) | intron (NM_153412, intron 12 of 17) | -20967 | NM_172511 | 213012 | Mm.247453 | NM_172511 | ENSMUSG00000033157 | **Abhd10** | - | abhydrolase domain containing 10 | protein-coding |

| | | | | | | | | | | | | | | | | | |
|---|---|---|---|---|---|---|---|---|---|---|---|---|---|---|---|---|---|
| chr2_150869424_150870314 | chr2 | 150869425 | 150870314 | + | -0.892774 | intron (NM_024465, intron 1 of 12) | intron (NM_024465, intron 1 of 12) | 34862 | NM_024465 | 76192 | Mm.112632 | NM_024465 | ENSMUSG00000032046 | **Abhd12** | 1500011G07Rik\|6330583M11Rik\|AI431047\|AW547313 | abhydrolase domain containing 12 | protein-coding |
| chr2_150869354_150870054 | chr2 | 150869355 | 150870054 | + | -0.955447 | intron (NM_024465, intron 1 of 12) | intron (NM_024465, intron 1 of 12) | 35027 | NM_024465 | 76192 | Mm.112632 | NM_024465 | ENSMUSG00000032046 | **Abhd12** | 1500011G07Rik\|6330583M11Rik\|AI431047\|AW547313 | abhydrolase domain containing 12 | protein-coding |
| chr2_150869422_150869907 | chr2 | 150869423 | 150869907 | + | -1.01697 | intron (NM_024465, intron 1 of 12) | intron (NM_024465, intron 1 of 12) | 35066 | NM_024465 | 76192 | Mm.112632 | NM_024465 | ENSMUSG00000032046 | **Abhd12** | 1500011G07Rik\|6330583M11Rik\|AI431047\|AW547313 | abhydrolase domain containing 12 | protein-coding |
| chr12_70135796_70136850 | chr12 | 70135797 | 70136850 | + | -0.693743 | Intergenic | Intergenic | -17848 | NM_001195033 | 100504285 | Mm.291105 | NM_001195033 | | **Abhd12b** | Gm804 | abhydrolase domain containing 12B | protein-coding |
| chr11_77518891_77519536 | chr11 | 77518892 | 77519536 | + | -0.757845 | 3' UTR (NM_026185, exon 2 of 2) | 3' UTR (NM_026185, exon 2 of 2) | 4097 | NM_026185 | 67477 | Mm.432526 | NM_026185 | ENSMUSG00000000686 | **Abhd15** | 1300007F04Rik | abhydrolase domain containing 15 | protein-coding |
| chr11_77521207_77521412 | chr11 | 77521208 | 77521412 | + | -1.09776 | TTS (NM_026185) | TTS (NM_026185) | 6193 | NM_026185 | 67477 | Mm.432526 | NM_026185 | ENSMUSG00000000686 | **Abhd15** | 1300007F04Rik | abhydrolase domain containing 15 | protein-coding |
| chr19_21713308_21714279 | chr19 | 21713309 | 21714279 | + | -0.639674 | Intergenic | RLTR20D\|LTR\|ERVK | 60485 | NM_146096 | 226016 | Mm.28796 | NM_146096 | ENSMUSG00000047368 | **Abhd17b** | 5730446C15Rik\|Cgi67\|Fam108b\|Fam108b1 | abhydrolase domain containing 17B | protein-coding |
| chr19_21713154_21714737 | chr19 | 21713155 | 21714737 | + | -0.660799 | Intergenic | RLTR20D\|LTR\|ERVK | 60637 | NM_146096 | 226016 | Mm.28796 | NM_146096 | ENSMUSG00000047368 | **Abhd17b** | 5730446C15Rik\|Cgi67\|Fam108b\|Fam108b1 | abhydrolase domain containing 17B | protein-coding |
| chr7_79300513_79301050 | chr7 | 79300514 | 79301050 | + | -0.891814 | intron (NM_018811, intron 3 of 10) | intron (NM_018811, intron 3 of 10) | 27516 | NM_018811 | 54608 | Mm.365490 | NM_018811 | | **Abhd2** | 2210009N18Rik\|LABH2\|Labh-2 | abhydrolase domain containing 2 | protein-coding |
| chr7_79300507_79301293 | chr7 | 79300508 | 79301293 | + | -0.926555 | intron (NM_018811, intron 3 of 10) | intron (NM_018811, intron 3 of 10) | 27634 | NM_018811 | 54608 | Mm.365490 | NM_018811 | | **Abhd2** | 2210009N18Rik\|LABH2\|Labh-2 | abhydrolase domain containing 2 | protein-coding |
| chr7_79293492_79294206 | chr7 | 79293493 | 79294206 | + | -0.929405 | intron (NM_018811, intron 1 of 10) | intron (NM_018811, intron 1 of 10) | 20583 | NM_018811 | 54608 | Mm.365490 | NM_018811 | | **Abhd2** | 2210009N18Rik\|LABH2\|Labh-2 | abhydrolase domain containing 2 | protein-coding |
| chr8_71462598_71463249 | chr8 | 71462599 | 71463249 | + | -0.674482 | intron (NM_022419, intron 1 of 4) | intron (NM_022419, intron 1 of 4) | 733 | NM_022419 | 64296 | Mm.276383 | NM_022419 | ENSMUSG00000007950 | **Abhd8** | 0910001L24Rik\|AB030191 | abhydrolase domain containing 8 | protein-coding |
| chr2_23002714_23003109 | chr2 | 23002715 | 23003109 | + | -0.686262 | intron (NM_145994, intron 1 of 9) | intron (NM_145994, intron 1 of 9) | 37329 | NM_001077192 | 11308 | Mm.205647 | NM_007380 | ENSMUSG00000058835 | **Abi1** | E3B1\|NAP1\|Ssh3bp1 | abl-interactor 1 | protein-coding |
| chr1_60482934_60483215 | chr1 | 60482935 | 60483215 | + | -0.774332 | TTS (NM_001045513) | TTS (NM_001045513) | 73456 | NM_001198570 | 329165 | Mm.212066 | NM_198127 | ENSMUSG00000026782 | **Abi2** | 8430425M24Rik\|AI839867\|C130078H13 | abl-interactor 2 | protein-coding |
| chr1_60460695_60460958 | chr1 | 60460696 | 60460958 | + | -0.82449 | intron (NM_198127, intron 7 of 8) | intron (NM_198127, intron 7 of 8) | 51208 | NM_001198570 | 329165 | Mm.212066 | NM_198127 | ENSMUSG00000026782 | **Abi2** | 8430425M24Rik\|AI839867\|C130078H13 | abl-interactor 2 | protein-coding |
| chr1_60460511_60461140 | chr1 | 60460512 | 60461140 | + | -0.852344 | intron (NM_198127, intron 7 of 8) | intron (NM_198127, intron 7 of 8) | 51207 | NM_001198570 | 329165 | Mm.212066 | NM_198127 | ENSMUSG00000026782 | **Abi2** | 8430425M24Rik\|AI839867\|C130078H13 | abl-interactor 2 | protein-coding |
| chr1_60416601_60417225 | chr1 | 60416602 | 60417225 | + | -1.06126 | intron (NM_198127, intron 1 of 8) | LTR75\|LTR\|ERVL | 7294 | NM_001198570 | 329165 | Mm.212066 | NM_198127 | ENSMUSG00000026782 | **Abi2** | 8430425M24Rik\|AI839867\|C130078H13 | abl-interactor 2 | protein-coding |
| chr2_31727031_31728168 | chr2 | 31727032 | 31728168 | + | -0.65758 | intron (NM_001283047, intron 2 of 11) | intron (NM_001283047, intron 2 of 11) | -32339 | NM_009594 | 11350 | Mm.1318 | NM_009594 | ENSMUSG00000026842 | **Abl1** | AI325092\|Abl\|E430008G22Rik\|c-Abl | c-abl oncogene 1, non-receptor tyrosine kinase | protein-coding |
| chr2_31727604_31728342 | chr2 | 31727605 | 31728342 | + | -0.698069 | intron (NM_001283047, intron 2 of 11) | intron (NM_001283047, intron 2 of 11) | -31966 | NM_009594 | 11350 | Mm.1318 | NM_009594 | ENSMUSG00000026842 | **Abl1** | AI325092\|Abl\|E430008G22Rik\|c-Abl | c-abl oncogene 1, non-receptor tyrosine kinase | protein-coding |
| chr1_156573939_156574476 | chr1 | 156573940 | 156574476 | + | -0.774731 | intron (NM_009595, intron 1 of 11) | RSINE1\|SINE\|B4 | 15421 | NM_001136104 | 11352 | Mm.329515 | NM_009595 | ENSMUSG00000026596 | **Abl2** | AA536808\|Abll\|Arg | v-abl Abelson murine leukemia viral oncogene 2 (arg, Abelson-related gene) | protein-coding |
| chr1_156574064_156574435 | chr1 | 156574065 | 156574435 | + | -0.785538 | intron (NM_009595, intron 1 of 11) | PB1D7\|SINE\|Alu | 15463 | NM_001136104 | 11352 | Mm.329515 | NM_009595 | ENSMUSG00000026596 | **Abl2** | AA536808\|Abll\|Arg | v-abl Abelson murine leukemia viral oncogene 2 (arg, Abelson-related gene) | protein-coding |
| chr19_57082881_57083458 | chr19 | 57082882 | 57083458 | + | -0.702384 | intron (NM_001290813, intron 7 of 20) | intron (NM_001290813, intron 7 of 20) | 35854 | NM_001103178 | 226251 | Mm.217161 | NM_178688 | ENSMUSG00000025085 | **Ablim1** | 2210411C18Rik\|2610209L21Rik\|4833406P10Rik\|9330196J19Rik\|AV079770\|AW060987\|AW215784\|Limab1\|mKIAA0059 | actin-binding LIM protein 1 | protein-coding |
| chr19_57093281_57093986 | chr19 | 57093282 | 57093986 | + | -0.76267 | intron (NM_001290813, intron 6 of 20) | L1ME1\|LINE\|L1 | 25390 | NM_001103178 | 226251 | Mm.217161 | NM_178688 | ENSMUSG00000025085 | **Ablim1** | 2210411C18Rik\|2610209L21Rik\|4833406P10Rik\|9330196J19Rik\|AV079770\|AW060987\|AW215784\|Limab1\|mKIAA0059 | actin-binding LIM protein 1 | protein-coding |
| chr19_57093651_57093927 | chr19 | 57093652 | 57093927 | + | -0.794154 | intron (NM_001290813, intron 6 of 20) | L1ME1\|LINE\|L1 | 25235 | NM_001103178 | 226251 | Mm.217161 | NM_178688 | ENSMUSG00000025085 | **Ablim1** | 2210411C18Rik\|2610209L21Rik\|4833406P10Rik\|9330196J19Rik\|AV079770\|AW060987\|AW215784\|Limab1\|mKIAA0059 | actin-binding LIM protein 1 | protein-coding |
| chr19_57082926_57083138 | chr19 | 57082927 | 57083138 | + | -0.848212 | intron (NM_001290813, intron 7 of 20) | intron (NM_001290813, intron 7 of 20) | 35992 | NM_001103178 | 226251 | Mm.217161 | NM_178688 | ENSMUSG00000025085 | **Ablim1** | 2210411C18Rik\|2610209L21Rik\|4833406P10Rik\|9330196J19Rik\|AV079770\|AW060987\|AW215784\|Limab1\|mKIAA0059 | actin-binding LIM protein 1 | protein-coding |
| chr19_57072795_57073287 | chr19 | 57072796 | 57073287 | + | -0.851197 | intron (NM_001290813, intron 8 of 20) | intron (NM_001290813, intron 8 of 20) | 45983 | NM_001103178 | 226251 | Mm.217161 | NM_178688 | ENSMUSG00000025085 | **Ablim1** | 2210411C18Rik\|2610209L21Rik\|4833406P10Rik\|9330196J19Rik\|AV079770\|AW060987\|AW215784\|Limab1\|mKIAA0059 | actin-binding LIM protein 1 | protein-coding |
| chr19_57115025_57115275 | chr19 | 57115026 | 57115275 | + | -0.9665 | intron (NM_001290813, intron 6 of 20) | intron (NM_001290813, intron 6 of 20) | 3874 | NM_001103178 | 226251 | Mm.217161 | NM_178688 | ENSMUSG00000025085 | **Ablim1** | 2210411C18Rik\|2610209L21Rik\|4833406P10Rik\|9330196J19Rik\|AV079770\|AW060987\|AW215784\|Limab1\|mKIAA0059 | actin-binding LIM protein 1 | protein-coding |
| chr2_26773383_26773669 | chr2 | 26773384 | 26773669 | + | -0.782924 | Intergenic | ORR1B1\|LTR\|ERVL-MaLR | 91469 | NM_030718 | 80908 | Mm.160386 | NM_030718 | ENSMUSG00000015787 | **Abo** | NAGAT | ABO blood group (transferase A, alpha 1-3-N-acetylgalactosaminyltransferase, transferase B, alpha 1-3-galactosyltransferase) | protein-coding |
| chr11_76496460_76496651 | chr11 | 76496461 | 76496651 | + | -0.870498 | intron (NM_198894, intron 2 of 22) | intron (NM_198894, intron 2 of 22) | 12863 | NM_198018 | 109934 | Mm.258939 | NM_198018 | ENSMUSG00000017631 | **Abr** | - | active BCR-related gene | protein-coding |
| chr2_103651946_103652526 | chr2 | 103651947 | 103652526 | + | -0.842716 | intron (NM_178890, intron 1 of 16) | ID4\|SINE\|ID | 85926 | NM_178890 | 99382 | Mm.35850 | NM_178890 | ENSMUSG00000032724 | **Abtb2** | AW539457\|BPOZ-2 | ankyrin repeat and BTB (POZ) domain containing 2 | protein-coding |
| chr2_103652081_103652486 | chr2 | 103652082 | 103652486 | + | -0.940992 | intron (NM_178890, intron 1 of 16) | intron (NM_178890, intron 1 of 16) | 85974 | NM_178890 | 99382 | Mm.35850 | NM_178890 | ENSMUSG00000032724 | **Abtb2** | AW539457\|BPOZ-2 | ankyrin repeat and BTB (POZ) domain containing 2 | protein-coding |
| chr18_74789182_74789991 | chr18 | 74789183 | 74789991 | + | -0.707507 | intron (NM_177470, intron 2 of 9) | intron (NM_177470, intron 2 of 9) | 10375 | NM_177470 | 52538 | Mm.245724 | NM_177470 | ENSMUSG00000036880 | **Acaa2** | 0610011L04Rik\|AI255831\|AI265397\|D18Ertd240e | acetyl-Coenzyme A acyltransferase 2 (mitochondrial 3-oxoacyl-Coenzyme A thiolase) | protein-coding |
| chr5_114199137_114199442 | chr5 | 114199138 | 114199442 | + | -0.654983 | intron (NM_133904, intron 12 of 51) | intron (NM_133904, intron 12 of 51) | 33772 | NM_133904 | 100705 | Mm.81793 | NM_133904 | ENSMUSG00000042010 | **Acacb** | AI597064\|AW743042\|Acc2\|Accb | acetyl-Coenzyme A carboxylase beta | protein-coding |
| chr5_114193909_114194571 | chr5 | 114193910 | 114194571 | + | -0.777011 | intron (NM_133904, intron 8 of 51) | B1F2\|SINE\|Alu | 28722 | NM_133904 | 100705 | Mm.81793 | NM_133904 | ENSMUSG00000042010 | **Acacb** | AI597064\|AW743042\|Acc2\|Accb | acetyl-Coenzyme A carboxylase beta | protein-coding |
| chr1_66874625_66875052 | chr1 | 66874626 | 66875052 | + | -0.679095 | Intergenic | Intergenic | -11530 | NM_007381 | 11363 | Mm.2445 | NM_007381 | ENSMUSG00000026003 | **Acadl** | AA960361\|AU018452\|C79855\|LCAD | acyl-Coenzyme A dehydrogenase, long-chain | protein-coding |
| chr16_31235950_31236309 | chr16 | 31235951 | 31236309 | + | -0.694438 | intron (NR_035427, intron 1 of 2) | intron (NR_035427, intron 1 of 2) | -34892 | NM_030138 | 78618 | Mm.274646 | NM_030138 | ENSMUSG00000049076 | **Acap2** | 4832442G16\|9530039J15Rik\|CNT-B2\|Centb2\|mKIAA0041 | ArfGAP with coiled-coil, ankyrin repeat and PH domains 2 | protein-coding |
| chr16_31191807_31192382 | chr16 | 31191808 | 31192382 | + | -0.7273 | intron (NM_030138, intron 1 of 22) | ORR1E\|LTR\|ERVL-MaLR | 9143 | NM_030138 | 78618 | Mm.274646 | NM_030138 | ENSMUSG00000049076 | **Acap2** | 4832442G16\|9530039J15Rik\|CNT-B2\|Centb2\|mKIAA0041 | ArfGAP with coiled-coil, ankyrin repeat and PH domains 2 | protein-coding |
| chr1_180723773_180724037 | chr1 | 180723774 | 180724037 | + | -0.940556 | Intergenic | MLT1A0\|LTR\|ERVL-MaLR | -2138 | NM_133225 | 170760 | Mm.272981 | NM_133225 | ENSMUSG00000026499 | **Acbd3** | 60kDa\|8430407O11Rik\|D1Ertd10e\|GCP60\|GOLPH1\|Gocap1\|Pap7 | acyl-Coenzyme A binding domain containing 3 | protein-coding |
| chr4_86885728_86886686 | chr4 | 86885729 | 86886686 | + | -1.20359 | exon (NM_001290541, exon 2 of 5) | exon (NM_001290541, exon 2 of 5) | 11793 | NM_139306 | 230379 | Mm.45019 | NM_139306 | ENSMUSG00000038007 | **Acer2** | 2410116I05Rik\|Asah3l\|CRG-L1\|maCER2 | alkaline ceramidase 2 | protein-coding |
| chr5_137268227_137268932 | chr5 | 137268228 | 137268932 | + | -0.683077 | Intergenic | Intergenic | -19674 | NM_009599 | 11423 | Mm.255464 | NM_009599 | ENSMUSG00000023328 | **Ache** | - | acetylcholinesterase | protein-coding |
| chr4_106794209_106795053 | chr4 | 106794210 | 106795053 | + | -0.883387 | intron (NM_025590, intron 1 of 15) | intron (NM_025590, intron 1 of 15) | 5200 | NM_025590 | 329910 | Mm.222956 | NM_025590 | ENSMUSG00000034853 | **Acot11** | 1110020M10Rik\|2010309H15Rik\|AW060409\|BFIT\|BFIT1\|Thea\|Them1\|mKIAA0707 | acyl-CoA thioesterase 11 | protein-coding |
| chr4_106794201_106794774 | chr4 | 106794202 | 106794774 | + | -0.924283 | intron (NM_025590, intron 1 of 15) | intron (NM_025590, intron 1 of 15) | 5343 | NM_025590 | 329910 | Mm.222956 | NM_025590 | ENSMUSG00000034853 | **Acot11** | 1110020M10Rik\|2010309H15Rik\|AW060409\|BFIT\|BFIT1\|Thea\|Them1\|mKIAA0707 | acyl-CoA thioesterase 11 | protein-coding |
| chr4_152189835_152190147 | chr4 | 152189836 | 152190147 | + | -0.644619 | intron (NM_001146057, intron 1 of 8) | intron (NM_001146057, intron 1 of 8) | 3857 | NM_001146057 | 70025 | Mm.296191 | NM_133348 | ENSMUSG00000028937 | **Acot7** | 2410041A17Rik\|Ach1\|Act\|Bach\|CTE-IIa\|Cte-II\|Lach1 | acyl-CoA thioesterase 7 | protein-coding |

| | | | | | | | | | | | | | | | | | |
|---|---|---|---|---|---|---|---|---|---|---|---|---|---|---|---|---|---|
| chr4_152188711_152188980 | chr4 | 152188712 | 152188980 | + | -0.699449 | intron (NM_001146057, intron 1 of 8) | ID_B1\|SINE\|B4 | 2712 | NM_001146057 | 70025 | Mm.296191 | NM_133348 | ENSMUSG00000028937 | **Acot7** | 2410041A17Rik\|Ach1\|Act\|Bach\|CTE-IIa\|Cte-II\|Lach1 | acyl-CoA thioesterase 7 | protein-coding |
| chr4_152201486_152201914 | chr4 | 152201487 | 152201914 | + | -0.815358 | intron (NM_001146058, intron 1 of 8) | intron (NM_001146058, intron 1 of 8) | 1856 | NM_001146058 | 70025 | Mm.296191 | NM_133348 | ENSMUSG00000028937 | **Acot7** | 2410041A17Rik\|Ach1\|Act\|Bach\|CTE-IIa\|Cte-II\|Lach1 | acyl-CoA thioesterase 7 | protein-coding |
| chr4_152197476_152198016 | chr4 | 152197477 | 152198016 | + | -1.01202 | intron (NM_001146057, intron 1 of 8) | intron (NM_001146057, intron 1 of 8) | -2098 | NM_001146058 | 70025 | Mm.296191 | NM_133348 | ENSMUSG00000028937 | **Acot7** | 2410041A17Rik\|Ach1\|Act\|Bach\|CTE-IIa\|Cte-II\|Lach1 | acyl-CoA thioesterase 7 | protein-coding |
| chr2_127881053_127881374 | chr2 | 127881054 | 127881374 | + | -0.784487 | intron (NM_028765, intron 7 of 17) | intron (NM_028765, intron 7 of 17) | 26586 | NM_028765 | 74121 | Mm.244835 | NM_028765 | ENSMUSG00000027380 | **Acoxl** | 1200014P05Rik\|AV025673 | acyl-Coenzyme A oxidase-like | protein-coding |
| chr9_22136007_22136414 | chr9 | 22136008 | 22136414 | + | -1.0328 | promoter-TSS (NM_001102405) | promoter-TSS (NM_001102405) | -465 | NM_001102405 | 11433 | Mm.46354 | NM_007388 | ENSMUSG00000001348 | **Acp5** | TRACP\|TRAP | acid phosphatase 5, tartrate resistant | protein-coding |
| chr9_22134286_22134621 | chr9 | 22134287 | 22134621 | + | -1.06676 | intron (NM_001102405, intron 1 of 4) | C-rich\|Low_complexity\|Low_complexity | 1292 | NM_001102405 | 11433 | Mm.46354 | NM_007388 | ENSMUSG00000001348 | **Acp5** | TRACP\|TRAP | acid phosphatase 5, tartrate resistant | protein-coding |
| chr9_104331255_104331789 | chr9 | 104331256 | 104331789 | + | -0.615402 | intron (NM_019807, intron 1 of 10) | intron (NM_019807, intron 1 of 10) | 6200 | NM_019807 | 56318 | Mm.19941 | NM_019807 | ENSMUSG00000032561 | **Acpp** | 5'-NT\|A030005E02Rik\|FRAP\|Lap\|PAP\|Ppal | acid phosphatase, prostate | protein-coding |
| chr9_104331269_104331599 | chr9 | 104331270 | 104331599 | + | -0.663488 | intron (NM_019807, intron 1 of 10) | intron (NM_019807, intron 1 of 10) | 6288 | NM_019807 | 56318 | Mm.19941 | NM_019807 | ENSMUSG00000032561 | **Acpp** | 5'-NT\|A030005E02Rik\|FRAP\|Lap\|PAP\|Ppal | acid phosphatase, prostate | protein-coding |
| chr11_94591271_94592165 | chr11 | 94591272 | 94592165 | + | -0.853873 | intron (NM_153807, intron 1 of 15) | Lx8\|LINE\|L1 | 10068 | NM_153807 | 264895 | Mm.386885 | NM_153807 | ENSMUSG00000076435 | **Acsf2** | - | acyl-CoA synthetase family member 2 | protein-coding |
| chrX_142469983_142470519 | chrX | 142469984 | 142470519 | + | -0.635377 | Intergenic | Intergenic | -79716 | NM_019477 | 50790 | Mm.391337 | NM_019477 | ENSMUSG00000031278 | **Acsl4** | 9430020A05Rik\|ACS4\|AU018108\|Facl4\|Lacs4 | acyl-CoA synthetase long-chain family member 4 | protein-coding |
| chr2_155554686_155554919 | chr2 | 155554687 | 155554919 | + | -0.939132 | intron (NR_040613, intron 2 of 3) | intron (NR_040613, intron 2 of 3) | 2127 | NR_040613 | 74945 | Mm.401956 | NR_040613 | ENSMUSG00000086529 | **Acss2os** | 4930471I20Rik | acyl-CoA synthetase short-chain family member 2, opposite strand | ncRNA |
| chr2_155552889_155553460 | chr2 | 155552890 | 155553460 | + | -1.01009 | intron (NR_040613, intron 2 of 3) | intron (NR_040613, intron 2 of 3) | 3755 | NR_040613 | 74945 | Mm.401956 | NR_040613 | ENSMUSG00000086529 | **Acss2os** | 4930471I20Rik | acyl-CoA synthetase short-chain family member 2, opposite strand | ncRNA |
| chr12_80210870_80211861 | chr12 | 80210871 | 80211861 | + | -0.61247 | intron (NM_134156, intron 1 of 20) | intron (NM_134156, intron 1 of 20) | 49005 | NM_134156 | 109711 | Mm.253564 | NM_134156 | ENSMUSG00000015143 | **Actn1** | 3110023F10Rik\|Actn1a | actinin, alpha 1 | protein-coding |
| chr12_80258480_80258755 | chr12 | 80258481 | 80258755 | + | -0.670051 | intron (NM_134156, intron 1 of 20) | intron (NM_134156, intron 1 of 20) | 1753 | NM_134156 | 109711 | Mm.253564 | NM_134156 | ENSMUSG00000015143 | **Actn1** | 3110023F10Rik\|Actn1a | actinin, alpha 1 | protein-coding |
| chr12_80272926_80273660 | chr12 | 80272927 | 80273660 | + | -0.82339 | Intergenic | Intergenic | -12922 | NM_134156 | 109711 | Mm.253564 | NM_134156 | ENSMUSG00000015143 | **Actn1** | 3110023F10Rik\|Actn1a | actinin, alpha 1 | protein-coding |
| chr13_12349228_12349797 | chr13 | 12349229 | 12349797 | + | -1.11575 | Intergenic | Intergenic | -8781 | NM_033268 | 11472 | Mm.37638 | NM_033268 | ENSMUSG00000052374 | **Actn2** | 1110008F24Rik | actinin alpha 2 | protein-coding |
| chr12_70881923_70882477 | chr12 | 70881924 | 70882477 | + | -0.711793 | intron (NM_028127, intron 6 of 13) | intron (NM_028127, intron 6 of 13) | -55657 | NM_019785 | 56444 | Mm.29317 | NM_019785 | ENSMUSG00000021076 | **Actr10** | Actr11\|Arp11 | ARP10 actin-related protein 10 | protein-coding |
| chr12_70881937_70882288 | chr12 | 70881938 | 70882288 | + | -0.726997 | intron (NM_028127, intron 6 of 13) | intron (NM_028127, intron 6 of 13) | -55744 | NM_019785 | 56444 | Mm.29317 | NM_019785 | ENSMUSG00000021076 | **Actr10** | Actr11\|Arp11 | ARP10 actin-related protein 10 | protein-coding |
| chr1_125464116_125464311 | chr1 | 125464117 | 125464311 | + | -0.649994 | Intergenic | Intergenic | -28487 | NM_001205386 | 74117 | Mm.183102 | NM_023735 | ENSMUSG00000026341 | **Actr3** | 1200003A09Rik\|Arp3 | ARP3 actin-related protein 3 | protein-coding |
| chr15_101180224_101180997 | chr15 | 101180225 | 101180997 | + | -0.784972 | intron (NM_007395, intron 1 of 8) | MLT1E1A\|LTR\|ERVL-MaLR | 6486 | NM_007395 | 11479 | Mm.308467 | NM_007395 | ENSMUSG00000000532 | **Acvr1b** | 6820432J04\|ActR-IB\|ActRIB\|Acvrlk4\|Alk4\|SKR2 | activin A receptor, type 1B | protein-coding |
| chr2_48469464_48469730 | chr2 | 48469465 | 48469730 | + | -0.82343 | Intergenic | Intergenic | -344512 | NM_007396 | 11480 | Mm.314338 | NM_007396 | ENSMUSG00000052155 | **Acvr2a** | ActrIIa\|Acvr2\|TactrII | activin receptor IIA | protein-coding |
| chr2_48473077_48473648 | chr2 | 48473078 | 48473648 | + | -0.843194 | Intergenic | Intergenic | -340746 | NM_007396 | 11480 | Mm.314338 | NM_007396 | ENSMUSG00000052155 | **Acvr2a** | ActrIIa\|Acvr2\|TactrII | activin receptor IIA | protein-coding |
| chr2_48332545_48333227 | chr2 | 48332546 | 48333227 | + | -0.849947 | Intergenic | Intergenic | -481223 | NM_007396 | 11480 | Mm.314338 | NM_007396 | ENSMUSG00000052155 | **Acvr2a** | ActrIIa\|Acvr2\|TactrII | activin receptor IIA | protein-coding |
| chr2_48349831_48350405 | chr2 | 48349832 | 48350405 | + | -1.34103 | Intergenic | URR1A\|DNA\|hAT-Charlie | -463991 | NM_007396 | 11480 | Mm.314338 | NM_007396 | ENSMUSG00000052155 | **Acvr2a** | ActrIIa\|Acvr2\|TactrII | activin receptor IIA | protein-coding |
| chr12_113994241_113994811 | chr12 | 113994242 | 113994811 | + | -0.762385 | Intergenic | Intergenic | 450618 | NM_174885 | 238406 | Mm.188683 | NM_174885 | ENSMUSG00000043945 | **Adam6a** | Adam6\|tMDCIV | a disintegrin and metallopeptidase domain 6A | protein-coding |
| chr12_113446087_113446553 | chr12 | 113446088 | 113446553 | + | -1.4929 | Intergenic | MTD\|LTR\|ERVL-MaLR | -43245 | NM_001009545 | 238405 | Mm.442216 | NM_001009545 | ENSMUSG00000051804 | **Adam6b** | 4930523C11Rik | a disintegrin and metallopeptidase domain 6B | protein-coding |
| chr10_61232875_61233569 | chr10 | 61232876 | 61233569 | + | -1.43721 | intron (NM_001081127, intron 4 of 21) | intron (NM_001081127, intron 4 of 21) | 40216 | NM_001081127 | 237360 | Mm.333252 | NM_001081127 | ENSMUSG00000059901 | **Adamts14** | Adamts-14\|TS14 | a disintegrin-like and metallopeptidase (reprolysin type) with thrombospondin type 1 motif, 14 | protein-coding |
| chr8_113827226_113827749 | chr8 | 113827227 | 113827749 | + | -0.927173 | intron (NM_172466, intron 3 of 22) | intron (NM_172466, intron 3 of 22) | 21351 | NM_172466 | 208936 | Mm.167471 | NM_172466 | ENSMUSG00000053399 | **Adamts18** | 9630038L21\|ADAMTS21\|E130314N14Rik | a disintegrin-like and metallopeptidase (reprolysin type) with thrombospondin type 1 motif, 18 | protein-coding |
| chr3_95709099_95709377 | chr3 | 95709100 | 95709377 | + | -0.660474 | Intergenic | Intergenic | -21380 | NM_144899 | 229595 | Mm.490574 | NM_144899 | ENSMUSG00000015850 | **Adamtsl4** | Tsrc1 | ADAMTS-like 4 | protein-coding |
| chr3_89672958_89673495 | chr3 | 89672959 | 89673495 | + | -0.644106 | TTS (NM_080466) | TTS (NM_080466) | -41795 | NM_001038587 | 56417 | Mm.316628 | NM_019655 | ENSMUSG00000027951 | **Adar** | AV242451\|Adar1\|Adar1p110\|Adar1p150\|mZaADAR | adenosine deaminase, RNA-specific | protein-coding |
| chr3_89672972_89673216 | chr3 | 89672973 | 89673216 | + | -0.65758 | TTS (NM_080466) | TTS (NM_080466) | -41928 | NM_001038587 | 56417 | Mm.316628 | NM_019655 | ENSMUSG00000027951 | **Adar** | AV242451\|Adar1\|Adar1p110\|Adar1p150\|mZaADAR | adenosine deaminase, RNA-specific | protein-coding |
| chr3_89672951_89673279 | chr3 | 89672952 | 89673279 | + | -0.661889 | TTS (NM_080466) | TTS (NM_080466) | -41907 | NM_001038587 | 56417 | Mm.316628 | NM_019655 | ENSMUSG00000027951 | **Adar** | AV242451\|Adar1\|Adar1p110\|Adar1p150\|mZaADAR | adenosine deaminase, RNA-specific | protein-coding |
| chr13_8456526_8457359 | chr13 | 8456527 | 8457359 | + | -0.665292 | intron (NM_001289530, intron 1 of 9) | MLT1C\|LTR\|ERVL-MaLR | 254077 | NM_052977 | 94191 | Mm.425794 | NM_052977 | ENSMUSG00000052551 | **Adarb2** | Adar3\|RED2 | adenosine deaminase, RNA-specific, B2 | protein-coding |
| chr13_8426435_8426944 | chr13 | 8426436 | 8426944 | + | -0.734792 | intron (NM_001289530, intron 1 of 9) | intron (NM_001289530, intron 1 of 9) | 223824 | NM_052977 | 94191 | Mm.425794 | NM_052977 | ENSMUSG00000052551 | **Adarb2** | Adar3\|RED2 | adenosine deaminase, RNA-specific, B2 | protein-coding |
| chr13_8400712_8401156 | chr13 | 8400713 | 8401156 | + | -0.818827 | intron (NM_001289530, intron 1 of 9) | MTE2b\|LTR\|ERVL-MaLR | 198068 | NM_052977 | 94191 | Mm.425794 | NM_052977 | ENSMUSG00000052551 | **Adarb2** | Adar3\|RED2 | adenosine deaminase, RNA-specific, B2 | protein-coding |
| chr13_8400188_8401083 | chr13 | 8400189 | 8401083 | + | -0.84959 | intron (NM_001289530, intron 1 of 9) | MTE2b\|LTR\|ERVL-MaLR | 197770 | NM_052977 | 94191 | Mm.425794 | NM_052977 | ENSMUSG00000052551 | **Adarb2** | Adar3\|RED2 | adenosine deaminase, RNA-specific, B2 | protein-coding |
| chr8_111976106_111976913 | chr8 | 111976107 | 111976913 | + | -0.647989 | intron (NM_013925, intron 8 of 9) | intron (NM_013925, intron 8 of 9) | 15792 | NM_013925 | 30947 | Mm.282162 | NM_013925 | ENSMUSG00000031949 | **Adat1** | MMADAT1\|mADAT1 | adenosine deaminase, tRNA-specific 1 | protein-coding |
| chr15_98609616_98610258 | chr15 | 98609617 | 98610258 | + | -0.671886 | Intergenic | CpG | -2304 | NM_007405 | 11512 | Mm.157091 | NM_007405 | ENSMUSG00000022994 | **Adcy6** | AC6\|mKIAA0422 | adenylate cyclase 6 | protein-coding |
| chr15_98609893_98610206 | chr15 | 98609894 | 98610206 | + | -0.733661 | Intergenic | CpG | -2417 | NM_007405 | 11512 | Mm.157091 | NM_007405 | ENSMUSG00000022994 | **Adcy6** | AC6\|mKIAA0422 | adenylate cyclase 6 | protein-coding |
| chr19_53184216_53184574 | chr19 | 53184217 | 53184574 | + | -0.691046 | intron (NM_013758, intron 1 of 15) | intron (NM_013758, intron 1 of 15) | 40398 | NM_013758 | 27360 | Mm.426080 | NM_013758 | ENSMUSG00000025026 | **Add3** | AI463285\|R75380 | adducin 3 (gamma) | protein-coding |
| chr19_53188291_53188511 | chr19 | 53188292 | 53188511 | + | -0.723873 | intron (NM_013758, intron 1 of 15) | intron (NM_013758, intron 1 of 15) | 44404 | NM_013758 | 27360 | Mm.426080 | NM_013758 | ENSMUSG00000025026 | **Add3** | AI463285\|R75380 | adducin 3 (gamma) | protein-coding |
| chr19_53189761_53190025 | chr19 | 53189762 | 53190025 | + | -0.893162 | intron (NM_013758, intron 1 of 15) | intron (NM_013758, intron 1 of 15) | 45896 | NM_013758 | 27360 | Mm.426080 | NM_013758 | ENSMUSG00000025026 | **Add3** | AI463285\|R75380 | adducin 3 (gamma) | protein-coding |
| chr10_10396118_10396436 | chr10 | 10396119 | 10396436 | + | -0.669425 | intron (NM_001127353, intron 21 of 36) | intron (NM_001127353, intron 21 of 36) | 76037 | NM_001127353 | 215772 | Mm.246573 | NM_172547 | ENSMUSG00000050994 | **Adgb** | 9130014G24Rik | androglobin | protein-coding |
| chr16_23165399_23165916 | chr16 | 23165400 | 23165916 | + | -0.741582 | Intergenic | Intergenic | 19122 | NM_009605 | 11450 | Mm.3969 | NM_009605 | ENSMUSG00000022878 | **Adipoq** | 30kDa\|APN\|Acdc\|Acrp30\|GBP28\|adipo\|apM1 | adiponectin, C1Q and collagen domain containing | protein-coding |
| chr16_23165236_23166024 | chr16 | 23165237 | 23166024 | + | -0.754487 | Intergenic | Intergenic | 19094 | NM_009605 | 11450 | Mm.3969 | NM_009605 | ENSMUSG00000022878 | **Adipoq** | 30kDa\|APN\|Acdc\|Acrp30\|GBP28\|adipo\|apM1 | adiponectin, C1Q and collagen domain containing | protein-coding |
| chr16_23151403_23151751 | chr16 | 23151404 | 23151751 | + | -0.778329 | intron (NM_009605, intron 1 of 2) | intron (NM_009605, intron 1 of 2) | 5041 | NM_009605 | 11450 | Mm.3969 | NM_009605 | ENSMUSG00000022878 | **Adipoq** | 30kDa\|APN\|Acdc\|Acrp30\|GBP28\|adipo\|apM1 | adiponectin, C1Q and collagen domain containing | protein-coding |
| chr16_23151403_23151886 | chr16 | 23151404 | 23151886 | + | -0.784819 | intron (NM_009605, intron 1 of 2) | intron (NM_009605, intron 1 of 2) | 5109 | NM_009605 | 11450 | Mm.3969 | NM_009605 | ENSMUSG00000022878 | **Adipoq** | 30kDa\|APN\|Acdc\|Acrp30\|GBP28\|adipo\|apM1 | adiponectin, C1Q and collagen domain containing | protein-coding |
| chr16_23174651_23175733 | chr16 | 23174652 | 23175733 | + | -0.885052 | Intergenic | Intergenic | 28656 | NM_009605 | 11450 | Mm.3969 | NM_009605 | ENSMUSG00000022878 | **Adipoq** | 30kDa\|APN\|Acdc\|Acrp30\|GBP28\|adipo\|apM1 | adiponectin, C1Q and collagen domain containing | protein-coding |
| chr16_23181487_23182025 | chr16 | 23181488 | 23182025 | + | -1.28521 | Intergenic | Intergenic | 35220 | NM_009605 | 11450 | Mm.3969 | NM_009605 | ENSMUSG00000022878 | **Adipoq** | 30kDa\|APN\|Acdc\|Acrp30\|GBP28\|adipo\|apM1 | adiponectin, C1Q and collagen domain containing | protein-coding |
| chr16_23170786_23171252 | chr16 | 23170787 | 23171252 | + | -1.39163 | Intergenic | RMER19C\|LTR\|ERVK | 24483 | NM_009605 | 11450 | Mm.3969 | NM_009605 | ENSMUSG00000022878 | **Adipoq** | 30kDa\|APN\|Acdc\|Acrp30\|GBP28\|adipo\|apM1 | adiponectin, C1Q and collagen domain containing | protein-coding |

| | | | | | | | | | | | | | | | | | |
|---|---|---|---|---|---|---|---|---|---|---|---|---|---|---|---|---|---|
| chr6_119377610_119378071 | chr6 | 119377611 | 119378071 | + | -0.845571 | intron (NM_197985, intron 1 of 7) | intron (NM_197985, intron 1 of 7) | 39642 | NM_197985 | 68465 | Mm.291826 | NM_197985 | ENSMUSG00000030168 | **Adipor2** | 1110001I14Rik\|ADCR2\|AI115388\|AW554121\|D6Ucla1e\|Paqr2 | adiponectin receptor 2 | protein-coding |
| chr10_67581874_67582263 | chr10 | 67581875 | 67582263 | + | -0.868694 | Intergenic | Intergenic | -33114 | NM_001005419 | 211488 | Mm.297250 | NM_001005419 | ENSMUSG00000057134 | **Ado** | Gm237 | 2-aminoethanethiol (cysteamine) dioxygenase | protein-coding |
| chr10_75295013_75295480 | chr10 | 75295014 | 75295480 | + | -0.700818 | intron (NM_153406, intron 13 of 15) | intron (NM_153406, intron 13 of 15) | -21630 | NM_009630 | 11540 | Mm.333734 | NM_009630 | ENSMUSG00000020178 | **Adora2a** | A2AAR\|A2aR\|AA2AR | adenosine A2a receptor | protein-coding |
| chr10_75294128_75294594 | chr10 | 75294129 | 75294594 | + | -0.7298 | intron (NM_153406, intron 13 of 15) | intron (NM_153406, intron 13 of 15) | -22516 | NM_009630 | 11540 | Mm.333734 | NM_009630 | ENSMUSG00000020178 | **Adora2a** | A2AAR\|A2aR\|AA2AR | adenosine A2a receptor | protein-coding |
| chr18_62125381_62125767 | chr18 | 62125382 | 62125767 | + | -0.702193 | Intergenic | Intergenic | 54407 | NM_007420 | 11555 | Mm.5598 | NM_007420 | ENSMUSG00000045730 | **Adrb2** | Adrb-2\|Badm\|Gpcr7 | adrenergic receptor, beta 2 | protein-coding |
| chr18_62156539_62156866 | chr18 | 62156540 | 62156866 | + | -0.8048 | Intergenic | MIR3\|SINE\|MIR | 23278 | NM_007420 | 11555 | Mm.5598 | NM_007420 | ENSMUSG00000045730 | **Adrb2** | Adrb-2\|Badm\|Gpcr7 | adrenergic receptor, beta 2 | protein-coding |
| chr18_62156413_62156840 | chr18 | 62156414 | 62156840 | + | -0.83032 | Intergenic | Intergenic | 23354 | NM_007420 | 11555 | Mm.5598 | NM_007420 | ENSMUSG00000045730 | **Adrb2** | Adrb-2\|Badm\|Gpcr7 | adrenergic receptor, beta 2 | protein-coding |
| chr18_62151652_62152082 | chr18 | 62151653 | 62152082 | + | -0.838691 | Intergenic | Intergenic | 28114 | NM_007420 | 11555 | Mm.5598 | NM_007420 | ENSMUSG00000045730 | **Adrb2** | Adrb-2\|Badm\|Gpcr7 | adrenergic receptor, beta 2 | protein-coding |
| chr19_4302289_4302607 | chr19 | 4302290 | 4302607 | + | -0.663626 | intron (NM_001290818, intron 1 of 20) | intron (NM_001290818, intron 1 of 20) | 3774 | NM_130863 | 110355 | Mm.254144 | NM_130863 | ENSMUSG00000024858 | **Adrbk1** | Adrbk-1\|Bark-1\|GRK2\|beta ARK\|betaARK1 | adrenergic receptor kinase, beta 1 | protein-coding |
| chr19_4299641_4299885 | chr19 | 4299642 | 4299885 | + | -1.0338 | intron (NM_001290818, intron 1 of 20) | intron (NM_001290818, intron 1 of 20) | 6459 | NM_130863 | 110355 | Mm.254144 | NM_130863 | ENSMUSG00000024858 | **Adrbk1** | Adrbk-1\|Bark-1\|GRK2\|beta ARK\|betaARK1 | adrenergic receptor kinase, beta 1 | protein-coding |
| chr5_113034903_113035576 | chr5 | 113034904 | 113035576 | + | -0.689855 | Intergenic | L1_Rod\|LINE\|L1 | -19702 | NM_177078 | 320129 | Mm.285619 | NM_177078 | ENSMUSG00000042249 | **Adrbk2** | 4833444A01Rik\|AI851927\|AW551196\|Adrbk-2\|Bark-2\|GRK3 | adrenergic receptor kinase, beta 2 | protein-coding |
| chr6_140707887_140708555 | chr6 | 140707888 | 140708555 | + | -0.736971 | Intergenic | Intergenic | 83923 | NM_009637 | 11569 | Mm.86453 | NM_009637 | ENSMUSG00000030232 | **Aebp2** | AU023766\|B230313N05Rik | AE binding protein 2 | protein-coding |
| chr6_140657395_140657989 | chr6 | 140657396 | 140657989 | + | -0.809422 | intron (NM_009637, intron 8 of 8) | intron (NM_009637, intron 8 of 8) | 33394 | NM_009637 | 11569 | Mm.86453 | NM_009637 | ENSMUSG00000030232 | **Aebp2** | AU023766\|B230313N05Rik | AE binding protein 2 | protein-coding |
| chr6_140546049_140546471 | chr6 | 140546050 | 140546471 | + | -0.83191 | intron (NM_144920, intron 11 of 30) | intron (NM_144920, intron 11 of 30) | -76403 | NM_178803 | 11569 | Mm.86453 | NM_009637 | ENSMUSG00000030232 | **Aebp2** | AU023766\|B230313N05Rik | AE binding protein 2 | protein-coding |
| chr6_140645912_140646239 | chr6 | 140645913 | 140646239 | + | -0.853732 | intron (NM_001005605, intron 5 of 7) | intron (NM_001005605, intron 5 of 7) | 21778 | NM_009637 | 11569 | Mm.86453 | NM_009637 | ENSMUSG00000030232 | **Aebp2** | AU023766\|B230313N05Rik | AE binding protein 2 | protein-coding |
| chr6_140657024_140657944 | chr6 | 140657025 | 140657944 | + | -0.98951 | intron (NM_009637, intron 8 of 8) | B2_Mm2\|SINE\|B2 | 33186 | NM_009637 | 11569 | Mm.86453 | NM_009637 | ENSMUSG00000030232 | **Aebp2** | AU023766\|B230313N05Rik | AE binding protein 2 | protein-coding |
| chr6_140656981_140657343 | chr6 | 140656982 | 140657343 | + | -1.04362 | intron (NM_009637, intron 8 of 8) | intron (NM_009637, intron 8 of 8) | 32864 | NM_009637 | 11569 | Mm.86453 | NM_009637 | ENSMUSG00000030232 | **Aebp2** | AU023766\|B230313N05Rik | AE binding protein 2 | protein-coding |
| chr6_140816877_140817496 | chr6 | 140816878 | 140817496 | + | -1.05885 | Intergenic | Intergenic | 192889 | NM_009637 | 11569 | Mm.86453 | NM_009637 | ENSMUSG00000030232 | **Aebp2** | AU023766\|B230313N05Rik | AE binding protein 2 | protein-coding |
| chr6_140819347_140819929 | chr6 | 140819348 | 140819929 | + | -1.17233 | Intergenic | ORR1D2\|LTR\|ERVL-MaLR | 195340 | NM_009637 | 11569 | Mm.86453 | NM_009637 | ENSMUSG00000030232 | **Aebp2** | AU023766\|B230313N05Rik | AE binding protein 2 | protein-coding |
| chr10_81560892_81561590 | chr10 | 81560893 | 81561590 | + | -0.722011 | exon (NM_010347, exon 2 of 7) | exon (NM_010347, exon 2 of 7) | 1797 | NM_001276288 | 14797 | Mm.180013 | NM_010347 | ENSMUSG00000054452 | **Aes** | AL024115\|Esp1\|Grg\|Grg-5\|Grg5\|Tle5 | amino-terminal enhancer of split | protein-coding |
| chr7_28355216_28355540 | chr7 | 28355217 | 28355540 | + | -0.831728 | Intergenic | Intergenic | 3838 | NR_028129 | 100302567 | | NR_028129 | | **AF357399** | MBI-142 | snoRNA AF357399 | snoRNA |
| chr5_36006034_36006475 | chr5 | 36006035 | 36006475 | + | -0.788183 | Intergenic | Intergenic | 112936 | NM_027373 | 70292 | Mm.125167 | NM_027373 | ENSMUSG00000029094 | **Afap1** | 2600003E23Rik\|9630044L16Rik\|AI848729\|Afap\|mKIAA3018 | actin filament associated protein 1 | protein-coding |
| chr11_53351474_53352040 | chr11 | 53351475 | 53352040 | + | -0.676484 | intron (NM_033565, intron 1 of 20) | intron (NM_033565, intron 1 of 20) | 990 | NM_033565 | 93736 | Mm.395281 | NM_033565 | ENSMUSG00000049470 | **Aff4** | AF5Q31\|Alf4\|Laf4l\|MCEF | AF4/FMR2 family, member 4 | protein-coding |
| chr11_53351715_53352000 | chr11 | 53351716 | 53352000 | + | -0.724756 | intron (NM_033565, intron 1 of 20) | intron (NM_033565, intron 1 of 20) | 1091 | NM_033565 | 93736 | Mm.395281 | NM_033565 | ENSMUSG00000049470 | **Aff4** | AF5Q31\|Alf4\|Laf4l\|MCEF | AF4/FMR2 family, member 4 | protein-coding |
| chr5_90492407_90493199 | chr5 | 90492408 | 90493199 | + | -0.675938 | intron (NM_007423, intron 3 of 14) | intron (NM_007423, intron 3 of 14) | 2089 | NM_007423 | 11576 | Mm.358570 | NM_007423 | ENSMUSG00000054932 | **Afp** | - | alpha fetoprotein | protein-coding |
| chr7_76176032_76176208 | chr7 | 76176033 | 76176208 | + | -0.728527 | Intergenic | Intergenic | -53767 | NM_001199224 | 244071 | Mm.322829 | NM_172902 | ENSMUSG00000025754 | **Agbl1** | Ccp4\|D430020F16\|EG244071\|Nna1-l1 | ATP/GTP binding protein-like 1 | protein-coding |
| chr7_76176319_76176566 | chr7 | 76176320 | 76176566 | + | -0.731679 | Intergenic | Intergenic | -53444 | NM_001199224 | 244071 | Mm.322829 | NM_172902 | ENSMUSG00000025754 | **Agbl1** | Ccp4\|D430020F16\|EG244071\|Nna1-l1 | ATP/GTP binding protein-like 1 | protein-coding |
| chr7_76175941_76176256 | chr7 | 76175942 | 76176256 | + | -0.740409 | Intergenic | Intergenic | -53788 | NM_001199224 | 244071 | Mm.322829 | NM_172902 | ENSMUSG00000025754 | **Agbl1** | Ccp4\|D430020F16\|EG244071\|Nna1-l1 | ATP/GTP binding protein-like 1 | protein-coding |
| chr7_76176302_76176856 | chr7 | 76176303 | 76176856 | + | -0.747465 | Intergenic | Intergenic | -53308 | NM_001199224 | 244071 | Mm.322829 | NM_172902 | ENSMUSG00000025754 | **Agbl1** | Ccp4\|D430020F16\|EG244071\|Nna1-l1 | ATP/GTP binding protein-like 1 | protein-coding |
| chr7_76231290_76231713 | chr7 | 76231291 | 76231713 | + | -0.914471 | intron (NM_001199224, intron 1 of 22) | intron (NM_001199224, intron 1 of 22) | 1615 | NM_001199224 | 244071 | Mm.322829 | NM_172902 | ENSMUSG00000025754 | **Agbl1** | Ccp4\|D430020F16\|EG244071\|Nna1-l1 | ATP/GTP binding protein-like 1 | protein-coding |
| chr7_76237493_76238003 | chr7 | 76237494 | 76238003 | + | -1.06419 | intron (NM_001199224, intron 1 of 22) | RMER19B2\|LTR\|ERVK | 7861 | NM_001199224 | 244071 | Mm.322829 | NM_172902 | ENSMUSG00000025754 | **Agbl1** | Ccp4\|D430020F16\|EG244071\|Nna1-l1 | ATP/GTP binding protein-like 1 | protein-coding |
| chr7_76237549_76237868 | chr7 | 76237550 | 76237868 | + | -1.08108 | intron (NM_001199224, intron 1 of 22) | RMER19B2\|LTR\|ERVK | 7822 | NM_001199224 | 244071 | Mm.322829 | NM_172902 | ENSMUSG00000025754 | **Agbl1** | Ccp4\|D430020F16\|EG244071\|Nna1-l1 | ATP/GTP binding protein-like 1 | protein-coding |
| chr1_82807105_82807601 | chr1 | 82807106 | 82807601 | + | -0.705209 | Intergenic | Intergenic | -32130 | NM_010472 | 15463 | Mm.392569 | NM_010472 | ENSMUSG00000026159 | **Agfg1** | AU045498\|C130049H11Rik\|C85612\|D730048C23Rik\|Hrb\|RAB\|Rip | ArfGAP with FG repeats 1 | protein-coding |
| chr1_82833575_82833876 | chr1 | 82833576 | 82833876 | + | -0.771812 | Intergenic | ORR1D2\|LTR\|ERVL-MaLR | -5757 | NM_010472 | 15463 | Mm.392569 | NM_010472 | ENSMUSG00000026159 | **Agfg1** | AU045498\|C130049H11Rik\|C85612\|D730048C23Rik\|Hrb\|RAB\|Rip | ArfGAP with FG repeats 1 | protein-coding |
| chr15_73175831_73176310 | chr15 | 73175832 | 73176310 | + | -0.618993 | intron (NM_153178, intron 1 of 18) | RMER4B\|LTR\|ERVK | 8876 | NM_153178 | 239528 | Mm.23095 | NM_153178 | | **Ago2** | 1110029L17Rik\|2310051F07Rik\|AI225898\|AL022874\|AW546247\|ENSMUSG00000072493\|Eif2c2\|Gerp95\|Gm10365\|mKIAA4215 | argonaute RISC catalytic subunit 2 | protein-coding |
| chr15_73156498_73156845 | chr15 | 73156499 | 73156845 | + | -0.6358 | intron (NM_153178, intron 1 of 18) | PB1D10\|SINE\|Alu | 28275 | NM_153178 | 239528 | Mm.23095 | NM_153178 | | **Ago2** | 1110029L17Rik\|2310051F07Rik\|AI225898\|AL022874\|AW546247\|ENSMUSG00000072493\|Eif2c2\|Gerp95\|Gm10365\|mKIAA4215 | argonaute RISC catalytic subunit 2 | protein-coding |
| chr15_73140151_73140711 | chr15 | 73140152 | 73140711 | + | -0.655342 | intron (NM_153178, intron 2 of 18) | intron (NM_153178, intron 2 of 18) | 44516 | NM_153178 | 239528 | Mm.23095 | NM_153178 | | **Ago2** | 1110029L17Rik\|2310051F07Rik\|AI225898\|AL022874\|AW546247\|ENSMUSG00000072493\|Eif2c2\|Gerp95\|Gm10365\|mKIAA4215 | argonaute RISC catalytic subunit 2 | protein-coding |
| chr15_73228463_73228657 | chr15 | 73228464 | 73228657 | + | -0.724273 | intron (NM_007982, intron 26 of 31) | intron (NM_007982, intron 26 of 31) | -43613 | NM_153178 | 239528 | Mm.23095 | NM_153178 | | **Ago2** | 1110029L17Rik\|2310051F07Rik\|AI225898\|AL022874\|AW546247\|ENSMUSG00000072493\|Eif2c2\|Gerp95\|Gm10365\|mKIAA4215 | argonaute RISC catalytic subunit 2 | protein-coding |
| chr15_73228437_73228833 | chr15 | 73228438 | 73228833 | + | -0.842614 | intron (NM_007982, intron 26 of 31) | intron (NM_007982, intron 26 of 31) | -43688 | NM_153178 | 239528 | Mm.23095 | NM_153178 | | **Ago2** | 1110029L17Rik\|2310051F07Rik\|AI225898\|AL022874\|AW546247\|ENSMUSG00000072493\|Eif2c2\|Gerp95\|Gm10365\|mKIAA4215 | argonaute RISC catalytic subunit 2 | protein-coding |
| chr15_73137469_73138256 | chr15 | 73137470 | 73138256 | + | -0.999168 | intron (NM_153178, intron 3 of 18) | intron (NM_153178, intron 3 of 18) | 47084 | NM_153178 | 239528 | Mm.23095 | NM_153178 | | **Ago2** | 1110029L17Rik\|2310051F07Rik\|AI225898\|AL022874\|AW546247\|ENSMUSG00000072493\|Eif2c2\|Gerp95\|Gm10365\|mKIAA4215 | argonaute RISC catalytic subunit 2 | protein-coding |
| chr15_73137732_73138206 | chr15 | 73137733 | 73138206 | + | -1.01209 | intron (NM_153178, intron 3 of 18) | intron (NM_153178, intron 3 of 18) | 46978 | NM_153178 | 239528 | Mm.23095 | NM_153178 | | **Ago2** | 1110029L17Rik\|2310051F07Rik\|AI225898\|AL022874\|AW546247\|ENSMUSG00000072493\|Eif2c2\|Gerp95\|Gm10365\|mKIAA4215 | argonaute RISC catalytic subunit 2 | protein-coding |
| chr5_100929643_100930264 | chr5 | 100929644 | 100930264 | + | -0.693179 | Intergenic | MIRb\|SINE\|MIR | 83725 | NM_172715 | 231510 | Mm.271911 | NM_172715 | ENSMUSG00000029314 | **Agpat9** | 1-AGPAT\|1-AGPAT 9\|4933407I02Rik\|4933408F15\|9\|A230097K15Rik\|GPAT-3\|mGPAT3 | 1-acylglycerol-3-phosphate O-acyltransferase 9 | protein-coding |
| chr5_100911356_100911730 | chr5 | 100911357 | 100911730 | + | -0.79753 | Intergenic | Intergenic | 65314 | NM_172715 | 231510 | Mm.271911 | NM_172715 | ENSMUSG00000029314 | **Agpat9** | 1-AGPAT\|1-AGPAT 9\|4933407I02Rik\|4933408F15\|9\|A230097K15Rik\|GPAT-3\|mGPAT3 | 1-acylglycerol-3-phosphate O-acyltransferase 9 | protein-coding |

| | | | | | | | | | | | | | | | | | |
|---|---|---|---|---|---|---|---|---|---|---|---|---|---|---|---|---|---|
| chr5_100912294_100912828 | chr5 | 100912295 | 100912828 | + | -1.01898 | Intergenic | MLT1F2\|LTR\|ERVL-MaLR | 66332 | NM_172715 | 231510 | Mm.271911 | NM_172715 | ENSMUSG00000029314 | **Agpat9** | 1-AGPAT\|1-AGPAT 9\|4933407I02Rik\|4933408F15\|9\|A230097K15Rik\|GPAT-3\|mGPAT3 | 1-acylglycerol-3-phosphate O-acyltransferase 9 | protein-coding |
| chr2_75825648_75825942 | chr2 | 75825649 | 75825942 | + | -0.697422 | Intergenic | MTD\|LTR\|ERVL-MaLR | -6382 | NM_172666 | 228061 | Mm.31227 | NM_172666 | ENSMUSG00000042410 | **Agps** | 5832437L22\|9930035G10Rik\|AW123847\|Adaps\|Adas\|Adhaps\|Adps\|Aldhpsy\|bs2 | alkylglycerone phosphate synthase | protein-coding |
| chr12_35983513_35983923 | chr12 | 35983514 | 35983923 | + | -0.688731 | Intergenic | Intergenic | -9207 | NM_011783 | 23795 | Mm.7244 | NM_011783 | ENSMUSG00000020581 | **Agr2** | Agr2h\|Gob-4\|HAG-2\|XAG-2\|mAG-2 | anterior gradient 2 | protein-coding |
| chr12_35915494_35916150 | chr12 | 35915495 | 35916150 | + | -0.647414 | Intergenic | RLTR14-int\|LTR\|ERV1 | -9799 | NM_207531 | 403205 | Mm.190508 | NM_207531 | ENSMUSG00000036231 | **Agr3** | BCMP11\|E030025L21Rik\|Gm888 | anterior gradient 3 | protein-coding |
| chr4_133057358_133057819 | chr4 | 133057359 | 133057819 | + | -0.726645 | intron (NM_146155, intron 3 of 6) | intron (NM_146155, intron 3 of 6) | 46083 | NM_146155 | 230793 | Mm.31816 | NM_146155 | ENSMUSG00000037692 | **Ahdc1** | D030015G18Rik | AT hook, DNA binding motif, containing 1 | protein-coding |
| chr4_133057830_133058393 | chr4 | 133057831 | 133058393 | + | -0.735614 | intron (NM_146155, intron 3 of 6) | B1_Mur2\|SINE\|Alu | 46606 | NM_146155 | 230793 | Mm.31816 | NM_146155 | ENSMUSG00000037692 | **Ahdc1** | D030015G18Rik | AT hook, DNA binding motif, containing 1 | protein-coding |
| chr10_20882527_20882971 | chr10 | 20882528 | 20882971 | + | -0.633404 | Intergenic | Intergenic | -69798 | NM_026203 | 52906 | Mm.253280 | NM_026203 | ENSMUSG00000019986 | **Ahi1** | 1700015F03Rik\|Ahi-1\|D10Bwg0629e | Abelson helper integration site 1 | protein-coding |
| chr10_20882505_20883259 | chr10 | 20882506 | 20883259 | + | -0.646239 | Intergenic | Intergenic | -69665 | NM_026203 | 52906 | Mm.253280 | NM_026203 | ENSMUSG00000019986 | **Ahi1** | 1700015F03Rik\|Ahi-1\|D10Bwg0629e | Abelson helper integration site 1 | protein-coding |
| chr10_21092371_21093482 | chr10 | 21092372 | 21093482 | + | -0.695795 | Intergenic | Intergenic | 52107 | NM_001177776 | 52906 | Mm.253280 | NM_026203 | ENSMUSG00000019986 | **Ahi1** | 1700015F03Rik\|Ahi-1\|D10Bwg0629e | Abelson helper integration site 1 | protein-coding |
| chr4_53277718_53278611 | chr4 | 53277719 | 53278611 | + | -0.85089 | Intergenic | Intergenic | -7933 | NR_033139 | 381524 | Mm.23351 | NM_001033454 | | **AI427809** | Gm1659 | expressed sequence AI427809 | ncRNA |
| chr4_53277985_53278577 | chr4 | 53277986 | 53278577 | + | -0.89653 | Intergenic | Intergenic | -8049 | NR_033139 | 381524 | Mm.23351 | NM_001033454 | | **AI427809** | Gm1659 | expressed sequence AI427809 | ncRNA |
| chr19_5432976_5433589 | chr19 | 5432977 | 5433589 | + | -0.640204 | Intergenic | (CA)n\|Simple_repeat\|Simple_repeat | 8139 | NM_001256515 | 107242 | Mm.329658 | NM_134149 | ENSMUSG00000047423 | **AI837181** | Bles03\|N28173 | expressed sequence AI837181 | protein-coding |
| chr19_5433050_5433350 | chr19 | 5433051 | 5433350 | + | -0.803104 | Intergenic | Intergenic | 8056 | NM_001256515 | 107242 | Mm.329658 | NM_134149 | ENSMUSG00000047423 | **AI837181** | Bles03\|N28173 | expressed sequence AI837181 | protein-coding |
| chr2_129183676_129184116 | chr2 | 129183677 | 129184116 | + | -0.672744 | Intergenic | Intergenic | -3223 | NR_045264 | 100048701 | Mm.428544 | NR_045264 | ENSMUSG00000084826 | **AI847159** | OTTMUSG00000015420 | expressed sequence AI847159 | ncRNA |
| chr2_129180399_129180622 | chr2 | 129180400 | 129180622 | + | -0.794139 | non-coding (NR_045264, exon 1 of 3) | non-coding (NR_045264, exon 1 of 3) | 162 | NR_045264 | 100048701 | Mm.428544 | NR_045264 | ENSMUSG00000084826 | **AI847159** | OTTMUSG00000015420 | expressed sequence AI847159 | ncRNA |
| chr2_129180005_129180291 | chr2 | 129180006 | 129180291 | + | -0.858409 | intron (NR_045264, intron 2 of 2) | intron (NR_045264, intron 2 of 2) | 525 | NR_045264 | 100048701 | Mm.428544 | NR_045264 | ENSMUSG00000084826 | **AI847159** | OTTMUSG00000015420 | expressed sequence AI847159 | ncRNA |
| chr2_129179983_129180676 | chr2 | 129179984 | 129180676 | + | -0.873185 | intron (NR_045264, intron 1 of 2) | intron (NR_045264, intron 1 of 2) | 343 | NR_045264 | 100048701 | Mm.428544 | NR_045264 | ENSMUSG00000084826 | **AI847159** | OTTMUSG00000015420 | expressed sequence AI847159 | ncRNA |
| chr2_31929695_31930073 | chr2 | 31929696 | 31930073 | + | -0.817917 | intron (NM_011836, intron 21 of 27) | ORR1E\|LTR\|ERVL-MaLR | -20419 | NM_145144 | 108897 | Mm.24838 | NM_145144 | ENSMUSG00000001864 | **Aif1l** | 2810003C17Rik\|AI043124\|C87647\|Iba2 | allograft inflammatory factor 1-like | protein-coding |
| chr10_13820269_13820701 | chr10 | 13820270 | 13820701 | + | -0.890905 | intron (NM_025446, intron 2 of 5) | RMER6A\|LTR\|ERVK | 48345 | NM_025446 | 66253 | Mm.45755 | NM_025446 | ENSMUSG00000019806 | **Aig1** | 1500031O19Rik\|AV064870\|AW413422\|CGI-103 | androgen-induced 1 | protein-coding |
| chr10_43983964_43984200 | chr10 | 43983965 | 43984200 | + | -0.950678 | intron (NM_172393, intron 6 of 19) | L2\|LINE\|L2 | 20764 | NM_172393 | 11630 | Mm.292082 | NM_172393 | ENSMUSG00000019866 | **Aim1** | AI462491\|AI463325 | absent in melanoma 1 | protein-coding |
| chr10_43979995_43980578 | chr10 | 43979996 | 43980578 | + | -1.05275 | intron (NM_172393, intron 6 of 19) | intron (NM_172393, intron 6 of 19) | 24559 | NM_172393 | 11630 | Mm.292082 | NM_172393 | ENSMUSG00000019866 | **Aim1** | AI462491\|AI463325 | absent in melanoma 1 | protein-coding |
| chr1_173432070_173433017 | chr1 | 173432071 | 173433017 | + | -0.614492 | intron (NM_001013779, intron 2 of 8) | intron (NM_001013779, intron 2 of 8) | 11940 | NM_001013779 | 383619 | Mm.131453 | NM_001013779 | ENSMUSG00000037860 | **Aim2** | Gm1313\|Ifi210 | absent in melanoma 2 | protein-coding |
| chr1_173437054_173437431 | chr1 | 173437055 | 173437431 | + | -0.783321 | intron (NM_001013779, intron 2 of 8) | intron (NM_001013779, intron 2 of 8) | 16639 | NM_001013779 | 383619 | Mm.131453 | NM_001013779 | ENSMUSG00000037860 | **Aim2** | Gm1313\|Ifi210 | absent in melanoma 2 | protein-coding |
| chr19_4119601_4120105 | chr19 | 4119602 | 4120105 | + | -1.38603 | intron (NM_001276284, intron 1 of 5) | intron (NM_001276284, intron 1 of 5) | 1722 | NM_001276284 | 11632 | Mm.10433 | NM_016666 | ENSMUSG00000024847 | **Aip** | AA408703\|AW476050\|Ara9\|D19Bwg1412e\|Fkbp16\|Xap2 | aryl-hydrocarbon receptor-interacting protein | protein-coding |
| chr17_12745742_12746200 | chr17 | 12745743 | 12746200 | + | -0.623793 | intron (NR_027773, intron 1 of 2) | intron (NR_027773, intron 1 of 2) | 4660 | NR_027784 | 104103 | Mm.26553 | NR_002853 | ENSMUSG00000078247 | **Airn** | 2810051F02Rik\|2810434M15Rik\|AI256653\|AI597500\|AW049873\|Air\|B930018I07Rik\|D17Ertd663e\|IGF2RAS | antisense Igf2r RNA | ncRNA |
| chr7_75649856_75650295 | chr7 | 75649857 | 75650295 | + | -0.683689 | intron (NM_029332, intron 8 of 36) | intron (NM_029332, intron 8 of 36) | 194542 | NM_029332 | 75547 | Mm.216107 | NM_029332 | ENSMUSG00000066406 | **Akap13** | 1700026G02Rik\|5730522G15Rik\|5830460E08Rik\|AKAP-Lbc\|BRX\|Ht31\|LBC\|PROTO-LB\|PROTO-LBC | A kinase (PRKA) anchor protein 13 | protein-coding |
| chr7_75384541_75385194 | chr7 | 75384542 | 75385194 | + | -0.784105 | Intergenic | Intergenic | -70666 | NM_029332 | 75547 | Mm.216107 | NM_029332 | ENSMUSG00000066406 | **Akap13** | 1700026G02Rik\|5730522G15Rik\|5830460E08Rik\|AKAP-Lbc\|BRX\|Ht31\|LBC\|PROTO-LB\|PROTO-LBC | A kinase (PRKA) anchor protein 13 | protein-coding |
| chr7_75384472_75385359 | chr7 | 75384473 | 75385359 | + | -0.796882 | Intergenic | Intergenic | -70618 | NM_029332 | 75547 | Mm.216107 | NM_029332 | ENSMUSG00000066406 | **Akap13** | 1700026G02Rik\|5730522G15Rik\|5830460E08Rik\|AKAP-Lbc\|BRX\|Ht31\|LBC\|PROTO-LB\|PROTO-LBC | A kinase (PRKA) anchor protein 13 | protein-coding |
| chr7_75496813_75497161 | chr7 | 75496814 | 75497161 | + | -0.849854 | intron (NM_029332, intron 1 of 36) | intron (NM_029332, intron 1 of 36) | 41453 | NM_029332 | 75547 | Mm.216107 | NM_029332 | ENSMUSG00000066406 | **Akap13** | 1700026G02Rik\|5730522G15Rik\|5830460E08Rik\|AKAP-Lbc\|BRX\|Ht31\|LBC\|PROTO-LB\|PROTO-LBC | A kinase (PRKA) anchor protein 13 | protein-coding |
| chr7_75600269_75601033 | chr7 | 75600270 | 75601033 | + | -0.856451 | intron (NM_029332, intron 5 of 36) | intron (NM_029332, intron 5 of 36) | 145117 | NM_029332 | 75547 | Mm.216107 | NM_029332 | ENSMUSG00000066406 | **Akap13** | 1700026G02Rik\|5730522G15Rik\|5830460E08Rik\|AKAP-Lbc\|BRX\|Ht31\|LBC\|PROTO-LB\|PROTO-LBC | A kinase (PRKA) anchor protein 13 | protein-coding |
| chr7_75578719_75578970 | chr7 | 75578720 | 75578970 | + | -0.864149 | intron (NM_029332, intron 3 of 36) | intron (NM_029332, intron 3 of 36) | 123311 | NM_029332 | 75547 | Mm.216107 | NM_029332 | ENSMUSG00000066406 | **Akap13** | 1700026G02Rik\|5730522G15Rik\|5830460E08Rik\|AKAP-Lbc\|BRX\|Ht31\|LBC\|PROTO-LB\|PROTO-LBC | A kinase (PRKA) anchor protein 13 | protein-coding |
| chr7_75578704_75579159 | chr7 | 75578705 | 75579159 | + | -0.880776 | intron (NM_029332, intron 3 of 36) | intron (NM_029332, intron 3 of 36) | 123398 | NM_029332 | 75547 | Mm.216107 | NM_029332 | ENSMUSG00000066406 | **Akap13** | 1700026G02Rik\|5730522G15Rik\|5830460E08Rik\|AKAP-Lbc\|BRX\|Ht31\|LBC\|PROTO-LB\|PROTO-LBC | A kinase (PRKA) anchor protein 13 | protein-coding |
| chr7_75600391_75600704 | chr7 | 75600392 | 75600704 | + | -0.901992 | intron (NM_029332, intron 5 of 36) | intron (NM_029332, intron 5 of 36) | 145014 | NM_029332 | 75547 | Mm.216107 | NM_029332 | ENSMUSG00000066406 | **Akap13** | 1700026G02Rik\|5730522G15Rik\|5830460E08Rik\|AKAP-Lbc\|BRX\|Ht31\|LBC\|PROTO-LB\|PROTO-LBC | A kinase (PRKA) anchor protein 13 | protein-coding |
| chr7_75611576_75611898 | chr7 | 75611577 | 75611898 | + | -0.962292 | intron (NM_029332, intron 7 of 36) | intron (NM_029332, intron 7 of 36) | 156203 | NM_029332 | 75547 | Mm.216107 | NM_029332 | ENSMUSG00000066406 | **Akap13** | 1700026G02Rik\|5730522G15Rik\|5830460E08Rik\|AKAP-Lbc\|BRX\|Ht31\|LBC\|PROTO-LB\|PROTO-LBC | A kinase (PRKA) anchor protein 13 | protein-coding |
| chr7_75495994_75496337 | chr7 | 75495995 | 75496337 | + | -1.28405 | intron (NM_029332, intron 1 of 36) | intron (NM_029332, intron 1 of 36) | 40632 | NM_029332 | 75547 | Mm.216107 | NM_029332 | ENSMUSG00000066406 | **Akap13** | 1700026G02Rik\|5730522G15Rik\|5830460E08Rik\|AKAP-Lbc\|BRX\|Ht31\|LBC\|PROTO-LB\|PROTO-LBC | A kinase (PRKA) anchor protein 13 | protein-coding |
| chrX_36689701_36690030 | chrX | 36689702 | 36690030 | + | -0.724983 | Intergenic | MamGypLTR3a\|LTR\|Gypsy | -44452 | NM_001081956 | 338351 | Mm.116704 | NM_001081956 | ENSMUSG00000059708 | **Akap17b** | AA407619\|AKAP-17B\|B230333C21Rik\|C230056F04Rik\|PRKA17B\|Sfrs17b\|Srsf17b\|Talia | A kinase (PRKA) anchor protein 17B | protein-coding |
| chrX_36689665_36690222 | chrX | 36689666 | 36690222 | + | -0.765577 | Intergenic | MamGypLTR3a\|LTR\|Gypsy | -44530 | NM_001081956 | 338351 | Mm.116704 | NM_001081956 | ENSMUSG00000059708 | **Akap17b** | AA407619\|AKAP-17B\|B230333C21Rik\|C230056F04Rik\|PRKA17B\|Sfrs17b\|Srsf17b\|Talia | A kinase (PRKA) anchor protein 17B | protein-coding |

| | | | | | | | | | | | | | | | | | |
|---|---|---|---|---|---|---|---|---|---|---|---|---|---|---|---|---|---|
| chr4_57866958_57868065 | chr4 | 57866959 | 57868065 | + | -0.633123 | intron (NM_009649, intron 2 of 2) | intron (NM_009649, intron 2 of 2) | 12950 | NM_001035533 | 11641 | Mm.467752 | NM_009649 | ENSMUSG00000038729 | **Akap2** | AA959716\|AI649048\|Akap-2\|Akap-kl\|B230340M18Rik\|Prka2 | A kinase (PRKA) anchor protein 2 | protein-coding |
| chr4_57813320_57814102 | chr4 | 57813321 | 57814102 | + | -0.898629 | Intergenic | HAL1\|LINE\|L1 | -31537 | NM_009649 | 11641 | Mm.467752 | NM_009649 | ENSMUSG00000038729 | **Akap2** | AA959716\|AI649048\|Akap-2\|Akap-kl\|B230340M18Rik\|Prka2 | A kinase (PRKA) anchor protein 2 | protein-coding |
| chr4_57833977_57834485 | chr4 | 57833978 | 57834485 | + | -0.901043 | Intergenic | Intergenic | -11017 | NM_009649 | 11641 | Mm.467752 | NM_009649 | ENSMUSG00000038729 | **Akap2** | AA959716\|AI649048\|Akap-2\|Akap-kl\|B230340M18Rik\|Prka2 | A kinase (PRKA) anchor protein 2 | protein-coding |
| chr4_57813582_57813996 | chr4 | 57813583 | 57813996 | + | -0.901424 | Intergenic | Intergenic | -31459 | NM_009649 | 11641 | Mm.467752 | NM_009649 | ENSMUSG00000038729 | **Akap2** | AA959716\|AI649048\|Akap-2\|Akap-kl\|B230340M18Rik\|Prka2 | A kinase (PRKA) anchor protein 2 | protein-coding |
| chr4_57833962_57834717 | chr4 | 57833963 | 57834717 | + | -0.902006 | Intergenic | Intergenic | -10908 | NM_009649 | 11641 | Mm.467752 | NM_009649 | ENSMUSG00000038729 | **Akap2** | AA959716\|AI649048\|Akap-2\|Akap-kl\|B230340M18Rik\|Prka2 | A kinase (PRKA) anchor protein 2 | protein-coding |
| chr4_57867500_57868078 | chr4 | 57867501 | 57868078 | + | -0.920709 | intron (NM_009649, intron 2 of 2) | intron (NM_009649, intron 2 of 2) | 13227 | NM_001035533 | 11641 | Mm.467752 | NM_009649 | ENSMUSG00000038729 | **Akap2** | AA959716\|AI649048\|Akap-2\|Akap-kl\|B230340M18Rik\|Prka2 | A kinase (PRKA) anchor protein 2 | protein-coding |
| chr4_57832330_57832673 | chr4 | 57832331 | 57832673 | + | -1.19962 | Intergenic | Intergenic | -12746 | NM_009649 | 11641 | Mm.467752 | NM_009649 | ENSMUSG00000038729 | **Akap2** | AA959716\|AI649048\|Akap-2\|Akap-kl\|B230340M18Rik\|Prka2 | A kinase (PRKA) anchor protein 2 | protein-coding |
| chr12_52729016_52730122 | chr12 | 52729017 | 52730122 | + | -0.73564 | intron (NM_198111, intron 1 of 13) | intron (NM_198111, intron 1 of 13) | 30186 | NM_198111 | 238161 | Mm.310822 | NM_198111 | ENSMUSG00000061603 | **Akap6** | AI482140\|Akapalpha\|Akapbeta\|PRKA | A kinase (PRKA) anchor protein 6 | protein-coding |
| chr12_52795881_52796862 | chr12 | 52795882 | 52796862 | + | -0.804989 | intron (NM_198111, intron 2 of 13) | intron (NM_198111, intron 2 of 13) | 96989 | NM_198111 | 238161 | Mm.310822 | NM_198111 | ENSMUSG00000061603 | **Akap6** | AI482140\|Akapalpha\|Akapbeta\|PRKA | A kinase (PRKA) anchor protein 6 | protein-coding |
| chr12_52796431_52796881 | chr12 | 52796432 | 52796881 | + | -1.03628 | intron (NM_198111, intron 2 of 13) | intron (NM_198111, intron 2 of 13) | 97273 | NM_198111 | 238161 | Mm.310822 | NM_198111 | ENSMUSG00000061603 | **Akap6** | AI482140\|Akapalpha\|Akapbeta\|PRKA | A kinase (PRKA) anchor protein 6 | protein-coding |
| chr17_32341068_32341456 | chr17 | 32341069 | 32341456 | + | -0.866641 | intron (NM_017476, intron 2 of 13) | intron (NM_017476, intron 2 of 13) | 9315 | NM_017476 | 54194 | Mm.281005 | NM_017476 | ENSMUSG00000002625 | **Akap8l** | HAP95\|Nakap95 | A kinase (PRKA) anchor protein 8-like | protein-coding |
| chr6_34364640_34365125 | chr6 | 34364641 | 34365125 | + | -0.85721 | intron (NM_008012, intron 7 of 9) | ORR1D1\|LTR\|ERVL-MaLR | 10719 | NM_008012 | 14187 | Mm.5378 | NM_008012 | ENSMUSG00000029762 | **Akr1b8** | FR-1\|Fgfrp\|Fgrp | aldo-keto reductase family 1, member B8 | protein-coding |
| chr6_37633943_37634344 | chr6 | 37633944 | 37634344 | + | -0.798014 | Intergenic | ORR1D2\|LTR\|ERVL-MaLR | 103971 | NM_145364 | 208665 | Mm.262635 | NM_145364 | ENSMUSG00000038641 | **Akr1d1** | - | aldo-keto reductase family 1, member D1 | protein-coding |
| chr6_37607471_37608227 | chr6 | 37607472 | 37608227 | + | -1.0875 | Intergenic | Intergenic | 77676 | NM_145364 | 208665 | Mm.262635 | NM_145364 | ENSMUSG00000038641 | **Akr1d1** | - | aldo-keto reductase family 1, member D1 | protein-coding |
| chr13_4942426_4942896 | chr13 | 4942427 | 4942896 | + | -0.705325 | Intergenic | IAPLTR2_Mm\|LTR\|ERVK | -333497 | NM_018859 | 56043 | Mm.251908 | NM_018859 | ENSMUSG00000045410 | **Akr1e1** | 1810061I10Rik\|Akr1e2 | aldo-keto reductase family 1, member E1 | protein-coding |
| chr10_21479245_21479860 | chr10 | 21479246 | 21479860 | + | -0.825201 | Intergenic | Intergenic | 102253 | NM_178713 | 237320 | Mm.90181 | NM_178713 | ENSMUSG00000037542 | **Aldh8a1** | Raldh4 | aldehyde dehydrogenase 8 family, member A1 | protein-coding |
| chr3_121344682_121345308 | chr3 | 121344683 | 121345308 | + | -0.658688 | intron (NM_024178, intron 3 of 3) | intron (NM_024178, intron 3 of 3) | 53178 | NM_024178 | 66789 | Mm.269881 | NM_024178 | ENSMUSG00000039887 | **Alg14** | 5430428G01Rik\|AI854024 | asparagine-linked glycosylation 14 | protein-coding |
| chr3_121312205_121312872 | chr3 | 121312206 | 121312872 | + | -0.700857 | intron (NM_024178, intron 2 of 3) | intron (NM_024178, intron 2 of 3) | 20722 | NM_024178 | 66789 | Mm.269881 | NM_024178 | ENSMUSG00000039887 | **Alg14** | 5430428G01Rik\|AI854024 | asparagine-linked glycosylation 14 | protein-coding |
| chr3_121312443_121312882 | chr3 | 121312444 | 121312882 | + | -0.773892 | intron (NM_024178, intron 2 of 3) | intron (NM_024178, intron 2 of 3) | 20846 | NM_024178 | 66789 | Mm.269881 | NM_024178 | ENSMUSG00000039887 | **Alg14** | 5430428G01Rik\|AI854024 | asparagine-linked glycosylation 14 | protein-coding |
| chr3_121343015_121343558 | chr3 | 121343016 | 121343558 | + | -0.876396 | intron (NM_024178, intron 3 of 3) | intron (NM_024178, intron 3 of 3) | 51470 | NM_024178 | 66789 | Mm.269881 | NM_024178 | ENSMUSG00000039887 | **Alg14** | 5430428G01Rik\|AI854024 | asparagine-linked glycosylation 14 | protein-coding |
| chr5_136146776_136147438 | chr5 | 136146777 | 136147438 | + | -0.758671 | TTS (NM_178751) | TTS (NM_178751) | 8326 | NM_028070 | 72041 | Mm.486605 | NM_028070 | ENSMUSG00000039754 | **Alkbh4** | 2010004B12Rik\|Abj4 | alkB, alkylation repair homolog 4 (E. coli) | protein-coding |
| chr11_70265170_70265601 | chr11 | 70265171 | 70265601 | + | -0.633841 | Intergenic | Intergenic | -10045 | NM_007440 | 11684 | Mm.12286 | NM_007440 | ENSMUSG00000000320 | **Alox12** | 9930022G08Rik\|Alox12p\|P-12LO | arachidonate 12-lipoxygenase | protein-coding |
| chr11_70265149_70265815 | chr11 | 70265150 | 70265815 | + | -0.659967 | Intergenic | Intergenic | -10141 | NM_007440 | 11684 | Mm.12286 | NM_007440 | ENSMUSG00000000320 | **Alox12** | 9930022G08Rik\|Alox12p\|P-12LO | arachidonate 12-lipoxygenase | protein-coding |
| chr11_70280488_70280862 | chr11 | 70280489 | 70280862 | + | -0.921262 | Intergenic | Intergenic | -25334 | NM_007440 | 11684 | Mm.12286 | NM_007440 | ENSMUSG00000000320 | **Alox12** | 9930022G08Rik\|Alox12p\|P-12LO | arachidonate 12-lipoxygenase | protein-coding |
| chr11_70291337_70291544 | chr11 | 70291338 | 70291544 | + | -0.610809 | Intergenic | LTR33A\|LTR\|ERVL | 31077 | NM_145684 | 11685 | Mm.274093 | NM_145684 | ENSMUSG00000018907 | **Alox12e** | 8-LOX\|AW259591\|Alox12-ps1\|Alox12-ps2\|Aloxe\|e-LOX1 | arachidonate lipoxygenase, epidermal | protein-coding |
| chr11_70291144_70291563 | chr11 | 70291145 | 70291563 | + | -0.714887 | Intergenic | LTR33A\|LTR\|ERVL | 31164 | NM_145684 | 11685 | Mm.274093 | NM_145684 | ENSMUSG00000018907 | **Alox12e** | 8-LOX\|AW259591\|Alox12-ps1\|Alox12-ps2\|Aloxe\|e-LOX1 | arachidonate lipoxygenase, epidermal | protein-coding |
| chr11_70297450_70298453 | chr11 | 70297451 | 70298453 | + | -0.835355 | Intergenic | Intergenic | 24566 | NM_145684 | 11685 | Mm.274093 | NM_145684 | ENSMUSG00000018907 | **Alox12e** | 8-LOX\|AW259591\|Alox12-ps1\|Alox12-ps2\|Aloxe\|e-LOX1 | arachidonate lipoxygenase, epidermal | protein-coding |
| chr11_70297922_70298482 | chr11 | 70297923 | 70298482 | + | -0.974 | Intergenic | Intergenic | 24316 | NM_145684 | 11685 | Mm.274093 | NM_145684 | ENSMUSG00000018907 | **Alox12e** | 8-LOX\|AW259591\|Alox12-ps1\|Alox12-ps2\|Aloxe\|e-LOX1 | arachidonate lipoxygenase, epidermal | protein-coding |
| chr5_149320498_149321057 | chr5 | 149320499 | 149321057 | + | -0.783992 | Intergenic | Intergenic | 55720 | NM_009663 | 11690 | Mm.19844 | NM_009663 | ENSMUSG00000060063 | **Alox5ap** | Flap | arachidonate 5-lipoxygenase activating protein | protein-coding |
| chr5_149245838_149246642 | chr5 | 149245839 | 149246642 | + | -0.797478 | Intergenic | Intergenic | -18818 | NM_009663 | 11690 | Mm.19844 | NM_009663 | ENSMUSG00000060063 | **Alox5ap** | Flap | arachidonate 5-lipoxygenase activating protein | protein-coding |
| chr5_149265480_149265809 | chr5 | 149265481 | 149265809 | + | -0.799139 | intron (NM_009663, intron 1 of 4) | intron (NM_009663, intron 1 of 4) | 587 | NM_009663 | 11690 | Mm.19844 | NM_009663 | ENSMUSG00000060063 | **Alox5ap** | Flap | arachidonate 5-lipoxygenase activating protein | protein-coding |
| chr5_149255564_149255869 | chr5 | 149255565 | 149255869 | + | -0.805159 | Intergenic | L1ME2\|LINE\|L1 | -9341 | NM_009663 | 11690 | Mm.19844 | NM_009663 | ENSMUSG00000060063 | **Alox5ap** | Flap | arachidonate 5-lipoxygenase activating protein | protein-coding |
| chr7_81075208_81075942 | chr7 | 81075209 | 81075942 | + | -0.664651 | intron (NM_054085, intron 3 of 13) | intron (NM_054085, intron 3 of 13) | 17975 | NM_054085 | 116904 | Mm.475768 | NM_054085 | ENSMUSG00000038763 | **Alpk3** | AW319487\|D330016D04\|MAK\|Midori\|mKIAA1330 | alpha-kinase 3 | protein-coding |
| chr7_81075302_81075850 | chr7 | 81075303 | 81075850 | + | -0.682124 | intron (NM_054085, intron 3 of 13) | intron (NM_054085, intron 3 of 13) | 17976 | NM_054085 | 116904 | Mm.475768 | NM_054085 | ENSMUSG00000038763 | **Alpk3** | AW319487\|D330016D04\|MAK\|Midori\|mKIAA1330 | alpha-kinase 3 | protein-coding |
| chr9_110891121_110891435 | chr9 | 110891122 | 110891435 | + | -0.794017 | intron (NM_001146060, intron 15 of 26) | intron (NM_001146060, intron 15 of 26) | 10956 | NM_001146060 | 235633 | Mm.86338 | NM_146228 | ENSMUSG00000044037 | **Als2cl** | 79H19D\|79H19E\|BC022645\|D930044G19Rik\|mRn.49018 | ALS2 C-terminal like | protein-coding |
| chr9_110896464_110897064 | chr9 | 110896465 | 110897064 | + | -0.96304 | intron (NM_001146060, intron 23 of 26) | intron (NM_001146060, intron 23 of 26) | 16442 | NM_001146060 | 235633 | Mm.86338 | NM_146228 | ENSMUSG00000044037 | **Als2cl** | 79H19D\|79H19E\|BC022645\|D930044G19Rik\|mRn.49018 | ALS2 C-terminal like | protein-coding |
| chr15_97420410_97420716 | chr15 | 97420411 | 97420716 | + | -0.709906 | Intergenic | RLTR19\|LTR\|ERVK | -173276 | NM_001164602 | 105827 | Mm.24005 | NM_178114 | ENSMUSG00000048218 | **Amigo2** | AI415330\|AMIGO-2\|AW208913\|Ali1 | adhesion molecule with Ig like domain 2 | protein-coding |
| chr15_97342809_97343665 | chr15 | 97342810 | 97343665 | + | -0.82424 | intron (NM_172293, intron 3 of 5) | intron (NM_172293, intron 3 of 5) | -95950 | NM_001164602 | 105827 | Mm.24005 | NM_178114 | ENSMUSG00000048218 | **Amigo2** | AI415330\|AMIGO-2\|AW208913\|Ali1 | adhesion molecule with Ig like domain 2 | protein-coding |
| chr15_97391486_97391909 | chr15 | 97391487 | 97391909 | + | -0.941166 | Intergenic | Intergenic | -144411 | NM_001164602 | 105827 | Mm.24005 | NM_178114 | ENSMUSG00000048218 | **Amigo2** | AI415330\|AMIGO-2\|AW208913\|Ali1 | adhesion molecule with Ig like domain 2 | protein-coding |
| chr15_97252446_97253036 | chr15 | 97252447 | 97253036 | + | -0.949753 | intron (NM_172293, intron 1 of 5) | ORR1E\|LTR\|ERVL-MaLR | -5454 | NM_001164602 | 105827 | Mm.24005 | NM_178114 | ENSMUSG00000048218 | **Amigo2** | AI415330\|AMIGO-2\|AW208913\|Ali1 | adhesion molecule with Ig like domain 2 | protein-coding |
| chr15_97390797_97391129 | chr15 | 97390798 | 97391129 | + | -0.988583 | Intergenic | ID_B1\|SINE\|B4 | -143676 | NM_001164602 | 105827 | Mm.24005 | NM_178114 | ENSMUSG00000048218 | **Amigo2** | AI415330\|AMIGO-2\|AW208913\|Ali1 | adhesion molecule with Ig like domain 2 | protein-coding |
| chr15_97389601_97390035 | chr15 | 97389602 | 97390035 | + | -1.09463 | Intergenic | Intergenic | -142531 | NM_001164602 | 105827 | Mm.24005 | NM_178114 | ENSMUSG00000048218 | **Amigo2** | AI415330\|AMIGO-2\|AW208913\|Ali1 | adhesion molecule with Ig like domain 2 | protein-coding |
| chr6_149183287_149183619 | chr6 | 149183288 | 149183619 | + | -0.95953 | exon (NM_001113424, exon 3 of 7) | exon (NM_001113424, exon 3 of 7) | 5259 | NM_001113424 | 232566 | Mm.209147 | NM_001113424 | ENSMUSG00000068250 | **Amn1** | 5830467E07Rik\|A230083D09\|C730024G19Rik | antagonist of mitotic exit network 1 | protein-coding |
| chr6_149183387_149183608 | chr6 | 149183388 | 149183608 | + | -0.961632 | exon (NM_001113424, exon 3 of 7) | exon (NM_001113424, exon 3 of 7) | 5214 | NM_001113424 | 232566 | Mm.209147 | NM_001113424 | ENSMUSG00000068250 | **Amn1** | 5830467E07Rik\|A230083D09\|C730024G19Rik | antagonist of mitotic exit network 1 | protein-coding |
| chrX_145433647_145434051 | chrX | 145433648 | 145434051 | + | -0.677819 | Intergenic | MLTR11B\|LTR\|ERVK | 53790 | NM_001290274 | 27494 | Mm.100068 | NM_153319 | ENSMUSG00000041688 | **Amot** | CAG-2\|D0Kist1\|Sii6 | angiomotin | protein-coding |
| chr3_103091714_103092449 | chr3 | 103091715 | 103092449 | + | -0.830187 | intron (NM_001033303, intron 9 of 15) | intron (NM_001033303, intron 9 of 15) | 18068 | NM_001033303 | 229665 | Mm.331272 | NM_001033303 | ENSMUSG00000070385 | **Ampd1** | AI553520\|Ampd-1 | adenosine monophosphate deaminase 1 | protein-coding |
| chr3_103091732_103092155 | chr3 | 103091733 | 103092155 | + | -0.876941 | intron (NM_001033303, intron 9 of 15) | intron (NM_001033303, intron 9 of 15) | 17930 | NM_001033303 | 229665 | Mm.331272 | NM_001033303 | ENSMUSG00000070385 | **Ampd1** | AI553520\|Ampd-1 | adenosine monophosphate deaminase 1 | protein-coding |
| chr13_19162120_19162511 | chr13 | 19162121 | 19162511 | + | -1.19289 | Intergenic | MMAR1\|DNA\|TcMar-Mariner | 213965 | NM_001289546 | 218038 | Mm.101650 | NM_175007 | ENSMUSG00000021314 | **Amph** | - | amphiphysin | protein-coding |
| chr11_109419880_109420947 | chr11 | 109419881 | 109420947 | + | -0.726379 | Intergenic | Intergenic | -5532 | NM_025275 | 13929 | Mm.211850 | NM_025275 | ENSMUSG00000020610 | **Amz2** | AA408420\|ESTM12\|X83328 | archaelysin family metallopeptidase 2 | protein-coding |
| chr5_122456467_122456736 | chr5 | 122456468 | 122456736 | + | -1.01878 | non-coding (NR_027838, exon 20 of 21) | non-coding (NR_027838, exon 20 of 21) | 34158 | NM_019805 | 56317 | Mm.37341 | NM_019805 | ENSMUSG00000029466 | **Anapc7** | APC7\|AW545589 | anaphase promoting complex subunit 7 | protein-coding |
| chr19_36242398_36242753 | chr19 | 36242399 | 36242753 | + | -1.23356 | Intergenic | Intergenic | -122732 | NM_013468 | 107765 | Mm.10279 | NM_013468 | ENSMUSG00000024803 | **Ankrd1** | Alrp\|CARP\|Crap\|MARP1 | ankyrin repeat domain 1 (cardiac muscle) | protein-coding |

| | | | | | | | | | | | | | | | | | |
|---|---|---|---|---|---|---|---|---|---|---|---|---|---|---|---|---|---|
| chr8_122986854_122987103 | chr8 | 122986855 | 122987103 | + | -0.674581 | intron (NR_037865, intron 1 of 13) | intron (NR_037865, intron 1 of 13) | 55305 | NR_037865 | 77087 | Mm.31567 | NM_001081379 | ENSMUSG00000035569 | **Ankrd11** | 2410104C19Rik\|3010027A04Rik\|6330578C09Rik\|9530048I21Rik\|AA930108\|Gm176\|Yod | ankyrin repeat domain 11 | protein-coding |
| chr8_122960940_122961427 | chr8 | 122960941 | 122961427 | + | -0.705102 | intron (NR_037865, intron 2 of 13) | ORR1D1\|LTR\|ERVL-MaLR | 81100 | NR_037865 | 77087 | Mm.31567 | NM_001081379 | ENSMUSG00000035569 | **Ankrd11** | 2410104C19Rik\|3010027A04Rik\|6330578C09Rik\|9530048I21Rik\|AA930108\|Gm176\|Yod | ankyrin repeat domain 11 | protein-coding |
| chr8_122967371_122967665 | chr8 | 122967372 | 122967665 | + | -0.72536 | intron (NR_037865, intron 2 of 13) | MT2A\|LTR\|ERVL | 74766 | NR_037865 | 77087 | Mm.31567 | NM_001081379 | ENSMUSG00000035569 | **Ankrd11** | 2410104C19Rik\|3010027A04Rik\|6330578C09Rik\|9530048I21Rik\|AA930108\|Gm176\|Yod | ankyrin repeat domain 11 | protein-coding |
| chr8_122953022_122953433 | chr8 | 122953023 | 122953433 | + | -0.854657 | intron (NR_037865, intron 2 of 13) | intron (NR_037865, intron 2 of 13) | 89056 | NR_037865 | 77087 | Mm.31567 | NM_001081379 | ENSMUSG00000035569 | **Ankrd11** | 2410104C19Rik\|3010027A04Rik\|6330578C09Rik\|9530048I21Rik\|AA930108\|Gm176\|Yod | ankyrin repeat domain 11 | protein-coding |
| chr8_123025764_123026455 | chr8 | 123025765 | 123026455 | + | -1.14075 | intron (NR_037865, intron 1 of 13) | intron (NR_037865, intron 1 of 13) | 16174 | NR_037865 | 77087 | Mm.31567 | NM_001081379 | ENSMUSG00000035569 | **Ankrd11** | 2410104C19Rik\|3010027A04Rik\|6330578C09Rik\|9530048I21Rik\|AA930108\|Gm176\|Yod | ankyrin repeat domain 11 | protein-coding |
| chr8_123025969_123026238 | chr8 | 123025970 | 123026238 | + | -1.24473 | intron (NR_037865, intron 1 of 13) | intron (NR_037865, intron 1 of 13) | 16180 | NR_037865 | 77087 | Mm.31567 | NM_001081379 | ENSMUSG00000035569 | **Ankrd11** | 2410104C19Rik\|3010027A04Rik\|6330578C09Rik\|9530048I21Rik\|AA930108\|Gm176\|Yod | ankyrin repeat domain 11 | protein-coding |
| chr8_123000248_123000708 | chr8 | 123000249 | 123000708 | + | -1.24766 | intron (NR_037865, intron 1 of 13) | intron (NR_037865, intron 1 of 13) | 41806 | NR_037865 | 77087 | Mm.31567 | NM_001081379 | ENSMUSG00000035569 | **Ankrd11** | 2410104C19Rik\|3010027A04Rik\|6330578C09Rik\|9530048I21Rik\|AA930108\|Gm176\|Yod | ankyrin repeat domain 11 | protein-coding |
| chr8_123000269_123000961 | chr8 | 123000270 | 123000961 | + | -1.25336 | intron (NR_037865, intron 1 of 13) | intron (NR_037865, intron 1 of 13) | 41669 | NR_037865 | 77087 | Mm.31567 | NM_001081379 | ENSMUSG00000035569 | **Ankrd11** | 2410104C19Rik\|3010027A04Rik\|6330578C09Rik\|9530048I21Rik\|AA930108\|Gm176\|Yod | ankyrin repeat domain 11 | protein-coding |
| chr19_42018957_42019626 | chr19 | 42018958 | 42019626 | + | -0.624325 | intron (NM_145500, intron 1 of 2) | intron (NM_145500, intron 1 of 2) | -16746 | NM_020033 | 56642 | Mm.143737 | NM_020033 | ENSMUSG00000025172 | **Ankrd2** | Arpp\|mArpp | ankyrin repeat domain 2 (stretch responsive muscle) | protein-coding |
| chr19_34168393_34168682 | chr19 | 34168394 | 34168682 | + | -0.639707 | Intergenic | Intergenic | -2497 | NM_024204 | 52024 | Mm.183030 | NM_024204 | ENSMUSG00000024774 | **Ankrd22** | 5430429D21Rik\|D19Ertd675e | ankyrin repeat domain 22 | protein-coding |
| chr19_34167745_34168277 | chr19 | 34167746 | 34168277 | + | -0.768016 | Intergenic | Intergenic | -1970 | NM_024204 | 52024 | Mm.183030 | NM_024204 | ENSMUSG00000024774 | **Ankrd22** | 5430429D21Rik\|D19Ertd675e | ankyrin repeat domain 22 | protein-coding |
| chr19_34167796_34168666 | chr19 | 34167797 | 34168666 | + | -0.847006 | Intergenic | Intergenic | -2190 | NM_024204 | 52024 | Mm.183030 | NM_024204 | ENSMUSG00000024774 | **Ankrd22** | 5430429D21Rik\|D19Ertd675e | ankyrin repeat domain 22 | protein-coding |
| chr15_101113264_101113551 | chr15 | 101113265 | 101113551 | + | -0.887826 | Intergenic | Intergenic | -2347 | NM_144790 | 208258 | Mm.152952 | NM_144790 | ENSMUSG00000047034 | **Ankrd33** | A930021G21Rik\|Panky\|Panky-a | ankyrin repeat domain 33 | protein-coding |
| chr15_101113131_101113588 | chr15 | 101113132 | 101113588 | + | -0.900926 | Intergenic | Intergenic | -2395 | NM_144790 | 208258 | Mm.152952 | NM_144790 | ENSMUSG00000047034 | **Ankrd33** | A930021G21Rik\|Panky\|Panky-a | ankyrin repeat domain 33 | protein-coding |
| chr1_54913364_54913883 | chr1 | 54913365 | 54913883 | + | -0.632315 | intron (NM_001081433, intron 1 of 27) | intron (NM_001081433, intron 1 of 27) | 12763 | NM_001081433 | 329154 | Mm.397910 | NM_001081433 | ENSMUSG00000052331 | **Ankrd44** | 4930444A19Rik\|A130096K20\|E130014H08Rik\|PP6-ARS-B | ankyrin repeat domain 44 | protein-coding |
| chr1_54914437_54914734 | chr1 | 54914438 | 54914734 | + | -0.640924 | intron (NM_001081433, intron 1 of 27) | intron (NM_001081433, intron 1 of 27) | 11801 | NM_001081433 | 329154 | Mm.397910 | NM_001081433 | ENSMUSG00000052331 | **Ankrd44** | 4930444A19Rik\|A130096K20\|E130014H08Rik\|PP6-ARS-B | ankyrin repeat domain 44 | protein-coding |
| chr1_54882656_54883439 | chr1 | 54882657 | 54883439 | + | -0.722608 | intron (NM_001081433, intron 1 of 27) | intron (NM_001081433, intron 1 of 27) | 43339 | NM_001081433 | 329154 | Mm.397910 | NM_001081433 | ENSMUSG00000052331 | **Ankrd44** | 4930444A19Rik\|A130096K20\|E130014H08Rik\|PP6-ARS-B | ankyrin repeat domain 44 | protein-coding |
| chr1_54882808_54883346 | chr1 | 54882809 | 54883346 | + | -0.770856 | intron (NM_001081433, intron 1 of 27) | intron (NM_001081433, intron 1 of 27) | 43310 | NM_001081433 | 329154 | Mm.397910 | NM_001081433 | ENSMUSG00000052331 | **Ankrd44** | 4930444A19Rik\|A130096K20\|E130014H08Rik\|PP6-ARS-B | ankyrin repeat domain 44 | protein-coding |
| chr1_54747725_54748017 | chr1 | 54747726 | 54748017 | + | -0.79535 | intron (NM_001081433, intron 9 of 27) | L1ME4a\|LINE\|L1 | 178516 | NM_001081433 | 329154 | Mm.397910 | NM_001081433 | ENSMUSG00000052331 | **Ankrd44** | 4930444A19Rik\|A130096K20\|E130014H08Rik\|PP6-ARS-B | ankyrin repeat domain 44 | protein-coding |
| chr1_54860374_54860722 | chr1 | 54860375 | 54860722 | + | -1.21195 | intron (NM_001081433, intron 1 of 27) | intron (NM_001081433, intron 1 of 27) | 65839 | NM_001081433 | 329154 | Mm.397910 | NM_001081433 | ENSMUSG00000052331 | **Ankrd44** | 4930444A19Rik\|A130096K20\|E130014H08Rik\|PP6-ARS-B | ankyrin repeat domain 44 | protein-coding |
| chr15_36521306_36521807 | chr15 | 36521307 | 36521807 | + | -0.750259 | intron (NM_025712, intron 11 of 13) | MTC\|LTR\|ERVL-MaLR | -24766 | NM_175134 | 68839 | Mm.396537 | NM_175134 | ENSMUSG00000048307 | **Ankrd46** | 1110054N06Rik\|AI314978\|AI987733 | ankyrin repeat domain 46 | protein-coding |
| chr15_36440028_36440292 | chr15 | 36440029 | 36440292 | + | -0.762629 | Intergenic | Intergenic | 56631 | NM_175134 | 68839 | Mm.396537 | NM_175134 | ENSMUSG00000048307 | **Ankrd46** | 1110054N06Rik\|AI314978\|AI987733 | ankyrin repeat domain 46 | protein-coding |
| chr3_38568496_38568979 | chr3 | 38568497 | 38568979 | + | -0.731232 | Intergenic | Intergenic | -83922 | NM_001167883 | 99696 | Mm.290937 | NM_001033198 | ENSMUSG00000044864 | **Ankrd50** | AI662170\|E430012K20Rik\|mKIAA1223\|mKIAA3019 | ankyrin repeat domain 50 | protein-coding |
| chr3_38440394_38441089 | chr3 | 38440395 | 38441089 | + | -1.32762 | Intergenic | Intergenic | 44074 | NM_001167883 | 99696 | Mm.290937 | NM_001033198 | ENSMUSG00000044864 | **Ankrd50** | AI662170\|E430012K20Rik\|mKIAA1223\|mKIAA3019 | ankyrin repeat domain 50 | protein-coding |
| chr13_112287942_112289004 | chr13 | 112287943 | 112289004 | + | -0.648832 | promoter-TSS (NM_001168405) | promoter-TSS (NM_001168405) | 22 | NM_001168405 | 77318 | Mm.159895 | NM_029898 | ENSMUSG00000049985 | **Ankrd55** | C030011J08Rik | ankyrin repeat domain 55 | protein-coding |
| chr13_112288255_112289014 | chr13 | 112288256 | 112289014 | + | -0.65829 | 5' UTR (NM_001168405, exon 1 of 6) | 5' UTR (NM_001168405, exon 1 of 6) | 117 | NM_001168404 | 77318 | Mm.159895 | NM_029898 | ENSMUSG00000049985 | **Ankrd55** | C030011J08Rik | ankyrin repeat domain 55 | protein-coding |
| chr13_112354627_112355333 | chr13 | 112354628 | 112355333 | + | -0.773129 | intron (NM_029898, intron 6 of 10) | intron (NM_029898, intron 6 of 10) | 66462 | NM_001168404 | 77318 | Mm.159895 | NM_029898 | ENSMUSG00000049985 | **Ankrd55** | C030011J08Rik | ankyrin repeat domain 55 | protein-coding |
| chr13_112354821_112355351 | chr13 | 112354822 | 112355351 | + | -0.837088 | intron (NM_029898, intron 6 of 10) | intron (NM_029898, intron 6 of 10) | 66568 | NM_001168404 | 77318 | Mm.159895 | NM_029898 | ENSMUSG00000049985 | **Ankrd55** | C030011J08Rik | ankyrin repeat domain 55 | protein-coding |
| chr13_112355045_112355303 | chr13 | 112355046 | 112355303 | + | -0.903985 | intron (NM_029898, intron 6 of 10) | intron (NM_029898, intron 6 of 10) | 66656 | NM_001168404 | 77318 | Mm.159895 | NM_029898 | ENSMUSG00000049985 | **Ankrd55** | C030011J08Rik | ankyrin repeat domain 55 | protein-coding |
| chr13_112063501_112064136 | chr13 | 112063502 | 112064136 | + | -1.04143 | Intergenic | MTEa\|LTR\|ERVL-MaLR | -224632 | NM_001168403 | 77318 | Mm.159895 | NM_029898 | ENSMUSG00000049985 | **Ankrd55** | C030011J08Rik | ankyrin repeat domain 55 | protein-coding |
| chr4_32874766_32875045 | chr4 | 32874767 | 32875045 | + | -0.656882 | intron (NM_001012450, intron 1 of 16) | intron (NM_001012450, intron 1 of 16) | 48599 | NM_080471 | 140577 | Mm.214766 | NM_080471 | ENSMUSG00000040183 | **Ankrd6** | - | ankyrin repeat domain 6 | protein-coding |
| chr9_62425216_62425565 | chr9 | 62425217 | 62425565 | + | -0.815198 | intron (NM_175484, intron 11 of 11) | intron (NM_175484, intron 11 of 11) | 84048 | NM_009672 | 11737 | Mm.269088 | NM_009672 | ENSMUSG00000032249 | **Anp32a** | Anp32\|I1PP2A\|LANP\|PHAP1\|W91701\|pp32 | acidic (leucine-rich) nuclear phosphoprotein 32 family, member A | protein-coding |
| chr5_98033016_98033425 | chr5 | 98033017 | 98033425 | + | -0.695773 | Intergenic | Intergenic | -2259 | NM_133738 | 71914 | Mm.24842 | NM_133738 | | **Antxr2** | 2310046B19Rik\|AW561899\|CMG-2\|CMG2\|cl-35 | anthrax toxin receptor 2 | protein-coding |
| chr5_98033181_98033386 | chr5 | 98033182 | 98033386 | + | -0.760433 | Intergenic | Intergenic | -2322 | NM_133738 | 71914 | Mm.24842 | NM_133738 | | **Antxr2** | 2310046B19Rik\|AW561899\|CMG-2\|CMG2\|cl-35 | anthrax toxin receptor 2 | protein-coding |
| chr1_58311153_58311891 | chr1 | 58311154 | 58311891 | + | -0.6891 | intron (NM_001008419, intron 18 of 35) | MLT1B\|LTR\|ERVL-MaLR | 33196 | NM_001008419 | 213043 | Mm.414292 | NM_001008419 | ENSMUSG00000079554 | **Aox2** | Aoh3\|Aox3l1 | aldehyde oxidase 2 | protein-coding |
| chr3_127860234_127860426 | chr3 | 127860235 | 127860426 | + | -0.955496 | Intergenic | MT2B1\|LTR\|ERVL | -22838 | NM_145964 | 211556 | Mm.403486 | NM_145964 | ENSMUSG00000074238 | **Ap1ar** | AA407621\|C78909\|Gadkin | adaptor-related protein complex 1 associated regulatory protein | protein-coding |
| chr3_127844958_127845507 | chr3 | 127844959 | 127845507 | + | -1.07835 | Intergenic | Intergenic | -7741 | NM_145964 | 211556 | Mm.403486 | NM_145964 | ENSMUSG00000074238 | **Ap1ar** | AA407621\|C78909\|Gadkin | adaptor-related protein complex 1 associated regulatory protein | protein-coding |
| chr3_127862421_127862967 | chr3 | 127862422 | 127862967 | + | -1.16236 | Intergenic | Intergenic | -25202 | NM_145964 | 211556 | Mm.403486 | NM_145964 | ENSMUSG00000074238 | **Ap1ar** | AA407621\|C78909\|Gadkin | adaptor-related protein complex 1 associated regulatory protein | protein-coding |
| chr11_4989262_4989536 | chr11 | 4989263 | 4989536 | + | -0.862081 | intron (NM_007454, intron 1 of 20) | intron (NM_007454, intron 1 of 20) | 2575 | NM_007454 | 11764 | Mm.274816 | NM_007454 | ENSMUSG00000009090 | **Ap1b1** | Adtb1\|b2b1660Clo | adaptor protein complex AP-1, beta 1 subunit | protein-coding |
| chr14_21034235_21034748 | chr14 | 21034236 | 21034748 | + | -0.955037 | TTS (NM_009502) | TTS (NM_009502) | 17950 | NM_018829 | 55946 | Mm.376269 | NM_018829 | | **Ap3m1** | 1200013D09Rik\|C78982\|R75378 | adaptor-related protein complex 3, mu 1 subunit | protein-coding |
| chr8_22807018_22807314 | chr8 | 22807019 | 22807314 | + | -0.673398 | Intergenic | Intergenic | -1512 | NM_001122820 | 64933 | Mm.390023 | NM_029505 | ENSMUSG00000031539 | **Ap3m2** | 5830445E16Rik\|AP-3B | adaptor-related protein complex 3, mu 2 subunit | protein-coding |
| chr18_46745768_46746031 | chr18 | 46745769 | 46746031 | + | -1.22322 | intron (NM_009681, intron 1 of 5) | intron (NM_009681, intron 1 of 5) | 3983 | NM_009681 | 11777 | Mm.312536 | NM_009681 | ENSMUSG00000024480 | **Ap3s1** | [s]3A | adaptor-related protein complex 3, sigma 1 subunit | protein-coding |
| chr18_46745686_46746123 | chr18 | 46745687 | 46746123 | + | -1.22791 | intron (NM_009681, intron 1 of 5) | intron (NM_009681, intron 1 of 5) | 3988 | NM_009681 | 11777 | Mm.312536 | NM_009681 | ENSMUSG00000024480 | **Ap3s1** | [s]3A | adaptor-related protein complex 3, sigma 1 subunit | protein-coding |
| chr2_22816277_22816541 | chr2 | 22816278 | 22816541 | + | -0.644583 | intron (NM_019456, intron 1 of 13) | ORR1E\|LTR\|ERVL-MaLR | 42082 | NM_019456 | 54519 | Mm.14255 | NM_019456 | ENSMUSG00000026786 | **Apbb1ip** | 9930118P07Rik\|Prp48 | amyloid beta (A4) precursor protein-binding, family B, member 1 interacting protein | protein-coding |
| chr2_22784039_22784631 | chr2 | 22784040 | 22784631 | + | -0.867538 | intron (NM_019456, intron 1 of 13) | intron (NM_019456, intron 1 of 13) | 10008 | NM_019456 | 54519 | Mm.14255 | NM_019456 | ENSMUSG00000026786 | **Apbb1ip** | 9930118P07Rik\|Prp48 | amyloid beta (A4) precursor protein-binding, family B, member 1 interacting protein | protein-coding |

| | | | | | | | | | | | | | | | | | |
|---|---|---|---|---|---|---|---|---|---|---|---|---|---|---|---|---|---|
| chr2_22784888_22785087 | chr2 | 22784889 | 22785087 | + | -0.884954 | intron (NM_019456, intron 1 of 13) | ORR1E\|LTR\|ERVL-MaLR | 10661 | NM_019456 | 54519 | Mm.14255 | NM_019456 | ENSMUSG00000026786 | **Apbb1ip** | 9930118P07Rik\|Prp48 | amyloid beta (A4) precursor protein-binding, family B, member 1 interacting protein | protein-coding |
| chr2_22796463_22796904 | chr2 | 22796464 | 22796904 | + | -1.29809 | intron (NM_019456, intron 1 of 13) | intron (NM_019456, intron 1 of 13) | 22357 | NM_019456 | 54519 | Mm.14255 | NM_019456 | ENSMUSG00000026786 | **Apbb1ip** | 9930118P07Rik\|Prp48 | amyloid beta (A4) precursor protein-binding, family B, member 1 interacting protein | protein-coding |
| chr9_66873517_66873948 | chr9 | 66873518 | 66873948 | + | -0.611089 | intron (NM_173413, intron 1 of 7) | intron (NM_173413, intron 1 of 7) | -39027 | NM_026674 | 68318 | Mm.284229 | NM_026674 | ENSMUSG00000053040 | **Aph1c** | 0610008A10Rik\|APH-1b\|Aph1b | anterior pharynx defective 1c homolog (C. elegans) | protein-coding |
| chr9_46175250_46175889 | chr9 | 46175251 | 46175889 | + | -0.613218 | intron (NM_027498, intron 4 of 24) | ORR1B2\|LTR\|ERVL-MaLR | -53060 | NM_009692 | 11806 | Mm.26743 | NM_009692 | ENSMUSG00000032083 | **Apoa1** | Alp-1\|Apoa-1\|Brp-14\|Ltw-1\|Lvtw-1\|Sep-1\|Sep-2\|Sep2\|apo-AI\|apoA-I | apolipoprotein A-I | protein-coding |
| chr9_46175302_46175756 | chr9 | 46175303 | 46175756 | + | -0.630147 | intron (NM_027498, intron 4 of 24) | ORR1B2\|LTR\|ERVL-MaLR | -53101 | NM_009692 | 11806 | Mm.26743 | NM_009692 | ENSMUSG00000032083 | **Apoa1** | Alp-1\|Apoa-1\|Brp-14\|Ltw-1\|Lvtw-1\|Sep-1\|Sep-2\|Sep2\|apo-AI\|apoA-I | apolipoprotein A-I | protein-coding |
| chr9_46176098_46176313 | chr9 | 46176099 | 46176313 | + | -1.18792 | intron (NM_027498, intron 4 of 24) | intron (NM_027498, intron 4 of 24) | -52424 | NM_009692 | 11806 | Mm.26743 | NM_009692 | ENSMUSG00000032083 | **Apoa1** | Alp-1\|Apoa-1\|Brp-14\|Ltw-1\|Lvtw-1\|Sep-1\|Sep-2\|Sep2\|apo-AI\|apoA-I | apolipoprotein A-I | protein-coding |
| chr9_46176010_46176434 | chr9 | 46176011 | 46176434 | + | -1.19613 | intron (NM_027498, intron 4 of 24) | intron (NM_027498, intron 4 of 24) | -52408 | NM_009692 | 11806 | Mm.26743 | NM_009692 | ENSMUSG00000032083 | **Apoa1** | Alp-1\|Apoa-1\|Brp-14\|Ltw-1\|Lvtw-1\|Sep-1\|Sep-2\|Sep2\|apo-AI\|apoA-I | apolipoprotein A-I | protein-coding |
| chr15_77705320_77705683 | chr15 | 77705321 | 77705683 | + | -1.09648 | intron (NM_001134802, intron 1 of 4) | RMER5\|LTR\|ERV1 | 6613 | NM_001134802 | 666348 | Mm.303207 | NM_001134802 | ENSMUSG00000071716 | **Apol7e** | ENSMUSG00000071716 | apolipoprotein L 7e | protein-coding |
| chr15_77788911_77789899 | chr15 | 77788912 | 77789899 | + | -0.819216 | intron (NM_022410, intron 10 of 40) | intron (NM_022410, intron 10 of 40) | -34176 | NM_001081970 | 239552 | Mm.125650 | NM_001081970 | ENSMUSG00000056656 | **Apol8** | 9830006J20Rik\|APOL-II\|Apol2 | apolipoprotein L 8 | protein-coding |
| chr15_77406472_77406817 | chr15 | 77406473 | 77406817 | + | -0.824558 | intron (NM_001162883, intron 1 of 2) | intron (NM_001162883, intron 1 of 2) | 4389 | NM_173786 | 223672 | Mm.243758 | NM_173786 | ENSMUSG00000057346 | **Apol9a** | - | apolipoprotein L 9a | protein-coding |
| chr15_77733256_77733803 | chr15 | 77733257 | 77733803 | + | -0.651362 | intron (NM_001168660, intron 1 of 2) | intron (NM_001168660, intron 1 of 2) | 4409 | NM_001168660 | 71898 | Mm.422856 | NM_173743 | ENSMUSG00000068246 | **Apol9b** | 2310016F22Rik | apolipoprotein L 9b | protein-coding |
| chr10_83663854_83664194 | chr10 | 83663855 | 83664194 | + | -1.01732 | Intergenic | Intergenic | -15360 | NM_145220 | 216190 | Mm.282985 | NM_145220 | ENSMUSG00000020263 | **Appl2** | Dip3b | adaptor protein, phosphotyrosine interaction, PH domain and leucine zipper containing 2 | protein-coding |
| chr18_15342152_15342589 | chr18 | 15342153 | 15342589 | + | -0.685767 | Intergenic | MERV1_I-int\|LTR\|ERV1 | 61313 | NM_009700 | 11829 | Mm.250786 | NM_009700 | ENSMUSG00000024411 | **Aqp4** | WCH4 | aquaporin 4 | protein-coding |
| chr9_71165153_71165594 | chr9 | 71165154 | 71165594 | + | -0.618243 | Intergenic | Intergenic | -2085 | NM_022026 | 64008 | Mm.449427 | NM_022026 | ENSMUSG00000032204 | **Aqp9** | 1700020I22Rik\|AI266899\|AQP-9 | aquaporin 9 | protein-coding |
| chr9_71162906_71163359 | chr9 | 71162907 | 71163359 | + | -0.968048 | promoter-TSS (NM_001271843) | promoter-TSS (NM_001271843) | 156 | NR_073483 | 64008 | Mm.449427 | NM_022026 | ENSMUSG00000032204 | **Aqp9** | 1700020I22Rik\|AI266899\|AQP-9 | aquaporin 9 | protein-coding |
| chrX_98259457_98260024 | chrX | 98259458 | 98260024 | + | -0.676489 | intron (NM_013476, intron 2 of 7) | intron (NM_013476, intron 2 of 7) | 109991 | NM_013476 | 11835 | Mm.39005 | NM_013476 | | **Ar** | AW320017\|Tfm | androgen receptor | protein-coding |
| chrX_98291775_98292065 | chrX | 98291776 | 98292065 | + | -0.89566 | intron (NM_013476, intron 3 of 7) | intron (NM_013476, intron 3 of 7) | 142170 | NM_013476 | 11835 | Mm.39005 | NM_013476 | | **Ar** | AW320017\|Tfm | androgen receptor | protein-coding |
| chr5_62659161_62659547 | chr5 | 62659162 | 62659547 | + | -0.713395 | intron (NM_178407, intron 19 of 32) | PB1D9\|SINE\|Alu | 106823 | NM_178407 | 212285 | Mm.244403 | NM_178407 | ENSMUSG00000037999 | **Arap2** | Centd1\|Gm148\|mKIAA0580 | ArfGAP with RhoGAP domain, ankyrin repeat and PH domain 2 | protein-coding |
| chr5_62771844_62772519 | chr5 | 62771845 | 62772519 | + | -0.775141 | Intergenic | Intergenic | -6005 | NM_178407 | 212285 | Mm.244403 | NM_178407 | ENSMUSG00000037999 | **Arap2** | Centd1\|Gm148\|mKIAA0580 | ArfGAP with RhoGAP domain, ankyrin repeat and PH domain 2 | protein-coding |
| chr5_62771861_62772256 | chr5 | 62771862 | 62772256 | + | -0.818099 | Intergenic | Intergenic | -5882 | NM_178407 | 212285 | Mm.244403 | NM_178407 | ENSMUSG00000037999 | **Arap2** | Centd1\|Gm148\|mKIAA0580 | ArfGAP with RhoGAP domain, ankyrin repeat and PH domain 2 | protein-coding |
| chr5_62752893_62753302 | chr5 | 62752894 | 62753302 | + | -1.05216 | intron (NM_178407, intron 1 of 32) | intron (NM_178407, intron 1 of 32) | 13079 | NM_178407 | 212285 | Mm.244403 | NM_178407 | ENSMUSG00000037999 | **Arap2** | Centd1\|Gm148\|mKIAA0580 | ArfGAP with RhoGAP domain, ankyrin repeat and PH domain 2 | protein-coding |
| chr5_62778476_62779150 | chr5 | 62778477 | 62779150 | + | -1.44854 | Intergenic | Intergenic | -12636 | NM_178407 | 212285 | Mm.244403 | NM_178407 | ENSMUSG00000037999 | **Arap2** | Centd1\|Gm148\|mKIAA0580 | ArfGAP with RhoGAP domain, ankyrin repeat and PH domain 2 | protein-coding |
| chr11_59224550_59224756 | chr11 | 59224551 | 59224756 | + | -1.14669 | intron (NM_007476, intron 1 of 4) | intron (NM_007476, intron 1 of 4) | 3509 | NM_001130408 | 11840 | Mm.371546 | NM_007476 | ENSMUSG00000048076 | **Arf1** | - | ADP-ribosylation factor 1 | protein-coding |
| chr8_8847643_8847957 | chr8 | 8847644 | 8847957 | + | -0.872265 | Intergenic | B4\|SINE\|B4 | -157263 | NM_176849 | 234023 | Mm.251655 | NM_176849 | ENSMUSG00000040459 | **Arglu1** | 9430010O03Rik\|C130008N12 | arginine and glutamate rich 1 | protein-coding |
| chr2_113866237_113866503 | chr2 | 113866238 | 113866503 | + | -0.683558 | Intergenic | Intergenic | -17709 | NM_181416 | 228482 | Mm.45753 | NM_181416 | ENSMUSG00000041219 | **Arhgap11a** | 6530401L14Rik\|mKIAA0013 | Rho GTPase activating protein 11A | protein-coding |
| chr2_113914295_113914888 | chr2 | 113914296 | 113914888 | + | -0.751308 | Intergenic | Intergenic | -65931 | NM_181416 | 228482 | Mm.45753 | NM_181416 | ENSMUSG00000041219 | **Arhgap11a** | 6530401L14Rik\|mKIAA0013 | Rho GTPase activating protein 11A | protein-coding |
| chr2_113914264_113914634 | chr2 | 113914265 | 113914634 | + | -0.802926 | Intergenic | Lx8b\|LINE\|L1 | -65788 | NM_181416 | 228482 | Mm.45753 | NM_181416 | ENSMUSG00000041219 | **Arhgap11a** | 6530401L14Rik\|mKIAA0013 | Rho GTPase activating protein 11A | protein-coding |
| chr2_113902972_113903543 | chr2 | 113902973 | 113903543 | + | -0.868297 | Intergenic | MLT1B\|LTR\|ERVL-MaLR | -54597 | NM_181416 | 228482 | Mm.45753 | NM_181416 | ENSMUSG00000041219 | **Arhgap11a** | 6530401L14Rik\|mKIAA0013 | Rho GTPase activating protein 11A | protein-coding |
| chr18_6106582_6107108 | chr18 | 6106583 | 6107108 | + | -0.826265 | intron (NM_001039692, intron 3 of 18) | intron (NM_001039692, intron 3 of 18) | 29253 | NM_029277 | 75415 | Mm.239675 | NM_029277 | ENSMUSG00000041225 | **Arhgap12** | 2810011M08Rik | Rho GTPase activating protein 12 | protein-coding |
| chr2_43756625_43757218 | chr2 | 43756626 | 43757218 | + | -0.631061 | intron (NM_153820, intron 1 of 13) | B4\|SINE\|B4 | 8095 | NM_001025377 | 76117 | Mm.217350 | NM_153820 | ENSMUSG00000049744 | **Arhgap15** | 5830480G12Rik | Rho GTPase activating protein 15 | protein-coding |
| chr2_43756686_43757036 | chr2 | 43756687 | 43757036 | + | -0.742376 | intron (NM_153820, intron 1 of 13) | B4\|SINE\|B4 | 8034 | NM_001025377 | 76117 | Mm.217350 | NM_153820 | ENSMUSG00000049744 | **Arhgap15** | 5830480G12Rik | Rho GTPase activating protein 15 | protein-coding |
| chr2_43727557_43728279 | chr2 | 43727558 | 43728279 | + | -0.929871 | Intergenic | Intergenic | -20906 | NM_153820 | 76117 | Mm.217350 | NM_153820 | ENSMUSG00000049744 | **Arhgap15** | 5830480G12Rik | Rho GTPase activating protein 15 | protein-coding |
| chr2_44159574_44159834 | chr2 | 44159575 | 44159834 | + | -0.832209 | intron (NM_153820, intron 10 of 13) | intron (NM_153820, intron 10 of 13) | -94380 | NR_040622 | 320872 | Mm.445023 | NR_040622 | ENSMUSG00000085089 | **Arhgap15os** | A430092G05Rik | Rho GTPase activating protein 15, opposite strand | ncRNA |
| chr2_44159371_44159864 | chr2 | 44159372 | 44159864 | + | -0.839095 | intron (NM_153820, intron 10 of 13) | intron (NM_153820, intron 10 of 13) | -94294 | NR_040622 | 320872 | Mm.445023 | NR_040622 | ENSMUSG00000085089 | **Arhgap15os** | A430092G05Rik | Rho GTPase activating protein 15, opposite strand | ncRNA |
| chr2_43992498_43992853 | chr2 | 43992499 | 43992853 | + | -0.972543 | intron (NM_153820, intron 6 of 13) | intron (NM_153820, intron 6 of 13) | 72648 | NR_040622 | 320872 | Mm.445023 | NR_040622 | ENSMUSG00000085089 | **Arhgap15os** | A430092G05Rik | Rho GTPase activating protein 15, opposite strand | ncRNA |
| chr7_123354542_123355240 | chr7 | 123354543 | 123355240 | + | -0.734193 | intron (NM_001122643, intron 1 of 19) | B1_Mur1\|SINE\|Alu | 15024 | NM_001122643 | 70497 | Mm.403318 | NM_144529 | ENSMUSG00000030766 | **Arhgap17** | 5730403H17Rik\|Nadrin\|Nadrin2\|Rich1\|Wbp15 | Rho GTPase activating protein 17 | protein-coding |
| chr7_123354811_123355249 | chr7 | 123354812 | 123355249 | + | -0.782172 | intron (NM_001122643, intron 1 of 19) | intron (NM_001122643, intron 1 of 19) | 14885 | NM_001122643 | 70497 | Mm.403318 | NM_144529 | ENSMUSG00000030766 | **Arhgap17** | 5730403H17Rik\|Nadrin\|Nadrin2\|Rich1\|Wbp15 | Rho GTPase activating protein 17 | protein-coding |
| chr5_102497264_102497557 | chr5 | 102497265 | 102497557 | + | -0.757465 | intron (NM_146161, intron 1 of 8) | intron (NM_146161, intron 1 of 8) | 16020 | NM_029270 | 231532 | Mm.440191 | NM_029270 | ENSMUSG00000057315 | **Arhgap24** | 0610025G21Rik | Rho GTPase activating protein 24 | protein-coding |
| chr6_87502863_87503288 | chr6 | 87502864 | 87503288 | + | -0.77813 | intron (NM_001037727, intron 1 of 10) | intron (NM_001037727, intron 1 of 10) | -6749 | NM_175476 | 232201 | Mm.119564 | NM_175476 | | **Arhgap25** | A130039I20Rik | Rho GTPase activating protein 25 | protein-coding |
| chr6_87502379_87503679 | chr6 | 87502380 | 87503679 | + | -0.784341 | intron (NM_001037727, intron 1 of 10) | intron (NM_001037727, intron 1 of 10) | -6702 | NM_175476 | 232201 | Mm.119564 | NM_175476 | | **Arhgap25** | A130039I20Rik | Rho GTPase activating protein 25 | protein-coding |
| chr6_87503971_87504276 | chr6 | 87503972 | 87504276 | + | -0.864411 | intron (NM_001037727, intron 1 of 10) | intron (NM_001037727, intron 1 of 10) | -7797 | NM_175476 | 232201 | Mm.119564 | NM_175476 | | **Arhgap25** | A130039I20Rik | Rho GTPase activating protein 25 | protein-coding |
| chr6_87502373_87502797 | chr6 | 87502374 | 87502797 | + | -0.884809 | intron (NM_001037727, intron 1 of 10) | intron (NM_001037727, intron 1 of 10) | -6258 | NM_175476 | 232201 | Mm.119564 | NM_175476 | | **Arhgap25** | A130039I20Rik | Rho GTPase activating protein 25 | protein-coding |
| chr18_39082113_39082390 | chr18 | 39082114 | 39082390 | + | -0.905063 | intron (NM_175164, intron 1 of 22) | intron (NM_175164, intron 1 of 22) | 89107 | NM_175164 | 71302 | Mm.329396 | NM_175164 | ENSMUSG00000036452 | **Arhgap26** | 1810044B20Rik\|2610010G17Rik\|4933432P15Rik\|AI853435\|mKIAA0621 | Rho GTPase activating protein 26 | protein-coding |
| chr11_103381771_103382544 | chr11 | 103381772 | 103382544 | + | -0.64295 | intron (NM_183034, intron 5 of 11) | intron (NM_183034, intron 5 of 11) | -18466 | NM_001205236 | 544817 | Mm.259113 | NM_133715 | ENSMUSG00000034255 | **Arhgap27** | 2310069I04Rik\|5730442P18Rik\|Gm21552\|Sh3d20 | Rho GTPase activating protein 27 | protein-coding |
| chr11_103307949_103308240 | chr11 | 103307950 | 103308240 | + | -0.761171 | Intergenic | RMER19B\|LTR\|ERVK | 36607 | NM_133715 | 544817 | Mm.259113 | NM_133715 | ENSMUSG00000034255 | **Arhgap27** | 2310069I04Rik\|5730442P18Rik\|Gm21552\|Sh3d20 | Rho GTPase activating protein 27 | protein-coding |
| chr11_103375091_103375640 | chr11 | 103375092 | 103375640 | + | -0.805689 | intron (NM_183034, intron 7 of 11) | intron (NM_183034, intron 7 of 11) | -11674 | NM_001205236 | 544817 | Mm.259113 | NM_133715 | ENSMUSG00000034255 | **Arhgap27** | 2310069I04Rik\|5730442P18Rik\|Gm21552\|Sh3d20 | Rho GTPase activating protein 27 | protein-coding |

| | | | | | | | | | | | | | | | | | |
|---|---|---|---|---|---|---|---|---|---|---|---|---|---|---|---|---|---|
| chr16_38714229_38714447 | chr16 | 38714230 | 38714447 | + | -0.848168 | Intergenic | MER102b\|DNA\|hAT-Charlie | -1303 | NM_020260 | 12549 | Mm.370848 | NM_020260 | ENSMUSG00000022799 | **Arhgap31** | 5830477L08Rik\|AU041750\|Cdgap\|D330026I07Rik | Rho GTPase activating protein 31 | protein-coding |
| chrX_73911698_73912003 | chrX | 73911699 | 73912003 | + | -0.687451 | promoter-TSS (NM_138630) | promoter-TSS (NM_138630) | -553 | NM_001162424 | 171207 | Mm.482337 | NM_138630 | ENSMUSG00000031389 | **Arhgap4** | A130009C12Rik\|A530015A20Rik\|Rgc1\|c1\|mKIAA0131 | Rho GTPase activating protein 4 | protein-coding |
| chr12_52524749_52525054 | chr12 | 52524750 | 52525054 | + | -0.710891 | intron (NM_009706, intron 1 of 5) | intron (NM_009706, intron 1 of 5) | 8825 | NM_009706 | 11855 | Mm.35059 | NM_009706 | ENSMUSG00000035133 | **Arhgap5** | p190-B\|p190B | Rho GTPase activating protein 5 | protein-coding |
| chr12_52552104_52552465 | chr12 | 52552105 | 52552465 | + | -1.15012 | intron (NM_009706, intron 2 of 5) | ORR1D2\|LTR\|ERVL-MaLR | 36208 | NM_009706 | 11855 | Mm.35059 | NM_009706 | ENSMUSG00000035133 | **Arhgap5** | p190-B\|p190B | Rho GTPase activating protein 5 | protein-coding |
| chr10_127321367_127322007 | chr10 | 127321368 | 127322007 | + | -0.853238 | promoter-TSS (NM_001285785) | promoter-TSS (NM_001285785) | -277 | NM_001285785 | 216445 | Mm.227198 | NM_146011 | ENSMUSG00000040345 | **Arhgap9** | AU043488 | Rho GTPase activating protein 9 | protein-coding |
| chr6_136944605_136944994 | chr6 | 136944606 | 136944994 | + | -0.68141 | Intergenic | Lx9\|LINE\|L1 | -2901 | NM_001301309 | 11857 | Mm.2241 | NM_007486 | ENSMUSG00000030220 | **Arhgdib** | D4\|Gdid4\|Ly-GDI | Rho, GDP dissociation inhibitor (GDI) beta | protein-coding |
| chr6_136940992_136941542 | chr6 | 136940993 | 136941542 | + | -0.728441 | intron (NM_001301301, intron 1 of 5) | intron (NM_001301301, intron 1 of 5) | 236 | NM_001301305 | 11857 | Mm.2241 | NM_007486 | ENSMUSG00000030220 | **Arhgdib** | D4\|Gdid4\|Ly-GDI | Rho, GDP dissociation inhibitor (GDI) beta | protein-coding |
| chr6_136944543_136944882 | chr6 | 136944544 | 136944882 | + | -0.769466 | Intergenic | Lx9\|LINE\|L1 | -2814 | NM_001301309 | 11857 | Mm.2241 | NM_007486 | ENSMUSG00000030220 | **Arhgdib** | D4\|Gdid4\|Ly-GDI | Rho, GDP dissociation inhibitor (GDI) beta | protein-coding |
| chr6_136944600_136944850 | chr6 | 136944601 | 136944850 | + | -0.784104 | Intergenic | Lx9\|LINE\|L1 | -2826 | NM_001301309 | 11857 | Mm.2241 | NM_007486 | ENSMUSG00000030220 | **Arhgdib** | D4\|Gdid4\|Ly-GDI | Rho, GDP dissociation inhibitor (GDI) beta | protein-coding |
| chr8_14940993_14941508 | chr8 | 14940994 | 14941508 | + | -0.711979 | intron (NM_001037736, intron 8 of 27) | intron (NM_001037736, intron 8 of 27) | 29588 | NM_001037736 | 234094 | Mm.240298 | NM_172751 | ENSMUSG00000071176 | **Arhgef10** | 6430549H08Rik\|mKIAA0294 | Rho guanine nucleotide exchange factor (GEF) 10 | protein-coding |
| chr4_140601064_140601578 | chr4 | 140601065 | 140601578 | + | -0.963829 | intron (NM_001112722, intron 4 of 27) | intron (NM_001112722, intron 4 of 27) | 47435 | NM_001112723 | 72754 | Mm.29329 | NM_172415 | ENSMUSG00000040964 | **Arhgef10l** | 2810441C07Rik | Rho guanine nucleotide exchange factor (GEF) 10-like | protein-coding |
| chr14_27345539_27345819 | chr14 | 27345540 | 27345819 | + | -0.640986 | intron (NM_001289687, intron 1 of 9) | intron (NM_001289687, intron 1 of 9) | 9642 | NM_001289687 | 71704 | Mm.248606 | NM_027871 | ENSMUSG00000021895 | **Arhgef3** | 1200004I24Rik\|9830169H03Rik\|C76747\|Xpln | Rho guanine nucleotide exchange factor (GEF) 3 | protein-coding |
| chr14_27229179_27230063 | chr14 | 27229180 | 27230063 | + | -0.728475 | intron (NM_001289686, intron 3 of 12) | intron (NM_001289686, intron 3 of 12) | -8389 | NM_027871 | 71704 | Mm.248606 | NM_027871 | ENSMUSG00000021895 | **Arhgef3** | 1200004I24Rik\|9830169H03Rik\|C76747\|Xpln | Rho guanine nucleotide exchange factor (GEF) 3 | protein-coding |
| chr17_80323079_80324235 | chr17 | 80323080 | 80324235 | + | -0.651475 | intron (NM_001145452, intron 1 of 17) | intron (NM_001145452, intron 1 of 17) | 16250 | NM_001145452 | 381112 | Mm.360004 | NM_001145452 | ENSMUSG00000054901 | **Arhgef33** | Gm941 | Rho guanine nucleotide exchange factor (GEF) 33 | protein-coding |
| chr17_80323463_80324239 | chr17 | 80323464 | 80324239 | + | -0.756539 | intron (NM_001145452, intron 1 of 17) | intron (NM_001145452, intron 1 of 17) | 16444 | NM_001145452 | 381112 | Mm.360004 | NM_001145452 | ENSMUSG00000054901 | **Arhgef33** | Gm941 | Rho guanine nucleotide exchange factor (GEF) 33 | protein-coding |
| chr17_5195412_5195918 | chr17 | 5195413 | 5195918 | + | -0.659369 | intron (NM_001085355, intron 4 of 19) | intron (NM_001085355, intron 4 of 19) | 200591 | NM_001085355 | 239985 | Mm.133401 | NM_001085355 | ENSMUSG00000069729 | **Arid1b** | 8030481M12\|9330189K18Rik\|AI836955\|Ardi1b\|B230217J03Rik\|mKIAA1235 | AT rich interactive domain 1B (SWI-like) | protein-coding |
| chr17_5195408_5195778 | chr17 | 5195409 | 5195778 | + | -0.688517 | intron (NM_001085355, intron 4 of 19) | intron (NM_001085355, intron 4 of 19) | 200519 | NM_001085355 | 239985 | Mm.133401 | NM_001085355 | ENSMUSG00000069729 | **Arid1b** | 8030481M12\|9330189K18Rik\|AI836955\|Ardi1b\|B230217J03Rik\|mKIAA1235 | AT rich interactive domain 1B (SWI-like) | protein-coding |
| chr17_5009840_5010243 | chr17 | 5009841 | 5010243 | + | -0.738595 | intron (NM_001085355, intron 1 of 19) | intron (NM_001085355, intron 1 of 19) | 14968 | NM_001085355 | 239985 | Mm.133401 | NM_001085355 | ENSMUSG00000069729 | **Arid1b** | 8030481M12\|9330189K18Rik\|AI836955\|Ardi1b\|B230217J03Rik\|mKIAA1235 | AT rich interactive domain 1B (SWI-like) | protein-coding |
| chr15_96297554_96298324 | chr15 | 96297555 | 96298324 | + | -0.935404 | intron (NM_175251, intron 3 of 20) | B4A\|SINE\|B4 | 10417 | NM_175251 | 77044 | Mm.17166 | NM_175251 | ENSMUSG00000033237 | **Arid2** | 1700124K17Rik\|4432409D24Rik\|zipzap/p200 | AT rich interactive domain 2 (ARID, RFX-like) | protein-coding |
| chr12_71048097_71048586 | chr12 | 71048098 | 71048586 | + | -0.674561 | intron (NM_001081195, intron 11 of 23) | intron (NM_001081195, intron 11 of 23) | 32375 | NM_001081195 | 238247 | Mm.241601 | NM_001081195 | ENSMUSG00000048118 | **Arid4a** | A630009N03\|A630067N03Rik\|MmRBBP1\|Rbbp1 | AT rich interactive domain 4A (RBP1-like) | protein-coding |
| chr12_71040271_71040891 | chr12 | 71040272 | 71040891 | + | -0.832196 | intron (NM_001081195, intron 8 of 23) | PB1\|SINE\|Alu | 24614 | NM_001081195 | 238247 | Mm.241601 | NM_001081195 | ENSMUSG00000048118 | **Arid4a** | A630009N03\|A630067N03Rik\|MmRBBP1\|Rbbp1 | AT rich interactive domain 4A (RBP1-like) | protein-coding |
| chr12_71040439_71040857 | chr12 | 71040440 | 71040857 | + | -0.833925 | intron (NM_001081195, intron 8 of 23) | B4A\|SINE\|B4 | 24681 | NM_001081195 | 238247 | Mm.241601 | NM_001081195 | ENSMUSG00000048118 | **Arid4a** | A630009N03\|A630067N03Rik\|MmRBBP1\|Rbbp1 | AT rich interactive domain 4A (RBP1-like) | protein-coding |
| chr13_14119481_14119777 | chr13 | 14119482 | 14119777 | + | -0.903213 | intron (NM_198122, intron 2 of 22) | intron (NM_198122, intron 2 of 22) | 55845 | NM_198122 | 94246 | Mm.439784 | NM_194262 | ENSMUSG00000039219 | **Arid4b** | BCAA\|BRCAA1\|RBBP1L1\|Rbp1l1\|SAP180 | AT rich interactive domain 4B (RBP1-like) | protein-coding |
| chr1_36309572_36310017 | chr1 | 36309573 | 36310017 | + | -1.00839 | intron (NM_001290727, intron 1 of 5) | intron (NM_001290727, intron 1 of 5) | 2062 | NM_001172205 | 214855 | Mm.34316 | NM_145996 | ENSMUSG00000037447 | **Arid5a** | D430024K22Rik\|Mrf1 | AT rich interactive domain 5A (MRF1-like) | protein-coding |
| chr10_68240654_68241449 | chr10 | 68240655 | 68241449 | + | -0.857688 | intron (NM_023598, intron 3 of 9) | intron (NM_023598, intron 3 of 9) | 37674 | NM_023598 | 71371 | Mm.440357 | NM_023598 | ENSMUSG00000019947 | **Arid5b** | 4930580B11\|5430435G07Rik\|AI467247\|Desrt\|Mrf2\|Mrf2alpha\|Mrf2beta | AT rich interactive domain 5B (MRF1-like) | protein-coding |
| chr10_68240889_68241241 | chr10 | 68240890 | 68241241 | + | -0.919583 | intron (NM_023598, intron 3 of 9) | intron (NM_023598, intron 3 of 9) | 37661 | NM_023598 | 71371 | Mm.440357 | NM_023598 | ENSMUSG00000019947 | **Arid5b** | 4930580B11\|5430435G07Rik\|AI467247\|Desrt\|Mrf2\|Mrf2alpha\|Mrf2beta | AT rich interactive domain 5B (MRF1-like) | protein-coding |
| chr3_69177243_69177892 | chr3 | 69177244 | 69177892 | + | -0.63398 | Intergenic | Intergenic | -44851 | NM_027843 | 71619 | Mm.98122 | NM_027843 | ENSMUSG00000098207 | **Arl14** | 9130014L17Rik\|Arf7 | ADP-ribosylation factor-like 14 | protein-coding |
| chr13_113870435_113870888 | chr13 | 113870436 | 113870888 | + | -0.702406 | intron (NM_172595, intron 1 of 4) | intron (NM_172595, intron 1 of 4) | 76154 | NM_172595 | 218639 | Mm.79837 | NM_172595 | ENSMUSG00000042348 | **Arl15** | A430036I03\|Arfrp2\|C230032K13Rik | ADP-ribosylation factor-like 15 | protein-coding |
| chr13_113870434_113870756 | chr13 | 113870435 | 113870756 | + | -0.712955 | intron (NM_172595, intron 1 of 4) | intron (NM_172595, intron 1 of 4) | 76087 | NM_172595 | 218639 | Mm.79837 | NM_172595 | ENSMUSG00000042348 | **Arl15** | A430036I03\|Arfrp2\|C230032K13Rik | ADP-ribosylation factor-like 15 | protein-coding |
| chr8_94634116_94634704 | chr8 | 94634117 | 94634704 | + | -1.04808 | intron (NM_026274, intron 6 of 14) | intron (NM_026274, intron 6 of 14) | -32190 | NM_024191 | 107566 | Mm.393247 | NM_024191 | ENSMUSG00000031776 | **Arl2bp** | 1700010P10Rik\|1700027H16Rik\|6330544B05Rik\|AI482273\|AI849834\|BART\|Bart1 | ADP-ribosylation factor-like 2 binding protein | protein-coding |
| chr1_88638725_88638977 | chr1 | 88638726 | 88638977 | + | -0.717715 | Intergenic | RMER17C\|LTR\|ERVK | 63340 | NM_177305 | 320982 | Mm.27968 | NM_177305 | ENSMUSG00000049866 | **Arl4c** | A630084M22Rik\|Arl7\|LAK | ADP-ribosylation factor-like 4C | protein-coding |
| chr1_88654636_88655454 | chr1 | 88654637 | 88655454 | + | -0.755518 | Intergenic | Intergenic | 47146 | NM_177305 | 320982 | Mm.27968 | NM_177305 | ENSMUSG00000049866 | **Arl4c** | A630084M22Rik\|Arl7\|LAK | ADP-ribosylation factor-like 4C | protein-coding |
| chr1_88638711_88639129 | chr1 | 88638712 | 88639129 | + | -0.760759 | Intergenic | RMER17C\|LTR\|ERVK | 63271 | NM_177305 | 320982 | Mm.27968 | NM_177305 | ENSMUSG00000049866 | **Arl4c** | A630084M22Rik\|Arl7\|LAK | ADP-ribosylation factor-like 4C | protein-coding |
| chr1_88720572_88721296 | chr1 | 88720573 | 88721296 | + | -0.828483 | Intergenic | Intergenic | -18743 | NM_177305 | 320982 | Mm.27968 | NM_177305 | ENSMUSG00000049866 | **Arl4c** | A630084M22Rik\|Arl7\|LAK | ADP-ribosylation factor-like 4C | protein-coding |
| chr1_88656098_88656445 | chr1 | 88656099 | 88656445 | + | -0.941688 | Intergenic | Intergenic | 45919 | NM_177305 | 320982 | Mm.27968 | NM_177305 | ENSMUSG00000049866 | **Arl4c** | A630084M22Rik\|Arl7\|LAK | ADP-ribosylation factor-like 4C | protein-coding |
| chr1_88708997_88709316 | chr1 | 88708998 | 88709316 | + | -1.00118 | Intergenic | Intergenic | -6966 | NM_177305 | 320982 | Mm.27968 | NM_177305 | ENSMUSG00000049866 | **Arl4c** | A630084M22Rik\|Arl7\|LAK | ADP-ribosylation factor-like 4C | protein-coding |
| chr1_88684705_88685234 | chr1 | 88684706 | 88685234 | + | -1.05582 | Intergenic | Intergenic | 17221 | NM_177305 | 320982 | Mm.27968 | NM_177305 | ENSMUSG00000049866 | **Arl4c** | A630084M22Rik\|Arl7\|LAK | ADP-ribosylation factor-like 4C | protein-coding |
| chr1_88644397_88644862 | chr1 | 88644398 | 88644862 | + | -1.0996 | Intergenic | Intergenic | 57561 | NM_177305 | 320982 | Mm.27968 | NM_177305 | ENSMUSG00000049866 | **Arl4c** | A630084M22Rik\|Arl7\|LAK | ADP-ribosylation factor-like 4C | protein-coding |
| chr11_101697204_101697425 | chr11 | 101697205 | 101697425 | + | -0.625243 | Intergenic | PB1D10\|SINE\|Alu | 31774 | NM_025404 | 80981 | Mm.266840 | NM_025404 | ENSMUSG00000034936 | **Arl4d** | 1110036H21Rik\|ARL6\|AW456149\|Arf4l\|Arfl4\|Arl5 | ADP-ribosylation factor-like 4D | protein-coding |
| chr11_98001300_98001758 | chr11 | 98001301 | 98001758 | + | -0.707912 | TTS (NM_145121) | TTS (NM_145121) | -5356 | NM_207231 | 217151 | Mm.277896 | NM_207231 | ENSMUSG00000038352 | **Arl5c** | Arl12\|BC065791 | ADP-ribosylation factor-like 5C | protein-coding |
| chr11_98001300_98001553 | chr11 | 98001301 | 98001553 | + | -0.791932 | TTS (NM_145121) | TTS (NM_145121) | -5254 | NM_207231 | 217151 | Mm.277896 | NM_207231 | ENSMUSG00000038352 | **Arl5c** | Arl12\|BC065791 | ADP-ribosylation factor-like 5C | protein-coding |
| chr11_98000384_98000766 | chr11 | 98000385 | 98000766 | + | -1.14855 | TTS (NM_145121) | TTS (NM_145121) | -4402 | NM_207231 | 217151 | Mm.277896 | NM_207231 | ENSMUSG00000038352 | **Arl5c** | Arl12\|BC065791 | ADP-ribosylation factor-like 5C | protein-coding |
| chr6_97216119_97216387 | chr6 | 97216120 | 97216387 | + | -0.841207 | intron (NM_022992, intron 1 of 2) | intron (NM_022992, intron 1 of 2) | 5461 | NM_022992 | 65106 | Mm.291014 | NM_022992 | ENSMUSG00000035199 | **Arl6ip5** | 5930404D22Rik\|AV001879\|Aip-5\|Gtrap3-18\|addiscin | ADP-ribosylation factor-like 6 interacting protein 5 | protein-coding |
| chr6_97215956_97216407 | chr6 | 97215957 | 97216407 | + | -0.936549 | intron (NM_022992, intron 1 of 2) | intron (NM_022992, intron 1 of 2) | 5390 | NM_022992 | 65106 | Mm.291014 | NM_022992 | ENSMUSG00000035199 | **Arl6ip5** | 5930404D22Rik\|AV001879\|Aip-5\|Gtrap3-18\|addiscin | ADP-ribosylation factor-like 6 interacting protein 5 | protein-coding |
| chr2_19098889_19099254 | chr2 | 19098890 | 19099254 | + | -0.726525 | Intergenic | L1ME1\|LINE\|L1 | -100046 | NM_001271563 | 70882 | Mm.276696 | NM_001081083 | ENSMUSG00000037683 | **Armc3** | 4921513G22Rik | armadillo repeat containing 3 | protein-coding |
| chr2_19098767_19099320 | chr2 | 19098768 | 19099320 | + | -0.75344 | Intergenic | L1ME1\|LINE\|L1 | -100074 | NM_001271563 | 70882 | Mm.276696 | NM_001081083 | ENSMUSG00000037683 | **Armc3** | 4921513G22Rik | armadillo repeat containing 3 | protein-coding |
| chr11_115477461_115477716 | chr11 | 115477462 | 115477716 | + | -0.638818 | intron (NM_177778, intron 2 of 2) | intron (NM_177778, intron 2 of 2) | 1912 | NM_177778 | 276905 | Mm.38019 | NM_177778 | ENSMUSG00000057219 | **Armc7** | A630084C13\|BC023126 | armadillo repeat containing 7 | protein-coding |
| chr11_115481461_115481892 | chr11 | 115481462 | 115481892 | + | -0.842264 | intron (NM_177778, intron 2 of 2) | intron (NM_177778, intron 2 of 2) | 6000 | NM_177778 | 276905 | Mm.38019 | NM_177778 | ENSMUSG00000057219 | **Armc7** | A630084C13\|BC023126 | armadillo repeat containing 7 | protein-coding |
| chr11_115481563_115481878 | chr11 | 115481564 | 115481878 | + | -0.843787 | intron (NM_177778, intron 2 of 2) | intron (NM_177778, intron 2 of 2) | 6044 | NM_177778 | 276905 | Mm.38019 | NM_177778 | ENSMUSG00000057219 | **Armc7** | A630084C13\|BC023126 | armadillo repeat containing 7 | protein-coding |
| chr1_86228719_86228994 | chr1 | 86228720 | 86228994 | + | -0.845921 | intron (NM_027456, intron 19 of 20) | B3\|SINE\|B2 | 74042 | NM_027456 | 78795 | Mm.379264 | NM_027456 | ENSMUSG00000062590 | **Armc9** | 3830422A13Rik\|4831423D23Rik\|4930438O05Rik\|5730415N24Rik | armadillo repeat containing 9 | protein-coding |
| chr1_86228693_86229098 | chr1 | 86228694 | 86229098 | + | -0.85418 | intron (NM_027456, intron 19 of 20) | B3\|SINE\|B2 | 74081 | NM_027456 | 78795 | Mm.379264 | NM_027456 | ENSMUSG00000062590 | **Armc9** | 3830422A13Rik\|4831423D23Rik\|4930438O05Rik\|5730415N24Rik | armadillo repeat containing 9 | protein-coding |

| | | | | | | | | | | | | | | | | | |
|---|---|---|---|---|---|---|---|---|---|---|---|---|---|---|---|---|---|
| chr7_99534858_99536697 | chr7 | 99534859 | 99536697 | + | -0.645408 | 5' UTR (NM_178220, exon 1 of 15) | 5' UTR (NM_178220, exon 1 of 15) | 292 | NM_177231 | 109689 | Mm.260193 | NM_177231 | ENSMUSG00000018909 | **Arrb1** | 1200006I17Rik\|AW208571\|G430100A01Rik | arrestin, beta 1 | protein-coding |
| chr7_99535804_99536596 | chr7 | 99535805 | 99536596 | + | -0.686985 | intron (NM_177231, intron 1 of 15) | intron (NM_177231, intron 1 of 15) | 714 | NM_177231 | 109689 | Mm.260193 | NM_177231 | ENSMUSG00000018909 | **Arrb1** | 1200006I17Rik\|AW208571\|G430100A01Rik | arrestin, beta 1 | protein-coding |
| chr13_93762784_93763270 | chr13 | 93762785 | 93763270 | + | -0.830562 | Intergenic | ORR1E\|LTR\|ERVL-MaLR | -8652 | NM_009712 | 11881 | Mm.300178 | NM_009712 | ENSMUSG00000042082 | **Arsb** | 1110007C02Rik\|AI480648\|As-1\|As-1r\|As-1s\|As-1t\|As1\|As1-r\|As1-s\|As1-t\|Asr-1\|Ast-1 | arylsulfatase B | protein-coding |
| chr13_93756548_93757113 | chr13 | 93756549 | 93757113 | + | -0.884127 | Intergenic | Intergenic | -14848 | NM_009712 | 11881 | Mm.300178 | NM_009712 | ENSMUSG00000042082 | **Arsb** | 1110007C02Rik\|AI480648\|As-1\|As-1r\|As-1s\|As-1t\|As1\|As1-r\|As1-s\|As1-t\|Asr-1\|Ast-1 | arylsulfatase B | protein-coding |
| chr13_93756609_93756994 | chr13 | 93756610 | 93756994 | + | -0.896307 | Intergenic | Intergenic | -14877 | NM_009712 | 11881 | Mm.300178 | NM_009712 | ENSMUSG00000042082 | **Arsb** | 1110007C02Rik\|AI480648\|As-1\|As-1r\|As-1s\|As-1t\|As1\|As1-r\|As1-s\|As1-t\|Asr-1\|Ast-1 | arylsulfatase B | protein-coding |
| chr11_109554012_109554282 | chr11 | 109554013 | 109554282 | + | -0.737674 | intron (NM_028710, intron 11 of 13) | B3A\|SINE\|B2 | 10436 | NM_001166177 | 74008 | Mm.482224 | NM_028710 | ENSMUSG00000020604 | **Arsg** | 6330406P08Rik\|AI846872 | arylsulfatase G | protein-coding |
| chr7_101538256_101538486 | chr7 | 101538257 | 101538486 | + | -0.627341 | Intergenic | Intergenic | 17431 | NM_007490 | 11871 | Mm.439746 | NM_007490 | ENSMUSG00000092517 | **Art2a-ps** | ARTC2\|Art2a\|Ly92a\|Rt-6\|Rt6\|Rt6-1 | ADP-ribosyltransferase 2a, pseudogene | protein-coding |
| chr7_101555543_101555804 | chr7 | 101555544 | 101555804 | + | -1.16293 | intron (NR_033804, intron 2 of 3) | intron (NR_033804, intron 2 of 3) | 128 | NM_007490 | 11871 | Mm.439746 | NM_007490 | ENSMUSG00000092517 | **Art2a-ps** | ARTC2\|Art2a\|Ly92a\|Rt-6\|Rt6\|Rt6-1 | ADP-ribosyltransferase 2a, pseudogene | protein-coding |
| chr7_101555495_101555942 | chr7 | 101555496 | 101555942 | + | -1.17187 | promoter-TSS (NM_007490) | promoter-TSS (NM_007490) | 83 | NM_007490 | 11871 | Mm.439746 | NM_007490 | ENSMUSG00000092517 | **Art2a-ps** | ARTC2\|Art2a\|Ly92a\|Rt-6\|Rt6\|Rt6-1 | ADP-ribosyltransferase 2a, pseudogene | protein-coding |
| chr7_101600238_101600966 | chr7 | 101600239 | 101600966 | + | -0.684268 | Intergenic | L1MC1\|LINE\|L1 | -19441 | NM_019915 | 11872 | Mm.57008 | NM_019915 | ENSMUSG00000030651 | **Art2b** | ART2.2\|ARTC2\|Art\|Ly92b\|Rt-6\|Rt6\|Rt6-2 | ADP-ribosyltransferase 2b | protein-coding |
| chr7_101600350_101600771 | chr7 | 101600351 | 101600771 | + | -0.710086 | Intergenic | L1MC1\|LINE\|L1 | -19400 | NM_019915 | 11872 | Mm.57008 | NM_019915 | ENSMUSG00000030651 | **Art2b** | ART2.2\|ARTC2\|Art\|Ly92b\|Rt-6\|Rt6\|Rt6-2 | ADP-ribosyltransferase 2b | protein-coding |
| chr6_136854282_136854933 | chr6 | 136854283 | 136854933 | + | -0.881501 | exon (NM_026639, exon 2 of 3) | exon (NM_026639, exon 2 of 3) | 2992 | NM_026639 | 109978 | Mm.31840 | NM_026639 | ENSMUSG00000030217 | **Art4** | 4432404K01Rik\|ARTC4\|DO\|DOK1 | ADP-ribosyltransferase 4 | protein-coding |
| chr19_32148775_32149115 | chr19 | 32148776 | 32149115 | + | -0.802692 | intron (NM_001168526, intron 2 of 5) | MLT1A0\|LTR\|ERVL-MaLR | -45805 | NM_018830 | 54447 | Mm.491229 | NM_018830 | ENSMUSG00000024887 | **Asah2** | AI585898 | N-acylsphingosine amidohydrolase 2 | protein-coding |
| chr15_64389565_64389928 | chr15 | 64389566 | 64389928 | + | -0.772095 | Intergenic | Intergenic | -6828 | NM_010026 | 13196 | Mm.277236 | NM_010026 | ENSMUSG00000022377 | **Asap1** | AV239055\|DEF-1\|Ddef1\|PAP\|mKIAA1249\|s19 | ArfGAP with SH3 domain, ankyrin repeat and PH domain1 | protein-coding |
| chr15_64278527_64279342 | chr15 | 64278528 | 64279342 | + | -0.794604 | intron (NM_001276462, intron 1 of 27) | intron (NM_001276462, intron 1 of 27) | 33738 | NM_001276462 | 13196 | Mm.277236 | NM_010026 | ENSMUSG00000022377 | **Asap1** | AV239055\|DEF-1\|Ddef1\|PAP\|mKIAA1249\|s19 | ArfGAP with SH3 domain, ankyrin repeat and PH domain1 | protein-coding |
| chr15_64213085_64213411 | chr15 | 64213086 | 64213411 | + | -0.800253 | intron (NM_001276462, intron 1 of 27) | intron (NM_001276462, intron 1 of 27) | 99425 | NM_001276462 | 13196 | Mm.277236 | NM_010026 | ENSMUSG00000022377 | **Asap1** | AV239055\|DEF-1\|Ddef1\|PAP\|mKIAA1249\|s19 | ArfGAP with SH3 domain, ankyrin repeat and PH domain1 | protein-coding |
| chr15_64213149_64213352 | chr15 | 64213150 | 64213352 | + | -0.800493 | intron (NM_001276462, intron 1 of 27) | intron (NM_001276462, intron 1 of 27) | 99422 | NM_001276462 | 13196 | Mm.277236 | NM_010026 | ENSMUSG00000022377 | **Asap1** | AV239055\|DEF-1\|Ddef1\|PAP\|mKIAA1249\|s19 | ArfGAP with SH3 domain, ankyrin repeat and PH domain1 | protein-coding |
| chr4_136189815_136190337 | chr4 | 136189816 | 136190337 | + | -1.21642 | intron (NM_177733, intron 6 of 6) | intron (NM_177733, intron 6 of 6) | -16289 | NM_001008232 | 230837 | Mm.77788 | NM_001008232 | ENSMUSG00000036995 | **Asap3** | 9430088F20Rik\|Ddefl1\|Gm140\|UPLC1 | ArfGAP with SH3 domain, ankyrin repeat and PH domain 3 | protein-coding |
| chr1_91625811_91626442 | chr1 | 91625812 | 91626442 | + | -0.894698 | Intergenic | Intergenic | 85047 | NM_023046 | 65247 | Mm.234152 | NM_023046 | ENSMUSG00000026311 | **Asb1** | 1700029O08Rik\|1700054C17Rik\|AI426616\|AI851378\|mKIAA1146 | ankyrin repeat and SOCS box-containing 1 | protein-coding |
| chr1_91626090_91626464 | chr1 | 91626091 | 91626464 | + | -0.961156 | Intergenic | Intergenic | 85197 | NM_023046 | 65247 | Mm.234152 | NM_023046 | ENSMUSG00000026311 | **Asb1** | 1700029O08Rik\|1700054C17Rik\|AI426616\|AI851378\|mKIAA1146 | ankyrin repeat and SOCS box-containing 1 | protein-coding |
| chr1_91621156_91621535 | chr1 | 91621157 | 91621535 | + | -1.1532 | Intergenic | ORR1F\|LTR\|ERVL-MaLR | 80266 | NM_023046 | 65247 | Mm.234152 | NM_023046 | ENSMUSG00000026311 | **Asb1** | 1700029O08Rik\|1700054C17Rik\|AI426616\|AI851378\|mKIAA1146 | ankyrin repeat and SOCS box-containing 1 | protein-coding |
| chr1_91621020_91621505 | chr1 | 91621021 | 91621505 | + | -1.18984 | Intergenic | ORR1F\|LTR\|ERVL-MaLR | 80183 | NM_023046 | 65247 | Mm.234152 | NM_023046 | ENSMUSG00000026311 | **Asb1** | 1700029O08Rik\|1700054C17Rik\|AI426616\|AI851378\|mKIAA1146 | ankyrin repeat and SOCS box-containing 1 | protein-coding |
| chr6_5422126_5422734 | chr6 | 5422127 | 5422734 | + | -0.626897 | intron (NM_023048, intron 3 of 5) | intron (NM_023048, intron 3 of 5) | 39044 | NM_023048 | 65255 | Mm.51340 | NM_023048 | ENSMUSG00000042607 | **Asb4** | 8430401O13Rik\|AV117194\|AW060255 | ankyrin repeat and SOCS box-containing 4 | protein-coding |
| chr6_5422427_5422725 | chr6 | 5422428 | 5422725 | + | -0.681256 | intron (NM_023048, intron 3 of 5) | B3A\|SINE\|B2 | 39190 | NM_023048 | 65255 | Mm.51340 | NM_023048 | ENSMUSG00000042607 | **Asb4** | 8430401O13Rik\|AV117194\|AW060255 | ankyrin repeat and SOCS box-containing 4 | protein-coding |
| chr10_50650525_50650991 | chr10 | 50650526 | 50650991 | + | -0.657431 | intron (NM_198007, intron 11 of 41) | intron (NM_198007, intron 11 of 41) | 58089 | NM_198007 | 77987 | Mm.222497 | NM_198007 | ENSMUSG00000038774 | **Ascc3** | ASC1p200\|B630009I04Rik\|BC023451\|D430001L07Rik\|D630041L21\|Helic1\|RNAH | activating signal cointegrator 1 complex subunit 3 | protein-coding |
| chr7_109741407_109742185 | chr7 | 109741408 | 109742185 | + | -0.79035 | intron (NM_020050, intron 4 of 4) | intron (NM_020050, intron 4 of 4) | -10064 | NM_020051 | 56787 | Mm.29275 | NM_020051 | ENSMUSG00000035951 | **Ascl3** | Sgn1\|bHLHa42 | achaete-scute complex homolog 3 (Drosophila) | protein-coding |
| chr7_109740944_109742195 | chr7 | 109740945 | 109742195 | + | -0.811256 | intron (NM_020050, intron 4 of 4) | B4\|SINE\|B4 | -9838 | NM_020051 | 56787 | Mm.29275 | NM_020051 | ENSMUSG00000035951 | **Ascl3** | Sgn1\|bHLHa42 | achaete-scute complex homolog 3 (Drosophila) | protein-coding |
| chr6_86593120_86593898 | chr6 | 86593121 | 86593898 | + | -0.649004 | Intergenic | L1M2\|LINE\|L1 | -34665 | NM_026414 | 67855 | Mm.183043 | NM_026414 | ENSMUSG00000033508 | **Asprv1** | 2300003P22Rik\|AA986851\|SASP\|SASPase\|Taps | aspartic peptidase, retroviral-like 1 | protein-coding |
| chr6_86607413_86608043 | chr6 | 86607414 | 86608043 | + | -0.659433 | Intergenic | Intergenic | -20446 | NM_026414 | 67855 | Mm.183043 | NM_026414 | ENSMUSG00000033508 | **Asprv1** | 2300003P22Rik\|AA986851\|SASP\|SASPase\|Taps | aspartic peptidase, retroviral-like 1 | protein-coding |
| chr12_3441342_3441748 | chr12 | 3441343 | 3441748 | + | -0.772419 | intron (NM_001270988, intron 1 of 11) | L1MD2\|LINE\|L1 | 14688 | NM_172421 | 75302 | Mm.120475 | NM_172421 | ENSMUSG00000037486 | **Asxl2** | 4930556B16Rik\|mKIAA1685 | additional sex combs like 2 (Drosophila) | protein-coding |
| chr10_81194063_81194682 | chr10 | 81194064 | 81194682 | + | -0.716137 | promoter-TSS (NR_015477) | promoter-TSS (NR_015477) | -318 | NR_015477 | 69678 | Mm.81562 | NR_015477 | | **Atcayos** | 2310050B05Rik | ataxia, cerebellar, Cayman type homolog (human), opposite strand | ncRNA |
| chr1_170862967_170863416 | chr1 | 170862968 | 170863416 | + | -0.732546 | intron (NM_001081304, intron 1 of 15) | L2\|LINE\|L2 | 4579 | NM_001081304 | 226641 | Mm.377046 | NM_001081304 | ENSMUSG00000026663 | **Atf6** | 9130025P16Rik\|9630036G24\|AA789574\|Atf6alpha\|ESTM49 | activating transcription factor 6 | protein-coding |
| chr6_136516682_136517088 | chr6 | 136516683 | 136517088 | + | -0.719005 | Intergenic | MTC-int\|LTR\|ERVL-MaLR | -1966 | NM_019426 | 54343 | Mm.173271 | NM_019426 | ENSMUSG00000030213 | **Atf7ip** | 2610204M12Rik\|AM\|Mcaf1 | activating transcription factor 7 interacting protein | protein-coding |
| chr6_136491077_136491982 | chr6 | 136491078 | 136491982 | + | -0.825214 | Intergenic | Intergenic | -27321 | NM_019426 | 54343 | Mm.173271 | NM_019426 | ENSMUSG00000030213 | **Atf7ip** | 2610204M12Rik\|AM\|Mcaf1 | activating transcription factor 7 interacting protein | protein-coding |
| chr6_136531825_136532278 | chr6 | 136531826 | 136532278 | + | -0.844396 | intron (NM_019426, intron 1 of 14) | intron (NM_019426, intron 1 of 14) | 13201 | NM_019426 | 54343 | Mm.173271 | NM_019426 | ENSMUSG00000030213 | **Atf7ip** | 2610204M12Rik\|AM\|Mcaf1 | activating transcription factor 7 interacting protein | protein-coding |
| chr6_136490991_136491498 | chr6 | 136490992 | 136491498 | + | -0.935221 | Intergenic | Intergenic | -27606 | NM_019426 | 54343 | Mm.173271 | NM_019426 | ENSMUSG00000030213 | **Atf7ip** | 2610204M12Rik\|AM\|Mcaf1 | activating transcription factor 7 interacting protein | protein-coding |
| chr6_136491171_136491430 | chr6 | 136491172 | 136491430 | + | -0.95354 | Intergenic | Intergenic | -27550 | NM_019426 | 54343 | Mm.173271 | NM_019426 | ENSMUSG00000030213 | **Atf7ip** | 2610204M12Rik\|AM\|Mcaf1 | activating transcription factor 7 interacting protein | protein-coding |
| chr19_6242164_6242678 | chr19 | 6242165 | 6242678 | + | -0.65327 | intron (NM_194348, intron 1 of 39) | intron (NM_194348, intron 1 of 39) | 753 | NM_194348 | 329015 | Mm.277384 | NM_194348 | ENSMUSG00000024773 | **Atg2a** | 1810013C15Rik\|A830054M12\|BC023754\|mKIAA0404 | autophagy related 2A | protein-coding |
| chr19_6242331_6242621 | chr19 | 6242332 | 6242621 | + | -0.67864 | intron (NM_194348, intron 1 of 39) | intron (NM_194348, intron 1 of 39) | 808 | NM_194348 | 329015 | Mm.277384 | NM_194348 | ENSMUSG00000024773 | **Atg2a** | 1810013C15Rik\|A830054M12\|BC023754\|mKIAA0404 | autophagy related 2A | protein-coding |
| chr9_21260718_21260993 | chr9 | 21260719 | 21260993 | + | -0.839265 | Intergenic | B3A\|SINE\|B2 | -4429 | NM_153583 | 235040 | Mm.440165 | NM_153583 | ENSMUSG00000002820 | **Atg4d** | 9830134P12Rik\|APG4-D\|Apg4d\|Atg4dl\|Autl4 | autophagy related 4D, cysteine peptidase | protein-coding |
| chr17_79916851_79918052 | chr17 | 79916852 | 79918052 | + | -0.687814 | Intergenic | Intergenic | -21329 | NM_001286647 | 56298 | Mm.175403 | NM_019717 | ENSMUSG00000059811 | **Atl2** | 2010110I21Rik\|AA407293\|AV334690\|Aip-2\|Arl6ip2 | atlastin GTPase 2 | protein-coding |
| chr17_79916842_79917504 | chr17 | 79916843 | 79917504 | + | -0.773432 | Intergenic | L2b\|LINE\|L2 | -21050 | NM_001286647 | 56298 | Mm.175403 | NM_019717 | ENSMUSG00000059811 | **Atl2** | 2010110I21Rik\|AA407293\|AV334690\|Aip-2\|Arl6ip2 | atlastin GTPase 2 | protein-coding |
| chr6_124750417_124750823 | chr6 | 124750418 | 124750823 | + | -0.855321 | intron (NM_007881, intron 1 of 9) | intron (NM_007881, intron 1 of 9) | 5867 | NM_007881 | 13498 | Mm.333380 | NM_007881 | ENSMUSG00000004263 | **Atn1** | Atr1\|Drpla\|atrophin-1 | atrophin 1 | protein-coding |
| chr10_63089186_63089438 | chr10 | 63089187 | 63089438 | + | -0.689345 | Intergenic | Intergenic | -10844 | NM_016864 | 53404 | Mm.228661 | NM_016864 | ENSMUSG00000036816 | **Atoh7** | Math5\|bHLHa13 | atonal homolog 7 (Drosophila) | protein-coding |
| chr6_72190077_72190496 | chr6 | 72190078 | 72190496 | + | -0.813993 | Intergenic | Intergenic | 45290 | NM_153778 | 71093 | Mm.87449 | NM_153778 | ENSMUSG00000037621 | **Atoh8** | 4933425C05Rik\|Hath6\|Math6\|bHLHa21\|okadin | atonal homolog 8 (Drosophila) | protein-coding |
| chr7_58568057_58569169 | chr7 | 58568058 | 58569169 | + | -0.784974 | Intergenic | Intergenic | -89589 | NM_009728 | 11982 | Mm.135129 | NM_009728 | ENSMUSG00000025324 | **Atp10a** | Atp10c\|pfatp | ATPase, class V, type 10A | protein-coding |
| chr7_58697099_58697460 | chr7 | 58697100 | 58697460 | + | -0.843956 | intron (NM_009728, intron 1 of 20) | intron (NM_009728, intron 1 of 20) | 39078 | NM_009728 | 11982 | Mm.135129 | NM_009728 | ENSMUSG00000025324 | **Atp10a** | Atp10c\|pfatp | ATPase, class V, type 10A | protein-coding |

| | | | | | | | | | | | | | | | | | |
|---|---|---|---|---|---|---|---|---|---|---|---|---|---|---|---|---|---|
| chr7_58568423_58569155 | chr7 | 58568424 | 58569155 | + | -0.846387 | Intergenic | Intergenic | -89413 | NM_009728 | 11982 | Mm.135129 | NM_009728 | ENSMUSG00000025324 | **Atp10a** | Atp10c\|pfatp | ATPase, class V, type 10A | protein-coding |
| chr7_58448558_58449583 | chr7 | 58448559 | 58449583 | + | -0.901668 | Intergenic | RLTR14_RN\|LTR\|ERV1 | -209131 | NM_009728 | 11982 | Mm.135129 | NM_009728 | ENSMUSG00000025324 | **Atp10a** | Atp10c\|pfatp | ATPase, class V, type 10A | protein-coding |
| chr7_58568659_58569155 | chr7 | 58568660 | 58569155 | + | -0.937119 | Intergenic | (CAAAA)n\|Simple_repeat\|Simple_repeat | -89295 | NM_009728 | 11982 | Mm.135129 | NM_009728 | ENSMUSG00000025324 | **Atp10a** | Atp10c\|pfatp | ATPase, class V, type 10A | protein-coding |
| chr7_58730902_58731613 | chr7 | 58730903 | 58731613 | + | -1.05981 | intron (NM_009728, intron 1 of 20) | intron (NM_009728, intron 1 of 20) | 73056 | NM_009728 | 11982 | Mm.135129 | NM_009728 | ENSMUSG00000025324 | **Atp10a** | Atp10c\|pfatp | ATPase, class V, type 10A | protein-coding |
| chr7_58730884_58731400 | chr7 | 58730885 | 58731400 | + | -1.06433 | intron (NM_009728, intron 1 of 20) | intron (NM_009728, intron 1 of 20) | 72940 | NM_009728 | 11982 | Mm.135129 | NM_009728 | ENSMUSG00000025324 | **Atp10a** | Atp10c\|pfatp | ATPase, class V, type 10A | protein-coding |
| chr7_58451959_58452345 | chr7 | 58451960 | 58452345 | + | -1.11551 | Intergenic | L1MA4\|LINE\|L1 | -206050 | NM_009728 | 11982 | Mm.135129 | NM_009728 | ENSMUSG00000025324 | **Atp10a** | Atp10c\|pfatp | ATPase, class V, type 10A | protein-coding |
| chr7_58693421_58693688 | chr7 | 58693422 | 58693688 | + | -1.17435 | intron (NM_009728, intron 1 of 20) | L2a\|LINE\|L2 | 35353 | NM_009728 | 11982 | Mm.135129 | NM_009728 | ENSMUSG00000025324 | **Atp10a** | Atp10c\|pfatp | ATPase, class V, type 10A | protein-coding |
| chr3_35723605_35723976 | chr3 | 35723606 | 35723976 | + | -0.655891 | Intergenic | RLTR9A\|LTR\|ERVK | -30347 | NM_029570 | 76295 | Mm.239416 | NM_029570 | ENSMUSG00000037400 | **Atp11b** | 1110019I14Rik\|ATPIF\|ATPIR\|mKIAA0956 | ATPase, class VI, type 11B | protein-coding |
| chr3_101701019_101701599 | chr3 | 101701020 | 101701599 | + | -0.634196 | Intergenic | MIRc\|SINE\|MIR | -96602 | NM_144900 | 11928 | Mm.280103 | NM_144900 | ENSMUSG00000033161 | **Atp1a1** | Atpa-1\|BC010319 | ATPase, Na+/K+ transporting, alpha 1 polypeptide | protein-coding |
| chr3_101673728_101674199 | chr3 | 101673729 | 101674199 | + | -0.651836 | Intergenic | Intergenic | -69257 | NM_144900 | 11928 | Mm.280103 | NM_144900 | ENSMUSG00000033161 | **Atp1a1** | Atpa-1\|BC010319 | ATPase, Na+/K+ transporting, alpha 1 polypeptide | protein-coding |
| chr3_101673737_101674070 | chr3 | 101673738 | 101674070 | + | -0.750388 | Intergenic | Intergenic | -69197 | NM_144900 | 11928 | Mm.280103 | NM_144900 | ENSMUSG00000033161 | **Atp1a1** | Atpa-1\|BC010319 | ATPase, Na+/K+ transporting, alpha 1 polypeptide | protein-coding |
| chr3_101701293_101701779 | chr3 | 101701294 | 101701779 | + | -0.758138 | Intergenic | Intergenic | -96829 | NM_144900 | 11928 | Mm.280103 | NM_144900 | ENSMUSG00000033161 | **Atp1a1** | Atpa-1\|BC010319 | ATPase, Na+/K+ transporting, alpha 1 polypeptide | protein-coding |
| chr1_164413597_164414157 | chr1 | 164413598 | 164414157 | + | -0.642956 | intron (NM_138314, intron 13 of 13) | intron (NM_138314, intron 13 of 13) | 44478 | NM_009721 | 11931 | Mm.4550 | NM_009721 | ENSMUSG00000026576 | **Atp1b1** | Atp4b\|Atpb\|Atpb-1\|NKbeta1 | ATPase, Na+/K+ transporting, beta 1 polypeptide | protein-coding |
| chr1_164600716_164600958 | chr1 | 164600717 | 164600958 | + | -0.817986 | Intergenic | ORR1F\|LTR\|ERVL-MaLR | -142482 | NM_009721 | 11931 | Mm.4550 | NM_009721 | ENSMUSG00000026576 | **Atp1b1** | Atp4b\|Atpb\|Atpb-1\|NKbeta1 | ATPase, Na+/K+ transporting, beta 1 polypeptide | protein-coding |
| chr1_164453711_164454116 | chr1 | 164453712 | 164454116 | + | -0.837095 | intron (NM_009721, intron 1 of 5) | intron (NM_009721, intron 1 of 5) | 4441 | NM_009721 | 11931 | Mm.4550 | NM_009721 | ENSMUSG00000026576 | **Atp1b1** | Atp4b\|Atpb\|Atpb-1\|NKbeta1 | ATPase, Na+/K+ transporting, beta 1 polypeptide | protein-coding |
| chr1_164600652_164601064 | chr1 | 164600653 | 164601064 | + | -0.84359 | Intergenic | ORR1F\|LTR\|ERVL-MaLR | -142503 | NM_009721 | 11931 | Mm.4550 | NM_009721 | ENSMUSG00000026576 | **Atp1b1** | Atp4b\|Atpb\|Atpb-1\|NKbeta1 | ATPase, Na+/K+ transporting, beta 1 polypeptide | protein-coding |
| chr1_164494771_164495076 | chr1 | 164494772 | 164495076 | + | -0.98129 | Intergenic | Intergenic | -36569 | NM_009721 | 11931 | Mm.4550 | NM_009721 | ENSMUSG00000026576 | **Atp1b1** | Atp4b\|Atpb\|Atpb-1\|NKbeta1 | ATPase, Na+/K+ transporting, beta 1 polypeptide | protein-coding |
| chr9_96353241_96353723 | chr9 | 96353242 | 96353723 | + | -0.627097 | intron (NM_007502, intron 1 of 6) | intron (NM_007502, intron 1 of 6) | 10817 | NM_007502 | 11933 | Mm.424 | NM_007502 | ENSMUSG00000032412 | **Atp1b3** | AA409958\|AI664000\|AW212096 | ATPase, Na+/K+ transporting, beta 3 polypeptide | protein-coding |
| chr9_96399310_96399901 | chr9 | 96399311 | 96399901 | + | -0.68111 | Intergenic | Intergenic | -35307 | NM_007502 | 11933 | Mm.424 | NM_007502 | ENSMUSG00000032412 | **Atp1b3** | AA409958\|AI664000\|AW212096 | ATPase, Na+/K+ transporting, beta 3 polypeptide | protein-coding |
| chr9_96347842_96348180 | chr9 | 96347843 | 96348180 | + | -0.732443 | intron (NM_007502, intron 1 of 6) | intron (NM_007502, intron 1 of 6) | 16288 | NM_007502 | 11933 | Mm.424 | NM_007502 | ENSMUSG00000032412 | **Atp1b3** | AA409958\|AI664000\|AW212096 | ATPase, Na+/K+ transporting, beta 3 polypeptide | protein-coding |
| chr9_96369550_96370300 | chr9 | 96369551 | 96370300 | + | -0.75982 | Intergenic | Lx8\|LINE\|L1 | -5626 | NM_007502 | 11933 | Mm.424 | NM_007502 | ENSMUSG00000032412 | **Atp1b3** | AA409958\|AI664000\|AW212096 | ATPase, Na+/K+ transporting, beta 3 polypeptide | protein-coding |
| chr9_96393896_96394142 | chr9 | 96393897 | 96394142 | + | -0.850706 | Intergenic | Lx8\|LINE\|L1 | -29720 | NM_007502 | 11933 | Mm.424 | NM_007502 | ENSMUSG00000032412 | **Atp1b3** | AA409958\|AI664000\|AW212096 | ATPase, Na+/K+ transporting, beta 3 polypeptide | protein-coding |
| chr9_96394784_96395284 | chr9 | 96394785 | 96395284 | + | -0.870512 | Intergenic | Intergenic | -30735 | NM_007502 | 11933 | Mm.424 | NM_007502 | ENSMUSG00000032412 | **Atp1b3** | AA409958\|AI664000\|AW212096 | ATPase, Na+/K+ transporting, beta 3 polypeptide | protein-coding |
| chr9_96350407_96350896 | chr9 | 96350408 | 96350896 | + | -1.0006 | intron (NM_007502, intron 1 of 6) | intron (NM_007502, intron 1 of 6) | 13647 | NM_007502 | 11933 | Mm.424 | NM_007502 | ENSMUSG00000032412 | **Atp1b3** | AA409958\|AI664000\|AW212096 | ATPase, Na+/K+ transporting, beta 3 polypeptide | protein-coding |
| chr9_96350554_96350879 | chr9 | 96350555 | 96350879 | + | -1.03195 | intron (NM_007502, intron 1 of 6) | intron (NM_007502, intron 1 of 6) | 13582 | NM_007502 | 11933 | Mm.424 | NM_007502 | ENSMUSG00000032412 | **Atp1b3** | AA409958\|AI664000\|AW212096 | ATPase, Na+/K+ transporting, beta 3 polypeptide | protein-coding |
| chr10_98781754_98782382 | chr10 | 98781755 | 98782382 | + | -0.789261 | Intergenic | Intergenic | -133084 | NM_026482 | 67972 | Mm.166944 | NM_026482 | ENSMUSG00000019943 | **Atp2b1** | 2810442I22Rik\|E130111D10Rik\|Pmca1 | ATPase, Ca++ transporting, plasma membrane 1 | protein-coding |
| chr10_98774829_98775200 | chr10 | 98774830 | 98775200 | + | -0.865897 | Intergenic | Intergenic | -140137 | NM_026482 | 67972 | Mm.166944 | NM_026482 | ENSMUSG00000019943 | **Atp2b1** | 2810442I22Rik\|E130111D10Rik\|Pmca1 | ATPase, Ca++ transporting, plasma membrane 1 | protein-coding |
| chr10_98946083_98946414 | chr10 | 98946084 | 98946414 | + | -0.907359 | intron (NM_026482, intron 1 of 20) | intron (NM_026482, intron 1 of 20) | 31097 | NM_026482 | 67972 | Mm.166944 | NM_026482 | ENSMUSG00000019943 | **Atp2b1** | 2810442I22Rik\|E130111D10Rik\|Pmca1 | ATPase, Ca++ transporting, plasma membrane 1 | protein-coding |
| chr10_98785450_98785708 | chr10 | 98785451 | 98785708 | + | -1.26728 | Intergenic | Intergenic | -129573 | NM_026482 | 67972 | Mm.166944 | NM_026482 | ENSMUSG00000019943 | **Atp2b1** | 2810442I22Rik\|E130111D10Rik\|Pmca1 | ATPase, Ca++ transporting, plasma membrane 1 | protein-coding |
| chr10_98785388_98785772 | chr10 | 98785389 | 98785772 | + | -1.29725 | Intergenic | Intergenic | -129572 | NM_026482 | 67972 | Mm.166944 | NM_026482 | ENSMUSG00000019943 | **Atp2b1** | 2810442I22Rik\|E130111D10Rik\|Pmca1 | ATPase, Ca++ transporting, plasma membrane 1 | protein-coding |
| chr1_133762852_133763673 | chr1 | 133762853 | 133763673 | + | -0.687652 | Intergenic | Intergenic | -9516 | NM_213616 | 381290 | Mm.188617 | NM_213616 | ENSMUSG00000026463 | **Atp2b4** | Pmca4 | ATPase, Ca++ transporting, plasma membrane 4 | protein-coding |
| chr1_133762997_133763604 | chr1 | 133762998 | 133763604 | + | -0.719381 | Intergenic | Intergenic | -9554 | NM_213616 | 381290 | Mm.188617 | NM_213616 | ENSMUSG00000026463 | **Atp2b4** | Pmca4 | ATPase, Ca++ transporting, plasma membrane 4 | protein-coding |
| chrX_12577900_12578268 | chrX | 12577901 | 12578268 | + | -0.817666 | Intergenic | Intergenic | -9675 | NM_027439 | 70495 | Mm.25148 | NM_027439 | ENSMUSG00000031007 | **Atp6ap2** | (P)RR\|5730403E06Rik\|APT6M8-9\|ATP6M8-9\|Atp6ip2\|M8-9 | ATPase, H+ transporting, lysosomal accessory protein 2 | protein-coding |
| chrX_12583356_12583586 | chrX | 12583357 | 12583586 | + | -1.06018 | Intergenic | Intergenic | -4288 | NM_027439 | 70495 | Mm.25148 | NM_027439 | ENSMUSG00000031007 | **Atp6ap2** | (P)RR\|5730403E06Rik\|APT6M8-9\|ATP6M8-9\|Atp6ip2\|M8-9 | ATPase, H+ transporting, lysosomal accessory protein 2 | protein-coding |
| chr11_101008132_101008479 | chr11 | 101008133 | 101008479 | + | -0.704405 | Intergenic | Intergenic | -1146 | NM_016920 | 11975 | Mm.340818 | NM_016920 | ENSMUSG00000019302 | **Atp6v0a1** | AA959968\|ATP6a1\|Atp6n1\|Atp6n1a\|Atpv0a1\|Vpp-1\|Vpp1 | ATPase, H+ transporting, lysosomal V0 subunit A1 | protein-coding |
| chr5_124614583_124615136 | chr5 | 124614584 | 124615136 | + | -0.864831 | intron (NM_026486, intron 9 of 17) | intron (NM_026486, intron 9 of 17) | -14192 | NM_011596 | 21871 | Mm.1158 | NM_011596 | | **Atp6v0a2** | 8430408C20Rik\|AI385560\|ATP6a2\|AW489264\|Atp6n1d\|Atp6n2\|C76904\|ISF\|J6B7\|SHIF\|Stv1\|TJ6M\|TJ6s\|Tj6\|V-ATPase 116 kDa\|V-ATPase a2 | ATPase, H+ transporting, lysosomal V0 subunit A2 | protein-coding |
| chr5_124633514_124633786 | chr5 | 124633515 | 124633786 | + | -1.01997 | intron (NM_011596, intron 2 of 19) | intron (NM_011596, intron 2 of 19) | 4598 | NM_011596 | 21871 | Mm.1158 | NM_011596 | | **Atp6v0a2** | 8430408C20Rik\|AI385560\|ATP6a2\|AW489264\|Atp6n1d\|Atp6n2\|C76904\|ISF\|J6B7\|SHIF\|Stv1\|TJ6M\|TJ6s\|Tj6\|V-ATPase 116 kDa\|V-ATPase a2 | ATPase, H+ transporting, lysosomal V0 subunit A2 | protein-coding |
| chr6_38120601_38121101 | chr6 | 38120602 | 38121101 | + | -1.03304 | intron (NM_080467, intron 1 of 20) | intron (NM_080467, intron 1 of 20) | 3735 | NM_080467 | 140494 | Mm.462308 | NM_080467 | ENSMUSG00000038600 | **Atp6v0a4** | Atp6n1b\|a4 | ATPase, H+ transporting, lysosomal V0 subunit A4 | protein-coding |
| chr4_19950363_19950594 | chr4 | 19950364 | 19950594 | + | -0.615297 | Intergenic | B4\|SINE\|B4 | -27913 | NM_175406 | 242341 | Mm.19298 | NM_175406 | ENSMUSG00000028238 | **Atp6v0d2** | 1620401A02Rik\|AI324824\|V-ATPase | ATPase, H+ transporting, lysosomal V0 subunit D2 | protein-coding |
| chr4_19926834_19927319 | chr4 | 19926835 | 19927319 | + | -0.688583 | Intergenic | Intergenic | -4511 | NM_175406 | 242341 | Mm.19298 | NM_175406 | ENSMUSG00000028238 | **Atp6v0d2** | 1620401A02Rik\|AI324824\|V-ATPase | ATPase, H+ transporting, lysosomal V0 subunit D2 | protein-coding |
| chr4_19923268_19923699 | chr4 | 19923269 | 19923699 | + | -1.03238 | promoter-TSS (NM_175406) | promoter-TSS (NM_175406) | -918 | NM_175406 | 242341 | Mm.19298 | NM_175406 | ENSMUSG00000028238 | **Atp6v0d2** | 1620401A02Rik\|AI324824\|V-ATPase | ATPase, H+ transporting, lysosomal V0 subunit D2 | protein-coding |
| chr4_19932046_19932380 | chr4 | 19932047 | 19932380 | + | -1.1071 | Intergenic | Intergenic | -9647 | NM_175406 | 242341 | Mm.19298 | NM_175406 | ENSMUSG00000028238 | **Atp6v0d2** | 1620401A02Rik\|AI324824\|V-ATPase | ATPase, H+ transporting, lysosomal V0 subunit D2 | protein-coding |
| chr4_19928272_19928588 | chr4 | 19928273 | 19928588 | + | -1.2624 | Intergenic | Intergenic | -5864 | NM_175406 | 242341 | Mm.19298 | NM_175406 | ENSMUSG00000028238 | **Atp6v0d2** | 1620401A02Rik\|AI324824\|V-ATPase | ATPase, H+ transporting, lysosomal V0 subunit D2 | protein-coding |
| chr17_26630834_26631264 | chr17 | 26630835 | 26631264 | + | -1.16628 | intron (NM_026170, intron 5 of 9) | intron (NM_026170, intron 5 of 9) | -45347 | NM_025272 | 11974 | Mm.22602 | NM_025272 | ENSMUSG00000015575 | **Atp6v0e** | Atp6k\|Atp6v0e1\|M9.2 | ATPase, H+ transporting, lysosomal V0 subunit E | protein-coding |
| chr1_138232926_138233423 | chr1 | 138232927 | 138233423 | + | -0.845269 | Intergenic | Intergenic | -40563 | NM_177397 | 338375 | Mm.114424 | NM_177397 | ENSMUSG00000026394 | **Atp6v1g3** | - | ATPase, H+ transporting, lysosomal V1 subunit G3 | protein-coding |
| chr1_138233068_138233468 | chr1 | 138233069 | 138233468 | + | -0.861504 | Intergenic | Intergenic | -40470 | NM_177397 | 338375 | Mm.114424 | NM_177397 | ENSMUSG00000026394 | **Atp6v1g3** | - | ATPase, H+ transporting, lysosomal V1 subunit G3 | protein-coding |
| chr1_138251040_138251313 | chr1 | 138251041 | 138251313 | + | -1.00905 | Intergenic | Intergenic | -22561 | NM_177397 | 338375 | Mm.114424 | NM_177397 | ENSMUSG00000026394 | **Atp6v1g3** | - | ATPase, H+ transporting, lysosomal V1 subunit G3 | protein-coding |
| chr1_138250937_138251313 | chr1 | 138250938 | 138251313 | + | -1.04901 | Intergenic | Intergenic | -22613 | NM_177397 | 338375 | Mm.114424 | NM_177397 | ENSMUSG00000026394 | **Atp6v1g3** | - | ATPase, H+ transporting, lysosomal V1 subunit G3 | protein-coding |
| chr1_138303949_138304947 | chr1 | 138303950 | 138304947 | + | -1.05479 | Intergenic | Intergenic | 30710 | NM_177397 | 338375 | Mm.114424 | NM_177397 | ENSMUSG00000026394 | **Atp6v1g3** | - | ATPase, H+ transporting, lysosomal V1 subunit G3 | protein-coding |

| | | | | | | | | | | | | | | | | | |
|---|---|---|---|---|---|---|---|---|---|---|---|---|---|---|---|---|---|
| chr1_138248010_138248421 | chr1 | 138248011 | 138248421 | + | -1.05661 | Intergenic | (GA)n\|Simple_repeat\|Simple_repeat | -25522 | NM_177397 | 338375 | Mm.114424 | NM_177397 | ENSMUSG00000026394 | **Atp6v1g3** | - | ATPase, H+ transporting, lysosomal V1 subunit G3 | protein-coding |
| chr1_138248135_138248435 | chr1 | 138248136 | 138248435 | + | -1.09832 | Intergenic | Intergenic | -25453 | NM_177397 | 338375 | Mm.114424 | NM_177397 | ENSMUSG00000026394 | **Atp6v1g3** | - | ATPase, H+ transporting, lysosomal V1 subunit G3 | protein-coding |
| chr5_67866138_67866575 | chr5 | 67866139 | 67866575 | + | -0.686905 | Intergenic | MTE-int\|LTR\|ERVL-MaLR | -18926 | NM_009727 | 11980 | Mm.153230 | NM_009727 | ENSMUSG00000037685 | **Atp8a1** | AI481521\|AI853962\|APLT\|AW743152\|AW822227\|Atp3a2\|B230107D19Rik\|ClassI | ATPase, aminophospholipid transporter (APLT), class I, type 8A, member 1 | protein-coding |
| chr14_60131350_60131933 | chr14 | 60131351 | 60131933 | + | -0.636262 | Intergenic | Intergenic | -44808 | NM_015803 | 50769 | Mm.319599 | NM_015803 | ENSMUSG00000021983 | **Atp8a2** | AI415030\|Atpc1b\|wl | ATPase, aminophospholipid transporter-like, class I, type 8A, member 2 | protein-coding |
| chr14_60129270_60130049 | chr14 | 60129271 | 60130049 | + | -0.816687 | Intergenic | Intergenic | -42826 | NM_015803 | 50769 | Mm.319599 | NM_015803 | ENSMUSG00000021983 | **Atp8a2** | AI415030\|Atpc1b\|wl | ATPase, aminophospholipid transporter-like, class I, type 8A, member 2 | protein-coding |
| chr14_60129537_60129877 | chr14 | 60129538 | 60129877 | + | -0.817539 | Intergenic | Intergenic | -42873 | NM_015803 | 50769 | Mm.319599 | NM_015803 | ENSMUSG00000021983 | **Atp8a2** | AI415030\|Atpc1b\|wl | ATPase, aminophospholipid transporter-like, class I, type 8A, member 2 | protein-coding |
| chr14_60129270_60130345 | chr14 | 60129271 | 60130345 | + | -0.822615 | Intergenic | Intergenic | -42974 | NM_015803 | 50769 | Mm.319599 | NM_015803 | ENSMUSG00000021983 | **Atp8a2** | AI415030\|Atpc1b\|wl | ATPase, aminophospholipid transporter-like, class I, type 8A, member 2 | protein-coding |
| chr14_60072398_60073002 | chr14 | 60072399 | 60073002 | + | -1.08425 | intron (NM_015803, intron 1 of 35) | intron (NM_015803, intron 1 of 35) | 14134 | NM_015803 | 50769 | Mm.319599 | NM_015803 | ENSMUSG00000021983 | **Atp8a2** | AI415030\|Atpc1b\|wl | ATPase, aminophospholipid transporter-like, class I, type 8A, member 2 | protein-coding |
| chr14_60117810_60118966 | chr14 | 60117811 | 60118966 | + | -1.30276 | Intergenic | Intergenic | -31554 | NM_015803 | 50769 | Mm.319599 | NM_015803 | ENSMUSG00000021983 | **Atp8a2** | AI415030\|Atpc1b\|wl | ATPase, aminophospholipid transporter-like, class I, type 8A, member 2 | protein-coding |
| chr2_126500565_126500815 | chr2 | 126500566 | 126500815 | + | -0.76807 | Intergenic | Intergenic | -9137 | NM_001080944 | 241633 | Mm.222235 | NM_001080944 | ENSMUSG00000060131 | **Atp8b4** | A530043E15\|Im | ATPase, class I, type 8B, member 4 | protein-coding |
| chr2_126495867_126496168 | chr2 | 126495868 | 126496168 | + | -0.805456 | Intergenic | Intergenic | -4465 | NM_001080944 | 241633 | Mm.222235 | NM_001080944 | ENSMUSG00000060131 | **Atp8b4** | A530043E15\|Im | ATPase, class I, type 8B, member 4 | protein-coding |
| chr2_126474018_126474747 | chr2 | 126474019 | 126474747 | + | -0.885891 | intron (NM_001080944, intron 3 of 27) | intron (NM_001080944, intron 3 of 27) | 17170 | NM_001080944 | 241633 | Mm.222235 | NM_001080944 | ENSMUSG00000060131 | **Atp8b4** | A530043E15\|Im | ATPase, class I, type 8B, member 4 | protein-coding |
| chr2_126474035_126474426 | chr2 | 126474036 | 126474426 | + | -1.17025 | intron (NM_001080944, intron 3 of 27) | intron (NM_001080944, intron 3 of 27) | 17322 | NM_001080944 | 241633 | Mm.222235 | NM_001080944 | ENSMUSG00000060131 | **Atp8b4** | A530043E15\|Im | ATPase, class I, type 8B, member 4 | protein-coding |
| chr2_130918373_130918710 | chr2 | 130918374 | 130918710 | + | -0.898023 | intron (NM_009730, intron 1 of 28) | ID_B1\|SINE\|B4 | 12046 | NM_009730 | 11990 | Mm.119936 | NM_009730 | ENSMUSG00000027312 | **Atrn** | AW558010\|Mgca\|mKIAA0548\|mahogany\|mg | attractin | protein-coding |
| chr19_57802774_57803532 | chr19 | 57802775 | 57803532 | + | -1.00272 | intron (NM_181415, intron 26 of 28) | intron (NM_181415, intron 26 of 28) | 192119 | NM_181415 | 226255 | Mm.245340 | NM_181415 | ENSMUSG00000054843 | **Atrnl1** | AI504415\|AW555641\|Alp\|Atrnl\|Atrnlp | attractin like 1 | protein-coding |
| chr12_33314887_33315607 | chr12 | 33314888 | 33315607 | + | -0.722865 | intron (NM_001033436, intron 1 of 7) | intron (NM_001033436, intron 1 of 7) | 12732 | NM_001033436 | 380753 | Mm.41540 | NM_028139 | ENSMUSG00000020564 | **Atxn7l1** | 2810423G08Rik\|Atxn7l4 | ataxin 7-like 1 | protein-coding |
| chr12_33315161_33315634 | chr12 | 33315162 | 33315634 | + | -0.783593 | intron (NM_001033436, intron 1 of 7) | intron (NM_001033436, intron 1 of 7) | 12883 | NM_001033436 | 380753 | Mm.41540 | NM_028139 | ENSMUSG00000020564 | **Atxn7l1** | 2810423G08Rik\|Atxn7l4 | ataxin 7-like 1 | protein-coding |
| chr11_102295244_102295607 | chr11 | 102295245 | 102295607 | + | -0.665988 | intron (NM_001098836, intron 1 of 12) | intron (NM_001098836, intron 1 of 12) | 1203 | NM_001098836 | 217218 | Mm.390381 | NM_001098836 | ENSMUSG00000059995 | **Atxn7l3** | E030022H21Rik | ataxin 7-like 3 | protein-coding |
| chrX_93905596_93905871 | chrX | 93905597 | 93905871 | + | -0.76484 | Intergenic | Intergenic | -62922 | NR_028320 | 385493 | Mm.328014 | NM_001033480 | ENSMUSG00000081044 | **AU015836** | - | expressed sequence AU015836 | ncRNA |
| chrX_93905549_93905930 | chrX | 93905550 | 93905930 | + | -0.782675 | Intergenic | Intergenic | -62916 | NR_028320 | 385493 | Mm.328014 | NM_001033480 | ENSMUSG00000081044 | **AU015836** | - | expressed sequence AU015836 | ncRNA |
| chr11_79460802_79461127 | chr11 | 79460803 | 79461127 | + | -0.968012 | intron (NM_010897, intron 30 of 57) | intron (NM_010897, intron 30 of 57) | 23066 | NR_045305 | 104522 | Mm.436987 | NR_045305 | ENSMUSG00000091523 | **AU040972** | AU040954 | expressed sequence AU040972 | ncRNA |
| chr11_79460793_79461357 | chr11 | 79460794 | 79461357 | + | -0.969462 | intron (NM_010897, intron 30 of 57) | intron (NM_010897, intron 30 of 57) | 22956 | NR_045305 | 104522 | Mm.436987 | NR_045305 | ENSMUSG00000091523 | **AU040972** | AU040954 | expressed sequence AU040972 | ncRNA |
| chr13_52921943_52922323 | chr13 | 52921944 | 52922323 | + | -0.950381 | intron (NM_016709, intron 1 of 9) | L1M4\|LINE\|L1 | 7544 | NM_016709 | 11992 | Mm.252034 | NM_016709 | ENSMUSG00000021460 | **Auh** | C77140\|W91705 | AU RNA binding protein/enoyl-coenzyme A hydratase | protein-coding |
| chr2_120887695_120887923 | chr2 | 120887696 | 120887923 | + | -0.720906 | intron (NM_009461, intron 37 of 46) | PB1D7\|SINE\|Alu | 37153 | NR_038350 | 100502700 | Mm.330516 | NR_038349 | ENSMUSG00000087519 | **AV039307** | - | expressed sequence AV039307 | ncRNA |
| chr2_112681815_112682109 | chr2 | 112681816 | 112682109 | + | -0.697688 | intron (NM_177652, intron 69 of 103) | intron (NM_177652, intron 69 of 103) | 122280 | NM_001165935 | 74268 | Mm.315125 | NM_028844 | | **Aven** | 1700013A01Rik\|1700056A21Rik\|mAven-L\|mAven-S | apoptosis, caspase activation inhibitor | protein-coding |
| chr2_112510132_112510536 | chr2 | 112510133 | 112510536 | + | -0.720609 | intron (NM_028844, intron 1 of 5) | PB1D10\|SINE\|Alu | 17370 | NM_028844 | 74268 | Mm.315125 | NM_028844 | | **Aven** | 1700013A01Rik\|1700056A21Rik\|mAven-L\|mAven-S | apoptosis, caspase activation inhibitor | protein-coding |
| chr8_57650273_57650531 | chr8 | 57650274 | 57650531 | + | -0.798528 | intron (NM_001167981, intron 1 of 11) | intron (NM_001167981, intron 1 of 11) | -1351 | NR_040698 | 100502619 | Mm.393851 | NR_040698 | | **AW046200** | - | expressed sequence AW046200 | ncRNA |
| chr19_11064845_11065801 | chr19 | 11064846 | 11065801 | + | -0.624563 | Intergenic | RMER1C\|Other\|Other | -14757 | NR_102366 | 107350 | Mm.6381 | NM_001177351 | ENSMUSG00000075010 | **AW112010** | - | expressed sequence AW112010 | ncRNA |
| chr18_25241644_25242000 | chr18 | 25241645 | 25242000 | + | -0.865746 | intron (NM_001033532, intron 5 of 10) | intron (NM_001033532, intron 5 of 10) | 72802 | NM_001033532 | 225289 | Mm.391081 | NM_001033532 | ENSMUSG00000033632 | **AW554918** | 4921538P22\|AI461742\|Kiaa1328\|mKIAA1328 | expressed sequence AW554918 | protein-coding |
| chr17_26167865_26168550 | chr17 | 26167866 | 26168550 | + | -0.957576 | intron (NM_001159598, intron 2 of 9) | ORR1B1\|LTR\|ERVL-MaLR | 29522 | NM_001159598 | 12005 | Mm.23684 | NM_009733 | ENSMUSG00000024182 | **Axin1** | AI316800\|Axin\|Fu\|Kb\|Ki\|fused\|kinky\|knobbly | axin 1 | protein-coding |
| chr11_108910671_108911326 | chr11 | 108910672 | 108911326 | + | -0.857849 | Intergenic | Intergenic | -9350 | NM_015732 | 12006 | Mm.71710 | NM_015732 | ENSMUSG00000000142 | **Axin2** | Axi1\|Axil\|Conductin | axin2 | protein-coding |
| chr6_95515746_95516113 | chr6 | 95515747 | 95516113 | + | -0.766161 | intron (NM_011507, intron 9 of 10) | intron (NM_011507, intron 9 of 10) | -181937 | NR_033559 | 414067 | Mm.355990 | NR_033559 | ENSMUSG00000068180 | **AY512915** | - | cDNA sequence AY512915 | ncRNA |
| chr6_95515557_95516174 | chr6 | 95515558 | 95516174 | + | -0.778441 | intron (NM_011507, intron 9 of 10) | intron (NM_011507, intron 9 of 10) | -181873 | NR_033559 | 414067 | Mm.355990 | NR_033559 | ENSMUSG00000068180 | **AY512915** | - | cDNA sequence AY512915 | ncRNA |
| chr19_57343496_57344302 | chr19 | 57343497 | 57344302 | + | -0.756137 | intron (NR_040316, intron 1 of 2) | intron (NR_040316, intron 1 of 2) | 17000 | NR_040316 | 320879 | Mm.375089 | NR_040316 | ENSMUSG00000097785 | **B230217O12Rik** | - | RIKEN cDNA B230217O12 gene | ncRNA |
| chr19_57355440_57356249 | chr19 | 57355441 | 57356249 | + | -0.80221 | intron (NR_040316, intron 1 of 2) | (TGG)n\|Simple_repeat\|Simple_repeat | 5054 | NR_040316 | 320879 | Mm.375089 | NR_040316 | ENSMUSG00000097785 | **B230217O12Rik** | - | RIKEN cDNA B230217O12 gene | ncRNA |
| chr2_68243338_68243730 | chr2 | 68243339 | 68243730 | + | -0.618664 | intron (NM_016866, intron 16 of 17) | intron (NM_016866, intron 16 of 17) | 138862 | NM_020283 | 26877 | Mm.226435 | NM_020283 | ENSMUSG00000034780 | **B3galt1** | beta3Gal-T1 | UDP-Gal:betaGlcNAc beta 1,3-galactosyltransferase, polypeptide 1 | protein-coding |
| chr17_33947680_33948154 | chr17 | 33947681 | 33948154 | + | -0.949967 | intron (NM_020603, intron 10 of 14) | intron (NM_020603, intron 10 of 14) | 3571 | NM_019420 | 54218 | Mm.11132 | NM_019420 | ENSMUSG00000067370 | **B3galt4** | Gal-T2\|Galt2 | UDP-Gal:betaGlcNAc beta 1,3-galactosyltransferase, polypeptide 4 | protein-coding |
| chr11_22853419_22854153 | chr11 | 22853420 | 22854153 | + | -0.657454 | intron (NM_001169114, intron 1 of 1) | intron (NM_001169114, intron 1 of 1) | 5949 | NM_016888 | 53625 | Mm.395495 | NM_016888 | ENSMUSG00000051650 | **B3gnt2** | AA408337\|AA409404\|AW260308\|B3Galt6\|B3gnt1\|BGnT-1\|BGnT-2\|BGnT1\|BGnT2\|Beta3gnt | UDP-GlcNAc:betaGal beta-1,3-N-acetylglucosaminyltransferase 2 | protein-coding |
| chr11_22853576_22854114 | chr11 | 22853577 | 22854114 | + | -0.764565 | intron (NM_001169114, intron 1 of 1) | intron (NM_001169114, intron 1 of 1) | 5890 | NM_016888 | 53625 | Mm.395495 | NM_016888 | ENSMUSG00000051650 | **B3gnt2** | AA408337\|AA409404\|AW260308\|B3Galt6\|B3gnt1\|BGnT-1\|BGnT-2\|BGnT1\|BGnT2\|Beta3gnt | UDP-GlcNAc:betaGal beta-1,3-N-acetylglucosaminyltransferase 2 | protein-coding |
| chr7_25636337_25636720 | chr7 | 25636338 | 25636720 | + | -0.810324 | intron (NM_007533, intron 4 of 8) | ID4_\|SINE\|ID | 8905 | NM_146184 | 232984 | Mm.192369 | NM_146184 | ENSMUSG00000059479 | **B3gnt8** | B3galt7\|B7galt7\|BC025206 | UDP-GlcNAc:betaGal beta-1,3-N-acetylglucosaminyltransferase 8 | protein-coding |
| chr17_48315953_48316385 | chr17 | 48315954 | 48316385 | + | -0.63786 | promoter-TSS (NM_177083) | promoter-TSS (NM_177083) | 7 | NM_177083 | 320148 | Mm.123455 | NM_177083 | ENSMUSG00000043740 | **B430306N03Rik** | Treml6 | RIKEN cDNA B430306N03 gene | protein-coding |
| chr17_48315972_48316224 | chr17 | 48315973 | 48316224 | + | -0.714647 | promoter-TSS (NM_177083) | promoter-TSS (NM_177083) | -64 | NM_177083 | 320148 | Mm.123455 | NM_177083 | ENSMUSG00000043740 | **B430306N03Rik** | Treml6 | RIKEN cDNA B430306N03 gene | protein-coding |
| chr7_141064138_141064484 | chr7 | 141064139 | 141064484 | + | -0.753295 | exon (NM_177897, exon 3 of 20) | exon (NM_177897, exon 3 of 20) | 3037 | NM_177897 | 330671 | Mm.133741 | NM_177897 | ENSMUSG00000055629 | **B4galnt4** | BC038881\|Gm1095\|beta4GalNAcT4 | beta-1,4-N-acetyl-galactosaminyl transferase 4 | protein-coding |
| chr4_40875615_40876306 | chr4 | 40875616 | 40876306 | + | -0.828245 | Intergenic | Intergenic | -21424 | NM_022305 | 14595 | Mm.15622 | NM_022305 | ENSMUSG00000028413 | **B4galt1** | B-1,4-GalT\|B-1,4-GalT1\|GalT\|Ggtb\|Ggtb2\|b4Gal-T1\|beta-1,4-GalT\|beta-1,4-GalT1\|beta4Gal-T1 | UDP-Gal:betaGlcNAc beta 1,4-galactosyltransferase, polypeptide 1 | protein-coding |
| chr1_171262495_171263057 | chr1 | 171262496 | 171263057 | + | -0.91135 | Intergenic | Intergenic | -7552 | NM_020579 | 57370 | Mm.274011 | NM_020579 | ENSMUSG00000052423 | **B4galt3** | 9530061M23Rik\|AA104562\|AW125175\|ESTM26\|ESTM6\|R74981 | UDP-Gal:betaGlcNAc beta 1,4-galactosyltransferase, polypeptide 3 | protein-coding |
| chr6_13650568_13651295 | chr6 | 13650569 | 13651295 | + | -1.20361 | intron (NM_175312, intron 2 of 4) | MER21B\|LTR\|ERVL | 27034 | NM_175312 | 101148 | Mm.354863 | NM_175312 | ENSMUSG00000042742 | **B630005N14Rik** | AI666701 | RIKEN cDNA B630005N14 gene | protein-coding |
| chr6_13650818_13651272 | chr6 | 13650819 | 13651272 | + | -1.24575 | intron (NM_175312, intron 2 of 4) | MER21B\|LTR\|ERVL | 26921 | NM_175312 | 101148 | Mm.354863 | NM_175312 | ENSMUSG00000042742 | **B630005N14Rik** | AI666701 | RIKEN cDNA B630005N14 gene | protein-coding |

| | | | | | | | | | | | | | | | | | |
|---|---|---|---|---|---|---|---|---|---|---|---|---|---|---|---|---|---|
| chr9_45851305_45851749 | chr9 | 45851306 | 45851749 | + | -0.625535 | intron (NM_001145947, intron 1 of 7) | ORR1F\|LTR\|ERVL-MaLR | 12998 | NM_011792 | 23821 | Mm.24044 | NM_011792 | ENSMUSG00000032086 | **Bace1** | C76936 | beta-site APP cleaving enzyme 1 | protein-coding |
| chr9_45853706_45854019 | chr9 | 45853707 | 45854019 | + | -1.05879 | intron (NM_001145947, intron 2 of 7) | intron (NM_001145947, intron 2 of 7) | 15334 | NM_011792 | 23821 | Mm.24044 | NM_011792 | ENSMUSG00000032086 | **Bace1** | C76936 | beta-site APP cleaving enzyme 1 | protein-coding |
| chr16_87709119_87709641 | chr16 | 87709120 | 87709641 | + | -0.711342 | intron (NM_007520, intron 1 of 4) | intron (NM_007520, intron 1 of 4) | 10426 | NM_007520 | 12013 | Mm.26147 | NM_007520 | ENSMUSG00000025612 | **Bach1** | 6230421P05Rik\|AI323795 | BTB and CNC homology 1 | protein-coding |
| chr16_87656797_87657160 | chr16 | 87656798 | 87657160 | + | -0.727905 | Intergenic | Intergenic | -41975 | NM_007520 | 12013 | Mm.26147 | NM_007520 | ENSMUSG00000025612 | **Bach1** | 6230421P05Rik\|AI323795 | BTB and CNC homology 1 | protein-coding |
| chr4_32387521_32388250 | chr4 | 32387522 | 32388250 | + | -0.632806 | Intergenic | Intergenic | -29549 | NM_001109661 | 12014 | Mm.491223 | NM_007521 | ENSMUSG00000040270 | **Bach2** | E030004N02Rik | BTB and CNC homology 2 | protein-coding |
| chr4_32387856_32388250 | chr4 | 32387857 | 32388250 | + | -0.665158 | Intergenic | (CA)n\|Simple_repeat\|Simple_repeat | -29382 | NM_001109661 | 12014 | Mm.491223 | NM_007521 | ENSMUSG00000040270 | **Bach2** | E030004N02Rik | BTB and CNC homology 2 | protein-coding |
| chr4_32351934_32352244 | chr4 | 32351935 | 32352244 | + | -0.666808 | Intergenic | MER119\|DNA\|hAT-Charlie | -65346 | NM_001109661 | 12014 | Mm.491223 | NM_007521 | ENSMUSG00000040270 | **Bach2** | E030004N02Rik | BTB and CNC homology 2 | protein-coding |
| chr4_32335479_32336086 | chr4 | 32335480 | 32336086 | + | -0.675412 | Intergenic | Intergenic | -81652 | NM_001109661 | 12014 | Mm.491223 | NM_007521 | ENSMUSG00000040270 | **Bach2** | E030004N02Rik | BTB and CNC homology 2 | protein-coding |
| chr4_32343650_32344700 | chr4 | 32343651 | 32344700 | + | -0.679589 | Intergenic | Intergenic | -73260 | NM_001109661 | 12014 | Mm.491223 | NM_007521 | ENSMUSG00000040270 | **Bach2** | E030004N02Rik | BTB and CNC homology 2 | protein-coding |
| chr4_32363349_32364219 | chr4 | 32363350 | 32364219 | + | -0.855806 | Intergenic | Intergenic | -53651 | NM_001109661 | 12014 | Mm.491223 | NM_007521 | ENSMUSG00000040270 | **Bach2** | E030004N02Rik | BTB and CNC homology 2 | protein-coding |
| chr4_32363583_32364191 | chr4 | 32363584 | 32364191 | + | -0.863012 | Intergenic | Intergenic | -53548 | NM_001109661 | 12014 | Mm.491223 | NM_007521 | ENSMUSG00000040270 | **Bach2** | E030004N02Rik | BTB and CNC homology 2 | protein-coding |
| chr4_32328587_32328957 | chr4 | 32328588 | 32328957 | + | -0.88677 | Intergenic | Intergenic | -88663 | NM_001109661 | 12014 | Mm.491223 | NM_007521 | ENSMUSG00000040270 | **Bach2** | E030004N02Rik | BTB and CNC homology 2 | protein-coding |
| chr4_32560270_32561189 | chr4 | 32560271 | 32561189 | + | -1.08128 | intron (NR_026843, intron 4 of 4) | intron (NR_026843, intron 4 of 4) | 10932 | NR_026843 | 414068 | Mm.389550 | NR_026843 | ENSMUSG00000086150 | **Bach2os** | - | BTB and CNC homology 2, opposite strand | ncRNA |
| chr4_32560258_32560772 | chr4 | 32560259 | 32560772 | + | -1.2023 | non-coding (NR_026843, exon 5 of 5) | non-coding (NR_026843, exon 5 of 5) | 11147 | NR_026843 | 414068 | Mm.389550 | NR_026843 | ENSMUSG00000086150 | **Bach2os** | - | BTB and CNC homology 2, opposite strand | ncRNA |
| chr2_118896216_118897104 | chr2 | 118896217 | 118897104 | + | -0.6063 | Intergenic | Intergenic | -4955 | NM_001045523 | 228536 | Mm.330487 | NM_001045523 | ENSMUSG00000040007 | **Bahd1** | AL022997\|AW541238\|Gm117\|mKIAA0945 | bromo adjacent homology domain containing 1 | protein-coding |
| chr11_119933789_119934508 | chr11 | 119933790 | 119934508 | + | -0.775802 | Intergenic | Intergenic | -8943 | NM_001037754 | 108100 | Mm.197534 | NM_130862 | ENSMUSG00000025372 | **Baiap2** | IRSp53\|R75030 | brain-specific angiogenesis inhibitor 1-associated protein 2 | protein-coding |
| chr2_122466889_122467193 | chr2 | 122466890 | 122467193 | + | -0.845323 | non-coding (NR_027919, exon 1 of 1) | non-coding (NR_027919, exon 1 of 1) | 756 | NR_027919 | 81913 | Mm.122640 | NR_027919 | | **Bambi-ps1** | - | BMP and activin membrane-bound inhibitor, pseudogene (Xenopus laevis) | pseudo |
| chr2_122469061_122469758 | chr2 | 122469062 | 122469758 | + | -0.916994 | Intergenic | RMER19B\|LTR\|ERVK | -1613 | NR_027919 | 81913 | Mm.122640 | NR_027919 | | **Bambi-ps1** | - | BMP and activin membrane-bound inhibitor, pseudogene (Xenopus laevis) | pseudo |
| chr1_71100087_71100421 | chr1 | 71100088 | 71100421 | + | -0.632894 | intron (NM_007525, intron 1 of 10) | intron (NM_007525, intron 1 of 10) | 2718 | NM_007525 | 12021 | Mm.10764 | NM_007525 | ENSMUSG00000026196 | **Bard1** | ENSMUSG00000060893\|ENSMUSG00000073653 | BRCA1 associated RING domain 1 | protein-coding |
| chr15_25216475_25216941 | chr15 | 25216476 | 25216941 | + | -0.612521 | Intergenic | Intergenic | 197056 | NM_027395 | 70350 | Mm.29586 | NM_027395 | ENSMUSG00000045763 | **Basp1** | 2610024P12Rik\|CAP-23\|CAP23\|Ckap3\|NAP-22\|NAP22 | brain abundant, membrane attached signal protein 1 | protein-coding |
| chr15_25216419_25216806 | chr15 | 25216420 | 25216806 | + | -0.665872 | Intergenic | Intergenic | 197151 | NM_027395 | 70350 | Mm.29586 | NM_027395 | ENSMUSG00000045763 | **Basp1** | 2610024P12Rik\|CAP-23\|CAP23\|Ckap3\|NAP-22\|NAP22 | brain abundant, membrane attached signal protein 1 | protein-coding |
| chr15_25227424_25228444 | chr15 | 25227425 | 25228444 | + | -0.698495 | Intergenic | Intergenic | 185830 | NM_027395 | 70350 | Mm.29586 | NM_027395 | ENSMUSG00000045763 | **Basp1** | 2610024P12Rik\|CAP-23\|CAP23\|Ckap3\|NAP-22\|NAP22 | brain abundant, membrane attached signal protein 1 | protein-coding |
| chr15_25405954_25406342 | chr15 | 25405955 | 25406342 | + | -0.812872 | intron (NM_027395, intron 1 of 1) | intron (NM_027395, intron 1 of 1) | 7616 | NM_027395 | 70350 | Mm.29586 | NM_027395 | ENSMUSG00000045763 | **Basp1** | 2610024P12Rik\|CAP-23\|CAP23\|Ckap3\|NAP-22\|NAP22 | brain abundant, membrane attached signal protein 1 | protein-coding |
| chr2_60133195_60133567 | chr2 | 60133196 | 60133567 | + | -0.621206 | Intergenic | ORR1E\|LTR\|ERVL-MaLR | -7641 | NM_001001182 | 407823 | Mm.228071 | NM_001001182 | ENSMUSG00000026987 | **Baz2b** | 5830435C13Rik\|BC053917\|D2Ertd794e | bromodomain adjacent to zinc finger domain, 2B | protein-coding |
| chr2_60018683_60019693 | chr2 | 60018684 | 60019693 | + | -0.705419 | intron (NM_001001182, intron 2 of 35) | intron (NM_001001182, intron 2 of 35) | 106552 | NM_001001182 | 407823 | Mm.228071 | NM_001001182 | ENSMUSG00000026987 | **Baz2b** | 5830435C13Rik\|BC053917\|D2Ertd794e | bromodomain adjacent to zinc finger domain, 2B | protein-coding |
| chr13_52800478_52801464 | chr13 | 52800479 | 52801464 | + | -0.841593 | Intergenic | Intergenic | -43766 | NR_027893 | 105404 | Mm.184173 | NR_027893 | | **BB123696** | - | expressed sequence BB123696 | ncRNA |
| chr9_22646480_22646789 | chr9 | 22646481 | 22646789 | + | -0.654192 | intron (NM_178415, intron 15 of 22) | intron (NM_178415, intron 15 of 22) | 170920 | NM_178415 | 319845 | Mm.176725 | NM_178415 | ENSMUSG00000035919 | **Bbs9** | E130103I17Rik | Bardet-Biedl syndrome 9 (human) | protein-coding |
| chr9_22672982_22673193 | chr9 | 22672983 | 22673193 | + | -0.744929 | intron (NM_178415, intron 18 of 22) | RSINE1\|SINE\|B4 | 197373 | NM_178415 | 319845 | Mm.176725 | NM_178415 | ENSMUSG00000035919 | **Bbs9** | E130103I17Rik | Bardet-Biedl syndrome 9 (human) | protein-coding |
| chr9_22657291_22657546 | chr9 | 22657292 | 22657546 | + | -1.11539 | intron (NM_178415, intron 16 of 22) | intron (NM_178415, intron 16 of 22) | 181704 | NM_178415 | 319845 | Mm.176725 | NM_178415 | ENSMUSG00000035919 | **Bbs9** | E130103I17Rik | Bardet-Biedl syndrome 9 (human) | protein-coding |
| chr16_50372461_50373092 | chr16 | 50372462 | 50373092 | + | -0.812217 | intron (NM_027444, intron 2 of 16) | intron (NM_027444, intron 2 of 16) | 59612 | NM_027444 | 70508 | Mm.28940 | NM_027444 | ENSMUSG00000022641 | **Bbx** | 5530401J07Rik\|5730403O13Rik | bobby sox homolog (Drosophila) | protein-coding |
| chr16_50315404_50315930 | chr16 | 50315405 | 50315930 | + | -0.832657 | intron (NM_027444, intron 3 of 16) | intron (NM_027444, intron 3 of 16) | 116722 | NM_027444 | 70508 | Mm.28940 | NM_027444 | ENSMUSG00000022641 | **Bbx** | 5530401J07Rik\|5730403O13Rik | bobby sox homolog (Drosophila) | protein-coding |
| chr1_150396130_150396364 | chr1 | 150396131 | 150396364 | + | -0.805914 | intron (NM_133780, intron 3 of 51) | intron (NM_133780, intron 3 of 51) | -3192 | NM_145511 | 226499 | Mm.222093 | NM_145511 | ENSMUSG00000006010 | **BC003331** | 1810053E15Rik\|Odr4\|odr-4 | cDNA sequence BC003331 | protein-coding |
| chr16_44691935_44692195 | chr16 | 44691936 | 44692195 | + | -0.70433 | Intergenic | Intergenic | -32236 | NM_145972 | 212547 | Mm.310573 | NM_145972 | ENSMUSG00000036208 | **BC027231** | Nepro | cDNA sequence BC027231 | protein-coding |
| chr16_38131548_38132170 | chr16 | 38131549 | 38132170 | + | -0.769002 | intron (NM_019827, intron 1 of 10) | intron (NM_019827, intron 1 of 10) | -42599 | NR_033221 | 414072 | Mm.483602 | NR_033221 | | **BC031361** | - | cDNA sequence BC031361 | ncRNA |
| chr15_4085465_4086598 | chr15 | 4085466 | 4086598 | + | -0.616498 | intron (NM_024188, intron 7 of 16) | MLTR25C\|LTR\|ERVK | -58626 | NR_028266 | 414066 | Mm.394461 | NR_028266 | | **BC037032** | - | cDNA Sequence BC037032 | ncRNA |
| chr15_4042601_4043969 | chr15 | 4042602 | 4043969 | + | -0.706506 | intron (NM_024188, intron 3 of 16) | intron (NM_024188, intron 3 of 16) | -15879 | NR_028266 | 414066 | Mm.394461 | NR_028266 | | **BC037032** | - | cDNA Sequence BC037032 | ncRNA |
| chr15_4085569_4085952 | chr15 | 4085570 | 4085952 | + | -0.742562 | intron (NM_024188, intron 7 of 16) | MLTR25C\|LTR\|ERVK | -58355 | NR_028266 | 414066 | Mm.394461 | NR_028266 | | **BC037032** | - | cDNA Sequence BC037032 | ncRNA |
| chr15_4124628_4125195 | chr15 | 4124629 | 4125195 | + | -1.12251 | intron (NM_024188, intron 13 of 16) | intron (NM_024188, intron 13 of 16) | -97506 | NR_028266 | 414066 | Mm.394461 | NR_028266 | | **BC037032** | - | cDNA Sequence BC037032 | ncRNA |
| chr10_121886618_121887321 | chr10 | 121886619 | 121887321 | + | -0.939118 | intron (NM_001081037, intron 4 of 21) | intron (NM_001081037, intron 4 of 21) | 147033 | NM_173022 | 270802 | Mm.209172 | NM_173022 | ENSMUSG00000053684 | **BC048403** | E030027L10 | cDNA sequence BC048403 | protein-coding |
| chr6_136831722_136832028 | chr6 | 136831723 | 136832028 | + | -0.627075 | 5' UTR (NM_001039558, exon 2 of 2) | 5' UTR (NM_001039558, exon 2 of 2) | 3032 | NM_178776 | 320135 | Mm.391527 | NM_178776 | ENSMUSG00000047515 | **BC049715** | - | cDNA sequence BC049715 | protein-coding |
| chr4_106590224_106591239 | chr4 | 106590225 | 106591239 | + | -0.720793 | TTS (NM_183182) | TTS (NM_183182) | 26506 | NM_183182 | 242602 | Mm.235591 | NM_183182 | | **BC055111** | - | cDNA sequence BC055111 | protein-coding |
| chr4_106590837_106591191 | chr4 | 106590838 | 106591191 | + | -0.90129 | 3' UTR (NM_183182, exon 10 of 10) | 3' UTR (NM_183182, exon 10 of 10) | 26224 | NM_183182 | 242602 | Mm.235591 | NM_183182 | | **BC055111** | - | cDNA sequence BC055111 | protein-coding |
| chr11_118348521_118349259 | chr11 | 118348522 | 118349259 | + | -0.667178 | intron (NM_011594, intron 1 of 4) | intron (NM_011594, intron 1 of 4) | -6356 | NM_021440 | 58251 | Mm.439815 | NM_021440 | ENSMUSG00000076433 | **BC100451** | Ddc8\|FS39 | cDNA sequence BC100451 | protein-coding |
| chr11_118348450_118348722 | chr11 | 118348451 | 118348722 | + | -0.762282 | intron (NM_011594, intron 1 of 4) | intron (NM_011594, intron 1 of 4) | -6052 | NM_021440 | 58251 | Mm.439815 | NM_021440 | ENSMUSG00000076433 | **BC100451** | Ddc8\|FS39 | cDNA sequence BC100451 | protein-coding |
| chr12_31621622_31621953 | chr12 | 31621623 | 31621953 | + | -0.760754 | intron (NM_001164090, intron 5 of 7) | intron (NM_001164090, intron 5 of 7) | 12809 | NM_007530 | 12033 | Mm.242379 | NM_007530 | ENSMUSG00000020650 | **Bcap29** | AW208404\|Bap29 | B cell receptor associated protein 29 | protein-coding |
| chr6_144922800_144923124 | chr6 | 144922801 | 144923124 | + | -0.661102 | Intergenic | Intergenic | 125850 | NM_001024468 | 12035 | Mm.4606 | NM_007532 | ENSMUSG00000030268 | **Bcat1** | BCATc\|Eca39 | branched chain aminotransferase 1, cytosolic | protein-coding |
| chr6_144922788_144923394 | chr6 | 144922789 | 144923394 | + | -0.726029 | Intergenic | Intergenic | 125721 | NM_001024468 | 12035 | Mm.4606 | NM_007532 | ENSMUSG00000030268 | **Bcat1** | BCATc\|Eca39 | branched chain aminotransferase 1, cytosolic | protein-coding |
| chr3_145925618_145925830 | chr3 | 145925619 | 145925830 | + | -0.862322 | intron (NM_009740, intron 1 of 2) | intron (NM_009740, intron 1 of 2) | 1462 | NM_009740 | 12042 | Mm.239141 | NM_009740 | ENSMUSG00000028191 | **Bcl10** | AI132454\|BCL-10\|C81403\|CARMEN\|CIPER\|CLAP\|ME10\|cE10 | B cell leukemia/lymphoma 10 | protein-coding |
| chr12_107927907_107928445 | chr12 | 107927908 | 107928445 | + | -0.644196 | intron (NM_001286343, intron 1 of 1) | intron (NM_001286343, intron 1 of 1) | 75238 | NM_001079883 | 58208 | Mm.392694 | NM_021399 | ENSMUSG00000048251 | **Bcl11b** | 9130430L19Rik\|AI604821\|B630002E05Rik\|BCL-11B\|Ctip2\|Rit1 | B cell leukemia/lymphoma 11B | protein-coding |
| chr12_107921137_107921598 | chr12 | 107921138 | 107921598 | + | -0.850776 | intron (NM_001286343, intron 1 of 1) | intron (NM_001286343, intron 1 of 1) | 82046 | NM_001079883 | 58208 | Mm.392694 | NM_021399 | ENSMUSG00000048251 | **Bcl11b** | 9130430L19Rik\|AI604821\|B630002E05Rik\|BCL-11B\|Ctip2\|Rit1 | B cell leukemia/lymphoma 11B | protein-coding |
| chr12_107940802_107941174 | chr12 | 107940803 | 107941174 | + | -0.87586 | intron (NM_001286343, intron 1 of 1) | intron (NM_001286343, intron 1 of 1) | 62426 | NM_001079883 | 58208 | Mm.392694 | NM_021399 | ENSMUSG00000048251 | **Bcl11b** | 9130430L19Rik\|AI604821\|B630002E05Rik\|BCL-11B\|Ctip2\|Rit1 | B cell leukemia/lymphoma 11B | protein-coding |

| | | | | | | | | | | | | | | | | | |
|---|---|---|---|---|---|---|---|---|---|---|---|---|---|---|---|---|---|
| chr12_107918783_107919165 | chr12 | 107918784 | 107919165 | + | -1.02644 | intron (NM_001286343, intron 1 of 1) | intron (NM_001286343, intron 1 of 1) | 84440 | NM_001079883 | 58208 | Mm.392694 | NM_021399 | ENSMUSG00000048251 | **Bcl11b** | 9130430L19Rik\|AI604821\|B630002E05Rik\|BCL-11B\|Ctip2\|Rit1 | B cell leukemia/lymphoma 11B | protein-coding |
| chr12_107996229_107996543 | chr12 | 107996230 | 107996543 | + | -1.20879 | intron (NM_001079883, intron 1 of 3) | intron (NM_001079883, intron 1 of 3) | 7028 | NM_001079883 | 58208 | Mm.392694 | NM_021399 | ENSMUSG00000048251 | **Bcl11b** | 9130430L19Rik\|AI604821\|B630002E05Rik\|BCL-11B\|Ctip2\|Rit1 | B cell leukemia/lymphoma 11B | protein-coding |
| chr1_106595306_106595625 | chr1 | 106595307 | 106595625 | + | -0.681844 | intron (NM_009741, intron 1 of 1) | intron (NM_009741, intron 1 of 1) | 118824 | NM_009741 | 12043 | Mm.257460 | NM_009741 | ENSMUSG00000057329 | **Bcl2** | AW986256\|Bcl-2\|C430015F12Rik\|D630044D05Rik\|D830018M01Rik | B cell leukemia/lymphoma 2 | protein-coding |
| chr1_106549111_106549591 | chr1 | 106549112 | 106549591 | + | -0.963311 | intron (NM_009741, intron 1 of 1) | intron (NM_009741, intron 1 of 1) | 164939 | NM_009741 | 12043 | Mm.257460 | NM_009741 | ENSMUSG00000057329 | **Bcl2** | AW986256\|Bcl-2\|C430015F12Rik\|D630044D05Rik\|D830018M01Rik | B cell leukemia/lymphoma 2 | protein-coding |
| chr1_106549093_106549412 | chr1 | 106549094 | 106549412 | + | -1.05743 | intron (NM_009741, intron 1 of 1) | intron (NM_009741, intron 1 of 1) | 165037 | NM_009741 | 12043 | Mm.257460 | NM_009741 | ENSMUSG00000057329 | **Bcl2** | AW986256\|Bcl-2\|C430015F12Rik\|D630044D05Rik\|D830018M01Rik | B cell leukemia/lymphoma 2 | protein-coding |
| chr2_128152312_128154047 | chr2 | 128152313 | 128154047 | + | -0.615181 | intron (NM_009754, intron 3 of 3) | intron (NM_009754, intron 3 of 3) | 25605 | NM_001284410 | 12125 | Mm.141083 | NM_009754 | ENSMUSG00000027381 | **Bcl2l11** | 1500006F24Rik\|Bim\|Bod\|bcl2-L-11 | BCL2-like 11 (apoptosis facilitator) | protein-coding |
| chr2_128027426_128028180 | chr2 | 128027427 | 128028180 | + | -0.627679 | intron (NM_028765, intron 12 of 17) | intron (NM_028765, intron 12 of 17) | -98235 | NM_207681 | 12125 | Mm.141083 | NM_009754 | ENSMUSG00000027381 | **Bcl2l11** | 1500006F24Rik\|Bim\|Bod\|bcl2-L-11 | BCL2-like 11 (apoptosis facilitator) | protein-coding |
| chr2_128152739_128153836 | chr2 | 128152740 | 128153836 | + | -0.68383 | intron (NM_009754, intron 3 of 3) | intron (NM_009754, intron 3 of 3) | 25713 | NM_001284410 | 12125 | Mm.141083 | NM_009754 | ENSMUSG00000027381 | **Bcl2l11** | 1500006F24Rik\|Bim\|Bod\|bcl2-L-11 | BCL2-like 11 (apoptosis facilitator) | protein-coding |
| chr2_128168012_128168459 | chr2 | 128168013 | 128168459 | + | -0.78769 | Intergenic | Intergenic | 40661 | NM_001284410 | 12125 | Mm.141083 | NM_009754 | ENSMUSG00000027381 | **Bcl2l11** | 1500006F24Rik\|Bim\|Bod\|bcl2-L-11 | BCL2-like 11 (apoptosis facilitator) | protein-coding |
| chr2_128156065_128156344 | chr2 | 128156066 | 128156344 | + | -0.815274 | intron (NM_009754, intron 3 of 3) | (CACAC)n\|Simple_repeat\|Simple_repeat | 28630 | NM_001284410 | 12125 | Mm.141083 | NM_009754 | ENSMUSG00000027381 | **Bcl2l11** | 1500006F24Rik\|Bim\|Bod\|bcl2-L-11 | BCL2-like 11 (apoptosis facilitator) | protein-coding |
| chr2_128159737_128160095 | chr2 | 128159738 | 128160095 | + | -0.882964 | 3' UTR (NM_207681, exon 5 of 5) | 3' UTR (NM_207681, exon 5 of 5) | 32341 | NM_001284410 | 12125 | Mm.141083 | NM_009754 | ENSMUSG00000027381 | **Bcl2l11** | 1500006F24Rik\|Bim\|Bod\|bcl2-L-11 | BCL2-like 11 (apoptosis facilitator) | protein-coding |
| chr6_134414034_134414797 | chr6 | 134414035 | 134414797 | + | -0.652578 | intron (NM_025778, intron 2 of 6) | intron (NM_025778, intron 2 of 6) | 18087 | NM_025778 | 66813 | Mm.474472 | NM_025778 | ENSMUSG00000030200 | **Bcl2l14** | 4930452K23Rik\|4933405K19Rik\|9030625M01Rik\|AI429211\|Bclg | BCL2-like 14 (apoptosis facilitator) | protein-coding |
| chr6_134436833_134437416 | chr6 | 134436834 | 134437416 | + | -0.693904 | intron (NM_025778, intron 6 of 6) | ORR1E\|LTR\|ERVL-MaLR | 40796 | NM_025778 | 66813 | Mm.474472 | NM_025778 | ENSMUSG00000030200 | **Bcl2l14** | 4930452K23Rik\|4933405K19Rik\|9030625M01Rik\|AI429211\|Bclg | BCL2-like 14 (apoptosis facilitator) | protein-coding |
| chr6_134436904_134437335 | chr6 | 134436905 | 134437335 | + | -0.6951 | intron (NM_025778, intron 6 of 6) | ORR1E\|LTR\|ERVL-MaLR | 40791 | NM_025778 | 66813 | Mm.474472 | NM_025778 | ENSMUSG00000030200 | **Bcl2l14** | 4930452K23Rik\|4933405K19Rik\|9030625M01Rik\|AI429211\|Bclg | BCL2-like 14 (apoptosis facilitator) | protein-coding |
| chr16_24050766_24051201 | chr16 | 24050767 | 24051201 | + | -0.656994 | Intergenic | Intergenic | -62372 | NM_009744 | 12053 | Mm.347398 | NM_009744 | ENSMUSG00000022508 | **Bcl6** | Bcl5 | B cell leukemia/lymphoma 6 | protein-coding |
| chr12_109090441_109091210 | chr12 | 109090442 | 109091210 | + | -0.670006 | Intergenic | Intergenic | -22609 | NM_001163175 | 380785 | Mm.342085 | NM_001163175 | ENSMUSG00000040867 | **Begain** | BM948371\|Gm897 | brain-enriched guanylate kinase-associated | protein-coding |
| chr12_109090576_109091139 | chr12 | 109090577 | 109091139 | + | -0.7311 | Intergenic | Intergenic | -22641 | NM_001163175 | 380785 | Mm.342085 | NM_001163175 | ENSMUSG00000040867 | **Begain** | BM948371\|Gm897 | brain-enriched guanylate kinase-associated | protein-coding |
| chr5_67404436_67405215 | chr5 | 67404437 | 67405215 | + | -0.645676 | intron (NM_001164806, intron 3 of 4) | intron (NM_001164806, intron 3 of 4) | 22973 | NM_001164806 | 666938 | Mm.101339 | NM_001164806 | ENSMUSG00000092060 | **Bend4** | D330027G24Rik\|EG666938 | BEN domain containing 4 | protein-coding |
| chr5_67418477_67420178 | chr5 | 67418478 | 67420178 | + | -0.645789 | intron (NM_001164806, intron 1 of 4) | RMER6A\|LTR\|ERVK | 8471 | NM_001164806 | 666938 | Mm.101339 | NM_001164806 | ENSMUSG00000092060 | **Bend4** | D330027G24Rik\|EG666938 | BEN domain containing 4 | protein-coding |
| chr5_67404292_67405654 | chr5 | 67404293 | 67405654 | + | -0.660559 | intron (NM_001164806, intron 3 of 4) | intron (NM_001164806, intron 3 of 4) | 22826 | NM_001164806 | 666938 | Mm.101339 | NM_001164806 | ENSMUSG00000092060 | **Bend4** | D330027G24Rik\|EG666938 | BEN domain containing 4 | protein-coding |
| chr5_67404373_67404938 | chr5 | 67404374 | 67404938 | + | -0.706406 | intron (NM_001164806, intron 3 of 4) | intron (NM_001164806, intron 3 of 4) | 23143 | NM_001164806 | 666938 | Mm.101339 | NM_001164806 | ENSMUSG00000092060 | **Bend4** | D330027G24Rik\|EG666938 | BEN domain containing 4 | protein-coding |
| chr5_67394730_67395861 | chr5 | 67394731 | 67395861 | + | -0.746777 | 3' UTR (NM_001164806, exon 5 of 5) | 3' UTR (NM_001164806, exon 5 of 5) | 32503 | NM_001164806 | 666938 | Mm.101339 | NM_001164806 | ENSMUSG00000092060 | **Bend4** | D330027G24Rik\|EG666938 | BEN domain containing 4 | protein-coding |
| chr5_67375492_67375887 | chr5 | 67375493 | 67375887 | + | -0.887347 | Intergenic | Intergenic | 52109 | NM_001164806 | 666938 | Mm.101339 | NM_001164806 | ENSMUSG00000092060 | **Bend4** | D330027G24Rik\|EG666938 | BEN domain containing 4 | protein-coding |
| chr5_67395197_67395896 | chr5 | 67395198 | 67395896 | + | -0.90561 | 3' UTR (NM_001164806, exon 5 of 5) | 3' UTR (NM_001164806, exon 5 of 5) | 32252 | NM_001164806 | 666938 | Mm.101339 | NM_001164806 | ENSMUSG00000092060 | **Bend4** | D330027G24Rik\|EG666938 | BEN domain containing 4 | protein-coding |
| chr5_67418441_67419147 | chr5 | 67418442 | 67419147 | + | -0.989095 | intron (NM_001164806, intron 1 of 4) | intron (NM_001164806, intron 1 of 4) | 9005 | NM_001164806 | 666938 | Mm.101339 | NM_001164806 | ENSMUSG00000092060 | **Bend4** | D330027G24Rik\|EG666938 | BEN domain containing 4 | protein-coding |
| chr5_67418512_67419023 | chr5 | 67418513 | 67419023 | + | -1.01589 | intron (NM_001164806, intron 1 of 4) | intron (NM_001164806, intron 1 of 4) | 9031 | NM_001164806 | 666938 | Mm.101339 | NM_001164806 | ENSMUSG00000092060 | **Bend4** | D330027G24Rik\|EG666938 | BEN domain containing 4 | protein-coding |
| chr4_111440243_111440684 | chr4 | 111440244 | 111440684 | + | -0.664628 | intron (NR_033793, intron 3 of 5) | intron (NR_033793, intron 3 of 5) | 25458 | NM_026279 | 67621 | Mm.45533 | NM_026279 | ENSMUSG00000028545 | **Bend5** | 2310026E23Rik | BEN domain containing 5 | protein-coding |
| chr1_33898837_33899254 | chr1 | 33898838 | 33899254 | + | -0.716802 | intron (NM_177235, intron 1 of 6) | intron (NM_177235, intron 1 of 6) | 8575 | NM_177235 | 320705 | Mm.336619 | NM_177235 | ENSMUSG00000042182 | **Bend6** | B230209C24Rik | BEN domain containing 6 | protein-coding |
| chr9_103489894_103490310 | chr9 | 103489895 | 103490310 | + | -0.653354 | intron (NM_178638, intron 4 of 5) | intron (NM_178638, intron 4 of 5) | -9774 | NM_001002896 | 107993 | Mm.335403 | NM_001002896 | ENSMUSG00000032556 | **Bfsp2** | AI448593\|CP49 | beaded filament structural protein 2, phakinin | protein-coding |
| chr5_144165783_144166346 | chr5 | 144165784 | 144166346 | + | -0.766669 | intron (NM_001081109, intron 8 of 13) | intron (NM_001081109, intron 8 of 13) | -24221 | NM_010800 | 17341 | Mm.386767 | NM_010800 | ENSMUSG00000052271 | **Bhlha15** | 1810009C13Rik\|Bhlhb8\|MIST-1\|Mist1 | basic helix-loop-helix family, member a15 | protein-coding |
| chr13_93592972_93593763 | chr13 | 93592973 | 93593763 | + | -1.29014 | Intergenic | MTC\|LTR\|ERVL-MaLR | 44390 | NM_016668 | 12116 | Mm.329582 | NM_016668 | ENSMUSG00000074768 | **Bhmt** | - | betaine-homocysteine methyltransferase | protein-coding |
| chr13_49371384_49371991 | chr13 | 49371385 | 49371991 | + | -0.718417 | intron (NM_001039180, intron 3 of 8) | intron (NM_001039180, intron 3 of 8) | 30139 | NM_001039180 | 76895 | Mm.197387 | NM_029791 | ENSMUSG00000037933 | **Bicd2** | - | bicaudal D homolog 2 (Drosophila) | protein-coding |
| chr18_32352486_32353023 | chr18 | 32352487 | 32353023 | + | -0.801198 | Intergenic | Lx9\|LINE\|L1 | -24462 | NM_001083334 | 30948 | Mm.4383 | NM_009668 | ENSMUSG00000024381 | **Bin1** | ALP-1\|Amphl\|BRAMP-2\|SH3P9 | bridging integrator 1 | protein-coding |
| chr15_100667859_100668477 | chr15 | 100667860 | 100668477 | + | -0.770242 | intron (NM_001270537, intron 1 of 10) | intron (NM_001270537, intron 1 of 10) | 1332 | NM_001270537 | 668218 | Mm.451225 | NM_001270537 | ENSMUSG00000098112 | **Bin2** | - | bridging integrator 2 | protein-coding |
| chr1_44122268_44122628 | chr1 | 44122269 | 44122628 | + | -0.933093 | intron (NM_144558, intron 1 of 9) | intron (NM_144558, intron 1 of 9) | 2480 | NM_144558 | 246229 | Mm.386883 | NM_144558 | ENSMUSG00000041684 | **Bivm** | AI854267 | basic, immunoglobulin-like variable motif containing | protein-coding |
| chr1_44122195_44122792 | chr1 | 44122196 | 44122792 | + | -0.947144 | intron (NM_144558, intron 1 of 9) | intron (NM_144558, intron 1 of 9) | 2526 | NM_144558 | 246229 | Mm.386883 | NM_144558 | ENSMUSG00000041684 | **Bivm** | AI854267 | basic, immunoglobulin-like variable motif containing | protein-coding |
| chr7_80554356_80555468 | chr7 | 80554357 | 80555468 | + | -0.774992 | Intergenic | Intergenic | -19793 | NM_001042527 | 12144 | Mm.12932 | NM_007550 | ENSMUSG00000030528 | **Blm** | - | Bloom syndrome, RecQ helicase-like | protein-coding |
| chr7_80554307_80555131 | chr7 | 80554308 | 80555131 | + | -0.895561 | Intergenic | Intergenic | -19600 | NM_001042527 | 12144 | Mm.12932 | NM_007550 | ENSMUSG00000030528 | **Blm** | - | Bloom syndrome, RecQ helicase-like | protein-coding |
| chr5_96996951_96997205 | chr5 | 96996952 | 96997205 | + | -0.685262 | promoter-TSS (NM_080708) | promoter-TSS (NM_080708) | -611 | NM_080708 | 140780 | Mm.281490 | NM_080708 | ENSMUSG00000034663 | **Bmp2k** | 4933417M22Rik\|AA673486\|AV128808\|BIKE | BMP2 inducible kinase | protein-coding |
| chr17_26786652_26786920 | chr17 | 26786653 | 26786920 | + | -0.77767 | exon (NM_172149, exon 3 of 6) | exon (NM_172149, exon 3 of 6) | 5707 | NM_172149 | 224630 | Mm.21795 | NM_172149 | ENSMUSG00000024191 | **Bnip1** | 2010005M06Rik\|5930429G21Rik\|Sec20 | BCL2/adenovirus E1B interacting protein 1 | protein-coding |
| chr17_26752245_26752813 | chr17 | 26752246 | 26752813 | + | -1.23638 | intron (NM_029870, intron 4 of 8) | intron (NM_029870, intron 4 of 8) | -28550 | NM_172149 | 224630 | Mm.21795 | NM_172149 | ENSMUSG00000024191 | **Bnip1** | 2010005M06Rik\|5930429G21Rik\|Sec20 | BCL2/adenovirus E1B interacting protein 1 | protein-coding |
| chr17_26752295_26752699 | chr17 | 26752296 | 26752699 | + | -1.24627 | intron (NM_029870, intron 4 of 8) | intron (NM_029870, intron 4 of 8) | -28582 | NM_172149 | 224630 | Mm.21795 | NM_172149 | ENSMUSG00000024191 | **Bnip1** | 2010005M06Rik\|5930429G21Rik\|Sec20 | BCL2/adenovirus E1B interacting protein 1 | protein-coding |
| chr5_41841796_41842076 | chr5 | 41841797 | 41842076 | + | -1.20659 | intron (NM_001081422, intron 1 of 26) | intron (NM_001081422, intron 1 of 26) | 2379 | NM_001081422 | 665775 | Mm.158726 | NM_001081422 | ENSMUSG00000061755 | **Bod1l** | A230054D04Rik\|AI853319\|FAM44A\|mKIAA1327 | biorientation of chromosomes in cell division 1-like | protein-coding |
| chr6_34494923_34495573 | chr6 | 34494924 | 34495573 | + | -0.888955 | intron (NM_007563, intron 2 of 2) | intron (NM_007563, intron 2 of 2) | 18892 | NM_007563 | 12183 | Mm.282863 | NM_007563 | ENSMUSG00000038871 | **Bpgm** | AI323730\|AL022789\|C86192 | 2,3-bisphosphoglycerate mutase | protein-coding |
| chr17_32277025_32277628 | chr17 | 32277026 | 32277628 | + | -1.2637 | intron (NM_001286630, intron 1 of 19) | PB1D10\|SINE\|Alu | 6796 | NM_198094 | 57261 | Mm.253518 | NM_020508 | ENSMUSG00000024002 | **Brd4** | Brd5\|HUNK1\|MCAP\|WI-11513 | bromodomain containing 4 | protein-coding |
| chr5_107353991_107354405 | chr5 | 107353992 | 107354405 | + | -0.904596 | intron (NM_054054, intron 7 of 18) | L1M3\|LINE\|L1 | 23004 | NM_054054 | 114642 | Mm.182836 | NM_054054 | ENSMUSG00000029279 | **Brdt** | 7420412D09Rik\|Brd6\|Fsrg3 | bromodomain, testis-specific | protein-coding |

| | | | | | | | | | | | | | | | | | |
|---|---|---|---|---|---|---|---|---|---|---|---|---|---|---|---|---|---|
| chr5_31831553_31832013 | chr5 | 31831554 | 31832013 | + | -0.774015 | intron (NM_144541, intron 6 of 11) | intron (NM_144541, intron 6 of 11) | 133733 | NM_181282 | 107976 | Mm.482126 | NM_144541 | ENSMUSG00000052139 | **Bre** | 6030405P19Rik\|AI429776\|B830038C02Rik | brain and reproductive organ-expressed protein | protein-coding |
| chr1_183265850_183266606 | chr1 | 183265851 | 183266606 | + | -0.673664 | Intergenic | ORR1E\|LTR\|ERVL-MaLR | 30780 | NM_027861 | 71678 | Mm.392270 | NM_027861 | ENSMUSG00000046836 | **Brox** | 0610010K06Rik | BRO1 domain and CAAX motif containing | protein-coding |
| chr19_8844747_8845034 | chr19 | 8844748 | 8845034 | + | -0.741678 | intron (NM_001290823, intron 3 of 10) | intron (NM_001290823, intron 3 of 10) | 4388 | NM_001290823 | 14705 | Mm.345134 | NM_008144 | ENSMUSG00000071657 | **Bscl2** | 2900097C17Rik\|AI046355\|Gng3lg | Berardinelli-Seip congenital lipodystrophy 2 homolog (seipin) | protein-coding |
| chr9_108150517_108150847 | chr9 | 108150518 | 108150847 | + | -0.690803 | intron (NM_007567, intron 1 of 11) | intron (NM_007567, intron 1 of 11) | 39701 | NM_007567 | 12217 | Mm.20425 | NM_007567 | ENSMUSG00000032589 | **Bsn** | - | bassoon | protein-coding |
| chr9_108150414_108151045 | chr9 | 108150415 | 108151045 | + | -0.709346 | intron (NM_007567, intron 1 of 11) | intron (NM_007567, intron 1 of 11) | 39653 | NM_007567 | 12217 | Mm.20425 | NM_007567 | ENSMUSG00000032589 | **Bsn** | - | bassoon | protein-coding |
| chr4_62475830_62476188 | chr4 | 62475831 | 62476188 | + | -0.718346 | Intergenic | Intergenic | -4058 | NM_138653 | 192120 | Mm.29808 | NM_138653 | ENSMUSG00000028392 | **Bspry** | - | B-box and SPRY domain containing | protein-coding |
| chr10_85567032_85567489 | chr10 | 85567033 | 85567489 | + | -0.803031 | intron (NM_028709, intron 3 of 16) | MTE2b\|LTR\|ERVL-MaLR | -31150 | NM_001017525 | 74007 | Mm.36544 | NM_028709 | ENSMUSG00000020042 | **Btbd11** | 6330404E16Rik | BTB (POZ) domain containing 11 | protein-coding |
| chr10_85411063_85411660 | chr10 | 85411064 | 85411660 | + | -0.951368 | intron (NM_028709, intron 1 of 16) | intron (NM_028709, intron 1 of 16) | 24548 | NM_028709 | 74007 | Mm.36544 | NM_028709 | ENSMUSG00000020042 | **Btbd11** | 6330404E16Rik | BTB (POZ) domain containing 11 | protein-coding |
| chr10_85359335_85359992 | chr10 | 85359336 | 85359992 | + | -1.10405 | Intergenic | B1_Mur2\|SINE\|Alu | -27150 | NM_028709 | 74007 | Mm.36544 | NM_028709 | ENSMUSG00000020042 | **Btbd11** | 6330404E16Rik | BTB (POZ) domain containing 11 | protein-coding |
| chr10_80658644_80659129 | chr10 | 80658645 | 80659129 | + | -0.672241 | Intergenic | Intergenic | -1816 | NM_145361 | 208198 | Mm.60720 | NM_145361 | ENSMUSG00000003344 | **Btbd2** | 2610037C03Rik\|4930512K17Rik | BTB (POZ) domain containing 2 | protein-coding |
| chr12_112959399_112959835 | chr12 | 112959400 | 112959835 | + | -0.758217 | TTS (NM_028193) | TTS (NM_028193) | -16865 | NM_001145900 | 399566 | Mm.440140 | NM_201646 | ENSMUSG00000002803 | **Btbd6** | - | BTB (POZ) domain containing 6 | protein-coding |
| chr12_112967821_112968140 | chr12 | 112967822 | 112968140 | + | -0.89536 | intron (NM_028193, intron 10 of 17) | intron (NM_028193, intron 10 of 17) | -8501 | NM_001145900 | 399566 | Mm.440140 | NM_201646 | ENSMUSG00000002803 | **Btbd6** | - | BTB (POZ) domain containing 6 | protein-coding |
| chr12_112967543_112968273 | chr12 | 112967544 | 112968273 | + | -1.01111 | intron (NM_028193, intron 10 of 17) | intron (NM_028193, intron 10 of 17) | -8574 | NM_001145900 | 399566 | Mm.440140 | NM_201646 | ENSMUSG00000002803 | **Btbd6** | - | BTB (POZ) domain containing 6 | protein-coding |
| chr17_30552284_30552582 | chr17 | 30552285 | 30552582 | + | -0.743085 | intron (NM_027060, intron 1 of 10) | intron (NM_027060, intron 1 of 10) | 23854 | NM_172618 | 224671 | Mm.41220 | NM_027060 | ENSMUSG00000062202 | **Btbd9** | 1700023F20Rik\|4930402L05\|AI464421 | BTB (POZ) domain containing 9 | protein-coding |
| chr10_96697641_96698312 | chr10 | 96697642 | 96698312 | + | -0.64791 | Intergenic | Intergenic | 80976 | NM_007569 | 12226 | Mm.23811 | NM_007569 | ENSMUSG00000036478 | **Btg1** | AI426953\|AW546738 | B cell translocation gene 1, anti-proliferative | protein-coding |
| chr10_96706427_96707087 | chr10 | 96706428 | 96707087 | + | -0.710561 | Intergenic | Intergenic | 89756 | NM_007569 | 12226 | Mm.23811 | NM_007569 | ENSMUSG00000036478 | **Btg1** | AI426953\|AW546738 | B cell translocation gene 1, anti-proliferative | protein-coding |
| chr10_96620970_96621302 | chr10 | 96620971 | 96621302 | + | -0.714422 | 3' UTR (NM_007569, exon 2 of 2) | 3' UTR (NM_007569, exon 2 of 2) | 4135 | NM_007569 | 12226 | Mm.23811 | NM_007569 | ENSMUSG00000036478 | **Btg1** | AI426953\|AW546738 | B cell translocation gene 1, anti-proliferative | protein-coding |
| chr10_96706644_96707070 | chr10 | 96706645 | 96707070 | + | -0.734804 | Intergenic | Intergenic | 89856 | NM_007569 | 12226 | Mm.23811 | NM_007569 | ENSMUSG00000036478 | **Btg1** | AI426953\|AW546738 | B cell translocation gene 1, anti-proliferative | protein-coding |
| chr10_96579517_96579780 | chr10 | 96579518 | 96579780 | + | -0.794567 | Intergenic | Intergenic | -37352 | NM_007569 | 12226 | Mm.23811 | NM_007569 | ENSMUSG00000036478 | **Btg1** | AI426953\|AW546738 | B cell translocation gene 1, anti-proliferative | protein-coding |
| chr10_96471439_96471862 | chr10 | 96471440 | 96471862 | + | -0.800402 | Intergenic | Intergenic | -145350 | NM_007569 | 12226 | Mm.23811 | NM_007569 | ENSMUSG00000036478 | **Btg1** | AI426953\|AW546738 | B cell translocation gene 1, anti-proliferative | protein-coding |
| chr10_96663482_96664551 | chr10 | 96663483 | 96664551 | + | -0.829737 | Intergenic | Intergenic | 47016 | NM_007569 | 12226 | Mm.23811 | NM_007569 | ENSMUSG00000036478 | **Btg1** | AI426953\|AW546738 | B cell translocation gene 1, anti-proliferative | protein-coding |
| chr10_96693846_96694147 | chr10 | 96693847 | 96694147 | + | -0.880148 | Intergenic | Intergenic | 76996 | NM_007569 | 12226 | Mm.23811 | NM_007569 | ENSMUSG00000036478 | **Btg1** | AI426953\|AW546738 | B cell translocation gene 1, anti-proliferative | protein-coding |
| chr10_96663967_96664500 | chr10 | 96663968 | 96664500 | + | -0.95468 | Intergenic | Intergenic | 47233 | NM_007569 | 12226 | Mm.23811 | NM_007569 | ENSMUSG00000036478 | **Btg1** | AI426953\|AW546738 | B cell translocation gene 1, anti-proliferative | protein-coding |
| chr10_96496350_96496861 | chr10 | 96496351 | 96496861 | + | -0.992407 | Intergenic | Intergenic | -120395 | NM_007569 | 12226 | Mm.23811 | NM_007569 | ENSMUSG00000036478 | **Btg1** | AI426953\|AW546738 | B cell translocation gene 1, anti-proliferative | protein-coding |
| chr10_96609302_96609922 | chr10 | 96609303 | 96609922 | + | -0.995334 | Intergenic | Intergenic | -7389 | NM_007569 | 12226 | Mm.23811 | NM_007569 | ENSMUSG00000036478 | **Btg1** | AI426953\|AW546738 | B cell translocation gene 1, anti-proliferative | protein-coding |
| chr10_96466749_96467231 | chr10 | 96466750 | 96467231 | + | -1.00284 | Intergenic | L2a\|LINE\|L2 | -150011 | NM_007569 | 12226 | Mm.23811 | NM_007569 | ENSMUSG00000036478 | **Btg1** | AI426953\|AW546738 | B cell translocation gene 1, anti-proliferative | protein-coding |
| chr10_96480602_96481813 | chr10 | 96480603 | 96481813 | + | -1.0189 | Intergenic | Intergenic | -135793 | NM_007569 | 12226 | Mm.23811 | NM_007569 | ENSMUSG00000036478 | **Btg1** | AI426953\|AW546738 | B cell translocation gene 1, anti-proliferative | protein-coding |
| chr10_96787002_96787577 | chr10 | 96787003 | 96787577 | + | -1.06121 | Intergenic | Intergenic | 170289 | NM_007569 | 12226 | Mm.23811 | NM_007569 | ENSMUSG00000036478 | **Btg1** | AI426953\|AW546738 | B cell translocation gene 1, anti-proliferative | protein-coding |
| chr10_96786995_96787938 | chr10 | 96786996 | 96787938 | + | -1.06873 | Intergenic | Intergenic | 170466 | NM_007569 | 12226 | Mm.23811 | NM_007569 | ENSMUSG00000036478 | **Btg1** | AI426953\|AW546738 | B cell translocation gene 1, anti-proliferative | protein-coding |
| chr10_96609359_96609640 | chr10 | 96609360 | 96609640 | + | -1.08537 | Intergenic | Intergenic | -7501 | NM_007569 | 12226 | Mm.23811 | NM_007569 | ENSMUSG00000036478 | **Btg1** | AI426953\|AW546738 | B cell translocation gene 1, anti-proliferative | protein-coding |
| chr10_96822318_96823010 | chr10 | 96822319 | 96823010 | + | -1.08601 | Intergenic | Intergenic | 205663 | NM_007569 | 12226 | Mm.23811 | NM_007569 | ENSMUSG00000036478 | **Btg1** | AI426953\|AW546738 | B cell translocation gene 1, anti-proliferative | protein-coding |
| chr10_96466874_96467191 | chr10 | 96466875 | 96467191 | + | -1.09349 | Intergenic | L2a\|LINE\|L2 | -149968 | NM_007569 | 12226 | Mm.23811 | NM_007569 | ENSMUSG00000036478 | **Btg1** | AI426953\|AW546738 | B cell translocation gene 1, anti-proliferative | protein-coding |
| chr10_96481156_96481809 | chr10 | 96481157 | 96481809 | + | -1.17184 | Intergenic | Intergenic | -135518 | NM_007569 | 12226 | Mm.23811 | NM_007569 | ENSMUSG00000036478 | **Btg1** | AI426953\|AW546738 | B cell translocation gene 1, anti-proliferative | protein-coding |
| chr10_96602239_96602755 | chr10 | 96602240 | 96602755 | + | -1.45609 | Intergenic | Intergenic | -14504 | NM_007569 | 12226 | Mm.23811 | NM_007569 | ENSMUSG00000036478 | **Btg1** | AI426953\|AW546738 | B cell translocation gene 1, anti-proliferative | protein-coding |
| chr1_134077662_134077876 | chr1 | 134077663 | 134077876 | + | -0.7842 | intron (NM_007570, intron 1 of 1) | intron (NM_007570, intron 1 of 1) | 1386 | NM_007570 | 12227 | Mm.392646 | NM_007570 | ENSMUSG00000020423 | **Btg2** | AA959598\|APRO1\|Pc3\|TIS21 | B cell translocation gene 2, anti-proliferative | protein-coding |
| chr1_134058883_134059281 | chr1 | 134058884 | 134059281 | + | -1.58595 | Intergenic | Intergenic | 20073 | NM_007570 | 12227 | Mm.392646 | NM_007570 | ENSMUSG00000020423 | **Btg2** | AA959598\|APRO1\|Pc3\|TIS21 | B cell translocation gene 2, anti-proliferative | protein-coding |
| chr11_58927727_58927934 | chr11 | 58927728 | 58927934 | + | -0.924482 | TTS (NM_138678) | TTS (NM_138678) | 9774 | NM_138678 | 192194 | Mm.22540 | NM_138678 | ENSMUSG00000020490 | **Btnl10** | BUTR-1\|Butr1 | butyrophilin-like 10 | protein-coding |
| chr17_34354943_34355319 | chr17 | 34354944 | 34355319 | + | -0.727942 | intron (NM_079835, intron 1 of 8) | intron (NM_079835, intron 1 of 8) | 309 | NM_079835 | 547431 | Mm.441007 | NM_079835 | ENSMUSG00000024340 | **Btnl2** | BTL-II\|BTLN2\|Btl2\|Gm315\|NG9 | butyrophilin-like 2 | protein-coding |
| chr12_36135363_36135767 | chr12 | 36135364 | 36135767 | + | -0.805838 | intron (NM_025840, intron 1 of 11) | intron (NM_025840, intron 1 of 11) | 21260 | NM_025840 | 66912 | Mm.288669 | NM_025840 | ENSMUSG00000020547 | **Bzw2** | 1110001I24Rik\|Bdm2\|HSPC028 | basic leucine zipper and W2 domains 2 | protein-coding |
| chr12_36136310_36136843 | chr12 | 36136311 | 36136843 | + | -0.931528 | intron (NM_025840, intron 1 of 11) | intron (NM_025840, intron 1 of 11) | 20248 | NM_025840 | 66912 | Mm.288669 | NM_025840 | ENSMUSG00000020547 | **Bzw2** | 1110001I24Rik\|Bdm2\|HSPC028 | basic leucine zipper and W2 domains 2 | protein-coding |
| chr12_36155430_36155703 | chr12 | 36155431 | 36155703 | + | -1.02627 | intron (NM_025840, intron 1 of 11) | intron (NM_025840, intron 1 of 11) | 1258 | NM_025840 | 66912 | Mm.288669 | NM_025840 | ENSMUSG00000020547 | **Bzw2** | 1110001I24Rik\|Bdm2\|HSPC028 | basic leucine zipper and W2 domains 2 | protein-coding |
| chr5_64478257_64478968 | chr5 | 64478258 | 64478968 | + | -0.74433 | intron (NR_045411, intron 2 of 2) | intron (NR_045411, intron 2 of 2) | 1605 | NR_045413 | 77382 | Mm.366742 | NR_045411 | | **C030018K13Rik** | - | RIKEN cDNA C030018K13 gene | ncRNA |
| chr5_64518862_64519395 | chr5 | 64518863 | 64519395 | + | -0.832222 | Intergenic | Intergenic | 42121 | NR_045413 | 77382 | Mm.366742 | NR_045411 | | **C030018K13Rik** | - | RIKEN cDNA C030018K13 gene | ncRNA |
| chr5_64478598_64479030 | chr5 | 64478599 | 64479030 | + | -0.842067 | intron (NR_045411, intron 2 of 2) | intron (NR_045411, intron 2 of 2) | 1806 | NR_045413 | 77382 | Mm.366742 | NR_045411 | | **C030018K13Rik** | - | RIKEN cDNA C030018K13 gene | ncRNA |
| chr5_64478662_64478946 | chr5 | 64478663 | 64478946 | + | -0.88455 | intron (NR_045411, intron 2 of 2) | intron (NR_045411, intron 2 of 2) | 1796 | NR_045413 | 77382 | Mm.366742 | NR_045411 | | **C030018K13Rik** | - | RIKEN cDNA C030018K13 gene | ncRNA |
| chr5_64490646_64491009 | chr5 | 64490647 | 64491009 | + | -0.99547 | Intergenic | Intergenic | 13820 | NR_045413 | 77382 | Mm.366742 | NR_045411 | | **C030018K13Rik** | - | RIKEN cDNA C030018K13 gene | ncRNA |
| chr5_64496627_64497343 | chr5 | 64496628 | 64497343 | + | -1.30245 | Intergenic | Intergenic | 19977 | NR_045413 | 77382 | Mm.366742 | NR_045411 | | **C030018K13Rik** | - | RIKEN cDNA C030018K13 gene | ncRNA |
| chr5_64498340_64498840 | chr5 | 64498341 | 64498840 | + | -1.38459 | Intergenic | Intergenic | 21582 | NR_045413 | 77382 | Mm.366742 | NR_045411 | | **C030018K13Rik** | - | RIKEN cDNA C030018K13 gene | ncRNA |
| chr17_69412940_69413162 | chr17 | 69412941 | 69413162 | + | -0.634545 | Intergenic | Intergenic | -3396 | NR_026848 | 77533 | Mm.266592 | NR_026848 | | **C030034I22Rik** | - | RIKEN cDNA C030034I22 gene | ncRNA |
| chr17_69422713_69422953 | chr17 | 69422714 | 69422953 | + | -0.814718 | Intergenic | MLT1B\|LTR\|ERVL-MaLR | 6386 | NR_026848 | 77533 | Mm.266592 | NR_026848 | | **C030034I22Rik** | - | RIKEN cDNA C030034I22 gene | ncRNA |
| chr17_69422664_69423089 | chr17 | 69422665 | 69423089 | + | -0.817797 | Intergenic | MLT1B\|LTR\|ERVL-MaLR | 6430 | NR_026848 | 77533 | Mm.266592 | NR_026848 | | **C030034I22Rik** | - | RIKEN cDNA C030034I22 gene | ncRNA |
| chr3_9402832_9403380 | chr3 | 9402833 | 9403380 | + | -0.872283 | promoter-TSS (NR_015499) | promoter-TSS (NR_015499) | 28 | NR_015499 | 320088 | Mm.212908 | NR_015499 | ENSMUSG00000097365 | **C030034L19Rik** | A130044O20Rik | RIKEN cDNA C030034L19 gene | ncRNA |
| chr3_9400896_9401160 | chr3 | 9400897 | 9401160 | + | -0.875599 | Intergenic | Intergenic | -2050 | NR_015499 | 320088 | Mm.212908 | NR_015499 | ENSMUSG00000097365 | **C030034L19Rik** | A130044O20Rik | RIKEN cDNA C030034L19 gene | ncRNA |
| chr6_7843740_7844067 | chr6 | 7843741 | 7844067 | + | -1.03151 | Intergenic | Intergenic | -1320 | NM_052993 | 94192 | Mm.102752 | NM_052993 | ENSMUSG00000042460 | **C1galt1** | 2210410E06Rik\|AV284120\|T-synthase | core 1 synthase, glycoprotein-N-acetylgalactosamine 3-beta-galactosyltransferase, 1 | protein-coding |
| chr2_13132433_13132788 | chr2 | 13132434 | 13132788 | + | -0.631271 | intron (NM_009105, intron 8 of 8) | intron (NM_009105, intron 8 of 8) | -121747 | NM_153155 | 227580 | Mm.229322 | NM_153155 | | **C1ql3** | 1110065A22Rik\|AI661623\|C1ql\|C1qtnf13\|CTRP13\|K100 | C1q-like 3 | protein-coding |
| chr15_78518253_78518642 | chr15 | 78518254 | 78518642 | + | -1.10038 | Intergenic | Intergenic | 11203 | NM_001204153 | 72709 | Mm.34776 | NM_028331 | ENSMUSG00000022440 | **C1qtnf6** | 2810036M19Rik\|CTRP6 | C1q and tumor necrosis factor related protein 6 | protein-coding |
| chr15_78515696_78516085 | chr15 | 78515697 | 78516085 | + | -1.1992 | Intergenic | (CACCC)n\|Simple_repeat\|Simple_repeat | 13760 | NM_001204153 | 72709 | Mm.34776 | NM_028331 | ENSMUSG00000022440 | **C1qtnf6** | 2810036M19Rik\|CTRP6 | C1q and tumor necrosis factor related protein 6 | protein-coding |
| chr14_60709044_60709602 | chr14 | 60709045 | 60709602 | + | -0.678235 | intron (NM_001033272, intron 3 of 12) | intron (NM_001033272, intron 3 of 12) | -58811 | NM_183175 | 239126 | Mm.319636 | NM_183175 | ENSMUSG00000071347 | **C1qtnf9** | 9130217G22Rik\|CTRP9\|Ciqtnf9 | C1q and tumor necrosis factor related protein 9 | protein-coding |
| chr14_60732146_60732455 | chr14 | 60732147 | 60732455 | + | -0.941978 | intron (NM_001033272, intron 4 of 12) | intron (NM_001033272, intron 4 of 12) | -35833 | NM_183175 | 239126 | Mm.319636 | NM_183175 | ENSMUSG00000071347 | **C1qtnf9** | 9130217G22Rik\|CTRP9\|Ciqtnf9 | C1q and tumor necrosis factor related protein 9 | protein-coding |
| chr14_60715950_60716349 | chr14 | 60715951 | 60716349 | + | -1.01081 | intron (NM_001033272, intron 4 of 12) | ORR1D2-int\|LTR\|ERVL-MaLR | -51984 | NM_183175 | 239126 | Mm.319636 | NM_183175 | ENSMUSG00000071347 | **C1qtnf9** | 9130217G22Rik\|CTRP9\|Ciqtnf9 | C1q and tumor necrosis factor related protein 9 | protein-coding |
| chr10_119001284_119002191 | chr10 | 119001285 | 119002191 | + | -0.617772 | Intergenic | MTD\|LTR\|ERVL-MaLR | -74595 | NR_110482 | 216377 | Mm.208936 | NR_110482 | | **C230029M16** | - | uncharacterized protein C230029M16 | ncRNA |
| chr10_118952785_118953092 | chr10 | 118952786 | 118953092 | + | -0.724098 | Intergenic | Intergenic | -25796 | NR_110482 | 216377 | Mm.208936 | NR_110482 | | **C230029M16** | - | uncharacterized protein C230029M16 | ncRNA |
| chr10_118997200_118997423 | chr10 | 118997201 | 118997423 | + | -1.10148 | Intergenic | Intergenic | -70169 | NR_110482 | 216377 | Mm.208936 | NR_110482 | | **C230029M16** | - | uncharacterized protein C230029M16 | ncRNA |
| chr10_118997167_118997512 | chr10 | 118997168 | 118997512 | + | -1.12426 | Intergenic | RLTR11A\|LTR\|ERVK | -70197 | NR_110482 | 216377 | Mm.208936 | NR_110482 | | **C230029M16** | - | uncharacterized protein C230029M16 | ncRNA |
| chr10_119001952_119002252 | chr10 | 119001953 | 119002252 | + | -1.1313 | Intergenic | Intergenic | -74959 | NR_110482 | 216377 | Mm.208936 | NR_110482 | | **C230029M16** | - | uncharacterized protein C230029M16 | ncRNA |
| chr10_119005414_119005999 | chr10 | 119005415 | 119005999 | + | -1.33408 | Intergenic | ORR1F\|LTR\|ERVL-MaLR | -78564 | NR_110482 | 216377 | Mm.208936 | NR_110482 | | **C230029M16** | - | uncharacterized protein C230029M16 | ncRNA |

| | | | | | | | | | | | | | | | | | |
|---|---|---|---|---|---|---|---|---|---|---|---|---|---|---|---|---|---|
| chr6_143115069_143115532 | chr6 | 143115070 | 143115532 | + | -0.768365 | Intergenic | Intergenic | -15149 | NM_001109688 | 74741 | Mm.130260 | NM_029081 | ENSMUSG00000030279 | **C2cd5** | 5730419I09Rik\|C030008B15Rik\|CDP138 | C2 calcium-dependent domain containing 5 | protein-coding |
| chr17_51409991_51410242 | chr17 | 51409992 | 51410242 | + | -0.743357 | intron (NR_037250, intron 3 of 4) | intron (NR_037250, intron 3 of 4) | 38152 | NR_046166 | 328837 | Mm.236143 | NR_046166 | | **C330011F03Rik** | C330011F03 | RIKEN cDNA C330011F03 gene | ncRNA |
| chr17_51408259_51409184 | chr17 | 51408260 | 51409184 | + | -1.05878 | intron (NR_037250, intron 3 of 4) | intron (NR_037250, intron 3 of 4) | 39547 | NR_046166 | 328837 | Mm.236143 | NR_046166 | | **C330011F03Rik** | C330011F03 | RIKEN cDNA C330011F03 gene | ncRNA |
| chr15_100604623_100605434 | chr15 | 100604624 | 100605434 | + | -1.2123 | Intergenic | Intergenic | 10081 | NR_045701 | 78445 | Mm.44383 | NR_045701 | ENSMUSG00000097093 | **C330013E15Rik** | - | RIKEN cDNA C330013E15 gene | ncRNA |
| chr9_96785261_96785601 | chr9 | 96785262 | 96785601 | + | -0.66683 | Intergenic | Intergenic | 19869 | NR_102296 | 319707 | Mm.392191 | NR_102293 | | **C430002N11Rik** | - | RIKEN cDNA C430002N11 gene | ncRNA |
| chr1_151095803_151096491 | chr1 | 151095804 | 151096491 | + | -0.675748 | Intergenic | Intergenic | -41887 | NR_038011 | 402734 | Mm.337666 | NR_038011 | | **C730036E19Rik** | - | RIKEN cDNA C730036E19 gene | ncRNA |
| chr1_151096036_151096496 | chr1 | 151096037 | 151096496 | + | -0.71768 | Intergenic | Intergenic | -41768 | NR_038011 | 402734 | Mm.337666 | NR_038011 | | **C730036E19Rik** | - | RIKEN cDNA C730036E19 gene | ncRNA |
| chr1_151083696_151084202 | chr1 | 151083697 | 151084202 | + | -0.730885 | Intergenic | ORR1D2\|LTR\|ERVL-MaLR | -54085 | NR_038011 | 402734 | Mm.337666 | NR_038011 | | **C730036E19Rik** | - | RIKEN cDNA C730036E19 gene | ncRNA |
| chr1_151177166_151178316 | chr1 | 151177167 | 151178316 | + | -0.768702 | Intergenic | L2\|LINE\|L2 | 39707 | NR_038011 | 402734 | Mm.337666 | NR_038011 | | **C730036E19Rik** | - | RIKEN cDNA C730036E19 gene | ncRNA |
| chr1_151177451_151178172 | chr1 | 151177452 | 151178172 | + | -0.825182 | Intergenic | Intergenic | 39778 | NR_038011 | 402734 | Mm.337666 | NR_038011 | | **C730036E19Rik** | - | RIKEN cDNA C730036E19 gene | ncRNA |
| chr1_151177687_151178192 | chr1 | 151177688 | 151178192 | + | -0.885148 | Intergenic | Intergenic | 39906 | NR_038011 | 402734 | Mm.337666 | NR_038011 | | **C730036E19Rik** | - | RIKEN cDNA C730036E19 gene | ncRNA |
| chr13_46616689_46617070 | chr13 | 46616690 | 46617070 | + | -0.64958 | intron (NM_026056, intron 7 of 12) | intron (NM_026056, intron 7 of 12) | -52642 | NM_001033192 | 97863 | Mm.275934 | NM_001033192 | ENSMUSG00000069237 | **C78339** | - | expressed sequence C78339 | protein-coding |
| chr9_78176868_78177172 | chr9 | 78176869 | 78177172 | + | -0.874841 | non-coding (NR_040401, exon 3 of 4) | non-coding (NR_040401, exon 3 of 4) | 1106 | NR_040401 | 320295 | Mm.131285 | NR_040401 | ENSMUSG00000097574 | **C920006O11Rik** | - | RIKEN cDNA C920006O11 gene | ncRNA |
| chr10_22299428_22299683 | chr10 | 22299429 | 22299683 | + | -0.993874 | intron (NM_009018, intron 5 of 5) | intron (NM_009018, intron 5 of 5) | -7164 | NR_015465 | 606736 | Mm.387042 | NR_015465 | ENSMUSG00000097352 | **C920009B18Rik** | - | RIKEN cDNA C920009B18 gene | ncRNA |
| chr10_22299319_22299716 | chr10 | 22299320 | 22299716 | + | -1.03597 | intron (NM_009018, intron 5 of 5) | intron (NM_009018, intron 5 of 5) | -7202 | NR_015465 | 606736 | Mm.387042 | NR_015465 | ENSMUSG00000097352 | **C920009B18Rik** | - | RIKEN cDNA C920009B18 gene | ncRNA |
| chr10_22301640_22301960 | chr10 | 22301641 | 22301960 | + | -1.19536 | intron (NM_009018, intron 5 of 5) | ORR1C1\|LTR\|ERVL-MaLR | -4920 | NR_015465 | 606736 | Mm.387042 | NR_015465 | ENSMUSG00000097352 | **C920009B18Rik** | - | RIKEN cDNA C920009B18 gene | ncRNA |
| chr10_75721693_75722306 | chr10 | 75721694 | 75722306 | + | -0.913905 | intron (NM_172549, intron 22 of 36) | intron (NM_172549, intron 22 of 36) | 42357 | NM_172549 | 104248 | Mm.257404 | NM_172549 | ENSMUSG00000020196 | **Cabin1** | A330070M20Rik\|Cain\|Ppp3in | calcineurin binding protein 1 | protein-coding |
| chr10_75721980_75722340 | chr10 | 75721981 | 75722340 | + | -0.946268 | intron (NM_172549, intron 22 of 36) | intron (NM_172549, intron 22 of 36) | 42197 | NM_172549 | 104248 | Mm.257404 | NM_172549 | ENSMUSG00000020196 | **Cabin1** | A330070M20Rik\|Cain\|Ppp3in | calcineurin binding protein 1 | protein-coding |
| chr2_180276757_180277299 | chr2 | 180276758 | 180277299 | + | -0.946791 | TTS (NM_173031) | TTS (NM_173031) | -3563 | NM_145851 | 252966 | Mm.386884 | NM_145851 | ENSMUSG00000038990 | **Cables2** | ik3-2 | CDK5 and Abl enzyme substrate 2 | protein-coding |
| chr14_30249596_30250001 | chr14 | 30249597 | 30250001 | + | -0.995106 | intron (NM_001083616, intron 3 of 46) | intron (NM_001083616, intron 3 of 46) | 103687 | NM_001083616 | 12289 | Mm.9772 | NM_028981 | ENSMUSG00000015968 | **Cacna1d** | 8430418G19Rik\|C79217\|Cacnl1a2\|Cchl1a\|Cchl1a2\|D-LTCC | calcium channel, voltage-dependent, L type, alpha 1D subunit | protein-coding |
| chr1_154794482_154794937 | chr1 | 154794483 | 154794937 | + | -0.728953 | Intergenic | L2\|LINE\|L2 | -68790 | NM_009782 | 12290 | Mm.267517 | NM_009782 | ENSMUSG00000004110 | **Cacna1e** | A430040I15\|BII\|Cach6\|Cacnl1a6\|Cav2.3\|Cchra1\|alpha1E | calcium channel, voltage-dependent, R type, alpha 1E subunit | protein-coding |
| chrX_7617828_7618855 | chrX | 7617829 | 7618855 | + | -0.759337 | intron (NM_019582, intron 17 of 47) | intron (NM_019582, intron 17 of 47) | 11239 | NM_019582 | 54652 | Mm.289647 | NM_019582 | ENSMUSG00000031142 | **Cacna1f** | A930034B14\|Cav1.4\|Sfc17\|nerg1\|nob2 | calcium channel, voltage-dependent, alpha 1F subunit | protein-coding |
| chrX_7618195_7618727 | chrX | 7618196 | 7618727 | + | -0.76632 | intron (NM_019582, intron 17 of 47) | intron (NM_019582, intron 17 of 47) | 11358 | NM_019582 | 54652 | Mm.289647 | NM_019582 | ENSMUSG00000031142 | **Cacna1f** | A930034B14\|Cav1.4\|Sfc17\|nerg1\|nob2 | calcium channel, voltage-dependent, alpha 1F subunit | protein-coding |
| chr11_98020180_98020557 | chr11 | 98020181 | 98020557 | + | -0.661089 | intron (NM_001282977, intron 1 of 13) | intron (NM_001282977, intron 1 of 13) | -1997 | NM_001159320 | 12295 | Mm.41252 | NM_031173 | ENSMUSG00000020882 | **Cacnb1** | CAB1\|Cchb1\|Cchlb1 | calcium channel, voltage-dependent, beta 1 subunit | protein-coding |
| chr2_14879719_14880232 | chr2 | 14879720 | 14880232 | + | -0.60125 | intron (NR_045533, intron 2 of 12) | HAL1\|LINE\|L1 | 6320 | NR_045533 | 12296 | Mm.313930 | NM_023116 | ENSMUSG00000057914 | **Cacnb2** | AW060387\|CAB2\|Cavbeta2\|Cchb2 | calcium channel, voltage-dependent, beta 2 subunit | protein-coding |
| chr2_84347369_84347783 | chr2 | 84347370 | 84347783 | + | -1.48147 | intron (NM_018782, intron 11 of 15) | intron (NM_018782, intron 11 of 15) | 77690 | NM_018782 | 54598 | Mm.75467 | NM_018782 | ENSMUSG00000059588 | **Calcrl** | AV071593\|CRLR | calcitonin receptor-like | protein-coding |
| chr12_100228123_100228839 | chr12 | 100228124 | 100228839 | + | -0.836509 | Intergenic | Intergenic | 28940 | NM_009790 | 12313 | Mm.285993 | NM_009790 | ENSMUSG00000001175 | **Calm1** | AI256814\|AI327027\|AI461935\|AL024000\|CaM\|Calm | calmodulin 1 | protein-coding |
| chr12_100228496_100228813 | chr12 | 100228497 | 100228813 | + | -0.967711 | Intergenic | Intergenic | 29114 | NM_009790 | 12313 | Mm.285993 | NM_009790 | ENSMUSG00000001175 | **Calm1** | AI256814\|AI327027\|AI461935\|AL024000\|CaM\|Calm | calmodulin 1 | protein-coding |
| chr2_5724941_5725255 | chr2 | 5724942 | 5725255 | + | -0.919709 | Intergenic | Intergenic | -10336 | NM_177343 | 227541 | Mm.191949 | NM_177343 | ENSMUSG00000039145 | **Camk1d** | A630059D12Rik\|CKLiK\|CaMKIdelta\|E030025C11Rik\|mCKLiK | calcium/calmodulin-dependent protein kinase ID | protein-coding |
| chr1_193408534_193408933 | chr1 | 193408535 | 193408933 | + | -0.66588 | Intergenic | ORR1E\|LTR\|ERVL-MaLR | -38452 | NM_144817 | 215303 | Mm.40329 | NM_144817 | ENSMUSG00000016179 | **Camk1g** | CLICK-III\|CaMKIgamma | calcium/calmodulin-dependent protein kinase I gamma | protein-coding |
| chr1_193345032_193345299 | chr1 | 193345033 | 193345299 | + | -0.695096 | Intergenic | Intergenic | 25116 | NM_144817 | 215303 | Mm.40329 | NM_144817 | ENSMUSG00000016179 | **Camk1g** | CLICK-III\|CaMKIgamma | calcium/calmodulin-dependent protein kinase I gamma | protein-coding |
| chr1_193408322_193408826 | chr1 | 193408323 | 193408826 | + | -0.729821 | Intergenic | Intergenic | -38292 | NM_144817 | 215303 | Mm.40329 | NM_144817 | ENSMUSG00000016179 | **Camk1g** | CLICK-III\|CaMKIgamma | calcium/calmodulin-dependent protein kinase I gamma | protein-coding |
| chr1_193421873_193422151 | chr1 | 193421874 | 193422151 | + | -0.843384 | Intergenic | Intergenic | -51730 | NM_144817 | 215303 | Mm.40329 | NM_144817 | ENSMUSG00000016179 | **Camk1g** | CLICK-III\|CaMKIgamma | calcium/calmodulin-dependent protein kinase I gamma | protein-coding |
| chr1_193349696_193350146 | chr1 | 193349697 | 193350146 | + | -0.946593 | intron (NM_144817, intron 9 of 12) | (TG)n\|Simple_repeat\|Simple_repeat | 20361 | NM_144817 | 215303 | Mm.40329 | NM_144817 | ENSMUSG00000016179 | **Camk1g** | CLICK-III\|CaMKIgamma | calcium/calmodulin-dependent protein kinase I gamma | protein-coding |
| chr1_193349723_193350054 | chr1 | 193349724 | 193350054 | + | -0.95272 | intron (NM_144817, intron 9 of 12) | intron (NM_144817, intron 9 of 12) | 20393 | NM_144817 | 215303 | Mm.40329 | NM_144817 | ENSMUSG00000016179 | **Camk1g** | CLICK-III\|CaMKIgamma | calcium/calmodulin-dependent protein kinase I gamma | protein-coding |
| chr1_193421369_193422156 | chr1 | 193421370 | 193422156 | + | -1.19332 | Intergenic | Intergenic | -51481 | NM_144817 | 215303 | Mm.40329 | NM_144817 | ENSMUSG00000016179 | **Camk1g** | CLICK-III\|CaMKIgamma | calcium/calmodulin-dependent protein kinase I gamma | protein-coding |
| chr1_193421372_193421798 | chr1 | 193421373 | 193421798 | + | -1.24902 | Intergenic | Intergenic | -51303 | NM_144817 | 215303 | Mm.40329 | NM_144817 | ENSMUSG00000016179 | **Camk1g** | CLICK-III\|CaMKIgamma | calcium/calmodulin-dependent protein kinase I gamma | protein-coding |
| chr3_126589595_126589918 | chr3 | 126589596 | 126589918 | + | -0.735119 | Intergenic | Intergenic | -7194 | NM_001293663 | 108058 | Mm.255822 | NM_023813 | ENSMUSG00000053819 | **Camk2d** | 2810011D23Rik\|8030469K03Rik\|CaMK II\|[d]-CaMKII | calcium/calmodulin-dependent protein kinase II, delta | protein-coding |
| chr18_32970174_32970666 | chr18 | 32970175 | 32970666 | + | -0.81634 | intron (NM_009793, intron 1 of 10) | intron (NM_009793, intron 1 of 10) | 31379 | NM_009793 | 12326 | Mm.222329 | NM_009793 | ENSMUSG00000038128 | **Camk4** | A430110E23Rik\|AI666733\|CaMKIV\|CaMKIV/Gr\|D18Bwg0362e | calcium/calmodulin-dependent protein kinase IV | protein-coding |
| chr18_32970174_32970909 | chr18 | 32970175 | 32970909 | + | -0.841773 | intron (NM_009793, intron 1 of 10) | intron (NM_009793, intron 1 of 10) | 31501 | NM_009793 | 12326 | Mm.222329 | NM_009793 | ENSMUSG00000038128 | **Camk4** | A430110E23Rik\|AI666733\|CaMKIV\|CaMKIV/Gr\|D18Bwg0362e | calcium/calmodulin-dependent protein kinase IV | protein-coding |
| chr18_32954689_32955700 | chr18 | 32954690 | 32955700 | + | -0.936523 | intron (NM_009793, intron 1 of 10) | intron (NM_009793, intron 1 of 10) | 16154 | NM_009793 | 12326 | Mm.222329 | NM_009793 | ENSMUSG00000038128 | **Camk4** | A430110E23Rik\|AI666733\|CaMKIV\|CaMKIV/Gr\|D18Bwg0362e | calcium/calmodulin-dependent protein kinase IV | protein-coding |
| chr18_32940801_32941121 | chr18 | 32940802 | 32941121 | + | -0.94276 | intron (NM_009793, intron 1 of 10) | intron (NM_009793, intron 1 of 10) | 1920 | NM_009793 | 12326 | Mm.222329 | NM_009793 | ENSMUSG00000038128 | **Camk4** | A430110E23Rik\|AI666733\|CaMKIV\|CaMKIV/Gr\|D18Bwg0362e | calcium/calmodulin-dependent protein kinase IV | protein-coding |
| chr18_32993947_32994500 | chr18 | 32993948 | 32994500 | + | -1.1261 | intron (NM_009793, intron 1 of 10) | intron (NM_009793, intron 1 of 10) | 55183 | NM_009793 | 12326 | Mm.222329 | NM_009793 | ENSMUSG00000038128 | **Camk4** | A430110E23Rik\|AI666733\|CaMKIV\|CaMKIV/Gr\|D18Bwg0362e | calcium/calmodulin-dependent protein kinase IV | protein-coding |
| chr18_32958142_32958609 | chr18 | 32958143 | 32958609 | + | -1.14935 | intron (NM_009793, intron 1 of 10) | L2\|LINE\|L2 | 19335 | NM_009793 | 12326 | Mm.222329 | NM_009793 | ENSMUSG00000038128 | **Camk4** | A430110E23Rik\|AI666733\|CaMKIV\|CaMKIV/Gr\|D18Bwg0362e | calcium/calmodulin-dependent protein kinase IV | protein-coding |
| chr18_32953941_32954445 | chr18 | 32953942 | 32954445 | + | -1.38542 | intron (NM_009793, intron 1 of 10) | intron (NM_009793, intron 1 of 10) | 15152 | NM_009793 | 12326 | Mm.222329 | NM_009793 | ENSMUSG00000038128 | **Camk4** | A430110E23Rik\|AI666733\|CaMKIV\|CaMKIV/Gr\|D18Bwg0362e | calcium/calmodulin-dependent protein kinase IV | protein-coding |
| chr18_32953954_32954283 | chr18 | 32953955 | 32954283 | + | -1.56285 | intron (NM_009793, intron 1 of 10) | intron (NM_009793, intron 1 of 10) | 15078 | NM_009793 | 12326 | Mm.222329 | NM_009793 | ENSMUSG00000038128 | **Camk4** | A430110E23Rik\|AI666733\|CaMKIV\|CaMKIV/Gr\|D18Bwg0362e | calcium/calmodulin-dependent protein kinase IV | protein-coding |
| chr11_73019344_73019929 | chr11 | 73019345 | 73019929 | + | -0.882888 | intron (NM_018883, intron 1 of 15) | CpG | 629 | NM_018883 | 55984 | Mm.9998 | NM_018883 | ENSMUSG00000020785 | **Camkk1** | AI846603\|CaMKKalpha\|Camkk | calcium/calmodulin-dependent protein kinase kinase 1, alpha | protein-coding |
| chr10_119218608_119218982 | chr10 | 119218609 | 119218982 | + | -0.631108 | intron (NM_027994, intron 3 of 14) | intron (NM_027994, intron 3 of 14) | 21260 | NM_027994 | 71902 | Mm.203965 | NM_027994 | ENSMUSG00000020114 | **Cand1** | 2310038O07Rik\|6330512O03Rik\|AI195005\|AI846556\|D10Ertd516e\|Tp120a\|mKIAA0829 | cullin associated and neddylation disassociated 1 | protein-coding |
| chr10_119218230_119219011 | chr10 | 119218231 | 119219011 | + | -0.711777 | intron (NM_027994, intron 3 of 14) | intron (NM_027994, intron 3 of 14) | 21434 | NM_027994 | 71902 | Mm.203965 | NM_027994 | ENSMUSG00000020114 | **Cand1** | 2310038O07Rik\|6330512O03Rik\|AI195005\|AI846556\|D10Ertd516e\|Tp120a\|mKIAA0829 | cullin associated and neddylation disassociated 1 | protein-coding |
| chr6_72550771_72551056 | chr6 | 72550772 | 72551056 | + | -0.670679 | intron (NM_001271395, intron 1 of 9) | intron (NM_001271395, intron 1 of 9) | 1267 | NM_001271395 | 12332 | Mm.18626 | NM_007599 | ENSMUSG00000056737 | **Capg** | gCap39\|mbh1 | capping protein (actin filament), gelsolin-like | protein-coding |
| chr17_73270927_73271366 | chr17 | 73270928 | 73271366 | + | -0.633292 | Intergenic | Intergenic | 128149 | NM_001033444 | 381122 | Mm.81496 | NM_001033444 | ENSMUSG00000043705 | **Capn13** | CANP 13\|Gm943 | calpain 13 | protein-coding |
| chr17_73270929_73271208 | chr17 | 73270930 | 73271208 | + | -0.635176 | Intergenic | Intergenic | 128227 | NM_001033444 | 381122 | Mm.81496 | NM_001033444 | ENSMUSG00000043705 | **Capn13** | CANP 13\|Gm943 | calpain 13 | protein-coding |
| chr17_73273799_73274209 | chr17 | 73273800 | 73274209 | + | -0.927153 | Intergenic | Intergenic | 125292 | NM_001033444 | 381122 | Mm.81496 | NM_001033444 | ENSMUSG00000043705 | **Capn13** | CANP 13\|Gm943 | calpain 13 | protein-coding |
| chr17_73273660_73274190 | chr17 | 73273661 | 73274190 | + | -0.945249 | Intergenic | Intergenic | 125371 | NM_001033444 | 381122 | Mm.81496 | NM_001033444 | ENSMUSG00000043705 | **Capn13** | CANP 13\|Gm943 | calpain 13 | protein-coding |

| | | | | | | | | | | | | | | | | | |
|---|---|---|---|---|---|---|---|---|---|---|---|---|---|---|---|---|---|
| chr1_182521159_182521443 | chr1 | 182521160 | 182521443 | + | -0.709239 | Intergenic | ORR1F\|LTR\|ERVL-MaLR | -3818 | NM_009794 | 12334 | Mm.19306 | NM_009794 | ENSMUSG00000026509 | **Capn2** | AI326419\|Capa-2\|Capa2\|m-calpain\|m-calpin | calpain 2 | protein-coding |
| chr9_66705825_66706257 | chr9 | 66705826 | 66706257 | + | -0.734496 | Intergenic | Intergenic | -7645 | NM_178396 | 76459 | Mm.277921 | NM_178396 | ENSMUSG00000032373 | **Car12** | 2310047E01Rik\|AI314958\|Ca12 | carbonic anyhydrase 12 | protein-coding |
| chr9_66680002_66680635 | chr9 | 66680003 | 66680635 | + | -1.00189 | Intergenic | Intergenic | -33367 | NM_178396 | 76459 | Mm.277921 | NM_178396 | ENSMUSG00000032373 | **Car12** | 2310047E01Rik\|AI314958\|Ca12 | carbonic anyhydrase 12 | protein-coding |
| chr3_14915363_14915765 | chr3 | 14915364 | 14915765 | + | -0.985892 | Intergenic | Intergenic | 29138 | NM_009801 | 12349 | Mm.1186 | NM_009801 | ENSMUSG00000027562 | **Car2** | AI131712\|CAII\|Ca2\|Car-2\|Ltw-5\|Lvtw-5 | carbonic anhydrase 2 | protein-coding |
| chr5_141044036_141044573 | chr5 | 141044037 | 141044573 | + | -0.689262 | Intergenic | Intergenic | -43709 | NM_175362 | 108723 | Mm.46187 | NM_175362 | ENSMUSG00000036526 | **Card11** | 0610008L17Rik\|2410011D02Rik\|BIMP3\|CARMA1 | caspase recruitment domain family, member 11 | protein-coding |
| chr5_140947892_140948613 | chr5 | 140947893 | 140948613 | + | -0.883662 | intron (NM_175362, intron 1 of 24) | intron (NM_175362, intron 1 of 24) | 52343 | NM_175362 | 108723 | Mm.46187 | NM_175362 | ENSMUSG00000036526 | **Card11** | 0610008L17Rik\|2410011D02Rik\|BIMP3\|CARMA1 | caspase recruitment domain family, member 11 | protein-coding |
| chr15_5107791_5109054 | chr15 | 5107792 | 5109054 | + | -0.790755 | intron (NM_001163138, intron 1 of 3) | intron (NM_001163138, intron 1 of 3) | 110 | NM_001163138 | 239319 | Mm.31252 | NM_001163138 | ENSMUSG00000041849 | **Card6** | D730008L15 | caspase recruitment domain family, member 6 | protein-coding |
| chr13_99932317_99932874 | chr13 | 99932318 | 99932874 | + | -0.659891 | Intergenic | Intergenic | -31913 | NM_001081493 | 27220 | Mm.75498 | NM_013732 | ENSMUSG00000021647 | **Cartpt** | Cart | CART prepropeptide | protein-coding |
| chr13_99935093_99935487 | chr13 | 99935094 | 99935487 | + | -0.837193 | Intergenic | Intergenic | -34607 | NM_001081493 | 27220 | Mm.75498 | NM_013732 | ENSMUSG00000021647 | **Cartpt** | Cart | CART prepropeptide | protein-coding |
| chr13_99935244_99935440 | chr13 | 99935245 | 99935440 | + | -0.853659 | Intergenic | Intergenic | -34659 | NM_001081493 | 27220 | Mm.75498 | NM_013732 | ENSMUSG00000021647 | **Cartpt** | Cart | CART prepropeptide | protein-coding |
| chr6_145193132_145193887 | chr6 | 145193133 | 145193887 | + | -0.684526 | intron (NM_177222, intron 7 of 15) | intron (NM_177222, intron 7 of 15) | 17460 | NM_177222 | 320662 | Mm.207779 | NM_177222 | ENSMUSG00000043541 | **Casc1** | A230084G12Rik\|Las1 | cancer susceptibility candidate 1 | protein-coding |
| chr2_121865951_121866292 | chr2 | 121865952 | 121866292 | + | -0.70983 | promoter-TSS (NM_199038) | promoter-TSS (NM_199038) | -848 | NM_199038 | 319996 | Mm.292109 | NM_177054 | ENSMUSG00000060227 | **Casc4** | D130060C09Rik\|VGFG2573 | cancer susceptibility candidate 4 | protein-coding |
| chr1_58778302_58778551 | chr1 | 58778303 | 58778551 | + | -0.666035 | Intergenic | Intergenic | -16947 | NM_009812 | 12370 | Mm.336851 | NM_009812 | ENSMUSG00000026029 | **Casp8** | CASP-8\|FLICE\|MACH\|Mch5 | caspase 8 | protein-coding |
| chr1_58778172_58778557 | chr1 | 58778173 | 58778557 | + | -0.730186 | Intergenic | L1MB1\|LINE\|L1 | -17009 | NM_009812 | 12370 | Mm.336851 | NM_009812 | ENSMUSG00000026029 | **Casp8** | CASP-8\|FLICE\|MACH\|Mch5 | caspase 8 | protein-coding |
| chr1_58806706_58806983 | chr1 | 58806707 | 58806983 | + | -0.869951 | intron (NM_001277926, intron 2 of 9) | intron (NM_001277926, intron 2 of 9) | 4343 | NM_001080126 | 12370 | Mm.336851 | NM_009812 | ENSMUSG00000026029 | **Casp8** | CASP-8\|FLICE\|MACH\|Mch5 | caspase 8 | protein-coding |
| chr13_74845717_74846596 | chr13 | 74845718 | 74846596 | + | -0.680104 | Intergenic | RMER2\|LTR\|ERV1 | -37397 | NM_001301153 | 12380 | Mm.29163 | NM_009817 | ENSMUSG00000021585 | **Cast** | - | calpastatin | protein-coding |
| chr13_74845898_74846316 | chr13 | 74845899 | 74846316 | + | -0.709286 | Intergenic | RMER2\|LTR\|ERV1 | -37347 | NM_001301153 | 12380 | Mm.29163 | NM_009817 | ENSMUSG00000021585 | **Cast** | - | calpastatin | protein-coding |
| chr8_122666748_122667284 | chr8 | 122666749 | 122667284 | + | -0.636199 | intron (NM_009824, intron 1 of 11) | intron (NM_009824, intron 1 of 11) | 11159 | NM_001109873 | 12398 | Mm.194339 | NM_009824 | ENSMUSG00000006362 | **Cbfa2t3** | A630044F12Rik\|AI465270\|AW229127\|Cbfa2t3h\|ETO-2\|Eto2\|MTGR2 | core-binding factor, runt domain, alpha subunit 2, translocated to, 3 (human) | protein-coding |
| chr8_122666556_122667297 | chr8 | 122666557 | 122667297 | + | -0.639619 | intron (NM_009824, intron 1 of 11) | intron (NM_009824, intron 1 of 11) | 11248 | NM_001109873 | 12398 | Mm.194339 | NM_009824 | ENSMUSG00000006362 | **Cbfa2t3** | A630044F12Rik\|AI465270\|AW229127\|Cbfa2t3h\|ETO-2\|Eto2\|MTGR2 | core-binding factor, runt domain, alpha subunit 2, translocated to, 3 (human) | protein-coding |
| chr16_52064404_52064959 | chr16 | 52064405 | 52064959 | + | -0.632349 | intron (NM_001033238, intron 3 of 17) | intron (NM_001033238, intron 3 of 17) | 33133 | NM_001033238 | 208650 | Mm.328206 | NM_001033238 | ENSMUSG00000022637 | **Cblb** | AI429560\|AI851073\|Cbl-b | Casitas B-lineage lymphoma b | protein-coding |
| chr16_52086612_52087125 | chr16 | 52086613 | 52087125 | + | -0.666678 | intron (NM_001033238, intron 3 of 17) | intron (NM_001033238, intron 3 of 17) | 55320 | NM_001033238 | 208650 | Mm.328206 | NM_001033238 | ENSMUSG00000022637 | **Cblb** | AI429560\|AI851073\|Cbl-b | Casitas B-lineage lymphoma b | protein-coding |
| chr16_52129403_52130059 | chr16 | 52129404 | 52130059 | + | -0.732706 | intron (NM_001033238, intron 4 of 17) | intron (NM_001033238, intron 4 of 17) | 98182 | NM_001033238 | 208650 | Mm.328206 | NM_001033238 | ENSMUSG00000022637 | **Cblb** | AI429560\|AI851073\|Cbl-b | Casitas B-lineage lymphoma b | protein-coding |
| chr16_51807218_51808005 | chr16 | 51807219 | 51808005 | + | -0.780637 | Intergenic | Intergenic | -223937 | NM_001033238 | 208650 | Mm.328206 | NM_001033238 | ENSMUSG00000022637 | **Cblb** | AI429560\|AI851073\|Cbl-b | Casitas B-lineage lymphoma b | protein-coding |
| chr16_52132759_52133430 | chr16 | 52132760 | 52133430 | + | -0.786492 | intron (NM_001033238, intron 5 of 17) | intron (NM_001033238, intron 5 of 17) | 101546 | NM_001033238 | 208650 | Mm.328206 | NM_001033238 | ENSMUSG00000022637 | **Cblb** | AI429560\|AI851073\|Cbl-b | Casitas B-lineage lymphoma b | protein-coding |
| chr16_52081839_52082161 | chr16 | 52081840 | 52082161 | + | -0.823362 | intron (NM_001033238, intron 3 of 17) | intron (NM_001033238, intron 3 of 17) | 50451 | NM_001033238 | 208650 | Mm.328206 | NM_001033238 | ENSMUSG00000022637 | **Cblb** | AI429560\|AI851073\|Cbl-b | Casitas B-lineage lymphoma b | protein-coding |
| chr16_52072013_52072398 | chr16 | 52072014 | 52072398 | + | -0.887918 | intron (NM_001033238, intron 3 of 17) | intron (NM_001033238, intron 3 of 17) | 40657 | NM_001033238 | 208650 | Mm.328206 | NM_001033238 | ENSMUSG00000022637 | **Cblb** | AI429560\|AI851073\|Cbl-b | Casitas B-lineage lymphoma b | protein-coding |
| chr16_52133128_52133441 | chr16 | 52133129 | 52133441 | + | -0.95247 | intron (NM_001033238, intron 5 of 17) | intron (NM_001033238, intron 5 of 17) | 101736 | NM_001033238 | 208650 | Mm.328206 | NM_001033238 | ENSMUSG00000022637 | **Cblb** | AI429560\|AI851073\|Cbl-b | Casitas B-lineage lymphoma b | protein-coding |
| chr16_52009444_52009804 | chr16 | 52009445 | 52009804 | + | -1.03997 | Intergenic | Intergenic | -21925 | NM_001033238 | 208650 | Mm.328206 | NM_001033238 | ENSMUSG00000022637 | **Cblb** | AI429560\|AI851073\|Cbl-b | Casitas B-lineage lymphoma b | protein-coding |
| chr16_52143084_52143560 | chr16 | 52143085 | 52143560 | + | -1.06201 | intron (NM_001033238, intron 9 of 17) | intron (NM_001033238, intron 9 of 17) | 111773 | NM_001033238 | 208650 | Mm.328206 | NM_001033238 | ENSMUSG00000022637 | **Cblb** | AI429560\|AI851073\|Cbl-b | Casitas B-lineage lymphoma b | protein-coding |
| chr16_52142791_52143602 | chr16 | 52142792 | 52143602 | + | -1.08787 | intron (NM_001033238, intron 9 of 17) | intron (NM_001033238, intron 9 of 17) | 111648 | NM_001033238 | 208650 | Mm.328206 | NM_001033238 | ENSMUSG00000022637 | **Cblb** | AI429560\|AI851073\|Cbl-b | Casitas B-lineage lymphoma b | protein-coding |
| chr16_52009478_52009695 | chr16 | 52009479 | 52009695 | + | -1.09477 | Intergenic | Intergenic | -21962 | NM_001033238 | 208650 | Mm.328206 | NM_001033238 | ENSMUSG00000022637 | **Cblb** | AI429560\|AI851073\|Cbl-b | Casitas B-lineage lymphoma b | protein-coding |
| chr16_52056208_52056572 | chr16 | 52056209 | 52056572 | + | -1.18598 | intron (NM_001033238, intron 3 of 17) | intron (NM_001033238, intron 3 of 17) | 24841 | NM_001033238 | 208650 | Mm.328206 | NM_001033238 | ENSMUSG00000022637 | **Cblb** | AI429560\|AI851073\|Cbl-b | Casitas B-lineage lymphoma b | protein-coding |
| chr16_93608859_93609053 | chr16 | 93608860 | 93609053 | + | -0.675797 | intron (NM_007620, intron 2 of 2) | intron (NM_007620, intron 2 of 2) | 1119 | NM_007620 | 12408 | Mm.26940 | NM_007620 | ENSMUSG00000051483 | **Cbr1** | AW261796\|CR\|Cbr | carbonyl reductase 1 | protein-coding |
| chr11_119075408_119076304 | chr11 | 119075409 | 119076304 | + | -1.11303 | Intergenic | MIRb\|SINE\|MIR | 10381 | NM_007625 | 12418 | Mm.268070 | NM_007625 | ENSMUSG00000039989 | **Cbx4** | MPc2\|PC2 | chromobox 4 | protein-coding |
| chr11_119075765_119076248 | chr11 | 119075766 | 119076248 | + | -1.23815 | Intergenic | Intergenic | 10230 | NM_007625 | 12418 | Mm.268070 | NM_007625 | ENSMUSG00000039989 | **Cbx4** | MPc2\|PC2 | chromobox 4 | protein-coding |
| chr15_79956510_79956855 | chr15 | 79956511 | 79956855 | + | -0.8581 | Intergenic | Intergenic | -24037 | NM_144811 | 52609 | Mm.323442 | NM_144811 | ENSMUSG00000053411 | **Cbx7** | 1600014J01Rik\|AI851678\|D15Ertd417e | chromobox 7 | protein-coding |
| chr11_119050482_119050689 | chr11 | 119050483 | 119050689 | + | -0.727498 | Intergenic | Intergenic | -9673 | NM_013926 | 30951 | Mm.99953 | NM_013926 | ENSMUSG00000025578 | **Cbx8** | Pc3 | chromobox 8 | protein-coding |
| chr4_108638584_108639054 | chr4 | 108638585 | 108639054 | + | -0.670087 | TTS (NM_183300) | TTS (NM_183300) | 18863 | NM_177045 | 319965 | Mm.151129 | NM_177045 | ENSMUSG00000028582 | **Cc2d1b** | A830039B04Rik\|Freud2 | coiled-coil and C2 domain containing 1B | protein-coding |
| chr6_49321371_49321711 | chr6 | 49321372 | 49321711 | + | -0.726988 | intron (NM_175098, intron 1 of 2) | intron (NM_175098, intron 1 of 2) | 2188 | NM_175098 | 57895 | Mm.32416 | NM_175098 | ENSMUSG00000050786 | **Ccdc126** | 6330407D12Rik\|AA675040\|AI465320\|AL022711 | coiled-coil domain containing 126 | protein-coding |
| chr6_83095649_83095896 | chr6 | 83095650 | 83095896 | + | -0.726954 | Intergenic | A-rich\|Low_complexity\|Low_complexity | -5743 | NM_001081266 | 243510 | Mm.436553 | NM_001081266 | ENSMUSG00000079511 | **Ccdc142** | A230058J24Rik | coiled-coil domain containing 142 | protein-coding |
| chr6_83095511_83096546 | chr6 | 83095512 | 83096546 | + | -1.10021 | Intergenic | Intergenic | -5487 | NM_001081266 | 243510 | Mm.436553 | NM_001081266 | ENSMUSG00000079511 | **Ccdc142** | A230058J24Rik | coiled-coil domain containing 142 | protein-coding |
| chr6_83095942_83096545 | chr6 | 83095943 | 83096545 | + | -1.14035 | Intergenic | RMER15\|LTR\|ERVL | -5272 | NM_001081266 | 243510 | Mm.436553 | NM_001081266 | ENSMUSG00000079511 | **Ccdc142** | A230058J24Rik | coiled-coil domain containing 142 | protein-coding |
| chr5_21410944_21411944 | chr5 | 21410945 | 21411944 | + | -0.614571 | intron (NM_029195, intron 1 of 18) | intron (NM_029195, intron 1 of 18) | 13233 | NM_029195 | 75172 | Mm.426695 | NM_029195 | ENSMUSG00000064280 | **Ccdc146** | 4930528G09Rik\|mKIAA1505 | coiled-coil domain containing 146 | protein-coding |
| chr5_21411017_21411699 | chr5 | 21411018 | 21411699 | + | -0.69848 | intron (NM_029195, intron 1 of 18) | intron (NM_029195, intron 1 of 18) | 13319 | NM_029195 | 75172 | Mm.426695 | NM_029195 | ENSMUSG00000064280 | **Ccdc146** | 4930528G09Rik\|mKIAA1505 | coiled-coil domain containing 146 | protein-coding |
| chr9_37310619_37311122 | chr9 | 37310620 | 37311122 | + | -0.910659 | intron (NM_001081429, intron 10 of 15) | intron (NM_001081429, intron 10 of 15) | 37521 | NM_001081429 | 245902 | Mm.390150 | NM_001081429 | ENSMUSG00000034303 | **Ccdc15** | A630039F14Rik\|D030074L19 | coiled-coil domain containing 15 | protein-coding |
| chr15_3309430_3310328 | chr15 | 3309431 | 3310328 | + | -1.04115 | Intergenic | Intergenic | -6353 | NM_001166063 | 100039139 | Mm.455610 | NM_001166063 | ENSMUSG00000091119 | **Ccdc152** | ENSMUSG00000071814 | coiled-coil domain containing 152 | protein-coding |
| chr15_3309841_3310360 | chr15 | 3309842 | 3310360 | + | -1.18613 | Intergenic | Tigger1\|DNA\|TcMar-Tigger | -6575 | NM_001166063 | 100039139 | Mm.455610 | NM_001166063 | ENSMUSG00000091119 | **Ccdc152** | ENSMUSG00000071814 | coiled-coil domain containing 152 | protein-coding |
| chr9_44238515_44238828 | chr9 | 44238516 | 44238828 | + | -0.939837 | Intergenic | Intergenic | -2005 | NM_001081369 | 270150 | Mm.347681 | NM_001081369 | ENSMUSG00000070306 | **Ccdc153** | - | coiled-coil domain containing 153 | protein-coding |
| chr10_41602653_41603028 | chr10 | 41602654 | 41603028 | + | -0.719496 | Intergenic | Intergenic | -15088 | NM_001177571 | 75973 | Mm.228517 | NM_029493 | ENSMUSG00000075225 | **Ccdc162** | 5033413D22Rik\|Ccdc162p\|Gm61 | coiled-coil domain containing 162 | protein-coding |
| chr10_41602736_41603013 | chr10 | 41602737 | 41603013 | + | -0.730849 | Intergenic | Intergenic | -15122 | NM_001177571 | 75973 | Mm.228517 | NM_029493 | ENSMUSG00000075225 | **Ccdc162** | 5033413D22Rik\|Ccdc162p\|Gm61 | coiled-coil domain containing 162 | protein-coding |
| chr10_41603269_41603923 | chr10 | 41603270 | 41603923 | + | -0.742566 | Intergenic | B4\|SINE\|B4 | -15843 | NM_001177571 | 75973 | Mm.228517 | NM_029493 | ENSMUSG00000075225 | **Ccdc162** | 5033413D22Rik\|Ccdc162p\|Gm61 | coiled-coil domain containing 162 | protein-coding |
| chr10_41615495_41615793 | chr10 | 41615496 | 41615793 | + | -0.752593 | Intergenic | Intergenic | -27891 | NM_001177571 | 75973 | Mm.228517 | NM_029493 | ENSMUSG00000075225 | **Ccdc162** | 5033413D22Rik\|Ccdc162p\|Gm61 | coiled-coil domain containing 162 | protein-coding |
| chr10_41615368_41616013 | chr10 | 41615369 | 41616013 | + | -0.803273 | Intergenic | Intergenic | -27938 | NM_001177571 | 75973 | Mm.228517 | NM_029493 | ENSMUSG00000075225 | **Ccdc162** | 5033413D22Rik\|Ccdc162p\|Gm61 | coiled-coil domain containing 162 | protein-coding |
| chr10_41595383_41596158 | chr10 | 41595384 | 41596158 | + | -0.85382 | Intergenic | Intergenic | -8018 | NM_001177571 | 75973 | Mm.228517 | NM_029493 | ENSMUSG00000075225 | **Ccdc162** | 5033413D22Rik\|Ccdc162p\|Gm61 | coiled-coil domain containing 162 | protein-coding |
| chr10_41603596_41603916 | chr10 | 41603597 | 41603916 | + | -0.895907 | Intergenic | B4\|SINE\|B4 | -16003 | NM_001177571 | 75973 | Mm.228517 | NM_029493 | ENSMUSG00000075225 | **Ccdc162** | 5033413D22Rik\|Ccdc162p\|Gm61 | coiled-coil domain containing 162 | protein-coding |
| chr10_41638801_41639101 | chr10 | 41638802 | 41639101 | + | -1.19683 | Intergenic | ORR1E\|LTR\|ERVL-MaLR | -51198 | NM_001177571 | 75973 | Mm.228517 | NM_029493 | ENSMUSG00000075225 | **Ccdc162** | 5033413D22Rik\|Ccdc162p\|Gm61 | coiled-coil domain containing 162 | protein-coding |

| | | | | | | | | | | | | | | | | | |
|---|---|---|---|---|---|---|---|---|---|---|---|---|---|---|---|---|---|
| chr10_41638809_41639202 | chr10 | 41638810 | 41639202 | + | -1.19722 | Intergenic | ORR1E\|LTR\|ERVL-MaLR | -51253 | NM_001177571 | 75973 | Mm.228517 | NM_029493 | ENSMUSG00000075225 | **Ccdc162** | 5033413D22Rik\|Ccdc162p\|Gm61 | coiled-coil domain containing 162 | protein-coding |
| chr10_41620020_41620362 | chr10 | 41620021 | 41620362 | + | -1.30332 | Intergenic | B3\|SINE\|B2 | -32438 | NM_001177571 | 75973 | Mm.228517 | NM_029493 | ENSMUSG00000075225 | **Ccdc162** | 5033413D22Rik\|Ccdc162p\|Gm61 | coiled-coil domain containing 162 | protein-coding |
| chr10_41619918_41620385 | chr10 | 41619919 | 41620385 | + | -1.31467 | Intergenic | B3\|SINE\|B2 | -32399 | NM_001177571 | 75973 | Mm.228517 | NM_029493 | ENSMUSG00000075225 | **Ccdc162** | 5033413D22Rik\|Ccdc162p\|Gm61 | coiled-coil domain containing 162 | protein-coding |
| chr4_83567654_83568046 | chr4 | 83567655 | 83568046 | + | -0.721017 | intron (NM_001081012, intron 6 of 25) | intron (NM_001081012, intron 6 of 25) | 42305 | NM_001081012 | 320226 | Mm.180032 | NM_029117 | ENSMUSG00000052407 | **Ccdc171** | 4930418J05Rik\|4930473A06Rik\|A330015D16Rik | coiled-coil domain containing 171 | protein-coding |
| chr9_58112190_58112759 | chr9 | 58112191 | 58112759 | + | -1.46934 | intron (NM_001166282, intron 3 of 19) | intron (NM_001166282, intron 3 of 19) | 6348 | NM_001166282 | 382077 | Mm.335193 | NM_029212 | ENSMUSG00000037716 | **Ccdc33** | 4930535E21Rik\|Gm1117\|Tzip1 | coiled-coil domain containing 33 | protein-coding |
| chr10_88204320_88204560 | chr10 | 88204321 | 88204560 | + | -0.896594 | intron (NM_027487, intron 2 of 5) | L1M3e\|LINE\|L1 | 3343 | NM_027487 | 67282 | Mm.208465 | NM_026070 | ENSMUSG00000020056 | **Ccdc53** | 2900091E11Rik\|5730495F03Rik | coiled-coil domain containing 53 | protein-coding |
| chr10_88203294_88203580 | chr10 | 88203295 | 88203580 | + | -0.933861 | intron (NM_027487, intron 2 of 5) | intron (NM_027487, intron 2 of 5) | 2340 | NM_027487 | 67282 | Mm.208465 | NM_026070 | ENSMUSG00000020056 | **Ccdc53** | 2900091E11Rik\|5730495F03Rik | coiled-coil domain containing 53 | protein-coding |
| chr10_70032765_70033421 | chr10 | 70032766 | 70033421 | + | -0.69185 | Intergenic | Intergenic | -64028 | NM_001111121 | 76551 | Mm.281741 | NM_001111121 | ENSMUSG00000048701 | **Ccdc6** | 2810012H18Rik\|AA536681\|AA960498\|AW061011 | coiled-coil domain containing 6 | protein-coding |
| chr5_115760432_115761502 | chr5 | 115760433 | 115761502 | + | -0.633771 | Intergenic | Intergenic | -29408 | NM_001080808 | 75665 | Mm.233914 | NM_001080808 | ENSMUSG00000041609 | **Ccdc64** | 2210403N09Rik\|BICDR-1 | coiled-coil domain containing 64 | protein-coding |
| chr5_115760827_115761499 | chr5 | 115760828 | 115761499 | + | -0.736512 | Intergenic | RMER15\|LTR\|ERVL | -29604 | NM_001080808 | 75665 | Mm.233914 | NM_001080808 | ENSMUSG00000041609 | **Ccdc64** | 2210403N09Rik\|BICDR-1 | coiled-coil domain containing 64 | protein-coding |
| chr5_115754687_115755473 | chr5 | 115754688 | 115755473 | + | -1.06119 | Intergenic | Intergenic | -23521 | NM_001080808 | 75665 | Mm.233914 | NM_001080808 | ENSMUSG00000041609 | **Ccdc64** | 2210403N09Rik\|BICDR-1 | coiled-coil domain containing 64 | protein-coding |
| chr11_55072504_55072774 | chr11 | 55072505 | 55072774 | + | -0.681638 | intron (NM_177471, intron 1 of 6) | intron (NM_177471, intron 1 of 6) | 5492 | NM_177471 | 52570 | Mm.22361 | NM_177471 | ENSMUSG00000049588 | **Ccdc69** | 2210021E03Rik\|D11Ertd461e | coiled-coil domain containing 69 | protein-coding |
| chr11_55072423_55072995 | chr11 | 55072424 | 55072995 | + | -0.722461 | intron (NM_177471, intron 1 of 6) | intron (NM_177471, intron 1 of 6) | 5422 | NM_177471 | 52570 | Mm.22361 | NM_177471 | ENSMUSG00000049588 | **Ccdc69** | 2210021E03Rik\|D11Ertd461e | coiled-coil domain containing 69 | protein-coding |
| chr7_17009377_17009634 | chr7 | 17009378 | 17009634 | + | -0.670896 | intron (NM_011155, intron 4 of 12) | intron (NM_011155, intron 4 of 12) | 14918 | NM_001101535 | 434130 | Mm.459162 | NM_001101535 | | **Ccdc8** | ENSMUSG00000041117 | coiled-coil domain containing 8 | protein-coding |
| chr16_45025321_45025781 | chr16 | 45025322 | 45025781 | + | -0.68907 | Intergenic | Intergenic | -68502 | NM_026439 | 67896 | Mm.181074 | NM_026439 | ENSMUSG00000022665 | **Ccdc80** | 2610001E17Rik\|DRO1\|Ssg1\|Urb | coiled-coil domain containing 80 | protein-coding |
| chr16_45021490_45021942 | chr16 | 45021491 | 45021942 | + | -0.693138 | Intergenic | Intergenic | -72337 | NM_026439 | 67896 | Mm.181074 | NM_026439 | ENSMUSG00000022665 | **Ccdc80** | 2610001E17Rik\|DRO1\|Ssg1\|Urb | coiled-coil domain containing 80 | protein-coding |
| chr16_45018812_45019216 | chr16 | 45018813 | 45019216 | + | -1.04467 | Intergenic | Intergenic | -75039 | NM_026439 | 67896 | Mm.181074 | NM_026439 | ENSMUSG00000022665 | **Ccdc80** | 2610001E17Rik\|DRO1\|Ssg1\|Urb | coiled-coil domain containing 80 | protein-coding |
| chr9_13305867_13306078 | chr9 | 13305868 | 13306078 | + | -0.740819 | Intergenic | Intergenic | 58994 | NM_025534 | 66396 | Mm.290620 | NM_025534 | | **Ccdc82** | 2310043N13Rik | coiled-coil domain containing 82 | protein-coding |
| chr9_13323586_13324006 | chr9 | 13323587 | 13324006 | + | -0.769442 | Intergenic | Intergenic | 76817 | NM_025534 | 66396 | Mm.290620 | NM_025534 | | **Ccdc82** | 2310043N13Rik | coiled-coil domain containing 82 | protein-coding |
| chr9_121454341_121455624 | chr9 | 121454342 | 121455624 | + | -0.695448 | intron (NM_175114, intron 13 of 15) | intron (NM_175114, intron 13 of 15) | 40711 | NM_001284508 | 12424 | Mm.2619 | NM_031161 | ENSMUSG00000032532 | **Cck** | - | cholecystokinin | protein-coding |
| chr11_82214337_82214915 | chr11 | 82214338 | 82214915 | + | -1.03502 | Intergenic | Intergenic | -34814 | NM_011329 | 20290 | Mm.1283 | NM_011329 | ENSMUSG00000020702 | **Ccl1** | BF534335\|I-309\|Scya1\|Tca-3 | chemokine (C-C motif) ligand 1 | protein-coding |
| chr13_119596926_119597156 | chr13 | 119596927 | 119597156 | + | -0.678154 | intron (NM_001039244, intron 2 of 2) | intron (NM_001039244, intron 2 of 2) | -26778 | NM_020279 | 56838 | Mm.143745 | NM_020279 | ENSMUSG00000074715 | **Ccl28** | CCK1\|MEC\|Scya28 | chemokine (C-C motif) ligand 28 | protein-coding |
| chr11_6560907_6561415 | chr11 | 6560908 | 6561415 | + | -0.643904 | 5' UTR (NM_001190344, exon 2 of 10) | 5' UTR (NM_001190344, exon 2 of 10) | 978 | NM_001190344 | 216527 | Mm.221271 | NM_146014 | ENSMUSG00000000378 | **Ccm2** | BC029157 | cerebral cavernous malformation 2 | protein-coding |
| chr11_6560918_6561228 | chr11 | 6560919 | 6561228 | + | -0.69476 | intron (NM_001190344, intron 1 of 9) | intron (NM_001190344, intron 1 of 9) | 890 | NM_001190344 | 216527 | Mm.221271 | NM_146014 | ENSMUSG00000000378 | **Ccm2** | BC029157 | cerebral cavernous malformation 2 | protein-coding |
| chr11_6567935_6568313 | chr11 | 6567936 | 6568313 | + | -0.964527 | intron (NM_001190344, intron 2 of 9) | intron (NM_001190344, intron 2 of 9) | 7941 | NM_001190344 | 216527 | Mm.221271 | NM_146014 | ENSMUSG00000000378 | **Ccm2** | BC029157 | cerebral cavernous malformation 2 | protein-coding |
| chr17_47600913_47601302 | chr17 | 47600914 | 47601302 | + | -0.732752 | 3' UTR (NM_016859, exon 7 of 7) | 3' UTR (NM_016859, exon 7 of 7) | 7641 | NM_007632 | 12445 | Mm.27291 | NM_007632 | ENSMUSG00000034165 | **Ccnd3** | 9230106B05Rik\|AA682053\|AL024085\|AW146355\|C78795 | cyclin D3 | protein-coding |
| chr17_47521416_47522206 | chr17 | 47521417 | 47522206 | + | -0.778238 | intron (NM_001081636, intron 1 of 6) | intron (NM_001081636, intron 1 of 6) | 16760 | NM_001081635 | 12445 | Mm.27291 | NM_007632 | ENSMUSG00000034165 | **Ccnd3** | 9230106B05Rik\|AA682053\|AL024085\|AW146355\|C78795 | cyclin D3 | protein-coding |
| chr7_38094253_38094833 | chr7 | 38094254 | 38094833 | + | -0.90403 | Intergenic | Intergenic | 12947 | NM_007633 | 12447 | Mm.16110 | NM_007633 | ENSMUSG00000002068 | **Ccne1** | AW538188\|CycE1 | cyclin E1 | protein-coding |
| chr11_40744350_40744811 | chr11 | 40744351 | 40744811 | + | -0.707401 | Intergenic | ID_B1\|SINE\|B4 | 10705 | NM_009831 | 12450 | Mm.2103 | NM_009831 | ENSMUSG00000020326 | **Ccng1** | AI314029 | cyclin G1 | protein-coding |
| chr11_40744196_40744780 | chr11 | 40744197 | 40744780 | + | -0.758907 | Intergenic | Intergenic | 10798 | NM_009831 | 12450 | Mm.2103 | NM_009831 | ENSMUSG00000020326 | **Ccng1** | AI314029 | cyclin G1 | protein-coding |
| chr13_85126783_85127564 | chr13 | 85126784 | 85127564 | + | -0.820457 | Intergenic | Intergenic | -62303 | NM_023243 | 66671 | Mm.18474 | NM_023243 | ENSMUSG00000021548 | **Ccnh** | 6330408H09Rik\|AI661354\|AV102684\|AW538719 | cyclin H | protein-coding |
| chr13_85126517_85127574 | chr13 | 85126518 | 85127574 | + | -0.88208 | Intergenic | Intergenic | -62431 | NM_023243 | 66671 | Mm.18474 | NM_023243 | ENSMUSG00000021548 | **Ccnh** | 6330408H09Rik\|AI661354\|AV102684\|AW538719 | cyclin H | protein-coding |
| chr1_127796323_127797116 | chr1 | 127796324 | 127797116 | + | -0.667368 | intron (NM_028399, intron 5 of 8) | intron (NM_028399, intron 5 of 8) | 22556 | NM_028399 | 72949 | Mm.268672 | NM_028399 | | **Ccnt2** | 2900041I18Rik\|C81304\|CycT2\|CycT2a\|CycT2b | cyclin T2 | protein-coding |
| chr9_123941395_123941707 | chr9 | 123941396 | 123941707 | + | -0.683863 | Intergenic | Intergenic | 27141 | NM_009912 | 12768 | Mm.274927 | NM_009912 | ENSMUSG00000025804 | **Ccr1** | Cmkbr1\|Mip-1a-R | chemokine (C-C motif) receptor 1 | protein-coding |
| chr9_123928061_123929273 | chr9 | 123928062 | 123929273 | + | -0.69336 | Intergenic | ORR1F\|LTR\|ERVL-MaLR | 40025 | NM_009912 | 12768 | Mm.274927 | NM_009912 | ENSMUSG00000025804 | **Ccr1** | Cmkbr1\|Mip-1a-R | chemokine (C-C motif) receptor 1 | protein-coding |
| chr9_123928103_123928356 | chr9 | 123928104 | 123928356 | + | -0.697731 | Intergenic | MLT1F\|LTR\|ERVL-MaLR | 40462 | NM_009912 | 12768 | Mm.274927 | NM_009912 | ENSMUSG00000025804 | **Ccr1** | Cmkbr1\|Mip-1a-R | chemokine (C-C motif) receptor 1 | protein-coding |
| chr9_123941356_123941775 | chr9 | 123941357 | 123941775 | + | -0.72192 | Intergenic | Intergenic | 27126 | NM_009912 | 12768 | Mm.274927 | NM_009912 | ENSMUSG00000025804 | **Ccr1** | Cmkbr1\|Mip-1a-R | chemokine (C-C motif) receptor 1 | protein-coding |
| chr9_123931687_123931972 | chr9 | 123931688 | 123931972 | + | -0.871322 | Intergenic | Intergenic | 36862 | NM_009912 | 12768 | Mm.274927 | NM_009912 | ENSMUSG00000025804 | **Ccr1** | Cmkbr1\|Mip-1a-R | chemokine (C-C motif) receptor 1 | protein-coding |
| chr9_123938010_123938358 | chr9 | 123938011 | 123938358 | + | -1.02715 | Intergenic | Intergenic | 30508 | NM_009912 | 12768 | Mm.274927 | NM_009912 | ENSMUSG00000025804 | **Ccr1** | Cmkbr1\|Mip-1a-R | chemokine (C-C motif) receptor 1 | protein-coding |
| chr9_124079597_124080423 | chr9 | 124079598 | 124080423 | + | -0.622241 | Intergenic | Intergenic | -22173 | NM_009915 | 12772 | Mm.6272 | NM_009915 | ENSMUSG00000049103 | **Ccr2** | Cc-ckr-2\|Ccr2a\|Ccr2b\|Ckr2\|Ckr2a\|Ckr2b\|Cmkbr2\|mJe-r | chemokine (C-C motif) receptor 2 | protein-coding |
| chr9_124100837_124101324 | chr9 | 124100838 | 124101324 | + | -0.791716 | Intergenic | (GGAGA)n\|Simple_repeat\|Simple_repeat | -1102 | NM_009915 | 12772 | Mm.6272 | NM_009915 | ENSMUSG00000049103 | **Ccr2** | Cc-ckr-2\|Ccr2a\|Ccr2b\|Ckr2\|Ckr2a\|Ckr2b\|Cmkbr2\|mJe-r | chemokine (C-C motif) receptor 2 | protein-coding |
| chr9_124100351_124100750 | chr9 | 124100352 | 124100750 | + | -0.799942 | Intergenic | Intergenic | -1632 | NM_009915 | 12772 | Mm.6272 | NM_009915 | ENSMUSG00000049103 | **Ccr2** | Cc-ckr-2\|Ccr2a\|Ccr2b\|Ckr2\|Ckr2a\|Ckr2b\|Cmkbr2\|mJe-r | chemokine (C-C motif) receptor 2 | protein-coding |
| chr9_124039005_124039330 | chr9 | 124039006 | 124039330 | + | -0.979635 | Intergenic | Intergenic | 17197 | NM_009914 | 12771 | Mm.57050 | NM_009914 | ENSMUSG00000035448 | **Ccr3** | CC-CKR3\|CKR3\|Cmkbr1l2\|Cmkbr3 | chemokine (C-C motif) receptor 3 | protein-coding |
| chr11_99130782_99131022 | chr11 | 99130783 | 99131022 | + | -0.664315 | Intergenic | Intergenic | 24175 | NM_007719 | 12775 | Mm.2932 | NM_007719 | ENSMUSG00000037944 | **Ccr7** | CD197\|Cdw197\|Cmkbr7\|EBI1\|Ebi1h | chemokine (C-C motif) receptor 7 | protein-coding |
| chr11_99147850_99148181 | chr11 | 99147851 | 99148181 | + | -0.710429 | intron (NM_007719, intron 2 of 2) | intron (NM_007719, intron 2 of 2) | 7061 | NM_007719 | 12775 | Mm.2932 | NM_007719 | ENSMUSG00000037944 | **Ccr7** | CD197\|Cdw197\|Cmkbr7\|EBI1\|Ebi1h | chemokine (C-C motif) receptor 7 | protein-coding |
| chr11_99166293_99166597 | chr11 | 99166294 | 99166597 | + | -0.785449 | Intergenic | Intergenic | -11368 | NM_007719 | 12775 | Mm.2932 | NM_007719 | ENSMUSG00000037944 | **Ccr7** | CD197\|Cdw197\|Cmkbr7\|EBI1\|Ebi1h | chemokine (C-C motif) receptor 7 | protein-coding |
| chr11_99136382_99136881 | chr11 | 99136383 | 99136881 | + | -1.21938 | Intergenic | Intergenic | 18445 | NM_007719 | 12775 | Mm.2932 | NM_007719 | ENSMUSG00000037944 | **Ccr7** | CD197\|Cdw197\|Cmkbr7\|EBI1\|Ebi1h | chemokine (C-C motif) receptor 7 | protein-coding |
| chr9_120095975_120096415 | chr9 | 120095976 | 120096415 | + | -0.811455 | Intergenic | Intergenic | 4062 | NM_007720 | 12776 | Mm.442098 | NM_007720 | ENSMUSG00000042262 | **Ccr8** | C-C\|C-C CKR-8\|CC-CKR-8\|CCR-8\|CKR-8\|Cmkbr8\|mCCR8 | chemokine (C-C motif) receptor 8 | protein-coding |
| chr9_120084476_120084767 | chr9 | 120084477 | 120084767 | + | -0.952223 | Intergenic | Intergenic | -7511 | NM_007720 | 12776 | Mm.442098 | NM_007720 | ENSMUSG00000042262 | **Ccr8** | C-C\|C-C CKR-8\|CC-CKR-8\|CCR-8\|CKR-8\|Cmkbr8\|mCCR8 | chemokine (C-C motif) receptor 8 | protein-coding |
| chr9_123767022_123767475 | chr9 | 123767023 | 123767475 | + | -0.759126 | promoter-TSS (NM_001166625) | promoter-TSS (NM_001166625) | 38 | NM_001166625 | 12769 | Mm.440604 | NM_009913 | | **Ccr9** | A130091K22Rik\|Cmkbr10\|GPR-9-6 | chemokine (C-C motif) receptor 9 | protein-coding |
| chr9_123754325_123754848 | chr9 | 123754326 | 123754848 | + | -0.802978 | Intergenic | Intergenic | -12624 | NM_009913 | 12769 | Mm.440604 | NM_009913 | | **Ccr9** | A130091K22Rik\|Cmkbr10\|GPR-9-6 | chemokine (C-C motif) receptor 9 | protein-coding |
| chr9_123756571_123757137 | chr9 | 123756572 | 123757137 | + | -1.11802 | Intergenic | Intergenic | -10357 | NM_009913 | 12769 | Mm.440604 | NM_009913 | | **Ccr9** | A130091K22Rik\|Cmkbr10\|GPR-9-6 | chemokine (C-C motif) receptor 9 | protein-coding |
| chr14_36953148_36953547 | chr14 | 36953149 | 36953547 | + | -0.761925 | intron (NM_027045, intron 1 of 10) | Lx9\|LINE\|L1 | 15416 | NM_027045 | 72972 | Mm.27621 | NM_027045 | ENSMUSG00000058690 | **Ccser2** | 1700012P13Rik\|2900054P12Rik\|AI035535\|AW108503\|Fam190b\|Gcap14 | coiled-coil serine rich 2 | protein-coding |
| chr5_129790608_129790947 | chr5 | 129790609 | 129790947 | + | -0.60944 | intron (NM_009838, intron 5 of 12) | intron (NM_009838, intron 5 of 12) | 3422 | NM_009838 | 12466 | Mm.153159 | NM_009838 | | **Cct6a** | CCT-zeta\|CCT-zeta-1\|Cct6\|Cctz-1\|TCP-1-zeta | chaperonin containing Tcp1, subunit 6a (zeta) | protein-coding |
| chr3_101059295_101059549 | chr3 | 101059296 | 101059549 | + | -0.874689 | intron (NM_011197, intron 5 of 8) | L2\|LINE\|L2 | -29927 | NM_001167906 | 630146 | Mm.190915 | NM_001099332 | ENSMUSG00000086564 | **Cd101** | Gm1016\|Gm734\|Igsf2 | CD101 antigen | protein-coding |

| | | | | | | | | | | | | | | | | | |
|---|---|---|---|---|---|---|---|---|---|---|---|---|---|---|---|---|---|
| chr3_101059185_101059582 | chr3 | 101059186 | 101059582 | + | -0.885877 | intron (NM_011197, intron 5 of 8) | L2\|LINE\|L2 | -29889 | NM_001167906 | 630146 | Mm.190915 | NM_001099332 | ENSMUSG00000086564 | **Cd101** | Gm1016\|Gm734\|Igsf2 | CD101 antigen | protein-coding |
| chr3_101066082_101066582 | chr3 | 101066083 | 101066582 | + | -1.27526 | intron (NM_011197, intron 4 of 8) | intron (NM_011197, intron 4 of 8) | -36837 | NM_001167906 | 630146 | Mm.190915 | NM_001099332 | ENSMUSG00000086564 | **Cd101** | Gm1016\|Gm734\|Igsf2 | CD101 antigen | protein-coding |
| chr3_101029212_101029798 | chr3 | 101029213 | 101029798 | + | -1.32887 | promoter-TSS (NM_001167906) | promoter-TSS (NM_001167906) | -10 | NM_001167906 | 630146 | Mm.190915 | NM_001099332 | ENSMUSG00000086564 | **Cd101** | Gm1016\|Gm734\|Igsf2 | CD101 antigen | protein-coding |
| chr3_96814245_96814795 | chr3 | 96814246 | 96814795 | + | -0.777132 | promoter-TSS (NM_018767) | promoter-TSS (NM_018767) | 1 | NM_018767 | 54215 | Mm.34693 | NM_018767 | ENSMUSG00000038304 | **Cd160** | AU045688\|By55 | CD160 antigen | protein-coding |
| chr7_140213676_140214258 | chr7 | 140213677 | 140214258 | + | -1.01892 | Intergenic | Intergenic | -4300 | NM_172909 | 244233 | Mm.19283 | NM_172909 | ENSMUSG00000025461 | **Cd163l1** | B430307C05\|E430002D04Rik\|Scart1 | CD163 molecule-like 1 | protein-coding |
| chr7_126416512_126416977 | chr7 | 126416513 | 126416977 | + | -0.659234 | Intergenic | Intergenic | -1875 | NM_009844 | 12478 | Mm.4360 | NM_009844 | ENSMUSG00000030724 | **Cd19** | AW495831 | CD19 antigen | protein-coding |
| chr3_101283867_101284167 | chr3 | 101283868 | 101284167 | + | -0.680434 | intron (NM_013486, intron 2 of 4) | intron (NM_013486, intron 2 of 4) | 3922 | NM_013486 | 12481 | Mm.22842 | NM_013486 | ENSMUSG00000027863 | **Cd2** | LFA-2\|Ly-37\|Ly37 | CD2 antigen | protein-coding |
| chr3_101274626_101275213 | chr3 | 101274627 | 101275213 | + | -0.695354 | TTS (NM_013486) | TTS (NM_013486) | 13019 | NM_013486 | 12481 | Mm.22842 | NM_013486 | ENSMUSG00000027863 | **Cd2** | LFA-2\|Ly-37\|Ly37 | CD2 antigen | protein-coding |
| chr3_101296420_101296725 | chr3 | 101296421 | 101296725 | + | -0.707275 | Intergenic | Intergenic | -8634 | NM_013486 | 12481 | Mm.22842 | NM_013486 | ENSMUSG00000027863 | **Cd2** | LFA-2\|Ly-37\|Ly37 | CD2 antigen | protein-coding |
| chr3_101276477_101276876 | chr3 | 101276478 | 101276876 | + | -0.746382 | intron (NM_013486, intron 4 of 4) | B4\|SINE\|B4 | 11262 | NM_013486 | 12481 | Mm.22842 | NM_013486 | ENSMUSG00000027863 | **Cd2** | LFA-2\|Ly-37\|Ly37 | CD2 antigen | protein-coding |
| chr16_44875160_44875561 | chr16 | 44875161 | 44875561 | + | -0.658432 | intron (NM_206535, intron 1 of 5) | Lx6\|LINE\|L1 | 8264 | NM_206535 | 271375 | Mm.441535 | NM_206535 | ENSMUSG00000090176 | **Cd200r2** | AY230198\|Cd200r1l | Cd200 receptor 2 | protein-coding |
| chr16_45016524_45016850 | chr16 | 45016525 | 45016850 | + | -0.749923 | Intergenic | Intergenic | 73009 | NM_001128132 | 74603 | Mm.297337 | NM_027578 | ENSMUSG00000036172 | **Cd200r3** | 4733401I18Rik\|4833409J19Rik\|AI505817\|BB106877\|mCD200RLb | CD200 receptor 3 | protein-coding |
| chr16_44962851_44963095 | chr16 | 44962852 | 44963095 | + | -0.967958 | intron (NM_029018, intron 5 of 5) | MIR3\|SINE\|MIR | 19295 | NM_001128132 | 74603 | Mm.297337 | NM_027578 | ENSMUSG00000036172 | **Cd200r3** | 4733401I18Rik\|4833409J19Rik\|AI505817\|BB106877\|mCD200RLb | CD200 receptor 3 | protein-coding |
| chr16_45006952_45007504 | chr16 | 45006953 | 45007504 | + | -1.06024 | Intergenic | Intergenic | 63550 | NM_001128132 | 74603 | Mm.297337 | NM_027578 | ENSMUSG00000036172 | **Cd200r3** | 4733401I18Rik\|4833409J19Rik\|AI505817\|BB106877\|mCD200RLb | CD200 receptor 3 | protein-coding |
| chr16_44811573_44811924 | chr16 | 44811574 | 44811924 | + | -0.806873 | Intergenic | Intergenic | -8979 | NM_207244 | 239849 | Mm.445045 | NM_207244 | ENSMUSG00000062082 | **Cd200r4** | F630107N04Rik\|MCD200RLa | CD200 receptor 4 | protein-coding |
| chr18_89340286_89341175 | chr18 | 89340287 | 89341175 | + | -0.644369 | intron (NM_001039173, intron 7 of 7) | intron (NM_001039173, intron 7 of 7) | 143304 | NM_001039149 | 225825 | Mm.139293 | NM_178687 | ENSMUSG00000034028 | **Cd226** | BC051526\|DNAM-1\|DNAM1\|Pta1\|TLiSA1 | CD226 antigen | protein-coding |
| chr18_89241254_89241787 | chr18 | 89241255 | 89241787 | + | -0.682615 | intron (NM_178687, intron 3 of 6) | intron (NM_178687, intron 3 of 6) | 44094 | NM_001039149 | 225825 | Mm.139293 | NM_178687 | ENSMUSG00000034028 | **Cd226** | BC051526\|DNAM-1\|DNAM1\|Pta1\|TLiSA1 | CD226 antigen | protein-coding |
| chr18_89126432_89126798 | chr18 | 89126433 | 89126798 | + | -0.696958 | intron (NM_175542, intron 46 of 48) | ORR1E\|LTR\|ERVL-MaLR | -70812 | NM_178687 | 225825 | Mm.139293 | NM_178687 | ENSMUSG00000034028 | **Cd226** | BC051526\|DNAM-1\|DNAM1\|Pta1\|TLiSA1 | CD226 antigen | protein-coding |
| chr18_89340303_89340883 | chr18 | 89340304 | 89340883 | + | -0.712402 | intron (NM_001039173, intron 7 of 7) | intron (NM_001039173, intron 7 of 7) | 143166 | NM_001039149 | 225825 | Mm.139293 | NM_178687 | ENSMUSG00000034028 | **Cd226** | BC051526\|DNAM-1\|DNAM1\|Pta1\|TLiSA1 | CD226 antigen | protein-coding |
| chr18_89325874_89326275 | chr18 | 89325875 | 89326275 | + | -0.765342 | intron (NM_001039173, intron 7 of 7) | B4\|SINE\|B4 | 128648 | NM_001039149 | 225825 | Mm.139293 | NM_178687 | ENSMUSG00000034028 | **Cd226** | BC051526\|DNAM-1\|DNAM1\|Pta1\|TLiSA1 | CD226 antigen | protein-coding |
| chr18_89301169_89301698 | chr18 | 89301170 | 89301698 | + | -0.802287 | intron (NM_001039173, intron 7 of 7) | intron (NM_001039173, intron 7 of 7) | 104007 | NM_001039149 | 225825 | Mm.139293 | NM_178687 | ENSMUSG00000034028 | **Cd226** | BC051526\|DNAM-1\|DNAM1\|Pta1\|TLiSA1 | CD226 antigen | protein-coding |
| chr18_89206326_89206729 | chr18 | 89206327 | 89206729 | + | -1.07507 | intron (NM_178687, intron 1 of 6) | intron (NM_178687, intron 1 of 6) | 9101 | NM_001039149 | 225825 | Mm.139293 | NM_178687 | ENSMUSG00000034028 | **Cd226** | BC051526\|DNAM-1\|DNAM1\|Pta1\|TLiSA1 | CD226 antigen | protein-coding |
| chr1_165816360_165817155 | chr1 | 165816361 | 165817155 | + | -0.624926 | intron (NM_031162, intron 1 of 7) | intron (NM_031162, intron 1 of 7) | 28077 | NM_001113392 | 12503 | Mm.217308 | NM_031162 | ENSMUSG00000005763 | **Cd247** | 4930549J05Rik\|A430104F18Rik\|AW552088\|Cd3\|Cd3-eta\|Cd3-zeta\|Cd3h\|Cd3z\|Cd3zeta\|T3z\|Tcrk\|Tcrz | CD247 antigen | protein-coding |
| chr1_165816360_165816916 | chr1 | 165816361 | 165816916 | + | -0.63242 | intron (NM_031162, intron 1 of 7) | intron (NM_031162, intron 1 of 7) | 27957 | NM_001113392 | 12503 | Mm.217308 | NM_031162 | ENSMUSG00000005763 | **Cd247** | 4930549J05Rik\|A430104F18Rik\|AW552088\|Cd3\|Cd3-eta\|Cd3-zeta\|Cd3h\|Cd3z\|Cd3zeta\|T3z\|Tcrk\|Tcrz | CD247 antigen | protein-coding |
| chr1_165788520_165788903 | chr1 | 165788521 | 165788903 | + | -0.64356 | promoter-TSS (NM_001113391) | promoter-TSS (NM_001113391) | 31 | NM_001113392 | 12503 | Mm.217308 | NM_031162 | ENSMUSG00000005763 | **Cd247** | 4930549J05Rik\|A430104F18Rik\|AW552088\|Cd3\|Cd3-eta\|Cd3-zeta\|Cd3h\|Cd3z\|Cd3zeta\|T3z\|Tcrk\|Tcrz | CD247 antigen | protein-coding |
| chr1_165810682_165810955 | chr1 | 165810683 | 165810955 | + | -0.688659 | intron (NM_031162, intron 1 of 7) | intron (NM_031162, intron 1 of 7) | 22138 | NM_001113392 | 12503 | Mm.217308 | NM_031162 | ENSMUSG00000005763 | **Cd247** | 4930549J05Rik\|A430104F18Rik\|AW552088\|Cd3\|Cd3-eta\|Cd3-zeta\|Cd3h\|Cd3z\|Cd3zeta\|T3z\|Tcrk\|Tcrz | CD247 antigen | protein-coding |
| chr1_165837471_165838002 | chr1 | 165837472 | 165838002 | + | -0.707121 | intron (NM_031162, intron 1 of 7) | intron (NM_031162, intron 1 of 7) | 49056 | NM_001113392 | 12503 | Mm.217308 | NM_031162 | ENSMUSG00000005763 | **Cd247** | 4930549J05Rik\|A430104F18Rik\|AW552088\|Cd3\|Cd3-eta\|Cd3-zeta\|Cd3h\|Cd3z\|Cd3zeta\|T3z\|Tcrk\|Tcrz | CD247 antigen | protein-coding |
| chr1_165837439_165838249 | chr1 | 165837440 | 165838249 | + | -0.7262 | intron (NM_031162, intron 1 of 7) | intron (NM_031162, intron 1 of 7) | 49163 | NM_001113392 | 12503 | Mm.217308 | NM_031162 | ENSMUSG00000005763 | **Cd247** | 4930549J05Rik\|A430104F18Rik\|AW552088\|Cd3\|Cd3-eta\|Cd3-zeta\|Cd3h\|Cd3z\|Cd3zeta\|T3z\|Tcrk\|Tcrz | CD247 antigen | protein-coding |
| chr1_165848978_165849337 | chr1 | 165848979 | 165849337 | + | -0.902643 | intron (NM_031162, intron 1 of 7) | intron (NM_031162, intron 1 of 7) | 60477 | NM_001113392 | 12503 | Mm.217308 | NM_031162 | ENSMUSG00000005763 | **Cd247** | 4930549J05Rik\|A430104F18Rik\|AW552088\|Cd3\|Cd3-eta\|Cd3-zeta\|Cd3h\|Cd3z\|Cd3zeta\|T3z\|Tcrk\|Tcrz | CD247 antigen | protein-coding |
| chr1_60781649_60782121 | chr1 | 60781650 | 60782121 | + | -0.764133 | Intergenic | Lx9\|LINE\|L1 | 35497 | NM_007642 | 12487 | Mm.255003 | NM_007642 | ENSMUSG00000026012 | **Cd28** | - | CD28 antigen | protein-coding |
| chr1_60755819_60756231 | chr1 | 60755820 | 60756231 | + | -0.872489 | intron (NM_007642, intron 1 of 3) | intron (NM_007642, intron 1 of 3) | 9637 | NM_007642 | 12487 | Mm.255003 | NM_007642 | ENSMUSG00000026012 | **Cd28** | - | CD28 antigen | protein-coding |
| chr1_60793735_60794157 | chr1 | 60793736 | 60794157 | + | -0.915308 | Intergenic | Intergenic | 47558 | NM_007642 | 12487 | Mm.255003 | NM_007642 | ENSMUSG00000026012 | **Cd28** | - | CD28 antigen | protein-coding |
| chr1_60813460_60813791 | chr1 | 60813461 | 60813791 | + | -1.24782 | Intergenic | MIRc\|SINE\|MIR | 67238 | NM_007642 | 12487 | Mm.255003 | NM_007642 | ENSMUSG00000026012 | **Cd28** | - | CD28 antigen | protein-coding |
| chr1_60758129_60758656 | chr1 | 60758130 | 60758656 | + | -1.34346 | intron (NM_007642, intron 1 of 3) | intron (NM_007642, intron 1 of 3) | 12005 | NM_007642 | 12487 | Mm.255003 | NM_007642 | ENSMUSG00000026012 | **Cd28** | - | CD28 antigen | protein-coding |
| chr17_42875401_42875703 | chr17 | 42875402 | 42875703 | + | -0.668453 | intron (NM_009847, intron 1 of 17) | intron (NM_009847, intron 1 of 17) | 872 | NM_009847 | 12488 | Mm.218637 | NM_009847 | ENSMUSG00000061665 | **Cd2ap** | AL024079\|C78928\|METS-1\|Mets1 | CD2-associated protein | protein-coding |
| chr7_127180755_127181209 | chr7 | 127180756 | 127181209 | + | -0.873444 | Intergenic | Intergenic | 15095 | NM_001285907 | 70233 | Mm.28050 | NM_027353 | ENSMUSG00000042502 | **Cd2bp2** | 1500011B02Rik\|2410024K20Rik\|AW048183 | CD2 antigen (cytoplasmic tail) binding protein 2 | protein-coding |
| chr11_114876140_114876566 | chr11 | 114876141 | 114876566 | + | -0.764839 | Intergenic | B3\|SINE\|B2 | -13688 | NM_170758 | 217303 | Mm.294439 | NM_170758 | ENSMUSG00000034652 | **Cd300a** | B230315M08Rik\|Clm8\|LMIR1\|MAIR-I\|MAIR-Ia\|MMAC8\|Pigr4\|mcpir1 | CD300A antigen | protein-coding |
| chr11_114876167_114876741 | chr11 | 114876168 | 114876741 | + | -0.891673 | Intergenic | B3\|SINE\|B2 | -13587 | NM_170758 | 217303 | Mm.294439 | NM_170758 | ENSMUSG00000034652 | **Cd300a** | B230315M08Rik\|Clm8\|LMIR1\|MAIR-I\|MAIR-Ia\|MMAC8\|Pigr4\|mcpir1 | CD300A antigen | protein-coding |
| chr11_114872943_114873753 | chr11 | 114872944 | 114873753 | + | -1.12865 | TTS (NM_001110337) | TTS (NM_001110337) | -16693 | NM_170758 | 217303 | Mm.294439 | NM_170758 | ENSMUSG00000034652 | **Cd300a** | B230315M08Rik\|Clm8\|LMIR1\|MAIR-I\|MAIR-Ia\|MMAC8\|Pigr4\|mcpir1 | CD300A antigen | protein-coding |
| chr2_60278595_60279076 | chr2 | 60278596 | 60279076 | + | -0.679565 | intron (NM_001290660, intron 1 of 5) | ORR1E\|LTR\|ERVL-MaLR | 5648 | NM_025422 | 66205 | Mm.227281 | NM_025422 | ENSMUSG00000060703 | **Cd302** | 1110055L24Rik\|AI159627 | CD302 antigen | protein-coding |
| chr9_45015798_45016360 | chr9 | 45015799 | 45016360 | + | -1.05275 | Intergenic | MLT2F\|LTR\|ERVL | -6489 | NM_007648 | 12501 | Mm.210361 | NM_007648 | ENSMUSG00000032093 | **Cd3e** | AI504783\|CD3\|CD3epsilon\|T3e | CD3 antigen, epsilon polypeptide | protein-coding |
| chr6_124885099_124885695 | chr6 | 124885100 | 124885695 | + | -0.808806 | intron (NM_013488, intron 1 of 9) | intron (NM_013488, intron 1 of 9) | 2812 | NM_013488 | 12504 | Mm.2209 | NM_013488 | ENSMUSG00000023274 | **Cd4** | L3T4\|Ly-4 | CD4 antigen | protein-coding |
| chr6_124885848_124886483 | chr6 | 124885849 | 124886483 | + | -1.3243 | intron (NM_013488, intron 1 of 9) | intron (NM_013488, intron 1 of 9) | 2043 | NM_013488 | 12504 | Mm.2209 | NM_013488 | ENSMUSG00000023274 | **Cd4** | L3T4\|Ly-4 | CD4 antigen | protein-coding |
| chrX_57221828_57222348 | chrX | 57221829 | 57222348 | + | -0.706006 | intron (NM_011616, intron 4 of 4) | intron (NM_011616, intron 4 of 4) | 9945 | NM_011616 | 21947 | Mm.4861 | NM_011616 | ENSMUSG00000031132 | **Cd40lg** | CD154\|CD40-L\|Cd40l\|HIGM1\|IGM\|IMD3\|Ly-62\|Ly62\|T-BAM\|TRAP\|Tnfsf5\|gp39 | CD40 ligand | protein-coding |

| | | | | | | | | | | | | | | | | | |
|---|---|---|---|---|---|---|---|---|---|---|---|---|---|---|---|---|---|
| chrX_57253339_57254104 | chrX | 57253340 | 57254104 | + | -0.757154 | intron (NM_152801, intron 10 of 21) | intron (NM_152801, intron 10 of 21) | 41579 | NM_011616 | 21947 | Mm.4861 | NM_011616 | ENSMUSG00000031132 | **Cd40lg** | CD154\|CD40-L\|Cd40l\|HIGM1\|IGM\|IMD3\|Ly-62\|Ly62\|T-BAM\|TRAP\|Tnfsf5\|gp39 | CD40 ligand | protein-coding |
| chr2_102912964_102913368 | chr2 | 102912965 | 102913368 | + | -0.656316 | Intergenic | Intergenic | -11501 | NM_001039150 | 12505 | Mm.423621 | NM_009851 | ENSMUSG00000005087 | **Cd44** | AU023126\|AW121933\|AW146109\|HERMES\|Ly-24\|Pgp-1 | CD44 antigen | protein-coding |
| chr2_102873690_102874488 | chr2 | 102873691 | 102874488 | + | -0.682578 | intron (NM_001177787, intron 1 of 10) | intron (NM_001177787, intron 1 of 10) | 27576 | NM_001177785 | 12505 | Mm.423621 | NM_009851 | ENSMUSG00000005087 | **Cd44** | AU023126\|AW121933\|AW146109\|HERMES\|Ly-24\|Pgp-1 | CD44 antigen | protein-coding |
| chr2_102886390_102887015 | chr2 | 102886391 | 102887015 | + | -1.03093 | intron (NM_001177787, intron 1 of 10) | intron (NM_001177787, intron 1 of 10) | 14962 | NM_001177785 | 12505 | Mm.423621 | NM_009851 | ENSMUSG00000005087 | **Cd44** | AU023126\|AW121933\|AW146109\|HERMES\|Ly-24\|Pgp-1 | CD44 antigen | protein-coding |
| chr2_102848968_102849273 | chr2 | 102848969 | 102849273 | + | -1.29146 | intron (NM_009851, intron 5 of 18) | intron (NM_009851, intron 5 of 18) | 52544 | NM_001177785 | 12505 | Mm.423621 | NM_009851 | ENSMUSG00000005087 | **Cd44** | AU023126\|AW121933\|AW146109\|HERMES\|Ly-24\|Pgp-1 | CD44 antigen | protein-coding |
| chr2_102848931_102849416 | chr2 | 102848932 | 102849416 | + | -1.29616 | intron (NM_009851, intron 5 of 18) | intron (NM_009851, intron 5 of 18) | 52491 | NM_001177785 | 12505 | Mm.423621 | NM_009851 | ENSMUSG00000005087 | **Cd44** | AU023126\|AW121933\|AW146109\|HERMES\|Ly-24\|Pgp-1 | CD44 antigen | protein-coding |
| chr16_49801411_49801847 | chr16 | 49801412 | 49801847 | + | -0.622901 | Intergenic | Intergenic | -54025 | NM_010581 | 16423 | Mm.31752 | NM_010581 | ENSMUSG00000055447 | **Cd47** | 9130415E20Rik\|AA407862\|AI848868\|AW108519\|B430305P08Rik\|IAP\|Itgp | CD47 antigen (Rh-related antigen, integrin-associated signal transducer) | protein-coding |
| chr16_49798684_49799215 | chr16 | 49798685 | 49799215 | + | -0.636717 | Intergenic | Intergenic | -56704 | NM_010581 | 16423 | Mm.31752 | NM_010581 | ENSMUSG00000055447 | **Cd47** | 9130415E20Rik\|AA407862\|AI848868\|AW108519\|B430305P08Rik\|IAP\|Itgp | CD47 antigen (Rh-related antigen, integrin-associated signal transducer) | protein-coding |
| chr16_49792406_49793000 | chr16 | 49792407 | 49793000 | + | -0.670943 | Intergenic | Intergenic | -62951 | NM_010581 | 16423 | Mm.31752 | NM_010581 | ENSMUSG00000055447 | **Cd47** | 9130415E20Rik\|AA407862\|AI848868\|AW108519\|B430305P08Rik\|IAP\|Itgp | CD47 antigen (Rh-related antigen, integrin-associated signal transducer) | protein-coding |
| chr16_49839603_49840580 | chr16 | 49839604 | 49840580 | + | -0.724598 | Intergenic | Intergenic | -15562 | NM_010581 | 16423 | Mm.31752 | NM_010581 | ENSMUSG00000055447 | **Cd47** | 9130415E20Rik\|AA407862\|AI848868\|AW108519\|B430305P08Rik\|IAP\|Itgp | CD47 antigen (Rh-related antigen, integrin-associated signal transducer) | protein-coding |
| chr16_49781624_49781925 | chr16 | 49781625 | 49781925 | + | -0.731913 | Intergenic | Intergenic | -73879 | NM_010581 | 16423 | Mm.31752 | NM_010581 | ENSMUSG00000055447 | **Cd47** | 9130415E20Rik\|AA407862\|AI848868\|AW108519\|B430305P08Rik\|IAP\|Itgp | CD47 antigen (Rh-related antigen, integrin-associated signal transducer) | protein-coding |
| chr16_49818585_49819096 | chr16 | 49818586 | 49819096 | + | -0.733644 | Intergenic | MTD\|LTR\|ERVL-MaLR | -36813 | NM_010581 | 16423 | Mm.31752 | NM_010581 | ENSMUSG00000055447 | **Cd47** | 9130415E20Rik\|AA407862\|AI848868\|AW108519\|B430305P08Rik\|IAP\|Itgp | CD47 antigen (Rh-related antigen, integrin-associated signal transducer) | protein-coding |
| chr16_49781594_49782267 | chr16 | 49781595 | 49782267 | + | -0.738654 | Intergenic | Intergenic | -73723 | NM_010581 | 16423 | Mm.31752 | NM_010581 | ENSMUSG00000055447 | **Cd47** | 9130415E20Rik\|AA407862\|AI848868\|AW108519\|B430305P08Rik\|IAP\|Itgp | CD47 antigen (Rh-related antigen, integrin-associated signal transducer) | protein-coding |
| chr4_134091220_134091469 | chr4 | 134091221 | 134091469 | + | -0.714582 | intron (NM_001162970, intron 18 of 19) | intron (NM_001162970, intron 18 of 19) | 3728 | NM_013706 | 23833 | Mm.24130 | NM_013706 | ENSMUSG00000000682 | **Cd52** | AI463198\|B7\|B7-Ag\|CAMPATH-1\|CLS1\|MB7 | CD52 antigen | protein-coding |
| chr3_106787686_106788916 | chr3 | 106787687 | 106788916 | + | -0.640941 | intron (NM_007651, intron 1 of 7) | intron (NM_007651, intron 1 of 7) | 1848 | NM_007651 | 12508 | Mm.316861 | NM_007651 | ENSMUSG00000040747 | **Cd53** | AI323659\|Ox-44\|Tspan25 | CD53 antigen | protein-coding |
| chr3_106788385_106788916 | chr3 | 106788386 | 106788916 | + | -0.908902 | intron (NM_007651, intron 1 of 7) | intron (NM_007651, intron 1 of 7) | 1498 | NM_007651 | 12508 | Mm.316861 | NM_007651 | ENSMUSG00000040747 | **Cd53** | AI323659\|Ox-44\|Tspan25 | CD53 antigen | protein-coding |
| chr1_130474879_130475312 | chr1 | 130474880 | 130475312 | + | -0.869022 | Intergenic | MER102a\|DNA\|hAT-Charlie | -12356 | NM_010016 | 13136 | Mm.101591 | NM_010016 | ENSMUSG00000026399 | **Cd55** | Daf\|Daf-GPI\|Daf1\|GPI-DAF | CD55 antigen | protein-coding |
| chr11_121038942_121039666 | chr11 | 121038943 | 121039666 | + | -0.746149 | exon (NM_009854, exon 1 of 4) | exon (NM_009854, exon 1 of 4) | 174 | NM_009854 | 12516 | Mm.4100 | NM_009854 | ENSMUSG00000025163 | **Cd7** | - | CD7 antigen | protein-coding |
| chr11_121039275_121039736 | chr11 | 121039276 | 121039736 | + | -0.770472 | promoter-TSS (NM_009854) | promoter-TSS (NM_009854) | -28 | NM_009854 | 12516 | Mm.4100 | NM_009854 | ENSMUSG00000025163 | **Cd7** | - | CD7 antigen | protein-coding |
| chr4_43456492_43456917 | chr4 | 43456493 | 43456917 | + | -0.665616 | Intergenic | Intergenic | -2079 | NM_007654 | 12517 | Mm.188157 | NM_007654 | ENSMUSG00000028459 | **Cd72** | CD72c\|Ly-19\|Ly-32\|Ly-m19\|Lyb-2 | CD72 antigen | protein-coding |
| chr18_60805745_60806030 | chr18 | 60805746 | 60806030 | + | -0.639038 | intron (NM_001042605, intron 1 of 8) | intron (NM_001042605, intron 1 of 8) | 2039 | NM_010545 | 16149 | Mm.439737 | NM_010545 | ENSMUSG00000024610 | **Cd74** | CLIP\|DHLAG\|HLADG\|Ia-GAMMA\|Ii | CD74 antigen (invariant polypeptide of major histocompatibility complex, class II antigen-associated) | protein-coding |
| chr11_106317761_106318135 | chr11 | 106317762 | 106318135 | + | -0.875983 | TTS (NM_133199) | TTS (NM_133199) | -3419 | NM_008339 | 15985 | Mm.2987 | NM_008339 | ENSMUSG00000040592 | **Cd79b** | B29\|Ig-beta\|Igb\|Igbeta | CD79B antigen | protein-coding |
| chr2_93458948_93459512 | chr2 | 93458949 | 93459512 | + | -0.706155 | intron (NR_040532, intron 2 of 4) | intron (NR_040532, intron 2 of 4) | 2621 | NM_001271461 | 12521 | Mm.4261 | NM_007656 | ENSMUSG00000027215 | **Cd82** | AA682076\|AL023070\|C33\|IA4\|Kai1\|Tspan27 | CD82 antigen | protein-coding |
| chr13_43906047_43906725 | chr13 | 43906048 | 43906725 | + | -1.01179 | Intergenic | Intergenic | 121274 | NM_001289915 | 12522 | Mm.57175 | NM_009856 | ENSMUSG00000015396 | **Cd83** | - | CD83 antigen | protein-coding |
| chr1_171845097_171845346 | chr1 | 171845098 | 171845346 | + | -0.701647 | intron (NM_001289470, intron 2 of 8) | intron (NM_001289470, intron 2 of 8) | 5525 | NM_013489 | 12523 | Mm.259115 | NM_013489 | ENSMUSG00000038147 | **Cd84** | A130013D22Rik\|CDw84\|SLAMF5 | CD84 antigen | protein-coding |
| chr6_71369269_71369854 | chr6 | 71369270 | 71369854 | + | -0.643321 | Intergenic | Intergenic | -3865 | NM_009857 | 12525 | Mm.1858 | NM_009857 | ENSMUSG00000053977 | **Cd8a** | BB154331\|Ly-2\|Ly-35\|Ly-B\|Lyt-2 | CD8 antigen, alpha chain | protein-coding |
| chr6_71376764_71377080 | chr6 | 71376765 | 71377080 | + | -0.911936 | 3' UTR (NM_001081110, exon 5 of 5) | 3' UTR (NM_001081110, exon 5 of 5) | 3495 | NM_001081110 | 12525 | Mm.1858 | NM_009857 | ENSMUSG00000053977 | **Cd8a** | BB154331\|Ly-2\|Ly-35\|Ly-B\|Lyt-2 | CD8 antigen, alpha chain | protein-coding |
| chr6_71333888_71334183 | chr6 | 71333889 | 71334183 | + | -0.760678 | intron (NM_009858, intron 4 of 5) | intron (NM_009858, intron 4 of 5) | 11224 | NM_009858 | 12526 | Mm.333148 | NM_009858 | ENSMUSG00000053044 | **Cd8b1** | Cd8b\|Ly-3\|Ly-C\|Lyt-3 | CD8 antigen, beta chain 1 | protein-coding |
| chr6_71307531_71308064 | chr6 | 71307532 | 71308064 | + | -0.852837 | Intergenic | Intergenic | -15014 | NM_009858 | 12526 | Mm.333148 | NM_009858 | ENSMUSG00000053044 | **Cd8b1** | Cd8b\|Ly-3\|Ly-C\|Lyt-3 | CD8 antigen, beta chain 1 | protein-coding |
| chr16_46107465_46107821 | chr16 | 46107466 | 46107821 | + | -0.658731 | intron (NM_032465, intron 3 of 14) | intron (NM_032465, intron 3 of 14) | 12605 | NM_032465 | 84544 | Mm.29204 | NM_032465 | ENSMUSG00000022657 | **Cd96** | 1700109I12Rik\|Tactile | CD96 antigen | protein-coding |
| chr16_46104465_46104971 | chr16 | 46104466 | 46104971 | + | -0.737402 | intron (NM_032465, intron 3 of 14) | intron (NM_032465, intron 3 of 14) | 15530 | NM_032465 | 84544 | Mm.29204 | NM_032465 | ENSMUSG00000022657 | **Cd96** | 1700109I12Rik\|Tactile | CD96 antigen | protein-coding |
| chr8_83736709_83737175 | chr8 | 83736710 | 83737175 | + | -0.987731 | intron (NM_011925, intron 1 of 19) | intron (NM_011925, intron 1 of 19) | 4369 | NM_001163029 | 26364 | Mm.334648 | NM_011925 | ENSMUSG00000002885 | **Cd97** | AA409984\|TM7LN1 | CD97 antigen | protein-coding |
| chr8_83773521_83773886 | chr8 | 83773522 | 83773886 | + | -1.08833 | Intergenic | Intergenic | -32393 | NM_001163031 | 26364 | Mm.334648 | NM_011925 | ENSMUSG00000002885 | **Cd97** | AA409984\|TM7LN1 | CD97 antigen | protein-coding |
| chr14_59555014_59555537 | chr14 | 59555015 | 59555537 | + | -0.750158 | Intergenic | ORR1E\|LTR\|ERVL-MaLR | 42683 | NM_001168537 | 71891 | Mm.439850 | NM_027986 | ENSMUSG00000021982 | **Cdadc1** | 2310010M10Rik\|AI449174\|AI463398\|BB130816\|NYD-SP15 | cytidine and dCMP deaminase domain containing 1 | protein-coding |
| chr14_59554963_59555351 | chr14 | 59554964 | 59555351 | + | -0.851832 | Intergenic | ORR1E\|LTR\|ERVL-MaLR | 42802 | NM_001168537 | 71891 | Mm.439850 | NM_027986 | ENSMUSG00000021982 | **Cdadc1** | 2310010M10Rik\|AI449174\|AI463398\|BB130816\|NYD-SP15 | cytidine and dCMP deaminase domain containing 1 | protein-coding |
| chr2_5801312_5801625 | chr2 | 5801313 | 5801625 | + | -0.734272 | intron (NM_133837, intron 10 of 12) | intron (NM_133837, intron 10 of 12) | 43491 | NM_133837 | 98828 | Mm.181490 | NM_133837 | ENSMUSG00000039128 | **Cdc123** | AA959893\|G431001I09Rik | cell division cycle 123 | protein-coding |
| chr3_116415255_116415661 | chr3 | 116415256 | 116415661 | + | -0.633367 | intron (NM_001173553, intron 2 of 13) | Lx8\|LINE\|L1 | 8574 | NM_001080818 | 229776 | Mm.17647 | NM_001080818 | ENSMUSG00000033502 | **Cdc14a** | A830059A17Rik\|CDC14A2\|CDC14a1\|Cdc14 | CDC14 cell division cycle 14A | protein-coding |
| chr3_116436163_116436392 | chr3 | 116436164 | 116436392 | + | -0.714843 | Intergenic | Intergenic | -12246 | NM_001173553 | 229776 | Mm.17647 | NM_001080818 | ENSMUSG00000033502 | **Cdc14a** | A830059A17Rik\|CDC14A2\|CDC14a1\|Cdc14 | CDC14 cell division cycle 14A | protein-coding |
| chr3_116374346_116374880 | chr3 | 116374347 | 116374880 | + | -0.927724 | intron (NM_001080818, intron 4 of 15) | intron (NM_001080818, intron 4 of 15) | 49419 | NM_001080818 | 229776 | Mm.17647 | NM_001080818 | ENSMUSG00000033502 | **Cdc14a** | A830059A17Rik\|CDC14A2\|CDC14a1\|Cdc14 | CDC14 cell division cycle 14A | protein-coding |
| chr2_131189920_131190258 | chr2 | 131189921 | 131190258 | + | -0.753752 | intron (NM_001111075, intron 3 of 14) | intron (NM_001111075, intron 3 of 14) | 3141 | NM_001111075 | 12531 | Mm.38444 | NM_023117 | ENSMUSG00000027330 | **Cdc25b** | AI604853 | cell division cycle 25B | protein-coding |
| chr4_137358348_137358631 | chr4 | 137358349 | 137358631 | + | -0.701601 | promoter-TSS (NM_001243769) | promoter-TSS (NM_001243769) | -731 | NM_001243769 | 12540 | Mm.1022 | NM_009861 | ENSMUSG00000006699 | **Cdc42** | AI747189\|AU018915 | cell division cycle 42 | protein-coding |
| chr4_137322363_137322720 | chr4 | 137322364 | 137322720 | + | -0.918345 | exon (NM_001243769, exon 6 of 6) | exon (NM_001243769, exon 6 of 6) | 35217 | NM_009861 | 12540 | Mm.1022 | NM_009861 | ENSMUSG00000006699 | **Cdc42** | AI747189\|AU018915 | cell division cycle 42 | protein-coding |
| chr19_6302520_6302795 | chr19 | 6302521 | 6302795 | + | -0.77961 | Intergenic | Intergenic | -3799 | NM_001033342 | 240505 | Mm.329622 | NM_001033342 | ENSMUSG00000024769 | **Cdc42bpg** | BC046418\|MRCKgamma | CDC42 binding protein kinase gamma (DMPK-like) | protein-coding |
| chr19_6302549_6302753 | chr19 | 6302550 | 6302753 | + | -0.819041 | Intergenic | Intergenic | -3806 | NM_001033342 | 240505 | Mm.329622 | NM_001033342 | ENSMUSG00000024769 | **Cdc42bpg** | BC046418\|MRCKgamma | CDC42 binding protein kinase gamma (DMPK-like) | protein-coding |

| | | | | | | | | | | | | | | | | | |
|---|---|---|---|---|---|---|---|---|---|---|---|---|---|---|---|---|---|
| chr19_6293433_6294026 | chr19 | 6293434 | 6294026 | + | -0.840379 | intron (NM_010119, intron 2 of 4) | MER58A\|DNA\|hAT-Charlie | -12727 | NM_001033342 | 240505 | Mm.329622 | NM_001033342 | ENSMUSG00000024769 | **Cdc42bpg** | BC046418\|MRCKgamma | CDC42 binding protein kinase gamma (DMPK-like) | protein-coding |
| chr19_6293428_6293798 | chr19 | 6293429 | 6293798 | + | -0.925147 | intron (NM_010119, intron 2 of 4) | MER58A\|DNA\|hAT-Charlie | -12844 | NM_001033342 | 240505 | Mm.329622 | NM_001033342 | ENSMUSG00000024769 | **Cdc42bpg** | BC046418\|MRCKgamma | CDC42 binding protein kinase gamma (DMPK-like) | protein-coding |
| chr15_78826133_78826534 | chr15 | 78826134 | 78826534 | + | -0.718494 | Intergenic | Intergenic | -16313 | NM_027219 | 104445 | Mm.307488 | NM_027219 | ENSMUSG00000049521 | **Cdc42ep1** | 1810058K22Rik\|AA980734\|Borg5\|CEP1\|MSE55 | CDC42 effector protein (Rho GTPase binding) 1 | protein-coding |
| chr17_79379162_79379721 | chr17 | 79379163 | 79379721 | + | -0.629018 | Intergenic | MTD-int\|LTR\|ERVL-MaLR | -24351 | NM_026514 | 260409 | Mm.140601 | NM_026514 | ENSMUSG00000036533 | **Cdc42ep3** | 3200001F04Rik\|AA986861\|Borg2\|Cep3\|UB1 | CDC42 effector protein (Rho GTPase binding) 3 | protein-coding |
| chr17_79347767_79348593 | chr17 | 79347768 | 79348593 | + | -0.840629 | intron (NM_026514, intron 1 of 1) | L3b\|LINE\|CR1 | 6911 | NM_026514 | 260409 | Mm.140601 | NM_026514 | ENSMUSG00000036533 | **Cdc42ep3** | 3200001F04Rik\|AA986861\|Borg2\|Cep3\|UB1 | CDC42 effector protein (Rho GTPase binding) 3 | protein-coding |
| chr11_113788486_113788758 | chr11 | 113788487 | 113788758 | + | -0.818669 | intron (NM_172800, intron 44 of 44) | intron (NM_172800, intron 44 of 44) | -36807 | NM_001163346 | 56699 | Mm.293378 | NM_020006 | ENSMUSG00000041598 | **Cdc42ep4** | 1500041M20Rik\|Borg4\|CEP4 | CDC42 effector protein (Rho GTPase binding) 4 | protein-coding |
| chr3_95231262_95231567 | chr3 | 95231263 | 95231567 | + | -1.03817 | intron (NM_001038708, intron 1 of 4) | intron (NM_001038708, intron 1 of 4) | 2472 | NM_001038708 | 57912 | Mm.28189 | NM_172395 | ENSMUSG00000046722 | **Cdc42se1** | 1300002M12Rik\|AW558204\|Cdcse1\|SCIP1\|Spec1 | CDC42 small effector 1 | protein-coding |
| chr11_54764043_54764390 | chr11 | 54764044 | 54764390 | + | -0.630336 | intron (NM_178626, intron 1 of 5) | intron (NM_178626, intron 1 of 5) | 23486 | NM_178626 | 72729 | Mm.29476 | NM_178626 | ENSMUSG00000052298 | **Cdc42se2** | 2810404F18Rik\|AA408783\|AA536669\|SPEC2 | CDC42 small effector 2 | protein-coding |
| chr11_54741237_54741742 | chr11 | 54741238 | 54741742 | + | -0.688326 | intron (NM_178626, intron 2 of 5) | intron (NM_178626, intron 2 of 5) | 46213 | NM_178626 | 72729 | Mm.29476 | NM_178626 | ENSMUSG00000052298 | **Cdc42se2** | 2810404F18Rik\|AA408783\|AA536669\|SPEC2 | CDC42 small effector 2 | protein-coding |
| chr11_54764822_54765141 | chr11 | 54764823 | 54765141 | + | -0.715109 | intron (NM_178626, intron 1 of 5) | ORR1B2\|LTR\|ERVL-MaLR | 22721 | NM_178626 | 72729 | Mm.29476 | NM_178626 | ENSMUSG00000052298 | **Cdc42se2** | 2810404F18Rik\|AA408783\|AA536669\|SPEC2 | CDC42 small effector 2 | protein-coding |
| chr11_54708217_54708450 | chr11 | 54708218 | 54708450 | + | -0.954641 | Intergenic | RLTR33\|LTR\|ERVK | 79369 | NM_178626 | 72729 | Mm.29476 | NM_178626 | ENSMUSG00000052298 | **Cdc42se2** | 2810404F18Rik\|AA408783\|AA536669\|SPEC2 | CDC42 small effector 2 | protein-coding |
| chr5_107008823_107009025 | chr5 | 107008824 | 107009025 | + | -0.621267 | Intergenic | L1VL4\|LINE\|L1 | 44363 | NM_001271566 | 12545 | Mm.20842 | NM_009863 | ENSMUSG00000029283 | **Cdc7** | AI597260\|Cdc7l1\|muCdc7 | cell division cycle 7 (S. cerevisiae) | protein-coding |
| chr2_72412179_72412633 | chr2 | 72412180 | 72412633 | + | -1.01257 | intron (NM_023057, intron 14 of 19) | intron (NM_023057, intron 14 of 19) | -63813 | NM_025866 | 66953 | Mm.270676 | NM_025866 | ENSMUSG00000055612 | **Cdca7** | 2310021G01Rik\|AU044109\|JPO1 | cell division cycle associated 7 | protein-coding |
| chr8_122917348_122918363 | chr8 | 122917349 | 122918363 | + | -0.710976 | intron (NR_037865, intron 2 of 13) | intron (NR_037865, intron 2 of 13) | 69482 | NM_007662 | 12555 | Mm.1976 | NM_007662 | ENSMUSG00000031962 | **Cdh15** | AI323380\|Cdh14\|Mcad | cadherin 15 | protein-coding |
| chr8_122918132_122918366 | chr8 | 122918133 | 122918366 | + | -0.871474 | intron (NR_037865, intron 2 of 13) | RMER4B\|LTR\|ERVK | 69875 | NM_007662 | 12555 | Mm.1976 | NM_007662 | ENSMUSG00000031962 | **Cdh15** | AI323380\|Cdh14\|Mcad | cadherin 15 | protein-coding |
| chr8_122937472_122937933 | chr8 | 122937473 | 122937933 | + | -0.965611 | intron (NR_037865, intron 2 of 13) | intron (NR_037865, intron 2 of 13) | 89329 | NM_007662 | 12555 | Mm.1976 | NM_007662 | ENSMUSG00000031962 | **Cdh15** | AI323380\|Cdh14\|Mcad | cadherin 15 | protein-coding |
| chr8_104625653_104626012 | chr8 | 104625654 | 104626012 | + | -0.785818 | Intergenic | Intergenic | -1437 | NM_001252628 | 12556 | Mm.19423 | NM_007663 | ENSMUSG00000031881 | **Cdh16** | - | cadherin 16 | protein-coding |
| chr12_33048399_33048935 | chr12 | 33048400 | 33048935 | + | -1.05899 | intron (NM_001024478, intron 12 of 19) | intron (NM_001024478, intron 12 of 19) | 44208 | NM_001024478 | 68764 | Mm.60083 | NM_001024478 | ENSMUSG00000035860 | **Cdhr3** | 1110049B09Rik\|BB172694\|Cdh28 | cadherin-related family member 3 | protein-coding |
| chr7_126978884_126979298 | chr7 | 126978885 | 126979298 | + | -0.78968 | intron (NM_026638, intron 4 of 5) | intron (NM_026638, intron 4 of 5) | 3177 | NM_026638 | 52858 | Mm.28219 | NM_026638 | ENSMUSG00000030682 | **Cdipt** | 9530042F15Rik\|D7Bwg0575e\|Pis\|Pis1 | CDP-diacylglycerol–inositol 3-phosphatidyltransferase (phosphatidylinositol synthase) | protein-coding |
| chr10_40400307_40400862 | chr10 | 40400308 | 40400862 | + | -0.661363 | intron (NM_198164, intron 2 of 11) | Lx7\|LINE\|L1 | 51277 | NM_001291816 | 78334 | Mm.200924 | NM_198164 | ENSMUSG00000038481 | **Cdk19** | 2700084L06Rik\|AI845279\|AW228747\|CDK11\|Cdc2l6\|mKIAA1028 | cyclin-dependent kinase 19 | protein-coding |
| chr10_40371522_40371823 | chr10 | 40371523 | 40371823 | + | -0.682611 | intron (NM_001291816, intron 1 of 11) | intron (NM_001291816, intron 1 of 11) | 22365 | NM_001291816 | 78334 | Mm.200924 | NM_198164 | ENSMUSG00000038481 | **Cdk19** | 2700084L06Rik\|AI845279\|AW228747\|CDK11\|Cdc2l6\|mKIAA1028 | cyclin-dependent kinase 19 | protein-coding |
| chr10_40381336_40381634 | chr10 | 40381337 | 40381634 | + | -0.798172 | intron (NM_001291816, intron 1 of 11) | intron (NM_001291816, intron 1 of 11) | 32177 | NM_001291816 | 78334 | Mm.200924 | NM_198164 | ENSMUSG00000038481 | **Cdk19** | 2700084L06Rik\|AI845279\|AW228747\|CDK11\|Cdc2l6\|mKIAA1028 | cyclin-dependent kinase 19 | protein-coding |
| chr10_40419852_40420379 | chr10 | 40419853 | 40420379 | + | -0.873633 | intron (NM_198164, intron 2 of 11) | MIR\|SINE\|MIR | 70808 | NM_001291816 | 78334 | Mm.200924 | NM_198164 | ENSMUSG00000038481 | **Cdk19** | 2700084L06Rik\|AI845279\|AW228747\|CDK11\|Cdc2l6\|mKIAA1028 | cyclin-dependent kinase 19 | protein-coding |
| chr10_40419839_40420210 | chr10 | 40419840 | 40420210 | + | -0.898181 | intron (NM_198164, intron 2 of 11) | intron (NM_198164, intron 2 of 11) | 70717 | NM_001291816 | 78334 | Mm.200924 | NM_198164 | ENSMUSG00000038481 | **Cdk19** | 2700084L06Rik\|AI845279\|AW228747\|CDK11\|Cdc2l6\|mKIAA1028 | cyclin-dependent kinase 19 | protein-coding |
| chr10_40387675_40387904 | chr10 | 40387676 | 40387904 | + | -0.914521 | intron (NM_001291816, intron 1 of 11) | intron (NM_001291816, intron 1 of 11) | 38482 | NM_001291816 | 78334 | Mm.200924 | NM_198164 | ENSMUSG00000038481 | **Cdk19** | 2700084L06Rik\|AI845279\|AW228747\|CDK11\|Cdc2l6\|mKIAA1028 | cyclin-dependent kinase 19 | protein-coding |
| chr10_40386096_40386652 | chr10 | 40386097 | 40386652 | + | -1.17186 | intron (NM_001291816, intron 1 of 11) | RMER19C\|LTR\|ERVK | 37066 | NM_001291816 | 78334 | Mm.200924 | NM_198164 | ENSMUSG00000038481 | **Cdk19** | 2700084L06Rik\|AI845279\|AW228747\|CDK11\|Cdc2l6\|mKIAA1028 | cyclin-dependent kinase 19 | protein-coding |
| chr10_40386096_40386477 | chr10 | 40386097 | 40386477 | + | -1.21692 | intron (NM_001291816, intron 1 of 11) | RMER19C\|LTR\|ERVK | 36979 | NM_001291816 | 78334 | Mm.200924 | NM_198164 | ENSMUSG00000038481 | **Cdk19** | 2700084L06Rik\|AI845279\|AW228747\|CDK11\|Cdc2l6\|mKIAA1028 | cyclin-dependent kinase 19 | protein-coding |
| chr13_29780724_29781357 | chr13 | 29780725 | 29781357 | + | -0.690913 | intron (NM_144536, intron 4 of 15) | intron (NM_144536, intron 4 of 15) | 74547 | NM_144536 | 68916 | Mm.212227 | NM_144536 | ENSMUSG00000006191 | **Cdkal1** | 1190005B03Rik\|6620401C13Rik | CDK5 regulatory subunit associated protein 1-like 1 | protein-coding |
| chr13_29780696_29781151 | chr13 | 29780697 | 29781151 | + | -0.692385 | intron (NM_144536, intron 4 of 15) | intron (NM_144536, intron 4 of 15) | 74664 | NM_144536 | 68916 | Mm.212227 | NM_144536 | ENSMUSG00000006191 | **Cdkal1** | 1190005B03Rik\|6620401C13Rik | CDK5 regulatory subunit associated protein 1-like 1 | protein-coding |
| chr11_52035924_52036311 | chr11 | 52035925 | 52036311 | + | -0.722221 | intron (NM_001166653, intron 13 of 13) | intron (NM_001166653, intron 13 of 13) | 31897 | NM_001166653 | 213084 | Mm.280557 | NM_153785 | ENSMUSG00000020389 | **Cdkl3** | AI852479\|B230379H01Rik | cyclin-dependent kinase-like 3 | protein-coding |
| chr17_80555916_80556548 | chr17 | 80555917 | 80556548 | + | -0.784846 | intron (NM_001033443, intron 3 of 9) | (CTGGGG)n\|Simple_repeat\|Simple_repeat | 7602 | NM_001033443 | 381113 | Mm.329216 | NM_001033443 | ENSMUSG00000033966 | **Cdkl4** | AU067824\|Gm942 | cyclin-dependent kinase-like 4 | protein-coding |
| chr17_80555958_80556296 | chr17 | 80555959 | 80556296 | + | -0.866021 | intron (NM_001033443, intron 3 of 9) | intron (NM_001033443, intron 3 of 9) | 7707 | NM_001033443 | 381113 | Mm.329216 | NM_001033443 | ENSMUSG00000033966 | **Cdkl4** | AU067824\|Gm942 | cyclin-dependent kinase-like 4 | protein-coding |
| chr6_134917137_134917416 | chr6 | 134917138 | 134917416 | + | -0.784751 | Intergenic | Intergenic | -3124 | NM_009875 | 12576 | Mm.2958 | NM_009875 | ENSMUSG00000003031 | **Cdkn1b** | AA408329\|AI843786\|Kip1\|p27\|p27Kip1 | cyclin-dependent kinase inhibitor 1B | protein-coding |
| chr6_134921416_134921786 | chr6 | 134921417 | 134921786 | + | -1.03175 | intron (NM_009875, intron 1 of 2) | intron (NM_009875, intron 1 of 2) | 1200 | NM_009875 | 12576 | Mm.2958 | NM_009875 | ENSMUSG00000003031 | **Cdkn1b** | AA408329\|AI843786\|Kip1\|p27\|p27Kip1 | cyclin-dependent kinase inhibitor 1B | protein-coding |
| chr11_51938796_51939291 | chr11 | 51938797 | 51939291 | + | -0.712094 | Intergenic | ID4\|SINE\|ID | -28587 | NM_029976 | 52626 | Mm.289456 | NM_029976 | ENSMUSG00000020392 | **Cdkn2aipnl** | A430101B06Rik\|AA409260\|D11Ertd497e\|MNCb-1520 | CDKN2A interacting protein N-terminal like | protein-coding |
| chr11_51961831_51962607 | chr11 | 51961832 | 51962607 | + | -0.715195 | Intergenic | Intergenic | -5412 | NM_029976 | 52626 | Mm.289456 | NM_029976 | ENSMUSG00000020392 | **Cdkn2aipnl** | A430101B06Rik\|AA409260\|D11Ertd497e\|MNCb-1520 | CDKN2A interacting protein N-terminal like | protein-coding |
| chr11_51938679_51939416 | chr11 | 51938680 | 51939416 | + | -0.728236 | Intergenic | ID4\|SINE\|ID | -28583 | NM_029976 | 52626 | Mm.289456 | NM_029976 | ENSMUSG00000020392 | **Cdkn2aipnl** | A430101B06Rik\|AA409260\|D11Ertd497e\|MNCb-1520 | CDKN2A interacting protein N-terminal like | protein-coding |
| chr11_51938791_51939063 | chr11 | 51938792 | 51939063 | + | -0.778712 | Intergenic | MLT1F\|LTR\|ERVL-MaLR | -28704 | NM_029976 | 52626 | Mm.289456 | NM_029976 | ENSMUSG00000020392 | **Cdkn2aipnl** | A430101B06Rik\|AA409260\|D11Ertd497e\|MNCb-1520 | CDKN2A interacting protein N-terminal like | protein-coding |
| chr11_51938622_51939194 | chr11 | 51938623 | 51939194 | + | -0.78149 | Intergenic | MLT1F\|LTR\|ERVL-MaLR | -28723 | NM_029976 | 52626 | Mm.289456 | NM_029976 | ENSMUSG00000020392 | **Cdkn2aipnl** | A430101B06Rik\|AA409260\|D11Ertd497e\|MNCb-1520 | CDKN2A interacting protein N-terminal like | protein-coding |
| chr11_51961595_51962354 | chr11 | 51961596 | 51962354 | + | -0.81479 | Intergenic | Intergenic | -5656 | NM_029976 | 52626 | Mm.289456 | NM_029976 | ENSMUSG00000020392 | **Cdkn2aipnl** | A430101B06Rik\|AA409260\|D11Ertd497e\|MNCb-1520 | CDKN2A interacting protein N-terminal like | protein-coding |
| chr8_122561140_122561567 | chr8 | 122561141 | 122561567 | + | -0.959793 | Intergenic | Intergenic | -6661 | NM_026014 | 67177 | Mm.21873 | NM_026014 | ENSMUSG00000006585 | **Cdt1** | 2610318F11Rik\|AW545653\|C76791\|Ris2 | chromatin licensing and DNA replication factor 1 | protein-coding |
| chr7_25575927_25576174 | chr7 | 25575928 | 25576174 | + | -0.683687 | Intergenic | ERVB7_3-LTR_MM\|LTR\|ERVK | -36047 | NM_001113369 | 26367 | Mm.439731 | NM_007543 | ENSMUSG00000054385 | **Ceacam2** | AI267129\|Bgp2\|Ceacam2-L | carcinoembryonic antigen-related cell adhesion molecule 2 | protein-coding |
| chr7_16716657_16717079 | chr7 | 16716658 | 16717079 | + | -0.716038 | Intergenic | Intergenic | -5061 | NM_011927 | 26368 | Mm.27096 | NM_011927 | ENSMUSG00000007209 | **Ceacam9** | AA410097\|AW545709\|Cea-5\|Cea5\|mmCGM8 | carcinoembryonic antigen-related cell adhesion molecule 9 | protein-coding |
| chr7_16716803_16717085 | chr7 | 16716804 | 16717085 | + | -0.73858 | Intergenic | Intergenic | -4985 | NM_011927 | 26368 | Mm.27096 | NM_011927 | ENSMUSG00000007209 | **Ceacam9** | AA410097\|AW545709\|Cea-5\|Cea5\|mmCGM8 | carcinoembryonic antigen-related cell adhesion molecule 9 | protein-coding |
| chr7_16716293_16717084 | chr7 | 16716294 | 16717084 | + | -0.753702 | Intergenic | Intergenic | -5240 | NM_011927 | 26368 | Mm.27096 | NM_011927 | ENSMUSG00000007209 | **Ceacam9** | AA410097\|AW545709\|Cea-5\|Cea5\|mmCGM8 | carcinoembryonic antigen-related cell adhesion molecule 9 | protein-coding |

| | | | | | | | | | | | | | | | | | |
|---|---|---|---|---|---|---|---|---|---|---|---|---|---|---|---|---|---|
| chr2_6672314_6672777 | chr2 | 6672315 | 6672777 | + | -0.63046 | intron (NM_001110232, intron 4 of 15) | intron (NM_001110232, intron 4 of 15) | 212450 | NM_010160 | 14007 | Mm.398543 | NM_010160 | ENSMUSG00000002107 | **Celf2** | B230218O03\|B230345P09Rik\|C88023\|CELF-2\|CUG-BP2\|Cugbp2\|D230046B21Rik\|Etr-3\|Napor\|Napor-2\|mETR-3 | CUGBP, Elav-like family member 2 | protein-coding |
| chr2_6705477_6706564 | chr2 | 6705478 | 6706564 | + | -0.663133 | intron (NM_001110232, intron 4 of 15) | intron (NM_001110232, intron 4 of 15) | 178975 | NM_010160 | 14007 | Mm.398543 | NM_010160 | ENSMUSG00000002107 | **Celf2** | B230218O03\|B230345P09Rik\|C88023\|CELF-2\|CUG-BP2\|Cugbp2\|D230046B21Rik\|Etr-3\|Napor\|Napor-2\|mETR-3 | CUGBP, Elav-like family member 2 | protein-coding |
| chr2_6707329_6707714 | chr2 | 6707330 | 6707714 | + | -0.676403 | intron (NM_001110232, intron 4 of 15) | intron (NM_001110232, intron 4 of 15) | 177474 | NM_010160 | 14007 | Mm.398543 | NM_010160 | ENSMUSG00000002107 | **Celf2** | B230218O03\|B230345P09Rik\|C88023\|CELF-2\|CUG-BP2\|Cugbp2\|D230046B21Rik\|Etr-3\|Napor\|Napor-2\|mETR-3 | CUGBP, Elav-like family member 2 | protein-coding |
| chr2_7269903_7270463 | chr2 | 7269904 | 7270463 | + | -0.695063 | intron (NM_001110232, intron 1 of 15) | intron (NM_001110232, intron 1 of 15) | 125817 | NM_001110232 | 14007 | Mm.398543 | NM_010160 | ENSMUSG00000002107 | **Celf2** | B230218O03\|B230345P09Rik\|C88023\|CELF-2\|CUG-BP2\|Cugbp2\|D230046B21Rik\|Etr-3\|Napor\|Napor-2\|mETR-3 | CUGBP, Elav-like family member 2 | protein-coding |
| chr2_6705962_6706564 | chr2 | 6705963 | 6706564 | + | -0.74673 | intron (NM_001110232, intron 4 of 15) | intron (NM_001110232, intron 4 of 15) | 178733 | NM_010160 | 14007 | Mm.398543 | NM_010160 | ENSMUSG00000002107 | **Celf2** | B230218O03\|B230345P09Rik\|C88023\|CELF-2\|CUG-BP2\|Cugbp2\|D230046B21Rik\|Etr-3\|Napor\|Napor-2\|mETR-3 | CUGBP, Elav-like family member 2 | protein-coding |
| chr15_85981746_85982061 | chr15 | 85981747 | 85982061 | + | -0.661998 | intron (NM_009886, intron 1 of 34) | intron (NM_009886, intron 1 of 34) | 51873 | NM_009886 | 12614 | Mm.22680 | NM_009886 | ENSMUSG00000016028 | **Celsr1** | Crsh\|Scy\|crash | cadherin, EGF LAG seven-pass G-type receptor 1 (flamingo homolog, Drosophila) | protein-coding |
| chr15_85977045_85977355 | chr15 | 85977046 | 85977355 | + | -0.733245 | intron (NM_009886, intron 2 of 34) | intron (NM_009886, intron 2 of 34) | 56577 | NM_009886 | 12614 | Mm.22680 | NM_009886 | ENSMUSG00000016028 | **Celsr1** | Crsh\|Scy\|crash | cadherin, EGF LAG seven-pass G-type receptor 1 (flamingo homolog, Drosophila) | protein-coding |
| chr15_85990833_85991637 | chr15 | 85990834 | 85991637 | + | -0.846352 | intron (NM_009886, intron 1 of 34) | intron (NM_009886, intron 1 of 34) | 42542 | NM_009886 | 12614 | Mm.22680 | NM_009886 | ENSMUSG00000016028 | **Celsr1** | Crsh\|Scy\|crash | cadherin, EGF LAG seven-pass G-type receptor 1 (flamingo homolog, Drosophila) | protein-coding |
| chr15_85991030_85991572 | chr15 | 85991031 | 85991572 | + | -0.892299 | intron (NM_009886, intron 1 of 34) | intron (NM_009886, intron 1 of 34) | 42476 | NM_009886 | 12614 | Mm.22680 | NM_009886 | ENSMUSG00000016028 | **Celsr1** | Crsh\|Scy\|crash | cadherin, EGF LAG seven-pass G-type receptor 1 (flamingo homolog, Drosophila) | protein-coding |
| chr5_30666405_30667223 | chr5 | 30666406 | 30667223 | + | -0.662677 | promoter-TSS (NM_007681) | promoter-TSS (NM_007681) | -88 | NM_007681 | 12615 | Mm.290563 | NM_007681 | ENSMUSG00000029177 | **Cenpa** | Cenp-A | centromere protein A | protein-coding |
| chr1_189602201_189602638 | chr1 | 189602202 | 189602638 | + | -0.764208 | Intergenic | Intergenic | 85666 | NM_001081363 | 108000 | Mm.129746 | NM_001081363 | ENSMUSG00000026605 | **Cenpf** | 6530404A22Rik\|AI325968\|CENF\|Lek1\|PRO1779\|mitosin | centromere protein F | protein-coding |
| chr1_189602341_189602647 | chr1 | 189602342 | 189602647 | + | -0.789268 | Intergenic | Intergenic | 85592 | NM_001081363 | 108000 | Mm.129746 | NM_001081363 | ENSMUSG00000026605 | **Cenpf** | 6530404A22Rik\|AI325968\|CENF\|Lek1\|PRO1779\|mitosin | centromere protein F | protein-coding |
| chr1_161070537_161071067 | chr1 | 161070538 | 161071067 | + | -0.640377 | promoter-TSS (NM_027429) | promoter-TSS (NM_027429) | -90 | NM_027429 | 70454 | Mm.248212 | NM_027429 | ENSMUSG00000026708 | **Cenpl** | 2610300B10Rik\|AW121806\|AW550697 | centromere protein L | protein-coding |
| chr8_46543109_46543500 | chr8 | 46543110 | 46543500 | + | -0.863982 | Intergenic | B4\|SINE\|B4 | -8764 | NM_027973 | 71876 | Mm.22108 | NM_027973 | ENSMUSG00000031629 | **Cenpu** | 1700029A22Rik\|Mlf1ip | centromere protein U | protein-coding |
| chr11_62524137_62524802 | chr11 | 62524138 | 62524802 | + | -1.13925 | TTS (NM_028448) | TTS (NM_028448) | 14791 | NM_028448 | 73139 | Mm.390409 | NM_028448 | ENSMUSG00000018509 | **Cenpv** | 3110013H01Rik\|AA671303\|Prr6 | centromere protein V | protein-coding |
| chr11_62507618_62508138 | chr11 | 62507619 | 62508138 | + | -1.40687 | intron (NM_001039536, intron 4 of 6) | intron (NM_001039536, intron 4 of 6) | 31383 | NM_028448 | 73139 | Mm.390409 | NM_028448 | ENSMUSG00000018509 | **Cenpv** | 3110013H01Rik\|AA671303\|Prr6 | centromere protein V | protein-coding |
| chr9_45807808_45808312 | chr9 | 45807809 | 45808312 | + | -0.906238 | intron (NM_001081373, intron 4 of 29) | intron (NM_001081373, intron 4 of 29) | 20578 | NM_001081373 | 214552 | Mm.260103 | NM_172531 | ENSMUSG00000043987 | **Cep164** | AI450905\|BC027092\|D030051D21\|mKIAA1052 | centrosomal protein 164 | protein-coding |
| chr9_45807650_45808171 | chr9 | 45807651 | 45808171 | + | -0.952624 | intron (NM_001081373, intron 4 of 29) | intron (NM_001081373, intron 4 of 29) | 20727 | NM_001081373 | 214552 | Mm.260103 | NM_172531 | ENSMUSG00000043987 | **Cep164** | AI450905\|BC027092\|D030051D21\|mKIAA1052 | centrosomal protein 164 | protein-coding |
| chr9_45807805_45808161 | chr9 | 45807806 | 45808161 | + | -0.983129 | intron (NM_001081373, intron 4 of 29) | intron (NM_001081373, intron 4 of 29) | 20655 | NM_001081373 | 214552 | Mm.260103 | NM_172531 | ENSMUSG00000043987 | **Cep164** | AI450905\|BC027092\|D030051D21\|mKIAA1052 | centrosomal protein 164 | protein-coding |
| chr1_176800823_176801297 | chr1 | 176800824 | 176801297 | + | -0.669748 | intron (NM_001099637, intron 1 of 19) | intron (NM_001099637, intron 1 of 19) | 6064 | NM_001099637 | 545389 | Mm.269991 | NM_001024722 | ENSMUSG00000057335 | **Cep170** | 4933426L22Rik\|A330004A13Rik\|AI195353 | centrosomal protein 170 | protein-coding |
| chr9_99240614_99240927 | chr9 | 99240615 | 99240927 | + | -0.70507 | Intergenic | Intergenic | -2697 | NM_023873 | 68121 | Mm.288739 | NM_023873 | ENSMUSG00000056267 | **Cep70** | 6720484E09Rik\|Bite\|C030018L16Rik | centrosomal protein 70 | protein-coding |
| chr16_55924053_55924355 | chr16 | 55924054 | 55924355 | + | -0.652904 | TTS (NM_001159365) | TTS (NM_001159365) | 10644 | NM_001159364 | 74201 | Mm.178427 | NM_028815 | ENSMUSG00000022604 | **Cep97** | 2810403B08Rik\|4932439K18Rik\|AA437768\|E130116N02Rik\|Lrriq2 | centrosomal protein 97 | protein-coding |
| chr16_55947726_55948283 | chr16 | 55947727 | 55948283 | + | -1.00885 | Intergenic | MTC\|LTR\|ERVL-MaLR | -13157 | NM_001159365 | 74201 | Mm.178427 | NM_028815 | ENSMUSG00000022604 | **Cep97** | 2810403B08Rik\|4932439K18Rik\|AA437768\|E130116N02Rik\|Lrriq2 | centrosomal protein 97 | protein-coding |
| chr15_86173214_86174328 | chr15 | 86173215 | 86174328 | + | -0.701793 | intron (NM_145475, intron 1 of 12) | intron (NM_145475, intron 1 of 12) | 12370 | NM_145475 | 223753 | Mm.222685 | NM_145475 | ENSMUSG00000035891 | **Cerk** | AI848610\|D330016D08Rik\|mKIAA1646 | ceramide kinase | protein-coding |
| chr15_86182599_86183237 | chr15 | 86182600 | 86183237 | + | -0.706369 | intron (NM_145475, intron 1 of 12) | intron (NM_145475, intron 1 of 12) | 3223 | NM_145475 | 223753 | Mm.222685 | NM_145475 | ENSMUSG00000035891 | **Cerk** | AI848610\|D330016D08Rik\|mKIAA1646 | ceramide kinase | protein-coding |
| chr15_86182647_86183014 | chr15 | 86182648 | 86183014 | + | -0.723344 | intron (NM_145475, intron 1 of 12) | intron (NM_145475, intron 1 of 12) | 3310 | NM_145475 | 223753 | Mm.222685 | NM_145475 | ENSMUSG00000035891 | **Cerk** | AI848610\|D330016D08Rik\|mKIAA1646 | ceramide kinase | protein-coding |
| chr15_86173221_86173977 | chr15 | 86173222 | 86173977 | + | -0.728071 | intron (NM_145475, intron 1 of 12) | intron (NM_145475, intron 1 of 12) | 12542 | NM_145475 | 223753 | Mm.222685 | NM_145475 | ENSMUSG00000035891 | **Cerk** | AI848610\|D330016D08Rik\|mKIAA1646 | ceramide kinase | protein-coding |
| chr15_86168039_86168518 | chr15 | 86168040 | 86168518 | + | -0.763538 | intron (NM_145475, intron 1 of 12) | intron (NM_145475, intron 1 of 12) | 17862 | NM_145475 | 223753 | Mm.222685 | NM_145475 | ENSMUSG00000035891 | **Cerk** | AI848610\|D330016D08Rik\|mKIAA1646 | ceramide kinase | protein-coding |
| chr2_68932274_68932587 | chr2 | 68932275 | 68932587 | + | -0.680949 | intron (NM_172856, intron 1 of 9) | intron (NM_172856, intron 1 of 9) | 70874 | NM_172856 | 241447 | Mm.222222 | NM_172856 | ENSMUSG00000027035 | **Cers6** | 4732462C07Rik\|AW544719\|Lass6\|T1L | ceramide synthase 6 | protein-coding |
| chr2_68932113_68932580 | chr2 | 68932114 | 68932580 | + | -0.683862 | intron (NM_172856, intron 1 of 9) | intron (NM_172856, intron 1 of 9) | 70790 | NM_172856 | 241447 | Mm.222222 | NM_172856 | ENSMUSG00000027035 | **Cers6** | 4732462C07Rik\|AW544719\|Lass6\|T1L | ceramide synthase 6 | protein-coding |
| chr2_68960037_68960493 | chr2 | 68960038 | 68960493 | + | -0.812179 | intron (NM_172856, intron 3 of 9) | intron (NM_172856, intron 3 of 9) | 98708 | NM_172856 | 241447 | Mm.222222 | NM_172856 | ENSMUSG00000027035 | **Cers6** | 4732462C07Rik\|AW544719\|Lass6\|T1L | ceramide synthase 6 | protein-coding |
| chr2_68959564_68959979 | chr2 | 68959565 | 68959979 | + | -1.08518 | intron (NM_172856, intron 3 of 9) | intron (NM_172856, intron 3 of 9) | 98215 | NM_172856 | 241447 | Mm.222222 | NM_172856 | ENSMUSG00000027035 | **Cers6** | 4732462C07Rik\|AW544719\|Lass6\|T1L | ceramide synthase 6 | protein-coding |
| chr13_81793593_81794259 | chr13 | 81793594 | 81794259 | + | -0.742079 | intron (NM_007684, intron 4 of 4) | intron (NM_007684, intron 4 of 4) | 10634 | NM_007684 | 12626 | Mm.12481 | NM_007684 | ENSMUSG00000021537 | **Cetn3** | MmCEN3 | centrin 3 | protein-coding |
| chr13_81793809_81794294 | chr13 | 81793810 | 81794294 | + | -0.775477 | intron (NM_007684, intron 4 of 4) | B4A\|SINE\|B4 | 10760 | NM_007684 | 12626 | Mm.12481 | NM_007684 | ENSMUSG00000021537 | **Cetn3** | MmCEN3 | centrin 3 | protein-coding |
| chr1_58721033_58721255 | chr1 | 58721034 | 58721255 | + | -0.687205 | intron (NM_207653, intron 1 of 9) | intron (NM_207653, intron 1 of 9) | 7858 | NM_207653 | 12633 | Mm.336848 | NM_009805 | ENSMUSG00000026031 | **Cflar** | 2310024N18Rik\|A430105C05Rik\|CLARP\|Cash\|Casper\|ENSMUSG00000072980\|FLAME\|FLAME-1\|Flip\|Gm9845\|I-FLICE\|MRIT\|c-Flip | CASP8 and FADD-like apoptosis regulator | protein-coding |
| chr1_58720842_58721301 | chr1 | 58720843 | 58721301 | + | -0.699061 | intron (NM_207653, intron 1 of 9) | intron (NM_207653, intron 1 of 9) | 7786 | NM_207653 | 12633 | Mm.336848 | NM_009805 | ENSMUSG00000026031 | **Cflar** | 2310024N18Rik\|A430105C05Rik\|CLARP\|Cash\|Casper\|ENSMUSG00000072980\|FLAME\|FLAME-1\|Flip\|Gm9845\|I-FLICE\|MRIT\|c-Flip | CASP8 and FADD-like apoptosis regulator | protein-coding |
| chr16_64836103_64836580 | chr16 | 64836104 | 64836580 | + | -0.749428 | Intergenic | Intergenic | -15743 | NM_178647 | 106143 | Mm.10160 | NM_178647 | ENSMUSG00000054604 | **Cggbp1** | AA960172\|AL023996\|AW045841 | CGG triplet repeat binding protein 1 | protein-coding |
| chr16_64941524_64942530 | chr16 | 64941525 | 64942530 | + | -0.922982 | intron (NM_008310, intron 2 of 2) | RLTR14\|LTR\|ERV1 | 89942 | NM_178647 | 106143 | Mm.10160 | NM_178647 | ENSMUSG00000054604 | **Cggbp1** | AA960172\|AL023996\|AW045841 | CGG triplet repeat binding protein 1 | protein-coding |
| chr11_94570659_94571129 | chr11 | 94570660 | 94571129 | + | -0.723867 | intron (NM_153807, intron 6 of 15) | intron (NM_153807, intron 6 of 15) | 5820 | NM_007689 | 12643 | Mm.440523 | NM_007689 | ENSMUSG00000039084 | **Chad** | SLRR4A | chondroadherin | protein-coding |
| chr6_33055850_33056108 | chr6 | 33055851 | 33056108 | + | -0.795695 | intron (NM_025336, intron 1 of 7) | intron (NM_025336, intron 1 of 7) | 4173 | NM_025336 | 66075 | Mm.21501 | NM_025336 | ENSMUSG00000053768 | **Chchd3** | 0610041L09Rik\|1700039J09Rik\|AW558177 | coiled-coil-helix-coiled-coil-helix domain containing 3 | protein-coding |
| chr17_15742981_15743280 | chr17 | 15742982 | 15743280 | + | -0.690803 | exon (NM_007690, exon 17 of 36) | exon (NM_007690, exon 17 of 36) | 38164 | NM_007690 | 12648 | Mm.8137 | NM_007690 | ENSMUSG00000023852 | **Chd1** | 4930525N21Rik\|AI851787\|AW555109 | chromodomain helicase DNA binding protein 1 | protein-coding |
| chr17_15742866_15743279 | chr17 | 15742867 | 15743279 | + | -0.742512 | intron (NM_007690, intron 16 of 35) | intron (NM_007690, intron 16 of 35) | 38106 | NM_007690 | 12648 | Mm.8137 | NM_007690 | ENSMUSG00000023852 | **Chd1** | 4930525N21Rik\|AI851787\|AW555109 | chromodomain helicase DNA binding protein 1 | protein-coding |
| chr17_15705983_15706394 | chr17 | 15705984 | 15706394 | + | -0.765337 | intron (NM_007690, intron 1 of 35) | intron (NM_007690, intron 1 of 35) | 1222 | NM_007690 | 12648 | Mm.8137 | NM_007690 | ENSMUSG00000023852 | **Chd1** | 4930525N21Rik\|AI851787\|AW555109 | chromodomain helicase DNA binding protein 1 | protein-coding |

| | | | | | | | | | | | | | | | | | |
|---|---|---|---|---|---|---|---|---|---|---|---|---|---|---|---|---|---|
| chr17_15743427_15743791 | chr17 | 15743428 | 15743791 | + | -0.920222 | intron (NM_007690, intron 17 of 35) | intron (NM_007690, intron 17 of 35) | 38642 | NM_007690 | 12648 | Mm.8137 | NM_007690 | ENSMUSG00000023852 | **Chd1** | 4930525N21Rik\|AI851787\|AW555109 | chromodomain helicase DNA binding protein 1 | protein-coding |
| chr7_73478295_73478530 | chr7 | 73478296 | 73478530 | + | -0.801267 | intron (NM_001081345, intron 20 of 38) | intron (NM_001081345, intron 20 of 38) | 63333 | NM_001081345 | 244059 | Mm.34539 | NM_001081345 | ENSMUSG00000078671 | **Chd2** | 2810013C04Rik\|2810040A01Rik\|5630401D06Rik\|AI851092\|BC029703\|CHD-2 | chromodomain helicase DNA binding protein 2 | protein-coding |
| chr7_73523287_73523501 | chr7 | 73523288 | 73523501 | + | -1.36287 | intron (NM_001081345, intron 2 of 38) | intron (NM_001081345, intron 2 of 38) | 18352 | NM_001081345 | 244059 | Mm.34539 | NM_001081345 | ENSMUSG00000078671 | **Chd2** | 2810013C04Rik\|2810040A01Rik\|5630401D06Rik\|AI851092\|BC029703\|CHD-2 | chromodomain helicase DNA binding protein 2 | protein-coding |
| chr7_73523174_73523601 | chr7 | 73523175 | 73523601 | + | -1.38718 | intron (NM_001081345, intron 2 of 38) | intron (NM_001081345, intron 2 of 38) | 18358 | NM_001081345 | 244059 | Mm.34539 | NM_001081345 | ENSMUSG00000078671 | **Chd2** | 2810013C04Rik\|2810040A01Rik\|5630401D06Rik\|AI851092\|BC029703\|CHD-2 | chromodomain helicase DNA binding protein 2 | protein-coding |
| chr2_161511886_161512571 | chr2 | 161511887 | 161512571 | + | -0.667529 | Intergenic | Intergenic | -403173 | NM_173368 | 71389 | Mm.122738 | NM_173368 | ENSMUSG00000057133 | **Chd6** | 5430439G14Rik\|6330406J24Rik\|CHD-6 | chromodomain helicase DNA binding protein 6 | protein-coding |
| chr2_161511871_161512261 | chr2 | 161511872 | 161512261 | + | -0.670691 | Intergenic | IAPLTR2_Mm\|LTR\|ERVK | -403010 | NM_173368 | 71389 | Mm.122738 | NM_173368 | ENSMUSG00000057133 | **Chd6** | 5430439G14Rik\|6330406J24Rik\|CHD-6 | chromodomain helicase DNA binding protein 6 | protein-coding |
| chr4_8799089_8799904 | chr4 | 8799090 | 8799904 | + | -0.685641 | intron (NM_001277149, intron 3 of 37) | intron (NM_001277149, intron 3 of 37) | 109091 | NM_001277149 | 320790 | Mm.138792 | NM_001033395 | ENSMUSG00000041235 | **Chd7** | A730019I05Rik\|Cycn\|Cyn\|Dz\|Edy\|Flo\|GENA 47\|GENA 60\|Gena 52\|Lda\|Mt\|Obt\|Todo\|WBE1\|Whi\|metis | chromodomain helicase DNA binding protein 7 | protein-coding |
| chr4_8728678_8729133 | chr4 | 8728679 | 8729133 | + | -0.687936 | intron (NM_001277149, intron 1 of 37) | intron (NM_001277149, intron 1 of 37) | 38500 | NM_001277149 | 320790 | Mm.138792 | NM_001033395 | ENSMUSG00000041235 | **Chd7** | A730019I05Rik\|Cycn\|Cyn\|Dz\|Edy\|Flo\|GENA 47\|GENA 60\|Gena 52\|Lda\|Mt\|Obt\|Todo\|WBE1\|Whi\|metis | chromodomain helicase DNA binding protein 7 | protein-coding |
| chr4_8720906_8721578 | chr4 | 8720907 | 8721578 | + | -0.697251 | intron (NM_001277149, intron 1 of 37) | L1MA9\|LINE\|L1 | 30836 | NM_001277149 | 320790 | Mm.138792 | NM_001033395 | ENSMUSG00000041235 | **Chd7** | A730019I05Rik\|Cycn\|Cyn\|Dz\|Edy\|Flo\|GENA 47\|GENA 60\|Gena 52\|Lda\|Mt\|Obt\|Todo\|WBE1\|Whi\|metis | chromodomain helicase DNA binding protein 7 | protein-coding |
| chr4_8720906_8721185 | chr4 | 8720907 | 8721185 | + | -0.732197 | intron (NM_001277149, intron 1 of 37) | intron (NM_001277149, intron 1 of 37) | 30640 | NM_001277149 | 320790 | Mm.138792 | NM_001033395 | ENSMUSG00000041235 | **Chd7** | A730019I05Rik\|Cycn\|Cyn\|Dz\|Edy\|Flo\|GENA 47\|GENA 60\|Gena 52\|Lda\|Mt\|Obt\|Todo\|WBE1\|Whi\|metis | chromodomain helicase DNA binding protein 7 | protein-coding |
| chr4_8954178_8954665 | chr4 | 8954179 | 8954665 | + | -0.733346 | Intergenic | Intergenic | 264016 | NM_001277149 | 320790 | Mm.138792 | NM_001033395 | ENSMUSG00000041235 | **Chd7** | A730019I05Rik\|Cycn\|Cyn\|Dz\|Edy\|Flo\|GENA 47\|GENA 60\|Gena 52\|Lda\|Mt\|Obt\|Todo\|WBE1\|Whi\|metis | chromodomain helicase DNA binding protein 7 | protein-coding |
| chr4_8799512_8799911 | chr4 | 8799513 | 8799911 | + | -0.842354 | intron (NM_001277149, intron 3 of 37) | intron (NM_001277149, intron 3 of 37) | 109306 | NM_001277149 | 320790 | Mm.138792 | NM_001033395 | ENSMUSG00000041235 | **Chd7** | A730019I05Rik\|Cycn\|Cyn\|Dz\|Edy\|Flo\|GENA 47\|GENA 60\|Gena 52\|Lda\|Mt\|Obt\|Todo\|WBE1\|Whi\|metis | chromodomain helicase DNA binding protein 7 | protein-coding |
| chr4_8913145_8913517 | chr4 | 8913146 | 8913517 | + | -0.932328 | Intergenic | Intergenic | 222925 | NM_001277149 | 320790 | Mm.138792 | NM_001033395 | ENSMUSG00000041235 | **Chd7** | A730019I05Rik\|Cycn\|Cyn\|Dz\|Edy\|Flo\|GENA 47\|GENA 60\|Gena 52\|Lda\|Mt\|Obt\|Todo\|WBE1\|Whi\|metis | chromodomain helicase DNA binding protein 7 | protein-coding |
| chr4_8783164_8783897 | chr4 | 8783165 | 8783897 | + | -1.05292 | intron (NM_001277149, intron 2 of 37) | intron (NM_001277149, intron 2 of 37) | 93125 | NM_001277149 | 320790 | Mm.138792 | NM_001033395 | ENSMUSG00000041235 | **Chd7** | A730019I05Rik\|Cycn\|Cyn\|Dz\|Edy\|Flo\|GENA 47\|GENA 60\|Gena 52\|Lda\|Mt\|Obt\|Todo\|WBE1\|Whi\|metis | chromodomain helicase DNA binding protein 7 | protein-coding |
| chr12_102612298_102612977 | chr12 | 102612299 | 102612977 | + | -0.870097 | intron (NM_172584, intron 3 of 9) | intron (NM_172584, intron 3 of 9) | 57669 | NM_007693 | 12652 | Mm.4137 | NM_007693 | ENSMUSG00000021194 | **Chga** | ChrA | chromogranin A | protein-coding |
| chr12_102605680_102605972 | chr12 | 102605681 | 102605972 | + | -1.02654 | intron (NM_172584, intron 4 of 9) | intron (NM_172584, intron 4 of 9) | 50857 | NM_007693 | 12652 | Mm.4137 | NM_007693 | ENSMUSG00000021194 | **Chga** | ChrA | chromogranin A | protein-coding |
| chr6_103564865_103565281 | chr6 | 103564866 | 103565281 | + | -0.713432 | intron (NM_007697, intron 2 of 26) | intron (NM_007697, intron 2 of 26) | 54197 | NM_007697 | 12661 | Mm.251288 | NM_007697 | ENSMUSG00000030077 | **Chl1** | A530023M13Rik\|AI465420\|CALL\|LICAM2 | cell adhesion molecule with homology to L1CAM | protein-coding |
| chr8_123203355_123203955 | chr8 | 123203356 | 123203955 | + | -0.617898 | TTS (NM_145606) | TTS (NM_145606) | 9133 | NM_145606 | 234852 | Mm.41596 | NM_145606 | ENSMUSG00000000743 | **Chmp1a** | 2900018H07Rik\|Pcoln3 | charged multivesicular body protein 1A | protein-coding |
| chr18_67212351_67213135 | chr18 | 67212352 | 67213135 | + | -0.760004 | intron (NM_010307, intron 7 of 11) | intron (NM_010307, intron 7 of 11) | 7384 | NM_024190 | 67064 | Mm.73777 | NM_024190 | | **Chmp1b** | 2810405I11Rik\|Chmp1b1 | charged multivesicular body protein 1B | protein-coding |
| chr11_119877147_119877600 | chr11 | 119877148 | 119877600 | + | -0.814943 | intron (NM_028898, intron 24 of 33) | intron (NM_028898, intron 24 of 33) | -36436 | NM_001085498 | 208092 | Mm.306673 | NM_001085498 | ENSMUSG00000025371 | **Chmp6** | 2400004G01Rik | charged multivesicular body protein 6 | protein-coding |
| chr7_122251224_122252196 | chr7 | 122251225 | 122252196 | + | -0.679478 | Intergenic | Intergenic | 32214 | NM_027363 | 70261 | Mm.96997 | NM_027363 | ENSMUSG00000030865 | **Chp2** | 2010110P09Rik\|Cbhp2 | calcineurin-like EF hand protein 2 | protein-coding |
| chr15_73137453_73137694 | chr15 | 73137454 | 73137694 | + | -0.85333 | intron (NM_153178, intron 3 of 18) | intron (NM_153178, intron 3 of 18) | 47162 | NM_053068 | 93696 | Mm.23095 | NM_053068 | ENSMUSG00000068391 | **Chrac1** | 2410152E03Rik\|2810406L04Rik\|YCL1 | chromatin accessibility complex 1 | protein-coding |
| chr13_10070914_10071876 | chr13 | 10070915 | 10071876 | + | -0.624715 | intron (NM_033269, intron 3 of 4) | intron (NM_033269, intron 3 of 4) | 289408 | NM_033269 | 12671 | Mm.342315 | NM_033269 | | **Chrm3** | Chrm-3\|M3\|M3R | cholinergic receptor, muscarinic 3, cardiac | protein-coding |
| chr13_10071178_10071864 | chr13 | 10071179 | 10071864 | + | -0.634946 | intron (NM_033269, intron 3 of 4) | intron (NM_033269, intron 3 of 4) | 289282 | NM_033269 | 12671 | Mm.342315 | NM_033269 | | **Chrm3** | Chrm-3\|M3\|M3R | cholinergic receptor, muscarinic 3, cardiac | protein-coding |
| chr13_10344359_10344760 | chr13 | 10344360 | 10344760 | + | -0.643662 | intron (NM_033269, intron 1 of 4) | intron (NM_033269, intron 1 of 4) | 16243 | NM_033269 | 12671 | Mm.342315 | NM_033269 | | **Chrm3** | Chrm-3\|M3\|M3R | cholinergic receptor, muscarinic 3, cardiac | protein-coding |
| chr13_10344374_10344663 | chr13 | 10344375 | 10344663 | + | -0.686305 | intron (NM_033269, intron 1 of 4) | intron (NM_033269, intron 1 of 4) | 16284 | NM_033269 | 12671 | Mm.342315 | NM_033269 | | **Chrm3** | Chrm-3\|M3\|M3R | cholinergic receptor, muscarinic 3, cardiac | protein-coding |
| chr13_10065797_10066188 | chr13 | 10065798 | 10066188 | + | -0.756862 | intron (NM_033269, intron 3 of 4) | intron (NM_033269, intron 3 of 4) | 294810 | NM_033269 | 12671 | Mm.342315 | NM_033269 | | **Chrm3** | Chrm-3\|M3\|M3R | cholinergic receptor, muscarinic 3, cardiac | protein-coding |
| chr5_65929055_65929735 | chr5 | 65929056 | 65929735 | + | -0.935183 | Intergenic | Intergenic | -37729 | NM_001081104 | 231252 | Mm.486671 | NM_001081104 | ENSMUSG00000029205 | **Chrna9** | 2410015I05Rik\|Acra9\|EG666827\|Gm8311 | cholinergic receptor, nicotinic, alpha polypeptide 9 | protein-coding |
| chr1_38885942_38886480 | chr1 | 38885943 | 38886480 | + | -0.663094 | intron (NM_145142, intron 2 of 6) | intron (NM_145142, intron 2 of 6) | 11949 | NM_145142 | 98388 | Mm.260054 | NM_145142 | ENSMUSG00000026080 | **Chst10** | AI507003\|AU041319\|Hnk-1st\|ST | carbohydrate sulfotransferase 10 | protein-coding |
| chr10_83153903_83154250 | chr10 | 83153904 | 83154250 | + | -0.962966 | intron (NM_021439, intron 2 of 2) | intron (NM_021439, intron 2 of 2) | 168580 | NM_021439 | 58250 | Mm.360747 | NM_021439 | ENSMUSG00000034612 | **Chst11** | 1110020P09Rik\|C4ST\|C4ST-1\|C4ST1\|C4s | carbohydrate sulfotransferase 11 | protein-coding |
| chr10_83153624_83154230 | chr10 | 83153625 | 83154230 | + | -1.02462 | intron (NM_021439, intron 2 of 2) | intron (NM_021439, intron 2 of 2) | 168430 | NM_021439 | 58250 | Mm.360747 | NM_021439 | ENSMUSG00000034612 | **Chst11** | 1110020P09Rik\|C4ST\|C4ST-1\|C4ST1\|C4s | carbohydrate sulfotransferase 11 | protein-coding |
| chr7_132323211_132323814 | chr7 | 132323212 | 132323814 | + | -1.00059 | Intergenic | MLT2B4\|LTR\|ERVL | -6358 | NM_029935 | 77590 | Mm.213582 | NM_029935 | ENSMUSG00000030930 | **Chst15** | 4631426J05Rik\|BRAG\|GalNAcS-6ST\|MAd5\|mKIAA0598 | carbohydrate (N-acetylgalactosamine 4-sulfate 6-O) sulfotransferase 15 | protein-coding |
| chr9_95314858_95315516 | chr9 | 95314859 | 95315516 | + | -0.789135 | Intergenic | Intergenic | 92083 | NM_018763 | 54371 | Mm.212446 | NM_018763 | ENSMUSG00000033350 | **Chst2** | AI428561\|AW121776\|C130041E03Rik\|Chts2\|GST-2\|GlcNAc6ST\|Gn6st | carbohydrate sulfotransferase 2 | protein-coding |
| chr9_95316694_95317181 | chr9 | 95316695 | 95317181 | + | -0.789666 | Intergenic | MLT1A0\|LTR\|ERVL-MaLR | 90332 | NM_018763 | 54371 | Mm.212446 | NM_018763 | ENSMUSG00000033350 | **Chst2** | AI428561\|AW121776\|C130041E03Rik\|Chts2\|GST-2\|GlcNAc6ST\|Gn6st | carbohydrate sulfotransferase 2 | protein-coding |
| chr9_95113255_95113827 | chr9 | 95113256 | 95113827 | + | -1.64062 | intron (NM_177909, intron 12 of 15) | intron (NM_177909, intron 12 of 15) | 293729 | NM_018763 | 54371 | Mm.212446 | NM_018763 | ENSMUSG00000033350 | **Chst2** | AI428561\|AW121776\|C130041E03Rik\|Chts2\|GST-2\|GlcNAc6ST\|Gn6st | carbohydrate sulfotransferase 2 | protein-coding |
| chr7_66175784_66176136 | chr7 | 66175785 | 66176136 | + | -0.810953 | Intergenic | Intergenic | 66445 | NM_001081163 | 269941 | Mm.333916 | NM_001081163 | ENSMUSG00000032640 | **Chsy1** | mKIAA0990 | chondroitin sulfate synthase 1 | protein-coding |
| chr7_66230136_66230810 | chr7 | 66230137 | 66230810 | + | -0.989359 | Intergenic | Intergenic | 120958 | NM_001081163 | 269941 | Mm.333916 | NM_001081163 | ENSMUSG00000032640 | **Chsy1** | mKIAA0990 | chondroitin sulfate synthase 1 | protein-coding |
| chr12_76778353_76779053 | chr12 | 76778354 | 76779053 | + | -0.824208 | intron (NM_206534, intron 3 of 3) | intron (NM_206534, intron 3 of 3) | 13130 | NM_206534 | 211151 | Mm.80970 | NM_206534 | ENSMUSG00000090258 | **Churc1** | 1700030P15Rik\|My015\|chch | churchill domain containing 1 | protein-coding |
| chr3_135417782_135418227 | chr3 | 135417783 | 135418227 | + | -0.844729 | intron (NM_025902, intron 1 of 2) | intron (NM_025902, intron 1 of 2) | 5428 | NM_025902 | 67006 | Mm.269993 | NM_025902 | ENSMUSG00000028165 | **Cisd2** | 1500009M05Rik\|1500026J14Rik\|1500031D15Rik\|AI848398\|B630006A20Rik\|Miner1\|Noxp70\|Zcd2 | CDGSH iron sulfur domain 2 | protein-coding |
| chr10_17565325_17565646 | chr10 | 17565326 | 17565646 | + | -0.630767 | Intergenic | Intergenic | -157742 | NM_010828 | 17684 | Mm.272321 | NM_010828 | ENSMUSG00000039910 | **Cited2** | AI835299\|ER154-like\|Mrg1\|Msg2\|p35srj | Cbp/p300-interacting transactivator, with Glu/Asp-rich carboxy-terminal domain, 2 | protein-coding |
| chr10_17656944_17657215 | chr10 | 17656945 | 17657215 | + | -0.69663 | Intergenic | L2\|LINE\|L2 | -66148 | NM_010828 | 17684 | Mm.272321 | NM_010828 | ENSMUSG00000039910 | **Cited2** | AI835299\|ER154-like\|Mrg1\|Msg2\|p35srj | Cbp/p300-interacting transactivator, with Glu/Asp-rich carboxy-terminal domain, 2 | protein-coding |

| | | | | | | | | | | | | | | | | | |
|---|---|---|---|---|---|---|---|---|---|---|---|---|---|---|---|---|---|
| chr10_17728790_17728999 | chr10 | 17728791 | 17728999 | + | -0.922586 | Intergenic | Intergenic | 5667 | NM_010828 | 17684 | Mm.272321 | NM_010828 | ENSMUSG00000039910 | **Cited2** | AI835299\|ER154-like\|Mrg1\|Msg2\|p35srj | Cbp/p300-interacting transactivator, with Glu/Asp-rich carboxy-terminal domain, 2 | protein-coding |
| chr9_113792145_113792926 | chr9 | 113792146 | 113792926 | + | -0.701108 | promoter-TSS (NM_001286603) | promoter-TSS (NM_001286603) | -794 | NM_001286603 | 76499 | Mm.222272 | NM_029633 | ENSMUSG00000033392 | **Clasp2** | 1500004F14Rik\|8030404L10Rik\|C77448\|CLASP2beta\|mKIAA0627 | CLIP associating protein 2 | protein-coding |
| chr9_113792555_113792943 | chr9 | 113792556 | 113792943 | + | -0.999386 | promoter-TSS (NM_001286603) | promoter-TSS (NM_001286603) | -581 | NM_001286603 | 76499 | Mm.222272 | NM_029633 | ENSMUSG00000033392 | **Clasp2** | 1500004F14Rik\|8030404L10Rik\|C77448\|CLASP2beta\|mKIAA0627 | CLIP associating protein 2 | protein-coding |
| chr19_4218901_4219126 | chr19 | 4218902 | 4219126 | + | -0.702046 | intron (NM_019952, intron 1 of 2) | intron (NM_019952, intron 1 of 2) | 4622 | NM_019952 | 56708 | Mm.347919 | NM_019952 | | **Clcf1** | Bsf3\|CLC | cardiotrophin-like cytokine factor 1 | protein-coding |
| chr14_118834196_118834756 | chr14 | 118834197 | 118834756 | + | -0.694992 | intron (NM_001160097, intron 1 of 3) | intron (NM_001160097, intron 1 of 3) | -20474 | NM_021386 | 58187 | Mm.390755 | NM_021386 | ENSMUSG00000022132 | **Cldn10** | 6720456I16Rik\|Cldn10a\|Cldn10b\|D14Ertd728e | claudin 10 | protein-coding |
| chr14_118864917_118865316 | chr14 | 118864918 | 118865316 | + | -0.9969 | intron (NM_001160097, intron 3 of 3) | intron (NM_001160097, intron 3 of 3) | 10167 | NM_001160099 | 58187 | Mm.390755 | NM_021386 | ENSMUSG00000022132 | **Cldn10** | 6720456I16Rik\|Cldn10a\|Cldn10b\|D14Ertd728e | claudin 10 | protein-coding |
| chr16_93907795_93908118 | chr16 | 93907796 | 93908118 | + | -0.953335 | Intergenic | RLTR9A\|LTR\|ERVK | 21610 | NM_001165925 | 56173 | Mm.328716 | NM_019500 | ENSMUSG00000047109 | **Cldn14** | AI851731 | claudin 14 | protein-coding |
| chrX_139797814_139798498 | chrX | 139797815 | 139798498 | + | -0.6694 | Intergenic | Intergenic | -2652 | NM_016675 | 12738 | Mm.117068 | NM_016675 | ENSMUSG00000047230 | **Cldn2** | AL022813 | claudin 2 | protein-coding |
| chrX_139797755_139798272 | chrX | 139797756 | 139798272 | + | -0.720181 | Intergenic | Intergenic | -2794 | NM_016675 | 12738 | Mm.117068 | NM_016675 | ENSMUSG00000047230 | **Cldn2** | AL022813 | claudin 2 | protein-coding |
| chr5_134960460_134960713 | chr5 | 134960461 | 134960713 | + | -0.745251 | Intergenic | Intergenic | -13653 | NM_009903 | 12740 | Mm.7339 | NM_009903 | ENSMUSG00000047501 | **Cldn4** | Cep-r\|Cpetr\|Cpetr1 | claudin 4 | protein-coding |
| chr6_129186041_129186402 | chr6 | 129186042 | 129186402 | + | -0.847912 | 3' UTR (NM_053109, exon 5 of 5) | 3' UTR (NM_053109, exon 5 of 5) | 5607 | NM_053109 | 93694 | Mm.197536 | NM_053109 | ENSMUSG00000030157 | **Clec2d** | Clr-b\|Clrb\|Ocil | C-type lectin domain family 2, member d | protein-coding |
| chr6_128887153_128887691 | chr6 | 128887154 | 128887691 | + | -0.642776 | promoter-TSS (NM_001289707) | promoter-TSS (NM_001289707) | -165 | NM_001289706 | 93675 | Mm.391513 | NM_020257 | ENSMUSG00000030365 | **Clec2i** | Clr-g\|Clrg\|Dcl1\|LCL-1\|OCILrP2 | C-type lectin domain family 2, member i | protein-coding |
| chr9_123152117_123152454 | chr9 | 123152118 | 123152454 | + | -0.862327 | intron (NM_011606, intron 1 of 2) | intron (NM_011606, intron 1 of 2) | 1340 | NM_011606 | 21922 | Mm.34588 | NM_011606 | ENSMUSG00000025784 | **Clec3b** | Tna | C-type lectin domain family 3, member b | protein-coding |
| chr8_3711576_3712020 | chr8 | 3711577 | 3712020 | + | -0.721965 | Intergenic | Intergenic | 8853 | NM_029465 | 75863 | Mm.109183 | NM_029465 | ENSMUSG00000074491 | **Clec4g** | 4930572L20Rik | C-type lectin domain family 4, member g | protein-coding |
| chr4_135290261_135290614 | chr4 | 135290262 | 135290614 | + | -0.988915 | Intergenic | Intergenic | -17678 | NM_013885 | 29876 | Mm.257765 | NM_013885 | ENSMUSG00000037242 | **Clic4** | D0Jmb3\|TU-74\|mc3s5\|mtCLIC | chloride intracellular channel 4 (mitochondrial) | protein-coding |
| chr4_135248165_135248869 | chr4 | 135248166 | 135248869 | + | -1.1947 | intron (NM_013885, intron 1 of 5) | intron (NM_013885, intron 1 of 5) | 24243 | NM_013885 | 29876 | Mm.257765 | NM_013885 | ENSMUSG00000037242 | **Clic4** | D0Jmb3\|TU-74\|mc3s5\|mtCLIC | chloride intracellular channel 4 (mitochondrial) | protein-coding |
| chr11_45875622_45875856 | chr11 | 45875623 | 45875856 | + | -0.772819 | intron (NM_001045520, intron 1 of 11) | intron (NM_001045520, intron 1 of 11) | 23775 | NM_001045520 | 216705 | Mm.169673 | NM_001045520 | ENSMUSG00000006169 | **Clint1** | AI642036\|AW537616\|C530049I24Rik\|Enth\|Epn4\|F630206G17Rik\|mKIAA0171 | clathrin interactor 1 | protein-coding |
| chr11_45893806_45894466 | chr11 | 45893807 | 45894466 | + | -0.791335 | intron (NM_001045520, intron 7 of 11) | intron (NM_001045520, intron 7 of 11) | 42172 | NM_001045520 | 216705 | Mm.169673 | NM_001045520 | ENSMUSG00000006169 | **Clint1** | AI642036\|AW537616\|C530049I24Rik\|Enth\|Epn4\|F630206G17Rik\|mKIAA0171 | clathrin interactor 1 | protein-coding |
| chr11_45872118_45872538 | chr11 | 45872119 | 45872538 | + | -0.801854 | intron (NM_001045520, intron 1 of 11) | intron (NM_001045520, intron 1 of 11) | 20364 | NM_001045520 | 216705 | Mm.169673 | NM_001045520 | ENSMUSG00000006169 | **Clint1** | AI642036\|AW537616\|C530049I24Rik\|Enth\|Epn4\|F630206G17Rik\|mKIAA0171 | clathrin interactor 1 | protein-coding |
| chr11_45893887_45894210 | chr11 | 45893888 | 45894210 | + | -0.813134 | exon (NM_001045520, exon 7 of 12) | exon (NM_001045520, exon 7 of 12) | 42085 | NM_001045520 | 216705 | Mm.169673 | NM_001045520 | ENSMUSG00000006169 | **Clint1** | AI642036\|AW537616\|C530049I24Rik\|Enth\|Epn4\|F630206G17Rik\|mKIAA0171 | clathrin interactor 1 | protein-coding |
| chr11_45852864_45853251 | chr11 | 45852865 | 45853251 | + | -0.832842 | intron (NM_001045520, intron 1 of 11) | intron (NM_001045520, intron 1 of 11) | 1094 | NM_001045520 | 216705 | Mm.169673 | NM_001045520 | ENSMUSG00000006169 | **Clint1** | AI642036\|AW537616\|C530049I24Rik\|Enth\|Epn4\|F630206G17Rik\|mKIAA0171 | clathrin interactor 1 | protein-coding |
| chr5_123682351_123682793 | chr5 | 123682352 | 123682793 | + | -0.722685 | intron (NM_001291229, intron 1 of 23) | intron (NM_001291229, intron 1 of 23) | 1789 | NM_001291229 | 56430 | Mm.241109 | NM_019765 | ENSMUSG00000049550 | **Clip1** | 1110007I12Rik\|4631429H07Rik\|AV017631\|C81039\|CLIP-170\|CLIP170\|CYLN1\|Clip50\|Rsn\|mKIAA4046\|restin | CAP-GLY domain containing linker protein 1 | protein-coding |
| chr9_57770431_57770649 | chr9 | 57770432 | 57770649 | + | -0.906548 | Intergenic | Intergenic | -4680 | NM_007713 | 102414 | Mm.25720 | NM_007713 | ENSMUSG00000032316 | **Clk3** | AI256811 | CDC-like kinase 3 | protein-coding |
| chr7_101673170_101673955 | chr7 | 101673171 | 101673955 | + | -0.650587 | intron (NM_009191, intron 2 of 15) | intron (NM_009191, intron 2 of 15) | 9795 | NM_009191 | 20480 | Mm.3990 | NM_009191 | ENSMUSG00000001829 | **Clpb** | AL118244\|Skd3 | ClpB caseinolytic peptidase B | protein-coding |
| chr7_101673408_101673955 | chr7 | 101673409 | 101673955 | + | -0.665677 | intron (NM_009191, intron 2 of 15) | intron (NM_009191, intron 2 of 15) | 9914 | NM_009191 | 20480 | Mm.3990 | NM_009191 | ENSMUSG00000001829 | **Clpb** | AL118244\|Skd3 | ClpB caseinolytic peptidase B | protein-coding |
| chr7_101635711_101636538 | chr7 | 101635712 | 101636538 | + | -0.713244 | Intergenic | MLT1J1\|LTR\|ERVL-MaLR | -27643 | NM_009191 | 20480 | Mm.3990 | NM_009191 | ENSMUSG00000001829 | **Clpb** | AL118244\|Skd3 | ClpB caseinolytic peptidase B | protein-coding |
| chr7_101636132_101636519 | chr7 | 101636133 | 101636519 | + | -0.905666 | Intergenic | ORR1B1\|LTR\|ERVL-MaLR | -27442 | NM_009191 | 20480 | Mm.3990 | NM_009191 | ENSMUSG00000001829 | **Clpb** | AL118244\|Skd3 | ClpB caseinolytic peptidase B | protein-coding |
| chr4_44022193_44023182 | chr4 | 44022194 | 44023182 | + | -0.997794 | intron (NM_016760, intron 3 of 5) | B1F2\|SINE\|Alu | 10038 | NM_001290470 | 12757 | Mm.298875 | NM_016760 | ENSMUSG00000028478 | **Clta** | AV026556\|Lca | clathrin, light polypeptide (Lca) | protein-coding |
| chr4_8987974_8988653 | chr4 | 8987975 | 8988653 | + | -0.679303 | Intergenic | Intergenic | -281003 | NM_028940 | 74438 | Mm.473368 | NM_028940 | ENSMUSG00000041216 | **Clvs1** | 4933402J24Rik\|Clvl1\|Rlbp1l1 | clavesin 1 | protein-coding |
| chr4_8988014_8988874 | chr4 | 8988015 | 8988874 | + | -0.731731 | Intergenic | Intergenic | -280873 | NM_028940 | 74438 | Mm.473368 | NM_028940 | ENSMUSG00000041216 | **Clvs1** | 4933402J24Rik\|Clvl1\|Rlbp1l1 | clavesin 1 | protein-coding |
| chr4_9337385_9338041 | chr4 | 9337386 | 9338041 | + | -0.74196 | intron (NM_028940, intron 2 of 5) | intron (NM_028940, intron 2 of 5) | 68396 | NM_028940 | 74438 | Mm.473368 | NM_028940 | ENSMUSG00000041216 | **Clvs1** | 4933402J24Rik\|Clvl1\|Rlbp1l1 | clavesin 1 | protein-coding |
| chr4_9009408_9010139 | chr4 | 9009409 | 9010139 | + | -0.838994 | Intergenic | Intergenic | -259543 | NM_028940 | 74438 | Mm.473368 | NM_028940 | ENSMUSG00000041216 | **Clvs1** | 4933402J24Rik\|Clvl1\|Rlbp1l1 | clavesin 1 | protein-coding |
| chr4_9033109_9033586 | chr4 | 9033110 | 9033586 | + | -0.925296 | Intergenic | Intergenic | -235969 | NM_028940 | 74438 | Mm.473368 | NM_028940 | ENSMUSG00000041216 | **Clvs1** | 4933402J24Rik\|Clvl1\|Rlbp1l1 | clavesin 1 | protein-coding |
| chr14_122186957_122187194 | chr14 | 122186958 | 122187194 | + | -0.869156 | intron (NM_029556, intron 1 of 7) | L1MB8\|LINE\|L1 | 5382 | NM_029556 | 69634 | Mm.34608 | NM_029556 | ENSMUSG00000025545 | **Clybl** | 0610033J05Rik\|2310014M14Rik\|AI256068\|Clb | citrate lyase beta like | protein-coding |
| chr14_122186892_122187309 | chr14 | 122186893 | 122187309 | + | -0.914755 | intron (NM_029556, intron 1 of 7) | L1MB8\|LINE\|L1 | 5407 | NM_029556 | 69634 | Mm.34608 | NM_029556 | ENSMUSG00000025545 | **Clybl** | 0610033J05Rik\|2310014M14Rik\|AI256068\|Clb | citrate lyase beta like | protein-coding |
| chr13_24375722_24376265 | chr13 | 24375723 | 24376265 | + | -0.609952 | intron (NM_001284519, intron 2 of 17) | intron (NM_001284519, intron 2 of 17) | -39128 | NM_007717 | 12763 | Mm.8396 | NM_007717 | ENSMUSG00000016756 | **Cmah** | - | cytidine monophospho-N-acetylneuraminic acid hydroxylase | protein-coding |
| chr13_24415259_24416182 | chr13 | 24415260 | 24416182 | + | -0.703768 | intron (NM_001284520, intron 1 of 14) | intron (NM_001284520, intron 1 of 14) | 599 | NM_001284520 | 12763 | Mm.8396 | NM_007717 | ENSMUSG00000016756 | **Cmah** | - | cytidine monophospho-N-acetylneuraminic acid hydroxylase | protein-coding |
| chr13_24419561_24419929 | chr13 | 24419562 | 24419929 | + | -0.887842 | intron (NM_001284520, intron 3 of 14) | intron (NM_001284520, intron 3 of 14) | 4623 | NM_001284520 | 12763 | Mm.8396 | NM_007717 | ENSMUSG00000016756 | **Cmah** | - | cytidine monophospho-N-acetylneuraminic acid hydroxylase | protein-coding |
| chr6_142757611_142757914 | chr6 | 142757612 | 142757914 | + | -1.14293 | intron (NM_009908, intron 1 of 7) | intron (NM_009908, intron 1 of 7) | 1077 | NM_009908 | 12764 | Mm.3820 | NM_009908 | ENSMUSG00000030282 | **Cmas** | AW208911\|CMPNeu5Ac\|D6Bwg0250e | cytidine monophospho-N-acetylneuraminic acid synthetase | protein-coding |
| chr8_117306793_117307178 | chr8 | 117306794 | 117307178 | + | -0.638152 | intron (NM_001163262, intron 1 of 20) | intron (NM_001163262, intron 1 of 20) | -42184 | NM_028941 | 74440 | Mm.233181 | NM_028941 | ENSMUSG00000034390 | **Cmip** | 4933407C03Rik\|5830471E12Rik\|AI314065\|AI464126 | c-Maf inducing protein | protein-coding |
| chr8_117278086_117278588 | chr8 | 117278087 | 117278588 | + | -0.757172 | intron (NM_001163262, intron 1 of 20) | intron (NM_001163262, intron 1 of 20) | 21318 | NM_001163262 | 74440 | Mm.233181 | NM_028941 | ENSMUSG00000034390 | **Cmip** | 4933407C03Rik\|5830471E12Rik\|AI314065\|AI464126 | c-Maf inducing protein | protein-coding |
| chr5_113670473_113671082 | chr5 | 113670474 | 113671082 | + | -0.632037 | Intergenic | L1_Mur3\|LINE\|L1 | -20388 | NM_008153 | 14747 | Mm.5196 | NM_008153 | ENSMUSG00000042190 | **Cmklr1** | ChemR23\|DEZ\|Gpcr27\|mcmklr1 | chemokine-like receptor 1 | protein-coding |
| chr5_113670651_113670972 | chr5 | 113670652 | 113670972 | + | -0.656519 | Intergenic | L1_Mur3\|LINE\|L1 | -20422 | NM_008153 | 14747 | Mm.5196 | NM_008153 | ENSMUSG00000042190 | **Cmklr1** | ChemR23\|DEZ\|Gpcr27\|mcmklr1 | chemokine-like receptor 1 | protein-coding |
| chr9_114742471_114743180 | chr9 | 114742472 | 114743180 | + | -1.29941 | intron (NM_026036, intron 3 of 3) | intron (NM_026036, intron 3 of 3) | 11623 | NM_026036 | 67213 | Mm.28858 | NM_026036 | ENSMUSG00000032434 | **Cmtm6** | 2810051A14Rik\|AA536733\|Cklfsf6 | CKLF-like MARVEL transmembrane domain containing 6 | protein-coding |
| chr9_114742712_114743100 | chr9 | 114742713 | 114743100 | + | -1.39277 | intron (NM_026036, intron 3 of 3) | intron (NM_026036, intron 3 of 3) | 11703 | NM_026036 | 67213 | Mm.28858 | NM_026036 | ENSMUSG00000032434 | **Cmtm6** | 2810051A14Rik\|AA536733\|Cklfsf6 | CKLF-like MARVEL transmembrane domain containing 6 | protein-coding |
| chr9_114787552_114788093 | chr9 | 114787553 | 114788093 | + | -0.947735 | Intergenic | Intergenic | -5830 | NM_001252479 | 102545 | Mm.35600 | NM_133978 | ENSMUSG00000032436 | **Cmtm7** | AI481279\|Cklfsf7\|LNV | CKLF-like MARVEL transmembrane domain containing 7 | protein-coding |
| chr2_156267462_156267981 | chr2 | 156267463 | 156267981 | + | -0.763531 | intron (NM_172674, intron 6 of 17) | intron (NM_172674, intron 6 of 17) | -44751 | NM_027585 | 70873 | Mm.475234 | NM_027585 | ENSMUSG00000038085 | **Cnbd2** | 4921517L17Rik\|5430421B09Rik\|Cris | cyclic nucleotide binding domain containing 2 | protein-coding |
| chr2_156267621_156268086 | chr2 | 156267622 | 156268086 | + | -0.784921 | intron (NM_172674, intron 6 of 17) | intron (NM_172674, intron 6 of 17) | -44619 | NM_027585 | 70873 | Mm.475234 | NM_027585 | ENSMUSG00000038085 | **Cnbd2** | 4921517L17Rik\|5430421B09Rik\|Cris | cyclic nucleotide binding domain containing 2 | protein-coding |
| chr5_72634660_72635298 | chr5 | 72634661 | 72635298 | + | -0.726116 | intron (NM_007723, intron 1 of 8) | intron (NM_007723, intron 1 of 8) | 7773 | NM_007723 | 12788 | Mm.436652 | NM_007723 | ENSMUSG00000067220 | **Cnga1** | Cncg | cyclic nucleotide gated channel alpha 1 | protein-coding |
| chr8_95245137_95245656 | chr8 | 95245138 | 95245656 | + | -0.714413 | intron (NM_001195413, intron 31 of 32) | intron (NM_001195413, intron 31 of 32) | 61188 | NM_145601 | 333329 | Mm.484049 | NM_145601 | ENSMUSG00000031789 | **Cngb1** | Cngb1b\|Gm1959 | cyclic nucleotide gated channel beta 1 | protein-coding |

| | | | | | | | | | | | | | | | | | |
|---|---|---|---|---|---|---|---|---|---|---|---|---|---|---|---|---|---|
| chr1_181335132_181335895 | chr1 | 181335133 | 181335895 | + | -0.864843 | Intergenic | Intergenic | -17114 | NM_028408 | 72978 | Mm.338083 | NM_028408 | ENSMUSG00000026514 | **Cnih3** | 2900075G08Rik\|CNIH-3 | cornichon homolog 3 (Drosophila) | protein-coding |
| chr1_181335371_181335920 | chr1 | 181335372 | 181335920 | + | -0.878573 | Intergenic | Intergenic | -16982 | NM_028408 | 72978 | Mm.338083 | NM_028408 | ENSMUSG00000026514 | **Cnih3** | 2900075G08Rik\|CNIH-3 | cornichon homolog 3 (Drosophila) | protein-coding |
| chr3_121369446_121369969 | chr3 | 121369447 | 121369969 | + | -0.764944 | Intergenic | ORR1D2\|LTR\|ERVL-MaLR | -56833 | NM_028044 | 71994 | Mm.275555 | NM_028044 | ENSMUSG00000053931 | **Cnn3** | 1600014M03Rik\|C85854\|Calpo3 | calponin 3, acidic | protein-coding |
| chr3_121407974_121408622 | chr3 | 121407975 | 121408622 | + | -1.09531 | Intergenic | Intergenic | -18243 | NM_028044 | 71994 | Mm.275555 | NM_028044 | ENSMUSG00000053931 | **Cnn3** | 1600014M03Rik\|C85854\|Calpo3 | calponin 3, acidic | protein-coding |
| chr8_95810196_95810417 | chr8 | 95810197 | 95810417 | + | -0.777669 | intron (NR_015574, intron 2 of 3) | intron (NR_015574, intron 2 of 3) | -2845 | NM_001205226 | 234594 | Mm.259220 | NM_153164 | ENSMUSG00000036550 | **Cnot1** | 6030411K04Rik\|AA815922\|D830048B13 | CCR4-NOT transcription complex, subunit 1 | protein-coding |
| chr9_114612972_114613618 | chr9 | 114612973 | 114613618 | + | -0.809181 | intron (NM_153585, intron 12 of 19) | intron (NM_153585, intron 12 of 19) | 26905 | NM_153585 | 78893 | Mm.283708 | NM_153585 | ENSMUSG00000056167 | **Cnot10** | 2600001P13Rik | CCR4-NOT transcription complex, subunit 10 | protein-coding |
| chr9_114612897_114613417 | chr9 | 114612898 | 114613417 | + | -0.836933 | intron (NM_153585, intron 12 of 19) | RLTR34C_MM\|LTR\|ERVK | 27043 | NM_153585 | 78893 | Mm.283708 | NM_153585 | ENSMUSG00000056167 | **Cnot10** | 2600001P13Rik | CCR4-NOT transcription complex, subunit 10 | protein-coding |
| chr5_96176073_96176455 | chr5 | 96176074 | 96176455 | + | -0.664625 | Intergenic | Intergenic | -14274 | NM_144910 | 231464 | Mm.28374 | NM_144910 | ENSMUSG00000034724 | **Cnot6l** | 4932442K20Rik | CCR4-NOT transcription complex, subunit 6-like | protein-coding |
| chr5_28269451_28270484 | chr5 | 28269452 | 28270484 | + | -0.867686 | Intergenic | ORR1E\|LTR\|ERVL-MaLR | -32095 | NM_175651 | 269637 | Mm.68286 | NM_175651 | ENSMUSG00000044681 | **Cnpy1** | 1500012D20Rik\|9630008K15Rik\|AI853839 | canopy 1 homolog (zebrafish) | protein-coding |
| chr4_135904773_135905328 | chr4 | 135904774 | 135905328 | + | -1.11016 | intron (NM_009924, intron 1 of 1) | intron (NM_009924, intron 1 of 1) | 9647 | NM_009924 | 12802 | Mm.297251 | NM_009924 | ENSMUSG00000062585 | **Cnr2** | CB-2\|CB2\|CB2-R | cannabinoid receptor 2 (macrophage) | protein-coding |
| chr1_179578634_179579230 | chr1 | 179578635 | 179579230 | + | -0.66721 | intron (NM_146105, intron 1 of 10) | intron (NM_146105, intron 1 of 10) | 32403 | NM_146105 | 226744 | Mm.87662 | NM_146105 | ENSMUSG00000038949 | **Cnst** | 9630058J23Rik | consortin, connexin sorting protein | protein-coding |
| chr1_179578713_179578970 | chr1 | 179578714 | 179578970 | + | -0.784522 | intron (NM_146105, intron 1 of 10) | intron (NM_146105, intron 1 of 10) | 32313 | NM_146105 | 226744 | Mm.87662 | NM_146105 | ENSMUSG00000038949 | **Cnst** | 9630058J23Rik | consortin, connexin sorting protein | protein-coding |
| chr1_179549409_179549748 | chr1 | 179549410 | 179549748 | + | -0.867539 | intron (NM_146105, intron 1 of 10) | intron (NM_146105, intron 1 of 10) | 3050 | NM_146105 | 226744 | Mm.87662 | NM_146105 | ENSMUSG00000038949 | **Cnst** | 9630058J23Rik | consortin, connexin sorting protein | protein-coding |
| chr4_84946446_84946801 | chr4 | 84946447 | 84946801 | + | -1.15841 | intron (NM_177385, intron 2 of 4) | L1_Rod\|LINE\|L1 | 62315 | NM_177385 | 338349 | Mm.440562 | NM_175275 | ENSMUSG00000038070 | **Cntln** | AI448092\|B430108F07Rik\|D530005L17Rik | centlein, centrosomal protein | protein-coding |
| chr9_11396701_11397491 | chr9 | 11396702 | 11397491 | + | -0.945661 | Intergenic | Intergenic | -492321 | NM_001170787 | 244682 | Mm.187021 | NM_001033359 | ENSMUSG00000039488 | **Cntn5** | 6720426O10Rik\|A830025P08Rik\|Gm507\|NB-2 | contactin 5 | protein-coding |
| chr9_11579046_11579423 | chr9 | 11579047 | 11579423 | + | -0.987411 | Intergenic | Intergenic | -674460 | NM_001170787 | 244682 | Mm.187021 | NM_001033359 | ENSMUSG00000039488 | **Cntn5** | 6720426O10Rik\|A830025P08Rik\|Gm507\|NB-2 | contactin 5 | protein-coding |
| chr9_11584459_11584859 | chr9 | 11584460 | 11584859 | + | -1.00828 | Intergenic | Intergenic | -679884 | NM_001170787 | 244682 | Mm.187021 | NM_001033359 | ENSMUSG00000039488 | **Cntn5** | 6720426O10Rik\|A830025P08Rik\|Gm507\|NB-2 | contactin 5 | protein-coding |
| chr9_11396810_11397127 | chr9 | 11396811 | 11397127 | + | -1.05273 | Intergenic | Intergenic | -492194 | NM_001170787 | 244682 | Mm.187021 | NM_001033359 | ENSMUSG00000039488 | **Cntn5** | 6720426O10Rik\|A830025P08Rik\|Gm507\|NB-2 | contactin 5 | protein-coding |
| chr4_108318638_108319125 | chr4 | 108318639 | 108319125 | + | -1.0202 | Intergenic | Intergenic | -9270 | NM_027250 | 69893 | Mm.28354 | NM_027250 | | **Coa7** | 2010305A19Rik\|AI451324\|D4Ertd796e\|Selrc1 | cytochrome c oxidase assembly factor 7 | protein-coding |
| chr2_65214220_65214650 | chr2 | 65214221 | 65214650 | + | -0.692891 | intron (NM_177025, intron 1 of 13) | intron (NM_177025, intron 1 of 13) | 24191 | NM_177025 | 319876 | Mm.34159 | NM_027225 | ENSMUSG00000034903 | **Cobll1** | 1810047P18Rik\|Coblr1\|D430044D16Rik | Cobl-like 1 | protein-coding |
| chr2_65188651_65189010 | chr2 | 65188652 | 65189010 | + | -0.738371 | intron (NM_177025, intron 1 of 13) | intron (NM_177025, intron 1 of 13) | 49795 | NM_177025 | 319876 | Mm.34159 | NM_027225 | ENSMUSG00000034903 | **Cobll1** | 1810047P18Rik\|Coblr1\|D430044D16Rik | Cobl-like 1 | protein-coding |
| chr8_124509084_124509583 | chr8 | 124509085 | 124509583 | + | -1.28557 | Intergenic | L2\|LINE\|L2 | -11433 | NM_029746 | 76332 | Mm.240108 | NM_029746 | ENSMUSG00000031979 | **Cog2** | 1190002B08Rik\|2700012E02Rik\|C86089\|Ldlc | component of oligomeric golgi complex 2 | protein-coding |
| chr13_96555138_96555675 | chr13 | 96555139 | 96555675 | + | -0.663287 | intron (NM_023420, intron 2 of 17) | Lx7\|LINE\|L1 | 12672 | NM_023420 | 68018 | Mm.24125 | NM_023420 | ENSMUSG00000021669 | **Col4a3bp** | 2810404O15Rik\|9230101K08Rik\|AU016711\|CERT\|GPBP | collagen, type IV, alpha 3 (Goodpasture antigen) binding protein | protein-coding |
| chr2_18609846_18610126 | chr2 | 18609847 | 18610126 | + | -0.763308 | Intergenic | Intergenic | -62476 | NM_147778 | 12238 | Mm.249586 | NM_147778 | ENSMUSG00000051154 | **Commd3** | AW550818\|Bup\|D2Ertd542e | COMM domain containing 3 | protein-coding |
| chr9_57157504_57157888 | chr9 | 57157505 | 57157888 | + | -0.897424 | intron (NM_025417, intron 1 of 7) | intron (NM_025417, intron 1 of 7) | 603 | NM_025417 | 66199 | Mm.41687 | NM_025417 | ENSMUSG00000032299 | **Commd4** | 1110039H05Rik | COMM domain containing 4 | protein-coding |
| chr5_72182483_72183249 | chr5 | 72182484 | 72183249 | + | -0.608944 | Intergenic | Intergenic | -14683 | NM_178599 | 27784 | Mm.44223 | NM_178599 | ENSMUSG00000029213 | **Commd8** | C77770\|D5Buc26e | COMM domain containing 8 | protein-coding |
| chr5_72182788_72183247 | chr5 | 72182789 | 72183247 | + | -0.673294 | Intergenic | RMER20C_Mm\|LTR\|ERVK | -14835 | NM_178599 | 27784 | Mm.44223 | NM_178599 | ENSMUSG00000029213 | **Commd8** | C77770\|D5Buc26e | COMM domain containing 8 | protein-coding |
| chr5_72159564_72159908 | chr5 | 72159565 | 72159908 | + | -1.10703 | 3' UTR (NM_178599, exon 5 of 5) | 3' UTR (NM_178599, exon 5 of 5) | 8447 | NM_178599 | 27784 | Mm.44223 | NM_178599 | ENSMUSG00000029213 | **Commd8** | C77770\|D5Buc26e | COMM domain containing 8 | protein-coding |
| chr2_101992735_101993564 | chr2 | 101992736 | 101993564 | + | -0.672156 | intron (NM_178886, intron 4 of 5) | intron (NM_178886, intron 4 of 5) | 106888 | NM_029635 | 76501 | Mm.330424 | NM_029635 | ENSMUSG00000027163 | **Commd9** | 1810029F08Rik\|AA409293 | COMM domain containing 9 | protein-coding |
| chr2_101992829_101993311 | chr2 | 101992830 | 101993311 | + | -0.71342 | intron (NM_178886, intron 4 of 5) | intron (NM_178886, intron 4 of 5) | 106808 | NM_029635 | 76501 | Mm.330424 | NM_029635 | ENSMUSG00000027163 | **Commd9** | 1810029F08Rik\|AA409293 | COMM domain containing 9 | protein-coding |
| chr2_101992963_101993340 | chr2 | 101992964 | 101993340 | + | -0.715956 | intron (NM_178886, intron 4 of 5) | intron (NM_178886, intron 4 of 5) | 106890 | NM_029635 | 76501 | Mm.330424 | NM_029635 | ENSMUSG00000027163 | **Commd9** | 1810029F08Rik\|AA409293 | COMM domain containing 9 | protein-coding |
| chr7_114209842_114210227 | chr7 | 114209843 | 114210227 | + | -0.613539 | Intergenic | Intergenic | 44645 | NM_033370 | 70349 | Mm.277024 | NM_033370 | ENSMUSG00000030754 | **Copb1** | 2610019B04Rik | coatomer protein complex, subunit beta 1 | protein-coding |
| chr7_114234052_114234443 | chr7 | 114234053 | 114234443 | + | -0.770554 | intron (NM_033370, intron 12 of 21) | Lx\|LINE\|L1 | 20432 | NM_033370 | 70349 | Mm.277024 | NM_033370 | ENSMUSG00000030754 | **Copb1** | 2610019B04Rik | coatomer protein complex, subunit beta 1 | protein-coding |
| chr7_114210514_114211105 | chr7 | 114210515 | 114211105 | + | -0.918032 | Intergenic | ID4\|SINE\|ID | 43870 | NM_033370 | 70349 | Mm.277024 | NM_033370 | ENSMUSG00000030754 | **Copb1** | 2610019B04Rik | coatomer protein complex, subunit beta 1 | protein-coding |
| chr5_100526154_100526683 | chr5 | 100526155 | 100526683 | + | -0.698557 | intron (NM_012001, intron 1 of 9) | intron (NM_012001, intron 1 of 9) | 8110 | NM_012001 | 26891 | Mm.957 | NM_012001 | ENSMUSG00000035297 | **Cops4** | AW208976\|D5Ertd774e | COP9 (constitutive photomorphogenic) homolog, subunit 4 (Arabidopsis thaliana) | protein-coding |
| chr7_126702332_126702722 | chr7 | 126702333 | 126702722 | + | -0.643786 | intron (NM_009898, intron 2 of 10) | intron (NM_009898, intron 2 of 10) | 1693 | NM_001301374 | 12721 | Mm.290482 | NM_009898 | ENSMUSG00000030707 | **Coro1a** | Clabp\|Lmb3\|TACO\|p57 | coronin, actin binding protein 1A | protein-coding |
| chr9_62510420_62510957 | chr9 | 62510421 | 62510957 | + | -0.636598 | intron (NM_175484, intron 1 of 11) | intron (NM_175484, intron 1 of 11) | 26355 | NM_175484 | 235431 | Mm.335229 | NM_175484 | ENSMUSG00000041729 | **Coro2b** | CLIPINC\|E130012P22Rik | coronin, actin binding protein, 2B | protein-coding |
| chr9_62488749_62489322 | chr9 | 62488750 | 62489322 | + | -0.691624 | intron (NM_175484, intron 2 of 11) | intron (NM_175484, intron 2 of 11) | 48008 | NM_175484 | 235431 | Mm.335229 | NM_175484 | ENSMUSG00000041729 | **Coro2b** | CLIPINC\|E130012P22Rik | coronin, actin binding protein, 2B | protein-coding |
| chr9_62488734_62489142 | chr9 | 62488735 | 62489142 | + | -0.715552 | intron (NM_175484, intron 2 of 11) | intron (NM_175484, intron 2 of 11) | 48106 | NM_175484 | 235431 | Mm.335229 | NM_175484 | ENSMUSG00000041729 | **Coro2b** | CLIPINC\|E130012P22Rik | coronin, actin binding protein, 2B | protein-coding |
| chr16_4673868_4674538 | chr16 | 4673869 | 4674538 | + | -0.708054 | intron (NM_030205, intron 3 of 27) | intron (NM_030205, intron 3 of 27) | 5517 | NM_030205 | 78885 | Mm.325512 | NM_030205 | ENSMUSG00000039637 | **Coro7** | 0610011B16Rik\|AW048373\|AW556392 | coronin 7 | protein-coding |
| chr8_119812942_119813240 | chr8 | 119812943 | 119813240 | + | -0.755006 | intron (NM_028071, intron 3 of 3) | intron (NM_028071, intron 3 of 3) | 27488 | NM_028071 | 72042 | Mm.393405 | NM_028071 | ENSMUSG00000031827 | **Cotl1** | 1810074P22Rik\|2010004C08Rik\|Clp | coactosin-like 1 (Dictyostelium) | protein-coding |
| chr11_64051492_64051901 | chr11 | 64051493 | 64051901 | + | -0.748808 | intron (NM_178379, intron 3 of 6) | intron (NM_178379, intron 3 of 6) | 27775 | NM_178379 | 70383 | Mm.340211 | NM_178379 | ENSMUSG00000042148 | **Cox10** | 2410004F01Rik\|AU042636 | cytochrome c oxidase assembly protein 10 | protein-coding |
| chr11_64047323_64047874 | chr11 | 64047324 | 64047874 | + | -0.94321 | intron (NM_178379, intron 3 of 6) | intron (NM_178379, intron 3 of 6) | 31873 | NM_178379 | 70383 | Mm.340211 | NM_178379 | ENSMUSG00000042148 | **Cox10** | 2410004F01Rik\|AU042636 | cytochrome c oxidase assembly protein 10 | protein-coding |
| chr2_152775345_152775682 | chr2 | 152775346 | 152775682 | + | -1.25403 | intron (NM_001289739, intron 2 of 4) | B3\|SINE\|B2 | 21341 | NM_053091 | 84682 | Mm.196668 | NM_053091 | ENSMUSG00000009876 | **Cox4i2** | Cox4b\|CoxIV-2 | cytochrome c oxidase subunit IV isoform 2 | protein-coding |
| chr2_152775330_152775811 | chr2 | 152775331 | 152775811 | + | -1.27836 | intron (NM_001289739, intron 2 of 4) | B3\|SINE\|B2 | 21398 | NM_053091 | 84682 | Mm.196668 | NM_053091 | ENSMUSG00000009876 | **Cox4i2** | Cox4b\|CoxIV-2 | cytochrome c oxidase subunit IV isoform 2 | protein-coding |
| chr9_57524491_57524796 | chr9 | 57524492 | 57524796 | + | -0.694035 | intron (NM_007747, intron 1 of 4) | intron (NM_007747, intron 1 of 4) | 3412 | NM_007747 | 12858 | Mm.273403 | NM_007747 | ENSMUSG00000000088 | **Cox5a** | AA959768\|CcOX | cytochrome c oxidase subunit Va | protein-coding |
| chr17_83453674_83453927 | chr17 | 83453675 | 83453927 | + | -1.20636 | intron (NM_001286567, intron 14 of 23) | intron (NM_001286567, intron 14 of 23) | 60532 | NM_009187 | 20463 | Mm.30072 | NM_009187 | ENSMUSG00000024248 | **Cox7a2l** | COX7AR\|COX7RP\|EB1\|SIG-81\|SIG81\|Silg81 | cytochrome c oxidase subunit VIIa polypeptide 2-like | protein-coding |
| chr17_83453442_83454006 | chr17 | 83453443 | 83454006 | + | -1.24157 | intron (NM_001286567, intron 14 of 23) | intron (NM_001286567, intron 14 of 23) | 60609 | NM_009187 | 20463 | Mm.30072 | NM_009187 | ENSMUSG00000024248 | **Cox7a2l** | COX7AR\|COX7RP\|EB1\|SIG-81\|SIG81\|Silg81 | cytochrome c oxidase subunit VIIa polypeptide 2-like | protein-coding |
| chr17_83559040_83559614 | chr17 | 83559041 | 83559614 | + | -1.2879 | Intergenic | Intergenic | -44994 | NM_001159529 | 20463 | Mm.30072 | NM_009187 | ENSMUSG00000024248 | **Cox7a2l** | COX7AR\|COX7RP\|EB1\|SIG-81\|SIG81\|Silg81 | cytochrome c oxidase subunit VIIa polypeptide 2-like | protein-coding |
| chr3_19945589_19946294 | chr3 | 19945590 | 19946294 | + | -0.947781 | Intergenic | ORR1B2-int\|LTR\|ERVL-MaLR | -11112 | NM_001276250 | 12870 | Mm.13787 | NM_007752 | ENSMUSG00000003617 | **Cp** | D3Ertd555e | ceruloplasmin | protein-coding |
| chr3_20304793_20305264 | chr3 | 20304794 | 20305264 | + | -0.691813 | Intergenic | Intergenic | -29300 | NM_029706 | 76703 | Mm.34692 | NM_029706 | | **Cpb1** | 0910001A18Rik\|1810063F02Rik\|2210008M23Rik\|AI504870 | carboxypeptidase B1 (tissue) | protein-coding |
| chr3_20263491_20263848 | chr3 | 20263492 | 20263848 | + | -0.710214 | intron (NM_029706, intron 7 of 10) | intron (NM_029706, intron 7 of 10) | 12059 | NM_029706 | 76703 | Mm.34692 | NM_029706 | | **Cpb1** | 0910001A18Rik\|1810063F02Rik\|2210008M23Rik\|AI504870 | carboxypeptidase B1 (tissue) | protein-coding |

| | | | | | | | | | | | | | | | | | |
|---|---|---|---|---|---|---|---|---|---|---|---|---|---|---|---|---|---|
| chr3_20304577_20305253 | chr3 | 20304578 | 20305253 | + | -0.717527 | Intergenic | Intergenic | -29186 | NM_029706 | 76703 | Mm.34692 | NM_029706 | | **Cpb1** | 0910001A18Rik\|1810063F02Rik\|2210008M23Rik\|AI504870 | carboxypeptidase B1 (tissue) | protein-coding |
| chr16_30176353_30176671 | chr16 | 30176354 | 30176671 | + | -0.648315 | Intergenic | MT2A\|LTR\|ERVL | 91020 | NM_027904 | 71756 | Mm.290614 | NM_027904 | ENSMUSG00000023176 | **Cpn2** | 1300018K11Rik | carboxypeptidase N, polypeptide 2 | protein-coding |
| chr16_30177730_30178098 | chr16 | 30177731 | 30178098 | + | -1.10447 | Intergenic | Tigger7\|DNA\|TcMar-Tigger | 89618 | NM_027904 | 71756 | Mm.290614 | NM_027904 | ENSMUSG00000023176 | **Cpn2** | 1300018K11Rik | carboxypeptidase N, polypeptide 2 | protein-coding |
| chr10_117434855_117435231 | chr10 | 117434856 | 117435231 | + | -0.855927 | Intergenic | Intergenic | -58070 | NM_001013391 | 432508 | Mm.478881 | NM_001013391 | ENSMUSG00000055531 | **Cpsf6** | 4733401N12Rik\|AI256641\|CFIM\|CFIM68\|HPBRII-4\|HPBRII-7 | cleavage and polyadenylation specific factor 6 | protein-coding |
| chr6_127581845_127583317 | chr6 | 127581846 | 127583317 | + | -0.68971 | intron (NM_001033464, intron 1 of 6) | intron (NM_001033464, intron 1 of 6) | 4606 | NM_001033464 | 381812 | Mm.296093 | NM_001033464 | ENSMUSG00000061414 | **Cracr2a** | Efcab4b\|Gm1073\|Gm462 | calcium release activated channel regulator 2A | protein-coding |
| chr6_127582340_127583359 | chr6 | 127582341 | 127583359 | + | -0.750569 | intron (NM_001033464, intron 1 of 6) | intron (NM_001033464, intron 1 of 6) | 4875 | NM_001033464 | 381812 | Mm.296093 | NM_001033464 | ENSMUSG00000061414 | **Cracr2a** | Efcab4b\|Gm1073\|Gm462 | calcium release activated channel regulator 2A | protein-coding |
| chr6_127617230_127617649 | chr6 | 127617231 | 127617649 | + | -0.7742 | intron (NM_001033464, intron 5 of 6) | intron (NM_001033464, intron 5 of 6) | 39465 | NM_001033464 | 381812 | Mm.296093 | NM_001033464 | ENSMUSG00000061414 | **Cracr2a** | Efcab4b\|Gm1073\|Gm462 | calcium release activated channel regulator 2A | protein-coding |
| chr10_95178831_95179187 | chr10 | 95178832 | 95179187 | + | -0.658326 | intron (NM_009950, intron 2 of 2) | L1MB3\|LINE\|L1 | 145088 | NM_009950 | 12905 | Mm.473739 | NM_009950 | ENSMUSG00000045867 | **Cradd** | RAIDD | CASP2 and RIPK1 domain containing adaptor with death domain | protein-coding |
| chr10_95262357_95262785 | chr10 | 95262358 | 95262785 | + | -0.668719 | intron (NM_009950, intron 2 of 2) | intron (NM_009950, intron 2 of 2) | 61526 | NM_009950 | 12905 | Mm.473739 | NM_009950 | ENSMUSG00000045867 | **Cradd** | RAIDD | CASP2 and RIPK1 domain containing adaptor with death domain | protein-coding |
| chr10_95195704_95196160 | chr10 | 95195705 | 95196160 | + | -0.70982 | intron (NM_009950, intron 2 of 2) | ORR1E\|LTR\|ERVL-MaLR | 128165 | NM_009950 | 12905 | Mm.473739 | NM_009950 | ENSMUSG00000045867 | **Cradd** | RAIDD | CASP2 and RIPK1 domain containing adaptor with death domain | protein-coding |
| chr6_106791471_106791768 | chr6 | 106791472 | 106791768 | + | -1.14496 | intron (NM_175357, intron 4 of 10) | intron (NM_175357, intron 4 of 10) | 8467 | NM_021449 | 58799 | Mm.290085 | NM_021449 | | **Crbn** | 2610203G15Rik\|2900045O07Rik\|AF229032\|AW108261\|piL | cereblon | protein-coding |
| chr6_106792891_106793895 | chr6 | 106792892 | 106793895 | + | -1.22638 | intron (NM_175357, intron 4 of 10) | intron (NM_175357, intron 4 of 10) | 6694 | NM_021449 | 58799 | Mm.290085 | NM_021449 | | **Crbn** | 2610203G15Rik\|2900045O07Rik\|AF229032\|AW108261\|piL | cereblon | protein-coding |
| chr6_106792919_106793563 | chr6 | 106792920 | 106793563 | + | -1.43559 | intron (NM_175357, intron 4 of 10) | intron (NM_175357, intron 4 of 10) | 6846 | NM_021449 | 58799 | Mm.290085 | NM_021449 | | **Crbn** | 2610203G15Rik\|2900045O07Rik\|AF229032\|AW108261\|piL | cereblon | protein-coding |
| chr1_64486263_64486955 | chr1 | 64486264 | 64486955 | + | -0.648692 | Intergenic | Intergenic | -46195 | NM_133828 | 12912 | Mm.422634 | NM_009952 | ENSMUSG00000025958 | **Creb1** | 2310001E10Rik\|3526402H21Rik\|AV083133\|Creb\|Creb-1 | cAMP responsive element binding protein 1 | protein-coding |
| chr1_64482104_64482520 | chr1 | 64482105 | 64482520 | + | -0.763166 | Intergenic | Tigger15a\|DNA\|TcMar-Tigger | -50492 | NM_133828 | 12912 | Mm.422634 | NM_009952 | ENSMUSG00000025958 | **Creb1** | 2310001E10Rik\|3526402H21Rik\|AV083133\|Creb\|Creb-1 | cAMP responsive element binding protein 1 | protein-coding |
| chr1_64527669_64527979 | chr1 | 64527670 | 64527979 | + | -1.16837 | Intergenic | Intergenic | -4980 | NM_133828 | 12912 | Mm.422634 | NM_009952 | ENSMUSG00000025958 | **Creb1** | 2310001E10Rik\|3526402H21Rik\|AV083133\|Creb\|Creb-1 | cAMP responsive element binding protein 1 | protein-coding |
| chr18_3270497_3271118 | chr18 | 3270498 | 3271118 | + | -1.13805 | intron (NM_001110854, intron 3 of 3) | intron (NM_001110854, intron 3 of 3) | 10939 | NM_001110854 | 12916 | Mm.5244 | NM_013498 | ENSMUSG00000063889 | **Crem** | ICER\|ICERI | cAMP responsive element modulator | protein-coding |
| chr11_80072399_80072638 | chr11 | 80072400 | 80072638 | + | -0.734708 | intron (NM_001277106, intron 1 of 6) | intron (NM_001277106, intron 1 of 6) | 8472 | NM_001277106 | 54394 | Mm.272093 | NM_018776 | ENSMUSG00000017561 | **Crlf3** | BB164954\|Creme9\|Cytor4 | cytokine receptor-like factor 3 | protein-coding |
| chr11_80067580_80067979 | chr11 | 80067581 | 80067979 | + | -0.887252 | intron (NM_001277106, intron 1 of 6) | intron (NM_001277106, intron 1 of 6) | 13211 | NM_001277106 | 54394 | Mm.272093 | NM_018776 | ENSMUSG00000017561 | **Crlf3** | BB164954\|Creme9\|Cytor4 | cytokine receptor-like factor 3 | protein-coding |
| chr11_80067408_80067950 | chr11 | 80067409 | 80067950 | + | -0.893748 | intron (NM_001277106, intron 1 of 6) | intron (NM_001277106, intron 1 of 6) | 13312 | NM_001277106 | 54394 | Mm.272093 | NM_018776 | ENSMUSG00000017561 | **Crlf3** | BB164954\|Creme9\|Cytor4 | cytokine receptor-like factor 3 | protein-coding |
| chr11_80067475_80067806 | chr11 | 80067476 | 80067806 | + | -0.900768 | intron (NM_001277106, intron 1 of 6) | intron (NM_001277106, intron 1 of 6) | 13350 | NM_001277106 | 54394 | Mm.272093 | NM_018776 | ENSMUSG00000017561 | **Crlf3** | BB164954\|Creme9\|Cytor4 | cytokine receptor-like factor 3 | protein-coding |
| chr11_80075610_80076029 | chr11 | 80075611 | 80076029 | + | -1.55045 | intron (NM_001277106, intron 1 of 6) | intron (NM_001277106, intron 1 of 6) | 5171 | NM_001277106 | 54394 | Mm.272093 | NM_018776 | ENSMUSG00000017561 | **Crlf3** | BB164954\|Creme9\|Cytor4 | cytokine receptor-like factor 3 | protein-coding |
| chr5_9014806_9015173 | chr5 | 9014807 | 9015173 | + | -0.87672 | Intergenic | Intergenic | -17844 | NM_023733 | 74114 | Mm.28197 | NM_023733 | ENSMUSG00000003623 | **Crot** | 1200003H03Rik | carnitine O-octanoyltransferase | protein-coding |
| chr9_40997128_40997673 | chr9 | 40997129 | 40997673 | + | -0.807428 | intron (NM_019465, intron 1 of 9) | intron (NM_019465, intron 1 of 9) | 7227 | NM_001281954 | 54698 | Mm.42791 | NM_019465 | ENSMUSG00000032021 | **Crtam** | - | cytotoxic and regulatory T cell molecule | protein-coding |
| chr7_80672308_80672801 | chr7 | 80672309 | 80672801 | + | -0.835021 | intron (NM_173863, intron 2 of 14) | L1MB2\|LINE\|L1 | 16322 | NM_173863 | 70461 | Mm.392175 | NM_173863 | ENSMUSG00000030527 | **Crtc3** | 2610312F20Rik\|6332415K15Rik\|AI429792\|BC011210\|TORC3 | CREB regulated transcription coactivator 3 | protein-coding |
| chr7_80696981_80697326 | chr7 | 80696982 | 80697326 | + | -0.865161 | Intergenic | Intergenic | -8277 | NM_173863 | 70461 | Mm.392175 | NM_173863 | ENSMUSG00000030527 | **Crtc3** | 2610312F20Rik\|6332415K15Rik\|AI429792\|BC011210\|TORC3 | CREB regulated transcription coactivator 3 | protein-coding |
| chr7_80718418_80718846 | chr7 | 80718419 | 80718846 | + | -0.883315 | intron (NM_016721, intron 36 of 37) | L1MB2\|LINE\|L1 | -29755 | NM_173863 | 70461 | Mm.392175 | NM_173863 | ENSMUSG00000030527 | **Crtc3** | 2610312F20Rik\|6332415K15Rik\|AI429792\|BC011210\|TORC3 | CREB regulated transcription coactivator 3 | protein-coding |
| chr10_85166173_85166587 | chr10 | 85166174 | 85166587 | + | -0.904436 | intron (NM_007771, intron 1 of 12) | intron (NM_007771, intron 1 of 12) | 18674 | NM_007771 | 12952 | Mm.26237 | NM_007771 | ENSMUSG00000020038 | **Cry1** | AU020726\|AU021000\|Phll1 | cryptochrome 1 (photolyase-like) | protein-coding |
| chr2_92442690_92442973 | chr2 | 92442691 | 92442973 | + | -0.861289 | Intergenic | L1MB7\|LINE\|L1 | -8763 | NM_009963 | 12953 | Mm.254181 | NM_009963 | ENSMUSG00000068742 | **Cry2** | AV006279\|D130054K12Rik | cryptochrome 2 (photolyase-like) | protein-coding |
| chr5_112253185_112253586 | chr5 | 112253186 | 112253586 | + | -0.642411 | promoter-TSS (NM_021351) | promoter-TSS (NM_021351) | -884 | NM_021351 | 12959 | Mm.40324 | NM_021351 | ENSMUSG00000066975 | **Cryba4** | - | crystallin, beta A4 | protein-coding |
| chr5_112242050_112242873 | chr5 | 112242051 | 112242873 | + | -0.649147 | Intergenic | Intergenic | 10040 | NM_021351 | 12959 | Mm.40324 | NM_021351 | ENSMUSG00000066975 | **Cryba4** | - | crystallin, beta A4 | protein-coding |
| chr5_112253294_112253530 | chr5 | 112253295 | 112253530 | + | -0.653487 | promoter-TSS (NM_021351) | promoter-TSS (NM_021351) | -910 | NM_021351 | 12959 | Mm.40324 | NM_021351 | ENSMUSG00000066975 | **Cryba4** | - | crystallin, beta A4 | protein-coding |
| chr6_118166704_118167540 | chr6 | 118166705 | 118167540 | + | -0.870158 | intron (NM_009050, intron 15 of 19) | intron (NM_009050, intron 15 of 19) | -27982 | NM_030165 | 78752 | Mm.300317 | NM_030165 | ENSMUSG00000042042 | **Csgalnact2** | 4632415D10Rik\|Galnact2 | chondroitin sulfate N-acetylgalactosaminyltransferase 2 | protein-coding |
| chr9_57647516_57647735 | chr9 | 57647517 | 57647735 | + | -0.929039 | Intergenic | MIRb\|SINE\|MIR | -2446 | NM_007783 | 12988 | Mm.21974 | NM_007783 | ENSMUSG00000032312 | **Csk** | AW212630 | c-src tyrosine kinase | protein-coding |
| chr10_80618272_80618596 | chr10 | 80618273 | 80618596 | + | -0.96817 | Intergenic | Intergenic | -4346 | NM_134002 | 103236 | Mm.29873 | NM_134002 | ENSMUSG00000003345 | **Csnk1g2** | 2810429I12Rik\|AI463719 | casein kinase 1, gamma 2 | protein-coding |
| chr18_53937287_53937746 | chr18 | 53937288 | 53937746 | + | -0.742297 | intron (NM_152809, intron 10 of 12) | intron (NM_152809, intron 10 of 12) | 75404 | NM_152809 | 70425 | Mm.368668 | NM_152809 | ENSMUSG00000073563 | **Csnk1g3** | 3300002K07Rik\|C330049O21Rik | casein kinase 1, gamma 3 | protein-coding |
| chr1_135659414_135660187 | chr1 | 135659415 | 135660187 | + | -0.699111 | Intergenic | Intergenic | -69396 | NM_007791 | 13007 | Mm.196484 | NM_007791 | ENSMUSG00000026421 | **Csrp1** | AA408841\|AA959891\|AW545626\|CRP1\|Csrp | cysteine and glycine-rich protein 1 | protein-coding |
| chr1_135725428_135725662 | chr1 | 135725429 | 135725662 | + | -1.08531 | Intergenic | Intergenic | -3652 | NM_007791 | 13007 | Mm.196484 | NM_007791 | ENSMUSG00000026421 | **Csrp1** | AA408841\|AA959891\|AW545626\|CRP1\|Csrp | cysteine and glycine-rich protein 1 | protein-coding |
| chr2_150578257_150579117 | chr2 | 150578258 | 150579117 | + | -0.940312 | 3' UTR (NM_009977, exon 4 of 4) | 3' UTR (NM_009977, exon 4 of 4) | 8272 | NM_009977 | 13011 | Mm.12965 | NM_009977 | ENSMUSG00000068129 | **Cst7** | Cmap | cystatin F (leukocystatin) | protein-coding |
| chr2_30636129_30637302 | chr2 | 30636130 | 30637302 | + | -0.651856 | Intergenic | Intergenic | 41672 | NM_030137 | 78617 | Mm.40413 | NM_030137 | ENSMUSG00000047363 | **Cstad** | 9530039B02Rik | CSA-conditional, T cell activation-dependent protein | protein-coding |
| chr2_30586991_30587690 | chr2 | 30586992 | 30587690 | + | -0.690381 | Intergenic | Intergenic | -7703 | NM_030137 | 78617 | Mm.40413 | NM_030137 | ENSMUSG00000047363 | **Cstad** | 9530039B02Rik | CSA-conditional, T cell activation-dependent protein | protein-coding |
| chr2_30547504_30548206 | chr2 | 30547505 | 30548206 | + | -0.783088 | Intergenic | (CGGGGG)n\|Simple_repeat\|Simple_repeat | -47189 | NM_030137 | 78617 | Mm.40413 | NM_030137 | ENSMUSG00000047363 | **Cstad** | 9530039B02Rik | CSA-conditional, T cell activation-dependent protein | protein-coding |
| chr2_30625214_30626629 | chr2 | 30625215 | 30626629 | + | -0.784857 | Intergenic | MTE2a\|LTR\|ERVL-MaLR | 30878 | NM_030137 | 78617 | Mm.40413 | NM_030137 | ENSMUSG00000047363 | **Cstad** | 9530039B02Rik | CSA-conditional, T cell activation-dependent protein | protein-coding |
| chr2_30547849_30548211 | chr2 | 30547850 | 30548211 | + | -0.913758 | Intergenic | Intergenic | -47014 | NM_030137 | 78617 | Mm.40413 | NM_030137 | ENSMUSG00000047363 | **Cstad** | 9530039B02Rik | CSA-conditional, T cell activation-dependent protein | protein-coding |
| chr2_30625873_30626611 | chr2 | 30625874 | 30626611 | + | -0.936115 | Intergenic | ORR1D2-int\|LTR\|ERVL-MaLR | 31198 | NM_030137 | 78617 | Mm.40413 | NM_030137 | ENSMUSG00000047363 | **Cstad** | 9530039B02Rik | CSA-conditional, T cell activation-dependent protein | protein-coding |
| chr12_59125351_59125800 | chr12 | 59125352 | 59125800 | + | -0.761206 | Intergenic | RLTR33\|LTR\|ERVK | -4170 | NM_146034 | 217615 | Mm.244118 | NM_146034 | ENSMUSG00000021000 | **Ctage5** | D12Bwg0579e\|Mea6\|Mgea\|Mgea6 | CTAGE family, member 5 | protein-coding |
| chr12_59169467_59170503 | chr12 | 59169468 | 59170503 | + | -0.78486 | intron (NM_001165254, intron 13 of 22) | PB1D10\|SINE\|Alu | 38532 | NM_001165253 | 217615 | Mm.244118 | NM_146034 | ENSMUSG00000021000 | **Ctage5** | D12Bwg0579e\|Mea6\|Mgea\|Mgea6 | CTAGE family, member 5 | protein-coding |
| chr2_173082800_173083020 | chr2 | 173082801 | 173083020 | + | -0.665886 | Intergenic | Intergenic | 36615 | NM_001081387 | 664799 | Mm.389396 | NM_001081387 | ENSMUSG00000070495 | **Ctcfl** | Boris\|OTTMUSG00000016680 | CCCTC-binding factor (zinc finger protein)-like | protein-coding |
| chr2_173083527_173084319 | chr2 | 173083528 | 173084319 | + | -0.744968 | Intergenic | ORR1D1\|LTR\|ERVL-MaLR | 35602 | NM_001081387 | 664799 | Mm.389396 | NM_001081387 | ENSMUSG00000070495 | **Ctcfl** | Boris\|OTTMUSG00000016680 | CCCTC-binding factor (zinc finger protein)-like | protein-coding |
| chr10_24640502_24640928 | chr10 | 24640503 | 24640928 | + | -1.16031 | TTS (NM_008813) | TTS (NM_008813) | 45273 | NM_010217 | 14219 | Mm.390287 | NM_010217 | ENSMUSG00000019997 | **Ctgf** | Ccn2\|Fisp12\|Hcs24\|fisp-12 | connective tissue growth factor | protein-coding |

| | | | | | | | | | | | | | | | | | |
|---|---|---|---|---|---|---|---|---|---|---|---|---|---|---|---|---|---|
| chr13_60936392_60937144 | chr13 | 60936393 | 60937144 | + | -0.783345 | promoter-TSS (NM_007796) | promoter-TSS (NM_007796) | -143 | NM_007796 | 13024 | Mm.482089 | NM_007796 | ENSMUSG00000044258 | **Ctla2a** | Ctla-2a | cytotoxic T lymphocyte-associated protein 2 alpha | protein-coding |
| chr1_60907547_60907812 | chr1 | 60907548 | 60907812 | + | -1.02846 | Intergenic | Intergenic | -1345 | NM_009843 | 12477 | Mm.390 | NM_009843 | ENSMUSG00000026011 | **Ctla4** | Cd152\|Ctla-4\|Ly-56 | cytotoxic T-lymphocyte-associated protein 4 | protein-coding |
| chr1_60907409_60907796 | chr1 | 60907410 | 60907796 | + | -1.05507 | Intergenic | Intergenic | -1422 | NM_009843 | 12477 | Mm.390 | NM_009843 | ENSMUSG00000026011 | **Ctla4** | Cd152\|Ctla-4\|Ly-56 | cytotoxic T-lymphocyte-associated protein 4 | protein-coding |
| chr1_60882829_60883151 | chr1 | 60882830 | 60883151 | + | -1.36442 | Intergenic | Intergenic | -26035 | NM_009843 | 12477 | Mm.390 | NM_009843 | ENSMUSG00000026011 | **Ctla4** | Cd152\|Ctla-4\|Ly-56 | cytotoxic T-lymphocyte-associated protein 4 | protein-coding |
| chr2_84582147_84582805 | chr2 | 84582148 | 84582805 | + | -1.18449 | Intergenic | Intergenic | 68034 | NM_001085453 | 12388 | Mm.35738 | NM_007615 | ENSMUSG00000034101 | **Ctnnd1** | AA409437\|AU019353\|Catns\|Ctnnd\|P120\|mKIAA0384 | catenin (cadherin associated protein), delta 1 | protein-coding |
| chr7_88206310_88207755 | chr7 | 88206311 | 88207755 | + | -0.794259 | Intergenic | Intergenic | -71060 | NM_009982 | 13032 | Mm.322945 | NM_009982 | ENSMUSG00000030560 | **Ctsc** | AI047818\|DPP1\|DPPI | cathepsin C | protein-coding |
| chr7_88206320_88207238 | chr7 | 88206321 | 88207238 | + | -0.907747 | Intergenic | Intergenic | -71314 | NM_009982 | 13032 | Mm.322945 | NM_009982 | ENSMUSG00000030560 | **Ctsc** | AI047818\|DPP1\|DPPI | cathepsin C | protein-coding |
| chr7_142381070_142381691 | chr7 | 142381071 | 142381691 | + | -0.660996 | intron (NM_009983, intron 4 of 8) | intron (NM_009983, intron 4 of 8) | 6489 | NM_009983 | 13033 | Mm.231395 | NM_009983 | ENSMUSG00000007891 | **Ctsd** | CD\|CatD | cathepsin D | protein-coding |
| chr1_131653079_131653762 | chr1 | 131653080 | 131653762 | + | -0.63304 | intron (NM_007799, intron 1 of 9) | intron (NM_007799, intron 1 of 9) | 15107 | NM_007799 | 13034 | Mm.230249 | NM_007799 | ENSMUSG00000004552 | **Ctse** | A430072O03Rik\|C920004C08Rik\|CE\|CatE | cathepsin E | protein-coding |
| chr1_131653078_131653416 | chr1 | 131653079 | 131653416 | + | -0.723969 | intron (NM_007799, intron 1 of 9) | intron (NM_007799, intron 1 of 9) | 14933 | NM_007799 | 13034 | Mm.230249 | NM_007799 | ENSMUSG00000004552 | **Ctse** | A430072O03Rik\|C920004C08Rik\|CE\|CatE | cathepsin E | protein-coding |
| chr1_131638095_131638644 | chr1 | 131638096 | 131638644 | + | -0.938433 | promoter-TSS (NM_007799) | promoter-TSS (NM_007799) | 56 | NM_007799 | 13034 | Mm.230249 | NM_007799 | ENSMUSG00000004552 | **Ctse** | A430072O03Rik\|C920004C08Rik\|CE\|CatE | cathepsin E | protein-coding |
| chr1_131637839_131638731 | chr1 | 131637840 | 131638731 | + | -0.984354 | promoter-TSS (NM_007799) | promoter-TSS (NM_007799) | -29 | NM_007799 | 13034 | Mm.230249 | NM_007799 | ENSMUSG00000004552 | **Ctse** | A430072O03Rik\|C920004C08Rik\|CE\|CatE | cathepsin E | protein-coding |
| chr13_60826035_60826570 | chr13 | 60826036 | 60826570 | + | -0.760266 | Intergenic | Intergenic | -23459 | NM_027344 | 70202 | Mm.158337 | NM_027344 | ENSMUSG00000056728 | **Ctsll3** | 2310051M13Rik | cathepsin L-like 3 | protein-coding |
| chr3_81489410_81489679 | chr3 | 81489411 | 81489679 | + | -0.685629 | Intergenic | Lx7\|LINE\|L1 | -443071 | NM_177662 | 229445 | Mm.254642 | NM_177662 | ENSMUSG00000028015 | **Ctso** | A330105D01Rik\|A430056C07\|AI118514 | cathepsin O | protein-coding |
| chr3_81788825_81789466 | chr3 | 81788826 | 81789466 | + | -0.904517 | Intergenic | L1MCa\|LINE\|L1 | -143470 | NM_177662 | 229445 | Mm.254642 | NM_177662 | ENSMUSG00000028015 | **Ctso** | A330105D01Rik\|A430056C07\|AI118514 | cathepsin O | protein-coding |
| chr2_13413463_13414107 | chr2 | 13413464 | 13414107 | + | -0.65461 | intron (NM_001081084, intron 27 of 66) | intron (NM_001081084, intron 27 of 66) | 78091 | NM_001081084 | 65969 | Mm.313915 | NM_001081084 | ENSMUSG00000026726 | **Cubn** | AA408369\|AL022750\|D2Wsu88e | cubilin (intrinsic factor-cobalamin receptor) | protein-coding |
| chr2_13390915_13391427 | chr2 | 13390916 | 13391427 | + | -0.78672 | intron (NM_001081084, intron 28 of 66) | intron (NM_001081084, intron 28 of 66) | 100705 | NM_001081084 | 65969 | Mm.313915 | NM_001081084 | ENSMUSG00000026726 | **Cubn** | AA408369\|AL022750\|D2Wsu88e | cubilin (intrinsic factor-cobalamin receptor) | protein-coding |
| chr2_13390899_13391599 | chr2 | 13390900 | 13391599 | + | -0.799338 | intron (NM_001081084, intron 28 of 66) | intron (NM_001081084, intron 28 of 66) | 100627 | NM_001081084 | 65969 | Mm.313915 | NM_001081084 | ENSMUSG00000026726 | **Cubn** | AA408369\|AL022750\|D2Wsu88e | cubilin (intrinsic factor-cobalamin receptor) | protein-coding |
| chr11_88144813_88146068 | chr11 | 88144814 | 88146068 | + | -0.91793 | intron (NM_198013, intron 1 of 10) | MER3\|DNA\|hAT-Charlie | -24124 | NM_001172099 | 103841 | Mm.374846 | NM_198013 | ENSMUSG00000018378 | **Cuedc1** | AI841487\|C330016O16Rik | CUE domain containing 1 | protein-coding |
| chr11_88145155_88146049 | chr11 | 88145156 | 88146049 | + | -0.98238 | intron (NM_198013, intron 1 of 10) | intron (NM_198013, intron 1 of 10) | -23963 | NM_001172099 | 103841 | Mm.374846 | NM_198013 | ENSMUSG00000018378 | **Cuedc1** | AI841487\|C330016O16Rik | CUE domain containing 1 | protein-coding |
| chr11_88145191_88145823 | chr11 | 88145192 | 88145823 | + | -1.06987 | intron (NM_198013, intron 1 of 10) | MER3\|DNA\|hAT-Charlie | -24058 | NM_001172099 | 103841 | Mm.374846 | NM_198013 | ENSMUSG00000018378 | **Cuedc1** | AI841487\|C330016O16Rik | CUE domain containing 1 | protein-coding |
| chr9_120078314_120078592 | chr9 | 120078315 | 120078592 | + | -0.939518 | Intergenic | Lx8\|LINE\|L1 | -10157 | NM_009987 | 13051 | Mm.44065 | NM_009987 | ENSMUSG00000052336 | **Cx3cr1** | - | chemokine (C-X3-C motif) receptor 1 | protein-coding |
| chr9_120078333_120078535 | chr9 | 120078334 | 120078535 | + | -0.954552 | Intergenic | Lx8\|LINE\|L1 | -10138 | NM_009987 | 13051 | Mm.44065 | NM_009987 | ENSMUSG00000052336 | **Cx3cr1** | - | chemokine (C-X3-C motif) receptor 1 | protein-coding |
| chr1_128874256_128874613 | chr1 | 128874257 | 128874613 | + | -0.625469 | Intergenic | Intergenic | -282136 | NM_009911 | 12767 | Mm.1401 | NM_009911 | ENSMUSG00000045382 | **Cxcr4** | CD184\|Cmkar4\|LESTR\|PB-CKR\|PBSF/SDF-1\|Sdf1r\|b2b220Clo | chemokine (C-X-C motif) receptor 4 | protein-coding |
| chr1_128704899_128705605 | chr1 | 128704900 | 128705605 | + | -0.83646 | Intergenic | Intergenic | -112953 | NM_009911 | 12767 | Mm.1401 | NM_009911 | ENSMUSG00000045382 | **Cxcr4** | CD184\|Cmkar4\|LESTR\|PB-CKR\|PBSF/SDF-1\|Sdf1r\|b2b220Clo | chemokine (C-X-C motif) receptor 4 | protein-coding |
| chr1_128667231_128667442 | chr1 | 128667232 | 128667442 | + | -0.873339 | Intergenic | Intergenic | -75038 | NM_009911 | 12767 | Mm.1401 | NM_009911 | ENSMUSG00000045382 | **Cxcr4** | CD184\|Cmkar4\|LESTR\|PB-CKR\|PBSF/SDF-1\|Sdf1r\|b2b220Clo | chemokine (C-X-C motif) receptor 4 | protein-coding |
| chr1_128667166_128667626 | chr1 | 128667167 | 128667626 | + | -0.93 | Intergenic | Intergenic | -75097 | NM_009911 | 12767 | Mm.1401 | NM_009911 | ENSMUSG00000045382 | **Cxcr4** | CD184\|Cmkar4\|LESTR\|PB-CKR\|PBSF/SDF-1\|Sdf1r\|b2b220Clo | chemokine (C-X-C motif) receptor 4 | protein-coding |
| chr1_128675543_128676027 | chr1 | 128675544 | 128676027 | + | -0.937357 | Intergenic | Intergenic | -83486 | NM_009911 | 12767 | Mm.1401 | NM_009911 | ENSMUSG00000045382 | **Cxcr4** | CD184\|Cmkar4\|LESTR\|PB-CKR\|PBSF/SDF-1\|Sdf1r\|b2b220Clo | chemokine (C-X-C motif) receptor 4 | protein-coding |
| chr1_128782319_128782602 | chr1 | 128782320 | 128782602 | + | -0.971711 | Intergenic | Intergenic | -190162 | NM_009911 | 12767 | Mm.1401 | NM_009911 | ENSMUSG00000045382 | **Cxcr4** | CD184\|Cmkar4\|LESTR\|PB-CKR\|PBSF/SDF-1\|Sdf1r\|b2b220Clo | chemokine (C-X-C motif) receptor 4 | protein-coding |
| chr1_128614497_128615304 | chr1 | 128614498 | 128615304 | + | -1.08814 | Intergenic | Intergenic | -22602 | NM_009911 | 12767 | Mm.1401 | NM_009911 | ENSMUSG00000045382 | **Cxcr4** | CD184\|Cmkar4\|LESTR\|PB-CKR\|PBSF/SDF-1\|Sdf1r\|b2b220Clo | chemokine (C-X-C motif) receptor 4 | protein-coding |
| chr1_128602491_128602698 | chr1 | 128602492 | 128602698 | + | -1.2931 | Intergenic | PB1D10\|SINE\|Alu | -10296 | NM_009911 | 12767 | Mm.1401 | NM_009911 | ENSMUSG00000045382 | **Cxcr4** | CD184\|Cmkar4\|LESTR\|PB-CKR\|PBSF/SDF-1\|Sdf1r\|b2b220Clo | chemokine (C-X-C motif) receptor 4 | protein-coding |
| chr1_128602384_128603329 | chr1 | 128602385 | 128603329 | + | -1.32366 | Intergenic | Intergenic | -10558 | NM_009911 | 12767 | Mm.1401 | NM_009911 | ENSMUSG00000045382 | **Cxcr4** | CD184\|Cmkar4\|LESTR\|PB-CKR\|PBSF/SDF-1\|Sdf1r\|b2b220Clo | chemokine (C-X-C motif) receptor 4 | protein-coding |
| chr1_128602384_128602932 | chr1 | 128602385 | 128602932 | + | -1.49024 | Intergenic | Intergenic | -10359 | NM_009911 | 12767 | Mm.1401 | NM_009911 | ENSMUSG00000045382 | **Cxcr4** | CD184\|Cmkar4\|LESTR\|PB-CKR\|PBSF/SDF-1\|Sdf1r\|b2b220Clo | chemokine (C-X-C motif) receptor 4 | protein-coding |
| chr9_44533006_44533755 | chr9 | 44533007 | 44533755 | + | -1.14889 | Intergenic | Intergenic | -6960 | NM_007551 | 12145 | Mm.6246 | NM_007551 | ENSMUSG00000047880 | **Cxcr5** | Blr1\|CXC-R5\|CXCR-5\|Gpcr6\|MDR15 | chemokine (C-X-C motif) receptor 5 | protein-coding |
| chr9_123811845_123812296 | chr9 | 123811846 | 123812296 | + | -0.734939 | TTS (NM_030712) | TTS (NM_030712) | 5594 | NM_030712 | 80901 | Mm.124289 | NM_030712 | ENSMUSG00000048521 | **Cxcr6** | BB217514\|BONZO\|STRL33 | chemokine (C-X-C motif) receptor 6 | protein-coding |
| chr9_123814143_123815209 | chr9 | 123814144 | 123815209 | + | -0.757492 | intron (NM_148925, intron 15 of 18) | intron (NM_148925, intron 15 of 18) | 8199 | NM_030712 | 80901 | Mm.124289 | NM_030712 | ENSMUSG00000048521 | **Cxcr6** | BB217514\|BONZO\|STRL33 | chemokine (C-X-C motif) receptor 6 | protein-coding |
| chr9_123814169_123814931 | chr9 | 123814170 | 123814931 | + | -0.85572 | intron (NM_148925, intron 15 of 18) | intron (NM_148925, intron 15 of 18) | 8073 | NM_030712 | 80901 | Mm.124289 | NM_030712 | ENSMUSG00000048521 | **Cxcr6** | BB217514\|BONZO\|STRL33 | chemokine (C-X-C motif) receptor 6 | protein-coding |
| chr8_107165751_107166038 | chr8 | 107165752 | 107166038 | + | -0.755706 | intron (NM_025558, intron 1 of 4) | intron (NM_025558, intron 1 of 4) | 15234 | NM_025558 | 66427 | Mm.20242 | NM_025558 | ENSMUSG00000031924 | **Cyb5b** | 1810044O22Rik\|AU015618\|Cyb5m | cytochrome b5 type B | protein-coding |
| chr8_107164006_107164779 | chr8 | 107164007 | 107164779 | + | -0.799265 | intron (NM_025558, intron 1 of 4) | intron (NM_025558, intron 1 of 4) | 13732 | NM_025558 | 66427 | Mm.20242 | NM_025558 | ENSMUSG00000031924 | **Cyb5b** | 1810044O22Rik\|AU015618\|Cyb5m | cytochrome b5 type B | protein-coding |
| chr8_107164308_107164977 | chr8 | 107164309 | 107164977 | + | -0.936296 | intron (NM_025558, intron 1 of 4) | intron (NM_025558, intron 1 of 4) | 13982 | NM_025558 | 66427 | Mm.20242 | NM_025558 | ENSMUSG00000031924 | **Cyb5b** | 1810044O22Rik\|AU015618\|Cyb5m | cytochrome b5 type B | protein-coding |
| chr8_107164394_107164829 | chr8 | 107164395 | 107164829 | + | -1.02842 | intron (NM_025558, intron 1 of 4) | intron (NM_025558, intron 1 of 4) | 13951 | NM_025558 | 66427 | Mm.20242 | NM_025558 | ENSMUSG00000031924 | **Cyb5b** | 1810044O22Rik\|AU015618\|Cyb5m | cytochrome b5 type B | protein-coding |
| chr7_107687399_107687939 | chr7 | 107687400 | 107687939 | + | -0.621331 | intron (NM_008905, intron 5 of 24) | intron (NM_008905, intron 5 of 24) | 70363 | NM_001205227 | 320635 | Mm.132828 | NM_177216 | ENSMUSG00000048065 | **Cyb5r2** | D630003K02Rik\|b5R.2 | cytochrome b5 reductase 2 | protein-coding |
| chr7_55889975_55890460 | chr7 | 55889976 | 55890460 | + | -0.666376 | intron (NM_001164661, intron 12 of 31) | intron (NM_001164661, intron 12 of 31) | 48147 | NM_001164661 | 20430 | Mm.37249 | NM_011370 | ENSMUSG00000030447 | **Cyfip1** | E030028J09Rik\|P140SRA-1\|P140sra1\|Shyc\|Sra-1\|Sra1\|l(7)1Rl\|l71Rl\|l7Rl1\|mKIAA0068\|pl-1 | cytoplasmic FMR1 interacting protein 1 | protein-coding |
| chr7_55889839_55890342 | chr7 | 55889840 | 55890342 | + | -0.695651 | intron (NM_001164661, intron 12 of 31) | intron (NM_001164661, intron 12 of 31) | 48020 | NM_001164661 | 20430 | Mm.37249 | NM_011370 | ENSMUSG00000030447 | **Cyfip1** | E030028J09Rik\|P140SRA-1\|P140sra1\|Shyc\|Sra-1\|Sra1\|l(7)1Rl\|l71Rl\|l7Rl1\|mKIAA0068\|pl-1 | cytoplasmic FMR1 interacting protein 1 | protein-coding |
| chr11_46275309_46275683 | chr11 | 46275310 | 46275683 | + | -0.701296 | intron (NM_001252460, intron 8 of 30) | intron (NM_001252460, intron 8 of 30) | 36862 | NM_001252460 | 76884 | Mm.154358 | NM_133769 | ENSMUSG00000020340 | **Cyfip2** | 1500004I01Rik\|6430511D02Rik\|AA930218\|AU022376\|Pir121\|mKIAA1168 | cytoplasmic FMR1 interacting protein 2 | protein-coding |
| chr11_46298902_46299427 | chr11 | 46298903 | 46299427 | + | -0.929271 | intron (NM_001252459, intron 1 of 30) | intron (NM_001252459, intron 1 of 30) | 13193 | NM_001252460 | 76884 | Mm.154358 | NM_133769 | ENSMUSG00000020340 | **Cyfip2** | 1500004I01Rik\|6430511D02Rik\|AA930218\|AU022376\|Pir121\|mKIAA1168 | cytoplasmic FMR1 interacting protein 2 | protein-coding |

| | | | | | | | | | | | | | | | | | |
|---|---|---|---|---|---|---|---|---|---|---|---|---|---|---|---|---|---|
| chr11_46298907_46299229 | chr11 | 46298908 | 46299229 | + | -1.05217 | intron (NM_001252459, intron 1 of 30) | intron (NM_001252459, intron 1 of 30) | 13290 | NM_001252460 | 76884 | Mm.154358 | NM_133769 | ENSMUSG00000020340 | **Cyfip2** | 1500004I01Rik\|6430511D02Rik\|AA930218\|AU022376\|Pir121\|mKIAA1168 | cytoplasmic FMR1 interacting protein 2 | protein-coding |
| chr11_46343728_46344710 | chr11 | 46343729 | 46344710 | + | -1.3416 | TTS (NM_001281967) | TTS (NM_001281967) | -31360 | NM_001252459 | 76884 | Mm.154358 | NM_133769 | ENSMUSG00000020340 | **Cyfip2** | 1500004I01Rik\|6430511D02Rik\|AA930218\|AU022376\|Pir121\|mKIAA1168 | cytoplasmic FMR1 interacting protein 2 | protein-coding |
| chr3_107206069_107206297 | chr3 | 107206070 | 107206297 | + | -0.619001 | Intergenic | Intergenic | 15549 | NM_001111143 | 229697 | Mm.297051 | NM_001111143 | ENSMUSG00000046213 | **Cym** | Gm131 | chymosin | protein-coding |
| chr19_46653477_46654155 | chr19 | 46653478 | 46654155 | + | -0.815591 | intron (NM_146099, intron 3 of 3) | intron (NM_146099, intron 3 of 3) | 19184 | NM_007809 | 13074 | Mm.1262 | NM_007809 | ENSMUSG00000003555 | **Cyp17a1** | Cyp17\|p450c17 | cytochrome P450, family 17, subfamily a, polypeptide 1 | protein-coding |
| chr9_57667187_57667609 | chr9 | 57667188 | 57667609 | + | -0.641037 | Intergenic | Intergenic | 16257 | NM_009993 | 13077 | Mm.15537 | NM_009993 | ENSMUSG00000032310 | **Cyp1a2** | CP12\|Cyp1a1\|P450-3 | cytochrome P450, family 1, subfamily a, polypeptide 2 | protein-coding |
| chr17_79787121_79787596 | chr17 | 79787122 | 79787596 | + | -0.677754 | Intergenic | Intergenic | -72318 | NM_009994 | 13078 | Mm.214016 | NM_009994 | ENSMUSG00000024087 | **Cyp1b1** | CP1B\|P4501b1 | cytochrome P450, family 1, subfamily b, polypeptide 1 | protein-coding |
| chr17_79787230_79787550 | chr17 | 79787231 | 79787550 | + | -0.753093 | Intergenic | Intergenic | -72349 | NM_009994 | 13078 | Mm.214016 | NM_009994 | ENSMUSG00000024087 | **Cyp1b1** | CP1B\|P4501b1 | cytochrome P450, family 1, subfamily b, polypeptide 1 | protein-coding |
| chr17_79775040_79775536 | chr17 | 79775041 | 79775536 | + | -0.758663 | Intergenic | Intergenic | -60247 | NM_009994 | 13078 | Mm.214016 | NM_009994 | ENSMUSG00000024087 | **Cyp1b1** | CP1B\|P4501b1 | cytochrome P450, family 1, subfamily b, polypeptide 1 | protein-coding |
| chr1_60355829_60356471 | chr1 | 60355830 | 60356471 | + | -0.695926 | intron (NM_030013, intron 4 of 12) | intron (NM_030013, intron 4 of 12) | 12778 | NM_030013 | 77951 | Mm.197640 | NM_030013 | | **Cyp20a1** | A930011N14Rik\|Cypm | cytochrome P450, family 20, subfamily a, polypeptide 1 | protein-coding |
| chr1_60352846_60353235 | chr1 | 60352847 | 60353235 | + | -0.751829 | exon (NM_030013, exon 3 of 13) | exon (NM_030013, exon 3 of 13) | 9669 | NM_030013 | 77951 | Mm.197640 | NM_030013 | | **Cyp20a1** | A930011N14Rik\|Cypm | cytochrome P450, family 20, subfamily a, polypeptide 1 | protein-coding |
| chr6_84616950_84617603 | chr6 | 84616951 | 84617603 | + | -0.683665 | Intergenic | Intergenic | -23369 | NM_001177713 | 232174 | Mm.255246 | NM_175475 | ENSMUSG00000063415 | **Cyp26b1** | CP26\|P450RAI-2 | cytochrome P450, family 26, subfamily b, polypeptide 1 | protein-coding |
| chr15_82417421_82417899 | chr15 | 82417422 | 82417899 | + | -0.70277 | Intergenic | Intergenic | -10466 | NM_010005 | 13101 | Mm.174372 | NM_010005 | ENSMUSG00000094806 | **Cyp2d10** | AI303445\|Cyp2d\|P450-2D | cytochrome P450, family 2, subfamily d, polypeptide 10 | protein-coding |
| chr7_114494055_114494463 | chr7 | 114494056 | 114494463 | + | -1.10674 | intron (NM_011055, intron 2 of 15) | intron (NM_011055, intron 2 of 15) | 68713 | NR_110407 | 244209 | Mm.108037 | NM_177382 | ENSMUSG00000030670 | **Cyp2r1** | - | cytochrome P450, family 2, subfamily r, polypeptide 1 | protein-coding |
| chr7_25826004_25826369 | chr7 | 25826005 | 25826369 | + | -0.642435 | Intergenic | Intergenic | -9657 | NM_028775 | 74134 | Mm.275188 | NM_028775 | ENSMUSG00000040703 | **Cyp2s1** | 1200011C15Rik\|AU041727\|C79779 | cytochrome P450, family 2, subfamily s, polypeptide 1 | protein-coding |
| chr7_25826893_25827530 | chr7 | 25826894 | 25827530 | + | -0.672283 | Intergenic | (CA)n\|Simple_repeat\|Simple_repeat | -10682 | NM_028775 | 74134 | Mm.275188 | NM_028775 | ENSMUSG00000040703 | **Cyp2s1** | 1200011C15Rik\|AU041727\|C79779 | cytochrome P450, family 2, subfamily s, polypeptide 1 | protein-coding |
| chr7_25835053_25835962 | chr7 | 25835054 | 25835962 | + | -0.741584 | Intergenic | MER21B\|LTR\|ERVL | -18978 | NM_028775 | 74134 | Mm.275188 | NM_028775 | ENSMUSG00000040703 | **Cyp2s1** | 1200011C15Rik\|AU041727\|C79779 | cytochrome P450, family 2, subfamily s, polypeptide 1 | protein-coding |
| chr7_25827233_25827586 | chr7 | 25827234 | 25827586 | + | -0.74405 | Intergenic | Intergenic | -10880 | NM_028775 | 74134 | Mm.275188 | NM_028775 | ENSMUSG00000040703 | **Cyp2s1** | 1200011C15Rik\|AU041727\|C79779 | cytochrome P450, family 2, subfamily s, polypeptide 1 | protein-coding |
| chr7_25835417_25835912 | chr7 | 25835418 | 25835912 | + | -0.793272 | Intergenic | Intergenic | -19135 | NM_028775 | 74134 | Mm.275188 | NM_028775 | ENSMUSG00000040703 | **Cyp2s1** | 1200011C15Rik\|AU041727\|C79779 | cytochrome P450, family 2, subfamily s, polypeptide 1 | protein-coding |
| chr17_32944459_32944681 | chr17 | 32944460 | 32944681 | + | -0.641602 | intron (NM_130882, intron 2 of 12) | B3A\|SINE\|B2 | 2791 | NM_130882 | 170716 | Mm.254838 | NM_130882 | ENSMUSG00000024055 | **Cyp4f13** | 0610030I10Rik\|Cypf13 | cytochrome P450, family 4, subfamily f, polypeptide 13 | protein-coding |
| chr14_73045310_73045528 | chr14 | 73045311 | 73045528 | + | -0.601969 | intron (NM_001162412, intron 2 of 4) | intron (NM_001162412, intron 2 of 4) | 3695 | NM_133720 | 70086 | Mm.158324 | NM_133720 | ENSMUSG00000033470 | **Cysltr2** | 2300001H05Rik\|CYSLT2R\|Cltr2\|Cyslt2 | cysteinyl leukotriene receptor 2 | protein-coding |
| chr14_73045139_73045546 | chr14 | 73045140 | 73045546 | + | -0.640967 | intron (NM_001162412, intron 2 of 4) | intron (NM_001162412, intron 2 of 4) | 3771 | NM_133720 | 70086 | Mm.158324 | NM_133720 | ENSMUSG00000033470 | **Cysltr2** | 2300001H05Rik\|CYSLT2R\|Cltr2\|Cyslt2 | cysteinyl leukotriene receptor 2 | protein-coding |
| chr11_118245151_118245434 | chr11 | 118245152 | 118245434 | + | -0.679005 | intron (NM_001112699, intron 1 of 12) | intron (NM_001112699, intron 1 of 12) | 3299 | NM_001112699 | 19157 | Mm.86413 | NM_011180 | ENSMUSG00000017132 | **Cyth1** | CLM1\|CTH-1\|CYTIP\|Pscd1\|mKIAA4240 | cytohesin 1 | protein-coding |
| chr11_118209532_118209889 | chr11 | 118209533 | 118209889 | + | -0.731236 | intron (NM_001112699, intron 1 of 12) | intron (NM_001112699, intron 1 of 12) | 3031 | NM_001112700 | 19157 | Mm.86413 | NM_011180 | ENSMUSG00000017132 | **Cyth1** | CLM1\|CTH-1\|CYTIP\|Pscd1\|mKIAA4240 | cytohesin 1 | protein-coding |
| chr11_118260001_118260731 | chr11 | 118260002 | 118260731 | + | -0.796435 | 3' UTR (NM_001033528, exon 21 of 21) | 3' UTR (NM_001033528, exon 21 of 21) | -11774 | NM_011180 | 19157 | Mm.86413 | NM_011180 | ENSMUSG00000017132 | **Cyth1** | CLM1\|CTH-1\|CYTIP\|Pscd1\|mKIAA4240 | cytohesin 1 | protein-coding |
| chr11_118266289_118266969 | chr11 | 118266290 | 118266969 | + | -0.833762 | intron (NM_001033528, intron 15 of 20) | B1_Mur3\|SINE\|Alu | -18037 | NM_011180 | 19157 | Mm.86413 | NM_011180 | ENSMUSG00000017132 | **Cyth1** | CLM1\|CTH-1\|CYTIP\|Pscd1\|mKIAA4240 | cytohesin 1 | protein-coding |
| chr11_118266666_118266973 | chr11 | 118266667 | 118266973 | + | -0.947558 | intron (NM_001033528, intron 15 of 20) | B3\|SINE\|B2 | -18228 | NM_011180 | 19157 | Mm.86413 | NM_011180 | ENSMUSG00000017132 | **Cyth1** | CLM1\|CTH-1\|CYTIP\|Pscd1\|mKIAA4240 | cytohesin 1 | protein-coding |
| chr11_118206460_118206656 | chr11 | 118206461 | 118206656 | + | -1.05726 | intron (NM_001112699, intron 1 of 12) | intron (NM_001112699, intron 1 of 12) | 6184 | NM_001112700 | 19157 | Mm.86413 | NM_011180 | ENSMUSG00000017132 | **Cyth1** | CLM1\|CTH-1\|CYTIP\|Pscd1\|mKIAA4240 | cytohesin 1 | protein-coding |
| chr15_78594782_78595208 | chr15 | 78594783 | 78595208 | + | -0.789056 | Intergenic | MTE2b\|LTR\|ERVL-MaLR | -2052 | NM_028195 | 72318 | Mm.32911 | NM_028195 | ENSMUSG00000018008 | **Cyth4** | 2510004M07Rik\|5830469K17Rik\|AI467541\|Pscd4\|mFLJ00017 | cytohesin 4 | protein-coding |
| chr2_58155643_58155927 | chr2 | 58155644 | 58155927 | + | -0.683477 | intron (NM_139200, intron 1 of 7) | intron (NM_139200, intron 1 of 7) | 4337 | NM_139200 | 227929 | Mm.273905 | NM_139200 | ENSMUSG00000026832 | **Cytip** | A130053M09Rik\|AI462064\|C80816\|Cbp\|Cybr\|Pscdbp | cytohesin 1 interacting protein | protein-coding |
| chr2_58137612_58138066 | chr2 | 58137613 | 58138066 | + | -0.692485 | intron (NM_139200, intron 7 of 7) | intron (NM_139200, intron 7 of 7) | 22283 | NM_139200 | 227929 | Mm.273905 | NM_139200 | ENSMUSG00000026832 | **Cytip** | A130053M09Rik\|AI462064\|C80816\|Cbp\|Cybr\|Pscdbp | cytohesin 1 interacting protein | protein-coding |
| chr2_58123661_58124061 | chr2 | 58123662 | 58124061 | + | -0.77081 | Intergenic | L1MCa\|LINE\|L1 | 36261 | NM_139200 | 227929 | Mm.273905 | NM_139200 | ENSMUSG00000026832 | **Cytip** | A130053M09Rik\|AI462064\|C80816\|Cbp\|Cybr\|Pscdbp | cytohesin 1 interacting protein | protein-coding |
| chr2_58150073_58150414 | chr2 | 58150074 | 58150414 | + | -0.85818 | intron (NM_139200, intron 3 of 7) | intron (NM_139200, intron 3 of 7) | 9878 | NM_139200 | 227929 | Mm.273905 | NM_139200 | ENSMUSG00000026832 | **Cytip** | A130053M09Rik\|AI462064\|C80816\|Cbp\|Cybr\|Pscdbp | cytohesin 1 interacting protein | protein-coding |
| chr15_61428691_61429173 | chr15 | 61428692 | 61429173 | + | -0.694758 | Intergenic | Intergenic | 345519 | NR_040350 | 432956 | Mm.404918 | NR_040350 | | **D030024E09Rik** | - | RIKEN cDNA D030024E09 gene | ncRNA |
| chr15_61582026_61582676 | chr15 | 61582027 | 61582676 | + | -1.01706 | Intergenic | Intergenic | 192100 | NR_040350 | 432956 | Mm.404918 | NR_040350 | | **D030024E09Rik** | - | RIKEN cDNA D030024E09 gene | ncRNA |
| chr15_61670369_61670978 | chr15 | 61670370 | 61670978 | + | -1.06617 | Intergenic | Intergenic | 103777 | NR_040350 | 432956 | Mm.404918 | NR_040350 | | **D030024E09Rik** | - | RIKEN cDNA D030024E09 gene | ncRNA |
| chr15_61422966_61423495 | chr15 | 61422967 | 61423495 | + | -1.07987 | Intergenic | Intergenic | 351220 | NR_040350 | 432956 | Mm.404918 | NR_040350 | | **D030024E09Rik** | - | RIKEN cDNA D030024E09 gene | ncRNA |
| chr15_61481915_61482302 | chr15 | 61481916 | 61482302 | + | -1.15003 | Intergenic | Intergenic | 292342 | NR_040350 | 432956 | Mm.404918 | NR_040350 | | **D030024E09Rik** | - | RIKEN cDNA D030024E09 gene | ncRNA |
| chr15_61481933_61482205 | chr15 | 61481934 | 61482205 | + | -1.16488 | Intergenic | Intergenic | 292382 | NR_040350 | 432956 | Mm.404918 | NR_040350 | | **D030024E09Rik** | - | RIKEN cDNA D030024E09 gene | ncRNA |
| chr15_61481772_61482300 | chr15 | 61481773 | 61482300 | + | -1.18161 | Intergenic | Intergenic | 292415 | NR_040350 | 432956 | Mm.404918 | NR_040350 | | **D030024E09Rik** | - | RIKEN cDNA D030024E09 gene | ncRNA |
| chr6_127336594_127337040 | chr6 | 127336595 | 127337040 | + | -0.627987 | Intergenic | Intergenic | -28850 | NR_073373 | 232386 | Mm.257367 | NR_073373 | | **D130058E03** | - | uncharacterized D130058E03 | ncRNA |
| chr15_58418444_58418897 | chr15 | 58418445 | 58418897 | + | -0.714934 | intron (NM_145959, intron 1 of 23) | intron (NM_145959, intron 1 of 23) | 3203 | NM_145959 | 210998 | Mm.3224 | NM_145959 | ENSMUSG00000037119 | **D15Ertd621e** | AV220772\|BC033609\|Fam91a1\|SKIN\|mKIAA0493 | DNA segment, Chr 15, ERATO Doi 621, expressed | protein-coding |
| chr16_78606049_78606719 | chr16 | 78606050 | 78606719 | + | -0.631755 | Intergenic | MTE2a\|LTR\|ERVL-MaLR | -29696 | NM_025967 | 67102 | Mm.37332 | NM_025967 | | **D16Ertd472e** | 1700010I10Rik\|2310009O17Rik\|E330003K22Rik\|Eurl | DNA segment, Chr 16, ERATO Doi 472, expressed | protein-coding |
| chr16_78606225_78606796 | chr16 | 78606226 | 78606796 | + | -0.664832 | Intergenic | MTE2a\|LTR\|ERVL-MaLR | -29823 | NM_025967 | 67102 | Mm.37332 | NM_025967 | | **D16Ertd472e** | 1700010I10Rik\|2310009O17Rik\|E330003K22Rik\|Eurl | DNA segment, Chr 16, ERATO Doi 472, expressed | protein-coding |
| chr16_78606219_78606631 | chr16 | 78606220 | 78606631 | + | -0.692752 | Intergenic | MTE2a\|LTR\|ERVL-MaLR | -29737 | NM_025967 | 67102 | Mm.37332 | NM_025967 | | **D16Ertd472e** | 1700010I10Rik\|2310009O17Rik\|E330003K22Rik\|Eurl | DNA segment, Chr 16, ERATO Doi 472, expressed | protein-coding |
| chr16_78595649_78596173 | chr16 | 78595650 | 78596173 | + | -1.04701 | Intergenic | RLTR9D\|LTR\|ERVK | -19223 | NM_025967 | 67102 | Mm.37332 | NM_025967 | | **D16Ertd472e** | 1700010I10Rik\|2310009O17Rik\|E330003K22Rik\|Eurl | DNA segment, Chr 16, ERATO Doi 472, expressed | protein-coding |
| chr17_27792497_27792914 | chr17 | 27792498 | 27792914 | + | -0.906511 | intron (NM_001033279, intron 2 of 4) | intron (NM_001033279, intron 2 of 4) | 27836 | NM_001033279 | 224647 | Mm.196577 | NM_001033279 | ENSMUSG00000056692 | **D17Wsu92e** | AU020189\|C030010A15\|C80239 | DNA segment, Chr 17, Wayne State University 92, expressed | protein-coding |
| chr1_97580249_97580761 | chr1 | 97580250 | 97580761 | + | -0.634371 | Intergenic | L1MC1\|LINE\|L1 | 81513 | NM_133825 | 52392 | Mm.12309 | NM_133825 | ENSMUSG00000044768 | **D1Ertd622e** | AI987691\|AW047450 | DNA segment, Chr 1, ERATO Doi 622, expressed | protein-coding |
| chr2_104385972_104386307 | chr2 | 104385973 | 104386307 | + | -0.981771 | intron (NM_001033347, intron 1 of 20) | intron (NM_001033347, intron 1 of 20) | 24194 | NM_001033347 | 241589 | Mm.358909 | NM_001033347 | ENSMUSG00000068373 | **D430041D05Rik** | 9630056G07Rik\|G2 | RIKEN cDNA D430041D05 gene | protein-coding |
| chr15_94254977_94255528 | chr15 | 94254978 | 94255528 | + | -0.826756 | intron (NR_045629, intron 3 of 3) | intron (NR_045629, intron 3 of 3) | 8051 | NR_045629 | 328594 | Mm.135523 | NR_045629 | | **D630010B17Rik** | Gm811 | RIKEN cDNA D630010B17 gene | ncRNA |

| | | | | | | | | | | | | | | | | | |
|---|---|---|---|---|---|---|---|---|---|---|---|---|---|---|---|---|---|
| chr15_94276426_94276935 | chr15 | 94276427 | 94276935 | + | -1.12198 | intron (NM_177431, intron 37 of 38) | intron (NM_177431, intron 37 of 38) | 29479 | NR_045629 | 328594 | Mm.135523 | NR_045629 | | **D630010B17Rik** | Gm811 | RIKEN cDNA D630010B17 gene | ncRNA |
| chr4_57896728_57897194 | chr4 | 57896729 | 57897194 | + | -0.834923 | TTS (NM_009649) | TTS (NM_009649) | 19303 | NM_178727 | 242484 | Mm.23214 | NM_178727 | ENSMUSG00000052117 | **D630039A03Rik** | - | RIKEN cDNA D630039A03 gene | protein-coding |
| chr4_57896738_57897000 | chr4 | 57896739 | 57897000 | + | -0.948479 | 3' UTR (NM_009649, exon 3 of 3) | 3' UTR (NM_009649, exon 3 of 3) | 19395 | NM_178727 | 242484 | Mm.23214 | NM_178727 | ENSMUSG00000052117 | **D630039A03Rik** | - | RIKEN cDNA D630039A03 gene | protein-coding |
| chr12_73383534_73384237 | chr12 | 73383535 | 73384237 | + | -0.77892 | intron (NR_046013, intron 2 of 3) | MLT1B\|LTR\|ERVL-MaLR | 25677 | NR_046013 | 380765 | Mm.490476 | NR_046013 | | **D830013O20Rik** | Gm891 | RIKEN cDNA D830013O20 gene | ncRNA |
| chr12_73383939_73384250 | chr12 | 73383940 | 73384250 | + | -0.882445 | intron (NR_046013, intron 2 of 3) | MLT1B\|LTR\|ERVL-MaLR | 25468 | NR_046013 | 380765 | Mm.490476 | NR_046013 | | **D830013O20Rik** | Gm891 | RIKEN cDNA D830013O20 gene | ncRNA |
| chr12_73382214_73383168 | chr12 | 73382215 | 73383168 | + | -1.04717 | intron (NR_046013, intron 2 of 3) | intron (NR_046013, intron 2 of 3) | 26872 | NR_046013 | 380765 | Mm.490476 | NR_046013 | | **D830013O20Rik** | Gm891 | RIKEN cDNA D830013O20 gene | ncRNA |
| chr4_123415762_123416221 | chr4 | 123415763 | 123416221 | + | -0.753663 | intron (NM_001199136, intron 57 of 97) | intron (NM_001199136, intron 57 of 97) | -4081 | NM_001167918 | 442834 | Mm.483803 | NM_001167918 | | **D830031N03Rik** | Kiaa0754\|mKIAA0754 | RIKEN cDNA D830031N03 gene | protein-coding |
| chr8_36122342_36123486 | chr8 | 36122343 | 36123486 | + | -0.648476 | intron (NM_172911, intron 3 of 5) | intron (NM_172911, intron 3 of 5) | 28086 | NM_172911 | 244418 | Mm.244971 | NM_172911 | ENSMUSG00000050271 | **D8Ertd82e** | 9830148H23\|Sgk223\|mFLJ00269 | DNA segment, Chr 8, ERATO Doi 82, expressed | protein-coding |
| chr8_36158036_36158364 | chr8 | 36158037 | 36158364 | + | -0.73119 | Intergenic | Intergenic | 63372 | NM_172911 | 244418 | Mm.244971 | NM_172911 | ENSMUSG00000050271 | **D8Ertd82e** | 9830148H23\|Sgk223\|mFLJ00269 | DNA segment, Chr 8, ERATO Doi 82, expressed | protein-coding |
| chr8_36041318_36041961 | chr8 | 36041319 | 36041961 | + | -0.849042 | Intergenic | Intergenic | -53188 | NM_172911 | 244418 | Mm.244971 | NM_172911 | ENSMUSG00000050271 | **D8Ertd82e** | 9830148H23\|Sgk223\|mFLJ00269 | DNA segment, Chr 8, ERATO Doi 82, expressed | protein-coding |
| chr8_36010474_36011147 | chr8 | 36010475 | 36011147 | + | -0.87631 | Intergenic | Intergenic | -84017 | NM_172911 | 244418 | Mm.244971 | NM_172911 | ENSMUSG00000050271 | **D8Ertd82e** | 9830148H23\|Sgk223\|mFLJ00269 | DNA segment, Chr 8, ERATO Doi 82, expressed | protein-coding |
| chr8_36010474_36010835 | chr8 | 36010475 | 36010835 | + | -0.898011 | Intergenic | Intergenic | -84173 | NM_172911 | 244418 | Mm.244971 | NM_172911 | ENSMUSG00000050271 | **D8Ertd82e** | 9830148H23\|Sgk223\|mFLJ00269 | DNA segment, Chr 8, ERATO Doi 82, expressed | protein-coding |
| chr8_36168540_36169041 | chr8 | 36168541 | 36169041 | + | -1.02282 | Intergenic | Intergenic | 73963 | NM_172911 | 244418 | Mm.244971 | NM_172911 | ENSMUSG00000050271 | **D8Ertd82e** | 9830148H23\|Sgk223\|mFLJ00269 | DNA segment, Chr 8, ERATO Doi 82, expressed | protein-coding |
| chr8_36168609_36168953 | chr8 | 36168610 | 36168953 | + | -1.03354 | Intergenic | Intergenic | 73953 | NM_172911 | 244418 | Mm.244971 | NM_172911 | ENSMUSG00000050271 | **D8Ertd82e** | 9830148H23\|Sgk223\|mFLJ00269 | DNA segment, Chr 8, ERATO Doi 82, expressed | protein-coding |
| chr3_84045979_84046746 | chr3 | 84045980 | 84046746 | + | -0.615516 | Intergenic | MTEa\|LTR\|ERVL-MaLR | -6202 | NM_172681 | 229473 | Mm.28838 | NM_172681 | ENSMUSG00000033767 | **D930015E06Rik** | Kiaa0922\|mKIAA0922 | RIKEN cDNA D930015E06 gene | protein-coding |
| chr3_84065749_84066298 | chr3 | 84065750 | 84066298 | + | -0.729934 | Intergenic | Intergenic | -25863 | NM_172681 | 229473 | Mm.28838 | NM_172681 | ENSMUSG00000033767 | **D930015E06Rik** | Kiaa0922\|mKIAA0922 | RIKEN cDNA D930015E06 gene | protein-coding |
| chr7_25155596_25156431 | chr7 | 25155597 | 25156431 | + | -0.725804 | intron (NR_045847, intron 2 of 2) | intron (NR_045847, intron 2 of 2) | 945 | NR_045848 | 434147 | Mm.59171 | NM_001033781 | ENSMUSG00000074274 | **D930028M14Rik** | EG434147 | RIKEN cDNA D930028M14 gene | ncRNA |
| chr19_28612256_28612846 | chr19 | 28612257 | 28612846 | + | -0.818976 | intron (NM_175459, intron 2 of 10) | intron (NM_175459, intron 2 of 10) | -65673 | NR_045330 | 320176 | Mm.406716 | NR_045330 | ENSMUSG00000097735 | **D930032P07Rik** | - | RIKEN cDNA D930032P07 gene | ncRNA |
| chr11_51644301_51644684 | chr11 | 51644302 | 51644684 | + | -0.656182 | exon (NM_145974, exon 4 of 5) | exon (NM_145974, exon 4 of 5) | -6461 | NR_027958 | 97775 | Mm.220299 | NM_175289 | ENSMUSG00000052563 | **D930048N14Rik** | AI315703\|R75254 | RIKEN cDNA D930048N14 gene | ncRNA |
| chr12_71932742_71933359 | chr12 | 71932743 | 71933359 | + | -0.937168 | intron (NM_001286452, intron 3 of 24) | intron (NM_001286452, intron 3 of 24) | 43508 | NM_001286452 | 208846 | Mm.87417 | NM_026102 | ENSMUSG00000034574 | **Daam1** | - | dishevelled associated activator of morphogenesis 1 | protein-coding |
| chr7_16873958_16874168 | chr7 | 16873959 | 16874168 | + | -1.0869 | Intergenic | Intergenic | -1254 | NM_001081655 | 629378 | Mm.491123 | NM_001081655 | ENSMUSG00000078794 | **Dact3** | EG666671\|naa-3 | dapper homolog 3, antagonist of beta-catenin (xenopus) | protein-coding |
| chr14_54110981_54111444 | chr14 | 54110982 | 54111444 | + | -0.625142 | Intergenic | Intergenic | 142716 | NM_001113358 | 13135 | Mm.319038 | NM_010015 | ENSMUSG00000022174 | **Dad1** | AI323713 | defender against cell death 1 | protein-coding |
| chr14_54137085_54137502 | chr14 | 54137086 | 54137502 | + | -0.694587 | Intergenic | Intergenic | 116635 | NM_001113358 | 13135 | Mm.319038 | NM_010015 | ENSMUSG00000022174 | **Dad1** | AI323713 | defender against cell death 1 | protein-coding |
| chr14_54111108_54111451 | chr14 | 54111109 | 54111451 | + | -0.698454 | Intergenic | Intergenic | 142649 | NM_001113358 | 13135 | Mm.319038 | NM_010015 | ENSMUSG00000022174 | **Dad1** | AI323713 | defender against cell death 1 | protein-coding |
| chr14_54181278_54181659 | chr14 | 54181279 | 54181659 | + | -0.767855 | Intergenic | Intergenic | 72460 | NM_001113358 | 13135 | Mm.319038 | NM_010015 | ENSMUSG00000022174 | **Dad1** | AI323713 | defender against cell death 1 | protein-coding |
| chr14_54231123_54231478 | chr14 | 54231124 | 54231478 | + | -0.793524 | Intergenic | RSINE1\|SINE\|B4 | 22628 | NM_001113358 | 13135 | Mm.319038 | NM_010015 | ENSMUSG00000022174 | **Dad1** | AI323713 | defender against cell death 1 | protein-coding |
| chr14_53961519_53961771 | chr14 | 53961520 | 53961771 | + | -0.8325 | Intergenic | Intergenic | 292284 | NM_001113358 | 13135 | Mm.319038 | NM_010015 | ENSMUSG00000022174 | **Dad1** | AI323713 | defender against cell death 1 | protein-coding |
| chr14_54042077_54042884 | chr14 | 54042078 | 54042884 | + | -1.05165 | Intergenic | RMER20C_Mm\|LTR\|ERVK | 211448 | NM_001113358 | 13135 | Mm.319038 | NM_010015 | ENSMUSG00000022174 | **Dad1** | AI323713 | defender against cell death 1 | protein-coding |
| chr14_54052995_54053305 | chr14 | 54052996 | 54053305 | + | -1.06578 | Intergenic | MLTR73\|LTR\|ERV1 | 200779 | NM_001113358 | 13135 | Mm.319038 | NM_010015 | ENSMUSG00000022174 | **Dad1** | AI323713 | defender against cell death 1 | protein-coding |
| chr14_54042144_54042564 | chr14 | 54042145 | 54042564 | + | -1.09869 | Intergenic | Intergenic | 211575 | NM_001113358 | 13135 | Mm.319038 | NM_010015 | ENSMUSG00000022174 | **Dad1** | AI323713 | defender against cell death 1 | protein-coding |
| chr14_53991234_53991771 | chr14 | 53991235 | 53991771 | + | -1.30908 | Intergenic | Intergenic | 262426 | NM_001113358 | 13135 | Mm.319038 | NM_010015 | ENSMUSG00000022174 | **Dad1** | AI323713 | defender against cell death 1 | protein-coding |
| chr15_31166049_31167389 | chr15 | 31166050 | 31167389 | + | -0.760221 | Intergenic | Intergenic | -57666 | NM_146057 | 223453 | Mm.222867 | NM_146057 | ENSMUSG00000039168 | **Dap** | 4921531N22Rik\|AI789753\|AU040360 | death-associated protein | protein-coding |
| chr15_31225609_31225873 | chr15 | 31225610 | 31225873 | + | -0.81632 | intron (NM_146057, intron 1 of 3) | Lx7\|LINE\|L1 | 1356 | NM_146057 | 223453 | Mm.222867 | NM_146057 | ENSMUSG00000039168 | **Dap** | 4921531N22Rik\|AI789753\|AU040360 | death-associated protein | protein-coding |
| chr2_59527695_59528048 | chr2 | 59527696 | 59528048 | + | -0.772877 | Intergenic | Intergenic | 43219 | NM_029723 | 76747 | Mm.275769 | NM_029723 | ENSMUSG00000026989 | **Dapl1** | 2310032F03Rik\|EEDA | death associated protein-like 1 | protein-coding |
| chr2_59467981_59468412 | chr2 | 59467982 | 59468412 | + | -0.810182 | Intergenic | Intergenic | -16456 | NM_029723 | 76747 | Mm.275769 | NM_029723 | ENSMUSG00000026989 | **Dapl1** | 2310032F03Rik\|EEDA | death associated protein-like 1 | protein-coding |
| chr2_59469686_59470343 | chr2 | 59469687 | 59470343 | + | -0.993018 | Intergenic | Intergenic | -14638 | NM_029723 | 76747 | Mm.275769 | NM_029723 | ENSMUSG00000026989 | **Dapl1** | 2310032F03Rik\|EEDA | death associated protein-like 1 | protein-coding |
| chr10_80271172_80271813 | chr10 | 80271173 | 80271813 | + | -0.659361 | intron (NM_001122604, intron 1 of 11) | intron (NM_001122604, intron 1 of 11) | 6502 | NM_133188 | 70248 | Mm.148693 | NM_133188 | ENSMUSG00000069565 | **Dazap1** | 2410042M16Rik\|mPrrp | DAZ associated protein 1 | protein-coding |
| chr17_50331826_50332104 | chr17 | 50331827 | 50332104 | + | -0.643971 | Intergenic | ORR1E\|LTR\|ERVL-MaLR | -38345 | NM_001277863 | 13164 | Mm.280641 | NM_010021 | ENSMUSG00000010592 | **Dazl** | Daz-like\|Dazh\|Dazl1\|Dazla\|Tpx-2\|Tpx2 | deleted in azoospermia-like | protein-coding |
| chr17_50388820_50389208 | chr17 | 50388821 | 50389208 | + | -0.785341 | Intergenic | Intergenic | -95394 | NM_001277863 | 13164 | Mm.280641 | NM_010021 | ENSMUSG00000010592 | **Dazl** | Daz-like\|Dazh\|Dazl1\|Dazla\|Tpx-2\|Tpx2 | deleted in azoospermia-like | protein-coding |
| chr17_50388694_50389108 | chr17 | 50388695 | 50389108 | + | -0.882583 | Intergenic | Intergenic | -95281 | NM_001277863 | 13164 | Mm.280641 | NM_010021 | ENSMUSG00000010592 | **Dazl** | Daz-like\|Dazh\|Dazl1\|Dazla\|Tpx-2\|Tpx2 | deleted in azoospermia-like | protein-coding |
| chr11_106026770_106028187 | chr11 | 106026771 | 106028187 | + | -0.89505 | intron (NM_001037712, intron 11 of 12) | intron (NM_001037712, intron 11 of 12) | -9393 | NM_027946 | 71833 | Mm.307455 | NM_027946 | ENSMUSG00000049354 | **Dcaf7** | 1700012F10Rik\|2610037L01Rik\|C86529\|HAN11\|Wdr68 | DDB1 and CUL4 associated factor 7 | protein-coding |
| chr11_106033501_106033721 | chr11 | 106033502 | 106033721 | + | -1.06689 | intron (NM_001037712, intron 12 of 12) | intron (NM_001037712, intron 12 of 12) | -3261 | NM_027946 | 71833 | Mm.307455 | NM_027946 | ENSMUSG00000049354 | **Dcaf7** | 1700012F10Rik\|2610037L01Rik\|C86529\|HAN11\|Wdr68 | DDB1 and CUL4 associated factor 7 | protein-coding |
| chr11_106033340_106033755 | chr11 | 106033341 | 106033755 | + | -1.09712 | intron (NM_001037712, intron 12 of 12) | IAPLTR1_Mm\|LTR\|ERVK | -3324 | NM_027946 | 71833 | Mm.307455 | NM_027946 | ENSMUSG00000049354 | **Dcaf7** | 1700012F10Rik\|2610037L01Rik\|C86529\|HAN11\|Wdr68 | DDB1 and CUL4 associated factor 7 | protein-coding |
| chr5_88766156_88766476 | chr5 | 88766157 | 88766476 | + | -0.749385 | intron (NM_007832, intron 1 of 6) | intron (NM_007832, intron 1 of 6) | 1303 | NM_007832 | 13178 | Mm.298892 | NM_007832 | ENSMUSG00000029366 | **Dck** | - | deoxycytidine kinase | protein-coding |
| chr18_44392317_44392892 | chr18 | 44392318 | 44392892 | + | -0.660436 | intron (NM_027490, intron 1 of 10) | intron (NM_027490, intron 1 of 10) | 12105 | NM_027490 | 70640 | Mm.87629 | NM_027490 | ENSMUSG00000024472 | **Dcp2** | 2410015D23Rik\|5730537H01Rik\|AL118268 | DCP2 decapping enzyme homolog (S. cerevisiae) | protein-coding |
| chr8_34045253_34046028 | chr8 | 34045254 | 34046028 | + | -0.651399 | Intergenic | ORR1A2\|LTR\|ERVL-MaLR | 62173 | NM_001293758 | 22428 | Mm.150373 | NM_011722 | ENSMUSG00000031516 | **Dctn6** | AU044699\|WS-3\|p27 | dynactin 6 | protein-coding |
| chr8_34045672_34046020 | chr8 | 34045673 | 34046020 | + | -0.802051 | Intergenic | L1M4a1\|LINE\|L1 | 61968 | NM_001293758 | 22428 | Mm.150373 | NM_011722 | ENSMUSG00000031516 | **Dctn6** | AU044699\|WS-3\|p27 | dynactin 6 | protein-coding |
| chr8_34046733_34047108 | chr8 | 34046734 | 34047108 | + | -1.17638 | Intergenic | L1M4a1\|LINE\|L1 | 60893 | NM_001293758 | 22428 | Mm.150373 | NM_011722 | ENSMUSG00000031516 | **Dctn6** | AU044699\|WS-3\|p27 | dynactin 6 | protein-coding |
| chr17_35731615_35732091 | chr17 | 35731616 | 35732091 | + | -0.737577 | Intergenic | Intergenic | -27714 | NM_172962 | 12305 | Mm.5021 | NM_007584 | ENSMUSG00000003534 | **Ddr1** | 6030432F18\|AI323681\|CD167a\|Cak\|Nep\|PTK3A | discoidin domain receptor family, member 1 | protein-coding |
| chr17_35734563_35734772 | chr17 | 35734564 | 35734772 | + | -1.29496 | Intergenic | RLTR14_RN\|LTR\|ERV1 | -30529 | NM_172962 | 12305 | Mm.5021 | NM_007584 | ENSMUSG00000003534 | **Ddr1** | 6030432F18\|AI323681\|CD167a\|Cak\|Nep\|PTK3A | discoidin domain receptor family, member 1 | protein-coding |
| chr17_35734498_35734863 | chr17 | 35734499 | 35734863 | + | -1.31845 | Intergenic | RLTR14_RN\|LTR\|ERV1 | -30542 | NM_172962 | 12305 | Mm.5021 | NM_007584 | ENSMUSG00000003534 | **Ddr1** | 6030432F18\|AI323681\|CD167a\|Cak\|Nep\|PTK3A | discoidin domain receptor family, member 1 | protein-coding |
| chr9_53271683_53271989 | chr9 | 53271684 | 53271989 | + | -1.02151 | Intergenic | Intergenic | -23724 | NM_029936 | 77591 | Mm.133560 | NM_029936 | ENSMUSG00000053289 | **Ddx10** | 4632415A01Rik\|AI646054 | DEAD (Asp-Glu-Ala-Asp) box polypeptide 10 | protein-coding |
| chr8_106017139_106017494 | chr8 | 106017140 | 106017494 | + | -0.864599 | intron (NM_025518, intron 2 of 15) | intron (NM_025518, intron 2 of 15) | -5831 | NM_028038 | 71986 | Mm.307515 | NM_028038 | ENSMUSG00000045538 | **Ddx28** | 2410004K13Rik\|AI449652\|Mddx28 | DEAD (Asp-Glu-Ala-Asp) box polypeptide 28 | protein-coding |
| chr8_106017139_106017623 | chr8 | 106017140 | 106017623 | + | -0.935749 | intron (NM_025518, intron 2 of 15) | intron (NM_025518, intron 2 of 15) | -5895 | NM_028038 | 71986 | Mm.307515 | NM_028038 | ENSMUSG00000045538 | **Ddx28** | 2410004K13Rik\|AI449652\|Mddx28 | DEAD (Asp-Glu-Ala-Asp) box polypeptide 28 | protein-coding |
| chr2_28863349_28863617 | chr2 | 28863350 | 28863617 | + | -1.01733 | intron (NM_001033294, intron 13 of 19) | intron (NM_001033294, intron 13 of 19) | 23077 | NM_001033294 | 227674 | Mm.44179 | NM_001033294 | ENSMUSG00000026806 | **Ddx31** | 5830444G11Rik\|Gm997 | DEAD/H (Asp-Glu-Ala-Asp/His) box polypeptide 31 | protein-coding |
| chrX_13370309_13370788 | chrX | 13370310 | 13370788 | + | -0.822226 | Intergenic | RLTR33\|LTR\|ERVK | 89527 | NM_010028 | 13205 | Mm.289662 | NM_010028 | ENSMUSG00000000787 | **Ddx3x** | D1Pas1-rs2\|Ddx3\|Fin14 | DEAD/H (Asp-Glu-Ala-Asp/His) box polypeptide 3, X-linked | protein-coding |

| | | | | | | | | | | | | | | | | |
|---|---|---|---|---|---|---|---|---|---|---|---|---|---|---|---|---|
| chr9_44579320_44579786 | chr9 | 44579321 | 44579786 | + | -0.721325 | Intergenic | Intergenic | -25339 | NM_001110826 | 13209 | Mm.267061 | NM_007841 | ENSMUSG00000032097 | **Ddx6** | 1110001P04Rik\|E230023J21Rik\|HLR2\|mRCK/P54\|p54\|rck | DEAD (Asp-Glu-Ala-Asp) box polypeptide 6 | protein-coding |
| chr9_44567988_44568425 | chr9 | 44567989 | 44568425 | + | -0.800797 | Intergenic | Intergenic | -36685 | NM_001110826 | 13209 | Mm.267061 | NM_007841 | ENSMUSG00000032097 | **Ddx6** | 1110001P04Rik\|E230023J21Rik\|HLR2\|mRCK/P54\|p54\|rck | DEAD (Asp-Glu-Ala-Asp) box polypeptide 6 | protein-coding |
| chr9_44568077_44568392 | chr9 | 44568078 | 44568392 | + | -0.832996 | Intergenic | Intergenic | -36657 | NM_001110826 | 13209 | Mm.267061 | NM_007841 | ENSMUSG00000032097 | **Ddx6** | 1110001P04Rik\|E230023J21Rik\|HLR2\|mRCK/P54\|p54\|rck | DEAD (Asp-Glu-Ala-Asp) box polypeptide 6 | protein-coding |
| chr17_28217259_28218188 | chr17 | 28217260 | 28218188 | + | -0.79436 | exon (NM_027185, exon 3 of 11) | exon (NM_027185, exon 3 of 11) | 9946 | NM_027185 | 23853 | Mm.204731 | NM_027185 | ENSMUSG00000002257 | **Def6** | 2410003F05Rik\|6430538D02Rik\|AV094905\|Ibp\|Slat\|Slat2\|Slat6 | differentially expressed in FDCP 6 | protein-coding |
| chr2_38068237_38068661 | chr2 | 38068238 | 38068661 | + | -0.650378 | intron (NM_146122, intron 5 of 21) | intron (NM_146122, intron 5 of 21) | 218935 | NM_146122 | 227801 | Mm.314245 | NM_146122 | ENSMUSG00000035392 | **Dennd1a** | 6030446I19Rik\|connecdenn | DENN/MADD domain containing 1A | protein-coding |
| chr2_38068399_38068786 | chr2 | 38068400 | 38068786 | + | -0.705284 | intron (NM_146122, intron 5 of 21) | intron (NM_146122, intron 5 of 21) | 218791 | NM_146122 | 227801 | Mm.314245 | NM_146122 | ENSMUSG00000035392 | **Dennd1a** | 6030446I19Rik\|connecdenn | DENN/MADD domain containing 1A | protein-coding |
| chr2_38205375_38205779 | chr2 | 38205376 | 38205779 | + | -0.740198 | intron (NM_146122, intron 2 of 21) | intron (NM_146122, intron 2 of 21) | 81807 | NM_146122 | 227801 | Mm.314245 | NM_146122 | ENSMUSG00000035392 | **Dennd1a** | 6030446I19Rik\|connecdenn | DENN/MADD domain containing 1A | protein-coding |
| chr2_38072014_38072452 | chr2 | 38072015 | 38072452 | + | -0.885877 | intron (NM_146122, intron 5 of 21) | intron (NM_146122, intron 5 of 21) | 215151 | NM_146122 | 227801 | Mm.314245 | NM_146122 | ENSMUSG00000035392 | **Dennd1a** | 6030446I19Rik\|connecdenn | DENN/MADD domain containing 1A | protein-coding |
| chr3_106493263_106493520 | chr3 | 106493264 | 106493520 | + | -0.695331 | intron (NM_001278941, intron 8 of 12) | intron (NM_001278941, intron 8 of 12) | 7352 | NM_028110 | 72121 | Mm.14816 | NM_028110 | ENSMUSG00000027901 | **Dennd2d** | 2010308M01Rik\|AI451421 | DENN/MADD domain containing 2D | protein-coding |
| chr3_106461849_106462549 | chr3 | 106461850 | 106462549 | + | -0.707027 | Intergenic | Intergenic | -20232 | NM_001093754 | 72121 | Mm.14816 | NM_028110 | ENSMUSG00000027901 | **Dennd2d** | 2010308M01Rik\|AI451421 | DENN/MADD domain containing 2D | protein-coding |
| chr3_106493203_106493636 | chr3 | 106493204 | 106493636 | + | -0.718748 | intron (NM_001278941, intron 8 of 12) | intron (NM_001278941, intron 8 of 12) | 7380 | NM_028110 | 72121 | Mm.14816 | NM_028110 | ENSMUSG00000027901 | **Dennd2d** | 2010308M01Rik\|AI451421 | DENN/MADD domain containing 2D | protein-coding |
| chr3_106461802_106462884 | chr3 | 106461803 | 106462884 | + | -0.807513 | Intergenic | Intergenic | -20088 | NM_001093754 | 72121 | Mm.14816 | NM_028110 | ENSMUSG00000027901 | **Dennd2d** | 2010308M01Rik\|AI451421 | DENN/MADD domain containing 2D | protein-coding |
| chr15_73506333_73506784 | chr15 | 73506334 | 73506784 | + | -0.684359 | Intergenic | Intergenic | -6001 | NM_001081066 | 105841 | Mm.322557 | NM_001081066 | ENSMUSG00000036661 | **Dennd3** | AI447457\|E030003N15Rik | DENN/MADD domain containing 3 | protein-coding |
| chr15_73522923_73523696 | chr15 | 73522924 | 73523696 | + | -0.942504 | intron (NM_001081066, intron 2 of 22) | intron (NM_001081066, intron 2 of 22) | 10750 | NM_001081066 | 105841 | Mm.322557 | NM_001081066 | ENSMUSG00000036661 | **Dennd3** | AI447457\|E030003N15Rik | DENN/MADD domain containing 3 | protein-coding |
| chr15_73523163_73523609 | chr15 | 73523164 | 73523609 | + | -0.986277 | intron (NM_001081066, intron 2 of 22) | intron (NM_001081066, intron 2 of 22) | 10826 | NM_001081066 | 105841 | Mm.322557 | NM_001081066 | ENSMUSG00000036661 | **Dennd3** | AI447457\|E030003N15Rik | DENN/MADD domain containing 3 | protein-coding |
| chr9_64824368_64825086 | chr9 | 64824369 | 64825086 | + | -1.50039 | intron (NM_001162917, intron 2 of 34) | intron (NM_001162917, intron 2 of 34) | 13716 | NM_001162917 | 102442 | Mm.222473 | NM_176935 | ENSMUSG00000053641 | **Dennd4a** | AI115600\|AI851943\|F730015K02Rik | DENN/MADD domain containing 4A | protein-coding |
| chr4_86721877_86722387 | chr4 | 86721878 | 86722387 | + | -1.03622 | Intergenic | Intergenic | -26423 | NM_184088 | 329877 | Mm.273718 | NM_184088 | ENSMUSG00000038024 | **Dennd4c** | 1700065A05Rik\|AA420392 | DENN/MADD domain containing 4C | protein-coding |
| chr4_86721582_86722392 | chr4 | 86721583 | 86722392 | + | -1.04576 | Intergenic | Intergenic | -26568 | NM_184088 | 329877 | Mm.273718 | NM_184088 | ENSMUSG00000038024 | **Dennd4c** | 1700065A05Rik\|AA420392 | DENN/MADD domain containing 4C | protein-coding |
| chr13_108297267_108298117 | chr13 | 108297268 | 108298117 | + | -0.671031 | Intergenic | Intergenic | -18645 | NM_178683 | 218581 | Mm.32318 | NM_178683 | ENSMUSG00000021697 | **Depdc1b** | 9830132O03\|AW260467\|XTP1 | DEP domain containing 1B | protein-coding |
| chr13_108426300_108426685 | chr13 | 108426301 | 108426685 | + | -0.857515 | Intergenic | Intergenic | 110156 | NM_178683 | 218581 | Mm.32318 | NM_178683 | ENSMUSG00000021697 | **Depdc1b** | 9830132O03\|AW260467\|XTP1 | DEP domain containing 1B | protein-coding |
| chr13_108426219_108426800 | chr13 | 108426220 | 108426800 | + | -0.876155 | Intergenic | Intergenic | 110173 | NM_178683 | 218581 | Mm.32318 | NM_178683 | ENSMUSG00000021697 | **Depdc1b** | 9830132O03\|AW260467\|XTP1 | DEP domain containing 1B | protein-coding |
| chr15_57878191_57878746 | chr15 | 57878192 | 57878746 | + | -0.90492 | intron (NM_024207, intron 5 of 7) | intron (NM_024207, intron 5 of 7) | 13949 | NM_024207 | 67819 | Mm.289387 | NM_024207 | ENSMUSG00000022365 | **Derl1** | 1110021N07Rik\|AI195141\|AW551338\|Derlin-1 | Der1-like domain family, member 1 | protein-coding |
| chr7_78847576_78847888 | chr7 | 78847577 | 78847888 | + | -0.756147 | promoter-TSS (NM_029585) | promoter-TSS (NM_029585) | -521 | NM_029585 | 76375 | Mm.250541 | NM_029585 | ENSMUSG00000030610 | **Det1** | 2610034H20Rik | de-etiolated homolog 1 (Arabidopsis) | protein-coding |
| chr6_50281653_50282097 | chr6 | 50281654 | 50282097 | + | -1.05278 | Intergenic | Intergenic | -20106 | NM_018769 | 54722 | Mm.248361 | NM_018769 | ENSMUSG00000029821 | **Dfna5** | 2310037D07Rik\|4932441K13Rik\|Dfna5h\|EG14210\|Fin15 | deafness, autosomal dominant 5 (human) | protein-coding |
| chr6_50252483_50253223 | chr6 | 50252484 | 50253223 | + | -1.33545 | intron (NM_018769, intron 1 of 9) | MLTR11B\|LTR\|ERVK | 8916 | NM_018769 | 54722 | Mm.248361 | NM_018769 | ENSMUSG00000029821 | **Dfna5** | 2310037D07Rik\|4932441K13Rik\|Dfna5h\|EG14210\|Fin15 | deafness, autosomal dominant 5 (human) | protein-coding |
| chr16_17911041_17911600 | chr16 | 17911042 | 17911600 | + | -0.669588 | promoter-TSS (NM_001081633) | promoter-TSS (NM_001081633) | 27 | NM_022408 | 27886 | Mm.256480 | NM_022408 | ENSMUSG00000003527 | **Dgcr14** | AI462402\|D16H22S1269E\|Dgcr1\|Dgsi\|ES2\|Es2el | DiGeorge syndrome critical region gene 14 | protein-coding |
| chr1_87874324_87874567 | chr1 | 87874325 | 87874567 | + | -0.641555 | intron (NM_177646, intron 1 of 29) | intron (NM_177646, intron 1 of 29) | 21159 | NM_177646 | 227333 | Mm.277217 | NM_177646 | ENSMUSG00000070738 | **Dgkd** | AI841987\|D330025K09\|DGKdelta\|dgkd-2 | diacylglycerol kinase, delta | protein-coding |
| chr1_87829621_87830237 | chr1 | 87829622 | 87830237 | + | -0.688763 | intron (NM_009118, intron 10 of 15) | intron (NM_009118, intron 10 of 15) | -23358 | NM_177646 | 227333 | Mm.277217 | NM_177646 | ENSMUSG00000070738 | **Dgkd** | AI841987\|D330025K09\|DGKdelta\|dgkd-2 | diacylglycerol kinase, delta | protein-coding |
| chr1_87829608_87830069 | chr1 | 87829609 | 87830069 | + | -0.725245 | intron (NM_009118, intron 10 of 15) | intron (NM_009118, intron 10 of 15) | -23448 | NM_177646 | 227333 | Mm.277217 | NM_177646 | ENSMUSG00000070738 | **Dgkd** | AI841987\|D330025K09\|DGKdelta\|dgkd-2 | diacylglycerol kinase, delta | protein-coding |
| chr1_87846541_87847590 | chr1 | 87846542 | 87847590 | + | -0.770874 | Intergenic | Intergenic | -6221 | NM_177646 | 227333 | Mm.277217 | NM_177646 | ENSMUSG00000070738 | **Dgkd** | AI841987\|D330025K09\|DGKdelta\|dgkd-2 | diacylglycerol kinase, delta | protein-coding |
| chr1_87847299_87847551 | chr1 | 87847300 | 87847551 | + | -0.936661 | Intergenic | Intergenic | -5862 | NM_177646 | 227333 | Mm.277217 | NM_177646 | ENSMUSG00000070738 | **Dgkd** | AI841987\|D330025K09\|DGKdelta\|dgkd-2 | diacylglycerol kinase, delta | protein-coding |
| chr14_78618638_78619343 | chr14 | 78618639 | 78619343 | + | -0.67235 | intron (NR_104044, intron 6 of 27) | intron (NR_104044, intron 6 of 27) | 10299 | NM_001253766 | 380921 | Mm.89591 | NM_001081336 | ENSMUSG00000034731 | **Dgkh** | 5930402B05Rik\|D130015C16 | diacylglycerol kinase, eta | protein-coding |
| chr14_78687693_78688060 | chr14 | 78687694 | 78688060 | + | -1.03685 | intron (NM_001081336, intron 1 of 28) | intron (NM_001081336, intron 1 of 28) | 37212 | NM_001081336 | 380921 | Mm.89591 | NM_001081336 | ENSMUSG00000034731 | **Dgkh** | 5930402B05Rik\|D130015C16 | diacylglycerol kinase, eta | protein-coding |
| chr13_92274553_92275035 | chr13 | 92274554 | 92275035 | + | -0.785804 | exon (NM_010829, exon 14 of 24) | exon (NM_010829, exon 14 of 24) | -79989 | NM_010049 | 13361 | Mm.23695 | NM_010049 | ENSMUSG00000021707 | **Dhfr** | 8430436I03Rik\|AA607882\|AI662710\|AW555094 | dihydrofolate reductase | protein-coding |
| chr13_92274542_92274894 | chr13 | 92274543 | 92274894 | + | -0.796041 | intron (NM_010829, intron 14 of 23) | intron (NM_010829, intron 14 of 23) | -80065 | NM_010049 | 13361 | Mm.23695 | NM_010049 | ENSMUSG00000021707 | **Dhfr** | 8430436I03Rik\|AA607882\|AI662710\|AW555094 | dihydrofolate reductase | protein-coding |
| chr13_92274349_92274953 | chr13 | 92274350 | 92274953 | + | -0.843653 | intron (NM_010829, intron 14 of 23) | intron (NM_010829, intron 14 of 23) | -80132 | NM_010049 | 13361 | Mm.23695 | NM_010049 | ENSMUSG00000021707 | **Dhfr** | 8430436I03Rik\|AA607882\|AI662710\|AW555094 | dihydrofolate reductase | protein-coding |
| chr13_92244680_92245139 | chr13 | 92244681 | 92245139 | + | -0.854805 | intron (NM_010829, intron 19 of 23) | intron (NM_010829, intron 19 of 23) | -109873 | NM_010049 | 13361 | Mm.23695 | NM_010049 | ENSMUSG00000021707 | **Dhfr** | 8430436I03Rik\|AA607882\|AI662710\|AW555094 | dihydrofolate reductase | protein-coding |
| chr4_145019152_145019862 | chr4 | 145019153 | 145019862 | + | -1.13012 | intron (NM_001276465, intron 66 of 69) | intron (NM_001276465, intron 66 of 69) | 126370 | NM_011303 | 20148 | Mm.14063 | NM_011303 | ENSMUSG00000066026 | **Dhrs3** | Rsdr1\|retSDR1 | dehydrogenase/reductase (SDR family) member 3 | protein-coding |
| chr11_86820821_86821120 | chr11 | 86820822 | 86821120 | + | -0.630246 | intron (NR_106195, intron 2 of 8) | ORR1A3\|LTR\|ERVL-MaLR | -13311 | NM_026191 | 67487 | Mm.260627 | NM_026191 | ENSMUSG00000018425 | **Dhx40** | 2410016C14Rik\|ARG147\|DDX40\|PAD | DEAH (Asp-Glu-Ala-His) box polypeptide 40 | protein-coding |
| chr11_86834006_86834462 | chr11 | 86834007 | 86834462 | + | -0.65687 | intron (NR_106195, intron 2 of 8) | intron (NR_106195, intron 2 of 8) | -26574 | NM_026191 | 67487 | Mm.260627 | NM_026191 | ENSMUSG00000018425 | **Dhx40** | 2410016C14Rik\|ARG147\|DDX40\|PAD | DEAH (Asp-Glu-Ala-His) box polypeptide 40 | protein-coding |
| chr11_86810271_86811754 | chr11 | 86810272 | 86811754 | + | -0.925187 | intron (NR_106195, intron 2 of 8) | intron (NR_106195, intron 2 of 8) | -3353 | NM_026191 | 67487 | Mm.260627 | NM_026191 | ENSMUSG00000018425 | **Dhx40** | 2410016C14Rik\|ARG147\|DDX40\|PAD | DEAH (Asp-Glu-Ala-His) box polypeptide 40 | protein-coding |
| chr11_86827475_86827876 | chr11 | 86827476 | 86827876 | + | -1.0257 | intron (NR_106195, intron 2 of 8) | B1_Mur4\|SINE\|Alu | -20016 | NM_026191 | 67487 | Mm.260627 | NM_026191 | ENSMUSG00000018425 | **Dhx40** | 2410016C14Rik\|ARG147\|DDX40\|PAD | DEAH (Asp-Glu-Ala-His) box polypeptide 40 | protein-coding |
| chr11_86827472_86828786 | chr11 | 86827473 | 86828786 | + | -1.1461 | intron (NR_106195, intron 2 of 8) | B4A\|SINE\|B4 | -20469 | NM_026191 | 67487 | Mm.260627 | NM_026191 | ENSMUSG00000018425 | **Dhx40** | 2410016C14Rik\|ARG147\|DDX40\|PAD | DEAH (Asp-Glu-Ala-His) box polypeptide 40 | protein-coding |
| chr11_86828070_86828764 | chr11 | 86828071 | 86828764 | + | -1.16839 | intron (NR_106195, intron 2 of 8) | PB1D10\|SINE\|Alu | -20757 | NM_026191 | 67487 | Mm.260627 | NM_026191 | ENSMUSG00000018425 | **Dhx40** | 2410016C14Rik\|ARG147\|DDX40\|PAD | DEAH (Asp-Glu-Ala-His) box polypeptide 40 | protein-coding |
| chr11_86828297_86828816 | chr11 | 86828298 | 86828816 | + | -1.24135 | intron (NR_106195, intron 2 of 8) | intron (NR_106195, intron 2 of 8) | -20897 | NM_026191 | 67487 | Mm.260627 | NM_026191 | ENSMUSG00000018425 | **Dhx40** | 2410016C14Rik\|ARG147\|DDX40\|PAD | DEAH (Asp-Glu-Ala-His) box polypeptide 40 | protein-coding |
| chr11_86822354_86823068 | chr11 | 86822355 | 86823068 | + | -1.58508 | intron (NR_106195, intron 2 of 8) | RMER19B\|LTR\|ERVK | -15051 | NM_026191 | 67487 | Mm.260627 | NM_026191 | ENSMUSG00000018425 | **Dhx40** | 2410016C14Rik\|ARG147\|DDX40\|PAD | DEAH (Asp-Glu-Ala-His) box polypeptide 40 | protein-coding |
| chr18_37919798_37920200 | chr18 | 37919799 | 37920200 | + | -0.879666 | intron (NM_007858, intron 1 of 26) | intron (NM_007858, intron 1 of 26) | 15412 | NM_007858 | 13367 | Mm.195916 | NM_007858 | ENSMUSG00000024456 | **Diap1** | D18Wsu154e\|Dia1\|Diaph1\|Drf1\|mKIAA4062\|p140mDia | diaphanous homolog 1 (Drosophila) | protein-coding |
| chrX_130480974_130481238 | chrX | 130480975 | 130481238 | + | -0.654874 | Intergenic | Intergenic | 731364 | NM_172493 | 54004 | Mm.10763 | NM_017398 | ENSMUSG00000034480 | **Diap2** | Dia3\|Diaph2\|Drf2\|E430022I22Rik | diaphanous homolog 2 (Drosophila) | protein-coding |
| chrX_130958665_130958891 | chrX | 130958666 | 130958891 | + | -0.674459 | Intergenic | Intergenic | 1209036 | NM_172493 | 54004 | Mm.10763 | NM_017398 | ENSMUSG00000034480 | **Diap2** | Dia3\|Diaph2\|Drf2\|E430022I22Rik | diaphanous homolog 2 (Drosophila) | protein-coding |
| chrX_130415449_130415808 | chrX | 130415450 | 130415808 | + | -0.75254 | intron (NM_172493, intron 26 of 26) | Lx9\|LINE\|L1 | 665887 | NM_172493 | 54004 | Mm.10763 | NM_017398 | ENSMUSG00000034480 | **Diap2** | Dia3\|Diaph2\|Drf2\|E430022I22Rik | diaphanous homolog 2 (Drosophila) | protein-coding |
| chrX_130602015_130602326 | chrX | 130602016 | 130602326 | + | -0.773933 | Intergenic | Intergenic | 852429 | NM_172493 | 54004 | Mm.10763 | NM_017398 | ENSMUSG00000034480 | **Diap2** | Dia3\|Diaph2\|Drf2\|E430022I22Rik | diaphanous homolog 2 (Drosophila) | protein-coding |
| chrX_130548985_130549909 | chrX | 130548986 | 130549909 | + | -0.86204 | Intergenic | Intergenic | 799705 | NM_172493 | 54004 | Mm.10763 | NM_017398 | ENSMUSG00000034480 | **Diap2** | Dia3\|Diaph2\|Drf2\|E430022I22Rik | diaphanous homolog 2 (Drosophila) | protein-coding |
| chrX_130772758_130773106 | chrX | 130772759 | 130773106 | + | -0.959697 | Intergenic | B3A\|SINE\|B2 | 1023190 | NM_172493 | 54004 | Mm.10763 | NM_017398 | ENSMUSG00000034480 | **Diap2** | Dia3\|Diaph2\|Drf2\|E430022I22Rik | diaphanous homolog 2 (Drosophila) | protein-coding |
| chrX_130564817_130565314 | chrX | 130564818 | 130565314 | + | -1.07498 | Intergenic | MLTR25C\|LTR\|ERVK | 815324 | NM_172493 | 54004 | Mm.10763 | NM_017398 | ENSMUSG00000034480 | **Diap2** | Dia3\|Diaph2\|Drf2\|E430022I22Rik | diaphanous homolog 2 (Drosophila) | protein-coding |
| chrX_130564936_130565192 | chrX | 130564937 | 130565192 | + | -1.14986 | Intergenic | MLTR25C\|LTR\|ERVK | 815322 | NM_172493 | 54004 | Mm.10763 | NM_017398 | ENSMUSG00000034480 | **Diap2** | Dia3\|Diaph2\|Drf2\|E430022I22Rik | diaphanous homolog 2 (Drosophila) | protein-coding |

| | | | | | | | | | | | | | | | | | |
|---|---|---|---|---|---|---|---|---|---|---|---|---|---|---|---|---|---|
| chr14_86744044_86744421 | chr14 | 86744045 | 86744421 | + | -0.68662 | intron (NM_019670, intron 27 of 27) | intron (NM_019670, intron 27 of 27) | 396881 | NM_019670 | 56419 | Mm.440585 | NM_019670 | ENSMUSG00000022021 | **Diap3** | 4930417P13Rik\|Dia2\|Diaph3\|Drf3\|mDia2\|mKIAA4117\|p134MDia2 | diaphanous homolog 3 (Drosophila) | protein-coding |
| chr14_86731236_86731531 | chr14 | 86731237 | 86731531 | + | -0.687867 | intron (NM_019670, intron 27 of 27) | intron (NM_019670, intron 27 of 27) | 409730 | NM_019670 | 56419 | Mm.440585 | NM_019670 | ENSMUSG00000022021 | **Diap3** | 4930417P13Rik\|Dia2\|Diaph3\|Drf3\|mDia2\|mKIAA4117\|p134MDia2 | diaphanous homolog 3 (Drosophila) | protein-coding |
| chr14_86720483_86720760 | chr14 | 86720484 | 86720760 | + | -0.906179 | intron (NM_019670, intron 27 of 27) | B3A\|SINE\|B2 | 420492 | NM_019670 | 56419 | Mm.440585 | NM_019670 | ENSMUSG00000022021 | **Diap3** | 4930417P13Rik\|Dia2\|Diaph3\|Drf3\|mDia2\|mKIAA4117\|p134MDia2 | diaphanous homolog 3 (Drosophila) | protein-coding |
| chr14_86736695_86737153 | chr14 | 86736696 | 86737153 | + | -1.21403 | intron (NM_019670, intron 27 of 27) | intron (NM_019670, intron 27 of 27) | 404190 | NM_019670 | 56419 | Mm.440585 | NM_019670 | ENSMUSG00000022021 | **Diap3** | 4930417P13Rik\|Dia2\|Diaph3\|Drf3\|mDia2\|mKIAA4117\|p134MDia2 | diaphanous homolog 3 (Drosophila) | protein-coding |
| chr2_180684122_180684396 | chr2 | 180684123 | 180684396 | + | -0.749926 | intron (NM_001291433, intron 5 of 14) | intron (NM_001291433, intron 5 of 14) | 7789 | NR_111965 | 23856 | Mm.253836 | NM_011805 | ENSMUSG00000038914 | **Dido1** | 6720461J16Rik\|D130048F08Rik\|DATF-1\|DIO-1\|Datf1 | death inducer-obliterator 1 | protein-coding |
| chr15_100119280_100119777 | chr15 | 100119281 | 100119777 | + | -0.675925 | intron (NM_001159361, intron 2 of 37) | intron (NM_001159361, intron 2 of 37) | -34867 | NM_172819 | 239667 | Mm.243658 | NM_172819 | ENSMUSG00000023026 | **Dip2b** | 4932422C22\|AI317237\|AI854602\|mKIAA1463 | DIP2 disco-interacting protein 2 homolog B (Drosophila) | protein-coding |
| chr15_100041318_100041590 | chr15 | 100041319 | 100041590 | + | -0.960937 | intron (NM_001159361, intron 1 of 37) | intron (NM_001159361, intron 1 of 37) | 2790 | NM_001159361 | 239667 | Mm.243658 | NM_172819 | ENSMUSG00000023026 | **Dip2b** | 4932422C22\|AI317237\|AI854602\|mKIAA1463 | DIP2 disco-interacting protein 2 homolog B (Drosophila) | protein-coding |
| chr15_100041368_100041561 | chr15 | 100041369 | 100041561 | + | -0.963645 | intron (NM_001159361, intron 1 of 37) | intron (NM_001159361, intron 1 of 37) | 2801 | NM_001159361 | 239667 | Mm.243658 | NM_172819 | ENSMUSG00000023026 | **Dip2b** | 4932422C22\|AI317237\|AI854602\|mKIAA1463 | DIP2 disco-interacting protein 2 homolog B (Drosophila) | protein-coding |
| chr15_100140301_100140789 | chr15 | 100140302 | 100140789 | + | -1.111 | intron (NM_001159361, intron 3 of 37) | RMER1C\|Other\|Other | -13851 | NM_172819 | 239667 | Mm.243658 | NM_172819 | ENSMUSG00000023026 | **Dip2b** | 4932422C22\|AI317237\|AI854602\|mKIAA1463 | DIP2 disco-interacting protein 2 homolog B (Drosophila) | protein-coding |
| chr16_35765444_35765983 | chr16 | 35765445 | 35765983 | + | -0.917137 | intron (NM_153550, intron 1 of 8) | intron (NM_153550, intron 1 of 8) | 3642 | NM_153550 | 224132 | Mm.24094 | NM_153550 | ENSMUSG00000022848 | **Dirc2** | RCC4 | disrupted in renal carcinoma 2 (human) | protein-coding |
| chr16_35765562_35765798 | chr16 | 35765563 | 35765798 | + | -0.94022 | intron (NM_153550, intron 1 of 8) | intron (NM_153550, intron 1 of 8) | 3676 | NM_153550 | 224132 | Mm.24094 | NM_153550 | ENSMUSG00000022848 | **Dirc2** | RCC4 | disrupted in renal carcinoma 2 (human) | protein-coding |
| chr14_61971087_61972092 | chr14 | 61971088 | 61972092 | + | -0.751748 | Intergenic | Intergenic | -289217 | NR_028264 | 668253 | Mm.32886 | NR_028264 | ENSMUSG00000097589 | **Dleu2** | 1810047A16Rik\|4932411A06Rik\|7730401J12Rik\|AI197277\|AI788824\|Alt1\|Gm9069\|Leu2 | deleted in lymphocytic leukemia, 2 | ncRNA |
| chr14_61971002_61971467 | chr14 | 61971003 | 61971467 | + | -1.03894 | Intergenic | Intergenic | -288862 | NR_028264 | 668253 | Mm.32886 | NR_028264 | ENSMUSG00000097589 | **Dleu2** | 1810047A16Rik\|4932411A06Rik\|7730401J12Rik\|AI197277\|AI788824\|Alt1\|Gm9069\|Leu2 | deleted in lymphocytic leukemia, 2 | ncRNA |
| chr16_31683411_31684060 | chr16 | 31683412 | 31684060 | + | -0.674665 | intron (NM_001252434, intron 3 of 24) | B3A\|SINE\|B2 | 19801 | NM_001252434 | 13383 | Mm.382 | NM_007862 | ENSMUSG00000022770 | **Dlg1** | B130052P05Rik\|Dlgh1\|E-dlg/SAP97\|SAP-97\|SAP97\|mKIAA4187 | discs, large homolog 1 (Drosophila) | protein-coding |
| chr2_71522796_71523437 | chr2 | 71522797 | 71523437 | + | -0.633094 | intron (NM_025633, intron 7 of 9) | intron (NM_025633, intron 7 of 9) | -6328 | NM_010053 | 13390 | Mm.4543 | NM_010053 | ENSMUSG00000041911 | **Dlx1** | Dlx\|Dlx-1 | distal-less homeobox 1 | protein-coding |
| chr7_131045378_131045846 | chr7 | 131045379 | 131045846 | + | -0.776851 | intron (NM_007769, intron 5 of 46) | intron (NM_007769, intron 5 of 46) | 13536 | NM_007769 | 12945 | Mm.4138 | NM_007769 | ENSMUSG00000047517 | **Dmbt1** | CRP\|CRP-[a]\|CRP-[b]\|Crpd\|DBMT1\|gp300\|p80 | deleted in malignant brain tumors 1 | protein-coding |
| chr4_89547807_89548080 | chr4 | 89547808 | 89548080 | + | -0.87691 | Intergenic | Intergenic | -140254 | NM_175647 | 242523 | Mm.130167 | NM_175647 | ENSMUSG00000043753 | **Dmrta1** | Dmrt4 | doublesex and mab-3 related transcription factor like family A1 | protein-coding |
| chr4_89550148_89551105 | chr4 | 89550149 | 89551105 | + | -1.05063 | Intergenic | Intergenic | -137571 | NM_175647 | 242523 | Mm.130167 | NM_175647 | ENSMUSG00000043753 | **Dmrta1** | Dmrt4 | doublesex and mab-3 related transcription factor like family A1 | protein-coding |
| chr4_109907374_109907797 | chr4 | 109907375 | 109907797 | + | -0.789132 | intron (NM_007983, intron 15 of 18) | intron (NM_007983, intron 15 of 18) | -70439 | NM_172296 | 242620 | Mm.32825 | NM_172296 | ENSMUSG00000047143 | **Dmrta2** | Dmrt5 | doublesex and mab-3 related transcription factor like family A2 | protein-coding |
| chr8_119585737_119586540 | chr8 | 119585738 | 119586540 | + | -0.763332 | intron (NM_026648, intron 5 of 11) | RLTR1B-int\|LTR\|ERV1 | 10904 | NM_026648 | 68270 | Mm.41977 | NM_026648 | ENSMUSG00000031831 | **Dnaaf1** | 4930457P18Rik\|Lrrc50 | dynein, axonemal assembly factor 1 | protein-coding |
| chr11_118152968_118153348 | chr11 | 118152969 | 118153348 | + | -0.675041 | Intergenic | MLT1B\|LTR\|ERVL-MaLR | -23939 | NM_001167746 | 69926 | Mm.473893 | NM_001167746 | ENSMUSG00000033987 | **Dnah17** | 2810003K23Rik\|Dnahc17\|Dnahcl1\|Gm1178\|mKIAA3028 | dynein, axonemal, heavy chain 17 | protein-coding |
| chr11_118145697_118146415 | chr11 | 118145698 | 118146415 | + | -0.851544 | Intergenic | Intergenic | -16837 | NM_001167746 | 69926 | Mm.473893 | NM_001167746 | ENSMUSG00000033987 | **Dnah17** | 2810003K23Rik\|Dnahc17\|Dnahcl1\|Gm1178\|mKIAA3028 | dynein, axonemal, heavy chain 17 | protein-coding |
| chr11_118086019_118086298 | chr11 | 118086020 | 118086298 | + | -0.928425 | exon (NM_001167746, exon 37 of 81) | exon (NM_001167746, exon 37 of 81) | 43060 | NM_001167746 | 69926 | Mm.473893 | NM_001167746 | ENSMUSG00000033987 | **Dnah17** | 2810003K23Rik\|Dnahc17\|Dnahcl1\|Gm1178\|mKIAA3028 | dynein, axonemal, heavy chain 17 | protein-coding |
| chr11_118143801_118144081 | chr11 | 118143802 | 118144081 | + | -1.32012 | Intergenic | RLTR14\|LTR\|ERV1 | -14722 | NM_001167746 | 69926 | Mm.473893 | NM_001167746 | ENSMUSG00000033987 | **Dnah17** | 2810003K23Rik\|Dnahc17\|Dnahcl1\|Gm1178\|mKIAA3028 | dynein, axonemal, heavy chain 17 | protein-coding |
| chr17_30634784_30635096 | chr17 | 30634785 | 30635096 | + | -0.846305 | intron (NM_013811, intron 1 of 92) | MTD\|LTR\|ERVL-MaLR | 8004 | NM_013811 | 13417 | Mm.426017 | NM_013811 | ENSMUSG00000033826 | **Dnah8** | ATPase\|Dnahc8\|Hst6.7b\|P1-Loop | dynein, axonemal, heavy chain 8 | protein-coding |
| chr8_85618461_85619531 | chr8 | 85618462 | 85619531 | + | -0.626638 | Intergenic | RMER13B\|LTR\|ERVK | -63725 | NM_019794 | 56445 | Mm.475573 | NM_019794 | ENSMUSG00000031701 | **Dnaja2** | 1500017M13Rik\|2010206B19Rik\|DNAJ\|DNJ3\|Dnaj3\|HIRIP4\|PRO3015\|mDj3 | DnaJ (Hsp40) homolog, subfamily A, member 2 | protein-coding |
| chr8_83628284_83628741 | chr8 | 83628285 | 83628741 | + | -1.00826 | Intergenic | Lx7\|LINE\|L1 | 20338 | NM_018808 | 81489 | Mm.282092 | NM_018808 | ENSMUSG00000005483 | **Dnajb1** | 0610007I11Rik\|HSPF1\|Hsp40 | DnaJ (Hsp40) homolog, subfamily B, member 1 | protein-coding |
| chr8_83611853_83612399 | chr8 | 83611854 | 83612399 | + | -1.26727 | TTS (NM_018808) | TTS (NM_018808) | 3951 | NM_018808 | 81489 | Mm.282092 | NM_018808 | ENSMUSG00000005483 | **Dnajb1** | 0610007I11Rik\|HSPF1\|Hsp40 | DnaJ (Hsp40) homolog, subfamily B, member 1 | protein-coding |
| chr2_18307483_18308035 | chr2 | 18307484 | 18308035 | + | -0.679323 | intron (NM_001190817, intron 5 of 12) | intron (NM_001190817, intron 5 of 12) | 85071 | NM_001190817 | 13418 | Mm.246674 | NM_007869 | ENSMUSG00000026740 | **Dnajc1** | 4733401K02Rik\|AA960110\|D230036H06Rik\|Dnajl1\|ERdj1\|ERj1p\|MTJ1 | DnaJ (Hsp40) homolog, subfamily C, member 1 | protein-coding |
| chr14_77836821_77837211 | chr14 | 77836822 | 77837211 | + | -0.807183 | intron (NM_025384, intron 5 of 5) | intron (NM_025384, intron 5 of 5) | 37901 | NM_025384 | 66148 | Mm.248046 | NM_025384 | ENSMUSG00000022013 | **Dnajc15** | 1110003P16Rik\|Dnajd1 | DnaJ (Hsp40) homolog, subfamily C, member 15 | protein-coding |
| chr2_181537531_181538147 | chr2 | 181537532 | 181538147 | + | -0.623364 | intron (NM_001271584, intron 2 of 5) | intron (NM_001271584, intron 2 of 5) | 17354 | NM_001271584 | 13002 | Mm.140761 | NM_016775 | ENSMUSG00000000826 | **Dnajc5** | 2610314I24Rik\|AU018536\|Csp | DnaJ (Hsp40) homolog, subfamily C, member 5 | protein-coding |
| chr3_19429509_19430667 | chr3 | 19429510 | 19430667 | + | -0.826001 | Intergenic | Intergenic | -78507 | NM_001163536 | 66326 | Mm.45615 | NM_025489 | ENSMUSG00000027606 | **Dnajc5b** | 1700008A05Rik | DnaJ (Hsp40) homolog, subfamily C, member 5 beta | protein-coding |
| chr3_19411186_19411543 | chr3 | 19411187 | 19411543 | + | -1.14872 | Intergenic | Intergenic | -97230 | NM_001163536 | 66326 | Mm.45615 | NM_025489 | ENSMUSG00000027606 | **Dnajc5b** | 1700008A05Rik | DnaJ (Hsp40) homolog, subfamily C, member 5 beta | protein-coding |
| chr4_101495790_101496157 | chr4 | 101495791 | 101496157 | + | -0.954793 | promoter-TSS (NM_198412) | promoter-TSS (NM_198412) | -674 | NM_198412 | 72685 | Mm.76494 | NM_198412 | ENSMUSG00000028528 | **Dnajc6** | 2810027M23Rik\|mKIAA0473 | DnaJ (Hsp40) homolog, subfamily C, member 6 | protein-coding |
| chr11_100610267_100610505 | chr11 | 100610268 | 100610505 | + | -0.719869 | intron (NM_019795, intron 1 of 13) | (CA)n\|Simple_repeat\|Simple_repeat | 9782 | NM_019795 | 56354 | Mm.402409 | NM_019795 | ENSMUSG00000014195 | **Dnajc7** | 2010003F24Rik\|2010004G07Rik\|CCRP\|Ttc2\|mDj11\|mTpr2 | DnaJ (Hsp40) homolog, subfamily C, member 7 | protein-coding |
| chr11_100609653_100610201 | chr11 | 100609654 | 100610201 | + | -0.790644 | intron (NM_019795, intron 1 of 13) | PB1D9\|SINE\|Alu | 10241 | NM_019795 | 56354 | Mm.402409 | NM_019795 | ENSMUSG00000014195 | **Dnajc7** | 2010003F24Rik\|2010004G07Rik\|CCRP\|Ttc2\|mDj11\|mTpr2 | DnaJ (Hsp40) homolog, subfamily C, member 7 | protein-coding |
| chr11_100609748_100610515 | chr11 | 100609749 | 100610515 | + | -0.870547 | intron (NM_019795, intron 1 of 13) | intron (NM_019795, intron 1 of 13) | 10036 | NM_019795 | 56354 | Mm.402409 | NM_019795 | ENSMUSG00000014195 | **Dnajc7** | 2010003F24Rik\|2010004G07Rik\|CCRP\|Ttc2\|mDj11\|mTpr2 | DnaJ (Hsp40) homolog, subfamily C, member 7 | protein-coding |
| chr9_21419520_21420122 | chr9 | 21419521 | 21420122 | + | -0.651754 | intron (NM_021888, intron 9 of 9) | intron (NM_021888, intron 9 of 9) | -5087 | NM_007871 | 13430 | Mm.433257 | NM_007871 | ENSMUSG00000033335 | **Dnm2** | Dyn2\|Udnm\|b2b2159Clo | dynamin 2 | protein-coding |
| chr9_21449724_21450828 | chr9 | 21449725 | 21450828 | + | -0.905115 | intron (NM_007871, intron 1 of 19) | intron (NM_007871, intron 1 of 19) | 25368 | NM_001039520 | 13430 | Mm.433257 | NM_007871 | ENSMUSG00000033335 | **Dnm2** | Dyn2\|Udnm\|b2b2159Clo | dynamin 2 | protein-coding |
| chr1_162207086_162207657 | chr1 | 162207087 | 162207657 | + | -0.640678 | intron (NM_001038619, intron 14 of 20) | intron (NM_001038619, intron 14 of 20) | -10251 | NR_002870 | 474332 | Mm.479755 | NR_002870 | ENSMUSG00000078190 | **Dnm3os** | 6030416H16Rik\|Hag2\|Npn1\|P/L01 | dynamin 3, opposite strand | ncRNA |
| chr1_162179335_162179851 | chr1 | 162179336 | 162179851 | + | -0.740635 | intron (NM_001038619, intron 14 of 20) | intron (NM_001038619, intron 14 of 20) | -38030 | NR_002870 | 474332 | Mm.479755 | NR_002870 | ENSMUSG00000078190 | **Dnm3os** | 6030416H16Rik\|Hag2\|Npn1\|P/L01 | dynamin 3, opposite strand | ncRNA |
| chr19_41039041_41039596 | chr19 | 41039042 | 41039596 | + | -0.674702 | intron (NM_009345, intron 3 of 11) | intron (NM_009345, intron 3 of 11) | 10044 | NM_009345 | 21673 | Mm.25620 | NM_009345 | ENSMUSG00000025014 | **Dntt** | Tdt | deoxynucleotidyltransferase, terminal | protein-coding |
| chr19_41022213_41022620 | chr19 | 41022214 | 41022620 | + | -0.851913 | Intergenic | Intergenic | -6858 | NM_001043228 | 21673 | Mm.25620 | NM_009345 | ENSMUSG00000025014 | **Dntt** | Tdt | deoxynucleotidyltransferase, terminal | protein-coding |
| chr1_80755346_80755629 | chr1 | 80755347 | 80755629 | + | -0.638861 | intron (NM_175291, intron 1 of 55) | intron (NM_175291, intron 1 of 55) | 3065 | NM_175291 | 210293 | Mm.133473 | NM_175291 | ENSMUSG00000038608 | **Dock10** | 9330153B10Rik\|A630054M16Rik\|Jr4\|Jr5\|R75174\|ZIZ3\|Zizimin3\|mKIAA0694 | dedicator of cytokinesis 10 | protein-coding |

| | | | | | | | | | | | | | | | | | |
|---|---|---|---|---|---|---|---|---|---|---|---|---|---|---|---|---|---|
| chr1_80703200_80703656 | chr1 | 80703201 | 80703656 | + | -0.686757 | intron (NM_175291, intron 1 of 55) | ORR1D2\|LTR\|ERVL-MaLR | -38076 | NM_001285927 | 210293 | Mm.133473 | NM_175291 | ENSMUSG00000038608 | **Dock10** | 9330153B10Rik\|A630054M16Rik\|Jr4\|Jr5\|R75174\|ZIZ3\|Zizimin3\|mKIAA0694 | dedicator of cytokinesis 10 | protein-coding |
| chr1_80765522_80766063 | chr1 | 80765523 | 80766063 | + | -0.710585 | Intergenic | Intergenic | -7240 | NM_175291 | 210293 | Mm.133473 | NM_175291 | ENSMUSG00000038608 | **Dock10** | 9330153B10Rik\|A630054M16Rik\|Jr4\|Jr5\|R75174\|ZIZ3\|Zizimin3\|mKIAA0694 | dedicator of cytokinesis 10 | protein-coding |
| chr1_80755697_80756095 | chr1 | 80755698 | 80756095 | + | -1.08996 | intron (NM_175291, intron 1 of 55) | intron (NM_175291, intron 1 of 55) | 2657 | NM_175291 | 210293 | Mm.133473 | NM_175291 | ENSMUSG00000038608 | **Dock10** | 9330153B10Rik\|A630054M16Rik\|Jr4\|Jr5\|R75174\|ZIZ3\|Zizimin3\|mKIAA0694 | dedicator of cytokinesis 10 | protein-coding |
| chr1_80755731_80756009 | chr1 | 80755732 | 80756009 | + | -1.11187 | intron (NM_175291, intron 1 of 55) | intron (NM_175291, intron 1 of 55) | 2683 | NM_175291 | 210293 | Mm.133473 | NM_175291 | ENSMUSG00000038608 | **Dock10** | 9330153B10Rik\|A630054M16Rik\|Jr4\|Jr5\|R75174\|ZIZ3\|Zizimin3\|mKIAA0694 | dedicator of cytokinesis 10 | protein-coding |
| chr11_34770599_34770853 | chr11 | 34770600 | 34770853 | + | -0.811339 | intron (NM_033374, intron 1 of 51) | intron (NM_033374, intron 1 of 51) | 13179 | NM_033374 | 94176 | Mm.380679 | NM_033374 | ENSMUSG00000020143 | **Dock2** | AI662014\|AW122239\|CED-5\|Hch\|MBC | dedicator of cyto-kinesis 2 | protein-coding |
| chr11_34770557_34770957 | chr11 | 34770558 | 34770957 | + | -0.828516 | intron (NM_033374, intron 1 of 51) | intron (NM_033374, intron 1 of 51) | 13148 | NM_033374 | 94176 | Mm.380679 | NM_033374 | ENSMUSG00000020143 | **Dock2** | AI662014\|AW122239\|CED-5\|Hch\|MBC | dedicator of cyto-kinesis 2 | protein-coding |
| chr12_40535354_40536629 | chr12 | 40535355 | 40536629 | + | -0.725293 | intron (NM_172803, intron 1 of 52) | MLTR11A\|LTR\|ERVK | 89939 | NM_172803 | 238130 | Mm.341423 | NM_172803 | ENSMUSG00000035954 | **Dock4** | 5330406C03\|6330411N01Rik\|AF263288\|C030023J22\|mKIAA0716 | dedicator of cytokinesis 4 | protein-coding |
| chr12_40580357_40580743 | chr12 | 40580358 | 40580743 | + | -0.771706 | intron (NM_172803, intron 1 of 52) | intron (NM_172803, intron 1 of 52) | 134497 | NM_172803 | 238130 | Mm.341423 | NM_172803 | ENSMUSG00000035954 | **Dock4** | 5330406C03\|6330411N01Rik\|AF263288\|C030023J22\|mKIAA0716 | dedicator of cytokinesis 4 | protein-coding |
| chr12_40529722_40530064 | chr12 | 40529723 | 40530064 | + | -1.05829 | intron (NM_172803, intron 1 of 52) | intron (NM_172803, intron 1 of 52) | 83840 | NM_172803 | 238130 | Mm.341423 | NM_172803 | ENSMUSG00000035954 | **Dock4** | 5330406C03\|6330411N01Rik\|AF263288\|C030023J22\|mKIAA0716 | dedicator of cytokinesis 4 | protein-coding |
| chr12_40529739_40530243 | chr12 | 40529740 | 40530243 | + | -1.09385 | intron (NM_172803, intron 1 of 52) | intron (NM_172803, intron 1 of 52) | 83938 | NM_172803 | 238130 | Mm.341423 | NM_172803 | ENSMUSG00000035954 | **Dock4** | 5330406C03\|6330411N01Rik\|AF263288\|C030023J22\|mKIAA0716 | dedicator of cytokinesis 4 | protein-coding |
| chr14_67951053_67951555 | chr14 | 67951054 | 67951555 | + | -0.656588 | Intergenic | L1MB1\|LINE\|L1 | -17732 | NM_177780 | 68813 | Mm.258155 | NM_177780 | ENSMUSG00000044447 | **Dock5** | 1110060D06Rik\|AI666732\|AI956923\|BC016533\|E130320D18\|lr2\|rlc | dedicator of cytokinesis 5 | protein-coding |
| chr14_67951284_67951545 | chr14 | 67951285 | 67951545 | + | -0.69169 | Intergenic | L1MB1\|LINE\|L1 | -17843 | NM_177780 | 68813 | Mm.258155 | NM_177780 | ENSMUSG00000044447 | **Dock5** | 1110060D06Rik\|AI666732\|AI956923\|BC016533\|E130320D18\|lr2\|rlc | dedicator of cytokinesis 5 | protein-coding |
| chr14_67951193_67951636 | chr14 | 67951194 | 67951636 | + | -0.757658 | Intergenic | L1MB1\|LINE\|L1 | -17843 | NM_177780 | 68813 | Mm.258155 | NM_177780 | ENSMUSG00000044447 | **Dock5** | 1110060D06Rik\|AI666732\|AI956923\|BC016533\|E130320D18\|lr2\|rlc | dedicator of cytokinesis 5 | protein-coding |
| chr19_25052268_25052851 | chr19 | 25052269 | 25052851 | + | -0.785424 | intron (NM_028785, intron 2 of 47) | intron (NM_028785, intron 2 of 47) | 53031 | NM_028785 | 76088 | Mm.102485 | NM_028785 | ENSMUSG00000052085 | **Dock8** | 1200017A24Rik\|5830472H07Rik\|A130095G14Rik\|AI461977 | dedicator of cytokinesis 8 | protein-coding |
| chr14_121696739_121697512 | chr14 | 121696740 | 121697512 | + | -0.614206 | intron (NM_134074, intron 1 of 52) | intron (NM_134074, intron 1 of 52) | 1291 | NM_001081039 | 105445 | Mm.441054 | NM_134074 | ENSMUSG00000025558 | **Dock9** | AA959601\|AW538057\|B230309H04Rik\|D14Wsu89e\|Zizimin1\|mKIAA1058 | dedicator of cytokinesis 9 | protein-coding |
| chr16_93766606_93766949 | chr16 | 93766607 | 93766949 | + | -0.65269 | intron (NM_027293, intron 18 of 36) | L1MA7\|LINE\|L1 | 54871 | NM_027293 | 70028 | Mm.23230 | NM_026700 | ENSMUSG00000022946 | **Dopey2** | 0610038M01Rik\|2610510B01Rik\|AI503246\|C21orf5\|mKIAA0933 | dopey family member 2 | protein-coding |
| chr3_115899665_115899898 | chr3 | 115899666 | 115899898 | + | -0.613586 | exon (NM_027193, exon 4 of 8) | exon (NM_027193, exon 4 of 8) | 11599 | NM_027193 | 69740 | Mm.5915 | NM_027193 | ENSMUSG00000033554 | **Dph5** | 2410012M04Rik\|AU045680\|C80186 | DPH5 homolog (S. cerevisiae) | protein-coding |
| chr3_115899587_115899919 | chr3 | 115899588 | 115899919 | + | -0.640629 | exon (NM_027193, exon 4 of 8) | exon (NM_027193, exon 4 of 8) | 11570 | NM_027193 | 69740 | Mm.5915 | NM_027193 | ENSMUSG00000033554 | **Dph5** | 2410012M04Rik\|AU045680\|C80186 | DPH5 homolog (S. cerevisiae) | protein-coding |
| chr3_115933323_115934055 | chr3 | 115933324 | 115934055 | + | -0.692117 | Intergenic | Intergenic | 45506 | NM_027193 | 69740 | Mm.5915 | NM_027193 | ENSMUSG00000033554 | **Dph5** | 2410012M04Rik\|AU045680\|C80186 | DPH5 homolog (S. cerevisiae) | protein-coding |
| chr2_114545773_114545994 | chr2 | 114545774 | 114545994 | + | -0.637345 | intron (NM_025675, intron 5 of 8) | intron (NM_025675, intron 5 of 8) | 109044 | NM_025675 | 66632 | Mm.117809 | NM_025675 | ENSMUSG00000057147 | **Dph6** | 5730421E18Rik\|Atpbd4 | diphthamine biosynthesis 6 | protein-coding |
| chr2_62352034_62352387 | chr2 | 62352035 | 62352387 | + | -1.42368 | exon (NM_010074, exon 20 of 26) | exon (NM_010074, exon 20 of 26) | 60020 | NM_010074 | 13482 | Mm.1151 | NM_010074 | ENSMUSG00000035000 | **Dpp4** | Cd26\|Dpp-4\|THAM | dipeptidylpeptidase 4 | protein-coding |
| chr9_24331668_24332057 | chr9 | 24331669 | 24332057 | + | -1.23594 | Intergenic | Lx8b\|LINE\|L1 | 171277 | NM_172920 | 244745 | Mm.38526 | NM_172920 | ENSMUSG00000043067 | **Dpy19l1** | 1100001I19Rik\|6030402K12\|AI225861\|AW558481 | dpy-19-like 1 (C. elegans) | protein-coding |
| chr14_66843026_66843343 | chr14 | 66843027 | 66843343 | + | -0.678129 | intron (NM_009955, intron 3 of 13) | B4\|SINE\|B4 | 25415 | NM_009955 | 12934 | Mm.352648 | NM_009955 | ENSMUSG00000022048 | **Dpysl2** | AI851130\|Crmp2\|DRP2\|Musunc33\|TOAD-64\|Ulip2 | dihydropyrimidinase-like 2 | protein-coding |
| chr7_141302697_141303414 | chr7 | 141302698 | 141303414 | + | -0.853218 | intron (NM_001282072, intron 11 of 12) | intron (NM_001282072, intron 11 of 12) | 11050 | NM_007878 | 13491 | Mm.41075 | NM_007878 | ENSMUSG00000025496 | **Drd4** | AW125663\|D4R\|Drd-4 | dopamine receptor D4 | protein-coding |
| chr15_12923836_12924336 | chr15 | 12923837 | 12924336 | + | -1.02044 | exon (NM_026799, exon 30 of 35) | exon (NM_026799, exon 30 of 35) | 99271 | NM_001130149 | 14000 | Mm.293142 | NM_026799 | ENSMUSG00000022191 | **Drosha** | 1110013A17Rik\|AI874853\|Etohi2\|Rn3\|Rnasen | drosha, ribonuclease type III | protein-coding |
| chr10_34185539_34186262 | chr10 | 34185540 | 34186262 | + | -0.615409 | intron (NM_172508, intron 1 of 5) | intron (NM_172508, intron 1 of 5) | 21650 | NM_172508 | 212898 | Mm.34557 | NM_172508 | ENSMUSG00000039497 | **Dse** | 6030499O08\|AI480506\|B130024B19Rik\|Sart2 | dermatan sulfate epimerase | protein-coding |
| chr10_34235312_34236050 | chr10 | 34235313 | 34236050 | + | -0.686731 | Intergenic | Intergenic | -28130 | NM_172508 | 212898 | Mm.34557 | NM_172508 | ENSMUSG00000039497 | **Dse** | 6030499O08\|AI480506\|B130024B19Rik\|Sart2 | dermatan sulfate epimerase | protein-coding |
| chr10_34171067_34171685 | chr10 | 34171068 | 34171685 | + | -0.740694 | intron (NM_172508, intron 2 of 5) | intron (NM_172508, intron 2 of 5) | 36175 | NM_172508 | 212898 | Mm.34557 | NM_172508 | ENSMUSG00000039497 | **Dse** | 6030499O08\|AI480506\|B130024B19Rik\|Sart2 | dermatan sulfate epimerase | protein-coding |
| chr1_34187502_34187905 | chr1 | 34187503 | 34187905 | + | -1.1072 | intron (NM_133833, intron 35 of 92) | intron (NM_133833, intron 35 of 92) | 27374 | NM_010081 | 13518 | Mm.336625 | NM_010081 | ENSMUSG00000026131 | **Dst** | 2310001O04Rik\|A830042E19Rik\|AW554249\|BP230\|BPAG1-n\|Bpag\|Bpag1\|Macf2\|ah\|athetoid\|dt\|mKIAA0728\|nmf203\|nmf339 | dystonin | protein-coding |
| chr5_62791220_62791570 | chr5 | 62791221 | 62791570 | + | -0.625351 | Intergenic | ORR1E\|LTR\|ERVL-MaLR | -22428 | NM_001170705 | 100322896 | Mm.479821 | NM_001170705 | ENSMUSG00000090326 | **Dthd1** | Gm17384 | death domain containing 1 | protein-coding |
| chr5_62858168_62858897 | chr5 | 62858169 | 62858897 | + | -0.865191 | intron (NM_001170705, intron 7 of 8) | intron (NM_001170705, intron 7 of 8) | 44710 | NM_001170705 | 100322896 | Mm.479821 | NM_001170705 | ENSMUSG00000090326 | **Dthd1** | Gm17384 | death domain containing 1 | protein-coding |
| chr12_3568783_3569085 | chr12 | 3568784 | 3569085 | + | -0.765241 | Intergenic | Intergenic | -3457 | NM_001162465 | 13528 | Mm.286202 | NM_007886 | ENSMUSG00000071454 | **Dtnb** | - | dystrobrevin, beta | protein-coding |
| chr12_3547442_3547887 | chr12 | 3547443 | 3547887 | + | -0.801555 | Intergenic | RLTR19A\|LTR\|ERVK | -24726 | NM_001162465 | 13528 | Mm.286202 | NM_007886 | ENSMUSG00000071454 | **Dtnb** | - | dystrobrevin, beta | protein-coding |
| chr12_3548465_3549031 | chr12 | 3548466 | 3549031 | + | -0.926064 | Intergenic | Intergenic | -23643 | NM_001162465 | 13528 | Mm.286202 | NM_007886 | ENSMUSG00000071454 | **Dtnb** | - | dystrobrevin, beta | protein-coding |
| chr18_49770675_49771551 | chr18 | 49770676 | 49771551 | + | -0.637386 | Intergenic | Intergenic | -15512 | NM_001170960 | 68857 | Mm.96010 | NM_026854 | ENSMUSG00000024505 | **Dtwd2** | 1190002H09Rik\|8030470C17Rik\|AI414673\|BB115449 | DTW domain containing 2 | protein-coding |
| chr5_120685657_120686263 | chr5 | 120685658 | 120686263 | + | -0.683166 | intron (NM_008052, intron 3 of 9) | intron (NM_008052, intron 3 of 9) | 25709 | NM_008052 | 14357 | Mm.1645 | NM_008052 | ENSMUSG00000029603 | **Dtx1** | Fxit1\|mKIAA4160 | deltex 1 homolog (Drosophila) | protein-coding |
| chr5_120688437_120688764 | chr5 | 120688438 | 120688764 | + | -0.817818 | intron (NM_008052, intron 3 of 9) | intron (NM_008052, intron 3 of 9) | 23068 | NM_008052 | 14357 | Mm.1645 | NM_008052 | ENSMUSG00000029603 | **Dtx1** | Fxit1\|mKIAA4160 | deltex 1 homolog (Drosophila) | protein-coding |
| chr5_120696840_120697404 | chr5 | 120696841 | 120697404 | + | -0.840269 | intron (NM_008052, intron 2 of 9) | intron (NM_008052, intron 2 of 9) | 14547 | NM_008052 | 14357 | Mm.1645 | NM_008052 | ENSMUSG00000029603 | **Dtx1** | Fxit1\|mKIAA4160 | deltex 1 homolog (Drosophila) | protein-coding |
| chr2_122306824_122307114 | chr2 | 122306825 | 122307114 | + | -0.68172 | intron (NM_145395, intron 3 of 8) | L2a\|LINE\|L2 | 6761 | NM_145395 | 213696 | Mm.247121 | NM_145395 | ENSMUSG00000027224 | **Duoxa1** | BC019755\|Nip\|Nip1 | dual oxidase maturation factor 1 | protein-coding |
| chr14_21653535_21654731 | chr14 | 21653536 | 21654731 | + | -0.602764 | intron (NM_017479, intron 12 of 16) | intron (NM_017479, intron 12 of 16) | 60443 | NM_001013826 | 435391 | Mm.295860 | NM_001013826 | ENSMUSG00000063821 | **Dupd1** | EG435391 | dual specificity phosphatase and pro isomerase domain containing 1 | protein-coding |
| chr14_21653510_21654220 | chr14 | 21653511 | 21654220 | + | -0.64653 | intron (NM_017479, intron 12 of 16) | B1_Mur2\|SINE\|Alu | 60711 | NM_001013826 | 435391 | Mm.295860 | NM_001013826 | ENSMUSG00000063821 | **Dupd1** | EG435391 | dual specificity phosphatase and pro isomerase domain containing 1 | protein-coding |
| chr14_21653717_21654392 | chr14 | 21653718 | 21654392 | + | -0.675828 | intron (NM_017479, intron 12 of 16) | intron (NM_017479, intron 12 of 16) | 60521 | NM_001013826 | 435391 | Mm.295860 | NM_001013826 | ENSMUSG00000063821 | **Dupd1** | EG435391 | dual specificity phosphatase and pro isomerase domain containing 1 | protein-coding |
| chr14_21653855_21654278 | chr14 | 21653856 | 21654278 | + | -0.703777 | intron (NM_017479, intron 12 of 16) | intron (NM_017479, intron 12 of 16) | 60509 | NM_001013826 | 435391 | Mm.295860 | NM_001013826 | ENSMUSG00000063821 | **Dupd1** | EG435391 | dual specificity phosphatase and pro isomerase domain containing 1 | protein-coding |
| chr17_26507518_26507762 | chr17 | 26507519 | 26507762 | + | -0.689532 | exon (NM_013642, exon 2 of 4) | exon (NM_013642, exon 2 of 4) | 832 | NM_013642 | 19252 | Mm.239041 | NM_013642 | ENSMUSG00000024190 | **Dusp1** | 3CH134\|MKP1\|Ptpn16\|U19515\|erp\|mkp-1 | dual specificity phosphatase 1 | protein-coding |
| chr17_26507540_26507839 | chr17 | 26507541 | 26507839 | + | -0.726327 | exon (NM_013642, exon 2 of 4) | exon (NM_013642, exon 2 of 4) | 782 | NM_013642 | 19252 | Mm.239041 | NM_013642 | ENSMUSG00000024190 | **Dusp1** | 3CH134\|MKP1\|Ptpn16\|U19515\|erp\|mkp-1 | dual specificity phosphatase 1 | protein-coding |
| chr1_184064234_184064761 | chr1 | 184064235 | 184064761 | + | -0.860438 | intron (NM_022019, intron 2 of 3) | intron (NM_022019, intron 2 of 3) | 30037 | NM_022019 | 63953 | Mm.404024 | NM_022019 | ENSMUSG00000039384 | **Dusp10** | 2610306G15Rik\|AI158871\|MKP-5\|MKP5 | dual specificity phosphatase 10 | protein-coding |

| | | | | | | | | | | | | | | | | | |
|---|---|---|---|---|---|---|---|---|---|---|---|---|---|---|---|---|---|
| chr1_184066590_184067145 | chr1 | 184066591 | 184067145 | + | -1.03776 | intron (NM_022019, intron 2 of 3) | intron (NM_022019, intron 2 of 3) | 32407 | NM_022019 | 63953 | Mm.404024 | NM_022019 | ENSMUSG00000039384 | **Dusp10** | 2610306G15Rik\|AI158871\|MKP-5\|MKP5 | dual specificity phosphatase 10 | protein-coding |
| chr13_30681848_30682472 | chr13 | 30681849 | 30682472 | + | -0.970788 | intron (NM_001037955, intron 3 of 6) | intron (NM_001037955, intron 3 of 6) | 22101 | NM_134068 | 105352 | Mm.289646 | NM_134068 | ENSMUSG00000069255 | **Dusp22** | JKAP\|JSP1\|LMW-DSP2\|MKPX | dual specificity phosphatase 22 | protein-coding |
| chr8_34735108_34735314 | chr8 | 34735109 | 34735314 | + | -0.624923 | Intergenic | B2_Mm2\|SINE\|B2 | -72399 | NM_176933 | 319520 | Mm.170276 | NM_176933 | ENSMUSG00000031530 | **Dusp4** | 2700078F24Rik\|AI844617\|BB104621\|E130306H24Rik\|MKP2 | dual specificity phosphatase 4 | protein-coding |
| chr8_34735076_34735453 | chr8 | 34735077 | 34735453 | + | -0.755123 | Intergenic | B2_Mm2\|SINE\|B2 | -72345 | NM_176933 | 319520 | Mm.170276 | NM_176933 | ENSMUSG00000031530 | **Dusp4** | 2700078F24Rik\|AI844617\|BB104621\|E130306H24Rik\|MKP2 | dual specificity phosphatase 4 | protein-coding |
| chr10_99393521_99393957 | chr10 | 99393522 | 99393957 | + | -0.622743 | Intergenic | Intergenic | 130508 | NM_026268 | 67603 | Mm.1791 | NM_026268 | ENSMUSG00000019960 | **Dusp6** | 1300019I03Rik\|MKP-3\|MKP3\|PYST1 | dual specificity phosphatase 6 | protein-coding |
| chr10_99268443_99268733 | chr10 | 99268444 | 99268733 | + | -0.724981 | Intergenic | Intergenic | 5357 | NM_026268 | 67603 | Mm.1791 | NM_026268 | ENSMUSG00000019960 | **Dusp6** | 1300019I03Rik\|MKP-3\|MKP3\|PYST1 | dual specificity phosphatase 6 | protein-coding |
| chr10_99385852_99386351 | chr10 | 99385853 | 99386351 | + | -1.23062 | Intergenic | Intergenic | 122871 | NM_026268 | 67603 | Mm.1791 | NM_026268 | ENSMUSG00000019960 | **Dusp6** | 1300019I03Rik\|MKP-3\|MKP3\|PYST1 | dual specificity phosphatase 6 | protein-coding |
| chr10_58224325_58224505 | chr10 | 58224326 | 58224505 | + | -0.6647 | Intergenic | Intergenic | 8260 | NM_001081954 | 664783 | Mm.482575 | NM_001081954 | | **Dux** | AW822073\|Dux4\|Duxbl\|EG664783 | double homeobox | protein-coding |
| chr10_58231797_58232263 | chr10 | 58231798 | 58232263 | + | -1.10754 | exon (NM_001081954, exon 1 of 1) | exon (NM_001081954, exon 1 of 1) | 645 | NM_001081954 | 664783 | Mm.482575 | NM_001081954 | | **Dux** | AW822073\|Dux4\|Duxbl\|EG664783 | double homeobox | protein-coding |
| chr11_70005508_70006205 | chr11 | 70005509 | 70006205 | + | -0.648948 | intron (NM_007888, intron 5 of 14) | C-rich\|Low_complexity\|Low_complexity | 5231 | NM_007888 | 13543 | Mm.5114 | NM_007888 | ENSMUSG00000020888 | **Dvl2** | - | dishevelled 2, dsh homolog (Drosophila) | protein-coding |
| chr14_41050537_41051068 | chr14 | 41050538 | 41051068 | + | -0.615529 | intron (NM_027717, intron 3 of 3) | intron (NM_027717, intron 3 of 3) | 18271 | NM_027717 | 71200 | Mm.443973 | NM_027717 | ENSMUSG00000021791 | **Dydc2** | 4933428D01Rik | DPY30 domain containing 2 | protein-coding |
| chr18_75031170_75031469 | chr18 | 75031171 | 75031469 | + | -0.970096 | intron (NM_027727, intron 1 of 16) | B3A\|SINE\|B2 | 12548 | NM_027727 | 69190 | Mm.173826 | NM_027727 | ENSMUSG00000035765 | **Dym** | 1810041M12Rik\|4933427L07Rik\|C030019K18Rik | dymeclin | protein-coding |
| chr18_75031197_75031406 | chr18 | 75031198 | 75031406 | + | -1.01286 | intron (NM_027727, intron 1 of 16) | B3A\|SINE\|B2 | 12530 | NM_027727 | 69190 | Mm.173826 | NM_027727 | ENSMUSG00000035765 | **Dym** | 1810041M12Rik\|4933427L07Rik\|C030019K18Rik | dymeclin | protein-coding |
| chr17_84631904_84632192 | chr17 | 84631905 | 84632192 | + | -0.68 | intron (NM_172256, intron 3 of 12) | intron (NM_172256, intron 3 of 12) | 5549 | NM_172256 | 213575 | Mm.246119 | NM_172256 | ENSMUSG00000024253 | **Dync2li1** | 4933404O11Rik\|CGI-60\|D2lic\|LIC3\|mD2LIC | dynein cytoplasmic 2 light intermediate chain 1 | protein-coding |
| chr17_6429755_6430374 | chr17 | 6429756 | 6430374 | + | -0.747219 | promoter-TSS (NM_009342) | promoter-TSS (NM_009342) | -47 | NM_009342 | 21648 | Mm.1948 | NM_009342 | ENSMUSG00000096255 | **Dynlt1b** | AGS2\|Dynlt1\|Tctex-1\|Tctex1 | dynein light chain Tctex-type 1B | protein-coding |
| chr16_94600408_94600648 | chr16 | 94600409 | 94600648 | + | -0.685736 | intron (NM_001113389, intron 1 of 11) | intron (NM_001113389, intron 1 of 11) | 29621 | NM_001113389 | 13548 | Mm.310973 | NM_007890 | ENSMUSG00000022897 | **Dyrk1a** | 2310043O08Rik\|D16Ertd272e\|D16Ertd493e\|Dyrk\|ENSMUSG00000074897\|Gm10783\|Mnbh\|Mp86\|mmb | dual-specificity tyrosine-(Y)-phosphorylation regulated kinase 1a | protein-coding |
| chr16_94637517_94637877 | chr16 | 94637518 | 94637877 | + | -0.818378 | intron (NM_007890, intron 2 of 11) | intron (NM_007890, intron 2 of 11) | 66790 | NM_001113389 | 13548 | Mm.310973 | NM_007890 | ENSMUSG00000022897 | **Dyrk1a** | 2310043O08Rik\|D16Ertd272e\|D16Ertd493e\|Dyrk\|ENSMUSG00000074897\|Gm10783\|Mnbh\|Mp86\|mmb | dual-specificity tyrosine-(Y)-phosphorylation regulated kinase 1a | protein-coding |
| chr10_118816480_118816928 | chr10 | 118816481 | 118816928 | + | -0.631638 | Intergenic | ORR1F\|LTR\|ERVL-MaLR | 52199 | NM_001014390 | 69181 | Mm.45565 | NM_001014390 | ENSMUSG00000028630 | **Dyrk2** | 1810038L18Rik | dual-specificity tyrosine-(Y)-phosphorylation regulated kinase 2 | protein-coding |
| chr10_118813616_118814103 | chr10 | 118813617 | 118814103 | + | -0.639414 | Intergenic | B1_Mur4\|SINE\|Alu | 55043 | NM_001014390 | 69181 | Mm.45565 | NM_001014390 | ENSMUSG00000028630 | **Dyrk2** | 1810038L18Rik | dual-specificity tyrosine-(Y)-phosphorylation regulated kinase 2 | protein-coding |
| chr10_118882336_118883463 | chr10 | 118882337 | 118883463 | + | -0.689397 | Intergenic | Intergenic | -13997 | NM_001014390 | 69181 | Mm.45565 | NM_001014390 | ENSMUSG00000028630 | **Dyrk2** | 1810038L18Rik | dual-specificity tyrosine-(Y)-phosphorylation regulated kinase 2 | protein-coding |
| chr10_118758691_118759131 | chr10 | 118758692 | 118759131 | + | -0.697244 | Intergenic | Intergenic | 109992 | NM_001014390 | 69181 | Mm.45565 | NM_001014390 | ENSMUSG00000028630 | **Dyrk2** | 1810038L18Rik | dual-specificity tyrosine-(Y)-phosphorylation regulated kinase 2 | protein-coding |
| chr10_118714040_118714879 | chr10 | 118714041 | 118714879 | + | -0.759183 | Intergenic | Intergenic | 154443 | NM_001014390 | 69181 | Mm.45565 | NM_001014390 | ENSMUSG00000028630 | **Dyrk2** | 1810038L18Rik | dual-specificity tyrosine-(Y)-phosphorylation regulated kinase 2 | protein-coding |
| chr10_118721923_118722247 | chr10 | 118721924 | 118722247 | + | -0.962501 | Intergenic | Intergenic | 146818 | NM_001014390 | 69181 | Mm.45565 | NM_001014390 | ENSMUSG00000028630 | **Dyrk2** | 1810038L18Rik | dual-specificity tyrosine-(Y)-phosphorylation regulated kinase 2 | protein-coding |
| chr10_118721789_118722254 | chr10 | 118721790 | 118722254 | + | -0.975183 | Intergenic | Intergenic | 146881 | NM_001014390 | 69181 | Mm.45565 | NM_001014390 | ENSMUSG00000028630 | **Dyrk2** | 1810038L18Rik | dual-specificity tyrosine-(Y)-phosphorylation regulated kinase 2 | protein-coding |
| chr10_118884070_118884446 | chr10 | 118884071 | 118884446 | + | -1.20061 | Intergenic | B4A\|SINE\|B4 | -15355 | NM_001014390 | 69181 | Mm.45565 | NM_001014390 | ENSMUSG00000028630 | **Dyrk2** | 1810038L18Rik | dual-specificity tyrosine-(Y)-phosphorylation regulated kinase 2 | protein-coding |
| chr10_118712838_118713331 | chr10 | 118712839 | 118713331 | + | -1.29434 | Intergenic | Intergenic | 155818 | NM_001014390 | 69181 | Mm.45565 | NM_001014390 | ENSMUSG00000028630 | **Dyrk2** | 1810038L18Rik | dual-specificity tyrosine-(Y)-phosphorylation regulated kinase 2 | protein-coding |
| chr10_118712984_118713288 | chr10 | 118712985 | 118713288 | + | -1.33912 | Intergenic | Intergenic | 155767 | NM_001014390 | 69181 | Mm.45565 | NM_001014390 | ENSMUSG00000028630 | **Dyrk2** | 1810038L18Rik | dual-specificity tyrosine-(Y)-phosphorylation regulated kinase 2 | protein-coding |
| chr9_72969800_72970370 | chr9 | 72969801 | 72970370 | + | -0.632249 | intron (NM_001163725, intron 5 of 6) | intron (NM_001163725, intron 5 of 6) | 11300 | NM_001163725 | 67685 | Mm.31817 | NM_026314 | ENSMUSG00000092192 | **Dyx1c1** | 1700010I24Rik\|EKN1\|b2b811Clo | dyslexia susceptibility 1 candidate 1 homolog (human) | protein-coding |
| chr14_118920702_118921025 | chr14 | 118920703 | 118921025 | + | -0.645559 | intron (NM_025943, intron 2 of 19) | intron (NM_025943, intron 2 of 19) | 2303 | NM_025943 | 66573 | Mm.87456 | NM_025943 | ENSMUSG00000042156 | **Dzip1** | 2510025K24Rik\|2810422M04Rik\|mKIAA0996 | DAZ interacting protein 1 | protein-coding |
| chr7_63815967_63817031 | chr7 | 63815968 | 63817031 | + | -0.61302 | Intergenic | Intergenic | -100358 | NM_001256311 | 381994 | Mm.425466 | NM_001256311 | ENSMUSG00000095061 | **E030018B13Rik** | Gm1697 | RIKEN cDNA E030018B13 gene | protein-coding |
| chr7_63925338_63926039 | chr7 | 63925339 | 63926039 | + | -0.625632 | intron (NM_021366, intron 1 of 1) | intron (NM_021366, intron 1 of 1) | 8832 | NM_001256311 | 381994 | Mm.425466 | NM_001256311 | ENSMUSG00000095061 | **E030018B13Rik** | Gm1697 | RIKEN cDNA E030018B13 gene | protein-coding |
| chr7_63921546_63922378 | chr7 | 63921547 | 63922378 | + | -0.663317 | intron (NM_021366, intron 1 of 1) | intron (NM_021366, intron 1 of 1) | 5105 | NM_001256311 | 381994 | Mm.425466 | NM_001256311 | ENSMUSG00000095061 | **E030018B13Rik** | Gm1697 | RIKEN cDNA E030018B13 gene | protein-coding |
| chr7_63921968_63922247 | chr7 | 63921969 | 63922247 | + | -0.969622 | intron (NM_021366, intron 1 of 1) | intron (NM_021366, intron 1 of 1) | 5251 | NM_001256311 | 381994 | Mm.425466 | NM_001256311 | ENSMUSG00000095061 | **E030018B13Rik** | Gm1697 | RIKEN cDNA E030018B13 gene | protein-coding |
| chr11_109218245_109219585 | chr11 | 109218246 | 109219585 | + | -0.899443 | Intergenic | ORR1E\|LTR\|ERVL-MaLR | -74546 | NR_037978 | 268498 | Mm.227970 | NM_001085512 | ENSMUSG00000078605 | **E030025P04Rik** | Gm664 | RIKEN cDNA E030025P04 gene | ncRNA |
| chr10_22150581_22150982 | chr10 | 22150582 | 22150982 | + | -0.654208 | Intergenic | CpG | -1512 | NM_001254745 | 319887 | Mm.177746 | NM_001254744 | ENSMUSG00000055657 | **E030030I06Rik** | - | RIKEN cDNA E030030I06 gene | protein-coding |
| chr4_97673595_97674525 | chr4 | 97673596 | 97674525 | + | -0.663735 | intron (NM_001122952, intron 1 of 10) | intron (NM_001122952, intron 1 of 10) | -89469 | NR_015513 | 319865 | Mm.373807 | NM_177023 | | **E130114P18Rik** | - | RIKEN cDNA E130114P18 gene | ncRNA |
| chr4_97657461_97657953 | chr4 | 97657462 | 97657953 | + | -0.976876 | intron (NM_001122952, intron 1 of 10) | intron (NM_001122952, intron 1 of 10) | -73116 | NR_015513 | 319865 | Mm.373807 | NM_177023 | | **E130114P18Rik** | - | RIKEN cDNA E130114P18 gene | ncRNA |
| chr2_150666789_150668640 | chr2 | 150666790 | 150668640 | + | -0.702386 | non-coding (NR_040331, exon 1 of 2) | non-coding (NR_040331, exon 1 of 2) | 221 | NR_040331 | 109344 | Mm.440012 | NR_040331 | ENSMUSG00000087406 | **E130215H24Rik** | - | RIKEN cDNA E130215H24 gene | ncRNA |
| chr2_150667031_150667781 | chr2 | 150667032 | 150667781 | + | -0.885171 | promoter-TSS (NR_040331) | promoter-TSS (NR_040331) | -88 | NR_040331 | 109344 | Mm.440012 | NR_040331 | ENSMUSG00000087406 | **E130215H24Rik** | - | RIKEN cDNA E130215H24 gene | ncRNA |
| chr7_48898438_48898676 | chr7 | 48898439 | 48898676 | + | -0.677273 | Intergenic | ORR1D2\|LTR\|ERVL-MaLR | -17516 | NM_001013368 | 108961 | Mm.240566 | NM_001013368 | ENSMUSG00000046179 | **E2f8** | 4432406C08Rik\|AA410048 | E2F transcription factor 8 | protein-coding |
| chr6_40430196_40430550 | chr6 | 40430197 | 40430550 | + | -0.763388 | intron (NM_175528, intron 1 of 7) | intron (NM_175528, intron 1 of 7) | 5760 | NM_175528 | 243780 | Mm.333006 | NM_175528 | ENSMUSG00000037172 | **E330009J07Rik** | AI841796\|Lchn\|mKIAA1147 | RIKEN cDNA E330009J07 gene | protein-coding |
| chr6_40430326_40430545 | chr6 | 40430327 | 40430545 | + | -0.79641 | intron (NM_175528, intron 1 of 7) | intron (NM_175528, intron 1 of 7) | 5697 | NM_175528 | 243780 | Mm.333006 | NM_175528 | ENSMUSG00000037172 | **E330009J07Rik** | AI841796\|Lchn\|mKIAA1147 | RIKEN cDNA E330009J07 gene | protein-coding |
| chr1_153379895_153380286 | chr1 | 153379896 | 153380286 | + | -0.674853 | Intergenic | Intergenic | 34142 | NR_033736 | 626058 | Mm.31556 | NM_001177359 | | **E330020D12Rik** | EG626058\|F430003O07Rik\|Gm6648 | Riken cDNA E330020D12 gene | ncRNA |
| chr1_153379706_153380149 | chr1 | 153379707 | 153380149 | + | -0.743936 | Intergenic | Intergenic | 34305 | NR_033736 | 626058 | Mm.31556 | NM_001177359 | | **E330020D12Rik** | EG626058\|F430003O07Rik\|Gm6648 | Riken cDNA E330020D12 gene | ncRNA |
| chr9_121757445_121757828 | chr9 | 121757446 | 121757828 | + | -0.794319 | TTS (NM_010918) | TTS (NM_010918) | 2306 | NR_033503 | 320301 | Mm.373883 | NR_033503 | | **E530011L22Rik** | - | RIKEN cDNA E530011L22 gene | ncRNA |
| chr9_121757494_121757726 | chr9 | 121757495 | 121757726 | + | -0.844028 | TTS (NM_010918) | TTS (NM_010918) | 2333 | NR_033503 | 320301 | Mm.373883 | NR_033503 | | **E530011L22Rik** | - | RIKEN cDNA E530011L22 gene | ncRNA |
| chr15_44620985_44621216 | chr15 | 44620986 | 44621216 | + | -0.638999 | intron (NM_019480, intron 1 of 6) | intron (NM_019480, intron 1 of 6) | 1460 | NM_019480 | 55960 | Mm.287896 | NM_019480 | ENSMUSG00000022339 | **Ebag9** | AI835379\|Rcas1 | estrogen receptor-binding fragment-associated gene 9 | protein-coding |

| | | | | | | | | | | | | | | | | | |
|---|---|---|---|---|---|---|---|---|---|---|---|---|---|---|---|---|---|
| chr2_6256113_6257092 | chr2 | 6256114 | 6257092 | + | -0.646697 | intron (NR_040333, intron 3 of 6) | intron (NR_040333, intron 3 of 6) | -43609 | NM_024208 | 67856 | Mm.38342 | NM_024208 | ENSMUSG00000039063 | **Echdc3** | 2310005D12Rik\|AI662097 | enoyl Coenzyme A hydratase domain containing 3 | protein-coding |
| chr13_34956968_34957320 | chr13 | 34956969 | 34957320 | + | -0.913892 | intron (NM_026947, intron 3 of 8) | intron (NM_026947, intron 3 of 8) | 6665 | NM_026947 | 69123 | Mm.482475 | NM_026947 | ENSMUSG00000021416 | **Eci3** | 1810022C23Rik\|Kseg2 | enoyl-Coenzyme A delta isomerase 3 | protein-coding |
| chr10_18197009_18197833 | chr10 | 18197010 | 18197833 | + | -0.781808 | intron (NM_001195036, intron 3 of 20) | intron (NM_001195036, intron 3 of 20) | 13469 | NM_001195036 | 100045792 | Mm.380617 | NM_001195036 | ENSMUSG00000071392 | **Ect2l** | C330021H03Rik\|ENSMUSG00000071392\|Gm10331 | epithelial cell transforming sequence 2 oncogene-like | protein-coding |
| chr10_18197009_18197618 | chr10 | 18197010 | 18197618 | + | -0.811675 | intron (NM_001195036, intron 3 of 20) | intron (NM_001195036, intron 3 of 20) | 13576 | NM_001195036 | 100045792 | Mm.380617 | NM_001195036 | ENSMUSG00000071392 | **Ect2l** | C330021H03Rik\|ENSMUSG00000071392\|Gm10331 | epithelial cell transforming sequence 2 oncogene-like | protein-coding |
| chr10_18193119_18193763 | chr10 | 18193120 | 18193763 | + | -1.48198 | intron (NM_001195036, intron 3 of 20) | intron (NM_001195036, intron 3 of 20) | 17449 | NM_001195036 | 100045792 | Mm.380617 | NM_001195036 | ENSMUSG00000071392 | **Ect2l** | C330021H03Rik\|ENSMUSG00000071392\|Gm10331 | epithelial cell transforming sequence 2 oncogene-like | protein-coding |
| chr13_12507179_12507833 | chr13 | 12507180 | 12507833 | + | -0.83247 | intron (NM_133643, intron 2 of 5) | intron (NM_133643, intron 2 of 5) | 12944 | NM_133643 | 171211 | Mm.159671 | NM_133643 | ENSMUSG00000095105 | **Edaradd** | 1810032E07Rik\|5830469M23Rik\|A630043P06\|cr | EDAR (ectodysplasin-A receptor)-associated death domain | protein-coding |
| chr6_108833071_108833277 | chr6 | 108833072 | 108833277 | + | -0.757964 | intron (NM_138677, intron 1 of 11) | ID_B1\|SINE\|B4 | 4533 | NM_138677 | 192193 | Mm.21596 | NM_138677 | ENSMUSG00000030104 | **Edem1** | A130059K23Rik\|EDEM\|mKIAA0212 | ER degradation enhancer, mannosidase alpha-like 1 | protein-coding |
| chr1_151682897_151683337 | chr1 | 151682898 | 151683337 | + | -0.792148 | intron (NM_022018, intron 5 of 13) | intron (NM_022018, intron 5 of 13) | -72257 | NM_001039644 | 66967 | Mm.337691 | NM_001039644 | ENSMUSG00000043019 | **Edem3** | 2310050N11Rik | ER degradation enhancer, mannosidase alpha-like 3 | protein-coding |
| chr10_95779644_95780191 | chr10 | 95779645 | 95780191 | + | -0.938065 | Intergenic | Intergenic | -160745 | NM_001001932 | 216238 | Mm.210035 | NM_001001932 | ENSMUSG00000036499 | **Eea1** | A430109M19Rik\|B230358H09Rik\|ZFYVE2 | early endosome antigen 1 | protein-coding |
| chr10_95779655_95780027 | chr10 | 95779656 | 95780027 | + | -0.962194 | Intergenic | Intergenic | -160822 | NM_001001932 | 216238 | Mm.210035 | NM_001001932 | ENSMUSG00000036499 | **Eea1** | A430109M19Rik\|B230358H09Rik\|ZFYVE2 | early endosome antigen 1 | protein-coding |
| chr6_88442423_88443083 | chr6 | 88442424 | 88443083 | + | -0.88213 | intron (NM_023060, intron 1 of 6) | intron (NM_023060, intron 1 of 6) | 3786 | NM_023060 | 65967 | Mm.333237 | NM_023060 | ENSMUSG00000033216 | **Eefsec** | Selb\|sec | eukaryotic elongation factor, selenocysteine-tRNA-specific | protein-coding |
| chr6_88442511_88442861 | chr6 | 88442512 | 88442861 | + | -1.16679 | intron (NM_023060, intron 1 of 6) | intron (NM_023060, intron 1 of 6) | 3853 | NM_023060 | 65967 | Mm.333237 | NM_023060 | ENSMUSG00000033216 | **Eefsec** | Selb\|sec | eukaryotic elongation factor, selenocysteine-tRNA-specific | protein-coding |
| chrX_17365126_17365712 | chrX | 17365127 | 17365712 | + | -0.810178 | Intergenic | Intergenic | -46051 | NM_028916 | 74405 | Mm.248531 | NM_028916 | ENSMUSG00000025038 | **Efhc2** | 4933407D04Rik\|mRib72-2 | EF-hand domain (C-terminal) containing 2 | protein-coding |
| chrX_17365096_17365509 | chrX | 17365097 | 17365509 | + | -0.8676 | Intergenic | Intergenic | -45935 | NM_028916 | 74405 | Mm.248531 | NM_028916 | ENSMUSG00000025038 | **Efhc2** | 4933407D04Rik\|mRib72-2 | EF-hand domain (C-terminal) containing 2 | protein-coding |
| chr15_65726844_65727542 | chr15 | 65726845 | 65727542 | + | -0.860634 | Intergenic | Intergenic | -59848 | NM_133766 | 76740 | Mm.260647 | NM_133766 | ENSMUSG00000015002 | **Efr3a** | A130089M23Rik\|BB071175\|C76891\|C920006C10Rik\|D030063F01Rik\|mKIAA0143 | EFR3 homolog A (S. cerevisiae) | protein-coding |
| chr15_65727071_65727538 | chr15 | 65727072 | 65727538 | + | -0.8779 | Intergenic | Intergenic | -59736 | NM_133766 | 76740 | Mm.260647 | NM_133766 | ENSMUSG00000015002 | **Efr3a** | A130089M23Rik\|BB071175\|C76891\|C920006C10Rik\|D030063F01Rik\|mKIAA0143 | EFR3 homolog A (S. cerevisiae) | protein-coding |
| chr15_65731273_65731669 | chr15 | 65731274 | 65731669 | + | -1.07561 | Intergenic | MLT1A\|LTR\|ERVL-MaLR | -55570 | NM_133766 | 76740 | Mm.260647 | NM_133766 | ENSMUSG00000015002 | **Efr3a** | A130089M23Rik\|BB071175\|C76891\|C920006C10Rik\|D030063F01Rik\|mKIAA0143 | EFR3 homolog A (S. cerevisiae) | protein-coding |
| chr14_70010640_70011087 | chr14 | 70010641 | 70011087 | + | -0.712359 | intron (NM_028560, intron 4 of 6) | intron (NM_028560, intron 4 of 6) | -66581 | NM_018781 | 13655 | Mm.103737 | NM_018781 | ENSMUSG00000033730 | **Egr3** | Pilot | early growth response 3 | protein-coding |
| chr17_73839088_73839369 | chr17 | 73839089 | 73839369 | + | -0.945812 | Intergenic | Intergenic | 34388 | NM_020578 | 57440 | Mm.18526 | NM_020578 | ENSMUSG00000024065 | **Ehd3** | Ehd2 | EH-domain containing 3 | protein-coding |
| chr10_82895308_82895766 | chr10 | 82895309 | 82895766 | + | -0.714833 | intron (NM_001042513, intron 14 of 14) | intron (NM_001042513, intron 14 of 14) | 28911 | NM_025499 | 66341 | Mm.443284 | NM_025499 | | **Eid3** | 1700027M21Rik\|NS4EB | EP300 interacting inhibitor of differentiation 3 | protein-coding |
| chr10_82890916_82891315 | chr10 | 82890917 | 82891315 | + | -0.895491 | intron (NM_001042513, intron 14 of 14) | intron (NM_001042513, intron 14 of 14) | 24490 | NM_025499 | 66341 | Mm.443284 | NM_025499 | | **Eid3** | 1700027M21Rik\|NS4EB | EP300 interacting inhibitor of differentiation 3 | protein-coding |
| chr10_82887322_82887744 | chr10 | 82887323 | 82887744 | + | -1.00125 | intron (NM_001042513, intron 13 of 14) | intron (NM_001042513, intron 13 of 14) | 20907 | NM_025499 | 66341 | Mm.443284 | NM_025499 | | **Eid3** | 1700027M21Rik\|NS4EB | EP300 interacting inhibitor of differentiation 3 | protein-coding |
| chr9_120476375_120476637 | chr9 | 120476376 | 120476637 | + | -0.76472 | Intergenic | Intergenic | -16100 | NM_026892 | 68969 | Mm.28753 | NM_026892 | ENSMUSG00000006941 | **Eif1b** | 1500010M16Rik\|3110001N14Rik\|Gc20 | eukaryotic translation initiation factor 1B | protein-coding |
| chr6_70824455_70825368 | chr6 | 70824456 | 70825368 | + | -0.67673 | Intergenic | Intergenic | -19615 | NM_010121 | 13666 | Mm.247167 | NM_010121 | ENSMUSG00000031668 | **Eif2ak3** | AI427929\|Pek\|Perk | eukaryotic translation initiation factor 2 alpha kinase 3 | protein-coding |
| chr15_51871069_51871461 | chr15 | 51871070 | 51871461 | + | -0.603896 | Intergenic | ORR1E\|LTR\|ERVL-MaLR | -5804 | NM_080635 | 68135 | Mm.289800 | NM_080635 | ENSMUSG00000022312 | **Eif3h** | 1110008A16Rik\|40kD\|9430017H16Rik\|EIF3-P40\|EIF3-gamma\|Eif3s3 | eukaryotic translation initiation factor 3, subunit H | protein-coding |
| chr15_51736659_51737326 | chr15 | 51736660 | 51737326 | + | -0.655251 | Intergenic | Intergenic | 128468 | NM_080635 | 68135 | Mm.289800 | NM_080635 | ENSMUSG00000022312 | **Eif3h** | 1110008A16Rik\|40kD\|9430017H16Rik\|EIF3-P40\|EIF3-gamma\|Eif3s3 | eukaryotic translation initiation factor 3, subunit H | protein-coding |
| chr11_69671126_69671783 | chr11 | 69671127 | 69671783 | + | -0.680368 | intron (NM_144958, intron 1 of 10) | CpG | 968 | NM_144958 | 13681 | Mm.279821 | NM_144958 | ENSMUSG00000059796 | **Eif4a1** | BM-010\|Ddx2a\|Eif4 | eukaryotic translation initiation factor 4A1 | protein-coding |
| chr6_99646455_99646690 | chr6 | 99646456 | 99646690 | + | -0.694294 | intron (NM_025829, intron 2 of 6) | intron (NM_025829, intron 2 of 6) | 20198 | NM_025829 | 66892 | Mm.32889 | NM_025829 | ENSMUSG00000093661 | **Eif4e3** | 1300018P11Rik\|AI451927\|eIF4E-3 | eukaryotic translation initiation factor 4E member 3 | protein-coding |
| chr10_61451513_61451896 | chr10 | 61451514 | 61451896 | + | -0.944302 | intron (NM_010124, intron 1 of 2) | intron (NM_010124, intron 1 of 2) | 964 | NM_010124 | 13688 | Mm.259516 | NM_010124 | ENSMUSG00000020091 | **Eif4ebp2** | 2810011I19Rik\|4E-BP2\|AA792569\|BC010348\|PHAS-II | eukaryotic translation initiation factor 4E binding protein 2 | protein-coding |
| chr10_61451615_61451861 | chr10 | 61451616 | 61451861 | + | -0.946616 | intron (NM_010124, intron 1 of 2) | intron (NM_010124, intron 1 of 2) | 931 | NM_010124 | 13688 | Mm.259516 | NM_010124 | ENSMUSG00000020091 | **Eif4ebp2** | 2810011I19Rik\|4E-BP2\|AA792569\|BC010348\|PHAS-II | eukaryotic translation initiation factor 4E binding protein 2 | protein-coding |
| chr8_4313406_4314025 | chr8 | 4313407 | 4314025 | + | -0.645271 | intron (NM_010485, intron 1 of 5) | intron (NM_010485, intron 1 of 5) | 11384 | NM_010485 | 15568 | Mm.119162 | NM_010485 | ENSMUSG00000040028 | **Elavl1** | 2410055N02Rik\|HUR\|Hua\|W91709 | ELAV (embryonic lethal, abnormal vision)-like 1 (Hu antigen R) | protein-coding |
| chr8_4313566_4314025 | chr8 | 4313567 | 4314025 | + | -0.667857 | intron (NM_010485, intron 1 of 5) | MER20\|DNA\|hAT-Charlie | 11304 | NM_010485 | 15568 | Mm.119162 | NM_010485 | ENSMUSG00000040028 | **Elavl1** | 2410055N02Rik\|HUR\|Hua\|W91709 | ELAV (embryonic lethal, abnormal vision)-like 1 (Hu antigen R) | protein-coding |
| chr14_79530048_79530760 | chr14 | 79530049 | 79530760 | + | -0.681631 | intron (NM_001286411, intron 1 of 8) | MIR3\|SINE\|MIR | 14730 | NM_001286411 | 13709 | Mm.20938 | NM_007920 | ENSMUSG00000036461 | **Elf1** | Elf-1\|Sts1\|mElf-1\|p70 | E74-like factor 1 | protein-coding |
| chr14_79503513_79504186 | chr14 | 79503514 | 79504186 | + | -0.711866 | intron (NM_007920, intron 1 of 8) | MTE2a\|LTR\|ERVL-MaLR | -11824 | NM_001286412 | 13709 | Mm.20938 | NM_007920 | ENSMUSG00000036461 | **Elf1** | Elf-1\|Sts1\|mElf-1\|p70 | E74-like factor 1 | protein-coding |
| chr14_79530052_79530466 | chr14 | 79530053 | 79530466 | + | -0.730759 | intron (NM_001286411, intron 1 of 8) | ID4_\|SINE\|ID | 14585 | NM_001286411 | 13709 | Mm.20938 | NM_007920 | ENSMUSG00000036461 | **Elf1** | Elf-1\|Sts1\|mElf-1\|p70 | E74-like factor 1 | protein-coding |
| chr14_79526136_79526408 | chr14 | 79526137 | 79526408 | + | -0.74912 | intron (NM_001286411, intron 1 of 8) | intron (NM_001286411, intron 1 of 8) | 10598 | NM_001286411 | 13709 | Mm.20938 | NM_007920 | ENSMUSG00000036461 | **Elf1** | Elf-1\|Sts1\|mElf-1\|p70 | E74-like factor 1 | protein-coding |
| chr14_79532031_79532472 | chr14 | 79532032 | 79532472 | + | -0.786603 | intron (NM_001286411, intron 1 of 8) | intron (NM_001286411, intron 1 of 8) | 16578 | NM_001286411 | 13709 | Mm.20938 | NM_007920 | ENSMUSG00000036461 | **Elf1** | Elf-1\|Sts1\|mElf-1\|p70 | E74-like factor 1 | protein-coding |
| chr14_79521794_79522156 | chr14 | 79521795 | 79522156 | + | -0.961594 | intron (NM_001286411, intron 1 of 8) | intron (NM_001286411, intron 1 of 8) | 6301 | NM_001286411 | 13709 | Mm.20938 | NM_007920 | ENSMUSG00000036461 | **Elf1** | Elf-1\|Sts1\|mElf-1\|p70 | E74-like factor 1 | protein-coding |
| chr14_79521862_79522072 | chr14 | 79521863 | 79522072 | + | -0.994238 | intron (NM_001286411, intron 1 of 8) | intron (NM_001286411, intron 1 of 8) | 6293 | NM_001286411 | 13709 | Mm.20938 | NM_007920 | ENSMUSG00000036461 | **Elf1** | Elf-1\|Sts1\|mElf-1\|p70 | E74-like factor 1 | protein-coding |
| chrX_48451899_48452215 | chrX | 48451900 | 48452215 | + | -0.770843 | intron (NM_019680, intron 1 of 8) | intron (NM_019680, intron 1 of 8) | 11075 | NM_019680 | 56501 | Mm.36808 | NM_019680 | ENSMUSG00000031103 | **Elf4** | AV314029\|BC042423\|ENSMUSG00000053512\|Gm9907\|Mef | E74-like factor 4 (ets domain transcription factor) | protein-coding |
| chr5_139930884_139931341 | chr5 | 139930885 | 139931341 | + | -0.634068 | intron (NM_175522, intron 1 of 1) | intron (NM_175522, intron 1 of 1) | 23170 | NM_175522 | 243312 | Mm.237102 | NM_175522 | ENSMUSG00000048988 | **Elfn1** | A930017N06Rik\|AI197272\|Ppp1r28 | leucine rich repeat and fibronectin type III, extracellular 1 | protein-coding |
| chr5_139931054_139931585 | chr5 | 139931055 | 139931585 | + | -0.742872 | intron (NM_175522, intron 1 of 1) | intron (NM_175522, intron 1 of 1) | 23377 | NM_175522 | 243312 | Mm.237102 | NM_175522 | ENSMUSG00000048988 | **Elfn1** | A930017N06Rik\|AI197272\|Ppp1r28 | leucine rich repeat and fibronectin type III, extracellular 1 | protein-coding |
| chr15_78683415_78684110 | chr15 | 78683416 | 78684110 | + | -1.1078 | intron (NM_183141, intron 1 of 1) | intron (NM_183141, intron 1 of 1) | 34350 | NM_183141 | 207393 | Mm.323188 | NM_183141 | ENSMUSG00000043460 | **Elfn2** | 6330514E13\|AW048948\|BC094219\|Lrrc62\|Ppp1r29 | leucine rich repeat and fibronectin type III, extracellular 2 | protein-coding |

| | | | | | | | | | | | | | | | | | |
|---|---|---|---|---|---|---|---|---|---|---|---|---|---|---|---|---|---|
| chr15_78683460_78683909 | chr15 | 78683461 | 78683909 | + | -1.1333 | intron (NM_183141, intron 1 of 1) | intron (NM_183141, intron 1 of 1) | 34428 | NM_183141 | 207393 | Mm.323188 | NM_183141 | ENSMUSG00000043460 | **Elfn2** | 6330514E13\|AW048948\|BC094219\|Lrrc62\|Ppp1r29 | leucine rich repeat and fibronectin type III, extracellular 2 | protein-coding |
| chr15_78683648_78683926 | chr15 | 78683649 | 78683926 | + | -1.17214 | intron (NM_183141, intron 1 of 1) | intron (NM_183141, intron 1 of 1) | 34326 | NM_183141 | 207393 | Mm.323188 | NM_183141 | ENSMUSG00000043460 | **Elfn2** | 6330514E13\|AW048948\|BC094219\|Lrrc62\|Ppp1r29 | leucine rich repeat and fibronectin type III, extracellular 2 | protein-coding |
| chr8_70565512_70565995 | chr8 | 70565513 | 70565995 | + | -0.635389 | intron (NM_007924, intron 1 of 11) | intron (NM_007924, intron 1 of 11) | 26079 | NM_007924 | 13716 | Mm.271973 | NM_007924 | ENSMUSG00000070002 | **Ell** | Ell1\|Men | elongation factor RNA polymerase II | protein-coding |
| chr13_20502539_20503076 | chr13 | 20502540 | 20503076 | + | -0.651417 | intron (NM_198093, intron 1 of 6) | intron (NM_198093, intron 1 of 6) | 29818 | NM_198093 | 140580 | Mm.342392 | NM_080288 | ENSMUSG00000041112 | **Elmo1** | 6330578D22Rik\|C230095H21Rik\|CED-12 | engulfment and cell motility 1 | protein-coding |
| chr13_20538232_20538886 | chr13 | 20538233 | 20538886 | + | -0.898176 | intron (NM_198093, intron 1 of 6) | (T)n\|Simple_repeat\|Simple_repeat | 65569 | NM_198093 | 140580 | Mm.342392 | NM_080288 | ENSMUSG00000041112 | **Elmo1** | 6330578D22Rik\|C230095H21Rik\|CED-12 | engulfment and cell motility 1 | protein-coding |
| chr12_84177312_84177617 | chr12 | 84177313 | 84177617 | + | -0.699694 | intron (NM_001163501, intron 1 of 12) | intron (NM_001163501, intron 1 of 12) | 16550 | NM_001163502 | 238317 | Mm.31256 | NM_001163501 | ENSMUSG00000042507 | **Elmsan1** | 9430029N19Rik\|C130039O16Rik\|Gm260 | ELM2 and Myb/SANT-like domain containing 1 | protein-coding |
| chr12_84177285_84177810 | chr12 | 84177286 | 84177810 | + | -0.703506 | intron (NM_001163501, intron 1 of 12) | intron (NM_001163501, intron 1 of 12) | 16467 | NM_001163502 | 238317 | Mm.31256 | NM_001163501 | ENSMUSG00000042507 | **Elmsan1** | 9430029N19Rik\|C130039O16Rik\|Gm260 | ELM2 and Myb/SANT-like domain containing 1 | protein-coding |
| chr12_84206494_84207000 | chr12 | 84206495 | 84207000 | + | -0.760634 | intron (NM_001163501, intron 1 of 12) | intron (NM_001163501, intron 1 of 12) | 12134 | NM_001163501 | 238317 | Mm.31256 | NM_001163501 | ENSMUSG00000042507 | **Elmsan1** | 9430029N19Rik\|C130039O16Rik\|Gm260 | ELM2 and Myb/SANT-like domain containing 1 | protein-coding |
| chr12_84209099_84209470 | chr12 | 84209100 | 84209470 | + | -0.859095 | intron (NM_001163501, intron 1 of 12) | intron (NM_001163501, intron 1 of 12) | 9596 | NM_001163501 | 238317 | Mm.31256 | NM_001163501 | ENSMUSG00000042507 | **Elmsan1** | 9430029N19Rik\|C130039O16Rik\|Gm260 | ELM2 and Myb/SANT-like domain containing 1 | protein-coding |
| chr12_84242973_84243732 | chr12 | 84242974 | 84243732 | + | -0.866255 | Intergenic | Intergenic | -24472 | NM_001163501 | 238317 | Mm.31256 | NM_001163501 | ENSMUSG00000042507 | **Elmsan1** | 9430029N19Rik\|C130039O16Rik\|Gm260 | ELM2 and Myb/SANT-like domain containing 1 | protein-coding |
| chr12_84222652_84222940 | chr12 | 84222653 | 84222940 | + | -0.957468 | Intergenic | Intergenic | -3915 | NM_001163501 | 238317 | Mm.31256 | NM_001163501 | ENSMUSG00000042507 | **Elmsan1** | 9430029N19Rik\|C130039O16Rik\|Gm260 | ELM2 and Myb/SANT-like domain containing 1 | protein-coding |
| chr12_84222652_84223096 | chr12 | 84222653 | 84223096 | + | -1.01341 | Intergenic | Intergenic | -3993 | NM_001163501 | 238317 | Mm.31256 | NM_001163501 | ENSMUSG00000042507 | **Elmsan1** | 9430029N19Rik\|C130039O16Rik\|Gm260 | ELM2 and Myb/SANT-like domain containing 1 | protein-coding |
| chr12_84221032_84221484 | chr12 | 84221033 | 84221484 | + | -1.24828 | Intergenic | MLT1E1A\|LTR\|ERVL-MaLR | -2377 | NM_001163501 | 238317 | Mm.31256 | NM_001163501 | ENSMUSG00000042507 | **Elmsan1** | 9430029N19Rik\|C130039O16Rik\|Gm260 | ELM2 and Myb/SANT-like domain containing 1 | protein-coding |
| chr9_77941203_77941414 | chr9 | 77941204 | 77941414 | + | -0.654526 | intron (NM_134255, intron 1 of 7) | intron (NM_134255, intron 1 of 7) | 23944 | NM_134255 | 68801 | Mm.430736 | NM_134255 | ENSMUSG00000032349 | **Elovl5** | 1110059L23Rik\|AI747313\|AU043003\|HELO1 | ELOVL family member 5, elongation of long chain fatty acids (yeast) | protein-coding |
| chr9_77918277_77918550 | chr9 | 77918278 | 77918550 | + | -0.752878 | intron (NM_134255, intron 1 of 7) | intron (NM_134255, intron 1 of 7) | 1049 | NM_134255 | 68801 | Mm.430736 | NM_134255 | ENSMUSG00000032349 | **Elovl5** | 1110059L23Rik\|AI747313\|AU043003\|HELO1 | ELOVL family member 5, elongation of long chain fatty acids (yeast) | protein-coding |
| chr9_77918331_77918505 | chr9 | 77918332 | 77918505 | + | -0.759381 | intron (NM_134255, intron 1 of 7) | intron (NM_134255, intron 1 of 7) | 1053 | NM_134255 | 68801 | Mm.430736 | NM_134255 | ENSMUSG00000032349 | **Elovl5** | 1110059L23Rik\|AI747313\|AU043003\|HELO1 | ELOVL family member 5, elongation of long chain fatty acids (yeast) | protein-coding |
| chr9_77961255_77961654 | chr9 | 77961256 | 77961654 | + | -0.768994 | intron (NM_134255, intron 3 of 7) | intron (NM_134255, intron 3 of 7) | 44090 | NM_134255 | 68801 | Mm.430736 | NM_134255 | ENSMUSG00000032349 | **Elovl5** | 1110059L23Rik\|AI747313\|AU043003\|HELO1 | ELOVL family member 5, elongation of long chain fatty acids (yeast) | protein-coding |
| chr3_129494105_129495016 | chr3 | 129494106 | 129495016 | + | -0.664276 | Intergenic | Intergenic | -37825 | NM_130450 | 170439 | Mm.314113 | NM_130450 | ENSMUSG00000041220 | **Elovl6** | C77826\|FAE\|LCE | ELOVL family member 6, elongation of long chain fatty acids (yeast) | protein-coding |
| chr3_129545274_129545598 | chr3 | 129545275 | 129545598 | + | -0.771471 | intron (NM_130450, intron 2 of 4) | B1_Mur3\|SINE\|Alu | 13050 | NM_130450 | 170439 | Mm.314113 | NM_130450 | ENSMUSG00000041220 | **Elovl6** | C77826\|FAE\|LCE | ELOVL family member 6, elongation of long chain fatty acids (yeast) | protein-coding |
| chr3_129553173_129553597 | chr3 | 129553174 | 129553597 | + | -0.802335 | intron (NM_130450, intron 2 of 4) | B4\|SINE\|B4 | 20999 | NM_130450 | 170439 | Mm.314113 | NM_130450 | ENSMUSG00000041220 | **Elovl6** | C77826\|FAE\|LCE | ELOVL family member 6, elongation of long chain fatty acids (yeast) | protein-coding |
| chr3_129601927_129602906 | chr3 | 129601928 | 129602906 | + | -0.931095 | intron (NM_130450, intron 2 of 4) | intron (NM_130450, intron 2 of 4) | 70031 | NM_130450 | 170439 | Mm.314113 | NM_130450 | ENSMUSG00000041220 | **Elovl6** | C77826\|FAE\|LCE | ELOVL family member 6, elongation of long chain fatty acids (yeast) | protein-coding |
| chr13_117242108_117243066 | chr13 | 117242109 | 117243066 | + | -0.729152 | intron (NM_010330, intron 2 of 8) | intron (NM_010330, intron 2 of 8) | 22014 | NM_010330 | 13723 | Mm.274926 | NM_010330 | ENSMUSG00000021728 | **Emb** | AL022799\|Gp70 | embigin | protein-coding |
| chr13_117215701_117216528 | chr13 | 117215702 | 117216528 | + | -0.835082 | Intergenic | Intergenic | -4458 | NM_010330 | 13723 | Mm.274926 | NM_010330 | ENSMUSG00000021728 | **Emb** | AL022799\|Gp70 | embigin | protein-coding |
| chr13_117216170_117216535 | chr13 | 117216171 | 117216535 | + | -0.848177 | Intergenic | Intergenic | -4220 | NM_010330 | 13723 | Mm.274926 | NM_010330 | ENSMUSG00000021728 | **Emb** | AL022799\|Gp70 | embigin | protein-coding |
| chr13_117217652_117217877 | chr13 | 117217653 | 117217877 | + | -0.910355 | Intergenic | MLTR14\|LTR\|ERV1 | -2808 | NM_010330 | 13723 | Mm.274926 | NM_010330 | ENSMUSG00000021728 | **Emb** | AL022799\|Gp70 | embigin | protein-coding |
| chr13_117217640_117218027 | chr13 | 117217641 | 117218027 | + | -1.01901 | Intergenic | Intergenic | -2739 | NM_010330 | 13723 | Mm.274926 | NM_010330 | ENSMUSG00000021728 | **Emb** | AL022799\|Gp70 | embigin | protein-coding |
| chr13_117242108_117242491 | chr13 | 117242109 | 117242491 | + | -1.02278 | intron (NM_010330, intron 2 of 8) | intron (NM_010330, intron 2 of 8) | 21727 | NM_010330 | 13723 | Mm.274926 | NM_010330 | ENSMUSG00000021728 | **Emb** | AL022799\|Gp70 | embigin | protein-coding |
| chr13_117283967_117284407 | chr13 | 117283968 | 117284407 | + | -1.08381 | Intergenic | Intergenic | 63614 | NM_010330 | 13723 | Mm.274926 | NM_010330 | ENSMUSG00000021728 | **Emb** | AL022799\|Gp70 | embigin | protein-coding |
| chr6_113517515_113518141 | chr6 | 113517516 | 113518141 | + | -0.79222 | intron (NM_175101, intron 6 of 7) | intron (NM_175101, intron 6 of 7) | 13810 | NM_175101 | 66087 | Mm.275989 | NM_175101 | ENSMUSG00000030286 | **Emc3** | 0610039A15Rik\|AI225901\|AW260416\|Pob\|Tmem111 | ER membrane protein complex subunit 3 | protein-coding |
| chr6_124693981_124694676 | chr6 | 124693982 | 124694676 | + | -0.703463 | intron (NM_145130, intron 1 of 12) | intron (NM_145130, intron 1 of 12) | 17849 | NM_013536 | 14791 | Mm.6390 | NM_013536 | ENSMUSG00000004268 | **Emg1** | C2f\|Grcc2f | EMG1 nucleolar protein homolog (S. cerevisiae) | protein-coding |
| chr11_5176626_5177633 | chr11 | 5176627 | 5177633 | + | -0.777619 | Intergenic | Intergenic | -24908 | NM_080595 | 140703 | Mm.840 | NM_080595 | ENSMUSG00000034164 | **Emid1** | AW122071\|CO-5\|Emu1 | EMI domain containing 1 | protein-coding |
| chr11_5177024_5177607 | chr11 | 5177025 | 5177607 | + | -0.845756 | Intergenic | LTR84b\|LTR\|ERVL | -25094 | NM_080595 | 140703 | Mm.840 | NM_080595 | ENSMUSG00000034164 | **Emid1** | AW122071\|CO-5\|Emu1 | EMI domain containing 1 | protein-coding |
| chr17_83301489_83301808 | chr17 | 83301490 | 83301808 | + | -0.655151 | Intergenic | Intergenic | -49282 | NM_001114361 | 78798 | Mm.295565 | NM_199466 | ENSMUSG00000032624 | **Eml4** | 4930443C24Rik\|AI644019 | echinoderm microtubule associated protein like 4 | protein-coding |
| chr12_98896711_98897030 | chr12 | 98896712 | 98897030 | + | -0.751462 | intron (NM_001081191, intron 1 of 43) | ORR1B2\|LTR\|ERVL-MaLR | 4613 | NM_001081191 | 319670 | Mm.390484 | NM_001081191 | ENSMUSG00000051166 | **Eml5** | BC027154\|C130068M19Rik | echinoderm microtubule associated protein like 5 | protein-coding |
| chr6_135450388_135451025 | chr6 | 135450389 | 135451025 | + | -0.692724 | Intergenic | ORR1E\|LTR\|ERVL-MaLR | 83214 | NM_001288627 | 13730 | Mm.182785 | NM_010128 | ENSMUSG00000030208 | **Emp1** | I-8-09\|TMP | epithelial membrane protein 1 | protein-coding |
| chr6_135433571_135434177 | chr6 | 135433572 | 135434177 | + | -0.813834 | Intergenic | Intergenic | 66381 | NM_001288627 | 13730 | Mm.182785 | NM_010128 | ENSMUSG00000030208 | **Emp1** | I-8-09\|TMP | epithelial membrane protein 1 | protein-coding |
| chr13_97285645_97285965 | chr13 | 97285646 | 97285965 | + | -0.95996 | Intergenic | Intergenic | 44700 | NM_007930 | 13803 | Mm.241073 | NM_007930 | ENSMUSG00000041773 | **Enc1** | Nrpb\|PIG10 | ectodermal-neural cortex 1 | protein-coding |
| chr13_97322608_97322882 | chr13 | 97322609 | 97322882 | + | -1.10046 | Intergenic | Intergenic | 81640 | NM_007930 | 13803 | Mm.241073 | NM_007930 | ENSMUSG00000041773 | **Enc1** | Nrpb\|PIG10 | ectodermal-neural cortex 1 | protein-coding |
| chr13_97322536_97322902 | chr13 | 97322537 | 97322902 | + | -1.13384 | Intergenic | Intergenic | 81614 | NM_007930 | 13803 | Mm.241073 | NM_007930 | ENSMUSG00000041773 | **Enc1** | Nrpb\|PIG10 | ectodermal-neural cortex 1 | protein-coding |
| chr13_97286965_97287501 | chr13 | 97286966 | 97287501 | + | -1.13802 | Intergenic | Intergenic | 46128 | NM_007930 | 13803 | Mm.241073 | NM_007930 | ENSMUSG00000041773 | **Enc1** | Nrpb\|PIG10 | ectodermal-neural cortex 1 | protein-coding |
| chr13_97322418_97323049 | chr13 | 97322419 | 97323049 | + | -1.17966 | Intergenic | Intergenic | 81629 | NM_007930 | 13803 | Mm.241073 | NM_007930 | ENSMUSG00000041773 | **Enc1** | Nrpb\|PIG10 | ectodermal-neural cortex 1 | protein-coding |
| chr13_97309423_97310026 | chr13 | 97309424 | 97310026 | + | -1.22258 | Intergenic | Intergenic | 68620 | NM_007930 | 13803 | Mm.241073 | NM_007930 | ENSMUSG00000041773 | **Enc1** | Nrpb\|PIG10 | ectodermal-neural cortex 1 | protein-coding |
| chr13_97287116_97287497 | chr13 | 97287117 | 97287497 | + | -1.2276 | Intergenic | Intergenic | 46202 | NM_007930 | 13803 | Mm.241073 | NM_007930 | ENSMUSG00000041773 | **Enc1** | Nrpb\|PIG10 | ectodermal-neural cortex 1 | protein-coding |
| chr13_97309414_97309817 | chr13 | 97309415 | 97309817 | + | -1.28591 | Intergenic | Intergenic | 68511 | NM_007930 | 13803 | Mm.241073 | NM_007930 | ENSMUSG00000041773 | **Enc1** | Nrpb\|PIG10 | ectodermal-neural cortex 1 | protein-coding |
| chr2_32637737_32638427 | chr2 | 32637738 | 32638427 | + | -0.6328 | Intergenic | Intergenic | -8513 | NM_007932 | 13805 | Mm.225297 | NM_007932 | ENSMUSG00000026814 | **Eng** | AI528660\|AI662476\|CD105\|Endo\|S-endoglin | endoglin | protein-coding |
| chr4_150242772_150243130 | chr4 | 150242773 | 150243130 | + | -0.685744 | intron (NM_001025388, intron 4 of 11) | intron (NM_001025388, intron 4 of 11) | 5627 | NM_001025388 | 433182 | Mm.70666 | NM_001025388 | ENSMUSG00000059040 | **Eno1b** | EG433182\|Eno1\|Gm5506 | enolase 1B, retrotransposed | protein-coding |
| chr11_118992719_118993222 | chr11 | 118992720 | 118993222 | + | -0.673371 | TTS (NM_001030291) | TTS (NM_001030291) | 4783 | NM_001030291 | 238011 | Mm.244114 | NM_001030291 | ENSMUSG00000046697 | **Enpp7** | Alk-SMase\|Gm254 | ectonucleotide pyrophosphatase/phosphodiesterase 7 | protein-coding |
| chr15_80544071_80544575 | chr15 | 80544072 | 80544575 | + | -0.836961 | intron (NM_001163189, intron 1 of 5) | LTR101_Mam\|LTR\|ERVL | 16147 | NM_001163189 | 383075 | Mm.297626 | NM_001163189 | ENSMUSG00000050439 | **Enthd1** | Gm1242\|Gm86 | ENTH domain containing 1 | protein-coding |
| chr15_80562444_80562786 | chr15 | 80562445 | 80562786 | + | -0.87143 | Intergenic | Intergenic | -2145 | NM_001163189 | 383075 | Mm.297626 | NM_001163189 | ENSMUSG00000050439 | **Enthd1** | Gm1242\|Gm86 | ENTH domain containing 1 | protein-coding |
| chr15_80555528_80555760 | chr15 | 80555529 | 80555760 | + | -0.875412 | intron (NM_001163189, intron 1 of 5) | Lx9\|LINE\|L1 | 4826 | NM_001163189 | 383075 | Mm.297626 | NM_001163189 | ENSMUSG00000050439 | **Enthd1** | Gm1242\|Gm86 | ENTH domain containing 1 | protein-coding |
| chr15_80554937_80555441 | chr15 | 80554938 | 80555441 | + | -0.967987 | intron (NM_001163189, intron 1 of 5) | Lx9\|LINE\|L1 | 5281 | NM_001163189 | 383075 | Mm.297626 | NM_001163189 | ENSMUSG00000050439 | **Enthd1** | Gm1242\|Gm86 | ENTH domain containing 1 | protein-coding |
| chr15_80538618_80539264 | chr15 | 80538619 | 80539264 | + | -1.23377 | intron (NM_001163189, intron 1 of 5) | intron (NM_001163189, intron 1 of 5) | 21529 | NM_001163189 | 383075 | Mm.297626 | NM_001163189 | ENSMUSG00000050439 | **Enthd1** | Gm1242\|Gm86 | ENTH domain containing 1 | protein-coding |
| chr15_80559640_80559964 | chr15 | 80559641 | 80559964 | + | -1.30683 | intron (NM_001163189, intron 1 of 5) | intron (NM_001163189, intron 1 of 5) | 668 | NM_001163189 | 383075 | Mm.297626 | NM_001163189 | ENSMUSG00000050439 | **Enthd1** | Gm1242\|Gm86 | ENTH domain containing 1 | protein-coding |

| | | | | | | | | | | | | | | | | | |
|---|---|---|---|---|---|---|---|---|---|---|---|---|---|---|---|---|---|
| chr15_80559545_80560047 | chr15 | 80559546 | 80560047 | + | -1.32751 | intron (NM_001163189, intron 1 of 5) | intron (NM_001163189, intron 1 of 5) | 674 | NM_001163189 | 383075 | Mm.297626 | NM_001163189 | ENSMUSG00000050439 | **Enthd1** | Gm1242\|Gm86 | ENTH domain containing 1 | protein-coding |
| chr19_40708101_40708856 | chr19 | 40708102 | 40708856 | + | -0.62268 | intron (NM_009848, intron 1 of 9) | intron (NM_009848, intron 1 of 9) | 48685 | NM_009848 | 12495 | Mm.2824 | NM_009848 | ENSMUSG00000048120 | **Entpd1** | 2610206B08Rik\|AA408691\|Cd39\|NTPDase-1 | ectonucleoside triphosphate diphosphohydrolase 1 | protein-coding |
| chr19_40708053_40708557 | chr19 | 40708054 | 40708557 | + | -0.651748 | intron (NM_009848, intron 1 of 9) | intron (NM_009848, intron 1 of 9) | 48511 | NM_009848 | 12495 | Mm.2824 | NM_009848 | ENSMUSG00000048120 | **Entpd1** | 2610206B08Rik\|AA408691\|Cd39\|NTPDase-1 | ectonucleoside triphosphate diphosphohydrolase 1 | protein-coding |
| chr2_150750621_150750917 | chr2 | 150750622 | 150750917 | + | -0.66236 | intron (NM_172117, intron 1 of 13) | intron (NM_172117, intron 1 of 13) | 1688 | NM_172117 | 12497 | Mm.275524 | NM_172117 | ENSMUSG00000033068 | **Entpd6** | 2700026H11Rik\|Cd39l2\|NTPDase-6 | ectonucleoside triphosphate diphosphohydrolase 6 | protein-coding |
| chr17_86801313_86802881 | chr17 | 86801314 | 86802881 | + | -0.701617 | intron (NM_010137, intron 2 of 15) | intron (NM_010137, intron 2 of 15) | 48233 | NM_010137 | 13819 | Mm.1415 | NM_010137 | ENSMUSG00000024140 | **Epas1** | HIF-2alpha\|HIF2A\|HLF\|HRF\|MOP2\|bHLHe73 | endothelial PAS domain protein 1 | protein-coding |
| chr17_86674199_86674998 | chr17 | 86674200 | 86674998 | + | -0.868757 | Intergenic | Intergenic | -79265 | NM_010137 | 13819 | Mm.1415 | NM_010137 | ENSMUSG00000024140 | **Epas1** | HIF-2alpha\|HIF2A\|HLF\|HRF\|MOP2\|bHLHe73 | endothelial PAS domain protein 1 | protein-coding |
| chr4_132049882_132050222 | chr4 | 132049883 | 132050222 | + | -0.923871 | promoter-TSS (NM_001128606) | promoter-TSS (NM_001128606) | -974 | NM_001128606 | 269587 | Mm.30038 | NM_183428 | ENSMUSG00000028906 | **Epb4.1** | 4.1R\|AI415518\|D4Ertd442e\|Elp-1\|Elp1\|Epb41\|mKIAA4056 | erythrocyte protein band 4.1 | protein-coding |
| chr4_132007857_132009093 | chr4 | 132007858 | 132009093 | + | -0.939243 | intron (NM_183428, intron 1 of 20) | intron (NM_183428, intron 1 of 20) | 40603 | NM_001128606 | 269587 | Mm.30038 | NM_183428 | ENSMUSG00000028906 | **Epb4.1** | 4.1R\|AI415518\|D4Ertd442e\|Elp-1\|Elp1\|Epb41\|mKIAA4056 | erythrocyte protein band 4.1 | protein-coding |
| chr4_132008054_132008655 | chr4 | 132008055 | 132008655 | + | -0.963276 | intron (NM_183428, intron 1 of 20) | intron (NM_183428, intron 1 of 20) | 40723 | NM_001128606 | 269587 | Mm.30038 | NM_183428 | ENSMUSG00000028906 | **Epb4.1** | 4.1R\|AI415518\|D4Ertd442e\|Elp-1\|Elp1\|Epb41\|mKIAA4056 | erythrocyte protein band 4.1 | protein-coding |
| chr4_132041655_132041873 | chr4 | 132041656 | 132041873 | + | -0.987485 | intron (NM_183428, intron 1 of 20) | intron (NM_183428, intron 1 of 20) | 7314 | NM_001128606 | 269587 | Mm.30038 | NM_183428 | ENSMUSG00000028906 | **Epb4.1** | 4.1R\|AI415518\|D4Ertd442e\|Elp-1\|Elp1\|Epb41\|mKIAA4056 | erythrocyte protein band 4.1 | protein-coding |
| chr4_132040282_132040629 | chr4 | 132040283 | 132040629 | + | -1.11612 | intron (NM_183428, intron 1 of 20) | intron (NM_183428, intron 1 of 20) | 8622 | NM_001128606 | 269587 | Mm.30038 | NM_183428 | ENSMUSG00000028906 | **Epb4.1** | 4.1R\|AI415518\|D4Ertd442e\|Elp-1\|Elp1\|Epb41\|mKIAA4056 | erythrocyte protein band 4.1 | protein-coding |
| chr4_132040172_132040844 | chr4 | 132040173 | 132040844 | + | -1.11627 | intron (NM_183428, intron 1 of 20) | intron (NM_183428, intron 1 of 20) | 8570 | NM_001128606 | 269587 | Mm.30038 | NM_183428 | ENSMUSG00000028906 | **Epb4.1** | 4.1R\|AI415518\|D4Ertd442e\|Elp-1\|Elp1\|Epb41\|mKIAA4056 | erythrocyte protein band 4.1 | protein-coding |
| chr4_136816564_136817092 | chr4 | 136816565 | 136817092 | + | -0.700305 | intron (NM_001290753, intron 1 of 15) | intron (NM_001290753, intron 1 of 15) | 19183 | NM_001290753 | 13844 | Mm.250981 | NM_010142 | ENSMUSG00000028664 | **Ephb2** | Cek5\|Drt\|ETECK\|Erk\|Hek5\|Nuk\|Prkm5\|Qek5\|Sek3\|Tyro5 | Eph receptor B2 | protein-coding |
| chr17_32201023_32201552 | chr17 | 32201024 | 32201552 | + | -0.796002 | exon (NM_020508, exon 15 of 20) | exon (NM_020508, exon 15 of 20) | -11825 | NM_001033163 | 71932 | Mm.176240 | NM_001033163 | ENSMUSG00000037577 | **Ephx3** | 2310063B19Rik\|Abhd9 | epoxide hydrolase 3 | protein-coding |
| chr4_109273658_109274225 | chr4 | 109273659 | 109274225 | + | -0.978379 | Intergenic | Intergenic | -6333 | NM_007943 | 13858 | Mm.318250 | NM_007943 | ENSMUSG00000028552 | **Eps15** | 2410112D09Rik | epidermal growth factor receptor pathway substrate 15 | protein-coding |
| chr7_4464150_4464863 | chr7 | 4464151 | 4464863 | + | -0.853864 | promoter-TSS (NM_001290416) | promoter-TSS (NM_001290416) | -228 | NM_026146 | 67425 | Mm.360392 | NM_026146 | ENSMUSG00000006154 | **Eps8l1** | 2310051G19Rik\|4632407K17Rik\|AW060268\|DRC3\|EPS8R1 | EPS8-like 1 | protein-coding |
| chr18_32292832_32293565 | chr18 | 32292833 | 32293565 | + | -0.769952 | Intergenic | Intergenic | 52868 | NM_133658 | 13872 | Mm.282335 | NM_133658 | ENSMUSG00000024382 | **Ercc3** | BTF2 p89\|Ercc-3\|XPB | excision repair cross-complementing rodent repair deficiency, complementation group 3 | protein-coding |
| chr18_32293056_32293523 | chr18 | 32293057 | 32293523 | + | -0.782851 | Intergenic | Intergenic | 52959 | NM_133658 | 13872 | Mm.282335 | NM_133658 | ENSMUSG00000024382 | **Ercc3** | BTF2 p89\|Ercc-3\|XPB | excision repair cross-complementing rodent repair deficiency, complementation group 3 | protein-coding |
| chr16_13158926_13159472 | chr16 | 13158927 | 13159472 | + | -0.826896 | Intergenic | Intergenic | 49463 | NM_015769 | 50505 | Mm.287837 | NM_015769 | | **Ercc4** | AI606920\|Xpf | excision repair cross-complementing rodent repair deficiency, complementation group 4 | protein-coding |
| chr13_108169991_108170449 | chr13 | 108169992 | 108170449 | + | -1.03405 | intron (NM_028042, intron 4 of 11) | intron (NM_028042, intron 4 of 11) | 11482 | NM_028042 | 71991 | Mm.212208 | NM_028042 | ENSMUSG00000021694 | **Ercc8** | 2410022P04Rik\|2810431L23Rik\|4631412O06Rik\|B130065P18Rik\|Ckn1\|Csa | excision repaiross-complementing rodent repair deficiency, complementation group 8 | protein-coding |
| chr11_106481779_106482106 | chr11 | 106481780 | 106482106 | + | -0.836761 | intron (NM_023913, intron 1 of 21) | intron (NM_023913, intron 1 of 21) | 5853 | NM_023913 | 78943 | Mm.340943 | NM_023913 | | **Ern1** | 9030414B18Rik\|AI225830\|C85377\|Ire1a\|Ire1alpha\|Ire1p | endoplasmic reticulum (ER) to nucleus signalling 1 | protein-coding |
| chr13_113242182_113242435 | chr13 | 113242183 | 113242435 | + | -0.593435 | Intergenic | Intergenic | 32650 | NM_023612 | 71690 | Mm.38929 | NM_023612 | ENSMUSG00000042379 | **Esm1** | 0610042H23Rik\|AV004503\|ESM-1 | endothelial cell-specific molecule 1 | protein-coding |
| chr4_11404050_11404349 | chr4 | 11404051 | 11404349 | + | -0.685705 | Intergenic | Intergenic | -17417 | NM_194055 | 207920 | Mm.387719 | NM_194055 | ENSMUSG00000040728 | **Esrp1** | 2210008M09Rik\|A630065D16\|BC031468\|Rbm35a | epithelial splicing regulatory protein 1 | protein-coding |
| chr12_116313891_116314171 | chr12 | 116313892 | 116314171 | + | -0.721016 | intron (NM_028731, intron 1 of 21) | intron (NM_028731, intron 1 of 21) | 32809 | NM_028731 | 52635 | Mm.273755 | NM_028731 | ENSMUSG00000021171 | **Esyt2** | 2410017M09Rik\|4921504I16Rik\|CHR2SYT\|D12Ertd551e\|Fam62b | extended synaptotagmin-like protein 2 | protein-coding |
| chr6_143172219_143172501 | chr6 | 143172220 | 143172501 | + | -1.09824 | intron (NM_029250, intron 1 of 7) | intron (NM_029250, intron 1 of 7) | 5130 | NM_029250 | 75320 | Mm.272548 | NM_029250 | ENSMUSG00000030275 | **Etnk1** | 1110061E11Rik\|4930555L11Rik\|AI195356\|AI841245\|C80956\|D6Ertd3e\|EKI1 | ethanolamine kinase 1 | protein-coding |
| chr9_32940352_32940793 | chr9 | 32940353 | 32940793 | + | -0.609039 | Intergenic | Intergenic | 244531 | NM_001038642 | 23871 | Mm.292415 | NM_011808 | ENSMUSG00000032035 | **Ets1** | AI196000\|AI448617\|D230050P06\|Ets-1\|Tpl1 | E26 avian leukemia oncogene 1, 5' domain | protein-coding |
| chr9_32901103_32901782 | chr9 | 32901104 | 32901782 | + | -0.627443 | Intergenic | Intergenic | 205401 | NM_001038642 | 23871 | Mm.292415 | NM_011808 | ENSMUSG00000032035 | **Ets1** | AI196000\|AI448617\|D230050P06\|Ets-1\|Tpl1 | E26 avian leukemia oncogene 1, 5' domain | protein-coding |
| chr9_32910376_32911171 | chr9 | 32910377 | 32911171 | + | -0.639125 | Intergenic | Intergenic | 214732 | NM_001038642 | 23871 | Mm.292415 | NM_011808 | ENSMUSG00000032035 | **Ets1** | AI196000\|AI448617\|D230050P06\|Ets-1\|Tpl1 | E26 avian leukemia oncogene 1, 5' domain | protein-coding |
| chr9_32705842_32706471 | chr9 | 32705843 | 32706471 | + | -0.640933 | intron (NM_001038642, intron 1 of 6) | intron (NM_001038642, intron 1 of 6) | 10115 | NM_001038642 | 23871 | Mm.292415 | NM_011808 | ENSMUSG00000032035 | **Ets1** | AI196000\|AI448617\|D230050P06\|Ets-1\|Tpl1 | E26 avian leukemia oncogene 1, 5' domain | protein-coding |
| chr9_32745866_32746565 | chr9 | 32745867 | 32746565 | + | -0.658438 | intron (NM_011808, intron 6 of 7) | intron (NM_011808, intron 6 of 7) | 50174 | NM_001038642 | 23871 | Mm.292415 | NM_011808 | ENSMUSG00000032035 | **Ets1** | AI196000\|AI448617\|D230050P06\|Ets-1\|Tpl1 | E26 avian leukemia oncogene 1, 5' domain | protein-coding |
| chr9_32815432_32815868 | chr9 | 32815433 | 32815868 | + | -0.696086 | Intergenic | Intergenic | 119608 | NM_001038642 | 23871 | Mm.292415 | NM_011808 | ENSMUSG00000032035 | **Ets1** | AI196000\|AI448617\|D230050P06\|Ets-1\|Tpl1 | E26 avian leukemia oncogene 1, 5' domain | protein-coding |
| chr9_32747720_32747933 | chr9 | 32747721 | 32747933 | + | -0.710778 | intron (NM_011808, intron 6 of 7) | intron (NM_011808, intron 6 of 7) | 51785 | NM_001038642 | 23871 | Mm.292415 | NM_011808 | ENSMUSG00000032035 | **Ets1** | AI196000\|AI448617\|D230050P06\|Ets-1\|Tpl1 | E26 avian leukemia oncogene 1, 5' domain | protein-coding |
| chr9_32731630_32732275 | chr9 | 32731631 | 32732275 | + | -0.713348 | intron (NM_001038642, intron 3 of 6) | intron (NM_001038642, intron 3 of 6) | 35911 | NM_001038642 | 23871 | Mm.292415 | NM_011808 | ENSMUSG00000032035 | **Ets1** | AI196000\|AI448617\|D230050P06\|Ets-1\|Tpl1 | E26 avian leukemia oncogene 1, 5' domain | protein-coding |
| chr9_32747405_32748017 | chr9 | 32747406 | 32748017 | + | -0.721671 | intron (NM_011808, intron 6 of 7) | intron (NM_011808, intron 6 of 7) | 51669 | NM_001038642 | 23871 | Mm.292415 | NM_011808 | ENSMUSG00000032035 | **Ets1** | AI196000\|AI448617\|D230050P06\|Ets-1\|Tpl1 | E26 avian leukemia oncogene 1, 5' domain | protein-coding |
| chr9_32731843_32732301 | chr9 | 32731844 | 32732301 | + | -0.735832 | intron (NM_001038642, intron 3 of 6) | intron (NM_001038642, intron 3 of 6) | 36030 | NM_001038642 | 23871 | Mm.292415 | NM_011808 | ENSMUSG00000032035 | **Ets1** | AI196000\|AI448617\|D230050P06\|Ets-1\|Tpl1 | E26 avian leukemia oncogene 1, 5' domain | protein-coding |
| chr9_32745866_32746323 | chr9 | 32745867 | 32746323 | + | -0.74894 | intron (NM_011808, intron 6 of 7) | intron (NM_011808, intron 6 of 7) | 50053 | NM_001038642 | 23871 | Mm.292415 | NM_011808 | ENSMUSG00000032035 | **Ets1** | AI196000\|AI448617\|D230050P06\|Ets-1\|Tpl1 | E26 avian leukemia oncogene 1, 5' domain | protein-coding |
| chr9_32706044_32706498 | chr9 | 32706045 | 32706498 | + | -0.75568 | intron (NM_001038642, intron 1 of 6) | intron (NM_001038642, intron 1 of 6) | 10229 | NM_001038642 | 23871 | Mm.292415 | NM_011808 | ENSMUSG00000032035 | **Ets1** | AI196000\|AI448617\|D230050P06\|Ets-1\|Tpl1 | E26 avian leukemia oncogene 1, 5' domain | protein-coding |
| chr9_32740866_32741047 | chr9 | 32740867 | 32741047 | + | -0.764594 | intron (NM_011808, intron 6 of 7) | intron (NM_011808, intron 6 of 7) | 44915 | NM_001038642 | 23871 | Mm.292415 | NM_011808 | ENSMUSG00000032035 | **Ets1** | AI196000\|AI448617\|D230050P06\|Ets-1\|Tpl1 | E26 avian leukemia oncogene 1, 5' domain | protein-coding |
| chr9_32925824_32926713 | chr9 | 32925825 | 32926713 | + | -0.785944 | Intergenic | Intergenic | 230227 | NM_001038642 | 23871 | Mm.292415 | NM_011808 | ENSMUSG00000032035 | **Ets1** | AI196000\|AI448617\|D230050P06\|Ets-1\|Tpl1 | E26 avian leukemia oncogene 1, 5' domain | protein-coding |
| chr9_32706037_32706654 | chr9 | 32706038 | 32706654 | + | -0.797573 | intron (NM_001038642, intron 1 of 6) | intron (NM_001038642, intron 1 of 6) | 10304 | NM_001038642 | 23871 | Mm.292415 | NM_011808 | ENSMUSG00000032035 | **Ets1** | AI196000\|AI448617\|D230050P06\|Ets-1\|Tpl1 | E26 avian leukemia oncogene 1, 5' domain | protein-coding |
| chr9_32693537_32693800 | chr9 | 32693538 | 32693800 | + | -0.806268 | Intergenic | Intergenic | -2373 | NM_011808 | 23871 | Mm.292415 | NM_011808 | ENSMUSG00000032035 | **Ets1** | AI196000\|AI448617\|D230050P06\|Ets-1\|Tpl1 | E26 avian leukemia oncogene 1, 5' domain | protein-coding |
| chr9_32620526_32620823 | chr9 | 32620527 | 32620823 | + | -0.818574 | Intergenic | Intergenic | -75367 | NM_011808 | 23871 | Mm.292415 | NM_011808 | ENSMUSG00000032035 | **Ets1** | AI196000\|AI448617\|D230050P06\|Ets-1\|Tpl1 | E26 avian leukemia oncogene 1, 5' domain | protein-coding |
| chr9_32938885_32939225 | chr9 | 32938886 | 32939225 | + | -0.837194 | Intergenic | Intergenic | 243013 | NM_001038642 | 23871 | Mm.292415 | NM_011808 | ENSMUSG00000032035 | **Ets1** | AI196000\|AI448617\|D230050P06\|Ets-1\|Tpl1 | E26 avian leukemia oncogene 1, 5' domain | protein-coding |
| chr9_32928658_32929015 | chr9 | 32928659 | 32929015 | + | -0.911847 | Intergenic | Intergenic | 232795 | NM_001038642 | 23871 | Mm.292415 | NM_011808 | ENSMUSG00000032035 | **Ets1** | AI196000\|AI448617\|D230050P06\|Ets-1\|Tpl1 | E26 avian leukemia oncogene 1, 5' domain | protein-coding |
| chr9_32960656_32961113 | chr9 | 32960657 | 32961113 | + | -1.02129 | Intergenic | Intergenic | 264843 | NM_001038642 | 23871 | Mm.292415 | NM_011808 | ENSMUSG00000032035 | **Ets1** | AI196000\|AI448617\|D230050P06\|Ets-1\|Tpl1 | E26 avian leukemia oncogene 1, 5' domain | protein-coding |

| | | | | | | | | | | | | | | | | | |
|---|---|---|---|---|---|---|---|---|---|---|---|---|---|---|---|---|---|
| chr9_32906299_32906819 | chr9 | 32906300 | 32906819 | + | -1.02265 | Intergenic | Intergenic | 210517 | NM_001038642 | 23871 | Mm.292415 | NM_011808 | ENSMUSG00000032035 | **Ets1** | AI196000\|AI448617\|D230050P06\|Ets-1\|Tpl1 | E26 avian leukemia oncogene 1, 5' domain | protein-coding |
| chr9_32740761_32741438 | chr9 | 32740762 | 32741438 | + | -1.04061 | intron (NM_011808, intron 6 of 7) | intron (NM_011808, intron 6 of 7) | 45058 | NM_001038642 | 23871 | Mm.292415 | NM_011808 | ENSMUSG00000032035 | **Ets1** | AI196000\|AI448617\|D230050P06\|Ets-1\|Tpl1 | E26 avian leukemia oncogene 1, 5' domain | protein-coding |
| chr9_32928301_32929031 | chr9 | 32928302 | 32929031 | + | -1.04558 | Intergenic | Intergenic | 232624 | NM_001038642 | 23871 | Mm.292415 | NM_011808 | ENSMUSG00000032035 | **Ets1** | AI196000\|AI448617\|D230050P06\|Ets-1\|Tpl1 | E26 avian leukemia oncogene 1, 5' domain | protein-coding |
| chr9_32928323_32928559 | chr9 | 32928324 | 32928559 | + | -1.14246 | Intergenic | Intergenic | 232399 | NM_001038642 | 23871 | Mm.292415 | NM_011808 | ENSMUSG00000032035 | **Ets1** | AI196000\|AI448617\|D230050P06\|Ets-1\|Tpl1 | E26 avian leukemia oncogene 1, 5' domain | protein-coding |
| chr16_22466371_22466791 | chr16 | 22466372 | 22466791 | + | -0.639786 | TTS (NM_138650) | TTS (NM_138650) | -27011 | NM_023794 | 104156 | Mm.155708 | NM_023794 | ENSMUSG00000013089 | **Etv5** | 1110005E01Rik\|8430401F14Rik\|ERM | ets variant 5 | protein-coding |
| chr11_79528057_79528409 | chr11 | 79528058 | 79528409 | + | -0.743751 | 5' UTR (NM_001033711, exon 2 of 2) | 5' UTR (NM_001033711, exon 2 of 2) | 2376 | NM_146023 | 101488212 | Mm.439665 | NM_146023 | ENSMUSG00000093938 | **Evi2a-evi2b** | - | Evi2a-Evi2b readthrough | protein-coding |
| chr11_79519951_79520322 | chr11 | 79519952 | 79520322 | + | -0.604569 | intron (NM_146023, intron 3 of 3) | intron (NM_146023, intron 3 of 3) | 3625 | NM_001077496 | 216984 | Mm.439665 | NM_001077496 | | **Evi2b** | Evi-2\|Evi2 | ecotropic viral integration site 2b | protein-coding |
| chr11_79524003_79524191 | chr11 | 79524004 | 79524191 | + | -0.617673 | promoter-TSS (NM_001077496) | promoter-TSS (NM_001077496) | -335 | NM_001077496 | 216984 | Mm.439665 | NM_001077496 | | **Evi2b** | Evi-2\|Evi2 | ecotropic viral integration site 2b | protein-coding |
| chr12_108649407_108649671 | chr12 | 108649408 | 108649671 | + | -0.811082 | intron (NM_007965, intron 2 of 12) | intron (NM_007965, intron 2 of 12) | 13786 | NM_001163396 | 14026 | Mm.238841 | NM_007965 | ENSMUSG00000021262 | **Evl** | AI528774 | Ena-vasodilator stimulated phosphoprotein | protein-coding |
| chr6_33618602_33619094 | chr6 | 33618603 | 33619094 | + | -0.77522 | intron (NM_009148, intron 10 of 17) | (TGG)n\|Simple_repeat\|Simple_repeat | 369698 | NM_009148 | 20336 | Mm.265512 | NM_009148 | | **Exoc4** | C78892\|Sec8\|Sec8l1 | exocyst complex component 4 | protein-coding |
| chr6_33261027_33261541 | chr6 | 33261028 | 33261541 | + | -1.14282 | intron (NM_009148, intron 1 of 17) | ID_B1\|SINE\|B4 | 12134 | NM_009148 | 20336 | Mm.265512 | NM_009148 | | **Exoc4** | C78892\|Sec8\|Sec8l1 | exocyst complex component 4 | protein-coding |
| chr15_76317254_76317634 | chr15 | 76317255 | 76317634 | + | -1.17485 | Intergenic | Intergenic | -9953 | NM_175399 | 109075 | Mm.322752 | NM_175399 | ENSMUSG00000034259 | **Exosc4** | 1110039I09Rik\|1500001N04Rik\|Rrp41 | exosome component 4 | protein-coding |
| chr15_53100534_53101059 | chr15 | 53100535 | 53101059 | + | -0.66437 | intron (NM_010162, intron 4 of 10) | intron (NM_010162, intron 4 of 10) | 245386 | NM_010162 | 14042 | Mm.309395 | NM_010162 | | **Ext1** | AA409028 | exostoses (multiple) 1 | protein-coding |
| chr15_53266062_53267985 | chr15 | 53266063 | 53267985 | + | -0.669741 | intron (NM_010162, intron 1 of 10) | intron (NM_010162, intron 1 of 10) | 79159 | NM_010162 | 14042 | Mm.309395 | NM_010162 | | **Ext1** | AA409028 | exostoses (multiple) 1 | protein-coding |
| chr15_53100730_53100972 | chr15 | 53100731 | 53100972 | + | -0.712455 | intron (NM_010162, intron 4 of 10) | intron (NM_010162, intron 4 of 10) | 245332 | NM_010162 | 14042 | Mm.309395 | NM_010162 | | **Ext1** | AA409028 | exostoses (multiple) 1 | protein-coding |
| chr15_53266783_53267985 | chr15 | 53266784 | 53267985 | + | -0.730272 | intron (NM_010162, intron 1 of 10) | PB1D7\|SINE\|Alu | 78799 | NM_010162 | 14042 | Mm.309395 | NM_010162 | | **Ext1** | AA409028 | exostoses (multiple) 1 | protein-coding |
| chr15_53140284_53140767 | chr15 | 53140285 | 53140767 | + | -0.74327 | intron (NM_010162, intron 1 of 10) | intron (NM_010162, intron 1 of 10) | 205657 | NM_010162 | 14042 | Mm.309395 | NM_010162 | | **Ext1** | AA409028 | exostoses (multiple) 1 | protein-coding |
| chr15_53216303_53217036 | chr15 | 53216304 | 53217036 | + | -0.833989 | intron (NM_010162, intron 1 of 10) | intron (NM_010162, intron 1 of 10) | 129513 | NM_010162 | 14042 | Mm.309395 | NM_010162 | | **Ext1** | AA409028 | exostoses (multiple) 1 | protein-coding |
| chr15_53324704_53325124 | chr15 | 53324705 | 53325124 | + | -1.08654 | intron (NM_010162, intron 1 of 10) | intron (NM_010162, intron 1 of 10) | 21269 | NM_010162 | 14042 | Mm.309395 | NM_010162 | | **Ext1** | AA409028 | exostoses (multiple) 1 | protein-coding |
| chr15_53328081_53328430 | chr15 | 53328082 | 53328430 | + | -1.32531 | intron (NM_010162, intron 1 of 10) | intron (NM_010162, intron 1 of 10) | 17927 | NM_010162 | 14042 | Mm.309395 | NM_010162 | | **Ext1** | AA409028 | exostoses (multiple) 1 | protein-coding |
| chr15_53115301_53115969 | chr15 | 53115302 | 53115969 | + | -1.33866 | intron (NM_010162, intron 1 of 10) | intron (NM_010162, intron 1 of 10) | 230548 | NM_010162 | 14042 | Mm.309395 | NM_010162 | | **Ext1** | AA409028 | exostoses (multiple) 1 | protein-coding |
| chr4_134361232_134361479 | chr4 | 134361233 | 134361479 | + | -1.2036 | intron (NM_019578, intron 6 of 10) | intron (NM_019578, intron 6 of 10) | 11191 | NM_019578 | 56219 | Mm.30978 | NM_019578 | ENSMUSG00000028838 | **Extl1** | D430033M16Rik | exostoses (multiple)-like 1 | protein-coding |
| chr4_134361206_134361659 | chr4 | 134361207 | 134361659 | + | -1.26618 | intron (NM_019578, intron 6 of 10) | intron (NM_019578, intron 6 of 10) | 11114 | NM_019578 | 56219 | Mm.30978 | NM_019578 | ENSMUSG00000028838 | **Extl1** | D430033M16Rik | exostoses (multiple)-like 1 | protein-coding |
| chr14_65044328_65045596 | chr14 | 65044329 | 65045596 | + | -0.685036 | Intergenic | Intergenic | 53144 | NM_018788 | 54616 | Mm.103748 | NM_018788 | ENSMUSG00000021978 | **Extl3** | 2900009G18Rik\|AU040105\|Ext1l\|mKIAA0519 | exostoses (multiple)-like 3 | protein-coding |
| chr14_65078110_65078401 | chr14 | 65078111 | 65078401 | + | -1.05067 | intron (NM_018788, intron 2 of 6) | intron (NM_018788, intron 2 of 6) | 19850 | NM_018788 | 54616 | Mm.103748 | NM_018788 | ENSMUSG00000021978 | **Extl3** | 2900009G18Rik\|AU040105\|Ext1l\|mKIAA0519 | exostoses (multiple)-like 3 | protein-coding |
| chr14_65078115_65078817 | chr14 | 65078116 | 65078817 | + | -1.07989 | intron (NM_018788, intron 2 of 6) | intron (NM_018788, intron 2 of 6) | 19640 | NM_018788 | 54616 | Mm.103748 | NM_018788 | ENSMUSG00000021978 | **Extl3** | 2900009G18Rik\|AU040105\|Ext1l\|mKIAA0519 | exostoses (multiple)-like 3 | protein-coding |
| chr14_65078091_65078578 | chr14 | 65078092 | 65078578 | + | -1.1048 | intron (NM_018788, intron 2 of 6) | intron (NM_018788, intron 2 of 6) | 19771 | NM_018788 | 54616 | Mm.103748 | NM_018788 | ENSMUSG00000021978 | **Extl3** | 2900009G18Rik\|AU040105\|Ext1l\|mKIAA0519 | exostoses (multiple)-like 3 | protein-coding |
| chr2_165639387_165639757 | chr2 | 165639388 | 165639757 | + | -0.639855 | intron (NM_001271962, intron 1 of 15) | intron (NM_001271962, intron 1 of 15) | 9227 | NM_001271962 | 14049 | Mm.282719 | NM_010165 | ENSMUSG00000017897 | **Eya2** | Eab1 | eyes absent 2 homolog (Drosophila) | protein-coding |
| chr2_165639211_165639795 | chr2 | 165639212 | 165639795 | + | -0.694139 | intron (NM_001271962, intron 1 of 15) | intron (NM_001271962, intron 1 of 15) | 9158 | NM_001271962 | 14049 | Mm.282719 | NM_010165 | ENSMUSG00000017897 | **Eya2** | Eab1 | eyes absent 2 homolog (Drosophila) | protein-coding |
| chr11_101218706_101219036 | chr11 | 101218707 | 101219036 | + | -0.987301 | intron (NM_007970, intron 1 of 20) | intron (NM_007970, intron 1 of 20) | 7579 | NM_007970 | 14055 | Mm.5027 | NM_007970 | ENSMUSG00000006920 | **Ezh1** | AW557915\|mKIAA0388 | enhancer of zeste homolog 1 (Drosophila) | protein-coding |
| chr11_101218576_101219088 | chr11 | 101218577 | 101219088 | + | -1.02912 | intron (NM_007970, intron 1 of 20) | intron (NM_007970, intron 1 of 20) | 7618 | NM_007970 | 14055 | Mm.5027 | NM_007970 | ENSMUSG00000006920 | **Ezh1** | AW557915\|mKIAA0388 | enhancer of zeste homolog 1 (Drosophila) | protein-coding |
| chr17_6769634_6769918 | chr17 | 6769635 | 6769918 | + | -0.659466 | intron (NM_009510, intron 1 of 12) | intron (NM_009510, intron 1 of 12) | 13004 | NM_009510 | 22350 | Mm.277812 | NM_009510 | ENSMUSG00000052397 | **Ezr** | AW146364\|R75297\|Vil2\|p81 | ezrin | protein-coding |
| chr13_55431715_55432722 | chr13 | 55431716 | 55432722 | + | -0.812239 | Intergenic | L1MA4\|LINE\|L1 | -5417 | NM_021489 | 58992 | Mm.42224 | NM_021489 | ENSMUSG00000021492 | **F12** | FXII\|HAF | coagulation factor XII (Hageman factor) | protein-coding |
| chr13_55432038_55432733 | chr13 | 55432039 | 55432733 | + | -0.90908 | Intergenic | L1MA4\|LINE\|L1 | -5584 | NM_021489 | 58992 | Mm.42224 | NM_021489 | ENSMUSG00000021492 | **F12** | FXII\|HAF | coagulation factor XII (Hageman factor) | protein-coding |
| chr2_91638837_91639308 | chr2 | 91638838 | 91639308 | + | -1.22348 | Intergenic | Intergenic | -2659 | NM_010168 | 14061 | Mm.89048 | NM_010168 | ENSMUSG00000027249 | **F2** | Cf-2\|Cf2\|FII\|thrombin | coagulation factor II | protein-coding |
| chr13_95617096_95617448 | chr13 | 95617097 | 95617448 | + | -0.682567 | intron (NM_010169, intron 1 of 1) | intron (NM_010169, intron 1 of 1) | 1161 | NM_010169 | 14062 | Mm.24816 | NM_010169 | ENSMUSG00000048376 | **F2r** | AI482343\|Cf2r\|Par1\|ThrR | coagulation factor II (thrombin) receptor | protein-coding |
| chr13_95521053_95521382 | chr13 | 95521054 | 95521382 | + | -0.749385 | intron (NM_007974, intron 1 of 1) | intron (NM_007974, intron 1 of 1) | 4022 | NM_007974 | 14063 | Mm.1614 | NM_007974 | ENSMUSG00000021678 | **F2rl1** | Gpcr11\|PAR-2\|Par2 | coagulation factor II (thrombin) receptor-like 1 | protein-coding |
| chr11_45794406_45795196 | chr11 | 45794407 | 45795196 | + | -0.748372 | Intergenic | Intergenic | -13282 | NR_045876 | 100009664 | Mm.415439 | NR_045876 | | **F630206G17Rik** | - | RIKEN cDNA F630206G17 gene | ncRNA |
| chr12_70232167_70232593 | chr12 | 70232168 | 70232593 | + | -0.630245 | intron (NR_045174, intron 2 of 4) | intron (NR_045174, intron 2 of 4) | 4539 | NR_045174 | 414124 | Mm.416164 | NR_045174 | ENSMUSG00000085298 | **F730035M05Rik** | - | RIKEN cDNA F730035M05 gene | ncRNA |
| chr12_70257759_70258467 | chr12 | 70257760 | 70258467 | + | -0.818489 | intron (NM_053167, intron 8 of 13) | intron (NM_053167, intron 8 of 13) | 30272 | NR_045174 | 414124 | Mm.416164 | NR_045174 | ENSMUSG00000085298 | **F730035M05Rik** | - | RIKEN cDNA F730035M05 gene | ncRNA |
| chr12_70257836_70258220 | chr12 | 70257837 | 70258220 | + | -0.920862 | intron (NM_053167, intron 8 of 13) | intron (NM_053167, intron 8 of 13) | 30187 | NR_045174 | 414124 | Mm.416164 | NR_045174 | ENSMUSG00000085298 | **F730035M05Rik** | - | RIKEN cDNA F730035M05 gene | ncRNA |
| chr12_33128131_33128475 | chr12 | 33128132 | 33128475 | + | -0.853296 | intron (NR_015602, intron 4 of 5) | intron (NR_015602, intron 4 of 5) | 19283 | NR_015602 | 320046 | Mm.208337 | NR_015602 | ENSMUSG00000052125 | **F730043M19Rik** | - | RIKEN cDNA F730043M19 gene | ncRNA |
| chr5_125016433_125017152 | chr5 | 125016434 | 125017152 | + | -0.647683 | TTS (NM_001253905) | TTS (NM_001253905) | 13318 | NM_028443 | 73121 | Mm.296138 | NM_028443 | ENSMUSG00000037962 | **Fam101a** | 3110032G18Rik\|cfm\|cfm2 | family with sequence similarity 101, member A | protein-coding |
| chr5_125016438_125016860 | chr5 | 125016439 | 125016860 | + | -0.669467 | TTS (NM_001253905) | TTS (NM_001253905) | 13174 | NM_028443 | 73121 | Mm.296138 | NM_028443 | ENSMUSG00000037962 | **Fam101a** | 3110032G18Rik\|cfm\|cfm2 | family with sequence similarity 101, member A | protein-coding |
| chr5_124935429_124936337 | chr5 | 124935430 | 124936337 | + | -0.807977 | Intergenic | Intergenic | -67592 | NM_028443 | 73121 | Mm.296138 | NM_028443 | ENSMUSG00000037962 | **Fam101a** | 3110032G18Rik\|cfm\|cfm2 | family with sequence similarity 101, member A | protein-coding |
| chr5_124954656_124955410 | chr5 | 124954657 | 124955410 | + | -1.21932 | Intergenic | Intergenic | -48442 | NM_028443 | 73121 | Mm.296138 | NM_028443 | ENSMUSG00000037962 | **Fam101a** | 3110032G18Rik\|cfm\|cfm2 | family with sequence similarity 101, member A | protein-coding |
| chr11_76015198_76015746 | chr11 | 76015199 | 76015746 | + | -0.653725 | Intergenic | Intergenic | 12310 | NM_029658 | 76566 | Mm.34131 | NM_029658 | ENSMUSG00000020846 | **Fam101b** | 1500005K14Rik\|RefilinB\|cfm | family with sequence similarity 101, member B | protein-coding |
| chr11_76040150_76040420 | chr11 | 76040151 | 76040420 | + | -0.655568 | Intergenic | Intergenic | -12503 | NM_029658 | 76566 | Mm.34131 | NM_029658 | ENSMUSG00000020846 | **Fam101b** | 1500005K14Rik\|RefilinB\|cfm | family with sequence similarity 101, member B | protein-coding |
| chr11_76026385_76026843 | chr11 | 76026386 | 76026843 | + | -0.830393 | intron (NM_029658, intron 1 of 2) | intron (NM_029658, intron 1 of 2) | 1168 | NM_029658 | 76566 | Mm.34131 | NM_029658 | ENSMUSG00000020846 | **Fam101b** | 1500005K14Rik\|RefilinB\|cfm | family with sequence similarity 101, member B | protein-coding |
| chr11_76026620_76026848 | chr11 | 76026621 | 76026848 | + | -0.884824 | intron (NM_029658, intron 1 of 2) | intron (NM_029658, intron 1 of 2) | 1048 | NM_029658 | 76566 | Mm.34131 | NM_029658 | ENSMUSG00000020846 | **Fam101b** | 1500005K14Rik\|RefilinB\|cfm | family with sequence similarity 101, member B | protein-coding |
| chr11_76037602_76037884 | chr11 | 76037603 | 76037884 | + | -1.07984 | Intergenic | PB1D11\|SINE\|Alu | -9961 | NM_029658 | 76566 | Mm.34131 | NM_029658 | ENSMUSG00000020846 | **Fam101b** | 1500005K14Rik\|RefilinB\|cfm | family with sequence similarity 101, member B | protein-coding |
| chr2_32541910_32542520 | chr2 | 32541911 | 32542520 | + | -0.666088 | intron (NM_153560, intron 1 of 10) | intron (NM_153560, intron 1 of 10) | 6856 | NM_153560 | 98952 | Mm.4065 | NM_153560 | ENSMUSG00000039157 | **Fam102a** | AI426465\|C230093N12Rik\|Eeig1 | family with sequence similarity 102, member A | protein-coding |
| chr2_32520306_32520581 | chr2 | 32520307 | 32520581 | + | -0.820303 | Intergenic | L1M3\|LINE\|L1 | -14915 | NM_153560 | 98952 | Mm.4065 | NM_153560 | ENSMUSG00000039157 | **Fam102a** | AI426465\|C230093N12Rik\|Eeig1 | family with sequence similarity 102, member A | protein-coding |
| chr2_32495170_32495820 | chr2 | 32495171 | 32495820 | + | -0.968989 | Intergenic | Intergenic | -39864 | NM_153560 | 98952 | Mm.4065 | NM_153560 | ENSMUSG00000039157 | **Fam102a** | AI426465\|C230093N12Rik\|Eeig1 | family with sequence similarity 102, member A | protein-coding |
| chr15_27724245_27724739 | chr15 | 27724246 | 27724739 | + | -0.610753 | Intergenic | Intergenic | -42950 | NM_001242423 | 223433 | Mm.266485 | NM_198301 | ENSMUSG00000056069 | **Fam105a** | 9830126M18 | family with sequence similarity 105, member A | protein-coding |
| chr15_27785227_27785728 | chr15 | 27785228 | 27785728 | + | -0.668378 | intron (NM_001081302, intron 34 of 56) | Lx8b\|LINE\|L1 | -103936 | NM_001242423 | 223433 | Mm.266485 | NM_198301 | ENSMUSG00000056069 | **Fam105a** | 9830126M18 | family with sequence similarity 105, member A | protein-coding |

| | | | | | | | | | | | | | | | | | |
|---|---|---|---|---|---|---|---|---|---|---|---|---|---|---|---|---|---|
| chr15_27800211_27800562 | chr15 | 27800212 | 27800562 | + | -0.671589 | intron (NM_001081302, intron 34 of 56) | L1MB2\|LINE\|L1 | -118845 | NM_001242423 | 223433 | Mm.266485 | NM_198301 | ENSMUSG00000056069 | **Fam105a** | 9830126M18 | family with sequence similarity 105, member A | protein-coding |
| chr15_27795950_27796445 | chr15 | 27795951 | 27796445 | + | -0.699798 | intron (NM_001081302, intron 34 of 56) | intron (NM_001081302, intron 34 of 56) | -114656 | NM_001242423 | 223433 | Mm.266485 | NM_198301 | ENSMUSG00000056069 | **Fam105a** | 9830126M18 | family with sequence similarity 105, member A | protein-coding |
| chr15_27796544_27797279 | chr15 | 27796545 | 27797279 | + | -0.836376 | intron (NM_001081302, intron 34 of 56) | intron (NM_001081302, intron 34 of 56) | -115370 | NM_001242423 | 223433 | Mm.266485 | NM_198301 | ENSMUSG00000056069 | **Fam105a** | 9830126M18 | family with sequence similarity 105, member A | protein-coding |
| chr15_27827874_27828343 | chr15 | 27827875 | 27828343 | + | -1.0781 | intron (NM_001081302, intron 28 of 56) | intron (NM_001081302, intron 28 of 56) | -146567 | NM_001242423 | 223433 | Mm.266485 | NM_198301 | ENSMUSG00000056069 | **Fam105a** | 9830126M18 | family with sequence similarity 105, member A | protein-coding |
| chr15_27796578_27796890 | chr15 | 27796579 | 27796890 | + | -1.10443 | intron (NM_001081302, intron 34 of 56) | intron (NM_001081302, intron 34 of 56) | -115192 | NM_001242423 | 223433 | Mm.266485 | NM_198301 | ENSMUSG00000056069 | **Fam105a** | 9830126M18 | family with sequence similarity 105, member A | protein-coding |
| chr2_3730819_3731050 | chr2 | 3730820 | 3731050 | + | -0.640646 | intron (NM_025626, intron 1 of 3) | intron (NM_025626, intron 1 of 3) | 17477 | NM_025626 | 66540 | Mm.277864 | NM_025626 | ENSMUSG00000026655 | **Fam107b** | 3110001A13Rik\|AA589595\|AA959793\|AI132312 | family with sequence similarity 107, member B | protein-coding |
| chr2_3710503_3710898 | chr2 | 3710504 | 3710898 | + | -0.866461 | Intergenic | MTEb\|LTR\|ERVL-MaLR | -2757 | NM_025626 | 66540 | Mm.277864 | NM_025626 | ENSMUSG00000026655 | **Fam107b** | 3110001A13Rik\|AA589595\|AA959793\|AI132312 | family with sequence similarity 107, member B | protein-coding |
| chr2_3922780_3923088 | chr2 | 3922781 | 3923088 | + | -1.01194 | Intergenic | ORR1B1\|LTR\|ERVL-MaLR | 209476 | NM_025626 | 66540 | Mm.277864 | NM_025626 | ENSMUSG00000026655 | **Fam107b** | 3110001A13Rik\|AA589595\|AA959793\|AI132312 | family with sequence similarity 107, member B | protein-coding |
| chr19_12541148_12542141 | chr19 | 12541149 | 12542141 | + | -0.839334 | Intergenic | Intergenic | -31880 | NM_026640 | 107373 | Mm.1643 | NM_026640 | ENSMUSG00000024691 | **Fam111a** | 4632417K18Rik\|AW413625 | family with sequence similarity 111, member A | protein-coding |
| chr19_12541532_12542135 | chr19 | 12541533 | 12542135 | + | -0.938363 | Intergenic | Intergenic | -31691 | NM_026640 | 107373 | Mm.1643 | NM_026640 | ENSMUSG00000024691 | **Fam111a** | 4632417K18Rik\|AW413625 | family with sequence similarity 111, member A | protein-coding |
| chr11_95350810_95351360 | chr11 | 95350811 | 95351360 | + | -0.653003 | intron (NM_172543, intron 1 of 7) | intron (NM_172543, intron 1 of 7) | 14067 | NM_172543 | 215512 | Mm.262113 | NM_172543 | ENSMUSG00000038893 | **Fam117a** | 5730593F17Rik\|AW548096 | family with sequence similarity 117, member A | protein-coding |
| chr11_95333281_95333554 | chr11 | 95333282 | 95333554 | + | -0.722568 | Intergenic | Intergenic | -3600 | NM_172543 | 215512 | Mm.262113 | NM_172543 | ENSMUSG00000038893 | **Fam117a** | 5730593F17Rik\|AW548096 | family with sequence similarity 117, member A | protein-coding |
| chr11_95357137_95357324 | chr11 | 95357138 | 95357324 | + | -0.803669 | intron (NM_172543, intron 1 of 7) | intron (NM_172543, intron 1 of 7) | 20213 | NM_172543 | 215512 | Mm.262113 | NM_172543 | ENSMUSG00000038893 | **Fam117a** | 5730593F17Rik\|AW548096 | family with sequence similarity 117, member A | protein-coding |
| chr11_95356291_95357043 | chr11 | 95356292 | 95357043 | + | -0.944436 | intron (NM_172543, intron 1 of 7) | intron (NM_172543, intron 1 of 7) | 19649 | NM_172543 | 215512 | Mm.262113 | NM_172543 | ENSMUSG00000038893 | **Fam117a** | 5730593F17Rik\|AW548096 | family with sequence similarity 117, member A | protein-coding |
| chr11_95356504_95357429 | chr11 | 95356505 | 95357429 | + | -1.02617 | intron (NM_172543, intron 1 of 7) | intron (NM_172543, intron 1 of 7) | 19949 | NM_172543 | 215512 | Mm.262113 | NM_172543 | ENSMUSG00000038893 | **Fam117a** | 5730593F17Rik\|AW548096 | family with sequence similarity 117, member A | protein-coding |
| chr1_59895506_59896154 | chr1 | 59895507 | 59896154 | + | -0.774614 | Intergenic | Intergenic | -17176 | NM_001037725 | 72750 | Mm.258250 | NM_001037725 | ENSMUSG00000041040 | **Fam117b** | 2810425F24Rik\|6330416D14Rik\|Als2cr13 | family with sequence similarity 117, member B | protein-coding |
| chr1_59895298_59895846 | chr1 | 59895299 | 59895846 | + | -0.878365 | Intergenic | Intergenic | -17434 | NM_001037725 | 72750 | Mm.258250 | NM_001037725 | ENSMUSG00000041040 | **Fam117b** | 2810425F24Rik\|6330416D14Rik\|Als2cr13 | family with sequence similarity 117, member B | protein-coding |
| chr15_85042279_85043031 | chr15 | 85042280 | 85043031 | + | -0.666188 | intron (NM_133750, intron 1 of 8) | intron (NM_133750, intron 1 of 8) | 5593 | NM_133750 | 73225 | Mm.247535 | NM_133750 | ENSMUSG00000022434 | **Fam118a** | - | family with sequence similarity 118, member A | protein-coding |
| chr13_48929620_48930130 | chr13 | 48929621 | 48930130 | + | -0.684042 | intron (NM_001033268, intron 5 of 17) | intron (NM_001033268, intron 5 of 17) | 37953 | NM_001033268 | 218236 | Mm.426571 | NM_001033268 | ENSMUSG00000038014 | **Fam120a** | C9orf10\|Ossa | family with sequence similarity 120, member A | protein-coding |
| chr13_48994925_48995303 | chr13 | 48994926 | 48995303 | + | -0.945193 | Intergenic | Intergenic | 27002 | NR_015601 | 68128 | Mm.426304 | NR_015601 | | **Fam120aos** | C030044B11Rik | family with sequence similarity 120A, opposite strand | ncRNA |
| chr2_32890259_32890771 | chr2 | 32890260 | 32890771 | + | -0.988893 | intron (NM_146119, intron 1 of 13) | intron (NM_146119, intron 1 of 13) | 14381 | NM_146119 | 227737 | Mm.301646 | NM_146119 | ENSMUSG00000026796 | **Fam129b** | 9130404D14Rik | family with sequence similarity 129, member B | protein-coding |
| chr15_25909925_25910681 | chr15 | 25909926 | 25910681 | + | -0.666384 | intron (NM_001034851, intron 3 of 8) | intron (NM_001034851, intron 3 of 8) | -23249 | NM_001277318 | 66270 | Mm.25311 | NM_025459 | ENSMUSG00000022270 | **Fam134b** | 1810015C04Rik\|AU015349 | family with sequence similarity 134, member B | protein-coding |
| chr15_25910229_25910988 | chr15 | 25910230 | 25910988 | + | -0.785287 | intron (NM_001034851, intron 3 of 8) | intron (NM_001034851, intron 3 of 8) | -22943 | NM_001277318 | 66270 | Mm.25311 | NM_025459 | ENSMUSG00000022270 | **Fam134b** | 1810015C04Rik\|AU015349 | family with sequence similarity 134, member B | protein-coding |
| chr7_105395151_105395486 | chr7 | 105395152 | 105395486 | + | -0.703859 | intron (NM_001242363, intron 1 of 12) | intron (NM_001242363, intron 1 of 12) | 4735 | NM_001242364 | 74349 | Mm.30254 | NM_199009 | ENSMUSG00000044465 | **Fam160a2** | 4632419K20Rik\|6530415H11Rik\|FHIP | family with sequence similarity 160, member A2 | protein-coding |
| chr1_34831079_34831744 | chr1 | 34831080 | 34831744 | + | -0.73761 | intron (NM_174997, intron 2 of 6) | intron (NM_174997, intron 2 of 6) | 11638 | NM_001160235 | 214469 | Mm.120151 | NM_174997 | ENSMUSG00000037503 | **Fam168b** | mKIAA4042 | family with sequence similarity 168, member B | protein-coding |
| chr1_34831298_34831765 | chr1 | 34831299 | 34831765 | + | -0.817129 | intron (NM_174997, intron 2 of 6) | intron (NM_174997, intron 2 of 6) | 11518 | NM_001160235 | 214469 | Mm.120151 | NM_174997 | ENSMUSG00000037503 | **Fam168b** | mKIAA4042 | family with sequence similarity 168, member B | protein-coding |
| chr15_31648901_31650371 | chr15 | 31648902 | 31650371 | + | -0.735775 | Intergenic | Intergenic | 47520 | NM_026546 | 68073 | Mm.119185 | NM_026546 | ENSMUSG00000039065 | **Fam173b** | A930016P21Rik\|AA987072 | family with sequence similarity 173, member B | protein-coding |
| chr15_31649114_31650146 | chr15 | 31649115 | 31650146 | + | -0.800095 | Intergenic | Intergenic | 47514 | NM_026546 | 68073 | Mm.119185 | NM_026546 | ENSMUSG00000039065 | **Fam173b** | A930016P21Rik\|AA987072 | family with sequence similarity 173, member B | protein-coding |
| chr15_31649493_31650167 | chr15 | 31649494 | 31650167 | + | -0.895039 | Intergenic | MTE2a\|LTR\|ERVL-MaLR | 47714 | NM_026546 | 68073 | Mm.119185 | NM_026546 | ENSMUSG00000039065 | **Fam173b** | A930016P21Rik\|AA987072 | family with sequence similarity 173, member B | protein-coding |
| chr15_31650941_31651145 | chr15 | 31650942 | 31651145 | + | -1.27648 | Intergenic | Intergenic | 48927 | NM_026546 | 68073 | Mm.119185 | NM_026546 | ENSMUSG00000039065 | **Fam173b** | A930016P21Rik\|AA987072 | family with sequence similarity 173, member B | protein-coding |
| chr15_31650818_31651403 | chr15 | 31650819 | 31651403 | + | -1.38847 | Intergenic | Intergenic | 48995 | NM_026546 | 68073 | Mm.119185 | NM_026546 | ENSMUSG00000039065 | **Fam173b** | A930016P21Rik\|AA987072 | family with sequence similarity 173, member B | protein-coding |
| chr1_95450942_95451338 | chr1 | 95450943 | 95451338 | + | -0.644474 | Intergenic | Intergenic | 137512 | NM_026321 | 67698 | Mm.272705 | NM_026321 | ENSMUSG00000051185 | **Fam174a** | 2310044D20Rik\|Fam174\|Naa6\|Tmem157 | family with sequence similarity 174, member A | protein-coding |
| chr1_95451109_95451333 | chr1 | 95451110 | 95451333 | + | -0.692034 | Intergenic | Intergenic | 137593 | NM_026321 | 67698 | Mm.272705 | NM_026321 | ENSMUSG00000051185 | **Fam174a** | 2310044D20Rik\|Fam174\|Naa6\|Tmem157 | family with sequence similarity 174, member A | protein-coding |
| chr1_95451056_95451365 | chr1 | 95451057 | 95451365 | + | -0.73304 | Intergenic | Intergenic | 137583 | NM_026321 | 67698 | Mm.272705 | NM_026321 | ENSMUSG00000051185 | **Fam174a** | 2310044D20Rik\|Fam174\|Naa6\|Tmem157 | family with sequence similarity 174, member A | protein-coding |
| chr15_99275984_99276808 | chr15 | 99275985 | 99276808 | + | -0.791491 | intron (NM_001081254, intron 5 of 7) | MLTR14\|LTR\|ERV1 | 10784 | NM_001081254 | 545136 | Mm.324745 | NM_001081254 | ENSMUSG00000078907 | **Fam186b** | EG545136 | family with sequence similarity 186, member B | protein-coding |
| chr3_89190428_89190785 | chr3 | 89190429 | 89190785 | + | -0.934082 | Intergenic | Intergenic | 7382 | NM_001014995 | 68521 | Mm.274708 | NM_001014995 | ENSMUSG00000032657 | **Fam189b** | 1110013L07Rik | family with sequence similarity 189, member B | protein-coding |
| chr11_34398594_34398888 | chr11 | 34398595 | 34398888 | + | -0.680653 | intron (NM_001025382, intron 1 of 3) | intron (NM_001025382, intron 1 of 3) | 83919 | NM_001025382 | 574403 | Mm.444796 | NM_001025382 | ENSMUSG00000069911 | **Fam196b** | EG574403\|Gm6041 | family with sequence similarity 196, member B | protein-coding |
| chr6_97015890_97016642 | chr6 | 97015891 | 97016642 | + | -0.761218 | intron (NM_177233, intron 2 of 5) | intron (NM_177233, intron 2 of 5) | 44145 | NM_177233 | 320701 | Mm.131700 | NM_177233 | ENSMUSG00000046500 | **Fam19a4** | C130034I18Rik\|Fam19a4 Tafa-4\|Tafa-4 | family with sequence similarity 19, member A4 | protein-coding |
| chr6_22402166_22402397 | chr6 | 22402167 | 22402397 | + | -0.626483 | Intergenic | Intergenic | -46201 | NM_138587 | 27999 | Mm.490307 | NM_138587 | ENSMUSG00000029672 | **Fam3c** | D6Wsu176e\|Ilei | family with sequence similarity 3, member C | protein-coding |
| chr6_22350049_22350522 | chr6 | 22350050 | 22350522 | + | -0.639687 | intron (NM_138587, intron 1 of 9) | ORR1C2\|LTR\|ERVL-MaLR | 5795 | NM_138587 | 27999 | Mm.490307 | NM_138587 | ENSMUSG00000029672 | **Fam3c** | D6Wsu176e\|Ilei | family with sequence similarity 3, member C | protein-coding |
| chr6_22402161_22402476 | chr6 | 22402162 | 22402476 | + | -0.656792 | Intergenic | Intergenic | -46238 | NM_138587 | 27999 | Mm.490307 | NM_138587 | ENSMUSG00000029672 | **Fam3c** | D6Wsu176e\|Ilei | family with sequence similarity 3, member C | protein-coding |
| chr6_22350210_22350548 | chr6 | 22350211 | 22350548 | + | -0.71242 | intron (NM_138587, intron 1 of 9) | ID_B1\|SINE\|B4 | 5702 | NM_138587 | 27999 | Mm.490307 | NM_138587 | ENSMUSG00000029672 | **Fam3c** | D6Wsu176e\|Ilei | family with sequence similarity 3, member C | protein-coding |
| chr6_22444097_22444810 | chr6 | 22444098 | 22444810 | + | -0.736384 | Intergenic | Intergenic | -88373 | NM_138587 | 27999 | Mm.490307 | NM_138587 | ENSMUSG00000029672 | **Fam3c** | D6Wsu176e\|Ilei | family with sequence similarity 3, member C | protein-coding |
| chr6_22349490_22349939 | chr6 | 22349491 | 22349939 | + | -0.958663 | intron (NM_138587, intron 1 of 9) | intron (NM_138587, intron 1 of 9) | 6366 | NM_138587 | 27999 | Mm.490307 | NM_138587 | ENSMUSG00000029672 | **Fam3c** | D6Wsu176e\|Ilei | family with sequence similarity 3, member C | protein-coding |
| chr6_22374907_22375142 | chr6 | 22374908 | 22375142 | + | -0.987181 | Intergenic | Intergenic | -18944 | NM_138587 | 27999 | Mm.490307 | NM_138587 | ENSMUSG00000029672 | **Fam3c** | D6Wsu176e\|Ilei | family with sequence similarity 3, member C | protein-coding |
| chr6_22374735_22375202 | chr6 | 22374736 | 22375202 | + | -1.0089 | Intergenic | Intergenic | -18888 | NM_138587 | 27999 | Mm.490307 | NM_138587 | ENSMUSG00000029672 | **Fam3c** | D6Wsu176e\|Ilei | family with sequence similarity 3, member C | protein-coding |
| chr16_30686507_30686890 | chr16 | 30686508 | 30686890 | + | -0.788379 | Intergenic | Intergenic | 86976 | NM_177632 | 224093 | Mm.28244 | NM_177632 | ENSMUSG00000046546 | **Fam43a** | Tuf1 | family with sequence similarity 43, member A | protein-coding |
| chr9_85123671_85124193 | chr9 | 85123672 | 85124193 | + | -0.623576 | Intergenic | Intergenic | 203192 | NM_001160378 | 212943 | Mm.74610 | NM_001160378 | ENSMUSG00000032265 | **Fam46a** | D930050G01Rik | family with sequence similarity 46, member A | protein-coding |
| chr9_85125795_85126169 | chr9 | 85125796 | 85126169 | + | -0.735056 | Intergenic | MLT1C\|LTR\|ERVL-MaLR | 201142 | NM_001160378 | 212943 | Mm.74610 | NM_001160378 | ENSMUSG00000032265 | **Fam46a** | D930050G01Rik | family with sequence similarity 46, member A | protein-coding |
| chr9_85120113_85120426 | chr9 | 85120114 | 85120426 | + | -1.0371 | Intergenic | ID_B1\|SINE\|B4 | 206854 | NM_001160378 | 212943 | Mm.74610 | NM_001160378 | ENSMUSG00000032265 | **Fam46a** | D930050G01Rik | family with sequence similarity 46, member A | protein-coding |
| chr3_100582776_100583149 | chr3 | 100582777 | 100583149 | + | -0.607287 | intron (NM_010763, intron 10 of 12) | B4A\|SINE\|B4 | -93771 | NM_001142952 | 74645 | Mm.226900 | NM_001142952 | | **Fam46c** | 4930431B09Rik\|AI449797 | family with sequence similarity 46, member C | protein-coding |
| chr3_100582813_100583060 | chr3 | 100582814 | 100583060 | + | -0.63961 | intron (NM_010763, intron 10 of 12) | B4A\|SINE\|B4 | -93745 | NM_001142952 | 74645 | Mm.226900 | NM_001142952 | | **Fam46c** | 4930431B09Rik\|AI449797 | family with sequence similarity 46, member C | protein-coding |
| chr3_100480252_100480994 | chr3 | 100480253 | 100480994 | + | -0.670488 | intron (NM_001142952, intron 1 of 1) | intron (NM_001142952, intron 1 of 1) | 8569 | NM_001142952 | 74645 | Mm.226900 | NM_001142952 | | **Fam46c** | 4930431B09Rik\|AI449797 | family with sequence similarity 46, member C | protein-coding |

| | | | | | | | | | | | | | | | | | |
|---|---|---|---|---|---|---|---|---|---|---|---|---|---|---|---|---|---|
| chr3_100558153_100558710 | chr3 | 100558154 | 100558710 | + | -0.850578 | Intergenic | L2\|LINE\|L2 | -69240 | NM_001142952 | 74645 | Mm.226900 | NM_001142952 | | **Fam46c** | 4930431B09Rik\|AI449797 | family with sequence similarity 46, member C | protein-coding |
| chr3_100562033_100562218 | chr3 | 100562034 | 100562218 | + | -0.972942 | TTS (NM_010763) | TTS (NM_010763) | -72934 | NM_001142952 | 74645 | Mm.226900 | NM_001142952 | | **Fam46c** | 4930431B09Rik\|AI449797 | family with sequence similarity 46, member C | protein-coding |
| chr15_64085864_64086114 | chr15 | 64085865 | 64086114 | + | -0.638821 | TTS (NM_001276467) | TTS (NM_001276467) | -25541 | NM_144846 | 223601 | Mm.246108 | NM_144846 | ENSMUSG00000022378 | **Fam49b** | 0910001A06Rik\|AW122079 | family with sequence similarity 49, member B | protein-coding |
| chr15_64085821_64086346 | chr15 | 64085822 | 64086346 | + | -0.671463 | TTS (NM_001276467) | TTS (NM_001276467) | -25636 | NM_144846 | 223601 | Mm.246108 | NM_144846 | ENSMUSG00000022378 | **Fam49b** | 0910001A06Rik\|AW122079 | family with sequence similarity 49, member B | protein-coding |
| chr15_64085856_64086195 | chr15 | 64085857 | 64086195 | + | -0.681956 | TTS (NM_001276467) | TTS (NM_001276467) | -25578 | NM_144846 | 223601 | Mm.246108 | NM_144846 | ENSMUSG00000022378 | **Fam49b** | 0910001A06Rik\|AW122079 | family with sequence similarity 49, member B | protein-coding |
| chr15_64041301_64041892 | chr15 | 64041302 | 64041892 | + | -0.703554 | intron (NM_144846, intron 1 of 11) | intron (NM_144846, intron 1 of 11) | 18851 | NM_144846 | 223601 | Mm.246108 | NM_144846 | ENSMUSG00000022378 | **Fam49b** | 0910001A06Rik\|AW122079 | family with sequence similarity 49, member B | protein-coding |
| chr15_64041313_64042153 | chr15 | 64041314 | 64042153 | + | -0.734863 | intron (NM_144846, intron 1 of 11) | intron (NM_144846, intron 1 of 11) | 18715 | NM_144846 | 223601 | Mm.246108 | NM_144846 | ENSMUSG00000022378 | **Fam49b** | 0910001A06Rik\|AW122079 | family with sequence similarity 49, member B | protein-coding |
| chr15_64041493_64041892 | chr15 | 64041494 | 64041892 | + | -0.829838 | intron (NM_144846, intron 1 of 11) | intron (NM_144846, intron 1 of 11) | 18755 | NM_144846 | 223601 | Mm.246108 | NM_144846 | ENSMUSG00000022378 | **Fam49b** | 0910001A06Rik\|AW122079 | family with sequence similarity 49, member B | protein-coding |
| chr15_64106938_64107278 | chr15 | 64106939 | 64107278 | + | -0.83521 | intron (NM_001276461, intron 24 of 28) | intron (NM_001276461, intron 24 of 28) | -46660 | NM_144846 | 223601 | Mm.246108 | NM_144846 | ENSMUSG00000022378 | **Fam49b** | 0910001A06Rik\|AW122079 | family with sequence similarity 49, member B | protein-coding |
| chr11_78780009_78780921 | chr11 | 78780010 | 78780921 | + | -0.714244 | Intergenic | Intergenic | -28736 | NM_197989 | 69109 | Mm.341420 | NM_197989 | ENSMUSG00000049489 | **Fam58b** | 1810009O10Rik\|Fam58a | family with sequence similarity 58, member B | protein-coding |
| chr11_78766010_78766818 | chr11 | 78766011 | 78766818 | + | -0.761544 | Intergenic | MTEa\|LTR\|ERVL-MaLR | -14685 | NM_197989 | 69109 | Mm.341420 | NM_197989 | ENSMUSG00000049489 | **Fam58b** | 1810009O10Rik\|Fam58a | family with sequence similarity 58, member B | protein-coding |
| chr11_78771648_78772409 | chr11 | 78771649 | 78772409 | + | -0.874339 | Intergenic | Intergenic | -20300 | NM_197989 | 69109 | Mm.341420 | NM_197989 | ENSMUSG00000049489 | **Fam58b** | 1810009O10Rik\|Fam58a | family with sequence similarity 58, member B | protein-coding |
| chr11_78780662_78780982 | chr11 | 78780663 | 78780982 | + | -0.976816 | Intergenic | L2b\|LINE\|L2 | -29093 | NM_197989 | 69109 | Mm.341420 | NM_197989 | ENSMUSG00000049489 | **Fam58b** | 1810009O10Rik\|Fam58a | family with sequence similarity 58, member B | protein-coding |
| chr13_24588624_24588896 | chr13 | 24588625 | 24588896 | + | -0.679564 | intron (NM_001286100, intron 1 of 22) | MTD\|LTR\|ERVL-MaLR | 6571 | NM_001286100 | 193385 | Mm.217319 | NM_029679 | ENSMUSG00000036006 | **Fam65b** | - | family with sequence similarity 65, member B | protein-coding |
| chr13_24617503_24618169 | chr13 | 24617504 | 24618169 | + | -0.704438 | intron (NM_178658, intron 1 of 13) | intron (NM_178658, intron 1 of 13) | 3230 | NM_178658 | 193385 | Mm.217319 | NM_029679 | ENSMUSG00000036006 | **Fam65b** | - | family with sequence similarity 65, member B | protein-coding |
| chr5_107946434_107946752 | chr5 | 107946435 | 107946752 | + | -0.675045 | intron (NM_026062, intron 1 of 4) | intron (NM_026062, intron 1 of 4) | 40484 | NM_026062 | 67266 | Mm.277897 | NM_026062 | ENSMUSG00000029270 | **Fam69a** | 2900024C23Rik\|AI315274 | family with sequence similarity 69, member A | protein-coding |
| chr5_107946457_107946913 | chr5 | 107946458 | 107946913 | + | -0.690164 | intron (NM_026062, intron 1 of 4) | intron (NM_026062, intron 1 of 4) | 40392 | NM_026062 | 67266 | Mm.277897 | NM_026062 | ENSMUSG00000029270 | **Fam69a** | 2900024C23Rik\|AI315274 | family with sequence similarity 69, member A | protein-coding |
| chr5_107966861_107967176 | chr5 | 107966862 | 107967176 | + | -0.757158 | intron (NM_026062, intron 1 of 4) | intron (NM_026062, intron 1 of 4) | 20058 | NM_026062 | 67266 | Mm.277897 | NM_026062 | ENSMUSG00000029270 | **Fam69a** | 2900024C23Rik\|AI315274 | family with sequence similarity 69, member A | protein-coding |
| chr5_107977590_107978985 | chr5 | 107977591 | 107978985 | + | -0.791314 | intron (NM_026062, intron 1 of 4) | L2c\|LINE\|L2 | 8789 | NM_026062 | 67266 | Mm.277897 | NM_026062 | ENSMUSG00000029270 | **Fam69a** | 2900024C23Rik\|AI315274 | family with sequence similarity 69, member A | protein-coding |
| chr5_107977975_107978744 | chr5 | 107977976 | 107978744 | + | -0.917301 | intron (NM_026062, intron 1 of 4) | intron (NM_026062, intron 1 of 4) | 8717 | NM_026062 | 67266 | Mm.277897 | NM_026062 | ENSMUSG00000029270 | **Fam69a** | 2900024C23Rik\|AI315274 | family with sequence similarity 69, member A | protein-coding |
| chr5_107978178_107978766 | chr5 | 107978179 | 107978766 | + | -1.01441 | intron (NM_026062, intron 1 of 4) | intron (NM_026062, intron 1 of 4) | 8605 | NM_026062 | 67266 | Mm.277897 | NM_026062 | ENSMUSG00000029270 | **Fam69a** | 2900024C23Rik\|AI315274 | family with sequence similarity 69, member A | protein-coding |
| chr2_26622961_26623378 | chr2 | 26622962 | 26623378 | + | -1.12087 | Intergenic | Intergenic | -5287 | NM_019833 | 56279 | Mm.279863 | NM_019833 | ENSMUSG00000036186 | **Fam69b** | AB030186\|AW209000\|B230317C12Rik\|PIP49 | family with sequence similarity 69, member B | protein-coding |
| chr6_29286940_29287209 | chr6 | 29286941 | 29287209 | + | -0.657466 | intron (NM_001101486, intron 3 of 4) | intron (NM_001101486, intron 3 of 4) | 5934 | NM_001101486 | 245884 | Mm.33683 | NM_001101486 | ENSMUSG00000079652 | **Fam71f2** | AI507611\|GARI | family with sequence similarity 71, member F2 | protein-coding |
| chr6_29286930_29287326 | chr6 | 29286931 | 29287326 | + | -0.67096 | intron (NM_001101486, intron 3 of 4) | intron (NM_001101486, intron 3 of 4) | 5987 | NM_001101486 | 245884 | Mm.33683 | NM_001101486 | ENSMUSG00000079652 | **Fam71f2** | AI507611\|GARI | family with sequence similarity 71, member F2 | protein-coding |
| chr6_29286790_29287203 | chr6 | 29286791 | 29287203 | + | -0.716137 | intron (NM_001101486, intron 3 of 4) | PB1D10\|SINE\|Alu | 5856 | NM_001101486 | 245884 | Mm.33683 | NM_001101486 | ENSMUSG00000079652 | **Fam71f2** | AI507611\|GARI | family with sequence similarity 71, member F2 | protein-coding |
| chr2_32077786_32078020 | chr2 | 32077787 | 32078020 | + | -0.626501 | intron (NM_175511, intron 1 of 1) | intron (NM_175511, intron 1 of 1) | 5802 | NM_175511 | 241303 | Mm.250441 | NM_175511 | ENSMUSG00000050592 | **Fam78a** | A130092J06Rik | family with sequence similarity 78, member A | protein-coding |
| chr2_32084596_32084987 | chr2 | 32084597 | 32084987 | + | -0.681598 | Intergenic | (TC)n\|Simple_repeat\|Simple_repeat | -1087 | NM_175511 | 241303 | Mm.250441 | NM_175511 | ENSMUSG00000050592 | **Fam78a** | A130092J06Rik | family with sequence similarity 78, member A | protein-coding |
| chr15_80684571_80684778 | chr15 | 80684572 | 80684778 | + | -0.63793 | intron (NM_145986, intron 1 of 4) | intron (NM_145986, intron 1 of 4) | 12828 | NM_145986 | 213956 | Mm.132439 | NM_145986 | ENSMUSG00000022408 | **Fam83f** | AW544981 | family with sequence similarity 83, member F | protein-coding |
| chr11_61669705_61670092 | chr11 | 61669706 | 61670092 | + | -0.610616 | exon (NM_027817, exon 4 of 5) | exon (NM_027817, exon 4 of 5) | -14511 | NM_178618 | 69640 | Mm.458559 | NM_178618 | ENSMUSG00000042377 | **Fam83g** | 2310040C09Rik\|AI506990\|wly | family with sequence similarity 83, member G | protein-coding |
| chr16_5452767_5453532 | chr16 | 5452768 | 5453532 | + | -0.642532 | Intergenic | Intergenic | -197194 | NM_027446 | 70511 | Mm.45780 | NM_027446 | ENSMUSG00000022544 | **Fam86** | 5730409G15Rik\|Fam86a | family with sequence similarity 86 | protein-coding |
| chr16_5445546_5446143 | chr16 | 5445547 | 5446143 | + | -0.915186 | Intergenic | Intergenic | -189889 | NM_027446 | 70511 | Mm.45780 | NM_027446 | ENSMUSG00000022544 | **Fam86** | 5730409G15Rik\|Fam86a | family with sequence similarity 86 | protein-coding |
| chr13_63410912_63411574 | chr13 | 63410913 | 63411574 | + | -0.609031 | intron (NM_001042673, intron 1 of 15) | L1MEi\|LINE\|L1 | 20502 | NM_007985 | 14088 | Mm.126106 | NM_007985 | ENSMUSG00000021461 | **Fancc** | Facc | Fanconi anemia, complementation group C | protein-coding |
| chr7_79401496_79401917 | chr7 | 79401497 | 79401917 | + | -1.26119 | intron (NM_145946, intron 3 of 37) | intron (NM_145946, intron 3 of 37) | 9369 | NM_145946 | 208836 | Mm.349493 | NM_145946 | ENSMUSG00000039187 | **Fanci** | - | Fanconi anemia, complementation group I | protein-coding |
| chr7_113572740_113573437 | chr7 | 113572741 | 113573437 | + | -0.791735 | Intergenic | Intergenic | 59255 | NM_027379 | 67420 | Mm.206919 | NM_026143 | ENSMUSG00000030759 | **Far1** | 2600011M19Rik\|2900034E22Rik\|3732409C05Rik\|AI850429\|Mlstd2 | fatty acyl CoA reductase 1 | protein-coding |
| chr7_113572768_113573107 | chr7 | 113572769 | 113573107 | + | -0.948315 | Intergenic | Intergenic | 59104 | NM_027379 | 67420 | Mm.206919 | NM_026143 | ENSMUSG00000030759 | **Far1** | 2600011M19Rik\|2900034E22Rik\|3732409C05Rik\|AI850429\|Mlstd2 | fatty acyl CoA reductase 1 | protein-coding |
| chr1_78485314_78485588 | chr1 | 78485315 | 78485588 | + | -1.14742 | intron (NR_102757, intron 1 of 16) | MLT1F2-int\|LTR\|ERVL-MaLR | 3446 | NR_102758 | 23874 | Mm.28922 | NM_011811 | ENSMUSG00000026245 | **Farsb** | C76708\|Farsa\|Farsl\|Farslb\|Frsb | phenylalanyl-tRNA synthetase, beta subunit | protein-coding |
| chr1_161712882_161713401 | chr1 | 161712883 | 161713401 | + | -0.641382 | Intergenic | Intergenic | 75353 | NM_001205243 | 14103 | Mm.3355 | NM_010177 | ENSMUSG00000000817 | **Fasl** | APT1LG1\|CD178\|CD95-L\|CD95L\|Fas-L\|Faslg\|Tnfsf6\|gld | Fas ligand (TNF superfamily, member 6) | protein-coding |
| chr1_161788001_161788836 | chr1 | 161788002 | 161788836 | + | -0.72361 | promoter-TSS (NM_010177) | promoter-TSS (NM_010177) | 76 | NM_001205243 | 14103 | Mm.3355 | NM_010177 | ENSMUSG00000000817 | **Fasl** | APT1LG1\|CD178\|CD95-L\|CD95L\|Fas-L\|Faslg\|Tnfsf6\|gld | Fas ligand (TNF superfamily, member 6) | protein-coding |
| chr1_161704513_161705291 | chr1 | 161704514 | 161705291 | + | -0.751095 | Intergenic | Intergenic | 83593 | NM_001205243 | 14103 | Mm.3355 | NM_010177 | ENSMUSG00000000817 | **Fasl** | APT1LG1\|CD178\|CD95-L\|CD95L\|Fas-L\|Faslg\|Tnfsf6\|gld | Fas ligand (TNF superfamily, member 6) | protein-coding |
| chr1_161801380_161801654 | chr1 | 161801381 | 161801654 | + | -0.818264 | Intergenic | Intergenic | -13022 | NM_010177 | 14103 | Mm.3355 | NM_010177 | ENSMUSG00000000817 | **Fasl** | APT1LG1\|CD178\|CD95-L\|CD95L\|Fas-L\|Faslg\|Tnfsf6\|gld | Fas ligand (TNF superfamily, member 6) | protein-coding |
| chr1_161660714_161661182 | chr1 | 161660715 | 161661182 | + | -0.915853 | Intergenic | RMER15\|LTR\|ERVL | 127547 | NM_001205243 | 14103 | Mm.3355 | NM_010177 | ENSMUSG00000000817 | **Fasl** | APT1LG1\|CD178\|CD95-L\|CD95L\|Fas-L\|Faslg\|Tnfsf6\|gld | Fas ligand (TNF superfamily, member 6) | protein-coding |
| chr1_161779053_161779504 | chr1 | 161779054 | 161779504 | + | -0.969053 | Intergenic | Intergenic | 9216 | NM_001205243 | 14103 | Mm.3355 | NM_010177 | ENSMUSG00000000817 | **Fasl** | APT1LG1\|CD178\|CD95-L\|CD95L\|Fas-L\|Faslg\|Tnfsf6\|gld | Fas ligand (TNF superfamily, member 6) | protein-coding |
| chr1_161765913_161766325 | chr1 | 161765914 | 161766325 | + | -1.21529 | Intergenic | Intergenic | 22376 | NM_001205243 | 14103 | Mm.3355 | NM_010177 | ENSMUSG00000000817 | **Fasl** | APT1LG1\|CD178\|CD95-L\|CD95L\|Fas-L\|Faslg\|Tnfsf6\|gld | Fas ligand (TNF superfamily, member 6) | protein-coding |
| chr15_85250392_85251023 | chr15 | 85250393 | 85251023 | + | -1.17472 | intron (NM_010180, intron 14 of 16) | intron (NM_010180, intron 14 of 16) | 44700 | NM_010180 | 14114 | Mm.297992 | NM_010180 | ENSMUSG00000006369 | **Fbln1** | - | fibulin 1 | protein-coding |
| chr15_85250376_85250852 | chr15 | 85250377 | 85250852 | + | -1.19242 | intron (NM_010180, intron 14 of 16) | intron (NM_010180, intron 14 of 16) | 44606 | NM_010180 | 14114 | Mm.297992 | NM_010180 | ENSMUSG00000006369 | **Fbln1** | - | fibulin 1 | protein-coding |
| chr6_91259609_91260174 | chr6 | 91259610 | 91260174 | + | -0.746286 | intron (NM_007992, intron 9 of 17) | intron (NM_007992, intron 9 of 17) | 47128 | NM_007992 | 14115 | Mm.249146 | NM_007992 | ENSMUSG00000064080 | **Fbln2** | 5730577E14Rik\|FIBL-2 | fibulin 2 | protein-coding |
| chr6_91259490_91259947 | chr6 | 91259491 | 91259947 | + | -0.910592 | exon (NM_007992, exon 9 of 18) | exon (NM_007992, exon 9 of 18) | 46955 | NM_007992 | 14115 | Mm.249146 | NM_007992 | ENSMUSG00000064080 | **Fbln2** | 5730577E14Rik\|FIBL-2 | fibulin 2 | protein-coding |
| chr13_62879717_62880673 | chr13 | 62879718 | 62880673 | + | -0.743969 | intron (NM_019395, intron 1 of 6) | intron (NM_019395, intron 1 of 6) | 8087 | NM_019395 | 14121 | Mm.423078 | NM_019395 | ENSMUSG00000069805 | **Fbp1** | Fbp-2\|Fbp2\|Fbp3 | fructose bisphosphatase 1 | protein-coding |
| chr9_20609824_20610257 | chr9 | 20609825 | 20610257 | + | -0.850419 | intron (NR_033729, intron 1 of 2) | PB1D10\|SINE\|Alu | 2566 | NR_033729 | 66662 | Mm.477847 | NM_175110 | ENSMUSG00000062470 | **Fbxl12os** | 5730577I03Rik | F-box and leucine-rich repeat protein 12, opposite strand | pseudo |
| chr6_119499919_119500377 | chr6 | 119499920 | 119500377 | + | -0.659426 | intron (NM_009525, intron 1 of 4) | intron (NM_009525, intron 1 of 4) | 20480 | NM_133940 | 101358 | Mm.125477 | NM_133940 | ENSMUSG00000030019 | **Fbxl14** | AW322056\|Fbx14l | F-box and leucine-rich repeat protein 14 | protein-coding |
| chr6_119499340_119499645 | chr6 | 119499341 | 119499645 | + | -0.696172 | intron (NM_009525, intron 1 of 4) | intron (NM_009525, intron 1 of 4) | 19825 | NM_133940 | 101358 | Mm.125477 | NM_133940 | ENSMUSG00000030019 | **Fbxl14** | AW322056\|Fbx14l | F-box and leucine-rich repeat protein 14 | protein-coding |
| chr6_119508238_119508504 | chr6 | 119508239 | 119508504 | + | -0.736926 | intron (NM_009525, intron 1 of 4) | ORR1B1\|LTR\|ERVL-MaLR | 28703 | NM_133940 | 101358 | Mm.125477 | NM_133940 | ENSMUSG00000030019 | **Fbxl14** | AW322056\|Fbx14l | F-box and leucine-rich repeat protein 14 | protein-coding |
| chr6_119500079_119500469 | chr6 | 119500080 | 119500469 | + | -0.79715 | intron (NM_009525, intron 1 of 4) | intron (NM_009525, intron 1 of 4) | 20606 | NM_133940 | 101358 | Mm.125477 | NM_133940 | ENSMUSG00000030019 | **Fbxl14** | AW322056\|Fbx14l | F-box and leucine-rich repeat protein 14 | protein-coding |

| | | | | | | | | | | | | | | | | | |
|---|---|---|---|---|---|---|---|---|---|---|---|---|---|---|---|---|---|
| chr6_119508597_119509534 | chr6 | 119508598 | 119509534 | + | -0.872398 | intron (NM_009525, intron 1 of 4) | L1MCa\|LINE\|L1 | 29398 | NM_133940 | 101358 | Mm.125477 | NM_133940 | ENSMUSG00000030019 | **Fbxl14** | AW322056\|Fbx14l | F-box and leucine-rich repeat protein 14 | protein-coding |
| chr17_63362109_63362576 | chr17 | 63362110 | 63362576 | + | -0.639227 | intron (NM_015794, intron 5 of 8) | RMER19A\|LTR\|ERVK | 138237 | NM_015794 | 50758 | Mm.136366 | NM_015794 | | **Fbxl17** | 6330576B01Rik\|AI452053\|BB073797\|C130023C01Rik\|Fbx13\|Fbxo13 | F-box and leucine-rich repeat protein 17 | protein-coding |
| chr17_63405140_63405870 | chr17 | 63405141 | 63405870 | + | -0.694211 | intron (NM_015794, intron 4 of 8) | intron (NM_015794, intron 4 of 8) | 95075 | NM_015794 | 50758 | Mm.136366 | NM_015794 | | **Fbxl17** | 6330576B01Rik\|AI452053\|BB073797\|C130023C01Rik\|Fbx13\|Fbxo13 | F-box and leucine-rich repeat protein 17 | protein-coding |
| chr17_63405363_63405890 | chr17 | 63405364 | 63405890 | + | -0.72693 | intron (NM_015794, intron 4 of 8) | intron (NM_015794, intron 4 of 8) | 94953 | NM_015794 | 50758 | Mm.136366 | NM_015794 | | **Fbxl17** | 6330576B01Rik\|AI452053\|BB073797\|C130023C01Rik\|Fbx13\|Fbxo13 | F-box and leucine-rich repeat protein 17 | protein-coding |
| chr17_63356264_63356821 | chr17 | 63356265 | 63356821 | + | -0.767097 | intron (NM_015794, intron 6 of 8) | intron (NM_015794, intron 6 of 8) | 144037 | NM_015794 | 50758 | Mm.136366 | NM_015794 | | **Fbxl17** | 6330576B01Rik\|AI452053\|BB073797\|C130023C01Rik\|Fbx13\|Fbxo13 | F-box and leucine-rich repeat protein 17 | protein-coding |
| chr17_63324727_63325114 | chr17 | 63324728 | 63325114 | + | -0.796193 | intron (NM_015794, intron 6 of 8) | intron (NM_015794, intron 6 of 8) | 175659 | NM_015794 | 50758 | Mm.136366 | NM_015794 | | **Fbxl17** | 6330576B01Rik\|AI452053\|BB073797\|C130023C01Rik\|Fbx13\|Fbxo13 | F-box and leucine-rich repeat protein 17 | protein-coding |
| chr17_63356036_63356718 | chr17 | 63356037 | 63356718 | + | -0.826983 | intron (NM_015794, intron 6 of 8) | intron (NM_015794, intron 6 of 8) | 144203 | NM_015794 | 50758 | Mm.136366 | NM_015794 | | **Fbxl17** | 6330576B01Rik\|AI452053\|BB073797\|C130023C01Rik\|Fbx13\|Fbxo13 | F-box and leucine-rich repeat protein 17 | protein-coding |
| chr5_142891431_142891659 | chr5 | 142891432 | 142891659 | + | -1.00509 | intron (NM_001033312, intron 1 of 4) | B4A\|SINE\|B4 | 3693 | NM_001033312 | 231863 | Mm.89753 | NM_001033312 | ENSMUSG00000066640 | **Fbxl18** | AI427077\|B130019G13Rik\|C330021B20Rik | F-box and leucine-rich repeat protein 18 | protein-coding |
| chr9_66505514_66505913 | chr9 | 66505515 | 66505913 | + | -0.687611 | intron (NM_145617, intron 77 of 77) | intron (NM_145617, intron 77 of 77) | 8879 | NM_175206 | 74165 | Mm.297998 | NM_175206 | ENSMUSG00000050503 | **Fbxl22** | 1110004B15Rik | F-box and leucine-rich repeat protein 22 | protein-coding |
| chr9_66505593_66505879 | chr9 | 66505594 | 66505879 | + | -0.689532 | intron (NM_145617, intron 77 of 77) | intron (NM_145617, intron 77 of 77) | 8857 | NM_175206 | 74165 | Mm.297998 | NM_175206 | ENSMUSG00000050503 | **Fbxl22** | 1110004B15Rik | F-box and leucine-rich repeat protein 22 | protein-coding |
| chr9_66492156_66492760 | chr9 | 66492157 | 66492760 | + | -0.997081 | intron (NM_145617, intron 69 of 77) | intron (NM_145617, intron 69 of 77) | 22135 | NM_175206 | 74165 | Mm.297998 | NM_175206 | ENSMUSG00000050503 | **Fbxl22** | 1110004B15Rik | F-box and leucine-rich repeat protein 22 | protein-coding |
| chr14_103095102_103095336 | chr14 | 103095103 | 103095336 | + | -1.02792 | exon (NM_015822, exon 2 of 5) | exon (NM_015822, exon 2 of 5) | 4290 | NM_015822 | 50789 | Mm.214746 | NM_015822 | ENSMUSG00000022124 | **Fbxl3** | AU041772\|AW212966\|FBK\|Fbl3a\|Fbxl3a\|Ovtm\|Play68 | F-box and leucine-rich repeat protein 3 | protein-coding |
| chr5_43771520_43771903 | chr5 | 43771521 | 43771903 | + | -1.07224 | intron (NM_178729, intron 2 of 8) | ORR1D2\|LTR\|ERVL-MaLR | 10437 | NM_001159963 | 242960 | Mm.25794 | NM_178729 | ENSMUSG00000039753 | **Fbxl5** | Fbl4\|Fir4 | F-box and leucine-rich repeat protein 5 | protein-coding |
| chr18_84911614_84912148 | chr18 | 84911615 | 84912148 | + | -1.43063 | Intergenic | Intergenic | -23144 | NM_015798 | 50764 | Mm.28369 | NM_015798 | ENSMUSG00000034391 | **Fbxo15** | AU019763\|Fbx15\|ecat3 | F-box protein 15 | protein-coding |
| chr5_117977705_117978144 | chr5 | 117977706 | 117978144 | + | -0.878327 | exon (NM_145564, exon 2 of 12) | exon (NM_145564, exon 2 of 12) | 1155 | NM_145564 | 231670 | Mm.21912 | NM_145564 | ENSMUSG00000032898 | **Fbxo21** | 2810425J22Rik\|AU016673\|mKIAA0875 | F-box protein 21 | protein-coding |
| chr7_28677188_28677490 | chr7 | 28677189 | 28677490 | + | -0.787859 | Intergenic | Intergenic | -15510 | NM_207238 | 233040 | Mm.116116 | NM_207238 | ENSMUSG00000037463 | **Fbxo27** | E130008B10Rik\|FBG5\|Gm161 | F-box protein 27 | protein-coding |
| chr7_28671263_28671494 | chr7 | 28671264 | 28671494 | + | -0.873107 | Intergenic | ORR1D2\|LTR\|ERVL-MaLR | -21470 | NM_207238 | 233040 | Mm.116116 | NM_207238 | ENSMUSG00000037463 | **Fbxo27** | E130008B10Rik\|FBG5\|Gm161 | F-box protein 27 | protein-coding |
| chr7_28671205_28671582 | chr7 | 28671206 | 28671582 | + | -0.948648 | Intergenic | ORR1D2\|LTR\|ERVL-MaLR | -21455 | NM_207238 | 233040 | Mm.116116 | NM_207238 | ENSMUSG00000037463 | **Fbxo27** | E130008B10Rik\|FBG5\|Gm161 | F-box protein 27 | protein-coding |
| chr1_182357934_182358569 | chr1 | 182357935 | 182358569 | + | -1.54533 | Intergenic | Intergenic | -16646 | NM_175127 | 67948 | Mm.30737 | NM_175127 | ENSMUSG00000047539 | **Fbxo28** | 4833428J17Rik\|5730505P19Rik\|D1Ertd578e\|Fbx28\|mKIAA0483 | F-box protein 28 | protein-coding |
| chr1_182358184_182358473 | chr1 | 182358185 | 182358473 | + | -1.56533 | Intergenic | Intergenic | -16723 | NM_175127 | 67948 | Mm.30737 | NM_175127 | ENSMUSG00000047539 | **Fbxo28** | 4833428J17Rik\|5730505P19Rik\|D1Ertd578e\|Fbx28\|mKIAA0483 | F-box protein 28 | protein-coding |
| chr12_59206149_59207453 | chr12 | 59206150 | 59207453 | + | -0.625189 | intron (NM_001033156, intron 1 of 3) | intron (NM_001033156, intron 1 of 3) | 12682 | NM_001033156 | 70611 | Mm.311026 | NM_001033156 | ENSMUSG00000035329 | **Fbxo33** | 5730501N20Rik\|AI642135 | F-box protein 33 | protein-coding |
| chr12_59206700_59207433 | chr12 | 59206701 | 59207433 | + | -0.930311 | intron (NM_001033156, intron 1 of 3) | B1F\|SINE\|Alu | 12416 | NM_001033156 | 70611 | Mm.311026 | NM_001033156 | ENSMUSG00000035329 | **Fbxo33** | 5730501N20Rik\|AI642135 | F-box protein 33 | protein-coding |
| chr12_59207070_59207460 | chr12 | 59207071 | 59207460 | + | -0.986593 | intron (NM_001033156, intron 1 of 3) | L1ME1\|LINE\|L1 | 12218 | NM_001033156 | 70611 | Mm.311026 | NM_001033156 | ENSMUSG00000035329 | **Fbxo33** | 5730501N20Rik\|AI642135 | F-box protein 33 | protein-coding |
| chr11_97905438_97906105 | chr11 | 97905439 | 97906105 | + | -1.51938 | Intergenic | Intergenic | -21618 | NM_001081435 | 72973 | Mm.245939 | NM_001081435 | | **Fbxo47** | 2900052P03Rik\|Gm1562 | F-box protein 47 | protein-coding |
| chr11_97905497_97905943 | chr11 | 97905498 | 97905943 | + | -1.56765 | Intergenic | Intergenic | -21566 | NM_001081435 | 72973 | Mm.245939 | NM_001081435 | | **Fbxo47** | 2900052P03Rik\|Gm1562 | F-box protein 47 | protein-coding |
| chr11_97905596_97905873 | chr11 | 97905597 | 97905873 | + | -1.624 | Intergenic | Intergenic | -21581 | NM_001081435 | 72973 | Mm.245939 | NM_001081435 | | **Fbxo47** | 2900052P03Rik\|Gm1562 | F-box protein 47 | protein-coding |
| chr3_84848654_84849172 | chr3 | 84848655 | 84849172 | + | -0.709401 | intron (NM_001177773, intron 1 of 13) | ID4\|SINE\|ID | -15232 | NM_001177774 | 50754 | Mm.196475 | NM_080428 | ENSMUSG00000028086 | **Fbxw7** | 1110001A17Rik\|AGO\|Cdc4\|Fbw7\|Fbwd6\|Fbx30\|Fbxo30\|Fbxw6\|SEL-10 | F-box and WD-40 domain protein 7 | protein-coding |
| chr1_171060288_171060834 | chr1 | 171060289 | 171060834 | + | -0.926378 | Intergenic | RLTR9B\|LTR\|ERVK | -1158 | NM_010188 | 14131 | Mm.22119 | NM_010188 | ENSMUSG00000059498 | **Fcgr3** | CD16 | Fc receptor, IgG, low affinity III | protein-coding |
| chr17_78370329_78370766 | chr17 | 78370330 | 78370766 | + | -0.928761 | 3' UTR (NM_001285949, exon 8 of 8) | 3' UTR (NM_001285949, exon 8 of 8) | 47604 | NM_001285949 | 225020 | Mm.236835 | NM_199448 | ENSMUSG00000056121 | **Fez2** | 9030616F16\|D17Ertd315e | fasciculation and elongation protein zeta 2 (zygin II) | protein-coding |
| chr17_78351607_78352846 | chr17 | 78351608 | 78352846 | + | -0.936412 | intron (NM_015800, intron 12 of 16) | intron (NM_015800, intron 12 of 16) | 65925 | NM_001285949 | 225020 | Mm.236835 | NM_199448 | ENSMUSG00000056121 | **Fez2** | 9030616F16\|D17Ertd315e | fasciculation and elongation protein zeta 2 (zygin II) | protein-coding |
| chr17_78370469_78370805 | chr17 | 78370470 | 78370805 | + | -0.941297 | 3' UTR (NM_001285949, exon 8 of 8) | 3' UTR (NM_001285949, exon 8 of 8) | 47515 | NM_001285949 | 225020 | Mm.236835 | NM_199448 | ENSMUSG00000056121 | **Fez2** | 9030616F16\|D17Ertd315e | fasciculation and elongation protein zeta 2 (zygin II) | protein-coding |
| chr17_78429076_78429471 | chr17 | 78429077 | 78429471 | + | -1.58264 | Intergenic | Lx7\|LINE\|L1 | -11122 | NM_001285940 | 225020 | Mm.236835 | NM_199448 | ENSMUSG00000056121 | **Fez2** | 9030616F16\|D17Ertd315e | fasciculation and elongation protein zeta 2 (zygin II) | protein-coding |
| chr13_49307167_49307989 | chr13 | 49307168 | 49307989 | + | -0.636616 | intron (NM_015759, intron 1 of 17) | intron (NM_015759, intron 1 of 17) | 1630 | NM_015759 | 30938 | Mm.291089 | NM_015759 | ENSMUSG00000037946 | **Fgd3** | 5830461L01Rik\|ZFYVE5 | FYVE, RhoGEF and PH domain containing 3 | protein-coding |
| chr8_25535133_25535732 | chr8 | 25535134 | 25535732 | + | -0.672111 | intron (NM_001079909, intron 2 of 16) | L1ME3\|LINE\|L1 | 16674 | NM_001079908 | 14182 | Mm.265716 | NM_010206 | ENSMUSG00000031565 | **Fgfr1** | AW208770\|Eask\|FLG\|Fgfr-1\|Flt-2\|Hspy\|MFR\|bFGF-R-1\|c-fgr | fibroblast growth factor receptor 1 | protein-coding |
| chr4_132975808_132976209 | chr4 | 132975809 | 132976209 | + | -0.707987 | intron (NM_010208, intron 1 of 12) | intron (NM_010208, intron 1 of 12) | 1914 | NM_010208 | 14191 | Mm.271665 | NM_010208 | ENSMUSG00000028874 | **Fgr** | - | Gardner-Rasheed feline sarcoma viral (Fgr) oncogene homolog | protein-coding |
| chr4_132977695_132977975 | chr4 | 132977696 | 132977975 | + | -0.710704 | intron (NM_010208, intron 1 of 12) | intron (NM_010208, intron 1 of 12) | 3740 | NM_010208 | 14191 | Mm.271665 | NM_010208 | ENSMUSG00000028874 | **Fgr** | - | Gardner-Rasheed feline sarcoma viral (Fgr) oncogene homolog | protein-coding |
| chr4_132981685_132982030 | chr4 | 132981686 | 132982030 | + | -0.919401 | intron (NM_010208, intron 1 of 12) | intron (NM_010208, intron 1 of 12) | 7763 | NM_010208 | 14191 | Mm.271665 | NM_010208 | ENSMUSG00000028874 | **Fgr** | - | Gardner-Rasheed feline sarcoma viral (Fgr) oncogene homolog | protein-coding |
| chr14_11160748_11161107 | chr14 | 11160749 | 11161107 | + | -0.59256 | intron (NM_010210, intron 1 of 8) | intron (NM_010210, intron 1 of 8) | 1104 | NM_010210 | 14198 | Mm.397619 | NM_010210 | ENSMUSG00000060579 | **Fhit** | AW045638\|Fra14A2 | fragile histidine triad gene | protein-coding |
| chr14_10593432_10593819 | chr14 | 10593433 | 10593819 | + | -0.651031 | intron (NM_010210, intron 3 of 8) | intron (NM_010210, intron 3 of 8) | 568406 | NM_010210 | 14198 | Mm.397619 | NM_010210 | ENSMUSG00000060579 | **Fhit** | AW045638\|Fra14A2 | fragile histidine triad gene | protein-coding |
| chr14_10526747_10527403 | chr14 | 10526748 | 10527403 | + | -0.776037 | intron (NM_010210, intron 3 of 8) | intron (NM_010210, intron 3 of 8) | 634957 | NM_010210 | 14198 | Mm.397619 | NM_010210 | ENSMUSG00000060579 | **Fhit** | AW045638\|Fra14A2 | fragile histidine triad gene | protein-coding |
| chr4_124703882_124704278 | chr4 | 124703883 | 124704278 | + | -0.633345 | intron (NM_010213, intron 1 of 5) | intron (NM_010213, intron 1 of 5) | 3381 | NM_010213 | 14201 | Mm.331856 | NM_010213 | ENSMUSG00000032643 | **Fhl3** | SLIM2 | four and a half LIM domains 3 | protein-coding |
| chr16_57465214_57465962 | chr16 | 57465215 | 57465962 | + | -0.630613 | intron (NM_001040397, intron 1 of 4) | intron (NM_001040397, intron 1 of 4) | -83654 | NM_001177871 | 78749 | Mm.328360 | NM_001040397 | ENSMUSG00000043336 | **Filip1l** | 4631422O05Rik\|Doc1 | filamin A interacting protein 1-like | protein-coding |
| chr16_57434429_57435022 | chr16 | 57434430 | 57435022 | + | -0.722777 | intron (NM_001040397, intron 1 of 4) | L2a\|LINE\|L2 | 81449 | NM_001040397 | 78749 | Mm.328360 | NM_001040397 | ENSMUSG00000043336 | **Filip1l** | 4631422O05Rik\|Doc1 | filamin A interacting protein 1-like | protein-coding |
| chr16_57434564_57434995 | chr16 | 57434565 | 57434995 | + | -0.744129 | intron (NM_001040397, intron 1 of 4) | L2a\|LINE\|L2 | 81503 | NM_001040397 | 78749 | Mm.328360 | NM_001040397 | ENSMUSG00000043336 | **Filip1l** | 4631422O05Rik\|Doc1 | filamin A interacting protein 1-like | protein-coding |
| chr16_57423464_57423719 | chr16 | 57423465 | 57423719 | + | -0.749178 | intron (NM_001040397, intron 1 of 4) | intron (NM_001040397, intron 1 of 4) | 70315 | NM_001040397 | 78749 | Mm.328360 | NM_001040397 | ENSMUSG00000043336 | **Filip1l** | 4631422O05Rik\|Doc1 | filamin A interacting protein 1-like | protein-coding |

| | | | | | | | | | | | | | | | | | |
|---|---|---|---|---|---|---|---|---|---|---|---|---|---|---|---|---|---|
| chr16_57434563_57435209 | chr16 | 57434564 | 57435209 | + | -0.770183 | intron (NM_001040397, intron 1 of 4) | L2a\|LINE\|L2 | 81609 | NM_001040397 | 78749 | Mm.328360 | NM_001040397 | ENSMUSG00000043336 | **Filip1l** | 4631422O05Rik\|Doc1 | filamin A interacting protein 1-like | protein-coding |
| chr16_57470935_57471488 | chr16 | 57470936 | 57471488 | + | -0.78794 | intron (NM_001040397, intron 1 of 4) | intron (NM_001040397, intron 1 of 4) | -78030 | NM_001177871 | 78749 | Mm.328360 | NM_001040397 | ENSMUSG00000043336 | **Filip1l** | 4631422O05Rik\|Doc1 | filamin A interacting protein 1-like | protein-coding |
| chr16_57422991_57423408 | chr16 | 57422992 | 57423408 | + | -0.826292 | intron (NM_001040397, intron 1 of 4) | RMER17C\|LTR\|ERVK | 69923 | NM_001040397 | 78749 | Mm.328360 | NM_001040397 | ENSMUSG00000043336 | **Filip1l** | 4631422O05Rik\|Doc1 | filamin A interacting protein 1-like | protein-coding |
| chr16_57423006_57423688 | chr16 | 57423007 | 57423688 | + | -0.83018 | intron (NM_001040397, intron 1 of 4) | intron (NM_001040397, intron 1 of 4) | 70070 | NM_001040397 | 78749 | Mm.328360 | NM_001040397 | ENSMUSG00000043336 | **Filip1l** | 4631422O05Rik\|Doc1 | filamin A interacting protein 1-like | protein-coding |
| chr16_57471188_57471469 | chr16 | 57471189 | 57471469 | + | -0.839016 | intron (NM_001040397, intron 1 of 4) | intron (NM_001040397, intron 1 of 4) | -77913 | NM_001177871 | 78749 | Mm.328360 | NM_001040397 | ENSMUSG00000043336 | **Filip1l** | 4631422O05Rik\|Doc1 | filamin A interacting protein 1-like | protein-coding |
| chr16_57465207_57465608 | chr16 | 57465208 | 57465608 | + | -0.841153 | intron (NM_001040397, intron 1 of 4) | intron (NM_001040397, intron 1 of 4) | -83834 | NM_001177871 | 78749 | Mm.328360 | NM_001040397 | ENSMUSG00000043336 | **Filip1l** | 4631422O05Rik\|Doc1 | filamin A interacting protein 1-like | protein-coding |
| chrX_50599810_50600218 | chrX | 50599811 | 50600218 | + | -0.659417 | intron (NR_026976, intron 9 of 23) | intron (NR_026976, intron 9 of 23) | 35244 | NR_015505 | 103012 | Mm.482523 | NR_015505 | | **Firre** | 5830467J12Rik\|6720401G13Rik\|AW048145 | functional intergenic repeating RNA element | ncRNA |
| chrX_50599813_50600087 | chrX | 50599814 | 50600087 | + | -0.666052 | intron (NR_026976, intron 9 of 23) | intron (NR_026976, intron 9 of 23) | 35308 | NR_015505 | 103012 | Mm.482523 | NR_015505 | | **Firre** | 5830467J12Rik\|6720401G13Rik\|AW048145 | functional intergenic repeating RNA element | ncRNA |
| chrX_50565036_50565487 | chrX | 50565037 | 50565487 | + | -0.938645 | intron (NR_026976, intron 19 of 23) | IAPEY3C_LTR\|LTR\|ERVK | 69996 | NR_015505 | 103012 | Mm.482523 | NR_015505 | | **Firre** | 5830467J12Rik\|6720401G13Rik\|AW048145 | functional intergenic repeating RNA element | ncRNA |
| chr12_65065134_65065712 | chr12 | 65065135 | 65065712 | + | -0.89539 | intron (NM_013902, intron 5 of 6) | ORR1C2\|LTR\|ERVL-MaLR | 8515 | NM_013902 | 30795 | Mm.28480 | NM_013902 | ENSMUSG00000020949 | **Fkbp3** | 25kDa\|FKBP-3\|FKBP25 | FK506 binding protein 3 | protein-coding |
| chr17_28467740_28468053 | chr17 | 28467741 | 28468053 | + | -0.701812 | intron (NM_010220, intron 1 of 10) | intron (NM_010220, intron 1 of 10) | 18252 | NM_010220 | 14229 | Mm.276405 | NM_010220 | ENSMUSG00000024222 | **Fkbp5** | D17Ertd592e\|Dit1\|FKBP-5\|FKBP51 | FK506 binding protein 5 | protein-coding |
| chr17_28442472_28443209 | chr17 | 28442473 | 28443209 | + | -0.766285 | intron (NM_010220, intron 1 of 10) | intron (NM_010220, intron 1 of 10) | 43308 | NM_010220 | 14229 | Mm.276405 | NM_010220 | ENSMUSG00000024222 | **Fkbp5** | D17Ertd592e\|Dit1\|FKBP-5\|FKBP51 | FK506 binding protein 5 | protein-coding |
| chr7_16827213_16827722 | chr7 | 16827214 | 16827722 | + | -0.654592 | intron (NM_001039878, intron 5 of 17) | L2\|LINE\|L2 | -10736 | NM_173430 | 243853 | Mm.39703 | NM_173430 | ENSMUSG00000048920 | **Fkrp** | A830029B19Rik\|AI842067\|AI847300\|LGMD1I\|MDC1C | fukutin related protein | protein-coding |
| chr9_32495406_32496108 | chr9 | 32495407 | 32496108 | + | -0.725746 | intron (NM_008026, intron 1 of 8) | intron (NM_008026, intron 1 of 8) | 45695 | NM_008026 | 14247 | Mm.258908 | NM_008026 | ENSMUSG00000016087 | **Fli1** | EWSR2\|Fli-1\|SIC-1\|Sic1 | Friend leukemia integration 1 | protein-coding |
| chr9_32615866_32616316 | chr9 | 32615867 | 32616316 | + | -0.757167 | Intergenic | Intergenic | -74639 | NM_008026 | 14247 | Mm.258908 | NM_008026 | ENSMUSG00000016087 | **Fli1** | EWSR2\|Fli-1\|SIC-1\|Sic1 | Friend leukemia integration 1 | protein-coding |
| chr9_32500101_32500399 | chr9 | 32500102 | 32500399 | + | -0.801231 | intron (NM_008026, intron 1 of 8) | intron (NM_008026, intron 1 of 8) | 41202 | NM_008026 | 14247 | Mm.258908 | NM_008026 | ENSMUSG00000016087 | **Fli1** | EWSR2\|Fli-1\|SIC-1\|Sic1 | Friend leukemia integration 1 | protein-coding |
| chr9_32499486_32499807 | chr9 | 32499487 | 32499807 | + | -0.821275 | intron (NM_008026, intron 1 of 8) | intron (NM_008026, intron 1 of 8) | 41805 | NM_008026 | 14247 | Mm.258908 | NM_008026 | ENSMUSG00000016087 | **Fli1** | EWSR2\|Fli-1\|SIC-1\|Sic1 | Friend leukemia integration 1 | protein-coding |
| chr9_32581620_32582464 | chr9 | 32581621 | 32582464 | + | -0.838028 | Intergenic | Intergenic | -40590 | NM_008026 | 14247 | Mm.258908 | NM_008026 | ENSMUSG00000016087 | **Fli1** | EWSR2\|Fli-1\|SIC-1\|Sic1 | Friend leukemia integration 1 | protein-coding |
| chr9_32538156_32538440 | chr9 | 32538157 | 32538440 | + | -0.856734 | intron (NM_008026, intron 1 of 8) | intron (NM_008026, intron 1 of 8) | 3154 | NM_008026 | 14247 | Mm.258908 | NM_008026 | ENSMUSG00000016087 | **Fli1** | EWSR2\|Fli-1\|SIC-1\|Sic1 | Friend leukemia integration 1 | protein-coding |
| chr9_32570635_32571087 | chr9 | 32570636 | 32571087 | + | -1.20231 | Intergenic | Intergenic | -29409 | NM_008026 | 14247 | Mm.258908 | NM_008026 | ENSMUSG00000016087 | **Fli1** | EWSR2\|Fli-1\|SIC-1\|Sic1 | Friend leukemia integration 1 | protein-coding |
| chr5_147366382_147366840 | chr5 | 147366383 | 147366840 | + | -0.705311 | intron (NM_010229, intron 9 of 23) | intron (NM_010229, intron 9 of 23) | 33878 | NM_010229 | 14255 | Mm.194 | NM_010229 | ENSMUSG00000042817 | **Flt3** | B230315G04\|CD135\|Flk-2\|Flk2\|Flt-3\|Ly72\|wmfl | FMS-like tyrosine kinase 3 | protein-coding |
| chr5_147363108_147363393 | chr5 | 147363109 | 147363393 | + | -0.873978 | intron (NM_010229, intron 9 of 23) | intron (NM_010229, intron 9 of 23) | 37238 | NM_010229 | 14255 | Mm.194 | NM_010229 | ENSMUSG00000042817 | **Flt3** | B230315G04\|CD135\|Flk-2\|Flk2\|Flt-3\|Ly72\|wmfl | FMS-like tyrosine kinase 3 | protein-coding |
| chr15_99341060_99341556 | chr15 | 99341061 | 99341556 | + | -0.754792 | intron (NM_011711, intron 1 of 26) | intron (NM_011711, intron 1 of 26) | 29158 | NM_011711 | 22379 | Mm.29043 | NM_011711 | ENSMUSG00000023008 | **Fmnl3** | 2700073B04Rik\|FBP11\|Wbp3\|mKIAA2014 | formin-like 3 | protein-coding |
| chr15_99335858_99336354 | chr15 | 99335859 | 99336354 | + | -1.0433 | intron (NM_011711, intron 2 of 26) | intron (NM_011711, intron 2 of 26) | 34360 | NM_011711 | 22379 | Mm.29043 | NM_011711 | ENSMUSG00000023008 | **Fmnl3** | 2700073B04Rik\|FBP11\|Wbp3\|mKIAA2014 | formin-like 3 | protein-coding |
| chr2_31087737_31088097 | chr2 | 31087738 | 31088097 | + | -0.630899 | intron (NM_001038700, intron 4 of 14) | intron (NM_001038700, intron 4 of 14) | 8583 | NM_001177650 | 14269 | Mm.52297 | NM_019406 | ENSMUSG00000075415 | **Fnbp1** | 1110057E06Rik\|2210010H06Rik\|FBP1\|Fbp17 | formin binding protein 1 | protein-coding |
| chr2_31130994_31131472 | chr2 | 31130995 | 31131472 | + | -0.760633 | intron (NM_001038700, intron 1 of 14) | intron (NM_001038700, intron 1 of 14) | 10775 | NM_001177649 | 14269 | Mm.52297 | NM_019406 | ENSMUSG00000075415 | **Fnbp1** | 1110057E06Rik\|2210010H06Rik\|FBP1\|Fbp17 | formin binding protein 1 | protein-coding |
| chr2_31108051_31108669 | chr2 | 31108052 | 31108669 | + | -1.21283 | intron (NM_001038700, intron 1 of 14) | MTD\|LTR\|ERVL-MaLR | -11860 | NM_001177650 | 14269 | Mm.52297 | NM_019406 | ENSMUSG00000075415 | **Fnbp1** | 1110057E06Rik\|2210010H06Rik\|FBP1\|Fbp17 | formin binding protein 1 | protein-coding |
| chr4_129146390_129146735 | chr4 | 129146391 | 129146735 | + | -1.02314 | Intergenic | Intergenic | 9503 | NM_027402 | 384061 | Mm.44075 | NM_027402 | ENSMUSG00000001334 | **Fndc5** | 1500001L03Rik\|AI836596\|C87088\|PeP\|Pxp | fibronectin type III domain containing 5 | protein-coding |
| chr3_108865272_108866106 | chr3 | 108865273 | 108866106 | + | -1.24365 | intron (NM_177091, intron 9 of 12) | intron (NM_177091, intron 9 of 12) | 24319 | NM_177091 | 320181 | Mm.244233 | NM_177091 | ENSMUSG00000045326 | **Fndc7** | E230011A21Rik | fibronectin type III domain containing 7 | protein-coding |
| chr3_108865503_108866183 | chr3 | 108865504 | 108866183 | + | -1.25786 | intron (NM_177091, intron 9 of 12) | intron (NM_177091, intron 9 of 12) | 24165 | NM_177091 | 320181 | Mm.244233 | NM_177091 | ENSMUSG00000045326 | **Fndc7** | E230011A21Rik | fibronectin type III domain containing 7 | protein-coding |
| chr3_79535163_79536092 | chr3 | 79535164 | 79536092 | + | -0.599765 | intron (NM_001162999, intron 1 of 16) | intron (NM_001162999, intron 1 of 16) | 32051 | NM_001162999 | 329679 | Mm.244005 | NM_001162999 | ENSMUSG00000061175 | **Fnip2** | D630023B12Rik\|mKIAA1450 | folliculin interacting protein 2 | protein-coding |
| chr3_79535694_79536133 | chr3 | 79535695 | 79536133 | + | -0.766115 | intron (NM_001162999, intron 1 of 16) | intron (NM_001162999, intron 1 of 16) | 31765 | NM_001162999 | 329679 | Mm.244005 | NM_001162999 | ENSMUSG00000061175 | **Fnip2** | D630023B12Rik\|mKIAA1450 | folliculin interacting protein 2 | protein-coding |
| chr12_76844723_76845048 | chr12 | 76844724 | 76845048 | + | -0.909097 | intron (NM_145927, intron 1 of 11) | intron (NM_145927, intron 1 of 11) | 7419 | NM_145927 | 110606 | Mm.151174 | NM_145927 | ENSMUSG00000033373 | **Fntb** | 2010013E13Rik\|AA409500 | farnesyltransferase, CAAX box, beta | protein-coding |
| chr9_14926471_14926990 | chr9 | 14926472 | 14926990 | + | -0.781534 | Intergenic | Intergenic | -22780 | NM_022888 | 64931 | Mm.86738 | NM_022888 | ENSMUSG00000031933 | **Folr4** | 0910001L11Rik\|C86255\|Folbp3\|Juno | folate receptor 4 (delta) | protein-coding |
| chr12_85468694_85469006 | chr12 | 85468695 | 85469006 | + | -0.967776 | Intergenic | Intergenic | -5051 | NM_010234 | 14281 | Mm.246513 | NM_010234 | ENSMUSG00000021250 | **Fos** | D12Rfj1\|c-fos\|cFos | FBJ osteosarcoma oncogene | protein-coding |
| chr5_31918424_31918927 | chr5 | 31918425 | 31918927 | + | -0.757357 | intron (NM_144541, intron 7 of 11) | intron (NM_144541, intron 7 of 11) | -217796 | NM_008037 | 14284 | Mm.24684 | NM_008037 | ENSMUSG00000029135 | **Fosl2** | Fra-2 | fos-like antigen 2 | protein-coding |
| chr5_32110759_32111070 | chr5 | 32110760 | 32111070 | + | -0.758067 | Intergenic | MIR3\|SINE\|MIR | -25557 | NM_008037 | 14284 | Mm.24684 | NM_008037 | ENSMUSG00000029135 | **Fosl2** | Fra-2 | fos-like antigen 2 | protein-coding |
| chr5_32148622_32148978 | chr5 | 32148623 | 32148978 | + | -0.95451 | intron (NM_008037, intron 2 of 3) | intron (NM_008037, intron 2 of 3) | 12328 | NM_008037 | 14284 | Mm.24684 | NM_008037 | ENSMUSG00000029135 | **Fosl2** | Fra-2 | fos-like antigen 2 | protein-coding |
| chr5_32148387_32148937 | chr5 | 32148388 | 32148937 | + | -0.959481 | intron (NM_008037, intron 2 of 3) | intron (NM_008037, intron 2 of 3) | 12190 | NM_008037 | 14284 | Mm.24684 | NM_008037 | ENSMUSG00000029135 | **Fosl2** | Fra-2 | fos-like antigen 2 | protein-coding |
| chr11_116354528_116354958 | chr11 | 116354529 | 116354958 | + | -0.781564 | exon (NM_027258, exon 10 of 19) | exon (NM_027258, exon 10 of 19) | -19389 | NM_008240 | 15223 | Mm.378938 | NM_008240 | ENSMUSG00000034227 | **Foxj1** | FKHL-13\|HFH-4\|Hfh4 | forkhead box J1 | protein-coding |
| chr11_116369643_116370161 | chr11 | 116369644 | 116370161 | + | -0.929124 | intron (NM_027258, intron 2 of 18) | BGLII_B\|LTR\|ERVK | -34548 | NM_008240 | 15223 | Mm.378938 | NM_008240 | ENSMUSG00000034227 | **Foxj1** | FKHL-13\|HFH-4\|Hfh4 | forkhead box J1 | protein-coding |
| chr11_116369637_116369995 | chr11 | 116369638 | 116369995 | + | -0.948346 | intron (NM_027258, intron 2 of 18) | BGLII_B\|LTR\|ERVK | -34462 | NM_008240 | 15223 | Mm.378938 | NM_008240 | ENSMUSG00000034227 | **Foxj1** | FKHL-13\|HFH-4\|Hfh4 | forkhead box J1 | protein-coding |
| chr5_142404246_142404633 | chr5 | 142404247 | 142404633 | + | -0.683619 | intron (NM_199068, intron 1 of 8) | B4A\|SINE\|B4 | 2943 | NM_199068 | 17425 | Mm.24214 | NM_010812 | ENSMUSG00000056493 | **Foxk1** | A630048H08Rik\|AI463295\|ENSMUSG00000075577\|Gm10868\|Mnf | forkhead box K1 | protein-coding |
| chr5_142018151_142018681 | chr5 | 142018152 | 142018681 | + | -0.71377 | intron (NM_177879, intron 17 of 44) | intron (NM_177879, intron 17 of 44) | -383081 | NM_199068 | 17425 | Mm.24214 | NM_010812 | ENSMUSG00000056493 | **Foxk1** | A630048H08Rik\|AI463295\|ENSMUSG00000075577\|Gm10868\|Mnf | forkhead box K1 | protein-coding |
| chr5_114274295_114274736 | chr5 | 114274296 | 114274736 | + | -0.747812 | promoter-TSS (NM_148935) | promoter-TSS (NM_148935) | -755 | NM_148935 | 116810 | Mm.17709 | NM_148935 | ENSMUSG00000042002 | **Foxn4** | - | forkhead box N4 | protein-coding |
| chr3_52262474_52263230 | chr3 | 52262475 | 52263230 | + | -0.655673 | Intergenic | MTB_Mm\|LTR\|ERVL-MaLR | -5485 | NM_019739 | 56458 | Mm.29891 | NM_019739 | ENSMUSG00000044167 | **Foxo1** | AI876417\|Afxh\|FKHR\|Fkhr1\|Foxo1a | forkhead box O1 | protein-coding |
| chr3_52343255_52344265 | chr3 | 52343256 | 52344265 | + | -0.838079 | intron (NM_019739, intron 1 of 2) | intron (NM_019739, intron 1 of 2) | 75423 | NM_019739 | 56458 | Mm.29891 | NM_019739 | ENSMUSG00000044167 | **Foxo1** | AI876417\|Afxh\|FKHR\|Fkhr1\|Foxo1a | forkhead box O1 | protein-coding |

| | | | | | | | | | | | | | | | | | |
|---|---|---|---|---|---|---|---|---|---|---|---|---|---|---|---|---|---|
| chr3_52343223_52343938 | chr3 | 52343224 | 52343938 | + | -0.856626 | intron (NM_019739, intron 1 of 2) | intron (NM_019739, intron 1 of 2) | 75244 | NM_019739 | 56458 | Mm.29891 | NM_019739 | ENSMUSG00000044167 | **Foxo1** | AI876417\|Afxh\|FKHR\|Fkhr1\|Foxo1a | forkhead box O1 | protein-coding |
| chr3_52262478_52262712 | chr3 | 52262479 | 52262712 | + | -0.868386 | Intergenic | B3\|SINE\|B2 | -5742 | NM_019739 | 56458 | Mm.29891 | NM_019739 | ENSMUSG00000044167 | **Foxo1** | AI876417\|Afxh\|FKHR\|Fkhr1\|Foxo1a | forkhead box O1 | protein-coding |
| chr3_52395971_52396809 | chr3 | 52395972 | 52396809 | + | -0.8978 | Intergenic | Intergenic | 128053 | NM_019739 | 56458 | Mm.29891 | NM_019739 | ENSMUSG00000044167 | **Foxo1** | AI876417\|Afxh\|FKHR\|Fkhr1\|Foxo1a | forkhead box O1 | protein-coding |
| chr3_52411262_52411795 | chr3 | 52411263 | 52411795 | + | -0.902076 | Intergenic | Intergenic | 143192 | NM_019739 | 56458 | Mm.29891 | NM_019739 | ENSMUSG00000044167 | **Foxo1** | AI876417\|Afxh\|FKHR\|Fkhr1\|Foxo1a | forkhead box O1 | protein-coding |
| chr10_42269124_42270007 | chr10 | 42269125 | 42270007 | + | -0.614787 | intron (NM_019740, intron 2 of 3) | intron (NM_019740, intron 2 of 3) | 7176 | NM_019740 | 56484 | Mm.338613 | NM_019740 | ENSMUSG00000048756 | **Foxo3** | 1110048B16Rik\|2010203A17Rik\|C76856\|FKHRL1\|Fkhr2\|Foxo3a | forkhead box O3 | protein-coding |
| chr10_42226073_42226709 | chr10 | 42226074 | 42226709 | + | -0.769544 | intron (NM_019740, intron 2 of 3) | intron (NM_019740, intron 2 of 3) | 50351 | NM_019740 | 56484 | Mm.338613 | NM_019740 | ENSMUSG00000048756 | **Foxo3** | 1110048B16Rik\|2010203A17Rik\|C76856\|FKHRL1\|Fkhr2\|Foxo3a | forkhead box O3 | protein-coding |
| chr6_98842897_98843395 | chr6 | 98842898 | 98843395 | + | -0.622684 | Intergenic | MER5A\|DNA\|hAT-Charlie | 185152 | NM_001197322 | 108655 | Mm.234965 | NM_053202 | ENSMUSG00000030067 | **Foxp1** | 3110052D19Rik\|4932443N09Rik\|AI461938\|AW494214 | forkhead box P1 | protein-coding |
| chr6_99388675_99389594 | chr6 | 99388676 | 99389594 | + | -0.665464 | intron (NM_053202, intron 1 of 18) | intron (NM_053202, intron 1 of 18) | 46210 | NM_053202 | 108655 | Mm.234965 | NM_053202 | ENSMUSG00000030067 | **Foxp1** | 3110052D19Rik\|4932443N09Rik\|AI461938\|AW494214 | forkhead box P1 | protein-coding |
| chr6_99380100_99380540 | chr6 | 99380101 | 99380540 | + | -0.679909 | intron (NM_053202, intron 1 of 18) | intron (NM_053202, intron 1 of 18) | 55025 | NM_053202 | 108655 | Mm.234965 | NM_053202 | ENSMUSG00000030067 | **Foxp1** | 3110052D19Rik\|4932443N09Rik\|AI461938\|AW494214 | forkhead box P1 | protein-coding |
| chr6_99303033_99303381 | chr6 | 99303034 | 99303381 | + | -0.687728 | intron (NM_053202, intron 2 of 18) | intron (NM_053202, intron 2 of 18) | -36692 | NM_001197321 | 108655 | Mm.234965 | NM_053202 | ENSMUSG00000030067 | **Foxp1** | 3110052D19Rik\|4932443N09Rik\|AI461938\|AW494214 | forkhead box P1 | protein-coding |
| chr6_99388692_99389375 | chr6 | 99388693 | 99389375 | + | -0.725845 | intron (NM_053202, intron 1 of 18) | intron (NM_053202, intron 1 of 18) | 46311 | NM_053202 | 108655 | Mm.234965 | NM_053202 | ENSMUSG00000030067 | **Foxp1** | 3110052D19Rik\|4932443N09Rik\|AI461938\|AW494214 | forkhead box P1 | protein-coding |
| chr6_99126680_99127109 | chr6 | 99126681 | 99127109 | + | -0.870179 | intron (NM_053202, intron 4 of 18) | ORR1D2\|LTR\|ERVL-MaLR | -98597 | NM_001197322 | 108655 | Mm.234965 | NM_053202 | ENSMUSG00000030067 | **Foxp1** | 3110052D19Rik\|4932443N09Rik\|AI461938\|AW494214 | forkhead box P1 | protein-coding |
| chr6_99403895_99404367 | chr6 | 99403896 | 99404367 | + | -0.923239 | intron (NM_053202, intron 1 of 18) | intron (NM_053202, intron 1 of 18) | 31214 | NM_053202 | 108655 | Mm.234965 | NM_053202 | ENSMUSG00000030067 | **Foxp1** | 3110052D19Rik\|4932443N09Rik\|AI461938\|AW494214 | forkhead box P1 | protein-coding |
| chr6_99013440_99013812 | chr6 | 99013441 | 99013812 | + | -0.969039 | intron (NM_001197322, intron 3 of 14) | intron (NM_001197322, intron 3 of 14) | 14672 | NM_001197322 | 108655 | Mm.234965 | NM_053202 | ENSMUSG00000030067 | **Foxp1** | 3110052D19Rik\|4932443N09Rik\|AI461938\|AW494214 | forkhead box P1 | protein-coding |
| chr6_99322286_99323019 | chr6 | 99322287 | 99323019 | + | -1.01212 | intron (NM_053202, intron 1 of 18) | intron (NM_053202, intron 1 of 18) | -56138 | NM_001197321 | 108655 | Mm.234965 | NM_053202 | ENSMUSG00000030067 | **Foxp1** | 3110052D19Rik\|4932443N09Rik\|AI461938\|AW494214 | forkhead box P1 | protein-coding |
| chr6_99395744_99396093 | chr6 | 99395745 | 99396093 | + | -1.01518 | intron (NM_053202, intron 1 of 18) | intron (NM_053202, intron 1 of 18) | 39426 | NM_053202 | 108655 | Mm.234965 | NM_053202 | ENSMUSG00000030067 | **Foxp1** | 3110052D19Rik\|4932443N09Rik\|AI461938\|AW494214 | forkhead box P1 | protein-coding |
| chr6_99394501_99395198 | chr6 | 99394502 | 99395198 | + | -1.10376 | intron (NM_053202, intron 1 of 18) | intron (NM_053202, intron 1 of 18) | 40495 | NM_053202 | 108655 | Mm.234965 | NM_053202 | ENSMUSG00000030067 | **Foxp1** | 3110052D19Rik\|4932443N09Rik\|AI461938\|AW494214 | forkhead box P1 | protein-coding |
| chr6_99023634_99023960 | chr6 | 99023635 | 99023960 | + | -1.34158 | intron (NM_053202, intron 5 of 18) | intron (NM_053202, intron 5 of 18) | 4501 | NM_001197322 | 108655 | Mm.234965 | NM_053202 | ENSMUSG00000030067 | **Foxp1** | 3110052D19Rik\|4932443N09Rik\|AI461938\|AW494214 | forkhead box P1 | protein-coding |
| chr13_31505385_31505720 | chr13 | 31505386 | 31505720 | + | -0.7131 | Intergenic | Intergenic | -52617 | NM_008239 | 15220 | Mm.44235 | NM_008239 | ENSMUSG00000038415 | **Foxq1** | HFH-1\|Hfh1\|Hfh1l\|sa | forkhead box Q1 | protein-coding |
| chr13_31505139_31505725 | chr13 | 31505140 | 31505725 | + | -0.723592 | Intergenic | Intergenic | -52738 | NM_008239 | 15220 | Mm.44235 | NM_008239 | ENSMUSG00000038415 | **Foxq1** | HFH-1\|Hfh1\|Hfh1l\|sa | forkhead box Q1 | protein-coding |
| chr13_31557219_31557690 | chr13 | 31557220 | 31557690 | + | -0.743303 | promoter-TSS (NM_008239) | promoter-TSS (NM_008239) | -715 | NM_008239 | 15220 | Mm.44235 | NM_008239 | ENSMUSG00000038415 | **Foxq1** | HFH-1\|Hfh1\|Hfh1l\|sa | forkhead box Q1 | protein-coding |
| chr13_31503196_31503546 | chr13 | 31503197 | 31503546 | + | -1.13608 | Intergenic | Intergenic | -54799 | NM_008239 | 15220 | Mm.44235 | NM_008239 | ENSMUSG00000038415 | **Foxq1** | HFH-1\|Hfh1\|Hfh1l\|sa | forkhead box Q1 | protein-coding |
| chr17_18004690_18005148 | chr17 | 18004691 | 18005148 | + | -1.18539 | Intergenic | ID_B1\|SINE\|B4 | -16814 | NM_008041 | 14291 | Mm.455331 | NM_008041 | ENSMUSG00000048062 | **Fpr-rs4** | - | formyl peptide receptor, related sequence 4 | protein-coding |
| chr17_18002972_18003419 | chr17 | 18002973 | 18003419 | + | -1.21824 | Intergenic | B3A\|SINE\|B2 | -18537 | NM_008041 | 14291 | Mm.455331 | NM_008041 | ENSMUSG00000048062 | **Fpr-rs4** | - | formyl peptide receptor, related sequence 4 | protein-coding |
| chr17_17952327_17952663 | chr17 | 17952328 | 17952663 | + | -0.658098 | Intergenic | Lx3B\|LINE\|L1 | -17963 | NM_008042 | 14294 | Mm.378918 | NM_008042 | ENSMUSG00000079700 | **Fpr3** | ALX\|Fpr-rs1\|Fpr-s1\|Fprl1\|LXA4-R\|Lxa4r | formyl peptide receptor 3 | protein-coding |
| chr2_4580383_4581155 | chr2 | 4580384 | 4581155 | + | -0.843821 | intron (NM_001177843, intron 14 of 23) | intron (NM_001177843, intron 14 of 23) | 21015 | NM_001177844 | 209630 | Mm.37932 | NM_172475 | ENSMUSG00000026657 | **Frmd4a** | 2700017I06Rik\|5330420J21\|AI594879\|C230040M21Rik\|Frmd4\|Gm13190\|OTTMUSG00000010806\|mKIAA1294 | FERM domain containing 4A | protein-coding |
| chr6_97530978_97531400 | chr6 | 97530979 | 97531400 | + | -0.645279 | intron (NM_145148, intron 1 of 23) | intron (NM_145148, intron 1 of 23) | 86468 | NM_145148 | 232288 | Mm.27789 | NM_145148 | ENSMUSG00000030064 | **Frmd4b** | 6030440G05Rik\|C87375\|GOBLIN\|GRSP1\|R74720\|mKIAA1013 | FERM domain containing 4B | protein-coding |
| chr6_97457902_97458390 | chr6 | 97457903 | 97458390 | + | -0.768779 | intron (NM_145148, intron 2 of 23) | intron (NM_145148, intron 2 of 23) | 159511 | NM_145148 | 232288 | Mm.27789 | NM_145148 | ENSMUSG00000030064 | **Frmd4b** | 6030440G05Rik\|C87375\|GOBLIN\|GRSP1\|R74720\|mKIAA1013 | FERM domain containing 4B | protein-coding |
| chr6_97473633_97474071 | chr6 | 97473634 | 97474071 | + | -0.773133 | intron (NM_145148, intron 2 of 23) | L2\|LINE\|L2 | 143805 | NM_145148 | 232288 | Mm.27789 | NM_145148 | ENSMUSG00000030064 | **Frmd4b** | 6030440G05Rik\|C87375\|GOBLIN\|GRSP1\|R74720\|mKIAA1013 | FERM domain containing 4B | protein-coding |
| chr6_97473580_97474249 | chr6 | 97473581 | 97474249 | + | -0.775057 | intron (NM_145148, intron 2 of 23) | intron (NM_145148, intron 2 of 23) | 143742 | NM_145148 | 232288 | Mm.27789 | NM_145148 | ENSMUSG00000030064 | **Frmd4b** | 6030440G05Rik\|C87375\|GOBLIN\|GRSP1\|R74720\|mKIAA1013 | FERM domain containing 4B | protein-coding |
| chr6_97472973_97474352 | chr6 | 97472974 | 97474352 | + | -0.842835 | intron (NM_145148, intron 2 of 23) | L2\|LINE\|L2 | 143994 | NM_145148 | 232288 | Mm.27789 | NM_145148 | ENSMUSG00000030064 | **Frmd4b** | 6030440G05Rik\|C87375\|GOBLIN\|GRSP1\|R74720\|mKIAA1013 | FERM domain containing 4B | protein-coding |
| chr6_97472978_97473537 | chr6 | 97472979 | 97473537 | + | -0.857281 | intron (NM_145148, intron 2 of 23) | intron (NM_145148, intron 2 of 23) | 144399 | NM_145148 | 232288 | Mm.27789 | NM_145148 | ENSMUSG00000030064 | **Frmd4b** | 6030440G05Rik\|C87375\|GOBLIN\|GRSP1\|R74720\|mKIAA1013 | FERM domain containing 4B | protein-coding |
| chr6_97472978_97474005 | chr6 | 97472979 | 97474005 | + | -0.871936 | intron (NM_145148, intron 2 of 23) | intron (NM_145148, intron 2 of 23) | 144165 | NM_145148 | 232288 | Mm.27789 | NM_145148 | ENSMUSG00000030064 | **Frmd4b** | 6030440G05Rik\|C87375\|GOBLIN\|GRSP1\|R74720\|mKIAA1013 | FERM domain containing 4B | protein-coding |
| chrX_50930289_50930883 | chrX | 50930290 | 50930883 | + | -0.705054 | intron (NM_001190332, intron 1 of 11) | intron (NM_001190332, intron 1 of 11) | 12124 | NM_001190332 | 385354 | Mm.393050 | NM_001190332 | | **Frmd7** | EG665849\|Gm1533 | FERM domain containing 7 | protein-coding |
| chr10_117119535_117119826 | chr10 | 117119536 | 117119826 | + | -0.684668 | intron (NM_177798, intron 1 of 9) | intron (NM_177798, intron 1 of 9) | 28793 | NM_177798 | 327826 | Mm.135965 | NM_177798 | ENSMUSG00000020170 | **Frs2** | 4732458E18\|C330018A15Rik\|Frs2alpha\|SNT1 | fibroblast growth factor receptor substrate 2 | protein-coding |
| chr10_117127526_117127789 | chr10 | 117127527 | 117127789 | + | -0.924905 | intron (NM_177798, intron 1 of 9) | MT2A\|LTR\|ERVL | 20816 | NM_177798 | 327826 | Mm.135965 | NM_177798 | ENSMUSG00000020170 | **Frs2** | 4732458E18\|C330018A15Rik\|Frs2alpha\|SNT1 | fibroblast growth factor receptor substrate 2 | protein-coding |
| chr5_73206283_73206609 | chr5 | 73206284 | 73206609 | + | -0.690189 | intron (NM_028194, intron 2 of 62) | intron (NM_028194, intron 2 of 62) | 50172 | NM_028194 | 72313 | Mm.260594 | NM_028194 | ENSMUSG00000070733 | **Fryl** | 2010313D22Rik\|2310004H21Rik\|2510002A14Rik\|9030227G01Rik\|mKIAA0826 | furry homolog-like (Drosophila) | protein-coding |
| chr5_73208550_73208969 | chr5 | 73208551 | 73208969 | + | -0.793351 | intron (NM_028194, intron 2 of 62) | ID_B1\|SINE\|B4 | 47858 | NM_028194 | 72313 | Mm.260594 | NM_028194 | ENSMUSG00000070733 | **Fryl** | 2010313D22Rik\|2310004H21Rik\|2510002A14Rik\|9030227G01Rik\|mKIAA0826 | furry homolog-like (Drosophila) | protein-coding |
| chr5_73257189_73257396 | chr5 | 73257190 | 73257396 | + | -0.853424 | promoter-TSS (NM_028194) | promoter-TSS (NM_028194) | -675 | NM_028194 | 72313 | Mm.260594 | NM_028194 | ENSMUSG00000070733 | **Fryl** | 2010313D22Rik\|2310004H21Rik\|2510002A14Rik\|9030227G01Rik\|mKIAA0826 | furry homolog-like (Drosophila) | protein-coding |
| chr5_73265322_73265878 | chr5 | 73265323 | 73265878 | + | -1.0106 | Intergenic | L2b\|LINE\|L2 | -8982 | NM_028194 | 72313 | Mm.260594 | NM_028194 | ENSMUSG00000070733 | **Fryl** | 2010313D22Rik\|2310004H21Rik\|2510002A14Rik\|9030227G01Rik\|mKIAA0826 | furry homolog-like (Drosophila) | protein-coding |
| chr4_11616026_11616564 | chr4 | 11616027 | 11616564 | + | -0.671576 | TTS (NM_001039556) | TTS (NM_001039556) | 36633 | NM_001256142 | 100503583 | Mm.249437 | NM_001256142 | ENSMUSG00000094595 | **Fsbp** | - | fibrinogen silencer binding protein | protein-coding |
| chr4_11616076_11616400 | chr4 | 11616077 | 11616400 | + | -0.747219 | TTS (NM_001039556) | TTS (NM_001039556) | 36576 | NM_001256142 | 100503583 | Mm.249437 | NM_001256142 | ENSMUSG00000094595 | **Fsbp** | - | fibrinogen silencer binding protein | protein-coding |
| chr4_11609946_11610385 | chr4 | 11609947 | 11610385 | + | -1.28124 | intron (NM_001039556, intron 11 of 14) | intron (NM_001039556, intron 11 of 14) | 30504 | NM_001256142 | 100503583 | Mm.249437 | NM_001256142 | ENSMUSG00000094595 | **Fsbp** | - | fibrinogen silencer binding protein | protein-coding |
| chr6_28427062_28427432 | chr6 | 28427063 | 28427432 | + | -0.794864 | promoter-TSS (NM_019569) | promoter-TSS (NM_019569) | -654 | NM_019569 | 56223 | Mm.31708 | NM_019569 | ENSMUSG00000029707 | **Fscn3** | - | fascin homolog 3, actin-bundling protein, testicular (Strongylocentrotus purpuratus) | protein-coding |
| chr3_152211869_152212284 | chr3 | 152211870 | 152212284 | + | -0.716665 | intron (NM_057172, intron 1 of 19) | intron (NM_057172, intron 1 of 19) | 1619 | NM_057172 | 51886 | Mm.278922 | NM_057172 | ENSMUSG00000028034 | **Fubp1** | 9530027K12Rik\|D3Ertd330e\|FBP\|Fubp\|Fubp4 | far upstream element (FUSE) binding protein 1 | protein-coding |
| chr7_80449723_80450637 | chr7 | 80449724 | 80450637 | + | -1.06627 | Intergenic | RMER4B\|LTR\|ERVK | -44749 | NM_001081454 | 18550 | Mm.5241 | NM_011046 | ENSMUSG00000030530 | **Furin** | 9130404I01Rik\|Fur\|PACE\|Pcsk3\|SPC1 | furin (paired basic amino acid cleaving enzyme) | protein-coding |
| chr7_80449818_80450242 | chr7 | 80449819 | 80450242 | + | -1.24555 | Intergenic | RMER4B\|LTR\|ERVK | -44599 | NM_001081454 | 18550 | Mm.5241 | NM_011046 | ENSMUSG00000030530 | **Furin** | 9130404I01Rik\|Fur\|PACE\|Pcsk3\|SPC1 | furin (paired basic amino acid cleaving enzyme) | protein-coding |
| chr7_127968514_127968772 | chr7 | 127968515 | 127968772 | + | -0.84774 | intron (NM_139149, intron 1 of 14) | intron (NM_139149, intron 1 of 14) | 1164 | NM_139149 | 233908 | Mm.277680 | NM_139149 | ENSMUSG00000030795 | **Fus** | D430004D17Rik\|D930039C12Rik\|Fus1\|Tls | fused in sarcoma | protein-coding |
| chr12_77124220_77124620 | chr12 | 77124221 | 77124620 | + | -0.850509 | Intergenic | Intergenic | -113684 | NM_001252614 | 53618 | Mm.35628 | NM_016893 | ENSMUSG00000021065 | **Fut8** | - | fucosyltransferase 8 | protein-coding |
| chr12_77177328_77177662 | chr12 | 77177329 | 77177662 | + | -0.853604 | Intergenic | Intergenic | -60609 | NM_001252614 | 53618 | Mm.35628 | NM_016893 | ENSMUSG00000021065 | **Fut8** | - | fucosyltransferase 8 | protein-coding |
| chr12_77193208_77193771 | chr12 | 77193209 | 77193771 | + | -0.854998 | Intergenic | Intergenic | -44614 | NM_001252614 | 53618 | Mm.35628 | NM_016893 | ENSMUSG00000021065 | **Fut8** | - | fucosyltransferase 8 | protein-coding |
| chr12_77788174_77788795 | chr12 | 77788175 | 77788795 | + | -0.985761 | Intergenic | Intergenic | 378344 | NM_001252616 | 53618 | Mm.35628 | NM_016893 | ENSMUSG00000021065 | **Fut8** | - | fucosyltransferase 8 | protein-coding |

| | | | | | | | | | | | | | | | | | |
|---|---|---|---|---|---|---|---|---|---|---|---|---|---|---|---|---|---|
| chr12_77535133_77535504 | chr12 | 77535134 | 77535504 | + | -1.0773 | Intergenic | Intergenic | 125178 | NM_001252616 | 53618 | Mm.35628 | NM_016893 | ENSMUSG00000021065 | **Fut8** | - | fucosyltransferase 8 | protein-coding |
| chr12_77113508_77114057 | chr12 | 77113509 | 77114057 | + | -1.27207 | Intergenic | Intergenic | -124321 | NM_001252614 | 53618 | Mm.35628 | NM_016893 | ENSMUSG00000021065 | **Fut8** | - | fucosyltransferase 8 | protein-coding |
| chr7_31087335_31087589 | chr7 | 31087336 | 31087589 | + | -0.792021 | Intergenic | Intergenic | -10765 | NM_008557 | 17178 | Mm.263847 | NM_008557 | | **Fxyd3** | Mat8 | FXYD domain-containing ion transport regulator 3 | protein-coding |
| chr7_31089709_31090218 | chr7 | 31089710 | 31090218 | + | -1.27025 | Intergenic | ORR1D2\|LTR\|ERVL-MaLR | -13267 | NM_008557 | 17178 | Mm.263847 | NM_008557 | | **Fxyd3** | Mat8 | FXYD domain-containing ion transport regulator 3 | protein-coding |
| chr15_6554654_6555117 | chr15 | 6554655 | 6555117 | + | -0.626738 | Intergenic | Intergenic | -24961 | NM_001278269 | 23880 | Mm.170905 | NM_011815 | | **Fyb** | ADAP\|B630013F22Rik\|FYB-120/130\|Slap130\|p120/130\|p120/p130 | FYN binding protein | protein-coding |
| chr15_6579675_6579956 | chr15 | 6579676 | 6579956 | + | -0.701977 | promoter-TSS (NM_001278269) | promoter-TSS (NM_001278269) | -31 | NM_001278269 | 23880 | Mm.170905 | NM_011815 | | **Fyb** | ADAP\|B630013F22Rik\|FYB-120/130\|Slap130\|p120/130\|p120/p130 | FYN binding protein | protein-coding |
| chr15_6592353_6592933 | chr15 | 6592354 | 6592933 | + | -0.84093 | intron (NM_011815, intron 1 of 17) | intron (NM_011815, intron 1 of 17) | 12796 | NM_011815 | 23880 | Mm.170905 | NM_011815 | | **Fyb** | ADAP\|B630013F22Rik\|FYB-120/130\|Slap130\|p120/130\|p120/p130 | FYN binding protein | protein-coding |
| chr15_6610748_6611063 | chr15 | 6610749 | 6611063 | + | -0.841563 | intron (NM_011815, intron 1 of 17) | intron (NM_011815, intron 1 of 17) | 31059 | NM_011815 | 23880 | Mm.170905 | NM_011815 | | **Fyb** | ADAP\|B630013F22Rik\|FYB-120/130\|Slap130\|p120/130\|p120/p130 | FYN binding protein | protein-coding |
| chr15_6578030_6578315 | chr15 | 6578031 | 6578315 | + | -0.891849 | Intergenic | Intergenic | -1674 | NM_001278269 | 23880 | Mm.170905 | NM_011815 | | **Fyb** | ADAP\|B630013F22Rik\|FYB-120/130\|Slap130\|p120/130\|p120/p130 | FYN binding protein | protein-coding |
| chr15_6592595_6592901 | chr15 | 6592596 | 6592901 | + | -0.95591 | intron (NM_011815, intron 1 of 17) | intron (NM_011815, intron 1 of 17) | 12901 | NM_011815 | 23880 | Mm.170905 | NM_011815 | | **Fyb** | ADAP\|B630013F22Rik\|FYB-120/130\|Slap130\|p120/130\|p120/p130 | FYN binding protein | protein-coding |
| chr15_6592556_6593127 | chr15 | 6592557 | 6593127 | + | -0.999991 | intron (NM_011815, intron 1 of 17) | intron (NM_011815, intron 1 of 17) | 12995 | NM_011815 | 23880 | Mm.170905 | NM_011815 | | **Fyb** | ADAP\|B630013F22Rik\|FYB-120/130\|Slap130\|p120/130\|p120/p130 | FYN binding protein | protein-coding |
| chr15_6591567_6591809 | chr15 | 6591568 | 6591809 | + | -1.01485 | intron (NM_011815, intron 1 of 17) | intron (NM_011815, intron 1 of 17) | 11841 | NM_011815 | 23880 | Mm.170905 | NM_011815 | | **Fyb** | ADAP\|B630013F22Rik\|FYB-120/130\|Slap130\|p120/130\|p120/p130 | FYN binding protein | protein-coding |
| chr15_6601317_6601732 | chr15 | 6601318 | 6601732 | + | -1.03985 | intron (NM_011815, intron 1 of 17) | intron (NM_011815, intron 1 of 17) | 21678 | NM_011815 | 23880 | Mm.170905 | NM_011815 | | **Fyb** | ADAP\|B630013F22Rik\|FYB-120/130\|Slap130\|p120/130\|p120/p130 | FYN binding protein | protein-coding |
| chr15_6525380_6525856 | chr15 | 6525381 | 6525856 | + | -1.0561 | Intergenic | ORR1E\|LTR\|ERVL-MaLR | -54229 | NM_001278269 | 23880 | Mm.170905 | NM_011815 | | **Fyb** | ADAP\|B630013F22Rik\|FYB-120/130\|Slap130\|p120/130\|p120/p130 | FYN binding protein | protein-coding |
| chr15_6525490_6525713 | chr15 | 6525491 | 6525713 | + | -1.12265 | Intergenic | ORR1E\|LTR\|ERVL-MaLR | -54245 | NM_001278269 | 23880 | Mm.170905 | NM_011815 | | **Fyb** | ADAP\|B630013F22Rik\|FYB-120/130\|Slap130\|p120/130\|p120/p130 | FYN binding protein | protein-coding |
| chr10_39339235_39339511 | chr10 | 39339236 | 39339511 | + | -0.658332 | Intergenic | ORR1C1\|LTR\|ERVL-MaLR | -30426 | NM_001122892 | 14360 | Mm.4848 | NM_008054 | ENSMUSG00000019843 | **Fyn** | AI448320\|AW552119 | Fyn proto-oncogene | protein-coding |
| chr10_39339774_39340154 | chr10 | 39339775 | 39340154 | + | -0.776698 | Intergenic | RLTR14\|LTR\|ERV1 | -29835 | NM_001122892 | 14360 | Mm.4848 | NM_008054 | ENSMUSG00000019843 | **Fyn** | AI448320\|AW552119 | Fyn proto-oncogene | protein-coding |
| chr10_39339651_39340216 | chr10 | 39339652 | 39340216 | + | -0.82288 | Intergenic | RLTR14\|LTR\|ERV1 | -29865 | NM_001122892 | 14360 | Mm.4848 | NM_008054 | ENSMUSG00000019843 | **Fyn** | AI448320\|AW552119 | Fyn proto-oncogene | protein-coding |
| chr10_39371554_39371860 | chr10 | 39371555 | 39371860 | + | -0.83187 | intron (NM_008054, intron 1 of 13) | intron (NM_008054, intron 1 of 13) | 1908 | NM_001122893 | 14360 | Mm.4848 | NM_008054 | ENSMUSG00000019843 | **Fyn** | AI448320\|AW552119 | Fyn proto-oncogene | protein-coding |
| chr1_193285672_193286446 | chr1 | 193285673 | 193286446 | + | -1.25755 | Intergenic | ORR1A2-int\|LTR\|ERVL-MaLR | -12871 | NM_008059 | 14373 | Mm.439721 | NM_008059 | ENSMUSG00000009633 | **G0s2** | AI255151\|AV006465 | G0/G1 switch gene 2 | protein-coding |
| chr1_193285649_193286070 | chr1 | 193285650 | 193286070 | + | -1.30948 | Intergenic | ORR1A2-int\|LTR\|ERVL-MaLR | -12672 | NM_008059 | 14373 | Mm.439721 | NM_008059 | ENSMUSG00000009633 | **G0s2** | AI255151\|AV006465 | G0/G1 switch gene 2 | protein-coding |
| chr5_92094117_92094345 | chr5 | 92094118 | 92094345 | + | -0.657655 | Intergenic | RSINE1\|SINE\|B4 | -10496 | NM_001080795 | 23881 | Mm.290530 | NM_011816 | ENSMUSG00000029405 | **G3bp2** | AA409541\|E430034L04Rik\|G3BP\|mKIAA0660 | GTPase activating protein (SH3 domain) binding protein 2 | protein-coding |
| chr5_92103035_92103439 | chr5 | 92103036 | 92103439 | + | -1.00189 | Intergenic | Intergenic | -19502 | NM_001080795 | 23881 | Mm.290530 | NM_011816 | ENSMUSG00000029405 | **G3bp2** | AA409541\|E430034L04Rik\|G3BP\|mKIAA0660 | GTPase activating protein (SH3 domain) binding protein 2 | protein-coding |
| chr11_101357543_101357841 | chr11 | 101357544 | 101357841 | + | -0.713187 | Intergenic | Intergenic | -10038 | NM_008061 | 14377 | Mm.18064 | NM_008061 | ENSMUSG00000078650 | **G6pc** | AW107337\|G6Pase\|G6pt\|Glc-6-Pase | glucose-6-phosphatase, catalytic | protein-coding |
| chr16_50554909_50555364 | chr16 | 50554910 | 50555364 | + | -0.73088 | intron (NR_040380, intron 4 of 4) | intron (NR_040380, intron 4 of 4) | 26635 | NR_040379 | 619312 | Mm.427671 | NR_040379 | | **G730013B05Rik** | - | RIKEN cDNA G730013B05 gene | ncRNA |
| chrX_75035254_75035651 | chrX | 75035255 | 75035651 | + | -0.774388 | intron (NM_181584, intron 1 of 9) | B2_Mm2\|SINE\|B2 | 49452 | NM_181584 | 210710 | Mm.267056 | NM_181584 | ENSMUSG00000032750 | **Gab3** | 5930433H21Rik | growth factor receptor bound protein 2-associated protein 3 | protein-coding |
| chr4_47010663_47011465 | chr4 | 47010664 | 47011465 | + | -0.770447 | Intergenic | Intergenic | -19350 | NM_001081141 | 242425 | Mm.101909 | NM_001081141 | | **Gabbr2** | GABABR2\|Gb2\|Gm425\|Gpr51 | gamma-aminobutyric acid (GABA) B receptor, 2 | protein-coding |
| chr2_126674392_126676390 | chr2 | 126674393 | 126676390 | + | -0.649465 | non-coding (NR_073183, exon 1 of 6) | non-coding (NR_073183, exon 1 of 6) | 201 | NM_001271469 | 14391 | Mm.293266 | NM_010249 | ENSMUSG00000027361 | **Gabpb1** | BABPB2\|E4TF1\|E4TF1-47\|E4TF1-53\|E4Tf1B\|GABPB\|GABPB-1\|GABPB-2\|GABPB1-1\|GABPB1-2\|NRF2B1\|NRF2B2 | GA repeat binding protein, beta 1 | protein-coding |
| chr2_126675055_126676434 | chr2 | 126675056 | 126676434 | + | -0.723594 | promoter-TSS (NM_207669) | promoter-TSS (NM_207669) | -153 | NR_073183 | 14391 | Mm.293266 | NM_010249 | ENSMUSG00000027361 | **Gabpb1** | BABPB2\|E4TF1\|E4TF1-47\|E4TF1-53\|E4Tf1B\|GABPB\|GABPB-1\|GABPB-2\|GABPB1-1\|GABPB1-2\|NRF2B1\|NRF2B2 | GA repeat binding protein, beta 1 | protein-coding |
| chr5_71903415_71904007 | chr5 | 71903416 | 71904007 | + | -1.17577 | intron (NM_008069, intron 4 of 8) | intron (NM_008069, intron 4 of 8) | 203695 | NM_008069 | 14400 | Mm.38567 | NM_008069 | | **Gabrb1** | AW061132\|B230208N19Rik\|Gabrb-1 | gamma-aminobutyric acid (GABA) A receptor, subunit beta 1 | protein-coding |
| chr5_71903243_71903772 | chr5 | 71903244 | 71903772 | + | -1.21927 | intron (NM_008069, intron 4 of 8) | L2a\|LINE\|L2 | 203492 | NM_008069 | 14400 | Mm.38567 | NM_008069 | | **Gabrb1** | AW061132\|B230208N19Rik\|Gabrb-1 | gamma-aminobutyric acid (GABA) A receptor, subunit beta 1 | protein-coding |
| chr5_71903447_71903749 | chr5 | 71903448 | 71903749 | + | -1.26915 | intron (NM_008069, intron 4 of 8) | L2a\|LINE\|L2 | 203582 | NM_008069 | 14400 | Mm.38567 | NM_008069 | | **Gabrb1** | AW061132\|B230208N19Rik\|Gabrb-1 | gamma-aminobutyric acid (GABA) A receptor, subunit beta 1 | protein-coding |
| chr4_33072817_33073254 | chr4 | 33072818 | 33073254 | + | -0.880728 | intron (NM_008076, intron 2 of 8) | intron (NM_008076, intron 2 of 8) | 9924 | NM_008076 | 14409 | Mm.6227 | NM_008076 | ENSMUSG00000023267 | **Gabrr2** | - | gamma-aminobutyric acid (GABA) C receptor, subunit rho 2 | protein-coding |
| chr4_33077693_33078159 | chr4 | 33077694 | 33078159 | + | -1.15661 | intron (NM_008076, intron 3 of 8) | intron (NM_008076, intron 3 of 8) | 14814 | NM_008076 | 14409 | Mm.6227 | NM_008076 | ENSMUSG00000023267 | **Gabrr2** | - | gamma-aminobutyric acid (GABA) C receptor, subunit rho 2 | protein-coding |
| chr2_22590193_22590548 | chr2 | 22590194 | 22590548 | + | -0.755981 | Intergenic | Intergenic | -31956 | NM_008078 | 14417 | Mm.4784 | NM_008078 | ENSMUSG00000026787 | **Gad2** | 6330404F12Rik\|GAD(65)\|GAD65\|Gad-2 | glutamic acid decarboxylase 2 | protein-coding |
| chr2_22588903_22589332 | chr2 | 22588904 | 22589332 | + | -0.816054 | Intergenic | Intergenic | -33209 | NM_008078 | 14417 | Mm.4784 | NM_008078 | ENSMUSG00000026787 | **Gad2** | 6330404F12Rik\|GAD(65)\|GAD65\|Gad-2 | glutamic acid decarboxylase 2 | protein-coding |
| chr2_22588103_22589707 | chr2 | 22588104 | 22589707 | + | -0.888395 | Intergenic | Intergenic | -33422 | NM_008078 | 14417 | Mm.4784 | NM_008078 | ENSMUSG00000026787 | **Gad2** | 6330404F12Rik\|GAD(65)\|GAD65\|Gad-2 | glutamic acid decarboxylase 2 | protein-coding |
| chr6_67055940_67056784 | chr6 | 67055941 | 67056784 | + | -0.818511 | intron (NR_040278, intron 1 of 6) | intron (NR_040278, intron 1 of 6) | -18955 | NM_007836 | 13197 | Mm.72235 | NM_007836 | ENSMUSG00000036390 | **Gadd45a** | AA545191\|Ddit1\|GADD45 | growth arrest and DNA-damage-inducible 45 alpha | protein-coding |
| chr6_67056151_67056766 | chr6 | 67056152 | 67056766 | + | -0.833161 | intron (NR_040278, intron 1 of 6) | intron (NR_040278, intron 1 of 6) | -19052 | NM_007836 | 13197 | Mm.72235 | NM_007836 | ENSMUSG00000036390 | **Gadd45a** | AA545191\|Ddit1\|GADD45 | growth arrest and DNA-damage-inducible 45 alpha | protein-coding |
| chr9_115703618_115704206 | chr9 | 115703619 | 115704206 | + | -0.616419 | Intergenic | Intergenic | -205543 | NM_028638 | 73748 | Mm.485018 | NM_028638 | ENSMUSG00000056880 | **Gadl1** | 1110027M19Rik\|ADC\|CSADC | glutamate decarboxylase-like 1 | protein-coding |
| chr9_116018337_116019033 | chr9 | 116018338 | 116019033 | + | -0.974145 | intron (NM_028638, intron 12 of 14) | intron (NM_028638, intron 12 of 14) | 109230 | NM_028638 | 73748 | Mm.485018 | NM_028638 | ENSMUSG00000056880 | **Gadl1** | 1110027M19Rik\|ADC\|CSADC | glutamate decarboxylase-like 1 | protein-coding |
| chr9_116018553_116018950 | chr9 | 116018554 | 116018950 | + | -1.08566 | intron (NM_028638, intron 12 of 14) | intron (NM_028638, intron 12 of 14) | 109297 | NM_028638 | 73748 | Mm.485018 | NM_028638 | ENSMUSG00000056880 | **Gadl1** | 1110027M19Rik\|ADC\|CSADC | glutamate decarboxylase-like 1 | protein-coding |
| chr9_115640634_115641118 | chr9 | 115640635 | 115641118 | + | -1.31376 | Intergenic | Intergenic | -268579 | NM_028638 | 73748 | Mm.485018 | NM_028638 | ENSMUSG00000056880 | **Gadl1** | 1110027M19Rik\|ADC\|CSADC | glutamate decarboxylase-like 1 | protein-coding |
| chr9_115640785_115641155 | chr9 | 115640786 | 115641155 | + | -1.38298 | Intergenic | RLTR14_RN\|LTR\|ERV1 | -268485 | NM_028638 | 73748 | Mm.485018 | NM_028638 | ENSMUSG00000056880 | **Gadl1** | 1110027M19Rik\|ADC\|CSADC | glutamate decarboxylase-like 1 | protein-coding |
| chr8_122595931_122596348 | chr8 | 122595932 | 122596348 | + | -0.655761 | intron (NM_001193645, intron 8 of 11) | intron (NM_001193645, intron 8 of 11) | 15347 | NM_001193645 | 50917 | Mm.34702 | NM_016722 | ENSMUSG00000015027 | **Galns** | mFLJ00319 | galactosamine (N-acetyl)-6-sulfate sulfatase | protein-coding |

| | | | | | | | | | | | | | | | | | |
|---|---|---|---|---|---|---|---|---|---|---|---|---|---|---|---|---|---|
| chr8_122595796_122596358 | chr8 | 122595797 | 122596358 | + | -0.675051 | intron (NM_001193645, intron 8 of 11) | intron (NM_001193645, intron 8 of 11) | 15410 | NM_001193645 | 50917 | Mm.34702 | NM_016722 | ENSMUSG00000015027 | **Galns** | mFLJ00319 | galactosamine (N-acetyl)-6-sulfate sulfatase | protein-coding |
| chr11_57655682_57655981 | chr11 | 57655683 | 57655981 | + | -0.665576 | intron (NM_134189, intron 1 of 11) | intron (NM_134189, intron 1 of 11) | 10390 | NM_134189 | 171212 | Mm.271670 | NM_134189 | ENSMUSG00000020520 | **Galnt10** | AU018154\|C330012K04Rik\|GalNAc-T10\|GalNAc-T9\|Galnt9\|ppGaNTase | UDP-N-acetyl-alpha-D-galactosamine:polypeptide N-acetylgalactosaminyltransferase 10 | protein-coding |
| chr11_57660688_57661311 | chr11 | 57660689 | 57661311 | + | -0.761399 | intron (NM_134189, intron 1 of 11) | intron (NM_134189, intron 1 of 11) | 15558 | NM_134189 | 171212 | Mm.271670 | NM_134189 | ENSMUSG00000020520 | **Galnt10** | AU018154\|C330012K04Rik\|GalNAc-T10\|GalNAc-T9\|Galnt9\|ppGaNTase | UDP-N-acetyl-alpha-D-galactosamine:polypeptide N-acetylgalactosaminyltransferase 10 | protein-coding |
| chr11_57660711_57661117 | chr11 | 57660712 | 57661117 | + | -0.774903 | intron (NM_134189, intron 1 of 11) | intron (NM_134189, intron 1 of 11) | 15472 | NM_134189 | 171212 | Mm.271670 | NM_134189 | ENSMUSG00000020520 | **Galnt10** | AU018154\|C330012K04Rik\|GalNAc-T10\|GalNAc-T9\|Galnt9\|ppGaNTase | UDP-N-acetyl-alpha-D-galactosamine:polypeptide N-acetylgalactosaminyltransferase 10 | protein-coding |
| chr11_57686137_57686388 | chr11 | 57686138 | 57686388 | + | -0.827755 | intron (NM_134189, intron 1 of 11) | intron (NM_134189, intron 1 of 11) | 40821 | NM_134189 | 171212 | Mm.271670 | NM_134189 | ENSMUSG00000020520 | **Galnt10** | AU018154\|C330012K04Rik\|GalNAc-T10\|GalNAc-T9\|Galnt9\|ppGaNTase | UDP-N-acetyl-alpha-D-galactosamine:polypeptide N-acetylgalactosaminyltransferase 10 | protein-coding |
| chr11_57660855_57661157 | chr11 | 57660856 | 57661157 | + | -0.833779 | intron (NM_134189, intron 1 of 11) | MTE2a\|LTR\|ERVL-MaLR | 15564 | NM_134189 | 171212 | Mm.271670 | NM_134189 | ENSMUSG00000020520 | **Galnt10** | AU018154\|C330012K04Rik\|GalNAc-T10\|GalNAc-T9\|Galnt9\|ppGaNTase | UDP-N-acetyl-alpha-D-galactosamine:polypeptide N-acetylgalactosaminyltransferase 10 | protein-coding |
| chr11_57664792_57665088 | chr11 | 57664793 | 57665088 | + | -0.984838 | intron (NM_134189, intron 1 of 11) | intron (NM_134189, intron 1 of 11) | 19498 | NM_134189 | 171212 | Mm.271670 | NM_134189 | ENSMUSG00000020520 | **Galnt10** | AU018154\|C330012K04Rik\|GalNAc-T10\|GalNAc-T9\|Galnt9\|ppGaNTase | UDP-N-acetyl-alpha-D-galactosamine:polypeptide N-acetylgalactosaminyltransferase 10 | protein-coding |
| chr11_57673082_57673416 | chr11 | 57673083 | 57673416 | + | -0.998805 | intron (NM_134189, intron 1 of 11) | intron (NM_134189, intron 1 of 11) | 27807 | NM_134189 | 171212 | Mm.271670 | NM_134189 | ENSMUSG00000020520 | **Galnt10** | AU018154\|C330012K04Rik\|GalNAc-T10\|GalNAc-T9\|Galnt9\|ppGaNTase | UDP-N-acetyl-alpha-D-galactosamine:polypeptide N-acetylgalactosaminyltransferase 10 | protein-coding |
| chr8_124275731_124276100 | chr8 | 124275732 | 124276100 | + | -0.629424 | intron (NM_139272, intron 1 of 15) | intron (NM_139272, intron 1 of 15) | 44522 | NM_139272 | 108148 | Mm.33808 | NM_139272 | ENSMUSG00000089704 | **Galnt2** | AI480629 | UDP-N-acetyl-alpha-D-galactosamine:polypeptide N-acetylgalactosaminyltransferase 2 | protein-coding |
| chr8_124282044_124282702 | chr8 | 124282045 | 124282702 | + | -0.634816 | intron (NM_139272, intron 1 of 15) | intron (NM_139272, intron 1 of 15) | 50979 | NM_139272 | 108148 | Mm.33808 | NM_139272 | ENSMUSG00000089704 | **Galnt2** | AI480629 | UDP-N-acetyl-alpha-D-galactosamine:polypeptide N-acetylgalactosaminyltransferase 2 | protein-coding |
| chr8_124273506_124274168 | chr8 | 124273507 | 124274168 | + | -0.690077 | intron (NM_139272, intron 1 of 15) | intron (NM_139272, intron 1 of 15) | 42443 | NM_139272 | 108148 | Mm.33808 | NM_139272 | ENSMUSG00000089704 | **Galnt2** | AI480629 | UDP-N-acetyl-alpha-D-galactosamine:polypeptide N-acetylgalactosaminyltransferase 2 | protein-coding |
| chr8_124242033_124242649 | chr8 | 124242034 | 124242649 | + | -0.899104 | intron (NM_139272, intron 1 of 15) | MIR\|SINE\|MIR | 10947 | NM_139272 | 108148 | Mm.33808 | NM_139272 | ENSMUSG00000089704 | **Galnt2** | AI480629 | UDP-N-acetyl-alpha-D-galactosamine:polypeptide N-acetylgalactosaminyltransferase 2 | protein-coding |
| chr8_124317797_124318293 | chr8 | 124317798 | 124318293 | + | -1.18783 | intron (NM_139272, intron 3 of 15) | L1MB1\|LINE\|L1 | 86651 | NM_139272 | 108148 | Mm.33808 | NM_139272 | ENSMUSG00000089704 | **Galnt2** | AI480629 | UDP-N-acetyl-alpha-D-galactosamine:polypeptide N-acetylgalactosaminyltransferase 2 | protein-coding |
| chr8_124274507_124274878 | chr8 | 124274508 | 124274878 | + | -1.2309 | intron (NM_139272, intron 1 of 15) | intron (NM_139272, intron 1 of 15) | 43299 | NM_139272 | 108148 | Mm.33808 | NM_139272 | ENSMUSG00000089704 | **Galnt2** | AI480629 | UDP-N-acetyl-alpha-D-galactosamine:polypeptide N-acetylgalactosaminyltransferase 2 | protein-coding |
| chr8_124274365_124274836 | chr8 | 124274366 | 124274836 | + | -1.24438 | intron (NM_139272, intron 1 of 15) | intron (NM_139272, intron 1 of 15) | 43207 | NM_139272 | 108148 | Mm.33808 | NM_139272 | ENSMUSG00000089704 | **Galnt2** | AI480629 | UDP-N-acetyl-alpha-D-galactosamine:polypeptide N-acetylgalactosaminyltransferase 2 | protein-coding |
| chr10_99181578_99182525 | chr10 | 99181579 | 99182525 | + | -0.732055 | intron (NM_027740, intron 10 of 11) | intron (NM_027740, intron 10 of 11) | 73917 | NM_015737 | 14426 | Mm.314 | NM_015737 | ENSMUSG00000090035 | **Galnt4** | AV011803 | UDP-N-acetyl-alpha-D-galactosamine:polypeptide N-acetylgalactosaminyltransferase 4 | protein-coding |
| chr10_99119533_99119933 | chr10 | 99119534 | 99119933 | + | -0.921554 | intron (NM_027740, intron 2 of 11) | intron (NM_027740, intron 2 of 11) | 11598 | NM_015737 | 14426 | Mm.314 | NM_015737 | ENSMUSG00000090035 | **Galnt4** | AV011803 | UDP-N-acetyl-alpha-D-galactosamine:polypeptide N-acetylgalactosaminyltransferase 4 | protein-coding |
| chr10_99119547_99119786 | chr10 | 99119548 | 99119786 | + | -0.933564 | intron (NM_027740, intron 2 of 11) | PB1D10\|SINE\|Alu | 11532 | NM_015737 | 14426 | Mm.314 | NM_015737 | ENSMUSG00000090035 | **Galnt4** | AV011803 | UDP-N-acetyl-alpha-D-galactosamine:polypeptide N-acetylgalactosaminyltransferase 4 | protein-coding |
| chr10_99117301_99117635 | chr10 | 99117302 | 99117635 | + | -1.04648 | intron (NM_027740, intron 2 of 11) | intron (NM_027740, intron 2 of 11) | 9333 | NM_015737 | 14426 | Mm.314 | NM_015737 | ENSMUSG00000090035 | **Galnt4** | AV011803 | UDP-N-acetyl-alpha-D-galactosamine:polypeptide N-acetylgalactosaminyltransferase 4 | protein-coding |
| chr10_99116768_99117619 | chr10 | 99116769 | 99117619 | + | -1.37771 | intron (NM_027740, intron 2 of 11) | intron (NM_027740, intron 2 of 11) | 9059 | NM_015737 | 14426 | Mm.314 | NM_015737 | ENSMUSG00000090035 | **Galnt4** | AV011803 | UDP-N-acetyl-alpha-D-galactosamine:polypeptide N-acetylgalactosaminyltransferase 4 | protein-coding |
| chr10_99116784_99117219 | chr10 | 99116785 | 99117219 | + | -1.48611 | intron (NM_027740, intron 2 of 11) | B4A\|SINE\|B4 | 8867 | NM_015737 | 14426 | Mm.314 | NM_015737 | ENSMUSG00000090035 | **Galnt4** | AV011803 | UDP-N-acetyl-alpha-D-galactosamine:polypeptide N-acetylgalactosaminyltransferase 4 | protein-coding |
| chr2_57903480_57903729 | chr2 | 57903481 | 57903729 | + | -0.666392 | Intergenic | Intergenic | -94550 | NM_172855 | 241391 | Mm.484118 | NM_172855 | ENSMUSG00000026828 | **Galnt5** | 4832424J23 | UDP-N-acetyl-alpha-D-galactosamine:polypeptide N-acetylgalactosaminyltransferase 5 | protein-coding |
| chr2_57903363_57903724 | chr2 | 57903364 | 57903724 | + | -0.673766 | Intergenic | Intergenic | -94611 | NM_172855 | 241391 | Mm.484118 | NM_172855 | ENSMUSG00000026828 | **Galnt5** | 4832424J23 | UDP-N-acetyl-alpha-D-galactosamine:polypeptide N-acetylgalactosaminyltransferase 5 | protein-coding |
| chr8_57758864_57759205 | chr8 | 57758865 | 57759205 | + | -0.758959 | Intergenic | MLT1A1\|LTR\|ERVL-MaLR | -105980 | NM_144731 | 108150 | Mm.62886 | NM_144731 | ENSMUSG00000031608 | **Galnt7** | AI225872 | UDP-N-acetyl-alpha-D-galactosamine: polypeptide N-acetylgalactosaminyltransferase 7 | protein-coding |
| chr8_57828125_57828773 | chr8 | 57828126 | 57828773 | + | -0.91419 | intron (NM_175032, intron 9 of 11) | ORR1B2\|LTR\|ERVL-MaLR | -175394 | NM_144731 | 108150 | Mm.62886 | NM_144731 | ENSMUSG00000031608 | **Galnt7** | AI225872 | UDP-N-acetyl-alpha-D-galactosamine: polypeptide N-acetylgalactosaminyltransferase 7 | protein-coding |
| chr18_82244543_82245107 | chr18 | 82244544 | 82245107 | + | -0.855194 | Intergenic | Intergenic | 161952 | NM_008082 | 14427 | Mm.6219 | NM_008082 | ENSMUSG00000024553 | **Galr1** | Galnr1 | galanin receptor 1 | protein-coding |
| chr2_34614801_34615310 | chr2 | 34614802 | 34615310 | + | -0.806687 | intron (NM_001290625, intron 10 of 11) | B1_Mus2\|SINE\|Alu | 140176 | NM_025709 | 66691 | Mm.156452 | NM_025709 | ENSMUSG00000026867 | **Gapvd1** | 2010005B09Rik\|4432404J10Rik\|AW108497\|Gapex-5\|RAP6\|RME-6\|mKIAA1521 | GTPase activating protein and VPS9 domains 1 | protein-coding |
| chr2_9691556_9691823 | chr2 | 9691557 | 9691823 | + | -0.782187 | Intergenic | Intergenic | 186910 | NM_008091 | 14462 | Mm.313866 | NM_008091 | ENSMUSG00000015619 | **Gata3** | Gata-3\|jal | GATA binding protein 3 | protein-coding |
| chr2_9593773_9594514 | chr2 | 9593774 | 9594514 | + | -0.867408 | Intergenic | Intergenic | 284456 | NM_008091 | 14462 | Mm.313866 | NM_008091 | ENSMUSG00000015619 | **Gata3** | Gata-3\|jal | GATA binding protein 3 | protein-coding |
| chr2_9595365_9595775 | chr2 | 9595366 | 9595775 | + | -0.889887 | Intergenic | Intergenic | 283030 | NM_008091 | 14462 | Mm.313866 | NM_008091 | ENSMUSG00000015619 | **Gata3** | Gata-3\|jal | GATA binding protein 3 | protein-coding |
| chr3_90323201_90323585 | chr3 | 90323202 | 90323585 | + | -0.615593 | Intergenic | Intergenic | -18261 | NM_139304 | 229542 | Mm.270999 | NM_139304 | ENSMUSG00000042390 | **Gatad2b** | AL118180\|C430014D17Rik\|P66beta\|mKIAA1150 | GATA zinc finger domain containing 2B | protein-coding |
| chr3_90359334_90359594 | chr3 | 90359335 | 90359594 | + | -0.784351 | Intergenic | Intergenic | 17810 | NM_139304 | 229542 | Mm.270999 | NM_139304 | ENSMUSG00000042390 | **Gatad2b** | AL118180\|C430014D17Rik\|P66beta\|mKIAA1150 | GATA zinc finger domain containing 2B | protein-coding |
| chr11_4206536_4206963 | chr11 | 4206537 | 4206963 | + | -0.652562 | intron (NM_134023, intron 2 of 8) | intron (NM_134023, intron 2 of 8) | -11501 | NM_028022 | 71962 | Mm.7852 | NM_028022 | ENSMUSG00000020424 | **Gatsl3** | 2410008K03Rik | GATS protein-like 3 | protein-coding |
| chr11_4206659_4206985 | chr11 | 4206660 | 4206985 | + | -0.704687 | intron (NM_134023, intron 2 of 8) | intron (NM_134023, intron 2 of 8) | -11429 | NM_028022 | 71962 | Mm.7852 | NM_028022 | ENSMUSG00000020424 | **Gatsl3** | 2410008K03Rik | GATS protein-like 3 | protein-coding |
| chr16_70326311_70327464 | chr16 | 70326312 | 70327464 | + | -0.611853 | intron (NM_028803, intron 1 of 15) | L1MD2\|LINE\|L1 | 12939 | NM_028803 | 74185 | Mm.396102 | NM_028803 | ENSMUSG00000022707 | **Gbe1** | 2310045H19Rik\|2810426P10Rik\|D16Ertd536e | glucan (1,4-alpha-), branching enzyme 1 | protein-coding |
| chr16_70395205_70395755 | chr16 | 70395206 | 70395755 | + | -0.672248 | intron (NM_028803, intron 2 of 15) | intron (NM_028803, intron 2 of 15) | 81531 | NM_028803 | 74185 | Mm.396102 | NM_028803 | ENSMUSG00000022707 | **Gbe1** | 2310045H19Rik\|2810426P10Rik\|D16Ertd536e | glucan (1,4-alpha-), branching enzyme 1 | protein-coding |
| chr16_70326336_70326987 | chr16 | 70326337 | 70326987 | + | -0.697784 | intron (NM_028803, intron 1 of 15) | intron (NM_028803, intron 1 of 15) | 12713 | NM_028803 | 74185 | Mm.396102 | NM_028803 | ENSMUSG00000022707 | **Gbe1** | 2310045H19Rik\|2810426P10Rik\|D16Ertd536e | glucan (1,4-alpha-), branching enzyme 1 | protein-coding |
| chr5_105068419_105068685 | chr5 | 105068420 | 105068685 | + | -0.623842 | Intergenic | Intergenic | -14991 | NM_029509 | 76074 | Mm.458519 | NM_029509 | ENSMUSG00000034438 | **Gbp8** | 5830443L24Rik\|mGBP8 | guanylate-binding protein 8 | protein-coding |
| chr6_28409914_28410519 | chr6 | 28409915 | 28410519 | + | -0.689186 | Intergenic | MTE2a\|LTR\|ERVL-MaLR | 11507 | NM_028900 | 74375 | Mm.332950 | NM_028900 | ENSMUSG00000029708 | **Gcc1** | 4932417P04Rik | golgi coiled coil 1 | protein-coding |
| chr6_28350957_28351546 | chr6 | 28350958 | 28351546 | + | -1.34439 | Intergenic | MIRc\|SINE\|MIR | 70472 | NM_028900 | 74375 | Mm.332950 | NM_028900 | ENSMUSG00000029708 | **Gcc1** | 4932417P04Rik | golgi coiled coil 1 | protein-coding |
| chr14_47222793_47223482 | chr14 | 47222794 | 47223482 | + | -1.13829 | Intergenic | B4A\|SINE\|B4 | -33736 | NM_008102 | 14528 | Mm.10651 | NM_008102 | ENSMUSG00000037580 | **Gch1** | GTP-CH\|GTPCH\|Gch | GTP cyclohydrolase 1 | protein-coding |

| | | | | | | | | | | | | | | | | | |
|---|---|---|---|---|---|---|---|---|---|---|---|---|---|---|---|---|---|
| chr14_47187779_47188311 | chr14 | 47187780 | 47188311 | + | -1.31118 | intron (NM_008102, intron 1 of 5) | intron (NM_008102, intron 1 of 5) | 1357 | NM_008102 | 14528 | Mm.10651 | NM_008102 | ENSMUSG00000037580 | **Gch1** | GTP-CH\|GTPCH\|Gch | GTP cyclohydrolase 1 | protein-coding |
| chr3_122202050_122203023 | chr3 | 122202051 | 122203023 | + | -0.749029 | Intergenic | Intergenic | -43020 | NM_008129 | 14630 | Mm.292676 | NM_008129 | | **Gclm** | AI649393\|Gcmc\|Glclr | glutamate-cysteine ligase, modifier subunit | protein-coding |
| chr9_78010722_78011203 | chr9 | 78010723 | 78011203 | + | -0.730533 | Intergenic | Intergenic | -40995 | NM_008103 | 14531 | Mm.1400 | NM_008103 | ENSMUSG00000023333 | **Gcm1** | GCMa\|Gcm-rs2\|Gcm1-rs1\|glide | glial cells missing homolog 1 (Drosophila) | protein-coding |
| chr9_78010151_78010492 | chr9 | 78010152 | 78010492 | + | -1.35002 | Intergenic | Intergenic | -41636 | NM_008103 | 14531 | Mm.1400 | NM_008103 | ENSMUSG00000023333 | **Gcm1** | GCMa\|Gcm-rs2\|Gcm1-rs1\|glide | glial cells missing homolog 1 (Drosophila) | protein-coding |
| chr13_41097406_41098017 | chr13 | 41097407 | 41098017 | + | -0.717621 | Intergenic | ORR1D2\|LTR\|ERVL-MaLR | 12276 | NM_008104 | 107889 | Mm.1399 | NM_008104 | ENSMUSG00000021362 | **Gcm2** | Gcm-rs1\|Gcm1\|Gcm1-rs2 | glial cells missing homolog 2 (Drosophila) | protein-coding |
| chr13_40922515_40922798 | chr13 | 40922516 | 40922798 | + | -0.600404 | intron (NM_133219, intron 1 of 2) | intron (NM_133219, intron 1 of 2) | 5023 | NM_133219 | 14538 | Mm.314757 | NM_008105 | ENSMUSG00000021360 | **Gcnt2** | 5330430K10Rik\|IGnT\|IGnTA\|IGnTB\|IGnTC | glucosaminyl (N-acetyl) transferase 2, I-branching enzyme | protein-coding |
| chr13_40922363_40922800 | chr13 | 40922364 | 40922800 | + | -0.635371 | intron (NM_133219, intron 1 of 2) | intron (NM_133219, intron 1 of 2) | 4948 | NM_133219 | 14538 | Mm.314757 | NM_008105 | ENSMUSG00000021360 | **Gcnt2** | 5330430K10Rik\|IGnT\|IGnTA\|IGnTB\|IGnTC | glucosaminyl (N-acetyl) transferase 2, I-branching enzyme | protein-coding |
| chr11_53441868_53442302 | chr11 | 53441869 | 53442302 | + | -0.737878 | Intergenic | Intergenic | 8799 | NM_008110 | 14566 | Mm.9714 | NM_008110 | ENSMUSG00000018238 | **Gdf9** | Gdf-9 | growth differentiation factor 9 | protein-coding |
| chr11_53442000_53442287 | chr11 | 53442001 | 53442287 | + | -0.754832 | Intergenic | Intergenic | 8858 | NM_008110 | 14566 | Mm.9714 | NM_008110 | ENSMUSG00000018238 | **Gdf9** | Gdf-9 | growth differentiation factor 9 | protein-coding |
| chr13_3526226_3526919 | chr13 | 3526227 | 3526919 | + | -0.695281 | Intergenic | Intergenic | -11502 | NM_008112 | 14569 | Mm.153226 | NM_008112 | ENSMUSG00000021218 | **Gdi2** | GDI-B\|GDIB\|Gdi3 | guanosine diphosphate (GDP) dissociation inhibitor 2 | protein-coding |
| chr15_7925207_7926006 | chr15 | 7925208 | 7926006 | + | -0.705987 | intron (NM_001081402, intron 10 of 17) | intron (NM_001081402, intron 10 of 17) | 110057 | NM_001301357 | 14573 | Mm.4679 | NM_010275 | ENSMUSG00000022144 | **Gdnf** | AI385739 | glial cell line derived neurotrophic factor | protein-coding |
| **PeakID (cmd=annotatePeaks.pl ../averagescore_closing_peaks_allCA.bed mm10 -go go_closing_peaks_allCA)** | **Chr** | **Start** | **End** | **Strand** | **Peak Score** | **Annotation** | **Detailed Annotation** | **Distance to TSS** | **Nearest PromoterID** | **Entrez ID** | **Nearest Unigene** | **Nearest Refseq** | **Nearest Ensembl** | **Gene Name** | **Gene Alias** | **Gene Description** | **Gene Type** |
| chr5_107739079_107739375 | chr5 | 107739080 | 107739375 | + | -0.840077 | Intergenic | Intergenic | -13422 | NM_001267621 | 14581 | Mm.2078 | NM_010278 | ENSMUSG00000029275 | **Gfi1** | AW495828\|Gfi-1\|Pal-1\|Pal1 | growth factor independent 1 | protein-coding |
| chr8_105746091_105746753 | chr8 | 105746092 | 105746753 | + | -0.966221 | intron (NM_027469, intron 1 of 2) | intron (NM_027469, intron 1 of 2) | 12185 | NM_027469 | 70575 | Mm.40954 | NM_027469 | | **Gfod2** | 5730466C23Rik | glucose-fructose oxidoreductase domain containing 2 | protein-coding |
| chr8_105788825_105789571 | chr8 | 105788826 | 105789571 | + | -1.35558 | intron (NM_145824, intron 3 of 13) | intron (NM_145824, intron 3 of 13) | -30591 | NM_027469 | 70575 | Mm.40954 | NM_027469 | | **Gfod2** | 5730466C23Rik | glucose-fructose oxidoreductase domain containing 2 | protein-coding |
| chr6_87040965_87041418 | chr6 | 87040966 | 87041418 | + | -1.15143 | Intergenic | Intergenic | -1654 | NM_013528 | 14583 | Mm.19893 | NM_013528 | ENSMUSG00000029992 | **Gfpt1** | 2810423A18Rik\|AI324119\|AI449986\|GFA\|GFAT\|GFAT1\|GFAT1m\|Gfpt | glutamine fructose-6-phosphate transaminase 1 | protein-coding |
| chr19_58376567_58377120 | chr19 | 58376568 | 58377120 | + | -0.72058 | intron (NM_001285457, intron 3 of 8) | intron (NM_001285457, intron 3 of 8) | 77750 | NM_001285457 | 14585 | Mm.88367 | NM_010279 | ENSMUSG00000025089 | **Gfra1** | AU042498 | glial cell line derived neurotrophic factor family receptor alpha 1 | protein-coding |
| chr14_70962011_70962348 | chr14 | 70962012 | 70962348 | + | -1.03954 | intron (NM_008115, intron 4 of 8) | MER5A\|DNA\|hAT-Charlie | 72050 | NM_008115 | 14586 | Mm.32619 | NM_008115 | ENSMUSG00000022103 | **Gfra2** | - | glial cell line derived neurotrophic factor family receptor alpha 2 | protein-coding |
| chr14_122908077_122908539 | chr14 | 122908078 | 122908539 | + | -0.667563 | intron (NM_145466, intron 1 of 2) | Lx6\|LINE\|L1 | 4857 | NM_145466 | 223267 | Mm.379358 | NM_145466 | ENSMUSG00000041625 | **Ggact** | A2ld1 | gamma-glutamylamine cyclotransferase | protein-coding |
| chr14_122902064_122902579 | chr14 | 122902065 | 122902579 | + | -1.0285 | intron (NM_145466, intron 1 of 2) | intron (NM_145466, intron 1 of 2) | 10843 | NM_145466 | 223267 | Mm.379358 | NM_145466 | ENSMUSG00000041625 | **Ggact** | A2ld1 | gamma-glutamylamine cyclotransferase | protein-coding |
| chr10_75591435_75592032 | chr10 | 75591436 | 75592032 | + | -0.629772 | intron (NM_011820, intron 1 of 11) | intron (NM_011820, intron 1 of 11) | 2353 | NM_011820 | 23887 | Mm.257927 | NM_011820 | ENSMUSG00000006344 | **Ggt5** | AI551243\|GGL\|GGT-REL\|Ggtla1 | gamma-glutamyltransferase 5 | protein-coding |
| chr2_35453605_35454386 | chr2 | 35453606 | 35454386 | + | -0.859105 | intron (NM_010283, intron 1 of 8) | intron (NM_010283, intron 1 of 8) | 7429 | NM_010283 | 14594 | Mm.281124 | NM_010283 | ENSMUSG00000035778 | **Ggta1** | AW108479\|GALT\|Gal\|Ggta\|Ggta-1\|alpha3GalT | glycoprotein galactosyltransferase alpha 1, 3 | protein-coding |
| chr2_35453617_35454153 | chr2 | 35453618 | 35454153 | + | -0.953803 | intron (NM_010283, intron 1 of 8) | intron (NM_010283, intron 1 of 8) | 7540 | NM_010283 | 14594 | Mm.281124 | NM_010283 | ENSMUSG00000035778 | **Ggta1** | AW108479\|GALT\|Gal\|Ggta\|Ggta-1\|alpha3GalT | glycoprotein galactosyltransferase alpha 1, 3 | protein-coding |
| chr6_48738880_48739461 | chr6 | 48738881 | 48739461 | + | -0.718987 | intron (NM_175860, intron 1 of 2) | intron (NM_175860, intron 1 of 2) | 124 | NM_008376 | 16205 | Mm.252599 | NM_008376 | ENSMUSG00000090019 | **Gimap1** | IAP38\|Imap38\|imap | GTPase, IMAP family member 1 | protein-coding |
| chr6_48770643_48771402 | chr6 | 48770644 | 48771402 | + | -0.717133 | promoter-TSS (NM_031247) | promoter-TSS (NM_031247) | -172 | NM_031247 | 83408 | Mm.333050 | NM_031247 | | **Gimap3** | 2010110D23Rik\|Gimap5\|Ian4 | GTPase, IMAP family member 3 | protein-coding |
| chr6_48770717_48771168 | chr6 | 48770718 | 48771168 | + | -0.818304 | promoter-TSS (NM_031247) | promoter-TSS (NM_031247) | -92 | NM_031247 | 83408 | Mm.333050 | NM_031247 | | **Gimap3** | 2010110D23Rik\|Gimap5\|Ian4 | GTPase, IMAP family member 3 | protein-coding |
| chr6_48680555_48680745 | chr6 | 48680556 | 48680745 | + | -0.747173 | Intergenic | Intergenic | -3928 | NM_001243200 | 107526 | Mm.333048 | NM_174990 | ENSMUSG00000054435 | **Gimap4** | AU019574\|E430007K16Rik\|Ian-1\|Ian1\|Imap4 | GTPase, IMAP family member 4 | protein-coding |
| chr6_48680411_48680779 | chr6 | 48680412 | 48680779 | + | -0.815201 | Intergenic | Intergenic | -3983 | NM_001243200 | 107526 | Mm.333048 | NM_174990 | ENSMUSG00000054435 | **Gimap4** | AU019574\|E430007K16Rik\|Ian-1\|Ian1\|Imap4 | GTPase, IMAP family member 4 | protein-coding |
| chr6_48752496_48752945 | chr6 | 48752497 | 48752945 | + | -1.24613 | exon (NM_175035, exon 3 of 3) | exon (NM_175035, exon 3 of 3) | 6524 | NM_175035 | 317757 | Mm.455917 | NM_175035 | ENSMUSG00000043505 | **Gimap5** | D630024P16\|E230026N22Rik | GTPase, IMAP family member 5 | protein-coding |
| chr6_48705955_48706590 | chr6 | 48705956 | 48706590 | + | -0.61854 | intron (NM_153175, intron 1 of 2) | intron (NM_153175, intron 1 of 2) | 1971 | NM_153175 | 231931 | Mm.321047 | NM_153175 | ENSMUSG00000047867 | **Gimap6** | 4833419H03Rik\|Ian6\|mFLJ00102 | GTPase, IMAP family member 6 | protein-coding |
| chr6_48707874_48708420 | chr6 | 48707875 | 48708420 | + | -0.78448 | promoter-TSS (NM_153175) | promoter-TSS (NM_153175) | 97 | NM_153175 | 231931 | Mm.321047 | NM_153175 | ENSMUSG00000047867 | **Gimap6** | 4833419H03Rik\|Ian6\|mFLJ00102 | GTPase, IMAP family member 6 | protein-coding |
| chr6_48708109_48708415 | chr6 | 48708110 | 48708415 | + | -0.829435 | promoter-TSS (NM_153175) | promoter-TSS (NM_153175) | -18 | NM_153175 | 231931 | Mm.321047 | NM_153175 | ENSMUSG00000047867 | **Gimap6** | 4833419H03Rik\|Ian6\|mFLJ00102 | GTPase, IMAP family member 6 | protein-coding |
| chr6_48727566_48728259 | chr6 | 48727567 | 48728259 | + | -0.731534 | Intergenic | Intergenic | 9292 | NM_146167 | 231932 | Mm.30479 | NM_146167 | ENSMUSG00000043931 | **Gimap7** | IAN7\|Ian3 | GTPase, IMAP family member 7 | protein-coding |
| chr6_48725470_48725888 | chr6 | 48725471 | 48725888 | + | -0.793243 | Intergenic | Intergenic | 7058 | NM_146167 | 231932 | Mm.30479 | NM_146167 | ENSMUSG00000043931 | **Gimap7** | IAN7\|Ian3 | GTPase, IMAP family member 7 | protein-coding |
| chr6_48669504_48669969 | chr6 | 48669505 | 48669969 | + | -1.22607 | Intergenic | Intergenic | -6398 | NM_174960 | 317758 | Mm.116971 | NM_174960 | ENSMUSG00000051124 | **Gimap9** | A630002K24\|BB145400 | GTPase, IMAP family member 9 | protein-coding |
| chr6_48669615_48669934 | chr6 | 48669616 | 48669934 | + | -1.29549 | Intergenic | Intergenic | -6360 | NM_174960 | 317758 | Mm.116971 | NM_174960 | ENSMUSG00000051124 | **Gimap9** | A630002K24\|BB145400 | GTPase, IMAP family member 9 | protein-coding |
| chr2_114032301_114032615 | chr2 | 114032302 | 114032615 | + | -0.630541 | TTS (NR_028423) | TTS (NR_028423) | -18839 | NM_010290 | 14617 | Mm.389394 | NM_010290 | ENSMUSG00000068615 | **Gjd2** | Gja9\|connexin36\|cx36 | gap junction protein, delta 2 | protein-coding |
| chr2_114032222_114032662 | chr2 | 114032223 | 114032662 | + | -0.637322 | TTS (NR_028423) | TTS (NR_028423) | -18823 | NM_010290 | 14617 | Mm.389394 | NM_010290 | ENSMUSG00000068615 | **Gjd2** | Gja9\|connexin36\|cx36 | gap junction protein, delta 2 | protein-coding |
| chr6_8556906_8557202 | chr6 | 8556907 | 8557202 | + | -0.649108 | intron (NM_001286729, intron 2 of 7) | intron (NM_001286729, intron 2 of 7) | 36996 | NM_001286729 | 170772 | Mm.210787 | NM_133236 | ENSMUSG00000029638 | **Glcci1** | 2310047L21Rik\|A130036A18Rik\|Fam117c\|Gig18\|Tssn1 | glucocorticoid induced transcript 1 | protein-coding |
| chr10_127337546_127337943 | chr10 | 127337547 | 127337943 | + | -0.775614 | intron (NM_010296, intron 4 of 12) | B3A\|SINE\|B2 | 3834 | NM_010296 | 14632 | Mm.391450 | NM_010296 | ENSMUSG00000025407 | **Gli1** | AV235269\|Zfp-5\|Zfp5 | GLI-Kruppel family member GLI1 | protein-coding |
| chr10_127337568_127337831 | chr10 | 127337569 | 127337831 | + | -0.821387 | intron (NM_010296, intron 4 of 12) | B3A\|SINE\|B2 | 3879 | NM_010296 | 14632 | Mm.391450 | NM_010296 | ENSMUSG00000025407 | **Gli1** | AV235269\|Zfp-5\|Zfp5 | GLI-Kruppel family member GLI1 | protein-coding |
| chr4_43965886_43966406 | chr4 | 43965887 | 43966406 | + | -0.650076 | intron (NM_027450, intron 1 of 4) | intron (NM_027450, intron 1 of 4) | 8444 | NM_027450 | 384009 | Mm.22213 | NM_027450 | ENSMUSG00000028480 | **Glipr2** | 5730414A08Rik\|C77180\|GAPR-1 | GLI pathogenesis-related 2 | protein-coding |
| chr4_43941388_43941651 | chr4 | 43941389 | 43941651 | + | -0.7499 | intron (NM_016678, intron 19 of 20) | B3A\|SINE\|B2 | -16182 | NM_027450 | 384009 | Mm.22213 | NM_027450 | ENSMUSG00000028480 | **Glipr2** | 5730414A08Rik\|C77180\|GAPR-1 | GLI pathogenesis-related 2 | protein-coding |
| chr5_107612227_107612900 | chr5 | 107612228 | 107612900 | + | -0.622776 | intron (NM_144911, intron 6 of 12) | intron (NM_144911, intron 6 of 12) | -14676 | NM_001161738 | 170823 | Mm.41417 | NM_133248 | ENSMUSG00000029276 | **Glmn** | 9330160J16Rik\|AW227515\|Fap48\|Fap68 | glomulin, FKBP associated protein | protein-coding |
| chr1_88475406_88475861 | chr1 | 88475407 | 88475861 | + | -0.812894 | Intergenic | Intergenic | 34432 | NM_008132 | 14659 | Mm.5187 | NM_008132 | | **Glrp1** | GRP-1 | glutamine repeat protein 1 | protein-coding |
| chr1_88529863_88530143 | chr1 | 88529864 | 88530143 | + | -0.893263 | Intergenic | Intergenic | -19937 | NM_008132 | 14659 | Mm.5187 | NM_008132 | | **Glrp1** | GRP-1 | glutamine repeat protein 1 | protein-coding |
| chr1_88506380_88506866 | chr1 | 88506381 | 88506866 | + | -1.17319 | intron (NM_008132, intron 1 of 2) | MIRb\|SINE\|MIR | 3443 | NM_008132 | 14659 | Mm.5187 | NM_008132 | | **Glrp1** | GRP-1 | glutamine repeat protein 1 | protein-coding |
| chr1_88506412_88506773 | chr1 | 88506413 | 88506773 | + | -1.20418 | intron (NM_008132, intron 1 of 2) | intron (NM_008132, intron 1 of 2) | 3473 | NM_008132 | 14659 | Mm.5187 | NM_008132 | | **Glrp1** | GRP-1 | glutamine repeat protein 1 | protein-coding |
| chr5_114680625_114681379 | chr5 | 114680626 | 114681379 | + | -0.674883 | intron (NM_019821, intron 1 of 4) | intron (NM_019821, intron 1 of 4) | 9933 | NM_019821 | 56356 | Mm.275766 | NM_019821 | ENSMUSG00000011884 | **Gltp** | 1110001F24Rik\|C76925\|C77564 | glycolipid transfer protein | protein-coding |
| chr5_114680880_114681318 | chr5 | 114680881 | 114681318 | + | -0.789777 | intron (NM_019821, intron 1 of 4) | intron (NM_019821, intron 1 of 4) | 9836 | NM_019821 | 56356 | Mm.275766 | NM_019821 | ENSMUSG00000011884 | **Gltp** | 1110001F24Rik\|C76925\|C77564 | glycolipid transfer protein | protein-coding |
| chr7_16001670_16002698 | chr7 | 16001671 | 16002698 | + | -0.922118 | Intergenic | ID4\|SINE\|ID | -2689 | NM_001081418 | 243842 | Mm.440249 | NM_001081418 | ENSMUSG00000070808 | **Gltscr1** | - | glioma tumor suppressor candidate region gene 1 | protein-coding |
| chr7_16018467_16018789 | chr7 | 16018468 | 16018789 | + | -0.922495 | Intergenic | L1MB8\|LINE\|L1 | -19133 | NM_001081418 | 243842 | Mm.440249 | NM_001081418 | ENSMUSG00000070808 | **Gltscr1** | - | glioma tumor suppressor candidate region gene 1 | protein-coding |
| chr7_16001671_16002183 | chr7 | 16001672 | 16002183 | + | -1.01766 | Intergenic | B4\|SINE\|B4 | -2432 | NM_001081418 | 243842 | Mm.440249 | NM_001081418 | ENSMUSG00000070808 | **Gltscr1** | - | glioma tumor suppressor candidate region gene 1 | protein-coding |
| chr17_46835155_46835835 | chr17 | 46835156 | 46835835 | + | -0.695855 | Intergenic | Intergenic | -4082 | NM_001100452 | 210982 | Mm.37642 | NM_001100452 | ENSMUSG00000036568 | **Gltscr1l** | mKIAA0240 | GLTSCR1-like | protein-coding |
| chr17_46828896_46829283 | chr17 | 46828897 | 46829283 | + | -0.8387 | intron (NM_001100452, intron 3 of 10) | MIRb\|SINE\|MIR | 2323 | NM_001100452 | 210982 | Mm.37642 | NM_001100452 | ENSMUSG00000036568 | **Gltscr1l** | mKIAA0240 | GLTSCR1-like | protein-coding |

| | | | | | | | | | | | | | | | | | |
|---|---|---|---|---|---|---|---|---|---|---|---|---|---|---|---|---|---|
| chr18_5607903_5608160 | chr18 | 5607904 | 5608160 | + | -0.819005 | intron (NM_011546, intron 1 of 7) | intron (NM_011546, intron 1 of 7) | -15595 | NR_033552 | 791318 | Mm.84339 | NR_033552 | ENSMUSG00000063087 | **Gm10125** | ENSMUSG00000063087\|Zeb1os | predicted gene 10125 | ncRNA |
| chr18_5592464_5592990 | chr18 | 5592465 | 5592990 | + | -0.873554 | promoter-TSS (NR_033552) | promoter-TSS (NR_033552) | -290 | NR_033552 | 791318 | Mm.84339 | NR_033552 | ENSMUSG00000063087 | **Gm10125** | ENSMUSG00000063087\|Zeb1os | predicted gene 10125 | ncRNA |
| chr18_5593691_5594020 | chr18 | 5593692 | 5594020 | + | -1.12208 | intron (NM_011546, intron 1 of 7) | intron (NM_011546, intron 1 of 7) | -1419 | NR_033552 | 791318 | Mm.84339 | NR_033552 | ENSMUSG00000063087 | **Gm10125** | ENSMUSG00000063087\|Zeb1os | predicted gene 10125 | ncRNA |
| chr18_5593653_5594312 | chr18 | 5593654 | 5594312 | + | -1.24742 | intron (NM_011546, intron 1 of 7) | intron (NM_011546, intron 1 of 7) | -1546 | NR_033552 | 791318 | Mm.84339 | NR_033552 | ENSMUSG00000063087 | **Gm10125** | ENSMUSG00000063087\|Zeb1os | predicted gene 10125 | ncRNA |
| chr17_80391524_80392346 | chr17 | 80391525 | 80392346 | + | -0.826663 | Intergenic | Intergenic | -18393 | NR_028385 | 791338 | Mm.204722 | NR_028385 | | **Gm10190** | ENSMUSG00000066938 | predicted gene 10190 | ncRNA |
| chr18_75478409_75479398 | chr18 | 75478410 | 75479398 | + | -0.727184 | intron (NM_201354, intron 9 of 11) | intron (NM_201354, intron 9 of 11) | -35741 | NR_045879 | 100038353 | Mm.440724 | NR_045879 | ENSMUSG00000097712 | **Gm10532** | ENSMUSG00000073534 | predicted gene 10532 | ncRNA |
| chr18_75471806_75472273 | chr18 | 75471807 | 75472273 | + | -0.778909 | intron (NM_201354, intron 9 of 11) | intron (NM_201354, intron 9 of 11) | -42605 | NR_045879 | 100038353 | Mm.440724 | NR_045879 | ENSMUSG00000097712 | **Gm10532** | ENSMUSG00000073534 | predicted gene 10532 | ncRNA |
| chr18_75540735_75541597 | chr18 | 75540736 | 75541597 | + | -0.785864 | intron (NM_201354, intron 7 of 11) | intron (NM_201354, intron 7 of 11) | 26521 | NR_045879 | 100038353 | Mm.440724 | NR_045879 | ENSMUSG00000097712 | **Gm10532** | ENSMUSG00000073534 | predicted gene 10532 | ncRNA |
| chr18_75489331_75489696 | chr18 | 75489332 | 75489696 | + | -0.816082 | intron (NM_201354, intron 9 of 11) | intron (NM_201354, intron 9 of 11) | -25131 | NR_045879 | 100038353 | Mm.440724 | NR_045879 | ENSMUSG00000097712 | **Gm10532** | ENSMUSG00000073534 | predicted gene 10532 | ncRNA |
| chr18_75489116_75489696 | chr18 | 75489117 | 75489696 | + | -0.84261 | intron (NM_201354, intron 9 of 11) | intron (NM_201354, intron 9 of 11) | -25239 | NR_045879 | 100038353 | Mm.440724 | NR_045879 | ENSMUSG00000097712 | **Gm10532** | ENSMUSG00000073534 | predicted gene 10532 | ncRNA |
| chr18_75540881_75541397 | chr18 | 75540882 | 75541397 | + | -0.879343 | intron (NM_201354, intron 7 of 11) | (CCTG)n\|Simple_repeat\|Simple_repeat | 26494 | NR_045879 | 100038353 | Mm.440724 | NR_045879 | ENSMUSG00000097712 | **Gm10532** | ENSMUSG00000073534 | predicted gene 10532 | ncRNA |
| chr8_86699281_86699893 | chr8 | 86699282 | 86699893 | + | -0.724364 | intron (NM_001168591, intron 5 of 8) | Lx8\|LINE\|L1 | -46112 | NR_027829 | 666945 | Mm.391806 | NR_027829 | | **Gm10638** | ENSMUSG00000074178 | predicted gene 10638 | ncRNA |
| chr8_86699464_86699874 | chr8 | 86699465 | 86699874 | + | -0.755519 | intron (NM_001168591, intron 5 of 8) | Lx8\|LINE\|L1 | -46030 | NR_027829 | 666945 | Mm.391806 | NR_027829 | | **Gm10638** | ENSMUSG00000074178 | predicted gene 10638 | ncRNA |
| chr8_86699483_86699780 | chr8 | 86699484 | 86699780 | + | -0.785414 | intron (NM_001168591, intron 5 of 8) | Lx8\|LINE\|L1 | -46067 | NR_027829 | 666945 | Mm.391806 | NR_027829 | | **Gm10638** | ENSMUSG00000074178 | predicted gene 10638 | ncRNA |
| chr8_86735352_86735958 | chr8 | 86735353 | 86735958 | + | -0.83263 | intron (NM_009172, intron 1 of 1) | intron (NM_009172, intron 1 of 1) | -10044 | NR_027829 | 666945 | Mm.391806 | NR_027829 | | **Gm10638** | ENSMUSG00000074178 | predicted gene 10638 | ncRNA |
| chr8_86707689_86708348 | chr8 | 86707690 | 86708348 | + | -0.867719 | intron (NM_001168591, intron 5 of 8) | MTD\|LTR\|ERVL-MaLR | -37680 | NR_027829 | 666945 | Mm.391806 | NR_027829 | | **Gm10638** | ENSMUSG00000074178 | predicted gene 10638 | ncRNA |
| chr8_86719429_86720066 | chr8 | 86719430 | 86720066 | + | -0.936068 | Intergenic | ORR1F\|LTR\|ERVL-MaLR | -25951 | NR_027829 | 666945 | Mm.391806 | NR_027829 | | **Gm10638** | ENSMUSG00000074178 | predicted gene 10638 | ncRNA |
| chr8_86719405_86719864 | chr8 | 86719406 | 86719864 | + | -1.05922 | Intergenic | Intergenic | -26064 | NR_027829 | 666945 | Mm.391806 | NR_027829 | | **Gm10638** | ENSMUSG00000074178 | predicted gene 10638 | ncRNA |
| chr9_57056245_57056583 | chr9 | 57056246 | 57056583 | + | -0.760195 | TTS (NR_045886) | TTS (NR_045886) | 15552 | NR_045886 | 100038566 | Mm.361900 | NR_045886 | | **Gm10658** | ENSMUSG00000074284 | predicted gene 10658 | ncRNA |
| chr13_41932310_41932729 | chr13 | 41932311 | 41932729 | + | -0.719534 | Intergenic | Intergenic | 66605 | NR_033545 | 100038577 | Mm.387093 | NR_033545 | | **Gm10790** | ENSMUSG00000074927 | predicted gene 10790 | ncRNA |
| chr13_41921221_41921518 | chr13 | 41921222 | 41921518 | + | -0.743554 | Intergenic | Intergenic | 55455 | NR_033545 | 100038577 | Mm.387093 | NR_033545 | | **Gm10790** | ENSMUSG00000074927 | predicted gene 10790 | ncRNA |
| chr13_41948266_41948628 | chr13 | 41948267 | 41948628 | + | -0.904268 | Intergenic | Intergenic | 82532 | NR_033545 | 100038577 | Mm.387093 | NR_033545 | | **Gm10790** | ENSMUSG00000074927 | predicted gene 10790 | ncRNA |
| chr13_41929778_41930107 | chr13 | 41929779 | 41930107 | + | -0.984523 | Intergenic | L1MD\|LINE\|L1 | 64028 | NR_033545 | 100038577 | Mm.387093 | NR_033545 | | **Gm10790** | ENSMUSG00000074927 | predicted gene 10790 | ncRNA |
| chr16_84995443_84995996 | chr16 | 84995444 | 84995996 | + | -0.685599 | intron (NM_001198826, intron 13 of 16) | intron (NM_001198826, intron 13 of 16) | 23509 | NR_045889 | 100038614 | Mm.441351 | NR_045889 | | **Gm10791** | ENSMUSG00000074933 | predicted gene 10791 | ncRNA |
| chr16_84995438_84995805 | chr16 | 84995439 | 84995805 | + | -0.713879 | intron (NM_001198826, intron 13 of 16) | intron (NM_001198826, intron 13 of 16) | 23411 | NR_045889 | 100038614 | Mm.441351 | NR_045889 | | **Gm10791** | ENSMUSG00000074933 | predicted gene 10791 | ncRNA |
| chr16_84982633_84982952 | chr16 | 84982634 | 84982952 | + | -1.24369 | intron (NM_001198826, intron 13 of 16) | intron (NM_001198826, intron 13 of 16) | 10582 | NR_045889 | 100038614 | Mm.441351 | NR_045889 | | **Gm10791** | ENSMUSG00000074933 | predicted gene 10791 | ncRNA |
| chr19_6011659_6011940 | chr19 | 6011660 | 6011940 | + | -0.750597 | promoter-TSS (NR_045783) | promoter-TSS (NR_045783) | -820 | NR_045783 | 100502985 | Mm.465338 | NR_045783 | ENSMUSG00000097404 | **Gm10814** | ENSMUSG00000075230 | predicted gene 10814 | ncRNA |
| chr13_24603162_24603356 | chr13 | 24603163 | 24603356 | + | -0.694786 | intron (NR_024599, intron 1 of 1) | intron (NR_024599, intron 1 of 1) | 1508 | NR_024599 | 76024 | Mm.440710 | NR_024599 | | **Gm11346** | 5830431A10Rik\|AI447617\|Gm11347\|OTTMUSG00000000601\|OTTMUSG00000000602 | X-linked lymphocyte-regulated 5 pseudogene | pseudo |
| chr13_24603102_24603339 | chr13 | 24603103 | 24603339 | + | -0.70729 | intron (NR_024599, intron 1 of 1) | intron (NR_024599, intron 1 of 1) | 1546 | NR_024599 | 76024 | Mm.440710 | NR_024599 | | **Gm11346** | 5830431A10Rik\|AI447617\|Gm11347\|OTTMUSG00000000601\|OTTMUSG00000000602 | X-linked lymphocyte-regulated 5 pseudogene | pseudo |
| chr13_24594356_24594696 | chr13 | 24594357 | 24594696 | + | -0.798741 | intron (NM_001286100, intron 1 of 22) | intron (NM_001286100, intron 1 of 22) | 10241 | NR_024599 | 76024 | Mm.440710 | NR_024599 | | **Gm11346** | 5830431A10Rik\|AI447617\|Gm11347\|OTTMUSG00000000601\|OTTMUSG00000000602 | X-linked lymphocyte-regulated 5 pseudogene | pseudo |
| chr13_24599398_24600094 | chr13 | 24599399 | 24600094 | + | -0.813559 | intron (NR_024599, intron 1 of 1) | intron (NR_024599, intron 1 of 1) | 5021 | NR_024599 | 76024 | Mm.440710 | NR_024599 | | **Gm11346** | 5830431A10Rik\|AI447617\|Gm11347\|OTTMUSG00000000601\|OTTMUSG00000000602 | X-linked lymphocyte-regulated 5 pseudogene | pseudo |
| chr11_96441332_96442124 | chr11 | 96441333 | 96442124 | + | -1.17081 | Intergenic | Intergenic | 22819 | NR_033524 | 100038600 | Mm.483549 | NR_033524 | | **Gm11529** | OTTMUSG00000001775 | predicted gene 11529 | ncRNA |
| chr11_94697208_94698546 | chr11 | 94697209 | 94698546 | + | -0.973593 | intron (NM_001007584, intron 1 of 1) | intron (NM_001007584, intron 1 of 1) | 6622 | NM_001007584 | 432589 | Mm.342314 | NM_001007584 | ENSMUSG00000056008 | **Gm11541** | OTTMUSG00000002038 | predicted gene 11541 | protein-coding |
| chr11_109375361_109375937 | chr11 | 109375362 | 109375937 | + | -0.800463 | intron (NM_010303, intron 2 of 3) | intron (NM_010303, intron 2 of 3) | -11995 | NR_038097 | 100036768 | Mm.467767 | NR_038097 | ENSMUSG00000056687 | **Gm11696** | OTTMUSG00000003456 | predicted gene 11696 | ncRNA |
| chr11_3309859_3310291 | chr11 | 3309860 | 3310291 | + | -0.815544 | TTS (NM_001253691) | TTS (NM_001253691) | 1075 | NR_045708 | 100302648 | Mm.458664 | NR_045708 | ENSMUSG00000084941 | **Gm11944** | OTTMUSG00000005017 | predicted gene 11944 | ncRNA |
| chr11_6679691_6680777 | chr11 | 6679692 | 6680777 | + | -1.0777 | Intergenic | RSINE1\|SINE\|B4 | -19998 | NR_028586 | 100041311 | Mm.388018 | NR_028586 | ENSMUSG00000086373 | **Gm11978** | OTTMUSG00000005125\|OTTMUSG00000005127 | predicted gene 11978 | pseudo |
| chr11_6704820_6705245 | chr11 | 6704821 | 6705245 | + | -1.38243 | Intergenic | Intergenic | 15004 | NR_046025 | 626419 | Mm.390334 | NR_046025 | ENSMUSG00000086808 | **Gm11981** | OTTMUSG00000005129 | predicted gene 11981 | ncRNA |
| chr11_48933409_48933933 | chr11 | 48933410 | 48933933 | + | -0.672102 | Intergenic | Intergenic | -6489 | NM_001045540 | 620913 | Mm.452174 | NM_001045540 | ENSMUSG00000048852 | **Gm12185** | OTTMUSG00000005523 | predicted gene 12185 | protein-coding |
| chr11_53841971_53842452 | chr11 | 53841972 | 53842452 | + | -0.628994 | intron (NR_033332, intron 2 of 3) | MIRb\|SINE\|MIR | 17044 | NR_033332 | 622459 | Mm.479656 | NR_033332 | ENSMUSG00000081769 | **Gm12216** | 602-19-115\|OTTMUSG00000005650 | predicted gene 12216 | ncRNA |
| chr4_55397756_55398727 | chr4 | 55397757 | 55398727 | + | -0.940607 | Intergenic | Intergenic | 20600 | NR_040674 | 100415914 | Mm.261765 | NR_040674 | ENSMUSG00000087070 | **Gm12505** | OTTMUSG00000007226 | predicted gene 12505 | ncRNA |
| chr4_140952012_140952847 | chr4 | 140952013 | 140952847 | + | -0.773098 | 3' UTR (NM_008812, exon 16 of 16) | 3' UTR (NM_008812, exon 16 of 16) | 5472 | NR_045911 | 100126227 | Mm.404073 | NR_045911 | ENSMUSG00000087698 | **Gm13031** | OTTMUSG00000010081 | predicted gene 13031 | ncRNA |
| chr4_140952324_140952878 | chr4 | 140952325 | 140952878 | + | -0.840757 | TTS (NM_008812) | TTS (NM_008812) | 5301 | NR_045911 | 100126227 | Mm.404073 | NR_045911 | ENSMUSG00000087698 | **Gm13031** | OTTMUSG00000010081 | predicted gene 13031 | ncRNA |
| chr4_146457043_146458099 | chr4 | 146457044 | 146458099 | + | -0.71008 | intron (NM_001083918, intron 4 of 6).2 | intron (NM_001083918, intron 4 of 6).2 | 8541 | NM_001085522 | 433791 | Mm.276881 | NM_001085522 | ENSMUSG00000070605 | **Gm13251** | OTTMUSG00000011097 | predicted gene 13251 | protein-coding |
| chr2_58178858_58179616 | chr2 | 58178859 | 58179616 | + | -0.65826 | Intergenic | Intergenic | 15263 | NR_045895 | 100042926 | Mm.446515 | NR_045895 | ENSMUSG00000086862 | **Gm13546** | OTTMUSG00000012675 | predicted gene 13546 | ncRNA |
| chr2_58179038_58179585 | chr2 | 58179039 | 58179585 | + | -0.682265 | Intergenic | Intergenic | 15338 | NR_045895 | 100042926 | Mm.446515 | NR_045895 | ENSMUSG00000086862 | **Gm13546** | OTTMUSG00000012675 | predicted gene 13546 | ncRNA |
| chr2_29724902_29725381 | chr2 | 29724903 | 29725381 | + | -1.04061 | intron (NM_001039086, intron 15 of 25) | intron (NM_001039086, intron 15 of 25) | -36386 | NM_001177392 | 433416 | Mm.330112 | NM_001034887 | ENSMUSG00000075425 | **Gm13547** | OTTMUSG00000012686 | predicted gene 13547 | protein-coding |
| chr2_29725010_29725593 | chr2 | 29725011 | 29725593 | + | -1.09634 | intron (NM_001039086, intron 15 of 25) | intron (NM_001039086, intron 15 of 25) | -36226 | NM_001177392 | 433416 | Mm.330112 | NM_001034887 | ENSMUSG00000075425 | **Gm13547** | OTTMUSG00000012686 | predicted gene 13547 | protein-coding |
| chr2_84504824_84505439 | chr2 | 84504825 | 84505439 | + | -0.829904 | intron (NR_046046, intron 1 of 2) | Lx\|LINE\|L1 | 1741 | NR_046046 | 672763 | Mm.491046 | NR_046046 | ENSMUSG00000087362 | **Gm13710** | OTTMUSG00000013441 | predicted gene 13710 | ncRNA |
| chr2_84498817_84499330 | chr2 | 84498818 | 84499330 | + | -1.09861 | Intergenic | ORR1A4-int\|LTR\|ERVL-MaLR | 7799 | NR_046046 | 672763 | Mm.491046 | NR_046046 | ENSMUSG00000087362 | **Gm13710** | OTTMUSG00000013441 | predicted gene 13710 | ncRNA |
| chr5_115083463_115083849 | chr5 | 115083464 | 115083849 | + | -0.969991 | intron (NM_029012, intron 5 of 10) | intron (NM_029012, intron 5 of 10) | 24306 | NM_001271591 | 100502825 | Mm.361657 | NM_001271590 | | **Gm13826** | OTTMUSG00000014624 | predicted gene 13826 | protein-coding |
| chr2_128318970_128319516 | chr2 | 128318971 | 128319516 | + | -0.622801 | intron (NR_028590, intron 2 of 2) | intron (NR_028590, intron 2 of 2) | 110108 | NR_028591 | 100043424 | Mm.297222 | NR_028589 | | **Gm14005** | OTTMUSG00000015351 | predicted gene 14005 | ncRNA |

| | | | | | | | | | | | | | | | | | |
|---|---|---|---|---|---|---|---|---|---|---|---|---|---|---|---|---|---|
| chr2_128470343_128470585 | chr2 | 128470344 | 128470585 | + | -0.665889 | Intergenic | Intergenic | -41113 | NR_028590 | 100043424 | Mm.297222 | NR_028589 | | **Gm14005** | OTTMUSG00000015351 | predicted gene 14005 | ncRNA |
| chr2_128470139_128470978 | chr2 | 128470140 | 128470978 | + | -0.669315 | Intergenic | Intergenic | -41208 | NR_028590 | 100043424 | Mm.297222 | NR_028589 | | **Gm14005** | OTTMUSG00000015351 | predicted gene 14005 | ncRNA |
| chr2_122485705_122485955 | chr2 | 122485706 | 122485955 | + | -0.692595 | 5' UTR (NM_001085518, exon 1 of 18) | 5' UTR (NM_001085518, exon 1 of 18) | 889 | NM_001085518 | 381417 | Mm.482295 | NM_001085518 | ENSMUSG00000079071 | **Gm14085** | OTTMUSG00000015615 | predicted gene 14085 | protein-coding |
| chr19_34599318_34599592 | chr19 | 34599319 | 34599592 | + | -0.669195 | promoter-TSS (NM_001110517) | promoter-TSS (NM_001110517) | -28 | NM_001110517 | 667373 | Mm.371956 | NM_001101605 | ENSMUSG00000079339 | **Gm14446** | OTTMUSG00000016644 | predicted gene 14446 | protein-coding |
| chr19_34601949_34602401 | chr19 | 34601950 | 34602401 | + | -0.742835 | promoter-TSS (NM_001101605) | promoter-TSS (NM_001101605) | -207 | NM_001101605 | 667373 | Mm.371956 | NM_001101605 | ENSMUSG00000079339 | **Gm14446** | OTTMUSG00000016644 | predicted gene 14446 | protein-coding |
| chrX_57189288_57189779 | chrX | 57189289 | 57189779 | + | -1.15418 | Intergenic | Intergenic | -11103 | NR_038463 | 100502842 | Mm.164733 | NR_038463 | ENSMUSG00000085808 | **Gm14718** | OTTMUSG00000017790 | predicted gene 14718 | ncRNA |
| chr7_104290512_104291010 | chr7 | 104290513 | 104291010 | + | -0.661803 | Intergenic | Intergenic | -24270 | NR_040749 | 622070 | Mm.486406 | NR_040749 | ENSMUSG00000087175 | **Gm15133** | OTTMUSG00000019186 | predicted gene 15133 | ncRNA |
| chr7_104291640_104292011 | chr7 | 104291641 | 104292011 | + | -0.753327 | Intergenic | RMER19B2\|LTR\|ERVK | -23205 | NR_040749 | 622070 | Mm.486406 | NR_040749 | ENSMUSG00000087175 | **Gm15133** | OTTMUSG00000019186 | predicted gene 15133 | ncRNA |
| chr7_104292064_104292421 | chr7 | 104292065 | 104292421 | + | -0.931519 | Intergenic | RMER19B2\|LTR\|ERVK | -22788 | NR_040749 | 622070 | Mm.486406 | NR_040749 | ENSMUSG00000087175 | **Gm15133** | OTTMUSG00000019186 | predicted gene 15133 | ncRNA |
| chr5_149017708_149017964 | chr5 | 149017709 | 149017964 | + | -1.06456 | Intergenic | RLTR33\|LTR\|ERVK | 10874 | NR_040429 | 100503307 | Mm.188008 | NR_040429 | ENSMUSG00000086610 | **Gm15408** | OTTMUSG00000021225 | predicted gene 15408 | ncRNA |
| chr3_96564662_96564994 | chr3 | 96564663 | 96564994 | + | -0.754794 | intron (NR_040409, intron 1 of 3) | intron (NR_040409, intron 1 of 3) | 1973 | NR_040409 | 100038464 | Mm.475126 | NR_040409 | ENSMUSG00000074398 | **Gm15441** | ENSMUSG00000074398\|OTTMUSG00000021867 | predicted gene 15441 | ncRNA |
| chr3_96570571_96571087 | chr3 | 96570572 | 96571087 | + | -0.886082 | Intergenic | Intergenic | -4028 | NR_040409 | 100038464 | Mm.475126 | NR_040409 | ENSMUSG00000074398 | **Gm15441** | ENSMUSG00000074398\|OTTMUSG00000021867 | predicted gene 15441 | ncRNA |
| chr6_145284215_145284822 | chr6 | 145284216 | 145284822 | + | -0.693057 | Intergenic | Intergenic | 33967 | NR_045598 | 100039495 | Mm.121014 | NR_045598 | ENSMUSG00000086013 | **Gm15706** | OTTMUSG00000025463 | predicted gene 15706 | ncRNA |
| chr6_145284353_145284589 | chr6 | 145284354 | 145284589 | + | -0.712596 | Intergenic | Intergenic | 33919 | NR_045598 | 100039495 | Mm.121014 | NR_045598 | ENSMUSG00000086013 | **Gm15706** | OTTMUSG00000025463 | predicted gene 15706 | ncRNA |
| chr6_145303403_145304102 | chr6 | 145303404 | 145304102 | + | -0.768833 | Intergenic | Intergenic | 53201 | NR_045598 | 100039495 | Mm.121014 | NR_045598 | ENSMUSG00000086013 | **Gm15706** | OTTMUSG00000025463 | predicted gene 15706 | ncRNA |
| chr6_145284164_145284588 | chr6 | 145284165 | 145284588 | + | -0.784456 | Intergenic | B3A\|SINE\|B2 | 33824 | NR_045598 | 100039495 | Mm.121014 | NR_045598 | ENSMUSG00000086013 | **Gm15706** | OTTMUSG00000025463 | predicted gene 15706 | ncRNA |
| chr6_145285525_145286029 | chr6 | 145285526 | 145286029 | + | -0.890104 | Intergenic | RMER6A\|LTR\|ERVK | 35225 | NR_045598 | 100039495 | Mm.121014 | NR_045598 | ENSMUSG00000086013 | **Gm15706** | OTTMUSG00000025463 | predicted gene 15706 | ncRNA |
| chr6_145306665_145307079 | chr6 | 145306666 | 145307079 | + | -0.961291 | Intergenic | Intergenic | 56320 | NR_045598 | 100039495 | Mm.121014 | NR_045598 | ENSMUSG00000086013 | **Gm15706** | OTTMUSG00000025463 | predicted gene 15706 | ncRNA |
| chr16_43376662_43377474 | chr16 | 43376663 | 43377474 | + | -0.835014 | intron (NM_181058, intron 2 of 6) | intron (NM_181058, intron 2 of 6) | 43128 | NR_046026 | 626693 | Mm.447083 | NR_046026 | ENSMUSG00000085499 | **Gm15713** | OTTMUSG00000025617 | predicted gene 15713 | ncRNA |
| chr1_136127962_136128153 | chr1 | 136127963 | 136128153 | + | -0.751137 | intron (NR_046167, intron 3 of 3) | intron (NR_046167, intron 3 of 3) | 3125 | NR_046168 | 329253 | Mm.487858 | NR_046167 | ENSMUSG00000086264 | **Gm15850** | 6530439I21\|OTTMUSG00000026868 | predicted gene 15850 | ncRNA |
| chr14_77835813_77836272 | chr14 | 77835814 | 77836272 | + | -0.704192 | intron (NM_025384, intron 5 of 5) | intron (NM_025384, intron 5 of 5) | -37075 | NM_001033440 | 380920 | Mm.297771 | NM_001033440 | | **Gm1587** | - | predicted gene 1587 | protein-coding |
| chr14_77772085_77772342 | chr14 | 77772086 | 77772342 | + | -0.711804 | Intergenic | ORR1F\|LTR\|ERVL-MaLR | 26754 | NM_001033440 | 380920 | Mm.297771 | NM_001033440 | | **Gm1587** | - | predicted gene 1587 | protein-coding |
| chr7_80634818_80635069 | chr7 | 80634819 | 80635069 | + | -0.662618 | intron (NM_173863, intron 2 of 14) | intron (NM_173863, intron 2 of 14) | -1073 | NR_040343 | 100504375 | Mm.322881 | NR_040343 | ENSMUSG00000084821 | **Gm15880** | OTTMUSG00000027171 | predicted gene 15880 | ncRNA |
| chr7_80631480_80631754 | chr7 | 80631481 | 80631754 | + | -0.713304 | intron (NM_173863, intron 2 of 14) | intron (NM_173863, intron 2 of 14) | -4400 | NR_040343 | 100504375 | Mm.322881 | NR_040343 | ENSMUSG00000084821 | **Gm15880** | OTTMUSG00000027171 | predicted gene 15880 | ncRNA |
| chr5_119633607_119634194 | chr5 | 119633608 | 119634194 | + | -0.945889 | non-coding (NR_046178, exon 4 of 4) | non-coding (NR_046178, exon 4 of 4) | 7959 | NR_046178 | 330183 | Mm.413133 | NR_046178 | ENSMUSG00000085708 | **Gm16063** | G630008E18\|OTTMUSG00000029664 | predicted gene 16063 | ncRNA |
| chr5_119633629_119634018 | chr5 | 119633630 | 119634018 | + | -0.965461 | non-coding (NR_046178, exon 4 of 4) | non-coding (NR_046178, exon 4 of 4) | 7882 | NR_046178 | 330183 | Mm.413133 | NR_046178 | ENSMUSG00000085708 | **Gm16063** | G630008E18\|OTTMUSG00000029664 | predicted gene 16063 | ncRNA |
| chr7_68285873_68286298 | chr7 | 68285874 | 68286298 | + | -0.678176 | intron (NM_001013811, intron 1 of 8) | intron (NM_001013811, intron 1 of 8) | -9492 | NR_040340 | 100504026 | Mm.426764 | NR_040340 | ENSMUSG00000086628 | **Gm16157** | OTTMUSG00000031023 | predicted gene 16157 | ncRNA |
| chr7_68323048_68323539 | chr7 | 68323049 | 68323539 | + | -0.70612 | intron (NM_001013811, intron 4 of 8) | intron (NM_001013811, intron 4 of 8) | -46700 | NR_040340 | 100504026 | Mm.426764 | NR_040340 | ENSMUSG00000086628 | **Gm16157** | OTTMUSG00000031023 | predicted gene 16157 | ncRNA |
| chr7_68322821_68323544 | chr7 | 68322822 | 68323544 | + | -0.717006 | intron (NM_001013811, intron 4 of 8) | intron (NM_001013811, intron 4 of 8) | -46589 | NR_040340 | 100504026 | Mm.426764 | NR_040340 | ENSMUSG00000086628 | **Gm16157** | OTTMUSG00000031023 | predicted gene 16157 | ncRNA |
| chr7_68286706_68287297 | chr7 | 68286707 | 68287297 | + | -0.745591 | intron (NM_001013811, intron 1 of 8) | intron (NM_001013811, intron 1 of 8) | -10408 | NR_040340 | 100504026 | Mm.426764 | NR_040340 | ENSMUSG00000086628 | **Gm16157** | OTTMUSG00000031023 | predicted gene 16157 | ncRNA |
| chr7_68320176_68320510 | chr7 | 68320177 | 68320510 | + | -0.751144 | intron (NM_001013811, intron 3 of 8) | intron (NM_001013811, intron 3 of 8) | -43749 | NR_040340 | 100504026 | Mm.426764 | NR_040340 | ENSMUSG00000086628 | **Gm16157** | OTTMUSG00000031023 | predicted gene 16157 | ncRNA |
| chr7_68298308_68298794 | chr7 | 68298309 | 68298794 | + | -0.780234 | intron (NM_001013811, intron 1 of 8) | intron (NM_001013811, intron 1 of 8) | -21957 | NR_040340 | 100504026 | Mm.426764 | NR_040340 | ENSMUSG00000086628 | **Gm16157** | OTTMUSG00000031023 | predicted gene 16157 | ncRNA |
| chr7_68324722_68324965 | chr7 | 68324723 | 68324965 | + | -0.791681 | intron (NM_001013811, intron 4 of 8) | ORR1E\|LTR\|ERVL-MaLR | -48250 | NR_040340 | 100504026 | Mm.426764 | NR_040340 | ENSMUSG00000086628 | **Gm16157** | OTTMUSG00000031023 | predicted gene 16157 | ncRNA |
| chr2_71667766_71668301 | chr2 | 71667767 | 71668301 | + | -0.702708 | Intergenic | Intergenic | -51383 | NR_037979 | 381371 | Mm.330284 | NM_201366 | ENSMUSG00000064263 | **Gm1631** | - | predicted gene 1631 | ncRNA |
| chr3_69156330_69157058 | chr3 | 69156331 | 69157058 | + | -0.974935 | intron (NM_001243000, intron 2 of 2) | intron (NM_001243000, intron 2 of 2) | 25461 | NM_001243000 | 381452 | Mm.331067 | NM_001243000 | ENSMUSG00000094763 | **Gm1647** | - | predicted gene 1647 | protein-coding |
| chr3_69156509_69157022 | chr3 | 69156510 | 69157022 | + | -1.0229 | intron (NM_001243000, intron 2 of 2) | intron (NM_001243000, intron 2 of 2) | 25533 | NM_001243000 | 381452 | Mm.331067 | NM_001243000 | ENSMUSG00000094763 | **Gm1647** | - | predicted gene 1647 | protein-coding |
| chr15_79749154_79749550 | chr15 | 79749155 | 79749550 | + | -0.635043 | intron (NR_045069, intron 3 of 3) | intron (NR_045069, intron 3 of 3) | 6654 | NR_045069 | 100504191 | Mm.421420 | NR_045069 | ENSMUSG00000087543 | **Gm16576** | - | predicted gene 16576 | ncRNA |
| chr4_117220489_117221080 | chr4 | 117220490 | 117221080 | + | -0.84293 | intron (NM_001033774, intron 7 of 7) | ORR1B1\|LTR\|ERVL-MaLR | 31132 | NM_001145637 | 381544 | Mm.23504 | NM_001033774 | ENSMUSG00000060268 | **Gm1661** | Gm1662\|Ncrna00082 | predicted gene 1661 | protein-coding |
| chr17_8388985_8390193 | chr17 | 8388986 | 8390193 | + | -0.635223 | promoter-TSS (NR_045795) | promoter-TSS (NR_045795) | -121 | NR_045795 | 100504601 | Mm.365131 | NR_045795 | ENSMUSG00000097796 | **Gm16702** | AU016609 | predicted gene, 16702 | ncRNA |
| chr17_8389344_8390113 | chr17 | 8389345 | 8390113 | + | -0.674 | promoter-TSS (NR_045795) | promoter-TSS (NR_045795) | -261 | NR_045795 | 100504601 | Mm.365131 | NR_045795 | ENSMUSG00000097796 | **Gm16702** | AU016609 | predicted gene, 16702 | ncRNA |
| chr17_8389045_8389945 | chr17 | 8389046 | 8389945 | + | -0.680872 | promoter-TSS (NR_045795) | promoter-TSS (NR_045795) | -27 | NR_045795 | 100504601 | Mm.365131 | NR_045795 | ENSMUSG00000097796 | **Gm16702** | AU016609 | predicted gene, 16702 | ncRNA |
| chr17_8389530_8389897 | chr17 | 8389531 | 8389897 | + | -0.781182 | promoter-TSS (NR_045795) | promoter-TSS (NR_045795) | -246 | NR_045795 | 100504601 | Mm.365131 | NR_045795 | ENSMUSG00000097796 | **Gm16702** | AU016609 | predicted gene, 16702 | ncRNA |
| chr8_35677733_35678186 | chr8 | 35677734 | 35678186 | + | -0.844481 | 3' UTR (NM_001081279, exon 3 of 3) | 3' UTR (NM_001081279, exon 3 of 3) | -88940 | NR_040376 | 100504714 | Mm.381744 | NR_040376 | ENSMUSG00000097357 | **Gm16793** | - | predicted gene, 16793 | ncRNA |
| chr8_35692609_35693315 | chr8 | 35692610 | 35693315 | + | -0.918906 | Intergenic | Intergenic | -103942 | NR_040376 | 100504714 | Mm.381744 | NR_040376 | ENSMUSG00000097357 | **Gm16793** | - | predicted gene, 16793 | ncRNA |
| chr8_35677900_35678237 | chr8 | 35677901 | 35678237 | + | -0.956133 | 3' UTR (NM_001081279, exon 3 of 3) | 3' UTR (NM_001081279, exon 3 of 3) | -89049 | NR_040376 | 100504714 | Mm.381744 | NR_040376 | ENSMUSG00000097357 | **Gm16793** | - | predicted gene, 16793 | ncRNA |
| chr8_35692961_35693296 | chr8 | 35692962 | 35693296 | + | -0.96525 | Intergenic | MLT1B\|LTR\|ERVL-MaLR | -104109 | NR_040376 | 100504714 | Mm.381744 | NR_040376 | ENSMUSG00000097357 | **Gm16793** | - | predicted gene, 16793 | ncRNA |
| chr8_35678774_35679367 | chr8 | 35678775 | 35679367 | + | -1.16035 | 3' UTR (NM_001081279, exon 3 of 3) | 3' UTR (NM_001081279, exon 3 of 3) | -90051 | NR_040376 | 100504714 | Mm.381744 | NR_040376 | ENSMUSG00000097357 | **Gm16793** | - | predicted gene, 16793 | ncRNA |
| chr8_35680627_35681055 | chr8 | 35680628 | 35681055 | + | -1.19983 | Intergenic | Intergenic | -91821 | NR_040376 | 100504714 | Mm.381744 | NR_040376 | ENSMUSG00000097357 | **Gm16793** | - | predicted gene, 16793 | ncRNA |
| chr13_63284814_63285672 | chr13 | 63284815 | 63285672 | + | -0.782603 | intron (NR_110520, intron 3 of 5) | intron (NR_110520, intron 3 of 5) | 11904 | NR_045794 | 100504114 | Mm.437070 | NR_045794 | ENSMUSG00000097242 | **Gm16907** | - | predicted gene, 16907 | ncRNA |
| chr10_60380935_60382085 | chr10 | 60380936 | 60382085 | + | -0.718024 | intron (NM_023370, intron 33 of 68) | intron (NM_023370, intron 33 of 68) | -18216 | NM_001164374 | 100302688 | Mm.444135 | NM_001164374 | ENSMUSG00000090439 | **Gm17455** | - | predicted gene, 17455 | protein-coding |

| | | | | | | | | | | | | | | | | | |
|---|---|---|---|---|---|---|---|---|---|---|---|---|---|---|---|---|---|
| chr12_25187299_25187485 | chr12 | 25187300 | 25187485 | + | -0.664376 | Intergenic | ORR1E\|LTR\|ERVL-MaLR | 58218 | NR_045844 | 432641 | Mm.112955 | NR_045844 | | **Gm17746** | - | predicted gene, 17746 | ncRNA |
| chr12_25316003_25316309 | chr12 | 25316004 | 25316309 | + | -0.847329 | Intergenic | ORR1E\|LTR\|ERVL-MaLR | 186982 | NR_045844 | 432641 | Mm.112955 | NR_045844 | | **Gm17746** | - | predicted gene, 17746 | ncRNA |
| chr19_41768715_41769290 | chr19 | 41768716 | 41769290 | + | -0.728491 | 3' UTR (NM_027667, exon 12 of 12) | 3' UTR (NM_027667, exon 12 of 12) | 22042 | NR_040320 | 100502871 | Mm.394485 | NR_040320 | | **Gm19424** | - | predicted gene, 19424 | ncRNA |
| chr19_41768739_41769066 | chr19 | 41768740 | 41769066 | + | -0.795267 | 3' UTR (NM_027667, exon 12 of 12) | 3' UTR (NM_027667, exon 12 of 12) | 21942 | NR_040320 | 100502871 | Mm.394485 | NR_040320 | | **Gm19424** | - | predicted gene, 19424 | ncRNA |
| chr19_47510796_47511130 | chr19 | 47510797 | 47511130 | + | -0.756319 | intron (NM_175360, intron 7 of 9) | intron (NM_175360, intron 7 of 9) | -2019 | NR_045322 | 100503104 | Mm.442735 | NR_045322 | ENSMUSG00000097990 | **Gm19557** | - | predicted gene, 19557 | ncRNA |
| chr19_47512763_47513353 | chr19 | 47512764 | 47513353 | + | -1.01705 | promoter-TSS (NR_045322) | promoter-TSS (NR_045322) | 76 | NR_045322 | 100503104 | Mm.442735 | NR_045322 | ENSMUSG00000097990 | **Gm19557** | - | predicted gene, 19557 | ncRNA |
| chr19_47511353_47511670 | chr19 | 47511354 | 47511670 | + | -1.04248 | intron (NM_175360, intron 7 of 9) | intron (NM_175360, intron 7 of 9) | -1470 | NR_045322 | 100503104 | Mm.442735 | NR_045322 | ENSMUSG00000097990 | **Gm19557** | - | predicted gene, 19557 | ncRNA |
| chr1_88610559_88611615 | chr1 | 88610560 | 88611615 | + | -0.631992 | Intergenic | Intergenic | 59570 | NR_037971 | 100503197 | Mm.99352 | NR_037971 | | **Gm19589** | - | predicted gene, 19589 | ncRNA |
| chr1_88555558_88555890 | chr1 | 88555559 | 88555890 | + | -0.647466 | intron (NR_037971, intron 2 of 5) | MTD\|LTR\|ERVL-MaLR | 4207 | NR_037971 | 100503197 | Mm.99352 | NR_037971 | | **Gm19589** | - | predicted gene, 19589 | ncRNA |
| chr1_88622201_88622842 | chr1 | 88622202 | 88622842 | + | -0.721871 | Intergenic | Intergenic | 71005 | NR_037971 | 100503197 | Mm.99352 | NR_037971 | | **Gm19589** | - | predicted gene, 19589 | ncRNA |
| chr1_88622456_88622790 | chr1 | 88622457 | 88622790 | + | -0.781963 | Intergenic | Intergenic | 71106 | NR_037971 | 100503197 | Mm.99352 | NR_037971 | | **Gm19589** | - | predicted gene, 19589 | ncRNA |
| chr1_136672428_136673016 | chr1 | 136672429 | 136673016 | + | -0.803047 | Intergenic | Intergenic | -10674 | NR_045324 | 100503460 | Mm.30630 | NR_045324 | ENSMUSG00000097113 | **Gm19705** | - | predicted gene, 19705 | ncRNA |
| chr7_28781259_28781690 | chr7 | 28781260 | 28781690 | + | -0.716467 | promoter-TSS (NR_040339) | promoter-TSS (NR_040339) | 39 | NR_040339 | 100503790 | Mm.446894 | NR_040339 | | **Gm19897** | - | predicted gene, 19897 | ncRNA |
| chr7_28781244_28781871 | chr7 | 28781245 | 28781871 | + | -0.718266 | promoter-TSS (NR_040339) | promoter-TSS (NR_040339) | -44 | NR_040339 | 100503790 | Mm.446894 | NR_040339 | | **Gm19897** | - | predicted gene, 19897 | ncRNA |
| chr17_51986256_51986611 | chr17 | 51986257 | 51986611 | + | -0.62128 | Intergenic | Intergenic | 103707 | NR_045095 | 100504169 | Mm.417731 | NR_045095 | | **Gm20098** | - | predicted gene, 20098 | ncRNA |
| chr17_52062112_52062921 | chr17 | 52062113 | 52062921 | + | -0.686794 | Intergenic | Intergenic | 179790 | NR_045095 | 100504169 | Mm.417731 | NR_045095 | | **Gm20098** | - | predicted gene, 20098 | ncRNA |
| chr17_51876272_51876878 | chr17 | 51876273 | 51876878 | + | -0.72479 | Intergenic | Intergenic | -6152 | NR_045095 | 100504169 | Mm.417731 | NR_045095 | | **Gm20098** | - | predicted gene, 20098 | ncRNA |
| chr17_51903825_51904308 | chr17 | 51903826 | 51904308 | + | -0.825803 | intron (NR_045095, intron 1 of 2) | intron (NR_045095, intron 1 of 2) | 21340 | NR_045095 | 100504169 | Mm.417731 | NR_045095 | | **Gm20098** | - | predicted gene, 20098 | ncRNA |
| chr17_51905788_51906733 | chr17 | 51905789 | 51906733 | + | -0.853533 | intron (NR_045095, intron 2 of 2) | intron (NR_045095, intron 2 of 2) | 23534 | NR_045095 | 100504169 | Mm.417731 | NR_045095 | | **Gm20098** | - | predicted gene, 20098 | ncRNA |
| chr17_52168131_52169101 | chr17 | 52168132 | 52169101 | + | -0.864191 | Intergenic | Intergenic | 285889 | NR_045095 | 100504169 | Mm.417731 | NR_045095 | | **Gm20098** | - | predicted gene, 20098 | ncRNA |
| chr17_51906116_51906956 | chr17 | 51906117 | 51906956 | + | -1.02895 | intron (NR_045095, intron 2 of 2) | intron (NR_045095, intron 2 of 2) | 23809 | NR_045095 | 100504169 | Mm.417731 | NR_045095 | | **Gm20098** | - | predicted gene, 20098 | ncRNA |
| chr17_51880218_51880597 | chr17 | 51880219 | 51880597 | + | -1.03569 | Intergenic | Intergenic | -2319 | NR_045095 | 100504169 | Mm.417731 | NR_045095 | | **Gm20098** | - | predicted gene, 20098 | ncRNA |
| chr17_52177915_52178461 | chr17 | 52177916 | 52178461 | + | -1.79319 | Intergenic | Intergenic | 295461 | NR_045095 | 100504169 | Mm.417731 | NR_045095 | | **Gm20098** | - | predicted gene, 20098 | ncRNA |
| chr10_99599412_99599799 | chr10 | 99599413 | 99599799 | + | -0.648095 | Intergenic | IAPLTR2_Mm\|LTR\|ERVK | -9565 | NR_038019 | 100504202 | Mm.403801 | NR_038019 | | **Gm20110** | - | predicted gene, 20110 | ncRNA |
| chr10_99441066_99441443 | chr10 | 99441067 | 99441443 | + | -1.01465 | Intergenic | Intergenic | -167916 | NR_038019 | 100504202 | Mm.403801 | NR_038019 | | **Gm20110** | - | predicted gene, 20110 | ncRNA |
| chr10_99440965_99441472 | chr10 | 99440966 | 99441472 | + | -1.05484 | Intergenic | Intergenic | -167952 | NR_038019 | 100504202 | Mm.403801 | NR_038019 | | **Gm20110** | - | predicted gene, 20110 | ncRNA |
| chr10_19138297_19138507 | chr10 | 19138298 | 19138507 | + | -0.714011 | Intergenic | Intergenic | -3347 | NR_038021 | 100504265 | Mm.383417 | NR_038021 | | **Gm20139** | - | predicted gene, 20139 | ncRNA |
| chr10_19122123_19122601 | chr10 | 19122124 | 19122601 | + | -0.754886 | Intergenic | Intergenic | -19387 | NR_038021 | 100504265 | Mm.383417 | NR_038021 | | **Gm20139** | - | predicted gene, 20139 | ncRNA |
| chr10_19127440_19128247 | chr10 | 19127441 | 19128247 | + | -0.895492 | Intergenic | Intergenic | -13905 | NR_038021 | 100504265 | Mm.383417 | NR_038021 | | **Gm20139** | - | predicted gene, 20139 | ncRNA |
| chr10_19188181_19188740 | chr10 | 19188182 | 19188740 | + | -1.03172 | intron (NR_038022, intron 1 of 1) | intron (NR_038022, intron 1 of 1) | 21223 | NR_038022 | 100504265 | Mm.383417 | NR_038021 | | **Gm20139** | - | predicted gene, 20139 | ncRNA |
| chr10_19184735_19185324 | chr10 | 19184736 | 19185324 | + | -1.10625 | intron (NR_038022, intron 1 of 1) | intron (NR_038022, intron 1 of 1) | 17792 | NR_038022 | 100504265 | Mm.383417 | NR_038021 | | **Gm20139** | - | predicted gene, 20139 | ncRNA |
| chr10_19104540_19105371 | chr10 | 19104541 | 19105371 | + | -1.12596 | Intergenic | Intergenic | -36793 | NR_038021 | 100504265 | Mm.383417 | NR_038021 | | **Gm20139** | - | predicted gene, 20139 | ncRNA |
| chr10_19188420_19188754 | chr10 | 19188421 | 19188754 | + | -1.14216 | intron (NR_038022, intron 1 of 1) | intron (NR_038022, intron 1 of 1) | 21349 | NR_038022 | 100504265 | Mm.383417 | NR_038021 | | **Gm20139** | - | predicted gene, 20139 | ncRNA |
| chr10_19185788_19186207 | chr10 | 19185789 | 19186207 | + | -1.18187 | intron (NR_038022, intron 1 of 1) | intron (NR_038022, intron 1 of 1) | 18760 | NR_038022 | 100504265 | Mm.383417 | NR_038021 | | **Gm20139** | - | predicted gene, 20139 | ncRNA |
| chr10_19116814_19117545 | chr10 | 19116815 | 19117545 | + | -1.27187 | Intergenic | Intergenic | -24569 | NR_038021 | 100504265 | Mm.383417 | NR_038021 | | **Gm20139** | - | predicted gene, 20139 | ncRNA |
| chr10_19185775_19186634 | chr10 | 19185776 | 19186634 | + | -1.28318 | intron (NR_038022, intron 1 of 1) | intron (NR_038022, intron 1 of 1) | 18967 | NR_038022 | 100504265 | Mm.383417 | NR_038021 | | **Gm20139** | - | predicted gene, 20139 | ncRNA |
| chr10_19171830_19172216 | chr10 | 19171831 | 19172216 | + | -1.36477 | intron (NR_038022, intron 1 of 1) | intron (NR_038022, intron 1 of 1) | 4785 | NR_038022 | 100504265 | Mm.383417 | NR_038021 | | **Gm20139** | - | predicted gene, 20139 | ncRNA |
| chr10_19171826_19172093 | chr10 | 19171827 | 19172093 | + | -1.37032 | intron (NR_038022, intron 1 of 1) | intron (NR_038022, intron 1 of 1) | 4722 | NR_038022 | 100504265 | Mm.383417 | NR_038021 | | **Gm20139** | - | predicted gene, 20139 | ncRNA |
| chr10_19171773_19172315 | chr10 | 19171774 | 19172315 | + | -1.37719 | intron (NR_038022, intron 1 of 1) | intron (NR_038022, intron 1 of 1) | 4806 | NR_038022 | 100504265 | Mm.383417 | NR_038021 | | **Gm20139** | - | predicted gene, 20139 | ncRNA |
| chr10_19192315_19192662 | chr10 | 19192316 | 19192662 | + | -1.43876 | intron (NR_038021, intron 3 of 3) | intron (NR_038021, intron 3 of 3) | 25251 | NR_038022 | 100504265 | Mm.383417 | NR_038021 | | **Gm20139** | - | predicted gene, 20139 | ncRNA |
| chr1_101902783_101903116 | chr1 | 101902784 | 101903116 | + | -0.654806 | Intergenic | IAPLTR4\|LTR\|ERVK | 747182 | NR_037989 | 100504525 | Mm.87065 | NR_037989 | | **Gm20268** | - | predicted gene, 20268 | ncRNA |
| chr1_101902105_101902699 | chr1 | 101902106 | 101902699 | + | -0.695392 | Intergenic | IAPEY4_I-int\|LTR\|ERVK | 746634 | NR_037989 | 100504525 | Mm.87065 | NR_037989 | | **Gm20268** | - | predicted gene, 20268 | ncRNA |
| chr1_101902095_101903118 | chr1 | 101902096 | 101903118 | + | -0.718374 | Intergenic | IAPEY4_I-int\|LTR\|ERVK | 746839 | NR_037989 | 100504525 | Mm.87065 | NR_037989 | | **Gm20268** | - | predicted gene, 20268 | ncRNA |
| chr12_80938655_80939514 | chr12 | 80938656 | 80939514 | + | -0.819808 | Intergenic | MLT1C\|LTR\|ERVL-MaLR | 6314 | NR_045057 | 100504661 | Mm.442322 | NR_045057 | | **Gm20337** | - | predicted gene, 20337 | ncRNA |
| chr12_80938673_80939079 | chr12 | 80938674 | 80939079 | + | -1.01739 | Intergenic | Intergenic | 6523 | NR_045057 | 100504661 | Mm.442322 | NR_045057 | | **Gm20337** | - | predicted gene, 20337 | ncRNA |
| chr6_79818026_79818383 | chr6 | 79818027 | 79818383 | + | -0.797324 | 5' UTR (NM_001190732, exon 1 of 1) | 5' UTR (NM_001190732, exon 1 of 1) | 174 | NM_001190732 | 100463512 | Mm.461086 | NM_001190732 | ENSMUSG00000096887 | **Gm20594** | Mtrnr2l | predicted gene, 20594 | protein-coding |
| chr5_137631183_137631493 | chr5 | 137631184 | 137631493 | + | -0.654805 | intron (NR_033148, intron 1 of 21) | intron (NR_033148, intron 1 of 21) | 2215 | NR_033148 | 100316903 | Mm.200763 | NR_033148 | | **Gm20605** | Lrch4-sap25 | predicted gene 20605 | ncRNA |
| chr3_52929844_52930149 | chr3 | 52929845 | 52930149 | + | -0.711526 | Intergenic | Intergenic | -2331 | NR_040555 | 622124 | Mm.389434 | NR_040555 | | **Gm20750** | - | predicted gene, 20750 | ncRNA |
| chr3_52929836_52930244 | chr3 | 52929837 | 52930244 | + | -0.716369 | Intergenic | Intergenic | -2374 | NR_040555 | 622124 | Mm.389434 | NR_040555 | | **Gm20750** | - | predicted gene, 20750 | ncRNA |
| chr3_52927334_52927956 | chr3 | 52927335 | 52927956 | + | -0.739967 | promoter-TSS (NR_040555) | promoter-TSS (NR_040555) | 21 | NR_040555 | 622124 | Mm.389434 | NR_040555 | | **Gm20750** | - | predicted gene, 20750 | ncRNA |
| chr3_52840174_52840546 | chr3 | 52840175 | 52840546 | + | -1.03867 | Intergenic | Intergenic | 87306 | NR_040555 | 622124 | Mm.389434 | NR_040555 | | **Gm20750** | - | predicted gene, 20750 | ncRNA |
| chr3_52878049_52878807 | chr3 | 52878050 | 52878807 | + | -1.06662 | Intergenic | Intergenic | 49238 | NR_040555 | 622124 | Mm.389434 | NR_040555 | | **Gm20750** | - | predicted gene, 20750 | ncRNA |
| chr18_80597465_80597857 | chr18 | 80597466 | 80597857 | + | -0.707206 | Intergenic | Intergenic | -53115 | NR_028424 | 100039348 | Mm.254722 | NR_028424 | | **Gm2176** | A430076E10Rik | glucosamine-6-phosphate deaminase 1 pseudogene | pseudo |
| chr3_52670506_52671283 | chr3 | 52670507 | 52671283 | + | -0.691997 | Intergenic | MTE-int\|LTR\|ERVL-MaLR | -68829 | NR_038079 | 329639 | Mm.131245 | NR_038079 | | **Gm2447** | 100039832\|A530017F20 | predicted gene 2447 | ncRNA |
| chr3_52695379_52695714 | chr3 | 52695380 | 52695714 | + | -0.780767 | Intergenic | Intergenic | -44177 | NR_038079 | 329639 | Mm.131245 | NR_038079 | | **Gm2447** | 100039832\|A530017F20 | predicted gene 2447 | ncRNA |
| chr3_52572238_52572999 | chr3 | 52572239 | 52572999 | + | -0.799077 | Intergenic | RLTR14\|LTR\|ERV1 | -167105 | NR_038079 | 329639 | Mm.131245 | NR_038079 | | **Gm2447** | 100039832\|A530017F20 | predicted gene 2447 | ncRNA |
| chr3_52618265_52619160 | chr3 | 52618266 | 52619160 | + | -0.832928 | Intergenic | Intergenic | -121011 | NR_038079 | 329639 | Mm.131245 | NR_038079 | | **Gm2447** | 100039832\|A530017F20 | predicted gene 2447 | ncRNA |
| chr3_52832461_52832918 | chr3 | 52832462 | 52832918 | + | -0.848995 | Intergenic | Intergenic | 92966 | NR_038079 | 329639 | Mm.131245 | NR_038079 | | **Gm2447** | 100039832\|A530017F20 | predicted gene 2447 | ncRNA |
| chr3_52643813_52644254 | chr3 | 52643814 | 52644254 | + | -0.860312 | Intergenic | Intergenic | -95690 | NR_038079 | 329639 | Mm.131245 | NR_038079 | | **Gm2447** | 100039832\|A530017F20 | predicted gene 2447 | ncRNA |
| chr3_52572494_52573014 | chr3 | 52572495 | 52573014 | + | -0.923081 | Intergenic | RLTR14\|LTR\|ERV1 | -166970 | NR_038079 | 329639 | Mm.131245 | NR_038079 | | **Gm2447** | 100039832\|A530017F20 | predicted gene 2447 | ncRNA |
| chr3_52618301_52618561 | chr3 | 52618302 | 52618561 | + | -1.02158 | Intergenic | Intergenic | -121293 | NR_038079 | 329639 | Mm.131245 | NR_038079 | | **Gm2447** | 100039832\|A530017F20 | predicted gene 2447 | ncRNA |
| chr11_55120580_55121513 | chr11 | 55120581 | 55121513 | + | -1.00709 | Intergenic | PB1\|SINE\|Alu | 23062 | NM_010299 | 14667 | Mm.287807 | NM_010299 | ENSMUSG00000000594 | **Gm2a** | AA408702\|AW215435\|GM2-AP\|SAP-3 | GM2 ganglioside activator protein | protein-coding |
| chr2_19621743_19622579 | chr2 | 19621744 | 19622579 | + | -0.919767 | Intergenic | L1_Mus3\|LINE\|L1 | 35736 | NR_033642 | 100041244 | Mm.407223 | NR_033642 | | **Gm3230** | - | predicted gene 3230 | ncRNA |
| chr2_19621960_19622569 | chr2 | 19621961 | 19622569 | + | -0.921399 | Intergenic | RLTR9A2\|LTR\|ERVK | 35632 | NR_033642 | 100041244 | Mm.407223 | NR_033642 | | **Gm3230** | - | predicted gene 3230 | ncRNA |
| chr5_64599550_64600361 | chr5 | 64599551 | 64600361 | + | -0.615979 | intron (NR_045078, intron 3 of 3) | intron (NR_045078, intron 3 of 3) | 10743 | NR_045078 | 100042198 | Mm.25191 | NR_045078 | | **Gm3716** | Gm1683 | predicted gene 3716 | ncRNA |
| chr5_64676278_64676932 | chr5 | 64676279 | 64676932 | + | -0.89405 | Intergenic | Intergenic | -65906 | NR_045078 | 100042198 | Mm.25191 | NR_045078 | | **Gm3716** | Gm1683 | predicted gene 3716 | ncRNA |
| chr5_64609411_64609787 | chr5 | 64609412 | 64609787 | + | -1.01457 | intron (NR_045078, intron 2 of 3) | intron (NR_045078, intron 2 of 3) | 1100 | NR_045078 | 100042198 | Mm.25191 | NR_045078 | | **Gm3716** | Gm1683 | predicted gene 3716 | ncRNA |

| | | | | | | | | | | | | | | | | | |
|---|---|---|---|---|---|---|---|---|---|---|---|---|---|---|---|---|---|
| chr5_64681208_64681485 | chr5 | 64681209 | 64681485 | + | -1.10889 | Intergenic | MLT1A\|LTR\|ERVL-MaLR | -70648 | NR_045078 | 100042198 | Mm.25191 | NR_045078 | | **Gm3716** | Gm1683 | predicted gene 3716 | ncRNA |
| chr5_64684153_64684889 | chr5 | 64684154 | 64684889 | + | -1.53425 | Intergenic | Intergenic | -73822 | NR_045078 | 100042198 | Mm.25191 | NR_045078 | | **Gm3716** | Gm1683 | predicted gene 3716 | ncRNA |
| chr14_75828910_75829395 | chr14 | 75828911 | 75829395 | + | -0.680831 | Intergenic | Intergenic | 15811 | NR_045294 | 100043189 | Mm.426329 | NR_045294 | | **Gm4285** | - | predicted gene 4285 | ncRNA |
| chr17_32240848_32241237 | chr17 | 32240849 | 32241237 | + | -0.735565 | promoter-TSS (NR_102329) | promoter-TSS (NR_102329) | -877 | NR_102329 | 100043434 | | NR_102329 | | **Gm4432** | - | predicted gene 4432 | pseudo |
| chr13_93072817_93073694 | chr13 | 93072818 | 93073694 | + | -0.737594 | promoter-TSS (NR_036451) | promoter-TSS (NR_036451) | -48 | NR_036451 | 100502942 | Mm.441946 | NM_177623 | | **Gm4814** | C920025J10\|EG218444 | predicted gene 4814 | ncRNA |
| chrX_11647048_11647421 | chrX | 11647049 | 11647421 | + | -0.830269 | Intergenic | ORR1D2\|LTR\|ERVL-MaLR | 319413 | NM_001114529 | 236663 | Mm.473680 | NM_001114529 | ENSMUSG00000069038 | **Gm4906** | EG236663 | predicted gene 4906 | protein-coding |
| chr10_58239454_58239635 | chr10 | 58239455 | 58239635 | + | -0.784881 | Intergenic | Intergenic | -2885 | NM_001034869 | 245263 | Mm.291161 | NM_001034869 | ENSMUSG00000075045 | **Gm4981** | EG245263 | predicted gene 4981 | protein-coding |
| chr1_180355841_180356632 | chr1 | 180355842 | 180356632 | + | -0.634003 | intron (NM_001081175, intron 2 of 7) | intron (NM_001081175, intron 2 of 7) | -25688 | NR_003623 | 277333 | Mm.389223 | NM_199472 | | **Gm5069** | EG277333 | glyceraldehyde-3-phosphate dehydrogenase pseudogene | pseudo |
| chr1_180356021_180356582 | chr1 | 180356022 | 180356582 | + | -0.64884 | intron (NM_001081175, intron 2 of 7) | intron (NM_001081175, intron 2 of 7) | -25753 | NR_003623 | 277333 | Mm.389223 | NM_199472 | | **Gm5069** | EG277333 | glyceraldehyde-3-phosphate dehydrogenase pseudogene | pseudo |
| chr1_180359353_180359735 | chr1 | 180359354 | 180359735 | + | -0.650547 | intron (NM_001081175, intron 2 of 7) | MLT1L\|LTR\|ERVL-MaLR | -28995 | NR_003623 | 277333 | Mm.389223 | NM_199472 | | **Gm5069** | EG277333 | glyceraldehyde-3-phosphate dehydrogenase pseudogene | pseudo |
| chr1_180335819_180336085 | chr1 | 180335820 | 180336085 | + | -0.706287 | intron (NM_001081175, intron 2 of 7) | intron (NM_001081175, intron 2 of 7) | -5403 | NR_003623 | 277333 | Mm.389223 | NM_199472 | | **Gm5069** | EG277333 | glyceraldehyde-3-phosphate dehydrogenase pseudogene | pseudo |
| chr1_180334575_180334838 | chr1 | 180334576 | 180334838 | + | -0.824197 | intron (NM_001081175, intron 2 of 7) | intron (NM_001081175, intron 2 of 7) | -4158 | NR_003623 | 277333 | Mm.389223 | NM_199472 | | **Gm5069** | EG277333 | glyceraldehyde-3-phosphate dehydrogenase pseudogene | pseudo |
| chr1_180338431_180338746 | chr1 | 180338432 | 180338746 | + | -1.10569 | intron (NM_001081175, intron 2 of 7) | intron (NM_001081175, intron 2 of 7) | -8040 | NR_003623 | 277333 | Mm.389223 | NM_199472 | | **Gm5069** | EG277333 | glyceraldehyde-3-phosphate dehydrogenase pseudogene | pseudo |
| chr1_180338510_180338694 | chr1 | 180338511 | 180338694 | + | -1.12121 | intron (NM_001081175, intron 2 of 7) | intron (NM_001081175, intron 2 of 7) | -8053 | NR_003623 | 277333 | Mm.389223 | NM_199472 | | **Gm5069** | EG277333 | glyceraldehyde-3-phosphate dehydrogenase pseudogene | pseudo |
| chr1_180397786_180398083 | chr1 | 180397787 | 180398083 | + | -1.12873 | intron (NM_001081175, intron 2 of 7) | intron (NM_001081175, intron 2 of 7) | -67386 | NR_003623 | 277333 | Mm.389223 | NM_199472 | | **Gm5069** | EG277333 | glyceraldehyde-3-phosphate dehydrogenase pseudogene | pseudo |
| chr1_180397846_180398026 | chr1 | 180397847 | 180398026 | + | -1.13987 | intron (NM_001081175, intron 2 of 7) | intron (NM_001081175, intron 2 of 7) | -67387 | NR_003623 | 277333 | Mm.389223 | NM_199472 | | **Gm5069** | EG277333 | glyceraldehyde-3-phosphate dehydrogenase pseudogene | pseudo |
| chr13_44065729_44066430 | chr13 | 44065730 | 44066430 | + | -0.631923 | Intergenic | Intergenic | -55087 | NR_045285 | 328235 | Mm.188356 | NR_045285 | ENSMUSG00000089815 | **Gm5083** | A730030A06 | predicted gene 5083 | ncRNA |
| chr13_44065650_44066182 | chr13 | 44065651 | 44066182 | + | -0.712105 | Intergenic | Intergenic | -55251 | NR_045285 | 328235 | Mm.188356 | NR_045285 | ENSMUSG00000089815 | **Gm5083** | A730030A06 | predicted gene 5083 | ncRNA |
| chr13_44097140_44097576 | chr13 | 44097141 | 44097576 | + | -0.810828 | Intergenic | Intergenic | -23809 | NR_045285 | 328235 | Mm.188356 | NR_045285 | ENSMUSG00000089815 | **Gm5083** | A730030A06 | predicted gene 5083 | ncRNA |
| chr13_97857741_97858292 | chr13 | 97857742 | 97858292 | + | -0.633437 | Intergenic | Intergenic | 298017 | NR_046157 | 328314 | Mm.138047 | NR_046157 | | **Gm5086** | B230311A21\|EG328314 | predicted gene 5086 | ncRNA |
| chr13_97857838_97858229 | chr13 | 97857839 | 97858229 | + | -0.646115 | Intergenic | Intergenic | 298034 | NR_046157 | 328314 | Mm.138047 | NR_046157 | | **Gm5086** | B230311A21\|EG328314 | predicted gene 5086 | ncRNA |
| chr13_97857868_97858691 | chr13 | 97857869 | 97858691 | + | -0.734049 | Intergenic | Intergenic | 298280 | NR_046157 | 328314 | Mm.138047 | NR_046157 | | **Gm5086** | B230311A21\|EG328314 | predicted gene 5086 | ncRNA |
| chr13_97762155_97762613 | chr13 | 97762156 | 97762613 | + | -0.862898 | Intergenic | Intergenic | 202384 | NR_046157 | 328314 | Mm.138047 | NR_046157 | | **Gm5086** | B230311A21\|EG328314 | predicted gene 5086 | ncRNA |
| chr13_97645153_97645448 | chr13 | 97645154 | 97645448 | + | -0.881031 | Intergenic | Intergenic | 85301 | NR_046157 | 328314 | Mm.138047 | NR_046157 | | **Gm5086** | B230311A21\|EG328314 | predicted gene 5086 | ncRNA |
| chr13_97762146_97762438 | chr13 | 97762147 | 97762438 | + | -0.922258 | Intergenic | Intergenic | 202292 | NR_046157 | 328314 | Mm.138047 | NR_046157 | | **Gm5086** | B230311A21\|EG328314 | predicted gene 5086 | ncRNA |
| chr13_97762012_97762491 | chr13 | 97762013 | 97762491 | + | -0.958098 | Intergenic | Intergenic | 202252 | NR_046157 | 328314 | Mm.138047 | NR_046157 | | **Gm5086** | B230311A21\|EG328314 | predicted gene 5086 | ncRNA |
| chr13_97789769_97790184 | chr13 | 97789770 | 97790184 | + | -0.999158 | Intergenic | Intergenic | 229977 | NR_046157 | 328314 | Mm.138047 | NR_046157 | | **Gm5086** | B230311A21\|EG328314 | predicted gene 5086 | ncRNA |
| chr13_97789683_97790424 | chr13 | 97789684 | 97790424 | + | -1.00215 | Intergenic | Intergenic | 230054 | NR_046157 | 328314 | Mm.138047 | NR_046157 | | **Gm5086** | B230311A21\|EG328314 | predicted gene 5086 | ncRNA |
| chr13_97789801_97790342 | chr13 | 97789802 | 97790342 | + | -1.01417 | Intergenic | Intergenic | 230072 | NR_046157 | 328314 | Mm.138047 | NR_046157 | | **Gm5086** | B230311A21\|EG328314 | predicted gene 5086 | ncRNA |
| chr9_61056442_61057288 | chr9 | 61056443 | 61057288 | + | -0.644859 | Intergenic | Intergenic | 39796 | NR_040767 | 330956 | Mm.207985 | NR_040767 | | **Gm5122** | A430102L10 | predicted gene 5122 | ncRNA |
| chr9_60991373_60992048 | chr9 | 60991374 | 60992048 | + | -0.788794 | Intergenic | Intergenic | -25358 | NR_040767 | 330956 | Mm.207985 | NR_040767 | | **Gm5122** | A430102L10 | predicted gene 5122 | ncRNA |
| chr9_61054346_61054967 | chr9 | 61054347 | 61054967 | + | -1.33423 | Intergenic | Intergenic | 37588 | NR_040767 | 330956 | Mm.207985 | NR_040767 | | **Gm5122** | A430102L10 | predicted gene 5122 | ncRNA |
| chr9_61054368_61054792 | chr9 | 61054369 | 61054792 | + | -1.37067 | Intergenic | Intergenic | 37511 | NR_040767 | 330956 | Mm.207985 | NR_040767 | | **Gm5122** | A430102L10 | predicted gene 5122 | ncRNA |
| chr3_37724164_37724981 | chr3 | 37724165 | 37724981 | + | -0.682163 | promoter-TSS (NM_198657) | promoter-TSS (NM_198657) | -213 | NM_198657 | 381438 | Mm.490549 | NM_198657 | ENSMUSG00000058174 | **Gm5148** | EG381438 | predicted gene 5148 | protein-coding |
| chr3_59703551_59703909 | chr3 | 59703552 | 59703909 | + | -0.598676 | Intergenic | RMER19C\|LTR\|ERVK | -26056 | NM_001101531 | 433597 | Mm.297801 | NM_001101531 | ENSMUSG00000090527 | **Gm5538** | - | predicted gene 5538 | protein-coding |
| chr3_59703426_59703857 | chr3 | 59703427 | 59703857 | + | -0.674746 | Intergenic | RMER19C\|LTR\|ERVK | -26144 | NM_001101531 | 433597 | Mm.297801 | NM_001101531 | ENSMUSG00000090527 | **Gm5538** | - | predicted gene 5538 | protein-coding |
| chr3_59702393_59703176 | chr3 | 59702394 | 59703176 | + | -0.69109 | Intergenic | Intergenic | -27001 | NM_001101531 | 433597 | Mm.297801 | NM_001101531 | ENSMUSG00000090527 | **Gm5538** | - | predicted gene 5538 | protein-coding |
| chr3_105815211_105815784 | chr3 | 105815212 | 105815784 | + | -0.923033 | promoter-TSS (NR_045845) | promoter-TSS (NR_045845) | -69 | NR_045845 | 433637 | Mm.331285 | NR_045845 | | **Gm5547** | EG433637 | predicted gene 5547 | ncRNA |
| chr9_48532040_48532600 | chr9 | 48532041 | 48532600 | + | -0.716297 | Intergenic | ORR1D2\|LTR\|ERVL-MaLR | 36977 | NM_001004191 | 434402 | Mm.386950 | NM_001004191 | ENSMUSG00000042293 | **Gm5617** | EG434402 | predicted gene 5617 | protein-coding |
| chr9_48549101_48549348 | chr9 | 48549102 | 48549348 | + | -0.719926 | Intergenic | L1MC2\|LINE\|L1 | 53882 | NM_001004191 | 434402 | Mm.386950 | NM_001004191 | ENSMUSG00000042293 | **Gm5617** | EG434402 | predicted gene 5617 | protein-coding |
| chr1_138666337_138666649 | chr1 | 138666338 | 138666649 | + | -0.793604 | intron (NR_040304, intron 2 of 4) | intron (NR_040304, intron 2 of 4) | 15582 | NR_040304 | 545363 | Mm.362817 | NR_040304 | | **Gm5833** | - | predicted gene 5833 | ncRNA |
| chr9_110920710_110921132 | chr9 | 110920711 | 110921132 | + | -0.734158 | Intergenic | Intergenic | -4060 | NM_001195437 | 235634 | Mm.31788 | NM_001195437 | ENSMUSG00000096393 | **Gm590** | - | predicted gene 590 | protein-coding |
| chr9_110920393_110921083 | chr9 | 110920394 | 110921083 | + | -0.80282 | Intergenic | Intergenic | -3877 | NM_001195437 | 235634 | Mm.31788 | NM_001195437 | ENSMUSG00000096393 | **Gm590** | - | predicted gene 590 | protein-coding |
| chr9_110927255_110927623 | chr9 | 110927256 | 110927623 | + | -0.960952 | Intergenic | Intergenic | -10578 | NM_001195437 | 235634 | Mm.31788 | NM_001195437 | ENSMUSG00000096393 | **Gm590** | - | predicted gene 590 | protein-coding |
| chr10_20219256_20220033 | chr10 | 20219257 | 20220033 | + | -0.674687 | intron (NM_001198635, intron 1 of 17) | ORR1F\|LTR\|ERVL-MaLR | 11206 | NM_001101561 | 621697 | Mm.440587 | NM_001101561 | | **Gm6251** | EG621697 | predicted gene 6251 | protein-coding |
| chr3_143585425_143585984 | chr3 | 143585426 | 143585984 | + | -0.650655 | Intergenic | Intergenic | 126438 | NR_040405 | 621809 | Mm.360134 | NR_040405 | | **Gm6260** | EG621809 | predicted gene 6260 | ncRNA |
| chrX_36474758_36475273 | chrX | 36474759 | 36475273 | + | -0.772521 | Intergenic | Intergenic | 71539 | NR_044989 | 621901 | Mm.491051 | NR_044989 | ENSMUSG00000079642 | **Gm6268** | EG621901 | predicted gene 6268 | ncRNA |
| chr4_40720940_40721374 | chr4 | 40720941 | 40721374 | + | -0.690546 | non-coding (NR_077226, exon 3 of 3) | non-coding (NR_077226, exon 3 of 3) | 1160 | NR_077221 | 622208 | Mm.471544 | NR_077221 | | **Gm6297** | - | predicted gene 6297 | ncRNA |
| chr17_30363398_30363687 | chr17 | 30363399 | 30363687 | + | -0.953712 | intron (NM_027060, intron 6 of 10) | intron (NM_027060, intron 6 of 10) | 32637 | NR_030688 | 623169 | Mm.440508 | NR_030688 | | **Gm6402** | EG623169 | ribosomal protein S17 pseudogene | pseudo |
| chr14_48225855_48226806 | chr14 | 48225856 | 48226806 | + | -0.864257 | intron (NR_003630, intron 2 of 2) | intron (NR_003630, intron 2 of 2) | 27615 | NR_003630 | 624367 | Mm.445185 | NM_001037921 | | **Gm6498** | EG624367 | glyceraldehyde-3-phosphate dehydrogenase pseudogene | pseudo |
| chr14_48271103_48271477 | chr14 | 48271104 | 48271477 | + | -0.873646 | Intergenic | Intergenic | 72574 | NR_003630 | 624367 | Mm.445185 | NM_001037921 | | **Gm6498** | EG624367 | glyceraldehyde-3-phosphate dehydrogenase pseudogene | pseudo |
| chr14_48270943_48271657 | chr14 | 48270944 | 48271657 | + | -0.92822 | Intergenic | Intergenic | 72584 | NR_003630 | 624367 | Mm.445185 | NM_001037921 | | **Gm6498** | EG624367 | glyceraldehyde-3-phosphate dehydrogenase pseudogene | pseudo |
| chr6_12315130_12315966 | chr6 | 12315131 | 12315966 | + | -0.697393 | 3' UTR (NM_001164805, exon 27 of 27) | 3' UTR (NM_001164805, exon 27 of 27) | -205968 | NR_003631 | 625347 | Mm.389084 | NM_001039041 | ENSMUSG00000057223 | **Gm6578** | EG625347 | mitochondrial ribosomal protein L32 pseudogene | pseudo |
| chr6_12059931_12060317 | chr6 | 12059932 | 12060317 | + | -0.782564 | intron (NM_001168382, intron 17 of 17) | PB1D9\|SINE\|Alu | 49456 | NR_003631 | 625347 | Mm.389084 | NM_001039041 | ENSMUSG00000057223 | **Gm6578** | EG625347 | mitochondrial ribosomal protein L32 pseudogene | pseudo |
| chr6_12038217_12038663 | chr6 | 12038218 | 12038663 | + | -0.879921 | intron (NM_001168382, intron 16 of 17) | intron (NM_001168382, intron 16 of 17) | 71140 | NR_003631 | 625347 | Mm.389084 | NM_001039041 | ENSMUSG00000057223 | **Gm6578** | EG625347 | mitochondrial ribosomal protein L32 pseudogene | pseudo |
| chr3_35306051_35306699 | chr3 | 35306052 | 35306699 | + | -0.927633 | Intergenic | Intergenic | 359788 | NR_040748 | 625963 | Mm.251506 | NR_040748 | | **Gm6639** | - | predicted gene 6639 | ncRNA |
| chr16_49962274_49962665 | chr16 | 49962275 | 49962665 | + | -0.754204 | Intergenic | Intergenic | -17990 | NR_045001 | 628951 | Mm.428032 | NR_045001 | | **Gm6936** | - | predicted gene 6936 | ncRNA |
| chr14_77518915_77519285 | chr14 | 77518916 | 77519285 | + | -0.8837 | intron (NM_172813, intron 7 of 18) | B4\|SINE\|B4 | -12232 | NR_033141 | 629678 | Mm.390721 | NM_001037935 | ENSMUSG00000067973 | **Gm6994** | EG629678 | predicted gene 6994 | ncRNA |
| chr9_31271148_31271692 | chr9 | 31271149 | 31271692 | + | -1.36643 | promoter-TSS (NM_001101597) | promoter-TSS (NM_001101597) | 19 | NM_001101597 | 638580 | Mm.373655 | NM_001101597 | | **Gm7244** | EG638580 | predicted gene 7244 | protein-coding |

| | | | | | | | | | | | | | | | | | |
|---|---|---|---|---|---|---|---|---|---|---|---|---|---|---|---|---|---|
| chr6_142869940_142870230 | chr6 | 142869941 | 142870230 | + | -0.712015 | intron (NM_011374, intron 3 of 4) | intron (NM_011374, intron 3 of 4) | 55854 | NR_045707 | 665037 | Mm.401493 | NR_045707 | ENSMUSG00000072683 | **Gm7457** | EG665037\|ENSMUSG00000072683 | predicted gene 7457 | ncRNA |
| chr6_142887419_142887776 | chr6 | 142887420 | 142887776 | + | -1.21329 | intron (NM_011374, intron 2 of 4) | MTEa\|LTR\|ERVL-MaLR | 73367 | NR_045707 | 665037 | Mm.401493 | NR_045707 | ENSMUSG00000072683 | **Gm7457** | EG665037\|ENSMUSG00000072683 | predicted gene 7457 | ncRNA |
| chr5_119359744_119360441 | chr5 | 119359745 | 119360441 | + | -0.705647 | Intergenic | Intergenic | -160075 | NR_046033 | 665194 | Mm.491047 | NR_046033 | | **Gm7538** | EG665194 | predicted gene 7538 | ncRNA |
| chr5_119348014_119348767 | chr5 | 119348015 | 119348767 | + | -0.818575 | Intergenic | Intergenic | -148373 | NR_046033 | 665194 | Mm.491047 | NR_046033 | | **Gm7538** | EG665194 | predicted gene 7538 | ncRNA |
| chr5_119348257_119348702 | chr5 | 119348258 | 119348702 | + | -0.855903 | Intergenic | Intergenic | -148462 | NR_046033 | 665194 | Mm.491047 | NR_046033 | | **Gm7538** | EG665194 | predicted gene 7538 | ncRNA |
| chr6_98608986_98609515 | chr6 | 98608987 | 98609515 | + | -0.612629 | Intergenic | Intergenic | -266497 | NM_001128092 | 330390 | Mm.484161 | NM_001128092 | ENSMUSG00000090667 | **Gm765** | - | predicted gene 765 | protein-coding |
| chr6_98629673_98629982 | chr6 | 98629674 | 98629982 | + | -0.708551 | Intergenic | LTR16C\|LTR\|ERVL | -287074 | NM_001128092 | 330390 | Mm.484161 | NM_001128092 | ENSMUSG00000090667 | **Gm765** | - | predicted gene 765 | protein-coding |
| chr6_98629684_98630099 | chr6 | 98629685 | 98630099 | + | -0.766875 | Intergenic | LTR16C\|LTR\|ERVL | -287138 | NM_001128092 | 330390 | Mm.484161 | NM_001128092 | ENSMUSG00000090667 | **Gm765** | - | predicted gene 765 | protein-coding |
| chr19_11497427_11497599 | chr19 | 11497428 | 11497599 | + | -0.608223 | intron (NM_001164202, intron 1 of 4) | intron (NM_001164202, intron 1 of 4) | 5475 | NM_001164202 | 666926 | Mm.486382 | NM_001164202 | ENSMUSG00000058470 | **Gm8369** | EG666926 | predicted gene 8369 | protein-coding |
| chr19_11497335_11497599 | chr19 | 11497336 | 11497599 | + | -0.652221 | intron (NM_001164202, intron 1 of 4) | intron (NM_001164202, intron 1 of 4) | 5429 | NM_001164202 | 666926 | Mm.486382 | NM_001164202 | ENSMUSG00000058470 | **Gm8369** | EG666926 | predicted gene 8369 | protein-coding |
| chr6_89272147_89272975 | chr6 | 89272148 | 89272975 | + | -0.641141 | Intergenic | Intergenic | -56382 | NR_033190 | 330379 | Mm.297741 | NM_201356 | | **Gm839** | - | predicted gene 839 | ncRNA |
| chr6_89288172_89288530 | chr6 | 89288173 | 89288530 | + | -0.690429 | Intergenic | Intergenic | -72172 | NR_033190 | 330379 | Mm.297741 | NM_201356 | | **Gm839** | - | predicted gene 839 | ncRNA |
| chr10_26585064_26585490 | chr10 | 26585065 | 26585490 | + | -0.703147 | Intergenic | RMER13A2\|LTR\|ERVK | 89374 | NR_033633 | 667572 | Mm.482728 | NR_033633 | ENSMUSG00000098179 | **Gm8709** | EG667572 | glyceraldehyde-3-phosphate dehydrogenase pseudogene | pseudo |
| chrX_167108377_167108741 | chrX | 167108378 | 167108741 | + | -0.64077 | Intergenic | Intergenic | -38189 | NM_001101606 | 667794 | Mm.302232 | NM_001101606 | | **Gm8817** | EG667794 | predicted gene 8817 | protein-coding |
| chrX_167063803_167064219 | chrX | 167063804 | 167064219 | + | -0.933392 | intron (NM_001101606, intron 4 of 4) | intron (NM_001101606, intron 4 of 4) | 6359 | NM_001101606 | 667794 | Mm.302232 | NM_001101606 | | **Gm8817** | EG667794 | predicted gene 8817 | protein-coding |
| chr9_48450025_48450485 | chr9 | 48450026 | 48450485 | + | -1.14842 | Intergenic | Intergenic | -4089 | NR_026561 | 667933 | Mm.491076 | NR_026561 | | **Gm8884** | EG667933 | predicted gene 8884 | ncRNA |
| chr7_106270937_106271374 | chr7 | 106270938 | 106271374 | + | -0.707431 | Intergenic | RLTR45\|LTR\|ERVK | 53372 | NR_030720 | 668128 | Mm.484022 | NR_030720 | | **Gm8989** | EG668128\|Gm21911 | very large inducible GTPase 1 pseudogene | pseudo |
| chr14_72985204_72985474 | chr14 | 72985205 | 72985474 | + | -0.788174 | Intergenic | Intergenic | -40264 | NR_027860 | 668489 | Mm.158324 | NR_027860 | | **Gm9199** | EG668489 | glycine cleavage system protein H (aminomethyl carrier) pseudogene | pseudo |
| chr5_90402871_90403581 | chr5 | 90402872 | 90403581 | + | -0.675655 | Intergenic | Intergenic | 36229 | NR_045618 | 791294 | Mm.30508 | NR_045618 | ENSMUSG00000054945 | **Gm9958** | ENSMUSG00000054945 | predicted gene 9958 | ncRNA |
| chr6_86754530_86755094 | chr6 | 86754531 | 86755094 | + | -0.933128 | intron (NM_013471, intron 5 of 12) | intron (NM_013471, intron 5 of 12) | -21434 | NM_011818 | 23885 | Mm.321452 | NM_011818 | ENSMUSG00000001157 | **Gmcl1** | 2810049L19Rik\|Gcl\|mglc-1 | germ cell-less homolog 1 (Drosophila) | protein-coding |
| chr13_32154446_32154739 | chr13 | 32154447 | 32154739 | + | -0.674923 | intron (NM_146041, intron 4 of 10) | B4\|SINE\|B4 | 183951 | NM_146041 | 218138 | Mm.247143 | NM_146041 | ENSMUSG00000038372 | **Gmds** | BC031788\|C87208 | GDP-mannose 4, 6-dehydratase | protein-coding |
| chr7_28440788_28441073 | chr7 | 28440789 | 28441073 | + | -0.86791 | promoter-TSS (NM_001039192) | promoter-TSS (NM_001039192) | -6 | NM_001039192 | 63986 | Mm.194536 | NM_022024 | | **Gmfg** | 0610039G16Rik\|2310057N07Rik\|AI324845 | glia maturation factor, gamma | protein-coding |
| chr13_24778538_24779487 | chr13 | 24778539 | 24779487 | + | -0.6325 | Intergenic | RMER6A\|LTR\|ERVK | -17076 | NM_020567 | 57441 | Mm.12239 | NM_020567 | ENSMUSG00000006715 | **Gmnn** | AW546347\|Gem | geminin | protein-coding |
| chr13_24778822_24779451 | chr13 | 24778823 | 24779451 | + | -0.766698 | Intergenic | RMER6A\|LTR\|ERVK | -17200 | NM_020567 | 57441 | Mm.12239 | NM_020567 | ENSMUSG00000006715 | **Gmnn** | AW546347\|Gem | geminin | protein-coding |
| chr13_45455264_45456112 | chr13 | 45455265 | 45456112 | + | -0.603988 | Intergenic | ORR1E\|LTR\|ERVL-MaLR | -51781 | NM_025508 | 66355 | Mm.290802 | NM_025508 | ENSMUSG00000000253 | **Gmpr** | 2310004P21Rik\|AV028449 | guanosine monophosphate reductase | protein-coding |
| chr5_140858220_140858827 | chr5 | 140858221 | 140858827 | + | -0.831689 | Intergenic | Intergenic | -28093 | NM_010302 | 14673 | Mm.370185 | NM_010302 | | **Gna12** | AI414047\|AI504261\|Galpha12 | guanine nucleotide binding protein, alpha 12 | protein-coding |
| chr5_140858259_140858687 | chr5 | 140858260 | 140858687 | + | -0.848744 | Intergenic | Intergenic | -28042 | NM_010302 | 14673 | Mm.370185 | NM_010302 | | **Gna12** | AI414047\|AI504261\|Galpha12 | guanine nucleotide binding protein, alpha 12 | protein-coding |
| chr5_140858035_140858734 | chr5 | 140858036 | 140858734 | + | -0.858578 | Intergenic | B1_Mur1\|SINE\|Alu | -27954 | NM_010302 | 14673 | Mm.370185 | NM_010302 | | **Gna12** | AI414047\|AI504261\|Galpha12 | guanine nucleotide binding protein, alpha 12 | protein-coding |
| chr2_174361087_174361503 | chr2 | 174361088 | 174361503 | + | -0.68411 | Intergenic | Intergenic | 31206 | NM_001077510 | 14683 | Mm.125770 | NM_010309 | ENSMUSG00000027523 | **Gnas** | 5530400H20Rik\|A930027G11Rik\|C130027O20Rik\|GPSA\|GSP\|Galphas\|Gnas1\|Gnasxl\|Gsa\|Nesp\|Nespl\|Oed-Sml\|Oedsml\|P1\|P2\|P3\|PHP1A\|PHP1B\|POH | GNAS (guanine nucleotide binding protein, alpha stimulating) complex locus | protein-coding |
| chr9_107675658_107676704 | chr9 | 107675659 | 107676704 | + | -0.925894 | intron (NM_008140, intron 7 of 8) | CpG | 3411 | NM_008140 | 14685 | Mm.284853 | NM_008140 | ENSMUSG00000034837 | **Gnat1** | Gnat-1\|Ird1\|Ird2\|Tralpha\|irdc\|irdr\|transducin | guanine nucleotide binding protein, alpha transducing 1 | protein-coding |
| chr16_18503531_18504019 | chr16 | 18503532 | 18504019 | + | -0.940129 | intron (NM_001285493, intron 2 of 8) | intron (NM_001285493, intron 2 of 8) | 4904 | NR_104347 | 13972 | Mm.236139 | NM_023120 | ENSMUSG00000000884 | **Gnb1l** | ESTM55\|Gm16314\|Me49f07\|OTTMUSG00000033458\|Wdr14\|Wdvcf | guanine nucleotide binding protein (G protein), beta polypeptide 1-like | protein-coding |
| chr16_18503561_18503925 | chr16 | 18503562 | 18503925 | + | -1.00654 | intron (NM_001285493, intron 2 of 8) | intron (NM_001285493, intron 2 of 8) | 4872 | NR_104347 | 13972 | Mm.236139 | NM_023120 | ENSMUSG00000000884 | **Gnb1l** | ESTM55\|Gm16314\|Me49f07\|OTTMUSG00000033458\|Wdr14\|Wdvcf | guanine nucleotide binding protein (G protein), beta polypeptide 1-like | protein-coding |
| chr4_44059649_44060216 | chr4 | 44059650 | 44060216 | + | -0.950222 | exon (NM_015828, exon 3 of 12) | exon (NM_015828, exon 3 of 12) | 12740 | NM_001190414 | 50798 | Mm.256718 | NM_015828 | ENSMUSG00000028479 | **Gne** | 2310066H07Rik\|DMRV\|GLCNE\|IBM2\|NM\|Uae1 | glucosamine (UDP-N-acetyl)-2-epimerase/N-acetylmannosamine kinase | protein-coding |
| chr4_44097644_44098389 | chr4 | 44097645 | 44098389 | + | -0.963903 | Intergenic | Intergenic | -13840 | NM_015828 | 50798 | Mm.256718 | NM_015828 | ENSMUSG00000028479 | **Gne** | 2310066H07Rik\|DMRV\|GLCNE\|IBM2\|NM\|Uae1 | glucosamine (UDP-N-acetyl)-2-epimerase/N-acetylmannosamine kinase | protein-coding |
| chr14_19961883_19962289 | chr14 | 19961884 | 19962289 | + | -0.760038 | intron (NM_001285908, intron 1 of 3) | ID4\|SINE\|ID | 14607 | NM_001285911 | 14702 | Mm.41737 | NM_010315 | ENSMUSG00000043004 | **Gng2** | 82 | guanine nucleotide binding protein (G protein), gamma 2 | protein-coding |
| chr11_95834022_95834742 | chr11 | 95834023 | 95834742 | + | -0.607927 | exon (NM_001163464, exon 4 of 8) | exon (NM_001163464, exon 4 of 8) | -7913 | NM_023121 | 14710 | Mm.46299 | NM_023121 | ENSMUSG00000038811 | **Gngt2** | AV096488 | guanine nucleotide binding protein (G protein), gamma transducing activity polypeptide 2 | protein-coding |
| chr11_95835211_95835613 | chr11 | 95835212 | 95835613 | + | -0.922625 | intron (NM_025659, intron 3 of 7) | intron (NM_025659, intron 3 of 7) | -6883 | NM_023121 | 14710 | Mm.46299 | NM_023121 | ENSMUSG00000038811 | **Gngt2** | AV096488 | guanine nucleotide binding protein (G protein), gamma transducing activity polypeptide 2 | protein-coding |
| chr10_121419806_121420684 | chr10 | 121419807 | 121420684 | + | -0.779963 | intron (NM_138956, intron 1 of 4) | intron (NM_138956, intron 1 of 4) | 55155 | NM_029364 | 75612 | Mm.207683 | NM_029364 | ENSMUSG00000034707 | **Gns** | 2610016K11Rik\|AU042285\|C87209\|G6S\|N28088 | glucosamine (N-acetyl)-6-sulfatase | protein-coding |
| chr13_59668378_59669115 | chr13 | 59668379 | 59669115 | + | -0.667982 | intron (NM_001035122, intron 1 of 9) | intron (NM_001035122, intron 1 of 9) | 7037 | NM_001035122 | 105348 | Mm.171335 | NM_027307 | ENSMUSG00000021556 | **Golm1** | 2310001L02Rik\|AW125446\|D030064E01Rik\|GP73\|Golph2\|PSEC0257 | golgi membrane protein 1 | protein-coding |
| chr13_59672293_59672663 | chr13 | 59672294 | 59672663 | + | -0.686636 | intron (NM_001035122, intron 1 of 9) | intron (NM_001035122, intron 1 of 9) | 3306 | NM_001035122 | 105348 | Mm.171335 | NM_027307 | ENSMUSG00000021556 | **Golm1** | 2310001L02Rik\|AW125446\|D030064E01Rik\|GP73\|Golph2\|PSEC0257 | golgi membrane protein 1 | protein-coding |
| chr13_59657194_59657468 | chr13 | 59657195 | 59657468 | + | -0.688258 | intron (NM_027307, intron 4 of 10) | intron (NM_027307, intron 4 of 10) | 18453 | NM_001035122 | 105348 | Mm.171335 | NM_027307 | ENSMUSG00000021556 | **Golm1** | 2310001L02Rik\|AW125446\|D030064E01Rik\|GP73\|Golph2\|PSEC0257 | golgi membrane protein 1 | protein-coding |
| chr15_12435964_12436722 | chr15 | 12435965 | 12436722 | + | -0.622647 | intron (NM_001081064, intron 5 of 23) | intron (NM_001081064, intron 5 of 23) | 114847 | NM_025673 | 66629 | Mm.157655 | NM_025673 | ENSMUSG00000022200 | **Golph3** | 4733401N08Rik\|5730410D03Rik\|AW413496 | golgi phosphoprotein 3 | protein-coding |
| chr15_12424092_12424719 | chr15 | 12424093 | 12424719 | + | -0.679621 | intron (NM_001081064, intron 5 of 23) | intron (NM_001081064, intron 5 of 23) | 102910 | NM_025673 | 66629 | Mm.157655 | NM_025673 | ENSMUSG00000022200 | **Golph3** | 4733401N08Rik\|5730410D03Rik\|AW413496 | golgi phosphoprotein 3 | protein-coding |
| chr15_12435844_12436393 | chr15 | 12435845 | 12436393 | + | -0.694153 | intron (NM_001081064, intron 5 of 23) | intron (NM_001081064, intron 5 of 23) | 114623 | NM_025673 | 66629 | Mm.157655 | NM_025673 | ENSMUSG00000022200 | **Golph3** | 4733401N08Rik\|5730410D03Rik\|AW413496 | golgi phosphoprotein 3 | protein-coding |
| chr15_12285199_12285618 | chr15 | 12285200 | 12285618 | + | -0.842042 | Intergenic | Intergenic | -36087 | NM_025673 | 66629 | Mm.157655 | NM_025673 | ENSMUSG00000022200 | **Golph3** | 4733401N08Rik\|5730410D03Rik\|AW413496 | golgi phosphoprotein 3 | protein-coding |
| chr11_76739015_76740005 | chr11 | 76739016 | 76740005 | + | -0.621455 | intron (NM_016810, intron 6 of 8) | intron (NM_016810, intron 6 of 8) | 24045 | NM_016810 | 53334 | Mm.20931 | NM_016810 | ENSMUSG00000010392 | **Gosr1** | AI414660\|AI426320\|BB145494\|GOS-28\|GOSRI\|GS28 | golgi SNAP receptor complex member 1 | protein-coding |
| chr11_76739747_76739991 | chr11 | 76739748 | 76739991 | + | -0.746715 | intron (NM_016810, intron 6 of 8) | intron (NM_016810, intron 6 of 8) | 23686 | NM_016810 | 53334 | Mm.20931 | NM_016810 | ENSMUSG00000010392 | **Gosr1** | AI414660\|AI426320\|BB145494\|GOS-28\|GOSRI\|GS28 | golgi SNAP receptor complex member 1 | protein-coding |
| chr19_43530859_43531340 | chr19 | 43530860 | 43531340 | + | -1.30088 | Intergenic | Intergenic | -6495 | NM_010324 | 14718 | Mm.19039 | NM_010324 | ENSMUSG00000025190 | **Got1** | AI789014\|Got-1\|cAspAT\|cCAT | glutamate oxaloacetate transaminase 1, soluble | protein-coding |
| chr11_70629483_70629847 | chr11 | 70629484 | 70629847 | + | -0.796054 | Intergenic | Lx10\|LINE\|L1 | -9457 | NM_010326 | 14723 | Mm.377085 | NM_010326 | ENSMUSG00000050675 | **Gp1ba** | GPIba\|GPIbalpha | glycoprotein 1b, alpha polypeptide | protein-coding |
| chr1_187308380_187309050 | chr1 | 187308381 | 187309050 | + | -0.658641 | intron (NM_026367, intron 6 of 9) | intron (NM_026367, intron 6 of 9) | 93204 | NM_026367 | 67769 | Mm.138321 | NM_026367 | ENSMUSG00000039210 | **Gpatch2** | 5830433G22Rik\|5830436K05Rik\|AI427714\|AI447508\|AW107440\|AW491060\|Gpatc2 | G patch domain containing 2 | protein-coding |

| | | | | | | | | | | | | | | | | | |
|---|---|---|---|---|---|---|---|---|---|---|---|---|---|---|---|---|---|
| chr1_187348164_187348751 | chr1 | 187348165 | 187348751 | + | -0.661126 | exon (NM_026367, exon 10 of 10) | exon (NM_026367, exon 10 of 10) | 132947 | NM_026367 | 67769 | Mm.138321 | NM_026367 | ENSMUSG00000039210 | **Gpatch2** | 5830433G22Rik\|5830436K05Rik\|AI427714\|AI447508\|AW107440\|AW491060\|Gpatc2 | G patch domain containing 2 | protein-coding |
| chr1_187348102_187349166 | chr1 | 187348103 | 187349166 | + | -0.754657 | 3' UTR (NM_026367, exon 10 of 10) | 3' UTR (NM_026367, exon 10 of 10) | 133123 | NM_026367 | 67769 | Mm.138321 | NM_026367 | ENSMUSG00000039210 | **Gpatch2** | 5830433G22Rik\|5830436K05Rik\|AI427714\|AI447508\|AW107440\|AW491060\|Gpatc2 | G patch domain containing 2 | protein-coding |
| chr1_187348416_187349168 | chr1 | 187348417 | 187349168 | + | -0.842103 | 3' UTR (NM_026367, exon 10 of 10) | 3' UTR (NM_026367, exon 10 of 10) | 133281 | NM_026367 | 67769 | Mm.138321 | NM_026367 | ENSMUSG00000039210 | **Gpatch2** | 5830433G22Rik\|5830436K05Rik\|AI427714\|AI447508\|AW107440\|AW491060\|Gpatc2 | G patch domain containing 2 | protein-coding |
| chr1_74262410_74262906 | chr1 | 74262411 | 74262906 | + | -1.38554 | intron (NM_029711, intron 7 of 9) | ID2\|SINE\|ID | -15942 | NM_174985 | 227289 | Mm.246587 | NM_174985 | ENSMUSG00000064272 | **Gpbar1** | BG37\|GPCR\|GPR131\|M-BAR\|TGR5 | G protein-coupled bile acid receptor 1 | protein-coding |
| chr1_74262508_74262868 | chr1 | 74262509 | 74262868 | + | -1.39235 | intron (NM_029711, intron 7 of 9) | intron (NM_029711, intron 7 of 9) | -15912 | NM_174985 | 227289 | Mm.246587 | NM_174985 | ENSMUSG00000064272 | **Gpbar1** | BG37\|GPCR\|GPR131\|M-BAR\|TGR5 | G protein-coupled bile acid receptor 1 | protein-coding |
| chr13_111480701_111481036 | chr13 | 111480702 | 111481036 | + | -0.751095 | intron (NM_028487, intron 2 of 12) | B1F2\|SINE\|Alu | 9172 | NM_028487 | 73274 | Mm.57286 | NM_028487 | ENSMUSG00000032745 | **Gpbp1** | 1700034P14Rik\|AU019836\|D230035M11Rik\|Gpbp\|mGPBP | GC-rich promoter binding protein 1 | protein-coding |
| chr8_55347913_55348138 | chr8 | 55347914 | 55348138 | + | -0.686573 | Intergenic | Intergenic | 393298 | NM_001253756 | 234267 | Mm.241700 | NM_153581 | ENSMUSG00000031517 | **Gpm6a** | Gpm6\|M6A | glycoprotein m6a | protein-coding |
| chr6_49051715_49052658 | chr6 | 49051716 | 49052658 | + | -0.673705 | intron (NM_053110, intron 9 of 10) | intron (NM_053110, intron 9 of 10) | 15669 | NM_053110 | 93695 | Mm.302602 | NM_053110 | ENSMUSG00000029816 | **Gpnmb** | DC-HIL\|Dchil\|ipd | glycoprotein (transmembrane) nmb | protein-coding |
| chr6_49013453_49013840 | chr6 | 49013454 | 49013840 | + | -0.679124 | Intergenic | Intergenic | -22871 | NM_053110 | 93695 | Mm.302602 | NM_053110 | ENSMUSG00000029816 | **Gpnmb** | DC-HIL\|Dchil\|ipd | glycoprotein (transmembrane) nmb | protein-coding |
| chr10_14328809_14329303 | chr10 | 14328810 | 14329303 | + | -0.77739 | Intergenic | Intergenic | 215980 | NM_001002268 | 215798 | Mm.440142 | NM_001002268 | ENSMUSG00000039116 | **Gpr126** | 1190004A11Rik\|AI449247\|AW045736\|DREG\|Gm222 | G protein-coupled receptor 126 | protein-coding |
| chr10_14314554_14315020 | chr10 | 14314555 | 14315020 | + | -0.859579 | Intergenic | Intergenic | 230249 | NM_001002268 | 215798 | Mm.440142 | NM_001002268 | ENSMUSG00000039116 | **Gpr126** | 1190004A11Rik\|AI449247\|AW045736\|DREG\|Gm222 | G protein-coupled receptor 126 | protein-coding |
| chr10_14326194_14326564 | chr10 | 14326195 | 14326564 | + | -0.863554 | Intergenic | Intergenic | 218657 | NM_001002268 | 215798 | Mm.440142 | NM_001002268 | ENSMUSG00000039116 | **Gpr126** | 1190004A11Rik\|AI449247\|AW045736\|DREG\|Gm222 | G protein-coupled receptor 126 | protein-coding |
| chr12_112879804_112880109 | chr12 | 112879805 | 112880109 | + | -0.618312 | Intergenic | Intergenic | -19041 | NM_019925 | 56696 | Mm.20455 | NM_019925 | ENSMUSG00000021298 | **Gpr132** | G2a | G protein-coupled receptor 132 | protein-coding |
| chr12_112868046_112868733 | chr12 | 112868047 | 112868733 | + | -0.762307 | Intergenic | Intergenic | -7474 | NM_019925 | 56696 | Mm.20455 | NM_019925 | ENSMUSG00000021298 | **Gpr132** | G2a | G protein-coupled receptor 132 | protein-coding |
| chr12_112854690_112855493 | chr12 | 112854691 | 112855493 | + | -1.55444 | intron (NM_019925, intron 2 of 2) | L1MB3\|LINE\|L1 | 5824 | NM_019925 | 56696 | Mm.20455 | NM_019925 | ENSMUSG00000021298 | **Gpr132** | G2a | G protein-coupled receptor 132 | protein-coding |
| chr12_112854986_112855390 | chr12 | 112854987 | 112855390 | + | -1.65861 | intron (NM_019925, intron 2 of 2) | intron (NM_019925, intron 2 of 2) | 5728 | NM_019925 | 56696 | Mm.20455 | NM_019925 | ENSMUSG00000021298 | **Gpr132** | G2a | G protein-coupled receptor 132 | protein-coding |
| chr5_139382681_139383213 | chr5 | 139382682 | 139383213 | + | -0.607788 | intron (NM_030258, intron 1 of 1) | intron (NM_030258, intron 1 of 1) | 2280 | NM_030258 | 80290 | Mm.270003 | NM_030258 | ENSMUSG00000044197 | **Gpr146** | BC003323\|PGR8 | G protein-coupled receptor 146 | protein-coding |
| chr18_42585626_42586144 | chr18 | 42585627 | 42586144 | + | -0.967277 | Intergenic | Intergenic | -6233 | NM_181543 | 240239 | Mm.186779 | NM_181543 | ENSMUSG00000042816 | **Gpr151** | C130082O03Rik\|GalRL\|PGR7\|nGPCR-2037 | G protein-coupled receptor 151 | protein-coding |
| chrX_107217648_107218148 | chrX | 107217649 | 107218148 | + | -0.637734 | Intergenic | Intergenic | -37980 | NM_001177781 | 213439 | Mm.383562 | NM_001033251 | ENSMUSG00000073008 | **Gpr174** | Gm376\|Lypsr3 | G protein-coupled receptor 174 | protein-coding |
| chrX_107224722_107225331 | chrX | 107224723 | 107225331 | + | -0.663012 | Intergenic | Intergenic | -30851 | NM_001177781 | 213439 | Mm.383562 | NM_001033251 | ENSMUSG00000073008 | **Gpr174** | Gm376\|Lypsr3 | G protein-coupled receptor 174 | protein-coding |
| chrX_107223284_107223545 | chrX | 107223285 | 107223545 | + | -0.869386 | Intergenic | L1M4b\|LINE\|L1 | -32463 | NM_001177781 | 213439 | Mm.383562 | NM_001033251 | ENSMUSG00000073008 | **Gpr174** | Gm376\|Lypsr3 | G protein-coupled receptor 174 | protein-coding |
| chr14_121898446_121898927 | chr14 | 121898447 | 121898927 | + | -0.877863 | intron (NM_026861, intron 1 of 8) | intron (NM_026861, intron 1 of 8) | 17087 | NM_182806 | 110168 | Mm.37405 | NM_182806 | ENSMUSG00000050350 | **Gpr18** | - | G protein-coupled receptor 18 | protein-coding |
| chr14_121929282_121929879 | chr14 | 121929283 | 121929879 | + | -1.20496 | intron (NM_026861, intron 4 of 8) | intron (NM_026861, intron 4 of 8) | -13807 | NM_182806 | 110168 | Mm.37405 | NM_182806 | ENSMUSG00000050350 | **Gpr18** | - | G protein-coupled receptor 18 | protein-coding |
| chr14_118173464_118174013 | chr14 | 118173465 | 118174013 | + | -0.784713 | Intergenic | L1M2\|LINE\|L1 | 36612 | NM_021434 | 58245 | Mm.275411 | NM_021434 | ENSMUSG00000022131 | **Gpr180** | AB041545\|E130016I23Rik\|Itr | G protein-coupled receptor 180 | protein-coding |
| chr14_121954633_121954966 | chr14 | 121954634 | 121954966 | + | -0.654456 | exon (NM_183031, exon 2 of 2) | exon (NM_183031, exon 2 of 2) | 10393 | NM_183031 | 321019 | Mm.265618 | NM_183031 | ENSMUSG00000051212 | **Gpr183** | Ebi2 | G protein-coupled receptor 183 | protein-coding |
| chr14_121992608_121992929 | chr14 | 121992609 | 121992929 | + | -0.752216 | intron (NM_026861, intron 6 of 8) | intron (NM_026861, intron 6 of 8) | -27576 | NM_183031 | 321019 | Mm.265618 | NM_183031 | ENSMUSG00000051212 | **Gpr183** | Ebi2 | G protein-coupled receptor 183 | protein-coding |
| chr14_121987668_121988037 | chr14 | 121987669 | 121988037 | + | -0.79288 | intron (NM_026861, intron 6 of 8) | intron (NM_026861, intron 6 of 8) | -22660 | NM_183031 | 321019 | Mm.265618 | NM_183031 | ENSMUSG00000051212 | **Gpr183** | Ebi2 | G protein-coupled receptor 183 | protein-coding |
| chr14_121994365_121994631 | chr14 | 121994366 | 121994631 | + | -0.96608 | intron (NM_026861, intron 7 of 8) | intron (NM_026861, intron 7 of 8) | -29305 | NM_183031 | 321019 | Mm.265618 | NM_183031 | ENSMUSG00000051212 | **Gpr183** | Ebi2 | G protein-coupled receptor 183 | protein-coding |
| chr14_121951444_121951709 | chr14 | 121951445 | 121951709 | + | -1.00058 | TTS (NM_183031) | TTS (NM_183031) | 13616 | NM_183031 | 321019 | Mm.265618 | NM_183031 | ENSMUSG00000051212 | **Gpr183** | Ebi2 | G protein-coupled receptor 183 | protein-coding |
| chr14_121995518_121995827 | chr14 | 121995519 | 121995827 | + | -1.03209 | intron (NM_026861, intron 7 of 8) | intron (NM_026861, intron 7 of 8) | -30480 | NM_183031 | 321019 | Mm.265618 | NM_183031 | ENSMUSG00000051212 | **Gpr183** | Ebi2 | G protein-coupled receptor 183 | protein-coding |
| chr6_134883728_134884392 | chr6 | 134883729 | 134884392 | + | -0.832457 | intron (NM_001167697, intron 1 of 2) | intron (NM_001167697, intron 1 of 2) | 3725 | NR_074084 | 14760 | Mm.4787 | NM_008157 | ENSMUSG00000032641 | **Gpr19** | - | G protein-coupled receptor 19 | protein-coding |
| chr2_37565626_37566316 | chr2 | 37565627 | 37566316 | + | -0.688907 | 3' UTR (NM_146121, exon 26 of 26) | 3' UTR (NM_146121, exon 26 of 26) | 49345 | NM_177383 | 338346 | Mm.445325 | NM_177383 | | **Gpr21** | C230004C13Rik | G protein-coupled receptor 21 | protein-coding |
| chr2_37565610_37565990 | chr2 | 37565611 | 37565990 | + | -0.754658 | 3' UTR (NM_146121, exon 26 of 26) | 3' UTR (NM_146121, exon 26 of 26) | 49174 | NM_177383 | 338346 | Mm.445325 | NM_177383 | | **Gpr21** | C230004C13Rik | G protein-coupled receptor 21 | protein-coding |
| chr12_31755920_31756289 | chr12 | 31755921 | 31756289 | + | -0.833243 | intron (NM_001163126, intron 6 of 22) | MTB_Mm\|LTR\|ERVL-MaLR | -42179 | NM_175191 | 73010 | Mm.441517 | NM_175191 | ENSMUSG00000044067 | **Gpr22** | 2900068K05Rik\|AW061316 | G protein-coupled receptor 22 | protein-coding |
| chr12_52034777_52035570 | chr12 | 52034778 | 52035570 | + | -0.88101 | Intergenic | Intergenic | -7111 | NM_008159 | 14762 | Mm.12890 | NM_008159 | ENSMUSG00000035148 | **Gpr33** | - | G protein-coupled receptor 33 | protein-coding |
| chrX_13607820_13608130 | chrX | 13607821 | 13608130 | + | -0.723507 | intron (NM_009806, intron 9 of 24) | intron (NM_009806, intron 9 of 24) | -24794 | NM_011823 | 23890 | Mm.487326 | NM_011823 | ENSMUSG00000040229 | **Gpr34** | Lypsr1 | G protein-coupled receptor 34 | protein-coding |
| chrX_13618557_13618845 | chrX | 13618558 | 13618845 | + | -0.742712 | intron (NM_009806, intron 7 of 24) | intron (NM_009806, intron 7 of 24) | -14068 | NM_011823 | 23890 | Mm.487326 | NM_011823 | ENSMUSG00000040229 | **Gpr34** | Lypsr1 | G protein-coupled receptor 34 | protein-coding |
| chr12_100856398_100856646 | chr12 | 100856399 | 100856646 | + | -0.636311 | intron (NM_145448, intron 9 of 12) | intron (NM_145448, intron 9 of 12) | 43002 | NM_001177674 | 238377 | Mm.32160 | NM_175493 | ENSMUSG00000047415 | **Gpr68** | BB131428\|Ogr1 | G protein-coupled receptor 68 | protein-coding |
| chr12_100856370_100856820 | chr12 | 100856371 | 100856820 | + | -0.647418 | intron (NM_145448, intron 9 of 12) | intron (NM_145448, intron 9 of 12) | 42929 | NM_001177674 | 238377 | Mm.32160 | NM_175493 | ENSMUSG00000047415 | **Gpr68** | BB131428\|Ogr1 | G protein-coupled receptor 68 | protein-coding |
| chr12_100901524_100901740 | chr12 | 100901525 | 100901740 | + | -0.784727 | intron (NM_175493, intron 1 of 2) | intron (NM_175493, intron 1 of 2) | -2108 | NM_001177674 | 238377 | Mm.32160 | NM_175493 | ENSMUSG00000047415 | **Gpr68** | BB131428\|Ogr1 | G protein-coupled receptor 68 | protein-coding |
| chr12_100890571_100891095 | chr12 | 100890572 | 100891095 | + | -0.798317 | intron (NM_175493, intron 1 of 2) | RMER4B\|LTR\|ERVK | 8691 | NM_001177674 | 238377 | Mm.32160 | NM_175493 | ENSMUSG00000047415 | **Gpr68** | BB131428\|Ogr1 | G protein-coupled receptor 68 | protein-coding |
| chr12_100890546_100890954 | chr12 | 100890547 | 100890954 | + | -0.814072 | intron (NM_175493, intron 1 of 2) | RMER4B\|LTR\|ERVK | 8774 | NM_001177674 | 238377 | Mm.32160 | NM_175493 | ENSMUSG00000047415 | **Gpr68** | BB131428\|Ogr1 | G protein-coupled receptor 68 | protein-coding |
| chr8_95020300_95021235 | chr8 | 95020301 | 95021235 | + | -0.734057 | intron (NM_173036, intron 1 of 11) | (CCCCCA)n\|Simple_repeat\|Simple_repeat | 3076 | NM_173036 | 54672 | Mm.27995 | NM_173036 | ENSMUSG00000060470 | **Gpr97** | A030001G24Rik\|AW214383\|Pb1\|Pb99 | G protein-coupled receptor 97 | protein-coding |
| chr13_81347282_81348028 | chr13 | 81347283 | 81348028 | + | -0.681423 | intron (NM_054053, intron 83 of 89) | intron (NM_054053, intron 83 of 89) | 285489 | NM_054053 | 110789 | Mm.288694 | NM_054053 | ENSMUSG00000069170 | **Gpr98** | Frings\|Mass1\|Mgr1\|VLGR1 | G protein-coupled receptor 98 | protein-coding |
| chr13_81352336_81352673 | chr13 | 81352337 | 81352673 | + | -0.703239 | intron (NM_054053, intron 83 of 89) | intron (NM_054053, intron 83 of 89) | 280639 | NM_054053 | 110789 | Mm.288694 | NM_054053 | ENSMUSG00000069170 | **Gpr98** | Frings\|Mass1\|Mgr1\|VLGR1 | G protein-coupled receptor 98 | protein-coding |
| chr13_81347555_81347989 | chr13 | 81347556 | 81347989 | + | -0.795079 | intron (NM_054053, intron 83 of 89) | intron (NM_054053, intron 83 of 89) | 285372 | NM_054053 | 110789 | Mm.288694 | NM_054053 | ENSMUSG00000069170 | **Gpr98** | Frings\|Mass1\|Mgr1\|VLGR1 | G protein-coupled receptor 98 | protein-coding |
| chr6_59324861_59325380 | chr6 | 59324862 | 59325380 | + | -0.779273 | Intergenic | Intergenic | 101169 | NM_183183 | 243385 | Mm.138080 | NM_183183 | ENSMUSG00000045441 | **Gprin3** | C030038J10Rik\|C730021L23 | GPRIN family member 3 | protein-coding |
| chr3_108713084_108713535 | chr3 | 108713085 | 108713535 | + | -0.760316 | intron (NM_029522, intron 1 of 14) | intron (NM_029522, intron 1 of 14) | 8989 | NM_029522 | 76123 | Mm.226941 | NM_029522 | ENSMUSG00000027883 | **Gpsm2** | 6230410J09Rik\|LGN\|Pins | G-protein signalling modulator 2 (AGS3-like, C. elegans) | protein-coding |
| chr9_40397028_40397463 | chr9 | 40397029 | 40397463 | + | -0.766902 | intron (NM_172768, intron 2 of 19) | (CAG)n\|Simple_repeat\|Simple_repeat | 58518 | NM_172768 | 235283 | Mm.138434 | NM_172768 | ENSMUSG00000040111 | **Gramd1b** | 3222402H23\|A930008A22Rik\|AI593249\|mKIAA1201 | GRAM domain containing 1B | protein-coding |
| chr18_56473267_56473479 | chr18 | 56473268 | 56473479 | + | -0.661109 | intron (NM_026240, intron 2 of 13) | intron (NM_026240, intron 2 of 13) | 41241 | NM_026240 | 107022 | Mm.24356 | NM_026240 | ENSMUSG00000001700 | **Gramd3** | 9030613F08Rik\|9130427A09Rik\|AW061306\|NS3TP2 | GRAM domain containing 3 | protein-coding |
| chr18_56455542_56456100 | chr18 | 56455543 | 56456100 | + | -0.663904 | intron (NM_026240, intron 1 of 13) | intron (NM_026240, intron 1 of 13) | 23689 | NM_026240 | 107022 | Mm.24356 | NM_026240 | ENSMUSG00000001700 | **Gramd3** | 9030613F08Rik\|9130427A09Rik\|AW061306\|NS3TP2 | GRAM domain containing 3 | protein-coding |
| chr18_56373914_56374291 | chr18 | 56373915 | 56374291 | + | -0.732989 | Intergenic | RLTR14\|LTR\|ERV1 | -58029 | NM_026240 | 107022 | Mm.24356 | NM_026240 | ENSMUSG00000001700 | **Gramd3** | 9030613F08Rik\|9130427A09Rik\|AW061306\|NS3TP2 | GRAM domain containing 3 | protein-coding |
| chr18_56431163_56431449 | chr18 | 56431164 | 56431449 | + | -0.754382 | promoter-TSS (NM_026240) | promoter-TSS (NM_026240) | -826 | NM_026240 | 107022 | Mm.24356 | NM_026240 | ENSMUSG00000001700 | **Gramd3** | 9030613F08Rik\|9130427A09Rik\|AW061306\|NS3TP2 | GRAM domain containing 3 | protein-coding |
| chr18_56469006_56469718 | chr18 | 56469007 | 56469718 | + | -0.876815 | intron (NM_026240, intron 1 of 13) | intron (NM_026240, intron 1 of 13) | 37230 | NM_026240 | 107022 | Mm.24356 | NM_026240 | ENSMUSG00000001700 | **Gramd3** | 9030613F08Rik\|9130427A09Rik\|AW061306\|NS3TP2 | GRAM domain containing 3 | protein-coding |

| | | | | | | | | | | | | | | | | | |
|---|---|---|---|---|---|---|---|---|---|---|---|---|---|---|---|---|---|
| chr18_56440912_56441389 | chr18 | 56440913 | 56441389 | + | -0.893519 | intron (NM_026240, intron 1 of 13) | intron (NM_026240, intron 1 of 13) | 9019 | NM_026240 | 107022 | Mm.24356 | NM_026240 | ENSMUSG00000001700 | **Gramd3** | 9030613F08Rik\|9130427A09Rik\|AW061306\|NS3TP2 | GRAM domain containing 3 | protein-coding |
| chr18_56468961_56469469 | chr18 | 56468962 | 56469469 | + | -0.900167 | intron (NM_026240, intron 1 of 13) | intron (NM_026240, intron 1 of 13) | 37083 | NM_026240 | 107022 | Mm.24356 | NM_026240 | ENSMUSG00000001700 | **Gramd3** | 9030613F08Rik\|9130427A09Rik\|AW061306\|NS3TP2 | GRAM domain containing 3 | protein-coding |
| chr18_56451780_56452268 | chr18 | 56451781 | 56452268 | + | -0.959521 | intron (NM_026240, intron 1 of 13) | intron (NM_026240, intron 1 of 13) | 19892 | NM_026240 | 107022 | Mm.24356 | NM_026240 | ENSMUSG00000001700 | **Gramd3** | 9030613F08Rik\|9130427A09Rik\|AW061306\|NS3TP2 | GRAM domain containing 3 | protein-coding |
| chr18_56437233_56437516 | chr18 | 56437234 | 56437516 | + | -1.11296 | intron (NM_026240, intron 1 of 13) | intron (NM_026240, intron 1 of 13) | 5243 | NM_026240 | 107022 | Mm.24356 | NM_026240 | ENSMUSG00000001700 | **Gramd3** | 9030613F08Rik\|9130427A09Rik\|AW061306\|NS3TP2 | GRAM domain containing 3 | protein-coding |
| chr18_56446887_56447540 | chr18 | 56446888 | 56447540 | + | -1.14527 | intron (NM_026240, intron 1 of 13) | intron (NM_026240, intron 1 of 13) | 15082 | NM_026240 | 107022 | Mm.24356 | NM_026240 | ENSMUSG00000001700 | **Gramd3** | 9030613F08Rik\|9130427A09Rik\|AW061306\|NS3TP2 | GRAM domain containing 3 | protein-coding |
| chr18_56412559_56413507 | chr18 | 56412560 | 56413507 | + | -1.33447 | Intergenic | Intergenic | -19099 | NM_026240 | 107022 | Mm.24356 | NM_026240 | ENSMUSG00000001700 | **Gramd3** | 9030613F08Rik\|9130427A09Rik\|AW061306\|NS3TP2 | GRAM domain containing 3 | protein-coding |
| chr18_56412789_56413403 | chr18 | 56412790 | 56413403 | + | -1.35298 | Intergenic | Intergenic | -19036 | NM_026240 | 107022 | Mm.24356 | NM_026240 | ENSMUSG00000001700 | **Gramd3** | 9030613F08Rik\|9130427A09Rik\|AW061306\|NS3TP2 | GRAM domain containing 3 | protein-coding |
| chr15_80602422_80603201 | chr15 | 80602423 | 80603201 | + | -0.638837 | intron (NM_001289442, intron 1 of 7) | intron (NM_001289442, intron 1 of 7) | -20671 | NM_010815 | 17444 | Mm.12947 | NM_010815 | ENSMUSG00000042351 | **Grap2** | GRAP-2\|GRB2L\|GRBLG\|GRID\|Gads\|GrbX\|Grf40\|GrpL\|Mona\|P38 | GRB2-related adaptor protein 2 | protein-coding |
| chr15_80599174_80599828 | chr15 | 80599175 | 80599828 | + | -0.666713 | intron (NM_001289442, intron 1 of 7) | intron (NM_001289442, intron 1 of 7) | -23982 | NM_010815 | 17444 | Mm.12947 | NM_010815 | ENSMUSG00000042351 | **Grap2** | GRAP-2\|GRB2L\|GRBLG\|GRID\|Gads\|GrbX\|Grf40\|GrpL\|Mona\|P38 | GRB2-related adaptor protein 2 | protein-coding |
| chr15_80572203_80572830 | chr15 | 80572204 | 80572830 | + | -0.674979 | promoter-TSS (NM_001289442) | promoter-TSS (NM_001289442) | -80 | NM_001289442 | 17444 | Mm.12947 | NM_010815 | ENSMUSG00000042351 | **Grap2** | GRAP-2\|GRB2L\|GRBLG\|GRID\|Gads\|GrbX\|Grf40\|GrpL\|Mona\|P38 | GRB2-related adaptor protein 2 | protein-coding |
| chr15_80599426_80599803 | chr15 | 80599427 | 80599803 | + | -0.779066 | intron (NM_001289442, intron 1 of 7) | intron (NM_001289442, intron 1 of 7) | -23868 | NM_010815 | 17444 | Mm.12947 | NM_010815 | ENSMUSG00000042351 | **Grap2** | GRAP-2\|GRB2L\|GRBLG\|GRID\|Gads\|GrbX\|Grf40\|GrpL\|Mona\|P38 | GRB2-related adaptor protein 2 | protein-coding |
| chr15_80567122_80567448 | chr15 | 80567123 | 80567448 | + | -0.872061 | Intergenic | Intergenic | -5312 | NM_001289442 | 17444 | Mm.12947 | NM_010815 | ENSMUSG00000042351 | **Grap2** | GRAP-2\|GRB2L\|GRBLG\|GRID\|Gads\|GrbX\|Grf40\|GrpL\|Mona\|P38 | GRB2-related adaptor protein 2 | protein-coding |
| chr15_80630215_80630470 | chr15 | 80630216 | 80630470 | + | -0.875613 | intron (NM_001289442, intron 2 of 7) | intron (NM_001289442, intron 2 of 7) | 6860 | NM_010815 | 17444 | Mm.12947 | NM_010815 | ENSMUSG00000042351 | **Grap2** | GRAP-2\|GRB2L\|GRBLG\|GRID\|Gads\|GrbX\|Grf40\|GrpL\|Mona\|P38 | GRB2-related adaptor protein 2 | protein-coding |
| chr15_80610377_80610677 | chr15 | 80610378 | 80610677 | + | -1.17364 | intron (NM_001289442, intron 1 of 7) | Lx5c\|LINE\|L1 | -12956 | NM_010815 | 17444 | Mm.12947 | NM_010815 | ENSMUSG00000042351 | **Grap2** | GRAP-2\|GRB2L\|GRBLG\|GRID\|Gads\|GrbX\|Grf40\|GrpL\|Mona\|P38 | GRB2-related adaptor protein 2 | protein-coding |
| chr15_80610434_80610612 | chr15 | 80610435 | 80610612 | + | -1.21243 | intron (NM_001289442, intron 1 of 7) | Lx5c\|LINE\|L1 | -12960 | NM_010815 | 17444 | Mm.12947 | NM_010815 | ENSMUSG00000042351 | **Grap2** | GRAP-2\|GRB2L\|GRBLG\|GRID\|Gads\|GrbX\|Grf40\|GrpL\|Mona\|P38 | GRB2-related adaptor protein 2 | protein-coding |
| chr15_80611548_80611910 | chr15 | 80611549 | 80611910 | + | -1.30704 | intron (NM_001289442, intron 1 of 7) | intron (NM_001289442, intron 1 of 7) | -11754 | NM_010815 | 17444 | Mm.12947 | NM_010815 | ENSMUSG00000042351 | **Grap2** | GRAP-2\|GRB2L\|GRBLG\|GRID\|Gads\|GrbX\|Grf40\|GrpL\|Mona\|P38 | GRB2-related adaptor protein 2 | protein-coding |
| chr11_115698961_115700317 | chr11 | 115698962 | 115700317 | + | -0.663591 | intron (NM_008163, intron 1 of 5) | intron (NM_008163, intron 1 of 5) | 8958 | NM_008163 | 14784 | Mm.439649 | NM_008163 | ENSMUSG00000059923 | **Grb2** | AA408164\|Ash | growth factor receptor bound protein 2 | protein-coding |
| chr11_115716724_115717187 | chr11 | 115716725 | 115717187 | + | -0.71449 | Intergenic | Intergenic | -8359 | NM_008163 | 14784 | Mm.439649 | NM_008163 | ENSMUSG00000059923 | **Grb2** | AA408164\|Ash | growth factor receptor bound protein 2 | protein-coding |
| chr11_115698938_115699800 | chr11 | 115698939 | 115699800 | + | -0.766377 | intron (NM_008163, intron 1 of 5) | intron (NM_008163, intron 1 of 5) | 9228 | NM_008163 | 14784 | Mm.439649 | NM_008163 | ENSMUSG00000059923 | **Grb2** | AA408164\|Ash | growth factor receptor bound protein 2 | protein-coding |
| chr11_98487550_98488154 | chr11 | 98487551 | 98488154 | + | -0.644946 | intron (NM_011771, intron 5 of 7) | intron (NM_011771, intron 5 of 7) | 41018 | NM_010346 | 14786 | Mm.276702 | NM_010346 | ENSMUSG00000019312 | **Grb7** | mKIAA4028 | growth factor receptor bound protein 7 | protein-coding |
| chr11_98464565_98464868 | chr11 | 98464566 | 98464868 | + | -0.690743 | TTS (NM_011771) | TTS (NM_011771) | 17883 | NM_010346 | 14786 | Mm.276702 | NM_010346 | ENSMUSG00000019312 | **Grb7** | mKIAA4028 | growth factor receptor bound protein 7 | protein-coding |
| chrX_41400016_41400933 | chrX | 41400017 | 41400933 | + | -1.2098 | promoter-TSS (NM_001281929) | promoter-TSS (NM_001281929) | -826 | NM_016886 | 53623 | Mm.327681 | NM_016886 | ENSMUSG00000001986 | **Gria3** | 2900064I19Rik\|GluA3\|GluR-C\|GluR-K3\|Glur-3\|Glur3\|Gluralpha3 | glutamate receptor, ionotropic, AMPA3 (alpha 3) | protein-coding |
| chr5_140575471_140576020 | chr5 | 140575472 | 140576020 | + | -0.82241 | Intergenic | Intergenic | -10679 | NM_030022 | 77998 | Mm.160040 | NM_030022 | ENSMUSG00000036586 | **Grifin** | E130101O17Rik | galectin-related inter-fiber protein | protein-coding |
| chr5_140575655_140575871 | chr5 | 140575656 | 140575871 | + | -0.854102 | Intergenic | Intergenic | -10696 | NM_030022 | 77998 | Mm.160040 | NM_030022 | ENSMUSG00000036586 | **Grifin** | E130101O17Rik | galectin-related inter-fiber protein | protein-coding |
| chr13_55443877_55444369 | chr13 | 55443878 | 55444369 | + | -0.841763 | promoter-TSS (NM_001286063) | promoter-TSS (NM_001286063) | -949 | NM_001286063 | 26385 | Mm.10193 | NM_011938 | ENSMUSG00000074886 | **Grk6** | Gprk6 | G protein-coupled receptor kinase 6 | protein-coding |
| chr13_55444147_55444326 | chr13 | 55444148 | 55444326 | + | -0.905915 | promoter-TSS (NM_001286063) | promoter-TSS (NM_001286063) | -835 | NM_001286063 | 26385 | Mm.10193 | NM_011938 | ENSMUSG00000074886 | **Grk6** | Gprk6 | G protein-coupled receptor kinase 6 | protein-coding |
| chr13_55437260_55437547 | chr13 | 55437261 | 55437547 | + | -1.0043 | Intergenic | IAPEY5_I-int\|LTR\|ERVK | -7668 | NM_001286063 | 26385 | Mm.10193 | NM_011938 | ENSMUSG00000074886 | **Grk6** | Gprk6 | G protein-coupled receptor kinase 6 | protein-coding |
| chr5_21213865_21214410 | chr5 | 21213866 | 21214410 | + | -0.652701 | intron (NM_175437, intron 5 of 30) | intron (NM_175437, intron 5 of 30) | 27871 | NM_175437 | 212167 | Mm.121705 | NM_175437 | ENSMUSG00000039934 | **Gsap** | A530088I07Rik\|Pion | gamma-secretase activating protein | protein-coding |
| chr8_120497672_120498047 | chr8 | 120497673 | 120498047 | + | -0.739879 | intron (NM_198671, intron 1 of 15) | intron (NM_198671, intron 1 of 15) | 8994 | NM_198671 | 382034 | Mm.334856 | NM_198671 | ENSMUSG00000031822 | **Gse1** | 2210013I18Rik\|mKIAA0182 | genetic suppressor element 1 | protein-coding |
| chr2_35259697_35260071 | chr2 | 35259698 | 35260071 | + | -0.66798 | intron (NM_001206368, intron 1 of 16) | intron (NM_001206368, intron 1 of 16) | 3525 | NM_001206368 | 227753 | Mm.21109 | NM_146120 | ENSMUSG00000026879 | **Gsn** | ADF | gelsolin | protein-coding |
| chr19_47888145_47888375 | chr19 | 47888146 | 47888375 | + | -0.707349 | Intergenic | B3\|SINE\|B2 | 22466 | NM_026619 | 68214 | Mm.63791 | NM_026619 | ENSMUSG00000025069 | **Gsto2** | 1700020F09Rik\|4930425C18Rik\|GSTO 2-2 | glutathione S-transferase omega 2 | protein-coding |
| chr19_47887959_47888363 | chr19 | 47887960 | 47888363 | + | -0.767259 | Intergenic | B3\|SINE\|B2 | 22367 | NM_026619 | 68214 | Mm.63791 | NM_026619 | ENSMUSG00000025069 | **Gsto2** | 1700020F09Rik\|4930425C18Rik\|GSTO 2-2 | glutathione S-transferase omega 2 | protein-coding |
| chr10_75836002_75836323 | chr10 | 75836003 | 75836323 | + | -0.682488 | Intergenic | ORR1E\|LTR\|ERVL-MaLR | -1282 | NM_010361 | 14872 | Mm.24118 | NM_010361 | ENSMUSG00000033318 | **Gstt2** | AI266894\|Yrs\|mGSTT2 | glutathione S-transferase, theta 2 | protein-coding |
| chr12_91523582_91524256 | chr12 | 91523583 | 91524256 | + | -0.873642 | intron (NM_011648, intron 8 of 9) | intron (NM_011648, intron 8 of 9) | 65730 | NM_031391 | 83602 | Mm.275728 | NM_031391 | ENSMUSG00000020962 | **Gtf2a1** | 6330549H03Rik\|AA536742\|AA959775\|AW060250\|TfIIAa/b\|Tfiia1 | general transcription factor II A, 1 | protein-coding |
| chr5_134272749_134273015 | chr5 | 134272750 | 134273015 | + | -0.695455 | intron (NM_010365, intron 10 of 33) | intron (NM_010365, intron 10 of 33) | 41878 | NM_001080748 | 14886 | Mm.261570 | NM_010365 | ENSMUSG00000060261 | **Gtf2i** | 6030441I21Rik\|BAP-135\|Diws1t\|GtfII-I\|Spin\|TFII-I\|WBSCR6 | general transcription factor II I | protein-coding |
| chr5_134282510_134283205 | chr5 | 134282511 | 134283205 | + | -0.73482 | intron (NM_001080749, intron 7 of 32) | intron (NM_001080749, intron 7 of 32) | 31902 | NM_001080748 | 14886 | Mm.261570 | NM_010365 | ENSMUSG00000060261 | **Gtf2i** | 6030441I21Rik\|BAP-135\|Diws1t\|GtfII-I\|Spin\|TFII-I\|WBSCR6 | general transcription factor II I | protein-coding |
| chr5_134275566_134275923 | chr5 | 134275567 | 134275923 | + | -0.82792 | intron (NM_001080749, intron 9 of 32) | intron (NM_001080749, intron 9 of 32) | 39015 | NM_001080748 | 14886 | Mm.261570 | NM_010365 | ENSMUSG00000060261 | **Gtf2i** | 6030441I21Rik\|BAP-135\|Diws1t\|GtfII-I\|Spin\|TFII-I\|WBSCR6 | general transcription factor II I | protein-coding |
| chr5_134370480_134371318 | chr5 | 134370481 | 134371318 | + | -0.839203 | intron (NM_001081467, intron 22 of 26) | intron (NM_001081467, intron 22 of 26) | -56139 | NM_010365 | 14886 | Mm.261570 | NM_010365 | ENSMUSG00000060261 | **Gtf2i** | 6030441I21Rik\|BAP-135\|Diws1t\|GtfII-I\|Spin\|TFII-I\|WBSCR6 | general transcription factor II I | protein-coding |
| chr5_134481995_134482210 | chr5 | 134481996 | 134482210 | + | -0.743299 | Intergenic | L1MB8\|LINE\|L1 | -25387 | NM_001081464 | 57080 | Mm.332735 | NM_020331 | ENSMUSG00000023079 | **Gtf2ird1** | 1700012P16Rik\|BEN\|Cream1\|ESTM9\|GTF3\|Gtf2il\|MusTRD1\|Tg(Alb1-Myc)166.8Sst\|WBSCR11\|X83320 | general transcription factor II I repeat domain-containing 1 | protein-coding |
| chr5_134482263_134482608 | chr5 | 134482264 | 134482608 | + | -0.802797 | Intergenic | Intergenic | -25720 | NM_001081464 | 57080 | Mm.332735 | NM_020331 | ENSMUSG00000023079 | **Gtf2ird1** | 1700012P16Rik\|BEN\|Cream1\|ESTM9\|GTF3\|Gtf2il\|MusTRD1\|Tg(Alb1-Myc)166.8Sst\|WBSCR11\|X83320 | general transcription factor II I repeat domain-containing 1 | protein-coding |
| chr5_134481933_134482727 | chr5 | 134481934 | 134482727 | + | -0.897203 | Intergenic | Intergenic | -25614 | NM_001081464 | 57080 | Mm.332735 | NM_020331 | ENSMUSG00000023079 | **Gtf2ird1** | 1700012P16Rik\|BEN\|Cream1\|ESTM9\|GTF3\|Gtf2il\|MusTRD1\|Tg(Alb1-Myc)166.8Sst\|WBSCR11\|X83320 | general transcription factor II I repeat domain-containing 1 | protein-coding |
| chr5_134188646_134188875 | chr5 | 134188647 | 134188875 | + | -0.796279 | intron (NM_053266, intron 1 of 15) | intron (NM_053266, intron 1 of 15) | 4723 | NM_053266 | 114674 | Mm.218744 | NM_053266 | ENSMUSG00000015942 | **Gtf2ird2** | 1700012P16Rik | GTF2I repeat domain containing 2 | protein-coding |

| | | | | | | | | | | | | | | | | | |
|---|---|---|---|---|---|---|---|---|---|---|---|---|---|---|---|---|---|
| chr1_54422252_54422533 | chr1 | 54422253 | 54422533 | + | -0.682281 | intron (NM_001033194, intron 8 of 17) | intron (NM_001033194, intron 8 of 17) | 16578 | NM_001033194 | 98488 | Mm.25856 | NM_001033194 | ENSMUSG00000041303 | **Gtf3c3** | AL022818\|TF3C-gamma\|TFIIIC102 | general transcription factor IIIC, polypeptide 3 | protein-coding |
| chr1_54492366_54492972 | chr1 | 54492367 | 54492972 | + | -1.06026 | intron (NM_001163314, intron 22 of 26) | intron (NM_001163314, intron 22 of 26) | -53698 | NM_001033194 | 98488 | Mm.25856 | NM_001033194 | ENSMUSG00000041303 | **Gtf3c3** | AL022818\|TF3C-gamma\|TFIIIC102 | general transcription factor IIIC, polypeptide 3 | protein-coding |
| chr1_54493536_54493867 | chr1 | 54493537 | 54493867 | + | -1.08415 | intron (NM_001163314, intron 21 of 26) | intron (NM_001163314, intron 21 of 26) | -54731 | NM_001033194 | 98488 | Mm.25856 | NM_001033194 | ENSMUSG00000041303 | **Gtf3c3** | AL022818\|TF3C-gamma\|TFIIIC102 | general transcription factor IIIC, polypeptide 3 | protein-coding |
| chr1_54421702_54422830 | chr1 | 54421703 | 54422830 | + | -1.24213 | intron (NM_001033194, intron 8 of 17) | ID_B1\|SINE\|B4 | 16705 | NM_001033194 | 98488 | Mm.25856 | NM_001033194 | ENSMUSG00000041303 | **Gtf3c3** | AL022818\|TF3C-gamma\|TFIIIC102 | general transcription factor IIIC, polypeptide 3 | protein-coding |
| chr1_54421664_54422499 | chr1 | 54421665 | 54422499 | + | -1.3344 | intron (NM_001033194, intron 8 of 17) | intron (NM_001033194, intron 8 of 17) | 16889 | NM_001033194 | 98488 | Mm.25856 | NM_001033194 | ENSMUSG00000041303 | **Gtf3c3** | AL022818\|TF3C-gamma\|TFIIIC102 | general transcription factor IIIC, polypeptide 3 | protein-coding |
| chr1_54421678_54422207 | chr1 | 54421679 | 54422207 | + | -1.42808 | intron (NM_001033194, intron 8 of 17) | intron (NM_001033194, intron 8 of 17) | 17028 | NM_001033194 | 98488 | Mm.25856 | NM_001033194 | ENSMUSG00000041303 | **Gtf3c3** | AL022818\|TF3C-gamma\|TFIIIC102 | general transcription factor IIIC, polypeptide 3 | protein-coding |
| chr15_79693767_79694236 | chr15 | 79693768 | 79694236 | + | -0.738513 | intron (NM_013818, intron 1 of 11) | intron (NM_013818, intron 1 of 11) | 3106 | NM_013818 | 14904 | Mm.19080 | NM_013818 | ENSMUSG00000042535 | **Gtpbp1** | AL022987\|AW045683\|C85212\|GP-1\|Gtpbp | GTP binding protein 1 | protein-coding |
| chr15_79693745_79694080 | chr15 | 79693746 | 79694080 | + | -0.772577 | intron (NM_013818, intron 1 of 11) | intron (NM_013818, intron 1 of 11) | 3017 | NM_013818 | 14904 | Mm.19080 | NM_013818 | ENSMUSG00000042535 | **Gtpbp1** | AL022987\|AW045683\|C85212\|GP-1\|Gtpbp | GTP binding protein 1 | protein-coding |
| chr17_46172486_46173179 | chr17 | 46172487 | 46173179 | + | -0.867615 | exon (NM_030715, exon 11 of 11) | exon (NM_030715, exon 11 of 11) | 11801 | NM_001145979 | 56055 | Mm.22147 | NM_019581 | ENSMUSG00000023952 | **Gtpbp2** | nmf205 | GTP binding protein 2 | protein-coding |
| chr13_9030413_9030838 | chr13 | 9030414 | 9030838 | + | -0.890309 | Intergenic | Lx2\|LINE\|L1 | -34614 | NM_027000 | 69237 | Mm.41800 | NM_027000 | ENSMUSG00000021149 | **Gtpbp4** | 2610028C09Rik\|Crfg\|Gtpbp3\|NGB\|Nog1 | GTP binding protein 4 | protein-coding |
| chr13_9030431_9030746 | chr13 | 9030432 | 9030746 | + | -0.915383 | Intergenic | Lx2\|LINE\|L1 | -34577 | NM_027000 | 69237 | Mm.41800 | NM_027000 | ENSMUSG00000021149 | **Gtpbp4** | 2610028C09Rik\|Crfg\|Gtpbp3\|NGB\|Nog1 | GTP binding protein 4 | protein-coding |
| chr16_44735665_44736416 | chr16 | 44735666 | 44736416 | + | -1.06265 | exon (NM_145972, exon 9 of 9) | exon (NM_145972, exon 9 of 9) | 10322 | NM_001159329 | 66067 | Mm.34520 | NM_025332 | | **Gtpbp8** | 0610037H22Rik | GTP-binding protein 8 (putative) | protein-coding |
| chr16_44735707_44736070 | chr16 | 44735708 | 44736070 | + | -1.10974 | exon (NM_145972, exon 9 of 9) | exon (NM_145972, exon 9 of 9) | 10474 | NM_001159329 | 66067 | Mm.34520 | NM_025332 | | **Gtpbp8** | 0610037H22Rik | GTP-binding protein 8 (putative) | protein-coding |
| chr6_136745628_136746027 | chr6 | 136745629 | 136746027 | + | -0.795752 | intron (NM_145067, intron 9 of 25) | MTC\|LTR\|ERVL-MaLR | 35914 | NM_145067 | 14917 | Mm.222729 | NM_145067 | ENSMUSG00000042638 | **Gucy2c** | AI893437\|GC-C\|Gcc | guanylate cyclase 2c | protein-coding |
| chr6_136735728_136736202 | chr6 | 136735729 | 136736202 | + | -0.826246 | intron (NM_145067, intron 12 of 25) | ORR1E\|LTR\|ERVL-MaLR | 45777 | NM_145067 | 14917 | Mm.222729 | NM_145067 | ENSMUSG00000042638 | **Gucy2c** | AI893437\|GC-C\|Gcc | guanylate cyclase 2c | protein-coding |
| chr6_136742308_136742630 | chr6 | 136742309 | 136742630 | + | -1.30809 | intron (NM_145067, intron 10 of 25) | intron (NM_145067, intron 10 of 25) | 39273 | NM_145067 | 14917 | Mm.222729 | NM_145067 | ENSMUSG00000042638 | **Gucy2c** | AI893437\|GC-C\|Gcc | guanylate cyclase 2c | protein-coding |
| chr6_136742252_136742694 | chr6 | 136742253 | 136742694 | + | -1.32817 | intron (NM_145067, intron 10 of 25) | intron (NM_145067, intron 10 of 25) | 39269 | NM_145067 | 14917 | Mm.222729 | NM_145067 | ENSMUSG00000042638 | **Gucy2c** | AI893437\|GC-C\|Gcc | guanylate cyclase 2c | protein-coding |
| chr6_136743701_136744184 | chr6 | 136743702 | 136744184 | + | -1.40206 | exon (NM_001127318, exon 10 of 27) | exon (NM_001127318, exon 10 of 27) | 37799 | NM_145067 | 14917 | Mm.222729 | NM_145067 | ENSMUSG00000042638 | **Gucy2c** | AI893437\|GC-C\|Gcc | guanylate cyclase 2c | protein-coding |
| chr11_69275511_69275984 | chr11 | 69275512 | 69275984 | + | -0.887791 | Intergenic | Intergenic | -38726 | NM_008192 | 14919 | Mm.291816 | NM_008192 | ENSMUSG00000020890 | **Gucy2e** | GC-E\|GC1\|ROS-GC1 | guanylate cyclase 2e | protein-coding |
| chr15_93298220_93298586 | chr15 | 93298221 | 93298586 | + | -0.749224 | intron (NM_024189, intron 2 of 3) | intron (NM_024189, intron 2 of 3) | -23319 | NM_001033275 | 223827 | Mm.22819 | NM_001033275 | ENSMUSG00000036197 | **Gxylt1** | Glt8d3\|Gm1228\|Gm87 | glucoside xylosyltransferase 1 | protein-coding |
| chr8_80393201_80393434 | chr8 | 80393202 | 80393434 | + | -0.905883 | Intergenic | Intergenic | -100727 | NM_010369 | 14934 | Mm.13123 | NM_010369 | ENSMUSG00000051839 | **Gypa** | AI853584\|CD235a\|GPA | glycophorin A | protein-coding |
| chr8_80393097_80393765 | chr8 | 80393098 | 80393765 | + | -0.947272 | Intergenic | Intergenic | -100614 | NM_010369 | 14934 | Mm.13123 | NM_010369 | ENSMUSG00000051839 | **Gypa** | AI853584\|CD235a\|GPA | glycophorin A | protein-coding |
| chr18_32591217_32591572 | chr18 | 32591218 | 32591572 | + | -0.724864 | Intergenic | Intergenic | -31361 | NM_001048207 | 71683 | Mm.292145 | NM_027863 | ENSMUSG00000090523 | **Gypc** | 0610037F22Rik\|Cd236r | glycophorin C | protein-coding |
| chr18_32597951_32598636 | chr18 | 32597952 | 32598636 | + | -1.26806 | Intergenic | B4A\|SINE\|B4 | -38260 | NM_001048207 | 71683 | Mm.292145 | NM_027863 | ENSMUSG00000090523 | **Gypc** | 0610037F22Rik\|Cd236r | glycophorin C | protein-coding |
| chr18_32598045_32598464 | chr18 | 32598046 | 32598464 | + | -1.28677 | Intergenic | Intergenic | -38221 | NM_001048207 | 71683 | Mm.292145 | NM_027863 | ENSMUSG00000090523 | **Gypc** | 0610037F22Rik\|Cd236r | glycophorin C | protein-coding |
| chr6_142453130_142453670 | chr6 | 142453131 | 142453670 | + | -0.83971 | intron (NM_145572, intron 7 of 16) | intron (NM_145572, intron 7 of 16) | 19709 | NM_145572 | 232493 | Mm.275975 | NM_145572 | ENSMUSG00000030244 | **Gys2** | BC021322\|LGS | glycogen synthase 2 | protein-coding |
| chr13_113087645_113087972 | chr13 | 113087646 | 113087972 | + | -0.708785 | intron (NM_001281487, intron 11 of 11) | intron (NM_001281487, intron 11 of 11) | 13172 | NM_010370 | 14938 | Mm.15510 | NM_010370 | ENSMUSG00000023132 | **Gzma** | AW494114\|Ctla-3\|Ctla3\|Hf\|SE1\|TSP-1\|TSP1 | granzyme A | protein-coding |
| chr17_34127551_34127816 | chr17 | 34127552 | 34127816 | + | -0.785542 | intron (NM_010386, intron 1 of 5) | intron (NM_010386, intron 1 of 5) | 4852 | NM_010386 | 14998 | Mm.16373 | NM_010386 | ENSMUSG00000037649 | **H2-DMa** | H-2Ma\|H2-Ma | histocompatibility 2, class II, locus DMa | protein-coding |
| chr17_34329663_34330310 | chr17 | 34329664 | 34330310 | + | -0.729498 | intron (NM_001033978, intron 1 of 4) | intron (NM_001033978, intron 1 of 4) | 4312 | NM_001033978 | 381091 | Mm.139204 | NM_001033978 | | **H2-Eb2** | A130038H09Rik\|H-2Eb2\|Ia-5\|Ia5\|Iebeta2 | histocompatibility 2, class II antigen E beta2 | protein-coding |
| chr17_34070212_34070454 | chr17 | 34070213 | 34070454 | + | -0.737193 | Intergenic | Intergenic | -22046 | NM_008206 | 15001 | Mm.116 | NM_008206 | ENSMUSG00000024334 | **H2-Oa** | H-2Oa | histocompatibility 2, O region alpha locus | protein-coding |
| chr17_34239429_34240003 | chr17 | 34239430 | 34240003 | + | -1.17692 | intron (NM_010389, intron 1 of 5) | intron (NM_010389, intron 1 of 5) | 811 | NM_010389 | 15002 | Mm.327358 | NM_010389 | ENSMUSG00000041538 | **H2-Ob** | A-beta-2\|A-beta2\|H-2Ab\|H-2I\|H-2Ob\|H2-Ab\|H2-Ab2\|H2-IAb2\|Ob | histocompatibility 2, O region beta locus | protein-coding |
| chr17_34239698_34239955 | chr17 | 34239699 | 34239955 | + | -1.27531 | intron (NM_010389, intron 1 of 5) | intron (NM_010389, intron 1 of 5) | 922 | NM_010389 | 15002 | Mm.327358 | NM_010389 | ENSMUSG00000041538 | **H2-Ob** | A-beta-2\|A-beta2\|H-2Ab\|H-2I\|H-2Ob\|H2-Ab\|H2-Ab2\|H2-IAb2\|Ob | histocompatibility 2, O region beta locus | protein-coding |
| chr17_35353337_35353582 | chr17 | 35353338 | 35353582 | + | -0.931639 | Intergenic | MYSERV6-int\|LTR\|ERVK | 11127 | NM_010392 | 15013 | Mm.427969 | NM_010392 | ENSMUSG00000091705 | **H2-Q2** | BE136769\|ENSMUSG00000079507\|Gm11132\|H-2Q2\|gs14-2 | histocompatibility 2, Q region locus 2 | protein-coding |
| chr15_62164626_62165188 | chr15 | 62164627 | 62165188 | + | -1.11417 | intron (NR_003368, intron 4 of 8) | RLTR51A_Mm\|LTR\|ERVK? | 54544 | NR_003523 | 67552 | Mm.484027 | NM_026230 | | **H2afy3** | 4933432H23Rik | H2A histone family, member Y3 | pseudo |
| chr17_83918789_83919111 | chr17 | 83918790 | 83919111 | + | -0.641395 | Intergenic | Intergenic | -72160 | NM_025325 | 107766 | Mm.30100 | NM_025325 | ENSMUSG00000000673 | **Haao** | 0610007K21Rik\|0610012J07Rik\|3-HAO\|3-HAOxase\|3HAO | 3-hydroxyanthranilate 3,4-dioxygenase | protein-coding |
| chr17_83902530_83903039 | chr17 | 83902531 | 83903039 | + | -0.720808 | Intergenic | Intergenic | -55995 | NM_025325 | 107766 | Mm.30100 | NM_025325 | ENSMUSG00000000673 | **Haao** | 0610007K21Rik\|0610012J07Rik\|3-HAO\|3-HAOxase\|3HAO | 3-hydroxyanthranilate 3,4-dioxygenase | protein-coding |
| chr17_83935239_83935546 | chr17 | 83935240 | 83935546 | + | -0.90605 | Intergenic | Intergenic | -88603 | NM_025325 | 107766 | Mm.30100 | NM_025325 | ENSMUSG00000000673 | **Haao** | 0610007K21Rik\|0610012J07Rik\|3-HAO\|3-HAOxase\|3HAO | 3-hydroxyanthranilate 3,4-dioxygenase | protein-coding |
| chr17_83935239_83935706 | chr17 | 83935240 | 83935706 | + | -0.94421 | Intergenic | Intergenic | -88683 | NM_025325 | 107766 | Mm.30100 | NM_025325 | ENSMUSG00000000673 | **Haao** | 0610007K21Rik\|0610012J07Rik\|3-HAO\|3-HAOxase\|3HAO | 3-hydroxyanthranilate 3,4-dioxygenase | protein-coding |
| chr17_83799407_83800232 | chr17 | 83799408 | 83800232 | + | -1.03108 | intron (NM_054082, intron 14 of 16) | intron (NM_054082, intron 14 of 16) | 46970 | NM_025325 | 107766 | Mm.30100 | NM_025325 | ENSMUSG00000000673 | **Haao** | 0610007K21Rik\|0610012J07Rik\|3-HAO\|3-HAOxase\|3HAO | 3-hydroxyanthranilate 3,4-dioxygenase | protein-coding |
| chr3_131199548_131200307 | chr3 | 131199549 | 131200307 | + | -0.672955 | intron (NM_001276402, intron 7 of 9) | intron (NM_001276402, intron 7 of 9) | 72173 | NM_008212 | 15107 | Mm.260164 | NM_008212 | ENSMUSG00000027984 | **Hadh** | AA409008\|AU019341\|AW742602\|HCDH\|Hadhsc\|Schad | hydroxyacyl-Coenzyme A dehydrogenase | protein-coding |
| chr7_30940090_30940409 | chr7 | 30940091 | 30940409 | + | -0.672342 | Intergenic | Intergenic | 3767 | NM_032541 | 84506 | Mm.439939 | NM_032541 | ENSMUSG00000050440 | **Hamp** | Hamp1\|Hepc\|Hepc1 | hepcidin antimicrobial peptide | protein-coding |
| chr7_30940118_30940705 | chr7 | 30940119 | 30940705 | + | -0.835269 | Intergenic | Intergenic | 3605 | NM_032541 | 84506 | Mm.439939 | NM_032541 | ENSMUSG00000050440 | **Hamp** | Hamp1\|Hepc\|Hepc1 | hepcidin antimicrobial peptide | protein-coding |
| chr13_89401057_89401863 | chr13 | 89401058 | 89401863 | + | -1.08336 | Intergenic | Intergenic | -139176 | NM_013500 | 12950 | Mm.266790 | NM_013500 | ENSMUSG00000021613 | **Hapln1** | BB099155\|CLP\|Crtl1\|Crtl1l\|LP\|LP-1 | hyaluronan and proteoglycan link protein 1 | protein-coding |
| chr11_46448145_46448516 | chr11 | 46448146 | 46448516 | + | -1.12725 | intron (NR_105955, intron 2 of 4) | L1MC5a\|LINE\|L1 | -6600 | NM_134250 | 171285 | Mm.72168 | NM_134250 | ENSMUSG00000020399 | **Havcr2** | TIM-3\|Tim3\|Timd3 | hepatitis A virus cellular receptor 2 | protein-coding |
| chr12_31950006_31950951 | chr12 | 31950007 | 31950951 | + | -0.666042 | promoter-TSS (NM_177993) | promoter-TSS (NM_177993) | 56 | NM_177993 | 73389 | Mm.390461 | NM_153198 | ENSMUSG00000002996 | **Hbp1** | 1700058O05Rik\|BE629963\|C330012F01Rik\|C86454 | high mobility group box transcription factor 1 | protein-coding |
| chr10_21254618_21255194 | chr10 | 21254619 | 21255194 | + | -0.66517 | Intergenic | Intergenic | -41073 | NM_001042593 | 56422 | Mm.25527 | NM_019702 | ENSMUSG00000019977 | **Hbs1l** | 2810035F15Rik\|AI326327\|eRFS | Hbs1-like (S. cerevisiae) | protein-coding |
| chr10_21235208_21235590 | chr10 | 21235209 | 21235590 | + | -0.688143 | Intergenic | Intergenic | -60580 | NM_001042593 | 56422 | Mm.25527 | NM_019702 | ENSMUSG00000019977 | **Hbs1l** | 2810035F15Rik\|AI326327\|eRFS | Hbs1-like (S. cerevisiae) | protein-coding |
| chr10_21235176_21235815 | chr10 | 21235177 | 21235815 | + | -0.731456 | Intergenic | Intergenic | -60483 | NM_001042593 | 56422 | Mm.25527 | NM_019702 | ENSMUSG00000019977 | **Hbs1l** | 2810035F15Rik\|AI326327\|eRFS | Hbs1-like (S. cerevisiae) | protein-coding |
| chr1_92178145_92178640 | chr1 | 92178146 | 92178640 | + | -0.614887 | Intergenic | Intergenic | -30000 | NM_207225 | 208727 | Mm.318567 | NM_207225 | ENSMUSG00000026313 | **Hdac4** | 4932408F19Rik\|AI047285 | histone deacetylase 4 | protein-coding |
| chr1_91984397_91984913 | chr1 | 91984398 | 91984913 | + | -0.633628 | intron (NM_207225, intron 9 of 25) | intron (NM_207225, intron 9 of 25) | 163738 | NM_207225 | 208727 | Mm.318567 | NM_207225 | ENSMUSG00000026313 | **Hdac4** | 4932408F19Rik\|AI047285 | histone deacetylase 4 | protein-coding |
| chr1_92058749_92059120 | chr1 | 92058750 | 92059120 | + | -0.760381 | intron (NM_207225, intron 1 of 25) | L1M5\|LINE\|L1 | 89458 | NM_207225 | 208727 | Mm.318567 | NM_207225 | ENSMUSG00000026313 | **Hdac4** | 4932408F19Rik\|AI047285 | histone deacetylase 4 | protein-coding |
| chr1_92006688_92007058 | chr1 | 92006689 | 92007058 | + | -0.795678 | intron (NM_207225, intron 4 of 25) | intron (NM_207225, intron 4 of 25) | 141520 | NM_207225 | 208727 | Mm.318567 | NM_207225 | ENSMUSG00000026313 | **Hdac4** | 4932408F19Rik\|AI047285 | histone deacetylase 4 | protein-coding |

| | | | | | | | | | | | | | | | | |
|---|---|---|---|---|---|---|---|---|---|---|---|---|---|---|---|---|
| chr1_92130393_92131017 | chr1 | 92130394 | 92131017 | + | -0.852883 | intron (NM_207225, intron 1 of 25) | intron (NM_207225, intron 1 of 25) | 17688 | NM_207225 | 208727 | Mm.318567 | NM_207225 | ENSMUSG00000026313 | **Hdac4** | 4932408F19Rik\|AI047285 | histone deacetylase 4 | protein-coding |
| chr1_92130097_92131018 | chr1 | 92130098 | 92131018 | + | -0.860605 | intron (NM_207225, intron 1 of 25) | intron (NM_207225, intron 1 of 25) | 17835 | NM_207225 | 208727 | Mm.318567 | NM_207225 | ENSMUSG00000026313 | **Hdac4** | 4932408F19Rik\|AI047285 | histone deacetylase 4 | protein-coding |
| chr1_92130393_92130837 | chr1 | 92130394 | 92130837 | + | -0.907249 | intron (NM_207225, intron 1 of 25) | intron (NM_207225, intron 1 of 25) | 17778 | NM_207225 | 208727 | Mm.318567 | NM_207225 | ENSMUSG00000026313 | **Hdac4** | 4932408F19Rik\|AI047285 | histone deacetylase 4 | protein-coding |
| chr1_92069287_92069865 | chr1 | 92069288 | 92069865 | + | -1.03893 | intron (NM_207225, intron 1 of 25) | intron (NM_207225, intron 1 of 25) | 78817 | NM_207225 | 208727 | Mm.318567 | NM_207225 | ENSMUSG00000026313 | **Hdac4** | 4932408F19Rik\|AI047285 | histone deacetylase 4 | protein-coding |
| chr1_92235577_92235975 | chr1 | 92235578 | 92235975 | + | -1.10483 | Intergenic | Intergenic | -87383 | NM_207225 | 208727 | Mm.318567 | NM_207225 | ENSMUSG00000026313 | **Hdac4** | 4932408F19Rik\|AI047285 | histone deacetylase 4 | protein-coding |
| chr1_92148460_92148902 | chr1 | 92148461 | 92148902 | + | -1.2288 | promoter-TSS (NM_207225) | promoter-TSS (NM_207225) | -288 | NM_207225 | 208727 | Mm.318567 | NM_207225 | ENSMUSG00000026313 | **Hdac4** | 4932408F19Rik\|AI047285 | histone deacetylase 4 | protein-coding |
| chr1_92148436_92149069 | chr1 | 92148437 | 92149069 | + | -1.23828 | promoter-TSS (NM_207225) | promoter-TSS (NM_207225) | -360 | NM_207225 | 208727 | Mm.318567 | NM_207225 | ENSMUSG00000026313 | **Hdac4** | 4932408F19Rik\|AI047285 | histone deacetylase 4 | protein-coding |
| chr1_92069488_92069896 | chr1 | 92069489 | 92069896 | + | -1.25806 | intron (NM_207225, intron 1 of 25) | intron (NM_207225, intron 1 of 25) | 78701 | NM_207225 | 208727 | Mm.318567 | NM_207225 | ENSMUSG00000026313 | **Hdac4** | 4932408F19Rik\|AI047285 | histone deacetylase 4 | protein-coding |
| chrX_102464072_102464539 | chrX | 102464073 | 102464539 | + | -0.683246 | intron (NM_027382, intron 4 of 10) | intron (NM_027382, intron 4 of 10) | 40842 | NM_027382 | 70315 | Mm.328128 | NM_027382 | ENSMUSG00000067567 | **Hdac8** | 2610007D20Rik | histone deacetylase 8 | protein-coding |
| chr2_126605984_126606692 | chr2 | 126605985 | 126606692 | + | -0.687053 | intron (NM_008230, intron 4 of 11) | intron (NM_008230, intron 4 of 11) | 12340 | NM_008230 | 15186 | Mm.18603 | NM_008230 | ENSMUSG00000027360 | **Hdc** | AW108189\|Hdc-a\|Hdc-c\|Hdc-e\|Hdc-s | histidine decarboxylase | protein-coding |
| chr7_81928951_81929469 | chr7 | 81928952 | 81929469 | + | -0.613216 | intron (NM_013886, intron 1 of 5) | intron (NM_013886, intron 1 of 5) | 5249 | NM_013886 | 29877 | Mm.392934 | NM_013886 | ENSMUSG00000025104 | **Hdgfrp3** | 2700022B06Rik\|HRP-3 | hepatoma-derived growth factor, related protein 3 | protein-coding |
| chr13_12379369_12379875 | chr13 | 12379370 | 12379875 | + | -0.987424 | Intergenic | Intergenic | -15753 | NM_144835 | 217995 | Mm.171186 | NM_144835 | ENSMUSG00000050244 | **Heatr1** | AA517551\|B130016L12Rik\|BC019693 | HEAT repeat containing 1 | protein-coding |
| chr5_139166722_139167410 | chr5 | 139166723 | 139167410 | + | -0.681545 | intron (NM_001081265, intron 7 of 12) | intron (NM_001081265, intron 7 of 12) | 16843 | NM_001081265 | 433956 | Mm.5547 | NM_001081265 | ENSMUSG00000025857 | **Heatr2** | BC053401\|C76907 | HEAT repeat containing 2 | protein-coding |
| chr5_139174887_139175463 | chr5 | 139174888 | 139175463 | + | -0.910941 | intron (NM_001081265, intron 9 of 12) | intron (NM_001081265, intron 9 of 12) | 24952 | NM_001081265 | 433956 | Mm.5547 | NM_001081265 | ENSMUSG00000025857 | **Heatr2** | BC053401\|C76907 | HEAT repeat containing 2 | protein-coding |
| chr12_51976238_51977031 | chr12 | 51976239 | 51977031 | + | -0.945344 | Intergenic | Intergenic | -5314 | NM_177171 | 320487 | Mm.131941 | NM_177171 | ENSMUSG00000035181 | **Heatr5a** | C230081H03Rik\|D930036F22Rik\|mKIAA1316 | HEAT repeat containing 5A | protein-coding |
| chr12_51856528_51857209 | chr12 | 51856529 | 51857209 | + | -0.751552 | Intergenic | Intergenic | -27333 | NM_144788 | 207304 | Mm.249391 | NM_144788 | ENSMUSG00000035247 | **Hectd1** | A630086P08Rik\|AI844876\|b2b327Clo\|opm | HECT domain containing 1 | protein-coding |
| chr12_51856838_51857200 | chr12 | 51856839 | 51857200 | + | -0.849872 | Intergenic | Intergenic | -27483 | NM_144788 | 207304 | Mm.249391 | NM_144788 | ENSMUSG00000035247 | **Hectd1** | A630086P08Rik\|AI844876\|b2b327Clo\|opm | HECT domain containing 1 | protein-coding |
| chr12_51886356_51886742 | chr12 | 51886357 | 51886742 | + | -0.944239 | intron (NM_177171, intron 32 of 35) | B4\|SINE\|B4 | -57013 | NM_144788 | 207304 | Mm.249391 | NM_144788 | ENSMUSG00000035247 | **Hectd1** | A630086P08Rik\|AI844876\|b2b327Clo\|opm | HECT domain containing 1 | protein-coding |
| chr12_51871482_51872117 | chr12 | 51871483 | 51872117 | + | -1.18972 | Intergenic | Intergenic | -42264 | NM_144788 | 207304 | Mm.249391 | NM_144788 | ENSMUSG00000035247 | **Hectd1** | A630086P08Rik\|AI844876\|b2b327Clo\|opm | HECT domain containing 1 | protein-coding |
| chr16_33687580_33688052 | chr16 | 33687581 | 33688052 | + | -0.600012 | intron (NM_175256, intron 1 of 14) | intron (NM_175256, intron 1 of 14) | 3350 | NM_175256 | 77446 | Mm.245741 | NM_172934 | ENSMUSG00000075254 | **Heg1** | 4632417D23Rik\|5530401I02Rik\|9530025L16Rik\|BC038614\|Gm629\|OTTMUSG00000027315 | HEG homolog 1 (zebrafish) | protein-coding |
| chr11_107529246_107529840 | chr11 | 107529247 | 107529840 | + | -0.690181 | Intergenic | Intergenic | -18417 | NM_198298 | 78455 | Mm.272047 | NM_198298 | ENSMUSG00000020721 | **Helz** | 3110078M01Rik\|9430093I07Rik\|9630002H22Rik\|AI851979\|DRHC | helicase with zinc finger domain | protein-coding |
| chr9_66274535_66275066 | chr9 | 66274536 | 66275066 | + | -0.697921 | Intergenic | Intergenic | -75649 | NM_145617 | 235439 | Mm.244179 | NM_145617 | ENSMUSG00000038664 | **Herc1** | 2810449H11Rik\|B230218H07\|D130015N03Rik\|tbl | hect (homologous to the E6-AP (UBE3A) carboxyl terminus) domain and RCC1 (CHC1)-like domain (RLD) 1 | protein-coding |
| chr9_66368898_66369233 | chr9 | 66368899 | 66369233 | + | -1.07441 | intron (NM_145617, intron 1 of 77) | MIR\|SINE\|MIR | 18616 | NM_145617 | 235439 | Mm.244179 | NM_145617 | ENSMUSG00000038664 | **Herc1** | 2810449H11Rik\|B230218H07\|D130015N03Rik\|tbl | hect (homologous to the E6-AP (UBE3A) carboxyl terminus) domain and RCC1 (CHC1)-like domain (RLD) 1 | protein-coding |
| chr6_58844140_58844834 | chr6 | 58844141 | 58844834 | + | -0.910477 | intron (NM_028705, intron 2 of 24) | (TG)n\|Simple_repeat\|Simple_repeat | 10787 | NM_028705 | 73998 | Mm.191892 | NM_028705 | ENSMUSG00000029804 | **Herc3** | 5730409F18Rik\|AI646603\|mKIAA0032 | hect domain and RLD 3 | protein-coding |
| chr10_63231501_63231993 | chr10 | 63231502 | 63231993 | + | -1.02753 | Intergenic | Intergenic | -12050 | NM_030114 | 67345 | Mm.234437 | NM_026101 | ENSMUSG00000020064 | **Herc4** | 1700056O17Rik\|4921531D01Rik\|9530080M15Rik\|mKIAA1593 | hect domain and RLD 4 | protein-coding |
| chr10_63231541_63232354 | chr10 | 63231542 | 63232354 | + | -1.02838 | Intergenic | Intergenic | -11849 | NM_030114 | 67345 | Mm.234437 | NM_026101 | ENSMUSG00000020064 | **Herc4** | 1700056O17Rik\|4921531D01Rik\|9530080M15Rik\|mKIAA1593 | hect domain and RLD 4 | protein-coding |
| chr10_63231686_63231981 | chr10 | 63231687 | 63231981 | + | -1.06867 | Intergenic | Intergenic | -11963 | NM_030114 | 67345 | Mm.234437 | NM_026101 | ENSMUSG00000020064 | **Herc4** | 1700056O17Rik\|4921531D01Rik\|9530080M15Rik\|mKIAA1593 | hect domain and RLD 4 | protein-coding |
| chr10_63231671_63232104 | chr10 | 63231672 | 63232104 | + | -1.06996 | Intergenic | Intergenic | -11909 | NM_030114 | 67345 | Mm.234437 | NM_026101 | ENSMUSG00000020064 | **Herc4** | 1700056O17Rik\|4921531D01Rik\|9530080M15Rik\|mKIAA1593 | hect domain and RLD 4 | protein-coding |
| chr8_94372872_94373218 | chr8 | 94372873 | 94373218 | + | -0.868777 | Intergenic | RLTR33\|LTR\|ERVK | -13455 | NM_022331 | 64209 | Mm.29151 | NM_022331 | ENSMUSG00000031770 | **Herpud1** | Herp\|Mifl\|SUP | homocysteine-inducible, endoplasmic reticulum stress-inducible, ubiquitin-like domain member 1 | protein-coding |
| chr8_94372887_94373519 | chr8 | 94372888 | 94373519 | + | -0.931393 | Intergenic | RLTR33\|LTR\|ERVK | -13297 | NM_022331 | 64209 | Mm.29151 | NM_022331 | ENSMUSG00000031770 | **Herpud1** | Herp\|Mifl\|SUP | homocysteine-inducible, endoplasmic reticulum stress-inducible, ubiquitin-like domain member 1 | protein-coding |
| chr9_25191697_25192117 | chr9 | 25191698 | 25192117 | + | -0.6322 | Intergenic | Intergenic | -40126 | NM_020586 | 80517 | Mm.142843 | NM_020586 | ENSMUSG00000008429 | **Herpud2** | 5031400M07Rik\|AB041580 | HERPUD family member 2 | protein-coding |
| chr9_25196005_25197202 | chr9 | 25196006 | 25197202 | + | -0.680744 | Intergenic | B4\|SINE\|B4 | -44823 | NM_020586 | 80517 | Mm.142843 | NM_020586 | ENSMUSG00000008429 | **Herpud2** | 5031400M07Rik\|AB041580 | HERPUD family member 2 | protein-coding |
| chr9_25196594_25197204 | chr9 | 25196595 | 25197204 | + | -0.837721 | Intergenic | Intergenic | -45118 | NM_020586 | 80517 | Mm.142843 | NM_020586 | ENSMUSG00000008429 | **Herpud2** | 5031400M07Rik\|AB041580 | HERPUD family member 2 | protein-coding |
| chr13_97208087_97208510 | chr13 | 97208088 | 97208510 | + | -0.91638 | Intergenic | Intergenic | -9942 | NM_010422 | 15212 | Mm.27816 | NM_010422 | ENSMUSG00000021665 | **Hexb** | - | hexosaminidase B | protein-coding |
| chr3_8718126_8718663 | chr3 | 8718127 | 8718663 | + | -0.807864 | Intergenic | Intergenic | -51357 | NM_010423 | 15213 | Mm.29581 | NM_010423 | ENSMUSG00000040289 | **Hey1** | AI316788\|AI414254\|CHF2\|HRT1\|Herp2\|Hesr1\|bHLHb31\|hesr-1 | hairy/enhancer-of-split related with YRPW motif 1 | protein-coding |
| chr3_8766590_8767289 | chr3 | 8766591 | 8767289 | + | -0.975177 | Intergenic | Intergenic | -99902 | NM_010423 | 15213 | Mm.29581 | NM_010423 | ENSMUSG00000040289 | **Hey1** | AI316788\|AI414254\|CHF2\|HRT1\|Herp2\|Hesr1\|bHLHb31\|hesr-1 | hairy/enhancer-of-split related with YRPW motif 1 | protein-coding |
| chr13_65101263_65101629 | chr13 | 65101264 | 65101629 | + | -0.654211 | intron (NM_133680, intron 1 of 11) | intron (NM_133680, intron 1 of 11) | 11536 | NM_001083901 | 66631 | Mm.239196 | NM_133680 | ENSMUSG00000038212 | **Hiatl1** | 5730414C17Rik\|AU045608\|AW539692 | hippocampus abundant transcript-like 1 | protein-coding |
| chr11_115366148_115366799 | chr11 | 115366149 | 115366799 | + | -0.691686 | intron (NM_175454, intron 1 of 18) | intron (NM_175454, intron 1 of 18) | 1245 | NM_175454 | 217310 | Mm.205707 | NM_175454 | ENSMUSG00000034586 | **Hid1** | C630004H02Rik | HID1 domain containing | protein-coding |
| chr11_115366054_115366447 | chr11 | 115366055 | 115366447 | + | -0.89511 | intron (NM_175454, intron 1 of 18) | intron (NM_175454, intron 1 of 18) | 1468 | NM_175454 | 217310 | Mm.205707 | NM_175454 | ENSMUSG00000034586 | **Hid1** | C630004H02Rik | HID1 domain containing | protein-coding |
| chr6_38834806_38835104 | chr6 | 38834807 | 38835104 | + | -0.645747 | intron (NM_001294144, intron 1 of 14) | intron (NM_001294144, intron 1 of 14) | 2356 | NM_001136065 | 15258 | Mm.23790 | NM_010433 | ENSMUSG00000061436 | **Hipk2** | 1110014O20Rik\|B230339E18Rik\|Stank | homeodomain interacting protein kinase 2 | protein-coding |
| chr6_38834678_38835097 | chr6 | 38834679 | 38835097 | + | -0.703077 | intron (NM_001294144, intron 1 of 14) | intron (NM_001294144, intron 1 of 14) | 2423 | NM_001136065 | 15258 | Mm.23790 | NM_010433 | ENSMUSG00000061436 | **Hipk2** | 1110014O20Rik\|B230339E18Rik\|Stank | homeodomain interacting protein kinase 2 | protein-coding |
| chr13_42048985_42049241 | chr13 | 42048986 | 42049241 | + | -0.668569 | Intergenic | Intergenic | -2908 | NM_007772 | 110521 | Mm.4808 | NM_007772 | ENSMUSG00000021366 | **Hivep1** | Cryabp1 | human immunodeficiency virus type I enhancer binding protein 1 | protein-coding |
| chr13_42048985_42049336 | chr13 | 42048986 | 42049336 | + | -0.687342 | Intergenic | Intergenic | -2860 | NM_007772 | 110521 | Mm.4808 | NM_007772 | ENSMUSG00000021366 | **Hivep1** | Cryabp1 | human immunodeficiency virus type I enhancer binding protein 1 | protein-coding |
| chr10_13993235_13993904 | chr10 | 13993236 | 13993904 | + | -0.695891 | intron (NM_010437, intron 1 of 8) | intron (NM_010437, intron 1 of 8) | 27191 | NM_010437 | 15273 | Mm.42157 | NM_010437 | ENSMUSG00000015501 | **Hivep2** | MIBP1\|Schnurri-2\|Shn-2 | human immunodeficiency virus type I enhancer binding protein 2 | protein-coding |
| chr10_13998853_13999325 | chr10 | 13998854 | 13999325 | + | -0.736498 | intron (NM_010437, intron 1 of 8) | intron (NM_010437, intron 1 of 8) | 32710 | NM_010437 | 15273 | Mm.42157 | NM_010437 | ENSMUSG00000015501 | **Hivep2** | MIBP1\|Schnurri-2\|Shn-2 | human immunodeficiency virus type I enhancer binding protein 2 | protein-coding |
| chr10_14014108_14014425 | chr10 | 14014109 | 14014425 | + | -0.751242 | intron (NM_010437, intron 1 of 8) | intron (NM_010437, intron 1 of 8) | 47888 | NM_010437 | 15273 | Mm.42157 | NM_010437 | ENSMUSG00000015501 | **Hivep2** | MIBP1\|Schnurri-2\|Shn-2 | human immunodeficiency virus type I enhancer binding protein 2 | protein-coding |
| chr10_14115711_14116282 | chr10 | 14115712 | 14116282 | + | -0.792478 | intron (NM_010437, intron 2 of 8) | intron (NM_010437, intron 2 of 8) | 149618 | NM_010437 | 15273 | Mm.42157 | NM_010437 | ENSMUSG00000015501 | **Hivep2** | MIBP1\|Schnurri-2\|Shn-2 | human immunodeficiency virus type I enhancer binding protein 2 | protein-coding |
| chr10_14045030_14045390 | chr10 | 14045031 | 14045390 | + | -0.909691 | intron (NM_010437, intron 1 of 8) | intron (NM_010437, intron 1 of 8) | 78831 | NM_010437 | 15273 | Mm.42157 | NM_010437 | ENSMUSG00000015501 | **Hivep2** | MIBP1\|Schnurri-2\|Shn-2 | human immunodeficiency virus type I enhancer binding protein 2 | protein-coding |
| chr6_82775477_82775727 | chr6 | 82775478 | 82775727 | + | -0.631804 | Intergenic | L2\|LINE\|L2 | -1148 | NM_013820 | 15277 | Mm.255848 | NM_013820 | ENSMUSG00000000628 | **Hk2** | AI642394\|HKII | hexokinase 2 | protein-coding |

| | | | | | | | | | | | | | | | | | |
|---|---|---|---|---|---|---|---|---|---|---|---|---|---|---|---|---|---|
| chr13_55018315_55018687 | chr13 | 55018316 | 55018687 | + | -0.900057 | intron (NM_001206392, intron 1 of 17) | (GA)n\|Simple_repeat\|Simple_repeat | 2318 | NM_001206390 | 212032 | Mm.267479 | NM_001033245 | ENSMUSG00000025877 | **Hk3** | HK III\|HK-III | hexokinase 3 | protein-coding |
| chr13_55017892_55018635 | chr13 | 55017893 | 55018635 | + | -0.980568 | intron (NM_001206392, intron 1 of 17) | intron (NM_001206392, intron 1 of 17) | 2555 | NM_001206390 | 212032 | Mm.267479 | NM_001033245 | ENSMUSG00000025877 | **Hk3** | HK III\|HK-III | hexokinase 3 | protein-coding |
| chr16_94239615_94239899 | chr16 | 94239616 | 94239899 | + | -0.748709 | intron (NM_139145, intron 4 of 9) | ORR1D2\|LTR\|ERVL-MaLR | 48099 | NM_139145 | 110948 | Mm.30921 | NM_139145 | ENSMUSG00000040820 | **Hlcs** | 410I21.SP6\|D16Jhu34 | holocarboxylase synthetase (biotin- [propriony-Coenzyme A-carboxylase (ATP-hydrolysing)] ligase) | protein-coding |
| chr16_94254356_94254967 | chr16 | 94254357 | 94254967 | + | -1.37248 | intron (NM_139145, intron 4 of 9) | intron (NM_139145, intron 4 of 9) | 33194 | NM_139145 | 110948 | Mm.30921 | NM_139145 | ENSMUSG00000040820 | **Hlcs** | 410I21.SP6\|D16Jhu34 | holocarboxylase synthetase (biotin- [propriony-Coenzyme A-carboxylase (ATP-hydrolysing)] ligase) | protein-coding |
| chr14_64884839_64885236 | chr14 | 64884840 | 64885236 | + | -0.719992 | intron (NM_177338, intron 4 of 9) | intron (NM_177338, intron 4 of 9) | 64809 | NM_177338 | 219150 | Mm.344074 | NM_177338 | ENSMUSG00000021972 | **Hmbox1** | AI451877\|AI604847\|F830020C16Rik | homeobox containing 1 | protein-coding |
| chr14_64884674_64885226 | chr14 | 64884675 | 64885226 | + | -0.755456 | intron (NM_177338, intron 4 of 9) | intron (NM_177338, intron 4 of 9) | 64897 | NM_177338 | 219150 | Mm.344074 | NM_177338 | ENSMUSG00000021972 | **Hmbox1** | AI451877\|AI604847\|F830020C16Rik | homeobox containing 1 | protein-coding |
| chr6_87913753_87914810 | chr6 | 87913754 | 87914810 | + | -0.610462 | TTS (NM_017477) | TTS (NM_017477) | 306 | NM_173737 | 232210 | Mm.246241 | NM_173737 | ENSMUSG00000030060 | **Hmces** | 8430410A17Rik\|C85376 | 5-hydroxymethylcytosine (hmC) binding, ES cell specific | protein-coding |
| chr6_87921281_87921565 | chr6 | 87921282 | 87921565 | + | -1.20318 | intron (NM_173737, intron 3 of 6) | ID2\|SINE\|ID | 7447 | NM_173737 | 232210 | Mm.246241 | NM_173737 | ENSMUSG00000030060 | **Hmces** | 8430410A17Rik\|C85376 | 5-hydroxymethylcytosine (hmC) binding, ES cell specific | protein-coding |
| chr6_87921080_87921619 | chr6 | 87921081 | 87921619 | + | -1.20428 | intron (NM_173737, intron 3 of 6) | intron (NM_173737, intron 3 of 6) | 7374 | NM_173737 | 232210 | Mm.246241 | NM_173737 | ENSMUSG00000030060 | **Hmces** | 8430410A17Rik\|C85376 | 5-hydroxymethylcytosine (hmC) binding, ES cell specific | protein-coding |
| chr1_151024765_151026213 | chr1 | 151024766 | 151026213 | + | -0.7864 | Intergenic | Intergenic | -32054 | NM_001024720 | 545370 | Mm.379425 | NM_001024720 | ENSMUSG00000066842 | **Hmcn1** | EG545370\|Gm201 | hemicentin 1 | protein-coding |
| chr9_56422082_56422297 | chr9 | 56422083 | 56422297 | + | -0.624589 | intron (NM_025812, intron 1 of 9) | intron (NM_025812, intron 1 of 9) | 3344 | NM_025812 | 66867 | Mm.150856 | NM_025812 | ENSMUSG00000032329 | **Hmg20a** | 1200004E06Rik\|5730490E10Rik\|Hmgxb1\|Ibraf | high mobility group 20A | protein-coding |
| chr9_56509231_56509836 | chr9 | 56509232 | 56509836 | + | -0.643002 | Intergenic | Intergenic | 90688 | NM_025812 | 66867 | Mm.150856 | NM_025812 | ENSMUSG00000032329 | **Hmg20a** | 1200004E06Rik\|5730490E10Rik\|Hmgxb1\|Ibraf | high mobility group 20A | protein-coding |
| chr9_56456497_56456764 | chr9 | 56456498 | 56456764 | + | -0.81786 | intron (NM_025812, intron 1 of 9) | intron (NM_025812, intron 1 of 9) | 37785 | NM_025812 | 66867 | Mm.150856 | NM_025812 | ENSMUSG00000032329 | **Hmg20a** | 1200004E06Rik\|5730490E10Rik\|Hmgxb1\|Ibraf | high mobility group 20A | protein-coding |
| chr5_149080204_149080951 | chr5 | 149080205 | 149080951 | + | -1.18992 | Intergenic | Intergenic | -27541 | NM_010439 | 15289 | Mm.207047 | NM_010439 | ENSMUSG00000066551 | **Hmgb1** | DEF\|HMG-1\|Hmg1\|SBP-1\|amphoterin\|p30 | high mobility group box 1 | protein-coding |
| chr18_61187840_61188655 | chr18 | 61187841 | 61188655 | + | -0.982082 | Intergenic | Intergenic | -11198 | NM_134134 | 106894 | Mm.22574 | NM_134134 | ENSMUSG00000024622 | **Hmgxb3** | 2510002C16Rik\|A630042L21Rik\|AI413166\|mKIAA0194 | HMG box domain containing 3 | protein-coding |
| chr10_80022144_80022532 | chr10 | 80022145 | 80022532 | + | -0.63265 | exon (NM_001142701, exon 10 of 23) | exon (NM_001142701, exon 10 of 23) | 2663 | NM_027521 | 70719 | Mm.243954 | NM_027521 | ENSMUSG00000035697 | **Hmha1** | 6330406L22Rik\|AW539505\|Ha-1 | histocompatibility (minor) HA-1 | protein-coding |
| chr10_80018812_80019022 | chr10 | 80018813 | 80019022 | + | -0.686342 | promoter-TSS (NM_027521) | promoter-TSS (NM_027521) | -758 | NM_027521 | 70719 | Mm.243954 | NM_027521 | ENSMUSG00000035697 | **Hmha1** | 6330406L22Rik\|AW539505\|Ha-1 | histocompatibility (minor) HA-1 | protein-coding |
| chr11_40718089_40718711 | chr11 | 40718090 | 40718711 | + | -0.686333 | intron (NM_013552, intron 7 of 17) | RMER6B\|LTR\|ERVK | 15037 | NM_013552 | 15366 | Mm.116997 | NM_013552 | ENSMUSG00000020330 | **Hmmr** | AA386826\|CD168\|Rhamm | hyaluronan mediated motility receptor (RHAMM) | protein-coding |
| chr6_117890237_117890678 | chr6 | 117890238 | 117890678 | + | -0.620149 | Intergenic | Intergenic | -9882 | NM_001166432 | 98758 | Mm.422979 | NM_133834 | ENSMUSG00000042079 | **Hnrnpf** | 4833420I20Rik\|AA407306\|Hnrpf | heterogeneous nuclear ribonucleoprotein F | protein-coding |
| chr6_117881995_117882275 | chr6 | 117881996 | 117882275 | + | -0.624671 | Intergenic | MER68\|LTR\|ERVL | -18205 | NM_001166432 | 98758 | Mm.422979 | NM_133834 | ENSMUSG00000042079 | **Hnrnpf** | 4833420I20Rik\|AA407306\|Hnrpf | heterogeneous nuclear ribonucleoprotein F | protein-coding |
| chr6_117881986_117882543 | chr6 | 117881987 | 117882543 | + | -0.734717 | Intergenic | MER68\|LTR\|ERVL | -18075 | NM_001166432 | 98758 | Mm.422979 | NM_133834 | ENSMUSG00000042079 | **Hnrnpf** | 4833420I20Rik\|AA407306\|Hnrpf | heterogeneous nuclear ribonucleoprotein F | protein-coding |
| chr7_28799944_28800410 | chr7 | 28799945 | 28800410 | + | -1.25632 | Intergenic | ORR1E\|LTR\|ERVL-MaLR | -10713 | NM_177301 | 15388 | Mm.9043 | NM_177301 | ENSMUSG00000015165 | **Hnrnpl** | C79783\|D830027H13Rik\|Hnrpl | heterogeneous nuclear ribonucleoprotein L | protein-coding |
| chr5_77134278_77134836 | chr5 | 77134279 | 77134836 | + | -0.668919 | Intergenic | Intergenic | -19434 | NM_175606 | 74318 | Mm.181852 | NM_175606 | ENSMUSG00000059325 | **Hopx** | 1110018K11Rik\|1200015P04Rik\|2300002F06Rik\|AI848177\|AW490897\|Cameo\|Hdop\|Hod\|Hop\|Ob1\|Obl\|Toto | HOP homeobox | protein-coding |
| chr5_77134466_77134831 | chr5 | 77134467 | 77134831 | + | -0.674319 | Intergenic | Intergenic | -19526 | NM_175606 | 74318 | Mm.181852 | NM_175606 | ENSMUSG00000059325 | **Hopx** | 1110018K11Rik\|1200015P04Rik\|2300002F06Rik\|AI848177\|AW490897\|Cameo\|Hdop\|Hod\|Hop\|Ob1\|Obl\|Toto | HOP homeobox | protein-coding |
| chr11_4423873_4424251 | chr11 | 4423874 | 4424251 | + | -1.20326 | intron (NM_029458, intron 5 of 10) | intron (NM_029458, intron 5 of 10) | 17020 | NM_029458 | 75828 | Mm.159592 | NM_029458 | ENSMUSG00000020419 | **Hormad2** | 4930529M09Rik | HORMA domain containing 2 | protein-coding |
| chr6_52217450_52217729 | chr6 | 52217451 | 52217729 | + | -0.697468 | 5' UTR (NM_010455, exon 1 of 2) | 5' UTR (NM_010455, exon 1 of 2) | 983 | NM_010455 | 15404 | Mm.294826 | NM_010455 | | **Hoxa7** | AV118143\|Hox-1.1 | homeobox A7 | protein-coding |
| chr6_52217418_52217840 | chr6 | 52217419 | 52217840 | + | -0.700972 | 5' UTR (NM_010455, exon 1 of 2) | 5' UTR (NM_010455, exon 1 of 2) | 944 | NM_010455 | 15404 | Mm.294826 | NM_010455 | | **Hoxa7** | AV118143\|Hox-1.1 | homeobox A7 | protein-coding |
| chr4_138223733_138224017 | chr4 | 138223734 | 138224017 | + | -0.650359 | intron (NM_001285479, intron 4 of 12) | intron (NM_001285479, intron 4 of 12) | 6748 | NM_001122897 | 15441 | Mm.348429 | NM_010470 | ENSMUSG00000028759 | **Hp1bp3** | Hp1bp74 | heterochromatin protein 1, binding protein 3 | protein-coding |
| chr4_138222382_138222724 | chr4 | 138222383 | 138222724 | + | -0.684518 | intron (NM_001285480, intron 2 of 11) | intron (NM_001285480, intron 2 of 11) | 5426 | NM_001122897 | 15441 | Mm.348429 | NM_010470 | ENSMUSG00000028759 | **Hp1bp3** | Hp1bp74 | heterochromatin protein 1, binding protein 3 | protein-coding |
| chr12_17716152_17716463 | chr12 | 17716153 | 17716463 | + | -0.724854 | intron (NM_016677, intron 1 of 3) | ORR1B2\|LTR\|ERVL-MaLR | 25494 | NM_016677 | 53602 | Mm.313619 | NM_016677 | ENSMUSG00000071379 | **Hpcal1** | NVP-3\|Nvp3\|VILIP3\|Vnsl3\|Vsnl3 | hippocalcin-like 1 | protein-coding |
| chr12_17715689_17716051 | chr12 | 17715690 | 17716051 | + | -0.778217 | intron (NM_016677, intron 1 of 3) | MIR3\|SINE\|MIR | 25056 | NM_016677 | 53602 | Mm.313619 | NM_016677 | ENSMUSG00000071379 | **Hpcal1** | NVP-3\|Nvp3\|VILIP3\|Vnsl3\|Vsnl3 | hippocalcin-like 1 | protein-coding |
| chr12_17715667_17716444 | chr12 | 17715668 | 17716444 | + | -0.7999 | intron (NM_016677, intron 1 of 3) | intron (NM_016677, intron 1 of 3) | 25242 | NM_016677 | 53602 | Mm.313619 | NM_016677 | ENSMUSG00000071379 | **Hpcal1** | NVP-3\|Nvp3\|VILIP3\|Vnsl3\|Vsnl3 | hippocalcin-like 1 | protein-coding |
| chr6_65144365_65145241 | chr6 | 65144366 | 65145241 | + | -1.09324 | promoter-TSS (NM_019455) | promoter-TSS (NM_019455) | -73 | NM_019455 | 54486 | Mm.410688 | NM_019455 | ENSMUSG00000029919 | **Hpgds** | H-PGDS\|Ptgds2 | hematopoietic prostaglandin D synthase | protein-coding |
| chr3_144548180_144548642 | chr3 | 144548181 | 144548642 | + | -0.880055 | intron (NM_011828, intron 1 of 6) | intron (NM_011828, intron 1 of 6) | 21805 | NM_011828 | 23908 | Mm.12863 | NM_011828 | ENSMUSG00000040151 | **Hs2st1** | AW214369\|Hs2st\|mKIAA0448 | heparan sulfate 2-O-sulfotransferase 1 | protein-coding |
| chr3_144412527_144413235 | chr3 | 144412528 | 144413235 | + | -0.98545 | Intergenic | Intergenic | 157335 | NM_011828 | 23908 | Mm.12863 | NM_011828 | ENSMUSG00000040151 | **Hs2st1** | AW214369\|Hs2st\|mKIAA0448 | heparan sulfate 2-O-sulfotransferase 1 | protein-coding |
| chr11_63975031_63975912 | chr11 | 63975032 | 63975912 | + | -0.660607 | intron (NM_178379, intron 6 of 6) | intron (NM_178379, intron 6 of 6) | -53188 | NM_018805 | 54710 | Mm.274426 | NM_018805 | ENSMUSG00000070407 | **Hs3st3b1** | 3-OST-3B\|3OST3B1\|3Ost3b\|AU041882\|AW536289\|Hs3st3b\|m3-OST-3B | heparan sulfate (glucosamine) 3-O-sulfotransferase 3B1 | protein-coding |
| chr11_63985531_63985944 | chr11 | 63985532 | 63985944 | + | -0.858064 | intron (NM_178379, intron 5 of 6) | intron (NM_178379, intron 5 of 6) | -63454 | NM_018805 | 54710 | Mm.274426 | NM_018805 | ENSMUSG00000070407 | **Hs3st3b1** | 3-OST-3B\|3OST3B1\|3Ost3b\|AU041882\|AW536289\|Hs3st3b\|m3-OST-3B | heparan sulfate (glucosamine) 3-O-sulfotransferase 3B1 | protein-coding |
| chr11_63991740_63992383 | chr11 | 63991741 | 63992383 | + | -0.884858 | intron (NM_178379, intron 5 of 6) | intron (NM_178379, intron 5 of 6) | -69778 | NM_018805 | 54710 | Mm.274426 | NM_018805 | ENSMUSG00000070407 | **Hs3st3b1** | 3-OST-3B\|3OST3B1\|3Ost3b\|AU041882\|AW536289\|Hs3st3b\|m3-OST-3B | heparan sulfate (glucosamine) 3-O-sulfotransferase 3B1 | protein-coding |
| chr11_63984250_63985352 | chr11 | 63984251 | 63985352 | + | -1.21864 | intron (NM_178379, intron 5 of 6) | ORR1D1\|LTR\|ERVL-MaLR | -62517 | NM_018805 | 54710 | Mm.274426 | NM_018805 | ENSMUSG00000070407 | **Hs3st3b1** | 3-OST-3B\|3OST3B1\|3Ost3b\|AU041882\|AW536289\|Hs3st3b\|m3-OST-3B | heparan sulfate (glucosamine) 3-O-sulfotransferase 3B1 | protein-coding |
| chr11_63989582_63989847 | chr11 | 63989583 | 63989847 | + | -1.3419 | intron (NM_178379, intron 5 of 6) | intron (NM_178379, intron 5 of 6) | -67431 | NM_018805 | 54710 | Mm.274426 | NM_018805 | ENSMUSG00000070407 | **Hs3st3b1** | 3-OST-3B\|3OST3B1\|3Ost3b\|AU041882\|AW536289\|Hs3st3b\|m3-OST-3B | heparan sulfate (glucosamine) 3-O-sulfotransferase 3B1 | protein-coding |
| chr11_63989478_63989873 | chr11 | 63989479 | 63989873 | + | -1.35052 | intron (NM_178379, intron 5 of 6) | intron (NM_178379, intron 5 of 6) | -67392 | NM_018805 | 54710 | Mm.274426 | NM_018805 | ENSMUSG00000070407 | **Hs3st3b1** | 3-OST-3B\|3OST3B1\|3Ost3b\|AU041882\|AW536289\|Hs3st3b\|m3-OST-3B | heparan sulfate (glucosamine) 3-O-sulfotransferase 3B1 | protein-coding |
| chr10_36158012_36158716 | chr10 | 36158013 | 36158716 | + | -1.43734 | Intergenic | Intergenic | -348443 | NM_001253355 | 319415 | Mm.332442 | NM_001081208 | ENSMUSG00000044499 | **Hs3st5** | D930005L05Rik\|Gm1151\|Hs3ost5 | heparan sulfate (glucosamine) 3-O-sulfotransferase 5 | protein-coding |
| chr1_35971431_35972716 | chr1 | 35971432 | 35972716 | + | -0.710046 | Intergenic | Intergenic | -96326 | NM_015818 | 50785 | Mm.213566 | NM_015818 | ENSMUSG00000045216 | **Hs6st1** | 6Ost1 | heparan sulfate 6-O-sulfotransferase 1 | protein-coding |
| chr1_35983343_35983773 | chr1 | 35983344 | 35983773 | + | -0.723614 | Intergenic | Intergenic | -84842 | NM_015818 | 50785 | Mm.213566 | NM_015818 | ENSMUSG00000045216 | **Hs6st1** | 6Ost1 | heparan sulfate 6-O-sulfotransferase 1 | protein-coding |
| chrX_51702493_51702937 | chrX | 51702494 | 51702937 | + | -0.8693 | Intergenic | Intergenic | -21113 | NM_001290467 | 50786 | Mm.252561 | NM_015819 | ENSMUSG00000062184 | **Hs6st2** | 6OST2 | heparan sulfate 6-O-sulfotransferase 2 | protein-coding |
| chr5_104007754_104008159 | chr5 | 104007755 | 104008159 | + | -0.883062 | intron (NM_053262, intron 4 of 6) | intron (NM_053262, intron 4 of 6) | 13839 | NM_053262 | 114664 | Mm.46019 | NM_053262 | ENSMUSG00000029311 | **Hsd17b11** | Dhrs8\|Pan1b\|SDR2\|retSDR2 | hydroxysteroid (17-beta) dehydrogenase 11 | protein-coding |

| | | | | | | | | | | | | | | | | | |
|---|---|---|---|---|---|---|---|---|---|---|---|---|---|---|---|---|---|
| chr8_119569856_119570444 | chr8 | 119569857 | 119570444 | + | -0.842992 | intron (NM_175185, intron 1 of 4) | intron (NM_175185, intron 1 of 4) | 5050 | NM_175185 | 72552 | Mm.36756 | NM_175185 | ENSMUSG00000034189 | **Hsdl1** | 2700067E09Rik | hydroxysteroid dehydrogenase like 1 | protein-coding |
| chr8_119569778_119570280 | chr8 | 119569779 | 119570280 | + | -0.884106 | intron (NM_175185, intron 1 of 4) | intron (NM_175185, intron 1 of 4) | 5171 | NM_175185 | 72552 | Mm.36756 | NM_175185 | ENSMUSG00000034189 | **Hsdl1** | 2700067E09Rik | hydroxysteroid dehydrogenase like 1 | protein-coding |
| chr17_45581532_45582004 | chr17 | 45581533 | 45582004 | + | -0.704541 | Intergenic | Intergenic | -8507 | NM_008302 | 15516 | Mm.2180 | NM_008302 | ENSMUSG00000023944 | **Hsp90ab1** | 90kDa\|AL022974\|C81438\|Hsp84\|Hsp84-1\|Hsp90\|Hspcb | heat shock protein 90 alpha (cytosolic), class B member 1 | protein-coding |
| chr17_45581319_45581917 | chr17 | 45581320 | 45581917 | + | -0.722457 | Intergenic | Intergenic | -8357 | NM_008302 | 15516 | Mm.2180 | NM_008302 | ENSMUSG00000023944 | **Hsp90ab1** | 90kDa\|AL022974\|C81438\|Hsp84\|Hsp84-1\|Hsp90\|Hspcb | heat shock protein 90 alpha (cytosolic), class B member 1 | protein-coding |
| chr16_75823201_75823786 | chr16 | 75823202 | 75823786 | + | -0.73958 | Intergenic | Intergenic | -56676 | NM_030201 | 110920 | Mm.26704 | NM_030201 | ENSMUSG00000032932 | **Hspa13** | 1600002I10Rik\|AV006182\|B230217N24Rik\|Stch | heat shock protein 70 family, member 13 | protein-coding |
| chr9_40781672_40782620 | chr9 | 40781673 | 40782620 | + | -0.731489 | intron (NM_133733, intron 6 of 6) | intron (NM_133733, intron 6 of 6) | -19127 | NM_031165 | 15481 | Mm.290774 | NM_031165 | ENSMUSG00000015656 | **Hspa8** | 2410008N15Rik\|Hsc70\|Hsc71\|Hsc73\|Hsp73\|Hspa10 | heat shock protein 8 | protein-coding |
| chr9_40781642_40782138 | chr9 | 40781643 | 40782138 | + | -0.785635 | intron (NM_133733, intron 6 of 6) | MTD\|LTR\|ERVL-MaLR | -19383 | NM_031165 | 15481 | Mm.290774 | NM_031165 | ENSMUSG00000015656 | **Hspa8** | 2410008N15Rik\|Hsc70\|Hsc71\|Hsc73\|Hsp73\|Hspa10 | heat shock protein 8 | protein-coding |
| chr9_40813796_40814459 | chr9 | 40813797 | 40814459 | + | -0.902042 | Intergenic | Intergenic | 12855 | NM_031165 | 15481 | Mm.290774 | NM_031165 | ENSMUSG00000015656 | **Hspa8** | 2410008N15Rik\|Hsc70\|Hsc71\|Hsc73\|Hsp73\|Hspa10 | heat shock protein 8 | protein-coding |
| chr9_40813962_40814453 | chr9 | 40813963 | 40814453 | + | -0.921497 | Intergenic | Intergenic | 12935 | NM_031165 | 15481 | Mm.290774 | NM_031165 | ENSMUSG00000015656 | **Hspa8** | 2410008N15Rik\|Hsc70\|Hsc71\|Hsc73\|Hsp73\|Hspa10 | heat shock protein 8 | protein-coding |
| chr4_107260003_107260410 | chr4 | 107260004 | 107260410 | + | -0.921276 | intron (NM_028394, intron 1 of 5) | intron (NM_028394, intron 1 of 5) | 6273 | NM_028394 | 72938 | Mm.21787 | NM_028394 | ENSMUSG00000063172 | **Hspb11** | 2900042B11Rik\|IFT25 | heat shock protein family B (small), member 11 | protein-coding |
| chr4_107259831_107260402 | chr4 | 107259832 | 107260402 | + | -0.938698 | intron (NM_028394, intron 1 of 5) | intron (NM_028394, intron 1 of 5) | 6183 | NM_028394 | 72938 | Mm.21787 | NM_028394 | ENSMUSG00000063172 | **Hspb11** | 2900042B11Rik\|IFT25 | heat shock protein family B (small), member 11 | protein-coding |
| chr5_116380384_116380789 | chr5 | 116380385 | 116380789 | + | -0.829822 | Intergenic | B3A\|SINE\|B2 | 42277 | NM_030704 | 80888 | Mm.21549 | NM_030704 | ENSMUSG00000041548 | **Hspb8** | AU018630\|AW413033\|Cryac\|D5Ucla4\|E2IG1\|H11\|H11K\|HSP20-like\|HSP22 | heat shock protein 8 | protein-coding |
| chr7_49741462_49742220 | chr7 | 49741463 | 49742220 | + | -0.642802 | Intergenic | Intergenic | -17265 | NM_001146053 | 53415 | Mm.20801 | NM_016865 | ENSMUSG00000039745 | **Htatip2** | AW111545\|Cc3\|Tip30 | HIV-1 tat interactive protein 2, homolog (human) | protein-coding |
| chr7_49741480_49741964 | chr7 | 49741481 | 49741964 | + | -0.659792 | Intergenic | Intergenic | -17384 | NM_001146053 | 53415 | Mm.20801 | NM_016865 | ENSMUSG00000039745 | **Htatip2** | AW111545\|Cc3\|Tip30 | HIV-1 tat interactive protein 2, homolog (human) | protein-coding |
| chr7_49741464_49741841 | chr7 | 49741465 | 49741841 | + | -0.694498 | Intergenic | Intergenic | -17453 | NM_001146053 | 53415 | Mm.20801 | NM_016865 | ENSMUSG00000039745 | **Htatip2** | AW111545\|Cc3\|Tip30 | HIV-1 tat interactive protein 2, homolog (human) | protein-coding |
| chr5_34822552_34823093 | chr5 | 34822553 | 34823093 | + | -1.16261 | intron (NM_010414, intron 19 of 66) | intron (NM_010414, intron 19 of 66) | 61083 | NM_010414 | 15194 | Mm.209071 | NM_010414 | ENSMUSG00000029104 | **Htt** | AI256365\|C430023I11Rik\|Hd\|Hdh\|IT15 | huntingtin | protein-coding |
| chr5_34822792_34823054 | chr5 | 34822793 | 34823054 | + | -1.21529 | intron (NM_010414, intron 19 of 66) | intron (NM_010414, intron 19 of 66) | 61183 | NM_010414 | 15194 | Mm.209071 | NM_010414 | ENSMUSG00000029104 | **Htt** | AI256365\|C430023I11Rik\|Hd\|Hdh\|IT15 | huntingtin | protein-coding |
| chr5_34822681_34823077 | chr5 | 34822682 | 34823077 | + | -1.22853 | intron (NM_010414, intron 19 of 66) | intron (NM_010414, intron 19 of 66) | 61139 | NM_010414 | 15194 | Mm.209071 | NM_010414 | ENSMUSG00000029104 | **Htt** | AI256365\|C430023I11Rik\|Hd\|Hdh\|IT15 | huntingtin | protein-coding |
| chr16_90475578_90476356 | chr16 | 90475579 | 90476356 | + | -0.697104 | exon (NM_015755, exon 7 of 11) | exon (NM_015755, exon 7 of 11) | 89570 | NM_015755 | 26559 | Mm.125874 | NM_015755 | ENSMUSG00000053414 | **Hunk** | Bstk1\|Mak-v | hormonally upregulated Neu-associated kinase | protein-coding |
| chr5_122184383_122184817 | chr5 | 122184384 | 122184817 | + | -0.814373 | Intergenic | Intergenic | -25129 | NM_001042489 | 74096 | Mm.28804 | NM_028752 | ENSMUSG00000064267 | **Hvcn1** | 0610039P13Rik\|AI450555\|BTS\|HV1\|Vsop.\|mVSOP | hydrogen voltage-gated channel 1 | protein-coding |
| chr5_122184435_122184970 | chr5 | 122184436 | 122184970 | + | -0.875165 | Intergenic | Intergenic | -25026 | NM_001042489 | 74096 | Mm.28804 | NM_028752 | ENSMUSG00000064267 | **Hvcn1** | 0610039P13Rik\|AI450555\|BTS\|HV1\|Vsop.\|mVSOP | hydrogen voltage-gated channel 1 | protein-coding |
| chr5_122219603_122220702 | chr5 | 122219604 | 122220702 | + | -0.987074 | intron (NM_001042489, intron 3 of 7) | intron (NM_001042489, intron 3 of 7) | 9866 | NM_028752 | 74096 | Mm.28804 | NM_028752 | ENSMUSG00000064267 | **Hvcn1** | 0610039P13Rik\|AI450555\|BTS\|HV1\|Vsop.\|mVSOP | hydrogen voltage-gated channel 1 | protein-coding |
| chr9_61168933_61169399 | chr9 | 61168934 | 61169399 | + | -0.680492 | Intergenic | Intergenic | 30651 | NR_045705 | 619328 | Mm.325722 | NR_045705 | | **I730028E13Rik** | - | RIKEN cDNA I730028E13 gene | ncRNA |
| chr15_100746997_100747290 | chr15 | 100746998 | 100747290 | + | -0.749439 | Intergenic | B4\|SINE\|B4 | -14407 | NR_045782 | 619313 | Mm.123436 | NR_045781 | | **I730030J21Rik** | - | RIKEN cDNA I730030J21 gene | ncRNA |
| chr19_34623309_34623714 | chr19 | 34623310 | 34623714 | + | -0.676839 | intron (NM_053217, intron 2 of 2) | intron (NM_053217, intron 2 of 2) | 15555 | NM_001005858 | 667370 | Mm.271850 | NM_001005858 | ENSMUSG00000062488 | **I830012O16Rik** | - | RIKEN cDNA I830012O16 gene | protein-coding |
| chr9_85671388_85672071 | chr9 | 85671389 | 85672071 | + | -0.76111 | Intergenic | RMER6A\|LTR\|ERVK | 77604 | NM_001081282 | 108837 | Mm.22315 | NM_175371 | ENSMUSG00000035941 | **Ibtk** | 5430411K16Rik\|AA409502\|BTKI\|mKIAA1417 | inhibitor of Bruton agammaglobulinemia tyrosine kinase | protein-coding |
| chr11_106387615_106388382 | chr11 | 106387616 | 106388382 | + | -0.675078 | Intergenic | Intergenic | -5358 | NM_010494 | 15896 | Mm.394 | NM_010494 | ENSMUSG00000001029 | **Icam2** | CD102\|Icam-2\|Ly-60 | intercellular adhesion molecule 2 | protein-coding |
| chr11_106430272_106430857 | chr11 | 106430273 | 106430857 | + | -1.04402 | intron (NM_023913, intron 4 of 21) | intron (NM_023913, intron 4 of 21) | -47924 | NM_010494 | 15896 | Mm.394 | NM_010494 | ENSMUSG00000001029 | **Icam2** | CD102\|Icam-2\|Ly-60 | intercellular adhesion molecule 2 | protein-coding |
| chr1_60977642_60978058 | chr1 | 60977643 | 60978058 | + | -0.64764 | promoter-TSS (NM_017480) | promoter-TSS (NM_017480) | -64 | NM_017480 | 54167 | Mm.42044 | NM_017480 | ENSMUSG00000026009 | **Icos** | AILIM\|CCLP\|CRP-1\|H4\|Ly115 | inducible T cell co-stimulator | protein-coding |
| chr1_60990046_60991075 | chr1 | 60990047 | 60991075 | + | -0.861158 | intron (NM_017480, intron 1 of 4) | intron (NM_017480, intron 1 of 4) | 12647 | NM_017480 | 54167 | Mm.42044 | NM_017480 | ENSMUSG00000026009 | **Icos** | AILIM\|CCLP\|CRP-1\|H4\|Ly115 | inducible T cell co-stimulator | protein-coding |
| chr1_61068193_61068456 | chr1 | 61068194 | 61068456 | + | -1.06254 | Intergenic | Intergenic | 90411 | NM_017480 | 54167 | Mm.42044 | NM_017480 | ENSMUSG00000026009 | **Icos** | AILIM\|CCLP\|CRP-1\|H4\|Ly115 | inducible T cell co-stimulator | protein-coding |
| chr1_61068079_61068515 | chr1 | 61068080 | 61068515 | + | -1.06768 | Intergenic | Intergenic | 90383 | NM_017480 | 54167 | Mm.42044 | NM_017480 | ENSMUSG00000026009 | **Icos** | AILIM\|CCLP\|CRP-1\|H4\|Ly115 | inducible T cell co-stimulator | protein-coding |
| chr10_78078326_78078610 | chr10 | 78078327 | 78078610 | + | -0.717775 | 3' UTR (NM_015790, exon 7 of 7) | 3' UTR (NM_015790, exon 7 of 7) | 9100 | NM_015790 | 50723 | Mm.17819 | NM_015790 | ENSMUSG00000000732 | **Icosl** | AU044799\|B7-H2\|B7RP-1\|B7h\|BG071784\|GI50\|GL50\|GL50-B\|ICOS-L\|Icoslg\|LICOS\|Ly115l\|mKIAA0653 | icos ligand | protein-coding |
| chr10_78078327_78078711 | chr10 | 78078328 | 78078711 | + | -0.718886 | 3' UTR (NM_015790, exon 7 of 7) | 3' UTR (NM_015790, exon 7 of 7) | 9151 | NM_015790 | 50723 | Mm.17819 | NM_015790 | ENSMUSG00000000732 | **Icosl** | AU044799\|B7-H2\|B7RP-1\|B7h\|BG071784\|GI50\|GL50\|GL50-B\|ICOS-L\|Icoslg\|LICOS\|Ly115l\|mKIAA0653 | icos ligand | protein-coding |
| chr11_115406810_115407445 | chr11 | 115406811 | 115407445 | + | -0.872935 | intron (NM_026729, intron 2 of 5) | intron (NM_026729, intron 2 of 5) | 3362 | NM_026729 | 68572 | Mm.271062 | NM_026729 | ENSMUSG00000018858 | **Ict1** | 1110001A02Rik\|1110002E03Rik\|MRP-L58 | immature colon carcinoma transcript 1 | protein-coding |
| chr7_80122956_80123486 | chr7 | 80122957 | 80123486 | + | -0.658556 | Intergenic | Intergenic | -7871 | NM_173011 | 269951 | Mm.246432 | NM_173011 | ENSMUSG00000030541 | **Idh2** | E430004F23\|IDPm\|Idh-2 | isocitrate dehydrogenase 2 (NADP+), mitochondrial | protein-coding |
| chr7_80122956_80123881 | chr7 | 80122957 | 80123881 | + | -0.668721 | Intergenic | Intergenic | -8069 | NM_173011 | 269951 | Mm.246432 | NM_173011 | ENSMUSG00000030541 | **Idh2** | E430004F23\|IDPm\|Idh-2 | isocitrate dehydrogenase 2 (NADP+), mitochondrial | protein-coding |
| chr7_80123174_80123810 | chr7 | 80123175 | 80123810 | + | -0.729311 | Intergenic | Intergenic | -8142 | NM_173011 | 269951 | Mm.246432 | NM_173011 | ENSMUSG00000030541 | **Idh2** | E430004F23\|IDPm\|Idh-2 | isocitrate dehydrogenase 2 (NADP+), mitochondrial | protein-coding |
| chr7_80123207_80123622 | chr7 | 80123208 | 80123622 | + | -0.764037 | Intergenic | Intergenic | -8065 | NM_173011 | 269951 | Mm.246432 | NM_173011 | ENSMUSG00000030541 | **Idh2** | E430004F23\|IDPm\|Idh-2 | isocitrate dehydrogenase 2 (NADP+), mitochondrial | protein-coding |
| chr1_155040898_155041442 | chr1 | 155040899 | 155041442 | + | -0.663945 | Intergenic | ORR1D2\|LTR\|ERVL-MaLR | 58466 | NM_010500 | 15939 | Mm.12246 | NM_010500 | ENSMUSG00000056708 | **Ier5** | - | immediate early response 5 | protein-coding |
| chr1_155041145_155041435 | chr1 | 155041146 | 155041435 | + | -0.850236 | Intergenic | ORR1D2\|LTR\|ERVL-MaLR | 58346 | NM_010500 | 15939 | Mm.12246 | NM_010500 | ENSMUSG00000056708 | **Ier5** | - | immediate early response 5 | protein-coding |
| chr1_173949099_173949425 | chr1 | 173949100 | 173949425 | + | -0.661546 | Intergenic | Lx5\|LINE\|L1 | -6770 | NM_001045481 | 15950 | Mm.463437 | NM_008328 | ENSMUSG00000039997 | **Ifi203** | - | interferon activated gene 203 | protein-coding |
| chr19_34543667_34543912 | chr19 | 34543668 | 34543912 | + | -0.621323 | Intergenic | L1MB7\|LINE\|L1 | -6904 | NM_008332 | 15958 | Mm.2036 | NM_008332 | ENSMUSG00000045932 | **Ifit2** | AV302338\|Ifi54\|P54 | interferon-induced protein with tetratricopeptide repeats 2 | protein-coding |
| chr19_34562537_34562804 | chr19 | 34562538 | 34562804 | + | -0.835049 | intron (NM_008332, intron 1 of 2) | intron (NM_008332, intron 1 of 2) | 11977 | NM_008332 | 15958 | Mm.2036 | NM_008332 | ENSMUSG00000045932 | **Ifit2** | AV302338\|Ifi54\|P54 | interferon-induced protein with tetratricopeptide repeats 2 | protein-coding |
| chr7_142353844_142354374 | chr7 | 142353845 | 142354374 | + | -1.09177 | intron (NM_177265, intron 2 of 3) | intron (NM_177265, intron 2 of 3) | 18150 | NM_177265 | 320802 | Mm.154486 | NM_177265 | ENSMUSG00000045777 | **Ifitm10** | 6330512M04Rik\|AI848576\|DSPA3 | interferon induced transmembrane protein 10 | protein-coding |
| chr7_141038520_141038968 | chr7 | 141038521 | 141038968 | + | -0.705717 | Intergenic | (C)n\|Simple_repeat\|Simple_repeat | -21852 | NM_001033632 | 213002 | Mm.276440 | NM_001033632 | ENSMUSG00000059108 | **Ifitm6** | A330075D07\|Dspa2f\|Fgls5\|fragilis5 | interferon induced transmembrane protein 6 | protein-coding |
| chr4_88867916_88868519 | chr4 | 88867917 | 88868519 | + | -0.627683 | Intergenic | CpG | 11983 | NM_177348 | 230405 | Mm.246593 | NM_177348 | ENSMUSG00000045364 | **Ifne** | Ifn-tau-1\|Ifne1\|Ifnt1\|Infe1 | interferon epsilon | protein-coding |
| chr10_118455807_118456121 | chr10 | 118455808 | 118456121 | + | -0.796091 | Intergenic | Intergenic | 14917 | NM_008337 | 15978 | Mm.240327 | NM_008337 | ENSMUSG00000055170 | **Ifng** | IFN-g\|Ifg | interferon gamma | protein-coding |
| chr10_19505956_19506530 | chr10 | 19505957 | 19506530 | + | -0.614921 | Intergenic | Intergenic | -85715 | NM_010511 | 15979 | Mm.549 | NM_010511 | ENSMUSG00000020009 | **Ifngr1** | CD119\|IFN-gammaR\|Ifgr\|Ifngr\|Nktar | interferon gamma receptor 1 | protein-coding |
| chr10_19584045_19584481 | chr10 | 19584046 | 19584481 | + | -0.717742 | Intergenic | Intergenic | -7695 | NM_010511 | 15979 | Mm.549 | NM_010511 | ENSMUSG00000020009 | **Ifngr1** | CD119\|IFN-gammaR\|Ifgr\|Ifngr\|Nktar | interferon gamma receptor 1 | protein-coding |
| chr10_19503883_19504145 | chr10 | 19503884 | 19504145 | + | -0.759883 | Intergenic | Intergenic | -87944 | NM_010511 | 15979 | Mm.549 | NM_010511 | ENSMUSG00000020009 | **Ifngr1** | CD119\|IFN-gammaR\|Ifgr\|Ifngr\|Nktar | interferon gamma receptor 1 | protein-coding |

| | | | | | | | | | | | | | | | | | |
|---|---|---|---|---|---|---|---|---|---|---|---|---|---|---|---|---|---|
| chr10_19521740_19522833 | chr10 | 19521741 | 19522833 | + | -0.773939 | Intergenic | Intergenic | -69671 | NM_010511 | 15979 | Mm.549 | NM_010511 | ENSMUSG00000020009 | **Ifngr1** | CD119\|IFN-gammaR\|Ifgr\|Ifngr\|Nktar | interferon gamma receptor 1 | protein-coding |
| chr10_19529990_19530659 | chr10 | 19529991 | 19530659 | + | -0.778721 | Intergenic | Intergenic | -61633 | NM_010511 | 15979 | Mm.549 | NM_010511 | ENSMUSG00000020009 | **Ifngr1** | CD119\|IFN-gammaR\|Ifgr\|Ifngr\|Nktar | interferon gamma receptor 1 | protein-coding |
| chr10_19594616_19594871 | chr10 | 19594617 | 19594871 | + | -0.795487 | intron (NM_010511, intron 1 of 6) | intron (NM_010511, intron 1 of 6) | 2786 | NM_010511 | 15979 | Mm.549 | NM_010511 | ENSMUSG00000020009 | **Ifngr1** | CD119\|IFN-gammaR\|Ifgr\|Ifngr\|Nktar | interferon gamma receptor 1 | protein-coding |
| chr10_19530101_19530536 | chr10 | 19530102 | 19530536 | + | -0.846414 | Intergenic | Intergenic | -61639 | NM_010511 | 15979 | Mm.549 | NM_010511 | ENSMUSG00000020009 | **Ifngr1** | CD119\|IFN-gammaR\|Ifgr\|Ifngr\|Nktar | interferon gamma receptor 1 | protein-coding |
| chr10_19577500_19578393 | chr10 | 19577501 | 19578393 | + | -0.872356 | Intergenic | Intergenic | -14011 | NM_010511 | 15979 | Mm.549 | NM_010511 | ENSMUSG00000020009 | **Ifngr1** | CD119\|IFN-gammaR\|Ifgr\|Ifngr\|Nktar | interferon gamma receptor 1 | protein-coding |
| chr10_19596844_19597210 | chr10 | 19596845 | 19597210 | + | -0.929956 | intron (NM_010511, intron 1 of 6) | URR1B\|DNA\|hAT-Charlie | 5069 | NM_010511 | 15979 | Mm.549 | NM_010511 | ENSMUSG00000020009 | **Ifngr1** | CD119\|IFN-gammaR\|Ifgr\|Ifngr\|Nktar | interferon gamma receptor 1 | protein-coding |
| chr10_19577498_19578966 | chr10 | 19577499 | 19578966 | + | -0.95431 | Intergenic | Intergenic | -13726 | NM_010511 | 15979 | Mm.549 | NM_010511 | ENSMUSG00000020009 | **Ifngr1** | CD119\|IFN-gammaR\|Ifgr\|Ifngr\|Nktar | interferon gamma receptor 1 | protein-coding |
| chr10_19582235_19582588 | chr10 | 19582236 | 19582588 | + | -0.982322 | Intergenic | Intergenic | -9546 | NM_010511 | 15979 | Mm.549 | NM_010511 | ENSMUSG00000020009 | **Ifngr1** | CD119\|IFN-gammaR\|Ifgr\|Ifngr\|Nktar | interferon gamma receptor 1 | protein-coding |
| chr10_19577966_19578924 | chr10 | 19577967 | 19578924 | + | -1.28023 | Intergenic | Intergenic | -13513 | NM_010511 | 15979 | Mm.549 | NM_010511 | ENSMUSG00000020009 | **Ifngr1** | CD119\|IFN-gammaR\|Ifgr\|Ifngr\|Nktar | interferon gamma receptor 1 | protein-coding |
| chr16_91538579_91538779 | chr16 | 91538580 | 91538779 | + | -0.654459 | Intergenic | Intergenic | -8415 | NM_008338 | 15980 | Mm.249364 | NM_008338 | | **Ifngr2** | Ifgr2\|Ifgt | interferon gamma receptor 2 | protein-coding |
| chr16_91545471_91546238 | chr16 | 91545472 | 91546238 | + | -0.754583 | Intergenic | Intergenic | -1239 | NM_008338 | 15980 | Mm.249364 | NM_008338 | | **Ifngr2** | Ifgr2\|Ifgt | interferon gamma receptor 2 | protein-coding |
| chr16_91538581_91538870 | chr16 | 91538582 | 91538870 | + | -0.757319 | Intergenic | Intergenic | -8368 | NM_008338 | 15980 | Mm.249364 | NM_008338 | | **Ifngr2** | Ifgr2\|Ifgt | interferon gamma receptor 2 | protein-coding |
| chr16_91545471_91545899 | chr16 | 91545472 | 91545899 | + | -0.787405 | Intergenic | Intergenic | -1409 | NM_008338 | 15980 | Mm.249364 | NM_008338 | | **Ifngr2** | Ifgr2\|Ifgt | interferon gamma receptor 2 | protein-coding |
| chr16_91538611_91538998 | chr16 | 91538612 | 91538998 | + | -0.876025 | Intergenic | Intergenic | -8289 | NM_008338 | 15980 | Mm.249364 | NM_008338 | | **Ifngr2** | Ifgr2\|Ifgt | interferon gamma receptor 2 | protein-coding |
| chr16_91538144_91538390 | chr16 | 91538145 | 91538390 | + | -0.947492 | Intergenic | Intergenic | -8827 | NM_008338 | 15980 | Mm.249364 | NM_008338 | | **Ifngr2** | Ifgr2\|Ifgt | interferon gamma receptor 2 | protein-coding |
| chr16_91553601_91554193 | chr16 | 91553602 | 91554193 | + | -1.01748 | intron (NM_008338, intron 1 of 6) | intron (NM_008338, intron 1 of 6) | 6803 | NM_008338 | 15980 | Mm.249364 | NM_008338 | | **Ifngr2** | Ifgr2\|Ifgt | interferon gamma receptor 2 | protein-coding |
| chr16_91553723_91554061 | chr16 | 91553724 | 91554061 | + | -1.04248 | intron (NM_008338, intron 1 of 6) | intron (NM_008338, intron 1 of 6) | 6798 | NM_008338 | 15980 | Mm.249364 | NM_008338 | | **Ifngr2** | Ifgr2\|Ifgt | interferon gamma receptor 2 | protein-coding |
| chr5_31279246_31280010 | chr5 | 31279247 | 31280010 | + | -0.883686 | intron (NM_026298, intron 14 of 47) | intron (NM_026298, intron 14 of 47) | 11486 | NM_026298 | 67661 | Mm.293023 | NM_026298 | ENSMUSG00000038564 | **Ift172** | 4930553F24Rik\|Slb\|avc1\|wim | intraflagellar transport 172 | protein-coding |
| chr3_68974732_68975054 | chr3 | 68974733 | 68975054 | + | -0.905827 | intron (NM_026641, intron 5 of 19) | intron (NM_026641, intron 5 of 19) | 29677 | NM_026641 | 68259 | Mm.389451 | NM_026641 | ENSMUSG00000027778 | **Ift80** | 4921524P20Rik\|Wdr56\|mKIAA1374 | intraflagellar transport 80 | protein-coding |
| chr7_68102515_68103447 | chr7 | 68102516 | 68103447 | + | -0.631562 | intron (NM_010513, intron 2 of 20) | intron (NM_010513, intron 2 of 20) | 150724 | NM_010513 | 16001 | Mm.275742 | NM_010513 | | **Igf1r** | A330103N21Rik\|CD221\|D930020L01\|IGF-1R\|hyft | insulin-like growth factor I receptor | protein-coding |
| chr7_67958014_67958431 | chr7 | 67958015 | 67958431 | + | -0.807614 | intron (NM_010513, intron 1 of 20) | intron (NM_010513, intron 1 of 20) | 5966 | NM_010513 | 16001 | Mm.275742 | NM_010513 | | **Igf1r** | A330103N21Rik\|CD221\|D930020L01\|IGF-1R\|hyft | insulin-like growth factor I receptor | protein-coding |
| chr7_68070614_68071395 | chr7 | 68070615 | 68071395 | + | -1.13266 | intron (NM_010513, intron 2 of 20) | L1MD\|LINE\|L1 | 118748 | NM_010513 | 16001 | Mm.275742 | NM_010513 | | **Igf1r** | A330103N21Rik\|CD221\|D930020L01\|IGF-1R\|hyft | insulin-like growth factor I receptor | protein-coding |
| chr7_68070680_68071057 | chr7 | 68070681 | 68071057 | + | -1.13704 | intron (NM_010513, intron 2 of 20) | L1MD\|LINE\|L1 | 118612 | NM_010513 | 16001 | Mm.275742 | NM_010513 | | **Igf1r** | A330103N21Rik\|CD221\|D930020L01\|IGF-1R\|hyft | insulin-like growth factor I receptor | protein-coding |
| chr17_12779429_12779868 | chr17 | 12779430 | 12779868 | + | -1.12896 | intron (NR_027784, intron 2 of 2) | intron (NR_027784, intron 2 of 2) | -9943 | NM_010515 | 16004 | Mm.26553 | NM_010515 | ENSMUSG00000023830 | **Igf2r** | AI661837\|CD222\|CI-MPR\|M6P/IGF2R\|Mpr300 | insulin-like growth factor 2 receptor | protein-coding |
| chr7_19945820_19946248 | chr7 | 19945821 | 19946248 | + | -0.685106 | intron (NM_027308, intron 1 of 4) | intron (NM_027308, intron 1 of 4) | 4709 | NM_027308 | 70080 | Mm.98583 | NM_027308 | ENSMUSG00000040498 | **Igsf23** | 2210010C17Rik | immunoglobulin superfamily, member 23 | protein-coding |
| chr7_19940217_19940683 | chr7 | 19940218 | 19940683 | + | -0.815018 | intron (NM_027308, intron 3 of 4) | RMER15\|LTR\|ERVL | 10293 | NM_027308 | 70080 | Mm.98583 | NM_027308 | ENSMUSG00000040498 | **Igsf23** | 2210010C17Rik | immunoglobulin superfamily, member 23 | protein-coding |
| chr3_101407482_101407999 | chr3 | 101407483 | 101407999 | + | -0.676976 | intron (NM_207205, intron 2 of 10) | intron (NM_207205, intron 2 of 10) | 30616 | NM_207205 | 78908 | Mm.257997 | NM_207205 | ENSMUSG00000042035 | **Igsf3** | 1700016K10Rik\|2810035F16Rik\|4833439O17Rik\|mKIAA0466 | immunoglobulin superfamily, member 3 | protein-coding |
| chr7_121064368_121064715 | chr7 | 121064369 | 121064715 | + | -0.672176 | 3' UTR (NM_030691, exon 6 of 6) | 3' UTR (NM_030691, exon 6 of 6) | 9988 | NM_030691 | 80719 | Mm.160384 | NM_030691 | ENSMUSG00000035004 | **Igsf6** | - | immunoglobulin superfamily, member 6 | protein-coding |
| chr7_121059127_121060068 | chr7 | 121059128 | 121060068 | + | -0.787624 | intron (NM_021554, intron 4 of 4) | intron (NM_021554, intron 4 of 4) | 14932 | NM_030691 | 80719 | Mm.160384 | NM_030691 | ENSMUSG00000035004 | **Igsf6** | - | immunoglobulin superfamily, member 6 | protein-coding |
| chr7_121059379_121060073 | chr7 | 121059380 | 121060073 | + | -0.821934 | intron (NM_021554, intron 4 of 4) | intron (NM_021554, intron 4 of 4) | 14804 | NM_030691 | 80719 | Mm.160384 | NM_030691 | ENSMUSG00000035004 | **Igsf6** | - | immunoglobulin superfamily, member 6 | protein-coding |
| chr8_22724099_22724538 | chr8 | 22724100 | 22724538 | + | -0.914016 | Intergenic | RLTR14\|LTR\|ERV1 | -17736 | NM_001159774 | 16150 | Mm.277886 | NM_010546 | ENSMUSG00000031537 | **Ikbkb** | AI132552\|IKK-2\|IKK-beta\|IKK2\|IKK[b]\|IKKbeta | inhibitor of kappaB kinase beta | protein-coding |
| chr1_131324820_131325127 | chr1 | 131324821 | 131325127 | + | -0.82888 | intron (NM_001081011, intron 14 of 22) | B4A\|SINE\|B4 | -45411 | NM_019777 | 56489 | Mm.386783 | NM_019777 | ENSMUSG00000042349 | **Ikbke** | AW558201\|IKK-i\|IKKepsilon\|Ikki | inhibitor of kappaB kinase epsilon | protein-coding |
| chr1_131324617_131325465 | chr1 | 131324618 | 131325465 | + | -0.950169 | intron (NM_001081011, intron 14 of 22) | intron (NM_001081011, intron 14 of 22) | -45478 | NM_019777 | 56489 | Mm.386783 | NM_019777 | ENSMUSG00000042349 | **Ikbke** | AW558201\|IKK-i\|IKKepsilon\|Ikki | inhibitor of kappaB kinase epsilon | protein-coding |
| chr1_131349502_131349951 | chr1 | 131349503 | 131349951 | + | -0.969306 | intron (NM_001081011, intron 7 of 22) | intron (NM_001081011, intron 7 of 22) | -70164 | NM_019777 | 56489 | Mm.386783 | NM_019777 | ENSMUSG00000042349 | **Ikbke** | AW558201\|IKK-i\|IKKepsilon\|Ikki | inhibitor of kappaB kinase epsilon | protein-coding |
| chr11_11719806_11720170 | chr11 | 11719807 | 11720170 | + | -0.763951 | intron (NM_009578, intron 3 of 6) | intron (NM_009578, intron 3 of 6) | 33775 | NM_001025597 | 22778 | Mm.103545 | NM_009578 | ENSMUSG00000018654 | **Ikzf1** | 5832432G11Rik\|Ikaros\|LyF-1\|Zfpn1a1\|Znfn1a1\|hlk-1\|mKIAA4227 | IKAROS family zinc finger 1 | protein-coding |
| chr11_11741560_11741982 | chr11 | 11741561 | 11741982 | + | -0.836278 | intron (NM_009578, intron 3 of 6) | intron (NM_009578, intron 3 of 6) | 55558 | NM_001025597 | 22778 | Mm.103545 | NM_009578 | ENSMUSG00000018654 | **Ikzf1** | 5832432G11Rik\|Ikaros\|LyF-1\|Zfpn1a1\|Znfn1a1\|hlk-1\|mKIAA4227 | IKAROS family zinc finger 1 | protein-coding |
| chr11_11725953_11726546 | chr11 | 11725954 | 11726546 | + | -0.856723 | intron (NM_009578, intron 3 of 6) | intron (NM_009578, intron 3 of 6) | 40037 | NM_001025597 | 22778 | Mm.103545 | NM_009578 | ENSMUSG00000018654 | **Ikzf1** | 5832432G11Rik\|Ikaros\|LyF-1\|Zfpn1a1\|Znfn1a1\|hlk-1\|mKIAA4227 | IKAROS family zinc finger 1 | protein-coding |
| chr11_11703843_11704145 | chr11 | 11703844 | 11704145 | + | -1.02698 | intron (NM_001025597, intron 2 of 7) | intron (NM_001025597, intron 2 of 7) | 17781 | NM_001025597 | 22778 | Mm.103545 | NM_009578 | ENSMUSG00000018654 | **Ikzf1** | 5832432G11Rik\|Ikaros\|LyF-1\|Zfpn1a1\|Znfn1a1\|hlk-1\|mKIAA4227 | IKAROS family zinc finger 1 | protein-coding |
| chr1_69752727_69753041 | chr1 | 69752728 | 69753041 | + | -0.858383 | Intergenic | Intergenic | -66924 | NM_011770 | 22779 | Mm.106343 | NM_011770 | ENSMUSG00000025997 | **Ikzf2** | A730095J18Rik\|Helios\|Zfpn1a2\|Znfn1a2 | IKAROS family zinc finger 2 | protein-coding |
| chr1_69752686_69753194 | chr1 | 69752687 | 69753194 | + | -0.901415 | Intergenic | Intergenic | -66980 | NM_011770 | 22779 | Mm.106343 | NM_011770 | ENSMUSG00000025997 | **Ikzf2** | A730095J18Rik\|Helios\|Zfpn1a2\|Znfn1a2 | IKAROS family zinc finger 2 | protein-coding |
| chr1_69681050_69681348 | chr1 | 69681051 | 69681348 | + | -0.976421 | intron (NM_011770, intron 3 of 7) | intron (NM_011770, intron 3 of 7) | 4761 | NM_011770 | 22779 | Mm.106343 | NM_011770 | ENSMUSG00000025997 | **Ikzf2** | A730095J18Rik\|Helios\|Zfpn1a2\|Znfn1a2 | IKAROS family zinc finger 2 | protein-coding |
| chr11_98526585_98527739 | chr11 | 98526586 | 98527739 | + | -0.624725 | intron (NM_011771, intron 1 of 7) | intron (NM_011771, intron 1 of 7) | 18869 | NM_011771 | 22780 | Mm.133367 | NM_011771 | ENSMUSG00000018168 | **Ikzf3** | 5830411O07Rik\|Aiolos\|Zfpn1a3\|Znfn1a3 | IKAROS family zinc finger 3 | protein-coding |
| chr11_98526768_98527585 | chr11 | 98526769 | 98527585 | + | -0.742588 | intron (NM_011771, intron 1 of 7) | intron (NM_011771, intron 1 of 7) | 18854 | NM_011771 | 22780 | Mm.133367 | NM_011771 | ENSMUSG00000018168 | **Ikzf3** | 5830411O07Rik\|Aiolos\|Zfpn1a3\|Znfn1a3 | IKAROS family zinc finger 3 | protein-coding |
| chr3_68758539_68758966 | chr3 | 68758540 | 68758966 | + | -0.992084 | Intergenic | Intergenic | 67329 | NM_008351 | 16159 | Mm.103783 | NM_008351 | ENSMUSG00000027776 | **Il12a** | IL-12p35\|Il-12a\|Ll12a\|p35 | interleukin 12a | protein-coding |
| chr6_67339530_67339826 | chr6 | 67339531 | 67339826 | + | -1.00903 | intron (NM_008354, intron 7 of 15) | MLT1A\|LTR\|ERVL-MaLR | 36459 | NM_008354 | 16162 | Mm.188337 | NM_008354 | ENSMUSG00000018341 | **Il12rb2** | A930027I18Rik\|IL-12RB2\|Ifnm | interleukin 12 receptor, beta 2 | protein-coding |
| chr6_67339358_67339882 | chr6 | 67339359 | 67339882 | + | -1.04243 | intron (NM_008354, intron 7 of 15) | MLT1A\|LTR\|ERVL-MaLR | 36517 | NM_008354 | 16162 | Mm.188337 | NM_008354 | ENSMUSG00000018341 | **Il12rb2** | A930027I18Rik\|IL-12RB2\|Ifnm | interleukin 12 receptor, beta 2 | protein-coding |
| chr2_11718225_11718632 | chr2 | 11718226 | 11718632 | + | -2.16432 | exon (NM_008358, exon 2 of 7) | exon (NM_008358, exon 2 of 7) | 12717 | NM_001271500 | 16169 | Mm.200196 | NM_008358 | ENSMUSG00000023206 | **Il15ra** | AA690181\|IL-15RA | interleukin 15 receptor, alpha chain | protein-coding |
| chr1_40479452_40479933 | chr1 | 40479453 | 40479933 | + | -0.704868 | intron (NM_008365, intron 4 of 10) | intron (NM_008365, intron 4 of 10) | 13687 | NM_008365 | 16182 | Mm.253664 | NM_008365 | ENSMUSG00000026070 | **Il18r1** | Il18ralpha\|Il1rrp | interleukin 18 receptor 1 | protein-coding |
| chr1_40473701_40474052 | chr1 | 40473702 | 40474052 | + | -0.827816 | intron (NM_008365, intron 2 of 10) | intron (NM_008365, intron 2 of 10) | 7871 | NM_008365 | 16182 | Mm.253664 | NM_008365 | ENSMUSG00000026070 | **Il18r1** | Il18ralpha\|Il1rrp | interleukin 18 receptor 1 | protein-coding |
| chr1_40475337_40476072 | chr1 | 40475338 | 40476072 | + | -0.947842 | intron (NM_008365, intron 3 of 10) | intron (NM_008365, intron 3 of 10) | 9699 | NM_008365 | 16182 | Mm.253664 | NM_008365 | ENSMUSG00000026070 | **Il18r1** | Il18ralpha\|Il1rrp | interleukin 18 receptor 1 | protein-coding |
| chr1_40475673_40476072 | chr1 | 40475674 | 40476072 | + | -1.1829 | intron (NM_008365, intron 3 of 10) | intron (NM_008365, intron 3 of 10) | 9867 | NM_008365 | 16182 | Mm.253664 | NM_008365 | ENSMUSG00000026070 | **Il18r1** | Il18ralpha\|Il1rrp | interleukin 18 receptor 1 | protein-coding |

| | | | | | | | | | | | | | | | | | |
|---|---|---|---|---|---|---|---|---|---|---|---|---|---|---|---|---|---|
| chr2_129313955_129314369 | chr2 | 129313956 | 129314369 | + | -0.981982 | Intergenic | MIRb\|SINE\|MIR | -4190 | NM_010554 | 16175 | Mm.15534 | NM_010554 | ENSMUSG00000027399 | **Il1a** | Il-1a | interleukin 1 alpha | protein-coding |
| chr2_129313995_129314254 | chr2 | 129313996 | 129314254 | + | -0.989593 | Intergenic | MIRb\|SINE\|MIR | -4153 | NM_010554 | 16175 | Mm.15534 | NM_010554 | ENSMUSG00000027399 | **Il1a** | Il-1a | interleukin 1 alpha | protein-coding |
| chr1_40271254_40271523 | chr1 | 40271255 | 40271523 | + | -0.720916 | intron (NM_001123382, intron 1 of 10) | intron (NM_001123382, intron 1 of 10) | 4803 | NM_001123382 | 16177 | Mm.896 | NM_008362 | ENSMUSG00000026072 | **Il1r1** | CD121a\|CD121b\|IL-iR\|Il1r-1 | interleukin 1 receptor, type I | protein-coding |
| chr1_40270983_40271522 | chr1 | 40270984 | 40271522 | + | -0.773481 | intron (NM_001123382, intron 1 of 10) | intron (NM_001123382, intron 1 of 10) | 4667 | NM_001123382 | 16177 | Mm.896 | NM_008362 | ENSMUSG00000026072 | **Il1r1** | CD121a\|CD121b\|IL-iR\|Il1r-1 | interleukin 1 receptor, type I | protein-coding |
| chr1_40380966_40381439 | chr1 | 40380967 | 40381439 | + | -0.783159 | Intergenic | Intergenic | -48367 | NM_001294171 | 17082 | Mm.289824 | NM_010743 | ENSMUSG00000026069 | **Il1rl1** | DER4\|Fit-1\|Ly84\|ST2L\|St2\|St2-rs1\|T1\|T1/ST2 | interleukin 1 receptor-like 1 | protein-coding |
| chr1_40324435_40325147 | chr1 | 40324436 | 40325147 | + | -0.646252 | 5' UTR (NM_133193, exon 1 of 11) | 5' UTR (NM_133193, exon 1 of 11) | 164 | NM_133193 | 107527 | Mm.155583 | NM_133193 | ENSMUSG00000070942 | **Il1rl2** | AI481289\|AW551444\|IL-1Rrp2 | interleukin 1 receptor-like 2 | protein-coding |
| chr3_37152237_37152697 | chr3 | 37152238 | 37152697 | + | -0.61094 | Intergenic | Intergenic | -26513 | NM_008366 | 16183 | Mm.14190 | NM_008366 | ENSMUSG00000027720 | **Il2** | Il-2 | interleukin 2 | protein-coding |
| chr10_19691786_19692576 | chr10 | 19691787 | 19692576 | + | -0.827409 | Intergenic | Intergenic | -20406 | NM_172786 | 237313 | Mm.234667 | NM_172786 | ENSMUSG00000020007 | **Il20ra** | E230031K19 | interleukin 20 receptor, alpha | protein-coding |
| chr10_19747591_19748109 | chr10 | 19747592 | 19748109 | + | -0.889299 | intron (NM_172786, intron 2 of 6) | intron (NM_172786, intron 2 of 6) | 35263 | NM_172786 | 237313 | Mm.234667 | NM_172786 | ENSMUSG00000020007 | **Il20ra** | E230031K19 | interleukin 20 receptor, alpha | protein-coding |
| chr9_100457986_100458316 | chr9 | 100457987 | 100458316 | + | -0.626364 | 3' UTR (NM_001033543, exon 7 of 7) | 3' UTR (NM_001033543, exon 7 of 7) | 28322 | NM_001033543 | 213208 | Mm.242896 | NM_001033543 | ENSMUSG00000044244 | **Il20rb** | AV228068\|Fndc6\|Gm186\|Il20R2 | interleukin 20 receptor beta | protein-coding |
| chr9_100459983_100460831 | chr9 | 100459984 | 100460831 | + | -0.746896 | intron (NM_001033543, intron 6 of 6) | (TG)n\|Simple_repeat\|Simple_repeat | 26066 | NM_001033543 | 213208 | Mm.242896 | NM_001033543 | ENSMUSG00000044244 | **Il20rb** | AV228068\|Fndc6\|Gm186\|Il20R2 | interleukin 20 receptor beta | protein-coding |
| chr3_37203981_37204491 | chr3 | 37203982 | 37204491 | + | -0.673407 | Intergenic | Intergenic | 28400 | NM_001291041 | 60505 | Mm.157689 | NM_021782 | ENSMUSG00000027718 | **Il21** | IL-21 | interleukin 21 | protein-coding |
| chr3_37205084_37205855 | chr3 | 37205085 | 37205855 | + | -1.01892 | Intergenic | Intergenic | 27166 | NM_001291041 | 60505 | Mm.157689 | NM_021782 | ENSMUSG00000027718 | **Il21** | IL-21 | interleukin 21 | protein-coding |
| chr7_125635401_125635838 | chr7 | 125635402 | 125635838 | + | -0.631896 | Intergenic | Intergenic | 32191 | NM_021887 | 60504 | Mm.155643 | NM_021887 | ENSMUSG00000030745 | **Il21r** | NILR | interleukin 21 receptor | protein-coding |
| chr7_125630188_125630721 | chr7 | 125630189 | 125630721 | + | -0.696783 | intron (NM_021887, intron 7 of 8) | intron (NM_021887, intron 7 of 8) | 27026 | NM_021887 | 60504 | Mm.155643 | NM_021887 | ENSMUSG00000030745 | **Il21r** | NILR | interleukin 21 receptor | protein-coding |
| chr7_125625231_125625571 | chr7 | 125625232 | 125625571 | + | -0.926495 | intron (NM_021887, intron 3 of 8) | intron (NM_021887, intron 3 of 8) | 21972 | NM_021887 | 60504 | Mm.155643 | NM_021887 | ENSMUSG00000030745 | **Il21r** | NILR | interleukin 21 receptor | protein-coding |
| chr10_19613009_19614332 | chr10 | 19613010 | 19614332 | + | -0.94435 | Intergenic | Intergenic | -8357 | NM_178258 | 237310 | Mm.331979 | NM_178258 | ENSMUSG00000039760 | **Il22ra2** | CRF2-10\|CRF2-s1\|CRF2X\|Il-22bp | interleukin 22 receptor, alpha 2 | protein-coding |
| chr10_19613052_19613784 | chr10 | 19613053 | 19613784 | + | -0.948932 | Intergenic | MTE-int\|LTR\|ERVL-MaLR | -8610 | NM_178258 | 237310 | Mm.331979 | NM_178258 | ENSMUSG00000039760 | **Il22ra2** | CRF2-10\|CRF2-s1\|CRF2X\|Il-22bp | interleukin 22 receptor, alpha 2 | protein-coding |
| chr10_19619643_19620109 | chr10 | 19619644 | 19620109 | + | -1.25664 | Intergenic | Intergenic | -2152 | NM_178258 | 237310 | Mm.331979 | NM_178258 | ENSMUSG00000039760 | **Il22ra2** | CRF2-10\|CRF2-s1\|CRF2X\|Il-22bp | interleukin 22 receptor, alpha 2 | protein-coding |
| chr10_19619756_19620066 | chr10 | 19619757 | 19620066 | + | -1.26302 | Intergenic | Intergenic | -2117 | NM_178258 | 237310 | Mm.331979 | NM_178258 | ENSMUSG00000039760 | **Il22ra2** | CRF2-10\|CRF2-s1\|CRF2X\|Il-22bp | interleukin 22 receptor, alpha 2 | protein-coding |
| chr7_125561208_125561659 | chr7 | 125561209 | 125561659 | + | -0.641953 | intron (NM_001008700, intron 1 of 10) | intron (NM_001008700, intron 1 of 10) | 9152 | NM_001008700 | 16190 | Mm.233802 | NM_010557 | ENSMUSG00000030748 | **Il4ra** | CD124\|Il4r | interleukin 4 receptor, alpha | protein-coding |
| chr7_125541860_125542344 | chr7 | 125541861 | 125542344 | + | -0.652566 | Intergenic | Intergenic | -10180 | NM_001008700 | 16190 | Mm.233802 | NM_010557 | ENSMUSG00000030748 | **Il4ra** | CD124\|Il4r | interleukin 4 receptor, alpha | protein-coding |
| chr3_89872183_89872657 | chr3 | 89872184 | 89872657 | + | -0.887817 | intron (NM_010559, intron 9 of 9) | intron (NM_010559, intron 9 of 9) | 40742 | NM_010559 | 16194 | Mm.2856 | NM_010559 | ENSMUSG00000027947 | **Il6ra** | CD126\|IL-6R\|Il6r | interleukin 6 receptor, alpha | protein-coding |
| chr13_112424567_112425072 | chr13 | 112424568 | 112425072 | + | -0.696563 | Intergenic | Intergenic | -39250 | NM_010560 | 16195 | Mm.4364 | NM_010560 | ENSMUSG00000021756 | **Il6st** | 5133400A03Rik\|AA389424\|BB405851\|CD130\|D13Ertd699e\|gp130 | interleukin 6 signal transducer | protein-coding |
| chr13_112507577_112508111 | chr13 | 112507578 | 112508111 | + | -0.70169 | TTS (NM_010560) | TTS (NM_010560) | 43774 | NM_010560 | 16195 | Mm.4364 | NM_010560 | ENSMUSG00000021756 | **Il6st** | 5133400A03Rik\|AA389424\|BB405851\|CD130\|D13Ertd699e\|gp130 | interleukin 6 signal transducer | protein-coding |
| chr13_112472264_112472709 | chr13 | 112472265 | 112472709 | + | -0.705694 | intron (NM_010560, intron 1 of 16) | Lx8\|LINE\|L1 | 8417 | NM_010560 | 16195 | Mm.4364 | NM_010560 | ENSMUSG00000021756 | **Il6st** | 5133400A03Rik\|AA389424\|BB405851\|CD130\|D13Ertd699e\|gp130 | interleukin 6 signal transducer | protein-coding |
| chr13_112472414_112472691 | chr13 | 112472415 | 112472691 | + | -0.763148 | intron (NM_010560, intron 1 of 16) | intron (NM_010560, intron 1 of 16) | 8483 | NM_010560 | 16195 | Mm.4364 | NM_010560 | ENSMUSG00000021756 | **Il6st** | 5133400A03Rik\|AA389424\|BB405851\|CD130\|D13Ertd699e\|gp130 | interleukin 6 signal transducer | protein-coding |
| chr13_112427556_112427943 | chr13 | 112427557 | 112427943 | + | -0.849274 | Intergenic | Intergenic | -36320 | NM_010560 | 16195 | Mm.4364 | NM_010560 | ENSMUSG00000021756 | **Il6st** | 5133400A03Rik\|AA389424\|BB405851\|CD130\|D13Ertd699e\|gp130 | interleukin 6 signal transducer | protein-coding |
| chr13_112437948_112438346 | chr13 | 112437949 | 112438346 | + | -0.85662 | Intergenic | RCHARR1\|DNA\|hAT-Charlie | -25923 | NM_010560 | 16195 | Mm.4364 | NM_010560 | ENSMUSG00000021756 | **Il6st** | 5133400A03Rik\|AA389424\|BB405851\|CD130\|D13Ertd699e\|gp130 | interleukin 6 signal transducer | protein-coding |
| chr13_112427470_112428001 | chr13 | 112427471 | 112428001 | + | -0.858776 | Intergenic | Intergenic | -36334 | NM_010560 | 16195 | Mm.4364 | NM_010560 | ENSMUSG00000021756 | **Il6st** | 5133400A03Rik\|AA389424\|BB405851\|CD130\|D13Ertd699e\|gp130 | interleukin 6 signal transducer | protein-coding |
| chr13_112432981_112433705 | chr13 | 112432982 | 112433705 | + | -0.905605 | Intergenic | Intergenic | -30727 | NM_010560 | 16195 | Mm.4364 | NM_010560 | ENSMUSG00000021756 | **Il6st** | 5133400A03Rik\|AA389424\|BB405851\|CD130\|D13Ertd699e\|gp130 | interleukin 6 signal transducer | protein-coding |
| chr13_112433186_112433672 | chr13 | 112433187 | 112433672 | + | -0.966836 | Intergenic | Intergenic | -30641 | NM_010560 | 16195 | Mm.4364 | NM_010560 | ENSMUSG00000021756 | **Il6st** | 5133400A03Rik\|AA389424\|BB405851\|CD130\|D13Ertd699e\|gp130 | interleukin 6 signal transducer | protein-coding |
| chr15_9581350_9581889 | chr15 | 9581351 | 9581889 | + | -0.691653 | Intergenic | Intergenic | -51744 | NM_008372 | 16197 | Mm.389 | NM_008372 | ENSMUSG00000003882 | **Il7r** | CD127\|IL-7Ralpha | interleukin 7 receptor | protein-coding |
| chr15_9600095_9600467 | chr15 | 9600096 | 9600467 | + | -0.793645 | Intergenic | Intergenic | -70405 | NM_008372 | 16197 | Mm.389 | NM_008372 | ENSMUSG00000003882 | **Il7r** | CD127\|IL-7Ralpha | interleukin 7 receptor | protein-coding |
| chr15_9564084_9564725 | chr15 | 9564085 | 9564725 | + | -0.824288 | Intergenic | Intergenic | -34529 | NM_008372 | 16197 | Mm.389 | NM_008372 | ENSMUSG00000003882 | **Il7r** | CD127\|IL-7Ralpha | interleukin 7 receptor | protein-coding |
| chr15_9580483_9581278 | chr15 | 9580484 | 9581278 | + | -0.829564 | Intergenic | Intergenic | -51005 | NM_008372 | 16197 | Mm.389 | NM_008372 | ENSMUSG00000003882 | **Il7r** | CD127\|IL-7Ralpha | interleukin 7 receptor | protein-coding |
| chr15_9564176_9564547 | chr15 | 9564177 | 9564547 | + | -0.864345 | Intergenic | Intergenic | -34486 | NM_008372 | 16197 | Mm.389 | NM_008372 | ENSMUSG00000003882 | **Il7r** | CD127\|IL-7Ralpha | interleukin 7 receptor | protein-coding |
| chr15_9533272_9533837 | chr15 | 9533273 | 9533837 | + | -0.930604 | Intergenic | Intergenic | -3679 | NM_008372 | 16197 | Mm.389 | NM_008372 | ENSMUSG00000003882 | **Il7r** | CD127\|IL-7Ralpha | interleukin 7 receptor | protein-coding |
| chr15_9533370_9533784 | chr15 | 9533371 | 9533784 | + | -0.954485 | Intergenic | Intergenic | -3701 | NM_008372 | 16197 | Mm.389 | NM_008372 | ENSMUSG00000003882 | **Il7r** | CD127\|IL-7Ralpha | interleukin 7 receptor | protein-coding |
| chr15_9574742_9575296 | chr15 | 9574743 | 9575296 | + | -1.33236 | Intergenic | Intergenic | -45143 | NM_008372 | 16197 | Mm.389 | NM_008372 | ENSMUSG00000003882 | **Il7r** | CD127\|IL-7Ralpha | interleukin 7 receptor | protein-coding |
| chr15_9574596_9575425 | chr15 | 9574597 | 9575425 | + | -1.33463 | Intergenic | Intergenic | -45135 | NM_008372 | 16197 | Mm.389 | NM_008372 | ENSMUSG00000003882 | **Il7r** | CD127\|IL-7Ralpha | interleukin 7 receptor | protein-coding |
| chr11_32120456_32120953 | chr11 | 32120457 | 32120953 | + | -0.759877 | Intergenic | Intergenic | 79574 | NM_001134458 | 16199 | Mm.384 | NM_008374 | ENSMUSG00000020279 | **Il9r** | - | interleukin 9 receptor | protein-coding |
| chr11_32120588_32120939 | chr11 | 32120589 | 32120939 | + | -0.766159 | Intergenic | Intergenic | 79515 | NM_001134458 | 16199 | Mm.384 | NM_008374 | ENSMUSG00000020279 | **Il9r** | - | interleukin 9 receptor | protein-coding |
| chr11_32125640_32127020 | chr11 | 32125641 | 32127020 | + | -0.908454 | Intergenic | Intergenic | 73949 | NM_001134458 | 16199 | Mm.384 | NM_008374 | ENSMUSG00000020279 | **Il9r** | - | interleukin 9 receptor | protein-coding |
| chr11_32125855_32126712 | chr11 | 32125856 | 32126712 | + | -1.01947 | Intergenic | Intergenic | 73995 | NM_001134458 | 16199 | Mm.384 | NM_008374 | ENSMUSG00000020279 | **Il9r** | - | interleukin 9 receptor | protein-coding |
| chr4_4790217_4790494 | chr4 | 4790218 | 4790494 | + | -0.681275 | intron (NM_177730, intron 1 of 4) | intron (NM_177730, intron 1 of 4) | 2950 | NM_177730 | 242291 | Mm.218889 | NM_177730 | | **Impad1** | 1110001C20Rik\|AA408880\|AI451589\|AL022796\|B230207P20\|Jaws\|gPAPP | inositol monophosphatase domain containing 1 | protein-coding |
| chr4_4790058_4790573 | chr4 | 4790059 | 4790573 | + | -0.704103 | intron (NM_177730, intron 1 of 4) | intron (NM_177730, intron 1 of 4) | 2990 | NM_177730 | 242291 | Mm.218889 | NM_177730 | | **Impad1** | 1110001C20Rik\|AA408880\|AI451589\|AL022796\|B230207P20\|Jaws\|gPAPP | inositol monophosphatase domain containing 1 | protein-coding |
| chr4_4790126_4790500 | chr4 | 4790127 | 4790500 | + | -0.713117 | intron (NM_177730, intron 1 of 4) | intron (NM_177730, intron 1 of 4) | 2993 | NM_177730 | 242291 | Mm.218889 | NM_177730 | | **Impad1** | 1110001C20Rik\|AA408880\|AI451589\|AL022796\|B230207P20\|Jaws\|gPAPP | inositol monophosphatase domain containing 1 | protein-coding |
| chr6_29232298_29232786 | chr6 | 29232299 | 29232786 | + | -0.65215 | Intergenic | Intergenic | -20271 | NM_011829 | 23917 | Mm.260707 | NM_011829 | ENSMUSG00000003500 | **Impdh1** | B930086D20Rik | inosine 5'-phosphate dehydrogenase 1 | protein-coding |
| chr9_108540857_108541242 | chr9 | 108540858 | 108541242 | + | -1.17978 | intron (NM_175143, intron 3 of 9) | intron (NM_175143, intron 3 of 9) | -19451 | NM_011830 | 23918 | Mm.6065 | NM_011830 | ENSMUSG00000062867 | **Impdh2** | IMPD | inosine 5'-phosphate dehydrogenase 2 | protein-coding |
| chr9_80317886_80318527 | chr9 | 80317887 | 80318527 | + | -0.814447 | intron (NM_022016, intron 15 of 16) | intron (NM_022016, intron 15 of 16) | 147231 | NM_022016 | 63859 | Mm.422780 | NM_022016 | ENSMUSG00000032343 | **Impg1** | A930015H12Rik\|IMP150\|SPACR | interphotoreceptor matrix proteoglycan 1 | protein-coding |
| chr4_98436596_98437365 | chr4 | 98436597 | 98437365 | + | -0.918464 | intron (NM_001005784, intron 11 of 19) | intron (NM_001005784, intron 11 of 19) | 41155 | NM_172696 | 12695 | Mm.90218 | NM_007704 | ENSMUSG00000061859 | **Inadl** | Cipp\|Patj | InaD-like (Drosophila) | protein-coding |
| chr8_81703615_81703860 | chr8 | 81703616 | 81703860 | + | -0.871783 | intron (NM_001297596, intron 2 of 24) | intron (NM_001297596, intron 2 of 24) | 361176 | NM_001297593 | 234515 | Mm.324468 | NM_001024617 | ENSMUSG00000037940 | **Inpp4b** | E130107I17Rik | inositol polyphosphate-4-phosphatase, type II | protein-coding |
| chr8_81761511_81761854 | chr8 | 81761512 | 81761854 | + | -0.970246 | intron (NM_001297593, intron 4 of 23) | intron (NM_001297593, intron 4 of 23) | 419121 | NM_001297593 | 234515 | Mm.324468 | NM_001024617 | ENSMUSG00000037940 | **Inpp4b** | E130107I17Rik | inositol polyphosphate-4-phosphatase, type II | protein-coding |
| chr8_81704019_81704315 | chr8 | 81704020 | 81704315 | + | -0.972559 | intron (NM_001297596, intron 2 of 24) | intron (NM_001297596, intron 2 of 24) | 361605 | NM_001297593 | 234515 | Mm.324468 | NM_001024617 | ENSMUSG00000037940 | **Inpp4b** | E130107I17Rik | inositol polyphosphate-4-phosphatase, type II | protein-coding |
| chr8_81782699_81783273 | chr8 | 81782700 | 81783273 | + | -1.07343 | intron (NM_001024617, intron 7 of 24) | RMER19A\|LTR\|ERVK | 440424 | NM_001297593 | 234515 | Mm.324468 | NM_001024617 | ENSMUSG00000037940 | **Inpp4b** | E130107I17Rik | inositol polyphosphate-4-phosphatase, type II | protein-coding |

| | | | | | | | | | | | | | | | | | |
|---|---|---|---|---|---|---|---|---|---|---|---|---|---|---|---|---|---|
| chr8_81491775_81492182 | chr8 | 81491776 | 81492182 | + | -1.46721 | intron (NM_001297596, intron 2 of 24) | intron (NM_001297596, intron 2 of 24) | 149417 | NM_001297593 | 234515 | Mm.324468 | NM_001024617 | ENSMUSG00000037940 | **Inpp4b** | E130107I17Rik | inositol polyphosphate-4-phosphatase, type II | protein-coding |
| chr8_81694919_81695407 | chr8 | 81694920 | 81695407 | + | -1.47852 | intron (NM_001297596, intron 2 of 24) | intron (NM_001297596, intron 2 of 24) | 352601 | NM_001297593 | 234515 | Mm.324468 | NM_001024617 | ENSMUSG00000037940 | **Inpp4b** | E130107I17Rik | inositol polyphosphate-4-phosphatase, type II | protein-coding |
| chr1_87626369_87626956 | chr1 | 87626370 | 87626956 | + | -0.715651 | intron (NR_122078, intron 1 of 27) | intron (NR_122078, intron 1 of 27) | 6351 | NM_001110193 | 16331 | Mm.15105 | NM_010566 | ENSMUSG00000026288 | **Inpp5d** | SHIP\|SHIP-1\|SHIP1\|SIP-145\|p150Ship\|s-SHIP | inositol polyphosphate-5-phosphatase D | protein-coding |
| chr1_87607325_87608118 | chr1 | 87607326 | 87608118 | + | -0.91558 | Intergenic | Intergenic | -12590 | NM_001110192 | 16331 | Mm.15105 | NM_010566 | ENSMUSG00000026288 | **Inpp5d** | SHIP\|SHIP-1\|SHIP1\|SIP-145\|p150Ship\|s-SHIP | inositol polyphosphate-5-phosphatase D | protein-coding |
| chr4_103039130_103040184 | chr4 | 103039131 | 103040184 | + | -0.924396 | intron (NM_146254, intron 16 of 16) | intron (NM_146254, intron 16 of 16) | -12815 | NM_001290648 | 23919 | Mm.100163 | NM_011831 | ENSMUSG00000066090 | **Insl5** | RIF2 | insulin-like 5 | protein-coding |
| chr4_103039120_103039833 | chr4 | 103039121 | 103039833 | + | -0.994399 | intron (NM_146254, intron 16 of 16) | intron (NM_146254, intron 16 of 16) | -12635 | NM_001290648 | 23919 | Mm.100163 | NM_011831 | ENSMUSG00000066090 | **Insl5** | RIF2 | insulin-like 5 | protein-coding |
| chr1_191599057_191599728 | chr1 | 191599058 | 191599728 | + | -0.850189 | intron (NM_178632, intron 7 of 19) | intron (NM_178632, intron 7 of 19) | 23659 | NM_178632 | 77065 | Mm.338175 | NM_178632 | ENSMUSG00000037461 | **Ints7** | 5930412E23Rik | integrator complex subunit 7 | protein-coding |
| chr10_6988520_6988929 | chr10 | 6988521 | 6988929 | + | -0.642201 | intron (NM_001039652, intron 4 of 5) | intron (NM_001039652, intron 4 of 5) | -8260 | NM_001170801 | 320495 | Mm.331674 | NM_001033391 | ENSMUSG00000064065 | **Ipcef1** | A130090K04Rik | interaction protein for cytohesin exchange factors 1 | protein-coding |
| chr10_7005129_7005613 | chr10 | 7005130 | 7005613 | + | -0.650135 | intron (NM_001033391, intron 1 of 7) | intron (NM_001033391, intron 1 of 7) | 17814 | NM_001033391 | 320495 | Mm.331674 | NM_001033391 | ENSMUSG00000064065 | **Ipcef1** | A130090K04Rik | interaction protein for cytohesin exchange factors 1 | protein-coding |
| chr10_6954562_6954952 | chr10 | 6954563 | 6954952 | + | -0.777204 | intron (NM_001170801, intron 3 of 6) | intron (NM_001170801, intron 3 of 6) | 25708 | NM_001170800 | 320495 | Mm.331674 | NM_001033391 | ENSMUSG00000064065 | **Ipcef1** | A130090K04Rik | interaction protein for cytohesin exchange factors 1 | protein-coding |
| chr10_6986301_6986635 | chr10 | 6986302 | 6986635 | + | -0.98954 | intron (NM_001039652, intron 4 of 5) | intron (NM_001039652, intron 4 of 5) | -6003 | NM_001170801 | 320495 | Mm.331674 | NM_001033391 | ENSMUSG00000064065 | **Ipcef1** | A130090K04Rik | interaction protein for cytohesin exchange factors 1 | protein-coding |
| chr13_49435817_49436095 | chr13 | 49435818 | 49436095 | + | -0.717841 | intron (NM_001276399, intron 5 of 12) | intron (NM_001276399, intron 5 of 12) | 14342 | NM_001276399 | 75678 | Mm.390586 | NM_199056 | ENSMUSG00000021385 | **Ippk** | 1810043M15Rik\|InsP6 | inositol 1,3,4,5,6-pentakisphosphate 2-kinase | protein-coding |
| chr16_33012914_33013352 | chr16 | 33012915 | 33013352 | + | -0.782124 | 3' UTR (NM_001081255, exon 21 of 21) | 3' UTR (NM_001081255, exon 21 of 21) | 43053 | NM_178378 | 69707 | Mm.227918 | NM_178378 | ENSMUSG00000035578 | **Iqcg** | 2400003L07Rik | IQ motif containing G | protein-coding |
| chr16_33012926_33013200 | chr16 | 33012927 | 33013200 | + | -0.823036 | 3' UTR (NM_001081255, exon 21 of 21) | 3' UTR (NM_001081255, exon 21 of 21) | 43123 | NM_178378 | 69707 | Mm.227918 | NM_178378 | ENSMUSG00000035578 | **Iqcg** | 2400003L07Rik | IQ motif containing G | protein-coding |
| chr7_80832228_80832550 | chr7 | 80832229 | 80832550 | + | -0.728634 | Intergenic | RLTR19-int\|LTR\|ERVK | -29058 | NM_016721 | 29875 | Mm.207619 | NM_016721 | ENSMUSG00000030536 | **Iqgap1** | AA682088\|D7Ertd237e\|D7Ertd257e\|mKIAA0051 | IQ motif containing GTPase activating protein 1 | protein-coding |
| chr7_80799183_80799755 | chr7 | 80799184 | 80799755 | + | -0.820079 | intron (NM_016721, intron 2 of 37) | intron (NM_016721, intron 2 of 37) | 3862 | NM_016721 | 29875 | Mm.207619 | NM_016721 | ENSMUSG00000030536 | **Iqgap1** | AA682088\|D7Ertd237e\|D7Ertd257e\|mKIAA0051 | IQ motif containing GTPase activating protein 1 | protein-coding |
| chr13_95891586_95892842 | chr13 | 95891587 | 95892842 | + | -0.635773 | promoter-TSS (NM_027711) | promoter-TSS (NM_027711) | -292 | NM_027711 | 544963 | Mm.38878 | NM_027711 | ENSMUSG00000021676 | **Iqgap2** | 4933417J23Rik\|A630053O10\|AI788777 | IQ motif containing GTPase activating protein 2 | protein-coding |
| chr13_95859323_95859663 | chr13 | 95859324 | 95859663 | + | -0.76566 | intron (NM_027711, intron 1 of 35) | intron (NM_027711, intron 1 of 35) | 32429 | NM_027711 | 544963 | Mm.38878 | NM_027711 | ENSMUSG00000021676 | **Iqgap2** | 4933417J23Rik\|A630053O10\|AI788777 | IQ motif containing GTPase activating protein 2 | protein-coding |
| chr13_95956352_95957188 | chr13 | 95956353 | 95957188 | + | -0.793915 | Intergenic | (TG)n\|Simple_repeat\|Simple_repeat | -64848 | NM_027711 | 544963 | Mm.38878 | NM_027711 | ENSMUSG00000021676 | **Iqgap2** | 4933417J23Rik\|A630053O10\|AI788777 | IQ motif containing GTPase activating protein 2 | protein-coding |
| chr13_95956303_95956875 | chr13 | 95956304 | 95956875 | + | -0.852929 | Intergenic | Intergenic | -64667 | NM_027711 | 544963 | Mm.38878 | NM_027711 | ENSMUSG00000021676 | **Iqgap2** | 4933417J23Rik\|A630053O10\|AI788777 | IQ motif containing GTPase activating protein 2 | protein-coding |
| chr13_95942606_95943256 | chr13 | 95942607 | 95943256 | + | -0.967345 | Intergenic | PB1D10\|SINE\|Alu | -51009 | NM_027711 | 544963 | Mm.38878 | NM_027711 | ENSMUSG00000021676 | **Iqgap2** | 4933417J23Rik\|A630053O10\|AI788777 | IQ motif containing GTPase activating protein 2 | protein-coding |
| chr13_95964223_95964667 | chr13 | 95964224 | 95964667 | + | -1.08146 | intron (NM_029210, intron 12 of 12) | intron (NM_029210, intron 12 of 12) | -72523 | NM_027711 | 544963 | Mm.38878 | NM_027711 | ENSMUSG00000021676 | **Iqgap2** | 4933417J23Rik\|A630053O10\|AI788777 | IQ motif containing GTPase activating protein 2 | protein-coding |
| chr6_90740718_90741187 | chr6 | 90740719 | 90741187 | + | -0.749808 | intron (NM_001134384, intron 1 of 13) | intron (NM_001134384, intron 1 of 13) | -24424 | NM_001134383 | 232227 | Mm.196943 | NM_182784 | ENSMUSG00000034312 | **Iqsec1** | AW561907\|BC026481\|BRAG2\|D6Ertd349e\|cl-43 | IQ motif and Sec7 domain 1 | protein-coding |
| chr6_90748221_90748637 | chr6 | 90748222 | 90748637 | + | -0.881099 | intron (NM_001134384, intron 1 of 13) | intron (NM_001134384, intron 1 of 13) | -31900 | NM_001134383 | 232227 | Mm.196943 | NM_182784 | ENSMUSG00000034312 | **Iqsec1** | AW561907\|BC026481\|BRAG2\|D6Ertd349e\|cl-43 | IQ motif and Sec7 domain 1 | protein-coding |
| chr6_90807547_90807889 | chr6 | 90807548 | 90807889 | + | -0.963799 | intron (NM_001134384, intron 1 of 13) | intron (NM_001134384, intron 1 of 13) | 2405 | NM_001134384 | 232227 | Mm.196943 | NM_182784 | ENSMUSG00000034312 | **Iqsec1** | AW561907\|BC026481\|BRAG2\|D6Ertd349e\|cl-43 | IQ motif and Sec7 domain 1 | protein-coding |
| chr10_120193040_120193614 | chr10 | 120193041 | 120193614 | + | -0.654651 | intron (NM_028679, intron 1 of 11) | intron (NM_028679, intron 1 of 11) | 8210 | NM_028679 | 73914 | Mm.146194 | NM_028679 | ENSMUSG00000020227 | **Irak3** | 4833428C18Rik\|AI563835\|IRAK-M | interleukin-1 receptor-associated kinase 3 | protein-coding |
| chr11_53803058_53803449 | chr11 | 53803059 | 53803449 | + | -0.68709 | intron (NR_033332, intron 3 of 3) | intron (NR_033332, intron 3 of 3) | 32781 | NM_001159396 | 16362 | Mm.105218 | NM_008390 | ENSMUSG00000018899 | **Irf1** | AU020929\|Irf-1 | interferon regulatory factor 1 | protein-coding |
| chr11_53785468_53785868 | chr11 | 53785469 | 53785868 | + | -0.758853 | non-coding (NR_033332, exon 4 of 4) | non-coding (NR_033332, exon 4 of 4) | 15195 | NM_001159396 | 16362 | Mm.105218 | NM_008390 | ENSMUSG00000018899 | **Irf1** | AU020929\|Irf-1 | interferon regulatory factor 1 | protein-coding |
| chr8_46755750_46756363 | chr8 | 46755751 | 46756363 | + | -0.683093 | intron (NM_008391, intron 1 of 8) | A-rich\|Low_complexity\|Low_complexity | 16312 | NM_008391 | 16363 | Mm.1149 | NM_008391 | ENSMUSG00000031627 | **Irf2** | 9830146E22Rik\|AI646973\|Irf-2 | interferon regulatory factor 2 | protein-coding |
| chr8_46762673_46763011 | chr8 | 46762674 | 46763011 | + | -0.771439 | intron (NM_008391, intron 1 of 8) | intron (NM_008391, intron 1 of 8) | 23097 | NM_008391 | 16363 | Mm.1149 | NM_008391 | ENSMUSG00000031627 | **Irf2** | 9830146E22Rik\|AI646973\|Irf-2 | interferon regulatory factor 2 | protein-coding |
| chr8_126614343_126614787 | chr8 | 126614344 | 126614787 | + | -0.727795 | Intergenic | Intergenic | -21129 | NM_001164598 | 270110 | Mm.334918 | NM_001164598 | ENSMUSG00000051495 | **Irf2bp2** | E130305N23Rik\|IRF-2BP2 | interferon regulatory factor 2 binding protein 2 | protein-coding |
| chr8_126628660_126628962 | chr8 | 126628661 | 126628962 | + | -0.72863 | Intergenic | Intergenic | -35375 | NM_001164598 | 270110 | Mm.334918 | NM_001164598 | ENSMUSG00000051495 | **Irf2bp2** | E130305N23Rik\|IRF-2BP2 | interferon regulatory factor 2 binding protein 2 | protein-coding |
| chr8_126626597_126627357 | chr8 | 126626598 | 126627357 | + | -0.791238 | Intergenic | Intergenic | -33541 | NM_001164598 | 270110 | Mm.334918 | NM_001164598 | ENSMUSG00000051495 | **Irf2bp2** | E130305N23Rik\|IRF-2BP2 | interferon regulatory factor 2 binding protein 2 | protein-coding |
| chr8_126626881_126627419 | chr8 | 126626882 | 126627419 | + | -0.889409 | Intergenic | Intergenic | -33714 | NM_001164598 | 270110 | Mm.334918 | NM_001164598 | ENSMUSG00000051495 | **Irf2bp2** | E130305N23Rik\|IRF-2BP2 | interferon regulatory factor 2 binding protein 2 | protein-coding |
| chr12_86894225_86894879 | chr12 | 86894226 | 86894879 | + | -0.644417 | Intergenic | Intergenic | -9738 | NM_145836 | 238330 | Mm.248358 | NM_145836 | ENSMUSG00000034168 | **Irf2bpl** | 6430527G18Rik\|Eap1 | interferon regulatory factor 2 binding protein-like | protein-coding |
| chr12_86860093_86860852 | chr12 | 86860094 | 86860852 | + | -0.655449 | Intergenic | Intergenic | 24341 | NM_145836 | 238330 | Mm.248358 | NM_145836 | ENSMUSG00000034168 | **Irf2bpl** | 6430527G18Rik\|Eap1 | interferon regulatory factor 2 binding protein-like | protein-coding |
| chr12_86860049_86860529 | chr12 | 86860050 | 86860529 | + | -0.836822 | Intergenic | Intergenic | 24525 | NM_145836 | 238330 | Mm.248358 | NM_145836 | ENSMUSG00000034168 | **Irf2bpl** | 6430527G18Rik\|Eap1 | interferon regulatory factor 2 binding protein-like | protein-coding |
| chr12_86810674_86811384 | chr12 | 86810675 | 86811384 | + | -1.00179 | Intergenic | Intergenic | 73785 | NM_145836 | 238330 | Mm.248358 | NM_145836 | ENSMUSG00000034168 | **Irf2bpl** | 6430527G18Rik\|Eap1 | interferon regulatory factor 2 binding protein-like | protein-coding |
| chr13_30733091_30733411 | chr13 | 30733092 | 30733411 | + | -0.761099 | Intergenic | Intergenic | -16007 | NM_013674 | 16364 | Mm.4677 | NM_013674 | ENSMUSG00000021356 | **Irf4** | AI385587\|IRF-4\|LSIRF\|Spip | interferon regulatory factor 4 | protein-coding |
| chr8_120752081_120752438 | chr8 | 120752082 | 120752438 | + | -0.869849 | intron (NM_008320, intron 6 of 8) | intron (NM_008320, intron 6 of 8) | 15884 | NM_008320 | 15900 | Mm.334861 | NM_008320 | ENSMUSG00000041515 | **Irf8** | AI893568\|ICSBP\|IRF-8\|Icsbp1\|Myls | interferon regulatory factor 8 | protein-coding |
| chr7_24446653_24446914 | chr7 | 24446654 | 24446914 | + | -0.643504 | Intergenic | Intergenic | -1102 | NM_199013 | 210145 | Mm.33811 | NM_199013 | ENSMUSG00000062028 | **Irgc1** | F630044M05Rik\|Gm1102\|Gm474\|Ifgge\|Iigp5\|Irgc | immunity-related GTPase family, cinema 1 | protein-coding |
| chr7_24431660_24432337 | chr7 | 24431661 | 24432337 | + | -0.668986 | TTS (NM_199013) | TTS (NM_199013) | 13683 | NM_199013 | 210145 | Mm.33811 | NM_199013 | ENSMUSG00000062028 | **Irgc1** | F630044M05Rik\|Gm1102\|Gm474\|Ifgge\|Iigp5\|Irgc | immunity-related GTPase family, cinema 1 | protein-coding |
| chr7_24431756_24432062 | chr7 | 24431757 | 24432062 | + | -0.83135 | TTS (NM_199013) | TTS (NM_199013) | 13773 | NM_199013 | 210145 | Mm.33811 | NM_199013 | ENSMUSG00000062028 | **Irgc1** | F630044M05Rik\|Gm1102\|Gm474\|Ifgge\|Iigp5\|Irgc | immunity-related GTPase family, cinema 1 | protein-coding |
| chr7_24430555_24431488 | chr7 | 24430556 | 24431488 | + | -0.867155 | TTS (NM_199013) | TTS (NM_199013) | 14660 | NM_199013 | 210145 | Mm.33811 | NM_199013 | ENSMUSG00000062028 | **Irgc1** | F630044M05Rik\|Gm1102\|Gm474\|Ifgge\|Iigp5\|Irgc | immunity-related GTPase family, cinema 1 | protein-coding |
| chr7_24445543_24445944 | chr7 | 24445544 | 24445944 | + | -1.03101 | promoter-TSS (NM_199013) | promoter-TSS (NM_199013) | -62 | NM_199013 | 210145 | Mm.33811 | NM_199013 | ENSMUSG00000062028 | **Irgc1** | F630044M05Rik\|Gm1102\|Gm474\|Ifgge\|Iigp5\|Irgc | immunity-related GTPase family, cinema 1 | protein-coding |
| chr7_24445606_24445860 | chr7 | 24445607 | 24445860 | + | -1.04743 | promoter-TSS (NM_199013) | promoter-TSS (NM_199013) | -51 | NM_199013 | 210145 | Mm.33811 | NM_199013 | ENSMUSG00000062028 | **Irgc1** | F630044M05Rik\|Gm1102\|Gm474\|Ifgge\|Iigp5\|Irgc | immunity-related GTPase family, cinema 1 | protein-coding |
| chr8_11106890_11107426 | chr8 | 11106891 | 11107426 | + | -0.687424 | Intergenic | Intergenic | -98728 | NM_001081212 | 384783 | Mm.407207 | NM_001081212 | ENSMUSG00000038894 | **Irs2** | Irs-2 | insulin receptor substrate 2 | protein-coding |

| | | | | | | | | | | | | | | | | | |
|---|---|---|---|---|---|---|---|---|---|---|---|---|---|---|---|---|---|
| chr8_11097791_11098810 | chr8 | 11097792 | 11098810 | + | -0.699187 | Intergenic | Intergenic | -89871 | NM_001081212 | 384783 | Mm.407207 | NM_001081212 | ENSMUSG00000038894 | **Irs2** | Irs-2 | insulin receptor substrate 2 | protein-coding |
| chr8_11107059_11107434 | chr8 | 11107060 | 11107434 | + | -0.817184 | Intergenic | Intergenic | -98817 | NM_001081212 | 384783 | Mm.407207 | NM_001081212 | ENSMUSG00000038894 | **Irs2** | Irs-2 | insulin receptor substrate 2 | protein-coding |
| chr8_11098272_11098892 | chr8 | 11098273 | 11098892 | + | -0.824199 | Intergenic | Intergenic | -90152 | NM_001081212 | 384783 | Mm.407207 | NM_001081212 | ENSMUSG00000038894 | **Irs2** | Irs-2 | insulin receptor substrate 2 | protein-coding |
| chr8_109663681_109663982 | chr8 | 109663682 | 109663982 | + | -0.839007 | intron (NM_001039700, intron 27 of 33) | URR1B\|DNA\|hAT-Charlie | 29462 | NM_028018 | 71955 | Mm.290036 | NM_028018 | ENSMUSG00000031729 | **Ist1** | 2400003C14Rik\|AW536298\|mKIAA0174 | increased sodium tolerance 1 homolog (yeast) | protein-coding |
| chr8_109662987_109663545 | chr8 | 109662988 | 109663545 | + | -0.941804 | intron (NM_001039700, intron 27 of 33) | intron (NM_001039700, intron 27 of 33) | 30028 | NM_028018 | 71955 | Mm.290036 | NM_028018 | ENSMUSG00000031729 | **Ist1** | 2400003C14Rik\|AW536298\|mKIAA0174 | increased sodium tolerance 1 homolog (yeast) | protein-coding |
| chr8_109663239_109663572 | chr8 | 109663240 | 109663572 | + | -1.08369 | intron (NM_001039700, intron 27 of 33) | intron (NM_001039700, intron 27 of 33) | 29888 | NM_028018 | 71955 | Mm.290036 | NM_028018 | ENSMUSG00000031729 | **Ist1** | 2400003C14Rik\|AW536298\|mKIAA0174 | increased sodium tolerance 1 homolog (yeast) | protein-coding |
| chr8_109663284_109663532 | chr8 | 109663285 | 109663532 | + | -1.10129 | intron (NM_001039700, intron 27 of 33) | intron (NM_001039700, intron 27 of 33) | 29886 | NM_028018 | 71955 | Mm.290036 | NM_028018 | ENSMUSG00000031729 | **Ist1** | 2400003C14Rik\|AW536298\|mKIAA0174 | increased sodium tolerance 1 homolog (yeast) | protein-coding |
| chr17_26227742_26228251 | chr17 | 26227743 | 26228251 | + | -0.924925 | intron (NM_001206335, intron 1 of 12) | intron (NM_001206335, intron 1 of 12) | 16245 | NM_207217 | 106581 | Mm.28844 | NM_207217 | ENSMUSG00000024187 | **Itfg3** | - | integrin alpha FG-GAP repeat containing 3 | protein-coding |
| chr17_26227835_26228191 | chr17 | 26227836 | 26228191 | + | -0.946356 | intron (NM_001206335, intron 1 of 12) | intron (NM_001206335, intron 1 of 12) | 16229 | NM_207217 | 106581 | Mm.28844 | NM_207217 | ENSMUSG00000024187 | **Itfg3** | - | integrin alpha FG-GAP repeat containing 3 | protein-coding |
| chr17_26234430_26234985 | chr17 | 26234431 | 26234985 | + | -1.03348 | intron (NM_001206335, intron 1 of 12) | B2_Mm2\|SINE\|B2 | 9534 | NM_207217 | 106581 | Mm.28844 | NM_207217 | ENSMUSG00000024187 | **Itfg3** | - | integrin alpha FG-GAP repeat containing 3 | protein-coding |
| chr17_26234486_26234798 | chr17 | 26234487 | 26234798 | + | -1.03359 | intron (NM_001206335, intron 1 of 12) | intron (NM_001206335, intron 1 of 12) | 9600 | NM_207217 | 106581 | Mm.28844 | NM_207217 | ENSMUSG00000024187 | **Itfg3** | - | integrin alpha FG-GAP repeat containing 3 | protein-coding |
| chr13_114915724_114916198 | chr13 | 114915725 | 114916198 | + | -0.699049 | intron (NM_008396, intron 1 of 29) | MER20\|DNA\|hAT-Charlie | 16080 | NM_008396 | 16398 | Mm.5007 | NM_008396 | ENSMUSG00000015533 | **Itga2** | CD49B\|DX5 | integrin alpha 2 | protein-coding |
| chr2_79259867_79260104 | chr2 | 79259868 | 79260104 | + | -0.708429 | intron (NM_010576, intron 2 of 27) | intron (NM_010576, intron 2 of 27) | 4560 | NM_010576 | 16401 | Mm.31903 | NM_010576 | ENSMUSG00000027009 | **Itga4** | CD49D\|Itga4B | integrin alpha 4 | protein-coding |
| chr2_79127362_79127804 | chr2 | 79127363 | 79127804 | + | -0.800177 | Intergenic | Intergenic | -127843 | NM_010576 | 16401 | Mm.31903 | NM_010576 | ENSMUSG00000027009 | **Itga4** | CD49D\|Itga4B | integrin alpha 4 | protein-coding |
| chr2_79127460_79127704 | chr2 | 79127461 | 79127704 | + | -0.843193 | Intergenic | Intergenic | -127844 | NM_010576 | 16401 | Mm.31903 | NM_010576 | ENSMUSG00000027009 | **Itga4** | CD49D\|Itga4B | integrin alpha 4 | protein-coding |
| chr2_79230973_79231496 | chr2 | 79230974 | 79231496 | + | -0.942994 | Intergenic | Intergenic | -24191 | NM_010576 | 16401 | Mm.31903 | NM_010576 | ENSMUSG00000027009 | **Itga4** | CD49D\|Itga4B | integrin alpha 4 | protein-coding |
| chr9_118678196_118678758 | chr9 | 118678197 | 118678758 | + | -0.658046 | intron (NM_133721, intron 11 of 27) | intron (NM_133721, intron 11 of 27) | -33215 | NM_001113514 | 104099 | Mm.335520 | NM_133721 | ENSMUSG00000039115 | **Itga9** | (alpha)9\|2610002H11Rik\|6720458D17Rik\|AI461869\|D130073C02\|D9Ertd428e | integrin alpha 9 | protein-coding |
| chr9_118715641_118716534 | chr9 | 118715642 | 118716534 | + | -0.808697 | intron (NM_001113514, intron 1 of 13) | intron (NM_001113514, intron 1 of 13) | 4396 | NM_001113514 | 104099 | Mm.335520 | NM_133721 | ENSMUSG00000039115 | **Itga9** | (alpha)9\|2610002H11Rik\|6720458D17Rik\|AI461869\|D130073C02\|D9Ertd428e | integrin alpha 9 | protein-coding |
| chr9_118725101_118725472 | chr9 | 118725102 | 118725472 | + | -0.882381 | intron (NM_001113514, intron 1 of 13) | intron (NM_001113514, intron 1 of 13) | 13595 | NM_001113514 | 104099 | Mm.335520 | NM_133721 | ENSMUSG00000039115 | **Itga9** | (alpha)9\|2610002H11Rik\|6720458D17Rik\|AI461869\|D130073C02\|D9Ertd428e | integrin alpha 9 | protein-coding |
| chr9_118686182_118686683 | chr9 | 118686183 | 118686683 | + | -1.02288 | intron (NM_133721, intron 13 of 27) | intron (NM_133721, intron 13 of 27) | -25259 | NM_001113514 | 104099 | Mm.335520 | NM_133721 | ENSMUSG00000039115 | **Itga9** | (alpha)9\|2610002H11Rik\|6720458D17Rik\|AI461869\|D130073C02\|D9Ertd428e | integrin alpha 9 | protein-coding |
| chr11_73094331_73094570 | chr11 | 73094332 | 73094570 | + | -0.619432 | intron (NM_008399, intron 1 of 30) | intron (NM_008399, intron 1 of 30) | 3856 | NM_008399 | 16407 | Mm.96 | NM_008399 | ENSMUSG00000005947 | **Itgae** | A530055J10\|CD103\|aM290\|alpha-E1\|alpha-M290 | integrin alpha E, epithelial-associated | protein-coding |
| chr7_127287943_127288324 | chr7 | 127287944 | 127288324 | + | -0.649443 | Intergenic | Intergenic | -8126 | NM_001253873 | 16408 | Mm.1618 | NM_008400 | ENSMUSG00000030830 | **Itgal** | (p180)\|Cd11a\|LFA-1\|LFA-1A\|Ly-15\|Ly-21 | integrin alpha L | protein-coding |
| chr7_127287602_127288344 | chr7 | 127287603 | 127288344 | + | -0.658965 | Intergenic | Intergenic | -8287 | NM_001253873 | 16408 | Mm.1618 | NM_008400 | ENSMUSG00000030830 | **Itgal** | (p180)\|Cd11a\|LFA-1\|LFA-1A\|Ly-15\|Ly-21 | integrin alpha L | protein-coding |
| chr7_127285061_127285506 | chr7 | 127285062 | 127285506 | + | -0.788399 | Intergenic | Intergenic | -10976 | NM_001253873 | 16408 | Mm.1618 | NM_008400 | ENSMUSG00000030830 | **Itgal** | (p180)\|Cd11a\|LFA-1\|LFA-1A\|Ly-15\|Ly-21 | integrin alpha L | protein-coding |
| chr7_127285190_127285609 | chr7 | 127285191 | 127285609 | + | -0.81484 | Intergenic | Intergenic | -10860 | NM_001253873 | 16408 | Mm.1618 | NM_008400 | ENSMUSG00000030830 | **Itgal** | (p180)\|Cd11a\|LFA-1\|LFA-1A\|Ly-15\|Ly-21 | integrin alpha L | protein-coding |
| chr7_128119643_128120064 | chr7 | 128119644 | 128120064 | + | -0.644869 | Intergenic | Intergenic | -9714 | NM_021334 | 16411 | Mm.22378 | NM_021334 | ENSMUSG00000030789 | **Itgax** | AI449405\|Cd11c\|Cr4\|N418 | integrin alpha X | protein-coding |
| chr7_128119593_128120459 | chr7 | 128119594 | 128120459 | + | -0.868098 | Intergenic | Intergenic | -9542 | NM_021334 | 16411 | Mm.22378 | NM_021334 | ENSMUSG00000030789 | **Itgax** | AI449405\|Cd11c\|Cr4\|N418 | integrin alpha X | protein-coding |
| chr10_77537267_77537589 | chr10 | 77537268 | 77537589 | + | -0.705373 | intron (NM_008404, intron 1 of 15) | intron (NM_008404, intron 1 of 15) | 7080 | NM_008404 | 16414 | Mm.1137 | NM_008404 | ENSMUSG00000000290 | **Itgb2** | 2E6\|AI528527\|Cd18\|LAD\|LCAMB\|Lfa1\|MF17 | integrin beta 2 | protein-coding |
| chr10_77537273_77537756 | chr10 | 77537274 | 77537756 | + | -0.709014 | intron (NM_008404, intron 1 of 15) | (TG)n\|Simple_repeat\|Simple_repeat | 7167 | NM_008404 | 16414 | Mm.1137 | NM_008404 | ENSMUSG00000000290 | **Itgb2** | 2E6\|AI528527\|Cd18\|LAD\|LCAMB\|Lfa1\|MF17 | integrin beta 2 | protein-coding |
| chr10_77537881_77538144 | chr10 | 77537882 | 77538144 | + | -0.734481 | intron (NM_008404, intron 1 of 15) | intron (NM_008404, intron 1 of 15) | 7665 | NM_008404 | 16414 | Mm.1137 | NM_008404 | ENSMUSG00000000290 | **Itgb2** | 2E6\|AI528527\|Cd18\|LAD\|LCAMB\|Lfa1\|MF17 | integrin beta 2 | protein-coding |
| chr11_104768148_104768572 | chr11 | 104768149 | 104768572 | + | -0.661485 | Intergenic | Intergenic | 160360 | NM_016780 | 16416 | Mm.87150 | NM_016780 | ENSMUSG00000020689 | **Itgb3** | CD61\|GP3A\|INGRB3 | integrin beta 3 | protein-coding |
| chr11_104661051_104662265 | chr11 | 104661052 | 104662265 | + | -0.85455 | intron (NM_016780, intron 12 of 14) | intron (NM_016780, intron 12 of 14) | 53658 | NM_016780 | 16416 | Mm.87150 | NM_016780 | ENSMUSG00000020689 | **Itgb3** | CD61\|GP3A\|INGRB3 | integrin beta 3 | protein-coding |
| chr11_104663885_104664180 | chr11 | 104663886 | 104664180 | + | -1.00089 | intron (NM_016780, intron 13 of 14) | intron (NM_016780, intron 13 of 14) | 56033 | NM_016780 | 16416 | Mm.87150 | NM_016780 | ENSMUSG00000020689 | **Itgb3** | CD61\|GP3A\|INGRB3 | integrin beta 3 | protein-coding |
| chr11_104663885_104664284 | chr11 | 104663886 | 104664284 | + | -1.01881 | intron (NM_016780, intron 13 of 14) | intron (NM_016780, intron 13 of 14) | 56085 | NM_016780 | 16416 | Mm.87150 | NM_016780 | ENSMUSG00000020689 | **Itgb3** | CD61\|GP3A\|INGRB3 | integrin beta 3 | protein-coding |
| chr11_104667297_104668011 | chr11 | 104667298 | 104668011 | + | -1.02915 | 3' UTR (NM_016780, exon 15 of 15) | 3' UTR (NM_016780, exon 15 of 15) | 59654 | NM_016780 | 16416 | Mm.87150 | NM_016780 | ENSMUSG00000020689 | **Itgb3** | CD61\|GP3A\|INGRB3 | integrin beta 3 | protein-coding |
| chr2_60757059_60757573 | chr2 | 60757060 | 60757573 | + | -0.975378 | intron (NM_020296, intron 10 of 13) | intron (NM_020296, intron 10 of 13) | -34713 | NM_001159564 | 16420 | Mm.98193 | NM_021359 | ENSMUSG00000026971 | **Itgb6** | 2210409C20Rik\|4831415H04Rik | integrin beta 6 | protein-coding |
| chr15_102225100_102225597 | chr15 | 102225101 | 102225597 | + | -0.664167 | intron (NM_013566, intron 2 of 14) | intron (NM_013566, intron 2 of 14) | 6586 | NM_013566 | 16421 | Mm.58 | NM_013566 | ENSMUSG00000001281 | **Itgb7** | Ly69 | integrin beta 7 | protein-coding |
| chr12_119157050_119157462 | chr12 | 119157051 | 119157462 | + | -0.794998 | Intergenic | Intergenic | 81020 | NM_177290 | 320910 | Mm.217000 | NM_177290 | ENSMUSG00000025321 | **Itgb8** | 4832412O06Rik | integrin beta 8 | protein-coding |
| chr11_46358429_46358764 | chr11 | 46358430 | 46358764 | + | -0.841242 | intron (NM_001281967, intron 5 of 8) | MTD\|LTR\|ERVL-MaLR | 30918 | NM_001281968 | 16428 | Mm.339927 | NM_010583 | ENSMUSG00000020395 | **Itk** | Emt\|Tcsk\|Tsk | IL2 inducible T cell kinase | protein-coding |
| chr11_46372367_46373443 | chr11 | 46372368 | 46373443 | + | -0.884812 | intron (NM_001281966, intron 1 of 16) | Lx8b\|LINE\|L1 | 16610 | NM_001281968 | 16428 | Mm.339927 | NM_010583 | ENSMUSG00000020395 | **Itk** | Emt\|Tcsk\|Tsk | IL2 inducible T cell kinase | protein-coding |
| chr11_46372236_46373030 | chr11 | 46372237 | 46373030 | + | -1.16511 | intron (NM_001281966, intron 1 of 16) | Lx8b\|LINE\|L1 | 16882 | NM_001281968 | 16428 | Mm.339927 | NM_010583 | ENSMUSG00000020395 | **Itk** | Emt\|Tcsk\|Tsk | IL2 inducible T cell kinase | protein-coding |
| chr11_46372236_46372775 | chr11 | 46372237 | 46372775 | + | -1.38679 | intron (NM_001281966, intron 1 of 16) | Lx8b\|LINE\|L1 | 17009 | NM_001281968 | 16428 | Mm.339927 | NM_010583 | ENSMUSG00000020395 | **Itk** | Emt\|Tcsk\|Tsk | IL2 inducible T cell kinase | protein-coding |
| chrX_107421331_107421669 | chrX | 107421332 | 107421669 | + | -1.11199 | Intergenic | Intergenic | -18140 | NM_008409 | 16431 | Mm.193 | NM_008409 | ENSMUSG00000031239 | **Itm2a** | AW108051\|Bricd2a\|E25\|Itm2 | integral membrane protein 2A | protein-coding |
| chr14_73436635_73436882 | chr14 | 73436636 | 73436882 | + | -0.730406 | Intergenic | Intergenic | -51488 | NM_008410 | 16432 | Mm.4266 | NM_008410 | ENSMUSG00000022108 | **Itm2b** | AI256040\|Bri\|Bri2\|Bricd2b\|D14Sel6\|E25BMM\|imBRI2 | integral membrane protein 2B | protein-coding |
| chr14_73389101_73389456 | chr14 | 73389102 | 73389456 | + | -1.00456 | Intergenic | Intergenic | -4008 | NM_008410 | 16432 | Mm.4266 | NM_008410 | ENSMUSG00000022108 | **Itm2b** | AI256040\|Bri\|Bri2\|Bricd2b\|D14Sel6\|E25BMM\|imBRI2 | integral membrane protein 2B | protein-coding |
| chr14_73391509_73391818 | chr14 | 73391510 | 73391818 | + | -1.27229 | Intergenic | MTE2a\|LTR\|ERVL-MaLR | -6393 | NM_008410 | 16432 | Mm.4266 | NM_008410 | ENSMUSG00000022108 | **Itm2b** | AI256040\|Bri\|Bri2\|Bricd2b\|D14Sel6\|E25BMM\|imBRI2 | integral membrane protein 2B | protein-coding |
| chr1_180295975_180296824 | chr1 | 180295976 | 180296824 | + | -0.687428 | Intergenic | MLT1J\|LTR\|ERVL-MaLR | -34076 | NM_001081175 | 320404 | Mm.132535 | NM_001081175 | ENSMUSG00000038855 | **Itpkb** | 1110033J02Rik\|E130307H12Rik | inositol 1,4,5-trisphosphate 3-kinase B | protein-coding |
| chr1_180313831_180314434 | chr1 | 180313832 | 180314434 | + | -0.831596 | Intergenic | Intergenic | -16343 | NM_001081175 | 320404 | Mm.132535 | NM_001081175 | ENSMUSG00000038855 | **Itpkb** | 1110033J02Rik\|E130307H12Rik | inositol 1,4,5-trisphosphate 3-kinase B | protein-coding |
| chr6_146489906_146490202 | chr6 | 146489907 | 146490202 | + | -0.617359 | intron (NM_019923, intron 1 of 56) | intron (NM_019923, intron 1 of 56) | 12169 | NM_010586 | 16439 | Mm.7800 | NM_010586 | ENSMUSG00000030287 | **Itpr2** | AI649341\|InsP3R-2\|InsP3R-5\|Ip3r2\|Itpr5\|insP3R2 | inositol 1,4,5-triphosphate receptor 2 | protein-coding |

| | | | | | | | | | | | | | | | | | |
|---|---|---|---|---|---|---|---|---|---|---|---|---|---|---|---|---|---|
| chr6_146489724_146490218 | chr6 | 146489725 | 146490218 | + | -0.626093 | intron (NM_019923, intron 1 of 56) | intron (NM_019923, intron 1 of 56) | 12252 | NM_010586 | 16439 | Mm.7800 | NM_010586 | ENSMUSG00000030287 | **Itpr2** | AI649341\|InsP3R-2\|InsP3R-5\|Ip3r2\|Itpr5\|insP3R2 | inositol 1,4,5-triphosphate receptor 2 | protein-coding |
| chr6_146481402_146481826 | chr6 | 146481403 | 146481826 | + | -0.761652 | intron (NM_019923, intron 1 of 56) | intron (NM_019923, intron 1 of 56) | 20609 | NM_010586 | 16439 | Mm.7800 | NM_010586 | ENSMUSG00000030287 | **Itpr2** | AI649341\|InsP3R-2\|InsP3R-5\|Ip3r2\|Itpr5\|insP3R2 | inositol 1,4,5-triphosphate receptor 2 | protein-coding |
| chr6_146271124_146271644 | chr6 | 146271125 | 146271644 | + | -0.964184 | intron (NM_019923, intron 37 of 56) | intron (NM_019923, intron 37 of 56) | 230839 | NM_010586 | 16439 | Mm.7800 | NM_010586 | ENSMUSG00000030287 | **Itpr2** | AI649341\|InsP3R-2\|InsP3R-5\|Ip3r2\|Itpr5\|insP3R2 | inositol 1,4,5-triphosphate receptor 2 | protein-coding |
| chr17_27056089_27056622 | chr17 | 27056090 | 27056622 | + | -0.659683 | promoter-TSS (NM_080553) | promoter-TSS (NM_080553) | -948 | NM_080553 | 16440 | Mm.458106 | NM_080553 | ENSMUSG00000042644 | **Itpr3** | Ip3r3\|Itpr-3 | inositol 1,4,5-triphosphate receptor 3 | protein-coding |
| chr19_47910033_47910534 | chr19 | 47910034 | 47910534 | + | -0.624933 | intron (NM_001001738, intron 1 of 1) | intron (NM_001001738, intron 1 of 1) | 9015 | NM_001001738 | 414801 | Mm.29457 | NM_001001738 | ENSMUSG00000079254 | **Itprip** | 4833424O12Rik\|DANGER\|mKIAA1754 | inositol 1,4,5-triphosphate receptor interacting protein | protein-coding |
| chr12_4595123_4595389 | chr12 | 4595124 | 4595389 | + | -0.694682 | intron (NM_011365, intron 1 of 38) | intron (NM_011365, intron 1 of 38) | 2248 | NM_001198968 | 20403 | Mm.341204 | NM_011365 | ENSMUSG00000020640 | **Itsn2** | AI327390\|Ese2\|Sh3d1B\|Sh3p18\|mKIAA1256 | intersectin 2 | protein-coding |
| chr12_4558682_4559273 | chr12 | 4558683 | 4559273 | + | -0.775247 | Intergenic | ORR1F\|LTR\|ERVL-MaLR | -34030 | NM_001198969 | 20403 | Mm.341204 | NM_011365 | ENSMUSG00000020640 | **Itsn2** | AI327390\|Ese2\|Sh3d1B\|Sh3p18\|mKIAA1256 | intersectin 2 | protein-coding |
| chr12_4623973_4624537 | chr12 | 4623974 | 4624537 | + | -0.878219 | intron (NM_011365, intron 2 of 38) | intron (NM_011365, intron 2 of 38) | 31247 | NM_001198968 | 20403 | Mm.341204 | NM_011365 | ENSMUSG00000020640 | **Itsn2** | AI327390\|Ese2\|Sh3d1B\|Sh3p18\|mKIAA1256 | intersectin 2 | protein-coding |
| chr12_4624218_4624613 | chr12 | 4624219 | 4624613 | + | -0.916766 | intron (NM_011365, intron 2 of 38) | intron (NM_011365, intron 2 of 38) | 31408 | NM_001198968 | 20403 | Mm.341204 | NM_011365 | ENSMUSG00000020640 | **Itsn2** | AI327390\|Ese2\|Sh3d1B\|Sh3p18\|mKIAA1256 | intersectin 2 | protein-coding |
| chr11_51883419_51884146 | chr11 | 51883420 | 51884146 | + | -0.962595 | Intergenic | Intergenic | -26302 | NM_199299 | 76901 | Mm.259996 | NM_199299 | ENSMUSG00000020387 | **Jade2** | 1200017K05Rik\|AI480685\|Phf15\|mKIAA0239 | jade family PHD finger 2 | protein-coding |
| chr4_101298158_101298925 | chr4 | 101298159 | 101298925 | + | -0.619071 | Intergenic | Intergenic | -33260 | NM_146145 | 16451 | Mm.289657 | NM_146145 | ENSMUSG00000028530 | **Jak1** | AA960307\|C130039L05Rik | Janus kinase 1 | protein-coding |
| chr4_101298362_101298923 | chr4 | 101298363 | 101298923 | + | -0.656349 | Intergenic | Intergenic | -33361 | NM_146145 | 16451 | Mm.289657 | NM_146145 | ENSMUSG00000028530 | **Jak1** | AA960307\|C130039L05Rik | Janus kinase 1 | protein-coding |
| chr4_101300770_101301357 | chr4 | 101300771 | 101301357 | + | -0.716731 | Intergenic | Intergenic | -35782 | NM_146145 | 16451 | Mm.289657 | NM_146145 | ENSMUSG00000028530 | **Jak1** | AA960307\|C130039L05Rik | Janus kinase 1 | protein-coding |
| chr4_101182562_101182829 | chr4 | 101182563 | 101182829 | + | -0.725912 | intron (NM_146145, intron 5 of 24) | intron (NM_146145, intron 5 of 24) | 82586 | NM_146145 | 16451 | Mm.289657 | NM_146145 | ENSMUSG00000028530 | **Jak1** | AA960307\|C130039L05Rik | Janus kinase 1 | protein-coding |
| chr4_101216938_101217539 | chr4 | 101216939 | 101217539 | + | -0.747613 | intron (NM_146145, intron 1 of 24) | intron (NM_146145, intron 1 of 24) | 48043 | NM_146145 | 16451 | Mm.289657 | NM_146145 | ENSMUSG00000028530 | **Jak1** | AA960307\|C130039L05Rik | Janus kinase 1 | protein-coding |
| chr4_101187140_101187797 | chr4 | 101187141 | 101187797 | + | -0.763462 | intron (NM_146145, intron 4 of 24) | intron (NM_146145, intron 4 of 24) | 77813 | NM_146145 | 16451 | Mm.289657 | NM_146145 | ENSMUSG00000028530 | **Jak1** | AA960307\|C130039L05Rik | Janus kinase 1 | protein-coding |
| chr4_101260836_101261138 | chr4 | 101260837 | 101261138 | + | -0.865442 | intron (NM_146145, intron 1 of 24) | intron (NM_146145, intron 1 of 24) | 4295 | NM_146145 | 16451 | Mm.289657 | NM_146145 | ENSMUSG00000028530 | **Jak1** | AA960307\|C130039L05Rik | Janus kinase 1 | protein-coding |
| chr4_101167753_101168713 | chr4 | 101167754 | 101168713 | + | -0.93382 | intron (NM_146145, intron 11 of 24) | intron (NM_146145, intron 11 of 24) | 97049 | NM_146145 | 16451 | Mm.289657 | NM_146145 | ENSMUSG00000028530 | **Jak1** | AA960307\|C130039L05Rik | Janus kinase 1 | protein-coding |
| chr4_101175068_101175658 | chr4 | 101175069 | 101175658 | + | -1.35137 | intron (NM_146145, intron 8 of 24) | intron (NM_146145, intron 8 of 24) | 89919 | NM_146145 | 16451 | Mm.289657 | NM_146145 | ENSMUSG00000028530 | **Jak1** | AA960307\|C130039L05Rik | Janus kinase 1 | protein-coding |
| chr4_101167777_101168120 | chr4 | 101167778 | 101168120 | + | -1.46706 | intron (NM_146145, intron 11 of 24) | intron (NM_146145, intron 11 of 24) | 97333 | NM_146145 | 16451 | Mm.289657 | NM_146145 | ENSMUSG00000028530 | **Jak1** | AA960307\|C130039L05Rik | Janus kinase 1 | protein-coding |
| chr4_101185286_101185604 | chr4 | 101185287 | 101185604 | + | -1.49388 | intron (NM_146145, intron 4 of 24) | intron (NM_146145, intron 4 of 24) | 79837 | NM_146145 | 16451 | Mm.289657 | NM_146145 | ENSMUSG00000028530 | **Jak1** | AA960307\|C130039L05Rik | Janus kinase 1 | protein-coding |
| chr5_37051095_37051837 | chr5 | 37051096 | 37051837 | + | -0.644765 | intron (NM_178394, intron 1 of 12) | intron (NM_178394, intron 1 of 12) | 609 | NM_178394 | 76071 | Mm.85280 | NM_178394 | ENSMUSG00000063646 | **Jakmip1** | 5830437M04Rik\|C330021K24Rik\|Gababrbp\|Marlin-1 | janus kinase and microtubule interacting protein 1 | protein-coding |
| chr5_37066716_37067915 | chr5 | 37066717 | 37067915 | + | -0.680878 | intron (NM_178394, intron 1 of 12) | intron (NM_178394, intron 1 of 12) | 16459 | NM_178394 | 76071 | Mm.85280 | NM_178394 | ENSMUSG00000063646 | **Jakmip1** | 5830437M04Rik\|C330021K24Rik\|Gababrbp\|Marlin-1 | janus kinase and microtubule interacting protein 1 | protein-coding |
| chr5_37066780_37067644 | chr5 | 37066781 | 37067644 | + | -0.714122 | intron (NM_178394, intron 1 of 12) | intron (NM_178394, intron 1 of 12) | 16355 | NM_178394 | 76071 | Mm.85280 | NM_178394 | ENSMUSG00000063646 | **Jakmip1** | 5830437M04Rik\|C330021K24Rik\|Gababrbp\|Marlin-1 | janus kinase and microtubule interacting protein 1 | protein-coding |
| chr5_37056121_37056866 | chr5 | 37056122 | 37056866 | + | -0.731046 | intron (NM_178394, intron 1 of 12) | intron (NM_178394, intron 1 of 12) | 5637 | NM_178394 | 76071 | Mm.85280 | NM_178394 | ENSMUSG00000063646 | **Jakmip1** | 5830437M04Rik\|C330021K24Rik\|Gababrbp\|Marlin-1 | janus kinase and microtubule interacting protein 1 | protein-coding |
| chr5_37047272_37047540 | chr5 | 37047273 | 37047540 | + | -0.840418 | Intergenic | Intergenic | -3451 | NM_178394 | 76071 | Mm.85280 | NM_178394 | ENSMUSG00000063646 | **Jakmip1** | 5830437M04Rik\|C330021K24Rik\|Gababrbp\|Marlin-1 | janus kinase and microtubule interacting protein 1 | protein-coding |
| chr5_37056568_37056847 | chr5 | 37056569 | 37056847 | + | -0.843731 | intron (NM_178394, intron 1 of 12) | intron (NM_178394, intron 1 of 12) | 5851 | NM_178394 | 76071 | Mm.85280 | NM_178394 | ENSMUSG00000063646 | **Jakmip1** | 5830437M04Rik\|C330021K24Rik\|Gababrbp\|Marlin-1 | janus kinase and microtubule interacting protein 1 | protein-coding |
| chr5_37044635_37045191 | chr5 | 37044636 | 37045191 | + | -1.19949 | Intergenic | Intergenic | -5944 | NM_178394 | 76071 | Mm.85280 | NM_178394 | ENSMUSG00000063646 | **Jakmip1** | 5830437M04Rik\|C330021K24Rik\|Gababrbp\|Marlin-1 | janus kinase and microtubule interacting protein 1 | protein-coding |
| chr13_44852009_44852357 | chr13 | 44852010 | 44852357 | + | -0.630278 | intron (NM_001205043, intron 3 of 17) | intron (NM_001205043, intron 3 of 17) | 120912 | NM_001205043 | 16468 | Mm.25059 | NM_021878 | ENSMUSG00000038518 | **Jarid2** | Jmj\|jumonji | jumonji, AT rich interactive domain 2 | protein-coding |
| chr13_44811059_44812033 | chr13 | 44811060 | 44812033 | + | -0.793757 | intron (NM_001205043, intron 1 of 17) | intron (NM_001205043, intron 1 of 17) | 80275 | NM_001205043 | 16468 | Mm.25059 | NM_021878 | ENSMUSG00000038518 | **Jarid2** | Jmj\|jumonji | jumonji, AT rich interactive domain 2 | protein-coding |
| chr13_44819358_44819742 | chr13 | 44819359 | 44819742 | + | -1.23132 | intron (NM_001205043, intron 1 of 17) | intron (NM_001205043, intron 1 of 17) | 88279 | NM_001205043 | 16468 | Mm.25059 | NM_021878 | ENSMUSG00000038518 | **Jarid2** | Jmj\|jumonji | jumonji, AT rich interactive domain 2 | protein-coding |
| chr12_85545645_85546419 | chr12 | 85545646 | 85546419 | + | -0.67105 | Intergenic | Intergenic | -53073 | NM_001205052 | 81703 | Mm.103560 | NM_030887 | ENSMUSG00000034271 | **Jdp2** | Jundm2\|Jundp2\|TIF | Jun dimerization protein 2 | protein-coding |
| chr13_93499099_93500175 | chr13 | 93499100 | 93500175 | + | -0.707656 | 5' UTR (NM_021310, exon 1 of 11) | 5' UTR (NM_021310, exon 1 of 11) | 171 | NM_021310 | 57748 | Mm.100273 | NM_021310 | ENSMUSG00000021690 | **Jmy** | - | junction-mediating and regulatory protein | protein-coding |
| chr13_93546094_93546592 | chr13 | 93546095 | 93546592 | + | -0.7355 | Intergenic | Intergenic | -46535 | NM_021310 | 57748 | Mm.100273 | NM_021310 | ENSMUSG00000021690 | **Jmy** | - | junction-mediating and regulatory protein | protein-coding |
| chr13_93545883_93546561 | chr13 | 93545884 | 93546561 | + | -0.775393 | Intergenic | MLT1E3\|LTR\|ERVL-MaLR | -46414 | NM_021310 | 57748 | Mm.100273 | NM_021310 | ENSMUSG00000021690 | **Jmy** | - | junction-mediating and regulatory protein | protein-coding |
| chr13_93553938_93554328 | chr13 | 93553939 | 93554328 | + | -0.7909 | Intergenic | Intergenic | -54325 | NM_021310 | 57748 | Mm.100273 | NM_021310 | ENSMUSG00000021690 | **Jmy** | - | junction-mediating and regulatory protein | protein-coding |
| chr13_93485775_93486034 | chr13 | 93485776 | 93486034 | + | -0.955237 | intron (NM_021310, intron 1 of 10) | intron (NM_021310, intron 1 of 10) | 13903 | NM_021310 | 57748 | Mm.100273 | NM_021310 | ENSMUSG00000021690 | **Jmy** | - | junction-mediating and regulatory protein | protein-coding |
| chr4_95067941_95068624 | chr4 | 95067942 | 95068624 | + | -0.624995 | Intergenic | Intergenic | -16061 | NM_010591 | 16476 | Mm.275071 | NM_010591 | ENSMUSG00000052684 | **Jun** | AP-1\|Junc\|c-jun | jun proto-oncogene | protein-coding |
| chr4_95068078_95068519 | chr4 | 95068079 | 95068519 | + | -0.635642 | Intergenic | MIRb\|SINE\|MIR | -16077 | NM_010591 | 16476 | Mm.275071 | NM_010591 | ENSMUSG00000052684 | **Jun** | AP-1\|Junc\|c-jun | jun proto-oncogene | protein-coding |
| chr4_95196252_95196540 | chr4 | 95196253 | 95196540 | + | -0.66699 | Intergenic | Intergenic | -144174 | NM_010591 | 16476 | Mm.275071 | NM_010591 | ENSMUSG00000052684 | **Jun** | AP-1\|Junc\|c-jun | jun proto-oncogene | protein-coding |
| chr4_95232541_95232941 | chr4 | 95232542 | 95232941 | + | -0.809749 | Intergenic | Intergenic | -180519 | NM_010591 | 16476 | Mm.275071 | NM_010591 | ENSMUSG00000052684 | **Jun** | AP-1\|Junc\|c-jun | jun proto-oncogene | protein-coding |
| chr11_104425019_104425362 | chr11 | 104425020 | 104425362 | + | -0.710523 | exon (NM_001081045, exon 2 of 14) | exon (NM_001081045, exon 2 of 14) | 17100 | NM_001081045 | 76719 | Mm.136418 | NM_001081045 | ENSMUSG00000018412 | **Kansl1** | 1700081L11Rik\|9430041J06Rik\|mKIAA1267 | KAT8 regulatory NSL complex subunit 1 | protein-coding |
| chr11_104424875_104425399 | chr11 | 104424876 | 104425399 | + | -0.715112 | exon (NM_001081045, exon 2 of 14) | exon (NM_001081045, exon 2 of 14) | 17154 | NM_001081045 | 76719 | Mm.136418 | NM_001081045 | ENSMUSG00000018412 | **Kansl1** | 1700081L11Rik\|9430041J06Rik\|mKIAA1267 | KAT8 regulatory NSL complex subunit 1 | protein-coding |
| chr11_104381915_104382198 | chr11 | 104381916 | 104382198 | + | -0.9808 | intron (NM_001081045, intron 2 of 13) | B1_Mur2\|SINE\|Alu | 60234 | NM_001081045 | 76719 | Mm.136418 | NM_001081045 | ENSMUSG00000018412 | **Kansl1** | 1700081L11Rik\|9430041J06Rik\|mKIAA1267 | KAT8 regulatory NSL complex subunit 1 | protein-coding |
| chr1_66815981_66816322 | chr1 | 66815982 | 66816322 | + | -0.651715 | intron (NM_001122738, intron 1 of 14) | MIR\|SINE\|MIR | 1443 | NM_177645 | 68691 | Mm.240912 | NM_177645 | ENSMUSG00000026004 | **Kansl1l** | 1110028C15Rik\|1190030G24\|AI427598\|C430010P07Rik\|mKIAA4189 | KAT8 regulatory NSL complex subunit 1-like | protein-coding |
| chr14_21539772_21540077 | chr14 | 21539773 | 21540077 | + | -0.877467 | intron (NM_017479, intron 2 of 16) | intron (NM_017479, intron 2 of 16) | 40140 | NM_017479 | 54169 | Mm.248967 | NM_017479 | ENSMUSG00000021767 | **Kat6b** | AI507552\|B130044K16Rik\|MYST-4\|Morf\|Myst4\|mKIAA0383\|qkf\|querkopf | K(lysine) acetyltransferase 6B | protein-coding |
| chr5_148853406_148853875 | chr5 | 148853407 | 148853875 | + | -0.689797 | Intergenic | Intergenic | 75006 | NM_153572 | 231912 | Mm.397405 | NM_153572 | | **Katnal1** | - | katanin p60 subunit A-like 1 | protein-coding |
| chr5_148910196_148910710 | chr5 | 148910197 | 148910710 | + | -0.72395 | intron (NM_153572, intron 3 of 10) | intron (NM_153572, intron 3 of 10) | 18194 | NM_153572 | 231912 | Mm.397405 | NM_153572 | | **Katnal1** | - | katanin p60 subunit A-like 1 | protein-coding |
| chr5_148887888_148888663 | chr5 | 148887889 | 148888663 | + | -0.760741 | intron (NM_153572, intron 9 of 10) | L1ME2\|LINE\|L1 | 40371 | NM_153572 | 231912 | Mm.397405 | NM_153572 | | **Katnal1** | - | katanin p60 subunit A-like 1 | protein-coding |
| chr5_148887874_148888401 | chr5 | 148887875 | 148888401 | + | -0.769408 | intron (NM_153572, intron 9 of 10) | intron (NM_153572, intron 9 of 10) | 40509 | NM_153572 | 231912 | Mm.397405 | NM_153572 | | **Katnal1** | - | katanin p60 subunit A-like 1 | protein-coding |
| chr5_148867713_148868422 | chr5 | 148867714 | 148868422 | + | -0.774927 | Intergenic | Intergenic | 60579 | NM_153572 | 231912 | Mm.397405 | NM_153572 | | **Katnal1** | - | katanin p60 subunit A-like 1 | protein-coding |
| chr5_148910119_148910527 | chr5 | 148910120 | 148910527 | + | -0.827135 | intron (NM_153572, intron 3 of 10) | intron (NM_153572, intron 3 of 10) | 18324 | NM_153572 | 231912 | Mm.397405 | NM_153572 | | **Katnal1** | - | katanin p60 subunit A-like 1 | protein-coding |
| chr5_148866461_148867037 | chr5 | 148866462 | 148867037 | + | -1.15893 | Intergenic | Intergenic | 61898 | NM_153572 | 231912 | Mm.397405 | NM_153572 | | **Katnal1** | - | katanin p60 subunit A-like 1 | protein-coding |

| | | | | | | | | | | | | | | | | | |
|---|---|---|---|---|---|---|---|---|---|---|---|---|---|---|---|---|---|
| chr8_14961375_14962219 | chr8 | 14961376 | 14962219 | + | -0.829506 | intron (NM_001037736, intron 15 of 27) | intron (NM_001037736, intron 15 of 27) | -49228 | NM_029116 | 74901 | Mm.46675 | NM_029116 | ENSMUSG00000055675 | **Kbtbd11** | 2900016B01Rik\|4930465M17Rik\|mKIAA0711 | kelch repeat and BTB (POZ) domain containing 11 | protein-coding |
| chr6_72876691_72876951 | chr6 | 72876692 | 72876951 | + | -0.856491 | intron (NM_019715, intron 1 of 6) | intron (NM_019715, intron 1 of 6) | 23158 | NM_019715 | 74287 | Mm.29194 | NM_019715 | ENSMUSG00000055239 | **Kcmf1** | 1700094M07Rik\|Debt91\|Pmcf | potassium channel modulatory factor 1 | protein-coding |
| chr3_107185535_107186074 | chr3 | 107185536 | 107186074 | + | -1.0714 | intron (NM_001081140, intron 1 of 1) | intron (NM_001081140, intron 1 of 1) | 2662 | NM_001081140 | 242151 | Mm.245155 | NM_001081140 | ENSMUSG00000042861 | **Kcna10** | Gm1962\|Kcna8\|Kv1.8 | potassium voltage-gated channel, shaker-related subfamily, member 10 | protein-coding |
| chr3_107185653_107185909 | chr3 | 107185654 | 107185909 | + | -1.13636 | intron (NM_001081140, intron 1 of 1) | intron (NM_001081140, intron 1 of 1) | 2638 | NM_001081140 | 242151 | Mm.245155 | NM_001081140 | ENSMUSG00000042861 | **Kcna10** | Gm1962\|Kcna8\|Kv1.8 | potassium voltage-gated channel, shaker-related subfamily, member 10 | protein-coding |
| chr3_106974178_106974773 | chr3 | 106974179 | 106974773 | + | -0.791265 | Intergenic | Intergenic | -61686 | NM_008418 | 16491 | Mm.30640 | NM_008418 | ENSMUSG00000047959 | **Kcna3** | Kca1-3\|Kv1.3\|Mk-3 | potassium voltage-gated channel, shaker-related subfamily, member 3 | protein-coding |
| chr7_46499576_46499959 | chr7 | 46499577 | 46499959 | + | -0.878841 | intron (NM_013789, intron 10 of 10) | intron (NM_013789, intron 10 of 10) | 103300 | NM_001112739 | 16502 | Mm.249386 | NM_008421 | ENSMUSG00000058975 | **Kcnc1** | C230009H10Rik\|KShIIIB\|KV4\|Kcr2-1\|Kv3.1\|NGK2\|Shaw | potassium voltage gated channel, Shaw-related subfamily, member 1 | protein-coding |
| chr7_46391592_46392071 | chr7 | 46391593 | 46392071 | + | -1.11038 | Intergenic | MLTR11B\|LTR\|ERVK | -4636 | NM_008421 | 16502 | Mm.249386 | NM_008421 | ENSMUSG00000058975 | **Kcnc1** | C230009H10Rik\|KShIIIB\|KV4\|Kcr2-1\|Kv3.1\|NGK2\|Shaw | potassium voltage gated channel, Shaw-related subfamily, member 1 | protein-coding |
| chr7_46389461_46390050 | chr7 | 46389462 | 46390050 | + | -1.1532 | Intergenic | Intergenic | -6712 | NM_008421 | 16502 | Mm.249386 | NM_008421 | ENSMUSG00000058975 | **Kcnc1** | C230009H10Rik\|KShIIIB\|KV4\|Kcr2-1\|Kv3.1\|NGK2\|Shaw | potassium voltage gated channel, Shaw-related subfamily, member 1 | protein-coding |
| chr7_46391794_46392049 | chr7 | 46391795 | 46392049 | + | -1.18153 | Intergenic | MLTR11B\|LTR\|ERVK | -4546 | NM_008421 | 16502 | Mm.249386 | NM_008421 | ENSMUSG00000058975 | **Kcnc1** | C230009H10Rik\|KShIIIB\|KV4\|Kcr2-1\|Kv3.1\|NGK2\|Shaw | potassium voltage gated channel, Shaw-related subfamily, member 1 | protein-coding |
| chr17_83646424_83647171 | chr17 | 83646425 | 83647171 | + | -0.711372 | Intergenic | ID_B1\|SINE\|B4 | -14903 | NM_153512 | 225030 | Mm.223506 | NM_153512 | ENSMUSG00000045053 | **Kcng3** | KV6.3\|Kv10.1a\|Kv10.1b | potassium voltage-gated channel, subfamily G, member 3 | protein-coding |
| chr17_83654460_83655191 | chr17 | 83654461 | 83655191 | + | -0.772416 | Intergenic | Intergenic | -22931 | NM_153512 | 225030 | Mm.223506 | NM_153512 | ENSMUSG00000045053 | **Kcng3** | KV6.3\|Kv10.1a\|Kv10.1b | potassium voltage-gated channel, subfamily G, member 3 | protein-coding |
| chr17_52282303_52282666 | chr17 | 52282304 | 52282666 | + | -0.808333 | Intergenic | Intergenic | -320224 | NM_001031811 | 211468 | Mm.399068 | NM_001031811 | ENSMUSG00000035580 | **Kcnh8** | C130090D05Rik\|ELK\|ELK1\|ELK3\|Kv12.1 | potassium voltage-gated channel, subfamily H (eag-related), member 8 | protein-coding |
| chr17_52506633_52507595 | chr17 | 52506634 | 52507595 | + | -0.818482 | Intergenic | Intergenic | -95595 | NM_001031811 | 211468 | Mm.399068 | NM_001031811 | ENSMUSG00000035580 | **Kcnh8** | C130090D05Rik\|ELK\|ELK1\|ELK3\|Kv12.1 | potassium voltage-gated channel, subfamily H (eag-related), member 8 | protein-coding |
| chr17_52506784_52507483 | chr17 | 52506785 | 52507483 | + | -0.821312 | Intergenic | Intergenic | -95575 | NM_001031811 | 211468 | Mm.399068 | NM_001031811 | ENSMUSG00000035580 | **Kcnh8** | C130090D05Rik\|ELK\|ELK1\|ELK3\|Kv12.1 | potassium voltage-gated channel, subfamily H (eag-related), member 8 | protein-coding |
| chr17_52506779_52507244 | chr17 | 52506780 | 52507244 | + | -0.900785 | Intergenic | MIR\|SINE\|MIR | -95697 | NM_001031811 | 211468 | Mm.399068 | NM_001031811 | ENSMUSG00000035580 | **Kcnh8** | C130090D05Rik\|ELK\|ELK1\|ELK3\|Kv12.1 | potassium voltage-gated channel, subfamily H (eag-related), member 8 | protein-coding |
| chr17_52506636_52507189 | chr17 | 52506637 | 52507189 | + | -0.907836 | Intergenic | MIR\|SINE\|MIR | -95796 | NM_001031811 | 211468 | Mm.399068 | NM_001031811 | ENSMUSG00000035580 | **Kcnh8** | C130090D05Rik\|ELK\|ELK1\|ELK3\|Kv12.1 | potassium voltage-gated channel, subfamily H (eag-related), member 8 | protein-coding |
| chr17_52312109_52312394 | chr17 | 52312110 | 52312394 | + | -0.991116 | Intergenic | Intergenic | -290457 | NM_001031811 | 211468 | Mm.399068 | NM_001031811 | ENSMUSG00000035580 | **Kcnh8** | C130090D05Rik\|ELK\|ELK1\|ELK3\|Kv12.1 | potassium voltage-gated channel, subfamily H (eag-related), member 8 | protein-coding |
| chr17_52468324_52469117 | chr17 | 52468325 | 52469117 | + | -1.14781 | Intergenic | Intergenic | -133988 | NM_001031811 | 211468 | Mm.399068 | NM_001031811 | ENSMUSG00000035580 | **Kcnh8** | C130090D05Rik\|ELK\|ELK1\|ELK3\|Kv12.1 | potassium voltage-gated channel, subfamily H (eag-related), member 8 | protein-coding |
| chr17_52448638_52449096 | chr17 | 52448639 | 52449096 | + | -1.61378 | Intergenic | Intergenic | -153842 | NM_001031811 | 211468 | Mm.399068 | NM_001031811 | ENSMUSG00000035580 | **Kcnh8** | C130090D05Rik\|ELK\|ELK1\|ELK3\|Kv12.1 | potassium voltage-gated channel, subfamily H (eag-related), member 8 | protein-coding |
| chr17_52448438_52449132 | chr17 | 52448439 | 52449132 | + | -1.61443 | Intergenic | Intergenic | -153924 | NM_001031811 | 211468 | Mm.399068 | NM_001031811 | ENSMUSG00000035580 | **Kcnh8** | C130090D05Rik\|ELK\|ELK1\|ELK3\|Kv12.1 | potassium voltage-gated channel, subfamily H (eag-related), member 8 | protein-coding |
| chr19_45837423_45838063 | chr19 | 45837424 | 45838063 | + | -0.767601 | intron (NM_198296, intron 24 of 25) | intron (NM_198296, intron 24 of 25) | -21680 | NM_145703 | 80906 | Mm.213204 | NM_030716 | ENSMUSG00000025221 | **Kcnip2** | KChIP2 | Kv channel-interacting protein 2 | protein-coding |
| chr19_45837640_45838026 | chr19 | 45837641 | 45838026 | + | -0.782738 | intron (NM_198296, intron 24 of 25) | intron (NM_198296, intron 24 of 25) | -21770 | NM_145703 | 80906 | Mm.213204 | NM_030716 | ENSMUSG00000025221 | **Kcnip2** | KChIP2 | Kv channel-interacting protein 2 | protein-coding |
| chr19_45843451_45843943 | chr19 | 45843452 | 45843943 | + | -0.884121 | intron (NM_198296, intron 24 of 25) | (CGAG)n\|Simple_repeat\|Simple_repeat | -27634 | NM_145703 | 80906 | Mm.213204 | NM_030716 | ENSMUSG00000025221 | **Kcnip2** | KChIP2 | Kv channel-interacting protein 2 | protein-coding |
| chr1_87378877_87379117 | chr1 | 87378878 | 87379117 | + | -0.770932 | exon (NM_001110212, exon 7 of 29) | exon (NM_001110212, exon 7 of 29) | 15732 | NM_001110227 | 100040591 | Mm.443539 | NM_001110227 | ENSMUSG00000079436 | **Kcnj13** | - | potassium inwardly-rectifying channel, subfamily J, member 13 | protein-coding |
| chr1_87378635_87379189 | chr1 | 87378636 | 87379189 | + | -0.801617 | intron (NM_146112, intron 6 of 28) | intron (NM_146112, intron 6 of 28) | 15817 | NM_001110227 | 100040591 | Mm.443539 | NM_001110227 | ENSMUSG00000079436 | **Kcnj13** | - | potassium inwardly-rectifying channel, subfamily J, member 13 | protein-coding |
| chr16_95303706_95304152 | chr16 | 95303707 | 95304152 | + | -0.713658 | Intergenic | Intergenic | 46371 | NM_001271690 | 16516 | Mm.272239 | NM_019664 | ENSMUSG00000062609 | **Kcnj15** | 4930414N08Rik\|AI182284\|AI267127\|IRKK\|Kir4.2 | potassium inwardly-rectifying channel, subfamily J, member 15 | protein-coding |
| chr16_95218814_95219091 | chr16 | 95218815 | 95219091 | + | -0.87316 | Intergenic | Intergenic | -38605 | NM_001271693 | 16516 | Mm.272239 | NM_019664 | ENSMUSG00000062609 | **Kcnj15** | 4930414N08Rik\|AI182284\|AI267127\|IRKK\|Kir4.2 | potassium inwardly-rectifying channel, subfamily J, member 15 | protein-coding |
| chr16_95283993_95284561 | chr16 | 95283994 | 95284561 | + | -0.903768 | intron (NM_001271687, intron 3 of 4) | intron (NM_001271687, intron 3 of 4) | 26719 | NM_001271690 | 16516 | Mm.272239 | NM_019664 | ENSMUSG00000062609 | **Kcnj15** | 4930414N08Rik\|AI182284\|AI267127\|IRKK\|Kir4.2 | potassium inwardly-rectifying channel, subfamily J, member 15 | protein-coding |
| chr11_110997217_110997672 | chr11 | 110997218 | 110997672 | + | -0.691082 | promoter-TSS (NM_001252209) | promoter-TSS (NM_001252209) | -42 | NM_001252209 | 16517 | Mm.30176 | NM_010604 | ENSMUSG00000051497 | **Kcnj16** | 6430410F18Rik\|AI132396\|Kir5.1 | potassium inwardly-rectifying channel, subfamily J, member 16 | protein-coding |
| chr11_110993653_110994042 | chr11 | 110993654 | 110994042 | + | -0.985007 | intron (NM_001252207, intron 2 of 2) | intron (NM_001252207, intron 2 of 2) | -3639 | NM_001252209 | 16517 | Mm.30176 | NM_010604 | ENSMUSG00000051497 | **Kcnj16** | 6430410F18Rik\|AI132396\|Kir5.1 | potassium inwardly-rectifying channel, subfamily J, member 16 | protein-coding |
| chr11_110993597_110994207 | chr11 | 110993598 | 110994207 | + | -0.988803 | intron (NM_001252207, intron 2 of 2) | intron (NM_001252207, intron 2 of 2) | -3585 | NM_001252209 | 16517 | Mm.30176 | NM_010604 | ENSMUSG00000051497 | **Kcnj16** | 6430410F18Rik\|AI132396\|Kir5.1 | potassium inwardly-rectifying channel, subfamily J, member 16 | protein-coding |
| chr11_111800801_111801514 | chr11 | 111800802 | 111801514 | + | -0.679234 | Intergenic | L1MC\|LINE\|L1 | 734994 | NM_008425 | 16518 | Mm.4951 | NM_008425 | ENSMUSG00000041695 | **Kcnj2** | IRK1\|Kcnf1\|Kir2.1 | potassium inwardly-rectifying channel, subfamily J, member 2 | protein-coding |
| chr6_142561039_142561629 | chr6 | 142561040 | 142561629 | + | -0.65165 | Intergenic | ORR1E\|LTR\|ERVL-MaLR | 10022 | NM_008428 | 16523 | Mm.1482 | NM_008428 | ENSMUSG00000030247 | **Kcnj8** | AI448900\|Kir6.1\|gnite\|slmbr\|sltr | potassium inwardly-rectifying channel, subfamily J, member 8 | protein-coding |
| chr11_33938669_33939820 | chr11 | 33938670 | 33939820 | + | -0.746768 | intron (NM_001190886, intron 1 of 7) | intron (NM_001190886, intron 1 of 7) | -23768 | NM_031169 | 16533 | Mm.6206 | NM_031169 | ENSMUSG00000020155 | **Kcnmb1** | BKbeta1 | potassium large conductance calcium-activated channel, subfamily M, beta member 1 | protein-coding |
| chr11_33938669_33939460 | chr11 | 33938670 | 33939460 | + | -0.753539 | intron (NM_001190886, intron 1 of 7) | intron (NM_001190886, intron 1 of 7) | -23948 | NM_031169 | 16533 | Mm.6206 | NM_031169 | ENSMUSG00000020155 | **Kcnmb1** | BKbeta1 | potassium large conductance calcium-activated channel, subfamily M, beta member 1 | protein-coding |
| chr11_33939576_33940132 | chr11 | 33939577 | 33940132 | + | -0.84022 | intron (NM_001190886, intron 1 of 7) | RLTR14_RN\|LTR\|ERV1 | -23159 | NM_031169 | 16533 | Mm.6206 | NM_031169 | ENSMUSG00000020155 | **Kcnmb1** | BKbeta1 | potassium large conductance calcium-activated channel, subfamily M, beta member 1 | protein-coding |
| chr11_33939886_33940128 | chr11 | 33939887 | 33940128 | + | -0.96582 | intron (NM_001190886, intron 1 of 7) | RLTR14_RN\|LTR\|ERV1 | -23006 | NM_031169 | 16533 | Mm.6206 | NM_031169 | ENSMUSG00000020155 | **Kcnmb1** | BKbeta1 | potassium large conductance calcium-activated channel, subfamily M, beta member 1 | protein-coding |
| chr7_24378335_24378741 | chr7 | 24378336 | 24378741 | + | -0.625421 | intron (NM_001163510, intron 3 of 8) | intron (NM_001163510, intron 3 of 8) | 8275 | NM_008433 | 16534 | Mm.9911 | NM_008433 | ENSMUSG00000054342 | **Kcnn4** | IK1\|IKCA1\|KCA4\|KCa3.1\|SK4\|SKCas\|mIKCa1 | potassium intermediate/small conductance calcium-activated channel, subfamily N, member 4 | protein-coding |
| chr5_114378166_114378515 | chr5 | 114378167 | 114378515 | + | -1.14434 | intron (NM_026145, intron 1 of 6) | intron (NM_026145, intron 1 of 6) | 2164 | NM_026145 | 330171 | Mm.238285 | NM_026145 | ENSMUSG00000001098 | **Kctd10** | AW536343\|C87062\|mBACURD3 | potassium channel tetramerisation domain containing 10 | protein-coding |
| chr5_114378238_114378452 | chr5 | 114378239 | 114378452 | + | -1.16373 | intron (NM_026145, intron 1 of 6) | intron (NM_026145, intron 1 of 6) | 2160 | NM_026145 | 330171 | Mm.238285 | NM_026145 | ENSMUSG00000001098 | **Kctd10** | AW536343\|C87062\|mBACURD3 | potassium channel tetramerisation domain containing 10 | protein-coding |
| chr11_115423164_115423412 | chr11 | 115423165 | 115423412 | + | -0.802209 | intron (NM_183285, intron 2 of 5) | intron (NM_183285, intron 2 of 5) | 3162 | NM_183285 | 70382 | Mm.276299 | NM_183285 | ENSMUSG00000016940 | **Kctd2** | 2310012I15Rik | potassium channel tetramerisation domain containing 2 | protein-coding |
| chr1_189085093_189085630 | chr1 | 189085094 | 189085630 | + | -0.773358 | Intergenic | Intergenic | -77522 | NM_172650 | 226823 | Mm.209880 | NM_172650 | ENSMUSG00000026608 | **Kctd3** | 4930438A20Rik\|9330185B06\|E330032J19Rik\|NY-REN-45 | potassium channel tetramerisation domain containing 3 | protein-coding |
| chr5_143409409_143409646 | chr5 | 143409410 | 143409646 | + | -0.619109 | intron (NM_025841, intron 1 of 4) | ID4\|SINE\|ID | 5708 | NM_025841 | 66913 | Mm.181880 | NM_025841 | ENSMUSG00000079111 | **Kdelr2** | 1110007A14Rik | KDEL (Lys-Asp-Glu-Leu) endoplasmic reticulum protein retention receptor 2 | protein-coding |
| chr5_122968155_122968684 | chr5 | 122968156 | 122968684 | + | -0.633767 | intron (NM_001003953, intron 4 of 22) | intron (NM_001003953, intron 4 of 22) | 20113 | NM_001005866 | 30841 | Mm.86406 | NM_013910 | ENSMUSG00000029475 | **Kdm2b** | Cxxc2\|E430001G17\|Fbl10\|Fbxl10\|Jhdm1b\|PCCX2\|mKIAA3014 | lysine (K)-specific demethylase 2B | protein-coding |

| | | | | | | | | | | | | | | | | | |
|---|---|---|---|---|---|---|---|---|---|---|---|---|---|---|---|---|---|
| chr11_69409325_69409778 | chr11 | 69409326 | 69409778 | + | -0.615453 | intron (NM_001017426, intron 1 of 22) | intron (NM_001017426, intron 1 of 22) | 4123 | NM_001017426 | 216850 | Mm.261201 | NM_001017426 | ENSMUSG00000018476 | **Kdm6b** | 1700064E03Rik\|BC038313\|Jmjd3\|mKIAA0346 | KDM1 lysine (K)-specific demethylase 6B | protein-coding |
| chr6_39275098_39275868 | chr6 | 39275099 | 39275868 | + | -0.954408 | Intergenic | RLTR20A4\|LTR\|ERVK | -68710 | NM_001033430 | 338523 | Mm.293175 | NM_001033430 | ENSMUSG00000042599 | **Kdm7a** | A630082K20Rik\|BB041802\|ENSMUSG00000073143\|Jhdm1d\|mKIAA1718 | lysine (K)-specific demethylase 7A | protein-coding |
| chr6_39275176_39275581 | chr6 | 39275177 | 39275581 | + | -0.970734 | Intergenic | PB1D7\|SINE\|Alu | -68606 | NM_001033430 | 338523 | Mm.293175 | NM_001033430 | ENSMUSG00000042599 | **Kdm7a** | A630082K20Rik\|BB041802\|ENSMUSG00000073143\|Jhdm1d\|mKIAA1718 | lysine (K)-specific demethylase 7A | protein-coding |
| chr6_39275198_39275478 | chr6 | 39275199 | 39275478 | + | -0.992528 | Intergenic | PB1D7\|SINE\|Alu | -68565 | NM_001033430 | 338523 | Mm.293175 | NM_001033430 | ENSMUSG00000042599 | **Kdm7a** | A630082K20Rik\|BB041802\|ENSMUSG00000073143\|Jhdm1d\|mKIAA1718 | lysine (K)-specific demethylase 7A | protein-coding |
| chr4_129761604_129762078 | chr4 | 129761605 | 129762078 | + | -0.599638 | Intergenic | Intergenic | -19538 | NR_045036 | 20218 | Mm.8256 | NM_011317 | ENSMUSG00000028790 | **Khdrbs1** | Sam68\|p62\|p68 | KH domain containing, RNA binding, signal transduction associated 1 | protein-coding |
| chr4_129750425_129750762 | chr4 | 129750426 | 129750762 | + | -0.774598 | Intergenic | Intergenic | -8291 | NR_045036 | 20218 | Mm.8256 | NM_011317 | ENSMUSG00000028790 | **Khdrbs1** | Sam68\|p62\|p68 | KH domain containing, RNA binding, signal transduction associated 1 | protein-coding |
| chr4_129754289_129755219 | chr4 | 129754290 | 129755219 | + | -0.784705 | Intergenic | IMPB_01\|Satellite\|Satellite | -12451 | NR_045036 | 20218 | Mm.8256 | NM_011317 | ENSMUSG00000028790 | **Khdrbs1** | Sam68\|p62\|p68 | KH domain containing, RNA binding, signal transduction associated 1 | protein-coding |
| chr4_129754692_129755227 | chr4 | 129754693 | 129755227 | + | -0.831362 | Intergenic | L2b\|LINE\|L2 | -12657 | NR_045036 | 20218 | Mm.8256 | NM_011317 | ENSMUSG00000028790 | **Khdrbs1** | Sam68\|p62\|p68 | KH domain containing, RNA binding, signal transduction associated 1 | protein-coding |
| chr4_129765404_129765689 | chr4 | 129765405 | 129765689 | + | -0.983605 | Intergenic | Intergenic | -23244 | NR_045036 | 20218 | Mm.8256 | NM_011317 | ENSMUSG00000028790 | **Khdrbs1** | Sam68\|p62\|p68 | KH domain containing, RNA binding, signal transduction associated 1 | protein-coding |
| chr15_68959336_68959640 | chr15 | 68959337 | 68959640 | + | -0.820739 | intron (NM_010158, intron 1 of 8) | intron (NM_010158, intron 1 of 8) | 31068 | NM_010158 | 13992 | Mm.17964 | NM_010158 | ENSMUSG00000022332 | **Khdrbs3** | Etle\|SLM-2\|Salp\|Slm2\|T-STAR | KH domain containing, RNA binding, signal transduction associated 3 | protein-coding |
| chr15_68961745_68962136 | chr15 | 68961746 | 68962136 | + | -0.907706 | intron (NM_010158, intron 1 of 8) | intron (NM_010158, intron 1 of 8) | 33521 | NM_010158 | 13992 | Mm.17964 | NM_010158 | ENSMUSG00000022332 | **Khdrbs3** | Etle\|SLM-2\|Salp\|Slm2\|T-STAR | KH domain containing, RNA binding, signal transduction associated 3 | protein-coding |
| chr12_25012994_25013626 | chr12 | 25012995 | 25013626 | + | -1.00547 | exon (NM_001081378, exon 21 of 32) | exon (NM_001081378, exon 21 of 32) | 38378 | NM_001081378 | 77480 | Mm.250641 | NM_001081378 | ENSMUSG00000036333 | **Kidins220** | 3110039L19Rik\|AI194387\|AI316525\|C330002I19Rik\|mKIAA1250 | kinase D-interacting substrate 220 | protein-coding |
| chr12_25010400_25010788 | chr12 | 25010401 | 25010788 | + | -1.49693 | intron (NM_001081378, intron 18 of 31) | intron (NM_001081378, intron 18 of 31) | 35662 | NM_001081378 | 77480 | Mm.250641 | NM_001081378 | ENSMUSG00000036333 | **Kidins220** | 3110039L19Rik\|AI194387\|AI316525\|C330002I19Rik\|mKIAA1250 | kinase D-interacting substrate 220 | protein-coding |
| chr4_138257069_138257602 | chr4 | 138257070 | 138257602 | + | -0.667124 | intron (NM_001099631, intron 7 of 9) | intron (NM_001099631, intron 7 of 9) | -4915 | NM_001190978 | 16559 | Mm.271936 | NM_010623 | ENSMUSG00000028758 | **Kif17** | 5930435E01Rik\|AW492270\|Kif17b\|mKIAA1405 | kinesin family member 17 | protein-coding |
| chr4_138256644_138257571 | chr4 | 138256645 | 138257571 | + | -0.686569 | intron (NM_001099631, intron 6 of 9) | intron (NM_001099631, intron 6 of 9) | -5143 | NM_001190978 | 16559 | Mm.271936 | NM_010623 | ENSMUSG00000028758 | **Kif17** | 5930435E01Rik\|AW492270\|Kif17b\|mKIAA1405 | kinesin family member 17 | protein-coding |
| chr4_149282159_149282952 | chr4 | 149282160 | 149282952 | + | -1.01338 | intron (NM_001290995, intron 2 of 48) | intron (NM_001290995, intron 2 of 48) | 25142 | NM_207682 | 16561 | Mm.402393 | NM_008441 | ENSMUSG00000063077 | **Kif1b** | A530096N05Rik\|AI448212\|AI506502\|D4Mil1e\|KIF1Bp130\|KIF1Bp204 | kinesin family member 1B | protein-coding |
| chr9_61983268_61983664 | chr9 | 61983269 | 61983664 | + | -0.912486 | intron (NM_028748, intron 2 of 7) | ORR1E\|LTR\|ERVL-MaLR | -36667 | NM_024245 | 71819 | Mm.259374 | NM_024245 | ENSMUSG00000032254 | **Kif23** | 3110001D19Rik\|C87313\|CHO1\|Knsl5\|MKLP-1\|MKLP1 | kinesin family member 23 | protein-coding |
| chr9_61985581_61985793 | chr9 | 61985582 | 61985793 | + | -1.23648 | intron (NM_028748, intron 2 of 7) | intron (NM_028748, intron 2 of 7) | -38888 | NM_024245 | 71819 | Mm.259374 | NM_024245 | ENSMUSG00000032254 | **Kif23** | 3110001D19Rik\|C87313\|CHO1\|Knsl5\|MKLP-1\|MKLP1 | kinesin family member 23 | protein-coding |
| chr9_61985526_61985910 | chr9 | 61985527 | 61985910 | + | -1.24077 | intron (NM_028748, intron 2 of 7) | intron (NM_028748, intron 2 of 7) | -38919 | NM_024245 | 71819 | Mm.259374 | NM_024245 | ENSMUSG00000032254 | **Kif23** | 3110001D19Rik\|C87313\|CHO1\|Knsl5\|MKLP-1\|MKLP1 | kinesin family member 23 | protein-coding |
| chr13_106998605_106999165 | chr13 | 106998606 | 106999165 | + | -0.98891 | intron (NM_008442, intron 1 of 21) | ORR1A2\|LTR\|ERVL-MaLR | 23229 | NM_001145779 | 16563 | Mm.355686 | NM_008442 | ENSMUSG00000021693 | **Kif2a** | C530030B14Rik\|Kif2\|Kns2\|M-kinesin | kinesin family member 2A | protein-coding |
| chrX_52757016_52757507 | chrX | 52757017 | 52757507 | + | -0.695315 | Intergenic | Intergenic | -12669 | NR_003188 | 751866 | Mm.381916 | NR_003188 | | **Kis2** | Kis2-T1\|Kis2-T2\|Kis2-T3\|Kis2-T4\|Kis2-T5\|Xpcl1 | Kaplan integration site 2 | ncRNA |
| chrX_52757149_52757528 | chrX | 52757150 | 52757528 | + | -0.715053 | Intergenic | Intergenic | -12746 | NR_003188 | 751866 | Mm.381916 | NR_003188 | | **Kis2** | Kis2-T1\|Kis2-T2\|Kis2-T3\|Kis2-T4\|Kis2-T5\|Xpcl1 | Kaplan integration site 2 | ncRNA |
| chr5_75645416_75645677 | chr5 | 75645417 | 75645677 | + | -0.92718 | intron (NM_021099, intron 13 of 20) | intron (NM_021099, intron 13 of 20) | 70556 | NM_021099 | 16590 | Mm.247073 | NM_021099 | ENSMUSG00000005672 | **Kit** | Bs\|CD117\|Fdc\|Gsfsco1\|Gsfsco5\|Gsfsow3\|SCO1\|SCO5\|SOW3\|Ssm\|Tr-kit\|W\|c-KIT | kit oncogene | protein-coding |
| chr7_64038011_64038263 | chr7 | 64038012 | 64038263 | + | -0.709126 | Intergenic | Intergenic | -99222 | NM_021366 | 50794 | Mm.240473 | NM_021366 | ENSMUSG00000052040 | **Klf13** | 0610043C13Rik\|9430029L20Rik\|Bteb3\|FKLF-2\|FKLF2\|NSLP1\|RFLAT-1\|RFLAT1 | Kruppel-like factor 13 | protein-coding |
| chr7_64042697_64043403 | chr7 | 64042698 | 64043403 | + | -0.716901 | Intergenic | Intergenic | -104135 | NM_021366 | 50794 | Mm.240473 | NM_021366 | ENSMUSG00000052040 | **Klf13** | 0610043C13Rik\|9430029L20Rik\|Bteb3\|FKLF-2\|FKLF2\|NSLP1\|RFLAT-1\|RFLAT1 | Kruppel-like factor 13 | protein-coding |
| chr8_72360336_72360682 | chr8 | 72360337 | 72360682 | + | -0.639264 | intron (NM_007944, intron 21 of 23) | B3A\|SINE\|B2 | 41447 | NM_008452 | 16598 | Mm.26938 | NM_008452 | ENSMUSG00000055148 | **Klf2** | Lklf | Kruppel-like factor 2 (lung) | protein-coding |
| chr8_72318665_72319575 | chr8 | 72318666 | 72319575 | + | -0.690248 | promoter-TSS (NM_008452) | promoter-TSS (NM_008452) | 58 | NM_008452 | 16598 | Mm.26938 | NM_008452 | ENSMUSG00000055148 | **Klf2** | Lklf | Kruppel-like factor 2 (lung) | protein-coding |
| chr8_72291075_72291521 | chr8 | 72291076 | 72291521 | + | -0.843942 | Intergenic | Intergenic | -27764 | NM_008452 | 16598 | Mm.26938 | NM_008452 | ENSMUSG00000055148 | **Klf2** | Lklf | Kruppel-like factor 2 (lung) | protein-coding |
| chr8_72354308_72354780 | chr8 | 72354309 | 72354780 | + | -0.855553 | intron (NM_007944, intron 22 of 23) | intron (NM_007944, intron 22 of 23) | 35482 | NM_008452 | 16598 | Mm.26938 | NM_008452 | ENSMUSG00000055148 | **Klf2** | Lklf | Kruppel-like factor 2 (lung) | protein-coding |
| chr8_72295545_72296217 | chr8 | 72295546 | 72296217 | + | -0.941589 | Intergenic | Intergenic | -23181 | NM_008452 | 16598 | Mm.26938 | NM_008452 | ENSMUSG00000055148 | **Klf2** | Lklf | Kruppel-like factor 2 (lung) | protein-coding |
| chr5_64819431_64819779 | chr5 | 64819432 | 64819779 | + | -0.850035 | intron (NM_008453, intron 2 of 5) | intron (NM_008453, intron 2 of 5) | 16082 | NM_008453 | 16599 | Mm.392759 | NM_008453 | ENSMUSG00000029178 | **Klf3** | 9930027G08Rik\|AL024007\|Bklf\|Tef-2 | Kruppel-like factor 3 (basic) | protein-coding |
| chr5_64780677_64781262 | chr5 | 64780678 | 64781262 | + | -0.922915 | Intergenic | Intergenic | -22553 | NM_008453 | 16599 | Mm.392759 | NM_008453 | ENSMUSG00000029178 | **Klf3** | 9930027G08Rik\|AL024007\|Bklf\|Tef-2 | Kruppel-like factor 3 (basic) | protein-coding |
| chr5_64806144_64806522 | chr5 | 64806145 | 64806522 | + | -0.990829 | intron (NM_008453, intron 1 of 5) | intron (NM_008453, intron 1 of 5) | 2810 | NM_008453 | 16599 | Mm.392759 | NM_008453 | ENSMUSG00000029178 | **Klf3** | 9930027G08Rik\|AL024007\|Bklf\|Tef-2 | Kruppel-like factor 3 (basic) | protein-coding |
| chr5_64806013_64806530 | chr5 | 64806014 | 64806530 | + | -0.99932 | intron (NM_008453, intron 1 of 5) | intron (NM_008453, intron 1 of 5) | 2749 | NM_008453 | 16599 | Mm.392759 | NM_008453 | ENSMUSG00000029178 | **Klf3** | 9930027G08Rik\|AL024007\|Bklf\|Tef-2 | Kruppel-like factor 3 (basic) | protein-coding |
| chr13_5860627_5860980 | chr13 | 5860628 | 5860980 | + | -0.824884 | promoter-TSS (NM_011803) | promoter-TSS (NM_011803) | -685 | NM_011803 | 23849 | Mm.275036 | NM_011803 | ENSMUSG00000000078 | **Klf6** | AI448727\|BCD1\|C86813\|CPBP\|Copeb\|FM2\|FM6\|Ierepo1\|Ierepo3\|R75280\|Zf9 | Kruppel-like factor 6 | protein-coding |
| chr13_5955937_5956583 | chr13 | 5955938 | 5956583 | + | -0.869666 | Intergenic | MTEb\|LTR\|ERVL-MaLR | 94771 | NM_011803 | 23849 | Mm.275036 | NM_011803 | ENSMUSG00000000078 | **Klf6** | AI448727\|BCD1\|C86813\|CPBP\|Copeb\|FM2\|FM6\|Ierepo1\|Ierepo3\|R75280\|Zf9 | Kruppel-like factor 6 | protein-coding |
| chr13_5956175_5956480 | chr13 | 5956176 | 5956480 | + | -0.924623 | Intergenic | MTEb\|LTR\|ERVL-MaLR | 94839 | NM_011803 | 23849 | Mm.275036 | NM_011803 | ENSMUSG00000000078 | **Klf6** | AI448727\|BCD1\|C86813\|CPBP\|Copeb\|FM2\|FM6\|Ierepo1\|Ierepo3\|R75280\|Zf9 | Kruppel-like factor 6 | protein-coding |
| chr1_64104667_64105133 | chr1 | 64104668 | 64105133 | + | -0.637048 | intron (NM_033563, intron 1 of 3) | Lx8\|LINE\|L1 | 16489 | NM_033563 | 93691 | Mm.29466 | NM_033563 | ENSMUSG00000025959 | **Klf7** | 9830124P08Rik | Kruppel-like factor 7 (ubiquitous) | protein-coding |
| chr1_64087911_64088466 | chr1 | 64087912 | 64088466 | + | -0.76169 | intron (NM_033563, intron 1 of 3) | intron (NM_033563, intron 1 of 3) | 33200 | NM_033563 | 93691 | Mm.29466 | NM_033563 | ENSMUSG00000025959 | **Klf7** | 9830124P08Rik | Kruppel-like factor 7 (ubiquitous) | protein-coding |
| chr1_64135954_64136865 | chr1 | 64135955 | 64136865 | + | -0.763395 | Intergenic | Intergenic | -15021 | NM_033563 | 93691 | Mm.29466 | NM_033563 | ENSMUSG00000025959 | **Klf7** | 9830124P08Rik | Kruppel-like factor 7 (ubiquitous) | protein-coding |
| chr1_64085908_64086341 | chr1 | 64085909 | 64086341 | + | -0.80713 | intron (NM_033563, intron 1 of 3) | intron (NM_033563, intron 1 of 3) | 35264 | NM_033563 | 93691 | Mm.29466 | NM_033563 | ENSMUSG00000025959 | **Klf7** | 9830124P08Rik | Kruppel-like factor 7 (ubiquitous) | protein-coding |
| chr1_64136166_64136840 | chr1 | 64136167 | 64136840 | + | -0.817576 | Intergenic | PB1D7\|SINE\|Alu | -15114 | NM_033563 | 93691 | Mm.29466 | NM_033563 | ENSMUSG00000025959 | **Klf7** | 9830124P08Rik | Kruppel-like factor 7 (ubiquitous) | protein-coding |
| chr1_64087903_64088288 | chr1 | 64087904 | 64088288 | + | -0.889837 | intron (NM_033563, intron 1 of 3) | intron (NM_033563, intron 1 of 3) | 33293 | NM_033563 | 93691 | Mm.29466 | NM_033563 | ENSMUSG00000025959 | **Klf7** | 9830124P08Rik | Kruppel-like factor 7 (ubiquitous) | protein-coding |
| chr12_69257833_69258287 | chr12 | 69257834 | 69258287 | + | -0.862692 | exon (NM_178253, exon 7 of 13) | exon (NM_178253, exon 7 of 13) | 16228 | NM_178253 | 271005 | Mm.180969 | NM_178253 | | **Klhdc1** | - | kelch domain containing 1 | protein-coding |

| | | | | | | | | | | | | | | | | | |
|---|---|---|---|---|---|---|---|---|---|---|---|---|---|---|---|---|---|
| chr12_69255967_69256325 | chr12 | 69255968 | 69256325 | + | -0.960822 | intron (NM_178253, intron 5 of 12) | intron (NM_178253, intron 5 of 12) | 14314 | NM_178253 | 271005 | Mm.180969 | NM_178253 | | **Klhdc1** | - | kelch domain containing 1 | protein-coding |
| chr8_64826176_64826893 | chr8 | 64826177 | 64826893 | + | -0.775741 | intron (NM_178633, intron 2 of 14) | intron (NM_178633, intron 2 of 14) | 23389 | NM_178633 | 77113 | Mm.350347 | NM_178633 | ENSMUSG00000031605 | **Klhl2** | 6030411N21Rik\|ABP-KELCH\|AU020744\|Mav | kelch-like 2, Mayven | protein-coding |
| chr7_75915579_75916010 | chr7 | 75915580 | 75916010 | + | -0.65004 | Intergenic | Intergenic | 67457 | NM_029652 | 207952 | Mm.206223 | NM_029652 | ENSMUSG00000055652 | **Klhl25** | 2810402K13Rik\|BC027373\|Enc-1 | kelch-like 25 | protein-coding |
| chr7_75879343_75879807 | chr7 | 75879344 | 75879807 | + | -0.742324 | Intergenic | Intergenic | 31237 | NM_029652 | 207952 | Mm.206223 | NM_029652 | ENSMUSG00000055652 | **Klhl25** | 2810402K13Rik\|BC027373\|Enc-1 | kelch-like 25 | protein-coding |
| chr7_75910026_75910964 | chr7 | 75910027 | 75910964 | + | -0.824709 | Intergenic | Intergenic | 62157 | NM_029652 | 207952 | Mm.206223 | NM_029652 | ENSMUSG00000055652 | **Klhl25** | 2810402K13Rik\|BC027373\|Enc-1 | kelch-like 25 | protein-coding |
| chr7_75701753_75702276 | chr7 | 75701754 | 75702276 | + | -0.906136 | intron (NM_029332, intron 16 of 36) | intron (NM_029332, intron 16 of 36) | -146323 | NM_001122780 | 207952 | Mm.206223 | NM_029652 | ENSMUSG00000055652 | **Klhl25** | 2810402K13Rik\|BC027373\|Enc-1 | kelch-like 25 | protein-coding |
| chr7_75701756_75702134 | chr7 | 75701757 | 75702134 | + | -0.922957 | intron (NM_029332, intron 16 of 36) | intron (NM_029332, intron 16 of 36) | -146393 | NM_001122780 | 207952 | Mm.206223 | NM_029652 | ENSMUSG00000055652 | **Klhl25** | 2810402K13Rik\|BC027373\|Enc-1 | kelch-like 25 | protein-coding |
| chr9_77683495_77684190 | chr9 | 77683496 | 77684190 | + | -0.712613 | Intergenic | MTD\|LTR\|ERVL-MaLR | 47111 | NM_172925 | 244923 | Mm.138238 | NM_172925 | ENSMUSG00000044938 | **Klhl31** | 9830147P19Rik\|D930047P17Rik\|Kbtbd1 | kelch-like 31 | protein-coding |
| chr9_77606958_77607629 | chr9 | 77606959 | 77607629 | + | -0.740755 | Intergenic | Lx10\|LINE\|L1 | -29438 | NM_172925 | 244923 | Mm.138238 | NM_172925 | ENSMUSG00000044938 | **Klhl31** | 9830147P19Rik\|D930047P17Rik\|Kbtbd1 | kelch-like 31 | protein-coding |
| chr9_77607268_77607632 | chr9 | 77607269 | 77607632 | + | -0.870945 | Intergenic | Lx10\|LINE\|L1 | -29282 | NM_172925 | 244923 | Mm.138238 | NM_172925 | ENSMUSG00000044938 | **Klhl31** | 9830147P19Rik\|D930047P17Rik\|Kbtbd1 | kelch-like 31 | protein-coding |
| chr9_77671054_77671701 | chr9 | 77671055 | 77671701 | + | -0.987744 | Intergenic | ORR1D2\|LTR\|ERVL-MaLR | 34646 | NM_172925 | 244923 | Mm.138238 | NM_172925 | ENSMUSG00000044938 | **Klhl31** | 9830147P19Rik\|D930047P17Rik\|Kbtbd1 | kelch-like 31 | protein-coding |
| chr9_77670932_77671405 | chr9 | 77670933 | 77671405 | + | -1.27184 | Intergenic | ORR1D2\|LTR\|ERVL-MaLR | 34437 | NM_172925 | 244923 | Mm.138238 | NM_172925 | ENSMUSG00000044938 | **Klhl31** | 9830147P19Rik\|D930047P17Rik\|Kbtbd1 | kelch-like 31 | protein-coding |
| chr6_147102680_147102890 | chr6 | 147102681 | 147102890 | + | -0.638303 | intron (NM_001081237, intron 2 of 2) | PB1D10\|SINE\|Alu | 11710 | NM_001081237 | 232539 | Mm.491206 | NM_001081237 | ENSMUSG00000040102 | **Klhl42** | A330079N23Rik\|AI450355\|C230080I20Rik\|Ctb9\|Klhdc5\|mKIAA1340 | kelch-like 42 | protein-coding |
| chr6_147098889_147099947 | chr6 | 147098890 | 147099947 | + | -0.742361 | intron (NM_001081237, intron 1 of 2) | intron (NM_001081237, intron 1 of 2) | 8343 | NM_001081237 | 232539 | Mm.491206 | NM_001081237 | ENSMUSG00000040102 | **Klhl42** | A330079N23Rik\|AI450355\|C230080I20Rik\|Ctb9\|Klhdc5\|mKIAA1340 | kelch-like 42 | protein-coding |
| chr6_147099155_147099952 | chr6 | 147099156 | 147099952 | + | -0.795648 | intron (NM_001081237, intron 1 of 2) | intron (NM_001081237, intron 1 of 2) | 8479 | NM_001081237 | 232539 | Mm.491206 | NM_001081237 | ENSMUSG00000040102 | **Klhl42** | A330079N23Rik\|AI450355\|C230080I20Rik\|Ctb9\|Klhdc5\|mKIAA1340 | kelch-like 42 | protein-coding |
| chr5_65113203_65113738 | chr5 | 65113204 | 65113738 | + | -0.934849 | Intergenic | Intergenic | -17760 | NM_175174 | 71778 | Mm.10281 | NM_175174 | ENSMUSG00000054920 | **Klhl5** | 1300013C10Rik | kelch-like 5 | protein-coding |
| chr16_19982749_19983286 | chr16 | 19982750 | 19983286 | + | -0.769604 | promoter-TSS (NM_183390) | promoter-TSS (NM_183390) | 31 | NM_183390 | 239743 | Mm.86699 | NM_183390 | ENSMUSG00000043008 | **Klhl6** | - | kelch-like 6 | protein-coding |
| chr16_20000327_20000731 | chr16 | 20000328 | 20000731 | + | -0.849455 | Intergenic | RLTR45\|LTR\|ERVK | -17480 | NM_183390 | 239743 | Mm.86699 | NM_183390 | ENSMUSG00000043008 | **Klhl6** | - | kelch-like 6 | protein-coding |
| chr16_19985006_19985221 | chr16 | 19985007 | 19985221 | + | -1.01347 | Intergenic | B4A\|SINE\|B4 | -2065 | NM_183390 | 239743 | Mm.86699 | NM_183390 | ENSMUSG00000043008 | **Klhl6** | - | kelch-like 6 | protein-coding |
| chr16_19984922_19985287 | chr16 | 19984923 | 19985287 | + | -1.05592 | Intergenic | B4A\|SINE\|B4 | -2056 | NM_183390 | 239743 | Mm.86699 | NM_183390 | ENSMUSG00000043008 | **Klhl6** | - | kelch-like 6 | protein-coding |
| chr6_128766549_128767183 | chr6 | 128766550 | 128767183 | + | -0.781203 | Intergenic | L1M2\|LINE\|L1 | 21775 | NM_008527 | 17059 | Mm.480764 | NM_008527 | ENSMUSG00000030325 | **Klrb1c** | AI462337\|CD161\|Ly-59\|Ly55c\|Ly59\|NK-RP1\|NK1.1\|NKR-P1.9\|NKRP1\|NKRP140\|Nk-1\|Nk-1.2\|Nk1\|Nk1.2\|Nkrp1c\|ly-55c | killer cell lectin-like receptor subfamily B member 1C | protein-coding |
| chr6_129591394_129592026 | chr6 | 129591395 | 129592026 | + | -1.08614 | promoter-TSS (NM_010654) | promoter-TSS (NM_010654) | -101 | NM_010654 | 16643 | Mm.391515 | NM_010654 | ENSMUSG00000030165 | **Klrd1** | CD94 | killer cell lectin-like receptor, subfamily D, member 1 | protein-coding |
| chr6_129606959_129607259 | chr6 | 129606960 | 129607259 | + | -1.32594 | Intergenic | RSINE1\|SINE\|B4 | 15298 | NM_010654 | 16643 | Mm.391515 | NM_010654 | ENSMUSG00000030165 | **Klrd1** | CD94 | killer cell lectin-like receptor, subfamily D, member 1 | protein-coding |
| chr6_129608198_129609153 | chr6 | 129608199 | 129609153 | + | -0.852856 | Intergenic | Intergenic | 14007 | NM_033078 | 27007 | Mm.8217 | NM_033078 | ENSMUSG00000030149 | **Klrk1** | D6H12S2489E\|NKG2-D\|Nkg2d | killer cell lectin-like receptor subfamily K, member 1 | protein-coding |
| chr6_129608229_129608832 | chr6 | 129608230 | 129608832 | + | -0.886301 | Intergenic | Intergenic | 14152 | NM_033078 | 27007 | Mm.8217 | NM_033078 | ENSMUSG00000030149 | **Klrk1** | D6H12S2489E\|NKG2-D\|Nkg2d | killer cell lectin-like receptor subfamily K, member 1 | protein-coding |
| chr5_23453651_23453982 | chr5 | 23453652 | 23453982 | + | -0.817472 | intron (NM_026984, intron 2 of 25) | B2_Mm2\|SINE\|B2 | 19388 | NM_026984 | 69188 | Mm.205190 | NM_026984 | ENSMUSG00000029004 | **Kmt2e** | 1810033J14Rik\|9530077A04Rik\|D230038D11Rik\|Mll5\|NKp44L | lysine (K)-specific methyltransferase 2E | protein-coding |
| chr5_23524564_23525141 | chr5 | 23524565 | 23525141 | + | -1.22879 | intron (NM_009274, intron 9 of 14) | intron (NM_009274, intron 9 of 14) | 90424 | NM_026984 | 69188 | Mm.205190 | NM_026984 | ENSMUSG00000029004 | **Kmt2e** | 1810033J14Rik\|9530077A04Rik\|D230038D11Rik\|Mll5\|NKp44L | lysine (K)-specific methyltransferase 2E | protein-coding |
| chr7_118895573_118895796 | chr7 | 118895574 | 118895796 | + | -0.714386 | intron (NM_001081446, intron 4 of 8) | intron (NM_001081446, intron 4 of 8) | -39687 | NM_023197 | 66356 | Mm.482278 | NM_023197 | ENSMUSG00000030980 | **Knop1** | 2310008H09Rik\|Tsg118 | lysine rich nucleolar protein 1 | protein-coding |
| chr7_118895389_118895905 | chr7 | 118895390 | 118895905 | + | -0.726956 | intron (NM_001081446, intron 4 of 8) | intron (NM_001081446, intron 4 of 8) | -39649 | NM_023197 | 66356 | Mm.482278 | NM_023197 | ENSMUSG00000030980 | **Knop1** | 2310008H09Rik\|Tsg118 | lysine rich nucleolar protein 1 | protein-coding |
| chr7_118895493_118895819 | chr7 | 118895494 | 118895819 | + | -0.72781 | intron (NM_001081446, intron 4 of 8) | intron (NM_001081446, intron 4 of 8) | -39658 | NM_023197 | 66356 | Mm.482278 | NM_023197 | ENSMUSG00000030980 | **Knop1** | 2310008H09Rik\|Tsg118 | lysine rich nucleolar protein 1 | protein-coding |
| chr7_118895215_118895814 | chr7 | 118895216 | 118895814 | + | -0.788928 | intron (NM_001081446, intron 4 of 8) | intron (NM_001081446, intron 4 of 8) | -39517 | NM_023197 | 66356 | Mm.482278 | NM_023197 | ENSMUSG00000030980 | **Knop1** | 2310008H09Rik\|Tsg118 | lysine rich nucleolar protein 1 | protein-coding |
| chr5_123780615_123781114 | chr5 | 123780616 | 123781114 | + | -0.955791 | intron (NM_001042421, intron 27 of 63) | intron (NM_001042421, intron 27 of 63) | 31139 | NM_001042421 | 208628 | Mm.441089 | NM_001042421 | ENSMUSG00000029414 | **Kntc1** | D330012D13 | kinetochore associated 1 | protein-coding |
| chr5_123781701_123782281 | chr5 | 123781702 | 123782281 | + | -1.02841 | intron (NM_001042421, intron 30 of 63) | intron (NM_001042421, intron 30 of 63) | 32265 | NM_001042421 | 208628 | Mm.441089 | NM_001042421 | ENSMUSG00000029414 | **Kntc1** | D330012D13 | kinetochore associated 1 | protein-coding |
| chr5_123765376_123766014 | chr5 | 123765377 | 123766014 | + | -1.13076 | intron (NM_001042421, intron 12 of 63) | intron (NM_001042421, intron 12 of 63) | 15969 | NM_001042421 | 208628 | Mm.441089 | NM_001042421 | ENSMUSG00000029414 | **Kntc1** | D330012D13 | kinetochore associated 1 | protein-coding |
| chr11_106961402_106962105 | chr11 | 106961403 | 106962105 | + | -0.847663 | Intergenic | Intergenic | 37771 | NM_010655 | 16647 | Mm.12508 | NM_010655 | ENSMUSG00000018362 | **Kpna2** | 2410044B12Rik\|IPOA1\|PTAC58\|Rch1 | karyopherin (importin) alpha 2 | protein-coding |
| chr11_99255290_99255675 | chr11 | 99255291 | 99255675 | + | -0.864291 | Intergenic | B4A\|SINE\|B4 | -11416 | NM_172946 | 268481 | Mm.133193 | NM_172946 | ENSMUSG00000035849 | **Krt222** | 6330509G02Rik | keratin 222 | protein-coding |
| chr11_99255268_99255804 | chr11 | 99255269 | 99255804 | + | -0.871774 | Intergenic | Intergenic | -11469 | NM_172946 | 268481 | Mm.133193 | NM_172946 | ENSMUSG00000035849 | **Krt222** | 6330509G02Rik | keratin 222 | protein-coding |
| chr11_99238267_99238940 | chr11 | 99238268 | 99238940 | + | -0.899661 | exon (NM_172946, exon 3 of 6) | exon (NM_172946, exon 3 of 6) | 5463 | NM_172946 | 268481 | Mm.133193 | NM_172946 | ENSMUSG00000035849 | **Krt222** | 6330509G02Rik | keratin 222 | protein-coding |
| chr10_26520685_26521785 | chr10 | 26520686 | 26521785 | + | -0.725459 | Intergenic | Intergenic | -146050 | NM_172787 | 237339 | Mm.338439 | NM_172787 | ENSMUSG00000039089 | **L3mbtl3** | AI481284\|MBT-1 | l(3)mbt-like 3 (Drosophila) | protein-coding |
| chr10_26325131_26325724 | chr10 | 26325132 | 26325724 | + | -0.955817 | intron (NM_172787, intron 12 of 21) | RLTR14_RN\|LTR\|ERV1 | 49757 | NM_172787 | 237339 | Mm.338439 | NM_172787 | ENSMUSG00000039089 | **L3mbtl3** | AI481284\|MBT-1 | l(3)mbt-like 3 (Drosophila) | protein-coding |
| chr4_136626416_136626921 | chr4 | 136626417 | 136626921 | + | -0.61798 | intron (NM_001243262, intron 2 of 7) | intron (NM_001243262, intron 2 of 7) | 4048 | NM_001243262 | 242707 | Mm.448979 | NM_001243262 | ENSMUSG00000070683 | **Lactbl1** | - | lactamase, beta-like 1 | protein-coding |
| chr10_39023519_39024224 | chr10 | 39023520 | 39024224 | + | -1.11587 | intron (NM_010681, intron 4 of 37) | intron (NM_010681, intron 4 of 37) | 58357 | NM_010681 | 16775 | Mm.258065 | NM_010681 | ENSMUSG00000019846 | **Lama4** | - | laminin, alpha 4 | protein-coding |
| chr1_153087787_153088124 | chr1 | 153087788 | 153088124 | + | -0.953756 | intron (NM_175460, intron 5 of 10) | ORR1F\|LTR\|ERVL-MaLR | 98491 | NM_008485 | 16782 | Mm.329272 | NM_008485 | ENSMUSG00000026479 | **Lamc2** | AA589349 | laminin, gamma 2 | protein-coding |
| chr1_153087866_153088107 | chr1 | 153087867 | 153088107 | + | -0.959781 | intron (NM_175460, intron 5 of 10) | ORR1F\|LTR\|ERVL-MaLR | 98460 | NM_008485 | 16782 | Mm.329272 | NM_008485 | ENSMUSG00000026479 | **Lamc2** | AA589349 | laminin, gamma 2 | protein-coding |
| chr4_130913072_130913417 | chr4 | 130913073 | 130913417 | + | -0.613169 | promoter-TSS (NM_010686) | promoter-TSS (NM_010686) | -89 | NM_010686 | 16792 | Mm.271868 | NM_010686 | ENSMUSG00000028581 | **Laptm5** | E3 | lysosomal-associated protein transmembrane 5 | protein-coding |
| chr4_130920984_130921667 | chr4 | 130920985 | 130921667 | + | -0.73347 | intron (NM_010686, intron 1 of 7) | intron (NM_010686, intron 1 of 7) | 7992 | NM_010686 | 16792 | Mm.271868 | NM_010686 | ENSMUSG00000028581 | **Laptm5** | E3 | lysosomal-associated protein transmembrane 5 | protein-coding |
| chr4_130920981_130921378 | chr4 | 130920982 | 130921378 | + | -0.739432 | intron (NM_010686, intron 1 of 7) | intron (NM_010686, intron 1 of 7) | 7846 | NM_010686 | 16792 | Mm.271868 | NM_010686 | ENSMUSG00000028581 | **Laptm5** | E3 | lysosomal-associated protein transmembrane 5 | protein-coding |
| chr11_97810248_97810860 | chr11 | 97810249 | 97810860 | + | -0.644373 | intron (NM_010688, intron 2 of 6) | intron (NM_010688, intron 2 of 6) | 10882 | NM_010688 | 16796 | Mm.271967 | NM_010688 | ENSMUSG00000038366 | **Lasp1** | AA408629\|Def-4\|SH3P6\|Tg(Col1a1-lacZ)1Ngma | LIM and SH3 protein 1 | protein-coding |
| chr17_72939387_72939863 | chr17 | 72939388 | 72939863 | + | -0.645179 | 3' UTR (NM_029999, exon 3 of 3) | 3' UTR (NM_029999, exon 3 of 3) | 21320 | NM_029999 | 77889 | Mm.154109 | NM_029999 | ENSMUSG00000024063 | **Lbh** | 1810009F10Rik\|6720416L16Rik | limb-bud and heart | protein-coding |

| | | | | | | | | | | | | | | | | | |
|---|---|---|---|---|---|---|---|---|---|---|---|---|---|---|---|---|---|
| chr17_73008858_73009308 | chr17 | 73008859 | 73009308 | + | -0.773453 | Intergenic | Intergenic | 90778 | NM_029999 | 77889 | Mm.154109 | NM_029999 | ENSMUSG00000024063 | **Lbh** | 1810009F10Rik\|6720416L16Rik | limb-bud and heart | protein-coding |
| chr17_73009959_73010509 | chr17 | 73009960 | 73010509 | + | -0.796514 | Intergenic | Intergenic | 91929 | NM_029999 | 77889 | Mm.154109 | NM_029999 | ENSMUSG00000024063 | **Lbh** | 1810009F10Rik\|6720416L16Rik | limb-bud and heart | protein-coding |
| chr17_73008821_73009190 | chr17 | 73008822 | 73009190 | + | -0.832939 | Intergenic | Intergenic | 90701 | NM_029999 | 77889 | Mm.154109 | NM_029999 | ENSMUSG00000024063 | **Lbh** | 1810009F10Rik\|6720416L16Rik | limb-bud and heart | protein-coding |
| chr17_72996299_72997066 | chr17 | 72996300 | 72997066 | + | -1.07857 | Intergenic | Intergenic | 78378 | NM_029999 | 77889 | Mm.154109 | NM_029999 | ENSMUSG00000024063 | **Lbh** | 1810009F10Rik\|6720416L16Rik | limb-bud and heart | protein-coding |
| chr2_158322708_158323665 | chr2 | 158322709 | 158323665 | + | -0.91614 | intron (NM_008489, intron 9 of 14) | intron (NM_008489, intron 9 of 14) | 16694 | NM_008489 | 16803 | Mm.218846 | NM_008489 | ENSMUSG00000016024 | **Lbp** | Bpifd2\|Ly88 | lipopolysaccharide binding protein | protein-coding |
| chr1_181852074_181853556 | chr1 | 181852075 | 181853556 | + | -0.628992 | Intergenic | Intergenic | -10414 | NM_133815 | 98386 | Mm.4538 | NM_133815 | ENSMUSG00000004880 | **Lbr** | AI505894\|ic | lamin B receptor | protein-coding |
| chr1_181853019_181853665 | chr1 | 181853020 | 181853665 | + | -0.722111 | Intergenic | Intergenic | -10941 | NM_133815 | 98386 | Mm.4538 | NM_133815 | ENSMUSG00000004880 | **Lbr** | AI505894\|ic | lamin B receptor | protein-coding |
| chr9_83437610_83438106 | chr9 | 83437611 | 83438106 | + | -0.834009 | intron (NR_102437, intron 1 of 2) | intron (NR_102437, intron 1 of 2) | 3241 | NM_027448 | 75782 | Mm.199698 | NM_027448 | ENSMUSG00000032258 | **Lca5** | 4930431B11Rik\|5730406O13Rik\|AV274874\|ORF64 | Leber congenital amaurosis 5 (human) | protein-coding |
| chr17_73232630_73232990 | chr17 | 73232631 | 73232990 | + | -0.684075 | intron (NM_001177967, intron 5 of 5) | B4A\|SINE\|B4 | 124825 | NM_001177967 | 225010 | Mm.329194 | NM_001081071 | ENSMUSG00000054469 | **Lclat1** | AI181996\|Agpat8\|Alcat1\|Gm91\|Lycat | lysocardiolipin acyltransferase 1 | protein-coding |
| chr17_73163542_73164082 | chr17 | 73163543 | 73164082 | + | -1.00783 | intron (NM_001177967, intron 2 of 5) | MTE-int\|LTR\|ERVL-MaLR | 55827 | NM_001177967 | 225010 | Mm.329194 | NM_001081071 | ENSMUSG00000054469 | **Lclat1** | AI181996\|Agpat8\|Alcat1\|Gm91\|Lycat | lysocardiolipin acyltransferase 1 | protein-coding |
| chr17_73087008_73087806 | chr17 | 73087009 | 73087806 | + | -1.04121 | Intergenic | Intergenic | -20578 | NM_001177968 | 225010 | Mm.329194 | NM_001081071 | ENSMUSG00000054469 | **Lclat1** | AI181996\|Agpat8\|Alcat1\|Gm91\|Lycat | lysocardiolipin acyltransferase 1 | protein-coding |
| chr17_73086912_73087496 | chr17 | 73086913 | 73087496 | + | -1.16228 | Intergenic | Intergenic | -20781 | NM_001177968 | 225010 | Mm.329194 | NM_001081071 | ENSMUSG00000054469 | **Lclat1** | AI181996\|Agpat8\|Alcat1\|Gm91\|Lycat | lysocardiolipin acyltransferase 1 | protein-coding |
| chr2_26668710_26669298 | chr2 | 26668711 | 26669298 | + | -0.937923 | intron (NM_010695, intron 4 of 6) | intron (NM_010695, intron 4 of 6) | 2278 | NM_010695 | 16821 | Mm.123925 | NM_010695 | ENSMUSG00000026919 | **Lcn4** | A630045M08Rik\|Vnsp2 | lipocalin 4 | protein-coding |
| chr2_25828364_25828932 | chr2 | 25828365 | 25828932 | + | -0.856043 | Intergenic | Intergenic | 5495 | NM_029959 | 77704 | Mm.159974 | NM_029959 | ENSMUSG00000023210 | **Lcn9** | 9230102I19Rik | lipocalin 9 | protein-coding |
| chr11_34063630_34064549 | chr11 | 34063631 | 34064549 | + | -0.688064 | intron (NM_010696, intron 3 of 20) | intron (NM_010696, intron 3 of 20) | 16889 | NM_010696 | 16822 | Mm.265350 | NM_010696 | ENSMUSG00000002699 | **Lcp2** | AI323664\|BB161688\|SLP-76\|SLP76\|twm | lymphocyte cytosolic protein 2 | protein-coding |
| chr17_5238495_5239218 | chr17 | 5238496 | 5239218 | + | -0.646509 | intron (NM_001085355, intron 4 of 19) | intron (NM_001085355, intron 4 of 19) | 179910 | NM_175349 | 106557 | Mm.432764 | NM_175349 | | **Ldhal6b** | 4933402O15Rik\|AI326310\|LDHL\|Ldhal | lactate dehydrogenase A-like 6B | protein-coding |
| chr17_5220508_5220843 | chr17 | 5220509 | 5220843 | + | -0.881115 | intron (NM_001085355, intron 4 of 19) | intron (NM_001085355, intron 4 of 19) | 198091 | NM_175349 | 106557 | Mm.432764 | NM_175349 | | **Ldhal6b** | 4933402O15Rik\|AI326310\|LDHL\|Ldhal | lactate dehydrogenase A-like 6B | protein-coding |
| chr6_142538527_142538948 | chr6 | 142538528 | 142538948 | + | -0.657726 | Intergenic | Intergenic | -30795 | NM_008492 | 16832 | Mm.9745 | NM_008492 | ENSMUSG00000030246 | **Ldhb** | AI790582\|H-Ldh\|LDH-B\|LDH-H\|Ldh-2\|Ldh2 | lactate dehydrogenase B | protein-coding |
| chr8_111595706_111596263 | chr8 | 111595707 | 111596263 | + | -0.949323 | intron (NM_001168623, intron 1 of 4) | intron (NM_001168623, intron 1 of 4) | 34337 | NM_027570 | 52815 | Mm.27589 | NM_027570 | ENSMUSG00000031958 | **Ldhd** | 4733401P21Rik\|D8Bwg1320e | lactate dehydrogenase D | protein-coding |
| chr4_107201785_107202200 | chr4 | 107201786 | 107202200 | + | -0.71256 | TTS (NM_029565) | TTS (NM_029565) | -7187 | NM_001081272 | 546840 | Mm.331755 | NM_001081272 | ENSMUSG00000070877 | **Ldlrad1** | OTTMUSG00000008594 | low density lipoprotein receptor class A domain containing 1 | protein-coding |
| chr4_107196490_107196853 | chr4 | 107196491 | 107196853 | + | -0.743853 | intron (NM_029565, intron 6 of 7) | RMER4A\|LTR\|ERVK | -12508 | NM_001081272 | 546840 | Mm.331755 | NM_001081272 | ENSMUSG00000070877 | **Ldlrad1** | OTTMUSG00000008594 | low density lipoprotein receptor class A domain containing 1 | protein-coding |
| chr2_102056936_102057866 | chr2 | 102056937 | 102057866 | + | -0.722911 | intron (NM_178886, intron 4 of 5) | L1ME4c\|LINE\|L1 | 129059 | NM_178886 | 241576 | Mm.489740 | NM_178886 | ENSMUSG00000048058 | **Ldlrad3** | 6430500P08\|AI194318\|Lrad3 | low density lipoprotein receptor class A domain containing 3 | protein-coding |
| chr2_102039165_102039872 | chr2 | 102039166 | 102039872 | + | -0.723431 | intron (NM_178886, intron 4 of 5) | intron (NM_178886, intron 4 of 5) | 146941 | NM_178886 | 241576 | Mm.489740 | NM_178886 | ENSMUSG00000048058 | **Ldlrad3** | 6430500P08\|AI194318\|Lrad3 | low density lipoprotein receptor class A domain containing 3 | protein-coding |
| chr2_102057336_102057789 | chr2 | 102057337 | 102057789 | + | -0.7421 | intron (NM_178886, intron 4 of 5) | intron (NM_178886, intron 4 of 5) | 128897 | NM_178886 | 241576 | Mm.489740 | NM_178886 | ENSMUSG00000048058 | **Ldlrad3** | 6430500P08\|AI194318\|Lrad3 | low density lipoprotein receptor class A domain containing 3 | protein-coding |
| chr2_102040153_102040447 | chr2 | 102040154 | 102040447 | + | -0.797882 | intron (NM_178886, intron 4 of 5) | intron (NM_178886, intron 4 of 5) | 146160 | NM_178886 | 241576 | Mm.489740 | NM_178886 | ENSMUSG00000048058 | **Ldlrad3** | 6430500P08\|AI194318\|Lrad3 | low density lipoprotein receptor class A domain containing 3 | protein-coding |
| chr18_67966569_67966959 | chr18 | 67966570 | 67966959 | + | -0.876906 | intron (NM_172631, intron 1 of 5) | ORR1E\|LTR\|ERVL-MaLR | 33507 | NM_172631 | 52662 | Mm.440183 | NM_172631 | ENSMUSG00000024544 | **Ldlrad4** | 8230401C20Rik\|A430083H02\|A430108L08Rik\|C18orf1\|D18Ertd653e\|D330030L18Rik | low density lipoprotein receptor class A domain containing 4 | protein-coding |
| chr4_134752162_134752429 | chr4 | 134752163 | 134752429 | + | -0.675375 | intron (NM_145554, intron 4 of 8) | intron (NM_145554, intron 4 of 8) | 15708 | NM_145554 | 100017 | Mm.482148 | NM_145554 | | **Ldlrap1** | AA691260\|ARH2\|Arh\|Arh1\|FHCB1\|FHCB2 | low density lipoprotein receptor adaptor protein 1 | protein-coding |
| chr4_134786855_134787394 | chr4 | 134786856 | 134787394 | + | -0.816627 | Intergenic | Intergenic | -19121 | NM_145554 | 100017 | Mm.482148 | NM_145554 | | **Ldlrap1** | AA691260\|ARH2\|Arh\|Arh1\|FHCB1\|FHCB2 | low density lipoprotein receptor adaptor protein 1 | protein-coding |
| chr4_134753857_134754134 | chr4 | 134753858 | 134754134 | + | -0.989345 | intron (NM_145554, intron 4 of 8) | ORR1C2\|LTR\|ERVL-MaLR | 14008 | NM_145554 | 100017 | Mm.482148 | NM_145554 | | **Ldlrap1** | AA691260\|ARH2\|Arh\|Arh1\|FHCB1\|FHCB2 | low density lipoprotein receptor adaptor protein 1 | protein-coding |
| chr4_134750048_134750756 | chr4 | 134750049 | 134750756 | + | -1.26692 | exon (NM_145554, exon 5 of 9) | exon (NM_145554, exon 5 of 9) | 17602 | NM_145554 | 100017 | Mm.482148 | NM_145554 | | **Ldlrap1** | AA691260\|ARH2\|Arh\|Arh1\|FHCB1\|FHCB2 | low density lipoprotein receptor adaptor protein 1 | protein-coding |
| chr4_134750237_134750741 | chr4 | 134750238 | 134750741 | + | -1.27801 | intron (NM_145554, intron 4 of 8) | intron (NM_145554, intron 4 of 8) | 17515 | NM_145554 | 100017 | Mm.482148 | NM_145554 | | **Ldlrap1** | AA691260\|ARH2\|Arh\|Arh1\|FHCB1\|FHCB2 | low density lipoprotein receptor adaptor protein 1 | protein-coding |
| chr3_131190490_131191356 | chr3 | 131190491 | 131191356 | + | -0.647243 | intron (NM_001276402, intron 5 of 9) | intron (NM_001276402, intron 5 of 9) | 78132 | NM_001276403 | 16842 | Mm.255219 | NM_010703 | ENSMUSG00000027985 | **Lef1** | 3000002B05\|AI451430\|Lef-1 | lymphoid enhancer binding factor 1 | protein-coding |
| chr3_131013981_131014797 | chr3 | 131013982 | 131014797 | + | -0.671555 | Intergenic | Intergenic | -95908 | NM_010703 | 16842 | Mm.255219 | NM_010703 | ENSMUSG00000027985 | **Lef1** | 3000002B05\|AI451430\|Lef-1 | lymphoid enhancer binding factor 1 | protein-coding |
| chr3_130932206_130932608 | chr3 | 130932207 | 130932608 | + | -0.680178 | Intergenic | B4\|SINE\|B4 | -177890 | NM_010703 | 16842 | Mm.255219 | NM_010703 | ENSMUSG00000027985 | **Lef1** | 3000002B05\|AI451430\|Lef-1 | lymphoid enhancer binding factor 1 | protein-coding |
| chr3_131170152_131170622 | chr3 | 131170153 | 131170622 | + | -0.74559 | intron (NM_001276402, intron 3 of 9) | RLTR51B_Mm\|LTR\|ERVK? | 57596 | NM_001276403 | 16842 | Mm.255219 | NM_010703 | ENSMUSG00000027985 | **Lef1** | 3000002B05\|AI451430\|Lef-1 | lymphoid enhancer binding factor 1 | protein-coding |
| chr3_130932283_130932578 | chr3 | 130932284 | 130932578 | + | -0.754897 | Intergenic | B4\|SINE\|B4 | -177866 | NM_010703 | 16842 | Mm.255219 | NM_010703 | ENSMUSG00000027985 | **Lef1** | 3000002B05\|AI451430\|Lef-1 | lymphoid enhancer binding factor 1 | protein-coding |
| chr3_130946727_130947682 | chr3 | 130946728 | 130947682 | + | -0.808343 | Intergenic | Intergenic | -163092 | NM_010703 | 16842 | Mm.255219 | NM_010703 | ENSMUSG00000027985 | **Lef1** | 3000002B05\|AI451430\|Lef-1 | lymphoid enhancer binding factor 1 | protein-coding |
| chr3_130946978_130947706 | chr3 | 130946979 | 130947706 | + | -0.871292 | Intergenic | Intergenic | -162955 | NM_010703 | 16842 | Mm.255219 | NM_010703 | ENSMUSG00000027985 | **Lef1** | 3000002B05\|AI451430\|Lef-1 | lymphoid enhancer binding factor 1 | protein-coding |
| chr3_131026241_131026718 | chr3 | 131026242 | 131026718 | + | -0.9519 | Intergenic | Intergenic | -83817 | NM_010703 | 16842 | Mm.255219 | NM_010703 | ENSMUSG00000027985 | **Lef1** | 3000002B05\|AI451430\|Lef-1 | lymphoid enhancer binding factor 1 | protein-coding |
| chr3_131157319_131157872 | chr3 | 131157320 | 131157872 | + | -0.995079 | intron (NM_001276402, intron 3 of 9) | intron (NM_001276402, intron 3 of 9) | 44805 | NM_001276403 | 16842 | Mm.255219 | NM_010703 | ENSMUSG00000027985 | **Lef1** | 3000002B05\|AI451430\|Lef-1 | lymphoid enhancer binding factor 1 | protein-coding |
| chr3_131138276_131138578 | chr3 | 131138277 | 131138578 | + | -1.02996 | intron (NM_001276402, intron 3 of 9) | intron (NM_001276402, intron 3 of 9) | 25636 | NM_001276403 | 16842 | Mm.255219 | NM_010703 | ENSMUSG00000027985 | **Lef1** | 3000002B05\|AI451430\|Lef-1 | lymphoid enhancer binding factor 1 | protein-coding |
| chr3_131160283_131160555 | chr3 | 131160284 | 131160555 | + | -1.03095 | intron (NM_001276402, intron 3 of 9) | intron (NM_001276402, intron 3 of 9) | 47628 | NM_001276403 | 16842 | Mm.255219 | NM_010703 | ENSMUSG00000027985 | **Lef1** | 3000002B05\|AI451430\|Lef-1 | lymphoid enhancer binding factor 1 | protein-coding |
| chr3_131171387_131171637 | chr3 | 131171388 | 131171637 | + | -1.03962 | intron (NM_001276402, intron 3 of 9) | intron (NM_001276402, intron 3 of 9) | 58721 | NM_001276403 | 16842 | Mm.255219 | NM_010703 | ENSMUSG00000027985 | **Lef1** | 3000002B05\|AI451430\|Lef-1 | lymphoid enhancer binding factor 1 | protein-coding |
| chr3_131057292_131057730 | chr3 | 131057293 | 131057730 | + | -1.16982 | Intergenic | ORR1B2\|LTR\|ERVL-MaLR | -52786 | NM_010703 | 16842 | Mm.255219 | NM_010703 | ENSMUSG00000027985 | **Lef1** | 3000002B05\|AI451430\|Lef-1 | lymphoid enhancer binding factor 1 | protein-coding |
| chr3_131167774_131168149 | chr3 | 131167775 | 131168149 | + | -1.17108 | intron (NM_001276402, intron 3 of 9) | B3\|SINE\|B2 | 55171 | NM_001276403 | 16842 | Mm.255219 | NM_010703 | ENSMUSG00000027985 | **Lef1** | 3000002B05\|AI451430\|Lef-1 | lymphoid enhancer binding factor 1 | protein-coding |
| chr3_131056927_131057745 | chr3 | 131056928 | 131057745 | + | -1.19059 | Intergenic | Intergenic | -52961 | NM_010703 | 16842 | Mm.255219 | NM_010703 | ENSMUSG00000027985 | **Lef1** | 3000002B05\|AI451430\|Lef-1 | lymphoid enhancer binding factor 1 | protein-coding |
| chr1_180921961_180922672 | chr1 | 180921962 | 180922672 | + | -0.826092 | Intergenic | Intergenic | -12722 | NM_010094 | 13590 | Mm.378911 | NM_010094 | ENSMUSG00000038793 | **Lefty1** | AI450052\|Ebaf\|Leftb\|Stra3\|Tgfb4\|lefty\|lefty-1 | left right determination factor 1 | protein-coding |
| chr3_65663039_65663559 | chr3 | 65663040 | 65663559 | + | -0.998529 | Intergenic | Intergenic | -2929 | NM_001037923 | 624866 | Mm.296317 | NM_001037923 | ENSMUSG00000074579 | **Lekr1** | EG546798\|EG624866\|Gm6534 | leucine, glutamate and lysine rich 1 | protein-coding |
| chr6_29042065_29042409 | chr6 | 29042066 | 29042409 | + | -0.663588 | Intergenic | Intergenic | -17984 | NM_008493 | 16846 | Mm.277072 | NM_008493 | ENSMUSG00000059201 | **Lep** | ob\|obese | leptin | protein-coding |
| chr8_25578223_25578608 | chr8 | 25578224 | 25578608 | + | -0.677393 | TTS (NM_173012) | TTS (NM_173012) | 19071 | NM_173012 | 270035 | Mm.268349 | NM_173012 | ENSMUSG00000037363 | **Letm2** | 6030453H13\|D030041N04Rik | leucine zipper-EF-hand containing transmembrane protein 2 | protein-coding |
| chr8_25578223_25578784 | chr8 | 25578224 | 25578784 | + | -0.680013 | TTS (NM_173012) | TTS (NM_173012) | 18983 | NM_173012 | 270035 | Mm.268349 | NM_173012 | ENSMUSG00000037363 | **Letm2** | 6030453H13\|D030041N04Rik | leucine zipper-EF-hand containing transmembrane protein 2 | protein-coding |
| chr15_100478281_100478780 | chr15 | 100478282 | 100478780 | + | -1.18845 | 3' UTR (NM_134093, exon 9 of 9) | 3' UTR (NM_134093, exon 9 of 9) | 9497 | NM_134093 | 68614 | Mm.275272 | NM_134093 | ENSMUSG00000037353 | **Letmd1** | 1110019O13Rik\|AI593524\|BB130465\|BB235638\|HCCR-2\|HCCR1\|MCC-32\|Mccr | LETM1 domain containing 1 | protein-coding |

| | | | | | | | | | | | | | | | | | |
|---|---|---|---|---|---|---|---|---|---|---|---|---|---|---|---|---|---|
| chr5_140596310_140597665 | chr5 | 140596311 | 140597665 | + | -0.617491 | Intergenic | Intergenic | -10353 | NM_008494 | 16848 | Mm.12834 | NM_008494 | ENSMUSG00000029570 | **Lfng** | AW061165 | LFNG O-fucosylpeptide 3-beta-N-acetylglucosaminyltransferase | protein-coding |
| chr5_140596409_140597179 | chr5 | 140596410 | 140597179 | + | -0.778306 | Intergenic | Intergenic | -10547 | NM_008494 | 16848 | Mm.12834 | NM_008494 | ENSMUSG00000029570 | **Lfng** | AW061165 | LFNG O-fucosylpeptide 3-beta-N-acetylglucosaminyltransferase | protein-coding |
| chr5_140586578_140587066 | chr5 | 140586579 | 140587066 | + | -0.821313 | Intergenic | Intergenic | -20519 | NM_008494 | 16848 | Mm.12834 | NM_008494 | ENSMUSG00000029570 | **Lfng** | AW061165 | LFNG O-fucosylpeptide 3-beta-N-acetylglucosaminyltransferase | protein-coding |
| chr11_118397771_118398193 | chr11 | 118397772 | 118398193 | + | -1.3473 | intron (NM_011150, intron 4 of 5) | intron (NM_011150, intron 4 of 5) | 3949 | NM_011150 | 19039 | Mm.3152 | NM_011150 | ENSMUSG00000033880 | **Lgals3bp** | 90K\|CyCAP\|MAC-2BP\|Ppicap\|Tango10b | lectin, galactoside-binding, soluble, 3 binding protein | protein-coding |
| chr19_38298100_38298544 | chr19 | 38298101 | 38298544 | + | -0.793176 | intron (NM_020278, intron 5 of 7) | intron (NM_020278, intron 5 of 7) | 33540 | NM_020278 | 56839 | Mm.298251 | NM_020278 | ENSMUSG00000067242 | **Lgi1** | BB130740 | leucine-rich repeat LGI family, member 1 | protein-coding |
| chr12_102370922_102371450 | chr12 | 102370923 | 102371450 | + | -0.910016 | intron (NM_177620, intron 6 of 9) | intron (NM_177620, intron 6 of 9) | 68511 | NM_011175 | 19141 | Mm.17185 | NM_011175 | ENSMUSG00000021190 | **Lgmn** | AEP\|AI746452\|AU022324\|Prsc1 | legumain | protein-coding |
| chr17_88757971_88758356 | chr17 | 88757972 | 88758356 | + | -0.678121 | intron (NM_013582, intron 7 of 10) | intron (NM_013582, intron 7 of 10) | 33812 | NM_013582 | 16867 | Mm.1644 | NM_013582 | ENSMUSG00000024107 | **Lhcgr** | Gpcr19-rs1\|LH-R\|Lhr | luteinizing hormone/choriogonadotropin receptor | protein-coding |
| chr3_53032453_53032752 | chr3 | 53032454 | 53032752 | + | -0.811351 | Intergenic | MLT1A0\|LTR\|ERVL-MaLR | -8944 | NM_175386 | 108927 | Mm.118617 | NM_175386 | ENSMUSG00000048332 | **Lhfp** | 2810489O06Rik\|AI194968 | lipoma HMGIC fusion partner | protein-coding |
| chr13_93989589_93991109 | chr13 | 93989590 | 93991109 | + | -0.699434 | Intergenic | Intergenic | -67447 | NM_172589 | 218454 | Mm.316553 | NM_172589 | ENSMUSG00000045312 | **Lhfpl2** | 6030465B15\|AI447312\|AW050335\|mKIAA0206\|vgim | lipoma HMGIC fusion partner-like 2 | protein-coding |
| chr13_93990133_93990835 | chr13 | 93990134 | 93990835 | + | -0.756848 | Intergenic | Intergenic | -67312 | NM_172589 | 218454 | Mm.316553 | NM_172589 | ENSMUSG00000045312 | **Lhfpl2** | 6030465B15\|AI447312\|AW050335\|mKIAA0206\|vgim | lipoma HMGIC fusion partner-like 2 | protein-coding |
| chr13_93990897_93991099 | chr13 | 93990898 | 93991099 | + | -0.808844 | Intergenic | Intergenic | -66798 | NM_172589 | 218454 | Mm.316553 | NM_172589 | ENSMUSG00000045312 | **Lhfpl2** | 6030465B15\|AI447312\|AW050335\|mKIAA0206\|vgim | lipoma HMGIC fusion partner-like 2 | protein-coding |
| chr2_38319110_38319754 | chr2 | 38319111 | 38319754 | + | -0.768986 | Intergenic | Intergenic | -19849 | NM_001290646 | 16870 | Mm.142856 | NM_010710 | ENSMUSG00000000247 | **Lhx2** | LH2A\|Lh-2\|Lim2\|ap\|apterous | LIM homeobox protein 2 | protein-coding |
| chr9_123497824_123498450 | chr9 | 123497825 | 123498450 | + | -0.934558 | intron (NM_013860, intron 1 of 7) | intron (NM_013860, intron 1 of 7) | 19436 | NM_013860 | 29806 | Mm.10987 | NM_013860 | ENSMUSG00000025239 | **Limd1** | AW822033\|D9Ertd192e | LIM domains containing 1 | protein-coding |
| chr11_106167167_106167916 | chr11 | 106167168 | 106167916 | + | -1.24783 | intron (NM_001252448, intron 10 of 12) | intron (NM_001252448, intron 10 of 12) | -7400 | NM_172397 | 67803 | Mm.21687 | NM_172397 | ENSMUSG00000040699 | **Limd2** | 0610025L06Rik\|AI413966 | LIM domain containing 2 | protein-coding |
| chr11_3344756_3345378 | chr11 | 3344757 | 3345378 | + | -0.916239 | 3' UTR (NM_001034030, exon 12 of 12) | 3' UTR (NM_001034030, exon 12 of 12) | 11074 | NM_001034030 | 16886 | Mm.124176 | NM_010718 | ENSMUSG00000020451 | **Limk2** | Limk-2 | LIM motif-containing protein kinase 2 | protein-coding |
| chr13_97408487_97408780 | chr13 | 97408488 | 97408780 | + | -0.661412 | Intergenic | Intergenic | -26336 | NR_110432 | 100039691 | Mm.219358 | NR_110430 | | **Lincenc1** | ENSMUSG00000078952\|Gm11019\|Gm2373\|linc-Enc1 | long intergenic non-coding RNA, embryonic stem cells expressed 1 | ncRNA |
| chr13_97406403_97407226 | chr13 | 97406404 | 97407226 | + | -0.87879 | Intergenic | Intergenic | -28155 | NR_110432 | 100039691 | Mm.219358 | NR_110430 | | **Lincenc1** | ENSMUSG00000078952\|Gm11019\|Gm2373\|linc-Enc1 | long intergenic non-coding RNA, embryonic stem cells expressed 1 | ncRNA |
| chr13_97442274_97442792 | chr13 | 97442275 | 97442792 | + | -0.918954 | intron (NR_110432, intron 1 of 1) | intron (NR_110432, intron 1 of 1) | 7563 | NR_110432 | 100039691 | Mm.219358 | NR_110430 | | **Lincenc1** | ENSMUSG00000078952\|Gm11019\|Gm2373\|linc-Enc1 | long intergenic non-coding RNA, embryonic stem cells expressed 1 | ncRNA |
| chr13_97498904_97499469 | chr13 | 97498905 | 97499469 | + | -1.28791 | Intergenic | Intergenic | 16486 | NR_110431 | 100039691 | Mm.219358 | NR_110430 | | **Lincenc1** | ENSMUSG00000078952\|Gm11019\|Gm2373\|linc-Enc1 | long intergenic non-coding RNA, embryonic stem cells expressed 1 | ncRNA |
| chr6_31201168_31201921 | chr6 | 31201169 | 31201921 | + | -0.713502 | intron (NR_110469, intron 1 of 3) | Lx8\|LINE\|L1 | 16929 | NR_110470 | 232685 | Mm.443019 | NM_144555 | | **Lincpint** | MNCb-1768\|Pint\|linc-Pint | long intergenic non-protein coding RNA, Trp53 induced transcript | ncRNA |
| chr6_31176349_31176827 | chr6 | 31176350 | 31176827 | + | -0.782308 | intron (NR_110470, intron 2 of 2) | intron (NR_110470, intron 2 of 2) | 41886 | NR_110470 | 232685 | Mm.443019 | NM_144555 | | **Lincpint** | MNCb-1768\|Pint\|linc-Pint | long intergenic non-protein coding RNA, Trp53 induced transcript | ncRNA |
| chr6_31216122_31217043 | chr6 | 31216123 | 31217043 | + | -0.798973 | intron (NR_110469, intron 1 of 3) | CpG | 1891 | NR_110470 | 232685 | Mm.443019 | NM_144555 | | **Lincpint** | MNCb-1768\|Pint\|linc-Pint | long intergenic non-protein coding RNA, Trp53 induced transcript | ncRNA |
| chr6_31191534_31192325 | chr6 | 31191535 | 31192325 | + | -0.845246 | intron (NR_110470, intron 1 of 2) | intron (NR_110470, intron 1 of 2) | 26544 | NR_110470 | 232685 | Mm.443019 | NM_144555 | | **Lincpint** | MNCb-1768\|Pint\|linc-Pint | long intergenic non-protein coding RNA, Trp53 induced transcript | ncRNA |
| chr6_31190673_31190932 | chr6 | 31190674 | 31190932 | + | -0.904976 | intron (NR_110470, intron 1 of 2) | intron (NR_110470, intron 1 of 2) | 27671 | NR_110470 | 232685 | Mm.443019 | NM_144555 | | **Lincpint** | MNCb-1768\|Pint\|linc-Pint | long intergenic non-protein coding RNA, Trp53 induced transcript | ncRNA |
| chr6_31191449_31191991 | chr6 | 31191450 | 31191991 | + | -0.96713 | intron (NR_110470, intron 1 of 2) | intron (NR_110470, intron 1 of 2) | 26754 | NR_110470 | 232685 | Mm.443019 | NM_144555 | | **Lincpint** | MNCb-1768\|Pint\|linc-Pint | long intergenic non-protein coding RNA, Trp53 induced transcript | ncRNA |
| chr6_31157069_31157399 | chr6 | 31157070 | 31157399 | + | -1.00478 | intron (NR_110471, intron 2 of 3) | intron (NR_110471, intron 2 of 3) | 61240 | NR_110470 | 232685 | Mm.443019 | NM_144555 | | **Lincpint** | MNCb-1768\|Pint\|linc-Pint | long intergenic non-protein coding RNA, Trp53 induced transcript | ncRNA |
| chr6_31184477_31184936 | chr6 | 31184478 | 31184936 | + | -1.27082 | intron (NR_110470, intron 1 of 2) | intron (NR_110470, intron 1 of 2) | 33767 | NR_110470 | 232685 | Mm.443019 | NM_144555 | | **Lincpint** | MNCb-1768\|Pint\|linc-Pint | long intergenic non-protein coding RNA, Trp53 induced transcript | ncRNA |
| chr19_34510677_34511251 | chr19 | 34510678 | 34511251 | + | -0.835479 | exon (NM_001111100, exon 4 of 10) | exon (NM_001111100, exon 4 of 10) | 16510 | NM_001111100 | 16889 | Mm.157545 | NM_021460 | ENSMUSG00000024781 | **Lipa** | AA960673\|Lal\|Lip-1\|Lip1 | lysosomal acid lipase A | protein-coding |
| chr18_74957165_74957461 | chr18 | 74957166 | 74957461 | + | -0.905598 | exon (NM_010720, exon 2 of 10) | exon (NM_010720, exon 2 of 10) | 3950 | NM_010720 | 16891 | Mm.299647 | NM_010720 | ENSMUSG00000053846 | **Lipg** | 3110013K01Rik\|EL\|lipase\|mEDL | lipase, endothelial | protein-coding |
| chr16_10968826_10969689 | chr16 | 10968827 | 10969689 | + | -1.17047 | intron (NM_019980, intron 1 of 3) | intron (NM_019980, intron 1 of 3) | 23863 | NM_019980 | 56722 | Mm.294753 | NM_019980 | ENSMUSG00000022500 | **Litaf** | 3222402J11Rik\|C85531\|N4WBP3\|TBX1 | LPS-induced TN factor | protein-coding |
| chr16_10969133_10969736 | chr16 | 10969134 | 10969736 | + | -1.20865 | intron (NM_019980, intron 1 of 3) | intron (NM_019980, intron 1 of 3) | 23686 | NM_019980 | 56722 | Mm.294753 | NM_019980 | ENSMUSG00000022500 | **Litaf** | 3222402J11Rik\|C85531\|N4WBP3\|TBX1 | LPS-induced TN factor | protein-coding |
| chr16_10969203_10969643 | chr16 | 10969204 | 10969643 | + | -1.21316 | intron (NM_019980, intron 1 of 3) | intron (NM_019980, intron 1 of 3) | 23698 | NM_019980 | 56722 | Mm.294753 | NM_019980 | ENSMUSG00000022500 | **Litaf** | 3222402J11Rik\|C85531\|N4WBP3\|TBX1 | LPS-induced TN factor | protein-coding |
| chr17_17444358_17444813 | chr17 | 17444359 | 17444813 | + | -0.717461 | intron (NM_025681, intron 3 of 5) | C-rich\|Low_complexity\|Low_complexity | 41900 | NM_025681 | 66643 | Mm.268018 | NM_025681 | ENSMUSG00000047786 | **Lix1** | 5730466L18Rik | limb expression 1 homolog (chicken) | protein-coding |
| chr17_17444340_17444971 | chr17 | 17444341 | 17444971 | + | -0.746237 | intron (NM_025681, intron 3 of 5) | B1_Mur2\|SINE\|Alu | 41970 | NM_025681 | 66643 | Mm.268018 | NM_025681 | ENSMUSG00000047786 | **Lix1** | 5730466L18Rik | limb expression 1 homolog (chicken) | protein-coding |
| chr17_17442190_17442504 | chr17 | 17442191 | 17442504 | + | -0.872141 | intron (NM_025681, intron 2 of 5) | intron (NM_025681, intron 2 of 5) | 39661 | NM_025681 | 66643 | Mm.268018 | NM_025681 | ENSMUSG00000047786 | **Lix1** | 5730466L18Rik | limb expression 1 homolog (chicken) | protein-coding |
| chr10_120219967_120220420 | chr10 | 120219968 | 120220420 | + | -0.772694 | intron (NM_026617, intron 3 of 6) | intron (NM_026617, intron 3 of 6) | -6866 | NM_025431 | 66225 | Mm.291755 | NM_025431 | ENSMUSG00000020224 | **Llph** | 1190005P17Rik | LLP homolog, long-term synaptic facilitation (Aplysia) | protein-coding |
| chr9_57620660_57621037 | chr9 | 57620661 | 57621037 | + | -0.860605 | promoter-TSS (NM_199222) | promoter-TSS (NM_199222) | -75 | NM_199222 | 235416 | Mm.295630 | NM_199222 | ENSMUSG00000056271 | **Lman1l** | BC020188\|slamp | lectin, mannose-binding 1 like | protein-coding |
| chr9_57620733_57621338 | chr9 | 57620734 | 57621338 | + | -0.917526 | promoter-TSS (NM_199222) | promoter-TSS (NM_199222) | -262 | NM_199222 | 235416 | Mm.295630 | NM_199222 | ENSMUSG00000056271 | **Lman1l** | BC020188\|slamp | lectin, mannose-binding 1 like | protein-coding |
| chr17_25583984_25584211 | chr17 | 25583985 | 25584211 | + | -0.657956 | intron (NR_045520, intron 1 of 8) | intron (NR_045520, intron 1 of 8) | 4924 | NR_045520 | 76483 | Mm.12787 | NM_029624 | ENSMUSG00000002279 | **Lmf1** | 2400010G15Rik\|AW822050\|Tmem112\|cld | lipase maturation factor 1 | protein-coding |
| chr3_144260572_144261296 | chr3 | 144260573 | 144261296 | + | -0.750366 | Intergenic | Intergenic | -55679 | NM_010723 | 16911 | Mm.29187 | NM_010723 | ENSMUSG00000028266 | **Lmo4** | A730077C12Rik\|Crp3\|Etohi4 | LIM domain only 4 | protein-coding |
| chr3_144252797_144253392 | chr3 | 144252798 | 144253392 | + | -0.972405 | Intergenic | Intergenic | -47840 | NM_010723 | 16911 | Mm.29187 | NM_010723 | ENSMUSG00000028266 | **Lmo4** | A730077C12Rik\|Crp3\|Etohi4 | LIM domain only 4 | protein-coding |
| chr3_144231477_144232308 | chr3 | 144231478 | 144232308 | + | -1.01312 | Intergenic | Intergenic | -26638 | NM_010723 | 16911 | Mm.29187 | NM_010723 | ENSMUSG00000028266 | **Lmo4** | A730077C12Rik\|Crp3\|Etohi4 | LIM domain only 4 | protein-coding |
| chr6_97235208_97235597 | chr6 | 97235209 | 97235597 | + | -0.640153 | Intergenic | MTE2b\|LTR\|ERVL-MaLR | 17377 | NM_001081157 | 320502 | Mm.68163 | NM_001081157 | ENSMUSG00000044086 | **Lmod3** | 5430424A14Rik | leiomodin 3 (fetal) | protein-coding |
| chr6_97317859_97318367 | chr6 | 97317860 | 97318367 | + | -1.14215 | intron (NM_145148, intron 12 of 23) | intron (NM_145148, intron 12 of 23) | -65333 | NM_001081157 | 320502 | Mm.68163 | NM_001081157 | ENSMUSG00000044086 | **Lmod3** | 5430424A14Rik | leiomodin 3 (fetal) | protein-coding |
| chr7_45778961_45779867 | chr7 | 45778962 | 45779867 | + | -0.703766 | Intergenic | L1MA8\|LINE\|L1 | -4533 | NM_001005511 | 381983 | Mm.44928 | NM_001005511 | ENSMUSG00000062044 | **Lmtk3** | BC059845\|aatyk3 | lemur tyrosine kinase 3 | protein-coding |
| chr17_17567140_17567389 | chr17 | 17567141 | 17567389 | + | -0.634671 | intron (NM_172827, intron 7 of 17) | intron (NM_172827, intron 7 of 17) | 57224 | NM_172827 | 240028 | Mm.328807 | NM_172827 | ENSMUSG00000023845 | **Lnpep** | 2010309L07Rik\|4732490P18Rik\|CAP\|IRAP\|PLAP\|gp160\|vp165 | leucyl/cystinyl aminopeptidase | protein-coding |
| chr17_17567150_17567575 | chr17 | 17567151 | 17567575 | + | -0.687609 | intron (NM_172827, intron 7 of 17) | intron (NM_172827, intron 7 of 17) | 57126 | NM_172827 | 240028 | Mm.328807 | NM_172827 | ENSMUSG00000023845 | **Lnpep** | 2010309L07Rik\|4732490P18Rik\|CAP\|IRAP\|PLAP\|gp160\|vp165 | leucyl/cystinyl aminopeptidase | protein-coding |
| chr17_17636042_17636643 | chr17 | 17636043 | 17636643 | + | -0.97857 | Intergenic | Intergenic | -11854 | NM_172827 | 240028 | Mm.328807 | NM_172827 | ENSMUSG00000023845 | **Lnpep** | 2010309L07Rik\|4732490P18Rik\|CAP\|IRAP\|PLAP\|gp160\|vp165 | leucyl/cystinyl aminopeptidase | protein-coding |
| chr4_59331358_59332124 | chr4 | 59331359 | 59332124 | + | -0.612961 | intron (NM_001163288, intron 13 of 16) | intron (NM_001163288, intron 13 of 16) | 71690 | NR_105031 | 101055769 | Mm.421401 | NR_105031 | | **LOC101055769** | - | uncharacterized LOC101055769 | ncRNA |
| chr4_59334246_59334583 | chr4 | 59334247 | 59334583 | + | -0.657243 | intron (NM_001163288, intron 12 of 16) | intron (NM_001163288, intron 12 of 16) | 74364 | NR_105031 | 101055769 | Mm.421401 | NR_105031 | | **LOC101055769** | - | uncharacterized LOC101055769 | ncRNA |

| | | | | | | | | | | | | | | | | | |
|---|---|---|---|---|---|---|---|---|---|---|---|---|---|---|---|---|---|
| chr4_59319416_59319959 | chr4 | 59319417 | 59319959 | + | -0.820606 | intron (NM_001163288, intron 15 of 16) | intron (NM_001163288, intron 15 of 16) | 59637 | NR_105031 | 101055769 | Mm.421401 | NR_105031 | | **LOC101055769** | - | uncharacterized LOC101055769 | ncRNA |
| chr4_59336080_59336343 | chr4 | 59336081 | 59336343 | + | -0.850067 | intron (NM_001163288, intron 12 of 16) | intron (NM_001163288, intron 12 of 16) | 76161 | NR_105031 | 101055769 | Mm.421401 | NR_105031 | | **LOC101055769** | - | uncharacterized LOC101055769 | ncRNA |
| chr4_59265096_59265629 | chr4 | 59265097 | 59265629 | + | -0.875075 | intron (NR_105031, intron 3 of 3) | intron (NR_105031, intron 3 of 3) | 5312 | NR_105031 | 101055769 | Mm.421401 | NR_105031 | | **LOC101055769** | - | uncharacterized LOC101055769 | ncRNA |
| chr4_59275483_59275910 | chr4 | 59275484 | 59275910 | + | -1.03343 | Intergenic | Intergenic | 15646 | NR_105031 | 101055769 | Mm.421401 | NR_105031 | | **LOC101055769** | - | uncharacterized LOC101055769 | ncRNA |
| chr4_59275584_59275879 | chr4 | 59275585 | 59275879 | + | -1.04727 | Intergenic | Intergenic | 15681 | NR_105031 | 101055769 | Mm.421401 | NR_105031 | | **LOC101055769** | - | uncharacterized LOC101055769 | ncRNA |
| chr8_86678194_86678511 | chr8 | 86678195 | 86678511 | + | -0.659207 | intron (NM_001168591, intron 5 of 8) | intron (NM_001168591, intron 5 of 8) | 26784 | NM_001168591 | 66887 | Mm.324550 | NM_025827 | ENSMUSG00000047866 | **Lonp2** | 1300002A08Rik\|AU015403\|Lonp | lon peptidase 2, peroxisomal | protein-coding |
| chr8_86686585_86687141 | chr8 | 86686586 | 86687141 | + | -1.10446 | intron (NM_001168591, intron 5 of 8) | intron (NM_001168591, intron 5 of 8) | 35294 | NM_001168591 | 66887 | Mm.324550 | NM_025827 | ENSMUSG00000047866 | **Lonp2** | 1300002A08Rik\|AU015403\|Lonp | lon peptidase 2, peroxisomal | protein-coding |
| chr8_36193907_36194469 | chr8 | 36193908 | 36194469 | + | -0.675004 | Intergenic | MTEa\|LTR\|ERVL-MaLR | 55328 | NM_001081150 | 244421 | Mm.324032 | NM_001081150 | ENSMUSG00000039633 | **Lonrf1** | - | LON peptidase N-terminal domain and ring finger 1 | protein-coding |
| chr8_36194064_36194571 | chr8 | 36194065 | 36194571 | + | -0.709585 | Intergenic | Intergenic | 55198 | NM_001081150 | 244421 | Mm.324032 | NM_001081150 | ENSMUSG00000039633 | **Lonrf1** | - | LON peptidase N-terminal domain and ring finger 1 | protein-coding |
| chr8_36178610_36178928 | chr8 | 36178611 | 36178928 | + | -0.742005 | Intergenic | Intergenic | 70747 | NM_001081150 | 244421 | Mm.324032 | NM_001081150 | ENSMUSG00000039633 | **Lonrf1** | - | LON peptidase N-terminal domain and ring finger 1 | protein-coding |
| chr6_125073956_125074208 | chr6 | 125073957 | 125074208 | + | -0.716921 | intron (NM_001163268, intron 1 of 1) | intron (NM_001163268, intron 1 of 1) | 2805 | NM_001163268 | 381810 | Mm.333386 | NM_001163268 | ENSMUSG00000067714 | **Lpar5** | GPR93\|Gm1072\|Gpr92\|LPA5 | lysophosphatidic acid receptor 5 | protein-coding |
| chr3_148895490_148895756 | chr3 | 148895491 | 148895756 | + | -0.628347 | intron (NM_001081298, intron 1 of 23) | intron (NM_001081298, intron 1 of 23) | 59012 | NM_001081298 | 99633 | Mm.9776 | NM_001081298 | ENSMUSG00000028184 | **Lphn2** | AI450192\|Gm619\|Lec1\|Lphh1\|mKIAA0786 | latrophilin 2 | protein-coding |
| chr3_148913570_148914032 | chr3 | 148913571 | 148914032 | + | -0.665374 | intron (NM_001081298, intron 1 of 23) | intron (NM_001081298, intron 1 of 23) | 40834 | NM_001081298 | 99633 | Mm.9776 | NM_001081298 | ENSMUSG00000028184 | **Lphn2** | AI450192\|Gm619\|Lec1\|Lphh1\|mKIAA0786 | latrophilin 2 | protein-coding |
| chr3_148807090_148807383 | chr3 | 148807091 | 148807383 | + | -0.697579 | Intergenic | Intergenic | 147398 | NM_001081298 | 99633 | Mm.9776 | NM_001081298 | ENSMUSG00000028184 | **Lphn2** | AI450192\|Gm619\|Lec1\|Lphh1\|mKIAA0786 | latrophilin 2 | protein-coding |
| chr3_148806799_148807435 | chr3 | 148806800 | 148807435 | + | -0.723061 | Intergenic | Intergenic | 147518 | NM_001081298 | 99633 | Mm.9776 | NM_001081298 | ENSMUSG00000028184 | **Lphn2** | AI450192\|Gm619\|Lec1\|Lphh1\|mKIAA0786 | latrophilin 2 | protein-coding |
| chr3_148867611_148868127 | chr3 | 148867612 | 148868127 | + | -0.854505 | intron (NM_001081298, intron 2 of 23) | B3A\|SINE\|B2 | 86766 | NM_001081298 | 99633 | Mm.9776 | NM_001081298 | ENSMUSG00000028184 | **Lphn2** | AI450192\|Gm619\|Lec1\|Lphh1\|mKIAA0786 | latrophilin 2 | protein-coding |
| chr3_148852568_148853022 | chr3 | 148852569 | 148853022 | + | -0.883855 | exon (NM_001081298, exon 6 of 24) | exon (NM_001081298, exon 6 of 24) | 101840 | NM_001081298 | 99633 | Mm.9776 | NM_001081298 | ENSMUSG00000028184 | **Lphn2** | AI450192\|Gm619\|Lec1\|Lphh1\|mKIAA0786 | latrophilin 2 | protein-coding |
| chr3_148850035_148851118 | chr3 | 148850036 | 148851118 | + | -1.03459 | intron (NM_001081298, intron 8 of 23) | intron (NM_001081298, intron 8 of 23) | 104058 | NM_001081298 | 99633 | Mm.9776 | NM_001081298 | ENSMUSG00000028184 | **Lphn2** | AI450192\|Gm619\|Lec1\|Lphh1\|mKIAA0786 | latrophilin 2 | protein-coding |
| chr3_148850680_148851352 | chr3 | 148850681 | 148851352 | + | -1.4014 | exon (NM_001081298, exon 8 of 24) | exon (NM_001081298, exon 8 of 24) | 103619 | NM_001081298 | 99633 | Mm.9776 | NM_001081298 | ENSMUSG00000028184 | **Lphn2** | AI450192\|Gm619\|Lec1\|Lphh1\|mKIAA0786 | latrophilin 2 | protein-coding |
| chr16_24391167_24391474 | chr16 | 24391168 | 24391474 | + | -0.878672 | Intergenic | Intergenic | -2029 | NM_178665 | 210126 | Mm.209385 | NM_178665 | ENSMUSG00000033306 | **Lpp** | 9430020K16Rik\|AA959454\|AU024130\|B130055L10Rik\|C79715\|D630048H16 | LIM domain containing preferred translocation partner in lipoma | protein-coding |
| chr3_86196093_86196809 | chr3 | 86196094 | 86196809 | + | -0.848621 | Intergenic | Intergenic | -28239 | NM_001077688 | 80877 | Mm.439825 | NM_030695 | ENSMUSG00000028080 | **Lrba** | C80285\|D3Ertd775e\|Lab300\|Lba | LPS-responsive beige-like anchor | protein-coding |
| chr3_86196427_86196849 | chr3 | 86196428 | 86196849 | + | -0.975067 | Intergenic | MER2\|DNA\|TcMar-Tigger | -28052 | NM_001077688 | 80877 | Mm.439825 | NM_030695 | ENSMUSG00000028080 | **Lrba** | C80285\|D3Ertd775e\|Lab300\|Lba | LPS-responsive beige-like anchor | protein-coding |
| chr14_74908263_74908657 | chr14 | 74908264 | 74908657 | + | -0.760725 | intron (NM_001252132, intron 1 of 18) | intron (NM_001252132, intron 1 of 18) | 39417 | NM_001252132 | 380916 | Mm.288868 | NM_001033439 | ENSMUSG00000068015 | **Lrch1** | 4832412D13Rik\|AI552548\|AW215503\|Chdc1\|mKIAA1016 | leucine-rich repeats and calponin homology (CH) domain containing 1 | protein-coding |
| chr16_32953831_32954199 | chr16 | 32953832 | 32954199 | + | -0.66932 | intron (NM_001081255, intron 3 of 20) | intron (NM_001081255, intron 3 of 20) | 39915 | NM_001081255 | 70144 | Mm.491178 | NM_001081255 | ENSMUSG00000022801 | **Lrch3** | 2210409B11Rik\|AW215594\|Gm1742 | leucine-rich repeats and calponin homology (CH) domain containing 3 | protein-coding |
| chr16_32972443_32973004 | chr16 | 32972444 | 32973004 | + | -0.998981 | intron (NM_001081255, intron 6 of 20) | intron (NM_001081255, intron 6 of 20) | 58624 | NM_001081255 | 70144 | Mm.491178 | NM_001081255 | ENSMUSG00000022801 | **Lrch3** | 2210409B11Rik\|AW215594\|Gm1742 | leucine-rich repeats and calponin homology (CH) domain containing 3 | protein-coding |
| chr6_94690308_94690596 | chr6 | 94690309 | 94690596 | + | -0.635741 | intron (NM_008377, intron 1 of 18) | intron (NM_008377, intron 1 of 18) | 9693 | NM_008377 | 16206 | Mm.245210 | NM_008377 | ENSMUSG00000030029 | **Lrig1** | D6Bwg0781e\|Img\|LIG-1 | leucine-rich repeats and immunoglobulin-like domains 1 | protein-coding |
| chr19_3668919_3669475 | chr19 | 3668920 | 3669475 | + | -0.754164 | intron (NM_008513, intron 1 of 22) | intron (NM_008513, intron 1 of 22) | 17367 | NM_008513 | 16973 | Mm.274581 | NM_008513 | ENSMUSG00000024913 | **Lrp5** | BMND1\|HBM\|LR3\|LRP7\|OPPG\|mKIAA4142 | low density lipoprotein receptor-related protein 5 | protein-coding |
| chr6_134533064_134533408 | chr6 | 134533065 | 134533408 | + | -0.717342 | intron (NM_008514, intron 2 of 22) | intron (NM_008514, intron 2 of 22) | 33677 | NM_008514 | 16974 | Mm.321990 | NM_008514 | ENSMUSG00000030201 | **Lrp6** | C030016K15Rik\|Cd\|Gw\|ska26\|ska<m26Jus>\|skax26 | low density lipoprotein receptor-related protein 6 | protein-coding |
| chr17_84844719_84845545 | chr17 | 84844720 | 84845545 | + | -1.13821 | Intergenic | Intergenic | -54346 | NM_028233 | 72416 | Mm.217027 | NM_028233 | ENSMUSG00000024120 | **Lrpprc** | 3110001K13Rik\|C76645\|Gp130\|Lrp130\|Lsfc | leucine-rich PPR-motif containing | protein-coding |
| chr9_77513951_77514352 | chr9 | 77513952 | 77514352 | + | -0.938172 | intron (NM_001146048, intron 1 of 13) | intron (NM_001146048, intron 1 of 13) | 29907 | NM_172528 | 214345 | Mm.28534 | NM_172528 | ENSMUSG00000032352 | **Lrrc1** | A430093J20Rik\|AU016091\|D030059I21\|mKIAA4018 | leucine rich repeat containing 1 | protein-coding |
| chr7_98468769_98469709 | chr7 | 98468770 | 98469709 | + | -0.699539 | intron (NM_001130693, intron 14 of 18) | intron (NM_001130693, intron 14 of 18) | -24983 | NM_001113379 | 434215 | Mm.323057 | NM_001113379 | ENSMUSG00000090958 | **Lrrc32** | AI426318\|D11S833Eh\|D7H11S833E\|EG434215\|Garp | leucine rich repeat containing 32 | protein-coding |
| chr7_98469192_98469786 | chr7 | 98469193 | 98469786 | + | -0.864774 | intron (NM_001130693, intron 14 of 18) | intron (NM_001130693, intron 14 of 18) | -24733 | NM_001113379 | 434215 | Mm.323057 | NM_001113379 | ENSMUSG00000090958 | **Lrrc32** | AI426318\|D11S833Eh\|D7H11S833E\|EG434215\|Garp | leucine rich repeat containing 32 | protein-coding |
| chr14_15681723_15682030 | chr14 | 15681724 | 15682030 | + | -0.682762 | Intergenic | Intergenic | -242890 | NM_146052 | 218763 | Mm.138039 | NM_146052 | ENSMUSG00000045201 | **Lrrc3b** | BC019794\|LRP15 | leucine rich repeat containing 3B | protein-coding |
| chr14_15746642_15747191 | chr14 | 15746643 | 15747191 | + | -0.693939 | Intergenic | Intergenic | -307930 | NM_146052 | 218763 | Mm.138039 | NM_146052 | ENSMUSG00000045201 | **Lrrc3b** | BC019794\|LRP15 | leucine rich repeat containing 3B | protein-coding |
| chr14_15272899_15273701 | chr14 | 15272900 | 15273701 | + | -0.852944 | Intergenic | Intergenic | 165687 | NM_146052 | 218763 | Mm.138039 | NM_146052 | ENSMUSG00000045201 | **Lrrc3b** | BC019794\|LRP15 | leucine rich repeat containing 3B | protein-coding |
| chr14_15272430_15272676 | chr14 | 15272431 | 15272676 | + | -0.906643 | Intergenic | ORR1E\|LTR\|ERVL-MaLR | 166434 | NM_146052 | 218763 | Mm.138039 | NM_146052 | ENSMUSG00000045201 | **Lrrc3b** | BC019794\|LRP15 | leucine rich repeat containing 3B | protein-coding |
| chr15_66445211_66445689 | chr15 | 66445212 | 66445689 | + | -0.755242 | intron (NM_019457, intron 7 of 11) | L2\|LINE\|L2 | 55460 | NM_019457 | 54562 | Mm.244890 | NM_019457 | ENSMUSG00000022375 | **Lrrc6** | LRTP | leucine rich repeat containing 6 (testis) | protein-coding |
| chr15_66512917_66513151 | chr15 | 66512918 | 66513151 | + | -0.769818 | Intergenic | MIRb\|SINE\|MIR | -12124 | NM_019457 | 54562 | Mm.244890 | NM_019457 | ENSMUSG00000022375 | **Lrrc6** | LRTP | leucine rich repeat containing 6 (testis) | protein-coding |
| chr15_66512806_66513171 | chr15 | 66512807 | 66513171 | + | -0.781195 | Intergenic | MIRb\|SINE\|MIR | -12079 | NM_019457 | 54562 | Mm.244890 | NM_019457 | ENSMUSG00000022375 | **Lrrc6** | LRTP | leucine rich repeat containing 6 (testis) | protein-coding |
| chr6_48554404_48555065 | chr6 | 48554405 | 48555065 | + | -0.758786 | promoter-TSS (NM_001110160) | promoter-TSS (NM_001110160) | -64 | NM_001110160 | 243371 | Mm.290764 | NM_177736 | ENSMUSG00000073096 | **Lrrc61** | BC027309 | leucine rich repeat containing 61 | protein-coding |
| chr6_48554352_48555582 | chr6 | 48554353 | 48555582 | + | -0.797518 | 5' UTR (NM_177736, exon 1 of 3) | 5' UTR (NM_177736, exon 1 of 3) | 168 | NM_177736 | 243371 | Mm.290764 | NM_177736 | ENSMUSG00000073096 | **Lrrc61** | BC027309 | leucine rich repeat containing 61 | protein-coding |
| chr14_75103016_75103357 | chr14 | 75103017 | 75103357 | + | -0.798845 | intron (NM_027581, intron 7 of 9) | Lx6\|LINE\|L1 | 27696 | NM_027581 | 70859 | Mm.320289 | NM_027581 | ENSMUSG00000021997 | **Lrrc63** | 4921509B22Rik | leucine rich repeat containing 63 | protein-coding |
| chr12_36190126_36190666 | chr12 | 36190127 | 36190666 | + | -0.68438 | intron (NM_146033, intron 7 of 9) | intron (NM_146033, intron 7 of 9) | 27806 | NM_027699 | 71156 | Mm.250837 | NM_027699 | ENSMUSG00000020545 | **Lrrc72** | 1700108M19Rik\|4933421E18Rik | leucine rich repeat containing 72 | protein-coding |
| chr12_86807321_86808096 | chr12 | 86807322 | 86808096 | + | -0.671015 | Intergenic | Intergenic | 73340 | NM_001195767 | 627607 | Mm.341880 | NM_001195767 | ENSMUSG00000059114 | **Lrrc74** | EG627607\|Gm6772 | leucine rich repeat containing 74 | protein-coding |
| chr12_86807874_86808067 | chr12 | 86807875 | 86808067 | + | -0.834938 | Intergenic | Intergenic | 73602 | NM_001195767 | 627607 | Mm.341880 | NM_001195767 | ENSMUSG00000059114 | **Lrrc74** | EG627607\|Gm6772 | leucine rich repeat containing 74 | protein-coding |
| chr10_75565129_75566147 | chr10 | 75565130 | 75566147 | + | -0.800119 | Intergenic | Intergenic | -5308 | NM_198860 | 192734 | Mm.259320 | NM_198860 | ENSMUSG00000046807 | **Lrrc75b** | AI646023\|Fam211b | leucine rich repeat containing 75B | protein-coding |
| chr5_105443156_105443912 | chr5 | 105443157 | 105443912 | + | -0.686738 | intron (NM_001033550, intron 1 of 4) | intron (NM_001033550, intron 1 of 4) | 27759 | NM_001033550 | 433926 | Mm.439781 | NM_001033550 | ENSMUSG00000070639 | **Lrrc8b** | R75581\|Ta-lrrp\|mKIAA0231 | leucine rich repeat containing 8 family, member B | protein-coding |

| | | | | | | | | | | | | | | | | | |
|---|---|---|---|---|---|---|---|---|---|---|---|---|---|---|---|---|---|
| chr5_105443456_105443855 | chr5 | 105443457 | 105443855 | + | -0.714606 | intron (NM_001033550, intron 1 of 4) | intron (NM_001033550, intron 1 of 4) | 27881 | NM_001033550 | 433926 | Mm.439781 | NM_001033550 | ENSMUSG00000070639 | **Lrrc8b** | R75581\|Ta-lrrp\|mKIAA0231 | leucine rich repeat containing 8 family, member B | protein-coding |
| chr5_105541071_105541429 | chr5 | 105541072 | 105541429 | + | -0.703601 | intron (NM_133897, intron 1 of 3) | RSINE1\|SINE\|B4 | 21779 | NM_133897 | 100604 | Mm.319847 | NM_133897 | ENSMUSG00000054720 | **Lrrc8c** | AD158\|AI326115\|E430036I04Rik\|fad158 | leucine rich repeat containing 8 family, member C | protein-coding |
| chr5_105598391_105599251 | chr5 | 105598392 | 105599251 | + | -0.703877 | intron (NM_133897, intron 3 of 3) | L1MC1\|LINE\|L1 | 79350 | NM_133897 | 100604 | Mm.319847 | NM_133897 | ENSMUSG00000054720 | **Lrrc8c** | AD158\|AI326115\|E430036I04Rik\|fad158 | leucine rich repeat containing 8 family, member C | protein-coding |
| chr5_105598742_105599251 | chr5 | 105598743 | 105599251 | + | -0.726254 | intron (NM_133897, intron 3 of 3) | intron (NM_133897, intron 3 of 3) | 79526 | NM_133897 | 100604 | Mm.319847 | NM_133897 | ENSMUSG00000054720 | **Lrrc8c** | AD158\|AI326115\|E430036I04Rik\|fad158 | leucine rich repeat containing 8 family, member C | protein-coding |
| chr5_105588980_105589508 | chr5 | 105588981 | 105589508 | + | -0.75754 | intron (NM_133897, intron 3 of 3) | intron (NM_133897, intron 3 of 3) | 69773 | NM_133897 | 100604 | Mm.319847 | NM_133897 | ENSMUSG00000054720 | **Lrrc8c** | AD158\|AI326115\|E430036I04Rik\|fad158 | leucine rich repeat containing 8 family, member C | protein-coding |
| chr5_105717851_105718475 | chr5 | 105717852 | 105718475 | + | -0.622841 | intron (NM_178701, intron 1 of 2) | intron (NM_178701, intron 1 of 2) | 17353 | NM_178701 | 231549 | Mm.23837 | NM_178701 | ENSMUSG00000046079 | **Lrrc8d** | 2810473G09Rik\|4930525N13Rik\|A930019F03\|Lrrc5 | leucine rich repeat containing 8D | protein-coding |
| chr5_105718092_105718770 | chr5 | 105718093 | 105718770 | + | -0.653061 | intron (NM_178701, intron 1 of 2) | intron (NM_178701, intron 1 of 2) | 17621 | NM_178701 | 231549 | Mm.23837 | NM_178701 | ENSMUSG00000046079 | **Lrrc8d** | 2810473G09Rik\|4930525N13Rik\|A930019F03\|Lrrc5 | leucine rich repeat containing 8D | protein-coding |
| chr5_105648586_105649335 | chr5 | 105648587 | 105649335 | + | -0.659514 | Intergenic | Intergenic | -51008 | NM_001122768 | 231549 | Mm.23837 | NM_178701 | ENSMUSG00000046079 | **Lrrc8d** | 2810473G09Rik\|4930525N13Rik\|A930019F03\|Lrrc5 | leucine rich repeat containing 8D | protein-coding |
| chr5_105635472_105635956 | chr5 | 105635473 | 105635956 | + | -0.677633 | Intergenic | Intergenic | -64255 | NM_001122768 | 231549 | Mm.23837 | NM_178701 | ENSMUSG00000046079 | **Lrrc8d** | 2810473G09Rik\|4930525N13Rik\|A930019F03\|Lrrc5 | leucine rich repeat containing 8D | protein-coding |
| chr5_105635599_105635952 | chr5 | 105635600 | 105635952 | + | -0.727526 | Intergenic | Intergenic | -64193 | NM_001122768 | 231549 | Mm.23837 | NM_178701 | ENSMUSG00000046079 | **Lrrc8d** | 2810473G09Rik\|4930525N13Rik\|A930019F03\|Lrrc5 | leucine rich repeat containing 8D | protein-coding |
| chr5_105648591_105649032 | chr5 | 105648592 | 105649032 | + | -0.745237 | Intergenic | Intergenic | -51157 | NM_001122768 | 231549 | Mm.23837 | NM_178701 | ENSMUSG00000046079 | **Lrrc8d** | 2810473G09Rik\|4930525N13Rik\|A930019F03\|Lrrc5 | leucine rich repeat containing 8D | protein-coding |
| chr5_105642290_105643108 | chr5 | 105642291 | 105643108 | + | -0.773872 | Intergenic | Intergenic | -57270 | NM_001122768 | 231549 | Mm.23837 | NM_178701 | ENSMUSG00000046079 | **Lrrc8d** | 2810473G09Rik\|4930525N13Rik\|A930019F03\|Lrrc5 | leucine rich repeat containing 8D | protein-coding |
| chr5_105643384_105643717 | chr5 | 105643385 | 105643717 | + | -0.891625 | Intergenic | Intergenic | -56418 | NM_001122768 | 231549 | Mm.23837 | NM_178701 | ENSMUSG00000046079 | **Lrrc8d** | 2810473G09Rik\|4930525N13Rik\|A930019F03\|Lrrc5 | leucine rich repeat containing 8D | protein-coding |
| chr1_91064838_91065516 | chr1 | 91064839 | 91065516 | + | -0.929084 | intron (NM_001111311, intron 1 of 7) | intron (NM_001111311, intron 1 of 7) | 11733 | NM_008515 | 16978 | Mm.45039 | NM_008515 | ENSMUSG00000026305 | **Lrrfip1** | AU024550\|Fliiap1 | leucine rich repeat (in FLII) interacting protein 1 | protein-coding |
| chr12_41325634_41326307 | chr12 | 41325635 | 41326307 | + | -0.673749 | intron (NM_053122, intron 4 of 6) | ORR1E\|LTR\|ERVL-MaLR | 160086 | NM_001271708 | 16981 | Mm.404786 | NM_010733 | ENSMUSG00000036295 | **Lrrn3** | NLRR-3 | leucine rich repeat protein 3, neuronal | protein-coding |
| chr12_41325800_41326088 | chr12 | 41325801 | 41326088 | + | -0.729733 | intron (NM_053122, intron 4 of 6) | ORR1E\|LTR\|ERVL-MaLR | 160113 | NM_001271708 | 16981 | Mm.404786 | NM_010733 | ENSMUSG00000036295 | **Lrrn3** | NLRR-3 | leucine rich repeat protein 3, neuronal | protein-coding |
| chr6_119336075_119336519 | chr6 | 119336076 | 119336519 | + | -0.672254 | intron (NM_001033382, intron 26 of 37) | intron (NM_001033382, intron 26 of 37) | -5531 | NM_172492 | 211187 | Mm.121498 | NM_172492 | ENSMUSG00000055003 | **Lrtm2** | A230084J22\|AI841794 | leucine-rich repeats and transmembrane domains 2 | protein-coding |
| chr4_126090011_126090895 | chr4 | 126090012 | 126090895 | + | -0.780328 | Intergenic | Intergenic | -6200 | NM_001163266 | 116748 | Mm.41865 | NM_138721 | ENSMUSG00000050188 | **Lsm10** | - | U7 snRNP-specific Sm-like protein LSM10 | protein-coding |
| chr8_70675404_70675658 | chr8 | 70675405 | 70675658 | + | -0.681749 | intron (NM_015816, intron 1 of 4) | intron (NM_015816, intron 1 of 4) | 2300 | NM_015816 | 50783 | Mm.248188 | NM_015816 | ENSMUSG00000031848 | **Lsm4** | - | LSM4 homolog, U6 small nuclear RNA associated (S. cerevisiae) | protein-coding |
| chr17_35190023_35190411 | chr17 | 35190024 | 35190411 | + | -0.727815 | Intergenic | Intergenic | -1777 | NM_010734 | 16988 | Mm.19379 | NM_010734 | ENSMUSG00000073412 | **Lst1** | B144 | leukocyte specific transcript 1 | protein-coding |
| chr17_35190098_35190580 | chr17 | 35190099 | 35190580 | + | -0.834527 | Intergenic | Intergenic | -1899 | NM_010734 | 16988 | Mm.19379 | NM_010734 | ENSMUSG00000073412 | **Lst1** | B144 | leukocyte specific transcript 1 | protein-coding |
| chr15_75052492_75052781 | chr15 | 75052493 | 75052781 | + | -0.846909 | Intergenic | Intergenic | -3800 | NM_001252056 | 17067 | Mm.1583 | NM_010741 | ENSMUSG00000079018 | **Ly6c1** | AA682074\|AA959465\|Ly-6C\|Ly-6C1\|Ly6c | lymphocyte antigen 6 complex, locus C1 | protein-coding |
| chr15_74961832_74962096 | chr15 | 74961833 | 74962096 | + | -0.676477 | Intergenic | B3\|SINE\|B2 | 6858 | NM_001164040 | 17069 | Mm.788 | NM_008529 | ENSMUSG00000022587 | **Ly6e** | 9804\|Ly67\|RIG-E\|Sca-2\|TSA-1\|Tsa1 | lymphocyte antigen 6 complex, locus E | protein-coding |
| chr15_74925996_74926216 | chr15 | 74925997 | 74926216 | + | -0.691081 | Intergenic | Intergenic | -28945 | NM_001164039 | 17069 | Mm.788 | NM_008529 | ENSMUSG00000022587 | **Ly6e** | 9804\|Ly67\|RIG-E\|Sca-2\|TSA-1\|Tsa1 | lymphocyte antigen 6 complex, locus E | protein-coding |
| chr15_74961816_74962178 | chr15 | 74961817 | 74962178 | + | -0.714671 | Intergenic | (CAGA)n\|Simple_repeat\|Simple_repeat | 6891 | NM_001164040 | 17069 | Mm.788 | NM_008529 | ENSMUSG00000022587 | **Ly6e** | 9804\|Ly67\|RIG-E\|Sca-2\|TSA-1\|Tsa1 | lymphocyte antigen 6 complex, locus E | protein-coding |
| chr2_60375180_60375820 | chr2 | 60375181 | 60375820 | + | -1.04027 | intron (NM_013825, intron 2 of 34) | MLTR11B\|LTR\|ERVK | 7731 | NM_013825 | 17076 | Mm.2074 | NM_013825 | ENSMUSG00000026980 | **Ly75** | CD205\|DEC-205\|DEC205 | lymphocyte antigen 75 | protein-coding |
| chr13_37562637_37562984 | chr13 | 37562638 | 37562984 | + | -0.755104 | Intergenic | Intergenic | 217466 | NM_010745 | 17084 | Mm.2639 | NM_010745 | ENSMUSG00000021423 | **Ly86** | MD-1\|MD1 | lymphocyte antigen 86 | protein-coding |
| chr13_37562462_37563010 | chr13 | 37562463 | 37563010 | + | -0.826746 | Intergenic | Intergenic | 217391 | NM_010745 | 17084 | Mm.2639 | NM_010745 | ENSMUSG00000021423 | **Ly86** | MD-1\|MD1 | lymphocyte antigen 86 | protein-coding |
| chr13_37537745_37538081 | chr13 | 37537746 | 37538081 | + | -1.30502 | Intergenic | Intergenic | 192568 | NM_010745 | 17084 | Mm.2639 | NM_010745 | ENSMUSG00000021423 | **Ly86** | MD-1\|MD1 | lymphocyte antigen 86 | protein-coding |
| chr13_37537787_37537986 | chr13 | 37537788 | 37537986 | + | -1.34376 | Intergenic | Intergenic | 192542 | NM_010745 | 17084 | Mm.2639 | NM_010745 | ENSMUSG00000021423 | **Ly86** | MD-1\|MD1 | lymphocyte antigen 86 | protein-coding |
| chr7_24646504_24646849 | chr7 | 24646505 | 24646849 | + | -0.923195 | Intergenic | Intergenic | 10107 | NM_133743 | 72434 | Mm.281093 | NM_133743 | ENSMUSG00000057454 | **Lypd3** | 2310061G07Rik\|C4.4a | Ly6/Plaur domain containing 3 | protein-coding |
| chr2_49847753_49848231 | chr2 | 49847754 | 49848231 | + | -0.698879 | intron (NM_027990, intron 1 of 9) | Lx8b\|LINE\|L1 | 60306 | NM_027990 | 71897 | Mm.41381 | NM_027990 | ENSMUSG00000026765 | **Lypd6b** | 2310010M24Rik\|AW049525 | LY6/PLAUR domain containing 6B | protein-coding |
| chr2_49846148_49846479 | chr2 | 49846149 | 49846479 | + | -0.928941 | intron (NM_027990, intron 1 of 9) | L1ME2\|LINE\|L1 | 58628 | NM_027990 | 71897 | Mm.41381 | NM_027990 | ENSMUSG00000026765 | **Lypd6b** | 2310010M24Rik\|AW049525 | LY6/PLAUR domain containing 6B | protein-coding |
| chr4_32768055_32768427 | chr4 | 32768056 | 32768427 | + | -1.22251 | intron (NM_001081392, intron 96 of 101) | intron (NM_001081392, intron 96 of 101) | -32018 | NM_175364 | 108755 | Mm.1182 | NM_175364 | | **Lyrm2** | 2610208E05Rik | LYR motif containing 2 | protein-coding |
| chr4_32767931_32768490 | chr4 | 32767932 | 32768490 | + | -1.22608 | intron (NM_001081392, intron 96 of 101) | (TG)n\|Simple_repeat\|Simple_repeat | -32048 | NM_175364 | 108755 | Mm.1182 | NM_175364 | | **Lyrm2** | 2610208E05Rik | LYR motif containing 2 | protein-coding |
| chr11_54850409_54850860 | chr11 | 54850410 | 54850860 | + | -0.639195 | intron (NM_029327, intron 2 of 4) | B1F1\|SINE\|Alu | 9956 | NM_029327 | 75530 | Mm.83578 | NM_029327 | | **Lyrm7** | 1700024C24Rik\|9330147L21Rik | LYR motif containing 7 | protein-coding |
| chr11_54844425_54844974 | chr11 | 54844426 | 54844974 | + | -1.12573 | intron (NM_029327, intron 4 of 4) | intron (NM_029327, intron 4 of 4) | 15891 | NM_029327 | 75530 | Mm.83578 | NM_029327 | | **Lyrm7** | 1700024C24Rik\|9330147L21Rik | LYR motif containing 7 | protein-coding |
| chr11_78792983_78793512 | chr11 | 78792984 | 78793512 | + | -0.707623 | Intergenic | Intergenic | -33347 | NM_001076681 | 66274 | Mm.258932 | NM_001076681 | | **Lyrm9** | 1810012P15Rik | LYR motif containing 9 | protein-coding |
| chr11_78830206_78830661 | chr11 | 78830207 | 78830661 | + | -0.779776 | intron (NM_001076681, intron 1 of 3) | intron (NM_001076681, intron 1 of 3) | 3839 | NM_001076681 | 66274 | Mm.258932 | NM_001076681 | | **Lyrm9** | 1810012P15Rik | LYR motif containing 9 | protein-coding |
| chr11_78823141_78823381 | chr11 | 78823142 | 78823381 | + | -0.835358 | Intergenic | Intergenic | -3334 | NM_001076681 | 66274 | Mm.258932 | NM_001076681 | | **Lyrm9** | 1810012P15Rik | LYR motif containing 9 | protein-coding |
| chr3_95135680_95136194 | chr3 | 95135681 | 95136194 | + | -0.844109 | intron (NM_153121, intron 1 of 2) | intron (NM_153121, intron 1 of 2) | 1849 | NM_153121 | 217779 | Mm.341945 | NM_153121 | ENSMUSG00000053769 | **Lysmd1** | 2610022K04Rik\|AI415311 | LysM, putative peptidoglycan-binding, domain containing 1 | protein-coding |
| chr13_13537483_13537985 | chr13 | 13537484 | 13537985 | + | -0.653084 | Intergenic | Intergenic | -52675 | NM_010748 | 17101 | Mm.342337 | NM_010748 | ENSMUSG00000019726 | **Lyst** | D13Sfk13\|beige\|bg | lysosomal trafficking regulator | protein-coding |
| chr13_13537629_13537944 | chr13 | 13537630 | 13537944 | + | -0.686555 | Intergenic | Intergenic | -52622 | NM_010748 | 17101 | Mm.342337 | NM_010748 | ENSMUSG00000019726 | **Lyst** | D13Sfk13\|beige\|bg | lysosomal trafficking regulator | protein-coding |
| chr13_13520476_13520766 | chr13 | 13520477 | 13520766 | + | -0.72713 | Intergenic | RLTR20A3_MM\|LTR\|ERVK | -69788 | NM_010748 | 17101 | Mm.342337 | NM_010748 | ENSMUSG00000019726 | **Lyst** | D13Sfk13\|beige\|bg | lysosomal trafficking regulator | protein-coding |
| chr13_13530884_13531266 | chr13 | 13530885 | 13531266 | + | -0.770358 | Intergenic | PB1D10\|SINE\|Alu | -59334 | NM_010748 | 17101 | Mm.342337 | NM_010748 | ENSMUSG00000019726 | **Lyst** | D13Sfk13\|beige\|bg | lysosomal trafficking regulator | protein-coding |
| chr13_13574638_13574920 | chr13 | 13574639 | 13574920 | + | -0.794568 | Intergenic | ID_B1\|SINE\|B4 | -15630 | NM_010748 | 17101 | Mm.342337 | NM_010748 | ENSMUSG00000019726 | **Lyst** | D13Sfk13\|beige\|bg | lysosomal trafficking regulator | protein-coding |
| chr13_13564220_13564943 | chr13 | 13564221 | 13564943 | + | -0.914634 | Intergenic | Intergenic | -25827 | NM_010748 | 17101 | Mm.342337 | NM_010748 | ENSMUSG00000019726 | **Lyst** | D13Sfk13\|beige\|bg | lysosomal trafficking regulator | protein-coding |
| chr13_13564464_13564938 | chr13 | 13564465 | 13564938 | + | -0.934253 | Intergenic | Intergenic | -25708 | NM_010748 | 17101 | Mm.342337 | NM_010748 | ENSMUSG00000019726 | **Lyst** | D13Sfk13\|beige\|bg | lysosomal trafficking regulator | protein-coding |
| chr13_13609695_13609980 | chr13 | 13609696 | 13609980 | + | -1.09917 | intron (NM_010748, intron 1 of 52) | ORR1F\|LTR\|ERVL-MaLR | 19429 | NM_010748 | 17101 | Mm.342337 | NM_010748 | ENSMUSG00000019726 | **Lyst** | D13Sfk13\|beige\|bg | lysosomal trafficking regulator | protein-coding |
| chr18_3984206_3984490 | chr18 | 3984207 | 3984490 | + | -0.624139 | Intergenic | ORR1F\|LTR\|ERVL-MaLR | -181484 | NM_026092 | 67328 | Mm.158148 | NM_026092 | ENSMUSG00000024233 | **Lyzl1** | 1700038F02Rik\|Lyc2 | lysozyme-like 1 | protein-coding |
| chr9_123723156_123723404 | chr9 | 123723157 | 123723404 | + | -0.781282 | Intergenic | Intergenic | -5723 | NM_033322 | 93730 | Mm.378263 | NM_033322 | ENSMUSG00000025245 | **Lztfl1** | 5530402H04Rik\|6130400H19Rik\|AI414725\|AW048545 | leucine zipper transcription factor-like 1 | protein-coding |
| chr3_55743758_55744252 | chr3 | 55743759 | 55744252 | + | -0.656945 | intron (NM_030595, intron 41 of 57) | intron (NM_030595, intron 41 of 57) | -38505 | NM_010750 | 17116 | Mm.440209 | NM_010750 | ENSMUSG00000056947 | **Mab21l1** | AW047968 | mab-21-like 1 (C. elegans) | protein-coding |
| chr3_86682731_86683138 | chr3 | 86682732 | 86683138 | + | -0.792511 | intron (NM_001077687, intron 47 of 50) | PB1\|SINE\|Alu | -134652 | NM_011839 | 23937 | Mm.389466 | NM_011839 | ENSMUSG00000057777 | **Mab21l2** | - | mab-21-like 2 (C. elegans) | protein-coding |
| chr3_86682804_86683098 | chr3 | 86682805 | 86683098 | + | -0.834636 | intron (NM_001077687, intron 47 of 50) | PB1\|SINE\|Alu | -134668 | NM_011839 | 23937 | Mm.389466 | NM_011839 | ENSMUSG00000057777 | **Mab21l2** | - | mab-21-like 2 (C. elegans) | protein-coding |

| | | | | | | | | | | | | | | | | | |
|---|---|---|---|---|---|---|---|---|---|---|---|---|---|---|---|---|---|
| chr3_86682826_86683031 | chr3 | 86682827 | 86683031 | + | -0.861703 | intron (NM_001077687, intron 47 of 50) | PB1\|SINE\|Alu | -134646 | NM_011839 | 23937 | Mm.389466 | NM_011839 | ENSMUSG00000057777 | **Mab21l2** | - | mab-21-like 2 (C. elegans) | protein-coding |
| chr3_101860654_101861142 | chr3 | 101860655 | 101861142 | + | -0.627437 | exon (NM_001039371, exon 11 of 12) | exon (NM_001039371, exon 11 of 12) | -24675 | NM_172295 | 242125 | Mm.32900 | NM_172295 | ENSMUSG00000044313 | **Mab21l3** | - | mab-21-like 3 (C. elegans) | protein-coding |
| chr3_101748151_101748574 | chr3 | 101748152 | 101748574 | + | -0.648459 | Intergenic | Intergenic | 87860 | NM_172295 | 242125 | Mm.32900 | NM_172295 | ENSMUSG00000044313 | **Mab21l3** | - | mab-21-like 3 (C. elegans) | protein-coding |
| chr3_101731159_101731448 | chr3 | 101731160 | 101731448 | + | -0.664047 | Intergenic | Intergenic | 104919 | NM_172295 | 242125 | Mm.32900 | NM_172295 | ENSMUSG00000044313 | **Mab21l3** | - | mab-21-like 3 (C. elegans) | protein-coding |
| chr3_101745966_101746338 | chr3 | 101745967 | 101746338 | + | -0.665194 | Intergenic | Intergenic | 90071 | NM_172295 | 242125 | Mm.32900 | NM_172295 | ENSMUSG00000044313 | **Mab21l3** | - | mab-21-like 3 (C. elegans) | protein-coding |
| chr3_101746050_101746313 | chr3 | 101746051 | 101746313 | + | -0.674941 | Intergenic | B3A\|SINE\|B2 | 90041 | NM_172295 | 242125 | Mm.32900 | NM_172295 | ENSMUSG00000044313 | **Mab21l3** | - | mab-21-like 3 (C. elegans) | protein-coding |
| chr5_140208516_140209439 | chr5 | 140208517 | 140209439 | + | -0.738361 | intron (NM_010752, intron 10 of 17) | intron (NM_010752, intron 10 of 17) | 112574 | NM_010752 | 17120 | Mm.27250 | NM_010752 | ENSMUSG00000029554 | **Mad1l1** | AI173502\|AW550425\|Mad1 | MAD1 mitotic arrest deficient 1-like 1 | protein-coding |
| chr5_140270928_140271877 | chr5 | 140270929 | 140271877 | + | -0.923506 | intron (NM_010752, intron 9 of 17) | B1F\|SINE\|Alu | 50149 | NM_010752 | 17120 | Mm.27250 | NM_010752 | ENSMUSG00000029554 | **Mad1l1** | AI173502\|AW550425\|Mad1 | MAD1 mitotic arrest deficient 1-like 1 | protein-coding |
| chr5_140282531_140283327 | chr5 | 140282532 | 140283327 | + | -0.977292 | intron (NM_010752, intron 9 of 17) | intron (NM_010752, intron 9 of 17) | 38623 | NM_010752 | 17120 | Mm.27250 | NM_010752 | ENSMUSG00000029554 | **Mad1l1** | AI173502\|AW550425\|Mad1 | MAD1 mitotic arrest deficient 1-like 1 | protein-coding |
| chr5_140282840_140283351 | chr5 | 140282841 | 140283351 | + | -1.09717 | intron (NM_010752, intron 9 of 17) | intron (NM_010752, intron 9 of 17) | 38456 | NM_010752 | 17120 | Mm.27250 | NM_010752 | ENSMUSG00000029554 | **Mad1l1** | AI173502\|AW550425\|Mad1 | MAD1 mitotic arrest deficient 1-like 1 | protein-coding |
| chr5_140215352_140216148 | chr5 | 140215353 | 140216148 | + | -1.3314 | intron (NM_010752, intron 10 of 17) | intron (NM_010752, intron 10 of 17) | 105802 | NM_010752 | 17120 | Mm.27250 | NM_010752 | ENSMUSG00000029554 | **Mad1l1** | AI173502\|AW550425\|Mad1 | MAD1 mitotic arrest deficient 1-like 1 | protein-coding |
| chr6_66412960_66413367 | chr6 | 66412961 | 66413367 | + | -0.902243 | Intergenic | Intergenic | -122304 | NM_019499 | 56150 | Mm.489653 | NM_019499 | ENSMUSG00000029910 | **Mad2l1** | AA673185\|MAD2 | MAD2 mitotic arrest deficient-like 1 | protein-coding |
| chr6_66493392_66494262 | chr6 | 66493393 | 66494262 | + | -1.23236 | Intergenic | Intergenic | -41641 | NM_019499 | 56150 | Mm.489653 | NM_019499 | ENSMUSG00000029910 | **Mad2l1** | AA673185\|MAD2 | MAD2 mitotic arrest deficient-like 1 | protein-coding |
| chr6_66493683_66494230 | chr6 | 66493684 | 66494230 | + | -1.25095 | Intergenic | Intergenic | -41511 | NM_019499 | 56150 | Mm.489653 | NM_019499 | ENSMUSG00000029910 | **Mad2l1** | AA673185\|MAD2 | MAD2 mitotic arrest deficient-like 1 | protein-coding |
| chr6_66493758_66494089 | chr6 | 66493759 | 66494089 | + | -1.26834 | Intergenic | Intergenic | -41544 | NM_019499 | 56150 | Mm.489653 | NM_019499 | ENSMUSG00000029910 | **Mad2l1** | AA673185\|MAD2 | MAD2 mitotic arrest deficient-like 1 | protein-coding |
| chr8_115514453_115514805 | chr8 | 115514454 | 115514805 | + | -0.678963 | Intergenic | Intergenic | 192265 | NM_001025577 | 17132 | Mm.275549 | NM_001025577 | | **Maf** | 2810401A20Rik\|A230108G15Rik\|AW047063\|c-maf | avian musculoaponeurotic fibrosarcoma (v-maf) AS42 oncogene homolog | protein-coding |
| chr8_115189766_115190273 | chr8 | 115189767 | 115190273 | + | -0.818657 | intron (NM_019573, intron 8 of 8) | intron (NM_019573, intron 8 of 8) | 516874 | NM_001025577 | 17132 | Mm.275549 | NM_001025577 | | **Maf** | 2810401A20Rik\|A230108G15Rik\|AW047063\|c-maf | avian musculoaponeurotic fibrosarcoma (v-maf) AS42 oncogene homolog | protein-coding |
| chr5_139791779_139792207 | chr5 | 139791780 | 139792207 | + | -0.714084 | intron (NM_010757, intron 1 of 2) | CpG | 457 | NM_010757 | 17135 | Mm.157313 | NM_010757 | ENSMUSG00000018143 | **Mafk** | AW061068\|NF-E2\|Nfe2u | v-maf musculoaponeurotic fibrosarcoma oncogene family, protein K (avian) | protein-coding |
| chr7_30909517_30910248 | chr7 | 30909518 | 30910248 | + | -0.687451 | intron (NM_010758, intron 5 of 12) | MTD\|LTR\|ERVL-MaLR | 4949 | NM_010758 | 17136 | Mm.241355 | NM_010758 | ENSMUSG00000036634 | **Mag** | Gma\|siglec-4a | myelin-associated glycoprotein | protein-coding |
| chr7_30909742_30910318 | chr7 | 30909743 | 30910318 | + | -0.692084 | intron (NM_010758, intron 5 of 12) | MTD\|LTR\|ERVL-MaLR | 4802 | NM_010758 | 17136 | Mm.241355 | NM_010758 | ENSMUSG00000036634 | **Mag** | Gma\|siglec-4a | myelin-associated glycoprotein | protein-coding |
| chrX_144525777_144526609 | chrX | 144525778 | 144526609 | + | -1.2978 | intron (NM_009428, intron 1 of 10) | MTC\|LTR\|ERVL-MaLR | 30046 | NR_033647 | 100039436 | Mm.483818 | NM_001111315 | | **Mageb16-ps1** | EG623523\|Gm15072\|Mageb16\|OTMUSG00000018935 | melanoma antigen family B, 16, pseudogene 1 | pseudo |
| chrX_144526013_144526455 | chrX | 144526014 | 144526455 | + | -1.342 | intron (NM_009428, intron 1 of 10) | intron (NM_009428, intron 1 of 10) | 30005 | NR_033647 | 100039436 | Mm.483818 | NM_001111315 | | **Mageb16-ps1** | EG623523\|Gm15072\|Mageb16\|OTMUSG00000018935 | melanoma antigen family B, 16, pseudogene 1 | pseudo |
| chr5_20332439_20333010 | chr5 | 20332440 | 20333010 | + | -1.01134 | intron (NM_001170746, intron 6 of 22) | intron (NM_001170746, intron 6 of 22) | 425207 | NM_015823 | 50791 | Mm.332231 | NM_015823 | ENSMUSG00000040003 | **Magi2** | AIP-1\|Acvri1\|Acvrinp1\|Acvrip1\|Magi-2\|S-SCAM\|mKIAA0705 | membrane associated guanylate kinase, WW and PDZ domain containing 2 | protein-coding |
| chr5_20299818_20300459 | chr5 | 20299819 | 20300459 | + | -1.31292 | intron (NM_001170746, intron 6 of 22) | intron (NM_001170746, intron 6 of 22) | 392621 | NM_015823 | 50791 | Mm.332231 | NM_015823 | ENSMUSG00000040003 | **Magi2** | AIP-1\|Acvri1\|Acvrinp1\|Acvrip1\|Magi-2\|S-SCAM\|mKIAA0705 | membrane associated guanylate kinase, WW and PDZ domain containing 2 | protein-coding |
| chr5_20299870_20300146 | chr5 | 20299871 | 20300146 | + | -1.33613 | intron (NM_001170746, intron 6 of 22) | intron (NM_001170746, intron 6 of 22) | 392490 | NM_015823 | 50791 | Mm.332231 | NM_015823 | ENSMUSG00000040003 | **Magi2** | AIP-1\|Acvri1\|Acvrinp1\|Acvrip1\|Magi-2\|S-SCAM\|mKIAA0705 | membrane associated guanylate kinase, WW and PDZ domain containing 2 | protein-coding |
| chr8_31175575_31175841 | chr8 | 31175576 | 31175841 | + | -0.893812 | Intergenic | MYSERV16_I-int\|LTR\|ERVK | -6984 | NM_026453 | 67920 | Mm.289818 | NM_026453 | ENSMUSG00000031578 | **Mak16** | 2600016B03Rik\|AI314911\|Rbm13 | MAK16 homolog (S. cerevisiae) | protein-coding |
| chr11_50286405_50286970 | chr11 | 50286406 | 50286970 | + | -0.680158 | intron (NM_175334, intron 1 of 4) | intron (NM_175334, intron 1 of 4) | 5648 | NM_175334 | 103806 | Mm.51116 | NM_175334 | ENSMUSG00000050567 | **Maml1** | AI644666\|AW743257\|D930008C07Rik\|Mam-1\|Mam1\|mKIAA0200 | mastermind like 1 (Drosophila) | protein-coding |
| chr9_13536663_13537278 | chr9 | 13536664 | 13537278 | + | -0.648827 | Intergenic | Intergenic | -83018 | NM_001013813 | 270118 | Mm.116898 | NM_173776 | ENSMUSG00000031925 | **Maml2** | 5930431H10\|BC032967 | mastermind like 2 (Drosophila) | protein-coding |
| chr9_13530379_13530922 | chr9 | 13530380 | 13530922 | + | -0.702609 | Intergenic | Intergenic | -89338 | NM_001013813 | 270118 | Mm.116898 | NM_173776 | ENSMUSG00000031925 | **Maml2** | 5930431H10\|BC032967 | mastermind like 2 (Drosophila) | protein-coding |
| chr9_13536663_13537067 | chr9 | 13536664 | 13537067 | + | -0.742807 | Intergenic | Intergenic | -83124 | NM_001013813 | 270118 | Mm.116898 | NM_173776 | ENSMUSG00000031925 | **Maml2** | 5930431H10\|BC032967 | mastermind like 2 (Drosophila) | protein-coding |
| chr9_13613680_13614333 | chr9 | 13613681 | 13614333 | + | -0.765194 | Intergenic | Intergenic | -5982 | NM_001013813 | 270118 | Mm.116898 | NM_173776 | ENSMUSG00000031925 | **Maml2** | 5930431H10\|BC032967 | mastermind like 2 (Drosophila) | protein-coding |
| chr9_13613883_13614217 | chr9 | 13613884 | 13614217 | + | -0.825499 | Intergenic | Intergenic | -5939 | NM_001013813 | 270118 | Mm.116898 | NM_173776 | ENSMUSG00000031925 | **Maml2** | 5930431H10\|BC032967 | mastermind like 2 (Drosophila) | protein-coding |
| chr10_54032280_54033071 | chr10 | 54032281 | 54033071 | + | -1.00785 | intron (NM_008548, intron 2 of 12) | intron (NM_008548, intron 2 of 12) | 43120 | NM_008548 | 17155 | Mm.117294 | NM_008548 | ENSMUSG00000003746 | **Man1a** | Man1a1\|PCR1 | mannosidase 1, alpha | protein-coding |
| chr10_54065164_54065479 | chr10 | 54065165 | 54065479 | + | -1.03182 | intron (NM_008548, intron 2 of 12) | intron (NM_008548, intron 2 of 12) | 10474 | NM_008548 | 17155 | Mm.117294 | NM_008548 | ENSMUSG00000003746 | **Man1a** | Man1a1\|PCR1 | mannosidase 1, alpha | protein-coding |
| chr4_134734177_134735141 | chr4 | 134734178 | 134735141 | + | -0.669781 | Intergenic | Intergenic | -30369 | NM_207237 | 230815 | Mm.18905 | NM_207237 | ENSMUSG00000037306 | **Man1c1** | AI593348 | mannosidase, alpha, class 1C, member 1 | protein-coding |
| chr17_64628853_64629226 | chr17 | 64628854 | 64629226 | + | -0.670275 | intron (NM_008549, intron 3 of 21) | intron (NM_008549, intron 3 of 21) | 27391 | NM_008549 | 17158 | Mm.2433 | NM_008549 | ENSMUSG00000024085 | **Man2a1** | Mana-2\|Mana2\|Map-2 | mannosidase 2, alpha 1 | protein-coding |
| chr17_64579753_64580332 | chr17 | 64579754 | 64580332 | + | -0.921738 | Intergenic | Intergenic | -21606 | NM_008549 | 17158 | Mm.2433 | NM_008549 | ENSMUSG00000024085 | **Man2a1** | Mana-2\|Mana2\|Map-2 | mannosidase 2, alpha 1 | protein-coding |
| chr7_80368768_80369177 | chr7 | 80368769 | 80369177 | + | -0.814534 | exon (NM_172903, exon 4 of 24) | exon (NM_172903, exon 4 of 24) | 2402 | NM_172903 | 140481 | Mm.269245 | NM_172903 | ENSMUSG00000038886 | **Man2a2** | 1700052O22Rik\|4931438M07Rik\|AI480988\|MX\|Man IIx | mannosidase 2, alpha 2 | protein-coding |
| chr3_135515286_135515543 | chr3 | 135515287 | 135515543 | + | -1.08406 | intron (NM_027288, intron 4 of 16) | intron (NM_027288, intron 4 of 16) | 29804 | NM_027288 | 110173 | Mm.280536 | NM_027288 | ENSMUSG00000028164 | **Manba** | 2410030O07Rik\|B930014J03Rik\|Bmn | mannosidase, beta A, lysosomal | protein-coding |
| chr4_26342397_26342624 | chr4 | 26342398 | 26342624 | + | -0.912858 | intron (NM_172865, intron 1 of 4) | intron (NM_172865, intron 1 of 4) | 4141 | NM_172865 | 242362 | Mm.245602 | NM_172865 | ENSMUSG00000040520 | **Manea** | 4932703L02Rik | mannosidase, endo-alpha | protein-coding |
| chr8_125667015_125667807 | chr8 | 125667016 | 125667807 | + | -1.3226 | Intergenic | Intergenic | -2407 | NM_028908 | 74393 | Mm.41709 | NM_028908 | ENSMUSG00000050930 | **Map10** | 4933403G14Rik\|AI835009\|mKIAA1383 | microtubule-associated protein 10 | protein-coding |
| chr13_111796367_111796901 | chr13 | 111796368 | 111796901 | + | -0.617896 | intron (NM_011945, intron 1 of 19) | intron (NM_011945, intron 1 of 19) | 12349 | NM_011945 | 26401 | Mm.15918 | NM_011945 | ENSMUSG00000021754 | **Map3k1** | MAPKKK1\|MEKK1\|Mekk | mitogen-activated protein kinase kinase kinase 1 | protein-coding |
| chr13_111867741_111868805 | chr13 | 111867742 | 111868805 | + | -0.622073 | Intergenic | Intergenic | -59290 | NM_011945 | 26401 | Mm.15918 | NM_011945 | ENSMUSG00000021754 | **Map3k1** | MAPKKK1\|MEKK1\|Mekk | mitogen-activated protein kinase kinase kinase 1 | protein-coding |
| chr13_111977705_111978338 | chr13 | 111977706 | 111978338 | + | -0.63012 | Intergenic | CpG | -169039 | NM_011945 | 26401 | Mm.15918 | NM_011945 | ENSMUSG00000021754 | **Map3k1** | MAPKKK1\|MEKK1\|Mekk | mitogen-activated protein kinase kinase kinase 1 | protein-coding |
| chr13_112002712_112003085 | chr13 | 112002713 | 112003085 | + | -0.643146 | Intergenic | Intergenic | -193916 | NM_011945 | 26401 | Mm.15918 | NM_011945 | ENSMUSG00000021754 | **Map3k1** | MAPKKK1\|MEKK1\|Mekk | mitogen-activated protein kinase kinase kinase 1 | protein-coding |
| chr13_111778496_111779178 | chr13 | 111778497 | 111779178 | + | -0.643547 | intron (NM_011945, intron 1 of 19) | intron (NM_011945, intron 1 of 19) | 30146 | NM_011945 | 26401 | Mm.15918 | NM_011945 | ENSMUSG00000021754 | **Map3k1** | MAPKKK1\|MEKK1\|Mekk | mitogen-activated protein kinase kinase kinase 1 | protein-coding |
| chr13_112002677_112002956 | chr13 | 112002678 | 112002956 | + | -0.692104 | Intergenic | Intergenic | -193834 | NM_011945 | 26401 | Mm.15918 | NM_011945 | ENSMUSG00000021754 | **Map3k1** | MAPKKK1\|MEKK1\|Mekk | mitogen-activated protein kinase kinase kinase 1 | protein-coding |
| chr13_111867685_111868380 | chr13 | 111867686 | 111868380 | + | -0.692802 | Intergenic | Intergenic | -59050 | NM_011945 | 26401 | Mm.15918 | NM_011945 | ENSMUSG00000021754 | **Map3k1** | MAPKKK1\|MEKK1\|Mekk | mitogen-activated protein kinase kinase kinase 1 | protein-coding |
| chr13_111807173_111807357 | chr13 | 111807174 | 111807357 | + | -0.701526 | intron (NM_011945, intron 1 of 19) | intron (NM_011945, intron 1 of 19) | 1718 | NM_011945 | 26401 | Mm.15918 | NM_011945 | ENSMUSG00000021754 | **Map3k1** | MAPKKK1\|MEKK1\|Mekk | mitogen-activated protein kinase kinase kinase 1 | protein-coding |
| chr13_111807162_111807954 | chr13 | 111807163 | 111807954 | + | -0.876984 | intron (NM_011945, intron 1 of 19) | intron (NM_011945, intron 1 of 19) | 1425 | NM_011945 | 26401 | Mm.15918 | NM_011945 | ENSMUSG00000021754 | **Map3k1** | MAPKKK1\|MEKK1\|Mekk | mitogen-activated protein kinase kinase kinase 1 | protein-coding |
| chr13_111807667_111807884 | chr13 | 111807668 | 111807884 | + | -0.883313 | intron (NM_011945, intron 1 of 19) | CpG | 1207 | NM_011945 | 26401 | Mm.15918 | NM_011945 | ENSMUSG00000021754 | **Map3k1** | MAPKKK1\|MEKK1\|Mekk | mitogen-activated protein kinase kinase kinase 1 | protein-coding |
| chr13_111807453_111807951 | chr13 | 111807454 | 111807951 | + | -0.885457 | intron (NM_011945, intron 1 of 19) | intron (NM_011945, intron 1 of 19) | 1281 | NM_011945 | 26401 | Mm.15918 | NM_011945 | ENSMUSG00000021754 | **Map3k1** | MAPKKK1\|MEKK1\|Mekk | mitogen-activated protein kinase kinase kinase 1 | protein-coding |
| chr13_111995494_111996127 | chr13 | 111995495 | 111996127 | + | -0.960105 | Intergenic | Intergenic | -186828 | NM_011945 | 26401 | Mm.15918 | NM_011945 | ENSMUSG00000021754 | **Map3k1** | MAPKKK1\|MEKK1\|Mekk | mitogen-activated protein kinase kinase kinase 1 | protein-coding |
| chr13_111984004_111984757 | chr13 | 111984005 | 111984757 | + | -1.01261 | Intergenic | Intergenic | -175398 | NM_011945 | 26401 | Mm.15918 | NM_011945 | ENSMUSG00000021754 | **Map3k1** | MAPKKK1\|MEKK1\|Mekk | mitogen-activated protein kinase kinase kinase 1 | protein-coding |
| chr11_103256264_103256580 | chr11 | 103256265 | 103256580 | + | -0.643252 | intron (NM_016896, intron 1 of 15) | intron (NM_016896, intron 1 of 15) | 10979 | NM_016896 | 53859 | Mm.158981 | NM_016896 | ENSMUSG00000020941 | **Map3k14** | Nik\|aly | mitogen-activated protein kinase kinase kinase 14 | protein-coding |
| chr11_103260831_103261164 | chr11 | 103260832 | 103261164 | + | -0.905525 | intron (NM_016896, intron 1 of 15) | intron (NM_016896, intron 1 of 15) | 6403 | NM_016896 | 53859 | Mm.158981 | NM_016896 | ENSMUSG00000020941 | **Map3k14** | Nik\|aly | mitogen-activated protein kinase kinase kinase 14 | protein-coding |

| | | | | | | | | | | | | | | | | | |
|---|---|---|---|---|---|---|---|---|---|---|---|---|---|---|---|---|---|
| chr1_127830135_127830904 | chr1 | 127830136 | 127830904 | + | -0.994763 | exon (NM_011737, exon 5 of 10) | exon (NM_011737, exon 5 of 10) | 9770 | NM_011737 | 22625 | Mm.246566 | NM_011737 | ENSMUSG00000051590 | **Map3k19** | Ysk4 | mitogen-activated protein kinase kinase kinase 19 | protein-coding |
| chr4_133237966_133238251 | chr4 | 133237967 | 133238251 | + | -0.813738 | Intergenic | Intergenic | -2709 | NM_016693 | 53608 | Mm.36640 | NM_016693 | ENSMUSG00000028862 | **Map3k6** | Ask2\|MAPKKK6\|MEKK6 | mitogen-activated protein kinase kinase kinase 6 | protein-coding |
| chr4_32166114_32166423 | chr4 | 32166115 | 32166423 | + | -0.825543 | Intergenic | Intergenic | 202162 | NM_172688 | 26409 | Mm.258589 | NM_009316 | ENSMUSG00000028284 | **Map3k7** | B430101B05\|C87327\|Tak1 | mitogen-activated protein kinase kinase kinase 7 | protein-coding |
| chr18_4333558_4334145 | chr18 | 4333559 | 4334145 | + | -0.877081 | exon (NM_007746, exon 7 of 8) | exon (NM_007746, exon 7 of 8) | 19101 | NM_007746 | 26410 | Mm.3275 | NM_007746 | ENSMUSG00000024235 | **Map3k8** | Cot\|Cot/Tpl2\|Est\|Estf\|Tpl-2\|Tpl2\|c-COT | mitogen-activated protein kinase kinase kinase 8 | protein-coding |
| chr17_80718264_80718569 | chr17 | 80718265 | 80718569 | + | -0.695465 | intron (NM_001290345, intron 1 of 33) | ORR1F\|LTR\|ERVL-MaLR | 9608 | NM_001290345 | 225028 | Mm.311948 | NM_001081357 | ENSMUSG00000024242 | **Map4k3** | 4833416M01Rik\|4833416M07Rik\|9530052P13Rik\|Glk\|MAPKKKK3\|MEKKK 3\|MEKKK3\|RAB8IPL1 | mitogen-activated protein kinase kinase kinase kinase 3 | protein-coding |
| chr17_80778718_80779207 | chr17 | 80778719 | 80779207 | + | -0.85866 | Intergenic | Intergenic | -50938 | NM_001290345 | 225028 | Mm.311948 | NM_001081357 | ENSMUSG00000024242 | **Map4k3** | 4833416M01Rik\|4833416M07Rik\|9530052P13Rik\|Glk\|MAPKKKK3\|MEKKK 3\|MEKKK3\|RAB8IPL1 | mitogen-activated protein kinase kinase kinase kinase 3 | protein-coding |
| chr1_39922701_39923102 | chr1 | 39922702 | 39923102 | + | -0.707078 | intron (NM_008696, intron 2 of 30) | intron (NM_008696, intron 2 of 30) | 21989 | NM_008696 | 26921 | Mm.491139 | NM_008696 | ENSMUSG00000026074 | **Map4k4** | 9430080K19Rik\|AU043147\|AU045934\|AW046177\|HGK\|Nik | mitogen-activated protein kinase kinase kinase kinase 4 | protein-coding |
| chr1_39917617_39918962 | chr1 | 39917618 | 39918962 | + | -0.77756 | intron (NM_008696, intron 2 of 30) | PB1D10\|SINE\|Alu | 17377 | NM_008696 | 26921 | Mm.491139 | NM_008696 | ENSMUSG00000026074 | **Map4k4** | 9430080K19Rik\|AU043147\|AU045934\|AW046177\|HGK\|Nik | mitogen-activated protein kinase kinase kinase kinase 4 | protein-coding |
| chr1_39917631_39918421 | chr1 | 39917632 | 39918421 | + | -0.865226 | intron (NM_008696, intron 2 of 30) | URR1B\|DNA\|hAT-Charlie | 17113 | NM_008696 | 26921 | Mm.491139 | NM_008696 | ENSMUSG00000026074 | **Map4k4** | 9430080K19Rik\|AU043147\|AU045934\|AW046177\|HGK\|Nik | mitogen-activated protein kinase kinase kinase kinase 4 | protein-coding |
| chr1_39980697_39981370 | chr1 | 39980698 | 39981370 | + | -1.07968 | intron (NM_008696, intron 6 of 30) | intron (NM_008696, intron 6 of 30) | 80121 | NM_008696 | 26921 | Mm.491139 | NM_008696 | ENSMUSG00000026074 | **Map4k4** | 9430080K19Rik\|AU043147\|AU045934\|AW046177\|HGK\|Nik | mitogen-activated protein kinase kinase kinase kinase 4 | protein-coding |
| chr12_69912777_69913080 | chr12 | 69912778 | 69913080 | + | -0.796862 | intron (NM_178628, intron 2 of 14) | ORR1E\|LTR\|ERVL-MaLR | -19766 | NM_201519 | 399510 | Mm.291936 | NM_201519 | ENSMUSG00000034761 | **Map4k5** | 4432415E19Rik\|GCKR\|KHS\|MAPKKKK5 | mitogen-activated protein kinase kinase kinase kinase 5 | protein-coding |
| chr4_126235254_126235788 | chr4 | 126235255 | 126235788 | + | -0.86769 | intron (NM_001145970, intron 8 of 16) | intron (NM_001145970, intron 8 of 16) | 20798 | NM_144941 | 245877 | Mm.266716 | NM_144941 | ENSMUSG00000028849 | **Map7d1** | AV028413\|BC019977\|Mtap7d1\|Parcc1\|Rprc1\|mKIAA1187 | MAP7 domain containing 1 | protein-coding |
| chr4_126235278_126235625 | chr4 | 126235279 | 126235625 | + | -0.878477 | intron (NM_001145970, intron 8 of 16) | intron (NM_001145970, intron 8 of 16) | 20867 | NM_144941 | 245877 | Mm.266716 | NM_144941 | ENSMUSG00000028849 | **Map7d1** | AV028413\|BC019977\|Mtap7d1\|Parcc1\|Rprc1\|mKIAA1187 | MAP7 domain containing 1 | protein-coding |
| chr11_49837489_49838255 | chr11 | 49837490 | 49838255 | + | -0.816078 | 3' UTR (NM_013529, exon 19 of 19) | 3' UTR (NM_013529, exon 19 of 19) | -8879 | NM_001163671 | 26420 | Mm.68933 | NM_016961 | ENSMUSG00000020366 | **Mapk9** | AI851083\|JNK2\|Prkm9\|p54aSAPK | mitogen-activated protein kinase 9 | protein-coding |
| chr11_49837689_49838208 | chr11 | 49837690 | 49838208 | + | -0.868958 | 3' UTR (NM_013529, exon 19 of 19) | 3' UTR (NM_013529, exon 19 of 19) | -8802 | NM_001163671 | 26420 | Mm.68933 | NM_016961 | ENSMUSG00000020366 | **Mapk9** | AI851083\|JNK2\|Prkm9\|p54aSAPK | mitogen-activated protein kinase 9 | protein-coding |
| chr18_23828549_23828866 | chr18 | 23828550 | 23828866 | + | -0.777033 | intron (NM_153058, intron 1 of 6) | intron (NM_153058, intron 1 of 6) | 24738 | NM_153058 | 212307 | Mm.132237 | NM_153058 | ENSMUSG00000024277 | **Mapre2** | AI314113\|C820009F03Rik\|D18Abb1e\|EB1\|EB2\|RP1 | microtubule-associated protein, RP/EB family, member 2 | protein-coding |
| chr1_184792873_184793912 | chr1 | 184792874 | 184793912 | + | -0.664684 | intron (NM_001290273, intron 6 of 6) | intron (NM_001290273, intron 6 of 6) | 17907 | NM_001290273 | 66112 | Mm.272457 | NM_001081361 | ENSMUSG00000026621 | **Marc1** | 1300013F15Rik\|Mosc1 | mitochondrial amidoxime reducing component 1 | protein-coding |
| chr11_105444937_105445313 | chr11 | 105444938 | 105445313 | + | -1.07711 | intron (NM_172568, intron 3 of 11) | intron (NM_172568, intron 3 of 11) | 11610 | NM_172568 | 632687 | Mm.121570 | NM_172568 | ENSMUSG00000078627 | **March10** | 4933417C16Rik\|AV039223\|OTTMUSG00000002847\|Rnf190 | membrane-associated ring finger (C3HC4) 10 | protein-coding |
| chr18_56888759_56889442 | chr18 | 56888760 | 56889442 | + | -1.12683 | intron (NM_177115, intron 1 of 4) | intron (NM_177115, intron 1 of 4) | 36447 | NM_177115 | 320253 | Mm.312623 | NM_177115 | ENSMUSG00000032656 | **March3** | 6330411I15Rik\|A530081L18Rik\|AI843639\|MARCH-III | membrane-associated ring finger (C3HC4) 3 | protein-coding |
| chr18_56888759_56889172 | chr18 | 56888760 | 56889172 | + | -1.18142 | intron (NM_177115, intron 1 of 4) | intron (NM_177115, intron 1 of 4) | 36582 | NM_177115 | 320253 | Mm.312623 | NM_177115 | ENSMUSG00000032656 | **March3** | 6330411I15Rik\|A530081L18Rik\|AI843639\|MARCH-III | membrane-associated ring finger (C3HC4) 3 | protein-coding |
| chr2_60209314_60210928 | chr2 | 60209315 | 60210928 | + | -0.64288 | intron (NM_020575, intron 1 of 9) | CpG | 185 | NM_020575 | 57438 | Mm.260635 | NM_020575 | ENSMUSG00000026977 | **March7** | Axo\|Axot\|Gtrgeo17 | membrane-associated ring finger (C3HC4) 7 | protein-coding |
| chr2_60216551_60216887 | chr2 | 60216552 | 60216887 | + | -0.851309 | intron (NM_020575, intron 1 of 9) | intron (NM_020575, intron 1 of 9) | 6783 | NM_020575 | 57438 | Mm.260635 | NM_020575 | ENSMUSG00000026977 | **March7** | Axo\|Axot\|Gtrgeo17 | membrane-associated ring finger (C3HC4) 7 | protein-coding |
| chr2_60200027_60200518 | chr2 | 60200028 | 60200518 | + | -0.863622 | Intergenic | Intergenic | -9663 | NM_020575 | 57438 | Mm.260635 | NM_020575 | ENSMUSG00000026977 | **March7** | Axo\|Axot\|Gtrgeo17 | membrane-associated ring finger (C3HC4) 7 | protein-coding |
| chr10_37192723_37193038 | chr10 | 37192724 | 37193038 | + | -0.749358 | Intergenic | Intergenic | -53955 | NM_008538 | 17118 | Mm.30059 | NM_008538 | ENSMUSG00000069662 | **Marcks** | Macs\|PKCSL | myristoylated alanine rich protein kinase C substrate | protein-coding |
| chr10_81247726_81248140 | chr10 | 81247727 | 81248140 | + | -0.791498 | intron (NM_001034895, intron 12 of 17) | intron (NM_001034895, intron 12 of 17) | -5002 | NM_001285855 | 17179 | Mm.2918 | NM_010768 | ENSMUSG00000004933 | **Matk** | CHK\|HYL\|Ntk | megakaryocyte-associated tyrosine kinase | protein-coding |
| chr10_81247715_81248648 | chr10 | 81247716 | 81248648 | + | -0.801911 | intron (NM_001034895, intron 12 of 17) | L1ME4b\|LINE\|L1 | -4753 | NM_001285855 | 17179 | Mm.2918 | NM_010768 | ENSMUSG00000004933 | **Matk** | CHK\|HYL\|Ntk | megakaryocyte-associated tyrosine kinase | protein-coding |
| chr10_81247778_81248428 | chr10 | 81247779 | 81248428 | + | -0.881256 | intron (NM_001034895, intron 12 of 17) | L1ME4b\|LINE\|L1 | -4832 | NM_001285855 | 17179 | Mm.2918 | NM_010768 | ENSMUSG00000004933 | **Matk** | CHK\|HYL\|Ntk | megakaryocyte-associated tyrosine kinase | protein-coding |
| chr4_130930663_130931305 | chr4 | 130930664 | 130931305 | + | -0.783062 | intron (NM_010686, intron 6 of 7) | intron (NM_010686, intron 6 of 7) | -13401 | NM_010769 | 17180 | Mm.271796 | NM_010769 | ENSMUSG00000040533 | **Matn1** | CMP\|Crtm\|Mat1\|matrilin-1 | matrilin 1, cartilage matrix protein | protein-coding |
| chr12_76923019_76923790 | chr12 | 76923020 | 76923790 | + | -0.942064 | Intergenic | Intergenic | 38843 | NM_008558 | 17187 | Mm.268548 | NM_008558 | ENSMUSG00000059436 | **Max** | AA960152\|AI875693\|bHLHd4\|bHLHd5\|bHLHd6\|bHLHd7\|bHLHd8 | Max protein | protein-coding |
| chr7_127024359_127024654 | chr7 | 127024360 | 127024654 | + | -0.66621 | exon (NM_010772, exon 4 of 5) | exon (NM_010772, exon 4 of 5) | 1972 | NM_010772 | 17188 | Mm.378964 | NM_010772 | ENSMUSG00000030678 | **Maz** | PUR1\|Pur-1\|SAF-1\|SAF-2 | MYC-associated zinc finger protein (purine-binding transcription factor) | protein-coding |
| chr7_127031442_127031674 | chr7 | 127031443 | 127031674 | + | -0.93198 | intron (NM_145588, intron 8 of 13) | B4A\|SINE\|B4 | -5079 | NM_010772 | 17188 | Mm.378964 | NM_010772 | ENSMUSG00000030678 | **Maz** | PUR1\|Pur-1\|SAF-1\|SAF-2 | MYC-associated zinc finger protein (purine-binding transcription factor) | protein-coding |
| chr18_70574508_70574759 | chr18 | 70574509 | 70574759 | + | -0.653194 | intron (NM_010773, intron 1 of 6) | intron (NM_010773, intron 1 of 6) | 6342 | NM_010773 | 17191 | Mm.322 | NM_010773 | ENSMUSG00000024513 | **Mbd2** | MBD2a | methyl-CpG binding domain protein 2 | protein-coding |
| chr18_70603217_70603593 | chr18 | 70603218 | 70603593 | + | -0.716237 | intron (NM_010773, intron 3 of 6) | L1M5\|LINE\|L1 | 35113 | NM_010773 | 17191 | Mm.322 | NM_010773 | ENSMUSG00000024513 | **Mbd2** | MBD2a | methyl-CpG binding domain protein 2 | protein-coding |
| chr18_70611616_70612401 | chr18 | 70611617 | 70612401 | + | -0.720965 | intron (NM_010773, intron 3 of 6) | intron (NM_010773, intron 3 of 6) | 43717 | NM_010773 | 17191 | Mm.322 | NM_010773 | ENSMUSG00000024513 | **Mbd2** | MBD2a | methyl-CpG binding domain protein 2 | protein-coding |
| chr18_70678239_70678925 | chr18 | 70678240 | 70678925 | + | -0.737433 | Intergenic | Intergenic | 110290 | NM_010773 | 17191 | Mm.322 | NM_010773 | ENSMUSG00000024513 | **Mbd2** | MBD2a | methyl-CpG binding domain protein 2 | protein-coding |
| chr18_70678430_70678925 | chr18 | 70678431 | 70678925 | + | -0.822867 | Intergenic | Intergenic | 110386 | NM_010773 | 17191 | Mm.322 | NM_010773 | ENSMUSG00000024513 | **Mbd2** | MBD2a | methyl-CpG binding domain protein 2 | protein-coding |
| chr18_70571230_70571498 | chr18 | 70571231 | 70571498 | + | -0.854119 | intron (NM_010773, intron 1 of 6) | RSINE1\|SINE\|B4 | 3072 | NM_010773 | 17191 | Mm.322 | NM_010773 | ENSMUSG00000024513 | **Mbd2** | MBD2a | methyl-CpG binding domain protein 2 | protein-coding |
| chr3_60515942_60516833 | chr3 | 60515943 | 60516833 | + | -0.603772 | intron (NM_001253709, intron 1 of 8) | intron (NM_001253709, intron 1 of 8) | 15136 | NM_001253708 | 56758 | Mm.255723 | NM_020007 | ENSMUSG00000027763 | **Mbnl1** | Mbnl\|mKIAA0428 | muscleblind-like 1 (Drosophila) | protein-coding |
| chr3_60516238_60516752 | chr3 | 60516239 | 60516752 | + | -0.642366 | intron (NM_001253709, intron 1 of 8) | intron (NM_001253709, intron 1 of 8) | 15243 | NM_001253708 | 56758 | Mm.255723 | NM_020007 | ENSMUSG00000027763 | **Mbnl1** | Mbnl\|mKIAA0428 | muscleblind-like 1 (Drosophila) | protein-coding |
| chr3_60319971_60320861 | chr3 | 60319972 | 60320861 | + | -0.675148 | Intergenic | ORR1B1\|LTR\|ERVL-MaLR | -152414 | NM_001253713 | 56758 | Mm.255723 | NM_020007 | ENSMUSG00000027763 | **Mbnl1** | Mbnl\|mKIAA0428 | muscleblind-like 1 (Drosophila) | protein-coding |
| chr3_60500716_60502073 | chr3 | 60500717 | 60502073 | + | -0.682121 | 5' UTR (NM_020007, exon 1 of 10) | 5' UTR (NM_020007, exon 1 of 10) | 143 | NM_001253708 | 56758 | Mm.255723 | NM_020007 | ENSMUSG00000027763 | **Mbnl1** | Mbnl\|mKIAA0428 | muscleblind-like 1 (Drosophila) | protein-coding |
| chr3_60496236_60496654 | chr3 | 60496237 | 60496654 | + | -0.685901 | intron (NM_001253713, intron 2 of 9) | intron (NM_001253713, intron 2 of 9) | -4807 | NM_001253711 | 56758 | Mm.255723 | NM_020007 | ENSMUSG00000027763 | **Mbnl1** | Mbnl\|mKIAA0428 | muscleblind-like 1 (Drosophila) | protein-coding |
| chr3_60413798_60414639 | chr3 | 60413799 | 60414639 | + | -0.694387 | Intergenic | PB1D9\|SINE\|Alu | -58611 | NM_001253713 | 56758 | Mm.255723 | NM_020007 | ENSMUSG00000027763 | **Mbnl1** | Mbnl\|mKIAA0428 | muscleblind-like 1 (Drosophila) | protein-coding |
| chr3_60413787_60414295 | chr3 | 60413788 | 60414295 | + | -0.827499 | Intergenic | Intergenic | -58789 | NM_001253713 | 56758 | Mm.255723 | NM_020007 | ENSMUSG00000027763 | **Mbnl1** | Mbnl\|mKIAA0428 | muscleblind-like 1 (Drosophila) | protein-coding |
| chr3_60519429_60519654 | chr3 | 60519430 | 60519654 | + | -0.843948 | intron (NM_001253709, intron 1 of 8) | intron (NM_001253709, intron 1 of 8) | 18290 | NM_001253708 | 56758 | Mm.255723 | NM_020007 | ENSMUSG00000027763 | **Mbnl1** | Mbnl\|mKIAA0428 | muscleblind-like 1 (Drosophila) | protein-coding |
| chr3_60413769_60414132 | chr3 | 60413770 | 60414132 | + | -0.865469 | Intergenic | Intergenic | -58879 | NM_001253713 | 56758 | Mm.255723 | NM_020007 | ENSMUSG00000027763 | **Mbnl1** | Mbnl\|mKIAA0428 | muscleblind-like 1 (Drosophila) | protein-coding |
| chr3_60638847_60639409 | chr3 | 60638848 | 60639409 | + | -0.884073 | Intergenic | Lx8\|LINE\|L1 | 137876 | NM_001253708 | 56758 | Mm.255723 | NM_020007 | ENSMUSG00000027763 | **Mbnl1** | Mbnl\|mKIAA0428 | muscleblind-like 1 (Drosophila) | protein-coding |
| chr3_60528592_60528959 | chr3 | 60528593 | 60528959 | + | -0.890927 | 5' UTR (NM_020007, exon 2 of 10) | 5' UTR (NM_020007, exon 2 of 10) | 27524 | NM_001253708 | 56758 | Mm.255723 | NM_020007 | ENSMUSG00000027763 | **Mbnl1** | Mbnl\|mKIAA0428 | muscleblind-like 1 (Drosophila) | protein-coding |

| | | | | | | | | | | | | | | | | | |
|---|---|---|---|---|---|---|---|---|---|---|---|---|---|---|---|---|---|
| chr3_60575960_60576270 | chr3 | 60575961 | 60576270 | + | -0.913756 | intron (NM_001253708, intron 2 of 8) | intron (NM_001253708, intron 2 of 8) | 74863 | NM_001253708 | 56758 | Mm.255723 | NM_020007 | ENSMUSG00000027763 | **Mbnl1** | Mbnl\|mKIAA0428 | muscleblind-like 1 (Drosophila) | protein-coding |
| chr3_60611853_60612230 | chr3 | 60611854 | 60612230 | + | -0.920571 | intron (NM_001253708, intron 4 of 8) | intron (NM_001253708, intron 4 of 8) | 110790 | NM_001253708 | 56758 | Mm.255723 | NM_020007 | ENSMUSG00000027763 | **Mbnl1** | Mbnl\|mKIAA0428 | muscleblind-like 1 (Drosophila) | protein-coding |
| chr3_60501635_60502133 | chr3 | 60501636 | 60502133 | + | -0.928857 | intron (NM_001253709, intron 1 of 8) | intron (NM_001253709, intron 1 of 8) | 632 | NM_001253708 | 56758 | Mm.255723 | NM_020007 | ENSMUSG00000027763 | **Mbnl1** | Mbnl\|mKIAA0428 | muscleblind-like 1 (Drosophila) | protein-coding |
| chr3_60419239_60419603 | chr3 | 60419240 | 60419603 | + | -1.22779 | Intergenic | Intergenic | -53409 | NM_001253713 | 56758 | Mm.255723 | NM_020007 | ENSMUSG00000027763 | **Mbnl1** | Mbnl\|mKIAA0428 | muscleblind-like 1 (Drosophila) | protein-coding |
| chr13_30192149_30192432 | chr13 | 30192150 | 30192432 | + | -0.612414 | intron (NM_153546, intron 1 of 12) | intron (NM_153546, intron 1 of 12) | 55801 | NM_153546 | 218121 | Mm.89682 | NM_153546 | ENSMUSG00000038732 | **Mboat1** | 9130215M02Rik\|BC023845\|LPEAT1\|Moact1\|Oact1 | membrane bound O-acyltransferase domain containing 1 | protein-coding |
| chr8_34120401_34121038 | chr8 | 34120402 | 34121038 | + | -0.769817 | intron (NM_001126314, intron 2 of 2) | intron (NM_001126314, intron 2 of 2) | 5690 | NM_001126314 | 234155 | Mm.324008 | NM_001126314 | ENSMUSG00000071113 | **Mboat4** | GOAT\|Gm171 | membrane bound O-acyltransferase domain containing 4 | protein-coding |
| chr18_82607345_82607922 | chr18 | 82607346 | 82607922 | + | -0.626382 | intron (NM_177832, intron 26 of 29) | intron (NM_177832, intron 26 of 29) | 53126 | NM_001025254 | 17196 | Mm.252063 | NM_010777 | ENSMUSG00000041607 | **Mbp** | C76307\|Hmbpr\|R75289\|golli-mbp\|mld\|shi | myelin basic protein | protein-coding |
| chr18_82525844_82526933 | chr18 | 82525845 | 82526933 | + | -0.813642 | intron (NM_001025245, intron 3 of 3) | intron (NM_001025245, intron 3 of 3) | -28074 | NM_001025256 | 17196 | Mm.252063 | NM_010777 | ENSMUSG00000041607 | **Mbp** | C76307\|Hmbpr\|R75289\|golli-mbp\|mld\|shi | myelin basic protein | protein-coding |
| chr18_82526556_82526875 | chr18 | 82526557 | 82526875 | + | -1.27511 | intron (NM_001025245, intron 3 of 3) | intron (NM_001025245, intron 3 of 3) | -27747 | NM_001025256 | 17196 | Mm.252063 | NM_010777 | ENSMUSG00000041607 | **Mbp** | C76307\|Hmbpr\|R75289\|golli-mbp\|mld\|shi | myelin basic protein | protein-coding |
| chr8_123404157_123405137 | chr8 | 123404158 | 123405137 | + | -0.893276 | TTS (NM_001286362) | TTS (NM_001286362) | -2435 | NM_008559 | 17199 | Mm.324942 | NM_008559 | ENSMUSG00000074037 | **Mc1r** | Mshra\|Tob\|e | melanocortin 1 receptor | protein-coding |
| chr9_44181596_44181881 | chr9 | 44181597 | 44181881 | + | -0.965862 | intron (NM_007619, intron 2 of 15) | intron (NM_007619, intron 2 of 15) | 47081 | NM_023061 | 84004 | Mm.275003 | NM_023061 | ENSMUSG00000032135 | **Mcam** | 1-gicerin\|AV025631\|CD146\|CD149\|Muc18\|s-endo\|s-gicerin | melanoma cell adhesion molecule | protein-coding |
| chr1_9882844_9883727 | chr1 | 9882845 | 9883727 | + | -0.63836 | intron (NM_177547, intron 10 of 16) | intron (NM_177547, intron 10 of 16) | -25352 | NM_177722 | 240697 | Mm.259800 | NM_177722 | ENSMUSG00000046101 | **Mcmdc2** | 6030422M02Rik | minichromosome maintenance domain containing 2 | protein-coding |
| chr1_9883051_9883691 | chr1 | 9883052 | 9883691 | + | -0.667932 | intron (NM_177547, intron 10 of 16) | intron (NM_177547, intron 10 of 16) | -25267 | NM_177722 | 240697 | Mm.259800 | NM_177722 | ENSMUSG00000046101 | **Mcmdc2** | 6030422M02Rik | minichromosome maintenance domain containing 2 | protein-coding |
| chr7_72313703_72315105 | chr7 | 72313704 | 72315105 | + | -0.619958 | Intergenic | Intergenic | -7809 | NM_001024703 | 244049 | Mm.217412 | NM_001024703 | ENSMUSG00000032776 | **Mctp2** | Gm489 | multiple C2 domains, transmembrane 2 | protein-coding |
| chr7_72314442_72315095 | chr7 | 72314443 | 72315095 | + | -0.762678 | Intergenic | Intergenic | -8174 | NM_001024703 | 244049 | Mm.217412 | NM_001024703 | ENSMUSG00000032776 | **Mctp2** | Gm489 | multiple C2 domains, transmembrane 2 | protein-coding |
| chr7_72308061_72308452 | chr7 | 72308062 | 72308452 | + | -0.776639 | Intergenic | Intergenic | -1662 | NM_001024703 | 244049 | Mm.217412 | NM_001024703 | ENSMUSG00000032776 | **Mctp2** | Gm489 | multiple C2 domains, transmembrane 2 | protein-coding |
| chr7_72260816_72261679 | chr7 | 72260817 | 72261679 | + | -1.56556 | intron (NM_001024703, intron 1 of 22) | Lx7\|LINE\|L1 | 45347 | NM_001024703 | 244049 | Mm.217412 | NM_001024703 | ENSMUSG00000032776 | **Mctp2** | Gm489 | multiple C2 domains, transmembrane 2 | protein-coding |
| chr13_43554804_43555227 | chr13 | 43554805 | 43555227 | + | -0.642643 | intron (NM_001081059, intron 1 of 8) | intron (NM_001081059, intron 1 of 8) | 5175 | NM_001081059 | 76137 | Mm.426290 | NM_001081059 | ENSMUSG00000021371 | **Mcur1** | 6230416A05Rik\|AU015498\|AV136929\|AW554392\|C88263\|Ccdc90a | mitochondrial calcium uniporter regulator 1 | protein-coding |
| chr6_15748289_15748606 | chr6 | 15748290 | 15748606 | + | -0.716445 | intron (NM_175088, intron 3 of 4) | intron (NM_175088, intron 3 of 4) | 27787 | NM_175088 | 16543 | Mm.1314 | NM_175088 | ENSMUSG00000041390 | **Mdfic** | Kdt1\|Mdfid | MyoD family inhibitor domain containing | protein-coding |
| chr10_118121628_118122320 | chr10 | 118121629 | 118122320 | + | -0.637286 | Intergenic | Intergenic | -19813 | NM_001162904 | 17245 | Mm.101191 | NM_010785 | ENSMUSG00000020212 | **Mdm1** | Arrd2\|Mdm-1 | transformed mouse 3T3 cell double minute 1 | protein-coding |
| chr10_118121945_118122267 | chr10 | 118121946 | 118122267 | + | -0.659144 | Intergenic | Intergenic | -19681 | NM_001162904 | 17245 | Mm.101191 | NM_010785 | ENSMUSG00000020212 | **Mdm1** | Arrd2\|Mdm-1 | transformed mouse 3T3 cell double minute 1 | protein-coding |
| chr10_118064571_118065303 | chr10 | 118064572 | 118065303 | + | -0.661221 | Intergenic | Intergenic | -76850 | NM_001162904 | 17245 | Mm.101191 | NM_010785 | ENSMUSG00000020212 | **Mdm1** | Arrd2\|Mdm-1 | transformed mouse 3T3 cell double minute 1 | protein-coding |
| chr10_118087625_118087857 | chr10 | 118087626 | 118087857 | + | -0.701355 | Intergenic | Intergenic | -54046 | NM_001162904 | 17245 | Mm.101191 | NM_010785 | ENSMUSG00000020212 | **Mdm1** | Arrd2\|Mdm-1 | transformed mouse 3T3 cell double minute 1 | protein-coding |
| chr10_118083013_118083960 | chr10 | 118083014 | 118083960 | + | -0.780128 | Intergenic | Intergenic | -58300 | NM_001162904 | 17245 | Mm.101191 | NM_010785 | ENSMUSG00000020212 | **Mdm1** | Arrd2\|Mdm-1 | transformed mouse 3T3 cell double minute 1 | protein-coding |
| chr10_118082865_118083656 | chr10 | 118082866 | 118083656 | + | -0.862368 | Intergenic | Intergenic | -58526 | NM_001162904 | 17245 | Mm.101191 | NM_010785 | ENSMUSG00000020212 | **Mdm1** | Arrd2\|Mdm-1 | transformed mouse 3T3 cell double minute 1 | protein-coding |
| chr10_118057458_118058057 | chr10 | 118057459 | 118058057 | + | -0.962282 | Intergenic | L2a\|LINE\|L2 | -84029 | NM_001162904 | 17245 | Mm.101191 | NM_010785 | ENSMUSG00000020212 | **Mdm1** | Arrd2\|Mdm-1 | transformed mouse 3T3 cell double minute 1 | protein-coding |
| chr18_73809872_73810289 | chr18 | 73809873 | 73810289 | + | -0.692369 | intron (NM_145494, intron 1 of 15) | MER21C\|LTR\|ERVL | 5311 | NM_145494 | 107029 | Mm.36817 | NM_145494 | ENSMUSG00000024556 | **Me2** | AW120568\|D030040L20Rik\|NAD-ME | malic enzyme 2, NAD(+)-dependent, mitochondrial | protein-coding |
| chr18_73810001_73810322 | chr18 | 73810002 | 73810322 | + | -0.742456 | intron (NM_145494, intron 1 of 15) | intron (NM_145494, intron 1 of 15) | 5230 | NM_145494 | 107029 | Mm.36817 | NM_145494 | ENSMUSG00000024556 | **Me2** | AW120568\|D030040L20Rik\|NAD-ME | malic enzyme 2, NAD(+)-dependent, mitochondrial | protein-coding |
| chr18_73834992_73836091 | chr18 | 73834993 | 73836091 | + | -0.820376 | Intergenic | Intergenic | -20150 | NM_145494 | 107029 | Mm.36817 | NM_145494 | ENSMUSG00000024556 | **Me2** | AW120568\|D030040L20Rik\|NAD-ME | malic enzyme 2, NAD(+)-dependent, mitochondrial | protein-coding |
| chrX_101273422_101273703 | chrX | 101273423 | 101273703 | + | -0.627189 | promoter-TSS (NM_021521) | promoter-TSS (NM_021521) | -528 | NM_021521 | 59024 | Mm.20873 | NM_021521 | ENSMUSG00000079487 | **Med12** | 230kDa\|Mopa\|OPA-1\|Tnrc11\|Trap230 | mediator complex subunit 12 | protein-coding |
| chr5_118709919_118710315 | chr5 | 118709920 | 118710315 | + | -0.725372 | intron (NM_172424, intron 4 of 30) | intron (NM_172424, intron 4 of 30) | 149398 | NM_172424 | 76199 | Mm.206238 | NM_172424 | ENSMUSG00000018076 | **Med13l** | 2210413I17Rik\|6330591G05Rik\|9030618F05Rik\|Thrap2\|Trap240L\|mKIAA1025 | mediator complex subunit 13-like | protein-coding |
| chr5_118601497_118601905 | chr5 | 118601498 | 118601905 | + | -0.970894 | intron (NM_172424, intron 2 of 30) | intron (NM_172424, intron 2 of 30) | 40982 | NM_172424 | 76199 | Mm.206238 | NM_172424 | ENSMUSG00000018076 | **Med13l** | 2210413I17Rik\|6330591G05Rik\|9030618F05Rik\|Thrap2\|Trap240L\|mKIAA1025 | mediator complex subunit 13-like | protein-coding |
| chr5_118614583_118614889 | chr5 | 118614584 | 118614889 | + | -1.11188 | intron (NM_172424, intron 2 of 30) | intron (NM_172424, intron 2 of 30) | 54017 | NM_172424 | 76199 | Mm.206238 | NM_172424 | ENSMUSG00000018076 | **Med13l** | 2210413I17Rik\|6330591G05Rik\|9030618F05Rik\|Thrap2\|Trap240L\|mKIAA1025 | mediator complex subunit 13-like | protein-coding |
| chr15_52705218_52705395 | chr15 | 52705219 | 52705395 | + | -0.745377 | Intergenic | Intergenic | -7138 | NM_027212 | 69790 | Mm.195562 | NM_027212 | ENSMUSG00000038622 | **Med30** | 1810038N03Rik\|2510044J04Rik\|TRAP25\|Thrap6 | mediator complex subunit 30 | protein-coding |
| chr15_52705183_52705421 | chr15 | 52705184 | 52705421 | + | -0.78163 | Intergenic | Intergenic | -7143 | NM_027212 | 69790 | Mm.195562 | NM_027212 | ENSMUSG00000038622 | **Med30** | 1810038N03Rik\|2510044J04Rik\|TRAP25\|Thrap6 | mediator complex subunit 30 | protein-coding |
| chr15_52753659_52754045 | chr15 | 52753660 | 52754045 | + | -0.795335 | Intergenic | Intergenic | 41407 | NM_027212 | 69790 | Mm.195562 | NM_027212 | ENSMUSG00000038622 | **Med30** | 1810038N03Rik\|2510044J04Rik\|TRAP25\|Thrap6 | mediator complex subunit 30 | protein-coding |
| chr15_52752874_52753608 | chr15 | 52752875 | 52753608 | + | -0.830302 | Intergenic | ORR1B1\|LTR\|ERVL-MaLR | 40796 | NM_027212 | 69790 | Mm.195562 | NM_027212 | ENSMUSG00000038622 | **Med30** | 1810038N03Rik\|2510044J04Rik\|TRAP25\|Thrap6 | mediator complex subunit 30 | protein-coding |
| chr15_52753306_52754045 | chr15 | 52753307 | 52754045 | + | -0.945442 | Intergenic | Intergenic | 41231 | NM_027212 | 69790 | Mm.195562 | NM_027212 | ENSMUSG00000038622 | **Med30** | 1810038N03Rik\|2510044J04Rik\|TRAP25\|Thrap6 | mediator complex subunit 30 | protein-coding |
| chr15_52753267_52753608 | chr15 | 52753268 | 52753608 | + | -0.95745 | Intergenic | ORR1B1\|LTR\|ERVL-MaLR | 40993 | NM_027212 | 69790 | Mm.195562 | NM_027212 | ENSMUSG00000038622 | **Med30** | 1810038N03Rik\|2510044J04Rik\|TRAP25\|Thrap6 | mediator complex subunit 30 | protein-coding |
| chr15_52749906_52750312 | chr15 | 52749907 | 52750312 | + | -1.06969 | Intergenic | Intergenic | 37664 | NM_027212 | 69790 | Mm.195562 | NM_027212 | ENSMUSG00000038622 | **Med30** | 1810038N03Rik\|2510044J04Rik\|TRAP25\|Thrap6 | mediator complex subunit 30 | protein-coding |
| chr11_46420866_46421937 | chr11 | 46420867 | 46421937 | + | -0.65078 | intron (NR_105955, intron 2 of 4) | intron (NR_105955, intron 2 of 4) | -15545 | NM_001104557 | 66213 | Mm.24159 | NM_025426 | ENSMUSG00000020397 | **Med7** | 1110063B05Rik\|Crsp33\|Crsp9 | mediator complex subunit 7 | protein-coding |
| chr7_67419588_67420095 | chr7 | 67419589 | 67420095 | + | -0.692431 | Intergenic | Intergenic | -46984 | NM_001291195 | 17258 | Mm.132788 | NM_001033713 | ENSMUSG00000030557 | **Mef2a** | A430079H05Rik | myocyte enhancer factor 2A | protein-coding |
| chr7_67407012_67407387 | chr7 | 67407013 | 67407387 | + | -0.907254 | Intergenic | ORR1D2\|LTR\|ERVL-MaLR | -34342 | NM_001291195 | 17258 | Mm.132788 | NM_001033713 | ENSMUSG00000030557 | **Mef2a** | A430079H05Rik | myocyte enhancer factor 2A | protein-coding |
| chr3_88138290_88139039 | chr3 | 88138291 | 88139039 | + | -0.604157 | Intergenic | Intergenic | -3730 | NM_133665 | 17261 | Mm.28184 | NM_133665 | ENSMUSG00000001419 | **Mef2d** | C80750 | myocyte enhancer factor 2D | protein-coding |
| chr3_88138671_88139039 | chr3 | 88138672 | 88139039 | + | -0.774667 | Intergenic | Intergenic | -3540 | NM_133665 | 17261 | Mm.28184 | NM_133665 | ENSMUSG00000001419 | **Mef2d** | C80750 | myocyte enhancer factor 2D | protein-coding |
| chr11_121697149_121697743 | chr11 | 121697150 | 121697743 | + | -0.997873 | Intergenic | Intergenic | -4981 | NM_144797 | 210029 | Mm.153566 | NM_144797 | | **Metrnl** | 9430048M07Rik\|BC019776 | meteorin, glial cell differentiation regulator-like | protein-coding |
| chr3_123334825_123335233 | chr3 | 123334826 | 123335233 | + | -0.811511 | intron (NM_027135, intron 10 of 22) | intron (NM_027135, intron 10 of 22) | 50961 | NM_201638 | 210529 | Mm.260039 | NM_201638 | ENSMUSG00000028114 | **Mettl14** | G430022H21Rik\|mKIAA1627 | methyltransferase like 14 | protein-coding |
| chr16_8455050_8455544 | chr16 | 8455051 | 8455544 | + | -0.673647 | Intergenic | Intergenic | -15516 | NM_146247 | 239706 | Mm.214953 | NM_146247 | | **Mettl22** | - | methyltransferase like 22 | protein-coding |
| chr2_71046839_71047375 | chr2 | 71046840 | 71047375 | + | -0.771589 | intron (NM_145524, intron 1 of 9) | intron (NM_145524, intron 1 of 9) | 8499 | NM_145524 | 228019 | Mm.490343 | NM_145524 | ENSMUSG00000041975 | **Mettl8** | BC004636\|Tip | methyltransferase like 8 | protein-coding |
| chr2_70930716_70931273 | chr2 | 70930717 | 70931273 | + | -0.790865 | Intergenic | Intergenic | 87379 | NM_001110512 | 228019 | Mm.490343 | NM_145524 | ENSMUSG00000041975 | **Mettl8** | BC004636\|Tip | methyltransferase like 8 | protein-coding |
| chr2_70967278_70967548 | chr2 | 70967279 | 70967548 | + | -0.825252 | intron (NM_001110512, intron 6 of 8) | intron (NM_001110512, intron 6 of 8) | 50961 | NM_001110512 | 228019 | Mm.490343 | NM_145524 | ENSMUSG00000041975 | **Mettl8** | BC004636\|Tip | methyltransferase like 8 | protein-coding |
| chr2_70931005_70931236 | chr2 | 70931006 | 70931236 | + | -0.829514 | Intergenic | Intergenic | 87253 | NM_001110512 | 228019 | Mm.490343 | NM_145524 | ENSMUSG00000041975 | **Mettl8** | BC004636\|Tip | methyltransferase like 8 | protein-coding |
| chr2_70967186_70967532 | chr2 | 70967187 | 70967532 | + | -0.843512 | intron (NM_001110512, intron 6 of 8) | intron (NM_001110512, intron 6 of 8) | 51015 | NM_001110512 | 228019 | Mm.490343 | NM_145524 | ENSMUSG00000041975 | **Mettl8** | BC004636\|Tip | methyltransferase like 8 | protein-coding |
| chr2_70922661_70923040 | chr2 | 70922662 | 70923040 | + | -0.923296 | Intergenic | Intergenic | 95523 | NM_001110512 | 228019 | Mm.490343 | NM_145524 | ENSMUSG00000041975 | **Mettl8** | BC004636\|Tip | methyltransferase like 8 | protein-coding |

| | | | | | | | | | | | | | | | | | |
|---|---|---|---|---|---|---|---|---|---|---|---|---|---|---|---|---|---|
| chr2_71040827_71041356 | chr2 | 71040828 | 71041356 | + | -1.01975 | intron (NM_145524, intron 1 of 9) | B3\|SINE\|B2 | 14514 | NM_145524 | 228019 | Mm.490343 | NM_145524 | ENSMUSG00000041975 | **Mettl8** | BC004636\|Tip | methyltransferase like 8 | protein-coding |
| chr2_71040775_71041581 | chr2 | 71040776 | 71041581 | + | -1.07656 | intron (NM_145524, intron 1 of 9) | intron (NM_145524, intron 1 of 9) | 14428 | NM_145524 | 228019 | Mm.490343 | NM_145524 | ENSMUSG00000041975 | **Mettl8** | BC004636\|Tip | methyltransferase like 8 | protein-coding |
| chr7_121043531_121043882 | chr7 | 121043532 | 121043882 | + | -1.04804 | intron (NM_021554, intron 1 of 4) | intron (NM_021554, intron 1 of 4) | 9262 | NM_021554 | 59052 | Mm.29122 | NM_021554 | ENSMUSG00000030876 | **Mettl9** | 0610012D09Rik\|AA517660\|Drev | methyltransferase like 9 | protein-coding |
| chr7_82890224_82890652 | chr7 | 82890225 | 82890652 | + | -0.615632 | Intergenic | ORR1D2\|LTR\|ERVL-MaLR | 23105 | NM_175366 | 108797 | Mm.306789 | NM_175366 | ENSMUSG00000057706 | **Mex3b** | 4931439A04Rik\|M-04814\|Rkhd3\|mKIAA2009 | mex3 homolog B (C. elegans) | protein-coding |
| chr7_82868761_82868945 | chr7 | 82868762 | 82868945 | + | -0.742363 | exon (NM_175366, exon 2 of 2) | exon (NM_175366, exon 2 of 2) | 1520 | NM_175366 | 108797 | Mm.306789 | NM_175366 | ENSMUSG00000057706 | **Mex3b** | 4931439A04Rik\|M-04814\|Rkhd3\|mKIAA2009 | mex3 homolog B (C. elegans) | protein-coding |
| chr7_79190591_79191099 | chr7 | 79190592 | 79191099 | + | -0.948782 | Intergenic | Intergenic | -41785 | NM_008594 | 17304 | Mm.1451 | NM_008594 | ENSMUSG00000030605 | **Mfge8** | AA408458\|AI325141\|MFG-E8\|MP47\|Mfgm\|P47\|SED1\|lactadherin | milk fat globule-EGF factor 8 protein | protein-coding |
| chr1_132038625_132039197 | chr1 | 132038626 | 132039197 | + | -0.887107 | 3' UTR (NM_001114662, exon 10 of 10) | 3' UTR (NM_001114662, exon 10 of 10) | 29151 | NM_172510 | 213006 | Mm.208445 | NM_172510 | ENSMUSG00000059149 | **Mfsd4** | A230072B04\|A930031D07Rik\|AI850289 | major facilitator superfamily domain containing 4 | protein-coding |
| chr1_52726309_52727815 | chr1 | 52726310 | 52727815 | + | -0.788185 | 5' UTR (NM_178081, exon 1 of 6) | 5' UTR (NM_178081, exon 1 of 6) | 256 | NM_178081 | 98682 | Mm.491173 | NM_133829 | ENSMUSG00000041439 | **Mfsd6** | 2210010L05Rik\|9630025I22Rik\|AI836200\|AW212394\|MMR2 | major facilitator superfamily domain containing 6 | protein-coding |
| chr1_52753769_52754625 | chr1 | 52753770 | 52754625 | + | -0.988796 | Intergenic | Intergenic | -26879 | NM_133829 | 98682 | Mm.491173 | NM_133829 | ENSMUSG00000041439 | **Mfsd6** | 2210010L05Rik\|9630025I22Rik\|AI836200\|AW212394\|MMR2 | major facilitator superfamily domain containing 6 | protein-coding |
| chr1_52753788_52754318 | chr1 | 52753789 | 52754318 | + | -1.00818 | Intergenic | Intergenic | -26735 | NM_133829 | 98682 | Mm.491173 | NM_133829 | ENSMUSG00000041439 | **Mfsd6** | 2210010L05Rik\|9630025I22Rik\|AI836200\|AW212394\|MMR2 | major facilitator superfamily domain containing 6 | protein-coding |
| chr15_80191901_80192726 | chr15 | 80191902 | 80192726 | + | -0.98815 | intron (NM_010795, intron 1 of 1) | intron (NM_010795, intron 1 of 1) | 18593 | NM_010795 | 17309 | Mm.299693 | NM_010795 | ENSMUSG00000042428 | **Mgat3** | 1110038J12Rik\|AI848272\|GnT-III | mannoside acetylglucosaminyltransferase 3 | protein-coding |
| chr1_127319881_127320367 | chr1 | 127319882 | 127320367 | + | -0.656879 | intron (NM_145128, intron 3 of 17) | B4A\|SINE\|B4 | 115138 | NM_145128 | 107895 | Mm.214682 | NM_145128 | ENSMUSG00000036155 | **Mgat5** | 4930471A21Rik\|5330407H02Rik\|AI480971\|GlcNAc-TV | mannoside acetylglucosaminyltransferase 5 | protein-coding |
| chr1_127230562_127230854 | chr1 | 127230563 | 127230854 | + | -0.72544 | intron (NM_145128, intron 1 of 17) | L1MB8\|LINE\|L1 | 25722 | NM_145128 | 107895 | Mm.214682 | NM_145128 | ENSMUSG00000036155 | **Mgat5** | 4930471A21Rik\|5330407H02Rik\|AI480971\|GlcNAc-TV | mannoside acetylglucosaminyltransferase 5 | protein-coding |
| chr1_127212252_127212615 | chr1 | 127212253 | 127212615 | + | -0.884366 | intron (NM_145128, intron 1 of 17) | ID_B1\|SINE\|B4 | 7448 | NM_145128 | 107895 | Mm.214682 | NM_145128 | ENSMUSG00000036155 | **Mgat5** | 4930471A21Rik\|5330407H02Rik\|AI480971\|GlcNAc-TV | mannoside acetylglucosaminyltransferase 5 | protein-coding |
| chr1_127322295_127322905 | chr1 | 127322296 | 127322905 | + | -1.1163 | intron (NM_145128, intron 4 of 17) | B4A\|SINE\|B4 | 117614 | NM_145128 | 107895 | Mm.214682 | NM_145128 | ENSMUSG00000036155 | **Mgat5** | 4930471A21Rik\|5330407H02Rik\|AI480971\|GlcNAc-TV | mannoside acetylglucosaminyltransferase 5 | protein-coding |
| chr1_127214674_127215247 | chr1 | 127214675 | 127215247 | + | -1.40597 | intron (NM_145128, intron 1 of 17) | intron (NM_145128, intron 1 of 17) | 9975 | NM_145128 | 107895 | Mm.214682 | NM_145128 | ENSMUSG00000036155 | **Mgat5** | 4930471A21Rik\|5330407H02Rik\|AI480971\|GlcNAc-TV | mannoside acetylglucosaminyltransferase 5 | protein-coding |
| chr1_127214952_127215112 | chr1 | 127214953 | 127215112 | + | -1.45218 | intron (NM_145128, intron 1 of 17) | intron (NM_145128, intron 1 of 17) | 10046 | NM_145128 | 107895 | Mm.214682 | NM_145128 | ENSMUSG00000036155 | **Mgat5** | 4930471A21Rik\|5330407H02Rik\|AI480971\|GlcNAc-TV | mannoside acetylglucosaminyltransferase 5 | protein-coding |
| chr19_45780511_45780970 | chr19 | 45780512 | 45780970 | + | -0.677881 | intron (NM_023799, intron 1 of 15) | B1_Mur3\|SINE\|Alu | 2550 | NM_023799 | 76055 | Mm.122725 | NM_023799 | ENSMUSG00000025220 | **Mgea5** | 2810009A20Rik\|4833427O07Rik\|5830447M11Rik\|AA408215\|Hy5\|Ncoat\|mKIAA0679 | meningioma expressed antigen 5 (hyaluronidase) | protein-coding |
| chr19_45795717_45797250 | chr19 | 45795718 | 45797250 | + | -0.859767 | intron (NM_145703, intron 2 of 9) | intron (NM_145703, intron 2 of 9) | -13193 | NM_023799 | 76055 | Mm.122725 | NM_023799 | ENSMUSG00000025220 | **Mgea5** | 2810009A20Rik\|4833427O07Rik\|5830447M11Rik\|AA408215\|Hy5\|Ncoat\|mKIAA0679 | meningioma expressed antigen 5 (hyaluronidase) | protein-coding |
| chr19_45795969_45796698 | chr19 | 45795970 | 45796698 | + | -1.00009 | intron (NM_145703, intron 2 of 9) | intron (NM_145703, intron 2 of 9) | -13043 | NM_023799 | 76055 | Mm.122725 | NM_023799 | ENSMUSG00000025220 | **Mgea5** | 2810009A20Rik\|4833427O07Rik\|5830447M11Rik\|AA408215\|Hy5\|Ncoat\|mKIAA0679 | meningioma expressed antigen 5 (hyaluronidase) | protein-coding |
| chr19_45796243_45796751 | chr19 | 45796244 | 45796751 | + | -1.09943 | intron (NM_145703, intron 2 of 9) | intron (NM_145703, intron 2 of 9) | -13206 | NM_023799 | 76055 | Mm.122725 | NM_023799 | ENSMUSG00000025220 | **Mgea5** | 2810009A20Rik\|4833427O07Rik\|5830447M11Rik\|AA408215\|Hy5\|Ncoat\|mKIAA0679 | meningioma expressed antigen 5 (hyaluronidase) | protein-coding |
| chr2_144275660_144276139 | chr2 | 144275661 | 144276139 | + | -1.29225 | intron (NM_001289631, intron 2 of 4) | B1_Mur2\|SINE\|Alu | 4996 | NM_001289630 | 74528 | Mm.386817 | NM_028984 | ENSMUSG00000027424 | **Mgme1** | 8430406I07Rik\|AI426476 | mitochondrial genome maintainance exonuclease 1 | protein-coding |
| chr3_51654691_51655049 | chr3 | 51654692 | 51655049 | + | -1.03008 | Intergenic | Intergenic | -6323 | NM_174995 | 211666 | Mm.24679 | NM_174995 | ENSMUSG00000074604 | **Mgst2** | GST2\|MGST-II | microsomal glutathione S-transferase 2 | protein-coding |
| chr10_41474088_41474305 | chr10 | 41474089 | 41474305 | + | -0.679281 | Intergenic | Intergenic | -2117 | NM_001164433 | 171580 | Mm.290431 | NM_138315 | ENSMUSG00000019823 | **Mical1** | MICAL\|MICAL-1\|Nical | microtubule associated monooxygenase, calponin and LIM domain containing 1 | protein-coding |
| chr6_121062771_121063487 | chr6 | 121062772 | 121063487 | + | -0.668691 | intron (NM_001270475, intron 1 of 31) | intron (NM_001270475, intron 1 of 31) | 18498 | NM_153396 | 194401 | Mm.122399 | NM_153396 | ENSMUSG00000003178 | **Mical3** | C130040D16Rik\|Mical-3\|mKIAA1364 | microtubule associated monooxygenase, calponin and LIM domain containing 3 | protein-coding |
| chr10_59692481_59692771 | chr10 | 59692482 | 59692771 | + | -0.788113 | Intergenic | Intergenic | -9937 | NM_001291442 | 216001 | Mm.27199 | NM_144822 | ENSMUSG00000020111 | **Micu1** | C730016L05Rik\|Calc\|Cbara1 | mitochondrial calcium uptake 1 | protein-coding |
| chr10_59698980_59699226 | chr10 | 59698981 | 59699226 | + | -0.794046 | Intergenic | Intergenic | -3460 | NM_001291442 | 216001 | Mm.27199 | NM_144822 | ENSMUSG00000020111 | **Micu1** | C730016L05Rik\|Calc\|Cbara1 | mitochondrial calcium uptake 1 | protein-coding |
| chr10_59748714_59749039 | chr10 | 59748715 | 59749039 | + | -0.800952 | intron (NM_001291443, intron 4 of 12) | L1MEf\|LINE\|L1 | 46314 | NM_001291443 | 216001 | Mm.27199 | NM_144822 | ENSMUSG00000020111 | **Micu1** | C730016L05Rik\|Calc\|Cbara1 | mitochondrial calcium uptake 1 | protein-coding |
| chr4_103120131_103120292 | chr4 | 103120132 | 103120292 | + | -0.667315 | intron (NM_001039081, intron 1 of 12) | ID4\|SINE\|ID | 822 | NM_001039081 | 71148 | Mm.288721 | NM_027696 | ENSMUSG00000028522 | **Mier1** | 4933425I22Rik\|5830411K19Rik\|er1 | mesoderm induction early response 1 homolog (Xenopus laevis | protein-coding |
| chr4_103120099_103120357 | chr4 | 103120100 | 103120357 | + | -0.681685 | intron (NM_001039081, intron 1 of 12) | ID4\|SINE\|ID | 838 | NM_001039081 | 71148 | Mm.288721 | NM_027696 | ENSMUSG00000028522 | **Mier1** | 4933425I22Rik\|5830411K19Rik\|er1 | mesoderm induction early response 1 homolog (Xenopus laevis | protein-coding |
| chr10_75848445_75848705 | chr10 | 75848446 | 75848705 | + | -0.679412 | Intergenic | RLTR9A\|LTR\|ERVK | 11675 | NM_010798 | 17319 | Mm.2326 | NM_010798 | ENSMUSG00000033307 | **Mif** | GIF\|Glif | macrophage migration inhibitory factor | protein-coding |
| chr11_106752079_106752598 | chr11 | 106752080 | 106752598 | + | -0.638486 | intron (NR_073165, intron 2 of 9) | intron (NR_073165, intron 2 of 9) | 1113 | NM_001033435 | 380732 | Mm.343952 | NM_001033435 | ENSMUSG00000040528 | **Milr1** | Allergin-1\|Gm885\|Mca32 | mast cell immunoglobulin like receptor 1 | protein-coding |
| chr11_106750027_106750813 | chr11 | 106750028 | 106750813 | + | -0.811553 | promoter-TSS (NM_001033435) | promoter-TSS (NM_001033435) | -806 | NM_001271375 | 380732 | Mm.343952 | NM_001033435 | ENSMUSG00000040528 | **Milr1** | Allergin-1\|Gm885\|Mca32 | mast cell immunoglobulin like receptor 1 | protein-coding |
| chr4_101320585_101321001 | chr4 | 101320586 | 101321001 | + | -0.704981 | Intergenic | B3\|SINE\|B2 | 26234 | NR_029537 | 387143 | | NR_029537 | miRBase:MI0000148 | **Mir101a** | Mirn101\|Mirn101a\|miR-101 | microRNA 101a | ncRNA |
| chr4_101328960_101329600 | chr4 | 101328961 | 101329600 | + | -1.1322 | Intergenic | MTD\|LTR\|ERVL-MaLR | 17747 | NR_029537 | 387143 | | NR_029537 | miRBase:MI0000148 | **Mir101a** | Mirn101\|Mirn101a\|miR-101 | microRNA 101a | ncRNA |
| chr18_65301515_65301879 | chr18 | 65301516 | 65301879 | + | -1.21524 | intron (NM_001037294, intron 5 of 12) | intron (NM_001037294, intron 5 of 12) | 52836 | NR_029600 | 387231 | | NR_029600 | miRBase:MI0000256 | **Mir122a** | Mirn122a\|mmu-mir-122 | microRNA 122a | ncRNA |
| chr18_65312136_65312860 | chr18 | 65312137 | 65312860 | + | -1.26467 | intron (NM_001037294, intron 4 of 12) | intron (NM_001037294, intron 4 of 12) | 63637 | NR_029600 | 387231 | | NR_029600 | miRBase:MI0000256 | **Mir122a** | Mirn122a\|mmu-mir-122 | microRNA 122a | ncRNA |
| chr18_56507397_56507750 | chr18 | 56507398 | 56507750 | + | -0.975022 | Intergenic | Intergenic | -30565 | NR_106160 | 102465790 | | NR_106160 | miRBase:MI0025040 | **Mir1258** | mmu-mir-1258 | microRNA mir-1258 | ncRNA |
| chr18_56505507_56505839 | chr18 | 56505508 | 56505839 | + | -1.13475 | Intergenic | Intergenic | -32466 | NR_106160 | 102465790 | | NR_106160 | miRBase:MI0025040 | **Mir1258** | mmu-mir-1258 | microRNA mir-1258 | ncRNA |
| chr18_56504283_56504776 | chr18 | 56504284 | 56504776 | + | -1.18475 | TTS (NM_026240) | TTS (NM_026240) | -33609 | NR_106160 | 102465790 | | NR_106160 | miRBase:MI0025040 | **Mir1258** | mmu-mir-1258 | microRNA mir-1258 | ncRNA |
| chr18_56489940_56490223 | chr18 | 56489941 | 56490223 | + | -1.25617 | intron (NM_026240, intron 10 of 13) | ORR1B1\|LTR\|ERVL-MaLR | -48057 | NR_106160 | 102465790 | | NR_106160 | miRBase:MI0025040 | **Mir1258** | mmu-mir-1258 | microRNA mir-1258 | ncRNA |
| chr18_56489797_56490223 | chr18 | 56489798 | 56490223 | + | -1.27487 | intron (NM_026240, intron 10 of 13) | ORR1B1\|LTR\|ERVL-MaLR | -48129 | NR_106160 | 102465790 | | NR_106160 | miRBase:MI0025040 | **Mir1258** | mmu-mir-1258 | microRNA mir-1258 | ncRNA |
| chr18_56504305_56505231 | chr18 | 56504306 | 56505231 | + | -1.31319 | TTS (NM_026240) | TTS (NM_026240) | -33371 | NR_106160 | 102465790 | | NR_106160 | miRBase:MI0025040 | **Mir1258** | mmu-mir-1258 | microRNA mir-1258 | ncRNA |
| chr9_112100877_112101246 | chr9 | 112100878 | 112101246 | + | -0.994516 | intron (NM_001177618, intron 17 of 18) | intron (NM_001177618, intron 17 of 18) | 17649 | NR_029823 | 723815 | | NR_029823 | | **Mir128-2** | Mirn128-2\|Mirn128b\|mir-128b | microRNA 128-2 | ncRNA |
| chr7_101466101_101467148 | chr7 | 101466102 | 101467148 | + | -0.970261 | intron (NM_001243757, intron 2 of 30) | intron (NM_001243757, intron 2 of 30) | -8751 | NR_029791 | 387157 | | NR_029791 | | **Mir139** | Mirn139\|mmu-mir-139 | microRNA 139 | ncRNA |
| chr7_101486885_101487456 | chr7 | 101486886 | 101487456 | + | -0.97933 | intron (NM_001243757, intron 4 of 30) | intron (NM_001243757, intron 4 of 30) | 11795 | NR_029791 | 387157 | | NR_029791 | | **Mir139** | Mirn139\|mmu-mir-139 | microRNA 139 | ncRNA |
| chr11_78061396_78061863 | chr11 | 78061397 | 78061863 | + | -1.19795 | Intergenic | PB1D7\|SINE\|Alu | -11375 | NR_029556 | 387162 | | NR_029556 | miRBase:MI0000168 | **Mir144** | Mirn144\|mmu-mir-144 | microRNA 144 | ncRNA |
| chr18_69051005_69051567 | chr18 | 69051006 | 69051567 | + | -1.05743 | Intergenic | Intergenic | -29013 | NR_105780 | 102465187 | | NR_105780 | miRBase:MI0021890 | **Mir145b** | mmu-mir-145b | microRNA mir-145b | ncRNA |
| chr7_45121196_45121854 | chr7 | 45121197 | 45121854 | + | -1.1138 | promoter-TSS (NR_029560) | promoter-TSS (NR_029560) | -232 | NR_029560 | 387168 | | NR_029560 | miRBase:MI0000172 | **Mir150** | Mirn150\|mmu-mir-150 | microRNA 150 | ncRNA |

| | | | | | | | | | | | | | | | | | |
|---|---|---|---|---|---|---|---|---|---|---|---|---|---|---|---|---|---|
| chr7_45121376_45121782 | chr7 | 45121377 | 45121782 | + | -1.15531 | promoter-TSS (NR_029560) | promoter-TSS (NR_029560) | -178 | NR_029560 | 387168 | | NR_029560 | miRBase:MI0000172 | **Mir150** | Mirn150\|mmu-mir-150 | microRNA 150 | ncRNA |
| chr1_137814665_137814926 | chr1 | 137814666 | 137814926 | + | -0.67805 | Intergenic | Intergenic | -151659 | NR_029795 | 735252 | | NR_029795 | miRBase:MI0000697 | **Mir181a-1** | Mirn181a-1\|Mirn213\|miR-213\|mmu-mir-213 | microRNA 181a-1 | ncRNA |
| chr18_6377493_6377946 | chr18 | 6377494 | 6377946 | + | -0.658999 | Intergenic | Intergenic | 112926 | NR_035446 | 100316773 | | NR_035446 | | **Mir1893** | Mirn1893\|mmu-mir-1893 | microRNA 1893 | ncRNA |
| chr18_6410994_6411364 | chr18 | 6410995 | 6411364 | + | -0.675788 | Intergenic | Intergenic | 79467 | NR_035446 | 100316773 | | NR_035446 | | **Mir1893** | Mirn1893\|mmu-mir-1893 | microRNA 1893 | ncRNA |
| chr18_6484549_6485131 | chr18 | 6484550 | 6485131 | + | -0.989954 | intron (NM_007935, intron 1 of 13) | intron (NM_007935, intron 1 of 13) | 5806 | NR_035446 | 100316773 | | NR_035446 | | **Mir1893** | Mirn1893\|mmu-mir-1893 | microRNA 1893 | ncRNA |
| chr18_11935794_11936337 | chr18 | 11935795 | 11936337 | + | -0.704337 | intron (NM_022021, intron 6 of 8) | intron (NM_022021, intron 6 of 8) | -95628 | NR_035447 | 100316686 | | NR_035447 | miRBase:MI0008328 | **Mir1901** | Mirn1901\|mmu-mir-1901 | microRNA 1901 | ncRNA |
| chr8_128433396_128433735 | chr8 | 128433397 | 128433735 | + | -0.757723 | intron (NM_008737, intron 5 of 16) | ID_B1\|SINE\|B4 | 74325 | NR_035436 | 100316680 | | NR_035436 | miRBase:MI0008317 | **Mir1903** | Mirn1903\|mmu-mir-1903 | microRNA 1903 | ncRNA |
| chr13_109907558_109907966 | chr13 | 109907559 | 109907966 | + | -0.815176 | intron (NM_011056, intron 8 of 16) | intron (NM_011056, intron 8 of 16) | 3953 | NR_035442 | 100316683 | | NR_035442 | | **Mir1904** | Mirn1904\|mmu-mir-1904 | microRNA 1904 | ncRNA |
| chr13_109824609_109824780 | chr13 | 109824610 | 109824780 | + | -1.10843 | intron (NM_011056, intron 7 of 16) | intron (NM_011056, intron 7 of 16) | -79114 | NR_035442 | 100316683 | | NR_035442 | | **Mir1904** | Mirn1904\|mmu-mir-1904 | microRNA 1904 | ncRNA |
| chr13_109820197_109820413 | chr13 | 109820198 | 109820413 | + | -1.19681 | intron (NM_011056, intron 7 of 16) | MIRb\|SINE\|MIR | -83504 | NR_035442 | 100316683 | | NR_035442 | | **Mir1904** | Mirn1904\|mmu-mir-1904 | microRNA 1904 | ncRNA |
| chr13_109837346_109838369 | chr13 | 109837347 | 109838369 | + | -1.29384 | intron (NM_011056, intron 7 of 16) | CT-rich\|Low_complexity\|Low_complexity | -65951 | NR_035442 | 100316683 | | NR_035442 | | **Mir1904** | Mirn1904\|mmu-mir-1904 | microRNA 1904 | ncRNA |
| chr15_50687231_50687622 | chr15 | 50687232 | 50687622 | + | -0.663642 | intron (NM_032000, intron 4 of 5) | intron (NM_032000, intron 4 of 5) | 201687 | NR_035444 | 100316685 | | NR_035444 | miRBase:MI0008325 | **Mir1907** | Mirn1907\|mmu-mir-1907 | microRNA 1907 | ncRNA |
| chr15_50402598_50402975 | chr15 | 50402599 | 50402975 | + | -0.830297 | Intergenic | Intergenic | 486327 | NR_035444 | 100316685 | | NR_035444 | miRBase:MI0008325 | **Mir1907** | Mirn1907\|mmu-mir-1907 | microRNA 1907 | ncRNA |
| chr15_50402471_50402953 | chr15 | 50402472 | 50402953 | + | -0.836536 | Intergenic | Intergenic | 486402 | NR_035444 | 100316685 | | NR_035444 | miRBase:MI0008325 | **Mir1907** | Mirn1907\|mmu-mir-1907 | microRNA 1907 | ncRNA |
| chr15_50836584_50837115 | chr15 | 50836585 | 50837115 | + | -0.907493 | intron (NM_032000, intron 2 of 5) | intron (NM_032000, intron 2 of 5) | 52264 | NR_035444 | 100316685 | | NR_035444 | miRBase:MI0008325 | **Mir1907** | Mirn1907\|mmu-mir-1907 | microRNA 1907 | ncRNA |
| chr15_50841687_50842060 | chr15 | 50841688 | 50842060 | + | -0.995932 | intron (NM_032000, intron 2 of 5) | intron (NM_032000, intron 2 of 5) | 47240 | NR_035444 | 100316685 | | NR_035444 | miRBase:MI0008325 | **Mir1907** | Mirn1907\|mmu-mir-1907 | microRNA 1907 | ncRNA |
| chr15_50841524_50841996 | chr15 | 50841525 | 50841996 | + | -1.05514 | intron (NM_032000, intron 2 of 5) | intron (NM_032000, intron 2 of 5) | 47354 | NR_035444 | 100316685 | | NR_035444 | miRBase:MI0008325 | **Mir1907** | Mirn1907\|mmu-mir-1907 | microRNA 1907 | ncRNA |
| chr15_49899872_49900170 | chr15 | 49899873 | 49900170 | + | -1.18671 | Intergenic | Intergenic | 989093 | NR_035444 | 100316685 | | NR_035444 | miRBase:MI0008325 | **Mir1907** | Mirn1907\|mmu-mir-1907 | microRNA 1907 | ncRNA |
| chr5_65795252_65795690 | chr5 | 65795253 | 65795690 | + | -1.21385 | intron (NM_001024917, intron 8 of 19) | intron (NM_001024917, intron 8 of 19) | -25230 | NR_035449 | 100316687 | | NR_035449 | miRBase:MI0009917 | **Mir1928** | mmu-mir-1928 | microRNA 1928 | ncRNA |
| chr10_93205310_93206078 | chr10 | 93205311 | 93206078 | + | -0.732675 | intron (NM_146239, intron 1 of 16) | intron (NM_146239, intron 1 of 16) | 42909 | NR_035452 | 100316689 | | NR_035452 | | **Mir1931** | mmu-mir-1931 | microRNA 1931 | ncRNA |
| chr10_93171175_93171527 | chr10 | 93171176 | 93171527 | + | -0.76065 | intron (NM_146239, intron 1 of 16) | ORR1E\|LTR\|ERVL-MaLR | 8566 | NR_035452 | 100316689 | | NR_035452 | | **Mir1931** | mmu-mir-1931 | microRNA 1931 | ncRNA |
| chr10_93171165_93171835 | chr10 | 93171166 | 93171835 | + | -0.781073 | intron (NM_146239, intron 1 of 16) | intron (NM_146239, intron 1 of 16) | 8715 | NR_035452 | 100316689 | | NR_035452 | | **Mir1931** | mmu-mir-1931 | microRNA 1931 | ncRNA |
| chr10_93205324_93205880 | chr10 | 93205325 | 93205880 | + | -0.816192 | intron (NM_146239, intron 1 of 16) | MER77\|LTR\|ERVL | 42817 | NR_035452 | 100316689 | | NR_035452 | | **Mir1931** | mmu-mir-1931 | microRNA 1931 | ncRNA |
| chr11_119384888_119385333 | chr11 | 119384889 | 119385333 | + | -0.661986 | Intergenic | MLTR13\|LTR\|ERVK | -5361 | NR_035453 | 100316690 | | NR_035453 | miRBase:MI0009921 | **Mir1932** | mmu-mir-1932 | microRNA 1932 | ncRNA |
| chr11_21303472_21303999 | chr11 | 21303473 | 21303999 | + | -0.968179 | intron (NM_001290628, intron 12 of 22) | MLT1J\|LTR\|ERVL-MaLR | 40939 | NR_035454 | 100316753 | | NR_035454 | | **Mir1933** | mmu-mir-1933 | microRNA 1933 | ncRNA |
| chr11_21303659_21303956 | chr11 | 21303660 | 21303956 | + | -0.99816 | intron (NM_001290628, intron 12 of 22) | intron (NM_001290628, intron 12 of 22) | 40867 | NR_035454 | 100316753 | | NR_035454 | | **Mir1933** | mmu-mir-1933 | microRNA 1933 | ncRNA |
| chr11_21299221_21299622 | chr11 | 21299222 | 21299622 | + | -1.22992 | intron (NM_001290628, intron 11 of 22) | L2b\|LINE\|L2 | 45253 | NR_035454 | 100316753 | | NR_035454 | | **Mir1933** | mmu-mir-1933 | microRNA 1933 | ncRNA |
| chr12_102643290_102643626 | chr12 | 102643291 | 102643626 | + | -0.631097 | intron (NM_172584, intron 2 of 9) | intron (NM_172584, intron 2 of 9) | 41562 | NR_035457 | 100316812 | | NR_035457 | | **Mir1936** | mmu-mir-1936 | microRNA 1936 | ncRNA |
| chr12_102643193_102643735 | chr12 | 102643194 | 102643735 | + | -0.652023 | intron (NM_172584, intron 2 of 9) | intron (NM_172584, intron 2 of 9) | 41556 | NR_035457 | 100316812 | | NR_035457 | | **Mir1936** | mmu-mir-1936 | microRNA 1936 | ncRNA |
| chr12_102651465_102651842 | chr12 | 102651466 | 102651842 | + | -0.684638 | intron (NM_172584, intron 2 of 9) | intron (NM_172584, intron 2 of 9) | 33366 | NR_035457 | 100316812 | | NR_035457 | | **Mir1936** | mmu-mir-1936 | microRNA 1936 | ncRNA |
| chr12_102651511_102651765 | chr12 | 102651512 | 102651765 | + | -0.685671 | intron (NM_172584, intron 2 of 9) | intron (NM_172584, intron 2 of 9) | 33382 | NR_035457 | 100316812 | | NR_035457 | | **Mir1936** | mmu-mir-1936 | microRNA 1936 | ncRNA |
| chr12_102651892_102652918 | chr12 | 102651893 | 102652918 | + | -0.851592 | intron (NM_172584, intron 2 of 9) | intron (NM_172584, intron 2 of 9) | 32615 | NR_035457 | 100316812 | | NR_035457 | | **Mir1936** | mmu-mir-1936 | microRNA 1936 | ncRNA |
| chr12_102619852_102620176 | chr12 | 102619853 | 102620176 | + | -0.971997 | intron (NM_172584, intron 3 of 9) | intron (NM_172584, intron 3 of 9) | 65006 | NR_035457 | 100316812 | | NR_035457 | | **Mir1936** | mmu-mir-1936 | microRNA 1936 | ncRNA |
| chr2_151943921_151944939 | chr2 | 151943922 | 151944939 | + | -0.635597 | intron (NM_009641, intron 8 of 8) | intron (NM_009641, intron 8 of 8) | 23187 | NR_035478 | 100316704 | | NR_035478 | miRBase:MI0009948 | **Mir1953** | mmu-mir-1953 | microRNA 1953 | ncRNA |
| chr2_92202365_92202625 | chr2 | 92202366 | 92202625 | + | -0.634225 | intron (NM_001109691, intron 2 of 18) | intron (NM_001109691, intron 2 of 18) | 10518 | NR_035480 | 100316756 | | NR_035480 | miRBase:MI0009950 | **Mir1955** | mmu-mir-1955 | microRNA 1955 | ncRNA |
| chr7_135598317_135598928 | chr7 | 135598318 | 135598928 | + | -1.09182 | intron (NM_011212, intron 1 of 20) | intron (NM_011212, intron 1 of 20) | 32461 | NR_035488 | 100316818 | | NR_035488 | | **Mir1962** | mmu-mir-1962 | microRNA 1962 | ncRNA |
| chr8_105623037_105623495 | chr8 | 105623038 | 105623495 | + | -0.769367 | Intergenic | Intergenic | 7800 | NR_035492 | 100316712 | | NR_035492 | | **Mir1966** | mmu-mir-1966 | microRNA 1966 | ncRNA |
| chr4_156076939_156077263 | chr4 | 156076940 | 156077263 | + | -0.647443 | Intergenic | Intergenic | -21351 | NR_029587 | 387243 | | NR_029587 | | **Mir200b** | Mirn200b | microRNA 200b | ncRNA |
| chr7_64195997_64196518 | chr7 | 64195998 | 64196518 | + | -0.63344 | intron (NM_018752, intron 1 of 13) | intron (NM_018752, intron 1 of 13) | -9548 | NR_029805 | 387207 | | NR_029805 | | **Mir211** | Mirn211\|mmu-mir-211 | microRNA 211 | ncRNA |
| chr11_35505315_35505781 | chr11 | 35505316 | 35505781 | + | -0.660549 | intron (NM_011412, intron 4 of 35) | intron (NM_011412, intron 4 of 35) | -111268 | NR_029799 | 723924 | | NR_029799 | | **Mir218-2** | Mirn218-2\|mir-218-2 | microRNA 218-2 | ncRNA |
| chr11_86578867_86579408 | chr11 | 86578868 | 86579408 | + | -1.02934 | Intergenic | Intergenic | 5020 | NR_029738 | 387140 | | NR_029738 | miRBase:MI0000569 | **Mir21a** | Mir21\|Mirn21\|mmu-mir-21\|mmu-mir-21a | microRNA 21a | ncRNA |
| chr8_128165624_128165937 | chr8 | 128165625 | 128165937 | + | -0.65223 | Intergenic | Intergenic | 112444 | NR_105822 | 102465212 | | NR_105822 | miRBase:MI0021932 | **Mir21c** | mmu-mir-21c | microRNA mir-21c | ncRNA |
| chr8_127966890_127967247 | chr8 | 127966891 | 127967247 | + | -0.692237 | Intergenic | Intergenic | 311156 | NR_105822 | 102465212 | | NR_105822 | miRBase:MI0021932 | **Mir21c** | mmu-mir-21c | microRNA mir-21c | ncRNA |
| chr8_127967987_127968536 | chr8 | 127967988 | 127968536 | + | -0.781373 | Intergenic | Intergenic | 309963 | NR_105822 | 102465212 | | NR_105822 | miRBase:MI0021932 | **Mir21c** | mmu-mir-21c | microRNA mir-21c | ncRNA |
| chr8_128223156_128223442 | chr8 | 128223157 | 128223442 | + | -0.791513 | Intergenic | L1MA4\|LINE\|L1 | 54926 | NR_105822 | 102465212 | | NR_105822 | miRBase:MI0021932 | **Mir21c** | mmu-mir-21c | microRNA mir-21c | ncRNA |
| chr6_31114467_31115262 | chr6 | 31114468 | 31115262 | + | -0.715732 | intron (NR_110471, intron 3 of 3) | intron (NR_110471, intron 3 of 3) | -51772 | NR_029532 | 387223 | | NR_029532 | miRBase:MI0000143 | **Mir29b-1** | Mirn29b\|Mirn29b-1 | microRNA 29b-1 | ncRNA |
| chr14_25522413_25522690 | chr14 | 25522414 | 25522690 | + | -0.676157 | intron (NM_183208, intron 2 of 23) | intron (NM_183208, intron 2 of 23) | -11887 | NR_037236 | 100526533 | | NR_037236 | miRBase:MI0014038 | **Mir3075** | mmu-mir-3075 | microRNA 3075 | ncRNA |
| chr14_25522403_25522866 | chr14 | 25522404 | 25522866 | + | -0.702271 | intron (NM_183208, intron 2 of 23) | intron (NM_183208, intron 2 of 23) | -11804 | NR_037236 | 100526533 | | NR_037236 | miRBase:MI0014038 | **Mir3075** | mmu-mir-3075 | microRNA 3075 | ncRNA |
| chr2_30727879_30728352 | chr2 | 30727880 | 30728352 | + | -1.19712 | Intergenic | MT2A\|LTR\|ERVL | 6906 | NR_037272 | 100526499 | | NR_037272 | miRBase:MI0014082 | **Mir3089** | mmu-mir-3089 | microRNA 3089 | ncRNA |
| chr2_131654751_131655119 | chr2 | 131654752 | 131655119 | + | -0.766145 | Intergenic | Intergenic | -50111 | NR_037281 | 100526526 | | NR_037281 | miRBase:MI0014091 | **Mir3098** | mmu-mir-3098 | microRNA 3098 | ncRNA |
| chr8_108847299_108847815 | chr8 | 108847300 | 108847815 | + | -0.734296 | intron (NM_007496, intron 3 of 9) | intron (NM_007496, intron 3 of 9) | -89303 | NR_037295 | 100526511 | | NR_037295 | | **Mir3108** | mmu-mir-3108 | microRNA 3108 | ncRNA |
| chr8_108879967_108880657 | chr8 | 108879968 | 108880657 | + | -0.992463 | intron (NM_007496, intron 4 of 9) | intron (NM_007496, intron 4 of 9) | -56548 | NR_037295 | 100526511 | | NR_037295 | | **Mir3108** | mmu-mir-3108 | microRNA 3108 | ncRNA |
| chr8_126829571_126830128 | chr8 | 126829572 | 126830128 | + | -0.841097 | Intergenic | Intergenic | -108050 | NR_037297 | 100526512 | | NR_037297 | miRBase:MI0014107 | **Mir3110** | mmu-mir-3110 | microRNA 3110 | ncRNA |
| chr8_126662742_126663233 | chr8 | 126662743 | 126663233 | + | -0.866396 | Intergenic | Intergenic | 58812 | NR_037297 | 100526512 | | NR_037297 | miRBase:MI0014107 | **Mir3110** | mmu-mir-3110 | microRNA 3110 | ncRNA |
| chr8_126829866_126830089 | chr8 | 126829867 | 126830089 | + | -0.990704 | Intergenic | Intergenic | -108178 | NR_037297 | 100526512 | | NR_037297 | miRBase:MI0014107 | **Mir3110** | mmu-mir-3110 | microRNA 3110 | ncRNA |
| chr8_126721575_126722054 | chr8 | 126721576 | 126722054 | + | -1.35432 | promoter-TSS (NR_037297).3 | promoter-TSS (NR_037297).3 | -15 | NR_037297 | 100526512 | | NR_037297 | miRBase:MI0014107 | **Mir3110** | mmu-mir-3110 | microRNA 3110 | ncRNA |

| | | | | | | | | | | | | | | | | | |
|---|---|---|---|---|---|---|---|---|---|---|---|---|---|---|---|---|---|
| chr4_56915783_56916450 | chr4 | 56915784 | 56916450 | + | -0.856108 | intron (NM_175518, intron 7 of 17) | L1M3\|LINE\|L1 | -20819 | NR_029789 | 723837 | | NR_029789 | | **Mir32** | Mirn32\|mmu-mir-32 | microRNA 32 | ncRNA |
| chr7_99572751_99573462 | chr7 | 99572752 | 99573462 | + | -0.698653 | intron (NM_177231, intron 1 of 15) | intron (NM_177231, intron 1 of 15) | 20838 | NR_029760 | 723840 | | NR_029760 | | **Mir326** | Mirn326\|mmu-mir-326 | microRNA 326 | ncRNA |
| chr7_99572812_99573740 | chr7 | 99572813 | 99573740 | + | -0.705458 | intron (NM_177231, intron 1 of 15) | intron (NM_177231, intron 1 of 15) | 21007 | NR_029760 | 723840 | | NR_029760 | | **Mir326** | Mirn326\|mmu-mir-326 | microRNA 326 | ncRNA |
| chr15_82186757_82187611 | chr15 | 82186758 | 82187611 | + | -1.09694 | intron (NM_033218, intron 11 of 18) | intron (NM_033218, intron 11 of 18) | -10938 | NR_029804 | 723897 | | NR_029804 | | **Mir33** | Mirn33\|mmu-mir-33 | microRNA 33 | ncRNA |
| chr15_100180251_100180568 | chr15 | 100180252 | 100180568 | + | -0.698292 | intron (NM_172819, intron 15 of 32) | intron (NM_172819, intron 15 of 32) | 7626 | NR_037311 | 100499528 | | NR_037311 | | **Mir3473a** | Mir3473\|mmu-mir-3473\|mmu-mir-3473a | microRNA 3473a | ncRNA |
| chr15_100181744_100182377 | chr15 | 100181745 | 100182377 | + | -1.07927 | intron (NM_172819, intron 16 of 32) | intron (NM_172819, intron 16 of 32) | 5975 | NR_037311 | 100499528 | | NR_037311 | | **Mir3473a** | Mir3473\|mmu-mir-3473\|mmu-mir-3473a | microRNA 3473a | ncRNA |
| chr15_100182653_100183245 | chr15 | 100182654 | 100183245 | + | -1.34785 | intron (NM_172819, intron 16 of 32) | intron (NM_172819, intron 16 of 32) | 5087 | NR_037311 | 100499528 | | NR_037311 | | **Mir3473a** | Mir3473\|mmu-mir-3473\|mmu-mir-3473a | microRNA 3473a | ncRNA |
| chr3_97294304_97294979 | chr3 | 97294305 | 97294979 | + | -0.662642 | Intergenic | Intergenic | -14614 | NR_105813 | 102465206 | | NR_105813 | miRBase:MI0021923 | **Mir378c** | mmu-mir-378c | microRNA mir-378c | ncRNA |
| chr3_97273871_97274678 | chr3 | 97273872 | 97274678 | + | -0.795956 | Intergenic | Intergenic | -34981 | NR_105813 | 102465206 | | NR_105813 | miRBase:MI0021923 | **Mir378c** | mmu-mir-378c | microRNA mir-378c | ncRNA |
| chr3_97273802_97274410 | chr3 | 97273803 | 97274410 | + | -0.866432 | Intergenic | MIRc\|SINE\|MIR | -35150 | NR_105813 | 102465206 | | NR_105813 | miRBase:MI0021923 | **Mir378c** | mmu-mir-378c | microRNA mir-378c | ncRNA |
| chr2_5486687_5487023 | chr2 | 5486688 | 5487023 | + | -0.623156 | intron (NM_177343, intron 2 of 10) | intron (NM_177343, intron 2 of 10) | 107465 | NR_030601 | 100124465 | | NR_030601 | | **Mir466d** | Mirn466d | microRNA 466d | ncRNA |
| chr2_5486705_5487336 | chr2 | 5486706 | 5487336 | + | -0.678769 | intron (NM_177343, intron 2 of 10) | intron (NM_177343, intron 2 of 10) | 107631 | NR_030601 | 100124465 | | NR_030601 | | **Mir466d** | Mirn466d | microRNA 466d | ncRNA |
| chr15_80896616_80897243 | chr15 | 80896617 | 80897243 | + | -0.830273 | intron (NM_177124, intron 11 of 21) | intron (NM_177124, intron 11 of 21) | -43109 | NR_039573 | 100628612 | | NR_039573 | miRBase:MI0018022 | **Mir5113** | mmu-mir-5113\|mu-mir-5113 | microRNA 5113 | ncRNA |
| chr15_80930755_80930932 | chr15 | 80930756 | 80930932 | + | -1.05225 | 3' UTR (NM_144812, exon 25 of 25) | 3' UTR (NM_144812, exon 25 of 25) | -9195 | NR_039573 | 100628612 | | NR_039573 | miRBase:MI0018022 | **Mir5113** | mmu-mir-5113\|mu-mir-5113 | microRNA 5113 | ncRNA |
| chr15_80930716_80931040 | chr15 | 80930717 | 80931040 | + | -1.09959 | 3' UTR (NM_144812, exon 25 of 25) | 3' UTR (NM_144812, exon 25 of 25) | -9161 | NR_039573 | 100628612 | | NR_039573 | miRBase:MI0018022 | **Mir5113** | mmu-mir-5113\|mu-mir-5113 | microRNA 5113 | ncRNA |
| chr15_80892349_80892738 | chr15 | 80892350 | 80892738 | + | -1.4916 | intron (NM_177124, intron 10 of 21) | intron (NM_177124, intron 10 of 21) | -47495 | NR_039573 | 100628612 | | NR_039573 | miRBase:MI0018022 | **Mir5113** | mmu-mir-5113\|mu-mir-5113 | microRNA 5113 | ncRNA |
| chr4_40833248_40833693 | chr4 | 40833249 | 40833693 | + | -0.685973 | intron (NM_022305, intron 1 of 5) | intron (NM_022305, intron 1 of 5) | 16667 | NR_039584 | 100628614 | | NR_039584 | | **Mir5123** | mmu-mir-5123\|mu-mir-5123 | microRNA 5123 | ncRNA |
| chr4_40839147_40839890 | chr4 | 40839148 | 40839890 | + | -0.710142 | intron (NM_022305, intron 1 of 5) | intron (NM_022305, intron 1 of 5) | 10619 | NR_039584 | 100628614 | | NR_039584 | | **Mir5123** | mmu-mir-5123\|mu-mir-5123 | microRNA 5123 | ncRNA |
| chr18_81863039_81864153 | chr18 | 81863040 | 81864153 | + | -0.778338 | Intergenic | Intergenic | 128484 | NR_039589 | 100628596 | | NR_039589 | miRBase:MI0018039 | **Mir5127** | mmu-mir-5127\|mu-mir-5127 | microRNA 5127 | ncRNA |
| chr18_81652776_81653050 | chr18 | 81652777 | 81653050 | + | -0.784033 | Intergenic | Intergenic | 339167 | NR_039589 | 100628596 | | NR_039589 | miRBase:MI0018039 | **Mir5127** | mmu-mir-5127\|mu-mir-5127 | microRNA 5127 | ncRNA |
| chr18_81863041_81863773 | chr18 | 81863042 | 81863773 | + | -0.901269 | Intergenic | Intergenic | 128673 | NR_039589 | 100628596 | | NR_039589 | miRBase:MI0018039 | **Mir5127** | mmu-mir-5127\|mu-mir-5127 | microRNA 5127 | ncRNA |
| chr18_81568615_81569189 | chr18 | 81568616 | 81569189 | + | -0.991354 | Intergenic | Intergenic | 423178 | NR_039589 | 100628596 | | NR_039589 | miRBase:MI0018039 | **Mir5127** | mmu-mir-5127\|mu-mir-5127 | microRNA 5127 | ncRNA |
| chr18_81520990_81521791 | chr18 | 81520991 | 81521791 | + | -1.29768 | Intergenic | MTEb\|LTR\|ERVL-MaLR | 470689 | NR_039589 | 100628596 | | NR_039589 | miRBase:MI0018039 | **Mir5127** | mmu-mir-5127\|mu-mir-5127 | microRNA 5127 | ncRNA |
| chr18_81521067_81521591 | chr18 | 81521068 | 81521591 | + | -1.40965 | Intergenic | MTEb\|LTR\|ERVL-MaLR | 470751 | NR_039589 | 100628596 | | NR_039589 | miRBase:MI0018039 | **Mir5127** | mmu-mir-5127\|mu-mir-5127 | microRNA 5127 | ncRNA |
| chr2_44960520_44960976 | chr2 | 44960521 | 44960976 | + | -0.756537 | Intergenic | Intergenic | 62429 | NR_039591 | 100628598 | | NR_039591 | | **Mir5129** | mmu-mir-5129\|mu-mir-5129 | microRNA 5129 | ncRNA |
| chr9_7775945_7776351 | chr9 | 7775946 | 7776351 | + | -0.974747 | intron (NM_133739, intron 2 of 4) | ORR1B2\|LTR\|ERVL-MaLR | -8242 | NR_049191 | 100885830 | | NR_049191 | | **Mir5618** | mmu-mir-5618 | microRNA mir-5618 | ncRNA |
| chr9_7780280_7780608 | chr9 | 7780281 | 7780608 | + | -1.2043 | intron (NM_133739, intron 2 of 4) | intron (NM_133739, intron 2 of 4) | -3946 | NR_049191 | 100885830 | | NR_049191 | | **Mir5618** | mmu-mir-5618 | microRNA mir-5618 | ncRNA |
| chr19_58143633_58144419 | chr19 | 58143634 | 58144419 | + | -0.806959 | Intergenic | Intergenic | 92859 | NR_049196 | 100885835 | | NR_049196 | | **Mir5623** | mmu-mir-5623 | microRNA mir-5623 | ncRNA |
| chr16_43624820_43625248 | chr16 | 43624821 | 43625248 | + | -1.01958 | Intergenic | Intergenic | -15621 | NR_030576 | 100124467 | | NR_030576 | miRBase:MI0005517 | **Mir568** | Mirn568\|mmu-mir-568 | microRNA 568 | ncRNA |
| chr5_65019053_65019603 | chr5 | 65019054 | 65019603 | + | -0.812849 | intron (NM_026667, intron 7 of 13) | intron (NM_026667, intron 7 of 13) | 49010 | NR_030577 | 100124451 | | NR_030577 | miRBase:MI0005518 | **Mir574** | Mirn574\|mmu-mir-574 | microRNA 574 | ncRNA |
| chr5_64989831_64990373 | chr5 | 64989832 | 64990373 | + | -1.07041 | intron (NM_026667, intron 2 of 13) | intron (NM_026667, intron 2 of 13) | 19784 | NR_030577 | 100124451 | | NR_030577 | miRBase:MI0005518 | **Mir574** | Mirn574\|mmu-mir-574 | microRNA 574 | ncRNA |
| chr13_109231896_109232547 | chr13 | 109231897 | 109232547 | + | -0.71236 | intron (NM_011056, intron 3 of 16) | intron (NM_011056, intron 3 of 16) | -92522 | NR_030644 | 100124489 | | NR_030644 | | **Mir582** | Mirn582\|mmu-mir-582 | microRNA 582 | ncRNA |
| chr13_109239237_109239717 | chr13 | 109239238 | 109239717 | + | -0.729345 | intron (NM_011056, intron 3 of 16) | intron (NM_011056, intron 3 of 16) | -85267 | NR_030644 | 100124489 | | NR_030644 | | **Mir582** | Mirn582\|mmu-mir-582 | microRNA 582 | ncRNA |
| chr1_39102150_39102486 | chr1 | 39102151 | 39102486 | + | -0.930321 | Intergenic | Intergenic | 84674 | NR_105767 | 102466631 | | NR_105767 | miRBase:MI0021877 | **Mir6349** | mmu-mir-6349 | microRNA mir-6349 | ncRNA |
| chr1_39102002_39102485 | chr1 | 39102003 | 39102485 | + | -0.989314 | Intergenic | Intergenic | 84748 | NR_105767 | 102466631 | | NR_105767 | miRBase:MI0021877 | **Mir6349** | mmu-mir-6349 | microRNA mir-6349 | ncRNA |
| chr1_39090993_39091557 | chr1 | 39090994 | 39091557 | + | -1.07571 | Intergenic | MT2A\|LTR\|ERVL | 95717 | NR_105767 | 102466631 | | NR_105767 | miRBase:MI0021877 | **Mir6349** | mmu-mir-6349 | microRNA mir-6349 | ncRNA |
| chr2_166630219_166630784 | chr2 | 166630220 | 166630784 | + | -0.605409 | intron (NM_177782, intron 5 of 39) | intron (NM_177782, intron 5 of 39) | -42216 | NR_105783 | 102465188 | | NR_105783 | | **Mir6364** | mmu-mir-6364 | microRNA mir-6364 | ncRNA |
| chr2_166672555_166673015 | chr2 | 166672556 | 166673015 | + | -0.752246 | promoter-TSS (NR_105783) | promoter-TSS (NR_105783) | 67 | NR_105783 | 102465188 | | NR_105783 | | **Mir6364** | mmu-mir-6364 | microRNA mir-6364 | ncRNA |
| chr2_166622529_166622935 | chr2 | 166622530 | 166622935 | + | -0.841082 | intron (NM_177782, intron 5 of 39) | intron (NM_177782, intron 5 of 39) | -49986 | NR_105783 | 102465188 | | NR_105783 | | **Mir6364** | mmu-mir-6364 | microRNA mir-6364 | ncRNA |
| chr2_166671508_166671973 | chr2 | 166671509 | 166671973 | + | -0.886938 | promoter-TSS (NR_105783) | promoter-TSS (NR_105783) | -977 | NR_105783 | 102465188 | | NR_105783 | | **Mir6364** | mmu-mir-6364 | microRNA mir-6364 | ncRNA |
| chr2_166620108_166621022 | chr2 | 166620109 | 166621022 | + | -0.907676 | intron (NM_177782, intron 6 of 39) | MTE2b\|LTR\|ERVL-MaLR | -52153 | NR_105783 | 102465188 | | NR_105783 | | **Mir6364** | mmu-mir-6364 | microRNA mir-6364 | ncRNA |
| chr2_166617722_166618054 | chr2 | 166617723 | 166618054 | + | -1.05393 | exon (NM_177782, exon 7 of 40) | exon (NM_177782, exon 7 of 40) | -54830 | NR_105783 | 102465188 | | NR_105783 | | **Mir6364** | mmu-mir-6364 | microRNA mir-6364 | ncRNA |
| chr19_9038141_9038596 | chr19 | 9038142 | 9038596 | + | -0.836641 | intron (NM_001039959, intron 5 of 5) | intron (NM_001039959, intron 5 of 5) | 20465 | NR_105786 | 102465190 | | NR_105786 | miRBase:MI0021896 | **Mir6367** | mmu-mir-6367 | microRNA mir-6367 | ncRNA |
| chr13_28757773_28758335 | chr13 | 28757774 | 28758335 | + | -0.638056 | intron (NR_045053, intron 3 of 8) | intron (NR_045053, intron 3 of 8) | -47181 | NR_105787 | 102466635 | | NR_105787 | miRBase:MI0021897 | **Mir6368** | mmu-mir-6368 | microRNA mir-6368 | ncRNA |
| chr13_28600463_28601295 | chr13 | 28600464 | 28601295 | + | -0.704603 | intron (NR_045053, intron 6 of 8) | intron (NR_045053, intron 6 of 8) | 109994 | NR_105787 | 102466635 | | NR_105787 | miRBase:MI0021897 | **Mir6368** | mmu-mir-6368 | microRNA mir-6368 | ncRNA |
| chr13_28720078_28720571 | chr13 | 28720079 | 28720571 | + | -0.712269 | intron (NR_045053, intron 3 of 8) | intron (NR_045053, intron 3 of 8) | -9452 | NR_105787 | 102466635 | | NR_105787 | miRBase:MI0021897 | **Mir6368** | mmu-mir-6368 | microRNA mir-6368 | ncRNA |
| chr13_28762620_28762918 | chr13 | 28762621 | 28762918 | + | -0.762052 | intron (NR_045053, intron 3 of 8) | ORR1B2\|LTR\|ERVL-MaLR | -51896 | NR_105787 | 102466635 | | NR_105787 | miRBase:MI0021897 | **Mir6368** | mmu-mir-6368 | microRNA mir-6368 | ncRNA |
| chr13_28512495_28513179 | chr13 | 28512496 | 28513179 | + | -0.86759 | intron (NR_045053, intron 7 of 8) | intron (NR_045053, intron 7 of 8) | 198036 | NR_105787 | 102466635 | | NR_105787 | miRBase:MI0021897 | **Mir6368** | mmu-mir-6368 | microRNA mir-6368 | ncRNA |
| chr13_28551484_28552042 | chr13 | 28551485 | 28552042 | + | -1.08093 | intron (NR_045053, intron 7 of 8) | intron (NR_045053, intron 7 of 8) | 159110 | NR_105787 | 102466635 | | NR_105787 | miRBase:MI0021897 | **Mir6368** | mmu-mir-6368 | microRNA mir-6368 | ncRNA |
| chr6_83411635_83411978 | chr6 | 83411636 | 83411978 | + | -0.65776 | intron (NM_183138, intron 2 of 10) | intron (NM_183138, intron 2 of 10) | -10705 | NR_105793 | 102465194 | | NR_105793 | | **Mir6374** | mmu-mir-6374 | microRNA mir-6374 | ncRNA |
| chr6_83381467_83382221 | chr6 | 83381468 | 83382221 | + | -0.719347 | intron (NM_183138, intron 4 of 10) | intron (NM_183138, intron 4 of 10) | 19258 | NR_105793 | 102465194 | | NR_105793 | | **Mir6374** | mmu-mir-6374 | microRNA mir-6374 | ncRNA |
| chr3_142626024_142626249 | chr3 | 142626025 | 142626249 | + | -0.649172 | intron (NM_010260, intron 4 of 10) | intron (NM_010260, intron 4 of 10) | -2191 | NR_105802 | 102465200 | | NR_105802 | miRBase:MI0021912 | **Mir6381** | mmu-mir-6381 | microRNA mir-6381 | ncRNA |
| chr12_15808488_15808803 | chr12 | 15808489 | 15808803 | + | -0.904512 | intron (NM_144551, intron 2 of 2) | intron (NM_144551, intron 2 of 2) | -7545 | NR_105809 | 102465204 | | NR_105809 | | **Mir6387** | mmu-mir-6387 | microRNA mir-6387 | ncRNA |
| chr12_15804758_15805273 | chr12 | 15804759 | 15805273 | + | -1.23795 | intron (NM_144551, intron 2 of 2) | intron (NM_144551, intron 2 of 2) | -3915 | NR_105809 | 102465204 | | NR_105809 | | **Mir6387** | mmu-mir-6387 | microRNA mir-6387 | ncRNA |
| chr12_115691686_115692271 | chr12 | 115691687 | 115692271 | + | -0.631872 | Intergenic | Intergenic | 3815 | NR_105810 | 102465205 | | NR_105810 | miRBase:MI0021920 | **Mir6388** | mmu-mir-6388 | microRNA mir-6388 | ncRNA |
| chr12_115165432_115166056 | chr12 | 115165433 | 115166056 | + | -0.719884 | Intergenic | Intergenic | 530050 | NR_105810 | 102465205 | | NR_105810 | miRBase:MI0021920 | **Mir6388** | mmu-mir-6388 | microRNA mir-6388 | ncRNA |
| chr12_115847738_115848176 | chr12 | 115847739 | 115848176 | + | -0.769078 | Intergenic | Intergenic | -152163 | NR_105810 | 102465205 | | NR_105810 | miRBase:MI0021920 | **Mir6388** | mmu-mir-6388 | microRNA mir-6388 | ncRNA |
| chr12_115847743_115848045 | chr12 | 115847744 | 115848045 | + | -0.793287 | Intergenic | Intergenic | -152100 | NR_105810 | 102465205 | | NR_105810 | miRBase:MI0021920 | **Mir6388** | mmu-mir-6388 | microRNA mir-6388 | ncRNA |
| chr8_34215214_34216548 | chr8 | 34215215 | 34216548 | + | -0.711029 | Intergenic | Intergenic | -49169 | NR_105820 | 102465211 | | NR_105820 | miRBase:MI0021930 | **Mir6395** | mmu-mir-6395 | microRNA mir-6395 | ncRNA |
| chr8_34215635_34216576 | chr8 | 34215636 | 34216576 | + | -0.77254 | Intergenic | Intergenic | -49394 | NR_105820 | 102465211 | | NR_105820 | miRBase:MI0021930 | **Mir6395** | mmu-mir-6395 | microRNA mir-6395 | ncRNA |

| | | | | | | | | | | | | | | | | | |
|---|---|---|---|---|---|---|---|---|---|---|---|---|---|---|---|---|---|
| chr8_34215973_34216573 | chr8 | 34215974 | 34216573 | + | -0.812274 | Intergenic | Intergenic | -49561 | NR_105820 | 102465211 | | NR_105820 | miRBase:MI0021930 | **Mir6395** | mmu-mir-6395 | microRNA mir-6395 | ncRNA |
| chr8_34315176_34315687 | chr8 | 34315177 | 34315687 | + | -0.891072 | Intergenic | MTEa\|LTR\|ERVL-MaLR | -148720 | NR_105820 | 102465211 | | NR_105820 | miRBase:MI0021930 | **Mir6395** | mmu-mir-6395 | microRNA mir-6395 | ncRNA |
| chr8_34166752_34167148 | chr8 | 34166753 | 34167148 | + | -1.1081 | promoter-TSS (NR_105820) | promoter-TSS (NR_105820) | -238 | NR_105820 | 102465211 | | NR_105820 | miRBase:MI0021930 | **Mir6395** | mmu-mir-6395 | microRNA mir-6395 | ncRNA |
| chr8_34166840_34167133 | chr8 | 34166841 | 34167133 | + | -1.16415 | promoter-TSS (NR_105820) | promoter-TSS (NR_105820) | -275 | NR_105820 | 102465211 | | NR_105820 | miRBase:MI0021930 | **Mir6395** | mmu-mir-6395 | microRNA mir-6395 | ncRNA |
| chr8_34163321_34163843 | chr8 | 34163322 | 34163843 | + | -1.46517 | intron (NM_026432, intron 2 of 5) | intron (NM_026432, intron 2 of 5) | 3130 | NR_105820 | 102465211 | | NR_105820 | miRBase:MI0021930 | **Mir6395** | mmu-mir-6395 | microRNA mir-6395 | ncRNA |
| chr4_108082030_108082786 | chr4 | 108082031 | 108082786 | + | -0.837466 | intron (NM_011327, intron 10 of 15) | intron (NM_011327, intron 10 of 15) | -4949 | NR_105823 | 102465213 | | NR_105823 | miRBase:MI0021933 | **Mir6397** | mmu-mir-6397 | microRNA mir-6397 | ncRNA |
| chr4_108082285_108082786 | chr4 | 108082286 | 108082786 | + | -0.913017 | intron (NM_011327, intron 10 of 15) | intron (NM_011327, intron 10 of 15) | -4821 | NR_105823 | 102465213 | | NR_105823 | miRBase:MI0021933 | **Mir6397** | mmu-mir-6397 | microRNA mir-6397 | ncRNA |
| chr4_134601282_134602314 | chr4 | 134601283 | 134602314 | + | -0.744956 | intron (NM_207237, intron 2 of 11) | RMER15\|LTR\|ERVL | -33947 | NR_105829 | 102465216 | | NR_105829 | | **Mir6403** | mmu-mir-6403 | microRNA mir-6403 | ncRNA |
| chr4_134601412_134601944 | chr4 | 134601413 | 134601944 | + | -0.782855 | intron (NM_207237, intron 2 of 11) | MIRb\|SINE\|MIR | -33827 | NR_105829 | 102465216 | | NR_105829 | | **Mir6403** | mmu-mir-6403 | microRNA mir-6403 | ncRNA |
| chr10_20257689_20258471 | chr10 | 20257690 | 20258471 | + | -0.706884 | intron (NM_008635, intron 7 of 17) | intron (NM_008635, intron 7 of 17) | 39608 | NR_105840 | 102466647 | | NR_105840 | miRBase:MI0021950 | **Mir6413** | mmu-mir-6413 | microRNA mir-6413 | ncRNA |
| chr10_20257635_20258247 | chr10 | 20257636 | 20258247 | + | -0.715793 | intron (NM_008635, intron 7 of 17) | intron (NM_008635, intron 7 of 17) | 39747 | NR_105840 | 102466647 | | NR_105840 | miRBase:MI0021950 | **Mir6413** | mmu-mir-6413 | microRNA mir-6413 | ncRNA |
| chr10_20262913_20263276 | chr10 | 20262914 | 20263276 | + | -0.887918 | intron (NM_001198635, intron 8 of 17) | intron (NM_001198635, intron 8 of 17) | 34593 | NR_105840 | 102466647 | | NR_105840 | miRBase:MI0021950 | **Mir6413** | mmu-mir-6413 | microRNA mir-6413 | ncRNA |
| chr10_20281513_20282399 | chr10 | 20281514 | 20282399 | + | -1.00401 | TTS (NM_008635) | TTS (NM_008635) | 15732 | NR_105840 | 102466647 | | NR_105840 | miRBase:MI0021950 | **Mir6413** | mmu-mir-6413 | microRNA mir-6413 | ncRNA |
| chr5_92465972_92466345 | chr5 | 92465973 | 92466345 | + | -0.818289 | intron (NM_007644, intron 2 of 11) | intron (NM_007644, intron 2 of 11) | 7602 | NR_105843 | 102466159 | | NR_105843 | miRBase:MI0021953 | **Mir6415** | mmu-mir-6415 | microRNA mir-6415 | ncRNA |
| chr5_92466018_92466279 | chr5 | 92466019 | 92466279 | + | -0.824434 | intron (NM_007644, intron 2 of 11) | intron (NM_007644, intron 2 of 11) | 7592 | NR_105843 | 102466159 | | NR_105843 | miRBase:MI0021953 | **Mir6415** | mmu-mir-6415 | microRNA mir-6415 | ncRNA |
| chr7_64248293_64248718 | chr7 | 64248294 | 64248718 | + | -1.12741 | intron (NM_001039104, intron 26 of 26) | intron (NM_001039104, intron 26 of 26) | 1593 | NR_105844 | 102465226 | | NR_105844 | miRBase:MI0021954 | **Mir6416** | mmu-mir-6416 | microRNA mir-6416 | ncRNA |
| chr12_26401640_26402070 | chr12 | 26401641 | 26402070 | + | -0.716447 | intron (NM_080563, intron 1 of 8) | intron (NM_080563, intron 1 of 8) | 13155 | NR_105856 | 102466676 | | NR_105856 | miRBase:MI0022354 | **Mir6538** | mmu-mir-6538 | microRNA mir-6538 | ncRNA |
| chr14_62862641_62862918 | chr14 | 62862642 | 62862918 | + | -0.920646 | intron (NM_175546, intron 1 of 11) | intron (NM_175546, intron 1 of 11) | -2762 | NR_105860 | 102465311 | | NR_105860 | | **Mir6541** | mmu-mir-6541 | microRNA mir-6541 | ncRNA |
| chrX_77469257_77469622 | chrX | 77469258 | 77469622 | + | -0.692577 | intron (NR_035475, intron 1 of 3) | intron (NR_035475, intron 1 of 3) | 6496 | NR_035475 | 100316834 | | NR_035475 | | **Mir669m-2** | mmu-mir-669m-2 | microRNA 669m-2 | ncRNA |
| chr11_75012821_75013475 | chr11 | 75012822 | 75013475 | + | -0.785044 | intron (NM_001002764, intron 10 of 18) | intron (NM_001002764, intron 10 of 18) | 52843 | NR_030454 | 735274 | | NR_030454 | | **Mir684-1** | Mirn684-1 | microRNA 684-1 | ncRNA |
| chr11_75032765_75033379 | chr11 | 75032766 | 75033379 | + | -0.839471 | intron (NM_001002764, intron 10 of 18) | intron (NM_001002764, intron 10 of 18) | 32919 | NR_030454 | 735274 | | NR_030454 | | **Mir684-1** | Mirn684-1 | microRNA 684-1 | ncRNA |
| chr4_11143242_11143487 | chr4 | 11143243 | 11143487 | + | -0.606278 | Intergenic | PB1D10\|SINE\|Alu | 9268 | NR_030455 | 735261 | | NR_030455 | | **Mir684-2** | Mirn684-2 | microRNA 684-2 | ncRNA |
| chr1_36131307_36131501 | chr1 | 36131308 | 36131501 | + | -0.645639 | Intergenic | MLTR14\|LTR\|ERV1 | 12909 | NR_105862 | 102465540 | | NR_105862 | | **Mir6897** | mmu-mir-6897 | microRNA mir-6897 | ncRNA |
| chr1_36148857_36149745 | chr1 | 36148858 | 36149745 | + | -0.704046 | intron (NM_198899, intron 38 of 40) | intron (NM_198899, intron 38 of 40) | -4988 | NR_105862 | 102465540 | | NR_105862 | | **Mir6897** | mmu-mir-6897 | microRNA mir-6897 | ncRNA |
| chr1_36148857_36149284 | chr1 | 36148858 | 36149284 | + | -1.07983 | intron (NM_198899, intron 38 of 40) | intron (NM_198899, intron 38 of 40) | -4758 | NR_105862 | 102465540 | | NR_105862 | | **Mir6897** | mmu-mir-6897 | microRNA mir-6897 | ncRNA |
| chr1_92468720_92469399 | chr1 | 92468721 | 92469399 | + | -1.18068 | intron (NM_024197, intron 3 of 9) | intron (NM_024197, intron 3 of 9) | -4518 | NR_105865 | 102465542 | | NR_105865 | | **Mir6900** | mmu-mir-6900 | microRNA mir-6900 | ncRNA |
| chr10_78399252_78399782 | chr10 | 78399253 | 78399782 | + | -0.828352 | TTS (NM_010925) | TTS (NM_010925) | 9028 | NR_105872 | 102465547 | | NR_105872 | | **Mir6907** | mmu-mir-6907 | microRNA mir-6907 | ncRNA |
| chr10_80610173_80610498 | chr10 | 80610174 | 80610498 | + | -1.10505 | intron (NM_019575, intron 4 of 6) | intron (NM_019575, intron 4 of 6) | 1076 | NR_105877 | 102465550 | | NR_105877 | | **Mir6912** | mmu-mir-6912 | microRNA mir-6912 | ncRNA |
| chr11_101046348_101046921 | chr11 | 101046349 | 101046921 | + | -0.801176 | intron (NM_001243051, intron 17 of 20) | intron (NM_001243051, intron 17 of 20) | 16295 | NR_105893 | 102466767 | | NR_105893 | | **Mir6928** | mmu-mir-6928 | microRNA mir-6928 | ncRNA |
| chr11_107668158_107668695 | chr11 | 107668159 | 107668695 | + | -1.32437 | intron (NM_198298, intron 27 of 31) | intron (NM_198298, intron 27 of 31) | 30705 | NR_105897 | 102465562 | | NR_105897 | | **Mir6932** | mmu-mir-6932 | microRNA mir-6932 | ncRNA |
| chr11_118014194_118015093 | chr11 | 118014195 | 118015093 | + | -0.623978 | exon (NM_133757, exon 8 of 10) | exon (NM_133757, exon 8 of 10) | 12374 | NR_105898 | 102466768 | | NR_105898 | | **Mir6933** | mmu-mir-6933 | microRNA mir-6933 | ncRNA |
| chr11_118014323_118014919 | chr11 | 118014324 | 118014919 | + | -0.689181 | exon (NM_133757, exon 8 of 10) | exon (NM_133757, exon 8 of 10) | 12351 | NR_105898 | 102466768 | | NR_105898 | | **Mir6933** | mmu-mir-6933 | microRNA mir-6933 | ncRNA |
| chr12_85257291_85257631 | chr12 | 85257292 | 85257631 | + | -0.675864 | intron (NM_175337, intron 5 of 11) | intron (NM_175337, intron 5 of 11) | -11476 | NR_105903 | 102466769 | | NR_105903 | | **Mir6938** | mmu-mir-6938 | microRNA mir-6938 | ncRNA |
| chr14_20682404_20682924 | chr14 | 20682405 | 20682924 | + | -0.790291 | intron (NM_172596, intron 3 of 22) | intron (NM_172596, intron 3 of 22) | -7971 | NR_105911 | 102465571 | | NR_105911 | | **Mir6946** | mmu-mir-6946 | microRNA mir-6946 | ncRNA |
| chr15_89282422_89283156 | chr15 | 89282423 | 89283156 | + | -0.659802 | intron (NM_026813, intron 22 of 24) | intron (NM_026813, intron 22 of 24) | 22943 | NR_105924 | 102465981 | | NR_105924 | | **Mir6959** | mmu-mir-6959 | microRNA mir-6959 | ncRNA |
| chr15_101190824_101191119 | chr15 | 101190825 | 101191119 | + | -0.713208 | intron (NM_007395, intron 1 of 8) | MIR3\|SINE\|MIR | -2897 | NR_105927 | 102466774 | | NR_105927 | | **Mir6962** | mmu-mir-6962 | microRNA mir-6962 | ncRNA |
| chr17_35245981_35246603 | chr17 | 35245982 | 35246603 | + | -0.722694 | intron (NM_001252457, intron 4 of 10) | intron (NM_001252457, intron 4 of 10) | 2217 | NR_105941 | 102465589 | | NR_105941 | | **Mir6975** | mmu-mir-6975 | microRNA mir-6975 | ncRNA |
| chr7_78868753_78869143 | chr7 | 78868754 | 78869143 | + | -1.05703 | Intergenic | MER47B\|DNA\|TcMar-Tigger | -19329 | NR_029826 | 723884 | | NR_029826 | miRBase:MI0000729 | **Mir7-2** | Mirn7-2\|mmu-mir-7-2 | microRNA 7-2 | ncRNA |
| chr2_114144761_114145824 | chr2 | 114144762 | 114145824 | + | -0.777426 | intron (NM_009702, intron 14 of 34) | intron (NM_009702, intron 14 of 34) | 5277 | NR_105968 | 102465982 | | NR_105968 | | **Mir7002** | mmu-mir-7002 | microRNA mir-7002 | ncRNA |
| chr2_114136761_114137358 | chr2 | 114136762 | 114137358 | + | -0.951717 | intron (NM_009702, intron 16 of 34) | intron (NM_009702, intron 16 of 34) | 13510 | NR_105968 | 102465982 | | NR_105968 | | **Mir7002** | mmu-mir-7002 | microRNA mir-7002 | ncRNA |
| chr2_114153318_114153953 | chr2 | 114153319 | 114153953 | + | -1.06998 | intron (NM_009702, intron 9 of 34) | MLTR11A\|LTR\|ERVK | -3066 | NR_105968 | 102465982 | | NR_105968 | | **Mir7002** | mmu-mir-7002 | microRNA mir-7002 | ncRNA |
| chr2_114153123_114153768 | chr2 | 114153124 | 114153768 | + | -1.11477 | intron (NM_009702, intron 9 of 34) | MLTR11A\|LTR\|ERVK | -2876 | NR_105968 | 102465982 | | NR_105968 | | **Mir7002** | mmu-mir-7002 | microRNA mir-7002 | ncRNA |
| chr2_114153315_114153669 | chr2 | 114153316 | 114153669 | + | -1.21216 | intron (NM_009702, intron 9 of 34) | MLTR11A\|LTR\|ERVK | -2922 | NR_105968 | 102465982 | | NR_105968 | | **Mir7002** | mmu-mir-7002 | microRNA mir-7002 | ncRNA |
| chr3_20222154_20222620 | chr3 | 20222155 | 20222620 | + | -0.846172 | promoter-TSS (NR_105974) | promoter-TSS (NR_105974) | -28 | NR_105974 | 102465609 | | NR_105974 | | **Mir7007** | mmu-mir-7007 | microRNA mir-7007 | ncRNA |
| chr5_111115723_111116002 | chr5 | 111115724 | 111116002 | + | -0.739073 | intron (NM_001267622, intron 5 of 22) | intron (NM_001267622, intron 5 of 22) | 111719 | NR_030482 | 735278 | | NR_030482 | | **Mir701** | Mirn701\|mmu-mir-701 | microRNA 701 | ncRNA |
| chr5_111039502_111040273 | chr5 | 111039503 | 111040273 | + | -0.768573 | intron (NM_001267622, intron 2 of 22) | intron (NM_001267622, intron 2 of 22) | 35744 | NR_030482 | 735278 | | NR_030482 | | **Mir701** | Mirn701\|mmu-mir-701 | microRNA 701 | ncRNA |
| chr5_111039676_111040238 | chr5 | 111039677 | 111040238 | + | -0.82068 | intron (NM_001267622, intron 2 of 22) | intron (NM_001267622, intron 2 of 22) | 35813 | NR_030482 | 735278 | | NR_030482 | | **Mir701** | Mirn701\|mmu-mir-701 | microRNA 701 | ncRNA |
| chr3_95720231_95720732 | chr3 | 95720232 | 95720732 | + | -0.729861 | Intergenic | Intergenic | 14491 | NR_105981 | 102466787 | | NR_105981 | | **Mir7014** | mmu-mir-7014 | microRNA mir-7014 | ncRNA |
| chr4_120982375_120982669 | chr4 | 120982376 | 120982669 | + | -1.09386 | intron (NM_133716, intron 3 of 9) | intron (NM_133716, intron 3 of 9) | -9200 | NR_105982 | 102465614 | | NR_105982 | | **Mir7015** | mmu-mir-7015 | microRNA mir-7015 | ncRNA |

| | | | | | | | | | | | | | | | | | |
|---|---|---|---|---|---|---|---|---|---|---|---|---|---|---|---|---|---|
| chr4_133176107_133176591 | chr4 | 133176108 | 133176591 | + | -0.61188 | intron (NM_153423, intron 1 of 8) | intron (NM_153423, intron 1 of 8) | -19306 | NR_105984 | 102466215 | | NR_105984 | | **Mir7017** | mmu-mir-7017 | microRNA mir-7017 | ncRNA |
| chr4_133169265_133170108 | chr4 | 133169266 | 133170108 | + | -0.729202 | intron (NM_153423, intron 1 of 8) | intron (NM_153423, intron 1 of 8) | -25968 | NR_105984 | 102466215 | | NR_105984 | | **Mir7017** | mmu-mir-7017 | microRNA mir-7017 | ncRNA |
| chr4_133174383_133174781 | chr4 | 133174384 | 133174781 | + | -1.1858 | intron (NM_153423, intron 1 of 8) | intron (NM_153423, intron 1 of 8) | -21073 | NR_105984 | 102466215 | | NR_105984 | | **Mir7017** | mmu-mir-7017 | microRNA mir-7017 | ncRNA |
| chr4_143154652_143154884 | chr4 | 143154653 | 143154884 | + | -0.691233 | intron (NM_001256380, intron 6 of 7) | intron (NM_001256380, intron 6 of 7) | -18606 | NR_105988 | 102465618 | | NR_105988 | | **Mir7021** | mmu-mir-7021 | microRNA mir-7021 | ncRNA |
| chr5_124011126_124011492 | chr5 | 124011127 | 124011492 | + | -1.0037 | intron (NM_177876, intron 1 of 3) | intron (NM_177876, intron 1 of 3) | 14457 | NR_105999 | 102465625 | | NR_105999 | | **Mir7032** | mmu-mir-7032 | microRNA mir-7032 | ncRNA |
| chr5_124010911_124011385 | chr5 | 124010912 | 124011385 | + | -1.08656 | intron (NM_177876, intron 1 of 3) | intron (NM_177876, intron 1 of 3) | 14296 | NR_105999 | 102465625 | | NR_105999 | | **Mir7032** | mmu-mir-7032 | microRNA mir-7032 | ncRNA |
| chr5_124003124_124003804 | chr5 | 124003125 | 124003804 | + | -1.37005 | TTS (NM_145070) | TTS (NM_145070) | 6612 | NR_105999 | 102465625 | | NR_105999 | | **Mir7032** | mmu-mir-7032 | microRNA mir-7032 | ncRNA |
| chr5_144919602_144920174 | chr5 | 144919603 | 144920174 | + | -0.625821 | intron (NM_029438, intron 1 of 17) | intron (NM_029438, intron 1 of 17) | -23072 | NR_106006 | 102465629 | | NR_106006 | | **Mir7039** | mmu-mir-7039 | microRNA mir-7039 | ncRNA |
| chr5_144925899_144926751 | chr5 | 144925900 | 144926751 | + | -0.676947 | intron (NM_029438, intron 1 of 17) | intron (NM_029438, intron 1 of 17) | -29509 | NR_106006 | 102465629 | | NR_106006 | | **Mir7039** | mmu-mir-7039 | microRNA mir-7039 | ncRNA |
| chr5_144926156_144926864 | chr5 | 144926157 | 144926864 | + | -0.787892 | intron (NM_029438, intron 1 of 17) | PB1D10\|SINE\|Alu | -29694 | NR_106006 | 102465629 | | NR_106006 | | **Mir7039** | mmu-mir-7039 | microRNA mir-7039 | ncRNA |
| chr7_28565423_28565845 | chr7 | 28565424 | 28565845 | + | -0.809113 | intron (NM_027470, intron 3 of 9) | intron (NM_027470, intron 3 of 9) | -4650 | NR_106016 | 102466795 | | NR_106016 | | **Mir7049** | mmu-mir-7049 | microRNA mir-7049 | ncRNA |
| chr7_31037812_31038055 | chr7 | 31037813 | 31038055 | + | -0.850665 | intron (NM_001287217, intron 4 of 8) | intron (NM_001287217, intron 4 of 8) | 2383 | NR_106017 | 102465635 | | NR_106017 | | **Mir7050** | mmu-mir-7050 | microRNA mir-7050 | ncRNA |
| chr7_31033683_31034173 | chr7 | 31033684 | 31034173 | + | -0.890072 | intron (NM_001287213, intron 8 of 8) | intron (NM_001287213, intron 8 of 8) | 6389 | NR_106017 | 102465635 | | NR_106017 | | **Mir7050** | mmu-mir-7050 | microRNA mir-7050 | ncRNA |
| chr7_31033461_31034211 | chr7 | 31033462 | 31034211 | + | -0.917649 | intron (NM_001287213, intron 8 of 8) | intron (NM_001287213, intron 8 of 8) | 6481 | NR_106017 | 102465635 | | NR_106017 | | **Mir7050** | mmu-mir-7050 | microRNA mir-7050 | ncRNA |
| chr7_66263461_66263803 | chr7 | 66263462 | 66263803 | + | -1.01588 | intron (NM_146191, intron 31 of 33) | intron (NM_146191, intron 31 of 33) | 118092 | NR_106024 | 102465640 | | NR_106024 | miRBase:MI0022906 | **Mir7057** | mmu-mir-7057 | microRNA mir-7057 | ncRNA |
| chr7_126329874_126330584 | chr7 | 126329875 | 126330584 | + | -0.888168 | Intergenic | Intergenic | 37816 | NR_106025 | 102466797 | | NR_106025 | | **Mir7058** | mmu-mir-7058 | microRNA mir-7058 | ncRNA |
| chr9_45944589_45945481 | chr9 | 45944590 | 45945481 | + | -0.699127 | intron (NM_172257, intron 15 of 25) | MIRb\|SINE\|MIR | -5445 | NR_106055 | 102465659 | | NR_106055 | | **Mir7087** | mmu-mir-7087 | microRNA mir-7087 | ncRNA |
| chr9_45945005_45945430 | chr9 | 45945006 | 45945430 | + | -0.731574 | intron (NM_172257, intron 14 of 25) | intron (NM_172257, intron 14 of 25) | -5628 | NR_106055 | 102465659 | | NR_106055 | | **Mir7087** | mmu-mir-7087 | microRNA mir-7087 | ncRNA |
| chr9_121019444_121019928 | chr9 | 121019445 | 121019928 | + | -0.887895 | intron (NM_177589, intron 35 of 36) | C-rich\|Low_complexity\|Low_complexity | 64600 | NR_106058 | 102465661 | | NR_106058 | | **Mir7090** | mmu-mir-7090 | microRNA mir-7090 | ncRNA |
| chr12_114059615_114060258 | chr12 | 114059616 | 114060258 | + | -0.799221 | Intergenic | Intergenic | 409529 | NR_106062 | 102465663 | | NR_106062 | | **Mir7094-1** | mmu-mir-7094-1 | microRNA mir-7094-1 | ncRNA |
| chr12_114299638_114300255 | chr12 | 114299639 | 114300255 | + | -0.828468 | Intergenic | Intergenic | 169519 | NR_106062 | 102465663 | | NR_106062 | | **Mir7094-1** | mmu-mir-7094-1 | microRNA mir-7094-1 | ncRNA |
| chr12_114612095_114612537 | chr12 | 114612096 | 114612537 | + | -0.632697 | Intergenic | Intergenic | -98622 | NR_106063 | 102465664 | | NR_106063 | miRBase:MI0022945 | **Mir7094-2** | mmu-mir-7094-2 | microRNA mir-7094-2 | ncRNA |
| chr12_114615619_114616196 | chr12 | 114615620 | 114616196 | + | -0.657372 | Intergenic | Intergenic | -102214 | NR_106063 | 102465664 | | NR_106063 | miRBase:MI0022945 | **Mir7094-2** | mmu-mir-7094-2 | microRNA mir-7094-2 | ncRNA |
| chr12_114936960_114937332 | chr12 | 114936961 | 114937332 | + | -0.6863 | Intergenic | Intergenic | -423452 | NR_106063 | 102465664 | | NR_106063 | miRBase:MI0022945 | **Mir7094-2** | mmu-mir-7094-2 | microRNA mir-7094-2 | ncRNA |
| chr12_114587181_114587589 | chr12 | 114587182 | 114587589 | + | -0.783281 | Intergenic | Intergenic | -73691 | NR_106063 | 102465664 | | NR_106063 | miRBase:MI0022945 | **Mir7094-2** | mmu-mir-7094-2 | microRNA mir-7094-2 | ncRNA |
| chr12_114597675_114598132 | chr12 | 114597676 | 114598132 | + | -0.823273 | Intergenic | Intergenic | -84210 | NR_106063 | 102465664 | | NR_106063 | miRBase:MI0022945 | **Mir7094-2** | mmu-mir-7094-2 | microRNA mir-7094-2 | ncRNA |
| chr12_114914417_114914903 | chr12 | 114914418 | 114914903 | + | -0.827542 | Intergenic | Intergenic | -400966 | NR_106063 | 102465664 | | NR_106063 | miRBase:MI0022945 | **Mir7094-2** | mmu-mir-7094-2 | microRNA mir-7094-2 | ncRNA |
| chr11_70435534_70435855 | chr11 | 70435535 | 70435855 | + | -0.986674 | intron (NM_001271360, intron 2 of 15) | intron (NM_001271360, intron 2 of 15) | -2560 | NR_106064 | 102465670 | | NR_106064 | | **Mir7115** | mmu-mir-7115 | microRNA mir-7115 | ncRNA |
| chr15_27596507_27597607 | chr15 | 27596508 | 27597607 | + | -0.805108 | Intergenic | Intergenic | 25563 | NR_106065 | 102466808 | | NR_106065 | | **Mir7117** | mmu-mir-7117 | microRNA mir-7117 | ncRNA |
| chr5_136449736_136450149 | chr5 | 136449737 | 136450149 | + | -1.21518 | intron (NM_001291239, intron 2 of 22) | ORR1E\|LTR\|ERVL-MaLR | -74140 | NR_030500 | 723975 | | NR_030500 | | **Mir721** | Mirn721\|miRY\|mir-721\|mmu-mir-721 | microRNA 721 | ncRNA |
| chr17_27322857_27323461 | chr17 | 27322858 | 27323461 | + | -0.757567 | Intergenic | RMER4A\|LTR\|ERVK | -5226 | NR_106075 | 102466818 | | NR_106075 | miRBase:MI0023711 | **Mir7216** | mmu-mir-7216 | microRNA mir-7216 | ncRNA |
| chr18_68203022_68203410 | chr18 | 68203023 | 68203410 | + | -0.649304 | intron (NM_172631, intron 3 of 5) | ORR1A3-int\|LTR\|ERVL-MaLR | 57757 | NR_106078 | 102465702 | | NR_106078 | miRBase:MI0023714 | **Mir7219** | mmu-mir-7219 | microRNA mir-7219 | ncRNA |
| chr18_68205031_68205315 | chr18 | 68205032 | 68205315 | + | -0.652517 | intron (NM_172631, intron 3 of 5) | MTD\|LTR\|ERVL-MaLR | 55800 | NR_106078 | 102465702 | | NR_106078 | miRBase:MI0023714 | **Mir7219** | mmu-mir-7219 | microRNA mir-7219 | ncRNA |
| chr18_68219063_68219478 | chr18 | 68219064 | 68219478 | + | -0.659182 | intron (NM_172631, intron 3 of 5) | B1_Mur1\|SINE\|Alu | 41702 | NR_106078 | 102465702 | | NR_106078 | miRBase:MI0023714 | **Mir7219** | mmu-mir-7219 | microRNA mir-7219 | ncRNA |
| chr18_68214471_68214806 | chr18 | 68214472 | 68214806 | + | -0.724082 | intron (NM_172631, intron 3 of 5) | intron (NM_172631, intron 3 of 5) | 46334 | NR_106078 | 102465702 | | NR_106078 | miRBase:MI0023714 | **Mir7219** | mmu-mir-7219 | microRNA mir-7219 | ncRNA |
| chr18_68131110_68131591 | chr18 | 68131111 | 68131591 | + | -0.809001 | intron (NM_172631, intron 3 of 5) | intron (NM_172631, intron 3 of 5) | 129622 | NR_106078 | 102465702 | | NR_106078 | miRBase:MI0023714 | **Mir7219** | mmu-mir-7219 | microRNA mir-7219 | ncRNA |
| chr18_68131141_68131828 | chr18 | 68131142 | 68131828 | + | -0.825249 | intron (NM_172631, intron 3 of 5) | intron (NM_172631, intron 3 of 5) | 129488 | NR_106078 | 102465702 | | NR_106078 | miRBase:MI0023714 | **Mir7219** | mmu-mir-7219 | microRNA mir-7219 | ncRNA |
| chr18_68227519_68228106 | chr18 | 68227520 | 68228106 | + | -0.850937 | intron (NM_172631, intron 3 of 5) | intron (NM_172631, intron 3 of 5) | 33160 | NR_106078 | 102465702 | | NR_106078 | miRBase:MI0023714 | **Mir7219** | mmu-mir-7219 | microRNA mir-7219 | ncRNA |
| chr18_68131280_68131576 | chr18 | 68131281 | 68131576 | + | -0.860056 | intron (NM_172631, intron 3 of 5) | intron (NM_172631, intron 3 of 5) | 129545 | NR_106078 | 102465702 | | NR_106078 | miRBase:MI0023714 | **Mir7219** | mmu-mir-7219 | microRNA mir-7219 | ncRNA |
| chr18_60952085_60952950 | chr18 | 60952086 | 60952950 | + | -0.838455 | intron (NM_177407, intron 6 of 17) | intron (NM_177407, intron 6 of 17) | -1355 | NR_106079 | 102465703 | | NR_106079 | | **Mir7220** | mmu-mir-7220 | microRNA mir-7220 | ncRNA |
| chr3_122286910_122287635 | chr3 | 122286911 | 122287635 | + | -0.813856 | Intergenic | Intergenic | 6430 | NR_030439 | 791077 | | NR_030439 | miRBase:MI0004605 | **Mir760** | Mirn760\|mir-760\|mmu-mir-760 | microRNA 760 | ncRNA |
| chr3_122287104_122287769 | chr3 | 122287105 | 122287769 | + | -0.839992 | Intergenic | RLTR14\|LTR\|ERV1 | 6266 | NR_030439 | 791077 | | NR_030439 | miRBase:MI0004605 | **Mir760** | Mirn760\|mir-760\|mmu-mir-760 | microRNA 760 | ncRNA |
| chr9_94945546_94946621 | chr9 | 94945547 | 94946621 | + | -0.646231 | intron (NM_177909, intron 8 of 15) | intron (NM_177909, intron 8 of 15) | 194255 | NR_106116 | 102465763 | | NR_106116 | | **Mir7656** | mmu-mir-7656 | microRNA mir-7656 | ncRNA |
| chr9_94993641_94994051 | chr9 | 94993642 | 94994051 | + | -0.654324 | intron (NM_177909, intron 9 of 15) | intron (NM_177909, intron 9 of 15) | 242017 | NR_106116 | 102465763 | | NR_106116 | | **Mir7656** | mmu-mir-7656 | microRNA mir-7656 | ncRNA |
| chr9_95072520_95073274 | chr9 | 95072521 | 95073274 | + | -0.930685 | intron (NM_177909, intron 12 of 15) | intron (NM_177909, intron 12 of 15) | 321068 | NR_106116 | 102465763 | | NR_106116 | | **Mir7656** | mmu-mir-7656 | microRNA mir-7656 | ncRNA |
| chr9_95072548_95073088 | chr9 | 95072549 | 95073088 | + | -1.00843 | intron (NM_177909, intron 12 of 15) | intron (NM_177909, intron 12 of 15) | 320989 | NR_106116 | 102465763 | | NR_106116 | | **Mir7656** | mmu-mir-7656 | microRNA mir-7656 | ncRNA |
| chr6_108526136_108526532 | chr6 | 108526137 | 108526532 | + | -0.712642 | intron (NM_010585, intron 60 of 61) | B4A\|SINE\|B4 | 36591 | NR_106121 | 102465766 | | NR_106121 | | **Mir7661** | mmu-mir-7661 | microRNA mir-7661 | ncRNA |
| chr6_108503662_108504100 | chr6 | 108503663 | 108504100 | + | -0.915858 | intron (NM_010585, intron 53 of 61) | L1ME1\|LINE\|L1 | 14138 | NR_106121 | 102465766 | | NR_106121 | | **Mir7661** | mmu-mir-7661 | microRNA mir-7661 | ncRNA |
| chr6_108503676_108504266 | chr6 | 108503677 | 108504266 | + | -0.922157 | intron (NM_010585, intron 53 of 61) | intron (NM_010585, intron 53 of 61) | 14228 | NR_106121 | 102465766 | | NR_106121 | | **Mir7661** | mmu-mir-7661 | microRNA mir-7661 | ncRNA |
| chr6_108465840_108466543 | chr6 | 108465841 | 108466543 | + | -1.07978 | intron (NM_010585, intron 44 of 61) | intron (NM_010585, intron 44 of 61) | -23551 | NR_106121 | 102465766 | | NR_106121 | | **Mir7661** | mmu-mir-7661 | microRNA mir-7661 | ncRNA |
| chr1_35709246_35709589 | chr1 | 35709247 | 35709589 | + | -0.761066 | Intergenic | ORR1B1\|LTR\|ERVL-MaLR | -186785 | NR_106129 | 102465771 | | NR_106129 | | **Mir7669** | mmu-mir-7669 | microRNA mir-7669 | ncRNA |
| chr1_35652922_35653882 | chr1 | 35652923 | 35653882 | + | -1.04925 | Intergenic | RLTR14\|LTR\|ERV1 | -130769 | NR_106129 | 102465771 | | NR_106129 | | **Mir7669** | mmu-mir-7669 | microRNA mir-7669 | ncRNA |
| chr1_35653228_35653893 | chr1 | 35653229 | 35653893 | + | -1.09795 | Intergenic | RLTR14\|LTR\|ERV1 | -130928 | NR_106129 | 102465771 | | NR_106129 | | **Mir7669** | mmu-mir-7669 | microRNA mir-7669 | ncRNA |
| chr1_35580474_35580878 | chr1 | 35580475 | 35580878 | + | -1.12974 | Intergenic | Intergenic | -58043 | NR_106129 | 102465771 | | NR_106129 | | **Mir7669** | mmu-mir-7669 | microRNA mir-7669 | ncRNA |
| chr2_32062973_32063485 | chr2 | 32062974 | 32063485 | + | -0.659804 | Intergenic | Intergenic | 12283 | NR_106135 | 102466844 | | NR_106135 | | **Mir7674** | mmu-mir-7674 | microRNA mir-7674 | ncRNA |
| chr2_32057332_32057752 | chr2 | 32057333 | 32057752 | + | -0.705888 | Intergenic | Intergenic | 6596 | NR_106135 | 102466844 | | NR_106135 | | **Mir7674** | mmu-mir-7674 | microRNA mir-7674 | ncRNA |
| chr2_32047035_32047759 | chr2 | 32047036 | 32047759 | + | -0.724136 | intron (NM_172268, intron 32 of 35) | intron (NM_172268, intron 32 of 35) | -3549 | NR_106135 | 102466844 | | NR_106135 | | **Mir7674** | mmu-mir-7674 | microRNA mir-7674 | ncRNA |

| | | | | | | | | | | | | | | | | | |
|---|---|---|---|---|---|---|---|---|---|---|---|---|---|---|---|---|---|
| chr2_32065266_32065855 | chr2 | 32065267 | 32065855 | + | -0.8211 | Intergenic | Intergenic | 14615 | NR_106135 | 102466844 | | NR_106135 | | **Mir7674** | mmu-mir-7674 | microRNA mir-7674 | ncRNA |
| chr2_32029597_32029920 | chr2 | 32029598 | 32029920 | + | -0.851052 | intron (NM_172268, intron 26 of 35) | RSINE1\|SINE\|B4 | -21187 | NR_106135 | 102466844 | | NR_106135 | | **Mir7674** | mmu-mir-7674 | microRNA mir-7674 | ncRNA |
| chr2_32062072_32062538 | chr2 | 32062073 | 32062538 | + | -1.24033 | Intergenic | RLTR8\|LTR\|ERVK | 11359 | NR_106135 | 102466844 | | NR_106135 | | **Mir7674** | mmu-mir-7674 | microRNA mir-7674 | ncRNA |
| chr2_32062178_32062496 | chr2 | 32062179 | 32062496 | + | -1.25814 | Intergenic | RLTR8\|LTR\|ERVK | 11391 | NR_106135 | 102466844 | | NR_106135 | | **Mir7674** | mmu-mir-7674 | microRNA mir-7674 | ncRNA |
| chr1_53906235_53906686 | chr1 | 53906236 | 53906686 | + | -0.677341 | intron (NM_001001883, intron 13 of 28) | intron (NM_001001883, intron 13 of 28) | 57037 | NR_106149 | 102465783 | | NR_106149 | miRBase:MI0025029 | **Mir7681** | mmu-mir-7681 | microRNA mir-7681 | ncRNA |
| chr1_53886696_53887127 | chr1 | 53886697 | 53887127 | + | -0.746298 | intron (NM_001001883, intron 16 of 28) | U1\|snRNA\|snRNA | 37488 | NR_106149 | 102465783 | | NR_106149 | miRBase:MI0025029 | **Mir7681** | mmu-mir-7681 | microRNA mir-7681 | ncRNA |
| chr1_53875972_53876210 | chr1 | 53875973 | 53876210 | + | -0.835216 | intron (NM_001001883, intron 17 of 28) | intron (NM_001001883, intron 17 of 28) | 26667 | NR_106149 | 102465783 | | NR_106149 | miRBase:MI0025029 | **Mir7681** | mmu-mir-7681 | microRNA mir-7681 | ncRNA |
| chr1_53875950_53876411 | chr1 | 53875951 | 53876411 | + | -0.88349 | intron (NM_001001883, intron 17 of 28) | intron (NM_001001883, intron 17 of 28) | 26757 | NR_106149 | 102465783 | | NR_106149 | miRBase:MI0025029 | **Mir7681** | mmu-mir-7681 | microRNA mir-7681 | ncRNA |
| chr1_53851355_53851729 | chr1 | 53851356 | 53851729 | + | -1.097 | intron (NM_001001883, intron 21 of 28) | intron (NM_001001883, intron 21 of 28) | 2118 | NR_106149 | 102465783 | | NR_106149 | miRBase:MI0025029 | **Mir7681** | mmu-mir-7681 | microRNA mir-7681 | ncRNA |
| chr1_53850849_53851293 | chr1 | 53850850 | 53851293 | + | -1.47493 | intron (NM_001001883, intron 21 of 28) | intron (NM_001001883, intron 21 of 28) | 1647 | NR_106149 | 102465783 | | NR_106149 | miRBase:MI0025029 | **Mir7681** | mmu-mir-7681 | microRNA mir-7681 | ncRNA |
| chr1_151456949_151457615 | chr1 | 151456950 | 151457615 | + | -0.955276 | intron (NM_026876, intron 13 of 14) | intron (NM_026876, intron 13 of 14) | 14889 | NR_106150 | 102466847 | | NR_106150 | | **Mir7682** | mmu-mir-7682 | microRNA mir-7682 | ncRNA |
| chr1_151456953_151457407 | chr1 | 151456954 | 151457407 | + | -0.956842 | intron (NM_026876, intron 13 of 14) | intron (NM_026876, intron 13 of 14) | 14787 | NR_106150 | 102466847 | | NR_106150 | | **Mir7682** | mmu-mir-7682 | microRNA mir-7682 | ncRNA |
| chr1_151457121_151457427 | chr1 | 151457122 | 151457427 | + | -0.985369 | intron (NM_026876, intron 13 of 14) | intron (NM_026876, intron 13 of 14) | 14881 | NR_106150 | 102466847 | | NR_106150 | | **Mir7682** | mmu-mir-7682 | microRNA mir-7682 | ncRNA |
| chr16_22497929_22498353 | chr16 | 22497930 | 22498353 | + | -0.807861 | intron (NM_138650, intron 22 of 24) | intron (NM_138650, intron 22 of 24) | 34043 | NR_106170 | 102465887 | | NR_106170 | | **Mir8095** | mmu-mir-8095 | microRNA mir-8095 | ncRNA |
| chr12_92976233_92976809 | chr12 | 92976234 | 92976809 | + | -0.714176 | Intergenic | LTR16C\|LTR\|ERVL | -950817 | NR_106201 | 102466888 | | NR_106201 | miRBase:MI0026053 | **Mir8099-2** | mmu-mir-8099-2 | microRNA mir-8099-2 | ncRNA |
| chr12_93019655_93020155 | chr12 | 93019656 | 93020155 | + | -0.881989 | Intergenic | Intergenic | -907433 | NR_106201 | 102466888 | | NR_106201 | miRBase:MI0026053 | **Mir8099-2** | mmu-mir-8099-2 | microRNA mir-8099-2 | ncRNA |
| chr12_93173798_93174106 | chr12 | 93173799 | 93174106 | + | -0.919958 | Intergenic | Intergenic | -753386 | NR_106201 | 102466888 | | NR_106201 | miRBase:MI0026053 | **Mir8099-2** | mmu-mir-8099-2 | microRNA mir-8099-2 | ncRNA |
| chr12_93173703_93174115 | chr12 | 93173704 | 93174115 | + | -0.931271 | Intergenic | Intergenic | -753429 | NR_106201 | 102466888 | | NR_106201 | miRBase:MI0026053 | **Mir8099-2** | mmu-mir-8099-2 | microRNA mir-8099-2 | ncRNA |
| chr12_93230223_93230917 | chr12 | 93230224 | 93230917 | + | -1.05029 | Intergenic | Intergenic | -696768 | NR_106201 | 102466888 | | NR_106201 | miRBase:MI0026053 | **Mir8099-2** | mmu-mir-8099-2 | microRNA mir-8099-2 | ncRNA |
| chr12_93230280_93230596 | chr12 | 93230281 | 93230596 | + | -1.11151 | Intergenic | RMER19B2\|LTR\|ERVK | -696900 | NR_106201 | 102466888 | | NR_106201 | miRBase:MI0026053 | **Mir8099-2** | mmu-mir-8099-2 | microRNA mir-8099-2 | ncRNA |
| chr10_122742116_122742566 | chr10 | 122742117 | 122742566 | + | -0.857557 | intron (NM_176919, intron 1 of 8) | intron (NM_176919, intron 1 of 8) | 63026 | NR_106182 | 102466884 | | NR_106182 | | **Mir8104** | mmu-mir-8104 | microRNA mir-8104 | ncRNA |
| chr3_65650415_65650996 | chr3 | 65650416 | 65650996 | + | -0.917996 | Intergenic | Intergenic | -8582 | NR_106200 | 102465905 | | NR_106200 | miRBase:MI0026052 | **Mir8120** | mmu-mir-8120 | microRNA mir-8120 | ncRNA |
| chr6_98113656_98114060 | chr6 | 98113657 | 98114060 | + | -0.712397 | Intergenic | Intergenic | 122075 | NM_008601 | 17342 | Mm.333284 | NM_008601 | ENSMUSG00000035158 | **Mitf** | BCC2\|Bhlhe32\|Gsfbcc2\|Vitiligo\|Wh\|bw\|mi\|vit | microphthalmia-associated transcription factor | protein-coding |
| chr15_81101418_81101705 | chr15 | 81101419 | 81101705 | + | -0.737074 | intron (NM_001082536, intron 2 of 13) | intron (NM_001082536, intron 2 of 13) | 3485 | NM_153049 | 223701 | Mm.439814 | NM_153049 | ENSMUSG00000042292 | **Mkl1** | AMKL\|Bsac\|Mal\|Mrtf-A | MKL (megakaryoblastic leukemia)/myocardin-like 1 | protein-coding |
| chr15_81097961_81098260 | chr15 | 81097962 | 81098260 | + | -0.749056 | intron (NM_001082536, intron 2 of 13) | intron (NM_001082536, intron 2 of 13) | 6936 | NM_153049 | 223701 | Mm.439814 | NM_153049 | ENSMUSG00000042292 | **Mkl1** | AMKL\|Bsac\|Mal\|Mrtf-A | MKL (megakaryoblastic leukemia)/myocardin-like 1 | protein-coding |
| chr15_81131113_81131519 | chr15 | 81131114 | 81131519 | + | -0.817596 | intron (NM_001082536, intron 1 of 13) | intron (NM_001082536, intron 1 of 13) | -26269 | NM_153049 | 223701 | Mm.439814 | NM_153049 | ENSMUSG00000042292 | **Mkl1** | AMKL\|Bsac\|Mal\|Mrtf-A | MKL (megakaryoblastic leukemia)/myocardin-like 1 | protein-coding |
| chr15_81166370_81166905 | chr15 | 81166371 | 81166905 | + | -0.935357 | intron (NM_001082536, intron 1 of 13) | intron (NM_001082536, intron 1 of 13) | 24119 | NM_001082536 | 223701 | Mm.439814 | NM_153049 | ENSMUSG00000042292 | **Mkl1** | AMKL\|Bsac\|Mal\|Mrtf-A | MKL (megakaryoblastic leukemia)/myocardin-like 1 | protein-coding |
| chr15_81166342_81166659 | chr15 | 81166343 | 81166659 | + | -1.02382 | intron (NM_001082536, intron 1 of 13) | PB1D9\|SINE\|Alu | 24256 | NM_001082536 | 223701 | Mm.439814 | NM_153049 | ENSMUSG00000042292 | **Mkl1** | AMKL\|Bsac\|Mal\|Mrtf-A | MKL (megakaryoblastic leukemia)/myocardin-like 1 | protein-coding |
| chr10_80678002_80678547 | chr10 | 80678003 | 80678547 | + | -0.673715 | Intergenic | Intergenic | -6299 | NM_021462 | 17347 | Mm.42126 | NM_021462 | ENSMUSG00000020190 | **Mknk2** | 2010016G11Rik\|Gprk7\|Mnk2 | MAP kinase-interacting serine/threonine kinase 2 | protein-coding |
| chr5_115167317_115167925 | chr5 | 115167318 | 115167925 | + | -1.0604 | Intergenic | Intergenic | -9445 | NM_175403 | 109154 | Mm.153963 | NM_175403 | ENSMUSG00000048578 | **Mlec** | 2410014A08Rik\|AI850530\|ESTM19\|X83352\|mKIAA0152 | malectin | protein-coding |
| chr5_115151005_115151200 | chr5 | 115151006 | 115151200 | + | -1.19725 | intron (NM_175403, intron 1 of 4) | intron (NM_175403, intron 1 of 4) | 7073 | NM_175403 | 109154 | Mm.153963 | NM_175403 | ENSMUSG00000048578 | **Mlec** | 2410014A08Rik\|AI850530\|ESTM19\|X83352\|mKIAA0152 | malectin | protein-coding |
| chr5_115150969_115151383 | chr5 | 115150970 | 115151383 | + | -1.24843 | intron (NM_175403, intron 1 of 4) | intron (NM_175403, intron 1 of 4) | 7000 | NM_175403 | 109154 | Mm.153963 | NM_175403 | ENSMUSG00000048578 | **Mlec** | 2410014A08Rik\|AI850530\|ESTM19\|X83352\|mKIAA0152 | malectin | protein-coding |
| chr17_56911489_56912064 | chr17 | 56911490 | 56912064 | + | -0.943254 | intron (NM_022328, intron 3 of 11) | L1MC\|LINE\|L1 | 23611 | NM_022328 | 64144 | Mm.148748 | NM_022328 | ENSMUSG00000024212 | **Mllt1** | AA407901\|BAM11\|ENL\|LTG19 | myeloid/lymphoid or mixed-lineage leukemia (trithorax homolog, Drosophila); translocated to, 1 | protein-coding |
| chr17_56911558_56911980 | chr17 | 56911559 | 56911980 | + | -0.947937 | intron (NM_022328, intron 3 of 11) | L1MC\|LINE\|L1 | 23619 | NM_022328 | 64144 | Mm.148748 | NM_022328 | ENSMUSG00000024212 | **Mllt1** | AA407901\|BAM11\|ENL\|LTG19 | myeloid/lymphoid or mixed-lineage leukemia (trithorax homolog, Drosophila); translocated to, 1 | protein-coding |
| chr11_97665040_97665200 | chr11 | 97665041 | 97665200 | + | -0.973112 | intron (NM_139311, intron 2 of 19) | intron (NM_139311, intron 2 of 19) | 1708 | NM_139311 | 246198 | Mm.23685 | NM_139311 | ENSMUSG00000038437 | **Mllt6** | AI315037\|Af17 | myeloid/lymphoid or mixed-lineage leukemia (trithorax homolog, Drosophila); translocated to, 6 | protein-coding |
| chr11_97664992_97665267 | chr11 | 97664993 | 97665267 | + | -0.992695 | intron (NM_139311, intron 2 of 19) | intron (NM_139311, intron 2 of 19) | 1718 | NM_139311 | 246198 | Mm.23685 | NM_139311 | ENSMUSG00000038437 | **Mllt6** | AI315037\|Af17 | myeloid/lymphoid or mixed-lineage leukemia (trithorax homolog, Drosophila); translocated to, 6 | protein-coding |
| chr11_90256901_90257918 | chr11 | 90256902 | 90257918 | + | -0.678956 | intron (NM_026178, intron 1 of 6) | intron (NM_026178, intron 1 of 6) | 7934 | NM_026178 | 67468 | Mm.277518 | NM_026178 | ENSMUSG00000003948 | **Mmd** | 1200017E07Rik\|1810073C06Rik\|AA690185 | monocyte to macrophage differentiation-associated | protein-coding |
| chr11_90256898_90257575 | chr11 | 90256899 | 90257575 | + | -0.86664 | intron (NM_026178, intron 1 of 6) | intron (NM_026178, intron 1 of 6) | 7761 | NM_026178 | 67468 | Mm.277518 | NM_026178 | ENSMUSG00000003948 | **Mmd** | 1200017E07Rik\|1810073C06Rik\|AA690185 | monocyte to macrophage differentiation-associated | protein-coding |
| chr8_95389981_95390808 | chr8 | 95389982 | 95390808 | + | -0.661321 | Intergenic | Intergenic | 38058 | NM_008609 | 17388 | Mm.217116 | NM_008609 | ENSMUSG00000031790 | **Mmp15** | AI503551\|MT2-MMP | matrix metallopeptidase 15 | protein-coding |
| chr2_164942740_164943320 | chr2 | 164942741 | 164943320 | + | -0.828031 | Intergenic | Intergenic | -5189 | NM_013599 | 17395 | Mm.4406 | NM_013599 | ENSMUSG00000017737 | **Mmp9** | AW743869\|B/MMP9\|Clg4b\|MMP-9\|pro-MMP-9 | matrix metallopeptidase 9 | protein-coding |
| chr2_164938670_164939116 | chr2 | 164938671 | 164939116 | + | -0.853596 | Intergenic | Intergenic | -9326 | NM_013599 | 17395 | Mm.4406 | NM_013599 | ENSMUSG00000017737 | **Mmp9** | AW743869\|B/MMP9\|Clg4b\|MMP-9\|pro-MMP-9 | matrix metallopeptidase 9 | protein-coding |
| chr14_10146400_10146837 | chr14 | 10146401 | 10146837 | + | -0.832452 | intron (NM_010210, intron 4 of 8) | HAL1\|LINE\|L1 | 262524 | NR_030680 | 100042415 | Mm.235938 | NR_030680 | | **Mnd1-ps** | Gm3833 | Mnd1 retrotransposed pseudogene | pseudo |
| chr14_10288890_10289566 | chr14 | 10288891 | 10289566 | + | -0.870528 | intron (NM_010210, intron 4 of 8) | intron (NM_010210, intron 4 of 8) | 405133 | NR_030680 | 100042415 | Mm.235938 | NR_030680 | | **Mnd1-ps** | Gm3833 | Mnd1 retrotransposed pseudogene | pseudo |
| chr14_10301479_10302068 | chr14 | 10301480 | 10302068 | + | -1.02712 | intron (NM_010210, intron 4 of 8) | intron (NM_010210, intron 4 of 8) | 417679 | NR_030680 | 100042415 | Mm.235938 | NR_030680 | | **Mnd1-ps** | Gm3833 | Mnd1 retrotransposed pseudogene | pseudo |
| chr7_142056627_142057230 | chr7 | 142056628 | 142057230 | + | -0.760292 | intron (NM_028308, intron 1 of 4) | intron (NM_028308, intron 1 of 4) | 4105 | NM_028308 | 101513 | Mm.53259 | NM_028308 | ENSMUSG00000025147 | **Mob2** | 1110017M21Rik\|2700078K21Rik\|AI256456\|Mmh | MOB kinase activator 2 | protein-coding |
| chr10_80700697_80701053 | chr10 | 80700698 | 80701053 | + | -0.786755 | intron (NM_172457, intron 1 of 3) | intron (NM_172457, intron 1 of 3) | 945 | NM_172457 | 208228 | Mm.49309 | NM_172457 | ENSMUSG00000003348 | **Mob3a** | 5330417K06Rik\|A630029F06\|AV218468\|Mobkl2a | MOB kinase activator 3A | protein-coding |
| chr10_80700804_80701017 | chr10 | 80700805 | 80701017 | + | -0.787001 | intron (NM_172457, intron 1 of 3) | intron (NM_172457, intron 1 of 3) | 909 | NM_172457 | 208228 | Mm.49309 | NM_172457 | ENSMUSG00000003348 | **Mob3a** | 5330417K06Rik\|A630029F06\|AV218468\|Mobkl2a | MOB kinase activator 3A | protein-coding |
| chr13_114845663_114846465 | chr13 | 114845664 | 114846465 | + | -0.815054 | intron (NM_008396, intron 25 of 29) | intron (NM_008396, intron 25 of 29) | 27827 | NM_001113374 | 17434 | Mm.19027 | NM_013826 | ENSMUSG00000015536 | **Mocs2** | AI415403 | molybdenum cofactor synthesis 2 | protein-coding |

| | | | | | | | | | | | | | | | | | |
|---|---|---|---|---|---|---|---|---|---|---|---|---|---|---|---|---|---|
| chr16_48594430_48594887 | chr16 | 48594431 | 48594887 | + | -0.714201 | intron (NM_010816, intron 20 of 26) | intron (NM_010816, intron 20 of 26) | 163422 | NM_010816 | 17450 | Mm.250060 | NM_010816 | ENSMUSG00000022652 | **Morc1** | Morc | microrchidia 1 | protein-coding |
| chr4_155129512_155129785 | chr4 | 155129513 | 155129785 | + | -0.802848 | intron (NM_001081100, intron 12 of 13) | intron (NM_001081100, intron 12 of 13) | 43072 | NM_001081100 | 76866 | Mm.200590 | NM_001081100 | ENSMUSG00000029049 | **Morn1** | 2900057D20Rik\|4930417P05Rik\|AU042805 | MORN repeat containing 1 | protein-coding |
| chr19_12476930_12477399 | chr19 | 12476931 | 12477399 | + | -0.877383 | intron (NM_172442, intron 7 of 8) | intron (NM_172442, intron 7 of 8) | 16386 | NM_010821 | 17476 | Mm.3999 | NM_010821 | ENSMUSG00000046805 | **Mpeg1** | MPS1\|Mpg-1 | macrophage expressed gene 1 | protein-coding |
| chr19_12476924_12477222 | chr19 | 12476925 | 12477222 | + | -0.890907 | intron (NM_172442, intron 7 of 8) | intron (NM_172442, intron 7 of 8) | 16294 | NM_010821 | 17476 | Mm.3999 | NM_010821 | ENSMUSG00000046805 | **Mpeg1** | MPS1\|Mpg-1 | macrophage expressed gene 1 | protein-coding |
| chr19_12476954_12477175 | chr19 | 12476955 | 12477175 | + | -0.905172 | intron (NM_172442, intron 7 of 8) | intron (NM_172442, intron 7 of 8) | 16286 | NM_010821 | 17476 | Mm.3999 | NM_010821 | ENSMUSG00000046805 | **Mpeg1** | MPS1\|Mpg-1 | macrophage expressed gene 1 | protein-coding |
| chr6_50008929_50009770 | chr6 | 50008930 | 50009770 | + | -0.782582 | Intergenic | Intergenic | -100891 | NM_019939 | 56524 | Mm.41288 | NM_008016 | ENSMUSG00000038388 | **Mpp6** | P55t\|Pals2 | membrane protein, palmitoylated 6 (MAGUK p55 subfamily member 6) | protein-coding |
| chr9_45062417_45063153 | chr9 | 45062418 | 45063153 | + | -0.809478 | intron (NM_001093749, intron 2 of 5) | intron (NM_001093749, intron 2 of 5) | 7604 | NM_001093749 | 319742 | Mm.49611 | NM_176993 | ENSMUSG00000070305 | **Mpzl3** | 5430427F17Rik\|A530065I17Rik\|rc | myelin protein zero-like 3 | protein-coding |
| chr9_45062744_45063048 | chr9 | 45062745 | 45063048 | + | -0.835718 | intron (NM_001093749, intron 2 of 5) | intron (NM_001093749, intron 2 of 5) | 7715 | NM_001093749 | 319742 | Mm.49611 | NM_176993 | ENSMUSG00000070305 | **Mpzl3** | 5430427F17Rik\|A530065I17Rik\|rc | myelin protein zero-like 3 | protein-coding |
| chr9_45062607_45063132 | chr9 | 45062608 | 45063132 | + | -0.854447 | intron (NM_001093749, intron 2 of 5) | intron (NM_001093749, intron 2 of 5) | 7689 | NM_001093749 | 319742 | Mm.49611 | NM_176993 | ENSMUSG00000070305 | **Mpzl3** | 5430427F17Rik\|A530065I17Rik\|rc | myelin protein zero-like 3 | protein-coding |
| chr5_96209388_96209836 | chr5 | 96209389 | 96209836 | + | -0.638652 | promoter-TSS (NM_053158) | promoter-TSS (NM_053158) | -503 | NM_001039084 | 94061 | Mm.295499 | NM_053158 | ENSMUSG00000029486 | **Mrpl1** | 2410002L03Rik\|5830418D04Rik\|AI462572\|L1mt | mitochondrial ribosomal protein L1 | protein-coding |
| chr5_31614710_31615005 | chr5 | 31614711 | 31615005 | + | -0.668283 | intron (NM_025796, intron 2 of 3) | intron (NM_025796, intron 2 of 3) | 907 | NM_025796 | 66845 | Mm.120763 | NM_025796 | ENSMUSG00000029142 | **Mrpl33** | 0610009M10Rik\|L33mt\|MRP-L33 | mitochondrial ribosomal protein L33 | protein-coding |
| chr11_88234011_88234383 | chr11 | 88234012 | 88234383 | + | -0.720005 | Intergenic | Intergenic | 29809 | NM_001291271 | 64656 | Mm.168680 | NM_024174 | ENSMUSG00000023723 | **Mrps23** | D11Bwg1153e\|MRP-S23\|Rpms23\|S23mt | mitochondrial ribosomal protein S23 | protein-coding |
| chr11_88194179_88194805 | chr11 | 88194180 | 88194805 | + | -0.857288 | TTS (NM_198013) | TTS (NM_198013) | 386 | NM_001291270 | 64656 | Mm.168680 | NM_024174 | ENSMUSG00000023723 | **Mrps23** | D11Bwg1153e\|MRP-S23\|Rpms23\|S23mt | mitochondrial ribosomal protein S23 | protein-coding |
| chr2_127618966_127619286 | chr2 | 127618967 | 127619286 | + | -0.662374 | Intergenic | L1M3\|LINE\|L1 | 31700 | NM_029963 | 77721 | Mm.274601 | NM_029963 | | **Mrps5** | 1620401I16Rik\|AI850294 | mitochondrial ribosomal protein S5 | protein-coding |
| chr13_24989635_24990113 | chr13 | 24989636 | 24990113 | + | -1.12679 | intron (NM_008156, intron 25 of 25) | intron (NM_008156, intron 25 of 25) | 30443 | NM_001013389 | 380836 | Mm.387097 | NM_001013389 | ENSMUSG00000021339 | **Mrs2** | Gm902\|HPT\|Mrs2l\|RPT | MRS2 magnesium homeostasis factor homolog (S. cerevisiae) | protein-coding |
| chr7_110977483_110978341 | chr7 | 110977484 | 110978341 | + | -0.89736 | intron (NM_194464, intron 1 of 20) | intron (NM_194464, intron 1 of 20) | 4549 | NM_194464 | 17540 | Mm.250023 | NM_010826 | ENSMUSG00000005611 | **Mrvi1** | BB115629\|Irag\|Ris1 | MRV integration site 1 | protein-coding |
| chr7_110977726_110978428 | chr7 | 110977727 | 110978428 | + | -0.974999 | intron (NM_194464, intron 1 of 20) | intron (NM_194464, intron 1 of 20) | 4384 | NM_194464 | 17540 | Mm.250023 | NM_010826 | ENSMUSG00000005611 | **Mrvi1** | BB115629\|Irag\|Ris1 | MRV integration site 1 | protein-coding |
| chr7_110965408_110965844 | chr7 | 110965409 | 110965844 | + | -1.14787 | intron (NM_194464, intron 1 of 20) | Lx9\|LINE\|L1 | 16835 | NM_194464 | 17540 | Mm.250023 | NM_010826 | ENSMUSG00000005611 | **Mrvi1** | BB115629\|Irag\|Ris1 | MRV integration site 1 | protein-coding |
| chr7_110965423_110966002 | chr7 | 110965424 | 110966002 | + | -1.16559 | intron (NM_194464, intron 1 of 20) | Lx9\|LINE\|L1 | 16748 | NM_194464 | 17540 | Mm.250023 | NM_010826 | ENSMUSG00000005611 | **Mrvi1** | BB115629\|Irag\|Ris1 | MRV integration site 1 | protein-coding |
| chr19_11444658_11444929 | chr19 | 11444659 | 11444929 | + | -0.611752 | intron (NM_021718, intron 1 of 6) | intron (NM_021718, intron 1 of 6) | 1236 | NM_021718 | 60361 | Mm.33957 | NM_021718 | ENSMUSG00000056290 | **Ms4a4b** | AI463180\|Chandra\|Ly116 | membrane-spanning 4-domains, subfamily A, member 4B | protein-coding |
| chr19_11449376_11449658 | chr19 | 11449377 | 11449658 | + | -0.751367 | intron (NM_021718, intron 1 of 6) | intron (NM_021718, intron 1 of 6) | 5959 | NM_021718 | 60361 | Mm.33957 | NM_021718 | ENSMUSG00000056290 | **Ms4a4b** | AI463180\|Chandra\|Ly116 | membrane-spanning 4-domains, subfamily A, member 4B | protein-coding |
| chr19_11434951_11435345 | chr19 | 11434952 | 11435345 | + | -1.13223 | Intergenic | L1VL4\|LINE\|L1 | -8410 | NM_021718 | 60361 | Mm.33957 | NM_021718 | ENSMUSG00000056290 | **Ms4a4b** | AI463180\|Chandra\|Ly116 | membrane-spanning 4-domains, subfamily A, member 4B | protein-coding |
| chr19_11414013_11414483 | chr19 | 11414014 | 11414483 | + | -0.671151 | intron (NM_029499, intron 1 of 6) | RMER4B\|LTR\|ERVK | 6587 | NM_029499 | 64380 | Mm.440715 | NM_029499 | ENSMUSG00000024675 | **Ms4a4c** | 5830413L19Rik\|AI449185\|Ms4a9 | membrane-spanning 4-domains, subfamily A, member 4C | protein-coding |
| chr19_11387747_11388057 | chr19 | 11387748 | 11388057 | + | -0.70088 | Intergenic | RMER19B2\|LTR\|ERVK | -19759 | NM_029499 | 64380 | Mm.440715 | NM_029499 | ENSMUSG00000024675 | **Ms4a4c** | 5830413L19Rik\|AI449185\|Ms4a9 | membrane-spanning 4-domains, subfamily A, member 4C | protein-coding |
| chr19_11544480_11544699 | chr19 | 11544481 | 11544699 | + | -0.641782 | intron (NM_025658, intron 1 of 6) | intron (NM_025658, intron 1 of 6) | 7741 | NM_025658 | 66607 | Mm.441056 | NM_025658 | ENSMUSG00000024678 | **Ms4a4d** | 2210417J23Rik | membrane-spanning 4-domains, subfamily A, member 4D | protein-coding |
| chr19_11547891_11548263 | chr19 | 11547892 | 11548263 | + | -1.04154 | intron (NM_025658, intron 1 of 6) | intron (NM_025658, intron 1 of 6) | 11228 | NM_025658 | 66607 | Mm.441056 | NM_025658 | ENSMUSG00000024678 | **Ms4a4d** | 2210417J23Rik | membrane-spanning 4-domains, subfamily A, member 4D | protein-coding |
| chr19_11516854_11517323 | chr19 | 11516855 | 11517323 | + | -0.671499 | Intergenic | MLTR18C_MM\|LTR\|ERVK | -1470 | NM_027209 | 69774 | Mm.436639 | NM_027209 | ENSMUSG00000024677 | **Ms4a6b** | 1810027D10Rik | membrane-spanning 4-domains, subfamily A, member 6B | protein-coding |
| chr19_11589229_11589606 | chr19 | 11589230 | 11589606 | + | -0.98834 | intron (NM_026835, intron 5 of 6) | intron (NM_026835, intron 5 of 6) | 15386 | NM_026835 | 68774 | Mm.290390 | NM_026835 | ENSMUSG00000024679 | **Ms4a6d** | 1110058E16Rik\|AI447446\|Ms4a11 | membrane-spanning 4-domains, subfamily A, member 6D | protein-coding |
| chr19_11355534_11356239 | chr19 | 11355535 | 11356239 | + | -0.794613 | Intergenic | Intergenic | -19741 | NM_027836 | 109225 | Mm.491251 | NM_027836 | ENSMUSG00000024672 | **Ms4a7** | 9130422I10Rik\|A430103C15Rik\|CD20l4\|CFFMA | membrane-spanning 4-domains, subfamily A, member 7 | protein-coding |
| chr19_11371201_11371512 | chr19 | 11371202 | 11371512 | + | -0.843676 | Intergenic | LTR65\|LTR\|ERV1 | -35211 | NM_027836 | 109225 | Mm.491251 | NM_027836 | ENSMUSG00000024672 | **Ms4a7** | 9130422I10Rik\|A430103C15Rik\|CD20l4\|CFFMA | membrane-spanning 4-domains, subfamily A, member 7 | protein-coding |
| chr12_11210421_11210756 | chr12 | 11210422 | 11210756 | + | -0.6559 | Intergenic | Intergenic | -1641 | NM_019544 | 56184 | Mm.117113 | NM_019544 | ENSMUSG00000047002 | **Msgn1** | pMsgn1 | mesogenin 1 | protein-coding |
| chr12_11214795_11215211 | chr12 | 11214796 | 11215211 | + | -0.679105 | Intergenic | Intergenic | -6055 | NM_019544 | 56184 | Mm.117113 | NM_019544 | ENSMUSG00000047002 | **Msgn1** | pMsgn1 | mesogenin 1 | protein-coding |
| chr12_11229868_11231188 | chr12 | 11229869 | 11231188 | + | -0.872338 | Intergenic | Intergenic | -21580 | NM_019544 | 56184 | Mm.117113 | NM_019544 | ENSMUSG00000047002 | **Msgn1** | pMsgn1 | mesogenin 1 | protein-coding |
| chr12_11229868_11230662 | chr12 | 11229869 | 11230662 | + | -0.955692 | Intergenic | ID_B1\|SINE\|B4 | -21317 | NM_019544 | 56184 | Mm.117113 | NM_019544 | ENSMUSG00000047002 | **Msgn1** | pMsgn1 | mesogenin 1 | protein-coding |
| chr9_101061019_101061608 | chr9 | 101061020 | 101061608 | + | -0.94034 | Intergenic | Intergenic | -13448 | NM_001100451 | 77853 | Mm.326206 | NM_001100451 | | **Msl2** | E130103E02Rik\|Msl2l1\|Rnf184 | male-specific lethal 2 homolog (Drosophila) | protein-coding |
| chr8_64723250_64723534 | chr8 | 64723251 | 64723534 | + | -0.830361 | intron (NM_025436, intron 3 of 5) | ID_B1\|SINE\|B4 | 10186 | NM_025436 | 66234 | Mm.30119 | NM_025436 | ENSMUSG00000031604 | **Msmo1** | 1500001G16Rik\|C78600\|DESP4\|ERG25\|Sc4mol | methylsterol monoxygenase 1 | protein-coding |
| chr8_64788624_64789133 | chr8 | 64788625 | 64789133 | + | -1.03614 | intron (NM_178633, intron 4 of 14) | intron (NM_178633, intron 4 of 14) | -55301 | NM_025436 | 66234 | Mm.30119 | NM_025436 | ENSMUSG00000031604 | **Msmo1** | 1500001G16Rik\|C78600\|DESP4\|ERG25\|Sc4mol | methylsterol monoxygenase 1 | protein-coding |
| chr14_64383022_64383256 | chr14 | 64383023 | 64383256 | + | -0.855481 | intron (NM_001253715, intron 1 of 4) | intron (NM_001253715, intron 1 of 4) | 33817 | NM_001253716 | 110265 | Mm.26713 | NM_026322 | ENSMUSG00000054733 | **Msra** | 2310045J23Rik\|6530413P12Rik\|MSR-A | methionine sulfoxide reductase A | protein-coding |
| chr14_64392492_64392962 | chr14 | 64392493 | 64392962 | + | -0.965209 | intron (NM_001253715, intron 1 of 4) | (CA)n\|Simple_repeat\|Simple_repeat | 24229 | NM_001253716 | 110265 | Mm.26713 | NM_026322 | ENSMUSG00000054733 | **Msra** | 2310045J23Rik\|6530413P12Rik\|MSR-A | methionine sulfoxide reductase A | protein-coding |
| chr14_64272166_64272505 | chr14 | 64272167 | 64272505 | + | -0.997369 | intron (NM_001253715, intron 2 of 4) | L1MC1\|LINE\|L1 | 144620 | NM_001253716 | 110265 | Mm.26713 | NM_026322 | ENSMUSG00000054733 | **Msra** | 2310045J23Rik\|6530413P12Rik\|MSR-A | methionine sulfoxide reductase A | protein-coding |
| chr14_64486632_64487257 | chr14 | 64486633 | 64487257 | + | -1.02789 | Intergenic | (TG)n\|Simple_repeat\|Simple_repeat | -31042 | NM_026322 | 110265 | Mm.26713 | NM_026322 | ENSMUSG00000054733 | **Msra** | 2310045J23Rik\|6530413P12Rik\|MSR-A | methionine sulfoxide reductase A | protein-coding |
| chr14_64486841_64487179 | chr14 | 64486842 | 64487179 | + | -1.03034 | Intergenic | Intergenic | -31107 | NM_026322 | 110265 | Mm.26713 | NM_026322 | ENSMUSG00000054733 | **Msra** | 2310045J23Rik\|6530413P12Rik\|MSR-A | methionine sulfoxide reductase A | protein-coding |
| chr10_120882166_120883286 | chr10 | 120882167 | 120883286 | + | -0.735768 | intron (NM_177092, intron 1 of 6) | intron (NM_177092, intron 1 of 6) | 16245 | NM_177092 | 320183 | Mm.169755 | NM_177092 | ENSMUSG00000051236 | **Msrb3** | D430026P16Rik\|I-8-23\|MsrB2 | methionine sulfoxide reductase B3 | protein-coding |
| chr10_120895714_120896348 | chr10 | 120895715 | 120896348 | + | -0.806492 | intron (NM_177092, intron 1 of 6) | intron (NM_177092, intron 1 of 6) | 2940 | NM_177092 | 320183 | Mm.169755 | NM_177092 | ENSMUSG00000051236 | **Msrb3** | D430026P16Rik\|I-8-23\|MsrB2 | methionine sulfoxide reductase B3 | protein-coding |
| chr10_120882220_120882894 | chr10 | 120882221 | 120882894 | + | -0.826359 | intron (NM_177092, intron 1 of 6) | intron (NM_177092, intron 1 of 6) | 16414 | NM_177092 | 320183 | Mm.169755 | NM_177092 | ENSMUSG00000051236 | **Msrb3** | D430026P16Rik\|I-8-23\|MsrB2 | methionine sulfoxide reductase B3 | protein-coding |
| chr17_83764626_83764903 | chr17 | 83764627 | 83764903 | + | -1.01294 | intron (NM_054082, intron 7 of 16) | intron (NM_054082, intron 7 of 16) | 58602 | NM_001171052 | 116871 | Mm.277668 | NM_054082 | ENSMUSG00000055817 | **Mta3** | 1110002J22Rik\|mKIAA1266 | metastasis associated 3 | protein-coding |
| chr15_34050225_34050530 | chr15 | 34050226 | 34050530 | + | -0.768751 | Intergenic | ORR1D1\|LTR\|ERVL-MaLR | -32341 | NM_026002 | 67154 | Mm.130883 | NM_026002 | ENSMUSG00000022255 | **Mtdh** | 2610103J23Rik\|3D3\|3D3/Lyric\|AEG-1\|AV353288\|D8Bwg1112e\|Lyric | metadherin | protein-coding |
| chr15_34050206_34050727 | chr15 | 34050207 | 34050727 | + | -0.769575 | Intergenic | Lx7\|LINE\|L1 | -32252 | NM_026002 | 67154 | Mm.130883 | NM_026002 | ENSMUSG00000022255 | **Mtdh** | 2610103J23Rik\|3D3\|3D3/Lyric\|AEG-1\|AV353288\|D8Bwg1112e\|Lyric | metadherin | protein-coding |
| chr3_19189663_19189905 | chr3 | 19189664 | 19189905 | + | -0.693873 | intron (NR_045575, intron 1 of 8) | intron (NR_045575, intron 1 of 8) | 1563 | NR_045575 | 67472 | Mm.45125 | NM_026182 | ENSMUSG00000027601 | **Mtfr1** | 1300002C08Rik\|1700080D04Rik\|2810026O10Rik\|4930579E05Rik\|Chppr\|Fam54a2 | mitochondrial fission regulator 1 | protein-coding |

| | | | | | | | | | | | | | | | | | |
|---|---|---|---|---|---|---|---|---|---|---|---|---|---|---|---|---|---|
| chr10_4099872_4100594 | chr10 | 4099873 | 4100594 | + | -0.856109 | intron (NM_001170785, intron 24 of 27) | intron (NM_001170785, intron 24 of 27) | 127158 | NM_172308 | 270685 | Mm.184752 | NM_172308 | ENSMUSG00000040675 | **Mthfd1l** | 2410004L15Rik\|AI647056\|Fthfsdc 1 | methylenetetrahydrofolate dehydrogenase (NADP+ dependent) 1-like | protein-coding |
| chr10_4091902_4092243 | chr10 | 4091903 | 4092243 | + | -0.881124 | intron (NM_001170785, intron 24 of 27) | intron (NM_001170785, intron 24 of 27) | 118998 | NM_172308 | 270685 | Mm.184752 | NM_172308 | ENSMUSG00000040675 | **Mthfd1l** | 2410004L15Rik\|AI647056\|Fthfsdc 1 | methylenetetrahydrofolate dehydrogenase (NADP+ dependent) 1-like | protein-coding |
| chr5_146973939_146974550 | chr5 | 146973940 | 146974550 | + | -0.639221 | Intergenic | Intergenic | -10448 | NM_001256102 | 76366 | Mm.486234 | NM_029581 | ENSMUSG00000016510 | **Mtif3** | 2810012L14Rik\|AI414549 | mitochondrial translational initiation factor 3 | protein-coding |
| chrX_71378498_71378868 | chrX | 71378499 | 71378868 | + | -0.871433 | intron (NM_016985, intron 2 of 14) | intron (NM_016985, intron 2 of 14) | 13923 | NM_016985 | 53332 | Mm.219672 | NM_016985 | ENSMUSG00000015214 | **Mtmr1** | AW049210 | myotubularin related protein 1 | protein-coding |
| chr15_12229607_12229878 | chr15 | 12229608 | 12229878 | + | -0.807499 | intron (NM_172958, intron 1 of 15) | B4A\|SINE\|B4 | 24649 | NM_172958 | 268783 | Mm.54460 | NM_172958 | ENSMUSG00000039458 | **Mtmr12** | 3Pap\|4932703C11\|C730015A02Ri k\|Pip3ap\|mKIAA1682 | myotubularin related protein 12 | protein-coding |
| chr15_12229597_12230145 | chr15 | 12229598 | 12230145 | + | -0.809073 | intron (NM_172958, intron 1 of 15) | ID\|SINE\|ID | 24777 | NM_172958 | 268783 | Mm.54460 | NM_172958 | ENSMUSG00000039458 | **Mtmr12** | 3Pap\|4932703C11\|C730015A02Ri k\|Pip3ap\|mKIAA1682 | myotubularin related protein 12 | protein-coding |
| chr15_12225061_12225471 | chr15 | 12225062 | 12225471 | + | -0.955707 | intron (NM_172958, intron 1 of 15) | intron (NM_172958, intron 1 of 15) | 20172 | NM_172958 | 268783 | Mm.54460 | NM_172958 | ENSMUSG00000039458 | **Mtmr12** | 3Pap\|4932703C11\|C730015A02Ri k\|Pip3ap\|mKIAA1682 | myotubularin related protein 12 | protein-coding |
| chr15_12225070_12225355 | chr15 | 12225071 | 12225355 | + | -1.06986 | intron (NM_172958, intron 1 of 15) | intron (NM_172958, intron 1 of 15) | 20119 | NM_172958 | 268783 | Mm.54460 | NM_172958 | ENSMUSG00000039458 | **Mtmr12** | 3Pap\|4932703C11\|C730015A02Ri k\|Pip3ap\|mKIAA1682 | myotubularin related protein 12 | protein-coding |
| chr11_4536720_4537286 | chr11 | 4536721 | 4537286 | + | -0.621888 | intron (NM_028860, intron 1 of 17) | intron (NM_028860, intron 1 of 17) | 57812 | NM_028860 | 74302 | Mm.425669 | NM_028860 | ENSMUSG00000034354 | **Mtmr3** | 1700092A20Rik\|AI255150\|AW557 713\|FYVE-DSP1\|ZFYVE10\|mKIAA0371 | myotubularin related protein 3 | protein-coding |
| chr11_4536648_4536986 | chr11 | 4536649 | 4536986 | + | -0.623386 | intron (NM_028860, intron 1 of 17) | PB1D10\|SINE\|Alu | 57998 | NM_028860 | 74302 | Mm.425669 | NM_028860 | ENSMUSG00000034354 | **Mtmr3** | 1700092A20Rik\|AI255150\|AW557 713\|FYVE-DSP1\|ZFYVE10\|mKIAA0371 | myotubularin related protein 3 | protein-coding |
| chr6_35462464_35463069 | chr6 | 35462465 | 35463069 | + | -0.977913 | Intergenic | Intergenic | 77121 | NM_008098 | 14489 | Mm.182746 | NM_008098 | ENSMUSG00000029840 | **Mtpn** | 5033418D15Rik\|Gcdp\|V1 | myotrophin | protein-coding |
| chr6_35462464_35462866 | chr6 | 35462465 | 35462866 | + | -0.999577 | Intergenic | Intergenic | 77223 | NM_008098 | 14489 | Mm.182746 | NM_008098 | ENSMUSG00000029840 | **Mtpn** | 5033418D15Rik\|Gcdp\|V1 | myotrophin | protein-coding |
| chr13_12273938_12274306 | chr13 | 12273939 | 12274306 | + | -0.714862 | intron (NM_033268, intron 18 of 20) | B1F2\|SINE\|Alu | -16009 | NM_001081128 | 238505 | Mm.40335 | NM_001081128 | ENSMUSG00000021311 | **Mtr** | AI894170\|D830038K18Rik\|MS | 5-methyltetrahydrofolate-homocysteine methyltransferase | protein-coding |
| chr13_12277896_12278293 | chr13 | 12277897 | 12278293 | + | -1.00981 | intron (NM_033268, intron 15 of 20) | Charlie4a\|DNA\|hAT-Charlie | -19982 | NM_001081128 | 238505 | Mm.40335 | NM_001081128 | ENSMUSG00000021311 | **Mtr** | AI894170\|D830038K18Rik\|MS | 5-methyltetrahydrofolate-homocysteine methyltransferase | protein-coding |
| chr15_59089601_59090164 | chr15 | 59089602 | 59090164 | + | -0.697044 | Intergenic | LTR33\|LTR\|ERVL | -7857 | NM_144800 | 211401 | Mm.215481 | NM_144800 | ENSMUSG00000022353 | **Mtss1** | 2310003N14Rik\|BC024131\|D1300 01D01Rik\|Mim\|mKIAA0429 | metastasis suppressor 1 | protein-coding |
| chr6_54715566_54716277 | chr6 | 54715567 | 54716277 | + | -0.672411 | Intergenic | Intergenic | 34298 | NM_001289740 | 68235 | Mm.233860 | NM_026629 | ENSMUSG00000038065 | **Mturn** | 2410066E13Rik\|AI448550\|B23021 2L03Rik | maturin, neural progenitor differentiation regulator homolog (Xenopus) | protein-coding |
| chr6_54659712_54659946 | chr6 | 54659713 | 54659946 | + | -0.681511 | Intergenic | MIR\|SINE\|MIR | -21795 | NM_001289741 | 68235 | Mm.233860 | NM_026629 | ENSMUSG00000038065 | **Mturn** | 2410066E13Rik\|AI448550\|B23021 2L03Rik | maturin, neural progenitor differentiation regulator homolog (Xenopus) | protein-coding |
| chr6_54715644_54716032 | chr6 | 54715645 | 54716032 | + | -0.692225 | Intergenic | Intergenic | 34214 | NM_001289740 | 68235 | Mm.233860 | NM_026629 | ENSMUSG00000038065 | **Mturn** | 2410066E13Rik\|AI448550\|B23021 2L03Rik | maturin, neural progenitor differentiation regulator homolog (Xenopus) | protein-coding |
| chr6_54685017_54685410 | chr6 | 54685018 | 54685410 | + | -0.737894 | intron (NM_001289741, intron 1 of 4) | MIRc\|SINE\|MIR | 3590 | NM_001289740 | 68235 | Mm.233860 | NM_026629 | ENSMUSG00000038065 | **Mturn** | 2410066E13Rik\|AI448550\|B23021 2L03Rik | maturin, neural progenitor differentiation regulator homolog (Xenopus) | protein-coding |
| chr2_75054572_75055206 | chr2 | 75054573 | 75055206 | + | -1.16863 | Intergenic | Intergenic | 229077 | NM_016804 | 53375 | Mm.292613 | NM_016804 | | **Mtx2** | 1500012G02Rik | metaxin 2 | protein-coding |
| chr2_75054811_75055412 | chr2 | 75054812 | 75055412 | + | -1.17793 | Intergenic | Intergenic | 229300 | NM_016804 | 53375 | Mm.292613 | NM_016804 | | **Mtx2** | 1500012G02Rik | metaxin 2 | protein-coding |
| chr2_75054804_75055194 | chr2 | 75054805 | 75055194 | + | -1.2273 | Intergenic | Intergenic | 229187 | NM_016804 | 53375 | Mm.292613 | NM_016804 | | **Mtx2** | 1500012G02Rik | metaxin 2 | protein-coding |
| chr2_75073211_75073588 | chr2 | 75073212 | 75073588 | + | -1.37623 | Intergenic | Intergenic | 247588 | NM_016804 | 53375 | Mm.292613 | NM_016804 | | **Mtx2** | 1500012G02Rik | metaxin 2 | protein-coding |
| chr10_80211746_80212156 | chr10 | 80211747 | 80212156 | + | -1.00313 | intron (NR_038088, intron 1 of 4) | intron (NR_038088, intron 1 of 4) | -14647 | NM_023431 | 68114 | Mm.273418 | NM_023431 | ENSMUSG00000020156 | **Mum1** | 2610019J09\|9430059D04Rik\|UBE-1C2 | melanoma associated antigen (mutated) 1 | protein-coding |
| chr2_34041991_34042709 | chr2 | 34041992 | 34042709 | + | -0.813761 | Intergenic | Intergenic | -154404 | NM_175184 | 72543 | Mm.254914 | NM_175184 | ENSMUSG00000038740 | **Mvb12b** | 2610200O14Rik\|2610528K11Rik\|A I414895\|Fam125b | multivesicular body subunit 12B | protein-coding |
| chr2_33908317_33908872 | chr2 | 33908318 | 33908872 | + | -1.18081 | Intergenic | Intergenic | -20649 | NM_175184 | 72543 | Mm.254914 | NM_175184 | ENSMUSG00000038740 | **Mvb12b** | 2610200O14Rik\|2610528K11Rik\|A I414895\|Fam125b | multivesicular body subunit 12B | protein-coding |
| chr2_34073753_34074463 | chr2 | 34073754 | 34074463 | + | -1.28504 | Intergenic | MTD-int\|LTR\|ERVL-MaLR | -186162 | NM_175184 | 72543 | Mm.254914 | NM_175184 | ENSMUSG00000038740 | **Mvb12b** | 2610200O14Rik\|2610528K11Rik\|A I414895\|Fam125b | multivesicular body subunit 12B | protein-coding |
| chr5_114505217_114506025 | chr5 | 114505218 | 114506025 | + | -1.03608 | Intergenic | ORR1D2\|LTR\|ERVL-MaLR | 61315 | NM_023556 | 17855 | Mm.28088 | NM_023556 | ENSMUSG00000041939 | **Mvk** | 2310010A05Rik\|AI256848\|AI4140 37\|MK | mevalonate kinase | protein-coding |
| chr5_114505263_114505794 | chr5 | 114505264 | 114505794 | + | -1.06231 | Intergenic | ORR1D2\|LTR\|ERVL-MaLR | 61223 | NM_023556 | 17855 | Mm.28088 | NM_023556 | ENSMUSG00000041939 | **Mvk** | 2310010A05Rik\|AI256848\|AI4140 37\|MK | mevalonate kinase | protein-coding |
| chr5_34198651_34199057 | chr5 | 34198652 | 34199057 | + | -1.07359 | exon (NM_001015039, exon 10 of 13) | exon (NM_001015039, exon 10 of 13) | -11144 | NM_010753 | 17122 | Mm.259260 | NM_010753 | ENSMUSG00000037235 | **Mxd4** | 2810410A03Rik\|Mad4\|bHLHc12 | Max dimerization protein 4 | protein-coding |
| chr19_53299833_53300164 | chr19 | 53299834 | 53300164 | + | -0.90504 | Intergenic | Intergenic | -10507 | NM_001008542 | 17859 | Mm.2154 | NM_010847 | ENSMUSG00000025025 | **Mxi1** | ENSMUSG00000067085\|Gm1019 7\|Mad2\|bHLHc11 | Max interacting protein 1 | protein-coding |
| chr15_77877267_77877711 | chr15 | 77877268 | 77877711 | + | -0.619258 | Intergenic | Intergenic | -35314 | NM_022410 | 17886 | Mm.29677 | NM_022410 | ENSMUSG00000022443 | **Myh9** | Fltn\|Myhn-1\|Myhn1\|NMHCIIA\|NMMHC-A\|NMMHC-IIA\|TU72.6 | myosin, heavy polypeptide 9, non-muscle | protein-coding |
| chr15_77799941_77800177 | chr15 | 77799942 | 77800177 | + | -0.677012 | intron (NM_022410, intron 3 of 40) | intron (NM_022410, intron 3 of 40) | 42116 | NM_022410 | 17886 | Mm.29677 | NM_022410 | ENSMUSG00000022443 | **Myh9** | Fltn\|Myhn-1\|Myhn1\|NMHCIIA\|NMMHC-A\|NMMHC-IIA\|TU72.6 | myosin, heavy polypeptide 9, non-muscle | protein-coding |
| chr15_77810454_77810954 | chr15 | 77810455 | 77810954 | + | -0.701042 | intron (NM_022410, intron 2 of 40) | intron (NM_022410, intron 2 of 40) | 31471 | NM_022410 | 17886 | Mm.29677 | NM_022410 | ENSMUSG00000022443 | **Myh9** | Fltn\|Myhn-1\|Myhn1\|NMHCIIA\|NMMHC-A\|NMMHC-IIA\|TU72.6 | myosin, heavy polypeptide 9, non-muscle | protein-coding |
| chr15_77849512_77850011 | chr15 | 77849513 | 77850011 | + | -0.764015 | Intergenic | Intergenic | -7587 | NM_022410 | 17886 | Mm.29677 | NM_022410 | ENSMUSG00000022443 | **Myh9** | Fltn\|Myhn-1\|Myhn1\|NMHCIIA\|NMMHC-A\|NMMHC-IIA\|TU72.6 | myosin, heavy polypeptide 9, non-muscle | protein-coding |
| chr15_77805671_77806170 | chr15 | 77805672 | 77806170 | + | -0.88449 | intron (NM_022410, intron 3 of 40) | intron (NM_022410, intron 3 of 40) | 36254 | NM_022410 | 17886 | Mm.29677 | NM_022410 | ENSMUSG00000022443 | **Myh9** | Fltn\|Myhn-1\|Myhn1\|NMHCIIA\|NMMHC-A\|NMMHC-IIA\|TU72.6 | myosin, heavy polypeptide 9, non-muscle | protein-coding |
| chr15_77834261_77834900 | chr15 | 77834262 | 77834900 | + | -0.99716 | intron (NM_022410, intron 1 of 40) | intron (NM_022410, intron 1 of 40) | 7594 | NM_022410 | 17886 | Mm.29677 | NM_022410 | ENSMUSG00000022443 | **Myh9** | Fltn\|Myhn-1\|Myhn1\|NMHCIIA\|NMMHC-A\|NMMHC-IIA\|TU72.6 | myosin, heavy polypeptide 9, non-muscle | protein-coding |
| chr15_77834299_77834697 | chr15 | 77834300 | 77834697 | + | -1.02402 | intron (NM_022410, intron 1 of 40) | intron (NM_022410, intron 1 of 40) | 7677 | NM_022410 | 17886 | Mm.29677 | NM_022410 | ENSMUSG00000022443 | **Myh9** | Fltn\|Myhn-1\|Myhn1\|NMHCIIA\|NMMHC-A\|NMMHC-IIA\|TU72.6 | myosin, heavy polypeptide 9, non-muscle | protein-coding |
| chr5_136646777_136647096 | chr5 | 136646778 | 136647096 | + | -0.802429 | Intergenic | Intergenic | -46209 | NM_021611 | 59310 | Mm.5178 | NM_021611 | ENSMUSG00000005474 | **Myl10** | 1700027I08Rik\|Mylc2pl\|PLRLC\|PL RLC-A\|PLRLC-B\|PLRLC-C | myosin, light chain 10, regulatory | protein-coding |
| chr5_136646704_136647136 | chr5 | 136646705 | 136647136 | + | -0.820701 | Intergenic | Intergenic | -46226 | NM_021611 | 59310 | Mm.5178 | NM_021611 | ENSMUSG00000005474 | **Myl10** | 1700027I08Rik\|Mylc2pl\|PLRLC\|PL RLC-A\|PLRLC-B\|PLRLC-C | myosin, light chain 10, regulatory | protein-coding |
| chr17_70985909_70986382 | chr17 | 70985910 | 70986382 | + | -0.750506 | intron (NM_023402, intron 1 of 3) | intron (NM_023402, intron 1 of 3) | 4370 | NM_023402 | 67938 | Mm.261329 | NM_023402 | | **Myl12b** | 1500001M02Rik\|C77744\|Mylc2b\|R LC-B | myosin, light chain 12B, regulatory | protein-coding |
| chr10_128504478_128505295 | chr10 | 128504479 | 128505295 | + | -0.627995 | Intergenic | Intergenic | -6202 | NM_172259 | 216459 | Mm.291688 | NM_172259 | ENSMUSG00000039824 | **Myl6b** | 5730437E04Rik\|BC037527\|MLC1S A | myosin, light polypeptide 6B | protein-coding |
| chr13_45391578_45391865 | chr13 | 45391579 | 45391865 | + | -0.684568 | intron (NM_153789, intron 2 of 6) | intron (NM_153789, intron 2 of 6) | 1980 | NM_153789 | 218203 | Mm.212855 | NM_153789 | ENSMUSG00000038175 | **Mylip** | 9430057C20Rik\|Mir | myosin regulatory light chain interacting protein | protein-coding |
| chr13_45371703_45372018 | chr13 | 45371704 | 45372018 | + | -0.978604 | Intergenic | Intergenic | -17881 | NM_153789 | 218203 | Mm.212855 | NM_153789 | ENSMUSG00000038175 | **Mylip** | 9430057C20Rik\|Mir | myosin regulatory light chain interacting protein | protein-coding |
| chr3_30610890_30611559 | chr3 | 30610891 | 30611559 | + | -0.704123 | intron (NM_030557, intron 4 of 7) | intron (NM_030557, intron 4 of 7) | 9138 | NM_001289621 | 80732 | Mm.200378 | NM_030557 | ENSMUSG00000037730 | **Mynn** | 2810011C24Rik\|AA415053\|AI6610 31\|AW049392\|Mynr\|SBBIZ1 | myoneurin | protein-coding |
| chr11_77791868_77792405 | chr11 | 77791869 | 77792405 | + | -0.863385 | intron (NM_011586, intron 1 of 39) | intron (NM_011586, intron 1 of 39) | -9161 | NM_001291214 | 360013 | Mm.341248 | NM_011586 | ENSMUSG00000000631 | **Myo18a** | MAJN\|MyoPDZ\|MysPDZ | myosin XVIIIA | protein-coding |
| chr1_51934174_51935155 | chr1 | 51934175 | 51935155 | + | -0.992963 | Intergenic | Intergenic | -18687 | NM_001161817 | 17912 | Mm.3390 | NM_010863 | ENSMUSG00000018417 | **Myo1b** | AA406997\|MIH-L\|MMIa | myosin IB | protein-coding |
| chr1_51929491_51930119 | chr1 | 51929492 | 51930119 | + | -1.37753 | Intergenic | Intergenic | -13827 | NM_001161817 | 17912 | Mm.3390 | NM_010863 | ENSMUSG00000018417 | **Myo1b** | AA406997\|MIH-L\|MMIa | myosin IB | protein-coding |
| chr9_80313565_80313961 | chr9 | 80313566 | 80313961 | + | -0.692639 | 3' UTR (NM_022016, exon 17 of 17) | 3' UTR (NM_022016, exon 17 of 17) | 148729 | NM_001039546 | 17920 | Mm.4040 | NM_008662 | ENSMUSG00000033577 | **Myo6** | BC029719\|Tlc\|sv | myosin VI | protein-coding |

| | | | | | | | | | | | | | | | | | |
|---|---|---|---|---|---|---|---|---|---|---|---|---|---|---|---|---|---|
| chr9_80313692_80313982 | chr9 | 80313693 | 80313982 | + | -0.720415 | 3' UTR (NM_022016, exon 17 of 17) | 3' UTR (NM_022016, exon 17 of 17) | 148803 | NM_001039546 | 17920 | Mm.4040 | NM_008662 | ENSMUSG00000033577 | **Myo6** | BC029719\|Tlc\|sv | myosin VI | protein-coding |
| chr1_162615498_162615923 | chr1 | 162615499 | 162615923 | + | -0.814076 | Intergenic | RMER17C-int\|LTR\|ERVK | -23439 | NM_010865 | 17926 | Mm.10694 | NM_010865 | ENSMUSG00000026697 | **Myoc** | AI957332\|GLC1A\|TIGR | myocilin | protein-coding |
| chr19_38030748_38031105 | chr19 | 38030749 | 38031105 | + | -0.85627 | intron (NM_001099634, intron 1 of 52) | RLTR26C_MM\|LTR\|ERVK | 12650 | NM_001099634 | 226101 | Mm.34674 | NM_177035 | ENSMUSG00000048612 | **Myof** | 2310004N10Rik\|2310051D19Rik\|E030042N20Rik\|Fer1l3 | myoferlin | protein-coding |
| chr10_116913590_116914265 | chr10 | 116913591 | 116914265 | + | -0.794914 | intron (NM_001003950, intron 7 of 9) | intron (NM_001003950, intron 7 of 9) | -17049 | NM_001033333 | 237558 | Mm.336386 | NM_001033333 | ENSMUSG00000034057 | **Myrfl** | Gm239 | myelin regulatory factor-like | protein-coding |
| chr9_120259931_120260211 | chr9 | 120259932 | 120260211 | + | -0.639752 | Intergenic | Intergenic | -44002 | NM_144557 | 245049 | Mm.100936 | NM_144557 | ENSMUSG00000041794 | **Myrip** | A230081N12Rik\|Slac2-c\|Slac2c | myosin VIIA and Rab interacting protein | protein-coding |
| chr9_120289278_120289570 | chr9 | 120289279 | 120289570 | + | -0.646489 | Intergenic | MTEb\|LTR\|ERVL-MaLR | -14649 | NM_144557 | 245049 | Mm.100936 | NM_144557 | ENSMUSG00000041794 | **Myrip** | A230081N12Rik\|Slac2-c\|Slac2c | myosin VIIA and Rab interacting protein | protein-coding |
| chr9_120289131_120289526 | chr9 | 120289132 | 120289526 | + | -0.708919 | Intergenic | MTEb\|LTR\|ERVL-MaLR | -14744 | NM_144557 | 245049 | Mm.100936 | NM_144557 | ENSMUSG00000041794 | **Myrip** | A230081N12Rik\|Slac2-c\|Slac2c | myosin VIIA and Rab interacting protein | protein-coding |
| chr9_120289252_120289683 | chr9 | 120289253 | 120289683 | + | -0.745359 | Intergenic | Intergenic | -14605 | NM_144557 | 245049 | Mm.100936 | NM_144557 | ENSMUSG00000041794 | **Myrip** | A230081N12Rik\|Slac2-c\|Slac2c | myosin VIIA and Rab interacting protein | protein-coding |
| chr14_99041392_99041614 | chr14 | 99041393 | 99041614 | + | -0.78637 | intron (NM_175245, intron 1 of 2) | Lx6\|LINE\|L1 | 4633 | NM_175245 | 76789 | Mm.38912 | NM_175245 | ENSMUSG00000033186 | **Mzt1** | 2410129H14Rik\|AI120158\|Mozart1 | mitotic spindle organizing protein 1 | protein-coding |
| chr16_87324480_87324938 | chr16 | 87324481 | 87324938 | + | -0.878955 | Intergenic | Intergenic | -29476 | NM_001159331 | 67768 | Mm.296532 | NM_026366 | ENSMUSG00000044442 | **N6amt1** | 5830445C04Rik\|Hemk2\|Pred28 | N-6 adenine-specific DNA methyltransferase 1 (putative) | protein-coding |
| chr14_57616449_57616972 | chr14 | 57616450 | 57616972 | + | -0.710644 | intron (NM_020506, intron 7 of 22) | intron (NM_020506, intron 7 of 22) | -45142 | NM_026526 | 68043 | Mm.45225 | NM_026526 | ENSMUSG00000021951 | **N6amt2** | 2510005D08Rik\|AW045965\|Ayu21-96\|Gt(pU21)96Imeg\|GtAyu21-96 | N-6 adenine-specific DNA methyltransferase 2 (putative) | protein-coding |
| chr14_57611101_57611493 | chr14 | 57611102 | 57611493 | + | -0.813647 | intron (NM_020506, intron 9 of 22) | intron (NM_020506, intron 9 of 22) | -39728 | NM_026526 | 68043 | Mm.45225 | NM_026526 | ENSMUSG00000021951 | **N6amt2** | 2510005D08Rik\|AW045965\|Ayu21-96\|Gt(pU21)96Imeg\|GtAyu21-96 | N-6 adenine-specific DNA methyltransferase 2 (putative) | protein-coding |
| chr1_51584657_51585516 | chr1 | 51584658 | 51585516 | + | -0.684531 | Intergenic | ORR1F\|LTR\|ERVL-MaLR | -106688 | NM_028696 | 109019 | Mm.196290 | NM_028696 | ENSMUSG00000026107 | **Nabp1** | 4930434H03Rik\|4930442A21Rik\|4930488J04Rik\|4933440J18Rik\|5830411E10Rik\|AI852561\|Nbp1\|Obfc2a\|Ssb2 | nucleic acid binding protein 1 | protein-coding |
| chr1_51577683_51578237 | chr1 | 51577684 | 51578237 | + | -0.687721 | Intergenic | MLT1A0\|LTR\|ERVL-MaLR | -99561 | NM_028696 | 109019 | Mm.196290 | NM_028696 | ENSMUSG00000026107 | **Nabp1** | 4930434H03Rik\|4930442A21Rik\|4930488J04Rik\|4933440J18Rik\|5830411E10Rik\|AI852561\|Nbp1\|Obfc2a\|Ssb2 | nucleic acid binding protein 1 | protein-coding |
| chr1_51626391_51626672 | chr1 | 51626392 | 51626672 | + | -0.722314 | Intergenic | Intergenic | -148133 | NM_028696 | 109019 | Mm.196290 | NM_028696 | ENSMUSG00000026107 | **Nabp1** | 4930434H03Rik\|4930442A21Rik\|4930488J04Rik\|4933440J18Rik\|5830411E10Rik\|AI852561\|Nbp1\|Obfc2a\|Ssb2 | nucleic acid binding protein 1 | protein-coding |
| chr1_51529680_51530153 | chr1 | 51529681 | 51530153 | + | -0.827753 | Intergenic | Intergenic | -51518 | NM_028696 | 109019 | Mm.196290 | NM_028696 | ENSMUSG00000026107 | **Nabp1** | 4930434H03Rik\|4930442A21Rik\|4930488J04Rik\|4933440J18Rik\|5830411E10Rik\|AI852561\|Nbp1\|Obfc2a\|Ssb2 | nucleic acid binding protein 1 | protein-coding |
| chr11_101063510_101064045 | chr11 | 101063511 | 101064045 | + | -0.620737 | TTS (NM_016920) | TTS (NM_016920) | -6316 | NM_013792 | 27419 | Mm.95452 | NM_013792 | ENSMUSG00000001751 | **Naglu** | - | alpha-N-acetylglucosaminidase (Sanfilippo disease IIIB) | protein-coding |
| chr11_101063512_101064398 | chr11 | 101063513 | 101064398 | + | -0.655761 | TTS (NM_016920) | TTS (NM_016920) | -6139 | NM_013792 | 27419 | Mm.95452 | NM_013792 | ENSMUSG00000001751 | **Naglu** | - | alpha-N-acetylglucosaminidase (Sanfilippo disease IIIB) | protein-coding |
| chr11_101063657_101064045 | chr11 | 101063658 | 101064045 | + | -0.664576 | TTS (NM_016920) | TTS (NM_016920) | -6243 | NM_013792 | 27419 | Mm.95452 | NM_013792 | ENSMUSG00000001751 | **Naglu** | - | alpha-N-acetylglucosaminidase (Sanfilippo disease IIIB) | protein-coding |
| chr2_32447682_32447984 | chr2 | 32447683 | 32447984 | + | -1.09804 | intron (NM_146118, intron 1 of 10) | intron (NM_146118, intron 1 of 10) | -2624 | NM_194335 | 71254 | Mm.102341 | NM_194335 | | **Naif1** | 2310007O20Rik\|4933440H19Rik | nuclear apoptosis inducing factor 1 | protein-coding |
| chr4_46478786_46479635 | chr4 | 46478787 | 46479635 | + | -0.682406 | Intergenic | ORR1B1\|LTR\|ERVL-MaLR | -10118 | NM_053179 | 94181 | Mm.249349 | NM_053179 | ENSMUSG00000028334 | **Nans** | 4632418E04Rik\|BB056474\|Sas | N-acetylneuraminic acid synthase (sialic acid synthase) | protein-coding |
| chr6_58880521_58881271 | chr6 | 58880522 | 58881271 | + | -0.701821 | intron (NM_028705, intron 15 of 24) | RMER1C\|Other\|Other | 26230 | NM_021432 | 58243 | Mm.389747 | NM_021432 | ENSMUSG00000055430 | **Nap1l5** | 1110020M21Rik | nucleosome assembly protein 1-like 5 | protein-coding |
| chr2_103672723_103673097 | chr2 | 103672724 | 103673097 | + | -0.626046 | intron (NM_178890, intron 1 of 16) | intron (NM_178890, intron 1 of 16) | 88340 | NM_153126 | 98956 | Mm.288005 | NM_153126 | ENSMUSG00000027185 | **Nat10** | AI429152 | N-acetyltransferase 10 | protein-coding |
| chr2_103703146_103704040 | chr2 | 103703147 | 103704040 | + | -0.626502 | intron (NM_178890, intron 5 of 16) | intron (NM_178890, intron 5 of 16) | 57657 | NM_153126 | 98956 | Mm.288005 | NM_153126 | ENSMUSG00000027185 | **Nat10** | AI429152 | N-acetyltransferase 10 | protein-coding |
| chr2_103682134_103682824 | chr2 | 103682135 | 103682824 | + | -0.740039 | intron (NM_178890, intron 2 of 16) | intron (NM_178890, intron 2 of 16) | 78771 | NM_153126 | 98956 | Mm.288005 | NM_153126 | ENSMUSG00000027185 | **Nat10** | AI429152 | N-acetyltransferase 10 | protein-coding |
| chr1_135631583_135631829 | chr1 | 135631584 | 135631829 | + | -0.716567 | Intergenic | Intergenic | -46351 | NM_173437 | 215690 | Mm.34977 | NM_173437 | ENSMUSG00000009418 | **Nav1** | 9530089B19\|9930003A20Rik\|C230080M11Rik\|Pomfil3\|Unc53h1\|mKIAA1151\|steerin-1 | neuron navigator 1 | protein-coding |
| chr1_135631383_135631971 | chr1 | 135631384 | 135631971 | + | -0.758175 | Intergenic | Intergenic | -46322 | NM_173437 | 215690 | Mm.34977 | NM_173437 | ENSMUSG00000009418 | **Nav1** | 9530089B19\|9930003A20Rik\|C230080M11Rik\|Pomfil3\|Unc53h1\|mKIAA1151\|steerin-1 | neuron navigator 1 | protein-coding |
| chr7_49364365_49365123 | chr7 | 49364366 | 49365123 | + | -0.62104 | intron (NM_001111016, intron 3 of 38) | B4A\|SINE\|B4 | 118555 | NM_175272 | 78286 | Mm.333881 | NM_175272 | ENSMUSG00000052512 | **Nav2** | 5330421F07Rik\|AU067780\|BQ960892\|HELAD1\|POMFIL2\|RAINB1\|RAINB2\|Unc53H2\|mKIAA3015 | neuron navigator 2 | protein-coding |
| chr7_48937452_48938151 | chr7 | 48937453 | 48938151 | + | -0.86111 | Intergenic | Intergenic | -21271 | NM_001111016 | 78286 | Mm.333881 | NM_175272 | ENSMUSG00000052512 | **Nav2** | 5330421F07Rik\|AU067780\|BQ960892\|HELAD1\|POMFIL2\|RAINB1\|RAINB2\|Unc53H2\|mKIAA3015 | neuron navigator 2 | protein-coding |
| chr12_13518121_13518620 | chr12 | 13518122 | 13518620 | + | -1.10043 | intron (NM_027706, intron 45 of 51) | intron (NM_027706, intron 45 of 51) | 249244 | NM_027706 | 71169 | Mm.177520 | NM_027706 | ENSMUSG00000020576 | **Nbas** | 4933425L03Rik\|Nag | neuroblastoma amplified sequence | protein-coding |
| chr12_13518126_13518471 | chr12 | 13518127 | 13518471 | + | -1.1281 | intron (NM_027706, intron 45 of 51) | intron (NM_027706, intron 45 of 51) | 249172 | NM_027706 | 71169 | Mm.177520 | NM_027706 | ENSMUSG00000020576 | **Nbas** | 4933425L03Rik\|Nag | neuroblastoma amplified sequence | protein-coding |
| chr4_46163962_46164393 | chr4 | 46163963 | 46164393 | + | -0.903109 | intron (NM_001033201, intron 15 of 22) | intron (NM_001033201, intron 15 of 22) | 25667 | NM_001033201 | 433702 | Mm.389536 | NM_001033201 | ENSMUSG00000028330 | **Ncbp1** | AU014645\|AW538051\|CBP80 | nuclear cap binding protein subunit 1 | protein-coding |
| chr4_46151753_46152424 | chr4 | 46151754 | 46152424 | + | -1.4543 | intron (NM_001033201, intron 7 of 22) | intron (NM_001033201, intron 7 of 22) | 13578 | NM_001033201 | 433702 | Mm.389536 | NM_001033201 | ENSMUSG00000028330 | **Ncbp1** | AU014645\|AW538051\|CBP80 | nuclear cap binding protein subunit 1 | protein-coding |
| chr5_134266288_134266850 | chr5 | 134266289 | 134266850 | + | -0.632233 | exon (NM_001080746, exon 13 of 35) | exon (NM_001080746, exon 13 of 35) | -36944 | NR_104383 | 17969 | Mm.425296 | NM_010876 | ENSMUSG00000015950 | **Ncf1** | NCF-47K\|NOXO2\|Ncf-1\|p47-phox\|p47<phox>\|p47phox | neutrophil cytosolic factor 1 | protein-coding |
| chr5_134236623_134237550 | chr5 | 134236624 | 134237550 | + | -0.798574 | TTS (NM_001080746) | TTS (NM_001080746) | -7462 | NR_104383 | 17969 | Mm.425296 | NM_010876 | ENSMUSG00000015950 | **Ncf1** | NCF-47K\|NOXO2\|Ncf-1\|p47-phox\|p47<phox>\|p47phox | neutrophil cytosolic factor 1 | protein-coding |
| chr5_134236625_134237315 | chr5 | 134236626 | 134237315 | + | -0.831029 | TTS (NM_001080746) | TTS (NM_001080746) | -7345 | NR_104383 | 17969 | Mm.425296 | NM_010876 | ENSMUSG00000015950 | **Ncf1** | NCF-47K\|NOXO2\|Ncf-1\|p47-phox\|p47<phox>\|p47phox | neutrophil cytosolic factor 1 | protein-coding |
| chr5_134236887_134237517 | chr5 | 134236888 | 134237517 | + | -0.839333 | TTS (NM_001080746) | TTS (NM_001080746) | -7577 | NR_104383 | 17969 | Mm.425296 | NM_010876 | ENSMUSG00000015950 | **Ncf1** | NCF-47K\|NOXO2\|Ncf-1\|p47-phox\|p47<phox>\|p47phox | neutrophil cytosolic factor 1 | protein-coding |
| chr5_134253254_134254125 | chr5 | 134253255 | 134254125 | + | -0.841034 | intron (NM_001080749, intron 17 of 32) | intron (NM_001080749, intron 17 of 32) | -24065 | NR_104383 | 17969 | Mm.425296 | NM_010876 | ENSMUSG00000015950 | **Ncf1** | NCF-47K\|NOXO2\|Ncf-1\|p47-phox\|p47<phox>\|p47phox | neutrophil cytosolic factor 1 | protein-coding |
| chr5_134236846_134237324 | chr5 | 134236847 | 134237324 | + | -0.867928 | TTS (NM_001080746) | TTS (NM_001080746) | -7460 | NR_104383 | 17969 | Mm.425296 | NM_010876 | ENSMUSG00000015950 | **Ncf1** | NCF-47K\|NOXO2\|Ncf-1\|p47-phox\|p47<phox>\|p47phox | neutrophil cytosolic factor 1 | protein-coding |
| chr5_134253247_134253795 | chr5 | 134253248 | 134253795 | + | -0.940265 | intron (NM_001080749, intron 17 of 32) | intron (NM_001080749, intron 17 of 32) | -23896 | NR_104383 | 17969 | Mm.425296 | NM_010876 | ENSMUSG00000015950 | **Ncf1** | NCF-47K\|NOXO2\|Ncf-1\|p47-phox\|p47<phox>\|p47phox | neutrophil cytosolic factor 1 | protein-coding |
| chr5_134269556_134270311 | chr5 | 134269557 | 134270311 | + | -1.00547 | intron (NM_001080749, intron 10 of 32) | intron (NM_001080749, intron 10 of 32) | -40309 | NR_104383 | 17969 | Mm.425296 | NM_010876 | ENSMUSG00000015950 | **Ncf1** | NCF-47K\|NOXO2\|Ncf-1\|p47-phox\|p47<phox>\|p47phox | neutrophil cytosolic factor 1 | protein-coding |
| chr9_100521991_100522407 | chr9 | 100521992 | 100522407 | + | -1.00665 | intron (NM_010878, intron 1 of 3) | intron (NM_010878, intron 1 of 3) | 23854 | NM_010878 | 17973 | Mm.181485 | NM_010878 | ENSMUSG00000032475 | **Nck1** | 6330586M15Rik\|D230010O13Rik\|Nck\|Nck-1 | non-catalytic region of tyrosine kinase adaptor protein 1 | protein-coding |

| | | | | | | | | | | | | | | | | | |
|---|---|---|---|---|---|---|---|---|---|---|---|---|---|---|---|---|---|
| chr9_100522146_100522407 | chr9 | 100522147 | 100522407 | + | -1.02177 | intron (NM_010878, intron 1 of 3) | intron (NM_010878, intron 1 of 3) | 23776 | NM_010878 | 17973 | Mm.181485 | NM_010878 | ENSMUSG00000032475 | **Nck1** | 6330586M15Rik\|D230010O13Rik\|Nck\|Nck-1 | non-catalytic region of tyrosine kinase adaptor protein 1 | protein-coding |
| chr1_43542280_43543057 | chr1 | 43542281 | 43543057 | + | -0.755704 | intron (NM_010879, intron 3 of 4) | intron (NM_010879, intron 3 of 4) | 96918 | NM_010879 | 17974 | Mm.389903 | NM_010879 | ENSMUSG00000066877 | **Nck2** | 4833426I10Rik\|Grb4\|NCKbeta\|Nck-2 | non-catalytic region of tyrosine kinase adaptor protein 2 | protein-coding |
| chr1_43542393_43542738 | chr1 | 43542394 | 43542738 | + | -0.792761 | intron (NM_010879, intron 3 of 4) | intron (NM_010879, intron 3 of 4) | 96815 | NM_010879 | 17974 | Mm.389903 | NM_010879 | ENSMUSG00000066877 | **Nck2** | 4833426I10Rik\|Grb4\|NCKbeta\|Nck-2 | non-catalytic region of tyrosine kinase adaptor protein 2 | protein-coding |
| chr1_43501165_43501495 | chr1 | 43501166 | 43501495 | + | -0.876174 | intron (NM_010879, intron 1 of 4) | intron (NM_010879, intron 1 of 4) | 55579 | NM_010879 | 17974 | Mm.389903 | NM_010879 | ENSMUSG00000066877 | **Nck2** | 4833426I10Rik\|Grb4\|NCKbeta\|Nck-2 | non-catalytic region of tyrosine kinase adaptor protein 2 | protein-coding |
| chr1_43457067_43457365 | chr1 | 43457068 | 43457365 | + | -0.936031 | intron (NM_010879, intron 1 of 4) | intron (NM_010879, intron 1 of 4) | 11465 | NM_010879 | 17974 | Mm.389903 | NM_010879 | ENSMUSG00000066877 | **Nck2** | 4833426I10Rik\|Grb4\|NCKbeta\|Nck-2 | non-catalytic region of tyrosine kinase adaptor protein 2 | protein-coding |
| chr1_43464987_43465375 | chr1 | 43464988 | 43465375 | + | -1.18976 | intron (NM_010879, intron 1 of 4) | intron (NM_010879, intron 1 of 4) | 19430 | NM_010879 | 17974 | Mm.389903 | NM_010879 | ENSMUSG00000066877 | **Nck2** | 4833426I10Rik\|Grb4\|NCKbeta\|Nck-2 | non-catalytic region of tyrosine kinase adaptor protein 2 | protein-coding |
| chr1_43403368_43403757 | chr1 | 43403369 | 43403757 | + | -1.4029 | Intergenic | Intergenic | -42188 | NM_010879 | 17974 | Mm.389903 | NM_010879 | ENSMUSG00000066877 | **Nck2** | 4833426I10Rik\|Grb4\|NCKbeta\|Nck-2 | non-catalytic region of tyrosine kinase adaptor protein 2 | protein-coding |
| chr1_13294693_13295332 | chr1 | 13294694 | 13295332 | + | -0.615012 | intron (NM_001077695, intron 1 of 21) | intron (NM_001077695, intron 1 of 21) | 79070 | NM_001077695 | 17978 | Mm.2537 | NM_008678 | ENSMUSG00000005886 | **Ncoa2** | 9530095N19\|D1Ertd433e\|GRIP-1\|Grip1\|KAT13C\|SRC-2\|TIF-2\|TIF2\|bHLHe75 | nuclear receptor coactivator 2 | protein-coding |
| chr1_13426684_13427360 | chr1 | 13426685 | 13427360 | + | -0.668352 | Intergenic | RSINE1\|SINE\|B4 | -52939 | NM_008678 | 17978 | Mm.2537 | NM_008678 | ENSMUSG00000005886 | **Ncoa2** | 9530095N19\|D1Ertd433e\|GRIP-1\|Grip1\|KAT13C\|SRC-2\|TIF-2\|TIF2\|bHLHe75 | nuclear receptor coactivator 2 | protein-coding |
| chr1_13294745_13295115 | chr1 | 13294746 | 13295115 | + | -0.68506 | intron (NM_001077695, intron 1 of 21) | intron (NM_001077695, intron 1 of 21) | 79153 | NM_001077695 | 17978 | Mm.2537 | NM_008678 | ENSMUSG00000005886 | **Ncoa2** | 9530095N19\|D1Ertd433e\|GRIP-1\|Grip1\|KAT13C\|SRC-2\|TIF-2\|TIF2\|bHLHe75 | nuclear receptor coactivator 2 | protein-coding |
| chr1_13267126_13267834 | chr1 | 13267127 | 13267834 | + | -0.715995 | intron (NM_001077695, intron 2 of 21) | intron (NM_001077695, intron 2 of 21) | 106603 | NM_001077695 | 17978 | Mm.2537 | NM_008678 | ENSMUSG00000005886 | **Ncoa2** | 9530095N19\|D1Ertd433e\|GRIP-1\|Grip1\|KAT13C\|SRC-2\|TIF-2\|TIF2\|bHLHe75 | nuclear receptor coactivator 2 | protein-coding |
| chr1_13311623_13312320 | chr1 | 13311624 | 13312320 | + | -0.884521 | intron (NM_001077695, intron 1 of 21) | intron (NM_001077695, intron 1 of 21) | 62111 | NM_001077695 | 17978 | Mm.2537 | NM_008678 | ENSMUSG00000005886 | **Ncoa2** | 9530095N19\|D1Ertd433e\|GRIP-1\|Grip1\|KAT13C\|SRC-2\|TIF-2\|TIF2\|bHLHe75 | nuclear receptor coactivator 2 | protein-coding |
| chr1_13311894_13312197 | chr1 | 13311895 | 13312197 | + | -0.894784 | intron (NM_001077695, intron 1 of 21) | intron (NM_001077695, intron 1 of 21) | 62037 | NM_001077695 | 17978 | Mm.2537 | NM_008678 | ENSMUSG00000005886 | **Ncoa2** | 9530095N19\|D1Ertd433e\|GRIP-1\|Grip1\|KAT13C\|SRC-2\|TIF-2\|TIF2\|bHLHe75 | nuclear receptor coactivator 2 | protein-coding |
| chr2_165951483_165951766 | chr2 | 165951484 | 165951766 | + | -0.690952 | Intergenic | Intergenic | -41012 | NM_008679 | 17979 | Mm.469663 | NM_008679 | ENSMUSG00000027678 | **Ncoa3** | 2010305B15Rik\|AW321064\|Actr\|Aib1\|KAT13B\|Rac3\|Src3\|Tram-1\|Tram1\|bHLHe42\|p/Cip\|pCip | nuclear receptor coactivator 3 | protein-coding |
| chr2_166022268_166022667 | chr2 | 166022269 | 166022667 | + | -1.02986 | intron (NM_008679, intron 1 of 22) | intron (NM_008679, intron 1 of 22) | 29831 | NM_008679 | 17979 | Mm.469663 | NM_008679 | ENSMUSG00000027678 | **Ncoa3** | 2010305B15Rik\|AW321064\|Actr\|Aib1\|KAT13B\|Rac3\|Src3\|Tram-1\|Tram1\|bHLHe42\|p/Cip\|pCip | nuclear receptor coactivator 3 | protein-coding |
| chr5_125197063_125197784 | chr5 | 125197064 | 125197784 | + | -0.875844 | Intergenic | (GA)n\|Simple_repeat\|Simple_repeat | -18210 | NM_011424 | 20602 | Mm.278646 | NM_011424 | ENSMUSG00000029478 | **Ncor2** | N-CoR\|SMRT\|SMRTe | nuclear receptor co-repressor 2 | protein-coding |
| chr2_156970159_156970781 | chr2 | 156970160 | 156970781 | + | -0.918073 | intron (NM_180956, intron 1 of 16) | MLT1F2\|LTR\|ERVL-MaLR | 21641 | NM_013865 | 29812 | Mm.279256 | NM_013865 | ENSMUSG00000027634 | **Ndrg3** | 4833415O14Rik\|AI747193\|Ndr3 | N-myc downstream regulated gene 3 | protein-coding |
| chr2_156974973_156975332 | chr2 | 156974974 | 156975332 | + | -1.09524 | intron (NM_180956, intron 1 of 16) | RMER1C\|Other\|Other | 16958 | NM_013865 | 29812 | Mm.279256 | NM_013865 | ENSMUSG00000027634 | **Ndrg3** | 4833415O14Rik\|AI747193\|Ndr3 | N-myc downstream regulated gene 3 | protein-coding |
| chr8_69937456_69937851 | chr8 | 69937457 | 69937851 | + | -0.700924 | intron (NM_001286450, intron 2 of 12) | intron (NM_001286450, intron 2 of 12) | -35096 | NM_023312 | 67184 | Mm.21162 | NM_023312 | ENSMUSG00000036199 | **Ndufa13** | 2700054G14Rik\|AU022060\|CDA016\|CGI-39\|GRIM-19\|Grim19 | NADH dehydrogenase (ubiquinone) 1 alpha subcomplex, 13 | protein-coding |
| chr6_11900967_11901769 | chr6 | 11900968 | 11901769 | + | -0.74348 | intron (NM_010886, intron 3 of 3) | intron (NM_010886, intron 3 of 3) | 6078 | NM_010886 | 17992 | Mm.415865 | NM_010886 | ENSMUSG00000029632 | **Ndufa4** | MLRQ | NADH dehydrogenase (ubiquinone) 1 alpha subcomplex, 4 | protein-coding |
| chr6_126811430_126812002 | chr6 | 126811431 | 126812002 | + | -1.23788 | Intergenic | Intergenic | 37428 | NM_025358 | 66108 | Mm.29939 | NM_025358 | ENSMUSG00000000399 | **Ndufa9** | 1010001N11Rik | NADH dehydrogenase (ubiquinone) 1 alpha subcomplex, 9 | protein-coding |
| chr6_126811498_126811909 | chr6 | 126811499 | 126811909 | + | -1.24062 | Intergenic | Intergenic | 37440 | NM_025358 | 66108 | Mm.29939 | NM_025358 | ENSMUSG00000000399 | **Ndufa9** | 1010001N11Rik | NADH dehydrogenase (ubiquinone) 1 alpha subcomplex, 9 | protein-coding |
| chr13_114387164_114387538 | chr13 | 114387165 | 114387538 | + | -0.82014 | intron (NM_010887, intron 1 of 4) | B1_Mur2\|SINE\|Alu | 743 | NM_010887 | 17993 | Mm.253142 | NM_010887 | ENSMUSG00000021764 | **Ndufs4** | 6720411N02Rik\|C1-18k | NADH dehydrogenase (ubiquinone) Fe-S protein 4 | protein-coding |
| chr18_64919376_64919562 | chr18 | 64919377 | 64919562 | + | -0.632224 | intron (NM_001114386, intron 1 of 30) | intron (NM_001114386, intron 1 of 30) | 31713 | NM_001114386 | 83814 | Mm.98668 | NM_031881 | ENSMUSG00000024589 | **Nedd4l** | 1300012C07Rik\|Nedd4-2\|Nedd4b | neural precursor cell expressed, developmentally down-regulated gene 4-like | protein-coding |
| chr9_57148014_57148797 | chr9 | 57148015 | 57148797 | + | -0.962278 | Intergenic | Intergenic | -1372 | NM_028347 | 72774 | Mm.35749 | NM_028347 | ENSMUSG00000032298 | **Neil1** | 2810450N13Rik\|Nei1 | nei endonuclease VIII-like 1 (E. coli) | protein-coding |
| chr14_15062548_15063623 | chr14 | 15062549 | 15063623 | + | -0.663974 | Intergenic | Intergenic | 242271 | NM_001195229 | 674895 | Mm.317563 | NM_001034865 | ENSMUSG00000042567 | **Nek10** | Gm282 | NIMA (never in mitosis gene a)- related kinase 10 | protein-coding |
| chr14_15062885_15063419 | chr14 | 15062886 | 15063419 | + | -0.670117 | Intergenic | Intergenic | 242337 | NM_001195229 | 674895 | Mm.317563 | NM_001034865 | ENSMUSG00000042567 | **Nek10** | Gm282 | NIMA (never in mitosis gene a)- related kinase 10 | protein-coding |
| chr14_15083458_15084224 | chr14 | 15083459 | 15084224 | + | -0.712401 | Intergenic | Intergenic | 263026 | NM_001195229 | 674895 | Mm.317563 | NM_001034865 | ENSMUSG00000042567 | **Nek10** | Gm282 | NIMA (never in mitosis gene a)- related kinase 10 | protein-coding |
| chr14_14800801_14801265 | chr14 | 14800802 | 14801265 | + | -0.735129 | Intergenic | Intergenic | -19782 | NM_001195229 | 674895 | Mm.317563 | NM_001034865 | ENSMUSG00000042567 | **Nek10** | Gm282 | NIMA (never in mitosis gene a)- related kinase 10 | protein-coding |
| chr14_15083830_15084215 | chr14 | 15083831 | 15084215 | + | -0.793788 | Intergenic | Intergenic | 263208 | NM_001195229 | 674895 | Mm.317563 | NM_001034865 | ENSMUSG00000042567 | **Nek10** | Gm282 | NIMA (never in mitosis gene a)- related kinase 10 | protein-coding |
| chr14_15099661_15100334 | chr14 | 15099662 | 15100334 | + | -1.09629 | Intergenic | MTEa\|LTR\|ERVL-MaLR | 279183 | NM_001195229 | 674895 | Mm.317563 | NM_001034865 | ENSMUSG00000042567 | **Nek10** | Gm282 | NIMA (never in mitosis gene a)- related kinase 10 | protein-coding |
| chr14_15099932_15100396 | chr14 | 15099933 | 15100396 | + | -1.14304 | Intergenic | Intergenic | 279349 | NM_001195229 | 674895 | Mm.317563 | NM_001034865 | ENSMUSG00000042567 | **Nek10** | Gm282 | NIMA (never in mitosis gene a)- related kinase 10 | protein-coding |
| chr1_138584628_138585086 | chr1 | 138584629 | 138585086 | + | -0.779206 | intron (NM_021605, intron 1 of 9) | intron (NM_021605, intron 1 of 9) | 34839 | NM_021605 | 59125 | Mm.143817 | NM_021605 | ENSMUSG00000026393 | **Nek7** | 2810460C19Rik\|AU020186 | NIMA (never in mitosis gene a)-related expressed kinase 7 | protein-coding |
| chr1_191259023_191259560 | chr1 | 191259024 | 191259560 | + | -0.768703 | Intergenic | ORR1E\|LTR\|ERVL-MaLR | 58826 | NM_025424 | 66208 | Mm.46444 | NM_025424 | ENSMUSG00000037499 | **Nenf** | 1110060M21Rik\|SCIRP10\|Spuf | neuron derived neurotrophic factor | protein-coding |
| chr8_85624165_85625018 | chr8 | 85624166 | 85625018 | + | -0.842222 | Intergenic | Intergenic | 66417 | NM_001081324 | 74513 | Mm.126079 | NM_001081324 | ENSMUSG00000036902 | **Neto2** | 5530601C23Rik\|AI854825\|BTCL2 | neuropilin (NRP) and tolloid (TLL)-like 2 | protein-coding |
| chr8_85623259_85623571 | chr8 | 85623260 | 85623571 | + | -1.529 | Intergenic | Intergenic | 67594 | NM_001081324 | 74513 | Mm.126079 | NM_001081324 | ENSMUSG00000036902 | **Neto2** | 5530601C23Rik\|AI854825\|BTCL2 | neuropilin (NRP) and tolloid (TLL)-like 2 | protein-coding |
| chr8_85623174_85623788 | chr8 | 85623175 | 85623788 | + | -1.58771 | Intergenic | Intergenic | 67528 | NM_001081324 | 74513 | Mm.126079 | NM_001081324 | ENSMUSG00000036902 | **Neto2** | 5530601C23Rik\|AI854825\|BTCL2 | neuropilin (NRP) and tolloid (TLL)-like 2 | protein-coding |
| chr17_26368992_26369254 | chr17 | 26368993 | 26369254 | + | -0.96029 | Intergenic | Intergenic | -45842 | NM_001081656 | 240055 | Mm.133211 | NM_001081656 | ENSMUSG00000034413 | **Neurl1b** | C230078M08Rik\|EG240055 | neuralized homolog 1b (Drosophila) | protein-coding |
| chr17_26370397_26371291 | chr17 | 26370398 | 26371291 | + | -1.2809 | Intergenic | Intergenic | -44121 | NM_001081656 | 240055 | Mm.133211 | NM_001081656 | ENSMUSG00000034413 | **Neurl1b** | C230078M08Rik\|EG240055 | neuralized homolog 1b (Drosophila) | protein-coding |
| chr1_36260567_36260819 | chr1 | 36260568 | 36260819 | + | -0.824995 | Intergenic | Intergenic | 12732 | NM_153408 | 214854 | Mm.389110 | NM_153408 | ENSMUSG00000047180 | **Neurl3** | 2010300P06Rik\|Lincr | neuralized homolog 3 homolog (Drosophila) | protein-coding |
| chr1_36272397_36272907 | chr1 | 36272398 | 36272907 | + | -0.868041 | intron (NM_153408, intron 1 of 3) | intron (NM_153408, intron 1 of 3) | 773 | NM_153408 | 214854 | Mm.389110 | NM_153408 | ENSMUSG00000047180 | **Neurl3** | 2010300P06Rik\|Lincr | neuralized homolog 3 homolog (Drosophila) | protein-coding |
| chr1_36263929_36264717 | chr1 | 36263930 | 36264717 | + | -0.889594 | TTS (NM_153408) | TTS (NM_153408) | 9102 | NM_153408 | 214854 | Mm.389110 | NM_153408 | ENSMUSG00000047180 | **Neurl3** | 2010300P06Rik\|Lincr | neuralized homolog 3 homolog (Drosophila) | protein-coding |
| chr1_36281837_36282554 | chr1 | 36281838 | 36282554 | + | -0.973627 | Intergenic | ORR1G\|LTR\|ERVL-MaLR | -8771 | NM_153408 | 214854 | Mm.389110 | NM_153408 | ENSMUSG00000047180 | **Neurl3** | 2010300P06Rik\|Lincr | neuralized homolog 3 homolog (Drosophila) | protein-coding |
| chr1_36281843_36282188 | chr1 | 36281844 | 36282188 | + | -0.981982 | Intergenic | B3\|SINE\|B2 | -8591 | NM_153408 | 214854 | Mm.389110 | NM_153408 | ENSMUSG00000047180 | **Neurl3** | 2010300P06Rik\|Lincr | neuralized homolog 3 homolog (Drosophila) | protein-coding |
| chr15_83042727_83043168 | chr15 | 83042728 | 83043168 | + | -0.663088 | Intergenic | Intergenic | -9551 | NM_001271411 | 74039 | Mm.32584 | NM_028728 | | **Nfam1** | 4921501M20Rik\|Cnaip | Nfat activating molecule with ITAM motif 1 | protein-coding |
| chr15_83030265_83030783 | chr15 | 83030266 | 83030783 | + | -0.736939 | promoter-TSS (NM_001271413) | promoter-TSS (NM_001271413) | -574 | NM_001271413 | 74039 | Mm.32584 | NM_028728 | | **Nfam1** | 4921501M20Rik\|Cnaip | Nfat activating molecule with ITAM motif 1 | protein-coding |
| chr18_80707043_80707362 | chr18 | 80707044 | 80707362 | + | -0.655908 | intron (NM_198429, intron 1 of 9) | intron (NM_198429, intron 1 of 9) | 970 | NM_001164111 | 18018 | Mm.329560 | NM_016791 | ENSMUSG00000033016 | **Nfatc1** | 2210017P03Rik\|AI449492\|AV076380\|NF-ATc\|NFAT2\|NFATc\|Nfatcb | nuclear factor of activated T cells, cytoplasmic, calcineurin dependent 1 | protein-coding |
| chr18_80706073_80706363 | chr18 | 80706074 | 80706363 | + | -0.905774 | intron (NM_198429, intron 1 of 9) | intron (NM_198429, intron 1 of 9) | 1955 | NM_001164111 | 18018 | Mm.329560 | NM_016791 | ENSMUSG00000033016 | **Nfatc1** | 2210017P03Rik\|AI449492\|AV076380\|NF-ATc\|NFAT2\|NFATc\|Nfatcb | nuclear factor of activated T cells, cytoplasmic, calcineurin dependent 1 | protein-coding |
| chr18_80738182_80738852 | chr18 | 80738183 | 80738852 | + | -1.02615 | intron (NM_001201569, intron 28 of 29) | intron (NM_001201569, intron 28 of 29) | -25446 | NM_001164109 | 18018 | Mm.329560 | NM_016791 | ENSMUSG00000033016 | **Nfatc1** | 2210017P03Rik\|AI449492\|AV076380\|NF-ATc\|NFAT2\|NFATc\|Nfatcb | nuclear factor of activated T cells, cytoplasmic, calcineurin dependent 1 | protein-coding |
| chr8_106054108_106054852 | chr8 | 106054109 | 106054852 | + | -0.944173 | TTS (NM_001301176) | TTS (NM_001301176) | -5123 | NM_010901 | 18021 | Mm.383185 | NM_010901 | ENSMUSG00000031902 | **Nfatc3** | C80703\|D8Ertd281e\|NFAT4\|NFATx | nuclear factor of activated T cells, cytoplasmic, calcineurin dependent 3 | protein-coding |

| | | | | | | | | | | | | | | | | | |
|---|---|---|---|---|---|---|---|---|---|---|---|---|---|---|---|---|---|
| chr4_98040019_98040666 | chr4 | 98040020 | 98040666 | + | -1.01291 | intron (NM_001122953, intron 6 of 9) | intron (NM_001122953, intron 6 of 9) | 262717 | NM_010905 | 18027 | Mm.31274 | NM_010905 | ENSMUSG00000028565 | **Nfia** | 1110047K16Rik\|9430022M17Rik\|CTF\|NF1-A\|NF1A | nuclear factor I/A | protein-coding |
| chr4_98040219_98040560 | chr4 | 98040220 | 98040560 | + | -1.0182 | intron (NM_001122953, intron 6 of 9) | intron (NM_001122953, intron 6 of 9) | 262764 | NM_010905 | 18027 | Mm.31274 | NM_010905 | ENSMUSG00000028565 | **Nfia** | 1110047K16Rik\|9430022M17Rik\|CTF\|NF1-A\|NF1A | nuclear factor I/A | protein-coding |
| chr10_81434054_81434352 | chr10 | 81434055 | 81434352 | + | -0.629185 | Intergenic | Intergenic | -7030 | NM_026756 | 18029 | Mm.426936 | NM_008688 | ENSMUSG00000055053 | **Nfic** | 1110019L22Rik\|1500041O16Rik\|AA589446\|AI746521\|NF1-C | nuclear factor I/C | protein-coding |
| chr3_135602777_135603172 | chr3 | 135602778 | 135603172 | + | -0.838246 | intron (NM_008689, intron 16 of 24) | intron (NM_008689, intron 16 of 24) | 88572 | NM_008689 | 18033 | Mm.256765 | NM_008689 | ENSMUSG00000028163 | **Nfkb1** | NF-KB1\|NF-kappaB\|NF-kappaB1\|p105\|p50\|p50/p105 | nuclear factor of kappa light polypeptide gene enhancer in B cells 1, p105 | protein-coding |
| chr3_135602843_135603097 | chr3 | 135602844 | 135603097 | + | -0.867706 | intron (NM_008689, intron 16 of 24) | intron (NM_008689, intron 16 of 24) | 88577 | NM_008689 | 18033 | Mm.256765 | NM_008689 | ENSMUSG00000028163 | **Nfkb1** | NF-KB1\|NF-kappaB\|NF-kappaB1\|p105\|p50\|p50/p105 | nuclear factor of kappa light polypeptide gene enhancer in B cells 1, p105 | protein-coding |
| chr12_55489112_55489395 | chr12 | 55489113 | 55489395 | + | -0.692615 | TTS (NM_010907) | TTS (NM_010907) | 3393 | NM_010907 | 18035 | Mm.170515 | NM_010907 | ENSMUSG00000021025 | **Nfkbia** | AI462015\|Nfkbi | nuclear factor of kappa light polypeptide gene enhancer in B cells inhibitor, alpha | protein-coding |
| chr7_30422705_30422935 | chr7 | 30422706 | 30422935 | + | -0.728663 | promoter-TSS (NM_172142) | promoter-TSS (NM_172142) | -484 | NM_172142 | 243910 | Mm.386878 | NM_172142 | ENSMUSG00000036931 | **Nfkbid** | IkappaBNS | nuclear factor of kappa light polypeptide gene enhancer in B cells inhibitor, delta | protein-coding |
| chr16_55820930_55821207 | chr16 | 55820931 | 55821207 | + | -0.609117 | intron (NM_001159395, intron 2 of 12) | intron (NM_001159395, intron 2 of 12) | 1069 | NM_001159394 | 80859 | Mm.247272 | NM_030612 | ENSMUSG00000035356 | **Nfkbiz** | AA408868\|INAP\|Mail | nuclear factor of kappa light polypeptide gene enhancer in B cells inhibitor, zeta | protein-coding |
| chr4_120815635_120816254 | chr4 | 120815636 | 120816254 | + | -0.672029 | intron (NM_001048168, intron 1 of 9) | intron (NM_001048168, intron 1 of 9) | 9796 | NM_001277095 | 18046 | Mm.435590 | NM_008692 | ENSMUSG00000032897 | **Nfyc** | - | nuclear transcription factor-Y gamma | protein-coding |
| chr14_16023788_16024396 | chr14 | 16023789 | 16024396 | + | -0.628504 | Intergenic | Intergenic | -225222 | NM_021504 | 59007 | Mm.258381 | NM_021504 | ENSMUSG00000021785 | **Ngly1** | 1110002C09Rik\|PNGase\|Png1 | N-glycanase 1 | protein-coding |
| chr14_16164591_16165248 | chr14 | 16164592 | 16165248 | + | -0.688467 | Intergenic | Lx6\|LINE\|L1 | -84394 | NM_021504 | 59007 | Mm.258381 | NM_021504 | ENSMUSG00000021785 | **Ngly1** | 1110002C09Rik\|PNGase\|Png1 | N-glycanase 1 | protein-coding |
| chr14_16172357_16172691 | chr14 | 16172358 | 16172691 | + | -0.75879 | Intergenic | Intergenic | -76790 | NM_021504 | 59007 | Mm.258381 | NM_021504 | ENSMUSG00000021785 | **Ngly1** | 1110002C09Rik\|PNGase\|Png1 | N-glycanase 1 | protein-coding |
| chr14_16151853_16152424 | chr14 | 16151854 | 16152424 | + | -0.802567 | Intergenic | Intergenic | -97175 | NM_021504 | 59007 | Mm.258381 | NM_021504 | ENSMUSG00000021785 | **Ngly1** | 1110002C09Rik\|PNGase\|Png1 | N-glycanase 1 | protein-coding |
| chr14_16151826_16152261 | chr14 | 16151827 | 16152261 | + | -0.827106 | Intergenic | Intergenic | -97270 | NM_021504 | 59007 | Mm.258381 | NM_021504 | ENSMUSG00000021785 | **Ngly1** | 1110002C09Rik\|PNGase\|Png1 | N-glycanase 1 | protein-coding |
| chr14_16170209_16170628 | chr14 | 16170210 | 16170628 | + | -1.02239 | Intergenic | L1M5\|LINE\|L1 | -78895 | NM_021504 | 59007 | Mm.258381 | NM_021504 | ENSMUSG00000021785 | **Ngly1** | 1110002C09Rik\|PNGase\|Png1 | N-glycanase 1 | protein-coding |
| chr14_16173249_16173914 | chr14 | 16173250 | 16173914 | + | -1.29999 | Intergenic | Intergenic | -75732 | NM_021504 | 59007 | Mm.258381 | NM_021504 | ENSMUSG00000021785 | **Ngly1** | 1110002C09Rik\|PNGase\|Png1 | N-glycanase 1 | protein-coding |
| chrX_101917659_101918235 | chrX | 101917660 | 101918235 | + | -0.91821 | intron (NM_001163610, intron 1 of 7) | ORR1D2\|LTR\|ERVL-MaLR | 68562 | NM_001163610 | 100042480 | Mm.232 | NM_001163610 | ENSMUSG00000079481 | **Nhsl2** | 1110062M06Rik\|6430511F03\|AI225852\|AW488865\|EG621083\|ENSMUSG00000073037\|Gm10456 | NHS-like 2 | protein-coding |
| chr1_58454179_58454820 | chr1 | 58454180 | 58454820 | + | -1.17243 | intron (NM_022988, intron 3 of 5) | intron (NM_022988, intron 3 of 5) | 6867 | NM_022988 | 65102 | Mm.340248 | NM_022988 | ENSMUSG00000026036 | **Nif3l1** | 1110030G24Rik | Ngg1 interacting factor 3-like 1 (S. pombe) | protein-coding |
| chr12_70104812_70105069 | chr12 | 70104813 | 70105069 | + | -0.808504 | intron (NM_001286079, intron 1 of 28) | intron (NM_001286079, intron 1 of 28) | -2087 | NM_001081453 | 18080 | Mm.244047 | NM_008697 | ENSMUSG00000021068 | **Nin** | 3110068G20Rik\|AI385615\|AU024711\|mKIAA1565 | ninein | protein-coding |
| chr12_70074964_70075530 | chr12 | 70074965 | 70075530 | + | -0.914924 | intron (NM_001286079, intron 4 of 28) | intron (NM_001286079, intron 4 of 28) | 27607 | NM_001081453 | 18080 | Mm.244047 | NM_008697 | ENSMUSG00000021068 | **Nin** | 3110068G20Rik\|AI385615\|AU024711\|mKIAA1565 | ninein | protein-coding |
| chr12_70075043_70075465 | chr12 | 70075044 | 70075465 | + | -0.927711 | intron (NM_001286079, intron 4 of 28) | intron (NM_001286079, intron 4 of 28) | 27600 | NM_001081453 | 18080 | Mm.244047 | NM_008697 | ENSMUSG00000021068 | **Nin** | 3110068G20Rik\|AI385615\|AU024711\|mKIAA1565 | ninein | protein-coding |
| chr12_70104607_70105278 | chr12 | 70104608 | 70105278 | + | -0.937341 | intron (NM_001286079, intron 1 of 28) | intron (NM_001286079, intron 1 of 28) | -2089 | NM_001081453 | 18080 | Mm.244047 | NM_008697 | ENSMUSG00000021068 | **Nin** | 3110068G20Rik\|AI385615\|AU024711\|mKIAA1565 | ninein | protein-coding |
| chr6_120065802_120066336 | chr6 | 120065803 | 120066336 | + | -1.1687 | Intergenic | Intergenic | -27311 | NM_016718 | 29862 | Mm.103668 | NM_016718 | ENSMUSG00000041377 | **Ninj2** | - | ninjurin 2 | protein-coding |
| chr15_8568310_8569593 | chr15 | 8568311 | 8569593 | + | -0.790017 | Intergenic | CpG | -124489 | NM_201232 | 71175 | Mm.240329 | NM_027707 | ENSMUSG00000022141 | **Nipbl** | Idn3 | Nipped-B homolog (Drosophila) | protein-coding |
| chr15_8451101_8451448 | chr15 | 8451102 | 8451448 | + | -0.893301 | Intergenic | Intergenic | -6812 | NM_201232 | 71175 | Mm.240329 | NM_027707 | ENSMUSG00000022141 | **Nipbl** | Idn3 | Nipped-B homolog (Drosophila) | protein-coding |
| chr15_8568310_8570091 | chr15 | 8568311 | 8570091 | + | -0.942935 | Intergenic | CpG | -124738 | NM_201232 | 71175 | Mm.240329 | NM_027707 | ENSMUSG00000022141 | **Nipbl** | Idn3 | Nipped-B homolog (Drosophila) | protein-coding |
| chr15_8569688_8570104 | chr15 | 8569689 | 8570104 | + | -1.14457 | Intergenic | Intergenic | -125433 | NM_201232 | 71175 | Mm.240329 | NM_027707 | ENSMUSG00000022141 | **Nipbl** | Idn3 | Nipped-B homolog (Drosophila) | protein-coding |
| chr14_31194478_31194786 | chr14 | 31194479 | 31194786 | + | -0.73463 | intron (NM_022656, intron 3 of 20) | B2_Mm1t\|SINE\|B2 | 12194 | NM_022656 | 64652 | Mm.298728 | NM_022656 | ENSMUSG00000021910 | **Nisch** | 1200007D05Rik\|3202002H23Rik\|AW494485\|I-1 | nischarin | protein-coding |
| chrX_37141696_37142178 | chrX | 37141697 | 37142178 | + | -1.2517 | intron (NM_025937, intron 7 of 8) | intron (NM_025937, intron 7 of 8) | 15174 | NM_025937 | 67050 | Mm.55422 | NM_025937 | ENSMUSG00000016409 | **Nkap** | 2610020O08Rik\|AA987160\|AI849147\|AL024236 | NFKB activating protein | protein-coding |
| chr7_43438484_43439036 | chr7 | 43438485 | 43439036 | + | -0.668974 | TTS (NM_024253) | TTS (NM_024253) | 1622 | NM_024253 | 72310 | Mm.34613 | NM_024253 | ENSMUSG00000004612 | **Nkg7** | 2500004F03Rik | natural killer cell group 7 sequence | protein-coding |
| chr7_43438704_43438983 | chr7 | 43438705 | 43438983 | + | -0.784431 | TTS (NM_024253) | TTS (NM_024253) | 1706 | NM_024253 | 72310 | Mm.34613 | NM_024253 | ENSMUSG00000004612 | **Nkg7** | 2500004F03Rik | natural killer cell group 7 sequence | protein-coding |
| chr5_33468829_33469149 | chr5 | 33468830 | 33469149 | + | -0.66668 | Intergenic | Intergenic | -34899 | NM_011320 | 672284 | | NM_011320 | | **Nkx1-1** | Nkx-1.1\|Sax2 | NK1 transcription factor related, locus 1 (Drosophila) | protein-coding |
| chr5_33468726_33469283 | chr5 | 33468727 | 33469283 | + | -0.682195 | Intergenic | Intergenic | -34915 | NM_011320 | 672284 | | NM_011320 | | **Nkx1-1** | Nkx-1.1\|Sax2 | NK1 transcription factor related, locus 1 (Drosophila) | protein-coding |
| chr11_78689179_78689612 | chr11 | 78689180 | 78689612 | + | -0.668633 | intron (NM_008702, intron 1 of 10) | intron (NM_008702, intron 1 of 10) | 8029 | NM_008702 | 18099 | Mm.9001 | NM_008702 | ENSMUSG00000017376 | **Nlk** | AI194375 | nemo like kinase | protein-coding |
| chr11_78690594_78690879 | chr11 | 78690595 | 78690879 | + | -0.676151 | intron (NM_008702, intron 1 of 10) | intron (NM_008702, intron 1 of 10) | 6688 | NM_008702 | 18099 | Mm.9001 | NM_008702 | ENSMUSG00000017376 | **Nlk** | AI194375 | nemo like kinase | protein-coding |
| chr11_78673579_78673994 | chr11 | 78673580 | 78673994 | + | -0.791007 | intron (NM_008702, intron 1 of 10) | B1_Mur1\|SINE\|Alu | 23638 | NM_008702 | 18099 | Mm.9001 | NM_008702 | ENSMUSG00000017376 | **Nlk** | AI194375 | nemo like kinase | protein-coding |
| chr11_78624440_78624757 | chr11 | 78624441 | 78624757 | + | -0.963868 | intron (NM_008702, intron 2 of 10) | intron (NM_008702, intron 2 of 10) | 72826 | NM_008702 | 18099 | Mm.9001 | NM_008702 | ENSMUSG00000017376 | **Nlk** | AI194375 | nemo like kinase | protein-coding |
| chr11_78678663_78679189 | chr11 | 78678664 | 78679189 | + | -1.21003 | intron (NM_008702, intron 1 of 10) | intron (NM_008702, intron 1 of 10) | 18499 | NM_008702 | 18099 | Mm.9001 | NM_008702 | ENSMUSG00000017376 | **Nlk** | AI194375 | nemo like kinase | protein-coding |
| chr11_78678663_78679522 | chr11 | 78678664 | 78679522 | + | -1.26257 | intron (NM_008702, intron 1 of 10) | intron (NM_008702, intron 1 of 10) | 18332 | NM_008702 | 18099 | Mm.9001 | NM_008702 | ENSMUSG00000017376 | **Nlk** | AI194375 | nemo like kinase | protein-coding |
| chr13_104042765_104043300 | chr13 | 104042766 | 104043300 | + | -0.74759 | intron (NM_029447, intron 9 of 12) | intron (NM_029447, intron 9 of 12) | 66581 | NM_029447 | 75805 | Mm.127692 | NM_029447 | ENSMUSG00000021710 | **Nln** | 4930472G13Rik\|C79345 | neurolysin (metallopeptidase M3 family) | protein-coding |
| chr2_51972111_51972457 | chr2 | 51972112 | 51972457 | + | -0.849042 | intron (NM_019401, intron 1 of 7) | intron (NM_019401, intron 1 of 7) | 924 | NM_001141949 | 64685 | Mm.7491 | NM_019401 | ENSMUSG00000026946 | **Nmi** | - | N-myc (and STAT) interactor | protein-coding |
| chr1_153058258_153058653 | chr1 | 153058259 | 153058653 | + | -0.798157 | intron (NM_175460, intron 1 of 10) | intron (NM_175460, intron 1 of 10) | 103355 | NM_175460 | 226518 | Mm.40548 | NM_175460 | ENSMUSG00000042751 | **Nmnat2** | AI843915\|D030041I09Rik\|PNAT1\|PNAT2 | nicotinamide nucleotide adenylyltransferase 2 | protein-coding |
| chr1_153058219_153058537 | chr1 | 153058220 | 153058537 | + | -0.812451 | intron (NM_175460, intron 1 of 10) | intron (NM_175460, intron 1 of 10) | 103277 | NM_175460 | 226518 | Mm.40548 | NM_175460 | ENSMUSG00000042751 | **Nmnat2** | AI843915\|D030041I09Rik\|PNAT1\|PNAT2 | nicotinamide nucleotide adenylyltransferase 2 | protein-coding |
| chr9_98333154_98333716 | chr9 | 98333155 | 98333716 | + | -0.894456 | intron (NM_144533, intron 1 of 4) | MTD\|LTR\|ERVL-MaLR | 36852 | NM_144533 | 74080 | Mm.294082 | NM_144533 | ENSMUSG00000032456 | **Nmnat3** | 4933408N02Rik\|PNAT3 | nicotinamide nucleotide adenylyltransferase 3 | protein-coding |
| chr9_48562433_48562719 | chr9 | 48562434 | 48562719 | + | -1.23052 | Intergenic | L1MC1\|LINE\|L1 | 42501 | NM_010924 | 18113 | Mm.8362 | NM_010924 | ENSMUSG00000032271 | **Nnmt** | - | nicotinamide N-methyltransferase | protein-coding |
| chr9_48562296_48562753 | chr9 | 48562297 | 48562753 | + | -1.25504 | Intergenic | L1MC1\|LINE\|L1 | 42552 | NM_010924 | 18113 | Mm.8362 | NM_010924 | ENSMUSG00000032271 | **Nnmt** | - | nicotinamide N-methyltransferase | protein-coding |
| chr6_54902586_54902933 | chr6 | 54902587 | 54902933 | + | -0.941531 | Intergenic | Intergenic | 68901 | NM_001171007 | 107607 | Mm.28498 | NM_172729 | ENSMUSG00000038058 | **Nod1** | C230079P11\|Card4\|F830007N14Rik\|Nlrc1 | nucleotide-binding oligomerization domain containing 1 | protein-coding |
| chr3_98027158_98027716 | chr3 | 98027159 | 98027716 | + | -0.682044 | intron (NM_010928, intron 1 of 33) | intron (NM_010928, intron 1 of 33) | 13899 | NM_010928 | 18129 | Mm.485843 | NM_010928 | ENSMUSG00000027878 | **Notch2** | AI853703\|N2 | notch 2 | protein-coding |
| chr15_102526054_102526542 | chr15 | 102526055 | 102526542 | + | -0.720329 | Intergenic | (TG)n\|Simple_repeat\|Simple_repeat | -1356 | NR_033409 | 54615 | Mm.445915 | NM_018787 | ENSMUSG00000023052 | **Npff** | - | neuropeptide FF-amide peptide precursor | protein-coding |
| chr15_102526077_102526953 | chr15 | 102526078 | 102526953 | + | -0.995587 | Intergenic | PB1D10\|SINE\|Alu | -1573 | NR_033409 | 54615 | Mm.445915 | NM_018787 | ENSMUSG00000023052 | **Npff** | - | neuropeptide FF-amide peptide precursor | protein-coding |
| chr15_102526675_102526955 | chr15 | 102526676 | 102526955 | + | -1.04175 | Intergenic | Intergenic | -1873 | NR_033409 | 54615 | Mm.445915 | NM_018787 | ENSMUSG00000023052 | **Npff** | - | neuropeptide FF-amide peptide precursor | protein-coding |
| chr10_61535950_61536482 | chr10 | 61535951 | 61536482 | + | -0.717901 | intron (NM_153542, intron 3 of 4) | intron (NM_153542, intron 3 of 4) | -59268 | NM_001177511 | 237362 | Mm.131297 | NM_001177511 | ENSMUSG00000020090 | **Npffr1** | Gm236\|Gpr147\|NPFF1\|OT7T022 | neuropeptide FF receptor 1 | protein-coding |
| chr6_49930120_49930562 | chr6 | 49930121 | 49930562 | + | -0.782766 | Intergenic | Intergenic | 107612 | NM_023456 | 109648 | Mm.154796 | NM_023456 | ENSMUSG00000029819 | **Npy** | 0710005A05Rik | neuropeptide Y | protein-coding |
| chr9_59941581_59942320 | chr9 | 59941582 | 59942320 | + | -0.765394 | TTS (NM_013708) | TTS (NM_013708) | 8128 | NM_013708 | 23958 | Mm.103641 | NM_013708 | ENSMUSG00000032292 | **Nr2e3** | A930035N01Rik\|PNR\|RNR\|rd7 | nuclear receptor subfamily 2, group E, member 3 | protein-coding |
| chr7_71242495_71242948 | chr7 | 71242496 | 71242948 | + | -0.937955 | Intergenic | Intergenic | -875976 | NM_183261 | 11819 | Mm.158143 | NM_009697 | ENSMUSG00000030551 | **Nr2f2** | 2700033K02Rik\|9430015G03Rik\|ARP-1\|Aporp1\|COUP-TF2\|COUP-TFII\|COUPTFB\|EAR3\|SVP40\|Tcfcoup2 | nuclear receptor subfamily 2, group F, member 2 | protein-coding |

| | | | | | | | | | | | | | | | | | |
|---|---|---|---|---|---|---|---|---|---|---|---|---|---|---|---|---|---|
| chr4_48076742_48076945 | chr4 | 48076743 | 48076945 | + | -0.64772 | intron (NM_015743, intron 5 of 5) | intron (NM_015743, intron 5 of 5) | 25596 | NM_015743 | 18124 | Mm.247261 | NM_015743 | ENSMUSG00000028341 | **Nr4a3** | AI573420\|CHN\|CSMF\|MINOR\|NOR-1\|Nor1\|TEC | nuclear receptor subfamily 4, group A, member 3 | protein-coding |
| chr4_48074572_48075289 | chr4 | 48074573 | 48075289 | + | -0.805062 | intron (NM_015743, intron 5 of 5) | intron (NM_015743, intron 5 of 5) | 23683 | NM_015743 | 18124 | Mm.247261 | NM_015743 | ENSMUSG00000028341 | **Nr4a3** | AI573420\|CHN\|CSMF\|MINOR\|NOR-1\|Nor1\|TEC | nuclear receptor subfamily 4, group A, member 3 | protein-coding |
| chr4_48074786_48075381 | chr4 | 48074787 | 48075381 | + | -0.870452 | intron (NM_015743, intron 5 of 5) | intron (NM_015743, intron 5 of 5) | 23836 | NM_015743 | 18124 | Mm.247261 | NM_015743 | ENSMUSG00000028341 | **Nr4a3** | AI573420\|CHN\|CSMF\|MINOR\|NOR-1\|Nor1\|TEC | nuclear receptor subfamily 4, group A, member 3 | protein-coding |
| chr2_25159084_25159423 | chr2 | 25159085 | 25159423 | + | -0.735777 | Intergenic | ID_B1\|SINE\|B4 | -21504 | NM_025980 | 67122 | Mm.46539 | NM_025980 | ENSMUSG00000078202 | **Nrarp** | 2700054M22Rik | Notch-regulated ankyrin repeat protein | protein-coding |
| chr4_108986941_108987876 | chr4 | 108986942 | 108987876 | + | -0.697117 | Intergenic | ID_B1\|SINE\|B4 | -13396 | NM_146150 | 230598 | Mm.274950 | NM_146150 | ENSMUSG00000053510 | **Nrd1** | 2600011I06Rik\|AI875733\|NRD-C | nardilysin, N-arginine dibasic convertase, NRD convertase 1 | protein-coding |
| chr4_108987315_108987871 | chr4 | 108987316 | 108987871 | + | -0.780276 | Intergenic | Intergenic | -13212 | NM_146150 | 230598 | Mm.274950 | NM_146150 | ENSMUSG00000053510 | **Nrd1** | 2600011I06Rik\|AI875733\|NRD-C | nardilysin, N-arginine dibasic convertase, NRD convertase 1 | protein-coding |
| chr6_30053153_30053483 | chr6 | 30053154 | 30053483 | + | -1.09497 | intron (NM_001164226, intron 1 of 11) | intron (NM_001164226, intron 1 of 11) | 5330 | NM_001164226 | 18181 | Mm.259258 | NM_010938 | ENSMUSG00000058440 | **Nrf1** | C87038\|D6Ertd415e | nuclear respiratory factor 1 | protein-coding |
| chr16_76370735_76370948 | chr16 | 76370736 | 76370948 | + | -0.616768 | intron (NM_173440, intron 1 of 3) | intron (NM_173440, intron 1 of 3) | 2207 | NM_173440 | 268903 | Mm.74711 | NM_173440 | ENSMUSG00000048490 | **Nrip1** | 6030458L20Rik\|8430438I05Rik\|9630050P12\|AA959574\|AW456757\|RIP140 | nuclear receptor interacting protein 1 | protein-coding |
| chr16_76339289_76339678 | chr16 | 76339290 | 76339678 | + | -0.655961 | intron (NM_173440, intron 2 of 3) | ORR1D2\|LTR\|ERVL-MaLR | 33565 | NM_173440 | 268903 | Mm.74711 | NM_173440 | ENSMUSG00000048490 | **Nrip1** | 6030458L20Rik\|8430438I05Rik\|9630050P12\|AA959574\|AW456757\|RIP140 | nuclear receptor interacting protein 1 | protein-coding |
| chr16_76283210_76284555 | chr16 | 76283211 | 76284555 | + | -0.689876 | Intergenic | Intergenic | 89166 | NM_173440 | 268903 | Mm.74711 | NM_173440 | ENSMUSG00000048490 | **Nrip1** | 6030458L20Rik\|8430438I05Rik\|9630050P12\|AA959574\|AW456757\|RIP140 | nuclear receptor interacting protein 1 | protein-coding |
| chr16_76323215_76323968 | chr16 | 76323216 | 76323968 | + | -0.703205 | intron (NM_173440, intron 3 of 3) | intron (NM_173440, intron 3 of 3) | 49457 | NM_173440 | 268903 | Mm.74711 | NM_173440 | ENSMUSG00000048490 | **Nrip1** | 6030458L20Rik\|8430438I05Rik\|9630050P12\|AA959574\|AW456757\|RIP140 | nuclear receptor interacting protein 1 | protein-coding |
| chr16_76321631_76322078 | chr16 | 76321632 | 76322078 | + | -0.723319 | intron (NM_173440, intron 3 of 3) | intron (NM_173440, intron 3 of 3) | 51194 | NM_173440 | 268903 | Mm.74711 | NM_173440 | ENSMUSG00000048490 | **Nrip1** | 6030458L20Rik\|8430438I05Rik\|9630050P12\|AA959574\|AW456757\|RIP140 | nuclear receptor interacting protein 1 | protein-coding |
| chr16_76323445_76324216 | chr16 | 76323446 | 76324216 | + | -0.761376 | intron (NM_173440, intron 3 of 3) | intron (NM_173440, intron 3 of 3) | 49218 | NM_173440 | 268903 | Mm.74711 | NM_173440 | ENSMUSG00000048490 | **Nrip1** | 6030458L20Rik\|8430438I05Rik\|9630050P12\|AA959574\|AW456757\|RIP140 | nuclear receptor interacting protein 1 | protein-coding |
| chr16_76341610_76342051 | chr16 | 76341611 | 76342051 | + | -0.78207 | intron (NM_173440, intron 2 of 3) | RMER15-int\|LTR\|ERVL | 31218 | NM_173440 | 268903 | Mm.74711 | NM_173440 | ENSMUSG00000048490 | **Nrip1** | 6030458L20Rik\|8430438I05Rik\|9630050P12\|AA959574\|AW456757\|RIP140 | nuclear receptor interacting protein 1 | protein-coding |
| chr16_76283581_76284398 | chr16 | 76283582 | 76284398 | + | -0.817578 | Intergenic | MTC\|LTR\|ERVL-MaLR | 89059 | NM_173440 | 268903 | Mm.74711 | NM_173440 | ENSMUSG00000048490 | **Nrip1** | 6030458L20Rik\|8430438I05Rik\|9630050P12\|AA959574\|AW456757\|RIP140 | nuclear receptor interacting protein 1 | protein-coding |
| chr16_76354295_76354705 | chr16 | 76354296 | 76354705 | + | -0.82795 | intron (NM_173440, intron 1 of 3) | intron (NM_173440, intron 1 of 3) | 18549 | NM_173440 | 268903 | Mm.74711 | NM_173440 | ENSMUSG00000048490 | **Nrip1** | 6030458L20Rik\|8430438I05Rik\|9630050P12\|AA959574\|AW456757\|RIP140 | nuclear receptor interacting protein 1 | protein-coding |
| chr16_76549581_76550352 | chr16 | 76549582 | 76550352 | + | -1.08127 | Intergenic | Intergenic | -176918 | NM_173440 | 268903 | Mm.74711 | NM_173440 | ENSMUSG00000048490 | **Nrip1** | 6030458L20Rik\|8430438I05Rik\|9630050P12\|AA959574\|AW456757\|RIP140 | nuclear receptor interacting protein 1 | protein-coding |
| chr16_76299477_76299948 | chr16 | 76299478 | 76299948 | + | -1.15509 | intron (NM_173440, intron 3 of 3) | intron (NM_173440, intron 3 of 3) | 73336 | NM_173440 | 268903 | Mm.74711 | NM_173440 | ENSMUSG00000048490 | **Nrip1** | 6030458L20Rik\|8430438I05Rik\|9630050P12\|AA959574\|AW456757\|RIP140 | nuclear receptor interacting protein 1 | protein-coding |
| chr16_76299575_76299837 | chr16 | 76299576 | 76299837 | + | -1.25097 | intron (NM_173440, intron 3 of 3) | intron (NM_173440, intron 3 of 3) | 73343 | NM_173440 | 268903 | Mm.74711 | NM_173440 | ENSMUSG00000048490 | **Nrip1** | 6030458L20Rik\|8430438I05Rik\|9630050P12\|AA959574\|AW456757\|RIP140 | nuclear receptor interacting protein 1 | protein-coding |
| chr16_76310317_76310679 | chr16 | 76310318 | 76310679 | + | -1.33664 | intron (NM_173440, intron 3 of 3) | ORR1A4-int\|LTR\|ERVL-MaLR | 62551 | NM_173440 | 268903 | Mm.74711 | NM_173440 | ENSMUSG00000048490 | **Nrip1** | 6030458L20Rik\|8430438I05Rik\|9630050P12\|AA959574\|AW456757\|RIP140 | nuclear receptor interacting protein 1 | protein-coding |
| chr16_76345951_76346481 | chr16 | 76345952 | 76346481 | + | -1.51158 | intron (NM_173440, intron 2 of 3) | intron (NM_173440, intron 2 of 3) | 26833 | NM_173440 | 268903 | Mm.74711 | NM_173440 | ENSMUSG00000048490 | **Nrip1** | 6030458L20Rik\|8430438I05Rik\|9630050P12\|AA959574\|AW456757\|RIP140 | nuclear receptor interacting protein 1 | protein-coding |
| chr7_109768672_109769290 | chr7 | 109768673 | 109769290 | + | -0.689748 | intron (NM_020610, intron 1 of 6) | ORR1C2\|LTR\|ERVL-MaLR | 12564 | NM_020610 | 78593 | Mm.275374 | NM_020610 | ENSMUSG00000034825 | **Nrip3** | A330103B05Rik\|D7H11orf14\|ICRFP703B1614Q5.2 | nuclear receptor interacting protein 3 | protein-coding |
| chr12_90236041_90236455 | chr12 | 90236042 | 90236455 | + | -0.635598 | intron (NM_172544, intron 18 of 19) | intron (NM_172544, intron 18 of 19) | 423765 | NM_001252074 | 18191 | Mm.425766 | NM_172544 | ENSMUSG00000066392 | **Nrxn3** | - | neurexin III | protein-coding |
| chr13_55255095_55255710 | chr13 | 55255096 | 55255710 | + | -0.878557 | intron (NM_008739, intron 5 of 22) | intron (NM_008739, intron 5 of 22) | 45621 | NM_008739 | 18193 | Mm.12964 | NM_008739 | ENSMUSG00000021488 | **Nsd1** | AI528500\|KMT3B | nuclear receptor-binding SET-domain protein 1 | protein-coding |
| chr11_31999494_32000197 | chr11 | 31999495 | 32000197 | + | -0.641069 | promoter-TSS (NM_008741) | promoter-TSS (NM_008741) | -573 | NM_008741 | 18197 | Mm.3304 | NM_008741 | ENSMUSG00000020297 | **Nsg2** | - | neuron specific gene family member 2 | protein-coding |
| chr11_32014969_32015220 | chr11 | 32014970 | 32015220 | + | -0.723837 | intron (NM_001290681, intron 2 of 4) | intron (NM_001290681, intron 2 of 4) | 13818 | NM_001290681 | 18197 | Mm.3304 | NM_008741 | ENSMUSG00000020297 | **Nsg2** | - | neuron specific gene family member 2 | protein-coding |
| chr11_32061627_32062547 | chr11 | 32061628 | 32062547 | + | -0.779525 | Intergenic | Intergenic | 60810 | NM_001290681 | 18197 | Mm.3304 | NM_008741 | ENSMUSG00000020297 | **Nsg2** | - | neuron specific gene family member 2 | protein-coding |
| chr11_31999839_32000197 | chr11 | 31999840 | 32000197 | + | -0.840057 | promoter-TSS (NM_008741) | promoter-TSS (NM_008741) | -401 | NM_008741 | 18197 | Mm.3304 | NM_008741 | ENSMUSG00000020297 | **Nsg2** | - | neuron specific gene family member 2 | protein-coding |
| chr7_125487248_125487471 | chr7 | 125487249 | 125487471 | + | -0.829206 | intron (NM_026330, intron 1 of 7) | intron (NM_026330, intron 1 of 7) | 4182 | NM_026330 | 67711 | Mm.4467 | NM_026330 | ENSMUSG00000030750 | **Nsmce1** | 2510027N19Rik | non-SMC element 1 homolog (S. cerevisiae) | protein-coding |
| chr7_125488639_125489226 | chr7 | 125488640 | 125489226 | + | -0.969663 | intron (NM_026330, intron 1 of 7) | L1ME3E\|LINE\|L1 | 2609 | NM_026330 | 67711 | Mm.4467 | NM_026330 | ENSMUSG00000030750 | **Nsmce1** | 2510027N19Rik | non-SMC element 1 homolog (S. cerevisiae) | protein-coding |
| chr7_125507192_125508055 | chr7 | 125507193 | 125508055 | + | -1.65317 | Intergenic | Intergenic | -16082 | NM_026330 | 67711 | Mm.4467 | NM_026330 | ENSMUSG00000030750 | **Nsmce1** | 2510027N19Rik | non-SMC element 1 homolog (S. cerevisiae) | protein-coding |
| chr7_125507320_125507623 | chr7 | 125507321 | 125507623 | + | -1.75427 | Intergenic | B1_Mm\|SINE\|Alu | -15930 | NM_026330 | 67711 | Mm.4467 | NM_026330 | ENSMUSG00000030750 | **Nsmce1** | 2510027N19Rik | non-SMC element 1 homolog (S. cerevisiae) | protein-coding |
| chr2_14992502_14993017 | chr2 | 14992503 | 14993017 | + | -0.724033 | Intergenic | ERVB2_1A-I_MM-int\|LTR\|ERVK | 56552 | NM_001165942 | 74455 | Mm.478923 | NM_028950 | ENSMUSG00000026707 | **Nsun6** | 4933403D21Rik\|4933414E04Rik\|NOPD1 | NOL1/NOP2/Sun domain family member 6 | protein-coding |
| chr6_56919946_56920202 | chr6 | 56919947 | 56920202 | + | -0.603836 | intron (NM_026004, intron 1 of 8) | intron (NM_026004, intron 1 of 8) | 3858 | NM_026004 | 107569 | Mm.158936 | NM_026004 | ENSMUSG00000029780 | **Nt5c3** | 1600024P05Rik\|2610206B05Rik\|3110004A18Rik\|AI842776\|Nt5c3a\|Nt5y\|PN-1\|PN-I\|PSN1\|UMPH\|Umph-1\|Umph1\|cN-III\|lupin\|p36 | 5'-nucleotidase, cytosolic III | protein-coding |
| chr6_56901773_56902337 | chr6 | 56901774 | 56902337 | + | -0.739081 | promoter-TSS (NM_001252374) | promoter-TSS (NM_001252374) | -169 | NM_001252374 | 107569 | Mm.158936 | NM_026004 | ENSMUSG00000029780 | **Nt5c3** | 1600024P05Rik\|2610206B05Rik\|3110004A18Rik\|AI842776\|Nt5c3a\|Nt5y\|PN-1\|PN-I\|PSN1\|UMPH\|Umph-1\|Umph1\|cN-III\|lupin\|p36 | 5'-nucleotidase, cytosolic III | protein-coding |
| chr9_88362147_88362480 | chr9 | 88362148 | 88362480 | + | -0.810479 | intron (NM_011851, intron 3 of 8) | intron (NM_011851, intron 3 of 8) | 34705 | NM_011851 | 23959 | Mm.244235 | NM_011851 | ENSMUSG00000032420 | **Nt5e** | 2210401F01Rik\|AI447961\|CD73\|NT\|Nt5\|eNT | 5' nucleotidase, ecto | protein-coding |
| chr8_125725017_125725876 | chr8 | 125725018 | 125725876 | + | -0.666033 | Intergenic | Intergenic | -8756 | NM_025636 | 66566 | Mm.25296 | NM_025636 | ENSMUSG00000031851 | **Ntpcr** | 2310079N02Rik\|AI449709 | nucleoside-triphosphatase, cancer-related | protein-coding |
| chr8_125725212_125725750 | chr8 | 125725213 | 125725750 | + | -0.678306 | Intergenic | Intergenic | -8722 | NM_025636 | 66566 | Mm.25296 | NM_025636 | ENSMUSG00000031851 | **Ntpcr** | 2310079N02Rik\|AI449709 | nucleoside-triphosphatase, cancer-related | protein-coding |
| chr7_78451551_78452261 | chr7 | 78451552 | 78452261 | + | -1.24981 | exon (NM_182809, exon 10 of 14) | exon (NM_182809, exon 10 of 14) | 125932 | NM_008746 | 18213 | Mm.33496 | NM_008746 | | **Ntrk3** | AW125844\|Ntrk3_tv3\|TrkC | neurotrophic tyrosine kinase, receptor, type 3 | protein-coding |
| chr1_132341605_132342469 | chr1 | 132341606 | 132342469 | + | -0.990452 | Intergenic | Intergenic | 25912 | NM_001195025 | 74137 | Mm.100666 | NM_028778 | ENSMUSG00000009772 | **Nuak2** | 1200013B22Rik\|Omphk2\|Snark\|mKIAA0537 | NUAK family, SNF1-like kinase, 2 | protein-coding |
| chr1_132341751_132342126 | chr1 | 132341752 | 132342126 | + | -0.992912 | Intergenic | Intergenic | 25814 | NM_001195025 | 74137 | Mm.100666 | NM_028778 | ENSMUSG00000009772 | **Nuak2** | 1200013B22Rik\|Omphk2\|Snark\|mKIAA0537 | NUAK family, SNF1-like kinase, 2 | protein-coding |
| chr7_116529866_116530095 | chr7 | 116529867 | 116530095 | + | -0.705397 | intron (NM_001130479, intron 10 of 13) | ORR1B2\|LTR\|ERVL-MaLR | 25612 | NM_001130479 | 53322 | Mm.9901 | NM_016773 | ENSMUSG00000030659 | **Nucb2** | AI607786\|Calnuc\|Nefa\|Nesfatin-1 | nucleobindin 2 | protein-coding |

| | | | | | | | | | | | | | | | | | |
|---|---|---|---|---|---|---|---|---|---|---|---|---|---|---|---|---|---|
| chr7_116529799_116530132 | chr7 | 116529800 | 116530132 | + | -0.735829 | intron (NM_001130479, intron 10 of 13) | ORR1B2\|LTR\|ERVL-MaLR | 25597 | NM_001130479 | 53322 | Mm.9901 | NM_016773 | ENSMUSG00000030659 | **Nucb2** | AI607786\|Calnuc\|Nefa\|Nesfatin-1 | nucleobindin 2 | protein-coding |
| chr9_105134514_105134794 | chr9 | 105134515 | 105134794 | + | -0.758526 | Intergenic | L1MC3\|LINE\|L1 | -2849 | NM_029385 | 75686 | Mm.305810 | NM_029385 | ENSMUSG00000032565 | **Nudt16** | 2310041H06Rik\|2810047L04Rik\|2900006H04Rik\|AI851783 | nudix (nucleoside diphosphate linked moiety X)-type motif 16 | protein-coding |
| chr3_96715913_96716174 | chr3 | 96715914 | 96716174 | + | -0.929455 | intron (NM_028925, intron 9 of 14) | intron (NM_028925, intron 9 of 14) | -7484 | NM_030094 | 78373 | Mm.423186 | NM_030094 | ENSMUSG00000028100 | **Nudt17** | 2410015C20Rik | nudix (nucleoside diphosphate linked moiety X)-type motif 17 | protein-coding |
| chr10_95690954_95691261 | chr10 | 95690955 | 95691261 | + | -0.855891 | Intergenic | Intergenic | -126941 | NM_027722 | 71207 | Mm.24397 | NM_027722 | ENSMUSG00000020029 | **Nudt4** | 4933436C10Rik\|DIPP2\|DIPP2alpha\|DIPP2beta\|HDCMB47P | nudix (nucleoside diphosphate linked moiety X)-type motif 4 | protein-coding |
| chr10_95698895_95699428 | chr10 | 95698896 | 95699428 | + | -0.915519 | Intergenic | (CA)n\|Simple_repeat\|Simple_repeat | -134995 | NM_027722 | 71207 | Mm.24397 | NM_027722 | ENSMUSG00000020029 | **Nudt4** | 4933436C10Rik\|DIPP2\|DIPP2alpha\|DIPP2beta\|HDCMB47P | nudix (nucleoside diphosphate linked moiety X)-type motif 4 | protein-coding |
| chr12_83929317_83929930 | chr12 | 83929318 | 83929930 | + | -0.919118 | Intergenic | Intergenic | -7690 | NR_073564 | 18222 | Mm.4390 | NM_010949 | ENSMUSG00000021224 | **Numb** | Nb | numb gene homolog (Drosophila) | protein-coding |
| chr10_117803980_117804207 | chr10 | 117803981 | 117804207 | + | -0.760772 | Intergenic | MTD\|LTR\|ERVL-MaLR | -11389 | NM_134010 | 103468 | Mm.12568 | NM_134010 | ENSMUSG00000052798 | **Nup107** | AW541137\|C76801 | nucleoporin 107 | protein-coding |
| chr10_117784681_117785135 | chr10 | 117784682 | 117785135 | + | -1.14902 | intron (NM_134010, intron 5 of 27) | B1_Mur1\|SINE\|Alu | 7797 | NM_134010 | 103468 | Mm.12568 | NM_134010 | ENSMUSG00000052798 | **Nup107** | AW541137\|C76801 | nucleoporin 107 | protein-coding |
| chr2_90667687_90668410 | chr2 | 90667688 | 90668410 | + | -0.645165 | Intergenic | Intergenic | -9166 | NM_021512 | 59015 | Mm.24532 | NM_021512 | ENSMUSG00000051329 | **Nup160** | 160kDa\|2810011M03Rik\|AA414952\|AU020188\|Gtl-13\|Gtl1-13\|Gtl13\|mKIAA0197 | nucleoporin 160 | protein-coding |
| chr2_80684716_80685034 | chr2 | 80684717 | 80685034 | + | -0.641727 | Intergenic | Lx9\|LINE\|L1 | 45611 | NM_001190179 | 69482 | Mm.29200 | NM_027091 | | **Nup35** | 2310006I24Rik\|35kDa\|5330402E05Rik\|MP44\|NO44 | nucleoporin 35 | protein-coding |
| chr1_181117126_181117637 | chr1 | 181117127 | 181117637 | + | -0.794134 | intron (NM_026171, intron 13 of 22) | intron (NM_026171, intron 13 of 22) | 26775 | NM_026171 | 67459 | Mm.263464 | NM_026171 | ENSMUSG00000026516 | **Nvl** | 1200009I24Rik | nuclear VCP-like | protein-coding |
| chrX_13458777_13459147 | chrX | 13458778 | 13459147 | + | -0.776486 | Intergenic | B4A\|SINE\|B4 | -8710 | NM_173415 | 236690 | Mm.90233 | NM_173415 | ENSMUSG00000051228 | **Nyx** | CLNP\|CSNB1\|CSNB4\|nob | nyctalopin | protein-coding |
| chrX_13431526_13431772 | chrX | 13431527 | 13431772 | + | -0.920654 | Intergenic | Intergenic | -36023 | NM_173415 | 236690 | Mm.90233 | NM_173415 | ENSMUSG00000051228 | **Nyx** | CLNP\|CSNB1\|CSNB4\|nob | nyctalopin | protein-coding |
| chrX_13431343_13431787 | chrX | 13431344 | 13431787 | + | -0.968727 | Intergenic | Intergenic | -36107 | NM_173415 | 236690 | Mm.90233 | NM_173415 | ENSMUSG00000051228 | **Nyx** | CLNP\|CSNB1\|CSNB4\|nob | nyctalopin | protein-coding |
| chrX_13516419_13517014 | chrX | 13516420 | 13517014 | + | -1.04564 | TTS (NM_001284504) | TTS (NM_001284504) | 49045 | NM_173415 | 236690 | Mm.90233 | NM_173415 | ENSMUSG00000051228 | **Nyx** | CLNP\|CSNB1\|CSNB4\|nob | nyctalopin | protein-coding |
| chrX_13516419_13516813 | chrX | 13516420 | 13516813 | + | -1.09364 | TTS (NM_001284504) | TTS (NM_001284504) | 48944 | NM_173415 | 236690 | Mm.90233 | NM_173415 | ENSMUSG00000051228 | **Nyx** | CLNP\|CSNB1\|CSNB4\|nob | nyctalopin | protein-coding |
| chr19_47531109_47531716 | chr19 | 47531110 | 47531716 | + | -0.859944 | intron (NM_175360, intron 2 of 9) | intron (NM_175360, intron 2 of 9) | 5607 | NM_175360 | 108689 | Mm.218048 | NM_175360 | ENSMUSG00000042694 | **Obfc1** | 0610009H20Rik\|2310057J23Rik\|AAF-44\|AAF44\|AI413458\|RPA-32 | oligonucleotide/oligosaccharide-binding fold containing 1 | protein-coding |
| chr19_47531189_47531518 | chr19 | 47531190 | 47531518 | + | -0.895647 | intron (NM_175360, intron 2 of 9) | intron (NM_175360, intron 2 of 9) | 5666 | NM_175360 | 108689 | Mm.218048 | NM_175360 | ENSMUSG00000042694 | **Obfc1** | 0610009H20Rik\|2310057J23Rik\|AAF-44\|AAF44\|AI413458\|RPA-32 | oligonucleotide/oligosaccharide-binding fold containing 1 | protein-coding |
| chr19_47524844_47525296 | chr19 | 47524845 | 47525296 | + | -1.00804 | intron (NM_175360, intron 3 of 9) | intron (NM_175360, intron 3 of 9) | 11950 | NM_175360 | 108689 | Mm.218048 | NM_175360 | ENSMUSG00000042694 | **Obfc1** | 0610009H20Rik\|2310057J23Rik\|AAF-44\|AAF44\|AI413458\|RPA-32 | oligonucleotide/oligosaccharide-binding fold containing 1 | protein-coding |
| chr13_100575969_100576391 | chr13 | 100575970 | 100576391 | + | -0.71005 | Intergenic | Intergenic | -23682 | NM_008756 | 18260 | Mm.4807 | NM_008756 | ENSMUSG00000021638 | **Ocln** | AI503564\|Ocl | occludin | protein-coding |
| chr13_100576058_100576282 | chr13 | 100576059 | 100576282 | + | -0.72849 | Intergenic | Intergenic | -23672 | NM_008756 | 18260 | Mm.4807 | NM_008756 | ENSMUSG00000021638 | **Ocln** | AI503564\|Ocl | occludin | protein-coding |
| chr12_17511014_17511343 | chr12 | 17511015 | 17511343 | + | -0.677457 | Intergenic | B1_Mur1\|SINE\|Alu | -33694 | NM_013614 | 18263 | Mm.34102 | NM_013614 | ENSMUSG00000011179 | **Odc1** | ODC | ornithine decarboxylase, structural 1 | protein-coding |
| chr12_17511014_17511639 | chr12 | 17511015 | 17511639 | + | -0.689198 | Intergenic | Intergenic | -33546 | NM_013614 | 18263 | Mm.34102 | NM_013614 | ENSMUSG00000011179 | **Odc1** | ODC | ornithine decarboxylase, structural 1 | protein-coding |
| chr11_6324293_6324677 | chr11 | 6324294 | 6324677 | + | -0.654848 | intron (NM_010956, intron 4 of 22) | intron (NM_010956, intron 4 of 22) | 31927 | NM_001252288 | 18293 | Mm.276348 | NM_010956 | ENSMUSG00000020456 | **Ogdh** | 2210403E04Rik\|2210412K19Rik\|AA409584\|d1401\|mKIAA4192 | oxoglutarate (alpha-ketoglutarate) dehydrogenase (lipoamide) | protein-coding |
| chr11_6332523_6332939 | chr11 | 6332524 | 6332939 | + | -0.689881 | intron (NM_001252283, intron 5 of 23) | intron (NM_001252283, intron 5 of 23) | 40173 | NM_001252288 | 18293 | Mm.276348 | NM_010956 | ENSMUSG00000020456 | **Ogdh** | 2210403E04Rik\|2210412K19Rik\|AA409584\|d1401\|mKIAA4192 | oxoglutarate (alpha-ketoglutarate) dehydrogenase (lipoamide) | protein-coding |
| chr11_6332648_6332934 | chr11 | 6332649 | 6332934 | + | -0.709922 | intron (NM_001252283, intron 5 of 23) | intron (NM_001252283, intron 5 of 23) | 40233 | NM_001252288 | 18293 | Mm.276348 | NM_010956 | ENSMUSG00000020456 | **Ogdh** | 2210403E04Rik\|2210412K19Rik\|AA409584\|d1401\|mKIAA4192 | oxoglutarate (alpha-ketoglutarate) dehydrogenase (lipoamide) | protein-coding |
| chr11_6323982_6324828 | chr11 | 6323983 | 6324828 | + | -0.737855 | intron (NM_010956, intron 4 of 22) | intron (NM_010956, intron 4 of 22) | 31847 | NM_001252288 | 18293 | Mm.276348 | NM_010956 | ENSMUSG00000020456 | **Ogdh** | 2210403E04Rik\|2210412K19Rik\|AA409584\|d1401\|mKIAA4192 | oxoglutarate (alpha-ketoglutarate) dehydrogenase (lipoamide) | protein-coding |
| chr9_20765018_20765827 | chr9 | 20765019 | 20765827 | + | -0.63908 | Intergenic | RLTR14\|LTR\|ERV1 | -37209 | NM_173777 | 244723 | Mm.293552 | NM_173777 | ENSMUSG00000032172 | **Olfm2** | A030009A06Rik | olfactomedin 2 | protein-coding |
| chr9_20765281_20765696 | chr9 | 20765282 | 20765696 | + | -0.672028 | Intergenic | Intergenic | -37275 | NM_173777 | 244723 | Mm.293552 | NM_173777 | ENSMUSG00000032172 | **Olfm2** | A030009A06Rik | olfactomedin 2 | protein-coding |
| chr11_49101324_49101896 | chr11 | 49101325 | 49101896 | + | -0.732413 | intron (NM_010999, intron 2 of 5) | RLTR12G\|LTR\|ERVK | 12147 | NM_146337 | 258334 | Mm.390388 | NM_146337 | ENSMUSG00000047511 | **Olfr1396** | MOR276-2 | olfactory receptor 1396 | protein-coding |
| chr19_12026638_12026960 | chr19 | 12026639 | 12026960 | + | -1.01053 | Intergenic | Intergenic | 9941 | NM_146680 | 258675 | Mm.271517 | NM_146680 | ENSMUSG00000067529 | **Olfr1423** | MOR239-3 | olfactory receptor 1423 | protein-coding |
| chr14_53073145_53073437 | chr14 | 53073146 | 53073437 | + | -0.660983 | Intergenic | Intergenic | -577596 | NM_001170918 | 57269 | Mm.88822 | NM_020512 | ENSMUSG00000059887 | **Olfr1507** | MOR244-1\|Mor28\|Or28 | olfactory receptor 1507 | protein-coding |
| chr14_52436821_52437172 | chr14 | 52436822 | 52437172 | + | -0.679918 | Intergenic | ORR1B2-int\|LTR\|ERVL-MaLR | -13396 | NM_020514 | 57271 | Mm.88824 | NM_020514 | ENSMUSG00000035626 | **Olfr1509** | MOR244-3\|Mor83\|Or83 | olfactory receptor 1509 | protein-coding |
| chr14_52427555_52428037 | chr14 | 52427556 | 52428037 | + | -1.11985 | Intergenic | Intergenic | -16869 | NM_146431 | 258423 | Mm.377492 | NM_146431 | ENSMUSG00000063106 | **Olfr1510** | MOR223-6 | olfactory receptor 1510 | protein-coding |
| chr4_43834692_43835223 | chr4 | 43834693 | 43835223 | + | -0.673465 | 3' UTR (NM_019475, exon 1 of 1) | 3' UTR (NM_019475, exon 1 of 1) | 1559 | NM_019475 | 100040268 | Mm.317638 | NM_019475 | ENSMUSG00000059101 | **Olfr157** | Mor262-12\|Olfr37c\|Or37c | olfactory receptor 157 | protein-coding |
| chr11_58788473_58788717 | chr11 | 58788474 | 58788717 | + | -0.931826 | Intergenic | Intergenic | 2456 | NM_001011760 | 257917 | Mm.377302 | NM_001011760 | ENSMUSG00000059504 | **Olfr314** | MOR278-2 | olfactory receptor 314 | protein-coding |
| chr8_72118688_72118954 | chr8 | 72118689 | 72118954 | + | -0.660317 | Intergenic | MMERVK9E_I-int\|LTR\|ERVK | 11781 | NM_146338 | 258335 | Mm.377439 | NM_146338 | ENSMUSG00000046881 | **Olfr374** | MOR130-1 | olfactory receptor 374 | protein-coding |
| chr11_87935140_87935863 | chr11 | 87935141 | 87935863 | + | -1.19037 | Intergenic | URR1A\|DNA\|hAT-Charlie | -20499 | NM_146412 | 258407 | Mm.271337 | NM_146412 | ENSMUSG00000060787 | **Olfr464** | MOR240-2 | olfactory receptor 464 | protein-coding |
| chr11_87935434_87935909 | chr11 | 87935435 | 87935909 | + | -1.23953 | Intergenic | Intergenic | -20669 | NM_146412 | 258407 | Mm.271337 | NM_146412 | ENSMUSG00000060787 | **Olfr464** | MOR240-2 | olfactory receptor 464 | protein-coding |
| chr7_106792908_106793346 | chr7 | 106792909 | 106793346 | + | -0.712101 | Intergenic | Intergenic | -2128 | NM_001011862 | 258180 | Mm.377371 | NM_001011862 | ENSMUSG00000096714 | **Olfr699** | MOR283-10P | olfactory receptor 699 | protein-coding |
| chr7_106793026_106793317 | chr7 | 106793027 | 106793317 | + | -0.717902 | Intergenic | Intergenic | -2173 | NM_001011862 | 258180 | Mm.377371 | NM_001011862 | ENSMUSG00000096714 | **Olfr699** | MOR283-10P | olfactory receptor 699 | protein-coding |
| chr7_106805496_106806431 | chr7 | 106805497 | 106806431 | + | -0.788554 | exon (NM_146600, exon 1 of 1) | exon (NM_146600, exon 1 of 1) | 496 | NM_146600 | 258593 | Mm.377583 | NM_146600 | ENSMUSG00000094493 | **Olfr700** | MOR283-4 | olfactory receptor 700 | protein-coding |
| chr7_106805521_106806003 | chr7 | 106805522 | 106806003 | + | -0.892256 | exon (NM_146600, exon 1 of 1) | exon (NM_146600, exon 1 of 1) | 698 | NM_146600 | 258593 | Mm.377583 | NM_146600 | ENSMUSG00000094493 | **Olfr700** | MOR283-4 | olfactory receptor 700 | protein-coding |
| chr10_128990381_128991564 | chr10 | 128990382 | 128991564 | + | -0.758225 | TTS (NM_146861) | TTS (NM_146861) | 1059 | NM_146861 | 18373 | Mm.377111 | NM_146861 | ENSMUSG00000069430 | **Olfr9** | MOR269-3 | olfactory receptor 9 | protein-coding |
| chr10_128990863_128991552 | chr10 | 128990864 | 128991552 | + | -0.838327 | TTS (NM_146861) | TTS (NM_146861) | 1294 | NM_146861 | 18373 | Mm.377111 | NM_146861 | ENSMUSG00000069430 | **Olfr9** | MOR269-3 | olfactory receptor 9 | protein-coding |
| chr10_19333442_19333730 | chr10 | 19333443 | 19333730 | + | -0.950436 | Intergenic | Intergenic | -22953 | NM_053008 | 94222 | Mm.156946 | NM_053008 | ENSMUSG00000045591 | **Olig3** | Bhlhb7\|bHLHe20 | oligodendrocyte transcription factor 3 | protein-coding |
| chr10_19333426_19333935 | chr10 | 19333427 | 19333935 | + | -0.960589 | Intergenic | (CA)n\|Simple_repeat\|Simple_repeat | -22858 | NM_053008 | 94222 | Mm.156946 | NM_053008 | ENSMUSG00000045591 | **Olig3** | Bhlhb7\|bHLHe20 | oligodendrocyte transcription factor 3 | protein-coding |
| chr10_19356164_19356565 | chr10 | 19356165 | 19356565 | + | -0.996687 | promoter-TSS (NM_053008) | promoter-TSS (NM_053008) | -174 | NM_053008 | 94222 | Mm.156946 | NM_053008 | ENSMUSG00000045591 | **Olig3** | Bhlhb7\|bHLHe20 | oligodendrocyte transcription factor 3 | protein-coding |
| chr10_19356236_19356522 | chr10 | 19356237 | 19356522 | + | -0.998891 | promoter-TSS (NM_053008) | promoter-TSS (NM_053008) | -160 | NM_053008 | 94222 | Mm.156946 | NM_053008 | ENSMUSG00000045591 | **Olig3** | Bhlhb7\|bHLHe20 | oligodendrocyte transcription factor 3 | protein-coding |
| chr10_19356221_19356704 | chr10 | 19356222 | 19356704 | + | -1.043 | promoter-TSS (NM_053008) | promoter-TSS (NM_053008) | -76 | NM_053008 | 94222 | Mm.156946 | NM_053008 | ENSMUSG00000045591 | **Olig3** | Bhlhb7\|bHLHe20 | oligodendrocyte transcription factor 3 | protein-coding |
| chr10_19321643_19322516 | chr10 | 19321644 | 19322516 | + | -1.28155 | Intergenic | Intergenic | -34459 | NM_053008 | 94222 | Mm.156946 | NM_053008 | ENSMUSG00000045591 | **Olig3** | Bhlhb7\|bHLHe20 | oligodendrocyte transcription factor 3 | protein-coding |
| chr15_76316783_76317215 | chr15 | 76316784 | 76317215 | + | -0.80211 | Intergenic | ORR1A3\|LTR\|ERVL-MaLR | -9754 | NM_153122 | 75475 | Mm.322738 | NM_153122 | ENSMUSG00000022562 | **Oplah** | 1700010G02Rik | 5-oxoprolinase (ATP-hydrolysing) | protein-coding |
| chr15_76312297_76312660 | chr15 | 76312298 | 76312660 | + | -0.843067 | Intergenic | (CA)n\|Simple_repeat\|Simple_repeat | -5234 | NM_153122 | 75475 | Mm.322738 | NM_153122 | ENSMUSG00000022562 | **Oplah** | 1700010G02Rik | 5-oxoprolinase (ATP-hydrolysing) | protein-coding |
| chr15_76316785_76317544 | chr15 | 76316786 | 76317544 | + | -1.0819 | Intergenic | L1M4b\|LINE\|L1 | -9920 | NM_153122 | 75475 | Mm.322738 | NM_153122 | ENSMUSG00000022562 | **Oplah** | 1700010G02Rik | 5-oxoprolinase (ATP-hydrolysing) | protein-coding |
| chr5_123021498_123021731 | chr5 | 123021499 | 123021731 | + | -0.892886 | intron (NM_175423, intron 1 of 1) | PB1D10\|SINE\|Alu | 6541 | NM_175423 | 109305 | Mm.218582 | NM_175423 | ENSMUSG00000049686 | **Orai1** | D730049H07Rik\|Tmem142a\|orai-1 | ORAI calcium release-activated calcium modulator 1 | protein-coding |
| chr5_136155050_136155803 | chr5 | 136155051 | 136155803 | + | -0.613763 | intron (NM_178751, intron 2 of 2) | intron (NM_178751, intron 2 of 2) | 15229 | NM_178751 | 269717 | Mm.308396 | NM_178751 | ENSMUSG00000039747 | **Orai2** | A730041O15Rik\|Tmem142b | ORAI calcium release-activated calcium modulator 2 | protein-coding |
| chr5_136179516_136180059 | chr5 | 136179517 | 136180059 | + | -0.735499 | TTS (NM_025774) | TTS (NM_025774) | -9132 | NM_178751 | 269717 | Mm.308396 | NM_178751 | ENSMUSG00000039747 | **Orai2** | A730041O15Rik\|Tmem142b | ORAI calcium release-activated calcium modulator 2 | protein-coding |
| chr5_136173608_136174062 | chr5 | 136173609 | 136174062 | + | -1.0754 | Intergenic | B3A\|SINE\|B2 | -3179 | NM_178751 | 269717 | Mm.308396 | NM_178751 | ENSMUSG00000039747 | **Orai2** | A730041O15Rik\|Tmem142b | ORAI calcium release-activated calcium modulator 2 | protein-coding |
| chr16_33203288_33203565 | chr16 | 33203289 | 33203565 | + | -0.734472 | intron (NM_176840, intron 2 of 12) | intron (NM_176840, intron 2 of 12) | 18356 | NM_176840 | 106326 | Mm.486405 | NM_176840 | ENSMUSG00000022807 | **Osbpl11** | 9430097N02Rik\|AI132306\|AU019795\|ORP-11 | oxysterol binding protein-like 11 | protein-coding |
| chr2_76429374_76429605 | chr2 | 76429375 | 76429605 | + | -0.667304 | intron (NM_001290734, intron 1 of 24) | intron (NM_001290734, intron 1 of 24) | 22982 | NM_001290733 | 99031 | Mm.240435 | NM_145525 | ENSMUSG00000042359 | **Osbpl6** | 1110062M20Rik\|AI596402\|ORP-6\|mKIAA4128 | oxysterol binding protein-like 6 | protein-coding |

| | | | | | | | | | | | | | | | | | |
|---|---|---|---|---|---|---|---|---|---|---|---|---|---|---|---|---|---|
| chr2_76421238_76421922 | chr2 | 76421239 | 76421922 | + | -0.97832 | intron (NM_001290734, intron 1 of 24) | B3\|SINE\|B2 | 15072 | NM_001290733 | 99031 | Mm.240435 | NM_145525 | ENSMUSG00000042359 | **Osbpl6** | 1110062M20Rik\|AI596402\|ORP-6\|mKIAA4128 | oxysterol binding protein-like 6 | protein-coding |
| chr4_109079662_109080292 | chr4 | 109079663 | 109080292 | + | -1.08994 | intron (NM_133885, intron 11 of 22) | intron (NM_133885, intron 11 of 22) | 38059 | NM_173350 | 100273 | Mm.366315 | NM_133885 | ENSMUSG00000028559 | **Osbpl9** | 2600011I06Rik\|AU015843\|Orp-9 | oxysterol binding protein-like 9 | protein-coding |
| chr4_109079657_109080102 | chr4 | 109079658 | 109080102 | + | -1.10383 | intron (NM_133885, intron 11 of 22) | intron (NM_133885, intron 11 of 22) | 38156 | NM_173350 | 100273 | Mm.366315 | NM_133885 | ENSMUSG00000028559 | **Osbpl9** | 2600011I06Rik\|AU015843\|Orp-9 | oxysterol binding protein-like 9 | protein-coding |
| chr8_119423622_119424309 | chr8 | 119423623 | 119424309 | + | -0.837334 | Intergenic | Intergenic | -13196 | NM_027950 | 71839 | Mm.272357 | NM_027950 | ENSMUSG00000074063 | **Osgin1** | 1700012B18Rik\|Okl38 | oxidative stress induced growth inhibitor 1 | protein-coding |
| chr19_18619125_18619470 | chr19 | 18619126 | 18619470 | + | -0.801687 | intron (NM_017375, intron 1 of 9) | intron (NM_017375, intron 1 of 9) | 12515 | NM_017375 | 20409 | Mm.172222 | NM_017375 | | **Ostf1** | C78236\|SH3P2\|Sh3d3 | osteoclast stimulating factor 1 | protein-coding |
| chr19_18618292_18618565 | chr19 | 18618293 | 18618565 | + | -0.848938 | intron (NM_017375, intron 1 of 9) | intron (NM_017375, intron 1 of 9) | 13384 | NM_017375 | 20409 | Mm.172222 | NM_017375 | | **Ostf1** | C78236\|SH3P2\|Sh3d3 | osteoclast stimulating factor 1 | protein-coding |
| chr19_18555649_18556018 | chr19 | 18555650 | 18556018 | + | -0.898325 | Intergenic | MTD\|LTR\|ERVL-MaLR | 75979 | NM_017375 | 20409 | Mm.172222 | NM_017375 | | **Ostf1** | C78236\|SH3P2\|Sh3d3 | osteoclast stimulating factor 1 | protein-coding |
| chr10_42699003_42699439 | chr10 | 42699004 | 42699439 | + | -0.836269 | intron (NM_172416, intron 5 of 5) | intron (NM_172416, intron 5 of 5) | 20305 | NM_172416 | 14628 | Mm.46636 | NM_172416 | ENSMUSG00000038280 | **Ostm1** | 1200002H13Rik\|HSPC019\|gl | osteopetrosis associated transmembrane protein 1 | protein-coding |
| chr10_107872592_107872980 | chr10 | 107872593 | 107872980 | + | -0.729084 | intron (NM_001177567, intron 17 of 57) | RCHARR1\|DNA\|hAT-Charlie | 39348 | NM_001177567 | 628870 | Mm.336090 | NM_001177567 | ENSMUSG00000091455 | **Otogl** | EG628870\|Gm6924 | otogelin-like | protein-coding |
| chr15_27605243_27605725 | chr15 | 27605244 | 27605725 | + | -1.1406 | TTS (NM_001013792) | TTS (NM_001013792) | 25209 | NM_001013792 | 432940 | Mm.309164 | NM_001013792 | ENSMUSG00000046034 | **Otulin** | C79097\|Fam105b\|m3Sapc\|m7-1Sapc | OTU deubiquitinase with linear linkage specificity | protein-coding |
| chr5_122731958_122732625 | chr5 | 122731959 | 122732625 | + | -0.805175 | 3' UTR (NM_001199676, exon 17 of 17) | 3' UTR (NM_001199676, exon 17 of 17) | 24735 | NM_011026 | 18438 | Mm.290884 | NM_011026 | ENSMUSG00000029470 | **P2rx4** | AI504491\|AW555605\|D5Ertd444e\|P2X4 | purinergic receptor P2X, ligand-gated ion channel 4 | protein-coding |
| chr5_122731995_122732239 | chr5 | 122731996 | 122732239 | + | -0.877841 | 3' UTR (NM_001199676, exon 17 of 17) | 3' UTR (NM_001199676, exon 17 of 17) | 24560 | NM_011026 | 18438 | Mm.290884 | NM_011026 | ENSMUSG00000029470 | **P2rx4** | AI504491\|AW555605\|D5Ertd444e\|P2X4 | purinergic receptor P2X, ligand-gated ion channel 4 | protein-coding |
| chr5_122718879_122719648 | chr5 | 122718880 | 122719648 | + | -1.22419 | intron (NM_011026, intron 5 of 11) | intron (NM_011026, intron 5 of 11) | 11707 | NM_011026 | 18438 | Mm.290884 | NM_011026 | ENSMUSG00000029470 | **P2rx4** | AI504491\|AW555605\|D5Ertd444e\|P2X4 | purinergic receptor P2X, ligand-gated ion channel 4 | protein-coding |
| chr5_122719103_122719577 | chr5 | 122719104 | 122719577 | + | -1.25266 | intron (NM_011026, intron 5 of 11) | intron (NM_011026, intron 5 of 11) | 11783 | NM_011026 | 18438 | Mm.290884 | NM_011026 | ENSMUSG00000029470 | **P2rx4** | AI504491\|AW555605\|D5Ertd444e\|P2X4 | purinergic receptor P2X, ligand-gated ion channel 4 | protein-coding |
| chr5_122650067_122650477 | chr5 | 122650068 | 122650477 | + | -0.689534 | promoter-TSS (NM_001284402) | promoter-TSS (NM_001284402) | -152 | NM_001284402 | 18439 | Mm.42026 | NM_011027 | ENSMUSG00000029468 | **P2rx7** | AI467586\|P2X(7)\|P2X7R | purinergic receptor P2X, ligand-gated ion channel, 7 | protein-coding |
| chr5_122650159_122650395 | chr5 | 122650160 | 122650395 | + | -0.76385 | promoter-TSS (NM_001284402) | promoter-TSS (NM_001284402) | -147 | NM_001284402 | 18439 | Mm.42026 | NM_011027 | ENSMUSG00000029468 | **P2rx7** | AI467586\|P2X(7)\|P2X7R | purinergic receptor P2X, ligand-gated ion channel, 7 | protein-coding |
| chr5_122650191_122650479 | chr5 | 122650192 | 122650479 | + | -0.813551 | promoter-TSS (NM_001284402) | promoter-TSS (NM_001284402) | -89 | NM_001284402 | 18439 | Mm.42026 | NM_011027 | ENSMUSG00000029468 | **P2rx7** | AI467586\|P2X(7)\|P2X7R | purinergic receptor P2X, ligand-gated ion channel, 7 | protein-coding |
| chr3_60954155_60954660 | chr3 | 60954156 | 60954660 | + | -0.901467 | Intergenic | MER34\|LTR\|ERV1 | -48377 | NM_008772 | 18441 | Mm.281452 | NM_008772 | ENSMUSG00000027765 | **P2ry1** | P2Y1 | purinergic receptor P2Y, G-protein coupled 1 | protein-coding |
| chr3_60954171_60954459 | chr3 | 60954172 | 60954459 | + | -0.922658 | Intergenic | MER34\|LTR\|ERV1 | -48470 | NM_008772 | 18441 | Mm.281452 | NM_008772 | ENSMUSG00000027765 | **P2ry1** | P2Y1 | purinergic receptor P2Y, G-protein coupled 1 | protein-coding |
| chr3_59260642_59261187 | chr3 | 59260643 | 59261187 | + | -0.663754 | intron (NM_177855, intron 32 of 43) | intron (NM_177855, intron 32 of 43) | 1916 | NM_027571 | 70839 | Mm.422675 | NM_027571 | ENSMUSG00000036353 | **P2ry12** | 2900079B22Rik\|4921504D23Rik\|P2Y12 | purinergic receptor P2Y, G-protein coupled 12 | protein-coding |
| chr3_59234756_59235270 | chr3 | 59234757 | 59235270 | + | -1.08679 | intron (NM_177855, intron 18 of 43) | intron (NM_177855, intron 18 of 43) | -24131 | NM_028808 | 74191 | Mm.90067 | NM_028808 | ENSMUSG00000036362 | **P2ry13** | 2010001L06Rik\|GPCR1\|GPR94\|Gpr86\|P2Y13\|P2yr13\|SP174 | purinergic receptor P2Y, G-protein coupled 13 | protein-coding |
| chr3_59130403_59130821 | chr3 | 59130404 | 59130821 | + | -0.645013 | 5' UTR (NM_133200, exon 1 of 2) | 5' UTR (NM_133200, exon 1 of 2) | 150 | NM_001008497 | 140795 | Mm.439021 | NM_133200 | ENSMUSG00000036381 | **P2ry14** | A330108O13Rik\|Gpr105\|P2Y14 | purinergic receptor P2Y, G-protein coupled, 14 | protein-coding |
| chr3_59130245_59130860 | chr3 | 59130246 | 59130860 | + | -0.720197 | 5' UTR (NM_133200, exon 1 of 2) | 5' UTR (NM_133200, exon 1 of 2) | 209 | NM_001008497 | 140795 | Mm.439021 | NM_133200 | ENSMUSG00000036381 | **P2ry14** | A330108O13Rik\|Gpr105\|P2Y14 | purinergic receptor P2Y, G-protein coupled, 14 | protein-coding |
| chr11_54105066_54105505 | chr11 | 54105067 | 54105505 | + | -0.834798 | intron (NM_001136076, intron 2 of 15) | MER44B\|DNA\|TcMar-Tigger | 4362 | NM_011031 | 18452 | Mm.3705 | NM_011031 | ENSMUSG00000018906 | **P4ha2** | AA407196\|C76437\|P4hl | procollagen-proline, 2-oxoglutarate 4-dioxygenase (proline 4-hydroxylase), alpha II polypeptide | protein-coding |
| chr9_108617230_108617659 | chr9 | 108617231 | 108617659 | + | -1.26106 | intron (NM_011790, intron 4 of 15) | intron (NM_011790, intron 4 of 15) | -19845 | NM_028944 | 74443 | Mm.226534 | NM_028944 | ENSMUSG00000006675 | **P4htm** | 4933406E20Rik\|AI853847\|BB128974\|Ph-4 | prolyl 4-hydroxylase, transmembrane (endoplasmic reticulum) | protein-coding |
| chr15_36640610_36641187 | chr15 | 36640611 | 36641187 | + | -0.785135 | Intergenic | Intergenic | -31926 | NM_008774 | 18458 | Mm.371570 | NM_008774 | ENSMUSG00000022283 | **Pabpc1** | PABP\|Pabp1\|Pabpl\|Pabpl1\|ePAB | poly(A) binding protein, cytoplasmic 1 | protein-coding |
| chr15_36640769_36641153 | chr15 | 36640770 | 36641153 | + | -0.786104 | Intergenic | Intergenic | -31988 | NM_008774 | 18458 | Mm.371570 | NM_008774 | ENSMUSG00000022283 | **Pabpc1** | PABP\|Pabp1\|Pabpl\|Pabpl1\|ePAB | poly(A) binding protein, cytoplasmic 1 | protein-coding |
| chr15_36640582_36641447 | chr15 | 36640583 | 36641447 | + | -0.786939 | Intergenic | Intergenic | -32042 | NM_008774 | 18458 | Mm.371570 | NM_008774 | ENSMUSG00000022283 | **Pabpc1** | PABP\|Pabp1\|Pabpl\|Pabpl1\|ePAB | poly(A) binding protein, cytoplasmic 1 | protein-coding |
| chr19_5257175_5257675 | chr19 | 5257176 | 5257675 | + | -1.22606 | intron (NM_153129, intron 1 of 23) | intron (NM_153129, intron 1 of 23) | 15694 | NM_153129 | 107975 | Mm.234923 | NM_153129 | ENSMUSG00000024855 | **Pacs1** | AI325977 | phosphofurin acidic cluster sorting protein 1 | protein-coding |
| chr19_5257216_5257515 | chr19 | 5257217 | 5257515 | + | -1.25064 | intron (NM_153129, intron 1 of 23) | intron (NM_153129, intron 1 of 23) | 15753 | NM_153129 | 107975 | Mm.234923 | NM_153129 | ENSMUSG00000024855 | **Pacs1** | AI325977 | phosphofurin acidic cluster sorting protein 1 | protein-coding |
| chr3_9895719_9896148 | chr3 | 9895720 | 9896148 | + | -0.614758 | Intergenic | Intergenic | -62255 | NM_001195031 | 94212 | Mm.23897 | NM_053182 | ENSMUSG00000027508 | **Pag1** | Cbp\|F730007C19Rik\|Pag | phosphoprotein associated with glycosphingolipid microdomains 1 | protein-coding |
| chr3_9865313_9866139 | chr3 | 9865314 | 9866139 | + | -0.674222 | Intergenic | Intergenic | -32047 | NM_001195031 | 94212 | Mm.23897 | NM_053182 | ENSMUSG00000027508 | **Pag1** | Cbp\|F730007C19Rik\|Pag | phosphoprotein associated with glycosphingolipid microdomains 1 | protein-coding |
| chr3_9828404_9829030 | chr3 | 9828405 | 9829030 | + | -0.735047 | intron (NM_001195031, intron 1 of 8) | intron (NM_001195031, intron 1 of 8) | 4962 | NM_053182 | 94212 | Mm.23897 | NM_053182 | ENSMUSG00000027508 | **Pag1** | Cbp\|F730007C19Rik\|Pag | phosphoprotein associated with glycosphingolipid microdomains 1 | protein-coding |
| chr3_9916731_9917088 | chr3 | 9916732 | 9917088 | + | -0.811657 | Intergenic | Intergenic | -83231 | NM_001195031 | 94212 | Mm.23897 | NM_053182 | ENSMUSG00000027508 | **Pag1** | Cbp\|F730007C19Rik\|Pag | phosphoprotein associated with glycosphingolipid microdomains 1 | protein-coding |
| chr3_9828713_9829029 | chr3 | 9828714 | 9829029 | + | -0.815299 | intron (NM_001195031, intron 1 of 8) | intron (NM_001195031, intron 1 of 8) | 4808 | NM_053182 | 94212 | Mm.23897 | NM_053182 | ENSMUSG00000027508 | **Pag1** | Cbp\|F730007C19Rik\|Pag | phosphoprotein associated with glycosphingolipid microdomains 1 | protein-coding |
| chr3_9828727_9828958 | chr3 | 9828728 | 9828958 | + | -0.82314 | intron (NM_001195031, intron 1 of 8) | intron (NM_001195031, intron 1 of 8) | 4836 | NM_053182 | 94212 | Mm.23897 | NM_053182 | ENSMUSG00000027508 | **Pag1** | Cbp\|F730007C19Rik\|Pag | phosphoprotein associated with glycosphingolipid microdomains 1 | protein-coding |
| chr7_97877867_97878220 | chr7 | 97877868 | 97878220 | + | -0.825544 | intron (NM_011035, intron 5 of 14) | ORR1D1\|LTR\|ERVL-MaLR | 35105 | NM_011035 | 18479 | Mm.260227 | NM_011035 | ENSMUSG00000030774 | **Pak1** | AW045634\|PAK-1\|Paka | p21 protein (Cdc42/Rac)-activated kinase 1 | protein-coding |
| chr16_32078097_32078287 | chr16 | 32078098 | 32078287 | + | -0.689234 | intron (NM_177326, intron 1 of 14) | intron (NM_177326, intron 1 of 14) | 1150 | NM_177326 | 224105 | Mm.234204 | NM_177326 | ENSMUSG00000022781 | **Pak2** | 5330420P17Rik\|A130002K10Rik\|AI836325\|D16Ertd269e\|PAK-2\|mKIAA4182 | p21 protein (Cdc42/Rac)-activated kinase 2 | protein-coding |
| chr16_32082069_32082401 | chr16 | 32082070 | 32082401 | + | -0.758407 | Intergenic | Intergenic | -2893 | NM_177326 | 224105 | Mm.234204 | NM_177326 | ENSMUSG00000022781 | **Pak2** | 5330420P17Rik\|A130002K10Rik\|AI836325\|D16Ertd269e\|PAK-2\|mKIAA4182 | p21 protein (Cdc42/Rac)-activated kinase 2 | protein-coding |
| chrX_143482946_143483177 | chrX | 143482947 | 143483177 | + | -0.787543 | Intergenic | SYNREP_MM\|Satellite\|Satellite | -35529 | NM_001195048 | 18481 | Mm.40035 | NM_008778 | ENSMUSG00000031284 | **Pak3** | PAK-3\|Pak65alpha\|Pak65beta\|Stk4 | p21 protein (Cdc42/Rac)-activated kinase 3 | protein-coding |
| chr2_118642413_118642964 | chr2 | 118642414 | 118642964 | + | -0.767661 | Intergenic | Intergenic | -20888 | NM_001033254 | 214230 | Mm.241857 | NM_001033254 | ENSMUSG00000074923 | **Pak6** | 4732456M09 | p21 protein (Cdc42/Rac)-activated kinase 6 | protein-coding |
| chr2_118643035_118643387 | chr2 | 118643036 | 118643387 | + | -0.987573 | Intergenic | Intergenic | -20366 | NM_001033254 | 214230 | Mm.241857 | NM_001033254 | ENSMUSG00000074923 | **Pak6** | 4732456M09 | p21 protein (Cdc42/Rac)-activated kinase 6 | protein-coding |
| chr4_57640025_57640735 | chr4 | 57640026 | 57640735 | + | -0.9642 | intron (NM_172868, intron 2 of 5) | intron (NM_172868, intron 2 of 5) | 72132 | NM_172868 | 242481 | Mm.331630 | NM_172868 | | **Palm2** | 53-H12\|B130066I24\|R75368\|pEN70 | paralemmin 2 | protein-coding |
| chr5_147448214_147448855 | chr5 | 147448215 | 147448855 | + | -0.897897 | intron (NM_028291, intron 1 of 18) | (TTTTG)n\|Simple_repeat\|Simple_repeat | 17955 | NM_028291 | 72587 | Mm.320469 | NM_028291 | ENSMUSG00000029647 | **Pan3** | 2700050F09Rik\|A430027N15Rik\|AU014670 | PAN3 polyA specific ribonuclease subunit homolog (S. cerevisiae) | protein-coding |
| chr5_147448508_147448871 | chr5 | 147448509 | 147448871 | + | -0.902857 | intron (NM_028291, intron 1 of 18) | intron (NM_028291, intron 1 of 18) | 18110 | NM_028291 | 72587 | Mm.320469 | NM_028291 | ENSMUSG00000029647 | **Pan3** | 2700050F09Rik\|A430027N15Rik\|AU014670 | PAN3 polyA specific ribonuclease subunit homolog (S. cerevisiae) | protein-coding |
| chr5_147475370_147475740 | chr5 | 147475371 | 147475740 | + | -0.954187 | intron (NM_028291, intron 4 of 18) | ORR1E\|LTR\|ERVL-MaLR | 44975 | NM_028291 | 72587 | Mm.320469 | NM_028291 | ENSMUSG00000029647 | **Pan3** | 2700050F09Rik\|A430027N15Rik\|AU014670 | PAN3 polyA specific ribonuclease subunit homolog (S. cerevisiae) | protein-coding |
| chr9_15059868_15060434 | chr9 | 15059869 | 15060434 | + | -0.69378 | intron (NM_001164797, intron 16 of 19) | intron (NM_001164797, intron 16 of 19) | -14673 | NM_019482 | 55991 | Mm.142253 | NM_019482 | | **Panx1** | AI847747 | pannexin 1 | protein-coding |
| chr12_83762844_83763107 | chr12 | 83762845 | 83763107 | + | -0.679116 | promoter-TSS (NM_001205343) | promoter-TSS (NM_001205343) | -658 | NM_001205343 | 170721 | Mm.133443 | NM_130887 | ENSMUSG00000021223 | **Papln** | E030033C16Rik | papilin, proteoglycan-like sulfated glycoprotein | protein-coding |
| chr12_83762734_83763174 | chr12 | 83762735 | 83763174 | + | -0.697688 | promoter-TSS (NM_001205343) | promoter-TSS (NM_001205343) | -680 | NM_001205343 | 170721 | Mm.133443 | NM_130887 | ENSMUSG00000021223 | **Papln** | E030033C16Rik | papilin, proteoglycan-like sulfated glycoprotein | protein-coding |
| chr19_32574543_32575286 | chr19 | 32574544 | 32575286 | + | -0.664258 | Intergenic | L1M4\|LINE\|L1 | -20800 | NM_001201470 | 23972 | Mm.203916 | NM_011864 | ENSMUSG00000024899 | **Papss2** | 1810018P12Rik\|AI159688\|AtpsU2\|Atpsk2\|Sk2\|bm | 3'-phosphoadenosine 5'-phosphosulfate synthase 2 | protein-coding |

| | | | | | | | | | | | | | | | | | |
|---|---|---|---|---|---|---|---|---|---|---|---|---|---|---|---|---|---|
| chr19_32574573_32574812 | chr19 | 32574574 | 32574812 | + | -0.783026 | Intergenic | Intergenic | -21022 | NM_001201470 | 23972 | Mm.203916 | NM_011864 | ENSMUSG00000024899 | **Papss2** | 1810018P12Rik\|AI159688\|AtpsU2\|Atpsk2\|Sk2\|bm | 3'-phosphoadenosine 5'-phosphosulfate synthase 2 | protein-coding |
| chr4_134494824_134495660 | chr4 | 134494825 | 134495660 | + | -1.04586 | Intergenic | Intergenic | -1519 | NM_001285845 | 71904 | Mm.142343 | NM_027995 | ENSMUSG00000037348 | **Paqr7** | 2310021M12Rik\|AV001994\|BB007237\|Mpra\|PGLP\|mPR | progestin and adipoQ receptor family member VII | protein-coding |
| chr1_20899351_20899854 | chr1 | 20899352 | 20899854 | + | -0.714287 | intron (NM_028829, intron 1 of 2) | B4A\|SINE\|B4 | 8981 | NM_028829 | 74229 | Mm.40780 | NM_028829 | ENSMUSG00000025931 | **Paqr8** | 1700019B16Rik\|3110001D06Rik | progestin and adipoQ receptor family member VIII | protein-coding |
| chr8_127499821_127500546 | chr8 | 127499822 | 127500546 | + | -0.630883 | intron (NM_033620, intron 22 of 24) | intron (NM_033620, intron 22 of 24) | 200532 | NM_001013580 | 93742 | Mm.299254 | NM_033620 | ENSMUSG00000025812 | **Pard3** | AA960621\|AI256638\|ASIP\|D8Ertd580e\|PAR-3\|Par3\|Pard3a | par-3 family cell polarity regulator | protein-coding |
| chr8_127501863_127502058 | chr8 | 127501864 | 127502058 | + | -0.75383 | intron (NM_033620, intron 22 of 24) | intron (NM_033620, intron 22 of 24) | 202309 | NM_001013580 | 93742 | Mm.299254 | NM_033620 | ENSMUSG00000025812 | **Pard3** | AA960621\|AI256638\|ASIP\|D8Ertd580e\|PAR-3\|Par3\|Pard3a | par-3 family cell polarity regulator | protein-coding |
| chr8_127501873_127502252 | chr8 | 127501874 | 127502252 | + | -0.795274 | intron (NM_033620, intron 22 of 24) | intron (NM_033620, intron 22 of 24) | 202411 | NM_001013580 | 93742 | Mm.299254 | NM_033620 | ENSMUSG00000025812 | **Pard3** | AA960621\|AI256638\|ASIP\|D8Ertd580e\|PAR-3\|Par3\|Pard3a | par-3 family cell polarity regulator | protein-coding |
| chr8_127324968_127325485 | chr8 | 127324969 | 127325485 | + | -0.851763 | intron (NM_001013580, intron 3 of 13) | intron (NM_001013580, intron 3 of 13) | 25575 | NM_001013580 | 93742 | Mm.299254 | NM_033620 | ENSMUSG00000025812 | **Pard3** | AA960621\|AI256638\|ASIP\|D8Ertd580e\|PAR-3\|Par3\|Pard3a | par-3 family cell polarity regulator | protein-coding |
| chr8_127515948_127516468 | chr8 | 127515949 | 127516468 | + | -1.18482 | intron (NM_033620, intron 22 of 24) | intron (NM_033620, intron 22 of 24) | 216556 | NM_001013580 | 93742 | Mm.299254 | NM_033620 | ENSMUSG00000025812 | **Pard3** | AA960621\|AI256638\|ASIP\|D8Ertd580e\|PAR-3\|Par3\|Pard3a | par-3 family cell polarity regulator | protein-coding |
| chr6_127473983_127474515 | chr6 | 127473984 | 127474515 | + | -0.690839 | exon (NM_181402, exon 5 of 9) | exon (NM_181402, exon 5 of 9) | 20526 | NM_181402 | 101187 | Mm.110818 | NM_181402 | ENSMUSG00000037997 | **Parp11** | 5330431N24Rik\|AI851877\|ARTD11\|HIN1L | poly (ADP-ribose) polymerase family, member 11 | protein-coding |
| chr6_127472009_127472592 | chr6 | 127472010 | 127472592 | + | -0.761775 | intron (NM_181402, intron 4 of 8) | intron (NM_181402, intron 4 of 8) | 18578 | NM_181402 | 101187 | Mm.110818 | NM_181402 | ENSMUSG00000037997 | **Parp11** | 5330431N24Rik\|AI851877\|ARTD11\|HIN1L | poly (ADP-ribose) polymerase family, member 11 | protein-coding |
| chr6_127473957_127474294 | chr6 | 127473958 | 127474294 | + | -0.820568 | intron (NM_181402, intron 4 of 8) | intron (NM_181402, intron 4 of 8) | 20403 | NM_181402 | 101187 | Mm.110818 | NM_181402 | ENSMUSG00000037997 | **Parp11** | 5330431N24Rik\|AI851877\|ARTD11\|HIN1L | poly (ADP-ribose) polymerase family, member 11 | protein-coding |
| chr6_127434063_127434522 | chr6 | 127434064 | 127434522 | + | -1.09053 | Intergenic | L3\|LINE\|CR1 | -19430 | NM_181402 | 101187 | Mm.110818 | NM_181402 | ENSMUSG00000037997 | **Parp11** | 5330431N24Rik\|AI851877\|ARTD11\|HIN1L | poly (ADP-ribose) polymerase family, member 11 | protein-coding |
| chr6_127433935_127434557 | chr6 | 127433936 | 127434557 | + | -1.12069 | Intergenic | Intergenic | -19477 | NM_181402 | 101187 | Mm.110818 | NM_181402 | ENSMUSG00000037997 | **Parp11** | 5330431N24Rik\|AI851877\|ARTD11\|HIN1L | poly (ADP-ribose) polymerase family, member 11 | protein-coding |
| chr13_117054776_117055185 | chr13 | 117054777 | 117055185 | + | -0.912934 | Intergenic | (CA)n\|Simple_repeat\|Simple_repeat | -29465 | NM_001081009 | 52552 | Mm.274466 | NM_027272 | ENSMUSG00000021725 | **Parp8** | 2810430O08Rik\|ARTD16\|D13Ertd275e | poly (ADP-ribose) polymerase family, member 8 | protein-coding |
| chr13_117051394_117051841 | chr13 | 117051395 | 117051841 | + | -1.10563 | Intergenic | Intergenic | -26102 | NM_001081009 | 52552 | Mm.274466 | NM_027272 | ENSMUSG00000021725 | **Parp8** | 2810430O08Rik\|ARTD16\|D13Ertd275e | poly (ADP-ribose) polymerase family, member 8 | protein-coding |
| chr13_117051460_117051754 | chr13 | 117051461 | 117051754 | + | -1.15992 | Intergenic | Intergenic | -26091 | NM_001081009 | 52552 | Mm.274466 | NM_027272 | ENSMUSG00000021725 | **Parp8** | 2810430O08Rik\|ARTD16\|D13Ertd275e | poly (ADP-ribose) polymerase family, member 8 | protein-coding |
| chr4_139813490_139814054 | chr4 | 139813491 | 139814054 | + | -0.645047 | intron (NM_011039, intron 4 of 8) | intron (NM_011039, intron 4 of 8) | 19196 | NM_011039 | 18509 | Mm.218760 | NM_011039 | ENSMUSG00000028736 | **Pax7** | Pax-7 | paired box 7 | protein-coding |
| chr18_37813669_37814340 | chr18 | 37813670 | 37814340 | + | -0.923168 | intron (NM_033581, intron 1 of 3) | intron (NM_033581, intron 1 of 3) | -1074 | NM_033582 | 93707 | Mm.247203 | NM_033582 | ENSMUSG00000023036 | **Pcdhgc4** | - | protocadherin gamma subfamily C, 4 | protein-coding |
| chr1_7095821_7096534 | chr1 | 7095822 | 7096534 | + | -0.749161 | intron (NM_183028, intron 1 of 5) | intron (NM_183028, intron 1 of 5) | 7258 | NM_183028 | 319263 | Mm.21539 | NM_183028 | ENSMUSG00000051285 | **Pcmtd1** | 8430411F12Rik\|A030012M09Rik | protein-L-isoaspartate (D-aspartate) O-methyltransferase domain containing 1 | protein-coding |
| chr13_75034853_75035757 | chr13 | 75034854 | 75035757 | + | -0.852497 | Intergenic | Intergenic | -54682 | NM_013628 | 18548 | Mm.1333 | NM_013628 | ENSMUSG00000021587 | **Pcsk1** | Nec-1\|Nec1\|PC1\|PC3\|Phpp-1\|SPC3 | proprotein convertase subtilisin/kexin type 1 | protein-coding |
| chr13_75034871_75035337 | chr13 | 75034872 | 75035337 | + | -1.00906 | Intergenic | RLTR8\|LTR\|ERVK | -54883 | NM_013628 | 18548 | Mm.1333 | NM_013628 | ENSMUSG00000021587 | **Pcsk1** | Nec-1\|Nec1\|PC1\|PC3\|Phpp-1\|SPC3 | proprotein convertase subtilisin/kexin type 1 | protein-coding |
| chr19_4590100_4590381 | chr19 | 4590101 | 4590381 | + | -0.724444 | intron (NM_001162946, intron 2 of 20) | intron (NM_001162946, intron 2 of 20) | -4078 | NM_008797 | 18563 | Mm.1845 | NM_008797 | ENSMUSG00000024892 | **Pcx** | Pc\|Pcb | pyruvate carboxylase | protein-coding |
| chr16_32434052_32434516 | chr16 | 32434053 | 32434516 | + | -0.796627 | intron (NM_001163160, intron 1 of 8) | intron (NM_001163160, intron 1 of 8) | 3054 | NM_001163160 | 13026 | Mm.98775 | NM_009981 | ENSMUSG00000005615 | **Pcyt1a** | CTalpha\|Cctalpha\|Ctpct\|Cttalpha | phosphate cytidylyltransferase 1, choline, alpha isoform | protein-coding |
| chr1_94091611_94092315 | chr1 | 94091612 | 94092315 | + | -1.2381 | Intergenic | Intergenic | -39410 | NM_008798 | 18566 | Mm.5024 | NM_008798 | ENSMUSG00000026285 | **Pdcd1** | Ly101\|PD-1\|Pdc1 | programmed cell death 1 | protein-coding |
| chr19_29418579_29419091 | chr19 | 29418580 | 29419091 | + | -1.03999 | intron (NM_021396, intron 1 of 5) | intron (NM_021396, intron 1 of 5) | 7916 | NM_021396 | 58205 | Mm.116737 | NM_021396 | | **Pdcd1lg2** | B7-DC\|Btdc\|F730015O22Rik\|PD-L2 | programmed cell death 1 ligand 2 | protein-coding |
| chr19_53893537_53894113 | chr19 | 53893538 | 53894113 | + | -0.771096 | intron (NM_011050, intron 1 of 11) | intron (NM_011050, intron 1 of 11) | 1594 | NM_011050 | 18569 | Mm.1605 | NM_011050 | ENSMUSG00000024975 | **Pdcd4** | D19Ucla1\|Ma3\|Tis | programmed cell death 4 | protein-coding |
| chr9_113699778_113700032 | chr9 | 113699779 | 113700032 | + | -0.930582 | intron (NM_001164677, intron 2 of 17) | intron (NM_001164677, intron 2 of 17) | 8354 | NM_001164678 | 18571 | Mm.29816 | NM_011052 | ENSMUSG00000032504 | **Pdcd6ip** | AI480591\|AW544830\|Aip1\|Alix\|C76364\|Eig2\|mKIAA1375 | programmed cell death 6 interacting protein | protein-coding |
| chr9_113706193_113706500 | chr9 | 113706194 | 113706500 | + | -1.04943 | intron (NM_001164677, intron 1 of 17) | RSINE1\|SINE\|B4 | 1912 | NM_001164678 | 18571 | Mm.29816 | NM_011052 | ENSMUSG00000032504 | **Pdcd6ip** | AI480591\|AW544830\|Aip1\|Alix\|C76364\|Eig2\|mKIAA1375 | programmed cell death 6 interacting protein | protein-coding |
| chr9_113706170_113706594 | chr9 | 113706171 | 113706594 | + | -1.06432 | intron (NM_001164677, intron 1 of 17) | intron (NM_001164677, intron 1 of 17) | 1877 | NM_001164678 | 18571 | Mm.29816 | NM_011052 | ENSMUSG00000032504 | **Pdcd6ip** | AI480591\|AW544830\|Aip1\|Alix\|C76364\|Eig2\|mKIAA1375 | programmed cell death 6 interacting protein | protein-coding |
| chr9_113706033_113706627 | chr9 | 113706034 | 113706627 | + | -1.0802 | intron (NM_001164677, intron 1 of 17) | RSINE1\|SINE\|B4 | 1929 | NM_001164678 | 18571 | Mm.29816 | NM_011052 | ENSMUSG00000032504 | **Pdcd6ip** | AI480591\|AW544830\|Aip1\|Alix\|C76364\|Eig2\|mKIAA1375 | programmed cell death 6 interacting protein | protein-coding |
| chr17_8562899_8563287 | chr17 | 8562900 | 8563287 | + | -0.868238 | intron (NM_001290707, intron 1 of 22) | intron (NM_001290707, intron 1 of 22) | 36292 | NM_001290707 | 23984 | Mm.87161 | NM_011866 | ENSMUSG00000023868 | **Pde10a** | - | phosphodiesterase 10A | protein-coding |
| chr7_101457725_101458217 | chr7 | 101457726 | 101458217 | + | -0.982832 | intron (NM_001243758, intron 2 of 30) | intron (NM_001243758, intron 2 of 30) | 6719 | NM_001243757 | 207728 | Mm.247564 | NM_001008548 | ENSMUSG00000030653 | **Pde2a** | CGS-PDE\|cGSPDE | phosphodiesterase 2A, cGMP-stimulated | protein-coding |
| chr9_21196141_21196393 | chr9 | 21196142 | 21196393 | + | -0.722214 | promoter-TSS (NM_019798) | promoter-TSS (NM_019798) | -438 | NM_019798 | 18577 | Mm.191749 | NM_019798 | ENSMUSG00000032177 | **Pde4a** | D9Ertd60e\|Dpde2 | phosphodiesterase 4A, cAMP specific | protein-coding |
| chr9_21196135_21196990 | chr9 | 21196136 | 21196990 | + | -0.757611 | promoter-TSS (NM_019798) | promoter-TSS (NM_019798) | -142 | NM_019798 | 18577 | Mm.191749 | NM_019798 | ENSMUSG00000032177 | **Pde4a** | D9Ertd60e\|Dpde2 | phosphodiesterase 4A, cAMP specific | protein-coding |
| chr4_102517000_102517253 | chr4 | 102517001 | 102517253 | + | -0.677975 | intron (NM_019840, intron 5 of 14) | RSINE1\|SINE\|B4 | -52933 | NM_001177980 | 18578 | Mm.20181 | NM_019840 | ENSMUSG00000028525 | **Pde4b** | Dpde4\|R74983\|dunce | phosphodiesterase 4B, cAMP specific | protein-coding |
| chr4_102506905_102507275 | chr4 | 102506906 | 102507275 | + | -0.780568 | intron (NM_019840, intron 5 of 14) | intron (NM_019840, intron 5 of 14) | -62970 | NM_001177980 | 18578 | Mm.20181 | NM_019840 | ENSMUSG00000028525 | **Pde4b** | Dpde4\|R74983\|dunce | phosphodiesterase 4B, cAMP specific | protein-coding |
| chr13_108596966_108597254 | chr13 | 108596967 | 108597254 | + | -0.658136 | Intergenic | Intergenic | -57067 | NM_011056 | 238871 | Mm.434429 | NM_011056 | | **Pde4d** | 9630011N22Rik\|Dpde3 | phosphodiesterase 4D, cAMP specific | protein-coding |
| chr13_108862106_108862483 | chr13 | 108862107 | 108862483 | + | -0.671433 | intron (NM_011056, intron 2 of 16) | MTEa\|LTR\|ERVL-MaLR | 208118 | NM_011056 | 238871 | Mm.434429 | NM_011056 | | **Pde4d** | 9630011N22Rik\|Dpde3 | phosphodiesterase 4D, cAMP specific | protein-coding |
| chr13_108667137_108667454 | chr13 | 108667138 | 108667454 | + | -0.68101 | intron (NM_011056, intron 1 of 16) | intron (NM_011056, intron 1 of 16) | 13119 | NM_011056 | 238871 | Mm.434429 | NM_011056 | | **Pde4d** | 9630011N22Rik\|Dpde3 | phosphodiesterase 4D, cAMP specific | protein-coding |
| chr13_108596867_108597544 | chr13 | 108596868 | 108597544 | + | -0.691156 | Intergenic | Intergenic | -56971 | NM_011056 | 238871 | Mm.434429 | NM_011056 | | **Pde4d** | 9630011N22Rik\|Dpde3 | phosphodiesterase 4D, cAMP specific | protein-coding |
| chr13_108596884_108597313 | chr13 | 108596885 | 108597313 | + | -0.696544 | Intergenic | Intergenic | -57078 | NM_011056 | 238871 | Mm.434429 | NM_011056 | | **Pde4d** | 9630011N22Rik\|Dpde3 | phosphodiesterase 4D, cAMP specific | protein-coding |
| chr3_19318966_19319454 | chr3 | 19318967 | 19319454 | + | -0.640683 | Intergenic | L1MB4\|LINE\|L1 | -7888 | NM_001122759 | 18583 | Mm.355614 | NM_008802 | ENSMUSG00000069094 | **Pde7a** | AU015378\|AW047537 | phosphodiesterase 7A | protein-coding |
| chr3_19332461_19333116 | chr3 | 19332462 | 19333116 | + | -0.679879 | Intergenic | RLTR34D_MM\|LTR\|ERVK | -21467 | NM_001122759 | 18583 | Mm.355614 | NM_008802 | ENSMUSG00000069094 | **Pde7a** | AU015378\|AW047537 | phosphodiesterase 7A | protein-coding |
| chr3_19268542_19268835 | chr3 | 19268543 | 19268835 | + | -0.705794 | intron (NM_001122759, intron 1 of 12) | RSINE1\|SINE\|B4 | -3681 | NM_008802 | 18583 | Mm.355614 | NM_008802 | ENSMUSG00000069094 | **Pde7a** | AU015378\|AW047537 | phosphodiesterase 7A | protein-coding |
| chr3_19239853_19240345 | chr3 | 19239854 | 19240345 | + | -0.966993 | intron (NM_008802, intron 5 of 12) | intron (NM_008802, intron 5 of 12) | 24909 | NM_008802 | 18583 | Mm.355614 | NM_008802 | ENSMUSG00000069094 | **Pde7a** | AU015378\|AW047537 | phosphodiesterase 7A | protein-coding |
| chr3_19258273_19258941 | chr3 | 19258274 | 19258941 | + | -1.22798 | intron (NM_008802, intron 2 of 12) | intron (NM_008802, intron 2 of 12) | 6401 | NM_008802 | 18583 | Mm.355614 | NM_008802 | ENSMUSG00000069094 | **Pde7a** | AU015378\|AW047537 | phosphodiesterase 7A | protein-coding |
| chr3_19258498_19258942 | chr3 | 19258499 | 19258942 | + | -1.23085 | intron (NM_008802, intron 2 of 12) | intron (NM_008802, intron 2 of 12) | 6288 | NM_008802 | 18583 | Mm.355614 | NM_008802 | ENSMUSG00000069094 | **Pde7a** | AU015378\|AW047537 | phosphodiesterase 7A | protein-coding |
| chr17_31439979_31440373 | chr17 | 31439980 | 31440373 | + | -0.83876 | intron (NM_001163748, intron 4 of 18) | MER54A\|LTR\|ERVL | 53942 | NM_001163748 | 18585 | Mm.10812 | NM_008804 | ENSMUSG00000041119 | **Pde9a** | PDE9A1 | phosphodiesterase 9A | protein-coding |

| | | | | | | | | | | | | | | | | | |
|---|---|---|---|---|---|---|---|---|---|---|---|---|---|---|---|---|---|
| chr15_80020539_80020921 | chr15 | 80020540 | 80020921 | + | -0.942426 | Intergenic | Intergenic | -5922 | NM_011057 | 18591 | Mm.144089 | NM_011057 | ENSMUSG00000000489 | **Pdgfb** | PDGF-B\|Sis | platelet derived growth factor, B polypeptide | protein-coding |
| chr4_134273119_134273768 | chr4 | 134273120 | 134273768 | + | -0.68064 | Intergenic | Intergenic | 13809 | NM_001163794 | 230809 | Mm.22778 | NM_146156 | ENSMUSG00000050890 | **Pdik1l** | BC027088\|stk35l2 | PDLIM1 interacting kinase 1 like | protein-coding |
| chr2_71863571_71863896 | chr2 | 71863572 | 71863896 | + | -0.774008 | Intergenic | Intergenic | -9490 | NM_172665 | 228026 | Mm.34411 | NM_172665 | ENSMUSG00000006494 | **Pdk1** | B830012B01\|D530020C15Rik | pyruvate dehydrogenase kinase, isoenzyme 1 | protein-coding |
| chr2_71866316_71867090 | chr2 | 71866317 | 71867090 | + | -0.8567 | Intergenic | PB1D9\|SINE\|Alu | -6521 | NM_172665 | 228026 | Mm.34411 | NM_172665 | ENSMUSG00000006494 | **Pdk1** | B830012B01\|D530020C15Rik | pyruvate dehydrogenase kinase, isoenzyme 1 | protein-coding |
| chr19_40238139_40238507 | chr19 | 40238140 | 40238507 | + | -0.663608 | intron (NM_016861, intron 4 of 6) | L1MB2\|LINE\|L1 | 33293 | NM_016861 | 54132 | Mm.5567 | NM_016861 | | **Pdlim1** | Clim1\|mClim1 | PDZ and LIM domain 1 (elfin) | protein-coding |
| chr19_40267156_40267796 | chr19 | 40267157 | 40267796 | + | -0.738473 | intron (NM_016861, intron 1 of 6) | intron (NM_016861, intron 1 of 6) | 4140 | NM_016861 | 54132 | Mm.5567 | NM_016861 | | **Pdlim1** | Clim1\|mClim1 | PDZ and LIM domain 1 (elfin) | protein-coding |
| chr19_40267381_40267771 | chr19 | 40267382 | 40267771 | + | -0.844925 | intron (NM_016861, intron 1 of 6) | intron (NM_016861, intron 1 of 6) | 4040 | NM_016861 | 54132 | Mm.5567 | NM_016861 | | **Pdlim1** | Clim1\|mClim1 | PDZ and LIM domain 1 (elfin) | protein-coding |
| chr11_54062614_54063864 | chr11 | 54062615 | 54063864 | + | -0.61283 | intron (NM_019417, intron 2 of 6) | intron (NM_019417, intron 2 of 6) | 5778 | NM_019417 | 30794 | Mm.21830 | NM_019417 | ENSMUSG00000020388 | **Pdlim4** | Ril | PDZ and LIM domain 4 | protein-coding |
| chr11_54082755_54083130 | chr11 | 54082756 | 54083130 | + | -0.67974 | Intergenic | Intergenic | -13926 | NM_019417 | 30794 | Mm.21830 | NM_019417 | ENSMUSG00000020388 | **Pdlim4** | Ril | PDZ and LIM domain 4 | protein-coding |
| chr11_54063115_54063783 | chr11 | 54063116 | 54063783 | + | -0.827318 | intron (NM_019417, intron 2 of 6) | intron (NM_019417, intron 2 of 6) | 5568 | NM_019417 | 30794 | Mm.21830 | NM_019417 | ENSMUSG00000020388 | **Pdlim4** | Ril | PDZ and LIM domain 4 | protein-coding |
| chr3_142281060_142281447 | chr3 | 142281061 | 142281447 | + | -0.724083 | intron (NM_019808, intron 7 of 12) | ORR1F\|LTR\|ERVL-MaLR | 110596 | NM_022554 | 56376 | Mm.117709 | NM_019808 | ENSMUSG00000028273 | **Pdlim5** | 1110001A05Rik\|AI987914\|C87059\|Enh\|Enh1\|Enh2\|Enh3\|LIM | PDZ and LIM domain 5 | protein-coding |
| chr3_142324643_142324976 | chr3 | 142324644 | 142324976 | + | -0.813392 | intron (NM_019808, intron 3 of 12) | intron (NM_019808, intron 3 of 12) | 67040 | NM_022554 | 56376 | Mm.117709 | NM_019808 | ENSMUSG00000028273 | **Pdlim5** | 1110001A05Rik\|AI987914\|C87059\|Enh\|Enh1\|Enh2\|Enh3\|LIM | PDZ and LIM domain 5 | protein-coding |
| chr10_43329937_43330345 | chr10 | 43329938 | 43330345 | + | -0.774084 | intron (NM_027772, intron 2 of 7) | intron (NM_027772, intron 2 of 7) | 108655 | NM_001168289 | 71365 | Mm.490430 | NM_027772 | ENSMUSG00000038240 | **Pdss2** | 5430420P03Rik\|Gm60\|Plmp\|kd\|mDLP1 | prenyl (solanesyl) diphosphate synthase, subunit 2 | protein-coding |
| chr19_59388800_59389674 | chr19 | 59388801 | 59389674 | + | -0.701303 | Intergenic | Intergenic | -43457 | NM_001033222 | 107368 | Mm.268797 | NM_001033222 | ENSMUSG00000074746 | **Pdzd8** | A630041P07Rik\|AV329468\|AW228944\|Pdzk8 | PDZ domain containing 8 | protein-coding |
| chr4_115075301_115076167 | chr4 | 115075302 | 115076167 | + | -0.90746 | Intergenic | Intergenic | -12974 | NM_001164558 | 67182 | Mm.30181 | NM_026018 | ENSMUSG00000028716 | **Pdzk1ip1** | 0610007F13Rik\|2700030M23Rik\|AU046042\|Map17 | PDZK1 interacting protein 1 | protein-coding |
| chr6_101359643_101360222 | chr6 | 101359644 | 101360222 | + | -0.660284 | intron (NM_018884, intron 2 of 9) | intron (NM_018884, intron 2 of 9) | 17964 | NM_018884 | 55983 | Mm.321654 | NM_018884 | ENSMUSG00000035357 | **Pdzrn3** | 1110020C07Rik\|AI429718\|AL023082\|LNX3\|SEMACAP3\|Semcap3 | PDZ domain containing RING finger 3 | protein-coding |
| chr6_101359660_101360066 | chr6 | 101359661 | 101360066 | + | -0.66147 | intron (NM_018884, intron 2 of 9) | intron (NM_018884, intron 2 of 9) | 18034 | NM_018884 | 55983 | Mm.321654 | NM_018884 | ENSMUSG00000035357 | **Pdzrn3** | 1110020C07Rik\|AI429718\|AL023082\|LNX3\|SEMACAP3\|Semcap3 | PDZ domain containing RING finger 3 | protein-coding |
| chr1_172197090_172197473 | chr1 | 172197091 | 172197473 | + | -1.10179 | TTS (NM_153555) | TTS (NM_153555) | 9499 | NM_011063 | 18611 | Mm.544 | NM_011063 | ENSMUSG00000013698 | **Pea15a** | Mat1\|PEA-15\|Pea15\|Pkcs15 | phosphoprotein enriched in astrocytes 15A | protein-coding |
| chr9_56292100_56292441 | chr9 | 56292101 | 56292441 | + | -0.686102 | intron (NM_172924, intron 3 of 6) | intron (NM_172924, intron 3 of 6) | 125779 | NM_172924 | 244895 | Mm.490331 | NM_172924 | ENSMUSG00000074305 | **Peak1** | 1110049L02Rik\|9530046P14\|AA690169\|BE132841\|C230081A13Rik\|SGK269\|mKIAA2002 | pseudopodium-enriched atypical kinase 1 | protein-coding |
| chr14_69830629_69831223 | chr14 | 69830630 | 69831223 | + | -0.652382 | Intergenic | Intergenic | -9481 | NM_028560 | 73523 | Mm.23509 | NM_028526 | ENSMUSG00000022085 | **Pebp4** | PEBP-4 | phosphatidylethanolamine binding protein 4 | protein-coding |
| chr11_106665355_106665849 | chr11 | 106665356 | 106665849 | + | -0.800527 | intron (NM_001032378, intron 13 of 13) | intron (NM_001032378, intron 13 of 13) | 49679 | NM_001032378 | 18613 | Mm.343951 | NM_008816 | ENSMUSG00000020717 | **Pecam1** | C85791\|Cd31\|PECAM-1\|Pecam | platelet/endothelial cell adhesion molecule 1 | protein-coding |
| chr15_72928691_72929827 | chr15 | 72928692 | 72929827 | + | -0.858428 | intron (NM_001164641, intron 15 of 17) | intron (NM_001164641, intron 15 of 17) | -118935 | NR_002864 | 353342 | Mm.440245 | NR_002864 | | **Peg13** | - | paternally expressed 13 | ncRNA |
| chr11_21151685_21152040 | chr11 | 21151686 | 21152040 | + | -0.717661 | Intergenic | Intergenic | 60539 | NM_023324 | 67245 | Mm.28957 | NM_023324 | ENSMUSG00000020134 | **Peli1** | 2810468L03Rik\|A930031K15Rik\|AA409794\|AI586297\|D11Ertd676e | pellino 1 | protein-coding |
| chr11_21141311_21141606 | chr11 | 21141312 | 21141606 | + | -0.743082 | intron (NM_023324, intron 3 of 6) | intron (NM_023324, intron 3 of 6) | 50135 | NM_023324 | 67245 | Mm.28957 | NM_023324 | ENSMUSG00000020134 | **Peli1** | 2810468L03Rik\|A930031K15Rik\|AA409794\|AI586297\|D11Ertd676e | pellino 1 | protein-coding |
| chr11_21156589_21156877 | chr11 | 21156590 | 21156877 | + | -0.754422 | Intergenic | Intergenic | 65409 | NM_023324 | 67245 | Mm.28957 | NM_023324 | ENSMUSG00000020134 | **Peli1** | 2810468L03Rik\|A930031K15Rik\|AA409794\|AI586297\|D11Ertd676e | pellino 1 | protein-coding |
| chr11_21150397_21150606 | chr11 | 21150398 | 21150606 | + | -0.766804 | TTS (NM_023324) | TTS (NM_023324) | 59178 | NM_023324 | 67245 | Mm.28957 | NM_023324 | ENSMUSG00000020134 | **Peli1** | 2810468L03Rik\|A930031K15Rik\|AA409794\|AI586297\|D11Ertd676e | pellino 1 | protein-coding |
| chr11_21103209_21103551 | chr11 | 21103210 | 21103551 | + | -1.23236 | intron (NM_023324, intron 1 of 6) | PB1D10\|SINE\|Alu | 12056 | NM_023324 | 67245 | Mm.28957 | NM_023324 | ENSMUSG00000020134 | **Peli1** | 2810468L03Rik\|A930031K15Rik\|AA409794\|AI586297\|D11Ertd676e | pellino 1 | protein-coding |
| chr11_60032540_60032907 | chr11 | 60032541 | 60032907 | + | -0.832813 | intron (NM_001290012, intron 1 of 6) | MER20\|DNA\|hAT-Charlie | 4281 | NM_008819 | 18618 | Mm.2731 | NM_008819 | ENSMUSG00000000301 | **Pemt** | PEAMT\|PEMT2\|Pempt\|Pempt2 | phosphatidylethanolamine N-methyltransferase | protein-coding |
| chr11_60032535_60033210 | chr11 | 60032536 | 60033210 | + | -0.841142 | intron (NM_001290012, intron 1 of 6) | intron (NM_001290012, intron 1 of 6) | 4132 | NM_008819 | 18618 | Mm.2731 | NM_008819 | ENSMUSG00000000301 | **Pemt** | PEAMT\|PEMT2\|Pempt\|Pempt2 | phosphatidylethanolamine N-methyltransferase | protein-coding |
| chr11_69099887_69100235 | chr11 | 69099888 | 69100235 | + | -0.643872 | promoter-TSS (NM_001159367) | promoter-TSS (NM_001159367) | -97 | NM_001159367 | 18626 | Mm.7373 | NM_011065 | ENSMUSG00000020893 | **Per1** | Per\|m-rigui\|mPer1 | period circadian clock 1 | protein-coding |
| chr4_155050080_155050513 | chr4 | 155050081 | 155050513 | + | -1.1966 | Intergenic | Intergenic | -16733 | NM_001042407 | 668173 | Mm.133114 | NM_001042407 | ENSMUSG00000029047 | **Pex10** | AV128229\|Gm142 | peroxisomal biogenesis factor 10 | protein-coding |
| chr10_19886299_19886567 | chr10 | 19886300 | 19886567 | + | -0.713365 | intron (NM_008822, intron 7 of 9) | intron (NM_008822, intron 7 of 9) | 21241 | NM_008822 | 18634 | Mm.338363 | NM_008822 | ENSMUSG00000020003 | **Pex7** | MmPEX7 | peroxisomal biogenesis factor 7 | protein-coding |
| chr2_11542157_11542502 | chr2 | 11542158 | 11542502 | + | -1.13117 | intron (NM_001177758, intron 1 of 14) | ORR1F\|LTR\|ERVL-MaLR | 11599 | NM_001177758 | 170768 | Mm.19669 | NM_133232 | ENSMUSG00000026773 | **Pfkfb3** | E330010H22Rik\|iPFK-2\|uPFK-2 | 6-phosphofructo-2-kinase/fructose-2,6-biphosphatase 3 | protein-coding |
| chr17_32423894_32424689 | chr17 | 32423895 | 32424689 | + | -0.935655 | promoter-TSS (NM_001271476) | promoter-TSS (NM_001271476) | -125 | NM_001271478 | 57757 | Mm.86752 | NM_021319 | | **Pglyrp2** | C730002N09Rik\|PGRP-L\|Pglyrpl\|tagL | peptidoglycan recognition protein 2 | protein-coding |
| chr7_68205812_68206325 | chr7 | 68205813 | 68206325 | + | -0.686105 | intron (NM_010513, intron 15 of 20) | intron (NM_010513, intron 15 of 20) | 58164 | NM_030101 | 78444 | Mm.425468 | NM_030101 | ENSMUSG00000030553 | **Pgpep1l** | C330024D12Rik | pyroglutamyl-peptidase I-like | protein-coding |
| chr7_68139337_68139891 | chr7 | 68139338 | 68139891 | + | -0.720264 | intron (NM_010513, intron 2 of 20) | intron (NM_010513, intron 2 of 20) | 124619 | NM_030101 | 78444 | Mm.425468 | NM_030101 | ENSMUSG00000030553 | **Pgpep1l** | C330024D12Rik | pyroglutamyl-peptidase I-like | protein-coding |
| chr7_68133719_68134051 | chr7 | 68133720 | 68134051 | + | -0.743147 | intron (NM_010513, intron 2 of 20) | intron (NM_010513, intron 2 of 20) | 130348 | NM_030101 | 78444 | Mm.425468 | NM_030101 | ENSMUSG00000030553 | **Pgpep1l** | C330024D12Rik | pyroglutamyl-peptidase I-like | protein-coding |
| chr7_68173824_68174150 | chr7 | 68173825 | 68174150 | + | -0.841294 | intron (NM_010513, intron 5 of 20) | ORR1D2\|LTR\|ERVL-MaLR | 90246 | NM_030101 | 78444 | Mm.425468 | NM_030101 | ENSMUSG00000030553 | **Pgpep1l** | C330024D12Rik | pyroglutamyl-peptidase I-like | protein-coding |
| chr7_68144234_68144718 | chr7 | 68144235 | 68144718 | + | -1.04133 | intron (NM_010513, intron 2 of 20) | intron (NM_010513, intron 2 of 20) | 119757 | NM_030101 | 78444 | Mm.425468 | NM_030101 | ENSMUSG00000030553 | **Pgpep1l** | C330024D12Rik | pyroglutamyl-peptidase I-like | protein-coding |
| chr7_68250357_68250887 | chr7 | 68250358 | 68250887 | + | -1.08329 | intron (NM_030101, intron 1 of 3) | RMER19B2\|LTR\|ERVK | 13611 | NM_030101 | 78444 | Mm.425468 | NM_030101 | ENSMUSG00000030553 | **Pgpep1l** | C330024D12Rik | pyroglutamyl-peptidase I-like | protein-coding |
| chrX_97156202_97156485 | chrX | 97156203 | 97156485 | + | -0.641168 | Intergenic | ORR1D2\|LTR\|ERVL-MaLR | 83748 | NM_001033361 | 245526 | Mm.336164 | NM_001033361 | ENSMUSG00000031212 | **Pgr15l** | - | G protein-coupled receptor 15-like | protein-coding |
| chr6_122340495_122340726 | chr6 | 122340496 | 122340726 | + | -0.608541 | promoter-TSS (NM_001042623) | promoter-TSS (NM_001042623) | -384 | NM_001271579 | 13619 | Mm.6822 | NM_007905 | ENSMUSG00000040669 | **Phc1** | AW557034\|Edr\|Edr1\|Mph1\|Rae-28\|rae28 | polyhomeotic-like 1 (Drosophila) | protein-coding |
| chr6_12000415_12000688 | chr6 | 12000416 | 12000688 | + | -0.694445 | intron (NM_029404, intron 14 of 16) | intron (NM_029404, intron 14 of 16) | 74671 | NM_001168382 | 75725 | Mm.212411 | NM_029404 | ENSMUSG00000029629 | **Phf14** | 1110001C23Rik\|4932409F11Rik\|5730446A07Rik\|AA623952\|AV297001\|mKIAA0783 | PHD finger protein 14 | protein-coding |
| chr2_92253762_92254116 | chr2 | 92253763 | 92254116 | + | -0.64962 | intron (NM_001109691, intron 7 of 18) | intron (NM_001109691, intron 7 of 18) | 32377 | NM_001109690 | 192285 | Mm.330408 | NM_138755 | ENSMUSG00000058318 | **Phf21a** | 80kDa\|Bhc\|Bhc80\|D030065N23Rik\|PFTF1 | PHD finger protein 21A | protein-coding |
| chr1_106318384_106318742 | chr1 | 106318385 | 106318742 | + | -1.01904 | intron (NM_133821, intron 3 of 16) | intron (NM_133821, intron 3 of 16) | 146694 | NM_133821 | 98432 | Mm.24115 | NM_133821 | ENSMUSG00000044340 | **Phlpp1** | AI836256\|Phlpp\|Plekhe1\|SCOP\|mKIAA0606 | PH domain and leucine rich repeat protein phosphatase 1 | protein-coding |
| chr1_106318359_106318594 | chr1 | 106318360 | 106318594 | + | -1.03075 | intron (NM_133821, intron 3 of 16) | intron (NM_133821, intron 3 of 16) | 146608 | NM_133821 | 98432 | Mm.24115 | NM_133821 | ENSMUSG00000044340 | **Phlpp1** | AI836256\|Phlpp\|Plekhe1\|SCOP\|mKIAA0606 | PH domain and leucine rich repeat protein phosphatase 1 | protein-coding |
| chr11_95805011_95805335 | chr11 | 95805012 | 95805335 | + | -0.667861 | Intergenic | Intergenic | -19327 | NM_153104 | 237928 | Mm.133075 | NM_153104 | ENSMUSG00000050860 | **Phospho1** | D11Moh36\|Phospo1 | phosphatase, orphan 1 | protein-coding |
| chr9_13443460_13443901 | chr9 | 13443461 | 13443901 | + | -0.670088 | Intergenic | Intergenic | 12320 | NR_028271 | 18689 | Mm.41972 | NM_008835 | | **Phxr4** | Sp5 | per-hexamer repeat gene 4 | ncRNA |
| chr9_13466397_13466760 | chr9 | 13466398 | 13466760 | + | -0.67636 | Intergenic | Intergenic | 35218 | NR_028271 | 18689 | Mm.41972 | NM_008835 | | **Phxr4** | Sp5 | per-hexamer repeat gene 4 | ncRNA |
| chr9_13443466_13444111 | chr9 | 13443467 | 13444111 | + | -0.687616 | Intergenic | Intergenic | 12428 | NR_028271 | 18689 | Mm.41972 | NM_008835 | | **Phxr4** | Sp5 | per-hexamer repeat gene 4 | ncRNA |
| chr9_13470061_13470439 | chr9 | 13470062 | 13470439 | + | -1.0251 | Intergenic | Intergenic | 38889 | NR_028271 | 18689 | Mm.41972 | NM_008835 | | **Phxr4** | Sp5 | per-hexamer repeat gene 4 | ncRNA |

| | | | | | | | | | | | | | | | | | |
|---|---|---|---|---|---|---|---|---|---|---|---|---|---|---|---|---|---|
| chr9_13422653_13422886 | chr9 | 13422654 | 13422886 | + | -1.4005 | Intergenic | Intergenic | -8591 | NR_028271 | 18689 | Mm.41972 | NM_008835 | | **Phxr4** | Sp5 | per-hexamer repeat gene 4 | ncRNA |
| chr9_13422595_13422988 | chr9 | 13422596 | 13422988 | + | -1.43067 | Intergenic | Intergenic | -8569 | NR_028271 | 18689 | Mm.41972 | NM_008835 | | **Phxr4** | Sp5 | per-hexamer repeat gene 4 | ncRNA |
| chr19_42089121_42089490 | chr19 | 42089122 | 42089490 | + | -0.652388 | Intergenic | Intergenic | -1129 | NM_145501 | 84095 | Mm.117037 | NM_145501 | ENSMUSG00000025178 | **Pi4k2a** | Pi4k2 | phosphatidylinositol 4-kinase type 2 alpha | protein-coding |
| chr10_81169705_81169953 | chr10 | 81169706 | 81169953 | + | -0.782793 | Intergenic | ID_B1\|SINE\|B4 | -2109 | NM_021501 | 59004 | Mm.34428 | NM_021501 | | **Pias4** | PIASY\|Pias-gamma\|Piasg | protein inhibitor of activated STAT 4 | protein-coding |
| chr10_81169635_81169972 | chr10 | 81169636 | 81169972 | + | -0.814661 | Intergenic | ID_B1\|SINE\|B4 | -2084 | NM_021501 | 59004 | Mm.34428 | NM_021501 | | **Pias4** | PIASY\|Pias-gamma\|Piasg | protein inhibitor of activated STAT 4 | protein-coding |
| chr15_79208556_79209572 | chr15 | 79208557 | 79209572 | + | -0.771887 | intron (NR_029470, intron 2 of 4) | intron (NR_029470, intron 2 of 4) | -20109 | NM_008837 | 18693 | Mm.259464 | NM_008837 | ENSMUSG00000068206 | **Pick1** | Prkcabp | protein interacting with C kinase 1 | protein-coding |
| chr15_79208951_79209535 | chr15 | 79208952 | 79209535 | + | -0.782037 | intron (NR_029470, intron 2 of 4) | intron (NR_029470, intron 2 of 4) | -19930 | NM_008837 | 18693 | Mm.259464 | NM_008837 | ENSMUSG00000068206 | **Pick1** | Prkcabp | protein interacting with C kinase 1 | protein-coding |
| chr15_79209087_79209482 | chr15 | 79209088 | 79209482 | + | -0.782514 | intron (NR_029470, intron 2 of 4) | intron (NR_029470, intron 2 of 4) | -19888 | NM_008837 | 18693 | Mm.259464 | NM_008837 | ENSMUSG00000068206 | **Pick1** | Prkcabp | protein interacting with C kinase 1 | protein-coding |
| chr1_84318597_84319022 | chr1 | 84318598 | 84319022 | + | -1.55316 | Intergenic | Intergenic | -34165 | NM_001003948 | 98496 | Mm.187470 | NM_001003948 | ENSMUSG00000045658 | **Pid1** | 5033414K04Rik\|NYGGF4 | phosphotyrosine interaction domain containing 1 | protein-coding |
| chr3_106032833_106033121 | chr3 | 106032834 | 106033121 | + | -0.921447 | Intergenic | Intergenic | -18331 | NM_001200028 | 100503311 | Mm.386825 | NM_029604 | ENSMUSG00000010136 | **Pifo** | 1700027A23Rik | primary cilia formation | protein-coding |
| chr3_106032651_106033139 | chr3 | 106032652 | 106033139 | + | -0.944204 | Intergenic | Intergenic | -18249 | NM_001200028 | 100503311 | Mm.386825 | NM_029604 | ENSMUSG00000010136 | **Pifo** | 1700027A23Rik | primary cilia formation | protein-coding |
| chr4_133666256_133666597 | chr4 | 133666257 | 133666597 | + | -0.689624 | intron (NM_178698, intron 2 of 3) | L2\|LINE\|L2 | 6220 | NM_001145955 | 230801 | Mm.217004 | NM_178698 | | **Pigv** | B330013B03\|D430024F16Rik | phosphatidylinositol glycan anchor biosynthesis, class V | protein-coding |
| chr19_41328708_41329284 | chr19 | 41328709 | 41329284 | + | -0.72157 | intron (NM_031376, intron 4 of 16) | intron (NM_031376, intron 4 of 16) | 56074 | NM_031376 | 83490 | Mm.222266 | NM_031376 | ENSMUSG00000025017 | **Pik3ap1** | 1810044J04Rik\|BCAP | phosphoinositide-3-kinase adaptor protein 1 | protein-coding |
| chr7_116392341_116392720 | chr7 | 116392342 | 116392720 | + | -0.847298 | intron (NM_011083, intron 4 of 32) | B1_Mur3\|SINE\|Alu | 50927 | NM_011083 | 18704 | Mm.3810 | NM_011083 | ENSMUSG00000030660 | **Pik3c2a** | Cpk-m\|PI3KC2 | phosphatidylinositol 3-kinase, C2 domain containing, alpha polypeptide | protein-coding |
| chr7_116392371_116392961 | chr7 | 116392372 | 116392961 | + | -0.949743 | intron (NM_011083, intron 4 of 32) | B2_Mm2\|SINE\|B2 | 50792 | NM_011083 | 18704 | Mm.3810 | NM_011083 | ENSMUSG00000030660 | **Pik3c2a** | Cpk-m\|PI3KC2 | phosphatidylinositol 3-kinase, C2 domain containing, alpha polypeptide | protein-coding |
| chr1_133073366_133073984 | chr1 | 133073367 | 133073984 | + | -1.04349 | intron (NM_001099276, intron 5 of 32) | intron (NM_001099276, intron 5 of 32) | 27663 | NM_001099276 | 240752 | Mm.387308 | NM_001099276 | ENSMUSG00000026447 | **Pik3c2b** | C330011J12Rik\|Gm199\|PI3K-C2beta\|PI3KC2beta | phosphoinositide-3-kinase, class 2, beta polypeptide | protein-coding |
| chr12_32208176_32208976 | chr12 | 32208177 | 32208976 | + | -0.638219 | promoter-TSS (NM_001146200) | promoter-TSS (NM_001146200) | 73 | NM_001146200 | 30955 | Mm.101369 | NM_020272 | ENSMUSG00000020573 | **Pik3cg** | 5830428L06Rik\|PI3Kgamma\|p110 gamma | phosphoinositide-3-kinase, catalytic, gamma polypeptide | protein-coding |
| chr12_32206772_32207151 | chr12 | 32206773 | 32207151 | + | -0.812457 | intron (NM_001146200, intron 1 of 10) | intron (NM_001146200, intron 1 of 10) | 1376 | NM_001146201 | 30955 | Mm.101369 | NM_020272 | ENSMUSG00000020573 | **Pik3cg** | 5830428L06Rik\|PI3Kgamma\|p110 gamma | phosphoinositide-3-kinase, catalytic, gamma polypeptide | protein-coding |
| chr12_32212765_32213072 | chr12 | 32212766 | 32213072 | + | -0.836007 | Intergenic | Intergenic | -4270 | NM_001146200 | 30955 | Mm.101369 | NM_020272 | ENSMUSG00000020573 | **Pik3cg** | 5830428L06Rik\|PI3Kgamma\|p110 gamma | phosphoinositide-3-kinase, catalytic, gamma polypeptide | protein-coding |
| chr11_3328340_3328720 | chr11 | 3328341 | 3328720 | + | -0.984693 | intron (NR_037301, intron 1 of 1) | intron (NR_037301, intron 1 of 1) | -2201 | NM_178149 | 216505 | Mm.390323 | NM_178149 | ENSMUSG00000034614 | **Pik3ip1** | 1500004A08Rik\|5830455E04Rik\|Crkd\|Hgfl | phosphoinositide-3-kinase interacting protein 1 | protein-coding |
| chr13_101702105_101702747 | chr13 | 101702106 | 101702747 | + | -0.77491 | intron (NM_001077495, intron 5 of 15) | intron (NM_001077495, intron 5 of 15) | -9796 | NM_001024955 | 18708 | Mm.259333 | NM_001024955 | ENSMUSG00000041417 | **Pik3r1** | PI3K\|p50alpha\|p55alpha\|p85alpha | phosphatidylinositol 3-kinase, regulatory subunit, polypeptide 1 (p85 alpha) | protein-coding |
| chr13_101701859_101702696 | chr13 | 101701860 | 101702696 | + | -0.791781 | intron (NM_001077495, intron 5 of 15) | intron (NM_001077495, intron 5 of 15) | -9648 | NM_001024955 | 18708 | Mm.259333 | NM_001024955 | ENSMUSG00000041417 | **Pik3r1** | PI3K\|p50alpha\|p55alpha\|p85alpha | phosphatidylinositol 3-kinase, regulatory subunit, polypeptide 1 (p85 alpha) | protein-coding |
| chr13_101711286_101711799 | chr13 | 101711287 | 101711799 | + | -0.792045 | intron (NM_001077495, intron 2 of 15) | intron (NM_001077495, intron 2 of 15) | -18913 | NM_001024955 | 18708 | Mm.259333 | NM_001024955 | ENSMUSG00000041417 | **Pik3r1** | PI3K\|p50alpha\|p55alpha\|p85alpha | phosphatidylinositol 3-kinase, regulatory subunit, polypeptide 1 (p85 alpha) | protein-coding |
| chr13_101589023_101589246 | chr13 | 101589024 | 101589246 | + | -0.99382 | Intergenic | Intergenic | 103495 | NM_001024955 | 18708 | Mm.259333 | NM_001024955 | ENSMUSG00000041417 | **Pik3r1** | PI3K\|p50alpha\|p55alpha\|p85alpha | phosphatidylinositol 3-kinase, regulatory subunit, polypeptide 1 (p85 alpha) | protein-coding |
| chr13_101588830_101589351 | chr13 | 101588831 | 101589351 | + | -1.05176 | Intergenic | Intergenic | 103539 | NM_001024955 | 18708 | Mm.259333 | NM_001024955 | ENSMUSG00000041417 | **Pik3r1** | PI3K\|p50alpha\|p55alpha\|p85alpha | phosphatidylinositol 3-kinase, regulatory subunit, polypeptide 1 (p85 alpha) | protein-coding |
| chr9_105689298_105689877 | chr9 | 105689299 | 105689877 | + | -1.17019 | exon (NM_001102607, exon 37 of 37) | exon (NM_001102607, exon 37 of 37) | 46593 | NM_001081309 | 75669 | Mm.274830 | NM_029380 | ENSMUSG00000032571 | **Pik3r4** | 2210010O15Rik\|C730038E05Rik\|C85833\|D9Ertd418e | phosphatidylinositol 3 kinase, regulatory subunit, polypeptide 4, p150 | protein-coding |
| chr11_68431704_68432613 | chr11 | 68431705 | 68432613 | + | -0.651202 | promoter-TSS (NM_177320) | promoter-TSS (NM_177320) | 34 | NM_177320 | 320207 | Mm.244960 | NM_177320 | ENSMUSG00000020901 | **Pik3r5** | AV230647\|F730038I15Rik | phosphoinositide-3-kinase, regulatory subunit 5, p101 | protein-coding |
| chr11_68417808_68418381 | chr11 | 68417809 | 68418381 | + | -0.689295 | Intergenic | MLTR11B\|LTR\|ERVK | -14030 | NM_177320 | 320207 | Mm.244960 | NM_177320 | ENSMUSG00000020901 | **Pik3r5** | AV230647\|F730038I15Rik | phosphoinositide-3-kinase, regulatory subunit 5, p101 | protein-coding |
| chr11_68417764_68418215 | chr11 | 68417765 | 68418215 | + | -0.711058 | Intergenic | MLTR11B\|LTR\|ERVK | -14135 | NM_177320 | 320207 | Mm.244960 | NM_177320 | ENSMUSG00000020901 | **Pik3r5** | AV230647\|F730038I15Rik | phosphoinositide-3-kinase, regulatory subunit 5, p101 | protein-coding |
| chr17_29430451_29430778 | chr17 | 29430452 | 29430778 | + | -0.696838 | Intergenic | Intergenic | -60430 | NM_008842 | 18712 | Mm.405293 | NM_008842 | | **Pim1** | Pim-1 | proviral integration site 1 | protein-coding |
| chr17_29459602_29460097 | chr17 | 29459603 | 29460097 | + | -0.738905 | Intergenic | Intergenic | -31195 | NM_008842 | 18712 | Mm.405293 | NM_008842 | | **Pim1** | Pim-1 | proviral integration site 1 | protein-coding |
| chr17_29431856_29432337 | chr17 | 29431857 | 29432337 | + | -0.795854 | Intergenic | MIRb\|SINE\|MIR | -58948 | NM_008842 | 18712 | Mm.405293 | NM_008842 | | **Pim1** | Pim-1 | proviral integration site 1 | protein-coding |
| chr17_29459193_29460125 | chr17 | 29459194 | 29460125 | + | -0.833798 | Intergenic | Intergenic | -31386 | NM_008842 | 18712 | Mm.405293 | NM_008842 | | **Pim1** | Pim-1 | proviral integration site 1 | protein-coding |
| chr17_29428805_29429276 | chr17 | 29428806 | 29429276 | + | -1.11484 | Intergenic | Charlie16a\|DNA\|hAT-Charlie | -62004 | NM_008842 | 18712 | Mm.405293 | NM_008842 | | **Pim1** | Pim-1 | proviral integration site 1 | protein-coding |
| chr2_18920491_18920876 | chr2 | 18920492 | 18920876 | + | -0.62072 | intron (NM_008845, intron 1 of 9) | RMER4B\|LTR\|ERVK | 77437 | NM_008845 | 18718 | Mm.313977 | NM_008845 | ENSMUSG00000026737 | **Pip4k2a** | AW742916\|Pip5k2a | phosphatidylinositol-5-phosphate 4-kinase, type II, alpha | protein-coding |
| chr2_18984007_18984373 | chr2 | 18984008 | 18984373 | + | -0.632097 | intron (NM_008845, intron 1 of 9) | intron (NM_008845, intron 1 of 9) | 13931 | NM_008845 | 18718 | Mm.313977 | NM_008845 | ENSMUSG00000026737 | **Pip4k2a** | AW742916\|Pip5k2a | phosphatidylinositol-5-phosphate 4-kinase, type II, alpha | protein-coding |
| chr2_18984003_18984891 | chr2 | 18984004 | 18984891 | + | -0.720754 | intron (NM_008845, intron 1 of 9) | intron (NM_008845, intron 1 of 9) | 13674 | NM_008845 | 18718 | Mm.313977 | NM_008845 | ENSMUSG00000026737 | **Pip4k2a** | AW742916\|Pip5k2a | phosphatidylinositol-5-phosphate 4-kinase, type II, alpha | protein-coding |
| chr2_18988844_18989338 | chr2 | 18988845 | 18989338 | + | -0.721505 | intron (NM_008845, intron 1 of 9) | MTEa\|LTR\|ERVL-MaLR | 9030 | NM_008845 | 18718 | Mm.313977 | NM_008845 | ENSMUSG00000026737 | **Pip4k2a** | AW742916\|Pip5k2a | phosphatidylinositol-5-phosphate 4-kinase, type II, alpha | protein-coding |
| chr2_18984420_18984887 | chr2 | 18984421 | 18984887 | + | -0.749572 | intron (NM_008845, intron 1 of 9) | intron (NM_008845, intron 1 of 9) | 13467 | NM_008845 | 18718 | Mm.313977 | NM_008845 | ENSMUSG00000026737 | **Pip4k2a** | AW742916\|Pip5k2a | phosphatidylinositol-5-phosphate 4-kinase, type II, alpha | protein-coding |
| chr2_18947033_18947315 | chr2 | 18947034 | 18947315 | + | -0.776155 | intron (NM_008845, intron 1 of 9) | intron (NM_008845, intron 1 of 9) | 50947 | NM_008845 | 18718 | Mm.313977 | NM_008845 | ENSMUSG00000026737 | **Pip4k2a** | AW742916\|Pip5k2a | phosphatidylinositol-5-phosphate 4-kinase, type II, alpha | protein-coding |
| chr2_18968921_18969382 | chr2 | 18968922 | 18969382 | + | -0.793964 | intron (NM_008845, intron 1 of 9) | intron (NM_008845, intron 1 of 9) | 28969 | NM_008845 | 18718 | Mm.313977 | NM_008845 | ENSMUSG00000026737 | **Pip4k2a** | AW742916\|Pip5k2a | phosphatidylinositol-5-phosphate 4-kinase, type II, alpha | protein-coding |
| chr2_18920231_18920705 | chr2 | 18920232 | 18920705 | + | -0.855144 | intron (NM_008845, intron 1 of 9) | intron (NM_008845, intron 1 of 9) | 77653 | NM_008845 | 18718 | Mm.313977 | NM_008845 | ENSMUSG00000026737 | **Pip4k2a** | AW742916\|Pip5k2a | phosphatidylinositol-5-phosphate 4-kinase, type II, alpha | protein-coding |
| chr2_18952104_18952676 | chr2 | 18952105 | 18952676 | + | -0.863881 | intron (NM_008845, intron 1 of 9) | intron (NM_008845, intron 1 of 9) | 45731 | NM_008845 | 18718 | Mm.313977 | NM_008845 | ENSMUSG00000026737 | **Pip4k2a** | AW742916\|Pip5k2a | phosphatidylinositol-5-phosphate 4-kinase, type II, alpha | protein-coding |
| chr2_18982589_18982923 | chr2 | 18982590 | 18982923 | + | -1.04515 | intron (NM_008845, intron 1 of 9) | ORR1E\|LTR\|ERVL-MaLR | 15365 | NM_008845 | 18718 | Mm.313977 | NM_008845 | ENSMUSG00000026737 | **Pip4k2a** | AW742916\|Pip5k2a | phosphatidylinositol-5-phosphate 4-kinase, type II, alpha | protein-coding |
| chr11_97728204_97729259 | chr11 | 97728205 | 97729259 | + | -0.877045 | intron (NM_054051, intron 3 of 9) | intron (NM_054051, intron 3 of 9) | 15972 | NM_054051 | 108083 | Mm.39700 | NM_054051 | ENSMUSG00000018547 | **Pip4k2b** | AI848124\|PI5P4Kbeta\|Pip5k2b\|c11 | phosphatidylinositol-5-phosphate 4-kinase, type II, beta | protein-coding |
| chr3_95100894_95101254 | chr3 | 95100895 | 95101254 | + | -0.644862 | intron (NM_001293707, intron 1 of 14) | RSINE1\|SINE\|B4 | 5856 | NM_001293707 | 18720 | Mm.296409 | NM_008847 | ENSMUSG00000028126 | **Pip5k1a** | PI4P5K-I[a]\|PIP5K1-alpha\|PIP5KIalpha\|Pipk5a | phosphatidylinositol-4-phosphate 5-kinase, type 1 alpha | protein-coding |
| chr10_81297943_81298475 | chr10 | 81297944 | 81298475 | + | -0.749406 | intron (NM_001146687, intron 1 of 16) | intron (NM_001146687, intron 1 of 16) | 5246 | NM_001146687 | 18717 | Mm.29836 | NM_008844 | ENSMUSG00000034902 | **Pip5k1c** | AI115456\|AI835305\|Pip5kIgamma | phosphatidylinositol-4-phosphate 5-kinase, type 1 gamma | protein-coding |
| chr10_81297943_81298258 | chr10 | 81297944 | 81298258 | + | -0.810043 | intron (NM_001146687, intron 1 of 16) | intron (NM_001146687, intron 1 of 16) | 5138 | NM_001146687 | 18717 | Mm.29836 | NM_008844 | ENSMUSG00000034902 | **Pip5k1c** | AI115456\|AI835305\|Pip5kIgamma | phosphatidylinositol-4-phosphate 5-kinase, type 1 gamma | protein-coding |
| chr5_111200162_111201136 | chr5 | 111200163 | 111201136 | + | -0.615372 | intron (NM_001267622, intron 6 of 22) | intron (NM_001267622, intron 6 of 22) | -130114 | NM_019640 | 56305 | Mm.200516 | NM_019640 | ENSMUSG00000050017 | **Pitpnb** | AI256223\|AU040890 | phosphatidylinositol transfer protein, beta | protein-coding |
| chr5_111200162_111200840 | chr5 | 111200163 | 111200840 | + | -0.631776 | intron (NM_001267622, intron 6 of 22) | intron (NM_001267622, intron 6 of 22) | -130262 | NM_019640 | 56305 | Mm.200516 | NM_019640 | ENSMUSG00000050017 | **Pitpnb** | AI256223\|AU040890 | phosphatidylinositol transfer protein, beta | protein-coding |

| | | | | | | | | | | | | | | | | | |
|---|---|---|---|---|---|---|---|---|---|---|---|---|---|---|---|---|---|
| chr5_111262108_111262531 | chr5 | 111262109 | 111262531 | + | -0.650391 | intron (NM_001267622, intron 12 of 22) | intron (NM_001267622, intron 12 of 22) | -68443 | NM_019640 | 56305 | Mm.200516 | NM_019640 | ENSMUSG00000050017 | **Pitpnb** | AI256223\|AU040890 | phosphatidylinositol transfer protein, beta | protein-coding |
| chr5_111262178_111262466 | chr5 | 111262179 | 111262466 | + | -0.70138 | intron (NM_001267622, intron 12 of 22) | intron (NM_001267622, intron 12 of 22) | -68441 | NM_019640 | 56305 | Mm.200516 | NM_019640 | ENSMUSG00000050017 | **Pitpnb** | AI256223\|AU040890 | phosphatidylinositol transfer protein, beta | protein-coding |
| chr5_111178161_111179025 | chr5 | 111178162 | 111179025 | + | -0.749813 | intron (NM_001267622, intron 5 of 22) | intron (NM_001267622, intron 5 of 22) | -152170 | NM_019640 | 56305 | Mm.200516 | NM_019640 | ENSMUSG00000050017 | **Pitpnb** | AI256223\|AU040890 | phosphatidylinositol transfer protein, beta | protein-coding |
| chr5_111262834_111263144 | chr5 | 111262835 | 111263144 | + | -0.76116 | intron (NM_001267622, intron 12 of 22) | intron (NM_001267622, intron 12 of 22) | -67774 | NM_019640 | 56305 | Mm.200516 | NM_019640 | ENSMUSG00000050017 | **Pitpnb** | AI256223\|AU040890 | phosphatidylinositol transfer protein, beta | protein-coding |
| chr5_111262757_111263244 | chr5 | 111262758 | 111263244 | + | -0.794309 | intron (NM_001267622, intron 12 of 22) | intron (NM_001267622, intron 12 of 22) | -67762 | NM_019640 | 56305 | Mm.200516 | NM_019640 | ENSMUSG00000050017 | **Pitpnb** | AI256223\|AU040890 | phosphatidylinositol transfer protein, beta | protein-coding |
| chr5_111356852_111357520 | chr5 | 111356853 | 111357520 | + | -0.806458 | intron (NM_019640, intron 7 of 11) | intron (NM_019640, intron 7 of 11) | 26423 | NM_019640 | 56305 | Mm.200516 | NM_019640 | ENSMUSG00000050017 | **Pitpnb** | AI256223\|AU040890 | phosphatidylinositol transfer protein, beta | protein-coding |
| chr11_107417009_107417708 | chr11 | 107417010 | 107417708 | + | -0.616096 | intron (NM_145823, intron 1 of 9) | intron (NM_145823, intron 1 of 9) | 53361 | NM_145823 | 71795 | Mm.397413 | NM_145823 | ENSMUSG00000040430 | **Pitpnc1** | 1110020B03Rik\|5830436L09Rik\|AI662802\|AI851387\|C330017I21Rik\|Dnr411\|RDGBB\|RDGBB1\|rdgB-beta | phosphatidylinositol transfer protein, cytoplasmic 1 | protein-coding |
| chr11_107454548_107454831 | chr11 | 107454549 | 107454831 | + | -0.7683 | intron (NM_145823, intron 1 of 9) | intron (NM_145823, intron 1 of 9) | 16030 | NM_145823 | 71795 | Mm.397413 | NM_145823 | ENSMUSG00000040430 | **Pitpnc1** | 1110020B03Rik\|5830436L09Rik\|AI662802\|AI851387\|C330017I21Rik\|Dnr411\|RDGBB\|RDGBB1\|rdgB-beta | phosphatidylinositol transfer protein, cytoplasmic 1 | protein-coding |
| chr11_107443560_107444164 | chr11 | 107443561 | 107444164 | + | -0.780193 | intron (NM_145823, intron 1 of 9) | intron (NM_145823, intron 1 of 9) | 26858 | NM_145823 | 71795 | Mm.397413 | NM_145823 | ENSMUSG00000040430 | **Pitpnc1** | 1110020B03Rik\|5830436L09Rik\|AI662802\|AI851387\|C330017I21Rik\|Dnr411\|RDGBB\|RDGBB1\|rdgB-beta | phosphatidylinositol transfer protein, cytoplasmic 1 | protein-coding |
| chr11_107447454_107447723 | chr11 | 107447455 | 107447723 | + | -0.892434 | intron (NM_145823, intron 1 of 9) | intron (NM_145823, intron 1 of 9) | 23131 | NM_145823 | 71795 | Mm.397413 | NM_145823 | ENSMUSG00000040430 | **Pitpnc1** | 1110020B03Rik\|5830436L09Rik\|AI662802\|AI851387\|C330017I21Rik\|Dnr411\|RDGBB\|RDGBB1\|rdgB-beta | phosphatidylinositol transfer protein, cytoplasmic 1 | protein-coding |
| chr11_107447488_107447659 | chr11 | 107447489 | 107447659 | + | -0.905436 | intron (NM_145823, intron 1 of 9) | B4A\|SINE\|B4 | 23146 | NM_145823 | 71795 | Mm.397413 | NM_145823 | ENSMUSG00000040430 | **Pitpnc1** | 1110020B03Rik\|5830436L09Rik\|AI662802\|AI851387\|C330017I21Rik\|Dnr411\|RDGBB\|RDGBB1\|rdgB-beta | phosphatidylinositol transfer protein, cytoplasmic 1 | protein-coding |
| chr5_124213694_124213932 | chr5 | 124213695 | 124213932 | + | -0.636756 | intron (NM_011256, intron 2 of 24) | intron (NM_011256, intron 2 of 24) | -15912 | NR_045369 | 75927 | Mm.410803 | NR_045369 | ENSMUSG00000090220 | **Pitpnm2os1** | 4930568K20Rik | phosphatidylinositol transfer protein, membrane-associated 2, opposite strand 1 | ncRNA |
| chr14_70402271_70402794 | chr14 | 70402272 | 70402794 | + | -0.654961 | intron (NM_021308, intron 10 of 22) | B3A\|SINE\|B2 | 26561 | NM_021308 | 57746 | Mm.85253 | NM_021308 | ENSMUSG00000033644 | **Piwil2** | Piwil1l\|mili | piwi-like RNA-mediated gene silencing 2 | protein-coding |
| chrX_99593018_99593666 | chrX | 99593019 | 99593666 | + | -0.752787 | Intergenic | Intergenic | -122607 | NM_008853 | 18744 | Mm.8211 | NM_008853 | ENSMUSG00000034403 | **Pja1** | praja-1 | praja ring finger 1, E3 ubiquitin protein ligase | protein-coding |
| chr17_64248240_64248616 | chr17 | 64248241 | 64248616 | + | -0.958144 | Intergenic | Intergenic | 83455 | NM_144859 | 224938 | Mm.489681 | NM_144859 | ENSMUSG00000024083 | **Pja2** | AI447901\|AL022700\|Neurodap1\|mKIAA0438 | praja 2, RING-H2 motif containing | protein-coding |
| chr2_163667013_163667354 | chr2 | 163667014 | 163667354 | + | -0.8765 | intron (NM_001039390, intron 2 of 4) | intron (NM_001039390, intron 2 of 4) | 8726 | NM_001164053 | 18769 | Mm.10091 | NM_011106 | ENSMUSG00000035268 | **Pkig** | PKIgamma | protein kinase inhibitor, gamma | protein-coding |
| chr7_141071918_141072747 | chr7 | 141071919 | 141072747 | + | -0.663227 | TTS (NM_177897) | TTS (NM_177897) | -5896 | NM_001162924 | 56460 | Mm.350037 | NM_019762 | ENSMUSG00000054065 | **Pkp3** | 2310056L12Rik | plakophilin 3 | protein-coding |
| chr5_100584807_100585644 | chr5 | 100584808 | 100585644 | + | -0.914922 | Intergenic | MLT1G\|LTR\|ERVL-MaLR | -13021 | NM_139198 | 231507 | Mm.34609 | NM_139198 | ENSMUSG00000029322 | **Plac8** | C15\|D5Wsu111e | placenta-specific 8 | protein-coding |
| chr8_22733463_22733772 | chr8 | 22733464 | 22733772 | + | -0.683086 | Intergenic | Intergenic | -24126 | NM_008872 | 18791 | Mm.154660 | NM_008872 | ENSMUSG00000031538 | **Plat** | AU020998\|AW212668\|D8Ertd2e\|tPA | plasminogen activator, tissue | protein-coding |
| chr8_22780460_22780958 | chr8 | 22780461 | 22780958 | + | -1.08594 | intron (NM_008872, intron 13 of 13) | intron (NM_008872, intron 13 of 13) | 22965 | NM_008872 | 18791 | Mm.154660 | NM_008872 | ENSMUSG00000031538 | **Plat** | AU020998\|AW212668\|D8Ertd2e\|tPA | plasminogen activator, tissue | protein-coding |
| chr14_20877385_20877690 | chr14 | 20877386 | 20877690 | + | -0.719708 | Intergenic | B1_Mur3\|SINE\|Alu | 40876 | NM_008873 | 18792 | Mm.4183 | NM_008873 | ENSMUSG00000021822 | **Plau** | u-PA\|uPA | plasminogen activator, urokinase | protein-coding |
| chr7_24457395_24458484 | chr7 | 24457396 | 24458484 | + | -0.837785 | Intergenic | Intergenic | -4560 | NM_011113 | 18793 | Mm.1359 | NM_011113 | ENSMUSG00000046223 | **Plaur** | Cd87\|u-PAR\|uPAR | plasminogen activator, urokinase receptor | protein-coding |
| chr7_24467882_24468302 | chr7 | 24467883 | 24468302 | + | -1.5222 | intron (NM_011113, intron 3 of 6) | intron (NM_011113, intron 3 of 6) | 5592 | NM_011113 | 18793 | Mm.1359 | NM_011113 | ENSMUSG00000046223 | **Plaur** | Cd87\|u-PAR\|uPAR | plasminogen activator, urokinase receptor | protein-coding |
| chr2_160701602_160702075 | chr2 | 160701603 | 160702075 | + | -1.07667 | intron (NM_009408, intron 9 of 20) | B4\|SINE\|B4 | -29471 | NM_021280 | 18803 | Mm.44463 | NM_021280 | ENSMUSG00000016933 | **Plcg1** | AI894140\|Cded\|Plc-1\|Plc-gamma1\|Plcg-1 | phospholipase C, gamma 1 | protein-coding |
| chr8_117548413_117548977 | chr8 | 117548414 | 117548977 | + | -0.828282 | intron (NM_172285, intron 2 of 32) | intron (NM_172285, intron 2 of 32) | 50404 | NM_172285 | 234779 | Mm.192699 | NM_172285 | ENSMUSG00000034330 | **Plcg2** | PLCgamma2\|Plcg-2 | phospholipase C, gamma 2 | protein-coding |
| chr8_117532387_117533133 | chr8 | 117532388 | 117533133 | + | -1.04255 | intron (NM_172285, intron 2 of 32) | intron (NM_172285, intron 2 of 32) | 34469 | NM_172285 | 234779 | Mm.192699 | NM_172285 | ENSMUSG00000034330 | **Plcg2** | PLCgamma2\|Plcg-2 | phospholipase C, gamma 2 | protein-coding |
| chr17_50416139_50416395 | chr17 | 50416140 | 50416395 | + | -0.641607 | Intergenic | Intergenic | -93280 | NM_013880 | 224860 | Mm.217362 | NM_013880 | ENSMUSG00000038910 | **Plcl2** | PLC-L2\|PRIP-2\|Plce2\|mKIAA1092 | phospholipase C-like 2 | protein-coding |
| chr17_50539515_50539745 | chr17 | 50539516 | 50539745 | + | -0.662591 | intron (NM_013880, intron 1 of 5) | B3A\|SINE\|B2 | 30083 | NM_013880 | 224860 | Mm.217362 | NM_013880 | ENSMUSG00000038910 | **Plcl2** | PLC-L2\|PRIP-2\|Plce2\|mKIAA1092 | phospholipase C-like 2 | protein-coding |
| chr17_50499409_50499753 | chr17 | 50499410 | 50499753 | + | -0.692738 | Intergenic | Intergenic | -9966 | NM_013880 | 224860 | Mm.217362 | NM_013880 | ENSMUSG00000038910 | **Plcl2** | PLC-L2\|PRIP-2\|Plce2\|mKIAA1092 | phospholipase C-like 2 | protein-coding |
| chr17_50484675_50485562 | chr17 | 50484676 | 50485562 | + | -0.773799 | Intergenic | Intergenic | -24428 | NM_013880 | 224860 | Mm.217362 | NM_013880 | ENSMUSG00000038910 | **Plcl2** | PLC-L2\|PRIP-2\|Plce2\|mKIAA1092 | phospholipase C-like 2 | protein-coding |
| chr16_46022051_46022917 | chr16 | 46022052 | 46022917 | + | -0.7669 | Intergenic | Intergenic | -12071 | NM_001134480 | 433022 | Mm.380993 | NM_001134480 | ENSMUSG00000087141 | **Plcxd2** | AW494639\|EG433022 | phosphatidylinositol-specific phospholipase C, X domain containing 2 | protein-coding |
| chr16_46027207_46027564 | chr16 | 46027208 | 46027564 | + | -0.785839 | Intergenic | L1MC4a\|LINE\|L1 | -16973 | NM_001134480 | 433022 | Mm.380993 | NM_001134480 | ENSMUSG00000087141 | **Plcxd2** | AW494639\|EG433022 | phosphatidylinositol-specific phospholipase C, X domain containing 2 | protein-coding |
| chr16_46010949_46011159 | chr16 | 46010950 | 46011159 | + | -0.860893 | promoter-TSS (NM_001134480) | promoter-TSS (NM_001134480) | -641 | NM_001134480 | 433022 | Mm.380993 | NM_001134480 | ENSMUSG00000087141 | **Plcxd2** | AW494639\|EG433022 | phosphatidylinositol-specific phospholipase C, X domain containing 2 | protein-coding |
| chr16_46001586_46002133 | chr16 | 46001587 | 46002133 | + | -0.862064 | intron (NM_001134480, intron 1 of 3) | intron (NM_001134480, intron 1 of 3) | 8553 | NM_001134480 | 433022 | Mm.380993 | NM_001134480 | ENSMUSG00000087141 | **Plcxd2** | AW494639\|EG433022 | phosphatidylinositol-specific phospholipase C, X domain containing 2 | protein-coding |
| chr16_46006245_46006655 | chr16 | 46006246 | 46006655 | + | -0.94317 | intron (NM_001134480, intron 1 of 3) | intron (NM_001134480, intron 1 of 3) | 3963 | NM_001134480 | 433022 | Mm.380993 | NM_001134480 | ENSMUSG00000087141 | **Plcxd2** | AW494639\|EG433022 | phosphatidylinositol-specific phospholipase C, X domain containing 2 | protein-coding |
| chr16_45942062_45942547 | chr16 | 45942063 | 45942547 | + | -1.10353 | Intergenic | Intergenic | 68108 | NM_001134480 | 433022 | Mm.380993 | NM_001134480 | ENSMUSG00000087141 | **Plcxd2** | AW494639\|EG433022 | phosphatidylinositol-specific phospholipase C, X domain containing 2 | protein-coding |
| chr16_46008787_46009128 | chr16 | 46008788 | 46009128 | + | -1.22724 | intron (NM_001134480, intron 1 of 3) | intron (NM_001134480, intron 1 of 3) | 1455 | NM_001134480 | 433022 | Mm.380993 | NM_001134480 | ENSMUSG00000087141 | **Plcxd2** | AW494639\|EG433022 | phosphatidylinositol-specific phospholipase C, X domain containing 2 | protein-coding |
| chr7_27544962_27545414 | chr7 | 27544963 | 27545414 | + | -0.778777 | 5' UTR (NM_011116, exon 2 of 14) | 5' UTR (NM_011116, exon 2 of 14) | 7924 | NM_011116 | 18807 | Mm.6483 | NM_011116 | ENSMUSG00000003363 | **Pld3** | Sam-9 | phospholipase D family, member 3 | protein-coding |
| chr7_27541290_27541980 | chr7 | 27541291 | 27541980 | + | -0.817452 | intron (NM_011116, intron 6 of 13) | intron (NM_011116, intron 6 of 13) | 11477 | NM_011116 | 18807 | Mm.6483 | NM_011116 | ENSMUSG00000003363 | **Pld3** | Sam-9 | phospholipase D family, member 3 | protein-coding |
| chr7_27541395_27541707 | chr7 | 27541396 | 27541707 | + | -0.856555 | intron (NM_011116, intron 6 of 13) | intron (NM_011116, intron 6 of 13) | 11561 | NM_011116 | 18807 | Mm.6483 | NM_011116 | ENSMUSG00000003363 | **Pld3** | Sam-9 | phospholipase D family, member 3 | protein-coding |
| chr6_140514738_140515170 | chr6 | 140514739 | 140515170 | + | -0.67707 | intron (NM_144920, intron 3 of 30) | intron (NM_144920, intron 3 of 30) | 90855 | NM_144920 | 109135 | Mm.247670 | NM_144920 | ENSMUSG00000030231 | **Plekha5** | 2810431N21Rik\|AI428202\|AK129423\|AMH-Cre\|Ayu21-9\|Gt(pU21)9Imeg\|Pepp2\|Tg(AMH-cre)1Flor | pleckstrin homology domain containing, family A member 5 | protein-coding |
| chr6_140313939_140314432 | chr6 | 140313940 | 140314432 | + | -0.79664 | Intergenic | Intergenic | -109913 | NM_144920 | 109135 | Mm.247670 | NM_144920 | ENSMUSG00000030231 | **Plekha5** | 2810431N21Rik\|AI428202\|AK129423\|AMH-Cre\|Ayu21-9\|Gt(pU21)9Imeg\|Pepp2\|Tg(AMH-cre)1Flor | pleckstrin homology domain containing, family A member 5 | protein-coding |
| chr1_133237668_133238359 | chr1 | 133237669 | 133238359 | + | -0.810932 | Intergenic | Intergenic | -8083 | NM_001160268 | 240753 | Mm.253559 | NM_182930 | ENSMUSG00000041757 | **Plekha6** | 9530028L01Rik\|AU042621\|Pepp3 | pleckstrin homology domain containing, family A member 6 | protein-coding |

| | | | | | | | | | | | | | | | | | |
|---|---|---|---|---|---|---|---|---|---|---|---|---|---|---|---|---|---|
| chr1_64970413_64971247 | chr1 | 64970414 | 64971247 | + | -1.05573 | Intergenic | Intergenic | -14006 | NM_001039493 | 241075 | Mm.239534 | NM_175509 | ENSMUSG00000051344 | **Plekhm3** | 9430067K14Rik\|A230102O09Rik\|AI449570\|DAPR\|Plekhm1l | pleckstrin homology domain containing, family M, member 3 | protein-coding |
| chr9_65578899_65579255 | chr9 | 65578900 | 65579255 | + | -0.707518 | intron (NM_153119, intron 1 of 5) | MIR3\|SINE\|MIR | 1010 | NM_153119 | 102595 | Mm.247303 | NM_153119 | | **Plekho2** | AI840980\|Plekhq1 | pleckstrin homology domain containing, family O member 2 | protein-coding |
| chr4_86683239_86683654 | chr4 | 86683240 | 86683654 | + | -1.30633 | Intergenic | B4A\|SINE\|B4 | -13388 | NM_007408 | 11520 | Mm.381 | NM_007408 | ENSMUSG00000028494 | **Plin2** | AA407157\|ADPH\|Adfp\|Adrp | perilipin 2 | protein-coding |
| chr2_164866696_164867383 | chr2 | 164866697 | 164867383 | + | -0.84042 | Intergenic | ID_B1\|SINE\|B4 | -9332 | NM_011125 | 18830 | Mm.6105 | NM_011125 | ENSMUSG00000017754 | **Pltp** | Bpife\|OD107 | phospholipid transfer protein | protein-coding |
| chr6_89344943_89345253 | chr6 | 89344944 | 89345253 | + | -0.6878 | intron (NM_008881, intron 3 of 31) | intron (NM_008881, intron 3 of 31) | 17515 | NM_008881 | 18844 | Mm.3789 | NM_008881 | ENSMUSG00000030084 | **Plxna1** | 2600013D04Rik\|NOV\|PlexA1\|Plxn1\|mKIAA4053 | plexin A1 | protein-coding |
| chr2_173349420_173350312 | chr2 | 173349421 | 173350312 | + | -0.716247 | Intergenic | Intergenic | -73333 | NM_022995 | 65112 | Mm.73682 | NM_022995 | ENSMUSG00000038400 | **Pmepa1** | 2210418I02Rik\|AW455466\|Erg1.2\|N4wbp4\|Stag1\|Tmepai | prostate transmembrane protein, androgen induced 1 | protein-coding |
| chr16_8639885_8640142 | chr16 | 8639886 | 8640142 | + | -0.957465 | intron (NM_016881, intron 1 of 8) | intron (NM_016881, intron 1 of 8) | 2307 | NM_016881 | 54128 | Mm.9699 | NM_016881 | ENSMUSG00000022711 | **Pmm2** | AI585868\|C86848 | phosphomannomutase 2 | protein-coding |
| chr12_84170634_84170986 | chr12 | 84170635 | 84170986 | + | -0.965427 | intron (NM_001163502, intron 3 of 12) | intron (NM_001163502, intron 3 of 12) | -22321 | NM_027438 | 70481 | Mm.444348 | NM_027438 | ENSMUSG00000054383 | **Pnma1** | 5730402C15Rik | paraneoplastic antigen MA1 | protein-coding |
| chr11_17192146_17192832 | chr11 | 17192147 | 17192832 | + | -0.801848 | intron (NM_024459, intron 2 of 5) | intron (NM_024459, intron 2 of 5) | 19100 | NM_025443 | 66249 | Mm.27831 | NM_025443 | ENSMUSG00000020116 | **Pno1** | 1810003N24Rik\|AA407421\|AU015468\|AU019495\|AU042323\|Emi3\|Imi3 | partner of NOB1 homolog (S. cerevisiae) | protein-coding |
| chr9_106312818_106313374 | chr9 | 106312819 | 106313374 | + | -0.912434 | intron (NM_027354, intron 9 of 10) | intron (NM_027354, intron 9 of 10) | 32035 | NM_027354 | 70235 | Mm.23054 | NM_027354 | ENSMUSG00000023345 | **Poc1a** | 2510040D07Rik\|Wdr51a | POC1 centriolar protein homolog A (Chlamydomonas) | protein-coding |
| chr19_4227505_4227756 | chr19 | 4227506 | 4227756 | + | -0.800598 | Intergenic | Intergenic | -4268 | NM_027196 | 69745 | Mm.32518 | NM_027196 | ENSMUSG00000024854 | **Pold4** | 2410012M21Rik\|AI463381\|AW060307\|Polds\|p12 | polymerase (DNA-directed), delta 4 | protein-coding |
| chr19_4226789_4227066 | chr19 | 4226790 | 4227066 | + | -0.83786 | Intergenic | Intergenic | -4971 | NM_027196 | 69745 | Mm.32518 | NM_027196 | ENSMUSG00000024854 | **Pold4** | 2410012M21Rik\|AI463381\|AW060307\|Polds\|p12 | polymerase (DNA-directed), delta 4 | protein-coding |
| chr7_79447524_79447987 | chr7 | 79447525 | 79447987 | + | -0.691454 | intron (NM_145946, intron 34 of 37) | intron (NM_145946, intron 34 of 37) | 18517 | NM_017462 | 18975 | Mm.3616 | NM_017462 | ENSMUSG00000039176 | **Polg** | AA409516\|PolgA | polymerase (DNA directed), gamma | protein-coding |
| chr18_70530936_70531285 | chr18 | 70530937 | 70531285 | + | -0.804254 | promoter-TSS (NM_001289515) | promoter-TSS (NM_001289515) | -790 | NM_001289515 | 26447 | Mm.244352 | NM_011972 | ENSMUSG00000038425 | **Poli** | Rad30b | polymerase (DNA directed), iota | protein-coding |
| chr19_45527626_45528513 | chr19 | 45527627 | 45528513 | + | -0.731186 | intron (NM_001286465, intron 14 of 14) | intron (NM_001286465, intron 14 of 14) | 32473 | NM_020032 | 56626 | Mm.46509 | NM_020032 | ENSMUSG00000025218 | **Poll** | 1110003P06Rik\|AV007317 | polymerase (DNA directed), lambda | protein-coding |
| chr6_71962185_71963208 | chr6 | 71962186 | 71963208 | + | -0.619661 | intron (NM_009088, intron 21 of 33) | intron (NM_009088, intron 21 of 33) | 53644 | NM_009088 | 20019 | Mm.135581 | NM_009088 | ENSMUSG00000049553 | **Polr1a** | 194kDa\|3010014K16Rik\|RPA194\|Rpo1-4\|mRPA1 | polymerase (RNA) I polypeptide A | protein-coding |
| chr6_71962623_71963118 | chr6 | 71962624 | 71963118 | + | -0.637624 | intron (NM_009088, intron 21 of 33) | intron (NM_009088, intron 21 of 33) | 53818 | NM_009088 | 20019 | Mm.135581 | NM_009088 | ENSMUSG00000049553 | **Polr1a** | 194kDa\|3010014K16Rik\|RPA194\|Rpo1-4\|mRPA1 | polymerase (RNA) I polypeptide A | protein-coding |
| chr3_54378007_54378459 | chr3 | 54378008 | 54378459 | + | -0.61248 | intron (NM_015784, intron 16 of 21) | intron (NM_015784, intron 16 of 21) | 17126 | NM_015784 | 50706 | Mm.236067 | NM_015784 | ENSMUSG00000027750 | **Postn** | A630052E07Rik\|AI747096\|OSF-2\|Osf2\|PLF\|PN\|peri | periostin, osteoblast specific factor | protein-coding |
| chr3_54399845_54400513 | chr3 | 54399846 | 54400513 | + | -0.668425 | Intergenic | Intergenic | 39072 | NM_015784 | 50706 | Mm.236067 | NM_015784 | ENSMUSG00000027750 | **Postn** | A630052E07Rik\|AI747096\|OSF-2\|Osf2\|PLF\|PN\|peri | periostin, osteoblast specific factor | protein-coding |
| chr1_166016316_166016811 | chr1 | 166016317 | 166016811 | + | -0.814285 | Intergenic | Intergenic | -13930 | NM_011137 | 18986 | Mm.245261 | NM_011137 | ENSMUSG00000026565 | **Pou2f1** | 2810482H01Rik\|NF-A1\|Oct-1\|Oct1\|Otf-1\|Otf1 | POU domain, class 2, transcription factor 1 | protein-coding |
| chr1_166016350_166016977 | chr1 | 166016351 | 166016977 | + | -0.824541 | Intergenic | B1_Mur2\|SINE\|Alu | -14030 | NM_011137 | 18986 | Mm.245261 | NM_011137 | ENSMUSG00000026565 | **Pou2f1** | 2810482H01Rik\|NF-A1\|Oct-1\|Oct1\|Otf-1\|Otf1 | POU domain, class 2, transcription factor 1 | protein-coding |
| chr15_100598198_100599200 | chr15 | 100598199 | 100599200 | + | -0.614506 | Intergenic | Intergenic | -12334 | NM_010127 | 19009 | Mm.28825 | NM_010127 | ENSMUSG00000009739 | **Pou6f1** | 2310038G18Rik\|Brn5.0\|C79593\|Emb\|cns-1 | POU domain, class 6, transcription factor 1 | protein-coding |
| chr15_100598239_100598692 | chr15 | 100598240 | 100598692 | + | -0.628458 | Intergenic | Intergenic | -12101 | NM_010127 | 19009 | Mm.28825 | NM_010127 | ENSMUSG00000009739 | **Pou6f1** | 2310038G18Rik\|Brn5.0\|C79593\|Emb\|cns-1 | POU domain, class 6, transcription factor 1 | protein-coding |
| chr15_100599835_100600613 | chr15 | 100599836 | 100600613 | + | -0.685139 | Intergenic | Intergenic | -13859 | NM_010127 | 19009 | Mm.28825 | NM_010127 | ENSMUSG00000009739 | **Pou6f1** | 2310038G18Rik\|Brn5.0\|C79593\|Emb\|cns-1 | POU domain, class 6, transcription factor 1 | protein-coding |
| chr3_133394555_133394892 | chr3 | 133394556 | 133394892 | + | -0.73875 | Intergenic | ORR1E\|LTR\|ERVL-MaLR | 84614 | NM_146141 | 74776 | Mm.210305 | NM_146141 | ENSMUSG00000028013 | **Ppa2** | 1110013G13Rik\|Sid6306 | pyrophosphatase (inorganic) 2 | protein-coding |
| chr8_25709200_25709514 | chr8 | 25709201 | 25709514 | + | -0.821114 | intron (NM_001081269, intron 20 of 22) | intron (NM_001081269, intron 20 of 22) | -10691 | NR_121642 | 71910 | Mm.385055 | NM_028000 | ENSMUSG00000031570 | **Ppapdc1b** | 1810019D05Rik\|2310022A04Rik\|AW045471\|Ppapdc1 | phosphatidic acid phosphatase type 2 domain containing 1B | protein-coding |
| chr8_25709083_25709622 | chr8 | 25709084 | 25709622 | + | -0.825457 | intron (NM_001081269, intron 20 of 22) | intron (NM_001081269, intron 20 of 22) | -10695 | NR_121642 | 71910 | Mm.385055 | NM_028000 | ENSMUSG00000031570 | **Ppapdc1b** | 1810019D05Rik\|2310022A04Rik\|AW045471\|Ppapdc1 | phosphatidic acid phosphatase type 2 domain containing 1B | protein-coding |
| chr18_61377723_61378066 | chr18 | 61377724 | 61378066 | + | -0.984498 | intron (NM_133249, intron 1 of 11) | intron (NM_133249, intron 1 of 11) | 22536 | NM_133249 | 170826 | Mm.415302 | NM_133249 | ENSMUSG00000033871 | **Ppargc1b** | 4631412G21Rik\|Perc | peroxisome proliferative activated receptor, gamma, coactivator 1 beta | protein-coding |
| chr5_92245323_92245993 | chr5 | 92245324 | 92245993 | + | -1.13683 | intron (NM_011148, intron 6 of 15) | intron (NM_011148, intron 6 of 15) | 7501 | NM_011148 | 19023 | Mm.87155 | NM_011148 | ENSMUSG00000029410 | **Ppef2** | - | protein phosphatase, EF hand calcium-binding domain 2 | protein-coding |
| chr7_107659053_107659592 | chr7 | 107659054 | 107659592 | + | -0.651251 | intron (NM_008905, intron 3 of 24) | intron (NM_008905, intron 3 of 24) | 52479 | NM_001163557 | 19024 | Mm.2817 | NM_008905 | ENSMUSG00000036528 | **Ppfibp2** | Cclp1 | PTPRF interacting protein, binding protein 2 (liprin beta 2) | protein-coding |
| chr11_6410168_6410590 | chr11 | 6410169 | 6410590 | + | -0.698779 | Intergenic | Intergenic | -5491 | NM_008907 | 268373 | Mm.5246 | NM_008907 | | **Ppia** | 2700098C05\|Cphn\|CyP-18\|CypA | peptidylprolyl isomerase A | protein-coding |
| chr1_58439925_58440240 | chr1 | 58439926 | 58440240 | + | -0.745479 | intron (NM_001285826, intron 4 of 6) | MTC\|LTR\|ERVL-MaLR | 5403 | NM_027374 | 70225 | Mm.340195 | NM_027351 | ENSMUSG00000026035 | **Ppil3** | 2310076N22Rik\|2510026K04Rik\|Cyp10l | peptidylprolyl isomerase (cyclophilin)-like 3 | protein-coding |
| chr17_84997775_84998055 | chr17 | 84997776 | 84998055 | + | -0.630121 | intron (NM_001159496, intron 2 of 5) | (CA)n\|Simple_repeat\|Simple_repeat | 23863 | NM_001159498 | 19043 | Mm.249695 | NM_011151 | ENSMUSG00000061130 | **Ppm1b** | PP2CB | protein phosphatase 1B, magnesium dependent, beta isoform | protein-coding |
| chr17_84944322_84944789 | chr17 | 84944323 | 84944789 | + | -0.769114 | intron (NR_027929, intron 2 of 3) | intron (NR_027929, intron 2 of 3) | -12185 | NM_001159497 | 19043 | Mm.249695 | NM_011151 | ENSMUSG00000061130 | **Ppm1b** | PP2CB | protein phosphatase 1B, magnesium dependent, beta isoform | protein-coding |
| chr17_84888413_84888971 | chr17 | 84888414 | 84888971 | + | -0.776986 | Intergenic | Intergenic | -68049 | NM_001159497 | 19043 | Mm.249695 | NM_011151 | ENSMUSG00000061130 | **Ppm1b** | PP2CB | protein phosphatase 1B, magnesium dependent, beta isoform | protein-coding |
| chr17_84937170_84937607 | chr17 | 84937171 | 84937607 | + | -1.0255 | intron (NR_027929, intron 2 of 3) | intron (NR_027929, intron 2 of 3) | -19352 | NM_001159497 | 19043 | Mm.249695 | NM_011151 | ENSMUSG00000061130 | **Ppm1b** | PP2CB | protein phosphatase 1B, magnesium dependent, beta isoform | protein-coding |
| chr5_31226676_31227130 | chr5 | 31226677 | 31227130 | + | -0.94132 | intron (NR_105788, intron 5 of 7) | MER74B\|LTR\|ERVL | -6358 | NM_008014 | 14208 | Mm.14501 | NM_008014 | ENSMUSG00000029147 | **Ppm1g** | AI385652\|Fin13 | protein phosphatase 1G (formerly 2C), magnesium-dependent, gamma isoform | protein-coding |
| chr10_122663070_122663510 | chr10 | 122663071 | 122663510 | + | -0.964237 | Intergenic | Intergenic | -15472 | NM_176919 | 319468 | Mm.489703 | NM_176919 | ENSMUSG00000034613 | **Ppm1h** | A430075L18Rik\|ARHCL1\|C030002B11Rik | protein phosphatase 1H (PP2C domain containing) | protein-coding |
| chr10_122651620_122652345 | chr10 | 122651621 | 122652345 | + | -1.03238 | Intergenic | Intergenic | -26779 | NM_176919 | 319468 | Mm.489703 | NM_176919 | ENSMUSG00000034613 | **Ppm1h** | A430075L18Rik\|ARHCL1\|C030002B11Rik | protein phosphatase 1H (PP2C domain containing) | protein-coding |
| chr5_32479055_32479385 | chr5 | 32479056 | 32479385 | + | -1.24082 | intron (NM_172707, intron 2 of 7) | intron (NM_172707, intron 2 of 7) | 20250 | NM_172707 | 19046 | Mm.241931 | NM_172707 | | **Ppp1cb** | 1200010B19 | protein phosphatase 1, catalytic subunit, beta isoform | protein-coding |
| chr5_122163142_122163640 | chr5 | 122163143 | 122163640 | + | -1.62058 | intron (NM_013636, intron 1 of 6) | intron (NM_013636, intron 1 of 6) | 5112 | NM_013636 | 19047 | Mm.280784 | NM_013636 | ENSMUSG00000004455 | **Ppp1cc** | PP-1G\|PP1\|dis2m1 | protein phosphatase 1, catalytic subunit, gamma isoform | protein-coding |
| chr7_4492665_4493259 | chr7 | 4492666 | 4493259 | + | -0.671877 | intron (NM_029834, intron 3 of 21) | intron (NM_029834, intron 3 of 21) | 8718 | NM_029834 | 232807 | Mm.360391 | NM_029834 | ENSMUSG00000019254 | **Ppp1r12c** | 2410197A17Rik\|AI839747\|B430101M08\|Mbs85 | protein phosphatase 1, regulatory (inhibitor) subunit 12C | protein-coding |
| chr12_111902514_111902834 | chr12 | 111902515 | 111902834 | + | -0.778248 | intron (NM_011625, intron 1 of 16) | intron (NM_011625, intron 1 of 16) | 5381 | NM_011625 | 21981 | Mm.313076 | NM_011625 | ENSMUSG00000021285 | **Ppp1r13b** | AI449786\|ASPP1\|AW545810\|Tp53bp2\|Trp53bp2\|p85 | protein phosphatase 1, regulatory (inhibitor) subunit 13B | protein-coding |
| chr12_111913806_111914449 | chr12 | 111913807 | 111914449 | + | -1.16612 | Intergenic | Intergenic | -6073 | NM_011625 | 21981 | Mm.313076 | NM_011625 | ENSMUSG00000021285 | **Ppp1r13b** | AI449786\|ASPP1\|AW545810\|Tp53bp2\|Trp53bp2\|p85 | protein phosphatase 1, regulatory (inhibitor) subunit 13B | protein-coding |
| chr12_111914047_111914407 | chr12 | 111914048 | 111914407 | + | -1.25906 | Intergenic | Intergenic | -6172 | NM_011625 | 21981 | Mm.313076 | NM_011625 | ENSMUSG00000021285 | **Ppp1r13b** | AI449786\|ASPP1\|AW545810\|Tp53bp2\|Trp53bp2\|p85 | protein phosphatase 1, regulatory (inhibitor) subunit 13B | protein-coding |
| chr7_45521791_45522234 | chr7 | 45521792 | 45522234 | + | -0.668186 | TTS (NM_008654) | TTS (NM_008654) | 4255 | NM_008654 | 17872 | Mm.4048 | NM_008654 | ENSMUSG00000040435 | **Ppp1r15a** | 9630030H21\|Gadd34\|Myd116 | protein phosphatase 1, regulatory (inhibitor) subunit 15A | protein-coding |
| chr1_133159039_133159655 | chr1 | 133159040 | 133159655 | + | -1.05117 | Intergenic | Intergenic | 28181 | NM_133819 | 108954 | Mm.293628 | NM_133819 | ENSMUSG00000046062 | **Ppp1r15b** | 1810033K10Rik\|AI606441\|C530022L24Rik\|CReP | protein phosphatase 1, regulatory (inhibitor) subunit 15b | protein-coding |
| chr1_133159293_133159683 | chr1 | 133159294 | 133159683 | + | -1.08488 | Intergenic | Intergenic | 28322 | NM_133819 | 108954 | Mm.293628 | NM_133819 | ENSMUSG00000046062 | **Ppp1r15b** | 1810033K10Rik\|AI606441\|C530022L24Rik\|CReP | protein phosphatase 1, regulatory (inhibitor) subunit 15b | protein-coding |
| chr8_35366449_35367023 | chr8 | 35366450 | 35367023 | + | -0.856287 | Intergenic | Intergenic | -9005 | NM_177741 | 244416 | Mm.247126 | NM_177741 | ENSMUSG00000046794 | **Ppp1r3b** | 6430576E21\|AW821953\|GL\|PTG | protein phosphatase 1, regulatory (inhibitor) subunit 3B | protein-coding |

| | | | | | | | | | | | | | | | | | |
|---|---|---|---|---|---|---|---|---|---|---|---|---|---|---|---|---|---|
| chr8_35348027_35348304 | chr8 | 35348028 | 35348304 | + | -1.01929 | Intergenic | Intergenic | -27575 | NM_177741 | 244416 | Mm.247126 | NM_177741 | ENSMUSG00000046794 | **Ppp1r3b** | 6430576E21\|AW821953\|GL\|PTG | protein phosphatase 1, regulatory (inhibitor) subunit 3B | protein-coding |
| chrX_7570677_7571232 | chrX | 7570678 | 7571232 | + | -0.964424 | intron (NM_001290574, intron 1 of 3) | intron (NM_001290574, intron 1 of 3) | -2645 | NR_029473 | 78185 | Mm.160055 | NR_029473 | ENSMUSG00000039545 | **Ppp1r3fos** | 4930524L23Rik | protein phosphatase 1, regulatory subunit 3F, opposite strand | ncRNA |
| chrX_7570753_7571150 | chrX | 7570754 | 7571150 | + | -1.00486 | intron (NM_001290574, intron 1 of 3) | intron (NM_001290574, intron 1 of 3) | -2648 | NR_029473 | 78185 | Mm.160055 | NR_029473 | ENSMUSG00000039545 | **Ppp1r3fos** | 4930524L23Rik | protein phosphatase 1, regulatory subunit 3F, opposite strand | ncRNA |
| chr11_94997698_94998527 | chr11 | 94997699 | 94998527 | + | -0.636561 | exon (NM_172261, exon 3 of 10) | exon (NM_172261, exon 3 of 10) | 6901 | NM_172261 | 217124 | Mm.229087 | NM_172261 | ENSMUSG00000038976 | **Ppp1r9b** | SPL\|Spn | protein phosphatase 1, regulatory subunit 9B | protein-coding |
| chr11_94998067_94998500 | chr11 | 94998068 | 94998500 | + | -0.822539 | intron (NM_172261, intron 3 of 9) | L2b\|LINE\|L2 | 7072 | NM_172261 | 217124 | Mm.229087 | NM_172261 | ENSMUSG00000038976 | **Ppp1r9b** | SPL\|Spn | protein phosphatase 1, regulatory subunit 9B | protein-coding |
| chr9_50840933_50841771 | chr9 | 50840934 | 50841771 | + | -0.618876 | intron (NR_045601, intron 13 of 13) | intron (NR_045601, intron 13 of 13) | -15572 | NM_001034085 | 73699 | Mm.7726 | NM_028614 | ENSMUSG00000032058 | **Ppp2r1b** | 2410091N08Rik\|AI790395 | protein phosphatase 2, regulatory subunit A, beta | protein-coding |
| chr9_50865845_50866230 | chr9 | 50865846 | 50866230 | + | -0.74806 | intron (NM_001286553, intron 5 of 16) | MTC\|LTR\|ERVL-MaLR | 9114 | NM_001286553 | 73699 | Mm.7726 | NM_028614 | ENSMUSG00000032058 | **Ppp2r1b** | 2410091N08Rik\|AI790395 | protein phosphatase 2, regulatory subunit A, beta | protein-coding |
| chr1_191390269_191390514 | chr1 | 191390270 | 191390514 | + | -0.661403 | intron (NM_144880, intron 1 of 12) | intron (NM_144880, intron 1 of 12) | 6649 | NM_144880 | 226849 | Mm.205569 | NM_144880 | ENSMUSG00000026626 | **Ppp2r5a** | PR61alpha | protein phosphatase 2, regulatory subunit B', alpha | protein-coding |
| chr1_191369159_191369490 | chr1 | 191369160 | 191369490 | + | -0.680188 | intron (NM_144880, intron 2 of 12) | intron (NM_144880, intron 2 of 12) | 27716 | NM_144880 | 226849 | Mm.205569 | NM_144880 | ENSMUSG00000026626 | **Ppp2r5a** | PR61alpha | protein phosphatase 2, regulatory subunit B', alpha | protein-coding |
| chr1_191420958_191421351 | chr1 | 191420959 | 191421351 | + | -0.697037 | Intergenic | Intergenic | -24114 | NM_144880 | 226849 | Mm.205569 | NM_144880 | ENSMUSG00000026626 | **Ppp2r5a** | PR61alpha | protein phosphatase 2, regulatory subunit B', alpha | protein-coding |
| chr1_191420743_191421394 | chr1 | 191420744 | 191421394 | + | -0.699849 | Intergenic | Intergenic | -24028 | NM_144880 | 226849 | Mm.205569 | NM_144880 | ENSMUSG00000026626 | **Ppp2r5a** | PR61alpha | protein phosphatase 2, regulatory subunit B', alpha | protein-coding |
| chr1_191422193_191422555 | chr1 | 191422194 | 191422555 | + | -0.721094 | Intergenic | Intergenic | -25333 | NM_144880 | 226849 | Mm.205569 | NM_144880 | ENSMUSG00000026626 | **Ppp2r5a** | PR61alpha | protein phosphatase 2, regulatory subunit B', alpha | protein-coding |
| chr1_191363219_191363439 | chr1 | 191363220 | 191363439 | + | -0.788006 | intron (NM_144880, intron 3 of 12) | intron (NM_144880, intron 3 of 12) | 33712 | NM_144880 | 226849 | Mm.205569 | NM_144880 | ENSMUSG00000026626 | **Ppp2r5a** | PR61alpha | protein phosphatase 2, regulatory subunit B', alpha | protein-coding |
| chr1_191363142_191363464 | chr1 | 191363143 | 191363464 | + | -0.818684 | intron (NM_144880, intron 3 of 12) | intron (NM_144880, intron 3 of 12) | 33738 | NM_144880 | 226849 | Mm.205569 | NM_144880 | ENSMUSG00000026626 | **Ppp2r5a** | PR61alpha | protein phosphatase 2, regulatory subunit B', alpha | protein-coding |
| chr1_191425930_191426225 | chr1 | 191425931 | 191426225 | + | -1.1721 | Intergenic | Intergenic | -29037 | NM_144880 | 226849 | Mm.205569 | NM_144880 | ENSMUSG00000026626 | **Ppp2r5a** | PR61alpha | protein phosphatase 2, regulatory subunit B', alpha | protein-coding |
| chr1_191425839_191426311 | chr1 | 191425840 | 191426311 | + | -1.1998 | Intergenic | Intergenic | -29034 | NM_144880 | 226849 | Mm.205569 | NM_144880 | ENSMUSG00000026626 | **Ppp2r5a** | PR61alpha | protein phosphatase 2, regulatory subunit B', alpha | protein-coding |
| chr12_110494939_110495207 | chr12 | 110494940 | 110495207 | + | -0.625741 | intron (NM_012023, intron 1 of 13) | (GA)n\|Simple_repeat\|Simple_repeat | 9334 | NM_012023 | 26931 | Mm.240396 | NM_012023 | ENSMUSG00000017843 | **Ppp2r5c** | 2610043M05Rik\|2700063L20Rik\|AI060890\|AW545884\|C85228\|D12Bwg0916e\|mKIAA0044 | protein phosphatase 2, regulatory subunit B', gamma | protein-coding |
| chr12_110506079_110506469 | chr12 | 110506080 | 110506469 | + | -0.686497 | intron (NM_012023, intron 1 of 13) | intron (NM_012023, intron 1 of 13) | 20535 | NM_012023 | 26931 | Mm.240396 | NM_012023 | ENSMUSG00000017843 | **Ppp2r5c** | 2610043M05Rik\|2700063L20Rik\|AI060890\|AW545884\|C85228\|D12Bwg0916e\|mKIAA0044 | protein phosphatase 2, regulatory subunit B', gamma | protein-coding |
| chr3_136611646_136612119 | chr3 | 136611647 | 136612119 | + | -0.792897 | Intergenic | Intergenic | -58183 | NM_008913 | 19055 | Mm.331389 | NM_008913 | ENSMUSG00000028161 | **Ppp3ca** | 2900074D19Rik\|CN\|Caln\|Calna\|CnA | protein phosphatase 3, catalytic subunit, alpha isoform | protein-coding |
| chr7_126794218_126794495 | chr7 | 126794219 | 126794495 | + | -0.768472 | TTS (NM_001177307) | TTS (NM_001177307) | -1886 | NM_019674 | 56420 | Mm.41998 | NM_019674 | ENSMUSG00000030697 | **Ppp4c** | 1110002D08Rik\|AU016079\|Ppx | protein phosphatase 4, catalytic subunit | protein-coding |
| chr7_4647314_4647819 | chr7 | 4647315 | 4647819 | + | -0.640136 | intron (NM_172894, intron 1 of 23) | intron (NM_172894, intron 1 of 23) | 11383 | NM_172894 | 243819 | Mm.258670 | NM_172894 | ENSMUSG00000052296 | **Ppp6r1** | 2010309P17Rik\|AI836219\|B430201G11Rik\|D030074N20Rik\|Pp6r1\|Saps1\|mKIAA1115 | protein phosphatase 6, regulatory subunit 1 | protein-coding |
| chr7_4655492_4655737 | chr7 | 4655493 | 4655737 | + | -0.697145 | intron (NM_172894, intron 1 of 23) | L2b\|LINE\|L2 | 3335 | NM_172894 | 243819 | Mm.258670 | NM_172894 | ENSMUSG00000052296 | **Ppp6r1** | 2010309P17Rik\|AI836219\|B430201G11Rik\|D030074N20Rik\|Pp6r1\|Saps1\|mKIAA1115 | protein phosphatase 6, regulatory subunit 1 | protein-coding |
| chr7_4653713_4654230 | chr7 | 4653714 | 4654230 | + | -0.74347 | intron (NM_172894, intron 1 of 23) | intron (NM_172894, intron 1 of 23) | 4978 | NM_172894 | 243819 | Mm.258670 | NM_172894 | ENSMUSG00000052296 | **Ppp6r1** | 2010309P17Rik\|AI836219\|B430201G11Rik\|D030074N20Rik\|Pp6r1\|Saps1\|mKIAA1115 | protein phosphatase 6, regulatory subunit 1 | protein-coding |
| chr7_4651554_4651960 | chr7 | 4651555 | 4651960 | + | -0.760294 | intron (NM_172894, intron 1 of 23) | intron (NM_172894, intron 1 of 23) | 7193 | NM_172894 | 243819 | Mm.258670 | NM_172894 | ENSMUSG00000052296 | **Ppp6r1** | 2010309P17Rik\|AI836219\|B430201G11Rik\|D030074N20Rik\|Pp6r1\|Saps1\|mKIAA1115 | protein phosphatase 6, regulatory subunit 1 | protein-coding |
| chr19_3556953_3557317 | chr19 | 3556954 | 3557317 | + | -0.943794 | intron (NM_029456, intron 1 of 22) | intron (NM_029456, intron 1 of 22) | 18614 | NM_001164159 | 52036 | Mm.284686 | NM_028999 | ENSMUSG00000024908 | **Ppp6r3** | 4930528G08Rik\|9130026N02Rik\|D19Bwg1430e\|D19Ertd703e\|Pp6r3\|Ppcs3\|Pptcs3\|Sapl\|Saps3\|mKIAA1558 | protein phosphatase 6, regulatory subunit 3 | protein-coding |
| chr5_122274635_122275274 | chr5 | 122274636 | 122275274 | + | -0.649887 | Intergenic | Intergenic | -9443 | NM_177242 | 320717 | Mm.489670 | NM_177242 | ENSMUSG00000038582 | **Pptc7** | 9130017A15Rik\|AA672638\|AI848390\|TA-PP2C | PTC7 protein phosphatase homolog (S. cerevisiae) | protein-coding |
| chr10_44443775_44444249 | chr10 | 44443776 | 44444249 | + | -0.65713 | intron (NM_007548, intron 4 of 6) | B1_Mur1\|SINE\|Alu | 14675 | NM_007548 | 12142 | Mm.4800 | NM_007548 | ENSMUSG00000038151 | **Prdm1** | Blimp-1\|Blimp1\|PRDI-BF1\|ZNFPR1A1\|b2b1765Clo | PR domain containing 1, with ZNF domain | protein-coding |
| chr10_44482960_44483770 | chr10 | 44482961 | 44483770 | + | -0.805443 | Intergenic | Intergenic | -24678 | NM_007548 | 12142 | Mm.4800 | NM_007548 | ENSMUSG00000038151 | **Prdm1** | Blimp-1\|Blimp1\|PRDI-BF1\|ZNFPR1A1\|b2b1765Clo | PR domain containing 1, with ZNF domain | protein-coding |
| chr10_44483007_44483488 | chr10 | 44483008 | 44483488 | + | -0.843729 | Intergenic | Intergenic | -24561 | NM_007548 | 12142 | Mm.4800 | NM_007548 | ENSMUSG00000038151 | **Prdm1** | Blimp-1\|Blimp1\|PRDI-BF1\|ZNFPR1A1\|b2b1765Clo | PR domain containing 1, with ZNF domain | protein-coding |
| chr10_44483178_44483612 | chr10 | 44483179 | 44483612 | + | -0.856183 | Intergenic | Intergenic | -24708 | NM_007548 | 12142 | Mm.4800 | NM_007548 | ENSMUSG00000038151 | **Prdm1** | Blimp-1\|Blimp1\|PRDI-BF1\|ZNFPR1A1\|b2b1765Clo | PR domain containing 1, with ZNF domain | protein-coding |
| chr10_44483170_44483488 | chr10 | 44483171 | 44483488 | + | -0.879526 | Intergenic | Intergenic | -24642 | NM_007548 | 12142 | Mm.4800 | NM_007548 | ENSMUSG00000038151 | **Prdm1** | Blimp-1\|Blimp1\|PRDI-BF1\|ZNFPR1A1\|b2b1765Clo | PR domain containing 1, with ZNF domain | protein-coding |
| chr9_31300975_31301546 | chr9 | 31300976 | 31301546 | + | -0.65516 | Intergenic | Intergenic | -13846 | NM_001080817 | 382066 | Mm.335039 | NM_199315 | ENSMUSG00000042496 | **Prdm10** | Gm1112\|tristanin | PR domain containing 10 | protein-coding |
| chr9_31300556_31300839 | chr9 | 31300557 | 31300839 | + | -0.791005 | Intergenic | PB1D9\|SINE\|Alu | -14409 | NM_001080817 | 382066 | Mm.335039 | NM_199315 | ENSMUSG00000042496 | **Prdm10** | Gm1112\|tristanin | PR domain containing 10 | protein-coding |
| chr9_31300946_31301964 | chr9 | 31300947 | 31301964 | + | -0.848194 | Intergenic | Intergenic | -13652 | NM_001080817 | 382066 | Mm.335039 | NM_199315 | ENSMUSG00000042496 | **Prdm10** | Gm1112\|tristanin | PR domain containing 10 | protein-coding |
| chr9_31301598_31301961 | chr9 | 31301599 | 31301961 | + | -1.12622 | Intergenic | (CAGG)n\|Simple_repeat\|Simple_repeat | -13327 | NM_001080817 | 382066 | Mm.335039 | NM_199315 | ENSMUSG00000042496 | **Prdm10** | Gm1112\|tristanin | PR domain containing 10 | protein-coding |
| chr2_93049475_93049733 | chr2 | 93049476 | 93049733 | + | -0.662187 | Intergenic | Intergenic | -3460 | NM_001177536 | 100042784 | Mm.102816 | NM_001177536 | ENSMUSG00000075028 | **Prdm11** | 8030443D09Rik\|AA163972\|Gm707 | PR domain containing 11 | protein-coding |
| chr4_143219241_143219813 | chr4 | 143219242 | 143219813 | + | -0.843922 | Intergenic | Intergenic | -6818 | NM_001256380 | 110593 | Mm.332020 | NM_001081355 | ENSMUSG00000057637 | **Prdm2** | 4833427P12Rik\|E330024L24Rik\|Gm1033\|Gm19732\|KMT8\|Riz\|Riz1\|Znfpr1c1 | PR domain containing 2, with ZNF domain | protein-coding |
| chr6_65818376_65818783 | chr6 | 65818377 | 65818783 | + | -0.68421 | intron (NM_027547, intron 2 of 14) | intron (NM_027547, intron 2 of 14) | 39618 | NM_027547 | 70779 | Mm.263355 | NM_027547 | ENSMUSG00000029913 | **Prdm5** | 4432417F03Rik\|6530401I24Rik\|AI197291\|E130112L17Rik\|PFM2 | PR domain containing 5 | protein-coding |
| chr5_30961563_30961950 | chr5 | 30961564 | 30961950 | + | -1.01259 | intron (NR_105788, intron 2 of 7) | intron (NR_105788, intron 2 of 7) | -1430 | NM_001294302 | 50907 | Mm.272414 | NM_016703 | ENSMUSG00000045302 | **Preb** | C85705 | prolactin regulatory element binding | protein-coding |
| chr2_166701421_166702370 | chr2 | 166701422 | 166702370 | + | -1.00004 | intron (NR_105783, intron 2 of 2) | intron (NR_105783, intron 2 of 2) | 11936 | NM_177782 | 277360 | Mm.34420 | NM_177782 | ENSMUSG00000039621 | **Prex1** | G630042G04\|P-REX1\|Setd6 | phosphatidylinositol-3,4,5-trisphosphate-dependent Rac exchange factor 1 | protein-coding |
| chr2_166701426_166702026 | chr2 | 166701427 | 166702026 | + | -1.01839 | intron (NR_105783, intron 2 of 2) | intron (NR_105783, intron 2 of 2) | 12106 | NM_177782 | 277360 | Mm.34420 | NM_177782 | ENSMUSG00000039621 | **Prex1** | G630042G04\|P-REX1\|Setd6 | phosphatidylinositol-3,4,5-trisphosphate-dependent Rac exchange factor 1 | protein-coding |
| chr10_61296833_61297888 | chr10 | 61296834 | 61297888 | + | -0.662914 | promoter-TSS (NM_011073) | promoter-TSS (NM_011073) | -475 | NM_011073 | 18646 | Mm.240313 | NM_011073 | ENSMUSG00000037202 | **Prf1** | Pfn\|Pfp\|Prf-1 | perforin 1 (pore forming protein) | protein-coding |
| chr10_61297509_61297915 | chr10 | 61297510 | 61297915 | + | -0.86192 | promoter-TSS (NM_011073) | promoter-TSS (NM_011073) | -124 | NM_011073 | 18646 | Mm.240313 | NM_011073 | ENSMUSG00000037202 | **Prf1** | Pfn\|Pfp\|Prf-1 | perforin 1 (pore forming protein) | protein-coding |
| chr10_61305879_61306616 | chr10 | 61305880 | 61306616 | + | -1.06767 | Intergenic | Intergenic | 8412 | NM_011073 | 18646 | Mm.240313 | NM_011073 | ENSMUSG00000037202 | **Prf1** | Pfn\|Pfp\|Prf-1 | perforin 1 (pore forming protein) | protein-coding |
| chr15_93563518_93564148 | chr15 | 93563519 | 93564148 | + | -0.667854 | intron (NM_001033217, intron 1 of 7) | intron (NM_001033217, intron 1 of 7) | 32058 | NM_001033217 | 106042 | Mm.150314 | NM_001033217 | ENSMUSG00000036158 | **Prickle1** | 1110058P22Rik\|AW215793\|Prickle\|b2b019Clo\|mpk1 | prickle homolog 1 (Drosophila) | protein-coding |
| chr1_33578423_33579111 | chr1 | 33578424 | 33579111 | + | -1.03949 | intron (NM_008922, intron 7 of 13) | intron (NM_008922, intron 7 of 13) | 91027 | NM_008922 | 19076 | Mm.27705 | NM_008922 | ENSMUSG00000026134 | **Prim2** | AI323589 | DNA primase, p58 subunit | protein-coding |
| chr5_116029115_116029711 | chr5 | 116029116 | 116029711 | + | -0.642197 | Intergenic | Intergenic | -4985 | NM_031869 | 19079 | Mm.458152 | NM_031869 | ENSMUSG00000029513 | **Prkab1** | 1300015D22Rik\|AU021155\|E430008F22 | protein kinase, AMP-activated, beta 1 non-catalytic subunit | protein-coding |

| | | | | | | | | | | | | | | | | | |
|---|---|---|---|---|---|---|---|---|---|---|---|---|---|---|---|---|---|
| chr5_116029097_116029550 | chr5 | 116029098 | 116029550 | + | -0.678477 | Intergenic | Intergenic | -4896 | NM_031869 | 19079 | Mm.458152 | NM_031869 | ENSMUSG00000029513 | **Prkab1** | 1300015D22Rik\|AU021155\|E430008F22 | protein kinase, AMP-activated, beta 1 non-catalytic subunit | protein-coding |
| chr12_32012234_32012462 | chr12 | 32012235 | 32012462 | + | -0.696906 | intron (NM_011158, intron 2 of 10) | L1M4a1\|LINE\|L1 | 48931 | NM_011158 | 19088 | Mm.25594 | NM_011158 | ENSMUSG00000002997 | **Prkar2b** | AI451071\|AW061005\|PKARIIbeta\|Pkarb2\|RII(beta) | protein kinase, cAMP dependent regulatory, type II beta | protein-coding |
| chr11_108216049_108216452 | chr11 | 108216050 | 108216452 | + | -0.684096 | intron (NM_011101, intron 2 of 16) | intron (NM_011101, intron 2 of 16) | 127637 | NM_011101 | 18750 | Mm.222178 | NM_011101 | ENSMUSG00000050965 | **Prkca** | AI875142\|Pkca | protein kinase C, alpha | protein-coding |
| chr11_108249428_108249998 | chr11 | 108249429 | 108249998 | + | -0.741103 | intron (NM_011101, intron 2 of 16) | intron (NM_011101, intron 2 of 16) | 94175 | NM_011101 | 18750 | Mm.222178 | NM_011101 | ENSMUSG00000050965 | **Prkca** | AI875142\|Pkca | protein kinase C, alpha | protein-coding |
| chr11_108249569_108249881 | chr11 | 108249570 | 108249881 | + | -0.780584 | intron (NM_011101, intron 2 of 16) | intron (NM_011101, intron 2 of 16) | 94163 | NM_011101 | 18750 | Mm.222178 | NM_011101 | ENSMUSG00000050965 | **Prkca** | AI875142\|Pkca | protein kinase C, alpha | protein-coding |
| chr11_108273579_108274081 | chr11 | 108273580 | 108274081 | + | -0.785986 | intron (NM_011101, intron 2 of 16) | intron (NM_011101, intron 2 of 16) | 70058 | NM_011101 | 18750 | Mm.222178 | NM_011101 | ENSMUSG00000050965 | **Prkca** | AI875142\|Pkca | protein kinase C, alpha | protein-coding |
| chr7_122414070_122414666 | chr7 | 122414071 | 122414666 | + | -0.637186 | intron (NM_008855, intron 2 of 16) | intron (NM_008855, intron 2 of 16) | 125243 | NM_008855 | 18751 | Mm.207496 | NM_008855 | ENSMUSG00000052889 | **Prkcb** | A130082F03Rik\|PKC-Beta\|Pkcb\|Prkcb1\|Prkcb2 | protein kinase C, beta | protein-coding |
| chr7_122465804_122466156 | chr7 | 122465805 | 122466156 | + | -0.640747 | intron (NM_008855, intron 5 of 16) | intron (NM_008855, intron 5 of 16) | 176855 | NM_008855 | 18751 | Mm.207496 | NM_008855 | ENSMUSG00000052889 | **Prkcb** | A130082F03Rik\|PKC-Beta\|Pkcb\|Prkcb1\|Prkcb2 | protein kinase C, beta | protein-coding |
| chr7_122410032_122410347 | chr7 | 122410033 | 122410347 | + | -0.727056 | intron (NM_008855, intron 2 of 16) | MTD\|LTR\|ERVL-MaLR | 121065 | NM_008855 | 18751 | Mm.207496 | NM_008855 | ENSMUSG00000052889 | **Prkcb** | A130082F03Rik\|PKC-Beta\|Pkcb\|Prkcb1\|Prkcb2 | protein kinase C, beta | protein-coding |
| chr7_122387862_122388305 | chr7 | 122387863 | 122388305 | + | -0.815267 | intron (NM_008855, intron 2 of 16) | intron (NM_008855, intron 2 of 16) | 98959 | NM_008855 | 18751 | Mm.207496 | NM_008855 | ENSMUSG00000052889 | **Prkcb** | A130082F03Rik\|PKC-Beta\|Pkcb\|Prkcb1\|Prkcb2 | protein kinase C, beta | protein-coding |
| chr7_122287241_122287503 | chr7 | 122287242 | 122287503 | + | -1.25626 | Intergenic | Intergenic | -1753 | NM_008855 | 18751 | Mm.207496 | NM_008855 | ENSMUSG00000052889 | **Prkcb** | A130082F03Rik\|PKC-Beta\|Pkcb\|Prkcb1\|Prkcb2 | protein kinase C, beta | protein-coding |
| chr17_86206168_86207147 | chr17 | 86206169 | 86207147 | + | -0.624778 | intron (NM_011104, intron 1 of 14) | intron (NM_011104, intron 1 of 14) | 38873 | NM_011104 | 18754 | Mm.24614 | NM_011104 | ENSMUSG00000045038 | **Prkce** | 5830406C15Rik\|PKC[e]\|PKCepsilon\|Pkce\|R75156 | protein kinase C, epsilon | protein-coding |
| chr17_86184748_86185256 | chr17 | 86184749 | 86185256 | + | -0.686538 | intron (NM_011104, intron 1 of 14) | intron (NM_011104, intron 1 of 14) | 17217 | NM_011104 | 18754 | Mm.24614 | NM_011104 | ENSMUSG00000045038 | **Prkce** | 5830406C15Rik\|PKC[e]\|PKCepsilon\|Pkce\|R75156 | protein kinase C, epsilon | protein-coding |
| chr12_73720398_73721130 | chr12 | 73720399 | 73721130 | + | -0.605124 | intron (NM_008856, intron 9 of 13) | intron (NM_008856, intron 9 of 13) | 135723 | NM_008856 | 18755 | Mm.341677 | NM_008856 | ENSMUSG00000021108 | **Prkch** | Pkch | protein kinase C, eta | protein-coding |
| chr12_73580900_73581190 | chr12 | 73580901 | 73581190 | + | -0.690999 | Intergenic | L2\|LINE\|L2 | -3996 | NM_008856 | 18755 | Mm.341677 | NM_008856 | ENSMUSG00000021108 | **Prkch** | Pkch | protein kinase C, eta | protein-coding |
| chr12_73609726_73610018 | chr12 | 73609727 | 73610018 | + | -0.741654 | intron (NM_008856, intron 1 of 13) | intron (NM_008856, intron 1 of 13) | 24831 | NM_008856 | 18755 | Mm.341677 | NM_008856 | ENSMUSG00000021108 | **Prkch** | Pkch | protein kinase C, eta | protein-coding |
| chr12_73609779_73609985 | chr12 | 73609780 | 73609985 | + | -0.786202 | intron (NM_008856, intron 1 of 13) | intron (NM_008856, intron 1 of 13) | 24841 | NM_008856 | 18755 | Mm.341677 | NM_008856 | ENSMUSG00000021108 | **Prkch** | Pkch | protein kinase C, eta | protein-coding |
| chr12_73625370_73625959 | chr12 | 73625371 | 73625959 | + | -0.816337 | intron (NM_008856, intron 1 of 13) | Lx9\|LINE\|L1 | 40624 | NM_008856 | 18755 | Mm.341677 | NM_008856 | ENSMUSG00000021108 | **Prkch** | Pkch | protein kinase C, eta | protein-coding |
| chr12_73625085_73626002 | chr12 | 73625086 | 73626002 | + | -0.816929 | intron (NM_008856, intron 1 of 13) | Lx9\|LINE\|L1 | 40503 | NM_008856 | 18755 | Mm.341677 | NM_008856 | ENSMUSG00000021108 | **Prkch** | Pkch | protein kinase C, eta | protein-coding |
| chr12_73679347_73679810 | chr12 | 73679348 | 73679810 | + | -0.961952 | intron (NM_008856, intron 2 of 13) | intron (NM_008856, intron 2 of 13) | 94538 | NM_008856 | 18755 | Mm.341677 | NM_008856 | ENSMUSG00000021108 | **Prkch** | Pkch | protein kinase C, eta | protein-coding |
| chr12_73611768_73612132 | chr12 | 73611769 | 73612132 | + | -1.01929 | intron (NM_008856, intron 1 of 13) | intron (NM_008856, intron 1 of 13) | 26909 | NM_008856 | 18755 | Mm.341677 | NM_008856 | ENSMUSG00000021108 | **Prkch** | Pkch | protein kinase C, eta | protein-coding |
| chr12_73686828_73687663 | chr12 | 73686829 | 73687663 | + | -1.03783 | intron (NM_008856, intron 2 of 13) | intron (NM_008856, intron 2 of 13) | 102205 | NM_008856 | 18755 | Mm.341677 | NM_008856 | ENSMUSG00000021108 | **Prkch** | Pkch | protein kinase C, eta | protein-coding |
| chr2_11135717_11136036 | chr2 | 11135718 | 11136036 | + | -0.681843 | Intergenic | ORR1F\|LTR\|ERVL-MaLR | -36505 | NM_008859 | 18761 | Mm.329993 | NM_008859 | ENSMUSG00000026778 | **Prkcq** | A130035A12Rik\|AW494342\|PKC-0\|PKC-theta\|PKCtheta\|Pkcq | protein kinase C, theta | protein-coding |
| chr2_11218080_11218543 | chr2 | 11218081 | 11218543 | + | -0.738876 | intron (NM_008859, intron 1 of 17) | intron (NM_008859, intron 1 of 17) | 45930 | NM_008859 | 18761 | Mm.329993 | NM_008859 | ENSMUSG00000026778 | **Prkcq** | A130035A12Rik\|AW494342\|PKC-0\|PKC-theta\|PKCtheta\|Pkcq | protein kinase C, theta | protein-coding |
| chr2_11204272_11204634 | chr2 | 11204273 | 11204634 | + | -0.773131 | intron (NM_008859, intron 1 of 17) | intron (NM_008859, intron 1 of 17) | 32071 | NM_008859 | 18761 | Mm.329993 | NM_008859 | ENSMUSG00000026778 | **Prkcq** | A130035A12Rik\|AW494342\|PKC-0\|PKC-theta\|PKCtheta\|Pkcq | protein kinase C, theta | protein-coding |
| chr2_11204012_11204634 | chr2 | 11204013 | 11204634 | + | -0.780462 | intron (NM_008859, intron 1 of 17) | intron (NM_008859, intron 1 of 17) | 31941 | NM_008859 | 18761 | Mm.329993 | NM_008859 | ENSMUSG00000026778 | **Prkcq** | A130035A12Rik\|AW494342\|PKC-0\|PKC-theta\|PKCtheta\|Pkcq | protein kinase C, theta | protein-coding |
| chr2_11239644_11239937 | chr2 | 11239645 | 11239937 | + | -0.791588 | intron (NM_008859, intron 4 of 17) | intron (NM_008859, intron 4 of 17) | 67409 | NM_008859 | 18761 | Mm.329993 | NM_008859 | ENSMUSG00000026778 | **Prkcq** | A130035A12Rik\|AW494342\|PKC-0\|PKC-theta\|PKCtheta\|Pkcq | protein kinase C, theta | protein-coding |
| chr2_11254425_11254900 | chr2 | 11254426 | 11254900 | + | -0.832928 | intron (NM_008859, intron 10 of 17) | intron (NM_008859, intron 10 of 17) | 82281 | NM_008859 | 18761 | Mm.329993 | NM_008859 | ENSMUSG00000026778 | **Prkcq** | A130035A12Rik\|AW494342\|PKC-0\|PKC-theta\|PKCtheta\|Pkcq | protein kinase C, theta | protein-coding |
| chr2_11039950_11040404 | chr2 | 11039951 | 11040404 | + | -0.945661 | Intergenic | MIR\|SINE\|MIR | -132205 | NM_008859 | 18761 | Mm.329993 | NM_008859 | ENSMUSG00000026778 | **Prkcq** | A130035A12Rik\|AW494342\|PKC-0\|PKC-theta\|PKCtheta\|Pkcq | protein kinase C, theta | protein-coding |
| chr2_11040048_11040297 | chr2 | 11040049 | 11040297 | + | -0.950502 | Intergenic | Intergenic | -132209 | NM_008859 | 18761 | Mm.329993 | NM_008859 | ENSMUSG00000026778 | **Prkcq** | A130035A12Rik\|AW494342\|PKC-0\|PKC-theta\|PKCtheta\|Pkcq | protein kinase C, theta | protein-coding |
| chr2_11159647_11160055 | chr2 | 11159648 | 11160055 | + | -1.03431 | Intergenic | Intergenic | -12531 | NM_008859 | 18761 | Mm.329993 | NM_008859 | ENSMUSG00000026778 | **Prkcq** | A130035A12Rik\|AW494342\|PKC-0\|PKC-theta\|PKCtheta\|Pkcq | protein kinase C, theta | protein-coding |
| chr2_11193170_11193574 | chr2 | 11193171 | 11193574 | + | -1.07471 | intron (NM_008859, intron 1 of 17) | intron (NM_008859, intron 1 of 17) | 20990 | NM_008859 | 18761 | Mm.329993 | NM_008859 | ENSMUSG00000026778 | **Prkcq** | A130035A12Rik\|AW494342\|PKC-0\|PKC-theta\|PKCtheta\|Pkcq | protein kinase C, theta | protein-coding |
| chr2_11118492_11119066 | chr2 | 11118493 | 11119066 | + | -1.12086 | Intergenic | MLT1F-int\|LTR\|ERVL-MaLR | -53603 | NM_008859 | 18761 | Mm.329993 | NM_008859 | ENSMUSG00000026778 | **Prkcq** | A130035A12Rik\|AW494342\|PKC-0\|PKC-theta\|PKCtheta\|Pkcq | protein kinase C, theta | protein-coding |
| chr2_11205789_11206157 | chr2 | 11205790 | 11206157 | + | -1.19771 | intron (NM_008859, intron 1 of 17) | intron (NM_008859, intron 1 of 17) | 33591 | NM_008859 | 18761 | Mm.329993 | NM_008859 | ENSMUSG00000026778 | **Prkcq** | A130035A12Rik\|AW494342\|PKC-0\|PKC-theta\|PKCtheta\|Pkcq | protein kinase C, theta | protein-coding |
| chr4_155354078_155354446 | chr4 | 155354079 | 155354446 | + | -0.67826 | intron (NM_008860, intron 4 of 17) | intron (NM_008860, intron 4 of 17) | 7166 | NM_008860 | 18762 | Mm.28561 | NM_008860 | ENSMUSG00000029053 | **Prkcz** | AI098070\|C80388\|Pkcz\|R74924\|aPKCzeta\|nPKC-zeta\|zetaPKC | protein kinase C, zeta | protein-coding |
| chr4_155353914_155354442 | chr4 | 155353915 | 155354442 | + | -0.687531 | intron (NM_008860, intron 4 of 17) | intron (NM_008860, intron 4 of 17) | 7250 | NM_008860 | 18762 | Mm.28561 | NM_008860 | ENSMUSG00000029053 | **Prkcz** | AI098070\|C80388\|Pkcz\|R74924\|aPKCzeta\|nPKC-zeta\|zetaPKC | protein kinase C, zeta | protein-coding |
| chr4_155352372_155353001 | chr4 | 155352373 | 155353001 | + | -0.834297 | intron (NM_008860, intron 4 of 17) | intron (NM_008860, intron 4 of 17) | -6898 | NM_001039079 | 18762 | Mm.28561 | NM_008860 | ENSMUSG00000029053 | **Prkcz** | AI098070\|C80388\|Pkcz\|R74924\|aPKCzeta\|nPKC-zeta\|zetaPKC | protein kinase C, zeta | protein-coding |
| chr5_136186729_136187070 | chr5 | 136186730 | 136187070 | + | -0.618604 | intron (NM_025774, intron 5 of 5) | intron (NM_025774, intron 5 of 5) | 12054 | NM_025774 | 66801 | Mm.425295 | NM_025774 | | **Prkrip1** | 8430424D23Rik\|C114 | Prkr interacting protein 1 (IL11 inducible) | protein-coding |
| chr7_98717456_98717927 | chr7 | 98717457 | 98717927 | + | -0.938142 | 3' UTR (NM_028410, exon 5 of 5) | 3' UTR (NM_028410, exon 5 of 5) | 14329 | NM_028410 | 72981 | Mm.4428 | NM_028410 | ENSMUSG00000030753 | **Prkrir** | 2900052B10Rik\|Dap4\|Rpkrir | protein-kinase, interferon-inducible double stranded RNA dependent inhibitor, repressor of (P58 repressor) | protein-coding |
| chr13_28417363_28418009 | chr13 | 28417364 | 28418009 | + | -0.701067 | Intergenic | Intergenic | 275202 | NM_023746 | 28078 | Mm.155058 | NM_023746 | ENSMUSG00000017064 | **Prl5a1** | 1600013P04Rik\|D13Wsu14e\|PLP-L\|Prlpl | prolactin family 5, subfamily a, member 1 | protein-coding |
| chr15_10035580_10035905 | chr15 | 10035581 | 10035905 | + | -0.725215 | Intergenic | Intergenic | -141495 | NM_011169 | 19116 | Mm.10516 | NM_011169 | ENSMUSG00000005268 | **Prlr** | AI987712\|Pr-1\|Pr-3\|Prlr-rs1 | prolactin receptor | protein-coding |
| chr15_10236527_10237181 | chr15 | 10236528 | 10237181 | + | -0.807838 | intron (NM_001253781, intron 1 of 9) | intron (NM_001253781, intron 1 of 9) | 12948 | NM_001253781 | 19116 | Mm.10516 | NM_011169 | ENSMUSG00000005268 | **Prlr** | AI987712\|Pr-1\|Pr-3\|Prlr-rs1 | prolactin receptor | protein-coding |
| chr15_10236720_10237251 | chr15 | 10236721 | 10237251 | + | -0.812662 | intron (NM_001253781, intron 1 of 9) | intron (NM_001253781, intron 1 of 9) | 13080 | NM_001253781 | 19116 | Mm.10516 | NM_011169 | ENSMUSG00000005268 | **Prlr** | AI987712\|Pr-1\|Pr-3\|Prlr-rs1 | prolactin receptor | protein-coding |
| chr15_10235978_10236530 | chr15 | 10235979 | 10236530 | + | -0.911037 | intron (NM_001253781, intron 1 of 9) | intron (NM_001253781, intron 1 of 9) | 12348 | NM_001253781 | 19116 | Mm.10516 | NM_011169 | ENSMUSG00000005268 | **Prlr** | AI987712\|Pr-1\|Pr-3\|Prlr-rs1 | prolactin receptor | protein-coding |
| chr14_54518593_54519057 | chr14 | 54518594 | 54519057 | + | -0.801603 | Intergenic | Intergenic | -1355 | NM_013768 | 27374 | Mm.196585 | NM_013768 | ENSMUSG00000023110 | **Prmt5** | Jbp1\|Skb1 | protein arginine N-methyltransferase 5 | protein-coding |
| chr11_78215544_78216267 | chr11 | 78215545 | 78216267 | + | -1.03257 | intron (NM_009297, intron 28 of 36) | intron (NM_009297, intron 28 of 36) | 22514 | NM_001045516 | 216974 | Mm.50974 | NM_001045516 | ENSMUSG00000044122 | **Proca1** | 4933404M19Rik | protein interacting with cyclin A1 | protein-coding |
| chr6_99785145_99785611 | chr6 | 99785146 | 99785611 | + | -0.691227 | Intergenic | Intergenic | -58986 | NM_001170419 | 50501 | Mm.87365 | NM_015768 | ENSMUSG00000030069 | **Prok2** | Bv8\|PK2\|Prok1 | prokineticin 2 | protein-coding |
| chr6_99784941_99785653 | chr6 | 99784942 | 99785653 | + | -0.691788 | Intergenic | Intergenic | -58905 | NM_001170419 | 50501 | Mm.87365 | NM_015768 | ENSMUSG00000030069 | **Prok2** | Bv8\|PK2\|Prok1 | prokineticin 2 | protein-coding |
| chr6_99785278_99785561 | chr6 | 99785279 | 99785561 | + | -0.786193 | Intergenic | Intergenic | -59028 | NM_001170419 | 50501 | Mm.87365 | NM_015768 | ENSMUSG00000030069 | **Prok2** | Bv8\|PK2\|Prok1 | prokineticin 2 | protein-coding |
| chr16_62940465_62940967 | chr16 | 62940466 | 62940967 | + | -0.693151 | Intergenic | Intergenic | 86382 | NM_011173 | 19128 | Mm.127156 | NM_011173 | ENSMUSG00000022912 | **Pros1** | AW214361 | protein S (alpha) | protein-coding |
| chr16_62895609_62896463 | chr16 | 62895610 | 62896463 | + | -0.726067 | intron (NM_011173, intron 3 of 14) | intron (NM_011173, intron 3 of 14) | 41702 | NM_011173 | 19128 | Mm.127156 | NM_011173 | ENSMUSG00000022912 | **Pros1** | AW214361 | protein S (alpha) | protein-coding |
| chr16_62895597_62896224 | chr16 | 62895598 | 62896224 | + | -0.788247 | intron (NM_011173, intron 3 of 14) | intron (NM_011173, intron 3 of 14) | 41577 | NM_011173 | 19128 | Mm.127156 | NM_011173 | ENSMUSG00000022912 | **Pros1** | AW214361 | protein S (alpha) | protein-coding |

| | | | | | | | | | | | | | | | | | |
|---|---|---|---|---|---|---|---|---|---|---|---|---|---|---|---|---|---|
| chr2_53051200_53051906 | chr2 | 53051201 | 53051906 | + | -0.730607 | intron (NM_172409, intron 5 of 26) | intron (NM_172409, intron 5 of 26) | 139634 | NM_018785 | 56194 | Mm.257474 | NM_018785 | ENSMUSG00000061136 | **Prpf40a** | 2810012K09Rik\|FBP11\|Fnbp11\|Fnbp3 | PRP40 pre-mRNA processing factor 40 homolog A (yeast) | protein-coding |
| chr2_53049468_53049726 | chr2 | 53049469 | 53049726 | + | -0.931607 | intron (NM_172409, intron 4 of 26) | intron (NM_172409, intron 4 of 26) | 141590 | NM_018785 | 56194 | Mm.257474 | NM_018785 | ENSMUSG00000061136 | **Prpf40a** | 2810012K09Rik\|FBP11\|Fnbp11\|Fnbp3 | PRP40 pre-mRNA processing factor 40 homolog A (yeast) | protein-coding |
| chr15_102458220_102458693 | chr15 | 102458221 | 102458693 | + | -0.781047 | promoter-TSS (NM_001170911) | promoter-TSS (NM_001170911) | -713 | NM_001170911 | 66151 | Mm.393955 | NM_025385 | ENSMUSG00000023048 | **Prr13** | 1110020C13Rik\|2010324E22Rik | proline rich 13 | protein-coding |
| chr18_57362631_57362868 | chr18 | 57362632 | 57362868 | + | -0.622455 | intron (NM_028447, intron 2 of 8) | intron (NM_028447, intron 2 of 8) | 8017 | NM_028447 | 73137 | Mm.29889 | NM_028447 | ENSMUSG00000024594 | **Prrc1** | 1190002C06Rik\|2310058D16Rik\|3110038B19Rik\|9430085A19Rik | proline-rich coiled-coil 1 | protein-coding |
| chr2_32167391_32168201 | chr2 | 32167392 | 32168201 | + | -0.609798 | intron (NM_001159634, intron 1 of 31) | intron (NM_001159634, intron 1 of 31) | 16648 | NM_001159634 | 227723 | Mm.25504 | NM_172661 | ENSMUSG00000039262 | **Prrc2b** | 5830434P21Rik\|AI173903\|Bat2l\|Bat2l1\|D430039P21\|mKIAA0515 | proline-rich coiled-coil 2B | protein-coding |
| chr2_32167829_32168216 | chr2 | 32167830 | 32168216 | + | -0.764664 | intron (NM_001159634, intron 1 of 31) | intron (NM_001159634, intron 1 of 31) | 16875 | NM_001159634 | 227723 | Mm.25504 | NM_172661 | ENSMUSG00000039262 | **Prrc2b** | 5830434P21Rik\|AI173903\|Bat2l\|Bat2l1\|D430039P21\|mKIAA0515 | proline-rich coiled-coil 2B | protein-coding |
| chr1_162752027_162752462 | chr1 | 162752028 | 162752462 | + | -0.716026 | Intergenic | MER57E1\|LTR\|ERV1 | -11689 | NM_001081290 | 226562 | Mm.245446 | NM_001081290 | ENSMUSG00000040225 | **Prrc2c** | 1810043M20Rik\|9630039I18Rik\|A630006J20\|Bat2d\|Bat2d1\|Bat2l2\|E130112L15Rik\|Prrc3\|mKIAA1096 | proline-rich coiled-coil 2C | protein-coding |
| chr1_162727999_162728447 | chr1 | 162728000 | 162728447 | + | -1.31901 | intron (NM_001081290, intron 1 of 34) | RMER12B\|LTR\|ERVK | 12333 | NM_001081290 | 226562 | Mm.245446 | NM_001081290 | ENSMUSG00000040225 | **Prrc2c** | 1810043M20Rik\|9630039I18Rik\|A630006J20\|Bat2d\|Bat2d1\|Bat2l2\|E130112L15Rik\|Prrc3\|mKIAA1096 | proline-rich coiled-coil 2C | protein-coding |
| chr3_123444598_123444772 | chr3 | 123444599 | 123444772 | + | -0.653128 | Intergenic | Intergenic | -2228 | NM_008939 | 19142 | Mm.9431 | NM_008939 | ENSMUSG00000027978 | **Prss12** | Bssp-3\|motopsin | protease, serine 12 neurotrypsin (motopsin) | protein-coding |
| chr3_123444561_123444819 | chr3 | 123444562 | 123444819 | + | -0.710472 | Intergenic | Intergenic | -2223 | NM_008939 | 19142 | Mm.9431 | NM_008939 | ENSMUSG00000027978 | **Prss12** | Bssp-3\|motopsin | protease, serine 12 neurotrypsin (motopsin) | protein-coding |
| chr3_123452981_123453301 | chr3 | 123452982 | 123453301 | + | -0.730692 | intron (NM_008939, intron 1 of 10) | intron (NM_008939, intron 1 of 10) | 6228 | NM_008939 | 19142 | Mm.9431 | NM_008939 | ENSMUSG00000027978 | **Prss12** | Bssp-3\|motopsin | protease, serine 12 neurotrypsin (motopsin) | protein-coding |
| chr3_123452964_123453674 | chr3 | 123452965 | 123453674 | + | -0.813372 | intron (NM_008939, intron 1 of 10) | intron (NM_008939, intron 1 of 10) | 6406 | NM_008939 | 19142 | Mm.9431 | NM_008939 | ENSMUSG00000027978 | **Prss12** | Bssp-3\|motopsin | protease, serine 12 neurotrypsin (motopsin) | protein-coding |
| chr3_123448180_123448428 | chr3 | 123448181 | 123448428 | + | -0.855886 | intron (NM_008939, intron 1 of 10) | intron (NM_008939, intron 1 of 10) | 1391 | NM_008939 | 19142 | Mm.9431 | NM_008939 | ENSMUSG00000027978 | **Prss12** | Bssp-3\|motopsin | protease, serine 12 neurotrypsin (motopsin) | protein-coding |
| chr6_41553352_41553680 | chr6 | 41553353 | 41553680 | + | -0.722136 | Intergenic | L3b\|LINE\|CR1 | 31740 | NM_009430 | 22072 | Mm.276926 | NM_009430 | ENSMUSG00000057163 | **Prss2** | TRY8\|TRYP\|Ta\|Tesp4\|Try2 | protease, serine 2 | protein-coding |
| chr17_23858541_23859170 | chr17 | 23858542 | 23859170 | + | -1.03666 | intron (NM_027220, intron 4 of 4) | intron (NM_027220, intron 4 of 4) | 5084 | NM_027220 | 69814 | Mm.244143 | NM_027220 | ENSMUSG00000048992 | **Prss32** | 2010001P08Rik\|mT5\|tryptase-5 | protease, serine 32 | protein-coding |
| chr1_34484376_34484821 | chr1 | 34484377 | 34484821 | + | -0.627843 | Intergenic | MTE2b\|LTR\|ERVL-MaLR | -13831 | NM_009355 | 21755 | Mm.10725 | NM_009355 | ENSMUSG00000026125 | **Prss39** | Tesp1 | protease, serine 39 | protein-coding |
| chr9_110825765_110826236 | chr9 | 110825766 | 110826236 | + | -0.955067 | promoter-TSS (NM_199471) | promoter-TSS (NM_199471) | -689 | NM_199471 | 272643 | Mm.458525 | NM_199471 | ENSMUSG00000058398 | **Prss43** | Gm661\|Tessp3 | protease, serine 43 | protein-coding |
| chr14_64212506_64213351 | chr14 | 64212507 | 64213351 | + | -0.668254 | intron (NM_001253716, intron 4 of 5) | intron (NM_001253716, intron 4 of 5) | 108606 | NM_028525 | 73382 | Mm.481055 | NM_028525 | ENSMUSG00000021966 | **Prss52** | 1700049K14Rik\|Tesp3 | protease, serine 52 | protein-coding |
| chr5_36141192_36141462 | chr5 | 36141193 | 36141462 | + | -0.66645 | intron (NM_030889, intron 3 of 26) | intron (NM_030889, intron 3 of 26) | -62694 | NM_175249 | 76943 | Mm.44242 | NM_175249 | ENSMUSG00000043430 | **Psapl1** | 2310020A21Rik | prosaposin-like 1 | protein-coding |
| chr18_35910087_35910512 | chr18 | 35910088 | 35910512 | + | -0.667501 | Intergenic | Intergenic | -54530 | NM_028707 | 74002 | Mm.196944 | NM_028707 | ENSMUSG00000024347 | **Psd2** | 6330404E20Rik\|AW125584\|EFA6C | pleckstrin and Sec7 domain containing 2 | protein-coding |
| chr1_180226227_180226742 | chr1 | 180226228 | 180226742 | + | -0.682689 | TTS (NM_001128605) | TTS (NM_001128605) | 19465 | NM_011183 | 19165 | Mm.330850 | NM_011183 | ENSMUSG00000010609 | **Psen2** | ALG-3\|Ad4h\|Alg3\|PS-2\|PS2\|Psnl2\|STM2 | presenilin 2 | protein-coding |
| chr8_105900970_105901335 | chr8 | 105900971 | 105901335 | + | -0.768582 | intron (NM_173432, intron 1 of 2) | intron (NM_173432, intron 1 of 2) | 679 | NM_173432 | 244631 | Mm.27627 | NM_173432 | ENSMUSG00000048310 | **Pskh1** | AW539964\|E130013P03Rik\|b2b1230Clo | protein serine kinase H1 | protein-coding |
| chr8_105901030_105901289 | chr8 | 105901031 | 105901289 | + | -0.788165 | intron (NM_173432, intron 1 of 2) | intron (NM_173432, intron 1 of 2) | 686 | NM_173432 | 244631 | Mm.27627 | NM_173432 | ENSMUSG00000048310 | **Pskh1** | AW539964\|E130013P03Rik\|b2b1230Clo | protein serine kinase H1 | protein-coding |
| chr7_114301400_114301895 | chr7 | 114301401 | 114301895 | + | -0.672059 | Intergenic | L4_A_Mam\|LINE\|RTE-X | -25532 | NM_011965 | 26440 | Mm.121265 | NM_011965 | ENSMUSG00000030751 | **Psma1** | C2\|HC2\|Pros-30\|alpha-type | proteasome (prosome, macropain) subunit, alpha type 1 | protein-coding |
| chr7_114317214_114317832 | chr7 | 114317215 | 114317832 | + | -0.796813 | intron (NR_029437, intron 3 of 3) | intron (NR_029437, intron 3 of 3) | -41407 | NM_011965 | 26440 | Mm.121265 | NM_011965 | ENSMUSG00000030751 | **Psma1** | C2\|HC2\|Pros-30\|alpha-type | proteasome (prosome, macropain) subunit, alpha type 1 | protein-coding |
| chr7_114284499_114284897 | chr7 | 114284500 | 114284897 | + | -1.04258 | Intergenic | RMER1C\|Other\|Other | -8582 | NM_011965 | 26440 | Mm.121265 | NM_011965 | ENSMUSG00000030751 | **Psma1** | C2\|HC2\|Pros-30\|alpha-type | proteasome (prosome, macropain) subunit, alpha type 1 | protein-coding |
| chr11_97711239_97711587 | chr11 | 97711240 | 97711587 | + | -0.728824 | intron (NM_011971, intron 4 of 5) | intron (NM_011971, intron 4 of 5) | 7979 | NM_011971 | 26446 | Mm.21874 | NM_011971 | ENSMUSG00000069744 | **Psmb3** | AL033320\|C10-II | proteasome (prosome, macropain) subunit, beta type 3 | protein-coding |
| chr17_34170647_34171511 | chr17 | 34170648 | 34171511 | + | -0.624406 | Intergenic | Intergenic | 16251 | NM_013585 | 16912 | Mm.390983 | NM_013585 | | **Psmb9** | Lmp-2\|Lmp2 | proteasome (prosome, macropain) subunit, beta type 9 (large multifunctional peptidase 2) | protein-coding |
| chr17_34171166_34171509 | chr17 | 34171167 | 34171509 | + | -0.751343 | Intergenic | MTD\|LTR\|ERVL-MaLR | 15992 | NM_013585 | 16912 | Mm.390983 | NM_013585 | | **Psmb9** | Lmp-2\|Lmp2 | proteasome (prosome, macropain) subunit, beta type 9 (large multifunctional peptidase 2) | protein-coding |
| chr14_14067611_14069060 | chr14 | 14067612 | 14069060 | + | -0.781907 | intron (NM_139227, intron 4 of 11) | intron (NM_139227, intron 4 of 11) | 52568 | NM_025550 | 66413 | Mm.27591 | NM_025550 | ENSMUSG00000021737 | **Psmd6** | 2400006A19Rik | proteasome (prosome, macropain) 26S subunit, non-ATPase, 6 | protein-coding |
| chr16_95999178_95999724 | chr16 | 95999179 | 95999724 | + | -0.665301 | intron (NM_145125, intron 41 of 41) | intron (NM_145125, intron 41 of 41) | -8548 | NM_019537 | 56088 | Mm.396248 | NM_019537 | ENSMUSG00000022913 | **Psmg1** | AW552102\|Dscr2 | proteasome (prosome, macropain) assembly chaperone 1 | protein-coding |
| chr9_56075914_56076419 | chr9 | 56075915 | 56076419 | + | -0.678902 | Intergenic | L2b\|LINE\|L2 | -13809 | NM_011193 | 19200 | Mm.2534 | NM_011193 | ENSMUSG00000032322 | **Pstpip1** | CD2BP1\|def-2 | proline-serine-threonine phosphatase-interacting protein 1 | protein-coding |
| chr9_56075826_56076535 | chr9 | 56075827 | 56076535 | + | -0.682521 | Intergenic | L2b\|LINE\|L2 | -13795 | NM_011193 | 19200 | Mm.2534 | NM_011193 | ENSMUSG00000032322 | **Pstpip1** | CD2BP1\|def-2 | proline-serine-threonine phosphatase-interacting protein 1 | protein-coding |
| chr9_56075957_56076250 | chr9 | 56075958 | 56076250 | + | -0.762273 | Intergenic | Intergenic | -13872 | NM_011193 | 19200 | Mm.2534 | NM_011193 | ENSMUSG00000032322 | **Pstpip1** | CD2BP1\|def-2 | proline-serine-threonine phosphatase-interacting protein 1 | protein-coding |
| chr9_56100158_56100343 | chr9 | 56100159 | 56100343 | + | -0.820743 | intron (NM_011193, intron 1 of 14) | ORR1F\|LTR\|ERVL-MaLR | 10275 | NM_011193 | 19200 | Mm.2534 | NM_011193 | ENSMUSG00000032322 | **Pstpip1** | CD2BP1\|def-2 | proline-serine-threonine phosphatase-interacting protein 1 | protein-coding |
| chr9_56100087_56100477 | chr9 | 56100088 | 56100477 | + | -0.828694 | intron (NM_011193, intron 1 of 14) | ORR1F\|LTR\|ERVL-MaLR | 10306 | NM_011193 | 19200 | Mm.2534 | NM_011193 | ENSMUSG00000032322 | **Pstpip1** | CD2BP1\|def-2 | proline-serine-threonine phosphatase-interacting protein 1 | protein-coding |
| chr19_32890795_32891308 | chr19 | 32890796 | 32891308 | + | -0.752314 | Intergenic | Intergenic | 133475 | NM_008960 | 19211 | Mm.245395 | NM_008960 | ENSMUSG00000013663 | **Pten** | 2310035O07Rik\|A130070J02Rik\|AI463227\|B430203M17Rik\|MMAC1\|TEP1 | phosphatase and tensin homolog | protein-coding |
| chr19_32905973_32906531 | chr19 | 32905974 | 32906531 | + | -0.7676 | Intergenic | UCON26\|Unknown\|Unknown | 148675 | NM_008960 | 19211 | Mm.245395 | NM_008960 | ENSMUSG00000013663 | **Pten** | 2310035O07Rik\|A130070J02Rik\|AI463227\|B430203M17Rik\|MMAC1\|TEP1 | phosphatase and tensin homolog | protein-coding |
| chr19_32963941_32964385 | chr19 | 32963942 | 32964385 | + | -0.903595 | Intergenic | RLTR9A3\|LTR\|ERVK | 206586 | NM_008960 | 19211 | Mm.245395 | NM_008960 | ENSMUSG00000013663 | **Pten** | 2310035O07Rik\|A130070J02Rik\|AI463227\|B430203M17Rik\|MMAC1\|TEP1 | phosphatase and tensin homolog | protein-coding |
| chr3_157616631_157617272 | chr3 | 157616632 | 157617272 | + | -0.629838 | intron (NM_011196, intron 2 of 2) | intron (NM_011196, intron 2 of 2) | 50060 | NM_011196 | 19218 | Mm.30424 | NM_011196 | ENSMUSG00000040016 | **Ptger3** | EP3\|Pgerep3\|Ptgerep3 | prostaglandin E receptor 3 (subtype EP3) | protein-coding |
| chr2_30896841_30897767 | chr2 | 30896842 | 30897767 | + | -0.597364 | intron (NM_022415, intron 2 of 2) | intron (NM_022415, intron 2 of 2) | 5993 | NM_022415 | 64292 | Mm.28768 | NM_022415 | ENSMUSG00000050737 | **Ptges** | 2410099E23Rik\|D2Ertd369e\|Pges\|mPGES\|mPGES-1 | prostaglandin E synthase | protein-coding |
| chr3_101072347_101073041 | chr3 | 101072348 | 101073041 | + | -0.734556 | intron (NM_011197, intron 4 of 8) | intron (NM_011197, intron 4 of 8) | 37472 | NM_011197 | 19221 | Mm.24807 | NM_011197 | ENSMUSG00000027864 | **Ptgfrn** | 4833445A08Rik\|AU042434\|CD9P-1\|FPRP\|Trim45 | prostaglandin F2 receptor negative regulator | protein-coding |
| chr3_101150040_101150359 | chr3 | 101150041 | 101150359 | + | -0.915196 | Intergenic | MLT1J\|LTR\|ERVL-MaLR | -40034 | NM_011197 | 19221 | Mm.24807 | NM_011197 | ENSMUSG00000027864 | **Ptgfrn** | 4833445A08Rik\|AU042434\|CD9P-1\|FPRP\|Trim45 | prostaglandin F2 receptor negative regulator | protein-coding |
| chr9_110734115_110734454 | chr9 | 110734116 | 110734454 | + | -0.80916 | intron (NM_001083935, intron 2 of 14) | intron (NM_001083935, intron 2 of 14) | 8325 | NM_001083936 | 19228 | Mm.3542 | NM_011199 | ENSMUSG00000032492 | **Pth1r** | PPR\|Pthr\|Pthr1 | parathyroid hormone 1 receptor | protein-coding |
| chr14_66218719_66219154 | chr14 | 66218720 | 66219154 | + | -0.72946 | intron (NM_001162365, intron 1 of 30) | intron (NM_001162365, intron 1 of 30) | -5385 | NM_001162366 | 19229 | Mm.21613 | NM_172498 | ENSMUSG00000059456 | **Ptk2b** | CADTK\|CAKB\|CAKbeta\|E430023O05Rik\|FADK2\|FAK2\|PYK2\|Raftk | PTK2 protein tyrosine kinase 2 beta | protein-coding |
| chr14_66218830_66219145 | chr14 | 66218831 | 66219145 | + | -0.754651 | intron (NM_001162365, intron 1 of 30) | intron (NM_001162365, intron 1 of 30) | -5436 | NM_001162366 | 19229 | Mm.21613 | NM_172498 | ENSMUSG00000059456 | **Ptk2b** | CADTK\|CAKB\|CAKbeta\|E430023O05Rik\|FADK2\|FAK2\|PYK2\|Raftk | PTK2 protein tyrosine kinase 2 beta | protein-coding |

| | | | | | | | | | | | | | | | | | |
|---|---|---|---|---|---|---|---|---|---|---|---|---|---|---|---|---|---|
| chr14_66232175_66232628 | chr14 | 66232176 | 66232628 | + | -0.782924 | intron (NM_001162365, intron 1 of 30) | intron (NM_001162365, intron 1 of 30) | -18850 | NM_001162366 | 19229 | Mm.21613 | NM_172498 | ENSMUSG00000059456 | **Ptk2b** | CADTK\|CAKB\|CAKbeta\|E430023O05Rik\|FADK2\|FAK2\|PYK2\|Raftk | PTK2 protein tyrosine kinase 2 beta | protein-coding |
| chr14_66274608_66275024 | chr14 | 66274609 | 66275024 | + | -0.823103 | intron (NM_001162365, intron 1 of 30) | intron (NM_001162365, intron 1 of 30) | 6236 | NM_172498 | 19229 | Mm.21613 | NM_172498 | ENSMUSG00000059456 | **Ptk2b** | CADTK\|CAKB\|CAKbeta\|E430023O05Rik\|FADK2\|FAK2\|PYK2\|Raftk | PTK2 protein tyrosine kinase 2 beta | protein-coding |
| chr14_66188935_66189624 | chr14 | 66188936 | 66189624 | + | -1.32817 | intron (NM_001162365, intron 4 of 30) | intron (NM_001162365, intron 4 of 30) | 24272 | NM_001162366 | 19229 | Mm.21613 | NM_172498 | ENSMUSG00000059456 | **Ptk2b** | CADTK\|CAKB\|CAKbeta\|E430023O05Rik\|FADK2\|FAK2\|PYK2\|Raftk | PTK2 protein tyrosine kinase 2 beta | protein-coding |
| chr14_66188946_66189373 | chr14 | 66188947 | 66189373 | + | -1.41914 | intron (NM_001162365, intron 4 of 30) | MER63A\|DNA\|hAT-Blackjack | 24392 | NM_001162366 | 19229 | Mm.21613 | NM_172498 | ENSMUSG00000059456 | **Ptk2b** | CADTK\|CAKB\|CAKbeta\|E430023O05Rik\|FADK2\|FAK2\|PYK2\|Raftk | PTK2 protein tyrosine kinase 2 beta | protein-coding |
| chr2_167840002_167840719 | chr2 | 167840003 | 167840719 | + | -0.736235 | Intergenic | Intergenic | -91966 | NM_011201 | 19246 | Mm.277916 | NM_011201 | ENSMUSG00000027540 | **Ptpn1** | PTP-1B\|PTP-HA2\|PTP1B | protein tyrosine phosphatase, non-receptor type 1 | protein-coding |
| chr2_167840405_167840720 | chr2 | 167840406 | 167840720 | + | -0.88327 | Intergenic | Intergenic | -91764 | NM_011201 | 19246 | Mm.277916 | NM_011201 | ENSMUSG00000027540 | **Ptpn1** | PTP-1B\|PTP-HA2\|PTP1B | protein tyrosine phosphatase, non-receptor type 1 | protein-coding |
| chr18_67733552_67733883 | chr18 | 67733553 | 67733883 | + | -0.744124 | Intergenic | Intergenic | -9097 | NM_008977 | 19255 | Mm.260433 | NM_008977 | ENSMUSG00000024539 | **Ptpn2** | AI325124\|Ptpt\|TC-PTP | protein tyrosine phosphatase, non-receptor type 2 | protein-coding |
| chr18_67724959_67725301 | chr18 | 67724960 | 67725301 | + | -0.984386 | promoter-TSS (NM_008977) | promoter-TSS (NM_008977) | -509 | NM_008977 | 19255 | Mm.260433 | NM_008977 | ENSMUSG00000024539 | **Ptpn2** | AI325124\|Ptpt\|TC-PTP | protein tyrosine phosphatase, non-receptor type 2 | protein-coding |
| chr3_103885355_103886364 | chr3 | 103885356 | 103886364 | + | -0.685486 | non-coding (NR_104070, exon 15 of 23) | non-coding (NR_104070, exon 15 of 23) | 25583 | NM_008979 | 19260 | Mm.395 | NM_008979 | ENSMUSG00000027843 | **Ptpn22** | 70zpep\|PEP\|Ptpn8 | protein tyrosine phosphatase, non-receptor type 22 (lymphoid) | protein-coding |
| chr3_103867525_103868463 | chr3 | 103867526 | 103868463 | + | -0.900717 | intron (NM_008979, intron 1 of 20) | intron (NM_008979, intron 1 of 20) | 7717 | NM_008979 | 19260 | Mm.395 | NM_008979 | ENSMUSG00000027843 | **Ptpn22** | 70zpep\|PEP\|Ptpn8 | protein tyrosine phosphatase, non-receptor type 22 (lymphoid) | protein-coding |
| chr3_103867769_103868458 | chr3 | 103867770 | 103868458 | + | -0.902963 | intron (NM_008979, intron 1 of 20) | intron (NM_008979, intron 1 of 20) | 7837 | NM_008979 | 19260 | Mm.395 | NM_008979 | ENSMUSG00000027843 | **Ptpn22** | 70zpep\|PEP\|Ptpn8 | protein tyrosine phosphatase, non-receptor type 22 (lymphoid) | protein-coding |
| chr3_103885929_103886381 | chr3 | 103885930 | 103886381 | + | -0.917478 | non-coding (NR_104070, exon 15 of 23) | non-coding (NR_104070, exon 15 of 23) | 25878 | NM_008979 | 19260 | Mm.395 | NM_008979 | ENSMUSG00000027843 | **Ptpn22** | 70zpep\|PEP\|Ptpn8 | protein tyrosine phosphatase, non-receptor type 22 (lymphoid) | protein-coding |
| chr2_90583590_90583995 | chr2 | 90583591 | 90583995 | + | -0.671518 | Intergenic | Intergenic | -3146 | NM_008982 | 19271 | Mm.330393 | NM_008982 | ENSMUSG00000025314 | **Ptprj** | AI450271\|BET\|Byp\|CD148\|DEP-1\|PTPbeta2\|Ptpb2\|RPTPJ\|Scc-1\|Scc1 | protein tyrosine phosphatase, receptor type, J | protein-coding |
| chr2_90583590_90583850 | chr2 | 90583591 | 90583850 | + | -0.731176 | Intergenic | Intergenic | -3073 | NM_008982 | 19271 | Mm.330393 | NM_008982 | ENSMUSG00000025314 | **Ptprj** | AI450271\|BET\|Byp\|CD148\|DEP-1\|PTPbeta2\|Ptpb2\|RPTPJ\|Scc-1\|Scc1 | protein tyrosine phosphatase, receptor type, J | protein-coding |
| chr12_8662162_8662634 | chr12 | 8662163 | 8662634 | + | -1.05048 | Intergenic | ORR1E\|LTR\|ERVL-MaLR | -11736 | NM_001160221 | 80913 | Mm.341243 | NM_030723 | ENSMUSG00000020594 | **Pum2** | 5730503J23Rik\|Pumm2\|mKIAA0235 | pumilio RNA-binding family member 2 | protein-coding |
| chr15_94478602_94479141 | chr15 | 94478603 | 94479141 | + | -0.662276 | Intergenic | Intergenic | 64635 | NM_172437 | 78895 | Mm.34344 | NM_172437 | ENSMUSG00000033356 | **Pus7l** | 3000003F02Rik | pseudouridylate synthase 7 homolog (S. cerevisiae)-like | protein-coding |
| chr15_62092389_62093205 | chr15 | 62092390 | 62093205 | + | -0.64055 | intron (NR_003368, intron 1 of 8) | intron (NR_003368, intron 1 of 8) | 54810 | NR_003368 | 19296 | Mm.4608 | NM_011222 | ENSMUSG00000097039 | **Pvt1** | Gt(pU21)84Imeg\|Mis-1\|Mlvi-1\|Pvt-1 | plasmacytoma variant translocation 1 | ncRNA |
| chr9_96953400_96954090 | chr9 | 96953401 | 96954090 | + | -0.738846 | intron (NM_145134, intron 2 of 2) | MIR\|SINE\|MIR | -64271 | NM_001289645 | 235534 | Mm.233031 | NM_153420 | ENSMUSG00000043587 | **Pxylp1** | 9430094M07Rik\|Acpl2\|BB177120\|C130099A20Rik | 2-phosphoxylose phosphatase 1 | protein-coding |
| chr9_96935576_96936328 | chr9 | 96935577 | 96936328 | + | -0.809993 | Intergenic | L1_Mur3\|LINE\|L1 | -46478 | NM_001289645 | 235534 | Mm.233031 | NM_153420 | ENSMUSG00000043587 | **Pxylp1** | 9430094M07Rik\|Acpl2\|BB177120\|C130099A20Rik | 2-phosphoxylose phosphatase 1 | protein-coding |
| chr9_96953447_96953877 | chr9 | 96953448 | 96953877 | + | -0.841038 | intron (NM_145134, intron 2 of 2) | intron (NM_145134, intron 2 of 2) | -64188 | NM_001289645 | 235534 | Mm.233031 | NM_153420 | ENSMUSG00000043587 | **Pxylp1** | 9430094M07Rik\|Acpl2\|BB177120\|C130099A20Rik | 2-phosphoxylose phosphatase 1 | protein-coding |
| chr9_96942913_96943362 | chr9 | 96942914 | 96943362 | + | -0.90227 | TTS (NM_145134) | TTS (NM_145134) | -53664 | NM_001289645 | 235534 | Mm.233031 | NM_153420 | ENSMUSG00000043587 | **Pxylp1** | 9430094M07Rik\|Acpl2\|BB177120\|C130099A20Rik | 2-phosphoxylose phosphatase 1 | protein-coding |
| chr9_96942644_96943312 | chr9 | 96942645 | 96943312 | + | -0.964703 | TTS (NM_145134) | TTS (NM_145134) | -53504 | NM_001289645 | 235534 | Mm.233031 | NM_153420 | ENSMUSG00000043587 | **Pxylp1** | 9430094M07Rik\|Acpl2\|BB177120\|C130099A20Rik | 2-phosphoxylose phosphatase 1 | protein-coding |
| chr1_180911699_180912363 | chr1 | 180911700 | 180912363 | + | -0.719215 | Intergenic | ORR1C2\|LTR\|ERVL-MaLR | 7757 | NM_133705 | 69051 | Mm.274180 | NM_133705 | ENSMUSG00000026520 | **Pycr2** | 1810018M05Rik\|P5cr2 | pyrroline-5-carboxylate reductase family, member 2 | protein-coding |
| chr1_180916362_180916735 | chr1 | 180916363 | 180916735 | + | -1.13155 | Intergenic | RMER19A\|LTR\|ERVK | 12275 | NM_133705 | 69051 | Mm.274180 | NM_133705 | ENSMUSG00000026520 | **Pycr2** | 1810018M05Rik\|P5cr2 | pyrroline-5-carboxylate reductase family, member 2 | protein-coding |
| chr1_180916406_180916918 | chr1 | 180916407 | 180916918 | + | -1.27026 | Intergenic | RMER19A\|LTR\|ERVK | 12388 | NM_133705 | 69051 | Mm.274180 | NM_133705 | ENSMUSG00000026520 | **Pycr2** | 1810018M05Rik\|P5cr2 | pyrroline-5-carboxylate reductase family, member 2 | protein-coding |
| chr1_173678673_173678956 | chr1 | 173678674 | 173678956 | + | -0.636679 | intron (NM_001162938, intron 2 of 6) | intron (NM_001162938, intron 2 of 6) | 5135 | NM_001162938 | 100033459 | Mm.303377 | NM_001162938 | ENSMUSG00000066677 | **Pydc3** | 9830148g24rik\|E430029J22Rik | pyrin domain containing 3 | protein-coding |
| chr12_70218892_70219626 | chr12 | 70218893 | 70219626 | + | -1.49504 | intron (NM_133198, intron 3 of 19) | intron (NM_133198, intron 3 of 19) | 8424 | NM_133198 | 110095 | Mm.256926 | NM_133198 | ENSMUSG00000021069 | **Pygl** | - | liver glycogen phosphorylase | protein-coding |
| chr12_70219122_70219644 | chr12 | 70219123 | 70219644 | + | -1.49635 | intron (NM_133198, intron 3 of 19) | intron (NM_133198, intron 3 of 19) | 8300 | NM_133198 | 110095 | Mm.256926 | NM_133198 | ENSMUSG00000021069 | **Pygl** | - | liver glycogen phosphorylase | protein-coding |
| chr9_72890591_72891216 | chr9 | 72890592 | 72891216 | + | -0.879688 | intron (NM_175485, intron 12 of 19) | intron (NM_175485, intron 12 of 19) | -34746 | NM_028116 | 72135 | Mm.273605 | NM_028116 | | **Pygo1** | 2600014C22Rik | pygopus 1 | protein-coding |
| chr17_10301053_10301485 | chr17 | 10301054 | 10301485 | + | -0.832622 | intron (NM_001159516, intron 1 of 6) | intron (NM_001159516, intron 1 of 6) | 18092 | NM_021881 | 19317 | Mm.130694 | NM_021881 | ENSMUSG00000062078 | **Qk** | 1110003F05Rik\|1500005P18\|Qkl\|l(17)-1Wis\|l17Wis1 | quaking | protein-coding |
| chr17_79043442_79043999 | chr17 | 79043443 | 79043999 | + | -1.06279 | Intergenic | Intergenic | -8185 | NM_027455 | 70536 | Mm.293870 | NM_027455 | ENSMUSG00000024084 | **Qpct** | 5730422A13Rik | glutaminyl-peptide cyclotransferase (glutaminyl cyclase) | protein-coding |
| chr2_31814707_31815014 | chr2 | 31814708 | 31815014 | + | -0.998156 | 3' UTR (NM_178887, exon 7 of 7) | 3' UTR (NM_178887, exon 7 of 7) | -4343 | NM_183424 | 227717 | Mm.185027 | NM_183424 | ENSMUSG00000043102 | **Qrfp** | P518 | pyroglutamylated RFamide peptide | protein-coding |
| chr2_31814589_31815047 | chr2 | 31814590 | 31815047 | + | -1.04518 | 3' UTR (NM_178887, exon 7 of 7) | 3' UTR (NM_178887, exon 7 of 7) | -4300 | NM_183424 | 227717 | Mm.185027 | NM_183424 | ENSMUSG00000043102 | **Qrfp** | P518 | pyroglutamylated RFamide peptide | protein-coding |
| chr11_79632091_79632327 | chr11 | 79632092 | 79632327 | + | -0.660166 | intron (NM_175543, intron 3 of 14) | intron (NM_175543, intron 3 of 14) | -9046 | NR_003283 | 790912 | Mm.447313 | NR_003283 | ENSMUSG00000087434 | **Rab11fip4os1** | Gm11202\|OTTMUSG00000000185 | RAB11 family interacting protein 4 (class II), opposite strand 1 | ncRNA |
| chr11_79614869_79615325 | chr11 | 79614870 | 79615325 | + | -0.915168 | intron (NR_003283, intron 2 of 2) | intron (NR_003283, intron 2 of 2) | 8066 | NR_003283 | 790912 | Mm.447313 | NR_003283 | ENSMUSG00000087434 | **Rab11fip4os1** | Gm11202\|OTTMUSG00000000185 | RAB11 family interacting protein 4 (class II), opposite strand 1 | ncRNA |
| chr11_79614938_79615176 | chr11 | 79614939 | 79615176 | + | -0.924139 | intron (NR_003283, intron 2 of 2) | intron (NR_003283, intron 2 of 2) | 8106 | NR_003283 | 790912 | Mm.447313 | NR_003283 | ENSMUSG00000087434 | **Rab11fip4os1** | Gm11202\|OTTMUSG00000000185 | RAB11 family interacting protein 4 (class II), opposite strand 1 | ncRNA |
| chr11_79640721_79641452 | chr11 | 79640722 | 79641452 | + | -0.926536 | intron (NM_175543, intron 3 of 14) | intron (NM_175543, intron 3 of 14) | -17924 | NR_003283 | 790912 | Mm.447313 | NR_003283 | ENSMUSG00000087434 | **Rab11fip4os1** | Gm11202\|OTTMUSG00000000185 | RAB11 family interacting protein 4 (class II), opposite strand 1 | ncRNA |
| chr2_35225177_35225567 | chr2 | 35225178 | 35225567 | + | -0.649726 | Intergenic | ORR1D2\|LTR\|ERVL-MaLR | -24252 | NM_026697 | 68365 | Mm.198264 | NM_026697 | ENSMUSG00000026878 | **Rab14** | 0610030G24Rik\|2810475J17Rik\|A830021G03Rik\|AI314285\|AI649155\|D030017L14Rik | RAB14, member RAS oncogene family | protein-coding |
| chr18_70062294_70062660 | chr18 | 70062295 | 70062660 | + | -0.765506 | intron (NM_001082553, intron 2 of 6) | intron (NM_001082553, intron 2 of 6) | -9108 | NM_030554 | 80718 | Mm.246753 | NM_030554 | ENSMUSG00000024511 | **Rab27b** | 2310021G14Rik\|B130064M09Rik | RAB27B, member RAS oncogene family | protein-coding |
| chr4_8418176_8418803 | chr4 | 8418177 | 8418803 | + | -0.870159 | Intergenic | Intergenic | -117154 | NM_021518 | 59021 | Mm.397471 | NM_021518 | ENSMUSG00000047187 | **Rab2a** | 9330148M11Rik\|C80220\|Rab2 | RAB2A, member RAS oncogene family | protein-coding |
| chr4_8418421_8418815 | chr4 | 8418422 | 8418815 | + | -0.995773 | Intergenic | Intergenic | -117026 | NM_021518 | 59021 | Mm.397471 | NM_021518 | ENSMUSG00000047187 | **Rab2a** | 9330148M11Rik\|C80220\|Rab2 | RAB2A, member RAS oncogene family | protein-coding |
| chr3_51495693_51496032 | chr3 | 51495694 | 51496032 | + | -0.657827 | 3' UTR (NM_016858, exon 2 of 2) | 3' UTR (NM_016858, exon 2 of 2) | 11897 | NM_016858 | 19338 | Mm.1664 | NM_016858 | ENSMUSG00000027739 | **Rab33b** | - | RAB33B, member RAS oncogene family | protein-coding |
| chr11_115157913_115158510 | chr11 | 115157914 | 115158510 | + | -0.957799 | intron (NM_021411, intron 3 of 8) | intron (NM_021411, intron 3 of 8) | 4034 | NM_021411 | 58222 | Mm.143789 | NM_021411 | ENSMUSG00000020732 | **Rab37** | B230331O03Rik\|B230354I04Rik | RAB37, member RAS oncogene family | protein-coding |
| chr11_115157926_115158176 | chr11 | 115157927 | 115158176 | + | -1.00673 | intron (NM_021411, intron 3 of 8) | intron (NM_021411, intron 3 of 8) | 3873 | NM_021411 | 58222 | Mm.143789 | NM_021411 | ENSMUSG00000020732 | **Rab37** | B230331O03Rik\|B230354I04Rik | RAB37, member RAS oncogene family | protein-coding |
| chr11_115157901_115158268 | chr11 | 115157902 | 115158268 | + | -1.02389 | intron (NM_021411, intron 3 of 8) | intron (NM_021411, intron 3 of 8) | 3907 | NM_021411 | 58222 | Mm.143789 | NM_021411 | ENSMUSG00000020732 | **Rab37** | B230331O03Rik\|B230354I04Rik | RAB37, member RAS oncogene family | protein-coding |
| chr1_127981864_127982954 | chr1 | 127981865 | 127982954 | + | -0.663413 | intron (NM_027678, intron 12 of 20) | intron (NM_027678, intron 12 of 20) | 113636 | NM_178690 | 226407 | Mm.489713 | NM_178690 | ENSMUSG00000036104 | **Rab3gap1** | 1700003B17Rik\|4732493F09Rik\|AL117896\|Rab3gap\|p130 | RAB3 GTPase activating protein subunit 1 | protein-coding |

| | | | | | | | | | | | | | | | | | |
|---|---|---|---|---|---|---|---|---|---|---|---|---|---|---|---|---|---|
| chr1_127978078_127978471 | chr1 | 127978079 | 127978471 | + | -0.667858 | intron (NM_027678, intron 12 of 20) | intron (NM_027678, intron 12 of 20) | 109502 | NM_178690 | 226407 | Mm.489713 | NM_178690 | ENSMUSG00000036104 | **Rab3gap1** | 1700003B17Rik\|4732493F09Rik\|AL117896\|Rab3gap\|p130 | RAB3 GTPase activating protein subunit 1 | protein-coding |
| chr1_127982021_127982636 | chr1 | 127982022 | 127982636 | + | -0.719753 | intron (NM_027678, intron 12 of 20) | intron (NM_027678, intron 12 of 20) | 113556 | NM_178690 | 226407 | Mm.489713 | NM_178690 | ENSMUSG00000036104 | **Rab3gap1** | 1700003B17Rik\|4732493F09Rik\|AL117896\|Rab3gap\|p130 | RAB3 GTPase activating protein subunit 1 | protein-coding |
| chr10_116925334_116925718 | chr10 | 116925335 | 116925718 | + | -0.978354 | intron (NM_001003950, intron 3 of 9) | L1M2\|LINE\|L1 | 24854 | NM_001003950 | 216363 | Mm.336394 | NM_001003950 | ENSMUSG00000064181 | **Rab3ip** | B230311A06\|Gtpat12\|Rabin3 | RAB3A interacting protein | protein-coding |
| chr17_29155512_29156231 | chr17 | 29155513 | 29156231 | + | -0.620809 | TTS (NM_153166) | TTS (NM_153166) | 20816 | NM_001002786 | 442827 | Mm.440341 | NM_001002786 | ENSMUSG00000064147 | **Rab44** | 9830134C10Rik | RAB44, member RAS oncogene family | protein-coding |
| chr17_29155698_29156228 | chr17 | 29155699 | 29156228 | + | -0.653777 | TTS (NM_153166) | TTS (NM_153166) | 20907 | NM_001002786 | 442827 | Mm.440341 | NM_001002786 | ENSMUSG00000064147 | **Rab44** | 9830134C10Rik | RAB44, member RAS oncogene family | protein-coding |
| chr17_29152561_29152906 | chr17 | 29152562 | 29152906 | + | -0.65579 | Intergenic | Intergenic | 17678 | NM_001002786 | 442827 | Mm.440341 | NM_001002786 | ENSMUSG00000064147 | **Rab44** | 9830134C10Rik | RAB44, member RAS oncogene family | protein-coding |
| chr17_29152596_29153055 | chr17 | 29152597 | 29153055 | + | -0.695687 | Intergenic | Intergenic | 17770 | NM_001002786 | 442827 | Mm.440341 | NM_001002786 | ENSMUSG00000064147 | **Rab44** | 9830134C10Rik | RAB44, member RAS oncogene family | protein-coding |
| chr17_29171957_29172517 | chr17 | 29171958 | 29172517 | + | -0.922669 | intron (NM_153166, intron 12 of 20) | intron (NM_153166, intron 12 of 20) | 37181 | NM_001002786 | 442827 | Mm.440341 | NM_001002786 | ENSMUSG00000064147 | **Rab44** | 9830134C10Rik | RAB44, member RAS oncogene family | protein-coding |
| chr8_123807014_123807371 | chr8 | 123807015 | 123807371 | + | -0.728315 | intron (NM_009003, intron 1 of 7) | intron (NM_009003, intron 1 of 7) | 1197 | NM_009003 | 19341 | Mm.9221 | NM_009003 | ENSMUSG00000019478 | **Rab4a** | AI848268\|Rab4 | RAB4A, member RAS oncogene family | protein-coding |
| chr9_103108315_103108775 | chr9 | 103108316 | 103108775 | + | -0.873587 | Intergenic | L1MD1\|LINE\|L1 | -3529 | NM_173781 | 270192 | Mm.490252 | NM_173781 | ENSMUSG00000032549 | **Rab6b** | 9630015C03\|AI844641\|C330006L04Rik\|D9Bwg0185e | RAB6B, member RAS oncogene family | protein-coding |
| chr9_103108330_103108977 | chr9 | 103108331 | 103108977 | + | -0.880252 | Intergenic | IAPLTR2_Mm\|LTR\|ERVK | -3420 | NM_173781 | 270192 | Mm.490252 | NM_173781 | ENSMUSG00000032549 | **Rab6b** | 9630015C03\|AI844641\|C330006L04Rik\|D9Bwg0185e | RAB6B, member RAS oncogene family | protein-coding |
| chr9_66888019_66888524 | chr9 | 66888020 | 66888524 | + | -0.624973 | intron (NM_173413, intron 1 of 7) | intron (NM_173413, intron 1 of 7) | 31433 | NM_173413 | 235442 | Mm.260376 | NM_173413 | ENSMUSG00000036943 | **Rab8b** | 5930437D16\|D330025I23Rik | RAB8B, member RAS oncogene family | protein-coding |
| chr9_66883441_66883922 | chr9 | 66883442 | 66883922 | + | -0.627718 | intron (NM_173413, intron 1 of 7) | intron (NM_173413, intron 1 of 7) | 36023 | NM_173413 | 235442 | Mm.260376 | NM_173413 | ENSMUSG00000036943 | **Rab8b** | 5930437D16\|D330025I23Rik | RAB8B, member RAS oncogene family | protein-coding |
| chr1_160306975_160307632 | chr1 | 160306976 | 160307632 | + | -0.656198 | intron (NM_013862, intron 19 of 20) | intron (NM_013862, intron 19 of 20) | 44267 | NM_001038621 | 29809 | Mm.25833 | NM_013862 | ENSMUSG00000026721 | **Rabgap1l** | 5830411O09Rik\|8430421H08Rik\|9630005B12Rik\|AW049894\|HHL\|Hh1\|mKIAA0471 | RAB GTPase activating protein 1-like | protein-coding |
| chr1_160786608_160786839 | chr1 | 160786609 | 160786839 | + | -0.749776 | intron (NM_013862, intron 1 of 20) | intron (NM_013862, intron 1 of 20) | 6214 | NM_013862 | 29809 | Mm.25833 | NM_013862 | ENSMUSG00000026721 | **Rabgap1l** | 5830411O09Rik\|8430421H08Rik\|9630005B12Rik\|AW049894\|HHL\|Hh1\|mKIAA0471 | RAB GTPase activating protein 1-like | protein-coding |
| chr15_78570226_78570528 | chr15 | 78570227 | 78570528 | + | -0.848072 | intron (NM_009008, intron 2 of 6) | intron (NM_009008, intron 2 of 6) | 2406 | NM_009008 | 19354 | Mm.1972 | NM_009008 | ENSMUSG00000033220 | **Rac2** | AI323801\|AI452260 | RAS-related C3 botulinum substrate 2 | protein-coding |
| chr15_78576382_78576805 | chr15 | 78576383 | 78576805 | + | -1.0289 | Intergenic | B4A\|SINE\|B4 | -3811 | NM_009008 | 19354 | Mm.1972 | NM_009008 | ENSMUSG00000033220 | **Rac2** | AI323801\|AI452260 | RAS-related C3 botulinum substrate 2 | protein-coding |
| chr15_99646386_99646610 | chr15 | 99646387 | 99646610 | + | -0.927212 | intron (NM_012025, intron 1 of 17) | intron (NM_012025, intron 1 of 17) | 5158 | NM_001253808 | 26934 | Mm.273804 | NM_012025 | ENSMUSG00000023015 | **Racgap1** | AI227039\|AI327394\|Band25\|GTPase\|MgcRacGAP\|gtl11\|mKIAA1478 | Rac GTPase-activating protein 1 | protein-coding |
| chr15_99646320_99646675 | chr15 | 99646321 | 99646675 | + | -0.932679 | intron (NM_012025, intron 1 of 17) | intron (NM_012025, intron 1 of 17) | 5158 | NM_001253808 | 26934 | Mm.273804 | NM_012025 | ENSMUSG00000023015 | **Racgap1** | AI227039\|AI327394\|Band25\|GTPase\|MgcRacGAP\|gtl11\|mKIAA1478 | Rac GTPase-activating protein 1 | protein-coding |
| chr19_4203993_4204320 | chr19 | 4203994 | 4204320 | + | -0.81019 | Intergenic | Intergenic | -2554 | NM_011237 | 19367 | Mm.277629 | NM_011237 | ENSMUSG00000024824 | **Rad9a** | Rad9 | RAD9 homolog A | protein-coding |
| chr10_22345653_22346120 | chr10 | 22345654 | 22346120 | + | -0.772349 | intron (NM_009018, intron 5 of 5) | intron (NM_009018, intron 5 of 5) | -16007 | NM_020030 | 56554 | Mm.458004 | NM_020030 | ENSMUSG00000078452 | **Raet1d** | RAE-1delta | retinoic acid early transcript delta | protein-coding |
| chr11_60130454_60130998 | chr11 | 60130455 | 60130998 | + | -0.827936 | intron (NM_009021, intron 1 of 5) | intron (NM_009021, intron 1 of 5) | -9356 | NM_001037764 | 19377 | Mm.296366 | NM_009021 | ENSMUSG00000062115 | **Rai1** | Gt1 | retinoic acid induced 1 | protein-coding |
| chr1_119476657_119477398 | chr1 | 119476658 | 119477398 | + | -0.703036 | intron (NM_022327, intron 3 of 4) | intron (NM_022327, intron 3 of 4) | 27754 | NM_022327 | 64143 | Mm.27832 | NM_022327 | ENSMUSG00000004451 | **Ralb** | 5730472O18Rik | v-ral simian leukemia viral oncogene homolog B (ras related) | protein-coding |
| chr1_119476857_119477285 | chr1 | 119476858 | 119477285 | + | -0.76814 | intron (NM_022327, intron 3 of 4) | intron (NM_022327, intron 3 of 4) | 27711 | NM_022327 | 64143 | Mm.27832 | NM_022327 | ENSMUSG00000004451 | **Ralb** | 5730472O18Rik | v-ral simian leukemia viral oncogene homolog B (ras related) | protein-coding |
| chr17_65906823_65907542 | chr17 | 65906824 | 65907542 | + | -0.661096 | Intergenic | ORR1F\|LTR\|ERVL-MaLR | -21428 | NM_009067 | 19765 | Mm.17009 | NM_009067 | ENSMUSG00000024096 | **Ralbp1** | RLIP76\|Rip1 | ralA binding protein 1 | protein-coding |
| chr8_105810216_105810854 | chr8 | 105810217 | 105810854 | + | -0.810789 | intron (NM_145824, intron 2 of 13) | L2c\|LINE\|L2 | 16815 | NM_145824 | 74334 | Mm.4206 | NM_145824 | ENSMUSG00000037415 | **Ranbp10** | 4432417N03Rik | RAN binding protein 10 | protein-coding |
| chr8_105825544_105825901 | chr8 | 105825545 | 105825901 | + | -0.975776 | intron (NM_145824, intron 2 of 13) | intron (NM_145824, intron 2 of 13) | 1627 | NM_145824 | 74334 | Mm.4206 | NM_145824 | ENSMUSG00000037415 | **Ranbp10** | 4432417N03Rik | RAN binding protein 10 | protein-coding |
| chr10_58464031_58464363 | chr10 | 58464032 | 58464363 | + | -0.668669 | exon (NM_011240, exon 11 of 29) | exon (NM_011240, exon 11 of 29) | 17345 | NM_011240 | 19386 | Mm.431695 | NM_011240 | ENSMUSG00000003226 | **Ranbp2** | A430087B05Rik\|AI256741\|NUP358 | RAN binding protein 2 | protein-coding |
| chr13_43450365_43450939 | chr13 | 43450366 | 43450939 | + | -0.692564 | intron (NM_019930, intron 2 of 13) | intron (NM_019930, intron 2 of 13) | 30321 | NM_019930 | 56705 | Mm.148781 | NM_019930 | ENSMUSG00000038546 | **Ranbp9** | IBAP-1\|Ibap1\|RanBPM | RAN binding protein 9 | protein-coding |
| chr13_43450411_43450702 | chr13 | 43450412 | 43450702 | + | -0.813057 | intron (NM_019930, intron 2 of 13) | ORR1D2\|LTR\|ERVL-MaLR | 30416 | NM_019930 | 56705 | Mm.148781 | NM_019930 | ENSMUSG00000038546 | **Ranbp9** | IBAP-1\|Ibap1\|RanBPM | RAN binding protein 9 | protein-coding |
| chr3_105756535_105756804 | chr3 | 105756536 | 105756804 | + | -0.66295 | intron (NM_145541, intron 1 of 7) | ID4_\|SINE\|ID | 44711 | NM_145541 | 109905 | Mm.101922 | NM_145541 | ENSMUSG00000068798 | **Rap1a** | AI848598\|G-22K\|Krev-1\|Rap1 | RAS-related protein-1a | protein-coding |
| chr3_105791121_105791490 | chr3 | 105791122 | 105791490 | + | -0.698699 | intron (NM_145541, intron 1 of 7) | intron (NM_145541, intron 1 of 7) | 10075 | NM_145541 | 109905 | Mm.101922 | NM_145541 | ENSMUSG00000068798 | **Rap1a** | AI848598\|G-22K\|Krev-1\|Rap1 | RAS-related protein-1a | protein-coding |
| chr10_117839850_117840052 | chr10 | 117839851 | 117840052 | + | -0.909378 | intron (NM_024457, intron 1 of 7) | intron (NM_024457, intron 1 of 7) | 6023 | NM_024457 | 215449 | Mm.233009 | NM_024457 | ENSMUSG00000052681 | **Rap1b** | 2810443E11Rik | RAS related protein 1b | protein-coding |
| chr4_137641455_137642276 | chr4 | 137641456 | 137642276 | + | -0.604092 | intron (NM_130879, intron 23 of 27) | intron (NM_130879, intron 23 of 27) | -22860 | NM_029563 | 110351 | Mm.180763 | NM_029563 | ENSMUSG00000041351 | **Rap1gap** | 1300019I11Rik\|2310004O14Rik\|AI427470\|ARPP-90\|Rap1GAP1\|Rap1ga1 | Rap1 GTPase-activating protein | protein-coding |
| chr14_120525691_120526090 | chr14 | 120525692 | 120526090 | + | -0.659108 | Intergenic | Intergenic | 47430 | NM_029519 | 76108 | Mm.261448 | NM_029519 | ENSMUSG00000051615 | **Rap2a** | 5830461H18Rik | RAS related protein 2a | protein-coding |
| chr3_61409564_61410151 | chr3 | 61409565 | 61410151 | + | -0.915007 | Intergenic | Intergenic | 45351 | NM_028712 | 74012 | Mm.422674 | NM_028712 | ENSMUSG00000036894 | **Rap2b** | 4021402C18Rik\|AA408554 | RAP2B, member of RAS oncogene family | protein-coding |
| chr2_29622253_29622865 | chr2 | 29622254 | 29622865 | + | -0.709151 | intron (NM_001039086, intron 1 of 25) | intron (NM_001039086, intron 1 of 25) | 2839 | NM_001039086 | 107746 | Mm.298274 | NM_054050 | ENSMUSG00000039844 | **Rapgef1** | 4932418O06Rik\|C3G\|Grf2 | Rap guanine nucleotide exchange factor (GEF) 1 | protein-coding |
| chr3_79100208_79100521 | chr3 | 79100209 | 79100521 | + | -0.711439 | intron (NM_001099624, intron 3 of 23) | intron (NM_001099624, intron 3 of 23) | 45510 | NM_001099624 | 76089 | Mm.31220 | NM_001099624 | ENSMUSG00000062232 | **Rapgef2** | 5830453M24Rik\|CNRasGEF\|Pdzgef1\|RA-GEF-1\|mKIAA0313\|nRapGEP | Rap guanine nucleotide exchange factor (GEF) 2 | protein-coding |
| chr3_79039715_79040065 | chr3 | 79039716 | 79040065 | + | -1.05148 | Intergenic | Intergenic | 105985 | NM_001099624 | 76089 | Mm.31220 | NM_001099624 | ENSMUSG00000062232 | **Rapgef2** | 5830453M24Rik\|CNRasGEF\|Pdzgef1\|RA-GEF-1\|mKIAA0313\|nRapGEP | Rap guanine nucleotide exchange factor (GEF) 2 | protein-coding |
| chr2_72146107_72146446 | chr2 | 72146108 | 72146446 | + | -0.748641 | intron (NM_019688, intron 6 of 29) | intron (NM_019688, intron 6 of 29) | 91660 | NM_001204166 | 56508 | Mm.196153 | NM_019688 | ENSMUSG00000049044 | **Rapgef4** | 1300003D15Rik\|5730402K07Rik\|6330581N18Rik\|EPAC 2\|Epac-2\|Epac2\|mKIAA4040 | Rap guanine nucleotide exchange factor (GEF) 4 | protein-coding |
| chr2_71941156_71942020 | chr2 | 71941157 | 71942020 | + | -0.779659 | Intergenic | RLTR14\|LTR\|ERV1 | -39627 | NM_001204167 | 56508 | Mm.196153 | NM_019688 | ENSMUSG00000049044 | **Rapgef4** | 1300003D15Rik\|5730402K07Rik\|6330581N18Rik\|EPAC 2\|Epac-2\|Epac2\|mKIAA4040 | Rap guanine nucleotide exchange factor (GEF) 4 | protein-coding |
| chr2_72136779_72137331 | chr2 | 72136780 | 72137331 | + | -1.09574 | intron (NM_001204166, intron 1 of 27) | intron (NM_001204166, intron 1 of 27) | 82438 | NM_001204166 | 56508 | Mm.196153 | NM_019688 | ENSMUSG00000049044 | **Rapgef4** | 1300003D15Rik\|5730402K07Rik\|6330581N18Rik\|EPAC 2\|Epac-2\|Epac2\|mKIAA4040 | Rap guanine nucleotide exchange factor (GEF) 4 | protein-coding |
| chr2_72117619_72118069 | chr2 | 72117620 | 72118069 | + | -1.13582 | intron (NM_001204166, intron 1 of 27) | intron (NM_001204166, intron 1 of 27) | 63227 | NM_001204166 | 56508 | Mm.196153 | NM_019688 | ENSMUSG00000049044 | **Rapgef4** | 1300003D15Rik\|5730402K07Rik\|6330581N18Rik\|EPAC 2\|Epac-2\|Epac2\|mKIAA4040 | Rap guanine nucleotide exchange factor (GEF) 4 | protein-coding |
| chr11_54533855_54534267 | chr11 | 54533856 | 54534267 | + | -0.73284 | intron (NM_001252496, intron 1 of 28) | RLTR14\|LTR\|ERV1 | 11216 | NM_001252496 | 192786 | Mm.254404 | NM_175258 | ENSMUSG00000037533 | **Rapgef6** | A530068K01\|AI844632\|BB085664\|C030018K18Rik\|PDZ-GEF2\|Pdzgef2\|RA-GEF-2\|Ragef2\|mKIAA4052 | Rap guanine nucleotide exchange factor (GEF) 6 | protein-coding |
| chr11_54567965_54568355 | chr11 | 54567966 | 54568355 | + | -0.875421 | intron (NM_001252496, intron 5 of 28) | intron (NM_001252496, intron 5 of 28) | -27986 | NM_001252498 | 192786 | Mm.254404 | NM_175258 | ENSMUSG00000037533 | **Rapgef6** | A530068K01\|AI844632\|BB085664\|C030018K18Rik\|PDZ-GEF2\|Pdzgef2\|RA-GEF-2\|Ragef2\|mKIAA4052 | Rap guanine nucleotide exchange factor (GEF) 6 | protein-coding |

| | | | | | | | | | | | | | | | | | |
|---|---|---|---|---|---|---|---|---|---|---|---|---|---|---|---|---|---|
| chr11_54621827_54622119 | chr11 | 54621828 | 54622119 | + | -1.32655 | intron (NM_001252496, intron 8 of 28) | intron (NM_001252496, intron 8 of 28) | 25827 | NM_001252497 | 192786 | Mm.254404 | NM_175258 | ENSMUSG00000037533 | **Rapgef6** | A530068K01\|AI844632\|BB085664\|C030018K18Rik\|PDZ-GEF2\|Pdzgef2\|RA-GEF-2\|Ragef2\|mKIAA4052 | Rap guanine nucleotide exchange factor (GEF) 6 | protein-coding |
| chr1_60652273_60653055 | chr1 | 60652274 | 60653055 | + | -0.635399 | Intergenic | Intergenic | -85899 | NM_001045513 | 77300 | Mm.440325 | NM_001045513 | ENSMUSG00000026014 | **Raph1** | 9430025M21Rik\|C730009O10Rik\|Lpd | Ras association (RalGDS/AF-6) and pleckstrin homology domains 1 | protein-coding |
| chr11_98959532_98959813 | chr11 | 98959533 | 98959813 | + | -0.66612 | promoter-TSS (NM_001176528) | promoter-TSS (NM_001176528) | -798 | NM_001176528 | 19401 | Mm.439744 | NM_009024 | ENSMUSG00000037992 | **Rara** | Nr1b1\|RAR\|RARalpha1 | retinoic acid receptor, alpha | protein-coding |
| chr11_98959433_98960017 | chr11 | 98959434 | 98960017 | + | -0.670082 | promoter-TSS (NM_001176528) | promoter-TSS (NM_001176528) | -746 | NM_001176528 | 19401 | Mm.439744 | NM_009024 | ENSMUSG00000037992 | **Rara** | Nr1b1\|RAR\|RARalpha1 | retinoic acid receptor, alpha | protein-coding |
| chr11_98939541_98939850 | chr11 | 98939542 | 98939850 | + | -0.692457 | promoter-TSS (NM_001177303) | promoter-TSS (NM_001177303) | -15 | NM_001177303 | 19401 | Mm.439744 | NM_009024 | ENSMUSG00000037992 | **Rara** | Nr1b1\|RAR\|RARalpha1 | retinoic acid receptor, alpha | protein-coding |
| chr11_98945400_98945731 | chr11 | 98945401 | 98945731 | + | -0.756728 | intron (NM_001177303, intron 1 of 8) | intron (NM_001177303, intron 1 of 8) | 5855 | NM_001177303 | 19401 | Mm.439744 | NM_009024 | ENSMUSG00000037992 | **Rara** | Nr1b1\|RAR\|RARalpha1 | retinoic acid receptor, alpha | protein-coding |
| chr11_98945460_98945643 | chr11 | 98945461 | 98945643 | + | -0.785968 | intron (NM_001177303, intron 1 of 8) | intron (NM_001177303, intron 1 of 8) | 5841 | NM_001177303 | 19401 | Mm.439744 | NM_009024 | ENSMUSG00000037992 | **Rara** | Nr1b1\|RAR\|RARalpha1 | retinoic acid receptor, alpha | protein-coding |
| chr13_85376342_85376703 | chr13 | 85376343 | 85376703 | + | -0.663651 | Intergenic | Intergenic | -87037 | NM_145452 | 218397 | Mm.259653 | NM_145452 | ENSMUSG00000021549 | **Rasa1** | Gap\|RasGAP\|Rasa | RAS p21 protein activator 1 | protein-coding |
| chr13_85479370_85479759 | chr13 | 85479371 | 85479759 | + | -0.741846 | Intergenic | B1F1\|SINE\|Alu | -190079 | NM_145452 | 218397 | Mm.259653 | NM_145452 | ENSMUSG00000021549 | **Rasa1** | Gap\|RasGAP\|Rasa | RAS p21 protein activator 1 | protein-coding |
| chr13_85509687_85510069 | chr13 | 85509688 | 85510069 | + | -0.846938 | Intergenic | Intergenic | -220392 | NM_145452 | 218397 | Mm.259653 | NM_145452 | ENSMUSG00000021549 | **Rasa1** | Gap\|RasGAP\|Rasa | RAS p21 protein activator 1 | protein-coding |
| chr8_13630690_13631267 | chr8 | 13630691 | 13631267 | + | -0.684557 | intron (NM_009025, intron 2 of 23) | intron (NM_009025, intron 2 of 23) | 46608 | NM_009025 | 19414 | Mm.18517 | NM_009025 | ENSMUSG00000031453 | **Rasa3** | AI326412\|C86362\|E130011G04\|GAPIII\|scat | RAS p21 protein activator 3 | protein-coding |
| chr17_32403413_32403961 | chr17 | 32403414 | 32403961 | + | -0.730477 | promoter-TSS (NM_178785) | promoter-TSS (NM_178785) | -106 | NM_178785 | 320484 | Mm.122284 | NM_178785 | ENSMUSG00000052142 | **Rasal3** | A430107D22Rik | RAS protein activator like 3 | protein-coding |
| chr17_32403430_32403799 | chr17 | 32403431 | 32403799 | + | -0.747809 | promoter-TSS (NM_178785) | promoter-TSS (NM_178785) | -34 | NM_178785 | 320484 | Mm.122284 | NM_178785 | ENSMUSG00000052142 | **Rasal3** | A430107D22Rik | RAS protein activator like 3 | protein-coding |
| chr2_117421523_117421902 | chr2 | 117421524 | 117421902 | + | -0.627487 | Intergenic | Intergenic | -78836 | NM_011246 | 19419 | Mm.42150 | NM_011246 | ENSMUSG00000027347 | **Rasgrp1** | Rasgrp | RAS guanyl releasing protein 1 | protein-coding |
| chr2_117322341_117322751 | chr2 | 117322342 | 117322751 | + | -0.721931 | intron (NM_011246, intron 2 of 16) | B4A\|SINE\|B4 | 20331 | NM_011246 | 19419 | Mm.42150 | NM_011246 | ENSMUSG00000027347 | **Rasgrp1** | Rasgrp | RAS guanyl releasing protein 1 | protein-coding |
| chr2_117408304_117408599 | chr2 | 117408305 | 117408599 | + | -0.778908 | Intergenic | Intergenic | -65575 | NM_011246 | 19419 | Mm.42150 | NM_011246 | ENSMUSG00000027347 | **Rasgrp1** | Rasgrp | RAS guanyl releasing protein 1 | protein-coding |
| chr2_117421486_117421744 | chr2 | 117421487 | 117421744 | + | -0.888293 | Intergenic | Intergenic | -78738 | NM_011246 | 19419 | Mm.42150 | NM_011246 | ENSMUSG00000027347 | **Rasgrp1** | Rasgrp | RAS guanyl releasing protein 1 | protein-coding |
| chr19_6398472_6400163 | chr19 | 6398473 | 6400163 | + | -0.634565 | TTS (NM_011224) | TTS (NM_011224) | -1265 | NM_011242 | 19395 | Mm.77017 | NM_011242 | ENSMUSG00000032946 | **Rasgrp2** | CDC25L\|Caldaggef1 | RAS, guanyl releasing protein 2 | protein-coding |
| chr19_6400327_6400732 | chr19 | 6400328 | 6400732 | + | -0.986645 | promoter-TSS (NM_011242) | promoter-TSS (NM_011242) | -53 | NM_011242 | 19395 | Mm.77017 | NM_011242 | ENSMUSG00000032946 | **Rasgrp2** | CDC25L\|Caldaggef1 | RAS, guanyl releasing protein 2 | protein-coding |
| chr19_6400392_6400679 | chr19 | 6400393 | 6400679 | + | -1.00679 | promoter-TSS (NM_011242) | promoter-TSS (NM_011242) | -47 | NM_011242 | 19395 | Mm.77017 | NM_011242 | ENSMUSG00000032946 | **Rasgrp2** | CDC25L\|Caldaggef1 | RAS, guanyl releasing protein 2 | protein-coding |
| chr7_29134691_29135308 | chr7 | 29134692 | 29135308 | + | -0.691105 | promoter-TSS (NM_001174155) | promoter-TSS (NM_001174155) | 67 | NR_045676 | 233046 | Mm.465324 | NM_145149 | ENSMUSG00000030589 | **Rasgrp4** | - | RAS guanyl releasing protein 4 | protein-coding |
| chr7_29134678_29135520 | chr7 | 29134679 | 29135520 | + | -0.702715 | intron (NM_145149, intron 1 of 16) | intron (NM_145149, intron 1 of 16) | 166 | NR_045676 | 233046 | Mm.465324 | NM_145149 | ENSMUSG00000030589 | **Rasgrp4** | - | RAS guanyl releasing protein 4 | protein-coding |
| chr7_29134678_29135043 | chr7 | 29134679 | 29135043 | + | -0.71139 | promoter-TSS (NM_001174155) | promoter-TSS (NM_001174155) | -72 | NM_001174155 | 233046 | Mm.465324 | NM_145149 | ENSMUSG00000030589 | **Rasgrp4** | - | RAS guanyl releasing protein 4 | protein-coding |
| chr7_29141447_29141827 | chr7 | 29141448 | 29141827 | + | -0.88379 | intron (NM_145149, intron 6 of 16) | PB1D10\|SINE\|Alu | 6704 | NR_045676 | 233046 | Mm.465324 | NM_145149 | ENSMUSG00000030589 | **Rasgrp4** | - | RAS guanyl releasing protein 4 | protein-coding |
| chr5_74200199_74201028 | chr5 | 74200200 | 74201028 | + | -0.646404 | Intergenic | Intergenic | 5288 | NM_026878 | 68939 | Mm.293316 | NM_026878 | ENSMUSG00000049907 | **Rasl11b** | 1190017B18Rik | RAS-like, family 11, member B | protein-coding |
| chr2_132022650_132022980 | chr2 | 132022651 | 132022980 | + | -0.684588 | intron (NM_175445, intron 1 of 10) | B1_Mur4\|SINE\|Alu | 7173 | NM_175445 | 215653 | Mm.204737 | NM_175445 | ENSMUSG00000027339 | **Rassf2** | 3830431H01Rik\|9030412M04Rik\|AI852669\|AW495050 | Ras association (RalGDS/AF-6) domain family member 2 | protein-coding |
| chr2_132051719_132052188 | chr2 | 132051720 | 132052188 | + | -0.818628 | TTS (NM_018824) | TTS (NM_018824) | -21966 | NM_175445 | 215653 | Mm.204737 | NM_175445 | ENSMUSG00000027339 | **Rassf2** | 3830431H01Rik\|9030412M04Rik\|AI852669\|AW495050 | Ras association (RalGDS/AF-6) domain family member 2 | protein-coding |
| chr2_132035443_132035910 | chr2 | 132035444 | 132035910 | + | -1.02723 | Intergenic | ORR1D1\|LTR\|ERVL-MaLR | -5689 | NM_175445 | 215653 | Mm.204737 | NM_175445 | ENSMUSG00000027339 | **Rassf2** | 3830431H01Rik\|9030412M04Rik\|AI852669\|AW495050 | Ras association (RalGDS/AF-6) domain family member 2 | protein-coding |
| chr2_131992509_131992849 | chr2 | 131992510 | 131992849 | + | -1.06787 | TTS (NM_175445) | TTS (NM_175445) | 37309 | NM_175445 | 215653 | Mm.204737 | NM_175445 | ENSMUSG00000027339 | **Rassf2** | 3830431H01Rik\|9030412M04Rik\|AI852669\|AW495050 | Ras association (RalGDS/AF-6) domain family member 2 | protein-coding |
| chr10_121473823_121474470 | chr10 | 121473824 | 121474470 | + | -0.724411 | intron (NM_138956, intron 1 of 4) | intron (NM_138956, intron 1 of 4) | 2103 | NM_138956 | 192678 | Mm.41265 | NM_138956 | ENSMUSG00000025795 | **Rassf3** | AW212023\|AW322379 | Ras association (RalGDS/AF-6) domain family member 3 | protein-coding |
| chr10_121473880_121474289 | chr10 | 121473881 | 121474289 | + | -0.743477 | intron (NM_138956, intron 1 of 4) | intron (NM_138956, intron 1 of 4) | 2165 | NM_138956 | 192678 | Mm.41265 | NM_138956 | ENSMUSG00000025795 | **Rassf3** | AW212023\|AW322379 | Ras association (RalGDS/AF-6) domain family member 3 | protein-coding |
| chr10_121474016_121474290 | chr10 | 121474017 | 121474290 | + | -0.771855 | intron (NM_138956, intron 1 of 4) | intron (NM_138956, intron 1 of 4) | 2097 | NM_138956 | 192678 | Mm.41265 | NM_138956 | ENSMUSG00000025795 | **Rassf3** | AW212023\|AW322379 | Ras association (RalGDS/AF-6) domain family member 3 | protein-coding |
| chr10_121493989_121494574 | chr10 | 121493990 | 121494574 | + | -1.12842 | Intergenic | Intergenic | -18032 | NM_138956 | 192678 | Mm.41265 | NM_138956 | ENSMUSG00000025795 | **Rassf3** | AW212023\|AW322379 | Ras association (RalGDS/AF-6) domain family member 3 | protein-coding |
| chr14_73349230_73349619 | chr14 | 73349231 | 73349619 | + | -1.12365 | Intergenic | Intergenic | -23634 | NM_009029 | 19645 | Mm.273862 | NM_009029 | ENSMUSG00000022105 | **Rb1** | Rb\|Rb-1\|pRb | retinoblastoma 1 | protein-coding |
| chr7_122972075_122972404 | chr7 | 122972076 | 122972404 | + | -0.659559 | intron (NM_011247, intron 1 of 17) | intron (NM_011247, intron 1 of 17) | 1676 | NM_175023 | 19647 | Mm.4480 | NM_011247 | ENSMUSG00000030779 | **Rbbp6** | 4933422O15Rik\|AI316869\|BB233631\|C030034J04Rik\|C77662\|P2P-R\|PACT | retinoblastoma binding protein 6 | protein-coding |
| chr18_80178281_80178721 | chr18 | 80178282 | 80178721 | + | -0.783271 | Intergenic | ORR1E\|LTR\|ERVL-MaLR | 22118 | NM_199197 | 68731 | Mm.186936 | NM_199197 | ENSMUSG00000024570 | **Rbfa** | 1110032A13Rik\|AI595940 | ribosome binding factor A | protein-coding |
| chr18_80178394_80178732 | chr18 | 80178395 | 80178732 | + | -0.811829 | Intergenic | ORR1E\|LTR\|ERVL-MaLR | 22056 | NM_199197 | 68731 | Mm.186936 | NM_199197 | ENSMUSG00000024570 | **Rbfa** | 1110032A13Rik\|AI595940 | ribosome binding factor A | protein-coding |
| chr5_120124863_120125498 | chr5 | 120124864 | 120125498 | + | -0.811354 | intron (NM_028762, intron 8 of 23) | intron (NM_028762, intron 8 of 23) | 8668 | NM_028762 | 74111 | Mm.41022 | NM_028762 | | **Rbm19** | 1200009A02Rik\|AV302714\|NPO | RNA binding motif protein 19 | protein-coding |
| chr14_105164835_105165143 | chr14 | 105164836 | 105165143 | + | -0.690937 | intron (NM_134077, intron 1 of 20) | MTEa\|LTR\|ERVL-MaLR | 12338 | NM_134077 | 74213 | Mm.474531 | NM_134077 | ENSMUSG00000022119 | **Rbm26** | 1700009P03Rik\|AI447318\|C230097K14Rik\|C86225\|MLZ-393\|Pro1777\|Se70-2 | RNA binding motif protein 26 | protein-coding |
| chr5_28382327_28382737 | chr5 | 28382328 | 28382737 | + | -0.686813 | intron (NM_028234, intron 10 of 18) | intron (NM_028234, intron 10 of 18) | 65343 | NM_028234 | 381626 | Mm.304075 | NM_028234 | ENSMUSG00000048271 | **Rbm33** | 3200001K10Rik\|6430512A10Rik\|AA386890\|Prr8 | RNA binding motif protein 33 | protein-coding |
| chr5_28382270_28382974 | chr5 | 28382271 | 28382974 | + | -0.703241 | intron (NM_028234, intron 10 of 18) | intron (NM_028234, intron 10 of 18) | 65433 | NM_028234 | 381626 | Mm.304075 | NM_028234 | ENSMUSG00000048271 | **Rbm33** | 3200001K10Rik\|6430512A10Rik\|AA386890\|Prr8 | RNA binding motif protein 33 | protein-coding |
| chr5_28317988_28318240 | chr5 | 28317989 | 28318240 | + | -0.757878 | intron (NM_028234, intron 1 of 18) | intron (NM_028234, intron 1 of 18) | 925 | NM_028234 | 381626 | Mm.304075 | NM_028234 | ENSMUSG00000048271 | **Rbm33** | 3200001K10Rik\|6430512A10Rik\|AA386890\|Prr8 | RNA binding motif protein 33 | protein-coding |
| chr2_173039127_173039341 | chr2 | 173039128 | 173039341 | + | -0.665009 | Intergenic | Intergenic | 17332 | NM_019547 | 56190 | Mm.485782 | NM_019547 | ENSMUSG00000027510 | **Rbm38** | Rnpc1\|Seb4\|Seb4l | RNA binding motif protein 38 | protein-coding |
| chr2_173026607_173027204 | chr2 | 173026608 | 173027204 | + | -0.665805 | intron (NM_019547, intron 3 of 3) | intron (NM_019547, intron 3 of 3) | 5004 | NM_019547 | 56190 | Mm.485782 | NM_019547 | ENSMUSG00000027510 | **Rbm38** | Rnpc1\|Seb4\|Seb4l | RNA binding motif protein 38 | protein-coding |
| chr2_173023560_173023986 | chr2 | 173023561 | 173023986 | + | -0.688926 | intron (NM_019547, intron 3 of 3) | intron (NM_019547, intron 3 of 3) | 1871 | NM_019547 | 56190 | Mm.485782 | NM_019547 | ENSMUSG00000027510 | **Rbm38** | Rnpc1\|Seb4\|Seb4l | RNA binding motif protein 38 | protein-coding |
| chr2_173063246_173063573 | chr2 | 173063247 | 173063573 | + | -0.909141 | Intergenic | Intergenic | 41508 | NM_019547 | 56190 | Mm.485782 | NM_019547 | ENSMUSG00000027510 | **Rbm38** | Rnpc1\|Seb4\|Seb4l | RNA binding motif protein 38 | protein-coding |
| chr2_173056394_173056648 | chr2 | 173056395 | 173056648 | + | -1.1093 | Intergenic | Intergenic | 34619 | NM_019547 | 56190 | Mm.485782 | NM_019547 | ENSMUSG00000027510 | **Rbm38** | Rnpc1\|Seb4\|Seb4l | RNA binding motif protein 38 | protein-coding |
| chr2_156179449_156179845 | chr2 | 156179450 | 156179845 | + | -0.6721 | promoter-TSS (NM_001291115) | promoter-TSS (NM_001291115) | -67 | NM_001291115 | 170791 | Mm.392436 | NM_133242 | ENSMUSG00000027620 | **Rbm39** | 1500012C14Rik\|2310040E03Rik\|B330012G18Rik\|C79248\|R75070\|Rnpc2\|caper | RNA binding motif protein 39 | protein-coding |
| chr2_156179413_156179718 | chr2 | 156179414 | 156179718 | + | -0.721838 | promoter-TSS (NM_001291115) | promoter-TSS (NM_001291115) | 14 | NM_001291115 | 170791 | Mm.392436 | NM_133242 | ENSMUSG00000027620 | **Rbm39** | 1500012C14Rik\|2310040E03Rik\|B330012G18Rik\|C79248\|R75070\|Rnpc2\|caper | RNA binding motif protein 39 | protein-coding |
| chr5_66074935_66075657 | chr5 | 66074936 | 66075657 | + | -0.808355 | intron (NM_139065, intron 2 of 6) | B3\|SINE\|B2 | 22883 | NM_139065 | 245945 | Mm.36863 | NM_139065 | ENSMUSG00000070780 | **Rbm47** | 9530077J19Rik | RNA binding motif protein 47 | protein-coding |
| chr2_60868061_60868454 | chr2 | 60868062 | 60868454 | + | -0.751326 | intron (NM_020296, intron 1 of 13) | intron (NM_020296, intron 1 of 13) | 13180 | NM_001141932 | 56878 | Mm.259667 | NM_020296 | ENSMUSG00000026970 | **Rbms1** | 2600014B10Rik\|AI255215\|MSSP-1\|MSSP-2\|MSSP-3\|YC1 | RNA binding motif, single stranded interacting protein 1 | protein-coding |
| chr2_60810654_60811010 | chr2 | 60810655 | 60811010 | + | -0.964886 | intron (NM_020296, intron 2 of 13) | intron (NM_020296, intron 2 of 13) | 70606 | NM_001141932 | 56878 | Mm.259667 | NM_020296 | ENSMUSG00000026970 | **Rbms1** | 2600014B10Rik\|AI255215\|MSSP-1\|MSSP-2\|MSSP-3\|YC1 | RNA binding motif, single stranded interacting protein 1 | protein-coding |
| chr2_60904177_60904511 | chr2 | 60904178 | 60904511 | + | -1.13109 | intron (NM_020296, intron 1 of 13) | intron (NM_020296, intron 1 of 13) | -22906 | NM_001141931 | 56878 | Mm.259667 | NM_020296 | ENSMUSG00000026970 | **Rbms1** | 2600014B10Rik\|AI255215\|MSSP-1\|MSSP-2\|MSSP-3\|YC1 | RNA binding motif, single stranded interacting protein 1 | protein-coding |

| | | | | | | | | | | | | | | | | |
|---|---|---|---|---|---|---|---|---|---|---|---|---|---|---|---|---|
| chr4_149469515_149470053 | chr4 | 149469516 | 149470053 | + | -1.2019 | intron (NM_133435, intron 3 of 4) | intron (NM_133435, intron 3 of 4) | -14816 | NM_022020 | 63954 | Mm.46023 | NM_022020 | ENSMUSG00000028996 | **Rbp7** | 1110002J23Rik\|CRBP-III | retinol binding protein 7, cellular | protein-coding |
| chr4_135436326_135436628 | chr4 | 135436327 | 135436628 | + | -0.71645 | Intergenic | Intergenic | -2672 | NM_022980 | 53902 | Mm.331970 | NM_022980 | ENSMUSG00000059713 | **Rcan3** | AU041093\|Csp3\|Dscr1l2 | regulator of calcineurin 3 | protein-coding |
| chr4_135428054_135428403 | chr4 | 135428055 | 135428403 | + | -1.20732 | intron (NM_022980, intron 1 of 4) | intron (NM_022980, intron 1 of 4) | 5576 | NM_022980 | 53902 | Mm.331970 | NM_022980 | ENSMUSG00000059713 | **Rcan3** | AU041093\|Csp3\|Dscr1l2 | regulator of calcineurin 3 | protein-coding |
| chr14_59201853_59202171 | chr14 | 59201854 | 59202171 | + | -0.865674 | intron (NM_027764, intron 1 of 11) | intron (NM_027764, intron 1 of 11) | 784 | NM_027764 | 71330 | Mm.234195 | NM_027764 | ENSMUSG00000035469 | **Rcbtb1** | 5430409I18Rik\|AW111883\|CLLD7\|CLLL7 | regulator of chromosome condensation (RCC1) and BTB (POZ) domain containing protein 1 | protein-coding |
| chr14_73123763_73124310 | chr14 | 73123764 | 73124310 | + | -0.727048 | Intergenic | Intergenic | -18473 | NM_001170694 | 105670 | Mm.280068 | NM_134083 | ENSMUSG00000022106 | **Rcbtb2** | 2610028E02Rik\|2810420M18Rik\|AI448345\|AW240694\|Chc1l\|RC/BTB2 | regulator of chromosome condensation (RCC1) and BTB (POZ) domain containing protein 2 | protein-coding |
| chr14_73138575_73138836 | chr14 | 73138576 | 73138836 | + | -0.850392 | Intergenic | Intergenic | -3804 | NM_001170694 | 105670 | Mm.280068 | NM_134083 | ENSMUSG00000022106 | **Rcbtb2** | 2610028E02Rik\|2810420M18Rik\|AI448345\|AW240694\|Chc1l\|RC/BTB2 | regulator of chromosome condensation (RCC1) and BTB (POZ) domain containing protein 2 | protein-coding |
| chr14_73161738_73162132 | chr14 | 73161739 | 73162132 | + | -0.854347 | intron (NR_033185, intron 5 of 15) | intron (NR_033185, intron 5 of 15) | 19150 | NR_033185 | 105670 | Mm.280068 | NM_134083 | ENSMUSG00000022106 | **Rcbtb2** | 2610028E02Rik\|2810420M18Rik\|AI448345\|AW240694\|Chc1l\|RC/BTB2 | regulator of chromosome condensation (RCC1) and BTB (POZ) domain containing protein 2 | protein-coding |
| chr5_91947125_91947338 | chr5 | 91947126 | 91947338 | + | -0.673035 | Intergenic | L1M3\|LINE\|L1 | 15324 | NR_073442 | 68098 | Mm.159453 | NM_026557 | ENSMUSG00000029397 | **Rchy1** | ARNIP\|CHIMP\|Pirh2\|Zfp363\|Znf363 | ring finger and CHY zinc finger domain containing 1 | protein-coding |
| chr1_165705830_165706344 | chr1 | 165705831 | 165706344 | + | -0.64063 | intron (NM_001038846, intron 1 of 5) | intron (NM_001038846, intron 1 of 5) | 2007 | NM_178593 | 226594 | Mm.265552 | NM_178593 | ENSMUSG00000040723 | **Rcsd1** | A430105K13Rik\|BC025872 | RCSD domain containing 1 | protein-coding |
| chr1_165705835_165706058 | chr1 | 165705836 | 165706058 | + | -0.712193 | intron (NM_001038846, intron 1 of 5) | intron (NM_001038846, intron 1 of 5) | 2147 | NM_178593 | 226594 | Mm.265552 | NM_178593 | ENSMUSG00000040723 | **Rcsd1** | A430105K13Rik\|BC025872 | RCSD domain containing 1 | protein-coding |
| chr18_34827605_34828119 | chr18 | 34827606 | 34828119 | + | -0.797257 | intron (NM_001081256, intron 16 of 23) | intron (NM_001081256, intron 16 of 23) | -12727 | NM_001204914 | 225362 | Mm.293588 | NM_144865 | ENSMUSG00000038555 | **Reep2** | - | receptor accessory protein 2 | protein-coding |
| chr18_34827308_34828102 | chr18 | 34827309 | 34828102 | + | -0.802961 | intron (NM_001081256, intron 16 of 23) | intron (NM_001081256, intron 16 of 23) | -12884 | NM_001204914 | 225362 | Mm.293588 | NM_144865 | ENSMUSG00000038555 | **Reep2** | - | receptor accessory protein 2 | protein-coding |
| chr11_23782098_23782383 | chr11 | 23782099 | 23782383 | + | -1.02682 | Intergenic | Intergenic | -11271 | NM_009044 | 19696 | Mm.4869 | NM_009044 | ENSMUSG00000020275 | **Rel** | c-Rel | reticuloendotheliosis oncogene | protein-coding |
| chr5_63967646_63968089 | chr5 | 63967647 | 63968089 | + | -0.612329 | intron (NM_145923, intron 1 of 6) | intron (NM_145923, intron 1 of 6) | 1029 | NM_145923 | 100532 | Mm.243632 | NM_145923 | ENSMUSG00000047881 | **Rell1** | AA536743 | RELT-like 1 | protein-coding |
| chr5_63967671_63967852 | chr5 | 63967672 | 63967852 | + | -0.647894 | intron (NM_145923, intron 1 of 6) | intron (NM_145923, intron 1 of 6) | 1135 | NM_145923 | 100532 | Mm.243632 | NM_145923 | ENSMUSG00000047881 | **Rell1** | AA536743 | RELT-like 1 | protein-coding |
| chr5_22360482_22360939 | chr5 | 22360483 | 22360939 | + | -0.941643 | Intergenic | Intergenic | -16006 | NM_011261 | 19699 | Mm.425236 | NM_011261 | ENSMUSG00000042453 | **Reln** | reeler\|rl | reelin | protein-coding |
| chr6_147032182_147032986 | chr6 | 147032183 | 147032986 | + | -0.618236 | promoter-TSS (NM_025620) | promoter-TSS (NM_025620) | 47 | NM_025620 | 66532 | Mm.45888 | NM_025620 | ENSMUSG00000040121 | **Rep15** | 2210417D09Rik | RAB15 effector protein | protein-coding |
| chr10_18070588_18071152 | chr10 | 18070589 | 18071152 | + | -0.70217 | intron (NM_009048, intron 1 of 19) | intron (NM_009048, intron 1 of 19) | 14930 | NM_001111065 | 19707 | Mm.4479 | NM_009048 | ENSMUSG00000019854 | **Reps1** | BB161292 | RalBP1 associated Eps domain containing protein | protein-coding |
| chr10_39793249_39794072 | chr10 | 39793250 | 39794072 | + | -0.598517 | intron (NM_011264, intron 3 of 31) | intron (NM_011264, intron 3 of 31) | 61501 | NM_011264 | 19714 | Mm.439723 | NM_011264 | ENSMUSG00000019841 | **Rev3l** | Rev\|Rev3\|Sez4 | REV3-like, catalytic subunit of DNA polymerase zeta RAD54 like (S. cerevisiae) | protein-coding |
| chr10_39793204_39793680 | chr10 | 39793205 | 39793680 | + | -0.74556 | intron (NM_011264, intron 3 of 31) | intron (NM_011264, intron 3 of 31) | 61282 | NM_011264 | 19714 | Mm.439723 | NM_011264 | ENSMUSG00000019841 | **Rev3l** | Rev\|Rev3\|Sez4 | REV3-like, catalytic subunit of DNA polymerase zeta RAD54 like (S. cerevisiae) | protein-coding |
| chr11_82869995_82870390 | chr11 | 82869996 | 82870390 | + | -0.807362 | intron (NM_001164569, intron 1 of 7) | intron (NM_001164569, intron 1 of 7) | 551 | NM_001164570 | 67338 | Mm.341608 | NM_026097 | ENSMUSG00000020696 | **Rffl** | 1700051E09Rik\|4930516L10Rik\|BG080975\|Carp2 | ring finger and FYVE like domain containing protein | protein-coding |
| chr14_30674991_30675462 | chr14 | 30674992 | 30675462 | + | -0.976498 | intron (NM_177815, intron 7 of 12) | intron (NM_177815, intron 7 of 12) | 20852 | NM_177815 | 328370 | Mm.259657 | NM_177815 | ENSMUSG00000052395 | **Rft1** | D930025H04 | RFT1 homolog (S. cerevisiae) | protein-coding |
| chr17_50093996_50094539 | chr17 | 50093997 | 50094539 | + | -0.638154 | intron (NM_181397, intron 3 of 9) | intron (NM_181397, intron 3 of 9) | 96229 | NM_181397 | 76438 | Mm.41854 | NM_181397 | | **Rftn1** | 2310015N21Rik\|AI853526 | raftlin lipid raft linker 1 | protein-coding |
| chr17_50149536_50150210 | chr17 | 50149537 | 50150210 | + | -0.692216 | intron (NM_181397, intron 2 of 9) | intron (NM_181397, intron 2 of 9) | 40624 | NM_181397 | 76438 | Mm.41854 | NM_181397 | | **Rftn1** | 2310015N21Rik\|AI853526 | raftlin lipid raft linker 1 | protein-coding |
| chr17_50142216_50142495 | chr17 | 50142217 | 50142495 | + | -0.83412 | intron (NM_181397, intron 2 of 9) | intron (NM_181397, intron 2 of 9) | 48141 | NM_181397 | 76438 | Mm.41854 | NM_181397 | | **Rftn1** | 2310015N21Rik\|AI853526 | raftlin lipid raft linker 1 | protein-coding |
| chr17_50135898_50136491 | chr17 | 50135899 | 50136491 | + | -0.879552 | intron (NM_181397, intron 2 of 9) | intron (NM_181397, intron 2 of 9) | 54302 | NM_181397 | 76438 | Mm.41854 | NM_181397 | | **Rftn1** | 2310015N21Rik\|AI853526 | raftlin lipid raft linker 1 | protein-coding |
| chr17_50125259_50125650 | chr17 | 50125260 | 50125650 | + | -0.925628 | intron (NM_181397, intron 2 of 9) | B4A\|SINE\|B4 | 65042 | NM_181397 | 76438 | Mm.41854 | NM_181397 | | **Rftn1** | 2310015N21Rik\|AI853526 | raftlin lipid raft linker 1 | protein-coding |
| chr17_50125014_50125633 | chr17 | 50125015 | 50125633 | + | -0.939462 | intron (NM_181397, intron 2 of 9) | B4A\|SINE\|B4 | 65173 | NM_181397 | 76438 | Mm.41854 | NM_181397 | | **Rftn1** | 2310015N21Rik\|AI853526 | raftlin lipid raft linker 1 | protein-coding |
| chr1_55208859_55209087 | chr1 | 55208860 | 55209087 | + | -0.674882 | intron (NM_028713, intron 3 of 8) | MER5A\|DNA\|hAT-Charlie | 17809 | NM_028713 | 74013 | Mm.292050 | NM_028713 | ENSMUSG00000025978 | **Rftn2** | 2700010E02Rik\|3222401M22Rik | raftlin family member 2 | protein-coding |
| chr8_84073635_84074189 | chr8 | 84073636 | 84074189 | + | -0.671738 | exon (NM_009055, exon 2 of 21) | exon (NM_009055, exon 2 of 21) | 7076 | NM_009055 | 19724 | Mm.7892 | NM_009055 | ENSMUSG00000031706 | **Rfx1** | AI047719\|AI385641 | regulatory factor X, 1 (influences HLA class II expression) | protein-coding |
| chr19_28009762_28010225 | chr19 | 28009763 | 28010225 | + | -0.790405 | intron (NM_011265, intron 1 of 16) | intron (NM_011265, intron 1 of 16) | 1172 | NM_011265 | 19726 | Mm.313321 | NM_011265 | ENSMUSG00000040929 | **Rfx3** | C230093O12Rik\|MRFX3 | regulatory factor X, 3 (influences HLA class II expression) | protein-coding |
| chr14_79284807_79285158 | chr14 | 79284808 | 79285158 | + | -0.616117 | Intergenic | RSINE1\|SINE\|B4 | 16652 | NM_025427 | 66214 | Mm.29811 | NM_025427 | ENSMUSG00000022018 | **Rgcc** | 1190002H23Rik\|Rgc-32\|Rgc32 | regulator of cell cycle | protein-coding |
| chr14_79315776_79316357 | chr14 | 79315777 | 79316357 | + | -1.09364 | Intergenic | Intergenic | -14432 | NM_025427 | 66214 | Mm.29811 | NM_025427 | ENSMUSG00000022018 | **Rgcc** | 1190002H23Rik\|Rgc-32\|Rgc32 | regulator of cell cycle | protein-coding |
| chr14_79315966_79316372 | chr14 | 79315967 | 79316372 | + | -1.1912 | Intergenic | Intergenic | -14534 | NM_025427 | 66214 | Mm.29811 | NM_025427 | ENSMUSG00000022018 | **Rgcc** | 1190002H23Rik\|Rgc-32\|Rgc32 | regulator of cell cycle | protein-coding |
| chr1_144330238_144331004 | chr1 | 144330239 | 144331004 | + | -0.775893 | Intergenic | Intergenic | -81517 | NM_015811 | 50778 | Mm.103701 | NM_015811 | ENSMUSG00000026358 | **Rgs1** | BL34 | regulator of G-protein signaling 1 | protein-coding |
| chr7_128383505_128384470 | chr7 | 128383506 | 128384470 | + | -0.658103 | intron (NM_026418, intron 4 of 4) | intron (NM_026418, intron 4 of 4) | 34184 | NM_026418 | 67865 | Mm.18635 | NM_026418 | ENSMUSG00000030844 | **Rgs10** | 2310010N19Rik | regulator of G-protein signalling 10 | protein-coding |
| chr7_128384179_128384470 | chr7 | 128384180 | 128384470 | + | -0.79527 | intron (NM_026418, intron 4 of 4) | intron (NM_026418, intron 4 of 4) | 33847 | NM_026418 | 67865 | Mm.18635 | NM_026418 | ENSMUSG00000030844 | **Rgs10** | 2310010N19Rik | regulator of G-protein signalling 10 | protein-coding |
| chr7_128373861_128374645 | chr7 | 128373862 | 128374645 | + | -0.8834 | intron (NM_026418, intron 4 of 4) | RSINE1\|SINE\|B4 | 43919 | NM_026418 | 67865 | Mm.18635 | NM_026418 | ENSMUSG00000030844 | **Rgs10** | 2310010N19Rik | regulator of G-protein signalling 10 | protein-coding |
| chr7_128374131_128374534 | chr7 | 128374132 | 128374534 | + | -0.995804 | intron (NM_026418, intron 4 of 4) | RSINE1\|SINE\|B4 | 43839 | NM_026418 | 67865 | Mm.18635 | NM_026418 | ENSMUSG00000030844 | **Rgs10** | 2310010N19Rik | regulator of G-protein signalling 10 | protein-coding |
| chr1_143907359_143907723 | chr1 | 143907360 | 143907723 | + | -1.08937 | Intergenic | ID_B1\|SINE\|B4 | 96608 | NM_009061 | 19735 | Mm.28262 | NM_009061 | ENSMUSG00000026360 | **Rgs2** | GOS8 | regulator of G-protein signaling 2 | protein-coding |
| chr1_143918906_143919352 | chr1 | 143918907 | 143919352 | + | -1.18936 | Intergenic | Intergenic | 85020 | NM_009061 | 19735 | Mm.28262 | NM_009061 | ENSMUSG00000026360 | **Rgs2** | GOS8 | regulator of G-protein signaling 2 | protein-coding |
| chr1_143993092_143993402 | chr1 | 143993093 | 143993402 | + | -1.19652 | Intergenic | MYSERV6-int\|LTR\|ERVK | 10902 | NM_009061 | 19735 | Mm.28262 | NM_009061 | ENSMUSG00000026360 | **Rgs2** | GOS8 | regulator of G-protein signaling 2 | protein-coding |
| chr4_62756572_62757485 | chr4 | 62756573 | 62757485 | + | -0.684771 | Intergenic | Intergenic | 137357 | NM_134257 | 50780 | Mm.286753 | NM_019492 | ENSMUSG00000059810 | **Rgs3** | 4930506N09Rik\|C2PA-RGS3\|C2pa\|PDZ-RGS3\|RGS3S | regulator of G-protein signaling 3 | protein-coding |
| chr4_62587142_62587409 | chr4 | 62587143 | 62587409 | + | -0.73756 | intron (NM_019492, intron 3 of 19) | intron (NM_019492, intron 3 of 19) | 27429 | NM_019492 | 50780 | Mm.286753 | NM_019492 | ENSMUSG00000059810 | **Rgs3** | 4930506N09Rik\|C2PA-RGS3\|C2pa\|PDZ-RGS3\|RGS3S | regulator of G-protein signaling 3 | protein-coding |
| chr4_62593495_62594236 | chr4 | 62593496 | 62594236 | + | -0.744375 | intron (NM_019492, intron 3 of 19) | intron (NM_019492, intron 3 of 19) | -25806 | NM_001081650 | 50780 | Mm.286753 | NM_019492 | ENSMUSG00000059810 | **Rgs3** | 4930506N09Rik\|C2PA-RGS3\|C2pa\|PDZ-RGS3\|RGS3S | regulator of G-protein signaling 3 | protein-coding |
| chr4_62593948_62594270 | chr4 | 62593949 | 62594270 | + | -0.767551 | intron (NM_019492, intron 3 of 19) | Lx8\|LINE\|L1 | -25563 | NM_001081650 | 50780 | Mm.286753 | NM_019492 | ENSMUSG00000059810 | **Rgs3** | 4930506N09Rik\|C2PA-RGS3\|C2pa\|PDZ-RGS3\|RGS3S | regulator of G-protein signaling 3 | protein-coding |
| chr4_62562036_62562390 | chr4 | 62562037 | 62562390 | + | -0.804961 | intron (NM_019492, intron 1 of 19) | intron (NM_019492, intron 1 of 19) | 2366 | NM_019492 | 50780 | Mm.286753 | NM_019492 | ENSMUSG00000059810 | **Rgs3** | 4930506N09Rik\|C2PA-RGS3\|C2pa\|PDZ-RGS3\|RGS3S | regulator of G-protein signaling 3 | protein-coding |
| chr4_62562031_62562512 | chr4 | 62562032 | 62562512 | + | -0.810375 | intron (NM_019492, intron 1 of 19) | RSINE1\|SINE\|B4 | 2425 | NM_019492 | 50780 | Mm.286753 | NM_019492 | ENSMUSG00000059810 | **Rgs3** | 4930506N09Rik\|C2PA-RGS3\|C2pa\|PDZ-RGS3\|RGS3S | regulator of G-protein signaling 3 | protein-coding |
| chr14_69806780_69807077 | chr14 | 69806781 | 69807077 | + | -1.00748 | Intergenic | Intergenic | -1384 | NM_153514 | 246710 | Mm.102367 | NM_153514 | ENSMUSG00000022075 | **Rhobtb2** | Dbc2\|E130206H14Rik\|mKIAA0717 | Rho-related BTB domain containing 2 | protein-coding |
| chr5_65888097_65888405 | chr5 | 65888098 | 65888405 | + | -0.760805 | intron (NM_001081105, intron 1 of 2) | intron (NM_001081105, intron 1 of 2) | 24682 | NM_001081105 | 74734 | Mm.20323 | NM_001081105 | ENSMUSG00000029204 | **Rhoh** | 5830400A04Rik\|AU019774\|Arhh | ras homolog gene family, member H | protein-coding |
| chr5_65904501_65904864 | chr5 | 65904502 | 65904864 | + | -0.935612 | Intergenic | Intergenic | 41114 | NM_001081105 | 74734 | Mm.20323 | NM_001081105 | ENSMUSG00000029204 | **Rhoh** | 5830400A04Rik\|AU019774\|Arhh | ras homolog gene family, member H | protein-coding |

| | | | | | | | | | | | | | | | | | |
|---|---|---|---|---|---|---|---|---|---|---|---|---|---|---|---|---|---|
| chr17_86957961_86958330 | chr17 | 86957962 | 86958330 | + | -0.789494 | Intergenic | Intergenic | -4965 | NM_145491 | 104215 | Mm.826 | NM_145491 | ENSMUSG00000024143 | **Rhoq** | Arhq\|TC10A\|Tc10 | ras homolog gene family, member Q | protein-coding |
| chr17_86957845_86958359 | chr17 | 86957846 | 86958359 | + | -0.803599 | Intergenic | Intergenic | -5009 | NM_145491 | 104215 | Mm.826 | NM_145491 | ENSMUSG00000024143 | **Rhoq** | Arhq\|TC10A\|Tc10 | ras homolog gene family, member Q | protein-coding |
| chr15_6708777_6709361 | chr15 | 6708778 | 6709361 | + | -0.813761 | intron (NM_030168, intron 2 of 37) | CpG-4767 | 688 | NM_030168 | 78757 | Mm.275811 | NM_030168 | ENSMUSG00000050310 | **Rictor** | 4921505C17Rik\|6030405M08Rik\|AVO3\|AW492497\|D530039E11Rik | RPTOR independent companion of MTOR, complex 2 | protein-coding |
| chr2_52176392_52176964 | chr2 | 52176393 | 52176964 | + | -0.735578 | exon (NM_010889, exon 130 of 157) | exon (NM_010889, exon 130 of 157) | 103841 | NM_175238 | 51869 | Mm.254530 | NM_175238 | ENSMUSG00000036202 | **Rif1** | 5730435J01Rik\|6530403D07Rik\|AU016181\|AW549474\|D2Ertd145e | Rap1 interacting factor 1 homolog (yeast) | protein-coding |
| chr2_145832012_145832904 | chr2 | 145832013 | 145832904 | + | -0.671784 | intron (NM_028724, intron 4 of 12) | intron (NM_028724, intron 4 of 12) | 46342 | NM_028724 | 74030 | Mm.228799 | NM_028724 | ENSMUSG00000001768 | **Rin2** | 2010003K16Rik\|4632403N06Rik\|AW821980\|RASSF4 | Ras and Rab interactor 2 | protein-coding |
| chr12_102308222_102308999 | chr12 | 102308223 | 102308999 | + | -0.643251 | intron (NM_001161365, intron 1 of 9) | intron (NM_001161365, intron 1 of 9) | 24970 | NM_001161365 | 217835 | Mm.24145 | NM_177620 | ENSMUSG00000044456 | **Rin3** | 6430500K07\|C86933 | Ras and Rab interactor 3 | protein-coding |
| chr13_38037844_38038300 | chr13 | 38037845 | 38038300 | + | -1.10096 | intron (NM_024242, intron 1 of 16) | intron (NM_024242, intron 1 of 16) | 1083 | NM_024242 | 71340 | Mm.297130 | NM_024242 | ENSMUSG00000021428 | **Riok1** | 3110046C13Rik\|5430416A05Rik\|Ad034 | RIO kinase 1 (yeast) | protein-coding |
| chr14_55788111_55789211 | chr14 | 55788112 | 55789211 | + | -0.637915 | 5' UTR (NM_001164108, exon 1 of 10) | 5' UTR (NM_001164108, exon 1 of 10) | 196 | NM_001164107 | 56532 | Mm.46612 | NM_019955 | ENSMUSG00000022221 | **Ripk3** | 2610528K09Rik\|AW107945\|Rip3 | receptor-interacting serine-threonine kinase 3 | protein-coding |
| chr4_121230933_121231670 | chr4 | 121230934 | 121231670 | + | -0.647932 | Intergenic | Intergenic | -42768 | NM_001081013 | 109263 | Mm.215745 | NM_001081013 | ENSMUSG00000049878 | **Rlf** | 9230110M18Rik\|AI195322\|MommeD8 | rearranged L-myc fusion sequence | protein-coding |
| chr16_10827141_10827494 | chr16 | 10827142 | 10827494 | + | -0.786415 | Intergenic | RLTR8\|LTR\|ERVK | -7741 | NM_001162932 | 223970 | Mm.325777 | NM_001033278 | ENSMUSG00000037991 | **Rmi2** | A630055G03Rik\|Gm537\|Gm929 | RMI2, RecQ mediated genome instability 2, homolog (S. cerevisiae) | protein-coding |
| chr14_62342570_62342796 | chr14 | 62342571 | 62342796 | + | -0.810279 | intron (NM_026001, intron 1 of 10) | intron (NM_026001, intron 1 of 10) | 10578 | NM_026001 | 67153 | Mm.26040 | NM_026001 | ENSMUSG00000021932 | **Rnaseh2b** | 1110019N06Rik\|2610207P08Rik\|Dleu8 | ribonuclease H2, subunit B | protein-coding |
| chr14_62342537_62343063 | chr14 | 62342538 | 62343063 | + | -0.839825 | intron (NM_026001, intron 1 of 10) | intron (NM_026001, intron 1 of 10) | 10695 | NM_026001 | 67153 | Mm.26040 | NM_026001 | ENSMUSG00000021932 | **Rnaseh2b** | 1110019N06Rik\|2610207P08Rik\|Dleu8 | ribonuclease H2, subunit B | protein-coding |
| chr1_153748620_153748865 | chr1 | 153748621 | 153748865 | + | -0.659584 | promoter-TSS (NM_011882) | promoter-TSS (NM_011882) | -683 | NM_011882 | 24014 | Mm.259254 | NM_011882 | ENSMUSG00000066800 | **Rnasel** | E230029I04Rik | ribonuclease L (2', 5'-oligoisoadenylate synthetase-dependent) | protein-coding |
| chr9_70506246_70506487 | chr9 | 70506247 | 70506487 | + | -0.646466 | Intergenic | Intergenic | -2642 | NM_033604 | 93836 | Mm.29783 | NM_033604 | ENSMUSG00000032217 | **Rnf111** | ARK\|Arkadia | ring finger 111 | protein-coding |
| chr9_70511258_70511497 | chr9 | 70511259 | 70511497 | + | -0.678343 | Intergenic | Intergenic | -7653 | NM_033604 | 93836 | Mm.29783 | NM_033604 | ENSMUSG00000032217 | **Rnf111** | ARK\|Arkadia | ring finger 111 | protein-coding |
| chr8_31114916_31115165 | chr8 | 31114917 | 31115165 | + | -0.689848 | intron (NM_175136, intron 1 of 5) | intron (NM_175136, intron 1 of 5) | 3195 | NM_175136 | 68867 | Mm.29532 | NM_175136 | ENSMUSG00000039328 | **Rnf122** | 1110063C11Rik | ring finger protein 122 | protein-coding |
| chr18_20931370_20931660 | chr18 | 20931371 | 20931660 | + | -0.656407 | Intergenic | Intergenic | -13110 | NM_026301 | 67664 | Mm.45980 | NM_026301 | ENSMUSG00000033107 | **Rnf125** | 4930553F04Rik | ring finger protein 125 | protein-coding |
| chr18_20962629_20963056 | chr18 | 20962630 | 20963056 | + | -0.955327 | intron (NM_026301, intron 1 of 5) | intron (NM_026301, intron 1 of 5) | 18218 | NM_026301 | 67664 | Mm.45980 | NM_026301 | ENSMUSG00000033107 | **Rnf125** | 4930553F04Rik | ring finger protein 125 | protein-coding |
| chr18_20962647_20962856 | chr18 | 20962648 | 20962856 | + | -1.00931 | intron (NM_026301, intron 1 of 5) | intron (NM_026301, intron 1 of 5) | 18127 | NM_026301 | 67664 | Mm.45980 | NM_026301 | ENSMUSG00000033107 | **Rnf125** | 4930553F04Rik | ring finger protein 125 | protein-coding |
| chr18_20945658_20946841 | chr18 | 20945659 | 20946841 | + | -1.04393 | intron (NM_026301, intron 1 of 5) | ID_B1\|SINE\|B4 | 1625 | NM_026301 | 67664 | Mm.45980 | NM_026301 | ENSMUSG00000033107 | **Rnf125** | 4930553F04Rik | ring finger protein 125 | protein-coding |
| chr18_20958376_20958973 | chr18 | 20958377 | 20958973 | + | -1.12788 | intron (NM_026301, intron 1 of 5) | intron (NM_026301, intron 1 of 5) | 14050 | NM_026301 | 67664 | Mm.45980 | NM_026301 | ENSMUSG00000033107 | **Rnf125** | 4930553F04Rik | ring finger protein 125 | protein-coding |
| chr18_20946090_20946811 | chr18 | 20946091 | 20946811 | + | -1.12988 | intron (NM_026301, intron 1 of 5) | intron (NM_026301, intron 1 of 5) | 1826 | NM_026301 | 67664 | Mm.45980 | NM_026301 | ENSMUSG00000033107 | **Rnf125** | 4930553F04Rik | ring finger protein 125 | protein-coding |
| chr11_44545152_44545633 | chr11 | 44545153 | 44545633 | + | -0.631954 | intron (NM_028862, intron 4 of 10) | intron (NM_028862, intron 4 of 10) | 26016 | NM_001166553 | 74315 | Mm.156771 | NM_028862 | ENSMUSG00000019189 | **Rnf145** | 3732413I11Rik\|TMRF1 | ring finger protein 145 | protein-coding |
| chr11_44528638_44529644 | chr11 | 44528639 | 44529644 | + | -0.670461 | intron (NM_028862, intron 2 of 10) | intron (NM_028862, intron 2 of 10) | 9764 | NM_001166553 | 74315 | Mm.156771 | NM_028862 | ENSMUSG00000019189 | **Rnf145** | 3732413I11Rik\|TMRF1 | ring finger protein 145 | protein-coding |
| chr11_44519762_44520734 | chr11 | 44519763 | 44520734 | + | -0.676397 | intron (NM_001166553, intron 1 of 5) | intron (NM_001166553, intron 1 of 5) | 871 | NM_001166553 | 74315 | Mm.156771 | NM_028862 | ENSMUSG00000019189 | **Rnf145** | 3732413I11Rik\|TMRF1 | ring finger protein 145 | protein-coding |
| chr11_44529190_44529636 | chr11 | 44529191 | 44529636 | + | -0.821653 | intron (NM_028862, intron 2 of 10) | intron (NM_028862, intron 2 of 10) | 10036 | NM_001166553 | 74315 | Mm.156771 | NM_028862 | ENSMUSG00000019189 | **Rnf145** | 3732413I11Rik\|TMRF1 | ring finger protein 145 | protein-coding |
| chr11_44529010_44529674 | chr11 | 44529011 | 44529674 | + | -0.835701 | intron (NM_028862, intron 2 of 10) | intron (NM_028862, intron 2 of 10) | 9965 | NM_001166553 | 74315 | Mm.156771 | NM_028862 | ENSMUSG00000019189 | **Rnf145** | 3732413I11Rik\|TMRF1 | ring finger protein 145 | protein-coding |
| chr11_44500968_44501413 | chr11 | 44500969 | 44501413 | + | -1.09708 | Intergenic | ORR1E\|LTR\|ERVL-MaLR | -17773 | NM_028862 | 74315 | Mm.156771 | NM_028862 | ENSMUSG00000019189 | **Rnf145** | 3732413I11Rik\|TMRF1 | ring finger protein 145 | protein-coding |
| chr8_122469960_122470296 | chr8 | 122469961 | 122470296 | + | -0.766962 | intron (NM_001033142, intron 2 of 5) | intron (NM_001033142, intron 2 of 5) | 5936 | NM_001033142 | 68718 | Mm.34356 | NM_001033142 | ENSMUSG00000014470 | **Rnf166** | 1110031E24Rik\|AW555115\|Zfp313l | ring finger protein 166 | protein-coding |
| chr11_70650336_70650696 | chr11 | 70650337 | 70650696 | + | -0.699974 | intron (NM_027445, intron 7 of 8) | intron (NM_027445, intron 7 of 8) | 2927 | NM_027445 | 70510 | Mm.261818 | NM_027445 | ENSMUSG00000040746 | **Rnf167** | 0610010G05Rik\|5730408C10Rik\|AU019305\|AV328608 | ring finger protein 167 | protein-coding |
| chr7_99954203_99954687 | chr7 | 99954204 | 99954687 | + | -0.600952 | intron (NM_175388, intron 2 of 5) | intron (NM_175388, intron 2 of 5) | 26013 | NM_175388 | 108937 | Mm.323087 | NM_175388 | ENSMUSG00000058761 | **Rnf169** | 2900057K09Rik | ring finger protein 169 | protein-coding |
| chr4_138962787_138963060 | chr4 | 138962788 | 138963060 | + | -0.775629 | Intergenic | Intergenic | -4188 | NM_025786 | 66825 | Mm.252312 | NM_025786 | ENSMUSG00000070661 | **Rnf186** | 9130020G10Rik | ring finger protein 186 | protein-coding |
| chr15_36336829_36337422 | chr15 | 36336830 | 36337422 | + | -1.00613 | Intergenic | B1F1\|SINE\|Alu | -53979 | NM_013923 | 30945 | Mm.5181 | NM_013923 | ENSMUSG00000022280 | **Rnf19a** | AA032313\|Dorfin\|Rnf19\|UIP117\|Ubce7ip2\|XYbp | ring finger protein 19A | protein-coding |
| chr15_36345986_36346685 | chr15 | 36345987 | 36346685 | + | -1.07776 | Intergenic | Intergenic | -63189 | NM_013923 | 30945 | Mm.5181 | NM_013923 | ENSMUSG00000022280 | **Rnf19a** | AA032313\|Dorfin\|Rnf19\|UIP117\|Ubce7ip2\|XYbp | ring finger protein 19A | protein-coding |
| chr5_143077411_143077650 | chr5 | 143077412 | 143077650 | + | -0.647358 | intron (NM_080561, intron 9 of 16) | intron (NM_080561, intron 9 of 16) | 35489 | NM_080561 | 108086 | Mm.381440 | NM_080561 | ENSMUSG00000045078 | **Rnf216** | 2810055G22Rik\|AI647468\|AU019462\|C86502\|F830018F18Rik\|TRIAD3\|UIP83\|Ubce7ip1 | ring finger protein 216 | protein-coding |
| chr2_131342938_131343331 | chr2 | 131342939 | 131343331 | + | -0.927746 | intron (NM_178607, intron 1 of 5) | intron (NM_178607, intron 1 of 5) | 9757 | NM_178607 | 51902 | Mm.477523 | NM_178607 | ENSMUSG00000048911 | **Rnf24** | 2810473M14Rik\|4930505A13Rik\|AI317164\|C86507\|D2Ertd504e | ring finger protein 24 | protein-coding |
| chr5_29205637_29206051 | chr5 | 29205638 | 29206051 | + | -0.638738 | intron (NR_110371, intron 5 of 7) | intron (NR_110371, intron 5 of 7) | 9851 | NM_001289757 | 56874 | Mm.87352 | NM_021470 | ENSMUSG00000029130 | **Rnf32** | 1700009J01Rik\|2700025B22Rik\|4930542N22Rik\|Lmbr2 | ring finger protein 32 | protein-coding |
| chr4_44159825_44160314 | chr4 | 44159826 | 44160314 | + | -0.868736 | intron (NM_001038993, intron 1 of 11) | intron (NM_001038993, intron 1 of 11) | 7468 | NM_175201 | 73469 | Mm.262859 | NM_175201 | ENSMUSG00000035696 | **Rnf38** | 1700065B19Rik\|2610202O07Rik\|AA673263\|Oip1 | ring finger protein 38 | protein-coding |
| chr9_96436963_96438965 | chr9 | 96436964 | 96438965 | + | -0.817812 | Intergenic | Intergenic | 40631 | NM_011279 | 19823 | Mm.28235 | NM_011279 | ENSMUSG00000051234 | **Rnf7** | Rbx2\|SAG | ring finger protein 7 | protein-coding |
| chr9_96437649_96438964 | chr9 | 96437650 | 96438964 | + | -0.911205 | Intergenic | Intergenic | 40288 | NM_011279 | 19823 | Mm.28235 | NM_011279 | ENSMUSG00000051234 | **Rnf7** | Rbx2\|SAG | ring finger protein 7 | protein-coding |
| chr4_33315720_33316198 | chr4 | 33315721 | 33316198 | + | -0.922818 | intron (NM_011884, intron 1 of 15) | intron (NM_011884, intron 1 of 15) | 5648 | NM_011884 | 24018 | Mm.240024 | NM_011884 | ENSMUSG00000028274 | **Rngtt** | AU020997 | RNA guanylyltransferase and 5'-phosphatase | protein-coding |
| chr19_33265217_33265592 | chr19 | 33265218 | 33265592 | + | -0.870586 | intron (NM_001167818, intron 4 of 6) | intron (NM_001167818, intron 4 of 6) | 126890 | NM_001167818 | 67795 | Mm.204591 | NM_001146342 | ENSMUSG00000071573 | **Rnls** | 6530404N21Rik\|AI452315\|AW060440\|C10orf59 | renalase, FAD-dependent amine oxidase | protein-coding |
| chr19_33429463_33429915 | chr19 | 33429464 | 33429915 | + | -1.11725 | Intergenic | Intergenic | -37394 | NM_001146342 | 67795 | Mm.204591 | NM_001146342 | ENSMUSG00000071573 | **Rnls** | 6530404N21Rik\|AI452315\|AW060440\|C10orf59 | renalase, FAD-dependent amine oxidase | protein-coding |
| chr19_14500892_14501136 | chr19 | 14500893 | 14501136 | + | -0.72335 | intron (NM_011600, intron 8 of 19) | intron (NM_011600, intron 8 of 19) | -62508 | NR_003027 | 19862 | | NR_003027 | ENSMUSG00000095132 | **Rnu6** | - | U6 small nuclear RNA | snRNA |
| chr12_16874180_16874737 | chr12 | 16874181 | 16874737 | + | -0.845954 | Intergenic | (CA)n\|Simple_repeat\|Simple_repeat | -20519 | NM_009072 | 19878 | Mm.276024 | NM_009072 | ENSMUSG00000020580 | **Rock2** | B230113H15Rik\|ROKalpha\|Rho-kinase\|Rock-II\|Rock2m\|mKIAA0619 | Rho-associated coiled-coil containing protein kinase 2 | protein-coding |
| chr9_69092114_69092884 | chr9 | 69092115 | 69092884 | + | -0.626817 | intron (NM_013646, intron 1 of 10) | intron (NM_013646, intron 1 of 10) | -197342 | NM_001289916 | 19883 | Mm.378450 | NM_013646 | ENSMUSG00000032238 | **Rora** | 9530021D13Rik\|Nr1f1\|ROR1\|ROR2\|ROR3\|nmf267\|sg\|staggerer\|tmgc26 | RAR-related orphan receptor alpha | protein-coding |
| chr9_69165199_69165405 | chr9 | 69165200 | 69165405 | + | -0.922538 | intron (NM_013646, intron 1 of 10) | intron (NM_013646, intron 1 of 10) | -124539 | NM_001289916 | 19883 | Mm.378450 | NM_013646 | ENSMUSG00000032238 | **Rora** | 9530021D13Rik\|Nr1f1\|ROR1\|ROR2\|ROR3\|nmf267\|sg\|staggerer\|tmgc26 | RAR-related orphan receptor alpha | protein-coding |
| chr9_69165137_69165415 | chr9 | 69165138 | 69165415 | + | -0.943722 | intron (NM_013646, intron 1 of 10) | intron (NM_013646, intron 1 of 10) | -124565 | NM_001289916 | 19883 | Mm.378450 | NM_013646 | ENSMUSG00000032238 | **Rora** | 9530021D13Rik\|Nr1f1\|ROR1\|ROR2\|ROR3\|nmf267\|sg\|staggerer\|tmgc26 | RAR-related orphan receptor alpha | protein-coding |

| | | | | | | | | | | | | | | | | | |
|---|---|---|---|---|---|---|---|---|---|---|---|---|---|---|---|---|---|
| chr11_75318402_75319302 | chr11 | 75318403 | 75319302 | + | -0.628284 | intron (NM_001164223, intron 6 of 17) | MER126\|DNA\|DNA | 29531 | NM_001164223 | 68275 | Mm.180734 | NM_026653 | ENSMUSG00000000751 | **Rpa1** | 5031405K23Rik\|70kDa\|AA589576\|AW557552\|RF-A\|RP-A\|Rpa | replication protein A1 | protein-coding |
| chr11_75336581_75337172 | chr11 | 75336582 | 75337172 | + | -0.762343 | intron (NM_026653, intron 4 of 16) | MTEa\|LTR\|ERVL-MaLR | 11506 | NM_001164223 | 68275 | Mm.180734 | NM_026653 | ENSMUSG00000000751 | **Rpa1** | 5031405K23Rik\|70kDa\|AA589576\|AW557552\|RF-A\|RP-A\|Rpa | replication protein A1 | protein-coding |
| chr11_75282404_75283106 | chr11 | 75282405 | 75283106 | + | -0.793394 | Intergenic | Intergenic | 65628 | NM_001164223 | 68275 | Mm.180734 | NM_026653 | ENSMUSG00000000751 | **Rpa1** | 5031405K23Rik\|70kDa\|AA589576\|AW557552\|RF-A\|RP-A\|Rpa | replication protein A1 | protein-coding |
| chr15_97520459_97520729 | chr15 | 97520460 | 97520729 | + | -0.659804 | Intergenic | ORR1E-int\|LTR\|ERVL-MaLR | 185228 | NM_028003 | 71919 | Mm.12255 | NM_028003 | ENSMUSG00000022466 | **Rpap3** | 2310042P20Rik\|D15Ertd682e | RNA polymerase II associated protein 3 | protein-coding |
| chr15_97520272_97520891 | chr15 | 97520273 | 97520891 | + | -0.696631 | Intergenic | ORR1E-int\|LTR\|ERVL-MaLR | 185240 | NM_028003 | 71919 | Mm.12255 | NM_028003 | ENSMUSG00000022466 | **Rpap3** | 2310042P20Rik\|D15Ertd682e | RNA polymerase II associated protein 3 | protein-coding |
| chr10_40232443_40232836 | chr10 | 40232444 | 40232836 | + | -0.751088 | intron (NM_023323, intron 7 of 9) | MTE-int\|LTR\|ERVL-MaLR | 14339 | NM_001042556 | 67239 | Mm.11480 | NM_023323 | ENSMUSG00000038510 | **Rpf2** | 2810470K21Rik\|AU040229\|AU043242\|Bxdc1 | ribosome production factor 2 homolog (S. cerevisiae) | protein-coding |
| chr8_91274796_91275272 | chr8 | 91274797 | 91275272 | + | -0.672668 | intron (NM_173431, intron 12 of 25) | intron (NM_173431, intron 12 of 25) | 38188 | NM_173431 | 244585 | Mm.324592 | NM_173431 | ENSMUSG00000033282 | **Rpgrip1l** | 1700047E16Rik\|4931437C01\|Ftm\|Nphp8 | Rpgrip1-like | protein-coding |
| chr8_91274793_91275068 | chr8 | 91274794 | 91275068 | + | -0.772181 | intron (NM_173431, intron 12 of 25) | intron (NM_173431, intron 12 of 25) | 38291 | NM_173431 | 244585 | Mm.324592 | NM_173431 | ENSMUSG00000033282 | **Rpgrip1l** | 1700047E16Rik\|4931437C01\|Ftm\|Nphp8 | Rpgrip1-like | protein-coding |
| chr11_75926349_75927233 | chr11 | 75926350 | 75927233 | + | -0.630961 | promoter-TSS (NM_001291159) | promoter-TSS (NM_001291159) | -900 | NM_001291159 | 380714 | Mm.425697 | NM_029548 | ENSMUSG00000020847 | **Rph3al** | Gm1753\|Noc2 | rabphilin 3A-like (without C2 domains) | protein-coding |
| chr11_75956347_75957268 | chr11 | 75956348 | 75957268 | + | -0.658322 | Intergenic | Intergenic | -30917 | NM_001291159 | 380714 | Mm.425697 | NM_029548 | ENSMUSG00000020847 | **Rph3al** | Gm1753\|Noc2 | rabphilin 3A-like (without C2 domains) | protein-coding |
| chr11_75935642_75936454 | chr11 | 75935643 | 75936454 | + | -0.749437 | Intergenic | Intergenic | -10157 | NM_001291159 | 380714 | Mm.425697 | NM_029548 | ENSMUSG00000020847 | **Rph3al** | Gm1753\|Noc2 | rabphilin 3A-like (without C2 domains) | protein-coding |
| chr11_75956466_75957124 | chr11 | 75956467 | 75957124 | + | -0.758738 | Intergenic | Intergenic | -30904 | NM_001291159 | 380714 | Mm.425697 | NM_029548 | ENSMUSG00000020847 | **Rph3al** | Gm1753\|Noc2 | rabphilin 3A-like (without C2 domains) | protein-coding |
| chr11_75935815_75936063 | chr11 | 75935816 | 75936063 | + | -0.764884 | Intergenic | Intergenic | -10048 | NM_001291159 | 380714 | Mm.425697 | NM_029548 | ENSMUSG00000020847 | **Rph3al** | Gm1753\|Noc2 | rabphilin 3A-like (without C2 domains) | protein-coding |
| chr11_75961799_75962331 | chr11 | 75961800 | 75962331 | + | -0.806481 | Intergenic | Intergenic | -36174 | NM_001291159 | 380714 | Mm.425697 | NM_029548 | ENSMUSG00000020847 | **Rph3al** | Gm1753\|Noc2 | rabphilin 3A-like (without C2 domains) | protein-coding |
| chr11_75935632_75936247 | chr11 | 75935633 | 75936247 | + | -0.808609 | Intergenic | Intergenic | -10049 | NM_001291159 | 380714 | Mm.425697 | NM_029548 | ENSMUSG00000020847 | **Rph3al** | Gm1753\|Noc2 | rabphilin 3A-like (without C2 domains) | protein-coding |
| chr11_75935805_75936269 | chr11 | 75935806 | 75936269 | + | -0.810009 | Intergenic | Intergenic | -10146 | NM_001291159 | 380714 | Mm.425697 | NM_029548 | ENSMUSG00000020847 | **Rph3al** | Gm1753\|Noc2 | rabphilin 3A-like (without C2 domains) | protein-coding |
| chr6_70813932_70814833 | chr6 | 70813933 | 70814833 | + | -0.620597 | Intergenic | B4A\|SINE\|B4 | -22208 | NM_009075 | 19895 | Mm.17905 | NM_009075 | ENSMUSG00000053604 | **Rpia** | RPI | ribose 5-phosphate isomerase A | protein-coding |
| chr6_70813932_70814413 | chr6 | 70813933 | 70814413 | + | -0.656636 | Intergenic | Intergenic | -21998 | NM_009075 | 19895 | Mm.17905 | NM_009075 | ENSMUSG00000053604 | **Rpia** | RPI | ribose 5-phosphate isomerase A | protein-coding |
| chr6_70815560_70815831 | chr6 | 70815561 | 70815831 | + | -0.954373 | Intergenic | Intergenic | -23521 | NM_009075 | 19895 | Mm.17905 | NM_009075 | ENSMUSG00000053604 | **Rpia** | RPI | ribose 5-phosphate isomerase A | protein-coding |
| chr16_55952876_55953085 | chr16 | 55952877 | 55953085 | + | -0.850156 | Intergenic | Intergenic | -13294 | NM_024218 | 68193 | Mm.29221 | NM_024218 | ENSMUSG00000098274 | **Rpl24** | 0610008L05Rik\|Bst | ribosomal protein L24 | protein-coding |
| chr5_107926520_107927201 | chr5 | 107926521 | 107927201 | + | -0.803057 | intron (NM_026062, intron 1 of 4) | L1MC5\|LINE\|L1 | 26333 | NM_016980 | 100503670 | Mm.277897 | NM_016980 | ENSMUSG00000058558 | **Rpl5** | U21RNA | ribosomal protein L5 | protein-coding |
| chr5_107932475_107933000 | chr5 | 107932476 | 107933000 | + | -0.947064 | intron (NM_026062, intron 1 of 4) | intron (NM_026062, intron 1 of 4) | 32210 | NM_016980 | 100503670 | Mm.277897 | NM_016980 | ENSMUSG00000058558 | **Rpl5** | U21RNA | ribosomal protein L5 | protein-coding |
| chr9_61682378_61682780 | chr9 | 61682379 | 61682780 | + | -0.62293 | Intergenic | MIR3\|SINE\|MIR | 231931 | NM_018853 | 56040 | Mm.3158 | NM_018853 | ENSMUSG00000007892 | **Rplp1** | 2410042H16Rik\|Arpp1\|C430017H15Rik | ribosomal protein, large, P1 | protein-coding |
| chr9_61666099_61666523 | chr9 | 61666100 | 61666523 | + | -0.668474 | Intergenic | Intergenic | 248199 | NM_018853 | 56040 | Mm.3158 | NM_018853 | ENSMUSG00000007892 | **Rplp1** | 2410042H16Rik\|Arpp1\|C430017H15Rik | ribosomal protein, large, P1 | protein-coding |
| chr9_61679628_61680436 | chr9 | 61679629 | 61680436 | + | -0.686875 | Intergenic | Intergenic | 234478 | NM_018853 | 56040 | Mm.3158 | NM_018853 | ENSMUSG00000007892 | **Rplp1** | 2410042H16Rik\|Arpp1\|C430017H15Rik | ribosomal protein, large, P1 | protein-coding |
| chr9_61859143_61860019 | chr9 | 61859144 | 61860019 | + | -0.703966 | Intergenic | Intergenic | 54929 | NM_018853 | 56040 | Mm.3158 | NM_018853 | ENSMUSG00000007892 | **Rplp1** | 2410042H16Rik\|Arpp1\|C430017H15Rik | ribosomal protein, large, P1 | protein-coding |
| chr9_61748867_61749584 | chr9 | 61748868 | 61749584 | + | -0.72907 | Intergenic | (TG)n\|Simple_repeat\|Simple_repeat | 165284 | NM_018853 | 56040 | Mm.3158 | NM_018853 | ENSMUSG00000007892 | **Rplp1** | 2410042H16Rik\|Arpp1\|C430017H15Rik | ribosomal protein, large, P1 | protein-coding |
| chr9_61859274_61859840 | chr9 | 61859275 | 61859840 | + | -0.759678 | Intergenic | Intergenic | 54953 | NM_018853 | 56040 | Mm.3158 | NM_018853 | ENSMUSG00000007892 | **Rplp1** | 2410042H16Rik\|Arpp1\|C430017H15Rik | ribosomal protein, large, P1 | protein-coding |
| chr9_61925225_61925594 | chr9 | 61925226 | 61925594 | + | -0.783361 | intron (NM_024245, intron 15 of 22) | intron (NM_024245, intron 15 of 22) | -10900 | NM_018853 | 56040 | Mm.3158 | NM_018853 | ENSMUSG00000007892 | **Rplp1** | 2410042H16Rik\|Arpp1\|C430017H15Rik | ribosomal protein, large, P1 | protein-coding |
| chr9_61679622_61680091 | chr9 | 61679623 | 61680091 | + | -0.788956 | Intergenic | Intergenic | 234653 | NM_018853 | 56040 | Mm.3158 | NM_018853 | ENSMUSG00000007892 | **Rplp1** | 2410042H16Rik\|Arpp1\|C430017H15Rik | ribosomal protein, large, P1 | protein-coding |
| chr9_61925285_61925764 | chr9 | 61925286 | 61925764 | + | -0.79012 | intron (NM_024245, intron 15 of 22) | intron (NM_024245, intron 15 of 22) | -11015 | NM_018853 | 56040 | Mm.3158 | NM_018853 | ENSMUSG00000007892 | **Rplp1** | 2410042H16Rik\|Arpp1\|C430017H15Rik | ribosomal protein, large, P1 | protein-coding |
| chr9_61790841_61791475 | chr9 | 61790842 | 61791475 | + | -0.79203 | Intergenic | Intergenic | 123352 | NM_018853 | 56040 | Mm.3158 | NM_018853 | ENSMUSG00000007892 | **Rplp1** | 2410042H16Rik\|Arpp1\|C430017H15Rik | ribosomal protein, large, P1 | protein-coding |
| chr9_61679726_61680075 | chr9 | 61679727 | 61680075 | + | -0.795029 | Intergenic | Intergenic | 234609 | NM_018853 | 56040 | Mm.3158 | NM_018853 | ENSMUSG00000007892 | **Rplp1** | 2410042H16Rik\|Arpp1\|C430017H15Rik | ribosomal protein, large, P1 | protein-coding |
| chr9_61925347_61925590 | chr9 | 61925348 | 61925590 | + | -0.795359 | intron (NM_024245, intron 15 of 22) | intron (NM_024245, intron 15 of 22) | -10959 | NM_018853 | 56040 | Mm.3158 | NM_018853 | ENSMUSG00000007892 | **Rplp1** | 2410042H16Rik\|Arpp1\|C430017H15Rik | ribosomal protein, large, P1 | protein-coding |
| chr9_61859560_61859876 | chr9 | 61859561 | 61859876 | + | -0.819481 | Intergenic | Intergenic | 54792 | NM_018853 | 56040 | Mm.3158 | NM_018853 | ENSMUSG00000007892 | **Rplp1** | 2410042H16Rik\|Arpp1\|C430017H15Rik | ribosomal protein, large, P1 | protein-coding |
| chr9_61680682_61681354 | chr9 | 61680683 | 61681354 | + | -0.820885 | Intergenic | Intergenic | 233492 | NM_018853 | 56040 | Mm.3158 | NM_018853 | ENSMUSG00000007892 | **Rplp1** | 2410042H16Rik\|Arpp1\|C430017H15Rik | ribosomal protein, large, P1 | protein-coding |
| chr9_61680984_61681364 | chr9 | 61680985 | 61681364 | + | -0.999679 | Intergenic | Intergenic | 233336 | NM_018853 | 56040 | Mm.3158 | NM_018853 | ENSMUSG00000007892 | **Rplp1** | 2410042H16Rik\|Arpp1\|C430017H15Rik | ribosomal protein, large, P1 | protein-coding |
| chr9_61668714_61669125 | chr9 | 61668715 | 61669125 | + | -1.12556 | Intergenic | Intergenic | 245590 | NM_018853 | 56040 | Mm.3158 | NM_018853 | ENSMUSG00000007892 | **Rplp1** | 2410042H16Rik\|Arpp1\|C430017H15Rik | ribosomal protein, large, P1 | protein-coding |
| chr3_95794336_95794642 | chr3 | 95794337 | 95794642 | + | -1.08876 | intron (NM_001081293, intron 1 of 10) | intron (NM_001081293, intron 1 of 10) | 24464 | NM_001081293 | 75137 | Mm.196275 | NM_028328 | ENSMUSG00000028106 | **Rprd2** | 2810036A19Rik\|4930535B03Rik\|6720469I21Rik\|AL022841\|AL022940\|AU021304\|BB077382\|mKIAA0460 | regulation of nuclear pre-mRNA domain containing 2 | protein-coding |
| chr7_116327947_116328250 | chr7 | 116327948 | 116328250 | + | -0.696408 | Intergenic | MER63D\|DNA\|hAT-Blackjack | 6091 | NM_026533 | 68052 | Mm.14798 | NM_026533 | ENSMUSG00000090862 | **Rps13** | 2700063M04Rik | ribosomal protein S13 | protein-coding |
| chr7_118119878_118120326 | chr7 | 118119879 | 118120326 | + | -0.839433 | 3' UTR (NM_019419, exon 6 of 6) | 3' UTR (NM_019419, exon 6 of 6) | -3955 | NM_170669 | 267019 | Mm.288212 | NM_170669 | ENSMUSG00000008683 | **Rps15a** | A630031B11Rik | ribosomal protein S15A | protein-coding |
| chr17_7147852_7148165 | chr17 | 7147853 | 7148165 | + | -0.796478 | Intergenic | Intergenic | -22106 | NM_011299 | 20112 | Mm.268383 | NM_011299 | ENSMUSG00000023809 | **Rps6ka2** | 90kDa\|D17Wsu134e\|Rps6ka-rs1\|Rsk3\|p90rsk\|pp90rsk | ribosomal protein S6 kinase, polypeptide 2 | protein-coding |
| chr17_7147847_7148436 | chr17 | 7147848 | 7148436 | + | -0.804024 | Intergenic | Intergenic | -21973 | NM_011299 | 20112 | Mm.268383 | NM_011299 | ENSMUSG00000023809 | **Rps6ka2** | 90kDa\|D17Wsu134e\|Rps6ka-rs1\|Rsk3\|p90rsk\|pp90rsk | ribosomal protein S6 kinase, polypeptide 2 | protein-coding |
| chrX_159210884_159211255 | chrX | 159210885 | 159211255 | + | -0.607271 | Intergenic | Intergenic | -44712 | NM_148945 | 110651 | Mm.328476 | NM_148945 | ENSMUSG00000031309 | **Rps6ka3** | C130006E23\|MAPKAPK-1b\|MPK-9\|Rsk2\|S6K-alpha3\|p90RSK3\|pp90RSK2 | ribosomal protein S6 kinase polypeptide 3 | protein-coding |
| chrX_159080350_159081053 | chrX | 159080351 | 159081053 | + | -0.907948 | Intergenic | Intergenic | -175080 | NM_148945 | 110651 | Mm.328476 | NM_148945 | ENSMUSG00000031309 | **Rps6ka3** | C130006E23\|MAPKAPK-1b\|MPK-9\|Rsk2\|S6K-alpha3\|p90RSK3\|pp90RSK2 | ribosomal protein S6 kinase polypeptide 3 | protein-coding |
| chr12_100750456_100750839 | chr12 | 100750457 | 100750839 | + | -0.78127 | Intergenic | Intergenic | -25620 | NM_153587 | 73086 | Mm.220417 | NM_153587 | ENSMUSG00000021180 | **Rps6ka5** | 3110005L17Rik\|6330404E13Rik\|AI854034\|MSK1\|MSPK1\|RLPK | ribosomal protein S6 kinase, polypeptide 5 | protein-coding |
| chr11_119670714_119671563 | chr11 | 119670715 | 119671563 | + | -0.844026 | intron (NM_028898, intron 2 of 33) | intron (NM_028898, intron 2 of 33) | 68144 | NM_028898 | 74370 | Mm.209933 | NM_028898 | ENSMUSG00000025583 | **Rptor** | 4932417H02Rik\|Rap\|Raptor\|mKIAA1303 | regulatory associated protein of MTOR, complex 1 | protein-coding |
| chr11_119670933_119671564 | chr11 | 119670934 | 119671564 | + | -0.862727 | intron (NM_028898, intron 2 of 33) | intron (NM_028898, intron 2 of 33) | 68254 | NM_028898 | 74370 | Mm.209933 | NM_028898 | ENSMUSG00000025583 | **Rptor** | 4932417H02Rik\|Rap\|Raptor\|mKIAA1303 | regulatory associated protein of MTOR, complex 1 | protein-coding |
| chr11_119847086_119847331 | chr11 | 119847087 | 119847331 | + | -0.685773 | intron (NM_028898, intron 16 of 33) | intron (NM_028898, intron 16 of 33) | -12844 | NR_045312 | 319454 | Mm.442144 | NR_045312 | ENSMUSG00000085081 | **Rptoros** | A430071A18Rik | regulatory associated protein of MTOR, complex 1, opposite strand | ncRNA |
| chr11_119846842_119847596 | chr11 | 119846843 | 119847596 | + | -0.858165 | intron (NM_028898, intron 16 of 33) | intron (NM_028898, intron 16 of 33) | -12854 | NR_045312 | 319454 | Mm.442144 | NR_045312 | ENSMUSG00000085081 | **Rptoros** | A430071A18Rik | regulatory associated protein of MTOR, complex 1, opposite strand | ncRNA |
| chr11_119847071_119847596 | chr11 | 119847072 | 119847596 | + | -0.861007 | intron (NM_028898, intron 16 of 33) | (TG)n\|Simple_repeat\|Simple_repeat | -12969 | NR_045312 | 319454 | Mm.442144 | NR_045312 | ENSMUSG00000085081 | **Rptoros** | A430071A18Rik | regulatory associated protein of MTOR, complex 1, opposite strand | ncRNA |
| chr11_119833896_119834474 | chr11 | 119833897 | 119834474 | + | -0.868173 | intron (NR_045312, intron 1 of 3) | intron (NR_045312, intron 1 of 3) | 180 | NR_045313 | 319454 | Mm.442144 | NR_045312 | ENSMUSG00000085081 | **Rptoros** | A430071A18Rik | regulatory associated protein of MTOR, complex 1, opposite strand | ncRNA |
| chr7_114161127_114161458 | chr7 | 114161128 | 114161458 | + | -1.00848 | Intergenic | Intergenic | -43512 | NM_025846 | 66922 | Mm.276572 | NM_025846 | ENSMUSG00000055723 | **Rras2** | 2610016H24Rik\|C86394\|TC21 | related RAS viral (r-ras) oncogene homolog 2 | protein-coding |
| chr7_114087896_114088519 | chr7 | 114087897 | 114088519 | + | -1.12687 | intron (NM_025846, intron 1 of 5) | intron (NM_025846, intron 1 of 5) | 29573 | NM_025846 | 66922 | Mm.276572 | NM_025846 | ENSMUSG00000055723 | **Rras2** | 2610016H24Rik\|C86394\|TC21 | related RAS viral (r-ras) oncogene homolog 2 | protein-coding |

| | | | | | | | | | | | | | | | | | |
|---|---|---|---|---|---|---|---|---|---|---|---|---|---|---|---|---|---|
| chr7_114069019_114069665 | chr7 | 114069020 | 114069665 | + | -1.13576 | intron (NM_025846, intron 1 of 5) | intron (NM_025846, intron 1 of 5) | 48439 | NM_025846 | 66922 | Mm.276572 | NM_025846 | ENSMUSG00000055723 | **Rras2** | 2610016H24Rik\|C86394\|TC21 | related RAS viral (r-ras) oncogene homolog 2 | protein-coding |
| chr7_114093619_114093886 | chr7 | 114093620 | 114093886 | + | -1.29088 | intron (NM_025846, intron 1 of 5) | (CA)n\|Simple_repeat\|Simple_repeat | 24028 | NM_025846 | 66922 | Mm.276572 | NM_025846 | ENSMUSG00000055723 | **Rras2** | 2610016H24Rik\|C86394\|TC21 | related RAS viral (r-ras) oncogene homolog 2 | protein-coding |
| chr7_114093540_114094123 | chr7 | 114093541 | 114094123 | + | -1.29232 | intron (NM_025846, intron 1 of 5) | intron (NM_025846, intron 1 of 5) | 23949 | NM_025846 | 66922 | Mm.276572 | NM_025846 | ENSMUSG00000055723 | **Rras2** | 2610016H24Rik\|C86394\|TC21 | related RAS viral (r-ras) oncogene homolog 2 | protein-coding |
| chr13_37751097_37751880 | chr13 | 37751098 | 37751880 | + | -0.975729 | Intergenic | (TCTG)n\|Simple_repeat\|Simple_repeat | -74549 | NM_026830 | 68750 | Mm.491109 | NM_026830 | ENSMUSG00000039087 | **Rreb1** | 1110037N09Rik\|AA414966\|B930013M22Rik | ras responsive element binding protein 1 | protein-coding |
| chr13_37870203_37870929 | chr13 | 37870204 | 37870929 | + | -0.985382 | intron (NM_001039188, intron 1 of 9) | intron (NM_001039188, intron 1 of 9) | -17868 | NR_033218 | 68750 | Mm.491109 | NM_026830 | ENSMUSG00000039087 | **Rreb1** | 1110037N09Rik\|AA414966\|B930013M22Rik | ras responsive element binding protein 1 | protein-coding |
| chr16_13796292_13797278 | chr16 | 13796293 | 13797278 | + | -0.898292 | intron (NM_001039521, intron 10 of 17) | intron (NM_001039521, intron 10 of 17) | 16086 | NM_001039521 | 106298 | Mm.490362 | NM_001039521 | ENSMUSG00000022682 | **Rrn3** | AL023001\|E130302O19Rik\|R75565\|TIF-1A\|Tif1a | RRN3 RNA polymerase I transcription factor homolog (yeast) | protein-coding |
| chr16_13796564_13797273 | chr16 | 13796565 | 13797273 | + | -0.943362 | intron (NM_001039521, intron 10 of 17) | B3\|SINE\|B2 | 16220 | NM_001039521 | 106298 | Mm.490362 | NM_001039521 | ENSMUSG00000022682 | **Rrn3** | AL023001\|E130302O19Rik\|R75565\|TIF-1A\|Tif1a | RRN3 RNA polymerase I transcription factor homolog (yeast) | protein-coding |
| chr19_41901643_41902075 | chr19 | 41901644 | 41902075 | + | -0.850804 | Intergenic | Intergenic | -5706 | NM_199447 | 107094 | Mm.276044 | NM_199447 | ENSMUSG00000035049 | **Rrp12** | AA408556\|AA536972\|mKIAA0690 | ribosomal RNA processing 12 homolog (S. cerevisiae) | protein-coding |
| chr19_41874238_41874919 | chr19 | 41874239 | 41874919 | + | -1.0233 | exon (NM_199447, exon 22 of 34) | exon (NM_199447, exon 22 of 34) | 21574 | NM_199447 | 107094 | Mm.276044 | NM_199447 | ENSMUSG00000035049 | **Rrp12** | AA408556\|AA536972\|mKIAA0690 | ribosomal RNA processing 12 homolog (S. cerevisiae) | protein-coding |
| chr5_20955451_20955931 | chr5 | 20955452 | 20955931 | + | -0.759648 | non-coding (NR_030699, exon 1 of 1) | non-coding (NR_030699, exon 1 of 1) | -3869 | NM_001080977 | 242860 | Mm.254286 | NM_001080977 | ENSMUSG00000039968 | **Rsbn1l** | 8430412F05Rik\|AI447441\|C330002G24Rik | round spermatid basic protein 1-like | protein-coding |
| chr7_97598593_97599002 | chr7 | 97598594 | 97599002 | + | -0.664974 | intron (NM_001081267, intron 1 of 15) | ORR1A4\|LTR\|ERVL-MaLR | 18902 | NM_001081267 | 233532 | Mm.43977 | NM_001081267 | ENSMUSG00000035623 | **Rsf1** | 4832420A03Rik\|C030033M12Rik\|Gm164\|Hbxap\|XAP8\|p325 | remodeling and spacing factor 1 | protein-coding |
| chr4_141210459_141211343 | chr4 | 141210460 | 141211343 | + | -0.66491 | Intergenic | Intergenic | -3055 | NM_001081174 | 76166 | Mm.271064 | NM_001081174 | ENSMUSG00000073733 | **Rsg1** | 6330545A04Rik\|Gm723 | REM2 and RAB-like small GTPase 1 | protein-coding |
| chr4_141210401_141210865 | chr4 | 141210402 | 141210865 | + | -0.671117 | Intergenic | RSINE1\|SINE\|B4 | -3323 | NM_001081174 | 76166 | Mm.271064 | NM_001081174 | ENSMUSG00000073733 | **Rsg1** | 6330545A04Rik\|Gm723 | REM2 and RAB-like small GTPase 1 | protein-coding |
| chr17_31275235_31275479 | chr17 | 31275236 | 31275479 | + | -0.993985 | intron (NM_025290, intron 1 of 8) | ORR1D2\|LTR\|ERVL-MaLR | 1999 | NM_025290 | 22092 | Mm.12743 | NM_025290 | ENSMUSG00000024033 | **Rsph1** | MCA\|Tsga2 | radial spoke head 1 homolog (Chlamydomonas) | protein-coding |
| chr17_31275147_31275523 | chr17 | 31275148 | 31275523 | + | -1.02385 | intron (NM_025290, intron 1 of 8) | ORR1D2\|LTR\|ERVL-MaLR | 2021 | NM_025290 | 22092 | Mm.12743 | NM_025290 | ENSMUSG00000024033 | **Rsph1** | MCA\|Tsga2 | radial spoke head 1 homolog (Chlamydomonas) | protein-coding |
| chr2_13233250_13233788 | chr2 | 13233251 | 13233788 | + | -0.752295 | intron (NM_009105, intron 3 of 8) | intron (NM_009105, intron 3 of 8) | 37896 | NM_009105 | 20163 | Mm.905 | NM_009105 | ENSMUSG00000026727 | **Rsu1** | RsuI\|rsp-1 | Ras suppressor protein 1 | protein-coding |
| chr2_13224219_13225089 | chr2 | 13224220 | 13225089 | + | -1.34148 | intron (NM_009105, intron 3 of 8) | intron (NM_009105, intron 3 of 8) | 46761 | NM_009105 | 20163 | Mm.905 | NM_009105 | ENSMUSG00000026727 | **Rsu1** | RsuI\|rsp-1 | Ras suppressor protein 1 | protein-coding |
| chr2_13224609_13225013 | chr2 | 13224610 | 13225013 | + | -1.45164 | intron (NM_009105, intron 3 of 8) | intron (NM_009105, intron 3 of 8) | 46604 | NM_009105 | 20163 | Mm.905 | NM_009105 | ENSMUSG00000026727 | **Rsu1** | RsuI\|rsp-1 | Ras suppressor protein 1 | protein-coding |
| chr10_68119220_68119625 | chr10 | 68119221 | 68119625 | + | -0.839401 | intron (NM_023598, intron 7 of 9) | intron (NM_023598, intron 7 of 9) | 139825 | NM_001081346 | 170799 | Mm.209844 | NM_133244 | ENSMUSG00000037846 | **Rtkn2** | B130039D23Rik\|Mbf\|Plekhk1 | rhotekin 2 | protein-coding |
| chr10_68105064_68105709 | chr10 | 68105065 | 68105709 | + | -0.977359 | intron (NM_023598, intron 8 of 9) | intron (NM_023598, intron 8 of 9) | 125789 | NM_001081346 | 170799 | Mm.209844 | NM_133244 | ENSMUSG00000037846 | **Rtkn2** | B130039D23Rik\|Mbf\|Plekhk1 | rhotekin 2 | protein-coding |
| chr10_68105258_68105694 | chr10 | 68105259 | 68105694 | + | -1.03464 | intron (NM_023598, intron 8 of 9) | intron (NM_023598, intron 8 of 9) | 125878 | NM_001081346 | 170799 | Mm.209844 | NM_133244 | ENSMUSG00000037846 | **Rtkn2** | B130039D23Rik\|Mbf\|Plekhk1 | rhotekin 2 | protein-coding |
| chr10_68081671_68082192 | chr10 | 68081672 | 68082192 | + | -1.16173 | Intergenic | Intergenic | 102334 | NM_001081346 | 170799 | Mm.209844 | NM_133244 | ENSMUSG00000037846 | **Rtkn2** | B130039D23Rik\|Mbf\|Plekhk1 | rhotekin 2 | protein-coding |
| chr11_75193578_75193942 | chr11 | 75193579 | 75193942 | + | -0.597954 | promoter-TSS (NM_177708) | promoter-TSS (NM_177708) | -233 | NM_177708 | 237847 | Mm.82661 | NM_177708 | ENSMUSG00000045287 | **Rtn4rl1** | Ngr3\|Ngrh2\|Ngrl2 | reticulon 4 receptor-like 1 | protein-coding |
| chr16_23374725_23375482 | chr16 | 23374726 | 23375482 | + | -0.915274 | Intergenic | RMER19C\|LTR\|ERVK | -54029 | NM_001004151 | 239766 | Mm.208751 | NM_001004151 | ENSMUSG00000033383 | **Rtp1** | Gm1766\|Gm604 | receptor transporter protein 1 | protein-coding |
| chr16_92927761_92928024 | chr16 | 92927762 | 92928024 | + | -0.763853 | Intergenic | Intergenic | -101819 | NM_001111022 | 12394 | Mm.4081 | NM_009821 | ENSMUSG00000022952 | **Runx1** | AML1\|CBF-alpha-2\|Cbfa2\|Pebp2a2\|Pebpa2b | runt related transcription factor 1 | protein-coding |
| chr16_92801476_92802224 | chr16 | 92801477 | 92802224 | + | -0.805478 | intron (NM_001111022, intron 2 of 6) | intron (NM_001111022, intron 2 of 6) | 24224 | NM_001111021 | 12394 | Mm.4081 | NM_009821 | ENSMUSG00000022952 | **Runx1** | AML1\|CBF-alpha-2\|Cbfa2\|Pebp2a2\|Pebpa2b | runt related transcription factor 1 | protein-coding |
| chr16_92821694_92822350 | chr16 | 92821695 | 92822350 | + | -0.87787 | intron (NM_001111022, intron 2 of 6) | intron (NM_001111022, intron 2 of 6) | 4052 | NM_001111021 | 12394 | Mm.4081 | NM_009821 | ENSMUSG00000022952 | **Runx1** | AML1\|CBF-alpha-2\|Cbfa2\|Pebp2a2\|Pebpa2b | runt related transcription factor 1 | protein-coding |
| chr16_92810498_92811293 | chr16 | 92810499 | 92811293 | + | -0.991466 | intron (NM_001111022, intron 2 of 6) | intron (NM_001111022, intron 2 of 6) | 15178 | NM_001111021 | 12394 | Mm.4081 | NM_009821 | ENSMUSG00000022952 | **Runx1** | AML1\|CBF-alpha-2\|Cbfa2\|Pebp2a2\|Pebpa2b | runt related transcription factor 1 | protein-coding |
| chr16_92810766_92811291 | chr16 | 92810767 | 92811291 | + | -1.03571 | intron (NM_001111022, intron 2 of 6) | intron (NM_001111022, intron 2 of 6) | 15045 | NM_001111021 | 12394 | Mm.4081 | NM_009821 | ENSMUSG00000022952 | **Runx1** | AML1\|CBF-alpha-2\|Cbfa2\|Pebp2a2\|Pebpa2b | runt related transcription factor 1 | protein-coding |
| chr16_92641024_92641491 | chr16 | 92641025 | 92641491 | + | -1.61011 | intron (NM_001111022, intron 5 of 6) | intron (NM_001111022, intron 5 of 6) | 56071 | NM_001111023 | 12394 | Mm.4081 | NM_009821 | ENSMUSG00000022952 | **Runx1** | AML1\|CBF-alpha-2\|Cbfa2\|Pebp2a2\|Pebpa2b | runt related transcription factor 1 | protein-coding |
| chr17_44690443_44690925 | chr17 | 44690444 | 44690925 | + | -0.719499 | intron (NR_073392, intron 4 of 7) | intron (NR_073392, intron 4 of 7) | 45492 | NR_073425 | 12393 | Mm.391013 | NM_009820 | ENSMUSG00000039153 | **Runx2** | AML3\|Cbf\|Cbfa-1\|Cbfa1\|LS3\|Osf2\|PEBP2aA\|Pebp2a1\|Pebpa2a | runt related transcription factor 2 | protein-coding |
| chr4_135155440_135155808 | chr4 | 135155441 | 135155808 | + | -0.986282 | intron (NM_019732, intron 3 of 5) | intron (NM_019732, intron 3 of 5) | 34979 | NM_019732 | 12399 | Mm.378894 | NM_019732 | ENSMUSG00000070691 | **Runx3** | AML2\|Cbfa3\|Pebp2a3 | runt related transcription factor 3 | protein-coding |
| chr6_100255299_100255749 | chr6 | 100255300 | 100255749 | + | -0.672651 | intron (NM_019743, intron 2 of 4) | B4A\|SINE\|B4 | 31834 | NM_019743 | 56353 | Mm.321633 | NM_019743 | ENSMUSG00000072872 | **Rybp** | 2410018J24Rik\|DEDAF\|YEAF1 | RING1 and YY1 binding protein | protein-coding |
| chr6_100011638_100012085 | chr6 | 100011639 | 100012085 | + | -0.710919 | Intergenic | Intergenic | 275496 | NM_019743 | 56353 | Mm.321633 | NM_019743 | ENSMUSG00000072872 | **Rybp** | 2410018J24Rik\|DEDAF\|YEAF1 | RING1 and YY1 binding protein | protein-coding |
| chr6_100255428_100255884 | chr6 | 100255429 | 100255884 | + | -0.806689 | intron (NM_019743, intron 2 of 4) | B4A\|SINE\|B4 | 31702 | NM_019743 | 56353 | Mm.321633 | NM_019743 | ENSMUSG00000072872 | **Rybp** | 2410018J24Rik\|DEDAF\|YEAF1 | RING1 and YY1 binding protein | protein-coding |
| chr13_11958305_11958629 | chr13 | 11958306 | 11958629 | + | -1.09526 | intron (NM_023868, intron 1 of 104) | intron (NM_023868, intron 1 of 104) | 148478 | NM_023868 | 20191 | Mm.239871 | NM_023868 | ENSMUSG00000021313 | **Ryr2** | 9330127I20Rik\|RYR-2 | ryanodine receptor 2, cardiac | protein-coding |
| chr13_11958353_11958579 | chr13 | 11958354 | 11958579 | + | -1.11876 | intron (NM_023868, intron 1 of 104) | intron (NM_023868, intron 1 of 104) | 148479 | NM_023868 | 20191 | Mm.239871 | NM_023868 | ENSMUSG00000021313 | **Ryr2** | 9330127I20Rik\|RYR-2 | ryanodine receptor 2, cardiac | protein-coding |
| chr3_93549905_93550373 | chr3 | 93549906 | 93550373 | + | -0.622366 | Intergenic | Intergenic | -4978 | NM_009112 | 20194 | Mm.1 | NM_009112 | ENSMUSG00000041959 | **S100a10** | 42C\|AA409961\|AL024248\|CAL12\|CLP11\|Cal1l\|p10\|p11 | S100 calcium binding protein A10 (calpactin) | protein-coding |
| chr10_76271278_76271785 | chr10 | 76271279 | 76271785 | + | -0.659113 | intron (NM_001081419, intron 32 of 37) | intron (NM_001081419, intron 32 of 37) | 17696 | NM_009115 | 20203 | Mm.235998 | NM_009115 | ENSMUSG00000033208 | **S100b** | AI850290\|Bpb | S100 protein, beta polypeptide, neural | protein-coding |
| chr13_95470121_95470529 | chr13 | 95470122 | 95470529 | + | -0.731757 | Intergenic | Intergenic | 8330 | NM_001081159 | 268686 | Mm.291249 | NM_001081159 | ENSMUSG00000021679 | **S100z** | Gm625 | S100 calcium binding protein, zeta | protein-coding |
| chr3_115669063_115669616 | chr3 | 115669064 | 115669616 | + | -0.63093 | Intergenic | ORR1D1\|LTR\|ERVL-MaLR | 45715 | NM_007901 | 13609 | Mm.982 | NM_007901 | ENSMUSG00000045092 | **S1pr1** | AI849002\|Edg1\|Lpb1\|S1p\|S1p1 | sphingosine-1-phosphate receptor 1 | protein-coding |
| chr3_115669271_115669616 | chr3 | 115669272 | 115669616 | + | -0.713678 | Intergenic | ORR1D1\|LTR\|ERVL-MaLR | 45611 | NM_007901 | 13609 | Mm.982 | NM_007901 | ENSMUSG00000045092 | **S1pr1** | AI849002\|Edg1\|Lpb1\|S1p\|S1p1 | sphingosine-1-phosphate receptor 1 | protein-coding |
| chr3_115704237_115705066 | chr3 | 115704238 | 115705066 | + | -0.716485 | Intergenic | Intergenic | 10403 | NM_007901 | 13609 | Mm.982 | NM_007901 | ENSMUSG00000045092 | **S1pr1** | AI849002\|Edg1\|Lpb1\|S1p\|S1p1 | sphingosine-1-phosphate receptor 1 | protein-coding |
| chr3_115800950_115801280 | chr3 | 115800951 | 115801280 | + | -0.784606 | Intergenic | Intergenic | -86060 | NM_007901 | 13609 | Mm.982 | NM_007901 | ENSMUSG00000045092 | **S1pr1** | AI849002\|Edg1\|Lpb1\|S1p\|S1p1 | sphingosine-1-phosphate receptor 1 | protein-coding |
| chr3_115702139_115702383 | chr3 | 115702140 | 115702383 | + | -0.786755 | Intergenic | Intergenic | 12794 | NM_007901 | 13609 | Mm.982 | NM_007901 | ENSMUSG00000045092 | **S1pr1** | AI849002\|Edg1\|Lpb1\|S1p\|S1p1 | sphingosine-1-phosphate receptor 1 | protein-coding |
| chr3_115706787_115707003 | chr3 | 115706788 | 115707003 | + | -0.790551 | Intergenic | ID_B1\|SINE\|B4 | 8160 | NM_007901 | 13609 | Mm.982 | NM_007901 | ENSMUSG00000045092 | **S1pr1** | AI849002\|Edg1\|Lpb1\|S1p\|S1p1 | sphingosine-1-phosphate receptor 1 | protein-coding |
| chr3_115793316_115794039 | chr3 | 115793317 | 115794039 | + | -1.01959 | Intergenic | Intergenic | -78623 | NM_007901 | 13609 | Mm.982 | NM_007901 | ENSMUSG00000045092 | **S1pr1** | AI849002\|Edg1\|Lpb1\|S1p\|S1p1 | sphingosine-1-phosphate receptor 1 | protein-coding |
| chr3_115682318_115683232 | chr3 | 115682319 | 115683232 | + | -1.18267 | Intergenic | Intergenic | 32280 | NM_007901 | 13609 | Mm.982 | NM_007901 | ENSMUSG00000045092 | **S1pr1** | AI849002\|Edg1\|Lpb1\|S1p\|S1p1 | sphingosine-1-phosphate receptor 1 | protein-coding |
| chr3_115673709_115674172 | chr3 | 115673710 | 115674172 | + | -1.29014 | Intergenic | Intergenic | 41114 | NM_007901 | 13609 | Mm.982 | NM_007901 | ENSMUSG00000045092 | **S1pr1** | AI849002\|Edg1\|Lpb1\|S1p\|S1p1 | sphingosine-1-phosphate receptor 1 | protein-coding |
| chr10_81499424_81499895 | chr10 | 81499425 | 81499895 | + | -0.662121 | 5' UTR (NM_010102, exon 1 of 1) | 5' UTR (NM_010102, exon 1 of 1) | 477 | NM_010102 | 13611 | Mm.33065 | NM_010102 | ENSMUSG00000044199 | **S1pr4** | Edg6\|Lpc1\|S1p4 | sphingosine-1-phosphate receptor 4 | protein-coding |
| chr10_81499551_81499893 | chr10 | 81499552 | 81499893 | + | -0.718873 | 5' UTR (NM_010102, exon 1 of 1) | 5' UTR (NM_010102, exon 1 of 1) | 415 | NM_010102 | 13611 | Mm.33065 | NM_010102 | ENSMUSG00000044199 | **S1pr4** | Edg6\|Lpc1\|S1p4 | sphingosine-1-phosphate receptor 4 | protein-coding |

| | | | | | | | | | | | | | | | | | |
|---|---|---|---|---|---|---|---|---|---|---|---|---|---|---|---|---|---|
| chr10_81507705_81508491 | chr10 | 81507706 | 81508491 | + | -0.777183 | intron (NM_010304, intron 5 of 6) | intron (NM_010304, intron 5 of 6) | -7961 | NM_010102 | 13611 | Mm.33065 | NM_010102 | ENSMUSG00000044199 | **S1pr4** | Edg6\|Lpc1\|S1p4 | sphingosine-1-phosphate receptor 4 | protein-coding |
| chr10_81510272_81510708 | chr10 | 81510273 | 81510708 | + | -0.80604 | intron (NM_010304, intron 4 of 6) | intron (NM_010304, intron 4 of 6) | -10353 | NM_010102 | 13611 | Mm.33065 | NM_010102 | ENSMUSG00000044199 | **S1pr4** | Edg6\|Lpc1\|S1p4 | sphingosine-1-phosphate receptor 4 | protein-coding |
| chr10_81507965_81508473 | chr10 | 81507966 | 81508473 | + | -0.833573 | intron (NM_010304, intron 5 of 6) | intron (NM_010304, intron 5 of 6) | -8082 | NM_010102 | 13611 | Mm.33065 | NM_010102 | ENSMUSG00000044199 | **S1pr4** | Edg6\|Lpc1\|S1p4 | sphingosine-1-phosphate receptor 4 | protein-coding |
| chr7_16364143_16364743 | chr7 | 16364144 | 16364743 | + | -0.883796 | intron (NM_019748, intron 5 of 8) | intron (NM_019748, intron 5 of 8) | 23453 | NM_001285892 | 56459 | Mm.258530 | NM_019748 | ENSMUSG00000052833 | **Sae1** | 2400010M20Rik\|2610044L12Rik\|AL033372\|AOS1\|AW743391\|D7Ertd177e\|HSPC140\|Sua1\|Uble1a | SUMO1 activating enzyme subunit 1 | protein-coding |
| chr1_87828125_87828690 | chr1 | 87828126 | 87828690 | + | -0.71737 | exon (NM_009118, exon 10 of 16) | exon (NM_009118, exon 10 of 16) | 24728 | NM_009118 | 20215 | Mm.1276 | NM_009118 | ENSMUSG00000056055 | **Sag** | A930001K18Rik\|Arr1\|Irbp\|arrestin | S-antigen, retina and pineal gland (arrestin) | protein-coding |
| chr18_81366760_81367059 | chr18 | 81366761 | 81367059 | + | -0.652577 | Intergenic | Intergenic | -380332 | NM_178280 | 20689 | Mm.215917 | NM_178280 | ENSMUSG00000024565 | **Sall3** | B130022O04Rik\|Msal\|Msal-1\|Sal\|Salt\|Spalt | sal-like 3 (Drosophila) | protein-coding |
| chr18_81176542_81176910 | chr18 | 81176543 | 81176910 | + | -0.770792 | Intergenic | ORR1E\|LTR\|ERVL-MaLR | -190148 | NM_178280 | 20689 | Mm.215917 | NM_178280 | ENSMUSG00000024565 | **Sall3** | B130022O04Rik\|Msal\|Msal-1\|Sal\|Salt\|Spalt | sal-like 3 (Drosophila) | protein-coding |
| chr18_81422434_81422737 | chr18 | 81422435 | 81422737 | + | -0.839516 | Intergenic | Intergenic | -436008 | NM_178280 | 20689 | Mm.215917 | NM_178280 | ENSMUSG00000024565 | **Sall3** | B130022O04Rik\|Msal\|Msal-1\|Sal\|Salt\|Spalt | sal-like 3 (Drosophila) | protein-coding |
| chr18_81360079_81360326 | chr18 | 81360080 | 81360326 | + | -0.857864 | Intergenic | Intergenic | -373625 | NM_178280 | 20689 | Mm.215917 | NM_178280 | ENSMUSG00000024565 | **Sall3** | B130022O04Rik\|Msal\|Msal-1\|Sal\|Salt\|Spalt | sal-like 3 (Drosophila) | protein-coding |
| chr18_81429860_81430195 | chr18 | 81429861 | 81430195 | + | -0.895136 | Intergenic | ORR1D2\|LTR\|ERVL-MaLR | -443450 | NM_178280 | 20689 | Mm.215917 | NM_178280 | ENSMUSG00000024565 | **Sall3** | B130022O04Rik\|Msal\|Msal-1\|Sal\|Salt\|Spalt | sal-like 3 (Drosophila) | protein-coding |
| chr18_81429852_81430411 | chr18 | 81429853 | 81430411 | + | -0.935463 | Intergenic | ORR1D2\|LTR\|ERVL-MaLR | -443554 | NM_178280 | 20689 | Mm.215917 | NM_178280 | ENSMUSG00000024565 | **Sall3** | B130022O04Rik\|Msal\|Msal-1\|Sal\|Salt\|Spalt | sal-like 3 (Drosophila) | protein-coding |
| chr18_81046771_81047214 | chr18 | 81046772 | 81047214 | + | -0.93594 | Intergenic | Intergenic | -60415 | NM_178280 | 20689 | Mm.215917 | NM_178280 | ENSMUSG00000024565 | **Sall3** | B130022O04Rik\|Msal\|Msal-1\|Sal\|Salt\|Spalt | sal-like 3 (Drosophila) | protein-coding |
| chr18_81404880_81405121 | chr18 | 81404881 | 81405121 | + | -0.980656 | Intergenic | ORR1D2\|LTR\|ERVL-MaLR | -418423 | NM_178280 | 20689 | Mm.215917 | NM_178280 | ENSMUSG00000024565 | **Sall3** | B130022O04Rik\|Msal\|Msal-1\|Sal\|Salt\|Spalt | sal-like 3 (Drosophila) | protein-coding |
| chr18_81046602_81047386 | chr18 | 81046603 | 81047386 | + | -0.995938 | Intergenic | Intergenic | -60416 | NM_178280 | 20689 | Mm.215917 | NM_178280 | ENSMUSG00000024565 | **Sall3** | B130022O04Rik\|Msal\|Msal-1\|Sal\|Salt\|Spalt | sal-like 3 (Drosophila) | protein-coding |
| chr18_81404831_81405184 | chr18 | 81404832 | 81405184 | + | -1.00465 | Intergenic | ORR1D2\|LTR\|ERVL-MaLR | -418430 | NM_178280 | 20689 | Mm.215917 | NM_178280 | ENSMUSG00000024565 | **Sall3** | B130022O04Rik\|Msal\|Msal-1\|Sal\|Salt\|Spalt | sal-like 3 (Drosophila) | protein-coding |
| chr18_81210582_81211469 | chr18 | 81210583 | 81211469 | + | -1.53525 | Intergenic | Intergenic | -224448 | NM_178280 | 20689 | Mm.215917 | NM_178280 | ENSMUSG00000024565 | **Sall3** | B130022O04Rik\|Msal\|Msal-1\|Sal\|Salt\|Spalt | sal-like 3 (Drosophila) | protein-coding |
| chr4_156254971_156255798 | chr4 | 156254972 | 156255798 | + | -0.616225 | promoter-TSS (NM_001110516) | promoter-TSS (NM_001110516) | -47 | NM_001110516 | 231004 | Mm.406833 | NM_173736 | ENSMUSG00000096351 | **Samd11** | BC026991\|mr-s | sterile alpha motif domain containing 11 | protein-coding |
| chr15_53724102_53724481 | chr15 | 53724103 | 53724481 | + | -0.779281 | intron (NM_177225, intron 2 of 4) | intron (NM_177225, intron 2 of 4) | 178144 | NM_177225 | 320679 | Mm.442455 | NM_177225 | ENSMUSG00000058656 | **Samd12** | A830094I09Rik | sterile alpha motif domain containing 12 | protein-coding |
| chr10_26259757_26260268 | chr10 | 26259758 | 26260268 | + | -0.616604 | 3' UTR (NM_001013766, exon 8 of 8) | 3' UTR (NM_001013766, exon 8 of 8) | 30306 | NM_001115154 | 268288 | Mm.213322 | NM_001013766 | ENSMUSG00000051354 | **Samd3** | Gm623 | sterile alpha motif domain containing 3 | protein-coding |
| chr14_21751294_21751616 | chr14 | 21751295 | 21751616 | + | -0.725756 | intron (NM_026283, intron 1 of 5) | intron (NM_026283, intron 1 of 5) | 924 | NM_026283 | 67630 | Mm.102765 | NM_026283 | ENSMUSG00000021770 | **Samd8** | 1110053F04Rik\|1700010P07Rik\|SMSr | sterile alpha motif domain containing 8 | protein-coding |
| chr6_3345890_3346301 | chr6 | 3345891 | 3346301 | + | -0.637726 | Intergenic | Intergenic | 53475 | NM_010156 | 209086 | Mm.196013 | NM_010156 | ENSMUSG00000047735 | **Samd9l** | AA175286\|ESTM25\|mKIAA2005 | sterile alpha motif domain containing 9-like | protein-coding |
| chr6_3399329_3399920 | chr6 | 3399330 | 3399920 | + | -0.672288 | promoter-TSS (NM_010156) | promoter-TSS (NM_010156) | -54 | NM_010156 | 209086 | Mm.196013 | NM_010156 | ENSMUSG00000047735 | **Samd9l** | AA175286\|ESTM25\|mKIAA2005 | sterile alpha motif domain containing 9-like | protein-coding |
| chr6_3399301_3399703 | chr6 | 3399302 | 3399703 | + | -0.696724 | promoter-TSS (NM_010156) | promoter-TSS (NM_010156) | 69 | NM_010156 | 209086 | Mm.196013 | NM_010156 | ENSMUSG00000047735 | **Samd9l** | AA175286\|ESTM25\|mKIAA2005 | sterile alpha motif domain containing 9-like | protein-coding |
| chr2_157147377_157148170 | chr2 | 157147378 | 157148170 | + | -0.668221 | intron (NM_011249, intron 21 of 21) | intron (NM_011249, intron 21 of 21) | -12552 | NM_018851 | 56045 | Mm.248478 | NM_018851 | ENSMUSG00000027639 | **Samhd1** | E330031J07Rik\|Mg11 | SAM domain and HD domain, 1 | protein-coding |
| chr2_157129107_157129586 | chr2 | 157129108 | 157129586 | + | -0.885069 | intron (NM_001139520, intron 2 of 16) | intron (NM_001139520, intron 2 of 16) | 5875 | NM_001139520 | 56045 | Mm.248478 | NM_018851 | ENSMUSG00000027639 | **Samhd1** | E330031J07Rik\|Mg11 | SAM domain and HD domain, 1 | protein-coding |
| chr2_157121611_157122189 | chr2 | 157121612 | 157122189 | + | -0.889168 | exon (NM_018851, exon 5 of 16) | exon (NM_018851, exon 5 of 16) | 13322 | NM_001139520 | 56045 | Mm.248478 | NM_018851 | ENSMUSG00000027639 | **Samhd1** | E330031J07Rik\|Mg11 | SAM domain and HD domain, 1 | protein-coding |
| chr19_5389624_5390504 | chr19 | 5389625 | 5390504 | + | -0.698558 | TTS (NR_033224) | TTS (NR_033224) | -1361 | NM_016882 | 20227 | Mm.34562 | NM_016882 | ENSMUSG00000039148 | **Sart1** | U5-110K | squamous cell carcinoma antigen recognized by T cells 1 | protein-coding |
| chr19_5389639_5390117 | chr19 | 5389640 | 5390117 | + | -0.767974 | non-coding (NR_033224, exon 1 of 1) | non-coding (NR_033224, exon 1 of 1) | -1175 | NM_016882 | 20227 | Mm.34562 | NM_016882 | ENSMUSG00000039148 | **Sart1** | U5-110K | squamous cell carcinoma antigen recognized by T cells 1 | protein-coding |
| chr17_51814224_51814772 | chr17 | 51814225 | 51814772 | + | -0.648755 | intron (NM_001163632, intron 1 of 10) | intron (NM_001163632, intron 1 of 10) | -2444 | NM_001163630 | 20230 | Mm.311655 | NM_009122 | ENSMUSG00000023927 | **Satb1** | 2610306G12Rik\|AW413156 | special AT-rich sequence binding protein 1 | protein-coding |
| chr17_51855126_51855641 | chr17 | 51855127 | 51855641 | + | -0.765606 | Intergenic | Intergenic | -22094 | NM_001163632 | 20230 | Mm.311655 | NM_009122 | ENSMUSG00000023927 | **Satb1** | 2610306G12Rik\|AW413156 | special AT-rich sequence binding protein 1 | protein-coding |
| chr17_51805241_51805958 | chr17 | 51805242 | 51805958 | + | -0.799538 | intron (NM_001163631, intron 2 of 10) | intron (NM_001163631, intron 2 of 10) | 6454 | NM_001163630 | 20230 | Mm.311655 | NM_009122 | ENSMUSG00000023927 | **Satb1** | 2610306G12Rik\|AW413156 | special AT-rich sequence binding protein 1 | protein-coding |
| chr17_51732523_51733031 | chr17 | 51732524 | 51733031 | + | -0.827895 | Intergenic | L1MD\|LINE\|L1 | 79277 | NM_001163630 | 20230 | Mm.311655 | NM_009122 | ENSMUSG00000023927 | **Satb1** | 2610306G12Rik\|AW413156 | special AT-rich sequence binding protein 1 | protein-coding |
| chr17_51805241_51805646 | chr17 | 51805242 | 51805646 | + | -0.840582 | intron (NM_001163631, intron 2 of 10) | intron (NM_001163631, intron 2 of 10) | 6610 | NM_001163630 | 20230 | Mm.311655 | NM_009122 | ENSMUSG00000023927 | **Satb1** | 2610306G12Rik\|AW413156 | special AT-rich sequence binding protein 1 | protein-coding |
| chr17_51855260_51855857 | chr17 | 51855261 | 51855857 | + | -0.843382 | Intergenic | Intergenic | -22269 | NM_001163632 | 20230 | Mm.311655 | NM_009122 | ENSMUSG00000023927 | **Satb1** | 2610306G12Rik\|AW413156 | special AT-rich sequence binding protein 1 | protein-coding |
| chr17_51735501_51736026 | chr17 | 51735502 | 51736026 | + | -0.879008 | TTS (NM_009122) | TTS (NM_009122) | 76290 | NM_001163630 | 20230 | Mm.311655 | NM_009122 | ENSMUSG00000023927 | **Satb1** | 2610306G12Rik\|AW413156 | special AT-rich sequence binding protein 1 | protein-coding |
| chr17_51857677_51857972 | chr17 | 51857678 | 51857972 | + | -0.952372 | Intergenic | Intergenic | -24535 | NM_001163632 | 20230 | Mm.311655 | NM_009122 | ENSMUSG00000023927 | **Satb1** | 2610306G12Rik\|AW413156 | special AT-rich sequence binding protein 1 | protein-coding |
| chr17_51857725_51857916 | chr17 | 51857726 | 51857916 | + | -0.953487 | Intergenic | Intergenic | -24531 | NM_001163632 | 20230 | Mm.311655 | NM_009122 | ENSMUSG00000023927 | **Satb1** | 2610306G12Rik\|AW413156 | special AT-rich sequence binding protein 1 | protein-coding |
| chr17_51735210_51736079 | chr17 | 51735211 | 51736079 | + | -0.959512 | TTS (NM_009122) | TTS (NM_009122) | 76409 | NM_001163630 | 20230 | Mm.311655 | NM_009122 | ENSMUSG00000023927 | **Satb1** | 2610306G12Rik\|AW413156 | special AT-rich sequence binding protein 1 | protein-coding |
| chr17_51800264_51800797 | chr17 | 51800265 | 51800797 | + | -1.23113 | intron (NM_001163631, intron 5 of 10) | intron (NM_001163631, intron 5 of 10) | 11523 | NM_001163630 | 20230 | Mm.311655 | NM_009122 | ENSMUSG00000023927 | **Satb1** | 2610306G12Rik\|AW413156 | special AT-rich sequence binding protein 1 | protein-coding |
| chr17_51800267_51800591 | chr17 | 51800268 | 51800591 | + | -1.28843 | intron (NM_001163631, intron 5 of 10) | intron (NM_001163631, intron 5 of 10) | 11625 | NM_001163630 | 20230 | Mm.311655 | NM_009122 | ENSMUSG00000023927 | **Satb1** | 2610306G12Rik\|AW413156 | special AT-rich sequence binding protein 1 | protein-coding |
| chr12_70018063_70018475 | chr12 | 70018064 | 70018475 | + | -0.799255 | TTS (NR_104397) | TTS (NR_104397) | -31267 | NM_022028 | 64010 | Mm.390480 | NM_022028 | ENSMUSG00000021067 | **Sav1** | 1700040G09Rik\|Salv\|Sav\|WW45\|Wwp3\|Wwp4 | salvador homolog 1 (Drosophila) | protein-coding |
| chr12_70018208_70018512 | chr12 | 70018209 | 70018512 | + | -0.799992 | TTS (NR_104397) | TTS (NR_104397) | -31358 | NM_022028 | 64010 | Mm.390480 | NM_022028 | ENSMUSG00000021067 | **Sav1** | 1700040G09Rik\|Salv\|Sav\|WW45\|Wwp3\|Wwp4 | salvador homolog 1 (Drosophila) | protein-coding |
| chr7_4948360_4948853 | chr7 | 4948361 | 4948853 | + | -0.771304 | Intergenic | Intergenic | 15741 | NM_001146329 | 381836 | Mm.297829 | NM_001146329 | ENSMUSG00000030433 | **Sbk2** | Gm1079\|Sgk069 | SH3-binding domain kinase family, member 2 | protein-coding |
| chr15_96543572_96543802 | chr15 | 96543573 | 96543802 | + | -0.682323 | Intergenic | Intergenic | -82844 | NM_028148 | 72193 | Mm.324474 | NM_028148 | ENSMUSG00000033228 | **Scaf11** | 1110061H03Rik\|2610510E10Rik\|AI462454\|CASP11\|SIP1\|SRRP129\|Sfrs2ip\|Srsf2ip\|mKIAA3013 | SR-related CTD-associated factor 11 | protein-coding |
| chr15_96497661_96498193 | chr15 | 96497662 | 96498193 | + | -1.11238 | Intergenic | ORR1D2\|LTR\|ERVL-MaLR | -37084 | NM_028148 | 72193 | Mm.324474 | NM_028148 | ENSMUSG00000033228 | **Scaf11** | 1110061H03Rik\|2610510E10Rik\|AI462454\|CASP11\|SIP1\|SRRP129\|Sfrs2ip\|Srsf2ip\|mKIAA3013 | SR-related CTD-associated factor 11 | protein-coding |
| chr15_96497757_96498058 | chr15 | 96497758 | 96498058 | + | -1.12793 | Intergenic | ORR1D2\|LTR\|ERVL-MaLR | -37065 | NM_028148 | 72193 | Mm.324474 | NM_028148 | ENSMUSG00000033228 | **Scaf11** | 1110061H03Rik\|2610510E10Rik\|AI462454\|CASP11\|SIP1\|SRRP129\|Sfrs2ip\|Srsf2ip\|mKIAA3013 | SR-related CTD-associated factor 11 | protein-coding |
| chr15_96497709_96498386 | chr15 | 96497710 | 96498386 | + | -1.1284 | Intergenic | Intergenic | -37205 | NM_028148 | 72193 | Mm.324474 | NM_028148 | ENSMUSG00000033228 | **Scaf11** | 1110061H03Rik\|2610510E10Rik\|AI462454\|CASP11\|SIP1\|SRRP129\|Sfrs2ip\|Srsf2ip\|mKIAA3013 | SR-related CTD-associated factor 11 | protein-coding |
| chr16_17807013_17807832 | chr16 | 17807014 | 17807832 | + | -0.596953 | exon (NM_153790, exon 11 of 11) | exon (NM_153790, exon 11 of 11) | 10141 | NM_153790 | 224024 | Mm.194950 | NM_153790 | ENSMUSG00000012017 | **Scarf2** | AI194363\|NSR1\|SREC-II\|SRECRP-1\|Scarf1\|Scatf1\|Srec2\|SrecII\|W53987\|W65560 | scavenger receptor class F, member 2 | protein-coding |

| | | | | | | | | | | | | | | | | | |
|---|---|---|---|---|---|---|---|---|---|---|---|---|---|---|---|---|---|
| chr18_74760302_74761097 | chr18 | 74760303 | 74761097 | + | -1.06431 | intron (NM_201600, intron 36 of 38) | intron (NM_201600, intron 36 of 38) | 17755 | NR_028560 | 100217466 | | NR_028560 | | **Scarna17** | MBII-119 | small Cajal body-specific RNA 17 | ncRNA |
| chr18_74760546_74761009 | chr18 | 74760547 | 74761009 | + | -1.19488 | intron (NM_201600, intron 36 of 38) | intron (NM_201600, intron 36 of 38) | 17677 | NR_028560 | 100217466 | | NR_028560 | | **Scarna17** | MBII-119 | small Cajal body-specific RNA 17 | ncRNA |
| chr5_74402621_74403224 | chr5 | 74402622 | 74403224 | + | -0.942378 | intron (NM_001286437, intron 4 of 7) | MTD\|LTR\|ERVL-MaLR | 128819 | NM_178672 | 212986 | Mm.134147 | NM_178672 | ENSMUSG00000062110 | **Scfd2** | E430013M20Rik\|STXBP1L1 | Sec1 family domain containing 2 | protein-coding |
| chr5_74404038_74404533 | chr5 | 74404039 | 74404533 | + | -1.12322 | intron (NM_001286437, intron 4 of 7) | ORR1E\|LTR\|ERVL-MaLR | 127456 | NM_178672 | 212986 | Mm.134147 | NM_178672 | ENSMUSG00000062110 | **Scfd2** | E430013M20Rik\|STXBP1L1 | Sec1 family domain containing 2 | protein-coding |
| chr2_113835105_113835802 | chr2 | 113835106 | 113835802 | + | -0.984024 | intron (NM_181416, intron 10 of 11) | L2\|LINE\|L2 | -6363 | NM_009162 | 20394 | Mm.4836 | NM_009162 | ENSMUSG00000023236 | **Scg5** | 7B2\|AI325031\|Sgne-1\|Sgne1 | secretogranin V | protein-coding |
| chr19_9065671_9066684 | chr19 | 9065672 | 9066684 | + | -0.614563 | intron (NM_001286518, intron 6 of 6) | MIR\|SINE\|MIR | 21778 | NM_011681 | 22287 | Mm.2258 | NM_011681 | ENSMUSG00000024653 | **Scgb1a1** | CC10\|CC16\|CCSP\|PCB-BP\|UG\|UGB\|Utg | secretoglobin, family 1A, member 1 (uteroglobin) | protein-coding |
| chr12_40089928_40090367 | chr12 | 40089929 | 40090367 | + | -0.81676 | intron (NM_009132, intron 4 of 13) | intron (NM_009132, intron 4 of 13) | 44080 | NM_009132 | 20259 | Mm.2416 | NM_009132 | ENSMUSG00000002565 | **Scin** | AW545522\|adseverin | scinderin | protein-coding |
| chr10_42888086_42888822 | chr10 | 42888087 | 42888822 | + | -0.678361 | intron (NM_172938, intron 1 of 8) | intron (NM_172938, intron 1 of 8) | 27942 | NM_172938 | 268297 | Mm.98731 | NM_172938 | ENSMUSG00000044770 | **Scml4** | 9330161D17\|mFLJ00197 | sex comb on midleg-like 4 (Drosophila) | protein-coding |
| chr10_42962183_42963010 | chr10 | 42962184 | 42963010 | + | -0.792784 | Intergenic | Intergenic | 102085 | NM_172938 | 268297 | Mm.98731 | NM_172938 | ENSMUSG00000044770 | **Scml4** | 9330161D17\|mFLJ00197 | sex comb on midleg-like 4 (Drosophila) | protein-coding |
| chr10_42843183_42843628 | chr10 | 42843184 | 42843628 | + | -0.837285 | Intergenic | Intergenic | -17106 | NM_172938 | 268297 | Mm.98731 | NM_172938 | ENSMUSG00000044770 | **Scml4** | 9330161D17\|mFLJ00197 | sex comb on midleg-like 4 (Drosophila) | protein-coding |
| chr10_42886949_42887414 | chr10 | 42886950 | 42887414 | + | -1.10918 | intron (NM_172938, intron 1 of 8) | intron (NM_172938, intron 1 of 8) | 26670 | NM_172938 | 268297 | Mm.98731 | NM_172938 | ENSMUSG00000044770 | **Scml4** | 9330161D17\|mFLJ00197 | sex comb on midleg-like 4 (Drosophila) | protein-coding |
| chr9_45098331_45099108 | chr9 | 45098332 | 45099108 | + | -0.711892 | intron (NM_001005421, intron 7 of 10) | intron (NM_001005421, intron 7 of 10) | -19156 | NM_001014761 | 72821 | Mm.229373 | NM_001014761 | ENSMUSG00000070304 | **Scn2b** | 2810451E09Rik\|AI840361\|Gm183 | sodium channel, voltage-gated, type II, beta | protein-coding |
| chr9_45098436_45098785 | chr9 | 45098437 | 45098785 | + | -0.738448 | intron (NM_001005421, intron 7 of 10) | intron (NM_001005421, intron 7 of 10) | -19265 | NM_001014761 | 72821 | Mm.229373 | NM_001014761 | ENSMUSG00000070304 | **Scn2b** | 2810451E09Rik\|AI840361\|Gm183 | sodium channel, voltage-gated, type II, beta | protein-coding |
| chr9_45116869_45117210 | chr9 | 45116870 | 45117210 | + | -0.936905 | promoter-TSS (NM_001014761) | promoter-TSS (NM_001014761) | -836 | NM_001014761 | 72821 | Mm.229373 | NM_001014761 | ENSMUSG00000070304 | **Scn2b** | 2810451E09Rik\|AI840361\|Gm183 | sodium channel, voltage-gated, type II, beta | protein-coding |
| chr9_45116634_45117260 | chr9 | 45116635 | 45117260 | + | -1.00519 | promoter-TSS (NM_001014761) | promoter-TSS (NM_001014761) | -929 | NM_001014761 | 72821 | Mm.229373 | NM_001014761 | ENSMUSG00000070304 | **Scn2b** | 2810451E09Rik\|AI840361\|Gm183 | sodium channel, voltage-gated, type II, beta | protein-coding |
| chr11_106351886_106352376 | chr11 | 106351887 | 106352376 | + | -0.84875 | Intergenic | Intergenic | -2741 | NM_133199 | 110880 | Mm.432528 | NM_133199 | ENSMUSG00000001027 | **Scn4a** | Nav1.4\|SkM1\|mH2 | sodium channel, voltage-gated, type IV, alpha | protein-coding |
| chr15_100844799_100845795 | chr15 | 100844800 | 100845795 | + | -0.621497 | Intergenic | Intergenic | -25348 | NM_001077499 | 20273 | Mm.385012 | NM_011323 | ENSMUSG00000023033 | **Scn8a** | AI853486\|C630029C19Rik\|NaCh6\|Nav1.6\|dmu\|med\|mnd-2\|mnd2\|nmf2\|nmf335\|nmf58\|nur14\|seal | sodium channel, voltage-gated, type VIII, alpha | protein-coding |
| chr15_100905899_100906212 | chr15 | 100905900 | 100906212 | + | -0.980814 | intron (NM_001077499, intron 1 of 26) | intron (NM_001077499, intron 1 of 26) | -30215 | NM_011323 | 20273 | Mm.385012 | NM_011323 | ENSMUSG00000023033 | **Scn8a** | AI853486\|C630029C19Rik\|NaCh6\|Nav1.6\|dmu\|med\|mnd-2\|mnd2\|nmf2\|nmf335\|nmf58\|nur14\|seal | sodium channel, voltage-gated, type VIII, alpha | protein-coding |
| chr15_100905049_100905419 | chr15 | 100905050 | 100905419 | + | -1.16113 | intron (NM_001077499, intron 1 of 26) | intron (NM_001077499, intron 1 of 26) | -31037 | NM_011323 | 20273 | Mm.385012 | NM_011323 | ENSMUSG00000023033 | **Scn8a** | AI853486\|C630029C19Rik\|NaCh6\|Nav1.6\|dmu\|med\|mnd-2\|mnd2\|nmf2\|nmf335\|nmf58\|nur14\|seal | sodium channel, voltage-gated, type VIII, alpha | protein-coding |
| chr10_89703134_89703472 | chr10 | 89703135 | 89703472 | + | -0.720389 | Intergenic | Intergenic | -17018 | NM_198021 | 213326 | Mm.27651 | NM_198021 | ENSMUSG00000069539 | **Scyl2** | AU015688\|AW536445\|BC030932\|CVAK104\|D10Ertd802e | SCY1-like 2 (S. cerevisiae) | protein-coding |
| chr4_130774063_130775210 | chr4 | 130774064 | 130775210 | + | -1.17276 | intron (NM_001159604, intron 21 of 21) | intron (NM_001159604, intron 21 of 21) | -17900 | NM_011520 | 20970 | Mm.206536 | NM_011520 | ENSMUSG00000025743 | **Sdc3** | Synd3\|mKIAA0468\|syn-3 | syndecan 3 | protein-coding |
| chr4_130774367_130775242 | chr4 | 130774368 | 130775242 | + | -1.27618 | intron (NM_001159604, intron 21 of 21) | intron (NM_001159604, intron 21 of 21) | -17732 | NM_011520 | 20970 | Mm.206536 | NM_011520 | ENSMUSG00000025743 | **Sdc3** | Synd3\|mKIAA0468\|syn-3 | syndecan 3 | protein-coding |
| chr7_80936682_80937274 | chr7 | 80936683 | 80937274 | + | -0.988486 | intron (NM_019951, intron 1 of 5) | intron (NM_019951, intron 1 of 5) | 10572 | NM_019951 | 56529 | Mm.43760 | NM_019951 | ENSMUSG00000025724 | **Sec11a** | 1810012E07Rik\|18kDa\|Sec11l1\|Sid2895p\|Spc18\|sid2895 | SEC11 homolog A (S. cerevisiae) | protein-coding |
| chr18_65818805_65819593 | chr18 | 65818806 | 65819593 | + | -0.968918 | Intergenic | RLTR40\|LTR\|ERVK | 18621 | NM_025468 | 66286 | Mm.27800 | NM_025468 | ENSMUSG00000024516 | **Sec11c** | 1810029G24Rik\|Sec11l3 | SEC11 homolog C (S. cerevisiae) | protein-coding |
| chr9_121682060_121682796 | chr9 | 121682061 | 121682796 | + | -0.680789 | 3' UTR (NM_001164562, exon 7 of 7) | 3' UTR (NM_001164562, exon 7 of 7) | 22601 | NM_001164562 | 215474 | Mm.31421 | NM_178677 | ENSMUSG00000061536 | **Sec22c** | 4932412K21\|5930407I15Rik\|C530046H07\|Sec22l3 | SEC22 vesicle trafficking protein homolog C (S. cerevisiae) | protein-coding |
| chr9_121677072_121677372 | chr9 | 121677073 | 121677372 | + | -0.734386 | Intergenic | Intergenic | 27807 | NM_001164562 | 215474 | Mm.31421 | NM_178677 | ENSMUSG00000061536 | **Sec22c** | 4932412K21\|5930407I15Rik\|C530046H07\|Sec22l3 | SEC22 vesicle trafficking protein homolog C (S. cerevisiae) | protein-coding |
| chr12_92775756_92776299 | chr12 | 92775757 | 92776299 | + | -0.680786 | Intergenic | Intergenic | -926871 | NM_011344 | 20338 | Mm.250605 | NM_011344 | ENSMUSG00000020964 | **Sel1l** | AW493766\|Sel1h\|mKIAA4137 | sel-1 suppressor of lin-12-like (C. elegans) | protein-coding |
| chr12_92775754_92776064 | chr12 | 92775755 | 92776064 | + | -0.707572 | Intergenic | Intergenic | -926752 | NM_011344 | 20338 | Mm.250605 | NM_011344 | ENSMUSG00000020964 | **Sel1l** | AW493766\|Sel1h\|mKIAA4137 | sel-1 suppressor of lin-12-like (C. elegans) | protein-coding |
| chr12_92775583_92776004 | chr12 | 92775584 | 92776004 | + | -0.749283 | Intergenic | Intergenic | -926637 | NM_011344 | 20338 | Mm.250605 | NM_011344 | ENSMUSG00000020964 | **Sel1l** | AW493766\|Sel1h\|mKIAA4137 | sel-1 suppressor of lin-12-like (C. elegans) | protein-coding |
| chr12_92823015_92823811 | chr12 | 92823016 | 92823811 | + | -0.754217 | Intergenic | Intergenic | -974256 | NM_011344 | 20338 | Mm.250605 | NM_011344 | ENSMUSG00000020964 | **Sel1l** | AW493766\|Sel1h\|mKIAA4137 | sel-1 suppressor of lin-12-like (C. elegans) | protein-coding |
| chr3_144591128_144591786 | chr3 | 144591129 | 144591786 | + | -1.05665 | intron (NM_053102, intron 3 of 4) | intron (NM_053102, intron 3 of 4) | 21030 | NM_053102 | 93684 | Mm.29812 | NM_053102 | ENSMUSG00000037072 | **Selenof** | 9430015P09Rik | selenoprotein | protein-coding |
| chr1_164063736_164064327 | chr1 | 164063737 | 164064327 | + | -0.608916 | intron (NM_011346, intron 2 of 8) | Lx8\|LINE\|L1 | 1956 | NM_001164059 | 20343 | Mm.1461 | NM_011346 | ENSMUSG00000026581 | **Sell** | AI528707\|CD62L\|L-selectin\|LECAM-1\|Lnhr\|Ly-22\|Ly-m22\|Lyam-1\|Lyam1 | selectin, lymphocyte | protein-coding |
| chr1_164063663_164064159 | chr1 | 164063664 | 164064159 | + | -0.657963 | intron (NM_011346, intron 2 of 8) | Lx8\|LINE\|L1 | 1835 | NM_001164059 | 20343 | Mm.1461 | NM_011346 | ENSMUSG00000026581 | **Sell** | AI528707\|CD62L\|L-selectin\|LECAM-1\|Lnhr\|Ly-22\|Ly-m22\|Lyam-1\|Lyam1 | selectin, lymphocyte | protein-coding |
| chr5_113841820_113842918 | chr5 | 113841821 | 113842918 | + | -0.602926 | TTS (NM_011779) | TTS (NM_011779) | -11868 | NM_009151 | 20345 | Mm.332590 | NM_009151 | ENSMUSG00000048163 | **Selplg** | CD162\|Psgl-1\|Psgl1\|Selp1\|Selpl | selectin, platelet (p-selectin) ligand | protein-coding |
| chr5_113830195_113830749 | chr5 | 113830196 | 113830749 | + | -0.610588 | promoter-TSS (NM_009151) | promoter-TSS (NM_009151) | 29 | NM_009151 | 20345 | Mm.332590 | NM_009151 | ENSMUSG00000048163 | **Selplg** | CD162\|Psgl-1\|Psgl1\|Selp1\|Selpl | selectin, platelet (p-selectin) ligand | protein-coding |
| chr5_113832345_113833080 | chr5 | 113832346 | 113833080 | + | -0.636188 | Intergenic | Intergenic | -2212 | NM_009151 | 20345 | Mm.332590 | NM_009151 | ENSMUSG00000048163 | **Selplg** | CD162\|Psgl-1\|Psgl1\|Selp1\|Selpl | selectin, platelet (p-selectin) ligand | protein-coding |
| chr5_113841839_113842248 | chr5 | 113841840 | 113842248 | + | -0.726345 | TTS (NM_011779) | TTS (NM_011779) | -11543 | NM_009151 | 20345 | Mm.332590 | NM_009151 | ENSMUSG00000048163 | **Selplg** | CD162\|Psgl-1\|Psgl1\|Selp1\|Selpl | selectin, platelet (p-selectin) ligand | protein-coding |
| chr5_113824283_113824769 | chr5 | 113824284 | 113824769 | + | -0.82157 | intron (NM_009151, intron 1 of 1) | intron (NM_009151, intron 1 of 1) | 5975 | NM_009151 | 20345 | Mm.332590 | NM_009151 | ENSMUSG00000048163 | **Selplg** | CD162\|Psgl-1\|Psgl1\|Selp1\|Selpl | selectin, platelet (p-selectin) ligand | protein-coding |
| chr5_113859001_113859271 | chr5 | 113859002 | 113859271 | + | -0.851289 | intron (NM_011779, intron 3 of 10) | intron (NM_011779, intron 3 of 10) | -28635 | NM_009151 | 20345 | Mm.332590 | NM_009151 | ENSMUSG00000048163 | **Selplg** | CD162\|Psgl-1\|Psgl1\|Selp1\|Selpl | selectin, platelet (p-selectin) ligand | protein-coding |
| chr5_113858996_113859624 | chr5 | 113858997 | 113859624 | + | -0.897293 | intron (NM_011779, intron 3 of 10) | intron (NM_011779, intron 3 of 10) | -28809 | NM_009151 | 20345 | Mm.332590 | NM_009151 | ENSMUSG00000048163 | **Selplg** | CD162\|Psgl-1\|Psgl1\|Selp1\|Selpl | selectin, platelet (p-selectin) ligand | protein-coding |
| chr5_113825607_113825919 | chr5 | 113825608 | 113825919 | + | -1.0844 | intron (NM_009151, intron 1 of 1) | intron (NM_009151, intron 1 of 1) | 4738 | NM_009151 | 20345 | Mm.332590 | NM_009151 | ENSMUSG00000048163 | **Selplg** | CD162\|Psgl-1\|Psgl1\|Selp1\|Selpl | selectin, platelet (p-selectin) ligand | protein-coding |
| chr5_113825654_113825885 | chr5 | 113825655 | 113825885 | + | -1.11103 | intron (NM_009151, intron 1 of 1) | intron (NM_009151, intron 1 of 1) | 4731 | NM_009151 | 20345 | Mm.332590 | NM_009151 | ENSMUSG00000048163 | **Selplg** | CD162\|Psgl-1\|Psgl1\|Selp1\|Selpl | selectin, platelet (p-selectin) ligand | protein-coding |
| chr3_58581288_58582162 | chr3 | 58581289 | 58582162 | + | -0.724022 | intron (NM_001040396, intron 1 of 5) | intron (NM_001040396, intron 1 of 5) | 5067 | NM_001040396 | 69227 | Mm.270950 | NM_001040396 | ENSMUSG00000075700 | **Selt** | 2810407C02Rik\|5730408P04Rik | selenoprotein T | protein-coding |
| chr3_58582885_58583398 | chr3 | 58582886 | 58583398 | + | -0.751316 | intron (NM_001040396, intron 1 of 5) | intron (NM_001040396, intron 1 of 5) | 6484 | NM_001040396 | 69227 | Mm.270950 | NM_001040396 | ENSMUSG00000075700 | **Selt** | 2810407C02Rik\|5730408P04Rik | selenoprotein T | protein-coding |
| chr3_58583115_58583386 | chr3 | 58583116 | 58583386 | + | -0.83213 | intron (NM_001040396, intron 1 of 5) | intron (NM_001040396, intron 1 of 5) | 6593 | NM_001040396 | 69227 | Mm.270950 | NM_001040396 | ENSMUSG00000075700 | **Selt** | 2810407C02Rik\|5730408P04Rik | selenoprotein T | protein-coding |
| chr3_58581498_58581942 | chr3 | 58581499 | 58581942 | + | -0.841432 | intron (NM_001040396, intron 1 of 5) | intron (NM_001040396, intron 1 of 5) | 5062 | NM_001040396 | 69227 | Mm.270950 | NM_001040396 | ENSMUSG00000075700 | **Selt** | 2810407C02Rik\|5730408P04Rik | selenoprotein T | protein-coding |

| | | | | | | | | | | | | | | | | | |
|---|---|---|---|---|---|---|---|---|---|---|---|---|---|---|---|---|---|
| chr3_58552051_58552413 | chr3 | 58552052 | 58552413 | + | -1.35099 | intron (NM_001005509, intron 10 of 13) | intron (NM_001005509, intron 10 of 13) | -24426 | NM_001040396 | 69227 | Mm.270950 | NM_001040396 | ENSMUSG00000075700 | **Selt** | 2810407C02Rik\|5730408P04Rik | selenoprotein T | protein-coding |
| chr3_58552029_58552871 | chr3 | 58552030 | 58552871 | + | -1.54567 | intron (NM_001005509, intron 10 of 13) | intron (NM_001005509, intron 10 of 13) | -24208 | NM_001040396 | 69227 | Mm.270950 | NM_001040396 | ENSMUSG00000075700 | **Selt** | 2810407C02Rik\|5730408P04Rik | selenoprotein T | protein-coding |
| chr9_107724130_107724532 | chr9 | 107724131 | 107724532 | + | -0.789978 | Intergenic | Intergenic | -13856 | NM_011349 | 20350 | Mm.12903 | NM_011349 | ENSMUSG00000034684 | **Sema3f** | Sema4\|Semak | sema domain, immunoglobulin domain (Ig), short basic domain, secreted, (semaphorin) 3F | protein-coding |
| chr3_88452138_88452538 | chr3 | 88452139 | 88452538 | + | -0.635745 | intron (NM_001163489, intron 3 of 14) | MIRb\|SINE\|MIR | 3423 | NM_013658 | 20351 | Mm.439752 | NM_013658 | ENSMUSG00000028064 | **Sema4a** | AI132332\|SemB\|Semab | sema domain, immunoglobulin domain (Ig), transmembrane domain (TM) and short cytoplasmic domain, (semaphorin) 4A | protein-coding |
| chr3_88453341_88453652 | chr3 | 88453342 | 88453652 | + | -0.697475 | intron (NM_001163489, intron 2 of 14) | intron (NM_001163489, intron 2 of 14) | 2264 | NM_013658 | 20351 | Mm.439752 | NM_013658 | ENSMUSG00000028064 | **Sema4a** | AI132332\|SemB\|Semab | sema domain, immunoglobulin domain (Ig), transmembrane domain (TM) and short cytoplasmic domain, (semaphorin) 4A | protein-coding |
| chr7_80204344_80204788 | chr7 | 80204345 | 80204788 | + | -0.701677 | intron (NM_013659, intron 2 of 14) | intron (NM_013659, intron 2 of 14) | 17725 | NM_013659 | 20352 | Mm.275909 | NM_013659 | ENSMUSG00000030539 | **Sema4b** | SemC\|Semac\|mKIAA1745 | sema domain, immunoglobulin domain (Ig), transmembrane domain (TM) and short cytoplasmic domain, (semaphorin) 4B | protein-coding |
| chr7_80204363_80204965 | chr7 | 80204364 | 80204965 | + | -0.761895 | intron (NM_013659, intron 2 of 14) | intron (NM_013659, intron 2 of 14) | 17823 | NM_013659 | 20352 | Mm.275909 | NM_013659 | ENSMUSG00000030539 | **Sema4b** | SemC\|Semac\|mKIAA1745 | sema domain, immunoglobulin domain (Ig), transmembrane domain (TM) and short cytoplasmic domain, (semaphorin) 4B | protein-coding |
| chr13_51767493_51767805 | chr13 | 51767494 | 51767805 | + | -0.630929 | intron (NM_013660, intron 1 of 15) | intron (NM_013660, intron 1 of 15) | 26098 | NM_001281880 | 20354 | Mm.33903 | NM_013660 | ENSMUSG00000021451 | **Sema4d** | CD100\|Semacl2\|Semaj\|Semcl2\|coll-4 | sema domain, immunoglobulin domain (Ig), transmembrane domain (TM) and short cytoplasmic domain, (semaphorin) 4D | protein-coding |
| chr13_51765432_51765638 | chr13 | 51765433 | 51765638 | + | -0.661699 | intron (NM_013660, intron 1 of 15) | intron (NM_013660, intron 1 of 15) | 28212 | NM_001281880 | 20354 | Mm.33903 | NM_013660 | ENSMUSG00000021451 | **Sema4d** | CD100\|Semacl2\|Semaj\|Semcl2\|coll-4 | sema domain, immunoglobulin domain (Ig), transmembrane domain (TM) and short cytoplasmic domain, (semaphorin) 4D | protein-coding |
| chr13_51763301_51763650 | chr13 | 51763302 | 51763650 | + | -0.789867 | intron (NM_013660, intron 1 of 15) | intron (NM_013660, intron 1 of 15) | 30271 | NM_001281880 | 20354 | Mm.33903 | NM_013660 | ENSMUSG00000021451 | **Sema4d** | CD100\|Semacl2\|Semaj\|Semcl2\|coll-4 | sema domain, immunoglobulin domain (Ig), transmembrane domain (TM) and short cytoplasmic domain, (semaphorin) 4D | protein-coding |
| chr6_82940785_82941072 | chr6 | 82940786 | 82941072 | + | -0.659116 | Intergenic | Intergenic | -1179 | NM_001113481 | 20355 | Mm.270543 | NM_011350 | ENSMUSG00000000627 | **Sema4f** | - | sema domain, immunoglobulin domain (Ig), TM domain, and short cytoplasmic domain | protein-coding |
| chr6_82921844_82922670 | chr6 | 82921845 | 82922670 | + | -0.818382 | intron (NM_011350, intron 5 of 13) | intron (NM_011350, intron 5 of 13) | 17493 | NM_011350 | 20355 | Mm.270543 | NM_011350 | ENSMUSG00000000627 | **Sema4f** | - | sema domain, immunoglobulin domain (Ig), TM domain, and short cytoplasmic domain | protein-coding |
| chr6_82921815_82922411 | chr6 | 82921816 | 82922411 | + | -0.95923 | intron (NM_011350, intron 5 of 13) | intron (NM_011350, intron 5 of 13) | 17637 | NM_011350 | 20355 | Mm.270543 | NM_011350 | ENSMUSG00000000627 | **Sema4f** | - | sema domain, immunoglobulin domain (Ig), TM domain, and short cytoplasmic domain | protein-coding |
| chr16_22062333_22062578 | chr16 | 22062334 | 22062578 | + | -0.667796 | intron (NM_183029, intron 14 of 15) | intron (NM_183029, intron 14 of 15) | 52972 | NR_027488 | 75826 | Mm.297431 | NM_029457 | ENSMUSG00000022855 | **Senp2** | 2310007L05Rik\|4930538C18Rik\|AI646780\|AW554757\|Smt3ip2\|SuPr-1\|mKIAA1331 | SUMO/sentrin specific peptidase 2 | protein-coding |
| chr16_22075010_22075601 | chr16 | 22075011 | 22075601 | + | -0.745003 | intron (NM_183029, intron 10 of 15) | intron (NM_183029, intron 10 of 15) | 65822 | NR_027488 | 75826 | Mm.297431 | NM_029457 | ENSMUSG00000022855 | **Senp2** | 2310007L05Rik\|4930538C18Rik\|AI646780\|AW554757\|Smt3ip2\|SuPr-1\|mKIAA1331 | SUMO/sentrin specific peptidase 2 | protein-coding |
| chr16_56068510_56069090 | chr16 | 56068511 | 56069090 | + | -0.773924 | Intergenic | Intergenic | -6609 | NM_025483 | 66315 | Mm.255784 | NM_025483 | ENSMUSG00000052917 | **Senp7** | 2410152H17Rik\|2810413I22Rik\|2900036C23Rik\|6030449K19Rik\|AI790676\|mKIAA1707 | SUMO1/sentrin specific peptidase 7 | protein-coding |
| chr16_56068598_56069023 | chr16 | 56068599 | 56069023 | + | -0.788578 | Intergenic | Intergenic | -6598 | NM_025483 | 66315 | Mm.255784 | NM_025483 | ENSMUSG00000052917 | **Senp7** | 2410152H17Rik\|2810413I22Rik\|2900036C23Rik\|6030449K19Rik\|AI790676\|mKIAA1707 | SUMO1/sentrin specific peptidase 7 | protein-coding |
| chr16_56068584_56068890 | chr16 | 56068585 | 56068890 | + | -0.875998 | Intergenic | Intergenic | -6672 | NM_025483 | 66315 | Mm.255784 | NM_025483 | ENSMUSG00000052917 | **Senp7** | 2410152H17Rik\|2810413I22Rik\|2900036C23Rik\|6030449K19Rik\|AI790676\|mKIAA1707 | SUMO1/sentrin specific peptidase 7 | protein-coding |
| chr2_4883347_4883654 | chr2 | 4883348 | 4883654 | + | -0.712621 | intron (NM_175400, intron 1 of 8) | intron (NM_175400, intron 1 of 8) | 1937 | NM_175400 | 109079 | Mm.34329 | NM_175400 | ENSMUSG00000026662 | **Sephs1** | 1110046B24Rik\|AA589574\|AI505014\|AW111620\|SELD\|SPS\|SPS1 | selenophosphate synthetase 1 | protein-coding |
| chr7_127220440_127220752 | chr7 | 127220441 | 127220752 | + | -0.80799 | Intergenic | Intergenic | -2151 | NM_017461 | 54204 | Mm.18 | NM_017461 | ENSMUSG00000000486 | **Sept1** | Diff6\|Pnutl3 | septin 1 | protein-coding |
| chrX_36980888_36981378 | chrX | 36980889 | 36981378 | + | -0.634444 | intron (NM_001177323, intron 1 of 9) | intron (NM_001177323, intron 1 of 9) | 8562 | NM_001177324 | 56526 | Mm.260036 | NM_019942 | ENSMUSG00000050379 | **Sept6** | 2810035H17Rik\|C920001C06Rik\|Sep6\|mKIAA0128 | septin 6 | protein-coding |
| chrX_36980637_36980845 | chrX | 36980638 | 36980845 | + | -0.663495 | intron (NM_001177323, intron 1 of 9) | intron (NM_001177323, intron 1 of 9) | 8954 | NM_001177324 | 56526 | Mm.260036 | NM_019942 | ENSMUSG00000050379 | **Sept6** | 2810035H17Rik\|C920001C06Rik\|Sep6\|mKIAA0128 | septin 6 | protein-coding |
| chrX_36980625_36981409 | chrX | 36980626 | 36981409 | + | -0.686704 | intron (NM_001177323, intron 1 of 9) | intron (NM_001177323, intron 1 of 9) | 8678 | NM_001177324 | 56526 | Mm.260036 | NM_019942 | ENSMUSG00000050379 | **Sept6** | 2810035H17Rik\|C920001C06Rik\|Sep6\|mKIAA0128 | septin 6 | protein-coding |
| chrX_36953417_36953715 | chrX | 36953418 | 36953715 | + | -1.00965 | intron (NM_001177324, intron 3 of 9) | intron (NM_001177324, intron 3 of 9) | 36129 | NM_001177324 | 56526 | Mm.260036 | NM_019942 | ENSMUSG00000050379 | **Sept6** | 2810035H17Rik\|C920001C06Rik\|Sep6\|mKIAA0128 | septin 6 | protein-coding |
| chr11_53516781_53517092 | chr11 | 53516782 | 53517092 | + | -0.828391 | Intergenic | ID4\|SINE\|ID | -2799 | NM_033144 | 20362 | Mm.274399 | NM_033144 | ENSMUSG00000018398 | **Sept8** | AW046166\|Sepl\|mKIAA0202 | septin 8 | protein-coding |
| chr6_67302492_67302846 | chr6 | 67302493 | 67302846 | + | -0.936955 | intron (NM_008354, intron 12 of 15) | intron (NM_008354, intron 12 of 15) | 35690 | NM_025814 | 66870 | Mm.240490 | NM_025814 | ENSMUSG00000036371 | **Serbp1** | 1200009K13Rik\|9330147J08Rik\|AL022786\|Pairbp1 | serpine1 mRNA binding protein 1 | protein-coding |
| chr6_67184522_67185070 | chr6 | 67184523 | 67185070 | + | -1.31635 | Intergenic | Lx7\|LINE\|L1 | -82183 | NM_001113564 | 66870 | Mm.240490 | NM_025814 | ENSMUSG00000036371 | **Serbp1** | 1200009K13Rik\|9330147J08Rik\|AL022786\|Pairbp1 | serpine1 mRNA binding protein 1 | protein-coding |
| chr6_67184766_67185060 | chr6 | 67184767 | 67185060 | + | -1.35968 | Intergenic | Lx7\|LINE\|L1 | -82066 | NM_001113564 | 66870 | Mm.240490 | NM_025814 | ENSMUSG00000036371 | **Serbp1** | 1200009K13Rik\|9330147J08Rik\|AL022786\|Pairbp1 | serpine1 mRNA binding protein 1 | protein-coding |
| chr15_83075696_83076578 | chr15 | 83075697 | 83076578 | + | -1.05826 | Intergenic | Intergenic | -24068 | NM_023475 | 68607 | Mm.30056 | NM_023475 | ENSMUSG00000058586 | **Serhl** | 0610008B10Rik\|1110019M09Rik\|AU019786 | serine hydrolase-like | protein-coding |
| chr14_76579690_76580243 | chr14 | 76579691 | 76580243 | + | -1.261 | Intergenic | Intergenic | -23078 | NR_027699 | 72661 | Mm.320384 | NM_001160326 | ENSMUSG00000052584 | **Serp2** | 2810032E02Rik | stress-associated endoplasmic reticulum protein family member 2 | protein-coding |
| chr11_75429055_75429374 | chr11 | 75429056 | 75429374 | + | -0.833464 | Intergenic | Intergenic | -6592 | NM_011340 | 20317 | Mm.2044 | NM_011340 | ENSMUSG00000000753 | **Serpinf1** | AI195227\|EPC-1\|Pedf\|Pedfl\|Sdf3 | serine (or cysteine) peptidase inhibitor, clade F, member 1 | protein-coding |
| chr11_75428891_75429596 | chr11 | 75428892 | 75429596 | + | -0.835267 | Intergenic | Intergenic | -6621 | NM_011340 | 20317 | Mm.2044 | NM_011340 | ENSMUSG00000000753 | **Serpinf1** | AI195227\|EPC-1\|Pedf\|Pedfl\|Sdf3 | serine (or cysteine) peptidase inhibitor, clade F, member 1 | protein-coding |
| chr10_41880810_41881128 | chr10 | 41880811 | 41881128 | + | -1.10441 | intron (NM_001162908, intron 1 of 9) | intron (NM_001162908, intron 1 of 9) | -6470 | NM_001013370 | 140742 | Mm.139418 | NM_001013370 | ENSMUSG00000038332 | **Sesn1** | 1110002G11Rik\|AU044290\|Pa26\|Sest1 | sestrin 1 | protein-coding |
| chr9_14295891_14296587 | chr9 | 14295892 | 14296587 | + | -0.641041 | intron (NM_030261, intron 1 of 9) | intron (NM_030261, intron 1 of 9) | 19938 | NM_030261 | 75747 | Mm.325126 | NM_030261 | ENSMUSG00000032009 | **Sesn3** | 5630400E15Rik\|BC003348\|SEST3 | sestrin 3 | protein-coding |
| chr9_14244465_14245078 | chr9 | 14244466 | 14245078 | + | -0.647982 | Intergenic | Intergenic | -31529 | NM_030261 | 75747 | Mm.325126 | NM_030261 | ENSMUSG00000032009 | **Sesn3** | 5630400E15Rik\|BC003348\|SEST3 | sestrin 3 | protein-coding |
| chr9_14092894_14093568 | chr9 | 14092895 | 14093568 | + | -0.75269 | Intergenic | Intergenic | -183070 | NM_030261 | 75747 | Mm.325126 | NM_030261 | ENSMUSG00000032009 | **Sesn3** | 5630400E15Rik\|BC003348\|SEST3 | sestrin 3 | protein-coding |
| chr18_79267721_79268164 | chr18 | 79267722 | 79268164 | + | -0.70681 | Intergenic | Intergenic | -158552 | NM_053099 | 240427 | Mm.312871 | NM_053099 | ENSMUSG00000024548 | **Setbp1** | C130092E12\|Seb\|mKIAA0437 | SET binding protein 1 | protein-coding |
| chr18_79234119_79234704 | chr18 | 79234120 | 79234704 | + | -0.864369 | Intergenic | Intergenic | -125021 | NM_053099 | 240427 | Mm.312871 | NM_053099 | ENSMUSG00000024548 | **Setbp1** | C130092E12\|Seb\|mKIAA0437 | SET binding protein 1 | protein-coding |
| chr18_79293603_79294176 | chr18 | 79293604 | 79294176 | + | -0.916175 | Intergenic | Intergenic | -184499 | NM_053099 | 240427 | Mm.312871 | NM_053099 | ENSMUSG00000024548 | **Setbp1** | C130092E12\|Seb\|mKIAA0437 | SET binding protein 1 | protein-coding |
| chr18_79234328_79234695 | chr18 | 79234329 | 79234695 | + | -0.939162 | Intergenic | Intergenic | -125121 | NM_053099 | 240427 | Mm.312871 | NM_053099 | ENSMUSG00000024548 | **Setbp1** | C130092E12\|Seb\|mKIAA0437 | SET binding protein 1 | protein-coding |
| chr18_79303300_79303965 | chr18 | 79303301 | 79303965 | + | -1.01342 | Intergenic | Intergenic | -194242 | NM_053099 | 240427 | Mm.312871 | NM_053099 | ENSMUSG00000024548 | **Setbp1** | C130092E12\|Seb\|mKIAA0437 | SET binding protein 1 | protein-coding |
| chr18_79472931_79473323 | chr18 | 79472932 | 79473323 | + | -1.02749 | Intergenic | Intergenic | -363736 | NM_053099 | 240427 | Mm.312871 | NM_053099 | ENSMUSG00000024548 | **Setbp1** | C130092E12\|Seb\|mKIAA0437 | SET binding protein 1 | protein-coding |
| chr16_93574382_93575239 | chr16 | 93574383 | 93575239 | + | -0.769773 | Intergenic | Intergenic | 29004 | NM_145482 | 224440 | Mm.10628 | NM_145482 | ENSMUSG00000022948 | **Setd4** | AY037804\|C21orf18\|ORF21 | SET domain containing 4 | protein-coding |
| chr16_93552579_93553086 | chr16 | 93552580 | 93553086 | + | -0.835238 | Intergenic | Intergenic | 50982 | NM_145482 | 224440 | Mm.10628 | NM_145482 | ENSMUSG00000022948 | **Setd4** | AY037804\|C21orf18\|ORF21 | SET domain containing 4 | protein-coding |
| chr16_93552556_93553462 | chr16 | 93552557 | 93553462 | + | -0.869932 | Intergenic | Intergenic | 50806 | NM_145482 | 224440 | Mm.10628 | NM_145482 | ENSMUSG00000022948 | **Setd4** | AY037804\|C21orf18\|ORF21 | SET domain containing 4 | protein-coding |
| chr16_93574368_93574930 | chr16 | 93574369 | 93574930 | + | -0.87879 | Intergenic | Intergenic | 29166 | NM_145482 | 224440 | Mm.10628 | NM_145482 | ENSMUSG00000022948 | **Setd4** | AY037804\|C21orf18\|ORF21 | SET domain containing 4 | protein-coding |
| chr16_93574479_93574851 | chr16 | 93574480 | 93574851 | + | -0.933953 | Intergenic | Intergenic | 29150 | NM_145482 | 224440 | Mm.10628 | NM_145482 | ENSMUSG00000022948 | **Setd4** | AY037804\|C21orf18\|ORF21 | SET domain containing 4 | protein-coding |
| chr16_93572261_93572854 | chr16 | 93572262 | 93572854 | + | -1.0438 | Intergenic | ORR1E\|LTR\|ERVL-MaLR | 31257 | NM_145482 | 224440 | Mm.10628 | NM_145482 | ENSMUSG00000022948 | **Setd4** | AY037804\|C21orf18\|ORF21 | SET domain containing 4 | protein-coding |

| | | | | | | | | | | | | | | | | | |
|---|---|---|---|---|---|---|---|---|---|---|---|---|---|---|---|---|---|
| chr3_51603821_51604225 | chr3 | 51603822 | 51604225 | + | -0.649987 | Intergenic | Intergenic | -43200 | NM_080793 | 73251 | Mm.192111 | NM_080793 | ENSMUSG00000037111 | **Setd7** | 1600028F23Rik\|H3K4MT\|KMT7\|Set7\|Set7/9\|mKIAA1717 | SET domain containing (lysine methyltransferase) 7 | protein-coding |
| chr3_51566345_51566761 | chr3 | 51566346 | 51566761 | + | -0.669078 | intron (NR_033624, intron 4 of 4) | intron (NR_033624, intron 4 of 4) | -5730 | NM_080793 | 73251 | Mm.192111 | NM_080793 | ENSMUSG00000037111 | **Setd7** | 1600028F23Rik\|H3K4MT\|KMT7\|Set7\|Set7/9\|mKIAA1717 | SET domain containing (lysine methyltransferase) 7 | protein-coding |
| chr2_29097508_29097855 | chr2 | 29097509 | 29097855 | + | -0.713724 | Intergenic | MT2A\|LTR\|ERVL | -27310 | NM_198033 | 269254 | Mm.41867 | NM_198033 | ENSMUSG00000043535 | **Setx** | A130090N03\|A930037J23Rik\|AOA2\|AW060766\|Als4\|SCAR1\|Sen1\|mKIAA0625 | senataxin | protein-coding |
| chr2_29111153_29111365 | chr2 | 29111154 | 29111365 | + | -0.968817 | Intergenic | ORR1A2-int\|LTR\|ERVL-MaLR | -13733 | NM_198033 | 269254 | Mm.41867 | NM_198033 | ENSMUSG00000043535 | **Setx** | A130090N03\|A930037J23Rik\|AOA2\|AW060766\|Als4\|SCAR1\|Sen1\|mKIAA0625 | senataxin | protein-coding |
| chr2_29111075_29111397 | chr2 | 29111076 | 29111397 | + | -0.980275 | Intergenic | ORR1A2-int\|LTR\|ERVL-MaLR | -13756 | NM_198033 | 269254 | Mm.41867 | NM_198033 | ENSMUSG00000043535 | **Setx** | A130090N03\|A930037J23Rik\|AOA2\|AW060766\|Als4\|SCAR1\|Sen1\|mKIAA0625 | senataxin | protein-coding |
| chr19_45057326_45057644 | chr19 | 45057327 | 45057644 | + | -0.612227 | Intergenic | Intergenic | 9909 | NM_001178013 | 94280 | Mm.36169 | NM_053197 | ENSMUSG00000025212 | **Sfxn3** | - | sideroflexin 3 | protein-coding |
| chr10_21911500_21912016 | chr10 | 21911501 | 21912016 | + | -0.643679 | intron (NM_001161845, intron 1 of 13) | intron (NM_001161845, intron 1 of 13) | 29574 | NM_001161845 | 20393 | Mm.28405 | NM_011361 | ENSMUSG00000019970 | **Sgk1** | Sgk | serum/glucocorticoid regulated kinase 1 | protein-coding |
| chr10_21908739_21909076 | chr10 | 21908740 | 21909076 | + | -0.686862 | intron (NM_001161845, intron 1 of 13) | MER5A\|DNA\|hAT-Charlie | 26724 | NM_001161845 | 20393 | Mm.28405 | NM_011361 | ENSMUSG00000019970 | **Sgk1** | Sgk | serum/glucocorticoid regulated kinase 1 | protein-coding |
| chr10_21975329_21975689 | chr10 | 21975330 | 21975689 | + | -0.736139 | intron (NM_001161845, intron 3 of 13) | Lx8\|LINE\|L1 | -3163 | NM_001161847 | 20393 | Mm.28405 | NM_011361 | ENSMUSG00000019970 | **Sgk1** | Sgk | serum/glucocorticoid regulated kinase 1 | protein-coding |
| chr10_21905664_21905946 | chr10 | 21905665 | 21905946 | + | -1.15693 | intron (NM_001161845, intron 1 of 13) | MTB_Mm\|LTR\|ERVL-MaLR | 23621 | NM_001161845 | 20393 | Mm.28405 | NM_011361 | ENSMUSG00000019970 | **Sgk1** | Sgk | serum/glucocorticoid regulated kinase 1 | protein-coding |
| chr10_21905562_21905978 | chr10 | 21905563 | 21905978 | + | -1.17845 | intron (NM_001161845, intron 1 of 13) | MTB_Mm\|LTR\|ERVL-MaLR | 23586 | NM_001161845 | 20393 | Mm.28405 | NM_011361 | ENSMUSG00000019970 | **Sgk1** | Sgk | serum/glucocorticoid regulated kinase 1 | protein-coding |
| chr1_9807591_9807945 | chr1 | 9807592 | 9807945 | + | -0.681106 | intron (NM_177547, intron 1 of 16) | B1_Mur4\|SINE\|Alu | 9638 | NM_177547 | 170755 | Mm.336410 | NM_133220 | ENSMUSG00000025915 | **Sgk3** | 2510015P22Rik\|A330005P07Rik\|Cisk\|fy\|fz | serum/glucocorticoid regulated kinase 3 | protein-coding |
| chr1_9807703_9808062 | chr1 | 9807704 | 9808062 | + | -0.802221 | intron (NM_177547, intron 1 of 16) | intron (NM_177547, intron 1 of 16) | 9753 | NM_177547 | 170755 | Mm.336410 | NM_133220 | ENSMUSG00000025915 | **Sgk3** | 2510015P22Rik\|A330005P07Rik\|Cisk\|fy\|fz | serum/glucocorticoid regulated kinase 3 | protein-coding |
| chr19_32266047_32266739 | chr19 | 32266048 | 32266739 | + | -0.603769 | intron (NM_001168525, intron 2 of 9) | L1MB3\|LINE\|L1 | -55380 | NM_001168526 | 208449 | Mm.329810 | NM_144792 | ENSMUSG00000040451 | **Sgms1** | 9530058O11Rik\|AI841905\|C80702\|Mob\|Sms1\|Sor1\|Tmem23 | sphingomyelin synthase 1 | protein-coding |
| chr19_32266457_32266746 | chr19 | 32266458 | 32266746 | + | -0.620908 | intron (NM_001168525, intron 2 of 9) | L1MB3\|LINE\|L1 | -55589 | NM_001168526 | 208449 | Mm.329810 | NM_144792 | ENSMUSG00000040451 | **Sgms1** | 9530058O11Rik\|AI841905\|C80702\|Mob\|Sms1\|Sor1\|Tmem23 | sphingomyelin synthase 1 | protein-coding |
| chr19_32292486_32293099 | chr19 | 32292487 | 32293099 | + | -0.622634 | intron (NM_144792, intron 2 of 9) | intron (NM_144792, intron 2 of 9) | 56523 | NM_001168525 | 208449 | Mm.329810 | NM_144792 | ENSMUSG00000040451 | **Sgms1** | 9530058O11Rik\|AI841905\|C80702\|Mob\|Sms1\|Sor1\|Tmem23 | sphingomyelin synthase 1 | protein-coding |
| chr19_32292519_32292838 | chr19 | 32292520 | 32292838 | + | -0.629584 | intron (NM_144792, intron 2 of 9) | intron (NM_144792, intron 2 of 9) | 56637 | NM_001168525 | 208449 | Mm.329810 | NM_144792 | ENSMUSG00000040451 | **Sgms1** | 9530058O11Rik\|AI841905\|C80702\|Mob\|Sms1\|Sor1\|Tmem23 | sphingomyelin synthase 1 | protein-coding |
| chr19_32344568_32345103 | chr19 | 32344569 | 32345103 | + | -0.744594 | intron (NM_144792, intron 2 of 9) | HAL1\|LINE\|L1 | 4480 | NM_001168525 | 208449 | Mm.329810 | NM_144792 | ENSMUSG00000040451 | **Sgms1** | 9530058O11Rik\|AI841905\|C80702\|Mob\|Sms1\|Sor1\|Tmem23 | sphingomyelin synthase 1 | protein-coding |
| chr19_32172448_32173026 | chr19 | 32172449 | 32173026 | + | -0.776157 | intron (NM_144792, intron 5 of 9) | ORR1C2\|LTR\|ERVL-MaLR | 38276 | NM_001168526 | 208449 | Mm.329810 | NM_144792 | ENSMUSG00000040451 | **Sgms1** | 9530058O11Rik\|AI841905\|C80702\|Mob\|Sms1\|Sor1\|Tmem23 | sphingomyelin synthase 1 | protein-coding |
| chr19_32302203_32302445 | chr19 | 32302204 | 32302445 | + | -0.786063 | intron (NM_144792, intron 2 of 9) | intron (NM_144792, intron 2 of 9) | 46992 | NM_001168525 | 208449 | Mm.329810 | NM_144792 | ENSMUSG00000040451 | **Sgms1** | 9530058O11Rik\|AI841905\|C80702\|Mob\|Sms1\|Sor1\|Tmem23 | sphingomyelin synthase 1 | protein-coding |
| chr19_32325146_32325403 | chr19 | 32325147 | 32325403 | + | -0.796085 | intron (NM_144792, intron 2 of 9) | intron (NM_144792, intron 2 of 9) | 24041 | NM_001168525 | 208449 | Mm.329810 | NM_144792 | ENSMUSG00000040451 | **Sgms1** | 9530058O11Rik\|AI841905\|C80702\|Mob\|Sms1\|Sor1\|Tmem23 | sphingomyelin synthase 1 | protein-coding |
| chr19_32196938_32197818 | chr19 | 32196939 | 32197818 | + | -0.858382 | intron (NM_144792, intron 5 of 9) | LTR41\|LTR\|ERVL | 13635 | NM_001168526 | 208449 | Mm.329810 | NM_144792 | ENSMUSG00000040451 | **Sgms1** | 9530058O11Rik\|AI841905\|C80702\|Mob\|Sms1\|Sor1\|Tmem23 | sphingomyelin synthase 1 | protein-coding |
| chr19_32258044_32258676 | chr19 | 32258045 | 32258676 | + | -0.860106 | intron (NM_144792, intron 3 of 9) | MLTR18B_MM\|LTR\|ERVK | -47347 | NM_001168526 | 208449 | Mm.329810 | NM_144792 | ENSMUSG00000040451 | **Sgms1** | 9530058O11Rik\|AI841905\|C80702\|Mob\|Sms1\|Sor1\|Tmem23 | sphingomyelin synthase 1 | protein-coding |
| chr19_32197037_32197580 | chr19 | 32197038 | 32197580 | + | -0.991782 | intron (NM_144792, intron 5 of 9) | LTR41\|LTR\|ERVL | 13704 | NM_001168526 | 208449 | Mm.329810 | NM_144792 | ENSMUSG00000040451 | **Sgms1** | 9530058O11Rik\|AI841905\|C80702\|Mob\|Sms1\|Sor1\|Tmem23 | sphingomyelin synthase 1 | protein-coding |
| chr19_32166905_32167223 | chr19 | 32166906 | 32167223 | + | -1.1137 | intron (NM_144792, intron 5 of 9) | intron (NM_144792, intron 5 of 9) | 43949 | NM_001168526 | 208449 | Mm.329810 | NM_144792 | ENSMUSG00000040451 | **Sgms1** | 9530058O11Rik\|AI841905\|C80702\|Mob\|Sms1\|Sor1\|Tmem23 | sphingomyelin synthase 1 | protein-coding |
| chr19_32166915_32167133 | chr19 | 32166916 | 32167133 | + | -1.11693 | intron (NM_144792, intron 5 of 9) | intron (NM_144792, intron 5 of 9) | 43989 | NM_001168526 | 208449 | Mm.329810 | NM_144792 | ENSMUSG00000040451 | **Sgms1** | 9530058O11Rik\|AI841905\|C80702\|Mob\|Sms1\|Sor1\|Tmem23 | sphingomyelin synthase 1 | protein-coding |
| chr19_32186433_32187032 | chr19 | 32186434 | 32187032 | + | -1.15362 | intron (NM_144792, intron 5 of 9) | intron (NM_144792, intron 5 of 9) | 24280 | NM_001168526 | 208449 | Mm.329810 | NM_144792 | ENSMUSG00000040451 | **Sgms1** | 9530058O11Rik\|AI841905\|C80702\|Mob\|Sms1\|Sor1\|Tmem23 | sphingomyelin synthase 1 | protein-coding |
| chr19_32182104_32182466 | chr19 | 32182105 | 32182466 | + | -1.4756 | intron (NM_144792, intron 5 of 9) | intron (NM_144792, intron 5 of 9) | 28728 | NM_001168526 | 208449 | Mm.329810 | NM_144792 | ENSMUSG00000040451 | **Sgms1** | 9530058O11Rik\|AI841905\|C80702\|Mob\|Sms1\|Sor1\|Tmem23 | sphingomyelin synthase 1 | protein-coding |
| chr12_75766827_75767541 | chr12 | 75766828 | 75767541 | + | -0.903927 | Intergenic | Intergenic | -31455 | NM_030750 | 81535 | Mm.280199 | NM_030750 | ENSMUSG00000021054 | **Sgpp1** | AI463453\|SPP\|SPP1\|Spph1 | sphingosine-1-phosphate phosphatase 1 | protein-coding |
| chr15_81035241_81036067 | chr15 | 81035242 | 81036067 | + | -0.752728 | intron (NM_001082536, intron 3 of 13) | B4\|SINE\|B4 | 57889 | NM_134091 | 105835 | Mm.274943 | NM_134091 | ENSMUSG00000042303 | **Sgsm3** | 1810012I01Rik\|AI428557\|BB175482\|Cip85\|R75178\|RUSC3\|Rutbc3 | small G protein signaling modulator 3 | protein-coding |
| chr15_81035279_81035811 | chr15 | 81035280 | 81035811 | + | -0.842862 | intron (NM_001082536, intron 3 of 13) | B4\|SINE\|B4 | 57780 | NM_134091 | 105835 | Mm.274943 | NM_134091 | ENSMUSG00000042303 | **Sgsm3** | 1810012I01Rik\|AI428557\|BB175482\|Cip85\|R75178\|RUSC3\|Rutbc3 | small G protein signaling modulator 3 | protein-coding |
| chr13_104112520_104112777 | chr13 | 104112521 | 104112777 | + | -1.04719 | intron (NM_144838, intron 2 of 10) | intron (NM_144838, intron 2 of 10) | 2859 | NM_144838 | 218544 | Mm.44416 | NM_144838 | ENSMUSG00000042743 | **Sgtb** | C630001O05Rik | small glutamine-rich tetratricopeptide repeat (TPR)-containing, beta | protein-coding |
| chr7_126470094_126470563 | chr7 | 126470095 | 126470563 | + | -0.892527 | intron (NR_110346, intron 4 of 9) | intron (NR_110346, intron 4 of 9) | 2765 | NR_110346 | 20399 | Mm.8538 | NM_011363 | ENSMUSG00000030733 | **Sh2b1** | AI425885\|C530001K22Rik\|Irip\|Psm\|SH2-B\|SH2-Bb\|Sh2bpsm1\|mKIAA1299 | SH2B adaptor protein 1 | protein-coding |
| chrX_42511152_42511372 | chrX | 42511153 | 42511372 | + | -0.853795 | intron (NM_011364, intron 1 of 3) | intron (NM_011364, intron 1 of 3) | 8654 | NM_011364 | 20400 | Mm.441197 | NM_011364 | ENSMUSG00000005696 | **Sh2d1a** | Gm686\|SAP | SH2 domain protein 1A | protein-coding |
| chrX_42511139_42511838 | chrX | 42511140 | 42511838 | + | -0.923162 | intron (NM_011364, intron 1 of 3) | intron (NM_011364, intron 1 of 3) | 8881 | NM_011364 | 20400 | Mm.441197 | NM_011364 | ENSMUSG00000005696 | **Sh2d1a** | Gm686\|SAP | SH2 domain protein 1A | protein-coding |
| chr2_32725279_32725918 | chr2 | 32725280 | 32725918 | + | -0.810272 | intron (NR_045535, intron 2 of 12) | intron (NR_045535, intron 2 of 12) | -2090 | NM_001252547 | 27387 | Mm.9593 | NM_013781 | ENSMUSG00000059013 | **Sh2d3c** | Chat\|Nsp3\|Shep1 | SH2 domain containing 3C | protein-coding |
| chr14_40924439_40925064 | chr14 | 40924440 | 40925064 | + | -0.630589 | intron (NM_145928, intron 2 of 8) | intron (NM_145928, intron 2 of 8) | -31787 | NM_177816 | 328381 | Mm.235694 | NM_177816 | ENSMUSG00000037833 | **Sh2d4b** | A430109M18Rik\|D030001E08 | SH2 domain containing 4B | protein-coding |
| chr4_134118372_134118755 | chr4 | 134118373 | 134118755 | + | -0.708369 | intron (NR_045571, intron 8 of 13) | intron (NR_045571, intron 8 of 13) | 10189 | NM_080559 | 73723 | Mm.22240 | NM_080559 | ENSMUSG00000028843 | **Sh3bgrl3** | 1110004L05Rik\|AU022188 | SH3 domain binding glutamic acid-rich protein-like 3 | protein-coding |
| chr4_134121662_134121918 | chr4 | 134121663 | 134121918 | + | -0.709373 | intron (NR_045571, intron 8 of 13) | intron (NR_045571, intron 8 of 13) | 6963 | NM_080559 | 73723 | Mm.22240 | NM_080559 | ENSMUSG00000028843 | **Sh3bgrl3** | 1110004L05Rik\|AU022188 | SH3 domain binding glutamic acid-rich protein-like 3 | protein-coding |
| chr4_134116898_134117407 | chr4 | 134116899 | 134117407 | + | -0.887983 | intron (NR_045571, intron 8 of 13) | intron (NR_045571, intron 8 of 13) | 11600 | NM_080559 | 73723 | Mm.22240 | NM_080559 | ENSMUSG00000028843 | **Sh3bgrl3** | 1110004L05Rik\|AU022188 | SH3 domain binding glutamic acid-rich protein-like 3 | protein-coding |
| chr14_31389828_31390149 | chr14 | 31389829 | 31390149 | + | -0.653139 | intron (NM_011894, intron 3 of 8) | intron (NM_011894, intron 3 of 8) | 46044 | NM_011894 | 24056 | Mm.383198 | NM_011894 | ENSMUSG00000021892 | **Sh3bp5** | AI606498\|SH3BP-5\|Sab | SH3-domain binding protein 5 (BTK-associated) | protein-coding |
| chr14_31433165_31433516 | chr14 | 31433166 | 31433516 | + | -0.698548 | intron (NM_011894, intron 2 of 8) | intron (NM_011894, intron 2 of 8) | 2692 | NM_011894 | 24056 | Mm.383198 | NM_011894 | ENSMUSG00000021892 | **Sh3bp5** | AI606498\|SH3BP-5\|Sab | SH3-domain binding protein 5 (BTK-associated) | protein-coding |
| chr14_31399265_31399697 | chr14 | 31399266 | 31399697 | + | -0.773477 | intron (NM_011894, intron 3 of 8) | intron (NM_011894, intron 3 of 8) | 36552 | NM_011894 | 24056 | Mm.383198 | NM_011894 | ENSMUSG00000021892 | **Sh3bp5** | AI606498\|SH3BP-5\|Sab | SH3-domain binding protein 5 (BTK-associated) | protein-coding |
| chr14_31414256_31414553 | chr14 | 31414257 | 31414553 | + | -0.85441 | intron (NM_011894, intron 2 of 8) | intron (NM_011894, intron 2 of 8) | 21628 | NM_011894 | 24056 | Mm.383198 | NM_011894 | ENSMUSG00000021892 | **Sh3bp5** | AI606498\|SH3BP-5\|Sab | SH3-domain binding protein 5 (BTK-associated) | protein-coding |
| chr14_31414301_31414519 | chr14 | 31414302 | 31414519 | + | -0.900639 | intron (NM_011894, intron 2 of 8) | intron (NM_011894, intron 2 of 8) | 21623 | NM_011894 | 24056 | Mm.383198 | NM_011894 | ENSMUSG00000021892 | **Sh3bp5** | AI606498\|SH3BP-5\|Sab | SH3-domain binding protein 5 (BTK-associated) | protein-coding |

| | | | | | | | | | | | | | | | | | |
|---|---|---|---|---|---|---|---|---|---|---|---|---|---|---|---|---|---|
| chrX_159757916_159758639 | chrX | 159757917 | 159758639 | + | -0.663675 | intron (NM_021389, intron 1 of 16) | intron (NM_021389, intron 1 of 16) | 49677 | NR_110970 | 58194 | Mm.286495 | NM_021389 | ENSMUSG00000040990 | **Sh3kbp1** | 1200007H22Rik\|1700125L08Rik\|5830464D22Rik\|AI447724\|IN85\|Ruk\|Seta | SH3-domain kinase binding protein 1 | protein-coding |
| chrX_159689830_159690122 | chrX | 159689831 | 159690122 | + | -0.698645 | intron (NM_001135727, intron 2 of 17) | MIRb\|SINE\|MIR | -18625 | NM_021389 | 58194 | Mm.286495 | NM_021389 | ENSMUSG00000040990 | **Sh3kbp1** | 1200007H22Rik\|1700125L08Rik\|5830464D22Rik\|AI447724\|IN85\|Ruk\|Seta | SH3-domain kinase binding protein 1 | protein-coding |
| chrX_159858130_159858442 | chrX | 159858131 | 159858442 | + | -0.973433 | intron (NM_001135728, intron 1 of 12) | intron (NM_001135728, intron 1 of 12) | 17818 | NM_001135728 | 58194 | Mm.286495 | NM_021389 | ENSMUSG00000040990 | **Sh3kbp1** | 1200007H22Rik\|1700125L08Rik\|5830464D22Rik\|AI447724\|IN85\|Ruk\|Seta | SH3-domain kinase binding protein 1 | protein-coding |
| chr19_47412875_47413366 | chr19 | 47412876 | 47413366 | + | -0.624217 | intron (NM_008018, intron 2 of 14) | intron (NM_008018, intron 2 of 14) | 51290 | NM_001164717 | 14218 | Mm.127560 | NM_008018 | ENSMUSG00000053617 | **Sh3pxd2a** | 2310014D11Rik\|AA589508\|AI256723\|AI413738\|C230050L11\|EG329070\|Fish\|Gm5098\|Sh3md1\|TSK5\|Tks5 | SH3 and PX domains 2A | protein-coding |
| chr13_51598100_51598538 | chr13 | 51598101 | 51598538 | + | -0.781577 | Intergenic | MER31A\|LTR\|ERV1 | -31235 | NM_009167 | 20418 | Mm.131870 | NM_009167 | ENSMUSG00000021448 | **Shc3** | N-Shc\|Rai\|ShcC | src homology 2 domain-containing transforming protein C3 | protein-coding |
| chr3_89863973_89864490 | chr3 | 89863974 | 89864490 | + | -0.699221 | Intergenic | PB1D10\|SINE\|Alu | 32862 | NM_172530 | 214547 | Mm.225 | NM_172530 | ENSMUSG00000046280 | **She** | 9430022A14 | src homology 2 domain-containing transforming protein E | protein-coding |
| chr16_44320095_44320742 | chr16 | 44320096 | 44320742 | + | -0.673759 | intron (NM_198034, intron 1 of 24) | intron (NM_198034, intron 1 of 24) | 12419 | NM_198034 | 320007 | Mm.242144 | NM_198034 | ENSMUSG00000022696 | **Sidt1** | AW045928\|B830021E24Rik | SID1 transmembrane family, member 1 | protein-coding |
| chr16_44320167_44320550 | chr16 | 44320168 | 44320550 | + | -0.693502 | intron (NM_198034, intron 1 of 24) | intron (NM_198034, intron 1 of 24) | 12479 | NM_198034 | 320007 | Mm.242144 | NM_198034 | ENSMUSG00000022696 | **Sidt1** | AW045928\|B830021E24Rik | SID1 transmembrane family, member 1 | protein-coding |
| chr16_44339173_44339477 | chr16 | 44339174 | 44339477 | + | -0.919857 | Intergenic | RMER19C\|LTR\|ERVK | -6487 | NM_001159419 | 320007 | Mm.242144 | NM_198034 | ENSMUSG00000022696 | **Sidt1** | AW045928\|B830021E24Rik | SID1 transmembrane family, member 1 | protein-coding |
| chr17_31821191_31821870 | chr17 | 31821192 | 31821870 | + | -0.68839 | Intergenic | Intergenic | 34261 | NM_010831 | 17691 | Mm.290941 | NM_010831 | ENSMUSG00000024042 | **Sik1** | Msk\|Sik\|Snf1lk | salt inducible kinase 1 | protein-coding |
| chr17_31840214_31840947 | chr17 | 31840215 | 31840947 | + | -0.884728 | Intergenic | L1ME1\|LINE\|L1 | 15211 | NM_010831 | 17691 | Mm.290941 | NM_010831 | ENSMUSG00000024042 | **Sik1** | Msk\|Sik\|Snf1lk | salt inducible kinase 1 | protein-coding |
| chr17_31840401_31840929 | chr17 | 31840402 | 31840929 | + | -0.896266 | Intergenic | L1ME1\|LINE\|L1 | 15127 | NM_010831 | 17691 | Mm.290941 | NM_010831 | ENSMUSG00000024042 | **Sik1** | Msk\|Sik\|Snf1lk | salt inducible kinase 1 | protein-coding |
| chr9_46019016_46019297 | chr9 | 46019017 | 46019297 | + | -0.793057 | intron (NM_027498, intron 1 of 24) | intron (NM_027498, intron 1 of 24) | 6337 | NM_027498 | 70661 | Mm.219459 | NM_027498 | ENSMUSG00000034135 | **Sik3** | 5730525O22Rik\|9030204A07Rik\|AI447252\|AI846254\|AI849445\|Qsk\|SIK-3\|mKIAA0999 | SIK family kinase 3 | protein-coding |
| chr18_35383186_35383805 | chr18 | 35383187 | 35383805 | + | -0.788473 | intron (NM_030749, intron 3 of 9) | intron (NM_030749, intron 3 of 9) | 115429 | NM_030749 | 81500 | Mm.291482 | NM_030749 | ENSMUSG00000024357 | **Sil1** | 1810057E01Rik\|AI042831\|wz | endoplasmic reticulum chaperone SIL1 homolog (S. cerevisiae) | protein-coding |
| chr16_94095284_94095581 | chr16 | 94095285 | 94095581 | + | -0.679195 | intron (NM_011377, intron 2 of 10) | intron (NM_011377, intron 2 of 10) | 10173 | NM_011377 | 20465 | Mm.4775 | NM_011377 | ENSMUSG00000062713 | **Sim2** | bHLHe15 | single-minded homolog 2 (Drosophila) | protein-coding |
| chr16_94067695_94068174 | chr16 | 94067696 | 94068174 | + | -0.875311 | Intergenic | Intergenic | -17325 | NM_011377 | 20465 | Mm.4775 | NM_011377 | ENSMUSG00000062713 | **Sim2** | bHLHe15 | single-minded homolog 2 (Drosophila) | protein-coding |
| chr16_94066822_94067500 | chr16 | 94066823 | 94067500 | + | -1.05836 | Intergenic | Intergenic | -18099 | NM_011377 | 20465 | Mm.4775 | NM_011377 | ENSMUSG00000062713 | **Sim2** | bHLHe15 | single-minded homolog 2 (Drosophila) | protein-coding |
| chr16_94185368_94185957 | chr16 | 94185369 | 94185957 | + | -1.10351 | intron (NM_139145, intron 5 of 9) | intron (NM_139145, intron 5 of 9) | 100403 | NM_011377 | 20465 | Mm.4775 | NM_011377 | ENSMUSG00000062713 | **Sim2** | bHLHe15 | single-minded homolog 2 (Drosophila) | protein-coding |
| chr12_82097656_82097952 | chr12 | 82097657 | 82097952 | + | -0.676504 | Intergenic | Intergenic | -72212 | NM_001167983 | 217692 | Mm.261333 | NM_172579 | ENSMUSG00000042700 | **Sipa1l1** | 4931426N11Rik\|AW213287\|Spar\|mKIAA0440 | signal-induced proliferation-associated 1 like 1 | protein-coding |
| chr12_82349434_82350476 | chr12 | 82349435 | 82350476 | + | -0.899191 | intron (NM_172579, intron 2 of 20) | intron (NM_172579, intron 2 of 20) | 38606 | NM_172579 | 217692 | Mm.261333 | NM_172579 | ENSMUSG00000042700 | **Sipa1l1** | 4931426N11Rik\|AW213287\|Spar\|mKIAA0440 | signal-induced proliferation-associated 1 like 1 | protein-coding |
| chr12_82166084_82166665 | chr12 | 82166085 | 82166665 | + | -0.920377 | Intergenic | Intergenic | -3641 | NM_001167983 | 217692 | Mm.261333 | NM_172579 | ENSMUSG00000042700 | **Sipa1l1** | 4931426N11Rik\|AW213287\|Spar\|mKIAA0440 | signal-induced proliferation-associated 1 like 1 | protein-coding |
| chr12_82264137_82264675 | chr12 | 82264138 | 82264675 | + | -0.980625 | intron (NM_001167983, intron 2 of 23) | intron (NM_001167983, intron 2 of 23) | -46943 | NM_172579 | 217692 | Mm.261333 | NM_172579 | ENSMUSG00000042700 | **Sipa1l1** | 4931426N11Rik\|AW213287\|Spar\|mKIAA0440 | signal-induced proliferation-associated 1 like 1 | protein-coding |
| chr12_82165645_82165940 | chr12 | 82165646 | 82165940 | + | -1.03689 | Intergenic | ORR1A4\|LTR\|ERVL-MaLR | -4223 | NM_001167983 | 217692 | Mm.261333 | NM_172579 | ENSMUSG00000042700 | **Sipa1l1** | 4931426N11Rik\|AW213287\|Spar\|mKIAA0440 | signal-induced proliferation-associated 1 like 1 | protein-coding |
| chr12_82264137_82264441 | chr12 | 82264138 | 82264441 | + | -1.05862 | intron (NM_001167983, intron 2 of 23) | intron (NM_001167983, intron 2 of 23) | -47060 | NM_172579 | 217692 | Mm.261333 | NM_172579 | ENSMUSG00000042700 | **Sipa1l1** | 4931426N11Rik\|AW213287\|Spar\|mKIAA0440 | signal-induced proliferation-associated 1 like 1 | protein-coding |
| chr12_82187118_82187565 | chr12 | 82187119 | 82187565 | + | -1.17866 | intron (NM_001167983, intron 1 of 23) | intron (NM_001167983, intron 1 of 23) | 17326 | NM_001167983 | 217692 | Mm.261333 | NM_172579 | ENSMUSG00000042700 | **Sipa1l1** | 4931426N11Rik\|AW213287\|Spar\|mKIAA0440 | signal-induced proliferation-associated 1 like 1 | protein-coding |
| chr12_82187226_82187465 | chr12 | 82187227 | 82187465 | + | -1.18093 | intron (NM_001167983, intron 1 of 23) | intron (NM_001167983, intron 1 of 23) | 17330 | NM_001167983 | 217692 | Mm.261333 | NM_172579 | ENSMUSG00000042700 | **Sipa1l1** | 4931426N11Rik\|AW213287\|Spar\|mKIAA0440 | signal-induced proliferation-associated 1 like 1 | protein-coding |
| chr12_82203193_82203581 | chr12 | 82203194 | 82203581 | + | -1.29341 | intron (NM_001167983, intron 1 of 23) | intron (NM_001167983, intron 1 of 23) | 33371 | NM_001167983 | 217692 | Mm.261333 | NM_172579 | ENSMUSG00000042700 | **Sipa1l1** | 4931426N11Rik\|AW213287\|Spar\|mKIAA0440 | signal-induced proliferation-associated 1 like 1 | protein-coding |
| chr12_82208640_82208987 | chr12 | 82208641 | 82208987 | + | -1.45204 | intron (NM_001167983, intron 1 of 23) | intron (NM_001167983, intron 1 of 23) | 38798 | NM_001167983 | 217692 | Mm.261333 | NM_172579 | ENSMUSG00000042700 | **Sipa1l1** | 4931426N11Rik\|AW213287\|Spar\|mKIAA0440 | signal-induced proliferation-associated 1 like 1 | protein-coding |
| chr8_125413020_125413370 | chr8 | 125413021 | 125413370 | + | -0.693137 | Intergenic | Intergenic | 79515 | NM_001081337 | 244668 | Mm.271668 | NM_001081337 | ENSMUSG00000001995 | **Sipa1l2** | BC058408\|mKIAA1389 | signal-induced proliferation-associated 1 like 2 | protein-coding |
| chr8_125337488_125337818 | chr8 | 125337489 | 125337818 | + | -0.967071 | Intergenic | Intergenic | 155057 | NM_001081337 | 244668 | Mm.271668 | NM_001081337 | ENSMUSG00000001995 | **Sipa1l2** | BC058408\|mKIAA1389 | signal-induced proliferation-associated 1 like 2 | protein-coding |
| chr8_125337541_125337775 | chr8 | 125337542 | 125337775 | + | -0.975479 | Intergenic | Intergenic | 155052 | NM_001081337 | 244668 | Mm.271668 | NM_001081337 | ENSMUSG00000001995 | **Sipa1l2** | BC058408\|mKIAA1389 | signal-induced proliferation-associated 1 like 2 | protein-coding |
| chr5_115488758_115489306 | chr5 | 115488759 | 115489306 | + | -0.815105 | Intergenic | Intergenic | -4548 | NM_133760 | 75387 | Mm.332616 | NM_133760 | ENSMUSG00000029524 | **Sirt4** | 4930596O17Rik | sirtuin 4 | protein-coding |
| chr4_43482122_43482379 | chr4 | 43482123 | 43482379 | + | -0.75567 | 3' UTR (NM_019436, exon 5 of 5) | 3' UTR (NM_019436, exon 5 of 5) | 1458 | NM_019436 | 54390 | Mm.31775 | NM_019436 | ENSMUSG00000028460 | **Sit1** | Sit | suppression inducing transmembrane adaptor 1 | protein-coding |
| chr4_155195219_155195592 | chr4 | 155195220 | 155195592 | + | -0.647994 | intron (NM_011385, intron 1 of 6) | intron (NM_011385, intron 1 of 6) | 27129 | NM_011385 | 20481 | Mm.28520 | NM_011385 | ENSMUSG00000029050 | **Ski** | 2310012I02Rik\|2610001A11Rik\|AA062172\|AA589460\|BC004088 | ski sarcoma viral oncogene homolog (avian) | protein-coding |
| chr4_155209528_155210009 | chr4 | 155209529 | 155210009 | + | -0.661076 | intron (NM_011385, intron 1 of 6) | B3A\|SINE\|B2 | 12766 | NM_011385 | 20481 | Mm.28520 | NM_011385 | ENSMUSG00000029050 | **Ski** | 2310012I02Rik\|2610001A11Rik\|AA062172\|AA589460\|BC004088 | ski sarcoma viral oncogene homolog (avian) | protein-coding |
| chr4_155197745_155198056 | chr4 | 155197746 | 155198056 | + | -0.683419 | intron (NM_011385, intron 1 of 6) | intron (NM_011385, intron 1 of 6) | 24634 | NM_011385 | 20481 | Mm.28520 | NM_011385 | ENSMUSG00000029050 | **Ski** | 2310012I02Rik\|2610001A11Rik\|AA062172\|AA589460\|BC004088 | ski sarcoma viral oncogene homolog (avian) | protein-coding |
| chr4_155178229_155179003 | chr4 | 155178230 | 155179003 | + | -0.68751 | intron (NM_011385, intron 1 of 6) | B3\|SINE\|B2 | 43919 | NM_011385 | 20481 | Mm.28520 | NM_011385 | ENSMUSG00000029050 | **Ski** | 2310012I02Rik\|2610001A11Rik\|AA062172\|AA589460\|BC004088 | ski sarcoma viral oncogene homolog (avian) | protein-coding |
| chr4_155195327_155195570 | chr4 | 155195328 | 155195570 | + | -0.707025 | intron (NM_011385, intron 1 of 6) | intron (NM_011385, intron 1 of 6) | 27086 | NM_011385 | 20481 | Mm.28520 | NM_011385 | ENSMUSG00000029050 | **Ski** | 2310012I02Rik\|2610001A11Rik\|AA062172\|AA589460\|BC004088 | ski sarcoma viral oncogene homolog (avian) | protein-coding |
| chr4_155178232_155178701 | chr4 | 155178233 | 155178701 | + | -0.727145 | intron (NM_011385, intron 1 of 6) | intron (NM_011385, intron 1 of 6) | 44068 | NM_011385 | 20481 | Mm.28520 | NM_011385 | ENSMUSG00000029050 | **Ski** | 2310012I02Rik\|2610001A11Rik\|AA062172\|AA589460\|BC004088 | ski sarcoma viral oncogene homolog (avian) | protein-coding |
| chr4_155168759_155168990 | chr4 | 155168760 | 155168990 | + | -0.990614 | intron (NM_011385, intron 1 of 6) | L1MB7\|LINE\|L1 | 53660 | NM_011385 | 20481 | Mm.28520 | NM_011385 | ENSMUSG00000029050 | **Ski** | 2310012I02Rik\|2610001A11Rik\|AA062172\|AA589460\|BC004088 | ski sarcoma viral oncogene homolog (avian) | protein-coding |
| chr4_155169046_155169577 | chr4 | 155169047 | 155169577 | + | -1.00317 | intron (NM_011385, intron 1 of 6) | intron (NM_011385, intron 1 of 6) | 53223 | NM_011385 | 20481 | Mm.28520 | NM_011385 | ENSMUSG00000029050 | **Ski** | 2310012I02Rik\|2610001A11Rik\|AA062172\|AA589460\|BC004088 | ski sarcoma viral oncogene homolog (avian) | protein-coding |
| chr4_155169048_155169438 | chr4 | 155169049 | 155169438 | + | -1.0512 | intron (NM_011385, intron 1 of 6) | intron (NM_011385, intron 1 of 6) | 53292 | NM_011385 | 20481 | Mm.28520 | NM_011385 | ENSMUSG00000029050 | **Ski** | 2310012I02Rik\|2610001A11Rik\|AA062172\|AA589460\|BC004088 | ski sarcoma viral oncogene homolog (avian) | protein-coding |
| chr4_155168612_155169528 | chr4 | 155168613 | 155169528 | + | -1.11312 | intron (NM_011385, intron 1 of 6) | intron (NM_011385, intron 1 of 6) | 53465 | NM_011385 | 20481 | Mm.28520 | NM_011385 | ENSMUSG00000029050 | **Ski** | 2310012I02Rik\|2610001A11Rik\|AA062172\|AA589460\|BC004088 | ski sarcoma viral oncogene homolog (avian) | protein-coding |
| chr3_31082866_31083186 | chr3 | 31082867 | 31083186 | + | -0.739215 | Intergenic | PB1\|SINE\|Alu | -12032 | NM_011386 | 20482 | Mm.15406 | NM_011386 | ENSMUSG00000027660 | **Skil** | Skir\|SnoN\|sno | SKI-like | protein-coding |
| chr9_63173919_63174343 | chr9 | 63173920 | 63174343 | + | -0.630362 | intron (NM_011840, intron 21 of 21) | intron (NM_011840, intron 21 of 21) | -25170 | NM_001163758 | 207667 | Mm.36349 | NM_172446 | ENSMUSG00000022245 | **Skor1** | AV273001\|C230094B15Rik\|Corl1\|Lbxcor1 | SKI family transcriptional corepressor 1 | protein-coding |
| chr9_63173704_63174347 | chr9 | 63173705 | 63174347 | + | -0.682526 | intron (NM_011840, intron 21 of 21) | ID_B1\|SINE\|B4 | -25065 | NM_001163758 | 207667 | Mm.36349 | NM_172446 | ENSMUSG00000022245 | **Skor1** | AV273001\|C230094B15Rik\|Corl1\|Lbxcor1 | SKI family transcriptional corepressor 1 | protein-coding |
| chr9_63086699_63087005 | chr9 | 63086700 | 63087005 | + | -1.20325 | Intergenic | HAL1\|LINE\|L1 | 60128 | NM_172446 | 207667 | Mm.36349 | NM_172446 | ENSMUSG00000022245 | **Skor1** | AV273001\|C230094B15Rik\|Corl1\|Lbxcor1 | SKI family transcriptional corepressor 1 | protein-coding |
| chr9_63089106_63089635 | chr9 | 63089107 | 63089635 | + | -1.22068 | Intergenic | B4\|SINE\|B4 | 57609 | NM_172446 | 207667 | Mm.36349 | NM_172446 | ENSMUSG00000022245 | **Skor1** | AV273001\|C230094B15Rik\|Corl1\|Lbxcor1 | SKI family transcriptional corepressor 1 | protein-coding |
| chr18_76892287_76892824 | chr18 | 76892288 | 76892824 | + | -1.2291 | intron (NM_001109743, intron 7 of 8) | intron (NM_001109743, intron 7 of 8) | 36151 | NM_001109743 | 664805 | Mm.329543 | NM_001109743 | ENSMUSG00000091519 | **Skor2** | Corl2\|EG664805\|Fussel18\|Gm7348 | SKI family transcriptional corepressor 2 | protein-coding |

| | | | | | | | | | | | | | | | | | |
|---|---|---|---|---|---|---|---|---|---|---|---|---|---|---|---|---|---|
| chr18_76892297_76892685 | chr18 | 76892298 | 76892685 | + | -1.2641 | intron (NM_001109743, intron 7 of 8) | intron (NM_001109743, intron 7 of 8) | 36086 | NM_001109743 | 664805 | Mm.329543 | NM_001109743 | ENSMUSG00000091519 | **Skor2** | Corl2\|EG664805\|Fussel18\|Gm7348 | SKI family transcriptional corepressor 2 | protein-coding |
| chr11_52240925_52241453 | chr11 | 52240926 | 52241453 | + | -0.816772 | intron (NM_011543, intron 2 of 5) | intron (NM_011543, intron 2 of 5) | 9194 | NM_011543 | 21402 | Mm.42944 | NM_011543 | ENSMUSG00000036309 | **Skp1a** | 15kDa\|2610043E24Rik\|2610206H23Rik\|EMC19\|OCP-II\|OCP2\|SKP1\|Tceb1l\|p19A\|p19Skp1 | S-phase kinase-associated protein 1A | protein-coding |
| chr11_52224377_52224802 | chr11 | 52224378 | 52224802 | + | -0.878091 | Intergenic | RLTR26C_MM\|LTR\|ERVK | -7405 | NM_011543 | 21402 | Mm.42944 | NM_011543 | ENSMUSG00000036309 | **Skp1a** | 15kDa\|2610043E24Rik\|2610206H23Rik\|EMC19\|OCP-II\|OCP2\|SKP1\|Tceb1l\|p19A\|p19Skp1 | S-phase kinase-associated protein 1A | protein-coding |
| chr11_52223976_52224950 | chr11 | 52223977 | 52224950 | + | -0.943488 | Intergenic | RLTR26C_MM\|LTR\|ERVK | -7532 | NM_011543 | 21402 | Mm.42944 | NM_011543 | ENSMUSG00000036309 | **Skp1a** | 15kDa\|2610043E24Rik\|2610206H23Rik\|EMC19\|OCP-II\|OCP2\|SKP1\|Tceb1l\|p19A\|p19Skp1 | S-phase kinase-associated protein 1A | protein-coding |
| chr15_66801442_66801932 | chr15 | 66801443 | 66801932 | + | -0.68743 | intron (NM_001029841, intron 1 of 7) | intron (NM_001029841, intron 1 of 7) | 30142 | NM_001029841 | 20491 | Mm.7601 | NM_009192 | ENSMUSG00000022372 | **Sla** | Slap\|Slap-1 | src-like adaptor | protein-coding |
| chr15_66755056_66755901 | chr15 | 66755057 | 66755901 | + | -0.724979 | intron (NM_009375, intron 35 of 47) | intron (NM_009375, intron 35 of 47) | 76350 | NM_001029841 | 20491 | Mm.7601 | NM_009192 | ENSMUSG00000022372 | **Sla** | Slap\|Slap-1 | src-like adaptor | protein-coding |
| chr15_66755055_66756499 | chr15 | 66755056 | 66756499 | + | -0.733565 | intron (NM_009375, intron 35 of 47) | intron (NM_009375, intron 35 of 47) | 76052 | NM_001029841 | 20491 | Mm.7601 | NM_009192 | ENSMUSG00000022372 | **Sla** | Slap\|Slap-1 | src-like adaptor | protein-coding |
| chr15_66789012_66789583 | chr15 | 66789013 | 66789583 | + | -0.92295 | intron (NM_001029841, intron 5 of 7) | intron (NM_001029841, intron 5 of 7) | 42531 | NM_001029841 | 20491 | Mm.7601 | NM_009192 | ENSMUSG00000022372 | **Sla** | Slap\|Slap-1 | src-like adaptor | protein-coding |
| chr15_66789010_66789409 | chr15 | 66789011 | 66789409 | + | -0.930324 | intron (NM_001029841, intron 5 of 7) | intron (NM_001029841, intron 5 of 7) | 42619 | NM_001029841 | 20491 | Mm.7601 | NM_009192 | ENSMUSG00000022372 | **Sla** | Slap\|Slap-1 | src-like adaptor | protein-coding |
| chr15_66793060_66793358 | chr15 | 66793061 | 66793358 | + | -1.15615 | intron (NM_001029841, intron 3 of 7) | intron (NM_001029841, intron 3 of 7) | 38620 | NM_001029841 | 20491 | Mm.7601 | NM_009192 | ENSMUSG00000022372 | **Sla** | Slap\|Slap-1 | src-like adaptor | protein-coding |
| chr2_156887013_156887433 | chr2 | 156887014 | 156887433 | + | -0.65561 | promoter-TSS (NM_029983) | promoter-TSS (NM_029983) | -145 | NM_029983 | 77799 | Mm.29264 | NM_029983 | ENSMUSG00000027636 | **Sla2** | A930009E21Rik\|AI430952\|SLAP-2\|SLAP2 | Src-like-adaptor 2 | protein-coding |
| chr2_156882376_156882904 | chr2 | 156882377 | 156882904 | + | -0.664205 | intron (NM_029983, intron 2 of 7) | intron (NM_029983, intron 2 of 7) | 4438 | NM_029983 | 77799 | Mm.29264 | NM_029983 | ENSMUSG00000027636 | **Sla2** | A930009E21Rik\|AI430952\|SLAP-2\|SLAP2 | Src-like-adaptor 2 | protein-coding |
| chr2_156918561_156919040 | chr2 | 156918562 | 156919040 | + | -0.782653 | Intergenic | Intergenic | -31723 | NM_029983 | 77799 | Mm.29264 | NM_029983 | ENSMUSG00000027636 | **Sla2** | A930009E21Rik\|AI430952\|SLAP-2\|SLAP2 | Src-like-adaptor 2 | protein-coding |
| chr14_103648994_103649225 | chr14 | 103648995 | 103649225 | + | -0.714232 | Intergenic | Intergenic | -1133 | NM_198014 | 105439 | Mm.27548 | NM_198014 | ENSMUSG00000055717 | **Slain1** | 9630044O09Rik\|AA675320\|AW742596 | SLAIN motif family, member 1 | protein-coding |
| chr5_72916796_72917142 | chr5 | 72916797 | 72917142 | + | -0.646268 | intron (NM_153567, intron 1 of 8) | intron (NM_153567, intron 1 of 8) | 2607 | NM_153567 | 75991 | Mm.121285 | NM_153567 | ENSMUSG00000036087 | **Slain2** | 5033405K12Rik\|8030444K12Rik\|AI596370 | SLAIN motif family, member 2 | protein-coding |
| chr1_171789436_171789876 | chr1 | 171789437 | 171789876 | + | -0.829025 | intron (NM_013730, intron 4 of 6) | intron (NM_013730, intron 4 of 6) | 22524 | NM_013730 | 27218 | Mm.103648 | NM_013730 | ENSMUSG00000015316 | **Slamf1** | 4933415F16\|AA177906\|CD150\|CDw150\|ESTM51\|IPO-3\|Slam | signaling lymphocytic activation molecule family member 1 | protein-coding |
| chr1_171955108_171955450 | chr1 | 171955109 | 171955450 | + | -0.715477 | Intergenic | Intergenic | 37742 | NM_030710 | 30925 | Mm.486404 | NM_030710 | ENSMUSG00000015314 | **Slamf6** | KAL1\|KAL1b\|Ly108\|NTB-A\|NTBA\|SF2000 | SLAM family member 6 | protein-coding |
| chr1_171963656_171963985 | chr1 | 171963657 | 171963985 | + | -0.760258 | Intergenic | Intergenic | 46284 | NM_030710 | 30925 | Mm.486404 | NM_030710 | ENSMUSG00000015314 | **Slamf6** | KAL1\|KAL1b\|Ly108\|NTB-A\|NTBA\|SF2000 | SLAM family member 6 | protein-coding |
| chr8_78622362_78622771 | chr8 | 78622363 | 78622771 | + | -0.771692 | intron (NM_029736, intron 5 of 11) | ORR1D2\|LTR\|ERVL-MaLR | 113239 | NM_001282108 | 76775 | Mm.334631 | NM_029736 | ENSMUSG00000031684 | **Slc10a7** | 2410193C02Rik\|P7 | solute carrier family 10 (sodium/bile acid cotransporter family), member 7 | protein-coding |
| chr8_94354197_94354881 | chr8 | 94354198 | 94354881 | + | -0.71784 | intron (NM_019415, intron 21 of 24) | intron (NM_019415, intron 21 of 24) | 25331 | NM_001205311 | 20497 | Mm.25804 | NM_019415 | ENSMUSG00000031766 | **Slc12a3** | AI035291\|NCC\|NCCT\|TSC | solute carrier family 12, member 3 | protein-coding |
| chr8_94354467_94354868 | chr8 | 94354468 | 94354868 | + | -0.804771 | intron (NM_019415, intron 21 of 24) | intron (NM_019415, intron 21 of 24) | 25460 | NM_001205311 | 20497 | Mm.25804 | NM_019415 | ENSMUSG00000031766 | **Slc12a3** | AI035291\|NCC\|NCCT\|TSC | solute carrier family 12, member 3 | protein-coding |
| chr13_73748981_73749478 | chr13 | 73748982 | 73749478 | + | -0.686896 | Intergenic | Intergenic | -14467 | NM_011390 | 20499 | Mm.275800 | NM_011390 | ENSMUSG00000017756 | **Slc12a7** | AA408796\|D13Ertd261e\|Kcc4 | solute carrier family 12, member 7 | protein-coding |
| chr13_73760107_73760501 | chr13 | 73760108 | 73760501 | + | -0.689613 | Intergenic | Intergenic | -3393 | NM_011390 | 20499 | Mm.275800 | NM_011390 | ENSMUSG00000017756 | **Slc12a7** | AA408796\|D13Ertd261e\|Kcc4 | solute carrier family 12, member 7 | protein-coding |
| chr13_73736889_73737515 | chr13 | 73736890 | 73737515 | + | -0.774532 | Intergenic | Intergenic | -26495 | NM_011390 | 20499 | Mm.275800 | NM_011390 | ENSMUSG00000017756 | **Slc12a7** | AA408796\|D13Ertd261e\|Kcc4 | solute carrier family 12, member 7 | protein-coding |
| chr13_73772603_73773023 | chr13 | 73772604 | 73773023 | + | -0.921786 | intron (NM_011390, intron 1 of 23) | intron (NM_011390, intron 1 of 23) | 9116 | NM_011390 | 20499 | Mm.275800 | NM_011390 | ENSMUSG00000017756 | **Slc12a7** | AA408796\|D13Ertd261e\|Kcc4 | solute carrier family 12, member 7 | protein-coding |
| chr13_73772566_73773203 | chr13 | 73772567 | 73773203 | + | -0.957496 | intron (NM_011390, intron 1 of 23) | intron (NM_011390, intron 1 of 23) | 9188 | NM_011390 | 20499 | Mm.275800 | NM_011390 | ENSMUSG00000017756 | **Slc12a7** | AA408796\|D13Ertd261e\|Kcc4 | solute carrier family 12, member 7 | protein-coding |
| chr2_165452827_165453790 | chr2 | 165452828 | 165453790 | + | -0.631147 | intron (NM_054055, intron 1 of 12) | intron (NM_054055, intron 1 of 12) | 19888 | NM_054055 | 114644 | Mm.250738 | NM_054055 | ENSMUSG00000018459 | **Slc13a3** | NaDC-3\|NaDC3\|SDCT2 | solute carrier family 13 (sodium-dependent dicarboxylate transporter), member 3 | protein-coding |
| chr14_121477909_121478757 | chr14 | 121477910 | 121478757 | + | -0.688219 | intron (NM_053079, intron 12 of 22) | intron (NM_053079, intron 12 of 22) | 26921 | NM_053079 | 56643 | Mm.155618 | NM_053079 | ENSMUSG00000025557 | **Slc15a1** | D630032F02\|PECT1\|PEPT1 | solute carrier family 15 (oligopeptide transporter), member 1 | protein-coding |
| chr14_121553753_121554323 | chr14 | 121553754 | 121554323 | + | -0.698552 | intron (NM_001081039, intron 50 of 53) | intron (NM_001081039, intron 50 of 53) | -48784 | NM_053079 | 56643 | Mm.155618 | NM_053079 | ENSMUSG00000025557 | **Slc15a1** | D630032F02\|PECT1\|PEPT1 | solute carrier family 15 (oligopeptide transporter), member 1 | protein-coding |
| chr14_121477870_121478307 | chr14 | 121477871 | 121478307 | + | -0.765904 | exon (NM_053079, exon 14 of 23) | exon (NM_053079, exon 14 of 23) | 27165 | NM_053079 | 56643 | Mm.155618 | NM_053079 | ENSMUSG00000025557 | **Slc15a1** | D630032F02\|PECT1\|PEPT1 | solute carrier family 15 (oligopeptide transporter), member 1 | protein-coding |
| chr14_121545239_121546463 | chr14 | 121545240 | 121546463 | + | -0.873029 | intron (NM_001128308, intron 52 of 53) | intron (NM_001128308, intron 52 of 53) | -40597 | NM_053079 | 56643 | Mm.155618 | NM_053079 | ENSMUSG00000025557 | **Slc15a1** | D630032F02\|PECT1\|PEPT1 | solute carrier family 15 (oligopeptide transporter), member 1 | protein-coding |
| chr14_121540945_121541206 | chr14 | 121540946 | 121541206 | + | -0.892048 | TTS (NM_001128307) | TTS (NM_001128307) | -35822 | NM_053079 | 56643 | Mm.155618 | NM_053079 | ENSMUSG00000025557 | **Slc15a1** | D630032F02\|PECT1\|PEPT1 | solute carrier family 15 (oligopeptide transporter), member 1 | protein-coding |
| chr14_121552373_121553133 | chr14 | 121552374 | 121553133 | + | -0.948554 | intron (NM_001081039, intron 50 of 53) | intron (NM_001081039, intron 50 of 53) | -47499 | NM_053079 | 56643 | Mm.155618 | NM_053079 | ENSMUSG00000025557 | **Slc15a1** | D630032F02\|PECT1\|PEPT1 | solute carrier family 15 (oligopeptide transporter), member 1 | protein-coding |
| chr14_121545465_121545921 | chr14 | 121545466 | 121545921 | + | -1.18985 | intron (NM_001128308, intron 52 of 53) | intron (NM_001128308, intron 52 of 53) | -40439 | NM_053079 | 56643 | Mm.155618 | NM_053079 | ENSMUSG00000025557 | **Slc15a1** | D630032F02\|PECT1\|PEPT1 | solute carrier family 15 (oligopeptide transporter), member 1 | protein-coding |
| chr5_127484817_127485767 | chr5 | 127484818 | 127485767 | + | -0.868832 | intron (NM_175432, intron 3 of 8) | intron (NM_175432, intron 3 of 8) | 132100 | NM_133895 | 100561 | Mm.28506 | NM_133895 | ENSMUSG00000029416 | **Slc15a4** | AA987064\|AW742963\|C130069N12Rik\|PHT1\|PTR4 | solute carrier family 15, member 4 | protein-coding |
| chr3_104657387_104657797 | chr3 | 104657388 | 104657797 | + | -0.766486 | 3' UTR (NM_009196, exon 5 of 5) | 3' UTR (NM_009196, exon 5 of 5) | 18928 | NM_009196 | 20501 | Mm.9086 | NM_009196 | ENSMUSG00000032902 | **Slc16a1** | AL022710\|Mct1 | solute carrier family 16 (monocarboxylic acid transporters), member 1 | protein-coding |
| chr3_104630738_104631413 | chr3 | 104630739 | 104631413 | + | -1.01052 | Intergenic | Intergenic | -7588 | NM_009196 | 20501 | Mm.9086 | NM_009196 | ENSMUSG00000032902 | **Slc16a1** | AL022710\|Mct1 | solute carrier family 16 (monocarboxylic acid transporters), member 1 | protein-coding |
| chr3_104630973_104631407 | chr3 | 104630974 | 104631407 | + | -1.09607 | Intergenic | B4A\|SINE\|B4 | -7474 | NM_009196 | 20501 | Mm.9086 | NM_009196 | ENSMUSG00000032902 | **Slc16a1** | AL022710\|Mct1 | solute carrier family 16 (monocarboxylic acid transporters), member 1 | protein-coding |
| chr10_40125180_40125602 | chr10 | 40125181 | 40125602 | + | -0.849907 | intron (NM_028247, intron 1 of 2) | intron (NM_028247, intron 1 of 2) | 16863 | NM_028247 | 72472 | Mm.186778 | NM_028247 | ENSMUSG00000019838 | **Slc16a10** | 2610103N14Rik\|9830169E08\|Mct10\|PRO0813\|TAT1 | solute carrier family 16 (monocarboxylic acid transporters), member 10 | protein-coding |
| chr10_40163340_40163742 | chr10 | 40163341 | 40163742 | + | -0.932208 | Intergenic | Intergenic | -21287 | NM_001114332 | 72472 | Mm.186778 | NM_028247 | ENSMUSG00000019838 | **Slc16a10** | 2610103N14Rik\|9830169E08\|Mct10\|PRO0813\|TAT1 | solute carrier family 16 (monocarboxylic acid transporters), member 10 | protein-coding |
| chr11_109470981_109471161 | chr11 | 109470982 | 109471161 | + | -1.03308 | intron (NM_001029842, intron 1 of 5) | B1F\|SINE\|Alu | 2158 | NM_134038 | 104681 | Mm.265874 | NM_134038 | ENSMUSG00000041920 | **Slc16a6** | AW743111\|ESTM12\|MCT 6\|MCT 7\|MCT6 | solute carrier family 16 (monocarboxylic acid transporters), member 6 | protein-coding |
| chr11_109470952_109471227 | chr11 | 109470953 | 109471227 | + | -1.04773 | intron (NM_001029842, intron 1 of 5) | B1F\|SINE\|Alu | 2139 | NM_134038 | 104681 | Mm.265874 | NM_134038 | ENSMUSG00000041920 | **Slc16a6** | AW743111\|ESTM12\|MCT 6\|MCT 7\|MCT6 | solute carrier family 16 (monocarboxylic acid transporters), member 6 | protein-coding |
| chr10_125290373_125290671 | chr10 | 125290374 | 125290671 | + | -0.677324 | intron (NM_011391, intron 4 of 6) | intron (NM_011391, intron 4 of 6) | 38013 | NM_011391 | 20503 | Mm.29161 | NM_011391 | ENSMUSG00000020102 | **Slc16a7** | 4921534N07Rik\|Mct2 | solute carrier family 16 (monocarboxylic acid transporters), member 7 | protein-coding |
| chr10_125370039_125370554 | chr10 | 125370040 | 125370554 | + | -0.688529 | Intergenic | ORR1D2\|LTR\|ERVL-MaLR | -41762 | NM_011391 | 20503 | Mm.29161 | NM_011391 | ENSMUSG00000020102 | **Slc16a7** | 4921534N07Rik\|Mct2 | solute carrier family 16 (monocarboxylic acid transporters), member 7 | protein-coding |

| | | | | | | | | | | | | | | | | | |
|---|---|---|---|---|---|---|---|---|---|---|---|---|---|---|---|---|---|
| chr10_89619455_89619987 | chr10 | 89619456 | 89619987 | + | -0.738202 | intron (NM_182959, intron 1 of 11) | intron (NM_182959, intron 1 of 11) | 1528 | NM_182959 | 216227 | Mm.233921 | NM_182959 | ENSMUSG00000019935 | **Slc17a8** | BC042593\|Vglut3 | solute carrier family 17 (sodium-dependent inorganic phosphate cotransporter), member 8 | protein-coding |
| chr1_82973933_82974309 | chr1 | 82973934 | 82974309 | + | -1.14095 | Intergenic | Intergenic | 64327 | NM_030556 | 80721 | Mm.261542 | NM_030556 | ENSMUSG00000038496 | **Slc19a3** | A230084E24Rik\|AI788884\|ThTr2 | solute carrier family 19, member 3 | protein-coding |
| chr10_40479202_40480099 | chr10 | 40479203 | 40480099 | + | -0.736493 | intron (NM_198164, intron 11 of 11) | intron (NM_198164, intron 11 of 11) | -90711 | NM_027572 | 70840 | Mm.332570 | NM_027572 | ENSMUSG00000019834 | **Slc22a16** | 4921504E14Rik\|CT2\|FLIPT2\|OCT6\|OKB1 | solute carrier family 22 (organic cation transporter), member 16 | protein-coding |
| chr10_40461957_40462695 | chr10 | 40461958 | 40462695 | + | -0.748559 | intron (NM_198164, intron 5 of 11) | intron (NM_198164, intron 5 of 11) | -108036 | NM_027572 | 70840 | Mm.332570 | NM_027572 | ENSMUSG00000019834 | **Slc22a16** | 4921504E14Rik\|CT2\|FLIPT2\|OCT6\|OKB1 | solute carrier family 22 (organic cation transporter), member 16 | protein-coding |
| chr10_40462166_40462682 | chr10 | 40462167 | 40462682 | + | -0.766776 | intron (NM_198164, intron 5 of 11) | intron (NM_198164, intron 5 of 11) | -107938 | NM_027572 | 70840 | Mm.332570 | NM_027572 | ENSMUSG00000019834 | **Slc22a16** | 4921504E14Rik\|CT2\|FLIPT2\|OCT6\|OKB1 | solute carrier family 22 (organic cation transporter), member 16 | protein-coding |
| chr10_40479693_40480149 | chr10 | 40479694 | 40480149 | + | -0.890533 | 3' UTR (NM_001291816, exon 12 of 12) | 3' UTR (NM_001291816, exon 12 of 12) | -90441 | NM_027572 | 70840 | Mm.332570 | NM_027572 | ENSMUSG00000019834 | **Slc22a16** | 4921504E14Rik\|CT2\|FLIPT2\|OCT6\|OKB1 | solute carrier family 22 (organic cation transporter), member 16 | protein-coding |
| chr10_40514871_40515737 | chr10 | 40514872 | 40515737 | + | -0.892859 | Intergenic | LTR33\|LTR\|ERVL | -55058 | NM_027572 | 70840 | Mm.332570 | NM_027572 | ENSMUSG00000019834 | **Slc22a16** | 4921504E14Rik\|CT2\|FLIPT2\|OCT6\|OKB1 | solute carrier family 22 (organic cation transporter), member 16 | protein-coding |
| chr10_40515101_40515962 | chr10 | 40515102 | 40515962 | + | -0.893501 | Intergenic | LTR33\|LTR\|ERVL | -54830 | NM_027572 | 70840 | Mm.332570 | NM_027572 | ENSMUSG00000019834 | **Slc22a16** | 4921504E14Rik\|CT2\|FLIPT2\|OCT6\|OKB1 | solute carrier family 22 (organic cation transporter), member 16 | protein-coding |
| chr10_40476915_40477764 | chr10 | 40476916 | 40477764 | + | -0.898474 | intron (NM_198164, intron 10 of 11) | intron (NM_198164, intron 10 of 11) | -93022 | NM_027572 | 70840 | Mm.332570 | NM_027572 | ENSMUSG00000019834 | **Slc22a16** | 4921504E14Rik\|CT2\|FLIPT2\|OCT6\|OKB1 | solute carrier family 22 (organic cation transporter), member 16 | protein-coding |
| chr10_40476915_40477345 | chr10 | 40476916 | 40477345 | + | -0.91526 | intron (NM_198164, intron 10 of 11) | intron (NM_198164, intron 10 of 11) | -93232 | NM_027572 | 70840 | Mm.332570 | NM_027572 | ENSMUSG00000019834 | **Slc22a16** | 4921504E14Rik\|CT2\|FLIPT2\|OCT6\|OKB1 | solute carrier family 22 (organic cation transporter), member 16 | protein-coding |
| chr2_71360183_71360571 | chr2 | 71360184 | 71360571 | + | -1.3935 | intron (NM_172436, intron 2 of 17) | B4A\|SINE\|B4 | 7177 | NM_172436 | 78830 | Mm.30928 | NM_172436 | | **Slc25a12** | 2610002D09Rik\|AI839531\|B230107K20Rik\|BB129864 | solute carrier family 25 (mitochondrial carrier, Aralar), member 12 | protein-coding |
| chr2_32423231_32423919 | chr2 | 32423232 | 32423919 | + | -0.7325 | intron (NM_146118, intron 1 of 10) | intron (NM_146118, intron 1 of 10) | 491 | NM_001290558 | 227731 | Mm.37395 | NM_146118 | ENSMUSG00000026819 | **Slc25a25** | 1110030N17Rik\|MCSC\|mKIAA1896 | solute carrier family 25 (mitochondrial carrier, phosphate carrier), member 25 | protein-coding |
| chr4_149856320_149856874 | chr4 | 149856321 | 149856874 | + | -0.879122 | Intergenic | Intergenic | -82330 | NM_027460 | 70556 | Mm.41877 | NM_027460 | ENSMUSG00000028982 | **Slc25a33** | 5730438N18Rik\|Pnc1 | solute carrier family 25, member 33 | protein-coding |
| chr5_8425975_8426224 | chr5 | 8425976 | 8426224 | + | -1.21981 | intron (NM_001289595, intron 1 of 10) | intron (NM_001289595, intron 1 of 10) | 3262 | NM_001289595 | 319653 | Mm.146262 | NM_178766 | ENSMUSG00000054099 | **Slc25a40** | B230315F11Rik | solute carrier family 25, member 40 | protein-coding |
| chr5_8425935_8426311 | chr5 | 8425936 | 8426311 | + | -1.23443 | intron (NM_001289595, intron 1 of 10) | intron (NM_001289595, intron 1 of 10) | 3285 | NM_001289595 | 319653 | Mm.146262 | NM_178766 | ENSMUSG00000054099 | **Slc25a40** | B230315F11Rik | solute carrier family 25, member 40 | protein-coding |
| chrX_36757703_36758415 | chrX | 36757704 | 36758415 | + | -0.832665 | intron (NR_075097, intron 2 of 3) | intron (NR_075097, intron 2 of 3) | 14427 | NR_075096 | 194744 | Mm.336014 | NM_001085497 | ENSMUSG00000037636 | **Slc25a43** | Gm8 | solute carrier family 25, member 43 | protein-coding |
| chr4_45402338_45403178 | chr4 | 45402339 | 45403178 | + | -0.771088 | intron (NM_001009949, intron 2 of 2) | intron (NM_001009949, intron 2 of 2) | 6008 | NM_001009949 | 230125 | Mm.260210 | NM_001009949 | ENSMUSG00000045973 | **Slc25a51** | 9130208E07Rik\|D130005A03Rik\|Gm138\|Mcart1 | solute carrier family 25, member 51 | protein-coding |
| chr4_45402376_45402639 | chr4 | 45402377 | 45402639 | + | -0.79705 | intron (NM_001009949, intron 2 of 2) | intron (NM_001009949, intron 2 of 2) | 6258 | NM_001009949 | 230125 | Mm.260210 | NM_001009949 | ENSMUSG00000045973 | **Slc25a51** | 9130208E07Rik\|D130005A03Rik\|Gm138\|Mcart1 | solute carrier family 25, member 51 | protein-coding |
| chr4_45402304_45402803 | chr4 | 45402305 | 45402803 | + | -0.800136 | intron (NM_001009949, intron 2 of 2) | URR1A\|DNA\|hAT-Charlie | 6212 | NM_001009949 | 230125 | Mm.260210 | NM_001009949 | ENSMUSG00000045973 | **Slc25a51** | 9130208E07Rik\|D130005A03Rik\|Gm138\|Mcart1 | solute carrier family 25, member 51 | protein-coding |
| chr18_58571637_58572250 | chr18 | 58571638 | 58572250 | + | -1.01927 | intron (NM_001081072, intron 1 of 9) | intron (NM_001081072, intron 1 of 9) | 15704 | NM_001081072 | 225579 | Mm.258517 | NM_001081072 | ENSMUSG00000024600 | **Slc27a6** | 4732438L20Rik\|FACVL2\|FATP6\|VLCS-H1 | solute carrier family 27 (fatty acid transporter), member 6 | protein-coding |
| chr4_119127250_119127675 | chr4 | 119127251 | 119127675 | + | -0.85972 | intron (NM_011400, intron 2 of 9) | intron (NM_011400, intron 2 of 9) | 18718 | NM_011400 | 20525 | Mm.21002 | NM_011400 | ENSMUSG00000028645 | **Slc2a1** | Glut-1\|Glut1 | solute carrier family 2 (facilitated glucose transporter), member 1 | protein-coding |
| chr4_119127018_119127755 | chr4 | 119127019 | 119127755 | + | -0.872756 | intron (NM_011400, intron 2 of 9) | intron (NM_011400, intron 2 of 9) | 18642 | NM_011400 | 20525 | Mm.21002 | NM_011400 | ENSMUSG00000028645 | **Slc2a1** | Glut-1\|Glut1 | solute carrier family 2 (facilitated glucose transporter), member 1 | protein-coding |
| chr3_28554849_28555353 | chr3 | 28554850 | 28555353 | + | -0.671399 | intron (NM_001163007, intron 8 of 31) | intron (NM_001163007, intron 8 of 31) | -142802 | NM_031197 | 20526 | Mm.18443 | NM_031197 | ENSMUSG00000027690 | **Slc2a2** | AI266973\|Glut-2\|Glut2 | solute carrier family 2 (facilitated glucose transporter), member 2 | protein-coding |
| chr3_115971926_115972595 | chr3 | 115971927 | 115972595 | + | -0.93271 | intron (NM_023214, intron 8 of 10) | intron (NM_023214, intron 8 of 10) | 35145 | NM_023214 | 66500 | Mm.34550 | NM_023214 | ENSMUSG00000054414 | **Slc30a7** | 1810059J10Rik\|2610034N15Rik\|4833428C12Rik\|AI467242\|ZnT-7\|ZnT7\|Zntl2 | solute carrier family 30 (zinc transporter), member 7 | protein-coding |
| chr5_67299890_67300812 | chr5 | 67299891 | 67300812 | + | -0.671632 | Intergenic | Intergenic | -6606 | NM_178651 | 109108 | Mm.234455 | NM_178651 | | **Slc30a9** | 2310024J23Rik\|AL024256\|GAC63\|HUEL | solute carrier family 30 (zinc transporter), member 9 | protein-coding |
| chr5_67360104_67360619 | chr5 | 67360105 | 67360619 | + | -1.11937 | Intergenic | Intergenic | 53405 | NM_178651 | 109108 | Mm.234455 | NM_178651 | | **Slc30a9** | 2310024J23Rik\|AL024256\|GAC63\|HUEL | solute carrier family 30 (zinc transporter), member 9 | protein-coding |
| chr11_95366616_95367715 | chr11 | 95366617 | 95367715 | + | -0.666927 | intron (NM_172543, intron 2 of 7) | intron (NM_172543, intron 2 of 7) | -17756 | NM_016752 | 110172 | Mm.4593 | NM_016752 | | **Slc35b1** | Ugalt2 | solute carrier family 35, member B1 | protein-coding |
| chr11_95366963_95367705 | chr11 | 95366964 | 95367705 | + | -0.7826 | intron (NM_172543, intron 2 of 7) | intron (NM_172543, intron 2 of 7) | -17588 | NM_016752 | 110172 | Mm.4593 | NM_016752 | | **Slc35b1** | Ugalt2 | solute carrier family 35, member B1 | protein-coding |
| chr13_38938355_38938802 | chr13 | 38938356 | 38938802 | + | -0.767039 | intron (NM_134060, intron 6 of 9) | intron (NM_134060, intron 6 of 9) | 21958 | NM_001170430 | 108652 | Mm.255506 | NM_134060 | ENSMUSG00000021432 | **Slc35b3** | 4921526O06Rik\|AI428480\|CGI-19\|PABST2\|PAPST2 | solute carrier family 35, member B3 | protein-coding |
| chr13_38938503_38938788 | chr13 | 38938504 | 38938788 | + | -0.800492 | intron (NM_134060, intron 6 of 9) | intron (NM_134060, intron 6 of 9) | 21891 | NM_001170430 | 108652 | Mm.255506 | NM_134060 | ENSMUSG00000021432 | **Slc35b3** | 4921526O06Rik\|AI428480\|CGI-19\|PABST2\|PAPST2 | solute carrier family 35, member B3 | protein-coding |
| chr6_34176096_34176379 | chr6 | 34176097 | 34176379 | + | -0.732994 | intron (NM_021435, intron 1 of 9) | intron (NM_021435, intron 1 of 9) | 816 | NM_021435 | 58246 | Mm.245527 | NM_021435 | | **Slc35b4** | 4930474D06Rik\|AB041549\|AW494345 | solute carrier family 35, member B4 | protein-coding |
| chr4_155603803_155604107 | chr4 | 155603804 | 155604107 | + | -1.2701 | intron (NM_177186, intron 1 of 8) | intron (NM_177186, intron 1 of 8) | 2539 | NM_177186 | 320541 | Mm.306018 | NM_177186 | ENSMUSG00000042202 | **Slc35e2** | A530082C11Rik\|AI957035 | solute carrier family 35, member E2 | protein-coding |
| chr19_38401144_38401401 | chr19 | 38401145 | 38401401 | + | -0.682483 | intron (NM_175507, intron 2 of 2) | intron (NM_175507, intron 2 of 2) | 5293 | NM_175507 | 240660 | Mm.262085 | NM_175507 | ENSMUSG00000044026 | **Slc35g1** | AA590464\|D330039I19Rik\|Tmem20 | solute carrier family 35, member G1 | protein-coding |
| chr19_38401449_38401816 | chr19 | 38401450 | 38401816 | + | -1.031 | intron (NM_175507, intron 2 of 2) | intron (NM_175507, intron 2 of 2) | 5653 | NM_175507 | 240660 | Mm.262085 | NM_175507 | ENSMUSG00000044026 | **Slc35g1** | AA590464\|D330039I19Rik\|Tmem20 | solute carrier family 35, member G1 | protein-coding |
| chr11_55221279_55221752 | chr11 | 55221280 | 55221752 | + | -0.67482 | intron (NM_153139, intron 5 of 10) | intron (NM_153139, intron 5 of 10) | 17176 | NM_153139 | 215335 | Mm.35732 | NM_153139 | ENSMUSG00000020261 | **Slc36a1** | 5830411H19Rik\|AI839897\|Pat1 | solute carrier family 36 (proton/amino acid symporter), member 1 | protein-coding |
| chr17_31325580_31325849 | chr17 | 31325581 | 31325849 | + | -0.704313 | intron (NM_153062, intron 9 of 19) | intron (NM_153062, intron 9 of 19) | 29491 | NM_001242427 | 224674 | Mm.311395 | NM_153062 | ENSMUSG00000024036 | **Slc37a1** | G3PP | solute carrier family 37 (glycerol-3-phosphate transporter), member 1 | protein-coding |
| chr17_31325920_31326809 | chr17 | 31325921 | 31326809 | + | -0.735804 | intron (NM_153062, intron 9 of 19) | intron (NM_153062, intron 9 of 19) | 30141 | NM_001242427 | 224674 | Mm.311395 | NM_153062 | ENSMUSG00000024036 | **Slc37a1** | G3PP | solute carrier family 37 (glycerol-3-phosphate transporter), member 1 | protein-coding |
| chr17_31325523_31326809 | chr17 | 31325524 | 31326809 | + | -0.755582 | intron (NM_153062, intron 9 of 19) | intron (NM_153062, intron 9 of 19) | 29942 | NM_001242427 | 224674 | Mm.311395 | NM_153062 | ENSMUSG00000024036 | **Slc37a1** | G3PP | solute carrier family 37 (glycerol-3-phosphate transporter), member 1 | protein-coding |
| chr6_39347935_39348759 | chr6 | 39347936 | 39348759 | + | -0.651275 | intron (NM_028123, intron 9 of 14) | intron (NM_028123, intron 9 of 14) | 29360 | NM_028123 | 72144 | Mm.482167 | NM_028123 | ENSMUSG00000029924 | **Slc37a3** | 2610507O21Rik\|AU044904 | solute carrier family 37 (glycerol-3-phosphate transporter), member 3 | protein-coding |
| chr6_39347694_39348453 | chr6 | 39347695 | 39348453 | + | -0.764434 | intron (NM_028123, intron 9 of 14) | intron (NM_028123, intron 9 of 14) | 29633 | NM_028123 | 72144 | Mm.482167 | NM_028123 | ENSMUSG00000029924 | **Slc37a3** | 2610507O21Rik\|AU044904 | solute carrier family 37 (glycerol-3-phosphate transporter), member 3 | protein-coding |
| chr6_39379209_39379673 | chr6 | 39379210 | 39379673 | + | -0.963179 | Intergenic | Intergenic | -1734 | NM_028123 | 72144 | Mm.482167 | NM_028123 | ENSMUSG00000029924 | **Slc37a3** | 2610507O21Rik\|AU044904 | solute carrier family 37 (glycerol-3-phosphate transporter), member 3 | protein-coding |
| chr6_39316191_39316744 | chr6 | 39316192 | 39316744 | + | -1.10936 | Intergenic | Intergenic | 61239 | NM_028123 | 72144 | Mm.482167 | NM_028123 | ENSMUSG00000029924 | **Slc37a3** | 2610507O21Rik\|AU044904 | solute carrier family 37 (glycerol-3-phosphate transporter), member 3 | protein-coding |
| chr6_39316131_39316502 | chr6 | 39316132 | 39316502 | + | -1.24916 | Intergenic | Intergenic | 61390 | NM_028123 | 72144 | Mm.482167 | NM_028123 | ENSMUSG00000029924 | **Slc37a3** | 2610507O21Rik\|AU044904 | solute carrier family 37 (glycerol-3-phosphate transporter), member 3 | protein-coding |
| chr15_96593001_96593750 | chr15 | 96593002 | 96593750 | + | -0.914548 | intron (NM_134086, intron 4 of 15) | intron (NM_134086, intron 4 of 15) | 48586 | NM_134086 | 105727 | Mm.103568 | NM_134086 | ENSMUSG00000023169 | **Slc38a1** | AA408026\|AA409865\|AL022800\|AU015942\|NAT2 | solute carrier family 38, member 1 | protein-coding |
| chr15_96598911_96599339 | chr15 | 96598912 | 96599339 | + | -1.03786 | intron (NM_134086, intron 4 of 15) | intron (NM_134086, intron 4 of 15) | 42837 | NM_134086 | 105727 | Mm.103568 | NM_134086 | ENSMUSG00000023169 | **Slc38a1** | AA408026\|AA409865\|AL022800\|AU015942\|NAT2 | solute carrier family 38, member 1 | protein-coding |
| chr11_120142586_120142958 | chr11 | 120142587 | 120142958 | + | -0.718408 | intron (NM_024249, intron 2 of 16) | intron (NM_024249, intron 2 of 16) | 8579 | NM_001164799 | 72055 | Mm.253403 | NM_024249 | ENSMUSG00000061306 | **Slc38a10** | 1810073N04Rik | solute carrier family 38, member 10 | protein-coding |
| chr11_120142632_120142858 | chr11 | 120142633 | 120142858 | + | -0.763497 | intron (NM_024249, intron 2 of 16) | intron (NM_024249, intron 2 of 16) | 8606 | NM_001164799 | 72055 | Mm.253403 | NM_024249 | ENSMUSG00000061306 | **Slc38a10** | 1810073N04Rik | solute carrier family 38, member 10 | protein-coding |

| | | | | | | | | | | | | | | | | | |
|---|---|---|---|---|---|---|---|---|---|---|---|---|---|---|---|---|---|
| chr2_65342025_65342483 | chr2 | 65342026 | 65342483 | + | -0.857154 | intron (NM_177074, intron 7 of 11) | intron (NM_177074, intron 7 of 11) | 21772 | NM_177074 | 320106 | Mm.227703 | NM_177074 | ENSMUSG00000061171 | **Slc38a11** | 9330158F14Rik | solute carrier family 38, member 11 | protein-coding |
| chr15_96778117_96778931 | chr15 | 96778118 | 96778931 | + | -0.607394 | Intergenic | Intergenic | -78826 | NM_175121 | 67760 | Mm.46754 | NM_175121 | ENSMUSG00000022462 | **Slc38a2** | 5033402L14Rik\|AI316867\|ATA2\|SAT2\|mKIAA1382 | solute carrier family 38, member 2 | protein-coding |
| chr15_96770783_96771132 | chr15 | 96770784 | 96771132 | + | -1.37011 | Intergenic | Intergenic | -71260 | NM_175121 | 67760 | Mm.46754 | NM_175121 | ENSMUSG00000022462 | **Slc38a2** | 5033402L14Rik\|AI316867\|ATA2\|SAT2\|mKIAA1382 | solute carrier family 38, member 2 | protein-coding |
| chr1_46874364_46874891 | chr1 | 46874365 | 46874891 | + | -0.665901 | Intergenic | L2a\|LINE\|L2 | -21119 | NM_172653 | 227059 | Mm.233889 | NM_172653 | ENSMUSG00000025986 | **Slc39a10** | 2900042E17Rik\|5430433I10\|mKIAA1265 | solute carrier family 39 (zinc transporter), member 10 | protein-coding |
| chr1_46894067_46895072 | chr1 | 46894068 | 46895072 | + | -1.01694 | Intergenic | ORR1D2\|LTR\|ERVL-MaLR | -41061 | NM_172653 | 227059 | Mm.233889 | NM_172653 | ENSMUSG00000025986 | **Slc39a10** | 2900042E17Rik\|5430433I10\|mKIAA1265 | solute carrier family 39 (zinc transporter), member 10 | protein-coding |
| chr1_46894120_46894802 | chr1 | 46894121 | 46894802 | + | -1.03077 | Intergenic | Intergenic | -40952 | NM_172653 | 227059 | Mm.233889 | NM_172653 | ENSMUSG00000025986 | **Slc39a10** | 2900042E17Rik\|5430433I10\|mKIAA1265 | solute carrier family 39 (zinc transporter), member 10 | protein-coding |
| chr1_46894365_46894985 | chr1 | 46894366 | 46894985 | + | -1.06704 | Intergenic | ORR1D2\|LTR\|ERVL-MaLR | -41166 | NM_172653 | 227059 | Mm.233889 | NM_172653 | ENSMUSG00000025986 | **Slc39a10** | 2900042E17Rik\|5430433I10\|mKIAA1265 | solute carrier family 39 (zinc transporter), member 10 | protein-coding |
| chr2_91080815_91081119 | chr2 | 91080816 | 91081119 | + | -0.88077 | Intergenic | Intergenic | -10652 | NM_001290765 | 68427 | Mm.192375 | NM_026721 | ENSMUSG00000002105 | **Slc39a13** | 1100001L14Rik\|AA387663\|ZIP-13\|ZIP13 | solute carrier family 39 (metal ion transporter), member 13 | protein-coding |
| chr14_70341008_70341872 | chr14 | 70341009 | 70341872 | + | -0.752603 | intron (NM_144808, intron 1 of 9) | intron (NM_144808, intron 1 of 9) | 6993 | NM_001135152 | 213053 | Mm.270647 | NM_144808 | ENSMUSG00000022094 | **Slc39a14** | FAD-123\|ZIP-14\|Zip14\|fad123 | solute carrier family 39 (zinc transporter), member 14 | protein-coding |
| chr18_24596775_24597089 | chr18 | 24596776 | 24597089 | + | -0.759557 | intron (NM_139143, intron 4 of 9) | intron (NM_139143, intron 4 of 9) | 6885 | NM_139143 | 106957 | Mm.21688 | NM_139143 | ENSMUSG00000024270 | **Slc39a6** | Ermelin | solute carrier family 39 (metal ion transporter), member 6 | protein-coding |
| chr17_85023277_85024125 | chr17 | 85023278 | 85024125 | + | -0.725451 | exon (NM_001159497, exon 6 of 6) | exon (NM_001159497, exon 6 of 6) | -4646 | NM_009205 | 20532 | Mm.227176 | NM_009205 | ENSMUSG00000024131 | **Slc3a1** | D2H\|NBAT\|NTAA\|RBAT | solute carrier family 3, member 1 | protein-coding |
| chr17_85023569_85024007 | chr17 | 85023570 | 85024007 | + | -0.751845 | 3' UTR (NM_001159498, exon 7 of 7) | 3' UTR (NM_001159498, exon 7 of 7) | -4559 | NM_009205 | 20532 | Mm.227176 | NM_009205 | ENSMUSG00000024131 | **Slc3a1** | D2H\|NBAT\|NTAA\|RBAT | solute carrier family 3, member 1 | protein-coding |
| chr10_83161729_83162318 | chr10 | 83161730 | 83162318 | + | -0.873582 | intron (NM_021439, intron 2 of 2) | intron (NM_021439, intron 2 of 2) | 175793 | NM_177388 | 338365 | Mm.67073 | NM_177388 | ENSMUSG00000034591 | **Slc41a2** | A230035L05Rik\|SLC41A1-L1 | solute carrier family 41, member 2 | protein-coding |
| chr10_83161722_83162152 | chr10 | 83161723 | 83162152 | + | -0.923517 | intron (NM_021439, intron 2 of 2) | intron (NM_021439, intron 2 of 2) | 175880 | NM_177388 | 338365 | Mm.67073 | NM_177388 | ENSMUSG00000034591 | **Slc41a2** | A230035L05Rik\|SLC41A1-L1 | solute carrier family 41, member 2 | protein-coding |
| chr10_83161732_83161954 | chr10 | 83161733 | 83161954 | + | -0.947973 | intron (NM_021439, intron 2 of 2) | intron (NM_021439, intron 2 of 2) | 175974 | NM_177388 | 338365 | Mm.67073 | NM_177388 | ENSMUSG00000034591 | **Slc41a2** | A230035L05Rik\|SLC41A1-L1 | solute carrier family 41, member 2 | protein-coding |
| chr6_90618131_90618653 | chr6 | 90618132 | 90618653 | + | -0.617703 | promoter-TSS (NM_001037493) | promoter-TSS (NM_001037493) | -854 | NM_001037493 | 71699 | Mm.272633 | NM_027868 | ENSMUSG00000030089 | **Slc41a3** | 1010001P06Rik\|AI480742\|SLC41A1-L2 | solute carrier family 41, member 3 | protein-coding |
| chr6_90618294_90618687 | chr6 | 90618295 | 90618687 | + | -0.67762 | promoter-TSS (NM_001037493) | promoter-TSS (NM_001037493) | -755 | NM_001037493 | 71699 | Mm.272633 | NM_027868 | ENSMUSG00000030089 | **Slc41a3** | 1010001P06Rik\|AI480742\|SLC41A1-L2 | solute carrier family 41, member 3 | protein-coding |
| chr11_75533070_75533304 | chr11 | 75533071 | 75533304 | + | -0.64243 | intron (NM_001199284, intron 2 of 13) | intron (NM_001199284, intron 2 of 13) | 1075 | NM_001199283 | 215113 | Mm.11186 | NM_173388 | ENSMUSG00000038178 | **Slc43a2** | 7630402D21Rik\|BC042513\|Lat4 | solute carrier family 43, member 2 | protein-coding |
| chr9_21332248_21332545 | chr9 | 21332249 | 21332545 | + | -0.76777 | intron (NM_001199186, intron 1 of 21) | RMER15\|LTR\|ERVL | -5431 | NM_152808 | 68682 | Mm.148425 | NM_152808 | ENSMUSG00000057193 | **Slc44a2** | 1110028E10Rik\|CTL2 | solute carrier family 44, member 2 | protein-coding |
| chr4_150518871_150519223 | chr4 | 150518872 | 150519223 | + | -0.625999 | intron (NM_001085492, intron 10 of 22) | intron (NM_001085492, intron 10 of 22) | 133127 | NM_173774 | 242773 | Mm.29280 | NM_173774 | | **Slc45a1** | C230078B22\|Dnb5 | solute carrier family 45, member 1 | protein-coding |
| chr4_150539834_150540217 | chr4 | 150539835 | 150540217 | + | -0.825559 | intron (NM_001085492, intron 11 of 22) | intron (NM_001085492, intron 11 of 22) | 112148 | NM_173774 | 242773 | Mm.29280 | NM_173774 | | **Slc45a1** | C230078B22\|Dnb5 | solute carrier family 45, member 1 | protein-coding |
| chr4_150556384_150556834 | chr4 | 150556385 | 150556834 | + | -0.837518 | intron (NM_001085492, intron 11 of 22) | intron (NM_001085492, intron 11 of 22) | 95565 | NM_173774 | 242773 | Mm.29280 | NM_173774 | | **Slc45a1** | C230078B22\|Dnb5 | solute carrier family 45, member 1 | protein-coding |
| chr4_150539954_150540287 | chr4 | 150539955 | 150540287 | + | -0.853238 | intron (NM_001085492, intron 11 of 22) | intron (NM_001085492, intron 11 of 22) | 112053 | NM_173774 | 242773 | Mm.29280 | NM_173774 | | **Slc45a1** | C230078B22\|Dnb5 | solute carrier family 45, member 1 | protein-coding |
| chr11_61309640_61310337 | chr11 | 61309641 | 61310337 | + | -0.701542 | exon (NM_001033542, exon 12 of 17) | exon (NM_001033542, exon 12 of 17) | 32871 | NM_001033542 | 380701 | Mm.340166 | NM_001033542 | ENSMUSG00000069855 | **Slc47a2** | 4933429E10Rik | solute carrier family 47, member 2 | protein-coding |
| chr5_89051622_89051975 | chr5 | 89051623 | 89051975 | + | -0.711477 | intron (NM_018760, intron 6 of 25) | intron (NM_018760, intron 6 of 25) | 164539 | NM_018760 | 54403 | Mm.41044 | NM_018760 | ENSMUSG00000060961 | **Slc4a4** | AI835705\|NBC\|NBC1 | solute carrier family 4 (anion exchanger), member 4 | protein-coding |
| chr5_89044584_89045212 | chr5 | 89044585 | 89045212 | + | -1.01194 | intron (NM_018760, intron 5 of 25) | B4A\|SINE\|B4 | 157638 | NM_018760 | 54403 | Mm.41044 | NM_018760 | ENSMUSG00000060961 | **Slc4a4** | AI835705\|NBC\|NBC1 | solute carrier family 4 (anion exchanger), member 4 | protein-coding |
| chr5_89090135_89090552 | chr5 | 89090136 | 89090552 | + | -1.07561 | intron (NM_018760, intron 7 of 25) | intron (NM_018760, intron 7 of 25) | 203084 | NM_018760 | 54403 | Mm.41044 | NM_018760 | ENSMUSG00000060961 | **Slc4a4** | AI835705\|NBC\|NBC1 | solute carrier family 4 (anion exchanger), member 4 | protein-coding |
| chr5_89044808_89045207 | chr5 | 89044809 | 89045207 | + | -1.09812 | intron (NM_018760, intron 5 of 25) | B4A\|SINE\|B4 | 157748 | NM_018760 | 54403 | Mm.41044 | NM_018760 | ENSMUSG00000060961 | **Slc4a4** | AI835705\|NBC\|NBC1 | solute carrier family 4 (anion exchanger), member 4 | protein-coding |
| chr5_89044900_89045168 | chr5 | 89044901 | 89045168 | + | -1.10797 | intron (NM_018760, intron 5 of 25) | B4A\|SINE\|B4 | 157774 | NM_018760 | 54403 | Mm.41044 | NM_018760 | ENSMUSG00000060961 | **Slc4a4** | AI835705\|NBC\|NBC1 | solute carrier family 4 (anion exchanger), member 4 | protein-coding |
| chr15_100747019_100747526 | chr15 | 100747020 | 100747526 | + | -0.71355 | Intergenic | C-rich\|Low_complexity\|Low_complexity | -14474 | NM_021530 | 59033 | Mm.209856 | NM_021530 | ENSMUSG00000023032 | **Slc4a8** | AW493845\|C230026C11\|NDCBE\|kNBC-3 | solute carrier family 4 (anion exchanger), member 8 | protein-coding |
| chr15_100747343_100747928 | chr15 | 100747344 | 100747928 | + | -0.876529 | Intergenic | L1M2\|LINE\|L1 | -14111 | NM_021530 | 59033 | Mm.209856 | NM_021530 | ENSMUSG00000023032 | **Slc4a8** | AW493845\|C230026C11\|NDCBE\|kNBC-3 | solute carrier family 4 (anion exchanger), member 8 | protein-coding |
| chr15_100750192_100750597 | chr15 | 100750193 | 100750597 | + | -0.891331 | Intergenic | Intergenic | -11352 | NM_021530 | 59033 | Mm.209856 | NM_021530 | ENSMUSG00000023032 | **Slc4a8** | AW493845\|C230026C11\|NDCBE\|kNBC-3 | solute carrier family 4 (anion exchanger), member 8 | protein-coding |
| chr15_100747572_100747956 | chr15 | 100747573 | 100747956 | + | -0.982803 | Intergenic | Intergenic | -13983 | NM_021530 | 59033 | Mm.209856 | NM_021530 | ENSMUSG00000023032 | **Slc4a8** | AW493845\|C230026C11\|NDCBE\|kNBC-3 | solute carrier family 4 (anion exchanger), member 8 | protein-coding |
| chr5_33142816_33143122 | chr5 | 33142817 | 33143122 | + | -0.642871 | intron (NM_019810, intron 5 of 14) | intron (NM_019810, intron 5 of 14) | 38750 | NM_019810 | 20537 | Mm.25237 | NM_019810 | ENSMUSG00000011034 | **Slc5a1** | Sglt1 | solute carrier family 5 (sodium/glucose cotransporter), member 1 | protein-coding |
| chr13_73725324_73725889 | chr13 | 73725325 | 73725889 | + | -0.841396 | Intergenic | RMER6A\|LTR\|ERVK | -24912 | NM_028878 | 74338 | Mm.271635 | NM_028878 | | **Slc6a19** | 4632401C08Rik\|B<0>AT1 | solute carrier family 6 (neurotransmitter transporter), member 19 | protein-coding |
| chr13_73702578_73703312 | chr13 | 73702579 | 73703312 | + | -1.002 | intron (NM_027780, intron 1 of 1) | intron (NM_027780, intron 1 of 1) | -2250 | NM_028878 | 74338 | Mm.271635 | NM_028878 | | **Slc6a19** | 4632401C08Rik\|B<0>AT1 | solute carrier family 6 (neurotransmitter transporter), member 19 | protein-coding |
| chr13_73702577_73703989 | chr13 | 73702578 | 73703989 | + | -1.16163 | intron (NM_027780, intron 1 of 1) | intron (NM_027780, intron 1 of 1) | -2588 | NM_028878 | 74338 | Mm.271635 | NM_028878 | | **Slc6a19** | 4632401C08Rik\|B<0>AT1 | solute carrier family 6 (neurotransmitter transporter), member 19 | protein-coding |
| chr13_73703383_73703976 | chr13 | 73703384 | 73703976 | + | -1.34467 | intron (NM_027780, intron 1 of 1) | RLTR8\|LTR\|ERVK | -2985 | NM_028878 | 74338 | Mm.271635 | NM_028878 | | **Slc6a19** | 4632401C08Rik\|B<0>AT1 | solute carrier family 6 (neurotransmitter transporter), member 19 | protein-coding |
| chr18_60991342_60991930 | chr18 | 60991343 | 60991930 | + | -0.846015 | Intergenic | ID_B1\|SINE\|B4 | 22563 | NM_201353 | 240332 | Mm.39846 | NM_201353 | ENSMUSG00000052026 | **Slc6a7** | AW455934\|PROT | solute carrier family 6 (neurotransmitter transporter, L-proline), member 7 | protein-coding |
| chr5_148428943_148429141 | chr5 | 148428944 | 148429141 | + | -0.646314 | Intergenic | Intergenic | -29138 | NM_007513 | 11987 | Mm.275489 | NM_007513 | ENSMUSG00000041313 | **Slc7a1** | 4831426K01Rik\|AI447493\|Atrc-1\|Atrc1\|Cat1\|Rec-1\|Rev-1\|mCAT-1 | solute carrier family 7 (cationic amino acid transporter, y+ system), member 1 | protein-coding |
| chr5_148388241_148388686 | chr5 | 148388242 | 148388686 | + | -0.723304 | intron (NM_007513, intron 1 of 12) | intron (NM_007513, intron 1 of 12) | 11440 | NM_007513 | 11987 | Mm.275489 | NM_007513 | ENSMUSG00000041313 | **Slc7a1** | 4831426K01Rik\|AI447493\|Atrc-1\|Atrc1\|Cat1\|Rec-1\|Rev-1\|mCAT-1 | solute carrier family 7 (cationic amino acid transporter, y+ system), member 1 | protein-coding |
| chr4_19844135_19844453 | chr4 | 19844136 | 19844453 | + | -0.881176 | Intergenic | Intergenic | 25567 | NM_028746 | 74087 | Mm.173833 | NM_028746 | ENSMUSG00000041052 | **Slc7a13** | 0610009O04Rik\|AGT-1\|AGT1\|XAT2 | solute carrier family 7, (cationic amino acid transporter, y+ system) member 13 | protein-coding |
| chrX_101147485_101147962 | chrX | 101147486 | 101147962 | + | -1.32074 | intron (NM_001110311, intron 4 of 5) | PB1\|SINE\|Alu | -62372 | NM_007515 | 11989 | Mm.5140 | NM_007515 | ENSMUSG00000031297 | **Slc7a3** | Atrc3\|CAT3\|SLC7A1\|SLC7A2 | solute carrier family 7 (cationic amino acid transporter, y+ system), member 3 | protein-coding |
| chr8_121906517_121906806 | chr8 | 121906518 | 121906806 | + | -0.620795 | intron (NM_011404, intron 1 of 9) | intron (NM_011404, intron 1 of 9) | 1024 | NM_011404 | 20539 | Mm.27943 | NM_011404 | ENSMUSG00000040010 | **Slc7a5** | 4F2LC\|D0H16S474E\|LAT1\|TA1 | solute carrier family 7 (cationic amino acid transporter, y+ system), member 5 | protein-coding |
| chr14_54748855_54749450 | chr14 | 54748856 | 54749450 | + | -1.07531 | intron (NM_016972, intron 3 of 10) | intron (NM_016972, intron 3 of 10) | 32733 | NM_016972 | 50934 | Mm.276831 | NM_016972 | ENSMUSG00000022180 | **Slc7a8** | AA408822\|LAT2 | solute carrier family 7 (cationic amino acid transporter, y+ system), member 8 | protein-coding |

| | | | | | | | | | | | | | | | | | |
|---|---|---|---|---|---|---|---|---|---|---|---|---|---|---|---|---|---|
| chr9_94691684_94692537 | chr9 | 94691685 | 94692537 | + | -0.636586 | intron (NM_177909, intron 2 of 15) | intron (NM_177909, intron 2 of 15) | 22219 | NM_177909 | 331004 | Mm.326125 | NM_177909 | ENSMUSG00000031129 | **Slc9a9** | 5730527A11Rik\|9930105B05\|AI854429\|Nhe9 | solute carrier family 9 (sodium/hydrogen exchanger), member 9 | protein-coding |
| chr9_94691679_94692195 | chr9 | 94691680 | 94692195 | + | -0.812921 | intron (NM_177909, intron 2 of 15) | intron (NM_177909, intron 2 of 15) | 22045 | NM_177909 | 331004 | Mm.326125 | NM_177909 | ENSMUSG00000031129 | **Slc9a9** | 5730527A11Rik\|9930105B05\|AI854429\|Nhe9 | solute carrier family 9 (sodium/hydrogen exchanger), member 9 | protein-coding |
| chr9_94653043_94653593 | chr9 | 94653044 | 94653593 | + | -0.858785 | Intergenic | Intergenic | -16574 | NM_177909 | 331004 | Mm.326125 | NM_177909 | ENSMUSG00000031129 | **Slc9a9** | 5730527A11Rik\|9930105B05\|AI854429\|Nhe9 | solute carrier family 9 (sodium/hydrogen exchanger), member 9 | protein-coding |
| chr7_74427175_74427491 | chr7 | 74427176 | 74427491 | + | -0.697216 | intron (NM_001038643, intron 2 of 10) | L2a\|LINE\|L2 | 127447 | NM_023908 | 108116 | Mm.425467 | NM_023908 | ENSMUSG00000025790 | **Slco3a1** | 5830414C08Rik\|Anr1\|MJAM\|OATP-D\|R75096\|Slc21a11 | solute carrier organic anion transporter family, member 3a1 | protein-coding |
| chr7_74690775_74691370 | chr7 | 74690776 | 74691370 | + | -0.783032 | Intergenic | Intergenic | -136293 | NM_001038643 | 108116 | Mm.425467 | NM_023908 | ENSMUSG00000025790 | **Slco3a1** | 5830414C08Rik\|Anr1\|MJAM\|OATP-D\|R75096\|Slc21a11 | solute carrier organic anion transporter family, member 3a1 | protein-coding |
| chr7_74690490_74691370 | chr7 | 74690491 | 74691370 | + | -0.876084 | Intergenic | Intergenic | -136150 | NM_001038643 | 108116 | Mm.425467 | NM_023908 | ENSMUSG00000025790 | **Slco3a1** | 5830414C08Rik\|Anr1\|MJAM\|OATP-D\|R75096\|Slc21a11 | solute carrier organic anion transporter family, member 3a1 | protein-coding |
| chr7_74690292_74691055 | chr7 | 74690293 | 74691055 | + | -0.934665 | Intergenic | Intergenic | -135894 | NM_001038643 | 108116 | Mm.425467 | NM_023908 | ENSMUSG00000025790 | **Slco3a1** | 5830414C08Rik\|Anr1\|MJAM\|OATP-D\|R75096\|Slc21a11 | solute carrier organic anion transporter family, member 3a1 | protein-coding |
| chr7_74663817_74664179 | chr7 | 74663818 | 74664179 | + | -1.18048 | Intergenic | Intergenic | -109218 | NM_001038643 | 108116 | Mm.425467 | NM_023908 | ENSMUSG00000025790 | **Slco3a1** | 5830414C08Rik\|Anr1\|MJAM\|OATP-D\|R75096\|Slc21a11 | solute carrier organic anion transporter family, member 3a1 | protein-coding |
| chr11_83067141_83067527 | chr11 | 83067142 | 83067527 | + | -0.756507 | intron (NM_011408, intron 1 of 1) | RLTR11A\|LTR\|ERVK | 2222 | NM_011408 | 20556 | Mm.278689 | NM_011408 | ENSMUSG00000072620 | **Slfn2** | Shlf2 | schlafen 2 | protein-coding |
| chr11_83079535_83080187 | chr11 | 83079536 | 83080187 | + | -1.19126 | Intergenic | RLTR8\|LTR\|ERVK | 14749 | NM_011408 | 20556 | Mm.278689 | NM_011408 | ENSMUSG00000072620 | **Slfn2** | Shlf2 | schlafen 2 | protein-coding |
| chr11_83079755_83080188 | chr11 | 83079756 | 83080188 | + | -1.26007 | Intergenic | RLTR8\|LTR\|ERVK | 14860 | NM_011408 | 20556 | Mm.278689 | NM_011408 | ENSMUSG00000072620 | **Slfn2** | Shlf2 | schlafen 2 | protein-coding |
| chr11_82952425_82953142 | chr11 | 82952426 | 82953142 | + | -0.681832 | intron (NM_183201, intron 1 of 5) | intron (NM_183201, intron 1 of 5) | 682 | NM_183201 | 327978 | Mm.42147 | NM_183201 | ENSMUSG00000054404 | **Slfn5** | AI956840 | schlafen 5 | protein-coding |
| chr11_82947320_82947897 | chr11 | 82947321 | 82947897 | + | -0.847071 | intron (NR_045932, intron 1 of 2) | intron (NR_045932, intron 1 of 2) | -4493 | NM_183201 | 327978 | Mm.42147 | NM_183201 | ENSMUSG00000054404 | **Slfn5** | AI956840 | schlafen 5 | protein-coding |
| chr11_35326322_35326999 | chr11 | 35326323 | 35326999 | + | -0.875281 | intron (NM_011412, intron 4 of 35) | intron (NM_011412, intron 4 of 35) | 205205 | NM_011412 | 20564 | Mm.478604 | NM_011412 | ENSMUSG00000056427 | **Slit3** | Slil2\|Slit1 | slit homolog 3 (Drosophila) | protein-coding |
| chr11_35326346_35326737 | chr11 | 35326347 | 35326737 | + | -0.939157 | intron (NM_011412, intron 4 of 35) | intron (NM_011412, intron 4 of 35) | 205086 | NM_011412 | 20564 | Mm.478604 | NM_011412 | ENSMUSG00000056427 | **Slit3** | Slil2\|Slit1 | slit homolog 3 (Drosophila) | protein-coding |
| chr2_174555241_174556440 | chr2 | 174555242 | 174556440 | + | -0.638908 | Intergenic | Intergenic | -82900 | NM_025531 | 66390 | Mm.182294 | NM_025531 | ENSMUSG00000016257 | **Slmo2** | 2310042G06Rik\|AA407013 | slowmo homolog 2 (Drosophila) | protein-coding |
| chr2_174555634_174556448 | chr2 | 174555635 | 174556448 | + | -0.688169 | Intergenic | Intergenic | -83100 | NM_025531 | 66390 | Mm.182294 | NM_025531 | ENSMUSG00000016257 | **Slmo2** | 2310042G06Rik\|AA407013 | slowmo homolog 2 (Drosophila) | protein-coding |
| chr2_174555690_174556184 | chr2 | 174555691 | 174556184 | + | -0.697838 | Intergenic | Intergenic | -82996 | NM_025531 | 66390 | Mm.182294 | NM_025531 | ENSMUSG00000016257 | **Slmo2** | 2310042G06Rik\|AA407013 | slowmo homolog 2 (Drosophila) | protein-coding |
| chr9_70524351_70524788 | chr9 | 70524352 | 70524788 | + | -0.889602 | Intergenic | Intergenic | -18208 | NM_026337 | 66660 | Mm.22379 | NM_025690 | ENSMUSG00000032212 | **Sltm** | 5730455C01Rik\|5730555F13Rik\|9130215G10Rik\|Met | SAFB-like, transcription modulator | protein-coding |
| chr9_63748962_63749687 | chr9 | 63748963 | 63749687 | + | -0.681076 | intron (NM_016769, intron 1 of 8) | intron (NM_016769, intron 1 of 8) | 8669 | NM_016769 | 17127 | Mm.7320 | NM_016769 | ENSMUSG00000032402 | **Smad3** | AU022421\|Madh3 | SMAD family member 3 | protein-coding |
| chr9_63756766_63756992 | chr9 | 63756767 | 63756992 | + | -0.687019 | intron (NM_016769, intron 1 of 8) | intron (NM_016769, intron 1 of 8) | 1115 | NM_016769 | 17127 | Mm.7320 | NM_016769 | ENSMUSG00000032402 | **Smad3** | AU022421\|Madh3 | SMAD family member 3 | protein-coding |
| chr9_63698005_63698852 | chr9 | 63698006 | 63698852 | + | -0.73206 | intron (NM_016769, intron 1 of 8) | intron (NM_016769, intron 1 of 8) | 59565 | NM_016769 | 17127 | Mm.7320 | NM_016769 | ENSMUSG00000032402 | **Smad3** | AU022421\|Madh3 | SMAD family member 3 | protein-coding |
| chr9_63749330_63749634 | chr9 | 63749331 | 63749634 | + | -0.807948 | intron (NM_016769, intron 1 of 8) | intron (NM_016769, intron 1 of 8) | 8512 | NM_016769 | 17127 | Mm.7320 | NM_016769 | ENSMUSG00000032402 | **Smad3** | AU022421\|Madh3 | SMAD family member 3 | protein-coding |
| chr9_63759060_63759572 | chr9 | 63759061 | 63759572 | + | -0.828746 | Intergenic | Intergenic | -1322 | NM_016769 | 17127 | Mm.7320 | NM_016769 | ENSMUSG00000032402 | **Smad3** | AU022421\|Madh3 | SMAD family member 3 | protein-coding |
| chr9_63740561_63740924 | chr9 | 63740562 | 63740924 | + | -0.828851 | intron (NM_016769, intron 1 of 8) | intron (NM_016769, intron 1 of 8) | 17251 | NM_016769 | 17127 | Mm.7320 | NM_016769 | ENSMUSG00000032402 | **Smad3** | AU022421\|Madh3 | SMAD family member 3 | protein-coding |
| chr9_63776810_63777129 | chr9 | 63776811 | 63777129 | + | -0.83808 | Intergenic | L1M5\|LINE\|L1 | -18976 | NM_016769 | 17127 | Mm.7320 | NM_016769 | ENSMUSG00000032402 | **Smad3** | AU022421\|Madh3 | SMAD family member 3 | protein-coding |
| chr9_63719794_63720161 | chr9 | 63719795 | 63720161 | + | -0.869939 | intron (NM_016769, intron 1 of 8) | intron (NM_016769, intron 1 of 8) | 38016 | NM_016769 | 17127 | Mm.7320 | NM_016769 | ENSMUSG00000032402 | **Smad3** | AU022421\|Madh3 | SMAD family member 3 | protein-coding |
| chr9_63720713_63721316 | chr9 | 63720714 | 63721316 | + | -0.916857 | intron (NM_016769, intron 1 of 8) | intron (NM_016769, intron 1 of 8) | 36979 | NM_016769 | 17127 | Mm.7320 | NM_016769 | ENSMUSG00000032402 | **Smad3** | AU022421\|Madh3 | SMAD family member 3 | protein-coding |
| chr18_73644783_73645199 | chr18 | 73644784 | 73645199 | + | -0.838005 | intron (NM_008540, intron 10 of 11) | Lx8\|LINE\|L1 | 58750 | NM_008540 | 17128 | Mm.100399 | NM_008540 | ENSMUSG00000024515 | **Smad4** | AW743858\|D18Wsu70e\|DPC4\|Madh4 | SMAD family member 4 | protein-coding |
| chr13_56703875_56704298 | chr13 | 56703876 | 56704298 | + | -0.899777 | intron (NM_001164042, intron 1 of 6) | intron (NM_001164042, intron 1 of 6) | 419 | NM_001164042 | 17129 | Mm.272920 | NM_008541 | ENSMUSG00000021540 | **Smad5** | 1110051M15Rik\|AI451355\|Dwf-C\|Madh5\|MusMLP | SMAD family member 5 | protein-coding |
| chr18_75387340_75387925 | chr18 | 75387341 | 75387925 | + | -0.678663 | intron (NM_001042660, intron 3 of 3) | intron (NM_001042660, intron 3 of 3) | 20268 | NM_001042660 | 17131 | Mm.34407 | NM_008543 | ENSMUSG00000025880 | **Smad7** | Madh7 | SMAD family member 7 | protein-coding |
| chr18_75387345_75387756 | chr18 | 75387346 | 75387756 | + | -0.760272 | intron (NM_001042660, intron 3 of 3) | (CAG)n\|Simple_repeat\|Simple_repeat | 20186 | NM_001042660 | 17131 | Mm.34407 | NM_008543 | ENSMUSG00000025880 | **Smad7** | Madh7 | SMAD family member 7 | protein-coding |
| chr18_75424415_75424877 | chr18 | 75424416 | 75424877 | + | -0.836055 | Intergenic | Intergenic | 57281 | NM_001042660 | 17131 | Mm.34407 | NM_008543 | ENSMUSG00000025880 | **Smad7** | Madh7 | SMAD family member 7 | protein-coding |
| chr18_75377123_75378283 | chr18 | 75377124 | 75378283 | + | -0.845472 | intron (NM_001042660, intron 3 of 3) | intron (NM_001042660, intron 3 of 3) | 10338 | NM_001042660 | 17131 | Mm.34407 | NM_008543 | ENSMUSG00000025880 | **Smad7** | Madh7 | SMAD family member 7 | protein-coding |
| chr18_75419032_75419860 | chr18 | 75419033 | 75419860 | + | -1.24906 | Intergenic | Intergenic | 52081 | NM_001042660 | 17131 | Mm.34407 | NM_008543 | ENSMUSG00000025880 | **Smad7** | Madh7 | SMAD family member 7 | protein-coding |
| chr18_75338816_75339153 | chr18 | 75338817 | 75339153 | + | -1.30672 | Intergenic | (CAAAA)n\|Simple_repeat\|Simple_repeat | -28380 | NM_001042660 | 17131 | Mm.34407 | NM_008543 | ENSMUSG00000025880 | **Smad7** | Madh7 | SMAD family member 7 | protein-coding |
| chr1_23882718_23883153 | chr1 | 23882719 | 23883153 | + | -0.788967 | intron (NM_001290683, intron 2 of 9) | intron (NM_001290683, intron 2 of 9) | 26800 | NR_110972 | 98366 | Mm.329963 | NM_028534 | ENSMUSG00000026155 | **Smap1** | 1700056O10Rik\|4921514B13Rik\|4921525H11Rik\|AI462175 | small ArfGAP 1 | protein-coding |
| chr19_26646059_26646639 | chr19 | 26646060 | 26646639 | + | -0.637064 | intron (NM_011416, intron 5 of 32) | intron (NM_011416, intron 5 of 32) | 41189 | NM_011416 | 67155 | Mm.491146 | NM_011416 | ENSMUSG00000024921 | **Smarca2** | 2610209L14Rik\|SNF2alpha\|Snf2l2\|brahma\|brm | SWI/SNF related, matrix associated, actin dependent regulator of chromatin, subfamily a, member 2 | protein-coding |
| chr19_26607107_26607573 | chr19 | 26607108 | 26607573 | + | -0.648751 | intron (NM_011416, intron 1 of 32) | CpG | 2180 | NM_011416 | 67155 | Mm.491146 | NM_011416 | ENSMUSG00000024921 | **Smarca2** | 2610209L14Rik\|SNF2alpha\|Snf2l2\|brahma\|brm | SWI/SNF related, matrix associated, actin dependent regulator of chromatin, subfamily a, member 2 | protein-coding |
| chr19_26707021_26707337 | chr19 | 26707022 | 26707337 | + | -0.660979 | intron (NM_011416, intron 24 of 32) | intron (NM_011416, intron 24 of 32) | -41224 | NM_026003 | 67155 | Mm.491146 | NM_011416 | ENSMUSG00000024921 | **Smarca2** | 2610209L14Rik\|SNF2alpha\|Snf2l2\|brahma\|brm | SWI/SNF related, matrix associated, actin dependent regulator of chromatin, subfamily a, member 2 | protein-coding |
| chr19_26540186_26540569 | chr19 | 26540187 | 26540569 | + | -0.735128 | Intergenic | Intergenic | -64782 | NM_011416 | 67155 | Mm.491146 | NM_011416 | ENSMUSG00000024921 | **Smarca2** | 2610209L14Rik\|SNF2alpha\|Snf2l2\|brahma\|brm | SWI/SNF related, matrix associated, actin dependent regulator of chromatin, subfamily a, member 2 | protein-coding |
| chr11_106269123_106269389 | chr11 | 106269124 | 106269389 | + | -0.812453 | intron (NM_001130187, intron 1 of 12) | intron (NM_001130187, intron 1 of 12) | -1624 | NM_031878 | 83796 | Mm.21772 | NM_031878 | ENSMUSG00000078619 | **Smarcd2** | AW322457\|Baf60b | SWI/SNF related, matrix associated, actin dependent regulator of chromatin, subfamily d, member 2 | protein-coding |
| chr11_106269084_106269466 | chr11 | 106269085 | 106269466 | + | -0.834157 | intron (NM_001130187, intron 1 of 12) | intron (NM_001130187, intron 1 of 12) | -1643 | NM_031878 | 83796 | Mm.21772 | NM_031878 | ENSMUSG00000078619 | **Smarcd2** | AW322457\|Baf60b | SWI/SNF related, matrix associated, actin dependent regulator of chromatin, subfamily d, member 2 | protein-coding |
| chr11_99203585_99204395 | chr11 | 99203586 | 99204395 | + | -0.817034 | Intergenic | Intergenic | 27027 | NM_020618 | 57376 | Mm.379086 | NM_020618 | ENSMUSG00000037935 | **Smarce1** | 2810417B20Rik\|5830412H02Rik\|9030408N19Rik\|Baf27 | SWI/SNF related, matrix associated, actin dependent regulator of chromatin, subfamily e, member 1 | protein-coding |
| chr11_99203842_99204380 | chr11 | 99203843 | 99204380 | + | -0.879962 | Intergenic | Intergenic | 26906 | NM_020618 | 57376 | Mm.379086 | NM_020618 | ENSMUSG00000037935 | **Smarce1** | 2810417B20Rik\|5830412H02Rik\|9030408N19Rik\|Baf27 | SWI/SNF related, matrix associated, actin dependent regulator of chromatin, subfamily e, member 1 | protein-coding |
| chr19_23322848_23323613 | chr19 | 23322849 | 23323613 | + | -1.2942 | intron (NM_174857, intron 11 of 13) | intron (NM_174857, intron 11 of 13) | -49334 | NM_001252684 | 226026 | Mm.23267 | NM_153808 | ENSMUSG00000024943 | **Smc5** | MLZ-453\|SMC-5\|Smc5l1\|mKIAA0594\|mSMC5 | structural maintenance of chromosomes 5 | protein-coding |
| chr19_23323058_23323575 | chr19 | 23323059 | 23323575 | + | -1.31972 | intron (NM_174857, intron 11 of 13) | intron (NM_174857, intron 11 of 13) | -49420 | NM_001252684 | 226026 | Mm.23267 | NM_153808 | ENSMUSG00000024943 | **Smc5** | MLZ-453\|SMC-5\|Smc5l1\|mKIAA0594\|mSMC5 | structural maintenance of chromosomes 5 | protein-coding |
| chr12_11279958_11280235 | chr12 | 11279959 | 11280235 | + | -0.96903 | intron (NM_025695, intron 5 of 26) | intron (NM_025695, intron 5 of 26) | 14211 | NM_025695 | 67241 | Mm.173953 | NM_025695 | ENSMUSG00000020608 | **Smc6** | 2810489L22Rik\|3830418C19Rik\|AA990493\|AU018782\|AW742439\|SMC-6\|Smc6l1\|mKIAA4103 | structural maintenance of chromosomes 6 | protein-coding |

| | | | | | | | | | | | | | | | | | |
|---|---|---|---|---|---|---|---|---|---|---|---|---|---|---|---|---|---|
| chr17_71490694_71491432 | chr17 | 71490695 | 71491432 | + | -0.766668 | Intergenic | Intergenic | -15720 | NM_028887 | 74355 | Mm.194450 | NM_028887 | ENSMUSG00000024054 | **Smchd1** | 4931400A14Rik\|AW554188\|MommeD1\|mKIAA0650 | SMC hinge domain containing 1 | protein-coding |
| chr17_71490963_71491356 | chr17 | 71490964 | 71491356 | + | -1.02527 | Intergenic | (CCCA)n\|Simple_repeat\|Simple_repeat | -15817 | NM_028887 | 74355 | Mm.194450 | NM_028887 | ENSMUSG00000024054 | **Smchd1** | 4931400A14Rik\|AW554188\|MommeD1\|mKIAA0650 | SMC hinge domain containing 1 | protein-coding |
| chr9_15550281_15551096 | chr9 | 15550282 | 15551096 | + | -1.2884 | Intergenic | Intergenic | 45194 | NM_133214 | 170748 | Mm.86572 | NM_133214 | ENSMUSG00000058173 | **Smco4** | Fn5 | single-pass membrane protein with coiled-coil domains 4 | protein-coding |
| chr11_74959824_74960568 | chr11 | 74959825 | 74960568 | + | -0.623063 | intron (NM_001002764, intron 8 of 18) | intron (NM_001002764, intron 8 of 18) | 34324 | NM_001002764 | 103677 | Mm.288460 | NM_001002764 | ENSMUSG00000038290 | **Smg6** | AI317223\|AU041178 | Smg-6 homolog, nonsense mediated mRNA decay factor (C. elegans) | protein-coding |
| chr7_24413556_24414037 | chr7 | 24413557 | 24414037 | + | -1.07759 | intron (NM_028047, intron 6 of 13) | intron (NM_028047, intron 6 of 13) | 14169 | NM_028047 | 71997 | Mm.150977 | NM_028047 | ENSMUSG00000002210 | **Smg9** | 1500002O20Rik\|AI429693\|N28092 | smg-9 homolog, nonsense mediated mRNA decay factor (C. elegans) | protein-coding |
| chr5_65500034_65500973 | chr5 | 65500035 | 65500973 | + | -0.822209 | Intergenic | RMER19B2\|LTR\|ERVK | -7669 | NM_133697 | 68552 | Mm.10709 | NM_133697 | ENSMUSG00000037822 | **Smim14** | 1110003E01Rik\|1700127H04Rik\|5430439C17Rik\|MAd4 | small integral membrane protein 14 | protein-coding |
| chr5_65500031_65500627 | chr5 | 65500032 | 65500627 | + | -1.00487 | Intergenic | RMER19B2\|LTR\|ERVK | -7494 | NM_133697 | 68552 | Mm.10709 | NM_133697 | ENSMUSG00000037822 | **Smim14** | 1110003E01Rik\|1700127H04Rik\|5430439C17Rik\|MAd4 | small integral membrane protein 14 | protein-coding |
| chr11_32768626_32769476 | chr11 | 32768627 | 32769476 | + | -0.716704 | Intergenic | ORR1E\|LTR\|ERVL-MaLR | 55543 | NM_027050 | 69351 | Mm.65488 | NM_027050 | ENSMUSG00000020270 | **Smim23** | 1700008A04Rik | small integral membrane protein 23 | protein-coding |
| chr2_131509652_131510440 | chr2 | 131509653 | 131510440 | + | -0.779987 | intron (NM_001177840, intron 1 of 3) | intron (NM_001177840, intron 1 of 3) | 18184 | NM_001177838 | 228608 | Mm.136586 | NM_145533 | ENSMUSG00000027333 | **Smox** | B130066H01Rik\|PAO\|PAOh1\|SMO | spermine oxidase | protein-coding |
| chr2_131480544_131481067 | chr2 | 131480545 | 131481067 | + | -1.28444 | intron (NR_037281, intron 1 of 2) | intron (NR_037281, intron 1 of 2) | -11056 | NM_001177834 | 228608 | Mm.136586 | NM_145533 | ENSMUSG00000027333 | **Smox** | B130066H01Rik\|PAO\|PAOh1\|SMO | spermine oxidase | protein-coding |
| chr2_131480738_131481014 | chr2 | 131480739 | 131481014 | + | -1.28992 | intron (NR_037281, intron 1 of 2) | intron (NR_037281, intron 1 of 2) | -10986 | NM_001177834 | 228608 | Mm.136586 | NM_145533 | ENSMUSG00000027333 | **Smox** | B130066H01Rik\|PAO\|PAOh1\|SMO | spermine oxidase | protein-coding |
| chr15_76299328_76300197 | chr15 | 76299329 | 76300197 | + | -1.0321 | intron (NM_153122, intron 19 of 26) | B3A\|SINE\|B2 | 5329 | NM_001195537 | 100503915 | Mm.393455 | NM_001195537 | ENSMUSG00000071724 | **Smpd5** | ENSMUSG00000071724\|Gm10345\|MA-nSMase | sphingomyelin phosphodiesterase 5 | protein-coding |
| chr4_40798620_40799142 | chr4 | 40798621 | 40799142 | + | -0.674443 | Intergenic | Intergenic | -40996 | NM_021535 | 74255 | Mm.289929 | NM_021535 | | **Smu1** | 2600001O03Rik\|2610203K23Rik\|AB044414\|AI845086\|AW556129\|Bwd | smu-1 suppressor of mec-8 and unc-52 homolog (C. elegans) | protein-coding |
| chr4_40781147_40781565 | chr4 | 40781148 | 40781565 | + | -0.740252 | Intergenic | Intergenic | -23471 | NM_021535 | 74255 | Mm.289929 | NM_021535 | | **Smu1** | 2600001O03Rik\|2610203K23Rik\|AB044414\|AI845086\|AW556129\|Bwd | smu-1 suppressor of mec-8 and unc-52 homolog (C. elegans) | protein-coding |
| chr5_144936992_144937730 | chr5 | 144936993 | 144937730 | + | -0.740774 | intron (NM_029438, intron 1 of 17) | intron (NM_029438, intron 1 of 17) | 28469 | NM_029438 | 75788 | Mm.27735 | NM_029438 | ENSMUSG00000038780 | **Smurf1** | 4930431E10Rik\|mKIAA1625 | SMAD specific E3 ubiquitin protein ligase 1 | protein-coding |
| chr5_144937259_144937976 | chr5 | 144937260 | 144937976 | + | -0.796796 | intron (NM_029438, intron 1 of 17) | intron (NM_029438, intron 1 of 17) | 28212 | NM_029438 | 75788 | Mm.27735 | NM_029438 | ENSMUSG00000038780 | **Smurf1** | 4930431E10Rik\|mKIAA1625 | SMAD specific E3 ubiquitin protein ligase 1 | protein-coding |
| chr5_144937247_144937678 | chr5 | 144937248 | 144937678 | + | -0.801493 | intron (NM_029438, intron 1 of 17) | MTE2b\|LTR\|ERVL-MaLR | 28367 | NM_029438 | 75788 | Mm.27735 | NM_029438 | ENSMUSG00000038780 | **Smurf1** | 4930431E10Rik\|mKIAA1625 | SMAD specific E3 ubiquitin protein ligase 1 | protein-coding |
| chr11_106937221_106937727 | chr11 | 106937222 | 106937727 | + | -0.768219 | Intergenic | ID_B1\|SINE\|B4 | -16759 | NM_025481 | 66313 | Mm.340955 | NM_025481 | ENSMUSG00000018363 | **Smurf2** | 2810411E22Rik\|AI558114\|AI649275 | SMAD specific E3 ubiquitin protein ligase 2 | protein-coding |
| chr11_106937385_106937781 | chr11 | 106937386 | 106937781 | + | -0.776088 | Intergenic | Intergenic | -16868 | NM_025481 | 66313 | Mm.340955 | NM_025481 | ENSMUSG00000018363 | **Smurf2** | 2810411E22Rik\|AI558114\|AI649275 | SMAD specific E3 ubiquitin protein ligase 2 | protein-coding |
| chr1_179240580_179241219 | chr1 | 179240581 | 179241219 | + | -0.695907 | intron (NM_027188, intron 5 of 11) | intron (NM_027188, intron 5 of 11) | 277103 | NM_027188 | 69726 | Mm.222338 | NM_027188 | | **Smyd3** | 2410008A19Rik\|Zmynd1 | SET and MYND domain containing 3 | protein-coding |
| chr8_122463798_122464687 | chr8 | 122463799 | 122464687 | + | -0.872635 | Intergenic | Intergenic | -3551 | NM_013914 | 30927 | Mm.103673 | NM_013914 | ENSMUSG00000006587 | **Snai3** | AI643946\|Smuc\|Zfp293 | snail family zinc finger 3 | protein-coding |
| chr11_59425648_59426353 | chr11 | 59425649 | 59426353 | + | -1.14269 | intron (NM_144521, intron 3 of 4) | intron (NM_144521, intron 3 of 4) | 23955 | NM_144521 | 67826 | Mm.26680 | NM_144521 | ENSMUSG00000009894 | **Snap47** | 1110031B06Rik\|SNAP-47 | synaptosomal-associated protein, 47 | protein-coding |
| chr6_28588557_28589167 | chr6 | 28588558 | 28589167 | + | -0.868747 | intron (NM_019776, intron 10 of 23) | intron (NM_019776, intron 10 of 23) | 108514 | NM_019776 | 56463 | Mm.439987 | NM_019776 | ENSMUSG00000001424 | **Snd1** | AL033314\|Tudor-SN | staphylococcal nuclease and tudor domain containing 1 | protein-coding |
| chr16_11073056_11073747 | chr16 | 11073057 | 11073747 | + | -0.768278 | 3' UTR (NM_009223, exon 2 of 2) | 3' UTR (NM_009223, exon 2 of 2) | 7104 | NM_009223 | 20621 | Mm.325800 | NM_009223 | ENSMUSG00000037972 | **Snn** | 2810407J07Rik\|AI848521\|AW547286 | stannin | protein-coding |
| chr16_11073527_11073745 | chr16 | 11073528 | 11073745 | + | -0.87248 | 3' UTR (NM_009223, exon 2 of 2) | 3' UTR (NM_009223, exon 2 of 2) | 7338 | NM_009223 | 20621 | Mm.325800 | NM_009223 | ENSMUSG00000037972 | **Snn** | 2810407J07Rik\|AI848521\|AW547286 | stannin | protein-coding |
| chr4_132309300_132309563 | chr4 | 132309301 | 132309563 | + | -0.655594 | promoter-TSS (NR_029412) | promoter-TSS (NR_029412) | -33 | NR_029412 | 100310813 | Mm.482271 | NR_029412 | ENSMUSG00000065097 | **Snora16a** | - | small nucleolar RNA, H/ACA box 16A | snoRNA |
| chr5_74132669_74133281 | chr5 | 74132670 | 74133281 | + | -0.745243 | Intergenic | Intergenic | 39445 | NR_031758 | 100313943 | Mm.46547 | NR_031758 | ENSMUSG00000093355 | **Snora26** | MBI-6 | small nucleolar RNA, H/ACA box 26 | snoRNA |
| chr5_74132669_74133038 | chr5 | 74132670 | 74133038 | + | -0.784821 | Intergenic | Intergenic | 39324 | NR_031758 | 100313943 | Mm.46547 | NR_031758 | ENSMUSG00000093355 | **Snora26** | MBI-6 | small nucleolar RNA, H/ACA box 26 | snoRNA |
| chr2_32962791_32963184 | chr2 | 32962792 | 32963184 | + | -1.10961 | promoter-TSS (NR_002898) | promoter-TSS (NR_002898) | -313 | NR_002898 | 104367 | Mm.250030 | NR_002898 | | **Snora65** | MBII-351\|Rnu65 | small nucleolar RNA, H/ACA box 65 | snoRNA |
| chr1_86344367_86344676 | chr1 | 86344368 | 86344676 | + | -0.719583 | TTS (NM_010880) | TTS (NM_010880) | 6763 | NR_028478 | 100303740 | Mm.154378 | NR_028478 | | **Snora75** | MBI-55 | small nucleolar RNA, H/ACA box 75 | snoRNA |
| chr1_86344423_86344641 | chr1 | 86344424 | 86344641 | + | -0.755481 | TTS (NM_010880) | TTS (NM_010880) | 6753 | NR_028478 | 100303740 | Mm.154378 | NR_028478 | | **Snora75** | MBI-55 | small nucleolar RNA, H/ACA box 75 | snoRNA |
| chr11_106535914_106536654 | chr11 | 106535915 | 106536654 | + | -1.11422 | intron (NM_198292, intron 5 of 11) | intron (NM_198292, intron 5 of 11) | 35293 | NR_030703 | 100216537 | | NR_030703 | ENSMUSG00000065126 | **Snord104** | - | small nucleolar RNA, C/D box 104 | snoRNA |
| chr2_167081378_167082163 | chr2 | 167081379 | 167082163 | + | -0.647578 | Intergenic | Intergenic | 16478 | NR_028540 | 100217443 | | NR_028540 | | **Snord12** | MBII-99 | small nucleolar RNA, C/D box 12 | snoRNA |
| chr2_167081538_167082019 | chr2 | 167081539 | 167082019 | + | -0.671764 | Intergenic | Intergenic | 16486 | NR_028540 | 100217443 | | NR_028540 | | **Snord12** | MBII-99 | small nucleolar RNA, C/D box 12 | snoRNA |
| chr2_167080827_167082036 | chr2 | 167080828 | 167082036 | + | -0.792748 | Intergenic | Intergenic | 16139 | NR_028540 | 100217443 | | NR_028540 | | **Snord12** | MBII-99 | small nucleolar RNA, C/D box 12 | snoRNA |
| chr2_144230998_144231525 | chr2 | 144230999 | 144231525 | + | -1.07432 | Intergenic | MTEa\|LTR\|ERVL-MaLR | 34940 | NR_030762 | 100313519 | Mm.453538 | NR_030762 | | **Snord17** | MBI-43 | small nucleolar RNA, C/D box 17 | snoRNA |
| chr2_144228498_144229189 | chr2 | 144228499 | 144229189 | + | -1.29419 | Intergenic | Intergenic | 37358 | NR_030762 | 100313519 | Mm.453538 | NR_030762 | | **Snord17** | MBI-43 | small nucleolar RNA, C/D box 17 | snoRNA |
| chr2_144228516_144228933 | chr2 | 144228517 | 144228933 | + | -1.31314 | Intergenic | (GGGGA)n\|Simple_repeat\|Simple_repeat | 37477 | NR_030762 | 100313519 | Mm.453538 | NR_030762 | | **Snord17** | MBI-43 | small nucleolar RNA, C/D box 17 | snoRNA |
| chr9_122193718_122194122 | chr9 | 122193719 | 122194122 | + | -0.609402 | intron (NM_133979, intron 12 of 12) | intron (NM_133979, intron 12 of 12) | 76654 | NM_133741 | 20623 | Mm.257989 | NM_133741 | ENSMUSG00000038145 | **Snrk** | 2010012F07Rik\|AI448042\|AW547029\|E030034B15\|R74830\|mKIAA0096 | SNF related kinase | protein-coding |
| chr9_122128620_122129212 | chr9 | 122128621 | 122129212 | + | -0.632575 | intron (NM_001164572, intron 1 of 5) | intron (NM_001164572, intron 1 of 5) | 11650 | NM_133741 | 20623 | Mm.257989 | NM_133741 | ENSMUSG00000038145 | **Snrk** | 2010012F07Rik\|AI448042\|AW547029\|E030034B15\|R74830\|mKIAA0096 | SNF related kinase | protein-coding |
| chr9_122188614_122189193 | chr9 | 122188615 | 122189193 | + | -0.695866 | intron (NM_133979, intron 12 of 12) | intron (NM_133979, intron 12 of 12) | 71638 | NM_133741 | 20623 | Mm.257989 | NM_133741 | ENSMUSG00000038145 | **Snrk** | 2010012F07Rik\|AI448042\|AW547029\|E030034B15\|R74830\|mKIAA0096 | SNF related kinase | protein-coding |
| chr5_124496622_124497443 | chr5 | 124496623 | 124497443 | + | -0.760551 | intron (NM_021430, intron 6 of 6) | intron (NM_021430, intron 6 of 6) | 13878 | NM_029532 | 76167 | Mm.156035 | NM_029532 | ENSMUSG00000029402 | **Snrnp35** | 6330548G22Rik | small nuclear ribonucleoprotein 35 (U11/U12) | protein-coding |
| chr5_124496804_124497201 | chr5 | 124496805 | 124497201 | + | -0.844477 | intron (NM_021430, intron 6 of 6) | intron (NM_021430, intron 6 of 6) | 13848 | NM_029532 | 76167 | Mm.156035 | NM_029532 | ENSMUSG00000029402 | **Snrnp35** | 6330548G22Rik | small nuclear ribonucleoprotein 35 (U11/U12) | protein-coding |
| chr10_75535602_75536609 | chr10 | 75535603 | 75536609 | + | -0.832729 | TTS (NM_026095) | TTS (NM_026095) | 18064 | NM_026095 | 67332 | Mm.45151 | NM_026095 | ENSMUSG00000020180 | **Snrpd3** | 1700043E15Rik\|2310009E13Rik\|AW046420\|SMD3 | small nuclear ribonucleoprotein D3 | protein-coding |
| chr10_75531618_75532069 | chr10 | 75531619 | 75532069 | + | -1.12627 | intron (NM_026095, intron 2 of 3) | intron (NM_026095, intron 2 of 3) | 13802 | NM_026095 | 67332 | Mm.45151 | NM_026095 | ENSMUSG00000020180 | **Snrpd3** | 1700043E15Rik\|2310009E13Rik\|AW046420\|SMD3 | small nuclear ribonucleoprotein D3 | protein-coding |
| chr10_75535983_75536625 | chr10 | 75535984 | 75536625 | + | -1.18039 | TTS (NM_026095) | TTS (NM_026095) | 18262 | NM_026095 | 67332 | Mm.45151 | NM_026095 | ENSMUSG00000020180 | **Snrpd3** | 1700043E15Rik\|2310009E13Rik\|AW046420\|SMD3 | small nuclear ribonucleoprotein D3 | protein-coding |
| chr10_75531735_75532103 | chr10 | 75531736 | 75532103 | + | -1.18478 | intron (NM_026095, intron 2 of 3) | intron (NM_026095, intron 2 of 3) | 13877 | NM_026095 | 67332 | Mm.45151 | NM_026095 | ENSMUSG00000020180 | **Snrpd3** | 1700043E15Rik\|2310009E13Rik\|AW046420\|SMD3 | small nuclear ribonucleoprotein D3 | protein-coding |
| chr1_133541798_133542695 | chr1 | 133541799 | 133542695 | + | -0.775242 | Intergenic | Intergenic | 68033 | NM_009227 | 20643 | Mm.249110 | NM_009227 | ENSMUSG00000090553 | **Snrpe** | AL022645\|C76690\|SME | small nuclear ribonucleoprotein E | protein-coding |
| chr1_133541955_133542398 | chr1 | 133541956 | 133542398 | + | -0.858445 | Intergenic | Intergenic | 68103 | NM_009227 | 20643 | Mm.249110 | NM_009227 | ENSMUSG00000090553 | **Snrpe** | AL022645\|C76690\|SME | small nuclear ribonucleoprotein E | protein-coding |
| chr1_133544860_133545403 | chr1 | 133544861 | 133545403 | + | -1.09664 | Intergenic | Intergenic | 65148 | NM_009227 | 20643 | Mm.249110 | NM_009227 | ENSMUSG00000090553 | **Snrpe** | AL022645\|C76690\|SME | small nuclear ribonucleoprotein E | protein-coding |
| chr15_55828552_55829431 | chr15 | 55828553 | 55829431 | + | -0.612755 | intron (NM_016667, intron 1 of 6) | intron (NM_016667, intron 1 of 6) | 77957 | NM_016667 | 20649 | Mm.309425 | NM_016667 | ENSMUSG00000060429 | **Sntb1** | - | syntrophin, basic 1 | protein-coding |
| chr15_55848316_55848632 | chr15 | 55848317 | 55848632 | + | -0.698645 | intron (NM_016667, intron 1 of 6) | intron (NM_016667, intron 1 of 6) | 58475 | NM_016667 | 20649 | Mm.309425 | NM_016667 | ENSMUSG00000060429 | **Sntb1** | - | syntrophin, basic 1 | protein-coding |
| chr15_55828981_55829383 | chr15 | 55828982 | 55829383 | + | -0.705524 | intron (NM_016667, intron 1 of 6) | intron (NM_016667, intron 1 of 6) | 77767 | NM_016667 | 20649 | Mm.309425 | NM_016667 | ENSMUSG00000060429 | **Sntb1** | - | syntrophin, basic 1 | protein-coding |

| | | | | | | | | | | | | | | | | | |
|---|---|---|---|---|---|---|---|---|---|---|---|---|---|---|---|---|---|
| chr15_55854578_55854990 | chr15 | 55854579 | 55854990 | + | -0.805363 | intron (NM_016667, intron 1 of 6) | intron (NM_016667, intron 1 of 6) | 52165 | NM_016667 | 20649 | Mm.309425 | NM_016667 | ENSMUSG00000060429 | **Sntb1** | - | syntrophin, basic 1 | protein-coding |
| chr15_55844596_55845142 | chr15 | 55844597 | 55845142 | + | -0.913124 | intron (NM_016667, intron 1 of 6) | intron (NM_016667, intron 1 of 6) | 62080 | NM_016667 | 20649 | Mm.309425 | NM_016667 | ENSMUSG00000060429 | **Sntb1** | - | syntrophin, basic 1 | protein-coding |
| chr15_55764274_55764514 | chr15 | 55764275 | 55764514 | + | -0.913549 | intron (NM_016667, intron 2 of 6) | intron (NM_016667, intron 2 of 6) | 142555 | NM_016667 | 20649 | Mm.309425 | NM_016667 | ENSMUSG00000060429 | **Sntb1** | - | syntrophin, basic 1 | protein-coding |
| chr8_106973606_106973900 | chr8 | 106973607 | 106973900 | + | -0.767855 | intron (NM_009229, intron 1 of 6) | ID_B1\|SINE\|B4 | 38003 | NM_009229 | 20650 | Mm.30228 | NM_009229 | ENSMUSG00000041308 | **Sntb2** | Snt2 | syntrophin, basic 2 | protein-coding |
| chr11_96666679_96667392 | chr11 | 96666680 | 96667392 | + | -0.804967 | intron (NM_001177898, intron 4 of 10) | (GTCTG)n\|Simple_repeat\|Simple_repeat | 110519 | NM_028965 | 74479 | Mm.270781 | NM_028965 | ENSMUSG00000020876 | **Snx11** | 4933439F10Rik\|A930041K09Rik\|AI117694 | sorting nexin 11 | protein-coding |
| chr12_35123504_35123895 | chr12 | 35123505 | 35123895 | + | -0.777674 | intron (NM_001014973, intron 19 of 25) | intron (NM_001014973, intron 19 of 25) | 76511 | NM_001014973 | 217463 | Mm.213991 | NM_001014973 | ENSMUSG00000020590 | **Snx13** | Rgs-px1\|mKIAA0713 | sorting nexin 13 | protein-coding |
| chr13_113574184_113574790 | chr13 | 113574185 | 113574790 | + | -0.665711 | Intergenic | Intergenic | 44077 | NM_130796 | 170625 | Mm.33721 | NM_130796 | ENSMUSG00000042364 | **Snx18** | Snag1 | sorting nexin 18 | protein-coding |
| chr13_113579926_113580700 | chr13 | 113579927 | 113580700 | + | -0.751752 | Intergenic | Intergenic | 38251 | NM_130796 | 170625 | Mm.33721 | NM_130796 | ENSMUSG00000042364 | **Snx18** | Snag1 | sorting nexin 18 | protein-coding |
| chr18_53173426_53173631 | chr18 | 53173427 | 53173631 | + | -0.606504 | Intergenic | Intergenic | -2836 | NM_026386 | 67804 | Mm.252171 | NM_026386 | ENSMUSG00000034484 | **Snx2** | 0610030A03Rik | sorting nexin 2 | protein-coding |
| chr16_11519202_11519504 | chr16 | 11519203 | 11519504 | + | -0.724283 | intron (NM_001290148, intron 7 of 13) | MTD\|LTR\|ERVL-MaLR | 113705 | NM_001290148 | 74478 | Mm.325831 | NM_028964 | ENSMUSG00000071669 | **Snx29** | 4933437K13Rik\|ENSMUSG00000079738\|Gm11170\|Gm1737\|Gm930\|Rundc2a | sorting nexin 29 | protein-coding |
| chr16_11519130_11519667 | chr16 | 11519131 | 11519667 | + | -0.725451 | intron (NM_001290148, intron 7 of 13) | MTD\|LTR\|ERVL-MaLR | 113751 | NM_001290148 | 74478 | Mm.325831 | NM_028964 | ENSMUSG00000071669 | **Snx29** | 4933437K13Rik\|ENSMUSG00000079738\|Gm11170\|Gm1737\|Gm930\|Rundc2a | sorting nexin 29 | protein-coding |
| chr16_11526147_11526849 | chr16 | 11526148 | 11526849 | + | -0.749781 | intron (NM_001290148, intron 7 of 13) | intron (NM_001290148, intron 7 of 13) | 120850 | NM_001290148 | 74478 | Mm.325831 | NM_028964 | ENSMUSG00000071669 | **Snx29** | 4933437K13Rik\|ENSMUSG00000079738\|Gm11170\|Gm1737\|Gm930\|Rundc2a | sorting nexin 29 | protein-coding |
| chr16_11461480_11462023 | chr16 | 11461481 | 11462023 | + | -0.870547 | intron (NM_001290148, intron 6 of 13) | intron (NM_001290148, intron 6 of 13) | 56104 | NM_001290148 | 74478 | Mm.325831 | NM_028964 | ENSMUSG00000071669 | **Snx29** | 4933437K13Rik\|ENSMUSG00000079738\|Gm11170\|Gm1737\|Gm930\|Rundc2a | sorting nexin 29 | protein-coding |
| chr15_36562707_36563240 | chr15 | 36562708 | 36563240 | + | -0.622061 | Intergenic | Intergenic | -7402 | NM_025712 | 66696 | Mm.253627 | NM_025712 | ENSMUSG00000013611 | **Snx31** | 4631426E05Rik | sorting nexin 31 | protein-coding |
| chr19_5532457_5532952 | chr19 | 5532458 | 5532952 | + | -0.935481 | Intergenic | Intergenic | -22216 | NM_001024560 | 225861 | Mm.20071 | NM_001024560 | ENSMUSG00000056185 | **Snx32** | B930037P14Rik\|Snx6b | sorting nexin 32 | protein-coding |
| chr19_5532572_5532829 | chr19 | 5532573 | 5532829 | + | -0.978076 | Intergenic | Intergenic | -22212 | NM_001024560 | 225861 | Mm.20071 | NM_001024560 | ENSMUSG00000056185 | **Snx32** | B930037P14Rik\|Snx6b | sorting nexin 32 | protein-coding |
| chr19_5532577_5532934 | chr19 | 5532578 | 5532934 | + | -0.978117 | Intergenic | Intergenic | -22267 | NM_001024560 | 225861 | Mm.20071 | NM_001024560 | ENSMUSG00000056185 | **Snx32** | B930037P14Rik\|Snx6b | sorting nexin 32 | protein-coding |
| chr16_33261256_33262123 | chr16 | 33261257 | 33262123 | + | -0.783585 | intron (NM_080557, intron 1 of 13) | RMER4A\|LTR\|ERVK | 10234 | NM_080557 | 69150 | Mm.28196 | NM_080557 | ENSMUSG00000022808 | **Snx4** | 1810036H14Rik\|AI037066 | sorting nexin 4 | protein-coding |
| chr1_156446434_156446989 | chr1 | 156446435 | 156446989 | + | -1.20499 | intron (NM_009230, intron 3 of 15) | intron (NM_009230, intron 3 of 15) | 27616 | NM_009230 | 20652 | Mm.28099 | NM_009230 | ENSMUSG00000026600 | **Soat1** | 8430426K15Rik\|ACAT-1\|AW550831\|Acact\|ald\|hid | sterol O-acyltransferase 1 | protein-coding |
| chr11_117952592_117952858 | chr11 | 117952593 | 117952858 | + | -0.726093 | Intergenic | (TG)n\|Simple_repeat\|Simple_repeat | 16641 | NM_007707 | 12702 | Mm.3468 | NM_007707 | ENSMUSG00000053113 | **Socs3** | Cis3\|Cish3\|EF-10\|Ef10\|SSI-3\|Ssi3 | suppressor of cytokine signaling 3 | protein-coding |
| chr11_117952055_117952524 | chr11 | 117952056 | 117952524 | + | -0.867223 | Intergenic | Intergenic | 17076 | NM_007707 | 12702 | Mm.3468 | NM_007707 | ENSMUSG00000053113 | **Socs3** | Cis3\|Cish3\|EF-10\|Ef10\|SSI-3\|Ssi3 | suppressor of cytokine signaling 3 | protein-coding |
| chr11_117952027_117952857 | chr11 | 117952028 | 117952857 | + | -0.871453 | Intergenic | Intergenic | 16924 | NM_007707 | 12702 | Mm.3468 | NM_007707 | ENSMUSG00000053113 | **Socs3** | Cis3\|Cish3\|EF-10\|Ef10\|SSI-3\|Ssi3 | suppressor of cytokine signaling 3 | protein-coding |
| chr11_117952218_117952518 | chr11 | 117952219 | 117952518 | + | -0.873057 | Intergenic | Intergenic | 16998 | NM_007707 | 12702 | Mm.3468 | NM_007707 | ENSMUSG00000053113 | **Socs3** | Cis3\|Cish3\|EF-10\|Ef10\|SSI-3\|Ssi3 | suppressor of cytokine signaling 3 | protein-coding |
| chr18_88924391_88924891 | chr18 | 88924392 | 88924891 | + | -0.669632 | Intergenic | Intergenic | -30434 | NM_018821 | 54607 | Mm.91920 | NM_018821 | ENSMUSG00000056153 | **Socs6** | 1500012M23Rik\|5830401B18Rik\|AI447482\|Cis4\|Cish4\|HSPC060\|SOCS-4\|SOCS-6\|SSI4\|STAI4\|STATI4\|Socs4 | suppressor of cytokine signaling 6 | protein-coding |
| chr16_90221557_90222012 | chr16 | 90221558 | 90222012 | + | -0.930708 | intron (NM_011434, intron 1 of 4) | intron (NM_011434, intron 1 of 4) | 1043 | NM_011434 | 20655 | Mm.276325 | NM_011434 | ENSMUSG00000022982 | **Sod1** | B430204E11Rik\|Cu/Zn-SOD\|CuZnSOD\|Ipo-1\|Ipo1\|SODC\|Sod-1 | superoxide dismutase 1, soluble | protein-coding |
| chr9_42096767_42098259 | chr9 | 42096768 | 42098259 | + | -0.782972 | intron (NM_011436, intron 3 of 47) | intron (NM_011436, intron 3 of 47) | 26776 | NM_011436 | 20660 | Mm.121920 | NM_011436 | | **Sorl1** | 2900010L19Rik\|AI596264\|AW261561\|LR11\|SorLA\|gp250\|mSorLA | sortilin-related receptor, LDLR class A repeats-containing | protein-coding |
| chr9_42032413_42032939 | chr9 | 42032414 | 42032939 | + | -0.861632 | intron (NM_011436, intron 19 of 47) | intron (NM_011436, intron 19 of 47) | 91613 | NM_011436 | 20660 | Mm.121920 | NM_011436 | | **Sorl1** | 2900010L19Rik\|AI596264\|AW261561\|LR11\|SorLA\|gp250\|mSorLA | sortilin-related receptor, LDLR class A repeats-containing | protein-coding |
| chr9_42076265_42076760 | chr9 | 42076266 | 42076760 | + | -0.949842 | intron (NM_011436, intron 6 of 47) | Lx8\|LINE\|L1 | 47776 | NM_011436 | 20660 | Mm.121920 | NM_011436 | | **Sorl1** | 2900010L19Rik\|AI596264\|AW261561\|LR11\|SorLA\|gp250\|mSorLA | sortilin-related receptor, LDLR class A repeats-containing | protein-coding |
| chr9_42141802_42142214 | chr9 | 42141803 | 42142214 | + | -1.12532 | Intergenic | Intergenic | -17719 | NM_011436 | 20660 | Mm.121920 | NM_011436 | | **Sorl1** | 2900010L19Rik\|AI596264\|AW261561\|LR11\|SorLA\|gp250\|mSorLA | sortilin-related receptor, LDLR class A repeats-containing | protein-coding |
| chr9_42158584_42159330 | chr9 | 42158585 | 42159330 | + | -1.18261 | Intergenic | Charlie24\|DNA\|hAT-Charlie | -34668 | NM_011436 | 20660 | Mm.121920 | NM_011436 | | **Sorl1** | 2900010L19Rik\|AI596264\|AW261561\|LR11\|SorLA\|gp250\|mSorLA | sortilin-related receptor, LDLR class A repeats-containing | protein-coding |
| chr17_80479143_80480054 | chr17 | 80479144 | 80480054 | + | -0.615022 | intron (NM_009231, intron 1 of 21) | CpG | 854 | NM_009231 | 20662 | Mm.60975 | NM_009231 | ENSMUSG00000024241 | **Sos1** | 4430401P03Rik\|9630010N06\|AI449023 | son of sevenless homolog 1 (Drosophila) | protein-coding |
| chr17_80441810_80442103 | chr17 | 80441811 | 80442103 | + | -0.684052 | intron (NM_009231, intron 8 of 21) | intron (NM_009231, intron 8 of 21) | 38496 | NM_009231 | 20662 | Mm.60975 | NM_009231 | ENSMUSG00000024241 | **Sos1** | 4430401P03Rik\|9630010N06\|AI449023 | son of sevenless homolog 1 (Drosophila) | protein-coding |
| chr17_80441341_80442124 | chr17 | 80441342 | 80442124 | + | -0.754397 | intron (NM_009231, intron 8 of 21) | intron (NM_009231, intron 8 of 21) | 38720 | NM_009231 | 20662 | Mm.60975 | NM_009231 | ENSMUSG00000024241 | **Sos1** | 4430401P03Rik\|9630010N06\|AI449023 | son of sevenless homolog 1 (Drosophila) | protein-coding |
| chr17_80521525_80521806 | chr17 | 80521526 | 80521806 | + | -0.798014 | Intergenic | Intergenic | -41213 | NM_009231 | 20662 | Mm.60975 | NM_009231 | ENSMUSG00000024241 | **Sos1** | 4430401P03Rik\|9630010N06\|AI449023 | son of sevenless homolog 1 (Drosophila) | protein-coding |
| chr6_144572999_144573265 | chr6 | 144573000 | 144573265 | + | -0.67638 | Intergenic | Intergenic | -99736 | NR_040519 | 66760 | Mm.233631 | NR_040519 | ENSMUSG00000085785 | **Sox5os3** | 4933425H06Rik | SRY (sex determining region Y)-box 5, opposite strand 3 | ncRNA |
| chr6_144677492_144678374 | chr6 | 144677493 | 144678374 | + | -0.711604 | intron (NR_040519, intron 1 of 2) | intron (NR_040519, intron 1 of 2) | 5065 | NR_040519 | 66760 | Mm.233631 | NR_040519 | ENSMUSG00000085785 | **Sox5os3** | 4933425H06Rik | SRY (sex determining region Y)-box 5, opposite strand 3 | ncRNA |
| chr6_144670405_144671209 | chr6 | 144670406 | 144671209 | + | -0.711708 | Intergenic | Intergenic | -2061 | NR_040519 | 66760 | Mm.233631 | NR_040519 | ENSMUSG00000085785 | **Sox5os3** | 4933425H06Rik | SRY (sex determining region Y)-box 5, opposite strand 3 | ncRNA |
| chr6_144677494_144678037 | chr6 | 144677495 | 144678037 | + | -0.793852 | intron (NR_040519, intron 1 of 2) | intron (NR_040519, intron 1 of 2) | 4898 | NR_040519 | 66760 | Mm.233631 | NR_040519 | ENSMUSG00000085785 | **Sox5os3** | 4933425H06Rik | SRY (sex determining region Y)-box 5, opposite strand 3 | ncRNA |
| chr6_144674025_144674246 | chr6 | 144674026 | 144674246 | + | -0.912975 | intron (NR_040519, intron 1 of 2) | intron (NR_040519, intron 1 of 2) | 1268 | NR_040519 | 66760 | Mm.233631 | NR_040519 | ENSMUSG00000085785 | **Sox5os3** | 4933425H06Rik | SRY (sex determining region Y)-box 5, opposite strand 3 | ncRNA |
| chr6_144678685_144678913 | chr6 | 144678686 | 144678913 | + | -0.964665 | intron (NR_040519, intron 1 of 2) | intron (NR_040519, intron 1 of 2) | 5931 | NR_040519 | 66760 | Mm.233631 | NR_040519 | ENSMUSG00000085785 | **Sox5os3** | 4933425H06Rik | SRY (sex determining region Y)-box 5, opposite strand 3 | ncRNA |
| chr1_85687347_85687893 | chr1 | 85687348 | 85687893 | + | -0.605177 | intron (NM_013673, intron 11 of 19) | intron (NM_013673, intron 11 of 19) | 37570 | NM_013673 | 20684 | Mm.290906 | NM_013673 | ENSMUSG00000026222 | **Sp100** | A430075G10Rik | nuclear antigen Sp100 | protein-coding |
| chr1_85686694_85687900 | chr1 | 85686695 | 85687900 | + | -0.632561 | intron (NM_013673, intron 11 of 19) | intron (NM_013673, intron 11 of 19) | 37247 | NM_013673 | 20684 | Mm.290906 | NM_013673 | ENSMUSG00000026222 | **Sp100** | A430075G10Rik | nuclear antigen Sp100 | protein-coding |
| chr1_85687119_85687766 | chr1 | 85687120 | 85687766 | + | -0.651387 | intron (NM_013673, intron 11 of 19) | intron (NM_013673, intron 11 of 19) | 37393 | NM_013673 | 20684 | Mm.290906 | NM_013673 | ENSMUSG00000026222 | **Sp100** | A430075G10Rik | nuclear antigen Sp100 | protein-coding |
| chr1_85651844_85652467 | chr1 | 85651845 | 85652467 | + | -0.689304 | intron (NM_013673, intron 1 of 19) | intron (NM_013673, intron 1 of 19) | 2106 | NM_013673 | 20684 | Mm.290906 | NM_013673 | ENSMUSG00000026222 | **Sp100** | A430075G10Rik | nuclear antigen Sp100 | protein-coding |
| chr1_85652208_85652467 | chr1 | 85652209 | 85652467 | + | -0.836609 | intron (NM_013673, intron 1 of 19) | intron (NM_013673, intron 1 of 19) | 2288 | NM_013673 | 20684 | Mm.290906 | NM_013673 | ENSMUSG00000026222 | **Sp100** | A430075G10Rik | nuclear antigen Sp100 | protein-coding |
| chr11_96993010_96993307 | chr11 | 96993011 | 96993307 | + | -1.00771 | Intergenic | Intergenic | -15471 | NM_001080964 | 78912 | Mm.491183 | NM_030220 | ENSMUSG00000018678 | **Sp2** | 4930480I16Rik\|mKIAA0048 | Sp2 transcription factor | protein-coding |
| chr2_73054058_73054506 | chr2 | 73054059 | 73054506 | + | -0.685684 | Intergenic | Intergenic | -73836 | NM_001018042 | 20687 | Mm.124328 | NM_011450 | ENSMUSG00000027109 | **Sp3** | D130027J01Rik | trans-acting transcription factor 3 | protein-coding |
| chr2_70425694_70425988 | chr2 | 70425695 | 70425988 | + | -0.989409 | intron (NM_177376, intron 33 of 34) | intron (NM_177376, intron 33 of 34) | -49082 | NM_022435 | 64406 | Mm.423397 | NM_022435 | ENSMUSG00000075304 | **Sp5** | - | trans-acting transcription factor 5 | protein-coding |
| chr11_97007439_97007723 | chr11 | 97007440 | 97007723 | + | -0.79366 | Intergenic | Intergenic | -5988 | NM_031183 | 83395 | Mm.156282 | NM_031183 | ENSMUSG00000038560 | **Sp6** | 1110025J03Rik\|AA591031\|AI592962\|Epfn\|Klf14 | trans-acting transcription factor 6 | protein-coding |
| chr4_34049884_34050416 | chr4 | 34049885 | 34050416 | + | -0.700824 | promoter-TSS (NM_001290443) | promoter-TSS (NM_001290443) | -83 | NM_026293 | 67652 | Mm.55848 | NM_026293 | ENSMUSG00000028264 | **Spaca1** | 1700124L11Rik\|4930540L03Rik | sperm acrosome associated 1 | protein-coding |
| chr4_33988505_33989663 | chr4 | 33988506 | 33989663 | + | -0.797186 | Intergenic | Intergenic | 60981 | NM_001290443 | 67652 | Mm.55848 | NM_026293 | ENSMUSG00000028264 | **Spaca1** | 1700124L11Rik\|4930540L03Rik | sperm acrosome associated 1 | protein-coding |
| chr4_33988573_33989063 | chr4 | 33988574 | 33989063 | + | -0.960622 | Intergenic | Lx8b\|LINE\|L1 | 61247 | NM_001290443 | 67652 | Mm.55848 | NM_026293 | ENSMUSG00000028264 | **Spaca1** | 1700124L11Rik\|4930540L03Rik | sperm acrosome associated 1 | protein-coding |
| chr7_45733328_45733852 | chr7 | 45733329 | 45733852 | + | -1.02369 | intron (NM_017465, intron 5 of 6) | intron (NM_017465, intron 5 of 6) | -7774 | NM_027055 | 69363 | Mm.45607 | NM_027055 | ENSMUSG00000070563 | **Spaca4** | 1700008E09Rik\|AV043694\|Samp14 | sperm acrosome associated 4 | protein-coding |

| | | | | | | | | | | | | | | | | | |
|---|---|---|---|---|---|---|---|---|---|---|---|---|---|---|---|---|---|
| chr7_45733267_45734025 | chr7 | 45733268 | 45734025 | + | -1.02432 | exon (NM_017465, exon 5 of 7) | exon (NM_017465, exon 5 of 7) | -7830 | NM_027055 | 69363 | Mm.45607 | NM_027055 | ENSMUSG00000070563 | **Spaca4** | 1700008E09Rik\|AV043694\|Samp14 | sperm acrosome associated 4 | protein-coding |
| chr1_69768553_69769400 | chr1 | 69768554 | 69769400 | + | -0.641685 | Intergenic | Intergenic | -57993 | NM_029160 | 66722 | Mm.440554 | NM_025728 | ENSMUSG00000053153 | **Spag16** | 4921511D23Rik\|4930524F24Rik\|4930585K05Rik\|AV261009\|Pf20\|Wdr29 | sperm associated antigen 16 | protein-coding |
| chr11_103217777_103218662 | chr11 | 103217778 | 103218662 | + | -0.781677 | intron (NM_177801, intron 1 of 4) | intron (NM_177801, intron 1 of 4) | 212 | NM_177801 | 328019 | Mm.189646 | NM_177801 | ENSMUSG00000044787 | **Spata32** | 4933400C05Rik\|VAD1.2 | spermatogenesis associated 32 | protein-coding |
| chr11_103218158_103218652 | chr11 | 103218159 | 103218652 | + | -0.848727 | promoter-TSS (NM_177801) | promoter-TSS (NM_177801) | 27 | NM_177801 | 328019 | Mm.189646 | NM_177801 | ENSMUSG00000044787 | **Spata32** | 4933400C05Rik\|VAD1.2 | spermatogenesis associated 32 | protein-coding |
| chr3_37483727_37484318 | chr3 | 37483728 | 37484318 | + | -0.790309 | intron (NM_001163511, intron 14 of 15) | intron (NM_001163511, intron 14 of 15) | 63743 | NM_021343 | 57815 | Mm.172679 | NM_021343 | ENSMUSG00000027722 | **Spata5** | 2510048F20Rik\|C78064\|Spaf | spermatogenesis associated 5 | protein-coding |
| chr4_111721106_111721410 | chr4 | 111721107 | 111721410 | + | -0.797675 | intron (NM_026470, intron 1 of 12) | intron (NM_026470, intron 1 of 12) | 1248 | NM_026470 | 67946 | Mm.275526 | NM_026470 | ENSMUSG00000034401 | **Spata6** | 1700062C23Rik\|AI790763\|Hash\|KRP\|Mash | spermatogenesis associated 6 | protein-coding |
| chr17_45488289_45489326 | chr17 | 45488290 | 45489326 | + | -0.829361 | Intergenic | L1MB7\|LINE\|L1 | -13870 | NM_027649 | 71020 | Mm.44257 | NM_027649 | ENSMUSG00000023935 | **Spats1** | 1700011H05Rik\|4933400B06Rik\|Daip2\|Ddip\|Srsp1 | spermatogenesis associated, serine-rich 1 | protein-coding |
| chr15_99153489_99153874 | chr15 | 99153490 | 99153874 | + | -0.834258 | intron (NM_139140, intron 2 of 12) | intron (NM_139140, intron 2 of 12) | 26837 | NM_139140 | 72572 | Mm.276650 | NM_139140 | ENSMUSG00000051934 | **Spats2** | 2700012F11Rik\|59kDa\|Scr59\|p59 | spermatogenesis associated, serine-rich 2 | protein-coding |
| chr11_62131580_62131954 | chr11 | 62131581 | 62131954 | + | -0.603375 | intron (NM_001281818, intron 5 of 14) | intron (NM_001281818, intron 5 of 14) | 54743 | NM_001029936 | 432572 | Mm.458025 | NM_001029936 | ENSMUSG00000042331 | **Specc1** | 2810012G08Rik\|B230396K10Rik\|Cytsb | sperm antigen with calponin homology and coiled-coil domains 1 | protein-coding |
| chr10_75137098_75137701 | chr10 | 75137099 | 75137701 | + | -0.746652 | intron (NM_001081412, intron 5 of 22) | intron (NM_001081412, intron 5 of 22) | -74673 | NM_153406 | 74392 | Mm.27716 | NM_153406 | ENSMUSG00000033444 | **Specc1l** | 4930470P14Rik\|4932439K10Rik\|9530057A13Rik\|AI317249\|Cytsa\|mKIAA0376 | sperm antigen with calponin homology and coiled-coil domains 1-like | protein-coding |
| chr16_69937586_69938130 | chr16 | 69937587 | 69938130 | + | -0.655202 | Intergenic | RMER1C\|Other\|Other | -74114 | NM_173069 | 224318 | Mm.212743 | NM_173069 | ENSMUSG00000063163 | **Speer2** | SPEER-2 | spermatogenesis associated glutamate (E)-rich protein 2 | protein-coding |
| chr16_70006834_70007290 | chr16 | 70006835 | 70007290 | + | -1.03014 | Intergenic | Lx8\|LINE\|L1 | -143318 | NM_173069 | 224318 | Mm.212743 | NM_173069 | ENSMUSG00000063163 | **Speer2** | SPEER-2 | spermatogenesis associated glutamate (E)-rich protein 2 | protein-coding |
| chr15_9701850_9702365 | chr15 | 9701851 | 9702365 | + | -1.07735 | intron (NM_177123, intron 9 of 17) | intron (NM_177123, intron 9 of 17) | 46698 | NM_177123 | 320277 | Mm.127822 | NM_177123 | ENSMUSG00000072663 | **Spef2** | C230086A09Rik | sperm flagellar 2 | protein-coding |
| chr2_122125686_122126227 | chr2 | 122125687 | 122126227 | + | -0.621154 | intron (NM_026251, intron 8 of 17) | intron (NM_026251, intron 8 of 17) | -7571 | NM_145531 | 214585 | Mm.222303 | NM_145531 | ENSMUSG00000033396 | **Spg11** | 6030465E24Rik\|A330015I11\|C530005A01Rik | spastic paraplegia 11 | protein-coding |
| chr2_122125666_122125985 | chr2 | 122125667 | 122125985 | + | -0.676613 | exon (NM_026251, exon 9 of 18) | exon (NM_026251, exon 9 of 18) | -7440 | NM_145531 | 214585 | Mm.222303 | NM_145531 | ENSMUSG00000033396 | **Spg11** | 6030465E24Rik\|A330015I11\|C530005A01Rik | spastic paraplegia 11 | protein-coding |
| chr16_44349889_44350168 | chr16 | 44349890 | 44350168 | + | -1.05957 | intron (NM_144550, intron 1 of 17) | intron (NM_144550, intron 1 of 17) | 2628 | NM_144550 | 212514 | Mm.24035 | NM_144550 | ENSMUSG00000043065 | **Spice1** | Ccdc52\|D16Ertd480e | spindle and centriole associated protein 1 | protein-coding |
| chr18_62726656_62727133 | chr18 | 62726657 | 62727133 | + | -0.977371 | intron (NM_001168423, intron 1 of 4) | MTD\|LTR\|ERVL-MaLR | 14492 | NM_001168423 | 100038417 | Mm.387557 | NM_001168423 | ENSMUSG00000073551 | **Spink13** | ENSMUSG00000073551\|Gm10534\|Spink5l3 | serine peptidase inhibitor, Kazal type 13 | protein-coding |
| chr18_62719660_62720429 | chr18 | 62719661 | 62720429 | + | -0.985428 | intron (NM_001168423, intron 1 of 4) | intron (NM_001168423, intron 1 of 4) | 21342 | NM_001168423 | 100038417 | Mm.387557 | NM_001168423 | ENSMUSG00000073551 | **Spink13** | ENSMUSG00000073551\|Gm10534\|Spink5l3 | serine peptidase inhibitor, Kazal type 13 | protein-coding |
| chr7_127144358_127144932 | chr7 | 127144359 | 127144932 | + | -0.731052 | Intergenic | Intergenic | -6822 | NM_001037810 | 20737 | Mm.283714 | NM_009259 | ENSMUSG00000051457 | **Spn** | A630014B01Rik\|Cd43\|Galgp\|Ly-48\|Ly48 | sialophorin | protein-coding |
| chr5_115049338_115049672 | chr5 | 115049339 | 115049672 | + | -0.620253 | intron (NM_029012, intron 1 of 10) | ID_B1\|SINE\|B4 | 37981 | NM_029012 | 74585 | Mm.355894 | NM_029012 | ENSMUSG00000029550 | **Sppl3** | 4833416I09Rik\|Psl4\|Usmg3 | signal peptide peptidase 3 | protein-coding |
| chr11_19924713_19924969 | chr11 | 19924714 | 19924969 | + | -0.722288 | 5' UTR (NM_033523, exon 1 of 6) | 5' UTR (NM_033523, exon 1 of 6) | 399 | NM_033523 | 114716 | Mm.266627 | NM_033523 | ENSMUSG00000045671 | **Spred2** | C79158 | sprouty-related, EVH1 domain containing 2 | protein-coding |
| chr11_19963554_19963981 | chr11 | 19963555 | 19963981 | + | -0.978207 | intron (NM_033523, intron 1 of 5) | intron (NM_033523, intron 1 of 5) | 39326 | NM_033523 | 114716 | Mm.266627 | NM_033523 | ENSMUSG00000045671 | **Spred2** | C79158 | sprouty-related, EVH1 domain containing 2 | protein-coding |
| chr11_19963685_19964094 | chr11 | 19963686 | 19964094 | + | -1.15479 | intron (NM_033523, intron 1 of 5) | intron (NM_033523, intron 1 of 5) | 39448 | NM_033523 | 114716 | Mm.266627 | NM_033523 | ENSMUSG00000045671 | **Spred2** | C79158 | sprouty-related, EVH1 domain containing 2 | protein-coding |
| chr18_38700508_38701259 | chr18 | 38700509 | 38701259 | + | -0.72688 | Intergenic | Intergenic | -99616 | NM_011898 | 24066 | Mm.42038 | NM_011898 | ENSMUSG00000024427 | **Spry4** | A030006O18Rik\|sprouty4 | sprouty homolog 4 (Drosophila) | protein-coding |
| chr18_38700796_38701205 | chr18 | 38700797 | 38701205 | + | -0.729921 | Intergenic | Intergenic | -99733 | NM_011898 | 24066 | Mm.42038 | NM_011898 | ENSMUSG00000024427 | **Spry4** | A030006O18Rik\|sprouty4 | sprouty homolog 4 (Drosophila) | protein-coding |
| chr4_149872324_149872543 | chr4 | 149872325 | 149872543 | + | -0.73104 | Intergenic | Intergenic | 82572 | NM_029035 | 74646 | Mm.30 | NM_029035 | ENSMUSG00000039911 | **Spsb1** | 1110014L01Rik\|4930422J18Rik\|AI596360\|AI854583\|SSB-1\|SSB1 | splA/ryanodine receptor domain and SOCS box containing 1 | protein-coding |
| chr4_149871623_149871949 | chr4 | 149871624 | 149871949 | + | -0.878271 | Intergenic | Intergenic | 83220 | NM_029035 | 74646 | Mm.30 | NM_029035 | ENSMUSG00000039911 | **Spsb1** | 1110014L01Rik\|4930422J18Rik\|AI596360\|AI854583\|SSB-1\|SSB1 | splA/ryanodine receptor domain and SOCS box containing 1 | protein-coding |
| chr4_149879922_149880543 | chr4 | 149879923 | 149880543 | + | -1.08708 | Intergenic | MLT1D\|LTR\|ERVL-MaLR | 74773 | NM_029035 | 74646 | Mm.30 | NM_029035 | ENSMUSG00000039911 | **Spsb1** | 1110014L01Rik\|4930422J18Rik\|AI596360\|AI854583\|SSB-1\|SSB1 | splA/ryanodine receptor domain and SOCS box containing 1 | protein-coding |
| chr9_96957249_96957809 | chr9 | 96957250 | 96957809 | + | -0.858415 | intron (NM_145134, intron 2 of 2) | MIR3\|SINE\|MIR | 60826 | NM_145134 | 211949 | Mm.491271 | NM_145134 | ENSMUSG00000046997 | **Spsb4** | D030068E18Rik\|SSB-4\|Ssb4 | splA/ryanodine receptor domain and SOCS box containing 4 | protein-coding |
| chr11_30126644_30127780 | chr11 | 30126645 | 30127780 | + | -0.686768 | intron (NM_175836, intron 17 of 34) | intron (NM_175836, intron 17 of 34) | 71045 | NM_009260 | 20742 | Mm.123110 | NM_009260 | ENSMUSG00000020315 | **Sptbn1** | 9930031C03Rik\|AL033301\|SPTB2\|Spnb-2\|Spnb2\|elf1\|elf3\|mKIAA4049 | spectrin beta, non-erythrocytic 1 | protein-coding |
| chr11_30126644_30127366 | chr11 | 30126645 | 30127366 | + | -0.712036 | intron (NM_175836, intron 17 of 34) | intron (NM_175836, intron 17 of 34) | 71252 | NM_009260 | 20742 | Mm.123110 | NM_009260 | ENSMUSG00000020315 | **Sptbn1** | 9930031C03Rik\|AL033301\|SPTB2\|Spnb-2\|Spnb2\|elf1\|elf3\|mKIAA4049 | spectrin beta, non-erythrocytic 1 | protein-coding |
| chr11_30126867_30127354 | chr11 | 30126868 | 30127354 | + | -0.7913 | intron (NM_175836, intron 17 of 34) | intron (NM_175836, intron 17 of 34) | 71146 | NM_009260 | 20742 | Mm.123110 | NM_009260 | ENSMUSG00000020315 | **Sptbn1** | 9930031C03Rik\|AL033301\|SPTB2\|Spnb-2\|Spnb2\|elf1\|elf3\|mKIAA4049 | spectrin beta, non-erythrocytic 1 | protein-coding |
| chr2_122789380_122789823 | chr2 | 122789381 | 122789823 | + | -0.675214 | intron (NR_027888, intron 3 of 9) | intron (NR_027888, intron 3 of 9) | 7603 | NM_021507 | 59010 | Mm.28986 | NM_021507 | ENSMUSG00000005803 | **Sqrdl** | 0610039J17Rik\|4930557M22Rik | sulfide quinone reductase-like (yeast) | protein-coding |
| chr15_82166215_82166669 | chr15 | 82166216 | 82166669 | + | -0.656059 | intron (NM_033218, intron 1 of 18) | intron (NM_033218, intron 1 of 18) | 19173 | NM_033218 | 20788 | Mm.38016 | NM_033218 | ENSMUSG00000022463 | **Srebf2** | AI608257\|SREBP-2\|SREBP2\|SREBP2gc\|bHLHd2\|lop13\|nuc | sterol regulatory element binding factor 2 | protein-coding |
| chr15_82162304_82162526 | chr15 | 82162305 | 82162526 | + | -0.712704 | intron (NM_033218, intron 1 of 18) | intron (NM_033218, intron 1 of 18) | 15146 | NM_033218 | 20788 | Mm.38016 | NM_033218 | ENSMUSG00000022463 | **Srebf2** | AI608257\|SREBP-2\|SREBP2\|SREBP2gc\|bHLHd2\|lop13\|nuc | sterol regulatory element binding factor 2 | protein-coding |
| chr15_82165992_82166670 | chr15 | 82165993 | 82166670 | + | -0.718464 | intron (NM_033218, intron 1 of 18) | intron (NM_033218, intron 1 of 18) | 19062 | NM_033218 | 20788 | Mm.38016 | NM_033218 | ENSMUSG00000022463 | **Srebf2** | AI608257\|SREBP-2\|SREBP2\|SREBP2gc\|bHLHd2\|lop13\|nuc | sterol regulatory element binding factor 2 | protein-coding |
| chr1_131427247_131427812 | chr1 | 131427248 | 131427812 | + | -0.682256 | intron (NM_001081011, intron 3 of 22) | intron (NM_001081011, intron 3 of 22) | 99831 | NM_001081011 | 14270 | Mm.276259 | NM_019520 | ENSMUSG00000026425 | **Srgap2** | 9930124L22Rik\|AI448945\|FBP2\|Fnbp2\|srGAP3 | SLIT-ROBO Rho GTPase activating protein 2 | protein-coding |
| chr10_62503408_62503667 | chr10 | 62503409 | 62503667 | + | -0.953478 | intron (NM_011157, intron 1 of 2) | intron (NM_011157, intron 1 of 2) | 4217 | NM_011157 | 19073 | Mm.338790 | NM_011157 | ENSMUSG00000020077 | **Srgn** | Prg\|Prg1\|Sgc | serglycin | protein-coding |
| chr17_28617730_28617985 | chr17 | 28617731 | 28617985 | + | -0.715441 | intron (NM_016795, intron 2 of 15) | intron (NM_016795, intron 2 of 15) | 4596 | NM_016795 | 20815 | Mm.15252 | NM_016795 | | **Srpk1** | AU017960 | serine/arginine-rich protein specific kinase 1 | protein-coding |
| chr17_28600846_28601141 | chr17 | 28600847 | 28601141 | + | -1.00686 | intron (NM_016795, intron 8 of 15) | intron (NM_016795, intron 8 of 15) | 21460 | NM_016795 | 20815 | Mm.15252 | NM_016795 | | **Srpk1** | AU017960 | serine/arginine-rich protein specific kinase 1 | protein-coding |
| chr17_28612590_28613123 | chr17 | 28612591 | 28613123 | + | -1.10882 | intron (NM_016795, intron 2 of 15) | intron (NM_016795, intron 2 of 15) | 9597 | NM_016795 | 20815 | Mm.15252 | NM_016795 | | **Srpk1** | AU017960 | serine/arginine-rich protein specific kinase 1 | protein-coding |
| chr5_23604407_23604855 | chr5 | 23604408 | 23604855 | + | -0.661804 | intron (NM_009274, intron 1 of 14) | intron (NM_009274, intron 1 of 14) | 11940 | NM_009274 | 20817 | Mm.288728 | NM_009274 | ENSMUSG00000062604 | **Srpk2** | AW226533\|AW492537\|AW547358\|Wbp6\|mSRPK2 | serine/arginine-rich protein specific kinase 2 | protein-coding |
| chr5_23631483_23631907 | chr5 | 23631484 | 23631907 | + | -0.761042 | Intergenic | Intergenic | -15124 | NM_009274 | 20817 | Mm.288728 | NM_009274 | ENSMUSG00000062604 | **Srpk2** | AW226533\|AW492537\|AW547358\|Wbp6\|mSRPK2 | serine/arginine-rich protein specific kinase 2 | protein-coding |
| chr5_23631520_23632076 | chr5 | 23631521 | 23632076 | + | -0.823585 | Intergenic | Intergenic | -15227 | NM_009274 | 20817 | Mm.288728 | NM_009274 | ENSMUSG00000062604 | **Srpk2** | AW226533\|AW492537\|AW547358\|Wbp6\|mSRPK2 | serine/arginine-rich protein specific kinase 2 | protein-coding |
| chr5_23637216_23637788 | chr5 | 23637217 | 23637788 | + | -0.851736 | Intergenic | Intergenic | -20931 | NM_009274 | 20817 | Mm.288728 | NM_009274 | ENSMUSG00000062604 | **Srpk2** | AW226533\|AW492537\|AW547358\|Wbp6\|mSRPK2 | serine/arginine-rich protein specific kinase 2 | protein-coding |

| | | | | | | | | | | | | | | | | | |
|---|---|---|---|---|---|---|---|---|---|---|---|---|---|---|---|---|---|
| chr4_33226643_33227508 | chr4 | 33226644 | 33227508 | + | -0.709243 | intron (NM_177774, intron 2 of 2) | B1F1\|SINE\|Alu | 18085 | NM_177774 | 272009 | Mm.156636 | NM_177774 | ENSMUSG00000054679 | **Srsf12** | B830039L16\|Sfrs13b\|Srrp\|Srsf13b | serine/arginine-rich splicing factor 12 | protein-coding |
| chr4_33226580_33226953 | chr4 | 33226581 | 33226953 | + | -0.85307 | intron (NM_177774, intron 2 of 2) | B4A\|SINE\|B4 | 17776 | NM_177774 | 272009 | Mm.156636 | NM_177774 | ENSMUSG00000054679 | **Srsf12** | B830039L16\|Sfrs13b\|Srrp\|Srsf13b | serine/arginine-rich splicing factor 12 | protein-coding |
| chr17_80169833_80170165 | chr17 | 80169834 | 80170165 | + | -0.978175 | intron (NM_176963, intron 4 of 6) | intron (NM_176963, intron 4 of 6) | 37306 | NM_001195486 | 225027 | Mm.292016 | NM_146083 | ENSMUSG00000024097 | **Srsf7** | 35kDa\|9430065L19Rik\|9G8\|NX-96\|Sfrs7 | serine/arginine-rich splicing factor 7 | protein-coding |
| chr18_14604531_14604812 | chr18 | 14604532 | 14604812 | + | -0.686406 | Intergenic | MTEb\|LTR\|ERVL-MaLR | 78242 | NM_001161369 | 268996 | Mm.31761 | NM_009280 | ENSMUSG00000037013 | **Ss18** | D130059H17\|Ssxt\|Syt | synovial sarcoma translocation, Chromosome 18 | protein-coding |
| chr18_14603209_14604013 | chr18 | 14603210 | 14604013 | + | -0.863312 | Intergenic | ORR1D2\|LTR\|ERVL-MaLR | 79303 | NM_001161369 | 268996 | Mm.31761 | NM_009280 | ENSMUSG00000037013 | **Ss18** | D130059H17\|Ssxt\|Syt | synovial sarcoma translocation, Chromosome 18 | protein-coding |
| chr18_14666958_14667581 | chr18 | 14666959 | 14667581 | + | -0.92291 | intron (NM_001161369, intron 3 of 9) | RMER15\|LTR\|ERVL | 15644 | NM_001161369 | 268996 | Mm.31761 | NM_009280 | ENSMUSG00000037013 | **Ss18** | D130059H17\|Ssxt\|Syt | synovial sarcoma translocation, Chromosome 18 | protein-coding |
| chr18_14656710_14657755 | chr18 | 14656711 | 14657755 | + | -1.06709 | intron (NM_001161369, intron 3 of 9) | intron (NM_001161369, intron 3 of 9) | 25681 | NM_001161369 | 268996 | Mm.31761 | NM_009280 | ENSMUSG00000037013 | **Ss18** | D130059H17\|Ssxt\|Syt | synovial sarcoma translocation, Chromosome 18 | protein-coding |
| chr18_14656705_14657281 | chr18 | 14656706 | 14657281 | + | -1.23914 | intron (NM_001161369, intron 3 of 9) | intron (NM_001161369, intron 3 of 9) | 25921 | NM_001161369 | 268996 | Mm.31761 | NM_009280 | ENSMUSG00000037013 | **Ss18** | D130059H17\|Ssxt\|Syt | synovial sarcoma translocation, Chromosome 18 | protein-coding |
| chr13_91540806_91541594 | chr13 | 91540807 | 91541594 | + | -0.62306 | intron (NM_024272, intron 3 of 16) | intron (NM_024272, intron 3 of 16) | 80103 | NM_024186 | 66970 | Mm.343095 | NM_024186 | ENSMUSG00000003992 | **Ssbp2** | 1500004K09Rik\|2310079I02Rik\|9330163K02Rik\|A830008M03Rik\|AU067692\|AW558684\|Hspc116\|SSDP2 | single-stranded DNA binding protein 2 | protein-coding |
| chr13_91484116_91484760 | chr13 | 91484117 | 91484760 | + | -0.626213 | intron (NM_024272, intron 1 of 16) | intron (NM_024272, intron 1 of 16) | 23341 | NM_024186 | 66970 | Mm.343095 | NM_024186 | ENSMUSG00000003992 | **Ssbp2** | 1500004K09Rik\|2310079I02Rik\|9330163K02Rik\|A830008M03Rik\|AU067692\|AW558684\|Hspc116\|SSDP2 | single-stranded DNA binding protein 2 | protein-coding |
| chr13_91453236_91453765 | chr13 | 91453237 | 91453765 | + | -0.925868 | Intergenic | Intergenic | -7596 | NM_024272 | 66970 | Mm.343095 | NM_024186 | ENSMUSG00000003992 | **Ssbp2** | 1500004K09Rik\|2310079I02Rik\|9330163K02Rik\|A830008M03Rik\|AU067692\|AW558684\|Hspc116\|SSDP2 | single-stranded DNA binding protein 2 | protein-coding |
| chr11_77348788_77349394 | chr11 | 77348789 | 77349394 | + | -0.960849 | intron (NM_001291190, intron 1 of 13) | intron (NM_001291190, intron 1 of 13) | 811 | NM_001291190 | 237860 | Mm.440381 | NM_177710 | ENSMUSG00000037926 | **Ssh2** | SSH-2\|SSH-2L\|mSSH-2L | slingshot homolog 2 (Drosophila) | protein-coding |
| chr11_77349005_77349383 | chr11 | 77349006 | 77349383 | + | -0.963005 | intron (NM_001291190, intron 1 of 13) | intron (NM_001291190, intron 1 of 13) | 914 | NM_001291190 | 237860 | Mm.440381 | NM_177710 | ENSMUSG00000037926 | **Ssh2** | SSH-2\|SSH-2L\|mSSH-2L | slingshot homolog 2 (Drosophila) | protein-coding |
| chr11_77355282_77355699 | chr11 | 77355283 | 77355699 | + | -1.0836 | intron (NM_001291190, intron 1 of 13) | intron (NM_001291190, intron 1 of 13) | 7211 | NM_001291190 | 237860 | Mm.440381 | NM_177710 | ENSMUSG00000037926 | **Ssh2** | SSH-2\|SSH-2L\|mSSH-2L | slingshot homolog 2 (Drosophila) | protein-coding |
| chr11_77386150_77386743 | chr11 | 77386151 | 77386743 | + | -1.09599 | intron (NM_001291190, intron 1 of 13) | intron (NM_001291190, intron 1 of 13) | 38167 | NM_001291190 | 237860 | Mm.440381 | NM_177710 | ENSMUSG00000037926 | **Ssh2** | SSH-2\|SSH-2L\|mSSH-2L | slingshot homolog 2 (Drosophila) | protein-coding |
| chr11_77372228_77372565 | chr11 | 77372229 | 77372565 | + | -1.30528 | intron (NM_001291190, intron 1 of 13) | intron (NM_001291190, intron 1 of 13) | 24117 | NM_001291190 | 237860 | Mm.440381 | NM_177710 | ENSMUSG00000037926 | **Ssh2** | SSH-2\|SSH-2L\|mSSH-2L | slingshot homolog 2 (Drosophila) | protein-coding |
| chr6_146107190_146107950 | chr6 | 146107191 | 146107950 | + | -0.757984 | TTS (NM_010586) | TTS (NM_010586) | 173423 | NM_010656 | 16651 | Mm.49689 | NM_010656 | ENSMUSG00000030255 | **Sspn** | Krag | sarcospan | protein-coding |
| chr6_146077189_146077540 | chr6 | 146077190 | 146077540 | + | -0.826352 | Intergenic | ORR1E\|LTR\|ERVL-MaLR | 143218 | NM_010656 | 16651 | Mm.49689 | NM_010656 | ENSMUSG00000030255 | **Sspn** | Krag | sarcospan | protein-coding |
| chr6_146077277_146077492 | chr6 | 146077278 | 146077492 | + | -0.84158 | Intergenic | ORR1E\|LTR\|ERVL-MaLR | 143238 | NM_010656 | 16651 | Mm.49689 | NM_010656 | ENSMUSG00000030255 | **Sspn** | Krag | sarcospan | protein-coding |
| chr6_146005210_146005760 | chr6 | 146005211 | 146005760 | + | -0.940061 | Intergenic | Intergenic | 71338 | NM_010656 | 16651 | Mm.49689 | NM_010656 | ENSMUSG00000030255 | **Sspn** | Krag | sarcospan | protein-coding |
| chr6_146114581_146115029 | chr6 | 146114582 | 146115029 | + | -1.08119 | intron (NM_019923, intron 55 of 56) | intron (NM_019923, intron 55 of 56) | 180658 | NM_010656 | 16651 | Mm.49689 | NM_010656 | ENSMUSG00000030255 | **Sspn** | Krag | sarcospan | protein-coding |
| chr15_78557732_78558606 | chr15 | 78557733 | 78558606 | + | -0.631422 | TTS (NM_009008) | TTS (NM_009008) | -13824 | NM_009218 | 20607 | Mm.211411 | NM_009218 | ENSMUSG00000044933 | **Sstr3** | Smstr-3\|Smstr3\|sst3 | somatostatin receptor 3 | protein-coding |
| chr16_23196659_23197396 | chr16 | 23196660 | 23197396 | + | -0.622575 | Intergenic | Intergenic | -27712 | NM_145933 | 20440 | Mm.149029 | NM_145933 | ENSMUSG00000022885 | **St6gal1** | AW742324\|Siat1\|St6Gal-I\|St6gal\|St6gall | beta galactoside alpha 2,6 sialyltransferase 1 | protein-coding |
| chr16_23195502_23196000 | chr16 | 23195503 | 23196000 | + | -0.771269 | Intergenic | MTEa\|LTR\|ERVL-MaLR | -28989 | NM_145933 | 20440 | Mm.149029 | NM_145933 | ENSMUSG00000022885 | **St6gal1** | AW742324\|Siat1\|St6Gal-I\|St6gal\|St6gall | beta galactoside alpha 2,6 sialyltransferase 1 | protein-coding |
| chr16_23286310_23286853 | chr16 | 23286311 | 23286853 | + | -0.777987 | intron (NM_001252505, intron 2 of 7) | PB1D10\|SINE\|Alu | -3888 | NM_001252506 | 20440 | Mm.149029 | NM_145933 | ENSMUSG00000022885 | **St6gal1** | AW742324\|Siat1\|St6Gal-I\|St6gal\|St6gall | beta galactoside alpha 2,6 sialyltransferase 1 | protein-coding |
| chr16_23225068_23225757 | chr16 | 23225069 | 23225757 | + | -0.799224 | promoter-TSS (NM_001252505) | promoter-TSS (NM_001252505) | -608 | NM_001252505 | 20440 | Mm.149029 | NM_145933 | ENSMUSG00000022885 | **St6gal1** | AW742324\|Siat1\|St6Gal-I\|St6gal\|St6gall | beta galactoside alpha 2,6 sialyltransferase 1 | protein-coding |
| chr16_23357399_23357802 | chr16 | 23357400 | 23357802 | + | -0.86026 | intron (NM_001252505, intron 7 of 7) | intron (NM_001252505, intron 7 of 7) | 67131 | NM_001252506 | 20440 | Mm.149029 | NM_145933 | ENSMUSG00000022885 | **St6gal1** | AW742324\|Siat1\|St6Gal-I\|St6gal\|St6gall | beta galactoside alpha 2,6 sialyltransferase 1 | protein-coding |
| chr16_23226656_23226931 | chr16 | 23226657 | 23226931 | + | -0.975854 | intron (NM_001252505, intron 1 of 7) | intron (NM_001252505, intron 1 of 7) | 773 | NM_001252505 | 20440 | Mm.149029 | NM_145933 | ENSMUSG00000022885 | **St6gal1** | AW742324\|Siat1\|St6Gal-I\|St6gal\|St6gall | beta galactoside alpha 2,6 sialyltransferase 1 | protein-coding |
| chr3_152941427_152942025 | chr3 | 152941428 | 152942025 | + | -0.781054 | intron (NM_012028, intron 2 of 4) | intron (NM_012028, intron 2 of 4) | 40481 | NM_012028 | 26938 | Mm.40915 | NM_012028 | | **St6galnac5** | AI851940\|Siat7e | ST6 (alpha-N-acetyl-neuraminyl-2,3-beta-galactosyl-1,3)-N-acetylgalactosaminide alpha-2,6-sialyltransferase 5 | protein-coding |
| chr3_152950140_152950687 | chr3 | 152950141 | 152950687 | + | -0.800018 | intron (NM_012028, intron 2 of 4) | intron (NM_012028, intron 2 of 4) | 31793 | NM_012028 | 26938 | Mm.40915 | NM_012028 | | **St6galnac5** | AI851940\|Siat7e | ST6 (alpha-N-acetyl-neuraminyl-2,3-beta-galactosyl-1,3)-N-acetylgalactosaminide alpha-2,6-sialyltransferase 5 | protein-coding |
| chr3_152926239_152926587 | chr3 | 152926240 | 152926587 | + | -0.843783 | intron (NM_012028, intron 2 of 4) | ORR1E\|LTR\|ERVL-MaLR | 55794 | NM_012028 | 26938 | Mm.40915 | NM_012028 | | **St6galnac5** | AI851940\|Siat7e | ST6 (alpha-N-acetyl-neuraminyl-2,3-beta-galactosyl-1,3)-N-acetylgalactosaminide alpha-2,6-sialyltransferase 5 | protein-coding |
| chr3_152941658_152942029 | chr3 | 152941659 | 152942029 | + | -0.941412 | intron (NM_012028, intron 2 of 4) | intron (NM_012028, intron 2 of 4) | 40363 | NM_012028 | 26938 | Mm.40915 | NM_012028 | | **St6galnac5** | AI851940\|Siat7e | ST6 (alpha-N-acetyl-neuraminyl-2,3-beta-galactosyl-1,3)-N-acetylgalactosaminide alpha-2,6-sialyltransferase 5 | protein-coding |
| chr6_142946101_142946411 | chr6 | 142946102 | 142946411 | + | -0.833555 | intron (NM_011374, intron 1 of 4) | B4A\|SINE\|B4 | 18196 | NM_011374 | 20449 | Mm.260838 | NM_011374 | ENSMUSG00000030283 | **St8sia1** | 9330109E03Rik\|GD3S\|Sia-T\|Siat8\|Siat8a | ST8 alpha-N-acetyl-neuraminide alpha-2,8-sialyltransferase 1 | protein-coding |
| chr6_142909489_142909884 | chr6 | 142909490 | 142909884 | + | -1.11729 | intron (NM_011374, intron 2 of 4) | intron (NM_011374, intron 2 of 4) | 54765 | NM_011374 | 20449 | Mm.260838 | NM_011374 | ENSMUSG00000030283 | **St8sia1** | 9330109E03Rik\|GD3S\|Sia-T\|Siat8\|Siat8a | ST8 alpha-N-acetyl-neuraminide alpha-2,8-sialyltransferase 1 | protein-coding |
| chr6_142915336_142915820 | chr6 | 142915337 | 142915820 | + | -1.18387 | intron (NM_011374, intron 1 of 4) | intron (NM_011374, intron 1 of 4) | 48874 | NM_011374 | 20449 | Mm.260838 | NM_011374 | ENSMUSG00000030283 | **St8sia1** | 9330109E03Rik\|GD3S\|Sia-T\|Siat8\|Siat8a | ST8 alpha-N-acetyl-neuraminide alpha-2,8-sialyltransferase 1 | protein-coding |
| chr1_95666183_95666461 | chr1 | 95666184 | 95666461 | + | -0.871037 | intron (NM_001159745, intron 1 of 3) | intron (NM_001159745, intron 1 of 3) | 1272 | NM_009183 | 20452 | Mm.306228 | NM_009183 | ENSMUSG00000040710 | **St8sia4** | PST\|PST-1\|SIAT8-D\|ST8SiaIV\|Siat8d | ST8 alpha-N-acetyl-neuraminide alpha-2,8-sialyltransferase 4 | protein-coding |
| chr1_95666274_95666446 | chr1 | 95666275 | 95666446 | + | -0.906021 | intron (NM_001159745, intron 1 of 3) | intron (NM_001159745, intron 1 of 3) | 1234 | NM_009183 | 20452 | Mm.306228 | NM_009183 | ENSMUSG00000040710 | **St8sia4** | PST\|PST-1\|SIAT8-D\|ST8SiaIV\|Siat8d | ST8 alpha-N-acetyl-neuraminide alpha-2,8-sialyltransferase 4 | protein-coding |
| chr2_13760363_13760826 | chr2 | 13760364 | 13760826 | + | -0.846179 | intron (NM_145838, intron 2 of 7) | L2a\|LINE\|L2 | 32925 | NM_145838 | 241230 | Mm.330004 | NM_145838 | ENSMUSG00000003418 | **St8sia6** | 1700007J08Rik\|AI314453\|AI875066\|ST8SiaVI\|Siat8f | ST8 alpha-N-acetyl-neuraminide alpha-2,8-sialyltransferase 6 | protein-coding |
| chr2_13760457_13760744 | chr2 | 13760458 | 13760744 | + | -0.854319 | intron (NM_145838, intron 2 of 7) | L2a\|LINE\|L2 | 32919 | NM_145838 | 241230 | Mm.330004 | NM_145838 | ENSMUSG00000003418 | **St8sia6** | 1700007J08Rik\|AI314453\|AI875066\|ST8SiaVI\|Siat8f | ST8 alpha-N-acetyl-neuraminide alpha-2,8-sialyltransferase 6 | protein-coding |
| chr2_13757414_13757842 | chr2 | 13757415 | 13757842 | + | -1.09771 | intron (NM_145838, intron 2 of 7) | intron (NM_145838, intron 2 of 7) | 35892 | NM_145838 | 241230 | Mm.330004 | NM_145838 | ENSMUSG00000003418 | **St8sia6** | 1700007J08Rik\|AI314453\|AI875066\|ST8SiaVI\|Siat8f | ST8 alpha-N-acetyl-neuraminide alpha-2,8-sialyltransferase 6 | protein-coding |
| chr9_100669898_100670571 | chr9 | 100669899 | 100670571 | + | -0.649581 | intron (NM_009282, intron 1 of 33) | intron (NM_009282, intron 1 of 33) | 26612 | NM_009282 | 20842 | Mm.42135 | NM_009282 | ENSMUSG00000037286 | **Stag1** | AU045003\|SA-1\|Scc3 | stromal antigen 1 | protein-coding |

| | | | | | | | | | | | | | | | | | |
|---|---|---|---|---|---|---|---|---|---|---|---|---|---|---|---|---|---|
| chr9_100670212_100670473 | chr9 | 100670213 | 100670473 | + | -0.898082 | intron (NM_009282, intron 1 of 33) | intron (NM_009282, intron 1 of 33) | 26720 | NM_009282 | 20842 | Mm.42135 | NM_009282 | ENSMUSG00000037286 | **Stag1** | AU045003\|SA-1\|Scc3 | stromal antigen 1 | protein-coding |
| chrX_42202437_42202873 | chrX | 42202438 | 42202873 | + | -1.07641 | intron (NM_021465, intron 2 of 32) | ORR1C1\|LTR\|ERVL-MaLR | 800 | NM_001290713 | 20843 | Mm.290422 | NM_021465 | ENSMUSG00000025862 | **Stag2** | 9230105L23Rik\|B230112I07Rik\|SA-2\|SAP2 | stromal antigen 2 | protein-coding |
| chr5_86069203_86069473 | chr5 | 86069204 | 86069473 | + | -0.634327 | Intergenic | B1_Mur1\|SINE\|Alu | -2490 | NM_019992 | 56792 | Mm.131237 | NM_019992 | ENSMUSG00000029254 | **Stap1** | AI586015\|Brdg1\|STAP-1 | signal transducing adaptor family member 1 | protein-coding |
| chr5_86086492_86087334 | chr5 | 86086493 | 86087334 | + | -0.660366 | intron (NM_019992, intron 5 of 8) | intron (NM_019992, intron 5 of 8) | 15085 | NM_019992 | 56792 | Mm.131237 | NM_019992 | ENSMUSG00000029254 | **Stap1** | AI586015\|Brdg1\|STAP-1 | signal transducing adaptor family member 1 | protein-coding |
| chr13_19190143_19190470 | chr13 | 19190144 | 19190470 | + | -0.600165 | Intergenic | Intergenic | 205445 | NM_024270 | 76205 | Mm.41940 | NM_024270 | ENSMUSG00000003062 | **Stard3nl** | 0610035N01Rik\|6530409L22Rik\|AW124774\|MENTHO | STARD3 N-terminal like | protein-coding |
| chr13_19182270_19182781 | chr13 | 19182271 | 19182781 | + | -0.823482 | Intergenic | Intergenic | 213226 | NM_024270 | 76205 | Mm.41940 | NM_024270 | ENSMUSG00000003062 | **Stard3nl** | 0610035N01Rik\|6530409L22Rik\|AW124774\|MENTHO | STARD3 N-terminal like | protein-coding |
| chr13_19190550_19191138 | chr13 | 19190551 | 19191138 | + | -0.895581 | Intergenic | Intergenic | 204908 | NM_024270 | 76205 | Mm.41940 | NM_024270 | ENSMUSG00000003062 | **Stard3nl** | 0610035N01Rik\|6530409L22Rik\|AW124774\|MENTHO | STARD3 N-terminal like | protein-coding |
| chr7_83609727_83610614 | chr7 | 83609728 | 83610614 | + | -0.795688 | intron (NM_177695, intron 14 of 22) | RMER15\|LTR\|ERVL | -21846 | NM_023377 | 170460 | Mm.357953 | NM_023377 | ENSMUSG00000046027 | **Stard5** | 2310058G22Rik\|AI607880\|D7Ertd152e | StAR-related lipid transfer (START) domain containing 5 | protein-coding |
| chr1_52100380_52101012 | chr1 | 52100381 | 52101012 | + | -0.666204 | intron (NM_011487, intron 17 of 23) | intron (NM_011487, intron 17 of 23) | -18742 | NM_009283 | 20846 | Mm.277406 | NM_009283 | ENSMUSG00000026104 | **Stat1** | 2010005J02Rik\|AA408197 | signal transducer and activator of transcription 1 | protein-coding |
| chr1_52011250_52011614 | chr1 | 52011251 | 52011614 | + | -0.899785 | intron (NM_011487, intron 1 of 23) | intron (NM_011487, intron 1 of 23) | 3188 | NM_011487 | 20849 | Mm.1550 | NM_011487 | ENSMUSG00000062939 | **Stat4** | - | signal transducer and activator of transcription 4 | protein-coding |
| chr11_100891553_100892005 | chr11 | 100891554 | 100892005 | + | -0.662756 | intron (NM_213660, intron 21 of 23) | intron (NM_213660, intron 21 of 23) | 31295 | NM_001164062 | 20850 | Mm.277403 | NM_011488 | ENSMUSG00000004043 | **Stat5a** | AA959963\|STAT5 | signal transducer and activator of transcription 5A | protein-coding |
| chr11_100818445_100818642 | chr11 | 100818446 | 100818642 | + | -0.743513 | intron (NM_001113563, intron 1 of 18) | intron (NM_001113563, intron 1 of 18) | 4026 | NM_011489 | 20851 | Mm.34064 | NM_011489 | ENSMUSG00000020919 | **Stat5b** | - | signal transducer and activator of transcription 5B | protein-coding |
| chr11_100822385_100822804 | chr11 | 100822386 | 100822804 | + | -0.839869 | promoter-TSS (NM_011489) | promoter-TSS (NM_011489) | -25 | NM_011489 | 20851 | Mm.34064 | NM_011489 | ENSMUSG00000020919 | **Stat5b** | - | signal transducer and activator of transcription 5B | protein-coding |
| chr11_100823931_100824413 | chr11 | 100823932 | 100824413 | + | -0.865838 | intron (NM_001113563, intron 1 of 18) | intron (NM_001113563, intron 1 of 18) | -1602 | NM_011489 | 20851 | Mm.34064 | NM_011489 | ENSMUSG00000020919 | **Stat5b** | - | signal transducer and activator of transcription 5B | protein-coding |
| chr11_100804184_100805018 | chr11 | 100804185 | 100805018 | + | -1.10882 | intron (NM_001113563, intron 3 of 18) | MTC\|LTR\|ERVL-MaLR | 17969 | NM_011489 | 20851 | Mm.34064 | NM_011489 | ENSMUSG00000020919 | **Stat5b** | - | signal transducer and activator of transcription 5B | protein-coding |
| chr11_100848081_100848593 | chr11 | 100848082 | 100848593 | + | -1.43076 | intron (NM_001113563, intron 1 of 18) | intron (NM_001113563, intron 1 of 18) | 2248 | NM_001113563 | 20851 | Mm.34064 | NM_011489 | ENSMUSG00000020919 | **Stat5b** | - | signal transducer and activator of transcription 5B | protein-coding |
| chr11_100845784_100846384 | chr11 | 100845785 | 100846384 | + | -2.0125 | intron (NM_001113563, intron 1 of 18) | intron (NM_001113563, intron 1 of 18) | 4501 | NM_001113563 | 20851 | Mm.34064 | NM_011489 | ENSMUSG00000020919 | **Stat5b** | - | signal transducer and activator of transcription 5B | protein-coding |
| chr1_120293631_120294163 | chr1 | 120293632 | 120294163 | + | -1.18582 | Intergenic | ORR1E\|LTR\|ERVL-MaLR | -22815 | NM_001085409 | 68428 | Mm.181033 | NM_133186 | ENSMUSG00000026389 | **Steap3** | 1010001D01Rik\|Tsap6\|pHyde | STEAP family member 3 | protein-coding |
| chr7_102337884_102338454 | chr7 | 102337885 | 102338454 | + | -0.66647 | intron (NM_009287, intron 1 of 11) | intron (NM_009287, intron 1 of 11) | 70345 | NM_009287 | 20866 | Mm.645 | NM_009287 | ENSMUSG00000030987 | **Stim1** | SIM | stromal interaction molecule 1 | protein-coding |
| chr7_102260587_102260859 | chr7 | 102260588 | 102260859 | + | -0.698768 | Intergenic | ID_B1\|SINE\|B4 | -7101 | NM_009287 | 20866 | Mm.645 | NM_009287 | ENSMUSG00000030987 | **Stim1** | SIM | stromal interaction molecule 1 | protein-coding |
| chr7_102326311_102326636 | chr7 | 102326312 | 102326636 | + | -0.715855 | intron (NM_009287, intron 1 of 11) | intron (NM_009287, intron 1 of 11) | 58650 | NM_009287 | 20866 | Mm.645 | NM_009287 | ENSMUSG00000030987 | **Stim1** | SIM | stromal interaction molecule 1 | protein-coding |
| chr5_53999973_54000223 | chr5 | 53999974 | 54000223 | + | -0.721366 | intron (NM_001081103, intron 1 of 11) | intron (NM_001081103, intron 1 of 11) | 1575 | NM_001081103 | 116873 | Mm.265985 | NM_054083 | ENSMUSG00000039156 | **Stim2** | - | stromal interaction molecule 2 | protein-coding |
| chr11_32582843_32583306 | chr11 | 32582844 | 32583306 | + | -1.02718 | intron (NM_009288, intron 4 of 18) | intron (NM_009288, intron 4 of 18) | 49809 | NM_009288 | 20868 | Mm.8235 | NM_009288 | ENSMUSG00000020272 | **Stk10** | Gek1\|Lok\|mKIAA4026 | serine/threonine kinase 10 | protein-coding |
| chr11_32535640_32536064 | chr11 | 32535641 | 32536064 | + | -1.33835 | intron (NM_009288, intron 1 of 18) | intron (NM_009288, intron 1 of 18) | 2586 | NM_009288 | 20868 | Mm.8235 | NM_009288 | ENSMUSG00000020272 | **Stk10** | Gek1\|Lok\|mKIAA4026 | serine/threonine kinase 10 | protein-coding |
| chr1_75213115_75213522 | chr1 | 75213116 | 75213522 | + | -0.699096 | intron (NM_001277992, intron 6 of 7) | intron (NM_001277992, intron 6 of 7) | 2490 | NM_011494 | 20872 | Mm.1155 | NM_011494 | ENSMUSG00000026201 | **Stk16** | EDPK\|Krct\|MPSK\|PKL12\|TSF-1 | serine/threonine kinase 16 | protein-coding |
| chr1_53773916_53774339 | chr1 | 53773917 | 53774339 | + | -0.69819 | intron (NM_133810, intron 2 of 7) | intron (NM_133810, intron 2 of 7) | 11087 | NM_133810 | 98267 | Mm.25559 | NM_133810 | ENSMUSG00000026094 | **Stk17b** | 3110009A03Rik\|AI120141\|Drak2 | serine/threonine kinase 17b (apoptosis-inducing) | protein-coding |
| chr1_53784212_53784516 | chr1 | 53784213 | 53784516 | + | -0.773585 | intron (NM_133810, intron 1 of 7) | intron (NM_133810, intron 1 of 7) | 851 | NM_133810 | 98267 | Mm.25559 | NM_133810 | ENSMUSG00000026094 | **Stk17b** | 3110009A03Rik\|AI120141\|Drak2 | serine/threonine kinase 17b (apoptosis-inducing) | protein-coding |
| chr1_53781333_53781662 | chr1 | 53781334 | 53781662 | + | -0.844837 | intron (NM_133810, intron 1 of 7) | L2b\|LINE\|L2 | 3717 | NM_133810 | 98267 | Mm.25559 | NM_133810 | ENSMUSG00000026094 | **Stk17b** | 3110009A03Rik\|AI120141\|Drak2 | serine/threonine kinase 17b (apoptosis-inducing) | protein-coding |
| chr1_53796918_53797457 | chr1 | 53796919 | 53797457 | + | -1.03199 | Intergenic | Intergenic | -11973 | NM_133810 | 98267 | Mm.25559 | NM_133810 | ENSMUSG00000026094 | **Stk17b** | 3110009A03Rik\|AI120141\|Drak2 | serine/threonine kinase 17b (apoptosis-inducing) | protein-coding |
| chr1_53776575_53776937 | chr1 | 53776576 | 53776937 | + | -1.09871 | intron (NM_133810, intron 1 of 7) | intron (NM_133810, intron 1 of 7) | 8459 | NM_133810 | 98267 | Mm.25559 | NM_133810 | ENSMUSG00000026094 | **Stk17b** | 3110009A03Rik\|AI120141\|Drak2 | serine/threonine kinase 17b (apoptosis-inducing) | protein-coding |
| chr14_121405137_121406053 | chr14 | 121405138 | 121406053 | + | -0.619197 | Intergenic | RMER15\|LTR\|ERVL | -26365 | NM_145465 | 223255 | Mm.390756 | NM_145465 | ENSMUSG00000063410 | **Stk24** | 1810013H02Rik\|C76483\|MST-3\|Mst3\|STE20 | serine/threonine kinase 24 | protein-coding |
| chr14_121340597_121341178 | chr14 | 121340598 | 121341178 | + | -0.680998 | intron (NM_145465, intron 1 of 10) | intron (NM_145465, intron 1 of 10) | 38342 | NM_145465 | 223255 | Mm.390756 | NM_145465 | ENSMUSG00000063410 | **Stk24** | 1810013H02Rik\|C76483\|MST-3\|Mst3\|STE20 | serine/threonine kinase 24 | protein-coding |
| chr14_121349562_121349822 | chr14 | 121349563 | 121349822 | + | -0.703443 | intron (NM_145465, intron 1 of 10) | intron (NM_145465, intron 1 of 10) | 29538 | NM_145465 | 223255 | Mm.390756 | NM_145465 | ENSMUSG00000063410 | **Stk24** | 1810013H02Rik\|C76483\|MST-3\|Mst3\|STE20 | serine/threonine kinase 24 | protein-coding |
| chr14_121340496_121341293 | chr14 | 121340497 | 121341293 | + | -0.705428 | intron (NM_145465, intron 1 of 10) | intron (NM_145465, intron 1 of 10) | 38335 | NM_145465 | 223255 | Mm.390756 | NM_145465 | ENSMUSG00000063410 | **Stk24** | 1810013H02Rik\|C76483\|MST-3\|Mst3\|STE20 | serine/threonine kinase 24 | protein-coding |
| chr14_121405194_121405543 | chr14 | 121405195 | 121405543 | + | -0.748258 | Intergenic | Intergenic | -26139 | NM_145465 | 223255 | Mm.390756 | NM_145465 | ENSMUSG00000063410 | **Stk24** | 1810013H02Rik\|C76483\|MST-3\|Mst3\|STE20 | serine/threonine kinase 24 | protein-coding |
| chr14_121349234_121349509 | chr14 | 121349235 | 121349509 | + | -0.76864 | intron (NM_145465, intron 1 of 10) | MT2B2\|LTR\|ERVL | 29858 | NM_145465 | 223255 | Mm.390756 | NM_145465 | ENSMUSG00000063410 | **Stk24** | 1810013H02Rik\|C76483\|MST-3\|Mst3\|STE20 | serine/threonine kinase 24 | protein-coding |
| chr14_121349292_121349824 | chr14 | 121349293 | 121349824 | + | -0.819718 | intron (NM_145465, intron 1 of 10) | intron (NM_145465, intron 1 of 10) | 29672 | NM_145465 | 223255 | Mm.390756 | NM_145465 | ENSMUSG00000063410 | **Stk24** | 1810013H02Rik\|C76483\|MST-3\|Mst3\|STE20 | serine/threonine kinase 24 | protein-coding |
| chr14_121421572_121421826 | chr14 | 121421573 | 121421826 | + | -0.876494 | Intergenic | Intergenic | -42469 | NM_145465 | 223255 | Mm.390756 | NM_145465 | ENSMUSG00000063410 | **Stk24** | 1810013H02Rik\|C76483\|MST-3\|Mst3\|STE20 | serine/threonine kinase 24 | protein-coding |
| chr14_121290754_121291345 | chr14 | 121290755 | 121291345 | + | -1.08987 | intron (NM_145465, intron 10 of 10) | RSINE1\|SINE\|B4 | 88180 | NM_145465 | 223255 | Mm.390756 | NM_145465 | ENSMUSG00000063410 | **Stk24** | 1810013H02Rik\|C76483\|MST-3\|Mst3\|STE20 | serine/threonine kinase 24 | protein-coding |
| chr17_28994677_28994980 | chr17 | 28994678 | 28994980 | + | -0.829994 | intron (NM_134115, intron 2 of 13) | intron (NM_134115, intron 2 of 13) | 13108 | NM_134115 | 106504 | Mm.439986 | NM_134115 | ENSMUSG00000024006 | **Stk38** | 5830476G13Rik\|9530097A09Rik\|AA617404\|Ndr1 | serine/threonine kinase 38 | protein-coding |
| chr2_68319564_68320286 | chr2 | 68319565 | 68320286 | + | -0.688086 | intron (NM_016866, intron 10 of 17) | intron (NM_016866, intron 10 of 17) | 152056 | NM_016866 | 53416 | Mm.198414 | NM_016866 | ENSMUSG00000027030 | **Stk39** | AW227544\|AW556857\|DCHT\|RF005\|Rnl5\|SPAK | serine/threonine kinase 39 | protein-coding |
| chr2_68319621_68320068 | chr2 | 68319622 | 68320068 | + | -0.750079 | intron (NM_016866, intron 10 of 17) | intron (NM_016866, intron 10 of 17) | 152136 | NM_016866 | 53416 | Mm.198414 | NM_016866 | ENSMUSG00000027030 | **Stk39** | AW227544\|AW556857\|DCHT\|RF005\|Rnl5\|SPAK | serine/threonine kinase 39 | protein-coding |
| chr2_68460833_68461367 | chr2 | 68460834 | 68461367 | + | -0.760109 | intron (NM_016866, intron 1 of 17) | MTEa\|LTR\|ERVL-MaLR | 10881 | NM_016866 | 53416 | Mm.198414 | NM_016866 | ENSMUSG00000027030 | **Stk39** | AW227544\|AW556857\|DCHT\|RF005\|Rnl5\|SPAK | serine/threonine kinase 39 | protein-coding |
| chr2_164084054_164084701 | chr2 | 164084055 | 164084701 | + | -0.724523 | intron (NM_021420, intron 3 of 10) | intron (NM_021420, intron 3 of 10) | 10200 | NM_021420 | 58231 | Mm.234472 | NM_021420 | ENSMUSG00000018209 | **Stk4** | AI447339\|AU020804\|Kas-2\|Mst1\|Ysk3 | serine/threonine kinase 4 | protein-coding |
| chr2_164075113_164075337 | chr2 | 164075114 | 164075337 | + | -0.800734 | intron (NM_021420, intron 1 of 10) | intron (NM_021420, intron 1 of 10) | 1047 | NM_021420 | 58231 | Mm.234472 | NM_021420 | ENSMUSG00000018209 | **Stk4** | AI447339\|AU020804\|Kas-2\|Mst1\|Ysk3 | serine/threonine kinase 4 | protein-coding |
| chr2_164079524_164079856 | chr2 | 164079525 | 164079856 | + | -0.897227 | intron (NM_021420, intron 2 of 10) | intron (NM_021420, intron 2 of 10) | 5512 | NM_021420 | 58231 | Mm.234472 | NM_021420 | ENSMUSG00000018209 | **Stk4** | AI447339\|AU020804\|Kas-2\|Mst1\|Ysk3 | serine/threonine kinase 4 | protein-coding |
| chr2_164079509_164079952 | chr2 | 164079510 | 164079952 | + | -0.927317 | intron (NM_021420, intron 2 of 10) | intron (NM_021420, intron 2 of 10) | 5553 | NM_021420 | 58231 | Mm.234472 | NM_021420 | ENSMUSG00000018209 | **Stk4** | AI447339\|AU020804\|Kas-2\|Mst1\|Ysk3 | serine/threonine kinase 4 | protein-coding |
| chr4_134460272_134460870 | chr4 | 134460273 | 134460870 | + | -0.750001 | Intergenic | RMER1C\|Other\|Other | -7749 | NM_019641 | 16765 | Mm.378957 | NM_019641 | ENSMUSG00000028832 | **Stmn1** | 19k\|Lag\|Lap18\|Op18\|P18\|P19\|Pig\|Pp17\|Pp18\|Pp19\|Pr22\|Smn | stathmin 1 | protein-coding |

| | | | | | | | | | | | | | | | | | |
|---|---|---|---|---|---|---|---|---|---|---|---|---|---|---|---|---|---|
| chr13_46339898_46340397 | chr13 | 46339899 | 46340397 | + | -0.70142 | Intergenic | ORR1D2\|LTR\|ERVL-MaLR | 66427 | NM_001005422 | 380842 | Mm.188108 | NM_001005422 | ENSMUSG00000063529 | **Stmnd1** | Gm1574 | stathmin domain containing 1 | protein-coding |
| chr17_88581393_88582122 | chr17 | 88581394 | 88582122 | + | -0.740944 | intron (NM_028658, intron 19 of 21) | intron (NM_028658, intron 19 of 21) | -44797 | NM_029858 | 77057 | Mm.482203 | NM_029858 | ENSMUSG00000033855 | **Ston1** | 4921524J06Rik\|C86231\|Salf\|Sblf | stonin 1 | protein-coding |
| chr3_139245267_139245783 | chr3 | 139245268 | 139245783 | + | -0.792539 | intron (NM_198659, intron 6 of 13) | Lx8\|LINE\|L1 | 39632 | NM_198659 | 381476 | Mm.188334 | NM_198659 | ENSMUSG00000047940 | **Stpg2** | B930007M17Rik\|Gm1017 | sperm tail PG rich repeat containing 2 | protein-coding |
| chr9_58139866_58140464 | chr9 | 58139867 | 58140464 | + | -0.691829 | intron (NM_009291, intron 4 of 18) | intron (NM_009291, intron 4 of 18) | 5500 | NM_001162479 | 20897 | Mm.10801 | NM_009291 | ENSMUSG00000032327 | **Stra6** | AI891933 | stimulated by retinoic acid gene 6 | protein-coding |
| chr4_48109076_48109344 | chr4 | 48109077 | 48109344 | + | -0.73814 | Intergenic | Intergenic | -15709 | NM_026343 | 67727 | Mm.171334 | NM_026343 | ENSMUSG00000061455 | **Stx17** | 4833418L03Rik\|6330411F21Rik\|9030425C21Rik\|AW048351 | syntaxin 17 | protein-coding |
| chr5_135024971_135025423 | chr5 | 135024972 | 135025423 | + | -0.693431 | intron (NM_016801, intron 1 of 9) | intron (NM_016801, intron 1 of 9) | 1625 | NM_016801 | 20907 | Mm.6225 | NM_016801 | ENSMUSG00000007207 | **Stx1a** | HPC-1 | syntaxin 1A (brain) | protein-coding |
| chr5_135024964_135025294 | chr5 | 135024965 | 135025294 | + | -0.709128 | intron (NM_016801, intron 1 of 9) | intron (NM_016801, intron 1 of 9) | 1557 | NM_016801 | 20907 | Mm.6225 | NM_016801 | ENSMUSG00000007207 | **Stx1a** | HPC-1 | syntaxin 1A (brain) | protein-coding |
| chr5_135024523_135025428 | chr5 | 135024524 | 135025428 | + | -0.92984 | intron (NM_016801, intron 1 of 9) | B1F\|SINE\|Alu | 1404 | NM_016801 | 20907 | Mm.6225 | NM_016801 | ENSMUSG00000007207 | **Stx1a** | HPC-1 | syntaxin 1A (brain) | protein-coding |
| chr5_128982879_128983783 | chr5 | 128982880 | 128983783 | + | -0.719753 | Intergenic | Intergenic | 25241 | NM_001286033 | 13852 | Mm.260716 | NM_007941 | ENSMUSG00000029428 | **Stx2** | AW538950\|Epim\|G1-536-1\|Syn-2\|repro34 | syntaxin 2 | protein-coding |
| chr19_11807281_11807774 | chr19 | 11807282 | 11807774 | + | -1.17563 | intron (NM_001286543, intron 1 of 11) | intron (NM_001286543, intron 1 of 11) | 11875 | NM_001286543 | 20908 | Mm.272264 | NM_011502 | ENSMUSG00000041488 | **Stx3** | Syn-3 | syntaxin 3 | protein-coding |
| chr2_32834516_32835021 | chr2 | 32834517 | 32835021 | + | -0.874008 | intron (NM_001113569, intron 1 of 19) | intron (NM_001113569, intron 1 of 19) | 12463 | NM_009295 | 20910 | Mm.278865 | NM_009295 | ENSMUSG00000026797 | **Stxbp1** | AI317162\|AI326233\|MMS10-G\|Ms10g\|Munc-18a\|Munc18-1\|N-sec1\|Rb-sec1\|Sxtbp1\|Unc18-1\|Unc18h\|nsec1 | syntaxin binding protein 1 | protein-coding |
| chr11_90538601_90538931 | chr11 | 90538602 | 90538931 | + | -0.738333 | intron (NM_011505, intron 14 of 17) | intron (NM_011505, intron 14 of 17) | 99342 | NM_011505 | 20913 | Mm.207203 | NM_011505 | ENSMUSG00000020546 | **Stxbp4** | 6030470M02Rik\|Synip | syntaxin binding protein 4 | protein-coding |
| chr10_10184352_10184759 | chr10 | 10184353 | 10184759 | + | -0.613905 | Intergenic | Intergenic | -283516 | NM_001081344 | 78808 | Mm.331751 | NM_030191 | ENSMUSG00000019790 | **Stxbp5** | 0710001E20Rik\|4930565N16Rik\|AW742610\|LGL3\|mKIAA1006 | syntaxin binding protein 5 (tomosyn) | protein-coding |
| chr15_12007927_12008421 | chr15 | 12007928 | 12008421 | + | -0.697008 | Intergenic | Intergenic | -12167 | NM_011294 | 20024 | Mm.41746 | NM_011294 | ENSMUSG00000022205 | **Sub1** | AI842364\|P15\|P9\|Pc4\|Rpo2tc1 | SUB1 homolog (S. cerevisiae) | protein-coding |
| chr5_117111745_117112102 | chr5 | 117111746 | 117112102 | + | -0.704282 | intron (NM_001122666, intron 2 of 12) | PB1D7\|SINE\|Alu | 4069 | NM_001122666 | 71954 | Mm.34456 | NM_178622 | ENSMUSG00000066900 | **Suds3** | 2400003N08Rik\|2410008L21Rik\|AU067672\|mSds3 | suppressor of defective silencing 3 homolog (S. cerevisiae) | protein-coding |
| chr14_79593970_79594396 | chr14 | 79593971 | 79594396 | + | -0.664285 | intron (NM_026474, intron 2 of 12) | intron (NM_026474, intron 2 of 12) | 6492 | NM_026474 | 67955 | Mm.18972 | NM_026474 | ENSMUSG00000022024 | **Sugt1** | 2410174K12Rik\|SGT1 | SGT1, suppressor of G2 allele of SKP1 (S. cerevisiae) | protein-coding |
| chr14_79558969_79559524 | chr14 | 79558970 | 79559524 | + | -0.855801 | intron (NM_007920, intron 2 of 8) | intron (NM_007920, intron 2 of 8) | -28444 | NM_026474 | 67955 | Mm.18972 | NM_026474 | ENSMUSG00000022024 | **Sugt1** | 2410174K12Rik\|SGT1 | SGT1, suppressor of G2 allele of SKP1 (S. cerevisiae) | protein-coding |
| chr14_79590844_79591423 | chr14 | 79590845 | 79591423 | + | -0.941152 | intron (NM_026474, intron 2 of 12) | intron (NM_026474, intron 2 of 12) | 3443 | NM_026474 | 67955 | Mm.18972 | NM_026474 | ENSMUSG00000022024 | **Sugt1** | 2410174K12Rik\|SGT1 | SGT1, suppressor of G2 allele of SKP1 (S. cerevisiae) | protein-coding |
| chr6_108168321_108168800 | chr6 | 108168322 | 108168800 | + | -0.703829 | intron (NM_145937, intron 3 of 8) | L1MB7\|LINE\|L1 | 17022 | NM_145937 | 58911 | Mm.439876 | NM_145937 | ENSMUSG00000030101 | **Sumf1** | AA543204\|AI463102\|AI851573\|FGE | sulfatase modifying factor 1 | protein-coding |
| chr1_59602249_59603116 | chr1 | 59602250 | 59603116 | + | -0.806728 | intron (NM_001013771, intron 12 of 19) | intron (NM_001013771, intron 12 of 19) | 68151 | NM_009460 | 22218 | Mm.362118 | NM_009460 | ENSMUSG00000026021 | **Sumo1** | GMP1\|PIC1\|SENTRIN\|SMT3\|SMT3H3\|SMTP3\|SUMO-1\|Smt3C\|Ubl1 | SMT3 suppressor of mif two 3 homolog 1 (yeast) | protein-coding |
| chr1_59602588_59603312 | chr1 | 59602589 | 59603312 | + | -0.870788 | intron (NM_001013771, intron 12 of 19) | intron (NM_001013771, intron 12 of 19) | 67884 | NM_009460 | 22218 | Mm.362118 | NM_009460 | ENSMUSG00000026021 | **Sumo1** | GMP1\|PIC1\|SENTRIN\|SMT3\|SMT3H3\|SMTP3\|SUMO-1\|Smt3C\|Ubl1 | SMT3 suppressor of mif two 3 homolog 1 (yeast) | protein-coding |
| chr15_79739387_79739934 | chr15 | 79739388 | 79739934 | + | -0.68027 | intron (NM_194342, intron 2 of 16) | intron (NM_194342, intron 2 of 16) | 2875 | NM_194342 | 223697 | Mm.202715 | NM_194342 | ENSMUSG00000042524 | **Sun2** | B230369L08Rik\|C030011B15\|Unc84b | Sad1 and UNC84 domain containing 2 | protein-coding |
| chr15_79739376_79739680 | chr15 | 79739377 | 79739680 | + | -0.773627 | intron (NM_194342, intron 2 of 16) | intron (NM_194342, intron 2 of 16) | 3008 | NM_194342 | 223697 | Mm.202715 | NM_194342 | ENSMUSG00000042524 | **Sun2** | B230369L08Rik\|C030011B15\|Unc84b | Sad1 and UNC84 domain containing 2 | protein-coding |
| chr14_52166585_52166962 | chr14 | 52166586 | 52166962 | + | -0.750107 | intron (NM_033618, intron 22 of 25) | intron (NM_033618, intron 22 of 25) | 30465 | NM_033618 | 114741 | Mm.21662 | NM_033618 | ENSMUSG00000035726 | **Supt16** | Cdc68\|Fact140\|Spt16\|Supt16h | suppressor of Ty 16 | protein-coding |
| chr3_54567136_54567689 | chr3 | 54567137 | 54567689 | + | -0.733352 | Intergenic | Intergenic | -125692 | NM_019995 | 56790 | Mm.486375 | NM_019995 | ENSMUSG00000027751 | **Supt20** | AA667204\|AI450544\|D3Ertd300e\|Fam48a\|Supt20h\|p38IP | suppressor of Ty 20 | protein-coding |
| chr11_87734465_87734834 | chr11 | 87734466 | 87734834 | + | -0.629644 | 3' UTR (NM_172448, exon 10 of 10) | 3' UTR (NM_172448, exon 10 of 10) | -2915 | NM_009296 | 20922 | Mm.622 | NM_009296 | ENSMUSG00000020485 | **Supt4a** | AL022777\|Supt4h\|Supt4h1\|Supt4h1a | suppressor of Ty 4A | protein-coding |
| chr13_49242450_49243175 | chr13 | 49242451 | 49243175 | + | -0.622682 | intron (NM_028340, intron 1 of 4) | intron (NM_028340, intron 1 of 4) | 5350 | NM_025491 | 66329 | Mm.34308 | NM_025491 | ENSMUSG00000021384 | **Susd3** | 1700017I11Rik\|2810440J20Rik | sushi domain containing 3 | protein-coding |
| chr13_49267682_49268250 | chr13 | 49267683 | 49268250 | + | -0.799704 | intron (NM_015759, intron 15 of 17) | ORR1E\|LTR\|ERVL-MaLR | -19260 | NM_028340 | 66329 | Mm.34308 | NM_025491 | ENSMUSG00000021384 | **Susd3** | 1700017I11Rik\|2810440J20Rik | sushi domain containing 3 | protein-coding |
| chr13_49267748_49267978 | chr13 | 49267749 | 49267978 | + | -0.8221 | intron (NM_015759, intron 15 of 17) | ORR1E\|LTR\|ERVL-MaLR | -19157 | NM_028340 | 66329 | Mm.34308 | NM_025491 | ENSMUSG00000021384 | **Susd3** | 1700017I11Rik\|2810440J20Rik | sushi domain containing 3 | protein-coding |
| chr10_78594596_78595032 | chr10 | 78594597 | 78595032 | + | -1.26744 | Intergenic | Intergenic | -2850 | NM_027875 | 71709 | Mm.86439 | NM_027875 | ENSMUSG00000032714 | **Syde1** | 1200008N06Rik\|BB043000\|mSYD1A | synapse defective 1, Rho GTPase, homolog 1 (C. elegans) | protein-coding |
| chr3_146004377_146005191 | chr3 | 146004378 | 146005191 | + | -0.68175 | intron (NM_001166064, intron 3 of 6) | intron (NM_001166064, intron 3 of 6) | 16914 | NM_001166064 | 214804 | Mm.268854 | NM_001166064 | ENSMUSG00000036863 | **Syde2** | C430017H16Rik | synapse defective 1, Rho GTPase, homolog 2 (C. elegans) | protein-coding |
| chr4_134998004_134998372 | chr4 | 134998005 | 134998372 | + | -0.876853 | Intergenic | ORR1E\|LTR\|ERVL-MaLR | 67208 | NM_026780 | 68592 | Mm.29989 | NM_026780 | ENSMUSG00000028821 | **Syf2** | 1110018L13Rik\|Cbpin\|D4Bwg1551e\|Gcipip\|Ntc31\|mp29\|p29 | SYF2 homolog, RNA splicing factor (S. cerevisiae) | protein-coding |
| chr10_5187840_5188397 | chr10 | 5187841 | 5188397 | + | -0.660926 | intron (NM_001079686, intron 2 of 54) | intron (NM_001079686, intron 2 of 54) | 6588 | NM_001079686 | 64009 | Mm.331626 | NM_022027 | ENSMUSG00000019769 | **Syne1** | 8B\|A330049M09Rik\|BE692247\|C130039F11Rik\|CPG2\|Myne1\|mKIAA1756 | spectrin repeat containing, nuclear envelope 1 | protein-coding |
| chr10_5079721_5080437 | chr10 | 5079722 | 5080437 | + | -0.775504 | intron (NM_001079686, intron 37 of 54) | intron (NM_001079686, intron 37 of 54) | -11016 | NM_022027 | 64009 | Mm.331626 | NM_022027 | ENSMUSG00000019769 | **Syne1** | 8B\|A330049M09Rik\|BE692247\|C130039F11Rik\|CPG2\|Myne1\|mKIAA1756 | spectrin repeat containing, nuclear envelope 1 | protein-coding |
| chr10_5448092_5448953 | chr10 | 5448093 | 5448953 | + | -0.92029 | intron (NM_153399, intron 3 of 31) | intron (NM_153399, intron 3 of 31) | 102169 | NM_153399 | 64009 | Mm.331626 | NM_022027 | ENSMUSG00000019769 | **Syne1** | 8B\|A330049M09Rik\|BE692247\|C130039F11Rik\|CPG2\|Myne1\|mKIAA1756 | spectrin repeat containing, nuclear envelope 1 | protein-coding |
| chr10_5384043_5384836 | chr10 | 5384044 | 5384836 | + | -1.05359 | intron (NM_153399, intron 13 of 31) | intron (NM_153399, intron 13 of 31) | 166252 | NM_153399 | 64009 | Mm.331626 | NM_022027 | ENSMUSG00000019769 | **Syne1** | 8B\|A330049M09Rik\|BE692247\|C130039F11Rik\|CPG2\|Myne1\|mKIAA1756 | spectrin repeat containing, nuclear envelope 1 | protein-coding |
| chr10_5197347_5197722 | chr10 | 5197348 | 5197722 | + | -1.20407 | Intergenic | Intergenic | -2828 | NM_001079686 | 64009 | Mm.331626 | NM_022027 | ENSMUSG00000019769 | **Syne1** | 8B\|A330049M09Rik\|BE692247\|C130039F11Rik\|CPG2\|Myne1\|mKIAA1756 | spectrin repeat containing, nuclear envelope 1 | protein-coding |
| chr7_90341533_90342103 | chr7 | 90341534 | 90342103 | + | -0.698981 | intron (NR_110348, intron 1 of 8) | intron (NR_110348, intron 1 of 8) | 39463 | NR_110348 | 83671 | Mm.26751 | NM_031394 | ENSMUSG00000030616 | **Sytl2** | AI266830\|Slp2\|mKIAA1597 | synaptotagmin-like 2 | protein-coding |
| chr7_90286155_90286483 | chr7 | 90286156 | 90286483 | + | -0.777968 | Intergenic | Intergenic | -16036 | NM_001289583 | 83671 | Mm.26751 | NM_031394 | ENSMUSG00000030616 | **Sytl2** | AI266830\|Slp2\|mKIAA1597 | synaptotagmin-like 2 | protein-coding |
| chr7_90341534_90341913 | chr7 | 90341535 | 90341913 | + | -0.787219 | intron (NR_110348, intron 1 of 8) | intron (NR_110348, intron 1 of 8) | 39369 | NR_110348 | 83671 | Mm.26751 | NM_031394 | ENSMUSG00000030616 | **Sytl2** | AI266830\|Slp2\|mKIAA1597 | synaptotagmin-like 2 | protein-coding |
| chr7_90302792_90303371 | chr7 | 90302793 | 90303371 | + | -0.91632 | intron (NR_110348, intron 1 of 8) | intron (NR_110348, intron 1 of 8) | 727 | NR_110348 | 83671 | Mm.26751 | NM_031394 | ENSMUSG00000030616 | **Sytl2** | AI266830\|Slp2\|mKIAA1597 | synaptotagmin-like 2 | protein-coding |
| chr7_90353643_90354251 | chr7 | 90353644 | 90354251 | + | -0.983609 | intron (NR_110348, intron 2 of 8) | intron (NR_110348, intron 2 of 8) | -33034 | NM_001040087 | 83671 | Mm.26751 | NM_031394 | ENSMUSG00000030616 | **Sytl2** | AI266830\|Slp2\|mKIAA1597 | synaptotagmin-like 2 | protein-coding |
| chr17_6686629_6686863 | chr17 | 6686630 | 6686863 | + | -0.948138 | intron (NM_031395, intron 2 of 14) | MTD\|LTR\|ERVL-MaLR | -2326 | NM_183370 | 83672 | Mm.442201 | NM_031395 | ENSMUSG00000041831 | **Sytl3** | A430092N21\|Slp3\|Slp3-a\|Slp3-b\|mFLJ00307 | synaptotagmin-like 3 | protein-coding |
| chr15_80149241_80149972 | chr15 | 80149242 | 80149972 | + | -0.72702 | intron (NM_025609, intron 3 of 10) | intron (NM_025609, intron 3 of 10) | 16453 | NM_025609 | 66513 | Mm.288245 | NM_025609 | ENSMUSG00000022414 | **Tab1** | 2310012M03Rik\|Map3k7ip1\|b2b449Clo | TGF-beta activated kinase 1/MAP3K7 binding protein 1 | protein-coding |
| chr10_8065716_8066144 | chr10 | 8065717 | 8066144 | + | -0.628528 | Intergenic | Intergenic | -109807 | NM_138667 | 68652 | Mm.193041 | NM_138667 | ENSMUSG00000015755 | **Tab2** | 1110030N06Rik\|A530078N03Rik\|Map3k7ip2\|mKIAA0733 | TGF-beta activated kinase 1/MAP3K7 binding protein 2 | protein-coding |
| chr10_7976383_7976894 | chr10 | 7976384 | 7976894 | + | -1.06384 | Intergenic | Intergenic | -20516 | NM_138667 | 68652 | Mm.193041 | NM_138667 | ENSMUSG00000015755 | **Tab2** | 1110030N06Rik\|A530078N03Rik\|Map3k7ip2\|mKIAA0733 | TGF-beta activated kinase 1/MAP3K7 binding protein 2 | protein-coding |

| | | | | | | | | | | | | | | | | | |
|---|---|---|---|---|---|---|---|---|---|---|---|---|---|---|---|---|---|
| chr11_106056611_106057465 | chr11 | 106056612 | 106057465 | + | -0.952259 | 3' UTR (NM_027946, exon 7 of 7) | 3' UTR (NM_027946, exon 7 of 7) | -9069 | NM_027346 | 70207 | Mm.343857 | NM_027346 | ENSMUSG00000001983 | **Taco1** | 2310066I18Rik\|Ccdc44 | translational activator of mitochondrially encoded cytochrome c oxidase I | protein-coding |
| chr11_106056867_106057494 | chr11 | 106056868 | 106057494 | + | -0.994192 | 3' UTR (NM_027946, exon 7 of 7) | 3' UTR (NM_027946, exon 7 of 7) | -8926 | NM_027346 | 70207 | Mm.343857 | NM_027346 | ENSMUSG00000001983 | **Taco1** | 2310066I18Rik\|Ccdc44 | translational activator of mitochondrially encoded cytochrome c oxidase I | protein-coding |
| chr2_9997549_9997959 | chr2 | 9997550 | 9997959 | + | -0.834936 | intron (NM_027748, intron 2 of 6) | intron (NM_027748, intron 2 of 6) | 50855 | NM_027748 | 209361 | Mm.86343 | NM_027748 | ENSMUSG00000025782 | **Taf3** | 140kDa\|4933439M23Rik\|AW539625\|TAF140\|TAFII-140\|TAFII140\|mTAFII140 | TAF3 RNA polymerase II, TATA box binding protein (TBP)-associated factor | protein-coding |
| chr17_7927593_7928144 | chr17 | 7927594 | 7928144 | + | -0.691467 | intron (NM_145968, intron 4 of 9) | intron (NM_145968, intron 4 of 9) | 1869 | NM_145968 | 72536 | Mm.119680 | NM_145968 | ENSMUSG00000033450 | **Tagap** | 2610315E15Rik | T cell activation Rho GTPase activating protein | protein-coding |
| chr17_7927868_7928143 | chr17 | 7927869 | 7928143 | + | -0.697889 | intron (NM_145968, intron 4 of 9) | intron (NM_145968, intron 4 of 9) | 2006 | NM_145968 | 72536 | Mm.119680 | NM_145968 | ENSMUSG00000033450 | **Tagap** | 2610315E15Rik | T cell activation Rho GTPase activating protein | protein-coding |
| chr17_7902393_7902718 | chr17 | 7902394 | 7902718 | + | -0.748993 | Intergenic | Intergenic | -23444 | NM_145968 | 72536 | Mm.119680 | NM_145968 | ENSMUSG00000033450 | **Tagap** | 2610315E15Rik | T cell activation Rho GTPase activating protein | protein-coding |
| chr17_7927386_7928151 | chr17 | 7927387 | 7928151 | + | -0.786506 | intron (NM_145968, intron 4 of 9) | intron (NM_145968, intron 4 of 9) | 1769 | NM_145968 | 72536 | Mm.119680 | NM_145968 | ENSMUSG00000033450 | **Tagap** | 2610315E15Rik | T cell activation Rho GTPase activating protein | protein-coding |
| chr17_7884574_7884886 | chr17 | 7884575 | 7884886 | + | -0.868241 | Intergenic | Intergenic | -41270 | NM_145968 | 72536 | Mm.119680 | NM_145968 | ENSMUSG00000033450 | **Tagap** | 2610315E15Rik | T cell activation Rho GTPase activating protein | protein-coding |
| chr17_7927325_7928600 | chr17 | 7927326 | 7928600 | + | -1.10005 | intron (NM_145968, intron 4 of 9) | intron (NM_145968, intron 4 of 9) | 1963 | NM_145968 | 72536 | Mm.119680 | NM_145968 | ENSMUSG00000033450 | **Tagap** | 2610315E15Rik | T cell activation Rho GTPase activating protein | protein-coding |
| chr17_7928246_7928465 | chr17 | 7928247 | 7928465 | + | -1.4498 | intron (NM_145968, intron 4 of 9) | intron (NM_145968, intron 4 of 9) | 2356 | NM_145968 | 72536 | Mm.119680 | NM_145968 | ENSMUSG00000033450 | **Tagap** | 2610315E15Rik | T cell activation Rho GTPase activating protein | protein-coding |
| chr2_59631197_59631791 | chr2 | 59631198 | 59631791 | + | -0.721888 | intron (NM_198294, intron 1 of 26) | MTD\|LTR\|ERVL-MaLR | -15275 | NM_001290659 | 66860 | Mm.27917 | NM_198294 | ENSMUSG00000035168 | **Tanc1** | 1200003E16Rik\|Tanc\|mKIAA1728 | tetratricopeptide repeat, ankyrin repeat and coiled-coil containing 1 | protein-coding |
| chr2_59716517_59716855 | chr2 | 59716518 | 59716855 | + | -0.727191 | intron (NM_001290659, intron 2 of 25) | intron (NM_001290659, intron 2 of 25) | 69917 | NM_001290659 | 66860 | Mm.27917 | NM_198294 | ENSMUSG00000035168 | **Tanc1** | 1200003E16Rik\|Tanc\|mKIAA1728 | tetratricopeptide repeat, ankyrin repeat and coiled-coil containing 1 | protein-coding |
| chr2_59716332_59716870 | chr2 | 59716333 | 59716870 | + | -0.749122 | intron (NM_001290659, intron 2 of 25) | MLT1C\|LTR\|ERVL-MaLR | 69832 | NM_001290659 | 66860 | Mm.27917 | NM_198294 | ENSMUSG00000035168 | **Tanc1** | 1200003E16Rik\|Tanc\|mKIAA1728 | tetratricopeptide repeat, ankyrin repeat and coiled-coil containing 1 | protein-coding |
| chr8_106643043_106643538 | chr8 | 106643044 | 106643538 | + | -1.25528 | intron (NM_009864, intron 2 of 15) | intron (NM_009864, intron 2 of 15) | -39777 | NM_173037 | 272538 | Mm.87151 | NM_173037 | ENSMUSG00000041949 | **Tango6** | AW413431\|E330010G16\|Tmco7 | transport and golgi organization 6 | protein-coding |
| chr2_61513985_61514841 | chr2 | 61513986 | 61514841 | + | -0.702386 | Intergenic | Intergenic | -64173 | NM_011529 | 21353 | Mm.244393 | NM_011529 | ENSMUSG00000064289 | **Tank** | C86182\|E430026L09Rik\|I-TRAF | TRAF family member-associated Nf-kappa B activator | protein-coding |
| chr2_61513971_61514559 | chr2 | 61513972 | 61514559 | + | -0.751426 | Intergenic | Intergenic | -64321 | NM_011529 | 21353 | Mm.244393 | NM_011529 | ENSMUSG00000064289 | **Tank** | C86182\|E430026L09Rik\|I-TRAF | TRAF family member-associated Nf-kappa B activator | protein-coding |
| chr17_34211801_34212179 | chr17 | 34211802 | 34212179 | + | -0.744148 | exon (NM_011530, exon 6 of 12) | exon (NM_011530, exon 6 of 12) | 7511 | NM_011530 | 21355 | Mm.14814 | NM_011530 | | **Tap2** | ABC18\|AI462429\|APT2\|Abcb3\|Ham-2\|Ham2\|MTP2\|PSF2\|RING11\|Tap-2\|Y1\|jas | transporter 2, ATP-binding cassette, sub-family B (MDR/TAP) | protein-coding |
| chr5_44239074_44239517 | chr5 | 44239075 | 44239517 | + | -0.623184 | Intergenic | Intergenic | -12690 | NM_173764 | 231225 | Mm.30688 | NM_173764 | ENSMUSG00000046985 | **Tapt1** | 4932414K18Rik | transmembrane anterior posterior transformation 1 | protein-coding |
| chr5_44239250_44239510 | chr5 | 44239251 | 44239510 | + | -0.691034 | Intergenic | Intergenic | -12774 | NM_173764 | 231225 | Mm.30688 | NM_173764 | ENSMUSG00000046985 | **Tapt1** | 4932414K18Rik | transmembrane anterior posterior transformation 1 | protein-coding |
| chr5_44220437_44220808 | chr5 | 44220438 | 44220808 | + | -0.745396 | intron (NM_173764, intron 1 of 13) | intron (NM_173764, intron 1 of 13) | 5983 | NM_173764 | 231225 | Mm.30688 | NM_173764 | ENSMUSG00000046985 | **Tapt1** | 4932414K18Rik | transmembrane anterior posterior transformation 1 | protein-coding |
| chr5_44180835_44181173 | chr5 | 44180836 | 44181173 | + | -0.746909 | intron (NM_173764, intron 12 of 13) | intron (NM_173764, intron 12 of 13) | 45602 | NM_173764 | 231225 | Mm.30688 | NM_173764 | ENSMUSG00000046985 | **Tapt1** | 4932414K18Rik | transmembrane anterior posterior transformation 1 | protein-coding |
| chr5_44223227_44223602 | chr5 | 44223228 | 44223602 | + | -0.770772 | intron (NM_173764, intron 1 of 13) | intron (NM_173764, intron 1 of 13) | 3191 | NM_173764 | 231225 | Mm.30688 | NM_173764 | ENSMUSG00000046985 | **Tapt1** | 4932414K18Rik | transmembrane anterior posterior transformation 1 | protein-coding |
| chr15_11532952_11533307 | chr15 | 11532953 | 11533307 | + | -0.705248 | Intergenic | Intergenic | -133472 | NM_033074 | 110960 | Mm.286061 | NM_033074 | ENSMUSG00000022241 | **Tars** | D15Wsu59e\|ThrRS | threonyl-tRNA synthetase | protein-coding |
| chr2_139934542_139935144 | chr2 | 139934543 | 139935144 | + | -0.85642 | intron (NR_110351, intron 10 of 13) | intron (NR_110351, intron 10 of 13) | 131962 | NM_001159640 | 75812 | Mm.293449 | NM_175225 | ENSMUSG00000039033 | **Tasp1** | 4930485D02Rik\|AW986064 | taspase, threonine aspartase 1 | protein-coding |
| chr2_139924885_139925416 | chr2 | 139924886 | 139925416 | + | -0.915067 | intron (NR_110351, intron 10 of 13) | intron (NR_110351, intron 10 of 13) | 141654 | NM_001159640 | 75812 | Mm.293449 | NM_175225 | ENSMUSG00000039033 | **Tasp1** | 4930485D02Rik\|AW986064 | taspase, threonine aspartase 1 | protein-coding |
| chr5_64342036_64342435 | chr5 | 64342037 | 64342435 | + | -0.74409 | intron (NM_019636, intron 18 of 19) | intron (NM_019636, intron 18 of 19) | 111879 | NM_001289514 | 57915 | Mm.286353 | NM_019636 | ENSMUSG00000029174 | **Tbc1d1** | 1110062G02Rik\|AI385682\|AW555803\|Tbc1\|mKIAA1108 | TBC1 domain family, member 1 | protein-coding |
| chr5_64342163_64342480 | chr5 | 64342164 | 64342480 | + | -0.75976 | intron (NM_019636, intron 18 of 19) | intron (NM_019636, intron 18 of 19) | 111965 | NM_001289514 | 57915 | Mm.286353 | NM_019636 | ENSMUSG00000029174 | **Tbc1d1** | 1110062G02Rik\|AI385682\|AW555803\|Tbc1\|mKIAA1108 | TBC1 domain family, member 1 | protein-coding |
| chr5_64149410_64150380 | chr5 | 64149411 | 64150380 | + | -0.768119 | Intergenic | Intergenic | -10365 | NM_019636 | 57915 | Mm.286353 | NM_019636 | ENSMUSG00000029174 | **Tbc1d1** | 1110062G02Rik\|AI385682\|AW555803\|Tbc1\|mKIAA1108 | TBC1 domain family, member 1 | protein-coding |
| chr5_64157991_64158262 | chr5 | 64157992 | 64158262 | + | -0.805256 | Intergenic | Intergenic | -2133 | NM_019636 | 57915 | Mm.286353 | NM_019636 | ENSMUSG00000029174 | **Tbc1d1** | 1110062G02Rik\|AI385682\|AW555803\|Tbc1\|mKIAA1108 | TBC1 domain family, member 1 | protein-coding |
| chr5_64342162_64342743 | chr5 | 64342163 | 64342743 | + | -0.875771 | intron (NM_019636, intron 18 of 19) | intron (NM_019636, intron 18 of 19) | 112096 | NM_001289514 | 57915 | Mm.286353 | NM_019636 | ENSMUSG00000029174 | **Tbc1d1** | 1110062G02Rik\|AI385682\|AW555803\|Tbc1\|mKIAA1108 | TBC1 domain family, member 1 | protein-coding |
| chr19_4184814_4185839 | chr19 | 4184815 | 4185839 | + | -0.868265 | intron (NM_178650, intron 9 of 9) | intron (NM_178650, intron 9 of 9) | 5720 | NM_178650 | 108995 | Mm.288312 | NM_178650 | ENSMUSG00000040247 | **Tbc1d10c** | 1810062O14Rik\|AI428527 | TBC1 domain family, member 10c | protein-coding |
| chr5_36538567_36539165 | chr5 | 36538568 | 36539165 | + | -0.611901 | intron (NM_001113362, intron 4 of 14) | L1MEf\|LINE\|L1 | -8243 | NM_001113364 | 100855 | Mm.486203 | NM_133910 | ENSMUSG00000029192 | **Tbc1d14** | 2810413P16Rik\|AU043625\|C86258\|D5Ertd110e\|mKIAA1322 | TBC1 domain family, member 14 | protein-coding |
| chr15_86342930_86343387 | chr15 | 86342931 | 86343387 | + | -0.787636 | intron (NM_145476, intron 8 of 12) | intron (NM_145476, intron 8 of 12) | 128700 | NM_145476 | 223754 | Mm.28904 | NM_145476 | ENSMUSG00000051864 | **Tbc1d22a** | BC023106\|D15Ertd781e | TBC1 domain family, member 22a | protein-coding |
| chr17_29565552_29565847 | chr17 | 29565553 | 29565847 | + | -0.907456 | intron (NM_198647, intron 2 of 12) | B3A\|SINE\|B2 | 15898 | NM_198647 | 381085 | Mm.328934 | NM_198647 | ENSMUSG00000042203 | **Tbc1d22b** | BC045600 | TBC1 domain family, member 22B | protein-coding |
| chr17_29565397_29565841 | chr17 | 29565398 | 29565841 | + | -0.920477 | intron (NM_198647, intron 2 of 12) | intron (NM_198647, intron 2 of 12) | 15817 | NM_198647 | 381085 | Mm.328934 | NM_198647 | ENSMUSG00000042203 | **Tbc1d22b** | BC045600 | TBC1 domain family, member 22B | protein-coding |
| chr16_57243186_57243864 | chr16 | 57243187 | 57243864 | + | -0.671216 | Intergenic | Intergenic | -12059 | NM_026254 | 67581 | Mm.257819 | NM_026254 | ENSMUSG00000022749 | **Tbc1d23** | 4930451A13Rik\|AU015720\|AU043671\|AU043778\|D030022P07Rik\|W51689 | TBC1 domain family, member 23 | protein-coding |
| chr15_57938878_57939320 | chr15 | 57938879 | 57939320 | + | -0.83837 | intron (NM_001081396, intron 7 of 20) | intron (NM_001081396, intron 7 of 20) | 26900 | NM_001167679 | 210544 | Mm.390835 | NM_199300 | ENSMUSG00000022364 | **Tbc1d31** | 4B3\|D330013L20Rik\|Gm85\|Wdr67 | TBC1 domain family, member 31 | protein-coding |
| chr14_101477798_101478203 | chr14 | 101477799 | 101478203 | + | -0.943697 | intron (NM_001081278, intron 9 of 18) | intron (NM_001081278, intron 9 of 18) | 131190 | NM_001081278 | 210789 | Mm.320639 | NM_173380 | ENSMUSG00000033083 | **Tbc1d4** | 5930406J04Rik\|A930035N22\|AV295684\|As160 | TBC1 domain family, member 4 | protein-coding |
| chr17_46870857_46871286 | chr17 | 46870858 | 46871286 | + | -0.759963 | Intergenic | (TGGG)n\|Simple_repeat\|Simple_repeat | -19549 | NM_178385 | 72726 | Mm.33780 | NM_178385 | ENSMUSG00000036430 | **Tbcc** | 2810055C19Rik | tubulin-specific chaperone C | protein-coding |
| chr17_46870258_46870814 | chr17 | 46870259 | 46870814 | + | -0.820389 | Intergenic | Intergenic | -20085 | NM_178385 | 72726 | Mm.33780 | NM_178385 | ENSMUSG00000036430 | **Tbcc** | 2810055C19Rik | tubulin-specific chaperone C | protein-coding |
| chr17_46870290_46871251 | chr17 | 46870291 | 46871251 | + | -0.843533 | Intergenic | Intergenic | -19850 | NM_178385 | 72726 | Mm.33780 | NM_178385 | ENSMUSG00000036430 | **Tbcc** | 2810055C19Rik | tubulin-specific chaperone C | protein-coding |
| chr9_42673437_42674400 | chr9 | 42673438 | 42674400 | + | -0.739607 | intron (NM_175481, intron 5 of 20) | intron (NM_175481, intron 5 of 20) | -201693 | NM_173038 | 272589 | Mm.34483 | NM_173038 | ENSMUSG00000037287 | **Tbcel** | D330014K23\|E130107N23Rik\|Lrrc35 | tubulin folding cofactor E-like | protein-coding |
| chr9_42469380_42469676 | chr9 | 42469381 | 42469676 | + | -0.846094 | intron (NM_173038, intron 1 of 7) | intron (NM_173038, intron 1 of 7) | 2698 | NM_173038 | 272589 | Mm.34483 | NM_173038 | ENSMUSG00000037287 | **Tbcel** | D330014K23\|E130107N23Rik\|Lrrc35 | tubulin folding cofactor E-like | protein-coding |
| chr9_42465478_42466222 | chr9 | 42465479 | 42466222 | + | -1.13819 | intron (NM_173038, intron 1 of 7) | intron (NM_173038, intron 1 of 7) | 6376 | NM_173038 | 272589 | Mm.34483 | NM_173038 | ENSMUSG00000037287 | **Tbcel** | D330014K23\|E130107N23Rik\|Lrrc35 | tubulin folding cofactor E-like | protein-coding |
| chr9_42488060_42488468 | chr9 | 42488061 | 42488468 | + | -1.42913 | Intergenic | ORR1E\|LTR\|ERVL-MaLR | -16038 | NM_173038 | 272589 | Mm.34483 | NM_173038 | ENSMUSG00000037287 | **Tbcel** | D330014K23\|E130107N23Rik\|Lrrc35 | tubulin folding cofactor E-like | protein-coding |
| chrX_77580017_77580318 | chrX | 77580018 | 77580318 | + | -0.843181 | intron (NM_020601, intron 1 of 15) | intron (NM_020601, intron 1 of 15) | 68941 | NM_020601 | 21372 | Mm.258476 | NM_020601 | ENSMUSG00000025246 | **Tbl1x** | 5330429M20\|Tbl1 | transducin (beta)-like 1 X-linked | protein-coding |
| chrX_77579864_77580346 | chrX | 77579865 | 77580346 | + | -0.866477 | intron (NM_020601, intron 1 of 15) | intron (NM_020601, intron 1 of 15) | 68878 | NM_020601 | 21372 | Mm.258476 | NM_020601 | ENSMUSG00000025246 | **Tbl1x** | 5330429M20\|Tbl1 | transducin (beta)-like 1 X-linked | protein-coding |

| | | | | | | | | | | | | | | | | | |
|---|---|---|---|---|---|---|---|---|---|---|---|---|---|---|---|---|---|
| chr3_21910899_21911497 | chr3 | 21910900 | 21911497 | + | -0.669923 | Intergenic | Intergenic | -165454 | NM_030732 | 81004 | Mm.202966 | NM_030732 | ENSMUSG00000027630 | **Tbl1xr1** | 8030499H02Rik\|A630076E03Rik\|AW539987\|C21\|C230089I12Rik\|DC42\|Ira1\|TBLR1 | transducin (beta)-like 1X-linked receptor 1 | protein-coding |
| chr3_21986231_21986449 | chr3 | 21986232 | 21986449 | + | -0.914987 | Intergenic | Intergenic | -90312 | NM_030732 | 81004 | Mm.202966 | NM_030732 | ENSMUSG00000027630 | **Tbl1xr1** | 8030499H02Rik\|A630076E03Rik\|AW539987\|C21\|C230089I12Rik\|DC42\|Ira1\|TBLR1 | transducin (beta)-like 1X-linked receptor 1 | protein-coding |
| chr3_22152847_22153099 | chr3 | 22152848 | 22153099 | + | -1.07804 | intron (NM_030732, intron 2 of 15) | intron (NM_030732, intron 2 of 15) | 76321 | NM_030732 | 81004 | Mm.202966 | NM_030732 | ENSMUSG00000027630 | **Tbl1xr1** | 8030499H02Rik\|A630076E03Rik\|AW539987\|C21\|C230089I12Rik\|DC42\|Ira1\|TBLR1 | transducin (beta)-like 1X-linked receptor 1 | protein-coding |
| chr3_22152748_22153107 | chr3 | 22152749 | 22153107 | + | -1.08909 | intron (NM_030732, intron 2 of 15) | intron (NM_030732, intron 2 of 15) | 76276 | NM_030732 | 81004 | Mm.202966 | NM_030732 | ENSMUSG00000027630 | **Tbl1xr1** | 8030499H02Rik\|A630076E03Rik\|AW539987\|C21\|C230089I12Rik\|DC42\|Ira1\|TBLR1 | transducin (beta)-like 1X-linked receptor 1 | protein-coding |
| chr11_6627854_6628211 | chr11 | 6627855 | 6628211 | + | -0.998025 | Intergenic | RMER6A\|LTR\|ERVK | -1966 | NM_134011 | 21379 | Mm.247761 | NM_134011 | ENSMUSG00000000384 | **Tbrg4** | 2310042P22Rik\|AA120735\|AA408001\|AI527316\|Cpr2\|R74877\|Tb-12\|mKIAA0948 | transforming growth factor beta regulated gene 4 | protein-coding |
| chr11_6627908_6628145 | chr11 | 6627909 | 6628145 | + | -1.01426 | Intergenic | RMER6A\|LTR\|ERVK | -1960 | NM_134011 | 21379 | Mm.247761 | NM_134011 | ENSMUSG00000000384 | **Tbrg4** | 2310042P22Rik\|AA120735\|AA408001\|AI527316\|Cpr2\|R74877\|Tb-12\|mKIAA0948 | transforming growth factor beta regulated gene 4 | protein-coding |
| chr1_165151193_165151852 | chr1 | 165151194 | 165151852 | + | -0.656555 | intron (NM_032005, intron 3 of 7) | ORR1G\|LTR\|ERVL-MaLR | 9250 | NM_032005 | 83993 | Mm.190649 | NM_032005 | ENSMUSG00000026572 | **Tbx19** | AU019250\|D1Ertd754e\|Tpit | T-box 19 | protein-coding |
| chr1_165151198_165151618 | chr1 | 165151199 | 165151618 | + | -0.674721 | intron (NM_032005, intron 3 of 7) | intron (NM_032005, intron 3 of 7) | 9365 | NM_032005 | 83993 | Mm.190649 | NM_032005 | ENSMUSG00000026572 | **Tbx19** | AU019250\|D1Ertd754e\|Tpit | T-box 19 | protein-coding |
| chrX_136615922_136616230 | chrX | 136615923 | 136616230 | + | -0.865418 | Intergenic | Intergenic | -50299 | NM_001029978 | 594844 | Mm.260515 | NM_001029978 | ENSMUSG00000044550 | **Tceal3** | 1100001D19Rik | transcription elongation factor A (SII)-like 3 | protein-coding |
| chr1_16641904_16642657 | chr1 | 16641905 | 16642657 | + | -0.930495 | TTS (NM_026456) | TTS (NM_026456) | 14584 | NM_026456 | 67923 | Mm.289248 | NM_026456 | ENSMUSG00000079658 | **Tceb1** | 2610043E24Rik\|2610301I15Rik\|AA407206\|AI987979\|AW049146 | transcription elongation factor B (SIII), polypeptide 1 | protein-coding |
| chr9_72092178_72092508 | chr9 | 72092179 | 72092508 | + | -0.881883 | intron (NM_001253862, intron 3 of 19) | MLT1A1\|LTR\|ERVL-MaLR | 18858 | NM_001253862 | 21406 | Mm.171615 | NM_011544 | ENSMUSG00000032228 | **Tcf12** | A130037E08Rik\|ALF1\|HEB\|HEBAlt\|HTF-4\|HTF4\|ME1\|REB\|bHLHb20 | transcription factor 12 | protein-coding |
| chr15_82914395_82914895 | chr15 | 82914396 | 82914895 | + | -1.02036 | Intergenic | MIR\|SINE\|MIR | -2511 | NM_001114140 | 21411 | Mm.252156 | NM_013836 | ENSMUSG00000041852 | **Tcf20** | 2810438H08Rik\|SPBP\|mKIAA0292 | transcription factor 20 | protein-coding |
| chr8_123384916_123385586 | chr8 | 123384917 | 123385586 | + | -1.18756 | intron (NM_001037878, intron 5 of 17) | intron (NM_001037878, intron 5 of 17) | 11540 | NM_001286363 | 66855 | Mm.178818 | NM_025804 | ENSMUSG00000001472 | **Tcf25** | 1100001J13Rik\|1810041K11Rik\|D8Ertd325e\|Nulp1\|mKIAA1049 | transcription factor 25 (basic helix-loop-helix) | protein-coding |
| chr18_69365400_69365705 | chr18 | 69365401 | 69365705 | + | -0.674367 | intron (NM_001083967, intron 3 of 19) | intron (NM_001083967, intron 3 of 19) | 19832 | NM_001083967 | 21413 | Mm.4269 | NM_013685 | ENSMUSG00000053477 | **Tcf4** | 5730422P05Rik\|ASP-I2\|E2-2\|E2.2\|ITF-2\|ITF-2b\|ITF2\|ME2\|MITF-2A\|MITF-2B\|SEF-2\|SEF2\|SEF2-1\|TFE\|Tcf-4\|bHLHb19 | transcription factor 4 | protein-coding |
| chr11_52321623_52321886 | chr11 | 52321624 | 52321886 | + | -0.703186 | Intergenic | MER5A\|DNA\|hAT-Charlie | -39184 | NM_009331 | 21414 | Mm.31630 | NM_009331 | ENSMUSG00000000782 | **Tcf7** | AI465550\|TCF-1\|Tcf1 | transcription factor 7, T cell specific | protein-coding |
| chr11_52313917_52314890 | chr11 | 52313918 | 52314890 | + | -0.807299 | Intergenic | CpG | -31833 | NM_009331 | 21414 | Mm.31630 | NM_009331 | ENSMUSG00000000782 | **Tcf7** | AI465550\|TCF-1\|Tcf1 | transcription factor 7, T cell specific | protein-coding |
| chr11_52268243_52268951 | chr11 | 52268244 | 52268951 | + | -0.865826 | intron (NM_009331, intron 2 of 9) | intron (NM_009331, intron 2 of 9) | 13974 | NM_009331 | 21414 | Mm.31630 | NM_009331 | ENSMUSG00000000782 | **Tcf7** | AI465550\|TCF-1\|Tcf1 | transcription factor 7, T cell specific | protein-coding |
| chr11_52268539_52268957 | chr11 | 52268540 | 52268957 | + | -0.89995 | intron (NM_009331, intron 2 of 9) | intron (NM_009331, intron 2 of 9) | 13823 | NM_009331 | 21414 | Mm.31630 | NM_009331 | ENSMUSG00000000782 | **Tcf7** | AI465550\|TCF-1\|Tcf1 | transcription factor 7, T cell specific | protein-coding |
| chr11_52268399_52268870 | chr11 | 52268400 | 52268870 | + | -0.945422 | intron (NM_009331, intron 2 of 9) | intron (NM_009331, intron 2 of 9) | 13936 | NM_009331 | 21414 | Mm.31630 | NM_009331 | ENSMUSG00000000782 | **Tcf7** | AI465550\|TCF-1\|Tcf1 | transcription factor 7, T cell specific | protein-coding |
| chr11_52275345_52276019 | chr11 | 52275346 | 52276019 | + | -1.04508 | intron (NM_009331, intron 2 of 9) | intron (NM_009331, intron 2 of 9) | 6889 | NM_009331 | 21414 | Mm.31630 | NM_009331 | ENSMUSG00000000782 | **Tcf7** | AI465550\|TCF-1\|Tcf1 | transcription factor 7, T cell specific | protein-coding |
| chr11_52309416_52309826 | chr11 | 52309417 | 52309826 | + | -1.13147 | Intergenic | Intergenic | -27050 | NM_009331 | 21414 | Mm.31630 | NM_009331 | ENSMUSG00000000782 | **Tcf7** | AI465550\|TCF-1\|Tcf1 | transcription factor 7, T cell specific | protein-coding |
| chr11_52275425_52275748 | chr11 | 52275426 | 52275748 | + | -1.13812 | intron (NM_009331, intron 2 of 9) | intron (NM_009331, intron 2 of 9) | 6984 | NM_009331 | 21414 | Mm.31630 | NM_009331 | ENSMUSG00000000782 | **Tcf7** | AI465550\|TCF-1\|Tcf1 | transcription factor 7, T cell specific | protein-coding |
| chr19_55914753_55915886 | chr19 | 55914754 | 55915886 | + | -0.947549 | intron (NM_009333, intron 7 of 12) | intron (NM_009333, intron 7 of 12) | 19999 | NM_001142924 | 21416 | Mm.139815 | NM_009333 | ENSMUSG00000024985 | **Tcf7l2** | TCF4B\|TCF4E\|Tcf-4\|Tcf4\|mTcf-4B\|mTcf-4E | transcription factor 7 like 2, T cell specific, HMG box | protein-coding |
| chr19_55915285_55915941 | chr19 | 55915286 | 55915941 | + | -1.13089 | intron (NM_009333, intron 7 of 12) | intron (NM_009333, intron 7 of 12) | 20292 | NM_001142924 | 21416 | Mm.139815 | NM_009333 | ENSMUSG00000024985 | **Tcf7l2** | TCF4B\|TCF4E\|Tcf-4\|Tcf4\|mTcf-4B\|mTcf-4E | transcription factor 7 like 2, T cell specific, HMG box | protein-coding |
| chr17_28080376_28080981 | chr17 | 28080377 | 28080981 | + | -0.688179 | promoter-TSS (NM_001085555) | promoter-TSS (NM_001085555) | -40 | NM_001085555 | 21463 | Mm.29758 | NM_013687 | ENSMUSG00000062859 | **Tcp11** | D17Ken1\|Tcp-11 | t-complex protein 11 | protein-coding |
| chr17_27997079_27997668 | chr17 | 27997080 | 27997668 | + | -1.04936 | intron (NM_001286040, intron 11 of 24) | intron (NM_001286040, intron 11 of 24) | 83210 | NM_013687 | 21463 | Mm.29758 | NM_013687 | ENSMUSG00000062859 | **Tcp11** | D17Ken1\|Tcp-11 | t-complex protein 11 | protein-coding |
| chr17_27997199_27997632 | chr17 | 27997200 | 27997632 | + | -1.08388 | intron (NM_001286040, intron 11 of 24) | intron (NM_001286040, intron 11 of 24) | 83168 | NM_013687 | 21463 | Mm.29758 | NM_013687 | ENSMUSG00000062859 | **Tcp11** | D17Ken1\|Tcp-11 | t-complex protein 11 | protein-coding |
| chr10_84576002_84577618 | chr10 | 84576003 | 84577618 | + | -0.684079 | promoter-TSS (NM_146008) | promoter-TSS (NM_146008) | -137 | NM_146008 | 216198 | Mm.296142 | NM_146008 | ENSMUSG00000020034 | **Tcp11l2** | E430026E19Rik | t-complex 11 (mouse) like 2 | protein-coding |
| chr4_102967676_102968473 | chr4 | 102967677 | 102968473 | + | -1.32348 | non-coding (NR_040444, exon 4 of 4) | non-coding (NR_040444, exon 4 of 4) | -18304 | NM_001163767 | 67344 | Mm.111346 | NM_026100 | ENSMUSG00000028523 | **Tctex1d1** | 1700055O19Rik | Tctex1 domain containing 1 | protein-coding |
| chr14_87713578_87713885 | chr14 | 87713579 | 87713885 | + | -0.775276 | Intergenic | Intergenic | 297149 | NM_001253755 | 219249 | Mm.35168 | NM_172605 | ENSMUSG00000022019 | **Tdrd3** | 4732418C03Rik\|6720468N18 | tudor domain containing 3 | protein-coding |
| chr14_87544863_87545116 | chr14 | 87544864 | 87545116 | + | -0.856935 | intron (NM_172605, intron 13 of 13) | B1_Mus1\|SINE\|Alu | 128407 | NM_001253755 | 219249 | Mm.35168 | NM_172605 | ENSMUSG00000022019 | **Tdrd3** | 4732418C03Rik\|6720468N18 | tudor domain containing 3 | protein-coding |
| chr14_87497680_87498866 | chr14 | 87497681 | 87498866 | + | -1.01863 | intron (NM_172605, intron 10 of 13) | Lx7\|LINE\|L1 | 81690 | NM_001253755 | 219249 | Mm.35168 | NM_172605 | ENSMUSG00000022019 | **Tdrd3** | 4732418C03Rik\|6720468N18 | tudor domain containing 3 | protein-coding |
| chr8_14009564_14009916 | chr8 | 14009565 | 14009916 | + | -0.684117 | Intergenic | Intergenic | -34963 | NM_173744 | 72148 | Mm.5727 | NM_173744 | ENSMUSG00000050052 | **Tdrp** | 2610019F03Rik | testis development related protein | protein-coding |
| chr8_13986488_13986766 | chr8 | 13986489 | 13986766 | + | -0.783059 | Intergenic | Intergenic | -11850 | NM_173744 | 72148 | Mm.5727 | NM_173744 | ENSMUSG00000050052 | **Tdrp** | 2610019F03Rik | testis development related protein | protein-coding |
| chr8_13979765_13980629 | chr8 | 13979766 | 13980629 | + | -0.803213 | Intergenic | MER31-int\|LTR\|ERV1 | -5420 | NM_173744 | 72148 | Mm.5727 | NM_173744 | ENSMUSG00000050052 | **Tdrp** | 2610019F03Rik | testis development related protein | protein-coding |
| chr8_13979859_13980106 | chr8 | 13979860 | 13980106 | + | -0.960667 | Intergenic | MER31-int\|LTR\|ERV1 | -5206 | NM_173744 | 72148 | Mm.5727 | NM_173744 | ENSMUSG00000050052 | **Tdrp** | 2610019F03Rik | testis development related protein | protein-coding |
| chr5_144222838_144223111 | chr5 | 144222839 | 144223111 | + | -0.722787 | intron (NM_027410, intron 1 of 24) | intron (NM_027410, intron 1 of 24) | 603 | NM_027410 | 70381 | Mm.239412 | NM_027410 | ENSMUSG00000066621 | **Tecpr1** | 2210010N04Rik\|mKIAA1358 | tectonin beta-propeller repeat containing 1 | protein-coding |
| chr9_42423685_42423959 | chr9 | 42423686 | 42423959 | + | -0.888485 | intron (NM_173038, intron 7 of 7) | intron (NM_173038, intron 7 of 7) | -23893 | NM_009347 | 21683 | Mm.42209 | NM_009347 | ENSMUSG00000037705 | **Tecta** | Tctna | tectorin alpha | protein-coding |
| chr11_80136362_80136642 | chr11 | 80136363 | 80136642 | + | -0.629028 | TTS (NM_183275) | TTS (NM_183275) | 5651 | NM_183275 | 68550 | Mm.18466 | NM_183275 | ENSMUSG00000046909 | **Tefm** | 1110002N22Rik | transcription elongation factor, mitochondrial | protein-coding |
| chr16_10392626_10392996 | chr16 | 10392627 | 10392996 | + | -0.674232 | intron (NM_001291001, intron 1 of 6) | ORR1C1\|LTR\|ERVL-MaLR | 2637 | NM_001099275 | 70426 | Mm.142181 | NM_001099275 | ENSMUSG00000039179 | **Tekt5** | 3300001K11Rik | tektin 5 | protein-coding |
| chr6_17066290_17066532 | chr6 | 17066291 | 17066532 | + | -0.72046 | intron (NM_207176, intron 1 of 6) | intron (NM_207176, intron 1 of 6) | 1262 | NM_207176 | 21753 | Mm.436548 | NM_011570 | ENSMUSG00000029552 | **Tes** | D6Ertd352e\|TESS\|Tes1\|Tes2\|testin2 | testis derived transcript | protein-coding |
| chr5_118081902_118082702 | chr5 | 118081903 | 118082702 | + | -0.677479 | intron (NM_172721, intron 7 of 10) | intron (NM_172721, intron 7 of 10) | 54478 | NM_021344 | 57816 | Mm.273285 | NM_021344 | ENSMUSG00000029359 | **Tesc** | 1010001A17Rik\|2410011K10Rik\|TE-1 | tescalcin | protein-coding |
| chr5_118081839_118082319 | chr5 | 118081840 | 118082319 | + | -0.768563 | intron (NM_172721, intron 7 of 10) | intron (NM_172721, intron 7 of 10) | 54255 | NM_021344 | 57816 | Mm.273285 | NM_021344 | ENSMUSG00000029359 | **Tesc** | 1010001A17Rik\|2410011K10Rik\|TE-1 | tescalcin | protein-coding |
| chr5_118046983_118047655 | chr5 | 118046984 | 118047655 | + | -0.872971 | intron (NM_021344, intron 2 of 7) | intron (NM_021344, intron 2 of 7) | 19495 | NM_021344 | 57816 | Mm.273285 | NM_021344 | ENSMUSG00000029359 | **Tesc** | 1010001A17Rik\|2410011K10Rik\|TE-1 | tescalcin | protein-coding |
| chr4_116734796_116735365 | chr4 | 116734797 | 116735365 | + | -0.893567 | intron (NM_146151, intron 1 of 10) | polypyrimidine\|Low_complexity\|Low_complexity | 14126 | NM_146151 | 230661 | Mm.482431 | NM_146151 | ENSMUSG00000033985 | **Tesk2** | - | testis-specific kinase 2 | protein-coding |
| chr4_116735043_116735323 | chr4 | 116735044 | 116735323 | + | -0.951348 | intron (NM_146151, intron 1 of 10) | ID_B1\|SINE\|B4 | 14228 | NM_146151 | 230661 | Mm.482431 | NM_146151 | ENSMUSG00000033985 | **Tesk2** | - | testis-specific kinase 2 | protein-coding |
| chr10_130341912_130342240 | chr10 | 130341913 | 130342240 | + | -0.698538 | intron (NM_183264, intron 1 of 8) | Lx\|LINE\|L1 | 19224 | NM_183264 | 67596 | Mm.131264 | NM_183264 | ENSMUSG00000034833 | **Tespa1** | 5830405N20Rik\|A430001F24Rik\|AI450549 | thymocyte expressed, positive selection associated 1 | protein-coding |

| | | | | | | | | | | | | | | | | | |
|---|---|---|---|---|---|---|---|---|---|---|---|---|---|---|---|---|---|
| chr10_130335291_130335538 | chr10 | 130335292 | 130335538 | + | -0.975785 | intron (NM_183264, intron 1 of 8) | intron (NM_183264, intron 1 of 8) | 12563 | NM_183264 | 67596 | Mm.131264 | NM_183264 | ENSMUSG00000034833 | **Tespa1** | 5830405N20Rik\|A430001F24Rik\|AI450549 | thymocyte expressed, positive selection associated 1 | protein-coding |
| chr10_62858981_62859381 | chr10 | 62858982 | 62859381 | + | -0.730356 | intron (NM_027384, intron 2 of 10) | intron (NM_027384, intron 2 of 10) | 20833 | NM_027384 | 52463 | Mm.17774 | NM_027384 | ENSMUSG00000047146 | **Tet1** | 2510010B09Rik\|AA517754\|BB001228\|Cxxc6\|D10Ertd17e\|LCX\|mKIAA1676 | tet methylcytosine dioxygenase 1 | protein-coding |
| chr6_83437306_83437569 | chr6 | 83437307 | 83437569 | + | -1.00321 | intron (NM_183138, intron 2 of 10) | intron (NM_183138, intron 2 of 10) | 4240 | NM_183138 | 194388 | Mm.211030 | NM_183138 | ENSMUSG00000034832 | **Tet3** | B430006D22Rik\|BC037432\|D230004J03Rik | tet methylcytosine dioxygenase 3 | protein-coding |
| chr11_106562733_106563058 | chr11 | 106562734 | 106563058 | + | -0.936181 | intron (NM_198292, intron 2 of 11) | MTEb\|LTR\|ERVL-MaLR | 50034 | NM_198292 | 21763 | Mm.102407 | NM_198292 | ENSMUSG00000040548 | **Tex2** | 4930568E07Rik\|AI553404\|Def-5\|Taz4 | testis expressed gene 2 | protein-coding |
| chr12_113046657_113047231 | chr12 | 113046658 | 113047231 | + | -0.635657 | intron (NM_001291444, intron 3 of 23) | intron (NM_001291444, intron 3 of 23) | -27558 | NM_029381 | 75671 | Mm.23377 | NM_029381 | ENSMUSG00000012211 | **Tex22** | 1700028O09Rik\|AV210422\|Tep22 | testis expressed gene 22 | protein-coding |
| chr12_113046870_113047249 | chr12 | 113046871 | 113047249 | + | -0.794153 | exon (NM_001081170, exon 4 of 24) | exon (NM_001081170, exon 4 of 24) | -27442 | NM_029381 | 75671 | Mm.23377 | NM_029381 | ENSMUSG00000012211 | **Tex22** | 1700028O09Rik\|AV210422\|Tep22 | testis expressed gene 22 | protein-coding |
| chr1_157063637_157064134 | chr1 | 157063638 | 157064134 | + | -0.794915 | Intergenic | Intergenic | 44759 | NM_028540 | 73435 | Mm.116705 | NM_028540 | ENSMUSG00000026592 | **Tex35** | 1700057K13Rik | testis expressed 35 | protein-coding |
| chr9_72493534_72493800 | chr9 | 72493535 | 72493800 | + | -0.6953 | Intergenic | L1MC\|LINE\|L1 | -1708 | NM_009359 | 21778 | Mm.14485 | NM_009359 | | **Tex9** | Gm19407\|tsec-1 | testis expressed gene 9 | protein-coding |
| chr9_72479344_72479795 | chr9 | 72479345 | 72479795 | + | -0.959397 | intron (NM_009359, intron 7 of 11) | intron (NM_009359, intron 7 of 11) | 12389 | NM_009359 | 21778 | Mm.14485 | NM_009359 | | **Tex9** | Gm19407\|tsec-1 | testis expressed gene 9 | protein-coding |
| chr9_72479344_72479681 | chr9 | 72479345 | 72479681 | + | -0.976382 | intron (NM_009359, intron 7 of 11) | intron (NM_009359, intron 7 of 11) | 12446 | NM_009359 | 21778 | Mm.14485 | NM_009359 | | **Tex9** | Gm19407\|tsec-1 | testis expressed gene 9 | protein-coding |
| chr9_72465697_72466313 | chr9 | 72465698 | 72466313 | + | -1.23592 | intron (NM_009359, intron 10 of 11) | intron (NM_009359, intron 10 of 11) | 25954 | NM_009359 | 21778 | Mm.14485 | NM_009359 | | **Tex9** | Gm19407\|tsec-1 | testis expressed gene 9 | protein-coding |
| chr10_71246976_71247436 | chr10 | 71246977 | 71247436 | + | -0.76369 | Intergenic | URR1B\|DNA\|hAT-Charlie | -9162 | NM_009360 | 21780 | Mm.229292 | NM_009360 | ENSMUSG00000003923 | **Tfam** | AI661103\|Hmgts\|mtTFA\|tsHMG | transcription factor A, mitochondrial | protein-coding |
| chr1_118569447_118569861 | chr1 | 118569448 | 118569861 | + | -0.785887 | intron (NM_001293300, intron 34 of 40) | ORR1D2\|LTR\|ERVL-MaLR | -58291 | NM_023755 | 81879 | Mm.24621 | NM_023755 | ENSMUSG00000026380 | **Tfcp2l1** | 1810030F05Rik\|4932442M07Rik\|AA575098\|Cp2l1\|Crtr-1\|D930018N21Rik\|LBP-9\|Tcfcp2l1 | transcription factor CP2-like 1 | protein-coding |
| chr1_118569575_118569835 | chr1 | 118569576 | 118569835 | + | -0.813289 | intron (NM_001293300, intron 34 of 40) | ORR1D2\|LTR\|ERVL-MaLR | -58240 | NM_023755 | 81879 | Mm.24621 | NM_023755 | ENSMUSG00000026380 | **Tfcp2l1** | 1810030F05Rik\|4932442M07Rik\|AA575098\|Cp2l1\|Crtr-1\|D930018N21Rik\|LBP-9\|Tcfcp2l1 | transcription factor CP2-like 1 | protein-coding |
| chr17_31176290_31176731 | chr17 | 31176291 | 31176731 | + | -0.719148 | Intergenic | Intergenic | -11458 | NM_009362 | 21784 | Mm.2854 | NM_009362 | ENSMUSG00000024032 | **Tff1** | Bcei\|PS2 | trefoil factor 1 | protein-coding |
| chr17_31110289_31110493 | chr17 | 31110290 | 31110493 | + | -0.667322 | intron (NM_009593, intron 11 of 14) | ORR1E\|LTR\|ERVL-MaLR | 19220 | NM_011575 | 21786 | Mm.4641 | NM_011575 | ENSMUSG00000024029 | **Tff3** | ITF\|mITF | trefoil factor 3, intestinal | protein-coding |
| chr2_84481900_84482225 | chr2 | 84481901 | 84482225 | + | -0.717651 | Intergenic | RLTR42\|LTR\|ERVK | -5288 | NM_001177320 | 21788 | Mm.124316 | NM_011576 | ENSMUSG00000027082 | **Tfpi** | A630013F22Rik\|AW552122\|EPI\|LACI | tissue factor pathway inhibitor | protein-coding |
| chr2_84481700_84482225 | chr2 | 84481701 | 84482225 | + | -0.730788 | Intergenic | MLTR18B_MM\|LTR\|ERVK | -5188 | NM_001177320 | 21788 | Mm.124316 | NM_011576 | ENSMUSG00000027082 | **Tfpi** | A630013F22Rik\|AW552122\|EPI\|LACI | tissue factor pathway inhibitor | protein-coding |
| chr12_86034122_86034446 | chr12 | 86034123 | 86034446 | + | -0.676268 | intron (NM_001081423, intron 33 of 33) | B4A\|SINE\|B4 | 44757 | NM_009368 | 21809 | Mm.3992 | NM_009368 | ENSMUSG00000021253 | **Tgfb3** | Tgfb-3 | transforming growth factor, beta 3 | protein-coding |
| chr12_85973504_85973982 | chr12 | 85973505 | 85973982 | + | -0.692914 | intron (NM_001081423, intron 29 of 33) | intron (NM_001081423, intron 29 of 33) | 105298 | NM_009368 | 21809 | Mm.3992 | NM_009368 | ENSMUSG00000021253 | **Tgfb3** | Tgfb-3 | transforming growth factor, beta 3 | protein-coding |
| chr12_86047405_86047754 | chr12 | 86047406 | 86047754 | + | -0.701938 | intron (NM_001081423, intron 33 of 33) | intron (NM_001081423, intron 33 of 33) | 31461 | NM_009368 | 21809 | Mm.3992 | NM_009368 | ENSMUSG00000021253 | **Tgfb3** | Tgfb-3 | transforming growth factor, beta 3 | protein-coding |
| chr12_86047334_86047930 | chr12 | 86047335 | 86047930 | + | -0.732583 | intron (NM_001081423, intron 33 of 33) | intron (NM_001081423, intron 33 of 33) | 31409 | NM_009368 | 21809 | Mm.3992 | NM_009368 | ENSMUSG00000021253 | **Tgfb3** | Tgfb-3 | transforming growth factor, beta 3 | protein-coding |
| chr12_86034071_86034344 | chr12 | 86034072 | 86034344 | + | -0.771149 | intron (NM_001081423, intron 33 of 33) | intron (NM_001081423, intron 33 of 33) | 44833 | NM_009368 | 21809 | Mm.3992 | NM_009368 | ENSMUSG00000021253 | **Tgfb3** | Tgfb-3 | transforming growth factor, beta 3 | protein-coding |
| chr12_85969524_85970476 | chr12 | 85969525 | 85970476 | + | -0.939865 | intron (NM_001081423, intron 29 of 33) | intron (NM_001081423, intron 29 of 33) | 109041 | NM_009368 | 21809 | Mm.3992 | NM_009368 | ENSMUSG00000021253 | **Tgfb3** | Tgfb-3 | transforming growth factor, beta 3 | protein-coding |
| chr12_85969493_85970219 | chr12 | 85969494 | 85970219 | + | -1.00312 | intron (NM_001081423, intron 29 of 33) | intron (NM_001081423, intron 29 of 33) | 109185 | NM_009368 | 21809 | Mm.3992 | NM_009368 | ENSMUSG00000021253 | **Tgfb3** | Tgfb-3 | transforming growth factor, beta 3 | protein-coding |
| chr12_85969519_85970011 | chr12 | 85969520 | 85970011 | + | -1.02201 | intron (NM_001081423, intron 29 of 33) | intron (NM_001081423, intron 29 of 33) | 109276 | NM_009368 | 21809 | Mm.3992 | NM_009368 | ENSMUSG00000021253 | **Tgfb3** | Tgfb-3 | transforming growth factor, beta 3 | protein-coding |
| chr4_47319502_47320442 | chr4 | 47319503 | 47320442 | + | -0.959323 | Intergenic | GA-rich\|Low_complexity\|Low_complexity | -33333 | NM_009370 | 21812 | Mm.197552 | NM_009370 | ENSMUSG00000007613 | **Tgfbr1** | ALK5\|AU017191\|Alk-5\|TbetaR-I\|TbetaRI | transforming growth factor, beta receptor I | protein-coding |
| chr4_47319578_47319972 | chr4 | 47319579 | 47319972 | + | -1.03689 | Intergenic | Intergenic | -33530 | NM_009370 | 21812 | Mm.197552 | NM_009370 | ENSMUSG00000007613 | **Tgfbr1** | ALK5\|AU017191\|Alk-5\|TbetaR-I\|TbetaRI | transforming growth factor, beta receptor I | protein-coding |
| chr9_116293762_116294863 | chr9 | 116293763 | 116294863 | + | -0.604834 | Intergenic | B4A\|SINE\|B4 | -118950 | NM_009371 | 21813 | Mm.172346 | NM_009371 | ENSMUSG00000032440 | **Tgfbr2** | 1110020H15Rik\|AU042018\|DNIIR\|RIIDN\|TBR-II\|TbetaR-II\|TbetaRII | transforming growth factor, beta receptor II | protein-coding |
| chr9_116175803_116176123 | chr9 | 116175804 | 116176123 | + | -0.612377 | promoter-TSS (NM_009371) | promoter-TSS (NM_009371) | -600 | NM_009371 | 21813 | Mm.172346 | NM_009371 | ENSMUSG00000032440 | **Tgfbr2** | 1110020H15Rik\|AU042018\|DNIIR\|RIIDN\|TBR-II\|TbetaR-II\|TbetaRII | transforming growth factor, beta receptor II | protein-coding |
| chr9_116181364_116182065 | chr9 | 116181365 | 116182065 | + | -0.661792 | Intergenic | Intergenic | -6352 | NM_009371 | 21813 | Mm.172346 | NM_009371 | ENSMUSG00000032440 | **Tgfbr2** | 1110020H15Rik\|AU042018\|DNIIR\|RIIDN\|TBR-II\|TbetaR-II\|TbetaRII | transforming growth factor, beta receptor II | protein-coding |
| chr9_116145202_116145769 | chr9 | 116145203 | 116145769 | + | -0.731613 | intron (NM_029575, intron 1 of 6) | intron (NM_029575, intron 1 of 6) | 29877 | NM_029575 | 21813 | Mm.172346 | NM_009371 | ENSMUSG00000032440 | **Tgfbr2** | 1110020H15Rik\|AU042018\|DNIIR\|RIIDN\|TBR-II\|TbetaR-II\|TbetaRII | transforming growth factor, beta receptor II | protein-coding |
| chr9_116230323_116230901 | chr9 | 116230324 | 116230901 | + | -0.746852 | Intergenic | Intergenic | -55249 | NM_009371 | 21813 | Mm.172346 | NM_009371 | ENSMUSG00000032440 | **Tgfbr2** | 1110020H15Rik\|AU042018\|DNIIR\|RIIDN\|TBR-II\|TbetaR-II\|TbetaRII | transforming growth factor, beta receptor II | protein-coding |
| chr9_116366121_116366573 | chr9 | 116366122 | 116366573 | + | -0.751643 | Intergenic | Intergenic | -190984 | NM_009371 | 21813 | Mm.172346 | NM_009371 | ENSMUSG00000032440 | **Tgfbr2** | 1110020H15Rik\|AU042018\|DNIIR\|RIIDN\|TBR-II\|TbetaR-II\|TbetaRII | transforming growth factor, beta receptor II | protein-coding |
| chr9_116365858_116366644 | chr9 | 116365859 | 116366644 | + | -0.756529 | Intergenic | Intergenic | -190888 | NM_009371 | 21813 | Mm.172346 | NM_009371 | ENSMUSG00000032440 | **Tgfbr2** | 1110020H15Rik\|AU042018\|DNIIR\|RIIDN\|TBR-II\|TbetaR-II\|TbetaRII | transforming growth factor, beta receptor II | protein-coding |
| chr9_116249790_116250243 | chr9 | 116249791 | 116250243 | + | -0.759323 | Intergenic | Intergenic | -74654 | NM_009371 | 21813 | Mm.172346 | NM_009371 | ENSMUSG00000032440 | **Tgfbr2** | 1110020H15Rik\|AU042018\|DNIIR\|RIIDN\|TBR-II\|TbetaR-II\|TbetaRII | transforming growth factor, beta receptor II | protein-coding |
| chr9_116214341_116214719 | chr9 | 116214342 | 116214719 | + | -0.764632 | Intergenic | Intergenic | -39167 | NM_009371 | 21813 | Mm.172346 | NM_009371 | ENSMUSG00000032440 | **Tgfbr2** | 1110020H15Rik\|AU042018\|DNIIR\|RIIDN\|TBR-II\|TbetaR-II\|TbetaRII | transforming growth factor, beta receptor II | protein-coding |
| chr9_116216748_116217086 | chr9 | 116216749 | 116217086 | + | -0.786345 | Intergenic | MIRc\|SINE\|MIR | -41554 | NM_009371 | 21813 | Mm.172346 | NM_009371 | ENSMUSG00000032440 | **Tgfbr2** | 1110020H15Rik\|AU042018\|DNIIR\|RIIDN\|TBR-II\|TbetaR-II\|TbetaRII | transforming growth factor, beta receptor II | protein-coding |
| chr9_116373487_116373717 | chr9 | 116373488 | 116373717 | + | -0.843011 | Intergenic | Intergenic | -198239 | NM_009371 | 21813 | Mm.172346 | NM_009371 | ENSMUSG00000032440 | **Tgfbr2** | 1110020H15Rik\|AU042018\|DNIIR\|RIIDN\|TBR-II\|TbetaR-II\|TbetaRII | transforming growth factor, beta receptor II | protein-coding |
| chr9_116160841_116161289 | chr9 | 116160842 | 116161289 | + | -0.98701 | intron (NM_009371, intron 1 of 7) | intron (NM_009371, intron 1 of 7) | 14298 | NM_029575 | 21813 | Mm.172346 | NM_009371 | ENSMUSG00000032440 | **Tgfbr2** | 1110020H15Rik\|AU042018\|DNIIR\|RIIDN\|TBR-II\|TbetaR-II\|TbetaRII | transforming growth factor, beta receptor II | protein-coding |
| chr9_116188768_116189512 | chr9 | 116188769 | 116189512 | + | -0.987655 | Intergenic | Intergenic | -13777 | NM_009371 | 21813 | Mm.172346 | NM_009371 | ENSMUSG00000032440 | **Tgfbr2** | 1110020H15Rik\|AU042018\|DNIIR\|RIIDN\|TBR-II\|TbetaR-II\|TbetaRII | transforming growth factor, beta receptor II | protein-coding |
| chr9_116101767_116102237 | chr9 | 116101768 | 116102237 | + | -1.01378 | intron (NM_029575, intron 5 of 6) | intron (NM_029575, intron 5 of 6) | 73361 | NM_029575 | 21813 | Mm.172346 | NM_009371 | ENSMUSG00000032440 | **Tgfbr2** | 1110020H15Rik\|AU042018\|DNIIR\|RIIDN\|TBR-II\|TbetaR-II\|TbetaRII | transforming growth factor, beta receptor II | protein-coding |
| chr9_116122651_116123519 | chr9 | 116122652 | 116123519 | + | -1.02068 | intron (NM_029575, intron 3 of 6) | intron (NM_029575, intron 3 of 6) | 52278 | NM_029575 | 21813 | Mm.172346 | NM_009371 | ENSMUSG00000032440 | **Tgfbr2** | 1110020H15Rik\|AU042018\|DNIIR\|RIIDN\|TBR-II\|TbetaR-II\|TbetaRII | transforming growth factor, beta receptor II | protein-coding |
| chr9_116122918_116123504 | chr9 | 116122919 | 116123504 | + | -1.03687 | intron (NM_029575, intron 3 of 6) | intron (NM_029575, intron 3 of 6) | 52152 | NM_029575 | 21813 | Mm.172346 | NM_009371 | ENSMUSG00000032440 | **Tgfbr2** | 1110020H15Rik\|AU042018\|DNIIR\|RIIDN\|TBR-II\|TbetaR-II\|TbetaRII | transforming growth factor, beta receptor II | protein-coding |
| chr9_116160974_116161287 | chr9 | 116160975 | 116161287 | + | -1.03745 | intron (NM_009371, intron 1 of 7) | intron (NM_009371, intron 1 of 7) | 14232 | NM_029575 | 21813 | Mm.172346 | NM_009371 | ENSMUSG00000032440 | **Tgfbr2** | 1110020H15Rik\|AU042018\|DNIIR\|RIIDN\|TBR-II\|TbetaR-II\|TbetaRII | transforming growth factor, beta receptor II | protein-coding |

| | | | | | | | | | | | | | | | | | |
|---|---|---|---|---|---|---|---|---|---|---|---|---|---|---|---|---|---|
| chr9_116084411_116085026 | chr9 | 116084412 | 116085026 | + | -1.16025 | Intergenic | Intergenic | 90644 | NM_029575 | 21813 | Mm.172346 | NM_009371 | ENSMUSG00000032440 | **Tgfbr2** | 1110020H15Rik\|AU042018\|DNIIR\|RIIDN\|TBR-II\|TbetaR-II\|TbetaRII | transforming growth factor, beta receptor II | protein-coding |
| chr9_116084183_116084953 | chr9 | 116084184 | 116084953 | + | -1.20447 | Intergenic | Intergenic | 90795 | NM_029575 | 21813 | Mm.172346 | NM_009371 | ENSMUSG00000032440 | **Tgfbr2** | 1110020H15Rik\|AU042018\|DNIIR\|RIIDN\|TBR-II\|TbetaR-II\|TbetaRII | transforming growth factor, beta receptor II | protein-coding |
| chr9_116337918_116338801 | chr9 | 116337919 | 116338801 | + | -1.23958 | Intergenic | Intergenic | -162997 | NM_009371 | 21813 | Mm.172346 | NM_009371 | ENSMUSG00000032440 | **Tgfbr2** | 1110020H15Rik\|AU042018\|DNIIR\|RIIDN\|TBR-II\|TbetaR-II\|TbetaRII | transforming growth factor, beta receptor II | protein-coding |
| chr9_116163871_116164319 | chr9 | 116163872 | 116164319 | + | -1.2787 | intron (NM_009371, intron 1 of 7) | intron (NM_009371, intron 1 of 7) | 11268 | NM_029575 | 21813 | Mm.172346 | NM_009371 | ENSMUSG00000032440 | **Tgfbr2** | 1110020H15Rik\|AU042018\|DNIIR\|RIIDN\|TBR-II\|TbetaR-II\|TbetaRII | transforming growth factor, beta receptor II | protein-coding |
| chr9_116338324_116338753 | chr9 | 116338325 | 116338753 | + | -1.29569 | Intergenic | Intergenic | -163176 | NM_009371 | 21813 | Mm.172346 | NM_009371 | ENSMUSG00000032440 | **Tgfbr2** | 1110020H15Rik\|AU042018\|DNIIR\|RIIDN\|TBR-II\|TbetaR-II\|TbetaRII | transforming growth factor, beta receptor II | protein-coding |
| chr9_116368911_116369674 | chr9 | 116368912 | 116369674 | + | -1.34159 | Intergenic | ORR1D2\|LTR\|ERVL-MaLR | -193930 | NM_009371 | 21813 | Mm.172346 | NM_009371 | ENSMUSG00000032440 | **Tgfbr2** | 1110020H15Rik\|AU042018\|DNIIR\|RIIDN\|TBR-II\|TbetaR-II\|TbetaRII | transforming growth factor, beta receptor II | protein-coding |
| chr9_116187401_116187876 | chr9 | 116187402 | 116187876 | + | -1.3758 | Intergenic | Intergenic | -12276 | NM_009371 | 21813 | Mm.172346 | NM_009371 | ENSMUSG00000032440 | **Tgfbr2** | 1110020H15Rik\|AU042018\|DNIIR\|RIIDN\|TBR-II\|TbetaR-II\|TbetaRII | transforming growth factor, beta receptor II | protein-coding |
| chr5_107201561_107202251 | chr5 | 107201562 | 107202251 | + | -0.602582 | intron (NM_011578, intron 3 of 16) | intron (NM_011578, intron 3 of 16) | 87689 | NM_011578 | 21814 | Mm.200775 | NM_011578 | ENSMUSG00000029287 | **Tgfbr3** | 1110036H20Rik\|AU015626\|AW215636\|TBRIII | transforming growth factor, beta receptor III | protein-coding |
| chr5_107177241_107177623 | chr5 | 107177242 | 107177623 | + | -0.68279 | intron (NM_011578, intron 4 of 16) | intron (NM_011578, intron 4 of 16) | 112163 | NM_011578 | 21814 | Mm.200775 | NM_011578 | ENSMUSG00000029287 | **Tgfbr3** | 1110036H20Rik\|AU015626\|AW215636\|TBRIII | transforming growth factor, beta receptor III | protein-coding |
| chr5_107147951_107148695 | chr5 | 107147952 | 107148695 | + | -0.746304 | intron (NM_011578, intron 6 of 16) | intron (NM_011578, intron 6 of 16) | 141272 | NM_011578 | 21814 | Mm.200775 | NM_011578 | ENSMUSG00000029287 | **Tgfbr3** | 1110036H20Rik\|AU015626\|AW215636\|TBRIII | transforming growth factor, beta receptor III | protein-coding |
| chr5_107148244_107148690 | chr5 | 107148245 | 107148690 | + | -0.781314 | intron (NM_011578, intron 6 of 16) | intron (NM_011578, intron 6 of 16) | 141128 | NM_011578 | 21814 | Mm.200775 | NM_011578 | ENSMUSG00000029287 | **Tgfbr3** | 1110036H20Rik\|AU015626\|AW215636\|TBRIII | transforming growth factor, beta receptor III | protein-coding |
| chr5_107223033_107223434 | chr5 | 107223034 | 107223434 | + | -0.835184 | intron (NM_011578, intron 2 of 16) | intron (NM_011578, intron 2 of 16) | 66361 | NM_011578 | 21814 | Mm.200775 | NM_011578 | ENSMUSG00000029287 | **Tgfbr3** | 1110036H20Rik\|AU015626\|AW215636\|TBRIII | transforming growth factor, beta receptor III | protein-coding |
| chr5_107201573_107201787 | chr5 | 107201574 | 107201787 | + | -0.841885 | intron (NM_011578, intron 3 of 16) | intron (NM_011578, intron 3 of 16) | 87915 | NM_011578 | 21814 | Mm.200775 | NM_011578 | ENSMUSG00000029287 | **Tgfbr3** | 1110036H20Rik\|AU015626\|AW215636\|TBRIII | transforming growth factor, beta receptor III | protein-coding |
| chr9_123058552_123059100 | chr9 | 123058553 | 123059100 | + | -0.883502 | intron (NM_177911, intron 10 of 13) | L1M4\|LINE\|L1 | 24085 | NM_177911 | 331046 | Mm.195309 | NM_177911 | ENSMUSG00000025787 | **Tgm4** | 9530008N10Rik\|Eapa1\|TGP | transglutaminase 4 (prostate) | protein-coding |
| chr11_117881853_117882231 | chr11 | 117881854 | 117882231 | + | -0.609354 | Intergenic | Intergenic | -8516 | NM_027919 | 71776 | Mm.3200 | NM_027919 | ENSMUSG00000017713 | **Tha1** | 1300017K07Rik\|GLY1 | threonine aldolase 1 | protein-coding |
| chr11_117878072_117878794 | chr11 | 117878073 | 117878794 | + | -0.693424 | Intergenic | Intergenic | -4907 | NM_027919 | 71776 | Mm.3200 | NM_027919 | ENSMUSG00000017713 | **Tha1** | 1300017K07Rik\|GLY1 | threonine aldolase 1 | protein-coding |
| chr11_117871422_117872658 | chr11 | 117871423 | 117872658 | + | -0.695084 | intron (NM_027919, intron 1 of 6) | intron (NM_027919, intron 1 of 6) | 1486 | NM_027919 | 71776 | Mm.3200 | NM_027919 | ENSMUSG00000017713 | **Tha1** | 1300017K07Rik\|GLY1 | threonine aldolase 1 | protein-coding |
| chr11_117891526_117892555 | chr11 | 117891527 | 117892555 | + | -0.720416 | Intergenic | Intergenic | -18515 | NM_027919 | 71776 | Mm.3200 | NM_027919 | ENSMUSG00000017713 | **Tha1** | 1300017K07Rik\|GLY1 | threonine aldolase 1 | protein-coding |
| chr11_117909388_117909814 | chr11 | 117909389 | 117909814 | + | -0.797057 | Intergenic | Intergenic | -36075 | NM_027919 | 71776 | Mm.3200 | NM_027919 | ENSMUSG00000017713 | **Tha1** | 1300017K07Rik\|GLY1 | threonine aldolase 1 | protein-coding |
| chr11_117891551_117892151 | chr11 | 117891552 | 117892151 | + | -1.08581 | Intergenic | Intergenic | -18325 | NM_027919 | 71776 | Mm.3200 | NM_027919 | ENSMUSG00000017713 | **Tha1** | 1300017K07Rik\|GLY1 | threonine aldolase 1 | protein-coding |
| chr17_84422953_84423279 | chr17 | 84422954 | 84423279 | + | -0.748265 | exon (NM_183021, exon 21 of 38) | exon (NM_183021, exon 21 of 38) | 43092 | NM_183021 | 240174 | Mm.246767 | NM_177826 | ENSMUSG00000024251 | **Thada** | 6530405K19\|A930021O22\|AU019899\|BC052885\|GITA | thyroid adenoma associated | protein-coding |
| chr17_84352152_84352509 | chr17 | 84352153 | 84352509 | + | -0.847154 | intron (NM_183021, intron 26 of 37) | RSINE1\|SINE\|B4 | 113877 | NM_183021 | 240174 | Mm.246767 | NM_177826 | ENSMUSG00000024251 | **Thada** | 6530405K19\|A930021O22\|AU019899\|BC052885\|GITA | thyroid adenoma associated | protein-coding |
| chr17_84407710_84408423 | chr17 | 84407711 | 84408423 | + | -0.936225 | intron (NM_183021, intron 22 of 37) | intron (NM_183021, intron 22 of 37) | 58141 | NM_183021 | 240174 | Mm.246767 | NM_177826 | ENSMUSG00000024251 | **Thada** | 6530405K19\|A930021O22\|AU019899\|BC052885\|GITA | thyroid adenoma associated | protein-coding |
| chr17_84408042_84408403 | chr17 | 84408043 | 84408403 | + | -0.944516 | intron (NM_183021, intron 22 of 37) | B1F2\|SINE\|Alu | 57985 | NM_183021 | 240174 | Mm.246767 | NM_177826 | ENSMUSG00000024251 | **Thada** | 6530405K19\|A930021O22\|AU019899\|BC052885\|GITA | thyroid adenoma associated | protein-coding |
| chr17_84408082_84408292 | chr17 | 84408083 | 84408292 | + | -1.00346 | intron (NM_183021, intron 22 of 37) | B1F2\|SINE\|Alu | 58021 | NM_183021 | 240174 | Mm.246767 | NM_177826 | ENSMUSG00000024251 | **Thada** | 6530405K19\|A930021O22\|AU019899\|BC052885\|GITA | thyroid adenoma associated | protein-coding |
| chr17_84344234_84344587 | chr17 | 84344235 | 84344587 | + | -1.12197 | intron (NM_183021, intron 26 of 37) | intron (NM_183021, intron 26 of 37) | 121797 | NM_183021 | 240174 | Mm.246767 | NM_177826 | ENSMUSG00000024251 | **Thada** | 6530405K19\|A930021O22\|AU019899\|BC052885\|GITA | thyroid adenoma associated | protein-coding |
| chr17_84330569_84330888 | chr17 | 84330570 | 84330888 | + | -1.3238 | intron (NM_183021, intron 28 of 37) | intron (NM_183021, intron 28 of 37) | 135479 | NM_183021 | 240174 | Mm.246767 | NM_177826 | ENSMUSG00000024251 | **Thada** | 6530405K19\|A930021O22\|AU019899\|BC052885\|GITA | thyroid adenoma associated | protein-coding |
| chr17_84330445_84331064 | chr17 | 84330446 | 84331064 | + | -1.33128 | intron (NM_183021, intron 28 of 37) | intron (NM_183021, intron 28 of 37) | 135453 | NM_183021 | 240174 | Mm.246767 | NM_177826 | ENSMUSG00000024251 | **Thada** | 6530405K19\|A930021O22\|AU019899\|BC052885\|GITA | thyroid adenoma associated | protein-coding |
| chr4_151986012_151986395 | chr4 | 151986013 | 151986395 | + | -0.761964 | intron (NM_001145929, intron 1 of 2) | intron (NM_001145929, intron 1 of 2) | 2782 | NM_175152 | 69876 | Mm.226820 | NM_175152 | ENSMUSG00000039759 | **Thap3** | 2010013E08Rik\|2210418H06Rik | THAP domain containing, apoptosis associated protein 3 | protein-coding |
| chr2_117996168_117996680 | chr2 | 117996169 | 117996680 | + | -0.672891 | Intergenic | Intergenic | -115498 | NM_011580 | 21825 | Mm.4159 | NM_011580 | ENSMUSG00000040152 | **Thbs1** | TSP-1\|TSP1\|Thbs-1\|tbsp1 | thrombospondin 1 | protein-coding |
| chr2_117996231_117996993 | chr2 | 117996232 | 117996993 | + | -0.69107 | Intergenic | Intergenic | -115310 | NM_011580 | 21825 | Mm.4159 | NM_011580 | ENSMUSG00000040152 | **Thbs1** | TSP-1\|TSP1\|Thbs-1\|tbsp1 | thrombospondin 1 | protein-coding |
| chr2_118007815_118008487 | chr2 | 118007816 | 118008487 | + | -0.784834 | Intergenic | Intergenic | -103771 | NM_011580 | 21825 | Mm.4159 | NM_011580 | ENSMUSG00000040152 | **Thbs1** | TSP-1\|TSP1\|Thbs-1\|tbsp1 | thrombospondin 1 | protein-coding |
| chr2_117988502_117989071 | chr2 | 117988503 | 117989071 | + | -0.991536 | Intergenic | Intergenic | -123135 | NM_011580 | 21825 | Mm.4159 | NM_011580 | ENSMUSG00000040152 | **Thbs1** | TSP-1\|TSP1\|Thbs-1\|tbsp1 | thrombospondin 1 | protein-coding |
| chr2_117988488_117988884 | chr2 | 117988489 | 117988884 | + | -0.992375 | Intergenic | Intergenic | -123236 | NM_011580 | 21825 | Mm.4159 | NM_011580 | ENSMUSG00000040152 | **Thbs1** | TSP-1\|TSP1\|Thbs-1\|tbsp1 | thrombospondin 1 | protein-coding |
| chr10_28619435_28619749 | chr10 | 28619436 | 28619749 | + | -0.80669 | Intergenic | Intergenic | -48803 | NM_178666 | 210757 | Mm.123021 | NM_178666 | ENSMUSG00000049109 | **Themis** | A430108D01\|BB160712\|E430004N04Rik\|Gasp\|Tsepa\|thylex | thymocyte selection associated | protein-coding |
| chr10_28596934_28597307 | chr10 | 28596935 | 28597307 | + | -1.03424 | 3' UTR (NM_008983, exon 32 of 32) | 3' UTR (NM_008983, exon 32 of 32) | -71274 | NM_178666 | 210757 | Mm.123021 | NM_178666 | ENSMUSG00000049109 | **Themis** | A430108D01\|BB160712\|E430004N04Rik\|Gasp\|Tsepa\|thylex | thymocyte selection associated | protein-coding |
| chr6_71154863_71155277 | chr6 | 71154864 | 71155277 | + | -1.27549 | Intergenic | Intergenic | -10690 | NM_178413 | 232078 | Mm.37547 | NM_178413 | ENSMUSG00000054474 | **Thnsl2** | BC051244\|TSH2 | threonine synthase-like 2 (bacterial) | protein-coding |
| chr4_126194252_126194600 | chr4 | 126194253 | 126194600 | + | -1.02215 | intron (NM_146153, intron 1 of 10) | intron (NM_146153, intron 1 of 10) | 8284 | NM_146153 | 230753 | Mm.29813 | NM_146153 | ENSMUSG00000043962 | **Thrap3** | 9330151F09Rik\|B230333E16Rik\|Trap150 | thyroid hormone receptor associated protein 3 | protein-coding |
| chr4_126211815_126212380 | chr4 | 126211816 | 126212380 | + | -1.05352 | Intergenic | Intergenic | -9388 | NM_146153 | 230753 | Mm.29813 | NM_146153 | ENSMUSG00000043962 | **Thrap3** | 9330151F09Rik\|B230333E16Rik\|Trap150 | thyroid hormone receptor associated protein 3 | protein-coding |
| chr4_126211843_126212212 | chr4 | 126211844 | 126212212 | + | -1.07943 | Intergenic | Intergenic | -9318 | NM_146153 | 230753 | Mm.29813 | NM_146153 | ENSMUSG00000043962 | **Thrap3** | 9330151F09Rik\|B230333E16Rik\|Trap150 | thyroid hormone receptor associated protein 3 | protein-coding |
| chr14_18021783_18022178 | chr14 | 18021784 | 18022178 | + | -0.688606 | intron (NM_001113417, intron 7 of 9) | ORR1E\|LTR\|ERVL-MaLR | 40337 | NM_009380 | 21834 | Mm.32563 | NM_009380 | ENSMUSG00000021779 | **Thrb** | Nr1a2\|T3R[b]\|T3Rbeta\|Thrb1\|Thrb2\|c-erbAbeta | thyroid hormone receptor beta | protein-coding |
| chr14_18003793_18004004 | chr14 | 18003794 | 18004004 | + | -0.814597 | intron (NM_001113417, intron 5 of 9) | intron (NM_001113417, intron 5 of 9) | 22255 | NM_009380 | 21834 | Mm.32563 | NM_009380 | ENSMUSG00000021779 | **Thrb** | Nr1a2\|T3R[b]\|T3Rbeta\|Thrb1\|Thrb2\|c-erbAbeta | thyroid hormone receptor beta | protein-coding |
| chr14_18001859_18002725 | chr14 | 18001860 | 18002725 | + | -1.0919 | intron (NM_009380, intron 1 of 6) | intron (NM_009380, intron 1 of 6) | 20648 | NM_009380 | 21834 | Mm.32563 | NM_009380 | ENSMUSG00000021779 | **Thrb** | Nr1a2\|T3R[b]\|T3Rbeta\|Thrb1\|Thrb2\|c-erbAbeta | thyroid hormone receptor beta | protein-coding |
| chr14_18001857_18002427 | chr14 | 18001858 | 18002427 | + | -1.16533 | intron (NM_009380, intron 1 of 6) | intron (NM_009380, intron 1 of 6) | 20498 | NM_009380 | 21834 | Mm.32563 | NM_009380 | ENSMUSG00000021779 | **Thrb** | Nr1a2\|T3R[b]\|T3Rbeta\|Thrb1\|Thrb2\|c-erbAbeta | thyroid hormone receptor beta | protein-coding |
| chr17_81045162_81045452 | chr17 | 81045163 | 81045452 | + | -0.708637 | intron (NM_028138, intron 5 of 9) | intron (NM_028138, intron 5 of 9) | 19778 | NM_028138 | 72167 | Mm.273257 | NM_028138 | ENSMUSG00000024246 | **Thumpd2** | 2810025A12Rik | THUMP domain containing 2 | protein-coding |
| chr9_44029694_44029941 | chr9 | 44029695 | 44029941 | + | -0.667967 | Intergenic | Intergenic | -13566 | NM_009382 | 21838 | Mm.3951 | NM_009382 | ENSMUSG00000032011 | **Thy1** | CD90\|T25\|Thy-1\|Thy-1.2\|Thy1.1\|Thy1.2 | thymus cell antigen 1, theta | protein-coding |
| chr9_44017919_44018203 | chr9 | 44017920 | 44018203 | + | -1.0093 | Intergenic | Intergenic | -25323 | NM_009382 | 21838 | Mm.3951 | NM_009382 | ENSMUSG00000032011 | **Thy1** | CD90\|T25\|Thy-1\|Thy-1.2\|Thy1.1\|Thy1.2 | thymus cell antigen 1, theta | protein-coding |
| chr16_90042399_90042768 | chr16 | 90042400 | 90042768 | + | -0.633593 | Intergenic | Intergenic | -67885 | NM_001145886 | 21844 | Mm.310902 | NM_009384 | ENSMUSG00000002489 | **Tiam1** | AI847750\|D16Ium10\|D16Ium10e | T cell lymphoma invasion and metastasis 1 | protein-coding |
| chr16_89938181_89938872 | chr16 | 89938182 | 89938872 | + | -1.36032 | intron (NM_001145886, intron 2 of 27) | intron (NM_001145886, intron 2 of 27) | 22300 | NM_009384 | 21844 | Mm.310902 | NM_009384 | ENSMUSG00000002489 | **Tiam1** | AI847750\|D16Ium10\|D16Ium10e | T cell lymphoma invasion and metastasis 1 | protein-coding |
| chr17_56269108_56269699 | chr17 | 56269109 | 56269699 | + | -0.659222 | TTS (NM_174989) | TTS (NM_174989) | 7363 | NM_174989 | 106759 | Mm.203952 | NM_174989 | ENSMUSG00000047123 | **Ticam1** | AW046014\|AW547018\|TICAM-1\|TRIF | toll-like receptor adaptor molecule 1 | protein-coding |
| chr13_56158988_56159654 | chr13 | 56158989 | 56159654 | + | -0.613645 | Intergenic | Intergenic | 19564 | NM_145976 | 212937 | Mm.116697 | NM_145976 | ENSMUSG00000049625 | **Tifab** | - | TRAF-interacting protein with forkhead-associated domain, family member B | protein-coding |
| chr16_43689990_43690757 | chr16 | 43689991 | 43690757 | + | -0.752533 | Intergenic | Intergenic | -26190 | NM_001146325 | 100043314 | Mm.390896 | NM_001146325 | ENSMUSG00000071552 | **Tigit** | ENSMUSG00000071552\|Vstm3 | T cell immunoreceptor with Ig and ITIM domains | protein-coding |
| chr14_122009454_122009920 | chr14 | 122009455 | 122009920 | + | -0.680246 | intron (NM_026861, intron 8 of 8) | ORR1D2\|LTR\|ERVL-MaLR | -24987 | NM_001037744 | 223262 | Mm.297637 | NM_001037744 | ENSMUSG00000071229 | **Timm8a2** | - | translocase of inner mitochondrial membrane 8A2 | protein-coding |
| chr14_122011819_122012584 | chr14 | 122011820 | 122012584 | + | -0.733171 | intron (NM_026861, intron 8 of 8) | intron (NM_026861, intron 8 of 8) | -22472 | NM_001037744 | 223262 | Mm.297637 | NM_001037744 | ENSMUSG00000071229 | **Timm8a2** | - | translocase of inner mitochondrial membrane 8A2 | protein-coding |
| chr14_122070008_122070277 | chr14 | 122070009 | 122070277 | + | -0.741567 | Intergenic | RLTR6-int\|LTR\|ERV1 | 35469 | NM_001037744 | 223262 | Mm.297637 | NM_001037744 | ENSMUSG00000071229 | **Timm8a2** | - | translocase of inner mitochondrial membrane 8A2 | protein-coding |

| | | | | | | | | | | | | | | | | | |
|---|---|---|---|---|---|---|---|---|---|---|---|---|---|---|---|---|---|
| chr14_122011808_122012214 | chr14 | 122011809 | 122012214 | + | -0.920123 | intron (NM_026861, intron 8 of 8) | intron (NM_026861, intron 8 of 8) | -22663 | NM_001037744 | 223262 | Mm.297637 | NM_001037744 | ENSMUSG00000071229 | **Timm8a2** | - | translocase of inner mitochondrial membrane 8A2 | protein-coding |
| chr14_122025894_122026197 | chr14 | 122025895 | 122026197 | + | -0.95954 | Intergenic | ORR1C2\|LTR\|ERVL-MaLR | -8628 | NM_001037744 | 223262 | Mm.297637 | NM_001037744 | ENSMUSG00000071229 | **Timm8a2** | - | translocase of inner mitochondrial membrane 8A2 | protein-coding |
| chr14_122062567_122063111 | chr14 | 122062568 | 122063111 | + | -1.08273 | Intergenic | B4A\|SINE\|B4 | 28165 | NM_001037744 | 223262 | Mm.297637 | NM_001037744 | ENSMUSG00000071229 | **Timm8a2** | - | translocase of inner mitochondrial membrane 8A2 | protein-coding |
| chr14_122066271_122066727 | chr14 | 122066272 | 122066727 | + | -1.11627 | Intergenic | ORR1E\|LTR\|ERVL-MaLR | 31825 | NM_001037744 | 223262 | Mm.297637 | NM_001037744 | ENSMUSG00000071229 | **Timm8a2** | - | translocase of inner mitochondrial membrane 8A2 | protein-coding |
| chr14_122063684_122063976 | chr14 | 122063685 | 122063976 | + | -1.17885 | Intergenic | MLT1B\|LTR\|ERVL-MaLR | 29156 | NM_001037744 | 223262 | Mm.297637 | NM_001037744 | ENSMUSG00000071229 | **Timm8a2** | - | translocase of inner mitochondrial membrane 8A2 | protein-coding |
| chr14_122062825_122063070 | chr14 | 122062826 | 122063070 | + | -1.34281 | Intergenic | MTD\|LTR\|ERVL-MaLR | 28274 | NM_001037744 | 223262 | Mm.297637 | NM_001037744 | ENSMUSG00000071229 | **Timm8a2** | - | translocase of inner mitochondrial membrane 8A2 | protein-coding |
| chr9_61505992_61506441 | chr9 | 61505993 | 61506441 | + | -0.76434 | Intergenic | Intergenic | 133851 | NM_001083927 | 21887 | Mm.24255 | NM_009389 | ENSMUSG00000032280 | **Tle3** | 2610103N05Rik\|ESG\|Grg3a\|Grg3b\|mKIAA1547 | transducin-like enhancer of split 3, homolog of Drosophila E(spl) | protein-coding |
| chr9_61380181_61380932 | chr9 | 61380182 | 61380932 | + | -0.790722 | intron (NM_009389, intron 4 of 19) | L2b\|LINE\|L2 | 8191 | NM_001083927 | 21887 | Mm.24255 | NM_009389 | ENSMUSG00000032280 | **Tle3** | 2610103N05Rik\|ESG\|Grg3a\|Grg3b\|mKIAA1547 | transducin-like enhancer of split 3, homolog of Drosophila E(spl) | protein-coding |
| chr9_61426986_61427441 | chr9 | 61426987 | 61427441 | + | -0.839828 | Intergenic | ORR1E\|LTR\|ERVL-MaLR | 54848 | NM_001083927 | 21887 | Mm.24255 | NM_009389 | ENSMUSG00000032280 | **Tle3** | 2610103N05Rik\|ESG\|Grg3a\|Grg3b\|mKIAA1547 | transducin-like enhancer of split 3, homolog of Drosophila E(spl) | protein-coding |
| chr9_61426733_61427464 | chr9 | 61426734 | 61427464 | + | -0.85627 | Intergenic | ORR1E\|LTR\|ERVL-MaLR | 54733 | NM_001083927 | 21887 | Mm.24255 | NM_009389 | ENSMUSG00000032280 | **Tle3** | 2610103N05Rik\|ESG\|Grg3a\|Grg3b\|mKIAA1547 | transducin-like enhancer of split 3, homolog of Drosophila E(spl) | protein-coding |
| chr9_61380175_61380521 | chr9 | 61380176 | 61380521 | + | -0.896025 | intron (NM_009389, intron 4 of 19) | intron (NM_009389, intron 4 of 19) | 7982 | NM_001083927 | 21887 | Mm.24255 | NM_009389 | ENSMUSG00000032280 | **Tle3** | 2610103N05Rik\|ESG\|Grg3a\|Grg3b\|mKIAA1547 | transducin-like enhancer of split 3, homolog of Drosophila E(spl) | protein-coding |
| chr19_14678485_14678997 | chr19 | 14678486 | 14678997 | + | -0.776821 | Intergenic | MLTR11A\|LTR\|ERVK | -80758 | NM_011600 | 21888 | Mm.103638 | NM_011600 | ENSMUSG00000024642 | **Tle4** | 5730411M05Rik\|AA792082\|Bce-1\|Bce1\|ESTM13\|ESTM14\|Grg4\|X83332\|X83333 | transducin-like enhancer of split 4, homolog of Drosophila E(spl) | protein-coding |
| chr19_14641120_14641949 | chr19 | 14641121 | 14641949 | + | -0.827853 | Intergenic | Intergenic | -43552 | NM_011600 | 21888 | Mm.103638 | NM_011600 | ENSMUSG00000024642 | **Tle4** | 5730411M05Rik\|AA792082\|Bce-1\|Bce1\|ESTM13\|ESTM14\|Grg4\|X83332\|X83333 | transducin-like enhancer of split 4, homolog of Drosophila E(spl) | protein-coding |
| chr19_14588110_14588736 | chr19 | 14588111 | 14588736 | + | -1.12228 | intron (NM_011600, intron 4 of 19) | intron (NM_011600, intron 4 of 19) | 9560 | NM_011600 | 21888 | Mm.103638 | NM_011600 | ENSMUSG00000024642 | **Tle4** | 5730411M05Rik\|AA792082\|Bce-1\|Bce1\|ESTM13\|ESTM14\|Grg4\|X83332\|X83333 | transducin-like enhancer of split 4, homolog of Drosophila E(spl) | protein-coding |
| chr19_14588274_14588720 | chr19 | 14588275 | 14588720 | + | -1.13414 | intron (NM_011600, intron 4 of 19) | intron (NM_011600, intron 4 of 19) | 9486 | NM_011600 | 21888 | Mm.103638 | NM_011600 | ENSMUSG00000024642 | **Tle4** | 5730411M05Rik\|AA792082\|Bce-1\|Bce1\|ESTM13\|ESTM14\|Grg4\|X83332\|X83333 | transducin-like enhancer of split 4, homolog of Drosophila E(spl) | protein-coding |
| chr19_14588175_14588566 | chr19 | 14588176 | 14588566 | + | -1.17446 | intron (NM_011600, intron 4 of 19) | intron (NM_011600, intron 4 of 19) | 9612 | NM_011600 | 21888 | Mm.103638 | NM_011600 | ENSMUSG00000024642 | **Tle4** | 5730411M05Rik\|AA792082\|Bce-1\|Bce1\|ESTM13\|ESTM14\|Grg4\|X83332\|X83333 | transducin-like enhancer of split 4, homolog of Drosophila E(spl) | protein-coding |
| chr19_14695274_14695884 | chr19 | 14695275 | 14695884 | + | -1.61796 | Intergenic | Intergenic | -97596 | NM_011600 | 21888 | Mm.103638 | NM_011600 | ENSMUSG00000024642 | **Tle4** | 5730411M05Rik\|AA792082\|Bce-1\|Bce1\|ESTM13\|ESTM14\|Grg4\|X83332\|X83333 | transducin-like enhancer of split 4, homolog of Drosophila E(spl) | protein-coding |
| chr2_70835722_70836010 | chr2 | 70835723 | 70836010 | + | -0.617194 | Intergenic | Intergenic | -10386 | NM_172664 | 228012 | Mm.136511 | NM_172664 | ENSMUSG00000041997 | **Tlk1** | 4930545J15Rik | tousled-like kinase 1 | protein-coding |
| chr2_70808412_70809119 | chr2 | 70808413 | 70809119 | + | -0.66394 | intron (NM_172664, intron 1 of 19) | L1M4b\|LINE\|L1 | 16714 | NM_172664 | 228012 | Mm.136511 | NM_172664 | ENSMUSG00000041997 | **Tlk1** | 4930545J15Rik | tousled-like kinase 1 | protein-coding |
| chr2_70809917_70810323 | chr2 | 70809918 | 70810323 | + | -1.20782 | intron (NM_172664, intron 1 of 19) | ID_B1\|SINE\|B4 | 15360 | NM_172664 | 228012 | Mm.136511 | NM_172664 | ENSMUSG00000041997 | **Tlk1** | 4930545J15Rik | tousled-like kinase 1 | protein-coding |
| chr5_64923151_64923429 | chr5 | 64923152 | 64923429 | + | -0.723164 | Intergenic | Intergenic | 9409 | NM_030682 | 21897 | Mm.273024 | NM_030682 | ENSMUSG00000044827 | **Tlr1** | - | toll-like receptor 1 | protein-coding |
| chr5_64922963_64923412 | chr5 | 64922964 | 64923412 | + | -0.765656 | Intergenic | Intergenic | 9511 | NM_030682 | 21897 | Mm.273024 | NM_030682 | ENSMUSG00000044827 | **Tlr1** | - | toll-like receptor 1 | protein-coding |
| chr5_64959558_64959913 | chr5 | 64959559 | 64959913 | + | -0.713498 | intron (NM_011604, intron 1 of 1) | intron (NM_011604, intron 1 of 1) | 298 | NM_011604 | 21899 | Mm.42146 | NM_011604 | ENSMUSG00000051498 | **Tlr6** | - | toll-like receptor 6 | protein-coding |
| chr9_106215131_106215746 | chr9 | 106215132 | 106215746 | + | -0.695031 | TTS (NM_011876) | TTS (NM_011876) | -7159 | NM_031178 | 81897 | Mm.44889 | NM_031178 | ENSMUSG00000045322 | **Tlr9** | - | toll-like receptor 9 | protein-coding |
| chr9_106215400_106215679 | chr9 | 106215401 | 106215679 | + | -0.775228 | TTS (NM_011876) | TTS (NM_011876) | -7058 | NM_031178 | 81897 | Mm.44889 | NM_031178 | ENSMUSG00000045322 | **Tlr9** | - | toll-like receptor 9 | protein-coding |
| chr4_98091235_98092324 | chr4 | 98091236 | 98092324 | + | -0.644519 | intron (NM_001122953, intron 9 of 9) | intron (NM_001122953, intron 9 of 9) | 291485 | NM_053157 | 94043 | Mm.331719 | NM_053157 | ENSMUSG00000028563 | **Tm2d1** | 2310026L18Rik\|AA990549\|Bbp | TM2 domain containing 1 | protein-coding |
| chr4_98082620_98082914 | chr4 | 98082621 | 98082914 | + | -0.645478 | intron (NM_001122953, intron 9 of 9) | intron (NM_001122953, intron 9 of 9) | 300498 | NM_053157 | 94043 | Mm.331719 | NM_053157 | ENSMUSG00000028563 | **Tm2d1** | 2310026L18Rik\|AA990549\|Bbp | TM2 domain containing 1 | protein-coding |
| chr4_98243540_98244301 | chr4 | 98243541 | 98244301 | + | -0.694238 | Intergenic | C-rich\|Low_complexity\|Low_complexity | 139344 | NM_053157 | 94043 | Mm.331719 | NM_053157 | ENSMUSG00000028563 | **Tm2d1** | 2310026L18Rik\|AA990549\|Bbp | TM2 domain containing 1 | protein-coding |
| chr4_98082360_98082850 | chr4 | 98082361 | 98082850 | + | -0.694945 | intron (NM_001122953, intron 9 of 9) | intron (NM_001122953, intron 9 of 9) | 300660 | NM_053157 | 94043 | Mm.331719 | NM_053157 | ENSMUSG00000028563 | **Tm2d1** | 2310026L18Rik\|AA990549\|Bbp | TM2 domain containing 1 | protein-coding |
| chr4_98091252_98091867 | chr4 | 98091253 | 98091867 | + | -0.815668 | intron (NM_001122953, intron 9 of 9) | intron (NM_001122953, intron 9 of 9) | 291705 | NM_053157 | 94043 | Mm.331719 | NM_053157 | ENSMUSG00000028563 | **Tm2d1** | 2310026L18Rik\|AA990549\|Bbp | TM2 domain containing 1 | protein-coding |
| chr4_98131580_98132032 | chr4 | 98131581 | 98132032 | + | -0.873115 | Intergenic | Intergenic | 251459 | NM_053157 | 94043 | Mm.331719 | NM_053157 | ENSMUSG00000028563 | **Tm2d1** | 2310026L18Rik\|AA990549\|Bbp | TM2 domain containing 1 | protein-coding |
| chr4_98131553_98131883 | chr4 | 98131554 | 98131883 | + | -0.923523 | Intergenic | Intergenic | 251547 | NM_053157 | 94043 | Mm.331719 | NM_053157 | ENSMUSG00000028563 | **Tm2d1** | 2310026L18Rik\|AA990549\|Bbp | TM2 domain containing 1 | protein-coding |
| chr4_98243918_98244316 | chr4 | 98243919 | 98244316 | + | -0.938704 | Intergenic | Intergenic | 139148 | NM_053157 | 94043 | Mm.331719 | NM_053157 | ENSMUSG00000028563 | **Tm2d1** | 2310026L18Rik\|AA990549\|Bbp | TM2 domain containing 1 | protein-coding |
| chr4_98097048_98097508 | chr4 | 98097049 | 98097508 | + | -1.05213 | intron (NM_001122953, intron 9 of 9) | intron (NM_001122953, intron 9 of 9) | 285987 | NM_053157 | 94043 | Mm.331719 | NM_053157 | ENSMUSG00000028563 | **Tm2d1** | 2310026L18Rik\|AA990549\|Bbp | TM2 domain containing 1 | protein-coding |
| chr4_98113657_98114037 | chr4 | 98113658 | 98114037 | + | -1.3051 | 3' UTR (NM_001122952, exon 11 of 11) | 3' UTR (NM_001122952, exon 11 of 11) | 269418 | NM_053157 | 94043 | Mm.331719 | NM_053157 | ENSMUSG00000028563 | **Tm2d1** | 2310026L18Rik\|AA990549\|Bbp | TM2 domain containing 1 | protein-coding |
| chr7_81863174_81863561 | chr7 | 81863175 | 81863561 | + | -0.658766 | intron (NR_111910, intron 2 of 7) | intron (NR_111910, intron 2 of 7) | 681 | NM_001291282 | 107769 | Mm.221412 | NM_145375 | ENSMUSG00000038623 | **Tm6sf1** | AI428514\|BC023123\|C630016D09Rik | transmembrane 6 superfamily member 1 | protein-coding |
| chr19_20875909_20876163 | chr19 | 20875910 | 20876163 | + | -0.877469 | intron (NM_028953, intron 5 of 20) | ORR1D2\|LTR\|ERVL-MaLR | 78166 | NM_028953 | 13409 | Mm.219585 | NM_028953 | ENSMUSG00000024749 | **Tmc1** | 4933416G09Rik\|Beethoven\|Bth\|CWEA1\|dn | transmembrane channel-like gene family 1 | protein-coding |
| chr19_21036514_21037017 | chr19 | 21036515 | 21037017 | + | -0.917236 | Intergenic | Intergenic | -82564 | NM_028953 | 13409 | Mm.219585 | NM_028953 | ENSMUSG00000024749 | **Tmc1** | 4933416G09Rik\|Beethoven\|Bth\|CWEA1\|dn | transmembrane channel-like gene family 1 | protein-coding |
| chr19_21036570_21036862 | chr19 | 21036571 | 21036862 | + | -0.938696 | Intergenic | Intergenic | -82514 | NM_028953 | 13409 | Mm.219585 | NM_028953 | ENSMUSG00000024749 | **Tmc1** | 4933416G09Rik\|Beethoven\|Bth\|CWEA1\|dn | transmembrane channel-like gene family 1 | protein-coding |
| chr19_21036347_21037024 | chr19 | 21036348 | 21037024 | + | -0.968332 | Intergenic | Intergenic | -82484 | NM_028953 | 13409 | Mm.219585 | NM_028953 | ENSMUSG00000024749 | **Tmc1** | 4933416G09Rik\|Beethoven\|Bth\|CWEA1\|dn | transmembrane channel-like gene family 1 | protein-coding |
| chr11_117724504_117725270 | chr11 | 117724505 | 117725270 | + | -0.827093 | intron (NM_198022, intron 5 of 21) | intron (NM_198022, intron 5 of 21) | 55734 | NM_181321 | 217353 | Mm.286963 | NM_145439 | ENSMUSG00000025572 | **Tmc6** | D11Ertd204e\|EVER1 | transmembrane channel-like gene family 6 | protein-coding |
| chr11_117777133_117777429 | chr11 | 117777134 | 117777429 | + | -0.864929 | intron (NM_181321, intron 5 of 7) | MTE2a\|LTR\|ERVL-MaLR | 3340 | NM_181321 | 217353 | Mm.286963 | NM_145439 | ENSMUSG00000025572 | **Tmc6** | D11Ertd204e\|EVER1 | transmembrane channel-like gene family 6 | protein-coding |
| chr11_117718845_117719357 | chr11 | 117718846 | 117719357 | + | -1.51269 | intron (NM_198022, intron 4 of 21) | intron (NM_198022, intron 4 of 21) | 61520 | NM_181321 | 217353 | Mm.286963 | NM_145439 | ENSMUSG00000025572 | **Tmc6** | D11Ertd204e\|EVER1 | transmembrane channel-like gene family 6 | protein-coding |
| chr12_85416159_85416957 | chr12 | 85416160 | 85416957 | + | -0.764013 | Intergenic | MLT1J2\|LTR\|ERVL-MaLR | -41841 | NM_026775 | 68581 | Mm.379159 | NM_026775 | ENSMUSG00000021248 | **Tmed10** | 1110014C03Rik\|Tmp21\|p23\|p24delta1 | transmembrane emp24-like trafficking protein 10 (yeast) | protein-coding |
| chr12_85416318_85416850 | chr12 | 85416319 | 85416850 | + | -0.799643 | Intergenic | PB1D10\|SINE\|Alu | -41867 | NM_026775 | 68581 | Mm.379159 | NM_026775 | ENSMUSG00000021248 | **Tmed10** | 1110014C03Rik\|Tmp21\|p23\|p24delta1 | transmembrane emp24-like trafficking protein 10 (yeast) | protein-coding |
| chr11_115226718_115227383 | chr11 | 115226719 | 115227383 | + | -0.794034 | intron (NM_001033393, intron 8 of 9) | intron (NM_001033393, intron 8 of 9) | 39564 | NM_001033393 | 320534 | Mm.260587 | NM_001033393 | ENSMUSG00000045980 | **Tmem104** | C630005D06Rik\|mFLJ00021 | transmembrane protein 104 | protein-coding |
| chr6_13078732_13078985 | chr6 | 13078733 | 13078985 | + | -0.667347 | intron (NM_027992, intron 5 of 8) | intron (NM_027992, intron 5 of 8) | 9100 | NM_027992 | 71900 | Mm.27742 | NM_027992 | ENSMUSG00000029571 | **Tmem106b** | 2310036D22Rik\|5830455K21Rik\|6430519M21Rik\|AI428776\|AI661344 | transmembrane protein 106B | protein-coding |

| | | | | | | | | | | | | | | | | | |
|---|---|---|---|---|---|---|---|---|---|---|---|---|---|---|---|---|---|
| chr4_118548134_118548352 | chr4 | 118548135 | 118548352 | + | -0.686724 | Intergenic | G-rich\|Low_complexity\|Low_complexity | -4517 | NM_172383 | 230678 | Mm.23639 | NM_172383 | ENSMUSG00000050854 | **Tmem125** | 6330530A05Rik\|AV223516 | transmembrane protein 125 | protein-coding |
| chr1_36960370_36961172 | chr1 | 36960371 | 36961172 | + | -0.63311 | Intergenic | Intergenic | -21244 | NM_018872 | 56030 | Mm.186943 | NM_018872 | ENSMUSG00000026116 | **Tmem131** | 2610524E03Rik\|CC28\|D1Bwg0491e\|Neg\|RW1\|YR-23\|mKIAA0257 | transmembrane protein 131 | protein-coding |
| chr1_36911597_36911977 | chr1 | 36911598 | 36911977 | + | -0.684367 | intron (NM_018872, intron 1 of 40) | intron (NM_018872, intron 1 of 40) | 27740 | NM_018872 | 56030 | Mm.186943 | NM_018872 | ENSMUSG00000026116 | **Tmem131** | 2610524E03Rik\|CC28\|D1Bwg0491e\|Neg\|RW1\|YR-23\|mKIAA0257 | transmembrane protein 131 | protein-coding |
| chr1_36921602_36922321 | chr1 | 36921603 | 36922321 | + | -0.888723 | intron (NM_018872, intron 1 of 40) | intron (NM_018872, intron 1 of 40) | 17565 | NM_018872 | 56030 | Mm.186943 | NM_018872 | ENSMUSG00000026116 | **Tmem131** | 2610524E03Rik\|CC28\|D1Bwg0491e\|Neg\|RW1\|YR-23\|mKIAA0257 | transmembrane protein 131 | protein-coding |
| chr7_89248096_89248799 | chr7 | 89248097 | 89248799 | + | -0.852786 | intron (NM_028343, intron 4 of 14) | intron (NM_028343, intron 4 of 14) | 90339 | NM_028343 | 72759 | Mm.208477 | NM_028343 | ENSMUSG00000039428 | **Tmem135** | 2810439K08Rik\|AW319712\|PMP52 | transmembrane protein 135 | protein-coding |
| chr7_89248062_89248517 | chr7 | 89248063 | 89248517 | + | -0.918701 | intron (NM_028343, intron 4 of 14) | intron (NM_028343, intron 4 of 14) | 90497 | NM_028343 | 72759 | Mm.208477 | NM_028343 | ENSMUSG00000039428 | **Tmem135** | 2810439K08Rik\|AW319712\|PMP52 | transmembrane protein 135 | protein-coding |
| chr7_89232565_89232841 | chr7 | 89232566 | 89232841 | + | -1.09993 | intron (NM_028343, intron 5 of 14) | ORR1B2\|LTR\|ERVL-MaLR | 106084 | NM_028343 | 72759 | Mm.208477 | NM_028343 | ENSMUSG00000039428 | **Tmem135** | 2810439K08Rik\|AW319712\|PMP52 | transmembrane protein 135 | protein-coding |
| chr7_89232432_89232883 | chr7 | 89232433 | 89232883 | + | -1.13477 | intron (NM_028343, intron 5 of 14) | ORR1B2\|LTR\|ERVL-MaLR | 106129 | NM_028343 | 72759 | Mm.208477 | NM_028343 | ENSMUSG00000039428 | **Tmem135** | 2810439K08Rik\|AW319712\|PMP52 | transmembrane protein 135 | protein-coding |
| chr2_25620284_25620835 | chr2 | 25620285 | 25620835 | + | -0.986564 | intron (NM_001040130, intron 4 of 4) | intron (NM_001040130, intron 4 of 4) | 1442 | NM_001040130 | 51875 | Mm.271147 | NM_001040130 | ENSMUSG00000026939 | **Tmem141** | 1110065P14Rik\|AI663999\|D2Ertd217e | transmembrane protein 141 | protein-coding |
| chr3_84681643_84682137 | chr3 | 84681644 | 84682137 | + | -0.746807 | intron (NM_177260, intron 1 of 6) | PB1D10\|SINE\|Alu | 15698 | NM_177260 | 320782 | Mm.107370 | NM_177260 | ENSMUSG00000056498 | **Tmem154** | 9930117H01Rik | transmembrane protein 154 | protein-coding |
| chr16_8579562_8579909 | chr16 | 8579563 | 8579909 | + | -0.689655 | intron (NM_172961, intron 3 of 15) | intron (NM_172961, intron 3 of 15) | 57965 | NM_025708 | 66690 | Mm.10068 | NM_025708 | ENSMUSG00000043140 | **Tmem186** | 2810440M13Rik\|4432406C05Rik\|AI648175\|AW210586 | transmembrane protein 186 | protein-coding |
| chr1_52578229_52578940 | chr1 | 52578230 | 52578940 | + | -0.616719 | Intergenic | Intergenic | -52120 | NR_024590 | 227094 | Mm.83541 | NM_172654 | ENSMUSG00000043015 | **Tmem194b** | 5330401P04Rik\|9830163O08 | transmembrane protein 194B | protein-coding |
| chr1_52607635_52608094 | chr1 | 52607636 | 52608094 | + | -0.727279 | Intergenic | Intergenic | -22840 | NR_024590 | 227094 | Mm.83541 | NM_172654 | ENSMUSG00000043015 | **Tmem194b** | 5330401P04Rik\|9830163O08 | transmembrane protein 194B | protein-coding |
| chr1_52607731_52608073 | chr1 | 52607732 | 52608073 | + | -0.732611 | Intergenic | Intergenic | -22803 | NR_024590 | 227094 | Mm.83541 | NM_172654 | ENSMUSG00000043015 | **Tmem194b** | 5330401P04Rik\|9830163O08 | transmembrane protein 194B | protein-coding |
| chr1_52650037_52650570 | chr1 | 52650038 | 52650570 | + | -1.12091 | non-coding (NR_024590, exon 9 of 9) | non-coding (NR_024590, exon 9 of 9) | 19599 | NM_001142647 | 227094 | Mm.83541 | NM_172654 | ENSMUSG00000043015 | **Tmem194b** | 5330401P04Rik\|9830163O08 | transmembrane protein 194B | protein-coding |
| chr1_52650097_52650421 | chr1 | 52650098 | 52650421 | + | -1.15263 | non-coding (NR_024590, exon 9 of 9) | non-coding (NR_024590, exon 9 of 9) | 19554 | NM_001142647 | 227094 | Mm.83541 | NM_172654 | ENSMUSG00000043015 | **Tmem194b** | 5330401P04Rik\|9830163O08 | transmembrane protein 194B | protein-coding |
| chr1_52653261_52653799 | chr1 | 52653262 | 52653799 | + | -1.2234 | Intergenic | Intergenic | 22825 | NM_001142647 | 227094 | Mm.83541 | NM_172654 | ENSMUSG00000043015 | **Tmem194b** | 5330401P04Rik\|9830163O08 | transmembrane protein 194B | protein-coding |
| chr6_30499098_30499384 | chr6 | 30499099 | 30499384 | + | -0.944804 | intron (NM_178625, intron 6 of 14) | intron (NM_178625, intron 6 of 14) | 10465 | NM_178625 | 72649 | Mm.45418 | NM_178625 | ENSMUSG00000029782 | **Tmem209** | 2700094F01Rik\|AI428435 | transmembrane protein 209 | protein-coding |
| chr17_29533905_29534196 | chr17 | 29533906 | 29534196 | + | -0.957592 | intron (NM_001162902, intron 1 of 1) | Lx7\|LINE\|L1 | 15542 | NM_001162901 | 71138 | Mm.158581 | NM_001162901 | ENSMUSG00000079580 | **Tmem217** | 4933413N12Rik\|EG622644 | transmembrane protein 217 | protein-coding |
| chr4_133298519_133298893 | chr4 | 133298520 | 133298893 | + | -1.04533 | intron (NM_199306, intron 12 of 15) | intron (NM_199306, intron 12 of 15) | -20916 | NR_110978 | 52174 | Mm.244748 | NM_025667 | ENSMUSG00000028857 | **Tmem222** | 5730406H10Rik\|AA409341\|D4Ertd196e | transmembrane protein 222 | protein-coding |
| chr17_65561316_65561818 | chr17 | 65561317 | 65561818 | + | -0.660552 | Intergenic | RCHARR1\|DNA\|hAT-Charlie | -20785 | NM_001289486 | 381107 | Mm.141275 | NM_001008973 | ENSMUSG00000045036 | **Tmem232** | E130009J12Rik\|Gm940\|Tes13-L\|Tes13-S | transmembrane protein 232 | protein-coding |
| chr17_65561396_65561660 | chr17 | 65561397 | 65561660 | + | -0.679911 | Intergenic | RCHARR1\|DNA\|hAT-Charlie | -20746 | NM_001289486 | 381107 | Mm.141275 | NM_001008973 | ENSMUSG00000045036 | **Tmem232** | E130009J12Rik\|Gm940\|Tes13-L\|Tes13-S | transmembrane protein 232 | protein-coding |
| chr5_9073256_9073933 | chr5 | 9073257 | 9073933 | + | -0.683885 | Intergenic | Intergenic | -27142 | NM_001081029 | 652925 | Mm.439954 | NM_001081029 | ENSMUSG00000079659 | **Tmem243** | 4930420K17Rik\|E030031F02Rik | transmembrane protein 243, mitochondrial | protein-coding |
| chr5_9073278_9073624 | chr5 | 9073279 | 9073624 | + | -0.713724 | Intergenic | Intergenic | -27286 | NM_001081029 | 652925 | Mm.439954 | NM_001081029 | ENSMUSG00000079659 | **Tmem243** | 4930420K17Rik\|E030031F02Rik | transmembrane protein 243, mitochondrial | protein-coding |
| chr5_9072978_9073744 | chr5 | 9072979 | 9073744 | + | -0.798539 | Intergenic | Intergenic | -27376 | NM_001081029 | 652925 | Mm.439954 | NM_001081029 | ENSMUSG00000079659 | **Tmem243** | 4930420K17Rik\|E030031F02Rik | transmembrane protein 243, mitochondrial | protein-coding |
| chr5_9086937_9087714 | chr5 | 9086938 | 9087714 | + | -0.821098 | Intergenic | MTD\|LTR\|ERVL-MaLR | -13411 | NM_001081029 | 652925 | Mm.439954 | NM_001081029 | ENSMUSG00000079659 | **Tmem243** | 4930420K17Rik\|E030031F02Rik | transmembrane protein 243, mitochondrial | protein-coding |
| chr5_9086622_9087624 | chr5 | 9086623 | 9087624 | + | -0.839001 | Intergenic | Intergenic | -13614 | NM_001081029 | 652925 | Mm.439954 | NM_001081029 | ENSMUSG00000079659 | **Tmem243** | 4930420K17Rik\|E030031F02Rik | transmembrane protein 243, mitochondrial | protein-coding |
| chr5_9087009_9087250 | chr5 | 9087010 | 9087250 | + | -1.13862 | Intergenic | Intergenic | -13607 | NM_001081029 | 652925 | Mm.439954 | NM_001081029 | ENSMUSG00000079659 | **Tmem243** | 4930420K17Rik\|E030031F02Rik | transmembrane protein 243, mitochondrial | protein-coding |
| chr4_56932035_56932459 | chr4 | 56932036 | 56932459 | + | -0.699304 | intron (NM_175518, intron 3 of 17) | RLTR26D_MM\|LTR\|ERVK | 15182 | NM_175518 | 242474 | Mm.296632 | NM_175518 | | **Tmem245** | A630051L19Rik\|AI957324\|D730040F13Rik | transmembrane protein 245 | protein-coding |
| chr17_86880443_86880911 | chr17 | 86880444 | 86880911 | + | -0.752613 | Intergenic | Intergenic | -36671 | NM_001277980 | 78469 | Mm.67234 | NM_030104 | ENSMUSG00000037689 | **Tmem247** | 1700090G07Rik | transmembrane protein 247 | protein-coding |
| chr10_68673475_68674148 | chr10 | 68673476 | 68674148 | + | -0.709919 | Intergenic | MLT1E2\|LTR\|ERVL-MaLR | -49934 | NM_177794 | 327766 | Mm.2479 | NM_177794 | ENSMUSG00000060044 | **Tmem26** | 3230401A06\|AI195372 | transmembrane protein 26 | protein-coding |
| chr4_75277826_75278697 | chr4 | 75277827 | 75278697 | + | -0.686699 | promoter-TSS (NM_025849) | promoter-TSS (NM_025849) | 24 | NM_025849 | 66928 | Mm.24332 | NM_025849 | ENSMUSG00000028398 | **Tmem261** | 1700027K24Rik\|3110001D03Rik | transmembrane protein 261 | protein-coding |
| chr4_75277989_75278509 | chr4 | 75277990 | 75278509 | + | -0.686866 | promoter-TSS (NM_025849) | promoter-TSS (NM_025849) | 37 | NM_025849 | 66928 | Mm.24332 | NM_025849 | ENSMUSG00000028398 | **Tmem261** | 1700027K24Rik\|3110001D03Rik | transmembrane protein 261 | protein-coding |
| chr4_75063347_75063597 | chr4 | 75063348 | 75063597 | + | -0.716502 | Intergenic | Intergenic | 214814 | NM_025849 | 66928 | Mm.24332 | NM_025849 | ENSMUSG00000028398 | **Tmem261** | 1700027K24Rik\|3110001D03Rik | transmembrane protein 261 | protein-coding |
| chr4_75267139_75267492 | chr4 | 75267140 | 75267492 | + | -0.797536 | Intergenic | Lx8\|LINE\|L1 | 10970 | NM_025849 | 66928 | Mm.24332 | NM_025849 | ENSMUSG00000028398 | **Tmem261** | 1700027K24Rik\|3110001D03Rik | transmembrane protein 261 | protein-coding |
| chr4_75266984_75267495 | chr4 | 75266985 | 75267495 | + | -0.798371 | Intergenic | Lx8\|LINE\|L1 | 11046 | NM_025849 | 66928 | Mm.24332 | NM_025849 | ENSMUSG00000028398 | **Tmem261** | 1700027K24Rik\|3110001D03Rik | transmembrane protein 261 | protein-coding |
| chr4_75266792_75267403 | chr4 | 75266793 | 75267403 | + | -0.838722 | Intergenic | Lx8\|LINE\|L1 | 11188 | NM_025849 | 66928 | Mm.24332 | NM_025849 | ENSMUSG00000028398 | **Tmem261** | 1700027K24Rik\|3110001D03Rik | transmembrane protein 261 | protein-coding |
| chr4_75278830_75279442 | chr4 | 75278831 | 75279442 | + | -1.09363 | promoter-TSS (NM_025849) | promoter-TSS (NM_025849) | -850 | NM_025849 | 66928 | Mm.24332 | NM_025849 | ENSMUSG00000028398 | **Tmem261** | 1700027K24Rik\|3110001D03Rik | transmembrane protein 261 | protein-coding |
| chr12_73491275_73491804 | chr12 | 73491276 | 73491804 | + | -0.78075 | Intergenic | Intergenic | 54855 | NM_178715 | 238257 | Mm.251196 | NM_178715 | ENSMUSG00000034435 | **Tmem30b** | 9130011B11Rik | transmembrane protein 30B | protein-coding |
| chr5_67232603_67232906 | chr5 | 67232604 | 67232906 | + | -0.920906 | Intergenic | Intergenic | -27810 | NM_001285452 | 67878 | Mm.23217 | NM_028975 | ENSMUSG00000037720 | **Tmem33** | 1110006G02Rik\|1600019D15Rik\|2410089A21Rik\|2700052H22Rik\|5430406L04Rik\|AA388285\|AI314185\|AI426615\|db83 | transmembrane protein 33 | protein-coding |
| chr1_120055386_120056062 | chr1 | 120055387 | 120056062 | + | -0.652491 | intron (NM_001012322, intron 11 of 13) | intron (NM_001012322, intron 11 of 13) | 18056 | NM_019432 | 170706 | Mm.24750 | NM_019432 | | **Tmem37** | AI173373\|AW540203\|Pr\|Pr1 | transmembrane protein 37 | protein-coding |
| chr4_53920625_53921006 | chr4 | 53920626 | 53921006 | + | -0.739704 | Intergenic | Intergenic | 94771 | NM_028053 | 52076 | Mm.27184 | NM_028053 | ENSMUSG00000028420 | **Tmem38b** | 1600017F22Rik\|AA437809\|AV051057\|D4Ertd89e\|TRIC-B\|mg33b | transmembrane protein 38B | protein-coding |
| chr16_38616019_38616280 | chr16 | 38616020 | 38616280 | + | -0.688702 | intron (NM_020260, intron 8 of 11) | intron (NM_020260, intron 8 of 11) | 53336 | NM_026407 | 67846 | Mm.310476 | NM_026407 | ENSMUSG00000002845 | **Tmem39a** | 2610033C09Rik | transmembrane protein 39a | protein-coding |
| chr4_134918347_134918783 | chr4 | 134918348 | 134918783 | + | -0.974316 | Intergenic | RMER6B\|LTR\|ERVK | -3649 | NM_027935 | 71817 | Mm.88349 | NM_027935 | ENSMUSG00000028822 | **Tmem50a** | 3200001F09Rik\|CAM\|Smp1 | transmembrane protein 50A | protein-coding |
| chr16_91596427_91596706 | chr16 | 91596428 | 91596706 | + | -0.630959 | intron (NM_030018, intron 1 of 6) | Tigger16a\|DNA\|TcMar-Tigger | 1113 | NM_030018 | 77975 | Mm.290341 | NM_030018 | ENSMUSG00000022964 | **Tmem50b** | AU015466\|AU019872\|B230114J08Rik | transmembrane protein 50B | protein-coding |
| chr4_107184231_107184502 | chr4 | 107184232 | 107184502 | + | -0.723535 | intron (NM_029565, intron 1 of 7) | B3A\|SINE\|B2 | 5737 | NM_029565 | 56374 | Mm.291192 | NM_029565 | ENSMUSG00000028618 | **Tmem59** | 1110001M20Rik\|3110046P06Rik\|AI256529\|D4Ertd20e\|MTDCF1\|ORF18 | transmembrane protein 59 | protein-coding |
| chr4_15273203_15273560 | chr4 | 15273204 | 15273560 | + | -1.40985 | intron (NM_181401, intron 1 of 2) | intron (NM_181401, intron 1 of 2) | 7562 | NM_181401 | 100201 | Mm.394111 | NM_181401 | ENSMUSG00000043252 | **Tmem64** | 9630015D15Rik\|AI790744\|AV300874 | transmembrane protein 64 | protein-coding |
| chr4_15273187_15273723 | chr4 | 15273188 | 15273723 | + | -1.43512 | intron (NM_181401, intron 1 of 2) | intron (NM_181401, intron 1 of 2) | 7635 | NM_181401 | 100201 | Mm.394111 | NM_181401 | ENSMUSG00000043252 | **Tmem64** | 9630015D15Rik\|AI790744\|AV300874 | transmembrane protein 64 | protein-coding |
| chr15_58697099_58697405 | chr15 | 58697100 | 58697405 | + | -0.637369 | Intergenic | IAPEY5_I-int\|LTR\|ERVK | 126175 | NM_175212 | 74868 | Mm.293754 | NM_175212 | ENSMUSG00000062373 | **Tmem65** | 2610029O13Rik\|4930438D12Rik | transmembrane protein 65 | protein-coding |
| chr4_116540107_116540447 | chr4 | 116540108 | 116540447 | + | -0.988743 | Intergenic | Intergenic | 15666 | NM_177670 | 230657 | Mm.291484 | NM_177670 | | **Tmem69** | A630048M13Rik | transmembrane protein 69 | protein-coding |
| chr11_78600012_78600458 | chr11 | 78600013 | 78600458 | + | -1.23795 | intron (NM_008702, intron 3 of 10) | intron (NM_008702, intron 3 of 10) | -49500 | NM_133706 | 69071 | Mm.29431 | NM_133706 | ENSMUSG00000037278 | **Tmem97** | 1810014L12Rik\|AI115531\|AL022956\|D11Bhm182e | transmembrane protein 97 | protein-coding |
| chr10_118599699_118599997 | chr10 | 118599700 | 118599997 | + | -0.654893 | Intergenic | L1MB7\|LINE\|L1 | -43323 | NR_104123 | 103214 | Mm.30529 | NR_104123 | | **Tmevpg1** | AI447422 | Theiler's murine encephalomyelitis virus persistence candidate gene 1 | ncRNA |
| chr10_118556337_118556908 | chr10 | 118556338 | 118556908 | + | -0.751909 | promoter-TSS (NR_104123) | promoter-TSS (NR_104123) | -98 | NR_104123 | 103214 | Mm.30529 | NR_104123 | | **Tmevpg1** | AI447422 | Theiler's murine encephalomyelitis virus persistence candidate gene 1 | ncRNA |
| chr10_118633378_118634010 | chr10 | 118633379 | 118634010 | + | -1.51187 | Intergenic | Intergenic | -77169 | NR_104123 | 103214 | Mm.30529 | NR_104123 | | **Tmevpg1** | AI447422 | Theiler's murine encephalomyelitis virus persistence candidate gene 1 | ncRNA |
| chr10_118599120_118599607 | chr10 | 118599121 | 118599607 | + | -1.60496 | Intergenic | Intergenic | -42839 | NR_104123 | 103214 | Mm.30529 | NR_104123 | | **Tmevpg1** | AI447422 | Theiler's murine encephalomyelitis virus persistence candidate gene 1 | ncRNA |

| | | | | | | | | | | | | | | | | | |
|---|---|---|---|---|---|---|---|---|---|---|---|---|---|---|---|---|---|
| chr9_75546854_75547177 | chr9 | 75546855 | 75547177 | + | -0.828352 | intron (NM_016963, intron 1 of 9) | intron (NM_016963, intron 1 of 9) | 12641 | NM_016963 | 50875 | Mm.38445 | NM_016963 | ENSMUSG00000058587 | **Tmod3** | U-Tmod\|UTMOD | tropomodulin 3 | protein-coding |
| chr9_75536990_75537315 | chr9 | 75536991 | 75537315 | + | -0.990013 | intron (NM_016963, intron 1 of 9) | intron (NM_016963, intron 1 of 9) | 22504 | NM_016963 | 50875 | Mm.38445 | NM_016963 | ENSMUSG00000058587 | **Tmod3** | U-Tmod\|UTMOD | tropomodulin 3 | protein-coding |
| chr9_114443281_114443658 | chr9 | 114443282 | 114443658 | + | -1.0262 | intron (NR_108101, intron 9 of 14) | intron (NR_108101, intron 9 of 14) | 42375 | NM_001200002 | 100504715 | Mm.290516 | NM_001200002 | ENSMUSG00000079260 | **Tmppe** | - | transmembrane protein with metallophosphoesterase domain | protein-coding |
| chr9_114443281_114443790 | chr9 | 114443282 | 114443790 | + | -1.0719 | intron (NR_108101, intron 9 of 14) | intron (NR_108101, intron 9 of 14) | 42441 | NM_001200002 | 100504715 | Mm.290516 | NM_001200002 | ENSMUSG00000079260 | **Tmppe** | - | transmembrane protein with metallophosphoesterase domain | protein-coding |
| chr9_114443412_114443657 | chr9 | 114443413 | 114443657 | + | -1.07837 | intron (NR_108101, intron 9 of 14) | intron (NR_108101, intron 9 of 14) | 42440 | NM_001200002 | 100504715 | Mm.290516 | NM_001200002 | ENSMUSG00000079260 | **Tmppe** | - | transmembrane protein with metallophosphoesterase domain | protein-coding |
| chr6_72993133_72993728 | chr6 | 72993134 | 72993728 | + | -0.863907 | Intergenic | Intergenic | -34683 | NM_001190327 | 19240 | Mm.3532 | NM_025284 | ENSMUSG00000079523 | **Tmsb10** | Ptmb10\|Tb10 | thymosin, beta 10 | protein-coding |
| chr6_72993133_72994025 | chr6 | 72993134 | 72994025 | + | -0.893571 | Intergenic | Intergenic | -34831 | NM_001190327 | 19240 | Mm.3532 | NM_025284 | ENSMUSG00000079523 | **Tmsb10** | Ptmb10\|Tb10 | thymosin, beta 10 | protein-coding |
| chr6_72993770_72994100 | chr6 | 72993771 | 72994100 | + | -0.946666 | Intergenic | ID_B1\|SINE\|B4 | -35187 | NM_001190327 | 19240 | Mm.3532 | NM_025284 | ENSMUSG00000079523 | **Tmsb10** | Ptmb10\|Tb10 | thymosin, beta 10 | protein-coding |
| chr10_19043826_19044285 | chr10 | 19043827 | 19044285 | + | -0.640375 | Intergenic | RLTR16\|LTR\|ERVK | -28646 | NM_009397 | 21929 | Mm.116683 | NM_009397 | ENSMUSG00000019850 | **Tnfaip3** | A20\|Tnfip3 | tumor necrosis factor, alpha-induced protein 3 | protein-coding |
| chr10_18999278_18999761 | chr10 | 18999279 | 18999761 | + | -0.757182 | Intergenic | Intergenic | 12443 | NM_001166402 | 21929 | Mm.116683 | NM_009397 | ENSMUSG00000019850 | **Tnfaip3** | A20\|Tnfip3 | tumor necrosis factor, alpha-induced protein 3 | protein-coding |
| chr10_19061928_19063110 | chr10 | 19061929 | 19063110 | + | -1.10644 | Intergenic | Intergenic | -47109 | NM_009397 | 21929 | Mm.116683 | NM_009397 | ENSMUSG00000019850 | **Tnfaip3** | A20\|Tnfip3 | tumor necrosis factor, alpha-induced protein 3 | protein-coding |
| chr14_69774338_69774605 | chr14 | 69774339 | 69774605 | + | -0.674832 | intron (NM_020275, intron 3 of 7) | MTE2a\|LTR\|ERVL-MaLR | 7000 | NM_020275 | 21933 | Mm.193430 | NM_020275 | ENSMUSG00000022074 | **Tnfrsf10b** | DR5\|KILLER\|Ly98\|MK\|TRAILR2\|TRICK2A\|TRICK2B\|TRICKB | tumor necrosis factor receptor superfamily, member 10b | protein-coding |
| chr14_69774113_69775273 | chr14 | 69774114 | 69775273 | + | -1.147 | intron (NM_020275, intron 3 of 7) | intron (NM_020275, intron 3 of 7) | 7221 | NM_020275 | 21933 | Mm.193430 | NM_020275 | ENSMUSG00000022074 | **Tnfrsf10b** | DR5\|KILLER\|Ly98\|MK\|TRAILR2\|TRICK2A\|TRICK2B\|TRICKB | tumor necrosis factor receptor superfamily, member 10b | protein-coding |
| chr14_69774654_69775275 | chr14 | 69774655 | 69775275 | + | -1.35999 | intron (NM_020275, intron 3 of 7) | intron (NM_020275, intron 3 of 7) | 7493 | NM_020275 | 21933 | Mm.193430 | NM_020275 | ENSMUSG00000022074 | **Tnfrsf10b** | DR5\|KILLER\|Ly98\|MK\|TRAILR2\|TRICK2A\|TRICK2B\|TRICKB | tumor necrosis factor receptor superfamily, member 10b | protein-coding |
| chr14_69780306_69780886 | chr14 | 69780307 | 69780886 | + | -1.42258 | intron (NM_020275, intron 7 of 7) | intron (NM_020275, intron 7 of 7) | 13124 | NM_020275 | 21933 | Mm.193430 | NM_020275 | ENSMUSG00000022074 | **Tnfrsf10b** | DR5\|KILLER\|Ly98\|MK\|TRAILR2\|TRICK2A\|TRICK2B\|TRICKB | tumor necrosis factor receptor superfamily, member 10b | protein-coding |
| chr1_105752248_105752587 | chr1 | 105752249 | 105752587 | + | -0.633043 | intron (NM_173187, intron 28 of 28) | intron (NM_173187, intron 28 of 28) | -28305 | NM_009399 | 21934 | Mm.6251 | NM_009399 | ENSMUSG00000026321 | **Tnfrsf11a** | Ly109\|ODFR\|OFE\|Rank\|TRANCE-R\|mRANK | tumor necrosis factor receptor superfamily, member 11a, NFKB activator | protein-coding |
| chr4_154936478_154937236 | chr4 | 154936479 | 154937236 | + | -1.15901 | Intergenic | Intergenic | -8780 | NM_178931 | 230979 | Mm.215147 | NM_178931 | ENSMUSG00000042333 | **Tnfrsf14** | Atar\|HveA\|Hvem\|Tnfrs14 | tumor necrosis factor receptor superfamily, member 14 (herpesvirus entry mediator) | protein-coding |
| chr6_125351328_125351741 | chr6 | 125351329 | 125351741 | + | -0.805945 | intron (NM_011609, intron 1 of 9) | intron (NM_011609, intron 1 of 9) | 1812 | NM_011609 | 21937 | Mm.1258 | NM_011609 | ENSMUSG00000030341 | **Tnfrsf1a** | CD120a\|FPF\|TNF-R\|TNF-R-I\|TNF-R1\|TNF-R55\|TNF-alphaR1\|TNFAR\|TNFR60\|TNFRI\|TNFRp55\|TNFalpha-R1\|Tnfr-2\|Tnfr1\|p55\|p55-R | tumor necrosis factor receptor superfamily, member 1a | protein-coding |
| chr4_145229634_145230079 | chr4 | 145229635 | 145230079 | + | -0.67742 | intron (NM_011610, intron 1 of 9) | intron (NM_011610, intron 1 of 9) | 17013 | NM_011610 | 21938 | Mm.235328 | NM_011610 | | **Tnfrsf1b** | CD120b\|TNF-R-II\|TNF-R2\|TNF-R75\|TNF-alphaR2\|TNFBR\|TNFR80\|TNFRII\|TNFalpha-R2\|Tnfr-1\|Tnfr2\|p75 | tumor necrosis factor receptor superfamily, member 1b | protein-coding |
| chr4_145228713_145229483 | chr4 | 145228714 | 145229483 | + | -0.708138 | exon (NM_011610, exon 2 of 10) | exon (NM_011610, exon 2 of 10) | 17772 | NM_011610 | 21938 | Mm.235328 | NM_011610 | | **Tnfrsf1b** | CD120b\|TNF-R-II\|TNF-R2\|TNF-R75\|TNF-alphaR2\|TNFBR\|TNFR80\|TNFRII\|TNFalpha-R2\|Tnfr-1\|Tnfr2\|p75 | tumor necrosis factor receptor superfamily, member 1b | protein-coding |
| chr4_145228780_145230225 | chr4 | 145228781 | 145230225 | + | -0.759841 | intron (NM_011610, intron 1 of 9) | intron (NM_011610, intron 1 of 9) | 17367 | NM_011610 | 21938 | Mm.235328 | NM_011610 | | **Tnfrsf1b** | CD120b\|TNF-R-II\|TNF-R2\|TNF-R75\|TNF-alphaR2\|TNFBR\|TNFR80\|TNFRII\|TNFalpha-R2\|Tnfr-1\|Tnfr2\|p75 | tumor necrosis factor receptor superfamily, member 1b | protein-coding |
| chr4_145224681_145224942 | chr4 | 145224682 | 145224942 | + | -0.760779 | exon (NM_011610, exon 5 of 10) | exon (NM_011610, exon 5 of 10) | 22058 | NM_011610 | 21938 | Mm.235328 | NM_011610 | | **Tnfrsf1b** | CD120b\|TNF-R-II\|TNF-R2\|TNF-R75\|TNF-alphaR2\|TNFBR\|TNFR80\|TNFRII\|TNFalpha-R2\|Tnfr-1\|Tnfr2\|p75 | tumor necrosis factor receptor superfamily, member 1b | protein-coding |
| chr4_145240725_145240978 | chr4 | 145240726 | 145240978 | + | -0.77488 | intron (NM_011610, intron 1 of 9) | intron (NM_011610, intron 1 of 9) | 6018 | NM_011610 | 21938 | Mm.235328 | NM_011610 | | **Tnfrsf1b** | CD120b\|TNF-R-II\|TNF-R2\|TNF-R75\|TNF-alphaR2\|TNFBR\|TNFR80\|TNFRII\|TNFalpha-R2\|Tnfr-1\|Tnfr2\|p75 | tumor necrosis factor receptor superfamily, member 1b | protein-coding |
| chr4_145240652_145241342 | chr4 | 145240653 | 145241342 | + | -0.828872 | intron (NM_011610, intron 1 of 9) | intron (NM_011610, intron 1 of 9) | 5873 | NM_011610 | 21938 | Mm.235328 | NM_011610 | | **Tnfrsf1b** | CD120b\|TNF-R-II\|TNF-R2\|TNF-R75\|TNF-alphaR2\|TNFBR\|TNFR80\|TNFRII\|TNFalpha-R2\|Tnfr-1\|Tnfr2\|p75 | tumor necrosis factor receptor superfamily, member 1b | protein-coding |
| chr4_145235823_145236538 | chr4 | 145235824 | 145236538 | + | -0.839887 | intron (NM_011610, intron 1 of 9) | intron (NM_011610, intron 1 of 9) | 10689 | NM_011610 | 21938 | Mm.235328 | NM_011610 | | **Tnfrsf1b** | CD120b\|TNF-R-II\|TNF-R2\|TNF-R75\|TNF-alphaR2\|TNFBR\|TNFR80\|TNFRII\|TNFalpha-R2\|Tnfr-1\|Tnfr2\|p75 | tumor necrosis factor receptor superfamily, member 1b | protein-coding |
| chr4_145241058_145241326 | chr4 | 145241059 | 145241326 | + | -0.865391 | intron (NM_011610, intron 1 of 9) | intron (NM_011610, intron 1 of 9) | 5678 | NM_011610 | 21938 | Mm.235328 | NM_011610 | | **Tnfrsf1b** | CD120b\|TNF-R-II\|TNF-R2\|TNF-R75\|TNF-alphaR2\|TNFBR\|TNFR80\|TNFRII\|TNFalpha-R2\|Tnfr-1\|Tnfr2\|p75 | tumor necrosis factor receptor superfamily, member 1b | protein-coding |
| chr4_156004055_156004639 | chr4 | 156004056 | 156004639 | + | -0.943348 | intron (NM_011341, intron 4 of 6) | intron (NM_011341, intron 4 of 6) | -9348 | NM_011659 | 22163 | Mm.13885 | NM_011659 | ENSMUSG00000029075 | **Tnfrsf4** | ACT35\|CD134\|Ly-70\|Ox40\|TXGP1L\|Txgp1 | tumor necrosis factor receptor superfamily, member 4 | protein-coding |
| chr4_156004040_156004481 | chr4 | 156004041 | 156004481 | + | -0.995295 | intron (NM_011341, intron 4 of 6) | ORR1D2-int\|LTR\|ERVL-MaLR | -9434 | NM_011659 | 22163 | Mm.13885 | NM_011659 | ENSMUSG00000029075 | **Tnfrsf4** | ACT35\|CD134\|Ly-70\|Ox40\|TXGP1L\|Txgp1 | tumor necrosis factor receptor superfamily, member 4 | protein-coding |
| chr4_156004156_156004454 | chr4 | 156004157 | 156004454 | + | -1.01511 | intron (NM_011341, intron 4 of 6) | intron (NM_011341, intron 4 of 6) | -9390 | NM_011659 | 22163 | Mm.13885 | NM_011659 | ENSMUSG00000029075 | **Tnfrsf4** | ACT35\|CD134\|Ly-70\|Ox40\|TXGP1L\|Txgp1 | tumor necrosis factor receptor superfamily, member 4 | protein-coding |
| chr4_63704204_63704523 | chr4 | 63704205 | 63704523 | + | -0.883594 | Intergenic | Intergenic | 40749 | NM_177371 | 326623 | Mm.208152 | NM_177371 | | **Tnfsf15** | Tl1\|Tl1a\|Vegi | tumor necrosis factor (ligand) superfamily, member 15 | protein-coding |
| chr4_63849122_63849536 | chr4 | 63849123 | 63849536 | + | -0.629526 | intron (NM_009403, intron 2 of 3) | intron (NM_009403, intron 2 of 3) | 11955 | NM_009403 | 21949 | Mm.4664 | NM_009403 | | **Tnfsf8** | CD153\|CD30LG\|Cd30l | tumor necrosis factor (ligand) superfamily, member 8 | protein-coding |
| chr3_28255729_28256144 | chr3 | 28255730 | 28256144 | + | -0.647036 | Intergenic | Lx3_Mus\|LINE\|L1 | -7277 | NM_001163007 | 665113 | Mm.126193 | NM_026910 | ENSMUSG00000027692 | **Tnik** | 1500031A17Rik\|4831440I19Rik\|AI451411\|C530008O15Rik\|C630040K21Rik | TRAF2 and NCK interacting kinase | protein-coding |
| chr3_28274858_28275130 | chr3 | 28274859 | 28275130 | + | -0.856401 | intron (NM_001163007, intron 1 of 31) | intron (NM_001163007, intron 1 of 31) | 11780 | NM_001163009 | 665113 | Mm.126193 | NM_026910 | ENSMUSG00000027692 | **Tnik** | 1500031A17Rik\|4831440I19Rik\|AI451411\|C530008O15Rik\|C630040K21Rik | TRAF2 and NCK interacting kinase | protein-coding |
| chr6_65611864_65612191 | chr6 | 65611865 | 65612191 | + | -0.620616 | intron (NM_001001495, intron 6 of 7) | intron (NM_001001495, intron 6 of 7) | 21630 | NM_001001495 | 414084 | Mm.117558 | NM_001001495 | ENSMUSG00000044162 | **Tnip3** | 9030611K07Rik\|ABIN-3\|LIND | TNFAIP3 interacting protein 3 | protein-coding |
| chr6_65599826_65600597 | chr6 | 65599827 | 65600597 | + | -0.65495 | intron (NM_001001495, intron 5 of 7) | (A)n\|Simple_repeat\|Simple_repeat | 9814 | NM_001001495 | 414084 | Mm.117558 | NM_001001495 | ENSMUSG00000044162 | **Tnip3** | 9030611K07Rik\|ABIN-3\|LIND | TNFAIP3 interacting protein 3 | protein-coding |
| chr6_65569780_65570128 | chr6 | 65569781 | 65570128 | + | -0.691394 | Intergenic | Intergenic | -20444 | NM_001001495 | 414084 | Mm.117558 | NM_001001495 | ENSMUSG00000044162 | **Tnip3** | 9030611K07Rik\|ABIN-3\|LIND | TNFAIP3 interacting protein 3 | protein-coding |
| chr6_65573273_65573842 | chr6 | 65573274 | 65573842 | + | -0.780626 | Intergenic | Intergenic | -16840 | NM_001001495 | 414084 | Mm.117558 | NM_001001495 | ENSMUSG00000044162 | **Tnip3** | 9030611K07Rik\|ABIN-3\|LIND | TNFAIP3 interacting protein 3 | protein-coding |
| chr6_65600199_65600522 | chr6 | 65600200 | 65600522 | + | -0.842347 | intron (NM_001001495, intron 5 of 7) | intron (NM_001001495, intron 5 of 7) | 9963 | NM_001001495 | 414084 | Mm.117558 | NM_001001495 | ENSMUSG00000044162 | **Tnip3** | 9030611K07Rik\|ABIN-3\|LIND | TNFAIP3 interacting protein 3 | protein-coding |
| chr16_32650840_32651616 | chr16 | 32650841 | 32651616 | + | -0.701993 | intron (NM_016788, intron 1 of 15) | intron (NM_016788, intron 1 of 15) | 6585 | NM_016788 | 51789 | Mm.251115 | NM_016788 | ENSMUSG00000022791 | **Tnk2** | Ack\|Ack-1\|Ack1\|Cdgip\|Pyk1 | tyrosine kinase, non-receptor, 2 | protein-coding |
| chr8_34970109_34970467 | chr8 | 34970110 | 34970467 | + | -0.696093 | Intergenic | RLTR9A\|LTR\|ERVK | -4598 | NM_175091 | 21951 | Mm.88364 | NM_175091 | ENSMUSG00000031529 | **Tnks** | 4930554K12Rik\|AI662855\|ARTD5\|C86528\|D130072O21Rik\|TANK1\|mTNKS1 | tankyrase, TRF1-interacting ankyrin-related ADP-ribose polymerase | protein-coding |
| chr8_34970139_34970381 | chr8 | 34970140 | 34970381 | + | -0.711444 | Intergenic | RLTR9A\|LTR\|ERVK | -4570 | NM_175091 | 21951 | Mm.88364 | NM_175091 | ENSMUSG00000031529 | **Tnks** | 4930554K12Rik\|AI662855\|ARTD5\|C86528\|D130072O21Rik\|TANK1\|mTNKS1 | tankyrase, TRF1-interacting ankyrin-related ADP-ribose polymerase | protein-coding |

| | | | | | | | | | | | | | | | | | |
|---|---|---|---|---|---|---|---|---|---|---|---|---|---|---|---|---|---|
| chr8_34938657_34939014 | chr8 | 34938658 | 34939014 | + | -0.912081 | intron (NM_175091, intron 2 of 26) | intron (NM_175091, intron 2 of 26) | 26854 | NM_175091 | 21951 | Mm.88364 | NM_175091 | ENSMUSG00000031529 | **Tnks** | 4930554K12Rik\|AI662855\|ARTD5\|C86528\|D130072O21Rik\|TANK1\|mTNKS1 | tankyrase, TRF1-interacting ankyrin-related ADP-ribose polymerase | protein-coding |
| chr8_34945009_34945463 | chr8 | 34945010 | 34945463 | + | -1.37585 | intron (NM_175091, intron 1 of 26) | MTD\|LTR\|ERVL-MaLR | 20454 | NM_175091 | 21951 | Mm.88364 | NM_175091 | ENSMUSG00000031529 | **Tnks** | 4930554K12Rik\|AI662855\|ARTD5\|C86528\|D130072O21Rik\|TANK1\|mTNKS1 | tankyrase, TRF1-interacting ankyrin-related ADP-ribose polymerase | protein-coding |
| chr1_135807366_135808188 | chr1 | 135807367 | 135808188 | + | -0.82862 | intron (NM_001112702, intron 6 of 8) | intron (NM_001112702, intron 6 of 8) | 8266 | NM_001112702 | 21952 | Mm.44379 | NM_021467 | ENSMUSG00000026418 | **Tnni1** | 2700018B22Rik\|AI747285\|ssTnI | troponin I, skeletal, slow 1 | protein-coding |
| chr7_4518501_4518940 | chr7 | 4518502 | 4518940 | + | -0.774971 | intron (NM_009406, intron 7 of 7) | intron (NM_009406, intron 7 of 7) | -2746 | NM_011618 | 21955 | Mm.358643 | NM_011618 | ENSMUSG00000064179 | **Tnnt1** | AW146156\|Tnt\|sTnT\|ssTnT | troponin T1, skeletal, slow | protein-coding |
| chr13_98896262_98897365 | chr13 | 98896263 | 98897365 | + | -0.633974 | intron (NM_178716, intron 1 of 23) | MER20B\|DNA\|hAT-Charlie | -5772 | NM_001048267 | 238799 | Mm.173286 | NM_178716 | ENSMUSG00000009470 | **Tnpo1** | AU021749\|D13Ertd688e\|IPO2\|Kpnb2\|MIP\|MIP1\|TRN | transportin 1 | protein-coding |
| chr13_98896483_98897188 | chr13 | 98896484 | 98897188 | + | -0.694141 | intron (NM_178716, intron 1 of 23) | MER20B\|DNA\|hAT-Charlie | -5794 | NM_001048267 | 238799 | Mm.173286 | NM_178716 | ENSMUSG00000009470 | **Tnpo1** | AU021749\|D13Ertd688e\|IPO2\|Kpnb2\|MIP\|MIP1\|TRN | transportin 1 | protein-coding |
| chr5_142794123_142795351 | chr5 | 142794124 | 142795351 | + | -0.605749 | intron (NM_001122730, intron 2 of 27) | intron (NM_001122730, intron 2 of 27) | 22650 | NM_178242 | 231861 | Mm.354649 | NM_178242 | ENSMUSG00000039477 | **Tnrc18** | BC025631\|EG381742\|Zfp469 | trinucleotide repeat containing 18 | protein-coding |
| chr7_123135676_123136046 | chr7 | 123135677 | 123136046 | + | -0.67867 | intron (NM_144925, intron 2 of 23) | intron (NM_144925, intron 2 of 23) | 11976 | NM_144925 | 233833 | Mm.102305 | NM_144925 | ENSMUSG00000052707 | **Tnrc6a** | 2010321I05Rik\|3110054G10Rik\|AW557223\|CAGH26\|D130023A07Rik\|GW182\|Tnrc6 | trinucleotide repeat containing 6a | protein-coding |
| chr15_80814615_80815207 | chr15 | 80814616 | 80815207 | + | -0.72699 | intron (NM_177124, intron 1 of 21) | intron (NM_177124, intron 1 of 21) | 16301 | NM_177124 | 213988 | Mm.131328 | NM_144812 | ENSMUSG00000047888 | **Tnrc6b** | 2700090M07Rik\|A730065C02Rik\|AI848765\|D230019K20Rik | trinucleotide repeat containing 6b | protein-coding |
| chr15_80741884_80742173 | chr15 | 80741885 | 80742173 | + | -1.22122 | intron (NM_144812, intron 1 of 24) | ID_B1\|SINE\|B4 | 30716 | NM_144812 | 213988 | Mm.131328 | NM_144812 | ENSMUSG00000047888 | **Tnrc6b** | 2700090M07Rik\|A730065C02Rik\|AI848765\|D230019K20Rik | trinucleotide repeat containing 6b | protein-coding |
| chr11_117638565_117638806 | chr11 | 117638566 | 117638806 | + | -0.739911 | Intergenic | Intergenic | -15603 | NM_198022 | 217351 | Mm.40272 | NM_198022 | ENSMUSG00000025571 | **Tnrc6c** | 9930033H14Rik\|mKIAA1582 | trinucleotide repeat containing 6C | protein-coding |
| chr11_117638925_117640069 | chr11 | 117638926 | 117640069 | + | -0.747688 | Intergenic | Intergenic | -14792 | NM_198022 | 217351 | Mm.40272 | NM_198022 | ENSMUSG00000025571 | **Tnrc6c** | 9930033H14Rik\|mKIAA1582 | trinucleotide repeat containing 6C | protein-coding |
| chr11_117638861_117639305 | chr11 | 117638862 | 117639305 | + | -0.985175 | Intergenic | (CCCCCA)n\|Simple_repeat\|Simple_repeat | -15206 | NM_198022 | 217351 | Mm.40272 | NM_198022 | ENSMUSG00000025571 | **Tnrc6c** | 9930033H14Rik\|mKIAA1582 | trinucleotide repeat containing 6C | protein-coding |
| chr11_117638916_117639144 | chr11 | 117638917 | 117639144 | + | -1.16053 | Intergenic | Intergenic | -15259 | NM_198022 | 217351 | Mm.40272 | NM_198022 | ENSMUSG00000025571 | **Tnrc6c** | 9930033H14Rik\|mKIAA1582 | trinucleotide repeat containing 6C | protein-coding |
| chr11_99104566_99105526 | chr11 | 99104567 | 99105526 | + | -0.645734 | Intergenic | L2b\|LINE\|L2 | -15740 | NM_172564 | 217169 | Mm.262067 | NM_172564 | ENSMUSG00000017607 | **Tns4** | 9930017A07Rik\|AA589547\|AU016405\|Cten | tensin 4 | protein-coding |
| chr11_99116908_99117172 | chr11 | 99116909 | 99117172 | + | -0.717405 | Intergenic | Intergenic | -27734 | NM_172564 | 217169 | Mm.262067 | NM_172564 | ENSMUSG00000017607 | **Tns4** | 9930017A07Rik\|AA589547\|AU016405\|Cten | tensin 4 | protein-coding |
| chr11_60334278_60334596 | chr11 | 60334279 | 60334596 | + | -0.638341 | intron (NM_001039092, intron 1 of 13) | L1ME2\|LINE\|L1 | 18468 | NM_153080 | 216810 | Mm.218875 | NM_153080 | ENSMUSG00000000538 | **Tom1l2** | 2900016I08Rik\|A730055F12Rik\|AU042072\|Srebf1 | target of myb1-like 2 (chicken) | protein-coding |
| chr11_60330433_60330926 | chr11 | 60330434 | 60330926 | + | -0.680396 | intron (NM_001039092, intron 1 of 13) | intron (NM_001039092, intron 1 of 13) | 22225 | NM_153080 | 216810 | Mm.218875 | NM_153080 | ENSMUSG00000000538 | **Tom1l2** | 2900016I08Rik\|A730055F12Rik\|AU042072\|Srebf1 | target of myb1-like 2 (chicken) | protein-coding |
| chr11_60330646_60330919 | chr11 | 60330647 | 60330919 | + | -0.734968 | intron (NM_001039092, intron 1 of 13) | intron (NM_001039092, intron 1 of 13) | 22122 | NM_153080 | 216810 | Mm.218875 | NM_153080 | ENSMUSG00000000538 | **Tom1l2** | 2900016I08Rik\|A730055F12Rik\|AU042072\|Srebf1 | target of myb1-like 2 (chicken) | protein-coding |
| chr14_16372769_16373242 | chr14 | 16372770 | 16373242 | + | -0.943231 | intron (NM_009409, intron 1 of 35) | intron (NM_009409, intron 1 of 35) | 7800 | NM_009409 | 21974 | Mm.130362 | NM_009409 | ENSMUSG00000017485 | **Top2b** | D230016L12Rik\|Top-2 | topoisomerase (DNA) II beta | protein-coding |
| chr14_16372624_16373263 | chr14 | 16372625 | 16373263 | + | -0.952917 | intron (NM_009409, intron 1 of 35) | intron (NM_009409, intron 1 of 35) | 7738 | NM_009409 | 21974 | Mm.130362 | NM_009409 | ENSMUSG00000017485 | **Top2b** | D230016L12Rik\|Top-2 | topoisomerase (DNA) II beta | protein-coding |
| chr1_156076227_156076937 | chr1 | 156076228 | 156076937 | + | -0.704026 | 3' UTR (NM_177838, exon 5 of 5) | 3' UTR (NM_177838, exon 5 of 5) | 35721 | NM_022329 | 240832 | Mm.253335 | NM_022329 | ENSMUSG00000050565 | **Tor1aip2** | 1110020D10Rik\|15kDa\|A130072J07\|AA103493\|AW060462\|AW610675\|C77739\|Ifrg15\|Lull1 | torsin A interacting protein 2 | protein-coding |
| chr4_7207046_7207324 | chr4 | 7207047 | 7207324 | + | -0.677388 | Intergenic | Intergenic | -216462 | NM_145711 | 252838 | Mm.435828 | NM_145711 | ENSMUSG00000041272 | **Tox** | 1700007F02Rik | thymocyte selection-associated high mobility group box | protein-coding |
| chr4_7206964_7207352 | chr4 | 7206965 | 7207352 | + | -0.693713 | Intergenic | Intergenic | -216435 | NM_145711 | 252838 | Mm.435828 | NM_145711 | ENSMUSG00000041272 | **Tox** | 1700007F02Rik | thymocyte selection-associated high mobility group box | protein-coding |
| chr4_7206791_7207330 | chr4 | 7206792 | 7207330 | + | -0.711201 | Intergenic | Intergenic | -216338 | NM_145711 | 252838 | Mm.435828 | NM_145711 | ENSMUSG00000041272 | **Tox** | 1700007F02Rik | thymocyte selection-associated high mobility group box | protein-coding |
| chr4_6950300_6950919 | chr4 | 6950301 | 6950919 | + | -1.02633 | intron (NM_145711, intron 1 of 8) | MTD\|LTR\|ERVL-MaLR | 40113 | NM_145711 | 252838 | Mm.435828 | NM_145711 | ENSMUSG00000041272 | **Tox** | 1700007F02Rik | thymocyte selection-associated high mobility group box | protein-coding |
| chr4_6950298_6950727 | chr4 | 6950299 | 6950727 | + | -1.04541 | intron (NM_145711, intron 1 of 8) | intron (NM_145711, intron 1 of 8) | 40210 | NM_145711 | 252838 | Mm.435828 | NM_145711 | ENSMUSG00000041272 | **Tox** | 1700007F02Rik | thymocyte selection-associated high mobility group box | protein-coding |
| chr4_6975792_6976215 | chr4 | 6975793 | 6976215 | + | -1.25077 | intron (NM_145711, intron 1 of 8) | intron (NM_145711, intron 1 of 8) | 14719 | NM_145711 | 252838 | Mm.435828 | NM_145711 | ENSMUSG00000041272 | **Tox** | 1700007F02Rik | thymocyte selection-associated high mobility group box | protein-coding |
| chr4_6970375_6970965 | chr4 | 6970376 | 6970965 | + | -1.31263 | intron (NM_145711, intron 1 of 8) | intron (NM_145711, intron 1 of 8) | 20053 | NM_145711 | 252838 | Mm.435828 | NM_145711 | ENSMUSG00000041272 | **Tox** | 1700007F02Rik | thymocyte selection-associated high mobility group box | protein-coding |
| chr5_120578799_120579053 | chr5 | 120578800 | 120579053 | + | -0.80583 | intron (NM_145853, intron 2 of 27) | intron (NM_145853, intron 2 of 27) | 9687 | NM_145853 | 252972 | Mm.114054 | NM_145853 | ENSMUSG00000032741 | **Tpcn1** | 5730403B01Rik\|Tpc1\|mKIAA1169 | two pore channel 1 | protein-coding |
| chr3_8971662_8972004 | chr3 | 8971663 | 8972004 | + | -0.908143 | intron (NM_009412, intron 1 of 5) | MLT1J1\|LTR\|ERVL-MaLR | -7779 | NM_001025261 | 21985 | Mm.371590 | NM_009412 | ENSMUSG00000027506 | **Tpd52** | AI043038\|mD52 | tumor protein D52 | protein-coding |
| chr6_43619236_43619683 | chr6 | 43619237 | 43619683 | + | -0.64049 | intron (NM_013861, intron 2 of 8) | intron (NM_013861, intron 2 of 8) | 46715 | NM_013861 | 29807 | Mm.320979 | NM_013861 | ENSMUSG00000029735 | **Tpk1** | - | thiamine pyrophosphokinase | protein-coding |
| chr6_43619276_43619598 | chr6 | 43619277 | 43619598 | + | -0.676519 | intron (NM_013861, intron 2 of 8) | intron (NM_013861, intron 2 of 8) | 46738 | NM_013861 | 29807 | Mm.320979 | NM_013861 | ENSMUSG00000029735 | **Tpk1** | - | thiamine pyrophosphokinase | protein-coding |
| chr8_72131859_72132226 | chr8 | 72131860 | 72132226 | + | -0.736013 | Intergenic | Intergenic | -3249 | NM_001001491 | 326618 | Mm.295124 | NM_001001491 | ENSMUSG00000031799 | **Tpm4** | 2610528G24Rik | tropomyosin 4 | protein-coding |
| chr1_43896954_43897735 | chr1 | 43896955 | 43897735 | + | -0.675722 | Intergenic | Intergenic | -36662 | NM_009418 | 22019 | Mm.401675 | NM_009418 | ENSMUSG00000041763 | **Tpp2** | TPP-2\|TppII | tripeptidyl peptidase II | protein-coding |
| chr1_43896970_43897547 | chr1 | 43896971 | 43897547 | + | -0.69998 | Intergenic | Intergenic | -36748 | NM_009418 | 22019 | Mm.401675 | NM_009418 | ENSMUSG00000041763 | **Tpp2** | TPP-2\|TppII | tripeptidyl peptidase II | protein-coding |
| chr5_112270934_112271661 | chr5 | 112270935 | 112271661 | + | -0.687352 | Intergenic | Intergenic | -5409 | NM_009419 | 22022 | Mm.250631 | NM_009419 | ENSMUSG00000029344 | **Tpst2** | AI448750\|D5Ucla3\|Tango13b\|grm\|grt | protein-tyrosine sulfotransferase 2 | protein-coding |
| chr5_112286640_112287620 | chr5 | 112286641 | 112287620 | + | -0.888384 | intron (NM_009419, intron 1 of 6) | intron (NM_009419, intron 1 of 6) | 10423 | NM_009419 | 22022 | Mm.250631 | NM_009419 | ENSMUSG00000029344 | **Tpst2** | AI448750\|D5Ucla3\|Tango13b\|grm\|grt | protein-tyrosine sulfotransferase 2 | protein-coding |
| chr5_112286983_112287490 | chr5 | 112286984 | 112287490 | + | -0.986666 | intron (NM_009419, intron 1 of 6) | intron (NM_009419, intron 1 of 6) | 10530 | NM_009419 | 22022 | Mm.250631 | NM_009419 | ENSMUSG00000029344 | **Tpst2** | AI448750\|D5Ucla3\|Tango13b\|grm\|grt | protein-tyrosine sulfotransferase 2 | protein-coding |
| chr10_39564474_39565091 | chr10 | 39564475 | 39565091 | + | -0.805138 | 3' UTR (NM_001122892, exon 13 of 13) | 3' UTR (NM_001122892, exon 13 of 13) | -48151 | NM_134000 | 103213 | Mm.436686 | NM_134000 | ENSMUSG00000019842 | **Traf3ip2** | AI429613\|Act1\|CIKS | TRAF3 interacting protein 2 | protein-coding |
| chr10_39522040_39522396 | chr10 | 39522041 | 39522396 | + | -0.82026 | intron (NM_008054, intron 5 of 13) | intron (NM_008054, intron 5 of 13) | -90716 | NM_134000 | 103213 | Mm.436686 | NM_134000 | ENSMUSG00000019842 | **Traf3ip2** | AI429613\|Act1\|CIKS | TRAF3 interacting protein 2 | protein-coding |
| chr1_192078221_192079187 | chr1 | 192078222 | 192079187 | + | -0.735514 | intron (NM_011633, intron 1 of 9) | intron (NM_011633, intron 1 of 9) | 13895 | NM_011633 | 22033 | Mm.380491 | NM_011633 | ENSMUSG00000026637 | **Traf5** | - | TNF receptor-associated factor 5 | protein-coding |
| chr5_121335168_121335828 | chr5 | 121335169 | 121335828 | + | -1.24475 | intron (NM_181421, intron 53 of 75) | intron (NM_181421, intron 53 of 75) | 50117 | NM_001163470 | 231712 | Mm.20488 | NM_172275 | ENSMUSG00000042726 | **Trafd1** | 1110008K06Rik\|Fln29 | TRAF type zinc finger domain containing 1 | protein-coding |
| chr5_121335535_121335802 | chr5 | 121335536 | 121335802 | + | -1.3414 | intron (NM_181421, intron 53 of 75) | L1MB7\|LINE\|L1 | 49946 | NM_001163470 | 231712 | Mm.20488 | NM_172275 | ENSMUSG00000042726 | **Trafd1** | 1110008K06Rik\|Fln29 | TRAF type zinc finger domain containing 1 | protein-coding |
| chr1_58956241_58956651 | chr1 | 58956242 | 58956651 | + | -0.838822 | intron (NM_172406, intron 1 of 15) | intron (NM_172406, intron 1 of 15) | 17036 | NM_172406 | 70827 | Mm.222887 | NM_172406 | ENSMUSG00000026028 | **Trak2** | 2900022D04Rik\|4733401O11Rik\|AI480836\|AW046271\|Als2cr3\|CALS-C\|GRIF-1\|GRIF1\|OIP98\|mKIAA0549 | trafficking protein, kinesin binding 2 | protein-coding |
| chr1_21051894_21052747 | chr1 | 21051895 | 21052747 | + | -0.624317 | intron (NM_177409, intron 1 of 10) | intron (NM_177409, intron 1 of 10) | 26904 | NM_177409 | 170829 | Mm.185680 | NM_177409 | | **Tram2** | C330003D03Rik | translocating chain-associating membrane protein 2 | protein-coding |
| chr12_28735238_28736135 | chr12 | 28735239 | 28736135 | + | -0.874531 | intron (NM_178811, intron 3 of 11) | intron (NM_178811, intron 3 of 11) | 14707 | NM_001161410 | 217449 | Mm.132006 | NM_178811 | ENSMUSG00000020628 | **Trappc12** | CGI-87\|D930014A20Rik\|Ttc15 | trafficking protein particle complex 12 | protein-coding |
| chr18_20869250_20869813 | chr18 | 20869251 | 20869813 | + | -0.745507 | intron (NM_177038, intron 5 of 28) | ORR1D2\|LTR\|ERVL-MaLR | 26546 | NM_177038 | 75964 | Mm.273769 | NM_029491 | ENSMUSG00000033382 | **Trappc8** | 5033403J15Rik\|AI845423\|D030074E01Rik\|Trs85\|mKIAA1012 | trafficking protein particle complex 8 | protein-coding |

| | | | | | | | | | | | | | | | | | |
|---|---|---|---|---|---|---|---|---|---|---|---|---|---|---|---|---|---|
| chr18_20869399_20869756 | chr18 | 20869400 | 20869756 | + | -0.770026 | intron (NM_177038, intron 5 of 28) | ORR1D2\|LTR\|ERVL-MaLR | 26500 | NM_177038 | 75964 | Mm.273769 | NM_029491 | ENSMUSG00000033382 | **Trappc8** | 5033403J15Rik\|AI845423\|D030074E01Rik\|Trs85\|mKIAA1012 | trafficking protein particle complex 8 | protein-coding |
| chr16_48786275_48786692 | chr16 | 48786276 | 48786692 | + | -1.16533 | Intergenic | Intergenic | -14528 | NM_198297 | 77647 | Mm.167298 | NM_198297 | ENSMUSG00000030775 | **Trat1** | C030046M14Rik\|Tcrim\|Trim | T cell receptor associated transmembrane adaptor 1 | protein-coding |
| chr9_44690693_44692129 | chr9 | 44690694 | 44692129 | + | -0.663412 | intron (NM_153537, intron 20 of 21) | intron (NM_153537, intron 20 of 21) | 18178 | NM_021481 | 58866 | Mm.45380 | NM_021481 | ENSMUSG00000032098 | **Treh** | 2210412M19Rik | trehalase (brush-border membrane glycoprotein) | protein-coding |
| chr9_44648872_44649150 | chr9 | 44648873 | 44649150 | + | -0.712669 | Intergenic | Intergenic | -24222 | NM_001277851 | 58866 | Mm.45380 | NM_021481 | ENSMUSG00000032098 | **Treh** | 2210412M19Rik | trehalase (brush-border membrane glycoprotein) | protein-coding |
| chr9_44648743_44649157 | chr9 | 44648744 | 44649157 | + | -0.739524 | Intergenic | Intergenic | -24283 | NM_001277851 | 58866 | Mm.45380 | NM_021481 | ENSMUSG00000032098 | **Treh** | 2210412M19Rik | trehalase (brush-border membrane glycoprotein) | protein-coding |
| chr9_44690705_44691228 | chr9 | 44690706 | 44691228 | + | -0.823891 | intron (NM_153537, intron 20 of 21) | G-rich\|Low_complexity\|Low_complexity | 17734 | NM_021481 | 58866 | Mm.45380 | NM_021481 | ENSMUSG00000032098 | **Treh** | 2210412M19Rik | trehalase (brush-border membrane glycoprotein) | protein-coding |
| chr12_16133233_16133910 | chr12 | 16133234 | 16133910 | + | -0.661806 | Intergenic | Intergenic | -316787 | NM_144551 | 217410 | Mm.491136 | NM_144551 | ENSMUSG00000020601 | **Trib2** | AW319517\|TRB-2\|TRB2 | tribbles homolog 2 (Drosophila) | protein-coding |
| chr12_16014646_16015284 | chr12 | 16014647 | 16015284 | + | -0.691208 | Intergenic | Intergenic | -198180 | NM_144551 | 217410 | Mm.491136 | NM_144551 | ENSMUSG00000020601 | **Trib2** | AW319517\|TRB-2\|TRB2 | tribbles homolog 2 (Drosophila) | protein-coding |
| chr12_16134573_16134804 | chr12 | 16134574 | 16134804 | + | -0.761933 | Intergenic | Intergenic | -317904 | NM_144551 | 217410 | Mm.491136 | NM_144551 | ENSMUSG00000020601 | **Trib2** | AW319517\|TRB-2\|TRB2 | tribbles homolog 2 (Drosophila) | protein-coding |
| chr12_16049379_16050229 | chr12 | 16049380 | 16050229 | + | -0.883751 | Intergenic | Intergenic | -233019 | NM_144551 | 217410 | Mm.491136 | NM_144551 | ENSMUSG00000020601 | **Trib2** | AW319517\|TRB-2\|TRB2 | tribbles homolog 2 (Drosophila) | protein-coding |
| chr12_16191848_16192409 | chr12 | 16191849 | 16192409 | + | -0.898543 | Intergenic | RLTR11A2\|LTR\|ERVK | -375344 | NM_144551 | 217410 | Mm.491136 | NM_144551 | ENSMUSG00000020601 | **Trib2** | AW319517\|TRB-2\|TRB2 | tribbles homolog 2 (Drosophila) | protein-coding |
| chr12_16049449_16050036 | chr12 | 16049450 | 16050036 | + | -0.949886 | Intergenic | Intergenic | -232958 | NM_144551 | 217410 | Mm.491136 | NM_144551 | ENSMUSG00000020601 | **Trib2** | AW319517\|TRB-2\|TRB2 | tribbles homolog 2 (Drosophila) | protein-coding |
| chr14_61601305_61601733 | chr14 | 61601306 | 61601733 | + | -1.35319 | intron (NM_023233, intron 2 of 2) | intron (NM_023233, intron 2 of 2) | 2007 | NM_023233 | 66597 | Mm.491258 | NM_023233 | ENSMUSG00000035235 | **Trim13** | 3110001L12Rik\|CAR\|LEU5\|RNF77\|Rfp2 | tripartite motif-containing 13 | protein-coding |
| chr4_46526149_46526589 | chr4 | 46526150 | 46526589 | + | -0.731961 | intron (NM_029077, intron 2 of 5) | RLTR14\|LTR\|ERV1 | 9772 | NM_029077 | 74735 | Mm.240252 | NM_029077 | ENSMUSG00000039853 | **Trim14** | 5830400N10Rik\|pub | tripartite motif-containing 14 | protein-coding |
| chr6_37714148_37714837 | chr6 | 37714149 | 37714837 | + | -0.856741 | Intergenic | Intergenic | -156318 | NM_001272064 | 21848 | Mm.41063 | NM_145076 | | **Trim24** | A130082H20Rik\|AI447469\|D430004I05Rik\|TIF1\|TIF1-alpha\|TIF1alpha\|Tif1a | tripartite motif-containing 24 | protein-coding |
| chr11_89024212_89024566 | chr11 | 89024213 | 89024566 | + | -0.639866 | Intergenic | Intergenic | 24986 | NM_009546 | 217069 | Mm.248445 | NM_009546 | ENSMUSG00000000275 | **Trim25** | AA960166\|AL022677\|EFP\|Zfp147 | tripartite motif-containing 25 | protein-coding |
| chr11_89024171_89024435 | chr11 | 89024172 | 89024435 | + | -0.644712 | Intergenic | Intergenic | 24900 | NM_009546 | 217069 | Mm.248445 | NM_009546 | ENSMUSG00000000275 | **Trim25** | AA960166\|AL022677\|EFP\|Zfp147 | tripartite motif-containing 25 | protein-coding |
| chr11_89024073_89024586 | chr11 | 89024074 | 89024586 | + | -0.663442 | Intergenic | Intergenic | 24927 | NM_009546 | 217069 | Mm.248445 | NM_009546 | ENSMUSG00000000275 | **Trim25** | AA960166\|AL022677\|EFP\|Zfp147 | tripartite motif-containing 25 | protein-coding |
| chr3_103397331_103397971 | chr3 | 103397332 | 103397971 | + | -0.648192 | intron (NR_106164, intron 2 of 4) | intron (NR_106164, intron 2 of 4) | 118358 | NM_001079830 | 94093 | Mm.195036 | NM_053170 | ENSMUSG00000033014 | **Trim33** | 8030451N04Rik\|AI413936\|Ecto\|Tif1g | tripartite motif-containing 33 | protein-coding |
| chr3_103384901_103386175 | chr3 | 103384902 | 103386175 | + | -0.747915 | intron (NR_106164, intron 2 of 4) | intron (NR_106164, intron 2 of 4) | 106245 | NM_001079830 | 94093 | Mm.195036 | NM_053170 | ENSMUSG00000033014 | **Trim33** | 8030451N04Rik\|AI413936\|Ecto\|Tif1g | tripartite motif-containing 33 | protein-coding |
| chr3_103385420_103386289 | chr3 | 103385421 | 103386289 | + | -0.887454 | intron (NR_106164, intron 2 of 4) | intron (NR_106164, intron 2 of 4) | 106562 | NM_001079830 | 94093 | Mm.195036 | NM_053170 | ENSMUSG00000033014 | **Trim33** | 8030451N04Rik\|AI413936\|Ecto\|Tif1g | tripartite motif-containing 33 | protein-coding |
| chr3_103385426_103385871 | chr3 | 103385427 | 103385871 | + | -1.01055 | intron (NR_106164, intron 2 of 4) | intron (NR_106164, intron 2 of 4) | 106356 | NM_001079830 | 94093 | Mm.195036 | NM_053170 | ENSMUSG00000033014 | **Trim33** | 8030451N04Rik\|AI413936\|Ecto\|Tif1g | tripartite motif-containing 33 | protein-coding |
| chr18_46175426_46176425 | chr18 | 46175427 | 46176425 | + | -0.74744 | exon (NM_178872, exon 6 of 10) | exon (NM_178872, exon 6 of 10) | 22892 | NM_001170855 | 28105 | Mm.277382 | NM_178872 | ENSMUSG00000033949 | **Trim36** | D18Wsu100e\|haprin | tripartite motif-containing 36 | protein-coding |
| chr18_46175675_46176421 | chr18 | 46175676 | 46176421 | + | -0.777551 | exon (NM_178872, exon 6 of 10) | exon (NM_178872, exon 6 of 10) | 22770 | NM_001170855 | 28105 | Mm.277382 | NM_178872 | ENSMUSG00000033949 | **Trim36** | D18Wsu100e\|haprin | tripartite motif-containing 36 | protein-coding |
| chr18_46175834_46176632 | chr18 | 46175835 | 46176632 | + | -0.87439 | intron (NM_001170855, intron 5 of 9) | intron (NM_001170855, intron 5 of 9) | 22585 | NM_001170855 | 28105 | Mm.277382 | NM_178872 | ENSMUSG00000033949 | **Trim36** | D18Wsu100e\|haprin | tripartite motif-containing 36 | protein-coding |
| chr11_87240082_87240727 | chr11 | 87240083 | 87240727 | + | -0.871311 | intron (NM_177167, intron 4 of 6) | intron (NM_177167, intron 4 of 6) | 113330 | NM_197987 | 68729 | Mm.17436 | NM_197987 | ENSMUSG00000018548 | **Trim37** | 1110032A10Rik\|2810004E07Rik\|AI848587\|AU043018\|MUL\|TEF3\|mKIAA0898 | tripartite motif-containing 37 | protein-coding |
| chr15_27925825_27926218 | chr15 | 27925826 | 27926218 | + | -0.989162 | intron (NM_001081302, intron 1 of 56) | intron (NM_001081302, intron 1 of 56) | 99826 | NM_001081302 | 223435 | Mm.485422 | NM_001081302 | ENSMUSG00000022263 | **Trio** | 6720464I07Rik\|AA408740\|Solo | triple functional domain (PTPRF interacting) | protein-coding |
| chr12_111691105_111691625 | chr12 | 111691106 | 111691625 | + | -0.818682 | Intergenic | Intergenic | 13260 | NM_177374 | 328162 | Mm.425747 | NM_177374 | ENSMUSG00000060950 | **Trmt61a** | 6720458F09Rik\|AI606093\|Gcd14\|Trm61 | tRNA methyltransferase 61A | protein-coding |
| chr15_85956181_85956627 | chr15 | 85956182 | 85956627 | + | -0.763089 | intron (NM_009886, intron 5 of 34) | intron (NM_009886, intron 5 of 34) | 77077 | NM_028063 | 72026 | Mm.158211 | NM_028063 | ENSMUSG00000022386 | **Trmu** | 1110005N20Rik\|1600025P05Rik\|AI314320\|Trmt1 | tRNA 5-methylaminomethyl-2-thiouridylate methyltransferase | protein-coding |
| chr4_133506421_133506867 | chr4 | 133506422 | 133506867 | + | -0.970055 | Intergenic | RLTR14_RN\|LTR\|ERV1 | -8094 | NM_001081156 | 69539 | Mm.151594 | NM_001081156 | ENSMUSG00000056596 | **Trnp1** | 2300002D11Rik\|Tmp | TMF1-regulated nuclear protein 1 | protein-coding |
| chr4_133506319_133507077 | chr4 | 133506320 | 133507077 | + | -0.991246 | Intergenic | RLTR14_RN\|LTR\|ERV1 | -8148 | NM_001081156 | 69539 | Mm.151594 | NM_001081156 | ENSMUSG00000056596 | **Trnp1** | 2300002D11Rik\|Tmp | TMF1-regulated nuclear protein 1 | protein-coding |
| chr4_133506552_133507214 | chr4 | 133506553 | 133507214 | + | -1.09274 | Intergenic | RLTR14_RN\|LTR\|ERV1 | -8333 | NM_001081156 | 69539 | Mm.151594 | NM_001081156 | ENSMUSG00000056596 | **Trnp1** | 2300002D11Rik\|Tmp | TMF1-regulated nuclear protein 1 | protein-coding |
| chr4_11157329_11157732 | chr4 | 11157330 | 11157732 | + | -0.64183 | intron (NM_021897, intron 1 of 3) | intron (NM_021897, intron 1 of 3) | 1090 | NM_021897 | 60599 | Mm.393018 | NM_021897 | ENSMUSG00000028211 | **Trp53inp1** | 2700057G22Rik\|SIP\|SIP18\|SIP27\|Stinp\|Teap | transformation related protein 53 inducible nuclear protein 1 | protein-coding |
| chr4_11157493_11157701 | chr4 | 11157494 | 11157701 | + | -0.647889 | intron (NM_021897, intron 1 of 3) | intron (NM_021897, intron 1 of 3) | 1156 | NM_021897 | 60599 | Mm.393018 | NM_021897 | ENSMUSG00000028211 | **Trp53inp1** | 2700057G22Rik\|SIP\|SIP18\|SIP27\|Stinp\|Teap | transformation related protein 53 inducible nuclear protein 1 | protein-coding |
| chr2_155679616_155680110 | chr2 | 155679617 | 155680110 | + | -0.764377 | intron (NM_019828, intron 1 of 18) | intron (NM_019828, intron 1 of 18) | 12521 | NM_019828 | 56407 | Mm.268304 | NM_019828 | ENSMUSG00000038324 | **Trpc4ap** | 4833429F06Rik\|D2Ertd113e\|Trrp4ap\|mFLJ00177\|truss | transient receptor potential cation channel, subfamily C, member 4 associated protein | protein-coding |
| chr7_64184928_64185961 | chr7 | 64184929 | 64185961 | + | -0.62217 | promoter-TSS (NM_018752) | promoter-TSS (NM_018752) | -14 | NM_018752 | 17364 | Mm.38875 | NM_018752 | ENSMUSG00000030523 | **Trpm1** | 4732499L03Rik\|AI606771\|Ltrpc1\|Mlsn1\|melastatin | transient receptor potential cation channel, subfamily M, member 1 | protein-coding |
| chr7_64101652_64102616 | chr7 | 64101653 | 64102616 | + | -0.848877 | Intergenic | L1MDb\|LINE\|L1 | -51701 | NM_001039104 | 17364 | Mm.38875 | NM_018752 | ENSMUSG00000030523 | **Trpm1** | 4732499L03Rik\|AI606771\|Ltrpc1\|Mlsn1\|melastatin | transient receptor potential cation channel, subfamily M, member 1 | protein-coding |
| chr7_64076995_64077402 | chr7 | 64076996 | 64077402 | + | -0.852176 | Intergenic | Intergenic | -76636 | NM_001039104 | 17364 | Mm.38875 | NM_018752 | ENSMUSG00000030523 | **Trpm1** | 4732499L03Rik\|AI606771\|Ltrpc1\|Mlsn1\|melastatin | transient receptor potential cation channel, subfamily M, member 1 | protein-coding |
| chr7_64101652_64102196 | chr7 | 64101653 | 64102196 | + | -0.901163 | Intergenic | L1MDb\|LINE\|L1 | -51911 | NM_001039104 | 17364 | Mm.38875 | NM_018752 | ENSMUSG00000030523 | **Trpm1** | 4732499L03Rik\|AI606771\|Ltrpc1\|Mlsn1\|melastatin | transient receptor potential cation channel, subfamily M, member 1 | protein-coding |
| chr15_50928208_50928513 | chr15 | 50928209 | 50928513 | + | -0.719524 | Intergenic | Intergenic | -38320 | NM_032000 | 83925 | Mm.30466 | NM_032000 | ENSMUSG00000038679 | **Trps1** | AI115454\|AI447310\|D15Ertd586e | trichorhinophalangeal syndrome I (human) | protein-coding |
| chr11_62574278_62574685 | chr11 | 62574279 | 62574685 | + | -0.779801 | promoter-TSS (NM_011706) | promoter-TSS (NM_011706) | -4 | NM_011706 | 22368 | Mm.288064 | NM_011706 | ENSMUSG00000018507 | **Trpv2** | GRC\|OTRPC2\|VRL-1\|Vrl1 | transient receptor potential cation channel, subfamily V, member 2 | protein-coding |
| chr19_57477291_57477668 | chr19 | 57477292 | 57477668 | + | -0.84895 | intron (NM_029839, intron 4 of 6) | intron (NM_029839, intron 4 of 6) | 24574 | NM_028115 | 72133 | Mm.258404 | NM_028115 | ENSMUSG00000025086 | **Trub1** | 2610009I02Rik\|9030425C13Rik\|AI448235\|BB118943\|C76870 | TruB pseudouridine (psi) synthase homolog 1 (E. coli) | protein-coding |
| chr6_41215565_41216262 | chr6 | 41215566 | 41216262 | + | -0.936287 | Intergenic | Lx\|LINE\|L1 | -86358 | NM_011646 | 22074 | Mm.383263 | NM_011646 | ENSMUSG00000054106 | **Try4** | 0910001B19Rik\|Td | trypsin 4 | protein-coding |
| chr1_152269696_152270723 | chr1 | 152269697 | 152270723 | + | -0.668879 | Intergenic | Intergenic | 116472 | NM_025677 | 66637 | Mm.379120 | NM_025677 | ENSMUSG00000014980 | **Tsen15** | 5730449L18Rik\|AL023077\|Sen15 | tRNA splicing endonuclease 15 homolog (S. cerevisiae) | protein-coding |
| chr6_115559444_115560546 | chr6 | 115559445 | 115560546 | + | -0.710218 | exon (NM_199033, exon 5 of 12) | exon (NM_199033, exon 5 of 12) | 15291 | NM_199033 | 381802 | Mm.291208 | NM_199033 | ENSMUSG00000042389 | **Tsen2** | AU067695 | tRNA splicing endonuclease 2 homolog (S. cerevisiae) | protein-coding |
| chr18_83993117_83993674 | chr18 | 83993118 | 83993674 | + | -1.39611 | Intergenic | ORR1A2-int\|LTR\|ERVL-MaLR | 93166 | NM_001081300 | 110796 | Mm.102136 | NM_001081300 | ENSMUSG00000046982 | **Tshz1** | 5730407I04Rik\|D18Bwg1409e\|Mtsh1\|NY-CO-33\|Sdccag33\|Tsh1\|mKIAA4206\|teashirt1 | teashirt zinc finger family member 1 | protein-coding |
| chr6_127991516_127992364 | chr6 | 127991517 | 127992364 | + | -0.891214 | intron (NM_175414, intron 3 of 8) | intron (NM_175414, intron 3 of 8) | 104318 | NM_026743 | 68498 | Mm.174368 | NM_026743 | ENSMUSG00000030351 | **Tspan11** | 1110014F12Rik\|AU067806 | tetraspanin 11 | protein-coding |
| chr12_36041718_36041953 | chr12 | 36041719 | 36041953 | + | -0.626701 | intron (NM_025359, intron 1 of 5) | intron (NM_025359, intron 1 of 5) | 642 | NM_025359 | 66109 | Mm.254663 | NM_025359 | ENSMUSG00000020577 | **Tspan13** | 1100001I23Rik\|NET-6\|Tm4sf13 | tetraspanin 13 | protein-coding |
| chr12_36057777_36058156 | chr12 | 36057778 | 36058156 | + | -0.646792 | Intergenic | Lx8\|LINE\|L1 | -15489 | NM_025359 | 66109 | Mm.254663 | NM_025359 | ENSMUSG00000020577 | **Tspan13** | 1100001I23Rik\|NET-6\|Tm4sf13 | tetraspanin 13 | protein-coding |
| chr12_36025547_36026025 | chr12 | 36025548 | 36026025 | + | -0.712693 | intron (NM_025359, intron 1 of 5) | intron (NM_025359, intron 1 of 5) | 16692 | NM_025359 | 66109 | Mm.254663 | NM_025359 | ENSMUSG00000020577 | **Tspan13** | 1100001I23Rik\|NET-6\|Tm4sf13 | tetraspanin 13 | protein-coding |
| chr14_40929895_40930485 | chr14 | 40929896 | 40930485 | + | -1.05659 | intron (NM_145928, intron 2 of 8) | L1MB2\|LINE\|L1 | 36617 | NM_145928 | 52588 | Mm.28876 | NM_145928 | ENSMUSG00000037824 | **Tspan14** | AA958793\|D14Ertd226e\|Tm4sf14 | tetraspanin 14 | protein-coding |
| chr9_56146341_56146850 | chr9 | 56146342 | 56146850 | + | -0.792893 | exon (NM_019793, exon 4 of 7) | exon (NM_019793, exon 4 of 7) | 14474 | NM_019793 | 56434 | Mm.28484 | NM_019793 | ENSMUSG00000032324 | **Tspan3** | 1700055K04Rik\|TM4-A\|Tm4sf8 | tetraspanin 3 | protein-coding |

| | | | | | | | | | | | | | | | | | |
|---|---|---|---|---|---|---|---|---|---|---|---|---|---|---|---|---|---|
| chr7_142994978_142995410 | chr7 | 142994979 | 142995410 | + | -1.21459 | Intergenic | Intergenic | -9852 | NM_001128080 | 27027 | Mm.28172 | NM_020286 | ENSMUSG00000000244 | **Tspan32** | AW208513\|Art-1\|BB235973\|D7Wsu37e\|Phemx\|Tssc6 | tetraspanin 32 | protein-coding |
| chr6_128025457_128026259 | chr6 | 128025458 | 128026259 | + | -0.831157 | intron (NM_175414, intron 3 of 8) | intron (NM_175414, intron 3 of 8) | 117720 | NM_175414 | 109246 | Mm.21814 | NM_175414 | ENSMUSG00000030352 | **Tspan9** | 6720474K14Rik\|9430079M16Rik\|AU018597 | tetraspanin 9 | protein-coding |
| chr2_153214756_153215301 | chr2 | 153214757 | 153215301 | + | -0.842988 | Intergenic | Intergenic | 10412 | NM_198617 | 241732 | Mm.277243 | NM_198617 | ENSMUSG00000074671 | **Tspyl3** | AW212607\|Gm36 | TSPY-like 3 | protein-coding |
| chr12_28807679_28807983 | chr12 | 28807680 | 28807983 | + | -1.07603 | intron (NM_201357, intron 3 of 8) | intron (NM_201357, intron 3 of 8) | 56003 | NM_201357 | 380752 | Mm.273724 | NM_201357 | ENSMUSG00000036613 | **Tssc1** | D12Ertd604e | tumor suppressing subtransferable candidate 1 | protein-coding |
| chr12_28807679_28808341 | chr12 | 28807680 | 28808341 | + | -1.09372 | intron (NM_201357, intron 3 of 8) | intron (NM_201357, intron 3 of 8) | 56182 | NM_201357 | 380752 | Mm.273724 | NM_201357 | ENSMUSG00000036613 | **Tssc1** | D12Ertd604e | tumor suppressing subtransferable candidate 1 | protein-coding |
| chr12_28808663_28809468 | chr12 | 28808664 | 28809468 | + | -1.12793 | intron (NM_201357, intron 3 of 8) | intron (NM_201357, intron 3 of 8) | 57238 | NM_201357 | 380752 | Mm.273724 | NM_201357 | ENSMUSG00000036613 | **Tssc1** | D12Ertd604e | tumor suppressing subtransferable candidate 1 | protein-coding |
| chr12_28834045_28834837 | chr12 | 28834046 | 28834837 | + | -1.15428 | intron (NM_201357, intron 4 of 8) | intron (NM_201357, intron 4 of 8) | 82613 | NM_201357 | 380752 | Mm.273724 | NM_201357 | ENSMUSG00000036613 | **Tssc1** | D12Ertd604e | tumor suppressing subtransferable candidate 1 | protein-coding |
| chr12_28834308_28834842 | chr12 | 28834309 | 28834842 | + | -1.17121 | intron (NM_201357, intron 4 of 8) | intron (NM_201357, intron 4 of 8) | 82747 | NM_201357 | 380752 | Mm.273724 | NM_201357 | ENSMUSG00000036613 | **Tssc1** | D12Ertd604e | tumor suppressing subtransferable candidate 1 | protein-coding |
| chr12_28834296_28834668 | chr12 | 28834297 | 28834668 | + | -1.21712 | intron (NM_201357, intron 4 of 8) | intron (NM_201357, intron 4 of 8) | 82654 | NM_201357 | 380752 | Mm.273724 | NM_201357 | ENSMUSG00000036613 | **Tssc1** | D12Ertd604e | tumor suppressing subtransferable candidate 1 | protein-coding |
| chr12_28807741_28808086 | chr12 | 28807742 | 28808086 | + | -1.22008 | intron (NM_201357, intron 3 of 8) | intron (NM_201357, intron 3 of 8) | 56086 | NM_201357 | 380752 | Mm.273724 | NM_201357 | ENSMUSG00000036613 | **Tssc1** | D12Ertd604e | tumor suppressing subtransferable candidate 1 | protein-coding |
| chr12_28845796_28846687 | chr12 | 28845797 | 28846687 | + | -1.35629 | intron (NM_201357, intron 4 of 8) | L1_Rod\|LINE\|L1 | 94414 | NM_201357 | 380752 | Mm.273724 | NM_201357 | ENSMUSG00000036613 | **Tssc1** | D12Ertd604e | tumor suppressing subtransferable candidate 1 | protein-coding |
| chr8_124700745_124701475 | chr8 | 124700746 | 124701475 | + | -0.69075 | intron (NM_145607, intron 4 of 17) | intron (NM_145607, intron 4 of 17) | 20865 | NM_145607 | 234875 | Mm.38436 | NM_145607 | ENSMUSG00000037300 | **Ttc13** | BC017545 | tetratricopeptide repeat domain 13 | protein-coding |
| chr2_75915070_75915443 | chr2 | 75915071 | 75915443 | + | -1.22888 | intron (NM_172666, intron 19 of 19) | intron (NM_172666, intron 19 of 19) | 23205 | NM_028235 | 72421 | Mm.275109 | NM_028235 | ENSMUSG00000075273 | **Ttc30b** | 2510042P03Rik\|Tcc30b | tetratricopeptide repeat domain 30B | protein-coding |
| chr17_87299001_87299358 | chr17 | 87299002 | 87299358 | + | -0.674221 | intron (NM_028639, intron 3 of 19) | intron (NM_028639, intron 3 of 19) | 16294 | NM_028639 | 225049 | Mm.77396 | NM_028639 | ENSMUSG00000036918 | **Ttc7** | 1110035E02Rik\|1700007L07Rik\|Ttc7a\|fsn\|hea | tetratricopeptide repeat domain 7 | protein-coding |
| chr17_87298691_87299347 | chr17 | 87298692 | 87299347 | + | -0.70017 | intron (NM_028639, intron 3 of 19) | intron (NM_028639, intron 3 of 19) | 16133 | NM_028639 | 225049 | Mm.77396 | NM_028639 | ENSMUSG00000036918 | **Ttc7** | 1110035E02Rik\|1700007L07Rik\|Ttc7a\|fsn\|hea | tetratricopeptide repeat domain 7 | protein-coding |
| chr12_100501159_100501749 | chr12 | 100501160 | 100501749 | + | -0.97537 | intron (NM_001033213, intron 1 of 19) | intron (NM_001033213, intron 1 of 19) | 19368 | NM_001033213 | 104718 | Mm.295441 | NM_001033213 | ENSMUSG00000033530 | **Ttc7b** | AA408451\|Ttc7l1 | tetratricopeptide repeat domain 7B | protein-coding |
| chr12_100543991_100544546 | chr12 | 100543992 | 100544546 | + | -1.4125 | Intergenic | L1MEg\|LINE\|L1 | -23447 | NM_001033213 | 104718 | Mm.295441 | NM_001033213 | ENSMUSG00000033530 | **Ttc7b** | AA408451\|Ttc7l1 | tetratricopeptide repeat domain 7B | protein-coding |
| chr12_100544179_100544538 | chr12 | 100544180 | 100544538 | + | -1.43336 | Intergenic | L1MEg\|LINE\|L1 | -23537 | NM_001033213 | 104718 | Mm.295441 | NM_001033213 | ENSMUSG00000033530 | **Ttc7b** | AA408451\|Ttc7l1 | tetratricopeptide repeat domain 7B | protein-coding |
| chr15_83602507_83603298 | chr15 | 83602508 | 83603298 | + | -0.890108 | 3' UTR (NM_001271473, exon 20 of 20) | 3' UTR (NM_001271473, exon 20 of 20) | -7746 | NM_183017 | 223723 | Mm.46778 | NM_183017 | ENSMUSG00000016757 | **Ttll12** | BC055368\|D430005B17 | tubulin tyrosine ligase-like family, member 12 | protein-coding |
| chr5_140636683_140637144 | chr5 | 140636684 | 140637144 | + | -0.864796 | intron (NM_175274, intron 1 of 13) | intron (NM_175274, intron 1 of 13) | 12117 | NM_175274 | 78339 | Mm.28947 | NM_175274 | ENSMUSG00000036565 | **Ttyh3** | 2900029G13Rik\|AI414930 | tweety homolog 3 (Drosophila) | protein-coding |
| chr5_140637877_140638395 | chr5 | 140637878 | 140638395 | + | -1.09546 | intron (NM_175274, intron 1 of 13) | intron (NM_175274, intron 1 of 13) | 10895 | NM_175274 | 78339 | Mm.28947 | NM_175274 | ENSMUSG00000036565 | **Ttyh3** | 2900029G13Rik\|AI414930 | tweety homolog 3 (Drosophila) | protein-coding |
| chr15_99008677_99009192 | chr15 | 99008678 | 99009192 | + | -0.844162 | Intergenic | Intergenic | -20956 | NM_009448 | 22146 | Mm.88212 | NM_009448 | ENSMUSG00000043091 | **Tuba1c** | M[a]6\|Tuba6 | tubulin, alpha 1C | protein-coding |
| chr13_34090211_34090869 | chr13 | 34090212 | 34090869 | + | -1.24168 | intron (NR_046184, intron 3 of 4) | RMER19A\|LTR\|ERVK | -12532 | NM_009450 | 22151 | Mm.422827 | NM_009450 | ENSMUSG00000058672 | **Tubb2a** | M(beta)2\|Tubb2 | tubulin, beta 2A class IIA | protein-coding |
| chr13_34090410_34090875 | chr13 | 34090411 | 34090875 | + | -1.34669 | intron (NR_046184, intron 3 of 4) | RMER19A\|LTR\|ERVK | -12635 | NM_009450 | 22151 | Mm.422827 | NM_009450 | ENSMUSG00000058672 | **Tubb2a** | M(beta)2\|Tubb2 | tubulin, beta 2A class IIA | protein-coding |
| chr17_28417835_28418081 | chr17 | 28417836 | 28418081 | + | -0.952211 | intron (NM_010220, intron 5 of 10) | intron (NM_010220, intron 5 of 10) | -52815 | NM_021478 | 22157 | Mm.42102 | NM_021478 | ENSMUSG00000037446 | **Tulp1** | Tulp1l | tubby like protein 1 | protein-coding |
| chr17_28417812_28418183 | chr17 | 28417813 | 28418183 | + | -0.982484 | intron (NM_010220, intron 5 of 10) | intron (NM_010220, intron 5 of 10) | -52855 | NM_021478 | 22157 | Mm.42102 | NM_021478 | ENSMUSG00000037446 | **Tulp1** | Tulp1l | tubby like protein 1 | protein-coding |
| chr8_39108401_39108865 | chr8 | 39108402 | 39108865 | + | -0.677666 | intron (NM_030254, intron 6 of 7) | intron (NM_030254, intron 6 of 7) | 102766 | NM_030254 | 80286 | Mm.272722 | NM_030254 | | **Tusc3** | AU022242\|BC003311\|N33 | tumor suppressor candidate 3 | protein-coding |
| chr8_39087971_39088309 | chr8 | 39087972 | 39088309 | + | -0.846036 | intron (NM_030254, intron 5 of 7) | intron (NM_030254, intron 5 of 7) | 82273 | NM_030254 | 80286 | Mm.272722 | NM_030254 | | **Tusc3** | AU022242\|BC003311\|N33 | tumor suppressor candidate 3 | protein-coding |
| chr9_106210935_106211170 | chr9 | 106210936 | 106211170 | + | -0.671698 | intron (NM_011876, intron 2 of 8) | intron (NM_011876, intron 2 of 8) | 7945 | NM_011876 | 23999 | Mm.274346 | NM_011876 | ENSMUSG00000023277 | **Twf2** | A6-related\|AU014993\|Ptk9l\|Ptk9r | twinfilin, actin-binding protein, homolog 2 (Drosophila) | protein-coding |
| chr17_65957553_65957985 | chr17 | 65957554 | 65957985 | + | -1.19749 | Intergenic | Intergenic | -6582 | NM_023053 | 65960 | Mm.10153 | NM_023053 | ENSMUSG00000024098 | **Twsg1** | 1810013J15Rik\|9030422N06Rik\|AW552143\|D17Ertd403e\|Tsg\|Twg | twisted gastrulation homolog 1 (Drosophila) | protein-coding |
| chr5_72763509_72764119 | chr5 | 72763510 | 72764119 | + | -0.748902 | intron (NM_013689, intron 13 of 17) | intron (NM_013689, intron 13 of 17) | -11037 | NM_001122754 | 22165 | Mm.3264 | NM_013698 | ENSMUSG00000054892 | **Txk** | A130089B16Rik\|Btkl\|PTK-RL-18\|PTK4\|Rlk | TXK tyrosine kinase | protein-coding |
| chr5_72761143_72761631 | chr5 | 72761144 | 72761631 | + | -0.925055 | intron (NM_013689, intron 14 of 17) | intron (NM_013689, intron 14 of 17) | -8610 | NM_001122754 | 22165 | Mm.3264 | NM_013698 | ENSMUSG00000054892 | **Txk** | A130089B16Rik\|Btkl\|PTK-RL-18\|PTK4\|Rlk | TXK tyrosine kinase | protein-coding |
| chr5_72752591_72753079 | chr5 | 72752592 | 72753079 | + | -1.0148 | promoter-TSS (NM_001122754) | promoter-TSS (NM_001122754) | -58 | NM_001122754 | 22165 | Mm.3264 | NM_013698 | ENSMUSG00000054892 | **Txk** | A130089B16Rik\|Btkl\|PTK-RL-18\|PTK4\|Rlk | TXK tyrosine kinase | protein-coding |
| chr5_72800026_72800354 | chr5 | 72800027 | 72800354 | + | -1.01713 | intron (NM_013689, intron 3 of 17) | intron (NM_013689, intron 3 of 17) | -47413 | NM_001122754 | 22165 | Mm.3264 | NM_013698 | ENSMUSG00000054892 | **Txk** | A130089B16Rik\|Btkl\|PTK-RL-18\|PTK4\|Rlk | TXK tyrosine kinase | protein-coding |
| chr5_72800000_72800267 | chr5 | 72800001 | 72800267 | + | -1.02198 | intron (NM_013689, intron 3 of 17) | intron (NM_013689, intron 3 of 17) | -47357 | NM_001122754 | 22165 | Mm.3264 | NM_013698 | ENSMUSG00000054892 | **Txk** | A130089B16Rik\|Btkl\|PTK-RL-18\|PTK4\|Rlk | TXK tyrosine kinase | protein-coding |
| chr4_57946554_57947013 | chr4 | 57946555 | 57947013 | + | -0.988701 | intron (NM_011660, intron 3 of 4) | MTD\|LTR\|ERVL-MaLR | 9627 | NM_011660 | 22166 | Mm.260618 | NM_011660 | ENSMUSG00000028367 | **Txn1** | ADF\|AW550880\|Trx1\|Txn | thioredoxin 1 | protein-coding |
| chr14_45155386_45156021 | chr14 | 45155387 | 45156021 | + | -1.13851 | intron (NM_172597, intron 14 of 20) | MT2A\|LTR\|ERVL | 63690 | NM_172597 | 70561 | Mm.28129 | NM_172597 | ENSMUSG00000021830 | **Txndc16** | 5730420B22Rik\|C77647 | thioredoxin domain containing 16 | protein-coding |
| chr14_45155555_45155929 | chr14 | 45155556 | 45155929 | + | -1.1578 | intron (NM_172597, intron 14 of 20) | MT2A\|LTR\|ERVL | 63652 | NM_172597 | 70561 | Mm.28129 | NM_172597 | ENSMUSG00000021830 | **Txndc16** | 5730420B22Rik\|C77647 | thioredoxin domain containing 16 | protein-coding |
| chr13_38503341_38503913 | chr13 | 38503342 | 38503913 | + | -0.729176 | intron (NM_001289598, intron 8 of 9) | intron (NM_001289598, intron 8 of 9) | 24287 | NM_001289599 | 105245 | Mm.28622 | NM_145367 | ENSMUSG00000038991 | **Txndc5** | AL022641\|ERp46\|PC-TRP | thioredoxin domain containing 5 | protein-coding |
| chr4_58081080_58081323 | chr4 | 58081081 | 58081323 | + | -0.718174 | intron (NM_022814, intron 36 of 47) | intron (NM_022814, intron 36 of 47) | -72078 | NM_026132 | 67402 | Mm.87357 | NM_026132 | ENSMUSG00000038709 | **Txndc8** | 4930429J24Rik\|Sptrx-3\|Sptrx3\|Trx6 | thioredoxin domain containing 8 | protein-coding |
| chr3_96557057_96557321 | chr3 | 96557058 | 96557321 | + | -0.873347 | promoter-TSS (NM_001009935) | promoter-TSS (NM_001009935) | -768 | NM_001009935 | 56338 | Mm.410189 | NM_023719 | ENSMUSG00000038393 | **Txnip** | 1200008J08Rik\|AA682105\|Hyplip1\|THIF\|Tbp-2\|VDUP1 | thioredoxin interacting protein | protein-coding |
| chr4_80003375_80003612 | chr4 | 80003376 | 80003612 | + | -0.678173 | Intergenic | Intergenic | -830719 | NM_031202 | 22178 | Mm.30438 | NM_031202 | ENSMUSG00000005994 | **Tyrp1** | Oca3\|TRP-1\|TRP1\|Tyrp\|b\|brown\|isa | tyrosinase-related protein 1 | protein-coding |
| chr4_80003375_80003840 | chr4 | 80003376 | 80003840 | + | -0.817141 | Intergenic | Intergenic | -830605 | NM_031202 | 22178 | Mm.30438 | NM_031202 | ENSMUSG00000005994 | **Tyrp1** | Oca3\|TRP-1\|TRP1\|Tyrp\|b\|brown\|isa | tyrosinase-related protein 1 | protein-coding |
| chr4_80003983_80005239 | chr4 | 80003984 | 80005239 | + | -0.905045 | Intergenic | Intergenic | -829602 | NM_031202 | 22178 | Mm.30438 | NM_031202 | ENSMUSG00000005994 | **Tyrp1** | Oca3\|TRP-1\|TRP1\|Tyrp\|b\|brown\|isa | tyrosinase-related protein 1 | protein-coding |
| chr4_80003347_80005234 | chr4 | 80003348 | 80005234 | + | -0.929365 | Intergenic | Intergenic | -829922 | NM_031202 | 22178 | Mm.30438 | NM_031202 | ENSMUSG00000005994 | **Tyrp1** | Oca3\|TRP-1\|TRP1\|Tyrp\|b\|brown\|isa | tyrosinase-related protein 1 | protein-coding |
| chr5_92204503_92205069 | chr5 | 92204504 | 92205069 | + | -0.872492 | Intergenic | B3\|SINE\|B2 | 10622 | NR_033483 | 57425 | Mm.2548 | NM_020562 | ENSMUSG00000029409 | **U90926** | - | cDNA sequence U90926 | ncRNA |
| chr9_60858684_60859405 | chr9 | 60858685 | 60859405 | + | -0.927774 | intron (NM_028283, intron 9 of 18) | URR1B\|DNA\|hAT-Charlie | 64497 | NM_028283 | 72565 | Mm.68819 | NM_028283 | ENSMUSG00000034485 | **Uaca** | 2700059D02Rik\|nucling | uveal autoantigen with coiled-coil domains and ankyrin repeats | protein-coding |
| chr9_60859018_60859422 | chr9 | 60859019 | 60859422 | + | -1.06153 | intron (NM_028283, intron 10 of 18) | intron (NM_028283, intron 10 of 18) | 64672 | NM_028283 | 72565 | Mm.68819 | NM_028283 | ENSMUSG00000034485 | **Uaca** | 2700059D02Rik\|nucling | uveal autoantigen with coiled-coil domains and ankyrin repeats | protein-coding |
| chr1_170147549_170148035 | chr1 | 170147550 | 170148035 | + | -0.82306 | intron (NM_133806, intron 9 of 10) | intron (NM_133806, intron 9 of 10) | 27154 | NM_133806 | 107652 | Mm.27969 | NM_133806 | ENSMUSG00000026670 | **Uap1** | AA420407\|AA437972\|AGX-1\|AGX-2\|AGX1\|AgX\|ESTM38\|SPAG2 | UDP-N-acetylglucosamine pyrophosphorylase 1 | protein-coding |
| chr14_121879351_121879995 | chr14 | 121879352 | 121879995 | + | -0.781531 | intron (NM_026861, intron 1 of 8) | intron (NM_026861, intron 1 of 8) | 1067 | NM_026861 | 68889 | Mm.441151 | NM_026861 | ENSMUSG00000041765 | **Ubac2** | 1190008A14Rik\|AW554959\|Phgdhl1 | ubiquitin associated domain containing 2 | protein-coding |
| chr14_121879453_121879826 | chr14 | 121879454 | 121879826 | + | -0.798196 | intron (NM_026861, intron 1 of 8) | intron (NM_026861, intron 1 of 8) | 1034 | NM_026861 | 68889 | Mm.441151 | NM_026861 | ENSMUSG00000041765 | **Ubac2** | 1190008A14Rik\|AW554959\|Phgdhl1 | ubiquitin associated domain containing 2 | protein-coding |

| | | | | | | | | | | | | | | | | | |
|---|---|---|---|---|---|---|---|---|---|---|---|---|---|---|---|---|---|
| chr14_121880877_121881539 | chr14 | 121880878 | 121881539 | + | -0.86012 | intron (NM_026861, intron 1 of 8) | intron (NM_026861, intron 1 of 8) | 2602 | NM_026861 | 68889 | Mm.441151 | NM_026861 | ENSMUSG00000041765 | **Ubac2** | 1190008A14Rik\|AW554959\|Phgdhl1 | ubiquitin associated domain containing 2 | protein-coding |
| chr14_121881363_121881539 | chr14 | 121881364 | 121881539 | + | -0.887415 | intron (NM_026861, intron 1 of 8) | intron (NM_026861, intron 1 of 8) | 2845 | NM_026861 | 68889 | Mm.441151 | NM_026861 | ENSMUSG00000041765 | **Ubac2** | 1190008A14Rik\|AW554959\|Phgdhl1 | ubiquitin associated domain containing 2 | protein-coding |
| chr14_121881179_121881550 | chr14 | 121881180 | 121881550 | + | -0.977588 | intron (NM_026861, intron 1 of 8) | intron (NM_026861, intron 1 of 8) | 2759 | NM_026861 | 68889 | Mm.441151 | NM_026861 | ENSMUSG00000041765 | **Ubac2** | 1190008A14Rik\|AW554959\|Phgdhl1 | ubiquitin associated domain containing 2 | protein-coding |
| chr14_121846069_121846422 | chr14 | 121846070 | 121846422 | + | -1.11387 | Intergenic | Intergenic | -32360 | NM_026861 | 68889 | Mm.441151 | NM_026861 | ENSMUSG00000041765 | **Ubac2** | 1190008A14Rik\|AW554959\|Phgdhl1 | ubiquitin associated domain containing 2 | protein-coding |
| chr16_4878977_4879679 | chr16 | 4878978 | 4879679 | + | -0.639789 | intron (NM_145359, intron 1 of 2) | CpG | 523 | NM_145359 | 207740 | Mm.268895 | NM_145359 | | **Ubald1** | 1500031H01Rik\|BC013706\|Fam100a | UBA-like domain containing 1 | protein-coding |
| chr16_4878154_4878521 | chr16 | 4878155 | 4878521 | + | -0.650358 | intron (NM_145359, intron 1 of 2) | intron (NM_145359, intron 1 of 2) | 1513 | NM_145359 | 207740 | Mm.268895 | NM_145359 | | **Ubald1** | 1500031H01Rik\|BC013706\|Fam100a | UBA-like domain containing 1 | protein-coding |
| chr16_4878960_4879500 | chr16 | 4878961 | 4879500 | + | -0.663147 | intron (NM_145359, intron 1 of 2) | intron (NM_145359, intron 1 of 2) | 621 | NM_145359 | 207740 | Mm.268895 | NM_145359 | | **Ubald1** | 1500031H01Rik\|BC013706\|Fam100a | UBA-like domain containing 1 | protein-coding |
| chr17_31210399_31210664 | chr17 | 31210400 | 31210664 | + | -0.718296 | intron (NM_177823, intron 2 of 13) | intron (NM_177823, intron 2 of 13) | 2466 | NM_177823 | 328795 | Mm.387930 | NM_177823 | ENSMUSG00000042345 | **Ubash3a** | 5830413C03Rik\|C330001M22\|Sts-2\|TULA\|TULA-1 | ubiquitin associated and SH3 domain containing, A | protein-coding |
| chr17_31210355_31210722 | chr17 | 31210356 | 31210722 | + | -0.719786 | intron (NM_177823, intron 2 of 13) | intron (NM_177823, intron 2 of 13) | 2473 | NM_177823 | 328795 | Mm.387930 | NM_177823 | ENSMUSG00000042345 | **Ubash3a** | 5830413C03Rik\|C330001M22\|Sts-2\|TULA\|TULA-1 | ubiquitin associated and SH3 domain containing, A | protein-coding |
| chr9_41115842_41116539 | chr9 | 41115843 | 41116539 | + | -0.656228 | intron (NM_176860, intron 1 of 13) | intron (NM_176860, intron 1 of 13) | 41303 | NM_176860 | 72828 | Mm.133615 | NM_176860 | ENSMUSG00000032020 | **Ubash3b** | 2810457I06Rik\|BB125008\|TULA-2\|p70 | ubiquitin associated and SH3 domain containing, B | protein-coding |
| chr9_41141058_41141615 | chr9 | 41141059 | 41141615 | + | -0.708005 | intron (NM_176860, intron 1 of 13) | intron (NM_176860, intron 1 of 13) | 16157 | NM_176860 | 72828 | Mm.133615 | NM_176860 | ENSMUSG00000032020 | **Ubash3b** | 2810457I06Rik\|BB125008\|TULA-2\|p70 | ubiquitin associated and SH3 domain containing, B | protein-coding |
| chr9_41117907_41118445 | chr9 | 41117908 | 41118445 | + | -0.752754 | intron (NM_176860, intron 1 of 13) | intron (NM_176860, intron 1 of 13) | 39318 | NM_176860 | 72828 | Mm.133615 | NM_176860 | ENSMUSG00000032020 | **Ubash3b** | 2810457I06Rik\|BB125008\|TULA-2\|p70 | ubiquitin associated and SH3 domain containing, B | protein-coding |
| chr9_41118047_41118442 | chr9 | 41118048 | 41118442 | + | -0.800195 | intron (NM_176860, intron 1 of 13) | intron (NM_176860, intron 1 of 13) | 39249 | NM_176860 | 72828 | Mm.133615 | NM_176860 | ENSMUSG00000032020 | **Ubash3b** | 2810457I06Rik\|BB125008\|TULA-2\|p70 | ubiquitin associated and SH3 domain containing, B | protein-coding |
| chr9_41181307_41181797 | chr9 | 41181308 | 41181797 | + | -0.918041 | Intergenic | Intergenic | -24058 | NM_176860 | 72828 | Mm.133615 | NM_176860 | ENSMUSG00000032020 | **Ubash3b** | 2810457I06Rik\|BB125008\|TULA-2\|p70 | ubiquitin associated and SH3 domain containing, B | protein-coding |
| chr9_41080874_41081292 | chr9 | 41080875 | 41081292 | + | -1.21972 | intron (NM_176860, intron 1 of 13) | intron (NM_176860, intron 1 of 13) | 76411 | NM_176860 | 72828 | Mm.133615 | NM_176860 | ENSMUSG00000032020 | **Ubash3b** | 2810457I06Rik\|BB125008\|TULA-2\|p70 | ubiquitin associated and SH3 domain containing, B | protein-coding |
| chr18_35776480_35776903 | chr18 | 35776481 | 35776903 | + | -1.20584 | intron (NM_019912, intron 1 of 6) | Tigger15a\|DNA\|TcMar-Tigger | 5133 | NM_019912 | 56550 | Mm.180052 | NM_019912 | ENSMUSG00000091896 | **Ube2d2a** | 1500034D03Rik\|Ubc2e\|Ube2d2\|ubc4 | ubiquitin-conjugating enzyme E2D 2A | protein-coding |
| chr5_107840528_107841139 | chr5 | 107840529 | 107841139 | + | -1.23735 | intron (NM_007964, intron 2 of 17) | MLTR12\|LTR\|ERV1 | 10672 | NM_001276397 | 73318 | Mm.441500 | NM_028502 | | **Ube2d2b** | 1700013N18Rik | ubiquitin-conjugating enzyme E2D 2B | protein-coding |
| chr14_18618734_18619146 | chr14 | 18618735 | 18619146 | + | -0.770222 | intron (NM_144839, intron 4 of 5) | MER20\|DNA\|hAT-Charlie | 274687 | NM_144839 | 218793 | Mm.23551 | NM_144839 | ENSMUSG00000058317 | **Ube2e2** | BC016265 | ubiquitin-conjugating enzyme E2E 2 | protein-coding |
| chr14_18665598_18666018 | chr14 | 18665599 | 18666018 | + | -0.840853 | intron (NM_144839, intron 3 of 5) | intron (NM_144839, intron 3 of 5) | 227819 | NM_144839 | 218793 | Mm.23551 | NM_144839 | ENSMUSG00000058317 | **Ube2e2** | BC016265 | ubiquitin-conjugating enzyme E2E 2 | protein-coding |
| chr14_18665332_18666019 | chr14 | 18665333 | 18666019 | + | -0.860995 | intron (NM_144839, intron 3 of 5) | intron (NM_144839, intron 3 of 5) | 227951 | NM_144839 | 218793 | Mm.23551 | NM_144839 | ENSMUSG00000058317 | **Ube2e2** | BC016265 | ubiquitin-conjugating enzyme E2E 2 | protein-coding |
| chr14_18756632_18756822 | chr14 | 18756633 | 18756822 | + | -1.09234 | intron (NM_144839, intron 3 of 5) | intron (NM_144839, intron 3 of 5) | 136900 | NM_144839 | 218793 | Mm.23551 | NM_144839 | ENSMUSG00000058317 | **Ube2e2** | BC016265 | ubiquitin-conjugating enzyme E2E 2 | protein-coding |
| chr14_18756487_18756836 | chr14 | 18756488 | 18756836 | + | -1.18518 | intron (NM_144839, intron 3 of 5) | intron (NM_144839, intron 3 of 5) | 136965 | NM_144839 | 218793 | Mm.23551 | NM_144839 | ENSMUSG00000058317 | **Ube2e2** | BC016265 | ubiquitin-conjugating enzyme E2E 2 | protein-coding |
| chr2_78872224_78872449 | chr2 | 78872225 | 78872449 | + | -0.941121 | intron (NM_009454, intron 2 of 5) | intron (NM_009454, intron 2 of 5) | 3290 | NM_009454 | 22193 | Mm.1485 | NM_009454 | | **Ube2e3** | Ubce4\|ubcM2 | ubiquitin-conjugating enzyme E2E 3 | protein-coding |
| chr6_30336858_30337414 | chr6 | 30336859 | 30337414 | + | -0.631216 | Intergenic | Intergenic | -32597 | NM_001169576 | 22214 | Mm.332967 | NM_009459 | ENSMUSG00000039159 | **Ube2h** | 1500009C23Rik\|AI181839\|AI326965\|AI462483\|AW227540\|E2-20K\|Gid3\|N28148\|Ubc8 | ubiquitin-conjugating enzyme E2H | protein-coding |
| chr6_30230304_30230643 | chr6 | 30230305 | 30230643 | + | -0.657791 | intron (NM_001169576, intron 3 of 4) | ORR1E\|LTR\|ERVL-MaLR | 13329 | NM_001169577 | 22214 | Mm.332967 | NM_009459 | ENSMUSG00000039159 | **Ube2h** | 1500009C23Rik\|AI181839\|AI326965\|AI462483\|AW227540\|E2-20K\|Gid3\|N28148\|Ubc8 | ubiquitin-conjugating enzyme E2H | protein-coding |
| chr6_30311901_30312433 | chr6 | 30311902 | 30312433 | + | -0.722158 | Intergenic | Intergenic | -7628 | NM_001169576 | 22214 | Mm.332967 | NM_009459 | ENSMUSG00000039159 | **Ube2h** | 1500009C23Rik\|AI181839\|AI326965\|AI462483\|AW227540\|E2-20K\|Gid3\|N28148\|Ubc8 | ubiquitin-conjugating enzyme E2H | protein-coding |
| chr6_30326087_30326692 | chr6 | 30326088 | 30326692 | + | -0.790963 | Intergenic | Intergenic | -21851 | NM_001169576 | 22214 | Mm.332967 | NM_009459 | ENSMUSG00000039159 | **Ube2h** | 1500009C23Rik\|AI181839\|AI326965\|AI462483\|AW227540\|E2-20K\|Gid3\|N28148\|Ubc8 | ubiquitin-conjugating enzyme E2H | protein-coding |
| chr6_30326141_30326474 | chr6 | 30326142 | 30326474 | + | -0.828508 | Intergenic | Intergenic | -21769 | NM_001169576 | 22214 | Mm.332967 | NM_009459 | ENSMUSG00000039159 | **Ube2h** | 1500009C23Rik\|AI181839\|AI326965\|AI462483\|AW227540\|E2-20K\|Gid3\|N28148\|Ubc8 | ubiquitin-conjugating enzyme E2H | protein-coding |
| chr6_30226975_30227647 | chr6 | 30226976 | 30227647 | + | -0.840773 | intron (NM_001169576, intron 3 of 4) | intron (NM_001169576, intron 3 of 4) | 16492 | NM_001169577 | 22214 | Mm.332967 | NM_009459 | ENSMUSG00000039159 | **Ube2h** | 1500009C23Rik\|AI181839\|AI326965\|AI462483\|AW227540\|E2-20K\|Gid3\|N28148\|Ubc8 | ubiquitin-conjugating enzyme E2H | protein-coding |
| chr6_30227386_30227635 | chr6 | 30227387 | 30227635 | + | -0.87391 | intron (NM_001169576, intron 3 of 4) | intron (NM_001169576, intron 3 of 4) | 16292 | NM_001169577 | 22214 | Mm.332967 | NM_009459 | ENSMUSG00000039159 | **Ube2h** | 1500009C23Rik\|AI181839\|AI326965\|AI462483\|AW227540\|E2-20K\|Gid3\|N28148\|Ubc8 | ubiquitin-conjugating enzyme E2H | protein-coding |
| chr6_30340209_30340795 | chr6 | 30340210 | 30340795 | + | -0.966882 | Intergenic | Intergenic | -35963 | NM_001169576 | 22214 | Mm.332967 | NM_009459 | ENSMUSG00000039159 | **Ube2h** | 1500009C23Rik\|AI181839\|AI326965\|AI462483\|AW227540\|E2-20K\|Gid3\|N28148\|Ubc8 | ubiquitin-conjugating enzyme E2H | protein-coding |
| chr6_30340263_30340562 | chr6 | 30340264 | 30340562 | + | -1.04817 | Intergenic | Intergenic | -35874 | NM_001169576 | 22214 | Mm.332967 | NM_009459 | ENSMUSG00000039159 | **Ube2h** | 1500009C23Rik\|AI181839\|AI326965\|AI462483\|AW227540\|E2-20K\|Gid3\|N28148\|Ubc8 | ubiquitin-conjugating enzyme E2H | protein-coding |
| chr10_95526843_95527139 | chr10 | 95526844 | 95527139 | + | -0.707137 | intron (NM_080560, intron 1 of 3) | intron (NM_080560, intron 1 of 3) | 11829 | NM_080560 | 93765 | Mm.371667 | NM_080560 | ENSMUSG00000074781 | **Ube2n** | 1500026J17Rik\|AL022654\|BB101821\|UBC13 | ubiquitin-conjugating enzyme E2N | protein-coding |
| chr9_44954372_44955161 | chr9 | 44954373 | 44955161 | + | -1.22318 | intron (NM_145400, intron 3 of 19) | ID_B1\|SINE\|B4 | 10833 | NM_145400 | 140630 | Mm.215069 | NM_145400 | ENSMUSG00000059890 | **Ube4a** | 4732444G18Rik\|9930123J21Rik\|UFD2b | ubiquitination factor E4A | protein-coding |
| chr9_44954375_44954887 | chr9 | 44954376 | 44954887 | + | -1.2636 | intron (NM_145400, intron 3 of 19) | ID\|SINE\|ID | 10969 | NM_145400 | 140630 | Mm.215069 | NM_145400 | ENSMUSG00000059890 | **Ube4a** | 4732444G18Rik\|9930123J21Rik\|UFD2b | ubiquitination factor E4A | protein-coding |
| chr5_148495146_148495700 | chr5 | 148495147 | 148495700 | + | -0.627863 | Intergenic | Intergenic | 57365 | NM_011908 | 24109 | Mm.21846 | NM_011908 | ENSMUSG00000001687 | **Ubl3** | AW108023\|HCG | ubiquitin-like 3 | protein-coding |
| chr5_148501206_148501623 | chr5 | 148501207 | 148501623 | + | -0.644843 | Intergenic | RLTR14\|LTR\|ERV1 | 51373 | NM_011908 | 24109 | Mm.21846 | NM_011908 | ENSMUSG00000001687 | **Ubl3** | AW108023\|HCG | ubiquitin-like 3 | protein-coding |
| chr5_148594891_148595368 | chr5 | 148594892 | 148595368 | + | -0.953244 | Intergenic | MLT1N2\|LTR\|ERVL-MaLR | -42342 | NM_011908 | 24109 | Mm.21846 | NM_011908 | ENSMUSG00000001687 | **Ubl3** | AW108023\|HCG | ubiquitin-like 3 | protein-coding |
| chr5_148595443_148595661 | chr5 | 148595444 | 148595661 | + | -0.994843 | Intergenic | Intergenic | -42764 | NM_011908 | 24109 | Mm.21846 | NM_011908 | ENSMUSG00000001687 | **Ubl3** | AW108023\|HCG | ubiquitin-like 3 | protein-coding |
| chr5_148595037_148595348 | chr5 | 148595038 | 148595348 | + | -1.00391 | Intergenic | MLT1N2\|LTR\|ERVL-MaLR | -42405 | NM_011908 | 24109 | Mm.21846 | NM_011908 | ENSMUSG00000001687 | **Ubl3** | AW108023\|HCG | ubiquitin-like 3 | protein-coding |
| chr5_148595053_148595686 | chr5 | 148595054 | 148595686 | + | -1.11492 | Intergenic | MLT1N2\|LTR\|ERVL-MaLR | -42582 | NM_011908 | 24109 | Mm.21846 | NM_011908 | ENSMUSG00000001687 | **Ubl3** | AW108023\|HCG | ubiquitin-like 3 | protein-coding |
| chr11_44469586_44470222 | chr11 | 44469587 | 44470222 | + | -0.64242 | intron (NM_024475, intron 1 of 10) | intron (NM_024475, intron 1 of 10) | 644 | NM_024475 | 79560 | Mm.338784 | NM_024475 | ENSMUSG00000041231 | **Ublcp1** | 4930527B16Rik\|8430435I17Rik\|BC002236 | ubiquitin-like domain containing CTD phosphatase 1 | protein-coding |
| chrX_153375137_153375534 | chrX | 153375138 | 153375534 | + | -0.953399 | intron (NM_173780, intron 3 of 6) | intron (NM_173780, intron 3 of 6) | -122896 | NM_018798 | 54609 | Mm.430772 | NM_018798 | ENSMUSG00000050148 | **Ubqln2** | Chap1\|Dsk2\|HRIHFB2157\|Plic-2\|Plic2 | ubiquilin 2 | protein-coding |
| chr11_32445650_32445994 | chr11 | 32445651 | 32445994 | + | -0.767917 | Intergenic | Intergenic | -9550 | NM_173784 | 327900 | Mm.260956 | NM_173784 | | **Ubtd2** | 4930571L24\|9630054F20Rik\|AI645720 | ubiquitin domain containing 2 | protein-coding |
| chr11_32486647_32487075 | chr11 | 32486648 | 32487075 | + | -0.803308 | intron (NM_173784, intron 1 of 2) | intron (NM_173784, intron 1 of 2) | 31489 | NM_173784 | 327900 | Mm.260956 | NM_173784 | | **Ubtd2** | 4930571L24\|9630054F20Rik\|AI645720 | ubiquitin domain containing 2 | protein-coding |
| chr11_32486700_32486934 | chr11 | 32486701 | 32486934 | + | -0.80798 | intron (NM_173784, intron 1 of 2) | intron (NM_173784, intron 1 of 2) | 31445 | NM_173784 | 327900 | Mm.260956 | NM_173784 | | **Ubtd2** | 4930571L24\|9630054F20Rik\|AI645720 | ubiquitin domain containing 2 | protein-coding |
| chr11_32474770_32475095 | chr11 | 32474771 | 32475095 | + | -0.885045 | intron (NM_173784, intron 1 of 2) | ORR1E\|LTR\|ERVL-MaLR | 19561 | NM_173784 | 327900 | Mm.260956 | NM_173784 | | **Ubtd2** | 4930571L24\|9630054F20Rik\|AI645720 | ubiquitin domain containing 2 | protein-coding |
| chr11_32474821_32475046 | chr11 | 32474822 | 32475046 | + | -0.961214 | intron (NM_173784, intron 1 of 2) | ORR1E\|LTR\|ERVL-MaLR | 19562 | NM_173784 | 327900 | Mm.260956 | NM_173784 | | **Ubtd2** | 4930571L24\|9630054F20Rik\|AI645720 | ubiquitin domain containing 2 | protein-coding |

| | | | | | | | | | | | | | | | | | |
|---|---|---|---|---|---|---|---|---|---|---|---|---|---|---|---|---|---|
| chr9_64227660_64228092 | chr9 | 64227661 | 64228092 | + | -0.635401 | intron (NM_008927, intron 1 of 10) | intron (NM_008927, intron 1 of 10) | -7325 | NM_033607 | 93841 | Mm.390180 | NM_033607 | ENSMUSG00000035337 | **Uchl4** | - | ubiquitin carboxyl-terminal esterase L4 | protein-coding |
| chr1_167280129_167280403 | chr1 | 167280130 | 167280403 | + | -0.69156 | intron (NM_030724, intron 1 of 6) | intron (NM_030724, intron 1 of 6) | 4861 | NM_030724 | 80914 | Mm.280895 | NM_030724 | ENSMUSG00000026558 | **Uck2** | AA407809\|AI481316\|AU018180\|AU020720\|TSA903\|UK\|UMK\|Umpk | uridine-cytidine kinase 2 | protein-coding |
| chr7_100495512_100496163 | chr7 | 100495513 | 100496163 | + | -0.74098 | intron (NM_011671, intron 2 of 7) | intron (NM_011671, intron 2 of 7) | 2493 | NM_011671 | 22228 | Mm.171378 | NM_011671 | ENSMUSG00000033685 | **Ucp2** | Slc25a8 | uncoupling protein 2 (mitochondrial, proton carrier) | protein-coding |
| chr4_59205820_59206087 | chr4 | 59205821 | 59206087 | + | -0.666588 | intron (NM_011673, intron 1 of 8) | intron (NM_011673, intron 1 of 8) | 16404 | NM_011673 | 22234 | Mm.198803 | NM_011673 | ENSMUSG00000028381 | **Ugcg** | AU043821\|C80537\|Epcs21\|GlcT-1\|Ugcgl | UDP-glucose ceramide glucosyltransferase | protein-coding |
| chr4_59167192_59168242 | chr4 | 59167193 | 59168242 | + | -1.26043 | Intergenic | ORR1A2-int\|LTR\|ERVL-MaLR | -21833 | NM_011673 | 22234 | Mm.198803 | NM_011673 | ENSMUSG00000028381 | **Ugcg** | AU043821\|C80537\|Epcs21\|GlcT-1\|Ugcgl | UDP-glucose ceramide glucosyltransferase | protein-coding |
| chr4_59167198_59167745 | chr4 | 59167199 | 59167745 | + | -1.32888 | Intergenic | ORR1A2-int\|LTR\|ERVL-MaLR | -22078 | NM_011673 | 22234 | Mm.198803 | NM_011673 | ENSMUSG00000028381 | **Ugcg** | AU043821\|C80537\|Epcs21\|GlcT-1\|Ugcgl | UDP-glucose ceramide glucosyltransferase | protein-coding |
| chr13_55062336_55062722 | chr13 | 55062337 | 55062722 | + | -0.810861 | intron (NR_121592, intron 6 of 13) | intron (NR_121592, intron 6 of 13) | 37766 | NM_011307 | 20184 | Mm.259301 | NM_011307 | ENSMUSG00000025878 | **Uimc1** | 9430016E08Rik\|D330018D10Rik\|D630032M02Rik\|RIP110\|Rxrip110 | ubiquitin interaction motif containing 1 | protein-coding |
| chr1_66552373_66553041 | chr1 | 66552374 | 66553041 | + | -1.20454 | intron (NM_175510, intron 21 of 64) | L3\|LINE\|CR1 | 84260 | NM_175510 | 329178 | Mm.44798 | NM_175510 | ENSMUSG00000055567 | **Unc80** | C030018G13Rik\|C230061B10Rik\|UNC-80 | unc-80 homolog (C. elegans) | protein-coding |
| chr17_25224562_25225106 | chr17 | 25224563 | 25225106 | + | -0.699973 | intron (NM_028789, intron 1 of 4) | RLTR14\|LTR\|ERV1 | 2252 | NM_028789 | 74154 | Mm.267353 | NM_028789 | ENSMUSG00000015127 | **Unkl** | 1300004G08Rik | unkempt-like (Drosophila) | protein-coding |
| chr17_25219452_25219829 | chr17 | 25219453 | 25219829 | + | -0.993086 | intron (NM_001197024, intron 10 of 14) | RMER4B\|LTR\|ERVK | -2941 | NM_028789 | 74154 | Mm.267353 | NM_028789 | ENSMUSG00000015127 | **Unkl** | 1300004G08Rik | unkempt-like (Drosophila) | protein-coding |
| chr17_25219984_25220236 | chr17 | 25219985 | 25220236 | + | -1.06784 | intron (NM_001197024, intron 10 of 14) | intron (NM_001197024, intron 10 of 14) | -2472 | NM_028789 | 74154 | Mm.267353 | NM_028789 | ENSMUSG00000015127 | **Unkl** | 1300004G08Rik | unkempt-like (Drosophila) | protein-coding |
| chr10_75405949_75406997 | chr10 | 75405950 | 75406997 | + | -0.63307 | promoter-TSS (NM_133995) | promoter-TSS (NM_133995) | -438 | NM_133995 | 103149 | Mm.441195 | NM_133995 | | **Upb1** | AI195023 | ureidopropionase, beta | protein-coding |
| chr10_75405947_75406460 | chr10 | 75405948 | 75406460 | + | -0.705963 | promoter-TSS (NM_133995) | promoter-TSS (NM_133995) | -707 | NM_133995 | 103149 | Mm.441195 | NM_133995 | | **Upb1** | AI195023 | ureidopropionase, beta | protein-coding |
| chr10_75417684_75418062 | chr10 | 75417685 | 75418062 | + | -1.1255 | intron (NM_133995, intron 3 of 9) | HAL1\|LINE\|L1 | 10962 | NM_133995 | 103149 | Mm.441195 | NM_133995 | | **Upb1** | AI195023 | ureidopropionase, beta | protein-coding |
| chr2_58844469_58844941 | chr2 | 58844470 | 58844941 | + | -0.626299 | intron (NM_001001178, intron 11 of 13) | intron (NM_001001178, intron 11 of 13) | 89521 | NM_029692 | 76654 | Mm.200370 | NM_029692 | ENSMUSG00000026839 | **Upp2** | 1700124F02Rik\|AI266885\|UDRPASE2\|UP2\|UPASE2 | uridine phosphorylase 2 | protein-coding |
| chrX_104485711_104485960 | chrX | 104485712 | 104485960 | + | -0.917663 | intron (NM_001081189, intron 1 of 6) | B4A\|SINE\|B4 | 3054 | NM_001081189 | 331487 | Mm.438275 | NM_001081189 | | **Uprt** | Gm774 | uracil phosphoribosyltransferase (FUR1) homolog (S. cerevisiae) | protein-coding |
| chr13_30570311_30570500 | chr13 | 30570312 | 30570500 | + | -0.63309 | Intergenic | Intergenic | -25090 | NM_025710 | 66694 | Mm.181933 | NM_025710 | ENSMUSG00000038462 | **Uqcrfs1** | 4430402G14Rik\|AI875505 | ubiquinol-cytochrome c reductase, Rieske iron-sulfur polypeptide 1 | protein-coding |
| chr5_147340568_147341652 | chr5 | 147340569 | 147341652 | + | -0.808366 | intron (NM_010229, intron 20 of 23) | intron (NM_010229, intron 20 of 23) | -18670 | NM_001039678 | 231903 | Mm.291286 | NM_001039678 | ENSMUSG00000075543 | **Urad** | EG231903\|Prhoxnb | ureidoimidazoline (2-oxo-4-hydroxy-4-carboxy-5) decarboxylase | protein-coding |
| chr8_124075889_124076445 | chr8 | 124075890 | 124076445 | + | -0.760074 | Intergenic | Intergenic | 52694 | NM_001029876 | 382038 | Mm.334891 | NM_001029876 | ENSMUSG00000031976 | **Urb2** | C77575\|mKIAA0133 | URB2 ribosome biogenesis 2 homolog (S. cerevisiae) | protein-coding |
| chr8_124076128_124076445 | chr8 | 124076129 | 124076445 | + | -0.922351 | Intergenic | Intergenic | 52814 | NM_001029876 | 382038 | Mm.334891 | NM_001029876 | ENSMUSG00000031976 | **Urb2** | C77575\|mKIAA0133 | URB2 ribosome biogenesis 2 homolog (S. cerevisiae) | protein-coding |
| chr7_37999713_38000230 | chr7 | 37999714 | 38000230 | + | -0.757327 | intron (NM_011274, intron 1 of 10) | intron (NM_011274, intron 1 of 10) | 19580 | NM_011274 | 19777 | Mm.387220 | NM_011274 | ENSMUSG00000030421 | **Uri1** | C80913\|NNX3\|Rmp | URI1, prefoldin-like chaperone | protein-coding |
| chr5_92165807_92166306 | chr5 | 92165808 | 92166306 | + | -1.09143 | intron (NM_019490, intron 4 of 23) | intron (NM_019490, intron 4 of 23) | 28119 | NM_019490 | 56041 | Mm.15868 | NM_019490 | ENSMUSG00000029407 | **Uso1** | 115kDa\|TAP\|Vdp | USO1 vesicle docking factor | protein-coding |
| chr5_92165970_92166334 | chr5 | 92165971 | 92166334 | + | -1.10907 | intron (NM_019490, intron 4 of 23) | intron (NM_019490, intron 4 of 23) | 28214 | NM_019490 | 56041 | Mm.15868 | NM_019490 | ENSMUSG00000029407 | **Uso1** | 115kDa\|TAP\|Vdp | USO1 vesicle docking factor | protein-coding |
| chr8_119926455_119926842 | chr8 | 119926456 | 119926842 | + | -0.665623 | intron (NM_009462, intron 1 of 13) | B1_Mur4\|SINE\|Alu | 15797 | NM_009462 | 22224 | Mm.256910 | NM_009462 | ENSMUSG00000031826 | **Usp10** | 2610014N07Rik\|UBPO\|Uchrp\|mKIAA0190 | ubiquitin specific peptidase 10 | protein-coding |
| chr8_119946088_119946784 | chr8 | 119946089 | 119946784 | + | -0.88693 | intron (NM_009462, intron 4 of 13) | intron (NM_009462, intron 4 of 13) | 35584 | NM_009462 | 22224 | Mm.256910 | NM_009462 | ENSMUSG00000031826 | **Usp10** | 2610014N07Rik\|UBPO\|Uchrp\|mKIAA0190 | ubiquitin specific peptidase 10 | protein-coding |
| chr5_146781625_146782313 | chr5 | 146781626 | 146782313 | + | -1.0419 | intron (NM_011669, intron 1 of 8) | L1M4\|LINE\|L1 | 12987 | NM_011669 | 22217 | Mm.167971 | NM_011669 | ENSMUSG00000029640 | **Usp12** | Ubh1 | ubiquitin specific peptidase 12 | protein-coding |
| chr5_146781636_146781993 | chr5 | 146781637 | 146781993 | + | -1.04964 | intron (NM_011669, intron 1 of 8) | intron (NM_011669, intron 1 of 8) | 13141 | NM_011669 | 22217 | Mm.167971 | NM_011669 | ENSMUSG00000029640 | **Usp12** | Ubh1 | ubiquitin specific peptidase 12 | protein-coding |
| chr5_146783863_146784410 | chr5 | 146783864 | 146784410 | + | -1.46899 | intron (NM_011669, intron 1 of 8) | intron (NM_011669, intron 1 of 8) | 10819 | NM_011669 | 22217 | Mm.167971 | NM_011669 | ENSMUSG00000029640 | **Usp12** | Ubh1 | ubiquitin specific peptidase 12 | protein-coding |
| chr5_146783794_146784225 | chr5 | 146783795 | 146784225 | + | -1.53882 | intron (NM_011669, intron 1 of 8) | intron (NM_011669, intron 1 of 8) | 10946 | NM_011669 | 22217 | Mm.167971 | NM_011669 | ENSMUSG00000029640 | **Usp12** | Ubh1 | ubiquitin specific peptidase 12 | protein-coding |
| chr5_146783929_146784222 | chr5 | 146783930 | 146784222 | + | -1.59406 | intron (NM_011669, intron 1 of 8) | intron (NM_011669, intron 1 of 8) | 10880 | NM_011669 | 22217 | Mm.167971 | NM_011669 | ENSMUSG00000029640 | **Usp12** | Ubh1 | ubiquitin specific peptidase 12 | protein-coding |
| chr10_123158991_123159588 | chr10 | 123158992 | 123159588 | + | -0.853716 | intron (NM_027604, intron 6 of 21) | intron (NM_027604, intron 6 of 21) | 37633 | NM_027604 | 14479 | Mm.244209 | NM_027604 | ENSMUSG00000020124 | **Usp15** | 4921514G19Rik\|AI327321\|E430033I05Rik\|Gcap18 | ubiquitin specific peptidase 15 | protein-coding |
| chr10_123158998_123159231 | chr10 | 123158999 | 123159231 | + | -0.989901 | intron (NM_027604, intron 6 of 21) | intron (NM_027604, intron 6 of 21) | 37808 | NM_027604 | 14479 | Mm.244209 | NM_027604 | ENSMUSG00000020124 | **Usp15** | 4921514G19Rik\|AI327321\|E430033I05Rik\|Gcap18 | ubiquitin specific peptidase 15 | protein-coding |
| chr4_106314225_106314571 | chr4 | 106314226 | 106314571 | + | -0.728368 | Intergenic | Intergenic | -1815 | NM_183225 | 329908 | Mm.234544 | NM_183225 | ENSMUSG00000028514 | **Usp24** | 2700066K03Rik\|2810030C21Rik\|AI414051\|B130021E18\|C79851 | ubiquitin specific peptidase 24 | protein-coding |
| chr4_106314320_106314569 | chr4 | 106314321 | 106314569 | + | -0.733312 | Intergenic | Intergenic | -1768 | NM_183225 | 329908 | Mm.234544 | NM_183225 | ENSMUSG00000028514 | **Usp24** | 2700066K03Rik\|2810030C21Rik\|AI414051\|B130021E18\|C79851 | ubiquitin specific peptidase 24 | protein-coding |
| chr9_48980401_48980738 | chr9 | 48980402 | 48980738 | + | -0.720067 | Intergenic | Intergenic | -4815 | NM_175482 | 235323 | Mm.21630 | NM_175482 | ENSMUSG00000032267 | **Usp28** | 9830148O20Rik\|AU022237\|mKIAA1515 | ubiquitin specific peptidase 28 | protein-coding |
| chr9_48994660_48995203 | chr9 | 48994661 | 48995203 | + | -0.7577 | intron (NM_175482, intron 1 of 23) | intron (NM_175482, intron 1 of 23) | 9547 | NM_175482 | 235323 | Mm.21630 | NM_175482 | ENSMUSG00000032267 | **Usp28** | 9830148O20Rik\|AU022237\|mKIAA1515 | ubiquitin specific peptidase 28 | protein-coding |
| chr9_66586488_66586874 | chr9 | 66586489 | 66586874 | + | -0.687892 | intron (NM_144937, intron 1 of 14) | intron (NM_144937, intron 1 of 14) | 6299 | NM_144937 | 235441 | Mm.38976 | NM_144937 | ENSMUSG00000032376 | **Usp3** | AA409661\|BC017156 | ubiquitin specific peptidase 3 | protein-coding |
| chr9_66598583_66599022 | chr9 | 66598584 | 66599022 | + | -0.784606 | Intergenic | Intergenic | -5823 | NM_144937 | 235441 | Mm.38976 | NM_144937 | ENSMUSG00000032376 | **Usp3** | AA409661\|BC017156 | ubiquitin specific peptidase 3 | protein-coding |
| chr9_66615081_66615633 | chr9 | 66615082 | 66615633 | + | -0.943489 | Intergenic | Intergenic | -22377 | NM_144937 | 235441 | Mm.38976 | NM_144937 | ENSMUSG00000032376 | **Usp3** | AA409661\|BC017156 | ubiquitin specific peptidase 3 | protein-coding |
| chr9_66615121_66615473 | chr9 | 66615122 | 66615473 | + | -0.986624 | Intergenic | Intergenic | -22317 | NM_144937 | 235441 | Mm.38976 | NM_144937 | ENSMUSG00000032376 | **Usp3** | AA409661\|BC017156 | ubiquitin specific peptidase 3 | protein-coding |
| chr4_137603842_137604124 | chr4 | 137603843 | 137604124 | + | -0.720085 | intron (NM_130879, intron 1 of 27) | intron (NM_130879, intron 1 of 27) | 9794 | NM_130879 | 170707 | Mm.218478 | NM_130879 | ENSMUSG00000043411 | **Usp48** | 2810449C13Rik\|AI115503\|BC021769\|D330022K21Rik\|Usp31 | ubiquitin specific peptidase 48 | protein-coding |
| chr4_137603580_137604114 | chr4 | 137603581 | 137604114 | + | -0.797989 | intron (NM_130879, intron 1 of 27) | intron (NM_130879, intron 1 of 27) | 9658 | NM_130879 | 170707 | Mm.218478 | NM_130879 | ENSMUSG00000043411 | **Usp48** | 2810449C13Rik\|AI115503\|BC021769\|D330022K21Rik\|Usp31 | ubiquitin specific peptidase 48 | protein-coding |
| chr2_6391277_6391599 | chr2 | 6391278 | 6391599 | + | -0.935828 | intron (NM_001080548, intron 3 of 14) | intron (NM_001080548, intron 3 of 14) | 38702 | NM_181399 | 98910 | Mm.40673 | NM_181399 | ENSMUSG00000039046 | **Usp6nl** | AI316785\|RNTRE\|TRE2NL\|mKIAA0019 | USP6 N-terminal like | protein-coding |
| chr2_6478111_6479186 | chr2 | 6478112 | 6479186 | + | -1.0738 | Intergenic | Intergenic | 125913 | NM_181399 | 98910 | Mm.40673 | NM_181399 | ENSMUSG00000039046 | **Usp6nl** | AI316785\|RNTRE\|TRE2NL\|mKIAA0019 | USP6 N-terminal like | protein-coding |
| chr2_6478111_6478914 | chr2 | 6478112 | 6478914 | + | -1.09751 | Intergenic | Intergenic | 125777 | NM_181399 | 98910 | Mm.40673 | NM_181399 | ENSMUSG00000039046 | **Usp6nl** | AI316785\|RNTRE\|TRE2NL\|mKIAA0019 | USP6 N-terminal like | protein-coding |
| chr16_8745747_8746222 | chr16 | 8745748 | 8746222 | + | -0.791225 | Intergenic | Intergenic | -7643 | NM_001003918 | 252870 | Mm.295330 | NM_001003918 | ENSMUSG00000022710 | **Usp7** | 2210010O09Rik\|AA409944\|AA617399\|AU019296\|AW548146\|C80752\|Hausp | ubiquitin specific peptidase 7 | protein-coding |
| chr16_8719244_8719780 | chr16 | 8719245 | 8719780 | + | -0.991796 | intron (NM_001003918, intron 1 of 30) | intron (NM_001003918, intron 1 of 30) | 18830 | NM_001003918 | 252870 | Mm.295330 | NM_001003918 | ENSMUSG00000022710 | **Usp7** | 2210010O09Rik\|AA409944\|AA617399\|AU019296\|AW548146\|C80752\|Hausp | ubiquitin specific peptidase 7 | protein-coding |
| chr16_8719263_8719577 | chr16 | 8719264 | 8719577 | + | -0.994874 | intron (NM_001003918, intron 1 of 30) | intron (NM_001003918, intron 1 of 30) | 18922 | NM_001003918 | 252870 | Mm.295330 | NM_001003918 | ENSMUSG00000022710 | **Usp7** | 2210010O09Rik\|AA409944\|AA617399\|AU019296\|AW548146\|C80752\|Hausp | ubiquitin specific peptidase 7 | protein-coding |

| | | | | | | | | | | | | | | | | | |
|---|---|---|---|---|---|---|---|---|---|---|---|---|---|---|---|---|---|
| chr10_8501122_8501354 | chr10 | 8501123 | 8501354 | + | -0.728388 | intron (NM_177387, intron 1 of 7) | intron (NM_177387, intron 1 of 7) | 17587 | NM_177387 | 338362 | Mm.41163 | NM_177387 | ENSMUSG00000047712 | **Ust** | D930010O20Rik\|UA2OST | uronyl-2-sulfotransferase | protein-coding |
| chr10_8531531_8532007 | chr10 | 8531532 | 8532007 | + | -0.778263 | Intergenic | Intergenic | -12944 | NM_177387 | 338362 | Mm.41163 | NM_177387 | ENSMUSG00000047712 | **Ust** | D930010O20Rik\|UA2OST | uronyl-2-sulfotransferase | protein-coding |
| chr10_8535443_8535906 | chr10 | 8535444 | 8535906 | + | -0.830415 | Intergenic | Intergenic | -16850 | NM_177387 | 338362 | Mm.41163 | NM_177387 | ENSMUSG00000047712 | **Ust** | D930010O20Rik\|UA2OST | uronyl-2-sulfotransferase | protein-coding |
| chr7_99102854_99103263 | chr7 | 99102855 | 99103263 | + | -0.629537 | intron (NM_178635, intron 3 of 14) | intron (NM_178635, intron 3 of 14) | 38085 | NM_178635 | 78610 | Mm.323072 | NM_178635 | ENSMUSG00000035354 | **Uvrag** | 9530039D02Rik\|AI648770\|BB124205\|Uvrag1\|Uvragl | UV radiation resistance associated gene | protein-coding |
| chr7_99120139_99120379 | chr7 | 99120140 | 99120379 | + | -0.646324 | intron (NM_178635, intron 1 of 14) | ORR1F\|LTR\|ERVL-MaLR | 20885 | NM_178635 | 78610 | Mm.323072 | NM_178635 | ENSMUSG00000035354 | **Uvrag** | 9530039D02Rik\|AI648770\|BB124205\|Uvrag1\|Uvragl | UV radiation resistance associated gene | protein-coding |
| chr7_99086563_99087465 | chr7 | 99086564 | 99087465 | + | -0.740175 | intron (NM_178635, intron 5 of 14) | intron (NM_178635, intron 5 of 14) | 54130 | NM_178635 | 78610 | Mm.323072 | NM_178635 | ENSMUSG00000035354 | **Uvrag** | 9530039D02Rik\|AI648770\|BB124205\|Uvrag1\|Uvragl | UV radiation resistance associated gene | protein-coding |
| chr7_99152469_99153160 | chr7 | 99152470 | 99153160 | + | -0.740315 | TTS (NM_026384) | TTS (NM_026384) | -11671 | NM_178635 | 78610 | Mm.323072 | NM_178635 | ENSMUSG00000035354 | **Uvrag** | 9530039D02Rik\|AI648770\|BB124205\|Uvrag1\|Uvragl | UV radiation resistance associated gene | protein-coding |
| chr7_99003126_99003753 | chr7 | 99003127 | 99003753 | + | -0.758774 | intron (NM_178635, intron 8 of 14) | intron (NM_178635, intron 8 of 14) | 137704 | NM_178635 | 78610 | Mm.323072 | NM_178635 | ENSMUSG00000035354 | **Uvrag** | 9530039D02Rik\|AI648770\|BB124205\|Uvrag1\|Uvragl | UV radiation resistance associated gene | protein-coding |
| chr7_98993244_98993700 | chr7 | 98993245 | 98993700 | + | -0.972087 | intron (NM_178635, intron 8 of 14) | PB1D9\|SINE\|Alu | 147672 | NM_178635 | 78610 | Mm.323072 | NM_178635 | ENSMUSG00000035354 | **Uvrag** | 9530039D02Rik\|AI648770\|BB124205\|Uvrag1\|Uvragl | UV radiation resistance associated gene | protein-coding |
| chr7_98993393_98993739 | chr7 | 98993394 | 98993739 | + | -1.05799 | intron (NM_178635, intron 8 of 14) | L1M4\|LINE\|L1 | 147578 | NM_178635 | 78610 | Mm.323072 | NM_178635 | ENSMUSG00000035354 | **Uvrag** | 9530039D02Rik\|AI648770\|BB124205\|Uvrag1\|Uvragl | UV radiation resistance associated gene | protein-coding |
| chr1_43880309_43881137 | chr1 | 43880310 | 43881137 | + | -0.860056 | Intergenic | Intergenic | -53015 | NM_026430 | 67883 | Mm.387202 | NM_026430 | | **Uxs1** | 1600025I13Rik\|AI451869\|AI649125\|AW550562 | UDP-glucuronate decarboxylase 1 | protein-coding |
| chr1_43880326_43880792 | chr1 | 43880327 | 43880792 | + | -0.932715 | Intergenic | Intergenic | -52851 | NM_026430 | 67883 | Mm.387202 | NM_026430 | | **Uxs1** | 1600025I13Rik\|AI451869\|AI649125\|AW550562 | UDP-glucuronate decarboxylase 1 | protein-coding |
| chr6_125221832_125222558 | chr6 | 125221833 | 125222558 | + | -0.94418 | 3' UTR (NM_009496, exon 5 of 5) | 3' UTR (NM_009496, exon 5 of 5) | 6614 | NM_001080557 | 22317 | Mm.32321 | NM_009496 | ENSMUSG00000030337 | **Vamp1** | Syb-1\|Syb1\|VAMP-1\|lew | vesicle-associated membrane protein 1 | protein-coding |
| chr1_162571181_162571726 | chr1 | 162571182 | 162571726 | + | -0.898828 | intron (NM_016796, intron 1 of 7) | CpG | 626 | NM_016796 | 53330 | Mm.10699 | NM_016796 | ENSMUSG00000026696 | **Vamp4** | D1Ertd147e | vesicle-associated membrane protein 4 | protein-coding |
| chr1_162571357_162571682 | chr1 | 162571358 | 162571682 | + | -0.947226 | intron (NM_016796, intron 1 of 7) | CpG | 692 | NM_016796 | 53330 | Mm.10699 | NM_016796 | ENSMUSG00000026696 | **Vamp4** | D1Ertd147e | vesicle-associated membrane protein 4 | protein-coding |
| chr7_19256633_19256919 | chr7 | 19256634 | 19256919 | + | -0.67507 | TTS (NM_009499) | TTS (NM_009499) | 15078 | NM_001282021 | 22323 | Mm.9684 | NM_009499 | ENSMUSG00000030403 | **Vasp** | - | vasodilator-stimulated phosphoprotein | protein-coding |
| chr17_57286725_57287019 | chr17 | 57286726 | 57287019 | + | -0.849506 | intron (NM_011691, intron 1 of 26) | intron (NM_011691, intron 1 of 26) | 7772 | NM_001163815 | 22324 | Mm.248172 | NM_011691 | | **Vav1** | Vav\|vav-T | vav 1 oncogene | protein-coding |
| chr3_109371598_109371937 | chr3 | 109371599 | 109371937 | + | -0.98944 | intron (NM_020505, intron 1 of 26) | intron (NM_020505, intron 1 of 26) | 31085 | NM_020505 | 57257 | Mm.282257 | NM_020505 | ENSMUSG00000033721 | **Vav3** | A530094I06Rik\|AA986410\|Idd18.1 | vav 3 oncogene | protein-coding |
| chr11_52340995_52341457 | chr11 | 52340996 | 52341457 | + | -0.643551 | Intergenic | Intergenic | -19889 | NM_011694 | 22333 | Mm.3555 | NM_011694 | ENSMUSG00000020402 | **Vdac1** | AL033343\|Vdac5 | voltage-dependent anion channel 1 | protein-coding |
| chr11_88073213_88073874 | chr11 | 88073214 | 88073874 | + | -0.872561 | exon (NM_016686, exon 2 of 6) | exon (NM_016686, exon 2 of 6) | 5265 | NM_016686 | 22344 | Mm.46628 | NM_016686 | ENSMUSG00000018377 | **Vezf1** | AI848691\|AW046909\|db1 | vascular endothelial zinc finger 1 | protein-coding |
| chr6_114920600_114920998 | chr6 | 114920601 | 114920998 | + | -0.770009 | intron (NM_177683, intron 1 of 4) | intron (NM_177683, intron 1 of 4) | 953 | NM_177683 | 232334 | Mm.268357 | NM_177683 | | **Vgll4** | BC048841\|VGL-4 | vestigial like 4 (Drosophila) | protein-coding |
| chr6_114894542_114894823 | chr6 | 114894543 | 114894823 | + | -0.836299 | intron (NM_177683, intron 1 of 4) | intron (NM_177683, intron 1 of 4) | 27069 | NM_177683 | 232334 | Mm.268357 | NM_177683 | | **Vgll4** | BC048841\|VGL-4 | vestigial like 4 (Drosophila) | protein-coding |
| chr6_114898914_114899269 | chr6 | 114898915 | 114899269 | + | -0.93194 | intron (NM_177683, intron 1 of 4) | intron (NM_177683, intron 1 of 4) | 22660 | NM_177683 | 232334 | Mm.268357 | NM_177683 | | **Vgll4** | BC048841\|VGL-4 | vestigial like 4 (Drosophila) | protein-coding |
| chr6_114936311_114936931 | chr6 | 114936312 | 114936931 | + | -1.00466 | Intergenic | Intergenic | -14869 | NM_177683 | 232334 | Mm.268357 | NM_177683 | | **Vgll4** | BC048841\|VGL-4 | vestigial like 4 (Drosophila) | protein-coding |
| chr9_121647001_121647342 | chr9 | 121647002 | 121647342 | + | -0.745981 | intron (NM_011703, intron 1 of 12) | intron (NM_011703, intron 1 of 12) | 4456 | NM_011703 | 22354 | Mm.282007 | NM_011703 | ENSMUSG00000032528 | **Vipr1** | AV071699\|VIP-R1\|VPAC1 | vasoactive intestinal peptide receptor 1 | protein-coding |
| chr9_121647093_121647335 | chr9 | 121647094 | 121647335 | + | -0.781667 | intron (NM_011703, intron 1 of 12) | intron (NM_011703, intron 1 of 12) | 4498 | NM_011703 | 22354 | Mm.282007 | NM_011703 | ENSMUSG00000032528 | **Vipr1** | AV071699\|VIP-R1\|VPAC1 | vasoactive intestinal peptide receptor 1 | protein-coding |
| chr1_182162752_182163402 | chr1 | 182162753 | 182163402 | + | -1.40632 | Intergenic | Intergenic | -4979 | NM_001166728 | 625823 | Mm.479992 | NM_001166728 | | **Vmn1r1** | EG625823\|Gm6628 | vomeronasal 1 receptor 1 | protein-coding |
| chr17_19444681_19445378 | chr17 | 19444682 | 19445378 | + | -0.915012 | Intergenic | RLTR50A\|LTR\|ERVK | -59781 | NM_001104562 | 627537 | Mm.469858 | NM_001104562 | ENSMUSG00000091859 | **Vmn2r100** | EG627537 | vomeronasal 2, receptor 100 | protein-coding |
| chr17_18458166_18458831 | chr17 | 18458167 | 18458831 | + | -1.17374 | Intergenic | RLTR50A\|LTR\|ERVK | 34395 | NM_001102581 | 328759 | Mm.469839 | NM_001102581 | ENSMUSG00000091631 | **Vmn2r95** | - | vomeronasal 2, receptor 95 | protein-coding |
| chr17_18597258_18597847 | chr17 | 18597259 | 18597847 | + | -0.708254 | exon (NM_001104547, exon 5 of 5) | exon (NM_001104547, exon 5 of 5) | 15826 | NM_001104547 | 433070 | Mm.215962 | NM_001104547 | ENSMUSG00000091679 | **Vmn2r96** | EG433070 | vomeronasal 2, receptor 96 | protein-coding |
| chr17_18597137_18597626 | chr17 | 18597138 | 18597626 | + | -0.756136 | exon (NM_001104547, exon 5 of 5) | exon (NM_001104547, exon 5 of 5) | 15655 | NM_001104547 | 433070 | Mm.215962 | NM_001104547 | ENSMUSG00000091679 | **Vmn2r96** | EG433070 | vomeronasal 2, receptor 96 | protein-coding |
| chr17_18615234_18615617 | chr17 | 18615235 | 18615617 | + | -0.788102 | Intergenic | RLTR50A\|LTR\|ERVK | 33699 | NM_001104547 | 433070 | Mm.215962 | NM_001104547 | ENSMUSG00000091679 | **Vmn2r96** | EG433070 | vomeronasal 2, receptor 96 | protein-coding |
| chr17_18558169_18559197 | chr17 | 18558170 | 18559197 | + | -0.813222 | Intergenic | IAPLTR3-int\|LTR\|ERVK | -23044 | NM_001104547 | 433070 | Mm.215962 | NM_001104547 | ENSMUSG00000091679 | **Vmn2r96** | EG433070 | vomeronasal 2, receptor 96 | protein-coding |
| chr17_18718206_18718764 | chr17 | 18718207 | 18718764 | + | -0.922239 | Intergenic | RLTR50A\|LTR\|ERVK | 136758 | NM_001104547 | 433070 | Mm.215962 | NM_001104547 | ENSMUSG00000091679 | **Vmn2r96** | EG433070 | vomeronasal 2, receptor 96 | protein-coding |
| chr17_18968784_18969400 | chr17 | 18968785 | 18969400 | + | -0.609792 | Intergenic | IAPLTR4\|LTR\|ERVK | 54770 | NM_001104549 | 627367 | Mm.384131 | NM_001104549 | ENSMUSG00000091491 | **Vmn2r97** | EG627367\|Vn2r97 | vomeronasal 2, receptor 97 | protein-coding |
| chr17_18968352_18968731 | chr17 | 18968353 | 18968731 | + | -0.630627 | Intergenic | IAPLTR3-int\|LTR\|ERVK | 54220 | NM_001104549 | 627367 | Mm.384131 | NM_001104549 | ENSMUSG00000091491 | **Vmn2r97** | EG627367\|Vn2r97 | vomeronasal 2, receptor 97 | protein-coding |
| chr17_18968342_18969487 | chr17 | 18968343 | 18969487 | + | -0.649665 | Intergenic | IAPLTR3-int\|LTR\|ERVK | 54593 | NM_001104549 | 627367 | Mm.384131 | NM_001104549 | ENSMUSG00000091491 | **Vmn2r97** | EG627367\|Vn2r97 | vomeronasal 2, receptor 97 | protein-coding |
| chr17_18947091_18947814 | chr17 | 18947092 | 18947814 | + | -0.748487 | exon (NM_001104549, exon 6 of 6) | exon (NM_001104549, exon 6 of 6) | 33131 | NM_001104549 | 627367 | Mm.384131 | NM_001104549 | ENSMUSG00000091491 | **Vmn2r97** | EG627367\|Vn2r97 | vomeronasal 2, receptor 97 | protein-coding |
| chr17_18947080_18947537 | chr17 | 18947081 | 18947537 | + | -0.831659 | exon (NM_001104549, exon 6 of 6) | exon (NM_001104549, exon 6 of 6) | 32987 | NM_001104549 | 627367 | Mm.384131 | NM_001104549 | ENSMUSG00000091491 | **Vmn2r97** | EG627367\|Vn2r97 | vomeronasal 2, receptor 97 | protein-coding |
| chr17_19430073_19430606 | chr17 | 19430074 | 19430606 | + | -0.820386 | Intergenic | L1_Mus3\|LINE\|L1 | 68205 | NM_001104551 | 665376 | Mm.469859 | NM_001104551 | ENSMUSG00000090304 | **Vmn2r99** | EG665376 | vomeronasal 2, receptor 99 | protein-coding |
| chr17_19430156_19430464 | chr17 | 19430157 | 19430464 | + | -0.929838 | Intergenic | L1_Mus3\|LINE\|L1 | 68175 | NM_001104551 | 665376 | Mm.469859 | NM_001104551 | ENSMUSG00000090304 | **Vmn2r99** | EG665376 | vomeronasal 2, receptor 99 | protein-coding |
| chr17_19303736_19304329 | chr17 | 19303737 | 19304329 | + | -1.1844 | Intergenic | Intergenic | -58102 | NM_001104551 | 665376 | Mm.469859 | NM_001104551 | ENSMUSG00000090304 | **Vmn2r99** | EG665376 | vomeronasal 2, receptor 99 | protein-coding |
| chr17_19303673_19304138 | chr17 | 19303674 | 19304138 | + | -1.23858 | Intergenic | Intergenic | -58229 | NM_001104551 | 665376 | Mm.469859 | NM_001104551 | ENSMUSG00000090304 | **Vmn2r99** | EG665376 | vomeronasal 2, receptor 99 | protein-coding |
| chr6_57823071_57823385 | chr6 | 57823072 | 57823385 | + | -0.699451 | intron (NM_146168, intron 1 of 4) | intron (NM_146168, intron 1 of 4) | 1897 | NM_146168 | 232023 | Mm.24642 | NM_146168 | ENSMUSG00000037788 | **Vopp1** | AW146242\|Ecop | vesicular, overexpressed in cancer, prosurvival protein 1 | protein-coding |
| chr6_57822871_57823337 | chr6 | 57822872 | 57823337 | + | -0.748779 | intron (NM_146168, intron 1 of 4) | intron (NM_146168, intron 1 of 4) | 2021 | NM_146168 | 232023 | Mm.24642 | NM_146168 | ENSMUSG00000037788 | **Vopp1** | AW146242\|Ecop | vesicular, overexpressed in cancer, prosurvival protein 1 | protein-coding |
| chr4_145099667_145100307 | chr4 | 145099668 | 145100307 | + | -0.849273 | exon (NM_001276502, exon 40 of 69) | exon (NM_001276502, exon 40 of 69) | 90679 | NM_001276502 | 230895 | Mm.240310 | NM_001033190 | ENSMUSG00000020220 | **Vps13d** | - | vacuolar protein sorting 13 D (yeast) | protein-coding |
| chr9_27023675_27024210 | chr9 | 27023676 | 27024210 | + | -0.828399 | intron (NM_178027, intron 1 of 5) | intron (NM_178027, intron 1 of 5) | 6151 | NM_178027 | 69091 | Mm.218717 | NM_178027 | ENSMUSG00000031988 | **Vps26b** | 1810012I05Rik\|2310075A12Rik\|AI848392 | vacuolar protein sorting 26 homolog B (yeast) | protein-coding |
| chr5_124035673_124036192 | chr5 | 124035674 | 124036192 | + | -0.643937 | Intergenic | B1F1\|SINE\|Alu | -3673 | NM_177876 | 330192 | Mm.4079 | NM_177876 | ENSMUSG00000066278 | **Vps37b** | 2300007F24Rik\|AI415429\|BC026744 | vacuolar protein sorting 37B (yeast) | protein-coding |
| chr5_124019329_124019704 | chr5 | 124019330 | 124019704 | + | -0.74151 | intron (NM_177876, intron 1 of 3) | intron (NM_177876, intron 1 of 3) | 12743 | NM_177876 | 330192 | Mm.4079 | NM_177876 | ENSMUSG00000066278 | **Vps37b** | 2300007F24Rik\|AI415429\|BC026744 | vacuolar protein sorting 37B (yeast) | protein-coding |
| chr5_124032034_124032501 | chr5 | 124032035 | 124032501 | + | -0.76526 | promoter-TSS (NM_177876) | promoter-TSS (NM_177876) | -8 | NM_177876 | 330192 | Mm.4079 | NM_177876 | ENSMUSG00000066278 | **Vps37b** | 2300007F24Rik\|AI415429\|BC026744 | vacuolar protein sorting 37B (yeast) | protein-coding |
| chr5_124034287_124034949 | chr5 | 124034288 | 124034949 | + | -0.790461 | Intergenic | Intergenic | -2358 | NM_177876 | 330192 | Mm.4079 | NM_177876 | ENSMUSG00000066278 | **Vps37b** | 2300007F24Rik\|AI415429\|BC026744 | vacuolar protein sorting 37B (yeast) | protein-coding |
| chr5_124028674_124028944 | chr5 | 124028675 | 124028944 | + | -0.902357 | intron (NM_177876, intron 1 of 3) | intron (NM_177876, intron 1 of 3) | 3451 | NM_177876 | 330192 | Mm.4079 | NM_177876 | ENSMUSG00000066278 | **Vps37b** | 2300007F24Rik\|AI415429\|BC026744 | vacuolar protein sorting 37B (yeast) | protein-coding |
| chr3_96040263_96041403 | chr3 | 96040264 | 96041403 | + | -0.630639 | intron (NM_013841, intron 10 of 14) | intron (NM_013841, intron 10 of 14) | 17622 | NM_013841 | 22365 | Mm.263185 | NM_013841 | ENSMUSG00000015747 | **Vps45** | AI462172\|AW554165\|mVps45 | vacuolar protein sorting 45 (yeast) | protein-coding |
| chr11_76133646_76134059 | chr11 | 76133647 | 76134059 | + | -0.705237 | intron (NM_026664, intron 7 of 21) | intron (NM_026664, intron 7 of 21) | 45777 | NM_026664 | 68299 | Mm.46397 | NM_026664 | ENSMUSG00000017288 | **Vps53** | 2010002A08Rik\|2310040I21Rik\|3100002B05Rik\|Hccs1 | vacuolar protein sorting 53 (yeast) | protein-coding |
| chr11_21166553_21167232 | chr11 | 21166554 | 21167232 | + | -0.801539 | Intergenic | MTD\|LTR\|ERVL-MaLR | -72139 | NM_139061 | 245944 | Mm.170103 | NM_139061 | ENSMUSG00000020128 | **Vps54** | Hcc8\|Vps54l\|mSLP8\|wr | vacuolar protein sorting 54 (yeast) | protein-coding |
| chr7_44772308_44773133 | chr7 | 44772309 | 44773133 | + | -0.808067 | intron (NM_133945, intron 13 of 14) | B3\|SINE\|B2 | 24092 | NM_133945 | 101568 | Mm.28816 | NM_133945 | ENSMUSG00000002205 | **Vrk3** | AI428238 | vaccinia related kinase 3 | protein-coding |

| | | | | | | | | | | | | | | | | | |
|---|---|---|---|---|---|---|---|---|---|---|---|---|---|---|---|---|---|
| chr7_44772571_44773156 | chr7 | 44772572 | 44773156 | + | -0.914377 | intron (NM_133945, intron 13 of 14) | intron (NM_133945, intron 13 of 14) | 24235 | NM_133945 | 101568 | Mm.28816 | NM_133945 | ENSMUSG00000002205 | **Vrk3** | AI428238 | vaccinia related kinase 3 | protein-coding |
| chr3_100811843_100812184 | chr3 | 100811844 | 100812184 | + | -0.7782 | Intergenic | B3A\|SINE\|B2 | -13445 | NM_178594 | 242122 | Mm.137467 | NM_178594 | | **Vtcn1** | B7h4\|B7s1\|B7x\|BC032925 | V-set domain containing T cell activation inhibitor 1 | protein-coding |
| chr19_55408442_55408949 | chr19 | 55408443 | 55408949 | + | -1.09884 | intron (NM_016862, intron 5 of 7) | intron (NM_016862, intron 5 of 7) | 92639 | NM_001293686 | 53611 | Mm.258637 | NM_016862 | ENSMUSG00000024983 | **Vti1a** | 1110014F16Rik\|1110018K19Rik\|4921537J05Rik\|MVti1\|MVti1a\|Vti1\|Vti1-rp2 | vesicle transport through interaction with t-SNAREs 1A | protein-coding |
| chr14_78908160_78908786 | chr14 | 78908161 | 78908786 | + | -0.794549 | intron (NM_027906, intron 3 of 44) | intron (NM_027906, intron 3 of 44) | 59295 | NM_173758 | 219189 | Mm.159651 | NM_027906 | ENSMUSG00000058997 | **Vwa8** | 1300010F03Rik\|4932416F07Rik\|AI957255\|Kiaa0564\|mKIAA0564 | von Willebrand factor A domain containing 8 | protein-coding |
| chr4_133073091_133073807 | chr4 | 133073092 | 133073807 | + | -0.725374 | intron (NM_146155, intron 6 of 6) | intron (NM_146155, intron 6 of 6) | -57184 | NM_153423 | 242687 | Mm.23566 | NM_153423 | ENSMUSG00000028868 | **Wasf2** | AW742646\|D4Ertd13e\|WAVE2 | WAS protein family, member 2 | protein-coding |
| chr4_133073256_133073528 | chr4 | 133073257 | 133073528 | + | -0.72754 | intron (NM_146155, intron 6 of 6) | intron (NM_146155, intron 6 of 6) | -57241 | NM_153423 | 242687 | Mm.23566 | NM_153423 | ENSMUSG00000028868 | **Wasf2** | AW742646\|D4Ertd13e\|WAVE2 | WAS protein family, member 2 | protein-coding |
| chr4_133085837_133086373 | chr4 | 133085838 | 133086373 | + | -0.756235 | Intergenic | Intergenic | -44528 | NM_153423 | 242687 | Mm.23566 | NM_153423 | ENSMUSG00000028868 | **Wasf2** | AW742646\|D4Ertd13e\|WAVE2 | WAS protein family, member 2 | protein-coding |
| chr4_133158723_133159449 | chr4 | 133158724 | 133159449 | + | -0.816984 | intron (NM_153423, intron 1 of 8) | intron (NM_153423, intron 1 of 8) | 28453 | NM_153423 | 242687 | Mm.23566 | NM_153423 | ENSMUSG00000028868 | **Wasf2** | AW742646\|D4Ertd13e\|WAVE2 | WAS protein family, member 2 | protein-coding |
| chr4_133086022_133086300 | chr4 | 133086023 | 133086300 | + | -0.816991 | Intergenic | Intergenic | -44472 | NM_153423 | 242687 | Mm.23566 | NM_153423 | ENSMUSG00000028868 | **Wasf2** | AW742646\|D4Ertd13e\|WAVE2 | WAS protein family, member 2 | protein-coding |
| chr4_133162175_133162677 | chr4 | 133162176 | 133162677 | + | -0.860655 | intron (NM_153423, intron 1 of 8) | intron (NM_153423, intron 1 of 8) | 31793 | NM_153423 | 242687 | Mm.23566 | NM_153423 | ENSMUSG00000028868 | **Wasf2** | AW742646\|D4Ertd13e\|WAVE2 | WAS protein family, member 2 | protein-coding |
| chr19_46641595_46642036 | chr19 | 46641596 | 46642036 | + | -0.715555 | intron (NM_001177813, intron 1 of 3) | intron (NM_001177813, intron 1 of 3) | 18415 | NM_001177813 | 226178 | Mm.329895 | NM_146099 | ENSMUSG00000047731 | **Wbp1l** | D19Wsu162e\|Opal1 | WW domain binding protein 1 like | protein-coding |
| chr5_134166585_134167082 | chr5 | 134166586 | 134167082 | + | -0.79541 | intron (NM_033572, intron 4 of 10) | intron (NM_033572, intron 4 of 10) | 9933 | NM_033572 | 94254 | Mm.29508 | NM_033572 | ENSMUSG00000061979 | **Wbscr16** | 5730496C04Rik\|AU019812 | Williams-Beuren syndrome chromosome region 16 homolog (human) | protein-coding |
| chr5_134932946_134933716 | chr5 | 134932947 | 134933716 | + | -0.632224 | intron (NM_024479, intron 2 of 5) | intron (NM_024479, intron 2 of 5) | 958 | NM_024479 | 79565 | Mm.34235 | NM_024479 | ENSMUSG00000040557 | **Wbscr27** | AW492986\|BC002286 | Williams Beuren syndrome chromosome region 27 (human) | protein-coding |
| chr5_134939003_134939432 | chr5 | 134939004 | 134939432 | + | -0.795112 | intron (NM_024479, intron 5 of 5) | B1F\|SINE\|Alu | 6845 | NM_024479 | 79565 | Mm.34235 | NM_024479 | ENSMUSG00000040557 | **Wbscr27** | AW492986\|BC002286 | Williams Beuren syndrome chromosome region 27 (human) | protein-coding |
| chr5_134933322_134933683 | chr5 | 134933323 | 134933683 | + | -0.820781 | exon (NM_024479, exon 3 of 6) | exon (NM_024479, exon 3 of 6) | 1130 | NM_024479 | 79565 | Mm.34235 | NM_024479 | ENSMUSG00000040557 | **Wbscr27** | AW492986\|BC002286 | Williams Beuren syndrome chromosome region 27 (human) | protein-coding |
| chr1_60070654_60071199 | chr1 | 60070655 | 60071199 | + | -0.870919 | Intergenic | Intergenic | 26980 | NM_021312 | 57750 | Mm.281079 | NM_021312 | ENSMUSG00000026019 | **Wdr12** | 4933402C23Rik\|Ytm1\|Ytm1p | WD repeat domain 12 | protein-coding |
| chr1_60074946_60075413 | chr1 | 60074947 | 60075413 | + | -0.892192 | Intergenic | RLTR8\|LTR\|ERVK | 22727 | NM_021312 | 57750 | Mm.281079 | NM_021312 | ENSMUSG00000026019 | **Wdr12** | 4933402C23Rik\|Ytm1\|Ytm1p | WD repeat domain 12 | protein-coding |
| chr11_67949380_67949825 | chr11 | 67949381 | 67949825 | + | -0.673619 | exon (NM_027963, exon 4 of 14) | exon (NM_027963, exon 4 of 14) | 16039 | NM_027963 | 71860 | Mm.441823 | NM_027963 | ENSMUSG00000020904 | **Wdr16** | 1700019F09Rik\|4933417B11Rik\|WDRPUH | WD repeat domain 16 | protein-coding |
| chr12_110758909_110759418 | chr12 | 110758910 | 110759418 | + | -0.697949 | intron (NM_027149, intron 1 of 2) | MER21B\|LTR\|ERVL | 21215 | NM_027149 | 69641 | Mm.491226 | NM_027149 | ENSMUSG00000037957 | **Wdr20** | 2310040A13Rik\|AI549910\|Wdr20a | WD repeat domain 20 | protein-coding |
| chr12_110750744_110751503 | chr12 | 110750745 | 110751503 | + | -0.711967 | intron (NM_027149, intron 1 of 2) | B1_Mus1\|SINE\|Alu | 13175 | NM_027149 | 69641 | Mm.491226 | NM_027149 | ENSMUSG00000037957 | **Wdr20** | 2310040A13Rik\|AI549910\|Wdr20a | WD repeat domain 20 | protein-coding |
| chr1_181227800_181228230 | chr1 | 181227801 | 181228230 | + | -0.976677 | Intergenic | Intergenic | -16037 | NM_145514 | 226757 | Mm.289082 | NM_145514 | ENSMUSG00000038733 | **Wdr26** | 1600024A01Rik\|AA693241\|AI447817\|AU044014\|C77982\|Gid7 | WD repeat domain 26 | protein-coding |
| chr17_14907097_14907865 | chr17 | 14907098 | 14907865 | + | -0.74293 | intron (NM_175173, intron 17 of 24) | intron (NM_175173, intron 17 of 24) | 35643 | NM_175173 | 71682 | Mm.122698 | NM_175173 | ENSMUSG00000046991 | **Wdr27** | 0610012K18Rik | WD repeat domain 27 | protein-coding |
| chr17_14907090_14907596 | chr17 | 14907091 | 14907596 | + | -0.815962 | intron (NM_175173, intron 17 of 24) | intron (NM_175173, intron 17 of 24) | 35781 | NM_175173 | 71682 | Mm.122698 | NM_175173 | ENSMUSG00000046991 | **Wdr27** | 0610012K18Rik | WD repeat domain 27 | protein-coding |
| chr18_31807162_31807512 | chr18 | 31807163 | 31807512 | + | -0.814456 | intron (NM_001170966, intron 1 of 5) | intron (NM_001170966, intron 1 of 5) | 3280 | NM_001170966 | 74320 | Mm.277705 | NM_028866 | ENSMUSG00000024400 | **Wdr33** | 1110001N06Rik\|2310011G05Rik\|2810021O11Rik\|8430413N20Rik\|WDC146 | WD repeat domain 33 | protein-coding |
| chr13_8870116_8870415 | chr13 | 8870117 | 8870415 | + | -0.636317 | intron (NM_001039388, intron 1 of 13) | intron (NM_001039388, intron 1 of 13) | 778 | NM_001039389 | 207615 | Mm.284654 | NM_172445 | ENSMUSG00000021147 | **Wdr37** | - | WD repeat domain 37 | protein-coding |
| chr13_8742667_8743289 | chr13 | 8742668 | 8743289 | + | -0.760383 | intron (NM_001289530, intron 8 of 9) | intron (NM_001289530, intron 8 of 9) | 128066 | NM_001039389 | 207615 | Mm.284654 | NM_172445 | ENSMUSG00000021147 | **Wdr37** | - | WD repeat domain 37 | protein-coding |
| chr13_8831438_8831911 | chr13 | 8831439 | 8831911 | + | -0.764975 | intron (NM_001039388, intron 11 of 13) | intron (NM_001039388, intron 11 of 13) | 39369 | NM_001039389 | 207615 | Mm.284654 | NM_172445 | ENSMUSG00000021147 | **Wdr37** | - | WD repeat domain 37 | protein-coding |
| chr13_8742673_8743132 | chr13 | 8742674 | 8743132 | + | -0.77396 | intron (NM_001289530, intron 8 of 9) | intron (NM_001289530, intron 8 of 9) | 128141 | NM_001039389 | 207615 | Mm.284654 | NM_172445 | ENSMUSG00000021147 | **Wdr37** | - | WD repeat domain 37 | protein-coding |
| chr13_8831403_8832172 | chr13 | 8831404 | 8832172 | + | -0.809837 | intron (NM_001039388, intron 11 of 13) | intron (NM_001039388, intron 11 of 13) | 39256 | NM_001039389 | 207615 | Mm.284654 | NM_172445 | ENSMUSG00000021147 | **Wdr37** | - | WD repeat domain 37 | protein-coding |
| chr13_8783685_8784724 | chr13 | 8783686 | 8784724 | + | -0.892543 | Intergenic | Intergenic | 86839 | NM_001039389 | 207615 | Mm.284654 | NM_172445 | ENSMUSG00000021147 | **Wdr37** | - | WD repeat domain 37 | protein-coding |
| chr13_8783567_8784192 | chr13 | 8783568 | 8784192 | + | -1.14365 | Intergenic | MTEa\|LTR\|ERVL-MaLR | 87164 | NM_001039389 | 207615 | Mm.284654 | NM_172445 | ENSMUSG00000021147 | **Wdr37** | - | WD repeat domain 37 | protein-coding |
| chr2_38987778_38988627 | chr2 | 38987779 | 38988627 | + | -1.00921 | Intergenic | Intergenic | -10106 | NM_029687 | 76646 | Mm.358777 | NM_029687 | ENSMUSG00000035295 | **Wdr38** | 1700123D08Rik | WD repeat domain 38 | protein-coding |
| chr8_111505794_111506096 | chr8 | 111505795 | 111506096 | + | -0.833565 | intron (NM_001170743, intron 1 of 26) | intron (NM_001170743, intron 1 of 26) | 16162 | NM_001170742 | 319481 | Mm.247294 | NM_176923 | ENSMUSG00000031959 | **Wdr59** | 5430401O09Rik\|Cdw12 | WD repeat domain 59 | protein-coding |
| chr8_111505683_111506168 | chr8 | 111505684 | 111506168 | + | -0.86502 | exon (NM_001170742, exon 2 of 26) | exon (NM_001170742, exon 2 of 26) | 16181 | NM_001170742 | 319481 | Mm.247294 | NM_176923 | ENSMUSG00000031959 | **Wdr59** | 5430401O09Rik\|Cdw12 | WD repeat domain 59 | protein-coding |
| chr3_146062909_146063657 | chr3 | 146062910 | 146063657 | + | -0.995332 | exon (NM_172864, exon 15 of 23) | exon (NM_172864, exon 15 of 23) | 44753 | NM_172864 | 242253 | Mm.235040 | NM_172864 | ENSMUSG00000043020 | **Wdr63** | 4931433A13Rik | WD repeat domain 63 | protein-coding |
| chr15_7978650_7978972 | chr15 | 7978651 | 7978972 | + | -0.701145 | intron (NM_001081402, intron 9 of 17) | (TTGG)n\|Simple_repeat\|Simple_repeat | 120398 | NM_001081402 | 545085 | Mm.321312 | NM_198413 | ENSMUSG00000039828 | **Wdr70** | 4833422F06Rik | WD repeat domain 70 | protein-coding |
| chr2_121511447_121511786 | chr2 | 121511448 | 121511786 | + | -0.993214 | intron (NM_001290986, intron 2 of 12) | RMER6A\|LTR\|ERVK | 4894 | NM_001290986 | 241627 | Mm.293284 | NM_030234 | ENSMUSG00000027242 | **Wdr76** | 5830411K18Rik\|6030457D09\|AA517836 | WD repeat domain 76 | protein-coding |
| chr4_133327935_133328511 | chr4 | 133327936 | 133328511 | + | -0.755147 | intron (NM_199306, intron 1 of 15) | RSINE1\|SINE\|B4 | 11092 | NM_199306 | 230796 | Mm.226576 | NM_199306 | ENSMUSG00000037622 | **Wdtc1** | Gm695\|adipose\|adp | WD and tetratricopeptide repeats 1 | protein-coding |
| chr4_133328125_133328490 | chr4 | 133328126 | 133328490 | + | -0.798309 | intron (NM_199306, intron 1 of 15) | RSINE1\|SINE\|B4 | 11007 | NM_199306 | 230796 | Mm.226576 | NM_199306 | ENSMUSG00000037622 | **Wdtc1** | Gm695\|adipose\|adp | WD and tetratricopeptide repeats 1 | protein-coding |
| chr4_133330388_133330664 | chr4 | 133330389 | 133330664 | + | -0.904428 | intron (NM_199306, intron 1 of 15) | intron (NM_199306, intron 1 of 15) | 8789 | NM_199306 | 230796 | Mm.226576 | NM_199306 | ENSMUSG00000037622 | **Wdtc1** | Gm695\|adipose\|adp | WD and tetratricopeptide repeats 1 | protein-coding |
| chr4_133330405_133330773 | chr4 | 133330406 | 133330773 | + | -0.918278 | intron (NM_199306, intron 1 of 15) | intron (NM_199306, intron 1 of 15) | 8726 | NM_199306 | 230796 | Mm.226576 | NM_199306 | ENSMUSG00000037622 | **Wdtc1** | Gm695\|adipose\|adp | WD and tetratricopeptide repeats 1 | protein-coding |
| chr15_58164696_58165033 | chr15 | 58164697 | 58165033 | + | -0.802441 | Intergenic | Intergenic | 23429 | NM_029734 | 76773 | Mm.26771 | NM_029734 | ENSMUSG00000022359 | **Wdyhv1** | 2410187C16Rik\|AU014961\|AW550036\|Ntaq1 | WDYHV motif containing 1 | protein-coding |
| chr15_58165135_58165664 | chr15 | 58165136 | 58165664 | + | -0.925838 | Intergenic | Intergenic | 23964 | NM_029734 | 76773 | Mm.26771 | NM_029734 | ENSMUSG00000022359 | **Wdyhv1** | 2410187C16Rik\|AU014961\|AW550036\|Ntaq1 | WDYHV motif containing 1 | protein-coding |
| chr15_58175063_58175391 | chr15 | 58175064 | 58175391 | + | -1.30425 | TTS (NM_026346) | TTS (NM_026346) | 33791 | NM_029734 | 76773 | Mm.26771 | NM_029734 | ENSMUSG00000022359 | **Wdyhv1** | 2410187C16Rik\|AU014961\|AW550036\|Ntaq1 | WDYHV motif containing 1 | protein-coding |
| chr15_58174916_58175556 | chr15 | 58174917 | 58175556 | + | -1.3063 | TTS (NM_026346) | TTS (NM_026346) | 33800 | NM_029734 | 76773 | Mm.26771 | NM_029734 | ENSMUSG00000022359 | **Wdyhv1** | 2410187C16Rik\|AU014961\|AW550036\|Ntaq1 | WDYHV motif containing 1 | protein-coding |
| chr2_164534318_164534744 | chr2 | 164534319 | 164534744 | + | -1.10201 | Intergenic | Intergenic | -28185 | NM_026323 | 67701 | Mm.27289 | NM_026323 | ENSMUSG00000017723 | **Wfdc2** | 1600023A02Rik\|HE4\|WAP5 | WAP four-disulfide core domain 2 | protein-coding |
| chr8_25617727_25618399 | chr8 | 25617728 | 25618399 | + | -0.759857 | intron (NM_001001735, intron 1 of 9) | intron (NM_001001735, intron 1 of 9) | 15757 | NM_001001735 | 234135 | Mm.217337 | NM_001001735 | ENSMUSG00000054823 | **Whsc1l1** | 6720429E03\|A530023P05\|AI528490\|NSD3\|WHISTLE | Wolf-Hirschhorn syndrome candidate 1-like 1 (human) | protein-coding |
| chr10_128746068_128746671 | chr10 | 128746069 | 128746671 | + | -0.645444 | Intergenic | Intergenic | -1509 | NM_030100 | 78428 | Mm.27358 | NM_030100 | ENSMUSG00000064030 | **Wibg** | A030010B05Rik\|Pym | within bgcn homolog (Drosophila) | protein-coding |

| | | | | | | | | | | | | | | | | | |
|---|---|---|---|---|---|---|---|---|---|---|---|---|---|---|---|---|---|
| chr11_98878587_98878824 | chr11 | 98878588 | 98878824 | + | -0.750319 | intron (NM_197940, intron 1 of 7) | intron (NM_197940, intron 1 of 7) | 15108 | NM_197940 | 68524 | Mm.28661 | NM_197940 | ENSMUSG00000038013 | **Wipf2** | 1110014J05Rik\|5730509C05Rik\|AA407487\|Gm1176\|Wich\|Wire | WAS/WASL interacting protein family, member 2 | protein-coding |
| chr11_109591567_109591887 | chr11 | 109591568 | 109591887 | + | -0.921426 | intron (NM_145940, intron 4 of 12) | intron (NM_145940, intron 4 of 12) | 19662 | NM_145940 | 52639 | Mm.35817 | NM_145940 | ENSMUSG00000041895 | **Wipi1** | 4930533H01Rik\|AW411817\|D11Ertd498e | WD repeat domain, phosphoinositide interacting 1 | protein-coding |
| chr11_109591389_109591947 | chr11 | 109591390 | 109591947 | + | -0.944903 | intron (NM_145940, intron 4 of 12) | intron (NM_145940, intron 4 of 12) | 19721 | NM_145940 | 52639 | Mm.35817 | NM_145940 | ENSMUSG00000041895 | **Wipi1** | 4930533H01Rik\|AW411817\|D11Ertd498e | WD repeat domain, phosphoinositide interacting 1 | protein-coding |
| chr10_39182872_39183204 | chr10 | 39182873 | 39183204 | + | -0.685013 | Intergenic | Intergenic | -19244 | NM_001127376 | 327743 | Mm.208111 | NM_001127376 | ENSMUSG00000062074 | **Wisp3** | CCN6\|ENSMUSG00000062074\|Gm735\|WISP-3 | WNT1 inducible signaling pathway protein 3 | protein-coding |
| chr10_39192642_39193192 | chr10 | 39192643 | 39193192 | + | -1.07142 | Intergenic | Intergenic | -29123 | NM_001127376 | 327743 | Mm.208111 | NM_001127376 | ENSMUSG00000062074 | **Wisp3** | CCN6\|ENSMUSG00000062074\|Gm735\|WISP-3 | WNT1 inducible signaling pathway protein 3 | protein-coding |
| chr6_120055729_120056512 | chr6 | 120055730 | 120056512 | + | -0.95078 | Intergenic | Intergenic | -17466 | NM_001185021 | 232341 | Mm.333349 | NM_198703 | ENSMUSG00000045962 | **Wnk1** | 6430573H23Rik\|EG406236\|Hsn2\|Prkwnk1\|mKIAA0344 | WNK lysine deficient protein kinase 1 | protein-coding |
| chr6_120065794_120066097 | chr6 | 120065795 | 120066097 | + | -1.24163 | Intergenic | Intergenic | -27291 | NM_001185021 | 232341 | Mm.333349 | NM_198703 | ENSMUSG00000045962 | **Wnk1** | 6430573H23Rik\|EG406236\|Hsn2\|Prkwnk1\|mKIAA0344 | WNK lysine deficient protein kinase 1 | protein-coding |
| chr13_49090154_49090530 | chr13 | 49090155 | 49090530 | + | -0.743624 | intron (NM_001290311, intron 7 of 28) | intron (NM_001290311, intron 7 of 28) | 57986 | NM_029361 | 75607 | Mm.342723 | NM_029361 | ENSMUSG00000037989 | **Wnk2** | 1810073P09Rik\|AW122246\|ESTM15\|X83337\|mKIAA1760 | WNK lysine deficient protein kinase 2 | protein-coding |
| chr13_49090172_49090675 | chr13 | 49090173 | 49090675 | + | -0.747947 | intron (NM_001290311, intron 7 of 28) | intron (NM_001290311, intron 7 of 28) | 57904 | NM_029361 | 75607 | Mm.342723 | NM_029361 | ENSMUSG00000037989 | **Wnk2** | 1810073P09Rik\|AW122246\|ESTM15\|X83337\|mKIAA1760 | WNK lysine deficient protein kinase 2 | protein-coding |
| chr6_119630526_119631092 | chr6 | 119630527 | 119631092 | + | -0.647547 | intron (NR_104484, intron 16 of 20) | intron (NR_104484, intron 16 of 20) | -86462 | NM_009525 | 22419 | Mm.321818 | NM_009525 | ENSMUSG00000030170 | **Wnt5b** | AW545702\|Wnt-5b | wingless-type MMTV integration site family, member 5B | protein-coding |
| chr8_114973000_114973511 | chr8 | 114973001 | 114973511 | + | -0.642857 | intron (NM_019573, intron 8 of 8) | intron (NM_019573, intron 8 of 8) | 533604 | NM_019573 | 80707 | Mm.440420 | NM_019573 | ENSMUSG00000004637 | **Wwox** | 5330426P09Rik\|9030416C10Rik\|WOX1 | WW domain-containing oxidoreductase | protein-coding |
| chr8_114827891_114828148 | chr8 | 114827892 | 114828148 | + | -0.702546 | intron (NM_019573, intron 8 of 8) | intron (NM_019573, intron 8 of 8) | 388368 | NM_019573 | 80707 | Mm.440420 | NM_019573 | ENSMUSG00000004637 | **Wwox** | 5330426P09Rik\|9030416C10Rik\|WOX1 | WW domain-containing oxidoreductase | protein-coding |
| chr8_114954524_114954804 | chr8 | 114954525 | 114954804 | + | -0.716405 | intron (NM_019573, intron 8 of 8) | intron (NM_019573, intron 8 of 8) | 515012 | NM_019573 | 80707 | Mm.440420 | NM_019573 | ENSMUSG00000004637 | **Wwox** | 5330426P09Rik\|9030416C10Rik\|WOX1 | WW domain-containing oxidoreductase | protein-coding |
| chr8_114816281_114817390 | chr8 | 114816282 | 114817390 | + | -0.764748 | intron (NM_019573, intron 8 of 8) | intron (NM_019573, intron 8 of 8) | 377184 | NM_019573 | 80707 | Mm.440420 | NM_019573 | ENSMUSG00000004637 | **Wwox** | 5330426P09Rik\|9030416C10Rik\|WOX1 | WW domain-containing oxidoreductase | protein-coding |
| chr8_114954540_114954979 | chr8 | 114954541 | 114954979 | + | -0.794448 | intron (NM_019573, intron 8 of 8) | intron (NM_019573, intron 8 of 8) | 515108 | NM_019573 | 80707 | Mm.440420 | NM_019573 | ENSMUSG00000004637 | **Wwox** | 5330426P09Rik\|9030416C10Rik\|WOX1 | WW domain-containing oxidoreductase | protein-coding |
| chr8_114888394_114888762 | chr8 | 114888395 | 114888762 | + | -0.837027 | intron (NM_019573, intron 8 of 8) | B3A\|SINE\|B2 | 448926 | NM_019573 | 80707 | Mm.440420 | NM_019573 | ENSMUSG00000004637 | **Wwox** | 5330426P09Rik\|9030416C10Rik\|WOX1 | WW domain-containing oxidoreductase | protein-coding |
| chr8_114816306_114816771 | chr8 | 114816307 | 114816771 | + | -0.879333 | intron (NM_019573, intron 8 of 8) | MLT1E2\|LTR\|ERVL-MaLR | 376887 | NM_019573 | 80707 | Mm.440420 | NM_019573 | ENSMUSG00000004637 | **Wwox** | 5330426P09Rik\|9030416C10Rik\|WOX1 | WW domain-containing oxidoreductase | protein-coding |
| chr8_114803112_114803622 | chr8 | 114803113 | 114803622 | + | -1.07056 | intron (NM_019573, intron 8 of 8) | intron (NM_019573, intron 8 of 8) | 363715 | NM_019573 | 80707 | Mm.440420 | NM_019573 | ENSMUSG00000004637 | **Wwox** | 5330426P09Rik\|9030416C10Rik\|WOX1 | WW domain-containing oxidoreductase | protein-coding |
| chr8_114880737_114881269 | chr8 | 114880738 | 114881269 | + | -1.17112 | intron (NM_019573, intron 8 of 8) | intron (NM_019573, intron 8 of 8) | 441351 | NM_019573 | 80707 | Mm.440420 | NM_019573 | ENSMUSG00000004637 | **Wwox** | 5330426P09Rik\|9030416C10Rik\|WOX1 | WW domain-containing oxidoreductase | protein-coding |
| chr8_114880778_114881013 | chr8 | 114880779 | 114881013 | + | -1.1882 | intron (NM_019573, intron 8 of 8) | intron (NM_019573, intron 8 of 8) | 441244 | NM_019573 | 80707 | Mm.440420 | NM_019573 | ENSMUSG00000004637 | **Wwox** | 5330426P09Rik\|9030416C10Rik\|WOX1 | WW domain-containing oxidoreductase | protein-coding |
| chr8_114801755_114802112 | chr8 | 114801756 | 114802112 | + | -1.19738 | intron (NM_019573, intron 8 of 8) | intron (NM_019573, intron 8 of 8) | 362282 | NM_019573 | 80707 | Mm.440420 | NM_019573 | ENSMUSG00000004637 | **Wwox** | 5330426P09Rik\|9030416C10Rik\|WOX1 | WW domain-containing oxidoreductase | protein-coding |
| chr8_114875212_114875514 | chr8 | 114875213 | 114875514 | + | -1.22391 | intron (NM_019573, intron 8 of 8) | intron (NM_019573, intron 8 of 8) | 435711 | NM_019573 | 80707 | Mm.440420 | NM_019573 | ENSMUSG00000004637 | **Wwox** | 5330426P09Rik\|9030416C10Rik\|WOX1 | WW domain-containing oxidoreductase | protein-coding |
| chr8_114815181_114815784 | chr8 | 114815182 | 114815784 | + | -1.34953 | intron (NM_019573, intron 8 of 8) | intron (NM_019573, intron 8 of 8) | 375831 | NM_019573 | 80707 | Mm.440420 | NM_019573 | ENSMUSG00000004637 | **Wwox** | 5330426P09Rik\|9030416C10Rik\|WOX1 | WW domain-containing oxidoreductase | protein-coding |
| chr8_114870366_114870824 | chr8 | 114870367 | 114870824 | + | -1.42147 | intron (NM_019573, intron 8 of 8) | MLT1E1\|LTR\|ERVL-MaLR | 430943 | NM_019573 | 80707 | Mm.440420 | NM_019573 | ENSMUSG00000004637 | **Wwox** | 5330426P09Rik\|9030416C10Rik\|WOX1 | WW domain-containing oxidoreductase | protein-coding |
| chr8_114869100_114869648 | chr8 | 114869101 | 114869648 | + | -1.61418 | intron (NM_019573, intron 8 of 8) | RMER19A\|LTR\|ERVK | 429722 | NM_019573 | 80707 | Mm.440420 | NM_019573 | ENSMUSG00000004637 | **Wwox** | 5330426P09Rik\|9030416C10Rik\|WOX1 | WW domain-containing oxidoreductase | protein-coding |
| chr8_114869251_114869565 | chr8 | 114869252 | 114869565 | + | -1.65461 | intron (NM_019573, intron 8 of 8) | RMER19A\|LTR\|ERVK | 429756 | NM_019573 | 80707 | Mm.440420 | NM_019573 | ENSMUSG00000004637 | **Wwox** | 5330426P09Rik\|9030416C10Rik\|WOX1 | WW domain-containing oxidoreductase | protein-coding |
| chr17_73893262_73893776 | chr17 | 73893263 | 73893776 | + | -0.742225 | intron (NM_011723, intron 32 of 35) | intron (NM_011723, intron 32 of 35) | 56677 | NM_011723 | 22436 | Mm.11223 | NM_011723 | ENSMUSG00000024066 | **Xdh** | XO\|Xor\|Xox-1\|Xox1 | xanthine dehydrogenase | protein-coding |
| chr17_73905077_73905689 | chr17 | 73905078 | 73905689 | + | -1.05118 | intron (NM_011723, intron 24 of 35) | intron (NM_011723, intron 24 of 35) | 44813 | NM_011723 | 22436 | Mm.11223 | NM_011723 | ENSMUSG00000024066 | **Xdh** | XO\|Xor\|Xox-1\|Xox1 | xanthine dehydrogenase | protein-coding |
| chr17_73902341_73902687 | chr17 | 73902342 | 73902687 | + | -1.05684 | intron (NM_011723, intron 24 of 35) | intron (NM_011723, intron 24 of 35) | 47682 | NM_011723 | 22436 | Mm.11223 | NM_011723 | ENSMUSG00000024066 | **Xdh** | XO\|Xor\|Xox-1\|Xox1 | xanthine dehydrogenase | protein-coding |
| chr17_73902422_73902658 | chr17 | 73902423 | 73902658 | + | -1.1208 | intron (NM_011723, intron 24 of 35) | intron (NM_011723, intron 24 of 35) | 47656 | NM_011723 | 22436 | Mm.11223 | NM_011723 | ENSMUSG00000024066 | **Xdh** | XO\|Xor\|Xox-1\|Xox1 | xanthine dehydrogenase | protein-coding |
| chrX_134158171_134158486 | chrX | 134158172 | 134158486 | + | -0.731373 | intron (NM_183319, intron 1 of 2) | intron (NM_183319, intron 1 of 2) | 3599 | NM_183319 | 331524 | Mm.235055 | NM_183319 | ENSMUSG00000031258 | **Xkrx** | A630026F04 | X Kell blood group precursor related X linked | protein-coding |
| chr6_91505957_91506823 | chr6 | 91505958 | 91506823 | + | -0.660419 | intron (NM_009531, intron 4 of 15) | intron (NM_009531, intron 4 of 15) | 9498 | NM_009531 | 22591 | Mm.2806 | NM_009531 | ENSMUSG00000030094 | **Xpc** | - | xeroderma pigmentosum, complementation group C | protein-coding |
| chr6_91506000_91506541 | chr6 | 91506001 | 91506541 | + | -0.789447 | intron (NM_009531, intron 4 of 15) | intron (NM_009531, intron 4 of 15) | 9617 | NM_009531 | 22591 | Mm.2806 | NM_009531 | ENSMUSG00000030094 | **Xpc** | - | xeroderma pigmentosum, complementation group C | protein-coding |
| chr6_91505898_91506349 | chr6 | 91505899 | 91506349 | + | -0.821508 | intron (NM_009531, intron 4 of 15) | intron (NM_009531, intron 4 of 15) | 9764 | NM_009531 | 22591 | Mm.2806 | NM_009531 | ENSMUSG00000030094 | **Xpc** | - | xeroderma pigmentosum, complementation group C | protein-coding |
| chrX_48098405_48099066 | chrX | 48098406 | 48099066 | + | -0.859871 | Intergenic | Intergenic | -9989 | NM_001289729 | 170745 | Mm.129279 | NM_133213 | ENSMUSG00000037005 | **Xpnpep2** | 9030008G12Rik\|mAPP | X-prolyl aminopeptidase (aminopeptidase P) 2, membrane-bound | protein-coding |
| chrX_48096979_48097439 | chrX | 48096980 | 48097439 | + | -0.914806 | Intergenic | Intergenic | -11516 | NM_001289729 | 170745 | Mm.129279 | NM_133213 | ENSMUSG00000037005 | **Xpnpep2** | 9030008G12Rik\|mAPP | X-prolyl aminopeptidase (aminopeptidase P) 2, membrane-bound | protein-coding |
| chrX_48098458_48098833 | chrX | 48098459 | 48098833 | + | -0.922304 | Intergenic | Intergenic | -10079 | NM_001289729 | 170745 | Mm.129279 | NM_133213 | ENSMUSG00000037005 | **Xpnpep2** | 9030008G12Rik\|mAPP | X-prolyl aminopeptidase (aminopeptidase P) 2, membrane-bound | protein-coding |
| chr16_30942556_30943284 | chr16 | 30942557 | 30943284 | + | -0.763933 | intron (NR_035427, intron 1 of 2) | intron (NR_035427, intron 1 of 2) | 138512 | NM_198626 | 268880 | Mm.286693 | NM_198626 | ENSMUSG00000047434 | **Xxylt1** | AI480653 | xyloside xylosyltransferase 1 | protein-coding |
| chr7_117521286_117521810 | chr7 | 117521287 | 117521810 | + | -0.756706 | intron (NM_175645, intron 2 of 11) | intron (NM_175645, intron 2 of 11) | 140569 | NM_175645 | 233781 | Mm.461427 | NM_175645 | ENSMUSG00000030657 | **Xylt1** | 8030490L12 | xylosyltransferase 1 | protein-coding |
| chr7_117424846_117425212 | chr7 | 117424847 | 117425212 | + | -0.760954 | intron (NM_175645, intron 1 of 11) | intron (NM_175645, intron 1 of 11) | 44050 | NM_175645 | 233781 | Mm.461427 | NM_175645 | ENSMUSG00000030657 | **Xylt1** | 8030490L12 | xylosyltransferase 1 | protein-coding |
| chr7_117613733_117614314 | chr7 | 117613734 | 117614314 | + | -1.09938 | intron (NM_175645, intron 5 of 11) | intron (NM_175645, intron 5 of 11) | 233045 | NM_175645 | 233781 | Mm.461427 | NM_175645 | ENSMUSG00000030657 | **Xylt1** | 8030490L12 | xylosyltransferase 1 | protein-coding |
| chr16_20144676_20145064 | chr16 | 20144677 | 20145064 | + | -0.664091 | intron (NM_001033237, intron 1 of 29) | PB1D7\|SINE\|Alu | 3807 | NM_001145930 | 208146 | Mm.390895 | NM_001033237 | ENSMUSG00000041215 | **Yeats2** | BC042768\|mKIAA1197 | YEATS domain containing 2 | protein-coding |
| chr10_117215791_117216077 | chr10 | 117215792 | 117216077 | + | -0.692874 | intron (NM_026570, intron 6 of 6) | intron (NM_026570, intron 6 of 6) | 8573 | NM_026570 | 64050 | Mm.233529 | NM_026570 | ENSMUSG00000020171 | **Yeats4** | 4930573H17Rik\|B230215M10Rik\|GAS41\|NuBI-1 | YEATS domain containing 4 | protein-coding |
| chr5_32610075_32610795 | chr5 | 32610076 | 32610795 | + | -0.735732 | promoter-TSS (NM_009535) | promoter-TSS (NM_009535) | -736 | NM_001205133 | 22612 | Mm.4558 | NM_009535 | ENSMUSG00000014932 | **Yes1** | Yes\|p61-Yes | Yamaguchi sarcoma viral (v-yes) oncogene homolog 1 | protein-coding |
| chr11_86971716_86972141 | chr11 | 86971717 | 86972141 | + | -0.685901 | 5' UTR (NM_001005341, exon 2 of 5) | 5' UTR (NM_001005341, exon 2 of 5) | 21833 | NM_001005341 | 77864 | Mm.191894 | NM_001005341 | ENSMUSG00000018427 | **Ypel2** | 6430570G24\|E130113K08Rik | yippee-like 2 (Drosophila) | protein-coding |
| chr5_86816828_86817313 | chr5 | 86816829 | 86817313 | + | -0.772942 | exon (NM_177680, exon 4 of 17) | exon (NM_177680, exon 4 of 17) | 12581 | NM_177680 | 231386 | Mm.196325 | NM_177680 | ENSMUSG00000035851 | **Ythdc1** | A730098D12Rik\|C80342\|mKIAA1966 | YTH domain containing 1 | protein-coding |
| chr11_75756733_75757038 | chr11 | 75756734 | 75757038 | + | -0.898682 | exon (NM_009536, exon 4 of 6) | exon (NM_009536, exon 4 of 6) | 23999 | NM_009536 | 22627 | Mm.234700 | NM_009536 | ENSMUSG00000020849 | **Ywhae** | AU019196 | tyrosine 3-monooxygenase/tryptophan 5-monooxygenase activation protein, epsilon polypeptide | protein-coding |
| chr5_32950660_32950965 | chr5 | 32950661 | 32950965 | + | -0.868774 | intron (NM_001025426, intron 32 of 42) | intron (NM_001025426, intron 32 of 42) | -68003 | NM_011738 | 22629 | Mm.332314 | NM_011738 | ENSMUSG00000018965 | **Ywhah** | - | tyrosine 3-monooxygenase/tryptophan 5-monooxygenase activation protein, eta polypeptide | protein-coding |

| | | | | | | | | | | | | | | | | | |
|---|---|---|---|---|---|---|---|---|---|---|---|---|---|---|---|---|---|
| chr5_32950538_32951109 | chr5 | 32950539 | 32951109 | + | -0.923679 | intron (NM_001025426, intron 32 of 42) | intron (NM_001025426, intron 32 of 42) | -67992 | NM_011738 | 22629 | Mm.332314 | NM_011738 | ENSMUSG00000018965 | **Ywhah** | - | tyrosine 3-monooxygenase/tryptophan 5-monooxygenase activation protein, eta polypeptide | protein-coding |
| chr12_21480024_21480818 | chr12 | 21480025 | 21480818 | + | -0.646075 | Intergenic | Intergenic | -62985 | NM_011739 | 22630 | Mm.289630 | NM_011739 | ENSMUSG00000076432 | **Ywhaq** | 2700028P07Rik\|AA409740\|AU021156\|R74690 | tyrosine 3-monooxygenase/tryptophan 5-monooxygenase activation protein, theta polypeptide | protein-coding |
| chr12_21459666_21460157 | chr12 | 21459667 | 21460157 | + | -0.733958 | Intergenic | L1MC1\|LINE\|L1 | -42476 | NM_011739 | 22630 | Mm.289630 | NM_011739 | ENSMUSG00000076432 | **Ywhaq** | 2700028P07Rik\|AA409740\|AU021156\|R74690 | tyrosine 3-monooxygenase/tryptophan 5-monooxygenase activation protein, theta polypeptide | protein-coding |
| chr15_36866151_36866560 | chr15 | 36866152 | 36866560 | + | -0.800197 | Intergenic | ORR1B1\|LTR\|ERVL-MaLR | -69427 | NM_001253806 | 22631 | Mm.3360 | NM_011740 | ENSMUSG00000022285 | **Ywhaz** | 1110013I11Rik\|14-3-3zeta\|AI596267\|AL022924\|AU020854 | tyrosine 3-monooxygenase/tryptophan 5-monooxygenase activation protein, zeta polypeptide | protein-coding |
| chr12_108765613_108766306 | chr12 | 108765614 | 108766306 | + | -0.633079 | Intergenic | Intergenic | -27351 | NM_009537 | 22632 | Mm.3868 | NM_009537 | ENSMUSG00000021264 | **Yy1** | AW488674\|NF-E1 | YY1 transcription factor | protein-coding |
| chr18_84159097_84159874 | chr18 | 84159098 | 84159874 | + | -0.69559 | Intergenic | RMER13A1\|LTR\|ERVK | 71328 | NM_146090 | 225791 | Mm.147052 | NM_146090 | ENSMUSG00000049090 | **Zadh2** | C530046K17Rik\|Pthr3 | zinc binding alcohol dehydrogenase, domain containing 2 | protein-coding |
| chr18_84091489_84091982 | chr18 | 84091490 | 84091982 | + | -0.766734 | intron (NM_146090, intron 1 of 1) | intron (NM_146090, intron 1 of 1) | 3578 | NM_146090 | 225791 | Mm.147052 | NM_146090 | ENSMUSG00000049090 | **Zadh2** | C530046K17Rik\|Pthr3 | zinc binding alcohol dehydrogenase, domain containing 2 | protein-coding |
| chr1_36797422_36798006 | chr1 | 36797423 | 36798006 | + | -0.895004 | intron (NM_018872, intron 35 of 40) | ORR1F\|LTR\|ERVL-MaLR | 18730 | NM_001289612 | 22637 | Mm.8038 | NM_009539 | ENSMUSG00000026117 | **Zap70** | Srk\|ZAP-70\|mrtle\|mur | zeta-chain (TCR) associated protein kinase | protein-coding |
| chr17_69395980_69396768 | chr17 | 69395981 | 69396768 | + | -0.681541 | Intergenic | Intergenic | 12396 | NM_009547 | 22666 | Mm.29434 | NM_009547 | ENSMUSG00000049672 | **Zbtb14** | ZF5\|Zfp161\|b2b1982Clo | zinc finger and BTB domain containing 14 | protein-coding |
| chr9_48740190_48740538 | chr9 | 48740191 | 48740538 | + | -0.709961 | intron (NM_001033324, intron 3 of 6) | intron (NM_001033324, intron 3 of 6) | 95581 | NM_001033324 | 235320 | Mm.457803 | NM_001033324 | ENSMUSG00000066687 | **Zbtb16** | AI467657\|PLZF\|Zfp145\|lu | zinc finger and BTB domain containing 16 | protein-coding |
| chr1_177532717_177532983 | chr1 | 177532718 | 177532983 | + | -0.68429 | Intergenic | Intergenic | 87029 | NM_013915 | 30928 | Mm.330700 | NM_013915 | ENSMUSG00000063659 | **Zbtb18** | RP58\|Zfp238\|Znf238 | zinc finger and BTB domain containing 18 | protein-coding |
| chr12_76367386_76367638 | chr12 | 76367387 | 76367638 | + | -0.702793 | intron (NM_028356, intron 1 of 2) | intron (NM_028356, intron 1 of 2) | 1952 | NM_001172104 | 109929 | Mm.41880 | NM_028356 | ENSMUSG00000056459 | **Zbtb25** | 2810462M08Rik\|2900064P18Rik\|AI842128\|Kup\|Zfp-50\|Zfp50 | zinc finger and BTB domain containing 25 | protein-coding |
| chr12_76362241_76362863 | chr12 | 76362242 | 76362863 | + | -0.756791 | intron (NM_028356, intron 1 of 2) | intron (NM_028356, intron 1 of 2) | 6912 | NM_001172104 | 109929 | Mm.41880 | NM_028356 | ENSMUSG00000056459 | **Zbtb25** | 2810462M08Rik\|2900064P18Rik\|AI842128\|Kup\|Zfp-50\|Zfp50 | zinc finger and BTB domain containing 25 | protein-coding |
| chr9_96705626_96706247 | chr9 | 96705627 | 96706247 | + | -0.727297 | intron (NM_175537, intron 2 of 2) | intron (NM_175537, intron 2 of 2) | 25738 | NM_175537 | 245007 | Mm.485954 | NM_175537 | ENSMUSG00000040433 | **Zbtb38** | A930014K01Rik\|CIBZ | zinc finger and BTB domain containing 38 | protein-coding |
| chr9_96705999_96706262 | chr9 | 96706000 | 96706262 | + | -0.839769 | intron (NM_175537, intron 2 of 2) | intron (NM_175537, intron 2 of 2) | 25544 | NM_175537 | 245007 | Mm.485954 | NM_175537 | ENSMUSG00000040433 | **Zbtb38** | A930014K01Rik\|CIBZ | zinc finger and BTB domain containing 38 | protein-coding |
| chr9_96696211_96696876 | chr9 | 96696212 | 96696876 | + | -0.925826 | intron (NM_175537, intron 2 of 2) | intron (NM_175537, intron 2 of 2) | 35131 | NM_175537 | 245007 | Mm.485954 | NM_175537 | ENSMUSG00000040433 | **Zbtb38** | A930014K01Rik\|CIBZ | zinc finger and BTB domain containing 38 | protein-coding |
| chr9_96696012_96696960 | chr9 | 96696013 | 96696960 | + | -0.938577 | intron (NM_175537, intron 2 of 2) | intron (NM_175537, intron 2 of 2) | 35189 | NM_175537 | 245007 | Mm.485954 | NM_175537 | ENSMUSG00000040433 | **Zbtb38** | A930014K01Rik\|CIBZ | zinc finger and BTB domain containing 38 | protein-coding |
| chr10_81127332_81127929 | chr10 | 81127333 | 81127929 | + | -0.850767 | Intergenic | Intergenic | -8640 | NM_010731 | 16969 | Mm.20920 | NM_010731 | ENSMUSG00000035011 | **Zbtb7a** | 9030619K07Rik\|9130006G12Rik\|AI452336\|FBI-1\|Lrf\|Pokemon\|Zbtb7 | zinc finger and BTB domain containing 7a | protein-coding |
| chr17_26980871_26981506 | chr17 | 26980872 | 26981506 | + | -1.13729 | Intergenic | L1MB3\|LINE\|L1 | 8010 | NM_001005916 | 474156 | Mm.328890 | NM_001005916 | ENSMUSG00000079605 | **Zbtb9** | 3930402F13Rik | zinc finger and BTB domain containing 9 | protein-coding |
| chr14_75318741_75319143 | chr14 | 75318742 | 75319143 | + | -1.08954 | intron (NM_026083, intron 8 of 18) | intron (NM_026083, intron 8 of 18) | 34569 | NM_026083 | 67302 | Mm.74704 | NM_026083 | ENSMUSG00000022000 | **Zc3h13** | 2600010B19Rik\|3110050K21Rik\|4930570G11Rik\|C87618 | zinc finger CCCH type containing 13 | protein-coding |
| chr16_11157590_11158361 | chr16 | 11157591 | 11158361 | + | -0.822659 | intron (NR_027502, intron 7 of 22) | B2_Mm1a\|SINE\|B2 | 18417 | NR_027502 | 106205 | Mm.226860 | NM_145931 | ENSMUSG00000037965 | **Zc3h7a** | A430104C18Rik\|AI447294\|AW556219\|HSPC055\|Zc3h7\|Zc3hdc7 | zinc finger CCCH type containing 7 A | protein-coding |
| chr16_11157752_11158073 | chr16 | 11157753 | 11158073 | + | -0.855606 | intron (NR_027502, intron 7 of 22) | intron (NR_027502, intron 7 of 22) | 18480 | NR_027502 | 106205 | Mm.226860 | NM_145931 | ENSMUSG00000037965 | **Zc3h7a** | A430104C18Rik\|AI447294\|AW556219\|HSPC055\|Zc3h7\|Zc3hdc7 | zinc finger CCCH type containing 7 A | protein-coding |
| chr6_38360083_38360569 | chr6 | 38360084 | 38360569 | + | -0.660732 | Intergenic | MTD\|LTR\|ERVL-MaLR | -5723 | NM_028421 | 78781 | Mm.25243 | NM_028421 | ENSMUSG00000029826 | **Zc3hav1** | 1200014N16Rik\|2900058M19Rik\|9130009D18Rik\|9830115L13Rik\|ARTD13\|D6Bwg1452e\|ZAP | zinc finger CCCH type, antiviral 1 | protein-coding |
| chr6_38351998_38352228 | chr6 | 38351999 | 38352228 | + | -0.855218 | intron (NM_028864, intron 1 of 8) | intron (NM_028864, intron 1 of 8) | 2490 | NM_028864 | 78781 | Mm.25243 | NM_028421 | ENSMUSG00000029826 | **Zc3hav1** | 1200014N16Rik\|2900058M19Rik\|9130009D18Rik\|9830115L13Rik\|ARTD13\|D6Bwg1452e\|ZAP | zinc finger CCCH type, antiviral 1 | protein-coding |
| chr6_38309676_38310146 | chr6 | 38309677 | 38310146 | + | -0.853797 | TTS (NM_028421) | TTS (NM_028421) | -10652 | NM_172467 | 209032 | Mm.101389 | NM_172467 | ENSMUSG00000047749 | **Zc3hav1l** | B130055L09Rik\|E430016P22Rik | zinc finger CCCH-type, antiviral 1-like | protein-coding |
| chr6_30354767_30355036 | chr6 | 30354768 | 30355036 | + | -0.650231 | Intergenic | Intergenic | 36108 | NM_172735 | 232679 | Mm.29780 | NM_172735 | ENSMUSG00000039130 | **Zc3hc1** | 1110054L24Rik\|AU018540\|AU019789\|HSPC216\|Nipa | zinc finger, C3HC type 1 | protein-coding |
| chr4_108460484_108461009 | chr4 | 108460485 | 108461009 | + | -0.648733 | intron (NM_175472, intron 2 of 30) | CpG | 1321 | NM_175472 | 230594 | Mm.25181 | NM_175472 | ENSMUSG00000034610 | **Zcchc11** | 6030404K05Rik\|9230115F04Rik\|PAPD3\|mKIAA0191 | zinc finger, CCHC domain containing 11 | protein-coding |
| chrX_36195681_36196194 | chrX | 36195682 | 36196194 | + | -0.615074 | promoter-TSS (NM_028325) | promoter-TSS (NM_028325) | 34 | NM_028325 | 72693 | Mm.297698 | NM_028325 | ENSMUSG00000036699 | **Zcchc12** | 2810028A01Rik\|AV136720\|Sizn1 | zinc finger, CCHC domain containing 12 | protein-coding |
| chr8_121650387_121650978 | chr8 | 121650388 | 121650978 | + | -0.909967 | intron (NM_080855, intron 1 of 12) | intron (NM_080855, intron 1 of 12) | 1250 | NM_080855 | 142682 | Mm.103202 | NM_080855 | ENSMUSG00000061410 | **Zcchc14** | AA792890\|BDG29 | zinc finger, CCHC domain containing 14 | protein-coding |
| chr1_105980711_105981620 | chr1 | 105980712 | 105981620 | + | -0.896929 | Intergenic | Intergenic | -9240 | NM_001122676 | 227449 | Mm.210188 | NM_001122675 | ENSMUSG00000038866 | **Zcchc2** | 9930114B20Rik\|AW212015 | zinc finger, CCHC domain containing 2 | protein-coding |
| chr13_91827906_91828271 | chr13 | 91827907 | 91828271 | + | -0.763169 | Intergenic | ORR1B2\|LTR\|ERVL-MaLR | -20393 | NM_145453 | 69085 | Mm.52356 | NM_145453 | ENSMUSG00000021621 | **Zcchc9** | 1810019C21Rik | zinc finger, CCHC domain containing 9 | protein-coding |
| chr13_91814661_91815150 | chr13 | 91814662 | 91815150 | + | -0.854634 | Intergenic | Intergenic | -7210 | NM_145453 | 69085 | Mm.52356 | NM_145453 | ENSMUSG00000021621 | **Zcchc9** | 1810019C21Rik | zinc finger, CCHC domain containing 9 | protein-coding |
| chr13_91814454_91815140 | chr13 | 91814455 | 91815140 | + | -0.888038 | Intergenic | Intergenic | -7101 | NM_145453 | 69085 | Mm.52356 | NM_145453 | ENSMUSG00000021621 | **Zcchc9** | 1810019C21Rik | zinc finger, CCHC domain containing 9 | protein-coding |
| chr4_133625509_133626474 | chr4 | 133625510 | 133626474 | + | -0.598037 | intron (NM_001017968, intron 2 of 7) | intron (NM_001017968, intron 2 of 7) | 7437 | NM_001017968 | 503610 | Mm.331948 | NM_001017968 | | **Zdhhc18** | - | zinc finger, DHHC domain containing 18 | protein-coding |
| chr4_133625544_133626101 | chr4 | 133625545 | 133626101 | + | -0.641881 | intron (NM_001017968, intron 2 of 7) | intron (NM_001017968, intron 2 of 7) | 7606 | NM_001017968 | 503610 | Mm.331948 | NM_001017968 | | **Zdhhc18** | - | zinc finger, DHHC domain containing 18 | protein-coding |
| chr4_133620637_133621054 | chr4 | 133620638 | 133621054 | + | -0.733411 | intron (NM_001017968, intron 2 of 7) | intron (NM_001017968, intron 2 of 7) | 12583 | NM_001017968 | 503610 | Mm.331948 | NM_001017968 | | **Zdhhc18** | - | zinc finger, DHHC domain containing 18 | protein-coding |
| chr4_133629220_133629601 | chr4 | 133629221 | 133629601 | + | -1.23191 | intron (NM_001017968, intron 1 of 7) | ID_B1\|SINE\|B4 | 4018 | NM_001017968 | 503610 | Mm.331948 | NM_001017968 | | **Zdhhc18** | - | zinc finger, DHHC domain containing 18 | protein-coding |
| chr14_57908365_57908913 | chr14 | 57908366 | 57908913 | + | -0.809139 | Intergenic | ORR1D1\|LTR\|ERVL-MaLR | -18377 | NM_029492 | 75965 | Mm.29044 | NM_029492 | ENSMUSG00000021969 | **Zdhhc20** | 5033406L14Rik\|AI448102\|B230110O18Rik\|HK11 | zinc finger, DHHC domain containing 20 | protein-coding |
| chr14_57865232_57865662 | chr14 | 57865233 | 57865662 | + | -0.896672 | intron (NM_029492, intron 4 of 12) | intron (NM_029492, intron 4 of 12) | 24815 | NM_029492 | 75965 | Mm.29044 | NM_029492 | ENSMUSG00000021969 | **Zdhhc20** | 5033406L14Rik\|AI448102\|B230110O18Rik\|HK11 | zinc finger, DHHC domain containing 20 | protein-coding |
| chr14_57908472_57908707 | chr14 | 57908473 | 57908707 | + | -0.935752 | Intergenic | Intergenic | -18328 | NM_029492 | 75965 | Mm.29044 | NM_029492 | ENSMUSG00000021969 | **Zdhhc20** | 5033406L14Rik\|AI448102\|B230110O18Rik\|HK11 | zinc finger, DHHC domain containing 20 | protein-coding |
| chr9_123087458_123088280 | chr9 | 123087459 | 123088280 | + | -0.993899 | intron (NM_026917, intron 3 of 6) | intron (NM_026917, intron 3 of 6) | 25336 | NM_026917 | 69035 | Mm.28300 | NM_026917 | | **Zdhhc3** | 1110020O22Rik\|1810006O10Rik\|2210017C02Rik\|DHHC-3\|GODZ\|Gramp1\|Zfp373 | zinc finger, DHHC domain containing 3 | protein-coding |
| chr9_123087504_123088052 | chr9 | 123087505 | 123088052 | + | -1.1178 | intron (NM_026917, intron 3 of 6) | intron (NM_026917, intron 3 of 6) | 25427 | NM_026917 | 69035 | Mm.28300 | NM_026917 | | **Zdhhc3** | 1110020O22Rik\|1810006O10Rik\|2210017C02Rik\|DHHC-3\|GODZ\|Gramp1\|Zfp373 | zinc finger, DHHC domain containing 3 | protein-coding |
| chr5_139483151_139483450 | chr5 | 139483152 | 139483450 | + | -1.25585 | intron (NM_001159908, intron 1 of 6) | intron (NM_001159908, intron 1 of 6) | 1190 | NM_133349 | 100494 | Mm.24521 | NM_133349 | ENSMUSG00000053581 | **Zfand2a** | AA407930\|AU016206\|Airap | zinc finger, AN1-type domain 2A | protein-coding |
| chr17_30046594_30047049 | chr17 | 30046595 | 30047049 | + | -0.885867 | intron (NM_148926, intron 1 of 4) | intron (NM_148926, intron 1 of 4) | 41735 | NM_148926 | 21769 | Mm.220972 | NM_148926 | ENSMUSG00000044477 | **Zfand3** | AW539211\|TEG-27\|Tex27 | zinc finger, AN1-type domain 3 | protein-coding |
| chr17_30046594_30047323 | chr17 | 30046595 | 30047323 | + | -0.894626 | intron (NM_148926, intron 1 of 4) | intron (NM_148926, intron 1 of 4) | 41872 | NM_148926 | 21769 | Mm.220972 | NM_148926 | ENSMUSG00000044477 | **Zfand3** | AW539211\|TEG-27\|Tex27 | zinc finger, AN1-type domain 3 | protein-coding |
| chr17_30035691_30035968 | chr17 | 30035692 | 30035968 | + | -0.953173 | intron (NM_148926, intron 1 of 4) | B1_Mur4\|SINE\|Alu | 30743 | NM_148926 | 21769 | Mm.220972 | NM_148926 | ENSMUSG00000044477 | **Zfand3** | AW539211\|TEG-27\|Tex27 | zinc finger, AN1-type domain 3 | protein-coding |
| chr17_30035511_30035968 | chr17 | 30035512 | 30035968 | + | -0.977934 | intron (NM_148926, intron 1 of 4) | L1M4\|LINE\|L1 | 30653 | NM_148926 | 21769 | Mm.220972 | NM_148926 | ENSMUSG00000044477 | **Zfand3** | AW539211\|TEG-27\|Tex27 | zinc finger, AN1-type domain 3 | protein-coding |

| | | | | | | | | | | | | | | | | | |
|---|---|---|---|---|---|---|---|---|---|---|---|---|---|---|---|---|---|
| chr8_108738152_108738622 | chr8 | 108738153 | 108738622 | + | -1.00563 | intron (NM_007496, intron 1 of 9) | intron (NM_007496, intron 1 of 9) | 23743 | NM_007496 | 11906 | Mm.416972 | NM_007496 | ENSMUSG00000038872 | **Zfhx3** | A230102L03Rik\|Atbf1\|WBP9\|mKIAA4228 | zinc finger homeobox 3 | protein-coding |
| chr2_150111163_150111860 | chr2 | 150111164 | 150111860 | + | -0.809627 | Intergenic | Intergenic | 25166 | NM_023266 | 104348 | Mm.485851 | NM_023266 | ENSMUSG00000068134 | **Zfp120** | AI315103\|E030042N05Rik\|F80\|MZF31\|Zfp844 | zinc finger protein 120 | protein-coding |
| chr7_30045712_30046185 | chr7 | 30045713 | 30046185 | + | -0.598349 | intron (NM_178733, intron 1 of 4) | intron (NM_178733, intron 1 of 4) | 5447 | NM_011748 | 243906 | Mm.347644 | NM_011748 | | **Zfp14** | 4732429I09Rik\|Krox-9\|Zfp-14\|mKIAA1559 | zinc finger protein 14 | protein-coding |
| chr2_170205068_170205404 | chr2 | 170205069 | 170205404 | + | -0.635418 | Intergenic | Intergenic | -62561 | NM_001159683 | 228913 | Mm.22858 | NM_001033299 | ENSMUSG00000052056 | **Zfp217** | 4933431C08Rik\|AW987152\|Gm562\|ZABC1\|Znf217 | zinc finger protein 217 | protein-coding |
| chr2_170168877_170169536 | chr2 | 170168878 | 170169536 | + | -0.907613 | Intergenic | Intergenic | -26532 | NM_001159683 | 228913 | Mm.22858 | NM_001033299 | ENSMUSG00000052056 | **Zfp217** | 4933431C08Rik\|AW987152\|Gm562\|ZABC1\|Znf217 | zinc finger protein 217 | protein-coding |
| chr4_34857492_34857870 | chr4 | 34857493 | 34857870 | + | -0.728371 | intron (NM_013889, intron 1 of 7) | ORR1C2\|LTR\|ERVL-MaLR | 25267 | NM_013889 | 30046 | Mm.38193 | NM_013889 | ENSMUSG00000039967 | **Zfp292** | 5730450D02Rik\|5830493J20Rik\|9430062L07Rik\|AI449016\|Krox-10\|Zfp-15\|Zfp15\|Zn-15\|Zn-16\|mKIAA0530 | zinc finger protein 292 | protein-coding |
| chr11_70762685_70763066 | chr11 | 70762686 | 70763066 | + | -0.944432 | Intergenic | ORR1D-int\|LTR\|ERVL-MaLR | -1571 | NM_177565 | 193043 | Mm.40966 | NM_177565 | ENSMUSG00000043602 | **Zfp3** | AI854635\|B230309J24\|Fnp-1\|Zfp-3 | zinc finger protein 3 | protein-coding |
| chr2_154615715_154616074 | chr2 | 154615716 | 154616074 | + | -1.32402 | intron (NM_199304, intron 1 of 14) | intron (NM_199304, intron 1 of 14) | 2527 | NM_199304 | 228807 | Mm.103555 | NM_199304 | ENSMUSG00000059842 | **Zfp341** | Znf341 | zinc finger protein 341 | protein-coding |
| chr10_67887730_67888429 | chr10 | 67887731 | 67888429 | + | -0.615636 | 3' UTR (NM_178679, exon 5 of 5) | 3' UTR (NM_178679, exon 5 of 5) | 24582 | NM_178679 | 216049 | Mm.268346 | NM_178679 | ENSMUSG00000037855 | **Zfp365** | AI839779\|AV340892\|DBZ\|Su48\|Znf365\|talanin | zinc finger protein 365 | protein-coding |
| chr17_84308468_84308697 | chr17 | 84308469 | 84308697 | + | -0.633218 | intron (NM_183021, intron 29 of 37) | intron (NM_183021, intron 29 of 37) | -120636 | NM_001001806 | 12193 | Mm.259321 | NM_001001806 | ENSMUSG00000045817 | **Zfp36l2** | Brf2\|ERF2\|Tis11d | zinc finger protein 36, C3H type-like 2 | protein-coding |
| chr17_84166492_84166835 | chr17 | 84166493 | 84166835 | + | -0.671942 | Intergenic | Intergenic | 21283 | NM_001001806 | 12193 | Mm.259321 | NM_001001806 | ENSMUSG00000045817 | **Zfp36l2** | Brf2\|ERF2\|Tis11d | zinc finger protein 36, C3H type-like 2 | protein-coding |
| chr17_84260939_84261353 | chr17 | 84260940 | 84261353 | + | -0.722289 | intron (NM_183021, intron 30 of 37) | intron (NM_183021, intron 30 of 37) | -73199 | NM_001001806 | 12193 | Mm.259321 | NM_001001806 | ENSMUSG00000045817 | **Zfp36l2** | Brf2\|ERF2\|Tis11d | zinc finger protein 36, C3H type-like 2 | protein-coding |
| chr17_84180663_84182055 | chr17 | 84180664 | 84182055 | + | -0.774429 | Intergenic | Intergenic | 6588 | NM_001001806 | 12193 | Mm.259321 | NM_001001806 | ENSMUSG00000045817 | **Zfp36l2** | Brf2\|ERF2\|Tis11d | zinc finger protein 36, C3H type-like 2 | protein-coding |
| chr17_84272599_84272851 | chr17 | 84272600 | 84272851 | + | -0.832984 | exon (NM_183021, exon 30 of 38) | exon (NM_183021, exon 30 of 38) | -84778 | NM_001001806 | 12193 | Mm.259321 | NM_001001806 | ENSMUSG00000045817 | **Zfp36l2** | Brf2\|ERF2\|Tis11d | zinc finger protein 36, C3H type-like 2 | protein-coding |
| chr17_84238274_84238680 | chr17 | 84238275 | 84238680 | + | -0.856866 | intron (NM_183021, intron 31 of 37) | intron (NM_183021, intron 31 of 37) | -50530 | NM_001001806 | 12193 | Mm.259321 | NM_001001806 | ENSMUSG00000045817 | **Zfp36l2** | Brf2\|ERF2\|Tis11d | zinc finger protein 36, C3H type-like 2 | protein-coding |
| chr17_84180647_84181281 | chr17 | 84180648 | 84181281 | + | -0.90051 | Intergenic | Intergenic | 6983 | NM_001001806 | 12193 | Mm.259321 | NM_001001806 | ENSMUSG00000045817 | **Zfp36l2** | Brf2\|ERF2\|Tis11d | zinc finger protein 36, C3H type-like 2 | protein-coding |
| chr17_84268287_84268725 | chr17 | 84268288 | 84268725 | + | -1.01946 | intron (NM_183021, intron 30 of 37) | intron (NM_183021, intron 30 of 37) | -80559 | NM_001001806 | 12193 | Mm.259321 | NM_001001806 | ENSMUSG00000045817 | **Zfp36l2** | Brf2\|ERF2\|Tis11d | zinc finger protein 36, C3H type-like 2 | protein-coding |
| chr17_84313250_84313947 | chr17 | 84313251 | 84313947 | + | -1.11915 | intron (NM_183021, intron 28 of 37) | intron (NM_183021, intron 28 of 37) | -125652 | NM_001001806 | 12193 | Mm.259321 | NM_001001806 | ENSMUSG00000045817 | **Zfp36l2** | Brf2\|ERF2\|Tis11d | zinc finger protein 36, C3H type-like 2 | protein-coding |
| chr17_84313543_84313954 | chr17 | 84313544 | 84313954 | + | -1.16139 | intron (NM_183021, intron 28 of 37) | intron (NM_183021, intron 28 of 37) | -125802 | NM_001001806 | 12193 | Mm.259321 | NM_001001806 | ENSMUSG00000045817 | **Zfp36l2** | Brf2\|ERF2\|Tis11d | zinc finger protein 36, C3H type-like 2 | protein-coding |
| chr17_84272594_84273424 | chr17 | 84272595 | 84273424 | + | -1.16223 | intron (NM_183021, intron 29 of 37) | intron (NM_183021, intron 29 of 37) | -85062 | NM_001001806 | 12193 | Mm.259321 | NM_001001806 | ENSMUSG00000045817 | **Zfp36l2** | Brf2\|ERF2\|Tis11d | zinc finger protein 36, C3H type-like 2 | protein-coding |
| chr17_84314467_84314664 | chr17 | 84314468 | 84314664 | + | -1.22996 | intron (NM_183021, intron 28 of 37) | intron (NM_183021, intron 28 of 37) | -126619 | NM_001001806 | 12193 | Mm.259321 | NM_001001806 | ENSMUSG00000045817 | **Zfp36l2** | Brf2\|ERF2\|Tis11d | zinc finger protein 36, C3H type-like 2 | protein-coding |
| chr17_84272898_84273343 | chr17 | 84272899 | 84273343 | + | -1.2643 | intron (NM_183021, intron 29 of 37) | intron (NM_183021, intron 29 of 37) | -85174 | NM_001001806 | 12193 | Mm.259321 | NM_001001806 | ENSMUSG00000045817 | **Zfp36l2** | Brf2\|ERF2\|Tis11d | zinc finger protein 36, C3H type-like 2 | protein-coding |
| chr17_84314384_84314690 | chr17 | 84314385 | 84314690 | + | -1.2693 | intron (NM_183021, intron 28 of 37) | intron (NM_183021, intron 28 of 37) | -126590 | NM_001001806 | 12193 | Mm.259321 | NM_001001806 | ENSMUSG00000045817 | **Zfp36l2** | Brf2\|ERF2\|Tis11d | zinc finger protein 36, C3H type-like 2 | protein-coding |
| chr17_84239706_84240195 | chr17 | 84239707 | 84240195 | + | -1.27548 | intron (NM_183021, intron 31 of 37) | intron (NM_183021, intron 31 of 37) | -52004 | NM_001001806 | 12193 | Mm.259321 | NM_001001806 | ENSMUSG00000045817 | **Zfp36l2** | Brf2\|ERF2\|Tis11d | zinc finger protein 36, C3H type-like 2 | protein-coding |
| chr12_115920140_115920736 | chr12 | 115920141 | 115920736 | + | -0.659148 | Intergenic | Intergenic | -127286 | NM_001004066 | 56220 | Mm.254997 | NM_019565 | ENSMUSG00000042063 | **Zfp386** | Kzf1\|mKIAA4205 | zinc finger protein 386 (Kruppel-like) | protein-coding |
| chr12_115999785_116000374 | chr12 | 115999786 | 116000374 | + | -0.756661 | Intergenic | Intergenic | -47644 | NM_001004066 | 56220 | Mm.254997 | NM_019565 | ENSMUSG00000042063 | **Zfp386** | Kzf1\|mKIAA4205 | zinc finger protein 386 (Kruppel-like) | protein-coding |
| chr12_115887956_115888558 | chr12 | 115887957 | 115888558 | + | -0.757542 | Intergenic | Intergenic | -159467 | NM_001004066 | 56220 | Mm.254997 | NM_019565 | ENSMUSG00000042063 | **Zfp386** | Kzf1\|mKIAA4205 | zinc finger protein 386 (Kruppel-like) | protein-coding |
| chr12_115952396_115953080 | chr12 | 115952397 | 115953080 | + | -0.81676 | Intergenic | Intergenic | -94986 | NM_001004066 | 56220 | Mm.254997 | NM_019565 | ENSMUSG00000042063 | **Zfp386** | Kzf1\|mKIAA4205 | zinc finger protein 386 (Kruppel-like) | protein-coding |
| chr12_115887964_115888265 | chr12 | 115887965 | 115888265 | + | -0.904302 | Intergenic | Intergenic | -159609 | NM_001004066 | 56220 | Mm.254997 | NM_019565 | ENSMUSG00000042063 | **Zfp386** | Kzf1\|mKIAA4205 | zinc finger protein 386 (Kruppel-like) | protein-coding |
| chr12_115952370_115952809 | chr12 | 115952371 | 115952809 | + | -0.933062 | Intergenic | Intergenic | -95134 | NM_001004066 | 56220 | Mm.254997 | NM_019565 | ENSMUSG00000042063 | **Zfp386** | Kzf1\|mKIAA4205 | zinc finger protein 386 (Kruppel-like) | protein-coding |
| chr12_116010650_116011199 | chr12 | 116010651 | 116011199 | + | -1.05647 | Intergenic | Intergenic | -36799 | NM_001004066 | 56220 | Mm.254997 | NM_019565 | ENSMUSG00000042063 | **Zfp386** | Kzf1\|mKIAA4205 | zinc finger protein 386 (Kruppel-like) | protein-coding |
| chr2_91645737_91646305 | chr2 | 91645738 | 91646305 | + | -0.855968 | exon (NM_001033451, exon 4 of 4) | exon (NM_001033451, exon 4 of 4) | 3770 | NM_001033451 | 381410 | Mm.330401 | NM_001033451 | ENSMUSG00000075040 | **Zfp408** | Gm1011 | zinc finger protein 408 | protein-coding |
| chr2_91645824_91646144 | chr2 | 91645825 | 91646144 | + | -0.900804 | exon (NM_001033451, exon 4 of 4) | exon (NM_001033451, exon 4 of 4) | 3807 | NM_001033451 | 381410 | Mm.330401 | NM_001033451 | ENSMUSG00000075040 | **Zfp408** | Gm1011 | zinc finger protein 408 | protein-coding |
| chr18_5378399_5378757 | chr18 | 5378400 | 5378757 | + | -0.722901 | Intergenic | Intergenic | -44139 | NM_178722 | 240186 | Mm.118257 | NM_178722 | ENSMUSG00000050945 | **Zfp438** | 9430031G09\|9430091M14Rik\|B830013J05Rik | zinc finger protein 438 | protein-coding |
| chr18_5424244_5424800 | chr18 | 5424245 | 5424800 | + | -0.889816 | Intergenic | MTD\|LTR\|ERVL-MaLR | -90083 | NM_178722 | 240186 | Mm.118257 | NM_178722 | ENSMUSG00000050945 | **Zfp438** | 9430031G09\|9430091M14Rik\|B830013J05Rik | zinc finger protein 438 | protein-coding |
| chr7_35991080_35991533 | chr7 | 35991081 | 35991533 | + | -0.748491 | Intergenic | ORR1F\|LTR\|ERVL-MaLR | -188318 | NM_177739 | 668501 | Mm.286296 | NM_177739 | ENSMUSG00000044452 | **Zfp507** | 1810022O10Rik\|A230056M16Rik\|AL024009\|AU021817\|BB105092\|Gm480\|Znf507\|mKIAA1084 | zinc finger protein 507 | protein-coding |
| chr17_21463394_21464054 | chr17 | 21463395 | 21464054 | + | -0.695933 | exon (NM_009558, exon 5 of 5) | exon (NM_009558, exon 5 of 5) | 13372 | NM_009558 | 22709 | Mm.293030 | NM_009558 | ENSMUSG00000023892 | **Zfp51** | Zfp-51\|mszf92\|zfec12 | zinc finger protein 51 | protein-coding |
| chr17_21469304_21469571 | chr17 | 21469305 | 21469571 | + | -0.830638 | Intergenic | ORR1A2\|LTR\|ERVL-MaLR | 19086 | NM_009558 | 22709 | Mm.293030 | NM_009558 | ENSMUSG00000023892 | **Zfp51** | Zfp-51\|mszf92\|zfec12 | zinc finger protein 51 | protein-coding |
| chr18_13693852_13694115 | chr18 | 13693853 | 13694115 | + | -0.822497 | intron (NM_145492, intron 7 of 7) | intron (NM_145492, intron 7 of 7) | 278749 | NM_145492 | 225207 | Mm.40325 | NM_145492 | ENSMUSG00000024420 | **Zfp521** | B930086A16Rik\|Evi3\|Znf521 | zinc finger protein 521 | protein-coding |
| chr17_21473491_21474369 | chr17 | 21473492 | 21474369 | + | -0.798168 | Intergenic | MMERGLN-int\|LTR\|ERV1 | -15058 | NM_013843 | 24132 | Mm.42140 | NM_013843 | | **Zfp53** | D030067O06Rik\|KRAZ1\|Zfp-53\|Zfp118\|zfas8 | zinc finger protein 53 | protein-coding |
| chr7_37490165_37491134 | chr7 | 37490166 | 37491134 | + | -0.809879 | intron (NM_172385, intron 5 of 5) | (CTG)n\|Simple_repeat\|Simple_repeat | 279102 | NM_172385 | 243937 | Mm.322505 | NM_172385 | ENSMUSG00000043456 | **Zfp536** | 9630010P11Rik\|Znf536\|mKIAA0390 | zinc finger protein 536 | protein-coding |
| chr17_21408288_21408532 | chr17 | 21408289 | 21408532 | + | -0.874316 | Intergenic | RLTR19\|LTR\|ERVK | -14817 | NM_011760 | 22712 | Mm.12916 | NM_011760 | ENSMUSG00000023882 | **Zfp54** | 2810034D10Rik\|KRAB10\|Zfp-54\|Zfp76\|mszf83 | zinc finger protein 54 | protein-coding |
| chr7_16046022_16046381 | chr7 | 16046023 | 16046381 | + | -0.647815 | Intergenic | Intergenic | -25740 | NM_001099277 | 666528 | Mm.290718 | NM_001099277 | ENSMUSG00000078796 | **Zfp541** | EG666528\|Znf541 | zinc finger protein 541 | protein-coding |
| chr7_80973639_80973926 | chr7 | 80973640 | 80973926 | + | -0.85732 | Intergenic | Intergenic | -19901 | NM_178707 | 233410 | Mm.29525 | NM_178707 | ENSMUSG00000005621 | **Zfp592** | 8430405N24\|A730014M16Rik\|Znf592\|mKIAA0211 | zinc finger protein 592 | protein-coding |
| chr7_81004492_81005242 | chr7 | 81004493 | 81005242 | + | -0.868679 | intron (NM_178707, intron 1 of 9) | intron (NM_178707, intron 1 of 9) | 11183 | NM_178707 | 233410 | Mm.29525 | NM_178707 | ENSMUSG00000005621 | **Zfp592** | 8430405N24\|A730014M16Rik\|Znf592\|mKIAA0211 | zinc finger protein 592 | protein-coding |
| chr18_55398020_55398681 | chr18 | 55398021 | 55398681 | + | -0.683277 | Intergenic | MER34\|LTR\|ERV1 | -408171 | NM_175751 | 269023 | Mm.192984 | NM_175751 | ENSMUSG00000052713 | **Zfp608** | 4932417D18Rik\|D430007A19Rik\|Znf608 | zinc finger protein 608 | protein-coding |
| chr18_55681410_55681846 | chr18 | 55681411 | 55681846 | + | -0.907235 | Intergenic | Intergenic | -691448 | NM_175751 | 269023 | Mm.192984 | NM_175751 | ENSMUSG00000052713 | **Zfp608** | 4932417D18Rik\|D430007A19Rik\|Znf608 | zinc finger protein 608 | protein-coding |
| chr18_55681183_55681865 | chr18 | 55681184 | 55681865 | + | -0.927489 | Intergenic | Intergenic | -691344 | NM_175751 | 269023 | Mm.192984 | NM_175751 | ENSMUSG00000052713 | **Zfp608** | 4932417D18Rik\|D430007A19Rik\|Znf608 | zinc finger protein 608 | protein-coding |
| chr7_127632448_127632819 | chr7 | 127632449 | 127632819 | + | -0.779465 | Intergenic | ID_B1\|SINE\|B4 | -18201 | NM_177226 | 320683 | Mm.156718 | NM_177226 | ENSMUSG00000045639 | **Zfp629** | 9330199A09Rik\|Znf629 | zinc finger protein 629 | protein-coding |
| chr5_106663246_106663484 | chr5 | 106663247 | 106663484 | + | -0.7882 | intron (NM_026856, intron 2 of 6) | PB1D7\|SINE\|Alu | 33465 | NM_026856 | 52397 | Mm.249474 | NM_026856 | ENSMUSG00000049606 | **Zfp644** | 1110068L01Rik\|AA408882\|BM-005\|D5Ertd689e\|Zep-2 | zinc finger protein 644 | protein-coding |
| chr5_106676770_106677314 | chr5 | 106676771 | 106677314 | + | -0.939533 | intron (NM_026856, intron 1 of 6) | intron (NM_026856, intron 1 of 6) | 19788 | NM_026856 | 52397 | Mm.249474 | NM_026856 | ENSMUSG00000049606 | **Zfp644** | 1110068L01Rik\|AA408882\|BM-005\|D5Ertd689e\|Zep-2 | zinc finger protein 644 | protein-coding |
| chr5_106649878_106650322 | chr5 | 106649879 | 106650322 | + | -1.17074 | intron (NM_026856, intron 2 of 6) | (A)n\|Simple_repeat\|Simple_repeat | 46730 | NM_026856 | 52397 | Mm.249474 | NM_026856 | ENSMUSG00000049606 | **Zfp644** | 1110068L01Rik\|AA408882\|BM-005\|D5Ertd689e\|Zep-2 | zinc finger protein 644 | protein-coding |
| chr11_95713779_95714247 | chr11 | 95713780 | 95714247 | + | -0.654504 | Intergenic | (CA)n\|Simple_repeat\|Simple_repeat | -1259 | NR_045780 | 432396 | Mm.426849 | NR_045780 | ENSMUSG00000086191 | **Zfp652os** | 1110035M17Rik | zinc finger protein 652, opposite strand | ncRNA |
| chr11_95714463_95714672 | chr11 | 95714464 | 95714672 | + | -0.707902 | Intergenic | Intergenic | -1814 | NR_045780 | 432396 | Mm.426849 | NR_045780 | ENSMUSG00000086191 | **Zfp652os** | 1110035M17Rik | zinc finger protein 652, opposite strand | ncRNA |

| | | | | | | | | | | | | | | | | | |
|---|---|---|---|---|---|---|---|---|---|---|---|---|---|---|---|---|---|
| chr5_124873120_124873468 | chr5 | 124873121 | 124873468 | + | -0.714138 | intron (NM_001081750, intron 2 of 3) | intron (NM_001081750, intron 2 of 3) | 10589 | NM_001081750 | 269704 | Mm.138617 | NM_001081750 | ENSMUSG00000079215 | **Zfp664** | AW060232\|D930038J03Rik\|ZFOC1\|Znf664 | zinc finger protein 664 | protein-coding |
| chr11_58282170_58282456 | chr11 | 58282171 | 58282456 | + | -0.746299 | Intergenic | Intergenic | -24756 | NM_182996 | 103836 | Mm.41440 | NM_182996 | ENSMUSG00000037243 | **Zfp692** | AI746306\|AI839920\|Zfp692-ps\|Znf692 | zinc finger protein 692 | protein-coding |
| chr3_9719052_9719346 | chr3 | 9719053 | 9719346 | + | -0.621297 | intron (NM_053182, intron 2 of 7) | intron (NM_053182, intron 2 of 7) | -109114 | NM_133218 | 170753 | Mm.212908 | NM_133218 | ENSMUSG00000040209 | **Zfp704** | C030026M03\|C030034L19Rik\|Gig1\|Znf704 | zinc finger protein 704 | protein-coding |
| chr3_9719007_9719527 | chr3 | 9719008 | 9719527 | + | -0.623575 | intron (NM_053182, intron 2 of 7) | intron (NM_053182, intron 2 of 7) | -109182 | NM_133218 | 170753 | Mm.212908 | NM_133218 | ENSMUSG00000040209 | **Zfp704** | C030026M03\|C030034L19Rik\|Gig1\|Znf704 | zinc finger protein 704 | protein-coding |
| chr3_9547243_9547628 | chr3 | 9547244 | 9547628 | + | -0.657031 | intron (NM_133218, intron 2 of 8) | intron (NM_133218, intron 2 of 8) | 62649 | NM_133218 | 170753 | Mm.212908 | NM_133218 | ENSMUSG00000040209 | **Zfp704** | C030026M03\|C030034L19Rik\|Gig1\|Znf704 | zinc finger protein 704 | protein-coding |
| chr3_9520027_9520967 | chr3 | 9520028 | 9520967 | + | -0.664847 | intron (NM_133218, intron 2 of 8) | intron (NM_133218, intron 2 of 8) | 89588 | NM_133218 | 170753 | Mm.212908 | NM_133218 | ENSMUSG00000040209 | **Zfp704** | C030026M03\|C030034L19Rik\|Gig1\|Znf704 | zinc finger protein 704 | protein-coding |
| chr3_9512151_9513013 | chr3 | 9512152 | 9513013 | + | -0.737765 | intron (NM_133218, intron 2 of 8) | RMER1B\|Other\|Other | 97503 | NM_133218 | 170753 | Mm.212908 | NM_133218 | ENSMUSG00000040209 | **Zfp704** | C030026M03\|C030034L19Rik\|Gig1\|Znf704 | zinc finger protein 704 | protein-coding |
| chr3_9633438_9634336 | chr3 | 9633439 | 9634336 | + | -0.886202 | Intergenic | Intergenic | -23802 | NM_133218 | 170753 | Mm.212908 | NM_133218 | ENSMUSG00000040209 | **Zfp704** | C030026M03\|C030034L19Rik\|Gig1\|Znf704 | zinc finger protein 704 | protein-coding |
| chr3_9633459_9633939 | chr3 | 9633460 | 9633939 | + | -0.915627 | Intergenic | Intergenic | -23614 | NM_133218 | 170753 | Mm.212908 | NM_133218 | ENSMUSG00000040209 | **Zfp704** | C030026M03\|C030034L19Rik\|Gig1\|Znf704 | zinc finger protein 704 | protein-coding |
| chr15_36965820_36966125 | chr15 | 36965821 | 36966125 | + | -0.684659 | Intergenic | MLT1J\|LTR\|ERVL-MaLR | 41429 | NM_026521 | 68036 | Mm.371631 | NM_026521 | ENSMUSG00000062397 | **Zfp706** | 3110006P09Rik\|AA793007\|AW536850\|C77232\|Znf706 | zinc finger protein 706 | protein-coding |
| chr15_36931434_36931813 | chr15 | 36931435 | 36931813 | + | -0.720926 | Intergenic | ORR1B2\|LTR\|ERVL-MaLR | 75778 | NM_026521 | 68036 | Mm.371631 | NM_026521 | ENSMUSG00000062397 | **Zfp706** | 3110006P09Rik\|AA793007\|AW536850\|C77232\|Znf706 | zinc finger protein 706 | protein-coding |
| chr15_37037743_37038072 | chr15 | 37037744 | 37038072 | + | -1.27588 | Intergenic | MTE2b\|LTR\|ERVL-MaLR | -30506 | NM_026521 | 68036 | Mm.371631 | NM_026521 | ENSMUSG00000062397 | **Zfp706** | 3110006P09Rik\|AA793007\|AW536850\|C77232\|Znf706 | zinc finger protein 706 | protein-coding |
| chr15_37037808_37037998 | chr15 | 37037809 | 37037998 | + | -1.27699 | Intergenic | MTE2b\|LTR\|ERVL-MaLR | -30501 | NM_026521 | 68036 | Mm.371631 | NM_026521 | ENSMUSG00000062397 | **Zfp706** | 3110006P09Rik\|AA793007\|AW536850\|C77232\|Znf706 | zinc finger protein 706 | protein-coding |
| chr11_121559544_121559980 | chr11 | 121559545 | 121559980 | + | -0.765985 | intron (NM_029878, intron 16 of 38) | intron (NM_029878, intron 16 of 38) | -40420 | NM_178763 | 319530 | Mm.11395 | NM_178763 | ENSMUSG00000039238 | **Zfp750** | A030007D23Rik\|Znf750 | zinc finger protein 750 | protein-coding |
| chr6_28318894_28319974 | chr6 | 28318895 | 28319974 | + | -0.808929 | Intergenic | Intergenic | -57833 | NM_001081678 | 627049 | Mm.84441 | NM_001081678 | ENSMUSG00000039841 | **Zfp800** | AA407452\|Znf800 | zinc finger protein 800 | protein-coding |
| chr6_28319002_28319509 | chr6 | 28319003 | 28319509 | + | -0.912277 | Intergenic | Intergenic | -57655 | NM_001081678 | 627049 | Mm.84441 | NM_001081678 | ENSMUSG00000039841 | **Zfp800** | AA407452\|Znf800 | zinc finger protein 800 | protein-coding |
| chr6_28319277_28319502 | chr6 | 28319278 | 28319502 | + | -1.00091 | Intergenic | Intergenic | -57789 | NM_001081678 | 627049 | Mm.84441 | NM_001081678 | ENSMUSG00000039841 | **Zfp800** | AA407452\|Znf800 | zinc finger protein 800 | protein-coding |
| chr6_28322455_28322785 | chr6 | 28322456 | 28322785 | + | -1.1724 | Intergenic | MLTR25A\|LTR\|ERVK | -61019 | NM_001081678 | 627049 | Mm.84441 | NM_001081678 | ENSMUSG00000039841 | **Zfp800** | AA407452\|Znf800 | zinc finger protein 800 | protein-coding |
| chr6_28322178_28322792 | chr6 | 28322179 | 28322792 | + | -1.2185 | Intergenic | Intergenic | -60884 | NM_001081678 | 627049 | Mm.84441 | NM_001081678 | ENSMUSG00000039841 | **Zfp800** | AA407452\|Znf800 | zinc finger protein 800 | protein-coding |
| chr8_109707988_109708286 | chr8 | 109707989 | 109708286 | + | -0.933291 | 5' UTR (NM_001286391, exon 3 of 9) | 5' UTR (NM_001286391, exon 3 of 9) | 2118 | NM_029468 | 75871 | Mm.180410 | NM_029468 | ENSMUSG00000031728 | **Zfp821** | 4930566A11Rik\|Znf821 | zinc finger protein 821 | protein-coding |
| chr2_174612563_174613199 | chr2 | 174612564 | 174613199 | + | -0.641726 | Intergenic | Intergenic | -30653 | NM_001099328 | 100043757 | Mm.48711 | NM_001099328 | ENSMUSG00000050600 | **Zfp831** | ENSMUSG00000050600\|OTTMUSG00000017459 | zinc finger protein 831 | protein-coding |
| chr17_32880947_32881229 | chr17 | 32880948 | 32881229 | + | -0.925461 | 3' UTR (NM_207245, exon 4 of 4) | 3' UTR (NM_207245, exon 4 of 4) | 10515 | NM_207245 | 240066 | Mm.26636 | NM_207245 | ENSMUSG00000095325 | **Zfp870** | AI173952 | zinc finger protein 870 | protein-coding |
| chr17_32880145_32880887 | chr17 | 32880146 | 32880887 | + | -1.09776 | 3' UTR (NM_207245, exon 4 of 4) | 3' UTR (NM_207245, exon 4 of 4) | 11087 | NM_207245 | 240066 | Mm.26636 | NM_207245 | ENSMUSG00000095325 | **Zfp870** | AI173952 | zinc finger protein 870 | protein-coding |
| chr17_32880349_32880798 | chr17 | 32880350 | 32880798 | + | -1.13938 | 3' UTR (NM_207245, exon 4 of 4) | 3' UTR (NM_207245, exon 4 of 4) | 11029 | NM_207245 | 240066 | Mm.26636 | NM_207245 | ENSMUSG00000095325 | **Zfp870** | AI173952 | zinc finger protein 870 | protein-coding |
| chr17_32880363_32881274 | chr17 | 32880364 | 32881274 | + | -1.17039 | 3' UTR (NM_207245, exon 4 of 4) | 3' UTR (NM_207245, exon 4 of 4) | 10784 | NM_207245 | 240066 | Mm.26636 | NM_207245 | ENSMUSG00000095325 | **Zfp870** | AI173952 | zinc finger protein 870 | protein-coding |
| chr11_50835744_50836165 | chr11 | 50835745 | 50836165 | + | -0.924719 | intron (NM_001290779, intron 3 of 3) | intron (NM_001290779, intron 3 of 3) | 5597 | NM_001290779 | 214779 | Mm.191846 | NM_173387 | ENSMUSG00000044296 | **Zfp879** | Znf879 | zinc finger protein 879 | protein-coding |
| chr14_79180406_79180797 | chr14 | 79180407 | 79180797 | + | -0.621305 | intron (NM_027906, intron 40 of 44) | intron (NM_027906, intron 40 of 44) | 66765 | NM_001033215 | 105590 | Mm.486536 | NM_001033215 | ENSMUSG00000071262 | **Zfp957** | AU017455 | zinc finger protein 957 | protein-coding |
| chr14_79255822_79256162 | chr14 | 79255823 | 79256162 | + | -0.933268 | Intergenic | Intergenic | -8625 | NM_001033215 | 105590 | Mm.486536 | NM_001033215 | ENSMUSG00000071262 | **Zfp957** | AU017455 | zinc finger protein 957 | protein-coding |
| chr17_17004585_17005629 | chr17 | 17004586 | 17005629 | + | -0.683063 | Intergenic | Intergenic | -58996 | NM_001005358 | 449000 | Mm.483718 | NM_001005358 | ENSMUSG00000096696 | **Zfp960** | - | zinc finger protein 960 | protein-coding |
| chr17_16951085_16951838 | chr17 | 16951086 | 16951838 | + | -0.685606 | Intergenic | Intergenic | -112641 | NM_001005358 | 449000 | Mm.483718 | NM_001005358 | ENSMUSG00000096696 | **Zfp960** | - | zinc finger protein 960 | protein-coding |
| chr17_17234346_17235374 | chr17 | 17234347 | 17235374 | + | -0.641847 | Intergenic | Intergenic | 113477 | NM_011765 | 22759 | Mm.436633 | NM_011765 | ENSMUSG00000095990 | **Zfp97** | NGD16-4 | zinc finger protein 97 | protein-coding |
| chr15_57792879_57793623 | chr15 | 57792880 | 57793623 | + | -0.699064 | intron (NM_199449, intron 2 of 3) | intron (NM_199449, intron 2 of 3) | 98584 | NM_199449 | 387609 | Mm.36557 | NM_199449 | ENSMUSG00000071757 | **Zhx2** | Afr-1\|Afr1\|Raf\|mKIAA0854 | zinc fingers and homeoboxes 2 | protein-coding |
| chr15_57793057_57793490 | chr15 | 57793058 | 57793490 | + | -0.711576 | intron (NM_199449, intron 2 of 3) | intron (NM_199449, intron 2 of 3) | 98607 | NM_199449 | 387609 | Mm.36557 | NM_199449 | ENSMUSG00000071757 | **Zhx2** | Afr-1\|Afr1\|Raf\|mKIAA0854 | zinc fingers and homeoboxes 2 | protein-coding |
| chr15_57772374_57773523 | chr15 | 57772375 | 57773523 | + | -1.14179 | intron (NM_199449, intron 2 of 3) | intron (NM_199449, intron 2 of 3) | 78282 | NM_199449 | 387609 | Mm.36557 | NM_199449 | ENSMUSG00000071757 | **Zhx2** | Afr-1\|Afr1\|Raf\|mKIAA0854 | zinc fingers and homeoboxes 2 | protein-coding |
| chr4_58944324_58944807 | chr4 | 58944325 | 58944807 | + | -0.826279 | intron (NM_001099323, intron 1 of 5) | (TGGGGG)n\|Simple_repeat\|Simple_repeat | 938 | NM_001099323 | 100041581 | Mm.484169 | NM_001099323 | ENSMUSG00000038630 | **Zkscan16** | XM_143800\|Zfp483 | zinc finger with KRAB and SCAN domains 16 | protein-coding |
| chr8_23819844_23820230 | chr8 | 23819845 | 23820230 | + | -0.828911 | intron (NM_177086, intron 3 of 6) | intron (NM_177086, intron 3 of 6) | 150376 | NM_177086 | 320158 | Mm.121274 | NM_177086 | ENSMUSG00000037492 | **Zmat4** | 9630048M01Rik | zinc finger, matrin type 4 | protein-coding |
| chr8_23886467_23887350 | chr8 | 23886468 | 23887350 | + | -1.11074 | intron (NM_001277239, intron 4 of 7) | L1MB8\|LINE\|L1 | 217248 | NM_177086 | 320158 | Mm.121274 | NM_177086 | ENSMUSG00000037492 | **Zmat4** | 9630048M01Rik | zinc finger, matrin type 4 | protein-coding |
| chr13_9775893_9776233 | chr13 | 9775894 | 9776233 | + | -0.658153 | Intergenic | Intergenic | -10749 | NM_144516 | 66505 | Mm.397307 | NM_144516 | ENSMUSG00000021156 | **Zmynd11** | 2210402G22Rik\|BS69 | zinc finger, MYND domain containing 11 | protein-coding |
| chr2_165890275_165890896 | chr2 | 165890276 | 165890896 | + | -0.611052 | Intergenic | Intergenic | -5748 | NM_001252587 | 228880 | Mm.227598 | NM_027230 | ENSMUSG00000039671 | **Zmynd8** | 1110013E22Rik\|2010005I16Rik\|3632413B07Rik\|AI316811\|AL024039\|Prkcbp1\|RACK7 | zinc finger, MYND-type containing 8 | protein-coding |
| chr2_165923614_165924141 | chr2 | 165923615 | 165924141 | + | -1.25998 | Intergenic | Intergenic | -39040 | NM_001252587 | 228880 | Mm.227598 | NM_027230 | ENSMUSG00000039671 | **Zmynd8** | 1110013E22Rik\|2010005I16Rik\|3632413B07Rik\|AI316811\|AL024039\|Prkcbp1\|RACK7 | zinc finger, MYND-type containing 8 | protein-coding |
| chr8_111560048_111560230 | chr8 | 111560049 | 111560230 | + | -0.629923 | intron (NM_001168623, intron 1 of 4) | L2\|LINE\|L2 | 21796 | NM_001168623 | 170737 | Mm.271578 | NM_133206 | ENSMUSG00000033545 | **Znrf1** | B830022L21Rik\|Rnf42\|Zrfp1\|nin283 | zinc and ring finger 1 | protein-coding |
| chr8_111563735_111564060 | chr8 | 111563736 | 111564060 | + | -1.11195 | intron (NM_001168623, intron 1 of 4) | MLT1H1\|LTR\|ERVL-MaLR | 25555 | NM_001168623 | 170737 | Mm.271578 | NM_133206 | ENSMUSG00000033545 | **Znrf1** | B830022L21Rik\|Rnf42\|Zrfp1\|nin283 | zinc and ring finger 1 | protein-coding |
| chr8_111582457_111583244 | chr8 | 111582458 | 111583244 | + | -1.1689 | intron (NM_001168623, intron 1 of 4) | intron (NM_001168623, intron 1 of 4) | 44508 | NM_001168623 | 170737 | Mm.271578 | NM_133206 | ENSMUSG00000033545 | **Znrf1** | B830022L21Rik\|Rnf42\|Zrfp1\|nin283 | zinc and ring finger 1 | protein-coding |
| chr8_111582291_111582849 | chr8 | 111582292 | 111582849 | + | -1.38824 | intron (NM_001168623, intron 1 of 4) | intron (NM_001168623, intron 1 of 4) | 44227 | NM_001168623 | 170737 | Mm.271578 | NM_133206 | ENSMUSG00000033545 | **Znrf1** | B830022L21Rik\|Rnf42\|Zrfp1\|nin283 | zinc and ring finger 1 | protein-coding |
| chr6_54759750_54760461 | chr6 | 54759751 | 54760461 | + | -0.620911 | Intergenic | Intergenic | -56810 | NM_199143 | 387524 | Mm.286149 | NM_199143 | | **Znrf2** | 1190002C14Rik\|AU044205\|D6Ertd365e | zinc and ring finger 2 | protein-coding |
| chr6_54849244_54849760 | chr6 | 54849245 | 54849760 | + | -0.709829 | intron (NM_199143, intron 1 of 4) | intron (NM_199143, intron 1 of 4) | 32586 | NM_199143 | 387524 | Mm.286149 | NM_199143 | | **Znrf2** | 1190002C14Rik\|AU044205\|D6Ertd365e | zinc and ring finger 2 | protein-coding |
| chr6_54804135_54804430 | chr6 | 54804136 | 54804430 | + | -0.746501 | Intergenic | Intergenic | -12633 | NM_199143 | 387524 | Mm.286149 | NM_199143 | | **Znrf2** | 1190002C14Rik\|AU044205\|D6Ertd365e | zinc and ring finger 2 | protein-coding |
| chr6_54766486_54767024 | chr6 | 54766487 | 54767024 | + | -0.92016 | Intergenic | Intergenic | -50161 | NM_199143 | 387524 | Mm.286149 | NM_199143 | | **Znrf2** | 1190002C14Rik\|AU044205\|D6Ertd365e | zinc and ring finger 2 | protein-coding |
| chr6_54766501_54766829 | chr6 | 54766502 | 54766829 | + | -1.21534 | Intergenic | Intergenic | -50251 | NM_199143 | 387524 | Mm.286149 | NM_199143 | | **Znrf2** | 1190002C14Rik\|AU044205\|D6Ertd365e | zinc and ring finger 2 | protein-coding |
| chr11_5429840_5430146 | chr11 | 5429841 | 5430146 | + | -0.867743 | intron (NM_001080924, intron 1 of 8) | intron (NM_001080924, intron 1 of 8) | 14854 | NM_001080924 | 407821 | Mm.216313 | NM_001080924 | ENSMUSG00000041961 | **Znrf3** | Gm1167 | zinc and ring finger 3 | protein-coding |

<table>
<tr><td>chr11_5429862_5430079</td><td>chr11</td><td>5429863</td><td>5430079</td><td>+</td><td>-0.868082</td><td>intron (NM_001080924, intron 1 of 8)</td><td>intron (NM_001080924, intron 1 of 8)</td><td>14876</td><td>NM_001080924</td><td>407821</td><td>Mm.216313</td><td>NM_001080924</td><td>ENSMUSG00000041961</td><td>Znrf3</td><td>Gm1167</td><td>zinc and ring finger 3</td><td>protein-coding</td></tr>
<tr><td>chr11_5369047_5369563</td><td>chr11</td><td>5369048</td><td>5369563</td><td>+</td><td>-0.912725</td><td>intron (NM_001080924, intron 1 of 8)</td><td>intron (NM_001080924, intron 1 of 8)</td><td>12435</td><td>NM_001290501</td><td>407821</td><td>Mm.216313</td><td>NM_001080924</td><td>ENSMUSG00000041961</td><td>Znrf3</td><td>Gm1167</td><td>zinc and ring finger 3</td><td>protein-coding</td></tr>
<tr><td>chr11_5435524_5435929</td><td>chr11</td><td>5435525</td><td>5435929</td><td>+</td><td>-0.927007</td><td>intron (NM_001080924, intron 1 of 8)</td><td>intron (NM_001080924, intron 1 of 8)</td><td>9120</td><td>NM_001080924</td><td>407821</td><td>Mm.216313</td><td>NM_001080924</td><td>ENSMUSG00000041961</td><td>Znrf3</td><td>Gm1167</td><td>zinc and ring finger 3</td><td>protein-coding</td></tr>
<tr><td>chr11_5411566_5412044</td><td>chr11</td><td>5411567</td><td>5412044</td><td>+</td><td>-1.11393</td><td>intron (NM_001080924, intron 1 of 8)</td><td>intron (NM_001080924, intron 1 of 8)</td><td>-30065</td><td>NM_001290501</td><td>407821</td><td>Mm.216313</td><td>NM_001080924</td><td>ENSMUSG00000041961</td><td>Znrf3</td><td>Gm1167</td><td>zinc and ring finger 3</td><td>protein-coding</td></tr>
<tr><td>chr11_5411656_5411979</td><td>chr11</td><td>5411657</td><td>5411979</td><td>+</td><td>-1.11839</td><td>intron (NM_001080924, intron 1 of 8)</td><td>intron (NM_001080924, intron 1 of 8)</td><td>-30078</td><td>NM_001290501</td><td>407821</td><td>Mm.216313</td><td>NM_001080924</td><td>ENSMUSG00000041961</td><td>Znrf3</td><td>Gm1167</td><td>zinc and ring finger 3</td><td>protein-coding</td></tr>
<tr><td>chr1_130629718_130630032</td><td>chr1</td><td>130629719</td><td>130630032</td><td>+</td><td>-0.614134</td><td>promoter-TSS (NM_009581)</td><td>promoter-TSS (NM_009581)</td><td>-269</td><td>NM_009581</td><td>22789</td><td>Mm.4644</td><td>NM_009581</td><td>ENSMUSG00000042554</td><td>Zp3r</td><td>AV264399|SP56</td><td>zona pellucida 3 receptor</td><td>protein-coding</td></tr>
<tr><td>chr9_46272013_46272378</td><td>chr9</td><td>46272014</td><td>46272378</td><td>+</td><td>-0.996616</td><td>promoter-TSS (NM_011752)</td><td>promoter-TSS (NM_011752)</td><td>-868</td><td>NM_011752</td><td>22687</td><td>Mm.17519</td><td>NM_011752</td><td>ENSMUSG00000032078</td><td>Zpr1</td><td>AI303781|Zfp259|Znf259</td><td>ZPR1 zinc finger</td><td>protein-coding</td></tr>
<tr><td>chr17_23617035_23617592</td><td>chr17</td><td>23617036</td><td>23617592</td><td>+</td><td>-1.23493</td><td>Intergenic</td><td>MLTR11B|LTR|ERVK</td><td>16488</td><td>NM_001289482</td><td>332221</td><td>Mm.270315</td><td>NM_001033425</td><td>ENSMUSG00000023902</td><td>Zscan10</td><td>BC068284|Zfp206|Zkscan10|Znf206</td><td>zinc finger and SCAN domain containing 10</td><td>protein-coding</td></tr>
<tr><td>chr17_23620278_23620796</td><td>chr17</td><td>23620279</td><td>23620796</td><td>+</td><td>-1.53384</td><td>Intergenic</td><td>Intergenic</td><td>19711</td><td>NM_001289482</td><td>332221</td><td>Mm.270315</td><td>NM_001033425</td><td>ENSMUSG00000023902</td><td>Zscan10</td><td>BC068284|Zfp206|Zkscan10|Znf206</td><td>zinc finger and SCAN domain containing 10</td><td>protein-coding</td></tr>
<tr><td>chr7_12768148_12768489</td><td>chr7</td><td>12768149</td><td>12768489</td><td>+</td><td>-1.05929</td><td>3' UTR (NM_001017955, exon 6 of 6)</td><td>3' UTR (NM_001017955, exon 6 of 6)</td><td>7339</td><td>NM_001017955</td><td>232875</td><td>Mm.333573</td><td>NM_001017955</td><td>ENSMUSG00000070822</td><td>Zscan18</td><td>EG232875</td><td>zinc finger and SCAN domain containing 18</td><td>protein-coding</td></tr>
<tr><td>chr7_80864770_80865237</td><td>chr7</td><td>80864771</td><td>80865237</td><td>+</td><td>-0.691601</td><td>intron (NM_009553, intron 2 of 2)</td><td>intron (NM_009553, intron 2 of 2)</td><td>2896</td><td>NM_009553</td><td>22691</td><td>Mm.358645</td><td>NM_009553</td><td>ENSMUSG00000038797</td><td>Zscan2</td><td>Zfp-29|Zfp29</td><td>zinc finger and SCAN domain containing 2</td><td>protein-coding</td></tr>
<tr><td>chr4_128594035_128594626</td><td>chr4</td><td>128594036</td><td>128594626</td><td>+</td><td>-0.659504</td><td>intron (NM_177758, intron 2 of 7)</td><td>B4A|SINE|B4</td><td>15767</td><td>NM_177758</td><td>269585</td><td>Mm.153291</td><td>NM_177758</td><td>ENSMUSG00000061894</td><td>Zscan20</td><td>C130001F22|Zfp31|Zkscan20</td><td>zinc finger and SCAN domains 20</td><td>protein-coding</td></tr>
<tr><td>chr4_128593824_128594499</td><td>chr4</td><td>128593825</td><td>128594499</td><td>+</td><td>-0.775142</td><td>intron (NM_177758, intron 2 of 7)</td><td>L1MCa|LINE|L1</td><td>15936</td><td>NM_177758</td><td>269585</td><td>Mm.153291</td><td>NM_177758</td><td>ENSMUSG00000061894</td><td>Zscan20</td><td>C130001F22|Zfp31|Zkscan20</td><td>zinc finger and SCAN domains 20</td><td>protein-coding</td></tr>
<tr><td>chr2_83929225_83929989</td><td>chr2</td><td>83929226</td><td>83929989</td><td>+</td><td>-0.755935</td><td>intron (NM_027964, intron 3 of 8)</td><td>ORR1C2|LTR|ERVL-MaLR</td><td>11619</td><td>NM_027964</td><td>71861</td><td>Mm.116957</td><td>NM_027964</td><td>ENSMUSG00000034552</td><td>Zswim2</td><td>1700025P14Rik|4933437F18Rik|MEX</td><td>zinc finger SWIM-type containing 2</td><td>protein-coding</td></tr>
<tr><td>chr2_83929365_83929856</td><td>chr2</td><td>83929366</td><td>83929856</td><td>+</td><td>-0.777053</td><td>intron (NM_027964, intron 3 of 8)</td><td>ORR1C2|LTR|ERVL-MaLR</td><td>11615</td><td>NM_027964</td><td>71861</td><td>Mm.116957</td><td>NM_027964</td><td>ENSMUSG00000034552</td><td>Zswim2</td><td>1700025P14Rik|4933437F18Rik|MEX</td><td>zinc finger SWIM-type containing 2</td><td>protein-coding</td></tr>
<tr><td>chr2_83993713_83994844</td><td>chr2</td><td>83993714</td><td>83994844</td><td>+</td><td>-0.906446</td><td>Intergenic</td><td>Intergenic</td><td>-53053</td><td>NM_027964</td><td>71861</td><td>Mm.116957</td><td>NM_027964</td><td>ENSMUSG00000034552</td><td>Zswim2</td><td>1700025P14Rik|4933437F18Rik|MEX</td><td>zinc finger SWIM-type containing 2</td><td>protein-coding</td></tr>
<tr><td>chr2_83993719_83994531</td><td>chr2</td><td>83993720</td><td>83994531</td><td>+</td><td>-0.964416</td><td>Intergenic</td><td>Intergenic</td><td>-52899</td><td>NM_027964</td><td>71861</td><td>Mm.116957</td><td>NM_027964</td><td>ENSMUSG00000034552</td><td>Zswim2</td><td>1700025P14Rik|4933437F18Rik|MEX</td><td>zinc finger SWIM-type containing 2</td><td>protein-coding</td></tr>
<tr><td>chr2_84016207_84016754</td><td>chr2</td><td>84016208</td><td>84016754</td><td>+</td><td>-1.37038</td><td>Intergenic</td><td>Intergenic</td><td>-75255</td><td>NM_027964</td><td>71861</td><td>Mm.116957</td><td>NM_027964</td><td>ENSMUSG00000034552</td><td>Zswim2</td><td>1700025P14Rik|4933437F18Rik|MEX</td><td>zinc finger SWIM-type containing 2</td><td>protein-coding</td></tr>
<tr><td>chr2_84015962_84016730</td><td>chr2</td><td>84015963</td><td>84016730</td><td>+</td><td>-1.37075</td><td>Intergenic</td><td>B2_Mm2|SINE|B2</td><td>-75120</td><td>NM_027964</td><td>71861</td><td>Mm.116957</td><td>NM_027964</td><td>ENSMUSG00000034552</td><td>Zswim2</td><td>1700025P14Rik|4933437F18Rik|MEX</td><td>zinc finger SWIM-type containing 2</td><td>protein-coding</td></tr>
<tr><td>chr13_107831127_107831575</td><td>chr13</td><td>107831128</td><td>107831575</td><td>+</td><td>-0.648756</td><td>intron (NM_145456, intron 2 of 14)</td><td>intron (NM_145456, intron 2 of 14)</td><td>58713</td><td>NM_145456</td><td>67263</td><td>Mm.490418</td><td>NM_145456</td><td>ENSMUSG00000032846</td><td>Zswim6</td><td>2900036G02Rik|mKIAA1577</td><td>zinc finger SWIM-type containing 6</td><td>protein-coding</td></tr>
<tr><td>chr13_107868918_107869265</td><td>chr13</td><td>107868919</td><td>107869265</td><td>+</td><td>-0.753315</td><td>intron (NM_145456, intron 2 of 14)</td><td>intron (NM_145456, intron 2 of 14)</td><td>20972</td><td>NM_145456</td><td>67263</td><td>Mm.490418</td><td>NM_145456</td><td>ENSMUSG00000032846</td><td>Zswim6</td><td>2900036G02Rik|mKIAA1577</td><td>zinc finger SWIM-type containing 6</td><td>protein-coding</td></tr>
<tr><td>chr13_107843822_107844443</td><td>chr13</td><td>107843823</td><td>107844443</td><td>+</td><td>-0.767013</td><td>intron (NM_145456, intron 2 of 14)</td><td>intron (NM_145456, intron 2 of 14)</td><td>45931</td><td>NM_145456</td><td>67263</td><td>Mm.490418</td><td>NM_145456</td><td>ENSMUSG00000032846</td><td>Zswim6</td><td>2900036G02Rik|mKIAA1577</td><td>zinc finger SWIM-type containing 6</td><td>protein-coding</td></tr>
<tr><td>chr13_107837156_107837890</td><td>chr13</td><td>107837157</td><td>107837890</td><td>+</td><td>-0.78341</td><td>intron (NM_145456, intron 2 of 14)</td><td>intron (NM_145456, intron 2 of 14)</td><td>52541</td><td>NM_145456</td><td>67263</td><td>Mm.490418</td><td>NM_145456</td><td>ENSMUSG00000032846</td><td>Zswim6</td><td>2900036G02Rik|mKIAA1577</td><td>zinc finger SWIM-type containing 6</td><td>protein-coding</td></tr>
<tr><td>chr13_107844116_107844480</td><td>chr13</td><td>107844117</td><td>107844480</td><td>+</td><td>-0.846732</td><td>intron (NM_145456, intron 2 of 14)</td><td>intron (NM_145456, intron 2 of 14)</td><td>45766</td><td>NM_145456</td><td>67263</td><td>Mm.490418</td><td>NM_145456</td><td>ENSMUSG00000032846</td><td>Zswim6</td><td>2900036G02Rik|mKIAA1577</td><td>zinc finger SWIM-type containing 6</td><td>protein-coding</td></tr>
<tr><td>chr13_107902771_107903974</td><td>chr13</td><td>107902772</td><td>107903974</td><td>+</td><td>-0.853511</td><td>Intergenic</td><td>Intergenic</td><td>-13309</td><td>NM_145456</td><td>67263</td><td>Mm.490418</td><td>NM_145456</td><td>ENSMUSG00000032846</td><td>Zswim6</td><td>2900036G02Rik|mKIAA1577</td><td>zinc finger SWIM-type containing 6</td><td>protein-coding</td></tr>
<tr><td>chr13_107837169_107837592</td><td>chr13</td><td>107837170</td><td>107837592</td><td>+</td><td>-0.909262</td><td>intron (NM_145456, intron 2 of 14)</td><td>intron (NM_145456, intron 2 of 14)</td><td>52683</td><td>NM_145456</td><td>67263</td><td>Mm.490418</td><td>NM_145456</td><td>ENSMUSG00000032846</td><td>Zswim6</td><td>2900036G02Rik|mKIAA1577</td><td>zinc finger SWIM-type containing 6</td><td>protein-coding</td></tr>
<tr><td>chr4_108310612_108311208</td><td>chr4</td><td>108310613</td><td>108311208</td><td>+</td><td>-1.28335</td><td>Intergenic</td><td>Intergenic</td><td>-9820</td><td>NM_001033634</td><td>414872</td><td>Mm.426241</td><td>NM_001033634</td><td>ENSMUSG00000034636</td><td>Zyg11b</td><td>1110046I03Rik|2810482G21Rik|D4Mgi23|Gm431|mKIAA1730</td><td>zyg-ll family member B, cell cycle regulator</td><td>protein-coding</td></tr>
<tr><td>chr6_42343242_42344178</td><td>chr6</td><td>42343243</td><td>42344178</td><td>+</td><td>-0.698469</td><td>Intergenic</td><td>Intergenic</td><td>-6118</td><td>NM_011777</td><td>22793</td><td>Mm.282303</td><td>NM_011777</td><td>ENSMUSG00000029860</td><td>Zyx</td><td>9530098H06Rik|R75157</td><td>zyxin</td><td>protein-coding</td></tr>
</table>

**Table S2C_HOMER Closing peaks in both Cytokine-primed and Ag-stimulated cells.**
Data is sorted alphabetically by the gene name; Only peaks above the peak score value of 0.5 are shown.

| PeakID (cmd=annotatePeaks.pl ../averagescore_closing_peaks_BallCA.bed mm10 -go go_closing_peaks_BallCvA) | Chr | Start | End | Strand | Peak Score | Annotation | Detailed Annotation | Distance to TSS | Nearest PromoterID | Entrez ID | Nearest Unigene | Nearest Refseq | Nearest Ensembl | Gene Name | Gene Alias | Gene Description | Gene Type |
|---|---|---|---|---|---|---|---|---|---|---|---|---|---|---|---|---|---|
| chr2_132691701_132692100 | chr2 | 132691702 | 132692100 | + | -0.925234 | intron (NM_028637, intron 1 of 2) | intron (NM_028637, intron 1 of 2) | 1618 | NM_028637 | 73747 | Mm.284114 | NM_028637 | ENSMUSG00000044991 | **1110034G24Rik** | - | RIKEN cDNA 1110034G24 gene | protein-coding |
| chr16_95899720_95900588 | chr16 | 95899721 | 95900588 | + | -0.663766 | intron (NR_040484, intron 1 of 3) | intron (NR_040484, intron 1 of 3) | -1441 | NR_040485 | 69776 | Mm.120470 | NR_040484 | ENSMUSG00000086899 | **1600002D24Rik** | - | RIKEN cDNA 1600002D24 gene | ncRNA |
| chr16_95901395_95901682 | chr16 | 95901396 | 95901682 | + | -0.814961 | intron (NR_040484, intron 1 of 3) | intron (NR_040484, intron 1 of 3) | -2826 | NR_040485 | 69776 | Mm.120470 | NR_040484 | ENSMUSG00000086899 | **1600002D24Rik** | - | RIKEN cDNA 1600002D24 gene | ncRNA |
| chr16_95860698_95861353 | chr16 | 95860699 | 95861353 | + | -0.846939 | intron (NR_040484, intron 1 of 3) | intron (NR_040484, intron 1 of 3) | 37687 | NR_040485 | 69776 | Mm.120470 | NR_040484 | ENSMUSG00000086899 | **1600002D24Rik** | - | RIKEN cDNA 1600002D24 gene | ncRNA |
| chr16_95860787_95861174 | chr16 | 95860788 | 95861174 | + | -0.862702 | intron (NR_040484, intron 1 of 3) | intron (NR_040484, intron 1 of 3) | 37732 | NR_040485 | 69776 | Mm.120470 | NR_040484 | ENSMUSG00000086899 | **1600002D24Rik** | - | RIKEN cDNA 1600002D24 gene | ncRNA |
| chr16_95900040_95900625 | chr16 | 95900041 | 95900625 | + | -0.864233 | intron (NR_040484, intron 1 of 3) | intron (NR_040484, intron 1 of 3) | -1620 | NR_040485 | 69776 | Mm.120470 | NR_040484 | ENSMUSG00000086899 | **1600002D24Rik** | - | RIKEN cDNA 1600002D24 gene | ncRNA |
| chr1_183987633_183988190 | chr1 | 183987634 | 183988190 | + | -0.87025 | Intergenic | Intergenic | 46086 | NM_028516 | 73363 | Mm.50115 | NM_028516 | ENSMUSG00000044854 | **1700056E22Rik** | - | RIKEN cDNA 1700056E22 gene | protein-coding |
| chr18_75323821_75324179 | chr18 | 75323822 | 75324179 | + | -0.824752 | Intergenic | Intergenic | -26671 | NR_045393 | 100503117 | Mm.250517 | NR_045393 | | **2010010A06Rik** | 5033420E22Rik | RIKEN cDNA 2010010A06 gene | ncRNA |
| chr18_75304375_75305489 | chr18 | 75304376 | 75305489 | + | -0.873443 | Intergenic | Intergenic | -7603 | NR_045393 | 100503117 | Mm.250517 | NR_045393 | | **2010010A06Rik** | 5033420E22Rik | RIKEN cDNA 2010010A06 gene | ncRNA |
| chr18_75304315_75305067 | chr18 | 75304316 | 75305067 | + | -0.959233 | Intergenic | Intergenic | -7362 | NR_045393 | 100503117 | Mm.250517 | NR_045393 | | **2010010A06Rik** | 5033420E22Rik | RIKEN cDNA 2010010A06 gene | ncRNA |
| chr6_31221602_31221903 | chr6 | 31221603 | 31221903 | + | -0.939769 | intron (NR_040260, intron 1 of 5) | intron (NR_040260, intron 1 of 5) | 1402 | NR_040261 | 73652 | Mm.441297 | NR_040257 | ENSMUSG00000087380 | **2210408F21Rik** | - | RIKEN cDNA 2210408F21 gene | ncRNA |
| chr19_5405985_5406340 | chr19 | 5405986 | 5406340 | + | -0.693402 | promoter-TSS (NR_015545) | promoter-TSS (NR_015545) | -652 | NR_015545 | 74931 | Mm.388511 | NR_015545 | ENSMUSG00000086938 | **4930481A15Rik** | - | RIKEN cDNA 4930481A15 gene | ncRNA |
| chr1_36723118_36723653 | chr1 | 36723119 | 36723653 | + | -0.705867 | non-coding (NR_040290, exon 3 of 9) | non-coding (NR_040290, exon 3 of 9) | 8848 | NR_040290 | 71166 | Mm.157049 | NR_040290 | | **4933424G06Rik** | 1700104I15Rik | RIKEN cDNA 4933424G06 gene | ncRNA |
| chr9_118390703_118391123 | chr9 | 118390704 | 118391123 | + | -1.04279 | Intergenic | Intergenic | -38926 | NR_040716 | 71196 | Mm.73218 | NR_040716 | | **4933432G23Rik** | - | RIKEN cDNA 4933432G23 gene | ncRNA |
| chr17_84006786_84007167 | chr17 | 84006787 | 84007167 | + | -0.742869 | Intergenic | L1MA10\|LINE\|L1 | -71683 | NR_045458 | 74473 | Mm.399575 | NR_045458 | | **4933433H22Rik** | - | RIKEN cDNA 4933433H22 gene | ncRNA |
| chr2_59338374_59338812 | chr2 | 59338375 | 59338812 | + | -0.890359 | intron (NM_026361, intron 13 of 21) | intron (NM_026361, intron 13 of 21) | 48089 | NR_040510 | 76676 | Mm.110038 | NR_040510 | ENSMUSG00000087455 | **5330411J11Rik** | - | RIKEN cDNA 5330411J11 gene | ncRNA |
| chr13_24921296_24921856 | chr13 | 24921297 | 24921856 | + | -1.12217 | intron (NM_172532, intron 4 of 9) | intron (NM_172532, intron 4 of 9) | -15825 | NR_102323 | 493800 | Mm.458692 | NR_102323 | | **9330162012Rik** | - | cDNA RIKEN 9330162012 gene | ncRNA |
| chr2_75458078_75458547 | chr2 | 75458079 | 75458547 | + | -1.05982 | Intergenic | Intergenic | -14134 | NR_040635 | 77269 | Mm.389278 | NR_040635 | | **9430019J16Rik** | - | RIKEN cDNA 9430019J16 gene | ncRNA |
| chr14_53878677_53878921 | chr14 | 53878678 | 53878921 | + | -1.01376 | Intergenic | RMER15-int\|LTR\|ERVL | 212859 | NM_001205242 | 100568452 | Mm.213248 | NM_001205242 | | **A430107P09Rik** | Va10 | RIKEN cDNA A430107P09 gene | protein-coding |
| chr2_167777342_167778653 | chr2 | 167777343 | 167778653 | + | -0.686111 | Intergenic | Intergenic | 86821 | NR_015500 | 329562 | Mm.123101 | NM_177851 | ENSMUSG00000006462 | **A530013C23Rik** | - | RIKEN cDNA A530013C23 gene | ncRNA |
| chr2_167777349_167778277 | chr2 | 167777350 | 167778277 | + | -0.704176 | Intergenic | Intergenic | 86636 | NR_015500 | 329562 | Mm.123101 | NM_177851 | ENSMUSG00000006462 | **A530013C23Rik** | - | RIKEN cDNA A530013C23 gene | ncRNA |
| chr2_167777689_167778551 | chr2 | 167777690 | 167778551 | + | -0.743664 | Intergenic | Intergenic | 86943 | NR_015500 | 329562 | Mm.123101 | NM_177851 | ENSMUSG00000006462 | **A530013C23Rik** | - | RIKEN cDNA A530013C23 gene | ncRNA |
| chr2_167777703_167778191 | chr2 | 167777704 | 167778191 | + | -0.751272 | Intergenic | Intergenic | 86770 | NR_015500 | 329562 | Mm.123101 | NM_177851 | ENSMUSG00000006462 | **A530013C23Rik** | - | RIKEN cDNA A530013C23 gene | ncRNA |
| chr5_122989013_122989663 | chr5 | 122989014 | 122989663 | + | -0.838047 | promoter-TSS (NR_045820) | promoter-TSS (NR_045820) | -16 | NR_045820 | 109202 | Mm.30769 | NM_175405 | ENSMUSG00000056735 | **A930024E05Rik** | - | RIKEN cDNA A930024E05 gene | ncRNA |
| chr5_122990470_122990823 | chr5 | 122990471 | 122990823 | + | -0.911817 | intron (NR_045820, intron 1 of 2) | intron (NR_045820, intron 1 of 2) | 1293 | NR_045820 | 109202 | Mm.30769 | NM_175405 | ENSMUSG00000056735 | **A930024E05Rik** | - | RIKEN cDNA A930024E05 gene | ncRNA |
| chr5_122990341_122990929 | chr5 | 122990342 | 122990929 | + | -0.931355 | intron (NR_045820, intron 1 of 2) | intron (NR_045820, intron 1 of 2) | 1281 | NR_045820 | 109202 | Mm.30769 | NM_175405 | ENSMUSG00000056735 | **A930024E05Rik** | - | RIKEN cDNA A930024E05 gene | ncRNA |
| chr5_8838752_8839476 | chr5 | 8838753 | 8839476 | + | -0.735718 | intron (NM_011075, intron 21 of 27) | intron (NM_011075, intron 21 of 27) | 40967 | NM_011075 | 18669 | Mm.146649 | NM_011075 | ENSMUSG00000028970 | **Abcb1b** | Abcb1\|Mdr1\|Mdr1b\|Pgy-1\|Pgy1\|mdr | ATP-binding cassette, sub-family B (MDR/TAP), member 1B | protein-coding |
| chr7_79300507_79301293 | chr7 | 79300508 | 79301293 | + | -0.801949 | intron (NM_018811, intron 3 of 10) | intron (NM_018811, intron 3 of 10) | 27634 | NM_018811 | 54608 | Mm.365490 | NM_018811 | | **Abhd2** | 2210009N18Rik\|LABH2\|Labh-2 | abhydrolase domain containing 2 | protein-coding |
| chr16_23170786_23171252 | chr16 | 23170787 | 23171252 | + | -0.989116 | Intergenic | RMER19C\|LTR\|ERVK | 24483 | NM_009605 | 11450 | Mm.3969 | NM_009605 | ENSMUSG00000022878 | **Adipoq** | 30kDa\|APN\|Acdc\|Acrp30\|GBP28\|adipo\|apM1 | adiponectin, C1Q and collagen domain containing | protein-coding |
| chr6_140816877_140817496 | chr6 | 140816878 | 140817496 | + | -0.835033 | Intergenic | Intergenic | 192889 | NM_009637 | 11569 | Mm.86453 | NM_009637 | ENSMUSG00000030232 | **Aebp2** | AU023766\|B230313N05Rik | AE binding protein 2 | protein-coding |
| chr6_140819347_140819929 | chr6 | 140819348 | 140819929 | + | -0.888626 | Intergenic | ORR1D2\|LTR\|ERVL-MaLR | 195340 | NM_009637 | 11569 | Mm.86453 | NM_009637 | ENSMUSG00000030232 | **Aebp2** | AU023766\|B230313N05Rik | AE binding protein 2 | protein-coding |
| chr11_53351474_53352040 | chr11 | 53351475 | 53352040 | + | -0.638728 | intron (NM_033565, intron 1 of 20) | intron (NM_033565, intron 1 of 20) | 990 | NM_033565 | 93736 | Mm.395281 | NM_033565 | ENSMUSG00000049470 | **Aff4** | AF5Q31\|Alf4\|Laf4l\|MCEF | AF4/FMR2 family, member 4 | protein-coding |
| chr11_53351715_53352000 | chr11 | 53351716 | 53352000 | + | -0.666513 | intron (NM_033565, intron 1 of 20) | intron (NM_033565, intron 1 of 20) | 1091 | NM_033565 | 93736 | Mm.395281 | NM_033565 | ENSMUSG00000049470 | **Aff4** | AF5Q31\|Alf4\|Laf4l\|MCEF | AF4/FMR2 family, member 4 | protein-coding |
| chr15_73137469_73138256 | chr15 | 73137470 | 73138256 | + | -0.812282 | intron (NM_153178, intron 3 of 18) | intron (NM_153178, intron 3 of 18) | 47084 | NM_153178 | 239528 | Mm.23095 | NM_153178 | | **Ago2** | 1110029L17Rik\|2310051F07Rik\|AI225898\|AL022874\|AW546247\|ENSMUSG00000072493\|Eif2c2\|Gerp95\|Gm10365\|mKIAA4215 | argonaute RISC catalytic subunit 2 | protein-coding |
| chr15_73137732_73138206 | chr15 | 73137733 | 73138206 | + | -0.903895 | intron (NM_153178, intron 3 of 18) | intron (NM_153178, intron 3 of 18) | 46978 | NM_153178 | 239528 | Mm.23095 | NM_153178 | | **Ago2** | 1110029L17Rik\|2310051F07Rik\|AI225898\|AL022874\|AW546247\|ENSMUSG00000072493\|Eif2c2\|Gerp95\|Gm10365\|mKIAA4215 | argonaute RISC catalytic subunit 2 | protein-coding |
| chr8_123000248_123000708 | chr8 | 123000249 | 123000708 | + | -0.917809 | intron (NR_037865, intron 1 of 13) | intron (NR_037865, intron 1 of 13) | 41806 | NR_037865 | 77087 | Mm.31567 | NM_001081379 | ENSMUSG00000035569 | **Ankrd11** | 2410104C19Rik\|3010027A04Rik\|6330578C09Rik\|9530048I21Rik\|AA930108\|Gm176\|Yod | ankyrin repeat domain 11 | protein-coding |
| chr2_43727557_43728279 | chr2 | 43727558 | 43728279 | + | -0.793302 | Intergenic | Intergenic | -20906 | NM_153820 | 76117 | Mm.217350 | NM_153820 | ENSMUSG00000049744 | **Arhgap15** | 5830480G12Rik | Rho GTPase activating protein 15 | protein-coding |
| chr6_87502863_87503288 | chr6 | 87502864 | 87503288 | + | -0.7349 | intron (NM_001037727, intron 1 of 10) | intron (NM_001037727, intron 1 of 10) | -6749 | NM_175476 | 232201 | Mm.119564 | NM_175476 | | **Arhgap25** | A130039I20Rik | Rho GTPase activating protein 25 | protein-coding |
| chr7_101538256_101538486 | chr7 | 101538257 | 101538486 | + | -0.71399 | Intergenic | Intergenic | 17431 | NM_007490 | 11871 | Mm.439746 | NM_007490 | ENSMUSG00000092517 | **Art2a-ps** | ARTC2\|Art2a\|Ly92a\|Rt-6\|Rt6\|Rt6-1 | ADP-ribosyltransferase 2a, pseudogene | protein-coding |
| chr10_81194063_81194682 | chr10 | 81194064 | 81194682 | + | -0.695613 | promoter-TSS (NR_015477) | promoter-TSS (NR_015477) | -318 | NR_015477 | 69678 | Mm.81562 | NR_015477 | | **Atcayos** | 2310050B05Rik | ataxia, cerebellar, Cayman type homolog (human), opposite strand | ncRNA |
| chr1_133762852_133763673 | chr1 | 133762853 | 133763673 | + | -0.650249 | Intergenic | Intergenic | -9516 | NM_213616 | 381290 | Mm.188617 | NM_213616 | ENSMUSG00000026463 | **Atp2b4** | Pmca4 | ATPase, Ca++ transporting, plasma membrane 4 | protein-coding |
| chr1_133762997_133763604 | chr1 | 133762998 | 133763604 | + | -0.666364 | Intergenic | Intergenic | -9554 | NM_213616 | 381290 | Mm.188617 | NM_213616 | ENSMUSG00000026463 | **Atp2b4** | Pmca4 | ATPase, Ca++ transporting, plasma membrane 4 | protein-coding |
| chr4_32351934_32352244 | chr4 | 32351935 | 32352244 | + | -0.63354 | Intergenic | MER119\|DNA\|hAT-Charlie | -65346 | NM_001109661 | 12014 | Mm.491223 | NM_007521 | ENSMUSG00000040270 | **Bach2** | E030004N02Rik | BTB and CNC homology 2 | protein-coding |
| chr4_32343650_32344700 | chr4 | 32343651 | 32344700 | + | -0.674521 | Intergenic | Intergenic | -73260 | NM_001109661 | 12014 | Mm.491223 | NM_007521 | ENSMUSG00000040270 | **Bach2** | E030004N02Rik | BTB and CNC homology 2 | protein-coding |
| chr4_32363583_32364191 | chr4 | 32363584 | 32364191 | + | -0.758217 | Intergenic | Intergenic | -53548 | NM_001109661 | 12014 | Mm.491223 | NM_007521 | ENSMUSG00000040270 | **Bach2** | E030004N02Rik | BTB and CNC homology 2 | protein-coding |
| chr4_32363349_32364219 | chr4 | 32363350 | 32364219 | + | -0.766204 | Intergenic | Intergenic | -53651 | NM_001109661 | 12014 | Mm.491223 | NM_007521 | ENSMUSG00000040270 | **Bach2** | E030004N02Rik | BTB and CNC homology 2 | protein-coding |
| chr3_145925618_145925830 | chr3 | 145925619 | 145925830 | + | -0.766093 | intron (NM_009740, intron 1 of 2) | intron (NM_009740, intron 1 of 2) | 1462 | NM_009740 | 12042 | Mm.239141 | NM_009740 | ENSMUSG00000028191 | **Bcl10** | AI132454\|BCL-10\|C81403\|CARMEN\|CIPER\|CLAP\|ME10\|cE10 | B cell leukemia/lymphoma 10 | protein-coding |
| chr5_67394730_67395861 | chr5 | 67394731 | 67395861 | + | -0.711189 | 3' UTR (NM_001164806, exon 5 of 5) | 3' UTR (NM_001164806, exon 5 of 5) | 32503 | NM_001164806 | 666938 | Mm.101339 | NM_001164806 | ENSMUSG00000092060 | **Bend4** | D330027G24Rik\|EG666938 | BEN domain containing 4 | protein-coding |
| chr5_67395197_67395896 | chr5 | 67395198 | 67395896 | + | -0.830771 | 3' UTR (NM_001164806, exon 5 of 5) | 3' UTR (NM_001164806, exon 5 of 5) | 32252 | NM_001164806 | 666938 | Mm.101339 | NM_001164806 | ENSMUSG00000092060 | **Bend4** | D330027G24Rik\|EG666938 | BEN domain containing 4 | protein-coding |
| chr1_134058883_134059281 | chr1 | 134058884 | 134059281 | + | -1.21351 | Intergenic | Intergenic | 20073 | NM_007570 | 12227 | Mm.392646 | NM_007570 | ENSMUSG00000020423 | **Btg2** | AA959598\|APRO1\|Pc3\|TIS21 | B cell translocation gene 2, anti-proliferative | protein-coding |
| chr5_64518862_64519395 | chr5 | 64518863 | 64519395 | + | -0.720473 | Intergenic | Intergenic | 42121 | NR_045413 | 77382 | Mm.366742 | NR_045411 | | **C030018K13Rik** | - | RIKEN cDNA C030018K13 gene | ncRNA |
| chr5_64498340_64498840 | chr5 | 64498341 | 64498840 | + | -1.01893 | Intergenic | Intergenic | 21582 | NR_045413 | 77382 | Mm.366742 | NR_045411 | | **C030018K13Rik** | - | RIKEN cDNA C030018K13 gene | ncRNA |
| chr15_100604623_100605434 | chr15 | 100604624 | 100605434 | + | -1.04493 | Intergenic | Intergenic | 10081 | NR_045701 | 78445 | Mm.44383 | NR_045701 | ENSMUSG00000097093 | **C330013E15Rik** | - | RIKEN cDNA C330013E15 gene | ncRNA |

| | | | | | | | | | | | | | | | | | |
|---|---|---|---|---|---|---|---|---|---|---|---|---|---|---|---|---|---|
| chr10_22299428_22299683 | chr10 | 22299429 | 22299683 | + | -0.831136 | intron (NM_009018, intron 5 of 5) | intron (NM_009018, intron 5 of 5) | -7164 | NR_015465 | 606736 | Mm.387042 | NR_015465 | ENSMUSG00000097352 | **C920009B18Rik** | - | RIKEN cDNA C920009B18 gene | ncRNA |
| chr10_22299319_22299716 | chr10 | 22299320 | 22299716 | + | -0.871535 | intron (NM_009018, intron 5 of 5) | intron (NM_009018, intron 5 of 5) | -7202 | NR_015465 | 606736 | Mm.387042 | NR_015465 | ENSMUSG00000097352 | **C920009B18Rik** | - | RIKEN cDNA C920009B18 gene | ncRNA |
| chr18_32970174_32970666 | chr18 | 32970175 | 32970666 | + | -0.819633 | intron (NM_009793, intron 1 of 10) | intron (NM_009793, intron 1 of 10) | 31379 | NM_009793 | 12326 | Mm.222329 | NM_009793 | ENSMUSG00000038128 | **Camk4** | A430110E23Rik\|AI666733\|CaMKIV\|CaMKIV/Gr\|D18Bwg0362e | calcium/calmodulin-dependent protein kinase IV | protein-coding |
| chr18_32970174_32970909 | chr18 | 32970175 | 32970909 | + | -0.845578 | intron (NM_009793, intron 1 of 10) | intron (NM_009793, intron 1 of 10) | 31501 | NM_009793 | 12326 | Mm.222329 | NM_009793 | ENSMUSG00000038128 | **Camk4** | A430110E23Rik\|AI666733\|CaMKIV\|CaMKIV/Gr\|D18Bwg0362e | calcium/calmodulin-dependent protein kinase IV | protein-coding |
| chr18_32958142_32958609 | chr18 | 32958143 | 32958609 | + | -1.06488 | intron (NM_009793, intron 1 of 10) | L2\|LINE\|L2 | 19335 | NM_009793 | 12326 | Mm.222329 | NM_009793 | ENSMUSG00000038128 | **Camk4** | A430110E23Rik\|AI666733\|CaMKIV\|CaMKIV/Gr\|D18Bwg0362e | calcium/calmodulin-dependent protein kinase IV | protein-coding |
| chr16_52143084_52143560 | chr16 | 52143085 | 52143560 | + | -0.964475 | intron (NM_001033238, intron 9 of 17) | intron (NM_001033238, intron 9 of 17) | 111773 | NM_001033238 | 208650 | Mm.328206 | NM_001033238 | ENSMUSG00000022637 | **Cblb** | AI429560\|AI851073\|Cbl-b | Casitas B-lineage lymphoma b | protein-coding |
| chr16_52142791_52143602 | chr16 | 52142792 | 52143602 | + | -0.975752 | intron (NM_001033238, intron 9 of 17) | intron (NM_001033238, intron 9 of 17) | 111648 | NM_001033238 | 208650 | Mm.328206 | NM_001033238 | ENSMUSG00000022637 | **Cblb** | AI429560\|AI851073\|Cbl-b | Casitas B-lineage lymphoma b | protein-coding |
| chr1_165837439_165838249 | chr1 | 165837440 | 165838249 | + | -0.659291 | intron (NM_031162, intron 1 of 7) | intron (NM_031162, intron 1 of 7) | 49163 | NM_001113392 | 12503 | Mm.217308 | NM_031162 | ENSMUSG00000005763 | **Cd247** | 4930549J05Rik\|A430104F18Rik\|AW552088\|Cd3\|Cd3-eta\|Cd3-zeta\|Cd3h\|Cd3z\|Cd3zeta\|T3z\|Tcrk\|Tcrz | CD247 antigen | protein-coding |
| chr1_60755819_60756231 | chr1 | 60755820 | 60756231 | + | -0.819622 | intron (NM_007642, intron 1 of 3) | intron (NM_007642, intron 1 of 3) | 9637 | NM_007642 | 12487 | Mm.255003 | NM_007642 | ENSMUSG00000026012 | **Cd28** | - | CD28 antigen | protein-coding |
| chr16_49792406_49793000 | chr16 | 49792407 | 49793000 | + | -0.733776 | Intergenic | Intergenic | -62951 | NM_010581 | 16423 | Mm.31752 | NM_010581 | ENSMUSG00000055447 | **Cd47** | 9130415E20Rik\|AA407862\|AI848868\|AW108519\|B430305P08Rik\|IAP\|Itgp | CD47 antigen (Rh-related antigen, integrin-associated signal transducer) | protein-coding |
| chr2_6705477_6706564 | chr2 | 6705478 | 6706564 | + | -0.637579 | intron (NM_001110232, intron 4 of 15) | intron (NM_001110232, intron 4 of 15) | 178975 | NM_010160 | 14007 | Mm.398543 | NM_010160 | ENSMUSG00000002107 | **Celf2** | B230218O03\|B230345P09Rik\|C88023\|CELF-2\|CUG-BP2\|Cugbp2\|D230046B21Rik\|Etr-3\|Napor\|Napor-2\|mETR-3 | CUGBP, Elav-like family member 2 | protein-coding |
| chr2_6705962_6706564 | chr2 | 6705963 | 6706564 | + | -0.71188 | intron (NM_001110232, intron 4 of 15) | intron (NM_001110232, intron 4 of 15) | 178733 | NM_010160 | 14007 | Mm.398543 | NM_010160 | ENSMUSG00000002107 | **Celf2** | B230218O03\|B230345P09Rik\|C88023\|CELF-2\|CUG-BP2\|Cugbp2\|D230046B21Rik\|Etr-3\|Napor\|Napor-2\|mETR-3 | CUGBP, Elav-like family member 2 | protein-coding |
| chr2_68959564_68959979 | chr2 | 68959565 | 68959979 | + | -0.85977 | intron (NM_172856, intron 3 of 9) | intron (NM_172856, intron 3 of 9) | 98215 | NM_172856 | 241447 | Mm.222222 | NM_172856 | ENSMUSG00000027035 | **Cers6** | 4732462C07Rik\|AW544719\|Lass6\|T1L | ceramide synthase 6 | protein-coding |
| chr7_73523287_73523501 | chr7 | 73523288 | 73523501 | + | -1.10838 | intron (NM_001081345, intron 2 of 38) | intron (NM_001081345, intron 2 of 38) | 18352 | NM_001081345 | 244059 | Mm.34539 | NM_001081345 | ENSMUSG00000078671 | **Chd2** | 2810013C04Rik\|2810040A01Rik\|5630401D06Rik\|AI851092\|BC029703\|CHD-2 | chromodomain helicase DNA binding protein 2 | protein-coding |
| chr7_73523174_73523601 | chr7 | 73523175 | 73523601 | + | -1.1566 | intron (NM_001081345, intron 2 of 38) | intron (NM_001081345, intron 2 of 38) | 18358 | NM_001081345 | 244059 | Mm.34539 | NM_001081345 | ENSMUSG00000078671 | **Chd2** | 2810013C04Rik\|2810040A01Rik\|5630401D06Rik\|AI851092\|BC029703\|CHD-2 | chromodomain helicase DNA binding protein 2 | protein-coding |
| chr9_95314858_95315516 | chr9 | 95314859 | 95315516 | + | -0.729452 | Intergenic | Intergenic | 92083 | NM_018763 | 54371 | Mm.212446 | NM_018763 | ENSMUSG00000033350 | **Chst2** | AI428561\|AW121776\|C130041E03Rik\|Chts2\|GST-2\|GlcNAc6ST\|Gn6st | carbohydrate sulfotransferase 2 | protein-coding |
| chr9_95113255_95113827 | chr9 | 95113256 | 95113827 | + | -1.19984 | intron (NM_177909, intron 12 of 15) | intron (NM_177909, intron 12 of 15) | 293729 | NM_018763 | 54371 | Mm.212446 | NM_018763 | ENSMUSG00000033350 | **Chst2** | AI428561\|AW121776\|C130041E03Rik\|Chts2\|GST-2\|GlcNAc6ST\|Gn6st | carbohydrate sulfotransferase 2 | protein-coding |
| chr13_24415259_24416182 | chr13 | 24415260 | 24416182 | + | -0.668079 | intron (NM_001284520, intron 1 of 14) | intron (NM_001284520, intron 1 of 14) | 599 | NM_001284520 | 12763 | Mm.8396 | NM_007717 | ENSMUSG00000016756 | **Cmah** | - | cytidine monophospho-N-acetylneuraminic acid hydroxylase | protein-coding |
| chr6_142757611_142757914 | chr6 | 142757612 | 142757914 | + | -0.887112 | intron (NM_009908, intron 1 of 7) | intron (NM_009908, intron 1 of 7) | 1077 | NM_009908 | 12764 | Mm.3820 | NM_009908 | ENSMUSG00000030282 | **Cmas** | AW208911\|CMPNeu5Ac\|D6Bwg0250e | cytidine monophospho-N-acetylneuraminic acid synthetase | protein-coding |
| chr1_179549409_179549748 | chr1 | 179549410 | 179549748 | + | -0.742201 | intron (NM_146105, intron 1 of 10) | intron (NM_146105, intron 1 of 10) | 3050 | NM_146105 | 226744 | Mm.87662 | NM_146105 | ENSMUSG00000038949 | **Cnst** | 9630058J23Rik | consortin, connexin sorting protein | protein-coding |
| chr6_106792891_106793895 | chr6 | 106792892 | 106793895 | + | -1.05497 | intron (NM_175357, intron 4 of 10) | intron (NM_175357, intron 4 of 10) | 6694 | NM_021449 | 58799 | Mm.290085 | NM_021449 | | **Crbn** | 2610203G15Rik\|2900045O07Rik\|AF229032\|AW108261\|piL | cereblon | protein-coding |
| chr6_106792919_106793563 | chr6 | 106792920 | 106793563 | + | -1.24143 | intron (NM_175357, intron 4 of 10) | intron (NM_175357, intron 4 of 10) | 6846 | NM_021449 | 58799 | Mm.290085 | NM_021449 | | **Crbn** | 2610203G15Rik\|2900045O07Rik\|AF229032\|AW108261\|piL | cereblon | protein-coding |
| chr2_30625214_30626629 | chr2 | 30625215 | 30626629 | + | -0.730023 | Intergenic | MTE2a\|LTR\|ERVL-MaLR | 30878 | NM_030137 | 78617 | Mm.40413 | NM_030137 | ENSMUSG00000047363 | **Cstad** | 9530039B02Rik | CSA-conditional, T cell activation-dependent protein | protein-coding |
| chr2_30625873_30626611 | chr2 | 30625874 | 30626611 | + | -0.827758 | Intergenic | ORR1D2-int\|LTR\|ERVL-MaLR | 31198 | NM_030137 | 78617 | Mm.40413 | NM_030137 | ENSMUSG00000047363 | **Cstad** | 9530039B02Rik | CSA-conditional, T cell activation-dependent protein | protein-coding |
| chr7_114494055_114494463 | chr7 | 114494056 | 114494463 | + | -0.867776 | intron (NM_011055, intron 2 of 15) | intron (NM_011055, intron 2 of 15) | 68713 | NR_110407 | 244209 | Mm.108037 | NM_177382 | ENSMUSG00000030670 | **Cyp2r1** | - | cytochrome P450, family 2, subfamily r, polypeptide 1 | protein-coding |
| chr15_78594782_78595208 | chr15 | 78594783 | 78595208 | + | -0.786908 | Intergenic | MTE2b\|LTR\|ERVL-MaLR | -2052 | NM_028195 | 72318 | Mm.32911 | NM_028195 | ENSMUSG00000018008 | **Cyth4** | 2510004M07Rik\|5830469K17Rik\|AI467541\|Pscd4\|mFLJ00017 | cytohesin 4 | protein-coding |
| chr14_54231123_54231478 | chr14 | 54231124 | 54231478 | + | -0.765612 | Intergenic | RSINE1\|SINE\|B4 | 22628 | NM_001113358 | 13135 | Mm.319038 | NM_010015 | ENSMUSG00000022174 | **Dad1** | AI323713 | defender against cell death 1 | protein-coding |
| chr11_106026770_106028187 | chr11 | 106026771 | 106028187 | + | -0.789504 | intron (NM_001037712, intron 11 of 12) | intron (NM_001037712, intron 11 of 12) | -9393 | NM_027946 | 71833 | Mm.307455 | NM_027946 | ENSMUSG00000049354 | **Dcaf7** | 1700012F10Rik\|2610037L01Rik\|C86529\|HAN11\|Wdr68 | DDB1 and CUL4 associated factor 7 | protein-coding |
| chr11_106033501_106033721 | chr11 | 106033502 | 106033721 | + | -0.921167 | intron (NM_001037712, intron 12 of 12) | intron (NM_001037712, intron 12 of 12) | -3261 | NM_027946 | 71833 | Mm.307455 | NM_027946 | ENSMUSG00000049354 | **Dcaf7** | 1700012F10Rik\|2610037L01Rik\|C86529\|HAN11\|Wdr68 | DDB1 and CUL4 associated factor 7 | protein-coding |
| chr11_106033340_106033755 | chr11 | 106033341 | 106033755 | + | -0.969903 | intron (NM_001037712, intron 12 of 12) | IAPLTR1_Mm\|LTR\|ERVK | -3324 | NM_027946 | 71833 | Mm.307455 | NM_027946 | ENSMUSG00000049354 | **Dcaf7** | 1700012F10Rik\|2610037L01Rik\|C86529\|HAN11\|Wdr68 | DDB1 and CUL4 associated factor 7 | protein-coding |
| chr9_44579320_44579786 | chr9 | 44579321 | 44579786 | + | -0.766074 | Intergenic | Intergenic | -25339 | NM_001110826 | 13209 | Mm.267061 | NM_007841 | ENSMUSG00000032097 | **Ddx6** | 1110001P04Rik\|E230023J21Rik\|HLR2\|mRCK/P54\|p54\|rck | DEAD (Asp-Glu-Ala-Asp) box polypeptide 6 | protein-coding |
| chr11_86810271_86811754 | chr11 | 86810272 | 86811754 | + | -0.902321 | intron (NR_106195, intron 2 of 8) | intron (NR_106195, intron 2 of 8) | -3353 | NM_026191 | 67487 | Mm.260627 | NM_026191 | ENSMUSG00000018425 | **Dhx40** | 2410016C14Rik\|ARG147\|DDX40\|PAD | DEAH (Asp-Glu-Ala-His) box polypeptide 40 | protein-coding |
| chr11_86822354_86823068 | chr11 | 86822355 | 86823068 | + | -1.17796 | intron (NR_106195, intron 2 of 8) | RMER19B\|LTR\|ERVK | -15051 | NM_026191 | 67487 | Mm.260627 | NM_026191 | ENSMUSG00000018425 | **Dhx40** | 2410016C14Rik\|ARG147\|DDX40\|PAD | DEAH (Asp-Glu-Ala-His) box polypeptide 40 | protein-coding |
| chr15_100041368_100041561 | chr15 | 100041369 | 100041561 | + | -0.774511 | intron (NM_001159361, intron 1 of 37) | intron (NM_001159361, intron 1 of 37) | 2801 | NM_001159361 | 239667 | Mm.243658 | NM_172819 | ENSMUSG00000023026 | **Dip2b** | 4932422C22\|AI317237\|AI854602\|mKIAA1463 | DIP2 disco-interacting protein 2 homolog B (Drosophila) | protein-coding |
| chr11_118145697_118146415 | chr11 | 118145698 | 118146415 | + | -0.740121 | Intergenic | Intergenic | -16837 | NM_001167746 | 69926 | Mm.473893 | NM_001167746 | ENSMUSG00000033987 | **Dnah17** | 2810003K23Rik\|Dnahc17\|Dnahcl1\|Gm1178\|mKIAA3028 | dynein, axonemal, heavy chain 17 | protein-coding |
| chr4_101495790_101496157 | chr4 | 101495791 | 101496157 | + | -0.865046 | promoter-TSS (NM_198412) | promoter-TSS (NM_198412) | -674 | NM_198412 | 72685 | Mm.76494 | NM_198412 | ENSMUSG00000028528 | **Dnajc6** | 2810027M23Rik\|mKIAA0473 | DnaJ (Hsp40) homolog, subfamily C, member 6 | protein-coding |
| chr1_80755697_80756095 | chr1 | 80755698 | 80756095 | + | -1.11556 | intron (NM_175291, intron 1 of 55) | intron (NM_175291, intron 1 of 55) | 2657 | NM_175291 | 210293 | Mm.133473 | NM_175291 | ENSMUSG00000038608 | **Dock10** | 9330153B10Rik\|A630054M16Rik\|Jr4\|Jr5\|R75174\|ZIZ3\|Zizimin3\|mKIAA0694 | dedicator of cytokinesis 10 | protein-coding |
| chr1_80755731_80756009 | chr1 | 80755732 | 80756009 | + | -1.14122 | intron (NM_175291, intron 1 of 55) | intron (NM_175291, intron 1 of 55) | 2683 | NM_175291 | 210293 | Mm.133473 | NM_175291 | ENSMUSG00000038608 | **Dock10** | 9330153B10Rik\|A630054M16Rik\|Jr4\|Jr5\|R75174\|ZIZ3\|Zizimin3\|mKIAA0694 | dedicator of cytokinesis 10 | protein-coding |
| chr1_184066590_184067145 | chr1 | 184066591 | 184067145 | + | -0.820024 | intron (NM_022019, intron 2 of 3) | intron (NM_022019, intron 2 of 3) | 32407 | NM_022019 | 63953 | Mm.404024 | NM_022019 | ENSMUSG00000039384 | **Dusp10** | 2610306G15Rik\|AI158871\|MKP-5\|MKP5 | dual specificity phosphatase 10 | protein-coding |
| chr10_118714040_118714879 | chr10 | 118714041 | 118714879 | + | -0.677194 | Intergenic | Intergenic | 154443 | NM_001014390 | 69181 | Mm.45565 | NM_001014390 | ENSMUSG00000028630 | **Dyrk2** | 1810038L18Rik | dual-specificity tyrosine-(Y)-phosphorylation regulated kinase 2 | protein-coding |
| chr10_18197009_18197833 | chr10 | 18197010 | 18197833 | + | -0.686433 | intron (NM_001195036, intron 3 of 20) | intron (NM_001195036, intron 3 of 20) | 13469 | NM_001195036 | 100045792 | Mm.380617 | NM_001195036 | ENSMUSG00000071392 | **Ect2l** | C330021H03Rik\|ENSMUSG00000071392\|Gm10331 | epithelial cell transforming sequence 2 oncogene-like | protein-coding |

| | | | | | | | | | | | | | | | | | |
|---|---|---|---|---|---|---|---|---|---|---|---|---|---|---|---|---|---|
| chr10_18197009_18197618 | chr10 | 18197010 | 18197618 | + | -0.711757 | intron (NM_001195036, intron 3 of 20) | intron (NM_001195036, intron 3 of 20) | 13576 | NM_001195036 | 100045792 | Mm.380617 | NM_001195036 | ENSMUSG00000071392 | **Ect2l** | C330021H03Rik\|ENSMUSG00000071392\|Gm10331 | epithelial cell transforming sequence 2 oncogene-like | protein-coding |
| chr14_79530048_79530760 | chr14 | 79530049 | 79530760 | + | -0.65173 | intron (NM_001286411, intron 1 of 8) | MIR3\|SINE\|MIR | 14730 | NM_001286411 | 13709 | Mm.20938 | NM_007920 | ENSMUSG00000036461 | **Elf1** | Elf-1\|Sts1\|mElf-1\|p70 | E74-like factor 1 | protein-coding |
| chr14_79530052_79530466 | chr14 | 79530053 | 79530466 | + | -0.659281 | intron (NM_001286411, intron 1 of 8) | ID4_\|SINE\|ID | 14585 | NM_001286411 | 13709 | Mm.20938 | NM_007920 | ENSMUSG00000036461 | **Elf1** | Elf-1\|Sts1\|mElf-1\|p70 | E74-like factor 1 | protein-coding |
| chr14_79526136_79526408 | chr14 | 79526137 | 79526408 | + | -0.701878 | intron (NM_001286411, intron 1 of 8) | intron (NM_001286411, intron 1 of 8) | 10598 | NM_001286411 | 13709 | Mm.20938 | NM_007920 | ENSMUSG00000036461 | **Elf1** | Elf-1\|Sts1\|mElf-1\|p70 | E74-like factor 1 | protein-coding |
| chr12_84221032_84221484 | chr12 | 84221033 | 84221484 | + | -1.04297 | Intergenic | MLT1E1A\|LTR\|ERVL-MaLR | -2377 | NM_001163501 | 238317 | Mm.31256 | NM_001163501 | ENSMUSG00000042507 | **Elmsan1** | 9430029N19Rik\|C130039O16Rik\|Gm260 | ELM2 and Myb/SANT-like domain containing 1 | protein-coding |
| chr15_80562444_80562786 | chr15 | 80562445 | 80562786 | + | -0.732834 | Intergenic | Intergenic | -2145 | NM_001163189 | 383075 | Mm.297626 | NM_001163189 | ENSMUSG00000050439 | **Enthd1** | Gm1242\|Gm86 | ENTH domain containing 1 | protein-coding |
| chr15_80559640_80559964 | chr15 | 80559641 | 80559964 | + | -1.13161 | intron (NM_001163189, intron 1 of 5) | intron (NM_001163189, intron 1 of 5) | 668 | NM_001163189 | 383075 | Mm.297626 | NM_001163189 | ENSMUSG00000050439 | **Enthd1** | Gm1242\|Gm86 | ENTH domain containing 1 | protein-coding |
| chr15_80559545_80560047 | chr15 | 80559546 | 80560047 | + | -1.13514 | intron (NM_001163189, intron 1 of 5) | intron (NM_001163189, intron 1 of 5) | 674 | NM_001163189 | 383075 | Mm.297626 | NM_001163189 | ENSMUSG00000050439 | **Enthd1** | Gm1242\|Gm86 | ENTH domain containing 1 | protein-coding |
| chr4_132007857_132009093 | chr4 | 132007858 | 132009093 | + | -0.850773 | intron (NM_183428, intron 1 of 20) | intron (NM_183428, intron 1 of 20) | 40603 | NM_001128606 | 269587 | Mm.30038 | NM_183428 | ENSMUSG00000028906 | **Epb4.1** | 4.1R\|AI415518\|D4Ertd442e\|Elp-1\|Elp1\|Epb41\|mKIAA4056 | erythrocyte protein band 4.1 | protein-coding |
| chr4_132008054_132008655 | chr4 | 132008055 | 132008655 | + | -0.885744 | intron (NM_183428, intron 1 of 20) | intron (NM_183428, intron 1 of 20) | 40723 | NM_001128606 | 269587 | Mm.30038 | NM_183428 | ENSMUSG00000028906 | **Epb4.1** | 4.1R\|AI415518\|D4Ertd442e\|Elp-1\|Elp1\|Epb41\|mKIAA4056 | erythrocyte protein band 4.1 | protein-coding |
| chr16_13158926_13159472 | chr16 | 13158927 | 13159472 | + | -0.756521 | Intergenic | Intergenic | 49463 | NM_015769 | 50505 | Mm.287837 | NM_015769 | | **Ercc4** | AI606920\|Xpf | excision repair cross-complementing rodent repair deficiency, complementation group 4 | protein-coding |
| chr9_32731843_32732301 | chr9 | 32731844 | 32732301 | + | -0.660697 | intron (NM_001038642, intron 3 of 6) | intron (NM_001038642, intron 3 of 6) | 36030 | NM_001038642 | 23871 | Mm.292415 | NM_011808 | ENSMUSG00000032035 | **Ets1** | AI196000\|AI448617\|D230050P06\|Ets-1\|Tpl1 | E26 avian leukemia oncogene 1, 5' domain | protein-coding |
| chr9_32906299_32906819 | chr9 | 32906300 | 32906819 | + | -0.851891 | Intergenic | Intergenic | 210517 | NM_001038642 | 23871 | Mm.292415 | NM_011808 | ENSMUSG00000032035 | **Ets1** | AI196000\|AI448617\|D230050P06\|Ets-1\|Tpl1 | E26 avian leukemia oncogene 1, 5' domain | protein-coding |
| chr2_91638837_91639308 | chr2 | 91638838 | 91639308 | + | -0.929036 | Intergenic | Intergenic | -2659 | NM_010168 | 14061 | Mm.89048 | NM_010168 | ENSMUSG00000027249 | **F2** | Cf-2\|Cf2\|FII\|thrombin | coagulation factor II | protein-coding |
| chr1_161704513_161705291 | chr1 | 161704514 | 161705291 | + | -0.692314 | Intergenic | Intergenic | 83593 | NM_001205243 | 14103 | Mm.3355 | NM_010177 | ENSMUSG00000000817 | **Fasl** | APT1LG1\|CD178\|CD95-L\|CD95L\|Fas-L\|Faslg\|Tnfsf6\|gld | Fas ligand (TNF superfamily, member 6) | protein-coding |
| chr1_161765913_161766325 | chr1 | 161765914 | 161766325 | + | -0.967243 | Intergenic | Intergenic | 22376 | NM_001205243 | 14103 | Mm.3355 | NM_010177 | ENSMUSG00000000817 | **Fasl** | APT1LG1\|CD178\|CD95-L\|CD95L\|Fas-L\|Faslg\|Tnfsf6\|gld | Fas ligand (TNF superfamily, member 6) | protein-coding |
| chr6_99023634_99023960 | chr6 | 99023635 | 99023960 | + | -1.00411 | intron (NM_053202, intron 5 of 18) | intron (NM_053202, intron 5 of 18) | 4501 | NM_001197322 | 108655 | Mm.234965 | NM_053202 | ENSMUSG00000030067 | **Foxp1** | 3110052D19Rik\|4932443N09Rik\|AI461938\|AW494214 | forkhead box P1 | protein-coding |
| chr5_73265322_73265878 | chr5 | 73265323 | 73265878 | + | -0.892622 | Intergenic | L2b\|LINE\|L2 | -8982 | NM_028194 | 72313 | Mm.260594 | NM_028194 | ENSMUSG00000070733 | **Fryl** | 2010313D22Rik\|2310004H21Rik\|2510002A14Rik\|9030227G01Rik\|mKIAA0826 | furry homolog-like (Drosophila) | protein-coding |
| chr15_6592556_6593127 | chr15 | 6592557 | 6593127 | + | -0.795518 | intron (NM_011815, intron 1 of 17) | intron (NM_011815, intron 1 of 17) | 12995 | NM_011815 | 23880 | Mm.170905 | NM_011815 | | **Fyb** | ADAP\|B630013F22Rik\|FYB-120/130\|Slap130\|p120/130\|p120/p130 | FYN binding protein | protein-coding |
| chr13_3526226_3526919 | chr13 | 3526227 | 3526919 | + | -0.644552 | Intergenic | Intergenic | -11502 | NM_008112 | 14569 | Mm.153226 | NM_008112 | ENSMUSG00000021218 | **Gdi2** | GDI-B\|GDIB\|Gdi3 | guanosine diphosphate (GDP) dissociation inhibitor 2 | protein-coding |
| chr18_5593691_5594020 | chr18 | 5593692 | 5594020 | + | -0.895606 | intron (NM_011546, intron 1 of 7) | intron (NM_011546, intron 1 of 7) | -1419 | NR_033552 | 791318 | Mm.84339 | NR_033552 | ENSMUSG00000063087 | **Gm10125** | ENSMUSG00000063087\|Zeb1os | predicted gene 10125 | ncRNA |
| chr18_5593653_5594312 | chr18 | 5593654 | 5594312 | + | -1.04135 | intron (NM_011546, intron 1 of 7) | intron (NM_011546, intron 1 of 7) | -1546 | NR_033552 | 791318 | Mm.84339 | NR_033552 | ENSMUSG00000063087 | **Gm10125** | ENSMUSG00000063087\|Zeb1os | predicted gene 10125 | ncRNA |
| chr11_6679691_6680777 | chr11 | 6679692 | 6680777 | + | -0.942802 | Intergenic | RSINE1\|SINE\|B4 | -19998 | NR_028586 | 100041311 | Mm.388018 | NR_028586 | ENSMUSG00000086373 | **Gm11978** | OTTMUSG00000005125\|OTTMUSG00000005127 | predicted gene 11978 | pseudo |
| chr6_145306665_145307079 | chr6 | 145306666 | 145307079 | + | -0.835972 | Intergenic | Intergenic | 56320 | NR_045598 | 100039495 | Mm.121014 | NR_045598 | ENSMUSG00000086013 | **Gm15706** | OTTMUSG00000025463 | predicted gene 15706 | ncRNA |
| chr8_35678774_35679367 | chr8 | 35678775 | 35679367 | + | -0.923007 | 3' UTR (NM_001081279, exon 3 of 3) | 3' UTR (NM_001081279, exon 3 of 3) | -90051 | NR_040376 | 100504714 | Mm.381744 | NR_040376 | ENSMUSG00000097357 | **Gm16793** | - | predicted gene, 16793 | ncRNA |
| chr19_47512763_47513353 | chr19 | 47512764 | 47513353 | + | -0.902978 | promoter-TSS (NR_045322) | promoter-TSS (NR_045322) | 76 | NR_045322 | 100503104 | Mm.442735 | NR_045322 | ENSMUSG00000097990 | **Gm19557** | - | predicted gene, 19557 | ncRNA |
| chr17_51903825_51904308 | chr17 | 51903826 | 51904308 | + | -0.707268 | intron (NR_045095, intron 1 of 2) | intron (NR_045095, intron 1 of 2) | 21340 | NR_045095 | 100504169 | Mm.417731 | NR_045095 | | **Gm20098** | - | predicted gene, 20098 | ncRNA |
| chr17_52168131_52169101 | chr17 | 52168132 | 52169101 | + | -0.888359 | Intergenic | Intergenic | 285889 | NR_045095 | 100504169 | Mm.417731 | NR_045095 | | **Gm20098** | - | predicted gene, 20098 | ncRNA |
| chr17_52177915_52178461 | chr17 | 52177916 | 52178461 | + | -1.32354 | Intergenic | Intergenic | 295461 | NR_045095 | 100504169 | Mm.417731 | NR_045095 | | **Gm20098** | - | predicted gene, 20098 | ncRNA |
| chr10_19122123_19122601 | chr10 | 19122124 | 19122601 | + | -0.808461 | Intergenic | Intergenic | -19387 | NR_038021 | 100504265 | Mm.383417 | NR_038021 | | **Gm20139** | - | predicted gene, 20139 | ncRNA |
| chr10_19104540_19105371 | chr10 | 19104541 | 19105371 | + | -0.86295 | Intergenic | Intergenic | -36793 | NR_038021 | 100504265 | Mm.383417 | NR_038021 | | **Gm20139** | - | predicted gene, 20139 | ncRNA |
| chr10_19171826_19172093 | chr10 | 19171827 | 19172093 | + | -1.02721 | intron (NR_038022, intron 1 of 1) | intron (NR_038022, intron 1 of 1) | 4722 | NR_038022 | 100504265 | Mm.383417 | NR_038021 | | **Gm20139** | - | predicted gene, 20139 | ncRNA |
| chr10_19185775_19186634 | chr10 | 19185776 | 19186634 | + | -1.03035 | intron (NR_038022, intron 1 of 1) | intron (NR_038022, intron 1 of 1) | 18967 | NR_038022 | 100504265 | Mm.383417 | NR_038021 | | **Gm20139** | - | predicted gene, 20139 | ncRNA |
| chr10_19171830_19172216 | chr10 | 19171831 | 19172216 | + | -1.06102 | intron (NR_038022, intron 1 of 1) | intron (NR_038022, intron 1 of 1) | 4785 | NR_038022 | 100504265 | Mm.383417 | NR_038021 | | **Gm20139** | - | predicted gene, 20139 | ncRNA |
| chr10_19171773_19172315 | chr10 | 19171774 | 19172315 | + | -1.07717 | intron (NR_038022, intron 1 of 1) | intron (NR_038022, intron 1 of 1) | 4806 | NR_038022 | 100504265 | Mm.383417 | NR_038021 | | **Gm20139** | - | predicted gene, 20139 | ncRNA |
| chr10_19116814_19117545 | chr10 | 19116815 | 19117545 | + | -1.11367 | Intergenic | Intergenic | -24569 | NR_038021 | 100504265 | Mm.383417 | NR_038021 | | **Gm20139** | - | predicted gene, 20139 | ncRNA |
| chr3_52927334_52927956 | chr3 | 52927335 | 52927956 | + | -0.697653 | promoter-TSS (NR_040555) | promoter-TSS (NR_040555) | 21 | NR_040555 | 622124 | Mm.389434 | NR_040555 | | **Gm20750** | - | predicted gene, 20750 | ncRNA |
| chr5_64684153_64684889 | chr5 | 64684154 | 64684889 | + | -1.12242 | Intergenic | Intergenic | -73822 | NR_045078 | 100042198 | Mm.25191 | NR_045078 | | **Gm3716** | Gm1683 | predicted gene 3716 | ncRNA |
| chr1_180359353_180359735 | chr1 | 180359354 | 180359735 | + | -0.680497 | intron (NM_001081175, intron 2 of 7) | MLT1L\|LTR\|ERVL-MaLR | -28995 | NR_003623 | 277333 | Mm.389223 | NM_199472 | | **Gm5069** | EG277333 | glyceraldehyde-3-phosphate dehydrogenase pseudogene | pseudo |
| chr9_107675658_107676704 | chr9 | 107675659 | 107676704 | + | -0.852529 | intron (NM_008140, intron 7 of 8) | CpG | 3411 | NM_008140 | 14685 | Mm.284853 | NM_008140 | ENSMUSG00000034837 | **Gnat1** | Gnat-1\|Ird1\|Ird2\|Tralpha\|irdc\|irdr\|transducin | guanine nucleotide binding protein, alpha transducing 1 | protein-coding |
| chr1_74262410_74262906 | chr1 | 74262411 | 74262906 | + | -1.12375 | intron (NM_029711, intron 7 of 9) | ID2\|SINE\|ID | -15942 | NM_174985 | 227289 | Mm.246587 | NM_174985 | ENSMUSG00000064272 | **Gpbar1** | BG37\|GPCR\|GPR131\|M-BAR\|TGR5 | G protein-coupled bile acid receptor 1 | protein-coding |
| chr1_74262508_74262868 | chr1 | 74262509 | 74262868 | + | -1.13878 | intron (NM_029711, intron 7 of 9) | intron (NM_029711, intron 7 of 9) | -15912 | NM_174985 | 227289 | Mm.246587 | NM_174985 | ENSMUSG00000064272 | **Gpbar1** | BG37\|GPCR\|GPR131\|M-BAR\|TGR5 | G protein-coupled bile acid receptor 1 | protein-coding |
| chr12_112854986_112855390 | chr12 | 112854987 | 112855390 | + | -1.13417 | intron (NM_019925, intron 2 of 2) | intron (NM_019925, intron 2 of 2) | 5728 | NM_019925 | 56696 | Mm.20455 | NM_019925 | ENSMUSG00000021298 | **Gpr132** | G2a | G protein-coupled receptor 132 | protein-coding |
| chr14_121929282_121929879 | chr14 | 121929283 | 121929879 | + | -1.0085 | intron (NM_026861, intron 4 of 8) | intron (NM_026861, intron 4 of 8) | -13807 | NM_182806 | 110168 | Mm.37405 | NM_182806 | ENSMUSG00000050350 | **Gpr18** | - | G protein-coupled receptor 18 | protein-coding |
| chr18_56468961_56469469 | chr18 | 56468962 | 56469469 | + | -0.747624 | intron (NM_026240, intron 1 of 13) | intron (NM_026240, intron 1 of 13) | 37083 | NM_026240 | 107022 | Mm.24356 | NM_026240 | ENSMUSG00000001700 | **Gramd3** | 9030613F08Rik\|9130427A09Rik\|AW061306\|NS3TP2 | GRAM domain containing 3 | protein-coding |
| chr5_134481933_134482727 | chr5 | 134481934 | 134482727 | + | -0.750761 | Intergenic | Intergenic | -25614 | NM_001081464 | 57080 | Mm.332735 | NM_020331 | ENSMUSG00000023079 | **Gtf2ird1** | 1700012P16Rik\|BEN\|Cream1\|ESTM9\|GTF3\|Gtf2il\|MusTRD1\|Tg(Alb1-Myc)166.8Sst\|WBSCR11\|X83320 | general transcription factor II I repeat domain-containing 1 | protein-coding |
| chr6_136742308_136742630 | chr6 | 136742309 | 136742630 | + | -1.10515 | intron (NM_145067, intron 10 of 25) | intron (NM_145067, intron 10 of 25) | 39273 | NM_145067 | 14917 | Mm.222729 | NM_145067 | ENSMUSG00000042638 | **Gucy2c** | AI893437\|GC-C\|Gcc | guanylate cyclase 2c | protein-coding |
| chr6_136743701_136744184 | chr6 | 136743702 | 136744184 | + | -1.1173 | exon (NM_001127318, exon 10 of 27) | exon (NM_001127318, exon 10 of 27) | 37799 | NM_145067 | 14917 | Mm.222729 | NM_145067 | ENSMUSG00000042638 | **Gucy2c** | AI893437\|GC-C\|Gcc | guanylate cyclase 2c | protein-coding |

| | | | | | | | | | | | | | | | | | |
|---|---|---|---|---|---|---|---|---|---|---|---|---|---|---|---|---|---|
| chr6_136742252_136742694 | chr6 | 136742253 | 136742694 | + | -1.13394 | intron (NM_145067, intron 10 of 25) | intron (NM_145067, intron 10 of 25) | 39269 | NM_145067 | 14917 | Mm.222729 | NM_145067 | ENSMUSG00000042638 | **Gucy2c** | AI893437\|GC-C\|Gcc | guanylate cyclase 2c | protein-coding |
| chr17_34239429_34240003 | chr17 | 34239430 | 34240003 | + | -0.921032 | intron (NM_010389, intron 1 of 5) | intron (NM_010389, intron 1 of 5) | 811 | NM_010389 | 15002 | Mm.327358 | NM_010389 | ENSMUSG00000041538 | **H2-Ob** | A-beta-2\|A-beta2\|H-2Ab\|H-2I\|H-2Ob\|H2-Ab\|H2-Ab2\|H2-IAb2\|Ob | histocompatibility 2, O region beta locus | protein-coding |
| chr10_21254618_21255194 | chr10 | 21254619 | 21255194 | + | -0.724798 | Intergenic | Intergenic | -41073 | NM_001042593 | 56422 | Mm.25527 | NM_019702 | ENSMUSG00000019977 | **Hbs1l** | 2810035F15Rik\|AI326327\|eRFS | Hbs1-like (S. cerevisiae) | protein-coding |
| chr3_8766590_8767289 | chr3 | 8766591 | 8767289 | + | -0.834162 | Intergenic | Intergenic | -99902 | NM_010423 | 15213 | Mm.29581 | NM_010423 | ENSMUSG00000040289 | **Hey1** | AI316788\|AI414254\|CHF2\|HRT1\|Herp2\|Hesr1\|bHLHb31\|hesr-1 | hairy/enhancer-of-split related with YRPW motif 1 | protein-coding |
| chr10_13993235_13993904 | chr10 | 13993236 | 13993904 | + | -0.698335 | intron (NM_010437, intron 1 of 8) | intron (NM_010437, intron 1 of 8) | 27191 | NM_010437 | 15273 | Mm.42157 | NM_010437 | ENSMUSG00000015501 | **Hivep2** | MIBP1\|Schnurri-2\|Shn-2 | human immunodeficiency virus type I enhancer binding protein 2 | protein-coding |
| chr16_91538144_91538390 | chr16 | 91538145 | 91538390 | + | -0.85621 | Intergenic | Intergenic | -8827 | NM_008338 | 15980 | Mm.249364 | NM_008338 | | **Ifngr2** | Ifgr2\|Ifgt | interferon gamma receptor 2 | protein-coding |
| chr2_11718225_11718632 | chr2 | 11718226 | 11718632 | + | -1.55987 | exon (NM_008358, exon 2 of 7) | exon (NM_008358, exon 2 of 7) | 12717 | NM_001271500 | 16169 | Mm.200196 | NM_008358 | ENSMUSG00000023206 | **Il15ra** | AA690181\|IL-15RA | interleukin 15 receptor, alpha chain | protein-coding |
| chr13_112427556_112427943 | chr13 | 112427557 | 112427943 | + | -0.717351 | Intergenic | Intergenic | -36320 | NM_010560 | 16195 | Mm.4364 | NM_010560 | ENSMUSG00000021756 | **Il6st** | 5133400A03Rik\|AA389424\|BB405851\|CD130\|D13Ertd699e\|gp130 | interleukin 6 signal transducer | protein-coding |
| chr13_112427470_112428001 | chr13 | 112427471 | 112428001 | + | -0.728225 | Intergenic | Intergenic | -36334 | NM_010560 | 16195 | Mm.4364 | NM_010560 | ENSMUSG00000021756 | **Il6st** | 5133400A03Rik\|AA389424\|BB405851\|CD130\|D13Ertd699e\|gp130 | interleukin 6 signal transducer | protein-coding |
| chr8_109662987_109663545 | chr8 | 109662988 | 109663545 | + | -0.777983 | intron (NM_001039700, intron 27 of 33) | intron (NM_001039700, intron 27 of 33) | 30028 | NM_028018 | 71955 | Mm.290036 | NM_028018 | ENSMUSG00000031729 | **Ist1** | 2400003C14Rik\|AW536298\|mKIAA0174 | increased sodium tolerance 1 homolog (yeast) | protein-coding |
| chr8_109663239_109663572 | chr8 | 109663240 | 109663572 | + | -0.887279 | intron (NM_001039700, intron 27 of 33) | intron (NM_001039700, intron 27 of 33) | 29888 | NM_028018 | 71955 | Mm.290036 | NM_028018 | ENSMUSG00000031729 | **Ist1** | 2400003C14Rik\|AW536298\|mKIAA0174 | increased sodium tolerance 1 homolog (yeast) | protein-coding |
| chr8_109663284_109663532 | chr8 | 109663285 | 109663532 | + | -0.891566 | intron (NM_001039700, intron 27 of 33) | intron (NM_001039700, intron 27 of 33) | 29886 | NM_028018 | 71955 | Mm.290036 | NM_028018 | ENSMUSG00000031729 | **Ist1** | 2400003C14Rik\|AW536298\|mKIAA0174 | increased sodium tolerance 1 homolog (yeast) | protein-coding |
| chr11_46372236_46373030 | chr11 | 46372237 | 46373030 | + | -0.903639 | intron (NM_001281966, intron 1 of 16) | Lx8b\|LINE\|L1 | 16882 | NM_001281968 | 16428 | Mm.339927 | NM_010583 | ENSMUSG00000020395 | **Itk** | Emt\|Tcsk\|Tsk | IL2 inducible T cell kinase | protein-coding |
| chr11_46372236_46372775 | chr11 | 46372237 | 46372775 | + | -1.00381 | intron (NM_001281966, intron 1 of 16) | Lx8b\|LINE\|L1 | 17009 | NM_001281968 | 16428 | Mm.339927 | NM_010583 | ENSMUSG00000020395 | **Itk** | Emt\|Tcsk\|Tsk | IL2 inducible T cell kinase | protein-coding |
| chr4_101185286_101185604 | chr4 | 101185287 | 101185604 | + | -1.05352 | intron (NM_146145, intron 4 of 24) | intron (NM_146145, intron 4 of 24) | 79837 | NM_146145 | 16451 | Mm.289657 | NM_146145 | ENSMUSG00000028530 | **Jak1** | AA960307\|C130039L05Rik | Janus kinase 1 | protein-coding |
| chr17_52506633_52507595 | chr17 | 52506634 | 52507595 | + | -0.707665 | Intergenic | Intergenic | -95595 | NM_001031811 | 211468 | Mm.399068 | NM_001031811 | ENSMUSG00000035580 | **Kcnh8** | C130090D05Rik\|ELK\|ELK1\|ELK3\|Kv12.1 | potassium voltage-gated channel, subfamily H (eag-related), member 8 | protein-coding |
| chr17_52506784_52507483 | chr17 | 52506785 | 52507483 | + | -0.73305 | Intergenic | Intergenic | -95575 | NM_001031811 | 211468 | Mm.399068 | NM_001031811 | ENSMUSG00000035580 | **Kcnh8** | C130090D05Rik\|ELK\|ELK1\|ELK3\|Kv12.1 | potassium voltage-gated channel, subfamily H (eag-related), member 8 | protein-coding |
| chr17_52506779_52507244 | chr17 | 52506780 | 52507244 | + | -0.791408 | Intergenic | MIR\|SINE\|MIR | -95697 | NM_001031811 | 211468 | Mm.399068 | NM_001031811 | ENSMUSG00000035580 | **Kcnh8** | C130090D05Rik\|ELK\|ELK1\|ELK3\|Kv12.1 | potassium voltage-gated channel, subfamily H (eag-related), member 8 | protein-coding |
| chr17_52506636_52507189 | chr17 | 52506637 | 52507189 | + | -0.794928 | Intergenic | MIR\|SINE\|MIR | -95796 | NM_001031811 | 211468 | Mm.399068 | NM_001031811 | ENSMUSG00000035580 | **Kcnh8** | C130090D05Rik\|ELK\|ELK1\|ELK3\|Kv12.1 | potassium voltage-gated channel, subfamily H (eag-related), member 8 | protein-coding |
| chr8_72295545_72296217 | chr8 | 72295546 | 72296217 | + | -0.790655 | Intergenic | Intergenic | -23181 | NM_008452 | 16598 | Mm.26938 | NM_008452 | ENSMUSG00000055148 | **Klf2** | Lklf | Kruppel-like factor 2 (lung) | protein-coding |
| chr5_64806144_64806522 | chr5 | 64806145 | 64806522 | + | -0.944479 | intron (NM_008453, intron 1 of 5) | intron (NM_008453, intron 1 of 5) | 2810 | NM_008453 | 16599 | Mm.392759 | NM_008453 | ENSMUSG00000029178 | **Klf3** | 9930027G08Rik\|AL024007\|Bklf\|Tef-2 | Kruppel-like factor 3 (basic) | protein-coding |
| chr5_64806013_64806530 | chr5 | 64806014 | 64806530 | + | -0.958987 | intron (NM_008453, intron 1 of 5) | intron (NM_008453, intron 1 of 5) | 2749 | NM_008453 | 16599 | Mm.392759 | NM_008453 | ENSMUSG00000029178 | **Klf3** | 9930027G08Rik\|AL024007\|Bklf\|Tef-2 | Kruppel-like factor 3 (basic) | protein-coding |
| chr12_69255967_69256325 | chr12 | 69255968 | 69256325 | + | -0.78393 | intron (NM_178253, intron 5 of 12) | intron (NM_178253, intron 5 of 12) | 14314 | NM_178253 | 271005 | Mm.180969 | NM_178253 | | **Klhdc1** | - | kelch domain containing 1 | protein-coding |
| chr3_131170152_131170622 | chr3 | 131170153 | 131170622 | + | -0.725453 | intron (NM_001276402, intron 3 of 9) | RLTR51B_Mm\|LTR\|ERVK? | 57596 | NM_001276403 | 16842 | Mm.255219 | NM_010703 | ENSMUSG00000027985 | **Lef1** | 3000002B05\|AI451430\|Lef-1 | lymphoid enhancer binding factor 1 | protein-coding |
| chr6_31201168_31201921 | chr6 | 31201169 | 31201921 | + | -0.658422 | intron (NR_110469, intron 1 of 3) | Lx8\|LINE\|L1 | 16929 | NR_110470 | 232685 | Mm.443019 | NM_144555 | | **Lincpint** | MNCb-1768\|Pint\|linc-Pint | long intergenic non-protein coding RNA, Trp53 induced transcript | ncRNA |
| chr7_45778961_45779867 | chr7 | 45778962 | 45779867 | + | -0.746247 | Intergenic | L1MA8\|LINE\|L1 | -4533 | NM_001005511 | 381983 | Mm.44928 | NM_001005511 | ENSMUSG00000062044 | **Lmtk3** | BC059845\|aatyk3 | lemur tyrosine kinase 3 | protein-coding |
| chr8_36194064_36194571 | chr8 | 36194065 | 36194571 | + | -0.673111 | Intergenic | Intergenic | 55198 | NM_001081150 | 244421 | Mm.324032 | NM_001081150 | ENSMUSG00000039633 | **Lonrf1** | - | LON peptidase N-terminal domain and ring finger 1 | protein-coding |
| chr5_105717851_105718475 | chr5 | 105717852 | 105718475 | + | -0.6946 | intron (NM_178701, intron 1 of 2) | intron (NM_178701, intron 1 of 2) | 17353 | NM_178701 | 231549 | Mm.23837 | NM_178701 | ENSMUSG00000046079 | **Lrrc8d** | 2810473G09Rik\|4930525N13Rik\|A930019F03\|Lrrc5 | leucine rich repeat containing 8D | protein-coding |
| chr5_105718092_105718770 | chr5 | 105718093 | 105718770 | + | -0.806217 | intron (NM_178701, intron 1 of 2) | intron (NM_178701, intron 1 of 2) | 17621 | NM_178701 | 231549 | Mm.23837 | NM_178701 | ENSMUSG00000046079 | **Lrrc8d** | 2810473G09Rik\|4930525N13Rik\|A930019F03\|Lrrc5 | leucine rich repeat containing 8D | protein-coding |
| chr9_13536663_13537278 | chr9 | 13536664 | 13537278 | + | -0.667246 | Intergenic | Intergenic | -83018 | NM_001013813 | 270118 | Mm.116898 | NM_173776 | ENSMUSG00000031925 | **Maml2** | 5930431H10\|BC032967 | mastermind like 2 (Drosophila) | protein-coding |
| chr9_13536663_13537067 | chr9 | 13536664 | 13537067 | + | -0.71801 | Intergenic | Intergenic | -83124 | NM_001013813 | 270118 | Mm.116898 | NM_173776 | ENSMUSG00000031925 | **Maml2** | 5930431H10\|BC032967 | mastermind like 2 (Drosophila) | protein-coding |
| chr13_111807667_111807884 | chr13 | 111807668 | 111807884 | + | -0.771359 | intron (NM_011945, intron 1 of 19) | CpG | 1207 | NM_011945 | 26401 | Mm.15918 | NM_011945 | ENSMUSG00000021754 | **Map3k1** | MAPKKK1\|MEKK1\|Mekk | mitogen-activated protein kinase kinase kinase 1 | protein-coding |
| chr13_111807162_111807954 | chr13 | 111807163 | 111807954 | + | -0.789627 | intron (NM_011945, intron 1 of 19) | intron (NM_011945, intron 1 of 19) | 1425 | NM_011945 | 26401 | Mm.15918 | NM_011945 | ENSMUSG00000021754 | **Map3k1** | MAPKKK1\|MEKK1\|Mekk | mitogen-activated protein kinase kinase kinase 1 | protein-coding |
| chr13_111807453_111807951 | chr13 | 111807454 | 111807951 | + | -0.825033 | intron (NM_011945, intron 1 of 19) | intron (NM_011945, intron 1 of 19) | 1281 | NM_011945 | 26401 | Mm.15918 | NM_011945 | ENSMUSG00000021754 | **Map3k1** | MAPKKK1\|MEKK1\|Mekk | mitogen-activated protein kinase kinase kinase 1 | protein-coding |
| chr11_49837689_49838208 | chr11 | 49837690 | 49838208 | + | -0.744301 | 3' UTR (NM_013529, exon 19 of 19) | 3' UTR (NM_013529, exon 19 of 19) | -8802 | NM_001163671 | 26420 | Mm.68933 | NM_016961 | ENSMUSG00000020366 | **Mapk9** | AI851083\|JNK2\|Prkm9\|p54aSAPK | mitogen-activated protein kinase 9 | protein-coding |
| chr18_70571230_70571498 | chr18 | 70571231 | 70571498 | + | -0.739351 | intron (NM_010773, intron 1 of 6) | RSINE1\|SINE\|B4 | 3072 | NM_010773 | 17191 | Mm.322 | NM_010773 | ENSMUSG00000024513 | **Mbd2** | MBD2a | methyl-CpG binding domain protein 2 | protein-coding |
| chr3_60501635_60502133 | chr3 | 60501636 | 60502133 | + | -0.780301 | intron (NM_001253709, intron 1 of 8) | intron (NM_001253709, intron 1 of 8) | 632 | NM_001253708 | 56758 | Mm.255723 | NM_020007 | ENSMUSG00000027763 | **Mbnl1** | Mbnl\|mKIAA0428 | muscleblind-like 1 (Drosophila) | protein-coding |
| chr2_144275660_144276139 | chr2 | 144275661 | 144276139 | + | -0.94397 | intron (NM_001289631, intron 2 of 4) | B1_Mur2\|SINE\|Alu | 4996 | NM_001289630 | 74528 | Mm.386817 | NM_028984 | ENSMUSG00000027424 | **Mgme1** | 8430406I07Rik\|AI426476 | mitochondrial genome maintainance exonuclease 1 | protein-coding |
| chr4_101328960_101329600 | chr4 | 101328961 | 101329600 | + | -0.902666 | Intergenic | MTD\|LTR\|ERVL-MaLR | 17747 | NR_029537 | 387143 | | NR_029537 | miRBase:MI0000148 | **Mir101a** | Mirn101\|Mirn101a\|miR-101 | microRNA 101a | ncRNA |
| chr18_56489940_56490223 | chr18 | 56489941 | 56490223 | + | -0.939509 | intron (NM_026240, intron 10 of 13) | ORR1B1\|LTR\|ERVL-MaLR | -48057 | NR_106160 | 102465790 | | NR_106160 | miRBase:MI0025040 | **Mir1258** | mmu-mir-1258 | microRNA mir-1258 | ncRNA |
| chr18_56489797_56490223 | chr18 | 56489798 | 56490223 | + | -0.959437 | intron (NM_026240, intron 10 of 13) | ORR1B1\|LTR\|ERVL-MaLR | -48129 | NR_106160 | 102465790 | | NR_106160 | miRBase:MI0025040 | **Mir1258** | mmu-mir-1258 | microRNA mir-1258 | ncRNA |
| chr18_56504283_56504776 | chr18 | 56504284 | 56504776 | + | -1.00242 | TTS (NM_026240) | TTS (NM_026240) | -33609 | NR_106160 | 102465790 | | NR_106160 | miRBase:MI0025040 | **Mir1258** | mmu-mir-1258 | microRNA mir-1258 | ncRNA |
| chr18_56504305_56505231 | chr18 | 56504306 | 56505231 | + | -1.01806 | TTS (NM_026240) | TTS (NM_026240) | -33371 | NR_106160 | 102465790 | | NR_106160 | miRBase:MI0025040 | **Mir1258** | mmu-mir-1258 | microRNA mir-1258 | ncRNA |
| chr7_101466101_101467148 | chr7 | 101466102 | 101467148 | + | -0.838744 | intron (NM_001243757, intron 2 of 30) | intron (NM_001243757, intron 2 of 30) | -8751 | NR_029791 | 387157 | | NR_029791 | | **Mir139** | Mirn139\|mmu-mir-139 | microRNA 139 | ncRNA |
| chr15_100181744_100182377 | chr15 | 100181745 | 100182377 | + | -0.902579 | intron (NM_172819, intron 16 of 32) | intron (NM_172819, intron 16 of 32) | 5975 | NR_037311 | 100499528 | | NR_037311 | | **Mir3473a** | Mir3473\|mmu-mir-3473\|mmu-mir-3473a | microRNA 3473a | ncRNA |
| chr15_100182653_100183245 | chr15 | 100182654 | 100183245 | + | -1.06174 | intron (NM_172819, intron 16 of 32) | intron (NM_172819, intron 16 of 32) | 5087 | NR_037311 | 100499528 | | NR_037311 | | **Mir3473a** | Mir3473\|mmu-mir-3473\|mmu-mir-3473a | microRNA 3473a | ncRNA |
| chr15_80930716_80931040 | chr15 | 80930717 | 80931040 | + | -0.851262 | 3' UTR (NM_144812, exon 25 of 25) | 3' UTR (NM_144812, exon 25 of 25) | -9161 | NR_039573 | 100628612 | | NR_039573 | miRBase:MI0018022 | **Mir5113** | mmu-mir-5113\|mu-mir-5113 | microRNA 5113 | ncRNA |
| chr2_166672555_166673015 | chr2 | 166672556 | 166673015 | + | -0.709027 | promoter-TSS (NR_105783) | promoter-TSS (NR_105783) | 67 | NR_105783 | 102465188 | | NR_105783 | | **Mir6364** | mmu-mir-6364 | microRNA mir-6364 | ncRNA |
| chr5_124003124_124003804 | chr5 | 124003125 | 124003804 | + | -1.17466 | TTS (NM_145070) | TTS (NM_145070) | 6612 | NR_105999 | 102465625 | | NR_105999 | | **Mir7032** | mmu-mir-7032 | microRNA mir-7032 | ncRNA |
| chr1_53850849_53851293 | chr1 | 53850850 | 53851293 | + | -1.18733 | intron (NM_001001883, intron 21 of 28) | intron (NM_001001883, intron 21 of 28) | 1647 | NR_106149 | 102465783 | | NR_106149 | miRBase:MI0025029 | **Mir7681** | mmu-mir-7681 | microRNA mir-7681 | ncRNA |
| chr10_80678002_80678547 | chr10 | 80678003 | 80678547 | + | -0.660859 | Intergenic | Intergenic | -6299 | NM_021462 | 17347 | Mm.42126 | NM_021462 | ENSMUSG00000020190 | **Mknk2** | 2010016G11Rik\|Gprk7\|Mnk2 | MAP kinase-interacting serine/threonine kinase 2 | protein-coding |

| | | | | | | | | | | | | | | | | | |
|---|---|---|---|---|---|---|---|---|---|---|---|---|---|---|---|---|---|
| chr10_120895714_120896348 | chr10 | 120895715 | 120896348 | + | -0.755792 | intron (NM_177092, intron 1 of 6) | intron (NM_177092, intron 1 of 6) | 2940 | NM_177092 | 320183 | Mm.169755 | NM_177092 | ENSMUSG00000051236 | **Msrb3** | D430026P16Rik\|I-8-23\|MsrB2 | methionine sulfoxide reductase B3 | protein-coding |
| chr15_12229607_12229878 | chr15 | 12229608 | 12229878 | + | -0.709837 | intron (NM_172958, intron 1 of 15) | B4A\|SINE\|B4 | 24649 | NM_172958 | 268783 | Mm.54460 | NM_172958 | ENSMUSG00000039458 | **Mtmr12** | 3Pap\|4932703C11\|C730015A02Rik\|Pip3ap\|mKIAA1682 | myotubularin related protein 12 | protein-coding |
| chr15_12229597_12230145 | chr15 | 12229598 | 12230145 | + | -0.737707 | intron (NM_172958, intron 1 of 15) | ID\|SINE\|ID | 24777 | NM_172958 | 268783 | Mm.54460 | NM_172958 | ENSMUSG00000039458 | **Mtmr12** | 3Pap\|4932703C11\|C730015A02Rik\|Pip3ap\|mKIAA1682 | myotubularin related protein 12 | protein-coding |
| chr1_51929491_51930119 | chr1 | 51929492 | 51930119 | + | -1.06132 | Intergenic | Intergenic | -13827 | NM_001161817 | 17912 | Mm.3390 | NM_010863 | ENSMUSG00000018417 | **Myo1b** | AA406997\|MIH-L\|MMIa | myosin IB | protein-coding |
| chr5_134269556_134270311 | chr5 | 134269557 | 134270311 | + | -0.805106 | intron (NM_001080749, intron 10 of 32) | intron (NM_001080749, intron 10 of 32) | -40309 | NR_104383 | 17969 | Mm.425296 | NM_010876 | ENSMUSG00000015950 | **Ncf1** | NCF-47K\|NOXO2\|Ncf-1\|p47-phox\|p47<phox>\|p47phox | neutrophil cytosolic factor 1 | protein-coding |
| chr8_85623259_85623571 | chr8 | 85623260 | 85623571 | + | -1.08617 | Intergenic | Intergenic | 67594 | NM_001081324 | 74513 | Mm.126079 | NM_001081324 | ENSMUSG00000036902 | **Neto2** | 5530601C23Rik\|AI854825\|BTCL2 | neuropilin (NRP) and tolloid (TLL)-like 2 | protein-coding |
| chr8_85623174_85623788 | chr8 | 85623175 | 85623788 | + | -1.14939 | Intergenic | Intergenic | 67528 | NM_001081324 | 74513 | Mm.126079 | NM_001081324 | ENSMUSG00000036902 | **Neto2** | 5530601C23Rik\|AI854825\|BTCL2 | neuropilin (NRP) and tolloid (TLL)-like 2 | protein-coding |
| chr17_26370397_26371291 | chr17 | 26370398 | 26371291 | + | -1.00123 | Intergenic | Intergenic | -44121 | NM_001081656 | 240055 | Mm.133211 | NM_001081656 | ENSMUSG00000034413 | **Neurl1b** | C230078M08Rik\|EG240055 | neuralized homolog 1b (Drosophila) | protein-coding |
| chr18_80706073_80706363 | chr18 | 80706074 | 80706363 | + | -0.766526 | intron (NM_198429, intron 1 of 9) | intron (NM_198429, intron 1 of 9) | 1955 | NM_001164111 | 18018 | Mm.329560 | NM_016791 | ENSMUSG00000033016 | **Nfatc1** | 2210017P03Rik\|AI449492\|AV076380\|NF-ATc\|NFAT2\|NFATc\|Nfatcb | nuclear factor of activated T cells, cytoplasmic, calcineurin dependent 1 | protein-coding |
| chr15_8569688_8570104 | chr15 | 8569689 | 8570104 | + | -0.947037 | Intergenic | Intergenic | -125433 | NM_201232 | 71175 | Mm.240329 | NM_027707 | ENSMUSG00000022141 | **Nipbl** | Idn3 | Nipped-B homolog (Drosophila) | protein-coding |
| chr16_76345951_76346481 | chr16 | 76345952 | 76346481 | + | -1.05998 | intron (NM_173440, intron 2 of 3) | intron (NM_173440, intron 2 of 3) | 26833 | NM_173440 | 268903 | Mm.74711 | NM_173440 | ENSMUSG00000048490 | **Nrip1** | 6030458L20Rik\|8430438I05Rik\|9630050P12\|AA959574\|AW456757\|RIP140 | nuclear receptor interacting protein 1 | protein-coding |
| chr7_125507192_125508055 | chr7 | 125507193 | 125508055 | + | -1.12634 | Intergenic | Intergenic | -16082 | NM_026330 | 67711 | Mm.4467 | NM_026330 | ENSMUSG00000030750 | **Nsmce1** | 2510027N19Rik | non-SMC element 1 homolog (S. cerevisiae) | protein-coding |
| chr15_27605243_27605725 | chr15 | 27605244 | 27605725 | + | -0.882321 | TTS (NM_001013792) | TTS (NM_001013792) | 25209 | NM_001013792 | 432940 | Mm.309164 | NM_001013792 | ENSMUSG00000046034 | **Otulin** | C79097\|Fam105b\|m3Sapc\|m7-1Sapc | OTU deubiquitinase with linear linkage specificity | protein-coding |
| chr5_122718879_122719648 | chr5 | 122718880 | 122719648 | + | -0.931745 | intron (NM_011026, intron 5 of 11) | intron (NM_011026, intron 5 of 11) | 11707 | NM_011026 | 18438 | Mm.290884 | NM_011026 | ENSMUSG00000029470 | **P2rx4** | AI504491\|AW555605\|D5Ertd444e\|P2X4 | purinergic receptor P2X, ligand-gated ion channel 4 | protein-coding |
| chr5_122719103_122719577 | chr5 | 122719104 | 122719577 | + | -0.947613 | intron (NM_011026, intron 5 of 11) | intron (NM_011026, intron 5 of 11) | 11783 | NM_011026 | 18438 | Mm.290884 | NM_011026 | ENSMUSG00000029470 | **P2rx4** | AI504491\|AW555605\|D5Ertd444e\|P2X4 | purinergic receptor P2X, ligand-gated ion channel 4 | protein-coding |
| chr2_18984007_18984373 | chr2 | 18984008 | 18984373 | + | -0.693123 | intron (NM_008845, intron 1 of 9) | intron (NM_008845, intron 1 of 9) | 13931 | NM_008845 | 18718 | Mm.313977 | NM_008845 | ENSMUSG00000026737 | **Pip4k2a** | AW742916\|Pip5k2a | phosphatidylinositol-5-phosphate 4-kinase, type II, alpha | protein-coding |
| chr16_46001586_46002133 | chr16 | 46001587 | 46002133 | + | -0.91135 | intron (NM_001134480, intron 1 of 3) | intron (NM_001134480, intron 1 of 3) | 8553 | NM_001134480 | 433022 | Mm.380993 | NM_001134480 | ENSMUSG00000087141 | **Plcxd2** | AW494639\|EG433022 | phosphatidylinositol-specific phospholipase C, X domain containing 2 | protein-coding |
| chr5_122163142_122163640 | chr5 | 122163143 | 122163640 | + | -1.29257 | intron (NM_013636, intron 1 of 6) | intron (NM_013636, intron 1 of 6) | 5112 | NM_013636 | 19047 | Mm.280784 | NM_013636 | ENSMUSG00000004455 | **Ppp1cc** | PP-1G\|PP1\|dis2m1 | protein phosphatase 1, catalytic subunit, gamma isoform | protein-coding |
| chr1_191425839_191426311 | chr1 | 191425840 | 191426311 | + | -1.00604 | Intergenic | Intergenic | -29034 | NM_144880 | 226849 | Mm.205569 | NM_144880 | ENSMUSG00000026626 | **Ppp2r5a** | PR61alpha | protein phosphatase 2, regulatory subunit B', alpha | protein-coding |
| chr1_191425930_191426225 | chr1 | 191425931 | 191426225 | + | -1.01557 | Intergenic | Intergenic | -29037 | NM_144880 | 226849 | Mm.205569 | NM_144880 | ENSMUSG00000026626 | **Ppp2r5a** | PR61alpha | protein phosphatase 2, regulatory subunit B', alpha | protein-coding |
| chr11_108273579_108274081 | chr11 | 108273580 | 108274081 | + | -0.704708 | intron (NM_011101, intron 2 of 16) | intron (NM_011101, intron 2 of 16) | 70058 | NM_011101 | 18750 | Mm.222178 | NM_011101 | ENSMUSG00000050965 | **Prkca** | AI875142\|Pkca | protein kinase C, alpha | protein-coding |
| chr13_28417363_28418009 | chr13 | 28417364 | 28418009 | + | -0.721755 | Intergenic | Intergenic | 275202 | NM_023746 | 28078 | Mm.155058 | NM_023746 | ENSMUSG00000017064 | **Prl5a1** | 1600013P04Rik\|D13Wsu14e\|PLP-L\|Prlpl | prolactin family 5, subfamily a, member 1 | protein-coding |
| chr2_32167829_32168216 | chr2 | 32167830 | 32168216 | + | -0.725539 | intron (NM_001159634, intron 1 of 31) | intron (NM_001159634, intron 1 of 31) | 16875 | NM_001159634 | 227723 | Mm.25504 | NM_172661 | ENSMUSG00000039262 | **Prrc2b** | 5830434P21Rik\|AI173903\|Bat2l\|Bat2l1\|D430039P21\|mKIAA0515 | proline-rich coiled-coil 2B | protein-coding |
| chr1_34484376_34484821 | chr1 | 34484377 | 34484821 | + | -0.633899 | Intergenic | MTE2b\|LTR\|ERVL-MaLR | -13831 | NM_009355 | 21755 | Mm.10725 | NM_009355 | ENSMUSG00000026125 | **Prss39** | Tesp1 | protease, serine 39 | protein-coding |
| chr11_98959532_98959813 | chr11 | 98959533 | 98959813 | + | -0.796253 | promoter-TSS (NM_001176528) | promoter-TSS (NM_001176528) | -798 | NM_001176528 | 19401 | Mm.439744 | NM_009024 | ENSMUSG00000037992 | **Rara** | Nr1b1\|RAR\|RARalpha1 | retinoic acid receptor, alpha | protein-coding |
| chr11_98959433_98960017 | chr11 | 98959434 | 98960017 | + | -0.797321 | promoter-TSS (NM_001176528) | promoter-TSS (NM_001176528) | -746 | NM_001176528 | 19401 | Mm.439744 | NM_009024 | ENSMUSG00000037992 | **Rara** | Nr1b1\|RAR\|RARalpha1 | retinoic acid receptor, alpha | protein-coding |
| chr19_6400327_6400732 | chr19 | 6400328 | 6400732 | + | -0.920049 | promoter-TSS (NM_011242) | promoter-TSS (NM_011242) | -53 | NM_011242 | 19395 | Mm.77017 | NM_011242 | ENSMUSG00000032946 | **Rasgrp2** | CDC25L\|Caldaggef1 | RAS, guanyl releasing protein 2 | protein-coding |
| chr19_6400392_6400679 | chr19 | 6400393 | 6400679 | + | -0.932381 | promoter-TSS (NM_011242) | promoter-TSS (NM_011242) | -47 | NM_011242 | 19395 | Mm.77017 | NM_011242 | ENSMUSG00000032946 | **Rasgrp2** | CDC25L\|Caldaggef1 | RAS, guanyl releasing protein 2 | protein-coding |
| chr7_128373861_128374645 | chr7 | 128373862 | 128374645 | + | -0.744538 | intron (NM_026418, intron 4 of 4) | RSINE1\|SINE\|B4 | 43919 | NM_026418 | 67865 | Mm.18635 | NM_026418 | ENSMUSG00000030844 | **Rgs10** | 2310010N19Rik | regulator of G-protein signalling 10 | protein-coding |
| chr7_128374131_128374534 | chr7 | 128374132 | 128374534 | + | -0.869328 | intron (NM_026418, intron 4 of 4) | RSINE1\|SINE\|B4 | 43839 | NM_026418 | 67865 | Mm.18635 | NM_026418 | ENSMUSG00000030844 | **Rgs10** | 2310010N19Rik | regulator of G-protein signalling 10 | protein-coding |
| chr13_38037844_38038300 | chr13 | 38037845 | 38038300 | + | -1.00741 | intron (NM_024242, intron 1 of 16) | intron (NM_024242, intron 1 of 16) | 1083 | NM_024242 | 71340 | Mm.297130 | NM_024242 | ENSMUSG00000021428 | **Riok1** | 3110046C13Rik\|5430416A05Rik\|Ad034 | RIO kinase 1 (yeast) | protein-coding |
| chr18_20962647_20962856 | chr18 | 20962648 | 20962856 | + | -0.867231 | intron (NM_026301, intron 1 of 5) | intron (NM_026301, intron 1 of 5) | 18127 | NM_026301 | 67664 | Mm.45980 | NM_026301 | ENSMUSG00000033107 | **Rnf125** | 4930553F04Rik | ring finger protein 125 | protein-coding |
| chr18_20962629_20963056 | chr18 | 20962630 | 20963056 | + | -0.895028 | intron (NM_026301, intron 1 of 5) | intron (NM_026301, intron 1 of 5) | 18218 | NM_026301 | 67664 | Mm.45980 | NM_026301 | ENSMUSG00000033107 | **Rnf125** | 4930553F04Rik | ring finger protein 125 | protein-coding |
| chr6_70813932_70814833 | chr6 | 70813933 | 70814833 | + | -0.663598 | Intergenic | B4A\|SINE\|B4 | -22208 | NM_009075 | 19895 | Mm.17905 | NM_009075 | ENSMUSG00000053604 | **Rpia** | RPI | ribose 5-phosphate isomerase A | protein-coding |
| chr6_70813932_70814413 | chr6 | 70813933 | 70814413 | + | -0.708573 | Intergenic | Intergenic | -21998 | NM_009075 | 19895 | Mm.17905 | NM_009075 | ENSMUSG00000053604 | **Rpia** | RPI | ribose 5-phosphate isomerase A | protein-coding |
| chr7_114069019_114069665 | chr7 | 114069020 | 114069665 | + | -0.976227 | intron (NM_025846, intron 1 of 5) | intron (NM_025846, intron 1 of 5) | 48439 | NM_025846 | 66922 | Mm.276572 | NM_025846 | ENSMUSG00000055723 | **Rras2** | 2610016H24Rik\|C86394\|TC21 | related RAS viral (r-ras) oncogene homolog 2 | protein-coding |
| chr7_114093540_114094123 | chr7 | 114093541 | 114094123 | + | -1.1227 | intron (NM_025846, intron 1 of 5) | intron (NM_025846, intron 1 of 5) | 23949 | NM_025846 | 66922 | Mm.276572 | NM_025846 | ENSMUSG00000055723 | **Rras2** | 2610016H24Rik\|C86394\|TC21 | related RAS viral (r-ras) oncogene homolog 2 | protein-coding |
| chr7_114093619_114093886 | chr7 | 114093620 | 114093886 | + | -1.1689 | intron (NM_025846, intron 1 of 5) | (CA)n\|Simple_repeat\|Simple_repeat | 24028 | NM_025846 | 66922 | Mm.276572 | NM_025846 | ENSMUSG00000055723 | **Rras2** | 2610016H24Rik\|C86394\|TC21 | related RAS viral (r-ras) oncogene homolog 2 | protein-coding |
| chr10_68081671_68082192 | chr10 | 68081672 | 68082192 | + | -0.878031 | Intergenic | Intergenic | 102334 | NM_001081346 | 170799 | Mm.209844 | NM_133244 | ENSMUSG00000037846 | **Rtkn2** | B130039D23Rik\|Mbf\|Plekhk1 | rhotekin 2 | protein-coding |
| chr3_115793316_115794039 | chr3 | 115793317 | 115794039 | + | -0.834832 | Intergenic | Intergenic | -78623 | NM_007901 | 13609 | Mm.982 | NM_007901 | ENSMUSG00000045092 | **S1pr1** | AI849002\|Edg1\|Lpb1\|S1p\|S1p1 | sphingosine-1-phosphate receptor 1 | protein-coding |
| chr3_58552051_58552413 | chr3 | 58552052 | 58552413 | + | -1.02595 | intron (NM_001005509, intron 10 of 13) | intron (NM_001005509, intron 10 of 13) | -24426 | NM_001040396 | 69227 | Mm.270950 | NM_001040396 | ENSMUSG00000075700 | **Selt** | 2810407C02Rik\|5730408P04Rik | selenoprotein T | protein-coding |
| chr19_32182104_32182466 | chr19 | 32182105 | 32182466 | + | -1.17131 | intron (NM_144792, intron 5 of 9) | intron (NM_144792, intron 5 of 9) | 28728 | NM_001168526 | 208449 | Mm.329810 | NM_144792 | ENSMUSG00000040451 | **Sgms1** | 9530058O11Rik\|AI841905\|C80702\|Mob\|Sms1\|Sor1\|Tmem23 | sphingomyelin synthase 1 | protein-coding |
| chr7_126470094_126470563 | chr7 | 126470095 | 126470563 | + | -0.762947 | intron (NR_110346, intron 4 of 9) | intron (NR_110346, intron 4 of 9) | 2765 | NR_110346 | 20399 | Mm.8538 | NM_011363 | ENSMUSG00000030733 | **Sh2b1** | AI425885\|C530001K22Rik\|Irip\|Psm\|SH2-B\|SH2-Bb\|Sh2bpsm1\|mKIAA1299 | SH2B adaptor protein 1 | protein-coding |
| chr5_115488758_115489306 | chr5 | 115488759 | 115489306 | + | -0.823117 | Intergenic | Intergenic | -4548 | NM_133760 | 75387 | Mm.332616 | NM_133760 | ENSMUSG00000029524 | **Sirt4** | 4930596O17Rik | sirtuin 4 | protein-coding |
| chr4_155168612_155169528 | chr4 | 155168613 | 155169528 | + | -0.867572 | intron (NM_011385, intron 1 of 6) | intron (NM_011385, intron 1 of 6) | 53465 | NM_011385 | 20481 | Mm.28520 | NM_011385 | ENSMUSG00000029050 | **Ski** | 2310012I02Rik\|2610001A11Rik\|AA062172\|AA589460\|BC004088 | ski sarcoma viral oncogene homolog (avian) | protein-coding |
| chr4_155169046_155169577 | chr4 | 155169047 | 155169577 | + | -0.867899 | intron (NM_011385, intron 1 of 6) | intron (NM_011385, intron 1 of 6) | 53223 | NM_011385 | 20481 | Mm.28520 | NM_011385 | ENSMUSG00000029050 | **Ski** | 2310012I02Rik\|2610001A11Rik\|AA062172\|AA589460\|BC004088 | ski sarcoma viral oncogene homolog (avian) | protein-coding |
| chr4_155169048_155169438 | chr4 | 155169049 | 155169438 | + | -0.895383 | intron (NM_011385, intron 1 of 6) | intron (NM_011385, intron 1 of 6) | 53292 | NM_011385 | 20481 | Mm.28520 | NM_011385 | ENSMUSG00000029050 | **Ski** | 2310012I02Rik\|2610001A11Rik\|AA062172\|AA589460\|BC004088 | ski sarcoma viral oncogene homolog (avian) | protein-coding |
| chr6_34176096_34176379 | chr6 | 34176097 | 34176379 | + | -0.66736 | intron (NM_021435, intron 1 of 9) | intron (NM_021435, intron 1 of 9) | 816 | NM_021435 | 58246 | Mm.245527 | NM_021435 | | **Slc35b4** | 4930474D06Rik\|AB041549\|AW494345 | solute carrier family 35, member B4 | protein-coding |
| chrX_101147485_101147962 | chrX | 101147486 | 101147962 | + | -1.08191 | intron (NM_001110311, intron 4 of 5) | PB1\|SINE\|Alu | -62372 | NM_007515 | 11989 | Mm.5140 | NM_007515 | ENSMUSG00000031297 | **Slc7a3** | Atrc3\|CAT3\|SLC7A1\|SLC7A2 | solute carrier family 7 (cationic amino acid transporter, y+ system), member 3 | protein-coding |
| chr11_99203585_99204395 | chr11 | 99203586 | 99204395 | + | -0.860703 | Intergenic | Intergenic | 27027 | NM_020618 | 57376 | Mm.379086 | NM_020618 | ENSMUSG00000037935 | **Smarce1** | 2810417B20Rik\|5830412H02Rik\|9030408N19Rik\|Baf27 | SWI/SNF related, matrix associated, actin dependent regulator of chromatin, subfamily e, member 1 | protein-coding |
| chr11_99203842_99204380 | chr11 | 99203843 | 99204380 | + | -0.885915 | Intergenic | Intergenic | 26906 | NM_020618 | 57376 | Mm.379086 | NM_020618 | ENSMUSG00000037935 | **Smarce1** | 2810417B20Rik\|5830412H02Rik\|9030408N19Rik\|Baf27 | SWI/SNF related, matrix associated, actin dependent regulator of chromatin, subfamily e, member 1 | protein-coding |

| | | | | | | | | | | | | | | | | | |
|---|---|---|---|---|---|---|---|---|---|---|---|---|---|---|---|---|---|
| chr2_131509652_131510440 | chr2 | 131509653 | 131510440 | + | -0.706561 | intron (NM_001177840, intron 1 of 3) | intron (NM_001177840, intron 1 of 3) | 18184 | NM_001177838 | 228608 | Mm.136586 | NM_145533 | ENSMUSG00000027333 | **Smox** | B130066H01Rik\|PAO\|PAOh1\|SMO | spermine oxidase | protein-coding |
| chr11_106535914_106536654 | chr11 | 106535915 | 106536654 | + | -0.94041 | intron (NM_198292, intron 5 of 11) | intron (NM_198292, intron 5 of 11) | 35293 | NR_030703 | 100216537 | | NR_030703 | ENSMUSG00000065126 | **Snord104** | - | small nucleolar RNA, C/D box 104 | snoRNA |
| chr13_113574184_113574790 | chr13 | 113574185 | 113574790 | + | -0.706501 | Intergenic | Intergenic | 44077 | NM_130796 | 170625 | Mm.33721 | NM_130796 | ENSMUSG00000042364 | **Snx18** | Snag1 | sorting nexin 18 | protein-coding |
| chr16_90221557_90222012 | chr16 | 90221558 | 90222012 | + | -0.845853 | intron (NM_011434, intron 1 of 4) | intron (NM_011434, intron 1 of 4) | 1043 | NM_011434 | 20655 | Mm.276325 | NM_011434 | ENSMUSG00000022982 | **Sod1** | B430204E11Rik\|Cu/Zn-SOD\|CuZnSOD\|Ipo-1\|Ipo1\|SODC\|Sod-1 | superoxide dismutase 1, soluble | protein-coding |
| chr18_14656710_14657755 | chr18 | 14656711 | 14657755 | + | -0.891031 | intron (NM_001161369, intron 3 of 9) | intron (NM_001161369, intron 3 of 9) | 25681 | NM_001161369 | 268996 | Mm.31761 | NM_009280 | ENSMUSG00000037013 | **Ss18** | D130059H17\|Ssxt\|Syt | synovial sarcoma translocation, Chromosome 18 | protein-coding |
| chr18_14656705_14657281 | chr18 | 14656706 | 14657281 | + | -1.03294 | intron (NM_001161369, intron 3 of 9) | intron (NM_001161369, intron 3 of 9) | 25921 | NM_001161369 | 268996 | Mm.31761 | NM_009280 | ENSMUSG00000037013 | **Ss18** | D130059H17\|Ssxt\|Syt | synovial sarcoma translocation, Chromosome 18 | protein-coding |
| chr6_142915336_142915820 | chr6 | 142915337 | 142915820 | + | -0.891461 | intron (NM_011374, intron 1 of 4) | intron (NM_011374, intron 1 of 4) | 48874 | NM_011374 | 20449 | Mm.260838 | NM_011374 | ENSMUSG00000030283 | **St8sia1** | 9330109E03Rik\|GD3S\|Sia-T\|Siat8\|Siat8a | ST8 alpha-N-acetyl-neuraminide alpha-2,8-sialyltransferase 1 | protein-coding |
| chr11_100848081_100848593 | chr11 | 100848082 | 100848593 | + | -1.09456 | intron (NM_001113563, intron 1 of 18) | intron (NM_001113563, intron 1 of 18) | 2248 | NM_001113563 | 20851 | Mm.34064 | NM_011489 | ENSMUSG00000020919 | **Stat5b** | - | signal transducer and activator of transcription 5B | protein-coding |
| chr11_100845784_100846384 | chr11 | 100845785 | 100846384 | + | -1.52961 | intron (NM_001113563, intron 1 of 18) | intron (NM_001113563, intron 1 of 18) | 4501 | NM_001113563 | 20851 | Mm.34064 | NM_011489 | ENSMUSG00000020919 | **Stat5b** | - | signal transducer and activator of transcription 5B | protein-coding |
| chr11_32535640_32536064 | chr11 | 32535641 | 32536064 | + | -1.09122 | intron (NM_009288, intron 1 of 18) | intron (NM_009288, intron 1 of 18) | 2586 | NM_009288 | 20868 | Mm.8235 | NM_009288 | ENSMUSG00000020272 | **Stk10** | Gek1\|Lok\|mKIAA4026 | serine/threonine kinase 10 | protein-coding |
| chr5_44223227_44223602 | chr5 | 44223228 | 44223602 | + | -0.693903 | intron (NM_173764, intron 1 of 13) | intron (NM_173764, intron 1 of 13) | 3191 | NM_173764 | 231225 | Mm.30688 | NM_173764 | ENSMUSG00000046985 | **Tapt1** | 4932414K18Rik | transmembrane anterior posterior transformation 1 | protein-coding |
| chr4_102967676_102968473 | chr4 | 102967677 | 102968473 | + | -1.10352 | non-coding (NR_040444, exon 4 of 4) | non-coding (NR_040444, exon 4 of 4) | -18304 | NM_001163767 | 67344 | Mm.111346 | NM_026100 | ENSMUSG00000028523 | **Tctex1d1** | 1700055O19Rik | Tctex1 domain containing 1 | protein-coding |
| chr9_116160841_116161289 | chr9 | 116160842 | 116161289 | + | -0.800979 | intron (NM_009371, intron 1 of 7) | intron (NM_009371, intron 1 of 7) | 14298 | NM_029575 | 21813 | Mm.172346 | NM_009371 | ENSMUSG00000032440 | **Tgfbr2** | 1110020H15Rik\|AU042018\|DNIIR\|RIIDN\|TBR-II\|TbetaR-II\|TbetaRII | transforming growth factor, beta receptor II | protein-coding |
| chr9_116160974_116161287 | chr9 | 116160975 | 116161287 | + | -0.819543 | intron (NM_009371, intron 1 of 7) | intron (NM_009371, intron 1 of 7) | 14232 | NM_029575 | 21813 | Mm.172346 | NM_009371 | ENSMUSG00000032440 | **Tgfbr2** | 1110020H15Rik\|AU042018\|DNIIR\|RIIDN\|TBR-II\|TbetaR-II\|TbetaRII | transforming growth factor, beta receptor II | protein-coding |
| chr9_116101767_116102237 | chr9 | 116101768 | 116102237 | + | -0.839267 | intron (NM_029575, intron 5 of 6) | intron (NM_029575, intron 5 of 6) | 73361 | NM_029575 | 21813 | Mm.172346 | NM_009371 | ENSMUSG00000032440 | **Tgfbr2** | 1110020H15Rik\|AU042018\|DNIIR\|RIIDN\|TBR-II\|TbetaR-II\|TbetaRII | transforming growth factor, beta receptor II | protein-coding |
| chr9_116163871_116164319 | chr9 | 116163872 | 116164319 | + | -0.955797 | intron (NM_009371, intron 1 of 7) | intron (NM_009371, intron 1 of 7) | 11268 | NM_029575 | 21813 | Mm.172346 | NM_009371 | ENSMUSG00000032440 | **Tgfbr2** | 1110020H15Rik\|AU042018\|DNIIR\|RIIDN\|TBR-II\|TbetaR-II\|TbetaRII | transforming growth factor, beta receptor II | protein-coding |
| chr9_116084411_116085026 | chr9 | 116084412 | 116085026 | + | -1.06072 | Intergenic | Intergenic | 90644 | NM_029575 | 21813 | Mm.172346 | NM_009371 | ENSMUSG00000032440 | **Tgfbr2** | 1110020H15Rik\|AU042018\|DNIIR\|RIIDN\|TBR-II\|TbetaR-II\|TbetaRII | transforming growth factor, beta receptor II | protein-coding |
| chr9_116084183_116084953 | chr9 | 116084184 | 116084953 | + | -1.07627 | Intergenic | Intergenic | 90795 | NM_029575 | 21813 | Mm.172346 | NM_009371 | ENSMUSG00000032440 | **Tgfbr2** | 1110020H15Rik\|AU042018\|DNIIR\|RIIDN\|TBR-II\|TbetaR-II\|TbetaRII | transforming growth factor, beta receptor II | protein-coding |
| chr9_116187401_116187876 | chr9 | 116187402 | 116187876 | + | -1.11587 | Intergenic | Intergenic | -12276 | NM_009371 | 21813 | Mm.172346 | NM_009371 | ENSMUSG00000032440 | **Tgfbr2** | 1110020H15Rik\|AU042018\|DNIIR\|RIIDN\|TBR-II\|TbetaR-II\|TbetaRII | transforming growth factor, beta receptor II | protein-coding |
| chr17_84352152_84352509 | chr17 | 84352153 | 84352509 | + | -0.722687 | intron (NM_183021, intron 26 of 37) | RSINE1\|SINE\|B4 | 113877 | NM_183021 | 240174 | Mm.246767 | NM_177826 | ENSMUSG00000024251 | **Thada** | 6530405K19\|A930021O22\|AU019899\|BC052885\|GITA | thyroid adenoma associated | protein-coding |
| chr14_122062567_122063111 | chr14 | 122062568 | 122063111 | + | -0.881238 | Intergenic | B4A\|SINE\|B4 | 28165 | NM_001037744 | 223262 | Mm.297637 | NM_001037744 | ENSMUSG00000071229 | **Timm8a2** | - | translocase of inner mitochondrial membrane 8A2 | protein-coding |
| chr14_122062825_122063070 | chr14 | 122062826 | 122063070 | + | -1.00259 | Intergenic | MTD\|LTR\|ERVL-MaLR | 28274 | NM_001037744 | 223262 | Mm.297637 | NM_001037744 | ENSMUSG00000071229 | **Timm8a2** | - | translocase of inner mitochondrial membrane 8A2 | protein-coding |
| chr1_52653261_52653799 | chr1 | 52653262 | 52653799 | + | -0.947484 | Intergenic | Intergenic | 22825 | NM_001142647 | 227094 | Mm.83541 | NM_172654 | ENSMUSG00000043015 | **Tmem194b** | 5330401P04Rik\|9830163O08 | transmembrane protein 194B | protein-coding |
| chr4_133298519_133298893 | chr4 | 133298520 | 133298893 | + | -0.825783 | intron (NM_199306, intron 12 of 15) | intron (NM_199306, intron 12 of 15) | -20916 | NR_110978 | 52174 | Mm.244748 | NM_025667 | ENSMUSG00000028857 | **Tmem222** | 5730406H10Rik\|AA409341\|D4Ertd196e | transmembrane protein 222 | protein-coding |
| chr4_15273203_15273560 | chr4 | 15273204 | 15273560 | + | -1.09697 | intron (NM_181401, intron 1 of 2) | intron (NM_181401, intron 1 of 2) | 7562 | NM_181401 | 100201 | Mm.394111 | NM_181401 | ENSMUSG00000043252 | **Tmem64** | 9630015D15Rik\|AI790744\|AV300874 | transmembrane protein 64 | protein-coding |
| chr4_15273187_15273723 | chr4 | 15273188 | 15273723 | + | -1.11375 | intron (NM_181401, intron 1 of 2) | intron (NM_181401, intron 1 of 2) | 7635 | NM_181401 | 100201 | Mm.394111 | NM_181401 | ENSMUSG00000043252 | **Tmem64** | 9630015D15Rik\|AI790744\|AV300874 | transmembrane protein 64 | protein-coding |
| chr10_118633378_118634010 | chr10 | 118633379 | 118634010 | + | -1.19793 | Intergenic | Intergenic | -77169 | NR_104123 | 103214 | Mm.30529 | NR_104123 | | **Tmevpg1** | AI447422 | Theiler's murine encephalomyelitis virus persistence candidate gene 1 | ncRNA |
| chr10_118599120_118599607 | chr10 | 118599121 | 118599607 | + | -1.26229 | Intergenic | Intergenic | -42839 | NR_104123 | 103214 | Mm.30529 | NR_104123 | | **Tmevpg1** | AI447422 | Theiler's murine encephalomyelitis virus persistence candidate gene 1 | ncRNA |
| chr10_18999278_18999761 | chr10 | 18999279 | 18999761 | + | -0.705099 | Intergenic | Intergenic | 12443 | NM_001166402 | 21929 | Mm.116683 | NM_009397 | ENSMUSG00000019850 | **Tnfaip3** | A20\|Tnfip3 | tumor necrosis factor, alpha-induced protein 3 | protein-coding |
| chr10_19061928_19063110 | chr10 | 19061929 | 19063110 | + | -0.950535 | Intergenic | Intergenic | -47109 | NM_009397 | 21929 | Mm.116683 | NM_009397 | ENSMUSG00000019850 | **Tnfaip3** | A20\|Tnfip3 | tumor necrosis factor, alpha-induced protein 3 | protein-coding |
| chr6_125351328_125351741 | chr6 | 125351329 | 125351741 | + | -0.846944 | intron (NM_011609, intron 1 of 9) | intron (NM_011609, intron 1 of 9) | 1812 | NM_011609 | 21937 | Mm.1258 | NM_011609 | ENSMUSG00000030341 | **Tnfrsf1a** | CD120a\|FPF\|TNF-R\|TNF-R-I\|TNF-R1\|TNF-R55\|TNF-alphaR1\|TNFAR\|TNFR60\|TNFRI\|TNFRp55\|TNFalpha-R1\|Tnfr-2\|Tnfr1\|p55\|p55-R | tumor necrosis factor receptor superfamily, member 1a | protein-coding |
| chr4_145240725_145240978 | chr4 | 145240726 | 145240978 | + | -0.724085 | intron (NM_011610, intron 1 of 9) | intron (NM_011610, intron 1 of 9) | 6018 | NM_011610 | 21938 | Mm.235328 | NM_011610 | | **Tnfrsf1b** | CD120b\|TNF-R-II\|TNF-R2\|TNF-R75\|TNF-alphaR2\|TNFBR\|TNFR80\|TNFRII\|TNFalpha-R2\|Tnfr-1\|Tnfr2\|p75 | tumor necrosis factor receptor superfamily, member 1b | protein-coding |
| chr4_145241058_145241326 | chr4 | 145241059 | 145241326 | + | -0.735141 | intron (NM_011610, intron 1 of 9) | intron (NM_011610, intron 1 of 9) | 5678 | NM_011610 | 21938 | Mm.235328 | NM_011610 | | **Tnfrsf1b** | CD120b\|TNF-R-II\|TNF-R2\|TNF-R75\|TNF-alphaR2\|TNFBR\|TNFR80\|TNFRII\|TNFalpha-R2\|Tnfr-1\|Tnfr2\|p75 | tumor necrosis factor receptor superfamily, member 1b | protein-coding |
| chr4_145240652_145241342 | chr4 | 145240653 | 145241342 | + | -0.760352 | intron (NM_011610, intron 1 of 9) | intron (NM_011610, intron 1 of 9) | 5873 | NM_011610 | 21938 | Mm.235328 | NM_011610 | | **Tnfrsf1b** | CD120b\|TNF-R-II\|TNF-R2\|TNF-R75\|TNF-alphaR2\|TNFBR\|TNFR80\|TNFRII\|TNFalpha-R2\|Tnfr-1\|Tnfr2\|p75 | tumor necrosis factor receptor superfamily, member 1b | protein-coding |
| chr11_89024212_89024566 | chr11 | 89024213 | 89024566 | + | -0.615715 | Intergenic | Intergenic | 24986 | NM_009546 | 217069 | Mm.248445 | NM_009546 | ENSMUSG00000000275 | **Trim25** | AA960166\|AL022677\|EFP\|Zfp147 | tripartite motif-containing 25 | protein-coding |
| chr14_121879351_121879995 | chr14 | 121879352 | 121879995 | + | -0.783376 | intron (NM_026861, intron 1 of 8) | intron (NM_026861, intron 1 of 8) | 1067 | NM_026861 | 68889 | Mm.441151 | NM_026861 | ENSMUSG00000041765 | **Ubac2** | 1190008A14Rik\|AW554959\|Phgdhl1 | ubiquitin associated domain containing 2 | protein-coding |
| chr14_121879453_121879826 | chr14 | 121879454 | 121879826 | + | -0.790566 | intron (NM_026861, intron 1 of 8) | intron (NM_026861, intron 1 of 8) | 1034 | NM_026861 | 68889 | Mm.441151 | NM_026861 | ENSMUSG00000041765 | **Ubac2** | 1190008A14Rik\|AW554959\|Phgdhl1 | ubiquitin associated domain containing 2 | protein-coding |
| chr16_4878977_4879679 | chr16 | 4878978 | 4879679 | + | -0.615417 | intron (NM_145359, intron 1 of 2) | CpG | 523 | NM_145359 | 207740 | Mm.268895 | NM_145359 | | **Ubald1** | 1500031H01Rik\|BC013706\|Fam100a | UBA-like domain containing 1 | protein-coding |
| chr16_4878960_4879500 | chr16 | 4878961 | 4879500 | + | -0.663507 | intron (NM_145359, intron 1 of 2) | intron (NM_145359, intron 1 of 2) | 621 | NM_145359 | 207740 | Mm.268895 | NM_145359 | | **Ubald1** | 1500031H01Rik\|BC013706\|Fam100a | UBA-like domain containing 1 | protein-coding |
| chr2_6478111_6479186 | chr2 | 6478112 | 6479186 | + | -0.950335 | Intergenic | Intergenic | 125913 | NM_181399 | 98910 | Mm.40673 | NM_181399 | ENSMUSG00000039046 | **Usp6nl** | AI316785\|RNTRE\|TRE2NL\|mKIAA0019 | USP6 N-terminal like | protein-coding |
| chr2_6478111_6478914 | chr2 | 6478112 | 6478914 | + | -0.969687 | Intergenic | Intergenic | 125777 | NM_181399 | 98910 | Mm.40673 | NM_181399 | ENSMUSG00000039046 | **Usp6nl** | AI316785\|RNTRE\|TRE2NL\|mKIAA0019 | USP6 N-terminal like | protein-coding |
| chr1_162571181_162571726 | chr1 | 162571182 | 162571726 | + | -0.844223 | intron (NM_016796, intron 1 of 7) | CpG | 626 | NM_016796 | 53330 | Mm.10699 | NM_016796 | ENSMUSG00000026696 | **Vamp4** | D1Ertd147e | vesicle-associated membrane protein 4 | protein-coding |
| chr1_162571357_162571682 | chr1 | 162571358 | 162571682 | + | -0.902363 | intron (NM_016796, intron 1 of 7) | CpG | 692 | NM_016796 | 53330 | Mm.10699 | NM_016796 | ENSMUSG00000026696 | **Vamp4** | D1Ertd147e | vesicle-associated membrane protein 4 | protein-coding |
| chr4_133162175_133162677 | chr4 | 133162176 | 133162677 | + | -0.807146 | intron (NM_153423, intron 1 of 8) | intron (NM_153423, intron 1 of 8) | 31793 | NM_153423 | 242687 | Mm.23566 | NM_153423 | ENSMUSG00000028868 | **Wasf2** | AW742646\|D4Ertd13e\|WAVE2 | WAS protein family, member 2 | protein-coding |
| chr1_181227800_181228230 | chr1 | 181227801 | 181228230 | + | -0.787435 | Intergenic | Intergenic | -16037 | NM_145514 | 226757 | Mm.289082 | NM_145514 | ENSMUSG00000038733 | **Wdr26** | 1600024A01Rik\|AA693241\|AI447817\|AU044014\|C77982\|Gid7 | WD repeat domain 26 | protein-coding |

<table>
<tr><td>chr15_58165135_58165664</td><td>chr15</td><td>58165136</td><td>58165664</td><td>+</td><td>-0.839641</td><td>Intergenic</td><td>Intergenic</td><td>23964</td><td>NM_029734</td><td>76773</td><td>Mm.26771</td><td>NM_029734</td><td>ENSMUSG00000022359</td><td>Wdyhv1</td><td>2410187C16Rik|AU014961|AW550036|Ntaq1</td><td>WDYHV motif containing 1</td><td>protein-coding</td></tr>
<tr><td>chr8_114973000_114973511</td><td>chr8</td><td>114973001</td><td>114973511</td><td>+</td><td>-0.781406</td><td>intron (NM_019573, intron 8 of 8)</td><td>intron (NM_019573, intron 8 of 8)</td><td>533604</td><td>NM_019573</td><td>80707</td><td>Mm.440420</td><td>NM_019573</td><td>ENSMUSG00000004637</td><td>Wwox</td><td>5330426P09Rik|9030416C10Rik|WOX1</td><td>WW domain-containing oxidoreductase</td><td>protein-coding</td></tr>
<tr><td>chr8_114803112_114803622</td><td>chr8</td><td>114803113</td><td>114803622</td><td>+</td><td>-0.839032</td><td>intron (NM_019573, intron 8 of 8)</td><td>intron (NM_019573, intron 8 of 8)</td><td>363715</td><td>NM_019573</td><td>80707</td><td>Mm.440420</td><td>NM_019573</td><td>ENSMUSG00000004637</td><td>Wwox</td><td>5330426P09Rik|9030416C10Rik|WOX1</td><td>WW domain-containing oxidoreductase</td><td>protein-coding</td></tr>
<tr><td>chr8_114801755_114802112</td><td>chr8</td><td>114801756</td><td>114802112</td><td>+</td><td>-0.915655</td><td>intron (NM_019573, intron 8 of 8)</td><td>intron (NM_019573, intron 8 of 8)</td><td>362282</td><td>NM_019573</td><td>80707</td><td>Mm.440420</td><td>NM_019573</td><td>ENSMUSG00000004637</td><td>Wwox</td><td>5330426P09Rik|9030416C10Rik|WOX1</td><td>WW domain-containing oxidoreductase</td><td>protein-coding</td></tr>
<tr><td>chr17_73893262_73893776</td><td>chr17</td><td>73893263</td><td>73893776</td><td>+</td><td>-0.877352</td><td>intron (NM_011723, intron 32 of 35)</td><td>intron (NM_011723, intron 32 of 35)</td><td>56677</td><td>NM_011723</td><td>22436</td><td>Mm.11223</td><td>NM_011723</td><td>ENSMUSG00000024066</td><td>Xdh</td><td>XO|Xor|Xox-1|Xox1</td><td>xanthine dehydrogenase</td><td>protein-coding</td></tr>
<tr><td>chrX_48098405_48099066</td><td>chrX</td><td>48098406</td><td>48099066</td><td>+</td><td>-0.779137</td><td>Intergenic</td><td>Intergenic</td><td>-9989</td><td>NM_001289729</td><td>170745</td><td>Mm.129279</td><td>NM_133213</td><td>ENSMUSG00000037005</td><td>Xpnpep2</td><td>9030008G12Rik|mAPP</td><td>X-prolyl aminopeptidase (aminopeptidase P) 2, membrane-bound</td><td>protein-coding</td></tr>
<tr><td>chrX_48096979_48097439</td><td>chrX</td><td>48096980</td><td>48097439</td><td>+</td><td>-0.781486</td><td>Intergenic</td><td>Intergenic</td><td>-11516</td><td>NM_001289729</td><td>170745</td><td>Mm.129279</td><td>NM_133213</td><td>ENSMUSG00000037005</td><td>Xpnpep2</td><td>9030008G12Rik|mAPP</td><td>X-prolyl aminopeptidase (aminopeptidase P) 2, membrane-bound</td><td>protein-coding</td></tr>
<tr><td>chrX_48098458_48098833</td><td>chrX</td><td>48098459</td><td>48098833</td><td>+</td><td>-0.789928</td><td>Intergenic</td><td>Intergenic</td><td>-10079</td><td>NM_001289729</td><td>170745</td><td>Mm.129279</td><td>NM_133213</td><td>ENSMUSG00000037005</td><td>Xpnpep2</td><td>9030008G12Rik|mAPP</td><td>X-prolyl aminopeptidase (aminopeptidase P) 2, membrane-bound</td><td>protein-coding</td></tr>
<tr><td>chr9_96696211_96696876</td><td>chr9</td><td>96696212</td><td>96696876</td><td>+</td><td>-0.854901</td><td>intron (NM_175537, intron 2 of 2)</td><td>intron (NM_175537, intron 2 of 2)</td><td>35131</td><td>NM_175537</td><td>245007</td><td>Mm.485954</td><td>NM_175537</td><td>ENSMUSG00000040433</td><td>Zbtb38</td><td>A930014K01Rik|CIBZ</td><td>zinc finger and BTB domain containing 38</td><td>protein-coding</td></tr>
<tr><td>chr9_96696012_96696960</td><td>chr9</td><td>96696013</td><td>96696960</td><td>+</td><td>-0.876567</td><td>intron (NM_175537, intron 2 of 2)</td><td>intron (NM_175537, intron 2 of 2)</td><td>35189</td><td>NM_175537</td><td>245007</td><td>Mm.485954</td><td>NM_175537</td><td>ENSMUSG00000040433</td><td>Zbtb38</td><td>A930014K01Rik|CIBZ</td><td>zinc finger and BTB domain containing 38</td><td>protein-coding</td></tr>
<tr><td>chr5_139483151_139483450</td><td>chr5</td><td>139483152</td><td>139483450</td><td>+</td><td>-0.952445</td><td>intron (NM_001159908, intron 1 of 6)</td><td>intron (NM_001159908, intron 1 of 6)</td><td>1190</td><td>NM_133349</td><td>100494</td><td>Mm.24521</td><td>NM_133349</td><td>ENSMUSG00000053581</td><td>Zfand2a</td><td>AA407930|AU016206|Airap</td><td>zinc finger, AN1-type domain 2A</td><td>protein-coding</td></tr>
<tr><td>chr11_70762685_70763066</td><td>chr11</td><td>70762686</td><td>70763066</td><td>+</td><td>-0.782151</td><td>Intergenic</td><td>ORR1D-int|LTR|ERVL-MaLR</td><td>-1571</td><td>NM_177565</td><td>193043</td><td>Mm.40966</td><td>NM_177565</td><td>ENSMUSG00000043602</td><td>Zfp3</td><td>AI854635|B230309J24|Fnp-1|Zfp-3</td><td>zinc finger protein 3</td><td>protein-coding</td></tr>
<tr><td>chr2_154615715_154616074</td><td>chr2</td><td>154615716</td><td>154616074</td><td>+</td><td>-1.06017</td><td>intron (NM_199304, intron 1 of 14)</td><td>intron (NM_199304, intron 1 of 14)</td><td>2527</td><td>NM_199304</td><td>228807</td><td>Mm.103555</td><td>NM_199304</td><td>ENSMUSG00000059842</td><td>Zfp341</td><td>Znf341</td><td>zinc finger protein 341</td><td>protein-coding</td></tr>
<tr><td>chr17_21473491_21474369</td><td>chr17</td><td>21473492</td><td>21474369</td><td>+</td><td>-0.69553</td><td>Intergenic</td><td>MMERGLN-int|LTR|ERV1</td><td>-15058</td><td>NM_013843</td><td>24132</td><td>Mm.42140</td><td>NM_013843</td><td></td><td>Zfp53</td><td>D030067O06Rik|KRAZ1|Zfp-53|Zfp118|zfas8</td><td>zinc finger protein 53</td><td>protein-coding</td></tr>
<tr><td>chr17_23617035_23617592</td><td>chr17</td><td>23617036</td><td>23617592</td><td>+</td><td>-0.99075</td><td>Intergenic</td><td>MLTR11B|LTR|ERVK</td><td>16488</td><td>NM_001289482</td><td>332221</td><td>Mm.270315</td><td>NM_001033425</td><td>ENSMUSG00000023902</td><td>Zscan10</td><td>BC068284|Zfp206|Zkscan10|Znf206</td><td>zinc finger and SCAN domain containing 10</td><td>protein-coding</td></tr>
<tr><td>chr17_23620278_23620796</td><td>chr17</td><td>23620279</td><td>23620796</td><td>+</td><td>-1.18578</td><td>Intergenic</td><td>Intergenic</td><td>19711</td><td>NM_001289482</td><td>332221</td><td>Mm.270315</td><td>NM_001033425</td><td>ENSMUSG00000023902</td><td>Zscan10</td><td>BC068284|Zfp206|Zkscan10|Znf206</td><td>zinc finger and SCAN domain containing 10</td><td>protein-coding</td></tr>
</table>

**Supplementary Table S3. GO enrichment analysis of genes near opening and closing peaks of CytP cells.**
Only GO terms with FDR value below 0.05 are shown. GO terms related to immune system and T cell activation are highlighted.

| #term ID | term description | observed gene count | background gene count | false discovery rate | matching proteins in your network (labels) |
|---|---|---|---|---|---|
| GO:0065007 | biological regulation | 242 | 10168 | 5.56E-13 | Abhd2,Adipoq,Aebp2,Aff4,Ago2,Agtr1b,Akt2,Ankrd11,Arhgap15,Arhgap25,Atp10a,Atp1a1,Atp2b4,Bach2,Bcl10,Btg1,Btg2,Btla,Camk4,Cblb,Ccdc80,Ccne1,Ccr1,Ccr4,Ccr7,Cd101,Cd247,Cd28,Cd300a,Cd47,Cd7,Cd97,Cdkn2b,Chd2,Chd7,Chst11,Chsy1,Clic4,Cnst,Crbn,Crem,Crim1,Cryaa,Cyfip2,Cyp2r1,Cyth4,Cytip,Dach2,Dad1,Ddx6,Dnajc6,Dock10,Dpp4,Dusp1,Dusp10,Dusp2,Dyrk2,E2f2,Eif4e3,Elf1,Elovl5,Epb4.1,Ercc4,Esrp1,Ets1,Ets2,F2,Fam20a,Fanci,Fasl,Fli1,Fntb,Fosl2,Foxn3,Foxp1,Fyb,Gadd45g,Gatsl3,Gdi2,Glrx,Glrx2,Gnat1,Gng2,Gpbar1,Gpr132,Gpr18,Gtf2ird1,Gucy2c,Gzmb,H2-Aa,Hectd1,Hey1,Hivep2,Igf2r,Il15ra,Il21r,Il6st,Inpp4b,Ints7,Ist1,Itga4,Itk,Jak1,Jarid2,Jun,Kcnh8,Kdm2b,Klf2,Klf3,Klf6,Lef1,Leprotl1,Lmtk3,Lonrf1,Lpcat3,Lrrc8d,Lrrn3,Lrtm2,Map3k1,Map3k8,Map4k1,Mapk14,Mapk9,Max,Mbd2,Mbnl1,Mknk2,Mospd1,Moxd1,Mtmr12,Nabp1,Ncf1,Neto2,Neurl1b,Nfatc1,Nfkbiz,Nipbl,Nrip1,Nrxn3,Nsg2,Nsmce1,Os9,Osm,Otulin,P2rx4,P2ry1,Pdcd4,Pde11a,Pde4a,Per1,Pim1,Pip4k2a,Pkp4,Plcxd2,Ppp1cc,Ppp2r5a,Prickle1,Prkca,Prkch,Prkcq,Prl5a1,Rara,Rasgrp2,Rftn1,Rgs10,Riok1,Rnf125,Rnf219,Rras2,Rtkn2,S1pr1,Sar1a,Scml4,Selt,Serpine2,Sgms1,Sh2b1,Shisa2,Sipa1l1,Sirt4,Ski,Sla,Slc30a4,Slc35b4,Slc35g1,Slc39a10,Slc7a3,Smarca2,Smarce1,Snx18,Sod1,Spag9,Spty2d1,Ss18,St8sia1,Stat4,Stat5b,Stim2,Stk10,Syne3,Tapt1,Tars2,Tax1bp1,Tbl1xr1,Tgfbr2,Thada,Tigit,Tmc1,Tmeff2,Tmem123,Tmem64,Tnfaip3,Tnfrsf1a,Tnfrsf1b,Tnfsf11,Tnfsf8,Tnrc6b,Trappc12,Trib1,Trim25,Txnl4b,Ubac2,Ubr2,Usp3,Usp6nl,Vamp4,Vgll4,Vps37b,Vsx2,Wasf2,Wwox,Xbp1,Xcr1,Xdh,Xylt1,Zfand2a,Zfp3,Zfp326,Zfp609,Zfp652,Zfp932,Zscan10 |
| GO:0050794 | regulation of cellular process | 222 | 9045 | 1.61E-12 | Abhd2,Adipoq,Aebp2,Aff4,Ago2,Agtr1b,Akt2,Arhgap15,Arhgap25,Atp1a1,Atp2b4,Bach2,Bcl10,Btg1,Btg2,Btla,Camk4,Cblb,Ccne1,Ccr1,Ccr4,Ccr7,Cd101,Cd247,Cd28,Cd300a,Cd47,Cd97,Cdkn2b,Chd2,Chd7,Chst11,Chsy1,Clic4,Cnst,Crbn,Crem,Crim1,Cryaa,Cyfip2,Cyth4,Dach2,Dad1,Ddx6,Dnajc6,Dock10,Dpp4,Dusp1,Dusp10,Dusp2,Dyrk2,E2f2,Eif4e3,Elf1,Elovl5,Epb4.1,Ercc4,Esrp1,Ets1,Ets2,F2,Fam20a,Fanci,Fasl,Fli1,Fntb,Fosl2,Foxn3,Foxp1,Fyb,Gadd45g,Gatsl3,Gdi2,Glrx,Glrx2,Gnat1,Gng2,Gpbar1,Gpr132,Gpr18,Gtf2ird1,Gucy2c,Gzmb,H2-Aa,Hey1,Hivep2,Igf2r,Il15ra,Il21r,Il6st,Inpp4b,Ints7,Ist1,Itga4,Itk,Jak1,Jarid2,Jun,Kcnh8,Kdm2b,Klf2,Klf3,Klf6,Lef1,Lmtk3,Lonrf1,Lrrc8d,Lrrn3,Lrtm2,Map3k1,Map3k8,Map4k1,Mapk14,Mapk9,Max,Mbd2,Mbnl1,Mknk2,Mospd1,Nabp1,Ncf1,Neto2,Neurl1b,Nfatc1,Nfkbiz,Nipbl,Nrip1,Nsg2,Nsmce1,Os9,Osm,Otulin,P2rx4,P2ry1,Pdcd4,Pde11a,Pde4a,Per1,Pim1,Pip4k2a,Pkp4,Plcxd2,Ppp1cc,Ppp2r5a,Prickle1,Prkca,Prkch,Prkcq,Prl5a1,Rara,Rasgrp2,Rftn1,Rgs10,Riok1,Rnf125,Rnf219,Rras2,Rtkn2,S1pr1,Sar1a,Scml4,Selt,Serpine2,Sgms1,Sh2b1,Shisa2,Sipa1l1,Sirt4,Ski,Sla,Slc30a4,Slc35b4,Slc39a10,Slc7a3,Smarca2,Smarce1,Sod1,Spag9,Spty2d1,Ss18,St8sia1,Stat4,Stat5b,Stk10,Syne3,Tapt1,Tax1bp1,Tbl1xr1,Tgfbr2,Tigit,Tmeff2,Tmem64,Tnfaip3,Tnfrsf1a,Tnfrsf1b,Tnfsf11,Tnfsf8,Tnrc6b,Trappc12,Trib1,Trim25,Txnl4b,Ubac2,Ubr2,Usp3,Usp6nl,Vamp4,Vgll4,Vsx2,Wasf2,Wwox,Xbp1,Xcr1,Xdh,Xylt1,Zfand2a,Zfp3,Zfp326,Zfp609,Zfp652,Zfp932,Zscan10 |
| GO:0050789 | regulation of biological process | 230 | 9594 | 2.36E-12 | Abhd2,Adipoq,Aebp2,Aff4,Ago2,Agtr1b,Akt2,Arhgap15,Arhgap25,Atp1a1,Atp2b4,Bach2,Bcl10,Btg1,Btg2,Btla,Camk4,Cblb,Ccdc80,Ccne1,Ccr1,Ccr4,Ccr7,Cd101,Cd247,Cd28,Cd300a,Cd47,Cd97,Cdkn2b,Chd2,Chd7,Chst11,Chsy1,Clic4,Cnst,Crbn,Crem,Crim1,Cryaa,Cyfip2,Cyth4,Cytip,Dach2,Dad1,Ddx6,Dnajc6,Dock10,Dpp4,Dusp1,Dusp10,Dusp2,Dyrk2,E2f2,Eif4e3,Elf1,Elovl5,Epb4.1,Ercc4,Esrp1,Ets1,Ets2,F2,Fam20a,Fanci,Fasl,Fli1,Fntb,Fosl2,Foxn3,Foxp1,Fyb,Gadd45g,Gatsl3,Gdi2,Glrx,Glrx2,Gnat1,Gng2,Gpbar1,Gpr132,Gpr18,Gtf2ird1,Gucy2c,Gzmb,H2-Aa,Hectd1,Hey1,Hivep2,Igf2r,Il15ra,Il21r,Il6st,Inpp4b,Ints7,Ist1,Itga4,Itk,Jak1,Jarid2,Jun,Kcnh8,Kdm2b,Klf2,Klf3,Klf6,Lef1,Leprotl1,Lmtk3,Lonrf1,Lpcat3,Lrrc8d,Lrrn3,Lrtm2,Map3k1,Map3k8,Map4k1,Mapk14,Mapk9,Max,Mbd2,Mbnl1,Mknk2,Mospd1,Nabp1,Ncf1,Neto2,Neurl1b,Nfatc1,Nfkbiz,Nipbl,Nrip1,Nsg2,Nsmce1,Os9,Osm,Otulin,P2rx4,P2ry1,Pdcd4,Pde11a,Pde4a,Per1,Pim1,Pip4k2a,Pkp4,Plcxd2,Ppp1cc,Ppp2r5a,Prickle1,Prkca,Prkch,Prkcq,Prl5a1,Rara,Rasgrp2,Rftn1,Rgs10,Riok1,Rnf125,Rnf219,Rras2,Rtkn2,S1pr1,Sar1a,Scml4,Selt,Serpine2,Sgms1,Sh2b1,Shisa2,Sipa1l1,Sirt4,Ski,Sla,Slc30a4,Slc35b4,Slc39a10,Slc7a3,Smarca2,Smarce1,Sod1,Spag9,Spty2d1,Ss18,St8sia1,Stat4,Stat5b,Stim2,Stk10,Syne3,Tapt1,Tax1bp1,Tbl1xr1,Tgfbr2,Tigit,Tmc1,Tmeff2,Tmem64,Tnfaip3,Tnfrsf1a,Tnfrsf1b,Tnfsf11,Tnfsf8,Tnrc6b,Trappc12,Trib1,Trim25,Txnl4b,Ubac2,Ubr2,Usp3,Usp6nl,Vamp4,Vgll4,Vps37b,Vsx2,Wasf2,Wwox,Xbp1,Xcr1,Xdh,Xylt1,Zfand2a,Zfp3,Zfp326,Zfp609,Zfp652,Zfp932,Zscan10 |
| GO:0050896 | response to stimulus | 177 | 6616 | 1.09E-11 | Abcb1b,Abhd2,Adipoq,Ago2,Agtr1b,Akt2,Arhgap15,Arhgap25,Atp1a1,Atp2b4,Bcl10,Btg1,Btg2,Btla,Camk4,Cblb,Ccdc80,Ccne1,Ccr1,Ccr4,Ccr7,Cd101,Cd247,Cd28,Cd300a,Cd47,Cd7,Cd97,Cdkn2b,Chd2,Chd7,Chst2,Chsy1,Clic4,Crem,Cryaa,Cyb5a,Cyp2r1,Dad1,Dock10,Dpp4,Dusp1,Dusp10,Dyrk2,E2f2,Ercc4,Ets1,Ext1,F2,Fanci,Fasl,Fkbp5,Fntb,Fosl2,Foxn3,Foxp1,Fyb,Gadd45g,Gatsl3,Gdi2,Glrx,Glrx2,Gnat1,Gng2,Gpbar1,Gpr132,Gpr18,Gucy2c,Gzmb,H2-Aa,Hey1,Hist1h2be,Hivep2,Igf2r,Il15ra,Il21r,Il6st,Ints7,Itga4,Itk,Jak1,Jarid2,Jun,Kcnh8,Klf2,Klf3,Klf6,Lef1,Lrrc8d,Lrtm2,Map3k1,Map3k8,Map4k1,Mapk14,Mapk9,Mbd2,Mgme1,Mknk2,Msrb3,Nabp1,Ncf1,Neurl1b,Nfatc1,Nfkbiz,Ninj2,Nipbl,Nrip1,Nsg2,Nsmce1,Os9,Osm,Otulin,P2rx4,P2ry1,Pdcd4,Pde11a,Pde4a,Per1,Pim1,Plcxd2,Polg2,Ppp1cc,Ppp2r5a,Prkca,Prkch,Prkcq,Rara,Rasgrp2,Rftn1,Rgs10,Rnf125,Rras2,Rtkn2,S1pr1,Scd2,Selt,Serpine2,Sgms1,Sh2b1,Sipa1l1,Sirt4,Ski,Sla,Slc30a4,Slc38a1,Slc3a2,Slc7a3,Sod1,Spaca4,Spag9,Ss18,St8sia1,Stat4,Stat5b,Stk10,Tbl1xr1,Tgfbr2,Tmc1,Tmeff2,Tmem64,Tnfaip3,Tnfrsf1a,Tnfrsf1b,Tnfsf11,Tnfsf8,Trib1,Trim25,Ubr2,Usp3,Vamp4,Wasf2,Wwox,Xbp1,Xcr1,Xdh,Xylt1,Zfand2a |
| GO:0048518 | positive regulation of biological process | 149 | 5340 | 2.84E-10 | Adipoq,Ago2,Agtr1b,Akt2,Atp1a1,Atp2b4,Bcl10,Btg1,Btg2,Camk4,Cblb,Ccdc80,Ccne1,Ccr1,Ccr4,Ccr7,Cd101,Cd247,Cd28,Cd300a,Cd47,Cd97,Cdkn2b,Chd7,Chsy1,Cnst,Crem,Cryaa,Cyfip2,Dpp4,Dusp1,Dusp10,Dyrk2,E2f2,Elf1,Elovl5,Epb4.1,Esrp1,Ets1,Ets2,F2,Fam20a,Fanci,Fasl,Fli1,Fntb,Fosl2,Foxn3,Foxp1,Gadd45g,Glrx,Gpr18,H2-Aa,Hey1,Hivep2,Igf2r,Il15ra,Il6st,Ist1,Itga4,Itk,Jak1,Jarid2,Jun,Kdm2b,Klf2,Klf6,Lef1,Lonrf1,Lrrn3,Lrtm2,Map3k1,Map3k8,Map4k1,Mapk14,Mapk9,Max,Mbd2,Mospd1,Ncf1,Nfatc1,Nfkbiz,Nipbl,Nrip1,Nsmce1,Osm,Otulin,P2rx4,P2ry1,Pdcd4,Per1,Pim1,Pip4k2a,Pkp4,Ppp1cc,Ppp2r5a,Prickle1,Prkca,Prkch,Prkcq,Rara,Rftn1,Riok1,Rnf125,Rnf219,Rras2,Rtkn2,S1pr1,Selt,Serpine2,Sgms1,Sh2b1,Sirt4,Ski,Sla,Slc39a10,Smarca2,Sod1,Spag9,Ss18,St8sia1,Stat4,Stat5b,Stim2,Stk10,Tapt1,Tbl1xr1,Tgfbr2,Tigit,Tmem64,Tnfaip3,Tnfrsf1a,Tnfrsf1b,Tnfsf11,Tnfsf8,Tnrc6b,Trappc12,Trib1,Trim25,Txnl4b,Vps37b,Vsx2,Wasf2,Wwox,Xbp1,Xdh,Zfand2a,Zfp326,Zfp609 |
| GO:0051716 | cellular response to stimulus | 141 | 5142 | 6.49E-09 | Abhd2,Adipoq,Ago2,Agtr1b,Akt2,Arhgap15,Arhgap25,Atp1a1,Atp2b4,Bcl10,Btg2,Btla,Camk4,Cblb,Ccne1,Ccr1,Ccr4,Ccr7,Cd101,Cd247,Cd28,Cd300a,Cd47,Cd97,Cdkn2b,Chd2,Clic4,Crem,Dock10,Dusp1,Dyrk2,E2f2,Ercc4,Ets1,F2,Fanci,Fasl,Foxn3,Foxp1,Fyb,Gatsl3,Gdi2,Glrx,Glrx2,Gnat1,Gng2,Gpbar1,Gpr132,Gpr18,Gucy2c,Gzmb,Hey1,Hivep2,Igf2r,Il15ra,Il21r,Il6st,Ints7,Itga4,Itk,Jak1,Jarid2,Jun,Kcnh8,Klf2,Klf3,Klf6,Lef1,Lrrc8d,Lrtm2,Map3k1,Map3k8,Map4k1,Mapk14,Mapk9,Mbd2,Mgme1,Mknk2,Nabp1,Ncf1,Neurl1b,Nfatc1,Nfkbiz,Nipbl,Nrip1,Nsg2,Nsmce1,Os9,Osm,Otulin,P2rx4,P2ry1,Pdcd4,Pde11a,Pde4a,Pim1,Plcxd2,Polg2,Ppp2r5a,Prkca,Prkch,Prkcq,Rara,Rasgrp2,Rftn1,Rgs10,Rnf125,Rras2,Rtkn2,S1pr1,Selt,Sgms1,Sh2b1,Sipa1l1,Sirt4,Ski,Sla,Sod1,Spag9,Ss18,St8sia1,Stat4,Stat5b,Stk10,Tgfbr2,Tmem64,Tnfaip3,Tnfrsf1a,Tnfrsf1b,Tnfsf11,Trib1,Trim25,Ubr2,Usp3,Vamp4,Wasf2,Wwox,Xbp1,Xcr1,Xylt1,Zfand2a |
| GO:0009987 | cellular process | 263 | 12459 | 7.44E-09 | Abcb1b,Abhd2,Adarb2,Adipoq,Aebp2,Aff4,Ago2,Agtr1b,Akt2,Arhgap15,Arhgap25,Atp10a,Atp1a1,Atp2b4,Bach2,Bcat1,Bcl10,Btg1,Btg2,Btla,Camk4,Cblb,Ccdc80,Ccne1,Ccr1,Ccr4,Ccr7,Cd101,Cd247,Cd28,Cd300a,Cd47,Cd97,Cdkn2b,Celf2,Cers6,Chd2,Chd7,Chst11,Chst2,Chsy1,Clic4,Clint1,Cmah,Cmas,Crbn,Crem,Cryaa,Cyb561,Cyb5a,Cyfip2,Cyp2r1,Dach2,Dad1,Dcaf7,Ddx6,Dhx40,Dnah17,Dnajc6,Dock10,Dpp4,Drc1,Dusp1,Dusp10,Dusp2,Dyrk2,E2f2,Eif4e3,Elf1,Elovl5,Epb4.1,Ercc4,Esrp1,Ets1,Ets2,Ext1,F2,Fanci,Fasl,Fbxl12,Fbxo33,Fkbp5,Fli1,Fntb,Fosl2,Foxn3,Foxp1,Fyb,Gadd45g,Gatsl3,Gdi2,Ggta1,Glrx,Glrx2,Gnat1,Gng2,Golga5,Gpbar1,Gpr132,Gpr18,Gtf2ird1,Gucy2c,Gzmb,Hbs1l,Hectd1,Hey1,Hist1h2be,Hivep2,Igf2r,Il15ra,Il21r,Il6st,Inpp4b,Ints7,Ip6k1,Ist1,Itga4,Itk,Jak1,Jarid2,Jun,Kcnh8,Kdm2b,Klf2,Klf3,Klf6,Lef1,Lmtk3,Lnx2,Lonrf1,Lpcat3,Lrrc8d,Lrtm2,Map3k1,Map3k8,Map4k1,Mapk14,Mapk9,Max,Mbd2,Mbip,Mbnl1,Mfsd2a,Mgme1,Mknk2,Mogat1,Mospd1,Moxd1,Msrb3,Myo1b,Nabp1,Ncf1,Neurl1b,Nfatc1,Nfkbiz,Nipbl,Nrip1,Nrxn3,Nsg2,Nsmce1,Os9,Osm,Otulin,P2rx4,P2ry1,Pdcd4,Pde11a,Pde4a,Per1,Pim1,Pip4k2a,Pkp4,Plcxd2,Polg2,Ppp1cc,Ppp2r5a,Prkca,Prkch,Prkcq,Prrc2b,Ptp4a2,Rara,Rasgrp2,Rftn1,Rgs10,Riok1,Rnf125,Rpia,Rras2,Rtkn2,S1pr1,Sar1a,Scd2,Scml4,Selt,Serpine2,Sgms1,Sgol1,Sh2b1,Sipa1l1,Sirt4,Ski,Sla,Slc35g1,Slc39a10,Smarca2,Smarce1,Smndc1,Smox,Snx18,Sod1,Spaca4,Spag9,Spty2d1,Ss18,St8sia1,Stat4,Stat5b,Stim2,Stk10,Syne3,Tapt1,Tars2,Tax1bp1,Tbl1xr1,Tgfbr2,Thada,Tigit,Tmc1,Tmeff2,Tmem123,Tmem64,Tnfaip3,Tnfrsf1a,Tnfrsf1b,Tnfsf11,Tnfsf8,Tnrc6b,Trappc12,Trib1,Trim25,Txnl4b,Ubr2,Usp3,Usp6nl,Vamp4,Vgll4,Vsx2,Wasf2,Wdyhv1,Wwox,Xbp1,Xcr1,Xdh,Xylt1,Zdhhc8,Zfand2a,Zfp3,Zfp326,Zfp609,Zfp652,Zfp932,Zscan10 |
| GO:0048522 | positive regulation of cellular process | 133 | 4827 | 2.13E-08 | Adipoq,Ago2,Akt2,Atp2b4,Bcl10,Btg1,Btg2,Camk4,Ccne1,Ccr1,Ccr7,Cd101,Cd28,Cd300a,Cd47,Cd97,Cdkn2b,Chd7,Chsy1,Cnst,Crem,Cryaa,Cyfip2,Dpp4,Dusp1,Dusp10,Dyrk2,E2f2,Elf1,Elovl5,Esrp1,Ets1,Ets2,F2,Fam20a,Fanci,Fasl,Fli1,Fntb,Fosl2,Foxn3,Foxp1,Gadd45g,Glrx,Gpr18,H2-Aa,Hey1,Hivep2,Igf2r,Il15ra,Il6st,Ist1,Itga4,Jak1,Jarid2,Jun,Kdm2b,Klf2,Klf6,Lef1,Lonrf1,Lrrn3,Lrtm2,Map3k1,Map3k8,Map4k1,Mapk14,Mapk9,Max,Mbd2,Mospd1,Ncf1,Nfatc1,Nfkbiz,Nipbl,Nrip1,Nsmce1,Osm,P2rx4,P2ry1,Pdcd4,Per1,Pim1,Pip4k2a,Pkp4,Ppp1cc,Ppp2r5a,Prickle1,Prkca,Prkch,Prkcq,Rara,Riok1,Rnf125,Rnf219,Rras2,Rtkn2,S1pr1,Selt,Serpine2,Sh2b1,Ski,Sla,Slc39a10,Smarca2,Sod1,Spag9,Ss18,St8sia1,Stat4,Stat5b,Stk10,Tapt1,Tbl1xr1,Tgfbr2,Tmem64,Tnfaip3,Tnfrsf1a,Tnfrsf1b,Tnfsf11,Tnfsf8,Tnrc6b,Trib1,Trim25,Txnl4b,Vsx2,Wasf2,Wwox,Xbp1,Xdh,Zfand2a,Zfp326,Zfp609 |
| GO:0042127 | regulation of cell population proliferation | 63 | 1594 | 3.70E-08 | Adipoq,Akt2,Btg1,Btg2,Btla,Cblb,Ccr7,Cd28,Cd300a,Cd47,Cdkn2b,Chst11,Dpp4,Dusp1,Dusp10,Esrp1,Ets1,F2,Fasl,Fntb,Fosl2,Foxp1,Gucy2c,H2-Aa,Hey1,Il6st,Itga4,Itk,Jak1,Jarid2,Jun,Kdm2b,Lef1,Mapk14,Mbd2,Nfatc1,Osm,Pdcd4,Pim1,Ppp1cc,Prkca,Prkch,Prkcq,Rara,Rtkn2,S1pr1,Serpine2,Ski,Sla,Slc39a10,Smarca2,St8sia1,Stat5b,Tgfbr2,Tnfaip3,Tnfrsf1a,Tnfrsf1b,Trib1,Txnl4b,Vsx2,Xbp1,Xdh,Zfp609 |
| GO:0022407 | regulation of cell-cell adhesion | 27 | 360 | 6.72E-08 | Adipoq,Bcl10,Btla,Cblb,Ccr7,Cd28,Cd300a,Cd47,Dpp4,Dusp10,Ets1,H2-Aa,Il6st,Itga4,Lef1,Nfkbiz,Prkca,Prkcq,Rara,Serpine2,Stat5b,Tgfbr2,Tigit,Tnfaip3,Tnfsf11,Xbp1,Zfp609 |
| GO:0031323 | regulation of cellular metabolic process | 140 | 5293 | 6.72E-08 | Adipoq,Aebp2,Aff4,Ago2,Akt2,Atp1a1,Atp2b4,Bach2,Bcl10,Btg2,Camk4,Cblb,Ccne1,Ccr1,Ccr7,Cd28,Cd300a,Cdkn2b,Chd2,Chd7,Cnst,Crem,Crim1,Cryaa,Cyfip2,Dach2,Ddx6,Dusp1,Dusp10,Dusp2,Dyrk2,E2f2,Eif4e3,Elf1,Elovl5,Ercc4,Esrp1,Ets1,Ets2,F2,Fam20a,Fanci,Fasl,Fli1,Fosl2,Foxn3,Foxp1,Gadd45g,Gtf2ird1,Hey1,Hivep2,Il6st,Ist1,Jarid2,Jun,Kcnh8,Kdm2b,Klf2,Klf3,Klf6,Lef1,Lmtk3,Lonrf1,Lrrn3,Lrtm2,Map3k1,Map3k8,Map4k1,Mapk14,Mapk9,Max,Mbd2,Mbnl1,Mknk2,Mospd1,Ncf1,Nfatc1,Nfkbiz,Nipbl,Nrip1,Os9,Osm,Otulin,P2ry1,Pdcd4,Per1,Pim1,Pip4k2a,Ppp2r5a,Prickle1,Prkca,Prkch,Prkcq,Rara,Riok1,Rnf125,Rnf219,S1pr1,Scml4,Serpine2,Sh2b1,Sirt4,Ski,Slc35b4,Slc39a10,Smarca2,Smarce1,Sod1,Spag9,Spty2d1,Ss18,Stat4,Stat5b,Stk10,Tax1bp1,Tbl1xr1,Tgfbr2,Tnfaip3,Tnfrsf1a,Tnfrsf1b,Tnfsf11,Tnfsf8,Tnrc6b,Trib1,Trim25,Ubac2,Ubr2,Usp3,Vgll4,Vsx2,Wwox,Xbp1,Xdh,Zfand2a,Zfp3,Zfp326,Zfp609,Zfp652,Zfp932,Zscan10 |
| GO:0048519 | negative regulation of biological process | 129 | 4771 | 1.12E-07 | Abhd2,Adipoq,Aebp2,Ago2,Akt2,Arhgap25,Atp1a1,Atp2b4,Bach2,Bcl10,Btg1,Btg2,Btla,Cblb,Ccne1,Ccr1,Ccr7,Cd28,Cd300a,Cdkn2b,Chst11,Chsy1,Clic4,Cnst,Crbn,Crem,Crim1,Cryaa,Dad1,Ddx6,Dpp4,Dusp1,Dusp10,Dusp2,Dyrk2,E2f2,Elf1,Ercc4,Ets1,Ets2,F2,Fanci,Fasl,Fntb,Foxn3,Foxp1,Gadd45g,Gatsl3,Glrx,Gpr132,Gpr18,H2-Aa,Hectd1,Hey1,Il15ra,Il6st,Inpp4b,Ints7,Jarid2,Jun,Kcnh8,Kdm2b,Klf2,Klf3,Lef1,Leprotl1,Lmtk3,Lrtm2,Mapk14,Max,Mbd2,Mospd1,Nabp1,Nfatc1,Nipbl,Nrip1,Os9,Osm,Otulin,P2rx4,P2ry1,Pdcd4,Pde11a,Per1,Pim1,Ppp2r5a,Prickle1,Prkca,Prkch,Prkcq,Rara,Rgs10,Rnf125,S1pr1,Sar1a,Serpine2,Sgms1,Shisa2,Sirt4,Ski,Slc39a10,Smarca2,Smarce1,Sod1,Stat5b,Tax1bp1,Tbl1xr1,Tgfbr2,Tigit,Tmeff2,Tmem64,Tnfaip3,Tnfrsf1a,Tnfrsf1b,Tnrc6b,Trib1,Trim25,Ubac2,Ubr2,Usp3,Vsx2,Wasf2,Wwox,Xbp1,Xcr1,Xdh,Xylt1,Zfp932,Zscan10 |
| GO:0060255 | regulation of macromolecule metabolic process | 138 | 5246 | 1.12E-07 | Adipoq,Aebp2,Aff4,Ago2,Akt2,Atp2b4,Bach2,Bcl10,Btg2,Camk4,Cblb,Ccne1,Ccr1,Ccr7,Cd28,Cd300a,Cdkn2b,Chd2,Chd7,Crem,Crim1,Cryaa,Cyfip2,Dach2,Ddx6,Dusp1,Dusp10,Dusp2,Dyrk2,E2f2,Eif4e3,Elf1,Ercc4,Esrp1,Ets1,Ets2,F2,Fam20a,Fanci,Fasl,Fli1,Fntb,Fosl2,Foxn3,Foxp1,Gadd45g,Gtf2ird1,Hey1,Hivep2,Il6st,Ist1,Jarid2,Jun,Kcnh8,Kdm2b,Klf2,Klf3,Klf6,Lef1,Lonrf1,Lrrn3,Lrtm2,Map3k1,Map3k8,Map4k1,Mapk14,Mapk9,Max,Mbd2,Mbnl1,Mknk2,Mospd1,Ncf1,Nfatc1,Nfkbiz,Nipbl,Nrip1,Os9,Osm,Otulin,P2ry1,Pdcd4,Per1,Pim1,Ppp1cc,Ppp2r5a,Prickle1,Prkca,Prkch,Prkcq,Rara,Riok1,Rnf125,Rnf219,S1pr1,Scml4,Serpine2,Sgms1,Sh2b1,Sirt4,Ski,Slc39a10,Smarca2,Smarce1,Sod1,Spag9,Spty2d1,Ss18,Stat4,Stat5b,Stk10,Tax1bp1,Tbl1xr1,Tgfbr2,Tmeff2,Tnfaip3,Tnfrsf1a,Tnfrsf1b,Tnfsf11,Tnfsf8,Tnrc6b,Trib1,Trim25,Ubac2,Ubr2,Usp3,Vgll4,Vsx2,Wwox,Xbp1,Xdh,Zfand2a,Zfp3,Zfp326,Zfp609,Zfp652,Zfp932,Zscan10 |
| GO:0080090 | regulation of primary metabolic process | 137 | 5184 | 1.12E-07 | Adipoq,Aebp2,Aff4,Ago2,Akt2,Atp1a1,Atp2b4,Bach2,Bcl10,Btg2,Camk4,Cblb,Ccne1,Ccr1,Ccr7,Cd28,Cd300a,Cdkn2b,Chd2,Chd7,Crem,Crim1,Cryaa,Cyfip2,Dach2,Ddx6,Dusp1,Dusp10,Dusp2,Dyrk2,E2f2,Eif4e3,Elf1,Elovl5,Ercc4,Esrp1,Ets1,Ets2,F2,Fam20a,Fanci,Fasl,Fli1,Fosl2,Foxn3,Foxp1,Gadd45g,Gtf2ird1,Hey1,Hivep2,Il6st,Ist1,Jarid2,Jun,Kcnh8,Kdm2b,Klf2,Klf3,Klf6,Lef1,Lonrf1,Lrrn3,Lrtm2,Map3k1,Map3k8,Map4k1,Mapk14,Mapk9,Max,Mbd2,Mbnl1,Mknk2,Mospd1,Ncf1,Nfatc1,Nfkbiz,Nipbl,Nrip1,Os9,Osm,Otulin,P2ry1,Pdcd4,Per1,Pim1,Ppp2r5a,Prickle1,Prkca,Prkch,Prkcq,Rara,Riok1,Rnf125,Rnf219,S1pr1,Scml4,Serpine2,Sh2b1,Sirt4,Ski,Slc35b4,Slc39a10,Smarca2,Smarce1,Sod1,Spag9,Spty2d1,Ss18,Stat4,Stat5b,Stk10,Tax1bp1,Tbl1xr1,Tgfbr2,Tnfaip3,Tnfrsf1a,Tnfrsf1b,Tnfsf11,Tnfsf8,Tnrc6b,Trib1,Trim25,Ubac2,Ubr2,Usp3,Vgll4,Vsx2,Wwox,Xbp1,Xdh,Zfand2a,Zfp3,Zfp326,Zfp609,Zfp652,Zfp932,Zscan10 |
| GO:1903708 | positive regulation of hemopoiesis | 20 | 199 | 1.12E-07 | Ccr1,Ccr7,Cd101,Dusp10,Ets1,H2-Aa,Il15ra,Jun,Lef1,Mapk14,Nfkbiz,Prkca,Rara,Stat5b,Tgfbr2,Tmem64,Tnfsf11,Trib1,Xbp1,Zfp609 |

| | | | | | |
|---|---|---|---|---|---|
| GO:0048523 | negative regulation of cellular process | 120 | 4336 | 1.36E-07 | Abhd2,Adipoq,Aebp2,Ago2,Akt2,Arhgap25,Atp1a1,Atp2b4,Bach2,Bcl10,Btg1,Btg2,Btla,Cblb,Ccne1,Ccr7,Cd300a,Cdkn2b,Chst11,Clic4,Cnst,Crbn,Crem,Crim1,Cryaa,Dad1,Ddx6,Dpp4,Dusp1,Dusp10,Dusp2,Dyrk2,Elf1,Ercc4,Ets1,Ets2,F2,Fanci,Fasl,Fntb,Foxn3,Foxp1,Gadd45g,Gatsl3,Glrx,Gpr132,Gpr18,H2-Aa,Hey1,Il15ra,Il6st,Inpp4b,Ints7,Jarid2,Jun,Kcnh8,Kdm2b,Klf2,Klf3,Lef1,Lmtk3,Lrtm2,Mapk14,Max,Mbd2,Mospd1,Nabp1,Nfatc1,Nipbl,Nrip1,Os9,Osm,P2ry1,Pdcd4,Pde11a,Per1,Pim1,Ppp2r5a,Prickle1,Prkca,Prkch,Prkcq,Rara,Rgs10,Rnf125,S1pr1,Sar1a,Serpine2,Sgms1,Shisa2,Sirt4,Ski,Slc39a10,Smarca2,Smarce1,Sod1,Stat5b,Tax1bp1,Tbl1xr1,Tgfbr2,Tigit,Tmeff2,Tmem64,Tnfaip3,Tnfrsf1a,Tnfrsf1b,Tnrc6b,Trib1,Ubac2,Ubr2,Usp3,Vsx2,Wasf2,Wwox,Xbp1,Xcr1,Xdh,Xylt1,Zfp932,Zscan10 |
| GO:0019222 | regulation of metabolic process | 144 | 5616 | 1.68E-07 | Adipoq,Aebp2,Aff4,Ago2,Akt2,Atp1a1,Atp2b4,Bach2,Bcl10,Btg2,Camk4,Cblb,Ccne1,Ccr1,Ccr7,Cd28,Cd300a,Cdkn2b,Chd2,Chd7,Cnst,Crem,Crim1,Cryaa,Cyfip2,Dach2,Ddx6,Dusp1,Dusp10,Dusp2,Dyrk2,E2f2,Eif4e3,Elf1,Elovl5,Ercc4,Esrp1,Ets1,Ets2,F2,Fam20a,Fanci,Fasl,Fli1,Fntb,Fosl2,Foxn3,Foxp1,Gadd45g,Gtf2ird1,Hey1,Hivep2,Il6st,Ist1,Jarid2,Jun,Kcnh8,Kdm2b,Klf2,Klf3,Klf6,Lef1,Lmtk3,Lonrf1,Lrrn3,Lrtm2,Map3k1,Map3k8,Map4k1,Mapk14,Mapk9,Max,Mbd2,Mbnl1,Mknk2,Mospd1,Ncf1,Nfatc1,Nfkbiz,Nipbl,Nrip1,Os9,Osm,Otulin,P2ry1,Pdcd4,Per1,Pim1,Pip4k2a,Ppp1cc,Ppp2r5a,Prickle1,Prkca,Prkch,Prkcq,Rara,Riok1,Rnf125,Rnf219,S1pr1,Scml4,Serpine2,Sgms1,Sh2b1,Sirt4,Ski,Slc35b4,Slc39a10,Smarca2,Smarce1,Sod1,Spag9,Spty2d1,Ss18,Stat4,Stat5b,Stk10,Tax1bp1,Tbl1xr1,Tgfbr2,Tmeff2,Tnfaip3,Tnfrsf1a,Tnfrsf1b,Tnfsf11,Tnfsf8,Tnrc6b,Trib1,Trim25,Ubac2,Ubr2,Usp3,Vgll4,Vsx2,Wwox,Xbp1,Xdh,Zfand2a,Zfp3,Zfp326,Zfp609,Zfp652,Zfp932,Zscan10 |
| GO:1902107 | positive regulation of leukocyte differentiation | 18 | 162 | 1.68E-07 | Ccr1,Ccr7,Cd101,Dusp10,H2-Aa,Il15ra,Jun,Lef1,Nfkbiz,Prkca,Rara,Stat5b,Tgfbr2,Tmem64,Tnfsf11,Trib1,Xbp1,Zfp609 |
| GO:0051171 | regulation of nitrogen compound metabolic process | 133 | 5052 | 2.02E-07 | Adipoq,Aebp2,Aff4,Ago2,Akt2,Atp2b4,Bach2,Bcl10,Btg2,Camk4,Cblb,Ccne1,Ccr1,Ccr7,Cd28,Cd300a,Cdkn2b,Chd2,Chd7,Crem,Crim1,Cryaa,Cyfip2,Dach2,Ddx6,Dusp1,Dusp10,Dusp2,E2f2,Eif4e3,Elf1,Ercc4,Esrp1,Ets1,Ets2,F2,Fam20a,Fanci,Fasl,Fli1,Fosl2,Foxn3,Foxp1,Gadd45g,Gtf2ird1,Hey1,Hivep2,Il6st,Ist1,Jarid2,Jun,Kcnh8,Kdm2b,Klf2,Klf3,Klf6,Lef1,Lonrf1,Lrrn3,Lrtm2,Map3k1,Map3k8,Map4k1,Mapk14,Mapk9,Max,Mbd2,Mbnl1,Mknk2,Mospd1,Ncf1,Nfatc1,Nfkbiz,Nipbl,Nrip1,Os9,Osm,Otulin,P2ry1,Pdcd4,Per1,Pim1,Ppp2r5a,Prickle1,Prkca,Prkch,Prkcq,Rara,Riok1,Rnf125,Rnf219,S1pr1,Scml4,Serpine2,Sh2b1,Sirt4,Ski,Slc39a10,Smarca2,Smarce1,Sod1,Spag9,Spty2d1,Ss18,Stat4,Stat5b,Stk10,Tax1bp1,Tbl1xr1,Tgfbr2,Tnfaip3,Tnfrsf1a,Tnfrsf1b,Tnfsf11,Tnfsf8,Tnrc6b,Trib1,Trim25,Ubac2,Ubr2,Usp3,Vgll4,Vsx2,Wwox,Xbp1,Xdh,Zfand2a,Zfp3,Zfp326,Zfp609,Zfp652,Zfp932,Zscan10 |
| GO:0002376 | immune system process | 63 | 1703 | 2.19E-07 | Bcl10,Btla,Camk4,Cblb,Ccr1,Ccr4,Ccr7,Cd247,Cd28,Cd300a,Cd47,Cd7,Cdkn2b,Chd2,Chd7,Dock10,Dpp4,Ets1,F2,Fasl,Fli1,Foxp1,Fyb,Gadd45g,Gpr18,Gzmb,H2-Aa,Hectd1,Hist1h2be,Itga4,Itk,Jak1,Jarid2,Jun,Klf2,Lef1,Map3k8,Mapk14,Mknk2,Ncf1,Nfatc1,Nfkbiz,Osm,Otulin,Pip4k2a,Prkca,Prkcq,Rftn1,Rnf125,Rtkn2,S1pr1,Sla,Sod1,Stat5b,Tgfbr2,Tnfaip3,Tnfrsf1a,Tnfrsf1b,Tnfsf11,Tnfsf8,Trim25,Vamp4,Wasf2 |
| GO:0010604 | positive regulation of macromolecule metabolic process | 92 | 3004 | 2.19E-07 | Adipoq,Ago2,Akt2,Atp2b4,Bcl10,Btg2,Camk4,Cblb,Ccr1,Ccr7,Cd28,Cd300a,Chd7,Crem,Cryaa,Cyfip2,Dyrk2,E2f2,Elf1,Ets1,Ets2,F2,Fam20a,Fanci,Fasl,Fli1,Fntb,Fosl2,Foxn3,Foxp1,Gadd45g,Hey1,Hivep2,Il6st,Ist1,Jarid2,Jun,Klf2,Klf6,Lef1,Lonrf1,Lrrn3,Map3k1,Map3k8,Map4k1,Mapk14,Mapk9,Max,Mospd1,Ncf1,Nfatc1,Nipbl,Nrip1,Osm,P2ry1,Per1,Pim1,Ppp2r5a,Prickle1,Prkca,Prkcq,Rara,Riok1,Rnf125,Rnf219,S1pr1,Sgms1,Ski,Slc39a10,Smarca2,Sod1,Spag9,Ss18,Stat4,Stat5b,Stk10,Tbl1xr1,Tgfbr2,Tnfaip3,Tnfrsf1a,Tnfrsf1b,Tnfsf11,Tnfsf8,Tnrc6b,Trib1,Vsx2,Wwox,Xbp1,Xdh,Zfand2a,Zfp326,Zfp609 |
| GO:0051173 | positive regulation of nitrogen compound metabolic process | 89 | 2866 | 2.19E-07 | Adipoq,Ago2,Akt2,Atp2b4,Bcl10,Btg2,Camk4,Cblb,Ccr1,Ccr7,Cd28,Cd300a,Chd7,Crem,Cryaa,Cyfip2,E2f2,Elf1,Ets1,Ets2,F2,Fam20a,Fanci,Fasl,Fli1,Fosl2,Foxn3,Foxp1,Gadd45g,Hey1,Hivep2,Il6st,Ist1,Jarid2,Jun,Klf2,Klf6,Lef1,Lonrf1,Lrrn3,Map3k1,Map3k8,Map4k1,Mapk14,Mapk9,Max,Mospd1,Ncf1,Nfatc1,Nipbl,Nrip1,Osm,P2ry1,Per1,Pim1,Ppp2r5a,Prickle1,Prkca,Prkcq,Rara,Riok1,Rnf125,Rnf219,S1pr1,Ski,Slc39a10,Smarca2,Sod1,Spag9,Ss18,Stat4,Stat5b,Stk10,Tbl1xr1,Tgfbr2,Tnfaip3,Tnfrsf1a,Tnfrsf1b,Tnfsf11,Tnfsf8,Tnrc6b,Trib1,Vsx2,Wwox,Xbp1,Xdh,Zfand2a,Zfp326,Zfp609 |
| GO:1903037 | regulation of leukocyte cell-cell adhesion | 22 | 264 | 2.52E-07 | Bcl10,Btla,Cblb,Ccr7,Cd28,Cd300a,Cd47,Dpp4,Dusp10,Ets1,H2-Aa,Il6st,Itga4,Nfkbiz,Prkcq,Rara,Stat5b,Tgfbr2,Tigit,Tnfsf11,Xbp1,Zfp609 |
| GO:0031325 | positive regulation of cellular metabolic process | 91 | 3004 | 4.08E-07 | Adipoq,Ago2,Akt2,Atp2b4,Bcl10,Btg2,Camk4,Ccr1,Ccr7,Cd28,Cd300a,Chd7,Crem,Cryaa,Cyfip2,Dyrk2,E2f2,Elf1,Elovl5,Ets1,Ets2,F2,Fam20a,Fanci,Fasl,Fli1,Fosl2,Foxn3,Foxp1,Gadd45g,Hey1,Hivep2,Il6st,Ist1,Jarid2,Jun,Klf2,Klf6,Lef1,Lonrf1,Lrrn3,Map3k1,Map3k8,Map4k1,Mapk14,Mapk9,Max,Mospd1,Ncf1,Nfatc1,Nipbl,Nrip1,Osm,P2ry1,Per1,Pim1,Pip4k2a,Ppp2r5a,Prickle1,Prkca,Prkcq,Rara,Riok1,Rnf125,Rnf219,S1pr1,Ski,Slc39a10,Smarca2,Sod1,Spag9,Ss18,Stat4,Stat5b,Stk10,Tbl1xr1,Tgfbr2,Tnfaip3,Tnfrsf1a,Tnfrsf1b,Tnfsf11,Tnfsf8,Tnrc6b,Trib1,Vsx2,Wwox,Xbp1,Xdh,Zfand2a,Zfp326,Zfp609 |
| GO:0048583 | regulation of response to stimulus | 102 | 3552 | 4.95E-07 | Adipoq,Akt2,Arhgap25,Atp2b4,Bcl10,Btla,Cblb,Ccr1,Ccr4,Ccr7,Cd247,Cd28,Cd300a,Cd47,Cdkn2b,Chst11,Chsy1,Cyfip2,Cyth4,Dpp4,Dusp1,Dusp10,Dusp2,Dyrk2,Elf1,Ercc4,Ets1,F2,Fasl,Foxp1,Gadd45g,Gatsl3,Glrx,Gpr18,Gtf2ird1,Hey1,Il6st,Inpp4b,Itk,Jak1,Jarid2,Jun,Lef1,Lrtm2,Map3k1,Map3k8,Map4k1,Mapk14,Mapk9,Mbd2,Ncf1,Neto2,Nfatc1,Nfkbiz,Nsmce1,Os9,Osm,Otulin,P2rx4,P2ry1,Pdcd4,Pde11a,Pde4a,Per1,Prickle1,Prkca,Prkch,Prkcq,Prl5a1,Rara,Rftn1,Rgs10,Rnf125,S1pr1,Serpine2,Sgms1,Sh2b1,Shisa2,Sipa1l1,Ski,Sla,Slc39a10,Slc7a3,Sod1,Spag9,Stat5b,Tbl1xr1,Tgfbr2,Tmem64,Tnfaip3,Tnfrsf1a,Tnfrsf1b,Tnfsf11,Tnfsf8,Trib1,Trim25,Ubac2,Ubr2,Wwox,Xbp1,Xdh,Xylt1 |
| GO:0009893 | positive regulation of metabolic process | 95 | 3215 | 5.12E-07 | Adipoq,Ago2,Akt2,Atp2b4,Bcl10,Btg2,Camk4,Cblb,Ccr1,Ccr7,Cd28,Cd300a,Chd7,Crem,Cryaa,Cyfip2,Dyrk2,E2f2,Elf1,Elovl5,Ets1,Ets2,F2,Fam20a,Fanci,Fasl,Fli1,Fntb,Fosl2,Foxn3,Foxp1,Gadd45g,Hey1,Hivep2,Il6st,Ist1,Jarid2,Jun,Klf2,Klf6,Lef1,Lonrf1,Lrrn3,Map3k1,Map3k8,Map4k1,Mapk14,Mapk9,Max,Mospd1,Ncf1,Nfatc1,Nipbl,Nrip1,Osm,P2ry1,Per1,Pim1,Pip4k2a,Ppp2r5a,Prickle1,Prkca,Prkcq,Rara,Riok1,Rnf125,Rnf219,S1pr1,Sgms1,Sirt4,Ski,Slc39a10,Smarca2,Sod1,Spag9,Ss18,Stat4,Stat5b,Stk10,Tbl1xr1,Tgfbr2,Tnfaip3,Tnfrsf1a,Tnfrsf1b,Tnfsf11,Tnfsf8,Tnrc6b,Trib1,Vsx2,Wwox,Xbp1,Xdh,Zfand2a,Zfp326,Zfp609 |
| GO:1902105 | regulation of leukocyte differentiation | 22 | 284 | 7.52E-07 | Adipoq,Ccr1,Ccr7,Cd101,Cd28,Dusp10,Foxp1,H2-Aa,Il15ra,Inpp4b,Jun,Lef1,Nfkbiz,Prkca,Rara,Stat5b,Tgfbr2,Tmem64,Tnfsf11,Trib1,Xbp1,Zfp609 |
| GO:0009889 | regulation of biosynthetic process | 102 | 3613 | 1.07E-06 | Adipoq,Aebp2,Aff4,Ago2,Akt2,Atp1a1,Atp2b4,Bach2,Bcl10,Btg2,Camk4,Ccne1,Ccr7,Cd28,Chd2,Chd7,Crem,Dach2,Ddx6,Dusp1,Dyrk2,E2f2,Eif4e3,Elf1,Elovl5,Ets1,Ets2,F2,Fasl,Fli1,Fntb,Fosl2,Foxn3,Foxp1,Gtf2ird1,Hey1,Hivep2,Jarid2,Jun,Kcnh8,Kdm2b,Klf2,Klf3,Klf6,Lef1,Mapk14,Mapk9,Max,Mbd2,Mknk2,Mospd1,Ncf1,Nfatc1,Nfkbiz,Nipbl,Nrip1,Osm,Otulin,P2ry1,Pdcd4,Per1,Pim1,Prickle1,Prkca,Prkch,Prkcq,Rara,S1pr1,Scml4,Sh2b1,Sirt4,Ski,Slc35b4,Smarca2,Smarce1,Sod1,Spty2d1,Ss18,Stat4,Stat5b,Tax1bp1,Tbl1xr1,Tmeff2,Tnfaip3,Tnfrsf1a,Tnfsf11,Tnfsf8,Tnrc6b,Trib1,Trim25,Ubr2,Usp3,Vgll4,Vsx2,Wwox,Xbp1,Zfp3,Zfp326,Zfp609,Zfp652,Zfp932,Zscan10 |
| GO:1903039 | positive regulation of leukocyte cell-cell adhesion | 18 | 190 | 1.07E-06 | Bcl10,Ccr7,Cd28,Cd47,Dpp4,Dusp10,Ets1,H2-Aa,Il6st,Itga4,Nfkbiz,Prkcq,Rara,Stat5b,Tgfbr2,Tnfsf11,Xbp1,Zfp609 |
| GO:0010468 | regulation of gene expression | 104 | 3720 | 1.12E-06 | Adipoq,Aebp2,Aff4,Ago2,Akt2,Atp2b4,Bach2,Bcl10,Btg2,Camk4,Ccne1,Ccr1,Cd28,Chd2,Chd7,Crem,Cryaa,Dach2,Ddx6,E2f2,Eif4e3,Elf1,Esrp1,Ets1,Ets2,F2,Fasl,Fli1,Fosl2,Foxn3,Foxp1,Gtf2ird1,Hey1,Hivep2,Jarid2,Jun,Kcnh8,Kdm2b,Klf2,Klf3,Klf6,Lef1,Mapk14,Mapk9,Max,Mbd2,Mbnl1,Mknk2,Mospd1,Ncf1,Nfatc1,Nfkbiz,Nipbl,Nrip1,Osm,Otulin,P2ry1,Pdcd4,Per1,Pim1,Ppp1cc,Prickle1,Prkca,Prkch,Prkcq,Rara,Riok1,Rnf219,S1pr1,Scml4,Serpine2,Sgms1,Sirt4,Ski,Smarca2,Smarce1,Spty2d1,Ss18,Stat4,Stat5b,Tax1bp1,Tbl1xr1,Tgfbr2,Tnfaip3,Tnfrsf1a,Tnfrsf1b,Tnfsf11,Tnfsf8,Tnrc6b,Trib1,Trim25,Ubr2,Usp3,Vgll4,Vsx2,Wwox,Xbp1,Xdh,Zfp3,Zfp326,Zfp609,Zfp652,Zfp932,Zscan10 |
| GO:1903706 | regulation of hemopoiesis | 25 | 377 | 1.12E-06 | Adipoq,Ccr1,Ccr7,Cd101,Cd28,Dusp10,Ets1,Foxp1,H2-Aa,Il15ra,Inpp4b,Jun,Lef1,Mapk14,Nfkbiz,Pim1,Prkca,Rara,Stat5b,Tgfbr2,Tmem64,Tnfsf11,Trib1,Xbp1,Zfp609 |
| GO:0022409 | positive regulation of cell-cell adhesion | 19 | 227 | 2.30E-06 | Bcl10,Ccr7,Cd28,Cd47,Dpp4,Dusp10,Ets1,H2-Aa,Il6st,Itga4,Lef1,Nfkbiz,Prkcq,Rara,Stat5b,Tgfbr2,Tnfsf11,Xbp1,Zfp609 |
| GO:0007165 | signal transduction | 100 | 3594 | 2.89E-06 | Abhd2,Agtr1b,Akt2,Arhgap15,Arhgap25,Bcl10,Btla,Camk4,Cblb,Ccne1,Ccr1,Ccr4,Ccr7,Cd101,Cd247,Cd28,Cd300a,Cd47,Cd97,Crem,Dock10,Dusp1,Dyrk2,E2f2,F2,Fasl,Fyb,Gdi2,Gnat1,Gng2,Gpbar1,Gpr132,Gpr18,Gucy2c,Gzmb,Hey1,Hivep2,Igf2r,Il15ra,Il21r,Il6st,Itga4,Itk,Jak1,Jun,Kcnh8,Klf6,Lef1,Lrrc8d,Lrtm2,Map3k1,Map3k8,Map4k1,Mapk14,Mapk9,Mknk2,Neurl1b,Nfatc1,Nfkbiz,Nsg2,Nsmce1,Osm,Otulin,P2rx4,P2ry1,Pdcd4,Pde11a,Pde4a,Pim1,Plcxd2,Ppp2r5a,Prkca,Prkch,Prkcq,Rara,Rasgrp2,Rftn1,Rgs10,Rras2,Rtkn2,S1pr1,Sh2b1,Sipa1l1,Ski,Sla,Spag9,Ss18,Stat4,Stat5b,Stk10,Tgfbr2,Tmem64,Tnfrsf1a,Tnfrsf1b,Tnfsf11,Trib1,Wasf2,Wwox,Xbp1,Xcr1 |
| GO:0051252 | regulation of RNA metabolic process | 91 | 3152 | 2.89E-06 | Adipoq,Aebp2,Aff4,Ago2,Akt2,Atp2b4,Bach2,Bcl10,Btg2,Camk4,Ccne1,Cd28,Chd2,Chd7,Crem,Dach2,E2f2,Elf1,Esrp1,Ets1,Ets2,Fasl,Fli1,Fosl2,Foxn3,Foxp1,Gtf2ird1,Hey1,Hivep2,Jarid2,Jun,Kcnh8,Kdm2b,Klf2,Klf3,Klf6,Lef1,Mapk14,Mapk9,Max,Mbd2,Mbnl1,Mospd1,Ncf1,Nfatc1,Nfkbiz,Nipbl,Nrip1,Osm,Otulin,P2ry1,Pdcd4,Per1,Pim1,Prickle1,Prkch,Prkcq,Rara,Riok1,Rnf219,S1pr1,Scml4,Ski,Smarca2,Smarce1,Spty2d1,Ss18,Stat4,Stat5b,Tax1bp1,Tbl1xr1,Tnfaip3,Tnfrsf1a,Tnfrsf1b,Tnfsf11,Tnfsf8,Tnrc6b,Trib1,Trim25,Ubr2,Usp3,Vgll4,Vsx2,Wwox,Xbp1,Zfp3,Zfp326,Zfp609,Zfp652,Zfp932,Zscan10 |
| GO:0002684 | positive regulation of immune system process | 36 | 771 | 3.46E-06 | Bcl10,Cblb,Ccr1,Ccr7,Cd101,Cd247,Cd28,Cd300a,Cd47,Dpp4,Dusp10,Ets1,Foxp1,H2-Aa,Il15ra,Il6st,Itga4,Itk,Jun,Lef1,Mapk14,Nfkbiz,Otulin,Prkca,Prkch,Prkcq,Rara,Rftn1,Slc39a10,Stat5b,Tgfbr2,Tmem64,Tnfsf11,Trib1,Xbp1,Zfp609 |
| GO:0010556 | regulation of macromolecule biosynthetic process | 96 | 3417 | 3.61E-06 | Adipoq,Aebp2,Aff4,Ago2,Akt2,Atp2b4,Bach2,Bcl10,Btg2,Camk4,Ccne1,Ccr7,Cd28,Chd2,Chd7,Crem,Dach2,Ddx6,Dusp1,Dyrk2,E2f2,Eif4e3,Elf1,Ets1,Ets2,Fasl,Fli1,Fntb,Fosl2,Foxn3,Foxp1,Gtf2ird1,Hey1,Hivep2,Jarid2,Jun,Kcnh8,Kdm2b,Klf2,Klf3,Klf6,Lef1,Mapk14,Mapk9,Max,Mbd2,Mknk2,Mospd1,Ncf1,Nfatc1,Nfkbiz,Nipbl,Nrip1,Osm,Otulin,P2ry1,Pdcd4,Per1,Pim1,Prickle1,Prkca,Prkch,Prkcq,Rara,S1pr1,Scml4,Sh2b1,Ski,Smarca2,Smarce1,Spty2d1,Ss18,Stat4,Stat5b,Tax1bp1,Tbl1xr1,Tmeff2,Tnfaip3,Tnfrsf1a,Tnfsf11,Tnfsf8,Tnrc6b,Trib1,Trim25,Ubr2,Usp3,Vgll4,Vsx2,Wwox,Xbp1,Zfp3,Zfp326,Zfp609,Zfp652,Zfp932,Zscan10 |
| GO:0044237 | cellular metabolic process | 169 | 7348 | 3.61E-06 | Abhd2,Adarb2,Adipoq,Aebp2,Aff4,Ago2,Akt2,Atp1a1,Bach2,Bcat1,Btg1,Btg2,Camk4,Cblb,Ccne1,Ccr7,Celf2,Cers6,Chd2,Chd7,Chst11,Chst2,Chsy1,Cmah,Cmas,Crbn,Crem,Cyb561,Cyb5a,Cyp2r1,Dach2,Dad1,Dcaf7,Ddx6,Dhx40,Dnajc6,Dusp1,Dusp10,Dusp2,Dyrk2,E2f2,Eif4e3,Elf1,Elovl5,Ercc4,Esrp1,Ets1,Ets2,Ext1,Fanci,Fasl,Fbxl12,Fbxo33,Fkbp5,Fli1,Fntb,Fosl2,Foxn3,Foxp1,Ggta1,Glrx,Glrx2,Gtf2ird1,Gucy2c,Hbs1l,Hectd1,Hey1,Hivep2,Il6st,Inpp4b,Ints7,Ip6k1,Itk,Jak1,Jarid2,Jun,Kcnh8,Kdm2b,Klf2,Klf3,Klf6,Lef1,Lmtk3,Lonrf1,Lpcat3,Map3k1,Map3k8,Map4k1,Mapk14,Mapk9,Max,Mbd2,Mbip,Mbnl1,Mgme1,Mknk2,Mogat1,Moxd1,Msrb3,Nabp1,Ncf1,Neurl1b,Nfatc1,Nfkbiz,Nipbl,Nrip1,Nsmce1,Os9,Osm,Otulin,P2rx4,Pde11a,Pde4a,Per1,Pim1,Pip4k2a,Polg2,Ppp1cc,Ppp2r5a,Prkca,Prkch,Prkcq,Ptp4a2,Rara,Riok1,Rnf125,Rpia,Scd2,Scml4,Sgms1,Sirt4,Ski,Sla,Smarca2,Smndc1,Smox,Sod1,Spaca4,Spag9,Spty2d1,Ss18,St8sia1,Stat4,Stat5b,Stk10,Tars2,Tbl1xr1,Tgfbr2,Tnfaip3,Tnfrsf1a,Tnfsf11,Trib1,Trim25,Txnl4b,Ubr2,Usp3,Vgll4,Vsx2,Wdyhv1,Xbp1,Xdh,Xylt1,Zdhhc8,Zfp3,Zfp326,Zfp609,Zfp652,Zfp932,Zscan10 |
| GO:2000112 | regulation of cellular macromolecule biosynthetic process | 94 | 3324 | 3.72E-06 | Adipoq,Aebp2,Aff4,Ago2,Akt2,Atp2b4,Bach2,Bcl10,Btg2,Camk4,Ccne1,Ccr7,Cd28,Chd2,Chd7,Crem,Dach2,Ddx6,Dusp1,Dyrk2,E2f2,Eif4e3,Elf1,Ets1,Ets2,Fasl,Fli1,Fosl2,Foxn3,Foxp1,Gtf2ird1,Hey1,Hivep2,Jarid2,Jun,Kcnh8,Kdm2b,Klf2,Klf3,Klf6,Lef1,Mapk14,Mapk9,Max,Mbd2,Mknk2,Mospd1,Ncf1,Nfatc1,Nfkbiz,Nipbl,Nrip1,Osm,Otulin,P2ry1,Pdcd4,Per1,Pim1,Prickle1,Prkca,Prkch,Prkcq,Rara,S1pr1,Scml4,Sh2b1,Ski,Smarca2,Smarce1,Spty2d1,Ss18,Stat4,Stat5b,Tax1bp1,Tbl1xr1,Tnfaip3,Tnfrsf1a,Tnfsf11,Tnfsf8,Tnrc6b,Trib1,Trim25,Ubr2,Usp3,Vgll4,Vsx2,Wwox,Xbp1,Zfp3,Zfp326,Zfp609,Zfp652,Zfp932,Zscan10 |
| GO:0044238 | primary metabolic process | 170 | 7426 | 4.21E-06 | Abhd2,Adarb2,Adipoq,Aebp2,Aff4,Ago2,Akt2,Bach2,Bcat1,Btg1,Btg2,Camk4,Cblb,Ccne1,Ccr7,Celf2,Cers6,Chd2,Chd7,Chst11,Chst2,Chsy1,Cmah,Cmas,Crbn,Crem,Cyb5a,Cyfip2,Cyp2r1,Dach2,Dad1,Dcaf7,Ddx6,Dhx40,Dnajc6,Dpp4,Dusp1,Dusp10,Dusp2,Dyrk2,E2f2,Eif4e3,Elf1,Elovl5,Ercc4,Esrp1,Ets1,Ets2,Ext1,F2,Fanci,Fasl,Fbxl12,Fbxo33,Fkbp5,Fli1,Fntb,Fosl2,Foxn3,Foxp1,Gadd45g,Ggta1,Gtf2ird1,Gucy2c,Gzmb,Hbs1l,Hectd1,Hey1,Hivep2,Il6st,Inpp4b,Ints7,Ip6k1,Itk,Jak1,Jarid2,Jun,Kcnh8,Kdm2b,Klf2,Klf3,Klf6,Lef1,Lmtk3,Lonrf1,Lpcat3,Map3k1,Map3k8,Map4k1,Mapk14,Mapk9,Max,Mbd2,Mbip,Mbnl1,Mgme1,Mknk2,Mogat1,Msrb3,Nabp1,Ncf1,Neurl1b,Nfatc1,Nfkbiz,Nipbl,Nrip1,Nsmce1,Os9,Osm,Otulin,Pde11a,Pde4a,Per1,Pim1,Pip4k2a,Plcxd2,Polg2,Ppp1cc,Ppp2r5a,Prkca,Prkch,Prkcq,Prss39,Ptp4a2,Rara,Riok1,Rnf125,Rpia,Scd2,Scml4,Sgms1,Sirt4,Ski,Sla,Slc3a2,Smarca2,Smndc1,Spag9,Spty2d1,Ss18,St8sia1,Stat4,Stat5b,Stk10,Tars2,Tbl1xr1,Tgfbr2,Tnfaip3,Tnfrsf1a,Tnfsf11,Trappc12,Trib1,Trim25,Txnl4b,Ubr2,Usp3,Vgll4,Vsx2,Wdyhv1,Xbp1,Xdh,Xpnpep2,Xylt1,Zdhhc8,Zfp3,Zfp326,Zfp609,Zfp652,Zfp932,Zscan10 |
| GO:0002682 | regulation of immune system process | 46 | 1165 | 4.37E-06 | Adipoq,Bcl10,Btla,Cblb,Ccr1,Ccr7,Cd101,Cd247,Cd28,Cd300a,Cd47,Dpp4,Dusp10,Elf1,Ets1,Foxp1,Gpr18,H2-Aa,Il15ra,Il6st,Inpp4b,Itga4,Itk,Jun,Lef1,Mapk14,Nfkbiz,Otulin,Pim1,Prkca,Prkch,Prkcq,Rara,Rftn1,Rnf125,Slc39a10,Stat5b,Stk10,Tgfbr2,Tigit,Tmem64,Tnfaip3,Tnfsf11,Trib1,Xbp1,Zfp609 |
| GO:0008152 | metabolic process | 182 | 8164 | 5.82E-06 | Abhd2,Adarb2,Adipoq,Aebp2,Aff4,Ago2,Akt2,Atp1a1,Bach2,Bcat1,Btg1,Btg2,Camk4,Cblb,Ccne1,Ccr7,Celf2,Cers6,Chd2,Chd7,Chst11,Chst2,Chsy1,Cmah,Cmas,Crbn,Crem,Cyb561,Cyb5a,Cyfip2,Cyp2r1,Dach2,Dad1,Dcaf7,Ddx6,Dhx40,Dip2b,Dnajc6,Dpp4,Dusp1,Dusp10,Dusp2,Dyrk2,E2f2,Eif4e3,Elf1,Elovl5,Ercc4,Esrp1,Ets1,Ets2,Ext1,F2,Fanci,Fasl,Fbxl12,Fbxo33,Fkbp5,Fli1,Fntb,Fosl2,Foxn3,Foxp1,Gadd45g,Ggta1,Glrx,Glrx2,Gtf2ird1,Gucy2c,Gzmb,Hbs1l,Hectd1,Hey1,Hivep2,Il6st,Inpp4b,Ints7,Ip6k1,Itk,Jak1,Jarid2,Jun,Kcnh8,Kdm2b,Klf2,Klf3,Klf6,Lef1,Lmtk3,Lonrf1,Lpcat3,Map3k1,Map3k8,Map4k1,Mapk14,Mapk9,Max,Mbd2,Mbip,Mbnl1,Mgme1,Mknk2,Mogat1,Moxd1,Msrb3,Nabp1,Ncf1,Neurl1b,Nfatc1,Nfkbiz,Nipbl,Nrip1,Nsmce1,Os9,Osm,Otulin,P2rx4,Pde11a,Pde4a,Per1,Pim1,Pip4k2a,Plcxd2,Polg2,Ppp1cc,Ppp2r5a,Prkca,Prkch,Prkcq,Prss39,Ptp4a2,Rara,Riok1,Rnf125,Rpia,Scd2,Scml4,Selt,Sgms1,Sirt4,Ski,Sla,Slc3a2,Smarca2,Smndc1,Smox,Sod1,Spaca4,Spag9,Spty2d1,Ss18,St8sia1,Stat4,Stat5b,Stk10,Tars2,Tbl1xr1,Tgfbr2,Tnfaip3,Tnfrsf1a,Tnfsf11,Trappc12,Trib1,Trim25,Txnl4b,Ubr2,Usp3,Vgll4,Vsx2,Wdyhv1,Wwox,Xbp1,Xdh,Xpnpep2,Xylt1,Zdhhc8,Zfp3,Zfp326,Zfp609,Zfp652,Zfp932,Zscan10 |

| | | | | | |
|---|---|---|---|---|---|
| GO:0042221 | response to chemical | 97 | 3532 | 7.63E-06 | Abcb1b,Abhd2,Adipoq,Ago2,Agtr1b,Akt2,Atp1a1,Atp2b4,Bcl10,Btg1,Btg2,Ccr1,Ccr4,Ccr7,Cdkn2b,Chd7,Clic4,Crem,Cryaa,Cyb5a,Cyp2r1,Dad1,Dusp1,Dusp10,Ets1,Ext1,Fasl,Fntb,Fosl2,Foxp1,Gatsl3,Glrx,Glrx2,Gnat1,Gucy2c,H2-Aa,Igf2r,Il21r,Il6st,Jak1,Jarid2,Jun,Klf2,Klf3,Klf6,Lef1,Lrtm2,Map3k1,Map4k1,Mapk14,Mapk9,Mbd2,Mknk2,Ncf1,Nrip1,Os9,Osm,P2rx4,P2ry1,Pdcd4,Pde4a,Per1,Pim1,Prkca,Prkcq,Rara,Rasgrp2,Rftn1,Rgs10,Rnf125,S1pr1,Scd2,Selt,Sgms1,Sirt4,Slc30a4,Slc38a1,Slc3a2,Slc7a3,Sod1,Ss18,Stat4,Stat5b,Tgfbr2,Tnfaip3,Tnfrsf1a,Tnfrsf1b,Tnfsf11,Trib1,Trim25,Ubr2,Vamp4,Wwox,Xbp1,Xcr1,Xdh,Zfand2a |
| GO:0071704 | organic substance metabolic process | 174 | 7733 | 8.09E-06 | Abhd2,Adarb2,Adipoq,Aebp2,Aff4,Ago2,Akt2,Bach2,Bcat1,Btg1,Btg2,Camk4,Cblb,Ccne1,Ccr7,Celf2,Cers6,Chd2,Chd7,Chst11,Chst2,Chsy1,Cmah,Cmas,Crbn,Crem,Cyb5a,Cyfip2,Cyp2r1,Dach2,Dad1,Dcaf7,Ddx6,Dhx40,Dnajc6,Dpp4,Dusp1,Dusp10,Dusp2,Dyrk2,E2f2,Eif4e3,Elf1,Elovl5,Ercc4,Esrp1,Ets1,Ets2,Ext1,F2,Fanci,Fasl,Fbxl12,Fbxo33,Fkbp5,Fli1,Fntb,Fosl2,Foxn3,Foxp1,Gadd45g,Ggta1,Gtf2ird1,Gucy2c,Gzmb,Hbs1l,Hectd1,Hey1,Hivep2,Il6st,Inpp4b,Ints7,Ip6k1,Itk,Jak1,Jarid2,Jun,Kcnh8,Kdm2b,Klf2,Klf3,Klf6,Lef1,Lmtk3,Lonrf1,Lpcat3,Map3k1,Map3k8,Map4k1,Mapk14,Mapk9,Max,Mbd2,Mbip,Mbnl1,Mgme1,Mknk2,Mogat1,Moxd1,Msrb3,Nabp1,Ncf1,Neurl1b,Nfatc1,Nfkbiz,Nipbl,Nrip1,Nsmce1,Os9,Osm,Otulin,Pde11a,Pde4a,Per1,Pim1,Pip4k2a,Plcxd2,Polg2,Ppp1cc,Ppp2r5a,Prkca,Prkch,Prkcq,Prss39,Ptp4a2,Rara,Riok1,Rnf125,Rpia,Scd2,Scml4,Sgms1,Sirt4,Ski,Sla,Slc3a2,Smarca2,Smndc1,Smox,Sod1,Spaca4,Spag9,Spty2d1,Ss18,St8sia1,Stat4,Stat5b,Stk10,Tars2,Tbl1xr1,Tgfbr2,Tnfaip3,Tnfrsf1a,Tnfsf11,Trappc12,Trib1,Trim25,Txnl4b,Ubr2,Usp3,Vgll4,Vsx2,Wdyhv1,Xbp1,Xdh,Xpnpep2,Xylt1,Zdhhc8,Zfp3,Zfp326,Zfp609,Zfp652,Zfp932,Zscan10 |
| GO:0006355 | regulation of transcription, DNA-templated | 85 | 2952 | 8.65E-06 | Adipoq,Aebp2,Aff4,Ago2,Akt2,Atp2b4,Bach2,Bcl10,Btg2,Camk4,Ccne1,Cd28,Chd2,Chd7,Crem,Dach2,E2f2,Elf1,Ets1,Ets2,Fasl,Fli1,Fosl2,Foxn3,Foxp1,Gtf2ird1,Hey1,Hivep2,Jarid2,Jun,Kcnh8,Kdm2b,Klf2,Klf3,Klf6,Lef1,Mapk14,Mapk9,Max,Mbd2,Mospd1,Ncf1,Nfatc1,Nfkbiz,Nipbl,Nrip1,Osm,Otulin,P2ry1,Pdcd4,Per1,Pim1,Prickle1,Prkch,Prkcq,Rara,S1pr1,Scml4,Ski,Smarca2,Smarce1,Spty2d1,Ss18,Stat4,Stat5b,Tax1bp1,Tbl1xr1,Tnfaip3,Tnfrsf1a,Tnfsf11,Tnfsf8,Trib1,Trim25,Ubr2,Usp3,Vgll4,Vsx2,Wwox,Xbp1,Zfp3,Zfp326,Zfp609,Zfp652,Zfp932,Zscan10 |
| GO:0023052 | signaling | 103 | 3859 | 9.69E-06 | Abhd2,Agtr1b,Akt2,Arhgap15,Arhgap25,Bcl10,Btla,Camk4,Cblb,Ccne1,Ccr1,Ccr4,Ccr7,Cd101,Cd247,Cd28,Cd300a,Cd47,Cd97,Crem,Dock10,Dusp1,Dyrk2,E2f2,F2,Fasl,Fyb,Gdi2,Gnat1,Gng2,Gpbar1,Gpr132,Gpr18,Gucy2c,Gzmb,Hey1,Hivep2,Igf2r,Il15ra,Il21r,Il6st,Itga4,Itk,Jak1,Jun,Kcnh8,Klf6,Lef1,Lrrc8d,Lrtm2,Map3k1,Map3k8,Map4k1,Mapk14,Mapk9,Mknk2,Neurl1b,Nfatc1,Nfkbiz,Nrxn3,Nsg2,Nsmce1,Osm,Otulin,P2rx4,P2ry1,Pdcd4,Pde11a,Pde4a,Pim1,Plcxd2,Ppp2r5a,Prkca,Prkch,Prkcq,Rara,Rasgrp2,Rftn1,Rgs10,Rras2,Rtkn2,S1pr1,Selt,Sh2b1,Sipa1l1,Ski,Sla,Sod1,Spag9,Ss18,Stat4,Stat5b,Stk10,Tgfbr2,Tmem64,Tnfrsf1a,Tnfrsf1b,Tnfsf11,Trib1,Wasf2,Wwox,Xbp1,Xcr1 |
| GO:0019219 | regulation of nucleobase-containing compound metabolic process | 94 | 3407 | 9.79E-06 | Adipoq,Aebp2,Aff4,Ago2,Akt2,Atp2b4,Bach2,Bcl10,Btg2,Camk4,Ccne1,Cd28,Chd2,Chd7,Crem,Dach2,Dusp1,E2f2,Elf1,Ercc4,Esrp1,Ets1,Ets2,Fasl,Fli1,Fosl2,Foxn3,Foxp1,Gtf2ird1,Hey1,Hivep2,Jarid2,Jun,Kcnh8,Kdm2b,Klf2,Klf3,Klf6,Lef1,Mapk14,Mapk9,Max,Mbd2,Mbnl1,Mospd1,Ncf1,Nfatc1,Nfkbiz,Nipbl,Nrip1,Osm,Otulin,P2ry1,Pdcd4,Per1,Pim1,Prickle1,Prkch,Prkcq,Rara,Riok1,Rnf219,S1pr1,Scml4,Sh2b1,Ski,Smarca2,Smarce1,Spty2d1,Ss18,Stat4,Stat5b,Tax1bp1,Tbl1xr1,Tnfaip3,Tnfrsf1a,Tnfrsf1b,Tnfsf11,Tnfsf8,Tnrc6b,Trib1,Trim25,Ubr2,Usp3,Vgll4,Vsx2,Wwox,Xbp1,Zfp3,Zfp326,Zfp609,Zfp652,Zfp932,Zscan10 |
| GO:0044260 | cellular macromolecule metabolic process | 129 | 5233 | 1.00E-05 | Aebp2,Aff4,Ago2,Akt2,Bach2,Btg1,Btg2,Camk4,Cblb,Ccne1,Ccr7,Chd2,Chd7,Chst11,Chsy1,Crbn,Crem,Dach2,Dad1,Dcaf7,Dnajc6,Dusp1,Dusp10,Dusp2,Dyrk2,E2f2,Eif4e3,Elf1,Ercc4,Ets1,Ets2,Ext1,Fanci,Fasl,Fbxl12,Fbxo33,Fkbp5,Fli1,Fntb,Fosl2,Foxn3,Foxp1,Ggta1,Gtf2ird1,Gucy2c,Hbs1l,Hectd1,Hey1,Hivep2,Il6st,Itk,Jak1,Jarid2,Jun,Kcnh8,Kdm2b,Klf2,Klf3,Klf6,Lef1,Lmtk3,Lonrf1,Map3k1,Map3k8,Map4k1,Mapk14,Mapk9,Max,Mbd2,Mbip,Mgme1,Mknk2,Msrb3,Nabp1,Neurl1b,Nfatc1,Nfkbiz,Nipbl,Nrip1,Nsmce1,Os9,Osm,Otulin,Per1,Pim1,Polg2,Ppp1cc,Ppp2r5a,Prkca,Prkch,Prkcq,Ptp4a2,Rara,Riok1,Rnf125,Scml4,Sirt4,Ski,Sla,Smarca2,Spaca4,Spag9,Spty2d1,Ss18,St8sia1,Stat4,Stat5b,Stk10,Tars2,Tbl1xr1,Tgfbr2,Tnfaip3,Tnfsf11,Trib1,Trim25,Ubr2,Usp3,Vgll4,Vsx2,Wdyhv1,Xbp1,Xylt1,Zdhhc8,Zfp3,Zfp326,Zfp609,Zfp652,Zfp932,Zscan10 |
| GO:0050870 | positive regulation of T cell activation | 16 | 180 | 1.00E-05 | Bcl10,Ccr7,Cd28,Cd47,Dpp4,Dusp10,H2-Aa,Il6st,Nfkbiz,Prkcq,Rara,Stat5b,Tgfbr2,Tnfsf11,Xbp1,Zfp609 |
| GO:0050863 | regulation of T cell activation | 20 | 287 | 1.05E-05 | Bcl10,Btla,Cblb,Ccr7,Cd28,Cd300a,Cd47,Dpp4,Dusp10,H2-Aa,Il6st,Nfkbiz,Prkcq,Rara,Stat5b,Tgfbr2,Tigit,Tnfsf11,Xbp1,Zfp609 |
| GO:0007154 | cell communication | 105 | 3984 | 1.12E-05 | Abhd2,Agtr1b,Akt2,Arhgap15,Arhgap25,Bcl10,Btla,Camk4,Cblb,Ccne1,Ccr1,Ccr4,Ccr7,Cd101,Cd247,Cd28,Cd300a,Cd47,Cd97,Cdkn2b,Crem,Dock10,Dusp1,Dyrk2,E2f2,F2,Fasl,Fyb,Gatsl3,Gdi2,Gnat1,Gng2,Gpbar1,Gpr132,Gpr18,Gucy2c,Gzmb,Hey1,Hivep2,Igf2r,Il15ra,Il21r,Il6st,Itga4,Itk,Jak1,Jun,Kcnh8,Klf6,Lef1,Lrrc8d,Lrtm2,Map3k1,Map3k8,Map4k1,Mapk14,Mapk9,Mknk2,Neurl1b,Nfatc1,Nfkbiz,Nrxn3,Nsg2,Nsmce1,Osm,Otulin,P2rx4,P2ry1,Pdcd4,Pde11a,Pde4a,Pim1,Plcxd2,Ppp2r5a,Prkca,Prkch,Prkcq,Rara,Rasgrp2,Rftn1,Rgs10,Rras2,Rtkn2,S1pr1,Selt,Sh2b1,Sipa1l1,Ski,Sla,Sod1,Spag9,Ss18,Stat4,Stat5b,Stk10,Tgfbr2,Tmem64,Tnfrsf1a,Tnfrsf1b,Tnfsf11,Trib1,Wasf2,Wwox,Xbp1,Xcr1 |
| GO:0051254 | positive regulation of RNA metabolic process | 54 | 1558 | 1.25E-05 | Ago2,Akt2,Bcl10,Btg2,Camk4,Cd28,Chd7,Crem,E2f2,Elf1,Ets1,Ets2,Fli1,Fosl2,Foxn3,Foxp1,Hey1,Hivep2,Jun,Klf2,Klf6,Lef1,Mapk14,Mapk9,Max,Mospd1,Ncf1,Nfatc1,Nipbl,Nrip1,Osm,P2ry1,Per1,Pim1,Rara,Riok1,Rnf219,S1pr1,Ski,Smarca2,Ss18,Stat4,Stat5b,Tbl1xr1,Tnfrsf1a,Tnfrsf1b,Tnfsf11,Tnfsf8,Tnrc6b,Vsx2,Wwox,Xbp1,Zfp326,Zfp609 |
| GO:0031326 | regulation of cellular biosynthetic process | 96 | 3549 | 1.45E-05 | Adipoq,Aebp2,Aff4,Ago2,Akt2,Atp1a1,Atp2b4,Bach2,Bcl10,Btg2,Camk4,Ccne1,Ccr7,Cd28,Chd2,Chd7,Crem,Dach2,Ddx6,Dusp1,Dyrk2,E2f2,Eif4e3,Elf1,Elovl5,Ets1,Ets2,Fasl,Fli1,Fosl2,Foxn3,Foxp1,Gtf2ird1,Hey1,Hivep2,Jarid2,Jun,Kcnh8,Kdm2b,Klf2,Klf3,Klf6,Lef1,Mapk14,Mapk9,Max,Mbd2,Mknk2,Mospd1,Ncf1,Nfatc1,Nfkbiz,Nipbl,Nrip1,Osm,Otulin,P2ry1,Pdcd4,Per1,Pim1,Prickle1,Prkca,Prkch,Prkcq,Rara,S1pr1,Scml4,Sh2b1,Ski,Smarca2,Smarce1,Spty2d1,Ss18,Stat4,Stat5b,Tax1bp1,Tbl1xr1,Tnfaip3,Tnfrsf1a,Tnfsf11,Tnfsf8,Tnrc6b,Trib1,Trim25,Ubr2,Usp3,Vgll4,Vsx2,Wwox,Xbp1,Zfp3,Zfp326,Zfp609,Zfp652,Zfp932,Zscan10 |
| GO:0050790 | regulation of catalytic activity | 65 | 2057 | 1.47E-05 | Adipoq,Akt2,Arhgap15,Arhgap25,Atp2b4,Bcl10,Btg1,Cblb,Ccne1,Ccr7,Cd300a,Cdkn2b,Cnst,Crim1,Cryaa,Cyfip2,Dock10,Dusp1,Dusp10,Dusp2,F2,Fam20a,Fasl,Gadd45g,Gdi2,Gnat1,Itk,Jun,Lef1,Lmtk3,Lrtm2,Map3k1,Map3k8,Map4k1,Mapk14,Mapk9,Mtmr12,Ncf1,Pdcd4,Pim1,Pkp4,Ppp2r5a,Prkca,Prkcq,Rara,Rasgrp2,Rgs10,S1pr1,Serpine2,Sipa1l1,Sirt4,Ski,Slc39a10,Snx18,Sod1,Spag9,Stk10,Tgfbr2,Tmem64,Tnfaip3,Tnfrsf1b,Tnfsf11,Trib1,Usp6nl,Xdh |
| GO:0043170 | macromolecule metabolic process | 146 | 6225 | 1.56E-05 | Adarb2,Aebp2,Aff4,Ago2,Akt2,Bach2,Btg1,Btg2,Camk4,Cblb,Ccne1,Ccr7,Celf2,Chd2,Chd7,Chst11,Chsy1,Crbn,Crem,Cyfip2,Dach2,Dad1,Dcaf7,Ddx6,Dhx40,Dnajc6,Dpp4,Dusp1,Dusp10,Dusp2,Dyrk2,E2f2,Eif4e3,Elf1,Ercc4,Esrp1,Ets1,Ets2,Ext1,F2,Fanci,Fasl,Fbxl12,Fbxo33,Fkbp5,Fli1,Fntb,Fosl2,Foxn3,Foxp1,Gadd45g,Ggta1,Gtf2ird1,Gucy2c,Gzmb,Hbs1l,Hectd1,Hey1,Hivep2,Il6st,Ints7,Itk,Jak1,Jarid2,Jun,Kcnh8,Kdm2b,Klf2,Klf3,Klf6,Lef1,Lmtk3,Lonrf1,Map3k1,Map3k8,Map4k1,Mapk14,Mapk9,Max,Mbd2,Mbip,Mbnl1,Mgme1,Mknk2,Msrb3,Nabp1,Neurl1b,Nfatc1,Nfkbiz,Nipbl,Nrip1,Nsmce1,Os9,Osm,Otulin,Per1,Pim1,Polg2,Ppp1cc,Ppp2r5a,Prkca,Prkch,Prkcq,Prss39,Ptp4a2,Rara,Riok1,Rnf125,Scml4,Sirt4,Ski,Sla,Smarca2,Smndc1,Spaca4,Spag9,Spty2d1,Ss18,St8sia1,Stat4,Stat5b,Stk10,Tars2,Tbl1xr1,Tgfbr2,Tnfaip3,Tnfsf11,Trappc12,Trib1,Trim25,Txnl4b,Ubr2,Usp3,Vgll4,Vsx2,Wdyhv1,Xbp1,Xpnpep2,Xylt1,Zdhhc8,Zfp3,Zfp326,Zfp609,Zfp652,Zfp932,Zscan10 |
| GO:0009966 | regulation of signal transduction | 81 | 2823 | 1.72E-05 | Adipoq,Akt2,Arhgap25,Atp2b4,Bcl10,Cblb,Ccr1,Ccr7,Cd28,Cd300a,Cdkn2b,Chst11,Chsy1,Cyfip2,Cyth4,Dusp1,Dusp10,Dusp2,Dyrk2,Elf1,F2,Fasl,Foxp1,Gadd45g,Gatsl3,Glrx,Gpr18,Hey1,Il6st,Inpp4b,Jak1,Jun,Lef1,Lrtm2,Map3k1,Map3k8,Map4k1,Mapk14,Mapk9,Mbd2,Ncf1,Neto2,Osm,Otulin,P2rx4,P2ry1,Pdcd4,Pde11a,Pde4a,Per1,Prickle1,Prkca,Prkch,Prkcq,Prl5a1,Rara,Rgs10,Rnf125,Serpine2,Sgms1,Sh2b1,Shisa2,Sipa1l1,Ski,Sla,Slc39a10,Slc7a3,Sod1,Spag9,Tbl1xr1,Tmem64,Tnfaip3,Tnfrsf1a,Tnfsf11,Tnfsf8,Trib1,Trim25,Ubr2,Wwox,Xbp1,Xdh |
| GO:0030155 | regulation of cell adhesion | 30 | 624 | 1.77E-05 | Adipoq,Bcl10,Btla,Cblb,Ccdc80,Ccr7,Cd28,Cd300a,Cd47,Cytip,Dpp4,Dusp10,Ets1,H2-Aa,Il6st,Itga4,Lef1,Nfkbiz,Prkca,Prkcq,Rara,S1pr1,Serpine2,Stat5b,Tgfbr2,Tigit,Tnfaip3,Tnfsf11,Xbp1,Zfp609 |
| GO:0006807 | nitrogen compound metabolic process | 159 | 6983 | 1.85E-05 | Adarb2,Aebp2,Aff4,Ago2,Akt2,Bach2,Bcat1,Btg1,Btg2,Camk4,Cblb,Ccne1,Ccr7,Celf2,Cers6,Chd2,Chd7,Chst11,Chsy1,Cmah,Cmas,Crbn,Crem,Cyfip2,Dach2,Dad1,Dcaf7,Ddx6,Dhx40,Dnajc6,Dpp4,Dusp1,Dusp10,Dusp2,Dyrk2,E2f2,Eif4e3,Elf1,Elovl5,Ercc4,Esrp1,Ets1,Ets2,Ext1,F2,Fanci,Fasl,Fbxl12,Fbxo33,Fkbp5,Fli1,Fntb,Fosl2,Foxn3,Foxp1,Gadd45g,Ggta1,Gtf2ird1,Gucy2c,Gzmb,Hbs1l,Hectd1,Hey1,Hivep2,Ints7,Itk,Jak1,Jarid2,Jun,Kcnh8,Kdm2b,Klf2,Klf3,Klf6,Lef1,Lmtk3,Lonrf1,Map3k1,Map3k8,Map4k1,Mapk14,Mapk9,Max,Mbd2,Mbip,Mbnl1,Mgme1,Mknk2,Moxd1,Msrb3,Nabp1,Ncf1,Neurl1b,Nfatc1,Nfkbiz,Nipbl,Nrip1,Nsmce1,Os9,Osm,Otulin,P2rx4,Pde11a,Pde4a,Per1,Pim1,Polg2,Ppp1cc,Ppp2r5a,Prkca,Prkch,Prkcq,Prss39,Ptp4a2,Rara,Riok1,Rnf125,Rpia,Scml4,Sgms1,Sirt4,Ski,Sla,Smarca2,Smndc1,Smox,Sod1,Spag9,Spty2d1,Ss18,St8sia1,Stat4,Stat5b,Stk10,Tars2,Tbl1xr1,Tgfbr2,Tnfaip3,Tnfsf11,Trappc12,Trib1,Trim25,Txnl4b,Ubr2,Usp3,Vgll4,Vsx2,Wdyhv1,Xbp1,Xdh,Xpnpep2,Xylt1,Zdhhc8,Zfp3,Zfp326,Zfp609,Zfp652,Zfp932,Zscan10 |
| GO:0065009 | regulation of molecular function | 86 | 3081 | 1.98E-05 | Adipoq,Akt2,Arhgap15,Arhgap25,Atp2b4,Bcl10,Btg1,Cblb,Ccne1,Ccr7,Cd300a,Cdkn2b,Cnst,Crbn,Crim1,Cryaa,Cyfip2,Cyth4,Dock10,Dusp1,Dusp10,Dusp2,Epb4.1,Ercc4,F2,Fam20a,Fasl,Gadd45g,Gdi2,Glrx,Gnat1,Hey1,Itga4,Itk,Jun,Lef1,Lmtk3,Lrtm2,Map3k1,Map3k8,Map4k1,Mapk14,Mapk9,Mbd2,Mtmr12,Ncf1,Neto2,Nipbl,Nrxn3,Osm,Otulin,P2ry1,Pdcd4,Pim1,Pkp4,Ppp2r5a,Prkca,Prkch,Prkcq,Prl5a1,Rara,Rasgrp2,Rgs10,S1pr1,Serpine2,Sipa1l1,Sirt4,Ski,Slc39a10,Snx18,Sod1,Spag9,Stim2,Stk10,Tax1bp1,Tgfbr2,Tmem64,Tnfaip3,Tnfrsf1b,Tnfsf11,Tnfsf8,Trib1,Trim25,Usp6nl,Xdh,Zfp932 |
| GO:0051249 | regulation of lymphocyte activation | 23 | 396 | 2.13E-05 | Bcl10,Btla,Cblb,Ccr7,Cd28,Cd300a,Cd47,Dpp4,Dusp10,H2-Aa,Il15ra,Il6st,Nfkbiz,Prkcq,Rara,Slc39a10,Stat5b,Tgfbr2,Tigit,Tnfaip3,Tnfsf11,Xbp1,Zfp609 |
| GO:0051247 | positive regulation of protein metabolic process | 53 | 1556 | 2.40E-05 | Adipoq,Akt2,Atp2b4,Bcl10,Cblb,Ccr1,Ccr7,Cd28,Cd300a,Cryaa,Cyfip2,F2,Fam20a,Fanci,Fasl,Gadd45g,Il6st,Ist1,Jarid2,Jun,Klf2,Lonrf1,Lrrn3,Map3k1,Map3k8,Map4k1,Mapk14,Mapk9,Ncf1,Nipbl,Osm,P2ry1,Pim1,Ppp2r5a,Prickle1,Prkca,Prkcq,Rara,Rnf125,Slc39a10,Sod1,Spag9,Stat5b,Stk10,Tgfbr2,Tnfaip3,Tnfrsf1a,Tnfrsf1b,Tnfsf11,Trib1,Xbp1,Xdh,Zfand2a |
| GO:0051251 | positive regulation of lymphocyte activation | 18 | 253 | 2.70E-05 | Bcl10,Ccr7,Cd28,Cd47,Dpp4,Dusp10,H2-Aa,Il15ra,Il6st,Nfkbiz,Prkcq,Rara,Slc39a10,Stat5b,Tgfbr2,Tnfsf11,Xbp1,Zfp609 |
| GO:0032502 | developmental process | 126 | 5213 | 3.50E-05 | Abcb1b,Abhd2,Adipoq,Aff4,Ago2,Agtr1b,Akt2,Ankrd11,Atp2b4,Bcl10,Btg2,Camk4,Ccr1,Ccr4,Ccr7,Cd28,Cdkn2b,Chd2,Chd7,Chst11,Chsy1,Clic4,Cryaa,Cyfip2,Dach2,Dad1,Dcaf7,Ddx6,Dock10,Drc1,Dusp1,Dusp10,Dusp2,Elf1,Esrp1,Ets1,Ets2,Ext1,F2,Fam20a,Fasl,Fli1,Fosl2,Foxn3,Foxp1,Gadd45g,Glrx2,Gnat1,Gpr18,Gtf2ird1,Hectd1,Hey1,Hivep2,Igf2r,Il15ra,Il6st,Ist1,Itga4,Itk,Jak1,Jarid2,Jun,Kdm2b,Klf2,Lef1,Map3k1,Mapk14,Mapk9,Mbd2,Mbnl1,Mctp2,Mfsd2a,Mknk2,Mospd1,Nfatc1,Ninj2,Nipbl,Nrip1,Nrxn3,Osm,Otulin,P2ry1,Pdcd4,Phc3,Pip4k2a,Polg2,Ppp1cc,Prickle1,Prkca,Prkch,Prrc2b,Rara,Rtkn2,S1pr1,Selt,Serpine2,Shisa2,Sipa1l1,Ski,Sla,Smarca2,Smarce1,Sod1,Spag9,Ss18,Stat5b,Tapt1,Tbl1xr1,Tgfbr2,Tmc1,Tmeff2,Tmem64,Tnfaip3,Tnfrsf1a,Tnfrsf1b,Tnfsf11,Tnfsf8,Vsx2,Wasf2,Wwox,Xbp1,Xdh,Xylt1,Zfp609,Zfp932,Zscan10 |
| GO:0010033 | response to organic substance | 74 | 2553 | 4.03E-05 | Abhd2,Adipoq,Ago2,Agtr1b,Akt2,Atp1a1,Atp2b4,Bcl10,Btg1,Btg2,Ccr1,Ccr4,Ccr7,Crem,Cryaa,Dusp1,Dusp10,Ets1,Fasl,Fntb,Fosl2,Foxp1,Gatsl3,Glrx,Glrx2,H2-Aa,Igf2r,Il21r,Il6st,Jak1,Jarid2,Jun,Klf2,Klf3,Klf6,Lef1,Lrtm2,Map3k1,Map4k1,Mapk14,Mapk9,Mbd2,Nrip1,Os9,Osm,P2rx4,P2ry1,Pdcd4,Per1,Pim1,Prkca,Rara,Rftn1,Rgs10,Rnf125,Scd2,Selt,Sgms1,Slc3a2,Sod1,Stat4,Stat5b,Tgfbr2,Tnfaip3,Tnfrsf1a,Tnfrsf1b,Tnfsf11,Trib1,Trim25,Ubr2,Vamp4,Wwox,Xbp1,Xcr1 |
| GO:0045893 | positive regulation of transcription, DNA-templated | 49 | 1413 | 4.04E-05 | Ago2,Akt2,Bcl10,Camk4,Cd28,Chd7,Crem,E2f2,Elf1,Ets1,Ets2,Fli1,Fosl2,Foxn3,Foxp1,Hey1,Hivep2,Jun,Klf2,Klf6,Lef1,Mapk14,Mapk9,Max,Mospd1,Ncf1,Nfatc1,Nipbl,Nrip1,Osm,P2ry1,Per1,Pim1,Rara,S1pr1,Ski,Smarca2,Ss18,Stat4,Stat5b,Tbl1xr1,Tnfrsf1a,Tnfsf11,Tnfsf8,Vsx2,Wwox,Xbp1,Zfp326,Zfp609 |
| GO:0006357 | regulation of transcription by RNA polymerase II | 59 | 1856 | 4.10E-05 | Aebp2,Ago2,Akt2,Atp2b4,Bach2,Btg2,Ccne1,Cd28,Chd7,Crem,Dach2,E2f2,Elf1,Ets1,Ets2,Fasl,Fli1,Fosl2,Foxp1,Gtf2ird1,Hey1,Hivep2,Jarid2,Jun,Kcnh8,Kdm2b,Klf2,Klf3,Klf6,Lef1,Mapk14,Max,Mbd2,Mospd1,Nfatc1,Nfkbiz,Nipbl,Nrip1,Osm,P2ry1,Pdcd4,Per1,Rara,S1pr1,Ski,Smarca2,Ss18,Stat4,Stat5b,Tbl1xr1,Tnfrsf1a,Tnfsf11,Tnfsf8,Usp3,Vsx2,Wwox,Xbp1,Zfp609,Zfp932 |
| GO:0045944 | positive regulation of transcription by RNA polymerase II | 42 | 1125 | 4.44E-05 | Ago2,Akt2,Cd28,Chd7,Crem,E2f2,Elf1,Ets1,Ets2,Fli1,Fosl2,Foxp1,Hey1,Hivep2,Jun,Klf2,Klf6,Lef1,Mapk14,Max,Mospd1,Nfatc1,Nipbl,Nrip1,Osm,P2ry1,Per1,Rara,S1pr1,Ski,Smarca2,Ss18,Stat4,Stat5b,Tbl1xr1,Tnfrsf1a,Tnfsf11,Tnfsf8,Vsx2,Wwox,Xbp1,Zfp609 |
| GO:1901700 | response to oxygen-containing compound | 49 | 1429 | 5.29E-05 | Abhd2,Adipoq,Agtr1b,Akt2,Atp2b4,Btg1,Btg2,Ccr7,Crem,Cryaa,Dusp1,Dusp10,Ets1,Fasl,Fosl2,Foxp1,Gatsl3,Glrx,Glrx2,Igf2r,Jun,Klf2,Klf3,Map4k1,Mapk14,Mapk9,Mbd2,Ncf1,Nrip1,P2rx4,Pdcd4,Per1,Pim1,Prkca,Rara,Rgs10,Scd2,Selt,Sgms1,Sod1,Stat5b,Tgfbr2,Tnfaip3,Tnfrsf1a,Tnfrsf1b,Trib1,Trim25,Ubr2,Xbp1 |
| GO:0009605 | response to external stimulus | 57 | 1794 | 6.34E-05 | Adipoq,Agtr1b,Atp1a1,Bcl10,Btg2,Ccdc80,Ccr1,Ccr4,Ccr7,Cd47,Cdkn2b,Chd7,Chsy1,Dad1,Dusp10,Ets1,Ext1,F2,Fasl,Fkbp5,Foxp1,Gatsl3,Gnat1,Hist1h2be,Jun,Lef1,Mapk14,Mapk9,Mbd2,Ncf1,P2rx4,P2ry1,Pdcd4,Per1,Pim1,Ppp1cc,Prkca,Prkcq,Rara,S1pr1,Serpine2,Sgms1,Sod1,Spaca4,Stat5b,Tbl1xr1,Tgfbr2,Tmc1,Tnfaip3,Tnfrsf1a,Tnfrsf1b,Tnfsf11,Tnfsf8,Trib1,Trim25,Xbp1,Xcr1 |
| GO:0045639 | positive regulation of myeloid cell differentiation | 11 | 95 | 6.34E-05 | Ccr1,Cd101,Ets1,Jun,Lef1,Mapk14,Prkca,Stat5b,Tmem64,Tnfsf11,Trib1 |
| GO:0030097 | hemopoiesis | 28 | 600 | 6.39E-05 | Camk4,Ccr1,Ccr7,Cd28,Cdkn2b,Chd2,Chd7,Dock10,Fli1,Foxp1,Gadd45g,Gpr18,Hectd1,Itk,Jun,Klf2,Lef1,Mapk14,Mknk2,Nfatc1,Pip4k2a,Rtkn2,Stat5b,Tgfbr2,Tnfaip3,Tnfsf11,Tnfsf8,Wasf2 |
| GO:0033993 | response to lipid | 34 | 825 | 6.52E-05 | Abhd2,Adipoq,Agtr1b,Atp1a1,Ccr7,Cryaa,Dusp1,Dusp10,Ets1,Fasl,Fosl2,Foxp1,Glrx,Igf2r,Jun,Lef1,Map4k1,Mapk14,Mbd2,Nrip1,Pdcd4,Pim1,Prkca,Rara,Scd2,Sgms1,Stat5b,Tgfbr2,Tnfaip3,Tnfrsf1a,Tnfrsf1b,Trib1,Trim25,Xbp1 |
| GO:0009617 | response to bacterium | 27 | 566 | 6.54E-05 | Adipoq,Bcl10,Ccdc80,Ccr4,Ccr7,Cd47,Chd7,Dusp10,F2,Fasl,Fkbp5,Foxp1,Hist1h2be,Jun,Mapk14,Ncf1,Pdcd4,Rara,Sgms1,Spaca4,Stat5b,Tnfaip3,Tnfrsf1a,Tnfrsf1b,Tnfsf8,Trib1,Xbp1 |
| GO:0032101 | regulation of response to external stimulus | 30 | 681 | 7.79E-05 | Adipoq,Akt2,Ccr1,Ccr4,Ccr7,Cd28,Cd47,Dusp10,Ets1,F2,Foxp1,Gpr18,Mapk14,Otulin,P2rx4,Pdcd4,Per1,Prkca,Prkcq,Rnf125,S1pr1,Serpine2,Stat5b,Tgfbr2,Tnfaip3,Tnfrsf1a,Tnfrsf1b,Tnfsf11,Trib1,Xylt1 |

| | | | | | |
|---|---|---|---|---|---|
| GO:0051246 | regulation of protein metabolic process | 74 | 2612 | 8.00E-05 | Adipoq,Ago2,Akt2,Atp2b4,Bcl10,Btg2,Cblb,Ccne1,Ccr1,Ccr7,Cd28,Cd300a,Cdkn2b,Crim1,Cryaa,Cyfip2,Ddx6,Dusp1,Dusp10,Dusp2,Eif4e3,F2,Fam20a,Fanci,Fasl,Gadd45g,Il6st,Ist1,Jarid2,Jun,Klf2,Lef1,Lonrf1,Lrrn3,Lrtm2,Map3k1,Map3k8,Map4k1,Mapk14,Mapk9,Mknk2,Ncf1,Nipbl,Os9,Osm,P2ry1,Pdcd4,Per1,Pim1,Ppp2r5a,Prickle1,Prkca,Prkcq,Rara,Rnf125,Serpine2,Sirt4,Ski,Slc39a10,Sod1,Spag9,Stat5b,Stk10,Tgfbr2,Tnfaip3,Tnfrsf1a,Tnfrsf1b,Tnfsf11,Tnrc6b,Trib1,Ubac2,Xbp1,Xdh,Zfand2a |
| GO:0044093 | positive regulation of molecular function | 52 | 1593 | 8.70E-05 | Adipoq,Akt2,Arhgap15,Arhgap25,Atp2b4,Bcl10,Btg1,Ccr7,Cd300a,Crbn,Cyfip2,Dock10,Epb4.1,F2,Fam20a,Fasl,Gadd45g,Gdi2,Glrx,Gnat1,Itk,Jun,Map3k1,Map3k8,Map4k1,Mapk14,Mapk9,Mbd2,Ncf1,Nipbl,Pim1,Pkp4,Prkch,Prkcq,Rara,Rasgrp2,Rgs10,S1pr1,Sipa1l1,Ski,Slc39a10,Snx18,Sod1,Spag9,Stim2,Stk10,Tgfbr2,Tnfsf11,Trib1,Trim25,Usp6nl,Xdh |
| GO:0002520 | immune system development | 30 | 687 | 8.89E-05 | Camk4,Ccr1,Ccr4,Ccr7,Cd28,Cdkn2b,Chd2,Chd7,Dock10,Fli1,Foxp1,Gadd45g,Gpr18,Hectd1,Itk,Jarid2,Jun,Klf2,Lef1,Mapk14,Mknk2,Nfatc1,Pip4k2a,Rtkn2,Stat5b,Tgfbr2,Tnfaip3,Tnfsf11,Tnfsf8,Wasf2 |
| GO:0070887 | cellular response to chemical stimulus | 67 | 2287 | 9.01E-05 | Abhd2,Adipoq,Ago2,Agtr1b,Akt2,Atp1a1,Atp2b4,Ccr1,Ccr4,Ccr7,Cdkn2b,Clic4,Dusp1,Ets1,Foxp1,Gatsl3,Glrx,Glrx2,Il21r,Il6st,Jak1,Jarid2,Jun,Klf2,Klf3,Klf6,Lef1,Lrtm2,Map3k1,Map4k1,Mapk14,Mapk9,Mbd2,Mknk2,Ncf1,Nrip1,Osm,P2rx4,P2ry1,Pdcd4,Pde4a,Pim1,Prkca,Prkcq,Rara,Rasgrp2,Rgs10,Rnf125,S1pr1,Selt,Sgms1,Sirt4,Sod1,Stat4,Stat5b,Tgfbr2,Tnfaip3,Tnfrsf1a,Tnfrsf1b,Tnfsf11,Trim25,Ubr2,Vamp4,Wwox,Xbp1,Xcr1,Zfand2a |
| GO:0045935 | positive regulation of nucleobase-containing compound metabolic process | 55 | 1732 | 9.08E-05 | Ago2,Akt2,Bcl10,Btg2,Camk4,Cd28,Chd7,Crem,E2f2,Elf1,Ets1,Ets2,Fli1,Fosl2,Foxn3,Foxp1,Hey1,Hivep2,Jun,Klf2,Klf6,Lef1,Mapk14,Mapk9,Max,Mospd1,Ncf1,Nfatc1,Nipbl,Nrip1,Osm,P2ry1,Per1,Pim1,Prkcq,Rara,Riok1,Rnf219,S1pr1,Ski,Smarca2,Ss18,Stat4,Stat5b,Tbl1xr1,Tnfrsf1a,Tnfrsf1b,Tnfsf11,Tnfsf8,Tnrc6b,Vsx2,Wwox,Xbp1,Zfp326,Zfp609 |
| GO:2000026 | regulation of multicellular organismal development | 60 | 1961 | 9.08E-05 | Adipoq,Ago2,Agtr1b,Btg1,Cblb,Ccr1,Ccr7,Cd101,Cd28,Cd97,Cdkn2b,Chd7,Ddx6,Dusp10,E2f2,Ets1,F2,Fasl,Foxp1,H2-Aa,Hey1,Il15ra,Il6st,Inpp4b,Ist1,Jak1,Jarid2,Jun,Klf2,Lef1,Lrrn3,Lrtm2,Mapk14,Nfatc1,Nfkbiz,Nipbl,Pdcd4,Pim1,Ppp1cc,Prickle1,Prkca,Prkch,Rara,S1pr1,Serpine2,Sipa1l1,Ski,Spag9,Stat5b,Tapt1,Tgfbr2,Tmem64,Tnfaip3,Tnfrsf1a,Tnfsf11,Trib1,Xbp1,Xdh,Xylt1,Zfp609 |
| GO:0032879 | regulation of localization | 73 | 2579 | 9.09E-05 | Abhd2,Adipoq,Ago2,Akt2,Atp1a1,Camk4,Ccne1,Ccr1,Ccr7,Cd247,Cd300a,Cd47,Chd7,Clic4,Cnst,Crbn,Dnajc6,Dock10,Dusp10,Epb4.1,Ets1,F2,Fasl,Foxp1,Glrx,Gpr18,Hectd1,Il6st,Inpp4b,Itga4,Jun,Kcnh8,Lef1,Leprotl1,Map3k1,Mapk14,Mapk9,Neto2,Nipbl,Os9,Osm,P2rx4,P2ry1,Per1,Ppp1cc,Ppp2r5a,Prkca,Prkcq,Rras2,S1pr1,Sar1a,Selt,Serpine2,Sirt4,Slc30a4,Spag9,Stat5b,Stim2,Stk10,Tgfbr2,Tmc1,Tmeff2,Tnfrsf1a,Tnfrsf1b,Tnfsf11,Trappc12,Trib1,Ubac2,Usp6nl,Vamp4,Xbp1,Xcr1,Zfp609 |
| GO:0051094 | positive regulation of developmental process | 48 | 1423 | 9.09E-05 | Adipoq,Ago2,Agtr1b,Btg1,Cblb,Ccne1,Ccr1,Ccr7,Cd101,Cd28,Cd97,Chd7,Dusp10,Ets1,Foxp1,H2-Aa,Hey1,Il15ra,Il6st,Ist1,Jak1,Jun,Kdm2b,Lef1,Lrrn3,Lrtm2,Mapk14,Mapk9,Nfkbiz,Nipbl,Pim1,Ppp1cc,Prkca,Prkch,Rara,Serpine2,Spag9,Stat5b,Tapt1,Tgfbr2,Tmem64,Tnfaip3,Tnfrsf1a,Tnfrsf1b,Tnfsf11,Trib1,Xbp1,Zfp609 |
| GO:1902531 | regulation of intracellular signal transduction | 52 | 1599 | 9.09E-05 | Adipoq,Arhgap25,Atp2b4,Bcl10,Ccr1,Ccr7,Cd28,Cd300a,Cyth4,Dusp1,Dusp10,Dusp2,Dyrk2,F2,Fasl,Gadd45g,Gatsl3,Glrx,Gpr18,Il6st,Inpp4b,Jak1,Jun,Lrtm2,Map3k1,Map3k8,Map4k1,Mapk9,Ncf1,Osm,P2rx4,P2ry1,Pdcd4,Pde11a,Pde4a,Per1,Prkca,Rara,Rnf125,Serpine2,Sipa1l1,Slc7a3,Sod1,Spag9,Tnfaip3,Tnfrsf1a,Tnfsf11,Trib1,Trim25,Ubr2,Xbp1,Xdh |
| GO:0045637 | regulation of myeloid cell differentiation | 15 | 199 | 9.63E-05 | Adipoq,Ccr1,Cd101,Ets1,Foxp1,Inpp4b,Jun,Lef1,Mapk14,Prkca,Rara,Stat5b,Tmem64,Tnfsf11,Trib1 |
| GO:0010646 | regulation of cell communication | 84 | 3130 | 9.70E-05 | Adipoq,Akt2,Arhgap25,Atp2b4,Bcl10,Cblb,Ccr1,Ccr7,Cd28,Cd300a,Cdkn2b,Chd7,Chst11,Chsy1,Cyfip2,Cyth4,Dusp1,Dusp10,Dusp2,Dyrk2,Elf1,F2,Fasl,Foxp1,Gadd45g,Gatsl3,Glrx,Gpr18,Hey1,Il6st,Inpp4b,Jak1,Jun,Lef1,Lrtm2,Map3k1,Map3k8,Map4k1,Mapk14,Mapk9,Mbd2,Ncf1,Neto2,Osm,Otulin,P2rx4,P2ry1,Pdcd4,Pde11a,Pde4a,Per1,Prickle1,Prkca,Prkch,Prkcq,Prl5a1,Rara,Rgs10,Rnf125,Selt,Serpine2,Sgms1,Sh2b1,Shisa2,Sipa1l1,Sirt4,Ski,Sla,Slc39a10,Slc7a3,Sod1,Spag9,Tbl1xr1,Tmem64,Tnfaip3,Tnfrsf1a,Tnfsf11,Tnfsf8,Trib1,Trim25,Ubr2,Wwox,Xbp1,Xdh |
| GO:0051240 | positive regulation of multicellular organismal process | 54 | 1695 | 9.70E-05 | Adipoq,Ago2,Agtr1b,Atp1a1,Bcl10,Btg1,Cblb,Ccr1,Ccr7,Cd101,Cd28,Cd97,Chd7,Dusp10,Ets1,F2,Foxp1,H2-Aa,Hey1,Il15ra,Il6st,Ist1,Jak1,Jun,Kdm2b,Lef1,Lrrn3,Lrtm2,Mapk14,Mapk9,Nfkbiz,Nipbl,P2rx4,P2ry1,Pim1,Ppp1cc,Prkca,Prkch,Prkcq,Rara,Serpine2,Sod1,Spag9,Stat5b,Tapt1,Tgfbr2,Tigit,Tmem64,Tnfaip3,Tnfrsf1a,Tnfsf11,Trib1,Xbp1,Zfp609 |
| GO:0032268 | regulation of cellular protein metabolic process | 70 | 2443 | 9.71E-05 | Adipoq,Ago2,Akt2,Atp2b4,Bcl10,Btg2,Cblb,Ccne1,Ccr1,Ccr7,Cd300a,Cdkn2b,Crim1,Cryaa,Cyfip2,Ddx6,Dusp1,Dusp10,Dusp2,Eif4e3,F2,Fam20a,Fanci,Fasl,Gadd45g,Il6st,Ist1,Jarid2,Jun,Lef1,Lonrf1,Lrrn3,Lrtm2,Map3k1,Map3k8,Map4k1,Mapk14,Mapk9,Mknk2,Ncf1,Nipbl,Os9,Osm,P2ry1,Pdcd4,Per1,Pim1,Ppp2r5a,Prickle1,Prkca,Rara,Rnf125,Serpine2,Sirt4,Ski,Slc39a10,Sod1,Spag9,Stk10,Tgfbr2,Tnfaip3,Tnfrsf1a,Tnfrsf1b,Tnfsf11,Tnrc6b,Trib1,Ubac2,Xbp1,Xdh,Zfand2a |
| GO:0002761 | regulation of myeloid leukocyte differentiation | 12 | 125 | 9.97E-05 | Adipoq,Ccr1,Cd101,Foxp1,Inpp4b,Jun,Lef1,Prkca,Rara,Tmem64,Tnfsf11,Trib1 |
| GO:0048534 | hematopoietic or lymphoid organ development | 29 | 659 | 0.0001 | Camk4,Ccr1,Ccr7,Cd28,Cdkn2b,Chd2,Chd7,Dock10,Fli1,Foxp1,Gadd45g,Gpr18,Hectd1,Itk,Jarid2,Jun,Klf2,Lef1,Mapk14,Mknk2,Nfatc1,Pip4k2a,Rtkn2,Stat5b,Tgfbr2,Tnfaip3,Tnfsf11,Tnfsf8,Wasf2 |
| GO:0048869 | cellular developmental process | 92 | 3545 | 0.0001 | Abcb1b,Abhd2,Adipoq,Aff4,Ago2,Akt2,Btg2,Camk4,Ccr1,Ccr4,Ccr7,Cd28,Cdkn2b,Chd2,Chd7,Chst11,Chsy1,Clic4,Cryaa,Cyfip2,Ddx6,Dock10,Dusp10,Elf1,Ets1,Ets2,Ext1,F2,Fasl,Fli1,Fosl2,Foxn3,Foxp1,Gadd45g,Gnat1,Gpr18,Hectd1,Hey1,Il15ra,Il6st,Ist1,Itga4,Itk,Jak1,Jarid2,Jun,Klf2,Lef1,Map3k1,Mapk14,Mapk9,Mbnl1,Mfsd2a,Mospd1,Nfatc1,Nipbl,P2ry1,Pdcd4,Pip4k2a,Polg2,Ppp1cc,Prkca,Prkch,Prrc2b,Rara,S1pr1,Serpine2,Sipa1l1,Ski,Sla,Smarca2,Smarce1,Sod1,Spag9,Ss18,Stat5b,Tapt1,Tbl1xr1,Tgfbr2,Tmc1,Tmem64,Tnfaip3,Tnfsf11,Tnfsf8,Vsx2,Wasf2,Wwox,Xbp1,Xylt1,Zfp609,Zfp932,Zscan10 |
| GO:0006950 | response to stress | 79 | 2899 | 0.00011 | Abhd2,Adipoq,Agtr1b,Atp2b4,Bcl10,Btg1,Btg2,Camk4,Ccr1,Ccr4,Ccr7,Cd300a,Chd2,Chst2,Cryaa,Dpp4,Dusp1,Dyrk2,Ercc4,Ets1,F2,Fanci,Fntb,Fosl2,Foxn3,Gadd45g,Glrx2,Gzmb,H2-Aa,Hist1h2be,Ints7,Itk,Jak1,Jun,Klf2,Map3k1,Map3k8,Map4k1,Mapk14,Mapk9,Mgme1,Msrb3,Nabp1,Ncf1,Nfkbiz,Ninj2,Nipbl,Nsmce1,Os9,Osm,Otulin,P2rx4,P2ry1,Polg2,Prkca,Prkcq,Serpine2,Sgms1,Sirt4,Sla,Sod1,Spaca4,Spag9,St8sia1,Stat5b,Stk10,Tgfbr2,Tmeff2,Tnfaip3,Tnfrsf1a,Tnfrsf1b,Tnfsf11,Tnfsf8,Trib1,Trim25,Usp3,Vamp4,Xbp1,Xylt1 |
| GO:0007275 | multicellular organism development | 112 | 4603 | 0.00011 | Abcb1b,Adipoq,Ago2,Agtr1b,Akt2,Ankrd11,Atp2b4,Bcl10,Btg2,Camk4,Ccr1,Ccr4,Ccr7,Cd28,Cdkn2b,Chd2,Chd7,Chst11,Chsy1,Clic4,Cryaa,Cyfip2,Dach2,Dad1,Dcaf7,Ddx6,Dock10,Drc1,Dusp1,Dusp10,Dusp2,Esrp1,Ets1,Ets2,Ext1,F2,Fam20a,Fasl,Fli1,Fosl2,Foxn3,Foxp1,Gadd45g,Gnat1,Gpr18,Gtf2ird1,Hectd1,Hey1,Hivep2,Igf2r,Il15ra,Il6st,Ist1,Itga4,Itk,Jarid2,Jun,Kdm2b,Klf2,Lef1,Map3k1,Mapk14,Mapk9,Mbd2,Mbnl1,Mctp2,Mfsd2a,Mknk2,Nfatc1,Nipbl,Nrip1,Nrxn3,Osm,Otulin,P2ry1,Pdcd4,Phc3,Pip4k2a,Polg2,Ppp1cc,Prickle1,Prkca,Prkch,Rara,Rtkn2,S1pr1,Selt,Serpine2,Shisa2,Sipa1l1,Ski,Smarca2,Smarce1,Sod1,Spag9,Stat5b,Tapt1,Tbl1xr1,Tgfbr2,Tmc1,Tnfaip3,Tnfrsf1a,Tnfrsf1b,Tnfsf11,Tnfsf8,Vsx2,Wasf2,Wwox,Xbp1,Xdh,Xylt1,Zfp609 |
| GO:0023051 | regulation of signaling | 84 | 3150 | 0.00011 | Adipoq,Akt2,Arhgap25,Atp2b4,Bcl10,Cblb,Ccr1,Ccr7,Cd28,Cd300a,Cdkn2b,Chd7,Chst11,Chsy1,Cyfip2,Cyth4,Dusp1,Dusp10,Dusp2,Dyrk2,Elf1,F2,Fasl,Foxp1,Gadd45g,Gatsl3,Glrx,Gpr18,Hey1,Il6st,Inpp4b,Jak1,Jun,Lef1,Lrtm2,Map3k1,Map3k8,Map4k1,Mapk14,Mapk9,Mbd2,Ncf1,Neto2,Osm,Otulin,P2rx4,P2ry1,Pdcd4,Pde11a,Pde4a,Per1,Prickle1,Prkca,Prkch,Prkcq,Prl5a1,Rara,Rgs10,Rnf125,Selt,Serpine2,Sgms1,Sh2b1,Shisa2,Sipa1l1,Sirt4,Ski,Sla,Slc39a10,Slc7a3,Sod1,Spag9,Tbl1xr1,Tmem64,Tnfaip3,Tnfrsf1a,Tnfsf11,Tnfsf8,Trib1,Trim25,Ubr2,Wwox,Xbp1,Xdh |
| GO:0045785 | positive regulation of cell adhesion | 21 | 383 | 0.00011 | Bcl10,Ccdc80,Ccr7,Cd28,Cd47,Dpp4,Dusp10,Ets1,H2-Aa,Il6st,Itga4,Lef1,Nfkbiz,Prkca,Prkcq,Rara,Stat5b,Tgfbr2,Tnfsf11,Xbp1,Zfp609 |
| GO:0050865 | regulation of cell activation | 25 | 520 | 0.00011 | Bcl10,Btla,Cblb,Ccr7,Cd28,Cd300a,Cd47,Dpp4,Dusp10,H2-Aa,Il15ra,Il6st,Nfkbiz,Prkca,Prkcq,Rara,Serpine2,Slc39a10,Stat5b,Tgfbr2,Tigit,Tnfaip3,Tnfsf11,Xbp1,Zfp609 |
| GO:0030154 | cell differentiation | 90 | 3462 | 0.00012 | Abcb1b,Abhd2,Adipoq,Aff4,Ago2,Akt2,Btg2,Camk4,Ccr1,Ccr4,Ccr7,Cd28,Cdkn2b,Chd2,Chd7,Chst11,Chsy1,Clic4,Cryaa,Cyfip2,Ddx6,Dock10,Dusp10,Elf1,Ets1,Ets2,Ext1,F2,Fasl,Fli1,Fosl2,Foxn3,Foxp1,Gadd45g,Gnat1,Gpr18,Hectd1,Hey1,Il15ra,Il6st,Ist1,Itga4,Itk,Jak1,Jarid2,Jun,Klf2,Lef1,Map3k1,Mapk14,Mapk9,Mbnl1,Mfsd2a,Mospd1,Nfatc1,Nipbl,P2ry1,Pdcd4,Pip4k2a,Ppp1cc,Prkca,Prkch,Prrc2b,Rara,S1pr1,Serpine2,Sipa1l1,Ski,Sla,Smarca2,Smarce1,Sod1,Spag9,Stat5b,Tapt1,Tbl1xr1,Tgfbr2,Tmc1,Tmem64,Tnfaip3,Tnfsf11,Tnfsf8,Vsx2,Wasf2,Wwox,Xbp1,Xylt1,Zfp609,Zfp932,Zscan10 |
| GO:0051239 | regulation of multicellular organismal process | 78 | 2858 | 0.00012 | Adipoq,Ago2,Agtr1b,Atp1a1,Atp2b4,Bcl10,Btg1,Cblb,Ccr1,Ccr7,Cd101,Cd28,Cd97,Cdkn2b,Chd7,Chsy1,Ddx6,Dusp10,E2f2,Elf1,Ets1,F2,Fasl,Foxp1,Gpr18,Gtf2ird1,H2-Aa,Hey1,Il15ra,Il6st,Inpp4b,Ist1,Jak1,Jarid2,Jun,Kdm2b,Klf2,Lef1,Lrrn3,Lrtm2,Mapk14,Mapk9,Nfatc1,Nfkbiz,Nipbl,P2rx4,P2ry1,Pdcd4,Per1,Pim1,Ppp1cc,Prickle1,Prkca,Prkch,Prkcq,Rara,Rnf125,S1pr1,Serpine2,Sipa1l1,Ski,Sod1,Spag9,Stat5b,Tapt1,Tgfbr2,Tigit,Tmem64,Tnfaip3,Tnfrsf1a,Tnfrsf1b,Tnfsf11,Trib1,Xbp1,Xdh,Xylt1,Zfp609,Zfp932 |
| GO:0006464 | cellular protein modification process | 70 | 2471 | 0.00013 | Akt2,Bach2,Btg1,Btg2,Camk4,Cblb,Ccne1,Ccr7,Crbn,Dad1,Dcaf7,Dnajc6,Dusp1,Dusp10,Dusp2,Dyrk2,Ext1,Fbxl12,Fbxo33,Fkbp5,Fntb,Ggta1,Gucy2c,Hectd1,Itk,Jak1,Kcnh8,Kdm2b,Lef1,Lmtk3,Lonrf1,Map3k1,Map3k8,Map4k1,Mapk14,Mapk9,Mbip,Mknk2,Neurl1b,Nsmce1,Os9,Osm,Otulin,Per1,Pim1,Ppp1cc,Ppp2r5a,Prkca,Prkch,Prkcq,Ptp4a2,Rara,Riok1,Rnf125,Sirt4,Sla,Spag9,St8sia1,Stat4,Stk10,Tbl1xr1,Tgfbr2,Tnfaip3,Tnfsf11,Trib1,Trim25,Ubr2,Usp3,Wdyhv1,Zdhhc8 |
| GO:0048585 | negative regulation of response to stimulus | 49 | 1496 | 0.00013 | Adipoq,Arhgap25,Atp2b4,Cblb,Ccr1,Cd300a,Chst11,Dusp1,Dusp10,Dusp2,Dyrk2,Elf1,Ercc4,Ets1,F2,Foxp1,Gatsl3,Glrx,Gpr18,Hey1,Il6st,Lef1,Lrtm2,Mapk14,Os9,Otulin,Pdcd4,Pde11a,Per1,Prickle1,Prkca,Prkcq,Rgs10,Rnf125,Serpine2,Sgms1,Shisa2,Ski,Tmem64,Tnfaip3,Tnfrsf1a,Tnfrsf1b,Trib1,Ubac2,Ubr2,Wwox,Xbp1,Xdh,Xylt1 |
| GO:0050793 | regulation of developmental process | 70 | 2475 | 0.00013 | Adipoq,Ago2,Agtr1b,Arhgap15,Btg1,Cblb,Ccne1,Ccr1,Ccr7,Cd101,Cd28,Cd97,Cdkn2b,Chd7,Ddx6,Dusp10,E2f2,Epb4.1,Ets1,F2,Fasl,Foxp1,H2-Aa,Hey1,Il15ra,Il6st,Inpp4b,Ist1,Jak1,Jarid2,Jun,Kdm2b,Klf2,Lef1,Lrrn3,Lrtm2,Mapk14,Mapk9,Nfatc1,Nfkbiz,Nipbl,P2ry1,Pdcd4,Pim1,Ppp1cc,Prickle1,Prkca,Prkch,Rara,S1pr1,Serpine2,Sipa1l1,Ski,Sod1,Spag9,Stat5b,Syne3,Tapt1,Tgfbr2,Tmem64,Tnfaip3,Tnfrsf1a,Tnfrsf1b,Tnfsf11,Trib1,Xbp1,Xdh,Xylt1,Zfp609,Zfp932 |
| GO:0032270 | positive regulation of cellular protein metabolic process | 48 | 1462 | 0.00014 | Adipoq,Akt2,Atp2b4,Bcl10,Ccr1,Ccr7,Cd300a,Cryaa,Cyfip2,F2,Fam20a,Fanci,Fasl,Gadd45g,Il6st,Ist1,Jarid2,Jun,Lonrf1,Lrrn3,Map3k1,Map3k8,Map4k1,Mapk14,Mapk9,Ncf1,Nipbl,Osm,P2ry1,Pim1,Ppp2r5a,Prickle1,Prkca,Rara,Rnf125,Slc39a10,Sod1,Spag9,Stk10,Tgfbr2,Tnfaip3,Tnfrsf1a,Tnfrsf1b,Tnfsf11,Trib1,Xbp1,Xdh,Zfand2a |
| GO:0034097 | response to cytokine | 32 | 792 | 0.00014 | Adipoq,Ccr1,Ccr4,Ccr7,Ets1,Fntb,Foxp1,H2-Aa,Il21r,Il6st,Jak1,Jarid2,Jun,Klf2,Klf6,Lef1,Lrtm2,Mapk14,Osm,Prkca,Rara,Rnf125,Sgms1,Stat4,Stat5b,Tnfrsf1a,Tnfrsf1b,Tnfsf11,Trim25,Vamp4,Xbp1,Xcr1 |
| GO:0006955 | immune response | 35 | 914 | 0.00015 | Bcl10,Btla,Camk4,Cblb,Ccr1,Ccr4,Ccr7,Cd247,Cd28,Cd300a,Cd7,Dock10,F2,Fasl,Foxp1,Fyb,Gadd45g,Gzmb,H2-Aa,Hist1h2be,Itk,Jak1,Lef1,Osm,Otulin,Rftn1,Rnf125,Sla,Tnfaip3,Tnfrsf1a,Tnfrsf1b,Tnfsf11,Tnfsf8,Trim25,Vamp4 |
| GO:0048856 | anatomical structure development | 117 | 4912 | 0.00015 | Abcb1b,Abhd2,Adipoq,Aff4,Ago2,Agtr1b,Akt2,Ankrd11,Atp2b4,Bcl10,Btg2,Camk4,Ccr1,Ccr4,Ccr7,Cd28,Cdkn2b,Chd2,Chd7,Chst11,Chsy1,Clic4,Cryaa,Cyfip2,Dach2,Dad1,Dcaf7,Ddx6,Dock10,Drc1,Dusp1,Dusp10,Dusp2,Esrp1,Ets1,Ets2,Ext1,F2,Fam20a,Fasl,Fli1,Fosl2,Foxn3,Foxp1,Gadd45g,Gnat1,Gpr18,Gtf2ird1,Hectd1,Hey1,Hivep2,Igf2r,Il15ra,Il6st,Ist1,Itga4,Itk,Jarid2,Jun,Kdm2b,Klf2,Lef1,Map3k1,Mapk14,Mapk9,Mbd2,Mbnl1,Mctp2,Mfsd2a,Mknk2,Nfatc1,Ninj2,Nipbl,Nrip1,Nrxn3,Osm,Otulin,P2ry1,Pdcd4,Phc3,Pip4k2a,Polg2,Ppp1cc,Prickle1,Prkca,Prkch,Rara,Rtkn2,S1pr1,Selt,Serpine2,Shisa2,Sipa1l1,Ski,Smarca2,Smarce1,Sod1,Spag9,Ss18,Stat5b,Tapt1,Tbl1xr1,Tgfbr2,Tmc1,Tmeff2,Tnfaip3,Tnfrsf1a,Tnfrsf1b,Tnfsf11,Tnfsf8,Vsx2,Wasf2,Wwox,Xbp1,Xdh,Xylt1,Zfp609 |
| GO:0080134 | regulation of response to stress | 41 | 1167 | 0.00016 | Adipoq,Atp2b4,Ccr1,Ccr7,Cd28,Cd300a,Cd47,Dusp10,Ercc4,Ets1,F2,Foxp1,Gadd45g,Glrx,Map3k1,Map4k1,Mapk14,Mapk9,Ncf1,Nsmce1,Os9,Otulin,Pdcd4,Per1,Prkca,Prkcq,Rftn1,Rnf125,Serpine2,Sod1,Spag9,Stat5b,Tgfbr2,Tnfaip3,Tnfrsf1a,Tnfrsf1b,Tnfsf11,Ubac2,Xbp1,Xdh,Xylt1 |
| GO:0072507 | divalent inorganic cation homeostasis | 22 | 435 | 0.00018 | Agtr1b,Atp2b4,Ccr1,F2,Fam20a,Fasl,Gpr18,Il6st,Inpp4b,P2rx4,P2ry1,Prkca,S1pr1,Selt,Slc30a4,Slc35g1,Slc39a10,Stim2,Thada,Tmem64,Tnfsf11,Xcr1 |
| GO:0009891 | positive regulation of biosynthetic process | 56 | 1847 | 0.0002 | Ago2,Akt2,Bcl10,Camk4,Ccr7,Cd28,Chd7,Crem,Dyrk2,E2f2,Elf1,Elovl5,Ets1,Ets2,F2,Fli1,Fntb,Fosl2,Foxn3,Foxp1,Hey1,Hivep2,Jun,Klf2,Klf6,Lef1,Mapk14,Mapk9,Max,Mospd1,Ncf1,Nfatc1,Nipbl,Nrip1,Osm,P2ry1,Per1,Pim1,Prkcq,Rara,S1pr1,Sirt4,Ski,Smarca2,Ss18,Stat4,Stat5b,Tbl1xr1,Tnfrsf1a,Tnfsf11,Tnfsf8,Vsx2,Wwox,Xbp1,Zfp326,Zfp609 |
| GO:0002521 | leukocyte differentiation | 19 | 340 | 0.00021 | Camk4,Ccr7,Cd28,Chd7,Dock10,Foxp1,Gadd45g,Gpr18,Hectd1,Itk,Jun,Lef1,Mapk14,Nfatc1,Stat5b,Tgfbr2,Tnfaip3,Tnfsf11,Tnfsf8 |
| GO:0010557 | positive regulation of macromolecule biosynthetic process | 53 | 1717 | 0.00022 | Ago2,Akt2,Bcl10,Camk4,Ccr7,Cd28,Chd7,Crem,Dyrk2,E2f2,Elf1,Ets1,Ets2,Fli1,Fntb,Fosl2,Foxn3,Foxp1,Hey1,Hivep2,Jun,Klf2,Klf6,Lef1,Mapk14,Mapk9,Max,Mospd1,Ncf1,Nfatc1,Nipbl,Nrip1,Osm,P2ry1,Per1,Pim1,Prkcq,Rara,S1pr1,Ski,Smarca2,Ss18,Stat4,Stat5b,Tbl1xr1,Tnfrsf1a,Tnfsf11,Tnfsf8,Vsx2,Wwox,Xbp1,Zfp326,Zfp609 |
| GO:0035556 | intracellular signal transduction | 43 | 1274 | 0.00023 | Akt2,Bcl10,Camk4,Cblb,Ccr7,Dock10,Dusp1,Dyrk2,E2f2,Gdi2,Gucy2c,Il15ra,Itk,Jak1,Jun,Kcnh8,Map3k1,Map3k8,Map4k1,Mapk14,Mapk9,Mknk2,Nfatc1,Nsmce1,Osm,Otulin,P2rx4,Prkca,Prkch,Prkcq,Rasgrp2,Rras2,Spag9,Ss18,Stat5b,Stk10,Tnfrsf1a,Tnfrsf1b,Tnfsf11,Trib1,Wasf2,Wwox,Xbp1 |
| GO:0043085 | positive regulation of catalytic activity | 42 | 1239 | 0.00027 | Adipoq,Akt2,Arhgap15,Arhgap25,Atp2b4,Bcl10,Btg1,Ccr7,Cd300a,Cyfip2,Dock10,F2,Fam20a,Fasl,Gadd45g,Gdi2,Gnat1,Itk,Jun,Map3k1,Map3k8,Map4k1,Mapk14,Mapk9,Ncf1,Pim1,Pkp4,Prkcq,Rasgrp2,Rgs10,S1pr1,Sipa1l1,Slc39a10,Snx18,Sod1,Spag9,Stk10,Tgfbr2,Tnfsf11,Trib1,Usp6nl,Xdh |
| GO:0048584 | positive regulation of response to stimulus | 57 | 1922 | 0.00029 | Adipoq,Akt2,Bcl10,Ccr1,Ccr4,Ccr7,Cd247,Cd28,Cd47,Cdkn2b,Chsy1,Cyfip2,Ets1,F2,Fasl,Foxp1,Gadd45g,Glrx,Gpr18,Il6st,Itk,Jak1,Jun,Map3k1,Map3k8,Map4k1,Mapk14,Mapk9,Mbd2,Ncf1,Nfkbiz,Nsmce1,Osm,Otulin,P2rx4,P2ry1,Pdcd4,Prkca,Prkch,Prkcq,Rara,Rftn1,S1pr1,Sh2b1,Sla,Slc39a10,Sod1,Spag9,Stat5b,Tbl1xr1,Tgfbr2,Tnfrsf1a,Tnfsf11,Trim25,Wwox,Xbp1,Xdh |

| | | | | | |
|---|---|---|---|---|---|
| GO:0008284 | positive regulation of cell population proliferation | 34 | 913 | 0.00033 | Akt2,Ccr7,Cd28,Cd47,Dpp4,Esrp1,Ets1,F2,Fasl,Fntb,Fosl2,Foxp1,Il6st,Itga4,Jun,Lef1,Mapk14,Osm,Pim1,Ppp1cc,Prkca,Prkch,Prkcq,Rara,Rtkn2,S1pr1,Slc39a10,St8sia1,Stat5b,Tgfbr2,Tnfaip3,Txnl4b,Vsx2,Xbp1 |
| GO:0030099 | myeloid cell differentiation | 15 | 229 | 0.00034 | Camk4,Ccr1,Ccr7,Cdkn2b,Fli1,Foxp1,Jun,Klf2,Lef1,Mapk14,Nfatc1,Pip4k2a,Tgfbr2,Tnfsf11,Wasf2 |
| GO:0014070 | response to organic cyclic compound | 33 | 883 | 0.00041 | Abhd2,Adipoq,Ago2,Agtr1b,Akt2,Atp1a1,Atp2b4,Btg2,Crem,Cryaa,Dusp1,Ets1,Fntb,Fosl2,Foxp1,Glrx,Jun,Klf2,Lef1,Mbd2,Nrip1,P2rx4,P2ry1,Per1,Pim1,Prkca,Rara,Rftn1,Slc3a2,Sod1,Stat5b,Tgfbr2,Trim25 |
| GO:0040012 | regulation of locomotion | 34 | 924 | 0.00041 | Abhd2,Adipoq,Ago2,Akt2,Ccr1,Ccr4,Ccr7,Cd300a,Clic4,Dock10,Dusp10,Ets1,Foxp1,Gpr18,Il6st,Itga4,Jun,Lef1,Map3k1,Mapk14,Nipbl,P2rx4,Prkca,Rras2,S1pr1,Spag9,Stk10,Tgfbr2,Tmeff2,Trib1,Trim25,Vps37b,Xbp1,Zfp609 |
| GO:0008285 | negative regulation of cell population proliferation | 27 | 648 | 0.00043 | Adipoq,Btg1,Btg2,Btla,Cblb,Cd300a,Cdkn2b,Dusp1,Dusp10,Fntb,H2-Aa,Jarid2,Jun,Kdm2b,Nfatc1,Osm,Pdcd4,Prkca,Rara,Serpine2,Ski,Smarca2,Tgfbr2,Tnfaip3,Trib1,Vsx2,Xdh |
| GO:0030334 | regulation of cell migration | 31 | 805 | 0.00043 | Abhd2,Adipoq,Ago2,Akt2,Ccr1,Ccr7,Cd300a,Clic4,Dock10,Dusp10,Ets1,Foxp1,Gpr18,Il6st,Itga4,Jun,Lef1,Map3k1,Mapk14,Nipbl,P2rx4,Prkca,Rras2,S1pr1,Spag9,Stk10,Tgfbr2,Tmeff2,Trib1,Xbp1,Zfp609 |
| GO:0071310 | cellular response to organic substance | 55 | 1858 | 0.00043 | Abhd2,Adipoq,Ago2,Agtr1b,Akt2,Atp1a1,Atp2b4,Ccr1,Ccr4,Ccr7,Dusp1,Foxp1,Gatsl3,Glrx,Il21r,Il6st,Jak1,Jarid2,Jun,Klf2,Klf3,Klf6,Lef1,Lrtm2,Map3k1,Map4k1,Mapk14,Mapk9,Mbd2,Nrip1,Osm,P2rx4,P2ry1,Pdcd4,Pim1,Prkca,Rara,Rgs10,Rnf125,Selt,Sgms1,Sod1,Stat4,Stat5b,Tgfbr2,Tnfaip3,Tnfrsf1a,Tnfrsf1b,Tnfsf11,Trim25,Ubr2,Vamp4,Wwox,Xbp1,Xcr1 |
| GO:0009892 | negative regulation of metabolic process | 70 | 2584 | 0.00044 | Adipoq,Aebp2,Ago2,Akt2,Atp1a1,Atp2b4,Bach2,Btg2,Cblb,Ccne1,Ccr1,Cd28,Cd300a,Cdkn2b,Cnst,Crem,Crim1,Cryaa,Dusp1,Dusp10,Dusp2,Ercc4,Ets2,F2,Fasl,Foxn3,Foxp1,Gadd45g,Hey1,Jarid2,Jun,Kdm2b,Klf2,Klf3,Lef1,Lmtk3,Lrtm2,Max,Mbd2,Mospd1,Nfatc1,Nipbl,Nrip1,Os9,Pdcd4,Per1,Ppp2r5a,Prickle1,Prkca,Rara,Serpine2,Sirt4,Ski,Smarca2,Smarce1,Sod1,Tbl1xr1,Tmeff2,Tnfaip3,Tnfrsf1a,Tnfrsf1b,Tnrc6b,Trib1,Ubac2,Ubr2,Usp3,Xbp1,Xdh,Zfp932,Zscan10 |
| GO:0048731 | system development | 99 | 4072 | 0.00044 | Abcb1b,Adipoq,Agtr1b,Akt2,Ankrd11,Atp2b4,Bcl10,Btg2,Camk4,Ccr1,Ccr4,Ccr7,Cd28,Cdkn2b,Chd2,Chd7,Chst11,Chsy1,Clic4,Cryaa,Cyfip2,Ddx6,Dock10,Drc1,Dusp10,Esrp1,Ets1,Ext1,F2,Fam20a,Fasl,Fli1,Fosl2,Foxn3,Foxp1,Gadd45g,Gnat1,Gpr18,Hectd1,Hey1,Igf2r,Il15ra,Il6st,Ist1,Itga4,Itk,Jarid2,Jun,Kdm2b,Klf2,Lef1,Map3k1,Mapk14,Mapk9,Mbd2,Mbnl1,Mfsd2a,Mknk2,Nfatc1,Nipbl,Nrip1,Nrxn3,Osm,Otulin,P2ry1,Pdcd4,Pip4k2a,Ppp1cc,Prickle1,Prkca,Prkch,Rara,Rtkn2,S1pr1,Selt,Serpine2,Sipa1l1,Ski,Smarca2,Smarce1,Sod1,Spag9,Stat5b,Tapt1,Tbl1xr1,Tgfbr2,Tmc1,Tnfaip3,Tnfrsf1a,Tnfrsf1b,Tnfsf11,Tnfsf8,Vsx2,Wasf2,Wwox,Xbp1,Xdh,Xylt1,Zfp609 |
| GO:0002763 | positive regulation of myeloid leukocyte differentiation | 8 | 61 | 0.00045 | Ccr1,Cd101,Jun,Lef1,Prkca,Tmem64,Tnfsf11,Trib1 |
| GO:0006915 | apoptotic process | 31 | 809 | 0.00045 | Akt2,Bcl10,Cd28,Cryaa,Cyfip2,Dad1,Dyrk2,E2f2,Fasl,Gadd45g,Gzmb,Jun,Lef1,Map3k1,Mapk14,Mapk9,Mknk2,Ncf1,P2rx4,Pdcd4,Pim1,Prkca,Sgms1,Smndc1,Tax1bp1,Tgfbr2,Tnfaip3,Tnfrsf1a,Tnfrsf1b,Wwox,Xbp1 |
| GO:0031401 | positive regulation of protein modification process | 39 | 1142 | 0.00045 | Adipoq,Akt2,Atp2b4,Bcl10,Ccr1,Ccr7,Cd300a,Cryaa,F2,Fam20a,Fanci,Gadd45g,Il6st,Jarid2,Jun,Lrrn3,Map3k1,Map3k8,Map4k1,Mapk9,Ncf1,Nipbl,Osm,P2ry1,Pim1,Ppp2r5a,Prickle1,Prkca,Rara,Slc39a10,Sod1,Spag9,Stk10,Tgfbr2,Tnfrsf1a,Tnfsf11,Trib1,Xbp1,Xdh |
| GO:0043412 | macromolecule modification | 71 | 2636 | 0.00045 | Akt2,Bach2,Btg1,Btg2,Camk4,Cblb,Ccne1,Ccr7,Crbn,Dad1,Dcaf7,Dnajc6,Dusp1,Dusp10,Dusp2,Dyrk2,Ext1,Fbxl12,Fbxo33,Fkbp5,Fntb,Ggta1,Gucy2c,Hectd1,Itk,Jak1,Kcnh8,Kdm2b,Lef1,Lmtk3,Lonrf1,Map3k1,Map3k8,Map4k1,Mapk14,Mapk9,Mbd2,Mbip,Mknk2,Neurl1b,Nsmce1,Os9,Osm,Otulin,Per1,Pim1,Ppp1cc,Ppp2r5a,Prkca,Prkch,Prkcq,Ptp4a2,Rara,Riok1,Rnf125,Sirt4,Sla,Spag9,St8sia1,Stat4,Stk10,Tbl1xr1,Tgfbr2,Tnfaip3,Tnfsf11,Trib1,Trim25,Ubr2,Usp3,Wdyhv1,Zdhhc8 |
| GO:0045621 | positive regulation of lymphocyte differentiation | 10 | 103 | 0.00045 | Ccr7,Dusp10,H2-Aa,Il15ra,Nfkbiz,Rara,Stat5b,Tgfbr2,Xbp1,Zfp609 |
| GO:0010614 | negative regulation of cardiac muscle hypertrophy | 6 | 28 | 0.00046 | Atp2b4,Foxp1,Jarid2,P2rx4,Tnfrsf1a,Tnfrsf1b |
| GO:0032501 | multicellular organismal process | 132 | 5888 | 0.00046 | Abcb1b,Abhd2,Adipoq,Aff4,Ago2,Agtr1b,Akt2,Ankrd11,Atp1a1,Atp2b4,Bcl10,Btg1,Btg2,Camk4,Ccr1,Ccr4,Ccr7,Cd247,Cd28,Cd300a,Cd7,Cdkn2b,Chd2,Chd7,Chst11,Chsy1,Clic4,Crem,Cryaa,Cyfip2,Dach2,Dad1,Dcaf7,Ddx6,Dock10,Dpp4,Drc1,Dusp1,Dusp10,Dusp2,Esrp1,Ets1,Ets2,Ext1,F2,Fam107b,Fam20a,Fasl,Fli1,Fosl2,Foxn3,Foxp1,Gadd45g,Gnat1,Gpr18,Grxcr2,Gtf2ird1,Hectd1,Hey1,Hivep2,Igf2r,Il15ra,Il6st,Ist1,Itga4,Itk,Jarid2,Jun,Kdm2b,Klf2,Lef1,Map3k1,Mapk14,Mapk9,Mbd2,Mbnl1,Mctp2,Mfsd2a,Mknk2,Nfatc1,Nipbl,Nrip1,Nrxn3,Osm,Otulin,P2rx4,P2ry1,Pdcd4,Pde4a,Per1,Phc3,Pip4k2a,Polg2,Ppp1cc,Prickle1,Prkca,Prkch,Rara,Rftn1,Rtkn2,S1pr1,Selt,Serpine2,Shisa2,Sipa1l1,Ski,Slc38a1,Smarca2,Smarce1,Sod1,Spag9,Ss18,Stat5b,Tapt1,Tbl1xr1,Tdrp,Tgfbr2,Tmc1,Tmem64,Tnfaip3,Tnfrsf1a,Tnfrsf1b,Tnfsf11,Tnfsf8,Ubr2,Vsx2,Wasf2,Wwox,Xbp1,Xdh,Xylt1,Zfp609 |
| GO:0007166 | cell surface receptor signaling pathway | 51 | 1683 | 0.00047 | Akt2,Bcl10,Btla,Cblb,Ccne1,Ccr1,Ccr4,Ccr7,Cd101,Cd247,Cd28,Cd97,Dyrk2,F2,Fasl,Hey1,Il15ra,Il21r,Il6st,Itga4,Itk,Jak1,Jun,Klf6,Lef1,Lrtm2,Map3k1,Mapk14,Mknk2,Neurl1b,Nfkbiz,Osm,Otulin,P2rx4,P2ry1,Pdcd4,Rftn1,Sipa1l1,Ski,Sla,Ss18,Stat4,Stat5b,Tgfbr2,Tmem64,Tnfrsf1a,Tnfrsf1b,Tnfsf11,Wwox,Xbp1,Xcr1 |
| GO:0050727 | regulation of inflammatory response | 17 | 301 | 0.00048 | Adipoq,Ccr7,Cd28,Cd47,Dusp10,Ets1,F2,Mapk14,Otulin,Pdcd4,Per1,Prkca,Stat5b,Tnfaip3,Tnfrsf1a,Tnfrsf1b,Tnfsf11 |
| GO:0051241 | negative regulation of multicellular organismal process | 40 | 1191 | 0.00048 | Adipoq,Atp1a1,Atp2b4,Ccr1,Cdkn2b,Chsy1,Ddx6,Dusp10,E2f2,F2,Fasl,Foxp1,Gpr18,Hey1,Il15ra,Inpp4b,Jarid2,Klf2,Lef1,Nfatc1,P2rx4,Pdcd4,Prickle1,Prkca,Rara,Rnf125,Serpine2,Ski,Sod1,Stat5b,Tgfbr2,Tigit,Tmem64,Tnfaip3,Tnfrsf1a,Tnfrsf1b,Trib1,Xdh,Xylt1,Zfp932 |
| GO:0071900 | regulation of protein serine/threonine kinase activity | 21 | 435 | 0.00048 | Adipoq,Atp2b4,Ccr7,Cd300a,Cdkn2b,Dusp1,Dusp10,Dusp2,Fam20a,Gadd45g,Map3k1,Map3k8,Map4k1,Pdcd4,Pim1,Prkca,Sod1,Spag9,Tnfaip3,Tnfsf11,Trib1 |
| GO:0031324 | negative regulation of cellular metabolic process | 65 | 2357 | 0.00051 | Adipoq,Aebp2,Ago2,Akt2,Atp1a1,Atp2b4,Bach2,Btg2,Cblb,Ccne1,Cd300a,Cdkn2b,Cnst,Crem,Crim1,Cryaa,Dusp1,Dusp10,Dusp2,Ercc4,Ets2,F2,Fasl,Foxn3,Foxp1,Gadd45g,Hey1,Jarid2,Jun,Kdm2b,Klf2,Klf3,Lef1,Lmtk3,Lrtm2,Max,Mbd2,Mospd1,Nfatc1,Nipbl,Nrip1,Os9,Pdcd4,Per1,Ppp2r5a,Prickle1,Prkca,Rara,Serpine2,Sirt4,Ski,Smarca2,Smarce1,Tbl1xr1,Tnfaip3,Tnfrsf1b,Tnrc6b,Trib1,Ubac2,Ubr2,Usp3,Xbp1,Xdh,Zfp932,Zscan10 |
| GO:0043549 | regulation of kinase activity | 30 | 780 | 0.00054 | Adipoq,Atp2b4,Cblb,Ccne1,Ccr7,Cd300a,Cdkn2b,Dusp1,Dusp10,Dusp2,F2,Fam20a,Gadd45g,Lrtm2,Map3k1,Map3k8,Map4k1,Ncf1,Pdcd4,Pim1,Ppp2r5a,Prkca,Rara,Sod1,Spag9,Stk10,Tgfbr2,Tnfaip3,Tnfsf11,Trib1 |
| GO:0055074 | calcium ion homeostasis | 20 | 405 | 0.00054 | Agtr1b,Atp2b4,Ccr1,F2,Fam20a,Fasl,Gpr18,Il6st,Inpp4b,P2rx4,P2ry1,Prkca,S1pr1,Selt,Slc35g1,Stim2,Thada,Tmem64,Tnfsf11,Xcr1 |
| GO:0010629 | negative regulation of gene expression | 47 | 1515 | 0.00057 | Adipoq,Aebp2,Ago2,Akt2,Bach2,Btg2,Ccne1,Ccr1,Cd28,Crem,Cryaa,Ets2,Fasl,Foxn3,Foxp1,Hey1,Jarid2,Jun,Kdm2b,Klf2,Klf3,Lef1,Max,Mbd2,Mospd1,Nfatc1,Nipbl,Nrip1,Pdcd4,Per1,Prickle1,Prkca,Rara,Serpine2,Sirt4,Ski,Smarca2,Smarce1,Tbl1xr1,Tnfrsf1a,Tnrc6b,Ubr2,Usp3,Xbp1,Xdh,Zfp932,Zscan10 |
| GO:0055065 | metal ion homeostasis | 24 | 555 | 0.00063 | Agtr1b,Atp1a1,Atp2b4,Ccr1,F2,Fam20a,Fasl,Gpr18,Il6st,Inpp4b,P2rx4,P2ry1,Prkca,S1pr1,Selt,Slc30a4,Slc35g1,Slc39a10,Sod1,Stim2,Thada,Tmem64,Tnfsf11,Xcr1 |
| GO:0006468 | protein phosphorylation | 30 | 793 | 0.0007 | Akt2,Camk4,Ccne1,Ccr7,Dyrk2,Gucy2c,Itk,Jak1,Kcnh8,Lmtk3,Map3k1,Map3k8,Map4k1,Mapk14,Mapk9,Mknk2,Osm,Pim1,Prkca,Prkch,Prkcq,Rara,Riok1,Sla,Spag9,Stat4,Stk10,Tgfbr2,Tnfsf11,Trib1 |
| GO:0031328 | positive regulation of cellular biosynthetic process | 53 | 1807 | 0.0007 | Ago2,Akt2,Bcl10,Camk4,Ccr7,Cd28,Chd7,Crem,Dyrk2,E2f2,Elf1,Elovl5,Ets1,Ets2,Fli1,Fosl2,Foxn3,Foxp1,Hey1,Hivep2,Jun,Klf2,Klf6,Lef1,Mapk14,Mapk9,Max,Mospd1,Ncf1,Nfatc1,Nipbl,Nrip1,Osm,P2ry1,Per1,Pim1,Prkcq,Rara,S1pr1,Ski,Smarca2,Ss18,Stat4,Stat5b,Tbl1xr1,Tnfrsf1a,Tnfsf11,Tnfsf8,Vsx2,Wwox,Xbp1,Zfp326,Zfp609 |
| GO:0008219 | cell death | 32 | 883 | 0.00082 | Akt2,Bcl10,Cd28,Cryaa,Cyfip2,Dad1,Dyrk2,E2f2,Fasl,Gadd45g,Gzmb,Jun,Lef1,Map3k1,Mapk14,Mapk9,Mknk2,Ncf1,P2rx4,Pdcd4,Pim1,Prkca,Sgms1,Smndc1,Tax1bp1,Tgfbr2,Tmem123,Tnfaip3,Tnfrsf1a,Tnfrsf1b,Wwox,Xbp1 |
| GO:0045582 | positive regulation of T cell differentiation | 9 | 90 | 0.00085 | Ccr7,Dusp10,H2-Aa,Nfkbiz,Rara,Stat5b,Tgfbr2,Xbp1,Zfp609 |
| GO:0051172 | negative regulation of nitrogen compound metabolic process | 61 | 2206 | 0.00085 | Adipoq,Aebp2,Ago2,Akt2,Atp2b4,Bach2,Btg2,Cblb,Ccne1,Cd300a,Cdkn2b,Crem,Crim1,Cryaa,Dusp1,Dusp10,Dusp2,Ercc4,Ets2,F2,Fasl,Foxn3,Foxp1,Gadd45g,Hey1,Jarid2,Jun,Kdm2b,Klf2,Klf3,Lef1,Lrtm2,Max,Mbd2,Mospd1,Nfatc1,Nipbl,Nrip1,Os9,Pdcd4,Per1,Prickle1,Prkca,Rara,Serpine2,Sirt4,Ski,Smarca2,Smarce1,Tbl1xr1,Tnfaip3,Tnfrsf1b,Tnrc6b,Trib1,Ubac2,Ubr2,Usp3,Xbp1,Xdh,Zfp932,Zscan10 |
| GO:0009890 | negative regulation of biosynthetic process | 44 | 1408 | 0.00086 | Adipoq,Aebp2,Ago2,Atp1a1,Atp2b4,Bach2,Btg2,Ccne1,Crem,Dusp1,Ets2,Fasl,Foxn3,Foxp1,Hey1,Jarid2,Jun,Kdm2b,Klf2,Klf3,Lef1,Max,Mbd2,Mospd1,Nfatc1,Nipbl,Nrip1,Pdcd4,Per1,Prickle1,Prkca,Rara,Ski,Smarca2,Smarce1,Sod1,Tbl1xr1,Tmeff2,Tnrc6b,Ubr2,Usp3,Xbp1,Zfp932,Zscan10 |
| GO:0051270 | regulation of cellular component movement | 33 | 928 | 0.00086 | Abhd2,Adipoq,Ago2,Akt2,Atp1a1,Ccr1,Ccr7,Cd300a,Clic4,Dock10,Dusp10,Ets1,Foxp1,Gpr18,Il6st,Itga4,Jun,Lef1,Map3k1,Mapk14,Nipbl,P2rx4,Prkca,Rras2,S1pr1,Spag9,Stat5b,Stk10,Tgfbr2,Tmeff2,Trib1,Xbp1,Zfp609 |
| GO:0009968 | negative regulation of signal transduction | 38 | 1147 | 0.00093 | Adipoq,Arhgap25,Atp2b4,Cblb,Cd300a,Chst11,Dusp1,Dusp10,Dusp2,Dyrk2,Elf1,Foxp1,Gatsl3,Glrx,Hey1,Il6st,Lef1,Lrtm2,Mapk14,Pdcd4,Pde11a,Per1,Prickle1,Prkca,Prkcq,Rgs10,Rnf125,Serpine2,Sgms1,Shisa2,Ski,Tmem64,Tnfaip3,Trib1,Ubr2,Wwox,Xbp1,Xdh |
| GO:0009628 | response to abiotic stimulus | 36 | 1063 | 0.00097 | Adipoq,Agtr1b,Akt2,Atp1a1,Atp2b4,Bcl10,Btg2,Ccr4,Cryaa,Cyp2r1,Dpp4,Dusp1,Ercc4,Ets1,Fosl2,Gnat1,Ints7,Jun,Mapk14,Mapk9,Mbd2,Nabp1,Nipbl,Osm,P2ry1,Per1,Ppp1cc,Serpine2,Sirt4,Sod1,St8sia1,Tgfbr2,Tmc1,Tnfrsf1a,Xbp1,Xylt1 |
| GO:0010605 | negative regulation of macromolecule metabolic process | 64 | 2373 | 0.001 | Adipoq,Aebp2,Ago2,Akt2,Bach2,Btg2,Cblb,Ccne1,Ccr1,Cd28,Cd300a,Cdkn2b,Crem,Crim1,Cryaa,Dusp1,Dusp10,Dusp2,Ercc4,Ets2,F2,Fasl,Foxn3,Foxp1,Gadd45g,Hey1,Jarid2,Jun,Kdm2b,Klf2,Klf3,Lef1,Lrtm2,Max,Mbd2,Mospd1,Nfatc1,Nipbl,Nrip1,Os9,Pdcd4,Per1,Prickle1,Prkca,Rara,Serpine2,Sirt4,Ski,Smarca2,Smarce1,Tbl1xr1,Tmeff2,Tnfaip3,Tnfrsf1a,Tnfrsf1b,Tnrc6b,Trib1,Ubac2,Ubr2,Usp3,Xbp1,Xdh,Zfp932,Zscan10 |
| GO:0031098 | stress-activated protein kinase signaling cascade | 10 | 117 | 0.001 | Ccr7,Map3k1,Map3k8,Map4k1,Mapk14,Mapk9,Spag9,Stk10,Tnfsf11,Trib1 |
| GO:0045321 | leukocyte activation | 21 | 464 | 0.001 | Bcl10,Camk4,Cblb,Ccr7,Cd28,Cd300a,Chd7,Dock10,Dpp4,Foxp1,Gadd45g,Gpr18,Hectd1,Itk,Jun,Lef1,Nfatc1,Stat5b,Tgfbr2,Tnfaip3,Tnfsf8 |
| GO:0050854 | regulation of antigen receptor-mediated signaling pathway | 7 | 52 | 0.0011 | Bcl10,Cblb,Ccr7,Cd300a,Elf1,Prkch,Slc39a10 |
| GO:0010628 | positive regulation of gene expression | 52 | 1811 | 0.0013 | Ago2,Akt2,Bcl10,Camk4,Cd28,Chd7,Crem,E2f2,Elf1,Ets1,Ets2,Fli1,Fosl2,Foxn3,Foxp1,Hey1,Hivep2,Jun,Klf2,Klf6,Lef1,Mapk14,Mapk9,Max,Mospd1,Ncf1,Nfatc1,Nipbl,Nrip1,Osm,P2ry1,Per1,Pim1,Rara,Riok1,Rnf219,S1pr1,Sgms1,Ski,Smarca2,Ss18,Stat4,Stat5b,Tbl1xr1,Tnfrsf1a,Tnfsf11,Tnfsf8,Vsx2,Wwox,Xbp1,Zfp326,Zfp609 |
| GO:0044249 | cellular biosynthetic process | 87 | 3571 | 0.0013 | Aebp2,Aff4,Ago2,Akt2,Bach2,Bcat1,Btg2,Ccne1,Cers6,Chd2,Chd7,Chst11,Chsy1,Cmas,Crem,Cyp2r1,Dach2,Dad1,E2f2,Eif4e3,Elf1,Elovl5,Ets1,Ets2,Ext1,Fasl,Fli1,Fosl2,Foxn3,Foxp1,Ggta1,Gtf2ird1,Gucy2c,Hbs1l,Hey1,Hivep2,Inpp4b,Jarid2,Jun,Kdm2b,Klf2,Klf3,Klf6,Lef1,Lpcat3,Mapk14,Max,Mbd2,Mbip,Mgme1,Mogat1,Moxd1,Ncf1,Nfatc1,Nfkbiz,Nipbl,Nrip1,P2rx4,Pde11a,Pde4a,Per1,Polg2,Rara,Scd2,Scml4,Sgms1,Ski,Smarca2,Sod1,Spty2d1,Ss18,St8sia1,Stat4,Stat5b,Tars2,Tbl1xr1,Vgll4,Vsx2,Xbp1,Xylt1,Zdhhc8,Zfp3,Zfp326,Zfp609,Zfp652,Zfp932,Zscan10 |
| GO:0044267 | cellular protein metabolic process | 76 | 2995 | 0.0013 | Akt2,Bach2,Btg1,Btg2,Camk4,Cblb,Ccne1,Ccr7,Crbn,Dad1,Dcaf7,Dnajc6,Dusp1,Dusp10,Dusp2,Dyrk2,Eif4e3,Ext1,Fbxl12,Fbxo33,Fkbp5,Fntb,Ggta1,Gucy2c,Hbs1l,Hectd1,Itk,Jak1,Kcnh8,Kdm2b,Lef1,Lmtk3,Lonrf1,Map3k1,Map3k8,Map4k1,Mapk14,Mapk9,Mbip,Mknk2,Msrb3,Neurl1b,Nsmce1,Os9,Osm,Otulin,Per1,Pim1,Polg2,Ppp1cc,Ppp2r5a,Prkca,Prkch,Prkcq,Ptp4a2,Rara,Riok1,Rnf125,Sirt4,Sla,Spag9,St8sia1,Stat4,Stk10,Tars2,Tbl1xr1,Tgfbr2,Tnfaip3,Tnfsf11,Trib1,Trim25,Ubr2,Usp3,Wdyhv1,Xbp1,Zdhhc8 |
| GO:0055080 | cation homeostasis | 25 | 626 | 0.0013 | Agtr1b,Atp1a1,Atp2b4,Ccr1,Clic4,F2,Fam20a,Fasl,Gpr18,Il6st,Inpp4b,P2rx4,P2ry1,Prkca,S1pr1,Selt,Slc30a4,Slc35g1,Slc39a10,Sod1,Stim2,Thada,Tmem64,Tnfsf11,Xcr1 |
| GO:1901701 | cellular response to oxygen-containing compound | 31 | 870 | 0.0013 | Adipoq,Agtr1b,Akt2,Atp2b4,Ets1,Foxp1,Gatsl3,Glrx,Glrx2,Jun,Klf2,Klf3,Map4k1,Mapk14,Mapk9,Ncf1,Nrip1,P2rx4,Pdcd4,Pim1,Prkca,Rara,Rgs10,Selt,Sgms1,Sod1,Stat5b,Tnfaip3,Tnfrsf1b,Ubr2,Xbp1 |
| GO:0001775 | cell activation | 23 | 552 | 0.0014 | Bcl10,Camk4,Cblb,Ccr7,Cd28,Cd300a,Chd7,Dock10,Dpp4,F2,Foxp1,Gadd45g,Gpr18,Hectd1,Itk,Jun,Lef1,Nfatc1,P2ry1,Stat5b,Tgfbr2,Tnfaip3,Tnfsf8 |
| GO:0019538 | protein metabolic process | 87 | 3580 | 0.0014 | Akt2,Bach2,Btg1,Btg2,Camk4,Cblb,Ccne1,Ccr7,Chst11,Chsy1,Crbn,Cyfip2,Dad1,Dcaf7,Dnajc6,Dpp4,Dusp1,Dusp10,Dusp2,Dyrk2,Eif4e3,Ext1,F2,Fbxl12,Fbxo33,Fkbp5,Fntb,Gadd45g,Ggta1,Gucy2c,Gzmb,Hbs1l,Hectd1,Itk,Jak1,Kcnh8,Kdm2b,Lef1,Lmtk3,Lonrf1,Map3k1,Map3k8,Map4k1,Mapk14,Mapk9,Mbip,Mknk2,Msrb3,Neurl1b,Nsmce1,Os9,Osm,Otulin,Per1,Pim1,Polg2,Ppp1cc,Ppp2r5a,Prkca,Prkch,Prkcq,Prss39,Ptp4a2,Rara,Riok1,Rnf125,Sirt4,Sla,Spag9,St8sia1,Stat4,Stk10,Tars2,Tbl1xr1,Tgfbr2,Tnfaip3,Tnfsf11,Trappc12,Trib1,Trim25,Ubr2,Usp3,Wdyhv1,Xbp1,Xpnpep2,Xylt1,Zdhhc8 |
| GO:0046634 | regulation of alpha-beta T cell activation | 9 | GO:0042102 | 0.0014 | Btla,Cblb,Ccr7,Cd28,Cd300a,Nfkbiz,Prkcq,Rara,Tgfbr2 |
| GO:0048513 | animal organ development | 76 | 3012 | 0.0014 | Adipoq,Agtr1b,Ankrd11,Atp2b4,Btg2,Camk4,Ccr1,Ccr7,Cd28,Cdkn2b,Chd2,Chd7,Chst11,Chsy1,Clic4,Cryaa,Dock10,Drc1,Esrp1,Ets1,Ext1,Fam20a,Fasl,Fli1,Fosl2,Foxn3,Foxp1,Gadd45g,Gnat1,Gpr18,Hectd1,Hey1,Igf2r,Itga4,Itk,Jarid2,Jun,Kdm2b,Klf2,Lef1,Map3k1,Mapk14,Mbd2,Mbnl1,Mfsd2a,Mknk2,Nfatc1,Nipbl,Nrip1,Pdcd4,Pip4k2a,Prickle1,Prkca,Rara,Rtkn2,S1pr1,Selt,Serpine2,Ski,Sod1,Stat5b,Tapt1,Tbl1xr1,Tgfbr2,Tmc1,Tnfaip3,Tnfrsf1a,Tnfrsf1b,Tnfsf11,Tnfsf8,Vsx2,Wasf2,Wwox,Xbp1,Xdh,Zfp609 |

| | | | | | |
|---|---|---|---|---|---|
| GO:0051338 | regulation of transferase activity | 31 | 873 | 0.0014 | Adipoq,Atp2b4,Cblb,Ccne1,Ccr7,Cd300a,Cdkn2b,Dusp1,Dusp10,Dusp2,F2,Fam20a,Gadd45g,Lrtm2,Map3k1,Map3k8,Map4k1,Ncf1,Pdcd4,Pim1,Ppp2r5a,Prkca,Prkcq,Rara,Sod1,Spag9,Stk10,Tgfbr2,Tnfaip3,Tnfsf11,Trib1 |
| GO:0001932 | regulation of protein phosphorylation | 42 | 1362 | 0.0015 | Adipoq,Akt2,Atp2b4,Cblb,Ccne1,Ccr1,Ccr7,Cd300a,Cdkn2b,Cryaa,Dusp1,Dusp10,Dusp2,F2,Fam20a,Gadd45g,Il6st,Jun,Lrrn3,Lrtm2,Map3k1,Map3k8,Map4k1,Mapk9,Ncf1,Osm,P2ry1,Pdcd4,Per1,Pim1,Prkca,Rara,Sod1,Spag9,Stk10,Tgfbr2,Tnfaip3,Tnfrsf1a,Tnfsf11,Trib1,Xbp1,Xdh |
| GO:0010648 | negative regulation of cell communication | 40 | 1269 | 0.0015 | Adipoq,Arhgap25,Atp2b4,Cblb,Cd300a,Chst11,Dusp1,Dusp10,Dusp2,Dyrk2,Elf1,Foxp1,Gatsl3,Glrx,Hey1,Il6st,Lef1,Lrtm2,Mapk14,Osm,Pdcd4,Pde11a,Per1,Prickle1,Prkca,Prkcq,Rgs10,Rnf125,Serpine2,Sgms1,Shisa2,Sirt4,Ski,Tmem64,Tnfaip3,Trib1,Ubr2,Wwox,Xbp1,Xdh |
| GO:0023057 | negative regulation of signaling | 40 | 1271 | 0.0015 | Adipoq,Arhgap25,Atp2b4,Cblb,Cd300a,Chst11,Dusp1,Dusp10,Dusp2,Dyrk2,Elf1,Foxp1,Gatsl3,Glrx,Hey1,Il6st,Lef1,Lrtm2,Mapk14,Osm,Pdcd4,Pde11a,Per1,Prickle1,Prkca,Prkcq,Rgs10,Rnf125,Serpine2,Sgms1,Shisa2,Sirt4,Ski,Tmem64,Tnfaip3,Trib1,Ubr2,Wwox,Xbp1,Xdh |
| GO:0031399 | regulation of protein modification process | 50 | 1733 | 0.0015 | Adipoq,Akt2,Atp2b4,Bcl10,Cblb,Ccne1,Ccr1,Ccr7,Cd300a,Cdkn2b,Cryaa,Dusp1,Dusp10,Dusp2,F2,Fam20a,Fanci,Gadd45g,Il6st,Jarid2,Jun,Lrrn3,Lrtm2,Map3k1,Map3k8,Map4k1,Mapk9,Ncf1,Nipbl,Osm,P2ry1,Pdcd4,Per1,Pim1,Ppp2r5a,Prickle1,Prkca,Rara,Ski,Slc39a10,Sod1,Spag9,Stk10,Tgfbr2,Tnfaip3,Tnfrsf1a,Tnfsf11,Trib1,Xbp1,Xdh |
| GO:0043067 | regulation of programmed cell death | 45 | 1495 | 0.0015 | Adipoq,Akt2,Bcl10,Btg1,Btg2,Ccr7,Chst11,Cryaa,Dad1,Dusp1,Ets1,Fasl,Foxp1,Gadd45g,Glrx,Igf2r,Itga4,Jun,Kcnh8,Kdm2b,Lef1,Map3k1,Map3k8,Map4k1,Mapk9,Osm,Pdcd4,Pim1,Prkca,Prkch,Prkcq,Rara,Sgms1,Sirt4,Slc39a10,Sod1,Stat5b,Stk10,Tax1bp1,Tnfaip3,Tnfrsf1a,Tnfrsf1b,Wwox,Xbp1,Xdh |
| GO:0043405 | regulation of MAP kinase activity | 16 | 304 | 0.0015 | Adipoq,Ccr7,Cd300a,Dusp1,Dusp10,Dusp2,Gadd45g,Map3k1,Map3k8,Map4k1,Pdcd4,Prkca,Sod1,Spag9,Tnfsf11,Trib1 |
| GO:0043502 | regulation of muscle adaptation | 9 | 99 | 0.0015 | Atp2b4,Foxp1,Gtf2ird1,Jarid2,Nfatc1,P2rx4,Prkca,Tnfrsf1a,Tnfrsf1b |
| GO:0045892 | negative regulation of transcription, DNA-templated | 36 | 1094 | 0.0015 | Adipoq,Aebp2,Bach2,Btg2,Ccne1,Crem,Ets2,Fasl,Foxn3,Foxp1,Hey1,Jarid2,Jun,Kdm2b,Klf2,Klf3,Lef1,Max,Mbd2,Mospd1,Nfatc1,Nipbl,Nrip1,Pdcd4,Per1,Prickle1,Rara,Ski,Smarca2,Smarce1,Tbl1xr1,Ubr2,Usp3,Xbp1,Zfp932,Zscan10 |
| GO:0072503 | cellular divalent inorganic cation homeostasis | 19 | 408 | 0.0015 | Agtr1b,Atp2b4,Ccr1,F2,Fasl,Gpr18,Il6st,Inpp4b,P2rx4,P2ry1,Prkca,S1pr1,Selt,Slc35g1,Slc39a10,Stim2,Thada,Tmem64,Xcr1 |
| GO:2000113 | negative regulation of cellular macromolecule biosynthetic process | 40 | 1269 | 0.0015 | Adipoq,Aebp2,Ago2,Bach2,Btg2,Ccne1,Crem,Dusp1,Ets2,Fasl,Foxn3,Foxp1,Hey1,Jarid2,Jun,Kdm2b,Klf2,Klf3,Lef1,Max,Mbd2,Mospd1,Nfatc1,Nipbl,Nrip1,Pdcd4,Per1,Prickle1,Prkca,Rara,Ski,Smarca2,Smarce1,Tbl1xr1,Tnrc6b,Ubr2,Usp3,Xbp1,Zfp932,Zscan10 |
| GO:0009058 | biosynthetic process | 89 | 3707 | 0.0016 | Aebp2,Aff4,Ago2,Akt2,Bach2,Bcat1,Btg2,Ccne1,Cers6,Chd2,Chd7,Chst11,Chsy1,Cmas,Crem,Cyp2r1,Dach2,Dad1,E2f2,Eif4e3,Elf1,Elovl5,Ets1,Ets2,Ext1,Fasl,Fli1,Fosl2,Foxn3,Foxp1,Gadd45g,Ggta1,Gtf2ird1,Gucy2c,Hbs1l,Hey1,Hivep2,Inpp4b,Ip6k1,Jarid2,Jun,Kdm2b,Klf2,Klf3,Klf6,Lef1,Lpcat3,Mapk14,Max,Mbd2,Mbip,Mgme1,Mogat1,Moxd1,Ncf1,Nfatc1,Nfkbiz,Nipbl,Nrip1,P2rx4,Pde11a,Pde4a,Per1,Polg2,Rara,Scd2,Scml4,Sgms1,Ski,Smarca2,Sod1,Spty2d1,Ss18,St8sia1,Stat4,Stat5b,Tars2,Tbl1xr1,Vgll4,Vsx2,Xbp1,Xylt1,Zdhhc8,Zfp3,Zfp326,Zfp609,Zfp652,Zfp932,Zscan10 |
| GO:0010558 | negative regulation of macromolecule biosynthetic process | 41 | 1322 | 0.0016 | Adipoq,Aebp2,Ago2,Bach2,Btg2,Ccne1,Crem,Dusp1,Ets2,Fasl,Foxn3,Foxp1,Hey1,Jarid2,Jun,Kdm2b,Klf2,Klf3,Lef1,Max,Mbd2,Mospd1,Nfatc1,Nipbl,Nrip1,Pdcd4,Per1,Prickle1,Prkca,Rara,Ski,Smarca2,Smarce1,Tbl1xr1,Tmeff2,Tnrc6b,Ubr2,Usp3,Xbp1,Zfp932,Zscan10 |
| GO:0030098 | lymphocyte differentiation | 14 | 243 | 0.0016 | Ccr7,Cd28,Chd7,Dock10,Foxp1,Gadd45g,Gpr18,Hectd1,Itk,Lef1,Nfatc1,Stat5b,Tnfaip3,Tnfsf8 |
| GO:0045859 | regulation of protein kinase activity | 27 | 717 | 0.0016 | Adipoq,Atp2b4,Cblb,Ccne1,Ccr7,Cd300a,Cdkn2b,Dusp1,Dusp10,Dusp2,Fam20a,Gadd45g,Lrtm2,Map3k1,Map3k8,Map4k1,Ncf1,Pdcd4,Pim1,Prkca,Sod1,Spag9,Stk10,Tgfbr2,Tnfaip3,Tnfsf11,Trib1 |
| GO:0042325 | regulation of phosphorylation | 44 | 1466 | 0.0017 | Adipoq,Akt2,Atp2b4,Bcl10,Cblb,Ccne1,Ccr1,Ccr7,Cd300a,Cdkn2b,Cryaa,Dusp1,Dusp10,Dusp2,F2,Fam20a,Gadd45g,Il6st,Jun,Lrrn3,Lrtm2,Map3k1,Map3k8,Map4k1,Mapk9,Ncf1,Osm,P2ry1,Pdcd4,Per1,Pim1,Ppp2r5a,Prkca,Rara,Sod1,Spag9,Stk10,Tgfbr2,Tnfaip3,Tnfrsf1a,Tnfsf11,Trib1,Xbp1,Xdh |
| GO:0051403 | stress-activated MAPK cascade | 8 | 79 | 0.0017 | Ccr7,Map3k1,Map4k1,Mapk14,Mapk9,Spag9,Tnfsf11,Trib1 |
| GO:0098771 | inorganic ion homeostasis | 25 | 642 | 0.0017 | Agtr1b,Atp1a1,Atp2b4,Ccr1,Clic4,F2,Fam20a,Fasl,Gpr18,Il6st,Inpp4b,P2rx4,P2ry1,Prkca,S1pr1,Selt,Slc30a4,Slc35g1,Slc39a10,Sod1,Stim2,Thada,Tmem64,Tnfsf11,Xcr1 |
| GO:0031327 | negative regulation of cellular biosynthetic process | 42 | 1375 | 0.0018 | Adipoq,Aebp2,Ago2,Atp1a1,Atp2b4,Bach2,Btg2,Ccne1,Crem,Dusp1,Ets2,Fasl,Foxn3,Foxp1,Hey1,Jarid2,Jun,Kdm2b,Klf2,Klf3,Lef1,Max,Mbd2,Mospd1,Nfatc1,Nipbl,Nrip1,Pdcd4,Per1,Prickle1,Prkca,Rara,Ski,Smarca2,Smarce1,Tbl1xr1,Tnrc6b,Ubr2,Usp3,Xbp1,Zfp932,Zscan10 |
| GO:0060324 | face development | 7 | 58 | 0.0018 | Ankrd11,Chd7,Itga4,Lef1,Nipbl,Rara,Ski |
| GO:0006875 | cellular metal ion homeostasis | 21 | 492 | 0.0019 | Agtr1b,Atp1a1,Atp2b4,Ccr1,F2,Fasl,Gpr18,Il6st,Inpp4b,P2rx4,P2ry1,Prkca,S1pr1,Selt,Slc35g1,Slc39a10,Sod1,Stim2,Thada,Tmem64,Xcr1 |
| GO:0002237 | response to molecule of bacterial origin | 16 | 315 | 0.002 | Bcl10,Ccr7,Dusp10,Fasl,Foxp1,Jun,Mapk14,Pdcd4,Rara,Sgms1,Stat5b,Tnfaip3,Tnfrsf1a,Tnfrsf1b,Trib1,Xbp1 |
| GO:0042981 | regulation of apoptotic process | 44 | 1476 | 0.002 | Adipoq,Akt2,Bcl10,Btg1,Btg2,Ccr7,Chst11,Cryaa,Dad1,Dusp1,Ets1,Fasl,Foxp1,Gadd45g,Igf2r,Itga4,Jun,Kcnh8,Kdm2b,Lef1,Map3k1,Map3k8,Map4k1,Mapk9,Osm,Pdcd4,Pim1,Prkca,Prkch,Prkcq,Rara,Sgms1,Sirt4,Slc39a10,Sod1,Stat5b,Stk10,Tax1bp1,Tnfaip3,Tnfrsf1a,Tnfrsf1b,Wwox,Xbp1,Xdh |
| GO:0009612 | response to mechanical stimulus | 13 | 219 | 0.0021 | Atp1a1,Bcl10,Btg2,Ets1,Jun,Mapk14,Mapk9,Mbd2,P2ry1,Serpine2,Tgfbr2,Tmc1,Tnfrsf1a |
| GO:0006351 | transcription, DNA-templated | 54 | 1960 | 0.0022 | Aebp2,Aff4,Ago2,Bach2,Btg2,Chd2,Chd7,Crem,Dach2,E2f2,Elf1,Ets1,Ets2,Fasl,Fli1,Fosl2,Foxn3,Foxp1,Gtf2ird1,Hey1,Hivep2,Jarid2,Jun,Kdm2b,Klf2,Klf3,Klf6,Lef1,Mapk14,Max,Mbd2,Nfatc1,Nfkbiz,Nipbl,Nrip1,Per1,Rara,Scml4,Ski,Smarca2,Spty2d1,Ss18,Stat4,Stat5b,Tbl1xr1,Vgll4,Vsx2,Xbp1,Zfp3,Zfp326,Zfp609,Zfp652,Zfp932,Zscan10 |
| GO:0006874 | cellular calcium ion homeostasis | 18 | 388 | 0.0022 | Agtr1b,Atp2b4,Ccr1,F2,Fasl,Gpr18,Il6st,Inpp4b,P2rx4,P2ry1,Prkca,S1pr1,Selt,Slc35g1,Stim2,Thada,Tmem64,Xcr1 |
| GO:0031347 | regulation of defense response | 22 | 538 | 0.0022 | Adipoq,Ccr1,Ccr7,Cd28,Cd300a,Cd47,Dusp10,Ets1,F2,Foxp1,Mapk14,Otulin,Pdcd4,Per1,Prkca,Rftn1,Rnf125,Stat5b,Tnfaip3,Tnfrsf1a,Tnfrsf1b,Tnfsf11 |
| GO:0032103 | positive regulation of response to external stimulus | 15 | 285 | 0.0022 | Akt2,Ccr1,Ccr4,Ccr7,Cd28,Cd47,Ets1,Mapk14,P2rx4,Pdcd4,Prkca,S1pr1,Stat5b,Tnfrsf1a,Tnfsf11 |
| GO:0051707 | response to other organism | 28 | 779 | 0.0022 | Adipoq,Bcl10,Ccdc80,Ccr4,Ccr7,Cd47,Chd7,Dusp10,F2,Fasl,Fkbp5,Foxp1,Hist1h2be,Jun,Mapk14,Ncf1,Pdcd4,Rara,Sgms1,Spaca4,Stat5b,Tnfaip3,Tnfrsf1a,Tnfrsf1b,Tnfsf8,Trib1,Trim25,Xbp1 |
| GO:0098751 | bone cell development | 5 | 25 | 0.0022 | Fli1,Foxp1,Pip4k2a,Tnfsf11,Wasf2 |
| GO:0006793 | phosphorus metabolic process | 50 | 1775 | 0.0023 | Akt2,Atp1a1,Camk4,Ccne1,Ccr7,Cmah,Cmas,Dnajc6,Dusp1,Dusp10,Dusp2,Dyrk2,Gucy2c,Inpp4b,Ip6k1,Itk,Jak1,Kcnh8,Lmtk3,Lpcat3,Map3k1,Map3k8,Map4k1,Mapk14,Mapk9,Mbip,Mknk2,Ncf1,Osm,Pde11a,Pde4a,Pim1,Pip4k2a,Ppp1cc,Ppp2r5a,Prkca,Prkch,Prkcq,Ptp4a2,Rara,Riok1,Rpia,Sgms1,Sla,Spag9,Stat4,Stk10,Tgfbr2,Tnfsf11,Trib1 |
| GO:1901576 | organic substance biosynthetic process | 87 | 3651 | 0.0023 | Aebp2,Aff4,Ago2,Akt2,Bach2,Bcat1,Btg2,Ccne1,Cers6,Chd2,Chd7,Chst11,Chsy1,Cmas,Crem,Cyp2r1,Dach2,Dad1,E2f2,Eif4e3,Elf1,Elovl5,Ets1,Ets2,Ext1,Fasl,Fli1,Fosl2,Foxn3,Foxp1,Gadd45g,Ggta1,Gtf2ird1,Gucy2c,Hbs1l,Hey1,Hivep2,Inpp4b,Ip6k1,Jarid2,Jun,Kdm2b,Klf2,Klf3,Klf6,Lef1,Lpcat3,Mapk14,Max,Mbd2,Mbip,Mgme1,Mogat1,Moxd1,Ncf1,Nfatc1,Nfkbiz,Nipbl,Nrip1,Pde11a,Pde4a,Per1,Polg2,Rara,Scd2,Scml4,Sgms1,Ski,Smarca2,Spty2d1,Ss18,St8sia1,Stat4,Stat5b,Tars2,Tbl1xr1,Vgll4,Vsx2,Xbp1,Xylt1,Zdhhc8,Zfp3,Zfp326,Zfp609,Zfp652,Zfp932,Zscan10 |
| GO:0050801 | ion homeostasis | 26 | 706 | 0.0026 | Agtr1b,Atp1a1,Atp2b4,Ccr1,Clic4,F2,Fam20a,Fasl,Gpr18,Il6st,Inpp4b,P2rx4,P2ry1,Prkca,S1pr1,Selt,Slc30a4,Slc35g1,Slc39a10,Sod1,Stim2,Thada,Tmem64,Tnfsf11,Xbp1,Xcr1 |
| GO:0033673 | negative regulation of kinase activity | 14 | 260 | 0.0028 | Adipoq,Cblb,Cd300a,Cdkn2b,Dusp1,Dusp10,Dusp2,Gadd45g,Lrtm2,Pdcd4,Ppp2r5a,Prkca,Tnfaip3,Trib1 |
| GO:0040017 | positive regulation of locomotion | 22 | 550 | 0.0028 | Ago2,Akt2,Ccr1,Ccr4,Ccr7,Ets1,Foxp1,Il6st,Itga4,Jun,Lef1,Mapk14,Nipbl,P2rx4,Prkca,Rras2,S1pr1,Spag9,Tgfbr2,Vps37b,Xbp1,Zfp609 |
| GO:0043408 | regulation of MAPK cascade | 26 | 709 | 0.0028 | Adipoq,Ccr1,Ccr7,Cd300a,Dusp1,Dusp10,Dusp2,Gadd45g,Jun,Map3k1,Map3k8,Map4k1,Mapk9,Ncf1,Osm,P2ry1,Pdcd4,Per1,Prkca,Rara,Sod1,Spag9,Tnfsf11,Trib1,Xbp1,Xdh |
| GO:0045937 | positive regulation of phosphate metabolic process | 34 | 1051 | 0.0028 | Adipoq,Akt2,Atp2b4,Bcl10,Ccr1,Ccr7,Cd300a,Cryaa,F2,Fam20a,Gadd45g,Il6st,Jun,Lrrn3,Map3k1,Map3k8,Map4k1,Mapk9,Ncf1,Osm,P2ry1,Pim1,Ppp2r5a,Prkca,Rara,Slc39a10,Sod1,Spag9,Stk10,Tgfbr2,Tnfrsf1a,Tnfsf11,Xbp1,Xdh |
| GO:0019220 | regulation of phosphate metabolic process | 47 | 1650 | 0.0029 | Adipoq,Akt2,Atp2b4,Bcl10,Cblb,Ccne1,Ccr1,Ccr7,Cd300a,Cdkn2b,Cnst,Cryaa,Dusp1,Dusp10,Dusp2,F2,Fam20a,Gadd45g,Il6st,Jun,Lmtk3,Lrrn3,Lrtm2,Map3k1,Map3k8,Map4k1,Mapk9,Ncf1,Osm,P2ry1,Pdcd4,Per1,Pim1,Ppp2r5a,Prkca,Rara,Slc39a10,Sod1,Spag9,Stk10,Tgfbr2,Tnfaip3,Tnfrsf1a,Tnfsf11,Trib1,Xbp1,Xdh |
| GO:0001817 | regulation of cytokine production | 23 | 592 | 0.003 | Adipoq,Bcl10,Ccr7,Cd28,Elf1,F2,Foxp1,Gpr18,Klf2,Lef1,Mapk14,Mapk9,Pdcd4,Per1,Prkcq,Rara,Rnf125,Sod1,Stat5b,Tigit,Tnfaip3,Tnfrsf1a,Xbp1 |
| GO:0002573 | myeloid leukocyte differentiation | 9 | 113 | 0.003 | Camk4,Ccr7,Foxp1,Jun,Lef1,Mapk14,Nfatc1,Tgfbr2,Tnfsf11 |
| GO:0007254 | JNK cascade | 7 | 65 | 0.003 | Ccr7,Map3k1,Map4k1,Mapk9,Spag9,Tnfsf11,Trib1 |
| GO:0007584 | response to nutrient | 12 | 199 | 0.003 | Adipoq,Agtr1b,Cdkn2b,Dad1,Gatsl3,P2rx4,Pim1,Rara,Sod1,Tgfbr2,Trim25,Xbp1 |
| GO:0030003 | cellular cation homeostasis | 22 | 554 | 0.003 | Agtr1b,Atp1a1,Atp2b4,Ccr1,Clic4,F2,Fasl,Gpr18,Il6st,Inpp4b,P2rx4,P2ry1,Prkca,S1pr1,Selt,Slc35g1,Slc39a10,Sod1,Stim2,Thada,Tmem64,Xcr1 |
| GO:0000302 | response to reactive oxygen species | 12 | 200 | 0.0031 | Ccr7,Cryaa,Dusp1,Ets1,Glrx2,Jun,Klf2,Mapk9,Ncf1,Prkca,Sod1,Tnfaip3 |
| GO:0032496 | response to lipopolysaccharide | 15 | 298 | 0.0031 | Ccr7,Dusp10,Fasl,Foxp1,Jun,Mapk14,Pdcd4,Rara,Sgms1,Stat5b,Tnfaip3,Tnfrsf1a,Tnfrsf1b,Trib1,Xbp1 |
| GO:0045580 | regulation of T cell differentiation | 10 | 141 | 0.0031 | Ccr7,Cd28,Dusp10,H2-Aa,Nfkbiz,Rara,Stat5b,Tgfbr2,Xbp1,Zfp609 |
| GO:0050776 | regulation of immune response | 24 | 635 | 0.0031 | Bcl10,Btla,Cblb,Ccr1,Ccr7,Cd247,Cd28,Cd300a,Dpp4,Dusp10,Elf1,Itk,Mapk14,Nfkbiz,Otulin,Prkch,Prkcq,Rara,Rftn1,Rnf125,Slc39a10,Stat5b,Tnfaip3,Xbp1 |
| GO:0071345 | cellular response to cytokine stimulus | 25 | 676 | 0.0031 | Ccr1,Ccr4,Ccr7,Foxp1,Il21r,Il6st,Jak1,Jarid2,Klf2,Klf6,Lef1,Lrtm2,Mapk14,Osm,Rnf125,Sgms1,Stat4,Stat5b,Tnfrsf1a,Tnfrsf1b,Tnfsf11,Trim25,Vamp4,Xbp1,Xcr1 |
| GO:0009059 | macromolecule biosynthetic process | 70 | 2809 | 0.0033 | Aebp2,Aff4,Ago2,Akt2,Bach2,Btg2,Ccne1,Chd2,Chd7,Chst11,Chsy1,Crem,Dach2,Dad1,E2f2,Eif4e3,Elf1,Ets1,Ets2,Ext1,Fasl,Fli1,Fosl2,Foxn3,Foxp1,Gadd45g,Ggta1,Gtf2ird1,Hbs1l,Hey1,Hivep2,Jarid2,Jun,Kdm2b,Klf2,Klf3,Klf6,Lef1,Mapk14,Max,Mbd2,Mgme1,Nfatc1,Nfkbiz,Nipbl,Nrip1,Per1,Polg2,Rara,Scml4,Ski,Smarca2,Spty2d1,Ss18,St8sia1,Stat4,Stat5b,Tars2,Tbl1xr1,Vgll4,Vsx2,Xbp1,Xylt1,Zdhhc8,Zfp3,Zfp326,Zfp609,Zfp652,Zfp932,Zscan10 |
| GO:0010611 | regulation of cardiac muscle hypertrophy | 7 | 67 | 0.0033 | Atp2b4,Foxp1,Jarid2,P2rx4,Prkca,Tnfrsf1a,Tnfrsf1b |
| GO:0032269 | negative regulation of cellular protein metabolic process | 33 | 1020 | 0.0033 | Adipoq,Ago2,Akt2,Btg2,Cblb,Cd300a,Cdkn2b,Crim1,Cryaa,Dusp1,Dusp10,Dusp2,F2,Gadd45g,Jarid2,Jun,Lef1,Lrtm2,Os9,Pdcd4,Per1,Prkca,Rara,Serpine2,Sirt4,Ski,Tnfaip3,Tnfrsf1b,Tnrc6b,Trib1,Ubac2,Xbp1,Xdh |
| GO:0043407 | negative regulation of MAP kinase activity | 7 | 67 | 0.0033 | Adipoq,Cd300a,Dusp1,Dusp10,Dusp2,Pdcd4,Prkca |
| GO:0045619 | regulation of lymphocyte differentiation | 11 | 171 | 0.0033 | Ccr7,Cd28,Dusp10,H2-Aa,Il15ra,Nfkbiz,Rara,Stat5b,Tgfbr2,Xbp1,Zfp609 |
| GO:0071901 | negative regulation of protein serine/threonine kinase activity | 9 | 115 | 0.0033 | Adipoq,Cd300a,Cdkn2b,Dusp1,Dusp10,Dusp2,Pdcd4,Prkca,Tnfaip3 |
| GO:0009967 | positive regulation of signal transduction | 42 | 1433 | 0.0034 | Adipoq,Akt2,Bcl10,Ccr1,Ccr7,Cd28,Cdkn2b,Chsy1,Cyfip2,F2,Fasl,Gadd45g,Glrx,Gpr18,Il6st,Jak1,Jun,Map3k1,Map3k8,Map4k1,Mapk9,Mbd2,Ncf1,Osm,P2rx4,P2ry1,Pdcd4,Prkca,Prkch,Rara,Sh2b1,Sla,Slc39a10,Sod1,Spag9,Tbl1xr1,Tnfrsf1a,Tnfsf11,Trim25,Wwox,Xbp1,Xdh |
| GO:0050729 | positive regulation of inflammatory response | 9 | 116 | 0.0034 | Ccr7,Cd28,Cd47,Ets1,Pdcd4,Prkca,Stat5b,Tnfrsf1a,Tnfsf11 |
| GO:0051253 | negative regulation of RNA metabolic process | 37 | 1204 | 0.0034 | Adipoq,Aebp2,Akt2,Bach2,Btg2,Ccne1,Crem,Ets2,Fasl,Foxn3,Foxp1,Hey1,Jarid2,Jun,Kdm2b,Klf2,Klf3,Lef1,Max,Mbd2,Mospd1,Nfatc1,Nipbl,Nrip1,Pdcd4,Per1,Prickle1,Rara,Ski,Smarca2,Smarce1,Tbl1xr1,Ubr2,Usp3,Xbp1,Zfp932,Zscan10 |
| GO:1903827 | regulation of cellular protein localization | 20 | 484 | 0.0034 | Adipoq,Akt2,Camk4,Ccne1,Cd247,Cnst,Epb4.1,F2,Hectd1,Leprotl1,Mapk14,Os9,Ppp2r5a,Prkcq,Sirt4,Tnfrsf1a,Trappc12,Ubac2,Vamp4,Xbp1 |
| GO:0046328 | regulation of JNK cascade | 11 | 174 | 0.0036 | Ccr7,Dusp10,Gadd45g,Map3k1,Map4k1,Mapk9,Ncf1,Pdcd4,Per1,Spag9,Tnfsf11 |
| GO:0034645 | cellular macromolecule biosynthetic process | 69 | 2778 | 0.0038 | Aebp2,Aff4,Ago2,Akt2,Bach2,Btg2,Ccne1,Chd2,Chd7,Chst11,Chsy1,Crem,Dach2,Dad1,E2f2,Eif4e3,Elf1,Ets1,Ets2,Ext1,Fasl,Fli1,Fosl2,Foxn3,Foxp1,Ggta1,Gtf2ird1,Hbs1l,Hey1,Hivep2,Jarid2,Jun,Kdm2b,Klf2,Klf3,Klf6,Lef1,Mapk14,Max,Mbd2,Mgme1,Nfatc1,Nfkbiz,Nipbl,Nrip1,Per1,Polg2,Rara,Scml4,Ski,Smarca2,Spty2d1,Ss18,St8sia1,Stat4,Stat5b,Tars2,Tbl1xr1,Vgll4,Vsx2,Xbp1,Xylt1,Zdhhc8,Zfp3,Zfp326,Zfp609,Zfp652,Zfp932,Zscan10 |
| GO:0043069 | negative regulation of programmed cell death | 30 | 901 | 0.0038 | Akt2,Bcl10,Btg2,Ccr7,Chst11,Cryaa,Dad1,Dusp1,Foxp1,Glrx,Jun,Kcnh8,Kdm2b,Lef1,Pdcd4,Pim1,Prkca,Prkch,Prkcq,Rara,Sgms1,Sirt4,Slc39a10,Sod1,Stat5b,Tax1bp1,Tnfaip3,Tnfrsf1a,Tnfrsf1b,Xbp1 |
| GO:1901564 | organonitrogen compound metabolic process | 101 | 4480 | 0.0038 | Akt2,Bach2,Bcat1,Btg1,Btg2,Camk4,Cblb,Ccne1,Ccr7,Cers6,Chst11,Chsy1,Crbn,Crem,Cyfip2,Dad1,Dcaf7,Dnajc6,Dpp4,Dusp1,Dusp10,Dusp2,Dyrk2,Eif4e3,Elovl5,Ext1,F2,Fbxl12,Fbxo33,Fkbp5,Fntb,Gadd45g,Ggta1,Gucy2c,Gzmb,Hbs1l,Hectd1,Itk,Jak1,Kcnh8,Kdm2b,Lef1,Lmtk3,Lonrf1,Map3k1,Map3k8,Map4k1,Mapk14,Mapk9,Mbip,Mknk2,Moxd1,Msrb3,Ncf1,Neurl1b,Nsmce1,Os9,Osm,Otulin,Pde11a,Pde4a,Per1,Pim1,Polg2,Ppp1cc,Ppp2r5a,Prkca,Prkch,Prkcq,Prss39,Ptp4a2,Rara,Riok1,Rnf125,Rpia,Sgms1,Sirt4,Sla,Smox,Sod1,Spag9,St8sia1,Stat4,Stat5b,Stk10,Tars2,Tbl1xr1,Tgfbr2,Tnfaip3,Tnfsf11,Trappc12,Trib1,Trim25,Ubr2,Usp3,Wdyhv1,Xbp1,Xdh,Xpnpep2,Xylt1,Zdhhc8 |

| | | | | | |
|---|---|---|---|---|---|
| GO:0000122 | negative regulation of transcription by RNA polymerase II | 28 | 817 | 0.0039 | Aebp2,Bach2,Btg2,Ccne1,Crem,Ets2,Fasl,Foxp1,Hey1,Jarid2,Jun,Kdm2b,Klf2,Klf3,Lef1,Max,Mbd2,Mospd1,Nfatc1,Nipbl,Nrip1,Per1,Rara,Ski,Tbl1xr1,Usp3,Xbp1,Zfp932 |
| GO:0016310 | phosphorylation | 33 | 1034 | 0.0039 | Akt2,Camk4,Ccne1,Ccr7,Dyrk2,Gucy2c,Ip6k1,Itk,Jak1,Kcnh8,Lmtk3,Map3k1,Map3k8,Map4k1,Mapk14,Mapk9,Mknk2,Osm,Pim1,Pip4k2a,Prkca,Prkch,Prkcq,Rara,Riok1,Sgms1,Sla,Spag9,Stat4,Stk10,Tgfbr2,Tnfsf11,Trib1 |
| GO:0044092 | negative regulation of molecular function | 34 | 1078 | 0.0039 | Adipoq,Akt2,Atp2b4,Cblb,Cd300a,Cdkn2b,Cnst,Crim1,Cryaa,Dusp1,Dusp10,Dusp2,Ercc4,Gadd45g,Hey1,Itga4,Jun,Lef1,Lmtk3,Lrtm2,Otulin,P2ry1,Pdcd4,Pim1,Ppp2r5a,Prkca,Serpine2,Ski,Spag9,Tax1bp1,Tnfaip3,Tnfrsf1b,Trib1,Zfp932 |
| GO:0045597 | positive regulation of cell differentiation | 33 | 1033 | 0.0039 | Adipoq,Btg1,Ccne1,Ccr1,Ccr7,Cd101,Dusp10,Ets1,Foxp1,H2-Aa,Hey1,Il15ra,Il6st,Ist1,Jun,Lef1,Mapk14,Mapk9,Nfkbiz,Nipbl,Ppp1cc,Prkca,Prkch,Rara,Serpine2,Spag9,Stat5b,Tgfbr2,Tmem64,Tnfsf11,Trib1,Xbp1,Zfp609 |
| GO:0060348 | bone development | 11 | 177 | 0.0039 | Ankrd11,Chsy1,Fli1,Foxn3,Foxp1,Pip4k2a,Rara,Ski,Tgfbr2,Tnfsf11,Wasf2 |
| GO:0097190 | apoptotic signaling pathway | 14 | 273 | 0.0039 | Cd28,Dyrk2,E2f2,Fasl,Gzmb,Jun,Mapk9,Mknk2,P2rx4,Prkca,Tnfrsf1a,Tnfrsf1b,Wwox,Xbp1 |
| GO:0001501 | skeletal system development | 19 | 454 | 0.004 | Ankrd11,Chd7,Chst11,Chsy1,Ext1,Fli1,Foxn3,Foxp1,Mapk14,Nipbl,Pip4k2a,Prkca,Rara,Ski,Tapt1,Tgfbr2,Tnfsf11,Wasf2,Wwox |
| GO:0006469 | negative regulation of protein kinase activity | 13 | 241 | 0.004 | Adipoq,Cblb,Cd300a,Cdkn2b,Dusp1,Dusp10,Dusp2,Gadd45g,Lrtm2,Pdcd4,Prkca,Tnfaip3,Trib1 |
| GO:0010941 | regulation of cell death | 46 | 1640 | 0.004 | Adipoq,Akt2,Bcl10,Btg1,Btg2,Ccr7,Chst11,Cryaa,Dad1,Dusp1,Ets1,Fasl,Foxp1,Gadd45g,Glrx,Igf2r,Itga4,Jun,Kcnh8,Kdm2b,Lef1,Map3k1,Map3k8,Map4k1,Mapk9,Osm,Pdcd4,Pim1,Prkca,Prkch,Prkcq,Rara,Rras2,Sgms1,Sirt4,Slc39a10,Sod1,Stat5b,Stk10,Tax1bp1,Tnfaip3,Tnfrsf1a,Tnfrsf1b,Wwox,Xbp1,Xdh |
| GO:0051345 | positive regulation of hydrolase activity | 24 | 652 | 0.004 | Akt2,Arhgap15,Arhgap25,Bcl10,Ccr7,Cd300a,Cyfip2,Dock10,Fasl,Gdi2,Gnat1,Itk,Jun,Mapk14,Mapk9,Pkp4,Rasgrp2,Rgs10,S1pr1,Sipa1l1,Slc39a10,Snx18,Usp6nl,Xdh |
| GO:0051272 | positive regulation of cellular component movement | 21 | 533 | 0.0041 | Ago2,Akt2,Ccr1,Ccr7,Ets1,Foxp1,Il6st,Itga4,Jun,Lef1,Mapk14,Nipbl,P2rx4,Prkca,Rras2,S1pr1,Spag9,Stat5b,Tgfbr2,Xbp1,Zfp609 |
| GO:0034654 | nucleobase-containing compound biosynthetic process | 60 | 2339 | 0.0045 | Aebp2,Aff4,Ago2,Bach2,Btg2,Chd2,Chd7,Cmas,Crem,Dach2,E2f2,Elf1,Ets1,Ets2,Fasl,Fli1,Fosl2,Foxn3,Foxp1,Gtf2ird1,Gucy2c,Hey1,Hivep2,Jarid2,Jun,Kdm2b,Klf2,Klf3,Klf6,Lef1,Mapk14,Max,Mbd2,Ncf1,Nfatc1,Nfkbiz,Nipbl,Nrip1,Pde11a,Pde4a,Per1,Polg2,Rara,Scml4,Ski,Smarca2,Spty2d1,Ss18,Stat4,Stat5b,Tbl1xr1,Vgll4,Vsx2,Xbp1,Zfp3,Zfp326,Zfp609,Zfp652,Zfp932,Zscan10 |
| GO:0019221 | cytokine-mediated signaling pathway | 15 | 313 | 0.0046 | Ccr1,Ccr4,Ccr7,Il21r,Il6st,Jak1,Klf6,Lrtm2,Osm,Stat4,Stat5b,Tnfrsf1a,Tnfrsf1b,Tnfsf11,Xcr1 |
| GO:0030335 | positive regulation of cell migration | 20 | 500 | 0.0047 | Ago2,Akt2,Ccr1,Ccr7,Ets1,Foxp1,Il6st,Itga4,Jun,Lef1,Mapk14,Nipbl,P2rx4,Prkca,Rras2,S1pr1,Spag9,Tgfbr2,Xbp1,Zfp609 |
| GO:0032872 | regulation of stress-activated MAPK cascade | 12 | 213 | 0.0047 | Ccr7,Dusp10,Gadd45g,Map3k1,Map4k1,Mapk9,Ncf1,Pdcd4,Per1,Spag9,Tnfsf11,Xdh |
| GO:0045595 | regulation of cell differentiation | 48 | 1751 | 0.0048 | Adipoq,Btg1,Ccne1,Ccr1,Ccr7,Cd101,Cd28,Cdkn2b,Chd7,Ddx6,Dusp10,Ets1,F2,Foxp1,H2-Aa,Hey1,Il15ra,Il6st,Inpp4b,Ist1,Jun,Lef1,Mapk14,Mapk9,Nfatc1,Nfkbiz,Nipbl,Pdcd4,Ppp1cc,Prickle1,Prkca,Prkch,Rara,Serpine2,Sipa1l1,Ski,Spag9,Stat5b,Tgfbr2,Tmem64,Tnfrsf1a,Tnfsf11,Trib1,Xbp1,Xdh,Xylt1,Zfp609,Zfp932 |
| GO:0045934 | negative regulation of nucleobase-containing compound metabolic process | 39 | 1326 | 0.0049 | Adipoq,Aebp2,Akt2,Bach2,Btg2,Ccne1,Crem,Dusp1,Ercc4,Ets2,Fasl,Foxn3,Foxp1,Hey1,Jarid2,Jun,Kdm2b,Klf2,Klf3,Lef1,Max,Mbd2,Mospd1,Nfatc1,Nipbl,Nrip1,Pdcd4,Per1,Prickle1,Rara,Ski,Smarca2,Smarce1,Tbl1xr1,Ubr2,Usp3,Xbp1,Zfp932,Zscan10 |
| GO:0006796 | phosphate-containing compound metabolic process | 48 | 1758 | 0.0051 | Akt2,Atp1a1,Camk4,Ccne1,Ccr7,Dnajc6,Dusp1,Dusp10,Dusp2,Dyrk2,Gucy2c,Inpp4b,Ip6k1,Itk,Jak1,Kcnh8,Lmtk3,Lpcat3,Map3k1,Map3k8,Map4k1,Mapk14,Mapk9,Mbip,Mknk2,Ncf1,Osm,Pde11a,Pde4a,Pim1,Pip4k2a,Ppp1cc,Ppp2r5a,Prkca,Prkch,Prkcq,Ptp4a2,Rara,Riok1,Rpia,Sgms1,Sla,Spag9,Stat4,Stk10,Tgfbr2,Tnfsf11,Trib1 |
| GO:0010035 | response to inorganic substance | 20 | 505 | 0.0051 | Atp2b4,Ccr7,Clic4,Cryaa,Cyb5a,Cyp2r1,Dusp1,Ets1,Fntb,Glrx2,Jun,Klf2,Mapk9,Ncf1,P2rx4,Rasgrp2,Slc30a4,Sod1,Tnfaip3,Xdh |
| GO:0018130 | heterocycle biosynthetic process | 61 | 2404 | 0.0051 | Aebp2,Aff4,Ago2,Bach2,Btg2,Chd2,Chd7,Cmas,Crem,Dach2,E2f2,Elf1,Ets1,Ets2,Fasl,Fli1,Fosl2,Foxn3,Foxp1,Gtf2ird1,Gucy2c,Hey1,Hivep2,Jarid2,Jun,Kdm2b,Klf2,Klf3,Klf6,Lef1,Mapk14,Max,Mbd2,Mbip,Ncf1,Nfatc1,Nfkbiz,Nipbl,Nrip1,Pde11a,Pde4a,Per1,Polg2,Rara,Scml4,Ski,Smarca2,Spty2d1,Ss18,Stat4,Stat5b,Tbl1xr1,Vgll4,Vsx2,Xbp1,Zfp3,Zfp326,Zfp609,Zfp652,Zfp932,Zscan10 |
| GO:0030217 | T cell differentiation | 10 | 154 | 0.0051 | Ccr7,Cd28,Chd7,Foxp1,Gadd45g,Gpr18,Itk,Lef1,Stat5b,Tnfsf8 |
| GO:0032102 | negative regulation of response to external stimulus | 14 | 282 | 0.0051 | Adipoq,Dusp10,Ets1,F2,Gpr18,Otulin,Pdcd4,Rnf125,Serpine2,Tnfaip3,Tnfrsf1a,Tnfrsf1b,Trib1,Xylt1 |
| GO:1901342 | regulation of vasculature development | 15 | 319 | 0.0053 | Ago2,Btg1,E2f2,Ets1,Fasl,Hey1,Jak1,Klf2,Nfatc1,Pdcd4,Prkca,Tgfbr2,Tnfrsf1a,Xbp1,Xdh |
| GO:0019438 | aromatic compound biosynthetic process | 61 | 2414 | 0.0055 | Aebp2,Aff4,Ago2,Bach2,Btg2,Chd2,Chd7,Cmas,Crem,Dach2,E2f2,Elf1,Ets1,Ets2,Fasl,Fli1,Fosl2,Foxn3,Foxp1,Gtf2ird1,Gucy2c,Hey1,Hivep2,Jarid2,Jun,Kdm2b,Klf2,Klf3,Klf6,Lef1,Mapk14,Max,Mbd2,Moxd1,Ncf1,Nfatc1,Nfkbiz,Nipbl,Nrip1,Pde11a,Pde4a,Per1,Polg2,Rara,Scml4,Ski,Smarca2,Spty2d1,Ss18,Stat4,Stat5b,Tbl1xr1,Vgll4,Vsx2,Xbp1,Zfp3,Zfp326,Zfp609,Zfp652,Zfp932,Zscan10 |
| GO:0031348 | negative regulation of defense response | 11 | 186 | 0.0055 | Adipoq,Ccr1,Dusp10,Ets1,F2,Otulin,Pdcd4,Rnf125,Tnfaip3,Tnfrsf1a,Tnfrsf1b |
| GO:0043066 | negative regulation of apoptotic process | 29 | 884 | 0.0055 | Akt2,Bcl10,Btg2,Ccr7,Chst11,Cryaa,Dad1,Dusp1,Foxp1,Jun,Kcnh8,Kdm2b,Lef1,Pdcd4,Pim1,Prkca,Prkch,Prkcq,Rara,Sgms1,Sirt4,Slc39a10,Sod1,Stat5b,Tax1bp1,Tnfaip3,Tnfrsf1a,Tnfrsf1b,Xbp1 |
| GO:1902533 | positive regulation of intracellular signal transduction | 30 | 927 | 0.0055 | Adipoq,Bcl10,Ccr1,Ccr7,Cd28,F2,Fasl,Gadd45g,Glrx,Gpr18,Il6st,Jak1,Jun,Map3k1,Map3k8,Map4k1,Ncf1,Osm,P2rx4,P2ry1,Pdcd4,Prkca,Rara,Sod1,Spag9,Tnfrsf1a,Tnfsf11,Trim25,Xbp1,Xdh |
| GO:1901362 | organic cyclic compound biosynthetic process | 63 | 2522 | 0.0058 | Aebp2,Aff4,Ago2,Bach2,Btg2,Chd2,Chd7,Cmas,Crem,Cyp2r1,Dach2,E2f2,Elf1,Ets1,Ets2,Fasl,Fli1,Fosl2,Foxn3,Foxp1,Gtf2ird1,Gucy2c,Hey1,Hivep2,Jarid2,Jun,Kdm2b,Klf2,Klf3,Klf6,Lef1,Mapk14,Max,Mbd2,Mbip,Moxd1,Ncf1,Nfatc1,Nfkbiz,Nipbl,Nrip1,Pde11a,Pde4a,Per1,Polg2,Rara,Scml4,Ski,Smarca2,Spty2d1,Ss18,Stat4,Stat5b,Tbl1xr1,Vgll4,Vsx2,Xbp1,Zfp3,Zfp326,Zfp609,Zfp652,Zfp932,Zscan10 |
| GO:2000379 | positive regulation of reactive oxygen species metabolic process | 8 | 101 | 0.0058 | Akt2,F2,Klf2,Mapk14,Mapk9,Sod1,Tgfbr2,Xdh |
| GO:0000188 | inactivation of MAPK activity | 4 | 18 | 0.0059 | Dusp1,Dusp10,Dusp2,Prkca |
| GO:0050728 | negative regulation of inflammatory response | 9 | 129 | 0.0059 | Adipoq,Dusp10,Ets1,F2,Otulin,Pdcd4,Tnfaip3,Tnfrsf1a,Tnfrsf1b |
| GO:2000516 | positive regulation of CD4-positive, alpha-beta T cell activation | 5 | 34 | 0.0059 | Ccr7,Nfkbiz,Prkcq,Rara,Tgfbr2 |
| GO:0001934 | positive regulation of protein phosphorylation | 30 | 937 | 0.0063 | Adipoq,Akt2,Atp2b4,Ccr1,Ccr7,Cryaa,F2,Fam20a,Gadd45g,Il6st,Jun,Lrrn3,Map3k1,Map3k8,Map4k1,Mapk9,Ncf1,Osm,P2ry1,Pim1,Prkca,Rara,Sod1,Spag9,Stk10,Tgfbr2,Tnfrsf1a,Tnfsf11,Xbp1,Xdh |
| GO:0007257 | activation of JUN kinase activity | 5 | 35 | 0.0065 | Ccr7,Map3k1,Map4k1,Spag9,Tnfsf11 |
| GO:0031670 | cellular response to nutrient | 6 | 55 | 0.0065 | Cdkn2b,Gatsl3,P2rx4,Pim1,Sod1,Xbp1 |
| GO:0042327 | positive regulation of phosphorylation | 31 | 983 | 0.0065 | Adipoq,Akt2,Atp2b4,Bcl10,Ccr1,Ccr7,Cryaa,F2,Fam20a,Gadd45g,Il6st,Jun,Lrrn3,Map3k1,Map3k8,Map4k1,Mapk9,Ncf1,Osm,P2ry1,Pim1,Prkca,Rara,Sod1,Spag9,Stk10,Tgfbr2,Tnfrsf1a,Tnfsf11,Xbp1,Xdh |
| GO:1900744 | regulation of p38MAPK cascade | 5 | 35 | 0.0065 | Dusp10,Gadd45g,Ncf1,Per1,Xdh |
| GO:0048660 | regulation of smooth muscle cell proliferation | 10 | 161 | 0.0067 | Adipoq,Foxp1,Jun,Pdcd4,Prkca,S1pr1,Stat5b,Tgfbr2,Tnfaip3,Trib1 |
| GO:0030852 | regulation of granulocyte differentiation | 4 | 19 | 0.0068 | Adipoq,Lef1,Rara,Trib1 |
| GO:0050852 | T cell receptor signaling pathway | 6 | 56 | 0.0069 | Bcl10,Cd247,Cd28,Itk,Nfkbiz,Rftn1 |
| GO:0032355 | response to estradiol | 9 | 133 | 0.007 | Dusp1,Ets1,Fosl2,Glrx,Mbd2,Nrip1,Prkca,Rara,Stat5b |
| GO:0051704 | multi-organism process | 49 | 1840 | 0.007 | Abhd2,Adipoq,Aff4,Atp2b4,Bcl10,Btg1,Ccdc80,Ccr4,Ccr7,Cd47,Chd7,Clic4,Crem,Ddx6,Dpp4,Dusp10,Ets1,F2,Fasl,Fkbp5,Foxp1,Hist1h2be,Igf2r,Ist1,Jun,Kdm2b,Lef1,Mapk14,Ncf1,Nrip1,Pdcd4,Rara,Serpine2,Sgms1,Slc38a1,Sod1,Spaca4,Stat5b,Tdrp,Tgfbr2,Tnfaip3,Tnfrsf1a,Tnfrsf1b,Tnfsf8,Trib1,Trim25,Ubr2,Usp6nl,Xbp1 |
| GO:1902532 | negative regulation of intracellular signal transduction | 20 | 521 | 0.007 | Adipoq,Arhgap25,Atp2b4,Cd300a,Dusp1,Dusp10,Dusp2,Dyrk2,Gatsl3,Lrtm2,Pdcd4,Pde11a,Per1,Prkca,Rnf125,Serpine2,Tnfaip3,Ubr2,Xbp1,Xdh |
| GO:0009888 | tissue development | 44 | 1599 | 0.0072 | Abcb1b,Bcl10,Btg2,Chd7,Chst11,Chsy1,Clic4,Cryaa,Dusp1,Dusp2,Esrp1,Ets2,Ext1,Fam20a,Fosl2,Foxp1,Hectd1,Hey1,Itga4,Jun,Kdm2b,Klf2,Lef1,Map3k1,Mapk14,Mbnl1,Nfatc1,Ninj2,Pdcd4,Prickle1,Prkca,Rara,S1pr1,Serpine2,Ski,Sod1,Stat5b,Tapt1,Tbl1xr1,Tgfbr2,Tmc1,Tmeff2,Tnfsf11,Xbp1 |
| GO:0051336 | regulation of hydrolase activity | 34 | 1128 | 0.0072 | Akt2,Arhgap15,Arhgap25,Bcl10,Ccr7,Cd300a,Cnst,Crim1,Cryaa,Cyfip2,Dock10,Fasl,Gdi2,Gnat1,Itk,Jun,Lef1,Lmtk3,Mapk14,Mapk9,Pkp4,Ppp2r5a,Rasgrp2,Rgs10,S1pr1,Serpine2,Sipa1l1,Slc39a10,Snx18,Sod1,Tmem64,Tnfrsf1b,Usp6nl,Xdh |
| GO:0023014 | signal transduction by protein phosphorylation | 11 | 195 | 0.0073 | Ccr7,Kcnh8,Map3k1,Map3k8,Map4k1,Mapk14,Mapk9,Spag9,Stk10,Tnfsf11,Trib1 |
| GO:0003170 | heart valve development | 6 | 57 | 0.0074 | Hectd1,Hey1,Nfatc1,Tgfbr2,Tnfrsf1a,Tnfrsf1b |
| GO:0003177 | pulmonary valve development | 4 | 20 | 0.0077 | Hey1,Nfatc1,Tnfrsf1a,Tnfrsf1b |
| GO:0019725 | cellular homeostasis | 26 | 777 | 0.0077 | Agtr1b,Atp1a1,Atp2b4,Ccr1,Clic4,F2,Fasl,Glrx,Glrx2,Gpr18,Il6st,Inpp4b,Lrrc8d,P2rx4,P2ry1,Prkca,S1pr1,Selt,Slc35g1,Slc39a10,Sod1,Stim2,Thada,Tmem64,Xbp1,Xcr1 |
| GO:2000353 | positive regulation of endothelial cell apoptotic process | 4 | 20 | 0.0077 | Fasl,Itga4,Pdcd4,Xbp1 |
| GO:0010647 | positive regulation of cell communication | 44 | 1608 | 0.0078 | Adipoq,Akt2,Bcl10,Ccr1,Ccr7,Cd28,Cdkn2b,Chsy1,Cyfip2,F2,Fasl,Gadd45g,Glrx,Gpr18,Il6st,Jak1,Jun,Map3k1,Map3k8,Map4k1,Mapk9,Mbd2,Ncf1,Osm,P2rx4,P2ry1,Pdcd4,Prkca,Prkch,Rara,Selt,Serpine2,Sh2b1,Sla,Slc39a10,Sod1,Spag9,Tbl1xr1,Tnfrsf1a,Tnfsf11,Trim25,Wwox,Xbp1,Xdh |
| GO:0051726 | regulation of cell cycle | 30 | 954 | 0.0078 | Atp2b4,Ccne1,Cd28,Cdkn2b,Dusp1,E2f2,Ets1,Fanci,Fntb,Foxn3,Gadd45g,Gpr132,Ints7,Jun,Lef1,Map3k1,Map3k8,Map4k1,Mapk14,Nabp1,Osm,Pim1,Pkp4,Prkca,Prkcq,Rara,Sh2b1,Stat5b,Stk10,Tnfaip3 |
| GO:0016070 | RNA metabolic process | 66 | 2725 | 0.0085 | Adarb2,Aebp2,Aff4,Ago2,Bach2,Btg2,Celf2,Chd2,Chd7,Crem,Dach2,Ddx6,Dhx40,E2f2,Elf1,Esrp1,Ets1,Ets2,Fasl,Fli1,Fosl2,Foxn3,Foxp1,Gtf2ird1,Hey1,Hivep2,Ints7,Jarid2,Jun,Kdm2b,Klf2,Klf3,Klf6,Lef1,Mapk14,Max,Mbd2,Mbnl1,Nfatc1,Nfkbiz,Nipbl,Nrip1,Per1,Polg2,Rara,Riok1,Scml4,Ski,Smarca2,Smndc1,Spty2d1,Ss18,Stat4,Stat5b,Tars2,Tbl1xr1,Txnl4b,Vgll4,Vsx2,Xbp1,Zfp3,Zfp326,Zfp609,Zfp652,Zfp932,Zscan10 |
| GO:0023056 | positive regulation of signaling | 44 | 1615 | 0.0085 | Adipoq,Akt2,Bcl10,Ccr1,Ccr7,Cd28,Cdkn2b,Chsy1,Cyfip2,F2,Fasl,Gadd45g,Glrx,Gpr18,Il6st,Jak1,Jun,Map3k1,Map3k8,Map4k1,Mapk9,Mbd2,Ncf1,Osm,P2rx4,P2ry1,Pdcd4,Prkca,Prkch,Rara,Selt,Serpine2,Sh2b1,Sla,Slc39a10,Sod1,Spag9,Tbl1xr1,Tnfrsf1a,Tnfsf11,Trim25,Wwox,Xbp1,Xdh |
| GO:0006979 | response to oxidative stress | 16 | 377 | 0.0088 | Adipoq,Btg1,Ccr7,Cryaa,Dusp1,Ets1,Glrx2,Jun,Klf2,Mapk9,Msrb3,Ncf1,Prkca,Sod1,Tnfaip3,Xbp1 |
| GO:0044271 | cellular nitrogen compound biosynthetic process | 67 | 2781 | 0.0088 | Aebp2,Aff4,Ago2,Bach2,Btg2,Cers6,Chd2,Chd7,Cmas,Crem,Dach2,E2f2,Eif4e3,Elf1,Ets1,Ets2,Fasl,Fli1,Fosl2,Foxn3,Foxp1,Gtf2ird1,Gucy2c,Hbs1l,Hey1,Hivep2,Jarid2,Jun,Kdm2b,Klf2,Klf3,Klf6,Lef1,Mapk14,Max,Mbd2,Moxd1,Ncf1,Nfatc1,Nfkbiz,Nipbl,Nrip1,P2rx4,Pde11a,Pde4a,Per1,Polg2,Rara,Scml4,Sgms1,Ski,Smarca2,Spty2d1,Ss18,Stat4,Stat5b,Tars2,Tbl1xr1,Vgll4,Vsx2,Xbp1,Zfp3,Zfp326,Zfp609,Zfp652,Zfp932,Zscan10 |
| GO:0046649 | lymphocyte activation | 16 | 378 | 0.009 | Cblb,Ccr7,Cd28,Chd7,Dock10,Dpp4,Foxp1,Gadd45g,Gpr18,Hectd1,Itk,Lef1,Nfatc1,Stat5b,Tnfaip3,Tnfsf8 |
| GO:0046632 | alpha-beta T cell differentiation | 6 | 60 | 0.0091 | Foxp1,Gadd45g,Gpr18,Itk,Lef1,Tnfsf8 |
| GO:0050856 | regulation of T cell receptor signaling pathway | 5 | 39 | 0.0093 | Bcl10,Cblb,Ccr7,Cd300a,Elf1 |
| GO:0055082 | cellular chemical homeostasis | 23 | 663 | 0.0097 | Agtr1b,Atp1a1,Atp2b4,Ccr1,Clic4,F2,Fasl,Gpr18,Il6st,Inpp4b,P2rx4,P2ry1,Prkca,S1pr1,Selt,Slc35g1,Slc39a10,Sod1,Stim2,Thada,Tmem64,Xbp1,Xcr1 |
| GO:0002683 | negative regulation of immune system process | 17 | 421 | 0.01 | Adipoq,Bcl10,Btla,Cblb,Ccr1,Cd300a,Dusp10,Elf1,Gpr18,H2-Aa,Inpp4b,Rara,Rnf125,Stat5b,Tigit,Tnfaip3,Trib1 |
| GO:0042493 | response to drug | 29 | 926 | 0.01 | Abcb1b,Abhd2,Adipoq,Agtr1b,Atp1a1,Ccr7,Cryaa,Dad1,Dusp1,Ets1,Gatsl3,Glrx2,Jun,Klf2,Map4k1,Mapk14,Mapk9,P2rx4,Pde4a,Prkca,Rara,Rgs10,Slc38a1,Slc7a3,Sod1,Ss18,Tgfbr2,Tnfaip3,Xbp1 |

| | | | | | |
|---|---|---|---|---|---|
| GO:0002028 | regulation of sodium ion transport | 7 | 86 | 0.0101 | Akt2,Atp1a1,Glrx,Neto2,P2rx4,Per1,Serpine2 |
| GO:0030278 | regulation of ossification | 11 | 205 | 0.0102 | Ccr1,Chsy1,Hey1,Il6st,Mapk14,Nfatc1,Nipbl,S1pr1,Ski,Tmem64,Zfp932 |
| GO:0090304 | nucleic acid metabolic process | 74 | 3169 | 0.0102 | Adarb2,Aebp2,Aff4,Ago2,Bach2,Btg2,Ccne1,Celf2,Chd2,Chd7,Crem,Dach2,Ddx6,Dhx40,E2f2,Elf1,Ercc4,Esrp1,Ets1,Ets2,Fanci,Fasl,Fli1,Fosl2,Foxn3,Foxp1,Gtf2ird1,Hey1,Hivep2,Ints7,Jarid2,Jun,Kdm2b,Klf2,Klf3,Klf6,Lef1,Mapk14,Max,Mbd2,Mbnl1,Mgme1,Nabp1,Nfatc1,Nfkbiz,Nipbl,Nrip1,Nsmce1,Per1,Polg2,Rara,Riok1,Scml4,Ski,Smarca2,Smndc1,Spty2d1,Ss18,Stat4,Stat5b,Tars2,Tbl1xr1,Txnl4b,Ubr2,Usp3,Vgll4,Vsx2,Xbp1,Zfp3,Zfp326,Zfp609,Zfp652,Zfp932,Zscan10 |
| GO:0061515 | myeloid cell development | 6 | 62 | 0.0104 | Fli1,Foxp1,Klf2,Pip4k2a,Tnfsf11,Wasf2 |
| GO:0043506 | regulation of JUN kinase activity | 7 | 87 | 0.0106 | Ccr7,Dusp10,Map3k1,Map4k1,Pdcd4,Spag9,Tnfsf11 |
| GO:0072359 | circulatory system development | 28 | 889 | 0.011 | Agtr1b,Chd7,Clic4,Drc1,Ets1,Foxp1,Hectd1,Hey1,Itga4,Jun,Lef1,Mapk14,Mbd2,Nfatc1,Nipbl,Nrxn3,Otulin,Pdcd4,Prickle1,Prkca,Rara,S1pr1,Smarca2,Tgfbr2,Tnfrsf1a,Tnfrsf1b,Wasf2,Xbp1 |
| GO:1903054 | negative regulation of extracellular matrix organization | 3 | 9 | 0.011 | Dpp4,Tnfrsf1a,Tnfrsf1b |
| GO:0001933 | negative regulation of protein phosphorylation | 17 | 426 | 0.0111 | Adipoq,Cblb,Cd300a,Cdkn2b,Dusp1,Dusp10,Dusp2,Gadd45g,Jun,Lrtm2,Pdcd4,Per1,Prkca,Tnfaip3,Trib1,Xbp1,Xdh |
| GO:0042326 | negative regulation of phosphorylation | 18 | 466 | 0.0112 | Adipoq,Cblb,Cd300a,Cdkn2b,Dusp1,Dusp10,Dusp2,Gadd45g,Jun,Lrtm2,Pdcd4,Per1,Ppp2r5a,Prkca,Tnfaip3,Trib1,Xbp1,Xdh |
| GO:0030219 | megakaryocyte differentiation | 4 | 23 | 0.0114 | Cdkn2b,Fli1,Pip4k2a,Wasf2 |
| GO:0072358 | cardiovascular system development | 20 | 548 | 0.0114 | Agtr1b,Chd7,Clic4,Ets1,Foxp1,Hectd1,Hey1,Jun,Lef1,Mapk14,Nfatc1,Nrxn3,Otulin,Prickle1,Prkca,S1pr1,Smarca2,Tgfbr2,Wasf2,Xbp1 |
| GO:0042110 | T cell activation | 12 | 244 | 0.0118 | Cblb,Ccr7,Cd28,Chd7,Dpp4,Foxp1,Gadd45g,Gpr18,Itk,Lef1,Stat5b,Tnfsf8 |
| GO:0046635 | positive regulation of alpha-beta T cell activation | 6 | 64 | 0.0118 | Ccr7,Cd28,Nfkbiz,Prkcq,Rara,Tgfbr2 |
| GO:0060341 | regulation of cellular localization | 26 | 811 | 0.0129 | Adipoq,Akt2,Camk4,Ccne1,Cd247,Cd300a,Chd7,Cnst,Epb4.1,F2,Hectd1,Inpp4b,Leprotl1,Mapk14,Os9,P2rx4,Ppp1cc,Ppp2r5a,Prkcq,Sar1a,Sirt4,Tnfrsf1a,Trappc12,Ubac2,Vamp4,Xbp1 |
| GO:0071407 | cellular response to organic cyclic compound | 18 | 475 | 0.0135 | Abhd2,Adipoq,Ago2,Agtr1b,Akt2,Atp1a1,Atp2b4,Foxp1,Glrx,Klf2,Lef1,Mbd2,Nrip1,P2rx4,P2ry1,Pim1,Rara,Sod1 |
| GO:0030853 | negative regulation of granulocyte differentiation | 3 | 10 | 0.0137 | Adipoq,Rara,Trib1 |
| GO:0033554 | cellular response to stress | 38 | 1370 | 0.0137 | Btg2,Ccr7,Chd2,Dyrk2,Ercc4,Ets1,Fanci,Foxn3,Glrx2,Ints7,Jun,Klf2,Map3k1,Map3k8,Map4k1,Mapk14,Mapk9,Mgme1,Nabp1,Ncf1,Nipbl,Nsmce1,Os9,Polg2,Sirt4,Sod1,Spag9,St8sia1,Stk10,Tnfaip3,Tnfrsf1a,Tnfrsf1b,Tnfsf11,Trib1,Trim25,Usp3,Xbp1,Xylt1 |
| GO:0042542 | response to hydrogen peroxide | 8 | 120 | 0.0139 | Cryaa,Dusp1,Ets1,Glrx2,Jun,Klf2,Sod1,Tnfaip3 |
| GO:0050670 | regulation of lymphocyte proliferation | 11 | 215 | 0.014 | Btla,Cblb,Ccr7,Cd28,Cd300a,H2-Aa,Il6st,Prkcq,Slc39a10,Stat5b,Tgfbr2 |
| GO:0006139 | nucleobase-containing compound metabolic process | 83 | 3702 | 0.0144 | Adarb2,Aebp2,Aff4,Ago2,Bach2,Btg2,Ccne1,Celf2,Chd2,Chd7,Cmah,Cmas,Crem,Dach2,Ddx6,Dhx40,E2f2,Elf1,Ercc4,Esrp1,Ets1,Ets2,Fanci,Fasl,Fli1,Fosl2,Foxn3,Foxp1,Gtf2ird1,Gucy2c,Hey1,Hivep2,Ints7,Jarid2,Jun,Kdm2b,Klf2,Klf3,Klf6,Lef1,Mapk14,Max,Mbd2,Mbip,Mbnl1,Mgme1,Nabp1,Ncf1,Nfatc1,Nfkbiz,Nipbl,Nrip1,Nsmce1,Pde11a,Pde4a,Per1,Polg2,Rara,Riok1,Rpia,Scml4,Ski,Smarca2,Smndc1,Spty2d1,Ss18,Stat4,Stat5b,Tars2,Tbl1xr1,Txnl4b,Ubr2,Usp3,Vgll4,Vsx2,Xbp1,Xdh,Zfp3,Zfp326,Zfp609,Zfp652,Zfp932,Zscan10 |
| GO:0018105 | peptidyl-serine phosphorylation | 10 | 183 | 0.0145 | Akt2,Camk4,Map4k1,Mapk14,Mapk9,Mknk2,Prkca,Prkch,Prkcq,Tgfbr2 |
| GO:0071276 | cellular response to cadmium ion | 4 | 25 | 0.0145 | Jun,Mapk9,Ncf1,Sod1 |
| GO:0080135 | regulation of cellular response to stress | 21 | 603 | 0.0145 | Ccr7,Dusp10,Ercc4,Foxp1,Gadd45g,Glrx,Map3k1,Map4k1,Mapk9,Ncf1,Nsmce1,Os9,Pdcd4,Per1,Sod1,Spag9,Tnfsf11,Ubac2,Xbp1,Xdh,Xylt1 |
| GO:0035239 | tube morphogenesis | 23 | 690 | 0.0149 | Bcl10,Chd7,Clic4,Esrp1,Ets1,Hectd1,Hey1,Jun,Kdm2b,Lef1,Mapk14,Nfatc1,Nipbl,Nrxn3,Otulin,Prickle1,Prkca,Rara,S1pr1,Ski,Tgfbr2,Wasf2,Xbp1 |
| GO:0032387 | negative regulation of intracellular transport | 6 | 68 | 0.0152 | Adipoq,Cd300a,Os9,Sar1a,Sirt4,Ubac2 |
| GO:0060043 | regulation of cardiac muscle cell proliferation | 5 | 45 | 0.0152 | Foxp1,Jarid2,Mapk14,Pim1,Tgfbr2 |
| GO:0045936 | negative regulation of phosphate metabolic process | 20 | 567 | 0.0161 | Adipoq,Cblb,Cd300a,Cdkn2b,Cnst,Dusp1,Dusp10,Dusp2,Gadd45g,Jun,Lmtk3,Lrtm2,Pdcd4,Per1,Ppp2r5a,Prkca,Tnfaip3,Trib1,Xbp1,Xdh |
| GO:0051128 | regulation of cellular component organization | 57 | 2337 | 0.0161 | Adipoq,Akt2,Arhgap15,Ccr7,Cd28,Cd300a,Cd47,Cd97,Crbn,Crim1,Cryaa,Dnajc6,Dpp4,Dusp1,Epb4.1,Ercc4,Ets1,F2,Foxp1,Gpbar1,Il15ra,Ist1,Jarid2,Jun,Kdm2b,Lef1,Lrrn3,Lrtm2,Map3k1,Mapk14,Mapk9,Nipbl,Osm,P2ry1,Pip4k2a,Prkca,Prkch,Prkcq,S1pr1,Sar1a,Serpine2,Sh2b1,Sipa1l1,Sirt4,Ski,Smarca2,Spty2d1,Syne3,Tapt1,Tmeff2,Tnfrsf1a,Tnfrsf1b,Trappc12,Usp6nl,Wasf2,Xbp1,Xylt1 |
| GO:0034616 | response to laminar fluid shear stress | 3 | 11 | 0.0165 | Ets1,Klf2,Xbp1 |
| GO:0045086 | positive regulation of interleukin-2 biosynthetic process | 3 | 11 | 0.0165 | Cd28,Prkcq,Stat5b |
| GO:0009653 | anatomical structure morphogenesis | 52 | 2089 | 0.0174 | Ankrd11,Bcl10,Chd7,Chst11,Chsy1,Clic4,Cryaa,Cyfip2,Dock10,Dusp1,Dusp2,Esrp1,Ets1,Ets2,Ext1,Fam20a,Fasl,Fli1,Foxn3,Foxp1,Gnat1,Hectd1,Hey1,Itga4,Jun,Kdm2b,Klf2,Lef1,Map3k1,Mapk14,Mbnl1,Nfatc1,Nipbl,Nrxn3,Otulin,Pdcd4,Polg2,Prickle1,Prkca,Rara,S1pr1,Serpine2,Ski,Sod1,Ss18,Tgfbr2,Tmeff2,Tnfsf11,Vsx2,Wasf2,Wwox,Xbp1 |
| GO:0051480 | regulation of cytosolic calcium ion concentration | 13 | 295 | 0.0178 | Agtr1b,Ccr1,F2,Fasl,Gpr18,Il6st,P2rx4,P2ry1,S1pr1,Selt,Slc35g1,Tmem64,Xcr1 |
| GO:0031331 | positive regulation of cellular catabolic process | 14 | 333 | 0.0181 | Ago2,Akt2,Btg2,Lonrf1,Mapk9,Pip4k2a,Prickle1,Rnf125,Tnfaip3,Tnfrsf1b,Tnrc6b,Trib1,Xbp1,Zfand2a |
| GO:0042592 | homeostatic process | 40 | 1494 | 0.0181 | Adipoq,Agtr1b,Ankrd11,Atp1a1,Atp2b4,Ccne1,Ccr1,Ccr4,Ccr7,Cd7,Clic4,Dock10,Ercc4,F2,Fam20a,Fasl,Glrx,Glrx2,Gpr18,Il6st,Inpp4b,Klf2,Lrrc8d,P2rx4,P2ry1,Prkca,S1pr1,Selt,Slc30a4,Slc35g1,Slc39a10,Sod1,Stat5b,Stim2,Thada,Tmem64,Tnfaip3,Tnfsf11,Xbp1,Xcr1 |
| GO:0001944 | vasculature development | 19 | 533 | 0.0182 | Agtr1b,Chd7,Clic4,Ets1,Hectd1,Hey1,Jun,Lef1,Mapk14,Nfatc1,Nrxn3,Otulin,Prickle1,Prkca,S1pr1,Smarca2,Tgfbr2,Wasf2,Xbp1 |
| GO:0034614 | cellular response to reactive oxygen species | 8 | 127 | 0.0183 | Ets1,Glrx2,Jun,Klf2,Mapk9,Ncf1,Sod1,Tnfaip3 |
| GO:0045862 | positive regulation of proteolysis | 14 | 334 | 0.0183 | Bcl10,Cyfip2,Fasl,Ist1,Lonrf1,Mapk14,Mapk9,Prickle1,Rnf125,Tnfrsf1b,Trib1,Xbp1,Xdh,Zfand2a |
| GO:0002649 | regulation of tolerance induction to self antigen | 2 | 2 | 0.0186 | Ccr7,Tgfbr2 |
| GO:0046381 | CMP-N-acetylneuraminate metabolic process | 2 | 2 | 0.0186 | Cmah,Cmas |
| GO:0006954 | inflammatory response | 17 | 454 | 0.0192 | Agtr1b,Camk4,Ccr1,Ccr4,Ccr7,Cd300a,Chst2,F2,Jun,Ncf1,Nfkbiz,Prkcq,Sgms1,Stat5b,Tnfaip3,Tnfrsf1a,Tnfrsf1b |
| GO:0030206 | chondroitin sulfate biosynthetic process | 3 | 12 | 0.0195 | Chst11,Chsy1,Xylt1 |
| GO:0060213 | positive regulation of nuclear-transcribed mRNA poly(A) tail shortening | 3 | 12 | 0.0195 | Ago2,Btg2,Tnrc6b |
| GO:0014013 | regulation of gliogenesis | 8 | 129 | 0.0196 | Cdkn2b,Dusp10,F2,Il6st,Ppp1cc,Prkch,Serpine2,Ski |
| GO:0032673 | regulation of interleukin-4 production | 4 | 28 | 0.0196 | Cd28,Lef1,Prkcq,Rara |
| GO:0050927 | positive regulation of positive chemotaxis | 4 | 28 | 0.0196 | Akt2,Ccr4,Prkca,S1pr1 |
| GO:0090257 | regulation of muscle system process | 11 | 228 | 0.0198 | Atp1a1,Atp2b4,Foxp1,Gtf2ird1,Jarid2,Nfatc1,P2rx4,Prkca,Sod1,Tnfrsf1a,Tnfrsf1b |
| GO:0043409 | negative regulation of MAPK cascade | 9 | 161 | 0.02 | Adipoq,Cd300a,Dusp1,Dusp10,Dusp2,Pdcd4,Per1,Prkca,Xbp1 |
| GO:0001818 | negative regulation of cytokine production | 11 | 229 | 0.0203 | Adipoq,F2,Gpr18,Klf2,Lef1,Pdcd4,Rara,Rnf125,Tigit,Tnfaip3,Tnfrsf1a |
| GO:0042129 | regulation of T cell proliferation | 9 | 162 | 0.0207 | Btla,Cblb,Ccr7,Cd28,H2-Aa,Il6st,Prkcq,Stat5b,Tgfbr2 |
| GO:0005975 | carbohydrate metabolic process | 15 | 381 | 0.0215 | Adipoq,Akt2,Chst11,Chst2,Crem,Ext1,Ggta1,Il6st,Inpp4b,Ip6k1,Mapk14,Mogat1,Ppp1cc,Rpia,Slc3a2 |
| GO:0030162 | regulation of proteolysis | 23 | 716 | 0.0215 | Akt2,Bcl10,Crim1,Cryaa,Cyfip2,F2,Fasl,Ist1,Lef1,Lonrf1,Mapk14,Mapk9,Os9,Prickle1,Rnf125,Serpine2,Sirt4,Tnfrsf1b,Trib1,Ubac2,Xbp1,Xdh,Zfand2a |
| GO:0031667 | response to nutrient levels | 17 | 461 | 0.0215 | Adipoq,Agtr1b,Bcl10,Cdkn2b,Chsy1,Dad1,Gatsl3,Jun,Mbd2,P2rx4,Pim1,Rara,Sod1,Tbl1xr1,Tgfbr2,Trim25,Xbp1 |
| GO:0034110 | regulation of homotypic cell-cell adhesion | 4 | 29 | 0.0215 | Prkca,Prkcq,Serpine2,Tnfsf11 |
| GO:0002903 | negative regulation of B cell apoptotic process | 3 | 13 | 0.0227 | Bcl10,Foxp1,Slc39a10 |
| GO:0043086 | negative regulation of catalytic activity | 24 | 765 | 0.023 | Adipoq,Akt2,Atp2b4,Cblb,Cd300a,Cdkn2b,Cnst,Crim1,Cryaa,Dusp1,Dusp10,Dusp2,Gadd45g,Lef1,Lmtk3,Lrtm2,Pdcd4,Ppp2r5a,Prkca,Serpine2,Ski,Tnfaip3,Tnfrsf1b,Trib1 |
| GO:2000351 | regulation of endothelial cell apoptotic process | 5 | 51 | 0.023 | Fasl,Itga4,Pdcd4,Tnfaip3,Xbp1 |
| GO:0009896 | positive regulation of catabolic process | 15 | 388 | 0.0252 | Ago2,Akt2,Btg2,Cblb,Lonrf1,Mapk9,Pip4k2a,Prickle1,Rnf125,Tnfaip3,Tnfrsf1b,Tnrc6b,Trib1,Xbp1,Zfand2a |
| GO:0022408 | negative regulation of cell-cell adhesion | 9 | 168 | 0.0255 | Adipoq,Btla,Cblb,Cd300a,H2-Aa,Lef1,Serpine2,Tigit,Tnfaip3 |
| GO:0009719 | response to endogenous stimulus | 35 | 1289 | 0.0265 | Abhd2,Adipoq,Agtr1b,Akt2,Atp1a1,Atp2b4,Btg1,Btg2,Ccr7,Cryaa,Dusp1,Ets1,Fosl2,Foxp1,Gatsl3,Glrx,Jun,Klf2,Klf3,Lef1,Map3k1,Mbd2,Nrip1,P2rx4,P2ry1,Pdcd4,Prkca,Rara,Rgs10,Sod1,Stat5b,Tgfbr2,Ubr2,Wwox,Xbp1 |
| GO:0046686 | response to cadmium ion | 5 | 53 | 0.0265 | Cyb5a,Jun,Mapk9,Ncf1,Sod1 |
| GO:0000165 | MAPK cascade | 9 | 170 | 0.0272 | Ccr7,Map3k1,Map3k8,Map4k1,Mapk14,Mapk9,Spag9,Tnfsf11,Trib1 |
| GO:0034641 | cellular nitrogen compound metabolic process | 91 | 4247 | 0.0272 | Adarb2,Aebp2,Aff4,Ago2,Bach2,Btg2,Ccne1,Celf2,Cers6,Chd2,Chd7,Cmah,Cmas,Crem,Dach2,Ddx6,Dhx40,E2f2,Eif4e3,Elf1,Ercc4,Esrp1,Ets1,Ets2,Fanci,Fasl,Fli1,Fosl2,Foxn3,Foxp1,Gtf2ird1,Gucy2c,Hbs1l,Hey1,Hivep2,Ints7,Jarid2,Jun,Kdm2b,Klf2,Klf3,Klf6,Lef1,Mapk14,Max,Mbd2,Mbip,Mbnl1,Mgme1,Moxd1,Nabp1,Ncf1,Nfatc1,Nfkbiz,Nipbl,Nrip1,Nsmce1,P2rx4,Pde11a,Pde4a,Per1,Polg2,Rara,Riok1,Rpia,Scml4,Sgms1,Ski,Smarca2,Smndc1,Smox,Sod1,Spty2d1,Ss18,Stat4,Stat5b,Tars2,Tbl1xr1,Txnl4b,Ubr2,Usp3,Vgll4,Vsx2,Xbp1,Xdh,Zfp3,Zfp326,Zfp609,Zfp652,Zfp932,Zscan10 |
| GO:0043547 | positive regulation of GTPase activity | 14 | 353 | 0.0275 | Akt2,Arhgap15,Arhgap25,Ccr7,Dock10,Gdi2,Jun,Pkp4,Rasgrp2,Rgs10,S1pr1,Sipa1l1,Snx18,Usp6nl |
| GO:0003332 | negative regulation of extracellular matrix constituent secretion | 2 | 3 | 0.0282 | Tnfrsf1a,Tnfrsf1b |
| GO:0034405 | response to fluid shear stress | 4 | 32 | 0.0285 | Ets1,Klf2,P2rx4,Xbp1 |
| GO:0060251 | regulation of glial cell proliferation | 4 | 32 | 0.0285 | Cdkn2b,Ppp1cc,Prkch,Ski |
| GO:0046677 | response to antibiotic | 13 | 317 | 0.0292 | Adipoq,Ccr4,Cryaa,Dusp1,Ets1,Glrx2,Jak1,Jun,Klf2,Prkca,Rara,Sod1,Tnfaip3 |
| GO:1901800 | positive regulation of proteasomal protein catabolic process | 7 | 109 | 0.0293 | Lonrf1,Mapk9,Prickle1,Rnf125,Trib1,Xbp1,Zfand2a |

| | | | | | |
|---|---|---|---|---|---|
| GO:0030856 | regulation of epithelial cell differentiation | 8 | 140 | 0.0294 | Adipoq,Btg1,Foxp1,Hey1,Prkch,Stat5b,Tnfrsf1a,Xdh |
| GO:0030279 | negative regulation of ossification | 6 | 81 | 0.0301 | Ccr1,Chsy1,Nfatc1,Ski,Tmem64,Zfp932 |
| GO:0045732 | positive regulation of protein catabolic process | 10 | 208 | 0.0303 | Cblb,Lonrf1,Mapk9,Prickle1,Rnf125,Tnfaip3,Tnfrsf1b,Trib1,Xbp1,Zfand2a |
| GO:0046777 | protein autophosphorylation | 10 | 208 | 0.0303 | Camk4,Itk,Jak1,Map4k1,Mapk14,Mknk2,Pim1,Prkca,Sla,Stk10 |
| GO:0006725 | cellular aromatic compound metabolic process | 84 | 3879 | 0.0304 | Adarb2,Aebp2,Aff4,Ago2,Bach2,Btg2,Ccne1,Celf2,Chd2,Chd7,Cmah,Cmas,Crem,Dach2,Ddx6,Dhx40,E2f2,Elf1,Ercc4,Esrp1,Ets1,Ets2,Fanci,Fasl,Fli1,Fosl2,Foxn3,Foxp1,Gtf2ird1,Gucy2c,Hey1,Hivep2,Ints7,Jarid2,Jun,Kdm2b,Klf2,Klf3,Klf6,Lef1,Mapk14,Max,Mbd2,Mbip,Mbnl1,Mgme1,Moxd1,Nabp1,Ncf1,Nfatc1,Nfkbiz,Nipbl,Nrip1,Nsmce1,Pde11a,Pde4a,Per1,Polg2,Rara,Riok1,Rpia,Scml4,Ski,Smarca2,Smndc1,Spty2d1,Ss18,Stat4,Stat5b,Tars2,Tbl1xr1,Txnl4b,Ubr2,Usp3,Vgll4,Vsx2,Xbp1,Xdh,Zfp3,Zfp326,Zfp609,Zfp652,Zfp932,Zscan10 |
| GO:1903364 | positive regulation of cellular protein catabolic process | 8 | 141 | 0.0304 | Lonrf1,Mapk9,Prickle1,Rnf125,Tnfaip3,Trib1,Xbp1,Zfand2a |
| GO:0050855 | regulation of B cell receptor signaling pathway | 3 | 15 | 0.0305 | Cd300a,Prkch,Slc39a10 |
| GO:0007259 | receptor signaling pathway via JAK-STAT | 4 | 33 | 0.0307 | Il15ra,Jak1,Osm,Stat5b |
| GO:1900015 | regulation of cytokine production involved in inflammatory response | 4 | 33 | 0.0307 | F2,Mapk14,Pdcd4,Per1 |
| GO:1901698 | response to nitrogen compound | 28 | 971 | 0.0307 | Adipoq,Ago2,Agtr1b,Akt2,Atp2b4,Btg1,Btg2,Ccr7,Crem,Dusp1,Gatsl3,Jun,Klf2,Klf3,Mapk14,Os9,P2rx4,P2ry1,Per1,Rftn1,Rgs10,Slc3a2,Sod1,Stat5b,Tnfaip3,Trim25,Ubr2,Xbp1 |
| GO:0050921 | positive regulation of chemotaxis | 8 | 142 | 0.031 | Akt2,Ccr1,Ccr4,Ccr7,Mapk14,P2rx4,Prkca,S1pr1 |
| GO:0010467 | gene expression | 68 | 3013 | 0.0314 | Adarb2,Aebp2,Aff4,Ago2,Bach2,Btg2,Celf2,Chd2,Chd7,Crem,Cyfip2,Dach2,Dhx40,E2f2,Eif4e3,Elf1,Esrp1,Ets1,Ets2,Fasl,Fli1,Fosl2,Foxn3,Foxp1,Gtf2ird1,Hbs1l,Hey1,Hivep2,Ints7,Jarid2,Jun,Kdm2b,Klf2,Klf3,Klf6,Lef1,Mapk14,Max,Mbd2,Mbnl1,Nfatc1,Nfkbiz,Nipbl,Nrip1,Per1,Polg2,Rara,Riok1,Scml4,Ski,Smarca2,Smndc1,Spty2d1,Ss18,Stat4,Stat5b,Tars2,Tbl1xr1,Txnl4b,Vgll4,Vsx2,Xbp1,Zfp3,Zfp326,Zfp609,Zfp652,Zfp932,Zscan10 |
| GO:0046640 | regulation of alpha-beta T cell proliferation | 4 | 34 | 0.0332 | Btla,Cblb,Cd28,Tgfbr2 |
| GO:0032386 | regulation of intracellular transport | 14 | 364 | 0.0337 | Adipoq,Akt2,Camk4,Cd300a,Inpp4b,Mapk14,Os9,Ppp1cc,Prkcq,Sar1a,Sirt4,Tnfrsf1a,Ubac2,Xbp1 |
| GO:0045927 | positive regulation of growth | 12 | 286 | 0.0337 | Chd7,Cryaa,F2,Ist1,Kdm2b,Lef1,Mapk14,Nipbl,Pim1,Rftn1,Stat5b,Tgfbr2 |
| GO:0051093 | negative regulation of developmental process | 28 | 979 | 0.0337 | Adipoq,Ccr1,Cdkn2b,Ddx6,Dusp10,E2f2,F2,Fasl,Foxp1,Hey1,Il15ra,Inpp4b,Jarid2,Klf2,Lef1,Nfatc1,Pdcd4,Prickle1,Rara,Ski,Stat5b,Tgfbr2,Tmem64,Trib1,Xbp1,Xdh,Xylt1,Zfp932 |
| GO:0046850 | regulation of bone remodeling | 5 | 58 | 0.0352 | Inpp4b,Prkca,S1pr1,Tmem64,Tnfsf11 |
| GO:0061136 | regulation of proteasomal protein catabolic process | 9 | 179 | 0.0352 | Lonrf1,Mapk9,Os9,Prickle1,Rnf125,Trib1,Ubac2,Xbp1,Zfand2a |
| GO:1903828 | negative regulation of cellular protein localization | 7 | 114 | 0.0352 | Adipoq,Hectd1,Leprotl1,Os9,Ppp2r5a,Sirt4,Ubac2 |
| GO:0007399 | nervous system development | 52 | 2181 | 0.0353 | Abcb1b,Akt2,Atp2b4,Bcl10,Btg2,Ccr1,Ccr4,Cdkn2b,Chd7,Cyfip2,Ddx6,Dock10,Dusp10,Ets1,Ext1,F2,Foxp1,Gnat1,Hectd1,Hey1,Il15ra,Il6st,Ist1,Jun,Kdm2b,Lef1,Mapk9,Mbnl1,Mfsd2a,Nipbl,Nrxn3,Osm,P2ry1,Ppp1cc,Prickle1,Prkca,Prkch,Rara,S1pr1,Serpine2,Sipa1l1,Ski,Smarca2,Smarce1,Sod1,Spag9,Tgfbr2,Tmc1,Vsx2,Xbp1,Xylt1,Zfp609 |
| GO:0006952 | defense response | 30 | 1079 | 0.0356 | Agtr1b,Bcl10,Camk4,Ccr1,Ccr4,Ccr7,Cd300a,Chst2,Dpp4,F2,Gzmb,H2-Aa,Hist1h2be,Itk,Jak1,Jun,Ncf1,Nfkbiz,Otulin,Prkcq,Sgms1,Sla,Spaca4,Stat5b,Tnfaip3,Tnfrsf1a,Tnfrsf1b,Tnfsf8,Trim25,Vamp4 |
| GO:0006024 | glycosaminoglycan biosynthetic process | 4 | 35 | 0.0357 | Chst11,Chsy1,Ext1,Xylt1 |
| GO:0043087 | regulation of GTPase activity | 15 | 410 | 0.0374 | Akt2,Arhgap15,Arhgap25,Ccr7,Dock10,Gdi2,Jun,Pkp4,Rasgrp2,Rgs10,S1pr1,Sipa1l1,Snx18,Sod1,Usp6nl |
| GO:0065008 | regulation of biological quality | 75 | 3420 | 0.0374 | Adipoq,Agtr1b,Akt2,Ankrd11,Arhgap15,Atp10a,Atp1a1,Atp2b4,Bcl10,Ccne1,Ccr1,Ccr4,Ccr7,Cd7,Cd97,Chd7,Clic4,Cyp2r1,Dock10,Epb4.1,Ercc4,Ets1,F2,Fam20a,Fasl,Foxp1,Glrx,Glrx2,Gpr18,Il6st,Inpp4b,Ist1,Jun,Kcnh8,Klf2,Lrrc8d,Lrrn3,Lrtm2,Map3k1,Mapk14,Mapk9,Moxd1,Nrip1,Nrxn3,Os9,Osm,P2rx4,P2ry1,Pim1,Prkca,Prkcq,Rara,S1pr1,Selt,Serpine2,Sipa1l1,Sirt4,Slc30a4,Slc35g1,Slc39a10,Sod1,Stat5b,Stim2,Syne3,Tars2,Thada,Tmem123,Tmem64,Tnfaip3,Tnfrsf1b,Tnfsf11,Usp3,Xbp1,Xcr1,Xdh |
| GO:0014074 | response to purine-containing compound | 8 | 148 | 0.0378 | Adipoq,Crem,Dusp1,Jun,P2rx4,P2ry1,Per1,Sod1 |
| GO:0038165 | oncostatin-M-mediated signaling pathway | 2 | 4 | 0.0383 | Il6st,Osm |
| GO:0061384 | heart trabecula morphogenesis | 4 | 36 | 0.0383 | Chd7,Hey1,Nfatc1,S1pr1 |
| GO:1990086 | lens fiber cell apoptotic process | 2 | 4 | 0.0383 | E2f2,Tgfbr2 |
| GO:2000481 | positive regulation of cAMP-dependent protein kinase activity | 2 | 4 | 0.0383 | Adipoq,Atp2b4 |
| GO:0032436 | positive regulation of proteasomal ubiquitin-dependent protein catabolic process | 6 | 87 | 0.0387 | Lonrf1,Mapk9,Prickle1,Rnf125,Trib1,Zfand2a |
| GO:2000377 | regulation of reactive oxygen species metabolic process | 9 | 184 | 0.0401 | Akt2,Atp2b4,F2,Klf2,Mapk14,Mapk9,Sod1,Tgfbr2,Xdh |
| GO:0002697 | regulation of immune effector process | 13 | 335 | 0.0414 | Ccr7,Cd28,Cd300a,Dpp4,Dusp10,Foxp1,Mapk14,Nfkbiz,Rara,Rnf125,Stat5b,Tnfaip3,Xbp1 |
| GO:2000107 | negative regulation of leukocyte apoptotic process | 5 | 61 | 0.0414 | Bcl10,Ccr7,Foxp1,Prkcq,Slc39a10 |
| GO:0001654 | eye development | 14 | 376 | 0.042 | Atp2b4,Chd7,Clic4,Cryaa,Fasl,Gnat1,Jun,Kdm2b,Map3k1,Nipbl,Rara,Ski,Tgfbr2,Vsx2 |
| GO:0051098 | regulation of binding | 14 | 376 | 0.042 | Adipoq,Crbn,Epb4.1,Ercc4,Hey1,Itga4,Jun,Lef1,Mbd2,Nipbl,P2ry1,Rara,Ski,Spag9 |
| GO:0002250 | adaptive immune response | 11 | 260 | 0.0435 | Bcl10,Btla,Camk4,Cd247,Cd7,Gadd45g,Gzmb,Itk,Lef1,Rftn1,Rnf125 |
| GO:0001568 | blood vessel development | 17 | 504 | 0.0436 | Agtr1b,Chd7,Clic4,Ets1,Hectd1,Hey1,Jun,Lef1,Mapk14,Nrxn3,Otulin,Prickle1,Prkca,S1pr1,Tgfbr2,Wasf2,Xbp1 |
| GO:0002643 | regulation of tolerance induction | 3 | 18 | 0.0436 | Cblb,Ccr7,Tgfbr2 |
| GO:0009887 | animal organ morphogenesis | 27 | 956 | 0.0436 | Ankrd11,Chd7,Chst11,Chsy1,Cryaa,Esrp1,Fam20a,Fasl,Fli1,Foxn3,Gnat1,Hey1,Jun,Kdm2b,Lef1,Mapk14,Nfatc1,Nipbl,Pdcd4,Rara,S1pr1,Ski,Sod1,Tgfbr2,Tnfsf11,Vsx2,Wwox |
| GO:0031664 | regulation of lipopolysaccharide-mediated signaling pathway | 3 | 18 | 0.0436 | Prkca,Tnfaip3,Trib1 |
| GO:0035855 | megakaryocyte development | 3 | 18 | 0.0436 | Fli1,Pip4k2a,Wasf2 |
| GO:0090075 | relaxation of muscle | 3 | 18 | 0.0436 | P2rx4,P2ry1,Sod1 |
| GO:0060325 | face morphogenesis | 4 | 38 | 0.0437 | Ankrd11,Lef1,Nipbl,Ski |
| GO:1903053 | regulation of extracellular matrix organization | 4 | 38 | 0.0437 | Dpp4,Ets1,Tnfrsf1a,Tnfrsf1b |
| GO:0033157 | regulation of intracellular protein transport | 10 | 224 | 0.044 | Adipoq,Akt2,Camk4,Mapk14,Os9,Prkcq,Sirt4,Tnfrsf1a,Ubac2,Xbp1 |
| GO:0035295 | tube development | 26 | 912 | 0.0444 | Bcl10,Chd7,Clic4,Esrp1,Ets1,Foxp1,Hectd1,Hey1,Jun,Kdm2b,Klf2,Lef1,Mapk14,Nfatc1,Nipbl,Nrxn3,Otulin,Prickle1,Prkca,Rara,S1pr1,Serpine2,Ski,Tgfbr2,Wasf2,Xbp1 |
| GO:0071396 | cellular response to lipid | 16 | 464 | 0.045 | Abhd2,Agtr1b,Atp1a1,Foxp1,Glrx,Lef1,Map4k1,Mapk14,Nrip1,Pdcd4,Pim1,Rara,Sgms1,Tnfaip3,Tnfrsf1b,Xbp1 |
| GO:0031400 | negative regulation of protein modification process | 19 | 595 | 0.0458 | Adipoq,Cblb,Cd300a,Cdkn2b,Dusp1,Dusp10,Dusp2,Gadd45g,Jarid2,Jun,Lrtm2,Pdcd4,Per1,Prkca,Ski,Tnfaip3,Trib1,Xbp1,Xdh |
| GO:0070229 | negative regulation of lymphocyte apoptotic process | 4 | 39 | 0.0468 | Bcl10,Foxp1,Prkcq,Slc39a10 |
| GO:0001525 | angiogenesis | 12 | 303 | 0.0469 | Clic4,Ets1,Jun,Lef1,Mapk14,Nrxn3,Otulin,Prkca,S1pr1,Tgfbr2,Wasf2,Xbp1 |
| GO:0040008 | regulation of growth | 21 | 686 | 0.0469 | Chd7,Crim1,Cryaa,Dusp10,F2,Foxp1,Ist1,Jarid2,Kdm2b,Lef1,Mapk14,Nipbl,Osm,Pim1,Rftn1,Serpine2,Smarca2,Sod1,Stat5b,Tgfbr2,Xbp1 |
| GO:0003176 | aortic valve development | 3 | 19 | 0.0479 | Nfatc1,Tnfrsf1a,Tnfrsf1b |
| GO:0010038 | response to metal ion | 13 | 344 | 0.0479 | Clic4,Cryaa,Cyb5a,Cyp2r1,Dusp1,Jun,Mapk9,Ncf1,P2rx4,Rasgrp2,Slc30a4,Sod1,Xdh |
| GO:0045649 | regulation of macrophage differentiation | 3 | 19 | 0.0479 | Adipoq,Prkca,Trib1 |
| GO:1903409 | reactive oxygen species biosynthetic process | 3 | 19 | 0.0479 | Ncf1,P2rx4,Sod1 |
| GO:0032874 | positive regulation of stress-activated MAPK cascade | 8 | 157 | 0.0484 | Ccr7,Gadd45g,Map3k1,Map4k1,Ncf1,Spag9,Tnfsf11,Xdh |
| GO:0044691 | tooth eruption | 2 | 5 | 0.0485 | Fam20a,Tnfsf11 |
| GO:0090074 | negative regulation of protein homodimerization activity | 2 | 5 | 0.0485 | Itga4,Spag9 |
| GO:0002062 | chondrocyte differentiation | 5 | 65 | 0.0494 | Chst11,Chsy1,Mapk14,Prkca,Tgfbr2 |
| GO:0002768 | immune response-regulating cell surface receptor signaling pathway | 7 | 125 | 0.0496 | Bcl10,Btla,Cd247,Cd28,Itk,Nfkbiz,Rftn1 |
| GO:0009636 | response to toxic substance | 16 | 471 | 0.0496 | Adipoq,Agtr1b,Cryaa,Dusp1,Ets1,Glrx2,Gucy2c,Jun,Klf2,Prkca,Rara,Rgs10,Selt,Slc30a4,Sod1,Tnfaip3 |
| GO:0009725 | response to hormone | 24 | 829 | 0.0496 | Abhd2,Adipoq,Agtr1b,Akt2,Atp1a1,Btg1,Btg2,Ccr7,Cryaa,Dusp1,Ets1,Fosl2,Foxp1,Glrx,Jun,Lef1,Mbd2,Nrip1,Prkca,Rara,Rgs10,Stat5b,Tgfbr2,Xbp1 |
| GO:0030166 | proteoglycan biosynthetic process | 4 | 40 | 0.0496 | Chst11,Chsy1,Ext1,Xylt1 |
| GO:0031349 | positive regulation of defense response | 11 | 268 | 0.0496 | Ccr7,Cd28,Cd47,Ets1,Otulin,Pdcd4,Prkca,Rftn1,Stat5b,Tnfrsf1a,Tnfsf11 |
| GO:0045923 | positive regulation of fatty acid metabolic process | 4 | 40 | 0.0496 | Adipoq,Akt2,Elovl5,Mapk9 |
| GO:0050864 | regulation of B cell activation | 7 | 125 | 0.0496 | Btla,Cd28,Cd300a,Slc39a10,Stat5b,Tnfaip3,Xbp1 |
| GO:0060429 | epithelium development | 28 | 1020 | 0.0496 | Abcb1b,Bcl10,Chd7,Clic4,Cryaa,Esrp1,Fosl2,Foxp1,Hectd1,Hey1,Itga4,Jun,Kdm2b,Klf2,Lef1,Map3k1,Nfatc1,Prickle1,Rara,S1pr1,Serpine2,Ski,Sod1,Tgfbr2,Tmc1,Tmeff2,Tnfsf11,Xbp1 |
| GO:0061041 | regulation of wound healing | 7 | 125 | 0.0496 | Dusp10,F2,Prkca,Prkcq,Serpine2,Tgfbr2,Xbp1 |
| GO:1903034 | regulation of response to wounding | 8 | 158 | 0.0496 | Dusp10,F2,Prkca,Prkcq,Serpine2,Tgfbr2,Xbp1,Xylt1 |

**Supplementary Table S4. GO enrichment analysis of genes near openeing and closing peaks of CytP and Ags Cells**

Only GO terms with FDR value below 0.05 are shown. GO terms related to immune system and T cell activation are highlighted.

| #term ID | term description | observed gene count | background gene count | false discovery rate | matching proteins in the GO network of CytP and AgS cells |
|---|---|---|---|---|---|
| GO:0050794 | regulation of cellular process | 137 | 9045 | 3.55E-11 | Abhd2,Adipoq,Aebp2,Aff4,Ago2,Arhgap15,Arhgap25,Atp2b4,Bach2,Bcl10,Btg2,Camk4,Cblb,Ccr4,Ccr7,Cd247,Cd28,Cd47,Cdkn2b,Chd2,Cnst,Crbn,Crim1,Cyth4,Dad1,Ddx6,Dnajc6,Dock10,Dpp4,Dusp10,Dyrk2,Elf1,Elovl5,Epb4.1,Ercc4,Ets1,F2,Fam20a,Fasl,Foxn3,Foxp1,Fyb,Gdi2,Gnat1,Gpbar1,Gpr132,Gpr18,Gtf2ird1,Gucy2c,Gzmb,Hey1,Hivep2,Il15ra,Il6st,Ist1,Itk,Jak1,Kcnh8,Klf2,Klf3,Lef1,Lmtk3,Lonrf1,Lrrc8d,Lrtm2,Map3k1,Map3k8,Mapk9,Mbd2,Mbnl1,Mknk2,Nabp1,Ncf1,Neto2,Neurl1b,Nfatc1,Nfkbiz,Nipbl,Nrip1,Nsmce1,Otulin,P2rx4,Pdcd4,Pim1,Pip4k2a,Pkp4,Plcxd2,Ppp1cc,Ppp2r5a,Prkca,Prkcq,Prl5a1,Rara,Rasgrp2,Rftn1,Rgs10,Riok1,Rnf125,Rras2,Rtkn2,S1pr1,Selt,Sgms1,Sh2b1,Sirt4,Ski,Slc35b4,Slc39a10,Slc7a3,Smarce1,Sod1,Ss18,St8sia1,Stat5b,Stk10,Syne3,Tapt1,Tax1bp1,Tbl1xr1,Tgfbr2,Tmem64,Tnfaip3,Tnfrsf1a,Tnfrsf1b,Tnfsf8,Trim25,Ubac2,Usp6nl,Vamp4,Wasf2,Wwox,Xdh,Zfand2a,Zfp3,Zfp609,Zfp652,Zscan10 |
| GO:0065007 | biological regulation | 146 | 10168 | 5.17E-11 | Abhd2,Adipoq,Aebp2,Aff4,Ago2,Ankrd11,Arhgap15,Arhgap25,Atp10a,Atp2b4,Bach2,Bcl10,Btg2,Camk4,Cblb,Ccdc80,Ccr4,Ccr7,Cd247,Cd28,Cd47,Cdkn2b,Chd2,Cnst,Crbn,Crim1,Cyp2r1,Cyth4,Dad1,Ddx6,Dnajc6,Dock10,Dpp4,Dusp10,Dyrk2,Elf1,Elovl5,Epb4.1,Ercc4,Ets1,F2,Fam20a,Fasl,Foxn3,Foxp1,Fyb,Gdi2,Gnat1,Gpbar1,Gpr132,Gpr18,Gtf2ird1,Gucy2c,Gzmb,Hey1,Hivep2,Il15ra,Il6st,Ist1,Itk,Jak1,Kcnh8,Klf2,Klf3,Lef1,Leprotl1,Lmtk3,Lonrf1,Lpcat3,Lrrc8d,Lrtm2,Map3k1,Map3k8,Mapk9,Mbd2,Mbnl1,Mknk2,Mtmr12,Nabp1,Ncf1,Neto2,Neurl1b,Nfatc1,Nfkbiz,Nipbl,Nrip1,Nsmce1,Otulin,P2rx4,Pdcd4,Pim1,Pip4k2a,Pkp4,Plcxd2,Ppp1cc,Ppp2r5a,Prkca,Prkcq,Prl5a1,Rara,Rasgrp2,Rftn1,Rgs10,Riok1,Rnf125,Rras2,Rtkn2,S1pr1,Selt,Sgms1,Sh2b1,Sirt4,Ski,Slc35b4,Slc39a10,Slc7a3,Smarce1,Snx18,Sod1,Ss18,St8sia1,Stat5b,Stk10,Syne3,Tapt1,Tax1bp1,Tbl1xr1,Tgfbr2,Thada,Tmem64,Tnfaip3,Tnfrsf1a,Tnfrsf1b,Tnfsf8,Trim25,Ubac2,Usp6nl,Vamp4,Wasf2,Wwox,Xdh,Zfand2a,Zfp3,Zfp609,Zfp652,Zscan10 |
| GO:0050896 | response to stimulus | 112 | 6616 | 5.34E-11 | Abcb1b,Abhd2,Adipoq,Ago2,Arhgap15,Arhgap25,Atp2b4,Bcl10,Btg2,Camk4,Cblb,Ccdc80,Ccr4,Ccr7,Cd247,Cd28,Cd47,Cdkn2b,Chd2,Chst2,Cyb5a,Cyp2r1,Dad1,Dock10,Dpp4,Dusp10,Dyrk2,Ercc4,Ets1,F2,Fasl,Fkbp5,Foxn3,Foxp1,Fyb,Gdi2,Gnat1,Gpbar1,Gpr132,Gpr18,Gucy2c,Gzmb,Hey1,Hist1h2be,Hivep2,Il15ra,Il6st,Itk,Jak1,Kcnh8,Klf2,Klf3,Lef1,Lrrc8d,Lrtm2,Map3k1,Map3k8,Mapk9,Mbd2,Mgme1,Mknk2,Msrb3,Nabp1,Ncf1,Neurl1b,Nfatc1,Nfkbiz,Ninj2,Nipbl,Nrip1,Nsmce1,Otulin,P2rx4,Pdcd4,Pim1,Plcxd2,Ppp1cc,Ppp2r5a,Prkca,Prkcq,Rara,Rasgrp2,Rftn1,Rgs10,Rnf125,Rras2,Rtkn2,S1pr1,Selt,Sgms1,Sh2b1,Sirt4,Ski,Slc7a3,Sod1,Ss18,St8sia1,Stat5b,Stk10,Tbl1xr1,Tgfbr2,Tmem64,Tnfaip3,Tnfrsf1a,Tnfrsf1b,Tnfsf8,Trim25,Vamp4,Wasf2,Wwox,Xdh,Zfand2a |
| GO:0050789 | regulation of biological process | 140 | 9594 | 7.59E-11 | Abhd2,Adipoq,Aebp2,Aff4,Ago2,Arhgap15,Arhgap25,Atp2b4,Bach2,Bcl10,Btg2,Camk4,Cblb,Ccdc80,Ccr4,Ccr7,Cd247,Cd28,Cd47,Cdkn2b,Chd2,Cnst,Crbn,Crim1,Cyth4,Dad1,Ddx6,Dnajc6,Dock10,Dpp4,Dusp10,Dyrk2,Elf1,Elovl5,Epb4.1,Ercc4,Ets1,F2,Fam20a,Fasl,Foxn3,Foxp1,Fyb,Gdi2,Gnat1,Gpbar1,Gpr132,Gpr18,Gtf2ird1,Gucy2c,Gzmb,Hey1,Hivep2,Il15ra,Il6st,Ist1,Itk,Jak1,Kcnh8,Klf2,Klf3,Lef1,Leprotl1,Lmtk3,Lonrf1,Lpcat3,Lrrc8d,Lrtm2,Map3k1,Map3k8,Mapk9,Mbd2,Mbnl1,Mknk2,Nabp1,Ncf1,Neto2,Neurl1b,Nfatc1,Nfkbiz,Nipbl,Nrip1,Nsmce1,Otulin,P2rx4,Pdcd4,Pim1,Pip4k2a,Pkp4,Plcxd2,Ppp1cc,Ppp2r5a,Prkca,Prkcq,Prl5a1,Rara,Rasgrp2,Rftn1,Rgs10,Riok1,Rnf125,Rras2,Rtkn2,S1pr1,Selt,Sgms1,Sh2b1,Sirt4,Ski,Slc35b4,Slc39a10,Slc7a3,Smarce1,Sod1,Ss18,St8sia1,Stat5b,Stk10,Syne3,Tapt1,Tax1bp1,Tbl1xr1,Tgfbr2,Tmem64,Tnfaip3,Tnfrsf1a,Tnfrsf1b,Tnfsf8,Trim25,Ubac2,Usp6nl,Vamp4,Wasf2,Wwox,Xdh,Zfand2a,Zfp3,Zfp609,Zfp652,Zscan10 |
| GO:0051716 | cellular response to stimulus | 95 | 5142 | 1.00E-10 | Abhd2,Adipoq,Ago2,Arhgap15,Arhgap25,Atp2b4,Bcl10,Btg2,Camk4,Cblb,Ccr4,Ccr7,Cd247,Cd28,Cd47,Cdkn2b,Chd2,Dock10,Dyrk2,Ercc4,Ets1,F2,Fasl,Foxn3,Foxp1,Fyb,Gdi2,Gnat1,Gpbar1,Gpr132,Gpr18,Gucy2c,Gzmb,Hey1,Hivep2,Il15ra,Il6st,Itk,Jak1,Kcnh8,Klf2,Klf3,Lef1,Lrrc8d,Lrtm2,Map3k1,Map3k8,Mapk9,Mbd2,Mgme1,Mknk2,Nabp1,Ncf1,Neurl1b,Nfatc1,Nfkbiz,Nipbl,Nrip1,Nsmce1,Otulin,P2rx4,Pdcd4,Pim1,Plcxd2,Ppp2r5a,Prkca,Prkcq,Rara,Rasgrp2,Rftn1,Rgs10,Rnf125,Rras2,Rtkn2,S1pr1,Selt,Sgms1,Sh2b1,Sirt4,Ski,Sod1,Ss18,St8sia1,Stat5b,Stk10,Tgfbr2,Tmem64,Tnfaip3,Tnfrsf1a,Tnfrsf1b,Trim25,Vamp4,Wasf2,Wwox,Zfand2a |
| GO:0002376 | immune system process | 47 | 1703 | 5.03E-09 | Bcl10,Camk4,Cblb,Ccr4,Ccr7,Cd247,Cd28,Cd47,Cdkn2b,Chd2,Dock10,Dpp4,Ets1,F2,Fasl,Foxp1,Fyb,Gpr18,Gzmb,Hist1h2be,Itk,Jak1,Klf2,Lef1,Map3k8,Mknk2,Ncf1,Nfatc1,Nfkbiz,Otulin,Pip4k2a,Prkca,Prkcq,Rftn1,Rnf125,Rtkn2,S1pr1,Sod1,Stat5b,Tgfbr2,Tnfaip3,Tnfrsf1a,Tnfrsf1b,Tnfsf8,Trim25,Vamp4,Wasf2 |
| GO:0048518 | positive regulation of biological process | 91 | 5340 | 3.92E-08 | Adipoq,Ago2,Atp2b4,Bcl10,Btg2,Camk4,Cblb,Ccdc80,Ccr4,Ccr7,Cd247,Cd28,Cd47,Cdkn2b,Cnst,Dpp4,Dusp10,Dyrk2,Elf1,Elovl5,Epb4.1,Ets1,F2,Fam20a,Fasl,Foxn3,Foxp1,Gpr18,Hey1,Hivep2,Il15ra,Il6st,Ist1,Itk,Jak1,Klf2,Lef1,Lonrf1,Lrtm2,Map3k1,Map3k8,Mapk9,Mbd2,Ncf1,Nfatc1,Nfkbiz,Nipbl,Nrip1,Nsmce1,Otulin,P2rx4,Pdcd4,Pim1,Pip4k2a,Pkp4,Ppp1cc,Ppp2r5a,Prkca,Prkcq,Rara,Rftn1,Riok1,Rnf125,Rras2,Rtkn2,S1pr1,Selt,Sgms1,Sh2b1,Sirt4,Ski,Slc39a10,Sod1,Ss18,St8sia1,Stat5b,Stk10,Tapt1,Tbl1xr1,Tgfbr2,Tmem64,Tnfaip3,Tnfrsf1a,Tnfrsf1b,Tnfsf8,Trim25,Wasf2,Wwox,Xdh,Zfand2a,Zfp609 |
| GO:0031323 | regulation of cellular metabolic process | 88 | 5293 | 3.46E-07 | Adipoq,Aebp2,Aff4,Ago2,Atp2b4,Bach2,Bcl10,Btg2,Camk4,Cblb,Ccr7,Cd28,Cdkn2b,Chd2,Cnst,Crim1,Ddx6,Dusp10,Dyrk2,Elf1,Elovl5,Ercc4,Ets1,F2,Fam20a,Fasl,Foxn3,Foxp1,Gtf2ird1,Hey1,Hivep2,Il6st,Ist1,Kcnh8,Klf2,Klf3,Lef1,Lmtk3,Lonrf1,Lrtm2,Map3k1,Map3k8,Mapk9,Mbd2,Mbnl1,Mknk2,Ncf1,Nfatc1,Nfkbiz,Nipbl,Nrip1,Otulin,Pdcd4,Pim1,Pip4k2a,Ppp2r5a,Prkca,Prkcq,Rara,Riok1,Rnf125,S1pr1,Sh2b1,Sirt4,Ski,Slc35b4,Slc39a10,Smarce1,Sod1,Ss18,Stat5b,Stk10,Tax1bp1,Tbl1xr1,Tgfbr2,Tnfaip3,Tnfrsf1a,Tnfrsf1b,Tnfsf8,Trim25,Ubac2,Wwox,Xdh,Zfand2a,Zfp3,Zfp609,Zfp652,Zscan10 |
| GO:0050790 | regulation of catalytic activity | 48 | 2057 | 5.24E-07 | Adipoq,Arhgap15,Arhgap25,Atp2b4,Bcl10,Cblb,Ccr7,Cdkn2b,Cnst,Crim1,Dock10,Dusp10,F2,Fam20a,Fasl,Gdi2,Gnat1,Itk,Lef1,Lmtk3,Lrtm2,Map3k1,Map3k8,Mapk9,Mtmr12,Ncf1,Pdcd4,Pim1,Pkp4,Ppp2r5a,Prkca,Prkcq,Rara,Rasgrp2,Rgs10,S1pr1,Sirt4,Ski,Slc39a10,Snx18,Sod1,Stk10,Tgfbr2,Tmem64,Tnfaip3,Tnfrsf1b,Usp6nl,Xdh |
| GO:0022407 | regulation of cell-cell adhesion | 19 | 360 | 6.66E-07 | Adipoq,Bcl10,Cblb,Ccr7,Cd28,Cd47,Dpp4,Dusp10,Ets1,Il6st,Lef1,Nfkbiz,Prkca,Prkcq,Rara,Stat5b,Tgfbr2,Tnfaip3,Zfp609 |
| GO:0048583 | regulation of response to stimulus | 67 | 3552 | 6.66E-07 | Adipoq,Arhgap25,Atp2b4,Bcl10,Cblb,Ccr4,Ccr7,Cd247,Cd28,Cd47,Cdkn2b,Cyth4,Dpp4,Dusp10,Dyrk2,Elf1,Ercc4,Ets1,F2,Fasl,Foxp1,Gpr18,Gtf2ird1,Hey1,Il6st,Itk,Jak1,Lef1,Lrtm2,Map3k1,Map3k8,Mapk9,Mbd2,Ncf1,Neto2,Nfatc1,Nfkbiz,Nsmce1,Otulin,P2rx4,Pdcd4,Prkca,Prkcq,Prl5a1,Rara,Rftn1,Rgs10,Rnf125,S1pr1,Sgms1,Sh2b1,Ski,Slc39a10,Slc7a3,Sod1,Stat5b,Tbl1xr1,Tgfbr2,Tmem64,Tnfaip3,Tnfrsf1a,Tnfrsf1b,Tnfsf8,Trim25,Ubac2,Wwox,Xdh |
| GO:0065009 | regulation of molecular function | 61 | 3081 | 6.93E-07 | Adipoq,Arhgap15,Arhgap25,Atp2b4,Bcl10,Cblb,Ccr7,Cdkn2b,Cnst,Crbn,Crim1,Cyth4,Dock10,Dusp10,Epb4.1,Ercc4,F2,Fam20a,Fasl,Gdi2,Gnat1,Hey1,Itk,Lef1,Lmtk3,Lrtm2,Map3k1,Map3k8,Mapk9,Mbd2,Mtmr12,Ncf1,Neto2,Nipbl,Otulin,Pdcd4,Pim1,Pkp4,Ppp2r5a,Prkca,Prkcq,Prl5a1,Rara,Rasgrp2,Rgs10,S1pr1,Sirt4,Ski,Slc39a10,Snx18,Sod1,Stk10,Tax1bp1,Tgfbr2,Tmem64,Tnfaip3,Tnfrsf1b,Tnfsf8,Trim25,Usp6nl,Xdh |
| GO:0009987 | cellular process | 154 | 12459 | 7.06E-07 | Abcb1b,Abhd2,Adipoq,Aebp2,Aff4,Ago2,Arhgap15,Arhgap25,Atp10a,Atp2b4,Bach2,Bcl10,Btg2,Camk4,Cblb,Ccdc80,Ccr4,Ccr7,Cd247,Cd28,Cd47,Cdkn2b,Celf2,Cers6,Chd2,Chst2,Cmah,Cmas,Crbn,Cyb5a,Cyp2r1,Dad1,Dcaf7,Ddx6,Dhx40,Dnah17,Dnajc6,Dock10,Dpp4,Dusp10,Dyrk2,Elf1,Elovl5,Epb4.1,Ercc4,Ets1,F2,Fasl,Fkbp5,Foxn3,Foxp1,Fyb,Gdi2,Gnat1,Gpbar1,Gpr132,Gpr18,Gtf2ird1,Gucy2c,Gzmb,Hbs1l,Hey1,Hist1h2be,Hivep2,Il15ra,Il6st,Ist1,Itk,Jak1,Kcnh8,Klf2,Klf3,Lef1,Lmtk3,Lonrf1,Lpcat3,Lrrc8d,Lrtm2,Map3k1,Map3k8,Mapk9,Mbd2,Mbnl1,Mgme1,Mknk2,Msrb3,Myo1b,Nabp1,Ncf1,Neurl1b,Nfatc1,Nfkbiz,Nipbl,Nrip1,Nsmce1,Otulin,P2rx4,Pdcd4,Pim1,Pip4k2a,Pkp4,Plcxd2,Ppp1cc,Ppp2r5a,Prkca,Prkcq,Prrc2b,Rara,Rasgrp2,Rftn1,Rgs10,Riok1,Rnf125,Rpia,Rras2,Rtkn2,S1pr1,Selt,Sgms1,Sh2b1,Sirt4,Ski,Slc39a10,Smarce1,Smox,Snx18,Sod1,Ss18,St8sia1,Stat5b,Stk10,Syne3,Tapt1,Tax1bp1,Tbl1xr1,Tgfbr2,Thada,Tmem64,Tnfaip3,Tnfrsf1a,Tnfrsf1b,Tnfsf8,Trim25,Usp6nl,Vamp4,Wasf2,Wdyhv1,Wwox,Xdh,Zfand2a,Zfp3,Zfp609,Zfp652,Zscan10 |
| GO:0019222 | regulation of metabolic process | 90 | 5616 | 7.48E-07 | Adipoq,Aebp2,Aff4,Ago2,Atp2b4,Bach2,Bcl10,Btg2,Camk4,Cblb,Ccr7,Cd28,Cdkn2b,Chd2,Cnst,Crim1,Ddx6,Dusp10,Dyrk2,Elf1,Elovl5,Ercc4,Ets1,F2,Fam20a,Fasl,Foxn3,Foxp1,Gtf2ird1,Hey1,Hivep2,Il6st,Ist1,Kcnh8,Klf2,Klf3,Lef1,Lmtk3,Lonrf1,Lrtm2,Map3k1,Map3k8,Mapk9,Mbd2,Mbnl1,Mknk2,Ncf1,Nfatc1,Nfkbiz,Nipbl,Nrip1,Otulin,Pdcd4,Pim1,Pip4k2a,Ppp1cc,Ppp2r5a,Prkca,Prkcq,Rara,Riok1,Rnf125,S1pr1,Sgms1,Sh2b1,Sirt4,Ski,Slc35b4,Slc39a10,Smarce1,Sod1,Ss18,Stat5b,Stk10,Tax1bp1,Tbl1xr1,Tgfbr2,Tnfaip3,Tnfrsf1a,Tnfrsf1b,Tnfsf8,Trim25,Ubac2,Wwox,Xdh,Zfand2a,Zfp3,Zfp609,Zfp652,Zscan10 |
| GO:0007165 | signal transduction | 67 | 3594 | 7.49E-07 | Abhd2,Arhgap15,Arhgap25,Bcl10,Camk4,Cblb,Ccr4,Ccr7,Cd247,Cd28,Cd47,Dock10,Dyrk2,F2,Fasl,Fyb,Gdi2,Gnat1,Gpbar1,Gpr132,Gpr18,Gucy2c,Gzmb,Hey1,Hivep2,Il15ra,Il6st,Itk,Jak1,Kcnh8,Lef1,Lrrc8d,Lrtm2,Map3k1,Map3k8,Mapk9,Mknk2,Neurl1b,Nfatc1,Nfkbiz,Nsmce1,Otulin,P2rx4,Pdcd4,Pim1,Plcxd2,Ppp2r5a,Prkca,Prkcq,Rara,Rasgrp2,Rftn1,Rgs10,Rras2,Rtkn2,S1pr1,Sh2b1,Ski,Ss18,Stat5b,Stk10,Tgfbr2,Tmem64,Tnfrsf1a,Tnfrsf1b,Wasf2,Wwox |
| GO:0048522 | positive regulation of cellular process | 81 | 4827 | 8.63E-07 | Adipoq,Ago2,Atp2b4,Bcl10,Btg2,Camk4,Ccr7,Cd28,Cd47,Cdkn2b,Cnst,Dpp4,Dusp10,Dyrk2,Elf1,Elovl5,Ets1,F2,Fam20a,Fasl,Foxn3,Foxp1,Gpr18,Hey1,Hivep2,Il15ra,Il6st,Ist1,Jak1,Klf2,Lef1,Lonrf1,Lrtm2,Map3k1,Map3k8,Mapk9,Mbd2,Ncf1,Nfatc1,Nfkbiz,Nipbl,Nrip1,Nsmce1,P2rx4,Pdcd4,Pim1,Pip4k2a,Pkp4,Ppp1cc,Ppp2r5a,Prkca,Prkcq,Rara,Riok1,Rnf125,Rras2,Rtkn2,S1pr1,Selt,Sh2b1,Ski,Slc39a10,Sod1,Ss18,St8sia1,Stat5b,Stk10,Tapt1,Tbl1xr1,Tgfbr2,Tmem64,Tnfaip3,Tnfrsf1a,Tnfrsf1b,Tnfsf8,Trim25,Wasf2,Wwox,Xdh,Zfand2a,Zfp609 |
| GO:0080090 | regulation of primary metabolic process | 85 | 5184 | 8.63E-07 | Adipoq,Aebp2,Aff4,Ago2,Atp2b4,Bach2,Bcl10,Btg2,Camk4,Cblb,Ccr7,Cd28,Cdkn2b,Chd2,Crim1,Ddx6,Dusp10,Dyrk2,Elf1,Elovl5,Ercc4,Ets1,F2,Fam20a,Fasl,Foxn3,Foxp1,Gtf2ird1,Hey1,Hivep2,Il6st,Ist1,Kcnh8,Klf2,Klf3,Lef1,Lonrf1,Lrtm2,Map3k1,Map3k8,Mapk9,Mbd2,Mbnl1,Mknk2,Ncf1,Nfatc1,Nfkbiz,Nipbl,Nrip1,Otulin,Pdcd4,Pim1,Ppp2r5a,Prkca,Prkcq,Rara,Riok1,Rnf125,S1pr1,Sh2b1,Sirt4,Ski,Slc35b4,Slc39a10,Smarce1,Sod1,Ss18,Stat5b,Stk10,Tax1bp1,Tbl1xr1,Tgfbr2,Tnfaip3,Tnfrsf1a,Tnfrsf1b,Tnfsf8,Trim25,Ubac2,Wwox,Xdh,Zfand2a,Zfp3,Zfp609,Zfp652,Zscan10 |

| | | | | | |
|---|---|---|---|---|---|
| GO:1903039 | positive regulation of leukocyte cell-cell adhesion | 14 | 190 | 1.07E-06 | Bcl10,Ccr7,Cd28,Cd47,Dpp4,Dusp10,Ets1,Il6st,Nfkbiz,Prkcq,Rara,Stat5b,Tgfbr2,Zfp609 |
| GO:0022409 | positive regulation of cell-cell adhesion | 15 | 227 | 1.14E-06 | Bcl10,Ccr7,Cd28,Cd47,Dpp4,Dusp10,Ets1,Il6st,Lef1,Nfkbiz,Prkcq,Rara,Stat5b,Tgfbr2,Zfp609 |
| GO:0060255 | regulation of macromolecule metabolic process | 85 | 5246 | 1.24E-06 | Adipoq,Aebp2,Aff4,Ago2,Atp2b4,Bach2,Bcl10,Btg2,Camk4,Cblb,Ccr7,Cd28,Cdkn2b,Chd2,Crim1,Ddx6,Dusp10,Dyrk2,Elf1,Ercc4,Ets1,F2,Fam20a,Fasl,Foxn3,Foxp1,Gtf2ird1,Hey1,Hivep2,Il6st,Ist1,Kcnh8,Klf2,Klf3,Lef1,Lonrf1,Lrtm2,Map3k1,Map3k8,Mapk9,Mbd2,Mbnl1,Mknk2,Ncf1,Nfatc1,Nfkbiz,Nipbl,Nrip1,Otulin,Pdcd4,Pim1,Ppp1cc,Ppp2r5a,Prkca,Prkcq,Rara,Riok1,Rnf125,S1pr1,Sgms1,Sh2b1,Sirt4,Ski,Slc39a10,Smarce1,Sod1,Ss18,Stat5b,Stk10,Tax1bp1,Tbl1xr1,Tgfbr2,Tnfaip3,Tnfrsf1a,Tnfrsf1b,Tnfsf8,Trim25,Ubac2,Wwox,Xdh,Zfand2a,Zfp3,Zfp609,Zfp652,Zscan10 |
| GO:0023052 | signaling | 69 | 3859 | 1.56E-06 | Abhd2,Arhgap15,Arhgap25,Bcl10,Camk4,Cblb,Ccr4,Ccr7,Cd247,Cd28,Cd47,Dock10,Dyrk2,F2,Fasl,Fyb,Gdi2,Gnat1,Gpbar1,Gpr132,Gpr18,Gucy2c,Gzmb,Hey1,Hivep2,Il15ra,Il6st,Itk,Jak1,Kcnh8,Lef1,Lrrc8d,Lrtm2,Map3k1,Map3k8,Mapk9,Mknk2,Neurl1b,Nfatc1,Nfkbiz,Nsmce1,Otulin,P2rx4,Pdcd4,Pim1,Plcxd2,Ppp2r5a,Prkca,Prkcq,Rara,Rasgrp2,Rftn1,Rgs10,Rras2,Rtkn2,S1pr1,Selt,Sh2b1,Ski,Sod1,Ss18,Stat5b,Stk10,Tgfbr2,Tmem64,Tnfrsf1a,Tnfrsf1b,Wasf2,Wwox |
| GO:0007154 | cell communication | 70 | 3984 | 2.26E-06 | Abhd2,Arhgap15,Arhgap25,Bcl10,Camk4,Cblb,Ccr4,Ccr7,Cd247,Cd28,Cd47,Cdkn2b,Dock10,Dyrk2,F2,Fasl,Fyb,Gdi2,Gnat1,Gpbar1,Gpr132,Gpr18,Gucy2c,Gzmb,Hey1,Hivep2,Il15ra,Il6st,Itk,Jak1,Kcnh8,Lef1,Lrrc8d,Lrtm2,Map3k1,Map3k8,Mapk9,Mknk2,Neurl1b,Nfatc1,Nfkbiz,Nsmce1,Otulin,P2rx4,Pdcd4,Pim1,Plcxd2,Ppp2r5a,Prkca,Prkcq,Rara,Rasgrp2,Rftn1,Rgs10,Rras2,Rtkn2,S1pr1,Selt,Sh2b1,Ski,Sod1,Ss18,Stat5b,Stk10,Tgfbr2,Tmem64,Tnfrsf1a,Tnfrsf1b,Wasf2,Wwox |
| GO:0051171 | regulation of nitrogen compound metabolic process | 82 | 5052 | 2.29E-06 | Adipoq,Aebp2,Aff4,Ago2,Atp2b4,Bach2,Bcl10,Btg2,Camk4,Cblb,Ccr7,Cd28,Cdkn2b,Chd2,Crim1,Ddx6,Dusp10,Elf1,Ercc4,Ets1,F2,Fam20a,Fasl,Foxn3,Foxp1,Gtf2ird1,Hey1,Hivep2,Il6st,Ist1,Kcnh8,Klf2,Klf3,Lef1,Lonrf1,Lrtm2,Map3k1,Map3k8,Mapk9,Mbd2,Mbnl1,Mknk2,Ncf1,Nfatc1,Nfkbiz,Nipbl,Nrip1,Otulin,Pdcd4,Pim1,Ppp2r5a,Prkca,Prkcq,Rara,Riok1,Rnf125,S1pr1,Sh2b1,Sirt4,Ski,Slc39a10,Smarce1,Sod1,Ss18,Stat5b,Stk10,Tax1bp1,Tbl1xr1,Tgfbr2,Tnfaip3,Tnfrsf1a,Tnfrsf1b,Tnfsf8,Trim25,Ubac2,Wwox,Xdh,Zfand2a,Zfp3,Zfp609,Zfp652,Zscan10 |
| GO:0042127 | regulation of cell population proliferation | 39 | 1594 | 2.85E-06 | Adipoq,Btg2,Cblb,Ccr7,Cd28,Cd47,Cdkn2b,Dpp4,Dusp10,Ets1,F2,Fasl,Foxp1,Gucy2c,Hey1,Il6st,Itk,Jak1,Lef1,Mbd2,Nfatc1,Pdcd4,Pim1,Ppp1cc,Prkca,Prkcq,Rara,Rtkn2,S1pr1,Ski,Slc39a10,St8sia1,Stat5b,Tgfbr2,Tnfaip3,Tnfrsf1a,Tnfrsf1b,Xdh,Zfp609 |
| GO:0050870 | positive regulation of T cell activation | 13 | 180 | 3.40E-06 | Bcl10,Ccr7,Cd28,Cd47,Dpp4,Dusp10,Il6st,Nfkbiz,Prkcq,Rara,Stat5b,Tgfbr2,Zfp609 |
| GO:0051251 | positive regulation of lymphocyte activation | 15 | 253 | 3.40E-06 | Bcl10,Ccr7,Cd28,Cd47,Dpp4,Dusp10,Il15ra,Il6st,Nfkbiz,Prkcq,Rara,Slc39a10,Stat5b,Tgfbr2,Zfp609 |
| GO:0002682 | regulation of immune system process | 32 | 1165 | 4.59E-06 | Adipoq,Bcl10,Cblb,Ccr7,Cd247,Cd28,Cd47,Dpp4,Dusp10,Elf1,Ets1,Foxp1,Gpr18,Il15ra,Il6st,Itk,Lef1,Nfkbiz,Otulin,Pim1,Prkca,Prkcq,Rara,Rftn1,Rnf125,Slc39a10,Stat5b,Stk10,Tgfbr2,Tmem64,Tnfaip3,Zfp609 |
| GO:1903037 | regulation of leukocyte cell-cell adhesion | 15 | 264 | 5.17E-06 | Bcl10,Cblb,Ccr7,Cd28,Cd47,Dpp4,Dusp10,Ets1,Il6st,Nfkbiz,Prkcq,Rara,Stat5b,Tgfbr2,Zfp609 |
| GO:0002684 | positive regulation of immune system process | 25 | 771 | 7.89E-06 | Bcl10,Cblb,Ccr7,Cd247,Cd28,Cd47,Dpp4,Dusp10,Ets1,Foxp1,Il15ra,Il6st,Itk,Lef1,Nfkbiz,Otulin,Prkca,Prkcq,Rara,Rftn1,Slc39a10,Stat5b,Tgfbr2,Tmem64,Zfp609 |
| GO:0009893 | positive regulation of metabolic process | 59 | 3215 | 8.78E-06 | Adipoq,Ago2,Atp2b4,Bcl10,Btg2,Camk4,Cblb,Ccr7,Cd28,Dyrk2,Elf1,Elovl5,Ets1,F2,Fam20a,Fasl,Foxn3,Foxp1,Hey1,Hivep2,Il6st,Ist1,Klf2,Lef1,Lonrf1,Map3k1,Map3k8,Mapk9,Ncf1,Nfatc1,Nipbl,Nrip1,Pim1,Pip4k2a,Ppp2r5a,Prkca,Prkcq,Rara,Riok1,Rnf125,S1pr1,Sgms1,Sirt4,Ski,Slc39a10,Sod1,Ss18,Stat5b,Stk10,Tbl1xr1,Tgfbr2,Tnfaip3,Tnfrsf1a,Tnfrsf1b,Tnfsf8,Wwox,Xdh,Zfand2a,Zfp609 |
| GO:0009617 | response to bacterium | 21 | 566 | 1.06E-05 | Adipoq,Bcl10,Ccdc80,Ccr4,Ccr7,Cd47,Dusp10,F2,Fasl,Fkbp5,Foxp1,Hist1h2be,Ncf1,Pdcd4,Rara,Sgms1,Stat5b,Tnfaip3,Tnfrsf1a,Tnfrsf1b,Tnfsf8 |
| GO:0010604 | positive regulation of macromolecule metabolic process | 56 | 3004 | 1.21E-05 | Adipoq,Ago2,Atp2b4,Bcl10,Btg2,Camk4,Cblb,Ccr7,Cd28,Dyrk2,Elf1,Ets1,F2,Fam20a,Fasl,Foxn3,Foxp1,Hey1,Hivep2,Il6st,Ist1,Klf2,Lef1,Lonrf1,Map3k1,Map3k8,Mapk9,Ncf1,Nfatc1,Nipbl,Nrip1,Pim1,Ppp2r5a,Prkca,Prkcq,Rara,Riok1,Rnf125,S1pr1,Sgms1,Ski,Slc39a10,Sod1,Ss18,Stat5b,Stk10,Tbl1xr1,Tgfbr2,Tnfaip3,Tnfrsf1a,Tnfrsf1b,Tnfsf8,Wwox,Xdh,Zfand2a,Zfp609 |
| GO:0031325 | positive regulation of cellular metabolic process | 56 | 3004 | 1.21E-05 | Adipoq,Ago2,Atp2b4,Bcl10,Btg2,Camk4,Ccr7,Cd28,Dyrk2,Elf1,Elovl5,Ets1,F2,Fam20a,Fasl,Foxn3,Foxp1,Hey1,Hivep2,Il6st,Ist1,Klf2,Lef1,Lonrf1,Map3k1,Map3k8,Mapk9,Ncf1,Nfatc1,Nipbl,Nrip1,Pim1,Pip4k2a,Ppp2r5a,Prkca,Prkcq,Rara,Riok1,Rnf125,S1pr1,Ski,Slc39a10,Sod1,Ss18,Stat5b,Stk10,Tbl1xr1,Tgfbr2,Tnfaip3,Tnfrsf1a,Tnfrsf1b,Tnfsf8,Wwox,Xdh,Zfand2a,Zfp609 |
| GO:0051173 | positive regulation of nitrogen compound metabolic process | 54 | 2866 | 1.48E-05 | Adipoq,Ago2,Atp2b4,Bcl10,Btg2,Camk4,Cblb,Ccr7,Cd28,Elf1,Ets1,F2,Fam20a,Fasl,Foxn3,Foxp1,Hey1,Hivep2,Il6st,Ist1,Klf2,Lef1,Lonrf1,Map3k1,Map3k8,Mapk9,Ncf1,Nfatc1,Nipbl,Nrip1,Pim1,Ppp2r5a,Prkca,Prkcq,Rara,Riok1,Rnf125,S1pr1,Ski,Slc39a10,Sod1,Ss18,Stat5b,Stk10,Tbl1xr1,Tgfbr2,Tnfaip3,Tnfrsf1a,Tnfrsf1b,Tnfsf8,Wwox,Xdh,Zfand2a,Zfp609 |
| GO:0045785 | positive regulation of cell adhesion | 17 | 383 | 1.54E-05 | Bcl10,Ccdc80,Ccr7,Cd28,Cd47,Dpp4,Dusp10,Ets1,Il6st,Lef1,Nfkbiz,Prkca,Prkcq,Rara,Stat5b,Tgfbr2,Zfp609 |
| GO:0048519 | negative regulation of biological process | 76 | 4771 | 1.61E-05 | Abhd2,Adipoq,Aebp2,Ago2,Arhgap25,Atp2b4,Bach2,Bcl10,Btg2,Cblb,Ccr7,Cd28,Cdkn2b,Cnst,Crbn,Crim1,Dad1,Ddx6,Dpp4,Dusp10,Dyrk2,Elf1,Ercc4,Ets1,F2,Fasl,Foxn3,Foxp1,Gpr132,Gpr18,Hey1,Il15ra,Il6st,Kcnh8,Klf2,Klf3,Lef1,Leprotl1,Lmtk3,Lrtm2,Mbd2,Nabp1,Nfatc1,Nipbl,Nrip1,Otulin,P2rx4,Pdcd4,Pim1,Ppp2r5a,Prkca,Prkcq,Rara,Rgs10,Rnf125,S1pr1,Sgms1,Sirt4,Ski,Slc39a10,Smarce1,Sod1,Stat5b,Tax1bp1,Tbl1xr1,Tgfbr2,Tmem64,Tnfaip3,Tnfrsf1a,Tnfrsf1b,Trim25,Ubac2,Wasf2,Wwox,Xdh,Zscan10 |
| GO:0044093 | positive regulation of molecular function | 37 | 1593 | 1.66E-05 | Adipoq,Arhgap15,Arhgap25,Atp2b4,Bcl10,Ccr7,Crbn,Dock10,Epb4.1,F2,Fam20a,Fasl,Gdi2,Gnat1,Itk,Map3k1,Map3k8,Mapk9,Mbd2,Ncf1,Nipbl,Pim1,Pkp4,Prkcq,Rara,Rasgrp2,Rgs10,S1pr1,Ski,Slc39a10,Snx18,Sod1,Stk10,Tgfbr2,Trim25,Usp6nl,Xdh |
| GO:0048523 | negative regulation of cellular process | 71 | 4336 | 1.73E-05 | Abhd2,Adipoq,Aebp2,Ago2,Arhgap25,Atp2b4,Bach2,Bcl10,Btg2,Cblb,Ccr7,Cdkn2b,Cnst,Crbn,Crim1,Dad1,Ddx6,Dpp4,Dusp10,Dyrk2,Elf1,Ercc4,Ets1,F2,Fasl,Foxn3,Foxp1,Gpr132,Gpr18,Hey1,Il15ra,Il6st,Kcnh8,Klf2,Klf3,Lef1,Lmtk3,Lrtm2,Mbd2,Nabp1,Nfatc1,Nipbl,Nrip1,Pdcd4,Pim1,Ppp2r5a,Prkca,Prkcq,Rara,Rgs10,Rnf125,S1pr1,Sgms1,Sirt4,Ski,Slc39a10,Smarce1,Sod1,Stat5b,Tax1bp1,Tbl1xr1,Tgfbr2,Tmem64,Tnfaip3,Tnfrsf1a,Tnfrsf1b,Ubac2,Wasf2,Wwox,Xdh,Zscan10 |
| GO:0051249 | regulation of lymphocyte activation | 17 | 396 | 2.11E-05 | Bcl10,Cblb,Ccr7,Cd28,Cd47,Dpp4,Dusp10,Il15ra,Il6st,Nfkbiz,Prkcq,Rara,Slc39a10,Stat5b,Tgfbr2,Tnfaip3,Zfp609 |
| GO:0048584 | positive regulation of response to stimulus | 41 | 1922 | 2.67E-05 | Adipoq,Bcl10,Ccr4,Ccr7,Cd247,Cd28,Cd47,Cdkn2b,Ets1,F2,Fasl,Foxp1,Gpr18,Il6st,Itk,Jak1,Map3k1,Map3k8,Mapk9,Mbd2,Ncf1,Nfkbiz,Nsmce1,Otulin,P2rx4,Pdcd4,Prkca,Prkcq,Rara,Rftn1,S1pr1,Sh2b1,Slc39a10,Sod1,Stat5b,Tbl1xr1,Tgfbr2,Tnfrsf1a,Trim25,Wwox,Xdh |
| GO:0009889 | regulation of biosynthetic process | 62 | 3613 | 2.82E-05 | Adipoq,Aebp2,Aff4,Ago2,Atp2b4,Bach2,Bcl10,Btg2,Camk4,Ccr7,Cd28,Chd2,Ddx6,Dyrk2,Elf1,Elovl5,Ets1,F2,Fasl,Foxn3,Foxp1,Gtf2ird1,Hey1,Hivep2,Kcnh8,Klf2,Klf3,Lef1,Mapk9,Mbd2,Mknk2,Ncf1,Nfatc1,Nfkbiz,Nipbl,Nrip1,Otulin,Pdcd4,Pim1,Prkca,Prkcq,Rara,S1pr1,Sh2b1,Sirt4,Ski,Slc35b4,Smarce1,Sod1,Ss18,Stat5b,Tax1bp1,Tbl1xr1,Tnfaip3,Tnfrsf1a,Tnfsf8,Trim25,Wwox,Zfp3,Zfp609,Zfp652,Zscan10 |
| GO:0006955 | immune response | 26 | 914 | 3.23E-05 | Bcl10,Camk4,Cblb,Ccr4,Ccr7,Cd247,Cd28,Dock10,F2,Fasl,Foxp1,Fyb,Gzmb,Hist1h2be,Itk,Jak1,Lef1,Otulin,Rftn1,Rnf125,Tnfaip3,Tnfrsf1a,Tnfrsf1b,Tnfsf8,Trim25,Vamp4 |
| GO:0009605 | response to external stimulus | 39 | 1794 | 3.23E-05 | Adipoq,Bcl10,Btg2,Ccdc80,Ccr4,Ccr7,Cd47,Cdkn2b,Dad1,Dusp10,Ets1,F2,Fasl,Fkbp5,Foxp1,Gnat1,Hist1h2be,Lef1,Mapk9,Mbd2,Ncf1,P2rx4,Pdcd4,Pim1,Ppp1cc,Prkca,Prkcq,Rara,S1pr1,Sgms1,Sod1,Stat5b,Tbl1xr1,Tgfbr2,Tnfaip3,Tnfrsf1a,Tnfrsf1b,Tnfsf8,Trim25 |
| GO:0010468 | regulation of gene expression | 63 | 3720 | 3.23E-05 | Adipoq,Aebp2,Aff4,Ago2,Atp2b4,Bach2,Bcl10,Btg2,Camk4,Cd28,Chd2,Ddx6,Elf1,Ets1,F2,Fasl,Foxn3,Foxp1,Gtf2ird1,Hey1,Hivep2,Kcnh8,Klf2,Klf3,Lef1,Mapk9,Mbd2,Mbnl1,Mknk2,Ncf1,Nfatc1,Nfkbiz,Nipbl,Nrip1,Otulin,Pdcd4,Pim1,Ppp1cc,Prkca,Prkcq,Rara,Riok1,S1pr1,Sgms1,Sirt4,Ski,Smarce1,Ss18,Stat5b,Tax1bp1,Tbl1xr1,Tgfbr2,Tnfaip3,Tnfrsf1a,Tnfrsf1b,Tnfsf8,Trim25,Wwox,Xdh,Zfp3,Zfp609,Zfp652,Zscan10 |
| GO:0030155 | regulation of cell adhesion | 21 | 624 | 3.32E-05 | Adipoq,Bcl10,Cblb,Ccdc80,Ccr7,Cd28,Cd47,Dpp4,Dusp10,Ets1,Il6st,Lef1,Nfkbiz,Prkca,Prkcq,Rara,S1pr1,Stat5b,Tgfbr2,Tnfaip3,Zfp609 |
| GO:0032101 | regulation of response to external stimulus | 22 | 681 | 3.32E-05 | Adipoq,Ccr4,Ccr7,Cd28,Cd47,Dusp10,Ets1,F2,Foxp1,Gpr18,Otulin,P2rx4,Pdcd4,Prkca,Prkcq,Rnf125,S1pr1,Stat5b,Tgfbr2,Tnfaip3,Tnfrsf1a,Tnfrsf1b |
| GO:1902107 | positive regulation of leukocyte differentiation | 11 | 162 | 3.83E-05 | Ccr7,Dusp10,Il15ra,Lef1,Nfkbiz,Prkca,Rara,Stat5b,Tgfbr2,Tmem64,Zfp609 |
| GO:1903708 | positive regulation of hemopoiesis | 12 | 199 | 3.83E-05 | Ccr7,Dusp10,Ets1,Il15ra,Lef1,Nfkbiz,Prkca,Rara,Stat5b,Tgfbr2,Tmem64,Zfp609 |
| GO:1902105 | regulation of leukocyte differentiation | 14 | 284 | 4.19E-05 | Adipoq,Ccr7,Cd28,Dusp10,Foxp1,Il15ra,Lef1,Nfkbiz,Prkca,Rara,Stat5b,Tgfbr2,Tmem64,Zfp609 |
| GO:1903706 | regulation of hemopoiesis | 16 | 377 | 4.31E-05 | Adipoq,Ccr7,Cd28,Dusp10,Ets1,Foxp1,Il15ra,Lef1,Nfkbiz,Pim1,Prkca,Rara,Stat5b,Tgfbr2,Tmem64,Zfp609 |
| GO:0050863 | regulation of T cell activation | 14 | 287 | 4.54E-05 | Bcl10,Cblb,Ccr7,Cd28,Cd47,Dpp4,Dusp10,Il6st,Nfkbiz,Prkcq,Rara,Stat5b,Tgfbr2,Zfp609 |
| GO:0035556 | intracellular signal transduction | 31 | 1274 | 5.11E-05 | Bcl10,Camk4,Cblb,Ccr7,Dock10,Dyrk2,Gdi2,Gucy2c,Il15ra,Itk,Jak1,Kcnh8,Map3k1,Map3k8,Mapk9,Mknk2,Nfatc1,Nsmce1,Otulin,P2rx4,Prkca,Prkcq,Rasgrp2,Rras2,Ss18,Stat5b,Stk10,Tnfrsf1a,Tnfrsf1b,Wasf2,Wwox |
| GO:0030097 | hemopoiesis | 20 | 600 | 6.26E-05 | Camk4,Ccr7,Cd28,Cdkn2b,Chd2,Dock10,Foxp1,Gpr18,Itk,Klf2,Lef1,Mknk2,Nfatc1,Pip4k2a,Rtkn2,Stat5b,Tgfbr2,Tnfaip3,Tnfsf8,Wasf2 |
| GO:0050727 | regulation of inflammatory response | 14 | 301 | 7.38E-05 | Adipoq,Ccr7,Cd28,Cd47,Dusp10,Ets1,F2,Otulin,Pdcd4,Prkca,Stat5b,Tnfaip3,Tnfrsf1a,Tnfrsf1b |
| GO:2000112 | regulation of cellular macromolecule biosynthetic process | 57 | 3324 | 7.84E-05 | Adipoq,Aebp2,Aff4,Ago2,Atp2b4,Bach2,Bcl10,Btg2,Camk4,Ccr7,Cd28,Chd2,Ddx6,Dyrk2,Elf1,Ets1,Fasl,Foxn3,Foxp1,Gtf2ird1,Hey1,Hivep2,Kcnh8,Klf2,Klf3,Lef1,Mapk9,Mbd2,Mknk2,Ncf1,Nfatc1,Nfkbiz,Nipbl,Nrip1,Otulin,Pdcd4,Pim1,Prkca,Prkcq,Rara,S1pr1,Sh2b1,Ski,Smarce1,Ss18,Stat5b,Tax1bp1,Tbl1xr1,Tnfaip3,Tnfrsf1a,Tnfsf8,Trim25,Wwox,Zfp3,Zfp609,Zfp652,Zscan10 |

| | | | | | |
|---|---|---|---|---|---|
| GO:0043085 | positive regulation of catalytic activity | 30 | 1239 | 8.10E-05 | Adipoq,Arhgap15,Arhgap25,Atp2b4,Bcl10,Ccr7,Dock10,F2,Fam20a,Fasl,Gdi2,Gnat1,Itk,Map3k1,Map3k8,Mapk9,Ncf1,Pim1,Pkp4,Prkcq,Rasgrp2,Rgs10,S1pr1,Slc39a10,Snx18,Sod1,Stk10,Tgfbr2,Usp6nl,Xdh |
| GO:0070887 | cellular response to chemical stimulus | 44 | 2287 | 0.0001 | Abhd2,Adipoq,Ago2,Atp2b4,Ccr4,Ccr7,Cdkn2b,Ets1,Foxp1,Il6st,Jak1,Klf2,Klf3,Lef1,Lrtm2,Map3k1,Mapk9,Mbd2,Mknk2,Ncf1,Nrip1,P2rx4,Pdcd4,Pim1,Prkca,Prkcq,Rara,Rasgrp2,Rgs10,Rnf125,S1pr1,Selt,Sgms1,Sirt4,Sod1,Stat5b,Tgfbr2,Tnfaip3,Tnfrsf1a,Tnfrsf1b,Trim25,Vamp4,Wwox,Zfand2a |
| GO:0002520 | immune system development | 21 | 687 | 0.00012 | Camk4,Ccr4,Ccr7,Cd28,Cdkn2b,Chd2,Dock10,Foxp1,Gpr18,Itk,Klf2,Lef1,Mknk2,Nfatc1,Pip4k2a,Rtkn2,Stat5b,Tgfbr2,Tnfaip3,Tnfsf8,Wasf2 |
| GO:0044238 | primary metabolic process | 100 | 7426 | 0.00012 | Abhd2,Adipoq,Aebp2,Aff4,Ago2,Bach2,Btg2,Camk4,Cblb,Ccr7,Celf2,Cers6,Chd2,Chst2,Cmah,Cmas,Crbn,Cyb5a,Cyp2r1,Dad1,Dcaf7,Ddx6,Dhx40,Dnajc6,Dpp4,Dusp10,Dyrk2,Elf1,Elovl5,Ercc4,Ets1,F2,Fasl,Fkbp5,Foxn3,Foxp1,Gtf2ird1,Gucy2c,Gzmb,Hbs1l,Hey1,Hivep2,Il6st,Itk,Jak1,Kcnh8,Klf2,Klf3,Lef1,Lmtk3,Lonrf1,Lpcat3,Map3k1,Map3k8,Mapk9,Mbd2,Mbnl1,Mgme1,Mknk2,Msrb3,Nabp1,Ncf1,Neurl1b,Nfatc1,Nfkbiz,Nipbl,Nrip1,Nsmce1,Otulin,Pim1,Pip4k2a,Plcxd2,Ppp1cc,Ppp2r5a,Prkca,Prkcq,Prss39,Rara,Riok1,Rnf125,Rpia,Sgms1,Sirt4,Ski,Ss18,St8sia1,Stat5b,Stk10,Tbl1xr1,Tgfbr2,Tnfaip3,Tnfrsf1a,Trim25,Wdyhv1,Xdh,Xpnpep2,Zfp3,Zfp609,Zfp652,Zscan10 |
| GO:0050865 | regulation of cell activation | 18 | 520 | 0.00012 | Bcl10,Cblb,Ccr7,Cd28,Cd47,Dpp4,Dusp10,Il15ra,Il6st,Nfkbiz,Prkca,Prkcq,Rara,Slc39a10,Stat5b,Tgfbr2,Tnfaip3,Zfp609 |
| GO:0051247 | positive regulation of protein metabolic process | 34 | 1556 | 0.00013 | Adipoq,Atp2b4,Bcl10,Cblb,Ccr7,Cd28,F2,Fam20a,Fasl,Il6st,Ist1,Klf2,Lonrf1,Map3k1,Map3k8,Mapk9,Ncf1,Nipbl,Pim1,Ppp2r5a,Prkca,Prkcq,Rara,Rnf125,Slc39a10,Sod1,Stat5b,Stk10,Tgfbr2,Tnfaip3,Tnfrsf1a,Tnfrsf1b,Xdh,Zfand2a |
| GO:0051252 | regulation of RNA metabolic process | 54 | 3152 | 0.00015 | Adipoq,Aebp2,Aff4,Ago2,Atp2b4,Bach2,Bcl10,Btg2,Camk4,Cd28,Chd2,Elf1,Ets1,Fasl,Foxn3,Foxp1,Gtf2ird1,Hey1,Hivep2,Kcnh8,Klf2,Klf3,Lef1,Mapk9,Mbd2,Mbnl1,Ncf1,Nfatc1,Nfkbiz,Nipbl,Nrip1,Otulin,Pdcd4,Pim1,Prkcq,Rara,Riok1,S1pr1,Ski,Smarce1,Ss18,Stat5b,Tax1bp1,Tbl1xr1,Tnfaip3,Tnfrsf1a,Tnfrsf1b,Tnfsf8,Trim25,Wwox,Zfp3,Zfp609,Zfp652,Zscan10 |
| GO:0001817 | regulation of cytokine production | 19 | 592 | 0.00016 | Adipoq,Bcl10,Ccr7,Cd28,Elf1,F2,Foxp1,Gpr18,Klf2,Lef1,Mapk9,Pdcd4,Prkcq,Rara,Rnf125,Sod1,Stat5b,Tnfaip3,Tnfrsf1a |
| GO:0009966 | regulation of signal transduction | 50 | 2823 | 0.00016 | Adipoq,Arhgap25,Atp2b4,Bcl10,Cblb,Ccr7,Cd28,Cdkn2b,Cyth4,Dusp10,Dyrk2,Elf1,F2,Fasl,Foxp1,Gpr18,Hey1,Il6st,Jak1,Lef1,Lrtm2,Map3k1,Map3k8,Mapk9,Mbd2,Ncf1,Neto2,Otulin,P2rx4,Pdcd4,Prkca,Prkcq,Prl5a1,Rara,Rgs10,Rnf125,Sgms1,Sh2b1,Ski,Slc39a10,Slc7a3,Sod1,Tbl1xr1,Tmem64,Tnfaip3,Tnfrsf1a,Tnfsf8,Trim25,Wwox,Xdh |
| GO:0042221 | response to chemical | 58 | 3532 | 0.00019 | Abcb1b,Abhd2,Adipoq,Ago2,Atp2b4,Bcl10,Btg2,Ccr4,Ccr7,Cdkn2b,Cyb5a,Cyp2r1,Dad1,Dusp10,Ets1,Fasl,Foxp1,Gnat1,Gucy2c,Il6st,Jak1,Klf2,Klf3,Lef1,Lrtm2,Map3k1,Mapk9,Mbd2,Mknk2,Ncf1,Nrip1,P2rx4,Pdcd4,Pim1,Prkca,Prkcq,Rara,Rasgrp2,Rftn1,Rgs10,Rnf125,S1pr1,Selt,Sgms1,Sirt4,Slc7a3,Sod1,Ss18,Stat5b,Tgfbr2,Tnfaip3,Tnfrsf1a,Tnfrsf1b,Trim25,Vamp4,Wwox,Xdh,Zfand2a |
| GO:0051707 | response to other organism | 22 | 779 | 0.00019 | Adipoq,Bcl10,Ccdc80,Ccr4,Ccr7,Cd47,Dusp10,F2,Fasl,Fkbp5,Foxp1,Hist1h2be,Ncf1,Pdcd4,Rara,Sgms1,Stat5b,Tnfaip3,Tnfrsf1a,Tnfrsf1b,Tnfsf8,Trim25 |
| GO:0080134 | regulation of response to stress | 28 | 1167 | 0.00019 | Adipoq,Atp2b4,Ccr7,Cd28,Cd47,Dusp10,Ercc4,Ets1,F2,Foxp1,Map3k1,Mapk9,Ncf1,Nsmce1,Otulin,Pdcd4,Prkca,Prkcq,Rftn1,Rnf125,Sod1,Stat5b,Tgfbr2,Tnfaip3,Tnfrsf1a,Tnfrsf1b,Ubac2,Xdh |
| GO:0048869 | cellular developmental process | 58 | 3545 | 0.0002 | Abcb1b,Abhd2,Adipoq,Aff4,Ago2,Btg2,Camk4,Ccr4,Ccr7,Cd28,Cdkn2b,Chd2,Ddx6,Dock10,Dusp10,Elf1,Ets1,F2,Fasl,Foxn3,Foxp1,Gnat1,Gpr18,Hey1,Il15ra,Il6st,Ist1,Itk,Jak1,Klf2,Lef1,Map3k1,Mapk9,Mbnl1,Nfatc1,Nipbl,Pdcd4,Pip4k2a,Ppp1cc,Prkca,Prrc2b,Rara,S1pr1,Ski,Smarce1,Sod1,Ss18,Stat5b,Tapt1,Tbl1xr1,Tgfbr2,Tmem64,Tnfaip3,Tnfsf8,Wasf2,Wwox,Zfp609,Zscan10 |
| GO:0051246 | regulation of protein metabolic process | 47 | 2612 | 0.0002 | Adipoq,Ago2,Atp2b4,Bcl10,Btg2,Cblb,Ccr7,Cd28,Cdkn2b,Crim1,Ddx6,Dusp10,F2,Fam20a,Fasl,Il6st,Ist1,Klf2,Lef1,Lonrf1,Lrtm2,Map3k1,Map3k8,Mapk9,Mknk2,Ncf1,Nipbl,Pdcd4,Pim1,Ppp2r5a,Prkca,Prkcq,Rara,Rnf125,Sirt4,Ski,Slc39a10,Sod1,Stat5b,Stk10,Tgfbr2,Tnfaip3,Tnfrsf1a,Tnfrsf1b,Ubac2,Xdh,Zfand2a |
| GO:0071704 | organic substance metabolic process | 102 | 7733 | 0.0002 | Abhd2,Adipoq,Aebp2,Aff4,Ago2,Bach2,Btg2,Camk4,Cblb,Ccr7,Celf2,Cers6,Chd2,Chst2,Cmah,Cmas,Crbn,Cyb5a,Cyp2r1,Dad1,Dcaf7,Ddx6,Dhx40,Dnajc6,Dpp4,Dusp10,Dyrk2,Elf1,Elovl5,Ercc4,Ets1,F2,Fasl,Fkbp5,Foxn3,Foxp1,Gtf2ird1,Gucy2c,Gzmb,Hbs1l,Hey1,Hivep2,Il6st,Itk,Jak1,Kcnh8,Klf2,Klf3,Lef1,Lmtk3,Lonrf1,Lpcat3,Map3k1,Map3k8,Mapk9,Mbd2,Mbnl1,Mgme1,Mknk2,Msrb3,Nabp1,Ncf1,Neurl1b,Nfatc1,Nfkbiz,Nipbl,Nrip1,Nsmce1,Otulin,Pim1,Pip4k2a,Plcxd2,Ppp1cc,Ppp2r5a,Prkca,Prkcq,Prss39,Rara,Riok1,Rnf125,Rpia,Sgms1,Sirt4,Ski,Smox,Sod1,Ss18,St8sia1,Stat5b,Stk10,Tbl1xr1,Tgfbr2,Tnfaip3,Tnfrsf1a,Trim25,Wdyhv1,Xdh,Xpnpep2,Zfp3,Zfp609,Zfp652,Zscan10 |
| GO:0006355 | regulation of transcription, DNA-templated | 51 | 2952 | 0.00021 | Adipoq,Aebp2,Aff4,Ago2,Atp2b4,Bach2,Bcl10,Btg2,Camk4,Cd28,Chd2,Elf1,Ets1,Fasl,Foxn3,Foxp1,Gtf2ird1,Hey1,Hivep2,Kcnh8,Klf2,Klf3,Lef1,Mapk9,Mbd2,Ncf1,Nfatc1,Nfkbiz,Nipbl,Nrip1,Otulin,Pdcd4,Pim1,Prkcq,Rara,S1pr1,Ski,Smarce1,Ss18,Stat5b,Tax1bp1,Tbl1xr1,Tnfaip3,Tnfrsf1a,Tnfsf8,Trim25,Wwox,Zfp3,Zfp609,Zfp652,Zscan10 |
| GO:0008152 | metabolic process | 106 | 8164 | 0.00021 | Abhd2,Adipoq,Aebp2,Aff4,Ago2,Bach2,Btg2,Camk4,Cblb,Ccr7,Celf2,Cers6,Chd2,Chst2,Cmah,Cmas,Crbn,Cyb5a,Cyp2r1,Dad1,Dcaf7,Ddx6,Dhx40,Dip2b,Dnajc6,Dpp4,Dusp10,Dyrk2,Elf1,Elovl5,Ercc4,Ets1,F2,Fasl,Fkbp5,Foxn3,Foxp1,Gtf2ird1,Gucy2c,Gzmb,Hbs1l,Hey1,Hivep2,Il6st,Itk,Jak1,Kcnh8,Klf2,Klf3,Lef1,Lmtk3,Lonrf1,Lpcat3,Map3k1,Map3k8,Mapk9,Mbd2,Mbnl1,Mgme1,Mknk2,Msrb3,Nabp1,Ncf1,Neurl1b,Nfatc1,Nfkbiz,Nipbl,Nrip1,Nsmce1,Otulin,P2rx4,Pim1,Pip4k2a,Plcxd2,Ppp1cc,Ppp2r5a,Prkca,Prkcq,Prss39,Rara,Riok1,Rnf125,Rpia,Selt,Sgms1,Sirt4,Ski,Smox,Sod1,Ss18,St8sia1,Stat5b,Stk10,Tbl1xr1,Tgfbr2,Tnfaip3,Tnfrsf1a,Trim25,Wdyhv1,Wwox,Xdh,Xpnpep2,Zfp3,Zfp609,Zfp652,Zscan10 |
| GO:0030154 | cell differentiation | 57 | 3462 | 0.00021 | Abcb1b,Abhd2,Adipoq,Aff4,Ago2,Btg2,Camk4,Ccr4,Ccr7,Cd28,Cdkn2b,Chd2,Ddx6,Dock10,Dusp10,Elf1,Ets1,F2,Fasl,Foxn3,Foxp1,Gnat1,Gpr18,Hey1,Il15ra,Il6st,Ist1,Itk,Jak1,Klf2,Lef1,Map3k1,Mapk9,Mbnl1,Nfatc1,Nipbl,Pdcd4,Pip4k2a,Ppp1cc,Prkca,Prrc2b,Rara,S1pr1,Ski,Smarce1,Sod1,Stat5b,Tapt1,Tbl1xr1,Tgfbr2,Tmem64,Tnfaip3,Tnfsf8,Wasf2,Wwox,Zfp609,Zscan10 |
| GO:0031326 | regulation of cellular biosynthetic process | 58 | 3549 | 0.00021 | Adipoq,Aebp2,Aff4,Ago2,Atp2b4,Bach2,Bcl10,Btg2,Camk4,Ccr7,Cd28,Chd2,Ddx6,Dyrk2,Elf1,Elovl5,Ets1,Fasl,Foxn3,Foxp1,Gtf2ird1,Hey1,Hivep2,Kcnh8,Klf2,Klf3,Lef1,Mapk9,Mbd2,Mknk2,Ncf1,Nfatc1,Nfkbiz,Nipbl,Nrip1,Otulin,Pdcd4,Pim1,Prkca,Prkcq,Rara,S1pr1,Sh2b1,Ski,Smarce1,Ss18,Stat5b,Tax1bp1,Tbl1xr1,Tnfaip3,Tnfrsf1a,Tnfsf8,Trim25,Wwox,Zfp3,Zfp609,Zfp652,Zscan10 |
| GO:0050728 | negative regulation of inflammatory response | 9 | 129 | 0.00021 | Adipoq,Dusp10,Ets1,F2,Otulin,Pdcd4,Tnfaip3,Tnfrsf1a,Tnfrsf1b |
| GO:0051241 | negative regulation of multicellular organismal process | 28 | 1191 | 0.00023 | Adipoq,Atp2b4,Cdkn2b,Ddx6,Dusp10,F2,Fasl,Foxp1,Gpr18,Hey1,Il15ra,Klf2,Lef1,Nfatc1,P2rx4,Pdcd4,Prkca,Rara,Rnf125,Ski,Sod1,Stat5b,Tgfbr2,Tmem64,Tnfaip3,Tnfrsf1a,Tnfrsf1b,Xdh |
| GO:0019219 | regulation of nucleobase-containing compound metabolic process | 56 | 3407 | 0.00024 | Adipoq,Aebp2,Aff4,Ago2,Atp2b4,Bach2,Bcl10,Btg2,Camk4,Cd28,Chd2,Elf1,Ercc4,Ets1,Fasl,Foxn3,Foxp1,Gtf2ird1,Hey1,Hivep2,Kcnh8,Klf2,Klf3,Lef1,Mapk9,Mbd2,Mbnl1,Ncf1,Nfatc1,Nfkbiz,Nipbl,Nrip1,Otulin,Pdcd4,Pim1,Prkcq,Rara,Riok1,S1pr1,Sh2b1,Ski,Smarce1,Ss18,Stat5b,Tax1bp1,Tbl1xr1,Tnfaip3,Tnfrsf1a,Tnfrsf1b,Tnfsf8,Trim25,Wwox,Zfp3,Zfp609,Zfp652,Zscan10 |
| GO:0032502 | developmental process | 76 | 5213 | 0.00024 | Abcb1b,Abhd2,Adipoq,Aff4,Ago2,Ankrd11,Atp2b4,Bcl10,Btg2,Camk4,Ccr4,Ccr7,Cd28,Cdkn2b,Chd2,Dad1,Dcaf7,Ddx6,Dock10,Dusp10,Elf1,Ets1,F2,Fam20a,Fasl,Foxn3,Foxp1,Gnat1,Gpr18,Gtf2ird1,Hey1,Hivep2,Il15ra,Il6st,Ist1,Itk,Jak1,Klf2,Lef1,Map3k1,Mapk9,Mbd2,Mbnl1,Mknk2,Nfatc1,Ninj2,Nipbl,Nrip1,Otulin,Pdcd4,Pip4k2a,Ppp1cc,Prkca,Prrc2b,Rara,Rtkn2,S1pr1,Selt,Ski,Smarce1,Sod1,Ss18,Stat5b,Tapt1,Tbl1xr1,Tgfbr2,Tmem64,Tnfaip3,Tnfrsf1a,Tnfrsf1b,Tnfsf8,Wasf2,Wwox,Xdh,Zfp609,Zscan10 |
| GO:0043502 | regulation of muscle adaptation | 8 | 99 | 0.00024 | Atp2b4,Foxp1,Gtf2ird1,Nfatc1,P2rx4,Prkca,Tnfrsf1a,Tnfrsf1b |
| GO:0006915 | apoptotic process | 22 | 809 | 0.00028 | Bcl10,Cd28,Dad1,Dyrk2,Fasl,Gzmb,Lef1,Map3k1,Mapk9,Mknk2,Ncf1,P2rx4,Pdcd4,Pim1,Prkca,Sgms1,Tax1bp1,Tgfbr2,Tnfaip3,Tnfrsf1a,Tnfrsf1b,Wwox |
| GO:0045621 | positive regulation of lymphocyte differentiation | 8 | 103 | 0.00031 | Ccr7,Dusp10,Il15ra,Nfkbiz,Rara,Stat5b,Tgfbr2,Zfp609 |
| GO:0045321 | leukocyte activation | 16 | 464 | 0.00032 | Bcl10,Camk4,Cblb,Ccr7,Cd28,Dock10,Dpp4,Foxp1,Gpr18,Itk,Lef1,Nfatc1,Stat5b,Tgfbr2,Tnfaip3,Tnfsf8 |
| GO:2000026 | regulation of multicellular organismal development | 38 | 1961 | 0.00033 | Adipoq,Ago2,Cblb,Ccr7,Cd28,Cdkn2b,Ddx6,Dusp10,Ets1,F2,Fasl,Foxp1,Hey1,Il15ra,Il6st,Ist1,Jak1,Klf2,Lef1,Lrtm2,Nfatc1,Nfkbiz,Nipbl,Pdcd4,Pim1,Ppp1cc,Prkca,Rara,S1pr1,Ski,Stat5b,Tapt1,Tgfbr2,Tmem64,Tnfaip3,Tnfrsf1a,Xdh,Zfp609 |
| GO:0044237 | cellular metabolic process | 97 | 7348 | 0.00035 | Abhd2,Adipoq,Aebp2,Aff4,Ago2,Bach2,Btg2,Camk4,Cblb,Ccr7,Celf2,Cers6,Chd2,Chst2,Cmah,Cmas,Crbn,Cyb5a,Cyp2r1,Dad1,Dcaf7,Ddx6,Dhx40,Dnajc6,Dusp10,Dyrk2,Elf1,Elovl5,Ercc4,Ets1,Fasl,Fkbp5,Foxn3,Foxp1,Gtf2ird1,Gucy2c,Hbs1l,Hey1,Hivep2,Il6st,Itk,Jak1,Kcnh8,Klf2,Klf3,Lef1,Lmtk3,Lonrf1,Lpcat3,Map3k1,Map3k8,Mapk9,Mbd2,Mbnl1,Mgme1,Mknk2,Msrb3,Nabp1,Ncf1,Neurl1b,Nfatc1,Nfkbiz,Nipbl,Nrip1,Nsmce1,Otulin,P2rx4,Pim1,Pip4k2a,Ppp1cc,Ppp2r5a,Prkca,Prkcq,Rara,Riok1,Rnf125,Rpia,Sgms1,Sirt4,Ski,Smox,Sod1,Ss18,St8sia1,Stat5b,Stk10,Tbl1xr1,Tgfbr2,Tnfaip3,Tnfrsf1a,Trim25,Wdyhv1,Xdh,Zfp3,Zfp609,Zfp652,Zscan10 |
| GO:0051239 | regulation of multicellular organismal process | 49 | 2858 | 0.00035 | Adipoq,Ago2,Atp2b4,Bcl10,Cblb,Ccr7,Cd28,Cdkn2b,Ddx6,Dusp10,Elf1,Ets1,F2,Fasl,Foxp1,Gpr18,Gtf2ird1,Hey1,Il15ra,Il6st,Ist1,Jak1,Klf2,Lef1,Lrtm2,Mapk9,Nfatc1,Nfkbiz,Nipbl,P2rx4,Pdcd4,Pim1,Ppp1cc,Prkca,Prkcq,Rara,Rnf125,S1pr1,Ski,Sod1,Stat5b,Tapt1,Tgfbr2,Tmem64,Tnfaip3,Tnfrsf1a,Tnfrsf1b,Xdh,Zfp609 |
| GO:0010646 | regulation of cell communication | 52 | 3130 | 0.0004 | Adipoq,Arhgap25,Atp2b4,Bcl10,Cblb,Ccr7,Cd28,Cdkn2b,Cyth4,Dusp10,Dyrk2,Elf1,F2,Fasl,Foxp1,Gpr18,Hey1,Il6st,Jak1,Lef1,Lrtm2,Map3k1,Map3k8,Mapk9,Mbd2,Ncf1,Neto2,Otulin,P2rx4,Pdcd4,Prkca,Prkcq,Prl5a1,Rara,Rgs10,Rnf125,Selt,Sgms1,Sh2b1,Sirt4,Ski,Slc39a10,Slc7a3,Sod1,Tbl1xr1,Tmem64,Tnfaip3,Tnfrsf1a,Tnfsf8,Trim25,Wwox,Xdh |
| GO:0010614 | negative regulation of cardiac muscle hypertrophy | 5 | 28 | 0.00041 | Atp2b4,Foxp1,P2rx4,Tnfrsf1a,Tnfrsf1b |

| | | | | | |
|---|---|---|---|---|---|
| GO:0006807 | nitrogen compound metabolic process | 93 | 6983 | 0.00044 | Aebp2,Aff4,Ago2,Bach2,Btg2,Camk4,Cblb,Ccr7,Celf2,Cers6,Chd2,Cmah,Cmas,Crbn,Dad1,Dcaf7,Ddx6,Dhx40,Dnajc6,Dpp4,Dusp10,Dyrk2,Elf1,Elovl5,Ercc4,Ets1,F2,Fasl,Fkbp5,Foxn3,Foxp1,Gtf2ird1,Gucy2c,Gzmb,Hbs1l,Hey1,Hivep2,Itk,Jak1,Kcnh8,Klf2,Klf3,Lef1,Lmtk3,Lonrf1,Map3k1,Map3k8,Mapk9,Mbd2,Mbnl1,Mgme1,Mknk2,Msrb3,Nabp1,Ncf1,Neurl1b,Nfatc1,Nfkbiz,Nipbl,Nrip1,Nsmce1,Otulin,P2rx4,Pim1,Ppp1cc,Ppp2r5a,Prkca,Prkcq,Prss39,Rara,Riok1,Rnf125,Rpia,Sgms1,Sirt4,Ski,Smox,Sod1,Ss18,St8sia1,Stat5b,Stk10,Tbl1xr1,Tgfbr2,Tnfaip3,Trim25,Wdyhv1,Xdh,Xpnpep2,Zfp3,Zfp609,Zfp652,Zscan10 |
| GO:0031348 | negative regulation of defense response | 10 | 186 | 0.00045 | Adipoq,Dusp10,Ets1,F2,Otulin,Pdcd4,Rnf125,Tnfaip3,Tnfrsf1a,Tnfrsf1b |
| GO:0050793 | regulation of developmental process | 44 | 2475 | 0.00045 | Adipoq,Ago2,Arhgap15,Cblb,Ccr7,Cd28,Cdkn2b,Ddx6,Dusp10,Epb4.1,Ets1,F2,Fasl,Foxp1,Hey1,Il15ra,Il6st,Ist1,Jak1,Klf2,Lef1,Lrtm2,Mapk9,Nfatc1,Nfkbiz,Nipbl,Pdcd4,Pim1,Ppp1cc,Prkca,Rara,S1pr1,Ski,Sod1,Stat5b,Syne3,Tapt1,Tgfbr2,Tmem64,Tnfaip3,Tnfrsf1a,Tnfrsf1b,Xdh,Zfp609 |
| GO:0006950 | response to stress | 49 | 2899 | 0.00046 | Abhd2,Adipoq,Atp2b4,Bcl10,Btg2,Camk4,Ccr4,Ccr7,Chd2,Chst2,Dpp4,Dyrk2,Ercc4,Ets1,F2,Foxn3,Gzmb,Hist1h2be,Itk,Jak1,Klf2,Map3k1,Map3k8,Mapk9,Mgme1,Msrb3,Nabp1,Ncf1,Nfkbiz,Ninj2,Nipbl,Nsmce1,Otulin,P2rx4,Prkca,Prkcq,Sgms1,Sirt4,Sod1,St8sia1,Stat5b,Stk10,Tgfbr2,Tnfaip3,Tnfrsf1a,Tnfrsf1b,Tnfsf8,Trim25,Vamp4 |
| GO:0023051 | regulation of signaling | 52 | 3150 | 0.00046 | Adipoq,Arhgap25,Atp2b4,Bcl10,Cblb,Ccr7,Cd28,Cdkn2b,Cyth4,Dusp10,Dyrk2,Elf1,F2,Fasl,Foxp1,Gpr18,Hey1,Il6st,Jak1,Lef1,Lrtm2,Map3k1,Map3k8,Mapk9,Mbd2,Ncf1,Neto2,Otulin,P2rx4,Pdcd4,Prkca,Prkcq,Prl5a1,Rara,Rgs10,Rnf125,Selt,Sgms1,Sh2b1,Sirt4,Ski,Slc39a10,Slc7a3,Sod1,Tbl1xr1,Tmem64,Tnfaip3,Tnfrsf1a,Tnfsf8,Trim25,Wwox,Xdh |
| GO:0031347 | regulation of defense response | 17 | 538 | 0.00046 | Adipoq,Ccr7,Cd28,Cd47,Dusp10,Ets1,F2,Foxp1,Otulin,Pdcd4,Prkca,Rftn1,Rnf125,Stat5b,Tnfaip3,Tnfrsf1a,Tnfrsf1b |
| GO:0043549 | regulation of kinase activity | 21 | 780 | 0.00046 | Adipoq,Atp2b4,Cblb,Ccr7,Cdkn2b,Dusp10,F2,Fam20a,Lrtm2,Map3k1,Map3k8,Ncf1,Pdcd4,Pim1,Ppp2r5a,Prkca,Rara,Sod1,Stk10,Tgfbr2,Tnfaip3 |
| GO:0051240 | positive regulation of multicellular organismal process | 34 | 1695 | 0.00047 | Adipoq,Ago2,Bcl10,Cblb,Ccr7,Cd28,Dusp10,Ets1,F2,Foxp1,Hey1,Il15ra,Il6st,Ist1,Jak1,Lef1,Lrtm2,Mapk9,Nfkbiz,Nipbl,P2rx4,Pim1,Ppp1cc,Prkca,Prkcq,Rara,Sod1,Stat5b,Tapt1,Tgfbr2,Tmem64,Tnfaip3,Tnfrsf1a,Zfp609 |
| GO:0051336 | regulation of hydrolase activity | 26 | 1128 | 0.00056 | Arhgap15,Arhgap25,Bcl10,Ccr7,Cnst,Crim1,Dock10,Fasl,Gdi2,Gnat1,Itk,Lef1,Lmtk3,Mapk9,Pkp4,Ppp2r5a,Rasgrp2,Rgs10,S1pr1,Slc39a10,Snx18,Sod1,Tmem64,Tnfrsf1b,Usp6nl,Xdh |
| GO:0001775 | cell activation | 17 | 552 | 0.00058 | Bcl10,Camk4,Cblb,Ccr7,Cd28,Dock10,Dpp4,F2,Foxp1,Gpr18,Itk,Lef1,Nfatc1,Stat5b,Tgfbr2,Tnfaip3,Tnfsf8 |
| GO:0048731 | system development | 62 | 4072 | 0.00058 | Abcb1b,Adipoq,Ankrd11,Atp2b4,Bcl10,Btg2,Camk4,Ccr4,Ccr7,Cd28,Cdkn2b,Chd2,Ddx6,Dock10,Dusp10,Ets1,F2,Fam20a,Fasl,Foxn3,Foxp1,Gnat1,Gpr18,Hey1,Il15ra,Il6st,Ist1,Itk,Klf2,Lef1,Map3k1,Mapk9,Mbd2,Mbnl1,Mknk2,Nfatc1,Nipbl,Nrip1,Otulin,Pdcd4,Pip4k2a,Ppp1cc,Prkca,Rara,Rtkn2,S1pr1,Selt,Ski,Smarce1,Sod1,Stat5b,Tapt1,Tbl1xr1,Tgfbr2,Tnfaip3,Tnfrsf1a,Tnfrsf1b,Tnfsf8,Wasf2,Wwox,Xdh,Zfp609 |
| GO:0050729 | positive regulation of inflammatory response | 8 | 116 | 0.00058 | Ccr7,Cd28,Cd47,Ets1,Pdcd4,Prkca,Stat5b,Tnfrsf1a |
| GO:0048585 | negative regulation of response to stimulus | 31 | 1496 | 0.00061 | Adipoq,Arhgap25,Atp2b4,Cblb,Dusp10,Dyrk2,Elf1,Ercc4,Ets1,F2,Foxp1,Gpr18,Hey1,Il6st,Lef1,Lrtm2,Otulin,Pdcd4,Prkca,Prkcq,Rgs10,Rnf125,Sgms1,Ski,Tmem64,Tnfaip3,Tnfrsf1a,Tnfrsf1b,Ubac2,Wwox,Xdh |
| GO:0048856 | anatomical structure development | 71 | 4912 | 0.00061 | Abcb1b,Abhd2,Adipoq,Aff4,Ago2,Ankrd11,Atp2b4,Bcl10,Btg2,Camk4,Ccr4,Ccr7,Cd28,Cdkn2b,Chd2,Dad1,Dcaf7,Ddx6,Dock10,Dusp10,Ets1,F2,Fam20a,Fasl,Foxn3,Foxp1,Gnat1,Gpr18,Gtf2ird1,Hey1,Hivep2,Il15ra,Il6st,Ist1,Itk,Klf2,Lef1,Map3k1,Mapk9,Mbd2,Mbnl1,Mknk2,Nfatc1,Ninj2,Nipbl,Nrip1,Otulin,Pdcd4,Pip4k2a,Ppp1cc,Prkca,Rara,Rtkn2,S1pr1,Selt,Ski,Smarce1,Sod1,Ss18,Stat5b,Tapt1,Tbl1xr1,Tgfbr2,Tnfaip3,Tnfrsf1a,Tnfrsf1b,Tnfsf8,Wasf2,Wwox,Xdh,Zfp609 |
| GO:0051094 | positive regulation of developmental process | 30 | 1423 | 0.00061 | Adipoq,Ago2,Cblb,Ccr7,Cd28,Dusp10,Ets1,Foxp1,Hey1,Il15ra,Il6st,Ist1,Jak1,Lef1,Lrtm2,Mapk9,Nfkbiz,Nipbl,Pim1,Ppp1cc,Prkca,Rara,Stat5b,Tapt1,Tgfbr2,Tmem64,Tnfaip3,Tnfrsf1a,Tnfrsf1b,Zfp609 |
| GO:0002521 | leukocyte differentiation | 13 | 340 | 0.00064 | Camk4,Ccr7,Cd28,Dock10,Foxp1,Gpr18,Itk,Lef1,Nfatc1,Stat5b,Tgfbr2,Tnfaip3,Tnfsf8 |
| GO:0030098 | lymphocyte differentiation | 11 | 243 | 0.00064 | Ccr7,Cd28,Dock10,Foxp1,Gpr18,Itk,Lef1,Nfatc1,Stat5b,Tnfaip3,Tnfsf8 |
| GO:0032268 | regulation of cellular protein metabolic process | 43 | 2443 | 0.00064 | Adipoq,Ago2,Atp2b4,Bcl10,Btg2,Cblb,Ccr7,Cdkn2b,Crim1,Ddx6,Dusp10,F2,Fam20a,Fasl,Il6st,Ist1,Lef1,Lonrf1,Lrtm2,Map3k1,Map3k8,Mapk9,Mknk2,Ncf1,Nipbl,Pdcd4,Pim1,Ppp2r5a,Prkca,Rara,Rnf125,Sirt4,Ski,Slc39a10,Sod1,Stk10,Tgfbr2,Tnfaip3,Tnfrsf1a,Tnfrsf1b,Ubac2,Xdh,Zfand2a |
| GO:0050852 | T cell receptor signaling pathway | 6 | 56 | 0.00064 | Bcl10,Cd247,Cd28,Itk,Nfkbiz,Rftn1 |
| GO:0051338 | regulation of transferase activity | 22 | 873 | 0.00064 | Adipoq,Atp2b4,Cblb,Ccr7,Cdkn2b,Dusp10,F2,Fam20a,Lrtm2,Map3k1,Map3k8,Ncf1,Pdcd4,Pim1,Ppp2r5a,Prkca,Prkcq,Rara,Sod1,Stk10,Tgfbr2,Tnfaip3 |
| GO:1901700 | response to oxygen-containing compound | 30 | 1429 | 0.00064 | Abhd2,Adipoq,Atp2b4,Btg2,Ccr7,Dusp10,Ets1,Fasl,Foxp1,Klf2,Klf3,Mapk9,Mbd2,Ncf1,Nrip1,P2rx4,Pdcd4,Pim1,Prkca,Rara,Rgs10,Selt,Sgms1,Sod1,Stat5b,Tgfbr2,Tnfaip3,Tnfrsf1a,Tnfrsf1b,Trim25 |
| GO:0044260 | cellular macromolecule metabolic process | 74 | 5233 | 0.00073 | Aebp2,Aff4,Ago2,Bach2,Btg2,Camk4,Cblb,Ccr7,Chd2,Crbn,Dad1,Dcaf7,Dnajc6,Dusp10,Dyrk2,Elf1,Ercc4,Ets1,Fasl,Fkbp5,Foxn3,Foxp1,Gtf2ird1,Gucy2c,Hbs1l,Hey1,Hivep2,Il6st,Itk,Jak1,Kcnh8,Klf2,Klf3,Lef1,Lmtk3,Lonrf1,Map3k1,Map3k8,Mapk9,Mbd2,Mgme1,Mknk2,Msrb3,Nabp1,Neurl1b,Nfatc1,Nfkbiz,Nipbl,Nrip1,Nsmce1,Otulin,Pim1,Ppp1cc,Ppp2r5a,Prkca,Prkcq,Rara,Riok1,Rnf125,Sirt4,Ski,Ss18,St8sia1,Stat5b,Stk10,Tbl1xr1,Tgfbr2,Tnfaip3,Trim25,Wdyhv1,Zfp3,Zfp609,Zfp652,Zscan10 |
| GO:2000516 | positive regulation of CD4-positive, alpha-beta T cell activation | 5 | 34 | 0.00076 | Ccr7,Nfkbiz,Prkcq,Rara,Tgfbr2 |
| GO:0033993 | response to lipid | 21 | 825 | 0.00083 | Abhd2,Adipoq,Ccr7,Dusp10,Ets1,Fasl,Foxp1,Lef1,Mbd2,Nrip1,Pdcd4,Pim1,Prkca,Rara,Sgms1,Stat5b,Tgfbr2,Tnfaip3,Tnfrsf1a,Tnfrsf1b,Trim25 |
| GO:0045582 | positive regulation of T cell differentiation | 7 | 90 | 0.00083 | Ccr7,Dusp10,Nfkbiz,Rara,Stat5b,Tgfbr2,Zfp609 |
| GO:0050776 | regulation of immune response | 18 | 635 | 0.00083 | Bcl10,Cblb,Ccr7,Cd247,Cd28,Dpp4,Dusp10,Elf1,Itk,Nfkbiz,Otulin,Prkcq,Rara,Rftn1,Rnf125,Slc39a10,Stat5b,Tnfaip3 |
| GO:0007275 | multicellular organism development | 67 | 4603 | 0.00086 | Abcb1b,Adipoq,Ago2,Ankrd11,Atp2b4,Bcl10,Btg2,Camk4,Ccr4,Ccr7,Cd28,Cdkn2b,Chd2,Dad1,Dcaf7,Ddx6,Dock10,Dusp10,Ets1,F2,Fam20a,Fasl,Foxn3,Foxp1,Gnat1,Gpr18,Gtf2ird1,Hey1,Hivep2,Il15ra,Il6st,Ist1,Itk,Klf2,Lef1,Map3k1,Mapk9,Mbd2,Mbnl1,Mknk2,Nfatc1,Nipbl,Nrip1,Otulin,Pdcd4,Pip4k2a,Ppp1cc,Prkca,Rara,Rtkn2,S1pr1,Selt,Ski,Smarce1,Sod1,Stat5b,Tapt1,Tbl1xr1,Tgfbr2,Tnfaip3,Tnfrsf1a,Tnfrsf1b,Tnfsf8,Wasf2,Wwox,Xdh,Zfp609 |
| GO:0032879 | regulation of localization | 44 | 2579 | 0.00094 | Abhd2,Adipoq,Ago2,Camk4,Ccr7,Cd247,Cd47,Cnst,Crbn,Dnajc6,Dock10,Dusp10,Epb4.1,Ets1,F2,Fasl,Foxp1,Gpr18,Il6st,Kcnh8,Lef1,Leprotl1,Map3k1,Mapk9,Neto2,Nipbl,P2rx4,Ppp1cc,Ppp2r5a,Prkca,Prkcq,Rras2,S1pr1,Selt,Sirt4,Stat5b,Stk10,Tgfbr2,Tnfrsf1a,Tnfrsf1b,Ubac2,Usp6nl,Vamp4,Zfp609 |
| GO:0042981 | regulation of apoptotic process | 30 | 1476 | 0.001 | Adipoq,Bcl10,Btg2,Ccr7,Dad1,Ets1,Fasl,Foxp1,Kcnh8,Lef1,Map3k1,Map3k8,Mapk9,Pdcd4,Pim1,Prkca,Prkcq,Rara,Sgms1,Sirt4,Slc39a10,Sod1,Stat5b,Stk10,Tax1bp1,Tnfaip3,Tnfrsf1a,Tnfrsf1b,Wwox,Xdh |
| GO:0048513 | animal organ development | 49 | 3012 | 0.001 | Adipoq,Ankrd11,Atp2b4,Btg2,Camk4,Ccr7,Cd28,Cdkn2b,Chd2,Dock10,Ets1,Fam20a,Fasl,Foxn3,Foxp1,Gnat1,Gpr18,Hey1,Itk,Klf2,Lef1,Map3k1,Mbd2,Mbnl1,Mknk2,Nfatc1,Nipbl,Nrip1,Pdcd4,Pip4k2a,Prkca,Rara,Rtkn2,S1pr1,Selt,Ski,Sod1,Stat5b,Tapt1,Tbl1xr1,Tgfbr2,Tnfaip3,Tnfrsf1a,Tnfrsf1b,Tnfsf8,Wasf2,Wwox,Xdh,Zfp609 |
| GO:0071310 | cellular response to organic substance | 35 | 1858 | 0.001 | Abhd2,Adipoq,Ago2,Atp2b4,Ccr4,Ccr7,Foxp1,Il6st,Jak1,Klf2,Klf3,Lef1,Lrtm2,Map3k1,Mapk9,Mbd2,Nrip1,P2rx4,Pdcd4,Pim1,Prkca,Rara,Rgs10,Rnf125,Selt,Sgms1,Sod1,Stat5b,Tgfbr2,Tnfaip3,Tnfrsf1a,Tnfrsf1b,Trim25,Vamp4,Wwox |
| GO:0008284 | positive regulation of cell population proliferation | 22 | 913 | 0.0011 | Ccr7,Cd28,Cd47,Dpp4,Ets1,F2,Fasl,Foxp1,Il6st,Lef1,Pim1,Ppp1cc,Prkca,Prkcq,Rara,Rtkn2,S1pr1,Slc39a10,St8sia1,Stat5b,Tgfbr2,Tnfaip3 |
| GO:0045619 | regulation of lymphocyte differentiation | 9 | 171 | 0.0011 | Ccr7,Cd28,Dusp10,Il15ra,Nfkbiz,Rara,Stat5b,Tgfbr2,Zfp609 |
| GO:0046635 | positive regulation of alpha-beta T cell activation | 6 | 64 | 0.0011 | Ccr7,Cd28,Nfkbiz,Prkcq,Rara,Tgfbr2 |
| GO:0051254 | positive regulation of RNA metabolic process | 31 | 1558 | 0.0011 | Ago2,Bcl10,Btg2,Camk4,Cd28,Elf1,Ets1,Foxn3,Foxp1,Hey1,Hivep2,Klf2,Lef1,Mapk9,Ncf1,Nfatc1,Nipbl,Nrip1,Pim1,Rara,Riok1,S1pr1,Ski,Ss18,Stat5b,Tbl1xr1,Tnfrsf1a,Tnfrsf1b,Tnfsf8,Wwox,Zfp609 |
| GO:0051345 | positive regulation of hydrolase activity | 18 | 652 | 0.0011 | Arhgap15,Arhgap25,Bcl10,Ccr7,Dock10,Fasl,Gdi2,Gnat1,Itk,Mapk9,Pkp4,Rasgrp2,Rgs10,S1pr1,Slc39a10,Snx18,Usp6nl,Xdh |
| GO:0002237 | response to molecule of bacterial origin | 12 | 315 | 0.0012 | Bcl10,Ccr7,Dusp10,Fasl,Foxp1,Pdcd4,Rara,Sgms1,Stat5b,Tnfaip3,Tnfrsf1a,Tnfrsf1b |
| GO:0046634 | regulation of alpha-beta T cell activation | 7 | 98 | 0.0012 | Cblb,Ccr7,Cd28,Nfkbiz,Prkcq,Rara,Tgfbr2 |
| GO:0010611 | regulation of cardiac muscle hypertrophy | 6 | 67 | 0.0013 | Atp2b4,Foxp1,P2rx4,Prkca,Tnfrsf1a,Tnfrsf1b |
| GO:0040012 | regulation of locomotion | 22 | 924 | 0.0013 | Abhd2,Adipoq,Ago2,Ccr4,Ccr7,Dock10,Dusp10,Ets1,Foxp1,Gpr18,Il6st,Lef1,Map3k1,Nipbl,P2rx4,Prkca,Rras2,S1pr1,Stk10,Tgfbr2,Trim25,Zfp609 |
| GO:0043170 | macromolecule metabolic process | 83 | 6225 | 0.0013 | Aebp2,Aff4,Ago2,Bach2,Btg2,Camk4,Cblb,Ccr7,Celf2,Chd2,Crbn,Dad1,Dcaf7,Ddx6,Dhx40,Dnajc6,Dpp4,Dusp10,Dyrk2,Elf1,Ercc4,Ets1,F2,Fasl,Fkbp5,Foxn3,Foxp1,Gtf2ird1,Gucy2c,Gzmb,Hbs1l,Hey1,Hivep2,Il6st,Itk,Jak1,Kcnh8,Klf2,Klf3,Lef1,Lmtk3,Lonrf1,Map3k1,Map3k8,Mapk9,Mbd2,Mbnl1,Mgme1,Mknk2,Msrb3,Nabp1,Neurl1b,Nfatc1,Nfkbiz,Nipbl,Nrip1,Nsmce1,Otulin,Pim1,Ppp1cc,Ppp2r5a,Prkca,Prkcq,Prss39,Rara,Riok1,Rnf125,Sirt4,Ski,Ss18,St8sia1,Stat5b,Stk10,Tbl1xr1,Tgfbr2,Tnfaip3,Trim25,Wdyhv1,Xpnpep2,Zfp3,Zfp609,Zfp652,Zscan10 |
| GO:1901342 | regulation of vasculature development | 12 | 319 | 0.0013 | Ago2,Ets1,Fasl,Hey1,Jak1,Klf2,Nfatc1,Pdcd4,Prkca,Tgfbr2,Tnfrsf1a,Xdh |
| GO:0006464 | cellular protein modification process | 42 | 2471 | 0.0014 | Bach2,Btg2,Camk4,Cblb,Ccr7,Crbn,Dad1,Dcaf7,Dnajc6,Dusp10,Dyrk2,Fkbp5,Gucy2c,Itk,Jak1,Kcnh8,Lef1,Lmtk3,Lonrf1,Map3k1,Map3k8,Mapk9,Mknk2,Neurl1b,Nsmce1,Otulin,Pim1,Ppp1cc,Ppp2r5a,Prkca,Prkcq,Rara,Riok1,Rnf125,Sirt4,St8sia1,Stk10,Tbl1xr1,Tgfbr2,Tnfaip3,Trim25,Wdyhv1 |

| | | | | | |
|---|---|---|---|---|---|
| GO:0097190 | apoptotic signaling pathway | 11 | 273 | 0.0014 | Cd28,Dyrk2,Fasl,Gzmb,Mapk9,Mknk2,P2rx4,Prkca,Tnfrsf1a,Tnfrsf1b,Wwox |
| GO:0003177 | pulmonary valve development | 4 | 20 | 0.0015 | Hey1,Nfatc1,Tnfrsf1a,Tnfrsf1b |
| GO:0030334 | regulation of cell migration | 20 | 805 | 0.0015 | Abhd2,Adipoq,Ago2,Ccr7,Dock10,Dusp10,Ets1,Foxp1,Gpr18,Il6st,Lef1,Map3k1,Nipbl,P2rx4,Prkca,Rras2,S1pr1,Stk10,Tgfbr2,Zfp609 |
| GO:0045580 | regulation of T cell differentiation | 8 | 141 | 0.0015 | Ccr7,Cd28,Dusp10,Nfkbiz,Rara,Stat5b,Tgfbr2,Zfp609 |
| GO:0046649 | lymphocyte activation | 13 | 378 | 0.0015 | Cblb,Ccr7,Cd28,Dock10,Dpp4,Foxp1,Gpr18,Itk,Lef1,Nfatc1,Stat5b,Tnfaip3,Tnfsf8 |
| GO:1901701 | cellular response to oxygen-containing compound | 21 | 870 | 0.0015 | Adipoq,Atp2b4,Ets1,Foxp1,Klf2,Klf3,Mapk9,Ncf1,Nrip1,P2rx4,Pdcd4,Pim1,Prkca,Rara,Rgs10,Selt,Sgms1,Sod1,Stat5b,Tnfaip3,Tnfrsf1b |
| GO:0001818 | negative regulation of cytokine production | 10 | 229 | 0.0016 | Adipoq,F2,Gpr18,Klf2,Lef1,Pdcd4,Rara,Rnf125,Tnfaip3,Tnfrsf1a |
| GO:0007166 | cell surface receptor signaling pathway | 32 | 1683 | 0.0016 | Bcl10,Cblb,Ccr4,Ccr7,Cd247,Cd28,Dyrk2,F2,Fasl,Hey1,Il15ra,Il6st,Itk,Jak1,Lef1,Lrtm2,Map3k1,Mknk2,Neurl1b,Nfkbiz,Otulin,P2rx4,Pdcd4,Rftn1,Ski,Ss18,Stat5b,Tgfbr2,Tmem64,Tnfrsf1a,Tnfrsf1b,Wwox |
| GO:0030099 | myeloid cell differentiation | 10 | 229 | 0.0016 | Camk4,Ccr7,Cdkn2b,Foxp1,Klf2,Lef1,Nfatc1,Pip4k2a,Tgfbr2,Wasf2 |
| GO:0051704 | multi-organism process | 34 | 1840 | 0.0016 | Abhd2,Adipoq,Aff4,Atp2b4,Bcl10,Ccdc80,Ccr4,Ccr7,Cd47,Ddx6,Dpp4,Dusp10,Ets1,F2,Fasl,Fkbp5,Foxp1,Hist1h2be,Ist1,Lef1,Ncf1,Nrip1,Pdcd4,Rara,Sgms1,Sod1,Stat5b,Tgfbr2,Tnfaip3,Tnfrsf1a,Tnfrsf1b,Tnfsf8,Trim25,Usp6nl |
| GO:0009891 | positive regulation of biosynthetic process | 34 | 1847 | 0.0017 | Ago2,Bcl10,Camk4,Ccr7,Cd28,Dyrk2,Elf1,Elovl5,Ets1,F2,Foxn3,Foxp1,Hey1,Hivep2,Klf2,Lef1,Mapk9,Ncf1,Nfatc1,Nipbl,Nrip1,Pim1,Prkcq,Rara,S1pr1,Sirt4,Ski,Ss18,Stat5b,Tbl1xr1,Tnfrsf1a,Tnfsf8,Wwox,Zfp609 |
| GO:0032102 | negative regulation of response to external stimulus | 11 | 282 | 0.0017 | Adipoq,Dusp10,Ets1,F2,Gpr18,Otulin,Pdcd4,Rnf125,Tnfaip3,Tnfrsf1a,Tnfrsf1b |
| GO:0032270 | positive regulation of cellular protein metabolic process | 29 | 1462 | 0.0017 | Adipoq,Atp2b4,Bcl10,Ccr7,F2,Fam20a,Fasl,Il6st,Ist1,Lonrf1,Map3k1,Map3k8,Mapk9,Ncf1,Nipbl,Pim1,Ppp2r5a,Prkca,Rara,Rnf125,Slc39a10,Sod1,Stk10,Tgfbr2,Tnfaip3,Tnfrsf1a,Tnfrsf1b,Xdh,Zfand2a |
| GO:0043066 | negative regulation of apoptotic process | 21 | 884 | 0.0017 | Bcl10,Btg2,Ccr7,Dad1,Foxp1,Kcnh8,Lef1,Pdcd4,Pim1,Prkca,Prkcq,Rara,Sgms1,Sirt4,Slc39a10,Sod1,Stat5b,Tax1bp1,Tnfaip3,Tnfrsf1a,Tnfrsf1b |
| GO:0032103 | positive regulation of response to external stimulus | 11 | 285 | 0.0019 | Ccr4,Ccr7,Cd28,Cd47,Ets1,P2rx4,Pdcd4,Prkca,S1pr1,Stat5b,Tnfrsf1a |
| GO:0010941 | regulation of cell death | 31 | 1640 | 0.0023 | Adipoq,Bcl10,Btg2,Ccr7,Dad1,Ets1,Fasl,Foxp1,Kcnh8,Lef1,Map3k1,Map3k8,Mapk9,Pdcd4,Pim1,Prkca,Prkcq,Rara,Rras2,Sgms1,Sirt4,Slc39a10,Sod1,Stat5b,Stk10,Tax1bp1,Tnfaip3,Tnfrsf1a,Tnfrsf1b,Wwox,Xdh |
| GO:0045893 | positive regulation of transcription, DNA-templated | 28 | 1413 | 0.0023 | Ago2,Bcl10,Camk4,Cd28,Elf1,Ets1,Foxn3,Foxp1,Hey1,Hivep2,Klf2,Lef1,Mapk9,Ncf1,Nfatc1,Nipbl,Nrip1,Pim1,Rara,S1pr1,Ski,Ss18,Stat5b,Tbl1xr1,Tnfrsf1a,Tnfsf8,Wwox,Zfp609 |
| GO:0042110 | T cell activation | 10 | 244 | 0.0024 | Cblb,Ccr7,Cd28,Dpp4,Foxp1,Gpr18,Itk,Lef1,Stat5b,Tnfsf8 |
| GO:0030217 | T cell differentiation | 8 | 154 | 0.0025 | Ccr7,Cd28,Foxp1,Gpr18,Itk,Lef1,Stat5b,Tnfsf8 |
| GO:0043412 | macromolecule modification | 43 | 2636 | 0.0025 | Bach2,Btg2,Camk4,Cblb,Ccr7,Crbn,Dad1,Dcaf7,Dnajc6,Dusp10,Dyrk2,Fkbp5,Gucy2c,Itk,Jak1,Kcnh8,Lef1,Lmtk3,Lonrf1,Map3k1,Map3k8,Mapk9,Mbd2,Mknk2,Neurl1b,Nsmce1,Otulin,Pim1,Ppp1cc,Ppp2r5a,Prkca,Prkcq,Rara,Riok1,Rnf125,Sirt4,St8sia1,Stk10,Tbl1xr1,Tgfbr2,Tnfaip3,Trim25,Wdyhv1 |
| GO:0045935 | positive regulation of nucleobase-containing compound metabolic process | 32 | 1732 | 0.0025 | Ago2,Bcl10,Btg2,Camk4,Cd28,Elf1,Ets1,Foxn3,Foxp1,Hey1,Hivep2,Klf2,Lef1,Mapk9,Ncf1,Nfatc1,Nipbl,Nrip1,Pim1,Prkcq,Rara,Riok1,S1pr1,Ski,Ss18,Stat5b,Tbl1xr1,Tnfrsf1a,Tnfrsf1b,Tnfsf8,Wwox,Zfp609 |
| GO:0007584 | response to nutrient | 9 | 199 | 0.0026 | Adipoq,Cdkn2b,Dad1,P2rx4,Pim1,Rara,Sod1,Tgfbr2,Trim25 |
| GO:0032496 | response to lipopolysaccharide | 11 | 298 | 0.0026 | Ccr7,Dusp10,Fasl,Foxp1,Pdcd4,Rara,Sgms1,Stat5b,Tnfaip3,Tnfrsf1a,Tnfrsf1b |
| GO:0045859 | regulation of protein kinase activity | 18 | 717 | 0.0027 | Adipoq,Atp2b4,Cblb,Ccr7,Cdkn2b,Dusp10,Fam20a,Lrtm2,Map3k1,Map3k8,Ncf1,Pdcd4,Pim1,Prkca,Sod1,Stk10,Tgfbr2,Tnfaip3 |
| GO:0006468 | protein phosphorylation | 19 | 793 | 0.0031 | Camk4,Ccr7,Dyrk2,Gucy2c,Itk,Jak1,Kcnh8,Lmtk3,Map3k1,Map3k8,Mapk9,Mknk2,Pim1,Prkca,Prkcq,Rara,Riok1,Stk10,Tgfbr2 |
| GO:0009892 | negative regulation of metabolic process | 42 | 2584 | 0.0031 | Adipoq,Aebp2,Ago2,Atp2b4,Bach2,Btg2,Cblb,Cd28,Cdkn2b,Cnst,Crim1,Dusp10,Ercc4,F2,Fasl,Foxn3,Foxp1,Hey1,Klf2,Klf3,Lef1,Lmtk3,Lrtm2,Mbd2,Nfatc1,Nipbl,Nrip1,Pdcd4,Ppp2r5a,Prkca,Rara,Sirt4,Ski,Smarce1,Sod1,Tbl1xr1,Tnfaip3,Tnfrsf1a,Tnfrsf1b,Ubac2,Xdh,Zscan10 |
| GO:0034097 | response to cytokine | 19 | 792 | 0.0031 | Adipoq,Ccr4,Ccr7,Ets1,Foxp1,Il6st,Jak1,Klf2,Lef1,Lrtm2,Prkca,Rara,Rnf125,Sgms1,Stat5b,Tnfrsf1a,Tnfrsf1b,Trim25,Vamp4 |
| GO:0048660 | regulation of smooth muscle cell proliferation | 8 | 161 | 0.0031 | Adipoq,Foxp1,Pdcd4,Prkca,S1pr1,Stat5b,Tgfbr2,Tnfaip3 |
| GO:0051270 | regulation of cellular component movement | 21 | 928 | 0.0031 | Abhd2,Adipoq,Ago2,Ccr7,Dock10,Dusp10,Ets1,Foxp1,Gpr18,Il6st,Lef1,Map3k1,Nipbl,P2rx4,Prkca,Rras2,S1pr1,Stat5b,Stk10,Tgfbr2,Zfp609 |
| GO:1902531 | regulation of intracellular signal transduction | 30 | 1599 | 0.0031 | Adipoq,Arhgap25,Atp2b4,Bcl10,Ccr7,Cd28,Cyth4,Dusp10,Dyrk2,F2,Fasl,Gpr18,Il6st,Jak1,Lrtm2,Map3k1,Map3k8,Mapk9,Ncf1,P2rx4,Pdcd4,Prkca,Rara,Rnf125,Slc7a3,Sod1,Tnfaip3,Tnfrsf1a,Trim25,Xdh |
| GO:1903054 | negative regulation of extracellular matrix organization | 3 | 9 | 0.0031 | Dpp4,Tnfrsf1a,Tnfrsf1b |
| GO:0006952 | defense response | 23 | 1079 | 0.0033 | Bcl10,Camk4,Ccr4,Ccr7,Chst2,Dpp4,F2,Gzmb,Hist1h2be,Itk,Jak1,Ncf1,Nfkbiz,Otulin,Prkcq,Sgms1,Stat5b,Tnfaip3,Tnfrsf1a,Tnfrsf1b,Tnfsf8,Trim25,Vamp4 |
| GO:0050854 | regulation of antigen receptor-mediated signaling pathway | 5 | 52 | 0.0033 | Bcl10,Cblb,Ccr7,Elf1,Slc39a10 |
| GO:0006357 | regulation of transcription by RNA polymerase II | 33 | 1856 | 0.0037 | Aebp2,Ago2,Atp2b4,Bach2,Btg2,Cd28,Elf1,Ets1,Fasl,Foxp1,Gtf2ird1,Hey1,Hivep2,Kcnh8,Klf2,Klf3,Lef1,Mbd2,Nfatc1,Nfkbiz,Nipbl,Nrip1,Pdcd4,Rara,S1pr1,Ski,Ss18,Stat5b,Tbl1xr1,Tnfrsf1a,Tnfsf8,Wwox,Zfp609 |
| GO:0032673 | regulation of interleukin-4 production | 4 | 28 | 0.0037 | Cd28,Lef1,Prkcq,Rara |
| GO:0031324 | negative regulation of cellular metabolic process | 39 | 2357 | 0.0038 | Adipoq,Aebp2,Ago2,Atp2b4,Bach2,Btg2,Cblb,Cdkn2b,Cnst,Crim1,Dusp10,Ercc4,F2,Fasl,Foxn3,Foxp1,Hey1,Klf2,Klf3,Lef1,Lmtk3,Lrtm2,Mbd2,Nfatc1,Nipbl,Nrip1,Pdcd4,Ppp2r5a,Prkca,Rara,Sirt4,Ski,Smarce1,Tbl1xr1,Tnfaip3,Tnfrsf1b,Ubac2,Xdh,Zscan10 |
| GO:0014070 | response to organic cyclic compound | 20 | 883 | 0.004 | Abhd2,Adipoq,Ago2,Atp2b4,Btg2,Ets1,Foxp1,Klf2,Lef1,Mbd2,Nrip1,P2rx4,Pim1,Prkca,Rara,Rftn1,Sod1,Stat5b,Tgfbr2,Trim25 |
| GO:0010557 | positive regulation of macromolecule biosynthetic process | 31 | 1717 | 0.0043 | Ago2,Bcl10,Camk4,Ccr7,Cd28,Dyrk2,Elf1,Ets1,Foxn3,Foxp1,Hey1,Hivep2,Klf2,Lef1,Mapk9,Ncf1,Nfatc1,Nipbl,Nrip1,Pim1,Prkcq,Rara,S1pr1,Ski,Ss18,Stat5b,Tbl1xr1,Tnfrsf1a,Tnfsf8,Wwox,Zfp609 |
| GO:0031349 | positive regulation of defense response | 10 | 268 | 0.0044 | Ccr7,Cd28,Cd47,Ets1,Otulin,Pdcd4,Prkca,Rftn1,Stat5b,Tnfrsf1a |
| GO:0071900 | regulation of protein serine/threonine kinase activity | 13 | 435 | 0.0044 | Adipoq,Atp2b4,Ccr7,Cdkn2b,Dusp10,Fam20a,Map3k1,Map3k8,Pdcd4,Pim1,Prkca,Sod1,Tnfaip3 |
| GO:0072507 | divalent inorganic cation homeostasis | 13 | 435 | 0.0044 | Atp2b4,F2,Fam20a,Fasl,Gpr18,Il6st,P2rx4,Prkca,S1pr1,Selt,Slc39a10,Thada,Tmem64 |
| GO:0003170 | heart valve development | 5 | 57 | 0.0045 | Hey1,Nfatc1,Tgfbr2,Tnfrsf1a,Tnfrsf1b |
| GO:0010033 | response to organic substance | 41 | 2553 | 0.0045 | Abhd2,Adipoq,Ago2,Atp2b4,Bcl10,Btg2,Ccr4,Ccr7,Dusp10,Ets1,Fasl,Foxp1,Il6st,Jak1,Klf2,Klf3,Lef1,Lrtm2,Map3k1,Mapk9,Mbd2,Nrip1,P2rx4,Pdcd4,Pim1,Prkca,Rara,Rftn1,Rgs10,Rnf125,Selt,Sgms1,Sod1,Stat5b,Tgfbr2,Tnfaip3,Tnfrsf1a,Tnfrsf1b,Trim25,Vamp4,Wwox |
| GO:0045086 | positive regulation of interleukin-2 biosynthetic process | 3 | 11 | 0.0045 | Cd28,Prkcq,Stat5b |
| GO:0031328 | positive regulation of cellular biosynthetic process | 32 | 1807 | 0.0046 | Ago2,Bcl10,Camk4,Ccr7,Cd28,Dyrk2,Elf1,Elovl5,Ets1,Foxn3,Foxp1,Hey1,Hivep2,Klf2,Lef1,Mapk9,Ncf1,Nfatc1,Nipbl,Nrip1,Pim1,Prkcq,Rara,S1pr1,Ski,Ss18,Stat5b,Tbl1xr1,Tnfrsf1a,Tnfsf8,Wwox,Zfp609 |
| GO:0060324 | face development | 5 | 58 | 0.0048 | Ankrd11,Lef1,Nipbl,Rara,Ski |
| GO:0060348 | bone development | 8 | 177 | 0.0052 | Ankrd11,Foxn3,Foxp1,Pip4k2a,Rara,Ski,Tgfbr2,Wasf2 |
| GO:0046632 | alpha-beta T cell differentiation | 5 | 60 | 0.0055 | Foxp1,Gpr18,Itk,Lef1,Tnfsf8 |
| GO:0009967 | positive regulation of signal transduction | 27 | 1433 | 0.0056 | Adipoq,Bcl10,Ccr7,Cd28,Cdkn2b,F2,Fasl,Gpr18,Il6st,Jak1,Map3k1,Map3k8,Mapk9,Mbd2,Ncf1,P2rx4,Pdcd4,Prkca,Rara,Sh2b1,Slc39a10,Sod1,Tbl1xr1,Tnfrsf1a,Trim25,Wwox,Xdh |
| GO:0090257 | regulation of muscle system process | 9 | 228 | 0.0057 | Atp2b4,Foxp1,Gtf2ird1,Nfatc1,P2rx4,Prkca,Sod1,Tnfrsf1a,Tnfrsf1b |
| GO:0032680 | regulation of tumor necrosis factor production | 7 | 136 | 0.0058 | Adipoq,Ccr7,Foxp1,Gpr18,Rara,Tnfaip3,Tnfrsf1a |
| GO:0050671 | positive regulation of lymphocyte proliferation | 7 | 136 | 0.0058 | Ccr7,Cd28,Il6st,Prkcq,Slc39a10,Stat5b,Tgfbr2 |
| GO:2000107 | negative regulation of leukocyte apoptotic process | 5 | 61 | 0.0058 | Bcl10,Ccr7,Foxp1,Prkcq,Slc39a10 |
| GO:0006954 | inflammatory response | 13 | 454 | 0.006 | Camk4,Ccr4,Ccr7,Chst2,F2,Ncf1,Nfkbiz,Prkcq,Sgms1,Stat5b,Tnfaip3,Tnfrsf1a,Tnfrsf1b |
| GO:0042102 | positive regulation of T cell proliferation | 6 | 98 | 0.0063 | Ccr7,Cd28,Il6st,Prkcq,Stat5b,Tgfbr2 |
| GO:0002903 | negative regulation of B cell apoptotic process | 3 | 13 | 0.0064 | Bcl10,Foxp1,Slc39a10 |
| GO:0009968 | negative regulation of signal transduction | 23 | 1147 | 0.0066 | Adipoq,Arhgap25,Atp2b4,Cblb,Dusp10,Dyrk2,Elf1,Foxp1,Hey1,Il6st,Lef1,Lrtm2,Pdcd4,Prkca,Prkcq,Rgs10,Rnf125,Sgms1,Ski,Tmem64,Tnfaip3,Wwox,Xdh |
| GO:1902533 | positive regulation of intracellular signal transduction | 20 | 927 | 0.0066 | Adipoq,Bcl10,Ccr7,Cd28,F2,Fasl,Gpr18,Il6st,Jak1,Map3k1,Map3k8,Ncf1,P2rx4,Pdcd4,Prkca,Rara,Sod1,Tnfrsf1a,Trim25,Xdh |
| GO:2000379 | positive regulation of reactive oxygen species metabolic process | 6 | 101 | 0.0071 | F2,Klf2,Mapk9,Sod1,Tgfbr2,Xdh |
| GO:0055074 | calcium ion homeostasis | 12 | 405 | 0.0072 | Atp2b4,F2,Fam20a,Fasl,Gpr18,Il6st,P2rx4,Prkca,S1pr1,Selt,Thada,Tmem64 |

| | | | | | |
|---|---|---|---|---|---|
| GO:0051172 | negative regulation of nitrogen compound metabolic process | 36 | 2206 | 0.0076 | Adipoq,Aebp2,Ago2,Atp2b4,Bach2,Btg2,Cblb,Cdkn2b,Crim1,Dusp10,Ercc4,F2,Fasl,Foxn3,Foxp1,Hey1,Klf2,Klf3,Lef1,Lrtm2,Mbd2,Nfatc1,Nipbl,Nrip1,Pdcd4,Prkca,Rara,Sirt4,Ski,Smarce1,Tbl1xr1,Tnfaip3,Tnfrsf1b,Ubac2,Xdh,Zscan10 |
| GO:0072503 | cellular divalent inorganic cation homeostasis | 12 | 408 | 0.0076 | Atp2b4,F2,Fasl,Gpr18,Il6st,P2rx4,Prkca,S1pr1,Selt,Slc39a10,Thada,Tmem64 |
| GO:0051272 | positive regulation of cellular component movement | 14 | 533 | 0.0078 | Ago2,Ccr7,Ets1,Foxp1,Il6st,Lef1,Nipbl,P2rx4,Prkca,Rras2,S1pr1,Stat5b,Tgfbr2,Zfp609 |
| GO:0002649 | regulation of tolerance induction to self antigen | 2 | 2 | 0.0079 | Ccr7,Tgfbr2 |
| GO:0043087 | regulation of GTPase activity | 12 | 410 | 0.0079 | Arhgap15,Arhgap25,Ccr7,Dock10,Gdi2,Pkp4,Rasgrp2,Rgs10,S1pr1,Snx18,Sod1,Usp6nl |
| GO:0046381 | CMP-N-acetylneuraminate metabolic process | 2 | 2 | 0.0079 | Cmah,Cmas |
| GO:0043547 | positive regulation of GTPase activity | 11 | 353 | 0.008 | Arhgap15,Arhgap25,Ccr7,Dock10,Gdi2,Pkp4,Rasgrp2,Rgs10,S1pr1,Snx18,Usp6nl |
| GO:0071407 | cellular response to organic cyclic compound | 13 | 475 | 0.0084 | Abhd2,Adipoq,Ago2,Atp2b4,Foxp1,Klf2,Lef1,Mbd2,Nrip1,P2rx4,Pim1,Rara,Sod1 |
| GO:0060325 | face morphogenesis | 4 | 38 | 0.0088 | Ankrd11,Lef1,Nipbl,Ski |
| GO:1903053 | regulation of extracellular matrix organization | 4 | 38 | 0.0088 | Dpp4,Ets1,Tnfrsf1a,Tnfrsf1b |
| GO:0019220 | regulation of phosphate metabolic process | 29 | 1650 | 0.009 | Adipoq,Atp2b4,Bcl10,Cblb,Ccr7,Cdkn2b,Cnst,Dusp10,F2,Fam20a,Il6st,Lmtk3,Lrtm2,Map3k1,Map3k8,Mapk9,Ncf1,Pdcd4,Pim1,Ppp2r5a,Prkca,Rara,Slc39a10,Sod1,Stk10,Tgfbr2,Tnfaip3,Tnfrsf1a,Xdh |
| GO:0016310 | phosphorylation | 21 | 1034 | 0.0094 | Camk4,Ccr7,Dyrk2,Gucy2c,Itk,Jak1,Kcnh8,Lmtk3,Map3k1,Map3k8,Mapk9,Mknk2,Pim1,Pip4k2a,Prkca,Prkcq,Rara,Riok1,Sgms1,Stk10,Tgfbr2 |
| GO:0045637 | regulation of myeloid cell differentiation | 8 | 199 | 0.0095 | Adipoq,Ets1,Foxp1,Lef1,Prkca,Rara,Stat5b,Tmem64 |
| GO:0050856 | regulation of T cell receptor signaling pathway | 4 | 39 | 0.0095 | Bcl10,Cblb,Ccr7,Elf1 |
| GO:0070229 | negative regulation of lymphocyte apoptotic process | 4 | 39 | 0.0095 | Bcl10,Foxp1,Prkcq,Slc39a10 |
| GO:1903827 | regulation of cellular protein localization | 13 | 484 | 0.0095 | Adipoq,Camk4,Cd247,Cnst,Epb4.1,F2,Leprotl1,Ppp2r5a,Prkcq,Sirt4,Tnfrsf1a,Ubac2,Vamp4 |
| GO:0000302 | response to reactive oxygen species | 8 | 200 | 0.0096 | Ccr7,Ets1,Klf2,Mapk9,Ncf1,Prkca,Sod1,Tnfaip3 |
| GO:0040017 | positive regulation of locomotion | 14 | 550 | 0.0098 | Ago2,Ccr4,Ccr7,Ets1,Foxp1,Il6st,Lef1,Nipbl,P2rx4,Prkca,Rras2,S1pr1,Tgfbr2,Zfp609 |
| GO:0010648 | negative regulation of cell communication | 24 | 1269 | 0.0099 | Adipoq,Arhgap25,Atp2b4,Cblb,Dusp10,Dyrk2,Elf1,Foxp1,Hey1,Il6st,Lef1,Lrtm2,Pdcd4,Prkca,Prkcq,Rgs10,Rnf125,Sgms1,Sirt4,Ski,Tmem64,Tnfaip3,Wwox,Xdh |
| GO:0023057 | negative regulation of signaling | 24 | 1271 | 0.0101 | Adipoq,Arhgap25,Atp2b4,Cblb,Dusp10,Dyrk2,Elf1,Foxp1,Hey1,Il6st,Lef1,Lrtm2,Pdcd4,Prkca,Prkcq,Rgs10,Rnf125,Sgms1,Sirt4,Ski,Tmem64,Tnfaip3,Wwox,Xdh |
| GO:0032501 | multicellular organismal process | 75 | 5888 | 0.0102 | Abcb1b,Abhd2,Adipoq,Aff4,Ago2,Ankrd11,Atp2b4,Bcl10,Btg2,Camk4,Ccr4,Ccr7,Cd247,Cd28,Cdkn2b,Chd2,Dad1,Dcaf7,Ddx6,Dock10,Dpp4,Dusp10,Ets1,F2,Fam20a,Fasl,Foxn3,Foxp1,Gnat1,Gpr18,Gtf2ird1,Hey1,Hivep2,Il15ra,Il6st,Ist1,Itk,Klf2,Lef1,Map3k1,Mapk9,Mbd2,Mbnl1,Mknk2,Nfatc1,Nipbl,Nrip1,Otulin,P2rx4,Pdcd4,Pip4k2a,Ppp1cc,Prkca,Rara,Rftn1,Rtkn2,S1pr1,Selt,Ski,Smarce1,Sod1,Ss18,Stat5b,Tapt1,Tbl1xr1,Tgfbr2,Tmem64,Tnfaip3,Tnfrsf1a,Tnfrsf1b,Tnfsf8,Wasf2,Wwox,Xdh,Zfp609 |
| GO:0055065 | metal ion homeostasis | 14 | 555 | 0.0104 | Atp2b4,F2,Fam20a,Fasl,Gpr18,Il6st,P2rx4,Prkca,S1pr1,Selt,Slc39a10,Sod1,Thada,Tmem64 |
| GO:0006875 | cellular metal ion homeostasis | 13 | 492 | 0.0106 | Atp2b4,F2,Fasl,Gpr18,Il6st,P2rx4,Prkca,S1pr1,Selt,Slc39a10,Sod1,Thada,Tmem64 |
| GO:0010629 | negative regulation of gene expression | 27 | 1515 | 0.0107 | Adipoq,Aebp2,Ago2,Bach2,Btg2,Cd28,Fasl,Foxn3,Foxp1,Hey1,Klf2,Klf3,Lef1,Mbd2,Nfatc1,Nipbl,Nrip1,Pdcd4,Prkca,Rara,Sirt4,Ski,Smarce1,Tbl1xr1,Tnfrsf1a,Xdh,Zscan10 |
| GO:0045944 | positive regulation of transcription by RNA polymerase II | 22 | 1125 | 0.0107 | Ago2,Cd28,Elf1,Ets1,Foxp1,Hey1,Hivep2,Klf2,Lef1,Nfatc1,Nipbl,Nrip1,Rara,S1pr1,Ski,Ss18,Stat5b,Tbl1xr1,Tnfrsf1a,Tnfsf8,Wwox,Zfp609 |
| GO:0045937 | positive regulation of phosphate metabolic process | 21 | 1051 | 0.0108 | Adipoq,Atp2b4,Bcl10,Ccr7,F2,Fam20a,Il6st,Map3k1,Map3k8,Mapk9,Ncf1,Pim1,Ppp2r5a,Prkca,Rara,Slc39a10,Sod1,Stk10,Tgfbr2,Tnfrsf1a,Xdh |
| GO:0002573 | myeloid leukocyte differentiation | 6 | 113 | 0.0109 | Camk4,Ccr7,Foxp1,Lef1,Nfatc1,Tgfbr2 |
| GO:0002819 | regulation of adaptive immune response | 7 | 159 | 0.0115 | Ccr7,Cd28,Dpp4,Dusp10,Nfkbiz,Prkcq,Tnfaip3 |
| GO:0003332 | negative regulation of extracellular matrix constituent secretion | 2 | 3 | 0.0115 | Tnfrsf1a,Tnfrsf1b |
| GO:0033673 | negative regulation of kinase activity | 9 | 260 | 0.0115 | Adipoq,Cblb,Cdkn2b,Dusp10,Lrtm2,Pdcd4,Ppp2r5a,Prkca,Tnfaip3 |
| GO:0010647 | positive regulation of cell communication | 28 | 1608 | 0.0117 | Adipoq,Bcl10,Ccr7,Cd28,Cdkn2b,F2,Fasl,Gpr18,Il6st,Jak1,Map3k1,Map3k8,Mapk9,Mbd2,Ncf1,P2rx4,Pdcd4,Prkca,Rara,Selt,Sh2b1,Slc39a10,Sod1,Tbl1xr1,Tnfrsf1a,Trim25,Wwox,Xdh |
| GO:0071901 | negative regulation of protein serine/threonine kinase activity | 6 | 115 | 0.0117 | Adipoq,Cdkn2b,Dusp10,Pdcd4,Prkca,Tnfaip3 |
| GO:0030335 | positive regulation of cell migration | 13 | 500 | 0.0118 | Ago2,Ccr7,Ets1,Foxp1,Il6st,Lef1,Nipbl,P2rx4,Prkca,Rras2,S1pr1,Tgfbr2,Zfp609 |
| GO:0006793 | phosphorus metabolic process | 30 | 1775 | 0.0119 | Camk4,Ccr7,Cmah,Cmas,Dnajc6,Dusp10,Dyrk2,Gucy2c,Itk,Jak1,Kcnh8,Lmtk3,Lpcat3,Map3k1,Map3k8,Mapk9,Mknk2,Ncf1,Pim1,Pip4k2a,Ppp1cc,Ppp2r5a,Prkca,Prkcq,Rara,Riok1,Rpia,Sgms1,Stk10,Tgfbr2 |
| GO:0050778 | positive regulation of immune response | 12 | 438 | 0.0119 | Bcl10,Ccr7,Cd247,Cd28,Itk,Nfkbiz,Otulin,Prkcq,Rara,Rftn1,Slc39a10,Stat5b |
| GO:0002643 | regulation of tolerance induction | 3 | 18 | 0.012 | Cblb,Ccr7,Tgfbr2 |
| GO:0010605 | negative regulation of macromolecule metabolic process | 37 | 2373 | 0.0122 | Adipoq,Aebp2,Ago2,Bach2,Btg2,Cblb,Cd28,Cdkn2b,Crim1,Dusp10,Ercc4,F2,Fasl,Foxn3,Foxp1,Hey1,Klf2,Klf3,Lef1,Lrtm2,Mbd2,Nfatc1,Nipbl,Nrip1,Pdcd4,Prkca,Rara,Sirt4,Ski,Smarce1,Tbl1xr1,Tnfaip3,Tnfrsf1a,Tnfrsf1b,Ubac2,Xdh,Zscan10 |
| GO:0023056 | positive regulation of signaling | 28 | 1615 | 0.0122 | Adipoq,Bcl10,Ccr7,Cd28,Cdkn2b,F2,Fasl,Gpr18,Il6st,Jak1,Map3k1,Map3k8,Mapk9,Mbd2,Ncf1,P2rx4,Pdcd4,Prkca,Rara,Selt,Sh2b1,Slc39a10,Sod1,Tbl1xr1,Tnfrsf1a,Trim25,Wwox,Xdh |
| GO:0031401 | positive regulation of protein modification process | 22 | 1142 | 0.0122 | Adipoq,Atp2b4,Bcl10,Ccr7,F2,Fam20a,Il6st,Map3k1,Map3k8,Mapk9,Ncf1,Nipbl,Pim1,Ppp2r5a,Prkca,Rara,Slc39a10,Sod1,Stk10,Tgfbr2,Tnfrsf1a,Xdh |
| GO:0042129 | regulation of T cell proliferation | 7 | 162 | 0.0122 | Cblb,Ccr7,Cd28,Il6st,Prkcq,Stat5b,Tgfbr2 |
| GO:0010035 | response to inorganic substance | 13 | 505 | 0.0124 | Atp2b4,Ccr7,Cyb5a,Cyp2r1,Ets1,Klf2,Mapk9,Ncf1,P2rx4,Rasgrp2,Sod1,Tnfaip3,Xdh |
| GO:0044267 | cellular protein metabolic process | 44 | 2995 | 0.0125 | Bach2,Btg2,Camk4,Cblb,Ccr7,Crbn,Dad1,Dcaf7,Dnajc6,Dusp10,Dyrk2,Fkbp5,Gucy2c,Hbs1l,Itk,Jak1,Kcnh8,Lef1,Lmtk3,Lonrf1,Map3k1,Map3k8,Mapk9,Mknk2,Msrb3,Neurl1b,Nsmce1,Otulin,Pim1,Ppp1cc,Ppp2r5a,Prkca,Prkcq,Rara,Riok1,Rnf125,Sirt4,St8sia1,Stk10,Tbl1xr1,Tgfbr2,Tnfaip3,Trim25,Wdyhv1 |
| GO:0007346 | regulation of mitotic cell cycle | 13 | 507 | 0.0127 | Cd28,Cdkn2b,Foxn3,Gpr132,Map3k1,Map3k8,Nabp1,Pim1,Prkca,Prkcq,Sh2b1,Stat5b,Stk10 |
| GO:0003176 | aortic valve development | 3 | 19 | 0.0133 | Nfatc1,Tnfrsf1a,Tnfrsf1b |
| GO:0030852 | regulation of granulocyte differentiation | 3 | 19 | 0.0133 | Adipoq,Lef1,Rara |
| GO:0042325 | regulation of phosphorylation | 26 | 1466 | 0.0133 | Adipoq,Atp2b4,Bcl10,Cblb,Ccr7,Cdkn2b,Dusp10,F2,Fam20a,Il6st,Lrtm2,Map3k1,Map3k8,Mapk9,Ncf1,Pdcd4,Pim1,Ppp2r5a,Prkca,Rara,Sod1,Stk10,Tgfbr2,Tnfaip3,Tnfrsf1a,Xdh |
| GO:0044092 | negative regulation of molecular function | 21 | 1078 | 0.0133 | Adipoq,Atp2b4,Cblb,Cdkn2b,Cnst,Crim1,Dusp10,Ercc4,Hey1,Lef1,Lmtk3,Lrtm2,Otulin,Pdcd4,Pim1,Ppp2r5a,Prkca,Ski,Tax1bp1,Tnfaip3,Tnfrsf1b |
| GO:0050670 | regulation of lymphocyte proliferation | 8 | 215 | 0.0133 | Cblb,Ccr7,Cd28,Il6st,Prkcq,Slc39a10,Stat5b,Tgfbr2 |
| GO:1903409 | reactive oxygen species biosynthetic process | 3 | 19 | 0.0133 | Ncf1,P2rx4,Sod1 |
| GO:0006874 | cellular calcium ion homeostasis | 11 | 388 | 0.0139 | Atp2b4,F2,Fasl,Gpr18,Il6st,P2rx4,Prkca,S1pr1,Selt,Thada,Tmem64 |
| GO:0019216 | regulation of lipid metabolic process | 10 | 329 | 0.0141 | Adipoq,Elovl5,F2,Mapk9,Ppp2r5a,Rara,Sirt4,Sod1,Stat5b,Tbl1xr1 |
| GO:0022603 | regulation of anatomical structure morphogenesis | 20 | 1009 | 0.0141 | Adipoq,Ago2,Arhgap15,Epb4.1,Ets1,F2,Fasl,Hey1,Ist1,Jak1,Klf2,Mapk9,Pim1,Prkca,Syne3,Tgfbr2,Tnfaip3,Tnfrsf1a,Tnfrsf1b,Xdh |
| GO:0002757 | immune response-activating signal transduction | 7 | 168 | 0.0142 | Bcl10,Cd247,Cd28,Itk,Nfkbiz,Otulin,Rftn1 |
| GO:0001501 | skeletal system development | 12 | 454 | 0.0147 | Ankrd11,Foxn3,Foxp1,Nipbl,Pip4k2a,Prkca,Rara,Ski,Tapt1,Tgfbr2,Wasf2,Wwox |
| GO:0010628 | positive regulation of gene expression | 30 | 1811 | 0.0148 | Ago2,Bcl10,Camk4,Cd28,Elf1,Ets1,Foxn3,Foxp1,Hey1,Hivep2,Klf2,Lef1,Mapk9,Ncf1,Nfatc1,Nipbl,Nrip1,Pim1,Rara,Riok1,S1pr1,Sgms1,Ski,Ss18,Stat5b,Tbl1xr1,Tnfrsf1a,Tnfsf8,Wwox,Zfp609 |
| GO:0046638 | positive regulation of alpha-beta T cell differentiation | 4 | 47 | 0.0148 | Ccr7,Nfkbiz,Rara,Tgfbr2 |
| GO:2000481 | positive regulation of cAMP-dependent protein kinase activity | 2 | 4 | 0.0151 | Adipoq,Atp2b4 |
| GO:0045892 | negative regulation of transcription, DNA-templated | 21 | 1094 | 0.0152 | Adipoq,Aebp2,Bach2,Btg2,Fasl,Foxn3,Foxp1,Hey1,Klf2,Klf3,Lef1,Mbd2,Nfatc1,Nipbl,Nrip1,Pdcd4,Rara,Ski,Smarce1,Tbl1xr1,Zscan10 |
| GO:0002252 | immune effector process | 11 | 395 | 0.0153 | Bcl10,Cd47,Dock10,F2,Foxp1,Gzmb,Lef1,Ncf1,Rftn1,Tnfaip3,Trim25 |
| GO:0002697 | regulation of immune effector process | 10 | 335 | 0.0153 | Ccr7,Cd28,Dpp4,Dusp10,Foxp1,Nfkbiz,Rara,Rnf125,Stat5b,Tnfaip3 |

| | | | | | |
|---|---|---|---|---|---|
| GO:0009890 | negative regulation of biosynthetic process | 25 | 1408 | 0.0153 | Adipoq,Aebp2,Ago2,Atp2b4,Bach2,Btg2,Fasl,Foxn3,Foxp1,Hey1,Klf2,Klf3,Lef1,Mbd2,Nfatc1,Nipbl,Nrip1,Pdcd4,Prkca,Rara,Ski,Smarce1,Sod1,Tbl1xr1,Zscan10 |
| GO:0031399 | regulation of protein modification process | 29 | 1733 | 0.0153 | Adipoq,Atp2b4,Bcl10,Cblb,Ccr7,Cdkn2b,Dusp10,F2,Fam20a,Il6st,Lrtm2,Map3k1,Map3k8,Mapk9,Ncf1,Nipbl,Pdcd4,Pim1,Ppp2r5a,Prkca,Rara,Ski,Slc39a10,Sod1,Stk10,Tgfbr2,Tnfaip3,Tnfrsf1a,Xdh |
| GO:0002761 | regulation of myeloid leukocyte differentiation | 6 | 125 | 0.0154 | Adipoq,Foxp1,Lef1,Prkca,Rara,Tmem64 |
| GO:0014014 | negative regulation of gliogenesis | 4 | 48 | 0.0154 | Cdkn2b,Dusp10,F2,Ski |
| GO:0031667 | response to nutrient levels | 12 | 461 | 0.0159 | Adipoq,Bcl10,Cdkn2b,Dad1,Mbd2,P2rx4,Pim1,Rara,Sod1,Tbl1xr1,Tgfbr2,Trim25 |
| GO:0034614 | cellular response to reactive oxygen species | 6 | 127 | 0.0164 | Ets1,Klf2,Mapk9,Ncf1,Sod1,Tnfaip3 |
| GO:0051726 | regulation of cell cycle | 19 | 954 | 0.0167 | Atp2b4,Cd28,Cdkn2b,Ets1,Foxn3,Gpr132,Lef1,Map3k1,Map3k8,Nabp1,Pim1,Pkp4,Prkca,Prkcq,Rara,Sh2b1,Stat5b,Stk10,Tnfaip3 |
| GO:0045595 | regulation of cell differentiation | 29 | 1751 | 0.0172 | Adipoq,Ccr7,Cd28,Cdkn2b,Ddx6,Dusp10,Ets1,F2,Foxp1,Hey1,Il15ra,Il6st,Ist1,Lef1,Mapk9,Nfatc1,Nfkbiz,Nipbl,Pdcd4,Ppp1cc,Prkca,Rara,Ski,Stat5b,Tgfbr2,Tmem64,Tnfrsf1a,Xdh,Zfp609 |
| GO:0045597 | positive regulation of cell differentiation | 20 | 1033 | 0.0173 | Adipoq,Ccr7,Dusp10,Ets1,Foxp1,Hey1,Il15ra,Il6st,Ist1,Lef1,Mapk9,Nfkbiz,Nipbl,Ppp1cc,Prkca,Rara,Stat5b,Tgfbr2,Tmem64,Zfp609 |
| GO:0014013 | regulation of gliogenesis | 6 | 129 | 0.0174 | Cdkn2b,Dusp10,F2,Il6st,Ppp1cc,Ski |
| GO:0045624 | positive regulation of T-helper cell differentiation | 3 | 22 | 0.0174 | Ccr7,Nfkbiz,Rara |
| GO:0060438 | trachea development | 3 | 22 | 0.0174 | Lef1,Rara,Tgfbr2 |
| GO:2000113 | negative regulation of cellular macromolecule biosynthetic process | 23 | 1269 | 0.0177 | Adipoq,Aebp2,Ago2,Bach2,Btg2,Fasl,Foxn3,Foxp1,Hey1,Klf2,Klf3,Lef1,Mbd2,Nfatc1,Nipbl,Nrip1,Pdcd4,Prkca,Rara,Ski,Smarce1,Tbl1xr1,Zscan10 |
| GO:0071345 | cellular response to cytokine stimulus | 15 | 676 | 0.019 | Ccr4,Ccr7,Foxp1,Il6st,Jak1,Klf2,Lef1,Lrtm2,Rnf125,Sgms1,Stat5b,Tnfrsf1a,Tnfrsf1b,Trim25,Vamp4 |
| GO:0030219 | megakaryocyte differentiation | 3 | 23 | 0.0191 | Cdkn2b,Pip4k2a,Wasf2 |
| GO:0032753 | positive regulation of interleukin-4 production | 3 | 23 | 0.0191 | Cd28,Prkcq,Rara |
| GO:0032720 | negative regulation of tumor necrosis factor production | 4 | 52 | 0.0192 | Adipoq,Gpr18,Rara,Tnfaip3 |
| GO:0032355 | response to estradiol | 6 | 133 | 0.0195 | Ets1,Mbd2,Nrip1,Prkca,Rara,Stat5b |
| GO:0001932 | regulation of protein phosphorylation | 24 | 1362 | 0.0196 | Adipoq,Atp2b4,Cblb,Ccr7,Cdkn2b,Dusp10,F2,Fam20a,Il6st,Lrtm2,Map3k1,Map3k8,Mapk9,Ncf1,Pdcd4,Pim1,Prkca,Rara,Sod1,Stk10,Tgfbr2,Tnfaip3,Tnfrsf1a,Xdh |
| GO:0046686 | response to cadmium ion | 4 | 53 | 0.0203 | Cyb5a,Mapk9,Ncf1,Sod1 |
| GO:0002335 | mature B cell differentiation | 3 | 24 | 0.0208 | Dock10,Nfatc1,Tnfaip3 |
| GO:0033554 | cellular response to stress | 24 | 1370 | 0.0208 | Btg2,Ccr7,Chd2,Dyrk2,Ercc4,Ets1,Foxn3,Klf2,Map3k1,Map3k8,Mapk9,Mgme1,Nabp1,Ncf1,Nipbl,Nsmce1,Sirt4,Sod1,St8sia1,Stk10,Tnfaip3,Tnfrsf1a,Tnfrsf1b,Trim25 |
| GO:0051093 | negative regulation of developmental process | 19 | 979 | 0.0208 | Adipoq,Cdkn2b,Ddx6,Dusp10,F2,Fasl,Foxp1,Hey1,Il15ra,Klf2,Lef1,Nfatc1,Pdcd4,Rara,Ski,Stat5b,Tgfbr2,Tmem64,Xdh |
| GO:2000377 | regulation of reactive oxygen species metabolic process | 7 | 184 | 0.0208 | Atp2b4,F2,Klf2,Mapk9,Sod1,Tgfbr2,Xdh |
| GO:0031327 | negative regulation of cellular biosynthetic process | 24 | 1375 | 0.0216 | Adipoq,Aebp2,Ago2,Atp2b4,Bach2,Btg2,Fasl,Foxn3,Foxp1,Hey1,Klf2,Klf3,Lef1,Mbd2,Nfatc1,Nipbl,Nrip1,Pdcd4,Prkca,Rara,Ski,Smarce1,Tbl1xr1,Zscan10 |
| GO:0042327 | positive regulation of phosphorylation | 19 | 983 | 0.0216 | Adipoq,Atp2b4,Bcl10,Ccr7,F2,Fam20a,Il6st,Map3k1,Map3k8,Mapk9,Ncf1,Pim1,Prkca,Rara,Sod1,Stk10,Tgfbr2,Tnfrsf1a,Xdh |
| GO:0006351 | transcription, DNA-templated | 31 | 1960 | 0.0221 | Aebp2,Aff4,Ago2,Bach2,Btg2,Chd2,Elf1,Ets1,Fasl,Foxn3,Foxp1,Gtf2ird1,Hey1,Hivep2,Klf2,Klf3,Lef1,Mbd2,Nfatc1,Nfkbiz,Nipbl,Nrip1,Rara,Ski,Ss18,Stat5b,Tbl1xr1,Zfp3,Zfp609,Zfp652,Zscan10 |
| GO:0006469 | negative regulation of protein kinase activity | 8 | 241 | 0.0222 | Adipoq,Cblb,Cdkn2b,Dusp10,Lrtm2,Pdcd4,Prkca,Tnfaip3 |
| GO:0009628 | response to abiotic stimulus | 20 | 1063 | 0.0222 | Adipoq,Atp2b4,Bcl10,Btg2,Ccr4,Cyp2r1,Dpp4,Ercc4,Ets1,Gnat1,Mapk9,Mbd2,Nabp1,Nipbl,Ppp1cc,Sirt4,Sod1,St8sia1,Tgfbr2,Tnfrsf1a |
| GO:0019538 | protein metabolic process | 49 | 3580 | 0.0222 | Bach2,Btg2,Camk4,Cblb,Ccr7,Crbn,Dad1,Dcaf7,Dnajc6,Dpp4,Dusp10,Dyrk2,F2,Fkbp5,Gucy2c,Gzmb,Hbs1l,Itk,Jak1,Kcnh8,Lef1,Lmtk3,Lonrf1,Map3k1,Map3k8,Mapk9,Mknk2,Msrb3,Neurl1b,Nsmce1,Otulin,Pim1,Ppp1cc,Ppp2r5a,Prkca,Prkcq,Prss39,Rara,Riok1,Rnf125,Sirt4,St8sia1,Stk10,Tbl1xr1,Tgfbr2,Tnfaip3,Trim25,Wdyhv1,Xpnpep2 |
| GO:0031670 | cellular response to nutrient | 4 | 55 | 0.0222 | Cdkn2b,P2rx4,Pim1,Sod1 |
| GO:0032663 | regulation of interleukin-2 production | 4 | 55 | 0.0222 | Cd28,Prkcq,Stat5b,Tnfaip3 |
| GO:0043086 | negative regulation of catalytic activity | 16 | 765 | 0.0222 | Adipoq,Atp2b4,Cblb,Cdkn2b,Cnst,Crim1,Dusp10,Lef1,Lmtk3,Lrtm2,Pdcd4,Ppp2r5a,Prkca,Ski,Tnfaip3,Tnfrsf1b |
| GO:0071276 | cellular response to cadmium ion | 3 | 25 | 0.0222 | Mapk9,Ncf1,Sod1 |
| GO:0098751 | bone cell development | 3 | 25 | 0.0222 | Foxp1,Pip4k2a,Wasf2 |
| GO:0030856 | regulation of epithelial cell differentiation | 6 | 140 | 0.0232 | Adipoq,Foxp1,Hey1,Stat5b,Tnfrsf1a,Xdh |
| GO:0045639 | positive regulation of myeloid cell differentiation | 5 | 95 | 0.0232 | Ets1,Lef1,Prkca,Stat5b,Tmem64 |
| GO:0000820 | regulation of glutamine family amino acid metabolic process | 2 | 6 | 0.0235 | Atp2b4,Sirt4 |
| GO:0002666 | positive regulation of T cell tolerance induction | 2 | 6 | 0.0235 | Cblb,Tgfbr2 |
| GO:0070669 | response to interleukin-2 | 2 | 6 | 0.0235 | Jak1,Stat5b |
| GO:0070944 | neutrophil mediated killing of bacterium | 2 | 6 | 0.0235 | F2,Ncf1 |
| GO:0051128 | regulation of cellular component organization | 35 | 2337 | 0.0256 | Adipoq,Arhgap15,Ccr7,Cd28,Cd47,Crbn,Crim1,Dnajc6,Dpp4,Epb4.1,Ercc4,Ets1,F2,Foxp1,Gpbar1,Il15ra,Ist1,Lef1,Lrtm2,Map3k1,Mapk9,Nipbl,Pip4k2a,Prkca,Prkcq,S1pr1,Sh2b1,Sirt4,Ski,Syne3,Tapt1,Tnfrsf1a,Tnfrsf1b,Usp6nl,Wasf2 |
| GO:0033674 | positive regulation of kinase activity | 12 | 499 | 0.0257 | Adipoq,Atp2b4,Ccr7,F2,Fam20a,Map3k1,Map3k8,Ncf1,Pim1,Sod1,Stk10,Tgfbr2 |
| GO:0051347 | positive regulation of transferase activity | 13 | 569 | 0.0263 | Adipoq,Atp2b4,Ccr7,F2,Fam20a,Map3k1,Map3k8,Ncf1,Pim1,Prkcq,Sod1,Stk10,Tgfbr2 |
| GO:1901564 | organonitrogen compound metabolic process | 58 | 4480 | 0.0266 | Bach2,Btg2,Camk4,Cblb,Ccr7,Cers6,Crbn,Dad1,Dcaf7,Dnajc6,Dpp4,Dusp10,Dyrk2,Elovl5,F2,Fkbp5,Gucy2c,Gzmb,Hbs1l,Itk,Jak1,Kcnh8,Lef1,Lmtk3,Lonrf1,Map3k1,Map3k8,Mapk9,Mknk2,Msrb3,Ncf1,Neurl1b,Nsmce1,Otulin,Pim1,Ppp1cc,Ppp2r5a,Prkca,Prkcq,Prss39,Rara,Riok1,Rnf125,Rpia,Sgms1,Sirt4,Smox,Sod1,St8sia1,Stat5b,Stk10,Tbl1xr1,Tgfbr2,Tnfaip3,Trim25,Wdyhv1,Xdh,Xpnpep2 |
| GO:0002822 | regulation of adaptive immune response based on somatic recombination of immune receptors built from immunoglobulin superfamily domains | 6 | 146 | 0.027 | Ccr7,Cd28,Dpp4,Nfkbiz,Prkcq,Tnfaip3 |
| GO:0001934 | positive regulation of protein phosphorylation | 18 | 937 | 0.0272 | Adipoq,Atp2b4,Ccr7,F2,Fam20a,Il6st,Map3k1,Map3k8,Mapk9,Ncf1,Pim1,Prkca,Rara,Sod1,Stk10,Tgfbr2,Tnfrsf1a,Xdh |
| GO:0050927 | positive regulation of positive chemotaxis | 3 | 28 | 0.0276 | Ccr4,Prkca,S1pr1 |
| GO:0051098 | regulation of binding | 10 | 376 | 0.0282 | Adipoq,Crbn,Epb4.1,Ercc4,Hey1,Lef1,Mbd2,Nipbl,Rara,Ski |
| GO:0050665 | hydrogen peroxide biosynthetic process | 2 | 7 | 0.0284 | Ncf1,Sod1 |
| GO:0006979 | response to oxidative stress | 10 | 377 | 0.0286 | Adipoq,Ccr7,Ets1,Klf2,Mapk9,Msrb3,Ncf1,Prkca,Sod1,Tnfaip3 |
| GO:0032269 | negative regulation of cellular protein metabolic process | 19 | 1020 | 0.0287 | Adipoq,Ago2,Btg2,Cblb,Cdkn2b,Crim1,Dusp10,F2,Lef1,Lrtm2,Pdcd4,Prkca,Rara,Sirt4,Ski,Tnfaip3,Tnfrsf1b,Ubac2,Xdh |
| GO:0046677 | response to antibiotic | 9 | 317 | 0.0296 | Adipoq,Ccr4,Ets1,Jak1,Klf2,Prkca,Rara,Sod1,Tnfaip3 |
| GO:0006796 | phosphate-containing compound metabolic process | 28 | 1758 | 0.0297 | Camk4,Ccr7,Dnajc6,Dusp10,Dyrk2,Gucy2c,Itk,Jak1,Kcnh8,Lmtk3,Lpcat3,Map3k1,Map3k8,Mapk9,Mknk2,Ncf1,Pim1,Pip4k2a,Ppp1cc,Ppp2r5a,Prkca,Prkcq,Rara,Riok1,Rpia,Sgms1,Stk10,Tgfbr2 |
| GO:0061515 | myeloid cell development | 4 | 62 | 0.0299 | Foxp1,Klf2,Pip4k2a,Wasf2 |
| GO:0045596 | negative regulation of cell differentiation | 15 | 724 | 0.03 | Adipoq,Cdkn2b,Ddx6,Dusp10,F2,Foxp1,Hey1,Il15ra,Nfatc1,Pdcd4,Rara,Ski,Stat5b,Tmem64,Xdh |
| GO:0002250 | adaptive immune response | 8 | 260 | 0.0308 | Bcl10,Camk4,Cd247,Gzmb,Itk,Lef1,Rftn1,Rnf125 |
| GO:0001819 | positive regulation of cytokine production | 10 | 383 | 0.0312 | Adipoq,Bcl10,Ccr7,Cd28,Mapk9,Prkcq,Rara,Sod1,Stat5b,Tnfrsf1a |
| GO:0042108 | positive regulation of cytokine biosynthetic process | 4 | 63 | 0.0313 | Bcl10,Cd28,Prkcq,Stat5b |
| GO:0030278 | regulation of ossification | 7 | 205 | 0.0317 | Hey1,Il6st,Nfatc1,Nipbl,S1pr1,Ski,Tmem64 |
| GO:0048661 | positive regulation of smooth muscle cell proliferation | 5 | 105 | 0.0318 | Foxp1,Prkca,S1pr1,Stat5b,Tgfbr2 |

| | | | | | |
|---|---|---|---|---|---|
| GO:0014068 | positive regulation of phosphatidylinositol 3-kinase signaling | 4 | 64 | 0.0326 | Cd28,F2,Ncf1,Rara |
| GO:0009896 | positive regulation of catabolic process | 10 | 388 | 0.0336 | Ago2,Btg2,Cblb,Lonrf1,Mapk9,Pip4k2a,Rnf125,Tnfaip3,Tnfrsf1b,Zfand2a |
| GO:0001837 | epithelial to mesenchymal transition | 4 | 65 | 0.0337 | Hey1,Lef1,Nfatc1,Pdcd4 |
| GO:0001909 | leukocyte mediated cytotoxicity | 3 | 31 | 0.0337 | F2,Gzmb,Ncf1 |
| GO:0016999 | antibiotic metabolic process | 5 | 107 | 0.0337 | Cmah,Cmas,Ncf1,Sod1,Stat5b |
| GO:0045732 | positive regulation of protein catabolic process | 7 | 208 | 0.0337 | Cblb,Lonrf1,Mapk9,Rnf125,Tnfaip3,Tnfrsf1b,Zfand2a |
| GO:0046777 | protein autophosphorylation | 7 | 208 | 0.0337 | Camk4,Itk,Jak1,Mknk2,Pim1,Prkca,Stk10 |
| GO:0071872 | cellular response to epinephrine stimulus | 2 | 8 | 0.0337 | Adipoq,Atp2b4 |
| GO:1902339 | positive regulation of apoptotic process involved in morphogenesis | 2 | 8 | 0.0337 | Tnfrsf1a,Tnfrsf1b |
| GO:1905064 | negative regulation of vascular smooth muscle cell differentiation | 2 | 8 | 0.0337 | Nfatc1,Pdcd4 |
| GO:2000318 | positive regulation of T-helper 17 type immune response | 2 | 8 | 0.0337 | Nfkbiz,Prkcq |
| GO:0051253 | negative regulation of RNA metabolic process | 21 | 1204 | 0.0347 | Adipoq,Aebp2,Bach2,Btg2,Fasl,Foxn3,Foxp1,Hey1,Klf2,Klf3,Lef1,Mbd2,Nfatc1,Nipbl,Nrip1,Pdcd4,Rara,Ski,Smarce1,Tbl1xr1,Zscan10 |
| GO:0000122 | negative regulation of transcription by RNA polymerase II | 16 | 817 | 0.0356 | Aebp2,Bach2,Btg2,Fasl,Foxp1,Hey1,Klf2,Klf3,Lef1,Mbd2,Nfatc1,Nipbl,Nrip1,Rara,Ski,Tbl1xr1 |
| GO:0009719 | response to endogenous stimulus | 22 | 1289 | 0.0358 | Abhd2,Adipoq,Atp2b4,Btg2,Ccr7,Ets1,Foxp1,Klf2,Klf3,Lef1,Map3k1,Mbd2,Nrip1,P2rx4,Pdcd4,Prkca,Rara,Rgs10,Sod1,Stat5b,Tgfbr2,Wwox |
| GO:0034405 | response to fluid shear stress | 3 | 32 | 0.0358 | Ets1,Klf2,P2rx4 |
| GO:0060251 | regulation of glial cell proliferation | 3 | 32 | 0.0358 | Cdkn2b,Ppp1cc,Ski |
| GO:0043407 | negative regulation of MAP kinase activity | 4 | 67 | 0.0363 | Adipoq,Dusp10,Pdcd4,Prkca |
| GO:0032872 | regulation of stress-activated MAPK cascade | 7 | 213 | 0.0364 | Ccr7,Dusp10,Map3k1,Mapk9,Ncf1,Pdcd4,Xdh |
| GO:0045860 | positive regulation of protein kinase activity | 11 | 462 | 0.0364 | Adipoq,Atp2b4,Ccr7,Fam20a,Map3k1,Map3k8,Ncf1,Pim1,Sod1,Stk10,Tgfbr2 |
| GO:0031331 | positive regulation of cellular catabolic process | 9 | 333 | 0.037 | Ago2,Btg2,Lonrf1,Mapk9,Pip4k2a,Rnf125,Tnfaip3,Tnfrsf1b,Zfand2a |
| GO:0045862 | positive regulation of proteolysis | 9 | 334 | 0.0376 | Bcl10,Fasl,Ist1,Lonrf1,Mapk9,Rnf125,Tnfrsf1b,Xdh,Zfand2a |
| GO:0007259 | receptor signaling pathway via JAK-STAT | 3 | 33 | 0.0378 | Il15ra,Jak1,Stat5b |
| GO:0043122 | regulation of I-kappaB kinase/NF-kappaB signaling | 6 | 161 | 0.0378 | Adipoq,Bcl10,Fasl,Tnfaip3,Tnfrsf1a,Trim25 |
| GO:0007517 | muscle organ development | 8 | 274 | 0.0382 | Btg2,Chd2,Foxp1,Hey1,Mbnl1,S1pr1,Ski,Zfp609 |
| GO:0021761 | limbic system development | 5 | 112 | 0.0382 | Atp2b4,Btg2,Ets1,Lef1,Rara |
| GO:0042996 | regulation of Golgi to plasma membrane protein transport | 2 | 9 | 0.0382 | Cnst,Vamp4 |
| GO:0030032 | lamellipodium assembly | 3 | 34 | 0.0402 | S1pr1,Sh2b1,Wasf2 |
| GO:0034654 | nucleobase-containing compound biosynthetic process | 34 | 2339 | 0.0402 | Aebp2,Aff4,Ago2,Bach2,Btg2,Chd2,Cmas,Elf1,Ets1,Fasl,Foxn3,Foxp1,Gtf2ird1,Gucy2c,Hey1,Hivep2,Klf2,Klf3,Lef1,Mbd2,Ncf1,Nfatc1,Nfkbiz,Nipbl,Nrip1,Rara, Ski,Ss18,Stat5b,Tbl1xr1,Zfp3,Zfp609,Zfp652,Zscan10 |
| GO:0046640 | regulation of alpha-beta T cell proliferation | 3 | 34 | 0.0402 | Cblb,Cd28,Tgfbr2 |
| GO:0097028 | dendritic cell differentiation | 3 | 34 | 0.0402 | Camk4,Ccr7,Tgfbr2 |
| GO:0009612 | response to mechanical stimulus | 7 | 219 | 0.0403 | Bcl10,Btg2,Ets1,Mapk9,Mbd2,Tgfbr2,Tnfrsf1a |
| GO:0010565 | regulation of cellular ketone metabolic process | 5 | 114 | 0.0403 | Adipoq,Atp2b4,Elovl5,Mapk9,Sirt4 |
| GO:1903828 | negative regulation of cellular protein localization | 5 | 114 | 0.0403 | Adipoq,Leprotl1,Ppp2r5a,Sirt4,Ubac2 |
| GO:0044403 | symbiont process | 7 | 220 | 0.0411 | Ddx6,Dpp4,F2,Ist1,Lef1,Ncf1,Usp6nl |
| GO:0043123 | positive regulation of I-kappaB kinase/NF-kappaB signaling | 5 | 115 | 0.0413 | Adipoq,Bcl10,Fasl,Tnfrsf1a,Trim25 |
| GO:1900744 | regulation of p38MAPK cascade | 3 | 35 | 0.0423 | Dusp10,Ncf1,Xdh |
| GO:0002315 | marginal zone B cell differentiation | 2 | 10 | 0.0439 | Dock10,Tnfaip3 |
| GO:0030853 | negative regulation of granulocyte differentiation | 2 | 10 | 0.0439 | Adipoq,Rara |
| GO:0031098 | stress-activated protein kinase signaling cascade | 5 | 117 | 0.0439 | Ccr7,Map3k1,Map3k8,Mapk9,Stk10 |
| GO:0034115 | negative regulation of heterotypic cell-cell adhesion | 2 | 10 | 0.0439 | Adipoq,Tnfaip3 |
| GO:0043374 | CD8-positive, alpha-beta T cell differentiation | 2 | 10 | 0.0439 | Gpr18,Tnfsf8 |
| GO:0045766 | positive regulation of angiogenesis | 6 | 168 | 0.0439 | Ago2,Ets1,Jak1,Prkca,Tgfbr2,Tnfrsf1a |
| GO:0061384 | heart trabecula morphogenesis | 3 | 36 | 0.0442 | Hey1,Nfatc1,S1pr1 |
| GO:0045934 | negative regulation of nucleobase-containing compound metabolic process | 22 | 1326 | 0.0448 | Adipoq,Aebp2,Bach2,Btg2,Ercc4,Fasl,Foxn3,Foxp1,Hey1,Klf2,Klf3,Lef1,Mbd2,Nfatc1,Nipbl,Nrip1,Pdcd4,Rara,Ski,Smarce1,Tbl1xr1,Zscan10 |
| GO:0042592 | homeostatic process | 24 | 1494 | 0.0452 | Adipoq,Ankrd11,Atp2b4,Ccr4,Ccr7,Dock10,Ercc4,F2,Fam20a,Fasl,Gpr18,Il6st,Klf2,Lrrc8d,P2rx4,Prkca,S1pr1,Selt,Slc39a10,Sod1,Stat5b,Thada,Tmem64,Tnfa ip3 |
| GO:0042493 | response to drug | 17 | 926 | 0.0455 | Abcb1b,Abhd2,Adipoq,Ccr7,Dad1,Ets1,Klf2,Mapk9,P2rx4,Prkca,Rara,Rgs10,Slc7a3,Sod1,Ss18,Tgfbr2,Tnfaip3 |
| GO:0045927 | positive regulation of growth | 8 | 286 | 0.0455 | F2,Ist1,Lef1,Nipbl,Pim1,Rftn1,Stat5b,Tgfbr2 |
| GO:0006919 | activation of cysteine-type endopeptidase activity involved in apoptotic process | 4 | 74 | 0.0458 | Bcl10,Fasl,Mapk9,Xdh |
| GO:0045765 | regulation of angiogenesis | 8 | 287 | 0.0462 | Ago2,Ets1,Fasl,Jak1,Klf2,Prkca,Tgfbr2,Tnfrsf1a |
| GO:0051101 | regulation of DNA binding | 5 | 120 | 0.0467 | Ercc4,Hey1,Lef1,Nipbl,Ski |
| GO:0097305 | response to alcohol | 7 | 228 | 0.0468 | Adipoq,Ccr7,Klf2,Prkca,Rara,Sod1,Tgfbr2 |
| GO:0002507 | tolerance induction | 2 | 11 | 0.0491 | Ccr4,Tnfaip3 |
| GO:0002683 | negative regulation of immune system process | 10 | 421 | 0.0491 | Adipoq,Bcl10,Cblb,Dusp10,Elf1,Gpr18,Rara,Rnf125,Stat5b,Tnfaip3 |
| GO:0003151 | outflow tract morphogenesis | 4 | 76 | 0.0491 | Hey1,Nipbl,Rara,Tgfbr2 |
| GO:0006054 | N-acetylneuraminate metabolic process | 2 | 11 | 0.0491 | Cmah,Cmas |
| GO:0034616 | response to laminar fluid shear stress | 2 | 11 | 0.0491 | Ets1,Klf2 |
| GO:0042554 | superoxide anion generation | 2 | 11 | 0.0491 | Ncf1,Sod1 |
| GO:0046328 | regulation of JNK cascade | 6 | 174 | 0.0491 | Ccr7,Dusp10,Map3k1,Mapk9,Ncf1,Pdcd4 |
| GO:0048705 | skeletal system morphogenesis | 7 | 231 | 0.0491 | Ankrd11,Foxn3,Nipbl,Rara,Ski,Tgfbr2,Wwox |
| GO:1900016 | negative regulation of cytokine production involved in inflammatory response | 2 | 11 | 0.0491 | F2,Pdcd4 |
| GO:1901343 | negative regulation of vasculature development | 5 | 122 | 0.0491 | Fasl,Klf2,Nfatc1,Pdcd4,Xdh |

**Table S5A. HOMER Known Motifs near the opening peaks of cytokine-primed cells**
Only motifs with q-value below 0.05 are shown.

| Motif Name | Consensus | P-value | Log P-value | q-value (Benjamini) | # of Target Sequences with Motif (of 195) | % of Target Sequences with Motif | # of Background Sequences with Motif (of 49494) | % of Background Sequences with Motif |
|---|---|---|---|---|---|---|---|---|
| STAT5(Stat)/mCD4+-Stat5-ChIP-Seq(GSE12346)/Homer | RTTTCTNAGAAA | 1.00E-53 | -1.23E+02 | 0 | 119 | 61.03% | 6617.3 | 13.37% |
| STAT1(Stat)/HelaS3-STAT1-ChIP-Seq(GSE12782)/Homer | NATTTCCNGGAAAT | 1.00E-48 | -1.11E+02 | 0 | 106 | 54.36% | 5562.7 | 11.24% |
| STAT4(Stat)/CD4-Stat4-ChIP-Seq(GSE22104)/Homer | NYTTCCWGGAAR | 1.00E-35 | -8.26E+01 | 0 | 152 | 77.95% | 16682.8 | 33.71% |
| Stat3+il21(Stat)/CD4-Stat3-ChIP-Seq(GSE19198)/Homer | SVYTTCCNGGAARB | 1.00E-31 | -7.24E+01 | 0 | 125 | 64.10% | 11944.4 | 24.13% |
| Stat3(Stat)/mES-Stat3-ChIP-Seq(GSE11431)/Homer | CTTCCGGGAA | 1.00E-20 | -4.63E+01 | 0 | 88 | 45.13% | 8153.6 | 16.47% |
| RUNX1(Runt)/Jurkat-RUNX1-ChIP-Seq(GSE29180)/Homer | AAACCACARM | 1.00E-15 | -3.58E+01 | 0 | 125 | 64.10% | 17478.3 | 35.31% |
| RUNX(Runt)/HPC7-Runx1-ChIP-Seq(GSE22178)/Homer | SAAACCACAG | 1.00E-15 | -3.56E+01 | 0 | 102 | 52.31% | 12391.7 | 25.04% |
| Bcl6(Zf)/Liver-Bcl6-ChIP-Seq(GSE31578)/Homer | NNNCTTTCCAGGAAA | 1.00E-14 | -3.33E+01 | 0 | 141 | 72.31% | 22032.5 | 44.52% |
| ETS:E-box(ETS,bHLH)/HPC7-Scl-ChIP-Seq(GSE22178)/Homer | AGGAARCAGCTG | 1.00E-08 | -2.00E+01 | 0 | 27 | 13.85% | 1757.9 | 3.55% |
| RUNX2(Runt)/PCa-RUNX2-ChIP-Seq(GSE33889)/Homer | NWAACCACADNN | 1.00E-08 | -1.85E+01 | 0 | 97 | 49.74% | 14963.1 | 30.23% |
| STAT6(Stat)/Macrophage-Stat6-ChIP-Seq(GSE38377)/Homer | TTCCKNAGAA | 1.00E-07 | -1.68E+01 | 0 | 74 | 37.95% | 10400.4 | 21.01% |
| RUNX-AML(Runt)/CD4+-PolII-ChIP-Seq(Barski et al.)/Homer | GCTGTGGTTW | 1.00E-07 | -1.63E+01 | 0 | 85 | 43.59% | 12871.8 | 26.01% |
| EWS:FLI1-fusion(ETS)/SK_N_MC-EWS:FLI1-ChIP-Seq(SRA014231)/Homer | VACAGGAAAT | 1.00E-05 | -1.37E+01 | 0 | 66 | 33.85% | 9546.9 | 19.29% |
| EWS:ERG-fusion(ETS)/CADO_ES1-EWS:ERG-ChIP-Seq(SRA014231)/Homer | ATTTCCTGTN | 1.00E-05 | -1.30E+01 | 0 | 81 | 41.54% | 12952 | 26.17% |
| Fli1(ETS)/CD8-FLI-ChIP-Seq(GSE20898)/Homer | NRYTTCCGGH | 1.00E-05 | -1.19E+01 | 0.0001 | 94 | 48.21% | 16292.2 | 32.92% |
| Ptf1a(bHLH)/Panc1-Ptf1a-ChIP-Seq(GSE47459)/Homer | ACAGCTGTTN | 1.00E-04 | -1.15E+01 | 0.0002 | 155 | 79.49% | 32310.8 | 65.28% |
| Ets1-distal(ETS)/CD4+-PolII-ChIP-Seq(Barski et al.)/Homer | MACAGGAAGT | 1.00E-04 | -1.13E+01 | 0.0002 | 45 | 23.08% | 5949.9 | 12.02% |
| Atoh1(bHLH)/Cerebellum-Atoh1-ChIP-Seq(GSE22111)/Homer | VNRVCAGCTGGY | 1.00E-04 | -1.00E+01 | 0.0006 | 94 | 48.21% | 16980.1 | 34.31% |
| PU.1-IRF(ETS:IRF)/Bcell-PU.1-ChIP-Seq(GSE21512)/Homer | MGGAAGTGAAAC | 1.00E-04 | -9.23E+00 | 0.0014 | 112 | 57.44% | 21718.2 | 43.88% |
| Ascl1(bHLH)/NeuralTubes-Ascl1-ChIP-Seq(GSE55840)/Homer | NNVVCAGCTGBN | 1.00E-04 | -9.22E+00 | 0.0014 | 111 | 56.92% | 21475.7 | 43.39% |
| ETS:RUNX(ETS,Runt)/Jurkat-RUNX1-ChIP-Seq(GSE17954)/Homer | RCAGGATGTGGT | 1.00E-03 | -8.55E+00 | 0.0024 | 15 | 7.69% | 1281.6 | 2.59% |
| USF1(bHLH)/GM12878-Usf1-ChIP-Seq(GSE32465)/Homer | SGTCACGTGR | 1.00E-03 | -8.46E+00 | 0.0025 | 41 | 21.03% | 5900.2 | 11.92% |
| ETS1(ETS)/Jurkat-ETS1-ChIP-Seq(GSE17954)/Homer | ACAGGAAGTG | 1.00E-03 | -7.90E+00 | 0.0043 | 88 | 45.13% | 16454.7 | 33.25% |
| KLF5(Zf)/LoVo-KLF5-ChIP-Seq(GSE49402)/Homer | DGGGYGKGGC | 1.00E-03 | -7.73E+00 | 0.0048 | 81 | 41.54% | 14881.9 | 30.07% |
| ERG(ETS)/VCaP-ERG-ChIP-Seq(GSE14097)/Homer | ACAGGAAGTG | 1.00E-03 | -7.63E+00 | 0.0051 | 121 | 62.05% | 24761.6 | 50.03% |
| SPDEF(ETS)/VCaP-SPDEF-ChIP-Seq(SRA014231)/Homer | ASWTCCTGBT | 1.00E-03 | -7.55E+00 | 0.0054 | 88 | 45.13% | 16615 | 33.57% |
| PAX3:FKHR-fusion(Paired,Homeobox)/Rh4-PAX3:FKHR-ChIP-Seq(GSE19063)/Homer | ACCRTGACTAATTNN | 1.00E-03 | -7.12E+00 | 0.0079 | 26 | 13.33% | 3366.3 | 6.80% |
| E2A(bHLH)/proBcell-E2A-ChIP-Seq(GSE21978)/Homer | DNRCAGCTGY | 1.00E-03 | -7.05E+00 | 0.0082 | 101 | 51.79% | 20009 | 40.43% |
| PU.1(ETS)/ThioMac-PU.1-ChIP-Seq(GSE21512)/Homer | AGAGGAAGTG | 1.00E-02 | -6.85E+00 | 0.0096 | 52 | 26.67% | 8714.7 | 17.61% |
| Nkx3.1(Homeobox)/LNCaP-Nkx3.1-ChIP-Seq(GSE28264)/Homer | AAGCACTTAA | 1.00E-02 | -6.76E+00 | 0.0103 | 149 | 76.41% | 32716.4 | 66.10% |
| Maz(Zf)/HepG2-Maz-ChIP-Seq(GSE31477)/Homer | GGGGGGGG | 1.00E-02 | -6.71E+00 | 0.0104 | 83 | 42.56% | 15824.1 | 31.97% |
| IRF2(IRF)/Erythroblas-IRF2-ChIP-Seq(GSE36985)/Homer | GAAASYGAAASY | 1.00E-02 | -6.55E+00 | 0.0119 | 16 | 8.21% | 1723.9 | 3.48% |
| T1ISRE(IRF)/ThioMac-Ifnb-Expression/Homer | ACTTTCGTTTCT | 1.00E-02 | -6.43E+00 | 0.0129 | 5 | 2.56% | 211.6 | 0.43% |
| MyoG(bHLH)/C2C12-MyoG-ChIP-Seq(GSE36024)/Homer | AACAGCTG | 1.00E-02 | -5.93E+00 | 0.0207 | 82 | 42.05% | 15982.9 | 32.29% |
| Reverb(NR),DR2/RAW-Reverba.biotin-ChIP-Seq(GSE45914)/Homer | GTRGGTCASTGGGTCA | 1.00E-02 | -5.79E+00 | 0.0232 | 18 | 9.23% | 2212 | 4.47% |
| Ap4(bHLH)/AML-Tfap4-ChIP-Seq(GSE45738)/Homer | NAHCAGCTGD | 1.00E-02 | -5.78E+00 | 0.0232 | 91 | 46.67% | 18227.9 | 36.83% |
| Fosl2(bZIP)/3T3L1-Fosl2-ChIP-Seq(GSE56872)/Homer | NATGASTCABNN | 1.00E-02 | -5.63E+00 | 0.0255 | 33 | 16.92% | 5148.5 | 10.40% |
| STAT6(Stat)/CD4-Stat6-ChIP-Seq(GSE22104)/Homer | ABTTCYYRRGAA | 1.00E-02 | -5.63E+00 | 0.0255 | 57 | 29.23% | 10329.3 | 20.87% |
| Jun-AP1(bZIP)/K562-cJun-ChIP-Seq(GSE31477)/Homer | GATGASTCATCN | 1.00E-02 | -5.59E+00 | 0.0255 | 25 | 12.82% | 3565.3 | 7.20% |
| Elk4(ETS)/Hela-Elk4-ChIP-Seq(GSE31477)/Homer | NRYTTCCGGY | 1.00E-02 | -5.33E+00 | 0.032 | 41 | 21.03% | 6938.5 | 14.02% |
| EHF(ETS)/LoVo-EHF-ChIP-Seq(GSE49402)/Homer | AVCAGGAAGT | 1.00E-02 | -5.26E+00 | 0.0336 | 101 | 51.79% | 20997.1 | 42.42% |
| REST-NRSF(Zf)/Jurkat-NRSF-ChIP-Seq/Homer | GGMGCTGTCCATGGTGCTGA | 1.00E-02 | -5.24E+00 | 0.0336 | 3 | 1.54% | 88.4 | 0.18% |
| GLI3(Zf)/Limb-GLI3-ChIP-Chip(GSE11077)/Homer | CGTGGGTGGTCC | 1.00E-02 | -5.22E+00 | 0.0336 | 15 | 7.69% | 1799.8 | 3.64% |
| Myf5(bHLH)/GM-Myf5-ChIP-Seq(GSE24852)/Homer | BAACAGCTGT | 1.00E-02 | -5.19E+00 | 0.0336 | 60 | 30.77% | 11221 | 22.67% |
| Nkx2.5(Homeobox)/HL1-Nkx2.5.biotin-ChIP-Seq(GSE21529)/Homer | RRSCACTYAA | 1.00E-02 | -5.16E+00 | 0.0338 | 141 | 72.31% | 31421.5 | 63.49% |
| BMAL1(bHLH)/Liver-Bmal1-ChIP-Seq(GSE39860)/Homer | GNCACGTG | 1.00E-02 | -5.12E+00 | 0.0345 | 119 | 61.03% | 25646.2 | 51.82% |

| | | | | | | | | |
|---|---|---|---|---|---|---|---|---|
| NeuroD1(bHLH)/Islet-NeuroD1-ChIP-Seq(GSE30298)/Homer | GCCATCTGTT | 1.00E-02 | -5.10E+00 | 0.0345 | 68 | 34.87% | 13117.1 | 26.50% |
| E-box(bHLH)/Promoter/Homer | SSGGTCACGTGA | 1.00E-02 | -5.08E+00 | 0.0345 | 9 | 4.62% | 833.8 | 1.68% |
| ISRE(IRF)/ThioMac-LPS-Expression(GSE23622)/Homer | AGTTTCASTTTC | 1.00E-02 | -5.04E+00 | 0.0349 | 10 | 5.13% | 994.1 | 2.01% |
| ETV1(ETS)/GIST48-ETV1-ChIP-Seq(GSE22441)/Homer | AACCGGAAGT | 1.00E-02 | -4.75E+00 | 0.0455 | 99 | 50.77% | 20810.1 | 42.05% |
| NF1-halfsite(CTF)/LNCaP-NF1-ChIP-Seq(Unpublished)/Homer | YTGCCAAG | 1.00E-02 | -4.73E+00 | 0.0458 | 114 | 58.46% | 24611.7 | 49.73% |
| SpiB(ETS)/OCILY3-SPIB-ChIP-Seq(GSE56857)/Homer | AAAGRGGAAGTG | 1.00E-02 | -4.71E+00 | 0.0458 | 27 | 13.85% | 4234.2 | 8.56% |
| Tcf4(HMG)/Hct116-Tcf4-ChIP-Seq(SRA012054)/Homer | ASATCAAAGGVA | 1.00E-01 | -4.60E+00 | 0.05 | 48 | 24.62% | 8796.5 | 17.77% |
| MyoD(bHLH)/Myotube-MyoD-ChIP-Seq(GSE21614)/Homer | RRCAGCTGYTSY | 1.00E-01 | -4.59E+00 | 0.05 | 61 | 31.28% | 11767.1 | 23.78% |
| Tcf12(bHLH)/GM12878-Tcf12-ChIP-Seq(GSE32465)/Homer | VCAGCTGYTG | 1.00E-01 | -4.58E+00 | 0.05 | 72 | 36.92% | 14352.8 | 29.00% |
| ERE(NR),IR3/MCF7-ERa-ChIP-Seq(Unpublished)/Homer | VAGGTCACNSTGACC | 1.00E-01 | -4.58E+00 | 0.05 | 27 | 13.85% | 4279.1 | 8.65% |

**Table S5B. HOMER Known Motifs near the opening peaks of Ag-stimulated cells**
Only motifs with q-value below 0.05 are shown.

| Motif Name | Consensus | P-value | Log P-value | q-value (Benjamini) | # of Target Sequences with | % of Target Sequences with | # of Background Sequences with | % of Background |
|---|---|---|---|---|---|---|---|---|
| Fra1(bZIP)/BT549-Fra1-ChIP-Seq(GSE46166)/Homer | NNATGASTCATH | 1e-1618 | -3.73E+03 | 0 | 5961 | 53.83% | 6813.1 | 17.53% |
| Fosl2(bZIP)/3T3L1-Fosl2-ChIP-Seq(GSE56872)/Homer | NATGASTCABNN | 1e-1574 | -3.63E+03 | 0 | 4523 | 40.84% | 3881.4 | 9.99% |
| Atf3(bZIP)/GBM-ATF3-ChIP-Seq(GSE33912)/Homer | DATGASTCATHN | 1e-1549 | -3.57E+03 | 0 | 6425 | 58.02% | 8204.2 | 21.11% |
| BATF(bZIP)/Th17-BATF-ChIP-Seq(GSE39756)/Homer | DATGASTCAT | 1e-1507 | -3.47E+03 | 0 | 6391 | 57.71% | 8268.5 | 21.28% |
| Jun-AP1(bZIP)/K562-cJun-ChIP-Seq(GSE31477)/Homer | GATGASTCATCN | 1e-1418 | -3.27E+03 | 0 | 3651 | 32.97% | 2691.1 | 6.92% |
| AP-1(bZIP)/ThioMac-PU.1-ChIP-Seq(GSE21512)/Homer | VTGACTCATC | 1e-1380 | -3.18E+03 | 0 | 6553 | 59.17% | 9201.6 | 23.68% |
| Bach2(bZIP)/OCILy7-Bach2-ChIP-Seq(GSE44420)/Homer | TGCTGAGTCA | 1e-492 | -1.14E+03 | 0 | 2090 | 18.87% | 2244.8 | 5.78% |
| IRF2(IRF)/Erythroblas-IRF2-ChIP-Seq(GSE36985)/Homer | GAAASYGAAASY | 1.00E-204 | -4.72E+02 | 0 | 1058 | 9.55% | 1260.1 | 3.24% |
| bZIP:IRF(bZIP,IRF)/Th17-BatF-ChIP-Seq(GSE39756)/Homer | NAGTTTCABTHTGACTNW | 1.00E-204 | -4.72E+02 | 0 | 3178 | 28.70% | 6594.6 | 16.97% |
| IRF1(IRF)/PBMC-IRF1-ChIP-Seq(GSE43036)/Homer | GAAAGTGAAAGT | 1.00E-185 | -4.27E+02 | 0 | 1286 | 11.61% | 1834.7 | 4.72% |
| RUNX1(Runt)/Jurkat-RUNX1-ChIP-Seq(GSE29180)/Homer | AAACCACARM | 1.00E-153 | -3.53E+02 | 0 | 5045 | 45.56% | 12989.1 | 33.42% |
| NFkB-p65(RHD)/GM12787-p65-ChIP-Seq(GSE19485)/Homer | WGGGGATTTCCC | 1.00E-153 | -3.53E+02 | 0 | 2811 | 25.38% | 6070.7 | 15.62% |
| ISRE(IRF)/ThioMac-LPS-Expression(GSE23622)/Homer | AGTTTCASTTTC | 1.00E-152 | -3.52E+02 | 0 | 693 | 6.26% | 748.5 | 1.93% |
| IRF4(IRF)/GM12878-IRF4-ChIP-Seq(GSE32465)/Homer | ACTGAAACCA | 1.00E-148 | -3.43E+02 | 0 | 2755 | 24.88% | 5955.4 | 15.32% |
| STAT5(Stat)/mCD4+-Stat5-ChIP-Seq(GSE12346)/Homer | RTTTCTNAGAAA | 1.00E-144 | -3.34E+02 | 0 | 2457 | 22.19% | 5149.6 | 13.25% |
| RUNX(Runt)/HPC7-Runx1-ChIP-Seq(GSE22178)/Homer | SAAACCACAG | 1.00E-137 | -3.16E+02 | 0 | 3825 | 34.54% | 9317.3 | 23.97% |
| RUNX2(Runt)/PCa-RUNX2-ChIP-Seq(GSE33889)/Homer | NWAACCACADNN | 1.00E-137 | -3.16E+02 | 0 | 4420 | 39.91% | 11202.8 | 28.83% |
| NF-E2(bZIP)/K562-NFE2-ChIP-Seq(GSE31477)/Homer | GATGACTCAGCA | 1.00E-123 | -2.83E+02 | 0 | 554 | 5.00% | 593.7 | 1.53% |
| NFAT:AP1(RHD,bZIP)/Jurkat-NFATC1-ChIP-Seq(Jolma et al.)/Homer | SARTGGAAAAWRTGAGTCAB | 1.00E-122 | -2.83E+02 | 0 | 1412 | 12.75% | 2539.2 | 6.53% |
| RUNX-AML(Runt)/CD4+-PolII-ChIP-Seq(Barski et al.)/Homer | GCTGTGGTTW | 1.00E-120 | -2.78E+02 | 0 | 3889 | 35.12% | 9757.8 | 25.11% |
| NFkB-p50,p52(RHD)/Monocyte-p50-ChIP-Chip(Schreiber et al.)/Homer | GGGGGAATCCCC | 1.00E-115 | -2.66E+02 | 0 | 679 | 6.13% | 869.5 | 2.24% |
| Nrf2(bZIP)/Lymphoblast-Nrf2-ChIP-Seq(GSE37589)/Homer | HTGCTGAGTCAT | 1.00E-102 | -2.37E+02 | 0 | 453 | 4.09% | 477.5 | 1.23% |
| STAT1(Stat)/HelaS3-STAT1-ChIP-Seq(GSE12782)/Homer | NATTTCCNGGAAAT | 1.00E-95 | -2.20E+02 | 0 | 1994 | 18.01% | 4388.3 | 11.29% |
| Bach1(bZIP)/K562-Bach1-ChIP-Seq(GSE31477)/Homer | AWWNTGCTGAGTCAT | 1.00E-93 | -2.15E+02 | 0 | 496 | 4.48% | 594.9 | 1.53% |
| MafK(bZIP)/C2C12-MafK-ChIP-Seq(GSE36030)/Homer | GCTGASTCAGCA | 1.00E-91 | -2.11E+02 | 0 | 1361 | 12.29% | 2677.6 | 6.89% |
| NFAT(RHD)/Jurkat-NFATC1-ChIP-Seq(Jolma et al.)/Homer | ATTTTCCATT | 1.00E-85 | -1.97E+02 | 0 | 4342 | 39.21% | 11820.2 | 30.42% |
| Egr1(Zf)/K562-Egr1-ChIP-Seq(GSE32465)/Homer | TGCGTGGGYG | 1.00E-84 | -1.94E+02 | 0 | 2457 | 22.19% | 5896.2 | 15.17% |
| NFkB-p65-Rel(RHD)/ThioMac-LPS-Expression(GSE23622)/Homer | GGAAATTCCC | 1.00E-81 | -1.87E+02 | 0 | 513 | 4.63% | 683.6 | 1.76% |
| STAT4(Stat)/CD4-Stat4-ChIP-Seq(GSE22104)/Homer | NYTTCCWGGAAR | 1.00E-77 | -1.78E+02 | 0 | 4623 | 41.75% | 12913.7 | 33.23% |
| PU.1-IRF(ETS:IRF)/Bcell-PU.1-ChIP-Seq(GSE21512)/Homer | MGGAAGTGAAAC | 1.00E-66 | -1.52E+02 | 0 | 5612 | 50.68% | 16528.8 | 42.53% |
| Nur77(NR)/K562-NR4A1-ChIP-Seq(GSE31363)/Homer | TGACCTTTNCNT | 1.00E-63 | -1.46E+02 | 0 | 1285 | 11.60% | 2773.9 | 7.14% |
| Egr2(Zf)/Thymocytes-Egr2-ChIP-Seq(GSE34254)/Homer | NGCGTGGGCGGR | 1.00E-61 | -1.41E+02 | 0 | 688 | 6.21% | 1210.9 | 3.12% |
| Atf7(bZIP)/3T3L1-Atf7-ChIP-Seq(GSE56872)/Homer | NGRTGACGTCAY | 1.00E-56 | -1.29E+02 | 0 | 2223 | 20.07% | 5639.7 | 14.51% |
| Stat3+il21(Stat)/CD4-Stat3-ChIP-Seq(GSE19198)/Homer | SVYTTCCNGGAARB | 1.00E-54 | -1.26E+02 | 0 | 3296 | 29.76% | 9051.5 | 23.29% |
| PRDM1(Zf)/Hela-PRDM1-ChIP-Seq(GSE31477)/Homer | ACTTTCACTTTC | 1.00E-54 | -1.26E+02 | 0 | 2884 | 26.04% | 7736 | 19.91% |
| c-Jun-CRE(bZIP)/K562-cJun-ChIP-Seq(GSE31477)/Homer | ATGACGTCATCY | 1.00E-51 | -1.19E+02 | 0 | 1543 | 13.93% | 3668.7 | 9.44% |
| Tbet(T-box)/CD8-Tbet-ChIP-Seq(GSE33802)/Homer | AGGTGTGAAM | 1.00E-49 | -1.14E+02 | 0 | 4979 | 44.96% | 14780.3 | 38.03% |
| Atf2(bZIP)/3T3L1-Atf2-ChIP-Seq(GSE56872)/Homer | NRRTGACGTCAT | 1.00E-46 | -1.06E+02 | 0 | 1607 | 14.51% | 3950.2 | 10.16% |
| Atf1(bZIP)/K562-ATF1-ChIP-Seq(GSE31477)/Homer | GATGACGTCA | 1.00E-43 | -1.01E+02 | 0 | 2882 | 26.02% | 7968.1 | 20.50% |
| Atf4(bZIP)/MEF-Atf4-ChIP-Seq(GSE35681)/Homer | MTGATGCAAT | 1.00E-36 | -8.50E+01 | 0 | 1403 | 12.67% | 3499 | 9.00% |
| Stat3(Stat)/mES-Stat3-ChIP-Seq(GSE11431)/Homer | CTTCCGGGAA | 1.00E-34 | -8.05E+01 | 0 | 2259 | 20.40% | 6190.8 | 15.93% |
| JunD(bZIP)/K562-JunD-ChIP-Seq/Homer | ATGACGTCATCN | 1.00E-34 | -7.99E+01 | 0 | 545 | 4.92% | 1081.9 | 2.78% |
| STAT6(Stat)/Macrophage-Stat6-ChIP-Seq(GSE38377)/Homer | TTCCKNAGAA | 1.00E-33 | -7.62E+01 | 0 | 2888 | 26.08% | 8264.4 | 21.27% |
| Chop(bZIP)/MEF-Chop-ChIP-Seq(GSE35681)/Homer | ATTGCATCAT | 1.00E-31 | -7.23E+01 | 0 | 1130 | 10.20% | 2781.5 | 7.16% |
| Tbx5(T-box)/HL1-Tbx5.biotin-ChIP-Seq(GSE21529)/Homer | AGGTGTCA | 1.00E-30 | -6.94E+01 | 0 | 8885 | 80.23% | 29398.3 | 75.65% |
| ETS1(ETS)/Jurkat-ETS1-ChIP-Seq(GSE17954)/Homer | ACAGGAAGTG | 1.00E-28 | -6.54E+01 | 0 | 4055 | 36.62% | 12291.4 | 31.63% |
| Bcl6(Zf)/Liver-Bcl6-ChIP-Seq(GSE31578)/Homer | NNNCTTTCCAGGAAA | 1.00E-28 | -6.52E+01 | 0 | 5350 | 48.31% | 16731.5 | 43.05% |
| EWS:ERG-fusion(ETS)/CADO_ES1-EWS:ERG-ChIP-Seq(SRA014231)/Homer | ATTTCCTGTN | 1.00E-26 | -6.20E+01 | 0 | 3345 | 30.21% | 9959.2 | 25.63% |
| Tcf4(HMG)/Hct116-Tcf4-ChIP-Seq(SRA012054)/Homer | ASATCAAAGGVA | 1.00E-26 | -6.18E+01 | 0 | 2382 | 21.51% | 6798.9 | 17.49% |
| KLF5(Zf)/LoVo-KLF5-ChIP-Seq(GSE49402)/Homer | DGGGYGKGGC | 1.00E-26 | -6.12E+01 | 0 | 3631 | 32.79% | 10926.9 | 28.12% |

| | | | | | | | | |
|---|---|---|---|---|---|---|---|---|
| Maz(Zf)/HepG2-Maz-ChIP-Seq(GSE31477)/Homer | GGGGGGGG | 1.00E-25 | -5.96E+01 | 0 | 3871 | 34.96% | 11759.4 | 30.26% |
| ERG(ETS)/VCaP-ERG-ChIP-Seq(GSE14097)/Homer | ACAGGAAGTG | 1.00E-25 | -5.95E+01 | 0 | 5796 | 52.34% | 18382.5 | 47.30% |
| MITF(bHLH)/MastCells-MITF-ChIP-Seq(GSE48085)/Homer | RTCATGTGAC | 1.00E-24 | -5.57E+01 | 0 | 3748 | 33.85% | 11405.8 | 29.35% |
| Eomes(T-box)/H9-Eomes-ChIP-Seq(GSE26097)/Homer | ATTAACACCT | 1.00E-23 | -5.51E+01 | 0 | 7449 | 67.27% | 24336.6 | 62.62% |
| Fli1(ETS)/CD8-FLI-ChIP-Seq(GSE20898)/Homer | NRYTTCCGGH | 1.00E-23 | -5.36E+01 | 0 | 3952 | 35.69% | 12129.4 | 31.21% |
| CEBP(bZIP)/ThioMac-CEBPb-ChIP-Seq(GSE21512)/Homer | ATTGCGCAAC | 1.00E-22 | -5.15E+01 | 0 | 2750 | 24.83% | 8141.8 | 20.95% |
| MafA(bZIP)/Islet-MafA-ChIP-Seq(GSE30298)/Homer | TGCTGACTCA | 1.00E-22 | -5.15E+01 | 0 | 3278 | 29.60% | 9895.3 | 25.46% |
| CEBP:AP1(bZIP)/ThioMac-CEBPb-ChIP-Seq(GSE21512)/Homer | DRTGTTGCAA | 1.00E-21 | -5.06E+01 | 0 | 3203 | 28.92% | 9659.6 | 24.86% |
| PU.1(ETS)/ThioMac-PU.1-ChIP-Seq(GSE21512)/Homer | AGAGGAAGTG | 1.00E-20 | -4.78E+01 | 0 | 2214 | 19.99% | 6437.3 | 16.56% |
| Tcf3(HMG)/mES-Tcf3-ChIP-Seq(GSE11724)/Homer | ASWTCAAAGG | 1.00E-20 | -4.63E+01 | 0 | 1385 | 12.51% | 3801.1 | 9.78% |
| Ets1-distal(ETS)/CD4+-PolII-ChIP-Seq(Barski et al.)/Homer | MACAGGAAGT | 1.00E-19 | -4.42E+01 | 0 | 1586 | 14.32% | 4462.2 | 11.48% |
| Pdx1(Homeobox)/Islet-Pdx1-ChIP-Seq(SRA008281)/Homer | YCATYAATCA | 1.00E-18 | -4.23E+01 | 0 | 4114 | 37.15% | 12883.7 | 33.15% |
| MYB(HTH)/ERMYB-Myb-ChIPSeq(GSE22095)/Homer | GGCVGTTR | 1.00E-17 | -3.96E+01 | 0 | 6067 | 54.79% | 19713.7 | 50.73% |
| Oct4(POU,Homeobox)/mES-Oct4-ChIP-Seq(GSE11431)/Homer | ATTTGCATAW | 1.00E-16 | -3.87E+01 | 0 | 2071 | 18.70% | 6104.9 | 15.71% |
| STAT6(Stat)/CD4-Stat6-ChIP-Seq(GSE22104)/Homer | ABTTCYYRRGAA | 1.00E-16 | -3.86E+01 | 0 | 2711 | 24.48% | 8218.5 | 21.15% |
| Esrrb(NR)/mES-Esrrb-ChIP-Seq(GSE11431)/Homer | KTGACCTTGA | 1.00E-16 | -3.75E+01 | 0 | 2735 | 24.70% | 8317.8 | 21.40% |
| Oct2(POU,Homeobox)/Bcell-Oct2-ChIP-Seq(GSE21512)/Homer | ATATGCAAAT | 1.00E-15 | -3.62E+01 | 0 | 1352 | 12.21% | 3818.4 | 9.83% |
| CEBP:CEBP(bZIP)/MEF-Chop-ChIP-Seq(GSE35681)/Homer | NTNATGCAAYMNNHTGMAAY | 1.00E-15 | -3.57E+01 | 0 | 720 | 6.50% | 1856 | 4.78% |
| SpiB(ETS)/OCILY3-SPIB-ChIP-Seq(GSE56857)/Homer | AAAGRGGAAGTG | 1.00E-14 | -3.44E+01 | 0 | 1130 | 10.20% | 3137.9 | 8.07% |
| Rfx2(HTH)/LoVo-RFX2-ChIP-Seq(GSE49402)/Homer | GTTGCCATGGCAACM | 1.00E-14 | -3.36E+01 | 0 | 446 | 4.03% | 1061.9 | 2.73% |
| BMAL1(bHLH)/Liver-Bmal1-ChIP-Seq(GSE39860)/Homer | GNCACGTG | 1.00E-14 | -3.32E+01 | 0 | 6121 | 55.27% | 20045.8 | 51.58% |
| ETV1(ETS)/GIST48-ETV1-ChIP-Seq(GSE22441)/Homer | AACCGGAAGT | 1.00E-14 | -3.31E+01 | 0 | 4848 | 43.78% | 15600.8 | 40.14% |
| Unknown-ESC-element(?)/mES-Nanog-ChIP-Seq(GSE11724)/Homer | CACAGCAGGGGG | 1.00E-13 | -3.10E+01 | 0 | 2222 | 20.07% | 6729.4 | 17.32% |
| Mef2a(MADS)/HL1-Mef2a.biotin-ChIP-Seq(GSE21529)/Homer | CYAAAAATAG | 1.00E-12 | -2.78E+01 | 0 | 1914 | 17.28% | 5769.4 | 14.85% |
| EHF(ETS)/LoVo-EHF-ChIP-Seq(GSE49402)/Homer | AVCAGGAAGT | 1.00E-11 | -2.74E+01 | 0 | 4879 | 44.06% | 15844.2 | 40.77% |
| RFX(HTH)/K562-RFX3-ChIP-Seq(SRA012198)/Homer | CGGTTGCCATGGCAAC | 1.00E-11 | -2.74E+01 | 0 | 372 | 3.36% | 891.1 | 2.29% |
| X-box(HTH)/NPC-H3K4me1-ChIP-Seq(GSE16256)/Homer | GGTTGCCATGGCAA | 1.00E-11 | -2.70E+01 | 0 | 556 | 5.02% | 1440.7 | 3.71% |
| USF1(bHLH)/GM12878-Usf1-ChIP-Seq(GSE32465)/Homer | SGTCACGTGR | 1.00E-11 | -2.69E+01 | 0 | 1592 | 14.38% | 4728.4 | 12.17% |
| EWS:FLI1-fusion(ETS)/SK_N_MC-EWS:FLI1-ChIP-Seq(SRA014231)/Homer | VACAGGAAAT | 1.00E-11 | -2.65E+01 | 0 | 2317 | 20.92% | 7131.6 | 18.35% |
| GABPA(ETS)/Jurkat-GABPa-ChIP-Seq(GSE17954)/Homer | RACCGGAAGT | 1.00E-11 | -2.60E+01 | 0 | 3327 | 30.04% | 10546.2 | 27.14% |
| CHR(?)/Hela-CellCycle-Expression/Homer | SRGTTTCAAA | 1.00E-10 | -2.47E+01 | 0 | 2973 | 26.85% | 9373.6 | 24.12% |
| Nkx2.1(Homeobox)/LungAC-Nkx2.1-ChIP-Seq(GSE43252)/Homer | RSCACTYRAG | 1.00E-10 | -2.47E+01 | 0 | 8276 | 74.73% | 27956.4 | 71.94% |
| Nanog(Homeobox)/mES-Nanog-ChIP-Seq(GSE11724)/Homer | RGCCATTAAC | 1.00E-10 | -2.44E+01 | 0 | 9787 | 88.38% | 33527.6 | 86.27% |
| Nkx3.1(Homeobox)/LNCaP-Nkx3.1-ChIP-Seq(GSE28264)/Homer | AAGCACTTAA | 1.00E-10 | -2.43E+01 | 0 | 7540 | 68.09% | 25314.8 | 65.14% |
| BMYB(HTH)/Hela-BMYB-ChIP-Seq(GSE27030)/Homer | NHAACBGYYV | 1.00E-10 | -2.38E+01 | 0 | 5542 | 50.05% | 18253.6 | 46.97% |
| CRE(bZIP)/Promoter/Homer | CSGTGACGTCAC | 1.00E-10 | -2.35E+01 | 0 | 782 | 7.06% | 2176.3 | 5.60% |
| Erra(NR)/HepG2-Erra-ChIP-Seq(GSE31477)/Homer | CAAAGGTCAG | 1.00E-10 | -2.33E+01 | 0 | 6479 | 58.51% | 21563.2 | 55.49% |
| Mef2c(MADS)/GM12878-Mef2c-ChIP-Seq(GSE32465)/Homer | DCYAAAAATAGM | 1.00E-09 | -2.28E+01 | 0 | 1976 | 17.84% | 6070 | 15.62% |
| Tcf12(bHLH)/GM12878-Tcf12-ChIP-Seq(GSE32465)/Homer | VCAGCTGYTG | 1.00E-09 | -2.27E+01 | 0 | 3260 | 29.44% | 10398.9 | 26.76% |
| Meis1(Homeobox)/MastCells-Meis1-ChIP-Seq(GSE48085)/Homer | VGCTGWCAVB | 1.00E-09 | -2.26E+01 | 0 | 6115 | 55.22% | 20299.1 | 52.23% |
| Rfx1(HTH)/NPC-H3K4me1-ChIP-Seq(GSE16256)/Homer | KGTTGCCATGGCAA | 1.00E-09 | -2.24E+01 | 0 | 866 | 7.82% | 2455.5 | 6.32% |
| Srebp1a(bHLH)/HepG2-Srebp1a-ChIP-Seq(GSE31477)/Homer | RTCACSCCAY | 1.00E-09 | -2.23E+01 | 0 | 887 | 8.01% | 2523.6 | 6.49% |
| T1ISRE(IRF)/ThioMac-Ifnb-Expression/Homer | ACTTTCGTTTCT | 1.00E-09 | -2.22E+01 | 0 | 98 | 0.88% | 170.8 | 0.44% |
| AMYB(HTH)/Testes-AMYB-ChIP-Seq(GSE44588)/Homer | TGGCAGTTGG | 1.00E-09 | -2.20E+01 | 0 | 5533 | 49.96% | 18271.7 | 47.02% |
| SCL(bHLH)/HPC7-Scl-ChIP-Seq(GSE13511)/Homer | AVCAGCTG | 1.00E-09 | -2.11E+01 | 0 | 9642 | 87.07% | 33053.2 | 85.05% |
| Nkx6.1(Homeobox)/Islet-Nkx6.1-ChIP-Seq(GSE40975)/Homer | GKTAATGR | 1.00E-09 | -2.08E+01 | 0 | 7985 | 72.11% | 27007.7 | 69.49% |
| Usf2(bHLH)/C2C12-Usf2-ChIP-Seq(GSE36030)/Homer | GTCACGTGGT | 1.00E-08 | -2.07E+01 | 0 | 1216 | 10.98% | 3607.7 | 9.28% |
| Nkx2.5(Homeobox)/HL1-Nkx2.5.biotin-ChIP-Seq(GSE21529)/Homer | RRSCACTYAA | 1.00E-08 | -2.06E+01 | 0 | 7153 | 64.59% | 24034.4 | 61.84% |
| Brachyury(T-box)/Mesoendoderm-Brachyury-ChIP-exo(GSE54963)/Homer | ANTTMRCASBNNNGTGYKAAN | 1.00E-08 | -2.05E+01 | 0 | 1282 | 11.58% | 3827 | 9.85% |
| Rfx5(HTH)/GM12878-Rfx5-ChIP-Seq(GSE31477)/Homer | SCCTAGCAACAG | 1.00E-08 | -2.02E+01 | 0 | 1331 | 12.02% | 3991.9 | 10.27% |
| Pitx1(Homeobox)/Chicken-Pitx1-ChIP-Seq(GSE38910)/Homer | TAATCCCN | 1.00E-08 | -1.96E+01 | 0 | 9823 | 88.70% | 33759.6 | 86.87% |
| Sox3(HMG)/NPC-Sox3-ChIP-Seq(GSE33059)/Homer | CCWTTGTY | 1.00E-08 | -1.96E+01 | 0 | 6358 | 57.41% | 21244.6 | 54.67% |
| NPAS2(bHLH)/Liver-NPAS2-ChIP-Seq(GSE39860)/Homer | KCCACGTGAC | 1.00E-08 | -1.94E+01 | 0 | 3848 | 34.75% | 12498.3 | 32.16% |
| Rbpj1(?)/Panc1-Rbpj1-ChIP-Seq(GSE47459)/Homer | HTTTCCCASG | 1.00E-08 | -1.94E+01 | 0 | 5336 | 48.18% | 17660.6 | 45.44% |

| | | | | | | | | |
|---|---|---|---|---|---|---|---|---|
| NF1-halfsite(CTF)/LNCaP-NF1-ChIP-Seq(Unpublished)/Homer | YTGCCAAG | 1.00E-08 | -1.91E+01 | 0 | 5433 | 49.06% | 18007.9 | 46.34% |
| Ascl1(bHLH)/NeuralTubes-Ascl1-ChIP-Seq(GSE55840)/Homer | NNVVCAGCTGBN | 1.00E-08 | -1.91E+01 | 0 | 4822 | 43.54% | 15878.6 | 40.86% |
| TCFL2(HMG)/K562-TCF7L2-ChIP-Seq(GSE29196)/Homer | ACWTCAAAGG | 1.00E-08 | -1.85E+01 | 0 | 460 | 4.15% | 1232.4 | 3.17% |
| PPARE(NR),DR1/3T3L1-Pparg-ChIP-Seq(GSE13511)/Homer | TGACCTTTGCCCCA | 1.00E-07 | -1.83E+01 | 0 | 3507 | 31.67% | 11360.5 | 29.23% |
| CLOCK(bHLH)/Liver-Clock-ChIP-Seq(GSE39860)/Homer | GHCACGTG | 1.00E-07 | -1.79E+01 | 0 | 1966 | 17.75% | 6142.8 | 15.81% |
| Klf4(Zf)/mES-Klf4-ChIP-Seq(GSE11431)/Homer | GCCACACCCA | 1.00E-07 | -1.62E+01 | 0 | 1208 | 10.91% | 3662.5 | 9.42% |
| Ap4(bHLH)/AML-Tfap4-ChIP-Seq(GSE45738)/Homer | NAHCAGCTGD | 1.00E-06 | -1.60E+01 | 0 | 4084 | 36.88% | 13416.2 | 34.52% |
| E2A(bHLH)/proBcell-E2A-ChIP-Seq(GSE21978)/Homer | DNRCAGCTGY | 1.00E-06 | -1.56E+01 | 0 | 4427 | 39.98% | 14619.2 | 37.62% |
| MyoD(bHLH)/Myotube-MyoD-ChIP-Seq(GSE21614)/Homer | RRCAGCTGYTSY | 1.00E-06 | -1.54E+01 | 0 | 2653 | 23.96% | 8527.9 | 21.94% |
| ELF5(ETS)/T47D-ELF5-ChIP-Seq(GSE30407)/Homer | ACVAGGAAGT | 1.00E-06 | -1.52E+01 | 0 | 2957 | 26.70% | 9569.7 | 24.62% |
| Reverb(NR),DR2/RAW-Reverba.biotin-ChIP-Seq(GSE45914)/Homer | GTRGGTCASTGGGTCA | 1.00E-06 | -1.50E+01 | 0 | 589 | 5.32% | 1679.9 | 4.32% |
| Myf5(bHLH)/GM-Myf5-ChIP-Seq(GSE24852)/Homer | BAACAGCTGT | 1.00E-06 | -1.49E+01 | 0 | 2550 | 23.03% | 8193 | 21.08% |
| Sp1(Zf)/Promoter/Homer | GGCCCCGCCCCC | 1.00E-06 | -1.45E+01 | 0 | 390 | 3.52% | 1060.9 | 2.73% |
| Sox10(HMG)/SciaticNerve-Sox3-ChIP-Seq(GSE35132)/Homer | CCWTTGTYYB | 1.00E-06 | -1.45E+01 | 0 | 5977 | 53.97% | 20073.3 | 51.65% |
| Max(bHLH)/K562-Max-ChIP-Seq(GSE31477)/Homer | RCCACGTGGYYN | 1.00E-06 | -1.44E+01 | 0 | 2350 | 21.22% | 7526.3 | 19.37% |
| Ptf1a(bHLH)/Panc1-Ptf1a-ChIP-Seq(GSE47459)/Homer | ACAGCTGTTN | 1.00E-06 | -1.43E+01 | 0 | 7246 | 65.43% | 24567.6 | 63.22% |
| MyoG(bHLH)/C2C12-MyoG-ChIP-Seq(GSE36024)/Homer | AACAGCTG | 1.00E-06 | -1.41E+01 | 0 | 3583 | 32.36% | 11751.1 | 30.24% |
| GATA(Zf),IR3/iTreg-Gata3-ChIP-Seq(GSE20898)/Homer | NNNNNBAGATAWYATCTVHN | 1.00E-05 | -1.35E+01 | 0 | 705 | 6.37% | 2073.5 | 5.34% |
| TATA-Box(TBP)/Promoter/Homer | CCTTTTAWAGSC | 1.00E-05 | -1.33E+01 | 0 | 5148 | 46.49% | 17211.3 | 44.29% |
| PAX3:FKHR-fusion(Paired,Homeobox)/Rh4-PAX3:FKHR-ChIP-Seq(GSE19063)/Homer | ACCRTGACTAATTNN | 1.00E-05 | -1.30E+01 | 0 | 908 | 8.20% | 2742.2 | 7.06% |
| SPDEF(ETS)/VCaP-SPDEF-ChIP-Seq(SRA014231)/Homer | ASWTCCTGBT | 1.00E-05 | -1.29E+01 | 0 | 3743 | 33.80% | 12346.1 | 31.77% |
| Six1(Homeobox)/Myoblast-Six1-ChIP-Chip(GSE20150)/Homer | GKVTCADRTTWC | 1.00E-05 | -1.29E+01 | 0 | 1132 | 10.22% | 3481.7 | 8.96% |
| c-Myc(bHLH)/LNCAP-cMyc-ChIP-Seq(Unpublished)/Homer | VCCACGTG | 1.00E-05 | -1.23E+01 | 0 | 1220 | 11.02% | 3785.7 | 9.74% |
| Hoxb4(Homeobox)/ES-Hoxb4-ChIP-Seq(GSE34014)/Homer | TGATTRATGGCY | 1.00E-05 | -1.20E+01 | 0 | 855 | 7.72% | 2587.6 | 6.66% |
| Elk1(ETS)/Hela-Elk1-ChIP-Seq(GSE31477)/Homer | HACTTCCGGY | 1.00E-05 | -1.18E+01 | 0 | 1677 | 15.14% | 5325 | 13.70% |
| Foxo1(Forkhead)/RAW-Foxo1-ChIP-Seq(Fan et al.)/Homer | CTGTTTAC | 1.00E-04 | -1.08E+01 | 0 | 6538 | 59.04% | 22193.7 | 57.11% |
| Elk4(ETS)/Hela-Elk4-ChIP-Seq(GSE31477)/Homer | NRYTTCCGGY | 1.00E-04 | -1.07E+01 | 0 | 1662 | 15.01% | 5308.1 | 13.66% |
| CRX(Homeobox)/Retina-Crx-ChIP-Seq(GSE20012)/Homer | GCTAATCC | 1.00E-04 | -1.05E+01 | 0.0001 | 7242 | 65.40% | 24700.7 | 63.56% |
| Pit1(Homeobox)/GCrat-Pit1-ChIP-Seq(GSE58009)/Homer | ATGMATATDC | 1.00E-04 | -1.03E+01 | 0.0001 | 4103 | 37.05% | 13692.6 | 35.23% |
| Atoh1(bHLH)/Cerebellum-Atoh1-ChIP-Seq(GSE22111)/Homer | VNRVCAGCTGGY | 1.00E-04 | -1.03E+01 | 0.0001 | 3778 | 34.12% | 12566.1 | 32.33% |
| ETS(ETS)/Promoter/Homer | AACCGGAAGT | 1.00E-04 | -1.01E+01 | 0.0001 | 1092 | 9.86% | 3411.4 | 8.78% |
| Lhx3(Homeobox)/Neuron-Lhx3-ChIP-Seq(GSE31456)/Homer | ADBTAATTAR | 1.00E-04 | -9.89E+00 | 0.0001 | 5853 | 52.85% | 19821 | 51.00% |
| AP-2gamma(AP2)/MCF7-TFAP2C-ChIP-Seq(GSE21234)/Homer | SCCTSAGGSCAW | 1.00E-04 | -9.83E+00 | 0.0001 | 2819 | 25.46% | 9277.2 | 23.87% |
| c-Myc(bHLH)/mES-cMyc-ChIP-Seq(GSE11431)/Homer | VVCCACGTGG | 1.00E-04 | -9.79E+00 | 0.0001 | 1636 | 14.77% | 5246.6 | 13.50% |
| Tbox:Smad(T-box,MAD)/ESCd5-Smad2_3-ChIP-Seq(GSE29422)/Homer | AGGTGHCAGACA | 1.00E-03 | -8.75E+00 | 0.0003 | 842 | 7.60% | 2614.9 | 6.73% |
| RBPJ:Ebox(?,bHLH)/Panc1-Rbpj1-ChIP-Seq(GSE47459)/Homer | GGGRAARRGRMCAGMTG | 1.00E-03 | -8.54E+00 | 0.0004 | 1265 | 11.42% | 4033.9 | 10.38% |
| ERE(NR),IR3/MCF7-ERa-ChIP-Seq(Unpublished)/Homer | VAGGTCACNSTGACC | 1.00E-03 | -8.39E+00 | 0.0004 | 1018 | 9.19% | 3209.6 | 8.26% |
| Tbx20(T-box)/Heart-Tbx20-ChIP-Seq(GSE29636)/Homer | GGTGYTGACAGS | 1.00E-03 | -8.35E+00 | 0.0004 | 941 | 8.50% | 2953 | 7.60% |
| NeuroD1(bHLH)/Islet-NeuroD1-ChIP-Seq(GSE30298)/Homer | GCCATCTGTT | 1.00E-03 | -8.18E+00 | 0.0005 | 2922 | 26.39% | 9698.2 | 24.95% |
| n-Myc(bHLH)/mES-nMyc-ChIP-Seq(GSE11431)/Homer | VRCCACGTGG | 1.00E-03 | -8.17E+00 | 0.0005 | 2184 | 19.72% | 7165.9 | 18.44% |
| Tlx?(NR)/NPC-H3K4me1-ChIP-Seq(GSE16256)/Homer | CTGGCAGSCTGCCA | 1.00E-03 | -8.16E+00 | 0.0005 | 1470 | 13.27% | 4737 | 12.19% |
| RXR(NR),DR1/3T3L1-RXR-ChIP-Seq(GSE13511)/Homer | TAGGGCAAAGGTCA | 1.00E-03 | -8.16E+00 | 0.0005 | 3895 | 35.17% | 13064.6 | 33.62% |
| Olig2(bHLH)/Neuron-Olig2-ChIP-Seq(GSE30882)/Homer | RCCATMTGTT | 1.00E-03 | -7.92E+00 | 0.0007 | 6340 | 57.25% | 21629 | 55.65% |
| Isl1(Homeobox)/Neuron-Isl1-ChIP-Seq(GSE31456)/Homer | CTAATKGV | 1.00E-03 | -7.65E+00 | 0.0008 | 6601 | 59.61% | 22562.3 | 58.06% |
| Pbx3(Homeobox)/GM12878-PBX3-ChIP-Seq(GSE32465)/Homer | SCTGTCAMTCAN | 1.00E-03 | -7.40E+00 | 0.0011 | 1098 | 9.92% | 3505.8 | 9.02% |
| AP-2alpha(AP2)/Hela-AP2alpha-ChIP-Seq(GSE31477)/Homer | ATGCCCTGAGGC | 1.00E-03 | -7.28E+00 | 0.0012 | 2125 | 19.19% | 6999 | 18.01% |
| RORgt(NR)/EL4-RORgt.Flag-ChIP-Seq(GSE56019)/Homer | AAYTAGGTCA | 1.00E-03 | -7.01E+00 | 0.0016 | 547 | 4.94% | 1679.1 | 4.32% |
| Smad3(MAD)/NPC-Smad3-ChIP-Seq(GSE36673)/Homer | TWGTCTGV | 1.00E-02 | -6.60E+00 | 0.0024 | 8082 | 72.98% | 27864.7 | 71.70% |
| HIF-1b(HLH)/T47D-HIF1b-ChIP-Seq(GSE59937)/Homer | RTACGTGC | 1.00E-02 | -6.57E+00 | 0.0024 | 3107 | 28.06% | 10411.1 | 26.79% |
| HRE(HSF)/Striatum-HSF1-ChIP-Seq(GSE38000)/Homer | TTCTAGAABNTTCTA | 1.00E-02 | -6.25E+00 | 0.0033 | 1053 | 9.51% | 3389.6 | 8.72% |
| PR(NR)/T47D-PR-ChIP-Seq(GSE31130)/Homer | VAGRACAKNCTGTBC | 1.00E-02 | -6.21E+00 | 0.0034 | 7151 | 64.57% | 24582.3 | 63.25% |
| HRE(HSF)/HepG2-HSF1-ChIP-Seq(GSE31477)/Homer | BSTTCTRGAABVTTCYAGAA | 1.00E-02 | -6.11E+00 | 0.0038 | 658 | 5.94% | 2068.7 | 5.32% |
| ELF1(ETS)/Jurkat-ELF1-ChIP-Seq(SRA014231)/Homer | AVCCGGAAGT | 1.00E-02 | -6.09E+00 | 0.0038 | 1544 | 13.94% | 5061.9 | 13.02% |
| GRE(NR),IR3/RAW264.7-GRE-ChIP-Seq(Unpublished)/Homer | VAGRACAKWCTGTYC | 1.00E-02 | -6.04E+00 | 0.004 | 1003 | 9.06% | 3227.2 | 8.30% |

| | | | | | | | | |
|---|---|---|---|---|---|---|---|---|
| HIF-1a(bHLH)/MCF7-HIF1a-ChIP-Seq(GSE28352)/Homer | TACGTGCV | 1.00E-02 | -6.04E+00 | 0.004 | 680 | 6.14% | 2143.4 | 5.52% |
| Srebp2(bHLH)/HepG2-Srebp2-ChIP-Seq(GSE31477)/Homer | CGGTCACSCCAC | 1.00E-02 | -6.01E+00 | 0.0041 | 499 | 4.51% | 1543.3 | 3.97% |
| EBF1(EBF)/Near-E2A-ChIP-Seq(GSE21512)/Homer | GTCCCCWGGGGA | 1.00E-02 | -5.87E+00 | 0.0046 | 3203 | 28.92% | 10779.5 | 27.74% |
| Lhx2(Homeobox)/HFSC-Lhx2-ChIP-Seq(GSE48068)/Homer | TAATTAGN | 1.00E-02 | -5.65E+00 | 0.0057 | 4131 | 37.30% | 14016.4 | 36.07% |
| EKLF(Zf)/Erythrocyte-Klf1-ChIP-Seq(GSE20478)/Homer | NWGGGTGTGGCY | 1.00E-02 | -5.53E+00 | 0.0064 | 746 | 6.74% | 2378.2 | 6.12% |
| HOXD13(Homeobox)/Chicken-Hoxd13-ChIP-Seq(GSE38910)/Homer | NCYAATAAAA | 1.00E-02 | -5.49E+00 | 0.0067 | 4306 | 38.88% | 14636.3 | 37.66% |
| HIF2a(bHLH)/785_O-HIF2a-ChIP-Seq(GSE34871)/Homer | GCACGTACCC | 1.00E-02 | -5.22E+00 | 0.0086 | 1022 | 9.23% | 3319.6 | 8.54% |
| E2A(bHLH),near_PU.1/Bcell-PU.1-ChIP-Seq(GSE21512)/Homer | NVCACCTGBN | 1.00E-02 | -5.22E+00 | 0.0086 | 4383 | 39.58% | 14921.8 | 38.40% |
| GATA:SCL(Zf,bHLH)/Ter119-SCL-ChIP-Seq(GSE18720)/Homer | CRGCTGBNGNSNNSAGATAA | 1.00E-02 | -5.18E+00 | 0.0089 | 503 | 4.54% | 1576.4 | 4.06% |
| Sox2(HMG)/mES-Sox2-ChIP-Seq(GSE11431)/Homer | BCCATTGTTC | 1.00E-02 | -5.14E+00 | 0.0092 | 3515 | 31.74% | 11903.8 | 30.63% |
| HNF4a(NR),DR1/HepG2-HNF4a-ChIP-Seq(GSE25021)/Homer | CARRGKBCAAAGTYCA | 1.00E-02 | -5.11E+00 | 0.0095 | 1884 | 17.01% | 6267.6 | 16.13% |
| CArG(MADS)/PUER-Srf-ChIP-Seq(Sullivan et al.)/Homer | CCATATATGGNM | 1.00E-02 | -4.87E+00 | 0.0119 | 1200 | 10.84% | 3937.9 | 10.13% |
| Pit1+1bp(Homeobox)/GCrat-Pit1-ChIP-Seq(GSE58009)/Homer | ATGCATAATTCA | 1.00E-02 | -4.64E+00 | 0.0149 | 1496 | 13.51% | 4958.3 | 12.76% |
| GATA3(Zf)/iTreg-Gata3-ChIP-Seq(GSE20898)/Homer | AGATAASR | 1.00E-02 | -4.62E+00 | 0.0151 | 5579 | 50.38% | 19146.7 | 49.27% |
| Arnt:Ahr(bHLH)/MCF7-Arnt-ChIP-Seq(Lo et al.)/Homer | TBGCACGCAA | 1.00E-01 | -4.60E+00 | 0.0154 | 1772 | 16.00% | 5907.9 | 15.20% |
| Sox6(HMG)/Myotubes-Sox6-ChIP-Seq(GSE32627)/Homer | CCATTGTTNY | 1.00E-01 | -4.57E+00 | 0.0158 | 5803 | 52.40% | 19936.4 | 51.30% |
| OCT4-SOX2-TCF-NANOG(POU,Homeobox,HMG)/mES-Oct4-ChIP-Seq(GSE11431)/Homer | ATTTGCATAACAATG | 1.00E-01 | -4.54E+00 | 0.016 | 807 | 7.29% | 2615.5 | 6.73% |
| ETS:E-box(ETS,bHLH)/HPC7-Scl-ChIP-Seq(GSE22178)/Homer | AGGAARCAGCTG | 1.00E-01 | -4.50E+00 | 0.0167 | 438 | 3.96% | 1377.9 | 3.55% |
| PAX5(Paired,Homeobox),condensed/GM12878-PAX5-ChIP-Seq(GSE32465)/Homer | GTCACGCTCSCTGM | 1.00E-01 | -4.32E+00 | 0.0199 | 384 | 3.47% | 1202.2 | 3.09% |
| Znf263(Zf)/K562-Znf263-ChIP-Seq(GSE31477)/Homer | CVGTSCTCCC | 1.00E-01 | -4.23E+00 | 0.0216 | 4959 | 44.78% | 17001.1 | 43.75% |
| AR-halfsite(NR)/LNCaP-AR-ChIP-Seq(GSE27824)/Homer | CCAGGAACAG | 1.00E-01 | -4.12E+00 | 0.024 | 9298 | 83.96% | 32334.6 | 83.20% |
| GATA(Zf),IR4/iTreg-Gata3-ChIP-Seq(GSE20898)/Homer | NAGATWNBNATCTNN | 1.00E-01 | -4.05E+00 | 0.0255 | 379 | 3.42% | 1192.3 | 3.07% |
| Oct4:Sox17(POU,Homeobox,HMG)/F9-Sox17-ChIP-Seq(GSE44553)/Homer | CCATTGTATGCAAAT | 1.00E-01 | -4.02E+00 | 0.0263 | 649 | 5.86% | 2099.8 | 5.40% |
| NFY(CCAAT)/Promoter/Homer | RGCCAATSRG | 1.00E-01 | -3.87E+00 | 0.0301 | 2670 | 24.11% | 9049.2 | 23.29% |
| E2F6(E2F)/Hela-E2F6-ChIP-Seq(GSE31477)/Homer | GGCGGGAARN | 1.00E-01 | -3.81E+00 | 0.0319 | 912 | 8.24% | 2999.7 | 7.72% |
| ETS:RUNX(ETS,Runt)/Jurkat-RUNX1-ChIP-Seq(GSE17954)/Homer | RCAGGATGTGGT | 1.00E-01 | -3.79E+00 | 0.0324 | 319 | 2.88% | 999.5 | 2.57% |
| p53(p53)/mES-cMyc-ChIP-Seq(GSE11431)/Homer | ACATGCCCGGGCAT | 1.00E-01 | -3.73E+00 | 0.0344 | 51 | 0.46% | 133.5 | 0.34% |
| ZNF711(Zf)/SHSY5Y-ZNF711-ChIP-Seq(GSE20673)/Homer | AGGCCTAG | 1.00E-01 | -3.64E+00 | 0.0374 | 4333 | 39.13% | 14856.2 | 38.23% |
| PAX5(Paired,Homeobox)/GM12878-PAX5-ChIP-Seq(GSE32465)/Homer | GCAGCCAAGCRTGACH | 1.00E-01 | -3.53E+00 | 0.0416 | 1136 | 10.26% | 3777.7 | 9.72% |
| FXR(NR),IR1/Liver-FXR-ChIP-Seq(Chong et al.)/Homer | AGGTCANTGACCTB | 1.00E-01 | -3.32E+00 | 0.051 | 1354 | 12.23% | 4536.1 | 11.67% |
| p63(p53)/Keratinocyte-p63-ChIP-Seq(GSE17611)/Homer | NNDRCATGYCYNRRCATGYH | 1.00E-01 | -3.30E+00 | 0.0514 | 1261 | 11.39% | 4217.8 | 10.85% |

**Table S5C. HOMER Known Motifs near the opening peaks of both cytokine-primed and Ag-stimulated cells.**
Only motifs with q-value below 0.05 are shown.

| Motif Name | Consensus | P-value | Log P-value | q-value (Benjamini) | # of Target Sequences with Motif(of 65) | % of Target Sequences with Motif | # of Background Sequences with Motif(of 49567) | % of Background Sequences with Motif |
|---|---|---|---|---|---|---|---|---|
| STAT5(Stat)/mCD4+-Stat5-ChIP-Seq(GSE12346)/Homer | RTTTCTNAGAAA | 1.00E-29 | -6.80E+01 | 0 | 49 | 75.38% | 6448.7 | 13.01% |
| STAT1(Stat)/HelaS3-STAT1-ChIP-Seq(GSE12782)/Homer | NATTTCCNGGAAAT | 1.00E-28 | -6.65E+01 | 0 | 46 | 70.77% | 5476.7 | 11.05% |
| STAT4(Stat)/CD4-Stat4-ChIP-Seq(GSE22104)/Homer | NYTTCCWGGAAR | 1.00E-21 | -4.96E+01 | 0 | 59 | 90.77% | 16334.9 | 32.95% |
| Stat3+il21(Stat)/CD4-Stat3-ChIP-Seq(GSE19198)/Homer | SVYTTCCNGGAARB | 1.00E-18 | -4.16E+01 | 0 | 49 | 75.38% | 11493.6 | 23.19% |
| Stat3(Stat)/mES-Stat3-ChIP-Seq(GSE11431)/Homer | CTTCCGGGAA | 1.00E-12 | -2.89E+01 | 0 | 36 | 55.38% | 7832.2 | 15.80% |
| Bcl6(Zf)/Liver-Bcl6-ChIP-Seq(GSE31578)/Homer | NNNCTTTCCAGGAAA | 1.00E-09 | -2.25E+01 | 0 | 53 | 81.54% | 21186.4 | 42.74% |
| RUNX(Runt)/HPC7-Runx1-ChIP-Seq(GSE22178)/Homer | SAAACCACAG | 1.00E-08 | -1.89E+01 | 0 | 38 | 58.46% | 12174.1 | 24.56% |
| RUNX1(Runt)/Jurkat-RUNX1-ChIP-Seq(GSE29180)/Homer | AAACCACARM | 1.00E-06 | -1.53E+01 | 0 | 43 | 66.15% | 17153.8 | 34.60% |
| STAT6(Stat)/Macrophage-Stat6-ChIP-Seq(GSE38377)/Homer | TTCCKNAGAA | 1.00E-06 | -1.50E+01 | 0 | 32 | 49.23% | 10270.6 | 20.72% |
| ETS:E-box(ETS,bHLH)/HPC7-Scl-ChIP-Seq(GSE22178)/Homer | AGGAARCAGCTG | 1.00E-04 | -9.80E+00 | 0.0015 | 10 | 15.38% | 1648 | 3.32% |
| ETS:RUNX(ETS,Runt)/Jurkat-RUNX1-ChIP-Seq(GSE17954)/Homer | RCAGGATGTGGT | 1.00E-03 | -8.55E+00 | 0.0047 | 8 | 12.31% | 1221 | 2.46% |
| STAT6(Stat)/CD4-Stat6-ChIP-Seq(GSE22104)/Homer | ABTTCYYRRGAA | 1.00E-03 | -8.08E+00 | 0.0068 | 26 | 40.00% | 10267.5 | 20.71% |
| Jun-AP1(bZIP)/K562-cJun-ChIP-Seq(GSE31477)/Homer | GATGASTCATCN | 1.00E-03 | -7.77E+00 | 0.0086 | 13 | 20.00% | 3410 | 6.88% |
| Max(bHLH)/K562-Max-ChIP-Seq(GSE31477)/Homer | RCCACGTGGYYN | 1.00E-03 | -7.00E+00 | 0.0172 | 24 | 36.92% | 9747.2 | 19.66% |
| Rbpj1(?)/Panc1-Rbpj1-ChIP-Seq(GSE47459)/Homer | HTTTCCCASG | 1.00E-03 | -6.93E+00 | 0.0173 | 42 | 64.62% | 22176.9 | 44.74% |
| Fosl2(bZIP)/3T3L1-Fosl2-ChIP-Seq(GSE56872)/Homer | NATGASTCABNN | 1.00E-02 | -6.66E+00 | 0.0211 | 15 | 23.08% | 4845.5 | 9.77% |
| GATA3(Zf)/iTreg-Gata3-ChIP-Seq(GSE20898)/Homer | AGATAASR | 1.00E-02 | -6.52E+00 | 0.023 | 44 | 67.69% | 24107.9 | 48.63% |
| Ptf1a(bHLH)/Panc1-Ptf1a-ChIP-Seq(GSE47459)/Homer | ACAGCTGTTN | 1.00E-02 | -6.50E+00 | 0.023 | 53 | 81.54% | 31653.5 | 63.85% |
| Atf7(bZIP)/3T3L1-Atf7-ChIP-Seq(GSE56872)/Homer | NGRTGACGTCAY | 1.00E-02 | -6.45E+00 | 0.023 | 19 | 29.23% | 7145.2 | 14.41% |
| PAX3:FKHR-fusion(Paired,Homeobox)/Rh4-PAX3:FKHR-ChIP-Seq(GSE19063)/Homer | ACCRTGACTAATTNN | 1.00E-02 | -6.43E+00 | 0.023 | 12 | 18.46% | 3451.6 | 6.96% |
| RUNX2(Runt)/PCa-RUNX2-ChIP-Seq(GSE33889)/Homer | NWAACCACADNN | 1.00E-02 | -6.37E+00 | 0.023 | 31 | 47.69% | 14736.3 | 29.73% |
| RUNX-AML(Runt)/CD4+-PolII-ChIP-Seq(Barski et al.)/Homer | GCTGTGGTTW | 1.00E-02 | -6.36E+00 | 0.023 | 28 | 43.08% | 12739.1 | 25.70% |
| NPAS2(bHLH)/Liver-NPAS2-ChIP-Seq(GSE39860)/Homer | KCCACGTGAC | 1.00E-02 | -6.28E+00 | 0.023 | 33 | 50.77% | 16185.8 | 32.65% |
| BMAL1(bHLH)/Liver-Bmal1-ChIP-Seq(GSE39860)/Homer | GNCACGTG | 1.00E-02 | -6.01E+00 | 0.0269 | 46 | 70.77% | 26193.2 | 52.84% |
| CRX(Homeobox)/Retina-Crx-ChIP-Seq(GSE20012)/Homer | GCTAATCC | 1.00E-02 | -5.89E+00 | 0.0293 | 52 | 80.00% | 31362.3 | 63.27% |
| PU.1(ETS)/ThioMac-PU.1-ChIP-Seq(GSE21512)/Homer | AGAGGAAGTG | 1.00E-02 | -5.72E+00 | 0.0333 | 20 | 30.77% | 8198.4 | 16.54% |
| USF1(bHLH)/GM12878-Usf1-ChIP-Seq(GSE32465)/Homer | SGTCACGTGR | 1.00E-02 | -5.53E+00 | 0.0389 | 16 | 24.62% | 5998.3 | 12.10% |

**Table S6A. HOMER Known Motifs near the closing peaks of cytokine-primed cells.**
Only motifs with q-value below 0.05 are shown.

| Motif Name | Consensus | P-value | Log P-value | q-value (Benjamini) | # of Target Sequences with Motif(of 468) | % of Target Sequences with Motif | # of Background Sequences with Motif(of 46808) | % of Background Sequences with Motif |
|---|---|---|---|---|---|---|---|---|
| Atf1(bZIP)/K562-ATF1-ChIP-Seq(GSE31477)/Homer | GATGACGTCA | 1.00E-38 | -8.94E+01 | 0 | 227 | 48.50% | 9857.7 | 21.06% |
| Ets1-distal(ETS)/CD4+-PolII-ChIP-Seq(Barski et al.)/Homer | MACAGGAAGT | 1.00E-24 | -5.61E+01 | 0 | 142 | 30.34% | 5750.6 | 12.28% |
| Atf7(bZIP)/3T3L1-Atf7-ChIP-Seq(GSE56872)/Homer | NGRTGACGTCAY | 1.00E-23 | -5.40E+01 | 0 | 158 | 33.76% | 7005.5 | 14.96% |
| GABPA(ETS)/Jurkat-GABPa-ChIP-Seq(GSE17954)/Homer | RACCGGAAGT | 1.00E-22 | -5.23E+01 | 0 | 250 | 53.42% | 14595.4 | 31.18% |
| CRE(bZIP)/Promoter/Homer | CSGTGACGTCAC | 1.00E-21 | -5.06E+01 | 0 | 99 | 21.15% | 3313.3 | 7.08% |
| Atf2(bZIP)/3T3L1-Atf2-ChIP-Seq(GSE56872)/Homer | NRRTGACGTCAT | 1.00E-20 | -4.81E+01 | 0 | 124 | 26.50% | 5018.6 | 10.72% |
| Fli1(ETS)/CD8-FLI-ChIP-Seq(GSE20898)/Homer | NRYTTCCGGH | 1.00E-20 | -4.80E+01 | 0 | 272 | 58.12% | 17057.4 | 36.43% |
| PU.1(ETS)/ThioMac-PU.1-ChIP-Seq(GSE21512)/Homer | AGAGGAAGTG | 1.00E-20 | -4.74E+01 | 0 | 170 | 36.32% | 8380 | 17.90% |
| EWS:FLI1-fusion(ETS)/SK_N_MC-EWS:FLI1-ChIP-Seq(SRA014231)/Homer | VACAGGAAAT | 1.00E-19 | -4.40E+01 | 0 | 182 | 38.89% | 9604.4 | 20.51% |
| ETV1(ETS)/GIST48-ETV1-ChIP-Seq(GSE22441)/Homer | AACCGGAAGT | 1.00E-18 | -4.24E+01 | 0 | 305 | 65.17% | 20920.5 | 44.69% |
| ETS1(ETS)/Jurkat-ETS1-ChIP-Seq(GSE17954)/Homer | ACAGGAAGTG | 1.00E-17 | -4.03E+01 | 0 | 257 | 54.91% | 16502.2 | 35.25% |
| Elk4(ETS)/Hela-Elk4-ChIP-Seq(GSE31477)/Homer | NRYTTCCGGY | 1.00E-16 | -3.73E+01 | 0 | 162 | 34.62% | 8616.5 | 18.40% |
| ETS(ETS)/Promoter/Homer | AACCGGAAGT | 1.00E-14 | -3.29E+01 | 0 | 115 | 24.57% | 5432.6 | 11.60% |
| ELF1(ETS)/Jurkat-ELF1-ChIP-Seq(SRA014231)/Homer | AVCCGGAAGT | 1.00E-14 | -3.23E+01 | 0 | 150 | 32.05% | 8136.5 | 17.38% |
| EWS:ERG-fusion(ETS)/CADO_ES1-EWS:ERG-ChIP-Seq(SRA014231)/Homer | ATTTCCTGTN | 1.00E-13 | -3.19E+01 | 0 | 198 | 42.31% | 12167.2 | 25.99% |
| c-Jun-CRE(bZIP)/K562-cJun-ChIP-Seq(GSE31477)/Homer | ATGACGTCATCY | 1.00E-13 | -3.18E+01 | 0 | 101 | 21.58% | 4527.5 | 9.67% |
| JunD(bZIP)/K562-JunD-ChIP-Seq/Homer | ATGACGTCATCN | 1.00E-13 | -3.14E+01 | 0 | 50 | 10.68% | 1410.4 | 3.01% |
| EHF(ETS)/LoVo-EHF-ChIP-Seq(GSE49402)/Homer | AVCAGGAAGT | 1.00E-11 | -2.72E+01 | 0 | 280 | 59.83% | 20436.7 | 43.65% |
| ETS:RUNX(ETS,Runt)/Jurkat-RUNX1-ChIP-Seq(GSE17954)/Homer | RCAGGATGTGGT | 1.00E-11 | -2.62E+01 | 0 | 46 | 9.83% | 1408.4 | 3.01% |
| RUNX1(Runt)/Jurkat-RUNX1-ChIP-Seq(GSE29180)/Homer | AAACCACARM | 1.00E-11 | -2.59E+01 | 0 | 235 | 50.21% | 16296.1 | 34.81% |
| ELF5(ETS)/T47D-ELF5-ChIP-Seq(GSE30407)/Homer | ACVAGGAAGT | 1.00E-10 | -2.49E+01 | 0 | 192 | 41.03% | 12520.8 | 26.74% |
| Elk1(ETS)/Hela-Elk1-ChIP-Seq(GSE31477)/Homer | HACTTCCGGY | 1.00E-10 | -2.41E+01 | 0 | 145 | 30.98% | 8604.5 | 18.38% |
| ERG(ETS)/VCaP-ERG-ChIP-Seq(GSE14097)/Homer | ACAGGAAGTG | 1.00E-10 | -2.39E+01 | 0 | 310 | 66.24% | 24011.9 | 51.29% |
| RUNX2(Runt)/PCa-RUNX2-ChIP-Seq(GSE33889)/Homer | NWAACCACADNN | 1.00E-09 | -2.29E+01 | 0 | 204 | 43.59% | 13880 | 29.65% |
| RUNX(Runt)/HPC7-Runx1-ChIP-Seq(GSE22178)/Homer | SAAACCACAG | 1.00E-09 | -2.22E+01 | 0 | 180 | 38.46% | 11825 | 25.26% |
| SpiB(ETS)/OCILY3-SPIB-ChIP-Seq(GSE56857)/Homer | AAAGRGGAAGTG | 1.00E-09 | -2.11E+01 | 0 | 83 | 17.74% | 4106.1 | 8.77% |
| SPDEF(ETS)/VCaP-SPDEF-ChIP-Seq(SRA014231)/Homer | ASWTCCTGBT | 1.00E-07 | -1.71E+01 | 0 | 220 | 47.01% | 16296.3 | 34.81% |
| Rfx2(HTH)/LoVo-RFX2-ChIP-Seq(GSE49402)/Homer | GTTGCCATGGCAACM | 1.00E-07 | -1.66E+01 | 0 | 40 | 8.55% | 1535.1 | 3.28% |
| Foxa2(Forkhead)/Liver-Foxa2-ChIP-Seq(GSE25694)/Homer | CYTGTTTACWYW | 1.00E-06 | -1.58E+01 | 0 | 183 | 39.10% | 13109.7 | 28.00% |
| Ascl1(bHLH)/NeuralTubes-Ascl1-ChIP-Seq(GSE55840)/Homer | NNVVCAGCTGBN | 1.00E-06 | -1.52E+01 | 0 | 268 | 57.26% | 21341.6 | 45.58% |
| Rfx5(HTH)/GM12878-Rfx5-ChIP-Seq(GSE31477)/Homer | SCCTAGCAACAG | 1.00E-06 | -1.48E+01 | 0 | 89 | 19.02% | 5213.8 | 11.14% |
| Atoh1(bHLH)/Cerebellum-Atoh1-ChIP-Seq(GSE22111)/Homer | VNRVCAGCTGGY | 1.00E-06 | -1.40E+01 | 0 | 216 | 46.15% | 16510.9 | 35.27% |
| Foxo1(Forkhead)/RAW-Foxo1-ChIP-Seq(Fan et al.)/Homer | CTGTTTAC | 1.00E-05 | -1.36E+01 | 0 | 319 | 68.16% | 26890 | 57.44% |
| Tbx5(T-box)/HL1-Tbx5.biotin-ChIP-Seq(GSE21529)/Homer | AGGTGTCA | 1.00E-05 | -1.31E+01 | 0 | 406 | 86.75% | 36677.8 | 78.34% |
| NPAS2(bHLH)/Liver-NPAS2-ChIP-Seq(GSE39860)/Homer | KCCACGTGAC | 1.00E-04 | -1.15E+01 | 0.0001 | 210 | 44.87% | 16489.1 | 35.22% |
| c-Myc(bHLH)/LNCAP-cMyc-ChIP-Seq(Unpublished)/Homer | VCCACGTG | 1.00E-04 | -1.12E+01 | 0.0001 | 95 | 20.30% | 6189.5 | 13.22% |
| NFkB-p65(RHD)/GM12787-p65-ChIP-Seq(GSE19485)/Homer | WGGGGATTTCCC | 1.00E-04 | -1.10E+01 | 0.0001 | 115 | 24.57% | 7917.4 | 16.91% |
| Meis1(Homeobox)/MastCells-Meis1-ChIP-Seq(GSE48085)/Homer | VGCTGWCAVB | 1.00E-04 | -1.09E+01 | 0.0001 | 300 | 64.10% | 25554.9 | 54.58% |
| Tcf4(HMG)/Hct116-Tcf4-ChIP-Seq(SRA012054)/Homer | ASATCAAAGGVA | 1.00E-04 | -1.07E+01 | 0.0002 | 115 | 24.57% | 7986.3 | 17.06% |
| CLOCK(bHLH)/Liver-Clock-ChIP-Seq(GSE39860)/Homer | GHCACGTG | 1.00E-04 | -1.05E+01 | 0.0002 | 120 | 25.64% | 8449.4 | 18.05% |
| BMAL1(bHLH)/Liver-Bmal1-ChIP-Seq(GSE39860)/Homer | GNCACGTG | 1.00E-04 | -1.05E+01 | 0.0002 | 294 | 62.82% | 25042.6 | 53.49% |
| Egr1(Zf)/K562-Egr1-ChIP-Seq(GSE32465)/Homer | TGCGTGGGYG | 1.00E-04 | -1.02E+01 | 0.0002 | 130 | 27.78% | 9382.1 | 20.04% |
| Atf4(bZIP)/MEF-Atf4-ChIP-Seq(GSE35681)/Homer | MTGATGCAAT | 1.00E-04 | -1.02E+01 | 0.0002 | 63 | 13.46% | 3743.2 | 8.00% |
| CEBP:AP1(bZIP)/ThioMac-CEBPb-ChIP-Seq(GSE21512)/Homer | DRTGTTGCAA | 1.00E-04 | -1.01E+01 | 0.0003 | 147 | 31.41% | 10945 | 23.38% |
| Usf2(bHLH)/C2C12-Usf2-ChIP-Seq(GSE36030)/Homer | GTCACGTGGT | 1.00E-04 | -1.00E+01 | 0.0003 | 76 | 16.24% | 4805.9 | 10.27% |
| bHLHE40(bHLH)/HepG2-BHLHE40-ChIP-Seq(GSE31477)/Homer | KCACGTGMCN | 1.00E-04 | -1.00E+01 | 0.0003 | 71 | 15.17% | 4401.7 | 9.40% |
| USF1(bHLH)/GM12878-Usf1-ChIP-Seq(GSE32465)/Homer | SGTCACGTGR | 1.00E-04 | -1.00E+01 | 0.0003 | 99 | 21.15% | 6726.8 | 14.37% |
| KLF5(Zf)/LoVo-KLF5-ChIP-Seq(GSE49402)/Homer | DGGGYGKGGC | 1.00E-04 | -9.66E+00 | 0.0003 | 206 | 44.02% | 16537.3 | 35.32% |

| | | | | | | | | |
|---|---|---|---|---|---|---|---|---|
| RUNX-AML(Runt)/CD4+-PolII-ChIP-Seq(Barski et al.)/Homer | GCTGTGGTTW | 1.00E-04 | -9.44E+00 | 0.0004 | 158 | 33.76% | 12083.4 | 25.81% |
| STAT6(Stat)/CD4-Stat6-ChIP-Seq(GSE22104)/Homer | ABTTCYYRRGAA | 1.00E-04 | -9.35E+00 | 0.0005 | 126 | 26.92% | 9204.1 | 19.66% |
| HIF-1b(HLH)/T47D-HIF1b-ChIP-Seq(GSE59937)/Homer | RTACGTGC | 1.00E-03 | -9.19E+00 | 0.0005 | 193 | 41.24% | 15417.9 | 32.93% |
| RFX(HTH)/K562-RFX3-ChIP-Seq(SRA012198)/Homer | CGGTTGCCATGGCAAC | 1.00E-03 | -8.96E+00 | 0.0007 | 29 | 6.20% | 1351.1 | 2.89% |
| STAT6(Stat)/Macrophage-Stat6-ChIP-Seq(GSE38377)/Homer | TTCCKNAGAA | 1.00E-03 | -8.74E+00 | 0.0008 | 126 | 26.92% | 9333.5 | 19.94% |
| Ptf1a(bHLH)/Panc1-Ptf1a-ChIP-Seq(GSE47459)/Homer | ACAGCTGTTN | 1.00E-03 | -8.67E+00 | 0.0008 | 353 | 75.43% | 31720.9 | 67.75% |
| Eomes(T-box)/H9-Eomes-ChIP-Seq(GSE26097)/Homer | ATTAACACCT | 1.00E-03 | -8.40E+00 | 0.0011 | 327 | 69.87% | 29030 | 62.01% |
| FOXP1(Forkhead)/H9-FOXP1-ChIP-Seq(GSE31006)/Homer | NYYTGTTTACHN | 1.00E-03 | -8.40E+00 | 0.0011 | 103 | 22.01% | 7369.6 | 15.74% |
| MITF(bHLH)/MastCells-MITF-ChIP-Seq(GSE48085)/Homer | RTCATGTGAC | 1.00E-03 | -8.27E+00 | 0.0012 | 181 | 38.68% | 14517.2 | 31.01% |
| AP-1(bZIP)/ThioMac-PU.1-ChIP-Seq(GSE21512)/Homer | VTGACTCATC | 1.00E-03 | -8.21E+00 | 0.0012 | 141 | 30.13% | 10812.6 | 23.10% |
| Jun-AP1(bZIP)/K562-cJun-ChIP-Seq(GSE31477)/Homer | GATGASTCATCN | 1.00E-03 | -8.03E+00 | 0.0015 | 54 | 11.54% | 3317.2 | 7.09% |
| Tcf3(HMG)/mES-Tcf3-ChIP-Seq(GSE11724)/Homer | ASWTCAAAGG | 1.00E-03 | -7.71E+00 | 0.002 | 67 | 14.32% | 4424.6 | 9.45% |
| PU.1-IRF(ETS:IRF)/Bcell-PU.1-ChIP-Seq(GSE21512)/Homer | MGGAAGTGAAAC | 1.00E-03 | -7.70E+00 | 0.002 | 243 | 51.92% | 20682 | 44.18% |
| BATF(bZIP)/Th17-BATF-ChIP-Seq(GSE39756)/Homer | DATGASTCAT | 1.00E-03 | -7.68E+00 | 0.002 | 125 | 26.71% | 9481.5 | 20.25% |
| SCL(bHLH)/HPC7-Scl-ChIP-Seq(GSE13511)/Homer | AVCAGCTG | 1.00E-03 | -7.56E+00 | 0.0022 | 435 | 92.95% | 41314.6 | 88.25% |
| Tbet(T-box)/CD8-Tbet-ChIP-Seq(GSE33802)/Homer | AGGTGTGAAM | 1.00E-03 | -7.41E+00 | 0.0025 | 211 | 45.09% | 17632 | 37.66% |
| E2A(bHLH)/proBcell-E2A-ChIP-Seq(GSE21978)/Homer | DNRCAGCTGY | 1.00E-03 | -7.30E+00 | 0.0028 | 235 | 50.21% | 20012.5 | 42.75% |
| FOXA1(Forkhead)/LNCAP-FOXA1-ChIP-Seq(GSE27824)/Homer | WAAGTAAACA | 1.00E-03 | -6.95E+00 | 0.0039 | 216 | 46.15% | 18258.6 | 39.00% |
| Olig2(bHLH)/Neuron-Olig2-ChIP-Seq(GSE30882)/Homer | RCCATMTGTT | 1.00E-02 | -6.83E+00 | 0.0042 | 300 | 64.10% | 26699.5 | 57.03% |
| ZNF143\|STAF(Zf)/CUTLL-ZNF143-ChIP-Seq(GSE29600)/Homer | ATTTCCCAGVAKSCY | 1.00E-02 | -6.41E+00 | 0.0064 | 77 | 16.45% | 5511.2 | 11.77% |
| NFkB-p50,p52(RHD)/Monocyte-p50-ChIP-Chip(Schreiber et al.)/Homer | GGGGGAATCCCC | 1.00E-02 | -6.21E+00 | 0.0077 | 26 | 5.56% | 1390.3 | 2.97% |
| n-Myc(bHLH)/mES-nMyc-ChIP-Seq(GSE11431)/Homer | VRCCACGTGG | 1.00E-02 | -6.17E+00 | 0.0079 | 125 | 26.71% | 9862.9 | 21.07% |
| Tcf12(bHLH)/GM12878-Tcf12-ChIP-Seq(GSE32465)/Homer | VCAGCTGYTG | 1.00E-02 | -6.12E+00 | 0.0082 | 170 | 36.32% | 14086.2 | 30.09% |
| Fosl2(bZIP)/3T3L1-Fosl2-ChIP-Seq(GSE56872)/Homer | NATGASTCABNN | 1.00E-02 | -6.10E+00 | 0.0082 | 68 | 14.53% | 4800 | 10.25% |
| MYB(HTH)/ERMYB-Myb-ChIPSeq(GSE22095)/Homer | GGCVGTTR | 1.00E-02 | -6.07E+00 | 0.0084 | 277 | 59.19% | 24611.3 | 52.57% |
| Atf3(bZIP)/GBM-ATF3-ChIP-Seq(GSE33912)/Homer | DATGASTCATHN | 1.00E-02 | -6.03E+00 | 0.0086 | 122 | 26.07% | 9625.5 | 20.56% |
| Rfx1(HTH)/NPC-H3K4me1-ChIP-Seq(GSE16256)/Homer | KGTTGCCATGGCAA | 1.00E-02 | -6.02E+00 | 0.0086 | 49 | 10.47% | 3217.5 | 6.87% |
| Tlx?(NR)/NPC-H3K4me1-ChIP-Seq(GSE16256)/Homer | CTGGCAGSCTGCCA | 1.00E-02 | -5.71E+00 | 0.0115 | 87 | 18.59% | 6543.1 | 13.98% |
| Arnt:Ahr(bHLH)/MCF7-Arnt-ChIP-Seq(Lo et al.)/Homer | TBGCACGCAA | 1.00E-02 | -5.70E+00 | 0.0115 | 113 | 24.15% | 8893 | 19.00% |
| Fra1(bZIP)/BT549-Fra1-ChIP-Seq(GSE46166)/Homer | NNATGASTCATH | 1.00E-02 | -5.65E+00 | 0.0119 | 103 | 22.01% | 7995.4 | 17.08% |
| IRF2(IRF)/Erythroblas-IRF2-ChIP-Seq(GSE36985)/Homer | GAAASYGAAASY | 1.00E-02 | -5.64E+00 | 0.0119 | 28 | 5.98% | 1603.8 | 3.43% |
| Chop(bZIP)/MEF-Chop-ChIP-Seq(GSE35681)/Homer | ATTGCATCAT | 1.00E-02 | -5.61E+00 | 0.0121 | 45 | 9.62% | 2956.6 | 6.32% |
| NeuroD1(bHLH)/Islet-NeuroD1-ChIP-Seq(GSE30298)/Homer | GCCATCTGTT | 1.00E-02 | -5.56E+00 | 0.0125 | 154 | 32.91% | 12745 | 27.22% |
| Klf4(Zf)/mES-Klf4-ChIP-Seq(GSE11431)/Homer | GCCACACCCA | 1.00E-02 | -5.56E+00 | 0.0125 | 75 | 16.03% | 5521 | 11.79% |
| NF1(CTF)/LNCAP-NF1-ChIP-Seq(Unpublished)/Homer | CYTGGCABNSTGCCAR | 1.00E-02 | -5.49E+00 | 0.0131 | 76 | 16.24% | 5623.4 | 12.01% |
| Max(bHLH)/K562-Max-ChIP-Seq(GSE31477)/Homer | RCCACGTGGYYN | 1.00E-02 | -5.41E+00 | 0.0141 | 124 | 26.50% | 9986.8 | 21.33% |
| Cdx2(Homeobox)/mES-Cdx2-ChIP-Seq(GSE14586)/Homer | GYMATAAAAH | 1.00E-02 | -5.33E+00 | 0.0151 | 127 | 27.14% | 10287.9 | 21.97% |
| TCFL2(HMG)/K562-TCF7L2-ChIP-Seq(GSE29196)/Homer | ACWTCAAAGG | 1.00E-02 | -5.28E+00 | 0.0157 | 25 | 5.34% | 1418.7 | 3.03% |
| Mef2c(MADS)/GM12878-Mef2c-ChIP-Seq(GSE32465)/Homer | DCYAAAAATAGM | 1.00E-02 | -5.24E+00 | 0.0162 | 82 | 17.52% | 6214.2 | 13.27% |
| Tbx20(T-box)/Heart-Tbx20-ChIP-Seq(GSE29636)/Homer | GGTGYTGACAGS | 1.00E-02 | -5.18E+00 | 0.0169 | 54 | 11.54% | 3786.5 | 8.09% |
| Nkx2.1(Homeobox)/LungAC-Nkx2.1-ChIP-Seq(GSE43252)/Homer | RSCACTYRAG | 1.00E-02 | -5.17E+00 | 0.0169 | 369 | 78.85% | 34496.6 | 73.68% |
| Ap4(bHLH)/AML-Tfap4-ChIP-Seq(GSE45738)/Homer | NAHCAGCTGD | 1.00E-02 | -4.87E+00 | 0.0224 | 201 | 42.95% | 17500 | 37.38% |
| X-box(HTH)/NPC-H3K4me1-ChIP-Seq(GSE16256)/Homer | GGTTGCCATGGCAA | 1.00E-02 | -4.86E+00 | 0.0225 | 30 | 6.41% | 1859.2 | 3.97% |
| Fox:Ebox(Forkhead,bHLH)/Panc1-Foxa2-ChIP-Seq(GSE47459)/Homer | NNNVCTGWGYAAACASN | 1.00E-02 | -4.72E+00 | 0.0256 | 194 | 41.45% | 16876.1 | 36.05% |
| ETS:E-box(ETS,bHLH)/HPC7-Scl-ChIP-Seq(GSE22178)/Homer | AGGAARCAGCTG | 1.00E-01 | -4.53E+00 | 0.0306 | 27 | 5.77% | 1668.4 | 3.56% |
| Egr2(Zf)/Thymocytes-Egr2-ChIP-Seq(GSE34254)/Homer | NGCGTGGGCGGR | 1.00E-01 | -4.52E+00 | 0.0306 | 33 | 7.05% | 2151.6 | 4.60% |
| IRF1(IRF)/PBMC-IRF1-ChIP-Seq(GSE43036)/Homer | GAAAGTGAAAGT | 1.00E-01 | -4.48E+00 | 0.0314 | 34 | 7.26% | 2239.2 | 4.78% |
| Nkx6.1(Homeobox)/Islet-Nkx6.1-ChIP-Seq(GSE40975)/Homer | GKTAATGR | 1.00E-01 | -4.48E+00 | 0.0314 | 329 | 70.30% | 30537.2 | 65.23% |
| FOXA1(Forkhead)/MCF7-FOXA1-ChIP-Seq(GSE26831)/Homer | WAAGTAAACA | 1.00E-01 | -4.41E+00 | 0.0332 | 177 | 37.82% | 15348.9 | 32.78% |
| Myf5(bHLH)/GM-Myf5-ChIP-Seq(GSE24852)/Homer | BAACAGCTGT | 1.00E-01 | -4.27E+00 | 0.0377 | 128 | 27.35% | 10724.3 | 22.91% |
| PRDM1(Zf)/Hela-PRDM1-ChIP-Seq(GSE31477)/Homer | ACTTTCACTTTC | 1.00E-01 | -4.27E+00 | 0.0377 | 110 | 23.50% | 9041.6 | 19.31% |
| AR-halfsite(NR)/LNCaP-AR-ChIP-Seq(GSE27824)/Homer | CCAGGAACAG | 1.00E-01 | -4.26E+00 | 0.0377 | 417 | 89.10% | 40044.1 | 85.53% |
| Sox2(HMG)/mES-Sox2-ChIP-Seq(GSE11431)/Homer | BCCATTGTTC | 1.00E-01 | -4.10E+00 | 0.0432 | 164 | 35.04% | 14213.5 | 30.36% |
| Unknown-ESC-element(?)/mES-Nanog-ChIP-Seq(GSE11724)/Homer | CACAGCAGGGGG | 1.00E-01 | -3.99E+00 | 0.048 | 110 | 23.50% | 9134.2 | 19.51% |

| | | | | | | | | |
|---|---|---|---|---|---|---|---|---|
| Tbox:Smad(T-box,MAD)/ESCd5-Smad2_3-ChIP-Seq(GSE29422)/Homer | AGGTGHCAGACA | 1.00E-01 | -3.94E+00 | 0.0501 | 46 | 9.83% | 3356.9 | 7.17% |

**Table S6B. HOMER Known Motifs near the closing peaks of Ag-stimulated cells**

Only motifs with q-value below 0.05 are shown.

| Motif Name | Consensus | P-value | Log P-value | q-value (Benjamini) | # of Target Sequences with Motif(of 6470) | % of Target Sequences with Motif | # of Background Sequences with Motif(of 43318) | % of Background Sequences with Motif |
|---|---|---|---|---|---|---|---|---|
| Fli1(ETS)/CD8-FLI-ChIP-Seq(GSE20898)/Homer | NRYTTCCGGH | 1e-648 | -1.49E+03 | 0 | 4297 | 66.41% | 14364.2 | 33.16% |
| ETS1(ETS)/Jurkat-ETS1-ChIP-Seq(GSE17954)/Homer | ACAGGAAGTG | 1e-627 | -1.45E+03 | 0 | 4249 | 65.67% | 14286.1 | 32.98% |
| Ets1-distal(ETS)/CD4+-PolII-ChIP-Seq(Barski et al.)/Homer | MACAGGAAGT | 1e-625 | -1.44E+03 | 0 | 2411 | 37.26% | 4990.4 | 11.52% |
| GABPA(ETS)/Jurkat-GABPa-ChIP-Seq(GSE17954)/Homer | RACCGGAAGT | 1e-578 | -1.33E+03 | 0 | 3826 | 59.13% | 12286.5 | 28.36% |
| EWS:FLI1-fusion(ETS)/SK_N_MC-EWS:FLI1-ChIP-Seq(SRA014231)/Homer | VACAGGAAAT | 1e-567 | -1.31E+03 | 0 | 3040 | 46.99% | 8215.3 | 18.96% |
| ETV1(ETS)/GIST48-ETV1-ChIP-Seq(GSE22441)/Homer | AACCGGAAGT | 1e-551 | -1.27E+03 | 0 | 4700 | 72.64% | 18084 | 41.74% |
| EWS:ERG-fusion(ETS)/CADO_ES1-EWS:ERG-ChIP-Seq(SRA014231)/Homer | ATTTCCTGTN | 1e-476 | -1.10E+03 | 0 | 3387 | 52.35% | 10887.8 | 25.13% |
| ERG(ETS)/VCaP-ERG-ChIP-Seq(GSE14097)/Homer | ACAGGAAGTG | 1e-463 | -1.07E+03 | 0 | 4981 | 76.99% | 21245.5 | 49.04% |
| ETS:RUNX(ETS,Runt)/Jurkat-RUNX1-ChIP-Seq(GSE17954)/Homer | RCAGGATGTGGT | 1e-385 | -8.87E+02 | 0 | 949 | 14.67% | 1169 | 2.70% |
| ETS(ETS)/Promoter/Homer | AACCGGAAGT | 1e-369 | -8.50E+02 | 0 | 1763 | 27.25% | 4065.6 | 9.38% |
| Elk1(ETS)/Hela-Elk1-ChIP-Seq(GSE31477)/Homer | HACTTCCGGY | 1e-366 | -8.43E+02 | 0 | 2291 | 35.41% | 6422.5 | 14.82% |
| ELF1(ETS)/Jurkat-ELF1-ChIP-Seq(SRA014231)/Homer | AVCCGGAAGT | 1e-360 | -8.30E+02 | 0 | 2216 | 34.25% | 6125.7 | 14.14% |
| Elk4(ETS)/Hela-Elk4-ChIP-Seq(GSE31477)/Homer | NRYTTCCGGY | 1e-346 | -7.98E+02 | 0 | 2252 | 34.81% | 6423.3 | 14.83% |
| PU.1(ETS)/ThioMac-PU.1-ChIP-Seq(GSE21512)/Homer | AGAGGAAGTG | 1e-326 | -7.51E+02 | 0 | 2396 | 37.03% | 7332 | 16.92% |
| EHF(ETS)/LoVo-EHF-ChIP-Seq(GSE49402)/Homer | AVCAGGAAGT | 1.00E-287 | -6.62E+02 | 0 | 4143 | 64.03% | 18033.3 | 41.63% |
| SPDEF(ETS)/VCaP-SPDEF-ChIP-Seq(SRA014231)/Homer | ASWTCCTGBT | 1.00E-242 | -5.59E+02 | 0 | 3447 | 53.28% | 14341.8 | 33.10% |
| ELF5(ETS)/T47D-ELF5-ChIP-Seq(GSE30407)/Homer | ACVAGGAAGT | 1.00E-208 | -4.81E+02 | 0 | 2755 | 42.58% | 10815.9 | 24.97% |
| RUNX1(Runt)/Jurkat-RUNX1-ChIP-Seq(GSE29180)/Homer | AAACCACARM | 1.00E-204 | -4.71E+02 | 0 | 3447 | 53.28% | 15028.5 | 34.69% |
| RUNX2(Runt)/PCa-RUNX2-ChIP-Seq(GSE33889)/Homer | NWAACCACADNN | 1.00E-165 | -3.82E+02 | 0 | 2950 | 45.60% | 12729.7 | 29.38% |
| RUNX-AML(Runt)/CD4+-PolII-ChIP-Seq(Barski et al.)/Homer | GCTGTGGTTW | 1.00E-165 | -3.80E+02 | 0 | 2675 | 41.34% | 11121.9 | 25.67% |
| RUNX(Runt)/HPC7-Runx1-ChIP-Seq(GSE22178)/Homer | SAAACCACAG | 1.00E-153 | -3.53E+02 | 0 | 2554 | 39.47% | 10639.3 | 24.56% |
| Tcf4(HMG)/Hct116-Tcf4-ChIP-Seq(SRA012054)/Homer | ASATCAAAGGVA | 1.00E-102 | -2.36E+02 | 0 | 1808 | 27.94% | 7425 | 17.14% |
| Tcf3(HMG)/mES-Tcf3-ChIP-Seq(GSE11724)/Homer | ASWTCAAAGG | 1.00E-87 | -2.02E+02 | 0 | 1158 | 17.90% | 4245.9 | 9.80% |
| SpiB(ETS)/OCILY3-SPIB-ChIP-Seq(GSE56857)/Homer | AAAGRGGAAGTG | 1.00E-84 | -1.94E+02 | 0 | 1000 | 15.46% | 3508.1 | 8.10% |
| TCFL2(HMG)/K562-TCF7L2-ChIP-Seq(GSE29196)/Homer | ACWTCAAAGG | 1.00E-45 | -1.05E+02 | 0 | 438 | 6.77% | 1383.5 | 3.19% |
| KLF5(Zf)/LoVo-KLF5-ChIP-Seq(GSE49402)/Homer | DGGGYGKGGC | 1.00E-35 | -8.06E+01 | 0 | 2491 | 38.50% | 13519 | 31.21% |
| Klf4(Zf)/mES-Klf4-ChIP-Seq(GSE11431)/Homer | GCCACACCCA | 1.00E-32 | -7.56E+01 | 0 | 993 | 15.35% | 4543.6 | 10.49% |
| Maz(Zf)/HepG2-Maz-ChIP-Seq(GSE31477)/Homer | GGGGGGGG | 1.00E-32 | -7.44E+01 | 0 | 2571 | 39.74% | 14153.8 | 32.67% |
| Foxo1(Forkhead)/RAW-Foxo1-ChIP-Seq(Fan et al.)/Homer | CTGTTTAC | 1.00E-30 | -7.02E+01 | 0 | 4189 | 64.74% | 24998.3 | 57.70% |
| PU.1-IRF(ETS:IRF)/Bcell-PU.1-ChIP-Seq(GSE21512)/Homer | MGGAAGTGAAAC | 1.00E-25 | -5.98E+01 | 0 | 3172 | 49.03% | 18390.7 | 42.45% |
| Gata2(Zf)/K562-GATA2-ChIP-Seq(GSE18829)/Homer | BBCTTATCTS | 1.00E-25 | -5.94E+01 | 0 | 1880 | 29.06% | 10109.4 | 23.34% |
| Gata4(Zf)/Heart-Gata4-ChIP-Seq(GSE35151)/Homer | NBWGATAAGR | 1.00E-24 | -5.66E+01 | 0 | 2586 | 39.97% | 14646.3 | 33.81% |
| ETS:E-box(ETS,bHLH)/HPC7-Scl-ChIP-Seq(GSE22178)/Homer | AGGAARCAGCTG | 1.00E-24 | -5.53E+01 | 0 | 379 | 5.86% | 1446 | 3.34% |
| Nkx3.1(Homeobox)/LNCaP-Nkx3.1-ChIP-Seq(GSE28264)/Homer | AAGCACTTAA | 1.00E-22 | -5.16E+01 | 0 | 4574 | 70.70% | 28131.9 | 64.94% |
| EKLF(Zf)/Erythrocyte-Klf1-ChIP-Seq(GSE20478)/Homer | NWGGGTGTGGCY | 1.00E-21 | -4.99E+01 | 0 | 651 | 10.06% | 2958.9 | 6.83% |
| Gata1(Zf)/K562-GATA1-ChIP-Seq(GSE18829)/Homer | SAGATAAGRV | 1.00E-19 | -4.43E+01 | 0 | 1649 | 25.49% | 9004.5 | 20.78% |
| GATA3(Zf)/iTreg-Gata3-ChIP-Seq(GSE20898)/Homer | AGATAASR | 1.00E-17 | -4.13E+01 | 0 | 3440 | 53.17% | 20678.3 | 47.73% |
| Atf1(bZIP)/K562-ATF1-ChIP-Seq(GSE31477)/Homer | GATGACGTCA | 1.00E-16 | -3.91E+01 | 0 | 1610 | 24.88% | 8887.8 | 20.52% |
| E2A(bHLH)/proBcell-E2A-ChIP-Seq(GSE21978)/Homer | DNRCAGCTGY | 1.00E-16 | -3.76E+01 | 0 | 2946 | 45.53% | 17514.1 | 40.43% |
| Ascl1(bHLH)/NeuralTubes-Ascl1-ChIP-Seq(GSE55840)/Homer | NNVVCAGCTGBN | 1.00E-15 | -3.56E+01 | 0 | 3140 | 48.53% | 18860.6 | 43.54% |
| Meis1(Homeobox)/MastCells-Meis1-ChIP-Seq(GSE48085)/Homer | VGCTGWCAVB | 1.00E-14 | -3.38E+01 | 0 | 3723 | 57.54% | 22819.9 | 52.67% |
| Myf5(bHLH)/GM-Myf5-ChIP-Seq(GSE24852)/Homer | BAACAGCTGT | 1.00E-14 | -3.27E+01 | 0 | 1713 | 26.48% | 9702.4 | 22.40% |
| MITF(bHLH)/MastCells-MITF-ChIP-Seq(GSE48085)/Homer | RTCATGTGAC | 1.00E-13 | -3.22E+01 | 0 | 2196 | 33.94% | 12797.4 | 29.54% |
| FOXP1(Forkhead)/H9-FOXP1-ChIP-Seq(GSE31006)/Homer | NYYTGTTTACHN | 1.00E-13 | -3.04E+01 | 0 | 1292 | 19.97% | 7129 | 16.46% |
| bHLHE40(bHLH)/HepG2-BHLHE40-ChIP-Seq(GSE31477)/Homer | KCACGTGMCN | 1.00E-13 | -3.01E+01 | 0 | 694 | 10.73% | 3512.9 | 8.11% |

| | | | | | | | | |
|---|---|---|---|---|---|---|---|---|
| FOXA1(Forkhead)/LNCAP-FOXA1-ChIP-Seq(GSE27824)/Homer | WAAGTAAACA | 1.00E-13 | -3.00E+01 | 0 | 2963 | 45.80% | 17877.3 | 41.27% |
| Foxa2(Forkhead)/Liver-Foxa2-ChIP-Seq(GSE25694)/Homer | CYTGTTTACWYW | 1.00E-12 | -2.92E+01 | 0 | 2149 | 33.21% | 12591 | 29.06% |
| Ptf1a(bHLH)/Panc1-Ptf1a-ChIP-Seq(GSE47459)/Homer | ACAGCTGTTN | 1.00E-12 | -2.85E+01 | 0 | 4507 | 69.66% | 28362.6 | 65.47% |
| MyoD(bHLH)/Myotube-MyoD-ChIP-Seq(GSE21614)/Homer | RRCAGCTGYTSY | 1.00E-12 | -2.83E+01 | 0 | 1770 | 27.36% | 10191.6 | 23.53% |
| USF1(bHLH)/GM12878-Usf1-ChIP-Seq(GSE32465)/Homer | SGTCACGTGR | 1.00E-12 | -2.76E+01 | 0 | 1031 | 15.94% | 5591.8 | 12.91% |
| Fox:Ebox(Forkhead,bHLH)/Panc1-Foxa2-ChIP-Seq(GSE47459)/Homer | NNNVCTGWGYAAACASN | 1.00E-11 | -2.75E+01 | 0 | 2655 | 41.04% | 15940.7 | 36.80% |
| SCL(bHLH)/HPC7-Scl-ChIP-Seq(GSE13511)/Homer | AVCAGCTG | 1.00E-11 | -2.73E+01 | 0 | 5810 | 89.80% | 37674.2 | 86.96% |
| Usf2(bHLH)/C2C12-Usf2-ChIP-Seq(GSE36030)/Homer | GTCACGTGGT | 1.00E-11 | -2.72E+01 | 0 | 783 | 12.10% | 4099.7 | 9.46% |
| CLOCK(bHLH)/Liver-Clock-ChIP-Seq(GSE39860)/Homer | GHCACGTG | 1.00E-11 | -2.61E+01 | 0 | 1269 | 19.61% | 7101.8 | 16.39% |
| FOXA1(Forkhead)/MCF7-FOXA1-ChIP-Seq(GSE26831)/Homer | WAAGTAAACA | 1.00E-11 | -2.57E+01 | 0 | 2507 | 38.75% | 15036.1 | 34.71% |
| Tcf12(bHLH)/GM12878-Tcf12-ChIP-Seq(GSE32465)/Homer | VCAGCTGYTG | 1.00E-11 | -2.54E+01 | 0 | 2126 | 32.86% | 12572.8 | 29.02% |
| TATA-Box(TBP)/Promoter/Homer | CCTTTTAWAGSC | 1.00E-10 | -2.37E+01 | 0 | 3065 | 47.37% | 18789.3 | 43.37% |
| MyoG(bHLH)/C2C12-MyoG-ChIP-Seq(GSE36024)/Homer | AACAGCTG | 1.00E-09 | -2.26E+01 | 0 | 2308 | 35.67% | 13853 | 31.98% |
| c-Myc(bHLH)/LNCAP-cMyc-ChIP-Seq(Unpublished)/Homer | VCCACGTG | 1.00E-09 | -2.12E+01 | 0 | 858 | 13.26% | 4694.6 | 10.84% |
| CRE(bZIP)/Promoter/Homer | CSGTGACGTCAC | 1.00E-09 | -2.09E+01 | 0 | 521 | 8.05% | 2671.2 | 6.17% |
| Sox10(HMG)/SciaticNerve-Sox3-ChIP-Seq(GSE35132)/Homer | CCWTTGTYYB | 1.00E-08 | -1.95E+01 | 0 | 3593 | 55.53% | 22498.9 | 51.93% |
| PR(NR)/T47D-PR-ChIP-Seq(GSE31130)/Homer | VAGRACAKNCTGTBC | 1.00E-07 | -1.83E+01 | 0 | 4304 | 66.52% | 27374.6 | 63.19% |
| Sox6(HMG)/Myotubes-Sox6-ChIP-Seq(GSE32627)/Homer | CCATTGTTNY | 1.00E-07 | -1.82E+01 | 0 | 3489 | 53.93% | 21860 | 50.46% |
| NPAS2(bHLH)/Liver-NPAS2-ChIP-Seq(GSE39860)/Homer | KCCACGTGAC | 1.00E-07 | -1.81E+01 | 0 | 2350 | 36.32% | 14315.1 | 33.04% |
| Ap4(bHLH)/AML-Tfap4-ChIP-Seq(GSE45738)/Homer | NAHCAGCTGD | 1.00E-07 | -1.78E+01 | 0 | 2575 | 39.80% | 15802 | 36.48% |
| Sox3(HMG)/NPC-Sox3-ChIP-Seq(GSE33059)/Homer | CCWTTGTY | 1.00E-07 | -1.77E+01 | 0 | 3762 | 58.15% | 23722 | 54.76% |
| Tbx5(T-box)/HL1-Tbx5.biotin-ChIP-Seq(GSE21529)/Homer | AGGTGTCA | 1.00E-07 | -1.77E+01 | 0 | 5148 | 79.57% | 33238.7 | 76.72% |
| MYB(HTH)/ERMYB-Myb-ChIPSeq(GSE22095)/Homer | GGCVGTTR | 1.00E-07 | -1.73E+01 | 0 | 3558 | 54.99% | 22366.9 | 51.63% |
| Tbx20(T-box)/Heart-Tbx20-ChIP-Seq(GSE29636)/Homer | GGTGYTGACAGS | 1.00E-07 | -1.71E+01 | 0 | 625 | 9.66% | 3378.3 | 7.80% |
| GATA(Zf),IR4/iTreg-Gata3-ChIP-Seq(GSE20898)/Homer | NAGATWNBNATCTNN | 1.00E-07 | -1.70E+01 | 0 | 248 | 3.83% | 1163.8 | 2.69% |
| CHR(?)/Hela-CellCycle-Expression/Homer | SRGTTTCAAA | 1.00E-07 | -1.65E+01 | 0 | 1642 | 25.38% | 9788.5 | 22.59% |
| HIF-1b(HLH)/T47D-HIF1b-ChIP-Seq(GSE59937)/Homer | RTACGTGC | 1.00E-07 | -1.64E+01 | 0 | 2054 | 31.75% | 12458.4 | 28.76% |
| Sox4(HMG)/proB-Sox4-ChIP-Seq(GSE50066)/Homer | YCTTTGTTCC | 1.00E-07 | -1.63E+01 | 0 | 2125 | 32.84% | 12925.8 | 29.84% |
| Atf7(bZIP)/3T3L1-Atf7-ChIP-Seq(GSE56872)/Homer | NGRTGACGTCAY | 1.00E-07 | -1.62E+01 | 0 | 1086 | 16.79% | 6260.8 | 14.45% |
| Atoh1(bHLH)/Cerebellum-Atoh1-ChIP-Seq(GSE22111)/Homer | VNRVCAGCTGGY | 1.00E-06 | -1.58E+01 | 0 | 2403 | 37.14% | 14767.5 | 34.09% |
| Tbet(T-box)/CD8-Tbet-ChIP-Seq(GSE33802)/Homer | AGGTGTGAAM | 1.00E-06 | -1.57E+01 | 0 | 2601 | 40.20% | 16075.9 | 37.11% |
| Tlx?(NR)/NPC-H3K4me1-ChIP-Seq(GSE16256)/Homer | CTGGCAGSCTGCCA | 1.00E-06 | -1.53E+01 | 0 | 998 | 15.43% | 5738.2 | 13.25% |
| Nkx2.1(Homeobox)/LungAC-Nkx2.1-ChIP-Seq(GSE43252)/Homer | RSCACTYRAG | 1.00E-06 | -1.48E+01 | 0 | 4867 | 75.22% | 31408.5 | 72.50% |
| BMAL1(bHLH)/Liver-Bmal1-ChIP-Seq(GSE39860)/Homer | GNCACGTG | 1.00E-06 | -1.42E+01 | 0 | 3526 | 54.50% | 22307.8 | 51.49% |
| Mef2c(MADS)/GM12878-Mef2c-ChIP-Seq(GSE32465)/Homer | DCYAAAAATAGM | 1.00E-06 | -1.41E+01 | 0 | 1080 | 16.69% | 6299 | 14.54% |
| Smad4(MAD)/ESC-SMAD4-ChIP-Seq(GSE29422)/Homer | VBSYGTCTGG | 1.00E-05 | -1.34E+01 | 0 | 3286 | 50.79% | 20744.5 | 47.88% |
| Eomes(T-box)/H9-Eomes-ChIP-Seq(GSE26097)/Homer | ATTAACACCT | 1.00E-05 | -1.32E+01 | 0 | 4161 | 64.31% | 26650.8 | 61.52% |
| Rfx5(HTH)/GM12878-Rfx5-ChIP-Seq(GSE31477)/Homer | SCCTAGCAACAG | 1.00E-05 | -1.29E+01 | 0 | 812 | 12.55% | 4654.1 | 10.74% |
| E-box(bHLH)/Promoter/Homer | SSGGTCACGTGA | 1.00E-05 | -1.29E+01 | 0 | 170 | 2.63% | 786.4 | 1.82% |
| E2A(bHLH),near_PU.1/Bcell-PU.1-ChIP-Seq(GSE21512)/Homer | NVCACCTGBN | 1.00E-05 | -1.28E+01 | 0 | 2849 | 44.03% | 17865.2 | 41.24% |
| IRF2(IRF)/Erythroblas-IRF2-ChIP-Seq(GSE36985)/Homer | GAAASYGAAASY | 1.00E-05 | -1.26E+01 | 0 | 279 | 4.31% | 1414.9 | 3.27% |
| CEBP:AP1(bZIP)/ThioMac-CEBPb-ChIP-Seq(GSE21512)/Homer | DRTGTTGCAA | 1.00E-05 | -1.24E+01 | 0 | 1735 | 26.82% | 10570 | 24.40% |
| BMYB(HTH)/Hela-BMYB-ChIP-Seq(GSE27030)/Homer | NHAACBGYYV | 1.00E-05 | -1.24E+01 | 0 | 3237 | 50.03% | 20470.8 | 47.25% |
| JunD(bZIP)/K562-JunD-ChIP-Seq/Homer | ATGACGTCATCN | 1.00E-05 | -1.19E+01 | 0 | 252 | 3.89% | 1271.9 | 2.94% |
| Gfi1b(Zf)/HPC7-Gfi1b-ChIP-Seq(GSE22178)/Homer | MAATCACTGC | 1.00E-05 | -1.16E+01 | 0 | 1444 | 22.32% | 8730.8 | 20.15% |
| Olig2(bHLH)/Neuron-Olig2-ChIP-Seq(GSE30882)/Homer | RCCATMTGTT | 1.00E-04 | -1.13E+01 | 0 | 3800 | 58.73% | 24320 | 56.14% |
| ISRE(IRF)/ThioMac-LPS-Expression(GSE23622)/Homer | AGTTTCASTTTC | 1.00E-04 | -1.05E+01 | 0.0001 | 178 | 2.75% | 868.2 | 2.00% |
| GATA(Zf),IR3/iTreg-Gata3-ChIP-Seq(GSE20898)/Homer | NNNNNBAGATAWYATCTVHN | 1.00E-04 | -1.02E+01 | 0.0001 | 380 | 5.87% | 2071.2 | 4.78% |
| Chop(bZIP)/MEF-Chop-ChIP-Seq(GSE35681)/Homer | ATTGCATCAT | 1.00E-04 | -1.00E+01 | 0.0001 | 519 | 8.02% | 2928.9 | 6.76% |
| n-Myc(bHLH)/mES-nMyc-ChIP-Seq(GSE11431)/Homer | VRCCACGTGG | 1.00E-04 | -9.96E+00 | 0.0001 | 1377 | 21.28% | 8377.7 | 19.34% |

| | | | | | | | | |
|---|---|---|---|---|---|---|---|---|
| Egr1(Zf)/K562-Egr1-ChIP-Seq(GSE32465)/Homer | TGCGTGGGYG | 1.00E-04 | -9.81E+00 | 0.0002 | 1207 | 18.66% | 7289.6 | 16.83% |
| Arnt:Ahr(bHLH)/MCF7-Arnt-ChIP-Seq(Lo et al.)/Homer | TBGCACGCAA | 1.00E-04 | -9.72E+00 | 0.0002 | 1199 | 18.53% | 7242.8 | 16.72% |
| IRF1(IRF)/PBMC-IRF1-ChIP-Seq(GSE43036)/Homer | GAAAGTGAAAGT | 1.00E-04 | -9.40E+00 | 0.0002 | 367 | 5.67% | 2014.8 | 4.65% |
| AR-halfsite(NR)/LNCaP-AR-ChIP-Seq(GSE27824)/Homer | CCAGGAACAG | 1.00E-03 | -9.20E+00 | 0.0003 | 5590 | 86.40% | 36717.1 | 84.75% |
| Stat3+il21(Stat)/CD4-Stat3-ChIP-Seq(GSE19198)/Homer | SVYTTCCNGGAARB | 1.00E-03 | -9.08E+00 | 0.0003 | 1651 | 25.52% | 10202.2 | 23.55% |
| NF1(CTF)/LNCAP-NF1-ChIP-Seq(Unpublished)/Homer | CYTGGCABNSTGCCAR | 1.00E-03 | -9.04E+00 | 0.0003 | 813 | 12.57% | 4808.7 | 11.10% |
| Max(bHLH)/K562-Max-ChIP-Seq(GSE31477)/Homer | RCCACGTGGYYN | 1.00E-03 | -8.81E+00 | 0.0004 | 1410 | 21.79% | 8651.6 | 19.97% |
| Smad3(MAD)/NPC-Smad3-ChIP-Seq(GSE36673)/Homer | TWGTCTGV | 1.00E-03 | -8.50E+00 | 0.0005 | 4850 | 74.96% | 31632.5 | 73.02% |
| Atf2(bZIP)/3T3L1-Atf2-ChIP-Seq(GSE56872)/Homer | NRRTGACGTCAT | 1.00E-03 | -8.48E+00 | 0.0005 | 758 | 11.72% | 4483.9 | 10.35% |
| CRX(Homeobox)/Retina-Crx-ChIP-Seq(GSE20012)/Homer | GCTAATCC | 1.00E-03 | -8.27E+00 | 0.0006 | 4213 | 65.12% | 27306.1 | 63.03% |
| Atf4(bZIP)/MEF-Atf4-ChIP-Seq(GSE35681)/Homer | MTGATGCAAT | 1.00E-03 | -8.21E+00 | 0.0007 | 645 | 9.97% | 3780.4 | 8.73% |
| Nkx2.5(Homeobox)/HL1-Nkx2.5.biotin-ChIP-Seq(GSE21529)/Homer | RRSCACTYAA | 1.00E-03 | -7.79E+00 | 0.001 | 4146 | 64.08% | 26888 | 62.06% |
| Mouse_Recombination_Hotspot(Zf)/Testis-DMC1-ChIP-Seq(GSE24438)/Homer | ACTYKNATTCGTGNTACTTC | 1.00E-03 | -7.71E+00 | 0.0011 | 190 | 2.94% | 991.8 | 2.29% |
| AMYB(HTH)/Testes-AMYB-ChIP-Seq(GSE44588)/Homer | TGGCAGTTGG | 1.00E-03 | -7.23E+00 | 0.0018 | 3202 | 49.49% | 20580.5 | 47.51% |
| PBX1(Homeobox)/MCF7-PBX1-ChIP-Seq(GSE28007)/Homer | GSCTGTCACTCA | 1.00E-03 | -7.18E+00 | 0.0018 | 244 | 3.77% | 1327.2 | 3.06% |
| RFX(HTH)/K562-RFX3-ChIP-Seq(SRA012198)/Homer | CGGTTGCCATGGCAAC | 1.00E-02 | -6.82E+00 | 0.0026 | 202 | 3.12% | 1083.6 | 2.50% |
| Mef2a(MADS)/HL1-Mef2a.biotin-ChIP-Seq(GSE21529)/Homer | CYAAAAATAG | 1.00E-02 | -6.71E+00 | 0.0029 | 976 | 15.09% | 5963.2 | 13.76% |
| Smad2(MAD)/ES-SMAD2-ChIP-Seq(GSE29422)/Homer | CTGTCTGG | 1.00E-02 | -6.66E+00 | 0.003 | 3142 | 48.56% | 20225 | 46.68% |
| c-Myc(bHLH)/mES-cMyc-ChIP-Seq(GSE11431)/Homer | VVCCACGTGG | 1.00E-02 | -6.14E+00 | 0.005 | 1015 | 15.69% | 6247 | 14.42% |
| Nanog(Homeobox)/mES-Nanog-ChIP-Seq(GSE11724)/Homer | RGCCATTAAC | 1.00E-02 | -6.09E+00 | 0.0052 | 5628 | 86.99% | 37152.2 | 85.76% |
| NF1-halfsite(CTF)/LNCaP-NF1-ChIP-Seq(Unpublished)/Homer | YTGCCAAG | 1.00E-02 | -6.09E+00 | 0.0052 | 3304 | 51.07% | 21355.5 | 49.29% |
| GATA:SCL(Zf,bHLH)/Ter119-SCL-ChIP-Seq(GSE18720)/Homer | CRGCTGBNGNSNNSAGATAA | 1.00E-02 | -6.05E+00 | 0.0053 | 312 | 4.82% | 1777.7 | 4.10% |
| STAT4(Stat)/CD4-Stat4-ChIP-Seq(GSE22104)/Homer | NYTTCCWGGAAR | 1.00E-02 | -5.98E+00 | 0.0057 | 2222 | 34.34% | 14164.3 | 32.70% |
| Rfx2(HTH)/LoVo-RFX2-ChIP-Seq(GSE49402)/Homer | GTTGCCATGGCAACM | 1.00E-02 | -5.65E+00 | 0.0078 | 229 | 3.54% | 1278.3 | 2.95% |
| Pbx3(Homeobox)/GM12878-PBX3-ChIP-Seq(GSE32465)/Homer | SCTGTCAMTCAN | 1.00E-02 | -5.59E+00 | 0.0082 | 653 | 10.09% | 3949.1 | 9.12% |
| GATA3(Zf),DR4/iTreg-Gata3-ChIP-Seq(GSE20898)/Homer | AGATGKDGAGATAAG | 1.00E-02 | -5.57E+00 | 0.0084 | 217 | 3.35% | 1207 | 2.79% |
| Pitx1(Homeobox)/Chicken-Pitx1-ChIP-Seq(GSE38910)/Homer | TAATCCCN | 1.00E-02 | -5.37E+00 | 0.0101 | 5652 | 87.36% | 37364.7 | 86.25% |
| Tbox:Smad(T-box,MAD)/ESCd5-Smad2_3-ChIP-Seq(GSE29422)/Homer | AGGTGHCAGACA | 1.00E-02 | -5.29E+00 | 0.0108 | 494 | 7.64% | 2950.4 | 6.81% |
| HOXA9(Homeobox)/HSC-Hoxa9-ChIP-Seq(GSE33509)/Homer | GGCCATAAATCA | 1.00E-02 | -5.11E+00 | 0.0129 | 1512 | 23.37% | 9557.3 | 22.06% |
| GRHL2(CP2)/HBE-GRHL2-ChIP-Seq(GSE46194)/Homer | AAACYKGTTWDACMRGTTTB | 1.00E-02 | -4.99E+00 | 0.0144 | 871 | 13.46% | 5386.8 | 12.43% |
| Rfx1(HTH)/NPC-H3K4me1-ChIP-Seq(GSE16256)/Homer | KGTTGCCATGGCAA | 1.00E-02 | -4.89E+00 | 0.0158 | 469 | 7.25% | 2810.4 | 6.49% |
| c-Jun-CRE(bZIP)/K562-cJun-ChIP-Seq(GSE31477)/Homer | ATGACGTCATCY | 1.00E-02 | -4.85E+00 | 0.0163 | 675 | 10.43% | 4130.1 | 9.53% |
| PRDM9(Zf)/Testis-DMC1-ChIP-Seq(GSE35498)/Homer | ADGGYAGYAGCATCT | 1.00E-02 | -4.63E+00 | 0.0201 | 962 | 14.87% | 6000.4 | 13.85% |
| HOXD13(Homeobox)/Chicken-Hoxd13-ChIP-Seq(GSE38910)/Homer | NCYAATAAAA | 1.00E-01 | -4.58E+00 | 0.0211 | 2344 | 36.23% | 15096.7 | 34.85% |
| OCT4-SOX2-TCF-NANOG(POU,Homeobox,HMG)/mES-Oct4-ChIP-Seq(GSE11431)/Homer | ATTTGCATAACAATG | 1.00E-01 | -4.51E+00 | 0.0223 | 425 | 6.57% | 2548.7 | 5.88% |
| Brachyury(T-box)/Mesoendoderm-Brachyury-ChIP-exo(GSE54963)/Homer | ANTTMRCASBNNNGTGYKAAN | 1.00E-01 | -4.26E+00 | 0.0284 | 674 | 10.42% | 4158.6 | 9.60% |
| Unknown-ESC-element(?)/mES-Nanog-ChIP-Seq(GSE11724)/Homer | CACAGCAGGGGG | 1.00E-01 | -4.14E+00 | 0.0319 | 1321 | 20.42% | 8383.6 | 19.35% |
| Srebp2(bHLH)/HepG2-Srebp2-ChIP-Seq(GSE31477)/Homer | CGGTCACSCCAC | 1.00E-01 | -4.10E+00 | 0.0328 | 299 | 4.62% | 1768.8 | 4.08% |
| NeuroD1(bHLH)/Islet-NeuroD1-ChIP-Seq(GSE30298)/Homer | GCCATCTGTT | 1.00E-01 | -4.04E+00 | 0.0348 | 1788 | 27.64% | 11467.2 | 26.47% |
| AP-2gamma(AP2)/MCF7-TFAP2C-ChIP-Seq(GSE21234)/Homer | SCCTSAGGSCAW | 1.00E-01 | -4.00E+00 | 0.0357 | 1811 | 27.99% | 11622.4 | 26.83% |
| RORgt(NR)/EL4-RORgt.Flag-ChIP-Seq(GSE56019)/Homer | AAYTAGGTCA | 1.00E-01 | -3.79E+00 | 0.0437 | 307 | 4.74% | 1832.9 | 4.23% |
| Sox2(HMG)/mES-Sox2-ChIP-Seq(GSE11431)/Homer | BCCATTGTTC | 1.00E-01 | -3.72E+00 | 0.0466 | 2045 | 31.61% | 13199.9 | 30.47% |
| FOXA1:AR(Forkhead,NR)/LNCAP-AR-ChIP-Seq(GSE27824)/Homer | AGTAAACAAAAAAGAACAND | 1.00E-01 | -3.72E+00 | 0.0466 | 195 | 3.01% | 1130.4 | 2.61% |

**Table S6C. HOMER Known Motifs near the closing peaks of both cytokine-primed and Ag-stimulated cells**

Only motifs with q-value below 0.05 are shown.

| Motif Name | Consensus | P-value | Log P-value | q-value (Benjamini) | # of Target Sequences with Motif (of 294) | % of Target Sequences with Motif | # of Background Sequences with Motif (of 47421) | % of Background Sequences with Motif |
|---|---|---|---|---|---|---|---|---|
| GABPA(ETS)/Jurkat-GABPa-ChIP-Seq(GSE17954)/Homer | RACCGGAAGT | 1.00E-25 | -5.83E+01 | 0 | 174 | 59.18% | 13994 | 29.51% |
| Atf1(bZIP)/K562-ATF1-ChIP-Seq(GSE31477)/Homer | GATGACGTCA | 1.00E-23 | -5.51E+01 | 0 | 143 | 48.64% | 10227 | 21.57% |
| Fli1(ETS)/CD8-FLI-ChIP-Seq(GSE20898)/Homer | NRYTTCCGGH | 1.00E-23 | -5.34E+01 | 0 | 188 | 63.95% | 16607.4 | 35.02% |
| EWS:FLI1-fusion(ETS)/SK_N_MC-EWS:FLI1-ChIP-Seq(SRA014231)/Homer | VACAGGAAAT | 1.00E-23 | -5.30E+01 | 0 | 134 | 45.58% | 9332.6 | 19.68% |
| Ets1-distal(ETS)/CD4+-PolII-ChIP-Seq(Barski et al.)/Homer | MACAGGAAGT | 1.00E-22 | -5.28E+01 | 0 | 101 | 34.35% | 5662.2 | 11.94% |
| ETV1(ETS)/GIST48-ETV1-ChIP-Seq(GSE22441)/Homer | AACCGGAAGT | 1.00E-20 | -4.64E+01 | 0 | 206 | 70.07% | 20409.9 | 43.04% |
| ETS1(ETS)/Jurkat-ETS1-ChIP-Seq(GSE17954)/Homer | ACAGGAAGTG | 1.00E-19 | -4.48E+01 | 0 | 176 | 59.86% | 15945.1 | 33.62% |
| PU.1(ETS)/ThioMac-PU.1-ChIP-Seq(GSE21512)/Homer | AGAGGAAGTG | 1.00E-19 | -4.42E+01 | 0 | 116 | 39.46% | 8058.2 | 16.99% |
| Elk4(ETS)/Hela-Elk4-ChIP-Seq(GSE31477)/Homer | NRYTTCCGGY | 1.00E-16 | -3.82E+01 | 0 | 111 | 37.76% | 8099.4 | 17.08% |
| ETS(ETS)/Promoter/Homer | AACCGGAAGT | 1.00E-15 | -3.68E+01 | 0 | 81 | 27.55% | 4887.2 | 10.31% |
| EWS:ERG-fusion(ETS)/CADO_ES1-EWS:ERG-ChIP-Seq(SRA014231)/Homer | ATTTCCTGTN | 1.00E-15 | -3.63E+01 | 0 | 140 | 47.62% | 12018.2 | 25.34% |
| ELF1(ETS)/Jurkat-ELF1-ChIP-Seq(SRA014231)/Homer | AVCCGGAAGT | 1.00E-13 | -3.19E+01 | 0 | 100 | 34.01% | 7502.1 | 15.82% |
| CRE(bZIP)/Promoter/Homer | CSGTGACGTCAC | 1.00E-13 | -3.13E+01 | 0 | 60 | 20.41% | 3242.2 | 6.84% |
| Atf7(bZIP)/3T3L1-Atf7-ChIP-Seq(GSE56872)/Homer | NGRTGACGTCAY | 1.00E-12 | -2.87E+01 | 0 | 94 | 31.97% | 7163 | 15.10% |
| Atf2(bZIP)/3T3L1-Atf2-ChIP-Seq(GSE56872)/Homer | NRRTGACGTCAT | 1.00E-11 | -2.64E+01 | 0 | 74 | 25.17% | 5125.5 | 10.81% |
| ETS:RUNX(ETS,Runt)/Jurkat-RUNX1-ChIP-Seq(GSE17954)/Homer | RCAGGATGTGGT | 1.00E-11 | -2.58E+01 | 0 | 34 | 11.56% | 1341.2 | 2.83% |
| Elk1(ETS)/Hela-Elk1-ChIP-Seq(GSE31477)/Homer | HACTTCCGGY | 1.00E-10 | -2.52E+01 | 0 | 97 | 32.99% | 7980.7 | 16.83% |
| ELF5(ETS)/T47D-ELF5-ChIP-Seq(GSE30407)/Homer | ACVAGGAAGT | 1.00E-10 | -2.47E+01 | 0 | 128 | 43.54% | 12131.2 | 25.58% |
| EHF(ETS)/LoVo-EHF-ChIP-Seq(GSE49402)/Homer | AVCAGGAAGT | 1.00E-10 | -2.47E+01 | 0 | 180 | 61.22% | 19864.7 | 41.89% |
| ERG(ETS)/VCaP-ERG-ChIP-Seq(GSE14097)/Homer | ACAGGAAGTG | 1.00E-10 | -2.33E+01 | 0 | 201 | 68.37% | 23594.7 | 49.75% |
| SPDEF(ETS)/VCaP-SPDEF-ChIP-Seq(SRA014231)/Homer | ASWTCCTGBT | 1.00E-08 | -2.06E+01 | 0 | 149 | 50.68% | 15929.4 | 33.59% |
| RUNX1(Runt)/Jurkat-RUNX1-ChIP-Seq(GSE29180)/Homer | AAACCACARM | 1.00E-08 | -1.86E+01 | 0 | 150 | 51.02% | 16506.8 | 34.81% |
| SpiB(ETS)/OCILY3-SPIB-ChIP-Seq(GSE56857)/Homer | AAAGRGGAAGTG | 1.00E-07 | -1.79E+01 | 0 | 55 | 18.71% | 3981.2 | 8.39% |
| RUNX2(Runt)/PCa-RUNX2-ChIP-Seq(GSE33889)/Homer | NWAACCACADNN | 1.00E-07 | -1.72E+01 | 0 | 132 | 44.90% | 14149.7 | 29.84% |
| JunD(bZIP)/K562-JunD-ChIP-Seq/Homer | ATGACGTCATCN | 1.00E-07 | -1.62E+01 | 0 | 29 | 9.86% | 1502.1 | 3.17% |
| c-Jun-CRE(bZIP)/K562-cJun-ChIP-Seq(GSE31477)/Homer | ATGACGTCATCY | 1.00E-06 | -1.53E+01 | 0 | 58 | 19.73% | 4656.6 | 9.82% |
| Atoh1(bHLH)/Cerebellum-Atoh1-ChIP-Seq(GSE22111)/Homer | VNRVCAGCTGGY | 1.00E-06 | -1.47E+01 | 0 | 142 | 48.30% | 16217.6 | 34.20% |
| RUNX(Runt)/HPC7-Runx1-ChIP-Seq(GSE22178)/Homer | SAAACCACAG | 1.00E-05 | -1.25E+01 | 0 | 109 | 37.07% | 11901.7 | 25.10% |
| Foxa2(Forkhead)/Liver-Foxa2-ChIP-Seq(GSE25694)/Homer | CYTGTTTACWYW | 1.00E-05 | -1.22E+01 | 0 | 119 | 40.48% | 13442 | 28.34% |
| Rfx5(HTH)/GM12878-Rfx5-ChIP-Seq(GSE31477)/Homer | SCCTAGCAACAG | 1.00E-04 | -1.09E+01 | 0.0002 | 55 | 18.71% | 4985.6 | 10.51% |
| Tbx5(T-box)/HL1-Tbx5.biotin-ChIP-Seq(GSE21529)/Homer | AGGTGTCA | 1.00E-04 | -1.07E+01 | 0.0002 | 255 | 86.73% | 36573.8 | 77.12% |
| Olig2(bHLH)/Neuron-Olig2-ChIP-Seq(GSE30882)/Homer | RCCATMTGTT | 1.00E-04 | -1.03E+01 | 0.0003 | 198 | 67.35% | 26458 | 55.79% |
| c-Myc(bHLH)/LNCAP-cMyc-ChIP-Seq(Unpublished)/Homer | VCCACGTG | 1.00E-04 | -9.64E+00 | 0.0005 | 62 | 21.09% | 6125.7 | 12.92% |
| bHLHE40(bHLH)/HepG2-BHLHE40-ChIP-Seq(GSE31477)/Homer | KCACGTGMCN | 1.00E-04 | -9.44E+00 | 0.0006 | 49 | 16.67% | 4500.1 | 9.49% |
| BMAL1(bHLH)/Liver-Bmal1-ChIP-Seq(GSE39860)/Homer | GNCACGTG | 1.00E-04 | -9.22E+00 | 0.0008 | 191 | 64.97% | 25648.1 | 54.08% |
| STAT6(Stat)/CD4-Stat6-ChIP-Seq(GSE22104)/Homer | ABTTCYYRRGAA | 1.00E-03 | -9.09E+00 | 0.0008 | 84 | 28.57% | 9241.4 | 19.49% |
| NPAS2(bHLH)/Liver-NPAS2-ChIP-Seq(GSE39860)/Homer | KCCACGTGAC | 1.00E-03 | -8.58E+00 | 0.0013 | 133 | 45.24% | 16602.5 | 35.01% |
| USF1(bHLH)/GM12878-Usf1-ChIP-Seq(GSE32465)/Homer | SGTCACGTGR | 1.00E-03 | -8.29E+00 | 0.0017 | 65 | 22.11% | 6831.4 | 14.40% |
| Ascl1(bHLH)/NeuralTubes-Ascl1-ChIP-Seq(GSE55840)/Homer | NNVVCAGCTGBN | 1.00E-03 | -8.05E+00 | 0.0022 | 160 | 54.42% | 21017.1 | 44.32% |
| Foxo1(Forkhead)/RAW-Foxo1-ChIP-Seq(Fan et al.)/Homer | CTGTTTAC | 1.00E-03 | -7.92E+00 | 0.0024 | 198 | 67.35% | 27293.7 | 57.55% |
| Meis1(Homeobox)/MastCells-Meis1-ChIP-Seq(GSE48085)/Homer | VGCTGWCAVB | 1.00E-03 | -7.73E+00 | 0.0028 | 186 | 63.27% | 25365.9 | 53.49% |
| Ptf1a(bHLH)/Panc1-Ptf1a-ChIP-Seq(GSE47459)/Homer | ACAGCTGTTN | 1.00E-03 | -7.53E+00 | 0.0034 | 223 | 75.85% | 31737 | 66.92% |
| CLOCK(bHLH)/Liver-Clock-ChIP-Seq(GSE39860)/Homer | GHCACGTG | 1.00E-03 | -7.11E+00 | 0.005 | 76 | 25.85% | 8671.7 | 18.29% |
| STAT6(Stat)/Macrophage-Stat6-ChIP-Seq(GSE38377)/Homer | TTCCKNAGAA | 1.00E-03 | -7.05E+00 | 0.0052 | 81 | 27.55% | 9401.3 | 19.82% |
| CEBP:AP1(bZIP)/ThioMac-CEBPb-ChIP-Seq(GSE21512)/Homer | DRTGTTGCAA | 1.00E-03 | -6.91E+00 | 0.0059 | 93 | 31.63% | 11182.2 | 23.58% |
| Nkx2.1(Homeobox)/LungAC-Nkx2.1-ChIP-Seq(GSE43252)/Homer | RSCACTYRAG | 1.00E-02 | -6.81E+00 | 0.0063 | 236 | 80.27% | 34312.9 | 72.35% |
| HIF-1b(HLH)/T47D-HIF1b-ChIP-Seq(GSE59937)/Homer | RTACGTGC | 1.00E-02 | -6.78E+00 | 0.0064 | 124 | 42.18% | 15872 | 33.47% |
| ZNF143\|STAF(Zf)/CUTLL-ZNF143-ChIP-Seq(GSE29600)/Homer | ATTTCCCAGVAKSCY | 1.00E-02 | -6.61E+00 | 0.0074 | 52 | 17.69% | 5497.4 | 11.59% |

| | | | | | | | | |
|---|---|---|---|---|---|---|---|---|
| Tcf3(HMG)/mES-Tcf3-ChIP-Seq(GSE11724)/Homer | ASWTCAAAGG | 1.00E-02 | -6.60E+00 | 0.0074 | 44 | 14.97% | 4441.7 | 9.37% |
| Tcf4(HMG)/Hct116-Tcf4-ChIP-Seq(SRA012054)/Homer | ASATCAAAGGVA | 1.00E-02 | -6.55E+00 | 0.0076 | 70 | 23.81% | 7993.4 | 16.86% |
| Eomes(T-box)/H9-Eomes-ChIP-Seq(GSE26097)/Homer | ATTAACACCT | 1.00E-02 | -6.54E+00 | 0.0076 | 206 | 70.07% | 29199.3 | 61.57% |
| AR-halfsite(NR)/LNCaP-AR-ChIP-Seq(GSE27824)/Homer | CCAGGAACAG | 1.00E-02 | -6.48E+00 | 0.0078 | 268 | 91.16% | 40402.4 | 85.19% |
| Atf4(bZIP)/MEF-Atf4-ChIP-Seq(GSE35681)/Homer | MTGATGCAAT | 1.00E-02 | -6.46E+00 | 0.0078 | 40 | 13.61% | 3952.5 | 8.33% |
| MYB(HTH)/ERMYB-Myb-ChIPSeq(GSE22095)/Homer | GGCVGTTR | 1.00E-02 | -6.44E+00 | 0.0078 | 179 | 60.88% | 24735.8 | 52.16% |
| Cdx2(Homeobox)/mES-Cdx2-ChIP-Seq(GSE14586)/Homer | GYMATAAAAH | 1.00E-02 | -6.27E+00 | 0.0091 | 88 | 29.93% | 10669.7 | 22.50% |
| RUNX-AML(Runt)/CD4+-PolII-ChIP-Seq(Barski et al.)/Homer | GCTGTGGTTW | 1.00E-02 | -6.15E+00 | 0.0101 | 99 | 33.67% | 12339.2 | 26.02% |
| SCL(bHLH)/HPC7-Scl-ChIP-Seq(GSE13511)/Homer | AVCAGCTG | 1.00E-02 | -5.84E+00 | 0.0134 | 272 | 92.52% | 41397.3 | 87.29% |
| n-Myc(bHLH)/mES-nMyc-ChIP-Seq(GSE11431)/Homer | VRCCACGTGG | 1.00E-02 | -5.83E+00 | 0.0134 | 83 | 28.23% | 10094.4 | 21.29% |
| NeuroD1(bHLH)/Islet-NeuroD1-ChIP-Seq(GSE30298)/Homer | GCCATCTGTT | 1.00E-02 | -5.78E+00 | 0.0139 | 99 | 33.67% | 12480.8 | 26.32% |
| MITF(bHLH)/MastCells-MITF-ChIP-Seq(GSE48085)/Homer | RTCATGTGAC | 1.00E-02 | -5.50E+00 | 0.0181 | 112 | 38.10% | 14558.9 | 30.70% |
| Usf2(bHLH)/C2C12-Usf2-ChIP-Seq(GSE36030)/Homer | GTCACGTGGT | 1.00E-02 | -5.49E+00 | 0.0181 | 46 | 15.65% | 4981.1 | 10.50% |
| STAT1(Stat)/HelaS3-STAT1-ChIP-Seq(GSE12782)/Homer | NATTTCCNGGAAAT | 1.00E-02 | -5.46E+00 | 0.0181 | 46 | 15.65% | 4989.4 | 10.52% |
| E-box(bHLH)/Promoter/Homer | SSGGTCACGTGA | 1.00E-02 | -5.41E+00 | 0.0188 | 14 | 4.76% | 1004.6 | 2.12% |
| Srebp2(bHLH)/HepG2-Srebp2-ChIP-Seq(GSE31477)/Homer | CGGTCACSCCAC | 1.00E-02 | -5.35E+00 | 0.0196 | 23 | 7.82% | 2044.4 | 4.31% |
| Tlx?(NR)/NPC-H3K4me1-ChIP-Seq(GSE16256)/Homer | CTGGCAGSCTGCCA | 1.00E-02 | -5.11E+00 | 0.0246 | 55 | 18.71% | 6333.7 | 13.36% |
| Tbet(T-box)/CD8-Tbet-ChIP-Seq(GSE33802)/Homer | AGGTGTGAAM | 1.00E-02 | -5.04E+00 | 0.0259 | 130 | 44.22% | 17540.3 | 36.99% |
| KLF5(Zf)/LoVo-KLF5-ChIP-Seq(GSE49402)/Homer | DGGGYGKGGC | 1.00E-02 | -4.96E+00 | 0.0277 | 118 | 40.14% | 15716.7 | 33.14% |
| PAX5(Paired,Homeobox),condensed/GM12878-PAX5-ChIP-Seq(GSE32465)/Homer | GTCACGCTCSCTGM | 1.00E-02 | -4.78E+00 | 0.0325 | 18 | 6.12% | 1542.7 | 3.25% |
| Max(bHLH)/K562-Max-ChIP-Seq(GSE31477)/Homer | RCCACGTGGYYN | 1.00E-02 | -4.73E+00 | 0.0336 | 81 | 27.55% | 10219.5 | 21.55% |
| FOXA1(Forkhead)/LNCAP-FOXA1-ChIP-Seq(GSE27824)/Homer | WAAGTAAACA | 1.00E-02 | -4.73E+00 | 0.0336 | 140 | 47.62% | 19262.8 | 40.62% |
| Chop(bZIP)/MEF-Chop-ChIP-Seq(GSE35681)/Homer | ATTGCATCAT | 1.00E-01 | -4.58E+00 | 0.0381 | 30 | 10.20% | 3084.9 | 6.50% |
| Rfx2(HTH)/LoVo-RFX2-ChIP-Seq(GSE49402)/Homer | GTTGCCATGGCAACM | 1.00E-01 | -4.41E+00 | 0.0445 | 17 | 5.78% | 1484.3 | 3.13% |
| NF1(CTF)/LNCAP-NF1-ChIP-Seq(Unpublished)/Homer | CYTGGCABNSTGCCAR | 1.00E-01 | -4.41E+00 | 0.0445 | 46 | 15.65% | 5298.8 | 11.17% |
| IRF2(IRF)/Erythroblas-IRF2-ChIP-Seq(GSE36985)/Homer | GAAASYGAAASY | 1.00E-01 | -4.39E+00 | 0.0445 | 18 | 6.12% | 1608.3 | 3.39% |
| c-Myc(bHLH)/mES-cMyc-ChIP-Seq(GSE11431)/Homer | VVCCACGTGG | 1.00E-01 | -4.27E+00 | 0.0491 | 61 | 20.75% | 7477.4 | 15.77% |
